# Evolutionary context can clarify teleosts gene names

**DOI:** 10.1101/2020.02.02.931493

**Authors:** E.V. Gasanov, J. Jędrychowska, J. Kuźnicki, V. Korzh

**Author notes:** equal contribution (E.V. Gasanov, J. Jędrychowska). Corresponding author: Vladimir Korzh.

## Abstract

The initial convention to name genes relied on historical precedent, order in the human genome or mutants in model systems. However, partial duplication of genes in teleosts required naming the duplicated genes, so ohnologs adopted the ‘a’ or ‘b’ extension. Rapid advances in deciphering the zebrafish genome in relation to the human genome instituted naming genes in all other fish genomes in the convention of zebrafish. Unfortunately, some ohnologs and their resembling orthologs suffered from incorrect nomenclature, which created confusion in particular instances like establishing disease models. We sought to overcome this barrier by establishing the *ex silico* evolutionary-based systematic approach to naming the ohnologs in teleosts and other fish. We compared gene synteny using the spotted gar genome as the reference, which represents the unduplicated ancestral state. Using new criteria, we identified several hundreds of potentially misnamed ohnologs and validated manually several ohnologs as a proof pf principle. This may help to establish a standard naming practice resulting in the improved evolutionary-based gene nomenclature. This approach may help to identify and rename ohnologs in all relevant EMBL-EBI and NCBI databases starting from the zebrafish genome to avoid further proliferation of misleading information.

## Gene naming problem

Using scientific terms suggests a certain level of precision in defining a subject. Gene names have quickly become a part of the scientific language in genomics and developmental biology. Their usage then spreads to related areas such as medicine and biotechnology. Correctly naming genes should create a comprehensive tool for communicating across the biological sciences and beyond. Ideally, gene names should utilize a system that reflects the relationship to homologous genes in other species to better define their evolutionary and functional relationship. However, this convention becomes constrained with duplicated genes or ohnologs (Wolfe, 2000) characteristic for teleosts. The existing convention of gene naming relies on historical precedent, an order of genes in the human genome or names of mutants in model systems. So, gene names provide insight into the history of a gene study rather than providing insight into their function or evolutionary history. Our aim is to propose an approach for naming duplicated genes based upon the natural process of genome development, i.e. evolution. Our approach can create a convincing system for gene naming that requires a relatively small effort and minor nomenclature changes.

The organized naming of genes is important for proper nomenclature of genes and gene families, synchronizing with mammalian counterparts and working with disease models using zebrafish (*Danio rerio*) and medaka (*Oryzias latipes*). Each animal taxon evolved with a continuous process of gene loss and gain, with the latter primarily through gene duplications. Meanwhile, some groups of animals passed through extremely rare but massive gene number changes like whole-genome duplications followed by periods of genome stabilization, i.e. partially duplicated gene inactivation and reduction (Glasauer & Neuhauss, 2014). The significant portion of genes in teleosts, the largest group of vertebrates, are duplicated as a result of whole-genome duplication that occured at the beginning of the taxon evolution (Glasauer & Neuhauss, 2014). Similar gene names in different species should suggest insights into the evolution of the genome.

An existent practice is to define ohnologs (two fish orthologs of the mammalian gene) by letters ‘a’ and ‘b’ following the gene’s name used in the human genome, e.g., *kcng4a* and *kcng4b,* the orthologs of human *KCNG4* (potassium voltage-gated channel subfamily G member 4 gene). This convention was established at the meeting in Ringberg, Germany, in March 1992 (ZFIN Database). However, this convention did not specify when to use the ‘a’ or ‘b’ to distinguish genes. The practice to name the second discovered ohnolog ‘b’ after adding ‘a’ to the first discovery reveals complications to follow the precedence of discovery and the independent evolution of ohnologs across teleost species. Some teleost species have both ohnologs, whereas others lost one of them. When a preserved singleton is discovered, corresponding to either ‘a’ or ‘b’ in related species, can easily result in incorrect nomenclature. Taken together, these examples reveal an unreliable gene naming system.

The insufficiency of the historical approach is clearly described by one example of the *tbx*-family (T-box family) genes. After cloning, the mammalian *Tbx2* zebrafish ohnologs were temporarily named *tbx-a* and *tbx-c* (Dheen et al., 1999). When described independently by other authors, *tbx-c* was named *tbx2* (Ruvinsky et al., 2000). *tbx-a* was later renamed for reasons unknown as *tbx2a*, whereas *tbx-c* became *tbx2b*. In this case, ‘a’ and ‘b’ carried no readily apparent information to link to mammalian orthologs. The naming of ohnologs in different teleost species has followed that of the zebrafish. This practice triggered a chain of incorrect naming events like the rather casual approach of naming of zebrafish Tbx2 genes. According to Ensembl GRCz11 (Ensembl Release 98), a bulk of Tbx2 genes in teleosts were named “*tbx2b*”, whereas “*tbx2a*” has been defined in addition to “*tbx2b*” in two species [channel catfish (*Ictalurus punctatus*) and northern pike (*Esox lucius*)] to date. Thus, this example represents a problem in gene naming in teleosts, which highlights the problem of establishing a unified criterion for naming ohnolog genes. In the article, we propose using an evolutionary-based systematic approach of naming genes.

## Sequence similarity criterion

We posit that a more reliable criterion should utilize sequence similarity to define ‘a’ and ‘b’ orthologs. Sequence similarity relies on evolutionary conservation of a gene function in teleosts compared to other vertebrates. The fish ohnolog gene most resembling the human ortholog (using the traditional gene naming process) or an ancestral gene (using the evolution-based systematic approach proposed here) would hold the name ‘a’ with the less resembling one holding ‘b’, respectively. If the fish species possesses only one gene according to this criterion, we propose a second criterion that compares an ortholog to ohnologs of other teleosts to determine a reliable ‘a’ or ‘b’ extension.

Unfortunately, additional problems arise to systematically assign gene names. One problem arises from uncertainty about which sequence, gene or protein, coding or/and non-coding should be utilized to systematically name genes. Returning to the *Tbx2* ohnologs, the genomic DNA similarity of zebrafish *tbx2a* to human *TBX2* is 48.1% and zebrafish *tbx2b* to human *TBX2* is 41.5%, but the coding sequence identity is 36.6% and 47.5%, respectively. So, the protein identity of zebrafish Tbx2a and Tbx2b to human TBX2 is 45.5% and 50.3%, respectively. Using a different sequence, studies defined the non-coding regulatory regions of zebrafish *tbx2a* and *tbx2b* (Chi et al., 2008; Kaaij et al., 2016). However, the DNA alignment demonstrated a much closer evolutionary relationship between the enhancers of mammalian *Tbx2* and zebrafish *tbx2b* compared to *tbx2a*. The latter contains a single nucleotide polymorphism (SNP) at the critical Foxn4-binding site that regulates zebrafish *tbx2a* cardiac expression, which is preserved in all other Tbx2 genes including zebrafish *tbx2b* (Chi et al., 2008). Taken together, these suppositions illustrate the controversy regarding assigning the ‘a’ and ‘b’ extensions to zebrafish *tbx2* ohnologs solely on the basis of DNA sequence comparison. More evidence suggests that the ‘b’ variant should be an ‘a’ one and *vice versa*. Indeed, despite the reliability of sequence similarity criterion, gene naming carries an enduring controversy that requires individual gene analysis and sometimes advanced knowledge about the gene function that yields a non-systematic and case-dependent application.

## Gene synteny criterion

We agree that a more convincing approach to gene naming is based on analyzing the syntenic relation of a gene in relation to its genomic environment (Force et al., 1999; Chong et al., 2001). Despite the significant evolutionary distance between the mammalian and fish genomes, many genes are organized in recognizable blocks with the same neighbors (Braasch et al., 2016) due to conserved genomic organization among all vertebrates. For comparing the ohnolog surroundings, i.e., synteny, where possible with the human genome or the fish ancestral genome, we propose choosing the most evolutionary conserved ohnolog as the ‘a’ variant. So, ‘b’ becomes the second gene independent of whether an ohnologs were preserved or lost in evolution.

Utilizing the *tbx2* example, we analyzed the synteny of *tbx2* ohnologs (Fig. 1). This analysis demonstrated an almost perfect preservation of syntenic relations between *TBX2* on human Chr. 17, where a pair of *TBX2-TBX4* is present, and *tbx2b* on zebrafish Chr. 15, where it exists in a pair with *tbx4.* The synteny encompasses several other genes. In contrast, an order of genes at the site of *tbx2a* in zebrafish Chr. 5 lacks not only an obligatory member, *tbx4,* but also all other orthologous genes present alongside *tbx2b* on Chr. 15. *Tbx2-Tbx4* and *Tbx3-Tbx5* gene pairs are tightly linked in evolution in invertebrates, as far as amphioxus, and vertebrates (Horton et al., 2008). By lacking the syntenic association to *tbx4* as well as several other genes, the genomic site harboring *tbx-a/tbx2a* varies significantly from *Tbx2* in all species studied. This suggests that based on genomic position and confirmed by the DNA/protein sequence and gene regulation that *tbx-c/tbx2b* is much more closely related to the mammalian *Tbx2* compared to *tbx-a/tbx2a*. According to our approach, the zebrafish *tbx2b* ohnolog should have ‘a’ extension, while the present ‘a’ variant should be renamed as ‘b’. In this case, the results of synteny and sequence analysis on the evolutionary position of *tbx2* ohnologs agree. This agreement reflects the reliability of both criteria representing different facets of an evolutionary process, namely, one gene sequence variation vs genomic rearrangements. We conclude that the synteny analysis could contribute to systematically renaming genes at the level of the higher taxa, and ultimately, in all vertebrates, independent of gene evolution in individual species.

**Figure 1.**
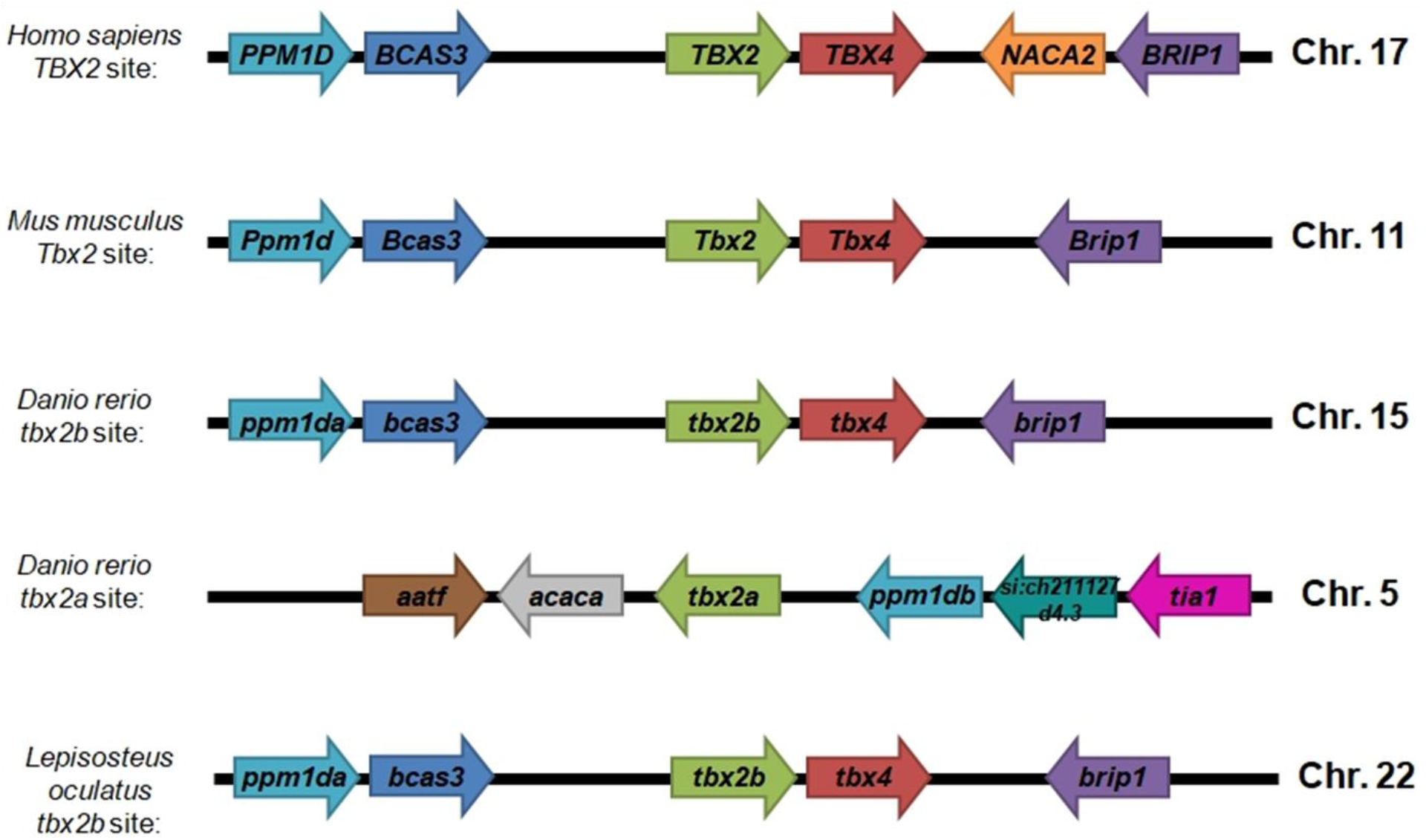
Renaming duplicated gene names *tbx2a* and *tbx2b* reciprocally revealed by synteny analysis. Since gene naming in teleosts uses the zebrafish genome as reference, synteny analysis indicates revising this practice for an eye on an ancient fish genome that did not undergo teleosts-specific gene duplication.

## Selecting the optimal reference genome

Usually, researchers use the well-studied human genome as a reference point to examine all other genomes. The anthropocentric view of the organic world presented as the tree of life has its roots in the several centuries-old cultural and religious dogmas that have very little in common with the theory of evolution. From the times of Ramon Llull (1232-1315) and Carl Linnaeus (1707-1778), science traditionally has considered the human and its beliefs as the final and most complex product of evolution. While these ideas were progressive in old days, this approach is not necessarily useful for naming genes that represent a specific problem within the context of evolution at the genomic level. So, it seems rather odd to take the most “evolutionarily advanced” genome as the framework for genomic organization and gene nomenclature. Humans like other mammals have their own long evolutionary history with genome rearrangements, including gene loss and duplications that differs not only from teleosts but also from many other vertebrate groups (Kapusta et al. 2017). In this respect, we suggest comparing the teleost genome, if not to the genome of an unknown teleosts ancestor or a genome of more primitive species, or as close to that state as possible. The genome most corresponding to these criteria is that of a spotted gar, *Lepisosteus oculatus* (Braasch et al., 2016). This species belongs to the basal Actinopterygian fish group, which represents the state of the genome before teleost whole-genome duplication. Indeed, the spotted gar, in terms of genome organization, seems closer to an ancestral state of tetrapods than any other known species (Braasch et al., 2016).

To avoid problems and misunderstanding in naming teleosts ohnologs, we propose an evolution-based systematic approach that utilizes the synteny with the spotted gar genome as a reference. However, this proposal requires adjusting the gene naming of the spotted gar first. Following the wrong naming in zebrafish, some spotted gar genes were named as ‘a’ and ‘b’ variants according to their similarity with zebrafish ohnologs. This practice occurred even though the spotted gar genome did not show teleost whole-genome duplication and these genes did not have a second copy in the gar genome. As a result of confusion in naming duplicated Tbx2 ohnologs in zebrafish, the only *tbx2* gene of spotted gar existing in its ancestral sole state (Fig. 1), i.e. the same as human, is irrationally annotated as the ‘b’-variant, *tbx2b* (Ensembl Release 98).

## Spotted gar duplicated genes

To test our proposed evolutionary-based systematic approach, we compared genes using the full genome data of human, zebrafish and spotted gar (Suppl. file 1). Using *ex silico* analysis and newly developed software for synteny analysis (Suppl. materials), we identified potentially duplicated genes, ‘a’ and ‘b’ variants, by comparing them to human genes by their names (Fig. 2, Suppl. file 2). Only a fraction of genes in the zebrafish and spotted gar exists in two copies, i.e. show true duplication (Table 1, Suppl. file 3). Genes present in the spotted gar genome as two copies and named as ohnologs (n=1006) could represent either individual gene duplications in the gar or gene losses in the human genome that happened during evolution when those two genomes diverged. We suggest analyzing the naming of these genes and their relation to teleost ohnologs not only by synteny but also by direct sequence comparison.

**Table 1.**

|  | **Zebrafish** | **Spotted gar** |
| --- | --- | --- |
| **Coding genes** | 25 592<sup>1</sup> | 18 341<sup>1</sup> |
| **Potentially duplicated\* genes comparing to human genes** | 7 231<sup>2</sup> | 3 093<sup>2</sup> |
| **True duplicated\* genes** | 6 760<sup>3</sup> | 1 006<sup>3</sup> |
| **Single genes with wrong names** | 471<sup>4</sup> | 2087<sup>4</sup> |
| **Duplicated\* genes with wrong names (synteny analysis)** | 335<sup>5</sup> | - |

\*Some genes exist in more than 2 copies only.
<sup>1</sup>Suppl. file 1
<sup>2</sup>Suppl. file 2
<sup>3</sup>Suppl. file 3
<sup>4</sup>Suppl. file 4
<sup>5</sup>Suppl. file 6

**Figure 2.**
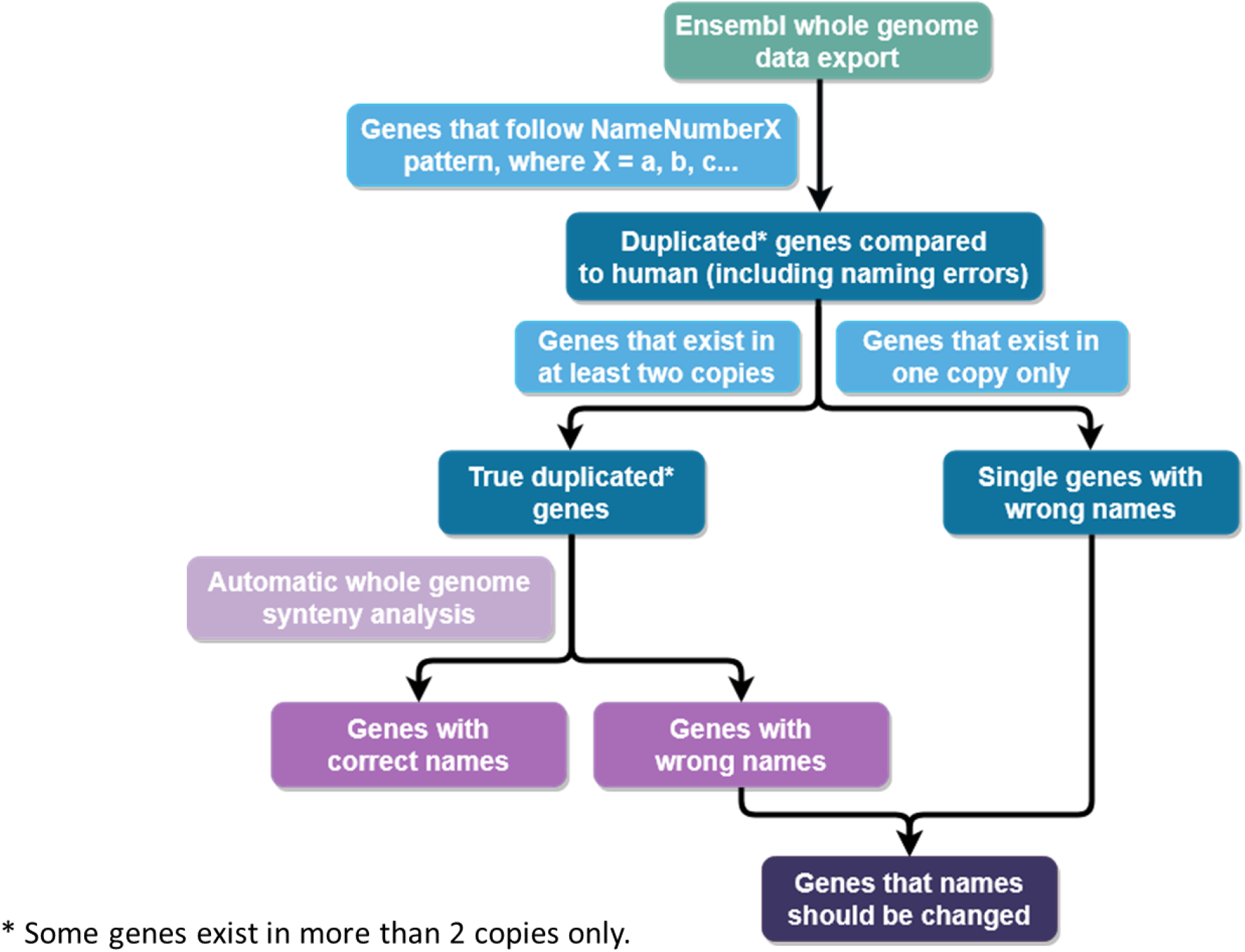
Diagram of the data analysis workflow. Human, zebrafish and spotted gar whole-genome databases were used for analysis. All zebrafish and spotted gar genes that follow the NameNumberX pattern where X = a, b, aa, bb, etc., were identified as duplicated genes. In this fraction, we identified single genes with wrong names and truly duplicated genes. For true duplicated genes, we applied automatic whole-genome synteny analysis, which generated a list of genes with correct and wrong names.

Another fraction of the zebrafish and spotted gar genes (n=2087) named as ‘a’ or ‘b’ variants exists in the genome as a single copy. These spotted gar genes do not represent ohnologs. They are the single orthologs of mammalian genes wrongly named in the spotted gar genome following the zebrafish nomenclature. The zebrafish genome also contains a fraction (n=471) of single genes named like ohnologs (Table 1, Suppl. file 4). They may represent a loss of one ohnolog in the gene pair. Alternatively, the naming could reflect the wrong naming when one known ohnolog had an individual name before the ‘a’ or ‘b’ extension convention. We again suggest that these names should be analyzed and corrected manually, as software cannot accurately determine these distinctions.

## Duplicated genes synteny analysis in zebrafish

The largest fraction of zebrafish genes classified as duplicated ones (n=6760) arose from the teleost genome duplication. These genes exist as two copies of the mammalian orthologs (Table 1). Using the Genome Synteny (GS) software for synteny analysis that we developed, we analyzed this group of genes by synteny in comparison to the human and spotted gar genomes.

We found a scored synteny similarity (Fig. 2, Suppl. file 5). Then, we manually selected the most unquestionable cases of inconsistency, where the zebrafish gene variant named as ‘b’ was flanked in the genome by at least two more homologous genes in human than the ‘a’ gene variant (Table 1; Suppl. file 6). According to this selection, 335 ohnolog pairs have the wrong ‘a’ and ‘b’ identifier, as the *tbx2* pair mentioned above and detected by GS software were taken as an internal positive control. Two examples of such genes, *gpa33* (glycoprotein A33 gene) and *fat1* (FAT atypical cadherin 1 gene) from the top of the list are represented graphically (Fig. 3).

**Figure 3.**
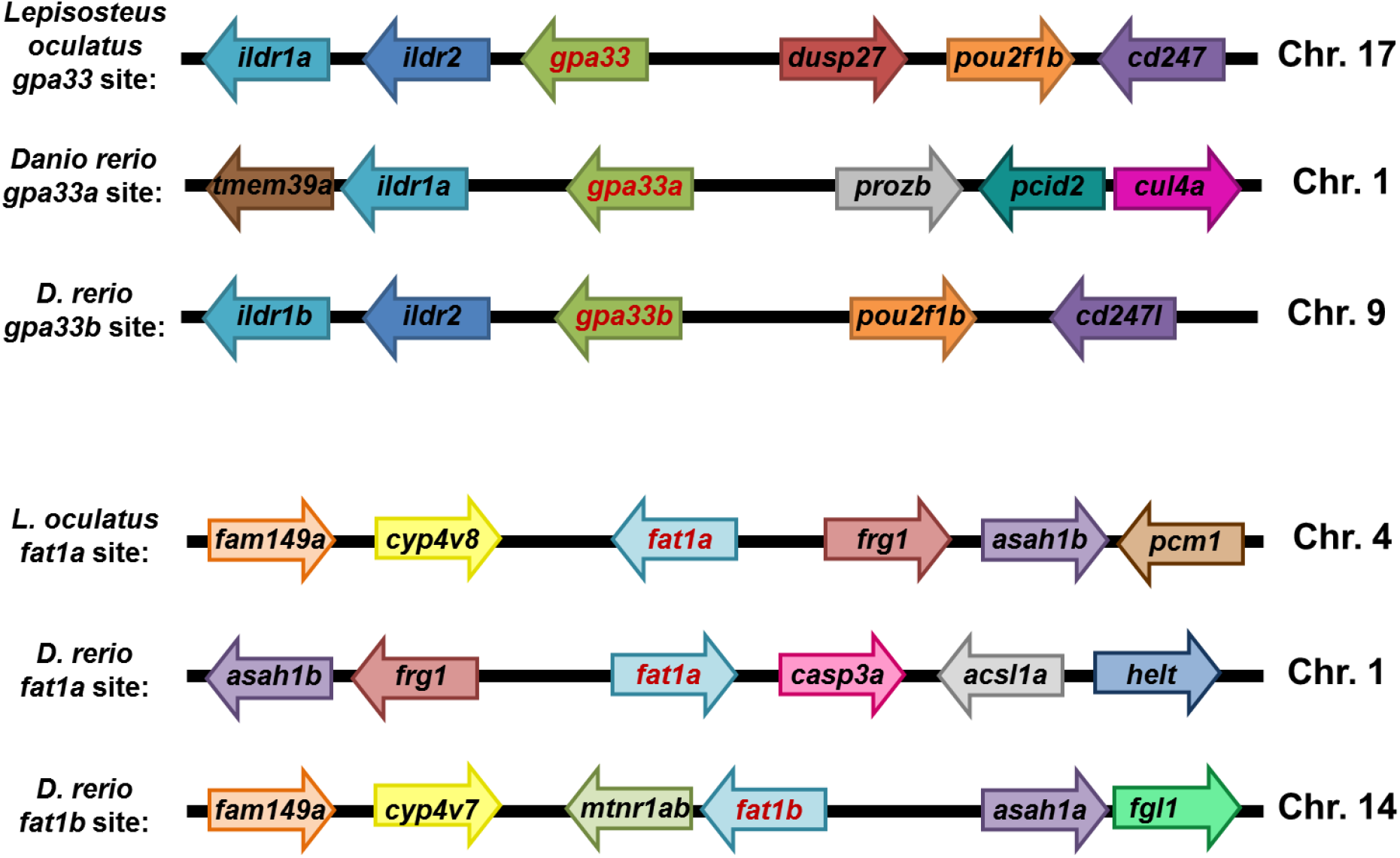
Two examples of gene renaming suggested by synteny analysis of the zebrafish (*D. rerio*) and spotted gar (*L. oculatus*) genomic regions. A – *gpa33a* and *gpa33b* genes should be renamed reciprocally; B – *fat1a* and *fat1b* genes should be renamed reciprocally.

Some ohnologs were preserved in evolution within the context of large, duplicated genomic fragments that remained relatively intact throughout evolution. This results in a close synteny scoring for such cases, so difficulties arise in distinguishing ‘a’ and ‘b’ variants. The same difficulties appear when, *vice versa*, the harboring ohnologs genome sites are strongly rearranged, and neighboring of both ohnologs does not correspond to the reference genome. We noticed such occurrences in the zebrafish genome with respect to *smcr8* (guanine nucleotide exchange protein 8), *pax3* (paired box proteins’ gene 3) or *ttyh3* (tweety family member 3 protein gene). To designate ‘a’ and ‘b’ variants correctly in such cases, gene sequence analysis or increased range of neighboring genes for synteny comparison will be useful. There are genes that possess extensions ‘a’ and ‘b’ that are not separated by digits in their names like *napaa* and *napab* (N-ethylmaleimide-sensitive factor attachment protein alpha gene). These genes were not analyzed given the GS software limitations.

It is unlikely that we detected all the cases of wrong gene naming in zebrafish. We aimed to propose the evolution-based systematic approach to gene naming based on synteny. The use of synteny criterion has limitations and in some cases also requires sequence comparisons and manual analysis. However, applying the evolution-based systematic approach referring to the spotted gar genome is an important filter that significantly reduces the number of questionable cases for further clarification by sequence similarity.

## Zebrafish ohnologs using synteny

To fit the synteny criterion and test our software, we conducted an initial random manual evaluation of a few examples of duplicated genes from the published literature for comparison with the results using the *ex silico* approach. Of the zebrafish *kcng4* genes (Shen et al., 2016), the *kcng4a* site, unlike *kcng4b* resembles its single-copy counterpart in gar given the presence of five common neighbors. This error was also picked up by the GS software (Suppl. file 4). Based on these results, we conclude that naming the single gene in gar *kcng4<u>a</u>* makes no sense. This conclusion also holds for the *ntrk2* (neurotrophic receptor tyrosine kinase 2) gene. According to synteny, the *ntrk2* gene has correctly named zebrafish ohnologs, ‘a’ and ‘b’ (Gasanov et al. 2015). The single spotted gar ortholog was initially incorrectly named *ntrk2<u>b</u>* (Braasch et al., 2016), and then wrongly renamed as *ntrk2<u>a</u>* (Ensembl Release 98). This error was also verified by the GS software (Suppl. file 4).

However, the gene *Pax6* highlights different challenges (Coutigno et al., 2011; Krauss et al., 1991). Comparing the gar and human *pax6* genes to the two genomic sites of the zebrafish *pax6-*s seemingly shows significant genomic rearrangements. The manual analysis suggested that the genomic neighborhood of the gar *pax6* site contains six common genes with the zebrafish *pax6a* and four in the vicinity of *pax6b*. Compared to the human genome, synteny returns scores of 5 and 4 for the ‘a’ and ‘b’ variants, respectively (Suppl. file 5). In the zebrafish *pax6b* site, one flank seems more divergent compared to the other one. This result suggests that some rearrangement in this area in the ancestral teleost genome after separation of the common ancestor of the gar-mammals and teleosts. Again, the naming of a single *pax6* of gar as *pax6<u>a</u>* makes no sense.

In the case of the *orai1* pair of genes (Wasilewska et al., 2018), the genomic neighborhood of the gar gene (*orai1b*) more closely resembles that of the zebrafish *orai1b* gene, even though the synteny between both zebrafish sites and the human and gar sites lies in only one flank. This result suggests chromosome fission in this region in the ancestors of teleosts unlike that in human and gar ancestors, that show more significant and present synteny in both flanks of *orai1*. Our synteny scoring for *orai1a* and *orai1b*, 3 and 6, respectively, suggests their reciprocal renaming as well as the renaming the single gar ortholog *orai1<u>b</u>* as *orai1* (Suppl. File 4, 6). In a pair of *stim1* genes (Wasilewska et al., 2018), the *stim1b* genomic site is more reminiscent of this site in both human and gar (scoring 3 vs 2, Suppl. File 6). We propose that this pair of ohnologs requires renaming as well. All our analyzed ohnolog cases using synteny and *ex silico* approaches revealed the same conclusion as the manual analysis.

We showed that gene naming requires more attention to minimize the number of genes across species that need renaming. We propose that the names of misnamed genes be corrected as exemplified by the *tbx-a/tbx2a* as *tbx2b* and *tbx-c/tbx2b* as *tbx2a* cases. While the wrongly named ohnolog pairs in zebrafish are relatively small problem, this problem will continue to increase and require additional diligence to resolve it when the problem becomes much more significant. The nonsystematic attempts to resolve this problem demonstrate the feasibility and precedents of such reciprocal renaming of ohnologs. In the case of the *six1a-six1b* pair (*sineoculis* homeobox homolog 1), “*six1b*” became “*six1a*” and *vice versa*. In addition the *crebbpa-crebbpb* pair (cyclic AMP response element-binding protein genes), *crebbpa* was previously named *crebbpb* and renamed to *crebbpa*. The somewhat different, but contextually similar renaming, occurred for the above-mentioned pair, *napaa* (previously α*-snap2*), and *napab* (previously known as α-*snap1 or napa*).

## Concluding remarks

The problem of incorrectly naming ohnologs is widespread and involves several hundreds of genes, just in case of the zebrafish genome alone (Table 1; Supp file 6). Moreover, these mistakes are repeated by incorrectly naming genes in other species. Thus, a large-scale effort to correctly rename genes using a standardized criteria with up-to-date technology must occur. The synteny analysis was not possible when genes were cloned by low stringency screening and mapped by radiation hybrid panels (Chong et al., 2001; Geisler et al., 1999; Kwok et al., 1998). We currently possess the tools to correct the errors and establish a logical ohnolog naming nomenclature, since teleosts represent approximately half of all vertebrate species on the planet. If we do not undertake this challenge, this problem will only grow in scale as more and more teleost genomes are being sequenced in a framework of Vertebrates Genomes Project [https://vertebrategenomesproject.org; (Braasch et al., 2016; Pasquier et al., 2016; Venkatesh et al., 2014)]. In conclusion, our proposed analysis represents an ideal method to detect the potential wrong naming. Our analysis and approach justify the suggested evolutionary approach to correct these mistakes.

## Author contribution

EG and JJ performed experiments with Tbx-a transgenics (in preparation) that led VK and JK to develop a concept for this paper, EG proposed and JJ developed software and performed automated synteny comparisons, EG, JJ, VK analyzed results, EG, JJ and VK wrote a paper.

## Acknowledgments

Authors appreciate the help of the team of Zebrafish Facility of the International Institute of Molecular and Cell Biology in Warsaw and Mr. J. Jędrychowski who helped to develop the software for *in silico* synteny analysis. We thank Dr. Brandi Mattson of Life Science Editors for editing assistance.

## Supplementary materials

### Data analysis workflow description

Whole-genome database of human (GRCh38.p13 (Genome Reference Consortium Human Build 38), INSDC Assembly GCA_000001405.28, Dec 2013, database version 98.38), zebrafish (GRCz11 (Genome Reference Consortium Zebrafish Build 11), INSDC Assembly GCA_000002035.4, May 2017, database version 98.11) and spotted gar (LepOcu1, INSDC Assembly GCA_000242695.1, Dec 2011, database version 98.1) were exported from Ensembl (http://ensembl.org/) using a dedicated MySQL server (Suppl. file 1). The lists of duplicated genes in zebrafish and spotted gar compared to human were exported by querying databases of genes that follow the NameNumberX pattern, where X = a, b, aa, ab, etc. (Suppl. file 2). Then the lists of genes were limited to new ones, which include only genes that physically occur in two copies (Supp. file 3). The remaining genes were collected in the lists of single genes with wrong names (Supp. file 4). The Ruby programming language was used to develop the Genome Synteny (GS) software for synteny analysis, which then was performed for all duplicated zebrafish genes. Each ohnolog was scored for the occurrence of homologous human genes in the vicinity of ten upstream and ten downstream genes from the analyzed gene. For the supporting results, the same analysis was performed for the spotted gar genes (Supp. file 5). Zebrafish genes with the highest score among a particular group of ohnologs named differently than ‘a’ were identified as genes incorrectly named (Supp. file 6). The full script and documentation for GS analysis were deposited as a GitHub repository (https://github.com/jjedrychowska/synteny-analysis).

### Legends for Supplementary files

File 1: Ensembl data for human, zebrafish and spotted gar genomes.

File 2: Complete sets of potentially duplicated genes in zebrafish and spotted gar comparing to human.

File 3: Complete sets of duplicated genes in zebrafish and spotted gar comparing to human.

File 4: Complete sets of single-copy genes named as duplicated ones in zebrafish and spotted gar.

File 5: The “in silico” synteny analysis of zebrafish and spotted gar genes compared to human. The program analyzes the neighboring genes and provides comparative scores for the ‘a’ and ‘b’ ohnologs.

File 6: Manually corrected results of “in silico” automatized synteny analysis of zebrafish and spotted gar genes comparing to human homolog genes from file 5. Genes with a score higher by 2 should be assigned the ‘a’ identity.

## References

1. Wolfe K., 2000. Robustness - It’s not where you think it is. Nat. Genet. 25(1):3–4. doi.org/10.1038/75560

2. Glasauer SM, Neuhauss SC. Whole-genome duplication in teleost fishes and its evolutionary consequences. Mol Genet Genomics. 2014; 289(6):1045–60. doi: 10.1007/s00438-014-0889-2.

3. ZFIN Database (http://www.zfin.org)

4. Dheen, T., Sleptsova-Friedrich, I., Xu, Y., Clark, M., Lehrach, H., Gong, Z., Korzh, V., 1999. Zebrafish tbx-c functions during formation of midline structures. Development 126, 2703–2713.

5. Ruvinsky, I., Oates, A.C., Silver, L.M., Ho, R.K., 2000. The evolution of paired appendages in vertebrates: T-box genes in the zebrafish. Dev. Genes Evol. 210(2): 82–91. doi.org/10.1007/s004270050014.

6. Ensembl Release 98. (http://www.ensembl.org/index.html)

7. Chi, N.C., Shaw, R.M., de Val, S., Kang, G., Jan, L.Y., Black, B.L., Stainier, D.Y.R., 2008. Foxn4 directly regulates tbx2b expression and atrioventricular canal formation. Genes Dev. 22(6): 734–739. doi.org/10.1101/gad.1629408.

8. Kaaij, L.J.T., Mokry, M., Zhou, M., Musheev, M., Geeven, G., Melquiond, A.S.J., de Jesus Domingues, A.M., de Laat, W., Niehrs, C., Smith, A.D., Ketting, R.F., 2016. Enhancers reside in a unique epigenetic environment during early zebrafish development. Genome Biol. 17(1): 146. doi.org/10.1186/s13059-016-1013-1.

9. Force A, Lynch M, Pickett FB, Amores A, Yan YL, Postlethwait J. Preservation of duplicate genes by complementary, degenerative mutations. Genetics. 1999; 151(4):1531–45.

10. Chong, S.W., Emelyanov, A., Gong, Z., Korzh, V., 2001. Expression pattern of two zebrafish genes, cxcr4a and cxcr4b. Mech. Dev. 109, 347–354. doi.org/10.1016/S0925-4773(01)00520-2.

11. Braasch, I., Gehrke, A.R., Smith, J.J., Kawasaki, K., Manousaki, T., Pasquier, J., Amores, A., Desvignes, T., Batzel, P., Catchen, J., Berlin, A.M., Campbell, M.S., Barrell, D., Martin, K.J., Mulley, J.F., Ravi, V., Lee, A.P., Nakamura, T., Chalopin, D., Fan, S., Wcisel, D., Caestro, C., Sydes, J., Beaudry, F.E.G., Sun, Y., Hertel, J., Beam, M.J., Fasold, M., Ishiyama, M., Johnson, J., Kehr, S., Lara, M., Letaw, J.H., Litman, G.W., Litman, R.T., Mikami, M., Ota, T., Saha, N.R., Williams, L., Stadler, P.F., Wang, H., Taylor, J.S., Fontenot, Q., Ferrara, A., Searle, S.M.J., Aken, B., Yandell, M., Schneider, I., Yoder, J.A., Volff, J.N., Meyer, A., Amemiya, C.T., Venkatesh, B., Holland, P.W.H., Guiguen, Y., Bobe, J., Shubin, N.H., Di Palma, F., Alföldi, J., Lindblad-Toh, K., Postlethwait, J.H., 2016. The spotted gar genome illuminates vertebrate evolution and facilitates human-teleost comparisons. Nat. Genet. 48(4): 427–437. doi.org/10.1038/ng.3526

12. Horton, A.C., Mahadevan, N.R., Minguillon, C., Osoegawa, K., Rokhsar, D.S., Ruvinsky, I., de Jong, P.J., Logan, M.P., Gibson-Brown, J.J., 2008. Conservation of linkage and evolution of developmental function within the Tbx2/3/4/5 subfamily of T-box genes: Implications for the origin of vertebrate limbs. Dev. Genes Evol. 218(11-12):613–628. doi.org/10.1007/s00427-008-0249-5.

13. Kapusta A, Suh A, and Feschotte C. Dynamics of genome size evolution in birds and mammals. 2017. PNAS, 114 (8): E1460–E1469. doi.org/10.1073/pnas.1616702114.

14. Shen H, Bocksteins E, Kondrychyn I, Snyders D, Korzh V. (2016) Functional antagonism of alpha-subunits of Kv channel in developing brain ventricular system. Development 143(22):4249–4260. doi: 10.1242/dev.140467.

15. Gasanov E.V., Rafieva L.M., Korzh V.P. BDNF-TrkB Axis Regulates Migration of the Lateral Line Primordium and Modulates the Maintenance of Mechanoreceptor Progenitors. // PLoS ONE. 2015; 10(3): e0119711. doi:10.1371/journal.pone.0119711.

16. Coutinho, P., Pavlou, S., Bhatia, S., Chalmers, K.J., Kleinjan, D.A., and van Heyningen, V. (2011) Discovery and assessment of conserved Pax6 target genes and enhancers. Genome research. 21(8):1349–59.

17. Krauss S., Johansen T., Korzh V., Moens U., Ericson J., A. Fjose. (1991) Zebrafish Pax(zf-a): a novel paired box-containing gene expressed in the neural tube. EMBO J., 10, 3609–3619.

18. Wasilewska I, Gupta RK, Palchevska O, Kuźnicki J. Identification of Zebrafish Calcium Toolkit Genes and their Expression in the Brain. Genes (Basel). 2019 Mar 18;10(3). pii: E230. doi: 10.3390/genes10030230.

19. Kwok, C., Korn, R.M., Davis, M.E., Burt, D.W., Critcher, R., McCarthy, L., Paw, B.H., Zon, L.I., Goodfellow, P.N., Schmitt, K., 1998. Characterization of whole genome radiation hybrid mapping resources for non-mammalian vertebrates. Nucleic Acids Res. 26(15):3562–3566. doi.org/10.1093/nar/26.15.3562.

20. Geisler, R., Rauch, G.J., Baier, H., Van Bebber, F., Broß, L., Dekens, M.P.S., Finger, K., Fricke, C., Gates, M.A., Geiger, H., Geiger-Rudolph, S., Gilmour, D., Glaser, S., Gnügge, L., Habeck, H., Hingst, K., Holley, S., Keenan, J., Kirn, A., Knaut, H., Lashkari, D., Maderspacher, F., Martyn, U., Neuhauss, S., Neumann, C., Nicolson, T., Pelegri, F., Ray, R., Rick, J.M., Roehl, H., Roeser, T., Schauerte, H.E., Schier, A.F., Schönberger, U., Schönthaler, H.B., Schulte-Merker, S., Seydler, C., Talbot, W.S., Weiler, C., Nüsslein-Volhard, C., Haffter, P., 1999. A radiation hybrid map of the zebrafish genome. Nat. Genet. 23(1): 86–89. doi.org/10.1038/12692.

21. Venkatesh, B., Lee, A.P., Ravi, V., Maurya, A.K., Lian, M.M., Swann, J.B., Ohta, Y., Flajnik, M.F., Sutoh, Y., Kasahara, M., Hoon, S., Gangu, V., Roy, S.W., Irimia, M., Korzh, V., Kondrychyn, I., Lim, Z.W., Tay, B.-H., Tohari, S., Kong, K.W., Ho, S., Lorente-Galdos, B., Quilez, J., Marques-Bonet, T., Raney, B.J., Ingham, P.W., Tay, A., Hillier, L.W., Minx, P., Boehm, T., Wilson, R.K., Brenner, S., Warren, W.C., 2014. Elephant shark genome provides unique insights into gnathostome evolution. Nature. 505(7482): 174–179. doi.org/10.1038/nature12826.

22. Pasquier, J., Cabau, C., Nguyen, T., Jouanno, E., Severac, D., Braasch, I., Journot, L., Pontarotti, P., Klopp, C., Postlethwait, J.H., Guiguen, Y., Bobe, J., 2016. Gene evolution and gene expression after whole genome duplication in fish: The PhyloFish database. BMC Genomics. 17(1): 368. doi.org/10.1186/s12864-016-2709-z.

