## Supplemental file 1 for "Evolutionary context can clarify teleosts gene names"

| name | chromosome | description |
|---|---|---|
| MUC6 | CHR_HG107_PATCH | mucin 6, oligomeric mucus/gel-forming [Sourc |
| LINC02688 | CHR_HG107_PATCH | long intergenic non-protein coding RNA 2688 |
| MUC2 | CHR_HG107_PATCH | mucin 2, oligomeric mucus/gel-forming [Sourc |
| MUC5AC | CHR_HG107_PATCH | mucin 5AC, oligomeric mucus/gel-forming [So |
| ZSWIM4 | CHR_HG109_PATCH | zinc finger SWIM-type containing 4 [Source:H( |
| AC020916.4 | CHR_HG109_PATCH | novel transcript, antisense to ZSWIM4 |
| RN7SL619P | CHR_HG109_PATCH | RNA, 7SL, cytoplasmic 619, pseudogene [Sour |
| AC020916.3 | CHR_HG109_PATCH | novel transcript |
| MIR24-2 | CHR_HG109_PATCH | microRNA 24-2 [Source:HGNC Symbol;Acc:HG |
| MIR27A | CHR_HG109_PATCH | microRNA 27a [Source:HGNC Symbol;Acc:HGN |
| MIR23A | CHR_HG109_PATCH | microRNA 23a [Source:HGNC Symbol;Acc:HGN |
| NANOS3 | CHR_HG109_PATCH | nanos C2HC-type zinc finger 3 [Source:HGNC S |
| MIR181C | CHR_HG109_PATCH | microRNA 181c [Source:HGNC Symbol;Acc:HG |
| MIR181D | CHR_HG109_PATCH | microRNA 181d [Source:HGNC Symbol;Acc:HG |
| C19orf57 | CHR_HG109_PATCH | chromosome 19 open reading frame 57 [Sour |
| CC2D1A | CHR_HG109_PATCH | coiled-coil and C2 domain containing 1A [Sour |
| PODNL1 | CHR_HG109_PATCH | podocan like 1 [Source:HGNC Symbol;Acc:HGI |
| DCAF15 | CHR_HG109_PATCH | DDB1 and CUL4 associated factor 15 [Source:I |
| RFX1 | CHR_HG109_PATCH | regulatory factor X1 [Source:HGNC Symbol;Ac |
| AC022098.6 | CHR_HG109_PATCH | novel transcript, antisense to RFX1 |
| RLN3 | CHR_HG109_PATCH | relaxin 3 [Source:HGNC Symbol;Acc:HGNC:17 |
| AC022098.7 | CHR_HG109_PATCH | novel transcript, antisense to RLN3 |
| IL27RA | CHR_HG109_PATCH | interleukin 27 receptor subunit alpha [Source: |
| PALM3 | CHR_HG109_PATCH | paralemmin 3 [Source:HGNC Symbol;Acc:HGN |
| EEF1DP1 | CHR_HG109_PATCH | eukaryotic translation elongation factor 1 delt |
| MISP3 | CHR_HG109_PATCH | MISP family member 3 [Source:HGNC Symbol; |
| MIR1199 | CHR_HG109_PATCH | microRNA 1199 [Source:HGNC Symbol;Acc:HG |
| C19orf67 | CHR_HG109_PATCH | chromosome 19 open reading frame 67 [Sour |
| NTIA01000060.1 | CHR_HG109_PATCH | NULL |
| PRKACA | CHR_HG109_PATCH | protein kinase cAMP-activated catalytic subur |
| AC022098.8 | CHR_HG109_PATCH | novel transcript, antisense to ASF1B |
| ASF1B | CHR_HG109_PATCH | anti-silencing function 1B histone chaperone [ |
| AC022098.5 | CHR_HG109_PATCH | novel transcript, antisense to LPHN1 |
| ADGRL1 | CHR_HG109_PATCH | adhesion G protein-coupled receptor L1 [Sour |
| RN7SL231P | CHR_HG109_PATCH | RNA, 7SL, cytoplasmic 231, pseudogene [Sour |
| AC011509.5 | CHR_HG109_PATCH | novel transcript |
| AC011509.4 | CHR_HG109_PATCH | novel transcript |
| AC011509.6 | CHR_HG109_PATCH | novel transcript |
| RYBP | CHR_HG126_PATCH | RING1 and YY1 binding protein [Source:NCBI g |
| AC255560.1 | CHR_HG126_PATCH | novel transcript |
| RNU1-62P | CHR_HG126_PATCH | RNA, U1 small nuclear 62, pseudogene [Sourc |
| AC097369.1 | CHR_HG126_PATCH | pseudogene similar to part of mitochondrially |
| RNA5SP136 | CHR_HG126_PATCH | RNA, 5S ribosomal pseudogene 136 [Source:H |
| FO681492.2 | CHR_HG1277_PATCH | synaptotagmin-15 [Source:NCBI gene;Acc:102 |
| LJII02000414.1 | CHR_HG1277_PATCH | NULL |
| AC245041.3 | CHR_HG1277_PATCH | novel transcript |
| NPY4R2 | CHR_HG1277_PATCH | neuropeptide Y receptor Y4-2 [Source:HGNC S |
| AC245041.4 | CHR_HG1277_PATCH | novel transcript |
| FAM25C | CHR_HG1277_PATCH | family with sequence similarity 25 member C |

| | | |
|---|---|---|
| AGAP12P | CHR_HG1277_PATCH | ArfGAP with GTPase domain, ankyrin repeat a |
| RNA5SP315 | CHR_HG1277_PATCH | RNA, 5S ribosomal pseudogene 315 [Source:H |
| BMS1P7 | CHR_HG1277_PATCH | BMS1 pseudogene 7 [Source:HGNC Symbol;Ac |
| PTPN20CP | CHR_HG1277_PATCH | protein tyrosine phosphatase non-receptor ty |
| AC209005.2 | CHR_HG1298_PATCH | uncharacterized LOC101928532 [Source:NCBI |
| HMX1 | CHR_HG1298_PATCH | H6 family homeobox 1 [Source:HGNC Symbol; |
| AC116612.2 | CHR_HG1298_PATCH | novel transcript, sense intronic to HMX1 |
| AC093627.26 | CHR_HG1309_PATCH | novel transcript |
| AC093627.24 | CHR_HG1309_PATCH | novel transcript |
| AC093627.23 | CHR_HG1309_PATCH | novel transcript |
| AC093627.29 | CHR_HG1309_PATCH | novel transcript |
| AC093627.27 | CHR_HG1309_PATCH | novel transcript |
| AC093627.30 | CHR_HG1309_PATCH | novel transcript |
| AC093627.25 | CHR_HG1309_PATCH | novel transcript |
| AC093627.28 | CHR_HG1309_PATCH | novel transcript |
| FAM20C | CHR_HG1309_PATCH | FAM20C golgi associated secretory pathway k |
| AC145676.2 | CHR_HG1309_PATCH | TEC |
| AC187652.1 | CHR_HG1309_PATCH | NULL |
| Y_RNA | CHR_HG1311_PATCH | Y RNA [Source:RFAM;Acc:RF00019] |
| AC000050.2 | CHR_HG1311_PATCH | novel transcript |
| RNU6-409P | CHR_HG1311_PATCH | RNA, U6 small nuclear 409, pseudogene [Sour |
| FO681507.1 | CHR_HG1311_PATCH | SH3 and multiple ankyrin repeat domains prot |
| ACR | CHR_HG1311_PATCH | acrosin [Source:HGNC Symbol;Acc:HGNC:126] |
| FO681507.2 | CHR_HG1311_PATCH | novel transcript |
| AC002056.3 | CHR_HG1311_PATCH | small nuclear ribonucleoprotein polypeptide A |
| RPL23AP82 | CHR_HG1311_PATCH | ribosomal protein L23a pseudogene 82 [Sourc |
| GCGR | CHR_HG1320_PATCH | glucagon receptor [Source:HGNC Symbol;Acc: |
| MCRIP1 | CHR_HG1320_PATCH | MAPK regulated corepressor interacting prote |
| PPP1R27 | CHR_HG1320_PATCH | protein phosphatase 1 regulatory subunit 27 [ |
| LINC01784 | CHR_HG1342_HG2282_PATC | long intergenic non-protein coding RNA 1784 |
| PRAMEF11 | CHR_HG1342_HG2282_PATC | PRAME family member 11 [Source:HGNC Sym |
| PRAMEF30P | CHR_HG1342_HG2282_PATC | PRAME family member 30, pseudogene [Sour |
| HNRNPCL1 | CHR_HG1342_HG2282_PATC | heterogeneous nuclear ribonucleoprotein C lil |
| PRAMEF2 | CHR_HG1342_HG2282_PATC | PRAME family member 2 [Source:HGNC Symb |
| PRAMEF4 | CHR_HG1342_HG2282_PATC | PRAME family member 4 [Source:HGNC Symb |
| PRAMEF10 | CHR_HG1342_HG2282_PATC | PRAME family member 10 [Source:HGNC Sym |
| PRAMEF7 | CHR_HG1342_HG2282_PATC | PRAME family member 7 [Source:HGNC Symb |
| RNU6-1072P | CHR_HG1342_HG2282_PATC | RNA, U6 small nuclear 1072, pseudogene [Sou |
| PRAMEF29P | CHR_HG1342_HG2282_PATC | PRAME family member 29, pseudogene [Sour |
| PRAMEF6 | CHR_HG1342_HG2282_PATC | PRAME family member 6 [Source:HGNC Symb |
| PRAMEF28P | CHR_HG1342_HG2282_PATC | PRAME family member 28, pseudogene [Sour |
| PRAMEF27 | CHR_HG1342_HG2282_PATC | PRAME family member 27 [Source:HGNC Sym |
| HNRNPCL3 | CHR_HG1342_HG2282_PATC | heterogeneous nuclear ribonucleoprotein C lil |
| PRAMEF25 | CHR_HG1342_HG2282_PATC | PRAME family member 25 [Source:HGNC Sym |
| PRAMEF34P | CHR_HG1342_HG2282_PATC | PRAME family member 34, pseudogene [Sour |
| PRAMEF35P | CHR_HG1342_HG2282_PATC | PRAME family member 35, pseudogene [Sour |
| HNRNPCL2 | CHR_HG1342_HG2282_PATC | heterogeneous nuclear ribonucleoprotein C lil |
| PRAMEF36P | CHR_HG1342_HG2282_PATC | PRAME family member 36, pseudogene [Sour |
| AC245056.2 | CHR_HG1342_HG2282_PATC | novel transcript |
| PRAMEF26 | CHR_HG1342_HG2282_PATC | PRAME family member 26 [Source:HGNC Sym |

| | | |
|---|---|---|
| HNRNPCL4 | CHR_HG1342_HG2282_PATC | heterogeneous nuclear ribonucleoprotein C lil |
| PRAMEF9 | CHR_HG1342_HG2282_PATC | PRAME family member 9 [Source:HGNC Symb |
| PRAMEF13 | CHR_HG1342_HG2282_PATC | PRAME family member 13 [Source:HGNC Sym |
| PRAMEF18 | CHR_HG1342_HG2282_PATC | PRAME family member 18 [Source:HGNC Sym |
| PRAMEF31P | CHR_HG1342_HG2282_PATC | PRAME family member 31, pseudogene [Sour |
| PRAMEF5 | CHR_HG1342_HG2282_PATC | PRAME family member 5 [Source:HGNC Symb |
| PRAMEF32P | CHR_HG1342_HG2282_PATC | PRAME family member 32, pseudogene [Sour |
| RNU6-771P | CHR_HG1342_HG2282_PATC | RNA, U6 small nuclear 771, pseudogene [Sour |
| PRAMEF8 | CHR_HG1342_HG2282_PATC | PRAME family member 8 [Source:HGNC Symb |
| PRAMEF33 | CHR_HG1342_HG2282_PATC | PRAME family member 33 [Source:HGNC Sym |
| BCL2L14 | CHR_HG1362_PATCH | BCL2 like 14 [Source:HGNC Symbol;Acc:HGNC |
| PTMAP9 | CHR_HG1362_PATCH | prothymosin alpha pseudogene 9 [Source:HGI |
| MIR1244-4 | CHR_HG1362_PATCH | microRNA 1244-4 [Source:HGNC Symbol;Acc:I |
| LRP6 | CHR_HG1362_PATCH | LDL receptor related protein 6 [Source:HGNC |
| AC007537.2 | CHR_HG1362_PATCH | ribosomal protein L21 (RPL21) pseudogene |
| RNU6-545P | CHR_HG1362_PATCH | RNA, U6 small nuclear 545, pseudogene [Sour |
| AC007621.2 | CHR_HG1362_PATCH | ubiquitin-conjugating enzyme E2 variant 2 (UE |
| RNU6-318P | CHR_HG1362_PATCH | RNA, U6 small nuclear 318, pseudogene [Sour |
| MANSC1 | CHR_HG1362_PATCH | MANSC domain containing 1 [Source:HGNC Sy |
| RPL23AP66 | CHR_HG1362_PATCH | ribosomal protein L23a pseudogene 66 [Sourc |
| LOH12CR2 | CHR_HG1362_PATCH | loss of heterozygosity, 12, chromosomal regic |
| BORCS5 | CHR_HG1362_PATCH | BLOC-1 related complex subunit 5 [Source:HG |
| AC007619.3 | CHR_HG1362_PATCH | adaptor-related protein complex 3, sigma 1 su |
| DUSP16 | CHR_HG1362_PATCH | dual specificity phosphatase 16 [Source:HGNC |
| AC007619.4 | CHR_HG1362_PATCH | novel transcript |
| AC092824.2 | CHR_HG1362_PATCH | ribosomal protein L19 (RPL19) pseudogene |
| AC130710.2 | CHR_HG1384_PATCH | novel transcript |
| AC174048.2 | CHR_HG1384_PATCH | polymerase (RNA) II (DNA directed) polypeptic |
| DNAJC18 | CHR_HG1395_PATCH | DnaJ heat shock protein family (Hsp40) memb |
| AC142391.2 | CHR_HG1395_PATCH | novel transcript |
| RNU5B-4P | CHR_HG1395_PATCH | RNA, U5B small nuclear 4, pseudogene [Sourc |
| ECSCR | CHR_HG1395_PATCH | endothelial cell surface expressed chemotaxis |
| SMIM33 | CHR_HG1395_PATCH | small integral membrane protein 33 [Source:H |
| TMEM173 | CHR_HG1395_PATCH | transmembrane protein 173 [Source:HGNC Sy |
| AC138517.2 | CHR_HG1395_PATCH | chromosome 1 open reading frame 57 pseudc |
| NCOA4P4 | CHR_HG1395_PATCH | nuclear receptor coactivator 4 pseudogene 4 |
| C1R | CHR_HG1398_PATCH | complement C1r [Source:HGNC Symbol;Acc:H |
| C1RL | CHR_HG1398_PATCH | complement C1r subcomponent like [Source:I |
| C1RL-AS1 | CHR_HG1398_PATCH | C1RL antisense RNA 1 [Source:HGNC Symbol;A |
| RBP5 | CHR_HG1398_PATCH | retinol binding protein 5 [Source:HGNC Symbc |
| RNU6-485P | CHR_HG1398_PATCH | RNA, U6 small nuclear 485, pseudogene [Sour |
| AC018653.7 | CHR_HG1398_PATCH | novel transcript |
| CLSTN3 | CHR_HG1398_PATCH | calsyntenin 3 [Source:HGNC Symbol;Acc:HGN( |
| AC018653.6 | CHR_HG1398_PATCH | novel transcript, antisense to CLSTN3 |
| AC018653.5 | CHR_HG1398_PATCH | novel transcript |
| PEX5 | CHR_HG1398_PATCH | peroxisomal biogenesis factor 5 [Source:HGN( |
| AC018653.8 | CHR_HG1398_PATCH | chromosome 10 open reading frame 129 (C1C |
| AC022882.2 | CHR_HG142_HG150_NOVEL_ | N(alpha)-acetyltransferase 50, NatE catalytic s |
| OR8J2 | CHR_HG142_HG150_NOVEL_ | olfactory receptor family 8 subfamily J membe |
| OR5T2 | CHR_HG142_HG150_NOVEL_ | olfactory receptor family 5 subfamily T memb |

| | | |
|---|---|---|
| OR5T3 | CHR_HG142_HG150_NOVEL_ | olfactory receptor family 5 subfamily T memb |
| OR5T1 | CHR_HG142_HG150_NOVEL_ | olfactory receptor family 5 subfamily T memb |
| OR8H1 | CHR_HG142_HG150_NOVEL_ | olfactory receptor family 8 subfamily H memb |
| OR8I1P | CHR_HG142_HG150_NOVEL_ | olfactory receptor family 8 subfamily I membe |
| OR8K3 | CHR_HG142_HG150_NOVEL_ | olfactory receptor family 8 subfamily K memb |
| FAM8A2P | CHR_HG142_HG150_NOVEL_ | family with sequence similarity 8 member A2, |
| OR8K2P | CHR_HG142_HG150_NOVEL_ | olfactory receptor family 8 subfamily K memb |
| OR8K1 | CHR_HG142_HG150_NOVEL_ | olfactory receptor family 8 subfamily K memb |
| RPL5P29 | CHR_HG142_HG150_NOVEL_ | ribosomal protein L5 pseudogene 29 [Source: |
| OR8J1 | CHR_HG142_HG150_NOVEL_ | olfactory receptor family 8 subfamily J membe |
| OR8U8 | CHR_HG142_HG150_NOVEL_ | olfactory receptor family 8 subfamily U memb |
| OR8U9 | CHR_HG142_HG150_NOVEL_ | olfactory receptor family 8 subfamily U memb |
| OR8L1P | CHR_HG142_HG150_NOVEL_ | olfactory receptor family 8 subfamily L memb |
| AP000676.6 | CHR_HG1445_PATCH | novel transcript |
| AP005436.4 | CHR_HG1445_PATCH | novel transcript |
| AP005436.5 | CHR_HG1445_PATCH | novel transcript |
| AP005436.6 | CHR_HG1445_PATCH | novel transcript |
| Y_RNA | CHR_HG1445_PATCH | Y RNA [Source:RFAM;Acc:RF00019] |
| AC092854.1 | CHR_HG1485_PATCH | NULL |
| U6 | CHR_HG1485_PATCH | U6 spliceosomal RNA [Source:RFAM;Acc:RF00 |
| ABCD1P4 | CHR_HG1485_PATCH | ATP binding cassette subfamily D member 1 p |
| AC137499.3 | CHR_HG1485_PATCH | zinc finger, CDGSH-type domain 2 (ZCD2) pseu |
| PABPC1P9 | CHR_HG1485_PATCH | poly(A) binding protein cytoplasmic 1 pseudog |
| SLC9B1P4 | CHR_HG1485_PATCH | solute carrier family 9 member B1 pseudogen |
| AC137499.4 | CHR_HG1485_PATCH | NULL |
| OR9G1 | CHR_HG151_NOVEL_TEST | olfactory receptor family 9 subfamily G memb |
| OR9G9 | CHR_HG151_NOVEL_TEST | olfactory receptor family 9 subfamily G memb |
| AP001803.2 | CHR_HG151_NOVEL_TEST | GNAS complex locus (GNAS) pseudogene |
| OR9G3P | CHR_HG151_NOVEL_TEST | olfactory receptor family 9 subfamily G memb |
| OR9G4 | CHR_HG151_NOVEL_TEST | olfactory receptor family 9 subfamily G memb |
| OR9G2P | CHR_HG151_NOVEL_TEST | olfactory receptor family 9 subfamily G memb |
| OR5G1P | CHR_HG151_NOVEL_TEST | olfactory receptor family 5 subfamily G memb |
| OR5G4P | CHR_HG151_NOVEL_TEST | olfactory receptor family 5 subfamily G memb |
| OR5G5P | CHR_HG151_NOVEL_TEST | olfactory receptor family 5 subfamily G memb |
| OR5G3 | CHR_HG151_NOVEL_TEST | olfactory receptor family 5 subfamily G memb |
| LINC02735 | CHR_HG151_NOVEL_TEST | long intergenic non-protein coding RNA 2735 |
| ATP4B | CHR_HG1524_PATCH | ATPase H+/K+ transporting subunit beta [Sour |
| GRK1 | CHR_HG1524_PATCH | G protein-coupled receptor kinase 1 [Source:H |
| LINC00552 | CHR_HG1524_PATCH | long intergenic non-protein coding RNA 552 [S |
| OFD1P3Y | CHR_HG1531_PATCH | OFD1 pseudogene 3 Y-linked [Source:HGNC Sy |
| TTTY13 | CHR_HG1535_PATCH | testis-specific transcript, Y-linked 13 [Source:H |
| ELOCP5 | CHR_HG1535_PATCH | elongin C pseudogene 5 [Source:HGNC Symbo |
| CDY10P | CHR_HG1535_PATCH | chromodomain Y-linked 10 pseudogene [Sour |
| USP9YP3 | CHR_HG1535_PATCH | USP9Y pseudogene 3 [Source:HGNC Symbol;A |
| USP9YP8 | CHR_HG1535_PATCH | USP9Y pseudogene 8 [Source:HGNC Symbol;A |
| CDY11P | CHR_HG1535_PATCH | chromodomain protein Y-linked 11 pseudoger |
| OFD1P16Y | CHR_HG1535_PATCH | OFD1 pseudogene 16 Y-linked [Source:HGNC S |
| AC007322.5 | CHR_HG1535_PATCH | novel transcript |
| AL356131.2 | CHR_HG1651_PATCH | proteasome subunit alpha type 4 (PROS29) ps |
| OR4C12 | CHR_HG1708_PATCH | olfactory receptor family 4 subfamily C memb |

| | | |
|---|---|---|
| OR4C48P | CHR_HG1708_PATCH | olfactory receptor family 4 subfamily C memb |
| OR4C49P | CHR_HG1708_PATCH | olfactory receptor family 4 subfamily C memb |
| MIR3649 | CHR_HG1815_PATCH | microRNA 3649 [Source:HGNC Symbol;Acc:HG |
| ADIPOR2 | CHR_HG1815_PATCH | adiponectin receptor 2 [Source:HGNC Symbol |
| AC005183.2 | CHR_HG1815_PATCH | SDA1 domain containing 1 (SDAD1) pseudoge |
| AC005343.2 | CHR_HG1815_PATCH | novel transcript, antisense to ADIPOR2 |
| RPS4XP14 | CHR_HG1815_PATCH | ribosomal protein S4X pseudogene 14 [Source |
| CACNA2D4 | CHR_HG1815_PATCH | calcium voltage-gated channel auxiliary subun |
| AC005343.3 | CHR_HG1815_PATCH | novel transcript, antisense to CACNA2D4 |
| LRTM2 | CHR_HG1815_PATCH | leucine rich repeats and transmembrane dom |
| AC005342.3 | CHR_HG1815_PATCH | novel transcript |
| LINC00940 | CHR_HG1815_PATCH | long intergenic non-protein coding RNA 940 [S |
| DCP1B | CHR_HG1815_PATCH | decapping mRNA 1B [Source:HGNC Symbol;Ac |
| CACNA1C | CHR_HG1815_PATCH | calcium voltage-gated channel subunit alpha1 |
| AC005342.4 | CHR_HG1815_PATCH | novel transcript |
| CACNA1C-IT1 | CHR_HG1815_PATCH | CACNA1C intronic transcript 1 [Source:HGNC S |
| CACNA1C-IT2 | CHR_HG1815_PATCH | CACNA1C intronic transcript 2 [Source:HGNC S |
| AC005344.2 | CHR_HG1815_PATCH | novel transcript, antisense to CACNA1C |
| CACNA1C-AS4 | CHR_HG1815_PATCH | CACNA1C antisense RNA 4 [Source:HGNC Sym |
| CACNA1C-IT3 | CHR_HG1815_PATCH | CACNA1C intronic transcript 3 [Source:HGNC S |
| AC005293.2 | CHR_HG1815_PATCH | novel transcript, antisense to CACNA1C |
| AC005414.2 | CHR_HG1815_PATCH | novel transcript, antisense to CACNA1C |
| CACNA1C-AS3 | CHR_HG1815_PATCH | CACNA1C antisense RNA 3 [Source:HGNC Sym |
| AC007618.2 | CHR_HG1815_PATCH | ribosomal protein S6 (RPS6) pseudogene |
| CACNA1C-AS2 | CHR_HG1815_PATCH | CACNA1C antisense RNA 2 [Source:HGNC Sym |
| CACNA1C-AS1 | CHR_HG1815_PATCH | CACNA1C antisense RNA 1 [Source:HGNC Sym |
| ITFG2-AS1 | CHR_HG1815_PATCH | ITFG2 antisense RNA 1 [Source:HGNC Symbol; |
| SERTAD4-AS1 | CHR_HG1832_PATCH | SERTAD4 antisense RNA 1 [Source:HGNC Symb |
| SERTAD4 | CHR_HG1832_PATCH | SERTA domain containing 4 [Source:HGNC Syr |
| ST13P19 | CHR_HG1832_PATCH | ST13, Hsp70 interacting protein pseudogene 1 |
| AL035414.2 | CHR_HG1832_PATCH | ribonuclease H1 (RNASEH1) pseudogene |
| HHAT | CHR_HG1832_PATCH | hedgehog acyltransferase [Source:HGNC Symb |
| AL034351.5 | CHR_HG1832_PATCH | novel transcript, antisense to HHAT |
| RNU5A-8P | CHR_HG1832_PATCH | RNA, U5A small nuclear 8, pseudogene [Sourc |
| AL034351.6 | CHR_HG1832_PATCH | novel transcript, antisense to HHAT |
| AL034351.2 | CHR_HG1832_PATCH | inositol monophosphatase domain containing |
| AL160237.5 | CHR_HG1_PATCH | novel transcript |
| BRD7P1 | CHR_HG1_PATCH | bromodomain containing 7 pseudogene 1 [So |
| AL160237.6 | CHR_HG1_PATCH | suppressor of zeste 12 homolog (Drosophila) ( |
| RN7SKP205 | CHR_HG1_PATCH | RN7SK pseudogene 205 [Source:HGNC Symbo |
| AL160237.4 | CHR_HG1_PATCH | dehydrogenase/reductase (SDR family) memb |
| LO018686.1 | CHR_HG1_PATCH | novel transcript |
| LINC00596 | CHR_HG1_PATCH | long intergenic non-protein coding RNA 596 [S |
| DHRS4-AS1 | CHR_HG1_PATCH | DHRS4 antisense RNA 1 [Source:HGNC Symbo |
| DHRS4 | CHR_HG1_PATCH | dehydrogenase/reductase 4 [Source:HGNC Sy |
| LO018686.2 | CHR_HG1_PATCH | novel transcript |
| AL136295.11 | CHR_HG1_PATCH | dehydrogenase/reductase 4 like 1 [Source:NC |
| DHRS4L1 | CHR_HG1_PATCH | dehydrogenase/reductase 4 like 1 [Source:HG |
| CARMIL3 | CHR_HG1_PATCH | capping protein regulator and myosin 1 linker |
| CPNE6 | CHR_HG1_PATCH | copine 6 [Source:HGNC Symbol;Acc:HGNC:23 |

| | | |
|---|---|---|
| NRL | CHR_HG1_PATCH | neural retina leucine zipper [Source:HGNC Syr |
| PCK2 | CHR_HG1_PATCH | phosphoenolpyruvate carboxykinase 2, mitocl |
| DCAF11 | CHR_HG1_PATCH | DDB1 and CUL4 associated factor 11 [Source:I |
| AL136295.9 | CHR_HG1_PATCH | novel protein |
| FITM1 | CHR_HG1_PATCH | fat storage inducing transmembrane protein 1 |
| PSME1 | CHR_HG1_PATCH | proteasome activator subunit 1 [Source:HGN( |
| EMC9 | CHR_HG1_PATCH | ER membrane protein complex subunit 9 [Sou |
| AL136295.13 | CHR_HG1_PATCH | novel transcript |
| PSME2 | CHR_HG1_PATCH | proteasome activator subunit 2 [Source:HGN( |
| MIR7703 | CHR_HG1_PATCH | microRNA 7703 [Source:HGNC Symbol;Acc:HG |
| RNF31 | CHR_HG1_PATCH | ring finger protein 31 [Source:HGNC Symbol;A |
| AL136295.10 | CHR_HG1_PATCH | novel transcript |
| RNA5SP383 | CHR_HG1_PATCH | RNA, 5S ribosomal pseudogene 383 [Source:H |
| IRF9 | CHR_HG1_PATCH | interferon regulatory factor 9 [Source:HGNC S |
| REC8 | CHR_HG1_PATCH | REC8 meiotic recombination protein [Source:I |
| IPO4 | CHR_HG1_PATCH | importin 4 [Source:HGNC Symbol;Acc:HGNC:1 |
| AL136295.12 | CHR_HG1_PATCH | novel protein |
| AL136295.16 | CHR_HG1_PATCH | novel transcript, antisense to TM9SF1 |
| TM9SF1 | CHR_HG1_PATCH | transmembrane 9 superfamily member 1 [Sou |
| AL136295.15 | CHR_HG1_PATCH | novel protein |
| AL136295.8 | CHR_HG1_PATCH | novel transcript |
| AL136295.14 | CHR_HG1_PATCH | novel transcript |
| TSSK4 | CHR_HG1_PATCH | testis specific serine kinase 4 [Source:HGNC Sy |
| CHMP4A | CHR_HG1_PATCH | charged multivesicular body protein 4A [Sourc |
| AL096870.3 | CHR_HG1_PATCH | novel transcript |
| MDP1 | CHR_HG1_PATCH | magnesium dependent phosphatase 1 [Source |
| NEDD8-MDP1 | CHR_HG1_PATCH | NEDD8-MDP1 readthrough [Source:HGNC Syn |
| NEDD8 | CHR_HG1_PATCH | NEDD8 ubiquitin like modifier [Source:HGNC S |
| GMPR2 | CHR_HG1_PATCH | guanosine monophosphate reductase 2 [Sour |
| TINF2 | CHR_HG1_PATCH | TERF1 interacting nuclear factor 2 [Source:HG |
| TGM1 | CHR_HG1_PATCH | transglutaminase 1 [Source:HGNC Symbol;Acc |
| RABGGTA | CHR_HG1_PATCH | Rab geranylgeranyltransferase subunit alpha [ |
| AL096870.7 | CHR_HG1_PATCH | novel transcript, antisense to RABGGTAand DI |
| AL096870.5 | CHR_HG1_PATCH | novel transcript, antisense to RABGGTA |
| DHRS1 | CHR_HG1_PATCH | dehydrogenase/reductase 1 [Source:HGNC Sy |
| NOP9 | CHR_HG1_PATCH | NOP9 nucleolar protein [Source:HGNC Symbo |
| CIDEB | CHR_HG1_PATCH | cell death inducing DFFA like effector b [Sourc |
| LTB4R2 | CHR_HG1_PATCH | leukotriene B4 receptor 2 [Source:HGNC Symb |
| LTB4R | CHR_HG1_PATCH | leukotriene B4 receptor [Source:HGNC Symbo |
| ADCY4 | CHR_HG1_PATCH | adenylate cyclase 4 [Source:HGNC Symbol;Acc |
| AL096870.4 | CHR_HG1_PATCH | novel transcript |
| RIPK3 | CHR_HG1_PATCH | receptor interacting serine/threonine kinase 3 |
| U6 | CHR_HG1_PATCH | U6 spliceosomal RNA [Source:RFAM;Acc:RF00 |
| NFATC4 | CHR_HG1_PATCH | nuclear factor of activated T cells 4 [Source:H( |
| AL139288.4 | CHR_HG2002_PATCH | novel transcript |
| TRIM17 | CHR_HG2002_PATCH | tripartite motif containing 17 [Source:HGNC S |
| HIST3H3 | CHR_HG2002_PATCH | histone cluster 3 H3 [Source:HGNC Symbol;Ac |
| RPL23AP15 | CHR_HG2002_PATCH | ribosomal protein L23a pseudogene 15 [Sourc |
| HIST3H2A | CHR_HG2002_PATCH | histone cluster 3 H2A [Source:HGNC Symbol;A |
| HIST3H2BB | CHR_HG2002_PATCH | histone cluster 3 H2B family member b [Sourc |

| | | |
|---|---|---|
| MIR4666A | CHR_HG2002_PATCH | microRNA 4666a [Source:HGNC Symbol;Acc:H |
| HIST3H2BA | CHR_HG2002_PATCH | histone cluster 3 H2B family member a (pseuc |
| AL139288.3 | CHR_HG2002_PATCH | novel transcript |
| RNF187 | CHR_HG2002_PATCH | ring finger protein 187 [Source:HGNC Symbol; |
| BTNL10 | CHR_HG2002_PATCH | butyrophilin like 10 [Source:HGNC Symbol;Acc |
| RNA5S1 | CHR_HG2002_PATCH | RNA, 5S ribosomal 1 [Source:HGNC Symbol;Ac |
| RNA5S2 | CHR_HG2002_PATCH | RNA, 5S ribosomal 2 [Source:HGNC Symbol;Ac |
| RNA5S3 | CHR_HG2002_PATCH | RNA, 5S ribosomal 3 [Source:HGNC Symbol;Ac |
| RNA5S16 | CHR_HG2002_PATCH | RNA, 5S ribosomal 16 [Source:HGNC Symbol;/ |
| RNA5S17 | CHR_HG2002_PATCH | RNA, 5S ribosomal 17 [Source:HGNC Symbol;/ |
| RNA5SP18 | CHR_HG2002_PATCH | RNA, 5S ribosomal pseudogene 18 [Source:HC |
| DUSP5P1 | CHR_HG2002_PATCH | dual specificity phosphatase 5 pseudogene 1 [ |
| AC275639.1 | CHR_HG2002_PATCH | NULL |
| FTH1P2 | CHR_HG2002_PATCH | ferritin heavy chain 1 pseudogene 2 [Source:H |
| RHOU | CHR_HG2002_PATCH | ras homolog family member U [Source:HGNC |
| LEUTX | CHR_HG2021_PATCH | leucine twenty homeobox [Source:HGNC Sym |
| AC005393.2 | CHR_HG2021_PATCH | novel transcript |
| DYRK1B | CHR_HG2021_PATCH | dual specificity tyrosine phosphorylation regu |
| MIR6719 | CHR_HG2021_PATCH | microRNA 6719 [Source:HGNC Symbol;Acc:HC |
| FBL | CHR_HG2021_PATCH | fibrillarin [Source:HGNC Symbol;Acc:HGNC:35 |
| FCGBP | CHR_HG2021_PATCH | Fc fragment of IgG binding protein [Source:HC |
| PRR13P5 | CHR_HG2021_PATCH | proline rich 13 pseudogene 5 [Source:HGNC S |
| PSMC4 | CHR_HG2021_PATCH | proteasome 26S subunit, ATPase 4 [Source:HC |
| ZNF546 | CHR_HG2021_PATCH | zinc finger protein 546 [Source:HGNC Symbol; |
| AC007842.2 | CHR_HG2021_PATCH | protein associated with topoisomerase II hom |
| ZNF780B | CHR_HG2021_PATCH | zinc finger protein 780B [Source:HGNC Symbo |
| AC005614.4 | CHR_HG2021_PATCH | novel transcript |
| ZNF780A | CHR_HG2021_PATCH | zinc finger protein 780A [Source:HGNC Symbo |
| AC005614.3 | CHR_HG2021_PATCH | novel transcript, antisense to ZNF780A |
| AL354926.1 | CHR_HG2022_PATCH | NULL |
| ABBA01000935.1 | CHR_HG2022_PATCH | primase, DNA, polypeptide 2 (58kDa) (PRIM2) |
| FRG1-DT | CHR_HG2023_PATCH | FRG1 divergent transcript [Source:HGNC Symb |
| FRG1 | CHR_HG2023_PATCH | FSHD region gene 1 [Source:HGNC Symbol;Acc |
| MLLT10P2 | CHR_HG2023_PATCH | MLLT10 pseudogene 2 [Source:HGNC Symbol; |
| AF146191.4 | CHR_HG2023_PATCH | family with sequence similarity 166, member |
| TUBB7P | CHR_HG2023_PATCH | tubulin beta 7 pseudogene [Source:HGNC Sym |
| RNA5SP174 | CHR_HG2023_PATCH | RNA, 5S ribosomal pseudogene 174 [Source:H |
| RNA5SP175 | CHR_HG2023_PATCH | RNA, 5S ribosomal pseudogene 175 [Source:H |
| DUX4L9 | CHR_HG2023_PATCH | double homeobox 4 like 9 (pseudogene) [Sour |
| FRG2 | CHR_HG2023_PATCH | FSHD region gene 2 [Source:HGNC Symbol;Acc |
| RARRES2P4 | CHR_HG2023_PATCH | retinoic acid receptor responder 2 pseudogen |
| DUX4L8 | CHR_HG2023_PATCH | double homeobox 4 like 8 (pseudogene) [Sour |
| CT476828.17 | CHR_HG2023_PATCH | NULL |
| CT476828.19 | CHR_HG2023_PATCH | NULL |
| CT476828.16 | CHR_HG2023_PATCH | NULL |
| CT476828.1 | CHR_HG2023_PATCH | NULL |
| CT476828.15 | CHR_HG2023_PATCH | NULL |
| CT476828.5 | CHR_HG2023_PATCH | NULL |
| CT476828.8 | CHR_HG2023_PATCH | NULL |
| CT476828.10 | CHR_HG2023_PATCH | NULL |

| | | |
|---|---|---|
| CT476828.4 | CHR_HG2023_PATCH | NULL |
| CT476828.14 | CHR_HG2023_PATCH | NULL |
| CT476828.2 | CHR_HG2023_PATCH | NULL |
| DUX4 | CHR_HG2023_PATCH | double homeobox 4 [Source:NCBI gene;Acc:1( |
| RPL23AP84 | CHR_HG2023_PATCH | ribosomal protein L23a pseudogene 84 [Sourc |
| AC215524.2 | CHR_HG2023_PATCH | major histocompatibility complex, class I, F (H |
| OBP2B | CHR_HG2030_PATCH | odorant binding protein 2B [Source:HGNC Syn |
| LCN1P1 | CHR_HG2030_PATCH | lipocalin 1 pseudogene 1 [Source:HGNC Symb |
| ABO | CHR_HG2030_PATCH | ABO, alpha 1-3-N-acetylgalactosaminyltransfe |
| Y_RNA | CHR_HG2030_PATCH | Y RNA [Source:RFAM;Acc:RF00019] |
| LCN1P2 | CHR_HG2030_PATCH | lipocalin 1 pseudogene 2 [Source:HGNC Symb |
| SURF6 | CHR_HG2030_PATCH | surfeit 6 [Source:HGNC Symbol;Acc:HGNC:114 |
| Y_RNA | CHR_HG2030_PATCH | Y RNA [Source:RFAM;Acc:RF00019] |
| MED22 | CHR_HG2030_PATCH | mediator complex subunit 22 [Source:HGNC S |
| AL772161.2 | CHR_HG2030_PATCH | ribosomal protein L21 (RPL21) pseudogene |
| RPL7A | CHR_HG2030_PATCH | ribosomal protein L7a [Source:HGNC Symbol;/ |
| SNORD24 | CHR_HG2030_PATCH | small nucleolar RNA, C/D box 24 [Source:HGN |
| SNORD36B | CHR_HG2030_PATCH | small nucleolar RNA, C/D box 36B [Source:HG |
| SNORD36A | CHR_HG2030_PATCH | small nucleolar RNA, C/D box 36A [Source:HG |
| AL593848.1 | CHR_HG2030_PATCH | NULL |
| SNORD36C | CHR_HG2030_PATCH | small nucleolar RNA, C/D box 36C [Source:HG |
| SURF1 | CHR_HG2030_PATCH | SURF1 cytochrome c oxidase assembly factor |
| SURF2 | CHR_HG2030_PATCH | surfeit 2 [Source:HGNC Symbol;Acc:HGNC:114 |
| SURF4 | CHR_HG2030_PATCH | surfeit 4 [Source:HGNC Symbol;Acc:HGNC:114 |
| STKLD1 | CHR_HG2030_PATCH | serine/threonine kinase like domain containin |
| REXO4 | CHR_HG2030_PATCH | REX4 homolog, 3'-5' exonuclease [Source:HGN |
| ADAMTS13 | CHR_HG2030_PATCH | ADAM metallopeptidase with thrombospondi |
| CACFD1 | CHR_HG2030_PATCH | calcium channel flower domain containing 1 [! |
| SLC2A6 | CHR_HG2030_PATCH | solute carrier family 2 member 6 [Source:HGN |
| TNK1 | CHR_HG2046_PATCH | tyrosine kinase non receptor 1 [Source:HGNC |
| PLSCR3 | CHR_HG2046_PATCH | phospholipid scramblase 3 [Source:HGNC Sym |
| TMEM256-PLSCR3 | CHR_HG2046_PATCH | TMEM256-PLSCR3 readthrough (NMD candida |
| TMEM256 | CHR_HG2046_PATCH | transmembrane protein 256 [Source:HGNC Sy |
| NLGN2 | CHR_HG2046_PATCH | neuroligin 2 [Source:HGNC Symbol;Acc:HGNC: |
| AC113189.5 | CHR_HG2046_PATCH | novel transcript |
| SPEM1 | CHR_HG2046_PATCH | spermatid maturation 1 [Source:HGNC Symbo |
| SPEM2 | CHR_HG2046_PATCH | SPEM family member 2 [Source:HGNC Symbo |
| SPEM3 | CHR_HG2046_PATCH | SPEM family member 3 [Source:HGNC Symbo |
| AC113189.10 | CHR_HG2046_PATCH | novel protein |
| TMEM102 | CHR_HG2046_PATCH | transmembrane protein 102 [Source:HGNC Sy |
| AC113189.6 | CHR_HG2046_PATCH | novel transcript, antisense to TMEM102 |
| FGF11 | CHR_HG2046_PATCH | fibroblast growth factor 11 [Source:HGNC Sym |
| AC113189.8 | CHR_HG2046_PATCH | novel transcript, antisense to FGF11 |
| AC113189.7 | CHR_HG2046_PATCH | novel transcript |
| CHRNB1 | CHR_HG2046_PATCH | cholinergic receptor nicotinic beta 1 subunit [! |
| ZBTB4 | CHR_HG2046_PATCH | zinc finger and BTB domain containing 4 [Sour |
| SLC35G6 | CHR_HG2046_PATCH | solute carrier family 35 member G6 [Source:H |
| POLR2A | CHR_HG2046_PATCH | RNA polymerase II subunit A [Source:HGNC Sy |
| RNU6-1004P | CHR_HG2047_PATCH | RNA, U6 small nuclear 1004, pseudogene [Sou |
| AC084018.3 | CHR_HG2047_PATCH | MORN repeat-containing protein 3 [Source:U |

| | | |
|---|---|---|
| GCNT2 | CHR_HG2057_PATCH | glucosaminyl (N-acetyl) transferase 2 (I blood |
| AL139039.4 | CHR_HG2057_PATCH | novel transcript, antisense to GCNT2 |
| AL139039.6 | CHR_HG2057_PATCH | glucosaminyl (N-acetyl) transferase 1, core 2 ( |
| AL139039.5 | CHR_HG2057_PATCH | ribosomal protein L21 (RPL21) pseudogene |
| GCNT2P1 | CHR_HG2057_PATCH | GCNT2 pseudogene 1 [Source:HGNC Symbol;A |
| AL358777.2 | CHR_HG2057_PATCH | novel transcript, antisense to C6orf52 |
| C6orf52 | CHR_HG2057_PATCH | chromosome 6 open reading frame 52 [Source |
| Y_RNA | CHR_HG2057_PATCH | Y RNA [Source:RFAM;Acc:RF00019] |
| PAK1IP1 | CHR_HG2057_PATCH | PAK1 interacting protein 1 [Source:HGNC Sym |
| TMEM14C | CHR_HG2057_PATCH | transmembrane protein 14C [Source:HGNC Sy |
| DPPA2P2 | CHR_HG2058_PATCH | developmental pluripotency associated 2 pseu |
| RPS6KA1 | CHR_HG2058_PATCH | ribosomal protein S6 kinase A1 [Source:HGNC |
| MIR1976 | CHR_HG2058_PATCH | microRNA 1976 [Source:HGNC Symbol;Acc:HG |
| RN7SL679P | CHR_HG2058_PATCH | RNA, 7SL, cytoplasmic 679, pseudogene [Sour |
| Y_RNA | CHR_HG2058_PATCH | Y RNA [Source:RFAM;Acc:RF00019] |
| RNU7-29P | CHR_HG2058_PATCH | RNA, U7 small nuclear 29 pseudogene [Source |
| AL627313.2 | CHR_HG2058_PATCH | chromosome 20 open reading frame 52 (C20o |
| AL627082.3 | CHR_HG2058_PATCH | NULL |
| AL627082.4 | CHR_HG2058_PATCH | chromosome 20 open reading frame 52 (C20o |
| AC136759.2 | CHR_HG2060_PATCH | glutamate receptor, metabotropic 5 (GRM5) p |
| AC130364.4 | CHR_HG2060_PATCH | novel pseudogene |
| AC130364.3 | CHR_HG2060_PATCH | tripartite motif-containing protein pseudogen |
| TRIM51FP | CHR_HG2060_PATCH | tripartite motif-containing 51F, pseudogene [S |
| CTBP2P1 | CHR_HG2062_PATCH | C-terminal binding protein 2 pseudogene 1 [So |
| MGAT4C | CHR_HG2063_PATCH | MGAT4 family member C [Source:HGNC Symb |
| AC010196.2 | CHR_HG2063_PATCH | novel transcript |
| AC079597.2 | CHR_HG2063_PATCH | novel transcript |
| ZNF445 | CHR_HG2066_PATCH | zinc finger protein 445 [Source:HGNC Symbol; |
| ZNF852 | CHR_HG2066_PATCH | zinc finger protein 852 [Source:HGNC Symbol; |
| EI24P3 | CHR_HG2066_PATCH | EI24 pseudogene 3 [Source:HGNC Symbol;Acc |
| AC099669.2 | CHR_HG2066_PATCH | zinc finger protein 347 (ZNF347) pseudogene |
| ZKSCAN7 | CHR_HG2066_PATCH | zinc finger with KRAB and SCAN domains 7 [Sc |
| ZKSCAN7-AS1 | CHR_HG2066_PATCH | ZKSCAN7 ZNF cluster antisense RNA 1 [Source |
| ZNF660 | CHR_HG2066_PATCH | zinc finger protein 660 [Source:HGNC Symbol; |
| MPRIPP1 | CHR_HG2066_PATCH | myosin phosphatase Rho interacting protein p |
| ZNF197 | CHR_HG2066_PATCH | zinc finger protein 197 [Source:HGNC Symbol; |
| ZNF197-AS1 | CHR_HG2066_PATCH | ZNF197 antisense RNA 1 [Source:HGNC Symbo |
| ZNF35 | CHR_HG2066_PATCH | zinc finger protein 35 [Source:HGNC Symbol;A |
| AC092046.1 | CHR_HG2066_PATCH | novel transcript |
| SOCS5P3 | CHR_HG2066_PATCH | suppressor of cytokine signaling 5 pseudogene |
| ZNF502 | CHR_HG2066_PATCH | zinc finger protein 502 [Source:HGNC Symbol; |
| ZNF501 | CHR_HG2066_PATCH | zinc finger protein 501 [Source:HGNC Symbol; |
| KIAA1143 | CHR_HG2066_PATCH | KIAA1143 [Source:HGNC Symbol;Acc:HGNC:29 |
| KIF15 | CHR_HG2066_PATCH | kinesin family member 15 [Source:HGNC Syml |
| MIR564 | CHR_HG2066_PATCH | microRNA 564 [Source:HGNC Symbol;Acc:HGN |
| TMEM42 | CHR_HG2066_PATCH | transmembrane protein 42 [Source:HGNC Syn |
| TGM4 | CHR_HG2066_PATCH | transglutaminase 4 [Source:HGNC Symbol;Acc |
| AC098649.2 | CHR_HG2066_PATCH | novel transcript |
| RNA5SP266 | CHR_HG2067_PATCH | RNA, 5S ribosomal pseudogene 266 [Source:H |
| LINC01606 | CHR_HG2067_PATCH | long intergenic non-protein coding RNA 1606 |

| | | |
|---|---|---|
| LINC02542 | CHR_HG2072_PATCH | long intergenic non-protein coding RNA 2542 |
| IBTK | CHR_HG2072_PATCH | inhibitor of Bruton tyrosine kinase [Source:HG |
| RNU6-130P | CHR_HG2072_PATCH | RNA, U6 small nuclear 130, pseudogene [Sour |
| AC209420.1 | CHR_HG2072_PATCH | single stranded DNA binding protein pseudoge |
| AL121977.3 | CHR_HG2072_PATCH | novel transcript, antisense to TPBG |
| TPBG | CHR_HG2072_PATCH | trophoblast glycoprotein [Source:HGNC Symb |
| CTDNEP1 | CHR_HG2087_PATCH | CTD nuclear envelope phosphatase 1 [Source: |
| AC003688.3 | CHR_HG2087_PATCH | novel protein |
| ELP5 | CHR_HG2087_PATCH | elongator acetyltransferase complex subunit ! |
| CLDN7 | CHR_HG2087_PATCH | claudin 7 [Source:HGNC Symbol;Acc:HGNC:20 |
| Y_RNA | CHR_HG2087_PATCH | Y RNA [Source:RFAM;Acc:RF00019] |
| SLC2A4 | CHR_HG2087_PATCH | solute carrier family 2 member 4 [Source:HGN |
| AC003688.4 | CHR_HG2087_PATCH | novel transcript, antisense to SLC2A4 |
| YBX2 | CHR_HG2087_PATCH | Y-box binding protein 2 [Source:HGNC Symbol |
| EIF5A | CHR_HG2087_PATCH | eukaryotic translation initiation factor 5A [Sou |
| GPS2 | CHR_HG2087_PATCH | G protein pathway suppressor 2 [Source:HGN |
| AC026954.6 | CHR_HG2087_PATCH | novel protein |
| NEURL4 | CHR_HG2087_PATCH | neuralized E3 ubiquitin protein ligase 4 [Sourc |
| AC026954.4 | CHR_HG2087_PATCH | novel protein phosphatase inhibitor 2 (IPP-2) |
| ACAP1 | CHR_HG2087_PATCH | ArfGAP with coiled-coil, ankyrin repeat and PH |
| KCTD11 | CHR_HG2087_PATCH | potassium channel tetramerization domain cc |
| AC026954.5 | CHR_HG2087_PATCH | novel transcript, antisense to KCTD11 |
| TMEM95 | CHR_HG2087_PATCH | transmembrane protein 95 [Source:HGNC Syn |
| SMURF1 | CHR_HG2088_PATCH | SMAD specific E3 ubiquitin protein ligase 1 [Sc |
| KPNA7 | CHR_HG2088_PATCH | karyopherin subunit alpha 7 [Source:HGNC Sy |
| LINC02783 | CHR_HG2095_PATCH | long intergenic non-protein coding RNA 2783 |
| AL590644.2 | CHR_HG2095_PATCH | novel transcript |
| PADI1 | CHR_HG2095_PATCH | peptidyl arginine deiminase 1 [Source:HGNC S |
| PADI3 | CHR_HG2095_PATCH | peptidyl arginine deiminase 3 [Source:HGNC S |
| MIR3972 | CHR_HG2095_PATCH | microRNA 3972 [Source:HGNC Symbol;Acc:HC |
| PADI4 | CHR_HG2095_PATCH | peptidyl arginine deiminase 4 [Source:HGNC S |
| PADI6 | CHR_HG2095_PATCH | peptidyl arginine deiminase 6 [Source:HGNC S |
| RCC2-AS1 | CHR_HG2095_PATCH | RCC2 antisense RNA 1 [Source:HGNC Symbol;/ |
| RCC2 | CHR_HG2095_PATCH | regulator of chromosome condensation 2 [Sou |
| SLC16A1 | CHR_HG2104_PATCH | solute carrier family 16 member 1 [Source:HG |
| AKR7A2P1 | CHR_HG2104_PATCH | aldo-keto reductase family 7 member A2 pseu |
| AL158844.1 | CHR_HG2104_PATCH | NULL |
| LRIG2-DT | CHR_HG2104_PATCH | LRIG2 divergent transcript [Source:HGNC Sym |
| ST13P5 | CHR_HG2111_PATCH | ST13, Hsp70 interacting protein pseudogene 5 |
| SAA1 | CHR_HG2111_PATCH | serum amyloid A1 [Source:HGNC Symbol;Acc: |
| RNA5SP334 | CHR_HG2111_PATCH | RNA, 5S ribosomal pseudogene 334 [Source:H |
| HPS5 | CHR_HG2111_PATCH | HPS5 biogenesis of lysosomal organelles comp |
| GTF2H1 | CHR_HG2111_PATCH | general transcription factor IIH subunit 1 [Sou |
| MIR3159 | CHR_HG2111_PATCH | microRNA 3159 [Source:HGNC Symbol;Acc:HC |
| LDHA | CHR_HG2111_PATCH | lactate dehydrogenase A [Source:HGNC Symb |
| AC084117.2 | CHR_HG2111_PATCH | novel transcript |
| NDUFS3 | CHR_HG2114_PATCH | NADH:ubiquinone oxidoreductase core subun |
| PTPMT1 | CHR_HG2114_PATCH | protein tyrosine phosphatase mitochondrial 1 |
| KBTBD4 | CHR_HG2114_PATCH | kelch repeat and BTB domain containing 4 [So |
| RNU5E-10P | CHR_HG2114_PATCH | RNA, U5E small nuclear 10, pseudogene [Sour |

| | | |
|---|---|---|
| KF459542.2 | CHR_HG2114_PATCH | novel transcript, antisense to KBTBD4 |
| FAM180B | CHR_HG2114_PATCH | family with sequence similarity 180 member E |
| C1QTNF4 | CHR_HG2114_PATCH | C1q and TNF related 4 [Source:HGNC Symbol;. |
| MTCH2 | CHR_HG2114_PATCH | mitochondrial carrier 2 [Source:HGNC Symbol |
| AGBL2 | CHR_HG2114_PATCH | ATP/GTP binding protein like 2 [Source:HGNC |
| FNBP4 | CHR_HG2114_PATCH | formin binding protein 4 [Source:HGNC Symb |
| Y_RNA | CHR_HG2114_PATCH | Y RNA [Source:RFAM;Acc:RF00019] |
| AC021443.2 | CHR_HG2114_PATCH | ribosomal protein L6 (RPL6) pseudogene |
| RNA5SP340 | CHR_HG2114_PATCH | RNA, 5S ribosomal pseudogene 340 [Source:H |
| Y_RNA | CHR_HG2114_PATCH | Y RNA [Source:RFAM;Acc:RF00019] |
| YPEL5P2 | CHR_HG2114_PATCH | YPEL5 pseudogene 2 [Source:HGNC Symbol;A |
| AP002336.5 | CHR_HG2115_PATCH | NULL |
| H2AFZP4 | CHR_HG2115_PATCH | H2A histone family member Z pseudogene 4 [ |
| AP002336.7 | CHR_HG2115_PATCH | novel transcript |
| MIR548K | CHR_HG2115_PATCH | microRNA 548k [Source:HGNC Symbol;Acc:HG |
| AP002336.6 | CHR_HG2115_PATCH | novel transcript |
| AP002336.4 | CHR_HG2115_PATCH | novel transcript |
| AP000487.4 | CHR_HG2115_PATCH | novel transcript |
| AP000487.3 | CHR_HG2115_PATCH | novel transcript |
| CTTN | CHR_HG2115_PATCH | cortactin [Source:HGNC Symbol;Acc:HGNC:33 |
| NDUFB11P1 | CHR_HG2116_PATCH | NADH:ubiquinone oxidoreductase subunit B1 |
| PGAM1P9 | CHR_HG2116_PATCH | phosphoglycerate mutase 1 pseudogene 9 [So |
| MDN1 | CHR_HG2121_PATCH | midasin AAA ATPase 1 [Source:HGNC Symbol; |
| AL353692.3 | CHR_HG2121_PATCH | family with sequence similarity 64, member A |
| CASP8AP2 | CHR_HG2121_PATCH | caspase 8 associated protein 2 [Source:HGNC |
| Y_RNA | CHR_HG2121_PATCH | Y RNA [Source:RFAM;Acc:RF00019] |
| RPL22P14 | CHR_HG2121_PATCH | ribosomal protein L22 pseudogene 14 [Source |
| AL353692.4 | CHR_HG2121_PATCH | G protein pathway suppressor 1 (GPS1) pseud |
| GJA10 | CHR_HG2121_PATCH | gap junction protein alpha 10 [Source:HGNC S |
| Y_RNA | CHR_HG2121_PATCH | Y RNA [Source:RFAM;Acc:RF00019] |
| RNU6-280P | CHR_HG2128_PATCH | RNA, U6 small nuclear 280, pseudogene [Sour |
| Y_RNA | CHR_HG2128_PATCH | Y RNA [Source:RFAM;Acc:RF00019] |
| ABBA01026962.1 | CHR_HG2133_PATCH | WD repeat domain 82 (WDR82) pseudogene |
| WDR82P1 | CHR_HG2133_PATCH | WD repeat domain 82 pseudogene 1 [Source: |
| LINC00879 | CHR_HG2133_PATCH | long intergenic non-protein coding RNA 879 [S |
| AL360225.1 | CHR_HG2191_PATCH | novel transcript |
| KAT6B | CHR_HG2191_PATCH | lysine acetyltransferase 6B [Source:HGNC Sym |
| AC018511.4 | CHR_HG2191_PATCH | novel transcript |
| AC018511.3 | CHR_HG2191_PATCH | novel transcript |
| DUPD1 | CHR_HG2191_PATCH | dual specificity phosphatase and pro isomeras |
| AC018511.6 | CHR_HG2191_PATCH | novel transcript, antisense to DUPD1 |
| AC010931.4 | CHR_HG2198_PATCH | novel transcript, antisense to ISLR |
| ISLR | CHR_HG2198_PATCH | immunoglobulin superfamily containing leucir |
| STRA6 | CHR_HG2198_PATCH | stimulated by retinoic acid 6 [Source:HGNC Sy |
| CCDC33 | CHR_HG2198_PATCH | coiled-coil domain containing 33 [Source:HGN |
| AC023300.5 | CHR_HG2198_PATCH | TEC |
| AC023300.6 | CHR_HG2198_PATCH | novel transcript |
| AC023300.4 | CHR_HG2198_PATCH | novel transcript, antisense to CCDC33 |
| CYP11A1 | CHR_HG2198_PATCH | cytochrome P450 family 11 subfamily A meml |
| PPIAP46 | CHR_HG2198_PATCH | peptidylprolyl isomerase A pseudogene 46 [So |

| | | |
|---|---|---|
| AC090826.4 | CHR_HG2198_PATCH | novel transcript, antisense to CYP11A1 |
| AC090826.5 | CHR_HG2198_PATCH | novel transcript |
| LINC02255 | CHR_HG2198_PATCH | long intergenic non-protein coding RNA 2255 |
| SEMA7A | CHR_HG2198_PATCH | semaphorin 7A (John Milton Hagen blood grou |
| MIR6881 | CHR_HG2198_PATCH | microRNA 6881 [Source:HGNC Symbol;Acc:HG |
| AC090826.6 | CHR_HG2198_PATCH | novel transcript, antisense to SEMA7A |
| UBL7 | CHR_HG2198_PATCH | ubiquitin like 7 [Source:HGNC Symbol;Acc:HG |
| UBL7-AS1 | CHR_HG2198_PATCH | UBL7 antisense RNA 1 (head to head) [Source: |
| AC012435.4 | CHR_HG2198_PATCH | coiled-coil-helix-coiled-coil-helix domain-conta |
| AC012435.5 | CHR_HG2198_PATCH | novel transcript |
| AC012435.6 | CHR_HG2198_PATCH | NULL |
| CTIF | CHR_HG2213_PATCH | cap binding complex dependent translation in |
| AC048380.4 | CHR_HG2213_PATCH | TEC |
| MIR4743 | CHR_HG2213_PATCH | microRNA 4743 [Source:HGNC Symbol;Acc:HG |
| AC022919.2 | CHR_HG2213_PATCH | novel transcript, antisense to CTIF |
| AC022919.3 | CHR_HG2213_PATCH | TEC |
| AC093567.2 | CHR_HG2213_PATCH | novel transcript, antisense to CTIF |
| FOXR1 | CHR_HG2217_PATCH | forkhead box R1 [Source:HGNC Symbol;Acc:H |
| Metazoa_SRP | CHR_HG2217_PATCH | Metazoan signal recognition particle RNA [Sou |
| Y_RNA | CHR_HG2217_PATCH | Y RNA [Source:RFAM;Acc:RF00019] |
| CCDC84-DT | CHR_HG2217_PATCH | CCDC84 divergent transcript [Source:HGNC Sy |
| CCDC84 | CHR_HG2217_PATCH | coiled-coil domain containing 84 [Source:HGN |
| RPL23AP64 | CHR_HG2217_PATCH | ribosomal protein L23a pseudogene 64 [Sourc |
| AP003392.6 | CHR_HG2217_PATCH | novel transcript, antisense to CCDC84 |
| AP003392.9 | CHR_HG2217_PATCH | potassium channel tetramerisation domain cc |
| RPS25 | CHR_HG2217_PATCH | ribosomal protein S25 [Source:HGNC Symbol; |
| TRAPPC4 | CHR_HG2217_PATCH | trafficking protein particle complex 4 [Source: |
| AP003392.12 | CHR_HG2217_PATCH | NULL |
| SLC37A4 | CHR_HG2217_PATCH | solute carrier family 37 member 4 [Source:HG |
| AP003392.10 | CHR_HG2217_PATCH | novel transcript, antisense to HYOU1 |
| HYOU1 | CHR_HG2217_PATCH | hypoxia up-regulated 1 [Source:HGNC Symbol |
| AP003392.8 | CHR_HG2217_PATCH | novel transcript |
| AP003392.7 | CHR_HG2217_PATCH | novel transcript, antisense to VPS11 |
| VPS11 | CHR_HG2217_PATCH | VPS11 core subunit of CORVET and HOPS com |
| INPP5D | CHR_HG2232_PATCH | inositol polyphosphate-5-phosphatase D [Sou |
| RN7SL32P | CHR_HG2232_PATCH | RNA, 7SL, cytoplasmic 32, pseudogene [Sourc |
| ATG16L1 | CHR_HG2232_PATCH | autophagy related 16 like 1 [Source:HGNC Syr |
| SCARNA5 | CHR_HG2232_PATCH | small Cajal body-specific RNA 5 [Source:HGNC |
| SCARNA6 | CHR_HG2232_PATCH | small Cajal body-specific RNA 6 [Source:HGNC |
| SAG | CHR_HG2232_PATCH | S-antigen visual arrestin [Source:HGNC Symbc |
| AC013726.2 | CHR_HG2232_PATCH | novel transcript, antisense to DGKD |
| DGKD | CHR_HG2232_PATCH | diacylglycerol kinase delta [Source:HGNC Sym |
| Y_RNA | CHR_HG2232_PATCH | Y RNA [Source:RFAM;Acc:RF00019] |
| AC093802.3 | CHR_HG2233_PATCH | novel transcript |
| AC093802.4 | CHR_HG2233_PATCH | novel transcript |
| MAGI1 | CHR_HG2235_PATCH | membrane associated guanylate kinase, WW |
| SLC25A26 | CHR_HG2235_PATCH | solute carrier family 25 member 26 [Source:H |
| RNU6-787P | CHR_HG2235_PATCH | RNA, U6 small nuclear 787, pseudogene [Sour |
| RN7SL482P | CHR_HG2235_PATCH | RNA, 7SL, cytoplasmic 482, pseudogene [Sour |
| AC091493.2 | CHR_HG2236_PATCH | novel transcript |

| | | |
|---|---|---|
| CDYLP1 | CHR_HG2236_PATCH | CDYL pseudogene 1 [Source:HGNC Symbol;Ac |
| PLCL2 | CHR_HG2236_PATCH | phospholipase C like 2 [Source:HGNC Symbol; |
| MIR3714 | CHR_HG2236_PATCH | microRNA 3714 [Source:HGNC Symbol;Acc:HG |
| PLCL2-AS1 | CHR_HG2236_PATCH | PLCL2 antisense RNA 1 [Source:HGNC Symbol, |
| AC024730.2 | CHR_HG2239_PATCH | novel transcript, antisense to DPP6 |
| EP400P1 | CHR_HG2246_HG2248_HG22 | EP400 pseudogene 1 [Source:HGNC Symbol;A |
| AC138466.10 | CHR_HG2246_HG2248_HG22 | novel transcript |
| AC138466.9 | CHR_HG2246_HG2248_HG22 | placenta-specific 8 (PLAC8) pseudogene |
| DDX51 | CHR_HG2246_HG2248_HG22 | DEAD-box helicase 51 [Source:HGNC Symbol;A |
| NOC4L | CHR_HG2246_HG2248_HG22 | nucleolar complex associated 4 homolog [Sou |
| AC138466.7 | CHR_HG2246_HG2248_HG22 | novel transcript |
| LINC02361 | CHR_HG2246_HG2248_HG22 | long intergenic non-protein coding RNA 2361 |
| AC138466.6 | CHR_HG2246_HG2248_HG22 | novel transcript |
| GALNT9 | CHR_HG2246_HG2248_HG22 | polypeptide N-acetylgalactosaminyltransferas |
| AC138466.8 | CHR_HG2246_HG2248_HG22 | novel transcript |
| AC148477.12 | CHR_HG2246_HG2248_HG22 | novel transcript |
| AC148477.19 | CHR_HG2246_HG2248_HG22 | novel transcript |
| AC148477.13 | CHR_HG2246_HG2248_HG22 | novel transcript |
| AC148477.16 | CHR_HG2246_HG2248_HG22 | novel transcript |
| AC148477.18 | CHR_HG2246_HG2248_HG22 | novel transcript |
| AC148477.15 | CHR_HG2246_HG2248_HG22 | novel transcript |
| AC148477.17 | CHR_HG2246_HG2248_HG22 | novel transcript |
| AC148477.20 | CHR_HG2246_HG2248_HG22 | novel transcript |
| AC148477.14 | CHR_HG2246_HG2248_HG22 | novel transcript |
| AC148477.11 | CHR_HG2246_HG2248_HG22 | novel transcript |
| BCL7A | CHR_HG2247_PATCH | BAF chromatin remodeling complex subunit B |
| AC156455.2 | CHR_HG2247_PATCH | novel transcript |
| MLXIP | CHR_HG2247_PATCH | MLX interacting protein [Source:HGNC Symbo |
| NALCN-AS1 | CHR_HG2249_PATCH | NALCN antisense RNA 1 [Source:HGNC Symbo |
| LINC00411 | CHR_HG2249_PATCH | long intergenic non-protein coding RNA 411 [S |
| AC109446.8 | CHR_HG2263_PATCH | novel transcript |
| AC109446.7 | CHR_HG2263_PATCH | transcription elongation regulator 1 (TCERG1) |
| AC109446.6 | CHR_HG2263_PATCH | LYR motif containing 2 (LYRM2) pseudogene |
| AC099494.7 | CHR_HG2263_PATCH | Xylosyltransferase 1 [Source:UniProtKB/Swiss |
| AC109446.5 | CHR_HG2263_PATCH | novel transcript, antisense to XYLT1 |
| AC099494.4 | CHR_HG2263_PATCH | novel transcript |
| AC099494.5 | CHR_HG2263_PATCH | novel transcript |
| AC099494.6 | CHR_HG2263_PATCH | novel transcript |
| AC009152.5 | CHR_HG2263_PATCH | novel transcript, antisense to XYLT1 |
| AC009152.2 | CHR_HG2263_PATCH | novel transcript, sense intronic to XYLT1 |
| AC009152.4 | CHR_HG2263_PATCH | novel transcript |
| PRKAR2B | CHR_HG2266_PATCH | protein kinase cAMP-dependent type II regula |
| HBP1 | CHR_HG2266_PATCH | HMG-box transcription factor 1 [Source:HGNC |
| AC004492.2 | CHR_HG2266_PATCH | novel transcript, antisense to HBP1 |
| COG5 | CHR_HG2266_PATCH | component of oligomeric golgi complex 5 [Sou |
| AC255406.1 | CHR_HG2266_PATCH | ribosomal protein L37a (RPL37A) pseudogene |
| GPR22 | CHR_HG2266_PATCH | G protein-coupled receptor 22 [Source:HGNC |
| DUS4L | CHR_HG2266_PATCH | dihydrouridine synthase 4 like [Source:HGNC S |
| AC004839.5 | CHR_HG2266_PATCH | novel transcript, antisense to BCAP29 |
| AC004839.3 | CHR_HG2266_PATCH | B cell receptor associated protein 29 [Source:I |

| | | |
|---|---|---|
| AC004839.4 | CHR_HG2266_PATCH | NULL |
| AC141424.2 | CHR_HG2285_HG106_HG225 | novel transcript, antisense to C17orf97 |
| C17orf97 | CHR_HG2285_HG106_HG225 | chromosome 17 open reading frame 97 [Sour |
| RFLNB | CHR_HG2285_HG106_HG225 | refilin B [Source:HGNC Symbol;Acc:HGNC:287 |
| AC015853.5 | CHR_HG2285_HG106_HG225 | novel transcript, antisense to VPS53 |
| VPS53 | CHR_HG2285_HG106_HG225 | VPS53 subunit of GARP complex [Source:HGN |
| AC015853.4 | CHR_HG2285_HG106_HG225 | novel transcript, antisense to VPS53 |
| AC015853.6 | CHR_HG2285_HG106_HG225 | novel transcript, antisense to VPS53 |
| AC027455.2 | CHR_HG2285_HG106_HG225 | BCL2-associated athanogene (BAG1) pseudog |
| RPS4XP17 | CHR_HG2285_HG106_HG225 | ribosomal protein S4X pseudogene 17 [Source |
| C13orf46 | CHR_HG2288_HG2289_PATC | chromosome 13 open reading frame 46 [Sour |
| RASA3 | CHR_HG2288_HG2289_PATC | RAS p21 protein activator 3 [Source:HGNC Syr |
| RASA3-IT1 | CHR_HG2288_HG2289_PATC | RASA3 intronic transcript 1 [Source:HGNC Sym |
| IGKV5-2 | CHR_HG2290_PATCH | immunoglobulin kappa variable 5-2 [Source:H |
| IGKV7-3 | CHR_HG2290_PATCH | immunoglobulin kappa variable 7-3 (pseudoge |
| PGBD4P5 | CHR_HG2290_PATCH | piggyBac transposable element derived 4 pseu |
| IGKV2-4 | CHR_HG2290_PATCH | immunoglobulin kappa variable 2-4 (pseudoge |
| IGKV1-5 | CHR_HG2290_PATCH | immunoglobulin kappa variable 1-5 [Source:H |
| IGKV1-6 | CHR_HG2290_PATCH | immunoglobulin kappa variable 1-6 [Source:H |
| IGKV3-7 | CHR_HG2290_PATCH | immunoglobulin kappa variable 3-7 (non-func |
| IGKV1-8 | CHR_HG2290_PATCH | immunoglobulin kappa variable 1-8 [Source:H |
| IGKV1-9 | CHR_HG2290_PATCH | immunoglobulin kappa variable 1-9 [Source:H |
| IGKV2-10 | CHR_HG2290_PATCH | immunoglobulin kappa variable 2-10 (pseudog |
| IGKV3-11 | CHR_HG2290_PATCH | immunoglobulin kappa variable 3-11 [Source: |
| IGKV1-12 | CHR_HG2290_PATCH | immunoglobulin kappa variable 1-12 [Source: |
| IGKV1-13 | CHR_HG2290_PATCH | immunoglobulin kappa variable 1-13 (gene/ps |
| IGKV2-14 | CHR_HG2290_PATCH | immunoglobulin kappa variable 2-14 (pseudog |
| IGKV3-15 | CHR_HG2290_PATCH | immunoglobulin kappa variable 3-15 [Source: |
| IGKV1-16 | CHR_HG2290_PATCH | immunoglobulin kappa variable 1-16 [Source: |
| IGKV1-17 | CHR_HG2290_PATCH | immunoglobulin kappa variable 1-17 [Source: |
| IGKV2-18 | CHR_HG2290_PATCH | immunoglobulin kappa variable 2-18 (pseudog |
| IGKV2-19 | CHR_HG2290_PATCH | immunoglobulin kappa variable 2-19 (pseudog |
| IGKV3-20 | CHR_HG2290_PATCH | immunoglobulin kappa variable 3-20 [Source: |
| IGKV6-21 | CHR_HG2290_PATCH | immunoglobulin kappa variable 6-21 (non-fun |
| IGKV1-22 | CHR_HG2290_PATCH | immunoglobulin kappa variable 1-22 (pseudog |
| IGKV2-23 | CHR_HG2290_PATCH | immunoglobulin kappa variable 2-23 (pseudog |
| IGKV2-24 | CHR_HG2290_PATCH | immunoglobulin kappa variable 2-24 [Source: |
| IGKV3-25 | CHR_HG2290_PATCH | immunoglobulin kappa variable 3-25 (pseudog |
| IGKV2-26 | CHR_HG2290_PATCH | immunoglobulin kappa variable 2-26 (pseudog |
| AC244255.2 | CHR_HG2290_PATCH | Ig kappa chain V-I region Lay pseudogene |
| IGKV1-27 | CHR_HG2290_PATCH | immunoglobulin kappa variable 1-27 [Source: |
| IGKV2-28 | CHR_HG2290_PATCH | immunoglobulin kappa variable 2-28 [Source: |
| IGKV2-29 | CHR_HG2290_PATCH | immunoglobulin kappa variable 2-29 (gene/ps |
| IGKV2-30 | CHR_HG2290_PATCH | immunoglobulin kappa variable 2-30 [Source: |
| IGKV3-31 | CHR_HG2290_PATCH | immunoglobulin kappa variable 3-31 (pseudog |
| IGKV1-32 | CHR_HG2290_PATCH | immunoglobulin kappa variable 1-32 (pseudog |
| IGKV1-33 | CHR_HG2290_PATCH | immunoglobulin kappa variable 1-33 [Source: |
| IGKV3-34 | CHR_HG2290_PATCH | immunoglobulin kappa variable 3-34 (pseudog |
| IGKV1-35 | CHR_HG2290_PATCH | immunoglobulin kappa variable 1-35 (pseudog |
| IGKV2-36 | CHR_HG2290_PATCH | immunoglobulin kappa variable 2-36 (pseudog |

| | | |
|---|---|---|
| IGKV1-37 | CHR_HG2290_PATCH | immunoglobulin kappa variable 1-37 (non-fun |
| IGKV2-38 | CHR_HG2290_PATCH | immunoglobulin kappa variable 2-38 (pseudog |
| IGKV1-39 | CHR_HG2290_PATCH | immunoglobulin kappa variable 1-39 (gene/ps |
| IGKV2-40 | CHR_HG2290_PATCH | immunoglobulin kappa variable 2-40 [Source:I |
| AL356585.7 | CHR_HG2291_PATCH | novel ankyrin repeat domain-containing prote |
| FAM230C | CHR_HG2291_PATCH | family with sequence similarity 230 member C |
| AL356585.6 | CHR_HG2291_PATCH | eukaryotic translation initiation factor 3 subur |
| AL356585.9 | CHR_HG2291_PATCH | NULL |
| AL356585.11 | CHR_HG2291_PATCH | gamma-glutamyltransferase 5 (GGT5) pseudog |
| GGT4P | CHR_HG2291_PATCH | gamma-glutamyltransferase 4 pseudogene [So |
| AL356585.5 | CHR_HG2291_PATCH | immunoglobulin superfamily, member 3 (IGSF |
| KMT5AP1 | CHR_HG2291_PATCH | KMT5A pseudogene 1 [Source:HGNC Symbol;A |
| AL356585.8 | CHR_HG2291_PATCH | lysine (K)-specific methyltransferase 2C (KMT2 |
| ATAD1 | CHR_HG2334_PATCH | ATPase family AAA domain containing 1 [Sour |
| CFL1P1 | CHR_HG2334_PATCH | cofilin 1 pseudogene 1 [Source:HGNC Symbol; |
| RN7SL78P | CHR_HG2334_PATCH | RNA, 7SL, cytoplasmic 78, pseudogene [Sourc |
| KLLN | CHR_HG2334_PATCH | killin, p53 regulated DNA replication inhibitor |
| PTEN | CHR_HG2334_PATCH | phosphatase and tensin homolog [Source:HGI |
| AC022016.1 | CHR_HG2334_PATCH | novel transcript, sense intronic to PTEN |
| RPL11P3 | CHR_HG2334_PATCH | ribosomal protein L11 pseudogene 3 [Source:I |
| AC063965.3 | CHR_HG2334_PATCH | NULL |
| MED6P1 | CHR_HG2334_PATCH | mediator complex subunit 6 pseudogene 1 [So |
| AC138701.3 | CHR_HG2365_PATCH | ankyrin repeat domain 30B (ANKRD30B) pseu |
| RNU6-978P | CHR_HG2365_PATCH | RNA, U6 small nuclear 978, pseudogene [Sour |
| AC138701.4 | CHR_HG2365_PATCH | connector enhancer of kinase suppressor of R |
| AC127381.3 | CHR_HG2365_PATCH | Ig heavy chain V-II pseudogene |
| AC275546.3 | CHR_HG2365_PATCH | NULL |
| AC275546.4 | CHR_HG2365_PATCH | NULL |
| AC275546.1 | CHR_HG2365_PATCH | NULL |
| AC275546.2 | CHR_HG2365_PATCH | NULL |
| AC270129.2 | CHR_HG2365_PATCH | NULL |
| AC270129.3 | CHR_HG2365_PATCH | NULL |
| AC270129.1 | CHR_HG2365_PATCH | NULL |
| AC270129.4 | CHR_HG2365_PATCH | NULL |
| AC275254.1 | CHR_HG2365_PATCH | NULL |
| AC256223.3 | CHR_HG2365_PATCH | NULL |
| AC256223.4 | CHR_HG2365_PATCH | NULL |
| AC256223.1 | CHR_HG2365_PATCH | NULL |
| AC256223.2 | CHR_HG2365_PATCH | NULL |
| AC275673.1 | CHR_HG2365_PATCH | NULL |
| AC255461.1 | CHR_HG2365_PATCH | NULL |
| AC255461.2 | CHR_HG2365_PATCH | NULL |
| AC270156.1 | CHR_HG2365_PATCH | NULL |
| AC270156.2 | CHR_HG2365_PATCH | NULL |
| AC270156.3 | CHR_HG2365_PATCH | NULL |
| AC275542.1 | CHR_HG2365_PATCH | NULL |
| AC256230.2 | CHR_HG2365_PATCH | NULL |
| AC256230.1 | CHR_HG2365_PATCH | NULL |
| AC275572.1 | CHR_HG2365_PATCH | NULL |
| AC270149.3 | CHR_HG2365_PATCH | NULL |

| | | |
|---|---|---|
| AC270149.2 | CHR_HG2365_PATCH | NULL |
| AC270149.1 | CHR_HG2365_PATCH | NULL |
| AC256234.2 | CHR_HG2365_PATCH | NULL |
| AC256234.1 | CHR_HG2365_PATCH | NULL |
| AC255360.1 | CHR_HG2365_PATCH | NULL |
| AC269752.2 | CHR_HG2365_PATCH | NULL |
| AC269752.1 | CHR_HG2365_PATCH | NULL |
| AC270091.2 | CHR_HG2365_PATCH | NULL |
| AC270091.1 | CHR_HG2365_PATCH | NULL |
| AC278124.1 | CHR_HG2365_PATCH | NULL |
| AC275614.1 | CHR_HG2365_PATCH | NULL |
| AC256231.1 | CHR_HG2365_PATCH | NULL |
| AC256231.2 | CHR_HG2365_PATCH | NULL |
| AC256231.3 | CHR_HG2365_PATCH | NULL |
| AC256221.1 | CHR_HG2365_PATCH | NULL |
| AC256221.2 | CHR_HG2365_PATCH | NULL |
| AC270239.2 | CHR_HG2365_PATCH | NULL |
| AC270239.1 | CHR_HG2365_PATCH | NULL |
| AC270126.1 | CHR_HG2365_PATCH | NULL |
| AC270230.1 | CHR_HG2365_PATCH | NULL |
| AC270170.1 | CHR_HG2365_PATCH | NULL |
| AC270313.1 | CHR_HG2365_PATCH | NULL |
| AC270313.2 | CHR_HG2365_PATCH | NULL |
| AC275284.3 | CHR_HG2365_PATCH | NULL |
| AC275284.2 | CHR_HG2365_PATCH | NULL |
| AC275284.4 | CHR_HG2365_PATCH | NULL |
| AC275284.1 | CHR_HG2365_PATCH | NULL |
| AC270153.2 | CHR_HG2365_PATCH | NULL |
| AC270153.1 | CHR_HG2365_PATCH | NULL |
| AC270095.1 | CHR_HG2365_PATCH | NULL |
| AC270095.2 | CHR_HG2365_PATCH | NULL |
| AC087463.4 | CHR_HG2365_PATCH | novel transcript |
| SLC14A2-AS1 | CHR_HG2412_PATCH | SLC14A2 antisense RNA 1 [Source:HGNC Symb |
| AC091151.2 | CHR_HG2412_PATCH | uncharacterized LOC112268408 [Source:NCBI |
| AC021517.3 | CHR_HG2412_PATCH | novel transcript |
| AC091151.3 | CHR_HG2412_PATCH | novel transcript |
| AC091151.4 | CHR_HG2412_PATCH | novel transcript, antisense to SLC14A2 |
| AC023421.3 | CHR_HG2412_PATCH | novel transcript, antisense to SLC14A1 and SL |
| BOP1 | CHR_HG2419_PATCH | BOP1 ribosomal biogenesis factor [Source:HG |
| HSF1 | CHR_HG2419_PATCH | heat shock transcription factor 1 [Source:HGN |
| DGAT1 | CHR_HG2419_PATCH | diacylglycerol O-acyltransferase 1 [Source:HG |
| AC233992.4 | CHR_HG2419_PATCH | novel transcript |
| MIR6848 | CHR_HG2419_PATCH | microRNA 6848 [Source:HGNC Symbol;Acc:HG |
| SCRT1 | CHR_HG2419_PATCH | scratch family transcriptional repressor 1 [Sou |
| TMEM249 | CHR_HG2419_PATCH | transmembrane protein 249 [Source:HGNC Sy |
| AC233992.6 | CHR_HG2419_PATCH | novel transcript |
| SLC52A2 | CHR_HG2419_PATCH | solute carrier family 52 member 2 [Source:HG |
| FBXL6 | CHR_HG2419_PATCH | F-box and leucine rich repeat protein 6 [Sourc |
| ADCK5 | CHR_HG2419_PATCH | aarF domain containing kinase 5 [Source:HGN |
| CPSF1 | CHR_HG2419_PATCH | cleavage and polyadenylation specific factor 1 |

| | | |
|---|---|---|
| MIR939 | CHR_HG2419_PATCH | microRNA 939 [Source:HGNC Symbol;Acc:HGN |
| MIR1234 | CHR_HG2419_PATCH | microRNA 1234 [Source:HGNC Symbol;Acc:HG |
| MIR6849 | CHR_HG2419_PATCH | microRNA 6849 [Source:HGNC Symbol;Acc:HG |
| AC233992.5 | CHR_HG2419_PATCH | novel transcript, antisense to SLC39A4 |
| SLC39A4 | CHR_HG2419_PATCH | solute carrier family 39 member 4 [Source:HG |
| VPS28 | CHR_HG2419_PATCH | VPS28 subunit of ESCRT-I [Source:HGNC Symb |
| RPS2P6 | CHR_HG2442_PATCH | ribosomal protein S2 pseudogene 6 [Source:H |
| AC091305.2 | CHR_HG2442_PATCH | TEC |
| AC091646.2 | CHR_HG2442_PATCH | novel transcript |
| TNRC6A | CHR_HG2471_PATCH | trinucleotide repeat containing adaptor 6A [So |
| AC008731.2 | CHR_HG2471_PATCH | novel transcript, antisense to TNRC6A |
| SLC5A11 | CHR_HG2471_PATCH | solute carrier family 5 member 11 [Source:HG |
| ARHGAP17 | CHR_HG2471_PATCH | Rho GTPase activating protein 17 [Source:HGN |
| AC140725.3 | CHR_HG2499_PATCH | olfactory receptor pseudogene |
| OR4F4 | CHR_HG2499_PATCH | olfactory receptor family 4 subfamily F memb |
| OR4G2P | CHR_HG2499_PATCH | olfactory receptor family 4 subfamily G memb |
| OR4G6P | CHR_HG2499_PATCH | olfactory receptor family 4 subfamily G memb |
| FAM138E | CHR_HG2499_PATCH | family with sequence similarity 138 member E |
| AC140725.4 | CHR_HG2499_PATCH | novel transcript |
| MIR1302-10 | CHR_HG2499_PATCH | microRNA 1302-10 [Source:HGNC Symbol;Acc |
| WASH3P | CHR_HG2499_PATCH | WASP family homolog 3, pseudogene [Source |
| MIR6859-3 | CHR_HG2499_PATCH | microRNA 6859-3 [Source:HGNC Symbol;Acc:H |
| DDX11L9 | CHR_HG2499_PATCH | DEAD/H-box helicase 11 like 9 [Source:HGNC S |
| AC207582.2 | CHR_HG2499_PATCH | NULL |
| AC207582.1 | CHR_HG2499_PATCH | NULL |
| MG910335.2 | CHR_HG2509_PATCH | NULL |
| MG910335.1 | CHR_HG2509_PATCH | NULL |
| MG910335.3 | CHR_HG2509_PATCH | NULL |
| MG910336.3 | CHR_HG2510_PATCH | NULL |
| MG910336.2 | CHR_HG2510_PATCH | NULL |
| MG910336.1 | CHR_HG2510_PATCH | NULL |
| MG910337.1 | CHR_HG2511_PATCH | NULL |
| MG910337.3 | CHR_HG2511_PATCH | NULL |
| MG910337.2 | CHR_HG2511_PATCH | NULL |
| MG910339.2 | CHR_HG2512_PATCH | NULL |
| MG910339.1 | CHR_HG2512_PATCH | NULL |
| MG910338.2 | CHR_HG2513_PATCH | NULL |
| MG910338.1 | CHR_HG2513_PATCH | NULL |
| MG910338.3 | CHR_HG2513_PATCH | NULL |
| MIR6724-3 | CHR_HG2513_PATCH | microRNA 6724-3 [Source:HGNC Symbol;Acc:H |
| FP236383.15 | CHR_HG2513_PATCH | NULL |
| FP236383.22 | CHR_HG2513_PATCH | novel transcript, similar to YY1 associated myo |
| RNA5-8SN3 | CHR_HG2513_PATCH | RNA, 5.8S ribosomal N3 [Source:HGNC Symbo |
| FP236383.24 | CHR_HG2513_PATCH | TEC |
| FP236383.21 | CHR_HG2513_PATCH | TEC |
| FP236383.16 | CHR_HG2513_PATCH | TEC |
| FP236383.23 | CHR_HG2513_PATCH | TEC |
| MIR6724-4 | CHR_HG2513_PATCH | microRNA 6724-4 [Source:HGNC Symbol;Acc:H |
| FP236383.19 | CHR_HG2513_PATCH | NULL |
| FP236383.17 | CHR_HG2513_PATCH | novel transcript, similar to YY1 associated myo |

| | | |
|---|---|---|
| RNA5-8SN1 | CHR_HG2513_PATCH | RNA, 5.8S ribosomal N1 [Source:HGNC Symbo |
| FP236383.14 | CHR_HG2513_PATCH | TEC |
| FP236383.20 | CHR_HG2513_PATCH | TEC |
| FP236383.25 | CHR_HG2513_PATCH | TEC |
| FP236383.18 | CHR_HG2513_PATCH | TEC |
| AC118282.5 | CHR_HG2525_PATCH | sorting nexin 18 (SNX18) pseudogene |
| AC118282.9 | CHR_HG2525_PATCH | ankyrin repeat domain 26 (ANKRD26) pseudo |
| AC118282.8 | CHR_HG2525_PATCH | tec |
| SNX18P23 | CHR_HG2525_PATCH | sorting nexin 18 pseudogene 23 [Source:HGN |
| AC118282.6 | CHR_HG2525_PATCH | ankyrin repeat domain 18B (ANKRD18B) pseu |
| MTCO3P39 | CHR_HG2525_PATCH | MT-CO3 pseudogene 39 [Source:HGNC Symbo |
| MTND3P22 | CHR_HG2525_PATCH | MT-ND3 pseudogene 22 [Source:HGNC Symbo |
| AC118282.7 | CHR_HG2525_PATCH | NULL |
| AC270106.1 | CHR_HG2525_PATCH | NULL |
| MAP4K1 | CHR_HG26_PATCH | mitogen-activated protein kinase kinase kinas |
| AC008649.4 | CHR_HG26_PATCH | novel transcript |
| EIF3K | CHR_HG26_PATCH | eukaryotic translation initiation factor 3 subur |
| ACTN4 | CHR_HG26_PATCH | actinin alpha 4 [Source:HGNC Symbol;Acc:HGI |
| AC008649.3 | CHR_HG26_PATCH | novel transcript, antisense to ACTN4 |
| AC256309.1 | CHR_HG26_PATCH | novel transcript, antisense to CAPN12 |
| LGALS7B | CHR_HG26_PATCH | galectin 7B [Source:HGNC Symbol;Acc:HGNC:3 |
| LGALS7 | CHR_HG26_PATCH | galectin 7 [Source:HGNC Symbol;Acc:HGNC:65 |
| RNU6-140P | CHR_HG26_PATCH | RNA, U6 small nuclear 140, pseudogene [Sour |
| LGALS4 | CHR_HG26_PATCH | galectin 4 [Source:HGNC Symbol;Acc:HGNC:65 |
| ECH1 | CHR_HG26_PATCH | enoyl-CoA hydratase 1 [Source:HGNC Symbol; |
| AC008982.3 | CHR_HG26_PATCH | novel protein |
| AC104534.2 | CHR_HG26_PATCH | novel transcript, antisense to ECH1 |
| HNRNPL | CHR_HG26_PATCH | heterogeneous nuclear ribonucleoprotein L [S |
| AC008982.4 | CHR_HG26_PATCH | novel transcript, sense intronic to HNRNPL |
| RINL | CHR_HG26_PATCH | Ras and Rab interactor like [Source:HGNC Sym |
| AC011455.4 | CHR_HG26_PATCH | novel transcript, antisense to RINL |
| SIRT2 | CHR_HG26_PATCH | sirtuin 2 [Source:HGNC Symbol;Acc:HGNC:108 |
| NFKBIB | CHR_HG26_PATCH | NFKB inhibitor beta [Source:HGNC Symbol;Acc |
| CCER2 | CHR_HG26_PATCH | coiled-coil glutamate rich protein 2 [Source:H |
| SARS2 | CHR_HG26_PATCH | seryl-tRNA synthetase 2, mitochondrial [Sourc |
| AC011455.3 | CHR_HG26_PATCH | novel protein |
| MRPS12 | CHR_HG26_PATCH | mitochondrial ribosomal protein S12 [Source: |
| ERVK9-11 | CHR_HG26_PATCH | endogenous retrovirus group K9, member 11 |
| FBXO17 | CHR_HG26_PATCH | F-box protein 17 [Source:HGNC Symbol;Acc:H |
| AC011455.5 | CHR_HG26_PATCH | novel transcript, antisense to FBXO17and FBX |
| SYT8 | CHR_HG28_PATCH | synaptotagmin 8 [Source:HGNC Symbol;Acc:H |
| TNNI2 | CHR_HG28_PATCH | troponin I2, fast skeletal type [Source:HGNC S |
| LSP1 | CHR_HG28_PATCH | lymphocyte specific protein 1 [Source:HGNC S |
| MIR4298 | CHR_HG28_PATCH | microRNA 4298 [Source:HGNC Symbol;Acc:HG |
| AC051649.6 | CHR_HG28_PATCH | novel transcript, sense overlapping LSP1 |
| MIR7847 | CHR_HG28_PATCH | microRNA 7847 [Source:HGNC Symbol;Acc:HG |
| PRR33 | CHR_HG28_PATCH | proline rich 33 [Source:HGNC Symbol;Acc:HGI |
| TNNT3 | CHR_HG28_PATCH | troponin T3, fast skeletal type [Source:HGNC S |
| LINC01150 | CHR_HG28_PATCH | long intergenic non-protein coding RNA 1150 |
| MRPL23 | CHR_HG28_PATCH | mitochondrial ribosomal protein L23 [Source: |

| | | |
|---|---|---|
| MRPL23-AS1 | CHR_HG28_PATCH | MRPL23 antisense RNA 1 [Source:HGNC Symb |
| LINC01219 | CHR_HG28_PATCH | long intergenic non-protein coding RNA 1219 |
| H19 | CHR_HG28_PATCH | H19 imprinted maternally expressed transcrip |
| AC051649.5 | CHR_HG28_PATCH | NULL |
| MIR675 | CHR_HG28_PATCH | microRNA 675 [Source:HGNC Symbol;Acc:HGI |
| AC051649.4 | CHR_HG28_PATCH | NULL |
| ADAMTS2 | CHR_HG30_PATCH | ADAM metallopeptidase with thrombospondi |
| AC109479.4 | CHR_HG30_PATCH | novel transcript |
| AC109479.5 | CHR_HG30_PATCH | novel transcript |
| AC109479.6 | CHR_HG30_PATCH | pseudogene similar to part of 5'-nucleotidase |
| AC270306.4 | CHR_HG30_PATCH | novel transcript |
| U4 | CHR_HG30_PATCH | U4 spliceosomal RNA [Source:RFAM;Acc:RF00 |
| AC270306.2 | CHR_HG30_PATCH | novel transcript |
| AC270306.5 | CHR_HG30_PATCH | novel pseudogene |
| AC270306.3 | CHR_HG30_PATCH | 6-pyruvoyltetrahydropterin synthase (PTS) ps |
| RUFY1 | CHR_HG30_PATCH | RUN and FYVE domain containing 1 [Source:H |
| PRDX2P3 | CHR_HG30_PATCH | peroxiredoxin 2 pseudogene 3 [Source:HGNC |
| AC270306.1 | CHR_HG30_PATCH | novel transcript |
| HNRNPH1 | CHR_HG30_PATCH | heterogeneous nuclear ribonucleoprotein H1 |
| Metazoa_SRP | CHR_HG30_PATCH | Metazoan signal recognition particle RNA [Sou |
| AC270227.2 | CHR_HG30_PATCH | chromosome 5 open reading frame 60 (C5orf6 |
| AC270227.1 | CHR_HG30_PATCH | Uncharacterized protein C5orf60 [Source:Uni |
| CBY3 | CHR_HG30_PATCH | chibby family member 3 [Source:HGNC Symbo |
| CANX | CHR_HG30_PATCH | calnexin [Source:HGNC Symbol;Acc:HGNC:147 |
| HMGB3P22 | CHR_HG30_PATCH | high mobility group box 3 pseudogene 22 [Sou |
| MAML1 | CHR_HG30_PATCH | mastermind like transcriptional coactivator 1 |
| LTC4S | CHR_HG30_PATCH | leukotriene C4 synthase [Source:HGNC Symbo |
| MGAT4B | CHR_HG30_PATCH | alpha-1,3-mannosyl-glycoprotein 4-beta-N-ac |
| MIR1229 | CHR_HG30_PATCH | microRNA 1229 [Source:HGNC Symbol;Acc:HG |
| AC008393.2 | CHR_HG30_PATCH | sequestosome 1 [Source:NCBI gene;Acc:8878] |
| MRNIP | CHR_HG30_PATCH | MRN complex interacting protein [Source:HGI |
| RN7SKP150 | CHR_HG30_PATCH | RN7SK pseudogene 150 [Source:HGNC Symbo |
| Y_RNA | CHR_HG30_PATCH | Y RNA [Source:RFAM;Acc:RF00019] |
| AC008393.3 | CHR_HG30_PATCH | novel transcript |
| TBC1D9B | CHR_HG30_PATCH | TBC1 domain family member 9B [Source:HGN |
| MCTS1 | CHR_HG439_PATCH | MCTS1 re-initiation and release factor [Source |
| C1GALT1C1 | CHR_HG439_PATCH | C1GALT1 specific chaperone 1 [Source:HGNC |
| AC011890.4 | CHR_HG439_PATCH | H3 histone, family 3B (H3.3B) (H3F3B) pseudo |
| AC011890.3 | CHR_HG439_PATCH | novel pseudogene |
| CT47B1 | CHR_HG439_PATCH | cancer/testis antigen family 47 member B1 [S |
| AC275592.3 | CHR_HG439_PATCH | coiled-coil domain containing 58 (CCDC58) ps |
| CT47A6 | CHR_HG439_PATCH | cancer/testis antigen family 47 member A6 [S |
| CT47A4 | CHR_HG439_PATCH | cancer/testis antigen family 47 member A4 [S |
| AC275592.2 | CHR_HG439_PATCH | coiled-coil domain containing 58 (CCDC58) ps |
| CT47A1 | CHR_HG439_PATCH | cancer/testis antigen family 47 member A1 [S |
| AC275592.1 | CHR_HG439_PATCH | coiled-coil domain containing 58 (CCDC58) ps |
| AC006144.4 | CHR_HG439_PATCH | E-cadherin binding protein E7 (HAKAI) pseudo |
| GLUD2 | CHR_HG439_PATCH | glutamate dehydrogenase 2 [Source:HGNC Sy |
| AC006144.3 | CHR_HG439_PATCH | novel transcript |
| HSD17B7P2 | CHR_HG545_PATCH | hydroxysteroid 17-beta dehydrogenase 7 pse |

| | | |
|---|---|---|
| SEPTIN7P9 | CHR_HG545_PATCH | septin 7 pseudogene 9 [Source:HGNC Symbol; |
| AL133216.4 | CHR_HG545_PATCH | general transcription factor IIi (GTF2I) pseudo |
| RNU6-1118P | CHR_HG545_PATCH | RNA, U6 small nuclear 1118, pseudogene [Sou |
| CICP9 | CHR_HG545_PATCH | capicua transcriptional repressor pseudogene |
| AL133216.3 | CHR_HG545_PATCH | novel transcript |
| ABCD1P2 | CHR_HG545_PATCH | ATP binding cassette subfamily D member 1 p |
| AL133173.3 | CHR_HG545_PATCH | CDGSH iron sulfur domain 2 (CISD2) pseudoge |
| PABPC1P12 | CHR_HG545_PATCH | poly(A) binding protein cytoplasmic 1 pseudo |
| SLC9B1P3 | CHR_HG545_PATCH | solute carrier family 9 member B1 pseudogen |
| ACTR3BP5 | CHR_HG545_PATCH | ACTR3B pseudogene 5 [Source:HGNC Symbol; |
| CHEK2P5 | CHR_HG545_PATCH | checkpoint kinase 2 pseudogene 5 [Source:H |
| AL133173.4 | CHR_HG545_PATCH | NULL |
| AL133173.5 | CHR_HG545_PATCH | ring finger protein 18 (RNF18) pseudogene |
| AC147067.4 | CHR_HG699_PATCH | novel transcript |
| AC147067.3 | CHR_HG699_PATCH | novel transcript |
| AC147067.5 | CHR_HG699_PATCH | NULL |
| AC017091.2 | CHR_HG705_PATCH | novel transcript |
| LINC02619 | CHR_HG705_PATCH | long intergenic non-protein coding RNA 2619 |
| LINC02429 | CHR_HG705_PATCH | long intergenic non-protein coding RNA 2429 |
| ZYX | CHR_HG708_PATCH | zyxin [Source:HGNC Symbol;Acc:HGNC:13200 |
| MIR6892 | CHR_HG708_PATCH | microRNA 6892 [Source:HGNC Symbol;Acc:H |
| EPHA1 | CHR_HG708_PATCH | EPH receptor A1 [Source:HGNC Symbol;Acc:H |
| EPHA1-AS1 | CHR_HG708_PATCH | EPHA1 antisense RNA 1 [Source:HGNC Symbo |
| TAS2R62P | CHR_HG708_PATCH | taste 2 receptor member 62 pseudogene [Sou |
| TAS2R60 | CHR_HG708_PATCH | taste 2 receptor member 60 [Source:HGNC Sy |
| TAS2R41 | CHR_HG708_PATCH | taste 2 receptor member 41 [Source:HGNC Sy |
| OR2R1P | CHR_HG708_PATCH | olfactory receptor family 2 subfamily R memb |
| OR10AC1 | CHR_HG708_PATCH | olfactory receptor family 10 subfamily AC mer |
| AC073264.4 | CHR_HG708_PATCH | ribosomal protein L26 (RPL26) pseudogene |
| PAICSP5 | CHR_HG708_PATCH | phosphoribosylaminoimidazole carboxylase, p |
| CTAGE15 | CHR_HG708_PATCH | CTAGE family member 15 [Source:HGNC Symb |
| RNU6-267P | CHR_HG708_PATCH | RNA, U6 small nuclear 267, pseudogene [Sour |
| CTAGE6 | CHR_HG708_PATCH | CTAGE family member 6 [Source:HGNC Symbo |
| AC254813.1 | CHR_HG708_PATCH | family with sequence similarity 115, member |
| AC254813.2 | CHR_HG708_PATCH | ribosomal protein L26 (RPL26) pseudogene |
| TCAF2P1 | CHR_HG708_PATCH | TRPM8 channel associated factor 2 pseudoge |
| OR2F2 | CHR_HG708_PATCH | olfactory receptor family 2 subfamily F memb |
| OR2F1 | CHR_HG708_PATCH | olfactory receptor family 2 subfamily F memb |
| AC004853.2 | CHR_HG708_PATCH | capping protein (actin filament) muscle Z-line, |
| OR2Q1P | CHR_HG708_PATCH | olfactory receptor family 2 subfamily Q memb |
| SLC16A1P1 | CHR_HG708_PATCH | SLC16A1 pseudogene 1 [Source:HGNC Symbol |
| OR6B1 | CHR_HG708_PATCH | olfactory receptor family 6 subfamily B memb |
| AC145625.4 | CHR_HG721_PATCH | novel transcript |
| AC145625.3 | CHR_HG721_PATCH | novel transcript |
| TWIST2 | CHR_HG721_PATCH | twist family bHLH transcription factor 2 [Sour |
| MCPH1 | CHR_HG76_PATCH | microcephalin 1 [Source:HGNC Symbol;Acc:H |
| AF287957.2 | CHR_HG76_PATCH | novel transcript |
| MCPH1-AS1 | CHR_HG76_PATCH | MCPH1 antisense RNA 1 [Source:HGNC Symbo |
| MIR8055 | CHR_HG76_PATCH | microRNA 8055 [Source:HGNC Symbol;Acc:H |
| AGPAT5 | CHR_HG76_PATCH | 1-acylglycerol-3-phosphate O-acyltransferase |

| | | |
|---|---|---|
| MIR4659A | CHR_HG76_PATCH | microRNA 4659a [Source:HGNC Symbol;Acc:H |
| MIR4659B | CHR_HG76_PATCH | microRNA 4659b [Source:HGNC Symbol;Acc:H |
| AC270252.1 | CHR_HG76_PATCH | novel transcript |
| XKR5 | CHR_HG76_PATCH | XK related 5 [Source:HGNC Symbol;Acc:HGNC |
| AC270297.1 | CHR_HG76_PATCH | novel transcript |
| DEFB1 | CHR_HG76_PATCH | defensin beta 1 [Source:HGNC Symbol;Acc:HG |
| DEFA6 | CHR_HG76_PATCH | defensin alpha 6 [Source:HGNC Symbol;Acc:H |
| AC270297.2 | CHR_HG76_PATCH | novel transcript |
| DEFA4 | CHR_HG76_PATCH | defensin alpha 4 [Source:HGNC Symbol;Acc:H |
| DEFA8P | CHR_HG76_PATCH | defensin alpha 8, pseudogene [Source:HGNC S |
| DEFA9P | CHR_HG76_PATCH | defensin alpha 9, pseudogene [Source:HGNC S |
| DEFA10P | CHR_HG76_PATCH | defensin alpha 10, pseudogene [Source:HGNC |
| DEFA1 | CHR_HG76_PATCH | defensin alpha 1 [Source:HGNC Symbol;Acc:H |
| AC270300.1 | CHR_HG76_PATCH | NULL |
| DEFA1B | CHR_HG76_PATCH | defensin alpha 1B [Source:HGNC Symbol;Acc:I |
| AC270300.2 | CHR_HG76_PATCH | NULL |
| DEFA3 | CHR_HG76_PATCH | defensin alpha 3 [Source:HGNC Symbol;Acc:H |
| DEFA11P | CHR_HG76_PATCH | defensin alpha 11, pseudogene [Source:HGNC |
| DEFA7P | CHR_HG76_PATCH | defensin alpha 7, pseudogene [Source:HGNC S |
| DEFA5 | CHR_HG76_PATCH | defensin alpha 5 [Source:HGNC Symbol;Acc:H |
| AC245425.1 | CHR_HG76_PATCH | chromosome 11 open reading frame 2 (C11or |
| AC270249.1 | CHR_HG76_PATCH | Ribosomal protein S24 pseudogene |
| RPS3AP30 | CHR_HG76_PATCH | ribosomal protein S3a pseudogene 30 [Source |
| SNRPCP6 | CHR_HG76_PATCH | small nuclear ribonucleoprotein polypeptide C |
| SNRPCP17 | CHR_HG76_PATCH | small nuclear ribonucleoprotein polypeptide C |
| RPS3AP33 | CHR_HG76_PATCH | ribosomal protein S3a pseudogene 33 [Source |
| AC270249.2 | CHR_HG76_PATCH | ribosomal protein S24 (RPS24) pseudogene |
| AC246795.1 | CHR_HG76_PATCH | RNA binding protein, fox-1 homolog (C. elegar |
| AC246795.3 | CHR_HG76_PATCH | RNA binding protein, fox-1 homolog (C. elegar |
| AC246795.2 | CHR_HG76_PATCH | chromosome 11 open reading frame 2 (C11or |
| OR7E154P | CHR_HG76_PATCH | olfactory receptor family 7 subfamily E memb |
| AC246795.7 | CHR_HG76_PATCH | family with sequence similarity 90, member A |
| AC246795.6 | CHR_HG76_PATCH | proline rich 23 domain containing 1 (PRR23D1 |
| PRR23D1 | CHR_HG76_PATCH | proline rich 23 domain containing 1 [Source:H |
| AC246795.5 | CHR_HG76_PATCH | proline rich 23 domain containing 1 (PRR23D1 |
| DEFB107B | CHR_HG76_PATCH | defensin beta 107B [Source:HGNC Symbol;Acc |
| DEFB105B | CHR_HG76_PATCH | defensin beta 105B [Source:HGNC Symbol;Acc |
| DEFB106A | CHR_HG76_PATCH | defensin beta 106A [Source:HGNC Symbol;Acc |
| DEFB104A | CHR_HG76_PATCH | defensin beta 104A [Source:HGNC Symbol;Acc |
| AC246795.4 | CHR_HG76_PATCH | sperm associated antigen 11A [Source:NCBI ge |
| DEFB103A | CHR_HG76_PATCH | defensin beta 103A [Source:HGNC Symbol;Acc |
| HSPD1P3 | CHR_HG76_PATCH | heat shock protein family D (Hsp60) member |
| DEFB4B | CHR_HG76_PATCH | defensin beta 4B [Source:HGNC Symbol;Acc:H |
| AC270258.1 | CHR_HG76_PATCH | Putative beta-defensin 108A [Source:UniProt |
| ZNF705B | CHR_HG76_PATCH | zinc finger protein 705B [Source:HGNC Symbo |
| AC275362.1 | CHR_HG76_PATCH | ubiquitin-specific protease (USP) pseudogene |
| USP17L4 | CHR_HG76_PATCH | ubiquitin specific peptidase 17 like family mer |
| USP17L3 | CHR_HG76_PATCH | ubiquitin specific peptidase 17 like family mer |
| DEFB109B | CHR_HG76_PATCH | defensin beta 109B [Source:HGNC Symbol;Acc |
| FAM90A20P | CHR_HG76_PATCH | family with sequence similarity 90 member A2 |

| | | |
|---|---|---|
| FAM90A3P | CHR_HG76_PATCH | family with sequence similarity 90 member A3 |
| AC275362.2 | CHR_HG76_PATCH | family with sequence similarity 90, member A |
| AC275401.1 | CHR_HG76_PATCH | NULL |
| ENPP7P12 | CHR_HG76_PATCH | ectonucleotide pyrophosphatase/phosphodie |
| AC275401.4 | CHR_HG76_PATCH | NULL |
| DEFB109A | CHR_HG76_PATCH | defensin beta 109A (pseudogene) [Source:HG |
| AC275401.2 | CHR_HG76_PATCH | ubiquitin specific peptidase 17-like 2 (USP17L2 |
| AC275401.3 | CHR_HG76_PATCH | ubiquitin specific peptidase 17-like 2 ( USP17L |
| AC270472.1 | CHR_HG76_PATCH | zinc finger protein 705D [Source:NCBI gene;Ac |
| DEFB108E | CHR_HG76_PATCH | defensin beta 108E (pseudogene) [Source:HGI |
| RNA5SP254 | CHR_HG76_PATCH | RNA, 5S ribosomal pseudogene 254 [Source:H |
| DEFB130A | CHR_HG76_PATCH | defensin beta 130A [Source:HGNC Symbol;Acc |
| DEFB131C | CHR_HG76_PATCH | defensin beta 131C (pseudogene) [Source:HG |
| AC270275.3 | CHR_HG76_PATCH | olfactory receptor, family 7, subfamily E, mem |
| AC270275.2 | CHR_HG76_PATCH | RNA binding protein, fox-1 homolog (C. elegar |
| DEFB131E | CHR_HG76_PATCH | defensin beta 131E (pseudogene) [Source:HGI |
| AC270275.1 | CHR_HG76_PATCH | Wolf-Hirschhorn syndrome candidate 1-like 1 |
| DEFB134 | CHR_HG76_PATCH | defensin beta 134 [Source:HGNC Symbol;Acc: |
| DEFB135 | CHR_HG76_PATCH | defensin beta 135 [Source:HGNC Symbol;Acc: |
| DEFB136 | CHR_HG76_PATCH | defensin beta 136 [Source:HGNC Symbol;Acc: |
| AC270275.5 | CHR_HG76_PATCH | novel transcript |
| AC270275.4 | CHR_HG76_PATCH | novel transcript |
| OR7E161P | CHR_HG76_PATCH | olfactory receptor family 7 subfamily E memb |
| AC270285.3 | CHR_HG76_PATCH | transmembrane protein 126A (TMEM126A) p |
| CTSB | CHR_HG76_PATCH | cathepsin B [Source:HGNC Symbol;Acc:HGNC: |
| AC270285.2 | CHR_HG76_PATCH | novel transcript, sense intronic to CTSB |
| FDFT1 | CHR_HG76_PATCH | farnesyl-diphosphate farnesyltransferase 1 [So |
| AC270285.4 | CHR_HG76_PATCH | novel transcript |
| SUB1P1 | CHR_HG76_PATCH | SUB1 pseudogene 1 [Source:HGNC Symbol;Ac |
| AC270285.1 | CHR_HG76_PATCH | NULL |
| C8orf49 | CHR_HG76_PATCH | chromosome 8 open reading frame 49 [Source |
| GATA4 | CHR_HG76_PATCH | GATA binding protein 4 [Source:HGNC Symbol |
| AC270284.1 | CHR_HG76_PATCH | novel transcript |
| AC270286.2 | CHR_HG76_PATCH | novel transcript, antisense to BLK |
| AC270286.1 | CHR_HG76_PATCH | BLK proto-oncogene, Src family tyrosine kinas |
| AC270272.3 | CHR_HG76_PATCH | novel transcript |
| AC270272.2 | CHR_HG76_PATCH | NULL |
| FAM167A-AS1 | CHR_HG76_PATCH | FAM167A antisense RNA 1 [Source:HGNC Sym |
| AC270272.1 | CHR_HG76_PATCH | NULL |
| RNU6-1084P | CHR_HG76_PATCH | RNA, U6 small nuclear 1084, pseudogene [Sou |
| RN7SL293P | CHR_HG76_PATCH | RNA, 7SL, cytoplasmic 293, pseudogene [Sour |
| TDH | CHR_HG76_PATCH | L-threonine dehydrogenase (pseudogene) [So |
| SLC35G5 | CHR_HG76_PATCH | solute carrier family 35 member G5 [Source:H |
| AC270277.3 | CHR_HG76_PATCH | uncharacterized LOC101929290 [Source:NCBI |
| MTMR9 | CHR_HG76_PATCH | myotubularin related protein 9 [Source:HGNC |
| AC270277.1 | CHR_HG76_PATCH | novel transcript, antisense to MTMR9 |
| AC270277.2 | CHR_HG76_PATCH | HERV-FRD provirus ancestral Env polyprotein |
| LINC00529 | CHR_HG76_PATCH | long intergenic non-protein coding RNA 529 [S |
| RPL19P13 | CHR_HG76_PATCH | ribosomal protein L19 pseudogene 13 [Source |
| AC270301.1 | CHR_HG76_PATCH | novel transcript |

| | | |
|---|---|---|
| AC270301.3 | CHR_HG76_PATCH | NULL |
| AC270301.5 | CHR_HG76_PATCH | ribosomal protein L17 (RPL17) pseudogene |
| AC270301.7 | CHR_HG76_PATCH | novel transcript |
| AC270301.4 | CHR_HG76_PATCH | novel transcript |
| AC270301.8 | CHR_HG76_PATCH | novel transcript |
| AC270301.2 | CHR_HG76_PATCH | novel transcript |
| AC270301.6 | CHR_HG76_PATCH | novel transcript |
| MIR598 | CHR_HG76_PATCH | microRNA 598 [Source:HGNC Symbol;Acc:HGI |
| AC270290.2 | CHR_HG76_PATCH | TEC |
| PINX1 | CHR_HG76_PATCH | PIN2 (TERF1) interacting telomerase inhibitor |
| MIR1322 | CHR_HG76_PATCH | microRNA 1322 [Source:HGNC Symbol;Acc:HG |
| AC270290.1 | CHR_HG76_PATCH | novel transcript, antisense to SOX7 and PINX1 |
| SOX7 | CHR_HG76_PATCH | SRY-box transcription factor 7 [Source:HGNC S |
| C8orf74 | CHR_HG76_PATCH | chromosome 8 open reading frame 74 [Source |
| RNA5SP252 | CHR_HG76_PATCH | RNA, 5S ribosomal pseudogene 252 [Source:H |
| MIR4286 | CHR_HG76_PATCH | microRNA 4286 [Source:HGNC Symbol;Acc:HG |
| PRSS55 | CHR_HG76_PATCH | serine protease 55 [Source:HGNC Symbol;Acc |
| AC270276.1 | CHR_HG76_PATCH | novel transcript |
| AC270296.3 | CHR_HG76_PATCH | novel transcript |
| AC270296.1 | CHR_HG76_PATCH | novel transcript |
| RNU6-729P | CHR_HG76_PATCH | RNA, U6 small nuclear 729, pseudogene [Sour |
| AC270296.2 | CHR_HG76_PATCH | novel protease, serine (PRSS) gene family pseu |
| MSRA | CHR_HG76_PATCH | methionine sulfoxide reductase A [Source:HGI |
| AC270296.4 | CHR_HG76_PATCH | transmembrane protein 126A (TMEM126A) ps |
| AC275404.1 | CHR_HG76_PATCH | novel transcript |
| AC270248.1 | CHR_HG76_PATCH | novel transcript |
| LINC00599 | CHR_HG76_PATCH | long intergenic non-protein coding RNA 599 [S |
| MIR124-1 | CHR_HG76_PATCH | microRNA 124-1 [Source:HGNC Symbol;Acc:H |
| TNKS | CHR_HG76_PATCH | tankyrase [Source:HGNC Symbol;Acc:HGNC:1 |
| MIR597 | CHR_HG76_PATCH | microRNA 597 [Source:HGNC Symbol;Acc:HGI |
| AC270292.1 | CHR_HG76_PATCH | microtubule associated serine/threonine kina |
| AC270292.2 | CHR_HG76_PATCH | DENN/MADD domain containing 4A (DENND4 |
| AC270263.2 | CHR_HG76_PATCH | novel transcript |
| AC270291.4 | CHR_HG76_PATCH | novel transcript |
| RNU6-526P | CHR_HG76_PATCH | RNA, U6 small nuclear 526, pseudogene [Sour |
| AC270263.1 | CHR_HG76_PATCH | novel transcript |
| RNU6-1151P | CHR_HG76_PATCH | RNA, U6 small nuclear 1151, pseudogene [Sou |
| AC270291.5 | CHR_HG76_PATCH | novel transcript |
| AC270291.3 | CHR_HG76_PATCH | ubiquitin-conjugating enzyme E2 variant 1 (UE |
| AC270291.2 | CHR_HG76_PATCH | novel transcript |
| AC270291.1 | CHR_HG76_PATCH | novel transcript |
| PPP1R3B | CHR_HG76_PATCH | protein phosphatase 1 regulatory subunit 3B [ |
| AC270254.1 | CHR_HG76_PATCH | novel transcript |
| Metazoa_SRP | CHR_HG76_PATCH | Metazoan signal recognition particle RNA [Sou |
| RNU7-55P | CHR_HG76_PATCH | RNA, U7 small nuclear 55 pseudogene [Source |
| MIR4660 | CHR_HG76_PATCH | microRNA 4660 [Source:HGNC Symbol;Acc:HG |
| RPL10P19 | CHR_HG76_PATCH | ribosomal protein L10 pseudogene 19 [Source |
| SNORA70 | CHR_HG76_PATCH | Small nucleolar RNA SNORA70 [Source:RFAM; |
| AC270253.1 | CHR_HG76_PATCH | mitochondrial ribosomal protein S18C (MRPS1 |
| RNU6-682P | CHR_HG76_PATCH | RNA, U6 small nuclear 682, pseudogene [Sour |

| | | |
|---|---|---|
| MFHAS1 | CHR_HG76_PATCH | malignant fibrous histiocytoma amplified sequ |
| AC270261.1 | CHR_HG76_PATCH | novel transcript |
| AC270261.2 | CHR_HG76_PATCH | novel lipoprotein receptor-related protein pse |
| CLDN23 | CHR_HG76_PATCH | claudin 23 [Source:HGNC Symbol;Acc:HGNC:1 |
| AC270261.3 | CHR_HG76_PATCH | novel transcript |
| RN7SL178P | CHR_HG76_PATCH | RNA, 7SL, cytoplasmic 178, pseudogene [Sour |
| AC270265.1 | CHR_HG76_PATCH | novel transcript |
| AC270265.2 | CHR_HG76_PATCH | novel transcript |
| AC270259.1 | CHR_HG76_PATCH | novel transcript |
| FAM86B3P | CHR_HG76_PATCH | family with sequence similarity 86 member B3 |
| ALG1L13P | CHR_HG76_PATCH | ALG1 like 13, pseudogene [Source:HGNC Syml |
| AC270267.1 | CHR_HG76_PATCH | novel transcript |
| AC270271.1 | CHR_HG76_PATCH | chromosome 11 open reading frame 2 (C11or |
| AC270271.2 | CHR_HG76_PATCH | FAM90A pseudogene [Source:NCBI gene;Acc:: |
| DEFB107A | CHR_HG76_PATCH | defensin beta 107A [Source:HGNC Symbol;Acc |
| DEFB105B | CHR_HG76_PATCH | defensin beta 105B [Source:NCBI gene;Acc:50 |
| DEFB106B | CHR_HG76_PATCH | defensin beta 106B [Source:NCBI gene;Acc:50 |
| DEFB104B | CHR_HG76_PATCH | defensin beta 104B [Source:NCBI gene;Acc:50 |
| SPAG11B | CHR_HG76_PATCH | sperm associated antigen 11B [Source:NCBI ge |
| DEFB103A | CHR_HG76_PATCH | defensin beta 103A [Source:HGNC Symbol;Acc |
| HSPD1P2 | CHR_HG76_PATCH | heat shock protein family D (Hsp60) member |
| DEFB4A | CHR_HG76_PATCH | defensin beta 4A [Source:HGNC Symbol;Acc:H |
| AC270273.1 | CHR_HG76_PATCH | NULL |
| ZNF705G | CHR_HG76_PATCH | zinc finger protein 705G [Source:NCBI gene;Ac |
| AC270273.2 | CHR_HG76_PATCH | deubiquitinating enzyme 3 pseudogene |
| USP17L7 | CHR_HG76_PATCH | ubiquitin specific peptidase 17 like family mer |
| USP17L2 | CHR_HG76_PATCH | ubiquitin specific peptidase 17 like family mer |
| DEFB109D | CHR_HG76_PATCH | defensin beta 109D (pseudogene) [Source:HG |
| FAM86B2 | CHR_HG76_PATCH | family with sequence similarity 86 member B2 |
| AC270293.1 | CHR_HG76_PATCH | NULL |
| RPS3AP34 | CHR_HG76_PATCH | ribosomal protein S3a pseudogene 34 [Source |
| AC275565.2 | CHR_HG76_PATCH | 40S ribosomal protein S24 (Rps24) pseudogen |
| AC275565.3 | CHR_HG76_PATCH | RNA binding protein, fox-1 homolog (C. elegar |
| AC275565.5 | CHR_HG76_PATCH | RNA binding protein, fox-1 homolog (C. elegar |
| RPS3AP35 | CHR_HG76_PATCH | ribosomal protein S3a pseudogene 35 [Source |
| AC275565.1 | CHR_HG76_PATCH | ribosomal protein S24 (Rps24) pseudogene |
| AC275565.4 | CHR_HG76_PATCH | RNA binding protein, fox-1 homolog (C. elegar |
| OR7E8P | CHR_HG76_PATCH | olfactory receptor family 7 subfamily E memb |
| OR7E15P | CHR_HG76_PATCH | olfactory receptor family 7 subfamily E memb |
| OR7E10P | CHR_HG76_PATCH | olfactory receptor family 7 subfamily E memb |
| MIR5692A2 | CHR_HG76_PATCH | microRNA 5692a-2 [Source:HGNC Symbol;Acc |
| LONRF1 | CHR_HG76_PATCH | LON peptidase N-terminal domain and ring fir |
| MIR3926-1 | CHR_HG76_PATCH | microRNA 3926-1 [Source:HGNC Symbol;Acc:I |
| MIR3926-2 | CHR_HG76_PATCH | microRNA 3926-2 [Source:HGNC Symbol;Acc:I |
| AC270298.1 | CHR_HG76_PATCH | novel transcript |
| LINC00681 | CHR_HG76_PATCH | long intergenic non-protein coding RNA 681 [S |
| AC270289.1 | CHR_HG76_PATCH | regulatory solute carrier protein, family 1, me |
| AC270289.2 | CHR_HG76_PATCH | tRNA methyltransferase 9B (putative) [Source |
| RNU6-842P | CHR_HG76_PATCH | RNA, U6 small nuclear 842, pseudogene [Sour |
| AC270289.3 | CHR_HG76_PATCH | DLC1 Rho GTPase activating protein [Source:N |

| | | |
|---|---|---|
| AC106845.2 | CHR_HG76_PATCH | novel transcript, antisense to DLC1 |
| Y_RNA | CHR_HG76_PATCH | Y RNA [Source:RFAM;Acc:RF00019] |
| TMEM159 | CHR_HG926_PATCH | transmembrane protein 159 [Source:HGNC Sy |
| AF001550.3 | CHR_HG926_PATCH | novel transcript, antisense to ZP2 |
| ZP2 | CHR_HG926_PATCH | zona pellucida glycoprotein 2 [Source:HGNC S |
| ANKS4B | CHR_HG926_PATCH | ankyrin repeat and sterile alpha motif domain |
| AC241640.1 | CHR_HG926_PATCH | NULL |
| AC241640.2 | CHR_HG926_PATCH | NULL |
| NPIPB5 | CHR_HG926_PATCH | nuclear pore complex interacting protein fami |
| AC241410.1 | CHR_HG926_PATCH | NULL |
| POLR3E | CHR_HG926_PATCH | RNA polymerase III subunit E [Source:HGNC Sy |
| EEF2K | CHR_HG926_PATCH | eukaryotic elongation factor 2 kinase [Source: |
| MOSMO | CHR_HG926_PATCH | modulator of smoothened [Source:HGNC Sym |
| PDZD9 | CHR_HG926_PATCH | PDZ domain containing 9 [Source:HGNC Symb |
| AC241401.3 | CHR_HG926_PATCH | NULL |
| AC241401.1 | CHR_HG926_PATCH | NULL |
| AC241401.2 | CHR_HG926_PATCH | nuclear pore complex interacting protein fami |
| AC241409.1 | CHR_HG926_PATCH | NULL |
| METTL9 | CHR_HG926_PATCH | methyltransferase like 9 [Source:HGNC Symbo |
| IGSF6 | CHR_HG926_PATCH | immunoglobulin superfamily member 6 [Sour |
| AC256236.1 | CHR_HG926_PATCH | NULL |
| AC256236.2 | CHR_HG926_PATCH | NULL |
| AC093151.9 | CHR_HG986_PATCH | novel transcript, antisense to SCMH1 |
| AC093151.5 | CHR_HG986_PATCH | novel transcript |
| AC093151.4 | CHR_HG986_PATCH | ribosomal protein L36a (RPL36A) pseudogene |
| AC093151.7 | CHR_HG986_PATCH | SET domain and mariner transposase fusion g |
| FOXO6 | CHR_HG986_PATCH | forkhead box O6 [Source:HGNC Symbol;Acc:H |
| FO681542.1 | CHR_HG986_PATCH | novel transcript |
| ARMC4P1 | CHR_HSCHR10_1_CTG1 | armadillo repeat containing 4 pseudogene 1 [ |
| U6 | CHR_HSCHR10_1_CTG1 | U6 spliceosomal RNA [Source:RFAM;Acc:RF00 |
| AL355493.2 | CHR_HSCHR10_1_CTG1 | pseudogene similar to part of breast cancer a |
| U6 | CHR_HSCHR10_1_CTG1 | U6 spliceosomal RNA [Source:RFAM;Acc:RF00 |
| FAM210CP | CHR_HSCHR10_1_CTG1 | family with sequence similarity 210 member ( |
| TRIAP1P1 | CHR_HSCHR10_1_CTG1 | TP53 regulated inhibitor of apoptosis 1 pseud |
| PTCHD3 | CHR_HSCHR10_1_CTG1 | patched domain containing 3 [Source:HGNC S |
| ANKRD30BP3 | CHR_HSCHR10_1_CTG2 | ankyrin repeat domain 30B pseudogene 3 [So |
| AL512324.2 | CHR_HSCHR10_1_CTG2 | novel transcript |
| AL512324.4 | CHR_HSCHR10_1_CTG2 | ankyrin repeat domain 30B-like (ANKRD30BL) |
| RNU6-1207P | CHR_HSCHR10_1_CTG2 | RNA, U6 small nuclear 1207, pseudogene [Sou |
| CUBNP2 | CHR_HSCHR10_1_CTG2 | cubilin pseudogene 2 [Source:HGNC Symbol;A |
| OR6D1P | CHR_HSCHR10_1_CTG2 | olfactory receptor family 6 subfamily D memb |
| OR13A1 | CHR_HSCHR10_1_CTG2 | olfactory receptor family 13 subfamily A mem |
| ALOX5 | CHR_HSCHR10_1_CTG2 | arachidonate 5-lipoxygenase [Source:HGNC Sy |
| AL731567.2 | CHR_HSCHR10_1_CTG2 | novel transcript |
| MARCH8 | CHR_HSCHR10_1_CTG2 | membrane associated ring-CH-type finger 8 [S |
| KSR1P1 | CHR_HSCHR10_1_CTG3 | kinase suppressor of ras 1 pseudogene 1 [Sou |
| IGKV1OR10-1 | CHR_HSCHR10_1_CTG3 | immunoglobulin kappa variable 1/OR10-1 (ps |
| AL031601.3 | CHR_HSCHR10_1_CTG3 | solute carrier family 9, subfamily B (NHA2, cat |
| PABPC1P8 | CHR_HSCHR10_1_CTG3 | poly(A) binding protein cytoplasmic 1 pseudo |
| RNU6-1266P | CHR_HSCHR10_1_CTG4 | RNA, U6 small nuclear 1266, pseudogene [Sou |

| | | |
|---|---|---|
| IMPDH1P5 | CHR_HSCHR10_1_CTG4 | inosine monophosphate dehydrogenase 1 pse |
| AL450306.2 | CHR_HSCHR10_1_CTG4 | novel transcript |
| DLG5 | CHR_HSCHR10_1_CTG4 | discs large MAGUK scaffold protein 5 [Source: |
| AL391421.2 | CHR_HSCHR10_1_CTG4 | novel transcript, antisense to DLG5 |
| RN7SL284P | CHR_HSCHR10_1_CTG4 | RNA, 7SL, cytoplasmic 284, pseudogene [Sour |
| DLG5-AS1 | CHR_HSCHR10_1_CTG4 | DLG5 antisense RNA 1 [Source:HGNC Symbol; |
| LINC01515 | CHR_HSCHR10_1_CTG6 | long intergenic non-protein coding RNA 1515 |
| AC021427.1 | CHR_HSCHR11_1_CTG1_2 | novel transcript, sense overlapping OR5P2, OF |
| AC044810.7 | CHR_HSCHR11_1_CTG1_2 | novel transcript, sense overlapping OR5P2, OF |
| AC044810.4 | CHR_HSCHR11_1_CTG1_2 | ribosomal protein L7a (RPL7A) pseudogene |
| AC044810.8 | CHR_HSCHR11_1_CTG1_2 | NULL |
| OR5P2 | CHR_HSCHR11_1_CTG1_2 | olfactory receptor family 5 subfamily P memb |
| OR5P3 | CHR_HSCHR11_1_CTG1_2 | olfactory receptor family 5 subfamily P memb |
| OR5E1P | CHR_HSCHR11_1_CTG1_2 | olfactory receptor family 5 subfamily E memb |
| RNU6-943P | CHR_HSCHR11_1_CTG1_2 | RNA, U6 small nuclear 943, pseudogene [Sour |
| OR10A6 | CHR_HSCHR11_1_CTG1_2 | olfactory receptor family 10 subfamily A mem |
| OR10A3 | CHR_HSCHR11_1_CTG1_2 | olfactory receptor family 10 subfamily A mem |
| NLRP10 | CHR_HSCHR11_1_CTG1_2 | NLR family pyrin domain containing 10 [Sourc |
| OOSP1 | CHR_HSCHR11_1_CTG3_1 | oocyte secreted protein 1 [Source:HGNC Syml |
| AP000790.3 | CHR_HSCHR11_1_CTG3_1 | novel transcript |
| OOSP4A | CHR_HSCHR11_1_CTG3_1 | oocyte secreted protein family member 4A [S |
| OOSP4B | CHR_HSCHR11_1_CTG3_1 | oocyte secreted protein family member 4B [S |
| OOSP2 | CHR_HSCHR11_1_CTG3_1 | oocyte secreted protein 2 [Source:HGNC Syml |
| AC226593.1 | CHR_HSCHR11_1_CTG3_1 | NULL |
| AC226593.2 | CHR_HSCHR11_1_CTG3_1 | tyrosine 3-monooxygenase/tryptophan 5-mor |
| AC044810.6 | CHR_HSCHR11_1_CTG5 | ribosomal protein L7a (RPL7A) pseudogene |
| OR5P2 | CHR_HSCHR11_1_CTG5 | olfactory receptor family 5 subfamily P memb |
| OR5P3 | CHR_HSCHR11_1_CTG5 | olfactory receptor family 5 subfamily P memb |
| OR5E1P | CHR_HSCHR11_1_CTG5 | olfactory receptor family 5 subfamily E memb |
| AC044810.5 | CHR_HSCHR11_1_CTG5 | novel transcript |
| U6 | CHR_HSCHR11_1_CTG5 | U6 spliceosomal RNA [Source:RFAM;Acc:RF00 |
| OR10A6 | CHR_HSCHR11_1_CTG5 | olfactory receptor family 10 subfamily A mem |
| OR10A3 | CHR_HSCHR11_1_CTG5 | olfactory receptor family 10 subfamily A mem |
| NLRP10 | CHR_HSCHR11_1_CTG5 | NLR family pyrin domain containing 10 [Sourc |
| DUSP8 | CHR_HSCHR11_1_CTG6 | dual specificity phosphatase 8 [Source:HGNC S |
| KRTAP5-AS1 | CHR_HSCHR11_1_CTG6 | KRTAP5-1/KRTAP5-2 antisense RNA 1 [Source |
| AP006285.2 | CHR_HSCHR11_1_CTG6 | NULL |
| KRTAP5-1 | CHR_HSCHR11_1_CTG6 | keratin associated protein 5-1 [Source:HGNC S |
| KRTAP5-2 | CHR_HSCHR11_1_CTG6 | keratin associated protein 5-2 [Source:HGNC S |
| KRTAP5-3 | CHR_HSCHR11_1_CTG6 | keratin associated protein 5-3 [Source:HGNC S |
| KRTAP5-4 | CHR_HSCHR11_1_CTG6 | keratin associated protein 5-4 [Source:HGNC S |
| KRTAP5-5 | CHR_HSCHR11_1_CTG6 | keratin associated protein 5-5 [Source:HGNC S |
| AP006285.3 | CHR_HSCHR11_1_CTG6 | novel transcript |
| FAM99A | CHR_HSCHR11_1_CTG6 | family with sequence similarity 99 member A |
| FAM99B | CHR_HSCHR11_1_CTG6 | family with sequence similarity 99 member B |
| LINC02708 | CHR_HSCHR11_1_CTG6 | long intergenic non-protein coding RNA 2708 |
| KCNQ1 | CHR_HSCHR11_1_CTG7 | potassium voltage-gated channel subfamily Q |
| COX6CP18 | CHR_HSCHR11_1_CTG7 | cytochrome c oxidase subunit 6C pseudogene |
| KCNQ1-AS1 | CHR_HSCHR11_1_CTG7 | KCNQ1 antisense RNA 1 [Source:HGNC Symbo |
| KCNQ1DN | CHR_HSCHR11_1_CTG7 | KCNQ1 downstream neighbor [Source:HGNC S |

| | | |
|---|---|---|
| AC001228.1 | CHR_HSCHR11_1_CTG7 | NULL |
| CDKN1C | CHR_HSCHR11_1_CTG7 | cyclin dependent kinase inhibitor 1C [Source:I |
| SLC22A18AS | CHR_HSCHR11_1_CTG7 | solute carrier family 22 member 18 antisense |
| SLC22A18 | CHR_HSCHR11_1_CTG7 | solute carrier family 22 member 18 [Source:H |
| PHLDA2 | CHR_HSCHR11_1_CTG7 | pleckstrin homology like domain family A mer |
| NAP1L4 | CHR_HSCHR11_1_CTG7 | nucleosome assembly protein 1 like 4 [Source |
| SNORA54 | CHR_HSCHR11_1_CTG7 | small nucleolar RNA, H/ACA box 54 [Source:H |
| AC001228.2 | CHR_HSCHR11_1_CTG7 | novel transcript, antisense to NAP1L4 |
| CARS | CHR_HSCHR11_1_CTG7 | cysteinyl-tRNA synthetase [Source:HGNC Sym |
| CARS-AS1 | CHR_HSCHR11_1_CTG7 | CARS antisense RNA 1 [Source:HGNC Symbol;/ |
| U1 | CHR_HSCHR11_1_CTG7 | U1 spliceosomal RNA [Source:RFAM;Acc:RF00 |
| PTDSS2 | CHR_HSCHR11_1_CTG8 | phosphatidylserine synthase 2 [Source:HGNC |
| RNH1 | CHR_HSCHR11_1_CTG8 | ribonuclease/angiogenin inhibitor 1 [Source:H |
| AC137894.2 | CHR_HSCHR11_1_CTG8 | artifact |
| HRAS | CHR_HSCHR11_1_CTG8 | HRas proto-oncogene, GTPase [Source:HGNC |
| LRRC56 | CHR_HSCHR11_1_CTG8 | leucine rich repeat containing 56 [Source:HGN |
| LMNTD2 | CHR_HSCHR11_1_CTG8 | lamin tail domain containing 2 [Source:HGNC |
| AC138374.1 | CHR_HSCHR11_1_CTG8 | novel transcript |
| RASSF7 | CHR_HSCHR11_1_CTG8 | Ras association domain family member 7 [Sou |
| MIR210HG | CHR_HSCHR11_1_CTG8 | MIR210 host gene [Source:HGNC Symbol;Acc: |
| PHRF1 | CHR_HSCHR11_1_CTG8 | PHD and ring finger domains 1 [Source:HGNC |
| IRF7 | CHR_HSCHR11_1_CTG8 | interferon regulatory factor 7 [Source:HGNC S |
| CDHR5 | CHR_HSCHR11_1_CTG8 | cadherin related family member 5 [Source:HG |
| SCT | CHR_HSCHR11_1_CTG8 | secretin [Source:HGNC Symbol;Acc:HGNC:106 |
| DRD4 | CHR_HSCHR11_1_CTG8 | dopamine receptor D4 [Source:HGNC Symbol; |
| DEAF1 | CHR_HSCHR11_1_CTG8 | DEAF1 transcription factor [Source:HGNC Sym |
| AC138374.2 | CHR_HSCHR11_1_CTG8 | novel transcript |
| AP2A2 | CHR_HSCHR11_2_CTG1 | adaptor related protein complex 2 subunit alp |
| RNU6-1025P | CHR_HSCHR11_2_CTG1 | RNA, U6 small nuclear 1025, pseudogene [Sou |
| MUC6 | CHR_HSCHR11_2_CTG1 | mucin 6, oligomeric mucus/gel-forming [Sour |
| DUSP8 | CHR_HSCHR11_2_CTG1_1 | dual specificity phosphatase 8 [Source:HGNC S |
| KRTAP5-AS1 | CHR_HSCHR11_2_CTG1_1 | KRTAP5-1/KRTAP5-2 antisense RNA 1 [Source |
| KRTAP5-1 | CHR_HSCHR11_2_CTG1_1 | keratin associated protein 5-1 [Source:HGNC S |
| KRTAP5-2 | CHR_HSCHR11_2_CTG1_1 | keratin associated protein 5-2 [Source:HGNC S |
| KRTAP5-3 | CHR_HSCHR11_2_CTG1_1 | keratin associated protein 5-3 [Source:HGNC S |
| KRTAP5-4 | CHR_HSCHR11_2_CTG1_1 | keratin associated protein 5-4 [Source:HGNC S |
| KRTAP5-5 | CHR_HSCHR11_2_CTG1_1 | keratin associated protein 5-5 [Source:HGNC S |
| AC130310.1 | CHR_HSCHR11_2_CTG1_1 | novel transcript |
| FAM99A | CHR_HSCHR11_2_CTG1_1 | family with sequence similarity 99 member A |
| FAM99B | CHR_HSCHR11_2_CTG1_1 | family with sequence similarity 99 member B |
| LINC02708 | CHR_HSCHR11_2_CTG1_1 | long intergenic non-protein coding RNA 2708 |
| KRTAP5-6 | CHR_HSCHR11_2_CTG1_1 | keratin associated protein 5-6 [Source:HGNC S |
| IFITM10 | CHR_HSCHR11_2_CTG1_1 | interferon induced transmembrane protein 1( |
| AP000648.5 | CHR_HSCHR11_2_CTG8 | tripartite motif-containing protein 43 (TRIM43 |
| AP000648.10 | CHR_HSCHR11_2_CTG8 | novel transcript |
| AP000648.6 | CHR_HSCHR11_2_CTG8 | NULL |
| AP000648.8 | CHR_HSCHR11_2_CTG8 | novel transcript |
| NAALAD2 | CHR_HSCHR11_2_CTG8 | N-acetylated alpha-linked acidic dipeptidase 2 |
| AP000648.7 | CHR_HSCHR11_2_CTG8 | novel transcript |
| CHORDC1 | CHR_HSCHR11_2_CTG8 | cysteine and histidine rich domain containing |

| | | |
|---|---|---|
| AP002364.2 | CHR_HSCHR11_2_CTG8 | novel transcript |
| TUBAP2 | CHR_HSCHR11_2_CTG8 | tubulin alpha pseudogene 2 [Source:HGNC Sy |
| AP2A2 | CHR_HSCHR11_3_CTG1 | adaptor related protein complex 2 subunit alp |
| RNU6-1025P | CHR_HSCHR11_3_CTG1 | RNA, U6 small nuclear 1025, pseudogene [Sou |
| MUC6 | CHR_HSCHR11_3_CTG1 | mucin 6, oligomeric mucus/gel-forming [Sour |
| LINC02688 | CHR_HSCHR11_3_CTG1 | long intergenic non-protein coding RNA 2688 |
| MUC2 | CHR_HSCHR11_3_CTG1 | mucin 2, oligomeric mucus/gel-forming [Sour |
| DDX11L8 | CHR_HSCHR12_1_CTG1 | DEAD/H-box helicase 11 like 8 [Source:HGNC |
| WASH8P | CHR_HSCHR12_1_CTG1 | WAS protein family homolog 8, pseudogene [ |
| FAM138D | CHR_HSCHR12_1_CTG1 | family with sequence similarity 138 member |
| AC026369.8 | CHR_HSCHR12_1_CTG1 | NULL |
| IQSEC3 | CHR_HSCHR12_1_CTG1 | IQ motif and Sec7 domain 3 [Source:HGNC Sy |
| AC026369.6 | CHR_HSCHR12_1_CTG1 | novel transcript |
| AC026369.7 | CHR_HSCHR12_1_CTG1 | novel transcript |
| AC026369.4 | CHR_HSCHR12_1_CTG1 | novel transcript |
| AC026369.5 | CHR_HSCHR12_1_CTG1 | novel transcript, antisense to IQSEC3 |
| GIHCG | CHR_HSCHR12_1_CTG2_1 | GIHCG inhibitor of miR-200b/200a/429 expre |
| ATP23 | CHR_HSCHR12_1_CTG2_1 | ATP23 metallopeptidase and ATP synthase as |
| AC084033.4 | CHR_HSCHR12_1_CTG2_1 | zinc finger protein pseudogene |
| RN7SKP65 | CHR_HSCHR12_1_CTG2_1 | RN7SK pseudogene 65 [Source:HGNC Symbol; |
| AC084033.3 | CHR_HSCHR12_1_CTG2_1 | suppression of tumorigenicity 13 (colon carcir |
| LINC02403 | CHR_HSCHR12_1_CTG2_1 | long intergenic non-protein coding RNA 2403 |
| TAS2R7 | CHR_HSCHR12_2_CTG2 | taste 2 receptor member 7 [Source:HGNC Syn |
| TAS2R8 | CHR_HSCHR12_2_CTG2 | taste 2 receptor member 8 [Source:HGNC Syn |
| TAS2R9 | CHR_HSCHR12_2_CTG2 | taste 2 receptor member 9 [Source:HGNC Syn |
| PRR4 | CHR_HSCHR12_2_CTG2 | proline rich 4 [Source:HGNC Symbol;Acc:HGN |
| TAS2R10 | CHR_HSCHR12_2_CTG2 | taste 2 receptor member 10 [Source:HGNC Sy |
| AC006518.3 | CHR_HSCHR12_2_CTG2 | taste receptor, type 2 pseudogene |
| AC006518.4 | CHR_HSCHR12_2_CTG2 | taste receptor, type 2, member 12 pseudoger |
| TAS2R13 | CHR_HSCHR12_2_CTG2 | taste 2 receptor member 13 [Source:HGNC Sy |
| PRH2 | CHR_HSCHR12_2_CTG2 | proline rich protein HaeIII subfamily 2 [Source |
| TAS2R14 | CHR_HSCHR12_2_CTG2 | taste 2 receptor member 14 [Source:HGNC Sy |
| TAS2R15P | CHR_HSCHR12_2_CTG2 | taste 2 receptor member 15 pseudogene [Sou |
| TAS2R50 | CHR_HSCHR12_2_CTG2 | taste 2 receptor member 50 [Source:HGNC Sy |
| TAS2R20 | CHR_HSCHR12_2_CTG2 | taste 2 receptor member 20 [Source:HGNC Sy |
| TAS2R19 | CHR_HSCHR12_2_CTG2 | taste 2 receptor member 19 [Source:HGNC Sy |
| TAS2R31 | CHR_HSCHR12_2_CTG2 | taste 2 receptor member 31 [Source:HGNC Sy |
| TAS2R63P | CHR_HSCHR12_2_CTG2 | taste 2 receptor member 63 pseudogene [Sou |
| TAS2R46 | CHR_HSCHR12_2_CTG2 | taste 2 receptor member 46 [Source:HGNC Sy |
| TAS2R64P | CHR_HSCHR12_2_CTG2 | taste 2 receptor member 64 pseudogene [Sou |
| TAS2R43 | CHR_HSCHR12_2_CTG2 | taste 2 receptor member 43 [Source:HGNC Sy |
| AC242308.1 | CHR_HSCHR12_2_CTG2 | iron-responsive element binding protein 2 (IR |
| AC242308.2 | CHR_HSCHR12_2_CTG2 | novel taste receptor, type 2 family pseudogen |
| TAS2R45 | CHR_HSCHR12_2_CTG2 | taste 2 receptor member 45 [Source:HGNC Sy |
| TAS2R30 | CHR_HSCHR12_2_CTG2 | taste 2 receptor member 30 [Source:HGNC Sy |
| AC134349.3 | CHR_HSCHR12_2_CTG2 | taste receptor, type 2, member 18 pseudoger |
| AC134349.4 | CHR_HSCHR12_2_CTG2 | novel transcript |
| SMIM10L1 | CHR_HSCHR12_2_CTG2 | small integral membrane protein 10 like 1 [So |
| RPS6P22 | CHR_HSCHR12_2_CTG2_1 | ribosomal protein S6 pseudogene 22 [Source: |
| AC068305.3 | CHR_HSCHR12_2_CTG2_1 | methyltransferase 5 domain containing 1 (ME |

| | | |
|---|---|---|
| AC068305.4 | CHR_HSCHR12_2_CTG2_1 | novel transcript, antisense to LRIG3 |
| TAS2R7 | CHR_HSCHR12_3_CTG2 | taste 2 receptor member 7 [Source:HGNC Syn |
| TAS2R8 | CHR_HSCHR12_3_CTG2 | taste 2 receptor member 8 [Source:HGNC Syn |
| TAS2R9 | CHR_HSCHR12_3_CTG2 | taste 2 receptor member 9 [Source:HGNC Syn |
| PRH1 | CHR_HSCHR12_3_CTG2 | proline rich protein HaeIII subfamily 1 [Source |
| TAS2R10 | CHR_HSCHR12_3_CTG2 | taste 2 receptor member 10 [Source:HGNC Sy |
| AC006518.5 | CHR_HSCHR12_3_CTG2 | taste receptor, type 2 pseudogene |
| PRR4 | CHR_HSCHR12_3_CTG2 | proline rich 4 [Source:HGNC Symbol;Acc:HGN |
| AC245078.1 | CHR_HSCHR12_3_CTG2 | PRH1-PRR4 readthrough |
| AC006518.7 | CHR_HSCHR12_3_CTG2 | NULL |
| AC006518.6 | CHR_HSCHR12_3_CTG2 | taste receptor, type 2, member 12 pseudogen |
| TAS2R13 | CHR_HSCHR12_3_CTG2 | taste 2 receptor member 13 [Source:HGNC Sy |
| PRH2 | CHR_HSCHR12_3_CTG2 | proline rich protein HaeIII subfamily 2 [Source |
| TAS2R14 | CHR_HSCHR12_3_CTG2 | taste 2 receptor member 14 [Source:HGNC Sy |
| TAS2R15P | CHR_HSCHR12_3_CTG2 | taste 2 receptor member 15 pseudogene [Sou |
| TAS2R50 | CHR_HSCHR12_3_CTG2 | taste 2 receptor member 50 [Source:HGNC Sy |
| TAS2R20 | CHR_HSCHR12_3_CTG2 | taste 2 receptor member 20 [Source:HGNC Sy |
| TAS2R19 | CHR_HSCHR12_3_CTG2 | taste 2 receptor member 19 [Source:HGNC Sy |
| TAS2R31 | CHR_HSCHR12_3_CTG2 | taste 2 receptor member 31 [Source:HGNC Sy |
| TAS2R63P | CHR_HSCHR12_3_CTG2 | taste 2 receptor member 63 pseudogene [Sou |
| TAS2R46 | CHR_HSCHR12_3_CTG2 | taste 2 receptor member 46 [Source:HGNC Sy |
| TAS2R64P | CHR_HSCHR12_3_CTG2 | taste 2 receptor member 64 pseudogene [Sou |
| TAS2R43 | CHR_HSCHR12_3_CTG2 | taste 2 receptor member 43 [Source:HGNC Sy |
| TAS2R30 | CHR_HSCHR12_3_CTG2 | taste 2 receptor member 30 [Source:HGNC Sy |
| AC245078.3 | CHR_HSCHR12_3_CTG2 | taste receptor, type 2, member 18 pseudogen |
| AC245078.2 | CHR_HSCHR12_3_CTG2 | novel transcript |
| SMIM10L1 | CHR_HSCHR12_3_CTG2 | small integral membrane protein 10 like 1 [So |
| TAS2R42 | CHR_HSCHR12_3_CTG2 | taste 2 receptor member 42 [Source:HGNC Sy |
| AC244131.3 | CHR_HSCHR12_3_CTG2 | taste receptor, type 2, member 7 pseudogene |
| AC244131.4 | CHR_HSCHR12_3_CTG2 | novel transcript |
| PRB3 | CHR_HSCHR12_3_CTG2 | proline rich protein BstNI subfamily 3 [Source: |
| PRB4 | CHR_HSCHR12_3_CTG2 | proline rich protein BstNI subfamily 4 [Source: |
| PRB1 | CHR_HSCHR12_3_CTG2 | proline rich protein BstNI subfamily 1 (gene/p |
| ANO4 | CHR_HSCHR12_3_CTG2_1 | anoctamin 4 [Source:HGNC Symbol;Acc:HGNC |
| SLC5A8 | CHR_HSCHR12_3_CTG2_1 | solute carrier family 5 member 8 [Source:HGN |
| U6 | CHR_HSCHR12_3_CTG2_1 | U6 spliceosomal RNA [Source:RFAM;Acc:RF00 |
| AC079953.2 | CHR_HSCHR12_3_CTG2_1 | ribosomal protein S6 (RPS6) pseudogene |
| AC024940.7 | CHR_HSCHR12_4_CTG2 | ovostatin 2 |
| AC024940.10 | CHR_HSCHR12_4_CTG2 | NADH dehydrogenase (ubiquinone) 1 alpha su |
| AC024940.9 | CHR_HSCHR12_4_CTG2 | ribosomal protein L13a (RPL13A) pseudogene |
| MREGP1 | CHR_HSCHR12_4_CTG2 | melanoregulin pseudogene 1 [Source:HGNC S |
| SINHCAF | CHR_HSCHR12_4_CTG2 | SIN3-HDAC complex associated factor [Source |
| AC024940.11 | CHR_HSCHR12_4_CTG2 | solute carrier family 25, member 13 (citrin) (S |
| AC024940.8 | CHR_HSCHR12_4_CTG2 | novel transcript, antisense to FAM60A |
| AC007368.2 | CHR_HSCHR12_4_CTG2_1 | novel transcript |
| AC007368.3 | CHR_HSCHR12_4_CTG2_1 | uncharacterized LOC101927531 [Source:NCBI |
| LINC02825 | CHR_HSCHR12_4_CTG2_1 | long intergenic non-protein coding RNA 2825 |
| PRB1 | CHR_HSCHR12_5_CTG2 | proline rich protein BstNI subfamily 1 (gene/p |
| PIWIL1 | CHR_HSCHR12_5_CTG2_1 | piwi like RNA-mediated gene silencing 1 [Sour |
| AC127071.3 | CHR_HSCHR12_5_CTG2_1 | novel transcript |

| | | |
|---|---|---|
| AC217627.1 | CHR_HSCHR12_5_CTG2_1 | novel transcript |
| DNAH10 | CHR_HSCHR12_6_CTG2_1 | dynein axonemal heavy chain 10 [Source:HGN |
| CCDC92 | CHR_HSCHR12_6_CTG2_1 | coiled-coil domain containing 92 [Source:HGN |
| AC208069.1 | CHR_HSCHR12_6_CTG2_1 | novel transcript, antisense to DNAH10 |
| MGAT4C | CHR_HSCHR12_8_CTG2_1 | MGAT4 family member C [Source:HGNC Symb |
| LINC02405 | CHR_HSCHR12_9_CTG2_1 | long intergenic non-protein coding RNA 2405 |
| AC079949.4 | CHR_HSCHR12_9_CTG2_1 | novel transcript |
| AC195745.1 | CHR_HSCHR12_9_CTG2_1 | novel transcript |
| AC195745.2 | CHR_HSCHR12_9_CTG2_1 | novel transcript |
| LINC01070 | CHR_HSCHR13_1_CTG1 | long intergenic non-protein coding RNA 1070 |
| LINC01043 | CHR_HSCHR13_1_CTG1 | long intergenic non-protein coding RNA 1043 |
| LINC01044 | CHR_HSCHR13_1_CTG1 | long intergenic non-protein coding RNA 1044 |
| SPACA7 | CHR_HSCHR13_1_CTG1 | sperm acrosome associated 7 [Source:HGNC S |
| TUBGCP3 | CHR_HSCHR13_1_CTG1 | tubulin gamma complex associated protein 3 |
| GXYLT1P1 | CHR_HSCHR13_1_CTG3 | glucoside xylosyltransferase 1 pseudogene 1 [ |
| CNN2P12 | CHR_HSCHR13_1_CTG3 | calponin 2 pseudogene 12 [Source:HGNC Sym |
| ZNF965P | CHR_HSCHR13_1_CTG3 | zinc finger protein 965, pseudogene [Source:H |
| CYP4F34P | CHR_HSCHR13_1_CTG3 | cytochrome P450 family 4 subfamily F membe |
| SNX18P26 | CHR_HSCHR13_1_CTG3 | sorting nexin 18 pseudogene 26 [Source:HGN |
| LINC00328-2P | CHR_HSCHR13_1_CTG3 | long intergenic non-protein coding RNA 328-2 |
| ANKRD20A9P | CHR_HSCHR13_1_CTG3 | ankyrin repeat domain 20 family member A9, |
| RNU6-55P | CHR_HSCHR13_1_CTG3 | RNA, U6 small nuclear 55, pseudogene [Sourc |
| U6 | CHR_HSCHR13_1_CTG3 | U6 spliceosomal RNA [Source:RFAM;Acc:RF00 |
| GRK1 | CHR_HSCHR13_1_CTG5 | G protein-coupled receptor kinase 1 [Source:H |
| LINC00552 | CHR_HSCHR13_1_CTG5 | long intergenic non-protein coding RNA 552 [S |
| TMEM255B | CHR_HSCHR13_1_CTG5 | transmembrane protein 255B [Source:HGNC S |
| AL133373.4 | CHR_HSCHR14_1_CTG1 | zinc finger protein 257 (ZNF257) pseudogene |
| AL133373.3 | CHR_HSCHR14_1_CTG1 | TEC |
| CATSPERB | CHR_HSCHR14_1_CTG1 | cation channel sperm associated auxiliary sub |
| POLR3GP1 | CHR_HSCHR14_1_CTG1 | RNA polymerase III subunit G pseudogene 1 [S |
| NANOGP7 | CHR_HSCHR14_1_CTG1 | Nanog homeobox pseudogene 7 [Source:HGN |
| AL121839.3 | CHR_HSCHR14_1_CTG1 | pyruvate dehydrogenase kinase, isozyme 3 (P |
| U3 | CHR_HSCHR14_1_CTG1 | Small nucleolar RNA U3 [Source:RFAM;Acc:RF |
| AL121839.4 | CHR_HSCHR14_1_CTG1 | novel transcript, intronic to CATSPERB |
| TC2N | CHR_HSCHR14_1_CTG1 | tandem C2 domains, nuclear [Source:HGNC Sy |
| TMEM179 | CHR_HSCHR14_2_CTG1 | transmembrane protein 179 [Source:HGNC Sy |
| C14orf180 | CHR_HSCHR14_2_CTG1 | chromosome 14 open reading frame 180 [Sou |
| BX927359.2 | CHR_HSCHR14_2_CTG1 | novel transcript, antisense to C14orf180 |
| ATP5MC1P1 | CHR_HSCHR14_3_CTG1 | ATP synthase membrane subunit c locus 1 pse |
| ELK2BP | CHR_HSCHR14_3_CTG1 | ETS transcription factor ELK2B, pseudogene [S |
| IGHA2 | CHR_HSCHR14_3_CTG1 | immunoglobulin heavy constant alpha 2 (A2m |
| IGHE | CHR_HSCHR14_3_CTG1 | immunoglobulin heavy constant epsilon [Sour |
| IGHG4 | CHR_HSCHR14_3_CTG1 | immunoglobulin heavy constant gamma 4 (G4 |
| IGHG2 | CHR_HSCHR14_3_CTG1 | immunoglobulin heavy constant gamma 2 (G2 |
| COPDA1 | CHR_HSCHR14_3_CTG1 | chronic obstructive pulmonary disease associa |
| IGHGP | CHR_HSCHR14_3_CTG1 | immunoglobulin heavy constant gamma P (no |
| ELK2AP | CHR_HSCHR14_3_CTG1 | ETS transcription factor ELK2A, pseudogene [S |
| AL928768.1 | CHR_HSCHR14_3_CTG1 | novel transcript |
| IGHA1 | CHR_HSCHR14_3_CTG1 | immunoglobulin heavy constant alpha 1 [Sour |
| IGHEP1 | CHR_HSCHR14_3_CTG1 | immunoglobulin heavy constant epsilon P1 (p |

| | | |
|---|---|---|
| IGHG1 | CHR_HSCHR14_3_CTG1 | immunoglobulin heavy constant gamma 1 (G1 |
| IGHG3 | CHR_HSCHR14_3_CTG1 | immunoglobulin heavy constant gamma 3 (G3 |
| AL122127.2 | CHR_HSCHR14_3_CTG1 | ATPase, H+ transporting, lysosomal 13kDa, V1 |
| IGHD | CHR_HSCHR14_3_CTG1 | immunoglobulin heavy constant delta [Source |
| IGHM | CHR_HSCHR14_3_CTG1 | immunoglobulin heavy constant mu [Source:H |
| IGHJ6 | CHR_HSCHR14_3_CTG1 | immunoglobulin heavy joining 6 [Source:HGN |
| IGHJ3P | CHR_HSCHR14_3_CTG1 | immunoglobulin heavy joining 3P (pseudogen |
| IGHJ5 | CHR_HSCHR14_3_CTG1 | immunoglobulin heavy joining 5 [Source:HGN |
| IGHJ4 | CHR_HSCHR14_3_CTG1 | immunoglobulin heavy joining 4 [Source:HGN |
| IGHJ3 | CHR_HSCHR14_3_CTG1 | immunoglobulin heavy joining 3 [Source:HGN |
| IGHJ2P | CHR_HSCHR14_3_CTG1 | immunoglobulin heavy joining 2P (pseudogen |
| IGHJ2 | CHR_HSCHR14_3_CTG1 | immunoglobulin heavy joining 2 [Source:HGN |
| IGHJ1 | CHR_HSCHR14_3_CTG1 | immunoglobulin heavy joining 1 [Source:HGN |
| IGHD7-27 | CHR_HSCHR14_3_CTG1 | immunoglobulin heavy diversity 7-27 [Source: |
| IGHJ1P | CHR_HSCHR14_3_CTG1 | immunoglobulin heavy joining 1P (pseudogen |
| IGHD1-26 | CHR_HSCHR14_3_CTG1 | immunoglobulin heavy diversity 1-26 [Source: |
| IGHD6-25 | CHR_HSCHR14_3_CTG1 | immunoglobulin heavy diversity 6-25 [Source: |
| IGHD5-24 | CHR_HSCHR14_3_CTG1 | immunoglobulin heavy diversity 5-24 (non-fur |
| IGHD4-23 | CHR_HSCHR14_3_CTG1 | immunoglobulin heavy diversity 4-23 (non-fur |
| IGHD3-22 | CHR_HSCHR14_3_CTG1 | immunoglobulin heavy diversity 3-22 [Source: |
| IGHD2-21 | CHR_HSCHR14_3_CTG1 | immunoglobulin heavy diversity 2-21 [Source: |
| AC246787.4 | CHR_HSCHR14_3_CTG1 | novel transcript |
| IGHD1-20 | CHR_HSCHR14_3_CTG1 | immunoglobulin heavy diversity 1-20 [Source: |
| IGHD6-19 | CHR_HSCHR14_3_CTG1 | immunoglobulin heavy diversity 6-19 [Source: |
| IGHD5-18 | CHR_HSCHR14_3_CTG1 | immunoglobulin heavy diversity 5-18 [Source: |
| IGHD4-17 | CHR_HSCHR14_3_CTG1 | immunoglobulin heavy diversity 4-17 [Source: |
| IGHD3-16 | CHR_HSCHR14_3_CTG1 | immunoglobulin heavy diversity 3-16 [Source: |
| IGHD2-15 | CHR_HSCHR14_3_CTG1 | immunoglobulin heavy diversity 2-15 [Source: |
| IGHD1-14 | CHR_HSCHR14_3_CTG1 | immunoglobulin heavy diversity 1-14 (non-fur |
| IGHD6-13 | CHR_HSCHR14_3_CTG1 | immunoglobulin heavy diversity 6-13 [Source: |
| IGHD5-12 | CHR_HSCHR14_3_CTG1 | immunoglobulin heavy diversity 5-12 [Source: |
| IGHD4-11 | CHR_HSCHR14_3_CTG1 | immunoglobulin heavy diversity 4-11 (non-fur |
| IGHD3-10 | CHR_HSCHR14_3_CTG1 | immunoglobulin heavy diversity 3-10 [Source: |
| IGHD3-9 | CHR_HSCHR14_3_CTG1 | immunoglobulin heavy diversity 3-9 [Source:H |
| IGHD2-8 | CHR_HSCHR14_3_CTG1 | immunoglobulin heavy diversity 2-8 [Source:H |
| IGHD1-7 | CHR_HSCHR14_3_CTG1 | immunoglobulin heavy diversity 1-7 [Source:H |
| IGHD6-6 | CHR_HSCHR14_3_CTG1 | immunoglobulin heavy diversity 6-6 [Source:H |
| IGHD5-5 | CHR_HSCHR14_3_CTG1 | immunoglobulin heavy diversity 5-5 [Source:H |
| IGHD4-4 | CHR_HSCHR14_3_CTG1 | immunoglobulin heavy diversity 4-4 [Source:H |
| IGHD3-3 | CHR_HSCHR14_3_CTG1 | immunoglobulin heavy diversity 3-3 [Source:H |
| IGHD2-2 | CHR_HSCHR14_3_CTG1 | immunoglobulin heavy diversity 2-2 [Source:H |
| FAM30A | CHR_HSCHR14_3_CTG1 | family with sequence similarity 30 member A |
| IGHD1-1 | CHR_HSCHR14_3_CTG1 | immunoglobulin heavy diversity 1-1 [Source:H |
| IGHV6-1 | CHR_HSCHR14_3_CTG1 | immunoglobulin heavy variable 6-1 [Source:H |
| IGHVII-1-1 | CHR_HSCHR14_3_CTG1 | immunoglobulin heavy variable (II)-1-1 (pseuc |
| ADAM6 | CHR_HSCHR14_3_CTG1 | ADAM metallopeptidase domain 6 (pseudoge |
| AC246787.3 | CHR_HSCHR14_3_CTG1 | ribosomal protein S8 (RPS8) pseudogene |
| IGHV1-2 | CHR_HSCHR14_3_CTG1 | immunoglobulin heavy variable 1-2 [Source:H |
| IGHVIII-2-1 | CHR_HSCHR14_3_CTG1 | immunoglobulin heavy variable (III)-2-1 (pseu |
| IGHV1-3 | CHR_HSCHR14_3_CTG1 | immunoglobulin heavy variable 1-3 [Source:H |

| | | |
|---|---|---|
| IGHV4-4 | CHR_HSCHR14_3_CTG1 | immunoglobulin heavy variable 4-4 [Source:H |
| AC244226.1 | CHR_HSCHR14_3_CTG1 | novel immunoglobulin heavy variable gene |
| IGHV2-5 | CHR_HSCHR14_3_CTG1 | immunoglobulin heavy variable 2-5 [Source:H |
| IGHVIII-5-1 | CHR_HSCHR14_3_CTG1 | immunoglobulin heavy variable (III)-5-1 (pseu |
| IGHVIII-5-2 | CHR_HSCHR14_3_CTG1 | immunoglobulin heavy variable (III)-5-2 (pseu |
| IGHV3-6 | CHR_HSCHR14_3_CTG1 | immunoglobulin heavy variable 3-6 (pseudoge |
| IGHV3-7 | CHR_HSCHR14_3_CTG1 | immunoglobulin heavy variable 3-7 [Source:H |
| AC247036.4 | CHR_HSCHR14_3_CTG1 | novel immunoglobulin heavy variable gene |
| AC247036.3 | CHR_HSCHR14_3_CTG1 | novel immunoglobulin heavy variable gene |
| IGHV3-11 | CHR_HSCHR14_3_CTG1 | immunoglobulin heavy variable 3-11 (gene/ps |
| IGHVIII-11-1 | CHR_HSCHR14_3_CTG1 | immunoglobulin heavy variable (III)-11-1 (pseu |
| IGHV1-12 | CHR_HSCHR14_3_CTG1 | immunoglobulin heavy variable 1-12 (pseudog |
| IGHV3-13 | CHR_HSCHR14_3_CTG1 | immunoglobulin heavy variable 3-13 [Source:I |
| IGHVIII-13-1 | CHR_HSCHR14_3_CTG1 | immunoglobulin heavy variable (III)-13-1 (pseu |
| IGHV1-14 | CHR_HSCHR14_3_CTG1 | immunoglobulin heavy variable 1-14 (pseudog |
| IGHV3-15 | CHR_HSCHR14_3_CTG1 | immunoglobulin heavy variable 3-15 [Source:I |
| IGHVII-15-1 | CHR_HSCHR14_3_CTG1 | immunoglobulin heavy variable (II)-15-1 (pseu |
| IGHV3-16 | CHR_HSCHR14_3_CTG1 | immunoglobulin heavy variable 3-16 (non-fun |
| IGHV1-17 | CHR_HSCHR14_3_CTG1 | immunoglobulin heavy variable 1-17 (pseudog |
| SLC20A1P2 | CHR_HSCHR14_3_CTG1 | solute carrier family 20 member 1 pseudogen |
| IGHV1-18 | CHR_HSCHR14_3_CTG1 | immunoglobulin heavy variable 1-18 [Source:I |
| IGHV3-19 | CHR_HSCHR14_3_CTG1 | immunoglobulin heavy variable 3-19 (pseudog |
| AC247036.2 | CHR_HSCHR14_3_CTG1 | novel transcript |
| IGHV3-20 | CHR_HSCHR14_3_CTG1 | immunoglobulin heavy variable 3-20 [Source:I |
| IGHV3-21 | CHR_HSCHR14_3_CTG1 | immunoglobulin heavy variable 3-21 [Source:I |
| IGHV3-22 | CHR_HSCHR14_3_CTG1 | immunoglobulin heavy variable 3-22 (pseudog |
| IGHVII-22-1 | CHR_HSCHR14_3_CTG1 | immunoglobulin heavy variable (II)-22-1 (pseu |
| IGHVIII-22-2 | CHR_HSCHR14_3_CTG1 | immunoglobulin heavy variable (III)-22-2 (pseu |
| IGHV3-23 | CHR_HSCHR14_3_CTG1 | immunoglobulin heavy variable 3-23 [Source:I |
| IGHV1-24 | CHR_HSCHR14_3_CTG1 | immunoglobulin heavy variable 1-24 [Source:I |
| HOMER2P2 | CHR_HSCHR14_3_CTG1 | HOMER2 pseudogene 2 [Source:HGNC Symbo |
| IGHV3-25 | CHR_HSCHR14_3_CTG1 | immunoglobulin heavy variable 3-25 (pseudog |
| IGHVIII-25-1 | CHR_HSCHR14_3_CTG1 | immunoglobulin heavy variable (III)-25-1 (pseu |
| IGHV2-26 | CHR_HSCHR14_3_CTG1 | immunoglobulin heavy variable 2-26 [Source:I |
| IGHVIII-26-1 | CHR_HSCHR14_3_CTG1 | immunoglobulin heavy variable (III)-26-1 (pseu |
| IGHVII-26-2 | CHR_HSCHR14_3_CTG1 | immunoglobulin heavy variable (II)-26-2 (pseu |
| IGHV7-27 | CHR_HSCHR14_3_CTG1 | immunoglobulin heavy variable 7-27 (pseudog |
| IGHV4-28 | CHR_HSCHR14_3_CTG1 | immunoglobulin heavy variable 4-28 [Source:I |
| AC245166.1 | CHR_HSCHR14_3_CTG1 | immunoglobulin heavy variable 3-29 (IGHV3-2 |
| IGHV3-30 | CHR_HSCHR14_3_CTG1 | immunoglobulin heavy variable 3-30 [Source:I |
| IGHVII-30-1 | CHR_HSCHR14_3_CTG1 | immunoglobulin heavy variable (II)-30-1 (pseu |
| IGHV3-30-2 | CHR_HSCHR14_3_CTG1 | immunoglobulin heavy variable 3-30-2 (pseud |
| AC234301.4 | CHR_HSCHR14_3_CTG1 | Immunoglobulin heavy variable 4-30-2 [Sourc |
| AC234301.6 | CHR_HSCHR14_3_CTG1 | immunoglobulin heavy variable (II)-31-1 (IGH\ |
| IGHV3-29 | CHR_HSCHR14_3_CTG1 | immunoglobulin heavy variable 3-29 (pseudog |
| IGHV3-33 | CHR_HSCHR14_3_CTG1 | immunoglobulin heavy variable 3-33 [Source:I |
| AC234301.2 | CHR_HSCHR14_3_CTG1 | novel immunoglobulin heavy variable pseudo |
| AC234301.1 | CHR_HSCHR14_3_CTG1 | novel immunoglobulin heavy variable gene |
| AC234301.5 | CHR_HSCHR14_3_CTG1 | novel immunoglobulin heavy variable pseudo |
| AC234301.3 | CHR_HSCHR14_3_CTG1 | novel immunoglobulin heavy variable gene |

| | | |
|---|---|---|
| IGHVII-33-1 | CHR_HSCHR14_3_CTG1 | immunoglobulin heavy variable (II)-33-1 (pseu |
| IGHV3-33-2 | CHR_HSCHR14_3_CTG1 | immunoglobulin heavy variable 3-33-2 (pseud |
| IGHV4-34 | CHR_HSCHR14_3_CTG1 | immunoglobulin heavy variable 4-34 [Source:I |
| IGHV7-34-1 | CHR_HSCHR14_3_CTG1 | immunoglobulin heavy variable 7-34-1 (pseud |
| IGHV3-35 | CHR_HSCHR14_3_CTG1 | immunoglobulin heavy variable 3-35 (non-fun |
| IGHV3-36 | CHR_HSCHR14_3_CTG1 | immunoglobulin heavy variable 3-36 (pseudog |
| IGHV3-37 | CHR_HSCHR14_3_CTG1 | immunoglobulin heavy variable 3-37 (pseudog |
| IGHV3-38 | CHR_HSCHR14_3_CTG1 | immunoglobulin heavy variable 3-38 (non-fun |
| IGHVIII-38-1 | CHR_HSCHR14_3_CTG1 | immunoglobulin heavy variable (III)-38-1 (pseu |
| IGHV4-39 | CHR_HSCHR14_3_CTG1 | immunoglobulin heavy variable 4-39 [Source:I |
| IGHV7-40 | CHR_HSCHR14_3_CTG1 | immunoglobulin heavy variable 7-40 (pseudog |
| AC244452.2 | CHR_HSCHR14_3_CTG1 | pseudogene similar to part of homer homolog |
| IGHVII-40-1 | CHR_HSCHR14_3_CTG1 | immunoglobulin heavy variable (II)-40-1 (pseu |
| IGHV3-41 | CHR_HSCHR14_3_CTG1 | immunoglobulin heavy variable 3-41 (pseudog |
| HOMER2P1 | CHR_HSCHR14_3_CTG1 | HOMER2 pseudogene 1 [Source:HGNC Symbo |
| IGHV3-42 | CHR_HSCHR14_3_CTG1 | immunoglobulin heavy variable 3-42 (pseudog |
| IGHV3-43 | CHR_HSCHR14_3_CTG1 | immunoglobulin heavy variable 3-43 [Source:I |
| IGHVIII-44 | CHR_HSCHR14_3_CTG1 | immunoglobulin heavy variable (III)-44 (pseud |
| IGHVIV-44-1 | CHR_HSCHR14_3_CTG1 | immunoglobulin heavy variable (IV)-44-1 (pse |
| IGHVII-44-2 | CHR_HSCHR14_3_CTG1 | immunoglobulin heavy variable (II)-44-2 (pseu |
| IGHV1-45 | CHR_HSCHR14_3_CTG1 | immunoglobulin heavy variable 1-45 [Source:I |
| IGHV1-46 | CHR_HSCHR14_3_CTG1 | immunoglobulin heavy variable 1-46 [Source:I |
| IGHVII-46-1 | CHR_HSCHR14_3_CTG1 | immunoglobulin heavy variable (II)-46-1 (pseu |
| IGHV3-47 | CHR_HSCHR14_3_CTG1 | immunoglobulin heavy variable 3-47 (pseudog |
| IGHVIII-47-1 | CHR_HSCHR14_3_CTG1 | immunoglobulin heavy variable (III)-47-1 (pseu |
| IGHV3-48 | CHR_HSCHR14_3_CTG1 | immunoglobulin heavy variable 3-48 [Source:I |
| IGHV3-49 | CHR_HSCHR14_3_CTG1 | immunoglobulin heavy variable 3-49 [Source:I |
| IGHVII-49-1 | CHR_HSCHR14_3_CTG1 | immunoglobulin heavy variable (II)-49-1 (pseu |
| IGHV3-50 | CHR_HSCHR14_3_CTG1 | immunoglobulin heavy variable 3-50 (pseudog |
| IGHV5-51 | CHR_HSCHR14_3_CTG1 | immunoglobulin heavy variable 5-51 [Source:I |
| IGHVIII-51-1 | CHR_HSCHR14_3_CTG1 | immunoglobulin heavy variable (III)-51-1 (pseu |
| IGHVII-51-2 | CHR_HSCHR14_3_CTG1 | immunoglobulin heavy variable (II)-51-2 (pseu |
| IGHV3-52 | CHR_HSCHR14_3_CTG1 | immunoglobulin heavy variable 3-52 (pseudog |
| IGHV3-53 | CHR_HSCHR14_3_CTG1 | immunoglobulin heavy variable 3-53 [Source:I |
| IGHVII-53-1 | CHR_HSCHR14_3_CTG1 | immunoglobulin heavy variable (II)-53-1 (pseu |
| IGHV3-54 | CHR_HSCHR14_3_CTG1 | immunoglobulin heavy variable 3-54 (pseudog |
| IGHV4-55 | CHR_HSCHR14_3_CTG1 | immunoglobulin heavy variable 4-55 (pseudog |
| IGHV7-56 | CHR_HSCHR14_3_CTG1 | immunoglobulin heavy variable 7-56 (pseudog |
| IGHV3-57 | CHR_HSCHR14_3_CTG1 | immunoglobulin heavy variable 3-57 (pseudog |
| IGHV1-58 | CHR_HSCHR14_3_CTG1 | immunoglobulin heavy variable 1-58 [Source:I |
| IGHV4-59 | CHR_HSCHR14_3_CTG1 | immunoglobulin heavy variable 4-59 [Source:I |
| IGHV3-60 | CHR_HSCHR14_3_CTG1 | immunoglobulin heavy variable 3-60 (pseudog |
| 5S_rRNA | CHR_HSCHR14_3_CTG1 | 5S ribosomal RNA [Source:RFAM;Acc:RF00001 |
| IGHVII-60-1 | CHR_HSCHR14_3_CTG1 | immunoglobulin heavy variable (II)-60-1 (pseu |
| IGHV4-61 | CHR_HSCHR14_3_CTG1 | immunoglobulin heavy variable 4-61 [Source:I |
| IGHV3-62 | CHR_HSCHR14_3_CTG1 | immunoglobulin heavy variable 3-62 (pseudog |
| IGHVII-62-1 | CHR_HSCHR14_3_CTG1 | immunoglobulin heavy variable (II)-62-1 (pseu |
| IGHV3-63 | CHR_HSCHR14_3_CTG1 | immunoglobulin heavy variable 3-63 (pseudog |
| IGHV3-64 | CHR_HSCHR14_3_CTG1 | immunoglobulin heavy variable 3-64 [Source:I |
| IGHV3-65 | CHR_HSCHR14_3_CTG1 | immunoglobulin heavy variable 3-65 (pseudog |

| | | |
|---|---|---|
| IGHVII-65-1 | CHR_HSCHR14_3_CTG1 | immunoglobulin heavy variable (II)-65-1 (pseu |
| IGHV3-66 | CHR_HSCHR14_3_CTG1 | immunoglobulin heavy variable 3-66 [Source:I |
| IGHV1-67 | CHR_HSCHR14_3_CTG1 | immunoglobulin heavy variable 1-67 (pseudog |
| SLC20A1P1 | CHR_HSCHR14_3_CTG1 | solute carrier family 20 member 1 pseudogen |
| IGHVII-67-1 | CHR_HSCHR14_3_CTG1 | immunoglobulin heavy variable (II)-67-1 (pseu |
| IGHVIII-67-2 | CHR_HSCHR14_3_CTG1 | immunoglobulin heavy variable (III)-67-2 (pse |
| IGHVIII-67-3 | CHR_HSCHR14_3_CTG1 | immunoglobulin heavy variable (III)-67-3 (pse |
| IGHVIII-67-4 | CHR_HSCHR14_3_CTG1 | immunoglobulin heavy variable (III)-67-4 (pse |
| IGHV1-68 | CHR_HSCHR14_3_CTG1 | immunoglobulin heavy variable 1-68 (pseudog |
| IGHV1-69 | CHR_HSCHR14_3_CTG1 | immunoglobulin heavy variable 1-69 [Source:I |
| IGHV2-70D | CHR_HSCHR14_3_CTG1 | immunoglobulin heavy variable 2-70D [Source |
| IGHV3-69-1 | CHR_HSCHR14_3_CTG1 | immunoglobulin heavy variable 3-69-1 (pseud |
| IGHV1-69-2 | CHR_HSCHR14_3_CTG1 | immunoglobulin heavy variable 1-69-2 [Sourc |
| AC242528.1 | CHR_HSCHR14_3_CTG1 | novel immunoglobulin heavy variable gene |
| IGHV1-69D | CHR_HSCHR14_3_CTG1 | immunoglobulin heavy variable 1-69D [Source |
| IGHV2-70 | CHR_HSCHR14_3_CTG1 | immunoglobulin heavy variable 2-70 [Source:I |
| IGHV3-71 | CHR_HSCHR14_3_CTG1 | immunoglobulin heavy variable 3-71 (pseudog |
| IGHV3-72 | CHR_HSCHR14_3_CTG1 | immunoglobulin heavy variable 3-72 [Source:I |
| IGHV3-73 | CHR_HSCHR14_3_CTG1 | immunoglobulin heavy variable 3-73 [Source:I |
| IGHV3-74 | CHR_HSCHR14_3_CTG1 | immunoglobulin heavy variable 3-74 [Source:I |
| IGHVII-74-1 | CHR_HSCHR14_3_CTG1 | immunoglobulin heavy variable (II)-74-1 (pseu |
| IGHV3-75 | CHR_HSCHR14_3_CTG1 | immunoglobulin heavy variable 3-75 (pseudog |
| IGHV3-76 | CHR_HSCHR14_3_CTG1 | immunoglobulin heavy variable 3-76 (pseudog |
| IGHVIII-76-1 | CHR_HSCHR14_3_CTG1 | immunoglobulin heavy variable (III)-76-1 (pse |
| IGHV5-78 | CHR_HSCHR14_3_CTG1 | immunoglobulin heavy variable 5-78 (pseudog |
| LINC02833 | CHR_HSCHR14_7_CTG1 | long intergenic non-protein coding RNA 2833 |
| LINC02287 | CHR_HSCHR14_7_CTG1 | long intergenic non-protein coding RNA 2287 |
| CHGA | CHR_HSCHR14_7_CTG1 | chromogranin A [Source:HGNC Symbol;Acc:H |
| ITPK1 | CHR_HSCHR14_7_CTG1 | inositol-tetrakisphosphate 1-kinase [Source:H |
| ITPK1-AS1 | CHR_HSCHR14_7_CTG1 | ITPK1 antisense RNA 1 [Source:HGNC Symbol; |
| AL110118.3 | CHR_HSCHR14_7_CTG1 | cytochrome b5 type B (outer mitochondrial m |
| MOAP1 | CHR_HSCHR14_7_CTG1 | modulator of apoptosis 1 [Source:HGNC Symb |
| TMEM251 | CHR_HSCHR14_7_CTG1 | transmembrane protein 251 [Source:HGNC Sy |
| AL110118.4 | CHR_HSCHR14_7_CTG1 | novel protein |
| GON7 | CHR_HSCHR14_7_CTG1 | GON7 subunit of KEOPS complex [Source:HGN |
| UBR7 | CHR_HSCHR14_7_CTG1 | ubiquitin protein ligase E3 component n-recog |
| AL132838.3 | CHR_HSCHR14_7_CTG1 | ribosomal protein L18a (RPL18A) pseudogene |
| BTBD7 | CHR_HSCHR14_7_CTG1 | BTB domain containing 7 [Source:HGNC Symb |
| AL132838.4 | CHR_HSCHR14_7_CTG1 | ribosomal protein L36a-like (RPL36AL) pseudo |
| UNC79 | CHR_HSCHR14_7_CTG1 | unc-79 homolog, NALCN channel complex sub |
| AL122023.2 | CHR_HSCHR14_7_CTG1 | novel transcript, sense intronic UNC79 |
| COX8C | CHR_HSCHR14_7_CTG1 | cytochrome c oxidase subunit 8C [Source:HGN |
| Y_RNA | CHR_HSCHR14_7_CTG1 | Y RNA [Source:RFAM;Acc:RF00019] |
| RNU6-1258P | CHR_HSCHR14_7_CTG1 | RNA, U6 small nuclear 1258, pseudogene [Sou |
| AL157858.1 | CHR_HSCHR14_7_CTG1 | NULL |
| PRIMA1 | CHR_HSCHR14_7_CTG1 | proline rich membrane anchor 1 [Source:HGN |
| AL132642.3 | CHR_HSCHR14_7_CTG1 | TEC |
| FAM181A-AS1 | CHR_HSCHR14_7_CTG1 | FAM181A antisense RNA 1 [Source:HGNC Sym |
| FAM181A | CHR_HSCHR14_7_CTG1 | family with sequence similarity 181 member A |
| ASB2 | CHR_HSCHR14_7_CTG1 | ankyrin repeat and SOCS box containing 2 [Sou |

| | | |
|---|---|---|
| AL132642.4 | CHR_HSCHR14_7_CTG1 | novel transcript, antisense to ASB2 |
| CCDC197 | CHR_HSCHR14_7_CTG1 | coiled-coil domain containing 197 [Source:HG |
| OTUB2 | CHR_HSCHR14_7_CTG1 | OTU deubiquitinase, ubiquitin aldehyde bindi |
| DDX24 | CHR_HSCHR14_7_CTG1 | DEAD-box helicase 24 [Source:HGNC Symbol; |
| IFI27L1 | CHR_HSCHR14_7_CTG1 | interferon alpha inducible protein 27 like 1 [S |
| IFI27 | CHR_HSCHR14_7_CTG1 | interferon alpha inducible protein 27 [Source: |
| IFI27L2 | CHR_HSCHR14_7_CTG1 | interferon alpha inducible protein 27 like 2 [S |
| PPP4R4 | CHR_HSCHR14_7_CTG1 | protein phosphatase 4 regulatory subunit 4 [S |
| SERPINA10 | CHR_HSCHR14_7_CTG1 | serpin family A member 10 [Source:HGNC Syn |
| SERPINA6 | CHR_HSCHR14_7_CTG1 | serpin family A member 6 [Source:HGNC Sym |
| SERPINA2 | CHR_HSCHR14_7_CTG1 | serpin family A member 2 (gene/pseudogene) |
| SERPINA1 | CHR_HSCHR14_7_CTG1 | serpin family A member 1 [Source:HGNC Sym |
| AL136038.6 | CHR_HSCHR14_8_CTG1 | novel transcript |
| SGPP1 | CHR_HSCHR14_8_CTG1 | sphingosine-1-phosphate phosphatase 1 [Sou |
| AL161670.3 | CHR_HSCHR14_8_CTG1 | eukaryotic translation initiation factor 2, subu |
| RNU7-116P | CHR_HSCHR14_8_CTG1 | RNA, U7 small nuclear 116 pseudogene [Sour |
| AL161670.4 | CHR_HSCHR14_8_CTG1 | RAP1B, member of RAS oncogene family (RAP |
| RPS28P1 | CHR_HSCHR14_8_CTG1 | ribosomal protein S28 pseudogene 1 [Source: |
| LINC00871 | CHR_HSCHR14_9_CTG1 | long intergenic non-protein coding RNA 871 [ |
| AC068446.4 | CHR_HSCHR15_1_CTG1 | neurobeachin (NBEA) pseudogene |
| AC136352.1 | CHR_HSCHR15_1_CTG1 | p21 protein (Cdc42/Rac)-activated kinase 2 (P |
| AC136352.10 | CHR_HSCHR15_1_CTG1 | novel transcript |
| U6 | CHR_HSCHR15_1_CTG1 | U6 spliceosomal RNA [Source:RFAM;Acc:RF00 |
| AC136352.8 | CHR_HSCHR15_1_CTG1 | novel transcript |
| AC136352.7 | CHR_HSCHR15_1_CTG1 | novel transcript |
| AC136352.6 | CHR_HSCHR15_1_CTG1 | LON peptidase N-terminal domain and ring fir |
| AC136352.9 | CHR_HSCHR15_1_CTG1 | novel transcript |
| CXADRP2 | CHR_HSCHR15_1_CTG1 | CXADR pseudogene 2 [Source:HGNC Symbol; |
| AC136352.5 | CHR_HSCHR15_1_CTG1 | GRAM domain containing 4 (GRAMD4) pseud |
| GRAMD4P6 | CHR_HSCHR15_1_CTG1 | GRAM domain containing 4 pseudogene 6 [So |
| POTEB3 | CHR_HSCHR15_1_CTG1 | POTE ankyrin domain family member B3 [Sou |
| U6 | CHR_HSCHR15_1_CTG1 | U6 spliceosomal RNA [Source:RFAM;Acc:RF00 |
| NF1P9 | CHR_HSCHR15_1_CTG1 | neurofibromin 1 pseudogene 9 [Source:HGNC |
| AC020679.1 | CHR_HSCHR15_1_CTG1 | novel transcript |
| AC020679.2 | CHR_HSCHR15_1_CTG1 | olfactory receptor, family 4 pseudogene |
| LINC02203 | CHR_HSCHR15_1_CTG1 | long intergenic non-protein coding RNA 2203 |
| AC171558.6 | CHR_HSCHR15_1_CTG1 | novel protein, similar to OR4M2 |
| AC171558.5 | CHR_HSCHR15_1_CTG1 | novel protein, similar to OR4N4 |
| AC171558.2 | CHR_HSCHR15_1_CTG1 | olfactory receptor, family 4, subfamily N, men |
| AC171558.7 | CHR_HSCHR15_1_CTG1 | ribosomal protein S8 (RPS8) pseudogene |
| CYFIP1 | CHR_HSCHR15_1_CTG3 | cytoplasmic FMR1 interacting protein 1 [Sour |
| TUBGCP5 | CHR_HSCHR15_1_CTG3 | tubulin gamma complex associated protein 5 |
| ELMO2P1 | CHR_HSCHR15_1_CTG3 | engulfment and cell motility 2 pseudogene 1 [ |
| GOLGA6L1 | CHR_HSCHR15_1_CTG3 | golgin A6 family like 1 [Source:HGNC Symbol; |
| AC116165.4 | CHR_HSCHR15_1_CTG3 | M-phase phosphoprotein 10 (U3 small nucleo |
| GOLGA8EP | CHR_HSCHR15_1_CTG3 | golgin A8 family member E, pseudogene [Sou |
| Metazoa_SRP | CHR_HSCHR15_1_CTG3 | Metazoan signal recognition particle RNA [Sou |
| ABCB10P1 | CHR_HSCHR15_1_CTG3 | ABCB10 pseudogene 1 [Source:HGNC Symbol; |
| AC116165.7 | CHR_HSCHR15_1_CTG3 | family with sequence similarity 208, member |
| AC116165.6 | CHR_HSCHR15_1_CTG3 | abhydrolase domain containing 17A (ABHD17 |

| | | |
|---|---|---|
| AC091304.7 | CHR_HSCHR15_1_CTG8 | NULL |
| HERC2 | CHR_HSCHR15_1_CTG8 | HECT and RLD domain containing E3 ubiquitin |
| AC091304.8 | CHR_HSCHR15_1_CTG8 | hect domain and RLD 2 (HERC2) pseudogene |
| AC138747.6 | CHR_HSCHR15_1_CTG8 | cholinergic receptor, nicotinic, beta 4 (CHRNB |
| AC138747.5 | CHR_HSCHR15_1_CTG8 | golgin A6 family-like [Source:NCBI gene;Acc:6 |
| AC138747.3 | CHR_HSCHR15_1_CTG8 | M-phase phosphoprotein 10 (U3 small nucleo |
| GOLGA8F | CHR_HSCHR15_1_CTG8 | golgin A8 family member F [Source:HGNC Sym |
| Metazoa_SRP | CHR_HSCHR15_1_CTG8 | Metazoan signal recognition particle RNA [Sou |
| AC138747.4 | CHR_HSCHR15_1_CTG8 | ATP-binding cassette, sub-family B (MDR/TAP |
| AC138747.10 | CHR_HSCHR15_1_CTG8 | family with sequence similarity 208, member |
| AC138747.7 | CHR_HSCHR15_1_CTG8 | ATP-binding cassette, sub-family B (MDR/TAP |
| AC138747.8 | CHR_HSCHR15_1_CTG8 | family with sequence similarity 208, member |
| AC138747.9 | CHR_HSCHR15_1_CTG8 | glypican 5 (GPC5) pseudogene |
| AC138747.1 | CHR_HSCHR15_1_CTG8 | family with sequence similarity 108, member |
| AC138747.2 | CHR_HSCHR15_1_CTG8 | glypican 5 (GPC5) pseudogene |
| AC138749.8 | CHR_HSCHR15_1_CTG8 | family with sequence similarity 208, member |
| ABCB10P4 | CHR_HSCHR15_1_CTG8 | ABCB10 pseudogene 4 [Source:HGNC Symbol; |
| GOLGA8G | CHR_HSCHR15_1_CTG8 | golgin A8 family member G [Source:HGNC Sym |
| Metazoa_SRP | CHR_HSCHR15_1_CTG8 | Metazoan signal recognition particle RNA [Sou |
| AC138749.9 | CHR_HSCHR15_1_CTG8 | M-phase phosphoprotein 10 (U3 small nucleo |
| AC138749.7 | CHR_HSCHR15_1_CTG8 | golgin subfamily A member 6-like protein 22 [ |
| AC138749.10 | CHR_HSCHR15_1_CTG8 | cholinergic receptor, nicotinic, beta 4 (CHRNB |
| HERC2P11 | CHR_HSCHR15_1_CTG8 | hect domain and RLD 2 pseudogene 11 [Sourc |
| AC138749.6 | CHR_HSCHR15_1_CTG8 | hect domain and RLD 2 (HERC2) pseudogene |
| TUBGCP5 | CHR_HSCHR15_2_CTG3 | tubulin gamma complex associated protein 5 |
| ELMO2P1 | CHR_HSCHR15_2_CTG3 | engulfment and cell motility 2 pseudogene 1 [ |
| GOLGA6L1 | CHR_HSCHR15_2_CTG3 | golgin A6 family like 1 [Source:HGNC Symbol;/ |
| AC116165.9 | CHR_HSCHR15_2_CTG3 | M-phase phosphoprotein 10 (U3 small nucleo |
| GOLGA8EP | CHR_HSCHR15_2_CTG3 | golgin A8 family member E, pseudogene [Sour |
| Metazoa_SRP | CHR_HSCHR15_2_CTG3 | Metazoan signal recognition particle RNA [Sou |
| ABCB10P1 | CHR_HSCHR15_2_CTG3 | ABCB10 pseudogene 1 [Source:HGNC Symbol; |
| AC116165.5 | CHR_HSCHR15_2_CTG3 | family with sequence similarity 208, member |
| AC116165.8 | CHR_HSCHR15_2_CTG3 | glypican 5 (GPC5) pseudogene |
| AC116165.10 | CHR_HSCHR15_2_CTG3 | abhydrolase domain containing 17A (ABHD17. |
| MEGF11 | CHR_HSCHR15_2_CTG8 | multiple EGF like domains 11 [Source:HGNC S |
| AC087382.2 | CHR_HSCHR15_2_CTG8 | novel transcript |
| AC087382.1 | CHR_HSCHR15_2_CTG8 | novel transcript, antisense to DIS3L |
| AC136687.2 | CHR_HSCHR15_3_CTG3 | Ig heavy chain V-I pseudogene |
| AC136687.3 | CHR_HSCHR15_3_CTG3 | ring finger protein 145 (Rnf145) pseudogene |
| SPATA31E3P | CHR_HSCHR15_3_CTG3 | SPATA31 subfamily E member 3, pseudogene |
| GOLGA8DP | CHR_HSCHR15_3_CTG3 | golgin A8 family member D, pseudogene [Sou |
| Metazoa_SRP | CHR_HSCHR15_3_CTG3 | Metazoan signal recognition particle RNA [Sou |
| AC136687.1 | CHR_HSCHR15_3_CTG3 | M-phase phosphoprotein 10 (U3 small nucleo |
| GOLGA6L22 | CHR_HSCHR15_3_CTG3 | golgin A6 family like 22 [Source:HGNC Symbol |
| HERC2P7 | CHR_HSCHR15_3_CTG3 | hect domain and RLD 2 pseudogene 7 [Source |
| HERC2P2 | CHR_HSCHR15_3_CTG3 | hect domain and RLD 2 pseudogene 2 [Source |
| GOLGA8IP | CHR_HSCHR15_3_CTG3 | golgin A8 family member I, pseudogene [Sour |
| Metazoa_SRP | CHR_HSCHR15_3_CTG3 | Metazoan signal recognition particle RNA [Sou |
| AC018901.1 | CHR_HSCHR15_3_CTG8 | TEC |
| Y_RNA | CHR_HSCHR15_3_CTG8 | Y RNA [Source:RFAM;Acc:RF00019] |

| | | |
|---|---|---|
| PATL2 | CHR_HSCHR15_3_CTG8 | PAT1 homolog 2 [Source:HGNC Symbol;Acc:H |
| B2M | CHR_HSCHR15_3_CTG8 | beta-2-microglobulin [Source:HGNC Symbol;A |
| TRIM69 | CHR_HSCHR15_3_CTG8 | tripartite motif containing 69 [Source:HGNC S |
| SORD2P | CHR_HSCHR15_3_CTG8 | sorbitol dehydrogenase 2, pseudogene [Sourc |
| Y_RNA | CHR_HSCHR15_3_CTG8 | Y RNA [Source:RFAM;Acc:RF00019] |
| U1 | CHR_HSCHR15_3_CTG8 | U1 spliceosomal RNA [Source:RFAM;Acc:RF00 |
| OCA2 | CHR_HSCHR15_4_CTG8 | OCA2 melanosomal transmembrane protein [ |
| RPL5P32 | CHR_HSCHR15_4_CTG8 | ribosomal protein L5 pseudogene 32 [Source: |
| HERC2 | CHR_HSCHR15_4_CTG8 | HECT and RLD domain containing E3 ubiquitin |
| RPL41P2 | CHR_HSCHR15_4_CTG8 | ribosomal protein L41 pseudogene 2 [Source: |
| AC243660.7 | CHR_HSCHR15_4_CTG8 | hect domain and RLD 2 (HERC2) pseudogene |
| AC243660.1 | CHR_HSCHR15_4_CTG8 | cholinergic receptor, nicotinic, beta 4 (CHRNB |
| AC243660.6 | CHR_HSCHR15_4_CTG8 | golgin A6 family-like [Source:NCBI gene;Acc:6 |
| AC243660.4 | CHR_HSCHR15_4_CTG8 | M-phase phosphoprotein 10 (U3 small nucleo |
| GOLGA8F | CHR_HSCHR15_4_CTG8 | golgin A8 family member F [Source:HGNC Syn |
| Metazoa_SRP | CHR_HSCHR15_4_CTG8 | Metazoan signal recognition particle RNA [Sou |
| ABCB10P3 | CHR_HSCHR15_4_CTG8 | ABCB10 pseudogene 3 [Source:HGNC Symbol; |
| AC243660.8 | CHR_HSCHR15_4_CTG8 | family with sequence similarity 208, member |
| AC243660.2 | CHR_HSCHR15_4_CTG8 | amily with sequence similarity 208, member A |
| AC243660.3 | CHR_HSCHR15_4_CTG8 | family with sequence similarity 208, member |
| AC243660.5 | CHR_HSCHR15_4_CTG8 | family with sequence similarity 208, member |
| AC243913.2 | CHR_HSCHR15_4_CTG8 | family with sequence similarity 208, member |
| AC243913.11 | CHR_HSCHR15_4_CTG8 | family with sequence similarity 208, member |
| AC243913.5 | CHR_HSCHR15_4_CTG8 | family with sequence similarity 208, member |
| AC243913.10 | CHR_HSCHR15_4_CTG8 | glypican 5 (GPC5) pseudogene |
| AC243913.6 | CHR_HSCHR15_4_CTG8 | family with sequence similarity 108, member |
| AC243913.3 | CHR_HSCHR15_4_CTG8 | glypican 5 (GPC5) pseudogene |
| AC243913.1 | CHR_HSCHR15_4_CTG8 | family with sequence similarity 208, member |
| ABCB10P4 | CHR_HSCHR15_4_CTG8 | ABCB10 pseudogene 4 [Source:HGNC Symbol; |
| GOLGA8G | CHR_HSCHR15_4_CTG8 | golgin A8 family member G [Source:HGNC Syn |
| Metazoa_SRP | CHR_HSCHR15_4_CTG8 | Metazoan signal recognition particle RNA [Sou |
| AC243913.4 | CHR_HSCHR15_4_CTG8 | Golgin subfamily A member 2 pseudogene |
| AC243913.8 | CHR_HSCHR15_4_CTG8 | golgin subfamily A member 6-like protein 22 [ |
| AC243913.7 | CHR_HSCHR15_4_CTG8 | cholinergic receptor, nicotinic, beta 4 (CHRNB |
| AC243913.9 | CHR_HSCHR15_4_CTG8 | hect domain and RLD 2 (HERC2) pseudogene |
| HERC2P9 | CHR_HSCHR15_4_CTG8 | hect domain and RLD 2 pseudogene 9 [Source |
| GOLGA8M | CHR_HSCHR15_4_CTG8 | golgin A8 family member M [Source:HGNC Syi |
| Metazoa_SRP | CHR_HSCHR15_4_CTG8 | Metazoan signal recognition particle RNA [Sou |
| WHAMMP2 | CHR_HSCHR15_4_CTG8 | WAS protein homolog associated with actin, g |
| AC243657.1 | CHR_HSCHR15_4_CTG8 | zinc finger, FYVE domain containing 9 (ZFYVE9 |
| AC243657.3 | CHR_HSCHR15_4_CTG8 | taspase, threonine aspartase, 1 (TASP1) pseuc |
| AC243657.2 | CHR_HSCHR15_4_CTG8 | dexamethasone-induced transcript (DEXI) pse |
| PDCD6IPP2 | CHR_HSCHR15_4_CTG8 | PDCD6IP pseudogene 2 [Source:HGNC Symbo |
| APBA2 | CHR_HSCHR15_4_CTG8 | amyloid beta precursor protein binding family |
| FAM189A1 | CHR_HSCHR15_4_CTG8 | family with sequence similarity 189 member A |
| AC243902.2 | CHR_HSCHR15_4_CTG8 | DDB1 and CUL4 associated factor 6 (DCAF6) p |
| AC243902.1 | CHR_HSCHR15_4_CTG8 | NULL |
| NSMCE3 | CHR_HSCHR15_4_CTG8 | NSE3 homolog, SMC5-SMC6 complex compon |
| AC243686.1 | CHR_HSCHR15_4_CTG8 | novel transcript |
| TJP1 | CHR_HSCHR15_4_CTG8 | tight junction protein 1 [Source:HGNC Symbol |

| | | |
|---|---|---|
| AC243587.1 | CHR_HSCHR15_4_CTG8 | NULL |
| GOLGA8J | CHR_HSCHR15_4_CTG8 | golgin A8 family member J [Source:HGNC Sym |
| Metazoa_SRP | CHR_HSCHR15_4_CTG8 | Metazoan signal recognition particle RNA [Sou |
| U8 | CHR_HSCHR15_4_CTG8 | U8 small nucleolar RNA [Source:RFAM;Acc:RF |
| Metazoa_SRP | CHR_HSCHR15_4_CTG8 | Metazoan signal recognition particle RNA [Sou |
| AC243790.1 | CHR_HSCHR15_4_CTG8 | NULL |
| U6 | CHR_HSCHR15_4_CTG8 | U6 spliceosomal RNA [Source:RFAM;Acc:RF00 |
| Y_RNA | CHR_HSCHR15_4_CTG8 | Y RNA [Source:RFAM;Acc:RF00019] |
| CHRFAM7A | CHR_HSCHR15_4_CTG8 | CHRNA7 (exons 5-10) and FAM7A (exons A-E) |
| U8 | CHR_HSCHR15_4_CTG8 | U8 small nucleolar RNA [Source:RFAM;Acc:RF |
| Metazoa_SRP | CHR_HSCHR15_4_CTG8 | Metazoan signal recognition particle RNA [Sou |
| Metazoa_SRP | CHR_HSCHR15_4_CTG8 | Metazoan signal recognition particle RNA [Sou |
| U8 | CHR_HSCHR15_4_CTG8 | U8 small nucleolar RNA [Source:RFAM;Acc:RF |
| Metazoa_SRP | CHR_HSCHR15_4_CTG8 | Metazoan signal recognition particle RNA [Sou |
| ARHGAP11B | CHR_HSCHR15_4_CTG8 | Rho GTPase activating protein 11B [Source:H |
| Y_RNA | CHR_HSCHR15_4_CTG8 | Y RNA [Source:RFAM;Acc:RF00019] |
| Metazoa_SRP | CHR_HSCHR15_4_CTG8 | Metazoan signal recognition particle RNA [Sou |
| FAN1 | CHR_HSCHR15_4_CTG8 | FANCD2 and FANCI associated nuclease 1 [Sou |
| MTMR10 | CHR_HSCHR15_4_CTG8 | myotubularin related protein 10 [Source:HGN |
| U6 | CHR_HSCHR15_4_CTG8 | U6 spliceosomal RNA [Source:RFAM;Acc:RF00 |
| TRPM1 | CHR_HSCHR15_4_CTG8 | transient receptor potential cation channel su |
| KLF13 | CHR_HSCHR15_4_CTG8 | Kruppel like factor 13 [Source:HGNC Symbol;A |
| AC243649.1 | CHR_HSCHR15_4_CTG8 | NULL |
| AC243734.1 | CHR_HSCHR15_4_CTG8 | cholinergic receptor nicotinic alpha 7 subunit |
| Y_RNA | CHR_HSCHR15_4_CTG8 | Y RNA [Source:RFAM;Acc:RF00019] |
| U6 | CHR_HSCHR15_4_CTG8 | U6 spliceosomal RNA [Source:RFAM;Acc:RF00 |
| Metazoa_SRP | CHR_HSCHR15_4_CTG8 | Metazoan signal recognition particle RNA [Sou |
| ARHGAP11A | CHR_HSCHR15_4_CTG8 | Rho GTPase activating protein 11A [Source:H |
| SCG5 | CHR_HSCHR15_4_CTG8 | secretogranin V [Source:HGNC Symbol;Acc:H |
| GREM1 | CHR_HSCHR15_4_CTG8 | gremlin 1, DAN family BMP antagonist [Source |
| ADAMTS7P1 | CHR_HSCHR15_5_CTG8 | ADAMTS7 pseudogene 1 [Source:HGNC Symb |
| GOLGA6L10 | CHR_HSCHR15_5_CTG8 | golgin A6 family like 10 [Source:NCBI gene;Ac |
| AC011295.3 | CHR_HSCHR15_5_CTG8 | chondroitin sulfate proteoglycan 4 (CSPG4) ps |
| AC011295.2 | CHR_HSCHR15_5_CTG8 | NULL |
| GOLGA6L10 | CHR_HSCHR15_5_CTG8 | golgin A6 family like 10 [Source:HGNC Symbol |
| UBE2Q2P2 | CHR_HSCHR15_5_CTG8 | ubiquitin conjugating enzyme E2 Q2 pseudoge |
| AC011295.1 | CHR_HSCHR15_5_CTG8 | ribosomal protein L9 pseudogene 9 |
| AC010724.3 | CHR_HSCHR15_5_CTG8 | novel transcript |
| GOLGA6L9 | CHR_HSCHR15_5_CTG8 | golgin A6 family like 9 [Source:HGNC Symbol;A |
| UBE2Q2P6 | CHR_HSCHR15_5_CTG8 | ubiquitin conjugating enzyme E2 Q2 pseudoge |
| CSPG4P10 | CHR_HSCHR15_5_CTG8 | chondroitin sulfate proteoglycan 4 pseudoger |
| AC010724.1 | CHR_HSCHR15_5_CTG8 | dynamin 1 (DNM1) pseudogene |
| GOLGA2P10 | CHR_HSCHR15_5_CTG8 | GOLGA2 pseudogene 10 [Source:HGNC Symbo |
| GOLGA6L17P | CHR_HSCHR15_5_CTG8 | golgin A6 family like 17, pseudogene [Source: |
| DNM1P38 | CHR_HSCHR15_5_CTG8 | dynamin 1 pseudogene 38 [Source:HGNC Sym |
| RPS17 | CHR_HSCHR15_5_CTG8 | ribosomal protein S17 [Source:HGNC Symbol; |
| AC010724.2 | CHR_HSCHR15_5_CTG8 | novel protein |
| CPEB1 | CHR_HSCHR15_5_CTG8 | cytoplasmic polyadenylation element binding |
| AC111152.4 | CHR_HSCHR15_6_CTG8 | novel transcript |
| AC254979.1 | CHR_HSCHR15_6_CTG8 | synaptogyrin 2 (SYNGR2) pseudogene |

| | | |
|---|---|---|
| AC254979.2 | CHR_HSCHR15_6_CTG8 | tight junction protein 1 (zona occludens 1) (TJ |
| AC254979.3 | CHR_HSCHR15_6_CTG8 | NULL |
| RN7SL673P | CHR_HSCHR15_6_CTG8 | RNA, 7SL, cytoplasmic 673, pseudogene [Sour |
| DNM1P28 | CHR_HSCHR15_6_CTG8 | dynamin 1 pseudogene 28 [Source:HGNC Sym |
| ULK4P3 | CHR_HSCHR15_6_CTG8 | ULK4 pseudogene 3 [Source:HGNC Symbol;Ac |
| U8 | CHR_HSCHR15_6_CTG8 | U8 small nucleolar RNA [Source:RFAM;Acc:RF |
| GOLGA8T | CHR_HSCHR15_6_CTG8 | golgin A8 family member T [Source:HGNC Sym |
| RN7SL469P | CHR_HSCHR15_6_CTG8 | RNA, 7SL, cytoplasmic 469, pseudogene [Sour |
| DNM1P30 | CHR_HSCHR15_6_CTG8 | dynamin 1 pseudogene 30 [Source:HGNC Sym |
| AC254962.2 | CHR_HSCHR15_6_CTG8 | synaptotagmin II (SYT2) pseudogene |
| AC254962.1 | CHR_HSCHR15_6_CTG8 | hyperpolarization activated cyclic nucleotide- |
| RNU6-17P | CHR_HSCHR15_6_CTG8 | RNA, U6 small nuclear 17, pseudogene [Sourc |
| AC254958.1 | CHR_HSCHR15_6_CTG8 | TEC |
| Y_RNA | CHR_HSCHR15_6_CTG8 | Y RNA [Source:RFAM;Acc:RF00019] |
| AC254952.1 | CHR_HSCHR15_6_CTG8 | cholinergic receptor nicotinic alpha 7 subunit |
| ARHGAP11B | CHR_HSCHR15_6_CTG8 | Rho GTPase activating protein 11B [Source:H |
| AC254944.2 | CHR_HSCHR15_6_CTG8 | novel transcript, antisense to ARHGAP11B |
| AC254944.3 | CHR_HSCHR15_6_CTG8 | novel transcript |
| RN7SL628P | CHR_HSCHR15_6_CTG8 | RNA, 7SL, cytoplasmic 628, pseudogene [Sour |
| AC254944.1 | CHR_HSCHR15_6_CTG8 | novel transcript, antisense to AC026150.11 |
| ULK4P2 | CHR_HSCHR15_6_CTG8 | ULK4 pseudogene 2 [Source:HGNC Symbol;Ac |
| U8 | CHR_HSCHR15_6_CTG8 | U8 small nucleolar RNA [Source:RFAM;Acc:RF |
| DNM1P50 | CHR_HSCHR15_6_CTG8 | dynamin 1 pseudogene 50 [Source:HGNC Sym |
| GOLGA8Q | CHR_HSCHR15_6_CTG8 | golgin A8 family member Q [Source:HGNC Syr |
| RN7SL796P | CHR_HSCHR15_6_CTG8 | RNA, 7SL, cytoplasmic 796, pseudogene [Sour |
| AC254960.1 | CHR_HSCHR15_6_CTG8 | novel transcript |
| AC254960.2 | CHR_HSCHR15_6_CTG8 | M-phase phosphoprotein 10 (U3 small nucleo |
| AC254945.1 | CHR_HSCHR15_6_CTG8 | novel pseudogene |
| GOLGA8N | CHR_HSCHR15_6_CTG8 | golgin A8 family member N [Source:HGNC Syr |
| RN7SL286P | CHR_HSCHR15_6_CTG8 | RNA, 7SL, cytoplasmic 286, pseudogene [Sour |
| AC254945.3 | CHR_HSCHR15_6_CTG8 | NULL |
| AC254945.4 | CHR_HSCHR15_6_CTG8 | novel transcript, antisense to ARHGAP11A |
| SCG5 | CHR_HSCHR15_6_CTG8 | secretogranin V [Source:HGNC Symbol;Acc:H |
| AC254945.2 | CHR_HSCHR15_6_CTG8 | novel transcript |
| AC090877.4 | CHR_HSCHR15_6_CTG8 | novel transcript |
| GREM1 | CHR_HSCHR15_6_CTG8 | gremlin 1, DAN family BMP antagonist [Source |
| AC090877.3 | CHR_HSCHR15_6_CTG8 | microtubule-associated protein 1 light chain 3 |
| PARN | CHR_HSCHR16_1_CTG1 | poly(A)-specific ribonuclease [Source:HGNC S |
| RN7SL274P | CHR_HSCHR16_1_CTG1 | RNA, 7SL, cytoplasmic 274, pseudogene [Sour |
| BFAR | CHR_HSCHR16_1_CTG1 | bifunctional apoptosis regulator [Source:HGN |
| U7 | CHR_HSCHR16_1_CTG1 | U7 small nuclear RNA [Source:RFAM;Acc:RF0 |
| PLA2G10 | CHR_HSCHR16_1_CTG1 | phospholipase A2 group X [Source:HGNC Sym |
| AC243753.1 | CHR_HSCHR16_1_CTG1 | novel transcript, antisense to a novel membe |
| NPIPA3 | CHR_HSCHR16_1_CTG1 | nuclear pore complex interacting protein fami |
| Y_RNA | CHR_HSCHR16_1_CTG1 | Y RNA [Source:RFAM;Acc:RF00019] |
| PKD1P6 | CHR_HSCHR16_1_CTG1 | polycystin 1, transient receptor potential char |
| AC243651.1 | CHR_HSCHR16_1_CTG1 | novel transcript, antisense to RRN3 and a nov |
| RRN3 | CHR_HSCHR16_1_CTG1 | RRN3 homolog, RNA polymerase I transcriptic |
| NTAN1 | CHR_HSCHR16_1_CTG1 | N-terminal asparagine amidase [Source:HGNC |
| PDXDC1 | CHR_HSCHR16_1_CTG1 | pyridoxal dependent decarboxylase domain c |

| | | |
|---|---|---|
| AC243651.2 | CHR_HSCHR16_1_CTG1 | novel transcript, antisense to PDXDC1 |
| AC243791.1 | CHR_HSCHR16_1_CTG1 | novel pseudogene KIAA2013 |
| NPIPA1 | CHR_HSCHR16_1_CTG1 | nuclear pore complex interacting protein fami |
| AC243791.3 | CHR_HSCHR16_1_CTG1 | novel transcript |
| AC243791.2 | CHR_HSCHR16_1_CTG1 | polycystic kidney disease 1 (autosomal domin |
| Y_RNA | CHR_HSCHR16_1_CTG1 | Y RNA [Source:RFAM;Acc:RF00019] |
| PDXDC1 | CHR_HSCHR16_1_CTG1 | pyridoxal dependent decarboxylase domain c |
| Y_RNA | CHR_HSCHR16_1_CTG1 | Y RNA [Source:RFAM;Acc:RF00019] |
| NOMO1 | CHR_HSCHR16_1_CTG1 | NODAL modulator 1 [Source:HGNC Symbol;Ac |
| AC245049.1 | CHR_HSCHR16_1_CTG1 | acid phosphatase 1, soluble (ACP1) pseudoger |
| ABCC6P2 | CHR_HSCHR16_1_CTG1 | ATP binding cassette subfamily C member 6 p |
| NPIPA5 | CHR_HSCHR16_1_CTG1 | nuclear pore complex interacting protein fami |
| MPV17L | CHR_HSCHR16_1_CTG1 | MPV17 mitochondrial inner membrane protei |
| AC243733.1 | CHR_HSCHR16_1_CTG1 | MPV17 mitochondrial inner membrane protei |
| BMERB1 | CHR_HSCHR16_1_CTG1 | bMERB domain containing 1 [Source:HGNC Sy |
| MARF1 | CHR_HSCHR16_1_CTG1 | meiosis regulator and mRNA stability factor 1 |
| AC243742.2 | CHR_HSCHR16_1_CTG1 | novel transcript |
| NDE1 | CHR_HSCHR16_1_CTG1 | nudE neurodevelopment protein 1 [Source:H( |
| AC243742.1 | CHR_HSCHR16_1_CTG1 | novel transcript, sense to NDE1 |
| AC243742.3 | CHR_HSCHR16_1_CTG1 | novel transcript |
| MYH11 | CHR_HSCHR16_1_CTG1 | myosin heavy chain 11 [Source:HGNC Symbol; |
| AC243629.2 | CHR_HSCHR16_1_CTG1 | novel transcript, antisense to MYH11 |
| AC243629.1 | CHR_HSCHR16_1_CTG1 | novel transcript, antisense to MYH11 |
| FOPNL | CHR_HSCHR16_1_CTG1 | FGFR1OP N-terminal like [Source:HGNC Symb |
| U6 | CHR_HSCHR16_1_CTG1 | U6 spliceosomal RNA [Source:RFAM;Acc:RF00 |
| AC243629.3 | CHR_HSCHR16_1_CTG1 | novel transcript, antisense to C16orf63 |
| RPL15P20 | CHR_HSCHR16_1_CTG1 | ribosomal protein L15 pseudogene 20 [Source |
| ABCC1 | CHR_HSCHR16_1_CTG1 | ATP binding cassette subfamily C member 1 [S |
| RPL17P40 | CHR_HSCHR16_1_CTG1 | ribosomal protein L17 pseudogene 40 [Source |
| ABCC6 | CHR_HSCHR16_1_CTG1 | ATP binding cassette subfamily C member 6 [S |
| AC243824.1 | CHR_HSCHR16_1_CTG1 | novel transcript, antisense to ABCC6 |
| NOMO3 | CHR_HSCHR16_1_CTG1 | NODAL modulator 3 [Source:HGNC Symbol;Ac |
| AC244489.4 | CHR_HSCHR16_1_CTG1 | acid phosphatase 1, soluble (ACP1) pseudoger |
| AC244489.3 | CHR_HSCHR16_1_CTG1 | novel transcript, antisense to NOMO3 |
| PKD1P1 | CHR_HSCHR16_1_CTG1 | polycystin 1, transient receptor potential char |
| Y_RNA | CHR_HSCHR16_1_CTG1 | Y RNA [Source:RFAM;Acc:RF00019] |
| AC244489.1 | CHR_HSCHR16_1_CTG1 | novel member of the nuclear pore complex in |
| AC244489.2 | CHR_HSCHR16_1_CTG1 | nuclear pore complex interacting protein fami |
| AC246789.1 | CHR_HSCHR16_1_CTG1 | geminin, DNA replication inhibitor (GMNN) ps |
| AC246789.2 | CHR_HSCHR16_1_CTG1 | novel transcript |
| AC243615.1 | CHR_HSCHR16_1_CTG1 | novel transcript |
| AC098965.2 | CHR_HSCHR16_1_CTG1 | chromosome 12 open reading frame 12 (C12c |
| CES1P1 | CHR_HSCHR16_1_CTG3_1 | carboxylesterase 1 pseudogene 1 [Source:HGI |
| CES1 | CHR_HSCHR16_1_CTG3_1 | carboxylesterase 1 [Source:HGNC Symbol;Acc |
| CES5A | CHR_HSCHR16_1_CTG3_1 | carboxylesterase 5A [Source:HGNC Symbol;Ac |
| UTP4 | CHR_HSCHR16_2_CTG3_1 | UTP4 small subunit processome component [S |
| U6 | CHR_HSCHR16_2_CTG3_1 | U6 spliceosomal RNA [Source:RFAM;Acc:RF00 |
| SNTB2 | CHR_HSCHR16_2_CTG3_1 | syntrophin beta 2 [Source:HGNC Symbol;Acc:I |
| Y_RNA | CHR_HSCHR16_2_CTG3_1 | Y RNA [Source:RFAM;Acc:RF00019] |
| GLIS2 | CHR_HSCHR16_3_CTG1 | GLIS family zinc finger 2 [Source:HGNC Symbo |

| | | |
|---|---|---|
| GLIS2-AS1 | CHR_HSCHR16_3_CTG1 | GLIS2 antisense RNA 1 [Source:HGNC Symbol; |
| PAM16 | CHR_HSCHR16_3_CTG1 | presequence translocase associated motor 16 |
| CORO7-PAM16 | CHR_HSCHR16_3_CTG1 | CORO7-PAM16 readthrough [Source:HGNC Sy |
| AC004789.1 | CHR_HSCHR16_3_CTG1 | tec |
| CORO7 | CHR_HSCHR16_3_CTG1 | coronin 7 [Source:HGNC Symbol;Acc:HGNC:26 |
| VASN | CHR_HSCHR16_3_CTG1 | vasorin [Source:HGNC Symbol;Acc:HGNC:185 |
| DNAJA3 | CHR_HSCHR16_3_CTG1 | DnaJ heat shock protein family (Hsp40) memb |
| AC012676.7 | CHR_HSCHR16_3_CTG1 | novel transcript, sense intronic with DNAJA3 |
| AC012676.8 | CHR_HSCHR16_3_CTG1 | novel transcript, sense intronic with DNAJA3 |
| AC012676.6 | CHR_HSCHR16_3_CTG1 | cytochrome c oxidase subunit VIIc (COX7C) ps |
| NMRAL1 | CHR_HSCHR16_3_CTG1 | NmrA like redox sensor 1 [Source:HGNC Symb |
| HMOX2 | CHR_HSCHR16_3_CTG1 | heme oxygenase 2 [Source:HGNC Symbol;Acc |
| CDIP1 | CHR_HSCHR16_3_CTG1 | cell death inducing p53 target 1 [Source:HGN |
| MEIOB | CHR_HSCHR16_4_CTG1 | meiosis specific with OB-fold [Source:HGNC Sy |
| LINC00254 | CHR_HSCHR16_4_CTG1 | long intergenic non-protein coding RNA 254 [S |
| LINC02124 | CHR_HSCHR16_4_CTG1 | long intergenic non-protein coding RNA 2124 |
| HS3ST6 | CHR_HSCHR16_4_CTG1 | heparan sulfate-glucosamine 3-sulfotransfera |
| HYDIN | CHR_HSCHR16_4_CTG3_1 | HYDIN axonemal central pair apparatus protei |
| AC009136.1 | CHR_HSCHR16_4_CTG3_1 | uncharacterized LOC102723786 [Source:NCBI |
| AC009136.2 | CHR_HSCHR16_4_CTG3_1 | NULL |
| CALB2 | CHR_HSCHR16_4_CTG3_1 | calbindin 2 [Source:HGNC Symbol;Acc:HGNC:1 |
| LINC02136 | CHR_HSCHR16_4_CTG3_1 | long intergenic non-protein coding RNA 2136 |
| PRSS21 | CHR_HSCHR16_5_CTG1 | serine protease 21 [Source:HGNC Symbol;Acc |
| EIF1P4 | CHR_HSCHR16_5_CTG1 | eukaryotic translation initiation factor 1 pseu |
| ZG16B | CHR_HSCHR16_5_CTG1 | zymogen granule protein 16B [Source:HGNC S |
| PRSS30P | CHR_HSCHR16_5_CTG1 | serine protease 30, pseudogene [Source:HGN |
| PRSS22 | CHR_HSCHR16_5_CTG1 | serine protease 22 [Source:HGNC Symbol;Acc |
| AC217628.1 | CHR_HSCHR16_5_CTG1 | novel transcript, antisense to PRSS22 |
| AC003965.3 | CHR_HSCHR16_5_CTG1 | novel transcript |
| ZFP1 | CHR_HSCHR16_5_CTG3_1 | ZFP1 zinc finger protein [Source:HGNC Symbo |
| FO681524.1 | CHR_HSCHR16_5_CTG3_1 | NULL |
| CTRB2 | CHR_HSCHR16_5_CTG3_1 | chymotrypsinogen B2 [Source:HGNC Symbol; |
| CTRB1 | CHR_HSCHR16_5_CTG3_1 | chymotrypsinogen B1 [Source:HGNC Symbol; |
| FO681525.1 | CHR_HSCHR16_5_CTG3_1 | chymotrypsinogen B (CTRB) pseudogene |
| BCAR1 | CHR_HSCHR16_5_CTG3_1 | BCAR1 scaffold protein, Cas family member [S |
| AC009078.4 | CHR_HSCHR16_5_CTG3_1 | TEC |
| DECR2 | CHR_HSCHR16_CTG2 | 2,4-dienoyl-CoA reductase 2 [Source:HGNC Sy |
| AL023881.2 | CHR_HSCHR16_CTG2 | novel transcript |
| RAB11FIP3 | CHR_HSCHR16_CTG2 | RAB11 family interacting protein 3 [Source:HG |
| Y_RNA | CHR_HSCHR16_CTG2 | Y RNA [Source:RFAM;Acc:RF00019] |
| AC208188.1 | CHR_HSCHR16_CTG2 | microtubule-associated protein 1 light chain 3 |
| LINC00235 | CHR_HSCHR16_CTG2 | long intergenic non-protein coding RNA 235 [S |
| CAPN15 | CHR_HSCHR16_CTG2 | calpain 15 [Source:HGNC Symbol;Acc:HGNC:1 |
| CCL18 | CHR_HSCHR17_10_CTG4 | C-C motif chemokine ligand 18 [Source:HGNC |
| AC243829.9 | CHR_HSCHR17_10_CTG4 | novel transcript, antisense CCL3 |
| CCL3 | CHR_HSCHR17_10_CTG4 | C-C motif chemokine ligand 3 [Source:HGNC S |
| CCL4 | CHR_HSCHR17_10_CTG4 | C-C motif chemokine ligand 4 [Source:HGNC S |
| RN7SL301P | CHR_HSCHR17_10_CTG4 | RNA, 7SL, cytoplasmic 301, pseudogene [Sour |
| AC243829.8 | CHR_HSCHR17_10_CTG4 | novel transcript, antisense an TBC1D3B pseud |
| AC131056.3 | CHR_HSCHR17_10_CTG4 | nitric oxide synthase 2, inducible (NOS2) pseu |

| | | |
|---|---|---|
| TBC1D3B | CHR_HSCHR17_10_CTG4 | TBC1 domain family member 3B [Source:HGN |
| AC131056.1 | CHR_HSCHR17_10_CTG4 | novel transcript, antisense to a novel gene |
| CCL3L3 | CHR_HSCHR17_10_CTG4 | C-C motif chemokine ligand 3 like 3 [Source:H |
| CCL4L2 | CHR_HSCHR17_10_CTG4 | C-C motif chemokine ligand 4 like 2 [Source:H |
| AC131056.5 | CHR_HSCHR17_10_CTG4 | nitric oxide synthase 2, inducible (NOS2) pseu |
| TBC1D3B | CHR_HSCHR17_10_CTG4 | TBC1 domain family member 3B [Source:HGN |
| AC131056.2 | CHR_HSCHR17_10_CTG4 | chemokine (C-C motif) ligand pseudogene |
| CCL3L3 | CHR_HSCHR17_10_CTG4 | C-C motif chemokine ligand 3 like 3 [Source:H |
| CCL4L2 | CHR_HSCHR17_10_CTG4 | C-C motif chemokine ligand 4 like 2 [Source:H |
| AC131056.6 | CHR_HSCHR17_10_CTG4 | novel transcript |
| AC131056.7 | CHR_HSCHR17_10_CTG4 | ribosomal protein S6 kinase, 70kDa, polypepti |
| AC131056.4 | CHR_HSCHR17_10_CTG4 | tyrosine 3-monooxygenase/tryptophan 5-mor |
| TBC1D3I | CHR_HSCHR17_10_CTG4 | TBC1 domain family member 3I [Source:HGN( |
| LINC01982 | CHR_HSCHR17_11_CTG4 | long intergenic non-protein coding RNA 1982 |
| AC011120.2 | CHR_HSCHR17_12_CTG4 | novel transcript, antisense to SLC39A11 |
| AC240565.3 | CHR_HSCHR17_1_CTG1 | novel transcript |
| U6 | CHR_HSCHR17_1_CTG1 | U6 spliceosomal RNA [Source:RFAM;Acc:RF00 |
| AC240565.4 | CHR_HSCHR17_1_CTG1 | novel transcript |
| SCGB1C2 | CHR_HSCHR17_1_CTG1 | secretoglobin family 1C member 2 [Source:HG |
| DOC2B | CHR_HSCHR17_1_CTG1 | double C2 domain beta [Source:HGNC Symbol |
| LINC02091 | CHR_HSCHR17_1_CTG1 | long intergenic non-protein coding RNA 2091 |
| RPH3AL | CHR_HSCHR17_1_CTG1 | rabphilin 3A like (without C2 domains) [Source |
| AC211856.1 | CHR_HSCHR17_1_CTG1 | novel transcript, antisense to RPH3AL |
| AC129507.5 | CHR_HSCHR17_1_CTG1 | uncharacterized LOC100506388 [Source:NCBI |
| AC243301.1 | CHR_HSCHR17_1_CTG1 | novel transcript, antisense to RPH3AL |
| AC243301.2 | CHR_HSCHR17_1_CTG1 | novel transcript, antisense to RPH3AL |
| SLC43A2 | CHR_HSCHR17_1_CTG2 | solute carrier family 43 member 2 [Source:HG |
| RN7SL105P | CHR_HSCHR17_1_CTG2 | RNA, 7SL, cytoplasmic 105, pseudogene [Sour |
| SCARF1 | CHR_HSCHR17_1_CTG2 | scavenger receptor class F member 1 [Source: |
| RILP | CHR_HSCHR17_1_CTG2 | Rab interacting lysosomal protein [Source:HG |
| PRPF8 | CHR_HSCHR17_1_CTG2 | pre-mRNA processing factor 8 [Source:HGNC S |
| TLCD2 | CHR_HSCHR17_1_CTG2 | TLC domain containing 2 [Source:HGNC Symb |
| MIR22HG | CHR_HSCHR17_1_CTG2 | MIR22 host gene [Source:HGNC Symbol;Acc:H |
| WDR81 | CHR_HSCHR17_1_CTG2 | WD repeat domain 81 [Source:HGNC Symbol; |
| SERPINF2 | CHR_HSCHR17_1_CTG2 | serpin family F member 2 [Source:HGNC Symb |
| SERPINF1 | CHR_HSCHR17_1_CTG2 | serpin family F member 1 [Source:HGNC Symb |
| KRTAP16-1 | CHR_HSCHR17_1_CTG4 | keratin associated protein 16-1 [Source:HGNC |
| KRTAP17-1 | CHR_HSCHR17_1_CTG4 | keratin associated protein 17-1 [Source:HGNC |
| TBC1D3P7 | CHR_HSCHR17_1_CTG4 | TBC1 domain family member 3 pseudogene 7 |
| AC003958.4 | CHR_HSCHR17_1_CTG4 | tumor necrosis factor receptor superfamily, m |
| KRT33A | CHR_HSCHR17_1_CTG4 | keratin 33A [Source:HGNC Symbol;Acc:HGNC: |
| KRT33B | CHR_HSCHR17_1_CTG4 | keratin 33B [Source:HGNC Symbol;Acc:HGNC: |
| KRT34 | CHR_HSCHR17_1_CTG4 | keratin 34 [Source:HGNC Symbol;Acc:HGNC:6 |
| AC207056.1 | CHR_HSCHR17_1_CTG4 | novel protein similar to keratin 34 KRT34 |
| KRT31 | CHR_HSCHR17_1_CTG4 | keratin 31 [Source:HGNC Symbol;Acc:HGNC:6 |
| AC003958.5 | CHR_HSCHR17_1_CTG4 | novel transcript |
| AC003958.3 | CHR_HSCHR17_1_CTG4 | novel keratin, hair, acidic (KRTHA) family pseu |
| KRT37 | CHR_HSCHR17_1_CTG4 | keratin 37 [Source:HGNC Symbol;Acc:HGNC:6 |
| AC003070.6 | CHR_HSCHR17_1_CTG5 | NULL |
| AC217778.1 | CHR_HSCHR17_1_CTG5 | novel transcript |

| | | |
|---|---|---|
| ARHGAP27 | CHR_HSCHR17_1_CTG5 | Rho GTPase activating protein 27 [Source:HGI |
| AC217778.2 | CHR_HSCHR17_1_CTG5 | novel transcript, antisense to ARHGAP27 |
| PLEKHM1 | CHR_HSCHR17_1_CTG5 | pleckstrin homology and RUN domain contain |
| LRRC37A2 | CHR_HSCHR17_1_CTG5 | leucine rich repeat containing 37 member A2 |
| Metazoa_SRP | CHR_HSCHR17_1_CTG5 | Metazoan signal recognition particle RNA [Sou |
| AC217769.1 | CHR_HSCHR17_1_CTG5 | RAD52 motif 1 (RDM1) pseudogene |
| NSFP1 | CHR_HSCHR17_1_CTG5 | N-ethylmaleimide-sensitive factor pseudogen |
| ARL17B | CHR_HSCHR17_1_CTG5 | ADP ribosylation factor like GTPase 17B [Sour |
| LRRC37A | CHR_HSCHR17_1_CTG5 | leucine rich repeat containing 37A [Source:HG |
| Metazoa_SRP | CHR_HSCHR17_1_CTG5 | Metazoan signal recognition particle RNA [Sou |
| AC127032.1 | CHR_HSCHR17_1_CTG5 | dead end homolog 1 (zebrafish) (DND1) pseuc |
| AC127032.2 | CHR_HSCHR17_1_CTG5 | mitogen-activated protein kinase 8 interacting |
| KANSL1 | CHR_HSCHR17_1_CTG5 | KAT8 regulatory NSL complex subunit 1 [Sourc |
| Y_RNA | CHR_HSCHR17_1_CTG5 | Y RNA [Source:RFAM;Acc:RF00019] |
| KANSL1-AS1 | CHR_HSCHR17_1_CTG5 | KANSL1 antisense RNA 1 [Source:HGNC Symb |
| AC217779.1 | CHR_HSCHR17_1_CTG5 | novel transcript, antisense to a novel protein |
| MAPT | CHR_HSCHR17_1_CTG5 | microtubule associated protein tau [Source:H |
| STH | CHR_HSCHR17_1_CTG5 | saitohin [Source:HGNC Symbol;Acc:HGNC:188 |
| Metazoa_SRP | CHR_HSCHR17_1_CTG5 | Metazoan signal recognition particle RNA [Sou |
| BX544879.1 | CHR_HSCHR17_1_CTG5 | novel transcript, antisense to MAPT |
| MAPT-IT1 | CHR_HSCHR17_1_CTG5 | MAPT intronic transcript 1 [Source:HGNC Sym |
| MAPT-AS1 | CHR_HSCHR17_1_CTG5 | MAPT antisense RNA 1 [Source:HGNC Symbol |
| SPPL2C | CHR_HSCHR17_1_CTG5 | signal peptide peptidase like 2C [Source:HGN |
| CRHR1 | CHR_HSCHR17_1_CTG5 | corticotropin releasing hormone receptor 1 [S |
| MAPK8IP1P1 | CHR_HSCHR17_1_CTG5 | mitogen-activated protein kinase 8 interacting |
| AC225613.1 | CHR_HSCHR17_1_CTG5 | C17orf58 pseudogene |
| Y_RNA | CHR_HSCHR17_1_CTG5 | Y RNA [Source:RFAM;Acc:RF00019] |
| AC139677.4 | CHR_HSCHR17_1_CTG5 | KAT8 regulatory NSL complex subunit 1 (KANS |
| AC139677.3 | CHR_HSCHR17_1_CTG5 | mitogen-activated protein kinase 8 interacting |
| AC139677.5 | CHR_HSCHR17_1_CTG5 | DND microRNA-mediated repression inhibitor |
| RDM1P2 | CHR_HSCHR17_1_CTG5 | RDM1 pseudogene 2 [Source:HGNC Symbol;A |
| AC139677.1 | CHR_HSCHR17_1_CTG5 | novel protein similar to leucine rich repeat co |
| AC139677.2 | CHR_HSCHR17_1_CTG5 | ADP-ribosylation factor-like protein 17 [Sourc |
| Metazoa_SRP | CHR_HSCHR17_1_CTG5 | Metazoan signal recognition particle RNA [Sou |
| AC217775.1 | CHR_HSCHR17_1_CTG5 | novel transcript |
| NSF | CHR_HSCHR17_1_CTG5 | N-ethylmaleimide sensitive factor, vesicle fusi |
| WNT3 | CHR_HSCHR17_1_CTG5 | Wnt family member 3 [Source:HGNC Symbol; |
| TBCD | CHR_HSCHR17_1_CTG9 | tubulin folding cofactor D [Source:HGNC Symb |
| B3GNTL1 | CHR_HSCHR17_1_CTG9 | UDP-GlcNAc:betaGal beta-1,3-N-acetylglucosa |
| AC127518.1 | CHR_HSCHR17_1_CTG9 | novel transcript, antisense to B3GNTL1 |
| METRNL | CHR_HSCHR17_1_CTG9 | meteorin like, glial cell differentiation regulat |
| RPH3AL | CHR_HSCHR17_2_CTG1 | rabphilin 3A like (without C2 domains) [Sourc |
| AC240137.1 | CHR_HSCHR17_2_CTG1 | novel transcript, antisense to RPH3AL |
| AC129507.6 | CHR_HSCHR17_2_CTG1 | uncharacterized LOC100506388 [Source:NCBI |
| AC129507.7 | CHR_HSCHR17_2_CTG1 | novel transcript, antisense to RPH3AL |
| AC129507.8 | CHR_HSCHR17_2_CTG1 | novel transcript, antisense to RPH3AL |
| NXN | CHR_HSCHR17_2_CTG2 | nucleoredoxin [Source:HGNC Symbol;Acc:HGI |
| TIMM22 | CHR_HSCHR17_2_CTG2 | translocase of inner mitochondrial membrane |
| ABR | CHR_HSCHR17_2_CTG2 | ABR activator of RhoGEF and GTPase [Source: |
| Metazoa_SRP | CHR_HSCHR17_2_CTG2 | Metazoan signal recognition particle RNA [Sou |

| | | |
|---|---|---|
| AC107911.1 | CHR_HSCHR17_2_CTG2 | mitochondrial ribosomal protein L14 (MRPL14 |
| AC107911.2 | CHR_HSCHR17_2_CTG2 | novel transcript, antisense to ABR |
| TRARG1 | CHR_HSCHR17_2_CTG2 | trafficking regulator of GLUT4 (SLC2A4) 1 [Sou |
| YWHAE | CHR_HSCHR17_2_CTG2 | tyrosine 3-monooxygenase/tryptophan 5-mor |
| AC207965.1 | CHR_HSCHR17_2_CTG4 | novel transcript |
| MAP3K14 | CHR_HSCHR17_2_CTG5 | mitogen-activated protein kinase kinase kinas |
| AC003070.3 | CHR_HSCHR17_2_CTG5 | NULL |
| AC003070.5 | CHR_HSCHR17_2_CTG5 | novel transcript |
| ARHGAP27 | CHR_HSCHR17_2_CTG5 | Rho GTPase activating protein 27 [Source:HGI |
| AC003070.4 | CHR_HSCHR17_2_CTG5 | novel transcript, antisense to ARHGAP27 |
| PLEKHM1 | CHR_HSCHR17_2_CTG5 | pleckstrin homology and RUN domain contain |
| AC244207.1 | CHR_HSCHR17_2_CTG5 | novel transcript, antisense to PLEKHM1 |
| LRRC37A4P | CHR_HSCHR17_2_CTG5 | leucine rich repeat containing 37 member A4, |
| Metazoa_SRP | CHR_HSCHR17_2_CTG5 | Metazoan signal recognition particle RNA [Sou |
| AC244207.2 | CHR_HSCHR17_2_CTG5 | novel transcript |
| AC243807.4 | CHR_HSCHR17_2_CTG5 | novel transcript |
| AC243807.1 | CHR_HSCHR17_2_CTG5 | novel transcript |
| DND1P1 | CHR_HSCHR17_2_CTG5 | DND microRNA-mediated repression inhibitor |
| MAPK8IP1P2 | CHR_HSCHR17_2_CTG5 | mitogen-activated protein kinase 8 interacting |
| RPS26P8 | CHR_HSCHR17_2_CTG5 | ribosomal protein S26 pseudogene 8 [Source: |
| AC243807.2 | CHR_HSCHR17_2_CTG5 | ADP-ribosylation factor 3 (ARF3) pseudogene |
| LINC02210-CRHR1 | CHR_HSCHR17_2_CTG5 | LINC02210-CRHR1 readthrough [Source:HGN( |
| AC243807.5 | CHR_HSCHR17_2_CTG5 | novel transcript |
| AC243807.3 | CHR_HSCHR17_2_CTG5 | novel transcript |
| CRHR1 | CHR_HSCHR17_2_CTG5 | corticotropin releasing hormone receptor 1 [S |
| MAPT-AS1 | CHR_HSCHR17_2_CTG5 | MAPT antisense RNA 1 [Source:HGNC Symbol |
| SPPL2C | CHR_HSCHR17_2_CTG5 | signal peptide peptidase like 2C [Source:HGN( |
| MAPT | CHR_HSCHR17_2_CTG5 | microtubule associated protein tau [Source:H |
| MAPT-IT1 | CHR_HSCHR17_2_CTG5 | MAPT intronic transcript 1 [Source:HGNC Sym |
| AC243621.1 | CHR_HSCHR17_2_CTG5 | novel transcript, antisense to MAPT |
| Metazoa_SRP | CHR_HSCHR17_2_CTG5 | Metazoan signal recognition particle RNA [Sou |
| KANSL1 | CHR_HSCHR17_2_CTG5 | KAT8 regulatory NSL complex subunit 1 [Sourc |
| AC243658.1 | CHR_HSCHR17_2_CTG5 | novel transcript, antisense to a novel protein |
| KANSL1-AS1 | CHR_HSCHR17_2_CTG5 | KANSL1 antisense RNA 1 [Source:HGNC Symb |
| Y_RNA | CHR_HSCHR17_2_CTG5 | Y RNA [Source:RFAM;Acc:RF00019] |
| MAPK8IP1P1 | CHR_HSCHR17_2_CTG5 | mitogen-activated protein kinase 8 interacting |
| AC243899.1 | CHR_HSCHR17_2_CTG5 | dead end homolog 1 (zebrafish) (DND1) pseuc |
| AC243899.2 | CHR_HSCHR17_2_CTG5 | C17orf58 pseudogene |
| ARL17B | CHR_HSCHR17_2_CTG5 | ADP ribosylation factor like GTPase 17B [Sour |
| LRRC37A2 | CHR_HSCHR17_2_CTG5 | leucine rich repeat containing 37 member A2 |
| Metazoa_SRP | CHR_HSCHR17_2_CTG5 | Metazoan signal recognition particle RNA [Sou |
| NSF | CHR_HSCHR17_2_CTG5 | N-ethylmaleimide sensitive factor, vesicle fusi |
| NSFP1 | CHR_HSCHR17_2_CTG5 | N-ethylmaleimide-sensitive factor pseudogen |
| RPS7P11 | CHR_HSCHR17_2_CTG5 | ribosomal protein S7 pseudogene 11 [Source: |
| WNT3 | CHR_HSCHR17_2_CTG5 | Wnt family member 3 [Source:HGNC Symbol; |
| WNT9B | CHR_HSCHR17_2_CTG5 | Wnt family member 9B [Source:HGNC Symbol |
| LLGL1 | CHR_HSCHR17_3_CTG1 | LLGL scribble cell polarity complex componen |
| FLII | CHR_HSCHR17_3_CTG1 | FLII actin remodeling protein [Source:HGNC Sy |
| MIEF2 | CHR_HSCHR17_3_CTG1 | mitochondrial elongation factor 2 [Source:HG |
| AC127537.2 | CHR_HSCHR17_3_CTG1 | novel transcript, antisense to TOP3A |

| | | |
|---|---|---|
| TOP3A | CHR_HSCHR17_3_CTG1 | DNA topoisomerase III alpha [Source:HGNC Sy |
| RPL7AP65 | CHR_HSCHR17_3_CTG1 | ribosomal protein L7a pseudogene 65 [Source |
| RPL21P121 | CHR_HSCHR17_3_CTG1 | ribosomal protein L21 pseudogene 121 [Sourc |
| SMCR8 | CHR_HSCHR17_3_CTG1 | SMCR8-C9orf72 complex subunit [Source:HGN |
| SHMT1 | CHR_HSCHR17_3_CTG1 | serine hydroxymethyltransferase 1 [Source:H( |
| MIR6778 | CHR_HSCHR17_3_CTG1 | microRNA 6778 [Source:HGNC Symbol;Acc:HG |
| EVPLL | CHR_HSCHR17_3_CTG1 | envoplakin like [Source:HGNC Symbol;Acc:HG |
| AL353997.10 | CHR_HSCHR17_3_CTG1 | novel transcript |
| AL353997.8 | CHR_HSCHR17_3_CTG1 | CMT1A duplicated region transcript 15-like 2 ( |
| AL353997.9 | CHR_HSCHR17_3_CTG1 | tumor necrosis factor receptor superfamily, m |
| TBC1D3P4 | CHR_HSCHR17_3_CTG1 | TBC1 domain family member 3 pseudogene 4 |
| LINC02076 | CHR_HSCHR17_3_CTG1 | long intergenic non-protein coding RNA 2076 |
| KRT17P5 | CHR_HSCHR17_3_CTG1 | keratin 17 pseudogene 5 [Source:HGNC Symb |
| YWHAEP2 | CHR_HSCHR17_3_CTG1 | tyrosine 3-monooxygenase/tryptophan 5-mor |
| KRT17P2 | CHR_HSCHR17_3_CTG1 | keratin 17 pseudogene 2 [Source:HGNC Symb |
| KRT16P1 | CHR_HSCHR17_3_CTG1 | keratin 16 pseudogene 1 [Source:HGNC Symb |
| KRT16P4 | CHR_HSCHR17_3_CTG1 | keratin 16 pseudogene 4 [Source:HGNC Symb |
| AL353997.7 | CHR_HSCHR17_3_CTG1 | tyrosine 3-monooxygenase/tryptophan 5-mor |
| AL353997.6 | CHR_HSCHR17_3_CTG1 | tyrosine 3-monooxygenase/tryptophan 5-mor |
| TNPO1P2 | CHR_HSCHR17_3_CTG1 | transportin 1 pseudogene 2 [Source:HGNC Syr |
| LGALS9C | CHR_HSCHR17_3_CTG1 | galectin 9C [Source:HGNC Symbol;Acc:HGNC:3 |
| NOS2P2 | CHR_HSCHR17_3_CTG1 | nitric oxide synthase 2 pseudogene 2 [Source: |
| NXN | CHR_HSCHR17_3_CTG2 | nucleoredoxin [Source:HGNC Symbol;Acc:HGN |
| TIMM22 | CHR_HSCHR17_3_CTG2 | translocase of inner mitochondrial membrane |
| ABR | CHR_HSCHR17_3_CTG2 | ABR activator of RhoGEF and GTPase [Source: |
| Metazoa_SRP | CHR_HSCHR17_3_CTG2 | Metazoan signal recognition particle RNA [Sou |
| SEPTIN9-DT | CHR_HSCHR17_3_CTG4 | SEPTIN9 divergent transcript [Source:HGNC Sy |
| AC068594.2 | CHR_HSCHR17_3_CTG4 | novel transcript |
| SEPTIN9 | CHR_HSCHR17_3_CTG4 | septin 9 [Source:HGNC Symbol;Acc:HGNC:732 |
| AC004231.2 | CHR_HSCHR17_4_CTG4 | novel transcript, antisense to KRT12 |
| KRT12 | CHR_HSCHR17_4_CTG4 | keratin 12 [Source:HGNC Symbol;Acc:HGNC:6 |
| KRT20 | CHR_HSCHR17_4_CTG4 | keratin 20 [Source:HGNC Symbol;Acc:HGNC:2 |
| AC037482.1 | CHR_HSCHR17_4_CTG4 | novel transcript |
| KRT23 | CHR_HSCHR17_4_CTG4 | keratin 23 [Source:HGNC Symbol;Acc:HGNC:6 |
| KRT39 | CHR_HSCHR17_4_CTG4 | keratin 39 [Source:HGNC Symbol;Acc:HGNC:3 |
| KRT40 | CHR_HSCHR17_4_CTG4 | keratin 40 [Source:HGNC Symbol;Acc:HGNC:2 |
| KRTAP3-3 | CHR_HSCHR17_4_CTG4 | keratin associated protein 3-3 [Source:HGNC S |
| KRTAP3-2 | CHR_HSCHR17_4_CTG4 | keratin associated protein 3-2 [Source:HGNC S |
| KRTAP3-4P | CHR_HSCHR17_4_CTG4 | keratin associated protein 3-4, pseudogene [S |
| KRTAP3-1 | CHR_HSCHR17_4_CTG4 | keratin associated protein 3-1 [Source:HGNC S |
| KRTAP1-5 | CHR_HSCHR17_4_CTG4 | keratin associated protein 1-5 [Source:HGNC S |
| KRTAP1-4 | CHR_HSCHR17_4_CTG4 | keratin associated protein 1-4 [Source:HGNC S |
| KRTAP1-3 | CHR_HSCHR17_4_CTG4 | keratin associated protein 1-3 [Source:HGNC S |
| KRTAP1-1 | CHR_HSCHR17_4_CTG4 | keratin associated protein 1-1 [Source:HGNC S |
| KRTAP2-1 | CHR_HSCHR17_4_CTG4 | keratin associated protein 2-1 [Source:HGNC S |
| AC037482.2 | CHR_HSCHR17_4_CTG4 | keratin associated protein 2-2 [Source:NCBI ge |
| KRTAP2-3 | CHR_HSCHR17_4_CTG4 | keratin associated protein 2-3 [Source:HGNC S |
| KRTAP2-4 | CHR_HSCHR17_4_CTG4 | keratin associated protein 2-4 [Source:HGNC S |
| KRTAP2-5P | CHR_HSCHR17_4_CTG4 | keratin associated protein 2-5, pseudogene [S |
| AC037482.3 | CHR_HSCHR17_4_CTG4 | keratin-associated protein 4-7 [Source:NCBI ge |

| | | |
|---|---|---|
| KRTAP4-8 | CHR_HSCHR17_4_CTG4 | keratin associated protein 4-8 [Source:NCBI ge |
| KRTAP4-16 | CHR_HSCHR17_4_CTG4 | keratin associated protein 4-16 [Source:HGNC |
| KRTAP4-9 | CHR_HSCHR17_4_CTG4 | keratin associated protein 4-9 [Source:HGNC S |
| KRTAP4-11 | CHR_HSCHR17_4_CTG4 | keratin associated protein 4-11 [Source:HGNC |
| KRTAP4-12 | CHR_HSCHR17_4_CTG4 | keratin associated protein 4-12 [Source:HGNC |
| KRTAP9-2 | CHR_HSCHR17_5_CTG4 | keratin associated protein 9-2 [Source:HGNC S |
| KRTAP9-3 | CHR_HSCHR17_5_CTG4 | keratin associated protein 9-3 [Source:HGNC S |
| AC209539.1 | CHR_HSCHR17_5_CTG4 | novel keratin associated protein 9 family pseu |
| KRTAP9-4 | CHR_HSCHR17_5_CTG4 | keratin associated protein 9-4 [Source:HGNC S |
| KRTAP9-9 | CHR_HSCHR17_5_CTG4 | keratin associated protein 9-9 [Source:NCBI ge |
| AC209539.2 | CHR_HSCHR17_5_CTG4 | keratin associated protein 9-6 [Source:NCBI ge |
| KRTAP9-10P | CHR_HSCHR17_5_CTG4 | keratin associated protein 9-10, pseudogene [ |
| KRTAP29-1 | CHR_HSCHR17_5_CTG4 | keratin associated like protein 29-1 [Source:H |
| KRTAP9-12P | CHR_HSCHR17_6_CTG4 | keratin associated protein 9-12, pseudogene [ |
| KRTAP9-2 | CHR_HSCHR17_6_CTG4 | keratin associated protein 9-2 [Source:HGNC S |
| KRTAP9-4 | CHR_HSCHR17_6_CTG4 | keratin associated protein 9-4 [Source:HGNC S |
| KRTAP9-9 | CHR_HSCHR17_6_CTG4 | keratin associated protein 9-9 [Source:HGNC S |
| KRTAP9-6 | CHR_HSCHR17_6_CTG4 | keratin associated protein 9-6 [Source:HGNC S |
| KRTAP9-11P | CHR_HSCHR17_6_CTG4 | keratin associated protein 9-11, pseudogene [ |
| KRTAP9-7 | CHR_HSCHR17_6_CTG4 | keratin associated protein 9-7 [Source:HGNC S |
| KRTAP9-10P | CHR_HSCHR17_6_CTG4 | keratin associated protein 9-10, pseudogene [ |
| KRTAP29-1 | CHR_HSCHR17_6_CTG4 | keratin associated like protein 29-1 [Source:H |
| MMP28 | CHR_HSCHR17_7_CTG4 | matrix metallopeptidase 28 [Source:HGNC Syr |
| AC015849.11 | CHR_HSCHR17_7_CTG4 | novel transcript |
| TAF15 | CHR_HSCHR17_7_CTG4 | TATA-box binding protein associated factor 15 |
| AC015849.10 | CHR_HSCHR17_7_CTG4 | novel transcript, antisense TAF15 |
| AC015849.7 | CHR_HSCHR17_7_CTG4 | novel transcript, sense overlapping TAF15 |
| HEATR9 | CHR_HSCHR17_7_CTG4 | HEAT repeat containing 9 [Source:HGNC Symb |
| AC015849.8 | CHR_HSCHR17_7_CTG4 | ribosomal L24 domain containing 1 (RSL24D1) |
| AC015849.12 | CHR_HSCHR17_7_CTG4 | novel transcript, antisense to CCL5 |
| CCL5 | CHR_HSCHR17_7_CTG4 | C-C motif chemokine ligand 5 [Source:HGNC S |
| 7SK | CHR_HSCHR17_7_CTG4 | 7SK RNA [Source:RFAM;Acc:RF00100] |
| AC015849.9 | CHR_HSCHR17_7_CTG4 | novel transcript |
| LRRC37A9P | CHR_HSCHR17_7_CTG4 | leucine rich repeat containing 37 member A9, |
| RDM1 | CHR_HSCHR17_7_CTG4 | RAD52 motif containing 1 [Source:HGNC Symb |
| LYZL6 | CHR_HSCHR17_7_CTG4 | lysozyme like 6 [Source:HGNC Symbol;Acc:HG |
| AC004675.2 | CHR_HSCHR17_7_CTG4 | tetratricopeptide repeat, ankyrin repeat and c |
| CCL16 | CHR_HSCHR17_7_CTG4 | C-C motif chemokine ligand 16 [Source:HGNC |
| CCL14 | CHR_HSCHR17_7_CTG4 | C-C motif chemokine ligand 14 [Source:HGNC |
| CCL15-CCL14 | CHR_HSCHR17_7_CTG4 | CCL15-CCL14 readthrough (NMD candidate) [S |
| AC004675.1 | CHR_HSCHR17_7_CTG4 | novel transcript, antisense to CCL14 |
| CCL15 | CHR_HSCHR17_7_CTG4 | C-C motif chemokine ligand 15 [Source:HGNC |
| AC069363.2 | CHR_HSCHR17_7_CTG4 | novel transcript, antisense CCL15 |
| AC069363.1 | CHR_HSCHR17_7_CTG4 | novel transcript |
| CCL23 | CHR_HSCHR17_7_CTG4 | C-C motif chemokine ligand 23 [Source:HGNC |
| CCL18 | CHR_HSCHR17_7_CTG4 | C-C motif chemokine ligand 18 [Source:HGNC |
| AC069363.3 | CHR_HSCHR17_7_CTG4 | novel transcript, antisense CCL3 |
| CCL3 | CHR_HSCHR17_7_CTG4 | C-C motif chemokine ligand 3 [Source:HGNC S |
| CCL4 | CHR_HSCHR17_7_CTG4 | C-C motif chemokine ligand 4 [Source:HGNC S |
| Metazoa_SRP | CHR_HSCHR17_7_CTG4 | Metazoan signal recognition particle RNA [Sou |

| | | |
|---|---|---|
| AC003976.2 | CHR_HSCHR17_7_CTG4 | novel transcript, antisense TBC1D3B |
| AC003976.1 | CHR_HSCHR17_7_CTG4 | nitric oxide synthase 2, inducible (NOS2) pseu |
| TBC1D3B | CHR_HSCHR17_7_CTG4 | TBC1 domain family member 3B [Source:HGN |
| AC233699.4 | CHR_HSCHR17_7_CTG4 | novel transcript, antisense CCL3L3 |
| CCL3L3 | CHR_HSCHR17_7_CTG4 | C-C motif chemokine ligand 3 like 3 [Source:H |
| CCL4L2 | CHR_HSCHR17_7_CTG4 | C-C motif chemokine ligand 4 like 2 [Source:H |
| AC233699.5 | CHR_HSCHR17_7_CTG4 | novel transcript |
| AC233699.2 | CHR_HSCHR17_7_CTG4 | ribosomal protein S6 kinase, 70kDa, polypepti |
| AC233699.3 | CHR_HSCHR17_7_CTG4 | tyrosine 3-monooxygenase/tryptophan 5-mor |
| TBC1D3I | CHR_HSCHR17_7_CTG4 | TBC1 domain family member 3I [Source:HGN( |
| AC233699.6 | CHR_HSCHR17_7_CTG4 | aminopeptidase puromycin sensitive (NPEPPS |
| Y_RNA | CHR_HSCHR17_7_CTG4 | Y RNA [Source:RFAM;Acc:RF00019] |
| TBC1D3G | CHR_HSCHR17_7_CTG4 | TBC1 domain family member 3G [Source:HGN |
| TBC1D3H | CHR_HSCHR17_7_CTG4 | TBC1 domain family member 3H [Source:HGN |
| U6 | CHR_HSCHR17_7_CTG4 | U6 spliceosomal RNA [Source:RFAM;Acc:RF00 |
| TBC1D3F | CHR_HSCHR17_7_CTG4 | TBC1 domain family member 3F [Source:HGN |
| TBC1D3J | CHR_HSCHR17_7_CTG4 | TBC1 domain family member 3J [Source:HGN( |
| ZNHIT3 | CHR_HSCHR17_7_CTG4 | zinc finger HIT-type containing 3 [Source:HGN |
| 5S_rRNA | CHR_HSCHR17_7_CTG4 | 5S ribosomal RNA [Source:RFAM;Acc:RF00001 |
| MYO19 | CHR_HSCHR17_7_CTG4 | myosin XIX [Source:HGNC Symbol;Acc:HGNC:2 |
| PIGW | CHR_HSCHR17_7_CTG4 | phosphatidylinositol glycan anchor biosynthes |
| GGNBP2 | CHR_HSCHR17_7_CTG4 | gametogenetin binding protein 2 [Source:HGN |
| AC003042.2 | CHR_HSCHR17_7_CTG4 | ribosomal protein S2 (RPS2) pseudogene |
| DHRS11 | CHR_HSCHR17_7_CTG4 | dehydrogenase/reductase 11 [Source:HGNC S |
| MRM1 | CHR_HSCHR17_7_CTG4 | mitochondrial rRNA methyltransferase 1 [Sou |
| AC003042.1 | CHR_HSCHR17_7_CTG4 | novel transcript |
| AC015938.1 | CHR_HSCHR17_7_CTG4 | novel transcript |
| AC015938.2 | CHR_HSCHR17_7_CTG4 | novel transcript |
| LHX1-DT | CHR_HSCHR17_7_CTG4 | LHX1 divergent transcript [Source:HGNC Symb |
| LHX1 | CHR_HSCHR17_7_CTG4 | LIM homeobox 1 [Source:HGNC Symbol;Acc:H |
| AC023315.1 | CHR_HSCHR17_7_CTG4 | novel transcript, antisense to LHX1 |
| AATF | CHR_HSCHR17_7_CTG4 | apoptosis antagonizing transcription factor [S |
| AC023315.2 | CHR_HSCHR17_7_CTG4 | novel transcript, sense intronic AATF |
| AC023315.3 | CHR_HSCHR17_7_CTG4 | novel transcript, sense intronic AATF |
| AC003103.1 | CHR_HSCHR17_7_CTG4 | novel transcript, sense intronic AATF |
| AC003103.2 | CHR_HSCHR17_7_CTG4 | novel transcript, antisense AATF |
| ACACA | CHR_HSCHR17_7_CTG4 | acetyl-CoA carboxylase alpha [Source:HGNC S |
| AC016482.2 | CHR_HSCHR17_7_CTG4 | ribosomal protein S2 (RPS2) pseudogene |
| HMGB1P24 | CHR_HSCHR17_7_CTG4 | high mobility group box 1 pseudogene 24 [Sou |
| AC016482.1 | CHR_HSCHR17_7_CTG4 | ribosomal protein L18a (RPL18A) pseudogene |
| C17orf78 | CHR_HSCHR17_7_CTG4 | chromosome 17 open reading frame 78 [Sour |
| AC068400.2 | CHR_HSCHR17_7_CTG4 | novel transcript, sense intronic to C17orf78 |
| TADA2A | CHR_HSCHR17_7_CTG4 | transcriptional adaptor 2A [Source:HGNC Sym |
| AC068400.1 | CHR_HSCHR17_7_CTG4 | novel transcript, antisense TADA2A |
| AC004099.1 | CHR_HSCHR17_7_CTG4 | ribosomal protein L24 (RPL24) pseudogene |
| DUSP14 | CHR_HSCHR17_7_CTG4 | dual specificity phosphatase 14 [Source:HGNC |
| SYNRG | CHR_HSCHR17_7_CTG4 | synergin gamma [Source:HGNC Symbol;Acc:H |
| DDX52 | CHR_HSCHR17_7_CTG4 | DExD-box helicase 52 [Source:HGNC Symbol;A |
| HNF1B | CHR_HSCHR17_7_CTG4 | HNF1 homeobox B [Source:HGNC Symbol;Acc |
| AC113211.1 | CHR_HSCHR17_7_CTG4 | novel transcript |

| | | |
|---|---|---|
| YWHAEP7 | CHR_HSCHR17_7_CTG4 | tyrosine 3-monooxygenase/tryptophan 5-mor |
| U6 | CHR_HSCHR17_7_CTG4 | U6 spliceosomal RNA [Source:RFAM;Acc:RF00 |
| TBC1D3K | CHR_HSCHR17_7_CTG4 | TBC1 domain family member 3K [Source:HGN |
| TBC1D3L | CHR_HSCHR17_7_CTG4 | TBC1 domain family member 3L [Source:HGN( |
| AC183084.1 | CHR_HSCHR17_7_CTG4 | aminopeptidase puromycin sensitive (NPEPPS |
| Y_RNA | CHR_HSCHR17_7_CTG4 | Y RNA [Source:RFAM;Acc:RF00019] |
| MRPL45 | CHR_HSCHR17_7_CTG4 | mitochondrial ribosomal protein L45 [Source:I |
| GPR179 | CHR_HSCHR17_7_CTG4 | G protein-coupled receptor 179 [Source:HGN( |
| SOCS7 | CHR_HSCHR17_7_CTG4 | suppressor of cytokine signaling 7 [Source:HG |
| ARHGAP23 | CHR_HSCHR17_7_CTG4 | Rho GTPase activating protein 23 [Source:HGI |
| AC115090.1 | CHR_HSCHR17_7_CTG4 | novel transcript, antisense to ARHGAP23 |
| SRCIN1 | CHR_HSCHR17_7_CTG4 | SRC kinase signaling inhibitor 1 [Source:HGNC |
| AC006449.8 | CHR_HSCHR17_7_CTG4 | novel transcript, antisense to SRCIN1 |
| EPOP | CHR_HSCHR17_7_CTG4 | elongin BC and polycomb repressive complex |
| AC006449.9 | CHR_HSCHR17_7_CTG4 | novel transcript |
| MLLT6 | CHR_HSCHR17_7_CTG4 | MLLT6, PHD finger containing [Source:HGNC S |
| AC006449.4 | CHR_HSCHR17_7_CTG4 | NULL |
| CISD3 | CHR_HSCHR17_7_CTG4 | CDGSH iron sulfur domain 3 [Source:HGNC Syı |
| 5S_rRNA | CHR_HSCHR17_7_CTG4 | 5S ribosomal RNA [Source:RFAM;Acc:RF00001 |
| PCGF2 | CHR_HSCHR17_7_CTG4 | polycomb group ring finger 2 [Source:HGNC S |
| AC006449.10 | CHR_HSCHR17_7_CTG4 | novel transcript, antisense to PCGF2 |
| PSMB3 | CHR_HSCHR17_7_CTG4 | proteasome subunit beta 3 [Source:HGNC Syn |
| U6 | CHR_HSCHR17_7_CTG4 | U6 spliceosomal RNA [Source:RFAM;Acc:RF00 |
| PIP4K2B | CHR_HSCHR17_7_CTG4 | phosphatidylinositol-5-phosphate 4-kinase typ |
| CWC25 | CHR_HSCHR17_7_CTG4 | CWC25 spliceosome associated protein homo |
| C17orf98 | CHR_HSCHR17_7_CTG4 | chromosome 17 open reading frame 98 [Sour |
| AC118653.2 | CHR_HSCHR17_8_CTG4 | novel transcript |
| MYL6P5 | CHR_HSCHR17_8_CTG4 | myosin light chain 6 pseudogene 5 [Source:HG |
| SLC39A11 | CHR_HSCHR17_9_CTG4 | solute carrier family 39 member 11 [Source:H |
| AC240577.1 | CHR_HSCHR17_9_CTG4 | novel transcript, antisense to SLC39A11 |
| AC019183.2 | CHR_HSCHR18_1_CTG1 | novel transcript |
| AP005057.2 | CHR_HSCHR18_1_CTG1 | novel transcript |
| AC105227.4 | CHR_HSCHR18_1_CTG1_1 | novel transcript |
| AC105227.3 | CHR_HSCHR18_1_CTG1_1 | ankyrin repeat domain 26 (ANKRD26) pseudog |
| RNA5SP458 | CHR_HSCHR18_1_CTG1_1 | RNA, 5S ribosomal pseudogene 458 [Source:H |
| SKA1 | CHR_HSCHR18_1_CTG1_1 | spindle and kinetochore associated complex s |
| AC013759.1 | CHR_HSCHR18_1_CTG1_1 | ribosomal protein, large, P0 (RPLP0) pseudoge |
| MAPK4 | CHR_HSCHR18_1_CTG1_1 | mitogen-activated protein kinase 4 [Source:H( |
| AC012572.2 | CHR_HSCHR18_1_CTG2_1 | novel transcript |
| Y_RNA | CHR_HSCHR18_1_CTG2_1 | Y RNA [Source:RFAM;Acc:RF00019] |
| AC208322.1 | CHR_HSCHR18_2_CTG1_1 | intraflagellar transport 57 homolog (Chlamydo |
| AC231988.1 | CHR_HSCHR18_2_CTG2 | TEC |
| AC110597.2 | CHR_HSCHR18_2_CTG2 | novel transcript |
| DSEL | CHR_HSCHR18_2_CTG2 | dermatan sulfate epimerase like [Source:HGN |
| AC110597.5 | CHR_HSCHR18_2_CTG2 | novel transcript, antisense to DSEL |
| AC110597.4 | CHR_HSCHR18_2_CTG2 | novel transcript |
| LINC01896 | CHR_HSCHR18_2_CTG2_1 | long intergenic non-protein coding RNA 1896 |
| SALL3 | CHR_HSCHR18_2_CTG2_1 | spalt like transcription factor 3 [Source:HGNC |
| ATP9B | CHR_HSCHR18_2_CTG2_1 | ATPase phospholipid transporting 9B (putative |
| AC068473.6 | CHR_HSCHR18_3_CTG2_1 | novel transcript |

| | | |
|---|---|---|
| AC215996.1 | CHR_HSCHR18_3_CTG2_1 | novel transcript |
| AC068473.9 | CHR_HSCHR18_3_CTG2_1 | novel transcript |
| AC068473.8 | CHR_HSCHR18_3_CTG2_1 | novel transcript |
| CTDP1 | CHR_HSCHR18_3_CTG2_1 | CTD phosphatase subunit 1 [Source:HGNC Syr |
| AC068473.7 | CHR_HSCHR18_3_CTG2_1 | novel transcript, antisense to CTDP1 |
| CTAGE1 | CHR_HSCHR18_5_CTG1_1 | cutaneous T cell lymphoma-associated antiger |
| AC099849.2 | CHR_HSCHR18_5_CTG1_1 | ATPase, Cu++ transporting, beta polypeptide, |
| AC012572.3 | CHR_HSCHR18_ALT21_CTG2_ | novel transcript |
| Y_RNA | CHR_HSCHR18_ALT21_CTG2_ | Y RNA [Source:RFAM;Acc:RF00019] |
| LINC01896 | CHR_HSCHR18_ALT2_CTG2_1 | long intergenic non-protein coding RNA 1896 |
| SALL3 | CHR_HSCHR18_ALT2_CTG2_1 | spalt like transcription factor 3 [Source:HGNC |
| ATP9B | CHR_HSCHR18_ALT2_CTG2_1 | ATPase phospholipid transporting 9B (putative |
| AC099689.2 | CHR_HSCHR18_ALT2_CTG2_1 | TEC |
| LILRP1 | CHR_HSCHR19KIR_0010-5217 | leukocyte immunoglobulin-like receptor pseu |
| AC245128.34 | CHR_HSCHR19KIR_0010-5217 | killer cell immunoglobulin-like receptor family |
| LILRP2 | CHR_HSCHR19KIR_0010-5217 | leukocyte immunoglobulin-like receptor pseu |
| KIR3DL3 | CHR_HSCHR19KIR_0010-5217 | killer cell immunoglobulin like receptor, three |
| KIR2DL3 | CHR_HSCHR19KIR_0010-5217 | killer cell immunoglobulin like receptor, two Ig |
| KIR2DP1 | CHR_HSCHR19KIR_0010-5217 | killer cell immunoglobulin like receptor, two Ig |
| KP420443.1 | CHR_HSCHR19KIR_0010-5217 | killer cell immunoglobulin like receptor, three |
| RNU6-222P | CHR_HSCHR19KIR_0010-5217 | RNA, U6 small nuclear 222, pseudogene [Sour |
| FCAR | CHR_HSCHR19KIR_0010-5217 | Fc fragment of IgA receptor [Source:HGNC Syr |
| LILRP1 | CHR_HSCHR19KIR_0019-4656 | leukocyte immunoglobulin-like receptor pseu |
| AC245128.20 | CHR_HSCHR19KIR_0019-4656 | killer cell immunoglobulin-like receptor family |
| LILRP2 | CHR_HSCHR19KIR_0019-4656 | leukocyte immunoglobulin-like receptor pseu |
| KIR3DL3 | CHR_HSCHR19KIR_0019-4656 | killer cell immunoglobulin like receptor, three |
| KP420439.1 | CHR_HSCHR19KIR_0019-4656 | novel transcript |
| KIR2DL3 | CHR_HSCHR19KIR_0019-4656 | killer cell immunoglobulin like receptor, two Ig |
| KIR2DL1 | CHR_HSCHR19KIR_0019-4656 | killer cell immunoglobulin like receptor, two Ig |
| KIR3DP1 | CHR_HSCHR19KIR_0019-4656 | killer cell immunoglobulin like receptor, three |
| KIR2DL4 | CHR_HSCHR19KIR_0019-4656 | killer cell immunoglobulin like receptor, two Ig |
| KIR3DL1 | CHR_HSCHR19KIR_0019-4656 | killer cell immunoglobulin like receptor, three |
| KIR2DS4 | CHR_HSCHR19KIR_0019-4656 | killer cell immunoglobulin like receptor, two Ig |
| KIR3DL2 | CHR_HSCHR19KIR_0019-4656 | killer cell immunoglobulin like receptor, three |
| LILRP1 | CHR_HSCHR19KIR_0019-4656 | leukocyte immunoglobulin-like receptor pseu |
| AC245128.38 | CHR_HSCHR19KIR_0019-4656 | killer cell immunoglobulin-like receptor family |
| LILRP2 | CHR_HSCHR19KIR_0019-4656 | leukocyte immunoglobulin-like receptor pseu |
| KP420440.3 | CHR_HSCHR19KIR_0019-4656 | Killer cell immunoglobulin-like receptor 2DS2 |
| KP420440.1 | CHR_HSCHR19KIR_0019-4656 | Killer cell immunoglobulin-like receptor 2DL2 |
| KIR2DL5B | CHR_HSCHR19KIR_0019-4656 | killer cell immunoglobulin like receptor, two Ig |
| KIR2DS3 | CHR_HSCHR19KIR_0019-4656 | killer cell immunoglobulin like receptor, two Ig |
| KIR2DP1 | CHR_HSCHR19KIR_0019-4656 | killer cell immunoglobulin like receptor, two Ig |
| KP420440.4 | CHR_HSCHR19KIR_0019-4656 | NULL |
| KIR3DP1 | CHR_HSCHR19KIR_0019-4656 | killer cell immunoglobulin like receptor, three |
| KIR2DL4 | CHR_HSCHR19KIR_0019-4656 | killer cell immunoglobulin like receptor, two Ig |
| KIR3DS1 | CHR_HSCHR19KIR_0019-4656 | killer cell immunoglobulin like receptor, three |
| KP420440.5 | CHR_HSCHR19KIR_0019-4656 | NULL |
| KIR2DS1 | CHR_HSCHR19KIR_0019-4656 | killer cell immunoglobulin like receptor, two Ig |
| KP420440.2 | CHR_HSCHR19KIR_0019-4656 | NULL |
| LILRP1 | CHR_HSCHR19KIR_50296000 | leukocyte immunoglobulin-like receptor pseu |

| | | |
|---|---|---|
| AC245128.37 | CHR_HSCHR19KIR_50296000 | killer cell immunoglobulin-like receptor family |
| LILRP2 | CHR_HSCHR19KIR_50296000 | leukocyte immunoglobulin-like receptor pseu |
| KIR3DL3 | CHR_HSCHR19KIR_50296000 | killer cell immunoglobulin like receptor, three |
| KP420438.1 | CHR_HSCHR19KIR_50296000 | novel transcript |
| KIR2DL3 | CHR_HSCHR19KIR_50296000 | killer cell immunoglobulin like receptor, two Ig |
| KIR2DP1 | CHR_HSCHR19KIR_50296000 | killer cell immunoglobulin like receptor, two Ig |
| KIR2DL1 | CHR_HSCHR19KIR_50296000 | killer cell immunoglobulin like receptor, two Ig |
| KIR3DP1 | CHR_HSCHR19KIR_50296000 | killer cell immunoglobulin like receptor, three |
| KIR2DL4 | CHR_HSCHR19KIR_50296000 | killer cell immunoglobulin like receptor, two Ig |
| KIR3DL1 | CHR_HSCHR19KIR_50296000 | killer cell immunoglobulin like receptor, three |
| KIR2DS4 | CHR_HSCHR19KIR_50296000 | killer cell immunoglobulin like receptor, two Ig |
| KIR3DL2 | CHR_HSCHR19KIR_50296000 | killer cell immunoglobulin like receptor, three |
| LILRP1 | CHR_HSCHR19KIR_50296000 | leukocyte immunoglobulin-like receptor pseu |
| AC245128.33 | CHR_HSCHR19KIR_50296000 | killer cell immunoglobulin-like receptor family |
| LILRP2 | CHR_HSCHR19KIR_50296000 | leukocyte immunoglobulin-like receptor pseu |
| AC245128.26 | CHR_HSCHR19KIR_50296000 | novel transcript |
| KIR2DP1 | CHR_HSCHR19KIR_50296000 | killer cell immunoglobulin like receptor, two Ig |
| KIR2DL1 | CHR_HSCHR19KIR_50296000 | killer cell immunoglobulin like receptor, two Ig |
| KIR3DP1 | CHR_HSCHR19KIR_50296000 | killer cell immunoglobulin like receptor, three |
| KIR2DL4 | CHR_HSCHR19KIR_50296000 | killer cell immunoglobulin like receptor, two Ig |
| KIR3DL1 | CHR_HSCHR19KIR_50296000 | killer cell immunoglobulin like receptor, three |
| KIR2DS4 | CHR_HSCHR19KIR_50296000 | killer cell immunoglobulin like receptor, two Ig |
| KIR3DL2 | CHR_HSCHR19KIR_50296000 | killer cell immunoglobulin like receptor, three |
| RNU6-222P | CHR_HSCHR19KIR_50296000 | RNA, U6 small nuclear 222, pseudogene [Sour |
| FCAR | CHR_HSCHR19KIR_50296000 | Fc fragment of IgA receptor [Source:HGNC Syr |
| LILRP1 | CHR_HSCHR19KIR_7191059-1 | leukocyte immunoglobulin-like receptor pseu |
| AC245128.30 | CHR_HSCHR19KIR_7191059-1 | killer cell immunoglobulin-like receptor family |
| LILRP2 | CHR_HSCHR19KIR_7191059-1 | leukocyte immunoglobulin-like receptor pseu |
| KIR3DL3 | CHR_HSCHR19KIR_7191059-1 | killer cell immunoglobulin like receptor, three |
| AC245128.35 | CHR_HSCHR19KIR_7191059-1 | novel transcript |
| KIR2DL3 | CHR_HSCHR19KIR_7191059-1 | killer cell immunoglobulin like receptor, two Ig |
| KIR2DP1 | CHR_HSCHR19KIR_7191059-1 | killer cell immunoglobulin like receptor, two Ig |
| KIR2DL1 | CHR_HSCHR19KIR_7191059-1 | killer cell immunoglobulin like receptor, two Ig |
| KIR3DP1 | CHR_HSCHR19KIR_7191059-1 | killer cell immunoglobulin like receptor, three |
| KIR2DL4 | CHR_HSCHR19KIR_7191059-1 | killer cell immunoglobulin like receptor, two Ig |
| KIR3DL1 | CHR_HSCHR19KIR_7191059-1 | killer cell immunoglobulin like receptor, three |
| KIR2DS4 | CHR_HSCHR19KIR_7191059-1 | killer cell immunoglobulin like receptor, two Ig |
| KIR3DL2 | CHR_HSCHR19KIR_7191059-1 | killer cell immunoglobulin like receptor, three |
| LILRP1 | CHR_HSCHR19KIR_7191059-2 | leukocyte immunoglobulin-like receptor pseu |
| AC245128.18 | CHR_HSCHR19KIR_7191059-2 | killer cell immunoglobulin-like receptor family |
| LILRP2 | CHR_HSCHR19KIR_7191059-2 | leukocyte immunoglobulin-like receptor pseu |
| KIR3DL3 | CHR_HSCHR19KIR_7191059-2 | killer cell immunoglobulin like receptor, three |
| AC245128.36 | CHR_HSCHR19KIR_7191059-2 | novel transcript |
| KIR2DL3 | CHR_HSCHR19KIR_7191059-2 | killer cell immunoglobulin like receptor, two Ig |
| KIR2DP1 | CHR_HSCHR19KIR_7191059-2 | killer cell immunoglobulin like receptor, two Ig |
| KP420446.1 | CHR_HSCHR19KIR_7191059-2 | NULL |
| KIR3DP1 | CHR_HSCHR19KIR_7191059-2 | killer cell immunoglobulin like receptor, three |
| KIR2DL4 | CHR_HSCHR19KIR_7191059-2 | killer cell immunoglobulin like receptor, two Ig |
| KIR3DL1 | CHR_HSCHR19KIR_7191059-2 | killer cell immunoglobulin like receptor, three |
| KIR2DS4 | CHR_HSCHR19KIR_7191059-2 | killer cell immunoglobulin like receptor, two Ig |

| | | |
|---|---|---|
| KIR3DL2 | CHR_HSCHR19KIR_7191059-2 | killer cell immunoglobulin like receptor, three |
| RNU6-222P | CHR_HSCHR19KIR_7191059-2 | RNA, U6 small nuclear 222, pseudogene [Sour |
| KIR3DL3 | CHR_HSCHR19KIR_ABC08_A1 | killer cell immunoglobulin like receptor, three |
| KIR2DL3 | CHR_HSCHR19KIR_ABC08_A1 | killer cell immunoglobulin like receptor, two Ig |
| KIR2DP1 | CHR_HSCHR19KIR_ABC08_A1 | killer cell immunoglobulin like receptor, two Ig |
| KIR2DL1 | CHR_HSCHR19KIR_ABC08_A1 | killer cell immunoglobulin like receptor, two Ig |
| GU182360.1 | CHR_HSCHR19KIR_ABC08_A1 | novel pseudogene |
| KIR2DL4 | CHR_HSCHR19KIR_ABC08_A1 | killer cell immunoglobulin like receptor, two Ig |
| KIR3DL1 | CHR_HSCHR19KIR_ABC08_A1 | killer cell immunoglobulin like receptor, three |
| KIR2DS4 | CHR_HSCHR19KIR_ABC08_A1 | killer cell immunoglobulin like receptor, two Ig |
| KIR3DL2 | CHR_HSCHR19KIR_ABC08_A1 | killer cell immunoglobulin like receptor, three |
| U6 | CHR_HSCHR19KIR_ABC08_A1 | U6 spliceosomal RNA [Source:RFAM;Acc:RF00 |
| FCAR | CHR_HSCHR19KIR_ABC08_A1 | Fc fragment of IgA receptor [Source:HGNC Syr |
| NCR1 | CHR_HSCHR19KIR_ABC08_A1 | natural cytotoxicity triggering receptor 1 [Sou |
| LILRP1 | CHR_HSCHR19KIR_ABC08_AE | leukocyte immunoglobulin-like receptor pseu |
| AC245128.8 | CHR_HSCHR19KIR_ABC08_AE | killer cell immunoglobulin-like receptor family |
| LILRP2 | CHR_HSCHR19KIR_ABC08_AE | leukocyte immunoglobulin-like receptor pseu |
| KIR3DL3 | CHR_HSCHR19KIR_ABC08_AE | killer cell immunoglobulin like receptor, three |
| KIR2DL3 | CHR_HSCHR19KIR_ABC08_AE | killer cell immunoglobulin like receptor, two Ig |
| KIR2DP1 | CHR_HSCHR19KIR_ABC08_AE | killer cell immunoglobulin like receptor, two Ig |
| KIR2DL1 | CHR_HSCHR19KIR_ABC08_AE | killer cell immunoglobulin like receptor, two Ig |
| KIR3DP1 | CHR_HSCHR19KIR_ABC08_AE | killer cell immunoglobulin like receptor, three |
| KIR2DL4 | CHR_HSCHR19KIR_ABC08_AE | killer cell immunoglobulin like receptor, two Ig |
| KIR3DL1 | CHR_HSCHR19KIR_ABC08_AE | killer cell immunoglobulin like receptor, three |
| LILRP1 | CHR_HSCHR19KIR_ABC08_AE | leukocyte immunoglobulin-like receptor pseu |
| AC245128.11 | CHR_HSCHR19KIR_ABC08_AE | killer cell immunoglobulin-like receptor family |
| LILRP2 | CHR_HSCHR19KIR_ABC08_AE | leukocyte immunoglobulin-like receptor pseu |
| KIR3DL3 | CHR_HSCHR19KIR_ABC08_AE | killer cell immunoglobulin like receptor, three |
| KIR2DL3 | CHR_HSCHR19KIR_ABC08_AE | killer cell immunoglobulin like receptor, two Ig |
| KIR2DP1 | CHR_HSCHR19KIR_ABC08_AE | killer cell immunoglobulin like receptor, two Ig |
| KIR2DL1 | CHR_HSCHR19KIR_ABC08_AE | killer cell immunoglobulin like receptor, two Ig |
| KIR3DP1 | CHR_HSCHR19KIR_ABC08_AE | killer cell immunoglobulin like receptor, three |
| KIR2DL4 | CHR_HSCHR19KIR_ABC08_AE | killer cell immunoglobulin like receptor, two Ig |
| KIR3DL1 | CHR_HSCHR19KIR_ABC08_AE | killer cell immunoglobulin like receptor, three |
| KIR2DS4 | CHR_HSCHR19KIR_ABC08_AE | killer cell immunoglobulin like receptor, two Ig |
| KIR3DL2 | CHR_HSCHR19KIR_ABC08_AE | killer cell immunoglobulin like receptor, three |
| U6 | CHR_HSCHR19KIR_ABC08_AE | U6 spliceosomal RNA [Source:RFAM;Acc:RF00 |
| LILRP1 | CHR_HSCHR19KIR_CA01-TA0 | leukocyte immunoglobulin-like receptor pseu |
| AC245128.22 | CHR_HSCHR19KIR_CA01-TA0 | killer cell immunoglobulin-like receptor family |
| LILRP2 | CHR_HSCHR19KIR_CA01-TA0 | leukocyte immunoglobulin-like receptor pseu |
| KIR3DL3 | CHR_HSCHR19KIR_CA01-TA0 | killer cell immunoglobulin like receptor, three |
| AC245128.21 | CHR_HSCHR19KIR_CA01-TA0 | novel transcript |
| KIR2DL3 | CHR_HSCHR19KIR_CA01-TA0 | killer cell immunoglobulin like receptor, two Ig |
| KIR2DP1 | CHR_HSCHR19KIR_CA01-TA0 | killer cell immunoglobulin like receptor, two Ig |
| KIR2DL1 | CHR_HSCHR19KIR_CA01-TA0 | killer cell immunoglobulin like receptor, two Ig |
| KIR3DP1 | CHR_HSCHR19KIR_CA01-TA0 | killer cell immunoglobulin like receptor, three |
| KIR2DL4 | CHR_HSCHR19KIR_CA01-TA0 | killer cell immunoglobulin like receptor, two Ig |
| KIR3DL1 | CHR_HSCHR19KIR_CA01-TA0 | killer cell immunoglobulin like receptor, three |
| KIR2DS4 | CHR_HSCHR19KIR_CA01-TA0 | killer cell immunoglobulin like receptor, two Ig |
| KIR3DL2 | CHR_HSCHR19KIR_CA01-TA0 | killer cell immunoglobulin like receptor, three |

| | | |
|---|---|---|
| LILRP1 | CHR_HSCHR19KIR_CA01-TA0 | leukocyte immunoglobulin-like receptor pseu |
| AC245128.32 | CHR_HSCHR19KIR_CA01-TA0 | killer cell immunoglobulin-like receptor family |
| LILRP2 | CHR_HSCHR19KIR_CA01-TA0 | leukocyte immunoglobulin-like receptor pseu |
| KIR3DL3 | CHR_HSCHR19KIR_CA01-TA0 | killer cell immunoglobulin like receptor, three |
| KU645195.1 | CHR_HSCHR19KIR_CA01-TA0 | novel transcript |
| KIR2DL3 | CHR_HSCHR19KIR_CA01-TA0 | killer cell immunoglobulin like receptor, two Ig |
| KIR2DP1 | CHR_HSCHR19KIR_CA01-TA0 | killer cell immunoglobulin like receptor, two Ig |
| KIR2DL1 | CHR_HSCHR19KIR_CA01-TA0 | killer cell immunoglobulin like receptor, two Ig |
| KIR3DP1 | CHR_HSCHR19KIR_CA01-TA0 | killer cell immunoglobulin like receptor, three |
| KIR2DL4 | CHR_HSCHR19KIR_CA01-TA0 | killer cell immunoglobulin like receptor, two Ig |
| KIR3DL1 | CHR_HSCHR19KIR_CA01-TA0 | killer cell immunoglobulin like receptor, three |
| KIR2DS4 | CHR_HSCHR19KIR_CA01-TA0 | killer cell immunoglobulin like receptor, two Ig |
| KIR3DL2 | CHR_HSCHR19KIR_CA01-TA0 | killer cell immunoglobulin like receptor, three |
| LILRP1 | CHR_HSCHR19KIR_CA01-TB0 | leukocyte immunoglobulin-like receptor pseu |
| AC245128.28 | CHR_HSCHR19KIR_CA01-TB0 | killer cell immunoglobulin-like receptor family |
| LILRP2 | CHR_HSCHR19KIR_CA01-TB0 | leukocyte immunoglobulin-like receptor pseu |
| AC245128.31 | CHR_HSCHR19KIR_CA01-TB0 | novel transcript |
| KIR2DL3 | CHR_HSCHR19KIR_CA01-TB0 | killer cell immunoglobulin like receptor, two Ig |
| KU645197.1 | CHR_HSCHR19KIR_CA01-TB0 | NULL |
| KIR2DL1 | CHR_HSCHR19KIR_CA01-TB0 | killer cell immunoglobulin like receptor, two Ig |
| KIR3DP1 | CHR_HSCHR19KIR_CA01-TB0 | killer cell immunoglobulin like receptor, three |
| KIR2DL4 | CHR_HSCHR19KIR_CA01-TB0 | killer cell immunoglobulin like receptor, two Ig |
| KIR3DS1 | CHR_HSCHR19KIR_CA01-TB0 | killer cell immunoglobulin like receptor, three |
| KIR2DS5 | CHR_HSCHR19KIR_CA01-TB0 | killer cell immunoglobulin like receptor, two Ig |
| KIR2DS1 | CHR_HSCHR19KIR_CA01-TB0 | killer cell immunoglobulin like receptor, two Ig |
| KIR3DL2 | CHR_HSCHR19KIR_CA01-TB0 | killer cell immunoglobulin like receptor, three |
| RNU6-222P | CHR_HSCHR19KIR_CA01-TB0 | RNA, U6 small nuclear 222, pseudogene [Sour |
| FCAR | CHR_HSCHR19KIR_CA01-TB0 | Fc fragment of IgA receptor [Source:HGNC Syr |
| AC245128.24 | CHR_HSCHR19KIR_CA01-TB0 | natural cytotoxicity triggering receptor 1 [Sou |
| LILRP1 | CHR_HSCHR19KIR_CA01-TB0 | leukocyte immunoglobulin-like receptor pseu |
| AC245128.27 | CHR_HSCHR19KIR_CA01-TB0 | killer cell immunoglobulin-like receptor family |
| LILRP2 | CHR_HSCHR19KIR_CA01-TB0 | leukocyte immunoglobulin-like receptor pseu |
| KIR3DL3 | CHR_HSCHR19KIR_CA01-TB0 | killer cell immunoglobulin like receptor, three |
| KIR2DL3 | CHR_HSCHR19KIR_CA01-TB0 | killer cell immunoglobulin like receptor, two Ig |
| KIR2DP1 | CHR_HSCHR19KIR_CA01-TB0 | killer cell immunoglobulin like receptor, two Ig |
| KU645196.5 | CHR_HSCHR19KIR_CA01-TB0 | NULL |
| KU645196.2 | CHR_HSCHR19KIR_CA01-TB0 | killer cell immunoglobulin-like receptor 2DL4- |
| KU645196.6 | CHR_HSCHR19KIR_CA01-TB0 | killer cell immunoglobulin-like receptor 3DS1- |
| KIR2DS3 | CHR_HSCHR19KIR_CA01-TB0 | killer cell immunoglobulin like receptor, two Ig |
| KU645196.3 | CHR_HSCHR19KIR_CA01-TB0 | killer cell immunoglobulin-like receptor 2DL1- |
| KIR3DP1 | CHR_HSCHR19KIR_CA01-TB0 | killer cell immunoglobulin like receptor, three |
| KU645196.1 | CHR_HSCHR19KIR_CA01-TB0 | NULL |
| KIR3DS1 | CHR_HSCHR19KIR_CA01-TB0 | killer cell immunoglobulin like receptor, three |
| KIR2DS5 | CHR_HSCHR19KIR_CA01-TB0 | killer cell immunoglobulin like receptor, two Ig |
| KIR2DS1 | CHR_HSCHR19KIR_CA01-TB0 | killer cell immunoglobulin like receptor, two Ig |
| KU645196.4 | CHR_HSCHR19KIR_CA01-TB0 | NULL |
| LILRP1 | CHR_HSCHR19KIR_CA04_CTG | leukocyte immunoglobulin-like receptor pseu |
| AC245128.25 | CHR_HSCHR19KIR_CA04_CTG | killer cell immunoglobulin-like receptor family |
| LILRP2 | CHR_HSCHR19KIR_CA04_CTG | leukocyte immunoglobulin-like receptor pseu |
| KIR2DL3 | CHR_HSCHR19KIR_CA04_CTG | killer cell immunoglobulin like receptor, two Ig |

| | | |
|---|---|---|
| KIR2DP1 | CHR_HSCHR19KIR_CA04_CTG | killer cell immunoglobulin like receptor, two Ig |
| KU645198.1 | CHR_HSCHR19KIR_CA04_CTG | NULL |
| RNU6-222P | CHR_HSCHR19KIR_CA04_CTG | RNA, U6 small nuclear 222, pseudogene [Sour |
| FCAR | CHR_HSCHR19KIR_CA04_CTG | Fc fragment of IgA receptor [Source:HGNC Syr |
| KIR3DL3 | CHR_HSCHR19KIR_FH05_A_H | killer cell immunoglobulin like receptor, three |
| KIR2DL3 | CHR_HSCHR19KIR_FH05_A_H | killer cell immunoglobulin like receptor, two Ig |
| KIR2DP1 | CHR_HSCHR19KIR_FH05_A_H | killer cell immunoglobulin like receptor, two Ig |
| KIR2DL1 | CHR_HSCHR19KIR_FH05_A_H | killer cell immunoglobulin like receptor, two Ig |
| GU182338.1 | CHR_HSCHR19KIR_FH05_A_H | killer cell immunoglobulin-like receptor family |
| KIR2DL4 | CHR_HSCHR19KIR_FH05_A_H | killer cell immunoglobulin like receptor, two Ig |
| KIR3DL1 | CHR_HSCHR19KIR_FH05_A_H | killer cell immunoglobulin like receptor, three |
| KIR2DS4 | CHR_HSCHR19KIR_FH05_A_H | killer cell immunoglobulin like receptor, two Ig |
| KIR3DL2 | CHR_HSCHR19KIR_FH05_A_H | killer cell immunoglobulin like receptor, three |
| U6 | CHR_HSCHR19KIR_FH05_A_H | U6 spliceosomal RNA [Source:RFAM;Acc:RF00 |
| FCAR | CHR_HSCHR19KIR_FH05_A_H | Fc fragment of IgA receptor [Source:HGNC Syr |
| NCR1 | CHR_HSCHR19KIR_FH05_A_H | natural cytotoxicity triggering receptor 1 [Sou |
| KIR3DL3 | CHR_HSCHR19KIR_FH05_B_H | killer cell immunoglobulin like receptor, three |
| KIR2DS2 | CHR_HSCHR19KIR_FH05_B_H | killer cell immunoglobulin like receptor, two Ig |
| KIR2DL2 | CHR_HSCHR19KIR_FH05_B_H | killer cell immunoglobulin like receptor, two Ig |
| KIR2DL5B | CHR_HSCHR19KIR_FH05_B_H | killer cell immunoglobulin like receptor, two Ig |
| KIR2DS3 | CHR_HSCHR19KIR_FH05_B_H | killer cell immunoglobulin like receptor, two Ig |
| GU182339.2 | CHR_HSCHR19KIR_FH05_B_H | killer cell immunoglobulin-like receptor, two c |
| KIR2DL1 | CHR_HSCHR19KIR_FH05_B_H | killer cell immunoglobulin like receptor, two Ig |
| KIR3DP1 | CHR_HSCHR19KIR_FH05_B_H | killer cell immunoglobulin like receptor, three |
| KIR2DL4 | CHR_HSCHR19KIR_FH05_B_H | killer cell immunoglobulin like receptor, two Ig |
| KIR3DS1 | CHR_HSCHR19KIR_FH05_B_H | killer cell immunoglobulin like receptor, three |
| KIR2DL5A | CHR_HSCHR19KIR_FH05_B_H | killer cell immunoglobulin like receptor, two Ig |
| GU182339.1 | CHR_HSCHR19KIR_FH05_B_H | killer cell immunoglobulin-like receptor 2DS3- |
| KIR2DS1 | CHR_HSCHR19KIR_FH05_B_H | killer cell immunoglobulin like receptor, two Ig |
| KIR3DL2 | CHR_HSCHR19KIR_FH05_B_H | killer cell immunoglobulin like receptor, three |
| U6 | CHR_HSCHR19KIR_FH05_B_H | U6 spliceosomal RNA [Source:RFAM;Acc:RF00 |
| FCAR | CHR_HSCHR19KIR_FH05_B_H | Fc fragment of IgA receptor [Source:HGNC Syr |
| NCR1 | CHR_HSCHR19KIR_FH05_B_H | natural cytotoxicity triggering receptor 1 [Sou |
| KIR3DL3 | CHR_HSCHR19KIR_FH06_A_H | killer cell immunoglobulin like receptor, three |
| KIR2DL3 | CHR_HSCHR19KIR_FH06_A_H | killer cell immunoglobulin like receptor, two Ig |
| KIR2DP1 | CHR_HSCHR19KIR_FH06_A_H | killer cell immunoglobulin like receptor, two Ig |
| KIR2DL1 | CHR_HSCHR19KIR_FH06_A_H | killer cell immunoglobulin like receptor, two Ig |
| KIR3DP1 | CHR_HSCHR19KIR_FH06_A_H | killer cell immunoglobulin like receptor, three |
| KIR2DL4 | CHR_HSCHR19KIR_FH06_A_H | killer cell immunoglobulin like receptor, two Ig |
| KIR3DL1 | CHR_HSCHR19KIR_FH06_A_H | killer cell immunoglobulin like receptor, three |
| KIR2DS4 | CHR_HSCHR19KIR_FH06_A_H | killer cell immunoglobulin like receptor, two Ig |
| KIR3DL2 | CHR_HSCHR19KIR_FH06_A_H | killer cell immunoglobulin like receptor, three |
| U6 | CHR_HSCHR19KIR_FH06_A_H | U6 spliceosomal RNA [Source:RFAM;Acc:RF00 |
| FCAR | CHR_HSCHR19KIR_FH06_A_H | Fc fragment of IgA receptor [Source:HGNC Syr |
| NCR1 | CHR_HSCHR19KIR_FH06_A_H | natural cytotoxicity triggering receptor 1 [Sou |
| LILRP2 | CHR_HSCHR19KIR_FH06_BA1 | leukocyte immunoglobulin-like receptor pseu |
| KIR3DL3 | CHR_HSCHR19KIR_FH06_BA1 | killer cell immunoglobulin like receptor, three |
| KIR2DS2 | CHR_HSCHR19KIR_FH06_BA1 | killer cell immunoglobulin like receptor, two Ig |
| KIR2DL2 | CHR_HSCHR19KIR_FH06_BA1 | killer cell immunoglobulin like receptor, two Ig |
| KIR3DP1 | CHR_HSCHR19KIR_FH06_BA1 | killer cell immunoglobulin like receptor, three |

| | | |
|---|---|---|
| KIR2DL4 | CHR_HSCHR19KIR_FH06_BA1 | killer cell immunoglobulin like receptor, two Ig |
| KIR3DL1 | CHR_HSCHR19KIR_FH06_BA1 | killer cell immunoglobulin like receptor, three |
| KIR2DS4 | CHR_HSCHR19KIR_FH06_BA1 | killer cell immunoglobulin like receptor, two Ig |
| KIR3DL2 | CHR_HSCHR19KIR_FH06_BA1 | killer cell immunoglobulin like receptor, three |
| U6 | CHR_HSCHR19KIR_FH06_BA1 | U6 spliceosomal RNA [Source:RFAM;Acc:RF00 |
| FCAR | CHR_HSCHR19KIR_FH06_BA1 | Fc fragment of IgA receptor [Source:HGNC Syr |
| NCR1 | CHR_HSCHR19KIR_FH06_BA1 | natural cytotoxicity triggering receptor 1 [Sou |
| KIR3DL3 | CHR_HSCHR19KIR_FH08_A_H | killer cell immunoglobulin like receptor, three |
| KIR2DL3 | CHR_HSCHR19KIR_FH08_A_H | killer cell immunoglobulin like receptor, two Ig |
| KIR2DP1 | CHR_HSCHR19KIR_FH08_A_H | killer cell immunoglobulin like receptor, two Ig |
| KIR2DL1 | CHR_HSCHR19KIR_FH08_A_H | killer cell immunoglobulin like receptor, two Ig |
| KIR3DP1 | CHR_HSCHR19KIR_FH08_A_H | killer cell immunoglobulin like receptor, three |
| KIR2DL4 | CHR_HSCHR19KIR_FH08_A_H | killer cell immunoglobulin like receptor, two Ig |
| KIR3DL1 | CHR_HSCHR19KIR_FH08_A_H | killer cell immunoglobulin like receptor, three |
| KIR2DS4 | CHR_HSCHR19KIR_FH08_A_H | killer cell immunoglobulin like receptor, two Ig |
| KIR3DL2 | CHR_HSCHR19KIR_FH08_A_H | killer cell immunoglobulin like receptor, three |
| U6 | CHR_HSCHR19KIR_FH08_A_H | U6 spliceosomal RNA [Source:RFAM;Acc:RF00 |
| FCAR | CHR_HSCHR19KIR_FH08_A_H | Fc fragment of IgA receptor [Source:HGNC Syr |
| NCR1 | CHR_HSCHR19KIR_FH08_A_H | natural cytotoxicity triggering receptor 1 [Sou |
| LILRP1 | CHR_HSCHR19KIR_FH08_BAX | leukocyte immunoglobulin-like receptor pseu |
| AC245128.14 | CHR_HSCHR19KIR_FH08_BAX | killer cell immunoglobulin-like receptor family |
| LILRP2 | CHR_HSCHR19KIR_FH08_BAX | leukocyte immunoglobulin-like receptor pseu |
| GU182343.4 | CHR_HSCHR19KIR_FH08_BAX | novel killer cell immunoglobulin-like receptor |
| KIR2DL3 | CHR_HSCHR19KIR_FH08_BAX | killer cell immunoglobulin like receptor, two Ig |
| GU182343.2 | CHR_HSCHR19KIR_FH08_BAX | killer cell immunoglobulin-like receptor, two c |
| GU182343.1 | CHR_HSCHR19KIR_FH08_BAX | killer cell immunoglobulin-like receptor, two c |
| GU182343.3 | CHR_HSCHR19KIR_FH08_BAX | novel killer cell immunoglobulin-like receptor |
| KIR2DL1 | CHR_HSCHR19KIR_FH08_BAX | killer cell immunoglobulin like receptor, two Ig |
| KIR3DL1 | CHR_HSCHR19KIR_FH08_BAX | killer cell immunoglobulin like receptor, three |
| GU182343.5 | CHR_HSCHR19KIR_FH08_BAX | novel killer cell immunoglobulin-like receptor |
| KIR2DL4 | CHR_HSCHR19KIR_FH08_BAX | killer cell immunoglobulin like receptor, two Ig |
| KIR2DS4 | CHR_HSCHR19KIR_FH08_BAX | killer cell immunoglobulin like receptor, two Ig |
| KIR3DL2 | CHR_HSCHR19KIR_FH08_BAX | killer cell immunoglobulin like receptor, three |
| U6 | CHR_HSCHR19KIR_FH08_BAX | U6 spliceosomal RNA [Source:RFAM;Acc:RF00 |
| LILRP1 | CHR_HSCHR19KIR_FH13_A_H | leukocyte immunoglobulin-like receptor pseu |
| AC245128.10 | CHR_HSCHR19KIR_FH13_A_H | killer cell immunoglobulin-like receptor family |
| LILRP2 | CHR_HSCHR19KIR_FH13_A_H | leukocyte immunoglobulin-like receptor pseu |
| KIR3DL3 | CHR_HSCHR19KIR_FH13_A_H | killer cell immunoglobulin like receptor, three |
| KIR2DL3 | CHR_HSCHR19KIR_FH13_A_H | killer cell immunoglobulin like receptor, two Ig |
| KIR2DP1 | CHR_HSCHR19KIR_FH13_A_H | killer cell immunoglobulin like receptor, two Ig |
| KIR2DL1 | CHR_HSCHR19KIR_FH13_A_H | killer cell immunoglobulin like receptor, two Ig |
| KIR3DP1 | CHR_HSCHR19KIR_FH13_A_H | killer cell immunoglobulin like receptor, three |
| KIR2DL4 | CHR_HSCHR19KIR_FH13_A_H | killer cell immunoglobulin like receptor, two Ig |
| KIR3DL1 | CHR_HSCHR19KIR_FH13_A_H | killer cell immunoglobulin like receptor, three |
| KIR2DS4 | CHR_HSCHR19KIR_FH13_A_H | killer cell immunoglobulin like receptor, two Ig |
| KIR3DL2 | CHR_HSCHR19KIR_FH13_A_H | killer cell immunoglobulin like receptor, three |
| U6 | CHR_HSCHR19KIR_FH13_A_H | U6 spliceosomal RNA [Source:RFAM;Acc:RF00 |
| LILRP1 | CHR_HSCHR19KIR_FH13_BA2 | leukocyte immunoglobulin-like receptor pseu |
| LILRP2 | CHR_HSCHR19KIR_FH13_BA2 | leukocyte immunoglobulin-like receptor pseu |
| KIR3DL3 | CHR_HSCHR19KIR_FH13_BA2 | killer cell immunoglobulin like receptor, three |

| | | |
|---|---|---|
| GU182345.3 | CHR_HSCHR19KIR_FH13_BA2 | Killer cell immunoglobulin-like receptor 2DS2 |
| GU182345.1 | CHR_HSCHR19KIR_FH13_BA2 | Killer cell immunoglobulin-like receptor 2DL2 |
| GU182345.2 | CHR_HSCHR19KIR_FH13_BA2 | killer cell immunoglobulin-like receptor, two d |
| GU182345.4 | CHR_HSCHR19KIR_FH13_BA2 | killer cell immunoglobulin-like receptor, two d |
| KIR2DP1 | CHR_HSCHR19KIR_FH13_BA2 | killer cell immunoglobulin like receptor, two Ig |
| KIR2DL1 | CHR_HSCHR19KIR_FH13_BA2 | killer cell immunoglobulin like receptor, two Ig |
| KIR3DP1 | CHR_HSCHR19KIR_FH13_BA2 | killer cell immunoglobulin like receptor, three |
| KIR2DL4 | CHR_HSCHR19KIR_FH13_BA2 | killer cell immunoglobulin like receptor, two Ig |
| KIR3DL1 | CHR_HSCHR19KIR_FH13_BA2 | killer cell immunoglobulin like receptor, three |
| KIR2DS4 | CHR_HSCHR19KIR_FH13_BA2 | killer cell immunoglobulin like receptor, two Ig |
| KIR3DL2 | CHR_HSCHR19KIR_FH13_BA2 | killer cell immunoglobulin like receptor, three |
| U6 | CHR_HSCHR19KIR_FH13_BA2 | U6 spliceosomal RNA [Source:RFAM;Acc:RF00 |
| LILRP1 | CHR_HSCHR19KIR_FH15_A_H | leukocyte immunoglobulin-like receptor pseu |
| AC245128.9 | CHR_HSCHR19KIR_FH15_A_H | killer cell immunoglobulin-like receptor family |
| LILRP2 | CHR_HSCHR19KIR_FH15_A_H | leukocyte immunoglobulin-like receptor pseu |
| KIR3DL3 | CHR_HSCHR19KIR_FH15_A_H | killer cell immunoglobulin like receptor, three |
| KIR2DL3 | CHR_HSCHR19KIR_FH15_A_H | killer cell immunoglobulin like receptor, two Ig |
| KIR2DP1 | CHR_HSCHR19KIR_FH15_A_H | killer cell immunoglobulin like receptor, two Ig |
| KIR2DL1 | CHR_HSCHR19KIR_FH15_A_H | killer cell immunoglobulin like receptor, two Ig |
| KIR3DP1 | CHR_HSCHR19KIR_FH15_A_H | killer cell immunoglobulin like receptor, three |
| KIR2DL4 | CHR_HSCHR19KIR_FH15_A_H | killer cell immunoglobulin like receptor, two Ig |
| KIR3DL1 | CHR_HSCHR19KIR_FH15_A_H | killer cell immunoglobulin like receptor, three |
| KIR2DS4 | CHR_HSCHR19KIR_FH15_A_H | killer cell immunoglobulin like receptor, two Ig |
| KIR3DL2 | CHR_HSCHR19KIR_FH15_A_H | killer cell immunoglobulin like receptor, three |
| U6 | CHR_HSCHR19KIR_FH15_A_H | U6 spliceosomal RNA [Source:RFAM;Acc:RF00 |
| LILRP1 | CHR_HSCHR19KIR_FH15_B_H | leukocyte immunoglobulin-like receptor pseu |
| AC245128.16 | CHR_HSCHR19KIR_FH15_B_H | killer cell immunoglobulin-like receptor family |
| LILRP2 | CHR_HSCHR19KIR_FH15_B_H | leukocyte immunoglobulin-like receptor pseu |
| KIR3DL3 | CHR_HSCHR19KIR_FH15_B_H | killer cell immunoglobulin like receptor, three |
| GU182347.3 | CHR_HSCHR19KIR_FH15_B_H | Killer cell immunoglobulin-like receptor 2DS2 |
| GU182347.5 | CHR_HSCHR19KIR_FH15_B_H | Killer cell immunoglobulin-like receptor 2DL2 |
| GU182347.8 | CHR_HSCHR19KIR_FH15_B_H | Killer cell immunoglobulin-like receptor 2DL5B |
| GU182347.6 | CHR_HSCHR19KIR_FH15_B_H | Killer cell immunoglobulin-like receptor 2DS3 |
| KIR2DP1 | CHR_HSCHR19KIR_FH15_B_H | killer cell immunoglobulin like receptor, two Ig |
| KIR2DL1 | CHR_HSCHR19KIR_FH15_B_H | killer cell immunoglobulin like receptor, two Ig |
| KIR3DP1 | CHR_HSCHR19KIR_FH15_B_H | killer cell immunoglobulin like receptor, three |
| KIR2DL4 | CHR_HSCHR19KIR_FH15_B_H | killer cell immunoglobulin like receptor, two Ig |
| GU182347.1 | CHR_HSCHR19KIR_FH15_B_H | killer cell immunoglobulin-like receptor, three |
| GU182347.4 | CHR_HSCHR19KIR_FH15_B_H | killer cell immunoglobulin-like receptor, two d |
| GU182347.7 | CHR_HSCHR19KIR_FH15_B_H | novel killer cell immunoglobulin-like receptor |
| GU182347.2 | CHR_HSCHR19KIR_FH15_B_H | killer cell immunoglobulin-like receptor, two d |
| KIR3DL2 | CHR_HSCHR19KIR_FH15_B_H | killer cell immunoglobulin like receptor, three |
| U6 | CHR_HSCHR19KIR_FH15_B_H | U6 spliceosomal RNA [Source:RFAM;Acc:RF00 |
| LILRP1 | CHR_HSCHR19KIR_G085_A_H | leukocyte immunoglobulin-like receptor pseu |
| AC245128.13 | CHR_HSCHR19KIR_G085_A_H | killer cell immunoglobulin-like receptor family |
| LILRP2 | CHR_HSCHR19KIR_G085_A_H | leukocyte immunoglobulin-like receptor pseu |
| KIR3DL3 | CHR_HSCHR19KIR_G085_A_H | killer cell immunoglobulin like receptor, three |
| KIR2DL3 | CHR_HSCHR19KIR_G085_A_H | killer cell immunoglobulin like receptor, two Ig |
| KIR2DP1 | CHR_HSCHR19KIR_G085_A_H | killer cell immunoglobulin like receptor, two Ig |
| KIR2DL1 | CHR_HSCHR19KIR_G085_A_H | killer cell immunoglobulin like receptor, two Ig |

| | | |
|---|---|---|
| KIR3DP1 | CHR_HSCHR19KIR_G085_A_H | killer cell immunoglobulin like receptor, three |
| KIR2DL4 | CHR_HSCHR19KIR_G085_A_H | killer cell immunoglobulin like receptor, two Ig |
| KIR3DL1 | CHR_HSCHR19KIR_G085_A_H | killer cell immunoglobulin like receptor, three |
| KIR2DS4 | CHR_HSCHR19KIR_G085_A_H | killer cell immunoglobulin like receptor, two Ig |
| KIR3DL2 | CHR_HSCHR19KIR_G085_A_H | killer cell immunoglobulin like receptor, three |
| LILRP1 | CHR_HSCHR19KIR_G085_BA1 | leukocyte immunoglobulin-like receptor pseu |
| AC245128.15 | CHR_HSCHR19KIR_G085_BA1 | killer cell immunoglobulin-like receptor family |
| LILRP2 | CHR_HSCHR19KIR_G085_BA1 | leukocyte immunoglobulin-like receptor pseu |
| KIR3DL3 | CHR_HSCHR19KIR_G085_BA1 | killer cell immunoglobulin like receptor, three |
| GU182349.2 | CHR_HSCHR19KIR_G085_BA1 | Killer cell immunoglobulin-like receptor 2DS2 |
| GU182349.1 | CHR_HSCHR19KIR_G085_BA1 | Killer cell immunoglobulin-like receptor 2DL2 |
| KIR3DP1 | CHR_HSCHR19KIR_G085_BA1 | killer cell immunoglobulin like receptor, three |
| KIR2DL4 | CHR_HSCHR19KIR_G085_BA1 | killer cell immunoglobulin like receptor, two Ig |
| KIR3DL1 | CHR_HSCHR19KIR_G085_BA1 | killer cell immunoglobulin like receptor, three |
| KIR2DS4 | CHR_HSCHR19KIR_G085_BA1 | killer cell immunoglobulin like receptor, two Ig |
| KIR3DL2 | CHR_HSCHR19KIR_G085_BA1 | killer cell immunoglobulin like receptor, three |
| U6 | CHR_HSCHR19KIR_G085_BA1 | U6 spliceosomal RNA [Source:RFAM;Acc:RF00 |
| LILRP1 | CHR_HSCHR19KIR_G248_A_H | leukocyte immunoglobulin-like receptor pseu |
| AC245128.4 | CHR_HSCHR19KIR_G248_A_H | killer cell immunoglobulin-like receptor family |
| LILRP2 | CHR_HSCHR19KIR_G248_A_H | leukocyte immunoglobulin-like receptor pseu |
| KIR3DL3 | CHR_HSCHR19KIR_G248_A_H | killer cell immunoglobulin like receptor, three |
| KIR2DL3 | CHR_HSCHR19KIR_G248_A_H | killer cell immunoglobulin like receptor, two Ig |
| KIR2DP1 | CHR_HSCHR19KIR_G248_A_H | killer cell immunoglobulin like receptor, two Ig |
| KIR2DL1 | CHR_HSCHR19KIR_G248_A_H | killer cell immunoglobulin like receptor, two Ig |
| KIR3DP1 | CHR_HSCHR19KIR_G248_A_H | killer cell immunoglobulin like receptor, three |
| KIR2DL4 | CHR_HSCHR19KIR_G248_A_H | killer cell immunoglobulin like receptor, two Ig |
| KIR3DL1 | CHR_HSCHR19KIR_G248_A_H | killer cell immunoglobulin like receptor, three |
| KIR2DS4 | CHR_HSCHR19KIR_G248_A_H | killer cell immunoglobulin like receptor, two Ig |
| KIR3DL2 | CHR_HSCHR19KIR_G248_A_H | killer cell immunoglobulin like receptor, three |
| U6 | CHR_HSCHR19KIR_G248_A_H | U6 spliceosomal RNA [Source:RFAM;Acc:RF00 |
| KIR3DL3 | CHR_HSCHR19KIR_G248_BA2 | killer cell immunoglobulin like receptor, three |
| GU182351.1 | CHR_HSCHR19KIR_G248_BA2 | Killer cell immunoglobulin-like receptor 2DS2 |
| GU182351.4 | CHR_HSCHR19KIR_G248_BA2 | Killer cell immunoglobulin-like receptor 2DL2 |
| GU182351.3 | CHR_HSCHR19KIR_G248_BA2 | Killer cell immunoglobulin-like receptor 2DL5B |
| GU182351.7 | CHR_HSCHR19KIR_G248_BA2 | Killer cell immunoglobulin-like receptor 2DS3 |
| GU182351.6 | CHR_HSCHR19KIR_G248_BA2 | killer cell immunoglobulin-like receptor, two c |
| KIR2DL1 | CHR_HSCHR19KIR_G248_BA2 | killer cell immunoglobulin like receptor, two Ig |
| GU182351.2 | CHR_HSCHR19KIR_G248_BA2 | KIR3DL3 killer cell immunoglobulin-like recept |
| KIR2DL4 | CHR_HSCHR19KIR_G248_BA2 | killer cell immunoglobulin like receptor, two Ig |
| KIR3DL1 | CHR_HSCHR19KIR_G248_BA2 | killer cell immunoglobulin like receptor, three |
| GU182351.5 | CHR_HSCHR19KIR_G248_BA2 | killer cell immunoglobulin-like receptor, two c |
| GU182352.5 | CHR_HSCHR19KIR_GRC212_A | killer cell immunoglobulin-like receptor, three |
| KIR3DL3 | CHR_HSCHR19KIR_GRC212_A | killer cell immunoglobulin like receptor, three |
| KIR2DL3 | CHR_HSCHR19KIR_GRC212_A | killer cell immunoglobulin like receptor, two Ig |
| GU182352.2 | CHR_HSCHR19KIR_GRC212_A | killer cell immunoglobulin-like receptor, two c |
| KIR2DL1 | CHR_HSCHR19KIR_GRC212_A | killer cell immunoglobulin like receptor, two Ig |
| GU182352.6 | CHR_HSCHR19KIR_GRC212_A | killer cell immunoglobulin-like receptor, three |
| KIR2DL4 | CHR_HSCHR19KIR_GRC212_A | killer cell immunoglobulin like receptor, two Ig |
| GU182352.1 | CHR_HSCHR19KIR_GRC212_A | killer cell immunoglobulin-like receptor, three |
| GU182352.4 | CHR_HSCHR19KIR_GRC212_A | killer cell immunoglobulin-like receptor, two c |

| | | |
|---|---|---|
| GU182352.3 | CHR_HSCHR19KIR_GRC212_A | killer cell immunoglobulin-like receptor, two c |
| GU182352.7 | CHR_HSCHR19KIR_GRC212_A | Killer cell immunoglobulin-like receptor 2DS1 |
| KIR3DL2 | CHR_HSCHR19KIR_GRC212_A | killer cell immunoglobulin like receptor, three |
| U6 | CHR_HSCHR19KIR_GRC212_A | U6 spliceosomal RNA [Source:RFAM;Acc:RF00 |
| GU182353.3 | CHR_HSCHR19KIR_GRC212_E | killer cell immunoglobulin-like receptor, three |
| KIR3DL3 | CHR_HSCHR19KIR_GRC212_E | killer cell immunoglobulin like receptor, three |
| GU182353.2 | CHR_HSCHR19KIR_GRC212_E | Killer cell immunoglobulin-like receptor 2DS2 |
| GU182353.4 | CHR_HSCHR19KIR_GRC212_E | killer cell immunoglobulin-like receptor, two c |
| GU182353.1 | CHR_HSCHR19KIR_GRC212_E | Novel priotein, similar to killer cell immunoglo |
| KIR2DL4 | CHR_HSCHR19KIR_GRC212_E | killer cell immunoglobulin like receptor, two Ig |
| KIR3DL1 | CHR_HSCHR19KIR_GRC212_E | killer cell immunoglobulin like receptor, three |
| KIR2DS4 | CHR_HSCHR19KIR_GRC212_E | killer cell immunoglobulin like receptor, two Ig |
| KIR3DL2 | CHR_HSCHR19KIR_GRC212_E | killer cell immunoglobulin like receptor, three |
| KP420441.2 | CHR_HSCHR19KIR_HG2393_C | NULL |
| KP420441.4 | CHR_HSCHR19KIR_HG2393_C | Killer cell immunoglobulin-like receptor 2DL2 |
| KP420441.1 | CHR_HSCHR19KIR_HG2393_C | NULL |
| KP420441.3 | CHR_HSCHR19KIR_HG2393_C | Killer cell immunoglobulin-like receptor 2DS5 |
| KP420441.5 | CHR_HSCHR19KIR_HG2393_C | Killer cell immunoglobulin-like receptor 2DS1 |
| KIR3DL2 | CHR_HSCHR19KIR_HG2393_C | killer cell immunoglobulin like receptor, three |
| RNU6-222P | CHR_HSCHR19KIR_HG2393_C | RNA, U6 small nuclear 222, pseudogene [Sour |
| FCAR | CHR_HSCHR19KIR_HG2393_C | Fc fragment of IgA receptor [Source:HGNC Syr |
| AC245128.23 | CHR_HSCHR19KIR_HG2393_C | novel transcript |
| AC245128.29 | CHR_HSCHR19KIR_HG2393_C | NULL |
| LILRP1 | CHR_HSCHR19KIR_HG2394_C | leukocyte immunoglobulin-like receptor pseu |
| AC245128.17 | CHR_HSCHR19KIR_HG2394_C | killer cell immunoglobulin-like receptor family |
| LILRP2 | CHR_HSCHR19KIR_HG2394_C | leukocyte immunoglobulin-like receptor pseu |
| KIR3DL3 | CHR_HSCHR19KIR_HG2394_C | killer cell immunoglobulin like receptor, three |
| KP420442.1 | CHR_HSCHR19KIR_HG2394_C | novel transcript |
| KIR2DL3 | CHR_HSCHR19KIR_HG2394_C | killer cell immunoglobulin like receptor, two Ig |
| KIR2DP1 | CHR_HSCHR19KIR_HG2394_C | killer cell immunoglobulin like receptor, two Ig |
| KIR2DL1 | CHR_HSCHR19KIR_HG2394_C | killer cell immunoglobulin like receptor, two Ig |
| KIR3DP1 | CHR_HSCHR19KIR_HG2394_C | killer cell immunoglobulin like receptor, three |
| KIR2DL4 | CHR_HSCHR19KIR_HG2394_C | killer cell immunoglobulin like receptor, two Ig |
| KIR3DL1 | CHR_HSCHR19KIR_HG2394_C | killer cell immunoglobulin like receptor, three |
| KIR2DS4 | CHR_HSCHR19KIR_HG2394_C | killer cell immunoglobulin like receptor, two Ig |
| KIR3DL2 | CHR_HSCHR19KIR_HG2394_C | killer cell immunoglobulin like receptor, three |
| LILRP1 | CHR_HSCHR19KIR_HG2396_C | leukocyte immunoglobulin-like receptor pseu |
| AC245128.19 | CHR_HSCHR19KIR_HG2396_C | killer cell immunoglobulin-like receptor family |
| LILRP2 | CHR_HSCHR19KIR_HG2396_C | leukocyte immunoglobulin-like receptor pseu |
| KP420444.1 | CHR_HSCHR19KIR_HG2396_C | novel transcript |
| KP420444.2 | CHR_HSCHR19KIR_HG2396_C | NULL |
| KIR2DL4 | CHR_HSCHR19KIR_HG2396_C | killer cell immunoglobulin like receptor, two Ig |
| KIR3DL1 | CHR_HSCHR19KIR_HG2396_C | killer cell immunoglobulin like receptor, three |
| KIR2DS4 | CHR_HSCHR19KIR_HG2396_C | killer cell immunoglobulin like receptor, two Ig |
| KIR3DL2 | CHR_HSCHR19KIR_HG2396_C | killer cell immunoglobulin like receptor, three |
| KIR3DL3 | CHR_HSCHR19KIR_LUCE_A_H | killer cell immunoglobulin like receptor, three |
| KIR2DL3 | CHR_HSCHR19KIR_LUCE_A_H | killer cell immunoglobulin like receptor, two Ig |
| GU182354.1 | CHR_HSCHR19KIR_LUCE_A_H | killer cell immunoglobulin-like receptor, two c |
| KIR2DL1 | CHR_HSCHR19KIR_LUCE_A_H | killer cell immunoglobulin like receptor, two Ig |
| GU182354.2 | CHR_HSCHR19KIR_LUCE_A_H | killer cell immunoglobulin-like receptor, three |

| | | |
|---|---|---|
| KIR2DL4 | CHR_HSCHR19KIR_LUCE_A_H | killer cell immunoglobulin like receptor, two Ig |
| KIR3DL1 | CHR_HSCHR19KIR_LUCE_A_H | killer cell immunoglobulin like receptor, three |
| KIR2DS4 | CHR_HSCHR19KIR_LUCE_A_H | killer cell immunoglobulin like receptor, two Ig |
| KIR3DL2 | CHR_HSCHR19KIR_LUCE_A_H | killer cell immunoglobulin like receptor, three |
| U6 | CHR_HSCHR19KIR_LUCE_A_H | U6 spliceosomal RNA [Source:RFAM;Acc:RF00 |
| LILRP1 | CHR_HSCHR19KIR_LUCE_BDE | leukocyte immunoglobulin-like receptor pseu |
| AC245128.12 | CHR_HSCHR19KIR_LUCE_BDE | killer cell immunoglobulin-like receptor family |
| LILRP2 | CHR_HSCHR19KIR_LUCE_BDE | leukocyte immunoglobulin-like receptor pseu |
| KIR3DL3 | CHR_HSCHR19KIR_LUCE_BDE | killer cell immunoglobulin like receptor, three |
| GU182355.5 | CHR_HSCHR19KIR_LUCE_BDE | Killer cell immunoglobulin-like receptor 2DS2 |
| GU182355.7 | CHR_HSCHR19KIR_LUCE_BDE | killer cell immunoglobulin-like receptor, two c |
| GU182355.6 | CHR_HSCHR19KIR_LUCE_BDE | novel killer cell immunoglobulin-like receptor |
| KIR2DL4 | CHR_HSCHR19KIR_LUCE_BDE | killer cell immunoglobulin like receptor, two Ig |
| GU182355.4 | CHR_HSCHR19KIR_LUCE_BDE | killer cell immunoglobulin-like receptor, three |
| GU182355.3 | CHR_HSCHR19KIR_LUCE_BDE | Killer cell immunoglobulin-like receptor 2DL5A |
| GU182355.2 | CHR_HSCHR19KIR_LUCE_BDE | Killer cell immunoglobulin-like receptor 2DS5 |
| GU182355.1 | CHR_HSCHR19KIR_LUCE_BDE | Killer cell immunoglobulin-like receptor 2DS1 |
| KIR3DL2 | CHR_HSCHR19KIR_LUCE_BDE | killer cell immunoglobulin like receptor, three |
| U6 | CHR_HSCHR19KIR_LUCE_BDE | U6 spliceosomal RNA [Source:RFAM;Acc:RF00 |
| LILRP1 | CHR_HSCHR19KIR_RP5_B_HA | leukocyte immunoglobulin-like receptor pseu |
| LILRP2 | CHR_HSCHR19KIR_RP5_B_HA | leukocyte immunoglobulin-like receptor pseu |
| KIR3DL3 | CHR_HSCHR19KIR_RP5_B_HA | killer cell immunoglobulin like receptor, three |
| AL133414.6 | CHR_HSCHR19KIR_RP5_B_HA | Killer cell immunoglobulin-like receptor 2DS2 |
| AL133414.2 | CHR_HSCHR19KIR_RP5_B_HA | killer cell immunoglobulin-like receptor, two c |
| KIR3DP1 | CHR_HSCHR19KIR_RP5_B_HA | killer cell immunoglobulin like receptor, three |
| KIR2DL4 | CHR_HSCHR19KIR_RP5_B_HA | killer cell immunoglobulin like receptor, two Ig |
| AL133414.4 | CHR_HSCHR19KIR_RP5_B_HA | killer cell immunoglobulin-like receptor, three |
| AL133414.3 | CHR_HSCHR19KIR_RP5_B_HA | Killer cell immunoglobulin-like receptor 2DL5A |
| AL133414.5 | CHR_HSCHR19KIR_RP5_B_HA | Killer cell immunoglobulin-like receptor 2DS5 |
| AL133414.1 | CHR_HSCHR19KIR_RP5_B_HA | Killer cell immunoglobulin-like receptor 2DS1 |
| KIR3DL2 | CHR_HSCHR19KIR_RP5_B_HA | killer cell immunoglobulin like receptor, three |
| LILRP1 | CHR_HSCHR19KIR_RSH_A_HA | leukocyte immunoglobulin-like receptor pseu |
| AC245128.5 | CHR_HSCHR19KIR_RSH_A_HA | killer cell immunoglobulin-like receptor family |
| LILRP2 | CHR_HSCHR19KIR_RSH_A_HA | leukocyte immunoglobulin-like receptor pseu |
| KIR3DL3 | CHR_HSCHR19KIR_RSH_A_HA | killer cell immunoglobulin like receptor, three |
| KIR2DL3 | CHR_HSCHR19KIR_RSH_A_HA | killer cell immunoglobulin like receptor, two Ig |
| KIR2DP1 | CHR_HSCHR19KIR_RSH_A_HA | killer cell immunoglobulin like receptor, two Ig |
| KIR2DL1 | CHR_HSCHR19KIR_RSH_A_HA | killer cell immunoglobulin like receptor, two Ig |
| KIR3DP1 | CHR_HSCHR19KIR_RSH_A_HA | killer cell immunoglobulin like receptor, three |
| KIR2DL4 | CHR_HSCHR19KIR_RSH_A_HA | killer cell immunoglobulin like receptor, two Ig |
| KIR3DL1 | CHR_HSCHR19KIR_RSH_A_HA | killer cell immunoglobulin like receptor, three |
| KIR2DS4 | CHR_HSCHR19KIR_RSH_A_HA | killer cell immunoglobulin like receptor, two Ig |
| KIR3DL2 | CHR_HSCHR19KIR_RSH_A_HA | killer cell immunoglobulin like receptor, three |
| RNU6-222P | CHR_HSCHR19KIR_RSH_A_HA | RNA, U6 small nuclear 222, pseudogene [Sour |
| LILRP1 | CHR_HSCHR19KIR_RSH_BA2_ | leukocyte immunoglobulin-like receptor pseu |
| AC245128.7 | CHR_HSCHR19KIR_RSH_BA2_ | killer cell immunoglobulin-like receptor family |
| LILRP2 | CHR_HSCHR19KIR_RSH_BA2_ | leukocyte immunoglobulin-like receptor pseu |
| KIR3DL3 | CHR_HSCHR19KIR_RSH_BA2_ | killer cell immunoglobulin like receptor, three |
| GU182357.2 | CHR_HSCHR19KIR_RSH_BA2_ | Killer cell immunoglobulin-like receptor 2DS2 |
| GU182357.5 | CHR_HSCHR19KIR_RSH_BA2_ | Killer cell immunoglobulin-like receptor 2DL2 |

| | | |
|---|---|---|
| GU182357.4 | CHR_HSCHR19KIR_RSH_BA2_ | killer cell immunoglobulin-like receptor, two c |
| GU182357.1 | CHR_HSCHR19KIR_RSH_BA2_ | novel killer cell immunoglobulin-like receptor |
| GU182357.3 | CHR_HSCHR19KIR_RSH_BA2_ | novel killer cell immunoglobulin-like receptor |
| KIR2DL1 | CHR_HSCHR19KIR_RSH_BA2_ | killer cell immunoglobulin like receptor, two Ig |
| KIR2DL4 | CHR_HSCHR19KIR_RSH_BA2_ | killer cell immunoglobulin like receptor, two Ig |
| KIR3DL1 | CHR_HSCHR19KIR_RSH_BA2_ | killer cell immunoglobulin like receptor, three |
| KIR2DS4 | CHR_HSCHR19KIR_RSH_BA2_ | killer cell immunoglobulin like receptor, two Ig |
| LILRP1 | CHR_HSCHR19KIR_T7526_A_ | leukocyte immunoglobulin-like receptor pseu |
| AC245128.6 | CHR_HSCHR19KIR_T7526_A_ | killer cell immunoglobulin-like receptor family |
| LILRP2 | CHR_HSCHR19KIR_T7526_A_ | leukocyte immunoglobulin-like receptor pseu |
| KIR3DL3 | CHR_HSCHR19KIR_T7526_A_ | killer cell immunoglobulin like receptor, three |
| KIR2DL3 | CHR_HSCHR19KIR_T7526_A_ | killer cell immunoglobulin like receptor, two Ig |
| GU182358.1 | CHR_HSCHR19KIR_T7526_A_ | novel killer cell immunoglobulin-like receptor |
| KIR2DL1 | CHR_HSCHR19KIR_T7526_A_ | killer cell immunoglobulin like receptor, two Ig |
| KIR3DP1 | CHR_HSCHR19KIR_T7526_A_ | killer cell immunoglobulin like receptor, three |
| KIR2DL4 | CHR_HSCHR19KIR_T7526_A_ | killer cell immunoglobulin like receptor, two Ig |
| KIR3DL1 | CHR_HSCHR19KIR_T7526_A_ | killer cell immunoglobulin like receptor, three |
| KIR2DS4 | CHR_HSCHR19KIR_T7526_A_ | killer cell immunoglobulin like receptor, two Ig |
| KIR3DL2 | CHR_HSCHR19KIR_T7526_A_ | killer cell immunoglobulin like receptor, three |
| U6 | CHR_HSCHR19KIR_T7526_A_ | U6 spliceosomal RNA [Source:RFAM;Acc:RF00 |
| LILRP1 | CHR_HSCHR19KIR_T7526_BD | leukocyte immunoglobulin-like receptor pseu |
| LILRP2 | CHR_HSCHR19KIR_T7526_BD | leukocyte immunoglobulin-like receptor pseu |
| KIR3DL3 | CHR_HSCHR19KIR_T7526_BD | killer cell immunoglobulin like receptor, three |
| GU182359.4 | CHR_HSCHR19KIR_T7526_BD | Killer cell immunoglobulin-like receptor 2DS2 |
| GU182359.1 | CHR_HSCHR19KIR_T7526_BD | killer cell immunoglobulin-like receptor, two c |
| KIR3DP1 | CHR_HSCHR19KIR_T7526_BD | killer cell immunoglobulin like receptor, three |
| KIR2DL4 | CHR_HSCHR19KIR_T7526_BD | killer cell immunoglobulin like receptor, two Ig |
| GU182359.5 | CHR_HSCHR19KIR_T7526_BD | killer cell immunoglobulin-like receptor, three |
| GU182359.3 | CHR_HSCHR19KIR_T7526_BD | killer cell immunoglobulin-like receptor, two c |
| GU182359.6 | CHR_HSCHR19KIR_T7526_BD | killer cell immunoglobulin-like receptor, two c |
| GU182359.2 | CHR_HSCHR19KIR_T7526_BD | Killer cell immunoglobulin-like receptor 2DS1 |
| KIR3DL2 | CHR_HSCHR19KIR_T7526_BD | killer cell immunoglobulin like receptor, three |
| U6 | CHR_HSCHR19KIR_T7526_BD | U6 spliceosomal RNA [Source:RFAM;Acc:RF00 |
| VSTM1 | CHR_HSCHR19LRC_COX1_CT | V-set and transmembrane domain containing |
| TARM1 | CHR_HSCHR19LRC_COX1_CT | T cell-interacting, activating receptor on myel |
| OSCAR | CHR_HSCHR19LRC_COX1_CT | osteoclast associated Ig-like receptor [Source: |
| AC012314.5 | CHR_HSCHR19LRC_COX1_CT | NADH:ubiquinone oxidoreductase subunit A3 |
| TFPT | CHR_HSCHR19LRC_COX1_CT | TCF3 fusion partner [Source:HGNC Symbol;Ac |
| AC012314.19 | CHR_HSCHR19LRC_COX1_CT | pre-mRNA processing factor 31 [Source:NCBI |
| CNOT3 | CHR_HSCHR19LRC_COX1_CT | CCR4-NOT transcription complex subunit 3 [So |
| LENG1 | CHR_HSCHR19LRC_COX1_CT | leukocyte receptor cluster member 1 [Source: |
| TMC4 | CHR_HSCHR19LRC_COX1_CT | transmembrane channel like 4 [Source:HGNC |
| MBOAT7 | CHR_HSCHR19LRC_COX1_CT | membrane bound O-acyltransferase domain c |
| TSEN34 | CHR_HSCHR19LRC_COX1_CT | tRNA splicing endonuclease subunit 34 [Sourc |
| RPS9 | CHR_HSCHR19LRC_COX1_CT | ribosomal protein S9 [Source:HGNC Symbol;A |
| LILRB3 | CHR_HSCHR19LRC_COX1_CT | leukocyte immunoglobulin like receptor B3 [S |
| CU457734.1 | CHR_HSCHR19LRC_COX1_CT | LSM14 homolog A (SCD6, S. cerevisiae) pseud |
| LILRA6 | CHR_HSCHR19LRC_COX1_CT | leukocyte immunoglobulin like receptor A6 [S |
| LILRB5 | CHR_HSCHR19LRC_COX1_CT | leukocyte immunoglobulin like receptor B5 [S |
| U6 | CHR_HSCHR19LRC_COX1_CT | U6 spliceosomal RNA [Source:RFAM;Acc:RF00 |

| | | |
|---|---|---|
| CU457734.2 | CHR_HSCHR19LRC_COX1_CT | YME1-like 1 (S. cerevisiae) pseudogene |
| LILRB2 | CHR_HSCHR19LRC_COX1_CT | leukocyte immunoglobulin like receptor B2 [S |
| LILRA3 | CHR_HSCHR19LRC_COX1_CT | leukocyte immunoglobulin like receptor A3 [S |
| LILRA5 | CHR_HSCHR19LRC_COX1_CT | leukocyte immunoglobulin like receptor A5 [S |
| TTYH1 | CHR_HSCHR19LRC_COX1_CT | tweety family member 1 [Source:HGNC Symb |
| CU467002.1 | CHR_HSCHR19LRC_COX1_CT | novel transcript |
| LENG8 | CHR_HSCHR19LRC_COX1_CT | leukocyte receptor cluster member 8 [Source: |
| LENG9 | CHR_HSCHR19LRC_COX1_CT | leukocyte receptor cluster member 9 [Source: |
| CDC42EP5 | CHR_HSCHR19LRC_COX1_CT | CDC42 effector protein 5 [Source:HGNC Symb |
| LAIR2 | CHR_HSCHR19LRC_COX1_CT | leukocyte associated immunoglobulin like rec |
| KIR3DX1 | CHR_HSCHR19LRC_COX1_CT | killer cell immunoglobulin like receptor, three |
| LILRA2 | CHR_HSCHR19LRC_COX1_CT | leukocyte immunoglobulin like receptor A2 [S |
| LILRA1 | CHR_HSCHR19LRC_COX1_CT | leukocyte immunoglobulin like receptor A1 [S |
| LILRB1 | CHR_HSCHR19LRC_COX1_CT | leukocyte immunoglobulin like receptor B1 [S |
| CU234142.1 | CHR_HSCHR19LRC_COX1_CT | vomeronasal 1 receptor 1 (VN1R1) pseudoger |
| LILRB4 | CHR_HSCHR19LRC_COX1_CT | leukocyte immunoglobulin like receptor B4 [S |
| LILRP1 | CHR_HSCHR19LRC_COX1_CT | leukocyte immunoglobulin-like receptor pseu |
| LILRP2 | CHR_HSCHR19LRC_COX1_CT | leukocyte immunoglobulin-like receptor pseu |
| KIR3DL3 | CHR_HSCHR19LRC_COX1_CT | killer cell immunoglobulin like receptor, three |
| KIR2DL3 | CHR_HSCHR19LRC_COX1_CT | killer cell immunoglobulin like receptor, two I |
| KIR2DP1 | CHR_HSCHR19LRC_COX1_CT | killer cell immunoglobulin like receptor, two I |
| KIR2DL1 | CHR_HSCHR19LRC_COX1_CT | killer cell immunoglobulin like receptor, two I |
| KIR3DP1 | CHR_HSCHR19LRC_COX1_CT | killer cell immunoglobulin like receptor, three |
| AC011476.9 | CHR_HSCHR19LRC_COX1_CT | natural cytotoxicity triggering receptor 1 [Sou |
| NLRP7 | CHR_HSCHR19LRC_COX1_CT | NLR family pyrin domain containing 7 [Source |
| NLRP2 | CHR_HSCHR19LRC_COX1_CT | NLR family pyrin domain containing 2 [Source |
| GP6 | CHR_HSCHR19LRC_COX1_CT | glycoprotein VI platelet [Source:NCBI gene;Ac |
| RDH13 | CHR_HSCHR19LRC_COX1_CT | retinol dehydrogenase 13 [Source:HGNC Symb |
| VSTM1 | CHR_HSCHR19LRC_COX2_CT | V-set and transmembrane domain containing |
| TARM1 | CHR_HSCHR19LRC_COX2_CT | T cell-interacting, activating receptor on myel |
| OSCAR | CHR_HSCHR19LRC_COX2_CT | osteoclast associated Ig-like receptor [Source: |
| AC012314.14 | CHR_HSCHR19LRC_COX2_CT | NADH:ubiquinone oxidoreductase subunit A3 |
| TFPT | CHR_HSCHR19LRC_COX2_CT | TCF3 fusion partner [Source:HGNC Symbol;Ac |
| AC012314.10 | CHR_HSCHR19LRC_COX2_CT | pre-mRNA processing factor 31 [Source:NCBI |
| CNOT3 | CHR_HSCHR19LRC_COX2_CT | CCR4-NOT transcription complex subunit 3 [S |
| LENG1 | CHR_HSCHR19LRC_COX2_CT | leukocyte receptor cluster member 1 [Source: |
| AC012314.15 | CHR_HSCHR19LRC_COX2_CT | transmembrane channel like 4 [Source:NCBI g |
| MBOAT7 | CHR_HSCHR19LRC_COX2_CT | membrane bound O-acyltransferase domain c |
| TSEN34 | CHR_HSCHR19LRC_COX2_CT | tRNA splicing endonuclease subunit 34 [Sourc |
| RPS9 | CHR_HSCHR19LRC_COX2_CT | ribosomal protein S9 [Source:HGNC Symbol;A |
| LILRA3 | CHR_HSCHR19LRC_COX2_CT | leukocyte immunoglobulin like receptor A3 [S |
| LILRA5 | CHR_HSCHR19LRC_COX2_CT | leukocyte immunoglobulin like receptor A5 [S |
| LILRA4 | CHR_HSCHR19LRC_COX2_CT | leukocyte immunoglobulin like receptor A4 [S |
| LAIR1 | CHR_HSCHR19LRC_COX2_CT | leukocyte associated immunoglobulin like rec |
| KIR2DL1 | CHR_HSCHR19LRC_COX2_CT | killer cell immunoglobulin like receptor, two I |
| KIR3DP1 | CHR_HSCHR19LRC_COX2_CT | killer cell immunoglobulin like receptor, three |
| KIR2DL4 | CHR_HSCHR19LRC_COX2_CT | killer cell immunoglobulin like receptor, two I |
| KIR3DS1 | CHR_HSCHR19LRC_COX2_CT | killer cell immunoglobulin like receptor, three |
| KIR2DL5A | CHR_HSCHR19LRC_COX2_CT | killer cell immunoglobulin like receptor, two I |
| KIR2DS5 | CHR_HSCHR19LRC_COX2_CT | killer cell immunoglobulin like receptor, two I |

| | | |
|---|---|---|
| KIR2DS1 | CHR_HSCHR19LRC_COX2_CT( | killer cell immunoglobulin like receptor, two Ig |
| KIR3DL2 | CHR_HSCHR19LRC_COX2_CT( | killer cell immunoglobulin like receptor, three |
| U6 | CHR_HSCHR19LRC_COX2_CT( | U6 spliceosomal RNA [Source:RFAM;Acc:RF00 |
| FCAR | CHR_HSCHR19LRC_COX2_CT( | Fc fragment of IgA receptor [Source:HGNC Syr |
| NCR1 | CHR_HSCHR19LRC_COX2_CT( | natural cytotoxicity triggering receptor 1 [Sou |
| NLRP7 | CHR_HSCHR19LRC_COX2_CT( | NLR family pyrin domain containing 7 [Source |
| NLRP2 | CHR_HSCHR19LRC_COX2_CT( | NLR family pyrin domain containing 2 [Source |
| GP6 | CHR_HSCHR19LRC_COX2_CT( | glycoprotein VI platelet [Source:NCBI gene;Ac |
| RDH13 | CHR_HSCHR19LRC_COX2_CT( | retinol dehydrogenase 13 [Source:HGNC Symb |
| VSTM1 | CHR_HSCHR19LRC_LRC_I_CT( | V-set and transmembrane domain containing |
| TARM1 | CHR_HSCHR19LRC_LRC_I_CT( | T cell-interacting, activating receptor on myel |
| OSCAR | CHR_HSCHR19LRC_LRC_I_CT( | osteoclast associated Ig-like receptor [Source: |
| AC012314.18 | CHR_HSCHR19LRC_LRC_I_CT( | NADH:ubiquinone oxidoreductase subunit A3 |
| TFPT | CHR_HSCHR19LRC_LRC_I_CT( | TCF3 fusion partner [Source:HGNC Symbol;Ac |
| AC012314.6 | CHR_HSCHR19LRC_LRC_I_CT( | pre-mRNA processing factor 31 [Source:NCBI g |
| CNOT3 | CHR_HSCHR19LRC_LRC_I_CT( | CCR4-NOT transcription complex subunit 3 [Sc |
| LENG1 | CHR_HSCHR19LRC_LRC_I_CT( | leukocyte receptor cluster member 1 [Source: |
| AC012314.9 | CHR_HSCHR19LRC_LRC_I_CT( | transmembrane channel like 4 [Source:NCBI g |
| MBOAT7 | CHR_HSCHR19LRC_LRC_I_CT( | membrane bound O-acyltransferase domain c |
| TSEN34 | CHR_HSCHR19LRC_LRC_I_CT( | tRNA splicing endonuclease subunit 34 [Sourc |
| RPS9 | CHR_HSCHR19LRC_LRC_I_CT( | ribosomal protein S9 [Source:HGNC Symbol;A |
| CU464060.1 | CHR_HSCHR19LRC_LRC_I_CT( | killer cell immunoglobulin like receptor, two Ig |
| KIR2DL2 | CHR_HSCHR19LRC_LRC_I_CT( | killer cell immunoglobulin like receptor, two Ig |
| KIR2DS4 | CHR_HSCHR19LRC_LRC_I_CT( | killer cell immunoglobulin like receptor, two Ig |
| KIR3DP1 | CHR_HSCHR19LRC_LRC_I_CT( | killer cell immunoglobulin like receptor, three |
| KIR3DL2 | CHR_HSCHR19LRC_LRC_I_CT( | killer cell immunoglobulin like receptor, three |
| AC011476.4 | CHR_HSCHR19LRC_LRC_I_CT( | natural cytotoxicity triggering receptor 1 [Sou |
| NLRP7 | CHR_HSCHR19LRC_LRC_I_CT( | NLR family pyrin domain containing 7 [Source |
| NLRP2 | CHR_HSCHR19LRC_LRC_I_CT( | NLR family pyrin domain containing 2 [Source |
| GP6 | CHR_HSCHR19LRC_LRC_I_CT( | glycoprotein VI platelet [Source:NCBI gene;Ac |
| RDH13 | CHR_HSCHR19LRC_LRC_I_CT( | retinol dehydrogenase 13 [Source:HGNC Symb |
| VSTM1 | CHR_HSCHR19LRC_LRC_J_CT( | V-set and transmembrane domain containing |
| TARM1 | CHR_HSCHR19LRC_LRC_J_CT( | T cell-interacting, activating receptor on myel |
| OSCAR | CHR_HSCHR19LRC_LRC_J_CT( | osteoclast associated Ig-like receptor [Source: |
| AC012314.1 | CHR_HSCHR19LRC_LRC_J_CT( | NADH:ubiquinone oxidoreductase subunit A3 |
| TFPT | CHR_HSCHR19LRC_LRC_J_CT( | TCF3 fusion partner [Source:HGNC Symbol;Ac |
| AC012314.8 | CHR_HSCHR19LRC_LRC_J_CT( | pre-mRNA processing factor 31 [Source:NCBI g |
| CNOT3 | CHR_HSCHR19LRC_LRC_J_CT( | CCR4-NOT transcription complex subunit 3 [Sc |
| LENG1 | CHR_HSCHR19LRC_LRC_J_CT( | leukocyte receptor cluster member 1 [Source: |
| AC012314.7 | CHR_HSCHR19LRC_LRC_J_CT( | transmembrane channel like 4 [Source:NCBI g |
| MBOAT7 | CHR_HSCHR19LRC_LRC_J_CT( | membrane bound O-acyltransferase domain c |
| TSEN34 | CHR_HSCHR19LRC_LRC_J_CT( | tRNA splicing endonuclease subunit 34 [Sourc |
| RPS9 | CHR_HSCHR19LRC_LRC_J_CT( | ribosomal protein S9 [Source:HGNC Symbol;A |
| KIR3DL3 | CHR_HSCHR19LRC_LRC_J_CT( | killer cell immunoglobulin like receptor, three |
| KIR2DP1 | CHR_HSCHR19LRC_LRC_J_CT( | killer cell immunoglobulin like receptor, two Ig |
| CU633846.1 | CHR_HSCHR19LRC_LRC_J_CT( | killer cell immunoglobulin like receptor, two Ig |
| KIR3DL2 | CHR_HSCHR19LRC_LRC_J_CT( | killer cell immunoglobulin like receptor, three |
| AC011476.10 | CHR_HSCHR19LRC_LRC_J_CT( | natural cytotoxicity triggering receptor 1 [Sou |
| NLRP7 | CHR_HSCHR19LRC_LRC_J_CT( | NLR family pyrin domain containing 7 [Source |
| NLRP2 | CHR_HSCHR19LRC_LRC_J_CT( | NLR family pyrin domain containing 2 [Source |

| | | |
|---|---|---|
| GP6 | CHR_HSCHR19LRC_LRC_J_CT | glycoprotein VI platelet [Source:NCBI gene;Ac |
| RDH13 | CHR_HSCHR19LRC_LRC_J_CT | retinol dehydrogenase 13 [Source:HGNC Sym |
| VSTM1 | CHR_HSCHR19LRC_LRC_S_CT | V-set and transmembrane domain containing |
| TARM1 | CHR_HSCHR19LRC_LRC_S_CT | T cell-interacting, activating receptor on myel |
| OSCAR | CHR_HSCHR19LRC_LRC_S_CT | osteoclast associated Ig-like receptor [Source: |
| AC012314.26 | CHR_HSCHR19LRC_LRC_S_CT | NADH:ubiquinone oxidoreductase subunit A3 |
| TFPT | CHR_HSCHR19LRC_LRC_S_CT | TCF3 fusion partner [Source:HGNC Symbol;Ac |
| AC012314.17 | CHR_HSCHR19LRC_LRC_S_CT | pre-mRNA processing factor 31 [Source:NCBI |
| CNOT3 | CHR_HSCHR19LRC_LRC_S_CT | CCR4-NOT transcription complex subunit 3 [S |
| LENG1 | CHR_HSCHR19LRC_LRC_S_CT | leukocyte receptor cluster member 1 [Source: |
| AC012314.24 | CHR_HSCHR19LRC_LRC_S_CT | transmembrane channel like 4 [Source:NCBI g |
| MBOAT7 | CHR_HSCHR19LRC_LRC_S_CT | membrane bound O-acyltransferase domain |
| TSEN34 | CHR_HSCHR19LRC_LRC_S_CT | tRNA splicing endonuclease subunit 34 [Sourc |
| RPS9 | CHR_HSCHR19LRC_LRC_S_CT | ribosomal protein S9 [Source:HGNC Symbol;A |
| KIR2DS4 | CHR_HSCHR19LRC_LRC_S_CT | killer cell immunoglobulin like receptor, two I |
| KIR3DL2 | CHR_HSCHR19LRC_LRC_S_CT | killer cell immunoglobulin like receptor, three |
| U6 | CHR_HSCHR19LRC_LRC_S_CT | U6 spliceosomal RNA [Source:RFAM;Acc:RF00 |
| AC011476.11 | CHR_HSCHR19LRC_LRC_S_CT | natural cytotoxicity triggering receptor 1 [Sou |
| NLRP7 | CHR_HSCHR19LRC_LRC_S_CT | NLR family pyrin domain containing 7 [Source |
| NLRP2 | CHR_HSCHR19LRC_LRC_S_CT | NLR family pyrin domain containing 2 [Source |
| GP6 | CHR_HSCHR19LRC_LRC_S_CT | glycoprotein VI platelet [Source:NCBI gene;Ac |
| RDH13 | CHR_HSCHR19LRC_LRC_S_CT | retinol dehydrogenase 13 [Source:HGNC Sym |
| VSTM1 | CHR_HSCHR19LRC_LRC_T_CT | V-set and transmembrane domain containing |
| TARM1 | CHR_HSCHR19LRC_LRC_T_CT | T cell-interacting, activating receptor on myel |
| OSCAR | CHR_HSCHR19LRC_LRC_T_CT | osteoclast associated Ig-like receptor [Source: |
| AC012314.12 | CHR_HSCHR19LRC_LRC_T_CT | NADH:ubiquinone oxidoreductase subunit A3 |
| TFPT | CHR_HSCHR19LRC_LRC_T_CT | TCF3 fusion partner [Source:HGNC Symbol;Ac |
| AC012314.23 | CHR_HSCHR19LRC_LRC_T_CT | pre-mRNA processing factor 31 [Source:NCBI |
| CNOT3 | CHR_HSCHR19LRC_LRC_T_CT | CCR4-NOT transcription complex subunit 3 [S |
| LENG1 | CHR_HSCHR19LRC_LRC_T_CT | leukocyte receptor cluster member 1 [Source: |
| AC012314.20 | CHR_HSCHR19LRC_LRC_T_CT | transmembrane channel like 4 [Source:NCBI g |
| MBOAT7 | CHR_HSCHR19LRC_LRC_T_CT | membrane bound O-acyltransferase domain |
| TSEN34 | CHR_HSCHR19LRC_LRC_T_CT | tRNA splicing endonuclease subunit 34 [Sourc |
| RPS9 | CHR_HSCHR19LRC_LRC_T_CT | ribosomal protein S9 [Source:HGNC Symbol;A |
| KIR3DL3 | CHR_HSCHR19LRC_LRC_T_CT | killer cell immunoglobulin like receptor, three |
| KIR2DL3 | CHR_HSCHR19LRC_LRC_T_CT | killer cell immunoglobulin like receptor, two I |
| KIR2DP1 | CHR_HSCHR19LRC_LRC_T_CT | killer cell immunoglobulin like receptor, two I |
| KIR2DS1 | CHR_HSCHR19LRC_LRC_T_CT | killer cell immunoglobulin like receptor, two I |
| KIR3DL2 | CHR_HSCHR19LRC_LRC_T_CT | killer cell immunoglobulin like receptor, three |
| U6 | CHR_HSCHR19LRC_LRC_T_CT | U6 spliceosomal RNA [Source:RFAM;Acc:RF00 |
| AC011476.5 | CHR_HSCHR19LRC_LRC_T_CT | natural cytotoxicity triggering receptor 1 [Sou |
| NLRP7 | CHR_HSCHR19LRC_LRC_T_CT | NLR family pyrin domain containing 7 [Source |
| NLRP2 | CHR_HSCHR19LRC_LRC_T_CT | NLR family pyrin domain containing 2 [Source |
| GP6 | CHR_HSCHR19LRC_LRC_T_CT | glycoprotein VI platelet [Source:NCBI gene;Ac |
| RDH13 | CHR_HSCHR19LRC_LRC_T_CT | retinol dehydrogenase 13 [Source:HGNC Sym |
| VSTM1 | CHR_HSCHR19LRC_PGF1_CT | V-set and transmembrane domain containing |
| TARM1 | CHR_HSCHR19LRC_PGF1_CT | T cell-interacting, activating receptor on myel |
| OSCAR | CHR_HSCHR19LRC_PGF1_CT | osteoclast associated Ig-like receptor [Source: |
| AC012314.11 | CHR_HSCHR19LRC_PGF1_CT | NADH:ubiquinone oxidoreductase subunit A3 |
| TFPT | CHR_HSCHR19LRC_PGF1_CT | TCF3 fusion partner [Source:HGNC Symbol;Ac |

| | | |
|---|---|---|
| AC012314.21 | CHR_HSCHR19LRC_PGF1_CT( | pre-mRNA processing factor 31 [Source:NCBI g |
| CNOT3 | CHR_HSCHR19LRC_PGF1_CT( | CCR4-NOT transcription complex subunit 3 [So |
| LENG1 | CHR_HSCHR19LRC_PGF1_CT( | leukocyte receptor cluster member 1 [Source: |
| TMC4 | CHR_HSCHR19LRC_PGF1_CT( | transmembrane channel like 4 [Source:HGNC |
| MBOAT7 | CHR_HSCHR19LRC_PGF1_CT( | membrane bound O-acyltransferase domain c |
| TSEN34 | CHR_HSCHR19LRC_PGF1_CT( | tRNA splicing endonuclease subunit 34 [Sourc |
| RPS9 | CHR_HSCHR19LRC_PGF1_CT( | ribosomal protein S9 [Source:HGNC Symbol;A |
| LILRB3 | CHR_HSCHR19LRC_PGF1_CT( | leukocyte immunoglobulin like receptor B3 [S |
| CU151838.1 | CHR_HSCHR19LRC_PGF1_CT( | LSM14 homolog A (SCD6, S. cerevisiae) pseud |
| LILRA6 | CHR_HSCHR19LRC_PGF1_CT( | leukocyte immunoglobulin like receptor A6 [S |
| LILRB5 | CHR_HSCHR19LRC_PGF1_CT( | leukocyte immunoglobulin like receptor B5 [S |
| U6 | CHR_HSCHR19LRC_PGF1_CT( | U6 spliceosomal RNA [Source:RFAM;Acc:RF00 |
| CU151838.2 | CHR_HSCHR19LRC_PGF1_CT( | YME1-like 1 (S. cerevisiae) pseudogene |
| LILRB2 | CHR_HSCHR19LRC_PGF1_CT( | leukocyte immunoglobulin like receptor B2 [S |
| LAIR1 | CHR_HSCHR19LRC_PGF1_CT( | leukocyte associated immunoglobulin like rec |
| TTYH1 | CHR_HSCHR19LRC_PGF1_CT( | tweety family member 1 [Source:HGNC Symb |
| CU207370.1 | CHR_HSCHR19LRC_PGF1_CT( | novel transcript |
| LENG8 | CHR_HSCHR19LRC_PGF1_CT( | leukocyte receptor cluster member 8 [Source: |
| LENG9 | CHR_HSCHR19LRC_PGF1_CT( | leukocyte receptor cluster member 9 [Source: |
| CDC42EP5 | CHR_HSCHR19LRC_PGF1_CT( | CDC42 effector protein 5 [Source:HGNC Symb |
| LAIR2 | CHR_HSCHR19LRC_PGF1_CT( | leukocyte associated immunoglobulin like rec |
| KIR3DX1 | CHR_HSCHR19LRC_PGF1_CT( | killer cell immunoglobulin like receptor, three |
| LILRA2 | CHR_HSCHR19LRC_PGF1_CT( | leukocyte immunoglobulin like receptor A2 [S |
| LILRA1 | CHR_HSCHR19LRC_PGF1_CT( | leukocyte immunoglobulin like receptor A1 [S |
| LILRB1 | CHR_HSCHR19LRC_PGF1_CT( | leukocyte immunoglobulin like receptor B1 [S |
| CU024913.1 | CHR_HSCHR19LRC_PGF1_CT( | vomeronasal 1 receptor 1 (VN1R1) pseudoger |
| LILRB4 | CHR_HSCHR19LRC_PGF1_CT( | leukocyte immunoglobulin like receptor B4 [S |
| LILRP1 | CHR_HSCHR19LRC_PGF1_CT( | leukocyte immunoglobulin-like receptor pseu |
| LILRP2 | CHR_HSCHR19LRC_PGF1_CT( | leukocyte immunoglobulin-like receptor pseu |
| KIR3DL3 | CHR_HSCHR19LRC_PGF1_CT( | killer cell immunoglobulin like receptor, three |
| KIR2DL3 | CHR_HSCHR19LRC_PGF1_CT( | killer cell immunoglobulin like receptor, two Ig |
| KIR2DP1 | CHR_HSCHR19LRC_PGF1_CT( | killer cell immunoglobulin like receptor, two Ig |
| KIR2DL1 | CHR_HSCHR19LRC_PGF1_CT( | killer cell immunoglobulin like receptor, two Ig |
| KIR3DP1 | CHR_HSCHR19LRC_PGF1_CT( | killer cell immunoglobulin like receptor, three |
| KIR2DL4 | CHR_HSCHR19LRC_PGF1_CT( | killer cell immunoglobulin like receptor, two Ig |
| KIR3DL1 | CHR_HSCHR19LRC_PGF1_CT( | killer cell immunoglobulin like receptor, three |
| KIR2DS4 | CHR_HSCHR19LRC_PGF1_CT( | killer cell immunoglobulin like receptor, two Ig |
| KIR3DL2 | CHR_HSCHR19LRC_PGF1_CT( | killer cell immunoglobulin like receptor, three |
| U6 | CHR_HSCHR19LRC_PGF1_CT( | U6 spliceosomal RNA [Source:RFAM;Acc:RF00 |
| FCAR | CHR_HSCHR19LRC_PGF1_CT( | Fc fragment of IgA receptor [Source:HGNC Syr |
| NCR1 | CHR_HSCHR19LRC_PGF1_CT( | natural cytotoxicity triggering receptor 1 [Sou |
| NLRP7 | CHR_HSCHR19LRC_PGF1_CT( | NLR family pyrin domain containing 7 [Source |
| NLRP2 | CHR_HSCHR19LRC_PGF1_CT( | NLR family pyrin domain containing 2 [Source |
| GP6 | CHR_HSCHR19LRC_PGF1_CT( | glycoprotein VI platelet [Source:NCBI gene;Ac |
| RDH13 | CHR_HSCHR19LRC_PGF1_CT( | retinol dehydrogenase 13 [Source:HGNC Symb |
| VSTM1 | CHR_HSCHR19LRC_PGF2_CT( | V-set and transmembrane domain containing |
| TARM1 | CHR_HSCHR19LRC_PGF2_CT( | T cell-interacting, activating receptor on myel |
| OSCAR | CHR_HSCHR19LRC_PGF2_CT( | osteoclast associated Ig-like receptor [Source: |
| AC012314.22 | CHR_HSCHR19LRC_PGF2_CT( | NADH:ubiquinone oxidoreductase subunit A3 |
| TFPT | CHR_HSCHR19LRC_PGF2_CT( | TCF3 fusion partner [Source:HGNC Symbol;Ac |

| | | |
|---|---|---|
| AC012314.4 | CHR_HSCHR19LRC_PGF2_CT( | pre-mRNA processing factor 31 [Source:NCBI | 
| CNOT3 | CHR_HSCHR19LRC_PGF2_CT( | CCR4-NOT transcription complex subunit 3 [So |
| LENG1 | CHR_HSCHR19LRC_PGF2_CT( | leukocyte receptor cluster member 1 [Source: |
| AC012314.25 | CHR_HSCHR19LRC_PGF2_CT( | transmembrane channel like 4 [Source:NCBI g |
| MBOAT7 | CHR_HSCHR19LRC_PGF2_CT( | membrane bound O-acyltransferase domain c |
| TSEN34 | CHR_HSCHR19LRC_PGF2_CT( | tRNA splicing endonuclease subunit 34 [Sourc |
| RPS9 | CHR_HSCHR19LRC_PGF2_CT( | ribosomal protein S9 [Source:HGNC Symbol;A |
| LILRA6 | CHR_HSCHR19LRC_PGF2_CT( | leukocyte immunoglobulin like receptor A6 [S |
| LILRB5 | CHR_HSCHR19LRC_PGF2_CT( | leukocyte immunoglobulin like receptor B5 [S |
| U6 | CHR_HSCHR19LRC_PGF2_CT( | U6 spliceosomal RNA [Source:RFAM;Acc:RF00 |
| CU104682.1 | CHR_HSCHR19LRC_PGF2_CT( | YME1-like 1 (S. cerevisiae) pseudogene |
| LILRB2 | CHR_HSCHR19LRC_PGF2_CT( | leukocyte immunoglobulin like receptor B2 [S |
| LILRA3 | CHR_HSCHR19LRC_PGF2_CT( | leukocyte immunoglobulin like receptor A3 [S |
| LILRA5 | CHR_HSCHR19LRC_PGF2_CT( | leukocyte immunoglobulin like receptor A5 [S |
| LILRA4 | CHR_HSCHR19LRC_PGF2_CT( | leukocyte immunoglobulin like receptor A4 [S |
| LAIR1 | CHR_HSCHR19LRC_PGF2_CT( | leukocyte associated immunoglobulin like rec |
| LILRB1 | CHR_HSCHR19LRC_PGF2_CT( | leukocyte immunoglobulin like receptor B1 [S |
| CU467612.1 | CHR_HSCHR19LRC_PGF2_CT( | vomeronasal 1 receptor 1 (VN1R1) pseudoger |
| LILRB4 | CHR_HSCHR19LRC_PGF2_CT( | leukocyte immunoglobulin like receptor B4 [S |
| LILRP1 | CHR_HSCHR19LRC_PGF2_CT( | leukocyte immunoglobulin-like receptor pseu |
| LILRP2 | CHR_HSCHR19LRC_PGF2_CT( | leukocyte immunoglobulin-like receptor pseu |
| KIR2DL2 | CHR_HSCHR19LRC_PGF2_CT( | killer cell immunoglobulin like receptor, two I |
| KIR3DL3 | CHR_HSCHR19LRC_PGF2_CT( | killer cell immunoglobulin like receptor, three |
| KIR2DS2 | CHR_HSCHR19LRC_PGF2_CT( | killer cell immunoglobulin like receptor, two I |
| KIR3DP1 | CHR_HSCHR19LRC_PGF2_CT( | killer cell immunoglobulin like receptor, three |
| AC011476.6 | CHR_HSCHR19LRC_PGF2_CT( | natural cytotoxicity triggering receptor 1 [Sou |
| NLRP7 | CHR_HSCHR19LRC_PGF2_CT( | NLR family pyrin domain containing 7 [Source |
| NLRP2 | CHR_HSCHR19LRC_PGF2_CT( | NLR family pyrin domain containing 2 [Source |
| GP6 | CHR_HSCHR19LRC_PGF2_CT( | glycoprotein VI platelet [Source:NCBI gene;Ac |
| RDH13 | CHR_HSCHR19LRC_PGF2_CT( | retinol dehydrogenase 13 [Source:HGNC Symb |
| AC010329.3 | CHR_HSCHR19_1_CTG2 | novel zinc finger pseudogene |
| AC010329.4 | CHR_HSCHR19_1_CTG2 | novel transcript |
| ZNF66 | CHR_HSCHR19_1_CTG2 | zinc finger protein 66 [Source:HGNC Symbol;A |
| BNIP3P24 | CHR_HSCHR19_1_CTG2 | BCL2 interacting protein 3 pseudogene 24 [So |
| AC008463.2 | CHR_HSCHR19_1_CTG2 | novel transcript |
| AC008463.1 | CHR_HSCHR19_1_CTG2 | novel zinc finger pseudogene |
| ZNF85 | CHR_HSCHR19_1_CTG2 | zinc finger protein 85 [Source:HGNC Symbol;A |
| AC008739.4 | CHR_HSCHR19_1_CTG2 | TEC |
| KRT18P40 | CHR_HSCHR19_1_CTG2 | keratin 18 pseudogene 40 [Source:HGNC Sym |
| AC008739.6 | CHR_HSCHR19_1_CTG2 | novel transcript |
| ZNF430 | CHR_HSCHR19_1_CTG2 | zinc finger protein 430 [Source:HGNC Symbol; |
| LSM14A | CHR_HSCHR19_1_CTG3_1 | LSM14A mRNA processing body assembly fact |
| KIAA0355 | CHR_HSCHR19_1_CTG3_1 | KIAA0355 [Source:HGNC Symbol;Acc:HGNC:2! |
| AC010614.1 | CHR_HSCHR19_1_CTG3_1 | ribosomal protein L29 (RPL29) pseudogene |
| AC226770.1 | CHR_HSCHR19_2_CTG2 | novel zinc finger protein pseudogene |
| MTDHP4 | CHR_HSCHR19_2_CTG2 | metadherin pseudogene 4 [Source:HGNC Sym |
| AC092364.3 | CHR_HSCHR19_2_CTG2 | metadherin (MTDH) pseudogene |
| VN1R84P | CHR_HSCHR19_2_CTG2 | vomeronasal 1 receptor 84 pseudogene [Sour |
| ZNF100 | CHR_HSCHR19_2_CTG2 | zinc finger protein 100 [Source:HGNC Symbol; |
| BRI3BPP1 | CHR_HSCHR19_2_CTG2 | BRI3 binding protein pseudogene 1 [Source:H( |

| | | |
|---|---|---|
| AC092364.5 | CHR_HSCHR19_2_CTG2 | novel transcript |
| AC092364.4 | CHR_HSCHR19_2_CTG2 | metadherin (MTDH) pseudogene |
| ZNF43 | CHR_HSCHR19_2_CTG2 | zinc finger protein 43 [Source:HGNC Symbol;A |
| KIAA0355 | CHR_HSCHR19_2_CTG3_1 | KIAA0355 [Source:HGNC Symbol;Acc:HGNC:2 |
| AC010504.2 | CHR_HSCHR19_2_CTG3_1 | novel transcript, antisense to KIAA0355 |
| GPI | CHR_HSCHR19_2_CTG3_1 | glucose-6-phosphate isomerase [Source:HGN |
| ZNF92P2 | CHR_HSCHR19_3_CTG2 | zinc finger protein 92 pseudogene 2 [Source:H |
| AC073539.4 | CHR_HSCHR19_3_CTG2 | ribosomal protein L17 (RPL17) pseudogene |
| PCGF7P | CHR_HSCHR19_3_CTG2 | polycomb group ring finger 7 pseudogene [So |
| MTDHP5 | CHR_HSCHR19_3_CTG2 | metadherin pseudogene 5 [Source:HGNC Sym |
| VN1R85P | CHR_HSCHR19_3_CTG2 | vomeronasal 1 receptor 85 pseudogene [Sour |
| ZNF676 | CHR_HSCHR19_3_CTG2 | zinc finger protein 676 [Source:HGNC Symbol; |
| BNIP3P30 | CHR_HSCHR19_3_CTG2 | BCL2 interacting protein 3 pseudogene 30 [So |
| AC073539.6 | CHR_HSCHR19_3_CTG2 | novel transcript |
| AC073539.5 | CHR_HSCHR19_3_CTG2 | novel transcript |
| ZNF729 | CHR_HSCHR19_3_CTG2 | zinc finger protein 729 [Source:HGNC Symbol; |
| BNIP3P31 | CHR_HSCHR19_3_CTG2 | BCL2 interacting protein 3 pseudogene 31 [So |
| AC243960.9 | CHR_HSCHR19_3_CTG3_1 | novel transcript (LOC100505495) |
| AC243960.6 | CHR_HSCHR19_3_CTG3_1 | tropomyosin pseudogene |
| PLEKHA3P1 | CHR_HSCHR19_3_CTG3_1 | pleckstrin homology domain containing A3 ps |
| AC236970.1 | CHR_HSCHR19_3_CTG3_1 | novel transcript |
| LINC01480 | CHR_HSCHR19_3_CTG3_1 | long intergenic non-protein coding RNA 1480 |
| CEACAM21 | CHR_HSCHR19_3_CTG3_1 | carcinoembryonic antigen related cell adhesic |
| AC243960.12 | CHR_HSCHR19_3_CTG3_1 | novel transcript, sense intronic to CEACAM21 |
| AC243960.10 | CHR_HSCHR19_3_CTG3_1 | TEC |
| CEACAMP3 | CHR_HSCHR19_3_CTG3_1 | carcinoembryonic antigen related cell adhesic |
| DNAJC19P2 | CHR_HSCHR19_3_CTG3_1 | DnaJ heat shock protein family (Hsp40) memb |
| CEACAM4 | CHR_HSCHR19_3_CTG3_1 | carcinoembryonic antigen related cell adhesic |
| DNAJC19P3 | CHR_HSCHR19_3_CTG3_1 | DnaJ heat shock protein family (Hsp40) memb |
| AC243960.8 | CHR_HSCHR19_3_CTG3_1 | pregnancy-specific glycoprotein pseudogene |
| AC243960.11 | CHR_HSCHR19_3_CTG3_1 | phosphodiesterase 7 pseudogene |
| CEACAM7 | CHR_HSCHR19_3_CTG3_1 | carcinoembryonic antigen related cell adhesic |
| SBNO2 | CHR_HSCHR19_4_CTG2 | strawberry notch homolog 2 [Source:HGNC Sy |
| VSTM1 | CHR_HSCHR19_4_CTG3_1 | V-set and transmembrane domain containing |
| TARM1 | CHR_HSCHR19_4_CTG3_1 | T cell-interacting, activating receptor on myel |
| OSCAR | CHR_HSCHR19_4_CTG3_1 | osteoclast associated Ig-like receptor [Source: |
| NDUFA3 | CHR_HSCHR19_4_CTG3_1 | NADH:ubiquinone oxidoreductase subunit A3 |
| TFPT | CHR_HSCHR19_4_CTG3_1 | TCF3 fusion partner [Source:HGNC Symbol;Ac |
| PRPF31 | CHR_HSCHR19_4_CTG3_1 | pre-mRNA processing factor 31 [Source:HGNC |
| AC012314.3 | CHR_HSCHR19_4_CTG3_1 | novel transcript |
| CNOT3 | CHR_HSCHR19_4_CTG3_1 | CCR4-NOT transcription complex subunit 3 [S |
| LENG1 | CHR_HSCHR19_4_CTG3_1 | leukocyte receptor cluster member 1 [Source: |
| TMC4 | CHR_HSCHR19_4_CTG3_1 | transmembrane channel like 4 [Source:HGNC |
| MBOAT7 | CHR_HSCHR19_4_CTG3_1 | membrane bound O-acyltransferase domain c |
| TSEN34 | CHR_HSCHR19_4_CTG3_1 | tRNA splicing endonuclease subunit 34 [Sourc |
| AC012314.16 | CHR_HSCHR19_4_CTG3_1 | novel transcript |
| RPS9 | CHR_HSCHR19_4_CTG3_1 | ribosomal protein S9 [Source:HGNC Symbol;A |
| AC012314.2 | CHR_HSCHR19_4_CTG3_1 | leukocyte immunoglobulin-like receptor, subf |
| LILRB3 | CHR_HSCHR19_4_CTG3_1 | leukocyte immunoglobulin like receptor B3 [S |
| AC010492.3 | CHR_HSCHR19_4_CTG3_1 | novel transcript |

| | | |
|---|---|---|
| AC010492.1 | CHR_HSCHR19_4_CTG3_1 | novel pseudogene |
| LILRA6 | CHR_HSCHR19_4_CTG3_1 | leukocyte immunoglobulin like receptor A6 [S |
| AC010492.2 | CHR_HSCHR19_4_CTG3_1 | pseudogene similar to part of leukocyte immu |
| LILRB5 | CHR_HSCHR19_4_CTG3_1 | leukocyte immunoglobulin like receptor B5 [S |
| U6 | CHR_HSCHR19_4_CTG3_1 | U6 spliceosomal RNA [Source:RFAM;Acc:RF00 |
| AC098789.1 | CHR_HSCHR19_4_CTG3_1 | YME1-like 1 (S. cerevisiae) (YME1L1) pseudog |
| LILRB2 | CHR_HSCHR19_4_CTG3_1 | leukocyte immunoglobulin like receptor B2 [S |
| LILRA3 | CHR_HSCHR19_4_CTG3_1 | leukocyte immunoglobulin like receptor A3 [S |
| AC010518.1 | CHR_HSCHR19_4_CTG3_1 | leukocyte immunoglobulin-like receptor, subf |
| AC008984.3 | CHR_HSCHR19_4_CTG3_1 | pseudogene similar to part of leukocyte immu |
| AC008984.5 | CHR_HSCHR19_4_CTG3_1 | leukocyte immunoglobulin-like receptor, subf |
| LILRA5 | CHR_HSCHR19_4_CTG3_1 | leukocyte immunoglobulin like receptor A5 [S |
| AC008984.2 | CHR_HSCHR19_4_CTG3_1 | novel transcript |
| AC008984.4 | CHR_HSCHR19_4_CTG3_1 | pseudogene similar to vomeronasal 1 recepto |
| AC008984.1 | CHR_HSCHR19_4_CTG3_1 | pseudogene similar to part of leukocyte immu |
| LILRA4 | CHR_HSCHR19_4_CTG3_1 | leukocyte immunoglobulin like receptor A4 [S |
| LAIR1 | CHR_HSCHR19_4_CTG3_1 | leukocyte associated immunoglobulin like rec |
| AC008746.5 | CHR_HSCHR19_4_CTG3_1 | mitochondrial ribosomal protein L40 (MRPL4C |
| TTYH1 | CHR_HSCHR19_4_CTG3_1 | tweety family member 1 [Source:HGNC Symb |
| AC008746.3 | CHR_HSCHR19_4_CTG3_1 | novel transcript, antisense to TTYH1 |
| AC008746.1 | CHR_HSCHR19_4_CTG3_1 | novel transcript |
| AC008746.4 | CHR_HSCHR19_4_CTG3_1 | novel transcript |
| LENG8-AS1 | CHR_HSCHR19_4_CTG3_1 | LENG8 antisense RNA 1 [Source:HGNC Symbo |
| LENG8 | CHR_HSCHR19_4_CTG3_1 | leukocyte receptor cluster member 8 [Source: |
| LENG9 | CHR_HSCHR19_4_CTG3_1 | leukocyte receptor cluster member 9 [Source: |
| CDC42EP5 | CHR_HSCHR19_4_CTG3_1 | CDC42 effector protein 5 [Source:HGNC Symb |
| LAIR2 | CHR_HSCHR19_4_CTG3_1 | leukocyte associated immunoglobulin like rec |
| AC008746.2 | CHR_HSCHR19_4_CTG3_1 | pseudogene similar to part of killer cell immu |
| KIR3DX1 | CHR_HSCHR19_4_CTG3_1 | killer cell immunoglobulin like receptor, three |
| AC009892.2 | CHR_HSCHR19_4_CTG3_1 | pseudogene similar to part of leukocyte immu |
| LILRA2 | CHR_HSCHR19_4_CTG3_1 | leukocyte immunoglobulin like receptor A2 [S |
| AC009892.1 | CHR_HSCHR19_4_CTG3_1 | novel transcript |
| LILRA1 | CHR_HSCHR19_4_CTG3_1 | leukocyte immunoglobulin like receptor A1 [S |
| AC009892.3 | CHR_HSCHR19_4_CTG3_1 | leukocyte immunoglobulin-like receptor prote |
| LILRB1 | CHR_HSCHR19_4_CTG3_1 | leukocyte immunoglobulin like receptor B1 [S |
| LILRB1-AS1 | CHR_HSCHR19_4_CTG3_1 | LILRB1 antisense RNA 1 [Source:HGNC Symbo |
| LILRB4 | CHR_HSCHR19_4_CTG3_1 | leukocyte immunoglobulin like receptor B4 [S |
| AC011515.1 | CHR_HSCHR19_4_CTG3_1 | pseudogene similar to leukocyte immunoglob |
| LILRP1 | CHR_HSCHR19_4_CTG3_1 | leukocyte immunoglobulin-like receptor pseu |
| AC098784.1 | CHR_HSCHR19_4_CTG3_1 | killer cell immunoglobulin-like receptor family |
| LILRP2 | CHR_HSCHR19_4_CTG3_1 | leukocyte immunoglobulin-like receptor pseu |
| KIR3DL3 | CHR_HSCHR19_4_CTG3_1 | killer cell immunoglobulin like receptor, three |
| KIR2DL3 | CHR_HSCHR19_4_CTG3_1 | killer cell immunoglobulin like receptor, two I |
| KIR2DP1 | CHR_HSCHR19_4_CTG3_1 | killer cell immunoglobulin like receptor, two I |
| KIR2DL1 | CHR_HSCHR19_4_CTG3_1 | killer cell immunoglobulin like receptor, two I |
| KIR3DP1 | CHR_HSCHR19_4_CTG3_1 | killer cell immunoglobulin like receptor, three |
| KIR2DL4 | CHR_HSCHR19_4_CTG3_1 | killer cell immunoglobulin like receptor, two I |
| KIR3DL1 | CHR_HSCHR19_4_CTG3_1 | killer cell immunoglobulin like receptor, three |
| KIR2DS4 | CHR_HSCHR19_4_CTG3_1 | killer cell immunoglobulin like receptor, two I |
| KIR3DL2 | CHR_HSCHR19_4_CTG3_1 | killer cell immunoglobulin like receptor, three |

| | | |
|---|---|---|
| U6 | CHR_HSCHR19_4_CTG3_1 | U6 spliceosomal RNA [Source:RFAM;Acc:RF00 |
| FCAR | CHR_HSCHR19_4_CTG3_1 | Fc fragment of IgA receptor [Source:HGNC Syr |
| NCR1 | CHR_HSCHR19_4_CTG3_1 | natural cytotoxicity triggering receptor 1 [Sou |
| NLRP7 | CHR_HSCHR19_4_CTG3_1 | NLR family pyrin domain containing 7 [Source |
| NLRP2 | CHR_HSCHR19_4_CTG3_1 | NLR family pyrin domain containing 2 [Source |
| GP6 | CHR_HSCHR19_4_CTG3_1 | glycoprotein VI platelet [Source:HGNC Symbo |
| AC011476.7 | CHR_HSCHR19_4_CTG3_1 | novel transcript |
| RDH13 | CHR_HSCHR19_4_CTG3_1 | retinol dehydrogenase 13 [Source:HGNC Symb |
| PLPPR3 | CHR_HSCHR19_5_CTG2 | phospholipid phosphatase related 3 [Source:H |
| AZU1 | CHR_HSCHR19_5_CTG2 | azurocidin 1 [Source:HGNC Symbol;Acc:HGNC |
| PRTN3 | CHR_HSCHR19_5_CTG2 | proteinase 3 [Source:HGNC Symbol;Acc:HGNC |
| ELANE | CHR_HSCHR19_5_CTG2 | elastase, neutrophil expressed [Source:HGNC |
| CFD | CHR_HSCHR19_5_CTG2 | complement factor D [Source:HGNC Symbol;A |
| MED16 | CHR_HSCHR19_5_CTG2 | mediator complex subunit 16 [Source:HGNC S |
| AC208814.1 | CHR_HSCHR1_1_CTG11 | novel transcript |
| AL161638.3 | CHR_HSCHR1_1_CTG11 | novel transcript |
| AL161638.4 | CHR_HSCHR1_1_CTG11 | novel transcript |
| PLCH2 | CHR_HSCHR1_1_CTG3 | phospholipase C eta 2 [Source:HGNC Symbol; |
| AL139246.7 | CHR_HSCHR1_1_CTG3 | novel transcript |
| AL139246.8 | CHR_HSCHR1_1_CTG3 | novel transcript |
| PANK4 | CHR_HSCHR1_1_CTG3 | pantothenate kinase 4 [Source:HGNC Symbol; |
| HES5 | CHR_HSCHR1_1_CTG3 | hes family bHLH transcription factor 5 [Source |
| AL139246.9 | CHR_HSCHR1_1_CTG3 | novel transcript |
| TNFRSF14-AS1 | CHR_HSCHR1_1_CTG3 | TNFRSF14 antisense RNA 1 [Source:HGNC Sym |
| TNFRSF14 | CHR_HSCHR1_1_CTG3 | TNF receptor superfamily member 14 [Source |
| AL139246.6 | CHR_HSCHR1_1_CTG3 | novel transcript |
| AL139246.10 | CHR_HSCHR1_1_CTG3 | novel transcript |
| PRXL2B | CHR_HSCHR1_1_CTG3 | peroxiredoxin like 2B [Source:HGNC Symbol;A |
| MMEL1 | CHR_HSCHR1_1_CTG3 | membrane metalloendopeptidase like 1 [Sour |
| AC234231.1 | CHR_HSCHR1_1_CTG3 | novel transcript |
| TTC34 | CHR_HSCHR1_1_CTG3 | tetratricopeptide repeat domain 34 [Source:H |
| Metazoa_SRP | CHR_HSCHR1_1_CTG31 | Metazoan signal recognition particle RNA [Sou |
| GEMIN2P1 | CHR_HSCHR1_1_CTG31 | gem nuclear organelle associated protein 2 ps |
| Metazoa_SRP | CHR_HSCHR1_1_CTG31 | Metazoan signal recognition particle RNA [Sou |
| AC217323.1 | CHR_HSCHR1_1_CTG31 | gem (nuclear organelle) associated protein 2 ( |
| Y_RNA | CHR_HSCHR1_1_CTG31 | Y RNA [Source:RFAM;Acc:RF00019] |
| INTS3 | CHR_HSCHR1_1_CTG31 | integrator complex subunit 3 [Source:HGNC S |
| AL513523.7 | CHR_HSCHR1_1_CTG31 | novel transcript |
| AL513523.8 | CHR_HSCHR1_1_CTG31 | novel transcript |
| AL513523.9 | CHR_HSCHR1_1_CTG31 | NULL |
| SLC27A3 | CHR_HSCHR1_1_CTG31 | solute carrier family 27 member 3 [Source:HG |
| Y_RNA | CHR_HSCHR1_1_CTG31 | Y RNA [Source:RFAM;Acc:RF00019] |
| AL513523.6 | CHR_HSCHR1_1_CTG31 | immunoglobulin superfamily, DCC subclass, m |
| GATAD2B | CHR_HSCHR1_1_CTG31 | GATA zinc finger domain containing 2B [Sourc |
| AL513523.5 | CHR_HSCHR1_1_CTG31 | adipose differentiation-related protein (ADFP) |
| KIF26B | CHR_HSCHR1_1_CTG32_1 | kinesin family member 26B [Source:HGNC Syr |
| CR854858.2 | CHR_HSCHR1_2_CTG3 | novel PRAME family member pseudogene |
| PRAMEF22 | CHR_HSCHR1_2_CTG3 | PRAME family member 22 [Source:HGNC Sym |
| CR854858.4 | CHR_HSCHR1_2_CTG3 | novel PRAME family pseudogene |
| CR854858.3 | CHR_HSCHR1_2_CTG3 | novel PRAME family member pseudogene |

| | | |
|---|---|---|
| HNRNPCL2 | CHR_HSCHR1_2_CTG3 | heterogeneous nuclear ribonucleoprotein C lil |
| CR854858.5 | CHR_HSCHR1_2_CTG3 | novel PRAME family pseudogene |
| PRAMEF25 | CHR_HSCHR1_2_CTG3 | PRAME family member 25 [Source:HGNC Sym |
| HNRNPCL4 | CHR_HSCHR1_2_CTG3 | heterogeneous nuclear ribonucleoprotein C lil |
| PRAMEF9 | CHR_HSCHR1_2_CTG3 | PRAME family member 9 [Source:NCBI gene;A |
| CR854858.1 | CHR_HSCHR1_2_CTG3 | novel PRAME family pseudogene |
| PRAMEF18 | CHR_HSCHR1_2_CTG3 | PRAME family member 18 [Source:HGNC Sym |
| AC244216.1 | CHR_HSCHR1_2_CTG3 | novel PRAME family member pseudogene |
| PRAMEF5 | CHR_HSCHR1_2_CTG3 | PRAME family member 5 [Source:HGNC Symb |
| AC244216.2 | CHR_HSCHR1_2_CTG3 | novel PRAME family member pseudogene |
| U6 | CHR_HSCHR1_2_CTG3 | U6 spliceosomal RNA [Source:RFAM;Acc:RF00 |
| PRAMEF8 | CHR_HSCHR1_2_CTG3 | PRAME family member 8 [Source:HGNC Symb |
| PRAMEF33 | CHR_HSCHR1_2_CTG3 | PRAME family member 33 [Source:NCBI gene; |
| AC217785.2 | CHR_HSCHR1_2_CTG31 | glucosidase, beta, acid (GBA) pseudogene |
| AC217785.1 | CHR_HSCHR1_2_CTG31 | metaxin 1 (MTX1) pseudogene |
| AC217785.3 | CHR_HSCHR1_2_CTG31 | glucosidase, beta, acid (GBA) pseudogene |
| MTX1P1 | CHR_HSCHR1_2_CTG31 | metaxin 1 pseudogene 1 [Source:HGNC Symb |
| GBA | CHR_HSCHR1_2_CTG31 | glucosylceramidase beta [Source:HGNC Symb |
| FAM189B | CHR_HSCHR1_2_CTG31 | family with sequence similarity 189 member E |
| SCAMP3 | CHR_HSCHR1_2_CTG31 | secretory carrier membrane protein 3 [Source |
| CLK2 | CHR_HSCHR1_2_CTG31 | CDC like kinase 2 [Source:HGNC Symbol;Acc:H |
| HCN3 | CHR_HSCHR1_2_CTG31 | hyperpolarization activated cyclic nucleotide g |
| PKLR | CHR_HSCHR1_2_CTG31 | pyruvate kinase L/R [Source:HGNC Symbol;Ac |
| OR2T4 | CHR_HSCHR1_2_CTG32_1 | olfactory receptor family 2 subfamily T memb |
| OR2T6 | CHR_HSCHR1_2_CTG32_1 | olfactory receptor family 2 subfamily T memb |
| OR2T1 | CHR_HSCHR1_2_CTG32_1 | olfactory receptor family 2 subfamily T memb |
| OR2T7 | CHR_HSCHR1_2_CTG32_1 | olfactory receptor family 2 subfamily T memb |
| OR2T2 | CHR_HSCHR1_2_CTG32_1 | olfactory receptor family 2 subfamily T memb |
| OR2T3 | CHR_HSCHR1_2_CTG32_1 | olfactory receptor family 2 subfamily T memb |
| OR2T5 | CHR_HSCHR1_2_CTG32_1 | olfactory receptor family 2 subfamily T memb |
| AC205875.1 | CHR_HSCHR1_3_CTG3 | adaptor-related protein complex 2, beta 1 sub |
| ATP6V1G3 | CHR_HSCHR1_3_CTG31 | ATPase H+ transporting V1 subunit G3 [Source |
| AL157402.4 | CHR_HSCHR1_3_CTG31 | novel transcript |
| PTPRC | CHR_HSCHR1_3_CTG31 | protein tyrosine phosphatase receptor type C |
| AL157402.3 | CHR_HSCHR1_3_CTG31 | novel transcript |
| PEBP1P3 | CHR_HSCHR1_3_CTG31 | phosphatidylethanolamine binding protein 1 p |
| AL606534.6 | CHR_HSCHR1_3_CTG32_1 | novel transcript |
| LINC01347 | CHR_HSCHR1_3_CTG32_1 | long intergenic non-protein coding RNA 1347 |
| AL606534.7 | CHR_HSCHR1_3_CTG32_1 | novel transcript |
| RNU6-747P | CHR_HSCHR1_3_CTG32_1 | RNA, U6 small nuclear 747, pseudogene [Sour |
| CEP170 | CHR_HSCHR1_3_CTG32_1 | centrosomal protein 170 [Source:HGNC Symb |
| SDCCAG8 | CHR_HSCHR1_3_CTG32_1 | serologically defined colon cancer antigen 8 [S |
| AKT3 | CHR_HSCHR1_3_CTG32_1 | AKT serine/threonine kinase 3 [Source:HGNC S |
| AC225099.1 | CHR_HSCHR1_3_CTG32_1 | novel transcript |
| AL109936.3 | CHR_HSCHR1_4_CTG3 | ribosomal protein L29 (RPL29) pseudogene |
| LINC01355 | CHR_HSCHR1_4_CTG3 | long intergenic non-protein coding RNA 1355 |
| AL109936.5 | CHR_HSCHR1_4_CTG3 | novel transcript |
| HNRNPR | CHR_HSCHR1_4_CTG3 | heterogeneous nuclear ribonucleoprotein R [S |
| ZNF436 | CHR_HSCHR1_4_CTG3 | zinc finger protein 436 [Source:HGNC Symbol; |
| ZNF436-AS1 | CHR_HSCHR1_4_CTG3 | ZNF436 antisense RNA 1 [Source:HGNC Symb |

| | | |
|---|---|---|
| Y_RNA | CHR_HSCHR1_4_CTG3 | Y RNA [Source:RFAM;Acc:RF00019] |
| AL109936.4 | CHR_HSCHR1_4_CTG3 | novel transcript |
| TCEA3 | CHR_HSCHR1_4_CTG3 | transcription elongation factor A3 [Source:HG |
| AL357134.2 | CHR_HSCHR1_4_CTG3 | single-stranded DNA binding protein (SSBP1) p |
| ASAP3 | CHR_HSCHR1_4_CTG3 | ArfGAP with SH3 domain, ankyrin repeat and |
| E2F2 | CHR_HSCHR1_4_CTG3 | E2F transcription factor 2 [Source:HGNC Symb |
| AL021154.2 | CHR_HSCHR1_4_CTG3 | novel transcript |
| ID3 | CHR_HSCHR1_4_CTG3 | inhibitor of DNA binding 3, HLH protein [Sourc |
| AL450043.2 | CHR_HSCHR1_4_CTG3 | novel transcript |
| U1 | CHR_HSCHR1_4_CTG31 | U1 spliceosomal RNA [Source:RFAM;Acc:RF00 |
| U1 | CHR_HSCHR1_4_CTG31 | U1 spliceosomal RNA [Source:RFAM;Acc:RF00 |
| U1 | CHR_HSCHR1_4_CTG31 | U1 spliceosomal RNA [Source:RFAM;Acc:RF00 |
| AC006359.4 | CHR_HSCHR1_4_CTG31 | novel transcript |
| AC006359.3 | CHR_HSCHR1_4_CTG31 | novel transcript |
| PPIAL4F | CHR_HSCHR1_4_CTG31 | peptidylprolyl isomerase A like 4F [Source:HG |
| AC006359.2 | CHR_HSCHR1_4_CTG31 | novel transcript |
| AC006359.1 | CHR_HSCHR1_4_CTG31 | novel transcript with similarity to B melanoma |
| PRAMEF28P | CHR_HSCHR1_5_CTG3 | PRAME family member 28, pseudogene [Sour |
| PRAMEF27 | CHR_HSCHR1_5_CTG3 | PRAME family member 27 [Source:HGNC Sym |
| HNRNPCL3 | CHR_HSCHR1_5_CTG3 | heterogeneous nuclear ribonucleoprotein C lil |
| PRAMEF25 | CHR_HSCHR1_5_CTG3 | PRAME family member 25 [Source:HGNC Sym |
| PRAMEF34P | CHR_HSCHR1_5_CTG3 | PRAME family member 34, pseudogene [Sour |
| PRAMEF35P | CHR_HSCHR1_5_CTG3 | PRAME family member 35, pseudogene [Sour |
| HNRNPCL2 | CHR_HSCHR1_5_CTG3 | heterogeneous nuclear ribonucleoprotein C lil |
| PRAMEF36P | CHR_HSCHR1_5_CTG3 | PRAME family member 36, pseudogene [Sour |
| AC245056.3 | CHR_HSCHR1_5_CTG3 | novel transcript |
| PRAMEF26 | CHR_HSCHR1_5_CTG3 | PRAME family member 26 [Source:HGNC Sym |
| HNRNPCL4 | CHR_HSCHR1_5_CTG3 | heterogeneous nuclear ribonucleoprotein C lil |
| PRAMEF9 | CHR_HSCHR1_5_CTG3 | PRAME family member 9 [Source:HGNC Symb |
| PRAMEF31P | CHR_HSCHR1_5_CTG3 | PRAME family member 31, pseudogene [Sour |
| PRAMEF5 | CHR_HSCHR1_5_CTG3 | PRAME family member 5 [Source:HGNC Symb |
| PRAMEF32P | CHR_HSCHR1_5_CTG3 | PRAME family member 32, pseudogene [Sour |
| RNU6-771P | CHR_HSCHR1_5_CTG3 | RNA, U6 small nuclear 771, pseudogene [Sour |
| PRAMEF8 | CHR_HSCHR1_5_CTG3 | PRAME family member 8 [Source:HGNC Symb |
| PRAMEF33 | CHR_HSCHR1_5_CTG3 | PRAME family member 33 [Source:HGNC Sym |
| TBCE | CHR_HSCHR1_5_CTG32_1 | tubulin folding cofactor E [Source:HGNC Symb |
| AL357556.2 | CHR_HSCHR1_5_CTG32_1 | dentatorubral-pallidoluysian atrophy (atrophi |
| AL357556.3 | CHR_HSCHR1_5_CTG32_1 | novel transcript, antisense to TBCE |
| RPS21P1 | CHR_HSCHR1_5_CTG32_1 | ribosomal protein S21 pseudogene 1 [Source: |
| B3GALNT2 | CHR_HSCHR1_5_CTG32_1 | beta-1,3-N-acetylgalactosaminyltransferase 2 |
| MTCYBP14 | CHR_HSCHR1_5_CTG32_1 | MT-CYB pseudogene 14 [Source:HGNC Symbo |
| MTND6P14 | CHR_HSCHR1_5_CTG32_1 | MT-ND6 pseudogene 14 [Source:HGNC Symbo |
| MTND5P19 | CHR_HSCHR1_5_CTG32_1 | MT-ND5 pseudogene 19 [Source:HGNC Symbo |
| MTND4P10 | CHR_HSCHR1_5_CTG32_1 | MT-ND4 pseudogene 10 [Source:HGNC Symbo |
| MTND4LP21 | CHR_HSCHR1_5_CTG32_1 | MT-ND4L pseudogene 21 [Source:HGNC Symb |
| MTND3P8 | CHR_HSCHR1_5_CTG32_1 | MT-ND3 pseudogene 8 [Source:HGNC Symbol |
| MTCO3P46 | CHR_HSCHR1_5_CTG32_1 | MT-CO3 pseudogene 46 [Source:HGNC Symbo |
| GNG4 | CHR_HSCHR1_5_CTG32_1 | G protein subunit gamma 4 [Source:HGNC Syr |
| FO393422.2 | CHR_HSCHR1_5_CTG32_1 | methylenetetrahydrofolate dehydrogenase (N |
| VAV3-AS1 | CHR_HSCHR1_6_CTG3 | VAV3 antisense RNA 1 [Source:HGNC Symbol; |

| | | |
|---|---|---|
| LINC02785 | CHR_HSCHR1_6_CTG3 | long intergenic non-protein coding RNA 2785 |
| SLC25A24 | CHR_HSCHR1_6_CTG3 | solute carrier family 25 member 24 [Source:H |
| AC275454.2 | CHR_HSCHR1_6_CTG3 | novel transcript, antisense to SLC25A24 |
| AC275454.1 | CHR_HSCHR1_6_CTG3 | TEC |
| AC275455.1 | CHR_HSCHR1_6_CTG3 | solute carrier family 25 (mitochondrial carrier |
| ST13P21 | CHR_HSCHR1_6_CTG3 | ST13, Hsp70 interacting protein pseudogene 2 |
| AL392088.3 | CHR_HSCHR1_6_CTG3 | ribosomal protein L7-like 1 (RPL7L1) pseudoge |
| DNAJC8 | CHR_HSCHR1_8_CTG3 | DnaJ heat shock protein family (Hsp40) memb |
| ATP5IF1 | CHR_HSCHR1_8_CTG3 | ATP synthase inhibitory factor subunit 1 [Sour |
| AL353622.4 | CHR_HSCHR1_8_CTG3 | novel transcript |
| AL353622.3 | CHR_HSCHR1_8_CTG3 | novel transcript |
| SESN2 | CHR_HSCHR1_8_CTG3 | sestrin 2 [Source:HGNC Symbol;Acc:HGNC:20 |
| MED18 | CHR_HSCHR1_8_CTG3 | mediator complex subunit 18 [Source:HGNC S |
| AL513166.2 | CHR_HSCHR1_9_CTG3 | novel transcript |
| RPL31P12 | CHR_HSCHR1_9_CTG3 | ribosomal protein L31 pseudogene 12 [Source |
| OR2T4 | CHR_HSCHR1_ALT2_1_CTG32 | olfactory receptor family 2 subfamily T memb |
| OR2T6 | CHR_HSCHR1_ALT2_1_CTG32 | olfactory receptor family 2 subfamily T memb |
| OR2T1 | CHR_HSCHR1_ALT2_1_CTG32 | olfactory receptor family 2 subfamily T memb |
| OR2T7 | CHR_HSCHR1_ALT2_1_CTG32 | olfactory receptor family 2 subfamily T memb |
| OR2T2 | CHR_HSCHR1_ALT2_1_CTG32 | olfactory receptor family 2 subfamily T memb |
| OR2T3 | CHR_HSCHR1_ALT2_1_CTG32 | olfactory receptor family 2 subfamily T memb |
| AC138089.2 | CHR_HSCHR1_ALT2_1_CTG32 | novel transcript |
| OR2T5 | CHR_HSCHR1_ALT2_1_CTG32 | olfactory receptor family 2 subfamily T memb |
| OR2AS2P | CHR_HSCHR1_ALT2_1_CTG32 | olfactory receptor family 2 subfamily AS mem |
| AL035045.2 | CHR_HSCHR20_1_CTG1 | novel transcript |
| AC207480.1 | CHR_HSCHR20_1_CTG1 | novel transcript |
| AL035045.3 | CHR_HSCHR20_1_CTG1 | novel transcript |
| CDH4 | CHR_HSCHR20_1_CTG2 | cadherin 4 [Source:HGNC Symbol;Acc:HGNC:1 |
| TAF4 | CHR_HSCHR20_1_CTG2 | TATA-box binding protein associated factor 4 |
| OPRL1 | CHR_HSCHR20_1_CTG3 | opioid related nociceptin receptor 1 [Source:H |
| NPBWR2 | CHR_HSCHR20_1_CTG3 | neuropeptides B and W receptor 2 [Source:HG |
| AC226131.1 | CHR_HSCHR20_1_CTG3 | novel transcript |
| MYT1 | CHR_HSCHR20_1_CTG3 | myelin transcription factor 1 [Source:HGNC Sy |
| AL121581.1 | CHR_HSCHR20_1_CTG3 | novel transcript |
| PCMTD2 | CHR_HSCHR20_1_CTG3 | protein-L-isoaspartate (D-aspartate) O-methyl |
| KCNQ2 | CHR_HSCHR20_1_CTG4 | potassium voltage-gated channel subfamily Q |
| AL353658.3 | CHR_HSCHR20_1_CTG4 | novel transcript, antisense to KCNQ2 |
| AC210433.1 | CHR_HSCHR20_1_CTG4 | novel transcript |
| AP000705.5 | CHR_HSCHR21_2_CTG1_1 | long intergenic non-protein coding RNA 308 |
| AP000705.6 | CHR_HSCHR21_2_CTG1_1 | phosphoglycerate kinase 1 (PGK1) pseudogen |
| AP000705.3 | CHR_HSCHR21_2_CTG1_1 | IBR domain containing 3 (IBRDC3) pseudogen |
| RNU4-45P | CHR_HSCHR21_2_CTG1_1 | RNA, U4 small nuclear 45, pseudogene [Sourc |
| AP000705.4 | CHR_HSCHR21_2_CTG1_1 | novel transcript |
| PAXBP1 | CHR_HSCHR21_3_CTG1_1 | PAX3 and PAX7 binding protein 1 [Source:HGN |
| C21orf62-AS1 | CHR_HSCHR21_3_CTG1_1 | C21orf62 antisense RNA 1 [Source:HGNC Sym |
| AP000280.2 | CHR_HSCHR21_3_CTG1_1 | TEC |
| C21orf62 | CHR_HSCHR21_3_CTG1_1 | chromosome 21 open reading frame 62 [Sour |
| IFNGR2 | CHR_HSCHR21_4_CTG1_1 | interferon gamma receptor 2 [Source:HGNC S |
| TMEM50B | CHR_HSCHR21_4_CTG1_1 | transmembrane protein 50B [Source:HGNC Sy |
| RPS5P3 | CHR_HSCHR21_4_CTG1_1 | ribosomal protein S5 pseudogene 3 [Source:H |

| | | |
|---|---|---|
| AC207981.1 | CHR_HSCHR21_4_CTG1_1 | novel transcript |
| DNAJC28 | CHR_HSCHR21_4_CTG1_1 | DnaJ heat shock protein family (Hsp40) memb |
| GART | CHR_HSCHR21_4_CTG1_1 | phosphoribosylglycinamide formyltransferase |
| FTCD | CHR_HSCHR21_5_CTG2 | formimidoyltransferase cyclodeaminase [Sour |
| FTCD-AS1 | CHR_HSCHR21_5_CTG2 | FTCD antisense RNA 1 [Source:HGNC Symbol; |
| SPATC1L | CHR_HSCHR21_5_CTG2 | spermatogenesis and centriole associated 1 lil |
| AP001468.2 | CHR_HSCHR21_5_CTG2 | novel transcript |
| LSS | CHR_HSCHR21_5_CTG2 | lanosterol synthase [Source:HGNC Symbol;Acc |
| BTG3 | CHR_HSCHR21_6_CTG1_1 | BTG anti-proliferation factor 3 [Source:HGNC |
| BTG3-AS1 | CHR_HSCHR21_6_CTG1_1 | BTG3 antisense RNA 1 [Source:HGNC Symbol; |
| AP000432.4 | CHR_HSCHR21_6_CTG1_1 | novel transcript |
| AP000432.3 | CHR_HSCHR21_6_CTG1_1 | TEC |
| AC208585.1 | CHR_HSCHR21_6_CTG1_1 | novel transcript |
| NEK4P1 | CHR_HSCHR21_8_CTG1_1 | NIMA-related kinase 4 pseudogene 1 [Source: |
| AP000457.2 | CHR_HSCHR21_8_CTG1_1 | G protein-coupled receptor 157 (GPR157) pse |
| SMDT1 | CHR_HSCHR22_1_CTG1 | single-pass membrane protein with aspartate |
| NDUFA6 | CHR_HSCHR22_1_CTG1 | NADH:ubiquinone oxidoreductase subunit A6 |
| NDUFA6-DT | CHR_HSCHR22_1_CTG1 | NDUFA6 divergent transcript [Source:HGNC Sy |
| OLA1P1 | CHR_HSCHR22_1_CTG1 | OLA1 pseudogene 1 [Source:HGNC Symbol;Ac |
| AL021878.1 | CHR_HSCHR22_1_CTG1 | novel transcript |
| CYP2D7 | CHR_HSCHR22_1_CTG1 | cytochrome P450 family 2 subfamily D memb |
| CYP2D8P | CHR_HSCHR22_1_CTG1 | ccytochrome P450 family 2 subfamily D meml |
| TCF20 | CHR_HSCHR22_1_CTG1 | transcription factor 20 [Source:HGNC Symbol; |
| AL022318.3 | CHR_HSCHR22_1_CTG2 | novel transcript |
| AC192820.1 | CHR_HSCHR22_1_CTG2 | novel pseudogene similar to apolipoprotein B |
| APOBEC3A | CHR_HSCHR22_1_CTG2 | apolipoprotein B mRNA editing enzyme cataly |
| APOBEC3B-AS1 | CHR_HSCHR22_1_CTG2 | APOBEC3B antisense RNA 1 [Source:HGNC Syr |
| IGLV5-45 | CHR_HSCHR22_1_CTG3 | immunoglobulin lambda variable 5-45 [Source |
| IGLV1-44 | CHR_HSCHR22_1_CTG3 | immunoglobulin lambda variable 1-44 [Source |
| IGLV7-43 | CHR_HSCHR22_1_CTG3 | immunoglobulin lambda variable 7-43 [Source |
| IGLVI-42 | CHR_HSCHR22_1_CTG3 | immunoglobulin lambda variable (I)-42 (pseuc |
| AC009286.3 | CHR_HSCHR22_1_CTG3 | Immunoglobulin lambda variable 5-39 [Sourc |
| IGLVVII-41-1 | CHR_HSCHR22_1_CTG3 | immunoglobulin lambda variable (VII)-41-1 (p: |
| IGLV1-41 | CHR_HSCHR22_1_CTG3 | immunoglobulin lambda variable 1-41 (pseud |
| IGLV1-40 | CHR_HSCHR22_1_CTG3 | immunoglobulin lambda variable 1-40 [Source |
| ASH2LP1 | CHR_HSCHR22_1_CTG3 | ASH2L pseudogene 1 [Source:HGNC Symbol;A |
| IGLVI-38 | CHR_HSCHR22_1_CTG3 | immunoglobulin lambda variable (I)-38 (pseuc |
| IGLV5-37 | CHR_HSCHR22_1_CTG3 | immunoglobulin lambda variable 5-37 [Source |
| IGLV1-36 | CHR_HSCHR22_1_CTG3 | immunoglobulin lambda variable 1-36 [Source |
| AC009286.1 | CHR_HSCHR22_1_CTG3 | ash2 (absent, small, or homeotic)-like (Drosop |
| IGLV7-35 | CHR_HSCHR22_1_CTG3 | immunoglobulin lambda variable 7-35 (pseud |
| AC009286.2 | CHR_HSCHR22_1_CTG3 | Ig lambda chain V-I region pseudogene |
| ZNF280B | CHR_HSCHR22_1_CTG3 | zinc finger protein 280B [Source:HGNC Symbo |
| ZNF280A | CHR_HSCHR22_1_CTG3 | zinc finger protein 280A [Source:HGNC Symbo |
| PRAME | CHR_HSCHR22_1_CTG3 | preferentially expressed antigen in melanoma |
| AC246793.2 | CHR_HSCHR22_1_CTG3 | novel transcript, antisense to PRAME |
| IGLV2-34 | CHR_HSCHR22_1_CTG3 | immunoglobulin lambda variable 2-34 (pseud |
| IGLV2-33 | CHR_HSCHR22_1_CTG3 | immunoglobulin lambda variable 2-33 (non-fu |
| IGLV3-32 | CHR_HSCHR22_1_CTG3 | immunoglobulin lambda variable 3-32 (non-fu |
| IGLV3-31 | CHR_HSCHR22_1_CTG3 | immunoglobulin lambda variable 3-31 (pseud |

| | | |
|---|---|---|
| IGLV3-30 | CHR_HSCHR22_1_CTG3 | immunoglobulin lambda variable 3-30 (pseud |
| AL049748.2 | CHR_HSCHR22_1_CTG4 | novel transcript |
| MRPS16P3 | CHR_HSCHR22_1_CTG4 | mitochondrial ribosomal protein S16 pseudog |
| APOL5 | CHR_HSCHR22_1_CTG4 | apolipoprotein L5 [Source:HGNC Symbol;Acc:I |
| RBFOX2 | CHR_HSCHR22_1_CTG4 | RNA binding fox-1 homolog 2 [Source:HGNC S |
| PVALB | CHR_HSCHR22_1_CTG5 | parvalbumin [Source:HGNC Symbol;Acc:HGNC |
| Z82185.2 | CHR_HSCHR22_1_CTG5 | novel transcript |
| NCF4-AS1 | CHR_HSCHR22_1_CTG5 | NCF4 antisense RNA 1 [Source:HGNC Symbol; |
| NCF4 | CHR_HSCHR22_1_CTG5 | neutrophil cytosolic factor 4 [Source:HGNC Sy |
| CES5AP1 | CHR_HSCHR22_1_CTG6 | carboxylesterase 5A pseudogene 1 [Source:H |
| ZDHHC8P1 | CHR_HSCHR22_1_CTG6 | zinc finger DHHC-type containing 8 pseudoger |
| AP000344.3 | CHR_HSCHR22_1_CTG6 | pseudogene similar to rat RGD1563782 protei |
| LINC01659 | CHR_HSCHR22_1_CTG6 | long intergenic non-protein coding RNA 1659 |
| AP000344.4 | CHR_HSCHR22_1_CTG6 | MAD1 mitotic arrest deficient-like 1 (yeast)(M |
| FAM230I | CHR_HSCHR22_1_CTG6 | family with sequence similarity 230 member I |
| LINC02557 | CHR_HSCHR22_1_CTG6 | long intergenic non-protein coding RNA 2557 |
| C22orf15 | CHR_HSCHR22_1_CTG7 | chromosome 22 open reading frame 15 [Sour |
| CHCHD10 | CHR_HSCHR22_1_CTG7 | coiled-coil-helix-coiled-coil-helix domain cont |
| MMP11 | CHR_HSCHR22_1_CTG7 | matrix metallopeptidase 11 [Source:HGNC Syr |
| AP000349.1 | CHR_HSCHR22_1_CTG7 | NULL |
| SMARCB1 | CHR_HSCHR22_1_CTG7 | SWI/SNF related, matrix associated, actin dep |
| DERL3 | CHR_HSCHR22_1_CTG7 | derlin 3 [Source:HGNC Symbol;Acc:HGNC:142 |
| AP000350.13 | CHR_HSCHR22_1_CTG7 | novel transcript, antisense to SLC2A11 |
| SLC2A11 | CHR_HSCHR22_1_CTG7 | solute carrier family 2 member 11 [Source:HG |
| AP000350.12 | CHR_HSCHR22_1_CTG7 | novel protein |
| AP000350.10 | CHR_HSCHR22_1_CTG7 | novel transcript, antisense to SLC2A11 |
| RN7SL268P | CHR_HSCHR22_1_CTG7 | RNA, 7SL, cytoplasmic 268, pseudogene [Sour |
| MIF | CHR_HSCHR22_1_CTG7 | macrophage migration inhibitory factor [Sour |
| MIF-AS1 | CHR_HSCHR22_1_CTG7 | MIF antisense RNA 1 [Source:HGNC Symbol;A |
| AP000350.14 | CHR_HSCHR22_1_CTG7 | novel transcript |
| AP000350.15 | CHR_HSCHR22_1_CTG7 | kelch-like 5 (Drosophila) (KLHL5) pseudogene |
| AP000350.9 | CHR_HSCHR22_1_CTG7 | glutathione S-transferase (GST) theta pseudog |
| AP000350.8 | CHR_HSCHR22_1_CTG7 | glutathione S-transferase (GST) theta pseudog |
| GSTT2B | CHR_HSCHR22_1_CTG7 | glutathione S-transferase theta 2B (gene/pseu |
| AP000350.11 | CHR_HSCHR22_1_CTG7 | novel pseudogene |
| DDTL | CHR_HSCHR22_1_CTG7 | D-dopachrome tautomerase like [Source:HGN |
| AP000351.8 | CHR_HSCHR22_1_CTG7 | novel transcript, sense intronic to DDTL |
| DDT | CHR_HSCHR22_1_CTG7 | D-dopachrome tautomerase [Source:HGNC Sy |
| AP000351.9 | CHR_HSCHR22_1_CTG7 | glutathione S-transferase (GST) theta pseudog |
| GSTT2 | CHR_HSCHR22_1_CTG7 | glutathione S-transferase theta 2 (gene/pseuc |
| AP000351.6 | CHR_HSCHR22_1_CTG7 | glutathione S-transferase (GST) theta pseudog |
| GSTT4 | CHR_HSCHR22_1_CTG7 | glutathione S-transferase theta 4 [Source:HGN |
| AP000351.4 | CHR_HSCHR22_1_CTG7 | glutathione S-transferase (GST) theta pseudog |
| AP000351.3 | CHR_HSCHR22_1_CTG7 | NULL |
| AP000351.1 | CHR_HSCHR22_1_CTG7 | Glutathione S-transferase pseudogene |
| AP000351.2 | CHR_HSCHR22_1_CTG7 | D-dopachrome tautomerase (DDT) pseudoger |
| GSTT1 | CHR_HSCHR22_1_CTG7 | glutathione S-transferase theta 1 [Source:HGN |
| AP000351.7 | CHR_HSCHR22_1_CTG7 | novel glutathione S-transferase theta family p |
| AP000351.5 | CHR_HSCHR22_1_CTG7 | Glutathione S-transferase pseudogene |
| CABIN1 | CHR_HSCHR22_1_CTG7 | calcineurin binding protein 1 [Source:HGNC Sy |

| | | |
|---|---|---|
| SMDT1 | CHR_HSCHR22_2_CTG1 | single-pass membrane protein with aspartate |
| NDUFA6 | CHR_HSCHR22_2_CTG1 | NADH:ubiquinone oxidoreductase subunit A6 |
| AL021878.3 | CHR_HSCHR22_2_CTG1 | novel transcript, sense intronic to NDUFA6 |
| NDUFA6-DT | CHR_HSCHR22_2_CTG1 | NDUFA6 divergent transcript [Source:HGNC Sy |
| OLA1P1 | CHR_HSCHR22_2_CTG1 | OLA1 pseudogene 1 [Source:HGNC Symbol;Ac |
| CYP2D6 | CHR_HSCHR22_2_CTG1 | cytochrome P450 family 2 subfamily D memb |
| AC236040.3 | CHR_HSCHR22_2_CTG1 | chromosome 6 open reading frame 108 (C6or |
| AC236040.1 | CHR_HSCHR22_2_CTG1 | NULL |
| AC236040.2 | CHR_HSCHR22_2_CTG1 | novel transcript |
| SMDT1 | CHR_HSCHR22_3_CTG1 | single-pass membrane protein with aspartate |
| NDUFA6 | CHR_HSCHR22_3_CTG1 | NADH:ubiquinone oxidoreductase subunit A6 |
| AL021878.4 | CHR_HSCHR22_3_CTG1 | novel transcript, sense intronic to NDUFA6 |
| NDUFA6-DT | CHR_HSCHR22_3_CTG1 | NDUFA6 divergent transcript [Source:HGNC Sy |
| OLA1P1 | CHR_HSCHR22_3_CTG1 | OLA1 pseudogene 1 [Source:HGNC Symbol;Ac |
| CYP2D6 | CHR_HSCHR22_3_CTG1 | cytochrome P450 family 2 subfamily D memb |
| BX247885.1 | CHR_HSCHR22_3_CTG1 | novel transcript |
| BX247885.2 | CHR_HSCHR22_3_CTG1 | novel transcript |
| CYP2D7 | CHR_HSCHR22_3_CTG1 | cytochrome P450 family 2 subfamily D memb |
| CYP2D8P | CHR_HSCHR22_3_CTG1 | ccytochrome P450 family 2 subfamily D meml |
| TCF20 | CHR_HSCHR22_3_CTG1 | transcription factor 20 [Source:HGNC Symbol; |
| NDUFA6 | CHR_HSCHR22_4_CTG1 | NADH:ubiquinone oxidoreductase subunit A6 |
| AC213220.2 | CHR_HSCHR22_4_CTG1 | novel transcript, sense intronic to NDUFA6 |
| OLA1P1 | CHR_HSCHR22_4_CTG1 | OLA1 pseudogene 1 [Source:HGNC Symbol;Ac |
| AC213220.1 | CHR_HSCHR22_4_CTG1 | novel transcript |
| AC213220.3 | CHR_HSCHR22_4_CTG1 | novel transcript |
| AC254560.1 | CHR_HSCHR22_4_CTG1 | NULL |
| CYP2D8P | CHR_HSCHR22_4_CTG1 | ccytochrome P450 family 2 subfamily D meml |
| TCF20 | CHR_HSCHR22_4_CTG1 | transcription factor 20 [Source:HGNC Symbol; |
| CYP2D6 | CHR_HSCHR22_5_CTG1 | cytochrome P450 family 2 subfamily D memb |
| AC208162.1 | CHR_HSCHR22_5_CTG1 | chromosome 6 open reading frame 108 (C6or |
| AC208162.2 | CHR_HSCHR22_5_CTG1 | novel transcript |
| AC208162.3 | CHR_HSCHR22_5_CTG1 | novel transcript |
| CYP2D8P | CHR_HSCHR22_5_CTG1 | ccytochrome P450 family 2 subfamily D meml |
| TCF20 | CHR_HSCHR22_5_CTG1 | transcription factor 20 [Source:HGNC Symbol; |
| OLA1P1 | CHR_HSCHR22_6_CTG1 | OLA1 pseudogene 1 [Source:HGNC Symbol;Ac |
| AC235563.2 | CHR_HSCHR22_6_CTG1 | novel transcript |
| AC235563.1 | CHR_HSCHR22_6_CTG1 | novel transcript |
| AC254788.1 | CHR_HSCHR22_6_CTG1 | NULL |
| CYP2D8P | CHR_HSCHR22_6_CTG1 | ccytochrome P450 family 2 subfamily D meml |
| TCF20 | CHR_HSCHR22_6_CTG1 | transcription factor 20 [Source:HGNC Symbol; |
| OLA1P1 | CHR_HSCHR22_7_CTG1 | OLA1 pseudogene 1 [Source:HGNC Symbol;Ac |
| CYP2D6 | CHR_HSCHR22_7_CTG1 | cytochrome P450 family 2 subfamily D memb |
| AC225828.1 | CHR_HSCHR22_7_CTG1 | chromosome 6 open reading frame 108 (C6or |
| AC254847.2 | CHR_HSCHR22_7_CTG1 | novel transcript |
| AC254847.1 | CHR_HSCHR22_7_CTG1 | novel transcript |
| CYP2D7 | CHR_HSCHR22_7_CTG1 | cytochrome P450 family 2 subfamily D memb |
| CYP2D8P | CHR_HSCHR22_7_CTG1 | ccytochrome P450 family 2 subfamily D meml |
| TCF20 | CHR_HSCHR22_7_CTG1 | transcription factor 20 [Source:HGNC Symbol; |
| CYP2D6 | CHR_HSCHR22_8_CTG1 | cytochrome P450 family 2 subfamily D memb |
| AC255093.3 | CHR_HSCHR22_8_CTG1 | chromosome 6 open reading frame 108 (C6or |

| | | |
|---|---|---|
| AC255093.1 | CHR_HSCHR22_8_CTG1 | novel transcript |
| AC255093.2 | CHR_HSCHR22_8_CTG1 | novel transcript |
| CYP2D7 | CHR_HSCHR22_8_CTG1 | cytochrome P450 family 2 subfamily D membe |
| CYP2D8P | CHR_HSCHR22_8_CTG1 | ccytochrome P450 family 2 subfamily D meml |
| TCF20 | CHR_HSCHR22_8_CTG1 | transcription factor 20 [Source:HGNC Symbol; |
| TRAPPC12 | CHR_HSCHR2_1_CTG1 | trafficking protein particle complex 12 [Source |
| AC216281.1 | CHR_HSCHR2_1_CTG1 | novel transcript sense intronic to TRAPPC12 |
| AC114810.2 | CHR_HSCHR2_1_CTG1 | novel transcript, antisense to ADI1 |
| ADI1 | CHR_HSCHR2_1_CTG1 | acireductone dioxygenase 1 [Source:HGNC Sy |
| LINC01238 | CHR_HSCHR2_1_CTG15 | long intergenic non-protein coding RNA 1238 |
| AC237191.1 | CHR_HSCHR2_1_CTG15 | novel transcript |
| AC237191.2 | CHR_HSCHR2_1_CTG15 | novel transcript |
| AC093642.4 | CHR_HSCHR2_1_CTG15 | novel transcript |
| AC093642.6 | CHR_HSCHR2_1_CTG15 | F-box protein 25 (FBXO25) pseudogene |
| AC009414.2 | CHR_HSCHR2_1_CTG5 | ribosomal protein L21 (RPL21) pseudogene |
| CRIM1-DT | CHR_HSCHR2_1_CTG5 | CRIM1 divergent transcript [Source:HGNC Syn |
| CRIM1 | CHR_HSCHR2_1_CTG5 | cysteine rich transmembrane BMP regulator 1 |
| AC233263.5 | CHR_HSCHR2_1_CTG7 | striatin, calmodulin binding protein (STRN) ps |
| IGKV1OR2-118 | CHR_HSCHR2_1_CTG7 | immunoglobulin kappa variable 1/OR2-118 (p |
| AC241851.1 | CHR_HSCHR2_1_CTG7 | pseudogene similar to part of kinase suppress |
| AC241851.2 | CHR_HSCHR2_1_CTG7 | novel transcript |
| NOSTRIN | CHR_HSCHR2_1_CTG7_2 | nitric oxide synthase trafficking [Source:HGNC |
| SPC25 | CHR_HSCHR2_1_CTG7_2 | SPC25 component of NDC80 kinetochore com |
| G6PC2 | CHR_HSCHR2_1_CTG7_2 | glucose-6-phosphatase catalytic subunit 2 [So |
| ABCB11 | CHR_HSCHR2_1_CTG7_2 | ATP binding cassette subfamily B member 11 |
| AC208876.2 | CHR_HSCHR2_2_CTG1 | novel transcript |
| AC208876.1 | CHR_HSCHR2_2_CTG1 | novel transcript |
| AC116609.4 | CHR_HSCHR2_2_CTG1 | novel transcript |
| LINC01238 | CHR_HSCHR2_2_CTG15 | long intergenic non-protein coding RNA 1238 |
| AC244167.1 | CHR_HSCHR2_2_CTG15 | novel transcript |
| AC244167.2 | CHR_HSCHR2_2_CTG15 | novel transcript |
| AC093642.3 | CHR_HSCHR2_2_CTG15 | novel transcript |
| AC093642.5 | CHR_HSCHR2_2_CTG15 | F-box protein 25 (FBXO25) pseudogene |
| IGKV1D-8 | CHR_HSCHR2_2_CTG7 | immunoglobulin kappa variable 1D-8 [Source: |
| AC145029.2 | CHR_HSCHR2_2_CTG7 | novel immunoglobulin kappa variable gene |
| IGKV3D-7 | CHR_HSCHR2_2_CTG7 | immunoglobulin kappa variable 3D-7 [Source: |
| AC145029.1 | CHR_HSCHR2_2_CTG7 | striatin, calmodulin binding protein (STRN) ps |
| IGKV1OR2-118 | CHR_HSCHR2_2_CTG7 | immunoglobulin kappa variable 1/OR2-118 (p |
| AC233263.4 | CHR_HSCHR2_2_CTG7 | pseudogene similar to part of kinase suppress |
| AC233263.3 | CHR_HSCHR2_2_CTG7 | novel transcript |
| KIF5C | CHR_HSCHR2_2_CTG7_2 | kinesin family member 5C [Source:HGNC Sym |
| SNTG2 | CHR_HSCHR2_3_CTG1 | syntrophin gamma 2 [Source:HGNC Symbol;A |
| GAL3ST2 | CHR_HSCHR2_3_CTG15 | galactose-3-O-sulfotransferase 2 [Source:HGN |
| AC131097.4 | CHR_HSCHR2_3_CTG15 | novel transcript |
| AC131097.5 | CHR_HSCHR2_3_CTG15 | novel transcript |
| NEU4 | CHR_HSCHR2_3_CTG15 | neuraminidase 4 [Source:HGNC Symbol;Acc:H |
| AC131097.6 | CHR_HSCHR2_3_CTG15 | novel transcript |
| PDCD1 | CHR_HSCHR2_3_CTG15 | programmed cell death 1 [Source:HGNC Symb |
| RTP5 | CHR_HSCHR2_3_CTG15 | receptor transporter protein 5 (putative) [Sou |
| LINC01237 | CHR_HSCHR2_3_CTG15 | long intergenic non-protein coding RNA 1237 |

| | | |
|---|---|---|
| FAM240C | CHR_HSCHR2_3_CTG15 | family with sequence similarity 240 member C |
| C2orf27A | CHR_HSCHR2_3_CTG7_2 | chromosome 2 open reading frame 27A [Sour |
| AC093724.2 | CHR_HSCHR2_3_CTG7_2 | translocase of outer mitochondrial membrane |
| C2orf27B | CHR_HSCHR2_3_CTG7_2 | chromosome 2 open reading frame 27B [Sour |
| SNTG2 | CHR_HSCHR2_4_CTG1 | syntrophin gamma 2 [Source:HGNC Symbol;Ac |
| AC108489.1 | CHR_HSCHR2_4_CTG1 | novel transcript |
| TPO | CHR_HSCHR2_4_CTG1 | thyroid peroxidase [Source:HGNC Symbol;Acc |
| RN7SKP178 | CHR_HSCHR2_6_CTG7_2 | RN7SK pseudogene 178 [Source:HGNC Symbo |
| PPIAP68 | CHR_HSCHR2_6_CTG7_2 | peptidylprolyl isomerase A pseudogene 68 [Sc |
| AC007679.3 | CHR_HSCHR2_6_CTG7_2 | novel transcript, antisense to INO80D |
| INO80D | CHR_HSCHR2_6_CTG7_2 | INO80 complex subunit D [Source:HGNC Symb |
| Vault | CHR_HSCHR2_6_CTG7_2 | Vault RNA [Source:RFAM;Acc:RF00006] |
| Y_RNA | CHR_HSCHR2_6_CTG7_2 | Y RNA [Source:RFAM;Acc:RF00019] |
| AC007679.4 | CHR_HSCHR2_6_CTG7_2 | ribosomal protein L27 (RPL27) pseudogene |
| AC007383.5 | CHR_HSCHR2_6_CTG7_2 | novel transcript |
| NDUFS1 | CHR_HSCHR2_6_CTG7_2 | NADH:ubiquinone oxidoreductase core subun |
| AC007383.7 | CHR_HSCHR2_6_CTG7_2 | novel transcript |
| GCSHP3 | CHR_HSCHR2_6_CTG7_2 | glycine cleavage system protein H pseudogene |
| EEF1B2 | CHR_HSCHR2_6_CTG7_2 | eukaryotic translation elongation factor 1 beta |
| AC007383.6 | CHR_HSCHR2_6_CTG7_2 | NULL |
| snoZ196 | CHR_HSCHR2_6_CTG7_2 | Small nucleolar RNA Z196/R39/R59 family [So |
| SNORD51 | CHR_HSCHR2_6_CTG7_2 | small nucleolar RNA, C/D box 51 [Source:HGN |
| SNORA41 | CHR_HSCHR2_6_CTG7_2 | small nucleolar RNA, H/ACA box 41 [Source:H |
| GPR1 | CHR_HSCHR2_6_CTG7_2 | G protein-coupled receptor 1 [Source:HGNC S |
| GPR1-AS | CHR_HSCHR2_6_CTG7_2 | GPR1 antisense RNA [Source:HGNC Symbol;Ac |
| RN7SKP200 | CHR_HSCHR2_6_CTG7_2 | RN7SK pseudogene 200 [Source:HGNC Symbo |
| HMGN1P6 | CHR_HSCHR2_6_CTG7_2 | high mobility group nucleosome binding doma |
| RN7SKP260 | CHR_HSCHR2_6_CTG7_2 | RN7SK pseudogene 260 [Source:HGNC Symbo |
| ZDBF2 | CHR_HSCHR2_6_CTG7_2 | zinc finger DBF-type containing 2 [Source:HGN |
| AC017081.2 | CHR_HSCHR2_6_CTG7_2 | N-acylsphingosine amidohydrolase 3-like (ASA |
| HNRNPA1P51 | CHR_HSCHR2_6_CTG7_2 | heterogeneous nuclear ribonucleoprotein A1 |
| AC142116.1 | CHR_HSCHR2_7_CTG7_2 | novel transcript |
| AC236964.1 | CHR_HSCHR2_7_CTG7_2 | solute carrier family 6 (amino acid transporter |
| PLEKHM3 | CHR_HSCHR2_8_CTG7_2 | pleckstrin homology domain containing M3 [S |
| RNU6-360P | CHR_HSCHR2_8_CTG7_2 | RNA, U6 small nuclear 360, pseudogene [Sour |
| AC093698.3 | CHR_HSCHR2_8_CTG7_2 | ribosomal protein L9 (RPL9) pseudogene |
| RNA5SP116 | CHR_HSCHR2_8_CTG7_2 | RNA, 5S ribosomal pseudogene 116 [Source:H |
| RPL12P17 | CHR_HSCHR2_8_CTG7_2 | ribosomal protein L12 pseudogene 17 [Source |
| CRYGEP | CHR_HSCHR2_8_CTG7_2 | crystallin gamma E, pseudogene [Source:HGN |
| CRYGD | CHR_HSCHR2_8_CTG7_2 | crystallin gamma D [Source:HGNC Symbol;Acc |
| CRYGC | CHR_HSCHR2_8_CTG7_2 | crystallin gamma C [Source:HGNC Symbol;Acc |
| TAMM41 | CHR_HSCHR3_1_CTG1 | TAM41 mitochondrial translocator assembly a |
| AC090958.3 | CHR_HSCHR3_1_CTG1 | NULL |
| AC226154.2 | CHR_HSCHR3_1_CTG1 | Fanconi anemia, complementation group D2 ( |
| AC226154.1 | CHR_HSCHR3_1_CTG1 | glypican 5 (CPG5) pseudogene |
| NUP210P2 | CHR_HSCHR3_1_CTG1 | nucleoporin 210 pseudogene 2 [Source:HGNC |
| MARK2P14 | CHR_HSCHR3_1_CTG1 | microtubule affinity regulating kinase 2 pseud |
| AC090958.2 | CHR_HSCHR3_1_CTG1 | novel pseudogene |
| Metazoa_SRP | CHR_HSCHR3_1_CTG1 | Metazoan signal recognition particle RNA [Sou |
| LINC02066 | CHR_HSCHR3_1_CTG2_1 | long intergenic non-protein coding RNA 2066 |

| | | |
|---|---|---|
| AADACL2 | CHR_HSCHR3_1_CTG2_1 | arylacetamide deacetylase like 2 [Source:HGN |
| AADACL2-AS1 | CHR_HSCHR3_1_CTG2_1 | AADACL2 antisense RNA 1 [Source:HGNC Sym |
| AC233280.10 | CHR_HSCHR3_1_CTG3 | novel transcript |
| AC233280.11 | CHR_HSCHR3_1_CTG3 | protein phosphatase 4, regulatory subunit 2, p |
| AC233280.12 | CHR_HSCHR3_1_CTG3 | novel transcript |
| MUC20-OT1 | CHR_HSCHR3_1_CTG3 | MUC20 overlapping transcript [Source:HGNC S |
| SDHAP2 | CHR_HSCHR3_1_CTG3 | succinate dehydrogenase complex flavoprotei |
| MUC20 | CHR_HSCHR3_1_CTG3 | mucin 20, cell surface associated [Source:HGN |
| MUC4 | CHR_HSCHR3_1_CTG3 | mucin 4, cell surface associated [Source:HGNC |
| ARHGEF26-AS1 | CHR_HSCHR3_2_CTG2_1 | ARHGEF26 antisense RNA 1 [Source:HGNC Syr |
| ARHGEF26 | CHR_HSCHR3_2_CTG2_1 | Rho guanine nucleotide exchange factor 26 [S |
| DHX36 | CHR_HSCHR3_2_CTG2_1 | DEAH-box helicase 36 [Source:HGNC Symbol;A |
| AC128709.4 | CHR_HSCHR3_2_CTG3 | novel transcript |
| AC231647.1 | CHR_HSCHR3_2_CTG3 | novel transcript |
| AC231647.2 | CHR_HSCHR3_2_CTG3 | novel transcript |
| BDH1 | CHR_HSCHR3_2_CTG3 | 3-hydroxybutyrate dehydrogenase 1 [Source:I |
| AC107622.1 | CHR_HSCHR3_3_CTG1 | novel transcript |
| AC233280.16 | CHR_HSCHR3_3_CTG3 | protein phosphatase 4, regulatory subunit 2, p |
| AC233280.5 | CHR_HSCHR3_3_CTG3 | novel transcript |
| MUC20-OT1 | CHR_HSCHR3_3_CTG3 | MUC20 overlapping transcript [Source:HGNC S |
| SDHAP2 | CHR_HSCHR3_3_CTG3 | succinate dehydrogenase complex flavoprotei |
| MUC20 | CHR_HSCHR3_3_CTG3 | mucin 20, cell surface associated [Source:HGN |
| MUC4 | CHR_HSCHR3_3_CTG3 | mucin 4, cell surface associated [Source:HGNC |
| LINC00971 | CHR_HSCHR3_4_CTG1 | long intergenic non-protein coding RNA 971 [S |
| LINC02025 | CHR_HSCHR3_4_CTG1 | long intergenic non-protein coding RNA 2025 |
| AC132660.1 | CHR_HSCHR3_4_CTG1 | novel transcript |
| AF186996.7 | CHR_HSCHR3_4_CTG2_1 | olfactory receptor pseudogene |
| AF186996.5 | CHR_HSCHR3_4_CTG2_1 | olfactory receptor pseudogene |
| AF186996.9 | CHR_HSCHR3_4_CTG2_1 | olfactory receptor pseudogene |
| AF186996.4 | CHR_HSCHR3_4_CTG2_1 | olfactory receptor pseudogene |
| AF186996.6 | CHR_HSCHR3_4_CTG2_1 | olfactory receptor pseudogene |
| AF186996.8 | CHR_HSCHR3_4_CTG2_1 | vacuolar protein sorting 51 homolog (S. cerev |
| AC092902.3 | CHR_HSCHR3_4_CTG2_1 | ribosomal protein S3A (RPS3A) pseudogene |
| AC233280.15 | CHR_HSCHR3_4_CTG3 | protein phosphatase 4, regulatory subunit 2, p |
| AC233280.9 | CHR_HSCHR3_4_CTG3 | novel transcript |
| MUC20-OT1 | CHR_HSCHR3_4_CTG3 | MUC20 overlapping transcript [Source:HGNC S |
| SDHAP2 | CHR_HSCHR3_4_CTG3 | succinate dehydrogenase complex flavoprotei |
| MUC20 | CHR_HSCHR3_4_CTG3 | mucin 20, cell surface associated [Source:HGN |
| MUC4 | CHR_HSCHR3_4_CTG3 | mucin 4, cell surface associated [Source:HGNC |
| OR7E55P | CHR_HSCHR3_5_CTG1 | olfactory receptor family 7 subfamily E memb |
| AC139453.3 | CHR_HSCHR3_5_CTG1 | chromosome 21 open reading frame 63 (C21o |
| LSP1P2 | CHR_HSCHR3_5_CTG1 | LSP1 pseudogene 2 [Source:HGNC Symbol;Acc |
| AC213033.1 | CHR_HSCHR3_5_CTG1 | pseudogene similar to part of ankyrin repeat d |
| ALG1L6P | CHR_HSCHR3_5_CTG1 | ALG1 like 6, pseudogene [Source:HGNC Symbo |
| AC213033.2 | CHR_HSCHR3_5_CTG1 | NULL |
| FAM86DP | CHR_HSCHR3_5_CTG1 | family with sequence similarity 86 member D, |
| LINC02018 | CHR_HSCHR3_5_CTG1 | long intergenic non-protein coding RNA 2018 |
| ENPP7P2 | CHR_HSCHR3_5_CTG1 | ectonucleotide pyrophosphatase/phosphodie |
| SNRPCP10 | CHR_HSCHR3_5_CTG1 | small nuclear ribonucleoprotein polypeptide C |
| AC133041.2 | CHR_HSCHR3_5_CTG1 | RNA binding protein, fox-1 homolog (C. elegar |

| | | |
|---|---|---|
| RPS3AP15 | CHR_HSCHR3_5_CTG1 | ribosomal protein S3a pseudogene 15 [Source |
| LINC02054 | CHR_HSCHR3_5_CTG2_1 | long intergenic non-protein coding RNA 2054 |
| LINC01839 | CHR_HSCHR3_5_CTG2_1 | long intergenic non-protein coding RNA 1839 |
| AC233280.17 | CHR_HSCHR3_5_CTG3 | novel transcript |
| AC233280.8 | CHR_HSCHR3_5_CTG3 | protein phosphatase 4, regulatory subunit 2, p |
| AC233280.3 | CHR_HSCHR3_5_CTG3 | novel transcript |
| MUC20-OT1 | CHR_HSCHR3_5_CTG3 | MUC20 overlapping transcript [Source:HGNC S |
| SDHAP2 | CHR_HSCHR3_5_CTG3 | succinate dehydrogenase complex flavoprotei |
| MUC20 | CHR_HSCHR3_5_CTG3 | mucin 20, cell surface associated [Source:HGN |
| MUC4 | CHR_HSCHR3_5_CTG3 | mucin 4, cell surface associated [Source:HGNC |
| C3orf52 | CHR_HSCHR3_6_CTG2_1 | chromosome 3 open reading frame 52 [Source |
| GCSAM | CHR_HSCHR3_6_CTG2_1 | germinal center associated signaling and moti |
| TBILA | CHR_HSCHR3_6_CTG2_1 | TGF-beta induced lncRNA [Source:HGNC Symb |
| SLC9C1 | CHR_HSCHR3_6_CTG2_1 | solute carrier family 9 member C1 [Source:HG |
| AC128688.2 | CHR_HSCHR3_6_CTG2_1 | chromosome 1 open reading frame 106 (C1or |
| AC233280.4 | CHR_HSCHR3_6_CTG3 | protein phosphatase 4, regulatory subunit 2, p |
| AC233280.6 | CHR_HSCHR3_6_CTG3 | novel transcript |
| MUC20-OT1 | CHR_HSCHR3_6_CTG3 | MUC20 overlapping transcript [Source:HGNC S |
| SDHAP2 | CHR_HSCHR3_6_CTG3 | succinate dehydrogenase complex flavoprotei |
| MUC20 | CHR_HSCHR3_6_CTG3 | mucin 20, cell surface associated [Source:HGN |
| MUC4 | CHR_HSCHR3_6_CTG3 | mucin 4, cell surface associated [Source:HGNC |
| AC233280.13 | CHR_HSCHR3_7_CTG3 | protein phosphatase 4, regulatory subunit 2, p |
| AC233280.14 | CHR_HSCHR3_7_CTG3 | novel transcript |
| MUC20-OT1 | CHR_HSCHR3_7_CTG3 | MUC20 overlapping transcript [Source:HGNC S |
| SDHAP2 | CHR_HSCHR3_7_CTG3 | succinate dehydrogenase complex flavoprotei |
| MUC20 | CHR_HSCHR3_7_CTG3 | mucin 20, cell surface associated [Source:HGN |
| MUC4 | CHR_HSCHR3_7_CTG3 | mucin 4, cell surface associated [Source:HGNC |
| MIR4789 | CHR_HSCHR3_8_CTG2_1 | microRNA 4789 [Source:HGNC Symbol;Acc:HG |
| RNU6-1233P | CHR_HSCHR3_8_CTG2_1 | RNA, U6 small nuclear 1233, pseudogene [Sou |
| AC008180.1 | CHR_HSCHR3_8_CTG2_1 | novel transcript, antisense to NAALADL2 |
| AC233280.18 | CHR_HSCHR3_8_CTG3 | protein phosphatase 4, regulatory subunit 2, p |
| AC233280.7 | CHR_HSCHR3_8_CTG3 | novel transcript |
| MUC20-OT1 | CHR_HSCHR3_8_CTG3 | MUC20 overlapping transcript [Source:HGNC S |
| SDHAP2 | CHR_HSCHR3_8_CTG3 | succinate dehydrogenase complex flavoprotei |
| MUC20 | CHR_HSCHR3_8_CTG3 | mucin 20, cell surface associated [Source:HGN |
| MUC4 | CHR_HSCHR3_8_CTG3 | mucin 4, cell surface associated [Source:HGNC |
| RUVBL1 | CHR_HSCHR3_9_CTG2_1 | RuvB like AAA ATPase 1 [Source:HGNC Symbo |
| RNU6-823P | CHR_HSCHR3_9_CTG2_1 | RNA, U6 small nuclear 823, pseudogene [Sour |
| EEFSEC | CHR_HSCHR3_9_CTG2_1 | eukaryotic elongation factor, selenocysteine-t |
| AC084784.1 | CHR_HSCHR3_9_CTG2_1 | Novel transcript, antisense to EEFSEC |
| AC073135.6 | CHR_HSCHR3_9_CTG3 | Ankyrin repeat domain 18A (ANKRD18A) pseu |
| RNU6-821P | CHR_HSCHR3_9_CTG3 | RNA, U6 small nuclear 821, pseudogene [Sour |
| AC240720.1 | CHR_HSCHR3_9_CTG3 | FSHD region gene 2 family pseudogene |
| AC240720.2 | CHR_HSCHR3_9_CTG3 | tubulin, beta 8 (TUBB8) pseudogene |
| AC240720.3 | CHR_HSCHR3_9_CTG3 | family with sequence similarity 166, member |
| AC073135.4 | CHR_HSCHR3_9_CTG3 | family with sequence similarity 157, member |
| AC073135.5 | CHR_HSCHR3_9_CTG3 | general transcription factor IIi (GTF2I) pseudo |
| U6 | CHR_HSCHR3_9_CTG3 | U6 spliceosomal RNA [Source:RFAM;Acc:RF00 |
| AC073135.3 | CHR_HSCHR3_9_CTG3 | capicua homolog (Drosophila) (CIC) pseudoge |
| AC073135.2 | CHR_HSCHR3_9_CTG3 | septin 14 (SEPT14) pseudogene |

| | | |
|---|---|---|
| FRG1-DT | CHR_HSCHR4_11_CTG12 | FRG1 divergent transcript [Source:HGNC Symb |
| FRG1 | CHR_HSCHR4_11_CTG12 | FSHD region gene 1 [Source:HGNC Symbol;Ac |
| MLLT10P2 | CHR_HSCHR4_11_CTG12 | MLLT10 pseudogene 2 [Source:HGNC Symbol; |
| AF146191.5 | CHR_HSCHR4_11_CTG12 | family with sequence similarity 166, member |
| TUBB7P | CHR_HSCHR4_11_CTG12 | tubulin beta 7 pseudogene [Source:HGNC Sym |
| RNA5SP174 | CHR_HSCHR4_11_CTG12 | RNA, 5S ribosomal pseudogene 174 [Source:H |
| RNA5SP175 | CHR_HSCHR4_11_CTG12 | RNA, 5S ribosomal pseudogene 175 [Source:H |
| DUX4L9 | CHR_HSCHR4_11_CTG12 | double homeobox 4 like 9 (pseudogene) [Sou |
| FRG2 | CHR_HSCHR4_11_CTG12 | FSHD region gene 2 [Source:HGNC Symbol;Ac |
| RARRES2P4 | CHR_HSCHR4_11_CTG12 | retinoic acid receptor responder 2 pseudogen |
| DUX4L8 | CHR_HSCHR4_11_CTG12 | double homeobox 4 like 8 (pseudogene) [Sou |
| CT476828.18 | CHR_HSCHR4_11_CTG12 | NULL |
| CT476828.9 | CHR_HSCHR4_11_CTG12 | NULL |
| CT476828.3 | CHR_HSCHR4_11_CTG12 | NULL |
| CT476828.7 | CHR_HSCHR4_11_CTG12 | NULL |
| CT476828.20 | CHR_HSCHR4_11_CTG12 | NULL |
| CT476828.6 | CHR_HSCHR4_11_CTG12 | NULL |
| CT476828.12 | CHR_HSCHR4_11_CTG12 | NULL |
| CT476828.21 | CHR_HSCHR4_11_CTG12 | NULL |
| CT476828.11 | CHR_HSCHR4_11_CTG12 | NULL |
| CT476828.13 | CHR_HSCHR4_11_CTG12 | NULL |
| DUX4L2 | CHR_HSCHR4_11_CTG12 | double homeobox 4 like 2 (pseudogene) [Sou |
| HM190170.1 | CHR_HSCHR4_11_CTG12 | NULL |
| AC104657.1 | CHR_HSCHR4_12_CTG12 | novel transcript, antisense to DCHS2 |
| DCHS2 | CHR_HSCHR4_12_CTG12 | dachsous cadherin-related 2 [Source:HGNC Sy |
| AC079298.5 | CHR_HSCHR4_12_CTG12 | novel transcript, antisense to DCHS2 |
| AC079298.4 | CHR_HSCHR4_12_CTG12 | novel transcript, antisense to DCHS2 |
| AC110775.2 | CHR_HSCHR4_12_CTG12 | kelch domain containing 4 pseudogene |
| AC093830.1 | CHR_HSCHR4_1_CTG12 | acid sensing ion channel subunit family memb |
| TDO2 | CHR_HSCHR4_1_CTG12 | tryptophan 2,3-dioxygenase [Source:HGNC Sy |
| CTSO | CHR_HSCHR4_1_CTG12 | cathepsin O [Source:HGNC Symbol;Acc:HGNC: |
| KCNIP4 | CHR_HSCHR4_1_CTG4 | potassium voltage-gated channel interacting |
| AC210756.1 | CHR_HSCHR4_1_CTG4 | novel transcript |
| AC096576.5 | CHR_HSCHR4_1_CTG4 | ribosomal protein L31 (RPL31) pseudogene |
| AC096576.4 | CHR_HSCHR4_1_CTG4 | novel transcript |
| AC093689.2 | CHR_HSCHR4_1_CTG6 | novel transcript |
| YTHDC1 | CHR_HSCHR4_1_CTG9 | YTH domain containing 1 [Source:HGNC Symb |
| MT2P1 | CHR_HSCHR4_1_CTG9 | metallothionein 2 pseudogene 1 [Source:HGN |
| TMPRSS11E | CHR_HSCHR4_1_CTG9 | transmembrane serine protease 11E [Source: |
| UGT2B29P | CHR_HSCHR4_1_CTG9 | UDP glucuronosyltransferase family 2 membe |
| UGT2B15 | CHR_HSCHR4_1_CTG9 | UDP glucuronosyltransferase family 2 membe |
| AC019173.1 | CHR_HSCHR4_1_CTG9 | pseudogene similar to part of UDP glucuronos |
| AC226496.4 | CHR_HSCHR4_1_CTG9 | pseudogene similar to part of UDP glucuronos |
| AC226496.2 | CHR_HSCHR4_1_CTG9 | pseudogene similar to part of UDP glucuronos |
| UGT2B10 | CHR_HSCHR4_1_CTG9 | UDP glucuronosyltransferase family 2 membe |
| AC226496.3 | CHR_HSCHR4_1_CTG9 | pseudogene similar to part of UDP glucuronos |
| AC226496.7 | CHR_HSCHR4_1_CTG9 | pseudogene similar to part of UDP glucuronos |
| AC226496.10 | CHR_HSCHR4_1_CTG9 | pseudogene similar to part of UDP glucuronos |
| AC226496.1 | CHR_HSCHR4_1_CTG9 | speckle-type POZ protein-like (SPOPL) pseudo |
| AC226496.5 | CHR_HSCHR4_1_CTG9 | pseudogene similar to part of UDP glucuronos |

| | | |
|---|---|---|
| AC226496.9 | CHR_HSCHR4_1_CTG9 | novel transcript |
| AC226496.6 | CHR_HSCHR4_1_CTG9 | pseudogene similar to part of UDP glucuronos |
| AC226496.8 | CHR_HSCHR4_1_CTG9 | pseudogene similar to part of UDP glucuronos |
| UGT2A3 | CHR_HSCHR4_1_CTG9 | UDP glucuronosyltransferase family 2 membe |
| AC021146.15 | CHR_HSCHR4_1_CTG9 | pseudogene similar to part of UDP glucuronos |
| AC021146.13 | CHR_HSCHR4_1_CTG9 | UDP glucuronosyltransferase 2 family pseudo |
| AC021146.14 | CHR_HSCHR4_1_CTG9 | speckle-type POZ protein-like (SPOPL) pseudo |
| UGT2B27P | CHR_HSCHR4_1_CTG9 | UDP glucuronosyltransferase family 2 membe |
| AC110772.3 | CHR_HSCHR4_2_CTG12 | novel transcript |
| CPEB2-DT | CHR_HSCHR4_2_CTG4 | CPEB2 divergent transcript [Source:HGNC Sym |
| AF250324.6 | CHR_HSCHR4_3_CTG12 | novel transcript |
| RNU1-51P | CHR_HSCHR4_3_CTG12 | RNA, U1 small nuclear 51, pseudogene [Sourc |
| FRG1-DT | CHR_HSCHR4_3_CTG12 | FRG1 divergent transcript [Source:HGNC Symb |
| AF250324.5 | CHR_HSCHR4_3_CTG12 | novel transcript |
| AC108866.1 | CHR_HSCHR4_4_CTG12 | novel transcript, antisense to MAD2L1 |
| AC093789.2 | CHR_HSCHR4_5_CTG12 | ribosomal protein L7a (RPL7A) pseudogene |
| ICE2P1 | CHR_HSCHR4_5_CTG12 | interactor of little elongation complex ELL sub |
| LINC01060 | CHR_HSCHR4_5_CTG12 | long intergenic non-protein coding RNA 1060 |
| AF250324.2 | CHR_HSCHR4_6_CTG12 | novel transcript |
| RNU1-51P | CHR_HSCHR4_6_CTG12 | RNA, U1 small nuclear 51, pseudogene [Sourc |
| FRG1-DT | CHR_HSCHR4_6_CTG12 | FRG1 divergent transcript [Source:HGNC Symb |
| AC226374.2 | CHR_HSCHR4_6_CTG12 | novel transcript |
| FRG1 | CHR_HSCHR4_6_CTG12 | FSHD region gene 1 [Source:HGNC Symbol;Ac |
| AC226374.1 | CHR_HSCHR4_6_CTG12 | myeloid/lymphoid or mixed-lineage leukemia |
| AF146191.2 | CHR_HSCHR4_6_CTG12 | family with sequence similarity 166, member |
| TUBB7P | CHR_HSCHR4_6_CTG12 | tubulin beta 7 pseudogene [Source:HGNC Sym |
| 5S_rRNA | CHR_HSCHR4_6_CTG12 | 5S ribosomal RNA [Source:RFAM;Acc:RF00001 |
| 5S_rRNA | CHR_HSCHR4_6_CTG12 | 5S ribosomal RNA [Source:RFAM;Acc:RF00001 |
| DUX4L9 | CHR_HSCHR4_6_CTG12 | double homeobox 4 like 9 (pseudogene) [Sour |
| FRG2 | CHR_HSCHR4_6_CTG12 | FSHD region gene 2 [Source:HGNC Symbol;Ac |
| AF146191.3 | CHR_HSCHR4_6_CTG12 | retinoic acid receptor responder (tazarotene i |
| HSP90AA4P | CHR_HSCHR4_7_CTG12 | heat shock protein 90 alpha family class A me |
| AC243969.1 | CHR_HSCHR4_7_CTG12 | novel transcript |
| AF250324.3 | CHR_HSCHR4_7_CTG12 | novel transcript |
| U1 | CHR_HSCHR4_7_CTG12 | U1 spliceosomal RNA [Source:RFAM;Acc:RF00 |
| FRG1-DT | CHR_HSCHR4_7_CTG12 | FRG1 divergent transcript [Source:HGNC Symb |
| AF250324.4 | CHR_HSCHR4_7_CTG12 | novel transcript |
| AC116049.3 | CHR_HSCHR4_8_CTG12 | novel transcript |
| AC096721.2 | CHR_HSCHR4_8_CTG12 | novel transcript |
| CSN2 | CHR_HSCHR4_9_CTG12 | casein beta [Source:HGNC Symbol;Acc:HGNC: |
| STATH | CHR_HSCHR4_9_CTG12 | statherin [Source:HGNC Symbol;Acc:HGNC:11 |
| HTN3 | CHR_HSCHR4_9_CTG12 | histatin 3 [Source:HGNC Symbol;Acc:HGNC:52 |
| HTN1 | CHR_HSCHR4_9_CTG12 | histatin 1 [Source:HGNC Symbol;Acc:HGNC:52 |
| CSN1S2AP | CHR_HSCHR4_9_CTG12 | casein alpha s2 like A, pseudogene [Source:HG |
| PRR27 | CHR_HSCHR4_9_CTG12 | proline rich 27 [Source:HGNC Symbol;Acc:HGI |
| LINC01194 | CHR_HSCHR5_1_CTG1 | long intergenic non-protein coding RNA 1194 |
| AC245690.1 | CHR_HSCHR5_1_CTG1_1 | glucuronidase, beta (GUSB) pseudogene |
| AC245690.2 | CHR_HSCHR5_1_CTG1_1 | glucuronidase, beta (GUSB) pseudogene |
| CDH12P2 | CHR_HSCHR5_1_CTG1_1 | cadherin 12 pseudogene 2 [Source:HGNC Sym |
| SERF1B | CHR_HSCHR5_1_CTG1_1 | small EDRK-rich factor 1B [Source:HGNC Symb |

| | | |
|---|---|---|
| SMN2 | CHR_HSCHR5_1_CTG1_1 | survival of motor neuron 2, centromeric [Sour |
| AC244635.3 | CHR_HSCHR5_1_CTG1_1 | novel transcript, antisense to SMN2 |
| AC244635.2 | CHR_HSCHR5_1_CTG1_1 | NLR family, apoptosis inhibitory protein pseuc |
| AC244635.1 | CHR_HSCHR5_1_CTG1_1 | cadherin 12 (N-cadherin 2) (CDH12) pseudoge |
| SERF1A | CHR_HSCHR5_1_CTG1_1 | small EDRK-rich factor 1A [Source:HGNC Symb |
| SMN1 | CHR_HSCHR5_1_CTG1_1 | survival of motor neuron 1, telomeric [Source |
| AC244260.2 | CHR_HSCHR5_1_CTG1_1 | novel transcript, antisense to SMN1 |
| NAIP | CHR_HSCHR5_1_CTG1_1 | NLR family apoptosis inhibitory protein [Sourc |
| GTF2H2 | CHR_HSCHR5_1_CTG1_1 | general transcription factor IIH subunit 2 [Sou |
| AC244260.1 | CHR_HSCHR5_1_CTG1_1 | NLR family, apoptosis inhibitory protein pseuc |
| AC243922.3 | CHR_HSCHR5_1_CTG1_1 | glucuronidase, beta (GUSB) pseudogene |
| AC243922.2 | CHR_HSCHR5_1_CTG1_1 | glucuronidase, beta (GUSB) pseudogene |
| AC243922.1 | CHR_HSCHR5_1_CTG1_1 | glucuronidase, beta (GUSB) pseudogene |
| LINC02197 | CHR_HSCHR5_1_CTG1_1 | long intergenic non-protein coding RNA 2197 |
| AC245931.2 | CHR_HSCHR5_1_CTG1_1 | novel transcript |
| PMCHL2 | CHR_HSCHR5_1_CTG1_1 | pro-melanin concentrating hormone like 2 (ps |
| AC245931.1 | CHR_HSCHR5_1_CTG1_1 | uncharacterized LOC107987434 [Source:NCBI |
| BDP1 | CHR_HSCHR5_1_CTG1_1 | B double prime 1, subunit of RNA polymerase |
| MCCC2 | CHR_HSCHR5_1_CTG1_1 | methylcrotonoyl-CoA carboxylase 2 [Source:H |
| AC091996.1 | CHR_HSCHR5_1_CTG5 | novel transcript |
| RNU6-164P | CHR_HSCHR5_1_CTG5 | RNA, U6 small nuclear 164, pseudogene [Sour |
| AC025771.2 | CHR_HSCHR5_2_CTG1 | NULL |
| AC025771.1 | CHR_HSCHR5_2_CTG1 | aldehyde dehydrogenase 6 family, member A1 |
| LINC02109 | CHR_HSCHR5_2_CTG1 | long intergenic non-protein coding RNA 2109 |
| AC112172.1 | CHR_HSCHR5_2_CTG1 | novel transcript |
| MRPS36 | CHR_HSCHR5_2_CTG1_1 | mitochondrial ribosomal protein S36 [Source: |
| CDK7 | CHR_HSCHR5_2_CTG1_1 | cyclin dependent kinase 7 [Source:HGNC Syml |
| CCDC125 | CHR_HSCHR5_2_CTG1_1 | coiled-coil domain containing 125 [Source:HG |
| CFL1P5 | CHR_HSCHR5_2_CTG1_1 | cofilin 1 pseudogene 5 [Source:HGNC Symbol; |
| CHCHD2P2 | CHR_HSCHR5_2_CTG1_1 | coiled-coil-helix-coiled-coil-helix domain conta |
| NDUFB9P1 | CHR_HSCHR5_2_CTG1_1 | NADH:ubiquinone oxidoreductase subunit B9 |
| AK6 | CHR_HSCHR5_2_CTG1_1 | adenylate kinase 6 [Source:HGNC Symbol;Acc |
| TAF9 | CHR_HSCHR5_2_CTG1_1 | TATA-box binding protein associated factor 9 |
| RAD17 | CHR_HSCHR5_2_CTG1_1 | RAD17 checkpoint clamp loader component [S |
| MARVELD2 | CHR_HSCHR5_2_CTG1_1 | MARVEL domain containing 2 [Source:HGNC S |
| RPS27P14 | CHR_HSCHR5_2_CTG1_1 | ribosomal protein S27 pseudogene 14 [Source |
| AC142525.2 | CHR_HSCHR5_2_CTG1_1 | novel transcript |
| Metazoa_SRP | CHR_HSCHR5_2_CTG1_1 | Metazoan signal recognition particle RNA [Sou |
| OCLN | CHR_HSCHR5_2_CTG1_1 | occludin [Source:HGNC Symbol;Acc:HGNC:810 |
| U6 | CHR_HSCHR5_2_CTG1_1 | U6 spliceosomal RNA [Source:RFAM;Acc:RF00 |
| AC142525.4 | CHR_HSCHR5_2_CTG1_1 | NULL |
| GTF2H2 | CHR_HSCHR5_2_CTG1_1 | general transcription factor IIH subunit 2 [Sou |
| AC142525.1 | CHR_HSCHR5_2_CTG1_1 | novel transcript |
| NAIP | CHR_HSCHR5_2_CTG1_1 | NLR family apoptosis inhibitory protein [Sourc |
| AC142525.3 | CHR_HSCHR5_2_CTG1_1 | TEC |
| AC139494.7 | CHR_HSCHR5_2_CTG1_1 | TEC |
| SMN2 | CHR_HSCHR5_2_CTG1_1 | survival of motor neuron 2, centromeric [Sour |
| AC139494.8 | CHR_HSCHR5_2_CTG1_1 | novel transcript, antisense to SMN2 |
| SERF1B | CHR_HSCHR5_2_CTG1_1 | small EDRK-rich factor 1B [Source:HGNC Symb |
| AC139494.1 | CHR_HSCHR5_2_CTG1_1 | glucuronidase, beta (GUSB) pseudogene |

| | | |
|---|---|---|
| AC139494.3 | CHR_HSCHR5_2_CTG1_1 | novel transcript |
| AC139494.6 | CHR_HSCHR5_2_CTG1_1 | novel transcript |
| AC139494.2 | CHR_HSCHR5_2_CTG1_1 | novel transcript |
| AC139494.5 | CHR_HSCHR5_2_CTG1_1 | cadherin 12 (N-cadherin 2) (CDH12) pseudoge |
| AC139494.4 | CHR_HSCHR5_2_CTG1_1 | novel transcript |
| AC118459.1 | CHR_HSCHR5_2_CTG1_1 | novel transcript |
| AC118459.2 | CHR_HSCHR5_2_CTG1_1 | glucuronidase, beta (GUSB) pseudogene |
| AC108108.1 | CHR_HSCHR5_2_CTG1_1 | glucuronidase, beta (GUSB) pseudogene |
| AC138827.1 | CHR_HSCHR5_2_CTG1_1 | NLR family, apoptosis inhibitory protein (NAIP |
| AC138827.2 | CHR_HSCHR5_2_CTG1_1 | novel transcript |
| GTF2H2C_2 | CHR_HSCHR5_2_CTG1_1 | GTF2H2 family member C, copy 2 [Source:HGI |
| AC138827.3 | CHR_HSCHR5_2_CTG1_1 | occludin (OCLN) pseudogene |
| AC146335.2 | CHR_HSCHR5_2_CTG1_1 | NLR family, apoptosis inhibitory protein (NAIP |
| CDH12P4 | CHR_HSCHR5_2_CTG1_1 | cadherin 12 pseudogene 4 [Source:HGNC Sym |
| AC146335.1 | CHR_HSCHR5_2_CTG1_1 | glucuronidase, beta (GUSB) pseudogene |
| LINC02197 | CHR_HSCHR5_2_CTG1_1 | long intergenic non-protein coding RNA 2197 |
| AC139463.1 | CHR_HSCHR5_2_CTG1_1 | novel transcript |
| PMCHL2 | CHR_HSCHR5_2_CTG1_1 | pro-melanin concentrating hormone like 2 (ps |
| AC138832.3 | CHR_HSCHR5_2_CTG1_1 | uncharacterized LOC107987434 [Source:NCBI |
| BDP1 | CHR_HSCHR5_2_CTG1_1 | B double prime 1, subunit of RNA polymerase |
| AC138832.2 | CHR_HSCHR5_2_CTG1_1 | sterol carrier protein (SCP2) pseudogene |
| HMGN1P12 | CHR_HSCHR5_2_CTG1_1 | high mobility group nucleosome binding doma |
| MCCC2 | CHR_HSCHR5_2_CTG1_1 | methylcrotonoyl-CoA carboxylase 2 [Source:H |
| AC106795.8 | CHR_HSCHR5_2_CTG5 | novel transcript |
| AC114936.1 | CHR_HSCHR5_2_CTG5 | THO complex 3 (THOC3) pseudogene |
| AC114936.3 | CHR_HSCHR5_2_CTG5 | novel transcript |
| AC114936.2 | CHR_HSCHR5_2_CTG5 | novel transcript, antisense to PROP1 |
| AC114936.4 | CHR_HSCHR5_2_CTG5 | novel transcript |
| SUDS3P1 | CHR_HSCHR5_2_CTG5 | SDS3 homolog, SIN3A corepressor complex cc |
| PROP1 | CHR_HSCHR5_2_CTG5 | PROP paired-like homeobox 1 [Source:HGNC S |
| CLPTM1L | CHR_HSCHR5_3_CTG1 | CLPTM1 like [Source:HGNC Symbol;Acc:HGNC |
| LINC01511 | CHR_HSCHR5_3_CTG1 | long intergenic non-protein coding RNA 1511 |
| MTCO2P32 | CHR_HSCHR5_3_CTG1 | MT-CO2 pseudogene 32 [Source:HGNC Symbo |
| AC026748.2 | CHR_HSCHR5_3_CTG1 | tec |
| SLC6A3 | CHR_HSCHR5_3_CTG1 | solute carrier family 6 member 3 [Source:HGN |
| LPCAT1 | CHR_HSCHR5_3_CTG1 | lysophosphatidylcholine acyltransferase 1 [So |
| AC106795.6 | CHR_HSCHR5_3_CTG5 | novel transcript |
| AC106795.4 | CHR_HSCHR5_3_CTG5 | THO complex 3 (THOC3) pseudogene |
| AC106795.7 | CHR_HSCHR5_3_CTG5 | novel transcript |
| AC106795.9 | CHR_HSCHR5_3_CTG5 | novel transcript, antisense to PROP1 |
| AC236592.1 | CHR_HSCHR5_3_CTG5 | novel transcript |
| SUDS3P1 | CHR_HSCHR5_3_CTG5 | SDS3 homolog, SIN3A corepressor complex cc |
| PROP1 | CHR_HSCHR5_3_CTG5 | PROP paired-like homeobox 1 [Source:HGNC S |
| AC116351.3 | CHR_HSCHR5_4_CTG1 | novel transcript |
| AC116351.4 | CHR_HSCHR5_4_CTG1 | novel transcript |
| NKD2 | CHR_HSCHR5_4_CTG1 | NKD inhibitor of WNT signaling pathway 2 [So |
| SLC12A7 | CHR_HSCHR5_4_CTG1 | solute carrier family 12 member 7 [Source:HG |
| AC109471.2 | CHR_HSCHR5_4_CTG1_1 | novel transcript |
| RN7SKP117 | CHR_HSCHR5_4_CTG1_1 | RN7SK pseudogene 117 [Source:HGNC Symbo |
| AC109471.3 | CHR_HSCHR5_4_CTG1_1 | novel transcript |

| | | |
|---|---|---|
| SLC9A3 | CHR_HSCHR5_5_CTG1 | solute carrier family 9 member A3 [Source:HG |
| AC106772.3 | CHR_HSCHR5_5_CTG1 | novel transcript |
| AC106772.4 | CHR_HSCHR5_5_CTG1 | novel transcript |
| CEP72 | CHR_HSCHR5_5_CTG1 | centrosomal protein 72 [Source:HGNC Symbo |
| ADAMTS12 | CHR_HSCHR5_6_CTG1 | ADAM metallopeptidase with thrombospondi |
| U6 | CHR_HSCHR5_6_CTG1 | U6 spliceosomal RNA [Source:RFAM;Acc:RF00 |
| RXFP3 | CHR_HSCHR5_6_CTG1 | relaxin family peptide receptor 3 [Source:HGN |
| SLC45A2 | CHR_HSCHR5_6_CTG1 | solute carrier family 45 member 2 [Source:HG |
| AC208067.1 | CHR_HSCHR5_7_CTG1 | stearoyl-CoA desaturase (delta-9-desaturase) |
| AC270107.1 | CHR_HSCHR5_8_CTG1 | NULL |
| AC270107.3 | CHR_HSCHR5_8_CTG1 | NULL |
| AC270107.8 | CHR_HSCHR5_8_CTG1 | TEC |
| AC270107.2 | CHR_HSCHR5_8_CTG1 | NULL |
| AC270107.7 | CHR_HSCHR5_8_CTG1 | NULL |
| AC270107.9 | CHR_HSCHR5_8_CTG1 | NULL |
| AC270107.6 | CHR_HSCHR5_8_CTG1 | NULL |
| AC270107.5 | CHR_HSCHR5_8_CTG1 | NULL |
| AC270107.4 | CHR_HSCHR5_8_CTG1 | novel transcript |
| AC091946.3 | CHR_HSCHR5_8_CTG1 | novel transcript |
| AL078601.5 | CHR_HSCHR6_1_CTG2 | high-mobility group (nonhistone chromosoma |
| AC209234.1 | CHR_HSCHR6_1_CTG2 | novel solute carrier family 25 family pseudoge |
| AL078601.6 | CHR_HSCHR6_1_CTG2 | novel solute carrier family 25 family pseudoge |
| AL078601.3 | CHR_HSCHR6_1_CTG2 | novel transcript |
| AL078601.4 | CHR_HSCHR6_1_CTG2 | diazepam binding inhibitor (DBI) pseudogene |
| AL513210.3 | CHR_HSCHR6_1_CTG3 | TEC |
| AL354892.5 | CHR_HSCHR6_1_CTG4 | novel transcript |
| AL354892.4 | CHR_HSCHR6_1_CTG4 | novel transcript |
| TCTE3 | CHR_HSCHR6_1_CTG4 | t-complex-associated-testis-expressed 3 [Sour |
| ERMARD | CHR_HSCHR6_1_CTG4 | ER membrane associated RNA degradation [So |
| AL354892.6 | CHR_HSCHR6_1_CTG4 | novel transcript |
| LINC00242 | CHR_HSCHR6_1_CTG4 | long intergenic non-protein coding RNA 242 [S |
| LINC00574 | CHR_HSCHR6_1_CTG4 | long intergenic non-protein coding RNA 574 [S |
| AL049612.2 | CHR_HSCHR6_1_CTG4 | novel transcript |
| LINC01624 | CHR_HSCHR6_1_CTG5 | long intergenic non-protein coding RNA 1624 |
| AL078605.2 | CHR_HSCHR6_1_CTG5 | novel transcript |
| DLL1 | CHR_HSCHR6_1_CTG5 | delta like canonical Notch ligand 1 [Source:HG |
| FAM120B | CHR_HSCHR6_1_CTG5 | family with sequence similarity 120B [Source: |
| AL078605.3 | CHR_HSCHR6_1_CTG5 | novel transcript |
| AL008628.2 | CHR_HSCHR6_1_CTG5 | novel transcript, overlapping to FAM120B |
| PSMB1 | CHR_HSCHR6_1_CTG5 | proteasome subunit beta 1 [Source:HGNC Syn |
| LINC02549 | CHR_HSCHR6_1_CTG6 | long intergenic non-protein coding RNA 2549 |
| SLC25A51P1 | CHR_HSCHR6_1_CTG7 | SLC25A51 pseudogene 1 [Source:HGNC Symb |
| ADH5P4 | CHR_HSCHR6_1_CTG7 | ADH5 pseudogene 4 [Source:HGNC Symbol;Ac |
| AL356432.5 | CHR_HSCHR6_1_CTG8 | transducin (beta)-like 1 X-linked receptor 1 (T |
| AL356432.4 | CHR_HSCHR6_1_CTG8 | transducin (beta)-like 1 X-linked receptor 1 (T |
| AL356432.6 | CHR_HSCHR6_1_CTG8 | transducin (beta)-like 1 X-linked receptor 1 (T |
| LINC02536 | CHR_HSCHR6_1_CTG8 | long intergenic non-protein coding RNA 2536 |
| THEMIS | CHR_HSCHR6_1_CTG8 | thymocyte selection associated [Source:HGN |
| MRPS17P5 | CHR_HSCHR6_1_CTG8 | mitochondrial ribosomal protein S17 pseudog |
| PTPRK | CHR_HSCHR6_1_CTG8 | protein tyrosine phosphatase receptor type K |

| | | |
|---|---|---|
| AL590006.2 | CHR_HSCHR6_1_CTG8 | novel transcript, antisense to PTPRK |
| AL034349.2 | CHR_HSCHR6_1_CTG8 | novel transcript, antisense to PTPRK |
| AL121956.3 | CHR_HSCHR6_1_CTG9 | heterogeneous nuclear ribonucleoprotein D (/ |
| AL121956.2 | CHR_HSCHR6_1_CTG9 | novel transcript |
| GNG5P1 | CHR_HSCHR6_1_CTG9 | G protein subunit gamma 5 pseudogene 1 [So |
| AC233714.1 | CHR_HSCHR6_1_CTG9 | novel transcript |
| AC226007.1 | CHR_HSCHR6_8_CTG1 | NULL |
| AL662796.1 | CHR_HSCHR6_8_CTG1 | NULL |
| NOP56P1 | CHR_HSCHR6_MHC_APD_CT( | NOP56 ribonucleoprotein pseudogene 1 [Sour |
| CR759954.1 | CHR_HSCHR6_MHC_APD_CT( | novel transcript |
| LINC01623 | CHR_HSCHR6_MHC_APD_CT( | long intergenic non-protein coding RNA 1623 |
| RPL13P | CHR_HSCHR6_MHC_APD_CT( | ribosomal protein L13 pseudogene [Source:H( |
| ZNF90P2 | CHR_HSCHR6_MHC_APD_CT( | zinc finger protein 90 pseudogene 2 [Source:H |
| HCG14 | CHR_HSCHR6_MHC_APD_CT( | HLA complex group 14 [Source:HGNC Symbol; |
| TRIM27 | CHR_HSCHR6_MHC_APD_CT( | tripartite motif containing 27 [Source:HGNC S |
| U6 | CHR_HSCHR6_MHC_APD_CT( | U6 spliceosomal RNA [Source:RFAM;Acc:RF00 |
| HCG15 | CHR_HSCHR6_MHC_APD_CT( | HLA complex group 15 [Source:HGNC Symbol; |
| CR759957.1 | CHR_HSCHR6_MHC_APD_CT( | HLA complex group 16 |
| ZNF311 | CHR_HSCHR6_MHC_APD_CT( | zinc finger protein 311 [Source:HGNC Symbol; |
| OR2AD1P | CHR_HSCHR6_MHC_APD_CT( | olfactory receptor family 2 subfamily AD mem |
| OR2W1 | CHR_HSCHR6_MHC_APD_CT( | olfactory receptor family 2 subfamily W meml |
| OR2B3 | CHR_HSCHR6_MHC_APD_CT( | olfactory receptor family 2 subfamily B memb |
| OR2J1 | CHR_HSCHR6_MHC_APD_CT( | olfactory receptor family 2 subfamily J membe |
| OR2J3 | CHR_HSCHR6_MHC_APD_CT( | olfactory receptor family 2 subfamily J membe |
| CR942175.1 | CHR_HSCHR6_MHC_APD_CT( | novel transcript |
| CR774181.1 | CHR_HSCHR6_MHC_APD_CT( | olfactory receptor, family 2, subfamily N, men |
| OR2J2 | CHR_HSCHR6_MHC_APD_CT( | olfactory receptor family 2 subfamily J membe |
| CR936918.1 | CHR_HSCHR6_MHC_APD_CT( | novel transcript |
| OR2U2P | CHR_HSCHR6_MHC_APD_CT( | olfactory receptor family 2 subfamily U memb |
| CR936918.2 | CHR_HSCHR6_MHC_APD_CT( | olfactory receptor, family 2, subfamily B, men |
| OR14J1 | CHR_HSCHR6_MHC_APD_CT( | olfactory receptor family 14 subfamily J meml |
| OR5V1 | CHR_HSCHR6_MHC_APD_CT( | olfactory receptor family 5 subfamily V memb |
| OR11A1 | CHR_HSCHR6_MHC_APD_CT( | olfactory receptor family 11 subfamily A mem |
| OR10C1 | CHR_HSCHR6_MHC_APD_CT( | olfactory receptor family 10 subfamily C mem |
| OR12D3 | CHR_HSCHR6_MHC_APD_CT( | olfactory receptor family 12 subfamily D mem |
| OR2H1 | CHR_HSCHR6_MHC_APD_CT( | olfactory receptor family 2 subfamily H memb |
| UBDP1 | CHR_HSCHR6_MHC_APD_CT( | ubiquitin D pseudogene 1 [Source:HGNC Symb |
| MAS1LP1 | CHR_HSCHR6_MHC_APD_CT( | MAS1L pseudogene 1 [Source:HGNC Symbol;/ |
| MAS1L | CHR_HSCHR6_MHC_APD_CT( | MAS1 proto-oncogene like, G protein-couplec |
| RPS17P1 | CHR_HSCHR6_MHC_APD_CT( | ribosomal protein S17 pseudogene 1 [Source: |
| LINC02829 | CHR_HSCHR6_MHC_APD_CT( | long intergenic non-protein coding RNA 2829 |
| LINC01015 | CHR_HSCHR6_MHC_APD_CT( | long intergenic non-protein coding RNA 1015 |
| GPR53P | CHR_HSCHR6_MHC_APD_CT( | G protein-coupled receptor 53, pseudogene [S |
| OR2I1P | CHR_HSCHR6_MHC_APD_CT( | olfactory receptor family 2 subfamily I membe |
| GABBR1 | CHR_HSCHR6_MHC_APD_CT( | gamma-aminobutyric acid type B receptor sub |
| UBD | CHR_HSCHR6_MHC_APD_CT( | ubiquitin D [Source:HGNC Symbol;Acc:HGNC:: |
| OR2H5P | CHR_HSCHR6_MHC_APD_CT( | olfactory receptor family 2 subfamily H memb |
| TMEM183AP1 | CHR_HSCHR6_MHC_APD_CT( | transmembrane protein 183A pseudogene 1 [ |
| SNORD32B | CHR_HSCHR6_MHC_APD_CT( | small nucleolar RNA, C/D box 32B [Source:HG |
| RPL13AP | CHR_HSCHR6_MHC_APD_CT( | ribosomal protein L13a pseudogene [Source:H |

| | | |
|---|---|---|
| OR2H2 | CHR_HSCHR6_MHC_APD_CT( | olfactory receptor family 2 subfamily H memb |
| SUMO2P1 | CHR_HSCHR6_MHC_APD_CT( | SUMO2 pseudogene 1 [Source:HGNC Symbol; |
| MOG | CHR_HSCHR6_MHC_APD_CT( | myelin oligodendrocyte glycoprotein [Source: |
| ZFP57 | CHR_HSCHR6_MHC_APD_CT( | ZFP57 zinc finger protein [Source:HGNC Symb |
| HLA-V | CHR_HSCHR6_MHC_APD_CT( | major histocompatibility complex, class I, V (p |
| HCG4 | CHR_HSCHR6_MHC_APD_CT( | HLA complex group 4 [Source:HGNC Symbol;A |
| HLA-P | CHR_HSCHR6_MHC_APD_CT( | major histocompatibility complex, class I, P (p |
| RPL7AP7 | CHR_HSCHR6_MHC_APD_CT( | ribosomal protein L7a pseudogene 7 [Source:I |
| MICG | CHR_HSCHR6_MHC_APD_CT( | MHC class I polypeptide-related sequence G (|
| HLA-G | CHR_HSCHR6_MHC_APD_CT( | major histocompatibility complex, class I, G [S |
| HCG4P8 | CHR_HSCHR6_MHC_APD_CT( | HLA complex group 4 pseudogene 8 [Source:H |
| CR925767.1 | CHR_HSCHR6_MHC_APD_CT( | MHC class I polypeptide-related sequence F p: |
| CT009496.1 | CHR_HSCHR6_MHC_APD_CT( | HLA complex group 4 pseudogene 7 |
| HLA-A | CHR_HSCHR6_MHC_APD_CT( | major histocompatibility complex, class I, A [S |
| HLA-H | CHR_HSCHR6_MHC_APD_CT( | major histocompatibility complex, class I, H (p |
| HLA-T | CHR_HSCHR6_MHC_APD_CT( | major histocompatibility complex, class I, T (p |
| DDX39BP1 | CHR_HSCHR6_MHC_APD_CT( | DEAD-box helicase 39B pseudogene 1 [Source |
| MCCD1P1 | CHR_HSCHR6_MHC_APD_CT( | mitochondrial coiled-coil domain 1 pseudoger |
| HCG4B | CHR_HSCHR6_MHC_APD_CT( | HLA complex group 4B [Source:HGNC Symbol; |
| HLA-K | CHR_HSCHR6_MHC_APD_CT( | major histocompatibility complex, class I, K (p |
| HLA-U | CHR_HSCHR6_MHC_APD_CT( | major histocompatibility complex, class I, U (p |
| CR759913.1 | CHR_HSCHR6_MHC_APD_CT( | HLA complex group 4 pseudogene 5 |
| HLA-W | CHR_HSCHR6_MHC_APD_CT( | major histocompatibility complex, class I, W (|
| DDX39BP2 | CHR_HSCHR6_MHC_APD_CT( | DEAD-box helicase 39B pseudogene 2 [Source |
| MCCD1P2 | CHR_HSCHR6_MHC_APD_CT( | mitochondrial coiled-coil domain 1 pseudoger |
| ZNRD1ASP | CHR_HSCHR6_MHC_APD_CT( | zinc ribbon domain containing 1 antisense, ps |
| CR759960.1 | CHR_HSCHR6_MHC_APD_CT( | HLA complex group 4 pseudogene 3 |
| HLA-J | CHR_HSCHR6_MHC_APD_CT( | major histocompatibility complex, class I, J (ps |
| ETF1P1 | CHR_HSCHR6_MHC_APD_CT( | eukaryotic translation termination factor 1 ps |
| CR759960.2 | CHR_HSCHR6_MHC_APD_CT( | NULL |
| CR759960.4 | CHR_HSCHR6_MHC_APD_CT( | NULL |
| ZNRD1 | CHR_HSCHR6_MHC_APD_CT( | zinc ribbon domain containing 1 [Source:HGN |
| CR759960.3 | CHR_HSCHR6_MHC_APD_CT( | NULL |
| PPP1R11 | CHR_HSCHR6_MHC_APD_CT( | protein phosphatase 1 regulatory inhibitor sul |
| RNF39 | CHR_HSCHR6_MHC_APD_CT( | ring finger protein 39 [Source:HGNC Symbol;A |
| TRIM31 | CHR_HSCHR6_MHC_APD_CT( | tripartite motif containing 31 [Source:HGNC S |
| TRIM31-AS1 | CHR_HSCHR6_MHC_APD_CT( | TRIM31 antisense RNA 1 [Source:HGNC Symb |
| CR925727.1 | CHR_HSCHR6_MHC_APD_CT( | NULL |
| TRIM40 | CHR_HSCHR6_MHC_APD_CT( | tripartite motif containing 40 [Source:HGNC S |
| TRIM10 | CHR_HSCHR6_MHC_APD_CT( | tripartite motif containing 10 [Source:HGNC S |
| TRIM15 | CHR_HSCHR6_MHC_APD_CT( | tripartite motif containing 15 [Source:HGNC S |
| TRIM26 | CHR_HSCHR6_MHC_APD_CT( | tripartite motif containing 26 [Source:HGNC S |
| PAIP1P1 | CHR_HSCHR6_MHC_APD_CT( | poly(A) binding protein interacting protein 1 p |
| HCG17 | CHR_HSCHR6_MHC_APD_CT( | HLA complex group 17 [Source:HGNC Symbol; |
| TRIM26BP | CHR_HSCHR6_MHC_APD_CT( | tripartite motif containing 26B, pseudogene [S |
| HCG18 | CHR_HSCHR6_MHC_APD_CT( | HLA complex group 18 [Source:HGNC Symbol; |
| TRIM39 | CHR_HSCHR6_MHC_APD_CT( | tripartite motif containing 39 [Source:HGNC S |
| RPP21 | CHR_HSCHR6_MHC_APD_CT( | ribonuclease P/MRP subunit p21 [Source:HGN |
| HLA-N | CHR_HSCHR6_MHC_APD_CT( | major histocompatibility complex, class I, N (p |
| CR759890.1 | CHR_HSCHR6_MHC_APD_CT( | HLA complex group 19 pseudogene |

| | | |
|---|---|---|
| UBQLN1P1 | CHR_HSCHR6_MHC_APD_CT( | ubiquilin 1 pseudogene 1 [Source:HGNC Symb |
| MICC | CHR_HSCHR6_MHC_APD_CT( | MHC class I polypeptide-related sequence C (p |
| LINC02569 | CHR_HSCHR6_MHC_APD_CT( | long intergenic non-protein coding RNA 2569 |
| GNL1 | CHR_HSCHR6_MHC_APD_CT( | G protein nucleolar 1 (putative) [Source:HGNC |
| DHX16 | CHR_HSCHR6_MHC_APD_CT( | DEAH-box helicase 16 [Source:HGNC Symbol;A |
| PPP1R18 | CHR_HSCHR6_MHC_APD_CT( | protein phosphatase 1 regulatory subunit 18 [ |
| NRM | CHR_HSCHR6_MHC_APD_CT( | nurim [Source:HGNC Symbol;Acc:HGNC:8003] |
| RPL7P4 | CHR_HSCHR6_MHC_APD_CT( | ribosomal protein L7 pseudogene 4 [Source:H |
| MDC1 | CHR_HSCHR6_MHC_APD_CT( | mediator of DNA damage checkpoint 1 [Sourc |
| MDC1-AS1 | CHR_HSCHR6_MHC_APD_CT( | MDC1 antisense RNA 1 [Source:HGNC Symbol |
| TUBB | CHR_HSCHR6_MHC_APD_CT( | tubulin beta class I [Source:HGNC Symbol;Acc |
| FLOT1 | CHR_HSCHR6_MHC_APD_CT( | flotillin 1 [Source:HGNC Symbol;Acc:HGNC:37 |
| DDR1 | CHR_HSCHR6_MHC_APD_CT( | discoidin domain receptor tyrosine kinase 1 [S |
| MUC21 | CHR_HSCHR6_MHC_APD_CT( | mucin 21, cell surface associated [Source:HGN |
| CR759964.2 | CHR_HSCHR6_MHC_APD_CT( | novel transcript |
| CR759964.1 | CHR_HSCHR6_MHC_APD_CT( | HLA complex group family pseudogene |
| MICB | CHR_HSCHR6_MHC_APD_CT( | MHC class I polypeptide-related sequence B [S |
| Y_RNA | CHR_HSCHR6_MHC_APD_CT( | Y RNA [Source:RFAM;Acc:RF00019] |
| PPIAP9 | CHR_HSCHR6_MHC_APD_CT( | peptidylprolyl isomerase A pseudogene 9 [Sou |
| RPL15P4 | CHR_HSCHR6_MHC_APD_CT( | ribosomal protein L15 pseudogene 4 [Source: |
| MCCD1 | CHR_HSCHR6_MHC_APD_CT( | mitochondrial coiled-coil domain 1 [Source:HC |
| DDX39B | CHR_HSCHR6_MHC_APD_CT( | DExD-box helicase 39B [Source:HGNC Symbol |
| SNORD117 | CHR_HSCHR6_MHC_APD_CT( | small nucleolar RNA, C/D box 117 [Source:HGI |
| TNF | CHR_HSCHR6_MHC_APD_CT( | tumor necrosis factor [Source:HGNC Symbol;A |
| LTB | CHR_HSCHR6_MHC_APD_CT( | lymphotoxin beta [Source:HGNC Symbol;Acc:I |
| LST1 | CHR_HSCHR6_MHC_APD_CT( | leukocyte specific transcript 1 [Source:HGNC S |
| NCR3 | CHR_HSCHR6_MHC_APD_CT( | natural cytotoxicity triggering receptor 3 [Sou |
| UQCRHP1 | CHR_HSCHR6_MHC_APD_CT( | ubiquinol-cytochrome c reductase hinge prote |
| MSH5 | CHR_HSCHR6_MHC_APD_CT( | mutS homolog 5 [Source:HGNC Symbol;Acc:H |
| U6 | CHR_HSCHR6_MHC_APD_CT( | U6 spliceosomal RNA [Source:RFAM;Acc:RF00 |
| MSH5-SAPCD1 | CHR_HSCHR6_MHC_APD_CT( | MSH5-SAPCD1 readthrough (NMD candidate) |
| SAPCD1 | CHR_HSCHR6_MHC_APD_CT( | suppressor APC domain containing 1 [Source:I |
| SAPCD1-AS1 | CHR_HSCHR6_MHC_APD_CT( | SAPCD1 antisense RNA 1 [Source:HGNC Symb |
| VWA7 | CHR_HSCHR6_MHC_APD_CT( | von Willebrand factor A domain containing 7 [ |
| VARS | CHR_HSCHR6_MHC_APD_CT( | valyl-tRNA synthetase [Source:HGNC Symbol;A |
| Y_RNA | CHR_HSCHR6_MHC_APD_CT( | Y RNA [Source:RFAM;Acc:RF00019] |
| LSM2 | CHR_HSCHR6_MHC_APD_CT( | LSM2 homolog, U6 small nuclear RNA and mR |
| HSPA1L | CHR_HSCHR6_MHC_APD_CT( | heat shock protein family A (Hsp70) member : |
| HSPA1A | CHR_HSCHR6_MHC_APD_CT( | heat shock protein family A (Hsp70) member : |
| HSPA1B | CHR_HSCHR6_MHC_APD_CT( | heat shock protein family A (Hsp70) member : |
| SNHG32 | CHR_HSCHR6_MHC_APD_CT( | small nucleolar RNA host gene 32 [Source:HGI |
| SNORD48 | CHR_HSCHR6_MHC_APD_CT( | small nucleolar RNA, C/D box 48 [Source:HGN |
| SNORD52 | CHR_HSCHR6_MHC_APD_CT( | small nucleolar RNA, C/D box 52 [Source:HGN |
| NEU1 | CHR_HSCHR6_MHC_APD_CT( | neuraminidase 1 [Source:HGNC Symbol;Acc:H |
| SLC44A4 | CHR_HSCHR6_MHC_APD_CT( | solute carrier family 44 member 4 [Source:HG |
| EHMT2 | CHR_HSCHR6_MHC_APD_CT( | euchromatic histone lysine methyltransferase |
| CR936237.1 | CHR_HSCHR6_MHC_APD_CT( | novel transcript |
| TNXB | CHR_HSCHR6_MHC_APD_CT( | tenascin XB [Source:HGNC Symbol;Acc:HGNC: |
| 5S_rRNA | CHR_HSCHR6_MHC_APD_CT( | 5S ribosomal RNA [Source:RFAM;Acc:RF00001 |
| PRRT1 | CHR_HSCHR6_MHC_APD_CT( | proline rich transmembrane protein 1 [Source |

| | | |
|---|---|---|
| PPT2 | CHR_HSCHR6_MHC_APD_CT( | palmitoyl-protein thioesterase 2 [Source:HGN |
| EGFL8 | CHR_HSCHR6_MHC_APD_CT( | EGF like domain multiple 8 [Source:HGNC Sym |
| AGPAT1 | CHR_HSCHR6_MHC_APD_CT( | 1-acylglycerol-3-phosphate O-acyltransferase |
| RNF5 | CHR_HSCHR6_MHC_APD_CT( | ring finger protein 5 [Source:HGNC Symbol;Ac |
| AGER | CHR_HSCHR6_MHC_APD_CT( | advanced glycosylation end-product specific r |
| PBX2 | CHR_HSCHR6_MHC_APD_CT( | PBX homeobox 2 [Source:HGNC Symbol;Acc:H |
| GPSM3 | CHR_HSCHR6_MHC_APD_CT( | G protein signaling modulator 3 [Source:HGN( |
| NOTCH4 | CHR_HSCHR6_MHC_APD_CT( | notch receptor 4 [Source:HGNC Symbol;Acc:H |
| TSBP1 | CHR_HSCHR6_MHC_APD_CT( | testis expressed basic protein 1 [Source:HGN( |
| HNRNPA1P2 | CHR_HSCHR6_MHC_APD_CT( | heterogeneous nuclear ribonucleoprotein A1 |
| U6 | CHR_HSCHR6_MHC_APD_CT( | U6 spliceosomal RNA [Source:RFAM;Acc:RF00 |
| TSBP1-AS1 | CHR_HSCHR6_MHC_APD_CT( | TSBP1 and BTNL2 antisense RNA 1 [Source:HG |
| BTNL2 | CHR_HSCHR6_MHC_APD_CT( | butyrophilin like 2 [Source:HGNC Symbol;Acc: |
| HLA-DRB9 | CHR_HSCHR6_MHC_APD_CT( | major histocompatibility complex, class II, DR |
| U1 | CHR_HSCHR6_MHC_APD_CT( | U1 spliceosomal RNA [Source:RFAM;Acc:RF00 |
| HLA-DRB3 | CHR_HSCHR6_MHC_APD_CT( | major histocompatibility complex, class II, DR |
| CR936921.1 | CHR_HSCHR6_MHC_APD_CT( | uncharacterized LOC102725019 [Source:NCBI |
| HLA-DQB3 | CHR_HSCHR6_MHC_APD_CT( | major histocompatibility complex, class II, DQ |
| HLA-DQA2 | CHR_HSCHR6_MHC_APD_CT( | major histocompatibility complex, class II, DQ |
| HLA-DQB2 | CHR_HSCHR6_MHC_APD_CT( | major histocompatibility complex, class II, DQ |
| HLA-DOB | CHR_HSCHR6_MHC_APD_CT( | major histocompatibility complex, class II, DO |
| TAP2 | CHR_HSCHR6_MHC_APD_CT( | transporter 2, ATP binding cassette subfamily |
| PSMB8 | CHR_HSCHR6_MHC_APD_CT( | proteasome subunit beta 8 [Source:HGNC Syn |
| PSMB8-AS1 | CHR_HSCHR6_MHC_APD_CT( | PSMB8 antisense RNA 1 (head to head) [Sourc |
| PSMB9 | CHR_HSCHR6_MHC_APD_CT( | proteasome subunit beta 9 [Source:HGNC Syn |
| TAP1 | CHR_HSCHR6_MHC_APD_CT( | transporter 1, ATP binding cassette subfamily |
| PPP1R2P1 | CHR_HSCHR6_MHC_APD_CT( | protein phosphatase 1 regulatory inhibitor sul |
| HLA-Z | CHR_HSCHR6_MHC_APD_CT( | major histocompatibility complex, class I, Z (p: |
| HLA-DMB | CHR_HSCHR6_MHC_APD_CT( | major histocompatibility complex, class II, DM |
| HLA-DMA | CHR_HSCHR6_MHC_APD_CT( | major histocompatibility complex, class II, DM |
| HLA-DPA1 | CHR_HSCHR6_MHC_APD_CT( | major histocompatibility complex, class II, DP |
| HLA-DPB1 | CHR_HSCHR6_MHC_APD_CT( | major histocompatibility complex, class II, DP |
| RPL32P1 | CHR_HSCHR6_MHC_APD_CT( | ribosomal protein L32 pseudogene 1 [Source: |
| HLA-DPA2 | CHR_HSCHR6_MHC_APD_CT( | major histocompatibility complex, class II, DP |
| COL11A2P1 | CHR_HSCHR6_MHC_APD_CT( | collagen type XI alpha 2 pseudogene 1 [Source |
| HLA-DPB2 | CHR_HSCHR6_MHC_APD_CT( | major histocompatibility complex, class II, DP |
| HLA-DPA3 | CHR_HSCHR6_MHC_APD_CT( | major histocompatibility complex, class II, DP |
| LYPLA2P1 | CHR_HSCHR6_MHC_APD_CT( | LYPLA2 pseudogene 1 [Source:HGNC Symbol;/ |
| GPX6 | CHR_HSCHR6_MHC_COX_CT( | glutathione peroxidase 6 [Source:NCBI gene;A |
| GPX5 | CHR_HSCHR6_MHC_COX_CT( | glutathione peroxidase 5 [Source:HGNC Symb |
| ZBED9 | CHR_HSCHR6_MHC_COX_CT( | zinc finger BED-type containing 9 [Source:HGN |
| AL671879.1 | CHR_HSCHR6_MHC_COX_CT( | novel transcript |
| LINC00533 | CHR_HSCHR6_MHC_COX_CT( | long intergenic non-protein coding RNA 533 [S |
| RPSAP2 | CHR_HSCHR6_MHC_COX_CT( | ribosomal protein SA pseudogene 2 [Source:H |
| NOP56P1 | CHR_HSCHR6_MHC_COX_CT( | NOP56 ribonucleoprotein pseudogene 1 [Sour |
| AL669978.1 | CHR_HSCHR6_MHC_COX_CT( | novel transcript |
| LINC01623 | CHR_HSCHR6_MHC_COX_CT( | long intergenic non-protein coding RNA 1623 |
| RPL13P | CHR_HSCHR6_MHC_COX_CT( | ribosomal protein L13 pseudogene [Source:H( |
| ZNF90P2 | CHR_HSCHR6_MHC_COX_CT( | zinc finger protein 90 pseudogene 2 [Source:H |
| HCG14 | CHR_HSCHR6_MHC_COX_CT( | HLA complex group 14 [Source:HGNC Symbol; |

| | | |
|---|---|---|
| TRIM27 | CHR_HSCHR6_MHC_COX_CT( | tripartite motif containing 27 [Source:HGNC S |
| U6 | CHR_HSCHR6_MHC_COX_CT( | U6 spliceosomal RNA [Source:RFAM;Acc:RF00 |
| LINC01556 | CHR_HSCHR6_MHC_COX_CT( | long intergenic non-protein coding RNA 1556 |
| KRT18P1 | CHR_HSCHR6_MHC_COX_CT( | keratin 18 pseudogene 1 [Source:HGNC Symb |
| Metazoa_SRP | CHR_HSCHR6_MHC_COX_CT( | Metazoan signal recognition particle RNA [Sou |
| HCG15 | CHR_HSCHR6_MHC_COX_CT( | HLA complex group 15 [Source:HGNC Symbol; |
| AL662865.1 | CHR_HSCHR6_MHC_COX_CT( | HLA complex group 16 |
| ZNF311 | CHR_HSCHR6_MHC_COX_CT( | zinc finger protein 311 [Source:HGNC Symbol; |
| OR2AD1P | CHR_HSCHR6_MHC_COX_CT( | olfactory receptor family 2 subfamily AD mem |
| OR2W1 | CHR_HSCHR6_MHC_COX_CT( | olfactory receptor family 2 subfamily W meml |
| AL662852.3 | CHR_HSCHR6_MHC_COX_CT( | NULL |
| OR2P1P | CHR_HSCHR6_MHC_COX_CT( | olfactory receptor family 2 subfamily P memb |
| SAR1AP1 | CHR_HSCHR6_MHC_COX_CT( | secretion associated Ras related GTPase 1A ps |
| OR2B3 | CHR_HSCHR6_MHC_COX_CT( | olfactory receptor family 2 subfamily B memb |
| OR2J1 | CHR_HSCHR6_MHC_COX_CT( | olfactory receptor family 2 subfamily J membe |
| OR2J3 | CHR_HSCHR6_MHC_COX_CT( | olfactory receptor family 2 subfamily J membe |
| AL662852.2 | CHR_HSCHR6_MHC_COX_CT( | novel transcript |
| AL662852.1 | CHR_HSCHR6_MHC_COX_CT( | olfactory receptor, family 2, subfamily N, men |
| OR2J2 | CHR_HSCHR6_MHC_COX_CT( | olfactory receptor family 2 subfamily J membe |
| OR2J4P | CHR_HSCHR6_MHC_COX_CT( | olfactory receptor family 2 subfamily J membe |
| AL662857.3 | CHR_HSCHR6_MHC_COX_CT( | olfactory receptor, family 2, subfamily H, men |
| AL662857.2 | CHR_HSCHR6_MHC_COX_CT( | novel transcript |
| OR2G1P | CHR_HSCHR6_MHC_COX_CT( | olfactory receptor family 2 subfamily G memb |
| OR2U1P | CHR_HSCHR6_MHC_COX_CT( | olfactory receptor family 2 subfamily U memb |
| OR2U2P | CHR_HSCHR6_MHC_COX_CT( | olfactory receptor family 2 subfamily U memb |
| AL662857.1 | CHR_HSCHR6_MHC_COX_CT( | olfactory receptor, family 2, subfamily B, men |
| OR14J1 | CHR_HSCHR6_MHC_COX_CT( | olfactory receptor family 14 subfamily J meml |
| DDX6P1 | CHR_HSCHR6_MHC_COX_CT( | DEAD-box helicase 6 pseudogene 1 [Source:H |
| OR5V1 | CHR_HSCHR6_MHC_COX_CT( | olfactory receptor family 5 subfamily V memb |
| OR12D3 | CHR_HSCHR6_MHC_COX_CT( | olfactory receptor family 12 subfamily D mem |
| OR12D2 | CHR_HSCHR6_MHC_COX_CT( | olfactory receptor family 12 subfamily D mem |
| OR12D1 | CHR_HSCHR6_MHC_COX_CT( | olfactory receptor family 12 subfamily D mem |
| OR11A1 | CHR_HSCHR6_MHC_COX_CT( | olfactory receptor family 11 subfamily A mem |
| OR10C1 | CHR_HSCHR6_MHC_COX_CT( | olfactory receptor family 10 subfamily C mem |
| OR2H1 | CHR_HSCHR6_MHC_COX_CT( | olfactory receptor family 2 subfamily H memb |
| UBDP1 | CHR_HSCHR6_MHC_COX_CT( | ubiquitin D pseudogene 1 [Source:HGNC Syml |
| MAS1LP1 | CHR_HSCHR6_MHC_COX_CT( | MAS1L pseudogene 1 [Source:HGNC Symbol;A |
| MAS1L | CHR_HSCHR6_MHC_COX_CT( | MAS1 proto-oncogene like, G protein-couplec |
| RPS17P1 | CHR_HSCHR6_MHC_COX_CT( | ribosomal protein S17 pseudogene 1 [Source: |
| LINC02829 | CHR_HSCHR6_MHC_COX_CT( | long intergenic non-protein coding RNA 2829 |
| LINC01015 | CHR_HSCHR6_MHC_COX_CT( | long intergenic non-protein coding RNA 1015 |
| GPR53P | CHR_HSCHR6_MHC_COX_CT( | G protein-coupled receptor 53, pseudogene [S |
| OR2I1P | CHR_HSCHR6_MHC_COX_CT( | olfactory receptor family 2 subfamily I membe |
| UBD | CHR_HSCHR6_MHC_COX_CT( | ubiquitin D [Source:HGNC Symbol;Acc:HGNC:1 |
| GABBR1 | CHR_HSCHR6_MHC_COX_CT( | gamma-aminobutyric acid type B receptor sub |
| OR2H5P | CHR_HSCHR6_MHC_COX_CT( | olfactory receptor family 2 subfamily H memb |
| TMEM183AP1 | CHR_HSCHR6_MHC_COX_CT( | transmembrane protein 183A pseudogene 1 [ |
| SNORD32B | CHR_HSCHR6_MHC_COX_CT( | small nucleolar RNA, C/D box 32B [Source:HG |
| RPL13AP | CHR_HSCHR6_MHC_COX_CT( | ribosomal protein L13a pseudogene [Source:H |
| OR2H2 | CHR_HSCHR6_MHC_COX_CT( | olfactory receptor family 2 subfamily H memb |

| | | |
|---|---|---|
| SUMO2P1 | CHR_HSCHR6_MHC_COX_CT( | SUMO2 pseudogene 1 [Source:HGNC Symbol; |
| MOG | CHR_HSCHR6_MHC_COX_CT( | myelin oligodendrocyte glycoprotein [Source: |
| ZFP57 | CHR_HSCHR6_MHC_COX_CT( | ZFP57 zinc finger protein [Source:HGNC Symb |
| ZDHHC20P1 | CHR_HSCHR6_MHC_COX_CT( | zinc finger DHHC-type containing 20 pseudoge |
| HLA-F | CHR_HSCHR6_MHC_COX_CT( | major histocompatibility complex, class I, F [S |
| AL669813.2 | CHR_HSCHR6_MHC_COX_CT( | HLA complex group 4 pseudogene 11 |
| HLA-F-AS1 | CHR_HSCHR6_MHC_COX_CT( | HLA-F antisense RNA 1 [Source:HGNC Symbol; |
| RPL23AP1 | CHR_HSCHR6_MHC_COX_CT( | ribosomal protein L23a pseudogene 1 [Source |
| MICE | CHR_HSCHR6_MHC_COX_CT( | MHC class I polypeptide-related sequence E (p |
| AL669813.1 | CHR_HSCHR6_MHC_COX_CT( | HLA complex group 9 pseudogene 5 |
| Y_RNA | CHR_HSCHR6_MHC_COX_CT( | Y RNA [Source:RFAM;Acc:RF00019] |
| IFITM4P | CHR_HSCHR6_MHC_COX_CT( | interferon induced transmembrane protein 4 |
| HLA-V | CHR_HSCHR6_MHC_COX_CT( | major histocompatibility complex, class I, V (p |
| HCG4 | CHR_HSCHR6_MHC_COX_CT( | HLA complex group 4 [Source:HGNC Symbol;A |
| HLA-P | CHR_HSCHR6_MHC_COX_CT( | major histocompatibility complex, class I, P (p |
| RPL7AP7 | CHR_HSCHR6_MHC_COX_CT( | ribosomal protein L7a pseudogene 7 [Source:I |
| MICG | CHR_HSCHR6_MHC_COX_CT( | MHC class I polypeptide-related sequence G (p |
| HLA-G | CHR_HSCHR6_MHC_COX_CT( | major histocompatibility complex, class I, G [S |
| HCG4P8 | CHR_HSCHR6_MHC_COX_CT( | HLA complex group 4 pseudogene 8 [Source:H |
| AL671561.1 | CHR_HSCHR6_MHC_COX_CT( | MHC class I polypeptide-related sequence F ps |
| AL671561.2 | CHR_HSCHR6_MHC_COX_CT( | HLA complex group 4 pseudogene 7 |
| HLA-A | CHR_HSCHR6_MHC_COX_CT( | major histocompatibility complex, class I, A [S |
| HLA-H | CHR_HSCHR6_MHC_COX_CT( | major histocompatibility complex, class I, H (p |
| HLA-T | CHR_HSCHR6_MHC_COX_CT( | major histocompatibility complex, class I, T (p |
| DDX39BP1 | CHR_HSCHR6_MHC_COX_CT( | DEAD-box helicase 39B pseudogene 1 [Source |
| MCCD1P1 | CHR_HSCHR6_MHC_COX_CT( | mitochondrial coiled-coil domain 1 pseudoger |
| HCG4B | CHR_HSCHR6_MHC_COX_CT( | HLA complex group 4B [Source:HGNC Symbol; |
| HLA-K | CHR_HSCHR6_MHC_COX_CT( | major histocompatibility complex, class I, K (p |
| HLA-U | CHR_HSCHR6_MHC_COX_CT( | major histocompatibility complex, class I, U (p |
| AL645935.1 | CHR_HSCHR6_MHC_COX_CT( | HLA complex group 4 pseudogene 5 |
| HLA-W | CHR_HSCHR6_MHC_COX_CT( | major histocompatibility complex, class I, W (p |
| MICD | CHR_HSCHR6_MHC_COX_CT( | MHC class I polypeptide-related sequence D (p |
| HCG9 | CHR_HSCHR6_MHC_COX_CT( | HLA complex group 9 [Source:HGNC Symbol;A |
| DDX39BP2 | CHR_HSCHR6_MHC_COX_CT( | DEAD-box helicase 39B pseudogene 2 [Source |
| MCCD1P2 | CHR_HSCHR6_MHC_COX_CT( | mitochondrial coiled-coil domain 1 pseudoger |
| ZNRD1ASP | CHR_HSCHR6_MHC_COX_CT( | zinc ribbon domain containing 1 antisense, ps |
| AL645935.2 | CHR_HSCHR6_MHC_COX_CT( | HLA complex group 4 pseudogene 3 |
| HLA-J | CHR_HSCHR6_MHC_COX_CT( | major histocompatibility complex, class I, J (ps |
| ETF1P1 | CHR_HSCHR6_MHC_COX_CT( | eukaryotic translation termination factor 1 ps |
| AL671859.4 | CHR_HSCHR6_MHC_COX_CT( | NULL |
| AL671859.2 | CHR_HSCHR6_MHC_COX_CT( | NULL |
| ZNRD1 | CHR_HSCHR6_MHC_COX_CT( | zinc ribbon domain containing 1 [Source:HGN |
| AL671859.3 | CHR_HSCHR6_MHC_COX_CT( | NULL |
| PPP1R11 | CHR_HSCHR6_MHC_COX_CT( | protein phosphatase 1 regulatory inhibitor sul |
| RNF39 | CHR_HSCHR6_MHC_COX_CT( | ring finger protein 39 [Source:HGNC Symbol;A |
| TRIM31 | CHR_HSCHR6_MHC_COX_CT( | tripartite motif containing 31 [Source:HGNC S |
| TRIM31-AS1 | CHR_HSCHR6_MHC_COX_CT( | TRIM31 antisense RNA 1 [Source:HGNC Symb |
| AL671859.1 | CHR_HSCHR6_MHC_COX_CT( | NULL |
| TRIM40 | CHR_HSCHR6_MHC_COX_CT( | tripartite motif containing 40 [Source:HGNC S |
| TRIM10 | CHR_HSCHR6_MHC_COX_CT( | tripartite motif containing 10 [Source:HGNC S |

| | | |
|---|---|---|
| TRIM15 | CHR_HSCHR6_MHC_COX_CT( | tripartite motif containing 15 [Source:HGNC S |
| TRIM26 | CHR_HSCHR6_MHC_COX_CT( | tripartite motif containing 26 [Source:HGNC S |
| PAIP1P1 | CHR_HSCHR6_MHC_COX_CT( | poly(A) binding protein interacting protein 1 p |
| HCG17 | CHR_HSCHR6_MHC_COX_CT( | HLA complex group 17 [Source:HGNC Symbol; |
| TRIM26BP | CHR_HSCHR6_MHC_COX_CT( | tripartite motif containing 26B, pseudogene [S |
| HLA-L | CHR_HSCHR6_MHC_COX_CT( | major histocompatibility complex, class I, L (ps |
| HCG18 | CHR_HSCHR6_MHC_COX_CT( | HLA complex group 18 [Source:HGNC Symbol; |
| TRIM39 | CHR_HSCHR6_MHC_COX_CT( | tripartite motif containing 39 [Source:HGNC S |
| RPP21 | CHR_HSCHR6_MHC_COX_CT( | ribonuclease P/MRP subunit p21 [Source:HGN |
| HLA-N | CHR_HSCHR6_MHC_COX_CT( | major histocompatibility complex, class I, N (p |
| AL662831.1 | CHR_HSCHR6_MHC_COX_CT( | HLA complex group 19 pseudogene |
| UBQLN1P1 | CHR_HSCHR6_MHC_COX_CT( | ubiquilin 1 pseudogene 1 [Source:HGNC Symb |
| MICC | CHR_HSCHR6_MHC_COX_CT( | MHC class I polypeptide-related sequence C (p |
| TMPOP1 | CHR_HSCHR6_MHC_COX_CT( | thymopoietin pseudogene 1 [Source:HGNC Sy |
| SUCLA2P1 | CHR_HSCHR6_MHC_COX_CT( | succinate-CoA ligase ADP-forming beta subun |
| RANP1 | CHR_HSCHR6_MHC_COX_CT( | RAN pseudogene 1 [Source:HGNC Symbol;Acc |
| HLA-E | CHR_HSCHR6_MHC_COX_CT( | major histocompatibility complex, class I, E [S |
| LINC02569 | CHR_HSCHR6_MHC_COX_CT( | long intergenic non-protein coding RNA 2569 |
| GNL1 | CHR_HSCHR6_MHC_COX_CT( | G protein nucleolar 1 (putative) [Source:HGNC |
| PRR3 | CHR_HSCHR6_MHC_COX_CT( | proline rich 3 [Source:HGNC Symbol;Acc:HGN |
| ABCF1 | CHR_HSCHR6_MHC_COX_CT( | ATP binding cassette subfamily F member 1 [S |
| PPP1R10 | CHR_HSCHR6_MHC_COX_CT( | protein phosphatase 1 regulatory subunit 10 [ |
| MRPS18B | CHR_HSCHR6_MHC_COX_CT( | mitochondrial ribosomal protein S18B [Source |
| ATAT1 | CHR_HSCHR6_MHC_COX_CT( | alpha tubulin acetyltransferase 1 [Source:HGN |
| PTMAP1 | CHR_HSCHR6_MHC_COX_CT( | prothymosin alpha pseudogene 1 [Source:HGI |
| C6orf136 | CHR_HSCHR6_MHC_COX_CT( | chromosome 6 open reading frame 136 [Sour |
| DHX16 | CHR_HSCHR6_MHC_COX_CT( | DEAH-box helicase 16 [Source:HGNC Symbol;/ |
| PPP1R18 | CHR_HSCHR6_MHC_COX_CT( | protein phosphatase 1 regulatory subunit 18 [ |
| NRM | CHR_HSCHR6_MHC_COX_CT( | nurim [Source:HGNC Symbol;Acc:HGNC:8003] |
| RPL7P4 | CHR_HSCHR6_MHC_COX_CT( | ribosomal protein L7 pseudogene 4 [Source:H |
| MDC1 | CHR_HSCHR6_MHC_COX_CT( | mediator of DNA damage checkpoint 1 [Sourc |
| MDC1-AS1 | CHR_HSCHR6_MHC_COX_CT( | MDC1 antisense RNA 1 [Source:HGNC Symbol |
| TUBB | CHR_HSCHR6_MHC_COX_CT( | tubulin beta class I [Source:HGNC Symbol;Acc |
| FLOT1 | CHR_HSCHR6_MHC_COX_CT( | flotillin 1 [Source:HGNC Symbol;Acc:HGNC:37 |
| Y_RNA | CHR_HSCHR6_MHC_COX_CT( | Y RNA [Source:RFAM;Acc:RF00019] |
| IER3 | CHR_HSCHR6_MHC_COX_CT( | immediate early response 3 [Source:HGNC Syi |
| Metazoa_SRP | CHR_HSCHR6_MHC_COX_CT( | Metazoan signal recognition particle RNA [Sou |
| HCG20 | CHR_HSCHR6_MHC_COX_CT( | HLA complex group 20 [Source:HGNC Symbol; |
| AL662848.1 | CHR_HSCHR6_MHC_COX_CT( | chromosome 6 open reading frame 214 (puta |
| LINC02570 | CHR_HSCHR6_MHC_COX_CT( | long intergenic non-protein coding RNA 2570 |
| 7SK | CHR_HSCHR6_MHC_COX_CT( | 7SK RNA [Source:RFAM;Acc:RF00100] |
| DDR1-DT | CHR_HSCHR6_MHC_COX_CT( | DDR1 divergent transcript [Source:HGNC Sym |
| DDR1 | CHR_HSCHR6_MHC_COX_CT( | discoidin domain receptor tyrosine kinase 1 [S |
| GTF2H4 | CHR_HSCHR6_MHC_COX_CT( | general transcription factor IIH subunit 4 [Sou |
| AL662854.1 | CHR_HSCHR6_MHC_COX_CT( | novel protein |
| VARS2 | CHR_HSCHR6_MHC_COX_CT( | valyl-tRNA synthetase 2, mitochondrial [Sourc |
| SFTA2 | CHR_HSCHR6_MHC_COX_CT( | surfactant associated 2 [Source:HGNC Symbol |
| Y_RNA | CHR_HSCHR6_MHC_COX_CT( | Y RNA [Source:RFAM;Acc:RF00019] |
| MUCL3 | CHR_HSCHR6_MHC_COX_CT( | mucin like 3 [Source:HGNC Symbol;Acc:HGNC |
| HCG21 | CHR_HSCHR6_MHC_COX_CT( | HLA complex group 21 [Source:HGNC Symbol; |

| | | |
|---|---|---|
| MUC21 | CHR_HSCHR6_MHC_COX_CT( | mucin 21, cell surface associated [Source:HGN |
| AL663093.1 | CHR_HSCHR6_MHC_COX_CT( | novel transcript |
| AL663093.2 | CHR_HSCHR6_MHC_COX_CT( | HLA complex group family pseudogene |
| U6 | CHR_HSCHR6_MHC_COX_CT( | U6 spliceosomal RNA [Source:RFAM;Acc:RF00 |
| C6orf15 | CHR_HSCHR6_MHC_COX_CT( | chromosome 6 open reading frame 15 [Source |
| PSORS1C1 | CHR_HSCHR6_MHC_COX_CT( | psoriasis susceptibility 1 candidate 1 [Source:I |
| CDSN | CHR_HSCHR6_MHC_COX_CT( | corneodesmosin [Source:HGNC Symbol;Acc:H |
| PSORS1C2 | CHR_HSCHR6_MHC_COX_CT( | psoriasis susceptibility 1 candidate 2 [Source:I |
| POLR2LP1 | CHR_HSCHR6_MHC_COX_CT( | RNA polymerase II subunit L pseudogene 1 [So |
| CCHCR1 | CHR_HSCHR6_MHC_COX_CT( | coiled-coil alpha-helical rod protein 1 [Source: |
| TCF19 | CHR_HSCHR6_MHC_COX_CT( | transcription factor 19 [Source:HGNC Symbol; |
| POU5F1 | CHR_HSCHR6_MHC_COX_CT( | POU class 5 homeobox 1 [Source:HGNC Symb |
| PSORS1C3 | CHR_HSCHR6_MHC_COX_CT( | psoriasis susceptibility 1 candidate 3 [Source:I |
| HCG27 | CHR_HSCHR6_MHC_COX_CT( | HLA complex group 27 [Source:HGNC Symbol; |
| AL662833.2 | CHR_HSCHR6_MHC_COX_CT( | novel transcript |
| HLA-C | CHR_HSCHR6_MHC_COX_CT( | major histocompatibility complex, class I, C [S |
| HLA-B | CHR_HSCHR6_MHC_COX_CT( | major histocompatibility complex, class I, B [S |
| USP8P1 | CHR_HSCHR6_MHC_COX_CT( | ubiquitin specific peptidase 8 pseudogene 1 [S |
| RPL3P2 | CHR_HSCHR6_MHC_COX_CT( | ribosomal protein L3 pseudogene 2 [Source:H |
| WASF5P | CHR_HSCHR6_MHC_COX_CT( | WASP family member 5, pseudogene [Source: |
| AL662833.1 | CHR_HSCHR6_MHC_COX_CT( | novel HLA class I pseudogene fragment |
| DHFRP2 | CHR_HSCHR6_MHC_COX_CT( | dihydrofolate reductase pseudogene 2 [Sourc |
| U6 | CHR_HSCHR6_MHC_COX_CT( | U6 spliceosomal RNA [Source:RFAM;Acc:RF00 |
| AL669854.1 | CHR_HSCHR6_MHC_COX_CT( | fibroblast growth factor receptor 3 pseudoger |
| ZDHHC20P2 | CHR_HSCHR6_MHC_COX_CT( | zinc finger DHHC-type containing 20 pseudoge |
| HLA-S | CHR_HSCHR6_MHC_COX_CT( | major histocompatibility complex, class I, S (p: |
| MICA | CHR_HSCHR6_MHC_COX_CT( | MHC class I polypeptide-related sequence A [S |
| Y_RNA | CHR_HSCHR6_MHC_COX_CT( | Y RNA [Source:RFAM;Acc:RF00019] |
| LINC01149 | CHR_HSCHR6_MHC_COX_CT( | long intergenic non-protein coding RNA 1149 |
| HCP5 | CHR_HSCHR6_MHC_COX_CT( | HLA complex P5 [Source:HGNC Symbol;Acc:H( |
| Y_RNA | CHR_HSCHR6_MHC_COX_CT( | Y RNA [Source:RFAM;Acc:RF00019] |
| MICB | CHR_HSCHR6_MHC_COX_CT( | MHC class I polypeptide-related sequence B [S |
| PPIAP9 | CHR_HSCHR6_MHC_COX_CT( | peptidylprolyl isomerase A pseudogene 9 [Sou |
| RPL15P4 | CHR_HSCHR6_MHC_COX_CT( | ribosomal protein L15 pseudogene 4 [Source: |
| MCCD1 | CHR_HSCHR6_MHC_COX_CT( | mitochondrial coiled-coil domain 1 [Source:H( |
| DDX39B | CHR_HSCHR6_MHC_COX_CT( | DExD-box helicase 39B [Source:HGNC Symbol; |
| SNORD117 | CHR_HSCHR6_MHC_COX_CT( | small nucleolar RNA, C/D box 117 [Source:HGI |
| SNORD84 | CHR_HSCHR6_MHC_COX_CT( | small nucleolar RNA, C/D box 84 [Source:HGN |
| DDX39B-AS1 | CHR_HSCHR6_MHC_COX_CT( | DDX39B antisense RNA 1 [Source:HGNC Symb |
| ATP6V1G2 | CHR_HSCHR6_MHC_COX_CT( | ATPase H+ transporting V1 subunit G2 [Source |
| NFKBIL1 | CHR_HSCHR6_MHC_COX_CT( | NFKB inhibitor like 1 [Source:HGNC Symbol;Ac |
| LTA | CHR_HSCHR6_MHC_COX_CT( | lymphotoxin alpha [Source:HGNC Symbol;Acc |
| TNF | CHR_HSCHR6_MHC_COX_CT( | tumor necrosis factor [Source:HGNC Symbol;A |
| LTB | CHR_HSCHR6_MHC_COX_CT( | lymphotoxin beta [Source:HGNC Symbol;Acc:I |
| LST1 | CHR_HSCHR6_MHC_COX_CT( | leukocyte specific transcript 1 [Source:HGNC S |
| NCR3 | CHR_HSCHR6_MHC_COX_CT( | natural cytotoxicity triggering receptor 3 [Sou |
| UQCRHP1 | CHR_HSCHR6_MHC_COX_CT( | ubiquinol-cytochrome c reductase hinge prote |
| AIF1 | CHR_HSCHR6_MHC_COX_CT( | allograft inflammatory factor 1 [Source:HGNC |
| PRRC2A | CHR_HSCHR6_MHC_COX_CT( | proline rich coiled-coil 2A [Source:HGNC Symb |
| SNORA38 | CHR_HSCHR6_MHC_COX_CT( | small nucleolar RNA, H/ACA box 38 [Source:H( |

| | | |
|---|---|---|
| BAG6 | CHR_HSCHR6_MHC_COX_CT( | BCL2 associated athanogene 6 [Source:HGNC |
| APOM | CHR_HSCHR6_MHC_COX_CT( | apolipoprotein M [Source:HGNC Symbol;Acc:H |
| C6orf47 | CHR_HSCHR6_MHC_COX_CT( | chromosome 6 open reading frame 47 [Source |
| C6orf47-AS1 | CHR_HSCHR6_MHC_COX_CT( | C6orf47 antisense RNA 1 [Source:HGNC Symb |
| GPANK1 | CHR_HSCHR6_MHC_COX_CT( | G-patch domain and ankyrin repeats 1 [Source |
| Y_RNA | CHR_HSCHR6_MHC_COX_CT( | Y RNA [Source:RFAM;Acc:RF00019] |
| CSNK2B | CHR_HSCHR6_MHC_COX_CT( | casein kinase 2 beta [Source:HGNC Symbol;Ac |
| LY6G5B | CHR_HSCHR6_MHC_COX_CT( | lymphocyte antigen 6 family member G5B [So |
| LY6G5C | CHR_HSCHR6_MHC_COX_CT( | lymphocyte antigen 6 family member G5C [So |
| ABHD16A | CHR_HSCHR6_MHC_COX_CT( | abhydrolase domain containing 16A [Source:H |
| LY6G6F | CHR_HSCHR6_MHC_COX_CT( | lymphocyte antigen 6 family member G6F [So |
| LY6G6D | CHR_HSCHR6_MHC_COX_CT( | lymphocyte antigen 6 family member G6D [Sc |
| LY6G6E | CHR_HSCHR6_MHC_COX_CT( | lymphocyte antigen 6 family member G6E [So |
| MPIG6B | CHR_HSCHR6_MHC_COX_CT( | megakaryocyte and platelet inhibitory recepto |
| LY6G6C | CHR_HSCHR6_MHC_COX_CT( | lymphocyte antigen 6 family member G6C [So |
| DDAH2 | CHR_HSCHR6_MHC_COX_CT( | dimethylarginine dimethylaminohydrolase 2 [ |
| CLIC1 | CHR_HSCHR6_MHC_COX_CT( | chloride intracellular channel 1 [Source:HGNC |
| MSH5 | CHR_HSCHR6_MHC_COX_CT( | mutS homolog 5 [Source:HGNC Symbol;Acc:H |
| MSH5-SAPCD1 | CHR_HSCHR6_MHC_COX_CT( | MSH5-SAPCD1 readthrough (NMD candidate) |
| U6 | CHR_HSCHR6_MHC_COX_CT( | U6 spliceosomal RNA [Source:RFAM;Acc:RF00 |
| SAPCD1 | CHR_HSCHR6_MHC_COX_CT( | suppressor APC domain containing 1 [Source:I |
| SAPCD1-AS1 | CHR_HSCHR6_MHC_COX_CT( | SAPCD1 antisense RNA 1 [Source:HGNC Symb |
| VWA7 | CHR_HSCHR6_MHC_COX_CT( | von Willebrand factor A domain containing 7 [ |
| VARS | CHR_HSCHR6_MHC_COX_CT( | valyl-tRNA synthetase [Source:HGNC Symbol;/ |
| Y_RNA | CHR_HSCHR6_MHC_COX_CT( | Y RNA [Source:RFAM;Acc:RF00019] |
| LSM2 | CHR_HSCHR6_MHC_COX_CT( | LSM2 homolog, U6 small nuclear RNA and mR |
| HSPA1L | CHR_HSCHR6_MHC_COX_CT( | heat shock protein family A (Hsp70) member : |
| HSPA1A | CHR_HSCHR6_MHC_COX_CT( | heat shock protein family A (Hsp70) member : |
| HSPA1B | CHR_HSCHR6_MHC_COX_CT( | heat shock protein family A (Hsp70) member : |
| SNHG32 | CHR_HSCHR6_MHC_COX_CT( | small nucleolar RNA host gene 32 [Source:HGI |
| SNORD48 | CHR_HSCHR6_MHC_COX_CT( | small nucleolar RNA, C/D box 48 [Source:HGN |
| SNORD52 | CHR_HSCHR6_MHC_COX_CT( | small nucleolar RNA, C/D box 52 [Source:HGN |
| NEU1 | CHR_HSCHR6_MHC_COX_CT( | neuraminidase 1 [Source:HGNC Symbol;Acc:H |
| SLC44A4 | CHR_HSCHR6_MHC_COX_CT( | solute carrier family 44 member 4 [Source:HG |
| EHMT2 | CHR_HSCHR6_MHC_COX_CT( | euchromatic histone lysine methyltransferase |
| AL662834.1 | CHR_HSCHR6_MHC_COX_CT( | novel transcript |
| C2 | CHR_HSCHR6_MHC_COX_CT( | complement C2 [Source:HGNC Symbol;Acc:HG |
| ZBTB12 | CHR_HSCHR6_MHC_COX_CT( | zinc finger and BTB domain containing 12 [Sou |
| C4B-AS1 | CHR_HSCHR6_MHC_COX_CT( | C4B antisense RNA 1 [Source:HGNC Symbol;A |
| CFB | CHR_HSCHR6_MHC_COX_CT( | complement factor B [Source:HGNC Symbol;A |
| NELFE | CHR_HSCHR6_MHC_COX_CT( | negative elongation factor complex member E |
| SKIV2L | CHR_HSCHR6_MHC_COX_CT( | Ski2 like RNA helicase [Source:HGNC Symbol;/ |
| DXO | CHR_HSCHR6_MHC_COX_CT( | decapping exoribonuclease [Source:HGNC Syr |
| STK19 | CHR_HSCHR6_MHC_COX_CT( | serine/threonine kinase 19 [Source:HGNC Sym |
| C4B | CHR_HSCHR6_MHC_COX_CT( | complement C4B (Chido blood group) [Source |
| AL662849.1 | CHR_HSCHR6_MHC_COX_CT( | novel transcript |
| CYP21A2 | CHR_HSCHR6_MHC_COX_CT( | cytochrome P450 family 21 subfamily A meml |
| TNXB | CHR_HSCHR6_MHC_COX_CT( | tenascin XB [Source:HGNC Symbol;Acc:HGNC: |
| 5S_rRNA | CHR_HSCHR6_MHC_COX_CT( | 5S ribosomal RNA [Source:RFAM;Acc:RF00001 |
| ATF6B | CHR_HSCHR6_MHC_COX_CT( | activating transcription factor 6 beta [Source:I |

| | | |
|---|---|---|
| FKBPL | CHR_HSCHR6_MHC_COX_CT( | FKBP prolyl isomerase like [Source:HGNC Sym |
| PRRT1 | CHR_HSCHR6_MHC_COX_CT( | proline rich transmembrane protein 1 [Source |
| PPT2 | CHR_HSCHR6_MHC_COX_CT( | palmitoyl-protein thioesterase 2 [Source:HGN |
| AL662828.1 | CHR_HSCHR6_MHC_COX_CT( | palmitoyl-protein thioesterase 2 (PPT2) and E( |
| EGFL8 | CHR_HSCHR6_MHC_COX_CT( | EGF like domain multiple 8 [Source:HGNC Sym |
| AGPAT1 | CHR_HSCHR6_MHC_COX_CT( | 1-acylglycerol-3-phosphate O-acyltransferase |
| RNF5 | CHR_HSCHR6_MHC_COX_CT( | ring finger protein 5 [Source:HGNC Symbol;Ac |
| AGER | CHR_HSCHR6_MHC_COX_CT( | advanced glycosylation end-product specific r |
| PBX2 | CHR_HSCHR6_MHC_COX_CT( | PBX homeobox 2 [Source:HGNC Symbol;Acc:H |
| GPSM3 | CHR_HSCHR6_MHC_COX_CT( | G protein signaling modulator 3 [Source:HGN( |
| NOTCH4 | CHR_HSCHR6_MHC_COX_CT( | notch receptor 4 [Source:HGNC Symbol;Acc:H |
| TSBP1 | CHR_HSCHR6_MHC_COX_CT( | testis expressed basic protein 1 [Source:HGN( |
| HNRNPA1P2 | CHR_HSCHR6_MHC_COX_CT( | heterogeneous nuclear ribonucleoprotein A1 |
| U6 | CHR_HSCHR6_MHC_COX_CT( | U6 spliceosomal RNA [Source:RFAM;Acc:RF00 |
| TSBP1-AS1 | CHR_HSCHR6_MHC_COX_CT( | TSBP1 and BTNL2 antisense RNA 1 [Source:HG |
| BTNL2 | CHR_HSCHR6_MHC_COX_CT( | butyrophilin like 2 [Source:HGNC Symbol;Acc: |
| HLA-DRA | CHR_HSCHR6_MHC_COX_CT( | major histocompatibility complex, class II, DR |
| HLA-DRB9 | CHR_HSCHR6_MHC_COX_CT( | major histocompatibility complex, class II, DR |
| RNU1-79P | CHR_HSCHR6_MHC_COX_CT( | RNA, U1 small nuclear 79, pseudogene [Sourc |
| HLA-DRB3 | CHR_HSCHR6_MHC_COX_CT( | major histocompatibility complex, class II, DR |
| U1 | CHR_HSCHR6_MHC_COX_CT( | U1 spliceosomal RNA [Source:RFAM;Acc:RF00 |
| AL662842.1 | CHR_HSCHR6_MHC_COX_CT( | major histocompatibility complex, class II, DR |
| RNU1-116P | CHR_HSCHR6_MHC_COX_CT( | RNA, U1 small nuclear 116, pseudogene [Sour |
| HLA-DRB1 | CHR_HSCHR6_MHC_COX_CT( | major histocompatibility complex, class II, DR |
| HLA-DQA2 | CHR_HSCHR6_MHC_COX_CT( | major histocompatibility complex, class II, DQ |
| HLA-DQB1 | CHR_HSCHR6_MHC_COX_CT( | major histocompatibility complex, class II, DQ |
| HLA-DQB1-AS1 | CHR_HSCHR6_MHC_COX_CT( | HLA-DQB1 antisense RNA 1 [Source:HGNC Syr |
| AL731683.2 | CHR_HSCHR6_MHC_COX_CT( | novel transcript |
| MTCO3P1 | CHR_HSCHR6_MHC_COX_CT( | MT-CO3 pseudogene 1 [Source:HGNC Symbol |
| AL731683.1 | CHR_HSCHR6_MHC_COX_CT( | uncharacterized LOC102725019 [Source:NCBI |
| HLA-DQB3 | CHR_HSCHR6_MHC_COX_CT( | major histocompatibility complex, class II, DQ |
| HLA-DQB2 | CHR_HSCHR6_MHC_COX_CT( | major histocompatibility complex, class II, DQ |
| HLA-DOB | CHR_HSCHR6_MHC_COX_CT( | major histocompatibility complex, class II, DO |
| TAP2 | CHR_HSCHR6_MHC_COX_CT( | transporter 2, ATP binding cassette subfamily |
| PSMB8 | CHR_HSCHR6_MHC_COX_CT( | proteasome subunit beta 8 [Source:HGNC Syn |
| PSMB8-AS1 | CHR_HSCHR6_MHC_COX_CT( | PSMB8 antisense RNA 1 (head to head) [Sourc |
| PSMB9 | CHR_HSCHR6_MHC_COX_CT( | proteasome subunit beta 9 [Source:HGNC Syn |
| TAP1 | CHR_HSCHR6_MHC_COX_CT( | transporter 1, ATP binding cassette subfamily |
| PPP1R2P1 | CHR_HSCHR6_MHC_COX_CT( | protein phosphatase 1 regulatory inhibitor sul |
| HLA-Z | CHR_HSCHR6_MHC_COX_CT( | major histocompatibility complex, class I, Z (p: |
| HLA-DMB | CHR_HSCHR6_MHC_COX_CT( | major histocompatibility complex, class II, DM |
| HLA-DMA | CHR_HSCHR6_MHC_COX_CT( | major histocompatibility complex, class II, DM |
| BRD2 | CHR_HSCHR6_MHC_COX_CT( | bromodomain containing 2 [Source:HGNC Syn |
| AL662845.1 | CHR_HSCHR6_MHC_COX_CT( | novel transcript |
| HLA-DOA | CHR_HSCHR6_MHC_COX_CT( | major histocompatibility complex, class II, DO |
| HLA-DPA1 | CHR_HSCHR6_MHC_COX_CT( | major histocompatibility complex, class II, DP |
| HLA-DPB1 | CHR_HSCHR6_MHC_COX_CT( | major histocompatibility complex, class II, DP |
| RPL32P1 | CHR_HSCHR6_MHC_COX_CT( | ribosomal protein L32 pseudogene 1 [Source: |
| HLA-DPA2 | CHR_HSCHR6_MHC_COX_CT( | major histocompatibility complex, class II, DP |
| COL11A2P1 | CHR_HSCHR6_MHC_COX_CT( | collagen type XI alpha 2 pseudogene 1 [Sourc |

| | | |
|---|---|---|
| HLA-DPB2 | CHR_HSCHR6_MHC_COX_CT( | major histocompatibility complex, class II, DP |
| HLA-DPA3 | CHR_HSCHR6_MHC_COX_CT( | major histocompatibility complex, class II, DP |
| HCG24 | CHR_HSCHR6_MHC_COX_CT( | HLA complex group 24 [Source:HGNC Symbol; |
| COL11A2 | CHR_HSCHR6_MHC_COX_CT( | collagen type XI alpha 2 chain [Source:HGNC S |
| RXRB | CHR_HSCHR6_MHC_COX_CT( | retinoid X receptor beta [Source:HGNC Symbo |
| Y_RNA | CHR_HSCHR6_MHC_COX_CT( | Y RNA [Source:RFAM;Acc:RF00019] |
| SLC39A7 | CHR_HSCHR6_MHC_COX_CT( | solute carrier family 39 member 7 [Source:HG |
| HSD17B8 | CHR_HSCHR6_MHC_COX_CT( | hydroxysteroid 17-beta dehydrogenase 8 [Sou |
| RING1 | CHR_HSCHR6_MHC_COX_CT( | ring finger protein 1 [Source:HGNC Symbol;Ac |
| ZNF70P1 | CHR_HSCHR6_MHC_COX_CT( | zinc finger protein 70 pseudogene 1 [Source:H |
| HTATSF1P1 | CHR_HSCHR6_MHC_COX_CT( | HIV-1 Tat specific factor 1 pseudogene 1 [Sour |
| HCG25 | CHR_HSCHR6_MHC_COX_CT( | HLA complex group 25 [Source:HGNC Symbol; |
| VPS52 | CHR_HSCHR6_MHC_COX_CT( | VPS52 subunit of GARP complex [Source:HGN |
| RPS18 | CHR_HSCHR6_MHC_COX_CT( | ribosomal protein S18 [Source:HGNC Symbol;/ |
| B3GALT4 | CHR_HSCHR6_MHC_COX_CT( | beta-1,3-galactosyltransferase 4 [Source:HGN |
| WDR46 | CHR_HSCHR6_MHC_COX_CT( | WD repeat domain 46 [Source:HGNC Symbol;/ |
| PFDN6 | CHR_HSCHR6_MHC_COX_CT( | prefoldin subunit 6 [Source:HGNC Symbol;Acc |
| RGL2 | CHR_HSCHR6_MHC_COX_CT( | ral guanine nucleotide dissociation stimulator |
| TAPBP | CHR_HSCHR6_MHC_COX_CT( | TAP binding protein [Source:HGNC Symbol;Ac |
| ZBTB22 | CHR_HSCHR6_MHC_COX_CT( | zinc finger and BTB domain containing 22 [Sou |
| DAXX | CHR_HSCHR6_MHC_COX_CT( | death domain associated protein [Source:HGN |
| MYL8P | CHR_HSCHR6_MHC_COX_CT( | myosin light chain 8, pseudogene [Source:HGI |
| LYPLA2P1 | CHR_HSCHR6_MHC_COX_CT( | LYPLA2 pseudogene 1 [Source:HGNC Symbol;/ |
| NOP56P1 | CHR_HSCHR6_MHC_DBB_CT( | NOP56 ribonucleoprotein pseudogene 1 [Sour |
| CR753651.1 | CHR_HSCHR6_MHC_DBB_CT( | novel transcript |
| LINC01623 | CHR_HSCHR6_MHC_DBB_CT( | long intergenic non-protein coding RNA 1623 |
| RPL13P | CHR_HSCHR6_MHC_DBB_CT( | ribosomal protein L13 pseudogene [Source:H( |
| Metazoa_SRP | CHR_HSCHR6_MHC_DBB_CT( | Metazoan signal recognition particle RNA [Sou |
| HCG15 | CHR_HSCHR6_MHC_DBB_CT( | HLA complex group 15 [Source:HGNC Symbol; |
| CR759835.1 | CHR_HSCHR6_MHC_DBB_CT( | HLA complex group 16 |
| ZNF311 | CHR_HSCHR6_MHC_DBB_CT( | zinc finger protein 311 [Source:HGNC Symbol; |
| OR2AD1P | CHR_HSCHR6_MHC_DBB_CT( | olfactory receptor family 2 subfamily AD mem |
| OR2W1 | CHR_HSCHR6_MHC_DBB_CT( | olfactory receptor family 2 subfamily W meml |
| CR388407.1 | CHR_HSCHR6_MHC_DBB_CT( | NULL |
| OR2P1P | CHR_HSCHR6_MHC_DBB_CT( | olfactory receptor family 2 subfamily P memb |
| SAR1AP1 | CHR_HSCHR6_MHC_DBB_CT( | secretion associated Ras related GTPase 1A ps |
| OR2B3 | CHR_HSCHR6_MHC_DBB_CT( | olfactory receptor family 2 subfamily B memb |
| OR2J1 | CHR_HSCHR6_MHC_DBB_CT( | olfactory receptor family 2 subfamily J membe |
| OR2J3 | CHR_HSCHR6_MHC_DBB_CT( | olfactory receptor family 2 subfamily J membe |
| CR388415.1 | CHR_HSCHR6_MHC_DBB_CT( | novel transcript |
| CR388415.2 | CHR_HSCHR6_MHC_DBB_CT( | olfactory receptor, family 2, subfamily N, men |
| OR2J2 | CHR_HSCHR6_MHC_DBB_CT( | olfactory receptor family 2 subfamily J membe |
| OR2J4P | CHR_HSCHR6_MHC_DBB_CT( | olfactory receptor family 2 subfamily J membe |
| CR933537.1 | CHR_HSCHR6_MHC_DBB_CT( | olfactory receptor, family 2, subfamily H, men |
| CR753509.2 | CHR_HSCHR6_MHC_DBB_CT( | novel transcript |
| OR2G1P | CHR_HSCHR6_MHC_DBB_CT( | olfactory receptor family 2 subfamily G memb |
| OR2U1P | CHR_HSCHR6_MHC_DBB_CT( | olfactory receptor family 2 subfamily U memb |
| OR2U2P | CHR_HSCHR6_MHC_DBB_CT( | olfactory receptor family 2 subfamily U memb |
| CR753509.1 | CHR_HSCHR6_MHC_DBB_CT( | olfactory receptor, family 2, subfamily B, men |
| OR14J1 | CHR_HSCHR6_MHC_DBB_CT( | olfactory receptor family 14 subfamily J meml |

| | | |
|---|---|---|
| DDX6P1 | CHR_HSCHR6_MHC_DBB_CT( | DEAD-box helicase 6 pseudogene 1 [Source:H( |
| OR12D3 | CHR_HSCHR6_MHC_DBB_CT( | olfactory receptor family 12 subfamily D mem |
| OR12D2 | CHR_HSCHR6_MHC_DBB_CT( | olfactory receptor family 12 subfamily D mem |
| OR12D1 | CHR_HSCHR6_MHC_DBB_CT( | olfactory receptor family 12 subfamily D mem |
| OR5V1 | CHR_HSCHR6_MHC_DBB_CT( | olfactory receptor family 5 subfamily V memb |
| OR11A1 | CHR_HSCHR6_MHC_DBB_CT( | olfactory receptor family 11 subfamily A mem |
| OR10C1 | CHR_HSCHR6_MHC_DBB_CT( | olfactory receptor family 10 subfamily C mem |
| OR2H1 | CHR_HSCHR6_MHC_DBB_CT( | olfactory receptor family 2 subfamily H memb |
| UBDP1 | CHR_HSCHR6_MHC_DBB_CT( | ubiquitin D pseudogene 1 [Source:HGNC Symb |
| MAS1LP1 | CHR_HSCHR6_MHC_DBB_CT( | MAS1L pseudogene 1 [Source:HGNC Symbol;A |
| MAS1L | CHR_HSCHR6_MHC_DBB_CT( | MAS1 proto-oncogene like, G protein-couplec |
| RPS17P1 | CHR_HSCHR6_MHC_DBB_CT( | ribosomal protein S17 pseudogene 1 [Source: |
| LINC02829 | CHR_HSCHR6_MHC_DBB_CT( | long intergenic non-protein coding RNA 2829 |
| LINC01015 | CHR_HSCHR6_MHC_DBB_CT( | long intergenic non-protein coding RNA 1015 |
| GPR53P | CHR_HSCHR6_MHC_DBB_CT( | G protein-coupled receptor 53, pseudogene [S |
| OR2I1P | CHR_HSCHR6_MHC_DBB_CT( | olfactory receptor family 2 subfamily I membe |
| UBD | CHR_HSCHR6_MHC_DBB_CT( | ubiquitin D [Source:HGNC Symbol;Acc:HGNC:1 |
| GABBR1 | CHR_HSCHR6_MHC_DBB_CT( | gamma-aminobutyric acid type B receptor sub |
| OR2H5P | CHR_HSCHR6_MHC_DBB_CT( | olfactory receptor family 2 subfamily H memb |
| TMEM183AP1 | CHR_HSCHR6_MHC_DBB_CT( | transmembrane protein 183A pseudogene 1 [ |
| SNORD32B | CHR_HSCHR6_MHC_DBB_CT( | small nucleolar RNA, C/D box 32B [Source:HG |
| RPL13AP | CHR_HSCHR6_MHC_DBB_CT( | ribosomal protein L13a pseudogene [Source:H |
| OR2H2 | CHR_HSCHR6_MHC_DBB_CT( | olfactory receptor family 2 subfamily H memb |
| SUMO2P1 | CHR_HSCHR6_MHC_DBB_CT( | SUMO2 pseudogene 1 [Source:HGNC Symbol; |
| MOG | CHR_HSCHR6_MHC_DBB_CT( | myelin oligodendrocyte glycoprotein [Source: |
| ZFP57 | CHR_HSCHR6_MHC_DBB_CT( | ZFP57 zinc finger protein [Source:HGNC Symb |
| ZDHHC20P1 | CHR_HSCHR6_MHC_DBB_CT( | zinc finger DHHC-type containing 20 pseudoge |
| HLA-F | CHR_HSCHR6_MHC_DBB_CT( | major histocompatibility complex, class I, F [S( |
| CR753818.1 | CHR_HSCHR6_MHC_DBB_CT( | HLA complex group 4 pseudogene 11 |
| HLA-F-AS1 | CHR_HSCHR6_MHC_DBB_CT( | HLA-F antisense RNA 1 [Source:HGNC Symbol; |
| RPL23AP1 | CHR_HSCHR6_MHC_DBB_CT( | ribosomal protein L23a pseudogene 1 [Source |
| MICE | CHR_HSCHR6_MHC_DBB_CT( | MHC class I polypeptide-related sequence E (p |
| CR753818.2 | CHR_HSCHR6_MHC_DBB_CT( | HLA complex group 9 pseudogene 5 |
| Y_RNA | CHR_HSCHR6_MHC_DBB_CT( | Y RNA [Source:RFAM;Acc:RF00019] |
| IFITM4P | CHR_HSCHR6_MHC_DBB_CT( | interferon induced transmembrane protein 4 |
| HLA-V | CHR_HSCHR6_MHC_DBB_CT( | major histocompatibility complex, class I, V (p |
| HCG4 | CHR_HSCHR6_MHC_DBB_CT( | HLA complex group 4 [Source:HGNC Symbol;A |
| HLA-P | CHR_HSCHR6_MHC_DBB_CT( | major histocompatibility complex, class I, P (p |
| RPL7AP7 | CHR_HSCHR6_MHC_DBB_CT( | ribosomal protein L7a pseudogene 7 [Source:I |
| MICG | CHR_HSCHR6_MHC_DBB_CT( | MHC class I polypeptide-related sequence G (p |
| HLA-G | CHR_HSCHR6_MHC_DBB_CT( | major histocompatibility complex, class I, G [S |
| HCG4P8 | CHR_HSCHR6_MHC_DBB_CT( | HLA complex group 4 pseudogene 8 [Source:H |
| CR388220.3 | CHR_HSCHR6_MHC_DBB_CT( | MHC class I polypeptide-related sequence F ps |
| CR388220.2 | CHR_HSCHR6_MHC_DBB_CT( | HLA complex group 4 pseudogene 7 |
| HLA-A | CHR_HSCHR6_MHC_DBB_CT( | major histocompatibility complex, class I, A [S |
| HLA-H | CHR_HSCHR6_MHC_DBB_CT( | major histocompatibility complex, class I, H (p |
| HLA-T | CHR_HSCHR6_MHC_DBB_CT( | major histocompatibility complex, class I, T (p |
| DDX39BP1 | CHR_HSCHR6_MHC_DBB_CT( | DEAD-box helicase 39B pseudogene 1 [Source |
| MCCD1P1 | CHR_HSCHR6_MHC_DBB_CT( | mitochondrial coiled-coil domain 1 pseudogen |
| HCG4B | CHR_HSCHR6_MHC_DBB_CT( | HLA complex group 4B [Source:HGNC Symbol; |

| | | |
|---|---|---|
| HLA-K | CHR_HSCHR6_MHC_DBB_CT( | major histocompatibility complex, class I, K (p |
| HLA-U | CHR_HSCHR6_MHC_DBB_CT( | major histocompatibility complex, class I, U (p |
| CR388220.1 | CHR_HSCHR6_MHC_DBB_CT( | HLA complex group 4 pseudogene 5 |
| HLA-W | CHR_HSCHR6_MHC_DBB_CT( | major histocompatibility complex, class I, W (p |
| MICD | CHR_HSCHR6_MHC_DBB_CT( | MHC class I polypeptide-related sequence D (p |
| HCG9 | CHR_HSCHR6_MHC_DBB_CT( | HLA complex group 9 [Source:HGNC Symbol;A |
| DDX39BP2 | CHR_HSCHR6_MHC_DBB_CT( | DEAD-box helicase 39B pseudogene 2 [Source |
| MCCD1P2 | CHR_HSCHR6_MHC_DBB_CT( | mitochondrial coiled-coil domain 1 pseudogen |
| ZNRD1ASP | CHR_HSCHR6_MHC_DBB_CT( | zinc ribbon domain containing 1 antisense, ps |
| CR388205.1 | CHR_HSCHR6_MHC_DBB_CT( | HLA complex group 4 pseudogene 3 |
| HLA-J | CHR_HSCHR6_MHC_DBB_CT( | major histocompatibility complex, class I, J (ps |
| ETF1P1 | CHR_HSCHR6_MHC_DBB_CT( | eukaryotic translation termination factor 1 ps |
| CR388205.4 | CHR_HSCHR6_MHC_DBB_CT( | NULL |
| CR388205.2 | CHR_HSCHR6_MHC_DBB_CT( | NULL |
| ZNRD1 | CHR_HSCHR6_MHC_DBB_CT( | zinc ribbon domain containing 1 [Source:HGN |
| CR388205.3 | CHR_HSCHR6_MHC_DBB_CT( | NULL |
| PPP1R11 | CHR_HSCHR6_MHC_DBB_CT( | protein phosphatase 1 regulatory inhibitor sul |
| RNF39 | CHR_HSCHR6_MHC_DBB_CT( | ring finger protein 39 [Source:HGNC Symbol;A |
| TRIM31 | CHR_HSCHR6_MHC_DBB_CT( | tripartite motif containing 31 [Source:HGNC S |
| TRIM31-AS1 | CHR_HSCHR6_MHC_DBB_CT( | TRIM31 antisense RNA 1 [Source:HGNC Symb |
| CR753815.1 | CHR_HSCHR6_MHC_DBB_CT( | NULL |
| TRIM40 | CHR_HSCHR6_MHC_DBB_CT( | tripartite motif containing 40 [Source:HGNC S |
| TRIM10 | CHR_HSCHR6_MHC_DBB_CT( | tripartite motif containing 10 [Source:HGNC S |
| TRIM15 | CHR_HSCHR6_MHC_DBB_CT( | tripartite motif containing 15 [Source:HGNC S |
| TRIM26 | CHR_HSCHR6_MHC_DBB_CT( | tripartite motif containing 26 [Source:HGNC S |
| PAIP1P1 | CHR_HSCHR6_MHC_DBB_CT( | poly(A) binding protein interacting protein 1 p |
| HCG17 | CHR_HSCHR6_MHC_DBB_CT( | HLA complex group 17 [Source:HGNC Symbol; |
| TRIM26BP | CHR_HSCHR6_MHC_DBB_CT( | tripartite motif containing 26B, pseudogene [S |
| HLA-L | CHR_HSCHR6_MHC_DBB_CT( | major histocompatibility complex, class I, L (ps |
| HCG18 | CHR_HSCHR6_MHC_DBB_CT( | HLA complex group 18 [Source:HGNC Symbol; |
| TRIM39 | CHR_HSCHR6_MHC_DBB_CT( | tripartite motif containing 39 [Source:HGNC S |
| RPP21 | CHR_HSCHR6_MHC_DBB_CT( | ribonuclease P/MRP subunit p21 [Source:HGN |
| HLA-N | CHR_HSCHR6_MHC_DBB_CT( | major histocompatibility complex, class I, N (p |
| CR759281.1 | CHR_HSCHR6_MHC_DBB_CT( | HLA complex group 19 pseudogene |
| UBQLN1P1 | CHR_HSCHR6_MHC_DBB_CT( | ubiquilin 1 pseudogene 1 [Source:HGNC Symb |
| MICC | CHR_HSCHR6_MHC_DBB_CT( | MHC class I polypeptide-related sequence C (p |
| TMPOP1 | CHR_HSCHR6_MHC_DBB_CT( | thymopoietin pseudogene 1 [Source:HGNC Sy |
| SUCLA2P1 | CHR_HSCHR6_MHC_DBB_CT( | succinate-CoA ligase ADP-forming beta subun |
| RANP1 | CHR_HSCHR6_MHC_DBB_CT( | RAN pseudogene 1 [Source:HGNC Symbol;Acc |
| HLA-E | CHR_HSCHR6_MHC_DBB_CT( | major histocompatibility complex, class I, E [S |
| LINC02569 | CHR_HSCHR6_MHC_DBB_CT( | long intergenic non-protein coding RNA 2569 |
| GNL1 | CHR_HSCHR6_MHC_DBB_CT( | G protein nucleolar 1 (putative) [Source:HGN |
| PRR3 | CHR_HSCHR6_MHC_DBB_CT( | proline rich 3 [Source:HGNC Symbol;Acc:HGN |
| ABCF1 | CHR_HSCHR6_MHC_DBB_CT( | ATP binding cassette subfamily F member 1 [S |
| PPP1R10 | CHR_HSCHR6_MHC_DBB_CT( | protein phosphatase 1 regulatory subunit 10 [ |
| MRPS18B | CHR_HSCHR6_MHC_DBB_CT( | mitochondrial ribosomal protein S18B [Source |
| ATAT1 | CHR_HSCHR6_MHC_DBB_CT( | alpha tubulin acetyltransferase 1 [Source:HGN |
| PTMAP1 | CHR_HSCHR6_MHC_DBB_CT( | prothymosin alpha pseudogene 1 [Source:HGI |
| C6orf136 | CHR_HSCHR6_MHC_DBB_CT( | chromosome 6 open reading frame 136 [Sour |
| DHX16 | CHR_HSCHR6_MHC_DBB_CT( | DEAH-box helicase 16 [Source:HGNC Symbol;/ |

| | | |
|---|---|---|
| PPP1R18 | CHR_HSCHR6_MHC_DBB_CT( | protein phosphatase 1 regulatory subunit 18 [ |
| NRM | CHR_HSCHR6_MHC_DBB_CT( | nurim [Source:HGNC Symbol;Acc:HGNC:8003] |
| RPL7P4 | CHR_HSCHR6_MHC_DBB_CT( | ribosomal protein L7 pseudogene 4 [Source:H |
| MDC1 | CHR_HSCHR6_MHC_DBB_CT( | mediator of DNA damage checkpoint 1 [Sourc |
| MDC1-AS1 | CHR_HSCHR6_MHC_DBB_CT( | MDC1 antisense RNA 1 [Source:HGNC Symbol |
| TUBB | CHR_HSCHR6_MHC_DBB_CT( | tubulin beta class I [Source:HGNC Symbol;Acc |
| FLOT1 | CHR_HSCHR6_MHC_DBB_CT( | flotillin 1 [Source:HGNC Symbol;Acc:HGNC:37 |
| Y_RNA | CHR_HSCHR6_MHC_DBB_CT( | Y RNA [Source:RFAM;Acc:RF00019] |
| IER3 | CHR_HSCHR6_MHC_DBB_CT( | immediate early response 3 [Source:HGNC Sy |
| Metazoa_SRP | CHR_HSCHR6_MHC_DBB_CT( | Metazoan signal recognition particle RNA [Sou |
| HCG20 | CHR_HSCHR6_MHC_DBB_CT( | HLA complex group 20 [Source:HGNC Symbol; |
| CR388215.1 | CHR_HSCHR6_MHC_DBB_CT( | chromosome 6 open reading frame 214 (puta |
| LINC02570 | CHR_HSCHR6_MHC_DBB_CT( | long intergenic non-protein coding RNA 2570 |
| 7SK | CHR_HSCHR6_MHC_DBB_CT( | 7SK RNA [Source:RFAM;Acc:RF00100] |
| DDR1-DT | CHR_HSCHR6_MHC_DBB_CT( | DDR1 divergent transcript [Source:HGNC Sym |
| DDR1 | CHR_HSCHR6_MHC_DBB_CT( | discoidin domain receptor tyrosine kinase 1 [S |
| GTF2H4 | CHR_HSCHR6_MHC_DBB_CT( | general transcription factor IIH subunit 4 [Sou |
| CR759747.1 | CHR_HSCHR6_MHC_DBB_CT( | novel protein |
| VARS2 | CHR_HSCHR6_MHC_DBB_CT( | valyl-tRNA synthetase 2, mitochondrial [Sourc |
| SFTA2 | CHR_HSCHR6_MHC_DBB_CT( | surfactant associated 2 [Source:HGNC Symbol |
| Y_RNA | CHR_HSCHR6_MHC_DBB_CT( | Y RNA [Source:RFAM;Acc:RF00019] |
| MUCL3 | CHR_HSCHR6_MHC_DBB_CT( | mucin like 3 [Source:HGNC Symbol;Acc:HGNC |
| HCG21 | CHR_HSCHR6_MHC_DBB_CT( | HLA complex group 21 [Source:HGNC Symbol; |
| MUC21 | CHR_HSCHR6_MHC_DBB_CT( | mucin 21, cell surface associated [Source:HGN |
| CR753812.2 | CHR_HSCHR6_MHC_DBB_CT( | novel transcript |
| CR753812.1 | CHR_HSCHR6_MHC_DBB_CT( | HLA complex group family pseudogene |
| U6 | CHR_HSCHR6_MHC_DBB_CT( | U6 spliceosomal RNA [Source:RFAM;Acc:RF00 |
| C6orf15 | CHR_HSCHR6_MHC_DBB_CT( | chromosome 6 open reading frame 15 [Sourc |
| PSORS1C1 | CHR_HSCHR6_MHC_DBB_CT( | psoriasis susceptibility 1 candidate 1 [Source:I |
| CDSN | CHR_HSCHR6_MHC_DBB_CT( | corneodesmosin [Source:HGNC Symbol;Acc:H |
| PSORS1C2 | CHR_HSCHR6_MHC_DBB_CT( | psoriasis susceptibility 1 candidate 2 [Source:I |
| POLR2LP1 | CHR_HSCHR6_MHC_DBB_CT( | RNA polymerase II subunit L pseudogene 1 [Sc |
| CCHCR1 | CHR_HSCHR6_MHC_DBB_CT( | coiled-coil alpha-helical rod protein 1 [Source: |
| TCF19 | CHR_HSCHR6_MHC_DBB_CT( | transcription factor 19 [Source:HGNC Symbol; |
| POU5F1 | CHR_HSCHR6_MHC_DBB_CT( | POU class 5 homeobox 1 [Source:HGNC Symb |
| PSORS1C3 | CHR_HSCHR6_MHC_DBB_CT( | psoriasis susceptibility 1 candidate 3 [Source:I |
| HCG27 | CHR_HSCHR6_MHC_DBB_CT( | HLA complex group 27 [Source:HGNC Symbol; |
| CR388229.1 | CHR_HSCHR6_MHC_DBB_CT( | novel transcript |
| HLA-C | CHR_HSCHR6_MHC_DBB_CT( | major histocompatibility complex, class I, C [S |
| USP8P1 | CHR_HSCHR6_MHC_DBB_CT( | ubiquitin specific peptidase 8 pseudogene 1 [S |
| RPL3P2 | CHR_HSCHR6_MHC_DBB_CT( | ribosomal protein L3 pseudogene 2 [Source:H |
| WASF5P | CHR_HSCHR6_MHC_DBB_CT( | WASP family member 5, pseudogene [Source: |
| CR759814.1 | CHR_HSCHR6_MHC_DBB_CT( | novel HLA class I pseudogene fragment |
| Y_RNA | CHR_HSCHR6_MHC_DBB_CT( | Y RNA [Source:RFAM;Acc:RF00019] |
| MICB | CHR_HSCHR6_MHC_DBB_CT( | MHC class I polypeptide-related sequence B [S |
| PPIAP9 | CHR_HSCHR6_MHC_DBB_CT( | peptidylprolyl isomerase A pseudogene 9 [Sou |
| RPL15P4 | CHR_HSCHR6_MHC_DBB_CT( | ribosomal protein L15 pseudogene 4 [Source: |
| MCCD1 | CHR_HSCHR6_MHC_DBB_CT( | mitochondrial coiled-coil domain 1 [Source:H( |
| DDX39B | CHR_HSCHR6_MHC_DBB_CT( | DExD-box helicase 39B [Source:HGNC Symbol, |
| SNORD117 | CHR_HSCHR6_MHC_DBB_CT( | small nucleolar RNA, C/D box 117 [Source:HG |

| | | |
|---|---|---|
| SNORD84 | CHR_HSCHR6_MHC_DBB_CT( | small nucleolar RNA, C/D box 84 [Source:HGN |
| DDX39B-AS1 | CHR_HSCHR6_MHC_DBB_CT( | DDX39B antisense RNA 1 [Source:HGNC Symb |
| ATP6V1G2 | CHR_HSCHR6_MHC_DBB_CT( | ATPase H+ transporting V1 subunit G2 [Source |
| NFKBIL1 | CHR_HSCHR6_MHC_DBB_CT( | NFKB inhibitor like 1 [Source:HGNC Symbol;Ac |
| LTA | CHR_HSCHR6_MHC_DBB_CT( | lymphotoxin alpha [Source:HGNC Symbol;Acc |
| TNF | CHR_HSCHR6_MHC_DBB_CT( | tumor necrosis factor [Source:HGNC Symbol;A |
| LTB | CHR_HSCHR6_MHC_DBB_CT( | lymphotoxin beta [Source:HGNC Symbol;Acc:I |
| LST1 | CHR_HSCHR6_MHC_DBB_CT( | leukocyte specific transcript 1 [Source:HGNC S |
| NCR3 | CHR_HSCHR6_MHC_DBB_CT( | natural cytotoxicity triggering receptor 3 [Sou |
| UQCRHP1 | CHR_HSCHR6_MHC_DBB_CT( | ubiquinol-cytochrome c reductase hinge prote |
| AIF1 | CHR_HSCHR6_MHC_DBB_CT( | allograft inflammatory factor 1 [Source:HGNC |
| PRRC2A | CHR_HSCHR6_MHC_DBB_CT( | proline rich coiled-coil 2A [Source:HGNC Symb |
| SNORA38 | CHR_HSCHR6_MHC_DBB_CT( | small nucleolar RNA, H/ACA box 38 [Source:H |
| BAG6 | CHR_HSCHR6_MHC_DBB_CT( | BCL2 associated athanogene 6 [Source:HGNC |
| APOM | CHR_HSCHR6_MHC_DBB_CT( | apolipoprotein M [Source:HGNC Symbol;Acc:H |
| C6orf47 | CHR_HSCHR6_MHC_DBB_CT( | chromosome 6 open reading frame 47 [Source |
| C6orf47-AS1 | CHR_HSCHR6_MHC_DBB_CT( | C6orf47 antisense RNA 1 [Source:HGNC Symb |
| GPANK1 | CHR_HSCHR6_MHC_DBB_CT( | G-patch domain and ankyrin repeats 1 [Source |
| Y_RNA | CHR_HSCHR6_MHC_DBB_CT( | Y RNA [Source:RFAM;Acc:RF00019] |
| CSNK2B | CHR_HSCHR6_MHC_DBB_CT( | casein kinase 2 beta [Source:HGNC Symbol;Ac |
| CR753842.1 | CHR_HSCHR6_MHC_DBB_CT( | novel protein |
| LY6G5B | CHR_HSCHR6_MHC_DBB_CT( | lymphocyte antigen 6 family member G5B [So |
| LY6G5C | CHR_HSCHR6_MHC_DBB_CT( | lymphocyte antigen 6 family member G5C [So |
| ABHD16A | CHR_HSCHR6_MHC_DBB_CT( | abhydrolase domain containing 16A [Source:H |
| LY6G6F | CHR_HSCHR6_MHC_DBB_CT( | lymphocyte antigen 6 family member G6F [So |
| LY6G6F-LY6G6D | CHR_HSCHR6_MHC_DBB_CT( | LY6G6F-LY6G6D readthrough [Source:NCBI ge |
| LY6G6E | CHR_HSCHR6_MHC_DBB_CT( | lymphocyte antigen 6 family member G6E [So |
| LY6G6D | CHR_HSCHR6_MHC_DBB_CT( | lymphocyte antigen 6 family member G6D [So |
| MPIG6B | CHR_HSCHR6_MHC_DBB_CT( | megakaryocyte and platelet inhibitory recepto |
| LY6G6C | CHR_HSCHR6_MHC_DBB_CT( | lymphocyte antigen 6 family member G6C [So |
| DDAH2 | CHR_HSCHR6_MHC_DBB_CT( | dimethylarginine dimethylaminohydrolase 2 [ |
| CLIC1 | CHR_HSCHR6_MHC_DBB_CT( | chloride intracellular channel 1 [Source:HGNC |
| MSH5 | CHR_HSCHR6_MHC_DBB_CT( | mutS homolog 5 [Source:HGNC Symbol;Acc:H |
| MSH5-SAPCD1 | CHR_HSCHR6_MHC_DBB_CT( | MSH5-SAPCD1 readthrough (NMD candidate) |
| U6 | CHR_HSCHR6_MHC_DBB_CT( | U6 spliceosomal RNA [Source:RFAM;Acc:RF00 |
| SAPCD1 | CHR_HSCHR6_MHC_DBB_CT( | suppressor APC domain containing 1 [Source:I |
| SAPCD1-AS1 | CHR_HSCHR6_MHC_DBB_CT( | SAPCD1 antisense RNA 1 [Source:HGNC Symb |
| VWA7 | CHR_HSCHR6_MHC_DBB_CT( | von Willebrand factor A domain containing 7 [ |
| VARS | CHR_HSCHR6_MHC_DBB_CT( | valyl-tRNA synthetase [Source:HGNC Symbol;A |
| Y_RNA | CHR_HSCHR6_MHC_DBB_CT( | Y RNA [Source:RFAM;Acc:RF00019] |
| LSM2 | CHR_HSCHR6_MHC_DBB_CT( | LSM2 homolog, U6 small nuclear RNA and mR |
| HSPA1L | CHR_HSCHR6_MHC_DBB_CT( | heat shock protein family A (Hsp70) member |
| HSPA1A | CHR_HSCHR6_MHC_DBB_CT( | heat shock protein family A (Hsp70) member |
| HSPA1B | CHR_HSCHR6_MHC_DBB_CT( | heat shock protein family A (Hsp70) member |
| SNHG32 | CHR_HSCHR6_MHC_DBB_CT( | small nucleolar RNA host gene 32 [Source:HGI |
| SNORD48 | CHR_HSCHR6_MHC_DBB_CT( | small nucleolar RNA, C/D box 48 [Source:HGN |
| SNORD52 | CHR_HSCHR6_MHC_DBB_CT( | small nucleolar RNA, C/D box 52 [Source:HGN |
| NEU1 | CHR_HSCHR6_MHC_DBB_CT( | neuraminidase 1 [Source:HGNC Symbol;Acc:H |
| SLC44A4 | CHR_HSCHR6_MHC_DBB_CT( | solute carrier family 44 member 4 [Source:HG |
| EHMT2 | CHR_HSCHR6_MHC_DBB_CT( | euchromatic histone lysine methyltransferase |

| | | |
|---|---|---|
| CR388202.1 | CHR_HSCHR6_MHC_DBB_CT( | novel transcript |
| C2 | CHR_HSCHR6_MHC_DBB_CT( | complement C2 [Source:HGNC Symbol;Acc:HG |
| ZBTB12 | CHR_HSCHR6_MHC_DBB_CT( | zinc finger and BTB domain containing 12 [Sou |
| C4B-AS1 | CHR_HSCHR6_MHC_DBB_CT( | C4B antisense RNA 1 [Source:HGNC Symbol;Ac |
| CFB | CHR_HSCHR6_MHC_DBB_CT( | complement factor B [Source:HGNC Symbol;A |
| NELFE | CHR_HSCHR6_MHC_DBB_CT( | negative elongation factor complex member E |
| SKIV2L | CHR_HSCHR6_MHC_DBB_CT( | Ski2 like RNA helicase [Source:HGNC Symbol;A |
| DXO | CHR_HSCHR6_MHC_DBB_CT( | decapping exoribonuclease [Source:HGNC Syr |
| STK19 | CHR_HSCHR6_MHC_DBB_CT( | serine/threonine kinase 19 [Source:HGNC Syn |
| C4A | CHR_HSCHR6_MHC_DBB_CT( | complement C4A (Rodgers blood group) [Sour |
| C4A-AS1 | CHR_HSCHR6_MHC_DBB_CT( | C4A antisense RNA 1 [Source:HGNC Symbol;A |
| CYP21A1P | CHR_HSCHR6_MHC_DBB_CT( | cytochrome P450 family 21 subfamily A meml |
| CR753845.2 | CHR_HSCHR6_MHC_DBB_CT( | NULL |
| TNXA | CHR_HSCHR6_MHC_DBB_CT( | tenascin XA (pseudogene) [Source:HGNC Sym |
| STK19B | CHR_HSCHR6_MHC_DBB_CT( | serine/threonine kinase 19B (pseudogene) [Sc |
| C4B | CHR_HSCHR6_MHC_DBB_CT( | complement C4B (Chido blood group) [Source |
| CR753845.1 | CHR_HSCHR6_MHC_DBB_CT( | novel transcript |
| CYP21A2 | CHR_HSCHR6_MHC_DBB_CT( | cytochrome P450 family 21 subfamily A meml |
| TNXB | CHR_HSCHR6_MHC_DBB_CT( | tenascin XB [Source:HGNC Symbol;Acc:HGNC: |
| 5S_rRNA | CHR_HSCHR6_MHC_DBB_CT( | 5S ribosomal RNA [Source:RFAM;Acc:RF00001 |
| ATF6B | CHR_HSCHR6_MHC_DBB_CT( | activating transcription factor 6 beta [Source:I |
| FKBPL | CHR_HSCHR6_MHC_DBB_CT( | FKBP prolyl isomerase like [Source:HGNC Sym |
| PRRT1 | CHR_HSCHR6_MHC_DBB_CT( | proline rich transmembrane protein 1 [Source |
| PPT2 | CHR_HSCHR6_MHC_DBB_CT( | palmitoyl-protein thioesterase 2 [Source:HGN |
| TSBP1 | CHR_HSCHR6_MHC_DBB_CT( | testis expressed basic protein 1 [Source:HGNC |
| HNRNPA1P2 | CHR_HSCHR6_MHC_DBB_CT( | heterogeneous nuclear ribonucleoprotein A1 |
| U6 | CHR_HSCHR6_MHC_DBB_CT( | U6 spliceosomal RNA [Source:RFAM;Acc:RF00 |
| TSBP1-AS1 | CHR_HSCHR6_MHC_DBB_CT( | TSBP1 and BTNL2 antisense RNA 1 [Source:HG |
| BTNL2 | CHR_HSCHR6_MHC_DBB_CT( | butyrophilin like 2 [Source:HGNC Symbol;Acc: |
| HLA-DRA | CHR_HSCHR6_MHC_DBB_CT( | major histocompatibility complex, class II, DR |
| HLA-DRB9 | CHR_HSCHR6_MHC_DBB_CT( | major histocompatibility complex, class II, DR |
| CR759837.1 | CHR_HSCHR6_MHC_DBB_CT( | major histocompatibility complex, class II, DR |
| CR753841.1 | CHR_HSCHR6_MHC_DBB_CT( | family with sequence similarity 136, member |
| CR753309.1 | CHR_HSCHR6_MHC_DBB_CT( | major histocompatibility complex, class II, DR |
| CR753841.2 | CHR_HSCHR6_MHC_DBB_CT( | family with sequence similarity 8, member A5 |
| U1 | CHR_HSCHR6_MHC_DBB_CT( | U1 spliceosomal RNA [Source:RFAM;Acc:RF00 |
| HLA-DRB1 | CHR_HSCHR6_MHC_DBB_CT( | major histocompatibility complex, class II, DR |
| HLA-DQA1 | CHR_HSCHR6_MHC_DBB_CT( | major histocompatibility complex, class II, DQ |
| HLA-DQB1 | CHR_HSCHR6_MHC_DBB_CT( | major histocompatibility complex, class II, DQ |
| HLA-DQB1-AS1 | CHR_HSCHR6_MHC_DBB_CT( | HLA-DQB1 antisense RNA 1 [Source:HGNC Syr |
| CR753846.2 | CHR_HSCHR6_MHC_DBB_CT( | novel transcript |
| MTCO3P1 | CHR_HSCHR6_MHC_DBB_CT( | MT-CO3 pseudogene 1 [Source:HGNC Symbol |
| CR753846.1 | CHR_HSCHR6_MHC_DBB_CT( | uncharacterized LOC102725019 [Source:NCBI |
| HLA-DQB3 | CHR_HSCHR6_MHC_DBB_CT( | major histocompatibility complex, class II, DQ |
| HLA-DQA2 | CHR_HSCHR6_MHC_DBB_CT( | major histocompatibility complex, class II, DQ |
| HLA-DQB2 | CHR_HSCHR6_MHC_DBB_CT( | major histocompatibility complex, class II, DQ |
| HLA-DOB | CHR_HSCHR6_MHC_DBB_CT( | major histocompatibility complex, class II, DO |
| TAP2 | CHR_HSCHR6_MHC_DBB_CT( | transporter 2, ATP binding cassette subfamily |
| PSMB8 | CHR_HSCHR6_MHC_DBB_CT( | proteasome subunit beta 8 [Source:HGNC Syn |
| PSMB8-AS1 | CHR_HSCHR6_MHC_DBB_CT( | PSMB8 antisense RNA 1 (head to head) [Sourc |

| | | |
|---|---|---|
| PSMB9 | CHR_HSCHR6_MHC_DBB_CT( | proteasome subunit beta 9 [Source:HGNC Syn |
| TAP1 | CHR_HSCHR6_MHC_DBB_CT( | transporter 1, ATP binding cassette subfamily |
| PPP1R2P1 | CHR_HSCHR6_MHC_DBB_CT( | protein phosphatase 1 regulatory inhibitor sul |
| HLA-Z | CHR_HSCHR6_MHC_DBB_CT( | major histocompatibility complex, class I, Z (p: |
| HLA-DMB | CHR_HSCHR6_MHC_DBB_CT( | major histocompatibility complex, class II, DM |
| HLA-DMA | CHR_HSCHR6_MHC_DBB_CT( | major histocompatibility complex, class II, DM |
| BRD2 | CHR_HSCHR6_MHC_DBB_CT( | bromodomain containing 2 [Source:HGNC Syn |
| HLA-DOA | CHR_HSCHR6_MHC_DBB_CT( | major histocompatibility complex, class II, DO |
| HLA-DPA1 | CHR_HSCHR6_MHC_DBB_CT( | major histocompatibility complex, class II, DP |
| HLA-DPB1 | CHR_HSCHR6_MHC_DBB_CT( | major histocompatibility complex, class II, DP |
| RPL32P1 | CHR_HSCHR6_MHC_DBB_CT( | ribosomal protein L32 pseudogene 1 [Source: |
| HLA-DPA2 | CHR_HSCHR6_MHC_DBB_CT( | major histocompatibility complex, class II, DP |
| COL11A2P1 | CHR_HSCHR6_MHC_DBB_CT( | collagen type XI alpha 2 pseudogene 1 [Source |
| HLA-DPB2 | CHR_HSCHR6_MHC_DBB_CT( | major histocompatibility complex, class II, DP |
| HLA-DPA3 | CHR_HSCHR6_MHC_DBB_CT( | major histocompatibility complex, class II, DP |
| HCG24 | CHR_HSCHR6_MHC_DBB_CT( | HLA complex group 24 [Source:HGNC Symbol; |
| COL11A2 | CHR_HSCHR6_MHC_DBB_CT( | collagen type XI alpha 2 chain [Source:HGNC S |
| RXRB | CHR_HSCHR6_MHC_DBB_CT( | retinoid X receptor beta [Source:HGNC Symbo |
| Y_RNA | CHR_HSCHR6_MHC_DBB_CT( | Y RNA [Source:RFAM;Acc:RF00019] |
| SLC39A7 | CHR_HSCHR6_MHC_DBB_CT( | solute carrier family 39 member 7 [Source:HG |
| HSD17B8 | CHR_HSCHR6_MHC_DBB_CT( | hydroxysteroid 17-beta dehydrogenase 8 [Sou |
| RING1 | CHR_HSCHR6_MHC_DBB_CT( | ring finger protein 1 [Source:HGNC Symbol;Ac |
| ZNF70P1 | CHR_HSCHR6_MHC_DBB_CT( | zinc finger protein 70 pseudogene 1 [Source:H |
| HTATSF1P1 | CHR_HSCHR6_MHC_DBB_CT( | HIV-1 Tat specific factor 1 pseudogene 1 [Sour |
| HCG25 | CHR_HSCHR6_MHC_DBB_CT( | HLA complex group 25 [Source:HGNC Symbol; |
| VPS52 | CHR_HSCHR6_MHC_DBB_CT( | VPS52 subunit of GARP complex [Source:HGN |
| RPS18 | CHR_HSCHR6_MHC_DBB_CT( | ribosomal protein S18 [Source:HGNC Symbol;. |
| B3GALT4 | CHR_HSCHR6_MHC_DBB_CT( | beta-1,3-galactosyltransferase 4 [Source:HGN |
| WDR46 | CHR_HSCHR6_MHC_DBB_CT( | WD repeat domain 46 [Source:HGNC Symbol;. |
| PFDN6 | CHR_HSCHR6_MHC_DBB_CT( | prefoldin subunit 6 [Source:HGNC Symbol;Acc |
| RGL2 | CHR_HSCHR6_MHC_DBB_CT( | ral guanine nucleotide dissociation stimulator |
| TAPBP | CHR_HSCHR6_MHC_DBB_CT( | TAP binding protein [Source:HGNC Symbol;Ac |
| ZBTB22 | CHR_HSCHR6_MHC_DBB_CT( | zinc finger and BTB domain containing 22 [Sou |
| DAXX | CHR_HSCHR6_MHC_DBB_CT( | death domain associated protein [Source:HGN |
| MYL8P | CHR_HSCHR6_MHC_DBB_CT( | myosin light chain 8, pseudogene [Source:HGI |
| NOP56P1 | CHR_HSCHR6_MHC_MANN_( | NOP56 ribonucleoprotein pseudogene 1 [Sour |
| BX908720.1 | CHR_HSCHR6_MHC_MANN_( | novel transcript |
| LINC01623 | CHR_HSCHR6_MHC_MANN_( | long intergenic non-protein coding RNA 1623 |
| RPL13P | CHR_HSCHR6_MHC_MANN_( | ribosomal protein L13 pseudogene [Source:H( |
| TRIM27 | CHR_HSCHR6_MHC_MANN_( | tripartite motif containing 27 [Source:HGNC S |
| U6 | CHR_HSCHR6_MHC_MANN_( | U6 spliceosomal RNA [Source:RFAM;Acc:RF00 |
| LINC01556 | CHR_HSCHR6_MHC_MANN_( | long intergenic non-protein coding RNA 1556 |
| KRT18P1 | CHR_HSCHR6_MHC_MANN_( | keratin 18 pseudogene 1 [Source:HGNC Symb |
| Metazoa_SRP | CHR_HSCHR6_MHC_MANN_( | Metazoan signal recognition particle RNA [Sou |
| HCG15 | CHR_HSCHR6_MHC_MANN_( | HLA complex group 15 [Source:HGNC Symbol; |
| BX927167.1 | CHR_HSCHR6_MHC_MANN_( | HLA complex group 16 |
| ZNF311 | CHR_HSCHR6_MHC_MANN_( | zinc finger protein 311 [Source:HGNC Symbol; |
| OR2AD1P | CHR_HSCHR6_MHC_MANN_( | olfactory receptor family 2 subfamily AD mem |
| OR2W1 | CHR_HSCHR6_MHC_MANN_( | olfactory receptor family 2 subfamily W meml |
| CR547123.1 | CHR_HSCHR6_MHC_MANN_( | NULL |

| | | |
|---|---|---|
| OR2P1P | CHR_HSCHR6_MHC_MANN_( | olfactory receptor family 2 subfamily P memb |
| SAR1AP1 | CHR_HSCHR6_MHC_MANN_( | secretion associated Ras related GTPase 1A ps |
| OR2B3 | CHR_HSCHR6_MHC_MANN_( | olfactory receptor family 2 subfamily B memb |
| OR2J1 | CHR_HSCHR6_MHC_MANN_( | olfactory receptor family 2 subfamily J membe |
| OR2J3 | CHR_HSCHR6_MHC_MANN_( | olfactory receptor family 2 subfamily J membe |
| BX927132.2 | CHR_HSCHR6_MHC_MANN_( | novel transcript |
| BX927132.1 | CHR_HSCHR6_MHC_MANN_( | olfactory receptor, family 2, subfamily N, men |
| OR2J2 | CHR_HSCHR6_MHC_MANN_( | olfactory receptor family 2 subfamily J membe |
| OR2J4P | CHR_HSCHR6_MHC_MANN_( | olfactory receptor family 2 subfamily J membe |
| BX908738.3 | CHR_HSCHR6_MHC_MANN_( | olfactory receptor, family 2, subfamily H, men |
| BX908738.1 | CHR_HSCHR6_MHC_MANN_( | novel transcript |
| OR2G1P | CHR_HSCHR6_MHC_MANN_( | olfactory receptor family 2 subfamily G memb |
| OR2U1P | CHR_HSCHR6_MHC_MANN_( | olfactory receptor family 2 subfamily U memb |
| OR2U2P | CHR_HSCHR6_MHC_MANN_( | olfactory receptor family 2 subfamily U memb |
| BX908738.2 | CHR_HSCHR6_MHC_MANN_( | olfactory receptor, family 2, subfamily B, men |
| OR14J1 | CHR_HSCHR6_MHC_MANN_( | olfactory receptor family 14 subfamily J meml |
| DDX6P1 | CHR_HSCHR6_MHC_MANN_( | DEAD-box helicase 6 pseudogene 1 [Source:H |
| OR5V1 | CHR_HSCHR6_MHC_MANN_( | olfactory receptor family 5 subfamily V memb |
| OR12D3 | CHR_HSCHR6_MHC_MANN_( | olfactory receptor family 12 subfamily D mem |
| OR12D2 | CHR_HSCHR6_MHC_MANN_( | olfactory receptor family 12 subfamily D mem |
| OR12D1 | CHR_HSCHR6_MHC_MANN_( | olfactory receptor family 12 subfamily D mem |
| OR11A1 | CHR_HSCHR6_MHC_MANN_( | olfactory receptor family 11 subfamily A mem |
| OR10C1 | CHR_HSCHR6_MHC_MANN_( | olfactory receptor family 10 subfamily C mem |
| OR2H1 | CHR_HSCHR6_MHC_MANN_( | olfactory receptor family 2 subfamily H memb |
| UBDP1 | CHR_HSCHR6_MHC_MANN_( | ubiquitin D pseudogene 1 [Source:HGNC Syml |
| MAS1LP1 | CHR_HSCHR6_MHC_MANN_( | MAS1L pseudogene 1 [Source:HGNC Symbol;/ |
| MAS1L | CHR_HSCHR6_MHC_MANN_( | MAS1 proto-oncogene like, G protein-couplec |
| RPS17P1 | CHR_HSCHR6_MHC_MANN_( | ribosomal protein S17 pseudogene 1 [Source: |
| LINC02829 | CHR_HSCHR6_MHC_MANN_( | long intergenic non-protein coding RNA 2829 |
| LINC01015 | CHR_HSCHR6_MHC_MANN_( | long intergenic non-protein coding RNA 1015 |
| GPR53P | CHR_HSCHR6_MHC_MANN_( | G protein-coupled receptor 53, pseudogene [S |
| OR2I1P | CHR_HSCHR6_MHC_MANN_( | olfactory receptor family 2 subfamily I membe |
| UBD | CHR_HSCHR6_MHC_MANN_( | ubiquitin D [Source:HGNC Symbol;Acc:HGNC:1 |
| GABBR1 | CHR_HSCHR6_MHC_MANN_( | gamma-aminobutyric acid type B receptor sul |
| OR2H5P | CHR_HSCHR6_MHC_MANN_( | olfactory receptor family 2 subfamily H memb |
| TMEM183AP1 | CHR_HSCHR6_MHC_MANN_( | transmembrane protein 183A pseudogene 1 [ |
| SNORD32B | CHR_HSCHR6_MHC_MANN_( | small nucleolar RNA, C/D box 32B [Source:HG |
| RPL13AP | CHR_HSCHR6_MHC_MANN_( | ribosomal protein L13a pseudogene [Source:H |
| OR2H2 | CHR_HSCHR6_MHC_MANN_( | olfactory receptor family 2 subfamily H memb |
| SUMO2P1 | CHR_HSCHR6_MHC_MANN_( | SUMO2 pseudogene 1 [Source:HGNC Symbol; |
| MOG | CHR_HSCHR6_MHC_MANN_( | myelin oligodendrocyte glycoprotein [Source: |
| ZFP57 | CHR_HSCHR6_MHC_MANN_( | ZFP57 zinc finger protein [Source:HGNC Symb |
| ZDHHC20P1 | CHR_HSCHR6_MHC_MANN_( | zinc finger DHHC-type containing 20 pseudoge |
| HLA-F | CHR_HSCHR6_MHC_MANN_( | major histocompatibility complex, class I, F [S |
| BX927250.1 | CHR_HSCHR6_MHC_MANN_( | HLA complex group 4 pseudogene 11 |
| HLA-F-AS1 | CHR_HSCHR6_MHC_MANN_( | HLA-F antisense RNA 1 [Source:HGNC Symbol; |
| RPL23AP1 | CHR_HSCHR6_MHC_MANN_( | ribosomal protein L23a pseudogene 1 [Source |
| MICE | CHR_HSCHR6_MHC_MANN_( | MHC class I polypeptide-related sequence E (p |
| BX927182.1 | CHR_HSCHR6_MHC_MANN_( | HLA complex group 9 pseudogene 5 |
| Y_RNA | CHR_HSCHR6_MHC_MANN_( | Y RNA [Source:RFAM;Acc:RF00019] |

| | | |
|---|---|---|
| IFITM4P | CHR_HSCHR6_MHC_MANN_( | interferon induced transmembrane protein 4 |
| HLA-V | CHR_HSCHR6_MHC_MANN_( | major histocompatibility complex, class I, V (p |
| HCG4 | CHR_HSCHR6_MHC_MANN_( | HLA complex group 4 [Source:HGNC Symbol;A |
| HLA-P | CHR_HSCHR6_MHC_MANN_( | major histocompatibility complex, class I, P (p |
| RPL7AP7 | CHR_HSCHR6_MHC_MANN_( | ribosomal protein L7a pseudogene 7 [Source:I |
| MICG | CHR_HSCHR6_MHC_MANN_( | MHC class I polypeptide-related sequence G (ı |
| HLA-G | CHR_HSCHR6_MHC_MANN_( | major histocompatibility complex, class I, G [S |
| HCG4P8 | CHR_HSCHR6_MHC_MANN_( | HLA complex group 4 pseudogene 8 [Source:H |
| BX927141.2 | CHR_HSCHR6_MHC_MANN_( | MHC class I polypeptide-related sequence F p: |
| BX927141.1 | CHR_HSCHR6_MHC_MANN_( | HLA complex group 4 pseudogene 7 |
| HLA-H | CHR_HSCHR6_MHC_MANN_( | major histocompatibility complex, class I, H (p |
| HLA-T | CHR_HSCHR6_MHC_MANN_( | major histocompatibility complex, class I, T (p |
| DDX39BP1 | CHR_HSCHR6_MHC_MANN_( | DEAD-box helicase 39B pseudogene 1 [Source |
| MCCD1P1 | CHR_HSCHR6_MHC_MANN_( | mitochondrial coiled-coil domain 1 pseudoger |
| HCG4B | CHR_HSCHR6_MHC_MANN_( | HLA complex group 4B [Source:HGNC Symbol; |
| HLA-K | CHR_HSCHR6_MHC_MANN_( | major histocompatibility complex, class I, K (p |
| HLA-U | CHR_HSCHR6_MHC_MANN_( | major histocompatibility complex, class I, U (p |
| HLA-A | CHR_HSCHR6_MHC_MANN_( | major histocompatibility complex, class I, A [S |
| CR382333.1 | CHR_HSCHR6_MHC_MANN_( | HLA complex group 4 pseudogene 5 |
| HLA-W | CHR_HSCHR6_MHC_MANN_( | major histocompatibility complex, class I, W (ı |
| MICD | CHR_HSCHR6_MHC_MANN_( | MHC class I polypeptide-related sequence D (ı |
| HCG9 | CHR_HSCHR6_MHC_MANN_( | HLA complex group 9 [Source:HGNC Symbol;A |
| DDX39BP2 | CHR_HSCHR6_MHC_MANN_( | DEAD-box helicase 39B pseudogene 2 [Source |
| MCCD1P2 | CHR_HSCHR6_MHC_MANN_( | mitochondrial coiled-coil domain 1 pseudoger |
| ZNRD1ASP | CHR_HSCHR6_MHC_MANN_( | zinc ribbon domain containing 1 antisense, ps |
| BX927229.1 | CHR_HSCHR6_MHC_MANN_( | HLA complex group 4 pseudogene 3 |
| HLA-J | CHR_HSCHR6_MHC_MANN_( | major histocompatibility complex, class I, J (ps |
| ETF1P1 | CHR_HSCHR6_MHC_MANN_( | eukaryotic translation termination factor 1 ps |
| BX927229.4 | CHR_HSCHR6_MHC_MANN_( | NULL |
| BX927229.3 | CHR_HSCHR6_MHC_MANN_( | NULL |
| ZNRD1 | CHR_HSCHR6_MHC_MANN_( | zinc ribbon domain containing 1 [Source:HGN |
| BX927229.2 | CHR_HSCHR6_MHC_MANN_( | NULL |
| PPP1R11 | CHR_HSCHR6_MHC_MANN_( | protein phosphatase 1 regulatory inhibitor sul |
| RNF39 | CHR_HSCHR6_MHC_MANN_( | ring finger protein 39 [Source:HGNC Symbol;A |
| TRIM31 | CHR_HSCHR6_MHC_MANN_( | tripartite motif containing 31 [Source:HGNC S |
| TRIM31-AS1 | CHR_HSCHR6_MHC_MANN_( | TRIM31 antisense RNA 1 [Source:HGNC Symb |
| BX927221.1 | CHR_HSCHR6_MHC_MANN_( | NULL |
| TRIM40 | CHR_HSCHR6_MHC_MANN_( | tripartite motif containing 40 [Source:HGNC S |
| TRIM10 | CHR_HSCHR6_MHC_MANN_( | tripartite motif containing 10 [Source:HGNC S |
| TRIM15 | CHR_HSCHR6_MHC_MANN_( | tripartite motif containing 15 [Source:HGNC S |
| TRIM26 | CHR_HSCHR6_MHC_MANN_( | tripartite motif containing 26 [Source:HGNC S |
| PAIP1P1 | CHR_HSCHR6_MHC_MANN_( | poly(A) binding protein interacting protein 1 p |
| HCG17 | CHR_HSCHR6_MHC_MANN_( | HLA complex group 17 [Source:HGNC Symbol; |
| TRIM26BP | CHR_HSCHR6_MHC_MANN_( | tripartite motif containing 26B, pseudogene [S |
| HLA-L | CHR_HSCHR6_MHC_MANN_( | major histocompatibility complex, class I, L (ps |
| HCG18 | CHR_HSCHR6_MHC_MANN_( | HLA complex group 18 [Source:HGNC Symbol; |
| TRIM39 | CHR_HSCHR6_MHC_MANN_( | tripartite motif containing 39 [Source:HGNC S |
| RPP21 | CHR_HSCHR6_MHC_MANN_( | ribonuclease P/MRP subunit p21 [Source:HGN |
| HLA-N | CHR_HSCHR6_MHC_MANN_( | major histocompatibility complex, class I, N (p |
| BX927214.1 | CHR_HSCHR6_MHC_MANN_( | HLA complex group 19 pseudogene |

| | | |
|---|---|---|
| UBQLN1P1 | CHR_HSCHR6_MHC_MANN_( | ubiquilin 1 pseudogene 1 [Source:HGNC Symb |
| MICC | CHR_HSCHR6_MHC_MANN_( | MHC class I polypeptide-related sequence C (p |
| TMPOP1 | CHR_HSCHR6_MHC_MANN_( | thymopoietin pseudogene 1 [Source:HGNC Sy |
| SUCLA2P1 | CHR_HSCHR6_MHC_MANN_( | succinate-CoA ligase ADP-forming beta subun |
| RANP1 | CHR_HSCHR6_MHC_MANN_( | RAN pseudogene 1 [Source:HGNC Symbol;Acc |
| HLA-E | CHR_HSCHR6_MHC_MANN_( | major histocompatibility complex, class I, E [S |
| LINC02569 | CHR_HSCHR6_MHC_MANN_( | long intergenic non-protein coding RNA 2569 |
| GNL1 | CHR_HSCHR6_MHC_MANN_( | G protein nucleolar 1 (putative) [Source:HGN( |
| PRR3 | CHR_HSCHR6_MHC_MANN_( | proline rich 3 [Source:HGNC Symbol;Acc:HGN |
| ABCF1 | CHR_HSCHR6_MHC_MANN_( | ATP binding cassette subfamily F member 1 [S |
| PPP1R10 | CHR_HSCHR6_MHC_MANN_( | protein phosphatase 1 regulatory subunit 10 [ |
| MRPS18B | CHR_HSCHR6_MHC_MANN_( | mitochondrial ribosomal protein S18B [Source |
| ATAT1 | CHR_HSCHR6_MHC_MANN_( | alpha tubulin acetyltransferase 1 [Source:HGN |
| PTMAP1 | CHR_HSCHR6_MHC_MANN_( | prothymosin alpha pseudogene 1 [Source:HGI |
| C6orf136 | CHR_HSCHR6_MHC_MANN_( | chromosome 6 open reading frame 136 [Sour |
| DHX16 | CHR_HSCHR6_MHC_MANN_( | DEAH-box helicase 16 [Source:HGNC Symbol;/ |
| PPP1R18 | CHR_HSCHR6_MHC_MANN_( | protein phosphatase 1 regulatory subunit 18 [ |
| NRM | CHR_HSCHR6_MHC_MANN_( | nurim [Source:HGNC Symbol;Acc:HGNC:8003] |
| RPL7P4 | CHR_HSCHR6_MHC_MANN_( | ribosomal protein L7 pseudogene 4 [Source:H |
| MDC1 | CHR_HSCHR6_MHC_MANN_( | mediator of DNA damage checkpoint 1 [Sourc |
| MDC1-AS1 | CHR_HSCHR6_MHC_MANN_( | MDC1 antisense RNA 1 [Source:HGNC Symbol |
| TUBB | CHR_HSCHR6_MHC_MANN_( | tubulin beta class I [Source:HGNC Symbol;Acc |
| FLOT1 | CHR_HSCHR6_MHC_MANN_( | flotillin 1 [Source:HGNC Symbol;Acc:HGNC:37 |
| IER3 | CHR_HSCHR6_MHC_MANN_( | immediate early response 3 [Source:HGNC Sy |
| Metazoa_SRP | CHR_HSCHR6_MHC_MANN_( | Metazoan signal recognition particle RNA [Sou |
| HCG20 | CHR_HSCHR6_MHC_MANN_( | HLA complex group 20 [Source:HGNC Symbol; |
| CR847861.1 | CHR_HSCHR6_MHC_MANN_( | chromosome 6 open reading frame 214 (puta |
| LINC02570 | CHR_HSCHR6_MHC_MANN_( | long intergenic non-protein coding RNA 2570 |
| 7SK | CHR_HSCHR6_MHC_MANN_( | 7SK RNA [Source:RFAM;Acc:RF00100] |
| DDR1-DT | CHR_HSCHR6_MHC_MANN_( | DDR1 divergent transcript [Source:HGNC Sym |
| DDR1 | CHR_HSCHR6_MHC_MANN_( | discoidin domain receptor tyrosine kinase 1 [S |
| GTF2H4 | CHR_HSCHR6_MHC_MANN_( | general transcription factor IIH subunit 4 [Sou |
| BX927194.1 | CHR_HSCHR6_MHC_MANN_( | novel protein |
| VARS2 | CHR_HSCHR6_MHC_MANN_( | valyl-tRNA synthetase 2, mitochondrial [Sourc |
| SFTA2 | CHR_HSCHR6_MHC_MANN_( | surfactant associated 2 [Source:HGNC Symbol |
| Y_RNA | CHR_HSCHR6_MHC_MANN_( | Y RNA [Source:RFAM;Acc:RF00019] |
| MUC21 | CHR_HSCHR6_MHC_MANN_( | mucin 21, cell surface associated [Source:NCB |
| BX927139.1 | CHR_HSCHR6_MHC_MANN_( | novel transcript |
| BX927139.2 | CHR_HSCHR6_MHC_MANN_( | HLA complex group family pseudogene |
| U6 | CHR_HSCHR6_MHC_MANN_( | U6 spliceosomal RNA [Source:RFAM;Acc:RF00 |
| C6orf15 | CHR_HSCHR6_MHC_MANN_( | chromosome 6 open reading frame 15 [Sourc |
| PSORS1C1 | CHR_HSCHR6_MHC_MANN_( | psoriasis susceptibility 1 candidate 1 [Source:I |
| CDSN | CHR_HSCHR6_MHC_MANN_( | corneodesmosin [Source:HGNC Symbol;Acc:H |
| PSORS1C2 | CHR_HSCHR6_MHC_MANN_( | psoriasis susceptibility 1 candidate 2 [Source:I |
| POLR2LP1 | CHR_HSCHR6_MHC_MANN_( | RNA polymerase II subunit L pseudogene 1 [Sc |
| CCHCR1 | CHR_HSCHR6_MHC_MANN_( | coiled-coil alpha-helical rod protein 1 [Source: |
| TCF19 | CHR_HSCHR6_MHC_MANN_( | transcription factor 19 [Source:HGNC Symbol; |
| POU5F1 | CHR_HSCHR6_MHC_MANN_( | POU class 5 homeobox 1 [Source:HGNC Symb |
| CR847794.2 | CHR_HSCHR6_MHC_MANN_( | NULL |
| PSORS1C3 | CHR_HSCHR6_MHC_MANN_( | psoriasis susceptibility 1 candidate 3 [Source:I |

| | | |
|---|---|---|
| HCG27 | CHR_HSCHR6_MHC_MANN_( | HLA complex group 27 [Source:HGNC Symbol; |
| CR847794.1 | CHR_HSCHR6_MHC_MANN_( | novel transcript |
| HLA-C | CHR_HSCHR6_MHC_MANN_( | major histocompatibility complex, class I, C [S |
| HLA-B | CHR_HSCHR6_MHC_MANN_( | major histocompatibility complex, class I, B [S |
| USP8P1 | CHR_HSCHR6_MHC_MANN_( | ubiquitin specific peptidase 8 pseudogene 1 [S |
| RPL3P2 | CHR_HSCHR6_MHC_MANN_( | ribosomal protein L3 pseudogene 2 [Source:H |
| WASF5P | CHR_HSCHR6_MHC_MANN_( | WASP family member 5, pseudogene [Source: |
| BX927178.1 | CHR_HSCHR6_MHC_MANN_( | novel HLA class I pseudogene fragment |
| DHFRP2 | CHR_HSCHR6_MHC_MANN_( | dihydrofolate reductase pseudogene 2 [Sourc |
| U6 | CHR_HSCHR6_MHC_MANN_( | U6 spliceosomal RNA [Source:RFAM;Acc:RF00 |
| CR318637.1 | CHR_HSCHR6_MHC_MANN_( | fibroblast growth factor receptor 3 pseudoger |
| ZDHHC20P2 | CHR_HSCHR6_MHC_MANN_( | zinc finger DHHC-type containing 20 pseudoge |
| HLA-S | CHR_HSCHR6_MHC_MANN_( | major histocompatibility complex, class I, S (p: |
| MICA | CHR_HSCHR6_MHC_MANN_( | MHC class I polypeptide-related sequence A [S |
| Y_RNA | CHR_HSCHR6_MHC_MANN_( | Y RNA [Source:RFAM;Acc:RF00019] |
| LINC01149 | CHR_HSCHR6_MHC_MANN_( | long intergenic non-protein coding RNA 1149 |
| MICB | CHR_HSCHR6_MHC_MANN_( | MHC class I polypeptide-related sequence B [S |
| PPIAP9 | CHR_HSCHR6_MHC_MANN_( | peptidylprolyl isomerase A pseudogene 9 [Sou |
| RPL15P4 | CHR_HSCHR6_MHC_MANN_( | ribosomal protein L15 pseudogene 4 [Source: |
| MCCD1 | CHR_HSCHR6_MHC_MANN_( | mitochondrial coiled-coil domain 1 [Source:H |
| DDX39B | CHR_HSCHR6_MHC_MANN_( | DExD-box helicase 39B [Source:HGNC Symbol |
| SNORD117 | CHR_HSCHR6_MHC_MANN_( | small nucleolar RNA, C/D box 117 [Source:HG |
| SNORD84 | CHR_HSCHR6_MHC_MANN_( | small nucleolar RNA, C/D box 84 [Source:HGN |
| DDX39B-AS1 | CHR_HSCHR6_MHC_MANN_( | DDX39B antisense RNA 1 [Source:HGNC Symb |
| ATP6V1G2 | CHR_HSCHR6_MHC_MANN_( | ATPase H+ transporting V1 subunit G2 [Source |
| NFKBIL1 | CHR_HSCHR6_MHC_MANN_( | NFKB inhibitor like 1 [Source:HGNC Symbol;Ac |
| LTA | CHR_HSCHR6_MHC_MANN_( | lymphotoxin alpha [Source:HGNC Symbol;Acc |
| TNF | CHR_HSCHR6_MHC_MANN_( | tumor necrosis factor [Source:HGNC Symbol;A |
| LTB | CHR_HSCHR6_MHC_MANN_( | lymphotoxin beta [Source:HGNC Symbol;Acc:I |
| LST1 | CHR_HSCHR6_MHC_MANN_( | leukocyte specific transcript 1 [Source:HGNC S |
| NCR3 | CHR_HSCHR6_MHC_MANN_( | natural cytotoxicity triggering receptor 3 [Sou |
| PRRC2A | CHR_HSCHR6_MHC_MANN_( | proline rich coiled-coil 2A [Source:HGNC Symb |
| SNORA38 | CHR_HSCHR6_MHC_MANN_( | small nucleolar RNA, H/ACA box 38 [Source:H |
| BAG6 | CHR_HSCHR6_MHC_MANN_( | BCL2 associated athanogene 6 [Source:HGNC |
| APOM | CHR_HSCHR6_MHC_MANN_( | apolipoprotein M [Source:HGNC Symbol;Acc:H |
| C6orf47 | CHR_HSCHR6_MHC_MANN_( | chromosome 6 open reading frame 47 [Source |
| C6orf47-AS1 | CHR_HSCHR6_MHC_MANN_( | C6orf47 antisense RNA 1 [Source:HGNC Symb |
| GPANK1 | CHR_HSCHR6_MHC_MANN_( | G-patch domain and ankyrin repeats 1 [Source |
| Y_RNA | CHR_HSCHR6_MHC_MANN_( | Y RNA [Source:RFAM;Acc:RF00019] |
| CSNK2B | CHR_HSCHR6_MHC_MANN_( | casein kinase 2 beta [Source:HGNC Symbol;Ac |
| CR354443.1 | CHR_HSCHR6_MHC_MANN_( | novel protein |
| LY6G5B | CHR_HSCHR6_MHC_MANN_( | lymphocyte antigen 6 family member G5B [So |
| LY6G5C | CHR_HSCHR6_MHC_MANN_( | lymphocyte antigen 6 family member G5C [So |
| ABHD16A | CHR_HSCHR6_MHC_MANN_( | abhydrolase domain containing 16A [Source:H |
| LY6G6F | CHR_HSCHR6_MHC_MANN_( | lymphocyte antigen 6 family member G6F [So |
| LY6G6F-LY6G6D | CHR_HSCHR6_MHC_MANN_( | LY6G6F-LY6G6D readthrough [Source:NCBI ge |
| LY6G6E | CHR_HSCHR6_MHC_MANN_( | lymphocyte antigen 6 family member G6E [So |
| LY6G6D | CHR_HSCHR6_MHC_MANN_( | lymphocyte antigen 6 family member G6D [Sc |
| MPIG6B | CHR_HSCHR6_MHC_MANN_( | megakaryocyte and platelet inhibitory recepto |
| LY6G6C | CHR_HSCHR6_MHC_MANN_( | lymphocyte antigen 6 family member G6C [So |

| | | |
|---|---|---|
| DDAH2 | CHR_HSCHR6_MHC_MANN_( | dimethylarginine dimethylaminohydrolase 2 [ |
| CLIC1 | CHR_HSCHR6_MHC_MANN_( | chloride intracellular channel 1 [Source:HGNC |
| MSH5 | CHR_HSCHR6_MHC_MANN_( | mutS homolog 5 [Source:HGNC Symbol;Acc:H |
| NEU1 | CHR_HSCHR6_MHC_MANN_( | neuraminidase 1 [Source:HGNC Symbol;Acc:H |
| SLC44A4 | CHR_HSCHR6_MHC_MANN_( | solute carrier family 44 member 4 [Source:HG |
| C2 | CHR_HSCHR6_MHC_MANN_( | complement C2 [Source:HGNC Symbol;Acc:HC |
| C4B-AS1 | CHR_HSCHR6_MHC_MANN_( | C4B antisense RNA 1 [Source:HGNC Symbol;A |
| CFB | CHR_HSCHR6_MHC_MANN_( | complement factor B [Source:HGNC Symbol;A |
| TNXB | CHR_HSCHR6_MHC_MANN_( | tenascin XB [Source:HGNC Symbol;Acc:HGNC: |
| 5S_rRNA | CHR_HSCHR6_MHC_MANN_( | 5S ribosomal RNA [Source:RFAM;Acc:RF00001 |
| PRRT1 | CHR_HSCHR6_MHC_MANN_( | proline rich transmembrane protein 1 [Source |
| PPT2 | CHR_HSCHR6_MHC_MANN_( | palmitoyl-protein thioesterase 2 [Source:HGN |
| EGFL8 | CHR_HSCHR6_MHC_MANN_( | EGF like domain multiple 8 [Source:HGNC Sym |
| AGPAT1 | CHR_HSCHR6_MHC_MANN_( | 1-acylglycerol-3-phosphate O-acyltransferase |
| RNF5 | CHR_HSCHR6_MHC_MANN_( | ring finger protein 5 [Source:HGNC Symbol;Ac |
| AGER | CHR_HSCHR6_MHC_MANN_( | advanced glycosylation end-product specific r |
| PBX2 | CHR_HSCHR6_MHC_MANN_( | PBX homeobox 2 [Source:HGNC Symbol;Acc:H |
| GPSM3 | CHR_HSCHR6_MHC_MANN_( | G protein signaling modulator 3 [Source:HGN( |
| NOTCH4 | CHR_HSCHR6_MHC_MANN_( | notch receptor 4 [Source:HGNC Symbol;Acc:H |
| TSBP1 | CHR_HSCHR6_MHC_MANN_( | testis expressed basic protein 1 [Source:HGN( |
| HNRNPA1P2 | CHR_HSCHR6_MHC_MANN_( | heterogeneous nuclear ribonucleoprotein A1 |
| U6 | CHR_HSCHR6_MHC_MANN_( | U6 spliceosomal RNA [Source:RFAM;Acc:RF00 |
| TSBP1-AS1 | CHR_HSCHR6_MHC_MANN_( | TSBP1 and BTNL2 antisense RNA 1 [Source:HG |
| BTNL2 | CHR_HSCHR6_MHC_MANN_( | butyrophilin like 2 [Source:HGNC Symbol;Acc: |
| HLA-DRA | CHR_HSCHR6_MHC_MANN_( | major histocompatibility complex, class II, DR |
| HLA-DRB9 | CHR_HSCHR6_MHC_MANN_( | major histocompatibility complex, class II, DR |
| CR354545.1 | CHR_HSCHR6_MHC_MANN_( | major histocompatibility complex, class II, DR |
| U1 | CHR_HSCHR6_MHC_MANN_( | U1 spliceosomal RNA [Source:RFAM;Acc:RF00 |
| BX927235.1 | CHR_HSCHR6_MHC_MANN_( | major histocompatibility complex, class II, DR |
| U1 | CHR_HSCHR6_MHC_MANN_( | U1 spliceosomal RNA [Source:RFAM;Acc:RF00 |
| HLA-DRB4 | CHR_HSCHR6_MHC_MANN_( | major histocompatibility complex, class II, DR |
| U1 | CHR_HSCHR6_MHC_MANN_( | U1 spliceosomal RNA [Source:RFAM;Acc:RF00 |
| BX927235.2 | CHR_HSCHR6_MHC_MANN_( | major histocompatibility complex, class II, DR |
| U1 | CHR_HSCHR6_MHC_MANN_( | U1 spliceosomal RNA [Source:RFAM;Acc:RF00 |
| CR933540.4 | CHR_HSCHR6_MHC_MANN_( | protein kinase interferon-inducible double str |
| CR933540.3 | CHR_HSCHR6_MHC_MANN_( | novel pseudogene |
| CR933540.2 | CHR_HSCHR6_MHC_MANN_( | major histocompatibility complex, class II, DR |
| CR933540.1 | CHR_HSCHR6_MHC_MANN_( | family with sequence similarity 8, member A5 |
| U1 | CHR_HSCHR6_MHC_MANN_( | U1 spliceosomal RNA [Source:RFAM;Acc:RF00 |
| HLA-DRB1 | CHR_HSCHR6_MHC_MANN_( | major histocompatibility complex, class II, DR |
| HLA-DQA1 | CHR_HSCHR6_MHC_MANN_( | major histocompatibility complex, class II, DQ |
| HLA-DQB1 | CHR_HSCHR6_MHC_MANN_( | major histocompatibility complex, class II, DQ |
| HLA-DQB1-AS1 | CHR_HSCHR6_MHC_MANN_( | HLA-DQB1 antisense RNA 1 [Source:HGNC Syr |
| BX927168.2 | CHR_HSCHR6_MHC_MANN_( | novel transcript |
| MTCO3P1 | CHR_HSCHR6_MHC_MANN_( | MT-CO3 pseudogene 1 [Source:HGNC Symbol |
| BX927168.1 | CHR_HSCHR6_MHC_MANN_( | uncharacterized LOC102725019 [Source:NCBI |
| HLA-DQB3 | CHR_HSCHR6_MHC_MANN_( | major histocompatibility complex, class II, DQ |
| HLA-DQA2 | CHR_HSCHR6_MHC_MANN_( | major histocompatibility complex, class II, DQ |
| HLA-DQB2 | CHR_HSCHR6_MHC_MANN_( | major histocompatibility complex, class II, DQ |
| TAP2 | CHR_HSCHR6_MHC_MANN_( | transporter 2, ATP binding cassette subfamily |

| | | |
|---|---|---|
| PSMB8 | CHR_HSCHR6_MHC_MANN_( | proteasome subunit beta 8 [Source:HGNC Syn |
| PSMB8-AS1 | CHR_HSCHR6_MHC_MANN_( | PSMB8 antisense RNA 1 (head to head) [Sourc |
| PSMB9 | CHR_HSCHR6_MHC_MANN_( | proteasome subunit beta 9 [Source:HGNC Syn |
| TAP1 | CHR_HSCHR6_MHC_MANN_( | transporter 1, ATP binding cassette subfamily |
| PPP1R2P1 | CHR_HSCHR6_MHC_MANN_( | protein phosphatase 1 regulatory inhibitor sul |
| HLA-Z | CHR_HSCHR6_MHC_MANN_( | major histocompatibility complex, class I, Z (p: |
| HLA-DMB | CHR_HSCHR6_MHC_MANN_( | major histocompatibility complex, class II, DM |
| HLA-DMA | CHR_HSCHR6_MHC_MANN_( | major histocompatibility complex, class II, DM |
| BRD2 | CHR_HSCHR6_MHC_MANN_( | bromodomain containing 2 [Source:HGNC Syn |
| BX908719.1 | CHR_HSCHR6_MHC_MANN_( | novel transcript |
| HLA-DOA | CHR_HSCHR6_MHC_MANN_( | major histocompatibility complex, class II, DO |
| HLA-DPA1 | CHR_HSCHR6_MHC_MANN_( | major histocompatibility complex, class II, DP |
| HLA-DPB1 | CHR_HSCHR6_MHC_MANN_( | major histocompatibility complex, class II, DP |
| RPL32P1 | CHR_HSCHR6_MHC_MANN_( | ribosomal protein L32 pseudogene 1 [Source: |
| HLA-DPA2 | CHR_HSCHR6_MHC_MANN_( | major histocompatibility complex, class II, DP |
| COL11A2P1 | CHR_HSCHR6_MHC_MANN_( | collagen type XI alpha 2 pseudogene 1 [Source |
| HLA-DPB2 | CHR_HSCHR6_MHC_MANN_( | major histocompatibility complex, class II, DP |
| HLA-DPA3 | CHR_HSCHR6_MHC_MANN_( | major histocompatibility complex, class II, DP |
| HCG24 | CHR_HSCHR6_MHC_MANN_( | HLA complex group 24 [Source:HGNC Symbol; |
| COL11A2 | CHR_HSCHR6_MHC_MANN_( | collagen type XI alpha 2 chain [Source:HGNC S |
| RXRB | CHR_HSCHR6_MHC_MANN_( | retinoid X receptor beta [Source:HGNC Symbc |
| Y_RNA | CHR_HSCHR6_MHC_MANN_( | Y RNA [Source:RFAM;Acc:RF00019] |
| SLC39A7 | CHR_HSCHR6_MHC_MANN_( | solute carrier family 39 member 7 [Source:HG |
| HSD17B8 | CHR_HSCHR6_MHC_MANN_( | hydroxysteroid 17-beta dehydrogenase 8 [Sou |
| RING1 | CHR_HSCHR6_MHC_MANN_( | ring finger protein 1 [Source:HGNC Symbol;Ac |
| ZNF70P1 | CHR_HSCHR6_MHC_MANN_( | zinc finger protein 70 pseudogene 1 [Source:H |
| HTATSF1P1 | CHR_HSCHR6_MHC_MANN_( | HIV-1 Tat specific factor 1 pseudogene 1 [Sour |
| HCG25 | CHR_HSCHR6_MHC_MANN_( | HLA complex group 25 [Source:HGNC Symbol; |
| VPS52 | CHR_HSCHR6_MHC_MANN_( | VPS52 subunit of GARP complex [Source:HGN |
| CR936879.1 | CHR_HSCHR6_MHC_MCF_CT | novel transcript |
| LINC01623 | CHR_HSCHR6_MHC_MCF_CT | long intergenic non-protein coding RNA 1623 |
| RPL13P | CHR_HSCHR6_MHC_MCF_CT | ribosomal protein L13 pseudogene [Source:H( |
| ZNF90P2 | CHR_HSCHR6_MHC_MCF_CT | zinc finger protein 90 pseudogene 2 [Source:H |
| HCG14 | CHR_HSCHR6_MHC_MCF_CT | HLA complex group 14 [Source:HGNC Symbol; |
| TRIM27 | CHR_HSCHR6_MHC_MCF_CT | tripartite motif containing 27 [Source:HGNC S |
| U6 | CHR_HSCHR6_MHC_MCF_CT | U6 spliceosomal RNA [Source:RFAM;Acc:RF00 |
| LINC01556 | CHR_HSCHR6_MHC_MCF_CT | long intergenic non-protein coding RNA 1556 |
| OR2AD1P | CHR_HSCHR6_MHC_MCF_CT | olfactory receptor family 2 subfamily AD mem |
| OR2W1 | CHR_HSCHR6_MHC_MCF_CT | olfactory receptor family 2 subfamily W meml |
| CR933783.1 | CHR_HSCHR6_MHC_MCF_CT | NULL |
| OR2P1P | CHR_HSCHR6_MHC_MCF_CT | olfactory receptor family 2 subfamily P memb |
| SAR1AP1 | CHR_HSCHR6_MHC_MCF_CT | secretion associated Ras related GTPase 1A ps |
| OR2B3 | CHR_HSCHR6_MHC_MCF_CT | olfactory receptor family 2 subfamily B memb |
| OR2J1 | CHR_HSCHR6_MHC_MCF_CT | olfactory receptor family 2 subfamily J membe |
| OR2J3 | CHR_HSCHR6_MHC_MCF_CT | olfactory receptor family 2 subfamily J membe |
| CR759834.1 | CHR_HSCHR6_MHC_MCF_CT | olfactory receptor, family 2, subfamily N, mem |
| OR2J2 | CHR_HSCHR6_MHC_MCF_CT | olfactory receptor family 2 subfamily J membe |
| OR2J4P | CHR_HSCHR6_MHC_MCF_CT | olfactory receptor family 2 subfamily J membe |
| CR759834.2 | CHR_HSCHR6_MHC_MCF_CT | olfactory receptor, family 2, subfamily H, mem |
| CR933842.1 | CHR_HSCHR6_MHC_MCF_CT | olfactory receptor, family 2, subfamily B, mem |

| | | |
|---|---|---|
| OR14J1 | CHR_HSCHR6_MHC_MCF_CT | olfactory receptor family 14 subfamily J meml |
| DDX6P1 | CHR_HSCHR6_MHC_MCF_CT | DEAD-box helicase 6 pseudogene 1 [Source:H |
| OR5V1 | CHR_HSCHR6_MHC_MCF_CT | olfactory receptor family 5 subfamily V memb |
| OR12D3 | CHR_HSCHR6_MHC_MCF_CT | olfactory receptor family 12 subfamily D mem |
| OR12D2 | CHR_HSCHR6_MHC_MCF_CT | olfactory receptor family 12 subfamily D mem |
| OR12D1 | CHR_HSCHR6_MHC_MCF_CT | olfactory receptor family 12 subfamily D mem |
| OR11A1 | CHR_HSCHR6_MHC_MCF_CT | olfactory receptor family 11 subfamily A mem |
| OR10C1 | CHR_HSCHR6_MHC_MCF_CT | olfactory receptor family 10 subfamily C mem |
| OR2H1 | CHR_HSCHR6_MHC_MCF_CT | olfactory receptor family 2 subfamily H memb |
| UBDP1 | CHR_HSCHR6_MHC_MCF_CT | ubiquitin D pseudogene 1 [Source:HGNC Syml |
| MAS1LP1 | CHR_HSCHR6_MHC_MCF_CT | MAS1L pseudogene 1 [Source:HGNC Symbol;/ |
| MAS1L | CHR_HSCHR6_MHC_MCF_CT | MAS1 proto-oncogene like, G protein-couplec |
| RPS17P1 | CHR_HSCHR6_MHC_MCF_CT | ribosomal protein S17 pseudogene 1 [Source: |
| LINC02829 | CHR_HSCHR6_MHC_MCF_CT | long intergenic non-protein coding RNA 2829 |
| LINC01015 | CHR_HSCHR6_MHC_MCF_CT | long intergenic non-protein coding RNA 1015 |
| GPR53P | CHR_HSCHR6_MHC_MCF_CT | G protein-coupled receptor 53, pseudogene [ |
| OR2I1P | CHR_HSCHR6_MHC_MCF_CT | olfactory receptor family 2 subfamily I membe |
| UBD | CHR_HSCHR6_MHC_MCF_CT | ubiquitin D [Source:HGNC Symbol;Acc:HGNC: |
| GABBR1 | CHR_HSCHR6_MHC_MCF_CT | gamma-aminobutyric acid type B receptor sul |
| OR2H5P | CHR_HSCHR6_MHC_MCF_CT | olfactory receptor family 2 subfamily H memb |
| TMEM183AP1 | CHR_HSCHR6_MHC_MCF_CT | transmembrane protein 183A pseudogene 1 [ |
| SNORD32B | CHR_HSCHR6_MHC_MCF_CT | small nucleolar RNA, C/D box 32B [Source:HG |
| RPL13AP | CHR_HSCHR6_MHC_MCF_CT | ribosomal protein L13a pseudogene [Source:H |
| OR2H2 | CHR_HSCHR6_MHC_MCF_CT | olfactory receptor family 2 subfamily H memb |
| SUMO2P1 | CHR_HSCHR6_MHC_MCF_CT | SUMO2 pseudogene 1 [Source:HGNC Symbol; |
| MOG | CHR_HSCHR6_MHC_MCF_CT | myelin oligodendrocyte glycoprotein [Source: |
| ZFP57 | CHR_HSCHR6_MHC_MCF_CT | ZFP57 zinc finger protein [Source:HGNC Symb |
| ZDHHC20P1 | CHR_HSCHR6_MHC_MCF_CT | zinc finger DHHC-type containing 20 pseudoge |
| HLA-F | CHR_HSCHR6_MHC_MCF_CT | major histocompatibility complex, class I, F [S |
| CR759790.1 | CHR_HSCHR6_MHC_MCF_CT | HLA complex group 4 pseudogene 11 |
| MICE | CHR_HSCHR6_MHC_MCF_CT | MHC class I polypeptide-related sequence E ( |
| CR788234.2 | CHR_HSCHR6_MHC_MCF_CT | novel transcript |
| CR788234.1 | CHR_HSCHR6_MHC_MCF_CT | HLA complex group 9 pseudogene 5 |
| Y_RNA | CHR_HSCHR6_MHC_MCF_CT | Y RNA [Source:RFAM;Acc:RF00019] |
| IFITM4P | CHR_HSCHR6_MHC_MCF_CT | interferon induced transmembrane protein 4 |
| HLA-V | CHR_HSCHR6_MHC_MCF_CT | major histocompatibility complex, class I, V (p |
| HCG4 | CHR_HSCHR6_MHC_MCF_CT | HLA complex group 4 [Source:HGNC Symbol;A |
| HLA-P | CHR_HSCHR6_MHC_MCF_CT | major histocompatibility complex, class I, P (p |
| RPL7AP7 | CHR_HSCHR6_MHC_MCF_CT | ribosomal protein L7a pseudogene 7 [Source: |
| MICG | CHR_HSCHR6_MHC_MCF_CT | MHC class I polypeptide-related sequence G ( |
| HLA-G | CHR_HSCHR6_MHC_MCF_CT | major histocompatibility complex, class I, G [S |
| HCG4P8 | CHR_HSCHR6_MHC_MCF_CT | HLA complex group 4 pseudogene 8 [Source:H |
| CT009517.2 | CHR_HSCHR6_MHC_MCF_CT | MHC class I polypeptide-related sequence F p |
| CT009517.1 | CHR_HSCHR6_MHC_MCF_CT | HLA complex group 4 pseudogene 7 |
| HLA-H | CHR_HSCHR6_MHC_MCF_CT | major histocompatibility complex, class I, H (p |
| HLA-T | CHR_HSCHR6_MHC_MCF_CT | major histocompatibility complex, class I, T (p |
| DDX39BP1 | CHR_HSCHR6_MHC_MCF_CT | DEAD-box helicase 39B pseudogene 1 [Source |
| MCCD1P1 | CHR_HSCHR6_MHC_MCF_CT | mitochondrial coiled-coil domain 1 pseudoger |
| HLA-A | CHR_HSCHR6_MHC_MCF_CT | major histocompatibility complex, class I, A [S |
| CR759762.1 | CHR_HSCHR6_MHC_MCF_CT | HLA complex group 4 pseudogene 5 |

| | | |
|---|---|---|
| HLA-W | CHR_HSCHR6_MHC_MCF_CT | major histocompatibility complex, class I, W (p |
| MICD | CHR_HSCHR6_MHC_MCF_CT | MHC class I polypeptide-related sequence D (p |
| HCG9 | CHR_HSCHR6_MHC_MCF_CT | HLA complex group 9 [Source:HGNC Symbol;A |
| DDX39BP2 | CHR_HSCHR6_MHC_MCF_CT | DEAD-box helicase 39B pseudogene 2 [Source |
| MCCD1P2 | CHR_HSCHR6_MHC_MCF_CT | mitochondrial coiled-coil domain 1 pseudogen |
| ZNRD1ASP | CHR_HSCHR6_MHC_MCF_CT | zinc ribbon domain containing 1 antisense, ps |
| CR759763.1 | CHR_HSCHR6_MHC_MCF_CT | HLA complex group 4 pseudogene 3 |
| HLA-J | CHR_HSCHR6_MHC_MCF_CT | major histocompatibility complex, class I, J (ps |
| ETF1P1 | CHR_HSCHR6_MHC_MCF_CT | eukaryotic translation termination factor 1 ps |
| CR759763.4 | CHR_HSCHR6_MHC_MCF_CT | NULL |
| CR759763.3 | CHR_HSCHR6_MHC_MCF_CT | NULL |
| ZNRD1 | CHR_HSCHR6_MHC_MCF_CT | zinc ribbon domain containing 1 [Source:HGN |
| CR759763.2 | CHR_HSCHR6_MHC_MCF_CT | NULL |
| PPP1R11 | CHR_HSCHR6_MHC_MCF_CT | protein phosphatase 1 regulatory inhibitor sul |
| RNF39 | CHR_HSCHR6_MHC_MCF_CT | ring finger protein 39 [Source:HGNC Symbol;A |
| TRIM31 | CHR_HSCHR6_MHC_MCF_CT | tripartite motif containing 31 [Source:HGNC S |
| TRIM31-AS1 | CHR_HSCHR6_MHC_MCF_CT | TRIM31 antisense RNA 1 [Source:HGNC Symb |
| CR759838.1 | CHR_HSCHR6_MHC_MCF_CT | NULL |
| TRIM40 | CHR_HSCHR6_MHC_MCF_CT | tripartite motif containing 40 [Source:HGNC S |
| TRIM10 | CHR_HSCHR6_MHC_MCF_CT | tripartite motif containing 10 [Source:HGNC S |
| TRIM15 | CHR_HSCHR6_MHC_MCF_CT | tripartite motif containing 15 [Source:HGNC S |
| TRIM26 | CHR_HSCHR6_MHC_MCF_CT | tripartite motif containing 26 [Source:HGNC S |
| PAIP1P1 | CHR_HSCHR6_MHC_MCF_CT | poly(A) binding protein interacting protein 1 p |
| MICC | CHR_HSCHR6_MHC_MCF_CT | MHC class I polypeptide-related sequence C (p |
| HLA-E | CHR_HSCHR6_MHC_MCF_CT | major histocompatibility complex, class I, E [S |
| LINC02569 | CHR_HSCHR6_MHC_MCF_CT | long intergenic non-protein coding RNA 2569 |
| GNL1 | CHR_HSCHR6_MHC_MCF_CT | G protein nucleolar 1 (putative) [Source:HGNC |
| PRR3 | CHR_HSCHR6_MHC_MCF_CT | proline rich 3 [Source:HGNC Symbol;Acc:HGN |
| ABCF1 | CHR_HSCHR6_MHC_MCF_CT | ATP binding cassette subfamily F member 1 [S |
| PPP1R10 | CHR_HSCHR6_MHC_MCF_CT | protein phosphatase 1 regulatory subunit 10 [ |
| MRPS18B | CHR_HSCHR6_MHC_MCF_CT | mitochondrial ribosomal protein S18B [Source |
| ATAT1 | CHR_HSCHR6_MHC_MCF_CT | alpha tubulin acetyltransferase 1 [Source:HGN |
| PTMAP1 | CHR_HSCHR6_MHC_MCF_CT | prothymosin alpha pseudogene 1 [Source:HGI |
| C6orf136 | CHR_HSCHR6_MHC_MCF_CT | chromosome 6 open reading frame 136 [Sour |
| DHX16 | CHR_HSCHR6_MHC_MCF_CT | DEAH-box helicase 16 [Source:HGNC Symbol;A |
| PPP1R18 | CHR_HSCHR6_MHC_MCF_CT | protein phosphatase 1 regulatory subunit 18 [ |
| NRM | CHR_HSCHR6_MHC_MCF_CT | nurim [Source:HGNC Symbol;Acc:HGNC:8003] |
| RPL7P4 | CHR_HSCHR6_MHC_MCF_CT | ribosomal protein L7 pseudogene 4 [Source:H |
| MDC1 | CHR_HSCHR6_MHC_MCF_CT | mediator of DNA damage checkpoint 1 [Sourc |
| MDC1-AS1 | CHR_HSCHR6_MHC_MCF_CT | MDC1 antisense RNA 1 [Source:HGNC Symbol |
| TUBB | CHR_HSCHR6_MHC_MCF_CT | tubulin beta class I [Source:HGNC Symbol;Acc |
| FLOT1 | CHR_HSCHR6_MHC_MCF_CT | flotillin 1 [Source:HGNC Symbol;Acc:HGNC:37 |
| Y_RNA | CHR_HSCHR6_MHC_MCF_CT | Y RNA [Source:RFAM;Acc:RF00019] |
| LINC02570 | CHR_HSCHR6_MHC_MCF_CT | long intergenic non-protein coding RNA 2570 |
| 7SK | CHR_HSCHR6_MHC_MCF_CT | 7SK RNA [Source:RFAM;Acc:RF00100] |
| DDR1-DT | CHR_HSCHR6_MHC_MCF_CT | DDR1 divergent transcript [Source:HGNC Sym |
| DDR1 | CHR_HSCHR6_MHC_MCF_CT | discoidin domain receptor tyrosine kinase 1 [S |
| GTF2H4 | CHR_HSCHR6_MHC_MCF_CT | general transcription factor IIH subunit 4 [Sou |
| CR936875.1 | CHR_HSCHR6_MHC_MCF_CT | novel protein |
| VARS2 | CHR_HSCHR6_MHC_MCF_CT | valyl-tRNA synthetase 2, mitochondrial [Sourc |

| | | |
|---|---|---|
| SFTA2 | CHR_HSCHR6_MHC_MCF_CT | surfactant associated 2 [Source:HGNC Symbol |
| Y_RNA | CHR_HSCHR6_MHC_MCF_CT | Y RNA [Source:RFAM;Acc:RF00019] |
| MUCL3 | CHR_HSCHR6_MHC_MCF_CT | mucin like 3 [Source:HGNC Symbol;Acc:HGNC |
| HCG21 | CHR_HSCHR6_MHC_MCF_CT | HLA complex group 21 [Source:HGNC Symbol; |
| MUC21 | CHR_HSCHR6_MHC_MCF_CT | mucin 21, cell surface associated [Source:HGN |
| CR759772.2 | CHR_HSCHR6_MHC_MCF_CT | novel transcript |
| CR759772.1 | CHR_HSCHR6_MHC_MCF_CT | HLA complex group family pseudogene |
| U6 | CHR_HSCHR6_MHC_MCF_CT | U6 spliceosomal RNA [Source:RFAM;Acc:RF00 |
| C6orf15 | CHR_HSCHR6_MHC_MCF_CT | chromosome 6 open reading frame 15 [Sourc |
| PSORS1C1 | CHR_HSCHR6_MHC_MCF_CT | psoriasis susceptibility 1 candidate 1 [Source: |
| CDSN | CHR_HSCHR6_MHC_MCF_CT | corneodesmosin [Source:HGNC Symbol;Acc:H |
| PSORS1C2 | CHR_HSCHR6_MHC_MCF_CT | psoriasis susceptibility 1 candidate 2 [Source: |
| POLR2LP1 | CHR_HSCHR6_MHC_MCF_CT | RNA polymerase II subunit L pseudogene 1 [S |
| CCHCR1 | CHR_HSCHR6_MHC_MCF_CT | coiled-coil alpha-helical rod protein 1 [Source: |
| TCF19 | CHR_HSCHR6_MHC_MCF_CT | transcription factor 19 [Source:HGNC Symbol; |
| POU5F1 | CHR_HSCHR6_MHC_MCF_CT | POU class 5 homeobox 1 [Source:HGNC Symb |
| CR759815.2 | CHR_HSCHR6_MHC_MCF_CT | NULL |
| PSORS1C3 | CHR_HSCHR6_MHC_MCF_CT | psoriasis susceptibility 1 candidate 3 [Source: |
| HCG27 | CHR_HSCHR6_MHC_MCF_CT | HLA complex group 27 [Source:HGNC Symbol; |
| CR759815.1 | CHR_HSCHR6_MHC_MCF_CT | novel transcript |
| HLA-C | CHR_HSCHR6_MHC_MCF_CT | major histocompatibility complex, class I, C [S |
| HLA-B | CHR_HSCHR6_MHC_MCF_CT | major histocompatibility complex, class I, B [S |
| USP8P1 | CHR_HSCHR6_MHC_MCF_CT | ubiquitin specific peptidase 8 pseudogene 1 [S |
| RPL3P2 | CHR_HSCHR6_MHC_MCF_CT | ribosomal protein L3 pseudogene 2 [Source:H |
| WASF5P | CHR_HSCHR6_MHC_MCF_CT | WASP family member 5, pseudogene [Source: |
| CR759828.1 | CHR_HSCHR6_MHC_MCF_CT | novel HLA class I pseudogene fragment |
| HCP5 | CHR_HSCHR6_MHC_MCF_CT | HLA complex P5 [Source:HGNC Symbol;Acc:H |
| Y_RNA | CHR_HSCHR6_MHC_MCF_CT | Y RNA [Source:RFAM;Acc:RF00019] |
| MICB | CHR_HSCHR6_MHC_MCF_CT | MHC class I polypeptide-related sequence B [S |
| PPIAP9 | CHR_HSCHR6_MHC_MCF_CT | peptidylprolyl isomerase A pseudogene 9 [Sou |
| RPL15P4 | CHR_HSCHR6_MHC_MCF_CT | ribosomal protein L15 pseudogene 4 [Source: |
| MCCD1 | CHR_HSCHR6_MHC_MCF_CT | mitochondrial coiled-coil domain 1 [Source:H |
| DDX39B | CHR_HSCHR6_MHC_MCF_CT | DExD-box helicase 39B [Source:HGNC Symbol |
| SNORD117 | CHR_HSCHR6_MHC_MCF_CT | small nucleolar RNA, C/D box 117 [Source:HG |
| SNORD84 | CHR_HSCHR6_MHC_MCF_CT | small nucleolar RNA, C/D box 84 [Source:HGN |
| DDX39B-AS1 | CHR_HSCHR6_MHC_MCF_CT | DDX39B antisense RNA 1 [Source:HGNC Symb |
| ATP6V1G2 | CHR_HSCHR6_MHC_MCF_CT | ATPase H+ transporting V1 subunit G2 [Source |
| NFKBIL1 | CHR_HSCHR6_MHC_MCF_CT | NFKB inhibitor like 1 [Source:HGNC Symbol;Ac |
| LTA | CHR_HSCHR6_MHC_MCF_CT | lymphotoxin alpha [Source:HGNC Symbol;Acc |
| TNF | CHR_HSCHR6_MHC_MCF_CT | tumor necrosis factor [Source:HGNC Symbol; |
| LTB | CHR_HSCHR6_MHC_MCF_CT | lymphotoxin beta [Source:HGNC Symbol;Acc: |
| LST1 | CHR_HSCHR6_MHC_MCF_CT | leukocyte specific transcript 1 [Source:HGNC S |
| NCR3 | CHR_HSCHR6_MHC_MCF_CT | natural cytotoxicity triggering receptor 3 [Sou |
| AIF1 | CHR_HSCHR6_MHC_MCF_CT | allograft inflammatory factor 1 [Source:HGNC |
| PRRC2A | CHR_HSCHR6_MHC_MCF_CT | proline rich coiled-coil 2A [Source:HGNC Symb |
| SNORA38 | CHR_HSCHR6_MHC_MCF_CT | small nucleolar RNA, H/ACA box 38 [Source:H |
| BAG6 | CHR_HSCHR6_MHC_MCF_CT | BCL2 associated athanogene 6 [Source:HGNC |
| APOM | CHR_HSCHR6_MHC_MCF_CT | apolipoprotein M [Source:HGNC Symbol;Acc: |
| C6orf47 | CHR_HSCHR6_MHC_MCF_CT | chromosome 6 open reading frame 47 [Sourc |
| C6orf47-AS1 | CHR_HSCHR6_MHC_MCF_CT | C6orf47 antisense RNA 1 [Source:HGNC Symb |

| | | |
|---|---|---|
| GPANK1 | CHR_HSCHR6_MHC_MCF_CT | G-patch domain and ankyrin repeats 1 [Source |
| Y_RNA | CHR_HSCHR6_MHC_MCF_CT | Y RNA [Source:RFAM;Acc:RF00019] |
| CSNK2B | CHR_HSCHR6_MHC_MCF_CT | casein kinase 2 beta [Source:HGNC Symbol;Ac |
| LY6G5B | CHR_HSCHR6_MHC_MCF_CT | lymphocyte antigen 6 family member G5B [So |
| LY6G5C | CHR_HSCHR6_MHC_MCF_CT | lymphocyte antigen 6 family member G5C [So |
| ABHD16A | CHR_HSCHR6_MHC_MCF_CT | abhydrolase domain containing 16A [Source:H |
| LY6G6F | CHR_HSCHR6_MHC_MCF_CT | lymphocyte antigen 6 family member G6F [So |
| LY6G6D | CHR_HSCHR6_MHC_MCF_CT | lymphocyte antigen 6 family member G6D [Sc |
| LY6G6E | CHR_HSCHR6_MHC_MCF_CT | lymphocyte antigen 6 family member G6E [So |
| MPIG6B | CHR_HSCHR6_MHC_MCF_CT | megakaryocyte and platelet inhibitory recepto |
| LY6G6C | CHR_HSCHR6_MHC_MCF_CT | lymphocyte antigen 6 family member G6C [So |
| DDAH2 | CHR_HSCHR6_MHC_MCF_CT | dimethylarginine dimethylaminohydrolase 2 [ |
| CLIC1 | CHR_HSCHR6_MHC_MCF_CT | chloride intracellular channel 1 [Source:HGNC |
| MSH5 | CHR_HSCHR6_MHC_MCF_CT | mutS homolog 5 [Source:HGNC Symbol;Acc:H |
| MSH5-SAPCD1 | CHR_HSCHR6_MHC_MCF_CT | MSH5-SAPCD1 readthrough (NMD candidate) |
| U6 | CHR_HSCHR6_MHC_MCF_CT | U6 spliceosomal RNA [Source:RFAM;Acc:RF00 |
| SAPCD1 | CHR_HSCHR6_MHC_MCF_CT | suppressor APC domain containing 1 [Source: |
| SAPCD1-AS1 | CHR_HSCHR6_MHC_MCF_CT | SAPCD1 antisense RNA 1 [Source:HGNC Symb |
| VWA7 | CHR_HSCHR6_MHC_MCF_CT | von Willebrand factor A domain containing 7 [ |
| VARS | CHR_HSCHR6_MHC_MCF_CT | valyl-tRNA synthetase [Source:HGNC Symbol;A |
| Y_RNA | CHR_HSCHR6_MHC_MCF_CT | Y RNA [Source:RFAM;Acc:RF00019] |
| LSM2 | CHR_HSCHR6_MHC_MCF_CT | LSM2 homolog, U6 small nuclear RNA and mR |
| NEU1 | CHR_HSCHR6_MHC_MCF_CT | neuraminidase 1 [Source:HGNC Symbol;Acc:H |
| SLC44A4 | CHR_HSCHR6_MHC_MCF_CT | solute carrier family 44 member 4 [Source:HG |
| EHMT2 | CHR_HSCHR6_MHC_MCF_CT | euchromatic histone lysine methyltransferase |
| CR759784.1 | CHR_HSCHR6_MHC_MCF_CT | novel transcript |
| C2 | CHR_HSCHR6_MHC_MCF_CT | complement C2 [Source:HGNC Symbol;Acc:HG |
| ZBTB12 | CHR_HSCHR6_MHC_MCF_CT | zinc finger and BTB domain containing 12 [Sou |
| C4B-AS1 | CHR_HSCHR6_MHC_MCF_CT | C4B antisense RNA 1 [Source:HGNC Symbol;A |
| CFB | CHR_HSCHR6_MHC_MCF_CT | complement factor B [Source:HGNC Symbol;A |
| NELFE | CHR_HSCHR6_MHC_MCF_CT | negative elongation factor complex member E |
| SKIV2L | CHR_HSCHR6_MHC_MCF_CT | Ski2 like RNA helicase [Source:HGNC Symbol;A |
| DXO | CHR_HSCHR6_MHC_MCF_CT | decapping exoribonuclease [Source:HGNC Syr |
| STK19 | CHR_HSCHR6_MHC_MCF_CT | serine/threonine kinase 19 [Source:HGNC Sym |
| C4B | CHR_HSCHR6_MHC_MCF_CT | complement C4B (Chido blood group) [Source |
| CYP21A1P | CHR_HSCHR6_MHC_MCF_CT | cytochrome P450 family 21 subfamily A meml |
| TNXA | CHR_HSCHR6_MHC_MCF_CT | tenascin XA (pseudogene) [Source:HGNC Sym |
| STK19B | CHR_HSCHR6_MHC_MCF_CT | serine/threonine kinase 19B (pseudogene) [Sc |
| C4A | CHR_HSCHR6_MHC_MCF_CT | complement C4A (Rodgers blood group) [Sour |
| C4A-AS1 | CHR_HSCHR6_MHC_MCF_CT | C4A antisense RNA 1 [Source:HGNC Symbol;A |
| CYP21A2 | CHR_HSCHR6_MHC_MCF_CT | cytochrome P450 family 21 subfamily A meml |
| TNXB | CHR_HSCHR6_MHC_MCF_CT | tenascin XB [Source:HGNC Symbol;Acc:HGNC: |
| 5S_rRNA | CHR_HSCHR6_MHC_MCF_CT | 5S ribosomal RNA [Source:RFAM;Acc:RF00001 |
| ATF6B | CHR_HSCHR6_MHC_MCF_CT | activating transcription factor 6 beta [Source: |
| FKBPL | CHR_HSCHR6_MHC_MCF_CT | FKBP prolyl isomerase like [Source:HGNC Sym |
| PRRT1 | CHR_HSCHR6_MHC_MCF_CT | proline rich transmembrane protein 1 [Source |
| PPT2 | CHR_HSCHR6_MHC_MCF_CT | palmitoyl-protein thioesterase 2 [Source:HGN |
| EGFL8 | CHR_HSCHR6_MHC_MCF_CT | EGF like domain multiple 8 [Source:HGNC Sym |
| AGPAT1 | CHR_HSCHR6_MHC_MCF_CT | 1-acylglycerol-3-phosphate O-acyltransferase |
| RNF5 | CHR_HSCHR6_MHC_MCF_CT | ring finger protein 5 [Source:HGNC Symbol;Ac |

| | | |
|---|---|---|
| AGER | CHR_HSCHR6_MHC_MCF_CT | advanced glycosylation end-product specific r |
| PBX2 | CHR_HSCHR6_MHC_MCF_CT | PBX homeobox 2 [Source:HGNC Symbol;Acc:H |
| GPSM3 | CHR_HSCHR6_MHC_MCF_CT | G protein signaling modulator 3 [Source:HGN( |
| NOTCH4 | CHR_HSCHR6_MHC_MCF_CT | notch receptor 4 [Source:HGNC Symbol;Acc:H |
| TSBP1 | CHR_HSCHR6_MHC_MCF_CT | testis expressed basic protein 1 [Source:HGN( |
| HNRNPA1P2 | CHR_HSCHR6_MHC_MCF_CT | heterogeneous nuclear ribonucleoprotein A1 |
| U6 | CHR_HSCHR6_MHC_MCF_CT | U6 spliceosomal RNA [Source:RFAM;Acc:RF00 |
| TSBP1-AS1 | CHR_HSCHR6_MHC_MCF_CT | TSBP1 and BTNL2 antisense RNA 1 [Source:HG |
| BTNL2 | CHR_HSCHR6_MHC_MCF_CT | butyrophilin like 2 [Source:HGNC Symbol;Acc: |
| HLA-DRA | CHR_HSCHR6_MHC_MCF_CT | major histocompatibility complex, class II, DR |
| HLA-DRB9 | CHR_HSCHR6_MHC_MCF_CT | major histocompatibility complex, class II, DR |
| CR788250.2 | CHR_HSCHR6_MHC_MCF_CT | major histocompatibility complex class II DR b |
| U1 | CHR_HSCHR6_MHC_MCF_CT | U1 spliceosomal RNA [Source:RFAM;Acc:RF00 |
| CR788250.3 | CHR_HSCHR6_MHC_MCF_CT | major histocompatibility complex class II DR b |
| U1 | CHR_HSCHR6_MHC_MCF_CT | U1 spliceosomal RNA [Source:RFAM;Acc:RF00 |
| CR788250.1 | CHR_HSCHR6_MHC_MCF_CT | major histocompatibility complex, class II, DR |
| HLA-DQA1 | CHR_HSCHR6_MHC_MCF_CT | major histocompatibility complex, class II, DQ |
| HLA-DQB1 | CHR_HSCHR6_MHC_MCF_CT | major histocompatibility complex, class II, DQ |
| HLA-DQB1-AS1 | CHR_HSCHR6_MHC_MCF_CT | HLA-DQB1 antisense RNA 1 [Source:HGNC Syr |
| CR933859.1 | CHR_HSCHR6_MHC_MCF_CT | novel transcript |
| MTCO3P1 | CHR_HSCHR6_MHC_MCF_CT | MT-CO3 pseudogene 1 [Source:HGNC Symbol |
| CR759848.1 | CHR_HSCHR6_MHC_MCF_CT | uncharacterized LOC102725019 [Source:NCBI |
| HLA-DQB3 | CHR_HSCHR6_MHC_MCF_CT | major histocompatibility complex, class II, DQ |
| HLA-DQA2 | CHR_HSCHR6_MHC_MCF_CT | major histocompatibility complex, class II, DQ |
| HLA-DQB2 | CHR_HSCHR6_MHC_MCF_CT | major histocompatibility complex, class II, DQ |
| HLA-DOB | CHR_HSCHR6_MHC_MCF_CT | major histocompatibility complex, class II, DO |
| TAP2 | CHR_HSCHR6_MHC_MCF_CT | transporter 2, ATP binding cassette subfamily |
| PSMB8 | CHR_HSCHR6_MHC_MCF_CT | proteasome subunit beta 8 [Source:HGNC Syn |
| PSMB8-AS1 | CHR_HSCHR6_MHC_MCF_CT | PSMB8 antisense RNA 1 (head to head) [Sourc |
| PSMB9 | CHR_HSCHR6_MHC_MCF_CT | proteasome subunit beta 9 [Source:HGNC Syn |
| TAP1 | CHR_HSCHR6_MHC_MCF_CT | transporter 1, ATP binding cassette subfamily |
| PPP1R2P1 | CHR_HSCHR6_MHC_MCF_CT | protein phosphatase 1 regulatory inhibitor sul |
| HLA-Z | CHR_HSCHR6_MHC_MCF_CT | major histocompatibility complex, class I, Z (p: |
| HLA-DMB | CHR_HSCHR6_MHC_MCF_CT | major histocompatibility complex, class II, DM |
| HLA-DMA | CHR_HSCHR6_MHC_MCF_CT | major histocompatibility complex, class II, DM |
| BRD2 | CHR_HSCHR6_MHC_MCF_CT | bromodomain containing 2 [Source:HGNC Syn |
| CR759798.1 | CHR_HSCHR6_MHC_MCF_CT | novel transcript |
| HLA-DOA | CHR_HSCHR6_MHC_MCF_CT | major histocompatibility complex, class II, DO |
| HLA-DPA1 | CHR_HSCHR6_MHC_MCF_CT | major histocompatibility complex, class II, DP |
| HLA-DPB1 | CHR_HSCHR6_MHC_MCF_CT | major histocompatibility complex, class II, DP |
| RPL32P1 | CHR_HSCHR6_MHC_MCF_CT | ribosomal protein L32 pseudogene 1 [Source: |
| HLA-DPA2 | CHR_HSCHR6_MHC_MCF_CT | major histocompatibility complex, class II, DP |
| COL11A2P1 | CHR_HSCHR6_MHC_MCF_CT | collagen type XI alpha 2 pseudogene 1 [Sourc |
| HLA-DPB2 | CHR_HSCHR6_MHC_MCF_CT | major histocompatibility complex, class II, DP |
| COL11A2 | CHR_HSCHR6_MHC_MCF_CT | collagen type XI alpha 2 chain [Source:HGNC S |
| RXRB | CHR_HSCHR6_MHC_MCF_CT | retinoid X receptor beta [Source:HGNC Symbo |
| Y_RNA | CHR_HSCHR6_MHC_MCF_CT | Y RNA [Source:RFAM;Acc:RF00019] |
| SLC39A7 | CHR_HSCHR6_MHC_MCF_CT | solute carrier family 39 member 7 [Source:HG |
| HSD17B8 | CHR_HSCHR6_MHC_MCF_CT | hydroxysteroid 17-beta dehydrogenase 8 [Sou |
| RING1 | CHR_HSCHR6_MHC_MCF_CT | ring finger protein 1 [Source:HGNC Symbol;Ac |

| | | |
|---|---|---|
| ZNF70P1 | CHR_HSCHR6_MHC_MCF_CT | zinc finger protein 70 pseudogene 1 [Source:H |
| HTATSF1P1 | CHR_HSCHR6_MHC_MCF_CT | HIV-1 Tat specific factor 1 pseudogene 1 [Sour |
| HCG25 | CHR_HSCHR6_MHC_MCF_CT | HLA complex group 25 [Source:HGNC Symbol; |
| VPS52 | CHR_HSCHR6_MHC_MCF_CT | VPS52 subunit of GARP complex [Source:HGN |
| RPS18 | CHR_HSCHR6_MHC_MCF_CT | ribosomal protein S18 [Source:HGNC Symbol;. |
| B3GALT4 | CHR_HSCHR6_MHC_MCF_CT | beta-1,3-galactosyltransferase 4 [Source:HGN |
| WDR46 | CHR_HSCHR6_MHC_MCF_CT | WD repeat domain 46 [Source:HGNC Symbol;. |
| PFDN6 | CHR_HSCHR6_MHC_MCF_CT | prefoldin subunit 6 [Source:HGNC Symbol;Acc |
| RGL2 | CHR_HSCHR6_MHC_MCF_CT | ral guanine nucleotide dissociation stimulator |
| TAPBP | CHR_HSCHR6_MHC_MCF_CT | TAP binding protein [Source:HGNC Symbol;Ac |
| ZBTB22 | CHR_HSCHR6_MHC_MCF_CT | zinc finger and BTB domain containing 22 [Sou |
| DAXX | CHR_HSCHR6_MHC_MCF_CT | death domain associated protein [Source:HGN |
| MYL8P | CHR_HSCHR6_MHC_MCF_CT | myosin light chain 8, pseudogene [Source:HGI |
| LYPLA2P1 | CHR_HSCHR6_MHC_MCF_CT | LYPLA2 pseudogene 1 [Source:HGNC Symbol;/ |
| RPL35AP4 | CHR_HSCHR6_MHC_MCF_CT | ribosomal protein L35a pseudogene 4 [Source |
| KIFC1 | CHR_HSCHR6_MHC_MCF_CT | kinesin family member C1 [Source:HGNC Sym |
| NOP56P1 | CHR_HSCHR6_MHC_QBL_CT( | NOP56 ribonucleoprotein pseudogene 1 [Sour |
| AL844539.1 | CHR_HSCHR6_MHC_QBL_CT( | novel transcript |
| LINC01623 | CHR_HSCHR6_MHC_QBL_CT( | long intergenic non-protein coding RNA 1623 |
| RPL13P | CHR_HSCHR6_MHC_QBL_CT( | ribosomal protein L13 pseudogene [Source:H( |
| ZNF90P2 | CHR_HSCHR6_MHC_QBL_CT( | zinc finger protein 90 pseudogene 2 [Source:H |
| HCG14 | CHR_HSCHR6_MHC_QBL_CT( | HLA complex group 14 [Source:HGNC Symbol; |
| TRIM27 | CHR_HSCHR6_MHC_QBL_CT( | tripartite motif containing 27 [Source:HGNC S |
| U6 | CHR_HSCHR6_MHC_QBL_CT( | U6 spliceosomal RNA [Source:RFAM;Acc:RF00 |
| LINC01556 | CHR_HSCHR6_MHC_QBL_CT( | long intergenic non-protein coding RNA 1556 |
| KRT18P1 | CHR_HSCHR6_MHC_QBL_CT( | keratin 18 pseudogene 1 [Source:HGNC Symb |
| Metazoa_SRP | CHR_HSCHR6_MHC_QBL_CT( | Metazoan signal recognition particle RNA [Sou |
| HCG15 | CHR_HSCHR6_MHC_QBL_CT( | HLA complex group 15 [Source:HGNC Symbol; |
| AL929561.3 | CHR_HSCHR6_MHC_QBL_CT( | HLA complex group 16 |
| ZNF311 | CHR_HSCHR6_MHC_QBL_CT( | zinc finger protein 311 [Source:HGNC Symbol; |
| OR2AD1P | CHR_HSCHR6_MHC_QBL_CT( | olfactory receptor family 2 subfamily AD mem |
| OR2W1 | CHR_HSCHR6_MHC_QBL_CT( | olfactory receptor family 2 subfamily W meml |
| AL929561.4 | CHR_HSCHR6_MHC_QBL_CT( | NULL |
| OR2P1P | CHR_HSCHR6_MHC_QBL_CT( | olfactory receptor family 2 subfamily P memb |
| SAR1AP1 | CHR_HSCHR6_MHC_QBL_CT( | secretion associated Ras related GTPase 1A ps |
| OR2B3 | CHR_HSCHR6_MHC_QBL_CT( | olfactory receptor family 2 subfamily B memb |
| OR2J1 | CHR_HSCHR6_MHC_QBL_CT( | olfactory receptor family 2 subfamily J membe |
| OR2J3 | CHR_HSCHR6_MHC_QBL_CT( | olfactory receptor family 2 subfamily J membe |
| AL929561.1 | CHR_HSCHR6_MHC_QBL_CT( | novel transcript |
| AL929561.2 | CHR_HSCHR6_MHC_QBL_CT( | olfactory receptor, family 2, subfamily N, men |
| OR2J2 | CHR_HSCHR6_MHC_QBL_CT( | olfactory receptor family 2 subfamily J membe |
| OR2J4P | CHR_HSCHR6_MHC_QBL_CT( | olfactory receptor family 2 subfamily J membe |
| AL935156.1 | CHR_HSCHR6_MHC_QBL_CT( | olfactory receptor, family 2, subfamily H, men |
| OR2G1P | CHR_HSCHR6_MHC_QBL_CT( | olfactory receptor family 2 subfamily G memb |
| OR12D1 | CHR_HSCHR6_MHC_QBL_CT( | olfactory receptor family 12 subfamily D mem |
| OR5V1 | CHR_HSCHR6_MHC_QBL_CT( | olfactory receptor family 5 subfamily V memb |
| OR11A1 | CHR_HSCHR6_MHC_QBL_CT( | olfactory receptor family 11 subfamily A mem |
| OR10C1 | CHR_HSCHR6_MHC_QBL_CT( | olfactory receptor family 10 subfamily C mem |
| OR2H1 | CHR_HSCHR6_MHC_QBL_CT( | olfactory receptor family 2 subfamily H memb |
| UBDP1 | CHR_HSCHR6_MHC_QBL_CT( | ubiquitin D pseudogene 1 [Source:HGNC Symb |

| | | |
|---|---|---|
| MAS1LP1 | CHR_HSCHR6_MHC_QBL_CT( | MAS1L pseudogene 1 [Source:HGNC Symbol;A |
| MAS1L | CHR_HSCHR6_MHC_QBL_CT( | MAS1 proto-oncogene like, G protein-couplec |
| RPS17P1 | CHR_HSCHR6_MHC_QBL_CT( | ribosomal protein S17 pseudogene 1 [Source: |
| LINC02829 | CHR_HSCHR6_MHC_QBL_CT( | long intergenic non-protein coding RNA 2829 |
| LINC01015 | CHR_HSCHR6_MHC_QBL_CT( | long intergenic non-protein coding RNA 1015 |
| GPR53P | CHR_HSCHR6_MHC_QBL_CT( | G protein-coupled receptor 53, pseudogene [S |
| OR2I1P | CHR_HSCHR6_MHC_QBL_CT( | olfactory receptor family 2 subfamily I membe |
| GABBR1 | CHR_HSCHR6_MHC_QBL_CT( | gamma-aminobutyric acid type B receptor sub |
| UBD | CHR_HSCHR6_MHC_QBL_CT( | ubiquitin D [Source:HGNC Symbol;Acc:HGNC:1 |
| OR2H5P | CHR_HSCHR6_MHC_QBL_CT( | olfactory receptor family 2 subfamily H memb |
| TMEM183AP1 | CHR_HSCHR6_MHC_QBL_CT( | transmembrane protein 183A pseudogene 1 [ |
| SNORD32B | CHR_HSCHR6_MHC_QBL_CT( | small nucleolar RNA, C/D box 32B [Source:HG |
| RPL13AP | CHR_HSCHR6_MHC_QBL_CT( | ribosomal protein L13a pseudogene [Source:H |
| OR2H2 | CHR_HSCHR6_MHC_QBL_CT( | olfactory receptor family 2 subfamily H memb |
| SUMO2P1 | CHR_HSCHR6_MHC_QBL_CT( | SUMO2 pseudogene 1 [Source:HGNC Symbol; |
| MOG | CHR_HSCHR6_MHC_QBL_CT( | myelin oligodendrocyte glycoprotein [Source: |
| ZFP57 | CHR_HSCHR6_MHC_QBL_CT( | ZFP57 zinc finger protein [Source:HGNC Symb |
| ZDHHC20P1 | CHR_HSCHR6_MHC_QBL_CT( | zinc finger DHHC-type containing 20 pseudoge |
| HLA-F | CHR_HSCHR6_MHC_QBL_CT( | major histocompatibility complex, class I, F [So |
| AL844851.1 | CHR_HSCHR6_MHC_QBL_CT( | HLA complex group 4 pseudogene 11 |
| HLA-F-AS1 | CHR_HSCHR6_MHC_QBL_CT( | HLA-F antisense RNA 1 [Source:HGNC Symbol; |
| RPL23AP1 | CHR_HSCHR6_MHC_QBL_CT( | ribosomal protein L23a pseudogene 1 [Source |
| MICE | CHR_HSCHR6_MHC_QBL_CT( | MHC class I polypeptide-related sequence E (p |
| AL844851.2 | CHR_HSCHR6_MHC_QBL_CT( | HLA complex group 9 pseudogene 5 |
| Y_RNA | CHR_HSCHR6_MHC_QBL_CT( | Y RNA [Source:RFAM;Acc:RF00019] |
| IFITM4P | CHR_HSCHR6_MHC_QBL_CT( | interferon induced transmembrane protein 4 |
| HLA-V | CHR_HSCHR6_MHC_QBL_CT( | major histocompatibility complex, class I, V (p |
| HCG4 | CHR_HSCHR6_MHC_QBL_CT( | HLA complex group 4 [Source:HGNC Symbol;A |
| HLA-P | CHR_HSCHR6_MHC_QBL_CT( | major histocompatibility complex, class I, P (p |
| RPL7AP7 | CHR_HSCHR6_MHC_QBL_CT( | ribosomal protein L7a pseudogene 7 [Source:I |
| MICG | CHR_HSCHR6_MHC_QBL_CT( | MHC class I polypeptide-related sequence G (p |
| HLA-G | CHR_HSCHR6_MHC_QBL_CT( | major histocompatibility complex, class I, G [S |
| HCG4P8 | CHR_HSCHR6_MHC_QBL_CT( | HLA complex group 4 pseudogene 8 [Source:H |
| BX001005.1 | CHR_HSCHR6_MHC_QBL_CT( | MHC class I polypeptide-related sequence F ps |
| AL845454.1 | CHR_HSCHR6_MHC_QBL_CT( | HLA complex group 4 pseudogene 7 |
| HLA-H | CHR_HSCHR6_MHC_QBL_CT( | major histocompatibility complex, class I, H (p |
| HLA-T | CHR_HSCHR6_MHC_QBL_CT( | major histocompatibility complex, class I, T (p |
| DDX39BP1 | CHR_HSCHR6_MHC_QBL_CT( | DEAD-box helicase 39B pseudogene 1 [Source |
| MCCD1P1 | CHR_HSCHR6_MHC_QBL_CT( | mitochondrial coiled-coil domain 1 pseudoger |
| HCG4B | CHR_HSCHR6_MHC_QBL_CT( | HLA complex group 4B [Source:HGNC Symbol; |
| HLA-K | CHR_HSCHR6_MHC_QBL_CT( | major histocompatibility complex, class I, K (p |
| HLA-U | CHR_HSCHR6_MHC_QBL_CT( | major histocompatibility complex, class I, U (p |
| HLA-A | CHR_HSCHR6_MHC_QBL_CT( | major histocompatibility complex, class I, A [S |
| AL845454.2 | CHR_HSCHR6_MHC_QBL_CT( | HLA complex group 4 pseudogene 5 |
| HLA-W | CHR_HSCHR6_MHC_QBL_CT( | major histocompatibility complex, class I, W (p |
| MICD | CHR_HSCHR6_MHC_QBL_CT( | MHC class I polypeptide-related sequence D (p |
| HCG9 | CHR_HSCHR6_MHC_QBL_CT( | HLA complex group 9 [Source:HGNC Symbol;A |
| DDX39BP2 | CHR_HSCHR6_MHC_QBL_CT( | DEAD-box helicase 39B pseudogene 2 [Source |
| MCCD1P2 | CHR_HSCHR6_MHC_QBL_CT( | mitochondrial coiled-coil domain 1 pseudoger |
| ZNRD1ASP | CHR_HSCHR6_MHC_QBL_CT( | zinc ribbon domain containing 1 antisense, ps |

| | | |
|---|---|---|
| AL845454.3 | CHR_HSCHR6_MHC_QBL_CT( | HLA complex group 4 pseudogene 3 |
| HLA-J | CHR_HSCHR6_MHC_QBL_CT( | major histocompatibility complex, class I, J (ps |
| ETF1P1 | CHR_HSCHR6_MHC_QBL_CT( | eukaryotic translation termination factor 1 ps |
| AL845439.3 | CHR_HSCHR6_MHC_QBL_CT( | NULL |
| AL845439.2 | CHR_HSCHR6_MHC_QBL_CT( | NULL |
| ZNRD1 | CHR_HSCHR6_MHC_QBL_CT( | zinc ribbon domain containing 1 [Source:HGN( |
| AL845439.1 | CHR_HSCHR6_MHC_QBL_CT( | NULL |
| PPP1R11 | CHR_HSCHR6_MHC_QBL_CT( | protein phosphatase 1 regulatory inhibitor sul |
| RNF39 | CHR_HSCHR6_MHC_QBL_CT( | ring finger protein 39 [Source:HGNC Symbol;A |
| TRIM31 | CHR_HSCHR6_MHC_QBL_CT( | tripartite motif containing 31 [Source:HGNC S |
| TRIM31-AS1 | CHR_HSCHR6_MHC_QBL_CT( | TRIM31 antisense RNA 1 [Source:HGNC Symb |
| BX322644.1 | CHR_HSCHR6_MHC_QBL_CT( | NULL |
| TRIM40 | CHR_HSCHR6_MHC_QBL_CT( | tripartite motif containing 40 [Source:HGNC S |
| TRIM10 | CHR_HSCHR6_MHC_QBL_CT( | tripartite motif containing 10 [Source:HGNC S |
| TRIM15 | CHR_HSCHR6_MHC_QBL_CT( | tripartite motif containing 15 [Source:HGNC S |
| TRIM26 | CHR_HSCHR6_MHC_QBL_CT( | tripartite motif containing 26 [Source:HGNC S |
| PAIP1P1 | CHR_HSCHR6_MHC_QBL_CT( | poly(A) binding protein interacting protein 1 p |
| HCG17 | CHR_HSCHR6_MHC_QBL_CT( | HLA complex group 17 [Source:HGNC Symbol; |
| TRIM26BP | CHR_HSCHR6_MHC_QBL_CT( | tripartite motif containing 26B, pseudogene [S |
| HLA-L | CHR_HSCHR6_MHC_QBL_CT( | major histocompatibility complex, class I, L (ps |
| HCG18 | CHR_HSCHR6_MHC_QBL_CT( | HLA complex group 18 [Source:HGNC Symbol; |
| TRIM39 | CHR_HSCHR6_MHC_QBL_CT( | tripartite motif containing 39 [Source:HGNC S |
| RPP21 | CHR_HSCHR6_MHC_QBL_CT( | ribonuclease P/MRP subunit p21 [Source:HGN |
| HLA-N | CHR_HSCHR6_MHC_QBL_CT( | major histocompatibility complex, class I, N (p |
| AL773535.1 | CHR_HSCHR6_MHC_QBL_CT( | HLA complex group 19 pseudogene |
| UBQLN1P1 | CHR_HSCHR6_MHC_QBL_CT( | ubiquilin 1 pseudogene 1 [Source:HGNC Symb |
| MICC | CHR_HSCHR6_MHC_QBL_CT( | MHC class I polypeptide-related sequence C (p |
| TMPOP1 | CHR_HSCHR6_MHC_QBL_CT( | thymopoietin pseudogene 1 [Source:HGNC Sy |
| SUCLA2P1 | CHR_HSCHR6_MHC_QBL_CT( | succinate-CoA ligase ADP-forming beta subun |
| RANP1 | CHR_HSCHR6_MHC_QBL_CT( | RAN pseudogene 1 [Source:HGNC Symbol;Acc |
| HLA-E | CHR_HSCHR6_MHC_QBL_CT( | major histocompatibility complex, class I, E [S |
| LINC02569 | CHR_HSCHR6_MHC_QBL_CT( | long intergenic non-protein coding RNA 2569 |
| GNL1 | CHR_HSCHR6_MHC_QBL_CT( | G protein nucleolar 1 (putative) [Source:HGN( |
| PRR3 | CHR_HSCHR6_MHC_QBL_CT( | proline rich 3 [Source:HGNC Symbol;Acc:HGN( |
| ABCF1 | CHR_HSCHR6_MHC_QBL_CT( | ATP binding cassette subfamily F member 1 [S |
| PPP1R10 | CHR_HSCHR6_MHC_QBL_CT( | protein phosphatase 1 regulatory subunit 10 [ |
| MRPS18B | CHR_HSCHR6_MHC_QBL_CT( | mitochondrial ribosomal protein S18B [Source |
| ATAT1 | CHR_HSCHR6_MHC_QBL_CT( | alpha tubulin acetyltransferase 1 [Source:HGN |
| PTMAP1 | CHR_HSCHR6_MHC_QBL_CT( | prothymosin alpha pseudogene 1 [Source:HGI |
| C6orf136 | CHR_HSCHR6_MHC_QBL_CT( | chromosome 6 open reading frame 136 [Sour |
| DHX16 | CHR_HSCHR6_MHC_QBL_CT( | DEAH-box helicase 16 [Source:HGNC Symbol;/ |
| PPP1R18 | CHR_HSCHR6_MHC_QBL_CT( | protein phosphatase 1 regulatory subunit 18 [ |
| NRM | CHR_HSCHR6_MHC_QBL_CT( | nurim [Source:HGNC Symbol;Acc:HGNC:8003] |
| RPL7P4 | CHR_HSCHR6_MHC_QBL_CT( | ribosomal protein L7 pseudogene 4 [Source:H |
| MDC1 | CHR_HSCHR6_MHC_QBL_CT( | mediator of DNA damage checkpoint 1 [Sourc |
| MDC1-AS1 | CHR_HSCHR6_MHC_QBL_CT( | MDC1 antisense RNA 1 [Source:HGNC Symbol |
| TUBB | CHR_HSCHR6_MHC_QBL_CT( | tubulin beta class I [Source:HGNC Symbol;Acc |
| FLOT1 | CHR_HSCHR6_MHC_QBL_CT( | flotillin 1 [Source:HGNC Symbol;Acc:HGNC:37. |
| Y_RNA | CHR_HSCHR6_MHC_QBL_CT( | Y RNA [Source:RFAM;Acc:RF00019] |
| IER3 | CHR_HSCHR6_MHC_QBL_CT( | immediate early response 3 [Source:HGNC Syı |

| | | |
|---|---|---|
| Metazoa_SRP | CHR_HSCHR6_MHC_QBL_CT( | Metazoan signal recognition particle RNA [Sou |
| HCG20 | CHR_HSCHR6_MHC_QBL_CT( | HLA complex group 20 [Source:HGNC Symbol; |
| AL773551.1 | CHR_HSCHR6_MHC_QBL_CT( | chromosome 6 open reading frame 214 (puta |
| LINC02570 | CHR_HSCHR6_MHC_QBL_CT( | long intergenic non-protein coding RNA 2570 |
| 7SK | CHR_HSCHR6_MHC_QBL_CT( | 7SK RNA [Source:RFAM;Acc:RF00100] |
| DDR1-DT | CHR_HSCHR6_MHC_QBL_CT( | DDR1 divergent transcript [Source:HGNC Sym |
| DDR1 | CHR_HSCHR6_MHC_QBL_CT( | discoidin domain receptor tyrosine kinase 1 [S |
| GTF2H4 | CHR_HSCHR6_MHC_QBL_CT( | general transcription factor IIH subunit 4 [Sou |
| AL773541.1 | CHR_HSCHR6_MHC_QBL_CT( | novel protein |
| VARS2 | CHR_HSCHR6_MHC_QBL_CT( | valyl-tRNA synthetase 2, mitochondrial [Sourc |
| SFTA2 | CHR_HSCHR6_MHC_QBL_CT( | surfactant associated 2 [Source:HGNC Symbol |
| Y_RNA | CHR_HSCHR6_MHC_QBL_CT( | Y RNA [Source:RFAM;Acc:RF00019] |
| MUCL3 | CHR_HSCHR6_MHC_QBL_CT( | mucin like 3 [Source:HGNC Symbol;Acc:HGNC |
| HCG21 | CHR_HSCHR6_MHC_QBL_CT( | HLA complex group 21 [Source:HGNC Symbol; |
| MUC21 | CHR_HSCHR6_MHC_QBL_CT( | mucin 21, cell surface associated [Source:NCB |
| AL805909.1 | CHR_HSCHR6_MHC_QBL_CT( | novel transcript |
| HCG22 | CHR_HSCHR6_MHC_QBL_CT( | HLA complex group 22 (gene/pseudogene) [So |
| U6 | CHR_HSCHR6_MHC_QBL_CT( | U6 spliceosomal RNA [Source:RFAM;Acc:RF00 |
| C6orf15 | CHR_HSCHR6_MHC_QBL_CT( | chromosome 6 open reading frame 15 [Sourc |
| PSORS1C1 | CHR_HSCHR6_MHC_QBL_CT( | psoriasis susceptibility 1 candidate 1 [Source:I |
| CDSN | CHR_HSCHR6_MHC_QBL_CT( | corneodesmosin [Source:HGNC Symbol;Acc:H |
| PSORS1C2 | CHR_HSCHR6_MHC_QBL_CT( | psoriasis susceptibility 1 candidate 2 [Source:I |
| POLR2LP1 | CHR_HSCHR6_MHC_QBL_CT( | RNA polymerase II subunit L pseudogene 1 [So |
| CCHCR1 | CHR_HSCHR6_MHC_QBL_CT( | coiled-coil alpha-helical rod protein 1 [Source: |
| TCF19 | CHR_HSCHR6_MHC_QBL_CT( | transcription factor 19 [Source:HGNC Symbol; |
| POU5F1 | CHR_HSCHR6_MHC_QBL_CT( | POU class 5 homeobox 1 [Source:HGNC Symb |
| PSORS1C3 | CHR_HSCHR6_MHC_QBL_CT( | psoriasis susceptibility 1 candidate 3 [Source:I |
| HCG27 | CHR_HSCHR6_MHC_QBL_CT( | HLA complex group 27 [Source:HGNC Symbol; |
| AL845458.1 | CHR_HSCHR6_MHC_QBL_CT( | novel transcript |
| HLA-C | CHR_HSCHR6_MHC_QBL_CT( | major histocompatibility complex, class I, C [S |
| HLA-B | CHR_HSCHR6_MHC_QBL_CT( | major histocompatibility complex, class I, B [S |
| USP8P1 | CHR_HSCHR6_MHC_QBL_CT( | ubiquitin specific peptidase 8 pseudogene 1 [S |
| RPL3P2 | CHR_HSCHR6_MHC_QBL_CT( | ribosomal protein L3 pseudogene 2 [Source:H |
| WASF5P | CHR_HSCHR6_MHC_QBL_CT( | WASP family member 5, pseudogene [Source: |
| AL845443.1 | CHR_HSCHR6_MHC_QBL_CT( | novel HLA class I pseudogene fragment |
| DHFRP2 | CHR_HSCHR6_MHC_QBL_CT( | dihydrofolate reductase pseudogene 2 [Sourc |
| U6 | CHR_HSCHR6_MHC_QBL_CT( | U6 spliceosomal RNA [Source:RFAM;Acc:RF00 |
| AL845443.2 | CHR_HSCHR6_MHC_QBL_CT( | fibroblast growth factor receptor 3 pseudoger |
| ZDHHC20P2 | CHR_HSCHR6_MHC_QBL_CT( | zinc finger DHHC-type containing 20 pseudoge |
| HLA-S | CHR_HSCHR6_MHC_QBL_CT( | major histocompatibility complex, class I, S (ps |
| MICA | CHR_HSCHR6_MHC_QBL_CT( | MHC class I polypeptide-related sequence A [S |
| Y_RNA | CHR_HSCHR6_MHC_QBL_CT( | Y RNA [Source:RFAM;Acc:RF00019] |
| Y_RNA | CHR_HSCHR6_MHC_QBL_CT( | Y RNA [Source:RFAM;Acc:RF00019] |
| MICB | CHR_HSCHR6_MHC_QBL_CT( | MHC class I polypeptide-related sequence B [S |
| PPIAP9 | CHR_HSCHR6_MHC_QBL_CT( | peptidylprolyl isomerase A pseudogene 9 [Sou |
| RPL15P4 | CHR_HSCHR6_MHC_QBL_CT( | ribosomal protein L15 pseudogene 4 [Source: |
| MCCD1 | CHR_HSCHR6_MHC_QBL_CT( | mitochondrial coiled-coil domain 1 [Source:H( |
| DDX39B | CHR_HSCHR6_MHC_QBL_CT( | DExD-box helicase 39B [Source:HGNC Symbol, |
| SNORD117 | CHR_HSCHR6_MHC_QBL_CT( | small nucleolar RNA, C/D box 117 [Source:HGI |
| SNORD84 | CHR_HSCHR6_MHC_QBL_CT( | small nucleolar RNA, C/D box 84 [Source:HGN |

| | | |
|---|---|---|
| DDX39B-AS1 | CHR_HSCHR6_MHC_QBL_CT( | DDX39B antisense RNA 1 [Source:HGNC Symb |
| ATP6V1G2 | CHR_HSCHR6_MHC_QBL_CT( | ATPase H+ transporting V1 subunit G2 [Source |
| NFKBIL1 | CHR_HSCHR6_MHC_QBL_CT( | NFKB inhibitor like 1 [Source:HGNC Symbol;Ac |
| LTA | CHR_HSCHR6_MHC_QBL_CT( | lymphotoxin alpha [Source:HGNC Symbol;Acc |
| TNF | CHR_HSCHR6_MHC_QBL_CT( | tumor necrosis factor [Source:HGNC Symbol;A |
| LTB | CHR_HSCHR6_MHC_QBL_CT( | lymphotoxin beta [Source:HGNC Symbol;Acc:I |
| LST1 | CHR_HSCHR6_MHC_QBL_CT( | leukocyte specific transcript 1 [Source:HGNC S |
| NCR3 | CHR_HSCHR6_MHC_QBL_CT( | natural cytotoxicity triggering receptor 3 [Sou |
| UQCRHP1 | CHR_HSCHR6_MHC_QBL_CT( | ubiquinol-cytochrome c reductase hinge prote |
| AIF1 | CHR_HSCHR6_MHC_QBL_CT( | allograft inflammatory factor 1 [Source:HGNC |
| PRRC2A | CHR_HSCHR6_MHC_QBL_CT( | proline rich coiled-coil 2A [Source:HGNC Symb |
| SNORA38 | CHR_HSCHR6_MHC_QBL_CT( | small nucleolar RNA, H/ACA box 38 [Source:H |
| BAG6 | CHR_HSCHR6_MHC_QBL_CT( | BCL2 associated athanogene 6 [Source:HGNC |
| APOM | CHR_HSCHR6_MHC_QBL_CT( | apolipoprotein M [Source:HGNC Symbol;Acc:H |
| C6orf47 | CHR_HSCHR6_MHC_QBL_CT( | chromosome 6 open reading frame 47 [Source |
| C6orf47-AS1 | CHR_HSCHR6_MHC_QBL_CT( | C6orf47 antisense RNA 1 [Source:HGNC Symb |
| GPANK1 | CHR_HSCHR6_MHC_QBL_CT( | G-patch domain and ankyrin repeats 1 [Source |
| Y_RNA | CHR_HSCHR6_MHC_QBL_CT( | Y RNA [Source:RFAM;Acc:RF00019] |
| CSNK2B | CHR_HSCHR6_MHC_QBL_CT( | casein kinase 2 beta [Source:HGNC Symbol;Ac |
| LY6G5B | CHR_HSCHR6_MHC_QBL_CT( | lymphocyte antigen 6 family member G5B [So |
| LY6G5C | CHR_HSCHR6_MHC_QBL_CT( | lymphocyte antigen 6 family member G5C [So |
| ABHD16A | CHR_HSCHR6_MHC_QBL_CT( | abhydrolase domain containing 16A [Source:H |
| LY6G6F | CHR_HSCHR6_MHC_QBL_CT( | lymphocyte antigen 6 family member G6F [So |
| LY6G6D | CHR_HSCHR6_MHC_QBL_CT( | lymphocyte antigen 6 family member G6D [So |
| LY6G6E | CHR_HSCHR6_MHC_QBL_CT( | lymphocyte antigen 6 family member G6E [So |
| MPIG6B | CHR_HSCHR6_MHC_QBL_CT( | megakaryocyte and platelet inhibitory recepto |
| LY6G6C | CHR_HSCHR6_MHC_QBL_CT( | lymphocyte antigen 6 family member G6C [So |
| DDAH2 | CHR_HSCHR6_MHC_QBL_CT( | dimethylarginine dimethylaminohydrolase 2 [ |
| CLIC1 | CHR_HSCHR6_MHC_QBL_CT( | chloride intracellular channel 1 [Source:HGNC |
| MSH5 | CHR_HSCHR6_MHC_QBL_CT( | mutS homolog 5 [Source:HGNC Symbol;Acc:H |
| U6 | CHR_HSCHR6_MHC_QBL_CT( | U6 spliceosomal RNA [Source:RFAM;Acc:RF00 |
| VARS | CHR_HSCHR6_MHC_QBL_CT( | valyl-tRNA synthetase [Source:HGNC Symbol;A |
| LSM2 | CHR_HSCHR6_MHC_QBL_CT( | LSM2 homolog, U6 small nuclear RNA and mR |
| HSPA1L | CHR_HSCHR6_MHC_QBL_CT( | heat shock protein family A (Hsp70) member 1 |
| HSPA1A | CHR_HSCHR6_MHC_QBL_CT( | heat shock protein family A (Hsp70) member 1 |
| HSPA1B | CHR_HSCHR6_MHC_QBL_CT( | heat shock protein family A (Hsp70) member 1 |
| SNHG32 | CHR_HSCHR6_MHC_QBL_CT( | small nucleolar RNA host gene 32 [Source:HGI |
| SNORD48 | CHR_HSCHR6_MHC_QBL_CT( | small nucleolar RNA, C/D box 48 [Source:HGN |
| SNORD52 | CHR_HSCHR6_MHC_QBL_CT( | small nucleolar RNA, C/D box 52 [Source:HGN |
| NEU1 | CHR_HSCHR6_MHC_QBL_CT( | neuraminidase 1 [Source:HGNC Symbol;Acc:H |
| CYP21A2 | CHR_HSCHR6_MHC_QBL_CT( | cytochrome P450 family 21 subfamily A meml |
| SLC44A4 | CHR_HSCHR6_MHC_QBL_CT( | solute carrier family 44 member 4 [Source:HG |
| EHMT2 | CHR_HSCHR6_MHC_QBL_CT( | euchromatic histone lysine methyltransferase |
| AL844853.1 | CHR_HSCHR6_MHC_QBL_CT( | novel transcript |
| C2 | CHR_HSCHR6_MHC_QBL_CT( | complement C2 [Source:HGNC Symbol;Acc:H( |
| ZBTB12 | CHR_HSCHR6_MHC_QBL_CT( | zinc finger and BTB domain containing 12 [Sou |
| C4B-AS1 | CHR_HSCHR6_MHC_QBL_CT( | C4B antisense RNA 1 [Source:HGNC Symbol;A |
| CFB | CHR_HSCHR6_MHC_QBL_CT( | complement factor B [Source:HGNC Symbol;A |
| NELFE | CHR_HSCHR6_MHC_QBL_CT( | negative elongation factor complex member E |
| SKIV2L | CHR_HSCHR6_MHC_QBL_CT( | Ski2 like RNA helicase [Source:HGNC Symbol;A |

| | | |
|---|---|---|
| DXO | CHR_HSCHR6_MHC_QBL_CT( | decapping exoribonuclease [Source:HGNC Syr |
| STK19 | CHR_HSCHR6_MHC_QBL_CT( | serine/threonine kinase 19 [Source:HGNC Syn |
| AL844853.2 | CHR_HSCHR6_MHC_QBL_CT( | novel transcript, STK19-C4A readthrough |
| C4A | CHR_HSCHR6_MHC_QBL_CT( | complement C4A (Rodgers blood group) [Sour |
| C4A-AS1 | CHR_HSCHR6_MHC_QBL_CT( | C4A antisense RNA 1 [Source:HGNC Symbol;A |
| TNXB | CHR_HSCHR6_MHC_QBL_CT( | tenascin XB [Source:HGNC Symbol;Acc:HGNC: |
| 5S_rRNA | CHR_HSCHR6_MHC_QBL_CT( | 5S ribosomal RNA [Source:RFAM;Acc:RF00001 |
| PRRT1 | CHR_HSCHR6_MHC_QBL_CT( | proline rich transmembrane protein 1 [Source |
| PPT2 | CHR_HSCHR6_MHC_QBL_CT( | palmitoyl-protein thioesterase 2 [Source:HGN |
| AL845464.1 | CHR_HSCHR6_MHC_QBL_CT( | novel transcript, PPT2-EGFL8 readthrough |
| EGFL8 | CHR_HSCHR6_MHC_QBL_CT( | EGF like domain multiple 8 [Source:HGNC Sym |
| AGPAT1 | CHR_HSCHR6_MHC_QBL_CT( | 1-acylglycerol-3-phosphate O-acyltransferase |
| RNF5 | CHR_HSCHR6_MHC_QBL_CT( | ring finger protein 5 [Source:HGNC Symbol;Ac |
| AGER | CHR_HSCHR6_MHC_QBL_CT( | advanced glycosylation end-product specific r |
| PBX2 | CHR_HSCHR6_MHC_QBL_CT( | PBX homeobox 2 [Source:HGNC Symbol;Acc:H |
| GPSM3 | CHR_HSCHR6_MHC_QBL_CT( | G protein signaling modulator 3 [Source:HGN( |
| NOTCH4 | CHR_HSCHR6_MHC_QBL_CT( | notch receptor 4 [Source:HGNC Symbol;Acc:H |
| TSBP1 | CHR_HSCHR6_MHC_QBL_CT( | testis expressed basic protein 1 [Source:HGN( |
| HNRNPA1P2 | CHR_HSCHR6_MHC_QBL_CT( | heterogeneous nuclear ribonucleoprotein A1 |
| U6 | CHR_HSCHR6_MHC_QBL_CT( | U6 spliceosomal RNA [Source:RFAM;Acc:RF00 |
| TSBP1-AS1 | CHR_HSCHR6_MHC_QBL_CT( | TSBP1 and BTNL2 antisense RNA 1 [Source:HG |
| BTNL2 | CHR_HSCHR6_MHC_QBL_CT( | butyrophilin like 2 [Source:HGNC Symbol;Acc: |
| HLA-DRA | CHR_HSCHR6_MHC_QBL_CT( | major histocompatibility complex, class II, DR |
| HLA-DRB9 | CHR_HSCHR6_MHC_QBL_CT( | major histocompatibility complex, class II, DR |
| U1 | CHR_HSCHR6_MHC_QBL_CT( | U1 spliceosomal RNA [Source:RFAM;Acc:RF00 |
| HLA-DRB3 | CHR_HSCHR6_MHC_QBL_CT( | major histocompatibility complex, class II, DR |
| U1 | CHR_HSCHR6_MHC_QBL_CT( | U1 spliceosomal RNA [Source:RFAM;Acc:RF00 |
| HLA-DRB2 | CHR_HSCHR6_MHC_QBL_CT( | major histocompatibility complex, class II, DR |
| U1 | CHR_HSCHR6_MHC_QBL_CT( | U1 spliceosomal RNA [Source:RFAM;Acc:RF00 |
| HLA-DRB1 | CHR_HSCHR6_MHC_QBL_CT( | major histocompatibility complex, class II, DR |
| HLA-DQA1 | CHR_HSCHR6_MHC_QBL_CT( | major histocompatibility complex, class II, DQ |
| HLA-DQB1 | CHR_HSCHR6_MHC_QBL_CT( | major histocompatibility complex, class II, DQ |
| HLA-DQB1-AS1 | CHR_HSCHR6_MHC_QBL_CT( | HLA-DQB1 antisense RNA 1 [Source:HGNC Syr |
| AL935026.1 | CHR_HSCHR6_MHC_QBL_CT( | novel transcript |
| MTCO3P1 | CHR_HSCHR6_MHC_QBL_CT( | MT-CO3 pseudogene 1 [Source:HGNC Symbol |
| AL773543.1 | CHR_HSCHR6_MHC_QBL_CT( | uncharacterized LOC102725019 [Source:NCBI |
| HLA-DQB3 | CHR_HSCHR6_MHC_QBL_CT( | major histocompatibility complex, class II, DQ |
| HLA-DQA2 | CHR_HSCHR6_MHC_QBL_CT( | major histocompatibility complex, class II, DQ |
| HLA-DQB2 | CHR_HSCHR6_MHC_QBL_CT( | major histocompatibility complex, class II, DQ |
| TAP2 | CHR_HSCHR6_MHC_QBL_CT( | transporter 2, ATP binding cassette subfamily |
| PSMB8 | CHR_HSCHR6_MHC_QBL_CT( | proteasome subunit beta 8 [Source:HGNC Syn |
| PSMB8-AS1 | CHR_HSCHR6_MHC_QBL_CT( | PSMB8 antisense RNA 1 (head to head) [Sourc |
| PSMB9 | CHR_HSCHR6_MHC_QBL_CT( | proteasome subunit beta 9 [Source:HGNC Syn |
| TAP1 | CHR_HSCHR6_MHC_QBL_CT( | transporter 1, ATP binding cassette subfamily |
| PPP1R2P1 | CHR_HSCHR6_MHC_QBL_CT( | protein phosphatase 1 regulatory inhibitor sul |
| HLA-Z | CHR_HSCHR6_MHC_QBL_CT( | major histocompatibility complex, class I, Z (p: |
| HLA-DMB | CHR_HSCHR6_MHC_QBL_CT( | major histocompatibility complex, class II, DM |
| HLA-DMA | CHR_HSCHR6_MHC_QBL_CT( | major histocompatibility complex, class II, DM |
| BRD2 | CHR_HSCHR6_MHC_QBL_CT( | bromodomain containing 2 [Source:HGNC Syn |
| AL935042.1 | CHR_HSCHR6_MHC_QBL_CT( | novel transcript |

| | | |
|---|---|---|
| HLA-DOA | CHR_HSCHR6_MHC_QBL_CT( | major histocompatibility complex, class II, DO |
| HLA-DPA1 | CHR_HSCHR6_MHC_QBL_CT( | major histocompatibility complex, class II, DP |
| HLA-DPB1 | CHR_HSCHR6_MHC_QBL_CT( | major histocompatibility complex, class II, DP |
| RPL32P1 | CHR_HSCHR6_MHC_QBL_CT( | ribosomal protein L32 pseudogene 1 [Source: |
| HLA-DPA2 | CHR_HSCHR6_MHC_QBL_CT( | major histocompatibility complex, class II, DP |
| COL11A2P1 | CHR_HSCHR6_MHC_QBL_CT( | collagen type XI alpha 2 pseudogene 1 [Source |
| HLA-DPB2 | CHR_HSCHR6_MHC_QBL_CT( | major histocompatibility complex, class II, DP |
| HLA-DPA3 | CHR_HSCHR6_MHC_QBL_CT( | major histocompatibility complex, class II, DP |
| HCG24 | CHR_HSCHR6_MHC_QBL_CT( | HLA complex group 24 [Source:HGNC Symbol; |
| COL11A2 | CHR_HSCHR6_MHC_QBL_CT( | collagen type XI alpha 2 chain [Source:HGNC S |
| RXRB | CHR_HSCHR6_MHC_QBL_CT( | retinoid X receptor beta [Source:HGNC Symbo |
| Y_RNA | CHR_HSCHR6_MHC_QBL_CT( | Y RNA [Source:RFAM;Acc:RF00019] |
| SLC39A7 | CHR_HSCHR6_MHC_QBL_CT( | solute carrier family 39 member 7 [Source:HG |
| HSD17B8 | CHR_HSCHR6_MHC_QBL_CT( | hydroxysteroid 17-beta dehydrogenase 8 [Sou |
| RING1 | CHR_HSCHR6_MHC_QBL_CT( | ring finger protein 1 [Source:HGNC Symbol;Ac |
| ZNF70P1 | CHR_HSCHR6_MHC_QBL_CT( | zinc finger protein 70 pseudogene 1 [Source:H |
| HTATSF1P1 | CHR_HSCHR6_MHC_QBL_CT( | HIV-1 Tat specific factor 1 pseudogene 1 [Sour |
| HCG25 | CHR_HSCHR6_MHC_QBL_CT( | HLA complex group 25 [Source:HGNC Symbol; |
| VPS52 | CHR_HSCHR6_MHC_QBL_CT( | VPS52 subunit of GARP complex [Source:HGN |
| RPS18 | CHR_HSCHR6_MHC_QBL_CT( | ribosomal protein S18 [Source:HGNC Symbol; |
| B3GALT4 | CHR_HSCHR6_MHC_QBL_CT( | beta-1,3-galactosyltransferase 4 [Source:HGN |
| WDR46 | CHR_HSCHR6_MHC_QBL_CT( | WD repeat domain 46 [Source:HGNC Symbol; |
| PFDN6 | CHR_HSCHR6_MHC_QBL_CT( | prefoldin subunit 6 [Source:HGNC Symbol;Acc |
| RGL2 | CHR_HSCHR6_MHC_QBL_CT( | ral guanine nucleotide dissociation stimulator |
| TAPBP | CHR_HSCHR6_MHC_QBL_CT( | TAP binding protein [Source:HGNC Symbol;Ac |
| ZBTB22 | CHR_HSCHR6_MHC_QBL_CT( | zinc finger and BTB domain containing 22 [Sou |
| DAXX | CHR_HSCHR6_MHC_QBL_CT( | death domain associated protein [Source:HGN |
| MYL8P | CHR_HSCHR6_MHC_QBL_CT( | myosin light chain 8, pseudogene [Source:HGI |
| LYPLA2P1 | CHR_HSCHR6_MHC_QBL_CT( | LYPLA2 pseudogene 1 [Source:HGNC Symbol;/ |
| RPL35AP4 | CHR_HSCHR6_MHC_QBL_CT( | ribosomal protein L35a pseudogene 4 [Source |
| KIFC1 | CHR_HSCHR6_MHC_QBL_CT( | kinesin family member C1 [Source:HGNC Sym |
| RPL12P1 | CHR_HSCHR6_MHC_QBL_CT( | ribosomal protein L12 pseudogene 1 [Source: |
| PHF1 | CHR_HSCHR6_MHC_QBL_CT( | PHD finger protein 1 [Source:HGNC Symbol;Ac |
| RPSAP2 | CHR_HSCHR6_MHC_SSTO_CT | ribosomal protein SA pseudogene 2 [Source:H |
| NOP56P1 | CHR_HSCHR6_MHC_SSTO_CT | NOP56 ribonucleoprotein pseudogene 1 [Sour |
| BX119924.1 | CHR_HSCHR6_MHC_SSTO_CT | novel transcript |
| LINC01623 | CHR_HSCHR6_MHC_SSTO_CT | long intergenic non-protein coding RNA 1623 |
| RPL13P | CHR_HSCHR6_MHC_SSTO_CT | ribosomal protein L13 pseudogene [Source:H( |
| ZNF90P2 | CHR_HSCHR6_MHC_SSTO_CT | zinc finger protein 90 pseudogene 2 [Source:H |
| HCG14 | CHR_HSCHR6_MHC_SSTO_CT | HLA complex group 14 [Source:HGNC Symbol; |
| TRIM27 | CHR_HSCHR6_MHC_SSTO_CT | tripartite motif containing 27 [Source:HGNC S |
| U6 | CHR_HSCHR6_MHC_SSTO_CT | U6 spliceosomal RNA [Source:RFAM;Acc:RF00 |
| LINC01556 | CHR_HSCHR6_MHC_SSTO_CT | long intergenic non-protein coding RNA 1556 |
| KRT18P1 | CHR_HSCHR6_MHC_SSTO_CT | keratin 18 pseudogene 1 [Source:HGNC Symb |
| Metazoa_SRP | CHR_HSCHR6_MHC_SSTO_CT | Metazoan signal recognition particle RNA [Sou |
| HCG15 | CHR_HSCHR6_MHC_SSTO_CT | HLA complex group 15 [Source:HGNC Symbol; |
| BX248109.1 | CHR_HSCHR6_MHC_SSTO_CT | HLA complex group 16 |
| ZNF311 | CHR_HSCHR6_MHC_SSTO_CT | zinc finger protein 311 [Source:HGNC Symbol; |
| OR2AD1P | CHR_HSCHR6_MHC_SSTO_CT | olfactory receptor family 2 subfamily AD mem |
| OR2W1 | CHR_HSCHR6_MHC_SSTO_CT | olfactory receptor family 2 subfamily W meml |

| | | |
|---|---|---|
| BX248413.1 | CHR_HSCHR6_MHC_SSTO_CT | NULL |
| OR2P1P | CHR_HSCHR6_MHC_SSTO_CT | olfactory receptor family 2 subfamily P memb |
| SAR1AP1 | CHR_HSCHR6_MHC_SSTO_CT | secretion associated Ras related GTPase 1A ps |
| OR2B3 | CHR_HSCHR6_MHC_SSTO_CT | olfactory receptor family 2 subfamily B memb |
| OR2J1 | CHR_HSCHR6_MHC_SSTO_CT | olfactory receptor family 2 subfamily J membe |
| OR2J3 | CHR_HSCHR6_MHC_SSTO_CT | olfactory receptor family 2 subfamily J membe |
| BX248093.2 | CHR_HSCHR6_MHC_SSTO_CT | novel transcript |
| BX248093.1 | CHR_HSCHR6_MHC_SSTO_CT | olfactory receptor family 2 subfamily N memb |
| OR2J2 | CHR_HSCHR6_MHC_SSTO_CT | olfactory receptor family 2 subfamily J membe |
| OR2J4P | CHR_HSCHR6_MHC_SSTO_CT | olfactory receptor family 2 subfamily J membe |
| BX248084.1 | CHR_HSCHR6_MHC_SSTO_CT | olfactory receptor family 2 subfamily H memb |
| BX248084.2 | CHR_HSCHR6_MHC_SSTO_CT | novel transcript |
| OR2G1P | CHR_HSCHR6_MHC_SSTO_CT | olfactory receptor family 2 subfamily G memb |
| OR2U1P | CHR_HSCHR6_MHC_SSTO_CT | olfactory receptor family 2 subfamily U memb |
| OR2U2P | CHR_HSCHR6_MHC_SSTO_CT | olfactory receptor family 2 subfamily U memb |
| BX005432.1 | CHR_HSCHR6_MHC_SSTO_CT | olfactory receptor family 2 subfamily B memb |
| OR14J1 | CHR_HSCHR6_MHC_SSTO_CT | olfactory receptor family 14 subfamily J meml |
| DDX6P1 | CHR_HSCHR6_MHC_SSTO_CT | DEAD-box helicase 6 pseudogene 1 [Source:H |
| OR5V1 | CHR_HSCHR6_MHC_SSTO_CT | olfactory receptor family 5 subfamily V memb |
| OR12D3 | CHR_HSCHR6_MHC_SSTO_CT | olfactory receptor family 12 subfamily D mem |
| OR12D2 | CHR_HSCHR6_MHC_SSTO_CT | olfactory receptor family 12 subfamily D mem |
| OR12D1 | CHR_HSCHR6_MHC_SSTO_CT | olfactory receptor family 12 subfamily D mem |
| OR11A1 | CHR_HSCHR6_MHC_SSTO_CT | olfactory receptor family 11 subfamily A mem |
| OR10C1 | CHR_HSCHR6_MHC_SSTO_CT | olfactory receptor family 10 subfamily C mem |
| OR2H1 | CHR_HSCHR6_MHC_SSTO_CT | olfactory receptor family 2 subfamily H memb |
| UBDP1 | CHR_HSCHR6_MHC_SSTO_CT | ubiquitin D pseudogene 1 [Source:HGNC Symb |
| MAS1LP1 | CHR_HSCHR6_MHC_SSTO_CT | MAS1L pseudogene 1 [Source:HGNC Symbol;A |
| MAS1L | CHR_HSCHR6_MHC_SSTO_CT | MAS1 proto-oncogene like, G protein-couplec |
| RPS17P1 | CHR_HSCHR6_MHC_SSTO_CT | ribosomal protein S17 pseudogene 1 [Source: |
| LINC02829 | CHR_HSCHR6_MHC_SSTO_CT | long intergenic non-protein coding RNA 2829 |
| LINC01015 | CHR_HSCHR6_MHC_SSTO_CT | long intergenic non-protein coding RNA 1015 |
| GPR53P | CHR_HSCHR6_MHC_SSTO_CT | G protein-coupled receptor 53, pseudogene [S |
| SUMO2P1 | CHR_HSCHR6_MHC_SSTO_CT | SUMO2 pseudogene 1 [Source:HGNC Symbol; |
| MOG | CHR_HSCHR6_MHC_SSTO_CT | myelin oligodendrocyte glycoprotein [Source: |
| ZFP57 | CHR_HSCHR6_MHC_SSTO_CT | ZFP57 zinc finger protein [Source:HGNC Symb |
| ZDHHC20P1 | CHR_HSCHR6_MHC_SSTO_CT | zinc finger DHHC-type containing 20 pseudoge |
| HLA-F | CHR_HSCHR6_MHC_SSTO_CT | major histocompatibility complex, class I, F [S |
| BX005428.2 | CHR_HSCHR6_MHC_SSTO_CT | HLA complex group 4 pseudogene 11 |
| HLA-F-AS1 | CHR_HSCHR6_MHC_SSTO_CT | HLA-F antisense RNA 1 [Source:HGNC Symbol; |
| RPL23AP1 | CHR_HSCHR6_MHC_SSTO_CT | ribosomal protein L23a pseudogene 1 [Source |
| MICE | CHR_HSCHR6_MHC_SSTO_CT | MHC class I polypeptide-related sequence E (p |
| BX005428.1 | CHR_HSCHR6_MHC_SSTO_CT | HLA complex group 9 pseudogene 5 |
| IFITM4P | CHR_HSCHR6_MHC_SSTO_CT | interferon induced transmembrane protein 4 |
| HLA-V | CHR_HSCHR6_MHC_SSTO_CT | major histocompatibility complex, class I, V (p |
| HCG4 | CHR_HSCHR6_MHC_SSTO_CT | HLA complex group 4 [Source:HGNC Symbol;A |
| HLA-P | CHR_HSCHR6_MHC_SSTO_CT | major histocompatibility complex, class I, P (p |
| RPL7AP7 | CHR_HSCHR6_MHC_SSTO_CT | ribosomal protein L7a pseudogene 7 [Source: |
| MICG | CHR_HSCHR6_MHC_SSTO_CT | MHC class I polypeptide-related sequence G ( |
| HLA-G | CHR_HSCHR6_MHC_SSTO_CT | major histocompatibility complex, class I, G [S |
| HCG4P8 | CHR_HSCHR6_MHC_SSTO_CT | HLA complex group 4 pseudogene 8 [Source:H |

| | | |
|---|---|---|
| BX247949.1 | CHR_HSCHR6_MHC_SSTO_CT | MHC class I polypeptide-related sequence F ps |
| BX284699.1 | CHR_HSCHR6_MHC_SSTO_CT | HLA complex group 4 pseudogene 7 |
| HLA-H | CHR_HSCHR6_MHC_SSTO_CT | major histocompatibility complex, class I, H (p |
| HLA-T | CHR_HSCHR6_MHC_SSTO_CT | major histocompatibility complex, class I, T (p |
| DDX39BP1 | CHR_HSCHR6_MHC_SSTO_CT | DEAD-box helicase 39B pseudogene 1 [Source |
| MCCD1P1 | CHR_HSCHR6_MHC_SSTO_CT | mitochondrial coiled-coil domain 1 pseudogen |
| HCG4B | CHR_HSCHR6_MHC_SSTO_CT | HLA complex group 4B [Source:HGNC Symbol; |
| HLA-K | CHR_HSCHR6_MHC_SSTO_CT | major histocompatibility complex, class I, K (p |
| HLA-U | CHR_HSCHR6_MHC_SSTO_CT | major histocompatibility complex, class I, U (p |
| HLA-A | CHR_HSCHR6_MHC_SSTO_CT | major histocompatibility complex, class I, A [S |
| BX005091.1 | CHR_HSCHR6_MHC_SSTO_CT | HLA complex group 4 pseudogene 5 |
| HLA-W | CHR_HSCHR6_MHC_SSTO_CT | major histocompatibility complex, class I, W (p |
| MICD | CHR_HSCHR6_MHC_SSTO_CT | MHC class I polypeptide-related sequence D (p |
| HCG9 | CHR_HSCHR6_MHC_SSTO_CT | HLA complex group 9 [Source:HGNC Symbol;A |
| DDX39BP2 | CHR_HSCHR6_MHC_SSTO_CT | DEAD-box helicase 39B pseudogene 2 [Source |
| MCCD1P2 | CHR_HSCHR6_MHC_SSTO_CT | mitochondrial coiled-coil domain 1 pseudogen |
| ZNRD1ASP | CHR_HSCHR6_MHC_SSTO_CT | zinc ribbon domain containing 1 antisense, ps |
| BX005091.2 | CHR_HSCHR6_MHC_SSTO_CT | HLA complex group 4 pseudogene 3 |
| HLA-J | CHR_HSCHR6_MHC_SSTO_CT | major histocompatibility complex, class I, J (ps |
| ETF1P1 | CHR_HSCHR6_MHC_SSTO_CT | eukaryotic translation termination factor 1 ps |
| BX088647.2 | CHR_HSCHR6_MHC_SSTO_CT | NULL |
| BX088647.1 | CHR_HSCHR6_MHC_SSTO_CT | NULL |
| ZNRD1 | CHR_HSCHR6_MHC_SSTO_CT | zinc ribbon domain containing 1 [Source:HGN |
| BX088647.3 | CHR_HSCHR6_MHC_SSTO_CT | NULL |
| PPP1R11 | CHR_HSCHR6_MHC_SSTO_CT | protein phosphatase 1 regulatory inhibitor sul |
| RNF39 | CHR_HSCHR6_MHC_SSTO_CT | ring finger protein 39 [Source:HGNC Symbol;A |
| TRIM31 | CHR_HSCHR6_MHC_SSTO_CT | tripartite motif containing 31 [Source:HGNC S |
| TRIM31-AS1 | CHR_HSCHR6_MHC_SSTO_CT | TRIM31 antisense RNA 1 [Source:HGNC Symb |
| BX005441.1 | CHR_HSCHR6_MHC_SSTO_CT | NULL |
| TRIM40 | CHR_HSCHR6_MHC_SSTO_CT | tripartite motif containing 40 [Source:HGNC S |
| TRIM10 | CHR_HSCHR6_MHC_SSTO_CT | tripartite motif containing 10 [Source:HGNC S |
| TRIM15 | CHR_HSCHR6_MHC_SSTO_CT | tripartite motif containing 15 [Source:HGNC S |
| TRIM26 | CHR_HSCHR6_MHC_SSTO_CT | tripartite motif containing 26 [Source:HGNC S |
| PAIP1P1 | CHR_HSCHR6_MHC_SSTO_CT | poly(A) binding protein interacting protein 1 p |
| HCG17 | CHR_HSCHR6_MHC_SSTO_CT | HLA complex group 17 [Source:HGNC Symbol; |
| TRIM26BP | CHR_HSCHR6_MHC_SSTO_CT | tripartite motif containing 26B, pseudogene [S |
| HLA-L | CHR_HSCHR6_MHC_SSTO_CT | major histocompatibility complex, class I, L (ps |
| HCG18 | CHR_HSCHR6_MHC_SSTO_CT | HLA complex group 18 [Source:HGNC Symbol; |
| TRIM39 | CHR_HSCHR6_MHC_SSTO_CT | tripartite motif containing 39 [Source:HGNC S |
| RPP21 | CHR_HSCHR6_MHC_SSTO_CT | ribonuclease P/MRP subunit p21 [Source:HGN |
| HLA-N | CHR_HSCHR6_MHC_SSTO_CT | major histocompatibility complex, class I, N (p |
| BX294158.1 | CHR_HSCHR6_MHC_SSTO_CT | HLA complex group 19 pseudogene |
| UBQLN1P1 | CHR_HSCHR6_MHC_SSTO_CT | ubiquilin 1 pseudogene 1 [Source:HGNC Symb |
| MICC | CHR_HSCHR6_MHC_SSTO_CT | MHC class I polypeptide-related sequence C (p |
| TMPOP1 | CHR_HSCHR6_MHC_SSTO_CT | thymopoietin pseudogene 1 [Source:HGNC Sy |
| SUCLA2P1 | CHR_HSCHR6_MHC_SSTO_CT | succinate-CoA ligase ADP-forming beta subun |
| RANP1 | CHR_HSCHR6_MHC_SSTO_CT | RAN pseudogene 1 [Source:HGNC Symbol;Acc |
| HLA-E | CHR_HSCHR6_MHC_SSTO_CT | major histocompatibility complex, class I, E [S |
| LINC02569 | CHR_HSCHR6_MHC_SSTO_CT | long intergenic non-protein coding RNA 2569 |
| GNL1 | CHR_HSCHR6_MHC_SSTO_CT | G protein nucleolar 1 (putative) [Source:HGN |

| | | |
|---|---|---|
| PRR3 | CHR_HSCHR6_MHC_SSTO_CT | proline rich 3 [Source:HGNC Symbol;Acc:HGN |
| ABCF1 | CHR_HSCHR6_MHC_SSTO_CT | ATP binding cassette subfamily F member 1 [S |
| PPP1R10 | CHR_HSCHR6_MHC_SSTO_CT | protein phosphatase 1 regulatory subunit 10 [ |
| MRPS18B | CHR_HSCHR6_MHC_SSTO_CT | mitochondrial ribosomal protein S18B [Source |
| ATAT1 | CHR_HSCHR6_MHC_SSTO_CT | alpha tubulin acetyltransferase 1 [Source:HGN |
| PTMAP1 | CHR_HSCHR6_MHC_SSTO_CT | prothymosin alpha pseudogene 1 [Source:HGI |
| C6orf136 | CHR_HSCHR6_MHC_SSTO_CT | chromosome 6 open reading frame 136 [Sour |
| DHX16 | CHR_HSCHR6_MHC_SSTO_CT | DEAH-box helicase 16 [Source:HGNC Symbol;/ |
| PPP1R18 | CHR_HSCHR6_MHC_SSTO_CT | protein phosphatase 1 regulatory subunit 18 [ |
| MDC1 | CHR_HSCHR6_MHC_SSTO_CT | mediator of DNA damage checkpoint 1 [Sourc |
| MDC1-AS1 | CHR_HSCHR6_MHC_SSTO_CT | MDC1 antisense RNA 1 [Source:HGNC Symbol |
| TUBB | CHR_HSCHR6_MHC_SSTO_CT | tubulin beta class I [Source:HGNC Symbol;Acc |
| FLOT1 | CHR_HSCHR6_MHC_SSTO_CT | flotillin 1 [Source:HGNC Symbol;Acc:HGNC:37 |
| Y_RNA | CHR_HSCHR6_MHC_SSTO_CT | Y RNA [Source:RFAM;Acc:RF00019] |
| IER3 | CHR_HSCHR6_MHC_SSTO_CT | immediate early response 3 [Source:HGNC Sy |
| Metazoa_SRP | CHR_HSCHR6_MHC_SSTO_CT | Metazoan signal recognition particle RNA [Sou |
| HCG20 | CHR_HSCHR6_MHC_SSTO_CT | HLA complex group 20 [Source:HGNC Symbol; |
| BX571801.1 | CHR_HSCHR6_MHC_SSTO_CT | chromosome 6 open reading frame 214 (puta |
| BX248321.1 | CHR_HSCHR6_MHC_SSTO_CT | novel transcript |
| HCG22 | CHR_HSCHR6_MHC_SSTO_CT | HLA complex group 22 (gene/pseudogene) [So |
| U6 | CHR_HSCHR6_MHC_SSTO_CT | U6 spliceosomal RNA [Source:RFAM;Acc:RF00 |
| TCF19 | CHR_HSCHR6_MHC_SSTO_CT | transcription factor 19 [Source:HGNC Symbol; |
| POU5F1 | CHR_HSCHR6_MHC_SSTO_CT | POU class 5 homeobox 1 [Source:HGNC Symb |
| PSORS1C3 | CHR_HSCHR6_MHC_SSTO_CT | psoriasis susceptibility 1 candidate 3 [Source:I |
| HCG27 | CHR_HSCHR6_MHC_SSTO_CT | HLA complex group 27 [Source:HGNC Symbol; |
| BX088580.1 | CHR_HSCHR6_MHC_SSTO_CT | novel transcript |
| HLA-C | CHR_HSCHR6_MHC_SSTO_CT | major histocompatibility complex, class I, C [S |
| HLA-B | CHR_HSCHR6_MHC_SSTO_CT | major histocompatibility complex, class I, B [S |
| USP8P1 | CHR_HSCHR6_MHC_SSTO_CT | ubiquitin specific peptidase 8 pseudogene 1 [S |
| RPL3P2 | CHR_HSCHR6_MHC_SSTO_CT | ribosomal protein L3 pseudogene 2 [Source:H |
| WASF5P | CHR_HSCHR6_MHC_SSTO_CT | WASP family member 5, pseudogene [Source: |
| BX248310.1 | CHR_HSCHR6_MHC_SSTO_CT | novel HLA class I pseudogene fragment |
| DHFRP2 | CHR_HSCHR6_MHC_SSTO_CT | dihydrofolate reductase pseudogene 2 [Sourc |
| U6 | CHR_HSCHR6_MHC_SSTO_CT | U6 spliceosomal RNA [Source:RFAM;Acc:RF00 |
| BX248310.2 | CHR_HSCHR6_MHC_SSTO_CT | fibroblast growth factor receptor 3 pseudoger |
| ZDHHC20P2 | CHR_HSCHR6_MHC_SSTO_CT | zinc finger DHHC-type containing 20 pseudoge |
| HLA-S | CHR_HSCHR6_MHC_SSTO_CT | major histocompatibility complex, class I, S (p: |
| MICA | CHR_HSCHR6_MHC_SSTO_CT | MHC class I polypeptide-related sequence A [S |
| Y_RNA | CHR_HSCHR6_MHC_SSTO_CT | Y RNA [Source:RFAM;Acc:RF00019] |
| LINC01149 | CHR_HSCHR6_MHC_SSTO_CT | long intergenic non-protein coding RNA 1149 |
| HCP5 | CHR_HSCHR6_MHC_SSTO_CT | HLA complex P5 [Source:HGNC Symbol;Acc:H( |
| Y_RNA | CHR_HSCHR6_MHC_SSTO_CT | Y RNA [Source:RFAM;Acc:RF00019] |
| MICB | CHR_HSCHR6_MHC_SSTO_CT | MHC class I polypeptide-related sequence B [S |
| PPIAP9 | CHR_HSCHR6_MHC_SSTO_CT | peptidylprolyl isomerase A pseudogene 9 [Sou |
| RPL15P4 | CHR_HSCHR6_MHC_SSTO_CT | ribosomal protein L15 pseudogene 4 [Source: |
| MCCD1 | CHR_HSCHR6_MHC_SSTO_CT | mitochondrial coiled-coil domain 1 [Source:H( |
| DDX39B | CHR_HSCHR6_MHC_SSTO_CT | DExD-box helicase 39B [Source:HGNC Symbol, |
| BX248516.1 | CHR_HSCHR6_MHC_SSTO_CT | ATPase H+ transporting lysosomal 13kDa V1 s |
| SNORD117 | CHR_HSCHR6_MHC_SSTO_CT | small nucleolar RNA, C/D box 117 [Source:HGI |
| SNORD84 | CHR_HSCHR6_MHC_SSTO_CT | small nucleolar RNA, C/D box 84 [Source:HGN |

| | | |
|---|---|---|
| DDX39B-AS1 | CHR_HSCHR6_MHC_SSTO_CT | DDX39B antisense RNA 1 [Source:HGNC Symb |
| ATP6V1G2 | CHR_HSCHR6_MHC_SSTO_CT | ATPase H+ transporting V1 subunit G2 [Source |
| NFKBIL1 | CHR_HSCHR6_MHC_SSTO_CT | NFKB inhibitor like 1 [Source:HGNC Symbol;Ac |
| LTA | CHR_HSCHR6_MHC_SSTO_CT | lymphotoxin alpha [Source:HGNC Symbol;Acc |
| TNF | CHR_HSCHR6_MHC_SSTO_CT | tumor necrosis factor [Source:HGNC Symbol;A |
| LTB | CHR_HSCHR6_MHC_SSTO_CT | lymphotoxin beta [Source:HGNC Symbol;Acc:H |
| LST1 | CHR_HSCHR6_MHC_SSTO_CT | leukocyte specific transcript 1 [Source:HGNC S |
| NCR3 | CHR_HSCHR6_MHC_SSTO_CT | natural cytotoxicity triggering receptor 3 [Sou |
| UQCRHP1 | CHR_HSCHR6_MHC_SSTO_CT | ubiquinol-cytochrome c reductase hinge prote |
| AIF1 | CHR_HSCHR6_MHC_SSTO_CT | allograft inflammatory factor 1 [Source:HGNC |
| PRRC2A | CHR_HSCHR6_MHC_SSTO_CT | proline rich coiled-coil 2A [Source:HGNC Symb |
| SNORA38 | CHR_HSCHR6_MHC_SSTO_CT | small nucleolar RNA, H/ACA box 38 [Source:H |
| BAG6 | CHR_HSCHR6_MHC_SSTO_CT | BCL2 associated athanogene 6 [Source:HGNC |
| APOM | CHR_HSCHR6_MHC_SSTO_CT | apolipoprotein M [Source:HGNC Symbol;Acc:H |
| C6orf47 | CHR_HSCHR6_MHC_SSTO_CT | chromosome 6 open reading frame 47 [Source |
| C6orf47-AS1 | CHR_HSCHR6_MHC_SSTO_CT | C6orf47 antisense RNA 1 [Source:HGNC Symb |
| GPANK1 | CHR_HSCHR6_MHC_SSTO_CT | G-patch domain and ankyrin repeats 1 [Source |
| Y_RNA | CHR_HSCHR6_MHC_SSTO_CT | Y RNA [Source:RFAM;Acc:RF00019] |
| CSNK2B | CHR_HSCHR6_MHC_SSTO_CT | casein kinase 2 beta [Source:HGNC Symbol;Ac |
| LY6G5B | CHR_HSCHR6_MHC_SSTO_CT | lymphocyte antigen 6 family member G5B [So |
| LY6G5C | CHR_HSCHR6_MHC_SSTO_CT | lymphocyte antigen 6 family member G5C [So |
| ABHD16A | CHR_HSCHR6_MHC_SSTO_CT | abhydrolase domain containing 16A [Source:H |
| LY6G6F | CHR_HSCHR6_MHC_SSTO_CT | lymphocyte antigen 6 family member G6F [So |
| LY6G6F-LY6G6D | CHR_HSCHR6_MHC_SSTO_CT | LY6G6F-LY6G6D readthrough [Source:NCBI ge |
| LY6G6E | CHR_HSCHR6_MHC_SSTO_CT | lymphocyte antigen 6 family member G6E [So |
| LY6G6D | CHR_HSCHR6_MHC_SSTO_CT | lymphocyte antigen 6 family member G6D [Sc |
| MPIG6B | CHR_HSCHR6_MHC_SSTO_CT | megakaryocyte and platelet inhibitory recepto |
| LY6G6C | CHR_HSCHR6_MHC_SSTO_CT | lymphocyte antigen 6 family member G6C [So |
| DDAH2 | CHR_HSCHR6_MHC_SSTO_CT | dimethylarginine dimethylaminohydrolase 2 [ |
| CLIC1 | CHR_HSCHR6_MHC_SSTO_CT | chloride intracellular channel 1 [Source:HGNC |
| MSH5 | CHR_HSCHR6_MHC_SSTO_CT | mutS homolog 5 [Source:HGNC Symbol;Acc:H |
| U6 | CHR_HSCHR6_MHC_SSTO_CT | U6 spliceosomal RNA [Source:RFAM;Acc:RF00 |
| MSH5-SAPCD1 | CHR_HSCHR6_MHC_SSTO_CT | MSH5-SAPCD1 readthrough (NMD candidate) |
| SAPCD1 | CHR_HSCHR6_MHC_SSTO_CT | suppressor APC domain containing 1 [Source:I |
| VWA7 | CHR_HSCHR6_MHC_SSTO_CT | von Willebrand factor A domain containing 7 [ |
| VARS | CHR_HSCHR6_MHC_SSTO_CT | valyl-tRNA synthetase [Source:HGNC Symbol;A |
| Y_RNA | CHR_HSCHR6_MHC_SSTO_CT | Y RNA [Source:RFAM;Acc:RF00019] |
| NEU1 | CHR_HSCHR6_MHC_SSTO_CT | neuraminidase 1 [Source:HGNC Symbol;Acc:H |
| SLC44A4 | CHR_HSCHR6_MHC_SSTO_CT | solute carrier family 44 member 4 [Source:HG |
| EHMT2 | CHR_HSCHR6_MHC_SSTO_CT | euchromatic histone lysine methyltransferase |
| BX005460.1 | CHR_HSCHR6_MHC_SSTO_CT | novel transcript |
| C2 | CHR_HSCHR6_MHC_SSTO_CT | complement C2 [Source:HGNC Symbol;Acc:HG |
| C4B-AS1 | CHR_HSCHR6_MHC_SSTO_CT | C4B antisense RNA 1 [Source:HGNC Symbol;Ac |
| CFB | CHR_HSCHR6_MHC_SSTO_CT | complement factor B [Source:HGNC Symbol;A |
| NELFE | CHR_HSCHR6_MHC_SSTO_CT | negative elongation factor complex member E |
| SKIV2L | CHR_HSCHR6_MHC_SSTO_CT | Ski2 like RNA helicase [Source:HGNC Symbol;A |
| DXO | CHR_HSCHR6_MHC_SSTO_CT | decapping exoribonuclease [Source:HGNC Syr |
| STK19 | CHR_HSCHR6_MHC_SSTO_CT | serine/threonine kinase 19 [Source:HGNC Sym |
| C4B_2 | CHR_HSCHR6_MHC_SSTO_CT | complement component 4B (Chido blood grou |
| BX679671.1 | CHR_HSCHR6_MHC_SSTO_CT | novel transcript |

| | | |
|---|---|---|
| CYP21A1P | CHR_HSCHR6_MHC_SSTO_CT | cytochrome P450 family 21 subfamily A meml |
| TNXA | CHR_HSCHR6_MHC_SSTO_CT | tenascin XA (pseudogene) [Source:HGNC Sym |
| STK19B | CHR_HSCHR6_MHC_SSTO_CT | serine/threonine kinase 19B (pseudogene) [Sc |
| C4B | CHR_HSCHR6_MHC_SSTO_CT | complement C4B (Chido blood group) [Source |
| C4A-AS1 | CHR_HSCHR6_MHC_SSTO_CT | C4A antisense RNA 1 [Source:HGNC Symbol;A |
| CYP21A2 | CHR_HSCHR6_MHC_SSTO_CT | cytochrome P450 family 21 subfamily A meml |
| TNXB | CHR_HSCHR6_MHC_SSTO_CT | tenascin XB [Source:HGNC Symbol;Acc:HGNC: |
| PRRT1 | CHR_HSCHR6_MHC_SSTO_CT | proline rich transmembrane protein 1 [Source |
| PPT2 | CHR_HSCHR6_MHC_SSTO_CT | palmitoyl-protein thioesterase 2 [Source:HGN |
| EGFL8 | CHR_HSCHR6_MHC_SSTO_CT | EGF like domain multiple 8 [Source:HGNC Syn |
| AGPAT1 | CHR_HSCHR6_MHC_SSTO_CT | 1-acylglycerol-3-phosphate O-acyltransferase |
| RNF5 | CHR_HSCHR6_MHC_SSTO_CT | ring finger protein 5 [Source:HGNC Symbol;Ac |
| AGER | CHR_HSCHR6_MHC_SSTO_CT | advanced glycosylation end-product specific r |
| PBX2 | CHR_HSCHR6_MHC_SSTO_CT | PBX homeobox 2 [Source:HGNC Symbol;Acc:H |
| GPSM3 | CHR_HSCHR6_MHC_SSTO_CT | G protein signaling modulator 3 [Source:HGN( |
| NOTCH4 | CHR_HSCHR6_MHC_SSTO_CT | notch receptor 4 [Source:HGNC Symbol;Acc:H |
| TSBP1 | CHR_HSCHR6_MHC_SSTO_CT | testis expressed basic protein 1 [Source:HGN( |
| HNRNPA1P2 | CHR_HSCHR6_MHC_SSTO_CT | heterogeneous nuclear ribonucleoprotein A1 |
| U6 | CHR_HSCHR6_MHC_SSTO_CT | U6 spliceosomal RNA [Source:RFAM;Acc:RF00 |
| TSBP1-AS1 | CHR_HSCHR6_MHC_SSTO_CT | TSBP1 and BTNL2 antisense RNA 1 [Source:HG |
| BTNL2 | CHR_HSCHR6_MHC_SSTO_CT | butyrophilin like 2 [Source:HGNC Symbol;Acc: |
| HLA-DRA | CHR_HSCHR6_MHC_SSTO_CT | major histocompatibility complex, class II, DR |
| HLA-DRB9 | CHR_HSCHR6_MHC_SSTO_CT | major histocompatibility complex, class II, DR |
| BX120007.2 | CHR_HSCHR6_MHC_SSTO_CT | major histocompatibility complex class II DR b |
| U1 | CHR_HSCHR6_MHC_SSTO_CT | U1 spliceosomal RNA [Source:RFAM;Acc:RF00 |
| BX120007.1 | CHR_HSCHR6_MHC_SSTO_CT | major histocompatibility complex class II DR b |
| U1 | CHR_HSCHR6_MHC_SSTO_CT | U1 spliceosomal RNA [Source:RFAM;Acc:RF00 |
| HLA-DRB4 | CHR_HSCHR6_MHC_SSTO_CT | major histocompatibility complex, class II, DR |
| U1 | CHR_HSCHR6_MHC_SSTO_CT | U1 spliceosomal RNA [Source:RFAM;Acc:RF00 |
| HLA-DRB8 | CHR_HSCHR6_MHC_SSTO_CT | major histocompatibility complex, class II, DR |
| U1 | CHR_HSCHR6_MHC_SSTO_CT | U1 spliceosomal RNA [Source:RFAM;Acc:RF00 |
| BX649210.2 | CHR_HSCHR6_MHC_SSTO_CT | protein kinase interferon-inducible double str |
| BX649210.1 | CHR_HSCHR6_MHC_SSTO_CT | novel pseudogene |
| HLA-DRB7 | CHR_HSCHR6_MHC_SSTO_CT | major histocompatibility complex, class II, DR |
| FAM8A5P | CHR_HSCHR6_MHC_SSTO_CT | family with sequence similarity 8 member A5, |
| U1 | CHR_HSCHR6_MHC_SSTO_CT | U1 spliceosomal RNA [Source:RFAM;Acc:RF00 |
| HLA-DRB1 | CHR_HSCHR6_MHC_SSTO_CT | major histocompatibility complex, class II, DR |
| HLA-DQA1 | CHR_HSCHR6_MHC_SSTO_CT | major histocompatibility complex, class II, DQ |
| HLA-DQB1 | CHR_HSCHR6_MHC_SSTO_CT | major histocompatibility complex, class II, DQ |
| HLA-DQB1-AS1 | CHR_HSCHR6_MHC_SSTO_CT | HLA-DQB1 antisense RNA 1 [Source:HGNC Syn |
| BX248406.2 | CHR_HSCHR6_MHC_SSTO_CT | novel transcript |
| MTCO3P1 | CHR_HSCHR6_MHC_SSTO_CT | MT-CO3 pseudogene 1 [Source:HGNC Symbol |
| BX248406.1 | CHR_HSCHR6_MHC_SSTO_CT | uncharacterized LOC102725019 [Source:NCBI |
| HLA-DQB3 | CHR_HSCHR6_MHC_SSTO_CT | major histocompatibility complex, class II, DQ |
| HLA-DQA2 | CHR_HSCHR6_MHC_SSTO_CT | major histocompatibility complex, class II, DQ |
| HLA-DQB2 | CHR_HSCHR6_MHC_SSTO_CT | major histocompatibility complex, class II, DQ |
| HLA-DOB | CHR_HSCHR6_MHC_SSTO_CT | major histocompatibility complex, class II, DO |
| TAP2 | CHR_HSCHR6_MHC_SSTO_CT | transporter 2, ATP binding cassette subfamily |
| PSMB8 | CHR_HSCHR6_MHC_SSTO_CT | proteasome subunit beta 8 [Source:HGNC Syn |
| PSMB8-AS1 | CHR_HSCHR6_MHC_SSTO_CT | PSMB8 antisense RNA 1 (head to head) [Sourc |

| | | |
|---|---|---|
| PSMB9 | CHR_HSCHR6_MHC_SSTO_CT | proteasome subunit beta 9 [Source:HGNC Sym |
| TAP1 | CHR_HSCHR6_MHC_SSTO_CT | transporter 1, ATP binding cassette subfamily |
| PPP1R2P1 | CHR_HSCHR6_MHC_SSTO_CT | protein phosphatase 1 regulatory inhibitor sul |
| HLA-Z | CHR_HSCHR6_MHC_SSTO_CT | major histocompatibility complex, class I, Z (p: |
| HLA-DMB | CHR_HSCHR6_MHC_SSTO_CT | major histocompatibility complex, class II, DM |
| HLA-DMA | CHR_HSCHR6_MHC_SSTO_CT | major histocompatibility complex, class II, DM |
| BRD2 | CHR_HSCHR6_MHC_SSTO_CT | bromodomain containing 2 [Source:HGNC Sym |
| BX664702.1 | CHR_HSCHR6_MHC_SSTO_CT | novel transcript |
| HLA-DOA | CHR_HSCHR6_MHC_SSTO_CT | major histocompatibility complex, class II, DO |
| HLA-DPA1 | CHR_HSCHR6_MHC_SSTO_CT | major histocompatibility complex, class II, DP |
| HLA-DPB1 | CHR_HSCHR6_MHC_SSTO_CT | major histocompatibility complex, class II, DP |
| RPL32P1 | CHR_HSCHR6_MHC_SSTO_CT | ribosomal protein L32 pseudogene 1 [Source: |
| HLA-DPA2 | CHR_HSCHR6_MHC_SSTO_CT | major histocompatibility complex, class II, DP |
| COL11A2P1 | CHR_HSCHR6_MHC_SSTO_CT | collagen type XI alpha 2 pseudogene 1 [Source |
| HLA-DPB2 | CHR_HSCHR6_MHC_SSTO_CT | major histocompatibility complex, class II, DP |
| HLA-DPA3 | CHR_HSCHR6_MHC_SSTO_CT | major histocompatibility complex, class II, DP |
| COL11A2 | CHR_HSCHR6_MHC_SSTO_CT | collagen type XI alpha 2 chain [Source:HGNC S |
| ZNF70P1 | CHR_HSCHR6_MHC_SSTO_CT | zinc finger protein 70 pseudogene 1 [Source:H |
| HTATSF1P1 | CHR_HSCHR6_MHC_SSTO_CT | HIV-1 Tat specific factor 1 pseudogene 1 [Sour |
| HCG25 | CHR_HSCHR6_MHC_SSTO_CT | HLA complex group 25 [Source:HGNC Symbol; |
| VPS52 | CHR_HSCHR6_MHC_SSTO_CT | VPS52 subunit of GARP complex [Source:HGN |
| RPS18 | CHR_HSCHR6_MHC_SSTO_CT | ribosomal protein S18 [Source:HGNC Symbol;, |
| DAXX | CHR_HSCHR6_MHC_SSTO_CT | death domain associated protein [Source:HGN |
| RPL35AP4 | CHR_HSCHR6_MHC_SSTO_CT | ribosomal protein L35a pseudogene 4 [Source |
| KIFC1 | CHR_HSCHR6_MHC_SSTO_CT | kinesin family member C1 [Source:HGNC Sym |
| RPL12P1 | CHR_HSCHR6_MHC_SSTO_CT | ribosomal protein L12 pseudogene 1 [Source: |
| PHF1 | CHR_HSCHR6_MHC_SSTO_CT | PHD finger protein 1 [Source:HGNC Symbol;Ac |
| CUTA | CHR_HSCHR6_MHC_SSTO_CT | cutA divalent cation tolerance homolog [Sourc |
| SYNGAP1 | CHR_HSCHR6_MHC_SSTO_CT | synaptic Ras GTPase activating protein 1 [Sour |
| ZBTB9 | CHR_HSCHR6_MHC_SSTO_CT | zinc finger and BTB domain containing 9 [Sour |
| AC093627.12 | CHR_HSCHR7_1_CTG1 | novel transcript |
| AC093627.13 | CHR_HSCHR7_1_CTG1 | novel transcript |
| AC093614.2 | CHR_HSCHR7_1_CTG1 | novel transcript |
| AC093614.1 | CHR_HSCHR7_1_CTG1 | novel transcript |
| AC093627.9 | CHR_HSCHR7_1_CTG1 | novel transcript |
| AC093627.8 | CHR_HSCHR7_1_CTG1 | novel transcript |
| AC093627.10 | CHR_HSCHR7_1_CTG1 | novel transcript |
| FAM20C | CHR_HSCHR7_1_CTG1 | FAM20C golgi associated secretory pathway k |
| ARMC10 | CHR_HSCHR7_1_CTG4_4 | armadillo repeat containing 10 [Source:HGNC |
| CRYZP1 | CHR_HSCHR7_1_CTG4_4 | crystallin zeta pseudogene 1 [Source:HGNC Sy |
| NAPEPLD | CHR_HSCHR7_1_CTG4_4 | N-acyl phosphatidylethanolamine phospholipa |
| AC216130.2 | CHR_HSCHR7_1_CTG4_4 | ribosomal protein L19 (RPL19) pseudogene |
| RPL23AP95 | CHR_HSCHR7_1_CTG4_4 | ribosomal protein L23a pseudogene 95 [Sourc |
| AC216130.1 | CHR_HSCHR7_1_CTG4_4 | tec |
| DPY19L2P2 | CHR_HSCHR7_1_CTG4_4 | DPY19L2 pseudogene 2 [Source:HGNC Symbo |
| AGK | CHR_HSCHR7_1_CTG6 | acylglycerol kinase [Source:HGNC Symbol;Acc |
| AC004918.3 | CHR_HSCHR7_1_CTG6 | novel transcript |
| DENND11 | CHR_HSCHR7_1_CTG6 | DENN domain containing 11 [Source:HGNC Sy |
| AC004918.2 | CHR_HSCHR7_1_CTG6 | novel transcript, antisense to WEE2 |
| WEE2 | CHR_HSCHR7_1_CTG6 | WEE1 homolog 2 [Source:HGNC Symbol;Acc:H |

| | | |
|---|---|---|
| U1 | CHR_HSCHR7_1_CTG6 | U1 spliceosomal RNA [Source:RFAM;Acc:RF00 |
| SSBP1 | CHR_HSCHR7_1_CTG6 | single stranded DNA binding protein 1 [Source |
| PTPRN2 | CHR_HSCHR7_1_CTG7 | protein tyrosine phosphatase receptor type N |
| AC093627.11 | CHR_HSCHR7_2_CTG1 | novel transcript similar to family member with |
| AC093627.15 | CHR_HSCHR7_2_CTG1 | von Hippel-Lindau binding protein 1 (VBP1) ps |
| AC093627.19 | CHR_HSCHR7_2_CTG1 | novel transcript |
| AC093627.17 | CHR_HSCHR7_2_CTG1 | novel transcript |
| AC093627.20 | CHR_HSCHR7_2_CTG1 | novel transcript |
| AC093627.16 | CHR_HSCHR7_2_CTG1 | novel transcript |
| AC093627.21 | CHR_HSCHR7_2_CTG1 | novel transcript |
| AC093627.14 | CHR_HSCHR7_2_CTG1 | novel transcript |
| AC093627.18 | CHR_HSCHR7_2_CTG1 | novel transcript |
| FAM20C | CHR_HSCHR7_2_CTG1 | FAM20C golgi associated secretory pathway k |
| DTX2 | CHR_HSCHR7_2_CTG4_4 | deltex E3 ubiquitin ligase 2 [Source:HGNC Sym |
| UPK3B | CHR_HSCHR7_2_CTG4_4 | uroplakin 3B [Source:HGNC Symbol;Acc:HGNC |
| AC237298.1 | CHR_HSCHR7_2_CTG4_4 | postmeiotic segregation increased 2-like 5 (PN |
| SPDYE16 | CHR_HSCHR7_2_CTG4_4 | speedy/RINGO cell cycle regulator family men |
| AC004980.5 | CHR_HSCHR7_2_CTG4_4 | pseudogene similar to part of postmeiotic seg |
| POMZP3 | CHR_HSCHR7_2_CTG4_4 | POM121 and ZP3 fusion [Source:HGNC Symbo |
| AC004980.6 | CHR_HSCHR7_2_CTG4_4 | pseudogene similar to part of RB-associated K |
| AC004980.7 | CHR_HSCHR7_2_CTG4_4 | novel transcript |
| MGAM | CHR_HSCHR7_2_CTG6 | maltase-glucoamylase [Source:HGNC Symbol; |
| MGAM2 | CHR_HSCHR7_2_CTG6 | maltase-glucoamylase 2 (putative) [Source:HG |
| MOXD2P | CHR_HSCHR7_2_CTG6 | monooxygenase DBH like 2, pseudogene [Sou |
| PRSS58 | CHR_HSCHR7_2_CTG6 | serine protease 58 [Source:HGNC Symbol;Acc |
| AC231381.1 | CHR_HSCHR7_2_CTG6 | protease, serine, 58 (PRSS58) pseudogene |
| AC231381.2 | CHR_HSCHR7_2_CTG6 | novel transcript |
| PRSS3P3 | CHR_HSCHR7_2_CTG6 | PRSS3 pseudogene 3 [Source:HGNC Symbol;A |
| TRBV1 | CHR_HSCHR7_2_CTG6 | T cell receptor beta variable 1 (pseudogene) [ |
| TRBV2 | CHR_HSCHR7_2_CTG6 | T cell receptor beta variable 2 [Source:HGNC S |
| TRBV3-1 | CHR_HSCHR7_2_CTG6 | T cell receptor beta variable 3-1 [Source:HGN |
| TRBV4-1 | CHR_HSCHR7_2_CTG6 | T cell receptor beta variable 4-1 [Source:HGN |
| TRBV5-1 | CHR_HSCHR7_2_CTG6 | T cell receptor beta variable 5-1 [Source:HGN |
| TRBV6-1 | CHR_HSCHR7_2_CTG6 | T cell receptor beta variable 6-1 [Source:HGN |
| TRBV7-1 | CHR_HSCHR7_2_CTG6 | T cell receptor beta variable 7-1 (non-function |
| TRBV4-2 | CHR_HSCHR7_2_CTG6 | T cell receptor beta variable 4-2 [Source:HGN |
| TRBV6-2 | CHR_HSCHR7_2_CTG6 | T cell receptor beta variable 6-2 (gene/pseudo |
| AC234635.2 | CHR_HSCHR7_2_CTG6 | T cell receptor beta variable 3-2 (pseudogene) |
| AC234635.3 | CHR_HSCHR7_2_CTG6 | T cell receptor beta variable 4-3 [Source:UniP |
| AC234635.1 | CHR_HSCHR7_2_CTG6 | T cell receptor beta variable 6-3 [Source:UniP |
| TRBV7-2 | CHR_HSCHR7_2_CTG6 | T cell receptor beta variable 7-2 [Source:HGN |
| TRBV8-1 | CHR_HSCHR7_2_CTG6 | T cell receptor beta variable 8-1 (pseudogene) |
| TRBV5-2 | CHR_HSCHR7_2_CTG6 | T cell receptor beta variable 5-2 (pseudogene) |
| TRBV6-4 | CHR_HSCHR7_2_CTG6 | T cell receptor beta variable 6-4 [Source:HGN |
| TRBV7-3 | CHR_HSCHR7_2_CTG6 | T cell receptor beta variable 7-3 [Source:HGN |
| TRBV8-2 | CHR_HSCHR7_2_CTG6 | T cell receptor beta variable 8-2 (pseudogene) |
| TRBV5-3 | CHR_HSCHR7_2_CTG6 | T cell receptor beta variable 5-3 (non-function |
| TRBV9 | CHR_HSCHR7_2_CTG6 | T cell receptor beta variable 9 [Source:HGNC S |
| TRBV10-1 | CHR_HSCHR7_2_CTG6 | T cell receptor beta variable 10-1(gene/pseud |
| TRBV11-1 | CHR_HSCHR7_2_CTG6 | T cell receptor beta variable 11-1 [Source:HGN |

| | | |
|---|---|---|
| TRBV12-1 | CHR_HSCHR7_2_CTG6 | T cell receptor beta variable 12-1 (pseudogene |
| TRBV10-2 | CHR_HSCHR7_2_CTG6 | T cell receptor beta variable 10-2 [Source:HGN |
| TRBV11-2 | CHR_HSCHR7_2_CTG6 | T cell receptor beta variable 11-2 [Source:HGN |
| TRBV12-2 | CHR_HSCHR7_2_CTG6 | T cell receptor beta variable 12-2 (pseudogene |
| TRBV6-5 | CHR_HSCHR7_2_CTG6 | T cell receptor beta variable 6-5 [Source:HGN( |
| TRBV7-4 | CHR_HSCHR7_2_CTG6 | T cell receptor beta variable 7-4 (gene/pseudo |
| TRBV5-4 | CHR_HSCHR7_2_CTG6 | T cell receptor beta variable 5-4 [Source:HGN( |
| TRBV6-6 | CHR_HSCHR7_2_CTG6 | T cell receptor beta variable 6-6 [Source:HGN( |
| TRBV7-5 | CHR_HSCHR7_2_CTG6 | T cell receptor beta variable 7-5 (pseudogene) |
| TRBV5-5 | CHR_HSCHR7_2_CTG6 | T cell receptor beta variable 5-5 [Source:HGN( |
| TRBV6-7 | CHR_HSCHR7_2_CTG6 | T cell receptor beta variable 6-7 (non-functior |
| TRBV7-6 | CHR_HSCHR7_2_CTG6 | T cell receptor beta variable 7-6 [Source:HGN( |
| TRBV5-6 | CHR_HSCHR7_2_CTG6 | T cell receptor beta variable 5-6 [Source:HGN( |
| TRBV6-8 | CHR_HSCHR7_2_CTG6 | T cell receptor beta variable 6-8 [Source:HGN( |
| TRBV7-7 | CHR_HSCHR7_2_CTG6 | T cell receptor beta variable 7-7 [Source:HGN( |
| TRBV5-7 | CHR_HSCHR7_2_CTG6 | T cell receptor beta variable 5-7 (non-functior |
| AC233282.2 | CHR_HSCHR7_2_CTG6 | T cell receptor beta variable 6-9 [Source:UniP |
| AC233282.1 | CHR_HSCHR7_2_CTG6 | T cell receptor beta variable 7-8 [Source:UniP |
| AC229888.1 | CHR_HSCHR7_2_CTG6 | T cell receptor beta variable 5-8 [Source:UniP |
| TRBV7-9 | CHR_HSCHR7_2_CTG6 | T cell receptor beta variable 7-9 [Source:HGN( |
| TRBV13 | CHR_HSCHR7_2_CTG6 | T cell receptor beta variable 13 [Source:HGNC |
| TRBV10-3 | CHR_HSCHR7_2_CTG6 | T cell receptor beta variable 10-3 [Source:HGN |
| TRBV11-3 | CHR_HSCHR7_2_CTG6 | T cell receptor beta variable 11-3 [Source:HGN |
| TRBV12-3 | CHR_HSCHR7_2_CTG6 | T cell receptor beta variable 12-3 [Source:HGN |
| TRBV12-4 | CHR_HSCHR7_2_CTG6 | T cell receptor beta variable 12-4 [Source:HGN |
| TRBV12-5 | CHR_HSCHR7_2_CTG6 | T cell receptor beta variable 12-5 [Source:HGN |
| TRBV14 | CHR_HSCHR7_2_CTG6 | T cell receptor beta variable 14 [Source:HGNC |
| TRBV15 | CHR_HSCHR7_2_CTG6 | T cell receptor beta variable 15 [Source:HGNC |
| TRBV16 | CHR_HSCHR7_2_CTG6 | T cell receptor beta variable 16 (gene/pseudo |
| TRBV17 | CHR_HSCHR7_2_CTG6 | T cell receptor beta variable 17 (non-functiona |
| TRBV18 | CHR_HSCHR7_2_CTG6 | T cell receptor beta variable 18 [Source:HGNC |
| TRBV19 | CHR_HSCHR7_2_CTG6 | T cell receptor beta variable 19 [Source:HGNC |
| TRBV20-1 | CHR_HSCHR7_2_CTG6 | T cell receptor beta variable 20-1 [Source:HGN |
| TRBV21-1 | CHR_HSCHR7_2_CTG6 | T cell receptor beta variable 21-1 (pseudogene |
| TRBV22-1 | CHR_HSCHR7_2_CTG6 | T cell receptor beta variable 22-1 (pseudogene |
| TRBV23-1 | CHR_HSCHR7_2_CTG6 | T cell receptor beta variable 23-1 (non-functic |
| TRBV24-1 | CHR_HSCHR7_2_CTG6 | T cell receptor beta variable 24-1 [Source:HGN |
| MTRNR2L6 | CHR_HSCHR7_2_CTG6 | MT-RNR2 like 6 [Source:HGNC Symbol;Acc:HG |
| TRBV25-1 | CHR_HSCHR7_2_CTG6 | T cell receptor beta variable 25-1 [Source:HGN |
| TRBVA | CHR_HSCHR7_2_CTG6 | T cell receptor beta variable A (pseudogene) [ |
| TRBV26 | CHR_HSCHR7_2_CTG6 | T cell receptor beta variable 26 (pseudogene) |
| TRBVB | CHR_HSCHR7_2_CTG6 | T cell receptor beta variable B (pseudogene) [ |
| TRBV27 | CHR_HSCHR7_2_CTG6 | T cell receptor beta variable 27 [Source:HGNC |
| TRBV28 | CHR_HSCHR7_2_CTG6 | T cell receptor beta variable 28 [Source:HGNC |
| PGBD4P1 | CHR_HSCHR7_2_CTG6 | piggyBac transposable element derived 4 pseu |
| TRBV29-1 | CHR_HSCHR7_2_CTG6 | T cell receptor beta variable 29-1 [Source:HGN |
| PRSS1 | CHR_HSCHR7_2_CTG6 | serine protease 1 [Source:HGNC Symbol;Acc:H |
| PRSS3P1 | CHR_HSCHR7_2_CTG6 | PRSS3 pseudogene 1 [Source:HGNC Symbol;A |
| PRSS3P2 | CHR_HSCHR7_2_CTG6 | PRSS3 pseudogene 2 [Source:HGNC Symbol;A |
| AC239618.2 | CHR_HSCHR7_2_CTG6 | trypsinogen D |

| | | |
|---|---|---|
| PRSS2 | CHR_HSCHR7_2_CTG6 | serine protease 2 [Source:HGNC Symbol;Acc:H |
| WBP1LP1 | CHR_HSCHR7_2_CTG6 | WBP1L pseudogene 1 [Source:HGNC Symbol;A |
| TRBD1 | CHR_HSCHR7_2_CTG6 | T cell receptor beta diversity 1 [Source:HGNC |
| TRBJ1-1 | CHR_HSCHR7_2_CTG6 | T cell receptor beta joining 1-1 [Source:HGNC |
| TRBJ1-2 | CHR_HSCHR7_2_CTG6 | T cell receptor beta joining 1-2 [Source:HGNC |
| TRBJ1-3 | CHR_HSCHR7_2_CTG6 | T cell receptor beta joining 1-3 [Source:HGNC |
| TRBJ1-4 | CHR_HSCHR7_2_CTG6 | T cell receptor beta joining 1-4 [Source:HGNC |
| TRBJ1-5 | CHR_HSCHR7_2_CTG6 | T cell receptor beta joining 1-5 [Source:HGNC |
| TRBJ1-6 | CHR_HSCHR7_2_CTG6 | T cell receptor beta joining 1-6 [Source:HGNC |
| TRBC1 | CHR_HSCHR7_2_CTG6 | T cell receptor beta constant 1 [Source:HGNC |
| TRBJ2-1 | CHR_HSCHR7_2_CTG6 | T cell receptor beta joining 2-1 [Source:HGNC |
| TRBJ2-2 | CHR_HSCHR7_2_CTG6 | T cell receptor beta joining 2-2 [Source:HGNC |
| TRBJ2-2P | CHR_HSCHR7_2_CTG6 | T cell receptor beta joining 2-2P (non-functior |
| TRBJ2-3 | CHR_HSCHR7_2_CTG6 | T cell receptor beta joining 2-3 [Source:HGNC |
| TRBJ2-4 | CHR_HSCHR7_2_CTG6 | T cell receptor beta joining 2-4 [Source:HGNC |
| TRBJ2-5 | CHR_HSCHR7_2_CTG6 | T cell receptor beta joining 2-5 [Source:HGNC |
| TRBJ2-6 | CHR_HSCHR7_2_CTG6 | T cell receptor beta joining 2-6 [Source:HGNC |
| TRBJ2-7 | CHR_HSCHR7_2_CTG6 | T cell receptor beta joining 2-7 [Source:HGNC |
| TRBC2 | CHR_HSCHR7_2_CTG6 | T cell receptor beta constant 2 [Source:HGNC |
| TRBV30 | CHR_HSCHR7_2_CTG6 | T cell receptor beta variable 30 (gene/pseudo |
| EPHB6 | CHR_HSCHR7_2_CTG6 | EPH receptor B6 [Source:HGNC Symbol;Acc:H |
| TRPV6 | CHR_HSCHR7_2_CTG6 | transient receptor potential cation channel su |
| AC239618.1 | CHR_HSCHR7_2_CTG6 | novel transcript, antisense to TRPV6 |
| TRPV5 | CHR_HSCHR7_2_CTG6 | transient receptor potential cation channel su |
| LLCFC1 | CHR_HSCHR7_2_CTG6 | LLLL and CFNLAS motif containing 1 [Source:H |
| KEL | CHR_HSCHR7_2_CTG6 | Kell metallo-endopeptidase (Kell blood group) |
| OR9A2 | CHR_HSCHR7_2_CTG6 | olfactory receptor family 9 subfamily A memb |
| OR9P1P | CHR_HSCHR7_2_CTG6 | olfactory receptor family 9 subfamily P memb |
| OR6V1 | CHR_HSCHR7_2_CTG6 | olfactory receptor family 6 subfamily V memb |
| OR6W1P | CHR_HSCHR7_2_CTG6 | olfactory receptor family 6 subfamily W meml |
| AC236756.1 | CHR_HSCHR7_2_CTG7 | novel transcript |
| AC004852.4 | CHR_HSCHR7_3_CTG1 | novel transcript |
| AC004852.6 | CHR_HSCHR7_3_CTG1 | LYR motif containing 9 (LYRM9) pseudogene |
| AC004852.5 | CHR_HSCHR7_3_CTG1 | ret finger protein-like 4A (RFPL4A) pseudogen |
| OR2A5 | CHR_HSCHR7_3_CTG4_4 | olfactory receptor family 2 subfamily A memb |
| OR2A25 | CHR_HSCHR7_3_CTG4_4 | olfactory receptor family 2 subfamily A memb |
| OR2A12 | CHR_HSCHR7_3_CTG4_4 | olfactory receptor family 2 subfamily A memb |
| OR2A2 | CHR_HSCHR7_3_CTG4_4 | olfactory receptor family 2 subfamily A memb |
| OR2A14 | CHR_HSCHR7_3_CTG4_4 | olfactory receptor family 2 subfamily A memb |
| CTAGE4 | CHR_HSCHR7_3_CTG4_4 | CTAGE family member 4 [Source:HGNC Symbo |
| ARHGEF35 | CHR_HSCHR7_3_CTG4_4 | Rho guanine nucleotide exchange factor 35 [S |
| OR2A42 | CHR_HSCHR7_3_CTG4_4 | olfactory receptor family 2 subfamily A memb |
| AC245175.1 | CHR_HSCHR7_3_CTG4_4 | NULL |
| OR2A7 | CHR_HSCHR7_3_CTG4_4 | olfactory receptor family 2 subfamily A memb |
| CTAGE8 | CHR_HSCHR7_3_CTG4_4 | CTAGE family member 8 [Source:NCBI gene;A |
| AC245175.2 | CHR_HSCHR7_3_CTG4_4 | NULL |
| AC256203.1 | CHR_HSCHR7_3_CTG4_4 | NULL |
| OR2A1 | CHR_HSCHR7_3_CTG4_4 | olfactory receptor family 2 subfamily A memb |
| AC256203.2 | CHR_HSCHR7_3_CTG4_4 | NULL |
| AC243837.2 | CHR_HSCHR7_3_CTG4_4 | NULL |

| | | |
|---|---|---|
| AC243837.1 | CHR_HSCHR7_3_CTG4_4 | olfactory receptor 2A7 [Source:NCBI gene;Acc |
| AC243837.3 | CHR_HSCHR7_3_CTG4_4 | NULL |
| AC243837.4 | CHR_HSCHR7_3_CTG4_4 | olfactory receptor 2A1/2A42-like [Source:NCB |
| ARHGEF5 | CHR_HSCHR7_3_CTG4_4 | Rho guanine nucleotide exchange factor 5 [So |
| NOBOX | CHR_HSCHR7_3_CTG4_4 | NOBOX oogenesis homeobox [Source:HGNC S |
| AC004864.1 | CHR_HSCHR7_3_CTG4_4 | NULL |
| CNTNAP2 | CHR_HSCHR7_3_CTG6 | contactin associated protein like 2 [Source:HG |
| U3 | CHR_HSCHR7_3_CTG6 | Small nucleolar RNA U3 [Source:RFAM;Acc:RF |
| Metazoa_SRP | CHR_HSCHR7_3_CTG6 | Metazoan signal recognition particle RNA [Sou |
| AC083849.3 | CHR_HSCHR7_3_CTG6 | novel transcript |
| AC083849.2 | CHR_HSCHR7_3_CTG6 | exosome component 6 (EXOSC6) pseudogene |
| AF067845.6 | CHR_HSCHR8_1_CTG1 | novel transcript |
| AF067845.3 | CHR_HSCHR8_1_CTG1 | novel transcript |
| AC131901.1 | CHR_HSCHR8_1_CTG1 | novel transcript |
| AC131901.2 | CHR_HSCHR8_1_CTG1 | novel transcript |
| DLGAP2 | CHR_HSCHR8_1_CTG1 | DLG associated protein 2 [Source:HGNC Symb |
| DLGAP2-AS1 | CHR_HSCHR8_1_CTG1 | DLGAP2 antisense RNA 1 [Source:HGNC Symb |
| LINC01606 | CHR_HSCHR8_1_CTG6 | long intergenic non-protein coding RNA 1606 |
| AC025674.4 | CHR_HSCHR8_1_CTG6 | ankyrin repeat domain 26 (ANKRD26) pseudo |
| AC025674.3 | CHR_HSCHR8_1_CTG6 | novel transcript |
| LINC00588 | CHR_HSCHR8_1_CTG6 | long intergenic non-protein coding RNA 588 [S |
| U6 | CHR_HSCHR8_1_CTG6 | U6 spliceosomal RNA [Source:RFAM;Acc:RF00 |
| LINC00824 | CHR_HSCHR8_1_CTG7 | long intergenic non-protein coding RNA 824 [S |
| AC068570.1 | CHR_HSCHR8_1_CTG7 | long intergenic non-protein coding RNA 976 |
| AC026950.1 | CHR_HSCHR8_2_CTG1 | ERICH1 antisense RNA 1 |
| AC129915.1 | CHR_HSCHR8_2_CTG1 | novel transcript |
| AC129915.2 | CHR_HSCHR8_2_CTG1 | novel transcript |
| AF186192.5 | CHR_HSCHR8_2_CTG7 | novel transcript |
| AF186192.6 | CHR_HSCHR8_2_CTG7 | zinc finger protein 154 (ZNF154) pseudogene |
| ZNF251 | CHR_HSCHR8_2_CTG7 | zinc finger protein 251 [Source:HGNC Symbol; |
| AF186192.4 | CHR_HSCHR8_2_CTG7 | novel transcript, antisense to ZNF251 |
| FAM66B | CHR_HSCHR8_3_CTG1 | family with sequence similarity 66 member B |
| AF252830.1 | CHR_HSCHR8_3_CTG1 | ubiquitin specific peptidase 17 like family mer |
| AF252830.2 | CHR_HSCHR8_3_CTG1 | ubiquitin-specific protease (USP) pseudogene |
| ZNF705G | CHR_HSCHR8_3_CTG1 | zinc finger protein 705G [Source:HGNC Symbo |
| DEFB108C | CHR_HSCHR8_3_CTG1 | defensin beta 108C [Source:HGNC Symbol;Acc |
| DEFB4B | CHR_HSCHR8_3_CTG1 | defensin beta 4B [Source:HGNC Symbol;Acc:H |
| HSPD1P3 | CHR_HSCHR8_3_CTG1 | heat shock protein family D (Hsp60) member |
| DEFB103B | CHR_HSCHR8_3_CTG1 | defensin beta 103B [Source:HGNC Symbol;Acc |
| SPAG11B | CHR_HSCHR8_3_CTG1 | sperm associated antigen 11B [Source:HGNC S |
| DEFB104B | CHR_HSCHR8_3_CTG1 | defensin beta 104B [Source:HGNC Symbol;Acc |
| DEFB106B | CHR_HSCHR8_3_CTG1 | defensin beta 106B [Source:HGNC Symbol;Acc |
| DEFB105B | CHR_HSCHR8_3_CTG1 | defensin beta 105B [Source:HGNC Symbol;Acc |
| DEFB107B | CHR_HSCHR8_3_CTG1 | defensin beta 107B [Source:HGNC Symbol;Acc |
| PRR23D1 | CHR_HSCHR8_3_CTG1 | proline rich 23 domain containing 1 [Source:H |
| FAM90A6P | CHR_HSCHR8_3_CTG1 | family with sequence similarity 90 member A6 |
| AC134684.3 | CHR_HSCHR8_3_CTG1 | family with sequence similarity 90, member A |
| FAM90A21P | CHR_HSCHR8_3_CTG1 | family with sequence similarity 90 member A2 |
| AC134684.4 | CHR_HSCHR8_3_CTG1 | family with sequence similarity 90, member A |
| AC134684.5 | CHR_HSCHR8_3_CTG1 | family with sequence similarity 90, member A |

| | | |
|---|---|---|
| OR7E157P | CHR_HSCHR8_3_CTG1 | olfactory receptor family 7 subfamily E memb |
| AC134684.2 | CHR_HSCHR8_3_CTG1 | chromosome 11 open reading frame 2 (C11or |
| ZC3H3 | CHR_HSCHR8_3_CTG7 | zinc finger CCCH-type containing 3 [Source:HG |
| RN7SKP175 | CHR_HSCHR8_3_CTG7 | RN7SK pseudogene 175 [Source:HGNC Symbo |
| GSDMD | CHR_HSCHR8_3_CTG7 | gasdermin D [Source:HGNC Symbol;Acc:HGNC |
| MROH6 | CHR_HSCHR8_3_CTG7 | maestro heat like repeat family member 6 [Sc |
| AC067930.7 | CHR_HSCHR8_3_CTG7 | novel transcript |
| NAPRT | CHR_HSCHR8_3_CTG7 | nicotinate phosphoribosyltransferase [Source |
| AC067930.6 | CHR_HSCHR8_3_CTG7 | novel transcript |
| EEF1D | CHR_HSCHR8_3_CTG7 | eukaryotic translation elongation factor 1 delt |
| TIGD5 | CHR_HSCHR8_3_CTG7 | tigger transposable element derived 5 [Source |
| PYCR3 | CHR_HSCHR8_3_CTG7 | pyrroline-5-carboxylate reductase 3 [Source:H |
| TSTA3 | CHR_HSCHR8_3_CTG7 | tissue specific transplantation antigen P35B [S |
| ZNF623 | CHR_HSCHR8_3_CTG7 | zinc finger protein 623 [Source:HGNC Symbol; |
| ZNF707 | CHR_HSCHR8_3_CTG7 | zinc finger protein 707 [Source:HGNC Symbol; |
| AC105219.5 | CHR_HSCHR8_3_CTG7 | breast cancer estrogen-induced apoptosis 2 (E |
| CCDC166 | CHR_HSCHR8_3_CTG7 | coiled-coil domain containing 166 [Source:HG |
| AC105219.7 | CHR_HSCHR8_3_CTG7 | novel transcript |
| MAPK15 | CHR_HSCHR8_3_CTG7 | mitogen-activated protein kinase 15 [Source:H |
| AC105219.8 | CHR_HSCHR8_3_CTG7 | novel transcript |
| FAM83H | CHR_HSCHR8_3_CTG7 | family with sequence similarity 83 member H |
| FAM83H-AS1 | CHR_HSCHR8_3_CTG7 | FAM83H antisense RNA 1 (head to head) [Sou |
| AC105219.6 | CHR_HSCHR8_3_CTG7 | novel transcript |
| SCRIB | CHR_HSCHR8_3_CTG7 | scribble planar cell polarity protein [Source:H( |
| PUF60 | CHR_HSCHR8_3_CTG7 | poly(U) binding splicing factor 60 [Source:HGN |
| AC100797.4 | CHR_HSCHR8_4_CTG1 | novel transcript |
| ERICH1 | CHR_HSCHR8_4_CTG1 | glutamate rich 1 [Source:HGNC Symbol;Acc:H( |
| AC100797.2 | CHR_HSCHR8_4_CTG1 | ERICH1 antisense RNA 1 |
| AC100797.3 | CHR_HSCHR8_4_CTG1 | novel transcript |
| LY6E-DT | CHR_HSCHR8_4_CTG7 | LY6E divergent transcript [Source:HGNC Symb |
| LY6E | CHR_HSCHR8_4_CTG7 | lymphocyte antigen 6 family member E [Sourc |
| C8orf31 | CHR_HSCHR8_4_CTG7 | chromosome 8 open reading frame 31 [Sourc |
| AK3P2 | CHR_HSCHR8_4_CTG7 | adenylate kinase 3 pseudogene 2 [Source:HGI |
| AC236721.1 | CHR_HSCHR8_4_CTG7 | ribosomal protein S26 (RPS26) pseudogene |
| LY6L | CHR_HSCHR8_4_CTG7 | lymphocyte antigen 6 family member L [Sourc |
| LY6H | CHR_HSCHR8_4_CTG7 | lymphocyte antigen 6 family member H [Sour |
| DLGAP2 | CHR_HSCHR8_5_CTG1 | DLG associated protein 2 [Source:HGNC Symb |
| DLGAP2-AS1 | CHR_HSCHR8_5_CTG1 | DLGAP2 antisense RNA 1 [Source:HGNC Symb |
| LINC01300 | CHR_HSCHR8_5_CTG7 | long intergenic non-protein coding RNA 1300 |
| AC100803.7 | CHR_HSCHR8_5_CTG7 | novel transcript |
| GPR20 | CHR_HSCHR8_5_CTG7 | G protein-coupled receptor 20 [Source:HGNC |
| PTP4A3 | CHR_HSCHR8_5_CTG7 | protein tyrosine phosphatase 4A3 [Source:HG |
| MROH5 | CHR_HSCHR8_5_CTG7 | maestro heat like repeat family member 5 (ge |
| AC100803.6 | CHR_HSCHR8_5_CTG7 | novel transcript, antisense to MROH5 |
| AC100803.5 | CHR_HSCHR8_5_CTG7 | NULL |
| HNRNPA1P38 | CHR_HSCHR8_5_CTG7 | heterogeneous nuclear ribonucleoprotein A1p |
| AC120035.1 | CHR_HSCHR8_6_CTG1 | novel transcript |
| AC242019.1 | CHR_HSCHR8_6_CTG1 | novel transcript |
| AF188030.1 | CHR_HSCHR8_7_CTG1 | DLGAP2 antisense RNA 1 |
| DLGAP2 | CHR_HSCHR8_7_CTG1 | DLG associated protein 2 [Source:HGNC Symb |

| | | |
|---|---|---|
| AF188030.2 | CHR_HSCHR8_7_CTG1 | DNA fragmentation factor, 45kDa, alpha polyp |
| CLN8 | CHR_HSCHR8_7_CTG1 | CLN8 transmembrane ER and ERGIC protein [S |
| AC100810.5 | CHR_HSCHR8_7_CTG1 | novel transcript |
| AC100810.4 | CHR_HSCHR8_7_CTG1 | novel transcript, antisense to CLN8 |
| MTRF1LP2 | CHR_HSCHR8_7_CTG7 | mitochondrial translational release factor 1 lik |
| GSDMC | CHR_HSCHR8_7_CTG7 | gasdermin C [Source:HGNC Symbol;Acc:HGNC |
| ARHGEF10 | CHR_HSCHR8_8_CTG1 | Rho guanine nucleotide exchange factor 10 [S |
| AC019257.4 | CHR_HSCHR8_8_CTG1 | novel transcript, antisense to ARHGEF10 |
| AC019257.7 | CHR_HSCHR8_8_CTG1 | novel transcript, antisense to ARHGEF10 |
| AC019257.6 | CHR_HSCHR8_8_CTG1 | novel transcript |
| AC019257.3 | CHR_HSCHR8_8_CTG1 | novel transcript, antisense to ARHGEF10 |
| KBTBD11 | CHR_HSCHR8_8_CTG1 | kelch repeat and BTB domain containing 11 [S |
| AC019257.5 | CHR_HSCHR8_8_CTG1 | novel transcript |
| AC125453.1 | CHR_HSCHR8_8_CTG1 | novel transcript |
| MYOM2 | CHR_HSCHR8_8_CTG1 | myomesin 2 [Source:HGNC Symbol;Acc:HGNC |
| AC022257.1 | CHR_HSCHR8_8_CTG1 | novel transcript |
| AC133633.2 | CHR_HSCHR8_8_CTG1 | novel transcript |
| AC133633.1 | CHR_HSCHR8_8_CTG1 | novel transcript |
| AC104021.1 | CHR_HSCHR8_8_CTG1 | novel transcript |
| AC246817.3 | CHR_HSCHR8_8_CTG1 | novel transcript |
| AC246817.4 | CHR_HSCHR8_8_CTG1 | novel transcript |
| AC246817.5 | CHR_HSCHR8_8_CTG1 | novel transcript |
| ADAM9 | CHR_HSCHR8_9_CTG1 | ADAM metallopeptidase domain 9 [Source:HG |
| ADAM32 | CHR_HSCHR8_9_CTG1 | ADAM metallopeptidase domain 32 [Source:H |
| RPL3P10 | CHR_HSCHR8_9_CTG1 | ribosomal protein L3 pseudogene 10 [Source: |
| AC245594.2 | CHR_HSCHR8_9_CTG1 | K(lysine) acetyltransferase 2B (KAT2B) pseudo |
| AC245594.1 | CHR_HSCHR8_9_CTG1 | ADAM metallopeptidase domain 5 pseudogen |
| ADAM18 | CHR_HSCHR8_9_CTG1 | ADAM metallopeptidase domain 18 [Source:H |
| ADAM2 | CHR_HSCHR8_9_CTG1 | ADAM metallopeptidase domain 2 [Source:HG |
| AP005902.4 | CHR_HSCHR8_9_CTG1 | ribosomal protein L7a (RPL7A) pseudogene |
| AP005902.3 | CHR_HSCHR8_9_CTG1 | Nipped-B homolog (Drosophila) (NIPBL) pseuc |
| RPL4P5 | CHR_HSCHR9_1_CTG1 | ribosomal protein L4 pseudogene 5 [Source:H |
| AC208386.1 | CHR_HSCHR9_1_CTG2 | novel pseudogene |
| APBA1 | CHR_HSCHR9_1_CTG2 | amyloid beta precursor protein binding family |
| AL355140.2 | CHR_HSCHR9_1_CTG2 | novel transcript |
| MAMDC2 | CHR_HSCHR9_1_CTG3 | MAM domain containing 2 [Source:HGNC Sym |
| MAMDC2-AS1 | CHR_HSCHR9_1_CTG3 | MAMDC2 antisense RNA 1 [Source:HGNC Sym |
| AC212493.1 | CHR_HSCHR9_1_CTG3 | novel transcript, antisense to MAMDC2 |
| U2 | CHR_HSCHR9_1_CTG3 | U2 spliceosomal RNA [Source:RFAM;Acc:RF00 |
| AL451142.3 | CHR_HSCHR9_1_CTG4 | novel pseudogene |
| AC215799.1 | CHR_HSCHR9_1_CTG4 | mitofusin 1 (MFN1) pseudogene |
| GAPDHP26 | CHR_HSCHR9_1_CTG5 | glyceraldehyde 3 phosphate dehydrogenase p |
| PLPPR1 | CHR_HSCHR9_1_CTG5 | phospholipid phosphatase related 1 [Source:H |
| AL161631.2 | CHR_HSCHR9_1_CTG5 | pseudogene similar to part of the transient re |
| AL359893.3 | CHR_HSCHR9_1_CTG5 | NULL |
| AC216089.2 | CHR_HSCHR9_1_CTG5 | bile acid Coenzyme A: amino acid N-acyltransf |
| TRMT112P4 | CHR_HSCHR9_1_CTG5 | tRNA methyltransferase subunit 11-2 pseudog |
| AC216089.1 | CHR_HSCHR9_1_CTG5 | zinc finger and BTB domain containing protein |
| BAAT | CHR_HSCHR9_1_CTG5 | bile acid-CoA:amino acid N-acyltransferase [So |
| FYTTD1P1 | CHR_HSCHR9_1_CTG5 | forty-two-three domain containing 1 pseudog |

| | | |
|---|---|---|
| PTPRD | CHR_HSCHR9_1_CTG6 | protein tyrosine phosphatase receptor type D |
| RPS26P3 | CHR_HSCHR9_1_CTG6 | ribosomal protein S26 pseudogene 3 [Source: |
| LINC00685 | CHR_HSCHRX_1_CTG3 | long intergenic non-protein coding RNA 685 [S |
| PPP2R3B | CHR_HSCHRX_1_CTG3 | protein phosphatase 2 regulatory subunit B''b |
| AL732314.3 | CHR_HSCHRX_1_CTG3 | novel transcript, antisense to PPP2R3B |
| Metazoa_SRP | CHR_HSCHRX_1_CTG3 | Metazoan signal recognition particle RNA [Sou |
| AL732314.2 | CHR_HSCHRX_1_CTG3 | novel transcript |
| FABP5P13 | CHR_HSCHRX_1_CTG3 | fatty acid binding protein 5 pseudogene 13 [S |
| KRT18P53 | CHR_HSCHRX_1_CTG3 | keratin 18 pseudogene 53 [Source:HGNC Sym |
| TBX22 | CHR_HSCHRX_2_CTG12 | T-box transcription factor 22 [Source:HGNC Sy |
| PPP2R3B | CHR_HSCHRX_2_CTG3 | protein phosphatase 2 regulatory subunit B''b |
| AL732314.1 | CHR_HSCHRX_2_CTG3 | novel transcript, antisense to PPP2R3B |
| AL732314.5 | CHR_HSCHRX_2_CTG3 | novel transcript |
| FABP5P13 | CHR_HSCHRX_2_CTG3 | fatty acid binding protein 5 pseudogene 13 [S |
| KRT18P53 | CHR_HSCHRX_2_CTG3 | keratin 18 pseudogene 53 [Source:HGNC Sym |
| BX004987.1 | GL000009.2 | NULL |
| AC145212.1 | GL000194.1 | MaFF-interacting protein [Source:UniProtKB/ |
| MAFIP | GL000194.1 | MAFF interacting protein (pseudogene) [Sourc |
| Y_RNA | GL000195.1 | Y RNA [Source:RFAM;Acc:RF00019] |
| AC011043.1 | GL000195.1 | uncharacterized LOC389831 [Source:NCBI ger |
| AC011043.2 | GL000195.1 | NULL |
| Y_RNA | GL000205.2 | Y RNA [Source:RFAM;Acc:RF00019] |
| AC011841.1 | GL000205.2 | NULL |
| BX072566.1 | GL000213.1 | POTE ankyrin domain family member D-like [S |
| U6 | GL000213.1 | U6 spliceosomal RNA [Source:RFAM;Acc:RF00 |
| U2 | GL000216.2 | U2 spliceosomal RNA [Source:RFAM;Acc:RF00 |
| AL354822.1 | GL000218.1 | NULL |
| AL592183.1 | GL000219.1 | NULL |
| AL592188.2 | GL000220.1 | NULL |
| RNA5-8SN5 | GL000220.1 | RNA, 5.8S ribosomal N5 [Source:HGNC Symbo |
| AL592188.1 | GL000220.1 | NULL |
| 5_8S_rRNA | GL000220.1 | 5.8S ribosomal RNA [Source:RFAM;Acc:RF000 |
| U2 | GL000225.1 | U2 spliceosomal RNA [Source:RFAM;Acc:RF00 |
| 5_8S_rRNA | KI270442.1 | 5.8S ribosomal RNA [Source:RFAM;Acc:RF000 |
| 5S_rRNA | KI270442.1 | 5S ribosomal RNA [Source:RFAM;Acc:RF00001 |
| AC240274.1 | KI270711.1 | neuroblastoma breakpoint family member 1 [ |
| U1 | KI270713.1 | U1 spliceosomal RNA [Source:RFAM;Acc:RF00 |
| U1 | KI270713.1 | U1 spliceosomal RNA [Source:RFAM;Acc:RF00 |
| AC213203.2 | KI270713.1 | NULL |
| AC213203.1 | KI270713.1 | FAM231A/C-like protein LOC102723383 [Sou |
| AC004556.3 | KI270721.1 | 39S ribosomal protein L23, mitochondrial [Sou |
| AC004556.1 | KI270721.1 | NULL |
| AC004556.4 | KI270721.1 | NULL |
| AC004556.2 | KI270721.1 | NULL |
| AC233755.2 | KI270726.1 | NULL |
| AC233755.1 | KI270726.1 | immunoglobulin heavy variable 4-38-2-like [S |
| AC136352.4 | KI270727.1 | NULL |
| U6 | KI270727.1 | U6 spliceosomal RNA [Source:RFAM;Acc:RF00 |
| AC136352.3 | KI270727.1 | NULL |
| AC136352.2 | KI270727.1 | POTE ankyrin domain family member B-like [S |

| | | |
|---|---|---|
| U6 | KI270727.1 | U6 spliceosomal RNA [Source:RFAM;Acc:RF00 |
| AC171558.3 | KI270727.1 | NULL |
| AC171558.1 | KI270727.1 | olfactory receptor 4N4-like [Source:NCBI gene |
| AC171558.4 | KI270727.1 | NULL |
| AC133551.1 | KI270728.1 | TP53-target gene 3 protein [Source:NCBI gene |
| AC009091.1 | KI270728.1 | NULL |
| AC136612.1 | KI270728.1 | TP53-target gene 3 protein [Source:NCBI gene |
| AC007908.1 | KI270728.1 | NULL |
| AC136616.1 | KI270728.1 | NULL |
| AC136616.3 | KI270728.1 | NULL |
| AC136616.2 | KI270728.1 | NULL |
| AC141272.1 | KI270728.1 | NULL |
| AC023491.2 | KI270731.1 | putative POM121-like protein 1 [Source:NCBI |
| AC023491.1 | KI270731.1 | NULL |
| AL353644.1 | KI270733.1 | NULL |
| RNA5-8SN4 | KI270733.1 | RNA, 5.8S ribosomal N4 [Source:HGNC Symbo |
| AL353644.2 | KI270733.1 | NULL |
| 5_8S_rRNA | KI270733.1 | 5.8S ribosomal RNA [Source:RFAM;Acc:RF000 |
| AC007325.3 | KI270734.1 | NULL |
| AC007325.1 | KI270734.1 | NULL |
| AC007325.4 | KI270734.1 | protein DGCR6 [Source:NCBI gene;Acc:102724 |
| AC007325.2 | KI270734.1 | proline dehydrogenase 1, mitochondrial [Sour |
| U6 | KI270744.1 | U6 spliceosomal RNA [Source:RFAM;Acc:RF00 |
| U1 | KI270750.1 | U1 spliceosomal RNA [Source:RFAM;Acc:RF00 |
| COL1A1 | LRG_1 | collagen type I alpha 1 chain [Source:HGNC Sy |
| PPIB | LRG_10 | peptidylprolyl isomerase B [Source:HGNC Sym |
| RASGRP2 | LRG_100 | RAS guanyl releasing protein 2 [Source:HGNC |
| HTR2A | LRG_1008 | 5-hydroxytryptamine receptor 2A [Source:HG |
| RFX5 | LRG_101 | regulatory factor X5 [Source:HGNC Symbol;Ac |
| COMT | LRG_1010 | catechol-O-methyltransferase [Source:HGNC S |
| GRIK4 | LRG_1012 | glutamate ionotropic receptor kainate type su |
| SRC | LRG_1018 | SRC proto-oncogene, non-receptor tyrosine ki |
| RFXANK | LRG_102 | regulatory factor X associated ankyrin contain |
| SLC13A5 | LRG_1020 | solute carrier family 13 member 5 [Source:HG |
| FGFR3 | LRG_1021 | fibroblast growth factor receptor 3 [Source:H( |
| SLCO1B1 | LRG_1022 | solute carrier organic anion transporter family |
| RNF43 | LRG_1026 | ring finger protein 43 [Source:HGNC Symbol;A |
| FLG | LRG_1028 | filaggrin [Source:HGNC Symbol;Acc:HGNC:374 |
| RFXAP | LRG_103 | regulatory factor X associated protein [Source |
| SBDS | LRG_104 | SBDS ribosome maturation factor [Source:HGI |
| SERPING1 | LRG_105 | serpin family G member 1 [Source:HGNC Sym |
| SH2D1A | LRG_106 | SH2 domain containing 1A [Source:HGNC Sym |
| SLC35C1 | LRG_107 | solute carrier family 35 member C1 [Source:H |
| HMBS | LRG_1076 | hydroxymethylbilane synthase [Source:HGNC |
| CPOX | LRG_1077 | coproporphyrinogen oxidase [Source:HGNC Sy |
| UROD | LRG_1079 | uroporphyrinogen decarboxylase [Source:HGN |
| SMARCAL1 | LRG_108 | SWI/SNF related, matrix associated, actin dep |
| SP110 | LRG_109 | SP110 nuclear body protein [Source:HGNC Syr |
| CDKN2A | LRG_11 | cyclin dependent kinase inhibitor 2A [Source:I |
| SPINK5 | LRG_110 | serine peptidase inhibitor, Kazal type 5 [Sourc |

| | | |
|---|---|---|
| STAT1 | LRG_111 | signal transducer and activator of transcriptio |
| STAT3 | LRG_112 | signal transducer and activator of transcriptio |
| RPL19 | LRG_1120 | ribosomal protein L19 [Source:HGNC Symbol; |
| NOTCH1 | LRG_1122 | notch receptor 1 [Source:HGNC Symbol;Acc:H |
| CTC1 | LRG_1124 | CST telomere replication complex component |
| RPL26 | LRG_1125 | ribosomal protein L26 [Source:HGNC Symbol; |
| ARHGAP26 | LRG_1127 | Rho GTPase activating protein 26 [Source:HGI |
| RPL35A | LRG_1129 | ribosomal protein L35a [Source:HGNC Symbol |
| STX11 | LRG_113 | syntaxin 11 [Source:HGNC Symbol;Acc:HGNC: |
| SPTB | LRG_1130 | spectrin beta, erythrocytic [Source:HGNC Sym |
| SPTA1 | LRG_1131 | spectrin alpha, erythrocytic 1 [Source:HGNC S |
| SLC2A1 | LRG_1132 | solute carrier family 2 member 1 [Source:HGN |
| SLC25A38 | LRG_1133 | solute carrier family 25 member 38 [Source:H |
| SEC23B | LRG_1134 | SEC23 homolog B, coat complex II component |
| PRKG1 | LRG_1135 | protein kinase cGMP-dependent 1 [Source:HG |
| PKLR | LRG_1136 | pyruvate kinase L/R [Source:HGNC Symbol;Ac |
| PIEZO1 | LRG_1137 | piezo type mechanosensitive ion channel com |
| RPS10 | LRG_1138 | ribosomal protein S10 [Source:HGNC Symbol; |
| TAPBP | LRG_114 | TAP binding protein [Source:HGNC Symbol;Ac |
| RPL11 | LRG_1140 | ribosomal protein L11 [Source:HGNC Symbol; |
| FBXW7 | LRG_1141 | F-box and WD repeat domain containing 7 [Sc |
| RPS17 | LRG_1143 | ribosomal protein S17 [Source:HGNC Symbol; |
| RPS26 | LRG_1146 | ribosomal protein S26 [Source:HGNC Symbol; |
| RPS7 | LRG_1148 | ribosomal protein S7 [Source:HGNC Symbol;A |
| TCIRG1 | LRG_115 | T cell immune regulator 1, ATPase H+ transpo |
| SETBP1 | LRG_1150 | SET binding protein 1 [Source:HGNC Symbol;A |
| SRP72 | LRG_1151 | signal recognition particle 72 [Source:HGNC S |
| RPL15 | LRG_1152 | ribosomal protein L15 [Source:HGNC Symbol; |
| ATRX | LRG_1153 | ATRX chromatin remodeler [Source:HGNC Syr |
| SH3BP1 | LRG_1154 | SH3 domain binding protein 1 [Source:HGNC S |
| RPL5 | LRG_1155 | ribosomal protein L5 [Source:HGNC Symbol;A |
| RPL27 | LRG_1156 | ribosomal protein L27 [Source:HGNC Symbol; |
| RPS27 | LRG_1158 | ribosomal protein S27 [Source:HGNC Symbol; |
| TCN2 | LRG_116 | transcobalamin 2 [Source:HGNC Symbol;Acc:H |
| CBLC | LRG_1161 | Cbl proto-oncogene C [Source:HGNC Symbol; |
| ABCB7 | LRG_1162 | ATP binding cassette subfamily B member 7 [S |
| ALAS2 | LRG_1163 | 5'-aminolevulinate synthase 2 [Source:HGNC S |
| CDAN1 | LRG_1164 | codanin 1 [Source:HGNC Symbol;Acc:HGNC:1 |
| ENO1 | LRG_1165 | enolase 1 [Source:HGNC Symbol;Acc:HGNC:33 |
| GCLC | LRG_1166 | glutamate-cysteine ligase catalytic subunit [Sc |
| GSS | LRG_1168 | glutathione synthetase [Source:HGNC Symbol |
| UBE2T | LRG_1169 | ubiquitin conjugating enzyme E2 T [Source:HG |
| TLR3 | LRG_117 | toll like receptor 3 [Source:HGNC Symbol;Acc: |
| VPS45 | LRG_1170 | vacuolar protein sorting 45 homolog [Source:I |
| EPB42 | LRG_1171 | erythrocyte membrane protein band 4.2 [Sou |
| KIF23 | LRG_1173 | kinesin family member 23 [Source:HGNC Syml |
| GPI | LRG_1178 | glucose-6-phosphate isomerase [Source:HGN( |
| CBLB | LRG_1179 | Cbl proto-oncogene B [Source:HGNC Symbol; |
| TMC6 | LRG_118 | transmembrane channel like 6 [Source:HGNC |
| ATP1A3 | LRG_1186 | ATPase Na+/K+ transporting subunit alpha 3 [S |

| | | |
|---|---|---|
| ARPC1B | LRG_1188 | actin related protein 2/3 complex subunit 1B [ |
| TMC8 | LRG_119 | transmembrane channel like 8 [Source:HGNC |
| FOXF2 | LRG_1190 | forkhead box F2 [Source:HGNC Symbol;Acc:H( |
| JAG1 | LRG_1191 | jagged canonical Notch ligand 1 [Source:HGN( |
| CYP2C9 | LRG_1195 | cytochrome P450 family 2 subfamily C membe |
| KDSR | LRG_1196 | 3-ketodihydrosphingosine reductase [Source:I |
| ABCB11 | LRG_1199 | ATP binding cassette subfamily B member 11 |
| FKBP10 | LRG_12 | FKBP prolyl isomerase 10 [Source:HGNC Symb |
| TNFRSF13B | LRG_120 | TNF receptor superfamily member 13B [Sourc |
| TJP2 | LRG_1201 | tight junction protein 2 [Source:HGNC Symbol |
| ADAM17 | LRG_1203 | ADAM metallopeptidase domain 17 [Source:H |
| ATP8B1 | LRG_1205 | ATPase phospholipid transporting 8B1 [Source |
| TBX4 | LRG_1206 | T-box transcription factor 4 [Source:HGNC Syr |
| AP3D1 | LRG_1207 | adaptor related protein complex 3 subunit de |
| ABCC2 | LRG_1208 | ATP binding cassette subfamily C member 2 [S |
| CCBE1 | LRG_1209 | collagen and calcium binding EGF domains 1 [ |
| TYK2 | LRG_121 | tyrosine kinase 2 [Source:HGNC Symbol;Acc:H |
| BCL10 | LRG_1210 | BCL10 immune signaling adaptor [Source:HGN |
| C8G | LRG_1211 | complement C8 gamma chain [Source:HGNC S |
| ADAR | LRG_1212 | adenosine deaminase RNA specific [Source:H( |
| VPS13D | LRG_1213 | vacuolar protein sorting 13 homolog D [Sourc |
| DNAJC21 | LRG_1214 | DnaJ heat shock protein family (Hsp40) memb |
| B2M | LRG_1215 | beta-2-microglobulin [Source:HGNC Symbol;A |
| CFHR2 | LRG_1216 | complement factor H related 2 [Source:HGNC |
| ADA2 | LRG_1217 | adenosine deaminase 2 [Source:HGNC Symbo |
| ACP5 | LRG_1218 | acid phosphatase 5, tartrate resistant [Source |
| DCLRE1B | LRG_1219 | DNA cross-link repair 1B [Source:HGNC Symbo |
| UNC13D | LRG_122 | unc-13 homolog D [Source:HGNC Symbol;Acc: |
| CTLA4 | LRG_1220 | cytotoxic T-lymphocyte associated protein 4 [ |
| MALT1 | LRG_1221 | MALT1 paracaspase [Source:HGNC Symbol;Ac |
| MAP3K14 | LRG_1222 | mitogen-activated protein kinase kinase kinas |
| IL17A | LRG_1223 | interleukin 17A [Source:HGNC Symbol;Acc:HG |
| CFHR4 | LRG_1224 | complement factor H related 4 [Source:HGNC |
| MOGS | LRG_1226 | mannosyl-oligosaccharide glucosidase [Source |
| DOCK2 | LRG_1227 | dedicator of cytokinesis 2 [Source:HGNC Symb |
| JAGN1 | LRG_1228 | jagunal homolog 1 [Source:HGNC Symbol;Acc: |
| CTPS1 | LRG_1229 | CTP synthase 1 [Source:HGNC Symbol;Acc:HG |
| UNC93B1 | LRG_123 | unc-93 homolog B1, TLR signaling regulator [S |
| IL10 | LRG_1230 | interleukin 10 [Source:HGNC Symbol;Acc:HGN |
| ISG15 | LRG_1231 | ISG15 ubiquitin like modifier [Source:HGNC Sy |
| INO80 | LRG_1233 | INO80 complex ATPase subunit [Source:HGNC |
| EPG5 | LRG_1234 | ectopic P-granules autophagy protein 5 homo |
| IFIH1 | LRG_1235 | interferon induced with helicase C domain 1 [ |
| ACD | LRG_1237 | ACD shelterin complex subunit and telomeras |
| FCGR3B | LRG_1238 | Fc fragment of IgG receptor IIIb [Source:HGN( |
| MCM4 | LRG_1239 | minichromosome maintenance complex comp |
| UNG | LRG_124 | uracil DNA glycosylase [Source:HGNC Symbol; |
| IKBKB | LRG_1241 | inhibitor of nuclear factor kappa B kinase subu |
| DSPP | LRG_1242 | dentin sialophosphoprotein [Source:HGNC Syr |
| MTHFD1 | LRG_1243 | methylenetetrahydrofolate dehydrogenase, c |

| | | |
|---|---|---|
| ALDOB | LRG_1244 | aldolase, fructose-bisphosphate B [Source:HG |
| COX6B1 | LRG_1246 | cytochrome c oxidase subunit 6B1 [Source:HG |
| AP5Z1 | LRG_1247 | adaptor related protein complex 5 subunit zet |
| IL22 | LRG_1248 | interleukin 22 [Source:HGNC Symbol;Acc:HGN |
| COL17A1 | LRG_1249 | collagen type XVII alpha 1 chain [Source:HGN( |
| WAS | LRG_125 | WASP actin nucleation promoting factor [Sour |
| COX7B | LRG_1250 | cytochrome c oxidase subunit 7B [Source:HGN |
| AGK | LRG_1251 | acylglycerol kinase [Source:HGNC Symbol;Acc |
| COL9A3 | LRG_1253 | collagen type IX alpha 3 chain [Source:HGNC S |
| CHRNE | LRG_1254 | cholinergic receptor nicotinic epsilon subunit |
| BBS10 | LRG_1255 | Bardet-Biedl syndrome 10 [Source:HGNC Sym |
| ALX4 | LRG_1256 | ALX homeobox 4 [Source:HGNC Symbol;Acc:H |
| ZAP70 | LRG_126 | zeta chain of T cell receptor associated protei |
| ALG6 | LRG_1260 | ALG6 alpha-1,3-glucosyltransferase [Source:H |
| CYP1A1 | LRG_1262 | cytochrome P450 family 1 subfamily A membe |
| IL21 | LRG_1263 | interleukin 21 [Source:HGNC Symbol;Acc:HGN |
| ALOX12B | LRG_1264 | arachidonate 12-lipoxygenase, 12R type [Sour |
| ALX3 | LRG_1265 | ALX homeobox 3 [Source:HGNC Symbol;Acc:H |
| CYP2B6 | LRG_1267 | cytochrome P450 family 2 subfamily B membe |
| CD55 | LRG_127 | CD55 molecule (Cromer blood group) [Source |
| CYP1A2 | LRG_1274 | cytochrome P450 family 1 subfamily A membe |
| BTK | LRG_128 | Bruton tyrosine kinase [Source:HGNC Symbol; |
| SART3 | LRG_1285 | spliceosome associated factor 3, U4/U6 recyc |
| PARN | LRG_1286 | poly(A)-specific ribonuclease [Source:HGNC Sy |
| SEMA3E | LRG_1287 | semaphorin 3E [Source:HGNC Symbol;Acc:HG |
| CFP | LRG_129 | complement factor properdin [Source:HGNC S |
| CALCA | LRG_13 | calcitonin related polypeptide alpha [Source:H |
| APC | LRG_130 | APC regulator of WNT signaling pathway [Sour |
| TBK1 | LRG_1306 | TANK binding kinase 1 [Source:HGNC Symbol; |
| TMEM173 | LRG_1308 | transmembrane protein 173 [Source:HGNC Sy |
| IL17RC | LRG_1311 | interleukin 17 receptor C [Source:HGNC Symb |
| PGM3 | LRG_1312 | phosphoglucomutase 3 [Source:HGNC Symbo |
| IRF7 | LRG_1313 | interferon regulatory factor 7 [Source:HGNC S |
| TRNT1 | LRG_1314 | tRNA nucleotidyl transferase 1 [Source:HGNC |
| RNF31 | LRG_1315 | ring finger protein 31 [Source:HGNC Symbol;A |
| NFKB1 | LRG_1316 | nuclear factor kappa B subunit 1 [Source:HGN |
| NLRC4 | LRG_1317 | NLR family CARD domain containing 4 [Source |
| SLC29A3 | LRG_1318 | solute carrier family 29 member 3 [Source:HG |
| TNFRSF4 | LRG_1319 | TNF receptor superfamily member 4 [Source:I |
| ACTB | LRG_132 | actin beta [Source:HGNC Symbol;Acc:HGNC:1 |
| TNFSF12 | LRG_1320 | TNF superfamily member 12 [Source:HGNC Sy |
| RORC | LRG_1322 | RAR related orphan receptor C [Source:HGNC |
| TTC7A | LRG_1323 | tetratricopeptide repeat domain 7A [Source:H |
| LRBA | LRG_1324 | LPS responsive beige-like anchor protein [Sou |
| TCF3 | LRG_1325 | transcription factor 3 [Source:HGNC Symbol;A |
| PRKCD | LRG_1327 | protein kinase C delta [Source:HGNC Symbol;A |
| PSMB8 | LRG_1328 | proteasome subunit beta 8 [Source:HGNC Sym |
| STAT2 | LRG_1329 | signal transducer and activator of transcriptio |
| AK2 | LRG_133 | adenylate kinase 2 [Source:HGNC Symbol;Acc |
| CARD14 | LRG_1330 | caspase recruitment domain family member 1 |

| | | |
|---|---|---|
| NFAT5 | LRG_1332 | nuclear factor of activated T cells 5 [Source:H( |
| ITGAM | LRG_1333 | integrin subunit alpha M [Source:HGNC Symb( |
| SH3BP2 | LRG_1334 | SH3 domain binding protein 2 [Source:HGNC S |
| COPA | LRG_1336 | coatomer protein complex subunit alpha [Sou |
| TRAF3IP2 | LRG_1337 | TRAF3 interacting protein 2 [Source:HGNC Syr |
| CLPB | LRG_1338 | ClpB homolog, mitochondrial AAA ATPase cha |
| FAS | LRG_134 | Fas cell surface death receptor [Source:HGNC |
| TPP2 | LRG_1341 | tripeptidyl peptidase 2 [Source:HGNC Symbol; |
| NFKB2 | LRG_1347 | nuclear factor kappa B subunit 2 [Source:HGN |
| ATM | LRG_135 | ATM serine/threonine kinase [Source:HGNC S |
| CFB | LRG_136 | complement factor B [Source:HGNC Symbol;A |
| CBFB | LRG_1364 | core-binding factor subunit beta [Source:HGN |
| GREM1 | LRG_1365 | gremlin 1, DAN family BMP antagonist [Source |
| NTHL1 | LRG_1366 | nth like DNA glycosylase 1 [Source:HGNC Sym |
| TACC3 | LRG_1367 | transforming acidic coiled-coil containing prot |
| TMPRSS2 | LRG_1368 | transmembrane serine protease 2 [Source:HG |
| STAT6 | LRG_1369 | signal transducer and activator of transcriptio |
| C4A | LRG_137 | complement C4A (Rodgers blood group) [Sour |
| RIT1 | LRG_1372 | Ras like without CAAX 1 [Source:HGNC Symbo |
| DEK | LRG_1375 | DEK proto-oncogene [Source:HGNC Symbol;A |
| C4B | LRG_138 | complement C4B (Chido blood group) [Source |
| NUP214 | LRG_1383 | nucleoporin 214 [Source:HGNC Symbol;Acc:H( |
| PAX5 | LRG_1384 | paired box 5 [Source:HGNC Symbol;Acc:HGNC |
| DDX41 | LRG_1386 | DEAD-box helicase 41 [Source:HGNC Symbol;/ |
| PTPRZ1 | LRG_1387 | protein tyrosine phosphatase receptor type Z: |
| SCN8A | LRG_1389 | sodium voltage-gated channel alpha subunit 8 |
| C8A | LRG_139 | complement C8 alpha chain [Source:HGNC Syı |
| DDR2 | LRG_1390 | discoidin domain receptor tyrosine kinase 2 [S |
| AKT2 | LRG_1391 | AKT serine/threonine kinase 2 [Source:HGNC : |
| PIK3R2 | LRG_1392 | phosphoinositide-3-kinase regulatory subunit |
| SMO | LRG_1393 | smoothened, frizzled class receptor [Source:H |
| AKT3 | LRG_1396 | AKT serine/threonine kinase 3 [Source:HGNC : |
| JAK1 | LRG_1398 | Janus kinase 1 [Source:HGNC Symbol;Acc:HGN |
| ETNK1 | LRG_1399 | ethanolamine kinase 1 [Source:HGNC Symbol; |
| MS4A1 | LRG_140 | membrane spanning 4-domains A1 [Source:H( |
| ACAT1 | LRG_1400 | acetyl-CoA acetyltransferase 1 [Source:HGNC |
| CD40LG | LRG_141 | CD40 ligand [Source:HGNC Symbol;Acc:HGNC |
| CD81 | LRG_142 | CD81 molecule [Source:HGNC Symbol;Acc:HG |
| LYST | LRG_143 | lysosomal trafficking regulator [Source:HGNC |
| CSF3R | LRG_144 | colony stimulating factor 3 receptor [Source:H |
| F12 | LRG_145 | coagulation factor XII [Source:HGNC Symbol;A |
| FPR1 | LRG_146 | formyl peptide receptor 1 [Source:HGNC Syml |
| G6PC | LRG_147 | glucose-6-phosphatase catalytic subunit [Sour |
| G6PD | LRG_148 | glucose-6-phosphate dehydrogenase [Source: |
| CFHR1 | LRG_149 | complement factor H related 1 [Source:HGNC |
| UBE3A | LRG_15 | ubiquitin protein ligase E3A [Source:HGNC Syr |
| IL2RG | LRG_150 | interleukin 2 receptor subunit gamma [Source |
| IL10RA | LRG_151 | interleukin 10 receptor subunit alpha [Source: |
| IL10RB | LRG_152 | interleukin 10 receptor subunit beta [Source:H |
| MBL2 | LRG_154 | mannose binding lectin 2 [Source:HGNC Symb |

| | | |
|---|---|---|
| CD46 | LRG_155 | CD46 molecule [Source:HGNC Symbol;Acc:HG |
| MVK | LRG_156 | mevalonate kinase [Source:HGNC Symbol;Acc |
| MYD88 | LRG_157 | MYD88 innate immune signal transduction ad |
| NBN | LRG_158 | nibrin [Source:HGNC Symbol;Acc:HGNC:7652] |
| NCF4 | LRG_159 | neutrophil cytosolic factor 4 [Source:HGNC Sy |
| ADA | LRG_16 | adenosine deaminase [Source:HGNC Symbol;/ |
| PIGA | LRG_160 | phosphatidylinositol glycan anchor biosynthes |
| PMS2 | LRG_161 | PMS1 homolog 2, mismatch repair system cor |
| PRKDC | LRG_162 | protein kinase, DNA-activated, catalytic subur |
| RMRP | LRG_163 | RNA component of mitochondrial RNA proces |
| STIM1 | LRG_164 | stromal interaction molecule 1 [Source:HGNC |
| STXBP2 | LRG_165 | syntaxin binding protein 2 [Source:HGNC Sym |
| THBD | LRG_168 | thrombomodulin [Source:HGNC Symbol;Acc:H |
| APOL1 | LRG_169 | apolipoprotein L1 [Source:HGNC Symbol;Acc:I |
| AICDA | LRG_17 | activation induced cytidine deaminase [Sourc |
| AP3B1 | LRG_170 | adaptor related protein complex 3 subunit be |
| FCN3 | LRG_171 | ficolin 3 [Source:HGNC Symbol;Acc:HGNC:362 |
| PSTPIP1 | LRG_172 | proline-serine-threonine phosphatase interac |
| TTC37 | LRG_173 | tetratricopeptide repeat domain 37 [Source:H |
| LPIN2 | LRG_174 | lipin 2 [Source:HGNC Symbol;Acc:HGNC:1445( |
| CFHR3 | LRG_175 | complement factor H related 3 [Source:HGNC |
| CHD7 | LRG_176 | chromodomain helicase DNA binding protein |
| NOD2 | LRG_177 | nucleotide binding oligomerization domain co |
| CARD9 | LRG_178 | caspase recruitment domain family member 9 |
| CLEC7A | LRG_179 | C-type lectin domain containing 7A [Source:H( |
| AIRE | LRG_18 | autoimmune regulator [Source:HGNC Symbol, |
| FERMT3 | LRG_180 | fermitin family member 3 [Source:HGNC Symb |
| G6PC3 | LRG_182 | glucose-6-phosphatase catalytic subunit 3 [So |
| SLC46A1 | LRG_183 | solute carrier family 46 member 1 [Source:HG |
| TNFRSF13C | LRG_184 | TNF receptor superfamily member 13C [Sourc |
| RNF168 | LRG_185 | ring finger protein 168 [Source:HGNC Symbol; |
| CSF2RA | LRG_186 | colony stimulating factor 2 receptor alpha sub |
| SLC37A4 | LRG_187 | solute carrier family 37 member 4 [Source:HG |
| IL1RN | LRG_188 | interleukin 1 receptor antagonist [Source:HGN |
| ITK | LRG_189 | IL2 inducible T cell kinase [Source:HGNC Symb |
| XIAP | LRG_19 | X-linked inhibitor of apoptosis [Source:HGNC S |
| MEFV | LRG_190 | MEFV innate immuity regulator, pyrin [Source |
| PIK3CD | LRG_191 | phosphatidylinositol-4,5-bisphosphate 3-kinas |
| STAT5B | LRG_192 | signal transducer and activator of transcriptio |
| TNFRSF1A | LRG_193 | TNF receptor superfamily member 1A [Source |
| TNFRSF11A | LRG_194 | TNF receptor superfamily member 11a [Sourc |
| CORO1A | LRG_195 | coronin 1A [Source:HGNC Symbol;Acc:HGNC:2 |
| DOCK8 | LRG_196 | dedicator of cytokinesis 8 [Source:HGNC Symb |
| NLRP3 | LRG_197 | NLR family pyrin domain containing 3 [Source |
| AGRN | LRG_198 | agrin [Source:HGNC Symbol;Acc:HGNC:329] |
| DMD | LRG_199 | dystrophin [Source:HGNC Symbol;Acc:HGNC:2 |
| COL1A2 | LRG_2 | collagen type I alpha 2 chain [Source:HGNC Sy |
| BLM | LRG_20 | BLM RecQ like helicase [Source:HGNC Symbol |
| MSTN | LRG_200 | myostatin [Source:HGNC Symbol;Acc:HGNC:4. |
| MYOT | LRG_201 | myotilin [Source:HGNC Symbol;Acc:HGNC:123 |

| | | |
|---|---|---|
| NEB | LRG_202 | nebulin [Source:HGNC Symbol;Acc:HGNC:772 |
| SGCA | LRG_203 | sarcoglycan alpha [Source:HGNC Symbol;Acc: |
| SGCB | LRG_204 | sarcoglycan beta [Source:HGNC Symbol;Acc:H |
| SGCD | LRG_205 | sarcoglycan delta [Source:HGNC Symbol;Acc: |
| SGCG | LRG_207 | sarcoglycan gamma [Source:HGNC Symbol;Ac |
| SGCZ | LRG_208 | sarcoglycan zeta [Source:HGNC Symbol;Acc:H |
| SSPN | LRG_209 | sarcospan [Source:HGNC Symbol;Acc:HGNC:1 |
| BLNK | LRG_21 | B cell linker [Source:HGNC Symbol;Acc:HGNC: |
| TCAP | LRG_210 | titin-cap [Source:HGNC Symbol;Acc:HGNC:11 |
| TRIM32 | LRG_211 | tripartite motif containing 32 [Source:HGNC S |
| ZMPSTE24 | LRG_212 | zinc metallopeptidase STE24 [Source:HGNC Sy |
| CFL2 | LRG_213 | cofilin 2 [Source:HGNC Symbol;Acc:HGNC:187 |
| NF1 | LRG_214 | neurofibromin 1 [Source:HGNC Symbol;Acc:H |
| EPCAM | LRG_215 | epithelial cell adhesion molecule [Source:HGN |
| MLH1 | LRG_216 | mutL homolog 1 [Source:HGNC Symbol;Acc:H |
| MLH3 | LRG_217 | mutL homolog 3 [Source:HGNC Symbol;Acc:H |
| MSH2 | LRG_218 | mutS homolog 2 [Source:HGNC Symbol;Acc:H |
| MSH6 | LRG_219 | mutS homolog 6 [Source:HGNC Symbol;Acc:H |
| C1QA | LRG_22 | complement C1q A chain [Source:HGNC Symb |
| MUTYH | LRG_220 | mutY DNA glycosylase [Source:HGNC Symbol; |
| PMS1 | LRG_221 | PMS1 homolog 1, mismatch repair system co |
| TBX1 | LRG_226 | T-box transcription factor 1 [Source:HGNC Sy |
| CFHR5 | LRG_227 | complement factor H related 5 [Source:HGNC |
| FADD | LRG_228 | Fas associated via death domain [Source:HGN |
| TRAF3 | LRG_229 | TNF receptor associated factor 3 [Source:HGN |
| C1QB | LRG_23 | complement C1q B chain [Source:HGNC Symb |
| COL4A3 | LRG_230 | collagen type IV alpha 3 chain [Source:HGNC S |
| COL4A4 | LRG_231 | collagen type IV alpha 4 chain [Source:HGNC S |
| ARHGEF10 | LRG_234 | Rho guanine nucleotide exchange factor 10 [S |
| BSCL2 | LRG_235 | BSCL2 lipid droplet biogenesis associated, sei |
| CTDP1 | LRG_236 | CTD phosphatase subunit 1 [Source:HGNC Sy |
| DCTN1 | LRG_237 | dynactin subunit 1 [Source:HGNC Symbol;Acc: |
| DNM2 | LRG_238 | dynamin 2 [Source:HGNC Symbol;Acc:HGNC:2 |
| EGR2 | LRG_239 | early growth response 2 [Source:HGNC Symbo |
| C1QC | LRG_24 | complement C1q C chain [Source:HGNC Symb |
| FGD4 | LRG_240 | FYVE, RhoGEF and PH domain containing 4 [S |
| FIG4 | LRG_241 | FIG4 phosphoinositide 5-phosphatase [Source |
| GAN | LRG_242 | gigaxonin [Source:HGNC Symbol;Acc:HGNC:41 |
| GARS | LRG_243 | glycyl-tRNA synthetase [Source:HGNC Symbol |
| GDAP1 | LRG_244 | ganglioside induced differentiation associated |
| GJB1 | LRG_245 | gap junction protein beta 1 [Source:HGNC Sy |
| HOXD10 | LRG_246 | homeobox D10 [Source:HGNC Symbol;Acc:HG |
| HSPB1 | LRG_248 | heat shock protein family B (small) member 1 |
| HSPB8 | LRG_249 | heat shock protein family B (small) member 8 |
| C1S | LRG_25 | complement C1s [Source:HGNC Symbol;Acc:H |
| IGHMBP2 | LRG_250 | immunoglobulin mu DNA binding protein 2 [S |
| ELP1 | LRG_251 | elongator complex protein 1 [Source:HGNC Sy |
| KIF1B | LRG_252 | kinesin family member 1B [Source:HGNC Sym |
| LITAF | LRG_253 | lipopolysaccharide induced TNF factor [Source |
| LMNA | LRG_254 | lamin A/C [Source:HGNC Symbol;Acc:HGNC:6 |

| | | |
|---|---|---|
| MFN2 | LRG_255 | mitofusin 2 [Source:HGNC Symbol;Acc:HGNC: |
| MPZ | LRG_256 | myelin protein zero [Source:HGNC Symbol;Acc |
| MTMR2 | LRG_257 | myotubularin related protein 2 [Source:HGNC |
| NDRG1 | LRG_258 | N-myc downstream regulated 1 [Source:HGN( |
| NEFL | LRG_259 | neurofilament light [Source:HGNC Symbol;Acc |
| C2 | LRG_26 | complement C2 [Source:HGNC Symbol;Acc:H( |
| NGF | LRG_260 | nerve growth factor [Source:HGNC Symbol;Ac |
| NTRK1 | LRG_261 | neurotrophic receptor tyrosine kinase 1 [Sour |
| PLEKHG5 | LRG_262 | pleckstrin homology and RhoGEF domain cont |
| PMP22 | LRG_263 | peripheral myelin protein 22 [Source:HGNC Sy |
| PRPS1 | LRG_264 | phosphoribosyl pyrophosphate synthetase 1 [ |
| PRX | LRG_265 | periaxin [Source:HGNC Symbol;Acc:HGNC:137 |
| RAB7A | LRG_266 | RAB7A, member RAS oncogene family [Source |
| SBF2 | LRG_267 | SET binding factor 2 [Source:HGNC Symbol;Ac |
| SETX | LRG_268 | senataxin [Source:HGNC Symbol;Acc:HGNC:44 |
| SH3TC2 | LRG_269 | SH3 domain and tetratricopeptide repeats 2 [ |
| C3 | LRG_27 | complement C3 [Source:HGNC Symbol;Acc:H( |
| SLC12A6 | LRG_270 | solute carrier family 12 member 6 [Source:HG |
| SOX10 | LRG_271 | SRY-box transcription factor 10 [Source:HGNC |
| SPTLC1 | LRG_272 | serine palmitoyltransferase long chain base su |
| YARS | LRG_273 | tyrosyl-tRNA synthetase [Source:HGNC Symbo |
| LDLR | LRG_274 | low density lipoprotein receptor [Source:HGN |
| PCSK9 | LRG_275 | proprotein convertase subtilisin/kexin type 9 |
| LDLRAP1 | LRG_276 | low density lipoprotein receptor adaptor prot |
| RNASEH2A | LRG_278 | ribonuclease H2 subunit A [Source:HGNC Sym |
| RNASEH2B | LRG_279 | ribonuclease H2 subunit B [Source:HGNC Sym |
| C5 | LRG_28 | complement C5 [Source:HGNC Symbol;Acc:H( |
| RNASEH2C | LRG_280 | ribonuclease H2 subunit C [Source:HGNC Sym |
| SAMHD1 | LRG_281 | SAM and HD domain containing deoxynucleos |
| LEPR | LRG_283 | leptin receptor [Source:HGNC Symbol;Acc:HG |
| ZNF778 | LRG_284 | zinc finger protein 778 [Source:HGNC Symbol; |
| FXR2 | LRG_285 | FMR1 autosomal homolog 2 [Source:HGNC Sy |
| COL7A1 | LRG_286 | collagen type VII alpha 1 chain [Source:HGNC |
| KCNQ1 | LRG_287 | potassium voltage-gated channel subfamily Q |
| KCNH2 | LRG_288 | potassium voltage-gated channel subfamily H |
| SCN5A | LRG_289 | sodium voltage-gated channel alpha subunit 5 |
| C6 | LRG_29 | complement C6 [Source:HGNC Symbol;Acc:H( |
| KCNE1 | LRG_290 | potassium voltage-gated channel subfamily E |
| KCNE2 | LRG_291 | potassium voltage-gated channel subfamily E |
| BRCA1 | LRG_292 | BRCA1 DNA repair associated [Source:HGNC S |
| BRCA2 | LRG_293 | BRCA2 DNA repair associated [Source:HGNC S |
| IRF8 | LRG_294 | interferon regulatory factor 8 [Source:HGNC S |
| GATA2 | LRG_295 | GATA binding protein 2 [Source:HGNC Symbol |
| AXIN2 | LRG_296 | axin 2 [Source:HGNC Symbol;Acc:HGNC:904] |
| BARD1 | LRG_297 | BRCA1 associated RING domain 1 [Source:HGI |
| BMPR1A | LRG_298 | bone morphogenetic protein receptor type 1/ |
| BRAF | LRG_299 | B-Raf proto-oncogene, serine/threonine kinas |
| COL3A1 | LRG_3 | collagen type III alpha 1 chain [Source:HGNC S |
| C7 | LRG_30 | complement C7 [Source:HGNC Symbol;Acc:H( |
| BRIP1 | LRG_300 | BRCA1 interacting protein C-terminal helicase |

| | | |
|---|---|---|
| CDH1 | LRG_301 | cadherin 1 [Source:HGNC Symbol;Acc:HGNC:1 |
| CYP2D6 | LRG_303 | cytochrome P450 family 2 subfamily D membe |
| EGFR | LRG_304 | epidermal growth factor receptor [Source:HG |
| FANCD2 | LRG_306 | FA complementation group D2 [Source:HGNC |
| KIT | LRG_307 | KIT proto-oncogene, receptor tyrosine kinase |
| PALB2 | LRG_308 | partner and localizer of BRCA2 [Source:HGNC |
| PDGFRA | LRG_309 | platelet derived growth factor receptor alpha |
| C8B | LRG_31 | complement C8 beta chain [Source:HGNC Sym |
| PIK3CA | LRG_310 | phosphatidylinositol-4,5-bisphosphate 3-kinas |
| PTEN | LRG_311 | phosphatase and tensin homolog [Source:HGI |
| RAD51C | LRG_314 | RAD51 paralog C [Source:HGNC Symbol;Acc:H |
| SDHB | LRG_316 | succinate dehydrogenase complex iron sulfur |
| SDHC | LRG_317 | succinate dehydrogenase complex subunit C [ |
| SMAD4 | LRG_318 | SMAD family member 4 [Source:HGNC Symbo |
| STK11 | LRG_319 | serine/threonine kinase 11 [Source:HGNC Sym |
| C9 | LRG_32 | complement C9 [Source:HGNC Symbol;Acc:HC |
| TP53 | LRG_321 | tumor protein p53 [Source:HGNC Symbol;Acc: |
| VHL | LRG_322 | von Hippel-Lindau tumor suppressor [Source:I |
| FLCN | LRG_325 | folliculin [Source:HGNC Symbol;Acc:HGNC:27 |
| ZBTB24 | LRG_326 | zinc finger and BTB domain containing 24 [Sou |
| ANK2 | LRG_327 | ankyrin 2 [Source:HGNC Symbol;Acc:HGNC:49 |
| KCNJ2 | LRG_328 | potassium voltage-gated channel subfamily J r |
| CAV3 | LRG_329 | caveolin 3 [Source:HGNC Symbol;Acc:HGNC:1 |
| CASP10 | LRG_33 | caspase 10 [Source:HGNC Symbol;Acc:HGNC: |
| SCN4B | LRG_330 | sodium voltage-gated channel beta subunit 4 |
| AKAP9 | LRG_331 | A-kinase anchoring protein 9 [Source:HGNC Sy |
| SNTA1 | LRG_332 | syntrophin alpha 1 [Source:HGNC Symbol;Acc |
| KCNJ5 | LRG_333 | potassium voltage-gated channel subfamily J r |
| CACNA1C | LRG_334 | calcium voltage-gated channel subunit alpha1 |
| ELF4 | LRG_335 | E74 like ETS transcription factor 4 [Source:HG |
| CPS1 | LRG_336 | carbamoyl-phosphate synthase 1 [Source:HGI |
| OPA1 | LRG_337 | OPA1 mitochondrial dynamin like GTPase [Sou |
| LCT | LRG_338 | lactase [Source:HGNC Symbol;Acc:HGNC:653C |
| CASP8 | LRG_34 | caspase 8 [Source:HGNC Symbol;Acc:HGNC:1 |
| DHCR7 | LRG_340 | 7-dehydrocholesterol reductase [Source:HGN |
| UNC119 | LRG_341 | unc-119 lipid binding chaperone [Source:HGN |
| TERT | LRG_343 | telomerase reverse transcriptase [Source:HGI |
| KRAS | LRG_344 | KRAS proto-oncogene, GTPase [Source:HGNC |
| NOP10 | LRG_345 | NOP10 ribonucleoprotein [Source:HGNC Symb |
| NHP2 | LRG_346 | NHP2 ribonucleoprotein [Source:HGNC Symbo |
| TERC | LRG_347 | telomerase RNA component [Source:HGNC Sy |
| CR2 | LRG_348 | complement C3d receptor 2 [Source:HGNC Sy |
| MASP1 | LRG_349 | mannan binding lectin serine peptidase 1 [Sou |
| CD19 | LRG_35 | CD19 molecule [Source:HGNC Symbol;Acc:HG |
| COLEC11 | LRG_350 | collectin subfamily member 11 [Source:HGNC |
| VPS13B | LRG_351 | vacuolar protein sorting 13 homolog B [Source |
| USB1 | LRG_352 | U6 snRNA biogenesis phosphodiesterase 1 [Sc |
| MAGT1 | LRG_353 | magnesium transporter 1 [Source:HGNC Symb |
| ITCH | LRG_354 | itchy E3 ubiquitin protein ligase [Source:HGNC |
| IL17RA | LRG_355 | interleukin 17 receptor A [Source:HGNC Symb |

| | | |
|---|---|---|
| IL17F | LRG_356 | interleukin 17F [Source:HGNC Symbol;Acc:HG |
| CD27 | LRG_357 | CD27 molecule [Source:HGNC Symbol;Acc:HG |
| AC027319.2 | LRG_358 | NULL |
| AARS | LRG_359 | alanyl-tRNA synthetase [Source:HGNC Symbol |
| CD247 | LRG_36 | CD247 molecule [Source:HGNC Symbol;Acc:H |
| ATL1 | LRG_360 | atlastin GTPase 1 [Source:HGNC Symbol;Acc:H |
| CCT5 | LRG_361 | chaperonin containing TCP1 subunit 5 [Source |
| DNMT1 | LRG_362 | DNA methyltransferase 1 [Source:HGNC Symb |
| RETREG1 | LRG_363 | reticulophagy regulator 1 [Source:HGNC Symb |
| FBLN5 | LRG_364 | fibulin 5 [Source:HGNC Symbol;Acc:HGNC:360 |
| HK1 | LRG_365 | hexokinase 1 [Source:HGNC Symbol;Acc:HGN |
| KARS | LRG_366 | lysyl-tRNA synthetase [Source:HGNC Symbol;A |
| KIF1A | LRG_367 | kinesin family member 1A [Source:HGNC Sym |
| MED25 | LRG_368 | mediator complex subunit 25 [Source:HGNC S |
| SCN9A | LRG_369 | sodium voltage-gated channel alpha subunit 9 |
| CD3D | LRG_37 | CD3d molecule [Source:HGNC Symbol;Acc:HG |
| SPTLC2 | LRG_371 | serine palmitoyltransferase long chain base su |
| TRPV4 | LRG_372 | transient receptor potential cation channel su |
| LRSAM1 | LRG_373 | leucine rich repeat and sterile alpha motif con |
| WIPF1 | LRG_374 | WAS/WASL interacting protein family membe |
| WRAP53 | LRG_375 | WD repeat containing antisense to TP53 [Sour |
| PLCG2 | LRG_376 | phospholipase C gamma 2 [Source:HGNC Sym |
| ABCC9 | LRG_377 | ATP binding cassette subfamily C member 9 [S |
| TNNC1 | LRG_378 | troponin C1, slow skeletal and cardiac type [So |
| ANKRD1 | LRG_379 | ankyrin repeat domain 1 [Source:HGNC Symbo |
| CD3E | LRG_38 | CD3e molecule [Source:HGNC Symbol;Acc:HG |
| DES | LRG_380 | desmin [Source:HGNC Symbol;Acc:HGNC:277 |
| RBM20 | LRG_382 | RNA binding motif protein 20 [Source:HGNC S |
| VCL | LRG_383 | vinculin [Source:HGNC Symbol;Acc:HGNC:126 |
| MYH7 | LRG_384 | myosin heavy chain 7 [Source:HGNC Symbol;A |
| LDB3 | LRG_385 | LIM domain binding 3 [Source:HGNC Symbol;A |
| MYBPC3 | LRG_386 | myosin binding protein C, cardiac [Source:HGI |
| ACTC1 | LRG_388 | actin alpha cardiac muscle 1 [Source:HGNC Sy |
| MYH6 | LRG_389 | myosin heavy chain 6 [Source:HGNC Symbol;A |
| CD3G | LRG_39 | CD3g molecule [Source:HGNC Symbol;Acc:HG |
| PLN | LRG_390 | phospholamban [Source:HGNC Symbol;Acc:H |
| TTN | LRG_391 | titin [Source:HGNC Symbol;Acc:HGNC:12403] |
| MYLK2 | LRG_392 | myosin light chain kinase 2 [Source:HGNC Sym |
| MYL2 | LRG_393 | myosin light chain 2 [Source:HGNC Symbol;Ac |
| JPH2 | LRG_394 | junctophilin 2 [Source:HGNC Symbol;Acc:HGN |
| MYL3 | LRG_395 | myosin light chain 3 [Source:HGNC Symbol;Ac |
| MYOZ2 | LRG_396 | myozenin 2 [Source:HGNC Symbol;Acc:HGNC: |
| DSG2 | LRG_397 | desmoglein 2 [Source:HGNC Symbol;Acc:HGN |
| PKP2 | LRG_398 | plakophilin 2 [Source:HGNC Symbol;Acc:HGN |
| TGFB3 | LRG_399 | transforming growth factor beta 3 [Source:HG |
| CRTAP | LRG_4 | cartilage associated protein [Source:HGNC Sy |
| CD40 | LRG_40 | CD40 molecule [Source:HGNC Symbol;Acc:HG |
| DSC2 | LRG_400 | desmocollin 2 [Source:HGNC Symbol;Acc:HGN |
| JUP | LRG_401 | junction plakoglobin [Source:HGNC Symbol;Ac |
| RYR2 | LRG_402 | ryanodine receptor 2 [Source:HGNC Symbol;A |

| | | |
|---|---|---|
| BMP10 | LRG_403 | bone morphogenetic protein 10 [Source:HGN[illegible] |
| CASQ2 | LRG_404 | calsequestrin 2 [Source:HGNC Symbol;Acc:HG[illegible] |
| CHRM2 | LRG_405 | cholinergic receptor muscarinic 2 [Source:HGN[illegible] |
| COX15 | LRG_406 | cytochrome c oxidase assembly homolog COX[illegible] |
| CTF1 | LRG_408 | cardiotrophin 1 [Source:HGNC Symbol;Acc:HG[illegible] |
| LAMA2 | LRG_409 | laminin subunit alpha 2 [Source:HGNC Symbol[illegible] |
| CD59 | LRG_41 | CD59 molecule (CD59 blood group) [Source:H[illegible] |
| MYPN | LRG_410 | myopalladin [Source:HGNC Symbol;Acc:HGNC[illegible] |
| NEBL | LRG_411 | nebulette [Source:HGNC Symbol;Acc:HGNC:1[illegible] |
| RAF1 | LRG_413 | Raf-1 proto-oncogene, serine/threonine kinas[illegible] |
| SRI | LRG_414 | sorcin [Source:HGNC Symbol;Acc:HGNC:1129[illegible] |
| SYNM | LRG_415 | synemin [Source:HGNC Symbol;Acc:HGNC:24[illegible] |
| TTR | LRG_416 | transthyretin [Source:HGNC Symbol;Acc:HGN[illegible] |
| TXNRD2 | LRG_417 | thioredoxin reductase 2 [Source:HGNC Symbo[illegible] |
| GPD1L | LRG_419 | glycerol-3-phosphate dehydrogenase 1 like [S[illegible] |
| CD79A | LRG_42 | CD79a molecule [Source:HGNC Symbol;Acc:H[illegible] |
| SCN3B | LRG_421 | sodium voltage-gated channel beta subunit 3 |
| CALR3 | LRG_422 | calreticulin 3 [Source:HGNC Symbol;Acc:HGN[illegible] |
| DSP | LRG_423 | desmoplakin [Source:HGNC Symbol;Acc:HGN[illegible] |
| ADORA3 | LRG_424 | adenosine A3 receptor [Source:HGNC Symbol[illegible] |
| MYOM1 | LRG_426 | myomesin 1 [Source:HGNC Symbol;Acc:HGNC[illegible] |
| ACTA1 | LRG_429 | actin alpha 1, skeletal muscle [Source:HGNC S[illegible] |
| CD79B | LRG_43 | CD79b molecule [Source:HGNC Symbol;Acc:H[illegible] |
| TNNI3 | LRG_432 | troponin I3, cardiac type [Source:HGNC Symb[illegible] |
| LAMA4 | LRG_433 | laminin subunit alpha 4 [Source:HGNC Symbol[illegible] |
| FKTN | LRG_434 | fukutin [Source:HGNC Symbol;Acc:HGNC:362[illegible] |
| TMEM43 | LRG_435 | transmembrane protein 43 [Source:HGNC Syn[illegible] |
| CACNA2D1 | LRG_437 | calcium voltage-gated channel auxiliary subun[illegible] |
| KCNE3 | LRG_439 | potassium voltage-gated channel subfamily E |
| CD8A | LRG_44 | CD8a molecule [Source:HGNC Symbol;Acc:HG[illegible] |
| CSRP3 | LRG_440 | cysteine and glycine rich protein 3 [Source:HG[illegible] |
| NEXN | LRG_442 | nexilin F-actin binding protein [Source:HGNC [illegible] |
| TMPO | LRG_443 | thymopoietin [Source:HGNC Symbol;Acc:HGN[illegible] |
| ILK | LRG_444 | integrin linked kinase [Source:HGNC Symbol;A[illegible] |
| KCND3 | LRG_445 | potassium voltage-gated channel subfamily D |
| TCF21 | LRG_446 | transcription factor 21 [Source:HGNC Symbol; |
| HMGCS2 | LRG_447 | 3-hydroxy-3-methylglutaryl-CoA synthase 2 [S[illegible] |
| ACAD8 | LRG_448 | acyl-CoA dehydrogenase family member 8 [S[illegible] |
| AUH | LRG_449 | AU RNA binding methylglutaconyl-CoA hydrat[illegible] |
| CEBPE | LRG_45 | CCAAT enhancer binding protein epsilon [Sour[illegible] |
| HSD17B10 | LRG_450 | hydroxysteroid 17-beta dehydrogenase 10 [S[illegible] |
| ACADSB | LRG_451 | acyl-CoA dehydrogenase short/branched chai[illegible] |
| FOXD4 | LRG_452 | forkhead box D4 [Source:HGNC Symbol;Acc:H[illegible] |
| FKBP14 | LRG_454 | FKBP prolyl isomerase 14 [Source:HGNC Symb[illegible] |
| ROGDI | LRG_455 | rogdi atypical leucine zipper [Source:HGNC Sy[illegible] |
| CEBPA | LRG_456 | CCAAT enhancer binding protein alpha [Sourc[illegible] |
| FLT3 | LRG_457 | fms related tyrosine kinase 3 [Source:HGNC S[illegible] |
| NPM1 | LRG_458 | nucleophosmin 1 [Source:HGNC Symbol;Acc:H[illegible] |
| DNMT3A | LRG_459 | DNA methyltransferase 3 alpha [Source:HGNC[illegible] |
| CFD | LRG_46 | complement factor D [Source:HGNC Symbol;A[illegible] |

| | | |
|---|---|---|
| AIP | LRG_460 | aryl hydrocarbon receptor interacting protein |
| ERCC2 | LRG_461 | ERCC excision repair 2, TFIIH core complex hel |
| ERCC3 | LRG_462 | ERCC excision repair 3, TFIIH core complex hel |
| ERCC4 | LRG_463 | ERCC excision repair 4, endonuclease catalytic |
| ERCC5 | LRG_464 | ERCC excision repair 5, endonuclease [Source: |
| ERCC6 | LRG_465 | ERCC excision repair 6, chromatin remodeling |
| ERCC8 | LRG_466 | ERCC excision repair 8, CSA ubiquitin ligase co |
| DDB2 | LRG_467 | damage specific DNA binding protein 2 [Sourc |
| GTF2H5 | LRG_469 | general transcription factor IIH subunit 5 [Sou |
| CFH | LRG_47 | complement factor H [Source:HGNC Symbol;A |
| POLH | LRG_470 | DNA polymerase eta [Source:HGNC Symbol;A |
| XPA | LRG_471 | XPA, DNA damage recognition and repair fact |
| XPC | LRG_472 | XPC complex subunit, DNA damage recognitio |
| COL6A3 | LRG_473 | collagen type VI alpha 3 chain [Source:HGNC S |
| DDB1 | LRG_474 | damage specific DNA binding protein 1 [Sourc |
| COL6A1 | LRG_475 | collagen type VI alpha 1 chain [Source:HGNC S |
| COL6A2 | LRG_476 | collagen type VI alpha 2 chain [Source:HGNC S |
| GP9 | LRG_477 | glycoprotein IX platelet [Source:HGNC Symbol |
| GP1BB | LRG_478 | glycoprotein Ib platelet subunit beta [Source:I |
| ITGA2B | LRG_479 | integrin subunit alpha 2b [Source:HGNC Symb |
| CFI | LRG_48 | complement factor I [Source:HGNC Symbol;A |
| ITGB3 | LRG_481 | integrin subunit beta 3 [Source:HGNC Symbol; |
| RUNX1 | LRG_482 | RUNX family transcription factor 1 [Source:HG |
| HNF4A | LRG_483 | hepatocyte nuclear factor 4 alpha [Source:HG |
| GLE1 | LRG_484 | GLE1 RNA export mediator [Source:HGNC Sym |
| CSTB | LRG_485 | cystatin B [Source:HGNC Symbol;Acc:HGNC:2 |
| TSC1 | LRG_486 | TSC complex subunit 1 [Source:HGNC Symbol; |
| TSC2 | LRG_487 | TSC complex subunit 2 [Source:HGNC Symbol; |
| ALK | LRG_488 | ALK receptor tyrosine kinase [Source:HGNC Sy |
| BUB1B | LRG_489 | BUB1 mitotic checkpoint serine/threonine kin |
| CIITA | LRG_49 | class II major histocompatibility complex trans |
| CDK4 | LRG_490 | cyclin dependent kinase 4 [Source:HGNC Sym |
| CYLD | LRG_491 | CYLD lysine 63 deubiquitinase [Source:HGNC S |
| DICER1 | LRG_492 | dicer 1, ribonuclease III [Source:HGNC Symbol |
| EXT1 | LRG_493 | exostosin glycosyltransferase 1 [Source:HGNC |
| EXT2 | LRG_494 | exostosin glycosyltransferase 2 [Source:HGNC |
| FANCA | LRG_495 | FA complementation group A [Source:HGNC S |
| FANCB | LRG_496 | FA complementation group B [Source:HGNC S |
| FANCC | LRG_497 | FA complementation group C [Source:HGNC S |
| FANCE | LRG_498 | FA complementation group E [Source:HGNC S |
| FANCG | LRG_499 | FA complementation group G [Source:HGNC S |
| P3H1 | LRG_5 | prolyl 3-hydroxylase 1 [Source:HGNC Symbol; |
| CTSC | LRG_50 | cathepsin C [Source:HGNC Symbol;Acc:HGNC: |
| FANCI | LRG_500 | FA complementation group I [Source:HGNC Sy |
| FANCL | LRG_501 | FA complementation group L [Source:HGNC S |
| FANCM | LRG_502 | FA complementation group M [Source:HGNC S |
| SLX4 | LRG_503 | SLX4 structure-specific endonuclease subunit |
| FH | LRG_504 | fumarate hydratase [Source:HGNC Symbol;Ac |
| GPC3 | LRG_505 | glypican 3 [Source:HGNC Symbol;Acc:HGNC:4 |
| CDC73 | LRG_507 | cell division cycle 73 [Source:HGNC Symbol;Ac |

| | | |
|---|---|---|
| MEN1 | LRG_509 | menin 1 [Source:HGNC Symbol;Acc:HGNC:701 |
| CXCR4 | LRG_51 | C-X-C motif chemokine receptor 4 [Source:HG |
| MPL | LRG_510 | MPL proto-oncogene, thrombopoietin recepto |
| NF2 | LRG_511 | neurofibromin 2 [Source:HGNC Symbol;Acc:H |
| NSD1 | LRG_512 | nuclear receptor binding SET domain protein 1 |
| PHOX2B | LRG_513 | paired like homeobox 2B [Source:HGNC Symb |
| PRKAR1A | LRG_514 | protein kinase cAMP-dependent type I regulat |
| PTCH1 | LRG_515 | patched 1 [Source:HGNC Symbol;Acc:HGNC:95 |
| RAD51D | LRG_516 | RAD51 paralog D [Source:HGNC Symbol;Acc:H |
| RB1 | LRG_517 | RB transcriptional corepressor 1 [Source:HGN |
| RET | LRG_518 | ret proto-oncogene [Source:HGNC Symbol;Ac |
| SDHAF2 | LRG_519 | succinate dehydrogenase complex assembly f |
| CYBA | LRG_52 | cytochrome b-245 alpha chain [Source:HGNC |
| SMARCB1 | LRG_520 | SWI/SNF related, matrix associated, actin dep |
| SUFU | LRG_521 | SUFU negative regulator of hedgehog signalin |
| HNF1A | LRG_522 | HNF1 homeobox A [Source:HGNC Symbol;Acc |
| TSHR | LRG_523 | thyroid stimulating hormone receptor [Source |
| WRN | LRG_524 | WRN RecQ like helicase [Source:HGNC Symbo |
| WT1 | LRG_525 | WT1 transcription factor [Source:HGNC Symb |
| CEP57 | LRG_526 | centrosomal protein 57 [Source:HGNC Symbo |
| FANCF | LRG_527 | FA complementation group F [Source:HGNC S |
| TMEM127 | LRG_528 | transmembrane protein 127 [Source:HGNC Sy |
| BAP1 | LRG_529 | BRCA1 associated protein 1 [Source:HGNC Syr |
| CYBB | LRG_53 | cytochrome b-245 beta chain [Source:HGNC S |
| MAX | LRG_530 | MYC associated factor X [Source:HGNC Symbo |
| EZH2 | LRG_531 | enhancer of zeste 2 polycomb repressive com |
| RHBDF2 | LRG_532 | rhomboid 5 homolog 2 [Source:HGNC Symbol |
| CDKN1C | LRG_533 | cyclin dependent kinase inhibitor 1C [Source:H |
| DIS3L2 | LRG_534 | DIS3 like 3'-5' exoribonuclease 2 [Source:HGN |
| STK4 | LRG_535 | serine/threonine kinase 4 [Source:HGNC Symb |
| FSHR | LRG_536 | follicle stimulating hormone receptor [Source |
| AMT | LRG_537 | aminomethyltransferase [Source:HGNC Symb |
| KERA | LRG_538 | keratocan [Source:HGNC Symbol;Acc:HGNC:6 |
| BCS1L | LRG_539 | BCS1 homolog, ubiquinol-cytochrome c reduc |
| DCLRE1C | LRG_54 | DNA cross-link repair 1C [Source:HGNC Symbo |
| CUBN | LRG_540 | cubilin [Source:HGNC Symbol;Acc:HGNC:2548 |
| GCSH | LRG_541 | glycine cleavage system protein H [Source:HG |
| ACVRL1 | LRG_543 | activin A receptor like type 1 [Source:HGNC Sy |
| ADAMTS13 | LRG_544 | ADAM metallopeptidase with thrombospondi |
| BLOC1S3 | LRG_546 | biogenesis of lysosomal organelles complex 1 |
| F10 | LRG_548 | coagulation factor X [Source:HGNC Symbol;Ac |
| F13A1 | LRG_549 | coagulation factor XIII A chain [Source:HGNC S |
| DKC1 | LRG_55 | dyskerin pseudouridine synthase 1 [Source:HG |
| F13B | LRG_550 | coagulation factor XIII B chain [Source:HGNC S |
| F2 | LRG_551 | coagulation factor II, thrombin [Source:HGNC |
| F5 | LRG_553 | coagulation factor V [Source:HGNC Symbol;Ac |
| F7 | LRG_554 | coagulation factor VII [Source:HGNC Symbol;A |
| F9 | LRG_556 | coagulation factor IX [Source:HGNC Symbol;A |
| FGA | LRG_557 | fibrinogen alpha chain [Source:HGNC Symbol; |
| FGB | LRG_558 | fibrinogen beta chain [Source:HGNC Symbol;A |

| | | |
|---|---|---|
| GATA1 | LRG_559 | GATA binding protein 1 [Source:HGNC Symbol |
| DNMT3B | LRG_56 | DNA methyltransferase 3 beta [Source:HGNC |
| GP6 | LRG_560 | glycoprotein VI platelet [Source:HGNC Symbo |
| HOXA11 | LRG_561 | homeobox A11 [Source:HGNC Symbol;Acc:HG |
| HPS6 | LRG_564 | HPS6 biogenesis of lysosomal organelles comp |
| MCFD2 | LRG_566 | multiple coagulation factor deficiency 2 [Sour |
| MYH9 | LRG_567 | myosin heavy chain 9 [Source:HGNC Symbol;A |
| NBEAL2 | LRG_568 | neurobeachin like 2 [Source:HGNC Symbol;Ac |
| P2RY12 | LRG_569 | purinergic receptor P2Y12 [Source:HGNC Sym |
| ELANE | LRG_57 | elastase, neutrophil expressed [Source:HGNC |
| PLAT | LRG_570 | plasminogen activator, tissue type [Source:HG |
| PLG | LRG_571 | plasminogen [Source:HGNC Symbol;Acc:HGNC |
| PROS1 | LRG_572 | protein S [Source:HGNC Symbol;Acc:HGNC:94 |
| RBM8A | LRG_574 | RNA binding motif protein 8A [Source:HGNC S |
| SERPINC1 | LRG_577 | serpin family C member 1 [Source:HGNC Symb |
| TBXA2R | LRG_578 | thromboxane A2 receptor [Source:HGNC Sym |
| TBXAS1 | LRG_579 | thromboxane A synthase 1 [Source:HGNC Sym |
| FASLG | LRG_58 | Fas ligand [Source:HGNC Symbol;Acc:HGNC:1 |
| THPO | LRG_580 | thrombopoietin [Source:HGNC Symbol;Acc:HG |
| TUBB1 | LRG_581 | tubulin beta 1 class VI [Source:HGNC Symbol;A |
| F11 | LRG_583 | coagulation factor XI [Source:HGNC Symbol;Ac |
| HPS5 | LRG_586 | HPS5 biogenesis of lysosomal organelles comp |
| DTNBP1 | LRG_588 | dystrobrevin binding protein 1 [Source:HGNC |
| ENG | LRG_589 | endoglin [Source:HGNC Symbol;Acc:HGNC:33 |
| FCGR1A | LRG_59 | Fc fragment of IgG receptor Ia [Source:HGNC S |
| HPS4 | LRG_590 | HPS4 biogenesis of lysosomal organelles comp |
| PLAU | LRG_593 | plasminogen activator, urokinase [Source:HGN |
| SERPINE1 | LRG_597 | serpin family E member 1 [Source:HGNC Symb |
| KNG1 | LRG_598 | kininogen 1 [Source:HGNC Symbol;Acc:HGNC: |
| PROC | LRG_599 | protein C, inactivator of coagulation factors V |
| ATP1A2 | LRG_6 | ATPase Na+/K+ transporting subunit alpha 2 [ |
| FCGR3A | LRG_60 | Fc fragment of IgG receptor IIIa [Source:HGNC |
| CHST14 | LRG_600 | carbohydrate sulfotransferase 14 [Source:HGN |
| HRG | LRG_601 | histidine rich glycoprotein [Source:HGNC Sym |
| ANKRD26 | LRG_605 | ankyrin repeat domain 26 [Source:HGNC Symb |
| TYROBP | LRG_607 | TYRO protein tyrosine kinase binding protein |
| CBL | LRG_608 | Cbl proto-oncogene [Source:HGNC Symbol;Ac |
| ETV6 | LRG_609 | ETS variant 6 [Source:HGNC Symbol;Acc:HGNC |
| FOXN1 | LRG_61 | forkhead box N1 [Source:HGNC Symbol;Acc:H |
| IDH1 | LRG_610 | isocitrate dehydrogenase (NADP(+)) 1, cytosol |
| IDH2 | LRG_611 | isocitrate dehydrogenase (NADP(+)) 2, mitoch |
| JAK2 | LRG_612 | Janus kinase 2 [Source:HGNC Symbol;Acc:HGN |
| KMT2A | LRG_613 | lysine methyltransferase 2A [Source:HGNC Sy |
| PTPN11 | LRG_614 | protein tyrosine phosphatase non-receptor ty |
| U2AF1 | LRG_615 | U2 small nuclear RNA auxiliary factor 1 [Sourc |
| KDM6A | LRG_616 | lysine demethylase 6A [Source:HGNC Symbol; |
| SF1 | LRG_617 | splicing factor 1 [Source:HGNC Symbol;Acc:HG |
| ZRSR2 | LRG_618 | zinc finger CCCH-type, RNA binding motif and |
| SOCS3 | LRG_619 | suppressor of cytokine signaling 3 [Source:HG |
| FOXP3 | LRG_62 | forkhead box P3 [Source:HGNC Symbol;Acc:H |

| | | |
|---|---|---|
| BCL2L11 | LRG_620 | BCL2 like 11 [Source:HGNC Symbol;Acc:HGNC |
| SH2B3 | LRG_621 | SH2B adaptor protein 3 [Source:HGNC Symbo |
| SF3A1 | LRG_622 | splicing factor 3a subunit 1 [Source:HGNC Syr |
| U2AF2 | LRG_623 | U2 small nuclear RNA auxiliary factor 2 [Sourc |
| SF3B1 | LRG_624 | splicing factor 3b subunit 1 [Source:HGNC Syr |
| PRPF40B | LRG_625 | pre-mRNA processing factor 40 homolog B [Sc |
| TET2 | LRG_626 | tet methylcytosine dioxygenase 2 [Source:HGI |
| BCOR | LRG_627 | BCL6 corepressor [Source:HGNC Symbol;Acc:H |
| BCORL1 | LRG_628 | BCL6 corepressor like 1 [Source:HGNC Symbol |
| PHF6 | LRG_629 | PHD finger protein 6 [Source:HGNC Symbol;Ac |
| GFI1 | LRG_63 | growth factor independent 1 transcriptional r |
| ASXL1 | LRG_630 | ASXL transcriptional regulator 1 [Source:HGN( |
| TREM2 | LRG_631 | triggering receptor expressed on myeloid cells |
| HAX1 | LRG_64 | HCLS1 associated protein X-1 [Source:HGNC S |
| SRSF2 | LRG_640 | serine and arginine rich splicing factor 2 [Sour |
| AMN | LRG_642 | amnion associated transmembrane protein [S |
| GLDC | LRG_643 | glycine decarboxylase [Source:HGNC Symbol;A |
| ICOS | LRG_65 | inducible T cell costimulator [Source:HGNC Sy |
| KRIT1 | LRG_650 | KRIT1 ankyrin repeat containing [Source:HGN( |
| SOD1 | LRG_652 | superoxide dismutase 1 [Source:HGNC Symbo |
| ANG | LRG_653 | angiogenin [Source:HGNC Symbol;Acc:HGNC:4 |
| FUS | LRG_655 | FUS RNA binding protein [Source:HGNC Symb |
| VAPB | LRG_656 | VAMP associated protein B and C [Source:HGI |
| VCP | LRG_657 | valosin containing protein [Source:HGNC Sym |
| TARDBP | LRG_659 | TAR DNA binding protein [Source:HGNC Symb |
| IFNGR1 | LRG_66 | interferon gamma receptor 1 [Source:HGNC S |
| GRN | LRG_661 | granulin precursor [Source:HGNC Symbol;Acc: |
| MET | LRG_662 | MET proto-oncogene, receptor tyrosine kinas |
| CFTR | LRG_663 | CF transmembrane conductance regulator [Sc |
| CCM2 | LRG_664 | CCM2 scaffold protein [Source:HGNC Symbol; |
| UBQLN2 | LRG_665 | ubiquilin 2 [Source:HGNC Symbol;Acc:HGNC:1 |
| AFG3L2 | LRG_666 | AFG3 like matrix AAA peptidase subunit 2 [Sou |
| PRKCG | LRG_669 | protein kinase C gamma [Source:HGNC Symbo |
| IFNGR2 | LRG_67 | interferon gamma receptor 2 [Source:HGNC S |
| TBX5 | LRG_670 | T-box transcription factor 5 [Source:HGNC Syr |
| GLA | LRG_672 | galactosidase alpha [Source:HGNC Symbol;Ac |
| GAA | LRG_673 | glucosidase alpha, acid [Source:HGNC Symbol |
| SALL1 | LRG_674 | spalt like transcription factor 1 [Source:HGNC |
| SALL4 | LRG_675 | spalt like transcription factor 4 [Source:HGNC |
| SMN1 | LRG_676 | survival of motor neuron 1, telomeric [Source |
| TPM2 | LRG_680 | tropomyosin 2 [Source:HGNC Symbol;Acc:HGI |
| KBTBD13 | LRG_682 | kelch repeat and BTB domain containing 13 [S |
| SLC26A3 | LRG_683 | solute carrier family 26 member 3 [Source:HG |
| SLC26A2 | LRG_684 | solute carrier family 26 member 2 [Source:HG |
| OAT | LRG_685 | ornithine aminotransferase [Source:HGNC Syr |
| MKS1 | LRG_687 | MKS transition zone complex subunit 1 [Sourc |
| TMEM67 | LRG_688 | transmembrane protein 67 [Source:HGNC Syn |
| CLN3 | LRG_689 | CLN3 lysosomal/endosomal transmembrane p |
| IGLL1 | LRG_69 | immunoglobulin lambda like polypeptide 1 [Sc |
| PPT1 | LRG_690 | palmitoyl-protein thioesterase 1 [Source:HGN |

| | | |
|---|---|---|
| CLN8 | LRG_691 | CLN8 transmembrane ER and ERGIC protein [S |
| CLN5 | LRG_692 | CLN5 intracellular trafficking protein [Source:H |
| NPHS1 | LRG_693 | NPHS1 adhesion molecule, nephrin [Source:H |
| SLC7A7 | LRG_695 | solute carrier family 7 member 7 [Source:HGN |
| CC2D2A | LRG_697 | coiled-coil and C2 domain containing 2A [Sour |
| CHM | LRG_699 | CHM Rab escort protein [Source:HGNC Symbo |
| CACNA1A | LRG_7 | calcium voltage-gated channel subunit alpha1 |
| IKBKG | LRG_70 | inhibitor of nuclear factor kappa B kinase regu |
| CLRN1 | LRG_700 | clarin 1 [Source:HGNC Symbol;Acc:HGNC:126 |
| RS1 | LRG_702 | retinoschisin 1 [Source:HGNC Symbol;Acc:HGI |
| SMAD9 | LRG_703 | SMAD family member 9 [Source:HGNC Symbo |
| IL12B | LRG_71 | interleukin 12B [Source:HGNC Symbol;Acc:HG |
| BMPR2 | LRG_712 | bone morphogenetic protein receptor type 2 |
| REEP1 | LRG_713 | receptor accessory protein 1 [Source:HGNC Sy |
| SPAST | LRG_714 | spastin [Source:HGNC Symbol;Acc:HGNC:112 |
| AVP | LRG_715 | arginine vasopressin [Source:HGNC Symbol;A |
| AQP2 | LRG_717 | aquaporin 2 [Source:HGNC Symbol;Acc:HGNC |
| SOX2 | LRG_719 | SRY-box transcription factor 2 [Source:HGNC S |
| IL12RB1 | LRG_72 | interleukin 12 receptor subunit beta 1 [Source |
| PAX6 | LRG_720 | paired box 6 [Source:HGNC Symbol;Acc:HGNC |
| AKT1 | LRG_721 | AKT serine/threonine kinase 1 [Source:HGNC S |
| DPYD | LRG_722 | dihydropyrimidine dehydrogenase [Source:HG |
| GSTP1 | LRG_723 | glutathione S-transferase pi 1 [Source:HGNC S |
| ERBB2 | LRG_724 | erb-b2 receptor tyrosine kinase 2 [Source:HGI |
| MAP2K1 | LRG_725 | mitogen-activated protein kinase kinase 1 [So |
| MTHFR | LRG_726 | methylenetetrahydrofolate reductase [Source |
| TYMP | LRG_727 | thymidine phosphorylase [Source:HGNC Symb |
| IL2RA | LRG_73 | interleukin 2 receptor subunit alpha [Source:H |
| IL36RN | LRG_730 | interleukin 36 receptor antagonist [Source:HG |
| UGT1A1 | LRG_733 | UDP glucuronosyltransferase family 1 membe |
| MTOR | LRG_734 | mechanistic target of rapamycin kinase [Sourc |
| COL5A1 | LRG_737 | collagen type V alpha 1 chain [Source:HGNC S |
| IL7R | LRG_74 | interleukin 7 receptor [Source:HGNC Symbol; |
| ALMS1 | LRG_741 | ALMS1 centrosome and basal body associated |
| BAG3 | LRG_742 | BCL2 associated athanogene 3 [Source:HGNC |
| DOLK | LRG_744 | dolichol kinase [Source:HGNC Symbol;Acc:HGI |
| EMD | LRG_745 | emerin [Source:HGNC Symbol;Acc:HGNC:333 |
| HFE | LRG_748 | homeostatic iron regulator [Source:HGNC Sym |
| LAMP2 | LRG_749 | lysosomal associated membrane protein 2 [Sc |
| IRAK4 | LRG_75 | interleukin 1 receptor associated kinase 4 [Sou |
| MAP2K2 | LRG_750 | mitogen-activated protein kinase kinase 2 [So |
| NPPA | LRG_751 | natriuretic peptide A [Source:HGNC Symbol;A |
| SHOC2 | LRG_753 | SHOC2 leucine rich repeat scaffold protein [Sc |
| SOS1 | LRG_754 | SOS Ras/Rac guanine nucleotide exchange fac |
| TBX20 | LRG_755 | T-box transcription factor 20 [Source:HGNC Sy |
| TRIM63 | LRG_757 | tripartite motif containing 63 [Source:HGNC S |
| MIB1 | LRG_759 | mindbomb E3 ubiquitin protein ligase 1 [Sourc |
| ITGB2 | LRG_76 | integrin subunit beta 2 [Source:HGNC Symbol; |
| CAVIN4 | LRG_760 | caveolae associated protein 4 [Source:HGNC S |
| FMR1 | LRG_762 | fragile X mental retardation 1 [Source:HGNC S |

| | | |
|---|---|---|
| HTT | LRG_763 | huntingtin [Source:HGNC Symbol;Acc:HGNC:4 |
| POLG | LRG_765 | DNA polymerase gamma, catalytic subunit [Sc |
| RYR1 | LRG_766 | ryanodine receptor 1 [Source:HGNC Symbol;A |
| APOA1 | LRG_767 | apolipoprotein A1 [Source:HGNC Symbol;Acc: |
| LYZ | LRG_768 | lysozyme [Source:HGNC Symbol;Acc:HGNC:67 |
| JAK3 | LRG_77 | Janus kinase 3 [Source:HGNC Symbol;Acc:HGN |
| PPM1D | LRG_770 | protein phosphatase, Mg2+/Mn2+ dependent |
| HOXB13 | LRG_771 | homeobox B13 [Source:HGNC Symbol;Acc:HG |
| RAD21 | LRG_772 | RAD21 cohesin complex component [Source:H |
| SMC1A | LRG_773 | structural maintenance of chromosomes 1A [S |
| SMC3 | LRG_774 | structural maintenance of chromosomes 3 [Sc |
| SETD2 | LRG_775 | SET domain containing 2, histone lysine methy |
| MITF | LRG_776 | melanocyte inducing transcription factor [Sou |
| CBS | LRG_777 | cystathionine-beta-synthase [Source:HGNC Sy |
| FBN1 | LRG_778 | fibrillin 1 [Source:HGNC Symbol;Acc:HGNC:36 |
| TGFBR2 | LRG_779 | transforming growth factor beta receptor 2 [S |
| LIG1 | LRG_78 | DNA ligase 1 [Source:HGNC Symbol;Acc:HGNC |
| EPHB2 | LRG_780 | EPH receptor B2 [Source:HGNC Symbol;Acc:H |
| ACTA2 | LRG_781 | actin alpha 2, smooth muscle [Source:HGNC S |
| STAG2 | LRG_782 | stromal antigen 2 [Source:HGNC Symbol;Acc:H |
| TYMS | LRG_783 | thymidylate synthetase [Source:HGNC Symbo |
| XRCC1 | LRG_784 | X-ray repair cross complementing 1 [Source:H |
| POLD1 | LRG_785 | DNA polymerase delta 1, catalytic subunit [So |
| MAPK1 | LRG_786 | mitogen-activated protein kinase 1 [Source:H |
| GFPT1 | LRG_787 | glutamine--fructose-6-phosphate transaminas |
| RRM2B | LRG_788 | ribonucleotide reductase regulatory TP53 indu |
| POLE | LRG_789 | DNA polymerase epsilon, catalytic subunit [So |
| LIG4 | LRG_79 | DNA ligase 4 [Source:HGNC Symbol;Acc:HGNC |
| BCAM | LRG_798 | basal cell adhesion molecule (Lutheran blood |
| SCN1A | LRG_8 | sodium voltage-gated channel alpha subunit 1 |
| LRRC8A | LRG_80 | leucine rich repeat containing 8 VRAC subunit |
| AQP1 | LRG_808 | aquaporin 1 (Colton blood group) [Source:HGI |
| LAMTOR2 | LRG_81 | late endosomal/lysosomal adaptor, MAPK and |
| SEMA7A | LRG_818 | semaphorin 7A (John Milton Hagen blood grou |
| RHAG | LRG_822 | Rh associated glycoprotein [Source:HGNC Sym |
| KLF1 | LRG_825 | Kruppel like factor 1 [Source:HGNC Symbol;Ac |
| CALR | LRG_828 | calreticulin [Source:HGNC Symbol;Acc:HGNC:1 |
| MLPH | LRG_83 | melanophilin [Source:HGNC Symbol;Acc:HGN |
| SLC40A1 | LRG_837 | solute carrier family 40 member 1 [Source:HG |
| ACADM | LRG_838 | acyl-CoA dehydrogenase medium chain [Sourc |
| MTM1 | LRG_839 | myotubularin 1 [Source:HGNC Symbol;Acc:HG |
| MPO | LRG_84 | myeloperoxidase [Source:HGNC Symbol;Acc:H |
| POMT2 | LRG_844 | protein O-mannosyltransferase 2 [Source:HGN |
| PIN1 | LRG_847 | peptidylprolyl cis/trans isomerase, NIMA-inter |
| MRE11 | LRG_85 | MRE11 homolog, double strand break repair r |
| CHKB | LRG_855 | choline kinase beta [Source:HGNC Symbol;Acc |
| MYO5A | LRG_86 | myosin VA [Source:HGNC Symbol;Acc:HGNC:7 |
| VMA21 | LRG_860 | vacuolar ATPase assembly factor VMA21 [Sou |
| TSPYL1 | LRG_862 | TSPY like 1 [Source:HGNC Symbol;Acc:HGNC:1 |
| ATXN1 | LRG_863 | ataxin 1 [Source:HGNC Symbol;Acc:HGNC:105 |

| | | |
|---|---|---|
| FRMD7 | LRG_867 | FERM domain containing 7 [Source:HGNC Sym |
| ANO5 | LRG_868 | anoctamin 5 [Source:HGNC Symbol;Acc:HGNC |
| FLNC | LRG_870 | filamin C [Source:HGNC Symbol;Acc:HGNC:37 |
| ARID1A | LRG_875 | AT-rich interaction domain 1A [Source:HGNC |
| CYCS | LRG_876 | cytochrome c, somatic [Source:HGNC Symbol; |
| GFI1B | LRG_879 | growth factor independent 1B transcriptional |
| NCF2 | LRG_88 | neutrophil cytosolic factor 2 [Source:HGNC Sy |
| EPPK1 | LRG_880 | epiplakin 1 [Source:HGNC Symbol;Acc:HGNC: |
| BLOC1S6 | LRG_883 | biogenesis of lysosomal organelles complex 1 |
| VPS33B | LRG_884 | VPS33B late endosome and lysosome associat |
| SERPINF2 | LRG_885 | serpin family F member 2 [Source:HGNC Symb |
| ACTN1 | LRG_886 | actinin alpha 1 [Source:HGNC Symbol;Acc:HG |
| NPHS2 | LRG_887 | NPHS2 stomatin family member, podocin [Sou |
| NFKBIA | LRG_89 | NFKB inhibitor alpha [Source:HGNC Symbol;A |
| NHEJ1 | LRG_90 | non-homologous end joining factor 1 [Source: |
| PNP | LRG_91 | purine nucleoside phosphorylase [Source:HGN |
| NRAS | LRG_92 | NRAS proto-oncogene, GTPase [Source:HGNC |
| ORAI1 | LRG_93 | ORAI calcium release-activated calcium modu |
| PRF1 | LRG_94 | perforin 1 [Source:HGNC Symbol;Acc:HGNC:9 |
| RAB27A | LRG_96 | RAB27A, member RAS oncogene family [Sourc |
| RAC2 | LRG_97 | Rac family small GTPase 2 [Source:HGNC Sym |
| RAG1 | LRG_98 | recombination activating 1 [Source:HGNC Sym |
| RAG2 | LRG_99 | recombination activating 2 [Source:HGNC Sym |
| CCND1 | LRG_990 | cyclin D1 [Source:HGNC Symbol;Acc:HGNC:15 |
| CDK6 | LRG_991 | cyclin dependent kinase 6 [Source:HGNC Sym |
| ESR1 | LRG_992 | estrogen receptor 1 [Source:HGNC Symbol;Ac |
| FGFR1 | LRG_993 | fibroblast growth factor receptor 1 [Source:H |
| FGFR2 | LRG_994 | fibroblast growth factor receptor 2 [Source:H |
| FUBP1 | LRG_995 | far upstream element binding protein 1 [Sour |
| ERBB3 | LRG_996 | erb-b2 receptor tyrosine kinase 3 [Source:HG |
| ROS1 | LRG_997 | ROS proto-oncogene 1, receptor tyrosine kina |
| CCND3 | LRG_998 | cyclin D3 [Source:HGNC Symbol;Acc:HGNC:15 |
| CIC | LRG_999 | capicua transcriptional repressor [Source:HGN |
| MT-TF | MT | mitochondrially encoded tRNA-Phe (UUU/C) [ |
| MT-RNR1 | MT | mitochondrially encoded 12S rRNA [Source:H |
| MT-TV | MT | mitochondrially encoded tRNA-Val (GUN) [Sou |
| MT-RNR2 | MT | mitochondrially encoded 16S rRNA [Source:H |
| MT-TL1 | MT | mitochondrially encoded tRNA-Leu (UUA/G) 1 |
| MT-ND1 | MT | mitochondrially encoded NADH:ubiquinone o |
| MT-TI | MT | mitochondrially encoded tRNA-Ile (AUU/C) [S |
| MT-TQ | MT | mitochondrially encoded tRNA-Gln (CAA/G) [S |
| MT-TM | MT | mitochondrially encoded tRNA-Met (AUA/G) [ |
| MT-ND2 | MT | mitochondrially encoded NADH:ubiquinone o |
| MT-TW | MT | mitochondrially encoded tRNA-Trp (UGA/G) [S |
| MT-TA | MT | mitochondrially encoded tRNA-Ala (GCN) [Sou |
| MT-TN | MT | mitochondrially encoded tRNA-Asn (AAU/C) [S |
| MT-TC | MT | mitochondrially encoded tRNA-Cys (UGU/C) [S |
| MT-TY | MT | mitochondrially encoded tRNA-Tyr (UAU/C) [S |
| MT-CO1 | MT | mitochondrially encoded cytochrome c oxidas |
| MT-TS1 | MT | mitochondrially encoded tRNA-Ser (UCN) 1 [S |

| | | |
|---|---|---|
| MT-TD | MT | mitochondrially encoded tRNA-Asp (GAU/C) [S |
| MT-CO2 | MT | mitochondrially encoded cytochrome c oxidas |
| MT-TK | MT | mitochondrially encoded tRNA-Lys (AAA/G) [S |
| MT-ATP8 | MT | mitochondrially encoded ATP synthase memb |
| MT-ATP6 | MT | mitochondrially encoded ATP synthase memb |
| MT-CO3 | MT | mitochondrially encoded cytochrome c oxidas |
| MT-TG | MT | mitochondrially encoded tRNA-Gly (GGN) [Sou |
| MT-ND3 | MT | mitochondrially encoded NADH:ubiquinone o |
| MT-TR | MT | mitochondrially encoded tRNA-Arg (CGN) [Sou |
| MT-ND4L | MT | mitochondrially encoded NADH:ubiquinone o |
| MT-ND4 | MT | mitochondrially encoded NADH:ubiquinone o |
| MT-TH | MT | mitochondrially encoded tRNA-His (CAU/C) [S |
| MT-TS2 | MT | mitochondrially encoded tRNA-Ser (AGU/C) 2 |
| MT-TL2 | MT | mitochondrially encoded tRNA-Leu (CUN) 2 [S |
| MT-ND5 | MT | mitochondrially encoded NADH:ubiquinone o |
| MT-ND6 | MT | mitochondrially encoded NADH:ubiquinone o |
| MT-TE | MT | mitochondrially encoded tRNA-Glu (GAA/G) [S |
| MT-CYB | MT | mitochondrially encoded cytochrome b [Sourc |
| MT-TT | MT | mitochondrially encoded tRNA-Thr (ACN) [Sou |
| MT-TP | MT | mitochondrially encoded tRNA-Pro (CCN) [Sou |
| AL954722.1 | X | regucalcin (RGN) pseudogene |
| PLCXD1 | X | phosphatidylinositol specific phospholipase C |
| GTPBP6 | X | GTP binding protein 6 (putative) [Source:HGN |
| LINC00685 | X | long intergenic non-protein coding RNA 685 [S |
| PPP2R3B | X | protein phosphatase 2 regulatory subunit B''b |
| AL732314.6 | X | novel transcript, antisense to PPP2R3B |
| Metazoa_SRP | X | Metazoan signal recognition particle RNA [Sou |
| AL732314.4 | X | novel transcript |
| FABP5P13 | X | fatty acid binding protein 5 pseudogene 13 [S |
| KRT18P53 | X | keratin 18 pseudogene 53 [Source:HGNC Sym |
| SHOX | X | short stature homeobox [Source:HGNC Symbo |
| AL672277.1 | X | novel transcript |
| RPL14P5 | X | ribosomal protein L14 pseudogene 5 [Source: |
| CRLF2 | X | cytokine receptor like factor 2 [Source:HGNC S |
| CSF2RA | X | colony stimulating factor 2 receptor alpha sub |
| MIR3690 | X | microRNA 3690 [Source:HGNC Symbol;Acc:HG |
| RNA5SP498 | X | RNA, 5S ribosomal pseudogene 498 [Source:H |
| IL3RA | X | interleukin 3 receptor subunit alpha [Source:H |
| SLC25A6 | X | solute carrier family 25 member 6 [Source:HG |
| LINC00106 | X | long intergenic non-protein coding RNA 106 [S |
| ASMTL-AS1 | X | ASMTL antisense RNA 1 [Source:HGNC Symbo |
| ASMTL | X | acetylserotonin O-methyltransferase like [Sou |
| P2RY8 | X | P2Y receptor family member 8 [Source:HGNC |
| AKAP17A | X | A-kinase anchoring protein 17A [Source:HGNC |
| ASMT | X | acetylserotonin O-methyltransferase [Source: |
| AL683807.1 | X | novel transcript |
| AL683807.2 | X | novel transcript |
| DHRSX | X | dehydrogenase/reductase X-linked [Source:HG |
| DHRSX-IT1 | X | DHRSX intronic transcript 1 [Source:HGNC Syn |
| ZBED1 | X | zinc finger BED-type containing 1 [Source:HGN |

| | | |
|---|---|---|
| MIR6089 | X | microRNA 6089 [Source:HGNC Symbol;Acc:HG |
| CD99P1 | X | CD99 molecule pseudogene 1 [Source:HGNC S |
| LINC00102 | X | long intergenic non-protein coding RNA 102 [S |
| CD99 | X | CD99 molecule (Xg blood group) [Source:HGN |
| XG | X | Xg glycoprotein (Xg blood group) [Source:HGN |
| GYG2 | X | glycogenin 2 [Source:HGNC Symbol;Acc:HGNC |
| GYG2-AS1 | X | GYG2 antisense RNA 1 [Source:HGNC Symbol; |
| ARSD | X | arylsulfatase D [Source:HGNC Symbol;Acc:HGI |
| ARSD-AS1 | X | ARSD antisense RNA 1 [Source:HGNC Symbol; |
| ARSE | X | arylsulfatase E [Source:HGNC Symbol;Acc:HGI |
| ARSH | X | arylsulfatase family member H [Source:HGNC |
| ARSF | X | arylsulfatase F [Source:HGNC Symbol;Acc:HGI |
| RN7SL578P | X | RNA, 7SL, cytoplasmic 578, pseudogene [Sour |
| AC112653.1 | X | ribosomal protein S24 (RPS24) pseudogene |
| LINC01546 | X | long intergenic non-protein coding RNA 1546 |
| MXRA5 | X | matrix remodeling associated 5 [Source:HGNC |
| ASS1P4 | X | argininosuccinate synthetase 1 pseudogene 4 |
| SNORA48B | X | small nucleolar RNA, H/ACA box 48B [Source:I |
| RNU6-114P | X | RNA, U6 small nuclear 114, pseudogene [Sour |
| PRKX | X | protein kinase X-linked [Source:HGNC Symbol |
| RNU6-146P | X | RNA, U6 small nuclear 146, pseudogene [Sour |
| Y_RNA | X | Y RNA [Source:RFAM;Acc:RF00019] |
| PRKX-AS1 | X | PRKX antisense RNA 1 [Source:HGNC Symbol;/ |
| AC007685.1 | X | ribosomal protein S27a (RPS27A) pseudogene |
| BX890604.2 | X | novel transcript |
| BX842570.1 | X | novel transcript |
| BX890604.1 | X | ribosomal protein L24 (RPL24) pseudogene |
| AC074035.1 | X | novel transcript |
| MTND6P12 | X | MT-ND6 pseudogene 12 [Source:HGNC Symbo |
| MTCYBP12 | X | MT-CYB pseudogene 12 [Source:HGNC Symbo |
| AC112656.1 | X | hydroxyacyl-Coenzyme A dehydrogenase\/3-k |
| AC095353.1 | X | NULL |
| AC110995.1 | X | novel transcript |
| NLGN4X | X | neuroligin 4 X-linked [Source:HGNC Symbol;A |
| AC079173.1 | X | NULL |
| MIR4770 | X | microRNA 4770 [Source:HGNC Symbol;Acc:HG |
| AC078991.1 | X | ribosomal protein S5 pseudogene |
| VCX3A | X | variable charge X-linked 3A [Source:HGNC Syn |
| PUDP | X | pseudouridine 5'-phosphatase [Source:HGNC |
| RPS27AP17 | X | ribosomal protein S27a pseudogene 17 [Sourc |
| AC073583.1 | X | ribosomal protein, large, P1 pseudogene |
| STS | X | steroid sulfatase [Source:HGNC Symbol;Acc:H |
| MIR4767 | X | microRNA 4767 [Source:HGNC Symbol;Acc:HG |
| AC120338.1 | X | TEC |
| VCX | X | variable charge X-linked [Source:HGNC Symbo |
| AC005296.1 | X | chromosome X open reading frame 40A (CXor |
| PNPLA4 | X | patatin like phospholipase domain containing |
| AC079142.1 | X | novel transcript |
| MIR651 | X | microRNA 651 [Source:HGNC Symbol;Acc:HGI |
| VCX2 | X | variable charge X-linked 2 [Source:HGNC Symb |

| | | |
|---|---|---|
| VCX3B | X | variable charge X-linked 3B [Source:HGNC Syn |
| AC006062.1 | X | novel transcript |
| ANOS1 | X | anosmin 1 [Source:HGNC Symbol;Acc:HGNC:6 |
| DRAXINP1 | X | dorsal inhibitory axon guidance protein pseud |
| FAM9A | X | family with sequence similarity 9 member A [S |
| AC003685.2 | X | novel pseudogene |
| AC003685.1 | X | novel transcript |
| AC074281.2 | X | novel transcript |
| AC074281.1 | X | nucleolar and coiled-body phosphoprotein 1 ( |
| FAM9B | X | family with sequence similarity 9 member B [S |
| RNA5SP499 | X | RNA, 5S ribosomal pseudogene 499 [Source:H |
| BRK1P1 | X | BRICK1 pseudogene 1 [Source:HGNC Symbol;A |
| AC003684.1 | X | novel transcript |
| AC073488.1 | X | chromodomain protein, Y chromosome-like (C |
| AC073488.7 | X | chromodomain protein, Y chromosome-like (C |
| AC073488.9 | X | chromodomain protein, Y chromosome-like (C |
| AC073488.8 | X | chromodomain protein, Y chromosome-like (C |
| AC073488.10 | X | chromodomain protein, Y chromosome-like (C |
| AC073488.5 | X | chromodomain protein, Y chromosome-like (C |
| AC073488.11 | X | chromodomain protein, Y chromosome-like (C |
| AC073488.4 | X | chromodomain protein, Y chromosome-like (C |
| AC073488.3 | X | chromodomain protein, Y chromosome-like (C |
| AC073488.2 | X | chromodomain protein, Y chromosome-like (C |
| AC073488.6 | X | chromodomain protein, Y chromosome-like (C |
| TBL1X | X | transducin beta like 1 X-linked [Source:HGNC S |
| GPR143 | X | G protein-coupled receptor 143 [Source:HGN( |
| SHROOM2 | X | shroom family member 2 [Source:HGNC Symb |
| EIF5P1 | X | eukaryotic translation initiation factor 5 pseuc |
| HMGN1P33 | X | high mobility group nucleosome binding doma |
| Y_RNA | X | Y RNA [Source:RFAM;Acc:RF00019] |
| CLDN34 | X | claudin 34 [Source:HGNC Symbol;Acc:HGNC:5 |
| WWC3 | X | WWC family member 3 [Source:HGNC Symbo |
| WWC3-AS1 | X | WWC3 antisense RNA 1 [Source:HGNC Symbo |
| Y_RNA | X | Y RNA [Source:RFAM;Acc:RF00019] |
| CLCN4 | X | chloride voltage-gated channel 4 [Source:HGN |
| AC003666.1 | X | novel transcript |
| MID1 | X | midline 1 [Source:HGNC Symbol;Acc:HGNC:70 |
| RNU6-800P | X | RNA, U6 small nuclear 800, pseudogene [Sour |
| AC073529.1 | X | novel transcript |
| HCCS | X | holocytochrome c synthase [Source:HGNC Syr |
| Y_RNA | X | Y RNA [Source:RFAM;Acc:RF00019] |
| ARHGAP6 | X | Rho GTPase activating protein 6 [Source:HGN( |
| AC003657.1 | X | NULL |
| AC004467.1 | X | family with sequence similarity 9, member C ( |
| AMELX | X | amelogenin X-linked [Source:HGNC Symbol;Ac |
| MIR548AX | X | microRNA 548ax [Source:HGNC Symbol;Acc:H |
| AC002366.1 | X | glutamic-oxaloacetic transaminase 2, mitocho |
| AC004554.2 | X | novel transcript |
| AC004554.1 | X | LIM domain kinase 2 (LIMK2) pseudogene |
| MSL3 | X | MSL complex subunit 3 [Source:HGNC Symbol |

| | | |
|---|---|---|
| FRMPD4 | X | FERM and PDZ domain containing 4 [Source:H |
| Y_RNA | X | Y RNA [Source:RFAM;Acc:RF00019] |
| Y_RNA | X | Y RNA [Source:RFAM;Acc:RF00019] |
| FRMPD4-AS1 | X | FRMPD4 antisense RNA 1 [Source:HGNC Symb |
| RPL17P49 | X | ribosomal protein L17 pseudogene 49 [Source |
| AC119619.1 | X | developmental pluripotency associated 5 (DPI |
| RN7SKP290 | X | RN7SK pseudogene 290 [Source:HGNC Symbo |
| PRPS2 | X | phosphoribosyl pyrophosphate synthetase 2 [ |
| PSMA6P2 | X | proteasome subunit alpha 6 pseudogene 2 [Sc |
| MRPL35P4 | X | mitochondrial ribosomal protein L35 pseudog |
| TLR7 | X | toll like receptor 7 [Source:HGNC Symbol;Acc: |
| TLR8-AS1 | X | TLR8 antisense RNA 1 [Source:HGNC Symbol;/ |
| TLR8 | X | toll like receptor 8 [Source:HGNC Symbol;Acc: |
| TMSB4X | X | thymosin beta 4 X-linked [Source:HGNC Symb |
| FAM9C | X | family with sequence similarity 9 member C [S |
| AC079171.1 | X | novel transcript |
| LINC02154 | X | long intergenic non-protein coding RNA 2154 |
| AC004674.1 | X | novel transcript |
| ATXN3L | X | ataxin 3 like [Source:HGNC Symbol;Acc:HGNC |
| AC004674.2 | X | novel pseudogene |
| LINC01203 | X | long intergenic non-protein coding RNA 1203 |
| GPX1P1 | X | glutathione peroxidase pseudogene 1 [Source |
| EGFL6 | X | EGF like domain multiple 6 [Source:HGNC Sym |
| RPL30P15 | X | ribosomal protein L30 pseudogene 15 [Source |
| MIR6086 | X | microRNA 6086 [Source:HGNC Symbol;Acc:HG |
| RN7SKP20 | X | RN7SK pseudogene 20 [Source:HGNC Symbol; |
| TCEANC | X | transcription elongation factor A N-terminal a |
| RAB9A | X | RAB9A, member RAS oncogene family [Source |
| TRAPPC2 | X | trafficking protein particle complex 2 [Source: |
| OFD1 | X | OFD1 centriole and centriolar satellite protein |
| GPM6B | X | glycoprotein M6B [Source:HGNC Symbol;Acc:I |
| AC003035.2 | X | novel transcript |
| AC003035.1 | X | novel transcript |
| GEMIN8 | X | gem nuclear organelle associated protein 8 [S |
| AC002524.1 | X | ubiquitin-conjugating enzyme E2E 3 (UBC4\/5 |
| GLRA2 | X | glycine receptor alpha 2 [Source:HGNC Symbo |
| NPM1P9 | X | nucleophosmin 1 pseudogene 9 [Source:HGN( |
| FANCB | X | FA complementation group B [Source:HGNC S |
| MOSPD2 | X | motile sperm domain containing 2 [Source:HG |
| TPT1P14 | X | tumor protein, translationally-controlled 1 pse |
| AC112498.1 | X | ribosomal protein L35a (RPL35A) pseudogene |
| ASB9 | X | ankyrin repeat and SOCS box containing 9 [So |
| ASB11 | X | ankyrin repeat and SOCS box containing 11 [S |
| PIGA | X | phosphatidylinositol glycan anchor biosynthes |
| VEGFD | X | vascular endothelial growth factor D [Source:I |
| PIR | X | pirin [Source:HGNC Symbol;Acc:HGNC:30048] |
| BMX | X | BMX non-receptor tyrosine kinase [Source:HG |
| ACE2 | X | angiotensin I converting enzyme 2 [Source:HG |
| AC097625.2 | X | novel protein |
| AC097625.1 | X | novel transcript |

| | | |
|---|---|---|
| CLTRN | X | collectrin, amino acid transport regulator [Sou |
| AC003669.1 | X | novel pseudogene |
| CA5BP1 | X | carbonic anhydrase 5B pseudogene 1 [Source: |
| CA5B | X | carbonic anhydrase 5B [Source:HGNC Symbol; |
| AC112497.2 | X | NULL |
| AC112497.1 | X | NULL |
| INE2 | X | inactivation escape 2 [Source:HGNC Symbol;A |
| ZRSR2 | X | zinc finger CCCH-type, RNA binding motif and |
| AP1S2 | X | adaptor related protein complex 1 subunit sig |
| RNU5F-7P | X | RNA, U5F small nuclear 7, pseudogene [Source |
| SETP15 | X | SET pseudogene 15 [Source:HGNC Symbol;Acc |
| GRPR | X | gastrin releasing peptide receptor [Source:HG |
| AC078993.1 | X | novel transcript, antisense to MAGEB17 |
| MAGEB17 | X | MAGE family member B17 [Source:HGNC Sym |
| RPL6P30 | X | ribosomal protein L6 pseudogene 30 [Source: |
| AC073909.1 | X | heat shock protein pseudogene |
| RN7SL658P | X | RNA, 7SL, cytoplasmic 658, pseudogene [Sour |
| CTPS2 | X | CTP synthase 2 [Source:HGNC Symbol;Acc:HG |
| MIR548AM | X | microRNA 548am [Source:HGNC Symbol;Acc:I |
| S100G | X | S100 calcium binding protein G [Source:HGNC |
| SYAP1 | X | synapse associated protein 1 [Source:HGNC Sy |
| RNU7-56P | X | RNA, U7 small nuclear 56 pseudogene [Source |
| TXLNG | X | taxilin gamma [Source:HGNC Symbol;Acc:HGN |
| RPL12P49 | X | ribosomal protein L12 pseudogene 49 [Source |
| RBBP7 | X | RB binding protein 7, chromatin remodeling fa |
| RNU4-6P | X | RNA, U4 small nuclear 6, pseudogene [Source |
| AL929302.1 | X | novel pseudogene |
| Y_RNA | X | Y RNA [Source:RFAM;Acc:RF00019] |
| REPS2 | X | RALBP1 associated Eps domain containing 2 [S |
| AL732371.1 | X | NULL |
| CBX1P2 | X | chromobox 1 pseudogene 2 [Source:HGNC Sy |
| CBX1P4 | X | chromobox 1 pseudogene 4 [Source:HGNC Sy |
| AL845433.1 | X | NULL |
| NHS | X | NHS actin remodeling regulator [Source:HGNC |
| MIR4768 | X | microRNA 4768 [Source:HGNC Symbol;Acc:HG |
| Z93022.1 | X | novel transcript |
| NHS-AS1 | X | NHS antisense RNA 1 [Source:HGNC Symbol;A |
| Z93242.1 | X | calcium binding protein P22 (CHP) pseudogen |
| SCML1 | X | Scm polycomb group protein like 1 [Source:HG |
| Z93242.2 | X | novel pseudogene |
| RAI2 | X | retinoic acid induced 2 [Source:HGNC Symbol; |
| Z93929.2 | X | zinc finger protein 831 (ZNF831) pseudogene |
| Z93929.1 | X | Mdm4, transformed 3T3 cell double minute 4, |
| LINC01456 | X | long intergenic non-protein coding RNA 1456 |
| BEND2 | X | BEN domain containing 2 [Source:HGNC Symb |
| SCML2 | X | Scm polycomb group protein like 2 [Source:HG |
| TMSB10P2 | X | thymosin beta 10 pseudogene 2 [Source:HGN |
| Y_RNA | X | Y RNA [Source:RFAM;Acc:RF00019] |
| CDKL5 | X | cyclin dependent kinase like 5 [Source:HGNC S |
| GJA6P | X | gap junction protein alpha 6, pseudogene [Sou |

| | | |
|---|---|---|
| RS1 | X | retinoschisin 1 [Source:HGNC Symbol;Acc:HGI |
| PPEF1 | X | protein phosphatase with EF-hand domain 1 [ |
| PPEF1-AS1 | X | PPEF1 antisense RNA 1 [Source:HGNC Symbol |
| AL096700.1 | X | pallidin homolog (mouse) (PLDN) pseudogene |
| HAUS1P2 | X | HAUS augmin like complex subunit 1 pseudog |
| PHKA2-AS1 | X | PHKA2 antisense RNA 1 [Source:HGNC Symbo |
| PHKA2 | X | phosphorylase kinase regulatory subunit alph |
| RN7SL48P | X | RNA, 7SL, cytoplasmic 48, pseudogene [Sourc |
| AL732509.1 | X | novel transcript, antisense to ADGRG2and PH |
| ADGRG2 | X | adhesion G protein-coupled receptor G2 [Sou |
| PDHA1 | X | pyruvate dehydrogenase E1 alpha 1 subunit [S |
| MAP3K15 | X | mitogen-activated protein kinase kinase kinas |
| Y_RNA | X | Y RNA [Source:RFAM;Acc:RF00019] |
| EIF5P2 | X | eukaryotic translation initiation factor 5 pseu |
| SH3KBP1 | X | SH3 domain containing kinase binding protein |
| BCLAF3 | X | BCLAF1 and THRAP3 family member 3 [Source |
| Y_RNA | X | Y RNA [Source:RFAM;Acc:RF00019] |
| MAP7D2 | X | MAP7 domain containing 2 [Source:HGNC Syn |
| MIR23C | X | microRNA 23c [Source:HGNC Symbol;Acc:HGN |
| EIF1AX | X | eukaryotic translation initiation factor 1A X-lir |
| AL732366.2 | X | NULL |
| AL732366.1 | X | NULL |
| EIF1AX-AS1 | X | EIF1AX antisense RNA 1 [Source:HGNC Symbo |
| RPS6KA3 | X | ribosomal protein S6 kinase A3 [Source:HGNC |
| AL928596.1 | X | novel transcript |
| AL807740.1 | X | novel transcript |
| RN7SKP183 | X | RN7SK pseudogene 183 [Source:HGNC Symbo |
| AL807742.1 | X | novel transcript |
| BX088723.1 | X | novel transcript |
| RNU6-133P | X | RNA, U6 small nuclear 133, pseudogene [Sour |
| CNKSR2 | X | connector enhancer of kinase suppressor of R |
| RARRES2P3 | X | retinoic acid receptor responder 2 pseudogen |
| KLHL34 | X | kelch like family member 34 [Source:HGNC Sy |
| SMPX | X | small muscle protein X-linked [Source:HGNC S |
| MBTPS2 | X | membrane bound transcription factor peptida |
| U73479.1 | X | LSM1 homolog, U6 small nuclear RNA associa |
| YY2 | X | YY2 transcription factor [Source:HGNC Symbo |
| Y_RNA | X | Y RNA [Source:RFAM;Acc:RF00019] |
| U73023.1 | X | cytohesin 1 (CYTH1) pseudogene |
| SMS | X | spermine synthase [Source:HGNC Symbol;Acc |
| PHEX | X | phosphate regulating endopeptidase homolog |
| PHEX-AS1 | X | PHEX antisense RNA 1 [Source:HGNC Symbol; |
| PTCHD1-AS | X | PTCHD1 antisense RNA (head to head) [Sourc |
| CBLL2 | X | Cbl proto-oncogene like 2 [Source:HGNC Syml |
| AC092832.2 | X | novel transcript |
| METTL15P3 | X | methyltransferase like 15 pseudogene 3 [Sour |
| RNU6-266P | X | RNA, U6 small nuclear 266, pseudogene [Sour |
| AC092832.1 | X | ribosomal protein L26 (RPL26) pseudogene |
| AC004470.1 | X | novel transcript |
| DDX53 | X | DEAD-box helicase 53 [Source:HGNC Symbol; |

| | | |
|---|---|---|
| FAM3C2 | X | family with sequence similarity 3 member C2 |
| PDCL2P1 | X | PDCL2 pseudogene 1 [Source:HGNC Symbol;A |
| PTCHD1 | X | patched domain containing 1 [Source:HGNC S |
| AC073910.1 | X | novel pseudogene |
| PRDX4 | X | peroxiredoxin 4 [Source:HGNC Symbol;Acc:HG |
| ACOT9 | X | acyl-CoA thioesterase 9 [Source:HGNC Symbo |
| AC131011.1 | X | novel transcript |
| Y_RNA | X | Y RNA [Source:RFAM;Acc:RF00019] |
| SAT1 | X | spermidine/spermine N1-acetyltransferase 1 [ |
| APOO | X | apolipoprotein O [Source:HGNC Symbol;Acc:H |
| RPL9P7 | X | ribosomal protein L9 pseudogene 7 [Source:H |
| CXorf58 | X | chromosome X open reading frame 58 [Sourc |
| KLHL15 | X | kelch like family member 15 [Source:HGNC Sy |
| EIF2S3 | X | eukaryotic translation initiation factor 2 subur |
| AC079169.1 | X | NULL |
| ZFX-AS1 | X | ZFX antisense RNA 1 [Source:HGNC Symbol;Ac |
| ZFX | X | zinc finger protein X-linked [Source:HGNC Sym |
| SUPT20HL2 | X | SUPT20H like 2 [Source:HGNC Symbol;Acc:HG |
| SUPT20HL1 | X | SUPT20H like 1 [Source:HGNC Symbol;Acc:HG |
| AC004552.1 | X | ribosomal protein S26 (RPS26) pseudogene |
| PDK3 | X | pyruvate dehydrogenase kinase 3 [Source:HG |
| SNRPEP9 | X | small nuclear ribonucleoprotein polypeptide E |
| PCYT1B | X | phosphate cytidylyltransferase 1, choline, bet |
| AC079168.1 | X | histone 1 H2bb (HIST1H2BB) pseudogene |
| PCYT1B-AS1 | X | PCYT1B antisense RNA 1 [Source:HGNC Symbo |
| AC079168.2 | X | ribosomal protein L23 (RPL23) pseudogene |
| POLA1 | X | DNA polymerase alpha 1, catalytic subunit [So |
| SCARNA23 | X | small Cajal body-specific RNA 23 [Source:HGN |
| EEF1B2P3 | X | eukaryotic translation elongation factor 1 bet |
| ARX | X | aristaless related homeobox [Source:HGNC Sy |
| AC002504.1 | X | platelet-activating factor acetylhydrolase, isof |
| RN7SL91P | X | RNA, 7SL, cytoplasmic 91, pseudogene [Sourc |
| AC112778.1 | X | methyltransferase-like 1 pseudogene |
| RPP40P1 | X | ribonuclease P/MRP subunit p40 pseudogene |
| AC097263.1 | X | RAN binding protein 1 pseudogene |
| AC121342.1 | X | novel transcript |
| MAGEB18 | X | MAGE family member B18 [Source:HGNC Sym |
| AC129850.1 | X | pseudogene similar to melanoma antigen fam |
| MAGEB6B | X | MAGE family member B6B [Source:HGNC Sym |
| AC005297.1 | X | pseudogene similar to part of melanoma antig |
| MAGEB6 | X | MAGE family member B6 [Source:HGNC Symb |
| AC005297.2 | X | pseudogene similar to part of melanoma antig |
| MAGEB5 | X | MAGE family member B5 [Source:HGNC Symb |
| AC005297.3 | X | pseudogene similar to part of melanoma antig |
| AC073614.1 | X | MAWD binding protein pseudogene |
| AC079178.2 | X | formin 2 (FMN2) pseudogene |
| AC079178.1 | X | VENT-like homeobox 2 pseudogene 1 |
| VENTXP1 | X | VENT homeobox pseudogene 1 [Source:HGNC |
| AC005345.1 | X | pseudogene similar to part of adaptor-related |
| RPL7P58 | X | ribosomal protein L7 pseudogene 58 [Source: |

| | | |
|---|---|---|
| HMGA1P1 | X | high mobility group AT-hook 1 pseudogene 1 |
| RNU1-142P | X | RNA, U1 small nuclear 142, pseudogene [Sour |
| AC112493.1 | X | novel transcript |
| AC107419.1 | X | novel transcript |
| AC097262.1 | X | novel pseudogene |
| Y_RNA | X | Y RNA [Source:RFAM;Acc:RF00019] |
| PTP4A1P5 | X | PTP4A1 pseudogene 5 [Source:HGNC Symbol; |
| PPP4R3C | X | protein phosphatase 4 regulatory subunit 3C [ |
| AC006210.1 | X | pseudogene similar to part of radixin (RDX) |
| AC006210.2 | X | pseudogene similar to part of radixin (RDX) |
| DCAF8L2 | X | DDB1 and CUL4 associated factor 8 like 2 [Sou |
| MAGEB10 | X | MAGE family member B10 [Source:HGNC Sym |
| AC107613.1 | X | MAGE family pseudogene |
| AC112492.2 | X | MAGE family pseudogene |
| AC112492.4 | X | pseudogene similar to part of adipose differer |
| AC112492.3 | X | MAGE family pseudogene |
| AC112492.5 | X | novel pseudogene |
| AC112492.1 | X | pseudogene similar to part of origin recognitio |
| DCAF8L1 | X | DDB1 and CUL4 associated factor 8 like 1 [Sou |
| MIR6134 | X | microRNA 6134 [Source:HGNC Symbol;Acc:HG |
| AC112495.1 | X | novel transcript |
| IL1RAPL1 | X | interleukin 1 receptor accessory protein like 1 |
| AC002367.1 | X | H3 histone, family 3B (H3.3B) (H3F3B) pseudo |
| AC121343.1 | X | NULL |
| AC003659.1 | X | novel transcript |
| RNA5SP500 | X | RNA, 5S ribosomal pseudogene 500 [Source:H |
| AL031575.1 | X | pseudogene similar to part of transcription fa |
| MIR4666B | X | microRNA 4666b [Source:HGNC Symbol;Acc:H |
| PIGFP3 | X | phosphatidylinositol glycan anchor biosynthes |
| AC005185.1 | X | melanoma antigen family A, 10 (MAGEA10) ps |
| MAGEB2 | X | MAGE family member B2 [Source:HGNC Symb |
| MAGEB3 | X | MAGE family member B3 [Source:HGNC Symb |
| MAGEB4 | X | MAGE family member B4 [Source:HGNC Symb |
| MAGEB1 | X | MAGE family member B1 [Source:HGNC Symb |
| PLCE1P1 | X | phospholipase C epsilon 1 pseudogene 1 [Sou |
| NR0B1 | X | nuclear receptor subfamily 0 group B member |
| CXorf21 | X | chromosome X open reading frame 21 [Sourc |
| CKS1BP6 | X | CDC28 protein kinase regulatory subunit 1B ps |
| FTLP2 | X | ferritin light chain pseudogene 2 [Source:HGN |
| GK | X | glycerol kinase [Source:HGNC Symbol;Acc:HGI |
| GK-IT1 | X | GK intronic transcript 1 [Source:HGNC Symbol |
| AC112496.1 | X | novel transcript, sense intronic to GK |
| GK-AS1 | X | GK antisense RNA 1 [Source:HGNC Symbol;Ac |
| AC108359.1 | X | myoneurin (MYNN) pseudogene |
| TAB3 | X | TGF-beta activated kinase 1 (MAP3K7) binding |
| TAB3-AS1 | X | TAB3 antisense RNA 1 [Source:HGNC Symbol;/ |
| TAB3-AS2 | X | TAB3 antisense RNA 2 [Source:HGNC Symbol;/ |
| FTHL17 | X | ferritin heavy chain like 17 [Source:HGNC Sym |
| DMD | X | dystrophin [Source:HGNC Symbol;Acc:HGNC:2 |
| RNU6-894P | X | RNA, U6 small nuclear 894, pseudogene [Sour |

| | | |
|---|---|---|
| AC079177.1 | X | novel transcript, antisense to DMD |
| RNA5SP501 | X | RNA, 5S ribosomal pseudogene 501 [Source:H |
| NPM1P8 | X | nucleophosmin 1 pseudogene 8 [Source:HGN( |
| MIR3915 | X | microRNA 3915 [Source:HGNC Symbol;Acc:HG |
| MIR548F5 | X | microRNA 548f-5 [Source:HGNC Symbol;Acc:H |
| DMD-AS3 | X | DMD antisense RNA 3 [Source:HGNC Symbol;/ |
| TBCAP1 | X | tubulin folding cofactor A pseudogene 1 [Sour |
| AL050305.1 | X | lysosomal-associated membrane protein 1 ps( |
| AL591378.1 | X | novel transcript |
| AL591501.1 | X | novel transcript |
| FAM47A | X | family with sequence similarity 47 member A |
| FTH1P14 | X | ferritin heavy chain 1 pseudogene 14 [Source: |
| AL592043.1 | X | novel transcript |
| AL592043.2 | X | serine/arginine-rich splicing factor 8 (SRSF8) p |
| AL592504.1 | X | Opa interacting protein 5 (OIP5) pseudogene |
| TMEM47 | X | transmembrane protein 47 [Source:HGNC Syn |
| AL606662.1 | X | pseudogene similar to part of poly(A) polymer |
| AL596245.1 | X | zinc finger protein 286B (ZNF286B) pseudoger |
| FAM47B | X | family with sequence similarity 47 member B |
| AL596268.1 | X | MAGE family pseudogene |
| RNU6-1087P | X | RNA, U6 small nuclear 1087, pseudogene [Sou |
| SIAH1P1 | X | siah E3 ubiquitin protein ligase 1 pseudogene |
| AL161722.1 | X | melanoma antigen pseudogene |
| AL161722.3 | X | CDC42 binding protein kinase beta (DMPK-like |
| AL161722.2 | X | nuclear transcription factor Y, gamma (NFYC) |
| MAGEB16 | X | MAGE family member B16 [Source:HGNC Sym |
| AL590065.1 | X | CDC42 binding protein kinase beta (DMPK-like |
| CFAP47 | X | cilia and flagella associated protein 47 [Source |
| RNU6-641P | X | RNA, U6 small nuclear 641, pseudogene [Sour |
| HMGB1P16 | X | high mobility group box 1 pseudogene 16 [Sou |
| AL606516.1 | X | novel transcript, antisense to CXorf30 |
| RPS15AP40 | X | ribosomal protein S15a pseudogene 40 [Sourc |
| AL592156.2 | X | novel transcript |
| AL592156.1 | X | H2A histone family pseudogene |
| AC006924.1 | X | nudix (nucleoside diphosphate linked moiety ) |
| BX842568.4 | X | transmembrane 4 superfamily member 10 (TN |
| BX842568.2 | X | transcription factor Dp-1 (TFDP1) pseudogene |
| BX842568.3 | X | novel pseudogene |
| FAM47C | X | family with sequence similarity 47 member C |
| FTHL18 | X | ferritin heavy chain like 18 [Source:HGNC Sym |
| BX842568.1 | X | ferritin, heavy polypeptide-like 17 (FTHL17) ps |
| MOB1AP2 | X | MOB kinase activator 1A pseudogene 2 [Sourc |
| PRRG1 | X | proline rich and Gla domain 1 [Source:HGNC S |
| AF241726.2 | X | novel proline rich Gla (G-carboxyglutamic acid |
| AC245096.1 | X | ferritin, heavy polypeptide-like 17 (FTHL17) ps |
| FTH1P19 | X | ferritin heavy chain 1 pseudogene 19 [Source: |
| FTH1P27 | X | ferritin heavy chain 1 pseudogene 27 [Source: |
| FAM47DP | X | family with sequence similarity 47 member D, |
| LANCL3 | X | LanC like 3 [Source:HGNC Symbol;Acc:HGNC:2 |
| XK | X | X-linked Kx blood group [Source:HGNC Symbo |

| | | |
|---|---|---|
| AC233297.1 | X | novel histone family pseudogene |
| CYBB | X | cytochrome b-245 beta chain [Source:HGNC S |
| DYNLT3 | X | dynein light chain Tctex-type 3 [Source:HGNC |
| RNU6-49P | X | RNA, U6 small nuclear 49, pseudogene [Sourc |
| AL121578.3 | X | novel transcript |
| AL121578.1 | X | sterol-C4-methyl oxidase-like (SC4MOL) pseuc |
| HYPM | X | huntingtin interacting protein M [Source:HGN |
| AL121578.2 | X | novel histone family protein |
| SYTL5 | X | synaptotagmin like 5 [Source:HGNC Symbol;A |
| MIR548AJ2 | X | microRNA 548aj-2 [Source:HGNC Symbol;Acc: |
| SRPX | X | sushi repeat containing protein X-linked [Sour |
| AL606748.1 | X | novel transcript |
| RPGR | X | retinitis pigmentosa GTPase regulator [Source |
| AL606748.2 | X | NULL |
| OTC | X | ornithine carbamoyltransferase [Source:HGN( |
| TDGF1P1 | X | teratocarcinoma-derived growth factor 1 pseu |
| AF241726.1 | X | Bcl2-associating athanogene (BAG1) pseudoge |
| FTLP16 | X | ferritin light chain pseudogene 16 [Source:HG |
| UBTFL11 | X | UBTF like 11 (pseudogene) [Source:HGNC Sym |
| TSPAN7 | X | tetraspanin 7 [Source:HGNC Symbol;Acc:HGN( |
| RNU7-7P | X | RNA, U7 small nuclear 7 pseudogene [Source: |
| AF241728.1 | X | novel transcript |
| AF241728.2 | X | novel transcript |
| MID1IP1 | X | MID1 interacting protein 1 [Source:HGNC Sym |
| MID1IP1-AS1 | X | MID1IP1 antisense RNA 1 [Source:HGNC Symb |
| RNU6-591P | X | RNA, U6 small nuclear 591, pseudogene [Sour |
| AC108879.1 | X | novel transcript |
| LINC01281 | X | long intergenic non-protein coding RNA 1281 |
| LINC01282 | X | long intergenic non-protein coding RNA 1282 |
| LINC01283 | X | long intergenic non-protein coding RNA 1283 |
| MIR3937 | X | microRNA 3937 [Source:HGNC Symbol;Acc:HG |
| RN7SL732P | X | RNA, 7SL, cytoplasmic 732, pseudogene [Sour |
| GAPDHP1 | X | glyceraldehyde-3-phosphate dehydrogenase p |
| AC091810.1 | X | oocyte expressed protein (OOEP) pseudogene |
| AL592164.1 | X | novel transcript |
| MIR1587 | X | microRNA 1587 [Source:HGNC Symbol;Acc:HG |
| RPS11P7 | X | ribosomal protein S11 pseudogene 7 [Source: |
| AL592164.2 | X | Novel transcript |
| AC092198.1 | X | novel transcript |
| BCOR | X | BCL6 corepressor [Source:HGNC Symbol;Acc:H |
| AC091806.1 | X | novel transcript |
| AC091807.1 | X | IMP (inosine monophosphate) dehydrogenase |
| Y_RNA | X | Y RNA [Source:RFAM;Acc:RF00019] |
| AC092473.1 | X | similar to part of ribosomal protein S20 (RPS2 |
| RNU7-164P | X | RNA, U7 small nuclear 164 pseudogene [Sourc |
| ATP6AP2 | X | ATPase H+ transporting accessory protein 2 [S |
| AC092473.2 | X | novel transcript |
| MPC1L | X | mitochondrial pyruvate carrier 1 like [Source:I |
| CXorf38 | X | chromosome X open reading frame 38 [Sourc |
| MED14 | X | mediator complex subunit 14 [Source:HGNC S |

| | | |
|---|---|---|
| MED14OS | X | MED14 opposite strand [Source:HGNC Symbo |
| DPRXP6 | X | divergent-paired related homeobox pseudoge |
| AC092474.1 | X | claudin-7 (CLDN7) pseudogene |
| AC092474.2 | X | fetal liver LKB1-interacting protein (FLIP1) pse |
| MKRN4P | X | makorin ring finger protein 4, pseudogene [So |
| SDCBPP3 | X | syndecan binding protein pseudogene 3 [Sour |
| AC092268.1 | X | Ribosomal protein L32 (RPL32) pseudogene |
| RPS2P55 | X | ribosomal protein S2 pseudogene 55 [Source: |
| AC092268.2 | X | novel transcript |
| CLIC4P3 | X | chloride intracellular channel 4 pseudogene 3 |
| USP9X | X | ubiquitin specific peptidase 9 X-linked [Source |
| LINC02601 | X | long intergenic non-protein coding RNA 2601 |
| RNA5SP502 | X | RNA, 5S ribosomal pseudogene 502 [Source:H |
| Y_RNA | X | Y RNA [Source:RFAM;Acc:RF00019] |
| AL391647.1 | X | novel pseudogene |
| DDX3X | X | DEAD-box helicase 3 X-linked [Source:HGNC S |
| RN7SL15P | X | RNA, 7SL, cytoplasmic 15, pseudogene [Sourc |
| NYX | X | nyctalopin [Source:HGNC Symbol;Acc:HGNC:8 |
| Z93015.1 | X | gem (nuclear organelle) associated protein 7 ( |
| CASK | X | calcium/calmodulin dependent serine protein |
| CASK-AS1 | X | CASK antisense RNA 1 [Source:HGNC Symbol;/ |
| RNU6-1321P | X | RNA, U6 small nuclear 1321, pseudogene [Sou |
| AL627402.1 | X | tyrosine 3-monooxygenase/tryptophan 5-mor |
| RN7SL144P | X | RNA, 7SL, cytoplasmic 144, pseudogene [Sour |
| GPR34 | X | G protein-coupled receptor 34 [Source:HGNC |
| GPR82 | X | G protein-coupled receptor 82 [Source:HGNC |
| AL353691.1 | X | ribosomal protein S15a (RPS15A) pseudogene |
| AL353691.2 | X | pseudogene similar to part of ribosomal prote |
| RNU6-202P | X | RNA, U6 small nuclear 202, pseudogene [Sour |
| AL133329.1 | X | novel transcript |
| ATP5MC2P4 | X | ATP synthase membrane subunit c locus 2 pse |
| RNU6-630P | X | RNA, U6 small nuclear 630, pseudogene [Sour |
| Z92545.1 | X | pseudogene similar to part of ribosomal prote |
| RNU6-1124P | X | RNA, U6 small nuclear 1124, pseudogene [Sou |
| Z93403.1 | X | novel transcript |
| AL023875.1 | X | TEC |
| Z94277.1 | X | novel transcript |
| PPP1R2C | X | PPP1R2C family member C [Source:HGNC Sym |
| AL451086.1 | X | small integral membrane protein 15 (SMIM15 |
| PINCR | X | p53-induced noncoding RNA [Source:HGNC Sy |
| IMPDH1P4 | X | inosine monophosphate dehydrogenase 1 pse |
| AL023574.1 | X | novel transcript |
| NANOGP10 | X | Nanog homeobox pseudogene 10 [Source:HG |
| MAOA | X | monoamine oxidase A [Source:HGNC Symbol; |
| MAOB | X | monoamine oxidase B [Source:HGNC Symbol; |
| NDP | X | norrin cystine knot growth factor NDP [Source |
| NDP-AS1 | X | NDP antisense RNA 1 [Source:HGNC Symbol;A |
| AL034370.1 | X | RNA-binding region (RNP1, RRM) containing 2 |
| EFHC2 | X | EF-hand domain containing 2 [Source:HGNC S |
| TATDN2P1 | X | TatD DNase domain containing 2 pseudogene |

| | | |
|---|---|---|
| RRM2P3 | X | ribonucleotide reductase M2 polypeptide pse |
| Y_RNA | X | Y RNA [Source:RFAM;Acc:RF00019] |
| AL022163.1 | X | ubinuclein 1 (UBN1) pseudogene |
| FDPSP5 | X | farnesyl diphosphate synthase pseudogene 5 |
| FUNDC1 | X | FUN14 domain containing 1 [Source:HGNC Sy |
| CHTF8P1 | X | chromosome transmission fidelity factor 8 ps |
| RPL19P20 | X | ribosomal protein L19 pseudogene 20 [Source |
| RPSAP61 | X | ribosomal protein SA pseudogene 61 [Source: |
| RN7SL291P | X | RNA, 7SL, cytoplasmic 291, pseudogene [Sour |
| DUSP21 | X | dual specificity phosphatase 21 [Source:HGNC |
| KDM6A | X | lysine demethylase 6A [Source:HGNC Symbol; |
| DIPK2B | X | divergent protein kinase domain 2B [Source:H |
| AC136489.1 | X | novel transcript |
| AL020989.1 | X | transmembrane 4 superfamily member 11 (pl |
| LINC01204 | X | long intergenic non-protein coding RNA 1204 |
| KRT8P14 | X | keratin 8 pseudogene 14 [Source:HGNC Symb |
| AC234772.1 | X | novel pseudogene (GL004) |
| MIR222HG | X | miR222/221 cluster host gene [Source:HGNC |
| MIR221 | X | microRNA 221 [Source:HGNC Symbol;Acc:HGI |
| MIR222 | X | microRNA 222 [Source:HGNC Symbol;Acc:HGI |
| AC234772.2 | X | novel transcript |
| LINC02595 | X | long intergenic non-protein coding RNA 2595 |
| AL031584.1 | X | NULL |
| AL031584.2 | X | pseudogene similar to part of splicing factor, |
| AL031584.3 | X | novel transcript |
| MED28P4 | X | mediator complex subunit 28 pseudogene 4 [ |
| KRT18P68 | X | keratin 18 pseudogene 68 [Source:HGNC Sym |
| PCNAP3 | X | proliferating cell nuclear antigen pseudogene |
| Metazoa_SRP | X | Metazoan signal recognition particle RNA [Sou |
| LINC01186 | X | long intergenic non-protein coding RNA 1186 |
| ACTBP1 | X | ACTB pseudogene 1 [Source:HGNC Symbol;Ac |
| AL139811.3 | X | novel transcript |
| AL139811.2 | X | catenin, beta like 1(CTNNBL1) pseudogene |
| GAPDHP65 | X | glyceraldehyde 3 phosphate dehydrogenase p |
| KRBOX4 | X | KRAB box domain containing 4 [Source:HGNC |
| AL139811.1 | X | transmembrane protein vezatin pseudogene |
| ZNF674 | X | zinc finger protein 674 [Source:HGNC Symbol; |
| RNU6-50P | X | RNA, U6 small nuclear 50, pseudogene [Sourc |
| ZNF674-AS1 | X | ZNF674 antisense RNA 1 (head to head) [Sour |
| Metazoa_SRP | X | Metazoan signal recognition particle RNA [Sou |
| MKI67P1 | X | marker of proliferation Ki-67 pseudogene 1 [S |
| CHST7 | X | carbohydrate sulfotransferase 7 [Source:HGN |
| SLC9A7 | X | solute carrier family 9 member A7 [Source:HG |
| PGAM1P7 | X | phosphoglycerate mutase 1 pseudogene 7 [Sc |
| YBX1P8 | X | Y-box binding protein 1 pseudogene 8 [Source |
| AL512633.1 | X | novel pseudogene (HSPC177) |
| RP2 | X | RP2 activator of ARL3 GTPase [Source:HGNC S |
| LINC01545 | X | long intergenic non-protein coding RNA 1545 |
| JADE3 | X | jade family PHD finger 3 [Source:HGNC Symbo |
| RGN | X | regucalcin [Source:HGNC Symbol;Acc:HGNC:9 |

| | | |
|---|---|---|
| RNU6-1189P | X | RNA, U6 small nuclear 1189, pseudogene [Sou |
| RNU12-2P | X | RNA, U12 small nuclear 2, pseudogene [Sourc |
| NDUFB11 | X | NADH:ubiquinone oxidoreductase subunit B1: |
| RBM10 | X | RNA binding motif protein 10 [Source:HGNC S |
| AL513366.1 | X | inositol 1,3,4-triphosphate 5/6 kinase (ITPK1) |
| UBA1 | X | ubiquitin like modifier activating enzyme 1 [Sc |
| INE1 | X | inactivation escape 1 [Source:HGNC Symbol;A |
| CDK16 | X | cyclin dependent kinase 16 [Source:HGNC Syn |
| USP11 | X | ubiquitin specific peptidase 11 [Source:HGNC |
| AL591503.3 | X | nicolin 1 (NICN1) pseudogene |
| RN7SL785P | X | RNA, 7SL, cytoplasmic 785, pseudogene [Sour |
| AL591503.1 | X | novel transcript |
| ZNF157 | X | zinc finger protein 157 [Source:HGNC Symbol; |
| SNORA11C | X | small nucleolar RNA, H/ACA box 11C [Source:I |
| AL591503.2 | X | UBX domain containing 2 (UBXD2) pseudogen |
| NPM1P49 | X | nucleophosmin 1 pseudogene 49 [Source:HGI |
| ZNF41 | X | zinc finger protein 41 [Source:HGNC Symbol;A |
| Metazoa_SRP | X | Metazoan signal recognition particle RNA [Sou |
| LINC01560 | X | long intergenic non-protein coding RNA 1560 |
| NUS1P1 | X | NUS1 pseudogene 1 [Source:HGNC Symbol;Ac |
| ARAF | X | A-Raf proto-oncogene, serine/threonine kinas |
| SYN1 | X | synapsin I [Source:HGNC Symbol;Acc:HGNC:1: |
| Z84466.1 | X | novel transcript |
| TIMP1 | X | TIMP metallopeptidase inhibitor 1 [Source:HG |
| MIR4769 | X | microRNA 4769 [Source:HGNC Symbol;Acc:HG |
| CFP | X | complement factor properdin [Source:HGNC S |
| ELK1 | X | ETS transcription factor ELK1 [Source:HGNC Sy |
| UXT | X | ubiquitously expressed prefoldin like chaperor |
| UXT-AS1 | X | UXT antisense RNA 1 [Source:HGNC Symbol;A |
| CXXC1P1 | X | CXXC finger protein 1 pseudogene 1 [Source:H |
| AL009172.1 | X | spermine synthase pseudogene |
| AL022578.1 | X | C2H2 type zinc finger pseudogene |
| WASF4P | X | WASP family member 4, pseudogene [Source: |
| ZNF81 | X | zinc finger protein 81 [Source:HGNC Symbol;A |
| RPL7P57 | X | ribosomal protein L7 pseudogene 57 [Source: |
| ZNF182 | X | zinc finger protein 182 [Source:HGNC Symbol; |
| ZNF630 | X | zinc finger protein 630 [Source:HGNC Symbol; |
| SPACA5 | X | sperm acrosome associated 5 [Source:HGNC S |
| ZNF630-AS1 | X | ZNF630 antisense RNA 1 [Source:HGNC Symbo |
| AC244636.3 | X | novel transcript, antisense to ZNF630 |
| SSX6P | X | SSX family member 6, pseudogene [Source:HC |
| SPACA5B | X | sperm acrosome associated 5B [Source:HGNC |
| AC244636.1 | X | zinc finger protein 21 (ZNF21) pseudogene |
| AC244636.2 | X | ornithine aminotransferase (gyrate atrophy) ( |
| RNU6-707P | X | RNA, U6 small nuclear 707, pseudogene [Sour |
| SSXP3 | X | SSX family pseudogene 3 [Source:HGNC Symb |
| AC245047.7 | X | S100 calcium binding protein A11 (calgizzarin) |
| SSX5 | X | SSX family member 5 [Source:HGNC Symbol;A |
| AC245047.5 | X | ornithine aminotransferase (gyrate atrophy)(C |
| RNA5SP503 | X | RNA, 5S ribosomal pseudogene 503 [Source:H |

| | | |
|---|---|---|
| AC245047.1 | X | SSX2 pseudogene (psiSSX2) |
| AC245047.3 | X | S100 calcium binding protein A11 (calgizzarin) |
| AC245047.2 | X | ornithine aminotransferase (gyrate atrophy)(( |
| SSX1 | X | SSX family member 1 [Source:HGNC Symbol;A |
| AC245047.4 | X | S100 calcium binding protein A11 (calgizzarin) |
| AC245047.8 | X | psiSSX8 pseudogene (psiSSX8) |
| AC245047.6 | X | ornithine aminotransferase (gyrate atrophy) p |
| SSX9P | X | SSX family member 9, pseudogene [Source:HC |
| AL606490.6 | X | ornithine aminotransferase (gyrate atrophy) ( |
| SSXP9 | X | SSX family pseudogene 9 [Source:HGNC Symb |
| AL606490.3 | X | novel transcript |
| AL606490.9 | X | S100 calcium binding protein A11 (S100A11) p |
| SSX3 | X | SSX family member 3 [Source:HGNC Symbol;A |
| AL606490.1 | X | ornithine aminotransferase (gyrate atrophy) ( |
| AL606490.7 | X | pseudogene similar to part of ornithine aminc |
| SSX4 | X | SSX family member 4 [Source:HGNC Symbol;A |
| SSX4B | X | SSX family member 4B [Source:HGNC Symbol; |
| AL606490.5 | X | ornithine aminotransferase (gyrate atrophy) ( |
| AL606490.4 | X | similar to part of the ornithine aminotransfera |
| AL606490.8 | X | SSX7 pseudogene (psiSSX7) |
| AL606490.2 | X | ornithine aminotransferase (gyrate atrophy) ( |
| SLC38A5 | X | solute carrier family 38 member 5 [Source:HG |
| FTSJ1 | X | FtsJ RNA 2'-O-methyltransferase 1 [Source:HG |
| AF196972.1 | X | novel transcript, antisense to PORCN, EBP |
| PORCN | X | porcupine O-acyltransferase [Source:HGNC Sy |
| EBP | X | EBP cholestenol delta-isomerase [Source:HGN |
| AF196969.1 | X | novel protein |
| Metazoa_SRP | X | Metazoan signal recognition particle RNA [Sou |
| TBC1D25 | X | TBC1 domain family member 25 [Source:HGN |
| AC115618.1 | X | novel transcript, antisense to RBM3 |
| RBM3 | X | RNA binding motif protein 3 [Source:HGNC Sy |
| AC115618.2 | X | novel transcript |
| AC115618.3 | X | TEC |
| MRPL32P1 | X | mitochondrial ribosomal protein L32 pseudog |
| WDR13 | X | WD repeat domain 13 [Source:HGNC Symbol; |
| VN1R110P | X | vomeronasal 1 receptor 110 pseudogene [Sou |
| WAS | X | WASP actin nucleation promoting factor [Sour |
| SUV39H1 | X | suppressor of variegation 3-9 homolog 1 [Sou |
| AC231533.1 | X | novel transcript |
| RNU6-1056P | X | RNA, U6 small nuclear 1056, pseudogene [Sou |
| GLOD5 | X | glyoxalase domain containing 5 [Source:HGNC |
| AC231533.2 | X | acetyl-Coenzyme A acyltransferase 2 (mitochc |
| RNU6-29P | X | RNA, U6 small nuclear 29, pseudogene [Sourc |
| GATA1 | X | GATA binding protein 1 [Source:HGNC Symbol |
| HDAC6 | X | histone deacetylase 6 [Source:HGNC Symbol; |
| ERAS | X | ES cell expressed Ras [Source:HGNC Symbol;A |
| PCSK1N | X | proprotein convertase subtilisin/kexin type 1 i |
| PQBP1 | X | polyglutamine binding protein 1 [Source:HGN |
| TIMM17B | X | translocase of inner mitochondrial membrane |
| SLC35A2 | X | solute carrier family 35 member A2 [Source:H |

| | | |
|---|---|---|
| PIM2 | X | Pim-2 proto-oncogene, serine/threonine kinas |
| OTUD5 | X | OTU deubiquitinase 5 [Source:HGNC Symbol;A |
| AC233300.1 | X | TEC |
| RNU6-722P | X | RNA, U6 small nuclear 722, pseudogene [Sour |
| KCND1 | X | potassium voltage-gated channel subfamily D |
| GRIPAP1 | X | GRIP1 associated protein 1 [Source:HGNC Sym |
| TFE3 | X | transcription factor binding to IGHM enhancer |
| CCDC120 | X | coiled-coil domain containing 120 [Source:HG |
| PRAF2 | X | PRA1 domain family member 2 [Source:HGNC |
| AC231657.3 | X | novel protein |
| WDR45 | X | WD repeat domain 45 [Source:HGNC Symbol;A |
| RNU4-52P | X | RNA, U4 small nuclear 52, pseudogene [Sourc |
| GPKOW | X | G-patch domain and KOW motifs [Source:HGN |
| RN7SL262P | X | RNA, 7SL, cytoplasmic 262, pseudogene [Sour |
| AC231657.2 | X | MAGI family member, X-linked (MAGIX) pseu |
| MAGIX | X | MAGI family member, X-linked [Source:HGNC |
| AC231657.1 | X | TEC |
| PLP2 | X | proteolipid protein 2 [Source:HGNC Symbol;A |
| PRICKLE3 | X | prickle planar cell polarity protein 3 [Source:H |
| SYP | X | synaptophysin [Source:HGNC Symbol;Acc:HGN |
| SYP-AS1 | X | SYP antisense RNA 1 [Source:HGNC Symbol;Ac |
| CACNA1F | X | calcium voltage-gated channel subunit alpha1 |
| HSPB1P2 | X | heat shock protein family B (small) member 1 |
| CCDC22 | X | coiled-coil domain containing 22 [Source:HGN |
| FOXP3 | X | forkhead box P3 [Source:HGNC Symbol;Acc:H |
| AC232271.3 | X | FOXP3 regulating long intergenic non-coding R |
| PPP1R3F | X | protein phosphatase 1 regulatory subunit 3F [ |
| AC232271.1 | X | novel transcript |
| AC232271.2 | X | novel transcript, antisense to PPP1R3F |
| GAGE10 | X | G antigen 10 [Source:HGNC Symbol;Acc:HGNC |
| GAGE12J | X | G antigen 12J [Source:HGNC Symbol;Acc:HGN |
| GAGE13 | X | G antigen 13 [Source:HGNC Symbol;Acc:HGNC |
| GAGE2E | X | G antigen 2E [Source:HGNC Symbol;Acc:HGNC |
| GAGE12B | X | G antigen 12B [Source:HGNC Symbol;Acc:HGN |
| GAGE12C | X | G antigen 12C [Source:HGNC Symbol;Acc:HGN |
| GAGE12D | X | G antigen 12D [Source:HGNC Symbol;Acc:HGN |
| GAGE12F | X | G antigen 12F [Source:HGNC Symbol;Acc:HGN |
| GAGE12E | X | G antigen 12E [Source:HGNC Symbol;Acc:HGN |
| GAGE12G | X | G antigen 12G [Source:HGNC Symbol;Acc:HGN |
| GAGE12H | X | G antigen 12H [Source:HGNC Symbol;Acc:HGN |
| GAGE1 | X | G antigen 1 [Source:HGNC Symbol;Acc:HGNC: |
| GAGE2A | X | G antigen 2A [Source:HGNC Symbol;Acc:HGNC |
| VDAC1P2 | X | voltage dependent anion channel 1 pseudoge |
| SALL1P1 | X | spalt like transcription factor 1 pseudogene 1 |
| PAGE1 | X | PAGE family member 1 [Source:HGNC Symbol |
| PAGE4 | X | PAGE family member 4 [Source:HGNC Symbol |
| USP27X-AS1 | X | USP27X antisense RNA 1 (head to head) [Sour |
| USP27X | X | ubiquitin specific peptidase 27 X-linked [Sourc |
| AC236430.1 | X | pseudogene similar to part of cleavage and po |
| CLCN5 | X | chloride voltage-gated channel 5 [Source:HGN |

| | | |
|---|---|---|
| RNU6-421P | X | RNA, U6 small nuclear 421, pseudogene [Sour |
| MIR532 | X | microRNA 532 [Source:HGNC Symbol;Acc:HGI |
| MIR188 | X | microRNA 188 [Source:HGNC Symbol;Acc:HGI |
| MIR500A | X | microRNA 500a [Source:HGNC Symbol;Acc:HG |
| MIR362 | X | microRNA 362 [Source:HGNC Symbol;Acc:HGI |
| MIR501 | X | microRNA 501 [Source:HGNC Symbol;Acc:HGI |
| MIR500B | X | microRNA 500b [Source:HGNC Symbol;Acc:HG |
| MIR660 | X | microRNA 660 [Source:HGNC Symbol;Acc:HGI |
| MIR502 | X | microRNA 502 [Source:HGNC Symbol;Acc:HGI |
| PYY3 | X | peptide YY 3 (pseudogene) [Source:HGNC Sym |
| Y_RNA | X | Y RNA [Source:RFAM;Acc:RF00019] |
| AKAP4 | X | A-kinase anchoring protein 4 [Source:HGNC Sy |
| CCNB3 | X | cyclin B3 [Source:HGNC Symbol;Acc:HGNC:18 |
| DGKK | X | diacylglycerol kinase kappa [Source:HGNC Syn |
| SHROOM4 | X | shroom family member 4 [Source:HGNC Symb |
| RNU6-935P | X | RNA, U6 small nuclear 935, pseudogene [Sour |
| H3F3AP5 | X | H3 histone, family 3A, pseudogene 5 [Source: |
| BMP15 | X | bone morphogenetic protein 15 [Source:HGN |
| HMGB1P15 | X | high mobility group box 1 pseudogene 15 [Sou |
| AC239367.1 | X | melanoma antigen pseudogene |
| AC239367.2 | X | pseudogene similar to part of SAH (SA hypert |
| LINC01284 | X | long intergenic non-protein coding RNA 1284 |
| AC239367.3 | X | melanoma antigen pseudogene |
| AC233976.1 | X | novel transcript |
| AC233976.2 | X | novel transcript |
| NUDT10 | X | nudix hydrolase 10 [Source:HGNC Symbol;Acc |
| AL158055.1 | X | novel transcript |
| EZHIP | X | EZH inhibitory protein [Source:HGNC Symbol; |
| NUDT11 | X | nudix hydrolase 11 [Source:HGNC Symbol;Acc |
| LINC01496 | X | long intergenic non-protein coding RNA 1496 |
| CENPVL3 | X | centromere protein V like 3 [Source:HGNC Syr |
| CENPVL2 | X | centromere protein V like 2 [Source:HGNC Syr |
| CENPVL1 | X | centromere protein V like 1 [Source:HGNC Syr |
| GSPT2 | X | G1 to S phase transition 2 [Source:HGNC Syml |
| MAGED1 | X | MAGE family member D1 [Source:HGNC Symb |
| AC241520.1 | X | zinc finger protein pseudogene |
| RNU6-504P | X | RNA, U6 small nuclear 504, pseudogene [Sour |
| AL929410.1 | X | pseudogene similar to part of PFKFB1 (6-phos |
| IPO7P1 | X | importin 7 pseudogene 1 [Source:HGNC Symb |
| AL929410.2 | X | ubiquinol-cytochrome c reductase complex (7 |
| TPMTP3 | X | thiopurine S-methyltransferase pseudogene 3 |
| AC239585.1 | X | novel transcript |
| AC239585.2 | X | novel transcript |
| MAGED4B | X | MAGE family member D4B [Source:HGNC Sym |
| SNORA11E | X | small nucleolar RNA, H/ACA box 11E [Source: |
| MAGED4 | X | MAGE family member D4 [Source:HGNC Symb |
| SNORA11D | X | small nucleolar RNA, H/ACA box 11D [Source: |
| AC231759.2 | X | novel transcript |
| AC245177.1 | X | novel transcript |
| AC231759.1 | X | family with sequence similarity 156, member |

| | | |
|---|---|---|
| MIR8088 | X | microRNA 8088 [Source:HGNC Symbol;Acc:HG |
| XAGE2 | X | X antigen family member 2 [Source:HGNC Syn |
| AC231532.1 | X | GAGE family member pseudogene |
| AC231532.2 | X | X antigen family, member 2 (XAGE2) pseudog |
| BX510359.1 | X | GAGE family member pseudogene |
| BX510359.5 | X | GAGE family member pseudogene |
| BX510359.8 | X | novel pseudogene |
| BX510359.7 | X | TBC1 domain family, member 3B (TBC1D3B) p |
| RBM22P6 | X | RNA binding motif protein 22 pseudogene 6 [S |
| XAGE1A | X | X antigen family member 1A [Source:HGNC Sy |
| RBM22P7 | X | RNA binding motif protein 22 pseudogene 7 [S |
| XAGE1B | X | X antigen family member 1B [Source:HGNC Sy |
| BX510359.6 | X | cytochrome P450, family 4, subfamily F, polyp |
| BX510359.2 | X | GAGE family member pseudogene |
| BX510359.4 | X | GAGE family member pseudogene |
| BX510359.3 | X | GAGE family member pseudogene |
| SSXP4 | X | SSX family pseudogene 4 [Source:HGNC Symb |
| SSXP1 | X | SSX family pseudogene 1 [Source:HGNC Symb |
| AL450023.2 | X | ornithine aminotransferase (gyrate atrophy) ( |
| SSX8P | X | SSX family member 8, pseudogene [Source:HG |
| SSX7 | X | SSX family member 7 [Source:HGNC Symbol;A |
| AL450023.1 | X | ornithine aminotransferase (gyrate atrophy) ( |
| RNA5SP504 | X | RNA, 5S ribosomal pseudogene 504 [Source:H |
| SSXP5 | X | SSX family pseudogene 5 [Source:HGNC Symb |
| AL450023.3 | X | S100 calcium binding protein A11 (calgizzarin) |
| SSX2 | X | SSX family member 2 [Source:HGNC Symbol;A |
| AC244505.2 | X | ornithine aminotransferase (gyrate atrophy) ( |
| AC244505.3 | X | pseudogene similar to part of ornithine aminc |
| AC244505.5 | X | pseudogene similar to part of ornithine aminc |
| SSX2B | X | SSX family member 2B [Source:HGNC Symbol; |
| AC244505.7 | X | S100 calcium binding protein A11 (S100A11) p |
| AC244505.4 | X | SSX family pseudogene |
| SPANXN5 | X | SPANX family member N5 [Source:HGNC Syml |
| AC244505.6 | X | X antigen family, member 3 (XAGE3) pseudog |
| XAGE5 | X | X antigen family member 5 [Source:HGNC Syn |
| AC244505.1 | X | GAGE family pseudogene |
| EIF4A2P4 | X | eukaryotic translation initiation factor 4A2 pse |
| XAGE3 | X | X antigen family member 3 [Source:HGNC Syn |
| FAM156B | X | family with sequence similarity 156 member E |
| AC234031.1 | X | novel transcript, antisense to FAM156A |
| FAM156A | X | family with sequence similarity 156 member A |
| GPR173 | X | G protein-coupled receptor 173 [Source:HGN( |
| BX322635.1 | X | ribosomal protein S3A (RPS3A) pseudogene |
| TSPYL2 | X | TSPY like 2 [Source:HGNC Symbol;Acc:HGNC:2 |
| AL591212.1 | X | novel transcript |
| KANTR | X | KDM5C adjacent transcript [Source:HGNC Syn |
| AL591212.2 | X | ribosomal protein L27 (RPL27) pseudogene |
| ACTG1P10 | X | actin gamma 1 pseudogene 10 [Source:HGNC |
| AL139396.1 | X | pseudogene similar to part of suppressor of va |
| KDM5C | X | lysine demethylase 5C [Source:HGNC Symbol; |

| | | |
|---|---|---|
| MIR6895 | X | microRNA 6895 [Source:HGNC Symbol;Acc:HG |
| MIR6894 | X | microRNA 6894 [Source:HGNC Symbol;Acc:HG |
| KDM5C-IT1 | X | KDM5C intronic transcript 1 [Source:HGNC Sy |
| IQSEC2 | X | IQ motif and Sec7 domain 2 [Source:HGNC Sy |
| RPSAP62 | X | ribosomal protein SA pseudogene 62 [Source: |
| Y_RNA | X | Y RNA [Source:RFAM;Acc:RF00019] |
| AC245102.2 | X | ribosomal protein S13 (RPS13) pseudogene |
| AC245102.1 | X | novel pseudogene |
| SMC1A | X | structural maintenance of chromosomes 1A [S |
| MIR6857 | X | microRNA 6857 [Source:HGNC Symbol;Acc:HG |
| RIBC1 | X | RIB43A domain with coiled-coils 1 [Source:HG |
| HSD17B10 | X | hydroxysteroid 17-beta dehydrogenase 10 [So |
| AC233728.1 | X | novel transcript |
| AC233728.2 | X | novel pseudogene (FLJ20516) |
| VTRNA3-1P | X | vault RNA 3-1, pseudogene [Source:HGNC Syn |
| HUWE1 | X | HECT, UBA and WWE domain containing E3 ul |
| MIR98 | X | microRNA 98 [Source:HGNC Symbol;Acc:HGN |
| MIRLET7F2 | X | microRNA let-7f-2 [Source:HGNC Symbol;Acc: |
| BX323845.1 | X | pseudogene similar to part of 6-phosphofruct |
| BX323845.2 | X | pseudogene similar to part of ribosomal prote |
| BX323845.3 | X | mitochondrial ribosomal protein L32 (MRPL32 |
| AC233279.1 | X | mitochondrial ribosomal protein S18C (MRPS1 |
| RNA5SP505 | X | RNA, 5S ribosomal pseudogene 505 [Source:H |
| PHF8 | X | PHD finger protein 8 [Source:HGNC Symbol;Ac |
| FAM120C | X | family with sequence similarity 120C [Source: |
| AL589872.1 | X | ribosomal protein L37 (RPL37) pseudogene |
| WNK3 | X | WNK lysine deficient protein kinase 3 [Source |
| RPL7AP71 | X | ribosomal protein L7a pseudogene 71 [Source |
| RNU6-434P | X | RNA, U6 small nuclear 434, pseudogene [Sour |
| TSR2 | X | TSR2 ribosome maturation factor [Source:HGI |
| FGD1 | X | FYVE, RhoGEF and PH domain containing 1 [So |
| GNL3L | X | G protein nucleolar 3 like [Source:HGNC Symb |
| PGAM4P1 | X | phosphoglycerate mutase family member 4 ps |
| ITIH6 | X | inter-alpha-trypsin inhibitor heavy chain famil |
| MAGED2 | X | MAGE family member D2 [Source:HGNC Symb |
| SNORA11 | X | small nucleolar RNA, H/ACA box 11 [Source:H |
| AL049732.1 | X | inter-alpha-trypsin inhibitor heavy chain famil |
| TRO | X | trophinin [Source:HGNC Symbol;Acc:HGNC:12 |
| SNORA11G | X | small nucleolar RNA, H/ACA box 11G [Source: |
| PFKFB1 | X | 6-phosphofructo-2-kinase/fructose-2,6-bipho |
| APEX2 | X | apurinic/apyrimidinic endodeoxyribonuclease |
| ALAS2 | X | 5'-aminolevulinate synthase 2 [Source:HGNC S |
| AL020991.1 | X | zinc finger protein 273 (ZNF273) pseudogene |
| AL590240.2 | X | likely ortholog of mouse hepatoma-derived gr |
| PAGE2B | X | PAGE family member 2B [Source:HGNC Symbo |
| PAGE2 | X | PAGE family member 2 [Source:HGNC Symbol |
| AL590240.4 | X | family with sequence similarity 156, member |
| AL590240.3 | X | component of oligomeric golgi complex 1 (CO |
| FAM104B | X | family with sequence similarity 104 member E |
| AL590240.1 | X | G antigen, family D pseudogene |

| | | |
|---|---|---|
| MTCO1P52 | X | MT-CO1 pseudogene 52 [Source:HGNC Symbo |
| MTND2P24 | X | MT-ND2 pseudogene 24 [Source:HGNC Symbo |
| MTND1P30 | X | MT-ND1 pseudogene 30 [Source:HGNC Symbo |
| MTRNR2L10 | X | MT-RNR2 like 10 [Source:HGNC Symbol;Acc:H |
| PAGE5 | X | PAGE family member 5 [Source:HGNC Symbol |
| PAGE3 | X | PAGE family member 3 [Source:HGNC Symbol |
| AL158819.1 | X | PAGE-5 protein (PAGE-5) pseudogene |
| MIR4536-2 | X | microRNA 4536-2 [Source:HGNC Symbol;Acc:I |
| MIR4536-1 | X | microRNA 4536-1 [Source:HGNC Symbol;Acc:I |
| MAGEH1 | X | MAGE family member H1 [Source:HGNC Symb |
| USP51 | X | ubiquitin specific peptidase 51 [Source:HGNC |
| AL590410.1 | X | melanoma antigen family E, 1 (MAGEE1) pseu |
| Y_RNA | X | Y RNA [Source:RFAM;Acc:RF00019] |
| AL627224.1 | X | proteasome (prosome, macropain) subunit al |
| FOXR2 | X | forkhead box R2 [Source:HGNC Symbol;Acc:H |
| AL159987.2 | X | X antigen family (XAGE) pseudogene |
| AL159987.1 | X | X antigen family, member 3 (XAGE3) pseudog |
| RRAGB | X | Ras related GTP binding B [Source:HGNC Symb |
| AL050309.1 | X | novel transcript |
| AL391419.1 | X | glutamic-oxaloacetic transaminase 2, mitocho |
| KLF8 | X | Kruppel like factor 8 [Source:HGNC Symbol;Ac |
| RPL23AP83 | X | ribosomal protein L23a pseudogene 83 [Sourc |
| UBQLN2 | X | ubiquilin 2 [Source:HGNC Symbol;Acc:HGNC:1 |
| AL354793.1 | X | novel transcript |
| NBDY | X | negative regulator of P-body association [Sour |
| UQCRBP1 | X | ubiquinol-cytochrome c reductase binding pro |
| SPIN3 | X | spindlin family member 3 [Source:HGNC Symb |
| AL139397.1 | X | novel transcript |
| AL139397.2 | X | spindlin-like protein 2 (SPIN2) pseudogne |
| SPIN2B | X | spindlin family member 2B [Source:HGNC Sym |
| AL022157.1 | X | novel transcript |
| SPIN2A | X | spindlin family member 2A [Source:HGNC Sym |
| Z83745.1 | X | novel transcript |
| PPP1R11P2 | X | protein phosphatase 1 regulatory inhibitor sul |
| FAAH2 | X | fatty acid amide hydrolase 2 [Source:HGNC Sy |
| MTHFD1P1 | X | methylenetetrahydrofolate dehydrogenase (N |
| ZXDB | X | zinc finger X-linked duplicated B [Source:HGN |
| NLRP2B | X | NLR family pyrin domain containing 2B [Sourc |
| Z93019.1 | X | malate dehydrogenase 1, NAD (soluble) (MDH |
| ZXDA | X | zinc finger X-linked duplicated A [Source:HGN |
| MYCLP2 | X | MYCL pseudogene 2 [Source:HGNC Symbol;Ac |
| KRT8P17 | X | keratin 8 pseudogene 17 [Source:HGNC Symb |
| BX322784.1 | X | Sjogren syndrome antigen B (autoantigen La) |
| MTND1P31 | X | MT-ND1 pseudogene 31 [Source:HGNC Symbo |
| MTND2P25 | X | MT-ND2 pseudogene 25 [Source:HGNC Symbo |
| ZFRP1 | X | zinc finger RNA binding protein pseudogene 1 |
| AL359854.1 | X | endogenous retrovirus group FRD, member 1 |
| LINC01278 | X | long intergenic non-protein coding RNA 1278 |
| CBX1P1 | X | chromobox 1 pseudogene 1 [Source:HGNC Sy |
| SPIN4 | X | spindlin family member 4 [Source:HGNC Symb |

| | | |
|---|---|---|
| SPIN4-AS1 | X | SPIN4 antisense RNA 1 [Source:HGNC Symbol; |
| ARHGEF9 | X | Cdc42 guanine nucleotide exchange factor 9 [ |
| ARHGEF9-IT1 | X | ARHGEF9 intronic transcript 1 [Source:HGNC S |
| BTF3P8 | X | basic transcription factor 3 pseudogene 8 [Sou |
| MIR1468 | X | microRNA 1468 [Source:HGNC Symbol;Acc:HG |
| HNRNPDP1 | X | heterogeneous nuclear ribonucleoprotein D p |
| AL355852.1 | X | zinc finger protein pseudogene |
| AMER1 | X | APC membrane recruitment protein 1 [Source |
| AL356317.2 | X | novel transcript |
| RN7SL799P | X | RNA, 7SL, cytoplasmic 799, pseudogene [Sour |
| ASB12 | X | ankyrin repeat and SOCS box containing 12 [S |
| MTMR8 | X | myotubularin related protein 8 [Source:HGNC |
| AL356317.1 | X | karyopherin alpha 4 (importin alpha 3) (KPNA |
| PFN5P | X | profilin 5, pseudogene [Source:HGNC Symbol; |
| SHC1P1 | X | SHC adaptor protein 1 pseudogene 1 [Source: |
| YWHAZP7 | X | tyrosine 3-monooxygenase/tryptophan 5-mor |
| KRT8P27 | X | keratin 8 pseudogene 27 [Source:HGNC Symb |
| GRPEL2P2 | X | GRPEL2 pseudogene 2 [Source:HGNC Symbol; |
| BLOC1S2P1 | X | BLOC1S2 pseudogene 1 [Source:HGNC Symbo |
| ZC4H2 | X | zinc finger C4H2-type containing [Source:HGN |
| AL445523.1 | X | karyopherin alpha 2 (RAG cohort 1, importin a |
| CCT4P2 | X | chaperonin containing TCP1 subunit 4 pseudo |
| MORF4L1P5 | X | mortality factor 4 like 1 pseudogene 5 [Source |
| AL441964.1 | X | novel pseudogene |
| ZC3H12B | X | zinc finger CCCH-type containing 12B [Source: |
| TLE1P1 | X | TLE family member 1 pseudogene 1 [Source:H |
| Z83841.1 | X | adaptor-related protein complex 1 (AP1M2) p |
| LAS1L | X | LAS1 like, ribosome biogenesis factor [Source: |
| FRMD8P1 | X | FERM domain containing 8 pseudogene 1 [Sou |
| MSN | X | moesin [Source:HGNC Symbol;Acc:HGNC:737 |
| NANOGP9 | X | Nanog homeobox pseudogene 9 [Source:HGN |
| AL450487.2 | X | family with sequence similarity 207, member |
| ETF1P3 | X | eukaryotic translation termination factor 1 ps |
| AL450487.1 | X | cyclin fold protein 1 (CFP1) pseudogene |
| AL034397.2 | X | novel transcript |
| AL034397.1 | X | Heterogenous nuclear ribonucleoprotein G ps |
| AL034397.3 | X | novel transcript, antisense VSIG4 |
| MIR223 | X | microRNA 223 [Source:HGNC Symbol;Acc:HGI |
| VSIG4 | X | V-set and immunoglobulin domain containing |
| AL139276.1 | X | novel pseudogene |
| EIF4BP9 | X | eukaryotic translation initiation factor 4B pseu |
| HEPH | X | hephaestin [Source:HGNC Symbol;Acc:HGNC: |
| AL157698.1 | X | likely ortholog of chicken chondrocyte proteir |
| AL592049.1 | X | novel pseudogene |
| PKMP2 | X | pyruvate kinase M1/2 pseudogene 2 [Source: |
| EDA2R | X | ectodysplasin A2 receptor [Source:HGNC Sym |
| RNU6-394P | X | RNA, U6 small nuclear 394, pseudogene [Sour |
| AL049641.1 | X | zinc finger protein pseudogene |
| AL049564.1 | X | poly(A) binding protein, nuclear 1 (PABPN1) p |
| AR | X | androgen receptor [Source:HGNC Symbol;Acc |

| | | |
|---|---|---|
| BMI1P1 | X | BMI1 proto-oncogene, polycomb ring finger p |
| AL157700.1 | X | novel transcript |
| OPHN1 | X | oligophrenin 1 [Source:HGNC Symbol;Acc:HGI |
| AL158201.1 | X | phosphoglycerate kinase 1 (PGK1) pseudogen |
| RNU6-1225P | X | RNA, U6 small nuclear 1225, pseudogene [Sou |
| AKIRIN1P2 | X | akirin 1 pseudogene 2 [Source:HGNC Symbol;/ |
| Y_RNA | X | Y RNA [Source:RFAM;Acc:RF00019] |
| AL672138.1 | X | nudE nuclear distribution gene E homolog like |
| YIPF6 | X | Yip1 domain family member 6 [Source:HGNC S |
| RNU6-245P | X | RNA, U6 small nuclear 245, pseudogene [Sour |
| RPL31P63 | X | ribosomal protein L31 pseudogene 63 [Source |
| COX6CP12 | X | cytochrome c oxidase subunit 6C pseudogene |
| STARD8 | X | StAR related lipid transfer domain containing |
| ACTR3P2 | X | ACTR3 pseudogene 2 [Source:HGNC Symbol;A |
| SERBP1P1 | X | SERPINE1 mRNA binding protein 1 pseudogen |
| EFNB1 | X | ephrin B1 [Source:HGNC Symbol;Acc:HGNC:32 |
| PJA1 | X | praja ring finger ubiquitin ligase 1 [Source:HGI |
| HMGN1P35 | X | high mobility group nucleosome binding dom |
| LINC00269 | X | long intergenic non-protein coding RNA 269 [S |
| CYCSP43 | X | CYCS pseudogene 43 [Source:HGNC Symbol;A |
| FAM155B | X | family with sequence similarity 155 member E |
| AL158069.1 | X | novel transcript |
| EDA | X | ectodysplasin A [Source:HGNC Symbol;Acc:HG |
| AL158069.2 | X | NULL |
| RNA5SP506 | X | RNA, 5S ribosomal pseudogene 506 [Source:H |
| U3 | X | Small nucleolar RNA U3 [Source:RFAM;Acc:RF |
| CNOT7P1 | X | CCR4-NOT transcription complex subunit 7 ps |
| MIR676 | X | microRNA 676 [Source:HGNC Symbol;Acc:HGI |
| AL158141.1 | X | novel transcript, antisense to EDA |
| AWAT2 | X | acyl-CoA wax alcohol acyltransferase 2 [Sourc |
| OTUD6A | X | OTU deubiquitinase 6A [Source:HGNC Symbol |
| MTCYBP31 | X | MT-CYB pseudogene 31 [Source:HGNC Symbo |
| MTND4P31 | X | MT-ND4 pseudogene 31 [Source:HGNC Symbo |
| IGBP1 | X | immunoglobulin binding protein 1 [Source:HG |
| IGBP1-AS2 | X | IGBP1 antisense RNA 2 [Source:HGNC Symbol |
| IGBP1-AS1 | X | IGBP1 antisense RNA 1 [Source:HGNC Symbol |
| DGAT2L6 | X | diacylglycerol O-acyltransferase 2 like 6 [Sour |
| RN7SL581P | X | RNA, 7SL, cytoplasmic 581, pseudogene [Sour |
| AWAT1 | X | acyl-CoA wax alcohol acyltransferase 1 [Sourc |
| RNA5SP507 | X | RNA, 5S ribosomal pseudogene 507 [Source:H |
| P2RY4 | X | pyrimidinergic receptor P2Y4 [Source:HGNC S |
| ARR3 | X | arrestin 3 [Source:HGNC Symbol;Acc:HGNC:71 |
| RAB41 | X | RAB41, member RAS oncogene family [Source |
| PDZD11 | X | PDZ domain containing 11 [Source:HGNC Sym |
| KIF4A | X | kinesin family member 4A [Source:HGNC Sym |
| AL357752.1 | X | novel pseudogene |
| RNY4P23 | X | RNY4 pseudogene 23 [Source:HGNC Symbol;A |
| GDPD2 | X | glycerophosphodiester phosphodiesterase do |
| AL139398.1 | X | uncharacterized LOC105373244 [Source:NCBI |
| DLG3 | X | discs large MAGUK scaffold protein 3 [Source: |

| | | |
|---|---|---|
| RNU4-81P | X | RNA, U4 small nuclear 81, pseudogene [Sourc |
| DLG3-AS1 | X | DLG3 antisense RNA 1 [Source:HGNC Symbol; |
| TEX11 | X | testis expressed 11 [Source:HGNC Symbol;Acc |
| AL139109.1 | X | ribosomal protein S26 (RPS26) pseudogene |
| AL627390.2 | X | NULL |
| SNORD3E | X | small nucleolar RNA, C/D box 3E [Source:HGN |
| AL627390.1 | X | novel pseudogene |
| SLC7A3 | X | solute carrier family 7 member 3 [Source:HGN |
| RPS23P8 | X | ribosomal protein S23 pseudogene 8 [Source: |
| NUTF2P7 | X | nuclear transport factor 2 pseudogene 7 [Sour |
| RNU1-56P | X | RNA, U1 small nuclear 56, pseudogene [Sourc |
| SOCS5P4 | X | suppressor of cytokine signaling 5 pseudogene |
| SNX12 | X | sorting nexin 12 [Source:HGNC Symbol;Acc:H |
| FOXO4 | X | forkhead box O4 [Source:HGNC Symbol;Acc:H |
| CXorf65 | X | chromosome X open reading frame 65 [Sourc |
| AL590764.2 | X | novel protein, IL2RG-CXorf65 readthrough |
| IL2RG | X | interleukin 2 receptor subunit gamma [Source |
| MED12 | X | mediator complex subunit 12 [Source:HGNC S |
| NLGN3 | X | neuroligin 3 [Source:HGNC Symbol;Acc:HGNC: |
| AL590764.1 | X | novel transcript |
| GJB1 | X | gap junction protein beta 1 [Source:HGNC Syn |
| ZMYM3 | X | zinc finger MYM-type containing 3 [Source:HG |
| NONO | X | non-POU domain containing octamer binding |
| ITGB1BP2 | X | integrin subunit beta 1 binding protein 2 [Sour |
| AL590762.3 | X | MADP-1 protein (MADP-1) pseudogene |
| AL590762.2 | X | ras homolog gene family, member G (rho G) ( |
| TAF1 | X | TATA-box binding protein associated factor 1 |
| Metazoa_SRP | X | Metazoan signal recognition particle RNA [Sou |
| AL590762.4 | X | pseudogene similar to part of ras-related C3 b |
| AL590762.1 | X | pseudogene similar to part of poly(A) binding |
| Y_RNA | X | Y RNA [Source:RFAM;Acc:RF00019] |
| INGX | X | inhibitor of growth family, X-linked (pseudoge |
| AL590763.1 | X | suppressor of cytokine signaling 4 (SOCS4) pse |
| OGT | X | O-linked N-acetylglucosamine (GlcNAc) transf |
| GCNA | X | germ cell nuclear acidic peptidase [Source:HG |
| CXCR3 | X | C-X-C motif chemokine receptor 3 [Source:HG |
| BX276092.7 | X | chromosome X open reading frame 49 (CXorf |
| BX276092.9 | X | chromosome X open reading frame 49-like [S |
| BX276092.4 | X | NADH dehydrogenase (ubiquinone) 1 alpha su |
| LINC00891 | X | long intergenic non-protein coding RNA 891 [ |
| CXorf49 | X | chromosome X open reading frame 49 [Sourc |
| BX276092.2 | X | chromosome X open reading frame 49 (CXorf |
| BX276092.6 | X | NADH dehydrogenase (ubiquinone) 1 alpha su |
| BX276092.8 | X | NADH dehydrogenase (ubiquinone) 1 alpha su |
| BX276092.5 | X | chromosome X open reading frame 49 (CXorf |
| CXorf49B | X | chromosome X open reading frame 49B [Sour |
| BX276092.1 | X | NADH dehydrogenase (ubiquinone) 1 alpha su |
| BX276092.3 | X | NADH dehydrogenase (ubiquinone) 1 alpha su |
| BX119927.1 | X | chromosome X open reading frame 49 (CXorf |
| NHSL2 | X | NHS like 2 [Source:HGNC Symbol;Acc:HGNC:3 |

| | | |
|---|---|---|
| RNU6-1078P | X | RNA, U6 small nuclear 1078, pseudogene [Sou |
| RPS26P11 | X | ribosomal protein S26 pseudogene 11 [Source |
| RTL5 | X | retrotransposon Gag like 5 [Source:HGNC Sym |
| BX119917.1 | X | tumor protein, translationally-controlled 1 (TF |
| PIN4 | X | peptidylprolyl cis/trans isomerase, NIMA-inter |
| RN7SL388P | X | RNA, 7SL, cytoplasmic 388, pseudogene [Sour |
| ERCC6L | X | ERCC excision repair 6 like, spindle assembly c |
| RPS4X | X | ribosomal protein S4 X-linked [Source:HGNC S |
| Y_RNA | X | Y RNA [Source:RFAM;Acc:RF00019] |
| CITED1 | X | Cbp/p300 interacting transactivator with Glu/ |
| AL133500.1 | X | novel protein |
| HDAC8 | X | histone deacetylase 8 [Source:HGNC Symbol;/ |
| RNU2-68P | X | RNA, U2 small nuclear 68, pseudogene [Sourc |
| PHKA1 | X | phosphorylase kinase regulatory subunit alph |
| BX295541.1 | X | matrin 3 (MATR3) pseudogene |
| PHKA1-AS1 | X | PHKA1 antisense RNA 1 [Source:HGNC Symbo |
| U3 | X | Small nucleolar RNA U3 [Source:RFAM;Acc:RF |
| AC240504.1 | X | capping protein (actin filament) muscle Z-line, |
| RNU1-112P | X | RNA, U1 small nuclear 112, pseudogene [Sour |
| DMRTC1B | X | DMRT like family C1B [Source:HGNC Symbol;/ |
| FAM226B | X | family with sequence similarity 226 member E |
| FAM236B | X | family with sequence similarity 236 member E |
| FAM236D | X | family with sequence similarity 236 member D |
| DMRTC1 | X | DMRT like family C1 [Source:HGNC Symbol;Ac |
| FAM236C | X | family with sequence similarity 236 member C |
| FAM236A | X | family with sequence similarity 235 member / |
| PABPC1L2B-AS1 | X | PABPC1L2B antisense RNA 1 (head to head) [S |
| PABPC1L2B | X | poly(A) binding protein cytoplasmic 1 like 2B [ |
| PABPC1L2A | X | poly(A) binding protein cytoplasmic 1 like 2A [ |
| AL662864.1 | X | novel transcript |
| NAP1L6 | X | nucleosome assembly protein 1 like 6 [Source |
| NAP1L2 | X | nucleosome assembly protein 1 like 2 [Source |
| TOMM20P4 | X | TOMM20 pseudogene 4 [Source:HGNC Symbo |
| AC004074.1 | X | leucine rich repeat (in FLII) interacting protein |
| YWHAZP8 | X | tyrosine 3-monooxygenase/tryptophan 5-mor |
| RNU6-1044P | X | RNA, U6 small nuclear 1044, pseudogene [Sou |
| CDX4 | X | caudal type homeobox 4 [Source:HGNC Symb |
| MORF4L1P6 | X | mortality factor 4 like 1 pseudogene 6 [Source |
| MAP2K4P1 | X | mitogen-activated protein kinase kinase 4 pse |
| RPL7P53 | X | ribosomal protein L7 pseudogene 53 [Source: |
| CHIC1 | X | cysteine rich hydrophobic domain 1 [Source:H |
| SEPHS1P4 | X | selenophosphate synthetase 1 pseudogene 4 |
| TSIX | X | TSIX transcript, XIST antisense RNA [Source:H |
| XIST | X | X inactive specific transcript [Source:HGNC Sy |
| AL353804.4 | X | NULL |
| AL353804.6 | X | NULL |
| AL353804.7 | X | NULL |
| FXYD6P3 | X | FXYD domain containing ion transport regulat |
| FTX | X | FTX transcript, XIST regulator [Source:HGNC S |
| JPX | X | JPX transcript, XIST activator [Source:HGNC Sy |

| | | |
|---|---|---|
| AL353804.3 | X | NULL |
| AL353804.5 | X | NULL |
| AL353804.1 | X | novel transcript |
| AL353804.2 | X | novel transcript, sense overlapping FTX |
| RPSAP14 | X | ribosomal protein SA pseudogene 14 [Source: |
| AL139400.1 | X | scotin (SCOTIN) pseudogene |
| AL627389.1 | X | novel alanyl-tRNA synthetase protein pseudog |
| UHRF2P1 | X | ubiquitin like with PHD and ring finger domair |
| DDX3P1 | X | DEAD-box helicase 3 pseudogene 1 [Source:H |
| AL359740.1 | X | TEC |
| DDX3P2 | X | DEAD-box helicase 3 pseudogene 2 [Source:H |
| MKRN5P | X | makorin ring finger protein 5, pseudogene [So |
| ATP5MDP1 | X | ATP synthase membrane subunit DAPIT pseuc |
| BMP2KL | X | BMP2 inducible kinase like (pseudogene) [Sou |
| RAB11FIP1P1 | X | RAB11 family interacting protein 1 pseudogen |
| Z83843.1 | X | novel transcript, sense intronic FTX |
| MIR421 | X | microRNA 421 [Source:HGNC Symbol;Acc:HGI |
| MIR374B | X | microRNA 374b [Source:HGNC Symbol;Acc:HG |
| MIR374C | X | microRNA 374c [Source:HGNC Symbol;Acc:HG |
| RN7SL648P | X | RNA, 7SL, cytoplasmic 648, pseudogene [Sour |
| AC004386.7 | X | NULL |
| AC004386.3 | X | NULL |
| AC004386.4 | X | NULL |
| AC004386.5 | X | NULL |
| MIR545 | X | microRNA 545 [Source:HGNC Symbol;Acc:HGI |
| MIR374A | X | microRNA 374a [Source:HGNC Symbol;Acc:HG |
| AC004386.6 | X | NULL |
| ZCCHC13 | X | zinc finger CCHC-type containing 13 [Source:H |
| AC004386.2 | X | ribosomal protein L39 (RPL39) pseudogene |
| AC004386.1 | X | ribosomal protein S6 (RPS6) pseudogene |
| RN7SL790P | X | RNA, 7SL, cytoplasmic 790, pseudogene [Sour |
| THAP12P1 | X | THAP domain containing 12 pseudogene 1 [Sc |
| RPS7P14 | X | ribosomal protein S7 pseudogene 14 [Source: |
| SLC16A2 | X | solute carrier family 16 member 2 [Source:HG |
| HNRNPA1P25 | X | heterogeneous nuclear ribonucleoprotein A1 |
| RLIM | X | ring finger protein, LIM domain interacting [Sc |
| PABPC1P3 | X | poly(A) binding protein cytoplasmic 1 pseudog |
| RNU6-330P | X | RNA, U6 small nuclear 330, pseudogene [Sour |
| NEXMIF | X | neurite extension and migration factor [Sourc |
| AL139395.1 | X | NLR family, pyrin domain containing 3 (NLRP3 |
| ABCB7 | X | ATP binding cassette subfamily B member 7 [S |
| UPRT | X | uracil phosphoribosyltransferase homolog [So |
| BUD31P2 | X | BUD31 homolog pseudogene 2 [Source:HGNC |
| RNU6-562P | X | RNA, U6 small nuclear 562, pseudogene [Sour |
| AL137013.1 | X | telomeric repeat binding factor (NIMA-interac |
| ZDHHC15 | X | zinc finger DHHC-type containing 15 [Source:H |
| RPL21P134 | X | ribosomal protein L21 pseudogene 134 [Sourc |
| PCNPP4 | X | PEST containing nuclear protein pseudogene 4 |
| BRAFP1 | X | BRAF pseudogene 1 [Source:HGNC Symbol;Ac |
| TTC3P1 | X | tetratricopeptide repeat domain 3 pseudogen |

| | | |
|---|---|---|
| MAGEE2 | X | MAGE family member E2 [Source:HGNC Symb |
| SAR1AP4 | X | secretion associated Ras related GTPase 1A ps |
| AL451105.1 | X | chromosome 14 open reading frame 116 (C14 |
| AL451105.2 | X | novel transcript |
| AC233982.1 | X | ADP-ribosylation factor-like 5 (ARL5) pseudog |
| PBDC1 | X | polysaccharide biosynthesis domain containin |
| AC233981.1 | X | novel pseudogene |
| RNU6-867P | X | RNA, U6 small nuclear 867, pseudogene [Sour |
| AC233981.2 | X | endogenous retrovirus group FRD, member 1 |
| LDHBP2 | X | lactate dehydrogenase B pseudogene 2 [Sourc |
| MAGEE1 | X | MAGE family member E1 [Source:HGNC Symb |
| RNA5SP508 | X | RNA, 5S ribosomal pseudogene 508 [Source:H |
| TRAPPC13P1 | X | trafficking protein particle complex 13 pseudo |
| AC233296.1 | X | MIR325 host gene (non-protein coding) |
| MIR384 | X | microRNA 384 [Source:HGNC Symbol;Acc:HGN |
| MIR325 | X | microRNA 325 [Source:HGNC Symbol;Acc:HGN |
| RNA5SP509 | X | RNA, 5S ribosomal pseudogene 509 [Source:H |
| SPRYD7P1 | X | SPRY domain containing 7 pseudogene 1 [Sou |
| FGF16 | X | fibroblast growth factor 16 [Source:HGNC Sym |
| ATRX | X | ATRX chromatin remodeler [Source:HGNC Syr |
| FABP5P15 | X | fatty acid binding protein 5 pseudogene 15 [S |
| MAGT1 | X | magnesium transporter 1 [Source:HGNC Symb |
| RNU6-854P | X | RNA, U6 small nuclear 854, pseudogene [Sour |
| RN7SL460P | X | RNA, 7SL, cytoplasmic 460, pseudogene [Sour |
| COX7B | X | cytochrome c oxidase subunit 7B [Source:HGN |
| ATP7A | X | ATPase copper transporting alpha [Source:HG |
| PGK1 | X | phosphoglycerate kinase 1 [Source:HGNC Sym |
| AL356235.1 | X | chromosome 4 open reading frame 46 (C4orf |
| PGAM4 | X | phosphoglycerate mutase family member 4 [S |
| TAF9B | X | TATA-box binding protein associated factor 9b |
| FNDC3CP | X | fibronectin type III domain containing 3C, pse |
| CYSLTR1 | X | cysteinyl leukotriene receptor 1 [Source:HGN |
| HMGN1P34 | X | high mobility group nucleosome binding dom |
| UBE2V1P7 | X | ubiquitin conjugating enzyme E2 V1 pseudoge |
| RTL3 | X | retrotransposon Gag like 3 [Source:HGNC Sym |
| PPATP2 | X | phosphoribosyl pyrophosphate amidotransfer |
| LPAR4 | X | lysophosphatidic acid receptor 4 [Source:HGN |
| RPL7P54 | X | ribosomal protein L7 pseudogene 54 [Source: |
| MIR4328 | X | microRNA 4328 [Source:HGNC Symbol;Acc:HG |
| P2RY10 | X | P2Y receptor family member 10 [Source:HGN |
| P2RY10BP | X | P2Y receptor family member 10B, pseudogen |
| GPR174 | X | G protein-coupled receptor 174 [Source:HGN |
| CTHRC1P1 | X | collagen triple helix repeat containing 1 pseu |
| KIF4CP | X | kinesin family member 4C, pseudogene [Sour |
| ITM2A | X | integral membrane protein 2A [Source:HGNC |
| AL009028.1 | X | coronin, actin binding protein, 1C (CORO1C) p |
| TBX22 | X | T-box transcription factor 22 [Source:HGNC S |
| CHMP1B2P | X | charged multivesicular body protein 1B2, pse |
| TENT5D | X | terminal nucleotidyltransferase 5D [Source:H |
| HNRNPH3P1 | X | heterogeneous nuclear ribonucleoprotein H3 |

| | | |
|---|---|---|
| AL512504.2 | X | WW domain binding protein 11 (WBP11) pseu |
| HK2P1 | X | hexokinase 2 pseudogene 1 [Source:HGNC Syr |
| BRWD3 | X | bromodomain and WD repeat domain contair |
| AL512504.1 | X | proteasome (prosome, macropain) subunit, al |
| AL590031.1 | X | NULL |
| RNU6-493P | X | RNA, U6 small nuclear 493, pseudogene [Sour |
| VDAC1P1 | X | voltage dependent anion channel 1 pseudoge |
| RNU6-995P | X | RNA, U6 small nuclear 995, pseudogene [Sour |
| AL391294.1 | X | novel transcript |
| HMGN5 | X | high mobility group nucleosome binding doma |
| SH3BGRL | X | SH3 domain binding glutamate rich protein lik |
| AL359885.1 | X | novel pseudogene |
| AL445213.1 | X | pseudogene similar to part of SLC9A2 (solute |
| AL445213.2 | X | novel transcript |
| AL691517.1 | X | pyruvate dehydrogenase kinase, isoenzyme 1 |
| RNU6-974P | X | RNA, U6 small nuclear 974, pseudogene [Sour |
| AL031313.1 | X | eukaryotic translation initiation factor 3 subur |
| RPL22P22 | X | ribosomal protein L22 pseudogene 22 [Source |
| AL109931.1 | X | NULL |
| AL008987.1 | X | dendritic cell protein (GA17) pseudogene |
| ATG4AP1 | X | autophagy related 4A cysteine peptidase pseu |
| Z82170.1 | X | TEC |
| POU3F4 | X | POU class 3 homeobox 4 [Source:HGNC Symb |
| AL121882.1 | X | novel pseudogene |
| TERF1P4 | X | telomeric repeat binding factor 1 pseudogene |
| CYLC1 | X | cylicin 1 [Source:HGNC Symbol;Acc:HGNC:258 |
| RPS6KA6 | X | ribosomal protein S6 kinase A6 [Source:HGNC |
| MIR548I4 | X | microRNA 548i-4 [Source:HGNC Symbol;Acc:H |
| HDX | X | highly divergent homeobox [Source:HGNC Syr |
| AL450471.1 | X | cytochrome b (CYTB) pseudogene |
| SETP4 | X | SET pseudogene 4 [Source:HGNC Symbol;Acc: |
| UBE2DNL | X | ubiquitin conjugating enzyme E2 D N-terminal |
| Z83820.1 | X | pseudogene similar to 4933403O08Rik |
| APOOL | X | apolipoprotein O like [Source:HGNC Symbol;A |
| SATL1 | X | spermidine/spermine N1-acetyl transferase lil |
| Z99571.1 | X | meprin A subunit beta (MEP1B) pseudogene |
| AC003001.2 | X | galactosidase, beta 1-like (GLB1L) pseudogene |
| AC003001.1 | X | novel transcript |
| ZNF711 | X | zinc finger protein 711 [Source:HGNC Symbol; |
| POF1B | X | POF1B actin binding protein [Source:HGNC Sy |
| MIR1321 | X | microRNA 1321 [Source:HGNC Symbol;Acc:HG |
| SFR1P2 | X | SFR1 pseudogene 2 [Source:HGNC Symbol;Acc |
| AL022401.1 | X | ring finger protein 34 (RNF34) pseudogene |
| CHM | X | CHM Rab escort protein [Source:HGNC Symbo |
| MIR361 | X | microRNA 361 [Source:HGNC Symbol;Acc:HGI |
| Y_RNA | X | Y RNA [Source:RFAM;Acc:RF00019] |
| NDUFA5P7 | X | NADH:ubiquinone oxidoreductase subunit A5 |
| STIP1P3 | X | stress induced phosphoprotein 1 pseudogene |
| TPMTP4 | X | thiopurine S-methyltransferase pseudogene 4 |
| DACH2 | X | dachshund family transcription factor 2 [Sourc |

| | | |
|---|---|---|
| EEF1A1P29 | X | eukaryotic translation elongation factor 1 alpl |
| GEMIN8P3 | X | gem nuclear organelle associated protein 8 ps |
| U6 | X | U6 spliceosomal RNA [Source:RFAM;Acc:RF00 |
| FCF1P9 | X | FCF1 pseudogene 9 [Source:HGNC Symbol;Acc |
| AL390058.1 | X | COP9 constitutive photomorphogenic homolo |
| Z82210.1 | X | ribosomal protein L7 (RPL7) pseudogene |
| AL590229.2 | X | general transcription factor IIE, polypeptide 2, |
| AL590229.1 | X | NULL |
| Z96811.1 | X | glyceraldehyde-3-phosphate dehydrogenase ( |
| KLHL4 | X | kelch like family member 4 [Source:HGNC Sym |
| RPSAP15 | X | ribosomal protein SA pseudogene 15 [Source: |
| Z83818.2 | X | novel transcript |
| Z83818.1 | X | zinc finger protein 486 (ZNF486) pseudogene |
| AL136363.1 | X | ribosomal protein L34 (RPL34) pseudogene |
| AL136363.2 | X | mitochondrial ribosomal protein S22 (MRPS22 |
| AL035246.1 | X | novel pseudogene |
| AL024493.1 | X | capping protein (actin filament) muscle Z-line, |
| AL024493.2 | X | NULL |
| AL929325.1 | X | platelet-activating factor acetylhydrolase 1b, ( |
| CPXCR1 | X | CPX chromosome region candidate 1 [Source: |
| AL022154.1 | X | adenosine deaminase, RNA-specific, B1 (ADAF |
| AL645638.1 | X | ubiquitin-conjugating enzyme E2D 2 (UBC4\\/ |
| AL731768.1 | X | riboflavin kinase (RFK) pseudogene |
| SRIP2 | X | sorcin pseudogene 2 [Source:HGNC Symbol;A |
| AL121872.1 | X | novel transcript |
| TGIF2LX | X | TGFB induced factor homeobox 2 like X-linkec |
| AL121877.1 | X | staufen RNA binding protein homolog 2 (Dros |
| USP12PX | X | ubiquitin specific peptidase 12 pseudogene X- |
| AL121873.1 | X | ring finger protein 19B (RNF19B) pseudogene |
| RNF19BPX | X | ring finger protein 19B pseudogene X-linked [ |
| UBE2V1P9 | X | ubiquitin conjugating enzyme E2 V1 pseudoge |
| RNU6-555P | X | RNA, U6 small nuclear 555, pseudogene [Sour |
| AL121823.1 | X | novel transcript |
| PABPC5-AS1 | X | PABPC5 antisense RNA 1 [Source:HGNC Symb |
| AL662874.1 | X | delta sleep inducing peptide, immunoreactor |
| PABPC5 | X | poly(A) binding protein cytoplasmic 5 [Source |
| SERBP1P4 | X | SERPINE1 mRNA binding protein 1 pseudogen |
| PCDH11X | X | protocadherin 11 X-linked [Source:HGNC Sym |
| RNU2-26P | X | RNA, U2 small nuclear 26, pseudogene [Sourc |
| VDAC1P3 | X | voltage dependent anion channel 1 pseudoge |
| EIF4A1P10 | X | eukaryotic translation initiation factor 4A1 ps |
| KRT18P11 | X | keratin 18 pseudogene 11 [Source:HGNC Sym |
| SNX3P1X | X | sorting nexin 3 pseudogene 1 X-linked [Source |
| AL121871.1 | X | ribosomal protein L26 (RPL26) pseudogene |
| TUSC2P2 | X | TUSC2 pseudogene 2 [Source:HGNC Symbol;A |
| AL158053.1 | X | discs, large homolog 7 (Drosophila) (DLG7) ps |
| AL359263.1 | X | adaptor-related protein complex 2, beta 1 sub |
| ST13P18 | X | ST13, Hsp70 interacting protein pseudogene 1 |
| AL023280.1 | X | 5'-nucleotidase, cytosolic II-like 1 (NT5C2L1) p |
| RPL7P55 | X | ribosomal protein L7 pseudogene 55 [Source: |

| | | |
|---|---|---|
| NAP1L3 | X | nucleosome assembly protein 1 like 3 [Source |
| FAM133A | X | family with sequence similarity 133 member A |
| Z94722.1 | X | ubiquitin specific peptidase 37 (USP37) pseud |
| RNU6-332P | X | RNA, U6 small nuclear 332, pseudogene [Sour |
| Y_RNA | X | Y RNA [Source:RFAM;Acc:RF00019] |
| AL390840.1 | X | tubulin, beta 8 class VIII (TUBB8) pseudogene |
| PAICSP7 | X | phosphoribosylaminoimidazole carboxylase, p |
| CCNB1IP1P3 | X | cyclin B1 interacting protein 1 pseudogene 3 [ |
| MIR548M | X | microRNA 548m [Source:HGNC Symbol;Acc:H |
| AL022161.1 | X | VAMP (vesicle-associated membrane protein) |
| CALM1P1 | X | calmodulin 1 pseudogene 1 [Source:HGNC Syr |
| RNA5SP510 | X | RNA, 5S ribosomal pseudogene 510 [Source:H |
| AL365400.1 | X | general transcription factor IIIC, polypeptide 6 |
| HNRNPDLP3 | X | HNRNPDL pseudogene 3 [Source:HGNC Symb |
| HNRNPDLP1 | X | HNRNPDL pseudogene 1 [Source:HGNC Symb |
| RPS7P13 | X | ribosomal protein S7 pseudogene 13 [Source: |
| AL365400.2 | X | ribosome binding factor A (putative) (RBFA) ps |
| AL137845.2 | X | MYST histone acetyltransferase 2 (MYST2) pse |
| RN7SL379P | X | RNA, 7SL, cytoplasmic 379, pseudogene [Sour |
| AL137845.1 | X | zinc finger protein 265 (ZNF265) pseudogene |
| AL022151.1 | X | novel transcript |
| RPS29P28 | X | ribosomal protein S29 pseudogene 28 [Source |
| BRDTP1 | X | bromodomain testis associated pseudogene 1 |
| RN7SKP194 | X | RN7SK pseudogene 194 [Source:HGNC Symbo |
| AL606530.1 | X | S-phase kinase-associated protein 2 (p45) (SKI |
| DIAPH2 | X | diaphanous related formin 2 [Source:HGNC Sy |
| Y_RNA | X | Y RNA [Source:RFAM;Acc:RF00019] |
| RPA4 | X | replication protein A4 [Source:HGNC Symbol;A |
| NDUFB5P2 | X | NADH:ubiquinone oxidoreductase subunit B5 |
| Metazoa_SRP | X | Metazoan signal recognition particle RNA [Sou |
| RN7SL74P | X | RNA, 7SL, cytoplasmic 74, pseudogene [Sourc |
| DIAPH2-AS1 | X | DIAPH2 antisense RNA 1 [Source:HGNC Symb |
| AL354685.1 | X | endogenous retroviral family W, env(C7), mer |
| AL354685.2 | X | pseudogene similar to part of NCK-associated |
| AL391666.1 | X | karyopherin (importin) beta 1 (KPNB1) pseud |
| RPL6P29 | X | ribosomal protein L6 pseudogene 29 [Source: |
| AL357312.1 | X | scavenger receptor class B, member 1 (SCARB |
| EEF1A1P15 | X | eukaryotic translation elongation factor 1 alph |
| HMGB1P32 | X | high mobility group box 1 pseudogene 32 [Sou |
| AL157778.1 | X | novel transcript |
| AL137843.1 | X | testis expressed 101 (TEX101) pseudogene |
| XRCC6P5 | X | X-ray repair cross complementing 6 pseudoge |
| B3GNT2P1 | X | B3GNT2 pseudogene 1 [Source:HGNC Symbol |
| AL359641.2 | X | NULL |
| AL359641.1 | X | novel pseudogene |
| RPSAP8 | X | ribosomal protein SA pseudogene 8 [Source:H |
| PCDH19 | X | protocadherin 19 [Source:HGNC Symbol;Acc:H |
| TNMD | X | tenomodulin [Source:HGNC Symbol;Acc:HGN |
| TSPAN6 | X | tetraspanin 6 [Source:HGNC Symbol;Acc:HGN |
| SRPX2 | X | sushi repeat containing protein X-linked 2 [Sou |

| | | |
|---|---|---|
| SYTL4 | X | synaptotagmin like 4 [Source:HGNC Symbol;A |
| PPIAP89 | X | peptidylprolyl isomerase A pseudogene 89 [Sc |
| Z95327.2 | X | RAD21 homolog (S. pombe) (LOC139319) pseu |
| Z95327.3 | X | tRNA methyltransferase 12 homolog (S. cerev |
| CSTF2 | X | cleavage stimulation factor subunit 2 [Source: |
| Z95327.1 | X | NULL |
| NOX1 | X | NADPH oxidase 1 [Source:HGNC Symbol;Acc:H |
| HNRNPA1P26 | X | heterogeneous nuclear ribonucleoprotein A1 |
| HNRNPA1P27 | X | heterogeneous nuclear ribonucleoprotein A1 |
| Z73417.1 | X | TGF beta-inducible nuclear protein 1 (TINP1) p |
| XKRX | X | XK related X-linked [Source:HGNC Symbol;Acc |
| Z97985.1 | X | chondroitin beta1,4 N-acetylgalactosaminyltra |
| ARL13A | X | ADP ribosylation factor like GTPase 13A [Sour |
| TRMT2B | X | tRNA methyltransferase 2 homolog B [Source: |
| TRMT2B-AS1 | X | TRMT2B antisense RNA 1 [Source:HGNC Symb |
| TMEM35A | X | transmembrane protein 35A [Source:HGNC Sy |
| CENPI | X | centromere protein I [Source:HGNC Symbol;A |
| Z70280.1 | X | tyrosine 3-monooxygenase/tryptophan 5-mor |
| DRP2 | X | dystrophin related protein 2 [Source:HGNC Sy |
| TAF7L | X | TATA-box binding protein associated factor 7 |
| RNU6-934P | X | RNA, U6 small nuclear 934, pseudogene [Sour |
| NANOGNBP3 | X | NANOGNB pseudogene 3 [Source:HGNC Symb |
| RPL21P132 | X | ribosomal protein L21 pseudogene 132 [Sourc |
| TIMM8A | X | translocase of inner mitochondrial membrane |
| BTK | X | Bruton tyrosine kinase [Source:HGNC Symbol; |
| RPL36A | X | ribosomal protein L36a [Source:HGNC Symbol |
| RPL36A-HNRNPH2 | X | RPL36A-HNRNPH2 readthrough [Source:HGN( |
| GLA | X | galactosidase alpha [Source:HGNC Symbol;Acc |
| HNRNPH2 | X | heterogeneous nuclear ribonucleoprotein H2 |
| ARMCX4 | X | armadillo repeat containing X-linked 4 [Source |
| PPIAP90 | X | peptidylprolyl isomerase A pseudogene 90 [Sc |
| AC234775.2 | X | protein kinase C, iota (PRKCI) pseudogene |
| ARMCX1 | X | armadillo repeat containing X-linked 1 [Source |
| ARMCX7P | X | armadillo repeat containing X-linked 7, pseud |
| ARMCX6 | X | armadillo repeat containing X-linked 6 [Source |
| ARMCX3 | X | armadillo repeat containing X-linked 3 [Source |
| ARMCX3-AS1 | X | ARMCX3 antisense RNA 1 [Source:HGNC Symb |
| AC234775.3 | X | novel transcript, overlapping to ARMCX3 |
| RNU6-30P | X | RNA, U6 small nuclear 30, pseudogene [Sourc |
| AC234775.4 | X | novel transcript |
| ARMCX2 | X | armadillo repeat containing X-linked 2 [Source |
| AC234775.1 | X | novel pseudogene |
| RNU6-587P | X | RNA, U6 small nuclear 587, pseudogene [Sour |
| FO082842.1 | X | novel zinc finger protein pseudogene |
| AL672207.1 | X | glycerol kinase (GK) pseudogene |
| NXF5 | X | nuclear RNA export factor 5 [Source:HGNC Sy |
| ZMAT1 | X | zinc finger matrin-type 1 [Source:HGNC Symb |
| Z68868.1 | X | zinc finger protein pseudogene |
| RNU6-345P | X | RNA, U6 small nuclear 345, pseudogene [Sour |
| Z80107.2 | X | histone H2B family pseudogene |

| | | |
|---|---|---|
| Z80107.1 | X | novel pseudogene |
| MTND6P13 | X | MT-ND6 pseudogene 13 [Source:HGNC Symbo |
| TCEAL2 | X | transcription elongation factor A like 2 [Sourc |
| TCEAL6 | X | transcription elongation factor A like 6 [Sourc |
| Z70719.1 | X | novel transcript, antisense to TCEAL6 |
| BEX5 | X | brain expressed X-linked 5 [Source:HGNC Sym |
| TCP11X3P | X | t-complex 11 family, X-linked 3, pseudogene [ |
| TCP11X1 | X | t-complex 11 family, X-linked 1 [Source:HGNC |
| NXF2 | X | nuclear RNA export factor 2 [Source:HGNC Sy |
| AC234791.1 | X | preferentially expressed antigen in melanoma |
| AC235565.2 | X | novel transcript |
| NXF2B | X | nuclear RNA export factor 2B [Source:HGNC S |
| AC235565.1 | X | preferentially expressed antigen in melanoma |
| TCP11X2 | X | t-complex 11 family, X-linked 2 [Source:HGNC |
| TMSB15A | X | thymosin beta 15a [Source:HGNC Symbol;Acc |
| FOXN3P1 | X | forkhead box N3 pseudogene 1 [Source:HGNC |
| NXF4 | X | nuclear RNA export factor 4 pseudogene [Sou |
| ARMCX5 | X | armadillo repeat containing X-linked 5 [Source |
| ARMCX5-GPRASP2 | X | ARMCX5-GPRASP2 readthrough [Source:HGN |
| AL035427.1 | X | novel transcript, antisense to ARMCX5 and GF |
| AL035551.1 | X | histone 3, H3 (HIST3H3) pseudogene |
| GPRASP1 | X | G protein-coupled receptor associated sorting |
| AL035427.2 | X | novel transcript |
| RNU6-589P | X | RNA, U6 small nuclear 589, pseudogene [Sour |
| BEND7P1 | X | BEN domain containing 7 pseudogene 1 [Sour |
| GPRASP2 | X | G protein-coupled receptor associated sorting |
| ARMCX5-GPRASP2 | X | ARMCX5-GPRASP2 readthrough [Source:NCBI |
| BHLHB9 | X | basic helix-loop-helix family member b9 [Sour |
| LINC00630 | X | long intergenic non-protein coding RNA 630 [S |
| MTND1P32 | X | MT-ND1 pseudogene 32 [Source:HGNC Symbo |
| MTND2P2 | X | MT-ND2 pseudogene 2 [Source:HGNC Symbol |
| MTCO1P19 | X | MT-CO1 pseudogene 19 [Source:HGNC Symbo |
| MTCO2P19 | X | MT-CO2 pseudogene 19 [Source:HGNC Symbo |
| MTATP6P19 | X | MT-ATP6 pseudogene 19 [Source:HGNC Symb |
| MTCO3P19 | X | MT-CO3 pseudogene 19 [Source:HGNC Symbo |
| MTND4P32 | X | MT-ND4 pseudogene 32 [Source:HGNC Symbo |
| MTND5P26 | X | MT-ND5 pseudogene 26 [Source:HGNC Symbo |
| MTND6P32 | X | MT-ND6 pseudogene 32 [Source:HGNC Symbo |
| MTCYBP32 | X | MT-CYB pseudogene 32 [Source:HGNC Symbo |
| Z68871.1 | X | novel transcript |
| RAB40AL | X | RAB40A like [Source:HGNC Symbol;Acc:HGNC |
| Z95624.1 | X | novel transcript |
| Z93943.1 | X | novel transcript |
| Z73361.1 | X | nudix (nucleoside diphosphate linked moiety X |
| BEX1 | X | brain expressed X-linked 1 [Source:HGNC Sym |
| NXF3 | X | nuclear RNA export factor 3 [Source:HGNC Sy |
| Z75746.1 | X | zinc finger protein 770 (ZNF770) pseudogene |
| RPSAP59 | X | ribosomal protein SA pseudogene 59 [Source: |
| BEX4 | X | brain expressed X-linked 4 [Source:HGNC Sym |
| TCEAL8 | X | transcription elongation factor A like 8 [Sourc |

| | | |
|---|---|---|
| Z68694.1 | X | guanosine monophosphate reductase (GMPR) |
| TCEAL5 | X | transcription elongation factor A like 5 [Source |
| BEX2 | X | brain expressed X-linked 2 [Source:HGNC Sym |
| TCEAL7 | X | transcription elongation factor A like 7 [Source |
| Z92846.1 | X | nerve growth factor receptor (TNFRSF16) asso |
| TCEAL9 | X | transcription elongation factor A like 9 [Source |
| BEX3 | X | brain expressed X-linked 3 [Source:HGNC Sym |
| AL117327.1 | X | novel transcript |
| Z69733.1 | X | novel transcript |
| RAB40A | X | RAB40A, member RAS oncogene family [Sourc |
| AL021308.1 | X | novel transcript |
| TCEAL4 | X | transcription elongation factor A like 4 [Source |
| Z73965.1 | X | TEC |
| TCEAL3 | X | transcription elongation factor A like 3 [Source |
| TCEAL3-AS1 | X | TCEAL3 antisense RNA 1 [Source:HGNC Symbo |
| TCEAL1 | X | transcription elongation factor A like 1 [Source |
| AL049610.1 | X | novel pseudogene |
| MORF4L2 | X | mortality factor 4 like 2 [Source:HGNC Symbo |
| MORF4L2-AS1 | X | MORF4L2 antisense RNA 1 [Source:HGNC Sym |
| GLRA4 | X | glycine receptor alpha 4 [Source:HGNC Symbo |
| TMEM31 | X | transmembrane protein 31 [Source:HGNC Syn |
| PLP1 | X | proteolipid protein 1 [Source:HGNC Symbol;A |
| RAB9B | X | RAB9B, member RAS oncogene family [Source |
| RNA5SP511 | X | RNA, 5S ribosomal pseudogene 511 [Source:H |
| TMSB15B-AS1 | X | TMSB15B antisense RNA 1 [Source:HGNC Sym |
| AC234782.2 | X | histone H2B pseudogene |
| AC234782.4 | X | HYDIN, axonemal central pair apparatus prote |
| AC234782.3 | X | E74-like factor 2 (ELF2) pseudogene |
| TMSB15B | X | thymosin beta 15B [Source:HGNC Symbol;Acc |
| AC234782.1 | X | solute carrier family 25 (mitochondrial oxodic |
| DPPA3P1 | X | developmental pluripotency associated 3 pseu |
| H2BFWT | X | H2B histone family member W, testis specific |
| H2BFM | X | H2B histone family member M [Source:HGNC |
| AC234783.1 | X | histone H2B pseudogene |
| TMSB15B | X | thymosin beta 15B [Source:NCBI gene;Acc:28 |
| SLC25A53 | X | solute carrier family 25 member 53 [Source:H |
| ZCCHC18 | X | zinc finger CCHC-type containing 18 [Source:H |
| FAM199X | X | family with sequence similarity 199, X-linked [ |
| ESX1 | X | ESX homeobox 1 [Source:HGNC Symbol;Acc:H |
| IL1RAPL2 | X | interleukin 1 receptor accessory protein like 2 |
| PHBP10 | X | prohibitin pseudogene 10 [Source:HGNC Symb |
| RPL18AP14 | X | ribosomal protein L18a pseudogene 14 [Sourc |
| TEX13A | X | testis expressed 13A [Source:HGNC Symbol;Ac |
| KCTD9P2 | X | potassium channel tetramerization domain co |
| RNU6-207P | X | RNA, U6 small nuclear 207, pseudogene [Sour |
| NRK | X | Nik related kinase [Source:HGNC Symbol;Acc: |
| SERPINA7 | X | serpin family A member 7 [Source:HGNC Symb |
| AL512661.1 | X | novel pseudogene |
| PWWP3B | X | PWWP domain containing 3B [Source:HGNC S |
| AL133271.1 | X | Chondroitin beta-1,4-N-acetylgalactosaminylt |

| | | |
|---|---|---|
| Y_RNA | X | Y RNA [Source:RFAM;Acc:RF00019] |
| NAP1L4P2 | X | nucleosome assembly protein 1 like 4 pseudo[illegible] |
| SERPINA7P1 | X | serpin family A member 7 pseudogene 1 [Sour[illegible] |
| AL590808.1 | X | novel zinc finger protein pseudogene |
| RADX | X | RPA1 related single stranded DNA binding pro[illegible] |
| MIR548AN | X | microRNA 548an [Source:HGNC Symbol;Acc:H[illegible] |
| RNF128 | X | ring finger protein 128 [Source:HGNC Symbol;[illegible] |
| TBC1D8B | X | TBC1 domain family member 8B [Source:HGN[illegible] |
| MORC4 | X | MORC family CW-type zinc finger 4 [Source:H[illegible] |
| RIPPLY1 | X | ripply transcriptional repressor 1 [Source:HGN[illegible] |
| CLDN2 | X | claudin 2 [Source:HGNC Symbol;Acc:HGNC:20[illegible] |
| EEF1A1P40 | X | eukaryotic translation elongation factor 1 alp[illegible] |
| RBM41 | X | RNA binding motif protein 41 [Source:HGNC S[illegible] |
| NUP62CL | X | nucleoporin 62 C-terminal like [Source:HGNC [illegible] |
| AL390039.1 | X | general transcription factor IIIC, polypeptide 6[illegible] |
| Y_RNA | X | Y RNA [Source:RFAM;Acc:RF00019] |
| PIH1D3 | X | PIH1 domain containing 3 [Source:HGNC Sym[illegible] |
| MYCLP1 | X | MYCL pseudogene 1 [Source:HGNC Symbol;Ac[illegible] |
| DNAJA1P3 | X | DnaJ heat shock protein family (Hsp40) memb[illegible] |
| KRT18P49 | X | keratin 18 pseudogene 49 [Source:HGNC Sym[illegible] |
| FRMPD3-AS1 | X | FRMPD3 antisense RNA 1 [Source:HGNC Symb[illegible] |
| FRMPD3 | X | FERM and PDZ domain containing 3 [Source:H[illegible] |
| PRPS1 | X | phosphoribosyl pyrophosphate synthetase 1 [[illegible] |
| TSC22D3 | X | TSC22 domain family member 3 [Source:HGN[illegible] |
| NCBP2L | X | nuclear cap binding protein subunit 2 like [Sou[illegible] |
| MID2 | X | midline 2 [Source:HGNC Symbol;Acc:HGNC:70[illegible] |
| AL109946.1 | X | novel transcript, antisense to MID2 |
| Y_RNA | X | Y RNA [Source:RFAM;Acc:RF00019] |
| AL034399.1 | X | novel pseudogene |
| NLRP3P1 | X | NLR family pyrin domain containing 3 pseudo[illegible] |
| TEX13B | X | testis expressed 13B [Source:HGNC Symbol;Ac[illegible] |
| VSIG1 | X | V-set and immunoglobulin domain containing [illegible] |
| PSMD10 | X | proteasome 26S subunit, non-ATPase 10 [Sou[illegible] |
| ATG4A | X | autophagy related 4A cysteine peptidase [Sou[illegible] |
| COL4A6 | X | collagen type IV alpha 6 chain [Source:HGNC [illegible] |
| AL034369.1 | X | pancreatic lipase-related protein 2 (PNLIPRP2[illegible] |
| COL4A5 | X | collagen type IV alpha 5 chain [Source:HGNC [illegible] |
| Y_RNA | X | Y RNA [Source:RFAM;Acc:RF00019] |
| AL035425.3 | X | novel transcript, |
| AL035425.2 | X | insulin receptor substrate 4 [Source:NCBI gen[illegible] |
| AL035425.4 | X | TEC |
| IRS4 | X | insulin receptor substrate 4 [Source:HGNC Sy[illegible] |
| AL035425.1 | X | novel transcript, antisense to IRS4 |
| RNU6-309P | X | RNA, U6 small nuclear 309, pseudogene [Sour[illegible] |
| AL928646.1 | X | NULL |
| AL034403.1 | X | glyceraldehyde-3-phosphate dehydrogenase ([illegible] |
| GUCY2F | X | guanylate cyclase 2F, retinal [Source:HGNC Sy[illegible] |
| NXT2 | X | nuclear transport factor 2 like export factor 2 [illegible] |
| AL118496.1 | X | mitofusin 1 (MFN1) pseudogene |
| KCNE5 | X | potassium voltage-gated channel subfamily E [illegible] |

| | | |
|---|---|---|
| ACSL4 | X | acyl-CoA synthetase long chain family membe |
| AL118496.2 | X | pseudogene similar to part of protein phospha |
| Y_RNA | X | Y RNA [Source:RFAM;Acc:RF00019] |
| RPS5P7 | X | ribosomal protein S5 pseudogene 7 [Source:H |
| TMEM164 | X | transmembrane protein 164 [Source:HGNC Sy |
| MIR652 | X | microRNA 652 [Source:HGNC Symbol;Acc:HGI |
| MIR3978 | X | microRNA 3978 [Source:HGNC Symbol;Acc:HG |
| AMMECR1 | X | Alport syndrome, mental retardation, midface |
| SNORD96B | X | small nucleolar RNA, C/D box 96B [Source:HG |
| AMMECR1-IT1 | X | AMMECR1 intronic transcript 1 [Source:HGNC |
| GNG5P2 | X | G protein subunit gamma 5 pseudogene 2 [So |
| RTL9 | X | retrotransposon Gag like 9 [Source:HGNC Sym |
| AC000113.1 | X | novel transcript |
| TDGF1P3 | X | teratocarcinoma-derived growth factor 1 pseu |
| AL512882.1 | X | mannose-6-phosphate receptor (cation deper |
| CHRDL1 | X | chordin like 1 [Source:HGNC Symbol;Acc:HGN |
| PAK3 | X | p21 (RAC1) activated kinase 3 [Source:HGNC S |
| FCF1P4 | X | FCF1 pseudogene 4 [Source:HGNC Symbol;Acc |
| GLUD1P9 | X | glutamate dehydrogenase 1 pseudogene 9 [Sc |
| CAPN6 | X | calpain 6 [Source:HGNC Symbol;Acc:HGNC:14 |
| DCX | X | doublecortin [Source:HGNC Symbol;Acc:HGN( |
| RNU6-496P | X | RNA, U6 small nuclear 496, pseudogene [Sour |
| HMGB1P12 | X | high mobility group box 1 pseudogene 12 [Sou |
| SERTM2 | X | serine rich and transmembrane domain conta |
| AL589880.1 | X | novel pseudogene (PRO1843) |
| EIF4BP7 | X | eukaryotic translation initiation factor 4B pseu |
| RPL18AP15 | X | ribosomal protein L18a pseudogene 15 [Sourc |
| ALG13 | X | ALG13 UDP-N-acetylglucosaminyltransferase s |
| RNA5SP512 | X | RNA, 5S ribosomal pseudogene 512 [Source:H |
| ALG13-AS1 | X | ALG13 antisense RNA 1 [Source:HGNC Symbol |
| TRPC5 | X | transient receptor potential cation channel su |
| TRPC5OS | X | TRPC5 opposite strand [Source:HGNC Symbol; |
| AC005191.1 | X | Finkel-Biskis-Reilly murine sarcoma virus (FBR- |
| DPRXP7 | X | divergent-paired related homeobox pseudoge |
| AL392112.1 | X | coiled-coil domain containing 121 (CCDC121) |
| RTL4 | X | retrotransposon Gag like 4 [Source:HGNC Sym |
| LHFPL1 | X | LHFPL tetraspan subfamily member 1 [Source |
| Y_RNA | X | Y RNA [Source:RFAM;Acc:RF00019] |
| HMGB3P30 | X | high mobility group box 3 pseudogene 30 [Sou |
| AMOT | X | angiomotin [Source:HGNC Symbol;Acc:HGNC: |
| MIR4329 | X | microRNA 4329 [Source:HGNC Symbol;Acc:HG |
| RN7SL266P | X | RNA, 7SL, cytoplasmic 266, pseudogene [Sour |
| AC002072.1 | X | novel transcript |
| RNU6-1015P | X | RNA, U6 small nuclear 1015, pseudogene [Sou |
| AL023877.1 | X | novel transcript |
| AL031183.1 | X | novel pseudogene |
| AC233289.1 | X | peptidyl-prolyl isomerase G (cyclophilin G) (PF |
| XACT | X | X active specific transcript [Source:HGNC Sym |
| QTRT1P1 | X | queuine tRNA-ribosyltransferase catalytic sub |
| U3 | X | Small nucleolar RNA U3 [Source:RFAM;Acc:RF |

| | | |
|---|---|---|
| RNU1-57P | X | RNA, U1 small nuclear 57, pseudogene [Sourc |
| RN7SL93P | X | RNA, 7SL, cytoplasmic 93, pseudogene [Sourc |
| SNORD30 | X | Small nucleolar RNA SNORD30 [Source:RFAM; |
| AL953862.1 | X | pseudogene similar to part of transportin 3 (T |
| AL953862.2 | X | transportin 3 (TNPO3) pseudogene |
| SSU72P1 | X | SSU72 pseudogene 1 [Source:HGNC Symbol;A |
| HTR2C | X | 5-hydroxytryptamine receptor 2C [Source:HG |
| RPL36AP53 | X | ribosomal protein L36a pseudogene 53 [Sourc |
| SNORA35 | X | small nucleolar RNA, H/ACA box 35 [Source:H |
| MIR764 | X | microRNA 764 [Source:HGNC Symbol;Acc:HGN |
| MIR1912 | X | microRNA 1912 [Source:HGNC Symbol;Acc:HG |
| MIR1264 | X | microRNA 1264 [Source:HGNC Symbol;Acc:HG |
| MIR1298 | X | microRNA 1298 [Source:HGNC Symbol;Acc:HG |
| MIR1911 | X | microRNA 1911 [Source:HGNC Symbol;Acc:HG |
| HSPA8P7 | X | heat shock protein family A (Hsp70) member |
| MIR448 | X | microRNA 448 [Source:HGNC Symbol;Acc:HGN |
| AL355812.1 | X | eukaryotic translation elongation factor 1 alp |
| IL13RA2 | X | interleukin 13 receptor subunit alpha 2 [Sourc |
| RNU6-648P | X | RNA, U6 small nuclear 648, pseudogene [Sour |
| AL121878.1 | X | zinc finger protein 33a (ZNF33A) pseudogene |
| YAP1P2 | X | Yes associated protein 1 pseudogene 2 [Sourc |
| LRCH2 | X | leucine rich repeats and calponin homology d |
| SNORA35B | X | small nucleolar RNA, H/ACA box 35B [Source: |
| Y_RNA | X | Y RNA [Source:RFAM;Acc:RF00019] |
| RBMXL3 | X | RBMX like 3 [Source:HGNC Symbol;Acc:HGNC |
| RPL36P18 | X | ribosomal protein L36 pseudogene 18 [Source |
| LUZP4 | X | leucine zipper protein 4 [Source:HGNC Symbo |
| AC239600.2 | X | alpha tubulin pseudogene |
| AC239600.1 | X | UDP-GlcNAc:betaGal beta-1,3-N-acetylglucosa |
| PLS3-AS1 | X | PLS3 antisense RNA 1 [Source:HGNC Symbol; |
| AL589842.1 | X | NULL |
| PLS3 | X | plastin 3 [Source:HGNC Symbol;Acc:HGNC:90 |
| AC005000.2 | X | PHD finger protein 5A (PHF5A) pseudogene |
| ASS1P5 | X | argininosuccinate synthetase 1 pseudogene 5 |
| DANT2 | X | DXZ4 associated non-coding transcript 2, dista |
| EEF1GP5 | X | eukaryotic translation elongation factor 1 gam |
| AC005000.1 | X | small ubiquitin-like modifier 2 (SUMO2) pseuc |
| DANT1 | X | DXZ4 associated non-coding transcript 1, prox |
| RN7SL712P | X | RNA, 7SL, cytoplasmic 712, pseudogene [Sour |
| AKR1B1P8 | X | aldo-keto reductase family 1 member B1 pseu |
| API5P1 | X | apoptosis inhibitor 5 pseudogene 1 [Source:H |
| AGTR2 | X | angiotensin II receptor type 2 [Source:HGNC S |
| RNU6-154P | X | RNA, U6 small nuclear 154, pseudogene [Sour |
| SLC6A14 | X | solute carrier family 6 member 14 [Source:HG |
| CT83 | X | cancer/testis antigen 83 [Source:HGNC Symbo |
| BX119904.3 | X | pseudogene similar to part of solute carrier fa |
| BX119904.4 | X | pseudogene similar to part of solute carrier fa |
| BX119904.2 | X | novel transcript |
| BX119904.1 | X | pseudogene similar to PRP4 pre-mRNA proces |
| RNU6-1323P | X | RNA, U6 small nuclear 1323, pseudogene [Sou |

| | | |
|---|---|---|
| SETP8 | X | SET pseudogene 8 [Source:HGNC Symbol;Acc: |
| AC005002.2 | X | novel pseudogene |
| AC005002.1 | X | pseudogene similar to part of TCERG1 (transcı |
| AC006963.1 | X | TEC |
| KLHL13 | X | kelch like family member 13 [Source:HGNC Sy |
| Y_RNA | X | Y RNA [Source:RFAM;Acc:RF00019] |
| TUBB4BP3 | X | tubulin beta 4B class IVb pseudogene 3 [Sourc |
| RPL12P43 | X | ribosomal protein L12 pseudogene 43 [Source |
| AL391358.1 | X | bromodomain adjacent to zinc finger domain, |
| WDR44 | X | WD repeat domain 44 [Source:HGNC Symbol; |
| MIR1277 | X | microRNA 1277 [Source:HGNC Symbol;Acc:HG |
| DOCK11 | X | dedicator of cytokinesis 11 [Source:HGNC Sym |
| Y_RNA | X | Y RNA [Source:RFAM;Acc:RF00019] |
| RBBP8P1 | X | RBBP8 pseudogene 1 [Source:HGNC Symbol;A |
| IL13RA1 | X | interleukin 13 receptor subunit alpha 1 [Sourc |
| AL391280.1 | X | NULL |
| AL391280.2 | X | novel pseudogene |
| ZCCHC12 | X | zinc finger CCHC-type containing 12 [Source:H |
| LINC01285 | X | long intergenic non-protein coding RNA 1285 |
| AC006965.1 | X | aldo-keto reductase family 7, member A2 (afla |
| AL772284.1 | X | pseudogene similar to part of KIAA0633 prote |
| LONRF3 | X | LON peptidase N-terminal domain and ring fir |
| AL772284.2 | X | novel cancer/testis antigen family protein |
| KIAA1210 | X | KIAA1210 [Source:HGNC Symbol;Acc:HGNC:2! |
| ARL5AP1 | X | ADP ribosylation factor like GTPase 5A pseudo |
| HNRNPA1P28 | X | heterogeneous nuclear ribonucleoprotein A1 |
| AC004835.2 | X | nudix (nucleoside diphosphate linked moiety ) |
| PGRMC1 | X | progesterone receptor membrane componen |
| AKAP17BP | X | A-kinase anchoring protein 17B, pseudogene |
| AC004835.1 | X | novel transcript |
| AC004973.1 | X | novel transcript |
| MRPS17P9 | X | mitochondrial ribosomal protein S17 pseudog |
| SLC25A43 | X | solute carrier family 25 member 43 [Source:H |
| AC004000.1 | X | novel transcript |
| RNU1-67P | X | RNA, U1 small nuclear 67, pseudogene [Sourc |
| RN7SL118P | X | RNA, 7SL, cytoplasmic 118, pseudogene [Sour |
| RNY3P16 | X | RNY3 pseudogene 16 [Source:HGNC Symbol;A |
| SLC25A5-AS1 | X | SLC25A5 antisense RNA 1 [Source:HGNC Symb |
| SLC25A5 | X | solute carrier family 25 member 5 [Source:HG |
| CXorf56 | X | chromosome X open reading frame 56 [Sourc |
| UBE2A | X | ubiquitin conjugating enzyme E2 A [Source:HG |
| NKRF | X | NFKB repressing factor [Source:HGNC Symbol |
| SEPTIN6 | X | septin 6 [Source:HGNC Symbol;Acc:HGNC:158 |
| MIR766 | X | microRNA 766 [Source:HGNC Symbol;Acc:HGI |
| AC005052.1 | X | TEC |
| SOWAHD | X | sosondowah ankyrin repeat domain family me |
| RPL39 | X | ribosomal protein L39 [Source:HGNC Symbol; |
| SNORA69 | X | small nucleolar RNA, H/ACA box 69 [Source:H |
| AC005052.2 | X | novel transcript, antisense to UPF3B |
| UPF3B | X | UPF3B regulator of nonsense mediated mRNA |

| | | |
|---|---|---|
| RNF113A | X | ring finger protein 113A [Source:HGNC Symbo |
| NDUFA1 | X | NADH:ubiquinone oxidoreductase subunit A1 |
| AKAP14 | X | A-kinase anchoring protein 14 [Source:HGNC S |
| NKAP | X | NFKB activating protein [Source:HGNC Symbo |
| RHOXF1P3 | X | Rhox homeobox family member 1 pseudogen |
| RHOXF1P1 | X | Rhox homeobox family member 1 pseudogen |
| RHOXF1P2 | X | Rhox homeobox family member 1 pseudogen |
| RHOXF1-AS1 | X | RHOXF1 antisense RNA 1 [Source:HGNC Symb |
| RHOXF2B | X | Rhox homeobox family member 2B [Source:H |
| RHOXF1 | X | Rhox homeobox family member 1 [Source:HG |
| LINC01402 | X | long intergenic non-protein coding RNA 1402 |
| NKAPP1 | X | NFKB activating protein pseudogene 1 [Source |
| RHOXF2 | X | Rhox homeobox family member 2 [Source:HG |
| SFR1P1 | X | SFR1 pseudogene 1 [Source:HGNC Symbol;Acc |
| EEF1A1P30 | X | eukaryotic translation elongation factor 1 alph |
| ZBTB33 | X | zinc finger and BTB domain containing 33 [Sou |
| TMEM255A | X | transmembrane protein 255A [Source:HGNC S |
| AL512286.1 | X | prefoldin 4 (PFDN4) pseudogene |
| ATP1B4 | X | ATPase Na+/K+ transporting family member b |
| LAMP2 | X | lysosomal associated membrane protein 2 [So |
| RNU7-37P | X | RNA, U7 small nuclear 37 pseudogene [Source |
| CUL4B | X | cullin 4B [Source:HGNC Symbol;Acc:HGNC:25 |
| MCTS1 | X | MCTS1 re-initiation and release factor [Source |
| C1GALT1C1 | X | C1GALT1 specific chaperone 1 [Source:HGNC S |
| AC011890.2 | X | H3 histone, family 3B (H3.3B) (H3F3B) pseudo |
| AC011890.1 | X | novel pseudogene |
| PA2G4P1 | X | proliferation-associated 2G4 pseudogene 1 [S |
| CT47B1 | X | cancer/testis antigen family 47 member B1 [S |
| AC008162.1 | X | coiled-coil domain containing 58 (CCDC58) ps |
| AC008162.2 | X | novel protein similar to cancer/testis antigen |
| CT47A12 | X | cancer/testis antigen family 47 member A12 [ |
| AL670379.6 | X | coiled-coil domain containing 58 (CCDC58) ps |
| CT47A11 | X | cancer/testis antigen family 47 member A11 [ |
| AL670379.4 | X | coiled-coil domain containing 58 (CCDC58) ps |
| CT47A10 | X | cancer/testis antigen family 47 member A10 [ |
| AL670379.10 | X | coiled-coil domain containing 58 (CCDC58) ps |
| CT47A9 | X | cancer/testis antigen family 47 member A9 [S |
| AL670379.9 | X | coiled-coil domain containing 58 (CCDC58) ps |
| CT47A8 | X | cancer/testis antigen family 47 member A8 [S |
| AL670379.2 | X | coiled-coil domain containing 58 (CCDC58) ps |
| CT47A7 | X | cancer/testis antigen family 47 member A7 [S |
| AL670379.12 | X | coiled-coil domain containing 58 (CCDC58) ps |
| CT47A6 | X | cancer/testis antigen family 47 member A6 [S |
| AL670379.11 | X | coiled-coil domain containing 58 (CCDC58) ps |
| CT47A5 | X | cancer/testis antigen family 47 member A5 [S |
| AL670379.7 | X | coiled-coil domain containing 58 (CCDC58) ps |
| CT47A4 | X | cancer/testis antigen family 47 member A4 [S |
| AL670379.8 | X | coiled-coil domain containing 58 (CCDC58) ps |
| CT47A3 | X | cancer/testis antigen family 47 member A3 [S |
| AL670379.5 | X | coiled-coil domain containing 58 (CCDC58) ps |

| | | |
|---|---|---|
| CT47A2 | X | cancer/testis antigen family 47 member A2 [S |
| AL670379.3 | X | coiled-coil domain containing 58 (CCDC58) ps |
| CT47A1 | X | cancer/testis antigen family 47 member A1 [S |
| AL670379.1 | X | coiled-coil domain containing 58 (CCDC58) ps |
| AC006144.1 | X | E-cadherin binding protein E7 (HAKAI) pseudo |
| GLUD2 | X | glutamate dehydrogenase 2 [Source:HGNC Sy |
| AC006144.2 | X | novel transcript |
| AC002377.1 | X | novel transcript |
| HSPA8P1 | X | heat shock protein family A (Hsp70) member |
| MIR3672 | X | microRNA 3672 [Source:HGNC Symbol;Acc:H |
| TPT1P13 | X | tumor protein, translationally-controlled 1 ps |
| AL359851.1 | X | novel transcript |
| AL441985.1 | X | pseudogene similar to part of ubiquitin-conju |
| AL357562.1 | X | TEC |
| AL513487.1 | X | novel transcript |
| RPL3P12 | X | ribosomal protein L3 pseudogene 12 [Source: |
| U3 | X | Small nucleolar RNA U3 [Source:RFAM;Acc:RF |
| MRRFP1 | X | mitochondrial ribosome recycling factor pseu |
| GRIA3 | X | glutamate ionotropic receptor AMPA type sub |
| AL590139.1 | X | H3 histone, family 3B (H3.3B) (H3F3B) pseudo |
| AL030996.1 | X | NADH dehydrogenase (ubiquinone) 1 alpha su |
| TUBB4AP1 | X | tubulin beta 4A class IVa pseudogene 1 [Sourc |
| THOC2 | X | THO complex 2 [Source:HGNC Symbol;Acc:HG |
| FERP1 | X | FER tyrosine kinase pseudogene 1 [Source:HG |
| MEMO1P4 | X | mediator of cell motility 1 pseudogene 4 [Sou |
| RNU7-69P | X | RNA, U7 small nuclear 69 pseudogene [Source |
| CHCHD2P1 | X | coiled-coil-helix-coiled-coil-helix domain conta |
| RN7SL190P | X | RNA, 7SL, cytoplasmic 190, pseudogene [Sour |
| Y_RNA | X | Y RNA [Source:RFAM;Acc:RF00019] |
| XIAP | X | X-linked inhibitor of apoptosis [Source:HGNC |
| XIAP-AS1 | X | XIAP antisense RNA 1 [Source:HGNC Symbol; |
| RNU6-122P | X | RNA, U6 small nuclear 122, pseudogene [Sour |
| AL121601.2 | X | par-6 partitioning defective 6 homolog beta ( |
| AL121601.1 | X | novel transcript |
| STAG2 | X | stromal antigen 2 [Source:HGNC Symbol;Acc: |
| ZIK1P1 | X | zinc finger protein interacting with K protein 1 |
| AL391241.1 | X | NULL |
| HSPA8P20 | X | heat shock protein family A (Hsp70) member |
| SH2D1A | X | SH2 domain containing 1A [Source:HGNC Sym |
| TEX13D | X | TEX13 family member D [Source:HGNC Symbo |
| NPM1P34 | X | nucleophosmin 1 pseudogene 34 [Source:HG |
| AL022718.1 | X | pseudogene similar to part of ribosomal prote |
| TENM1 | X | teneurin transmembrane protein 1 [Source:H |
| Z85995.1 | X | ribosomal protein (RPS26) pseudogene |
| AL008707.1 | X | odz, odd Oz\/ten-m homolog 1(Drosophila) ( |
| TEX13C | X | TEX13 family member C [Source:HGNC Symbo |
| AL627231.1 | X | filamin-binding LIM protein-1 (FBLP-1) pseudo |
| AL645820.1 | X | TEC |
| AL445072.1 | X | novel transcript |
| DCAF12L2 | X | DDB1 and CUL4 associated factor 12 like 2 [So |

| | | |
|---|---|---|
| MTCO1P53 | X | MT-CO1 pseudogene 53 [Source:HGNC Symbo |
| MTND4LP1 | X | MT-ND4L pseudogene 1 [Source:HGNC Symbo |
| MTND4P24 | X | MT-ND4 pseudogene 24 [Source:HGNC Symbo |
| Z69908.1 | X | olfactory receptor family 4 subfamily W meml |
| DCAF12L1 | X | DDB1 and CUL4 associated factor 12 like 1 [So |
| MTCYBP38 | X | MT-CYB pseudogene 38 [Source:HGNC Symbo |
| PRR32 | X | proline rich 32 [Source:HGNC Symbol;Acc:HGI |
| Z82209.1 | X | intracellular membrane-associated calcium-in |
| Z82209.2 | X | TEC |
| AL662814.1 | X | IMP1 inner mitochondrial membrane peptida: |
| AL591643.1 | X | novel transcript |
| Z75741.2 | X | kinesin heavy chain member 2 (KIF2) pseudog |
| Z75741.1 | X | Huntingtin interacting protein K (HYPK) pseud |
| AL008633.1 | X | novel transcript |
| ACTRT1 | X | actin related protein T1 [Source:HGNC Symbo |
| Z74696.2 | X | novel pseudogene |
| BTG3P1 | X | BTG anti-proliferation factor 3 pseudogene 1 |
| Z74696.1 | X | actin-related protein pseudogene |
| RPL7AP72 | X | ribosomal protein L7a pseudogene 72 [Source |
| AL442647.1 | X | RuvB-like 2 (E. coli) (RUVBL2) pseudogene |
| KRT18P44 | X | keratin 18 pseudogene 44 [Source:HGNC Sym |
| RPL32P35 | X | ribosomal protein L32 pseudogene 35 [Source |
| AL359845.1 | X | tight junction protein 4 (peripheral) (TJP4) pse |
| RPS26P56 | X | ribosomal protein S26 pseudogene 56 [Source |
| RNA5SP513 | X | RNA, 5S ribosomal pseudogene 513 [Source:H |
| SMARCA1 | X | SWI/SNF related, matrix associated, actin dep |
| AL138745.1 | X | NULL |
| OCRL | X | OCRL inositol polyphosphate-5-phosphatase [ |
| APLN | X | apelin [Source:HGNC Symbol;Acc:HGNC:1666! |
| AL022162.1 | X | novel transcript |
| XPNPEP2 | X | X-prolyl aminopeptidase 2 [Source:HGNC Sym |
| SASH3 | X | SAM and SH3 domain containing 3 [Source:H( |
| AL023653.1 | X | novel transcript |
| Metazoa_SRP | X | Metazoan signal recognition particle RNA [Sou |
| ZDHHC9 | X | zinc finger DHHC-type containing 9 [Source:H( |
| AL034405.1 | X | novel transcript |
| SALL4P2 | X | spalt like transcription factor 4 pseudogene 2 |
| UTP14A | X | UTP14A small subunit processome componen |
| BCORL1 | X | BCL6 corepressor like 1 [Source:HGNC Symbol |
| ELF4 | X | E74 like ETS transcription factor 4 [Source:HG |
| AIFM1 | X | apoptosis inducing factor mitochondria associ |
| AL139234.1 | X | novel transcript |
| RAB33A | X | RAB33A, member RAS oncogene family [Sourc |
| ZNF280C | X | zinc finger protein 280C [Source:HGNC Symbo |
| RNU6-1130P | X | RNA, U6 small nuclear 1130, pseudogene [Sou |
| AL035423.1 | X | SNRPN upstream reading frame pseudogene ( |
| SLC25A14 | X | solute carrier family 25 member 14 [Source:H |
| GPR119 | X | G protein-coupled receptor 119 [Source:HGN( |
| RBMX2 | X | RNA binding motif protein X-linked 2 [Source: |
| Z82195.2 | X | fibrous sheath interacting protein 2 (FSIP2) ps |

| | | |
|---|---|---|
| DENND10P1 | X | DENND10 pseudogene 1 [Source:HGNC Symb |
| Z82195.3 | X | novel transcript |
| Z82195.1 | X | myotubularin related protein 8 (MTMR8) pseu |
| ENOX2 | X | ecto-NOX disulfide-thiol exchanger 2 [Source: |
| AL049733.1 | X | novel pseudogene |
| LINC01201 | X | long intergenic non-protein coding RNA 1201 |
| ARHGAP36 | X | Rho GTPase activating protein 36 [Source:HGI |
| RN7SL191P | X | RNA, 7SL, cytoplasmic 191, pseudogene [Sour |
| OR11Q1P | X | olfactory receptor family 11 subfamily Q mem |
| AL603758.1 | X | novel olfactory receptor pseudogene |
| IGSF1 | X | immunoglobulin superfamily member 1 [Sour |
| OR11N1P | X | olfactory receptor family 11 subfamily N mem |
| OR5AW1P | X | olfactory receptor family 5 subfamily AW mer |
| AL135784.1 | X | novel transcript, antisense to IGSF1 |
| OR5BH1P | X | olfactory receptor family 5 subfamily BH mem |
| OR1AA1P | X | olfactory receptor family 1 subfamily AA mem |
| AL049734.2 | X | pseudogene similar to part of seven transmen |
| AL049734.1 | X | pseudogene similar to part of PYRIN-containir |
| OR13H1 | X | olfactory receptor family 13 subfamily H mem |
| OR13K1P | X | olfactory receptor family 13 subfamily K mem |
| AL365256.1 | X | novel transcript, antisense to IGSF1 |
| OR7L1P | X | olfactory receptor family 7 subfamily L memb |
| OR2AF1P | X | olfactory receptor family 2 subfamily AF mem |
| HDGFP1 | X | heparin binding growth factor pseudogene 1 [ |
| FIRRE | X | firre intergenic repeating RNA element [Sourc |
| MCRIP2P1 | X | MAPK regulated corepressor interacting prote |
| PNKDP1 | X | PNKD pseudogene 1 [Source:HGNC Symbol;Ac |
| AL008629.1 | X | pseudogene similar to part of SMT3 suppresso |
| RNA5SP514 | X | RNA, 5S ribosomal pseudogene 514 [Source:H |
| STK26 | X | serine/threonine kinase 26 [Source:HGNC Syn |
| FRMD7 | X | FERM domain containing 7 [Source:HGNC Sym |
| AL049792.1 | X | acylglycerol kinase, pseudogene |
| RAP2C | X | RAP2C, member of RAS oncogene family [Sou |
| RAP2C-AS1 | X | RAP2C antisense RNA 1 [Source:HGNC Symbo |
| MBNL3 | X | muscleblind like splicing regulator 3 [Source:H |
| RNU6-98P | X | RNA, U6 small nuclear 98, pseudogene [Sourc |
| HS6ST2 | X | heparan sulfate 6-O-sulfotransferase 2 [Sourc |
| HS6ST2-AS1 | X | HS6ST2 antisense RNA 1 [Source:HGNC Symbo |
| Z82205.1 | X | N-acetyltransferase 5 (ARD1 homolog, S. cere |
| USP26 | X | ubiquitin specific peptidase 26 [Source:HGNC |
| Z77249.2 | X | DEAD (Asp-Glu-Ala-Asp) box polypeptide 53 (I |
| Z77249.1 | X | NULL |
| TFDP3 | X | transcription factor Dp family member 3 [Sour |
| RNU1-115P | X | RNA, U1 small nuclear 115, pseudogene [Sour |
| GPC4 | X | glypican 4 [Source:HGNC Symbol;Acc:HGNC:4 |
| RNU6-203P | X | RNA, U6 small nuclear 203, pseudogene [Sour |
| GPC3 | X | glypican 3 [Source:HGNC Symbol;Acc:HGNC:4 |
| AL009174.1 | X | ribosomal protein S24 (RPS24) pseudogene |
| RPSAP63 | X | ribosomal protein SA pseudogene 63 [Source: |
| GPC3-AS1 | X | GPC3 antisense RNA 1 [Source:HGNC Symbol; |

| | | |
|---|---|---|
| AF003529.1 | X | novel pseudogene |
| Y_RNA | X | Y RNA [Source:RFAM;Acc:RF00019] |
| TIMM8BP2 | X | translocase of inner mitochondrial membrane |
| AC002407.1 | X | novel transcript |
| MIR363 | X | microRNA 363 [Source:HGNC Symbol;Acc:HGI |
| MIR92A2 | X | microRNA 92a-2 [Source:HGNC Symbol;Acc:H |
| MIR19B2 | X | microRNA 19b-2 [Source:HGNC Symbol;Acc:H |
| MIR20B | X | microRNA 20b [Source:HGNC Symbol;Acc:HGI |
| MIR18B | X | microRNA 18b [Source:HGNC Symbol;Acc:HGI |
| MIR106A | X | microRNA 106a [Source:HGNC Symbol;Acc:HG |
| LARP1BP3 | X | La ribonucleoprotein domain family member |
| CCDC160 | X | coiled-coil domain containing 160 [Source:HG |
| FO393409.1 | X | novel pseudogene |
| PHF6 | X | PHD finger protein 6 [Source:HGNC Symbol;Ac |
| HPRT1 | X | hypoxanthine phosphoribosyltransferase 1 [So |
| AC004383.1 | X | ribosomal protein L36a (RPL36A) pseudogene |
| MIR450B | X | microRNA 450b [Source:HGNC Symbol;Acc:HG |
| MIR450A1 | X | microRNA 450a-1 [Source:HGNC Symbol;Acc:I |
| MIR450A2 | X | microRNA 450a-2 [Source:HGNC Symbol;Acc:I |
| MIR542 | X | microRNA 542 [Source:HGNC Symbol;Acc:HGI |
| MIR503HG | X | MIR503 host gene [Source:HGNC Symbol;Acc: |
| MIR503 | X | microRNA 503 [Source:HGNC Symbol;Acc:HGI |
| MIR424 | X | microRNA 424 [Source:HGNC Symbol;Acc:HGI |
| LINC00629 | X | long intergenic non-protein coding RNA 629 [S |
| PLAC1 | X | placenta enriched 1 [Source:HGNC Symbol;Ac |
| AL672032.1 | X | novel transcript |
| RPL21P133 | X | ribosomal protein L21 pseudogene 133 [Sourc |
| RPS7P12 | X | ribosomal protein S7 pseudogene 12 [Source: |
| RNU4-44P | X | RNA, U4 small nuclear 44, pseudogene [Sourc |
| FAM122B | X | family with sequence similarity 122B [Source: |
| FAM122C | X | family with sequence similarity 122C [Source:I |
| AL691477.1 | X | NULL |
| MOSPD1 | X | motile sperm domain containing 1 [Source:HG |
| RNU6-616P | X | RNA, U6 small nuclear 616, pseudogene [Sour |
| LINC02243 | X | long intergenic non-protein coding RNA 2243 |
| HMGB3P31 | X | high mobility group box 3 pseudogene 31 [Sou |
| SMIM10 | X | small integral membrane protein 10 [Source:H |
| RTL8B | X | retrotransposon Gag like 8B [Source:HGNC Sy |
| RTL8C | X | retrotransposon Gag like 8C [Source:HGNC Sy |
| AL136169.1 | X | novel transcript |
| RTL8A | X | retrotransposon Gag like 8A [Source:HGNC Sy |
| SMIM10L2B-AS1 | X | SMIM10L2B antisense RNA 1 [Source:HGNC Sy |
| NCLP2 | X | nucleolin pseudogene 2 [Source:HGNC Symbo |
| SMIM10L2B | X | small integral membrane protein 10 like 2B [S |
| ETDB | X | embryonic testis differentiation homolog B [S |
| AC234771.1 | X | novel transcript |
| AC234771.3 | X | zinc finger protein 449, pseudogene |
| CT55 | X | cancer/testis antigen 55 [Source:HGNC Symbo |
| AC234771.4 | X | novel pseudogene |
| AC234771.2 | X | novel pseudogene |

| | | |
|---|---|---|
| AC234771.5 | X | novel zinc finger protein pseudogene |
| ZNF75D | X | zinc finger protein 75D [Source:HGNC Symbol; |
| ETDA | X | embryonic testis differentiation homolog A [S |
| AL590282.1 | X | tigger transposable element derived 7 (TIGD7 |
| ETDC | X | embryonic testis differentiation homolog C [S |
| AL590282.2 | X | novel transcript |
| ZNF449 | X | zinc finger protein 449 [Source:HGNC Symbol; |
| AL450472.1 | X | novel pseudogene similar to embryonic testis |
| SMIM10L2A | X | small integral membrane protein 10 like 2A [S |
| RNA5SP515 | X | RNA, 5S ribosomal pseudogene 515 [Source:H |
| CROCCP1 | X | CROCC pseudogene 1 [Source:HGNC Symbol;A |
| AL450472.3 | X | chromosome X open reading frame 48 (CXorf4 |
| AL450472.2 | X | WD repeat domain 4, pseudogene |
| INTS6L-AS1 | X | INTS6L antisense RNA 1 [Source:HGNC Symbo |
| INTS6L | X | integrator complex subunit 6 like [Source:HGN |
| AL391380.1 | X | v-ets erythroblastosis virus E26 oncogene hon |
| SAGE4P | X | sarcoma antigen 4, pseudogene [Source:HGN( |
| SAGE3P | X | sarcoma antigen 3, pseudogene [Source:HGN( |
| SAGE2P | X | sarcoma antigen 2, pseudogene [Source:HGN( |
| AC240442.1 | X | mannosyl (alpha-1,6-)-glycoprotein beta-1,2-N |
| CT45A1 | X | cancer/testis antigen family 45 member A1 [S |
| MGAT2P1 | X | MGAT2 pseudogene 1 [Source:HGNC Symbol; |
| MGAT2P2 | X | MGAT2 pseudogene 2 [Source:HGNC Symbol; |
| CT45A3 | X | cancer/testis antigen family 45 member A3 [S |
| CT45A5 | X | cancer/testis antigen family 45 member A5 [S |
| CT45A6 | X | cancer/testis antigen family 45 member A6 [S |
| CT45A2 | X | cancer/testis antigen family 45 member A2 [S |
| CT45A7 | X | cancer/testis antigen family 45 member A7 [S |
| CT45A8 | X | cancer/testis antigen family 45 member A8 [S |
| CT45A9 | X | cancer/testis antigen family 45 member A9 [S |
| CT45A10 | X | cancer/testis antigen family 45 member A10 [ |
| SAGE1 | X | sarcoma antigen 1 [Source:HGNC Symbol;Acc: |
| CT45A11P | X | cancer/testis antigen family 45 member A11, |
| MMGT1 | X | membrane magnesium transporter 1 [Source: |
| SLC9A6 | X | solute carrier family 9 member A6 [Source:HG |
| GAPDHP67 | X | glyceraldehyde 3 phosphate dehydrogenase p |
| E2F6P4 | X | E2F transcription factor 6 pseudogene 4 [Sour |
| AL807743.1 | X | NULL |
| FHL1 | X | four and a half LIM domains 1 [Source:HGNC S |
| MAP7D3 | X | MAP7 domain containing 3 [Source:HGNC Syn |
| STK24P1 | X | serine/threonine kinase 24 pseudogene 1 [Sou |
| ADGRG4 | X | adhesion G protein-coupled receptor G4 [Sour |
| BRS3 | X | bombesin receptor subtype 3 [Source:HGNC S |
| HTATSF1 | X | HIV-1 Tat specific factor 1 [Source:HGNC Symb |
| VGLL1 | X | vestigial like family member 1 [Source:HGNC S |
| MIR934 | X | microRNA 934 [Source:HGNC Symbol;Acc:HGN |
| Y_RNA | X | Y RNA [Source:RFAM;Acc:RF00019] |
| RNU6-320P | X | RNA, U6 small nuclear 320, pseudogene [Sour |
| LINC00892 | X | long intergenic non-protein coding RNA 892 [S |
| CD40LG | X | CD40 ligand [Source:HGNC Symbol;Acc:HGNC |

| | | |
|---|---|---|
| ARHGEF6 | X | Rac/Cdc42 guanine nucleotide exchange facto |
| RNU6-972P | X | RNA, U6 small nuclear 972, pseudogene [Sour |
| AL683813.2 | X | solute carrier family 25 (mitochondrial carrier |
| RPL7P56 | X | ribosomal protein L7 pseudogene 56 [Source: |
| RANP4 | X | RAN pseudogene 4 [Source:HGNC Symbol;Acc |
| Y_RNA | X | Y RNA [Source:RFAM;Acc:RF00019] |
| AL683813.1 | X | novel transcript |
| RAB28P5 | X | RAB28, member RAS oncogene family pseudo |
| RBMX | X | RNA binding motif protein X-linked [Source:H |
| SNORD61 | X | small nucleolar RNA, C/D box 61 [Source:HGN |
| AL390879.1 | X | transmembrane 9 superfamily member 2 (TM |
| RPL22P23 | X | ribosomal protein L22 pseudogene 23 [Source |
| SRRM1P3 | X | serine/arginine repetitive matrix 1 pseudogen |
| GPR101 | X | G protein-coupled receptor 101 [Source:HGN |
| AL355309.1 | X | ribosomal protein L28 pseudogene |
| RAC1P4 | X | Rac family small GTPase 1 pseudogene 4 [Sou |
| AL035443.1 | X | novel transcript |
| ZIC3 | X | Zic family member 3 [Source:HGNC Symbol;Ac |
| Z96074.1 | X | novel transcript |
| RN7SL325P | X | RNA, 7SL, cytoplasmic 325, pseudogene [Sour |
| ZFYVE9P1 | X | zinc finger FYVE-type containing 9 pseudogen |
| RNU6-985P | X | RNA, U6 small nuclear 985, pseudogene [Sour |
| RN7SKP31 | X | RN7SK pseudogene 31 [Source:HGNC Symbol; |
| MAGEA13P | X | MAGE family member A13, pseudogene [Sour |
| KRT8P6 | X | keratin 8 pseudogene 6 [Source:HGNC Symbo |
| AL357123.1 | X | histone protein pseudogene |
| FGF13 | X | fibroblast growth factor 13 [Source:HGNC Syn |
| AL031386.1 | X | novel transcript |
| MIR504 | X | microRNA 504 [Source:HGNC Symbol;Acc:HGN |
| FGF13-AS1 | X | FGF13 antisense RNA 1 [Source:HGNC Symbol |
| Z83313.1 | X | tropomyosin 2 (beta) (TPM2) pseudogene |
| SNURFL | X | SNRPN upstream reading frame like (pseudog |
| SRD5A1P1 | X | steroid 5 alpha-reductase 1 pseudogene 1 [So |
| F9 | X | coagulation factor IX [Source:HGNC Symbol;A |
| MCF2 | X | MCF.2 cell line derived transforming sequence |
| AL161777.1 | X | NULL |
| ATP11C | X | ATPase phospholipid transporting 11C [Source |
| RNU6ATAC23P | X | RNA, U6atac small nuclear 23, pseudogene [S |
| AL356785.1 | X | NULL |
| MIR505 | X | microRNA 505 [Source:HGNC Symbol;Acc:HGN |
| CXorf66 | X | chromosome X open reading frame 66 [Sourc |
| AL589987.2 | X | novel transcript |
| HNRNPA3P3 | X | heterogeneous nuclear ribonucleoprotein A3 |
| RN7SL727P | X | RNA, 7SL, cytoplasmic 727, pseudogene [Sour |
| AL589987.1 | X | novel pseudogene |
| FO393408.1 | X | novel transcript |
| AL137014.1 | X | embryonic ectoderm development (EED) pseu |
| U7 | X | U7 small nuclear RNA [Source:RFAM;Acc:RF0 |
| RPS17P17 | X | ribosomal protein S17 pseudogene 17 [Source |
| SOX3 | X | SRY-box transcription factor 3 [Source:HGNC |

| | | |
|---|---|---|
| AL844175.1 | X | novel transcript |
| LINC00632 | X | long intergenic non-protein coding RNA 632 [S |
| CDR1 | X | cerebellar degeneration related protein 1 [Sou |
| MIR320D2 | X | microRNA 320d-2 [Source:HGNC Symbol;Acc:I |
| AL451048.1 | X | novel transcript |
| SPANXB1 | X | SPANX family member B1 [Source:HGNC Symb |
| LDOC1 | X | LDOC1 regulator of NFKB signaling [Source:HG |
| RNU6-3P | X | RNA, U6 small nuclear 3, pseudogene [Source |
| AC234778.1 | X | ribosomal protein L36 (RPL36) pseudogene |
| SPANXA2-OT1 | X | SPANXA2 overlapping transcript 1 [Source:HG |
| SPANXC | X | SPANX family member C [Source:HGNC Symbo |
| RBMX2P2 | X | RBMX2 pseudogene 2 [Source:HGNC Symbol;A |
| AC234779.1 | X | NADH dehydrogenase (ubiquinone) 1 beta sub |
| SPANXA1 | X | sperm protein associated with the nucleus, X- |
| SPANXA2 | X | SPANX family member A2 [Source:HGNC Symb |
| AC235097.1 | X | uncharacterized LOC645188 [Source:NCBI ger |
| SPANXD | X | SPANX family member D [Source:HGNC Symbo |
| MAGEC3 | X | MAGE family member C3 [Source:HGNC Symb |
| MAGEC1 | X | MAGE family member C1 [Source:HGNC Symb |
| AL031073.2 | X | novel transcript |
| AL023773.2 | X | MAGE (Melanoma Associated Antigen) family |
| AL023773.1 | X | MAGE (Melanoma Associated Antigen) family |
| AL031073.1 | X | melanoma antigen, family E, 1, cancer/testis s |
| MAGEC2 | X | MAGE family member C2 [Source:HGNC Symb |
| AL109620.1 | X | zinc finger, MYM-type 1 (ZMYM1) pseudogene |
| RNA5SP516 | X | RNA, 5S ribosomal pseudogene 516 [Source:H |
| SPANXN4 | X | SPANX family member N4 [Source:HGNC Syml |
| RN7SKP81 | X | RN7SK pseudogene 81 [Source:HGNC Symbol; |
| RN7SKP149 | X | RN7SK pseudogene 149 [Source:HGNC Symbo |
| AC239727.1 | X | novel transcript, antisense to SPANXN3 |
| PGBD4P6 | X | piggyBac transposable element derived 4 pseu |
| MTND1P33 | X | MT-ND1 pseudogene 33 [Source:HGNC Symbo |
| MTND2P39 | X | MT-ND2 pseudogene 39 [Source:HGNC Symbo |
| SPANXN3 | X | SPANX family member N3 [Source:HGNC Syml |
| SLITRK4 | X | SLIT and NTRK like family member 4 [Source:H |
| HNRNPH1P2 | X | heterogeneous nuclear ribonucleoprotein H1 |
| SPANXN2 | X | SPANX family member N2 [Source:HGNC Syml |
| AL500522.1 | X | novel transcript |
| AL500522.2 | X | pseudogene similar to part of ribosomal prote |
| UBE2NL | X | ubiquitin conjugating enzyme E2 N like (gene/ |
| AC234064.1 | X | pseudogene similar to part of ribonucleotide r |
| RRM2P4 | X | ribonucleotide reductase M2 polypeptide pse |
| HNRNPCP10 | X | heterogeneous nuclear ribonucleoprotein C p |
| RN7SKP189 | X | RN7SK pseudogene 189 [Source:HGNC Symbo |
| CYCSP44 | X | CYCS pseudogene 44 [Source:HGNC Symbol;A |
| AL713923.1 | X | novel transcript |
| SPANXN1 | X | SPANX family member N1 [Source:HGNC Syml |
| AL391256.1 | X | novel pseudogene |
| AL109653.3 | X | novel transcript, antisense to SLITRK2 |
| AL109653.1 | X | ubiquitin-fold modifier 1 (UFM1) pseudogene |

| | | |
|---|---|---|
| AL109653.2 | X | TEC |
| SLITRK2 | X | SLIT and NTRK like family member 2 [Source:H |
| MIR892C | X | microRNA 892c [Source:HGNC Symbol;Acc:HG |
| MIR890 | X | microRNA 890 [Source:HGNC Symbol;Acc:HGI |
| MIR888 | X | microRNA 888 [Source:HGNC Symbol;Acc:HGI |
| MIR892A | X | microRNA 892a [Source:HGNC Symbol;Acc:HG |
| MIR892B | X | microRNA 892b [Source:HGNC Symbol;Acc:HG |
| MIR891B | X | microRNA 891b [Source:HGNC Symbol;Acc:HG |
| MIR891A | X | microRNA 891a [Source:HGNC Symbol;Acc:HG |
| RNA5SP517 | X | RNA, 5S ribosomal pseudogene 517 [Source:H |
| AL109913.1 | X | ADP-ribosylation factor-like 5 (ARL5) pseudog |
| ELL2P4 | X | elongation factor for RNA polymerase II 2 pse |
| Z97180.1 | X | novel pseudogene |
| CXorf51B | X | chromosome X open reading frame 51B [Sour |
| CXorf51A | X | chromosome X open reading frame 51A [Sour |
| Z99497.1 | X | pseudogene similar to part of a putative dime |
| Z99497.2 | X | kinesin family member 23 (KIF23) pseudogene |
| AL589669.1 | X | periphilin-1-like [Source:NCBI gene;Acc:10537 |
| MIR513C | X | microRNA 513c [Source:HGNC Symbol;Acc:HG |
| MIR513B | X | microRNA 513b [Source:HGNC Symbol;Acc:HG |
| MIR513A1 | X | microRNA 513a-1 [Source:HGNC Symbol;Acc:I |
| MIR513A2 | X | microRNA 513a-2 [Source:HGNC Symbol;Acc:I |
| MIR506 | X | microRNA 506 [Source:HGNC Symbol;Acc:HGI |
| MIR507 | X | microRNA 507 [Source:HGNC Symbol;Acc:HGI |
| MIR508 | X | microRNA 508 [Source:HGNC Symbol;Acc:HGI |
| MIR514B | X | microRNA 514b [Source:HGNC Symbol;Acc:HG |
| MIR509-2 | X | microRNA 509-2 [Source:HGNC Symbol;Acc:H |
| MIR509-3 | X | microRNA 509-3 [Source:HGNC Symbol;Acc:H |
| MIR509-1 | X | microRNA 509-1 [Source:HGNC Symbol;Acc:H |
| MIR510 | X | microRNA 510 [Source:HGNC Symbol;Acc:HGI |
| MIR514A1 | X | microRNA 514a-1 [Source:HGNC Symbol;Acc:I |
| AL591022.1 | X | novel transcript |
| MIR514A2 | X | microRNA 514a-2 [Source:HGNC Symbol;Acc:I |
| MIR514A3 | X | microRNA 514a-3 [Source:HGNC Symbol;Acc:I |
| AL096861.1 | X | ELOVL fatty acid elongase 6 (ELOVL6) pseudog |
| AL009048.1 | X | SMT3 suppressor of mif two 3 homolog 1 (yea |
| AC016925.1 | X | SRA stem-loop interacting RNA binding protei |
| AC016925.2 | X | cyclin-dependent kinase 16 (CDK16) pseudoge |
| RNU6-382P | X | RNA, U6 small nuclear 382, pseudogene [Sour |
| AC016925.3 | X | novel transcript |
| L29074.1 | X | novel transcript |
| FMR1-AS1 | X | FMR1 antisense RNA 1 [Source:HGNC Symbol; |
| L29074.2 | X | NULL |
| FMR1 | X | fragile X mental retardation 1 [Source:HGNC S |
| FMR1-IT1 | X | FMR1 intronic transcript 1 [Source:HGNC Sym |
| FMR1NB | X | FMR1 neighbor [Source:HGNC Symbol;Acc:HG |
| RNA5SP524 | X | RNA, 5S ribosomal pseudogene 524 [Source:H |
| FTH1P8 | X | ferritin heavy chain 1 pseudogene 8 [Source:H |
| U40455.1 | X | HS1 binding protein (HAX1) pseudogene |
| RPL7L1P11 | X | ribosomal protein L7 like 1 pseudogene 11 [So |

| | | |
|---|---|---|
| AC002368.1 | X | novel transcript |
| AFF2 | X | AF4/FMR2 family member 2 [Source:HGNC Sy |
| AFF2-IT1 | X | AFF2 intronic transcript 1 [Source:HGNC Symb |
| RN7SKP267 | X | RN7SK pseudogene 267 [Source:HGNC Symbo |
| AC231760.1 | X | pseudogene similar to part of taxol resistance |
| AC231656.1 | X | novel protein, similar to CXorf51A |
| IDS | X | iduronate 2-sulfatase [Source:HGNC Symbol;A |
| AC244197.3 | X | novel protein |
| AC244197.2 | X | novel transcript |
| IDSP1 | X | iduronate 2-sulfatase pseudogene 1 [Source:H |
| LINC00893 | X | long intergenic non-protein coding RNA 893 [S |
| CXorf40A | X | chromosome X open reading frame 40A [Sour |
| HSFX3 | X | heat shock transcription factor family, X-linke |
| MAGEA9B | X | MAGE family member A9B [Source:HGNC Sym |
| HSFX2 | X | heat shock transcription factor family, X-linke |
| TMEM185A | X | transmembrane protein 185A [Source:HGNC S |
| AC244197.1 | X | melanoma antigen family A pseudogene |
| MAGEA11 | X | MAGE family member A11 [Source:HGNC Sym |
| AC244098.2 | X | melanoma antigen family A, 11 (MAGEA11) ps |
| AC244098.1 | X | melanoma antigen family A, 11 (MAGEA11) ps |
| AC244098.3 | X | melanoma antigen family A, 11 (MAGEA11) ps |
| TMEM185AP1 | X | transmembrane protein 185A pseudogene 1 [ |
| HSFX1 | X | heat shock transcription factor family, X-linke |
| MAGEA9 | X | MAGE family member A9 [Source:HGNC Symb |
| MAGEA7P | X | MAGE family member A7, pseudogene [Sourc |
| DUTP4 | X | deoxyuridine triphosphatase pseudogene 4 [S |
| LINC00850 | X | long intergenic non-protein coding RNA 850 [S |
| MAGEA8-AS1 | X | MAGEA8 antisense RNA 1 [Source:HGNC Symb |
| MAGEA8 | X | MAGE family member A8 [Source:HGNC Symb |
| CXorf40B | X | chromosome X open reading frame 40B [Sour |
| HSFX4 | X | heat shock transcription factor family, X-linke |
| LINC00894 | X | long intergenic non-protein coding RNA 894 [S |
| BX322650.1 | X | nucleolar protein 11, pseudogene |
| MIR2114 | X | microRNA 2114 [Source:HGNC Symbol;Acc:HG |
| XRCC6P2 | X | X-ray repair cross complementing 6 pseudoge |
| MAMLD1 | X | mastermind like domain containing 1 [Source: |
| MTM1 | X | myotubularin 1 [Source:HGNC Symbol;Acc:HG |
| MTMR1 | X | myotubularin related protein 1 [Source:HGNC |
| CD99L2 | X | CD99 molecule like 2 [Source:HGNC Symbol;A |
| PPIAP91 | X | peptidylprolyl isomerase A pseudogene 91 [So |
| AF274573.1 | X | chromosome X novel transcript |
| RNU6-383P | X | RNA, U6 small nuclear 383, pseudogene [Sour |
| HMGB3 | X | high mobility group box 3 [Source:HGNC Symb |
| RPL19P21 | X | ribosomal protein L19 pseudogene 21 [Source |
| RPL12P50 | X | ribosomal protein L12 pseudogene 50 [Source |
| RNA5SP525 | X | RNA, 5S ribosomal pseudogene 525 [Source:H |
| MIR4330 | X | microRNA 4330 [Source:HGNC Symbol;Acc:HG |
| GPR50-AS1 | X | GPR50 antisense RNA 1 [Source:HGNC Symbo |
| GPR50 | X | G protein-coupled receptor 50 [Source:HGNC |
| KC877982.1 | X | novel transcript |

| | | |
|---|---|---|
| AF013593.1 | X | NGFI-A binding protein 1 (EGR1 binding protei |
| Metazoa_SRP | X | Metazoan signal recognition particle RNA [Sou |
| AF274853.1 | X | novel transcript, antisense to VMA21 |
| VMA21 | X | vacuolar ATPase assembly factor VMA21 [Sou |
| BX546450.2 | X | novel transcript |
| BX546450.1 | X | novel transcript |
| PASD1 | X | PAS domain containing repressor 1 [Source:H |
| PRRG3 | X | proline rich and Gla domain 3 [Source:HGNC S |
| FATE1 | X | fetal and adult testis expressed 1 [Source:HGN |
| CNGA2 | X | cyclic nucleotide gated channel subunit alpha |
| MAGEA4-AS1 | X | MAGEA4 antisense RNA 1 [Source:HGNC Syml |
| MAGEA4 | X | MAGE family member A4 [Source:HGNC Symb |
| GABRE | X | gamma-aminobutyric acid type A receptor ep |
| MIR224 | X | microRNA 224 [Source:HGNC Symbol;Acc:HGN |
| MIR452 | X | microRNA 452 [Source:HGNC Symbol;Acc:HGN |
| RPSAP60 | X | ribosomal protein SA pseudogene 60 [Source: |
| MAGEA5 | X | MAGE family member A5 [Source:HGNC Symb |
| MAGEA10-MAGEA5 | X | MAGEA10-MAGEA5 readthrough [Source:NCB |
| MAGEA10 | X | MAGE family member A10 [Source:HGNC Sym |
| AC116666.1 | X | novel transcript |
| GABRA3 | X | gamma-aminobutyric acid type A receptor alp |
| RNU6-764P | X | RNA, U6 small nuclear 764, pseudogene [Sour |
| MIR105-1 | X | microRNA 105-1 [Source:HGNC Symbol;Acc:H |
| MIR767 | X | microRNA 767 [Source:HGNC Symbol;Acc:HGN |
| MIR105-2 | X | microRNA 105-2 [Source:HGNC Symbol;Acc:H |
| KRT8P8 | X | keratin 8 pseudogene 8 [Source:HGNC Symbo |
| AC244102.4 | X | novel transcript, antisense to GABRQ |
| GABRQ | X | gamma-aminobutyric acid type A receptor the |
| AC244102.2 | X | melanoma antigen family A, 11 (MAGEA11) ps |
| MAGEA3 | X | MAGE family member A3 [Source:HGNC Symb |
| CSAG2 | X | CSAG family member 2 [Source:HGNC Symbol |
| MAGEA2B | X | MAGE family member A2B [Source:HGNC Sym |
| CSAG1 | X | chondrosarcoma associated gene 1 [Source:H |
| MAGEA12 | X | MAGE family member A12 [Source:HGNC Sym |
| CSAG4 | X | CSAG family member 4 (pseudogene) [Source |
| AC244102.1 | X | novel transcript |
| MAGEA2 | X | MAGE family member A2 [Source:HGNC Symb |
| CSAG3 | X | CSAG family member 3 [Source:HGNC Symbol |
| MAGEA6 | X | MAGE family member A6 [Source:HGNC Symb |
| AC244102.3 | X | novel transcript |
| CETN2 | X | centrin 2 [Source:HGNC Symbol;Acc:HGNC:18 |
| NSDHL | X | NAD(P) dependent steroid dehydrogenase-lik |
| ZNF185 | X | zinc finger protein 185 with LIM domain [Sour |
| U82671.1 | X | novel transcript, antisense to ZNF185 |
| PNMA5 | X | PNMA family member 5 [Source:HGNC Symbo |
| AC243591.1 | X | putative paraneoplastic antigen-like protein 6 |
| AC243428.1 | X | high-mobility group nucleosomal binding dom |
| PNMA3 | X | PNMA family member 3 [Source:HGNC Symbo |
| PNMA6A | X | PNMA family member 6A [Source:HGNC Symb |
| PNMA6B | X | PNMA family member 6B (pseudogene) [Sour |

| | | |
|---|---|---|
| AC152010.1 | X | melanoma associated antigen (mutated) 1-lik |
| MAGEA1 | X | MAGE family member A1 [Source:HGNC Symb |
| AC236972.2 | X | family with sequence similarity 9 (FAM9) pseu |
| AC236972.3 | X | uncharacterized LOC105373373 [Source:NCBI |
| AC236972.1 | X | zinc finger and BTB domain containing 20 (ZB |
| AC236972.4 | X | novel protein similar to melanoma associated |
| PNMA6F | X | PNMA family member 6F [Source:HGNC Symb |
| ZNF275 | X | zinc finger protein 275 [Source:HGNC Symbol; |
| AC152007.1 | X | renin binding protein (RENBP) pseudogene |
| PNMA6E | X | PNMA family member 6E [Source:HGNC Symb |
| RN7SL667P | X | RNA, 7SL, cytoplasmic 667, pseudogene [Sour |
| ZFP92 | X | ZFP92 zinc finger protein [Source:HGNC Symb |
| TREX2 | X | three prime repair exonuclease 2 [Source:HGI |
| HAUS7 | X | HAUS augmin like complex subunit 7 [Source: |
| U82695.1 | X | novel pseudogene |
| BGN | X | biglycan [Source:HGNC Symbol;Acc:HGNC:104 |
| ATP2B3 | X | ATPase plasma membrane Ca2+ transporting |
| CCNQ | X | cyclin Q [Source:HGNC Symbol;Acc:HGNC:284 |
| AF274858.1 | X | Morf4 family associated protein 1 (MRFAP1) p |
| KRT18P48 | X | keratin 18 pseudogene 48 [Source:HGNC Sym |
| DUSP9 | X | dual specificity phosphatase 9 [Source:HGNC S |
| RPL18AP16 | X | ribosomal protein L18a pseudogene 16 [Sourc |
| RN7SL687P | X | RNA, 7SL, cytoplasmic 687, pseudogene [Sour |
| PNCK | X | pregnancy up-regulated nonubiquitous CaM k |
| SLC6A8 | X | solute carrier family 6 member 8 [Source:HGN |
| BCAP31 | X | B cell receptor associated protein 31 [Source: |
| ABCD1 | X | ATP binding cassette subfamily D member 1 [S |
| U52111.1 | X | novel transcript |
| PLXNB3 | X | plexin B3 [Source:HGNC Symbol;Acc:HGNC:91 |
| SRPK3 | X | SRSF protein kinase 3 [Source:HGNC Symbol;A |
| IDH3G | X | isocitrate dehydrogenase (NAD(+)) 3 gamma [ |
| SSR4 | X | signal sequence receptor subunit 4 [Source:H |
| PDZD4 | X | PDZ domain containing 4 [Source:HGNC Symb |
| CYCSP45 | X | CYCS pseudogene 45 [Source:HGNC Symbol;A |
| L1CAM | X | L1 cell adhesion molecule [Source:HGNC Symb |
| U52112.2 | X | novel transcript |
| U52112.1 | X | lung carcinoma-associated 10 |
| AVPR2 | X | arginine vasopressin receptor 2 [Source:HGN |
| ARHGAP4 | X | Rho GTPase activating protein 4 [Source:HGN |
| NAA10 | X | N(alpha)-acetyltransferase 10, NatA catalytic s |
| RENBP | X | renin binding protein [Source:HGNC Symbol;A |
| HCFC1 | X | host cell factor C1 [Source:HGNC Symbol;Acc: |
| HCFC1-AS1 | X | HCFC1 antisense RNA 1 [Source:HGNC Symbol |
| TMEM187 | X | transmembrane protein 187 [Source:HGNC Sy |
| MIR3202-1 | X | microRNA 3202-1 [Source:HGNC Symbol;Acc: |
| MIR3202-2 | X | microRNA 3202-2 [Source:HGNC Symbol;Acc: |
| IRAK1 | X | interleukin 1 receptor associated kinase 1 [So |
| MIR718 | X | microRNA 718 [Source:HGNC Symbol;Acc:HGI |
| MECP2 | X | methyl-CpG binding protein 2 [Source:HGNC S |
| OPN1LW | X | opsin 1, long wave sensitive [Source:HGNC Sy |

| | | |
|---|---|---|
| TEX28P2 | X | testis expressed 28 pseudogene 2 [Source:HG |
| OPN1MW | X | opsin 1, medium wave sensitive [Source:HGN( |
| TEX28P1 | X | testis expressed 28 pseudogene 1 [Source:HG |
| OPN1MW2 | X | opsin 1, medium wave sensitive 2 [Source:HG |
| AC244097.1 | X | testis expressed 28 (TEX28) pseudogene |
| OPN1MW3 | X | opsin 1, medium wave sensitive 3 [Source:HG |
| TEX28 | X | testis expressed 28 [Source:HGNC Symbol;Acc |
| TKTL1 | X | transketolase like 1 [Source:HGNC Symbol;Acc |
| AC245140.1 | X | novel transcript |
| FLNA | X | filamin A [Source:HGNC Symbol;Acc:HGNC:37 |
| AC245140.3 | X | novel transcript, antisense to FLNA |
| EMD | X | emerin [Source:HGNC Symbol;Acc:HGNC:3331 |
| RPL10 | X | ribosomal protein L10 [Source:HGNC Symbol;/ |
| AC245140.2 | X | novel transcript, antisense to RPL10 |
| SNORA70 | X | small nucleolar RNA, H/ACA box 70 [Source:H |
| DNASE1L1 | X | deoxyribonuclease 1 like 1 [Source:HGNC Sym |
| TAZ | X | tafazzin [Source:HGNC Symbol;Acc:HGNC:115 |
| AC244090.1 | X | novel transcript |
| ATP6AP1 | X | ATPase H+ transporting accessory protein 1 [S |
| GDI1 | X | GDP dissociation inhibitor 1 [Source:HGNC Syı |
| FAM50A | X | family with sequence similarity 50 member A |
| MIR6858 | X | microRNA 6858 [Source:HGNC Symbol;Acc:HG |
| PLXNA3 | X | plexin A3 [Source:HGNC Symbol;Acc:HGNC:91 |
| LAGE3 | X | L antigen family member 3 [Source:HGNC Sym |
| UBL4A | X | ubiquitin like 4A [Source:HGNC Symbol;Acc:H( |
| SLC10A3 | X | solute carrier family 10 member 3 [Source:HG |
| RN7SL697P | X | RNA, 7SL, cytoplasmic 697, pseudogene [Sour |
| FAM3A | X | family with sequence similarity 3 member A [S |
| AC244090.3 | X | novel transcript, overlapping to FAM3A |
| G6PD | X | glucose-6-phosphate dehydrogenase [Source: |
| IKBKG | X | inhibitor of nuclear factor kappa B kinase regu |
| FAM223A | X | family with sequence similarity 223 member / |
| ATF4P2 | X | activating transcription factor 4 pseudogene 2 |
| CTAG1A | X | cancer/testis antigen 1A [Source:HGNC Symbo |
| AC244090.2 | X | coatomer protein complex, subunit zeta 1 (CC |
| CTAG1B | X | cancer/testis antigen 1B [Source:HGNC Symbo |
| AC244107.2 | X | solute carrier family 6 (neurotransmitter trans |
| ATF4P1 | X | activating transcription factor 4 pseudogene 1 |
| FAM223B | X | family with sequence similarity 223 member E |
| IKBKGP1 | X | inhibitor of nuclear factor kappa B kinase subu |
| CTAG2 | X | cancer/testis antigen 2 [Source:HGNC Symbol |
| AC244107.1 | X | similar to sodium- and chloride-dependent cre |
| OR3B1P | X | olfactory receptor family 3 subfamily B memb |
| GAB3 | X | GRB2 associated binding protein 3 [Source:HG |
| DKC1 | X | dyskerin pseudouridine synthase 1 [Source:H( |
| SNORA36A | X | small nucleolar RNA, H/ACA box 36A [Source:I |
| MIR664B | X | microRNA 664b [Source:HGNC Symbol;Acc:HG |
| SNORA56 | X | small nucleolar RNA, H/ACA box 56 [Source:H |
| MPP1 | X | membrane palmitoylated protein 1 [Source:H |
| HMGN1P37 | X | high mobility group nucleosome binding doma |

| | | |
|---|---|---|
| SMIM9 | X | small integral membrane protein 9 [Source:H( |
| F8 | X | coagulation factor VIII [Source:HGNC Symbol;/ |
| H2AFB1 | X | H2A histone family member B1 [Source:HGNC |
| F8A1 | X | coagulation factor VIII associated 1 [Source:H( |
| MIR1184-1 | X | microRNA 1184-1 [Source:HGNC Symbol;Acc:I |
| BX842559.1 | X | ClpX caseinolytic peptidase X homolog (E. coli |
| EEF1A1P31 | X | eukaryotic translation elongation factor 1 alpl |
| BX842559.2 | X | zinc finger-like protein 9 (ZPR) pseudogene |
| FUNDC2 | X | FUN14 domain containing 2 [Source:HGNC Syı |
| BX470111.1 | X | novel protein |
| CMC4 | X | C-X9-C motif containing 4 [Source:HGNC Symk |
| MTCP1 | X | mature T cell proliferation 1 [Source:HGNC Sy |
| BRCC3 | X | BRCA1/BRCA2-containing complex subunit 3 [ |
| BX293995.1 | X | phosphatidylinositol binding clathrin assembly |
| VBP1 | X | VHL binding protein 1 [Source:HGNC Symbol;/ |
| BX682237.1 | X | pseudogene similar to part of CD84 antigen (l( |
| RAB39B | X | RAB39B, member RAS oncogene family [Sourc |
| CLIC2 | X | chloride intracellular channel 2 [Source:HGNC |
| AC234781.2 | X | Twinfilin-1, pseudogene |
| AC234781.3 | X | PHD finger protein pseudogene |
| AC234781.4 | X | pseudogene similar to part of steroid-5-alpha- |
| AC234781.1 | X | novel transcript |
| AC234781.5 | X | trimethyllysine hydroxylase, epsilon (TMLHE) |
| H2AFB2 | X | H2A histone family member B2 [Source:HGNC |
| F8A2 | X | coagulation factor VIII associated 2 [Source:H( |
| MIR1184-2 | X | microRNA 1184-2 [Source:HGNC Symbol;Acc:I |
| F8A3 | X | coagulation factor VIII associated 3 [Source:H( |
| MIR1184-3 | X | microRNA 1184-3 [Source:HGNC Symbol;Acc:I |
| H2AFB3 | X | H2A histone family member B3 [Source:HGNC |
| TMLHE-AS1 | X | TMLHE antisense RNA 1 [Source:HGNC Symbc |
| BX571846.1 | X | novel transcript |
| TMLHE | X | trimethyllysine hydroxylase, epsilon [Source:H |
| SPRY3 | X | sprouty RTK signaling antagonist 3 [Source:HG |
| AMD1P2 | X | adenosylmethionine decarboxylase 1 pseudog |
| DPH3P2 | X | diphthamide biosynthesis 3 pseudogene 2 [So |
| VAMP7 | X | vesicle associated membrane protein 7 [Sourc |
| ELOCP24 | X | elongin C pseudogene 24 [Source:HGNC Symb |
| TRPC6P | X | transient receptor potential cation channel su |
| IL9R | X | interleukin 9 receptor [Source:HGNC Symbol;/ |
| AJ271736.1 | X | novel transcript |
| WASIR1 | X | WASH and IL9R antisense RNA 1 [Source:HGN |
| WASH6P | X | WASP family homolog 6, pseudogene [Source |
| DDX11L16 | X | DEAD/H-box helicase 11 like 16 [Source:HGN( |
| RNU6-1334P | Y | RNA, U6 small nuclear 1334, pseudogene [Sou |
| SRY | Y | sex determining region Y [Source:HGNC Symb |
| RNASEH2CP1 | Y | ribonuclease H2 subunit C pseudogene 1 [Sou |
| TOMM22P2 | Y | TOMM22 pseudogene 2 [Source:HGNC Symbc |
| AC006040.1 | Y | novel transcript |
| RPS4Y1 | Y | ribosomal protein S4 Y-linked 1 [Source:HGN( |
| HSFY3P | Y | heat shock transcription factor Y-linked 3, pse |

| | | |
|---|---|---|
| NAP1L1P2 | Y | nucleosome assembly protein 1 like 1 pseudo |
| AC006157.1 | Y | novel transcript |
| ZFY | Y | zinc finger protein Y-linked [Source:HGNC Sym |
| ZFY-AS1 | Y | ZFY antisense RNA 1 [Source:HGNC Symbol;Ac |
| EEF1A1P41 | Y | eukaryotic translation elongation factor 1 alph |
| LINC00278 | Y | long intergenic non-protein coding RNA 278 [S |
| AGPAT5P1 | Y | 1-acylglycerol-3-phosphate O-acyltransferase |
| PRRC2CP1 | Y | proline rich coiled-coil 2C pseudogene 1 [Sour |
| TGIF2LY | Y | TGFB induced factor homeobox 2 like Y-linked |
| USP12PY | Y | ubiquitin specific peptidase 12 pseudogene Y- |
| RNF19BPY | Y | ring finger protein 19B pseudogene Y-linked [S |
| AC012078.2 | Y | novel transcript |
| AC012078.1 | Y | pseudogene similar to primate FRD envelope |
| UBE2V1P3 | Y | ubiquitin conjugating enzyme E2 V1 pseudoge |
| AC010737.1 | Y | novel transcript |
| RNU6-303P | Y | RNA, U6 small nuclear 303, pseudogene [Sour |
| SERBP1P2 | Y | SERPINE1 mRNA binding protein 1 pseudogen |
| AC010722.1 | Y | novel transcript |
| PCDH11Y | Y | protocadherin 11 Y-linked [Source:HGNC Sym |
| RNU2-57P | Y | RNA, U2 small nuclear 57, pseudogene [Sourc |
| VDAC1P6 | Y | voltage dependent anion channel 1 pseudoge |
| EIF4A1P2 | Y | eukaryotic translation initiation factor 4A1 ps |
| KRT18P10 | Y | keratin 18 pseudogene 10 [Source:HGNC Sym |
| SNX3P1Y | Y | sorting nexin 3 pseudogene 1 Y-linked [Source |
| RPL26P37 | Y | ribosomal protein L26 pseudogene 37 [Source |
| TUSC2P1 | Y | TUSC2 pseudogene 1 [Source:HGNC Symbol;A |
| DLGAP5P1 | Y | discs large homolog associated protein 5 pseu |
| TTTY23B | Y | testis-specific transcript, Y-linked 23B [Source |
| TSPY2 | Y | testis specific protein Y-linked 2 [Source:HGN |
| FAM197Y9 | Y | family with sequence similarity 197 Y-linked m |
| TSPY11P | Y | testis specific protein Y-linked 11, pseudogene |
| AC006335.2 | Y | testis specific protein, Y-linked family pseudog |
| TSPY19P | Y | testis specific protein Y-linked 19, pseudogene |
| AC006335.1 | Y | testis specific protein, Y-linked family pseudog |
| RBMY2GP | Y | RNA binding motif protein, Y-linked, family 2, |
| LINC00280 | Y | long intergenic non-protein coding RNA 280 [S |
| TTTY1B | Y | testis-specific transcript, Y-linked 1B [Source:H |
| TTTY2B | Y | testis-specific transcript, Y-linked 2B [Source:H |
| TTTY21B | Y | testis-specific transcript, Y-linked 21B [Source |
| TTTY7 | Y | testis-specific transcript, Y-linked 7 [Source:H |
| TTTY8B | Y | testis-specific transcript, Y-linked 8B [Source:H |
| AC010154.1 | Y | RNA binding motif protein, Y-linked, family 1, |
| TSPY17P | Y | testis specific protein Y-linked 17, pseudogene |
| SRIP3 | Y | sorcin pseudogene 3 [Source:HGNC Symbol;A |
| AC013412.1 | Y | WWC family member 3 (WWC3) pseudogene |
| GOT2P5 | Y | glutamic-oxaloacetic transaminase 2 pseudog |
| AMELY | Y | amelogenin Y-linked [Source:HGNC Symbol;Ac |
| ATP5PFP1 | Y | ATP synthase peripheral stalk subunit F6 pseu |
| TBL1Y | Y | transducin beta like 1 Y-linked [Source:HGNC S |
| AC011297.1 | Y | novel transcript, sense intronic to TBL1Y |

| | | |
|---|---|---|
| GPR143P | Y | G protein-coupled receptor 143 pseudogene [ |
| PRKY | Y | protein kinase Y-linked (pseudogene) [Source: |
| RN7SKP282 | Y | RN7SK pseudogene 282 [Source:HGNC Symbo |
| Y_RNA | Y | Y RNA [Source:RFAM;Acc:RF00019] |
| RNU6-941P | Y | RNA, U6 small nuclear 941, pseudogene [Sour |
| RNU6-521P | Y | RNA, U6 small nuclear 521, pseudogene [Sour |
| AC007274.3 | Y | testis specific protein, Y-linked 1 (TSPY1) pseu |
| RBMY2HP | Y | RNA binding motif protein, Y-linked, family 2, |
| AC007274.1 | Y | testis specific protein, Y-linked 1 (TSPY1) pseu |
| TSPY12P | Y | testis specific protein Y-linked 12, pseudogene |
| AC007274.2 | Y | testis specific protein, Y-linked 1 (TSPY1) pseu |
| RFTN1P1 | Y | raftlin, lipid raft linker 1 pseudogene 1 [Source |
| TTTY12 | Y | testis-specific transcript, Y-linked 12 [Source:H |
| AC007275.1 | Y | tripartite motif containing 60 (TRIM60) pseud |
| ZNF92P1Y | Y | zinc finger protein 92 pseudogene 1, Y-linked |
| AC010678.1 | Y | tripartite motif containing 60 (TRIM60) pseud |
| ZNF736P8Y | Y | zinc finger protein 736 pseudogene 8, Y-linked |
| BPY2DP | Y | basic charge Y-linked 2D, pseudogene [Source |
| ZNF736P7Y | Y | zinc finger protein 736 pseudogene 7, Y-linked |
| ZNF736P9Y | Y | zinc finger protein 736 pseudogene 9, Y-linked |
| RBMY2JP | Y | RNA binding motif protein, Y-linked, family 2, |
| RBMY2KP | Y | RNA binding motif protein, Y-linked, family 2, |
| TSPY24P | Y | testis specific protein Y-linked 24, pseudogene |
| ZNF736P6Y | Y | zinc finger protein 736 pseudogene 6, Y-linked |
| MTND6P1 | Y | MT-ND6 pseudogene 1 [Source:HGNC Symbol |
| MTCYBP1 | Y | MT-CYB pseudogene 1 [Source:HGNC Symbol; |
| MTND1P1 | Y | MT-ND1 pseudogene 1 [Source:HGNC Symbol |
| MTND2P3 | Y | MT-ND2 pseudogene 3 [Source:HGNC Symbol |
| TRIM60P3Y | Y | tripartite motif containing 60 pseudogene 3, Y |
| ZNF736P10Y | Y | zinc finger protein 736 pseudogene 10, Y-linke |
| AC016991.1 | Y | testis specific protein, Y-linked 1 (TSPY1) pseu |
| AC064829.1 | Y | novel transcript |
| TTTY18 | Y | testis-specific transcript, Y-linked 18 [Source:H |
| TTTY19 | Y | testis-specific transcript, Y-linked 19 [Source:H |
| TTTY11 | Y | testis-specific transcript, Y-linked 11 [Source:H |
| AC007967.2 | Y | testis specific protein, Y-linked 1 (TSPY1) pseu |
| AC007967.1 | Y | RNA binding motif protein, Y-linked, family 1, |
| AC007967.4 | Y | testis specific protein, Y-linked 1 (TSPY1) pseu |
| AC007967.3 | Y | testis specific protein, Y-linked 1 (TSPY1) pseu |
| OFD1P3Y | Y | OFD1 pseudogene 3 Y-linked [Source:HGNC Sy |
| CDY3P | Y | chromodomain Y-linked 3 pseudogene [Source |
| USP9YP22 | Y | USP9Y pseudogene 22 [Source:HGNC Symbol; |
| USP9YP4 | Y | USP9Y pseudogene 4 [Source:HGNC Symbol;A |
| ELOCP4 | Y | elongin C pseudogene 4 [Source:HGNC Symbo |
| RBMY1A3P | Y | RNA binding motif protein, Y-linked, family 1, |
| AC006158.1 | Y | testis specific protein, Y-linked 1 (TSPY1) pse |
| TTTY20 | Y | testis-specific transcript, Y-linked 20 [Source:H |
| TSPY4 | Y | testis specific protein Y-linked 4 [Source:HGN( |
| FAM197Y8 | Y | family with sequence similarity 197 Y-linked m |
| TSPY8 | Y | testis specific protein Y-linked 8 [Source:HGN( |

| | | |
|---|---|---|
| FAM197Y7 | Y | family with sequence similarity 197 Y-linked m |
| AC006158.2 | Y | testis specific protein, Y-linked family pseudog |
| TSPY7P | Y | testis specific protein Y-linked 7, pseudogene |
| FAM197Y6 | Y | family with sequence similarity 197 Y-linked m |
| AC006158.3 | Y | testis specific protein, Y-linked family pseudog |
| TSPY3 | Y | testis specific protein Y-linked 3 [Source:HGN( |
| FAM197Y5 | Y | family with sequence similarity 197 Y-linked m |
| TSPY1 | Y | testis specific protein Y-linked 1 [Source:HGN( |
| FAM197Y4 | Y | family with sequence similarity 197 Y-linked m |
| TSPY9P | Y | testis specific protein Y-linked 9, pseudogene |
| FAM197Y3 | Y | family with sequence similarity 197 Y-linked m |
| TSPY6P | Y | testis specific protein Y-linked 6, pseudogene |
| FAM197Y2 | Y | family with sequence similarity 197 Y-linked m |
| TSPY10 | Y | testis specific protein Y-linked 10 [Source:HGN |
| FAM197Y1 | Y | family with sequence similarity 197 Y-linked m |
| TSPY15P | Y | testis specific protein Y-linked 15, pseudogene |
| RBMY3AP | Y | RNA binding motif protein, Y-linked, family 3, |
| TSPY25P | Y | testis specific protein Y-linked 25, pseudogene |
| TSPY16P | Y | testis specific protein Y-linked 16, pseudogene |
| AC017019.1 | Y | pseudogene similar to part of RNA binding mo |
| TTTY8 | Y | testis-specific transcript, Y-linked 8 [Source:H( |
| TTTY7B | Y | testis-specific transcript, Y-linked 7B [Source:H |
| TTTY21 | Y | testis-specific transcript, Y-linked 21 [Source:H |
| TTTY2 | Y | testis-specific transcript, Y-linked 2 [Source:H( |
| TTTY1 | Y | testis-specific transcript, Y-linked 1 [Source:H( |
| TTTY22 | Y | testis-specific transcript, Y-linked 22 [Source:H |
| AC010891.1 | Y | novel transcript |
| RBMY2NP | Y | RNA binding motif protein, Y-linked, family 2, |
| AC006986.1 | Y | testis specific protein, Y-linked 1 (TSPY1) pseu |
| TSPY18P | Y | testis specific protein Y-linked 18, pseudogene |
| TSPY13P | Y | testis specific protein Y-linked 13, pseudogene |
| TTTY23 | Y | testis-specific transcript, Y-linked 23 [Source:H |
| RBMY2OP | Y | RNA binding motif protein, Y-linked, family 2, |
| RBMY2QP | Y | RNA binding motif protein, Y-linked, family 2, |
| TSPY20P | Y | testis specific protein Y-linked 20, pseudogene |
| TSPY5P | Y | testis specific protein Y-linked 5, pseudogene |
| AC006987.1 | Y | testis specific protein, Y-linked 1 (TSPY1) pseu |
| RNA5SP518 | Y | RNA, 5S ribosomal pseudogene 518 [Source:H |
| RNA5SP519 | Y | RNA, 5S ribosomal pseudogene 519 [Source:H |
| DUX4L31 | Y | double homeobox 4 like 31 (pseudogene) [Sou |
| PCMTD1P1 | Y | protein-L-isoaspartate (D-aspartate) O-methyl |
| CDC27P2 | Y | cell division cycle 27 pseudogene 2 [Source:H( |
| AC010970.1 | Y | RNA, 18S ribosomal pseudogene |
| RNA5-8SP6 | Y | RNA, 5.8S ribosomal pseudogene 6 [Source:H( |
| AC140113.4 | Y | SH3 and multiple ankyrin repeat domains 2 (S |
| C2orf27AP1 | Y | chromosome 2 open reading frame 27A pseu( |
| AC140113.3 | Y | family with sequence similarity 201, member |
| AC140113.2 | Y | family with sequence similarity 201, member |
| AC140113.1 | Y | family with sequence similarity 201, member |
| AC134878.1 | Y | asparagine synthetase (glutamine-hydrolyzing |

| | | |
|---|---|---|
| AC134878.2 | Y | MAFF interacting protein (pseudogene) |
| ANKRD20A6P | Y | ankyrin repeat domain 20 family member A6, |
| SNX18P1Y | Y | sorting nexin 18 pseudogene 1, Y-linked [Sour |
| DUX4L16 | Y | double homeobox 4 like 16 (pseudogene) [Sou |
| DUX4L17 | Y | double homeobox 4 like 17 (pseudogene) [Sou |
| DUX4L18 | Y | double homeobox 4 like 18 (pseudogene) [Sou |
| DUX4L19 | Y | double homeobox 4 like 19 (pseudogene) [Sou |
| PABPC1P5 | Y | poly(A) binding protein cytoplasmic 1 pseudog |
| SLC9B1P1 | Y | solute carrier family 9 member B1 pseudogen |
| ACTR3BP1 | Y | ACTR3B pseudogene 1 [Source:HGNC Symbol; |
| AC134879.3 | Y | checkpoint kinase 2 (CHEK2) pseudogene |
| MTND1P12 | Y | MT-ND1 pseudogene 12 [Source:HGNC Symbo |
| AC134879.1 | Y | tripartite motif containing 43 (TRIM43) pseud |
| AC134879.2 | Y | immunoglobulin kappa variable 1D-39 (IGKV1 |
| RCC2P1 | Y | regulator of chromosome condensation 2 pse |
| ASS1P6 | Y | argininosuccinate synthetase 1 pseudogene 6 |
| MXRA5Y | Y | matrix remodeling associated 5 Y-linked (pseu |
| GYG2P1 | Y | glycogenin 2 pseudogene 1 [Source:HGNC Syn |
| RPS24P1 | Y | ribosomal protein S24 pseudogene 1 [Source: |
| ARSFP1 | Y | arylsulfatase F pseudogene 1 [Source:HGNC S |
| RN7SL702P | Y | RNA, 7SL, cytoplasmic 702, pseudogene [Sour |
| FAM8A4P | Y | family with sequence similarity 8 member A4, |
| ARSEP1 | Y | arylsulfatase E pseudogene 1 [Source:HGNC S |
| ARSDP1 | Y | arylsulfatase D pseudogene 1 [Source:HGNC S |
| XGY1 | Y | XG pseudogene, Y-linked 1 [Source:HGNC Sym |
| USP9Y | Y | ubiquitin specific peptidase 9 Y-linked [Source |
| SHROOM2P1 | Y | shroom family member 2 pseudogene 1 [Sour |
| MED14P1 | Y | mediator complex subunit 14 pseudogene 1 [S |
| AC244213.1 | Y | novel transcript, antisense to USP9Y |
| CDY4P | Y | chromodomain Y-linked 4 pseudogene [Sourc |
| DDX3Y | Y | DEAD-box helicase 3 Y-linked [Source:HGNC S |
| CASKP1 | Y | calcium/calmodulin dependent serine protein |
| SFPQP1 | Y | splicing factor proline and glutamine rich pseu |
| TAB3P1 | Y | TGF-beta activated kinase 1/MAP3K7 binding |
| DPPA2P1 | Y | developmental pluripotency associated 2 pseu |
| UTY | Y | ubiquitously transcribed tetratricopeptide rep |
| PSMA6P1 | Y | proteasome subunit alpha 6 pseudogene 1 [So |
| TMSB4Y | Y | thymosin beta 4 Y-linked [Source:HGNC Symb |
| ANOS2P | Y | anosmin 2, pseudogene [Source:HGNC Symbo |
| VCY | Y | variable charge Y-linked [Source:HGNC Symbo |
| VCY1B | Y | variable charge Y-linked 1B [Source:HGNC Syn |
| PNPLA4P1 | Y | patatin like phospholipase domain containing |
| AC010723.1 | Y | novel transcript |
| NLGN4Y | Y | neuroligin 4 Y-linked [Source:HGNC Symbol;A |
| AGKP1 | Y | acylglycerol kinase pseudogene 1 [Source:HGI |
| NLGN4Y-AS1 | Y | NLGN4Y antisense RNA 1 [Source:HGNC Symb |
| MED13P1 | Y | mediator complex subunit 13 pseudogene 1 [S |
| CYCSP46 | Y | CYCS pseudogene 46 [Source:HGNC Symbol;A |
| AC006989.1 | Y | cytochrome c, somatic (CYCS) pseudogene |
| PUDPP1 | Y | pseudouridine 5'-phosphatase pseudogene 1 |

| | | |
|---|---|---|
| STSP1 | Y | steroid sulfatase (microsomal) pseudogene 1 |
| AC011749.1 | Y | NULL |
| RNU6-109P | Y | RNA, U6 small nuclear 109, pseudogene [Sour |
| RNU6-184P | Y | RNA, U6 small nuclear 184, pseudogene [Sour |
| AC011751.1 | Y | novel transcript |
| SURF6P1 | Y | surfeit 6 pseudogene 1 [Source:HGNC Symbol |
| ELOCP35 | Y | elongin C pseudogene 35 [Source:HGNC Symb |
| FAM41AY1 | Y | family with sequence similarity 41 member A, |
| AC015978.2 | Y | family with sequence similarity 166, member |
| TUBB1P2 | Y | tubulin beta 1 class VI pseudogene 2 [Source:I |
| FAM224B | Y | family with sequence similarity 224 member E |
| RNA5SP520 | Y | RNA, 5S ribosomal pseudogene 520 [Source:H |
| RNA5SP521 | Y | RNA, 5S ribosomal pseudogene 521 [Source:H |
| AC015978.1 | Y | NULL |
| CLUHP1 | Y | clustered mitochondria homolog pseudogene |
| TAF9P1 | Y | TATA-box binding protein associated factor 9 |
| AC007742.2 | Y | ubiquitin specific peptidase 9, Y-linked (USP9Y |
| CDY5P | Y | chromodomain Y-linked 5 pseudogene [Sourc |
| AC007742.1 | Y | transcription elongation factor B (SIII), polype |
| PRYP1 | Y | PTPN13 like Y-linked pseudogene 1 [Source:H |
| ACTG1P2 | Y | actin gamma 1 pseudogene 2 [Source:HGNC S |
| XKRY | Y | XK related, Y-linked [Source:HGNC Symbol;Ac |
| USP9YP23 | Y | USP9Y pseudogene 23 [Source:HGNC Symbol; |
| USP9YP27 | Y | USP9Y pseudogene 27 [Source:HGNC Symbol; |
| RNU1-128P | Y | RNA, U1 small nuclear 128, pseudogene [Sour |
| TRAPPC2P3 | Y | trafficking protein particle complex 2 pseudog |
| OFD1P1Y | Y | OFD1 pseudogene 1 Y-linked [Source:HGNC Sy |
| ELOCP6 | Y | elongin C pseudogene 6 [Source:HGNC Symbo |
| CDY2B | Y | chromodomain Y-linked 2B [Source:HGNC Syn |
| CDY6P | Y | chromodomain Y-linked 6 pseudogene [Sourc |
| USP9YP7 | Y | USP9Y pseudogene 7 [Source:HGNC Symbol;A |
| USP9YP6 | Y | USP9Y pseudogene 6 [Source:HGNC Symbol;A |
| CDY7P | Y | chromodomain Y-linked 7 pseudogene [Sourc |
| USP9YP34 | Y | USP9Y pseudogene 34 [Source:HGNC Symbol; |
| USP9YP32 | Y | USP9Y pseudogene 32 [Source:HGNC Symbol; |
| CDY8P | Y | chromodomain Y-linked 8 pseudogene [Sourc |
| CDY2A | Y | chromodomain Y-linked 2A [Source:HGNC Syn |
| ELOCP12 | Y | elongin C pseudogene 12 [Source:HGNC Symb |
| OFD1P2Y | Y | OFD1 pseudogene 2 Y-linked [Source:HGNC Sy |
| TRAPPC2P8 | Y | trafficking protein particle complex 2 pseudog |
| USP9YP15 | Y | USP9Y pseudogene 15 [Source:HGNC Symbol; |
| RNU1-95P | Y | RNA, U1 small nuclear 95, pseudogene [Sourc |
| USP9YP16 | Y | USP9Y pseudogene 16 [Source:HGNC Symbol; |
| XKRY2 | Y | XK related, Y-linked 2 [Source:HGNC Symbol;A |
| ACTG1P11 | Y | actin gamma 1 pseudogene 11 [Source:HGNC |
| PRYP2 | Y | PTPN13 like Y-linked pseudogene 2 [Source:H |
| ELOCP26 | Y | elongin C pseudogene 26 [Source:HGNC Symb |
| CDY9P | Y | chromodomain Y-linked 9 pseudogene [Sourc |
| TAF9P2 | Y | TATA-box binding protein associated factor 9 |
| CLUHP2 | Y | clustered mitochondria homolog pseudogene |

| | | |
|---|---|---|
| FAM224A | Y | family with sequence similarity 224 member A |
| AC007241.1 | Y | NULL |
| RNA5SP522 | Y | RNA, 5S ribosomal pseudogene 522 [Source:H |
| RNA5SP523 | Y | RNA, 5S ribosomal pseudogene 523 [Source:H |
| TUBB1P1 | Y | tubulin beta 1 class VI pseudogene 1 [Source:I |
| AC007241.2 | Y | family with sequence similarity 166, member |
| FAM41AY2 | Y | family with sequence similarity 41 member A, |
| ELOCP13 | Y | elongin C pseudogene 13 [Source:HGNC Symb |
| OFD1P4Y | Y | OFD1 pseudogene 4 Y-linked [Source:HGNC Sy |
| USP9YP14 | Y | USP9Y pseudogene 14 [Source:HGNC Symbol; |
| RNU1-48P | Y | RNA, U1 small nuclear 48, pseudogene [Sourc |
| USP9YP5 | Y | USP9Y pseudogene 5 [Source:HGNC Symbol;A |
| AC022486.1 | Y | novel transcript |
| XKRYP1 | Y | XK related, Y-linked pseudogene 1 [Source:HG |
| PRYP5 | Y | PTPN13 like Y-linked pseudogene 5 [Source:H |
| ELOCP7 | Y | elongin C pseudogene 7 [Source:HGNC Symbo |
| USP9YP1 | Y | USP9Y pseudogene 1 [Source:HGNC Symbol;A |
| HSFY1 | Y | heat shock transcription factor Y-linked 1 [Sou |
| GPM6BP1 | Y | glycoprotein M6B pseudogene 1 [Source:HGN |
| TTTY9B | Y | testis-specific transcript, Y-linked 9B [Source:H |
| OFD1P5Y | Y | OFD1 pseudogene 5 Y-linked [Source:HGNC Sy |
| RAB9AP4 | Y | RAB9A, member RAS oncogene family pseudo |
| TRAPPC2P7 | Y | trafficking protein particle complex 2 pseudog |
| OFD1P6Y | Y | OFD1 pseudogene 6 Y-linked [Source:HGNC Sy |
| HSFY2 | Y | heat shock transcription factor Y-linked 2 [Sou |
| GPM6BP2 | Y | glycoprotein M6B pseudogene 2 [Source:HGN |
| TTTY14 | Y | testis-specific transcript, Y-linked 14 [Source:H |
| USP9YP2 | Y | USP9Y pseudogene 2 [Source:HGNC Symbol;A |
| ELOCP14 | Y | elongin C pseudogene 14 [Source:HGNC Symb |
| PRYP6 | Y | PTPN13 like Y-linked pseudogene 6 [Source:H |
| XKRYP2 | Y | XK related, Y-linked pseudogene 2 [Source:HG |
| USP9YP10 | Y | USP9Y pseudogene 10 [Source:HGNC Symbol; |
| USP9YP28 | Y | USP9Y pseudogene 28 [Source:HGNC Symbol; |
| RNU1-41P | Y | RNA, U1 small nuclear 41, pseudogene [Sourc |
| OFD1P7Y | Y | OFD1 pseudogene 7 Y-linked [Source:HGNC Sy |
| MTCYBP2 | Y | MT-CYB pseudogene 2 [Source:HGNC Symbol; |
| ZNF839P1 | Y | zinc finger protein 839 pseudogene 1 [Source: |
| CD24P4 | Y | CD24 molecule pseudogene 4 [Source:HGNC S |
| RNU6-255P | Y | RNA, U6 small nuclear 255, pseudogene [Sour |
| AC007244.1 | Y | NULL |
| AC010133.1 | Y | endogenous retrovirus group FRD, member 1 |
| GAPDHP19 | Y | glyceraldehyde 3 phosphate dehydrogenase p |
| BCORP1 | Y | BCL6 corepressor pseudogene 1 [Source:HGN |
| TXLNGY | Y | taxilin gamma pseudogene, Y-linked [Source:H |
| AC009977.1 | Y | ADAM metallopeptidase domain (ADAM) pseu |
| AC010889.1 | Y | novel transcript |
| KDM5D | Y | lysine demethylase 5D [Source:HGNC Symbol; |
| AC010889.2 | Y | uncharacterized LOC105377224 [Source:NCBI |
| AC009233.1 | Y | heat shock transcription factor, Y-linked 1 (HS |
| RCC2P2 | Y | regulator of chromosome condensation 2 pse |

| | | |
|---|---|---|
| ZNF886P | Y | zinc finger protein 886, pseudogene [Source:H |
| ZNF885P | Y | zinc finger protein 885, pseudogene [Source:H |
| TTTY10 | Y | testis-specific transcript, Y-linked 10 [Source:H |
| KDM5DP1 | Y | lysine demethylase 5D pseudogene 1 [Source: |
| EIF1AY | Y | eukaryotic translation initiation factor 1A Y-lir |
| AC009494.2 | Y | novel transcript |
| TBL1YP1 | Y | transducin beta like 1 Y-linked pseudogene 1 [ |
| RPS4Y2 | Y | ribosomal protein S4 Y-linked 2 [Source:HGNC |
| HSFY4P | Y | heat shock transcription factor Y-linked 4, pse |
| AC009494.1 | Y | novel pseudogene |
| GAPDHP17 | Y | glyceraldehyde-3-phosphate dehydrogenase p |
| TMEM167AP1 | Y | transmembrane protein 167A pseudogene 1 [ |
| AC007876.1 | Y | novel transcript |
| TOMM22P1 | Y | TOMM22 pseudogene 1 [Source:HGNC Symbo |
| NEFLP1 | Y | neurofilament light pseudogene 1 [Source:HG |
| AC010086.1 | Y | RNA binding motif protein, Y-linked, family 1, |
| PRORY | Y | proline rich Y-linked [Source:HGNC Symbol;Ac |
| RBMY2EP | Y | RNA binding motif protein, Y-linked, family 2, |
| AC010086.2 | Y | pseudogene similar to part of testis specific pr |
| AC010086.3 | Y | novel transcript |
| RBMY2TP | Y | RNA binding motif protein, Y-linked, family 2, |
| TSPY14P | Y | testis specific protein Y-linked 14, pseudogene |
| RBMY1HP | Y | RNA binding motif protein, Y-linked, family 1, |
| AC010141.2 | Y | testis specific protein, Y-linked 1 (TSPY1) pseu |
| RBMY1B | Y | RNA binding motif protein, Y-linked, family 1, |
| RBMY1A1 | Y | RNA binding motif protein, Y-linked, family 1, |
| AC010141.1 | Y | testis specific protein, Y-linked 1 (TSPY1) pseu |
| TTTY13 | Y | testis-specific transcript, Y-linked 13 [Source:H |
| ELOCP5 | Y | elongin C pseudogene 5 [Source:HGNC Symbo |
| CDY10P | Y | chromodomain Y-linked 10 pseudogene [Sour |
| USP9YP3 | Y | USP9Y pseudogene 3 [Source:HGNC Symbol;A |
| USP9YP8 | Y | USP9Y pseudogene 8 [Source:HGNC Symbol;A |
| CDY11P | Y | chromodomain protein Y-linked 11 pseudoger |
| OFD1P16Y | Y | OFD1 pseudogene 16 Y-linked [Source:HGNC S |
| AC024236.1 | Y | novel transcript |
| AC007322.2 | Y | testis specific protein, Y-linked 1 (TSPY1) pseu |
| RBMY1D | Y | RNA binding motif protein, Y-linked, family 1, |
| AC007322.4 | Y | testis specific protein, Y-linked 1 (TSPY1) pseu |
| RBMY1E | Y | RNA binding motif protein, Y-linked, family 1, |
| AC007322.1 | Y | testis specific protein, Y-linked 1 (TSPY1) pseu |
| RBMY2AP | Y | RNA binding motif protein, Y-linked, family 2, |
| AC007322.3 | Y | testis specific protein, Y-linked 1 (TSPY1) pseu |
| OFD1P8Y | Y | OFD1 pseudogene 8 Y-linked [Source:HGNC Sy |
| HSFY5P | Y | heat shock transcription factor Y-linked 5, pse |
| USP9YP17 | Y | USP9Y pseudogene 17 [Source:HGNC Symbol; |
| CDY12P | Y | chromodomain Y-linked 12 pseudogene [Sour |
| ELOCP15 | Y | elongin C pseudogene 15 [Source:HGNC Symb |
| PRY2 | Y | PTPN13 like Y-linked 2 [Source:HGNC Symbol; |
| AC007359.1 | Y | uncharacterized LOC102725532 [Source:NCBI |
| TTTY6B | Y | testis-specific transcript, Y-linked 6B [Source:H |

| | | |
|---|---|---|
| RBMY1F | Y | RNA binding motif protein, Y-linked, family 1, |
| TSPY23P | Y | testis specific protein Y-linked 23, pseudogene |
| RBMY2UP | Y | RNA binding motif protein, Y-linked, family 2, |
| RBMY1KP | Y | RNA binding motif protein, Y-linked, family 1, |
| TTTY5 | Y | testis-specific transcript, Y-linked 5 [Source:H( |
| TSPY22P | Y | testis specific protein Y-linked 22, pseudogene |
| RBMY2FP | Y | RNA binding motif protein, Y-linked, family 2, |
| TTTY25P | Y | testis-specific transcript, Y-linked 25, pseudog |
| TSPY21P | Y | testis specific protein Y-linked 21, pseudogene |
| RBMY1J | Y | RNA binding motif protein, Y-linked, family 1, |
| AC008175.1 | Y | uncharacterized LOC102725532 [Source:NCBI |
| TTTY6 | Y | testis-specific transcript, Y-linked 6 [Source:H( |
| PRY | Y | PTPN13 like Y-linked [Source:HGNC Symbol;A( |
| ELOCP8 | Y | elongin C pseudogene 8 [Source:HGNC Symbo |
| CDY13P | Y | chromodomain Y-linked 13 pseudogene [Sour |
| USP9YP24 | Y | USP9Y pseudogene 24 [Source:HGNC Symbol; |
| HSFY7P | Y | heat shock transcription factor Y-linked 7, pse |
| OFD1P9Y | Y | OFD1 pseudogene 9 Y-linked [Source:HGNC Sy |
| AC008175.2 | Y | testis specific protein, Y-linked 1 (TSPY1) pseu |
| RBMY2BP | Y | RNA binding motif protein, Y-linked, family 2, |
| AC016694.1 | Y | testis specific protein, Y-linked 1 (TSPY1) pseu |
| RBMY2WP | Y | RNA binding motif protein, Y-linked, family 2, |
| TTTY17A | Y | testis-specific transcript, Y-linked 17A [Source |
| AC016911.1 | Y | tripartite motif containing 60 (TRIM60) pseud |
| ZNF736P11Y | Y | zinc finger protein 736 pseudogene 11, Y-linke |
| TTTY4 | Y | testis-specific transcript, Y-linked 4 [Source:H( |
| BPY2 | Y | basic charge Y-linked 2 [Source:HGNC Symbol; |
| AC006366.1 | Y | zinc finger protein pseudogene |
| TRIM60P8Y | Y | tripartite motif containing 60 pseudogene 8, Y |
| ZNF736P3Y | Y | zinc finger protein 736 pseudogene 3, Y-linkec |
| TRIM60P9Y | Y | tripartite motif containing 60 pseudogene 9, Y |
| DAZ1 | Y | deleted in azoospermia 1 [Source:HGNC Symb |
| DAZ2 | Y | deleted in azoospermia 2 [Source:HGNC Symb |
| PPP1R12BP2 | Y | protein phosphatase 1 regulatory subunit 12B |
| SNORA70 | Y | Small nucleolar RNA SNORA70 [Source:RFAM; |
| AC009947.3 | Y | pseudogene similar to part of protein phospha |
| REREP1Y | Y | arginine-glutamic acid dipeptide repeats pseu |
| AC009947.1 | Y | testis specific protein, Y-linked 1 (TSPY1) pseu |
| RBMY2CP | Y | RNA binding motif protein, Y-linked, family 2, |
| AC009947.2 | Y | testis specific protein, Y-linked 1 (TSPY1) pseu |
| OFD1P10Y | Y | OFD1 pseudogene 10 Y-linked [Source:HGNC S |
| HSFY6P | Y | heat shock transcription factor Y-linked 6, pse |
| USP9YP18 | Y | USP9Y pseudogene 18 [Source:HGNC Symbol; |
| CDY14P | Y | chromodomain protein Y-linked 14 pseudoger |
| ELOCP16 | Y | elongin C pseudogene 16 [Source:HGNC Symb |
| PRYP3 | Y | PTPN13 like Y-linked pseudogene 3 [Source:H( |
| XKRYP3 | Y | XK related, Y-linked pseudogene 3 [Source:HG |
| USP9YP25 | Y | USP9Y pseudogene 25 [Source:HGNC Symbol; |
| USP9YP29 | Y | USP9Y pseudogene 29 [Source:HGNC Symbol; |
| RNU1-97P | Y | RNA, U1 small nuclear 97, pseudogene [Sourc |

| | | |
|---|---|---|
| RAB9AP1 | Y | RAB9A, member RAS oncogene family pseudo |
| TRAPPC2P9 | Y | trafficking protein particle complex 2 pseudog |
| OFD1P11Y | Y | OFD1 pseudogene 11 Y-linked [Source:HGNC S |
| ELOCP9 | Y | elongin C pseudogene 9 [Source:HGNC Symbo |
| CDY15P | Y | chromodomain Y-linked 15 pseudogene [Sour |
| XKRYP4 | Y | XK related, Y-linked pseudogene 4 [Source:HG |
| USP9YP13 | Y | USP9Y pseudogene 13 [Source:HGNC Symbol; |
| USP9YP11 | Y | USP9Y pseudogene 11 [Source:HGNC Symbol; |
| TTTY3B | Y | testis-specific transcript, Y-linked 3B [Source:H |
| USP9YP12 | Y | USP9Y pseudogene 12 [Source:HGNC Symbol; |
| RNU1-86P | Y | RNA, U1 small nuclear 86, pseudogene [Sourc |
| RAB9AP5 | Y | RAB9A, member RAS oncogene family pseudo |
| TRAPPC2P10 | Y | trafficking protein particle complex 2 pseudog |
| OFD1P12Y | Y | OFD1 pseudogene 12 Y-linked [Source:HGNC S |
| ELOCP10 | Y | elongin C pseudogene 10 [Source:HGNC Symb |
| CDY1B | Y | chromodomain Y-linked 1B [Source:HGNC Syn |
| CDY17P | Y | chromodomain Y-linked 17 pseudogene [Sour |
| USP9YP35 | Y | USP9Y pseudogene 35 [Source:HGNC Symbol; |
| USP9YP31 | Y | USP9Y pseudogene 31 [Source:HGNC Symbol; |
| CDY18P | Y | chromodomain Y-linked 18 pseudogene [Sour |
| AC023274.2 | Y | ubiquitin-conjugating enzyme E2 (UBE2Q2) ps |
| GOLGA6L11P | Y | golgin A6 family like 11, pseudogene [Source:I |
| DNM1P24 | Y | dynamin 1 pseudogene 24 [Source:HGNC Sym |
| AC023274.1 | Y | novel pseudogene |
| CSPG4P2Y | Y | CSPG4 pseudogene 2 Y-linked [Source:HGNC S |
| CSPG4P3Y | Y | CSPG4 pseudogene 3 Y-linked [Source:HGNC S |
| AC012005.2 | Y | ubiquitin conjugating enzyme E2 Q2 like ( UBE |
| GOLGA2P2Y | Y | GOLGA2 pseudogene 2 Y-linked [Source:HGN( |
| RN7SL818P | Y | RNA, 7SL, cytoplasmic 818, pseudogene [Sour |
| UBE2Q2P4Y | Y | ubiquitin conjugating enzyme E2 Q2 pseudoge |
| LINC00265-2P | Y | long intergenic non-protein coding RNA 265-2 |
| CICP2 | Y | capicua transcriptional repressor pseudogene |
| LINC00266-2P | Y | long intergenic non-protein coding RNA 266-2 |
| AC012005.1 | Y | ribosomal protein L41 (RPL41) pseudogene |
| RBMY2XP | Y | RNA binding motif protein, Y-linked, family 2, |
| TTTY17B | Y | testis-specific transcript, Y-linked 17B [Source |
| TRIM60P5Y | Y | tripartite motif containing 60 pseudogene 5, Y |
| ZNF736P12Y | Y | zinc finger protein 736 pseudogene 12, Y-linke |
| TTTY4B | Y | testis-specific transcript, Y-linked 4B [Source:H |
| BPY2B | Y | basic charge Y-linked 2B [Source:HGNC Symbo |
| AC010153.1 | Y | zinc finger protein pseudogene |
| TRIM60P10Y | Y | tripartite motif containing 60 pseudogene 10, |
| ZNF736P2Y | Y | zinc finger protein 736 pseudogene 2, Y-linkec |
| TRIM60P11Y | Y | tripartite motif containing 60 pseudogene 11, |
| DAZ3 | Y | deleted in azoospermia 3 [Source:HGNC Symb |
| DAZ4 | Y | deleted in azoospermia 4 [Source:HGNC Symb |
| AC006338.2 | Y | tripartite motif containing 60 (TRIM60) pseud |
| ZNF736P1Y | Y | zinc finger protein 736 pseudogene 1, Y-linkec |
| AC006338.1 | Y | tripartite motif containing 60 (TRIM60) pseud |
| ZNF736P5Y | Y | zinc finger protein 736 pseudogene 5, Y-linkec |

| | | |
|---|---|---|
| BPY2C | Y | basic charge Y-linked 2C [Source:HGNC Symbo |
| TTTY4C | Y | testis-specific transcript, Y-linked 4C [Source:H |
| ZNF736P4Y | Y | zinc finger protein 736 pseudogene 4, Y-linked |
| AC006386.1 | Y | tripartite motif containing 60 (TRIM60) pseud |
| TTTY17C | Y | testis-specific transcript, Y-linked 17C [Source |
| RBMY2YP | Y | RNA binding motif protein, Y-linked, family 2, |
| AC006386.2 | Y | ribosomal protein L41 (RPL41) pseudogene |
| LINC00266-4P | Y | long intergenic non-protein coding RNA 266-4 |
| CICP1 | Y | capicua transcriptional repressor pseudogene |
| LINC00265-3P | Y | long intergenic non-protein coding RNA 265-3 |
| UBE2Q2P5Y | Y | ubiquitin conjugating enzyme E2 Q2 pseudoge |
| GOLGA2P3Y | Y | GOLGA2 pseudogene 3 Y-linked [Source:HGNC |
| RN7SL725P | Y | RNA, 7SL, cytoplasmic 725, pseudogene [Sour |
| AC006328.2 | Y | ubiquitin conjugating enzyme E2 Q2 like (UBE |
| CSPG4P4Y | Y | CSPG4 pseudogene 4 Y-linked [Source:HGNC S |
| CSPG4P1Y | Y | CSPG4 pseudogene 1 Y-linked [Source:HGNC S |
| DNM1P48 | Y | dynamin 1 pseudogene 48 [Source:HGNC Sym |
| AC006328.1 | Y | novel pseudogene |
| GOLGA6L16P | Y | golgin A6 family like 16, pseudogene [Source: |
| AC007562.1 | Y | ubiquitin-conjugating enzyme pseudogene |
| CDY19P | Y | chromodomain Y-linked 19 pseudogene [Sour |
| USP9YP36 | Y | USP9Y pseudogene 36 [Source:HGNC Symbol; |
| USP9YP33 | Y | USP9Y pseudogene 33 [Source:HGNC Symbol; |
| CDY20P | Y | chromodomain Y-linked 20 pseudogene [Sour |
| CDY1 | Y | chromodomain Y-linked 1 [Source:HGNC Syml |
| ELOCP34 | Y | elongin C pseudogene 34 [Source:HGNC Symb |
| OFD1P13Y | Y | OFD1 pseudogene 13 Y-linked [Source:HGNC S |
| TRAPPC2P5 | Y | trafficking protein particle complex 2 pseudog |
| RAB9AP2 | Y | RAB9A, member RAS oncogene family pseudo |
| USP9YP30 | Y | USP9Y pseudogene 30 [Source:HGNC Symbol; |
| RNU1-107P | Y | RNA, U1 small nuclear 107, pseudogene [Sour |
| TTTY3 | Y | testis-specific transcript, Y-linked 3 [Source:H |
| USP9YP9 | Y | USP9Y pseudogene 9 [Source:HGNC Symbol;A |
| USP9YP19 | Y | USP9Y pseudogene 19 [Source:HGNC Symbol; |
| XKRYP5 | Y | XK related, Y-linked pseudogene 5 [Source:HG |
| CDY22P | Y | chromodomain Y-linked 22 pseudogene [Sour |
| ELOCP17 | Y | elongin C pseudogene 17 [Source:HGNC Symb |
| OFD1P18Y | Y | OFD1 pseudogene 18 Y-linked [Source:HGNC S |
| TRAPPC2P4 | Y | trafficking protein particle complex 2 pseudog |
| RAB9AP3 | Y | RAB9A, member RAS oncogene family pseudo |
| USP9YP20 | Y | USP9Y pseudogene 20 [Source:HGNC Symbol; |
| RNU1-40P | Y | RNA, U1 small nuclear 40, pseudogene [Sourc |
| USP9YP21 | Y | USP9Y pseudogene 21 [Source:HGNC Symbol; |
| XKRYP6 | Y | XK related, Y-linked pseudogene 6 [Source:HG |
| PRYP4 | Y | PTPN13 like Y-linked pseudogene 4 [Source:H |
| ELOCP11 | Y | elongin C pseudogene 11 [Source:HGNC Symb |
| CDY23P | Y | chromodomain Y-linked 23 pseudogene [Sour |
| USP9YP26 | Y | USP9Y pseudogene 26 [Source:HGNC Symbol; |
| HSFY8P | Y | heat shock transcription factor Y-linked 8, pse |
| OFD1P15Y | Y | OFD1 pseudogene 15 Y-linked [Source:HGNC S |

| | | | |
|---|---|---|---|
| RBMY2DP | Y | | RNA binding motif protein, Y-linked, family 2, |
| REREP2Y | Y | | arginine-glutamic acid dipeptide repeats pseu |
| AC024067.1 | Y | | novel pseudogene |
| SNORA70 | Y | | Small nucleolar RNA SNORA70 [Source:RFAM; |
| PPP1R12BP1 | Y | | protein phosphatase 1 regulatory subunit 12B |
| RNU6-1314P | Y | | RNA, U6 small nuclear 1314, pseudogene [Sou |
| CYCSP48 | Y | | CYCS pseudogene 48 [Source:HGNC Symbol;A |
| ANKRD36P1 | Y | | ankyrin repeat domain 36 pseudogene 1 [Sour |
| AC013734.1 | Y | | striatin, calmodulin binding protein 4 (STRN4) |
| TPTE2P4 | Y | | transmembrane phosphoinositide 3-phosphat |
| CYCSP49 | Y | | CYCS pseudogene 49 [Source:HGNC Symbol;A |
| SLC25A15P1 | Y | | solute carrier family 25 member 15 pseudoge |
| PARP4P1 | Y | | poly(ADP-ribose) polymerase family member |
| CCNQP2 | Y | | CCNQ pseudogene 2 [Source:HGNC Symbol;Ac |
| CTBP2P1 | Y | | C-terminal binding protein 2 pseudogene 1 [S |
| DDX11L1 | | 1 | DEAD/H-box helicase 11 like 1 [Source:HGNC |
| WASH7P | | 1 | WASP family homolog 7, pseudogene [Source |
| MIR6859-1 | | 1 | microRNA 6859-1 [Source:HGNC Symbol;Acc: |
| MIR1302-2HG | | 1 | MIR1302-2 host gene [Source:HGNC Symbol; |
| MIR1302-2 | | 1 | microRNA 1302-2 [Source:HGNC Symbol;Acc: |
| FAM138A | | 1 | family with sequence similarity 138 member |
| OR4G4P | | 1 | olfactory receptor family 4 subfamily G memb |
| OR4G11P | | 1 | olfactory receptor family 4 subfamily G memb |
| OR4F5 | | 1 | olfactory receptor family 4 subfamily F memb |
| AL627309.1 | | 1 | novel transcript |
| AL627309.3 | | 1 | novel transcript |
| CICP27 | | 1 | capicua transcriptional repressor pseudogene |
| AL627309.6 | | 1 | novel pseudogene |
| AL627309.7 | | 1 | novel pseudogene |
| AL627309.2 | | 1 | novel transcript |
| AL627309.5 | | 1 | novel transcript |
| RNU6-1100P | | 1 | RNA, U6 small nuclear 1100, pseudogene [Sou |
| AL627309.4 | | 1 | novel transcript |
| FO538757.1 | | 1 | DEAD/H-box helicase 11 (DDX11) pseudogene |
| WASH9P | | 1 | WAS protein family homolog 9, pseudogene [ |
| MIR6859-2 | | 1 | microRNA 6859-2 [Source:HGNC Symbol;Acc: |
| AP006222.1 | | 1 | ribosomal protein L23a (RPL23A) pseudogene |
| AP006222.2 | | 1 | novel transcript |
| RPL23AP24 | | 1 | ribosomal protein L23a pseudogene 24 [Sourc |
| AL732372.1 | | 1 | novel transcript |
| AL732372.2 | | 1 | pseudogene similar to part of septin 14 SEPT1 |
| WBP1LP7 | | 1 | WBP1L pseudogene 7 [Source:HGNC Symbol; |
| OR4F29 | | 1 | olfactory receptor family 4 subfamily F memb |
| CICP7 | | 1 | capicua transcriptional repressor pseudogene |
| AL732372.3 | | 1 | novel pseudogene |
| U6 | | 1 | U6 spliceosomal RNA [Source:RFAM;Acc:RF00 |
| AL669831.3 | | 1 | septin 14 (SEPT14) pseudogene |
| AC114498.1 | | 1 | novel transcript |
| MTND1P23 | | 1 | MT-ND1 pseudogene 23 [Source:HGNC Symbo |
| MTND2P28 | | 1 | MT-ND2 pseudogene 28 [Source:HGNC Symbo |

| | | |
|---|---|---|
| MTCO1P12 | 1 | MT-CO1 pseudogene 12 [Source:HGNC Symbo |
| AC114498.2 | 1 | NULL |
| MTCO2P12 | 1 | MT-CO2 pseudogene 12 [Source:HGNC Symbo |
| MTATP8P1 | 1 | MT-ATP8 pseudogene 1 [Source:HGNC Symbo |
| MTATP6P1 | 1 | MT-ATP6 pseudogene 1 [Source:HGNC Symbo |
| MTCO3P12 | 1 | MT-CO3 pseudogene 12 [Source:HGNC Symbo |
| WBP1LP6 | 1 | WBP1L pseudogene 6 [Source:HGNC Symbol;A |
| OR4F16 | 1 | olfactory receptor family 4 subfamily F memb |
| CICP3 | 1 | capicua transcriptional repressor pseudogene |
| AL669831.1 | 1 | general transcription factor IIi (GTF2I) pseudo |
| RNU6-1199P | 1 | RNA, U6 small nuclear 1199, pseudogene [Sou |
| AL669831.2 | 1 | novel transcript |
| LINC01409 | 1 | long intergenic non-protein coding RNA 1409 |
| AL669831.6 | 1 | novel septin family pseudogene |
| AL669831.4 | 1 | F-box only protein 25 (FBXO25) pseudogene |
| FAM87B | 1 | family with sequence similarity 87 member B |
| LINC01128 | 1 | long intergenic non-protein coding RNA 1128 |
| LINC00115 | 1 | long intergenic non-protein coding RNA 115 [S |
| FAM41C | 1 | family with sequence similarity 41 member C |
| TUBB8P11 | 1 | tubulin beta 8 class VIII pseudogene 11 [Sourc |
| AL669831.5 | 1 | FAM166A, pseudogene P3 |
| AL645608.6 | 1 | novel transcript |
| AL645608.2 | 1 | novel transcript |
| AL645608.4 | 1 | novel transcript |
| LINC02593 | 1 | long intergenic non-protein coding RNA 2593 |
| SAMD11 | 1 | sterile alpha motif domain containing 11 [Sou |
| NOC2L | 1 | NOC2 like nucleolar associated transcriptional |
| KLHL17 | 1 | kelch like family member 17 [Source:HGNC Sy |
| PLEKHN1 | 1 | pleckstrin homology domain containing N1 [S |
| PERM1 | 1 | PPARGC1 and ESRR induced regulator, muscle |
| AL645608.7 | 1 | novel transcript |
| HES4 | 1 | hes family bHLH transcription factor 4 [Source |
| ISG15 | 1 | ISG15 ubiquitin like modifier [Source:HGNC Sy |
| AL645608.3 | 1 | ribosomal protein L39 (RPL39) pseudogene |
| AL645608.1 | 1 | novel transcript |
| AGRN | 1 | agrin [Source:HGNC Symbol;Acc:HGNC:329] |
| AL645608.5 | 1 | novel transcript |
| AL390719.1 | 1 | protein tyrosine phosphatase family pseudoge |
| AL645608.8 | 1 | novel transcript |
| RNF223 | 1 | ring finger protein 223 [Source:HGNC Symbol; |
| C1orf159 | 1 | chromosome 1 open reading frame 159 [Sour |
| AL390719.3 | 1 | novel transcript, antisense to C1orf159 |
| LINC01342 | 1 | long intergenic non-protein coding RNA 1342 |
| MIR200B | 1 | microRNA 200b [Source:HGNC Symbol;Acc:HG |
| MIR200A | 1 | microRNA 200a [Source:HGNC Symbol;Acc:HG |
| MIR429 | 1 | microRNA 429 [Source:HGNC Symbol;Acc:HGN |
| AL390719.2 | 1 | novel transcript |
| TTLL10-AS1 | 1 | TTLL10 antisense RNA 1 [Source:HGNC Symbo |
| TTLL10 | 1 | tubulin tyrosine ligase like 10 [Source:HGNC S |
| TNFRSF18 | 1 | TNF receptor superfamily member 18 [Source |

| | | |
|---|---|---|
| TNFRSF4 | 1 | TNF receptor superfamily member 4 [Source: |
| SDF4 | 1 | stromal cell derived factor 4 [Source:HGNC S |
| B3GALT6 | 1 | beta-1,3-galactosyltransferase 6 [Source:HGN |
| C1QTNF12 | 1 | C1q and TNF related 12 [Source:HGNC Symbo |
| AL162741.1 | 1 | novel transcript |
| UBE2J2 | 1 | ubiquitin conjugating enzyme E2 J2 [Source:H |
| LINC01786 | 1 | long intergenic non-protein coding RNA 1786 |
| SCNN1D | 1 | sodium channel epithelial 1 delta subunit [Sou |
| ACAP3 | 1 | ArfGAP with coiled-coil, ankyrin repeat and P |
| MIR6726 | 1 | microRNA 6726 [Source:HGNC Symbol;Acc:H |
| PUSL1 | 1 | pseudouridine synthase like 1 [Source:HGNC |
| INTS11 | 1 | integrator complex subunit 11 [Source:HGNC |
| MIR6727 | 1 | microRNA 6727 [Source:HGNC Symbol;Acc:H |
| AL139287.1 | 1 | novel transcript |
| CPTP | 1 | ceramide-1-phosphate transfer protein [Sour |
| TAS1R3 | 1 | taste 1 receptor member 3 [Source:HGNC Sym |
| DVL1 | 1 | dishevelled segment polarity protein 1 [Sourc |
| MIR6808 | 1 | microRNA 6808 [Source:HGNC Symbol;Acc:H |
| MXRA8 | 1 | matrix remodeling associated 8 [Source:HGN |
| AURKAIP1 | 1 | aurora kinase A interacting protein 1 [Source: |
| NDUFB4P8 | 1 | NADH:ubiquinone oxidoreductase subunit B4 |
| CCNL2 | 1 | cyclin L2 [Source:HGNC Symbol;Acc:HGNC:20 |
| MRPL20-AS1 | 1 | MRPL20 antisense RNA 1 [Source:HGNC Symb |
| MRPL20 | 1 | mitochondrial ribosomal protein L20 [Source: |
| RN7SL657P | 1 | RNA, 7SL, cytoplasmic 657, pseudogene [Sour |
| AL391244.2 | 1 | novel transcript |
| ANKRD65 | 1 | ankyrin repeat domain 65 [Source:HGNC Sym |
| AL391244.1 | 1 | novel transcript |
| TMEM88B | 1 | transmembrane protein 88B [Source:HGNC S |
| LINC01770 | 1 | long intergenic non-protein coding RNA 1770 |
| VWA1 | 1 | von Willebrand factor A domain containing 1 |
| ATAD3C | 1 | ATPase family AAA domain containing 3C [Sou |
| ATAD3B | 1 | ATPase family AAA domain containing 3B [Sou |
| AL645728.2 | 1 | ATPase family, AAA domain containing 3C (AT |
| ATAD3A | 1 | ATPase family AAA domain containing 3A [Sou |
| TMEM240 | 1 | transmembrane protein 240 [Source:HGNC Sy |
| SSU72 | 1 | SSU72 homolog, RNA polymerase II CTD phos |
| AL645728.1 | 1 | novel transcript |
| AL691432.3 | 1 | sulfiredoxin 1 (SRXN1) pseudogene |
| FNDC10 | 1 | fibronectin type III domain containing 10 [Sou |
| AL691432.4 | 1 | novel transcript |
| AL691432.2 | 1 | artifact |
| MIB2 | 1 | mindbomb E3 ubiquitin protein ligase 2 [Sour |
| MMP23B | 1 | matrix metallopeptidase 23B [Source:HGNC S |
| CDK11B | 1 | cyclin dependent kinase 11B [Source:HGNC Sy |
| FO704657.1 | 1 | novel transcript, antisense to SLC35E2B |
| SLC35E2B | 1 | solute carrier family 35 member E2B [Source: |
| AL691432.1 | 1 | ribosomal protein S7 (RPS7) pseudogene |
| MMP23A | 1 | matrix metallopeptidase 23A (pseudogene) [S |
| CDK11A | 1 | cyclin dependent kinase 11A [Source:HGNC Sy |

| | | |
|---|---|---|
| AL031282.2 | 1 | novel transcript, readthrough between SLC35I |
| AL031282.1 | 1 | ribosomal protein S7 (RPS7) pseudogene |
| SLC35E2A | 1 | solute carrier family 35 member E2A [Source:I |
| NADK | 1 | NAD kinase [Source:HGNC Symbol;Acc:HGNC: |
| GNB1 | 1 | G protein subunit beta 1 [Source:HGNC Symb |
| AL109917.1 | 1 | novel transcript |
| CALML6 | 1 | calmodulin like 6 [Source:HGNC Symbol;Acc:H |
| TMEM52 | 1 | transmembrane protein 52 [Source:HGNC Syn |
| CFAP74 | 1 | cilia and flagella associated protein 74 [Source |
| AL391845.2 | 1 | novel transcript |
| GABRD | 1 | gamma-aminobutyric acid type A receptor del |
| AL391845.1 | 1 | novel transcript |
| PRKCZ | 1 | protein kinase C zeta [Source:HGNC Symbol;A |
| AL590822.2 | 1 | novel transcript, antisense PRKCZ |
| PRKCZ-AS1 | 1 | PRKCZ antisense RNA 1 [Source:HGNC Symbol |
| FAAP20 | 1 | FA core complex associated protein 20 [Sourc |
| AL590822.1 | 1 | novel transcript |
| SKI | 1 | SKI proto-oncogene [Source:HGNC Symbol;Ac |
| AL590822.3 | 1 | novel transcript, antisense to MORN1 |
| MORN1 | 1 | MORN repeat containing 1 [Source:HGNC Syn |
| AL589739.1 | 1 | novel transcript, sense intronic to MORN1 |
| AL513477.1 | 1 | small nuclear ribonucleoprotein N (SNRPN) ps |
| AL513477.2 | 1 | novel transcript, sense intronic to MORN1 |
| RER1 | 1 | retention in endoplasmic reticulum sorting re |
| PEX10 | 1 | peroxisomal biogenesis factor 10 [Source:HGI |
| PLCH2 | 1 | phospholipase C eta 2 [Source:HGNC Symbol; |
| AL139246.1 | 1 | novel transcript |
| AL139246.4 | 1 | novel transcript |
| PANK4 | 1 | pantothenate kinase 4 [Source:HGNC Symbol; |
| HES5 | 1 | hes family bHLH transcription factor 5 [Source |
| AL139246.5 | 1 | novel transcript |
| TNFRSF14-AS1 | 1 | TNFRSF14 antisense RNA 1 [Source:HGNC Syn |
| TNFRSF14 | 1 | TNF receptor superfamily member 14 [Source |
| AL139246.2 | 1 | novel transcript |
| AL139246.3 | 1 | novel transcript |
| PRXL2B | 1 | peroxiredoxin like 2B [Source:HGNC Symbol;A |
| MMEL1 | 1 | membrane metalloendopeptidase like 1 [Sour |
| AL831784.1 | 1 | novel transcript |
| TTC34 | 1 | tetratricopeptide repeat domain 34 [Source:H |
| AC242022.2 | 1 | novel transcript, antisense to TTC34 |
| AC242022.1 | 1 | novel transcript, antisense to TTC34 |
| AL592464.2 | 1 | novel transcript |
| AL592464.3 | 1 | novel transcript |
| AL592464.1 | 1 | novel transcript |
| AL589702.1 | 1 | novel transcript |
| ACTRT2 | 1 | actin related protein T2 [Source:HGNC Symbo |
| PRDM16-DT | 1 | PRDM16 divergent transcript [Source:HGNC S |
| PRDM16 | 1 | PR/SET domain 16 [Source:HGNC Symbol;Acc: |
| MIR4251 | 1 | microRNA 4251 [Source:HGNC Symbol;Acc:HG |
| AL008733.1 | 1 | novel transcript |

| | | |
|---|---|---|
| AL512383.1 | 1 | TEC |
| AL590438.1 | 1 | novel transcript |
| AL354743.2 | 1 | novel transcript, antisense to PRDM16 |
| AL354743.1 | 1 | novel transcript, antisense to PRDM16 |
| ARHGEF16 | 1 | Rho guanine nucleotide exchange factor 16 [S |
| AL512413.1 | 1 | novel transcript |
| MEGF6 | 1 | multiple EGF like domains 6 [Source:HGNC Sy |
| MIR551A | 1 | microRNA 551a [Source:HGNC Symbol;Acc:HG |
| AL513320.1 | 1 | novel transcript |
| TPRG1L | 1 | tumor protein p63 regulated 1 like [Source:HG |
| WRAP73 | 1 | WD repeat containing, antisense to TP73 [Sou |
| TP73 | 1 | tumor protein p73 [Source:HGNC Symbol;Acc: |
| AL136528.1 | 1 | novel transcript |
| AL136528.2 | 1 | novel transcript |
| TP73-AS1 | 1 | TP73 antisense RNA 1 [Source:HGNC Symbol;A |
| AL136528.3 | 1 | NULL |
| CCDC27 | 1 | coiled-coil domain containing 27 [Source:HGN |
| SMIM1 | 1 | small integral membrane protein 1 (Vel blood |
| LRRC47 | 1 | leucine rich repeat containing 47 [Source:HGN |
| RN7SL574P | 1 | RNA, 7SL, cytoplasmic 574, pseudogene [Sour |
| AL365330.1 | 1 | novel transcript, antisense to LRRC47 |
| CEP104 | 1 | centrosomal protein 104 [Source:HGNC Symb |
| DFFB | 1 | DNA fragmentation factor subunit beta [Sourc |
| C1orf174 | 1 | chromosome 1 open reading frame 174 [Sour |
| LINC01134 | 1 | long intergenic non-protein coding RNA 1134 |
| LINC01346 | 1 | long intergenic non-protein coding RNA 1346 |
| LINC01345 | 1 | long intergenic non-protein coding RNA 1345 |
| LINC02780 | 1 | long intergenic non-protein coding RNA 2780 |
| AL805961.1 | 1 | novel transcript |
| EEF1DP6 | 1 | eukaryotic translation elongation factor 1 delt |
| LINC01777 | 1 | long intergenic non-protein coding RNA 1777 |
| AL355602.1 | 1 | novel transcript |
| Z98747.1 | 1 | novel transcript |
| LINC01646 | 1 | long intergenic non-protein coding RNA 1646 |
| AJAP1 | 1 | adherens junctions associated protein 1 [Sour |
| Z98886.1 | 1 | novel transcript, antisense to AJAP1 |
| BX005132.1 | 1 | novel transcript |
| LINC02781 | 1 | long intergenic non-protein coding RNA 2781 |
| LINC02782 | 1 | long intergenic non-protein coding RNA 2782 |
| AL139823.1 | 1 | novel transcript |
| Z98259.3 | 1 | novel transcript |
| Z98259.2 | 1 | novel transcript |
| Z98259.1 | 1 | novel transcript |
| Z97988.2 | 1 | mitochondrially encoded cytochrome c oxidas |
| AL365255.1 | 1 | novel transcript |
| Z97988.1 | 1 | NULL |
| MIR4689 | 1 | microRNA 4689 [Source:HGNC Symbol;Acc:HG |
| NPHP4 | 1 | nephrocystin 4 [Source:HGNC Symbol;Acc:HG |
| KCNAB2 | 1 | potassium voltage-gated channel subfamily A |
| CHD5 | 1 | chromodomain helicase DNA binding protein |

| | | |
|---|---|---|
| AL031847.2 | 1 | novel transcript |
| RPL22 | 1 | ribosomal protein L22 [Source:HGNC Symbol;/ |
| AL031847.1 | 1 | novel transcript |
| RNF207 | 1 | ring finger protein 207 [Source:HGNC Symbol; |
| ICMT | 1 | isoprenylcysteine carboxyl methyltransferase |
| LINC00337 | 1 | long intergenic non-protein coding RNA 337 [S |
| HES3 | 1 | hes family bHLH transcription factor 3 [Source |
| GPR153 | 1 | G protein-coupled receptor 153 [Source:HGN( |
| ACOT7 | 1 | acyl-CoA thioesterase 7 [Source:HGNC Symbo |
| AL031848.2 | 1 | novel transcript, antisense to ACOT7 |
| HES2 | 1 | hes family bHLH transcription factor 2 [Source |
| ESPN | 1 | espin [Source:HGNC Symbol;Acc:HGNC:13281 |
| MIR4252 | 1 | microRNA 4252 [Source:HGNC Symbol;Acc:HC |
| AL031848.1 | 1 | novel transcript, antisense to ESPN |
| TNFRSF25 | 1 | TNF receptor superfamily member 25 [Source |
| PLEKHG5 | 1 | pleckstrin homology and RhoGEF domain cont |
| NOL9 | 1 | nucleolar protein 9 [Source:HGNC Symbol;Acc |
| RNU6-731P | 1 | RNA, U6 small nuclear 731, pseudogene [Sour |
| AL591866.1 | 1 | ribosomal protein L34 (RPL34) pseudogene |
| TAS1R1 | 1 | taste 1 receptor member 1 [Source:HGNC Syn |
| ZBTB48 | 1 | zinc finger and BTB domain containing 48 [Sou |
| KLHL21 | 1 | kelch like family member 21 [Source:HGNC Sy |
| PHF13 | 1 | PHD finger protein 13 [Source:HGNC Symbol;/ |
| THAP3 | 1 | THAP domain containing 3 [Source:HGNC Sym |
| DNAJC11 | 1 | DnaJ heat shock protein family (Hsp40) memb |
| LINC01672 | 1 | long intergenic non-protein coding RNA 1672 |
| AL591163.1 | 1 | novel transcript |
| CAMTA1-DT | 1 | CAMTA1 divergent transcript [Source:HGNC S |
| CAMTA1 | 1 | calmodulin binding transcription activator 1 [S |
| AL590128.1 | 1 | ribosomal protein L37 (RPL37) pseudogene |
| AL512330.1 | 1 | novel transcript, antisense to CAMTA1 |
| RNU1-8P | 1 | RNA, U1 small nuclear 8, pseudogene [Source |
| CAMTA1-IT1 | 1 | CAMTA1 intronic transcript 1 [Source:HGNC S |
| Z97987.1 | 1 | novel transcript |
| AL365194.1 | 1 | novel transcript |
| AL359881.3 | 1 | novel transcript |
| AL359881.2 | 1 | novel transcript |
| AL359881.1 | 1 | novel transcript |
| VAMP3 | 1 | vesicle associated membrane protein 3 [Sourc |
| Z98884.2 | 1 | novel transcript, sense intronic to VAMP3 |
| PER3 | 1 | period circadian regulator 3 [Source:HGNC Syı |
| Z98884.1 | 1 | novel transcript |
| UTS2 | 1 | urotensin 2 [Source:HGNC Symbol;Acc:HGNC: |
| TNFRSF9 | 1 | TNF receptor superfamily member 9 [Source:I |
| AL009183.1 | 1 | small EDRK-rich factor 2 (SERF2) pseudogene |
| PARK7 | 1 | Parkinsonism associated deglycase [Source:H( |
| Y_RNA | 1 | Y RNA [Source:RFAM;Acc:RF00019] |
| AL034417.4 | 1 | novel transcript, antisense to ERRFI1 |
| AL034417.3 | 1 | novel transcript |
| ERRFI1 | 1 | ERBB receptor feedback inhibitor 1 [Source:H( |

| | | |
|---|---|---|
| AL034417.2 | 1 | novel transcript |
| AL034417.1 | 1 | ribosomal protein L7a (RPL7A) pseudogene |
| AL358876.1 | 1 | 52kDa eukaryotic translation initiation factor : |
| LINC01714 | 1 | long intergenic non-protein coding RNA 1714 |
| RNU1-7P | 1 | RNA, U1 small nuclear 7, pseudogene [Source |
| AL358876.2 | 1 | novel transcript |
| RNU6-991P | 1 | RNA, U6 small nuclear 991, pseudogene [Sour |
| SLC45A1 | 1 | solute carrier family 45 member 1 [Source:HG |
| Y_RNA | 1 | Y RNA [Source:RFAM;Acc:RF00019] |
| RERE | 1 | arginine-glutamic acid dipeptide repeats [Sou |
| RERE-AS1 | 1 | RERE antisense RNA 1 [Source:HGNC Symbol;/ |
| AL096855.1 | 1 | NULL |
| AL096855.2 | 1 | ferritin, light polypeptide (FTL) pseudogene |
| RPL7P11 | 1 | ribosomal protein L7 pseudogene 11 [Source: |
| RPL7P7 | 1 | ribosomal protein L7 pseudogene 7 [Source:H |
| Y_RNA | 1 | Y RNA [Source:RFAM;Acc:RF00019] |
| AL357552.1 | 1 | ribosomal protein L27 (RPL27) pseudogene |
| AL357552.2 | 1 | novel transcript |
| RPL23AP19 | 1 | ribosomal protein L23a pseudogene 19 [Sourc |
| ENO1 | 1 | enolase 1 [Source:HGNC Symbol;Acc:HGNC:33 |
| MIR6728 | 1 | microRNA 6728 [Source:HGNC Symbol;Acc:HG |
| ENO1-AS1 | 1 | ENO1 antisense RNA 1 [Source:HGNC Symbol; |
| RNU6-304P | 1 | RNA, U6 small nuclear 304, pseudogene [Sour |
| HMGN2P17 | 1 | high mobility group nucleosomal binding dom |
| AL139415.1 | 1 | zinc finger protein 271 (ZNF) pseudogene |
| AL139415.2 | 1 | pseudogene similar to part of cytochrome oxi |
| CA6 | 1 | carbonic anhydrase 6 [Source:HGNC Symbol;A |
| RN7SL451P | 1 | RNA, 7SL, cytoplasmic 451, pseudogene [Sour |
| SLC2A7 | 1 | solute carrier family 2 member 7 [Source:HGN |
| SLC2A5 | 1 | solute carrier family 2 member 5 [Source:HGN |
| AL158048.1 | 1 | NULL |
| GPR157 | 1 | G protein-coupled receptor 157 [Source:HGN( |
| MIR34AHG | 1 | MIR34A host gene [Source:HGNC Symbol;Acc: |
| MIR34A | 1 | microRNA 34a [Source:HGNC Symbol;Acc:HGN |
| LNCTAM34A | 1 | long non coding transcriptional activator of m |
| H6PD | 1 | hexose-6-phosphate dehydrogenase/glucose |
| SPSB1 | 1 | splA/ryanodine receptor domain and SOCS bo |
| BX323043.1 | 1 | novel transcript |
| LINC02606 | 1 | long intergenic non-protein coding RNA 2606 |
| RNA5SP40 | 1 | RNA, 5S ribosomal pseudogene 40 [Source:HG |
| AL928921.1 | 1 | novel transcript |
| SLC25A33 | 1 | solute carrier family 25 member 33 [Source:H |
| AL954705.1 | 1 | ribosomal protein L9 (RPL9) pseudogene |
| TMEM201 | 1 | transmembrane protein 201 [Source:HGNC Sy |
| PIK3CD | 1 | phosphatidylinositol-4,5-bisphosphate 3-kinas |
| PIK3CD-AS1 | 1 | PIK3CD antisense RNA 1 [Source:HGNC Symbo |
| AL691449.1 | 1 | ribosomal protein L26 pseudogene (RPL26) |
| PIK3CD-AS2 | 1 | PIK3CD antisense RNA 2 [Source:HGNC Symbo |
| CLSTN1 | 1 | calsyntenin 1 [Source:HGNC Symbol;Acc:HGN( |
| MZT1P1 | 1 | mitotic spindle organizing protein 1 pseudoge |

| | | |
|---|---|---|
| AL357140.3 | 1 | TEC |
| CTNNBIP1 | 1 | catenin beta interacting protein 1 [Source:HG |
| AL357140.1 | 1 | novel transcript |
| AL357140.4 | 1 | novel transcript, antisense to CTNNBIP1 |
| LZIC | 1 | leucine zipper and CTNNBIP1 domain containi |
| AL357140.2 | 1 | novel transcript |
| NMNAT1 | 1 | nicotinamide nucleotide adenylyltransferase 1 |
| RN7SKP269 | 1 | RN7SK pseudogene 269 [Source:HGNC Symbo |
| TMEM274P | 1 | transmembrane protein 274, pseudogene [So |
| MIR5697 | 1 | microRNA 5697 [Source:HGNC Symbol;Acc:H( |
| AL603962.1 | 1 | novel transcript |
| RBP7 | 1 | retinol binding protein 7 [Source:HGNC Symbo |
| UBE4B | 1 | ubiquitination factor E4B [Source:HGNC Symb |
| AL590639.1 | 1 | pseudogene similar to part of ribosomal prote |
| PGAM1P11 | 1 | phosphoglycerate mutase 1 pseudogene 11 [S |
| RNU6-828P | 1 | RNA, U6 small nuclear 828, pseudogene [Sour |
| KIF1B | 1 | kinesin family member 1B [Source:HGNC Sym |
| AL358013.1 | 1 | NULL |
| RNU6-37P | 1 | RNA, U6 small nuclear 37, pseudogene [Sourc |
| RN7SL731P | 1 | RNA, 7SL, cytoplasmic 731, pseudogene [Sour |
| AL139424.3 | 1 | novel transcript |
| AL139424.2 | 1 | novel transcript |
| PGD | 1 | phosphogluconate dehydrogenase [Source:H( |
| AL139424.1 | 1 | novel transcript, antisense to APITD1-CORT |
| CENPS-CORT | 1 | CENPS-CORT readthrough [Source:HGNC Syml |
| CENPS | 1 | centromere protein S [Source:HGNC Symbol;A |
| CORT | 1 | cortistatin [Source:HGNC Symbol;Acc:HGNC:2 |
| DFFA | 1 | DNA fragmentation factor subunit alpha [Sour |
| AL354956.1 | 1 | novel transcript, antisense to DFFA |
| PEX14 | 1 | peroxisomal biogenesis factor 14 [Source:HGN |
| AL139423.2 | 1 | novel transcript, antisense to PEX14 |
| RN7SL614P | 1 | RNA, 7SL, cytoplasmic 614, pseudogene [Sour |
| CASZ1 | 1 | castor zinc finger 1 [Source:HGNC Symbol;Acc |
| AL139423.1 | 1 | novel transcript, antisense to CASZ1 |
| HSPE1P24 | 1 | heat shock protein family E (Hsp10) member 1 |
| C1orf127 | 1 | chromosome 1 open reading frame 127 [Sour |
| CFL1P6 | 1 | cofilin 1 pseudogene 6 [Source:HGNC Symbol; |
| TARDBP | 1 | TAR DNA binding protein [Source:HGNC Symb |
| AL109811.3 | 1 | novel protein |
| MASP2 | 1 | mannan binding lectin serine peptidase 2 [Sou |
| AL109811.2 | 1 | novel transcript, antisense to MASP2 |
| SRM | 1 | spermidine synthase [Source:HGNC Symbol;A |
| EXOSC10 | 1 | exosome component 10 [Source:HGNC Symbo |
| AL109811.1 | 1 | novel transcript |
| EXOSC10-AS1 | 1 | EXOSC10 antisense RNA 1 [Source:HGNC Syml |
| MTOR | 1 | mechanistic target of rapamycin kinase [Sourc |
| MTOR-AS1 | 1 | MTOR antisense RNA 1 [Source:HGNC Symbol |
| RNU6-537P | 1 | RNA, U6 small nuclear 537, pseudogene [Sour |
| ANGPTL7 | 1 | angiopoietin like 7 [Source:HGNC Symbol;Acc: |
| RNU6-291P | 1 | RNA, U6 small nuclear 291, pseudogene [Sour |

| | | |
|---|---|---|
| RPL39P6 | 1 | ribosomal protein L39 pseudogene 6 [Source:[illegible] |
| UBIAD1 | 1 | UbiA prenyltransferase domain containing 1 [[illegible] |
| UBE2V2P3 | 1 | ubiquitin conjugating enzyme E2 V2 pseudoge[illegible] |
| AL031291.1 | 1 | novel transcript |
| MTCYBP45 | 1 | MT-CYB pseudogene 45 [Source:HGNC Symbo[illegible] |
| DISP3 | 1 | dispatched RND transporter family member 3 |
| AL590989.1 | 1 | novel transcript, antisense to DISP3 |
| LINC01647 | 1 | long intergenic non-protein coding RNA 1647 |
| AL031731.1 | 1 | novel transcript |
| FBXO2 | 1 | F-box protein 2 [Source:HGNC Symbol;Acc:HG[illegible] |
| FBXO44 | 1 | F-box protein 44 [Source:HGNC Symbol;Acc:H[illegible] |
| FBXO6 | 1 | F-box protein 6 [Source:HGNC Symbol;Acc:HG[illegible] |
| MAD2L2 | 1 | mitotic arrest deficient 2 like 2 [Source:HGNC |
| DRAXIN | 1 | dorsal inhibitory axon guidance protein [Sourc[illegible] |
| AGTRAP | 1 | angiotensin II receptor associated protein [Sou[illegible] |
| C1orf167 | 1 | chromosome 1 open reading frame 167 [Sour[illegible] |
| AL953897.1 | 1 | novel transcript |
| MTHFR | 1 | methylenetetrahydrofolate reductase [Source[illegible] |
| CLCN6 | 1 | chloride voltage-gated channel 6 [Source:HGN[illegible] |
| AL021155.3 | 1 | NULL |
| AL021155.4 | 1 | NULL |
| NPPA | 1 | natriuretic peptide A [Source:HGNC Symbol;A[illegible] |
| AL021155.2 | 1 | NULL |
| NPPB | 1 | natriuretic peptide B [Source:HGNC Symbol;A[illegible] |
| AL021155.1 | 1 | pseudogene similar to part of SET binding fact[illegible] |
| AL021155.5 | 1 | novel transcript |
| RNU5E-1 | 1 | RNA, U5E small nuclear 1 [Source:HGNC Symb[illegible] |
| RNU5E-4P | 1 | RNA, U5E small nuclear 4, pseudogene [Sourc[illegible] |
| KIAA2013 | 1 | KIAA2013 [Source:HGNC Symbol;Acc:HGNC:2[illegible] |
| PLOD1 | 1 | procollagen-lysine,2-oxoglutarate 5-dioxygen[illegible] |
| AL096840.2 | 1 | novel transcript |
| MFN2 | 1 | mitofusin 2 [Source:HGNC Symbol;Acc:HGNC:[illegible] |
| AL096840.1 | 1 | inositol(myo)-1(or 4)-monophosphatase 1 (IM[illegible] |
| MIIP | 1 | migration and invasion inhibitory protein [Sou[illegible] |
| Y_RNA | 1 | Y RNA [Source:RFAM;Acc:RF00019] |
| MIR6729 | 1 | microRNA 6729 [Source:HGNC Symbol;Acc:H[illegible] |
| RN7SL649P | 1 | RNA, 7SL, cytoplasmic 649, pseudogene [Sour[illegible] |
| TNFRSF8 | 1 | TNF receptor superfamily member 8 [Source:[illegible] |
| RNU6-777P | 1 | RNA, U6 small nuclear 777, pseudogene [Sour[illegible] |
| RPL23AP89 | 1 | ribosomal protein L23a pseudogene 89 [Sourc[illegible] |
| AL357835.1 | 1 | novel transcript, antisense to TNFRSF8 |
| MIR7846 | 1 | microRNA 7846 [Source:HGNC Symbol;Acc:H[illegible] |
| TNFRSF1B | 1 | TNF receptor superfamily member 1B [Source[illegible] |
| MIR4632 | 1 | microRNA 4632 [Source:HGNC Symbol;Acc:H[illegible] |
| RPL10P17 | 1 | ribosomal protein L10 pseudogene 17 [Source[illegible] |
| SNORA70 | 1 | Small nucleolar RNA SNORA70 [Source:RFAM;[illegible] |
| VPS13D | 1 | vacuolar protein sorting 13 homolog D [Sourc[illegible] |
| SNORA59A | 1 | small nucleolar RNA, H/ACA box 59A [Source:[illegible] |
| LINC02766 | 1 | long intergenic non-protein coding RNA 2766 |
| DHRS3 | 1 | dehydrogenase/reductase 3 [Source:HGNC Sy[illegible] |

| | | |
|---|---|---|
| RNU6ATAC18P | 1 | RNA, U6atac small nuclear 18, pseudogene [So |
| MIR6730 | 1 | microRNA 6730 [Source:HGNC Symbol;Acc:HG |
| AC254633.1 | 1 | novel transcript |
| AADACL4 | 1 | arylacetamide deacetylase like 4 [Source:HGN |
| AC244670.1 | 1 | ribosomal protein L23a (RPL23A) pseudogene |
| AADACL3 | 1 | arylacetamide deacetylase like 3 [Source:HGN |
| C1orf158 | 1 | chromosome 1 open reading frame 158 [Sour |
| PRAMEF12 | 1 | PRAME family member 12 [Source:HGNC Sym |
| PRAMEF1 | 1 | PRAME family member 1 [Source:HGNC Symb |
| LINC01784 | 1 | long intergenic non-protein coding RNA 1784 |
| PRAMEF11 | 1 | PRAME family member 11 [Source:HGNC Sym |
| PRAMEF30P | 1 | PRAME family member 30, pseudogene [Sour |
| HNRNPCL1 | 1 | heterogeneous nuclear ribonucleoprotein C lil |
| PRAMEF2 | 1 | PRAME family member 2 [Source:HGNC Symb |
| PRAMEF4 | 1 | PRAME family member 4 [Source:HGNC Symb |
| PRAMEF10 | 1 | PRAME family member 10 [Source:HGNC Sym |
| PRAMEF7 | 1 | PRAME family member 7 [Source:HGNC Symb |
| RNU6-1072P | 1 | RNA, U6 small nuclear 1072, pseudogene [Sou |
| PRAMEF29P | 1 | PRAME family member 29, pseudogene [Sour |
| PRAMEF6 | 1 | PRAME family member 6 [Source:HGNC Symb |
| PRAMEF28P | 1 | PRAME family member 28, pseudogene [Sour |
| PRAMEF27 | 1 | PRAME family member 27 [Source:HGNC Sym |
| HNRNPCL3 | 1 | heterogeneous nuclear ribonucleoprotein C lil |
| PRAMEF25 | 1 | PRAME family member 25 [Source:HGNC Sym |
| PRAMEF34P | 1 | PRAME family member 34, pseudogene [Sour |
| PRAMEF35P | 1 | PRAME family member 35, pseudogene [Sour |
| HNRNPCL2 | 1 | heterogeneous nuclear ribonucleoprotein C lil |
| PRAMEF36P | 1 | PRAME family member 36, pseudogene [Sour |
| AC245056.1 | 1 | novel transcript |
| PRAMEF26 | 1 | PRAME family member 26 [Source:HGNC Sym |
| HNRNPCL4 | 1 | heterogeneous nuclear ribonucleoprotein C lil |
| PRAMEF9 | 1 | PRAME family member 9 [Source:HGNC Symb |
| PRAMEF13 | 1 | PRAME family member 13 [Source:HGNC Sym |
| PRAMEF18 | 1 | PRAME family member 18 [Source:HGNC Sym |
| PRAMEF31P | 1 | PRAME family member 31, pseudogene [Sour |
| PRAMEF5 | 1 | PRAME family member 5 [Source:HGNC Symb |
| PRAMEF32P | 1 | PRAME family member 32, pseudogene [Sour |
| RNU6-771P | 1 | RNA, U6 small nuclear 771, pseudogene [Sour |
| PRAMEF8 | 1 | PRAME family member 8 [Source:HGNC Symb |
| PRAMEF33 | 1 | PRAME family member 33 [Source:HGNC Sym |
| PRAMEF15 | 1 | PRAME family member 15 [Source:HGNC Sym |
| AC243961.1 | 1 | novel transcript |
| PRAMEF14 | 1 | PRAME family member 14 [Source:HGNC Sym |
| PRAMEF19 | 1 | PRAME family member 19 [Source:HGNC Sym |
| PRAMEF17 | 1 | PRAME family member 17 [Source:HGNC Sym |
| PRAMEF20 | 1 | PRAME family member 20 [Source:HGNC Sym |
| LRRC38 | 1 | leucine rich repeat containing 38 [Source:HGN |
| AL354712.1 | 1 | novel transcript |
| BRWD1P1 | 1 | bromodomain and WD repeat domain contair |
| PDPN | 1 | podoplanin [Source:HGNC Symbol;Acc:HGNC: |

| | | |
|---|---|---|
| RNA5SP41 | 1 | RNA, 5S ribosomal pseudogene 41 [Source:HG |
| AL359771.1 | 1 | novel transcript |
| AL359771.2 | 1 | NULL |
| PRDM2 | 1 | PR/SET domain 2 [Source:HGNC Symbol;Acc:H |
| KAZN | 1 | kazrin, periplakin interacting protein [Source:H |
| RNU6-1265P | 1 | RNA, U6 small nuclear 1265, pseudogene [Sou |
| AL357873.1 | 1 | novel transcript |
| KAZN-AS1 | 1 | KAZN antisense RNA 1 [Source:HGNC Symbol; |
| TBCAP2 | 1 | tubulin folding cofactor A pseudogene 2 [Sour |
| AL031293.1 | 1 | novel transcript, antisense to KAZN |
| TMEM51-AS1 | 1 | TMEM51 antisense RNA 1 [Source:HGNC Sym |
| TMEM51 | 1 | transmembrane protein 51 [Source:HGNC Syn |
| C1orf195 | 1 | chromosome 1 open reading frame 195 [Sour |
| AL035405.1 | 1 | novel pseudogene |
| AL391094.1 | 1 | zinc finger and BTB domain containing 2 (ZBTE |
| FHAD1 | 1 | forkhead associated phosphopeptide binding |
| AL031283.2 | 1 | novel transcript |
| AL031283.3 | 1 | novel transcript |
| AL031283.1 | 1 | novel transcript |
| EFHD2 | 1 | EF-hand domain family member D2 [Source:H |
| CTRC | 1 | chymotrypsin C [Source:HGNC Symbol;Acc:HG |
| CELA2A | 1 | chymotrypsin like elastase 2A [Source:HGNC S |
| CELA2B | 1 | chymotrypsin like elastase 2B [Source:HGNC S |
| CASP9 | 1 | caspase 9 [Source:HGNC Symbol;Acc:HGNC:15 |
| DNAJC16 | 1 | DnaJ heat shock protein family (Hsp40) memb |
| SCARNA21B | 1 | small Cajal body-specific RNA 21B [Source:HG |
| AL121992.3 | 1 | novel transcript, antisense to DNAJC16 |
| AGMAT | 1 | agmatinase [Source:HGNC Symbol;Acc:HGNC: |
| AL121992.1 | 1 | novel transcript |
| CHCHD2P6 | 1 | coiled-coil-helix-coiled-coil-helix domain conta |
| RNU7-179P | 1 | RNA, U7 small nuclear 179 pseudogene [Sourc |
| CD24P1 | 1 | CD24 molecule pseudogene 1 [Source:HGNC S |
| DDI2 | 1 | DNA damage inducible 1 homolog 2 [Source:H |
| RSC1A1 | 1 | regulator of solute carriers 1 [Source:HGNC Sy |
| AL121992.2 | 1 | novel transcript |
| PLEKHM2 | 1 | pleckstrin homology and RUN domain contain |
| AL450998.3 | 1 | novel transcript, antisense to SLC25A34 and P |
| SLC25A34 | 1 | solute carrier family 25 member 34 [Source:H |
| SLC25A34-AS1 | 1 | SLC25A34 and TMEM82 antisense RNA 1 [Sou |
| TMEM82 | 1 | transmembrane protein 82 [Source:HGNC Syn |
| FBLIM1 | 1 | filamin binding LIM protein 1 [Source:HGNC Sy |
| RPL12P14 | 1 | ribosomal protein L12 pseudogene 14 [Source |
| UQCRHL | 1 | ubiquinol-cytochrome c reductase hinge prote |
| AL450998.1 | 1 | ribosomal protein S16 (RPS16) pseudogene |
| AL450998.2 | 1 | novel transcript |
| SPEN | 1 | spen family transcriptional repressor [Source: |
| AL034555.1 | 1 | cytochrome c oxidase subunit VIc (COX6C) pse |
| ZBTB17 | 1 | zinc finger and BTB domain containing 17 [Sou |
| AL355994.4 | 1 | chromosome 16 open reading frame 52 (C16c |
| AL355994.1 | 1 | pseudogene similar to part of carboxypeptida |

| | | |
|---|---|---|
| AL355994.5 | 1 | tumor necrosis factor receptor superfamily, m |
| TBC1D3P6 | 1 | TBC1 domain family member 3 pseudogene 6 |
| SRARP | 1 | steroid receptor associated and regulated pro |
| AL355994.3 | 1 | novel transcript |
| HSPB7 | 1 | heat shock protein family B (small) member 7 |
| CLCNKA | 1 | chloride voltage-gated channel Ka [Source:HG |
| AL355994.2 | 1 | pseudogene similar to part of novel protein FL |
| CLCNKB | 1 | chloride voltage-gated channel Kb [Source:HG |
| FAM131C | 1 | family with sequence similarity 131 member C |
| EPHA2 | 1 | EPH receptor A2 [Source:HGNC Symbol;Acc:H |
| AL451042.2 | 1 | novel transcript |
| AL451042.1 | 1 | novel transcript |
| ARHGEF19-AS1 | 1 | ARHGEF19 antisense RNA 1 [Source:HGNC Syr |
| ARHGEF19 | 1 | Rho guanine nucleotide exchange factor 19 [S |
| ANO7L1 | 1 | anoctamin 7 like 1 (pseudogene) [Source:HGN |
| AL109627.1 | 1 | novel transcript |
| CPLANE2 | 1 | ciliogenesis and planar polarity effector 2 [Sou |
| MT1XP1 | 1 | metallothionein 1X pseudogene 1 [Source:HG |
| FBXO42 | 1 | F-box protein 42 [Source:HGNC Symbol;Acc:H |
| SZRD1 | 1 | SUZ RNA binding domain containing 1 [Source |
| RPL22P3 | 1 | ribosomal protein L22 pseudogene 3 [Source: |
| SPATA21 | 1 | spermatogenesis associated 21 [Source:HGNC |
| NECAP2 | 1 | NECAP endocytosis associated 2 [Source:HGN |
| LINC01772 | 1 | long intergenic non-protein coding RNA 1772 |
| CROCCP3 | 1 | CROCC pseudogene 3 [Source:HGNC Symbol;A |
| RNU1-1 | 1 | RNA, U1 small nuclear 1 [Source:HGNC Symbo |
| AL137802.2 | 1 | novel transcript |
| AL137802.3 | 1 | novel transcript |
| AL137802.1 | 1 | novel transcript |
| LINC01783 | 1 | long intergenic non-protein coding RNA 1783 |
| RNU1-6P | 1 | RNA, U1 small nuclear 6, pseudogene [Source |
| AL355149.1 | 1 | phosphodiesterase 4D interacting protein (my |
| NBPF1 | 1 | NBPF member 1 [Source:HGNC Symbol;Acc:H |
| AL137798.1 | 1 | novel transcript |
| CROCCP2 | 1 | CROCC pseudogene 2 [Source:HGNC Symbol;A |
| AL137798.3 | 1 | ring finger protein 123 (RNF123) pseudogene |
| MST1P2 | 1 | macrophage stimulating 1 pseudogene 2 [Sou |
| AL137798.2 | 1 | novel transcript |
| RNU1-3 | 1 | RNA, U1 small nuclear 3 [Source:HGNC Symbo |
| AL021920.2 | 1 | phosphodiesterase 4D interacting protein (my |
| EIF1AXP1 | 1 | EIF1AX pseudogene 1 [Source:HGNC Symbol;A |
| ESPNP | 1 | espin pseudogene [Source:HGNC Symbol;Acc: |
| AL021920.1 | 1 | novel transcript |
| AL021920.3 | 1 | novel transcript |
| CROCC | 1 | ciliary rootlet coiled-coil, rootletin [Source:HG |
| CROCCP4 | 1 | CROCC pseudogene 4 [Source:HGNC Symbol;A |
| RNU1-4 | 1 | RNA, U1 small nuclear 4 [Source:HGNC Symbo |
| MST1L | 1 | macrophage stimulating 1 like [Source:HGNC S |
| BX284668.1 | 1 | novel transcript |
| BX284668.3 | 1 | phosphodiesterase 4D interacting protein (PD |

| | | |
|---|---|---|
| MIR3675 | 1 | microRNA 3675 [Source:HGNC Symbol;Acc:HG |
| BX284668.2 | 1 | novel transcript |
| RNU1-5P | 1 | RNA, U1 small nuclear 5, pseudogene [Source |
| BX284668.5 | 1 | novel transcript |
| BX284668.4 | 1 | cyclin-dependent kinase 5 (CDK5) pseudogene |
| RNU1-2 | 1 | RNA, U1 small nuclear 2 [Source:HGNC Symbo |
| BX284668.6 | 1 | novel transcript |
| MFAP2 | 1 | microfibril associated protein 2 [Source:HGNC |
| AL049569.2 | 1 | novel transcript, antisense to MFAP2 |
| AL049569.1 | 1 | novel transcript |
| ATP13A2 | 1 | ATPase cation transporting 13A2 [Source:HGN |
| SDHB | 1 | succinate dehydrogenase complex iron sulfur |
| PADI2 | 1 | peptidyl arginine deiminase 2 [Source:HGNC S |
| LINC02783 | 1 | long intergenic non-protein coding RNA 2783 |
| AL590644.1 | 1 | novel transcript |
| PADI1 | 1 | peptidyl arginine deiminase 1 [Source:HGNC S |
| PADI3 | 1 | peptidyl arginine deiminase 3 [Source:HGNC S |
| MIR3972 | 1 | microRNA 3972 [Source:HGNC Symbol;Acc:HG |
| PADI4 | 1 | peptidyl arginine deiminase 4 [Source:HGNC S |
| PADI6 | 1 | peptidyl arginine deiminase 6 [Source:HGNC S |
| RCC2 | 1 | regulator of chromosome condensation 2 [Sou |
| RCC2-AS1 | 1 | RCC2 antisense RNA 1 [Source:HGNC Symbol; |
| ARHGEF10L | 1 | Rho guanine nucleotide exchange factor 10 lik |
| LINC02810 | 1 | long intergenic non-protein coding RNA 2810 |
| ACTL8 | 1 | actin like 8 [Source:HGNC Symbol;Acc:HGNC:2 |
| AL357509.1 | 1 | novel transcript |
| LINC01654 | 1 | long intergenic non-protein coding RNA 1654 |
| IGSF21 | 1 | immunoglobin superfamily member 21 [Sourc |
| AL365209.1 | 1 | TEC |
| IGSF21-AS1 | 1 | IGSF21 antisense RNA 1 [Source:HGNC Symbo |
| AL591896.1 | 1 | novel transcript |
| KLHDC7A | 1 | kelch domain containing 7A [Source:HGNC Syr |
| DYNLL1P3 | 1 | dynein light chain LC8-type 1 pseudogene 3 [S |
| AL031737.1 | 1 | MT-ND1 (MT-ND1) pseudogene |
| PAX7 | 1 | paired box 7 [Source:HGNC Symbol;Acc:HGNC |
| TAS1R2 | 1 | taste 1 receptor member 2 [Source:HGNC Sym |
| ALDH4A1 | 1 | aldehyde dehydrogenase 4 family member A1 |
| MIR4695 | 1 | microRNA 4695 [Source:HGNC Symbol;Acc:HG |
| MIR1290 | 1 | microRNA 1290 [Source:HGNC Symbol;Acc:HG |
| IFFO2 | 1 | intermediate filament family orphan 2 [Source |
| AL137127.1 | 1 | novel transcript |
| UBR4 | 1 | ubiquitin protein ligase E3 component n-recog |
| EMC1-AS1 | 1 | EMC1 antisense RNA 1 [Source:HGNC Symbol; |
| EMC1 | 1 | ER membrane protein complex subunit 1 [Sou |
| MRTO4 | 1 | MRT4 homolog, ribosome maturation factor [ |
| AL035413.2 | 1 | aldo-keto reductase family pseudogene |
| AKR7L | 1 | aldo-keto reductase family 7 like (gene/pseud |
| AKR7A3 | 1 | aldo-keto reductase family 7 member A3 [Sou |
| AL035413.1 | 1 | synaptonemal complex protein 3 (SYCP3) pseu |
| AKR7A2 | 1 | aldo-keto reductase family 7 member A2 [Sou |

| | | |
|---|---|---|
| RNU6-1099P | 1 | RNA, U6 small nuclear 1099, pseudogene [Sou |
| SLC66A1 | 1 | solute carrier family 66 member 1 [Source:HG |
| CAPZB | 1 | capping actin protein of muscle Z-line subunit |
| RN7SL277P | 1 | RNA, 7SL, cytoplasmic 277, pseudogene [Sour |
| MICOS10 | 1 | mitochondrial contact site and cristae organiz |
| RNU4-28P | 1 | RNA, U4 small nuclear 28, pseudogene [Sourc |
| AL031727.2 | 1 | novel transcript |
| NBL1 | 1 | NBL1, DAN family BMP antagonist [Source:HG |
| MICOS10-NBL1 | 1 | MICOS10-NBL1 readthrough [Source:HGNC Sy |
| AL031727.1 | 1 | ribosomal protein S14 (RPS14) pseudogene |
| HTR6 | 1 | 5-hydroxytryptamine receptor 6 [Source:HGN |
| TMCO4 | 1 | transmembrane and coiled-coil domains 4 [So |
| RNF186 | 1 | ring finger protein 186 [Source:HGNC Symbol; |
| AL391883.1 | 1 | novel transcript |
| OTUD3 | 1 | OTU deubiquitinase 3 [Source:HGNC Symbol;/ |
| PLA2G2E | 1 | phospholipase A2 group IIE [Source:HGNC Syn |
| RN7SL304P | 1 | RNA, 7SL, cytoplasmic 304, pseudogene [Sour |
| PLA2G2A | 1 | phospholipase A2 group IIA [Source:HGNC Syr |
| PLA2G5 | 1 | phospholipase A2 group V [Source:HGNC Sym |
| PLA2G2D | 1 | phospholipase A2 group IID [Source:HGNC Syr |
| PLA2G2F | 1 | phospholipase A2 group IIF [Source:HGNC Syn |
| Z98257.1 | 1 | novel transcript |
| PLA2G2C | 1 | phospholipase A2 group IIC [Source:HGNC Syr |
| UBXN10-AS1 | 1 | UBXN10 antisense RNA 1 [Source:HGNC Symb |
| UBXN10 | 1 | UBX domain protein 10 [Source:HGNC Symbo |
| LINC01757 | 1 | long intergenic non-protein coding RNA 1757 |
| Z98257.2 | 1 | novel transcript |
| VWA5B1 | 1 | von Willebrand factor A domain containing 5E |
| AL020998.1 | 1 | novel transcript |
| LINC01141 | 1 | long intergenic non-protein coding RNA 1141 |
| AL139254.1 | 1 | novel transcript |
| AL139254.2 | 1 | novel transcript |
| AL139254.3 | 1 | novel transcript |
| CAMK2N1 | 1 | calcium/calmodulin dependent protein kinase |
| MUL1 | 1 | mitochondrial E3 ubiquitin protein ligase 1 [Sc |
| RPS4XP4 | 1 | ribosomal protein S4X pseudogene 4 [Source: |
| FAM43B | 1 | family with sequence similarity 43 member B |
| CDA | 1 | cytidine deaminase [Source:HGNC Symbol;Acc |
| PINK1 | 1 | PTEN induced kinase 1 [Source:HGNC Symbol; |
| MIR6084 | 1 | microRNA 6084 [Source:HGNC Symbol;Acc:HG |
| PINK1-AS | 1 | PINK1 antisense RNA [Source:HGNC Symbol;A |
| DDOST | 1 | dolichyl-diphosphooligosaccharide--protein gl |
| KIF17 | 1 | kinesin family member 17 [Source:HGNC Syml |
| AL663074.2 | 1 | nucleolar protein 5A 56kDa with KKE/D repea |
| SH2D5 | 1 | SH2 domain containing 5 [Source:HGNC Symb |
| AL663074.1 | 1 | novel transcript |
| HP1BP3 | 1 | heterochromatin protein 1 binding protein 3 [ |
| EIF4G3 | 1 | eukaryotic translation initiation factor 4 gamn |
| RNU7-200P | 1 | RNA, U7 small nuclear 200 pseudogene [Sourc |
| MIR1256 | 1 | microRNA 1256 [Source:HGNC Symbol;Acc:HG |

| | | |
|---|---|---|
| RPS15AP6 | 1 | ribosomal protein S15a pseudogene 6 [Source |
| AL627311.1 | 1 | ribosomal protein L34 (RPL34) pseudogene |
| HSPE1P27 | 1 | heat shock protein family E (Hsp10) member : |
| AL031005.2 | 1 | novel transcript |
| ECE1 | 1 | endothelin converting enzyme 1 [Source:HGN |
| AL031005.1 | 1 | novel transcript |
| AL031728.1 | 1 | novel transcript |
| PPP1R11P1 | 1 | protein phosphatase 1 regulatory inhibitor sul |
| AL592309.1 | 1 | phosphodiesterase 4D interacting protein (my |
| AL592309.2 | 1 | novel transcript |
| NBPF2P | 1 | NBPF member 2, pseudogene [Source:HGNC S |
| HS6ST1P1 | 1 | heparan sulfate 6-O-sulfotransferase 1 pseudo |
| CROCCP5 | 1 | CROCC pseudogene 5 [Source:HGNC Symbol;A |
| NBPF3 | 1 | NBPF member 3 [Source:HGNC Symbol;Acc:H( |
| PFN1P10 | 1 | profilin 1 pseudogene 10 [Source:HGNC Symb |
| ALPL | 1 | alkaline phosphatase, biomineralization assoc |
| LINC02596 | 1 | long intergenic non-protein coding RNA 2596 |
| RAP1GAP | 1 | RAP1 GTPase activating protein [Source:HGN( |
| USP48 | 1 | ubiquitin specific peptidase 48 [Source:HGNC |
| AL590103.1 | 1 | S100 calcium binding protein A11 (calgizzarin) |
| LDLRAD2 | 1 | low density lipoprotein receptor class A doma |
| HSPG2 | 1 | heparan sulfate proteoglycan 2 [Source:HGNC |
| AL590556.1 | 1 | ribosomal protein L21 (RPL21) pseudogene |
| AL590556.2 | 1 | novel pseudogene |
| CELA3B | 1 | chymotrypsin like elastase 3B [Source:HGNC S |
| AL590556.3 | 1 | novel transcript |
| RN7SL386P | 1 | RNA, 7SL, cytoplasmic 386, pseudogene [Sour |
| RNU6-1022P | 1 | RNA, U6 small nuclear 1022, pseudogene [Sou |
| CELA3A | 1 | chymotrypsin like elastase 3A [Source:HGNC S |
| RN7SL186P | 1 | RNA, 7SL, cytoplasmic 186, pseudogene [Sour |
| RNU6-776P | 1 | RNA, U6 small nuclear 776, pseudogene [Sour |
| LINC01635 | 1 | long intergenic non-protein coding RNA 1635 |
| LINC00339 | 1 | long intergenic non-protein coding RNA 339 [S |
| CDC42 | 1 | cell division cycle 42 [Source:HGNC Symbol;Ac |
| CDC42-AS1 | 1 | CDC42 antisense RNA 1 [Source:HGNC Symbo |
| AL031281.2 | 1 | SCAN domain containing 1 (SCAND1) pseudog |
| CDC42-IT1 | 1 | CDC42 intronic transcript 1 [Source:HGNC Syn |
| AL031281.1 | 1 | M-phase phosphoprotein 6 (MPHOSPH6) pseu |
| WNT4 | 1 | Wnt family member 4 [Source:HGNC Symbol;/ |
| AL445253.1 | 1 | novel transcript |
| MIR4418 | 1 | microRNA 4418 [Source:HGNC Symbol;Acc:HG |
| PPIAP34 | 1 | peptidylprolyl isomerase A pseudogene 34 [Sc |
| AL591122.1 | 1 | TEC |
| ZBTB40 | 1 | zinc finger and BTB domain containing 40 [Sou |
| ZBTB40-IT1 | 1 | ZBTB40 intronic transcript 1 [Source:HGNC Sy |
| EPHA8 | 1 | EPH receptor A8 [Source:HGNC Symbol;Acc:H |
| MIR6127 | 1 | microRNA 6127 [Source:HGNC Symbol;Acc:HG |
| C1QA | 1 | complement C1q A chain [Source:HGNC Symb |
| C1QC | 1 | complement C1q C chain [Source:HGNC Symb |
| C1QB | 1 | complement C1q B chain [Source:HGNC Symb |

| | | |
|---|---|---|
| EPHB2 | 1 | EPH receptor B2 [Source:HGNC Symbol;Acc:H |
| MIR4684 | 1 | microRNA 4684 [Source:HGNC Symbol;Acc:HG |
| AL512444.1 | 1 | novel transcript |
| MIR4253 | 1 | microRNA 4253 [Source:HGNC Symbol;Acc:HG |
| LACTBL1 | 1 | lactamase beta like 1 [Source:HGNC Symbol;A |
| TEX46 | 1 | testis expressed 46 [Source:HGNC Symbol;Acc |
| KDM1A | 1 | lysine demethylase 1A [Source:HGNC Symbol; |
| AL031428.1 | 1 | novel transcript |
| MIR3115 | 1 | microRNA 3115 [Source:HGNC Symbol;Acc:HG |
| AL031428.2 | 1 | NULL |
| LUZP1 | 1 | leucine zipper protein 1 [Source:HGNC Symbo |
| AL161672.1 | 1 | novel pseudogene |
| RNU6-514P | 1 | RNA, U6 small nuclear 514, pseudogene [Sour |
| RNU6-135P | 1 | RNA, U6 small nuclear 135, pseudogene [Sour |
| HTR1D | 1 | 5-hydroxytryptamine receptor 1D [Source:HG |
| AL109936.1 | 1 | ribosomal protein L29 (RPL29) pseudogene |
| LINC01355 | 1 | long intergenic non-protein coding RNA 1355 |
| AL109936.6 | 1 | novel transcript |
| HNRNPR | 1 | heterogeneous nuclear ribonucleoprotein R [S |
| ZNF436 | 1 | zinc finger protein 436 [Source:HGNC Symbol; |
| ZNF436-AS1 | 1 | ZNF436 antisense RNA 1 [Source:HGNC Symbo |
| Y_RNA | 1 | Y RNA [Source:RFAM;Acc:RF00019] |
| AL109936.2 | 1 | novel transcript |
| TCEA3 | 1 | transcription elongation factor A3 [Source:HG |
| AL357134.1 | 1 | single-stranded DNA binding protein (SSBP1) p |
| ASAP3 | 1 | ArfGAP with SH3 domain, ankyrin repeat and |
| E2F2 | 1 | E2F transcription factor 2 [Source:HGNC Symb |
| AL021154.1 | 1 | novel transcript |
| ID3 | 1 | inhibitor of DNA binding 3, HLH protein [Sourc |
| AL450043.1 | 1 | novel transcript |
| MDS2 | 1 | myelodysplastic syndrome 2 translocation ass |
| AL451000.1 | 1 | eukaryotic translation elongation factor 1 alph |
| RPL11 | 1 | ribosomal protein L11 [Source:HGNC Symbol; |
| ELOA-AS1 | 1 | ELOA antisense RNA 1 [Source:HGNC Symbol; |
| ELOA | 1 | elongin A [Source:HGNC Symbol;Acc:HGNC:11 |
| PITHD1 | 1 | PITH domain containing 1 [Source:HGNC Symb |
| LYPLA2 | 1 | lysophospholipase 2 [Source:HGNC Symbol;Ac |
| GALE | 1 | UDP-galactose-4-epimerase [Source:HGNC Sy |
| HMGCL | 1 | 3-hydroxy-3-methylglutaryl-CoA lyase [Source |
| FUCA1 | 1 | alpha-L-fucosidase 1 [Source:HGNC Symbol;Ac |
| CNR2 | 1 | cannabinoid receptor 2 [Source:HGNC Symbol |
| BTBD6P1 | 1 | BTB domain containing 6 pseudogene 1 [Sourc |
| AL590609.3 | 1 | novel transcript, antisense to CNR2 |
| MIR378F | 1 | microRNA 378f [Source:HGNC Symbol;Acc:HG |
| AL590609.2 | 1 | H3 histone, family 3A (H3F3A) pseudogene |
| PNRC2 | 1 | proline rich nuclear receptor coactivator 2 [So |
| SRSF10 | 1 | serine and arginine rich splicing factor 10 [Sou |
| AL590609.1 | 1 | ribosomal protein L36 (RPL36) pseudogene |
| AL591178.1 | 1 | novel transcript, antisense to MYOM3 |
| MYOM3 | 1 | myomesin 3 [Source:HGNC Symbol;Acc:HGNC |

| | | |
|---|---|---|
| AL591178.2 | 1 | novel transcript, antisense to MYOM3 |
| IL22RA1 | 1 | interleukin 22 receptor subunit alpha 1 [Sourc |
| IFNLR1 | 1 | interferon lambda receptor 1 [Source:HGNC S |
| LINC02800 | 1 | long intergenic non-protein coding RNA 2800 |
| AL138902.1 | 1 | novel transcript |
| GRHL3 | 1 | grainyhead like transcription factor 3 [Source: |
| STPG1 | 1 | sperm tail PG-rich repeat containing 1 [Source |
| NIPAL3 | 1 | NIPA like domain containing 3 [Source:HGNC S |
| RCAN3AS | 1 | RCAN3 antisense RNA [Source:HGNC Symbol;A |
| RCAN3 | 1 | RCAN family member 3 [Source:HGNC Symbol |
| NCMAP-DT | 1 | NCMAP divergent transcript [Source:HGNC Sy |
| NCMAP | 1 | non-compact myelin associated protein [Sour |
| AL445686.1 | 1 | ribosomal protein L26 (RPL26) pseudogene |
| Y_RNA | 1 | Y RNA [Source:RFAM;Acc:RF00019] |
| SRRM1 | 1 | serine and arginine repetitive matrix 1 [Source |
| AL445648.1 | 1 | novel transcript |
| CLIC4 | 1 | chloride intracellular channel 4 [Source:HGNC |
| RNU6-1208P | 1 | RNA, U6 small nuclear 1208, pseudogene [Sou |
| RUNX3 | 1 | RUNX family transcription factor 3 [Source:HG |
| MIR6731 | 1 | microRNA 6731 [Source:HGNC Symbol;Acc:HG |
| AL445471.1 | 1 | novel transcript |
| AL445471.2 | 1 | novel transcript |
| MIR4425 | 1 | microRNA 4425 [Source:HGNC Symbol;Acc:HG |
| LINC02793 | 1 | long intergenic non-protein coding RNA 2793 |
| AL050344.1 | 1 | novel transcript |
| AL121994.1 | 1 | interferon induced transmembrane protein ps |
| AL031432.1 | 1 | novel transcript |
| SYF2 | 1 | SYF2 pre-mRNA splicing factor [Source:HGNC |
| AL031432.4 | 1 | novel transcript |
| AL031432.5 | 1 | novel transcript |
| RSRP1 | 1 | arginine and serine rich protein 1 [Source:HGN |
| AL031432.3 | 1 | novel transcript |
| AL031432.2 | 1 | novel pseudogene |
| RHD | 1 | Rh blood group D antigen [Source:HGNC Symb |
| SDHDP6 | 1 | succinate dehydrogenase complex subunit D p |
| AL928711.1 | 1 | novel pseudogene |
| TMEM50A | 1 | transmembrane protein 50A [Source:HGNC Sy |
| RHCE | 1 | Rh blood group CcEe antigens [Source:HGNC S |
| AL031284.1 | 1 | succinate dehydrogenase complex, subunit D, |
| MACO1 | 1 | macoilin 1 [Source:HGNC Symbol;Acc:HGNC:2 |
| LDLRAP1 | 1 | low density lipoprotein receptor adaptor prot |
| AL606491.1 | 1 | novel transcript |
| MAN1C1 | 1 | mannosidase alpha class 1C member 1 [Sourc |
| AL031280.1 | 1 | novel transcript |
| SELENON | 1 | selenoprotein N [Source:HGNC Symbol;Acc:HG |
| AL020996.2 | 1 | novel protein |
| AL020996.1 | 1 | novel transcript |
| MTFR1L | 1 | mitochondrial fission regulator 1 like [Source: |
| AL020996.3 | 1 | novel transcript, antisense to MTFR1L |
| AUNIP | 1 | aurora kinase A and ninein interacting protein |

| | | |
|---|---|---|
| AL033528.2 | 1 | novel transcript |
| PAQR7 | 1 | progestin and adipoQ receptor family membe |
| Y_RNA | 1 | Y RNA [Source:RFAM;Acc:RF00019] |
| STMN1 | 1 | stathmin 1 [Source:HGNC Symbol;Acc:HGNC:6 |
| SNRPFP2 | 1 | small nuclear ribonucleoprotein polypeptide F |
| AL033528.1 | 1 | ribosomal protein L39 (RPL39) pseudogene |
| MIR3917 | 1 | microRNA 3917 [Source:HGNC Symbol;Acc:HG |
| PAFAH2 | 1 | platelet activating factor acetylhydrolase 2 [So |
| RNU6-110P | 1 | RNA, U6 small nuclear 110, pseudogene [Sour |
| AL592064.1 | 1 | NULL |
| AL592064.2 | 1 | NULL |
| EXTL1 | 1 | exostosin like glycosyltransferase 1 [Source:H( |
| SLC30A2 | 1 | solute carrier family 30 member 2 [Source:HG |
| AL391650.2 | 1 | novel transcript |
| TRIM63 | 1 | tripartite motif containing 63 [Source:HGNC S |
| PDIK1L | 1 | PDLIM1 interacting kinase 1 like [Source:HGN( |
| FAM110D | 1 | family with sequence similarity 110 member D |
| C1orf232 | 1 | chromosome 1 open reading frame 230 [Sour |
| AL391650.1 | 1 | novel protein |
| ZNF593 | 1 | zinc finger protein 593 [Source:HGNC Symbol; |
| CNKSR1 | 1 | connector enhancer of kinase suppressor of R |
| CATSPER4 | 1 | cation channel sperm associated 4 [Source:HG |
| AL355877.1 | 1 | novel pseudogene |
| AL355877.3 | 1 | ADP-ribosylation factor-like 16 (ARL16) pseud |
| CEP85 | 1 | centrosomal protein 85 [Source:HGNC Symbo |
| AL355877.2 | 1 | mitochondrial ribosomal protein S6 (MRPS6) p |
| SH3BGRL3 | 1 | SH3 domain binding glutamate rich protein lik |
| UBXN11 | 1 | UBX domain protein 11 [Source:HGNC Symbo |
| CD52 | 1 | CD52 molecule [Source:HGNC Symbol;Acc:HG |
| CRYBG2 | 1 | crystallin beta-gamma domain containing 2 [S |
| AL451139.1 | 1 | ribosomal protein L34 (RPL34) pseudogene |
| ZNF683 | 1 | zinc finger protein 683 [Source:HGNC Symbol; |
| LIN28A | 1 | lin-28 homolog A [Source:HGNC Symbol;Acc:H |
| DHDDS | 1 | dehydrodolichyl diphosphate synthase subuni |
| AL513365.1 | 1 | ribosomal protein L17 (RPL17) pseudogene |
| AL513365.2 | 1 | novel transcript |
| HMGN2 | 1 | high mobility group nucleosomal binding dom |
| DPPA2P2 | 1 | developmental pluripotency associated 2 pseu |
| RPS6KA1 | 1 | ribosomal protein S6 kinase A1 [Source:HGNC |
| MIR1976 | 1 | microRNA 1976 [Source:HGNC Symbol;Acc:HG |
| RN7SL679P | 1 | RNA, 7SL, cytoplasmic 679, pseudogene [Sour |
| Y_RNA | 1 | Y RNA [Source:RFAM;Acc:RF00019] |
| RNU7-29P | 1 | RNA, U7 small nuclear 29 pseudogene [Source |
| AL627313.1 | 1 | chromosome 20 open reading frame 52 (C20o |
| AL627082.2 | 1 | NULL |
| AL627082.1 | 1 | chromosome 20 open reading frame 52 (C20o |
| AL512408.1 | 1 | novel transcript, antisense to ARID1A |
| ARID1A | 1 | AT-rich interaction domain 1A [Source:HGNC S |
| PIGV | 1 | phosphatidylinositol glycan anchor biosynthes |
| RN7SL165P | 1 | RNA, 7SL, cytoplasmic 165, pseudogene [Sour |

| | | |
|---|---|---|
| AL034380.2 | 1 | microtubule associated serine/threonine kina |
| ZDHHC18 | 1 | zinc finger DHHC-type containing 18 [Source:H |
| SFN | 1 | stratifin [Source:HGNC Symbol;Acc:HGNC:107 |
| GPN2 | 1 | GPN-loop GTPase 2 [Source:HGNC Symbol;Acc |
| AL034380.1 | 1 | novel transcript |
| GPATCH3 | 1 | G-patch domain containing 3 [Source:HGNC Sy |
| NUDC | 1 | nuclear distribution C, dynein complex regulat |
| NR0B2 | 1 | nuclear receptor subfamily 0 group B member |
| KDF1 | 1 | keratinocyte differentiation factor 1 [Source:H |
| RPL12P13 | 1 | ribosomal protein L12 pseudogene 13 [Source |
| AL356390.1 | 1 | ribosomal protein L32 (RPL32) pseudogene |
| OSTCP2 | 1 | oligosaccharyltransferase complex subunit ps |
| TRNP1 | 1 | TMF1 regulated nuclear protein 1 [Source:HG |
| TENT5B | 1 | terminal nucleotidyltransferase 5B [Source:HG |
| SLC9A1 | 1 | solute carrier family 9 member A1 [Source:HG |
| AL590640.2 | 1 | ribosomal protein L18a (RPL18A) pseudogene |
| CHCHD3P3 | 1 | coiled-coil-helix-coiled-coil-helix domain conta |
| NPM1P39 | 1 | nucleophosmin 1 pseudogene 39 [Source:HGN |
| SNRPEP7 | 1 | small nuclear ribonucleoprotein polypeptide E |
| AL590640.1 | 1 | novel transcript |
| WDTC1 | 1 | WD and tetratricopeptide repeats 1 [Source:H |
| Y_RNA | 1 | Y RNA [Source:RFAM;Acc:RF00019] |
| FO393419.3 | 1 | SMT3 suppressor of mif two 3 homolog 2 (yea |
| TMEM222 | 1 | transmembrane protein 222 [Source:HGNC Sy |
| RNU6-48P | 1 | RNA, U6 small nuclear 48, pseudogene [Sourc |
| ACTG1P20 | 1 | actin gamma 1 pseudogene 20 [Source:HGNC |
| SYTL1 | 1 | synaptotagmin like 1 [Source:HGNC Symbol;A |
| MAP3K6 | 1 | mitogen-activated protein kinase kinase kinas |
| FCN3 | 1 | ficolin 3 [Source:HGNC Symbol;Acc:HGNC:362 |
| CD164L2 | 1 | CD164 molecule like 2 [Source:HGNC Symbol; |
| FO393419.1 | 1 | ribosomal protein S24 (RPS24) pseudogene |
| GPR3 | 1 | G protein-coupled receptor 3 [Source:HGNC S |
| WASF2 | 1 | WASP family member 2 [Source:HGNC Symbo |
| FO393419.2 | 1 | novel transcript |
| BX293535.1 | 1 | novel transcript |
| AHDC1 | 1 | AT-hook DNA binding motif containing 1 [Sour |
| FGR | 1 | FGR proto-oncogene, Src family tyrosine kinas |
| AL031729.1 | 1 | novel pseudogene |
| LINC02574 | 1 | long intergenic non-protein coding RNA 2574 |
| IFI6 | 1 | interferon alpha inducible protein 6 [Source:H |
| AL445490.1 | 1 | novel transcript, antisense to IFI6 |
| RNU6-949P | 1 | RNA, U6 small nuclear 949, pseudogene [Sour |
| CHMP1AP1 | 1 | charged multivesicular body protein 1A pseud |
| RNU6-424P | 1 | RNA, U6 small nuclear 424, pseudogene [Sour |
| AL020997.5 | 1 | novel transcript |
| FAM76A | 1 | family with sequence similarity 76 member A |
| AL020997.1 | 1 | novel pseudogene |
| RPEP3 | 1 | ribulose-5-phosphate-3-epimerase pseudoger |
| STX12 | 1 | syntaxin 12 [Source:HGNC Symbol;Acc:HGNC: |
| AL020997.2 | 1 | novel transcript, sense intronic to STX12 |

| | | |
|---|---|---|
| AL020997.3 | 1 | novel transcript, sense intronic to STX12 |
| RNU6-1245P | 1 | RNA, U6 small nuclear 1245, pseudogene [Sou |
| AL020997.4 | 1 | novel transcript, antisense to PPP1R8 |
| PPP1R8 | 1 | protein phosphatase 1 regulatory subunit 8 [S |
| SCARNA1 | 1 | small Cajal body-specific RNA 1 [Source:HGNC |
| THEMIS2 | 1 | thymocyte selection associated family membe |
| RPA2 | 1 | replication protein A2 [Source:HGNC Symbol;A |
| SMPDL3B | 1 | sphingomyelin phosphodiesterase acid like 3B |
| AL512288.1 | 1 | novel transcript |
| XKR8 | 1 | XK related 8 [Source:HGNC Symbol;Acc:HGNC |
| EYA3 | 1 | EYA transcriptional coactivator and phosphata |
| AL512288.2 | 1 | interferon-induced transmembrane protein ps |
| RN7SL559P | 1 | RNA, 7SL, cytoplasmic 559, pseudogene [Sour |
| SPCS2P4 | 1 | signal peptidase complex subunit 2 pseudogen |
| AL137792.2 | 1 | NADH dehydrogenase (ubiquinone) 1 alpha su |
| AL137792.1 | 1 | novel ADP-ribosylation factor family domain c |
| RNU6-176P | 1 | RNA, U6 small nuclear 176, pseudogene [Sour |
| PTAFR | 1 | platelet activating factor receptor [Source:HG |
| DNAJC8 | 1 | DnaJ heat shock protein family (Hsp40) memb |
| ATP5IF1 | 1 | ATP synthase inhibitory factor subunit 1 [Sour |
| AL353622.1 | 1 | novel transcript |
| AL353622.2 | 1 | novel transcript |
| SESN2 | 1 | sestrin 2 [Source:HGNC Symbol;Acc:HGNC:20 |
| MED18 | 1 | mediator complex subunit 18 [Source:HGNC S |
| PHACTR4 | 1 | phosphatase and actin regulator 4 [Source:HG |
| Y_RNA | 1 | Y RNA [Source:RFAM;Acc:RF00019] |
| CR391992.1 | 1 | DnaJ (Hsp40) homolog, subfamily C member 8 |
| RNU6ATAC27P | 1 | RNA, U6atac small nuclear 27, pseudogene [So |
| RCC1 | 1 | regulator of chromosome condensation 1 [Sou |
| SNHG3 | 1 | small nucleolar RNA host gene 3 [Source:HGN |
| SNORA73A | 1 | small nucleolar RNA, H/ACA box 73A [Source:I |
| SNORA73B | 1 | small nucleolar RNA, H/ACA box 73B [Source:I |
| PRDX3P2 | 1 | peroxiredoxin 3 pseudogene 2 [Source:HGNC |
| AL513497.1 | 1 | TEC |
| TRNAU1AP | 1 | tRNA selenocysteine 1 associated protein 1 [S |
| SNHG12 | 1 | small nucleolar RNA host gene 12 [Source:HGI |
| SNORD99 | 1 | small nucleolar RNA, C/D box 99 [Source:HGN |
| SNORA61 | 1 | small nucleolar RNA, H/ACA box 61 [Source:H |
| SNORA44 | 1 | small nucleolar RNA, H/ACA box 44 [Source:H |
| SNORA16A | 1 | small nucleolar RNA, H/ACA box 16A [Source:I |
| SNORA16A | 1 | small nucleolar RNA, H/ACA box 16A [Source:I |
| TAF12 | 1 | TATA-box binding protein associated factor 12 |
| RAB42 | 1 | RAB42, member RAS oncogene family [Source |
| LINC01715 | 1 | long intergenic non-protein coding RNA 1715 |
| RNU11 | 1 | RNA, U11 small nuclear [Source:HGNC Symbo |
| AL360012.1 | 1 | RNA, U11 small nuclear |
| GMEB1 | 1 | glucocorticoid modulatory element binding pr |
| AL645729.1 | 1 | NULL |
| YTHDF2 | 1 | YTH N6-methyladenosine RNA binding proteir |
| OPRD1 | 1 | opioid receptor delta 1 [Source:HGNC Symbol |

| | | |
|---|---|---|
| AL009181.1 | 1 | novel transcript |
| Y_RNA | 1 | Y RNA [Source:RFAM;Acc:RF00019] |
| EPB41 | 1 | erythrocyte membrane protein band 4.1 [Sou |
| AL138785.1 | 1 | ribosomal protein L27 (RPL27) pseudogene |
| Y_RNA | 1 | Y RNA [Source:RFAM;Acc:RF00019] |
| Metazoa_SRP | 1 | Metazoan signal recognition particle RNA [Sou |
| TMEM200B | 1 | transmembrane protein 200B [Source:HGNC S |
| AL357500.1 | 1 | novel transcript |
| SRSF4 | 1 | serine and arginine rich splicing factor 4 [Sour |
| AL357500.2 | 1 | NULL |
| MECR | 1 | mitochondrial trans-2-enoyl-CoA reductase [S |
| AL590729.1 | 1 | novel transcript |
| PTPRU | 1 | protein tyrosine phosphatase receptor type U |
| LINC01756 | 1 | long intergenic non-protein coding RNA 1756 |
| AL671862.1 | 1 | protein tyrosine phosphatase, receptor type, |
| AC092265.1 | 1 | novel transcript |
| AL645944.2 | 1 | novel transcript |
| AL645944.1 | 1 | N(alpha)-acetyltransferase 20, NatB catalytic s |
| LINC01648 | 1 | long intergenic non-protein coding RNA 1648 |
| AL137076.1 | 1 | novel transcript |
| AL021921.1 | 1 | family with sequence similarity 32, member A |
| AL161638.2 | 1 | novel transcript |
| AL161638.1 | 1 | novel transcript |
| MATN1 | 1 | matrilin 1 [Source:HGNC Symbol;Acc:HGNC:69 |
| MATN1-AS1 | 1 | MATN1 antisense RNA 1 [Source:HGNC Symbo |
| AL137857.1 | 1 | novel transcript, antisense to LAPTM5 |
| LAPTM5 | 1 | lysosomal protein transmembrane 5 [Source:H |
| MIR4420 | 1 | microRNA 4420 [Source:HGNC Symbol;Acc:HG |
| AL137027.1 | 1 | novel transcript |
| LINC01778 | 1 | long intergenic non-protein coding RNA 1778 |
| RN7SKP91 | 1 | RN7SK pseudogene 91 [Source:HGNC Symbol; |
| AL445235.1 | 1 | novel transcript |
| SDC3 | 1 | syndecan 3 [Source:HGNC Symbol;Acc:HGNC: |
| RN7SL371P | 1 | RNA, 7SL, cytoplasmic 371, pseudogene [Sour |
| PUM1 | 1 | pumilio RNA binding family member 1 [Source |
| AL356320.2 | 1 | TAF9 RNA polymerase II, TATA box binding pro |
| AL356320.1 | 1 | ribosomal protein S29 (RPS29) pseudogene |
| SELENOWP1 | 1 | selenoprotein W pseudogene 1 [Source:HGNC |
| AC114495.1 | 1 | ribosomal protein L21 (RPL21) pseudogene |
| NKAIN1 | 1 | sodium/potassium transporting ATPase intera |
| SNRNP40 | 1 | small nuclear ribonucleoprotein U5 subunit 40 |
| AC114495.2 | 1 | general transcription factor IIA, 2, 12kDa (GTF |
| ZCCHC17 | 1 | zinc finger CCHC-type containing 17 [Source:H |
| AL451070.1 | 1 | novel transcript |
| FABP3 | 1 | fatty acid binding protein 3 [Source:HGNC Syn |
| SERINC2 | 1 | serine incorporator 2 [Source:HGNC Symbol;A |
| AC209007.1 | 1 | eukaryotic translation elongation factor 1 alph |
| RNU6-40P | 1 | RNA, U6 small nuclear 40, pseudogene [Sourc |
| LDC1P | 1 | leucine decarboxylase 1, pseudogene [Source |
| Y_RNA | 1 | Y RNA [Source:RFAM;Acc:RF00019] |

| | | |
|---|---|---|
| LINC01226 | 1 | long intergenic non-protein coding RNA 1226 |
| AC114488.1 | 1 | novel transcript |
| TINAGL1 | 1 | tubulointerstitial nephritis antigen like 1 [Sour |
| HCRTR1 | 1 | hypocretin receptor 1 [Source:HGNC Symbol;A |
| PEF1 | 1 | penta-EF-hand domain containing 1 [Source:H |
| AC114488.2 | 1 | novel transcript, antisense to COL16A1 |
| AC114488.3 | 1 | novel transcript, antisense to PEF1 |
| COL16A1 | 1 | collagen type XVI alpha 1 chain [Source:HGNC |
| ADGRB2 | 1 | adhesion G protein-coupled receptor B2 [Sour |
| MIR4254 | 1 | microRNA 4254 [Source:HGNC Symbol;Acc:HG |
| AL354919.2 | 1 | novel transcript |
| SPOCD1 | 1 | SPOC domain containing 1 [Source:HGNC Sym |
| AL354919.1 | 1 | novel transcript |
| AL136115.2 | 1 | novel transcript |
| PTP4A2 | 1 | protein tyrosine phosphatase 4A2 [Source:HG |
| AL136115.1 | 1 | novel transcript |
| AL136115.3 | 1 | novel transcript |
| KHDRBS1 | 1 | KH RNA binding domain containing, signal tran |
| AL445248.1 | 1 | novel transcript |
| TMEM39B | 1 | transmembrane protein 39B [Source:HGNC Sy |
| MIR5585 | 1 | microRNA 5585 [Source:HGNC Symbol;Acc:HG |
| Metazoa_SRP | 1 | Metazoan signal recognition particle RNA [Sou |
| KPNA6 | 1 | karyopherin subunit alpha 6 [Source:HGNC Sy |
| AL049795.2 | 1 | novel transcript |
| TXLNA | 1 | taxilin alpha [Source:HGNC Symbol;Acc:HGNC |
| CCDC28B | 1 | coiled-coil domain containing 28B [Source:HG |
| AL049795.1 | 1 | novel transcript |
| IQCC | 1 | IQ motif containing C [Source:HGNC Symbol;A |
| DCDC2B | 1 | doublecortin domain containing 2B [Source:H |
| TMEM234 | 1 | transmembrane protein 234 [Source:HGNC Sy |
| EIF3I | 1 | eukaryotic translation initiation factor 3 subur |
| MTMR9LP | 1 | myotubularin related protein 9 like, pseudoge |
| FAM167B | 1 | family with sequence similarity 167 member B |
| LCK | 1 | LCK proto-oncogene, Src family tyrosine kinas |
| Metazoa_SRP | 1 | Metazoan signal recognition particle RNA [Sou |
| Y_RNA | 1 | Y RNA [Source:RFAM;Acc:RF00019] |
| HDAC1 | 1 | histone deacetylase 1 [Source:HGNC Symbol;A |
| MARCKSL1 | 1 | MARCKS like 1 [Source:HGNC Symbol;Acc:HGN |
| AL109945.1 | 1 | novel transcript |
| TSSK3 | 1 | testis specific serine kinase 3 [Source:HGNC Sy |
| FAM229A | 1 | family with sequence similarity 229 member A |
| BSDC1 | 1 | BSD domain containing 1 [Source:HGNC Symb |
| GAPDHP20 | 1 | glyceraldehyde 3 phosphate dehydrogenase p |
| AL356986.1 | 1 | golgi SNAP receptor complex member 2 (GOS |
| LRRC37A12P | 1 | leucine rich repeat containing 37 member A12 |
| RN7SL122P | 1 | RNA, 7SL, cytoplasmic 122, pseudogene [Sour |
| AL033529.1 | 1 | novel transcript |
| ZBTB8B | 1 | zinc finger and BTB domain containing 8B [Sou |
| ZBTB8A | 1 | zinc finger and BTB domain containing 8A [Sou |
| ZBTB8OS | 1 | zinc finger and BTB domain containing 8 oppo |

| | | |
|---|---|---|
| Y_RNA | 1 | Y RNA [Source:RFAM;Acc:RF00019] |
| RBBP4 | 1 | RB binding protein 4, chromatin remodeling fa |
| SYNC | 1 | syncoilin, intermediate filament protein [Sour |
| AC114489.1 | 1 | novel transcript |
| KIAA1522 | 1 | KIAA1522 [Source:HGNC Symbol;Acc:HGNC:2 |
| YARS | 1 | tyrosyl-tRNA synthetase [Source:HGNC Symbo |
| S100PBP | 1 | S100P binding protein [Source:HGNC Symbol; |
| FNDC5 | 1 | fibronectin type III domain containing 5 [Sour |
| HPCA | 1 | hippocalcin [Source:HGNC Symbol;Acc:HGNC: |
| TMEM54 | 1 | transmembrane protein 54 [Source:HGNC Syn |
| AL031602.2 | 1 | novel transcript, antisense to RNF19B |
| RNF19B | 1 | ring finger protein 19B [Source:HGNC Symbol; |
| AL031602.1 | 1 | TEC |
| AL355864.1 | 1 | ribosomal protein L18a (RPL18A) pseudogene |
| AL355864.2 | 1 | ribosomal protein S2 (RPS2) pseudogene |
| AL020995.1 | 1 | novel transcript |
| AK2 | 1 | adenylate kinase 2 [Source:HGNC Symbol;Acc |
| Metazoa_SRP | 1 | Metazoan signal recognition particle RNA [Sou |
| AZIN2 | 1 | antizyme inhibitor 2 [Source:HGNC Symbol;Ac |
| AL662907.1 | 1 | TEC |
| TRIM62 | 1 | tripartite motif containing 62 [Source:HGNC S |
| AL662907.2 | 1 | TEC |
| AL662907.3 | 1 | novel transcript |
| ZNF362 | 1 | zinc finger protein 362 [Source:HGNC Symbol; |
| AL513327.3 | 1 | novel transcript |
| A3GALT2 | 1 | alpha 1,3-galactosyltransferase 2 [Source:HGN |
| AL513327.1 | 1 | novel transcript |
| PHC2 | 1 | polyhomeotic homolog 2 [Source:HGNC Symb |
| MIR3605 | 1 | microRNA 3605 [Source:HGNC Symbol;Acc:HG |
| RN7SKP16 | 1 | RN7SK pseudogene 16 [Source:HGNC Symbol; |
| AL513327.2 | 1 | novel transcript |
| TLR12P | 1 | toll like receptor 12, pseudogene [Source:HGN |
| ZSCAN20 | 1 | zinc finger and SCAN domain containing 20 [So |
| AC115285.1 | 1 | proteasome (prosome, macropain) subunit, al |
| CSMD2 | 1 | CUB and Sushi multiple domains 2 [Source:HG |
| HSPD1P14 | 1 | heat shock protein family D (Hsp60) member |
| HMGB4 | 1 | high mobility group box 4 [Source:HGNC Symb |
| CSMD2-AS1 | 1 | CSMD2 antisense RNA 1 [Source:HGNC Symbo |
| RNA5SP42 | 1 | RNA, 5S ribosomal pseudogene 42 [Source:HG |
| C1orf94 | 1 | chromosome 1 open reading frame 94 [Sourc |
| AC115286.1 | 1 | ferritin, heavy polypeptide 1 (FTH1) pseudoge |
| AC099565.1 | 1 | novel transcript |
| MIR552 | 1 | microRNA 552 [Source:HGNC Symbol;Acc:HGN |
| SMIM12 | 1 | small integral membrane protein 12 [Source:H |
| GJB5 | 1 | gap junction protein beta 5 [Source:HGNC Syn |
| GJB4 | 1 | gap junction protein beta 4 [Source:HGNC Syn |
| AL121988.1 | 1 | novel transcript, antisense to GJB4 and GJB3 |
| GJB3 | 1 | gap junction protein beta 3 [Source:HGNC Syn |
| GJA4 | 1 | gap junction protein alpha 4 [Source:HGNC Sy |
| AL122010.1 | 1 | novel transcript |

| | | |
|---|---|---|
| DLGAP3 | 1 | DLG associated protein 3 [Source:HGNC Symb |
| AC114490.3 | 1 | novel protein |
| AC114490.1 | 1 | novel platelet-activating factor receptor famil |
| AC114490.2 | 1 | novel transcript, ZMYM6-ZMYM6NB readthro |
| TMEM35B | 1 | transmembrane protein 35B [Source:HGNC Sy |
| ZMYM6 | 1 | zinc finger MYM-type containing 6 [Source:HG |
| ZMYM1 | 1 | zinc finger MYM-type containing 1 [Source:HG |
| RPL12P45 | 1 | ribosomal protein L12 pseudogene 45 [Source |
| EFCAB14P1 | 1 | EF-hand calcium binding domain 14 pseudoge |
| AL590434.1 | 1 | novel transcript |
| SFPQ | 1 | splicing factor proline and glutamine rich [Sou |
| Y_RNA | 1 | Y RNA [Source:RFAM;Acc:RF00019] |
| ZMYM4 | 1 | zinc finger MYM-type containing 4 [Source:HG |
| RN7SL503P | 1 | RNA, 7SL, cytoplasmic 503, pseudogene [Sour |
| SNORA62 | 1 | Small nucleolar RNA SNORA62/SNORA6 family |
| RPL5P4 | 1 | ribosomal protein L5 pseudogene 4 [Source:H |
| ZMYM4-AS1 | 1 | ZMYM4 antisense RNA 1 [Source:HGNC Symb |
| Metazoa_SRP | 1 | Metazoan signal recognition particle RNA [Sou |
| KIAA0319L | 1 | KIAA0319 like [Source:HGNC Symbol;Acc:HGN |
| AC004865.1 | 1 | zinc finger protein pseudogene |
| NCDN | 1 | neurochondrin [Source:HGNC Symbol;Acc:HG |
| AC004865.2 | 1 | novel transcript |
| TFAP2E | 1 | transcription factor AP-2 epsilon [Source:HGN |
| PSMB2 | 1 | proteasome subunit beta 2 [Source:HGNC Syn |
| C1orf216 | 1 | chromosome 1 open reading frame 216 [Sour |
| CLSPN | 1 | claspin [Source:HGNC Symbol;Acc:HGNC:1971 |
| AL354864.1 | 1 | novel transcript |
| AGO4 | 1 | argonaute RISC component 4 [Source:HGNC S |
| AGO1 | 1 | argonaute RISC component 1 [Source:HGNC S |
| AL139286.1 | 1 | novel pseudogene |
| AL139286.3 | 1 | novel transcript, antisense to AGO1 |
| AL139286.2 | 1 | novel transcript |
| AGO3 | 1 | argonaute RISC catalytic component 3 [Source |
| AL138787.2 | 1 | novel transcript, antisense to EIF2C3 |
| AL138787.1 | 1 | implantation-associated protein (DKFZp564K1 |
| TEKT2 | 1 | tektin 2 [Source:HGNC Symbol;Acc:HGNC:117 |
| ADPRHL2 | 1 | ADP-ribosylhydrolase like 2 [Source:HGNC Syr |
| COL8A2 | 1 | collagen type VIII alpha 2 chain [Source:HGNC |
| TRAPPC3 | 1 | trafficking protein particle complex 3 [Source: |
| MAP7D1 | 1 | MAP7 domain containing 1 [Source:HGNC Syn |
| RN7SL131P | 1 | RNA, 7SL, cytoplasmic 131, pseudogene [Sour |
| THRAP3 | 1 | thyroid hormone receptor associated protein |
| UBE2V2P4 | 1 | ubiquitin conjugating enzyme E2 V2 pseudoge |
| SH3D21 | 1 | SH3 domain containing 21 [Source:HGNC Sym |
| EVA1B | 1 | eva-1 homolog B [Source:HGNC Symbol;Acc:H |
| AL591845.1 | 1 | novel transcript |
| STK40 | 1 | serine/threonine kinase 40 [Source:HGNC Sym |
| LSM10 | 1 | LSM10, U7 small nuclear RNA associated [Sou |
| RNU4-27P | 1 | RNA, U4 small nuclear 27, pseudogene [Sourc |
| OSCP1 | 1 | organic solute carrier partner 1 [Source:HGNC |

| | | |
|---|---|---|
| SNORA63C | 1 | small nucleolar RNA, H/ACA box 63C [Source:I |
| MRPS15 | 1 | mitochondrial ribosomal protein S15 [Source: |
| CSF3R | 1 | colony stimulating factor 3 receptor [Source:H |
| FTLP18 | 1 | ferritin light chain pseudogene 18 [Source:HG |
| AL596257.1 | 1 | novel transcript |
| AC117945.2 | 1 | novel transcript |
| AC117945.1 | 1 | novel transcript |
| GRIK3 | 1 | glutamate ionotropic receptor kainate type su |
| AL031430.1 | 1 | novel transcript |
| AL353604.1 | 1 | novel transcript |
| MIR4255 | 1 | microRNA 4255 [Source:HGNC Symbol;Acc:HG |
| RNU6-636P | 1 | RNA, U6 small nuclear 636, pseudogene [Sour |
| RNA5SP43 | 1 | RNA, 5S ribosomal pseudogene 43 [Source:HG |
| RPS29P6 | 1 | ribosomal protein S29 pseudogene 6 [Source: |
| LINC01137 | 1 | long intergenic non-protein coding RNA 1137 |
| ZC3H12A | 1 | zinc finger CCCH-type containing 12A [Source: |
| MIR6732 | 1 | microRNA 6732 [Source:HGNC Symbol;Acc:HG |
| MEAF6 | 1 | MYST/Esa1 associated factor 6 [Source:HGNC |
| MIR5581 | 1 | microRNA 5581 [Source:HGNC Symbol;Acc:HG |
| SNIP1 | 1 | Smad nuclear interacting protein 1 [Source:HG |
| FTH1P1 | 1 | ferritin heavy chain 1 pseudogene 1 [Source:H |
| AL034379.1 | 1 | ribosomal protein S27 (metallopanstimulin 1) |
| DNALI1 | 1 | dynein axonemal light intermediate chain 1 [S |
| GNL2 | 1 | G protein nucleolar 2 [Source:HGNC Symbol;A |
| AL513220.1 | 1 | novel transcript |
| RSPO1 | 1 | R-spondin 1 [Source:HGNC Symbol;Acc:HGNC: |
| C1orf109 | 1 | chromosome 1 open reading frame 109 [Sour |
| CDCA8 | 1 | cell division cycle associated 8 [Source:HGNC S |
| EPHA10 | 1 | EPH receptor A10 [Source:HGNC Symbol;Acc:I |
| Y_RNA | 1 | Y RNA [Source:RFAM;Acc:RF00019] |
| ACTN4P2 | 1 | actinin alpha 4 pseudogene 2 [Source:HGNC S |
| AL929472.1 | 1 | novel pseudogene |
| MANEAL | 1 | mannosidase endo-alpha like [Source:HGNC S |
| AL929472.3 | 1 | novel transcript |
| YRDC | 1 | yrdC N6-threonylcarbamoyltransferase domai |
| C1orf122 | 1 | chromosome 1 open reading frame 122 [Sour |
| MTF1 | 1 | metal regulatory transcription factor 1 [Sourc |
| AL929472.4 | 1 | ribosomal protein S2 (RPS2) pseudogene |
| INPP5B | 1 | inositol polyphosphate-5-phosphatase B [Sour |
| AL929472.2 | 1 | novel transcript |
| SNORA63 | 1 | Small nucleolar RNA SNORA63 [Source:RFAM; |
| RNU6-584P | 1 | RNA, U6 small nuclear 584, pseudogene [Sour |
| Metazoa_SRP | 1 | Metazoan signal recognition particle RNA [Sou |
| SF3A3 | 1 | splicing factor 3a subunit 3 [Source:HGNC Sym |
| RNU6-510P | 1 | RNA, U6 small nuclear 510, pseudogene [Sour |
| FHL3 | 1 | four and a half LIM domains 3 [Source:HGNC S |
| UTP11 | 1 | UTP11 small subunit processome component |
| POU3F1 | 1 | POU class 3 homeobox 1 [Source:HGNC Symb |
| MIR3659HG | 1 | MIR3659 host gene [Source:HGNC Symbol;Acc |
| AL139158.1 | 1 | FK506 binding protein 3, 25kDa (FKBP3) pseuc |

| | | |
|---|---|---|
| MIR3659 | 1 | microRNA 3659 [Source:HGNC Symbol;Acc:HG |
| LINC02786 | 1 | long intergenic non-protein coding RNA 2786 |
| AL390839.1 | 1 | novel transcript |
| AL390839.2 | 1 | novel transcript |
| LINC01343 | 1 | long intergenic non-protein coding RNA 1343 |
| AL513479.1 | 1 | tubulin, beta 3 class III (TUBB3) pseudogene |
| RNU6-753P | 1 | RNA, U6 small nuclear 753, pseudogene [Sour |
| LINC01685 | 1 | long intergenic non-protein coding RNA 1685 |
| AL354702.1 | 1 | heat shock 70kDa protein 5 (glucose-regulate |
| AL354702.2 | 1 | novel transcript |
| RRAGC | 1 | Ras related GTP binding C [Source:HGNC Symb |
| AL139260.2 | 1 | novel transcript |
| AL139260.1 | 1 | novel transcript |
| MYCBP | 1 | MYC binding protein [Source:HGNC Symbol;Ac |
| AL139260.3 | 1 | novel protein |
| GJA9 | 1 | gap junction protein alpha 9 [Source:HGNC Sy |
| RHBDL2 | 1 | rhomboid like 2 [Source:HGNC Symbol;Acc:HG |
| RNU6-605P | 1 | RNA, U6 small nuclear 605, pseudogene [Sour |
| Y_RNA | 1 | Y RNA [Source:RFAM;Acc:RF00019] |
| EIF1P2 | 1 | eukaryotic translation initiation factor 1 pseu |
| AKIRIN1 | 1 | akirin 1 [Source:HGNC Symbol;Acc:HGNC:257 |
| NDUFS5 | 1 | NADH:ubiquinone oxidoreductase subunit S5 |
| MACF1 | 1 | microtubule actin crosslinking factor 1 [Source |
| RNU6-608P | 1 | RNA, U6 small nuclear 608, pseudogene [Sour |
| RNA5SP44 | 1 | RNA, 5S ribosomal pseudogene 44 [Source:HG |
| AL356055.1 | 1 | novel transcript, intronic to MACF1 |
| AL442071.2 | 1 | novel transcript, antisense to MACF1 |
| AL442071.1 | 1 | novel transcript |
| HSPE1P8 | 1 | heat shock protein family E (Hsp10) member 1 |
| BMP8A | 1 | bone morphogenetic protein 8a [Source:HGN |
| OXCT2P1 | 1 | 3-oxoacid CoA-transferase 2 pseudogene 1 [S |
| PPIEL | 1 | peptidylprolyl isomerase E like pseudogene [S |
| PABPC4 | 1 | poly(A) binding protein cytoplasmic 4 [Source |
| PABPC4-AS1 | 1 | PABPC4 antisense RNA 1 [Source:HGNC Symb |
| SNORA55 | 1 | small nucleolar RNA, H/ACA box 55 [Source:H |
| HEYL | 1 | hes related family bHLH transcription factor w |
| AL035404.2 | 1 | novel transcript |
| NT5C1A | 1 | 5'-nucleotidase, cytosolic IA [Source:HGNC Sy |
| HPCAL4 | 1 | hippocalcin like 4 [Source:HGNC Symbol;Acc:H |
| PPIE | 1 | peptidylprolyl isomerase E [Source:HGNC Sym |
| AL035404.1 | 1 | novel pseudogene |
| RNU7-121P | 1 | RNA, U7 small nuclear 121 pseudogene [Sourc |
| BMP8B | 1 | bone morphogenetic protein 8b [Source:HGN |
| OXCT2 | 1 | 3-oxoacid CoA-transferase 2 [Source:HGNC Sy |
| BMP8B-AS1 | 1 | BMP8B antisense RNA 1 [Source:HGNC Symbo |
| AL033527.3 | 1 | novel transcript, antisense to BMP8B |
| AL033527.1 | 1 | microtubule-actin crosslinking factor 1 (MACF |
| AL033527.4 | 1 | novel transcript |
| LINC02811 | 1 | long intergenic non-protein coding RNA 2811 |
| TRIT1 | 1 | tRNA isopentenyltransferase 1 [Source:HGNC |

| | | |
|---|---|---|
| Y_RNA | 1 | Y RNA [Source:RFAM;Acc:RF00019] |
| MYCL | 1 | MYCL proto-oncogene, bHLH transcription fac |
| AL033527.2 | 1 | novel transcript |
| Y_RNA | 1 | Y RNA [Source:RFAM;Acc:RF00019] |
| MFSD2A | 1 | major facilitator superfamily domain containir |
| AL663070.1 | 1 | ribosomal protein S2 (RPS2) pseudogene |
| CAP1 | 1 | cyclase associated actin cytoskeleton regulatc |
| PPT1 | 1 | palmitoyl-protein thioesterase 1 [Source:HGN |
| AL512599.1 | 1 | ornithine decarboxylase antizyme 1 (OAZ1) ps |
| RLF | 1 | rearranged L-myc fusion [Source:HGNC Symbo |
| RNU6-1237P | 1 | RNA, U6 small nuclear 1237, pseudogene [Sou |
| TMCO2 | 1 | transmembrane and coiled-coil domains 2 [So |
| AL050341.2 | 1 | novel transcript, antisense to ZMPSTE24 |
| ZMPSTE24 | 1 | zinc metallopeptidase STE24 [Source:HGNC Sy |
| AL050341.1 | 1 | pseudogene similar to part of RIO kinase 2 (ye |
| COL9A2 | 1 | collagen type IX alpha 2 chain [Source:HGNC S |
| AL031985.2 | 1 | ribosomal protein L21 (RPL21) |
| SMAP2 | 1 | small ArfGAP2 [Source:HGNC Symbol;Acc:HGN |
| AL031985.1 | 1 | ribosomal protein L10a (RPL10A) |
| AL031985.4 | 1 | novel transcript |
| ZFP69B | 1 | ZFP69 zinc finger protein B [Source:HGNC Syn |
| AL031985.3 | 1 | novel transcript, sense overlapping with ZNF6 |
| AL603839.4 | 1 | TEC |
| ZFP69 | 1 | ZFP69 zinc finger protein [Source:HGNC Symb |
| AL603839.3 | 1 | novel transcript |
| EXO5 | 1 | exonuclease 5 [Source:HGNC Symbol;Acc:HGN |
| AL603839.1 | 1 | novel transcript |
| AL603839.2 | 1 | novel transcript |
| ZNF684 | 1 | zinc finger protein 684 [Source:HGNC Symbol; |
| AL356379.2 | 1 | novel transcript |
| AL356379.1 | 1 | novel pseudogene |
| GTF2F2P2 | 1 | general transcription factor IIF subunit 2 pseu |
| RIMS3 | 1 | regulating synaptic membrane exocytosis 3 [S |
| AL031289.2 | 1 | novel transcript, antisense to RIMS3 |
| AL031289.1 | 1 | novel transcript |
| NFYC-AS1 | 1 | NFYC antisense RNA 1 [Source:HGNC Symbol;/ |
| NFYC | 1 | nuclear transcription factor Y subunit gamma |
| MIR30E | 1 | microRNA 30e [Source:HGNC Symbol;Acc:HGN |
| MIR30C1 | 1 | microRNA 30c-1 [Source:HGNC Symbol;Acc:H( |
| KCNQ4 | 1 | potassium voltage-gated channel subfamily Q |
| RN7SL326P | 1 | RNA, 7SL, cytoplasmic 326, pseudogene [Sour |
| CITED4 | 1 | Cbp/p300 interacting transactivator with Glu/ |
| AC119677.1 | 1 | novel transcript |
| AL391730.1 | 1 | protein phosphatase 2 (formerly 2A), catalytic |
| AL391730.2 | 1 | novel transcript |
| UBE2V1P8 | 1 | ubiquitin conjugating enzyme E2 V1 pseudoge |
| CTPS1 | 1 | CTP synthase 1 [Source:HGNC Symbol;Acc:HG |
| SLFNL1-AS1 | 1 | SLFNL1 antisense RNA 1 [Source:HGNC Symbc |
| SLFNL1 | 1 | schlafen like 1 [Source:HGNC Symbol;Acc:HGN |
| SCMH1 | 1 | Scm polycomb group protein homolog 1 [Sour |

| | | |
|---|---|---|
| RPL23AP17 | 1 | ribosomal protein L23a pseudogene 17 [Sourc |
| AC093151.8 | 1 | novel transcript, antisense to SCMH1 |
| AC093151.3 | 1 | novel transcript |
| AC093151.1 | 1 | ribosomal protein L36a (RPL36A) pseudogene |
| AC093151.6 | 1 | SET domain and mariner transposase fusion g |
| FOXO6 | 1 | forkhead box O6 [Source:HGNC Symbol;Acc:H |
| AC093151.2 | 1 | novel transcript |
| RNA5SP45 | 1 | RNA, 5S ribosomal pseudogene 45 [Source:HG |
| EDN2 | 1 | endothelin 2 [Source:HGNC Symbol;Acc:HGNC |
| HIVEP3 | 1 | human immunodeficiency virus type I enhanc |
| AL445933.2 | 1 | novel transcript |
| AL445933.1 | 1 | novel pseudogene |
| AC119676.1 | 1 | novel protein |
| AL158216.1 | 1 | novel transcript |
| HNRNPFP1 | 1 | heterogeneous nuclear ribonucleoprotein F ps |
| AC114492.1 | 1 | ribosomal protein L21 (RPL21) pseudogene |
| GUCA2B | 1 | guanylate cyclase activator 2B [Source:HGNC |
| GUCA2A | 1 | guanylate cyclase activator 2A [Source:HGNC |
| FOXJ3 | 1 | forkhead box J3 [Source:HGNC Symbol;Acc:HG |
| AC096540.1 | 1 | novel transcript |
| RIMKLA | 1 | ribosomal modification protein rimK like famil |
| AL513331.1 | 1 | ribosomal protein S15 (RPS15) pseudogene |
| ZMYND12 | 1 | zinc finger MYND-type containing 12 [Source: |
| PPCS | 1 | phosphopantothenoylcysteine synthetase [So |
| CCDC30 | 1 | coiled-coil domain containing 30 [Source:HGN |
| AL445669.1 | 1 | ribosomal protein S3a (RPS3A) pseudogene |
| TMSB4XP1 | 1 | TMSB4X pseudogene 1 [Source:HGNC Symbol |
| RNU6-536P | 1 | RNA, U6 small nuclear 536, pseudogene [Sour |
| AL445669.2 | 1 | novel pseudogene |
| PPIH | 1 | peptidylprolyl isomerase H [Source:HGNC Sym |
| AC098484.4 | 1 | novel transcript, antisense to PPIH |
| AC098484.2 | 1 | novel transcript |
| YBX1 | 1 | Y-box binding protein 1 [Source:HGNC Symbol |
| CLDN19 | 1 | claudin 19 [Source:HGNC Symbol;Acc:HGNC:2 |
| P3H1 | 1 | prolyl 3-hydroxylase 1 [Source:HGNC Symbol; |
| C1orf50 | 1 | chromosome 1 open reading frame 50 [Sourc |
| AC098484.3 | 1 | novel protein |
| AC098484.1 | 1 | novel transcript |
| TMEM269 | 1 | transmembrane protein 269 [Source:HGNC Sy |
| SVBP | 1 | small vasohibin binding protein [Source:HGNC |
| ERMAP | 1 | erythroblast membrane associated protein (S |
| AL512353.1 | 1 | novel transcript, antisense to ERMAP |
| ZNF691 | 1 | zinc finger protein 691 [Source:HGNC Symbol; |
| AL512353.2 | 1 | epidermodysplasia verruciformis 1 (EVER1) ps |
| AL512353.3 | 1 | makorin, ring finger protein, 1 (MKRN1) pseuc |
| ATP6V1E1P1 | 1 | ATPase H+ transporting V1 subunit E1 pseudo |
| SLC2A1 | 1 | solute carrier family 2 member 1 [Source:HGN |
| ATP6V0CP4 | 1 | ATPase H+ transporting V0 subunit c pseudog |
| SLC2A1-AS1 | 1 | SLC2A1 antisense RNA 1 [Source:HGNC Symbo |
| AC099795.1 | 1 | novel transcript |

| | | |
|---|---|---|
| RNU6-880P | 1 | RNA, U6 small nuclear 880, pseudogene [Sour |
| U6 | 1 | U6 spliceosomal RNA [Source:RFAM;Acc:RF00 |
| AL161637.1 | 1 | zinc finger and SCAN domain containing 2 (ZS( |
| FAM183A | 1 | family with sequence similarity 183 member / |
| EBNA1BP2 | 1 | EBNA1 binding protein 2 [Source:HGNC Symb( |
| MIR6733 | 1 | microRNA 6733 [Source:HGNC Symbol;Acc:HG |
| CFAP57 | 1 | cilia and flagella associated protein 57 [Source |
| RNA5SP46 | 1 | RNA, 5S ribosomal pseudogene 46 [Source:HG |
| TMEM125 | 1 | transmembrane protein 125 [Source:HGNC Sy |
| C1orf210 | 1 | chromosome 1 open reading frame 210 [Sour |
| TIE1 | 1 | tyrosine kinase with immunoglobulin like and |
| MPL | 1 | MPL proto-oncogene, thrombopoietin recept( |
| AL139289.3 | 1 | novel transcript, antisense to MPL |
| AL139289.2 | 1 | novel transcript |
| CDC20 | 1 | cell division cycle 20 [Source:HGNC Symbol;Ac |
| ELOVL1 | 1 | ELOVL fatty acid elongase 1 [Source:HGNC Syr |
| MIR6734 | 1 | microRNA 6734 [Source:HGNC Symbol;Acc:HG |
| MED8 | 1 | mediator complex subunit 8 [Source:HGNC Sy |
| AL139289.1 | 1 | novel transcript |
| SZT2 | 1 | SZT2 subunit of KICSTOR complex [Source:HGI |
| SZT2-AS1 | 1 | SZT2 antisense RNA 1 [Source:HGNC Symbol;/ |
| MIR6735 | 1 | microRNA 6735 [Source:HGNC Symbol;Acc:HG |
| HYI | 1 | hydroxypyruvate isomerase (putative) [Source |
| HYI-AS1 | 1 | HYI antisense RNA 1 [Source:HGNC Symbol;Ac |
| PTPRF | 1 | protein tyrosine phosphatase receptor type F |
| AL451062.3 | 1 | novel transcript |
| KDM4A | 1 | lysine demethylase 4A [Source:HGNC Symbol; |
| KDM4A-AS1 | 1 | KDM4A antisense RNA 1 [Source:HGNC Symb( |
| ST3GAL3 | 1 | ST3 beta-galactoside alpha-2,3-sialyltransfera: |
| AL451062.1 | 1 | novel transcript |
| U6 | 1 | U6 spliceosomal RNA [Source:RFAM;Acc:RF00 |
| AL451062.2 | 1 | SMT3 suppressor of mif two 3 (yeast) family p |
| MIR6079 | 1 | microRNA 6079 [Source:HGNC Symbol;Acc:HG |
| SHMT1P1 | 1 | serine hydroxymethyltransferase 1 (soluble) p |
| ARTN | 1 | artemin [Source:HGNC Symbol;Acc:HGNC:727 |
| AL357079.1 | 1 | novel transcript |
| IPO13 | 1 | importin 13 [Source:HGNC Symbol;Acc:HGNC: |
| AL357079.2 | 1 | novel transcript, antisense to DPH2 and ATP6' |
| DPH2 | 1 | diphthamide biosynthesis 2 [Source:HGNC Syr |
| ATP6V0B | 1 | ATPase H+ transporting V0 subunit b [Source: |
| B4GALT2 | 1 | beta-1,4-galactosyltransferase 2 [Source:HGN |
| CCDC24 | 1 | coiled-coil domain containing 24 [Source:HGN |
| SLC6A9 | 1 | solute carrier family 6 member 9 [Source:HGN |
| AL139220.2 | 1 | novel transcript |
| AL139220.1 | 1 | pseudogene similar to part of ATP synthase, H |
| KRT8P47 | 1 | keratin 8 pseudogene 47 [Source:HGNC Symb |
| RN7SL479P | 1 | RNA, 7SL, cytoplasmic 479, pseudogene [Sour |
| KLF17 | 1 | Kruppel like factor 17 [Source:HGNC Symbol;/ |
| AL356653.1 | 1 | small nuclear ribonucleoprotein D1 polypeptic |
| KLF18 | 1 | Kruppel like factor 18 [Source:HGNC Symbol;/ |

| | | |
|---|---|---|
| AL359839.1 | 1 | zinc finger protein 146 (ZNF146) pseudogene |
| Y_RNA | 1 | Y RNA [Source:RFAM;Acc:RF00019] |
| OOSP1P1 | 1 | oocyte secreted protein 1 pseudogene 1 [Sour |
| AL035417.1 | 1 | pseudogene similar to part of novel protein D |
| AL035417.2 | 1 | metaxin 3 (MTX3) pseudogene |
| DMAP1 | 1 | DNA methyltransferase 1 associated protein 1 |
| ERI3 | 1 | ERI1 exoribonuclease family member 3 [Sourc |
| ERI3-IT1 | 1 | ERI3 intronic transcript 1 [Source:HGNC Symb |
| RNU6-369P | 1 | RNA, U6 small nuclear 369, pseudogene [Sour |
| RNF220 | 1 | ring finger protein 220 [Source:HGNC Symbol; |
| MIR5584 | 1 | microRNA 5584 [Source:HGNC Symbol;Acc:HG |
| TMEM53 | 1 | transmembrane protein 53 [Source:HGNC Sym |
| ARMH1 | 1 | armadillo like helical domain containing 1 [Sou |
| RNU5F-1 | 1 | RNA, U5F small nuclear 1 [Source:HGNC Symb |
| RNU5D-1 | 1 | RNA, U5D small nuclear 1 [Source:HGNC Symb |
| KIF2C | 1 | kinesin family member 2C [Source:HGNC Sym |
| AL592166.1 | 1 | novel transcript |
| RPS8 | 1 | ribosomal protein S8 [Source:HGNC Symbol;A |
| SNORD55 | 1 | small nucleolar RNA, C/D box 55 [Source:HGN |
| SNORD46 | 1 | small nucleolar RNA, C/D box 46 [Source:HGN |
| SNORD38A | 1 | small nucleolar RNA, C/D box 38A [Source:HG |
| SNORD38B | 1 | small nucleolar RNA, C/D box 38B [Source:HG |
| SNORD38B | 1 | small nucleolar RNA, C/D box 38B [Source:HG |
| RPS15AP11 | 1 | ribosomal protein S15a pseudogene 11 [Sourc |
| BEST4 | 1 | bestrophin 4 [Source:HGNC Symbol;Acc:HGNC |
| PLK3 | 1 | polo like kinase 3 [Source:HGNC Symbol;Acc:H |
| TCTEX1D4 | 1 | Tctex1 domain containing 4 [Source:HGNC Syr |
| BTBD19 | 1 | BTB domain containing 19 [Source:HGNC Sym |
| PTCH2 | 1 | patched 2 [Source:HGNC Symbol;Acc:HGNC:9 |
| RNU5E-6P | 1 | RNA, U5E small nuclear 6, pseudogene [Sourc |
| AL136380.1 | 1 | ribosomal protein S2 (RPS2) pseudogene |
| EIF2B3 | 1 | eukaryotic translation initiation factor 2B sub |
| RNA5SP47 | 1 | RNA, 5S ribosomal pseudogene 47 [Source:HG |
| CCNB1IP1P1 | 1 | cyclin B1 interacting protein 1 pseudogene 1 [ |
| PPIAP35 | 1 | peptidylprolyl isomerase A pseudogene 35 [Sc |
| MRPS17P1 | 1 | mitochondrial ribosomal protein S17 pseudog |
| HECTD3 | 1 | HECT domain E3 ubiquitin protein ligase 3 [So |
| UROD | 1 | uroporphyrinogen decarboxylase [Source:HGN |
| ZSWIM5 | 1 | zinc finger SWIM-type containing 5 [Source:H |
| OSTCP5 | 1 | oligosaccharyltransferase complex subunit ps |
| LINC01144 | 1 | long intergenic non-protein coding RNA 1144 |
| HPDL | 1 | 4-hydroxyphenylpyruvate dioxygenase like [Sc |
| MUTYH | 1 | mutY DNA glycosylase [Source:HGNC Symbol; |
| AL451136.1 | 1 | novel protein |
| TOE1 | 1 | target of EGR1, exonuclease [Source:HGNC Sy |
| TESK2 | 1 | testis associated actin remodelling kinase 2 [S |
| PPIAP36 | 1 | peptidylprolyl isomerase A pseudogene 36 [Sc |
| CCDC163 | 1 | coiled-coil domain containing 163 [Source:HG |
| MMACHC | 1 | metabolism of cobalamin associated C [Sourc |
| PRDX1 | 1 | peroxiredoxin 1 [Source:HGNC Symbol;Acc:HG |

| | | |
|---|---|---|
| HMGB1P48 | 1 | high mobility group box 1 pseudogene 48 [Sou |
| AKR1A1 | 1 | aldo-keto reductase family 1 member A1 [Sou |
| AL355480.3 | 1 | NULL |
| AL355480.1 | 1 | NULL |
| NASP | 1 | nuclear autoantigenic sperm protein [Source:I |
| AL355480.2 | 1 | NULL |
| AL355480.4 | 1 | NULL |
| CCDC17 | 1 | coiled-coil domain containing 17 [Source:HGN |
| GPBP1L1 | 1 | GC-rich promoter binding protein 1 like 1 [Sou |
| RPS15AP10 | 1 | ribosomal protein S15a pseudogene 10 [Sourc |
| AL604028.2 | 1 | ribosomal protein L7a (RPL7A) pseudogene |
| TMEM69 | 1 | transmembrane protein 69 [Source:HGNC Syn |
| IPP | 1 | intracisternal A particle-promoted polypeptid |
| AL604028.1 | 1 | novel transcript |
| AL603882.1 | 1 | ribosomal protein L6 (RPL6) pseudogene |
| MAST2 | 1 | microtubule associated serine/threonine kina |
| TMA16P2 | 1 | translation machinery associated 16 homolog |
| PIK3R3 | 1 | phosphoinositide-3-kinase regulatory subunit |
| AL358075.4 | 1 | -PIK3R3 readthrough [Source:NCBI gene;Acc:1 |
| AL358075.1 | 1 | novel transcript |
| AL358075.3 | 1 | cytochrome c oxidase subunit VIIb (COX7B) ps |
| AL358075.2 | 1 | novel transcript |
| TSPAN1 | 1 | tetraspanin 1 [Source:HGNC Symbol;Acc:HGN |
| P3R3URF | 1 | PIK3R3 upstream reading frame [Source:HGN |
| POMGNT1 | 1 | protein O-linked mannose N-acetylglucosamir |
| LURAP1 | 1 | leucine rich adaptor protein 1 [Source:HGNC S |
| RAD54L | 1 | RAD54 like [Source:HGNC Symbol;Acc:HGNC:9 |
| LRRC41 | 1 | leucine rich repeat containing 41 [Source:HGN |
| UQCRH | 1 | ubiquinol-cytochrome c reductase hinge prote |
| NSUN4 | 1 | NOP2/Sun RNA methyltransferase 4 [Source:H |
| AL122001.1 | 1 | novel pseudogene |
| FAAH | 1 | fatty acid amide hydrolase [Source:HGNC Sym |
| FAAHP1 | 1 | fatty acid amide hydrolase pseudogene 1 [Sou |
| LINC01398 | 1 | long intergenic non-protein coding RNA 1398 |
| DMBX1 | 1 | diencephalon/mesencephalon homeobox 1 [S |
| TMEM275 | 1 | transmembrane protein 275 [Source:HGNC Sy |
| MKNK1-AS1 | 1 | MKNK1 antisense RNA 1 [Source:HGNC Symbo |
| KNCN | 1 | kinocilin [Source:HGNC Symbol;Acc:HGNC:264 |
| MKNK1 | 1 | MAPK interacting serine/threonine kinase 1 [S |
| MOB3C | 1 | MOB kinase activator 3C [Source:HGNC Symb |
| ATPAF1 | 1 | ATP synthase mitochondrial F1 complex asser |
| NENFP1 | 1 | neudesin neurotrophic factor pseudogene 1 [S |
| TEX38 | 1 | testis expressed 38 [Source:HGNC Symbol;Acc |
| EFCAB14-AS1 | 1 | EFCAB14 antisense RNA 1 [Source:HGNC Symb |
| EFCAB14 | 1 | EF-hand calcium binding domain 14 [Source:H |
| AL593856.1 | 1 | novel pseudogene |
| CYP4B1 | 1 | cytochrome P450 family 4 subfamily B membe |
| CYP4Z2P | 1 | cytochrome P450 family 4 subfamily Z membe |
| AL731892.1 | 1 | tubulin alpha pseudogene |
| CYP4A11 | 1 | cytochrome P450 family 4 subfamily A membe |

| | | |
|---|---|---|
| CYP4A26P | 1 | cytochrome P450 family 4 subfamily A membe |
| CYP4A43P | 1 | cytochrome P450 family 4 subfamily A membe |
| CYP4A27P | 1 | cytochrome P450 family 4 subfamily A membe |
| CYP4A44P | 1 | cytochrome P450 family 4 subfamily A membe |
| CYP4X1 | 1 | cytochrome P450 family 4 subfamily X membe |
| CYP4Z1 | 1 | cytochrome P450 family 4 subfamily Z membe |
| AL450996.1 | 1 | tubulin alpha pseudogene |
| CYP4A22-AS1 | 1 | CYP4A22 antisense RNA 1 [Source:HGNC Syml |
| CYP4A22 | 1 | cytochrome P450 family 4 subfamily A membe |
| MTND1P34 | 1 | MT-ND1 pseudogene 34 [Source:HGNC Symbo |
| LINC00853 | 1 | long intergenic non-protein coding RNA 853 [S |
| PDZK1IP1 | 1 | PDZK1 interacting protein 1 [Source:HGNC Syr |
| TAL1 | 1 | TAL bHLH transcription factor 1, erythroid diff |
| AL135960.1 | 1 | novel transcript |
| STIL | 1 | STIL centriolar assembly protein [Source:HGN |
| CMPK1 | 1 | cytidine/uridine monophosphate kinase 1 [Sou |
| LINC01389 | 1 | long intergenic non-protein coding RNA 1389 |
| FOXE3 | 1 | forkhead box E3 [Source:HGNC Symbol;Acc:H |
| FOXD2-AS1 | 1 | FOXD2 adjacent opposite strand RNA 1 [Sourc |
| FOXD2 | 1 | forkhead box D2 [Source:HGNC Symbol;Acc:H |
| AL356458.1 | 1 | myotubularin related protein 14 (MTMR14) ps |
| RPL21P24 | 1 | ribosomal protein L21 pseudogene 24 [Source |
| ATP6V0E1P4 | 1 | ATPase H+ transporting V0 subunit e1 pseudo |
| LINC01738 | 1 | long intergenic non-protein coding RNA 1738 |
| TRABD2B | 1 | TraB domain containing 2B [Source:HGNC Syn |
| AL691459.1 | 1 | novel transcript |
| AC096541.1 | 1 | novel transcript |
| LINC02794 | 1 | long intergenic non-protein coding RNA 2794 |
| AL356289.1 | 1 | novel transcript |
| CYP46A4P | 1 | cytochrome P450 family 46 subfamily A meml |
| SKINT1L | 1 | Skint1 like (pseudogene) [Source:HGNC Symbo |
| AL356289.2 | 1 | TEC |
| AL109659.1 | 1 | novel transcript |
| SLC5A9 | 1 | solute carrier family 5 member 9 [Source:HGN |
| AL109659.2 | 1 | novel transcript, antisense to SLC5A9 |
| AL109659.3 | 1 | TEC |
| SPATA6 | 1 | spermatogenesis associated 6 [Source:HGNC S |
| PPP1R8P1 | 1 | protein phosphatase 1 regulatory subunit 8 ps |
| RNU6-723P | 1 | RNA, U6 small nuclear 723, pseudogene [Sour |
| AL356968.2 | 1 | amyotrophic lateral sclerosis 2 (juvenile) chro |
| RNU4-61P | 1 | RNA, U4 small nuclear 61, pseudogene [Sourc |
| AL356968.1 | 1 | ribosomal protein L21 (RPL21) pseudogene |
| AGBL4 | 1 | ATP/GTP binding protein like 4 [Source:HGNC |
| AL391844.1 | 1 | novel transcript, antisense to AGBL4 |
| BEND5 | 1 | BEN domain containing 5 [Source:HGNC Symb |
| AL645507.1 | 1 | novel transcript, antisense to AGBL4 |
| AC099788.1 | 1 | novel transcript |
| AL590432.1 | 1 | novel transcript |
| AGBL4-IT1 | 1 | AGBL4 intronic transcript 1 [Source:HGNC Syn |
| AL390023.1 | 1 | pseudogene similar to part of ribosomal prote |

| | | |
|---|---|---|
| ZNF859P | 1 | zinc finger protein 859, pseudogene [Source:H |
| AL645730.1 | 1 | novel transcript |
| MTND2P29 | 1 | MT-ND2 pseudogene 29 [Source:HGNC Symbo |
| ELAVL4 | 1 | ELAV like RNA binding protein 4 [Source:HGN( |
| AL583843.1 | 1 | novel pseudogene |
| AL592182.2 | 1 | novel transcript |
| AL592182.1 | 1 | novel transcript |
| LINC02808 | 1 | long intergenic non-protein coding RNA 2808 |
| AL592182.3 | 1 | novel transcript |
| AL596275.1 | 1 | ribosomal protein S2 (RPS2) pseudogene |
| HMGB1P45 | 1 | high mobility group box 1 pseudogene 45 [Sou |
| FCF1P6 | 1 | FCF1 pseudogene 6 [Source:HGNC Symbol;Acc |
| DMRTA2 | 1 | DMRT like family A2 [Source:HGNC Symbol;Ac |
| AL049637.2 | 1 | novel transcript |
| FAF1 | 1 | Fas associated factor 1 [Source:HGNC Symbol; |
| AL049637.1 | 1 | novel transcript |
| RNU6-1026P | 1 | RNA, U6 small nuclear 1026, pseudogene [Sou |
| PHBP12 | 1 | prohibitin pseudogene 12 [Source:HGNC Symb |
| MRPS6P2 | 1 | mitochondrial ribosomal protein S6 pseudoge |
| CDKN2C | 1 | cyclin dependent kinase inhibitor 2C [Source:H |
| MIR4421 | 1 | microRNA 4421 [Source:HGNC Symbol;Acc:HG |
| MIR6500 | 1 | microRNA 6500 [Source:HGNC Symbol;Acc:HG |
| C1orf185 | 1 | chromosome 1 open reading frame 185 [Sour |
| Y_RNA | 1 | Y RNA [Source:RFAM;Acc:RF00019] |
| CFL1P2 | 1 | cofilin 1 pseudogene 2 [Source:HGNC Symbol; |
| LINC01562 | 1 | long intergenic non-protein coding RNA 1562 |
| RNF11 | 1 | ring finger protein 11 [Source:HGNC Symbol;A |
| AL162430.1 | 1 | ribosomal protein S2 (RPS2) pseudogene |
| AL162430.2 | 1 | novel transcript |
| TTC39A | 1 | tetratricopeptide repeat domain 39A [Source: |
| TTC39A-AS1 | 1 | TTC39A antisense RNA 1 [Source:HGNC Symbo |
| EPS15 | 1 | epidermal growth factor receptor pathway su |
| AL671986.1 | 1 | proteasome (prosome, macropain) 26S subun |
| RNU6-877P | 1 | RNA, U6 small nuclear 877, pseudogene [Sour |
| AC104170.2 | 1 | novel transcript |
| AC104170.1 | 1 | novel transcript |
| RNU6-1281P | 1 | RNA, U6 small nuclear 1281, pseudogene [Sou |
| CALR4P | 1 | calreticulin 4, pseudogene [Source:HGNC Sym |
| OSBPL9 | 1 | oxysterol binding protein like 9 [Source:HGNC |
| GAPDHP51 | 1 | glyceraldehyde 3 phosphate dehydrogenase p |
| SLC25A6P3 | 1 | solute carrier family 25 member 6 pseudogen |
| AL050343.1 | 1 | NULL |
| NRDC | 1 | nardilysin convertase [Source:HGNC Symbol;A |
| AL050343.2 | 1 | novel transcript, antisense to NRD1 |
| AL050343.3 | 1 | novel transcript, antisense to NRD1 |
| MIR761 | 1 | microRNA 761 [Source:HGNC Symbol;Acc:HGI |
| TSEN15P2 | 1 | tRNA splicing endonuclease subunit 15 pseud |
| RAB3B | 1 | RAB3B, member RAS oncogene family [Source |
| RNA5SP48 | 1 | RNA, 5S ribosomal pseudogene 48 [Source:HG |
| AL445685.2 | 1 | 40s ribosomal protein S26 (RPS26) pseudogen |

| | | |
|---|---|---|
| RN7SL290P | 1 | RNA, 7SL, cytoplasmic 290, pseudogene [Sour |
| TXNDC12 | 1 | thioredoxin domain containing 12 [Source:HG |
| AL445685.3 | 1 | novel transcript |
| KTI12 | 1 | KTI12 chromatin associated homolog [Source: |
| AL445685.1 | 1 | novel transcript |
| TXNDC12-AS1 | 1 | TXNDC12 antisense RNA 1 [Source:HGNC Sym |
| BTF3L4 | 1 | basic transcription factor 3 like 4 [Source:HGN |
| ZFYVE9 | 1 | zinc finger FYVE-type containing 9 [Source:HG |
| RN7SL788P | 1 | RNA, 7SL, cytoplasmic 788, pseudogene [Sour |
| AL139156.2 | 1 | pseudogene similar to part of alcohol dehydro |
| AL139156.1 | 1 | transformer-2 alpha (TRA2A) pseudogene |
| PDCL3P6 | 1 | PDCL3 pseudogene 6 [Source:HGNC Symbol;A |
| AL139156.3 | 1 | pseudogene similar to part of ribosomal prote |
| DNAJC19P7 | 1 | DnaJ heat shock protein family (Hsp40) memb |
| ANAPC10P1 | 1 | anaphase promoting complex subunit 10 pseu |
| CC2D1B | 1 | coiled-coil and C2 domain containing 1B [Sour |
| AL513218.1 | 1 | novel transcript, antisense to CC2D1B |
| AL513218.2 | 1 | novel transcript, antisense to CC2D1B |
| PLA2G12AP1 | 1 | phospholipase A2 group XIIA pseudogene 1 [S |
| ORC1 | 1 | origin recognition complex subunit 1 [Source: |
| PRPF38A | 1 | pre-mRNA processing factor 38A [Source:HGN |
| TUT4 | 1 | terminal uridylyl transferase 4 [Source:HGNC S |
| AL591167.1 | 1 | novel transcript |
| EEF1GP7 | 1 | eukaryotic translation elongation factor 1 gam |
| GPX7 | 1 | glutathione peroxidase 7 [Source:HGNC Symb |
| SHISAL2A | 1 | shisa like 2A [Source:HGNC Symbol;Acc:HGNC |
| COA7 | 1 | cytochrome c oxidase assembly factor 7 (puta |
| NDUFS5P3 | 1 | NADH:ubiquinone oxidoreductase subunit S5 |
| RN7SL62P | 1 | RNA, 7SL, cytoplasmic 62, pseudogene [Sourc |
| ZYG11B | 1 | zyg-11 family member B, cell cycle regulator [S |
| RNU2-30P | 1 | RNA, U2 small nuclear 30, pseudogene [Sourc |
| RPS13P2 | 1 | ribosomal protein S13 pseudogene 2 [Source: |
| RNU6-969P | 1 | RNA, U6 small nuclear 969, pseudogene [Sour |
| ZYG11A | 1 | zyg-11 family member A, cell cycle regulator [ |
| AC099677.4 | 1 | novel transcript, antisense to ZYG11A |
| ECHDC2 | 1 | enoyl-CoA hydratase domain containing 2 [So |
| AC099677.1 | 1 | translational machinery associated 7 homolog |
| AC099677.2 | 1 | chromosome 14 open reading frame 2 (C14or |
| SCP2 | 1 | sterol carrier protein 2 [Source:HGNC Symbol; |
| AC099677.5 | 1 | NULL |
| AC099677.7 | 1 | NULL |
| AC099677.6 | 1 | NULL |
| AC099677.3 | 1 | H3 histone family pseudogene |
| AL445183.3 | 1 | pseudogene similar to part of related RAS vira |
| TUBBP10 | 1 | tubulin beta class I pseudogene 10 [Source:HG |
| PODN | 1 | podocan [Source:HGNC Symbol;Acc:HGNC:23 |
| AL445183.1 | 1 | novel transcript |
| HIGD1AP11 | 1 | HIG1 hypoxia inducible domain family membe |
| SLC1A7 | 1 | solute carrier family 1 member 7 [Source:HGN |
| AL445183.2 | 1 | novel transcript |

| | | |
|---|---|---|
| AL606760.4 | 1 | carnitine palmitoyltransferase 2 (CPT2) pseud |
| CPT2 | 1 | carnitine palmitoyltransferase 2 [Source:HGN |
| AL606760.2 | 1 | novel transcript, antisense to CPT2 |
| CZIB | 1 | CXXC motif containing zinc binding protein [Sc |
| AL606760.3 | 1 | novel transcript, antisense to C1orf123 & MA |
| MAGOH | 1 | mago homolog, exon junction complex subun |
| AL606760.1 | 1 | novel transcript, antisense to MAGOH |
| LRP8 | 1 | LDL receptor related protein 8 [Source:HGNC |
| AL355483.3 | 1 | novel transcript |
| AL355483.1 | 1 | novel transcript |
| AL355483.2 | 1 | novel transcript |
| LINC01771 | 1 | long intergenic non-protein coding RNA 1771 |
| AC119428.2 | 1 | novel transcript |
| AC119428.1 | 1 | novel transcript |
| LINC02812 | 1 | long intergenic non-protein coding RNA 2812 |
| SLC25A3P1 | 1 | solute carrier family 25 member 3 pseudogen |
| DMRTB1 | 1 | DMRT like family B with proline rich C-termina |
| GLIS1 | 1 | GLIS family zinc finger 1 [Source:HGNC Symbo |
| RNU7-95P | 1 | RNA, U7 small nuclear 95 pseudogene [Source |
| NDC1 | 1 | NDC1 transmembrane nucleoporin [Source:H |
| AL049745.1 | 1 | NULL |
| AL049745.2 | 1 | ribosomal protein L37 (RPL37) pseudogene |
| YIPF1 | 1 | Yip1 domain family member 1 [Source:HGNC |
| DIO1 | 1 | iodothyronine deiodinase 1 [Source:HGNC Syr |
| HSPB11 | 1 | heat shock protein family B (small) member 1 |
| LRRC42 | 1 | leucine rich repeat containing 42 [Source:HGN |
| HNRNPA3P12 | 1 | heterogeneous nuclear ribonucleoprotein A3 |
| LDLRAD1 | 1 | low density lipoprotein receptor class A doma |
| TMEM59 | 1 | transmembrane protein 59 [Source:HGNC Syn |
| AL353898.3 | 1 | TEC |
| TCEANC2 | 1 | transcription elongation factor A N-terminal a |
| MIR4781 | 1 | microRNA 4781 [Source:HGNC Symbol;Acc:HG |
| AL353898.1 | 1 | cytochrome c oxidase subunit VIIb (COX7B) ps |
| AL353898.2 | 1 | NULL |
| AL357673.1 | 1 | novel transcript |
| CDCP2 | 1 | CUB domain containing protein 2 [Source:HGI |
| AL357673.2 | 1 | novel transcript, antisense to CDCP2 |
| CYB5RL | 1 | cytochrome b5 reductase like [Source:HGNC S |
| MRPL37 | 1 | mitochondrial ribosomal protein L37 [Source: |
| SSBP3 | 1 | single stranded DNA binding protein 3 [Source |
| SSBP3-AS1 | 1 | SSBP3 antisense RNA 1 [Source:HGNC Symbol |
| AL161644.1 | 1 | novel transcript |
| AL035415.1 | 1 | novel transcript |
| AC099796.2 | 1 | novel transcript |
| LINC02784 | 1 | long intergenic non-protein coding RNA 2784 |
| AC099796.1 | 1 | transaldolase 1 (TALDO1) pseudogene |
| HNRNPA1P63 | 1 | heterogeneous nuclear ribonucleoprotein A1 |
| ACOT11 | 1 | acyl-CoA thioesterase 11 [Source:HGNC Symb |
| FAM151A | 1 | family with sequence similarity 151 member |
| AL590093.1 | 1 | novel transcript |

| | | |
|---|---|---|
| MROH7 | 1 | maestro heat like repeat family member 7 [Sc |
| MROH7-TTC4 | 1 | MROH7-TTC4 readthrough (NMD candidate) [ |
| TTC4 | 1 | tetratricopeptide repeat domain 4 [Source:HG |
| PARS2 | 1 | prolyl-tRNA synthetase 2, mitochondrial [Sour |
| TTC22 | 1 | tetratricopeptide repeat domain 22 [Source:H |
| AC096536.1 | 1 | novel transcript, antisense to TTC22 |
| LEXM | 1 | lymphocyte expansion molecule [Source:HGN |
| DHCR24 | 1 | 24-dehydrocholesterol reductase [Source:HGI |
| AC096536.3 | 1 | novel transcript, antisense to DHCR24 |
| AC096536.2 | 1 | novel transcript |
| DHCR24-DT | 1 | DHCR24 divergent transcript [Source:HGNC Sy |
| AL590440.2 | 1 | novel transcript |
| TMEM61 | 1 | transmembrane protein 61 [Source:HGNC Syn |
| AL590440.1 | 1 | novel transcript |
| BSND | 1 | barttin CLCNK type accessory beta subunit [Sc |
| PCSK9 | 1 | proprotein convertase subtilisin/kexin type 9 |
| USP24 | 1 | ubiquitin specific peptidase 24 [Source:HGNC |
| MIR4422HG | 1 | MIR4422 host gene [Source:HGNC Symbol;Acc |
| GYG1P3 | 1 | glycogenin 1 pseudogene 3 [Source:HGNC Syn |
| MIR4422 | 1 | microRNA 4422 [Source:HGNC Symbol;Acc:HG |
| AL603840.1 | 1 | novel transcript |
| GOT2P1 | 1 | glutamic-oxaloacetic transaminase 2 pseudog |
| MTCO2P34 | 1 | MT-CO2 pseudogene 34 [Source:HGNC Symbo |
| RN7SKP291 | 1 | RN7SK pseudogene 291 [Source:HGNC Symbo |
| RNU6-830P | 1 | RNA, U6 small nuclear 830, pseudogene [Sour |
| Y_RNA | 1 | Y RNA [Source:RFAM;Acc:RF00019] |
| LINC01755 | 1 | long intergenic non-protein coding RNA 1755 |
| LINC01753 | 1 | long intergenic non-protein coding RNA 1753 |
| PIGQP1 | 1 | phosphatidylinositol glycan anchor biosynthes |
| AL353681.1 | 1 | novel transcript |
| AC119674.1 | 1 | novel transcript |
| AC119674.2 | 1 | novel transcript |
| RPSAP20 | 1 | ribosomal protein SA pseudogene 20 [Source: |
| LINC01767 | 1 | long intergenic non-protein coding RNA 1767 |
| PLPP3 | 1 | phospholipid phosphatase 3 [Source:HGNC Sy |
| RPL21P23 | 1 | ribosomal protein L21 pseudogene 23 [Source |
| RPL23AP85 | 1 | ribosomal protein L23a pseudogene 85 [Sourc |
| AC099789.1 | 1 | riribosomal protein S2 (RPS2) pseudogene |
| PRKAA2 | 1 | protein kinase AMP-activated catalytic subuni |
| FYB2 | 1 | FYN binding protein 2 [Source:HGNC Symbol;/ |
| AL360295.1 | 1 | novel transcript |
| C8A | 1 | complement C8 alpha chain [Source:HGNC Syr |
| C8B | 1 | complement C8 beta chain [Source:HGNC Sym |
| AL161740.1 | 1 | novel transcript |
| DAB1 | 1 | DAB adaptor protein 1 [Source:HGNC Symbol; |
| AL390243.1 | 1 | novel transcript |
| RPS20P5 | 1 | ribosomal protein S20 pseudogene 5 [Source: |
| DAB1-AS1 | 1 | DAB1 antisense RNA 1 [Source:HGNC Symbol; |
| HNRNPA1P6 | 1 | heterogeneous nuclear ribonucleoprotein A1 |
| RPS26P15 | 1 | ribosomal protein S26 pseudogene 15 [Source |

| | | |
|---|---|---|
| AL445193.2 | 1 | novel transcript |
| AL445193.1 | 1 | pseudogene similar to part of WAP four-disulf |
| AL357373.1 | 1 | novel pseudogene |
| OMA1 | 1 | OMA1 zinc metallopeptidase [Source:HGNC Sy |
| AL035411.3 | 1 | novel transcript, antisense to OMA1 |
| AL035411.2 | 1 | zinc finger protein 280 pseudogene |
| RN7SL713P | 1 | RNA, 7SL, cytoplasmic 713, pseudogene [Sour |
| TACSTD2 | 1 | tumor associated calcium signal transducer 2 |
| AL035411.1 | 1 | adenylate kinase 2 (AK2) pseudogene |
| MYSM1 | 1 | Myb like, SWIRM and MPN domains 1 [Source |
| AL136985.3 | 1 | novel transcript |
| JUN | 1 | Jun proto-oncogene, AP-1 transcription factor |
| LINC01135 | 1 | long intergenic non-protein coding RNA 1135 |
| AL136985.2 | 1 | novel transcript |
| AL136985.1 | 1 | novel transcript |
| LINC02777 | 1 | long intergenic non-protein coding RNA 2777 |
| LINC01358 | 1 | long intergenic non-protein coding RNA 1358 |
| PHBP3 | 1 | prohibitin pseudogene 3 [Source:HGNC Symbo |
| AL592431.2 | 1 | novel transcript |
| AL592431.1 | 1 | novel transcript |
| AC093425.1 | 1 | novel transcript |
| AC093424.1 | 1 | novel transcript |
| FGGY | 1 | FGGY carbohydrate kinase domain containing |
| MIR4711 | 1 | microRNA 4711 [Source:HGNC Symbol;Acc:HG |
| AL035416.1 | 1 | novel transcript |
| HOOK1 | 1 | hook microtubule tethering protein 1 [Source |
| CYP2J2 | 1 | cytochrome P450 family 2 subfamily J membe |
| RN7SL475P | 1 | RNA, 7SL, cytoplasmic 475, pseudogene [Sour |
| C1orf87 | 1 | chromosome 1 open reading frame 87 [Source |
| PGBD4P8 | 1 | piggyBac transposable element derived 4 pseu |
| LINC02778 | 1 | long intergenic non-protein coding RNA 2778 |
| LINC01748 | 1 | long intergenic non-protein coding RNA 1748 |
| AC099792.1 | 1 | novel transcript |
| NFIA | 1 | nuclear factor I A [Source:HGNC Symbol;Acc:H |
| NFIA-AS2 | 1 | NFIA antisense RNA 2 [Source:HGNC Symbol;A |
| NFIA-AS1 | 1 | NFIA antisense RNA 1 [Source:HGNC Symbol;A |
| AC099791.3 | 1 | novel transcript |
| AC099791.4 | 1 | novel transcript |
| AC099791.1 | 1 | sepiapterin reductase (7,8-dihydrobiopterin:N |
| TM2D1 | 1 | TM2 domain containing 1 [Source:HGNC Symb |
| AC099791.2 | 1 | novel transcript |
| PATJ | 1 | PATJ crumbs cell polarity complex component |
| RNU6-414P | 1 | RNA, U6 small nuclear 414, pseudogene [Sour |
| RNU6-1177P | 1 | RNA, U6 small nuclear 1177, pseudogene [Sou |
| LAMTOR5P1 | 1 | late endosomal/lysosomal adaptor, MAPK anc |
| RN7SL180P | 1 | RNA, 7SL, cytoplasmic 180, pseudogene [Sour |
| MIR3116-1 | 1 | microRNA 3116-1 [Source:HGNC Symbol;Acc:I |
| MIR3116-2 | 1 | microRNA 3116-2 [Source:HGNC Symbol;Acc:I |
| PIGPP2 | 1 | phosphatidylinositol glycan anchor biosynthes |
| RPS15AP7 | 1 | ribosomal protein S15a pseudogene 7 [Source |

| | | |
|---|---|---|
| L1TD1 | 1 | LINE1 type transposase domain containing 1 [ |
| AL162739.1 | 1 | ribosomal protein L36 (RPL36) pseudogene |
| Y_RNA | 1 | Y RNA [Source:RFAM;Acc:RF00019] |
| KANK4 | 1 | KN motif and ankyrin repeat domains 4 [Sourc |
| RNU6-371P | 1 | RNA, U6 small nuclear 371, pseudogene [Sour |
| USP1 | 1 | ubiquitin specific peptidase 1 [Source:HGNC S |
| DOCK7 | 1 | dedicator of cytokinesis 7 [Source:HGNC Symb |
| AL451044.1 | 1 | ubiquitin A-52 residue ribosomal protein fusic |
| ANGPTL3 | 1 | angiopoietin like 3 [Source:HGNC Symbol;Acc: |
| AL138847.2 | 1 | novel transcript |
| AL138847.1 | 1 | ribosomal protein L13a (RPL13A) pseudogene |
| AC103923.1 | 1 | novel transcript |
| ATG4C | 1 | autophagy related 4C cysteine peptidase [Sou |
| AC099794.1 | 1 | novel transcript |
| AC099794.2 | 1 | novel pseudogene |
| LINC01739 | 1 | long intergenic non-protein coding RNA 1739 |
| AL162400.3 | 1 | novel transcript |
| AL162400.1 | 1 | novel transcript |
| AL162400.2 | 1 | novel transcript |
| AC096543.1 | 1 | novel transcript |
| LINC00466 | 1 | long intergenic non-protein coding RNA 466 [S |
| RNA5SP49 | 1 | RNA, 5S ribosomal pseudogene 49 [Source:HG |
| AC096543.2 | 1 | novel transcript |
| RPSAP65 | 1 | ribosomal protein SA pseudogene 65 [Source: |
| FOXD3-AS1 | 1 | FOXD3 antisense RNA 1 [Source:HGNC Symbo |
| FOXD3 | 1 | forkhead box D3 [Source:HGNC Symbol;Acc:H |
| MIR6068 | 1 | microRNA 6068 [Source:HGNC Symbol;Acc:HG |
| Y_RNA | 1 | Y RNA [Source:RFAM;Acc:RF00019] |
| AL049636.1 | 1 | DnaJ (Hsp40) homolog, subfamily C, member |
| ALG6 | 1 | ALG6 alpha-1,3-glucosyltransferase [Source:H |
| ITGB3BP | 1 | integrin subunit beta 3 binding protein [Sourc |
| BX004807.1 | 1 | novel transcript, antisense to ITGB3BP |
| EFCAB7 | 1 | EF-hand calcium binding domain 7 [Source:HG |
| RN7SL488P | 1 | RNA, 7SL, cytoplasmic 488, pseudogene [Sour |
| RNU7-123P | 1 | RNA, U7 small nuclear 123 pseudogene [Sourc |
| DLEU2L | 1 | deleted in lymphocytic leukemia 2 like [Source |
| AL109925.3 | 1 | NULL |
| AL109925.6 | 1 | NULL |
| AL109925.4 | 1 | NULL |
| AL109925.2 | 1 | NULL |
| AL109925.5 | 1 | NULL |
| AL109925.1 | 1 | NULL |
| PGM1 | 1 | phosphoglucomutase 1 [Source:HGNC Symbo |
| RN7SL130P | 1 | RNA, 7SL, cytoplasmic 130, pseudogene [Sour |
| ROR1 | 1 | receptor tyrosine kinase like orphan receptor |
| AL161742.1 | 1 | ribosomal protein L19 (RPL19) pseudogene |
| CFL1P3 | 1 | cofilin 1 pseudogene 3 [Source:HGNC Symbol; |
| RNU6-809P | 1 | RNA, U6 small nuclear 809, pseudogene [Sour |
| Y_RNA | 1 | Y RNA [Source:RFAM;Acc:RF00019] |
| ROR1-AS1 | 1 | ROR1 antisense RNA 1 [Source:HGNC Symbol; |

| | | |
|---|---|---|
| AL445205.1 | 1 | novel transcript |
| UBE2U | 1 | ubiquitin conjugating enzyme E2 U [Source:H( |
| RNU7-62P | 1 | RNA, U7 small nuclear 62 pseudogene [Source |
| CACHD1 | 1 | cache domain containing 1 [Source:HGNC Syn |
| MIR4794 | 1 | microRNA 4794 [Source:HGNC Symbol;Acc:H( |
| RAVER2 | 1 | ribonucleoprotein, PTB binding 2 [Source:HGN |
| JAK1 | 1 | Janus kinase 1 [Source:HGNC Symbol;Acc:HGN |
| AL606517.2 | 1 | NADH dehydrogenase (ubiquinone) 1 alpha su |
| AL606517.1 | 1 | SIPL protein (SIPL) pseudogene |
| LINC01359 | 1 | long intergenic non-protein coding RNA 1359 |
| SLC2A3P2 | 1 | solute carrier family 2 member 3 pseudogene |
| AL357078.2 | 1 | novel transcript |
| RNU6-1176P | 1 | RNA, U6 small nuclear 1176, pseudogene [Sou |
| MIR3671 | 1 | microRNA 3671 [Source:HGNC Symbol;Acc:H( |
| MIR101-1 | 1 | microRNA 101-1 [Source:HGNC Symbol;Acc:H |
| AL357078.1 | 1 | ribosomal protein S24 (RPS24) pseudogene |
| MRPS21P1 | 1 | mitochondrial ribosomal protein S21 pseudog |
| AK4 | 1 | adenylate kinase 4 [Source:HGNC Symbol;Acc |
| RPS29P7 | 1 | ribosomal protein S29 pseudogene 7 [Source: |
| DNAJC6 | 1 | DnaJ heat shock protein family (Hsp40) memb |
| AL139294.1 | 1 | novel transcript |
| COX6CP13 | 1 | cytochrome c oxidase subunit 6C pseudogene |
| RNU2-15P | 1 | RNA, U2 small nuclear 15, pseudogene [Sourc |
| LEPROT | 1 | leptin receptor overlapping transcript [Source |
| LEPR | 1 | leptin receptor [Source:HGNC Symbol;Acc:HG |
| AC119800.1 | 1 | novel transcript |
| AC097063.1 | 1 | UDP-N-acetyl-alpha-D-galactosamine:polypep |
| AL513493.1 | 1 | novel transcript |
| RN7SL854P | 1 | RNA, 7SL, cytoplasmic 854, pseudogene [Sour |
| PDE4B | 1 | phosphodiesterase 4B [Source:HGNC Symbol; |
| AL590783.1 | 1 | novel transcript |
| RNU4-88P | 1 | RNA, U4 small nuclear 88, pseudogene [Sourc |
| SGIP1 | 1 | SH3 domain GRB2 like endophilin interacting |
| MIR3117 | 1 | microRNA 3117 [Source:HGNC Symbol;Acc:H( |
| AL139147.1 | 1 | novel transcript, antisense to SGIP1 |
| TCTEX1D1 | 1 | Tctex1 domain containing 1 [Source:HGNC Syı |
| INSL5 | 1 | insulin like 5 [Source:HGNC Symbol;Acc:HGNC |
| WDR78 | 1 | WD repeat domain 78 [Source:HGNC Symbol; |
| AL592161.1 | 1 | novel transcript |
| MIER1 | 1 | MIER1 transcriptional regulator [Source:HGN( |
| SLC35D1 | 1 | solute carrier family 35 member D1 [Source:H |
| C1orf141 | 1 | chromosome 1 open reading frame 141 [Sour |
| AL133320.1 | 1 | NULL |
| AL133320.2 | 1 | novel transcript, sense intronic to C1orf141 |
| IL23R | 1 | interleukin 23 receptor [Source:HGNC Symbol |
| RNU6-586P | 1 | RNA, U6 small nuclear 586, pseudogene [Sour |
| RNU4ATAC4P | 1 | RNA, U4atac small nuclear 4, pseudogene [So |
| AL389925.1 | 1 | DnaJ (Hsp40) homolog, subfamily B, member |
| IL12RB2 | 1 | interleukin 12 receptor subunit beta 2 [Source |
| SERBP1 | 1 | SERPINE1 mRNA binding protein 1 [Source:HG |

| | | |
|---|---|---|
| RNU6-387P | 1 | RNA, U6 small nuclear 387, pseudogene [Sour |
| LINC01702 | 1 | long intergenic non-protein coding RNA 1702 |
| RNU6-1031P | 1 | RNA, U6 small nuclear 1031, pseudogene [Sou |
| AL590559.1 | 1 | poly(A) binding protein, cytoplasmic 4 (PABPC |
| RN7SL392P | 1 | RNA, 7SL, cytoplasmic 392, pseudogene [Sour |
| HNRNPCP9 | 1 | heterogeneous nuclear ribonucleoprotein C p |
| GADD45A | 1 | growth arrest and DNA damage inducible alph |
| GNG12 | 1 | G protein subunit gamma 12 [Source:HGNC Sy |
| RNU7-80P | 1 | RNA, U7 small nuclear 80 pseudogene [Source |
| GNG12-AS1 | 1 | GNG12, DIRAS3 and WLS antisense RNA 1 [Sou |
| AL157407.1 | 1 | sirtuin (silent mating type information regulat |
| DIRAS3 | 1 | DIRAS family GTPase 3 [Source:HGNC Symbol; |
| ARL5AP3 | 1 | ADP ribosylation factor like GTPase 5A pseudo |
| WLS | 1 | Wnt ligand secretion mediator [Source:HGNC |
| CTBP2P8 | 1 | C-terminal binding protein 2 pseudogene 8 [S |
| MIR1262 | 1 | microRNA 1262 [Source:HGNC Symbol;Acc:HG |
| RPS7P4 | 1 | ribosomal protein S7 pseudogene 4 [Source:H |
| COX6B1P7 | 1 | cytochrome c oxidase subunit 6B1 pseudogen |
| ELOCP18 | 1 | elongin C pseudogene 18 [Source:HGNC Symb |
| AL139413.1 | 1 | pseudogene similar to part of BET1 homolog ( |
| RPE65 | 1 | retinoid isomerohydrolase RPE65 [Source:HGI |
| DEPDC1 | 1 | DEP domain containing 1 [Source:HGNC Symb |
| AL138789.1 | 1 | novel transcript |
| DEPDC1-AS1 | 1 | DEPDC1 antisense RNA 1 [Source:HGNC Symb |
| TXNP2 | 1 | thioredoxin pseudogene 2 [Source:HGNC Sym |
| AL035412.1 | 1 | novel transcript |
| AL033530.1 | 1 | novel transcript |
| AL691520.1 | 1 | novel transcript |
| LINC01707 | 1 | long intergenic non-protein coding RNA 1707 |
| LINC02791 | 1 | long intergenic non-protein coding RNA 2791 |
| LINC01758 | 1 | long intergenic non-protein coding RNA 1758 |
| AL359894.1 | 1 | novel transcript |
| LRRC7 | 1 | leucine rich repeat containing 7 [Source:HGN( |
| SGO1P1 | 1 | shugoshin 1 pseudogene 1 [Source:HGNC Sym |
| AL117353.1 | 1 | novel transcript, antisense to LRRC7 |
| RN7SL538P | 1 | RNA, 7SL, cytoplasmic 538, pseudogene [Sour |
| PIN1P1 | 1 | peptidylprolyl cis/trans isomerase, NIMA-inte |
| AL158840.1 | 1 | novel transcript |
| LRRC40 | 1 | leucine rich repeat containing 40 [Source:HGN |
| RN7SL242P | 1 | RNA, 7SL, cytoplasmic 242, pseudogene [Sour |
| SRSF11 | 1 | serine and arginine rich splicing factor 11 [Sou |
| AL353771.1 | 1 | novel transcript |
| ANKRD13C | 1 | ankyrin repeat domain 13C [Source:HGNC Syn |
| HHLA3 | 1 | HERV-H LTR-associating 3 [Source:HGNC Symb |
| AL158839.1 | 1 | novel transcript |
| CTH | 1 | cystathionine gamma-lyase [Source:HGNC Syr |
| Metazoa_SRP | 1 | Metazoan signal recognition particle RNA [Sou |
| AL354872.2 | 1 | novel transcript |
| AL354872.1 | 1 | cysteine and histidine-rich domain (CHORD) c |
| CASP3P1 | 1 | caspase 3 pseudogene 1 [Source:HGNC Symbo |

| | | |
|---|---|---|
| LINC01788 | 1 | long intergenic non-protein coding RNA 1788 |
| AL513285.1 | 1 | novel transcript |
| PTGER3 | 1 | prostaglandin E receptor 3 [Source:HGNC Sym |
| AL031429.1 | 1 | novel transcript |
| AL031429.2 | 1 | novel transcript, sense intronic to PTGER3 |
| ZRANB2-AS1 | 1 | ZRANB2 antisense RNA 1 [Source:HGNC Symb |
| ZRANB2 | 1 | zinc finger RANBP2-type containing 2 [Source: |
| MIR186 | 1 | microRNA 186 [Source:HGNC Symbol;Acc:HGI |
| ZRANB2-AS2 | 1 | ZRANB2 antisense RNA 2 (head to head) [Sour |
| AL358453.1 | 1 | cytochrome c oxidase subunit VIa polypeptide |
| NEGR1 | 1 | neuronal growth regulator 1 [Source:HGNC Sy |
| AL354949.1 | 1 | novel transcript |
| AL359821.1 | 1 | kinesin family member 2C (KIF2C) pseudogene |
| NEGR1-IT1 | 1 | NEGR1 intronic transcript 1 [Source:HGNC Syr |
| GDI2P2 | 1 | GDP dissociation inhibitor 2 pseudogene 2 [Sc |
| AL513166.1 | 1 | novel transcript |
| RPL31P12 | 1 | ribosomal protein L31 pseudogene 12 [Source |
| AL583808.1 | 1 | novel transcript |
| RNU6-1246P | 1 | RNA, U6 small nuclear 1246, pseudogene [Sou |
| LINC02796 | 1 | long intergenic non-protein coding RNA 2796 |
| LINC02797 | 1 | long intergenic non-protein coding RNA 2797 |
| AL732618.1 | 1 | TEC |
| KRT8P21 | 1 | keratin 8 pseudogene 21 [Source:HGNC Symb |
| RN7SKP19 | 1 | RN7SK pseudogene 19 [Source:HGNC Symbol; |
| LINC01360 | 1 | long intergenic non-protein coding RNA 1360 |
| LINC02238 | 1 | long intergenic non-protein coding RNA 2238 |
| RNA5SP50 | 1 | RNA, 5S ribosomal pseudogene 50 [Source:HG |
| AL591463.1 | 1 | novel transcript |
| RNU4ATAC8P | 1 | RNA, U4atac small nuclear 8, pseudogene [Sou |
| LRRIQ3 | 1 | leucine rich repeats and IQ motif containing 3 |
| FPGT | 1 | fucose-1-phosphate guanylyltransferase [Sour |
| FPGT-TNNI3K | 1 | FPGT-TNNI3K readthrough [Source:HGNC Sym |
| TNNI3K | 1 | TNNI3 interacting kinase [Source:HGNC Symb |
| AC093158.1 | 1 | novel transcript |
| AC105271.1 | 1 | novel transcript |
| LRRC53 | 1 | leucine rich repeat containing 53 [Source:HGN |
| ERICH3 | 1 | glutamate rich 3 [Source:HGNC Symbol;Acc:H |
| ERICH3-AS1 | 1 | ERICH3 antisense RNA 1 [Source:HGNC Symbo |
| AC135803.1 | 1 | novel transcript |
| CRYZ | 1 | crystallin zeta [Source:HGNC Symbol;Acc:HGN |
| TYW3 | 1 | tRNA-yW synthesizing protein 3 homolog [Sou |
| AC133865.1 | 1 | novel transcript |
| AC099786.3 | 1 | novel transcript |
| AC099786.1 | 1 | novel transcript, antisense to LHX8 |
| LHX8 | 1 | LIM homeobox 8 [Source:HGNC Symbol;Acc:H |
| AC099786.2 | 1 | novel transcript, antisense to LHX8 |
| RNU6-622P | 1 | RNA, U6 small nuclear 622, pseudogene [Sour |
| SLC44A5 | 1 | solute carrier family 44 member 5 [Source:HG |
| AC093156.1 | 1 | pseudogene similar to part of nucleostemin (N |
| RNU6-503P | 1 | RNA, U6 small nuclear 503, pseudogene [Sour |

| | | |
|---|---|---|
| AL096829.1 | 1 | ribosomal protein L29 (RPL29) pseudogene |
| AL137803.1 | 1 | histidine acid phosphatase domain containing |
| ACADM | 1 | acyl-CoA dehydrogenase medium chain [Sour |
| DLSTP1 | 1 | dihydrolipoamide S-succinyltransferase pseud |
| RABGGTB | 1 | Rab geranylgeranyltransferase subunit beta [S |
| SNORD45C | 1 | small nucleolar RNA, C/D box 45C [Source:HG |
| SNORD45A | 1 | small nucleolar RNA, C/D box 45A [Source:HG |
| SNORD45B | 1 | small nucleolar RNA, C/D box 45B [Source:HG |
| MSH4 | 1 | mutS homolog 4 [Source:HGNC Symbol;Acc:H |
| ASB17 | 1 | ankyrin repeat and SOCS box containing 17 [S |
| AC092813.1 | 1 | novel transcript |
| AC092813.2 | 1 | novel transcript |
| ST6GALNAC3 | 1 | ST6 N-acetylgalactosaminide alpha-2,6-sialyltr |
| AC103592.1 | 1 | peroxiredoxin 5 (PRDX5) pseudogene |
| AC104458.1 | 1 | novel transcript |
| TPI1P1 | 1 | triosephosphate isomerase 1 pseudogene 1 [S |
| RNU6-161P | 1 | RNA, U6 small nuclear 161, pseudogene [Sour |
| LINC02567 | 1 | long intergenic non-protein coding RNA 2567 |
| ST6GALNAC5 | 1 | ST6 N-acetylgalactosaminide alpha-2,6-sialyltr |
| MIR7156 | 1 | microRNA 7156 [Source:HGNC Symbol;Acc:HG |
| AL035409.1 | 1 | novel transcript |
| PIGK | 1 | phosphatidylinositol glycan anchor biosynthes |
| AC113935.1 | 1 | ribosomal protein L17 (RPL17) pseudogene |
| AC093433.1 | 1 | novel pseudogene |
| AC093433.2 | 1 | novel transcript |
| AK5 | 1 | adenylate kinase 5 [Source:HGNC Symbol;Acc |
| AC095030.1 | 1 | novel transcript |
| RNU7-8P | 1 | RNA, U7 small nuclear 8 pseudogene [Source: |
| AC093575.1 | 1 | novel transcript, antisense to AK5 |
| AC118549.1 | 1 | zinc finger ZZ-type containing 3 [Source:NCBI |
| RNA5SP20 | 1 | RNA, 5S ribosomal pseudogene 20 [Source:HG |
| RN7SL370P | 1 | RNA, 7SL, cytoplasmic 370, pseudogene [Sour |
| USP33 | 1 | ubiquitin specific peptidase 33 [Source:HGNC |
| ACTG1P21 | 1 | actin gamma 1 pseudogene 21 [Source:HGNC |
| MIGA1 | 1 | mitoguardin 1 [Source:HGNC Symbol;Acc:HGN |
| RNA5SP21 | 1 | RNA, 5S ribosomal pseudogene 21 [Source:HG |
| AC138392.1 | 1 | novel pseudogene |
| NSRP1P1 | 1 | nuclear speckle splicing regulatory protein 1 p |
| HSPE1P25 | 1 | heat shock protein family E (Hsp10) member |
| NEXN-AS1 | 1 | NEXN antisense RNA 1 [Source:HGNC Symbol; |
| NEXN | 1 | nexilin F-actin binding protein [Source:HGNC S |
| FUBP1 | 1 | far upstream element binding protein 1 [Sour |
| DNAJB4 | 1 | DnaJ heat shock protein family (Hsp40) memb |
| GIPC2 | 1 | GIPC PDZ domain containing family member 2 |
| AC103591.3 | 1 | novel transcript, antisense to DNAJB4 |
| AC103591.4 | 1 | novel transcript |
| AC103591.2 | 1 | Nit protein 2 (NIT2) pseudogene |
| RNU6-1102P | 1 | RNA, U6 small nuclear 1102, pseudogene [Sou |
| AC103591.1 | 1 | signal sequence receptor, gamma (translocon |
| RNA5SP22 | 1 | RNA, 5S ribosomal pseudogene 22 [Source:HG |

| | | |
|---|---|---|
| RNFT1P2 | 1 | ring finger protein, transmembrane 1 pseudog |
| AC096531.2 | 1 | novel transcript |
| PTGFR | 1 | prostaglandin F receptor [Source:HGNC Symb |
| AC096531.1 | 1 | pseudogene fragment similar to putative dim |
| RNA5SP23 | 1 | RNA, 5S ribosomal pseudogene 23 [Source:HG |
| IFI44L | 1 | interferon induced protein 44 like [Source:HG |
| IFI44 | 1 | interferon induced protein 44 [Source:HGNC S |
| AC104837.2 | 1 | ribosomal protein L23 (RPL23) pseudogene |
| AC104837.1 | 1 | SKI-interacting protein (SNW1) pseudogene |
| ADGRL4 | 1 | adhesion G protein-coupled receptor L4 [Sour |
| PSAT1P3 | 1 | phosphoserine aminotransferase 1 pseudoger |
| AL353651.2 | 1 | novel transcript |
| AL353651.1 | 1 | novel transcript |
| LINC02792 | 1 | long intergenic non-protein coding RNA 2792 |
| ADH5P2 | 1 | ADH5 pseudogene 2 [Source:HGNC Symbol;Ac |
| AC099671.1 | 1 | novel transcript |
| AC098657.1 | 1 | PAI-1 mRNA-binding protein (PAI-RBP1) (CGI-! |
| AC098657.2 | 1 | novel transcript |
| AC098657.3 | 1 | succinate-CoA ligase, ADP-forming, beta subu |
| HMGB1P18 | 1 | high mobility group box 1 pseudogene 18 [Sou |
| AL606519.1 | 1 | novel transcript |
| HNRNPA1P64 | 1 | heterogeneous nuclear ribonucleoprotein A1 |
| AL596276.1 | 1 | ubiquitin-conjugating enzyme E2 variant 1 (UE |
| AL596276.2 | 1 | cytochrome c oxidase subunit V1a polypeptid |
| AL136234.1 | 1 | novel transcript |
| LINC01781 | 1 | long intergenic non-protein coding RNA 1781 |
| MTND2P30 | 1 | MT-ND2 pseudogene 30 [Source:HGNC Symbo |
| RPL7P10 | 1 | ribosomal protein L7 pseudogene 10 [Source: |
| AL357632.1 | 1 | ribosomal protein L10a (RPL10A) pseudogene |
| AC093154.1 | 1 | novel transcript |
| RN7SKP247 | 1 | RN7SK pseudogene 247 [Source:HGNC Symbo |
| ADGRL2 | 1 | adhesion G protein-coupled receptor L2 [Sour |
| HNRNPA3P14 | 1 | heterogeneous nuclear ribonucleoprotein A3 |
| AL138799.1 | 1 | dead ringer (Drosophila)-like 2 (bright and dea |
| AL138799.2 | 1 | novel transcript |
| AL138799.4 | 1 | novel transcript |
| MED28P8 | 1 | mediator complex subunit 28 pseudogene 8 [! |
| AC117944.1 | 1 | novel transcript |
| AL138799.3 | 1 | ribosomal protein S20 (RPS20) pseudogene |
| ST13P20 | 1 | ST13, Hsp70 interacting protein pseudogene 2 |
| AL157944.1 | 1 | novel transcript |
| LINC01362 | 1 | long intergenic non-protein coding RNA 1362 |
| LINC01361 | 1 | long intergenic non-protein coding RNA 1361 |
| AC116099.1 | 1 | novel transcript |
| LINC01712 | 1 | long intergenic non-protein coding RNA 1712 |
| LINC01725 | 1 | long intergenic non-protein coding RNA 1725 |
| AL035706.1 | 1 | novel transcript |
| TTLL7 | 1 | tubulin tyrosine ligase like 7 [Source:HGNC Sy |
| TTLL7-IT1 | 1 | TTLL7 intronic transcript 1 [Source:HGNC Sym |
| AC104454.1 | 1 | nuclear factor (eythroid-derived 2)-like 2 (NFE |

| | | |
|---|---|---|
| AC104454.2 | 1 | novel transcript |
| AL359504.2 | 1 | novel transcript |
| PRKACB | 1 | protein kinase cAMP-activated catalytic subur |
| AL359504.1 | 1 | thioredoxin 2 (TXN2) pseudogene |
| AL450063.1 | 1 | neural precursor cell expressed, development |
| AL359273.1 | 1 | NULL |
| SAMD13 | 1 | sterile alpha motif domain containing 13 [Sou |
| AL359273.2 | 1 | novel pseudogene |
| UOX | 1 | urate oxidase (pseudogene) [Source:HGNC Syı |
| DNASE2B | 1 | deoxyribonuclease 2 beta [Source:HGNC Symb |
| AL844170.1 | 1 | novel transcript |
| RPF1 | 1 | ribosome production factor 1 homolog [Sourc |
| GNG5 | 1 | G protein subunit gamma 5 [Source:HGNC Syr |
| AL359762.3 | 1 | novel transcript |
| SPATA1 | 1 | spermatogenesis associated 1 [Source:HGNC S |
| CTBS | 1 | chitobiase [Source:HGNC Symbol;Acc:HGNC:2 |
| AL359762.1 | 1 | novel transcript |
| AL359762.2 | 1 | novel transcript |
| LINC01555 | 1 | long intergenic non-protein coding RNA 1555 |
| AC114482.1 | 1 | ribosomal protein S27-like (RPS27L) pseudoge |
| SSX2IP | 1 | SSX family member 2 interacting protein [Sou |
| AC104169.1 | 1 | novel transcript |
| LPAR3 | 1 | lysophosphatidic acid receptor 3 [Source:HGN |
| MCOLN2 | 1 | mucolipin 2 [Source:HGNC Symbol;Acc:HGNC: |
| WDR63 | 1 | WD repeat domain 63 [Source:HGNC Symbol;A |
| MCOLN3 | 1 | mucolipin 3 [Source:HGNC Symbol;Acc:HGNC: |
| MIR4423 | 1 | microRNA 4423 [Source:HGNC Symbol;Acc:HG |
| AL603750.1 | 1 | TEC |
| SYDE2 | 1 | synapse defective Rho GTPase homolog 2 [Sou |
| C1orf52 | 1 | chromosome 1 open reading frame 52 [Source |
| Y_RNA | 1 | Y RNA [Source:RFAM;Acc:RF00019] |
| BCL10 | 1 | BCL10 immune signaling adaptor [Source:HGN |
| AL078459.1 | 1 | novel transcript |
| DDAH1 | 1 | dimethylarginine dimethylaminohydrolase 1 [ |
| Y_RNA | 1 | Y RNA [Source:RFAM;Acc:RF00019] |
| AL360219.1 | 1 | novel transcript |
| AC092807.3 | 1 | novel transcript |
| AC092807.2 | 1 | novel transcript |
| CCN1 | 1 | cellular communication network factor 1 [Sou |
| SNORD81 | 1 | Small nucleolar RNA SNORD81 [Source:RFAM; |
| AC092807.1 | 1 | novel transcript |
| ZNHIT6 | 1 | zinc finger HIT-type containing 6 [Source:HGN |
| COL24A1 | 1 | collagen type XXIV alpha 1 chain [Source:HGN |
| RNA5SP51 | 1 | RNA, 5S ribosomal pseudogene 51 [Source:HG |
| AC104455.1 | 1 | novel pseudogene |
| LINC02795 | 1 | long intergenic non-protein coding RNA 2795 |
| ODF2L | 1 | outer dense fiber of sperm tails 2 like [Source |
| MIR7856 | 1 | microRNA 7856 [Source:HGNC Symbol;Acc:HG |
| AC119749.1 | 1 | ribosomal protein L7 (RPL7) pseudogene |
| CLCA2 | 1 | chloride channel accessory 2 [Source:HGNC Sy |

| | | |
|---|---|---|
| CLCA1 | 1 | chloride channel accessory 1 [Source:HGNC Sy |
| CLCA4 | 1 | chloride channel accessory 4 [Source:HGNC Sy |
| CDCA4P2 | 1 | cell division cycle associated 4 pseudogene 2 [ |
| CLCA4-AS1 | 1 | CLCA4 antisense RNA 1 [Source:HGNC Symbol |
| CLCA3P | 1 | chloride channel accessory 3, pseudogene [So |
| AL049597.2 | 1 | novel transcript, antisense to SH3GLB1 |
| SH3GLB1 | 1 | SH3 domain containing GRB2 like, endophilin |
| AL049597.1 | 1 | ribosomal protein L17 (RPL17) pseudogene |
| AL355981.1 | 1 | novel transcript |
| SELENOF | 1 | selenoprotein F [Source:HGNC Symbol;Acc:HG |
| HS2ST1 | 1 | heparan sulfate 2-O-sulfotransferase 1 [Sourc |
| AL121989.1 | 1 | novel transcript, antisense to HS2ST1 |
| AC093155.3 | 1 | novel protein |
| AC093155.2 | 1 | pseudogene similar to part of SUMO-1 activat |
| AC093155.1 | 1 | pseudogene similar to part of SUMO-1 activat |
| LINC01140 | 1 | long intergenic non-protein coding RNA 1140 |
| LINC02801 | 1 | long intergenic non-protein coding RNA 2801 |
| LMO4 | 1 | LIM domain only 4 [Source:HGNC Symbol;Acc |
| LINC01364 | 1 | long intergenic non-protein coding RNA 1364 |
| RNA5SP52 | 1 | RNA, 5S ribosomal pseudogene 52 [Source:HG |
| PKN2-AS1 | 1 | PKN2 antisense RNA 1 [Source:HGNC Symbol; |
| AL445437.1 | 1 | TEC |
| AL049861.1 | 1 | transmembrane protein 241 (TMEM241) pseu |
| AL445438.1 | 1 | novel transcript |
| RN7SL583P | 1 | RNA, 7SL, cytoplasmic 583, pseudogene [Sour |
| AC093578.1 | 1 | chromosome 11 open reading frame 23 (C11c |
| AC093578.2 | 1 | NULL |
| RPL36AP10 | 1 | ribosomal protein L36a pseudogene 10 [Sourc |
| PKN2 | 1 | protein kinase N2 [Source:HGNC Symbol;Acc:I |
| RNU6-125P | 1 | RNA, U6 small nuclear 125, pseudogene [Sour |
| ELOCP19 | 1 | elongin C pseudogene 19 [Source:HGNC Symb |
| GTF2B | 1 | general transcription factor IIB [Source:HGNC |
| AL445991.1 | 1 | novel pseudogene |
| KYAT3 | 1 | kynurenine aminotransferase 3 [Source:HGNC |
| RBMXL1 | 1 | RBMX like 1 [Source:HGNC Symbol;Acc:HGNC |
| GBP3 | 1 | guanylate binding protein 3 [Source:HGNC Syr |
| Y_RNA | 1 | Y RNA [Source:RFAM;Acc:RF00019] |
| GBP1 | 1 | guanylate binding protein 1 [Source:HGNC Syr |
| PTGES3P1 | 1 | prostaglandin E synthase 3 pseudogene 1 [Sou |
| GBP2 | 1 | guanylate binding protein 2 [Source:HGNC Syr |
| AC104459.1 | 1 | novel transcript, antisense to GBP7and GBP2 |
| GBP7 | 1 | guanylate binding protein 7 [Source:HGNC Syr |
| GBP4 | 1 | guanylate binding protein 4 [Source:HGNC Syr |
| AC099063.4 | 1 | novel transcript, antisense to GBP4 |
| AC099063.3 | 1 | guanylate binding protein 4 (GBP4) pseudoger |
| GBP5 | 1 | guanylate binding protein 5 [Source:HGNC Syr |
| AC099063.1 | 1 | novel transcript |
| AC099063.2 | 1 | chromosome 11 open reading frame 74 (C11c |
| BX119321.1 | 1 | guanylate binding protein pseudogene |
| GBP6 | 1 | guanylate binding protein family member 6 [S |

| | | |
|---|---|---|
| CAPNS1P1 | 1 | calpain small subunit 1 pseudogene 1 [Source |
| GBP1P1 | 1 | guanylate binding protein 1 pseudogene 1 [So |
| AL596214.1 | 1 | novel transcript, antisense to LRRC8B |
| LRRC8B | 1 | leucine rich repeat containing 8 VRAC subunit |
| LRRC8C-DT | 1 | LRRC8C divergent transcript [Source:HGNC Sy |
| AC093423.2 | 1 | novel transcript |
| LRRC8C | 1 | leucine rich repeat containing 8 VRAC subunit |
| AC093423.3 | 1 | novel transcript, LRRC8C-LRRC8D readthrough |
| AC093423.1 | 1 | NADH dehydrogenase (ubiquinone) 1 alpha su |
| AC099568.1 | 1 | novel transcript |
| AC099568.2 | 1 | novel transcript |
| LRRC8D | 1 | leucine rich repeat containing 8 VRAC subunit |
| AL391497.1 | 1 | novel transcript |
| RN7SKP272 | 1 | RN7SK pseudogene 272 [Source:HGNC Symbo |
| GEMIN8P4 | 1 | gem nuclear organelle associated protein 8 ps |
| ZNF326 | 1 | zinc finger protein 326 [Source:HGNC Symbol; |
| AL161797.1 | 1 | novel transcript |
| AC098656.1 | 1 | novel transcript |
| RNU6-695P | 1 | RNA, U6 small nuclear 695, pseudogene [Sour |
| AL627316.1 | 1 | novel transcript |
| LINC02787 | 1 | long intergenic non-protein coding RNA 2787 |
| SNORD3G | 1 | small nucleolar RNA, C/D box 3G [Source:HGN |
| BARHL2 | 1 | BarH like homeobox 2 [Source:HGNC Symbol; |
| AC092805.1 | 1 | novel transcript |
| LINC02609 | 1 | long intergenic non-protein coding RNA 2609 |
| LINC02788 | 1 | long intergenic non-protein coding RNA 2788 |
| LINC01763 | 1 | long intergenic non-protein coding RNA 1763 |
| AC091614.1 | 1 | novel transcript |
| PHKA1P1 | 1 | phosphorylase kinase regulatory subunit alph |
| ZNF644 | 1 | zinc finger protein 644 [Source:HGNC Symbol; |
| RPL5P6 | 1 | ribosomal protein L5 pseudogene 6 [Source:H |
| HFM1 | 1 | helicase for meiosis 1 [Source:HGNC Symbol; |
| Y_RNA | 1 | Y RNA [Source:RFAM;Acc:RF00019] |
| FEN1P1 | 1 | flap structure-specific endonuclease 1 pseudo |
| CDC7 | 1 | cell division cycle 7 [Source:HGNC Symbol;Acc |
| WDR82P2 | 1 | WD repeat domain 82 pseudogene 2 [Source: |
| AL714022.1 | 1 | ribosomal protein L39 (RPL39) pseudogene |
| HSP90B3P | 1 | heat shock protein 90 beta family member 3, |
| TGFBR3 | 1 | transforming growth factor beta receptor 3 [S |
| RN7SL653P | 1 | RNA, 7SL, cytoplasmic 653, pseudogene [Sour |
| BRDT | 1 | bromodomain testis associated [Source:HGNC |
| EPHX4 | 1 | epoxide hydrolase 4 [Source:HGNC Symbol;Ac |
| LPCAT2BP | 1 | lysophosphatidylcholine acyltransferase 2b, p |
| SETSIP | 1 | SET like protein [Source:HGNC Symbol;Acc:HG |
| BTBD8 | 1 | BTB domain containing 8 [Source:HGNC Symb |
| GAPDHP46 | 1 | glyceraldehyde 3 phosphate dehydrogenase p |
| PRKAR1AP1 | 1 | protein kinase cAMP-dependent type I regula |
| AC104836.1 | 1 | novel transcript |
| AL663109.1 | 1 | zinc finger pseudogene |
| C1orf146 | 1 | chromosome 1 open reading frame 146 [Sour |

| | | |
|---|---|---|
| ACTBP12 | 1 | ACTB pseudogene 12 [Source:HGNC Symbol;A |
| GLMN | 1 | glomulin, FKBP associated protein [Source:HG |
| RPAP2 | 1 | RNA polymerase II associated protein 2 [Sourc |
| RN7SL824P | 1 | RNA, 7SL, cytoplasmic 824, pseudogene [Sour |
| GFI1 | 1 | growth factor independent 1 transcriptional r |
| EVI5 | 1 | ecotropic viral integration site 5 [Source:HGN |
| AL354890.1 | 1 | ACN9 homolog (S. cerevisiae) pseudogene |
| HMGB3P9 | 1 | high mobility group box 3 pseudogene 9 [Sour |
| RNU4-59P | 1 | RNA, U4 small nuclear 59, pseudogene [Sourc |
| AC104332.1 | 1 | novel pseudogene |
| AC104332.2 | 1 | H3 histone, family 3A (H3F3A) pseudogene |
| CCNJP2 | 1 | cyclin J pseudogene 2 [Source:HGNC Symbol;A |
| RPL5 | 1 | ribosomal protein L5 [Source:HGNC Symbol;A |
| DIPK1A | 1 | divergent protein kinase domain 1A [Source:H |
| SNORD21 | 1 | small nucleolar RNA, C/D box 21 [Source:HGN |
| SNORA66 | 1 | small nucleolar RNA, H/ACA box 66 [Source:H |
| AL162740.1 | 1 | NULL |
| AL449283.1 | 1 | GAJ protein pseudogene |
| RNU6-970P | 1 | RNA, U6 small nuclear 970, pseudogene [Sour |
| RN7SL692P | 1 | RNA, 7SL, cytoplasmic 692, pseudogene [Sour |
| AC093577.1 | 1 | novel transcript |
| RNU6-210P | 1 | RNA, U6 small nuclear 210, pseudogene [Sour |
| RN7SKP123 | 1 | RN7SK pseudogene 123 [Source:HGNC Symbo |
| Y_RNA | 1 | Y RNA [Source:RFAM;Acc:RF00019] |
| MTF2 | 1 | metal response element binding transcription |
| TMED5 | 1 | transmembrane p24 trafficking protein 5 [Sou |
| CCDC18 | 1 | coiled-coil domain containing 18 [Source:HGN |
| AC126124.2 | 1 | ribosomal protein L36a (RPL36A) pseudogene |
| AC126124.1 | 1 | pseudogene similar to part of aldo-keto reduc |
| CCDC18-AS1 | 1 | CCDC18 antisense RNA 1 [Source:HGNC Symb |
| AL139421.1 | 1 | ribosomal protein L39 (RPL39) pseudogene |
| DR1 | 1 | down-regulator of transcription 1 [Source:HG |
| AL137159.1 | 1 | pseudogene similar to part of F-box and WD-4 |
| Y_RNA | 1 | Y RNA [Source:RFAM;Acc:RF00019] |
| FNBP1L | 1 | formin binding protein 1 like [Source:HGNC Sy |
| RNA5SP53 | 1 | RNA, 5S ribosomal pseudogene 53 [Source:HG |
| BCAR3 | 1 | BCAR3 adaptor protein, NSP family member [ |
| AL109613.1 | 1 | novel transcript |
| MIR760 | 1 | microRNA 760 [Source:HGNC Symbol;Acc:HGN |
| AL049796.1 | 1 | novel transcript |
| DNTTIP2 | 1 | deoxynucleotidyltransferase terminal interact |
| GCLM | 1 | glutamate-cysteine ligase modifier subunit [Sc |
| CHCHD2P5 | 1 | coiled-coil-helix-coiled-coil-helix domain conta |
| MTND4P11 | 1 | MT-ND4 pseudogene 11 [Source:HGNC Symbo |
| MTND3P21 | 1 | MT-ND3 pseudogene 21 [Source:HGNC Symbo |
| MTCO3P21 | 1 | MT-CO3 pseudogene 21 [Source:HGNC Symbo |
| MTATP6P13 | 1 | MT-ATP6 pseudogene 13 [Source:HGNC Symb |
| AL117351.2 | 1 | mitochondrially encoded cytochrome c oxidas |
| MTCO2P21 | 1 | MT-CO2 pseudogene 21 [Source:HGNC Symbo |
| MTCO1P21 | 1 | MT-CO1 pseudogene 21 [Source:HGNC Symbo |

| | | |
|---|---|---|
| AL117351.1 | 1 | pseudogene similar to part of NADH dehydrog |
| ABCA4 | 1 | ATP binding cassette subfamily A member 4 [S |
| AC093579.1 | 1 | novel transcript |
| ARHGAP29 | 1 | Rho GTPase activating protein 29 [Source:HGI |
| RN7SL440P | 1 | RNA, 7SL, cytoplasmic 440, pseudogene [Sour |
| ARHGAP29-AS1 | 1 | ARHGAP29 antisense RNA 1 [Source:HGNC Syı |
| GAPDHP29 | 1 | glyceraldehyde 3 phosphate dehydrogenase p |
| AC097059.1 | 1 | novel transcript |
| AC118469.1 | 1 | ribosomal protein L21 (RPL21) pseudogene |
| AC118469.2 | 1 | novel transcript |
| ABCD3 | 1 | ATP binding cassette subfamily D member 3 [S |
| F3 | 1 | coagulation factor III, tissue factor [Source:HG |
| AC093117.1 | 1 | novel transcript |
| SLC44A3-AS1 | 1 | SLC44A3 antisense RNA 1 [Source:HGNC Symb |
| KATNBL1P2 | 1 | katanin regulatory subunit B1 like 1 pseudoge |
| MIR378G | 1 | microRNA 378g [Source:HGNC Symbol;Acc:HG |
| SLC44A3 | 1 | solute carrier family 44 member 3 [Source:HG |
| CNN3 | 1 | calponin 3 [Source:HGNC Symbol;Acc:HGNC:2 |
| AC105942.1 | 1 | novel transcript |
| ALG14 | 1 | ALG14 UDP-N-acetylglucosaminyltransferase : |
| AC095033.1 | 1 | novel transcript |
| TLCD4 | 1 | TLC domain containing 4 [Source:HGNC Symb |
| TLCD4-RWDD3 | 1 | TLCD4-RWDD3 readthrough [Source:HGNC Sy |
| AC092802.3 | 1 | novel transcript |
| Y_RNA | 1 | Y RNA [Source:RFAM;Acc:RF00019] |
| AC092802.1 | 1 | novel transcript |
| RWDD3 | 1 | RWD domain containing 3 [Source:HGNC Sym |
| AC092802.2 | 1 | novel transcript, antisense to RWDD3 |
| AC092802.4 | 1 | novel transcript |
| LINC01760 | 1 | long intergenic non-protein coding RNA 1760 |
| LINC01650 | 1 | long intergenic non-protein coding RNA 1650 |
| AL445218.1 | 1 | novel transcript |
| LINC01761 | 1 | long intergenic non-protein coding RNA 1761 |
| LINC02607 | 1 | long intergenic non-protein coding RNA 2607 |
| AL683887.1 | 1 | novel transcript |
| LINC02790 | 1 | long intergenic non-protein coding RNA 2790 |
| RNU1-130P | 1 | RNA, U1 small nuclear 130, pseudogene [Sour |
| LINC01787 | 1 | long intergenic non-protein coding RNA 1787 |
| AC092393.1 | 1 | secretory carrier membrane protein 1 (SCAMI |
| UBE2WP1 | 1 | ubiquitin conjugating enzyme E2 W pseudoge |
| EEF1A1P11 | 1 | eukaryotic translation elongation factor 1 alpl |
| RN7SL831P | 1 | RNA, 7SL, cytoplasmic 831, pseudogene [Sour |
| NDUFS5P2 | 1 | NADH:ubiquinone oxidoreductase subunit S5 |
| RPL7P9 | 1 | ribosomal protein L7 pseudogene 9 [Source:H |
| RN7SKP270 | 1 | RN7SK pseudogene 270 [Source:HGNC Symbo |
| PTBP2 | 1 | polypyrimidine tract binding protein 2 [Source |
| DPYD | 1 | dihydropyrimidine dehydrogenase [Source:HG |
| DPYD-AS1 | 1 | DPYD antisense RNA 1 [Source:HGNC Symbol; |
| DPYD-IT1 | 1 | DPYD intronic transcript 1 [Source:HGNC Syml |
| SEC63P1 | 1 | SEC63 homolog, protein translocation regulat |

| | | |
|---|---|---|
| RPL26P9 | 1 | ribosomal protein L26 pseudogene 9 [Source:I |
| AC114878.1 | 1 | pseudogene similar to part of zinc finger prote |
| DPYD-AS2 | 1 | DPYD antisense RNA 2 [Source:HGNC Symbol; |
| AC114878.2 | 1 | CTAGE family, member 5 (CTAGE5) pseudoger |
| MIR137HG | 1 | MIR137 host gene [Source:HGNC Symbol;Acc: |
| BX005019.1 | 1 | novel transcript |
| MIR2682 | 1 | microRNA 2682 [Source:HGNC Symbol;Acc:HG |
| MIR137 | 1 | microRNA 137 [Source:HGNC Symbol;Acc:HGI |
| AC104453.1 | 1 | novel transcript |
| NFU1P2 | 1 | NFU1 iron-sulfur cluster scaffold pseudogene |
| LINC01776 | 1 | long intergenic non-protein coding RNA 1776 |
| AC095031.1 | 1 | novel transcript |
| SNX7 | 1 | sorting nexin 7 [Source:HGNC Symbol;Acc:HGI |
| PLPPR5 | 1 | phospholipid phosphatase related 5 [Source:H |
| AL445433.2 | 1 | novel transcript, antisense to PLPPR5 |
| AL445433.1 | 1 | ribosomal protein L5 (RPL5) pseudogene |
| PLPPR4 | 1 | phospholipid phosphatase related 4 [Source:H |
| AL365220.1 | 1 | chromosome 1 open reading frame 123 (C1or |
| LINC01708 | 1 | long intergenic non-protein coding RNA 1708 |
| PALMD | 1 | palmdelphin [Source:HGNC Symbol;Acc:HGNC |
| HMGB3P10 | 1 | high mobility group box 3 pseudogene 10 [Sou |
| FRRS1 | 1 | ferric chelate reductase 1 [Source:HGNC Symb |
| RNU4-75P | 1 | RNA, U4 small nuclear 75, pseudogene [Sourc |
| Y_RNA | 1 | Y RNA [Source:RFAM;Acc:RF00019] |
| AC096949.1 | 1 | ribosomal protein L39-like (RPL39L) pseudoge |
| AGL | 1 | amylo-alpha-1, 6-glucosidase, 4-alpha-glucanc |
| AC118553.1 | 1 | novel transcript |
| SLC35A3 | 1 | solute carrier family 35 member A3 [Source:H |
| AC118553.2 | 1 | novel protein |
| RNU6-750P | 1 | RNA, U6 small nuclear 750, pseudogene [Sour |
| RNU6-1318P | 1 | RNA, U6 small nuclear 1318, pseudogene [Sou |
| MFSD14A | 1 | major facilitator superfamily domain containi |
| AC093019.2 | 1 | novel transcript |
| SASS6 | 1 | SAS-6 centriolar assembly protein [Source:HG |
| AC093019.1 | 1 | novel pseudogene |
| TRMT13 | 1 | tRNA methyltransferase 13 homolog [Source:I |
| LRRC39 | 1 | leucine rich repeat containing 39 [Source:HGN |
| DBT | 1 | dihydrolipoamide branched chain transacylas |
| RPL23AP90 | 1 | ribosomal protein L23a pseudogene 90 [Sourc |
| BRI3P1 | 1 | brain protein I3 pseudogene 1 [Source:HGNC |
| AL445928.2 | 1 | novel transcript, antisense to DBT |
| AL445928.1 | 1 | novel pseudogene |
| RTCA-AS1 | 1 | RTCA antisense RNA 1 [Source:HGNC Symbol; |
| RTCA | 1 | RNA 3'-terminal phosphate cyclase [Source:H( |
| MIR553 | 1 | microRNA 553 [Source:HGNC Symbol;Acc:HGI |
| AC104457.1 | 1 | novel pseudogene |
| AC104457.2 | 1 | novel transcript |
| CDC14A | 1 | cell division cycle 14A [Source:HGNC Symbol; |
| BCAS2P2 | 1 | BCAS2 pseudogene 2 [Source:HGNC Symbol;A |
| AL589990.1 | 1 | novel transcript |

| | | |
|---|---|---|
| GPR88 | 1 | G protein-coupled receptor 88 [Source:HGNC |
| AC099670.1 | 1 | ribosomal protein L7a (RPL7A) pseudogene |
| LINC01349 | 1 | long intergenic non-protein coding RNA 1349 |
| AC099670.3 | 1 | novel transcript |
| AC099670.2 | 1 | ribosomal protein L36a (RPL36A) pseudogene |
| VCAM1 | 1 | vascular cell adhesion molecule 1 [Source:HGI |
| EXTL2 | 1 | exostosin like glycosyltransferase 2 [Source:H |
| AC104506.1 | 1 | novel transcript, antisense to EXTL2 |
| SLC30A7 | 1 | solute carrier family 30 member 7 [Source:HG |
| HNRNPA1P68 | 1 | heterogeneous nuclear ribonucleoprotein A1 |
| DPH5 | 1 | diphthamide biosynthesis 5 [Source:HGNC Syr |
| AC093157.2 | 1 | novel transcript |
| AC093157.1 | 1 | novel transcript |
| AC093157.3 | 1 | NULL |
| AC119501.1 | 1 | chromosome X open reading frame 40A (CXor |
| AL109741.1 | 1 | novel transcript |
| S1PR1 | 1 | sphingosine-1-phosphate receptor 1 [Source:I |
| AL109741.2 | 1 | ribosomal protein S20 (RPS20) pseudogene |
| PPIAP7 | 1 | peptidylprolyl isomerase A pseudogene 7 [Sou |
| LINC01307 | 1 | long intergenic non-protein coding RNA 1307 |
| LINC01709 | 1 | long intergenic non-protein coding RNA 1709 |
| RNU6-965P | 1 | RNA, U6 small nuclear 965, pseudogene [Sour |
| RPSAP19 | 1 | ribosomal protein SA pseudogene 19 [Source: |
| OLFM3 | 1 | olfactomedin 3 [Source:HGNC Symbol;Acc:HG |
| RNU6-352P | 1 | RNA, U6 small nuclear 352, pseudogene [Sour |
| AL356280.1 | 1 | phosphoglycerate mutase 1 (brain) (PGAM1) p |
| DNAJA1P5 | 1 | DnaJ heat shock protein family (Hsp40) memb |
| AC114485.1 | 1 | novel transcript |
| AC099567.1 | 1 | novel transcript |
| COL11A1 | 1 | collagen type XI alpha 1 chain [Source:HGNC S |
| SOD2P1 | 1 | superoxide dismutase 2 pseudogene 1 [Source |
| AC095032.2 | 1 | novel transcript |
| AC095032.1 | 1 | uncharacterized LOC101928436 [Source:NCBI |
| RN7SKP285 | 1 | RN7SK pseudogene 285 [Source:HGNC Symbo |
| RNPC3 | 1 | RNA binding region (RNP1, RRM) containing 3 |
| AMY2B | 1 | amylase alpha 2B (pancreatic) [Source:HGNC S |
| ACTG1P4 | 1 | actin gamma 1 pseudogene 4 [Source:HGNC S |
| AMY2A | 1 | amylase alpha 2A (pancreatic) [Source:HGNC S |
| AMY1A | 1 | amylase alpha 1A (salivary) [Source:HGNC Syn |
| AC105272.1 | 1 | pseudogene similar to part of alpha amylase |
| AMY1B | 1 | amylase alpha 1B (salivary) [Source:HGNC Syn |
| AMYP1 | 1 | amylase alpha pseudogene 1 [Source:HGNC Sy |
| AMY1C | 1 | amylase alpha 1C (salivary) [Source:HGNC Syn |
| AL136455.1 | 1 | adaptor-related protein complex 2, beta 1 sub |
| AC092506.1 | 1 | THAP domain containing, apoptosis associated |
| FTLP17 | 1 | ferritin light chain pseudogene 17 [Source:HG |
| AL591888.1 | 1 | Novel transcript |
| CDK4P1 | 1 | cyclin dependent kinase 4 pseudogene 1 [Sour |
| LINC01676 | 1 | long intergenic non-protein coding RNA 1676 |
| SEPTIN2P1 | 1 | septin 2 pseudogene 1 [Source:HGNC Symbol; |

| | | |
|---|---|---|
| AL512844.2 | 1 | matrin 3 (MATR3) pseudogene |
| AL512844.1 | 1 | matrin 3 (MATR3) pseudogene |
| LINC01677 | 1 | long intergenic non-protein coding RNA 1677 |
| AL355306.2 | 1 | novel transcript |
| AL355306.1 | 1 | tight junction protein 3 (zona occludens 3) (TJ |
| AL499605.1 | 1 | oocyte expressed protein (OOEP) pseudogene |
| AL596327.1 | 1 | ubiquitin C (UBC) pseudogene |
| MTATP6P14 | 1 | MT-ATP6 pseudogene 14 [Source:HGNC Symb |
| MTCO1P14 | 1 | MT-CO1 pseudogene 14 [Source:HGNC Symbo |
| LINC01661 | 1 | long intergenic non-protein coding RNA 1661 |
| PRMT6 | 1 | protein arginine methyltransferase 6 [Source: |
| NTNG1 | 1 | netrin G1 [Source:HGNC Symbol;Acc:HGNC:23 |
| AC114491.1 | 1 | NADH dehydrogenase (ubiquinone) 1 alpha su |
| VAV3 | 1 | vav guanine nucleotide exchange factor 3 [Sou |
| MIR7852 | 1 | microRNA 7852 [Source:HGNC Symbol;Acc:HG |
| VAV3-AS1 | 1 | VAV3 antisense RNA 1 [Source:HGNC Symbol; |
| LINC02785 | 1 | long intergenic non-protein coding RNA 2785 |
| SLC25A24 | 1 | solute carrier family 25 member 24 [Source:H |
| AL359258.2 | 1 | novel transcript, antisense to SLC25A24 |
| AL359258.3 | 1 | TEC |
| NBPF4 | 1 | NBPF member 4 [Source:HGNC Symbol;Acc:H |
| AL359258.1 | 1 | novel transcript |
| SLC25A24P1 | 1 | SLC25A24 pseudogene 1 [Source:HGNC Symb |
| NBPF5P | 1 | NBPF member 5, pseudogene [Source:HGNC S |
| SLC25A24P2 | 1 | SLC25A24 pseudogene 2 [Source:HGNC Symb |
| AL390038.1 | 1 | novel transcript |
| NBPF6 | 1 | NBPF member 6 [Source:HGNC Symbol;Acc:H |
| AL392088.2 | 1 | solute carrier family 25 (mitochondrial carrier |
| ST13P21 | 1 | ST13, Hsp70 interacting protein pseudogene 2 |
| AL392088.1 | 1 | ribosomal protein L7-like 1 (RPL7L1) pseudoge |
| FAM102B | 1 | family with sequence similarity 102 member B |
| HENMT1 | 1 | HEN methyltransferase 1 [Source:HGNC Symb |
| AL160171.1 | 1 | Novel transcript |
| PRPF38B | 1 | pre-mRNA processing factor 38B [Source:HGN |
| FNDC7 | 1 | fibronectin type III domain containing 7 [Sour |
| AL591719.2 | 1 | novel transcript, antisense to FNDC7 |
| STXBP3 | 1 | syntaxin binding protein 3 [Source:HGNC Sym |
| AL591719.1 | 1 | chromosome 20 open reading frame 147 (C20 |
| AKNAD1 | 1 | AKNA domain containing 1 [Source:HGNC Sym |
| SPATA42 | 1 | spermatogenesis associated 42 [Source:HGNC |
| GPSM2 | 1 | G protein signaling modulator 2 [Source:HGN |
| CLCC1 | 1 | chloride channel CLIC like 1 [Source:HGNC Syr |
| WDR47 | 1 | WD repeat domain 47 [Source:HGNC Symbol; |
| BX679664.2 | 1 | NADH dehydrogenase (ubiquinone) 1 beta sub |
| BX679664.3 | 1 | ribosomal protein L17 (RPL17) pseudogene |
| BX679664.1 | 1 | ribosomal protein S27 (metallopanstimulin 1) |
| RANP5 | 1 | RAN pseudogene 5 [Source:HGNC Symbol;Acc |
| TAF13 | 1 | TATA-box binding protein associated factor 13 |
| AL356488.3 | 1 | novel transcript, antisense to TMEM167B |
| TMEM167B | 1 | transmembrane protein 167B [Source:HGNC S |

| | | |
|---|---|---|
| SCARNA2 | 1 | small Cajal body-specific RNA 2 [Source:HGNC |
| AL356488.2 | 1 | small Cajal body-specific RNA 2 |
| AL356488.1 | 1 | poly(rC) binding protein 1 (PCBP1) pseudogen |
| C1orf194 | 1 | chromosome 1 open reading frame 194 [Sour |
| KIAA1324 | 1 | KIAA1324 [Source:HGNC Symbol;Acc:HGNC:29 |
| SARS | 1 | seryl-tRNA synthetase [Source:HGNC Symbol;. |
| CELSR2 | 1 | cadherin EGF LAG seven-pass G-type receptor |
| PSRC1 | 1 | proline and serine rich coiled-coil 1 [Source:H( |
| MYBPHL | 1 | myosin binding protein H like [Source:HGNC S |
| SORT1 | 1 | sortilin 1 [Source:HGNC Symbol;Acc:HGNC:11 |
| PSMA5 | 1 | proteasome subunit alpha 5 [Source:HGNC Sy |
| SYPL2 | 1 | synaptophysin like 2 [Source:HGNC Symbol;Ac |
| ATXN7L2 | 1 | ataxin 7 like 2 [Source:HGNC Symbol;Acc:HGN |
| CYB561D1 | 1 | cytochrome b561 family member D1 [Source: |
| AMIGO1 | 1 | adhesion molecule with Ig like domain 1 [Sour |
| GPR61 | 1 | G protein-coupled receptor 61 [Source:HGNC |
| AL355310.3 | 1 | novel transcript |
| GNAI3 | 1 | G protein subunit alpha i3 [Source:HGNC Sym |
| RNU6V | 1 | RNA, U6 small nuclear variant sequence with : |
| AL355310.1 | 1 | novel transcript |
| MIR197 | 1 | microRNA 197 [Source:HGNC Symbol;Acc:HGN |
| GNAT2 | 1 | G protein subunit alpha transducin 2 [Source:I |
| AMPD2 | 1 | adenosine monophosphate deaminase 2 [Sou |
| AL355310.2 | 1 | novel transcript |
| RPL7P8 | 1 | ribosomal protein L7 pseudogene 8 [Source:H |
| GSTM4 | 1 | glutathione S-transferase mu 4 [Source:HGNC |
| GSTM2 | 1 | glutathione S-transferase mu 2 [Source:HGNC |
| GSTM1 | 1 | glutathione S-transferase mu 1 [Source:HGNC |
| AC000032.1 | 1 | novel transcript, antisense to GSTM1 |
| GSTM5 | 1 | glutathione S-transferase mu 5 [Source:HGNC |
| AL158847.1 | 1 | novel transcript |
| GSTM3 | 1 | glutathione S-transferase mu 3 [Source:HGNC |
| EPS8L3 | 1 | EPS8 like 3 [Source:HGNC Symbol;Acc:HGNC:2 |
| NDUFA5P10 | 1 | NADH:ubiquinone oxidoreductase subunit A5 |
| LINC01768 | 1 | long intergenic non-protein coding RNA 1768 |
| AL450468.1 | 1 | novel transcript |
| AL450468.2 | 1 | novel transcript |
| CSF1 | 1 | colony stimulating factor 1 [Source:HGNC Sym |
| AHCYL1 | 1 | adenosylhomocysteinase like 1 [Source:HGNC |
| STRIP1 | 1 | striatin interacting protein 1 [Source:HGNC Sy |
| AL160006.1 | 1 | novel transcript, antisense to ALX3 |
| ALX3 | 1 | ALX homeobox 3 [Source:HGNC Symbol;Acc:H |
| LINC01397 | 1 | long intergenic non-protein coding RNA 1397 |
| UBL4B | 1 | ubiquitin like 4B [Source:HGNC Symbol;Acc:H( |
| SLC6A17 | 1 | solute carrier family 6 member 17 [Source:HG |
| AL355990.1 | 1 | novel transcript |
| AL355990.2 | 1 | novel transcript |
| LINC02586 | 1 | long intergenic non-protein coding RNA 2586 |
| KCNC4 | 1 | potassium voltage-gated channel subfamily C |
| AL137790.2 | 1 | cytochrome c oxidase assembly factor 5 (COA |

| | | |
|---|---|---|
| AL137790.1 | 1 | NULL |
| RBM15-AS1 | 1 | RBM15 antisense RNA 1 [Source:HGNC Symbo |
| RBM15 | 1 | RNA binding motif protein 15 [Source:HGNC S |
| LAMTOR5-AS1 | 1 | LAMTOR5 antisense RNA 1 [Source:HGNC Sym |
| SLC16A4 | 1 | solute carrier family 16 member 4 [Source:HG |
| AL355488.1 | 1 | novel transcript, antisense to SLC16A4 |
| LAMTOR5 | 1 | late endosomal/lysosomal adaptor, MAPK and |
| PROK1 | 1 | prokineticin 1 [Source:HGNC Symbol;Acc:HGN |
| AL358215.1 | 1 | novel transcript, antisense to PROK1 |
| AL358215.2 | 1 | novel transcript |
| AL358215.3 | 1 | novel transcript |
| CYMP | 1 | chymosin pseudogene [Source:HGNC Symbol; |
| CYMP-AS1 | 1 | CYMP antisense RNA 1 [Source:HGNC Symbol; |
| KCNA10 | 1 | potassium voltage-gated channel subfamily A |
| KCNA2 | 1 | potassium voltage-gated channel subfamily A |
| AL365361.1 | 1 | novel transcript |
| KCNA3 | 1 | potassium voltage-gated channel subfamily A |
| Y_RNA | 1 | Y RNA [Source:RFAM;Acc:RF00019] |
| NRBF2P3 | 1 | nuclear receptor binding factor 2 pseudogene |
| OR11I1P | 1 | olfactory receptor family 11 subfamily I memb |
| CD53 | 1 | CD53 molecule [Source:HGNC Symbol;Acc:HG |
| AL360270.2 | 1 | novel transcript, sense overlapping C1orf103 |
| AL360270.1 | 1 | novel transcript |
| LRIF1 | 1 | ligand dependent nuclear receptor interacting |
| AL360270.3 | 1 | novel transcript, antisense to LRIF1 |
| CCNT2P1 | 1 | cyclin T2 pseudogene 1 [Source:HGNC Symbol |
| RNA5SP54 | 1 | RNA, 5S ribosomal pseudogene 54 [Source:HG |
| DRAM2 | 1 | DNA damage regulated autophagy modulator |
| CEPT1 | 1 | choline/ethanolamine phosphotransferase 1 [ |
| AL355816.1 | 1 | novel transcript, antisense to CEPT1 |
| AL355816.2 | 1 | novel transcript, antisense to CEPT1 |
| DENND2D | 1 | DENN domain containing 2D [Source:HGNC Sy |
| CHI3L2 | 1 | chitinase 3 like 2 [Source:HGNC Symbol;Acc:H |
| CHIAP1 | 1 | chitinase, acidic pseudogene 1 [Source:HGNC |
| CHIAP2 | 1 | chitinase, acidic pseudogene 2 [Source:HGNC |
| CHIA | 1 | chitinase acidic [Source:HGNC Symbol;Acc:HG |
| AL356387.1 | 1 | novel transcript |
| AL356387.2 | 1 | novel pseudogene |
| PIFO | 1 | primary cilia formation [Source:HGNC Symbol |
| CHIAP3 | 1 | chitinase, acidic pseudogene 3 [Source:HGNC |
| HIGD1AP12 | 1 | HIG1 hypoxia inducible domain family membe |
| PGCP1 | 1 | progastricsin pseudogene 1 [Source:HGNC Syr |
| OVGP1 | 1 | oviductal glycoprotein 1 [Source:HGNC Symbo |
| AL390195.2 | 1 | novel transcript, sense overlapping WDR77 & |
| UBE2FP3 | 1 | UBE2F pseudogene 3 [Source:HGNC Symbol;A |
| AL390195.1 | 1 | novel transcript |
| WDR77 | 1 | WD repeat domain 77 [Source:HGNC Symbol;. |
| Y_RNA | 1 | Y RNA [Source:RFAM;Acc:RF00019] |
| ATP5PB | 1 | ATP synthase peripheral stalk-membrane subu |
| C1orf162 | 1 | chromosome 1 open reading frame 162 [Sour |

| | | |
|---|---|---|
| TMIGD3 | 1 | transmembrane and immunoglobulin domain |
| RNU6-792P | 1 | RNA, U6 small nuclear 792, pseudogene [Sour |
| ADORA3 | 1 | adenosine A3 receptor [Source:HGNC Symbol; |
| RAP1A | 1 | RAP1A, member of RAS oncogene family [Sou |
| LINC01160 | 1 | long intergenic non-protein coding RNA 1160 |
| KRT18P57 | 1 | keratin 18 pseudogene 57 [Source:HGNC Sym |
| RNU6-151P | 1 | RNA, U6 small nuclear 151, pseudogene [Sour |
| INKA2 | 1 | inka box actin regulator 2 [Source:HGNC Symb |
| INKA2-AS1 | 1 | INKA2 antisense RNA 1 [Source:HGNC Symbol |
| AL049557.1 | 1 | novel transcript |
| DDX20 | 1 | DEAD-box helicase 20 [Source:HGNC Symbol;A |
| KCND3 | 1 | potassium voltage-gated channel subfamily D |
| KCND3-IT1 | 1 | KCND3 intronic transcript 1 [Source:HGNC Syr |
| KCND3-AS1 | 1 | KCND3 antisense RNA 1 [Source:HGNC Symbo |
| LINC01750 | 1 | long intergenic non-protein coding RNA 1750 |
| AL445426.1 | 1 | novel transcript |
| TXNP3 | 1 | thioredoxin pseudogene 3 [Source:HGNC Sym |
| AL591521.1 | 1 | NULL |
| CTTNBP2NL | 1 | CTTNBP2 N-terminal like [Source:HGNC Symb |
| MIR4256 | 1 | microRNA 4256 [Source:HGNC Symbol;Acc:HG |
| WNT2B | 1 | Wnt family member 2B [Source:HGNC Symbol |
| AL354760.1 | 1 | novel transcript, antisense to WNT2B |
| ST7L | 1 | suppression of tumorigenicity 7 like [Source:H |
| CAPZA1 | 1 | capping actin protein of muscle Z-line subunit |
| MRPL53P1 | 1 | mitochondrial ribosomal protein L53 pseudog |
| RNU7-70P | 1 | RNA, U7 small nuclear 70 pseudogene [Source |
| MOV10 | 1 | Mov10 RISC complex RNA helicase [Source:HG |
| AL603832.1 | 1 | novel transcript |
| RHOC | 1 | ras homolog family member C [Source:HGNC |
| AL603832.3 | 1 | novel protein, PPM1J-RHOC readthrough |
| PPM1J | 1 | protein phosphatase, Mg2+/Mn2+ dependent |
| AL603832.2 | 1 | novel transcript, antisense to PPM1J |
| TAFA3 | 1 | TAFA chemokine like family member 3 [Sourc |
| NUTF2P4 | 1 | nuclear transport factor 2 pseudogene 4 [Sour |
| LINC01356 | 1 | long intergenic non-protein coding RNA 1356 |
| LINC01357 | 1 | long intergenic non-protein coding RNA 1357 |
| AL390729.1 | 1 | ribosomal protein L39 (RPL39) pseudogene |
| SLC16A1 | 1 | solute carrier family 16 member 1 [Source:HG |
| AKR7A2P1 | 1 | aldo-keto reductase family 7 member A2 pseu |
| SLC16A1-AS1 | 1 | SLC16A1 antisense RNA 1 [Source:HGNC Symb |
| AL390242.1 | 1 | novel transcript |
| LRIG2-DT | 1 | LRIG2 divergent transcript [Source:HGNC Sym |
| LRIG2 | 1 | leucine rich repeats and immunoglobulin like |
| RLIMP2 | 1 | ring finger protein, LIM domain interacting ps |
| AL357055.1 | 1 | ribosomal protein S19 (RPS19) pseudogene |
| AL357055.2 | 1 | ribosomal protein S15 (RPS15) pseudogene |
| MAGI3 | 1 | membrane associated guanylate kinase, WW |
| AL391058.1 | 1 | heterogeneous nuclear ribonucleoprotein A3 |
| MTND5P20 | 1 | MT-ND5 pseudogene 20 [Source:HGNC Symbo |
| PHTF1 | 1 | putative homeodomain transcription factor 1 |

| | | |
|---|---|---|
| AL133517.1 | 1 | ribosomal protein S2 (RPS2) pseudogene |
| RSBN1 | 1 | round spermatid basic protein 1 [Source:HGN |
| AL137856.1 | 1 | novel transcript, antisense to PTPN22 |
| PTPN22 | 1 | protein tyrosine phosphatase non-receptor ty |
| AP4B1-AS1 | 1 | AP4B1 antisense RNA 1 [Source:HGNC Symbo |
| BCL2L15 | 1 | BCL2 like 15 [Source:HGNC Symbol;Acc:HGNC |
| AP4B1 | 1 | adaptor related protein complex 4 subunit be |
| DCLRE1B | 1 | DNA cross-link repair 1B [Source:HGNC Symbo |
| HIPK1-AS1 | 1 | HIPK1 antisense RNA 1 [Source:HGNC Symbol |
| HIPK1 | 1 | homeodomain interacting protein kinase 1 [S |
| OLFML3 | 1 | olfactomedin like 3 [Source:HGNC Symbol;Acc |
| AL591742.1 | 1 | ribosomal protein L13a (RPL13A) pseudogene |
| AL591742.2 | 1 | novel transcript, sense intronic to OLFML3 |
| SYT6 | 1 | synaptotagmin 6 [Source:HGNC Symbol;Acc:H |
| AL121999.1 | 1 | novel transcript |
| MRPL57P1 | 1 | mitochondrial ribosomal protein L57 pseudog |
| AL035410.2 | 1 | trafficking protein, kinesin binding 2 (TRAK2) p |
| TRIM33 | 1 | tripartite motif containing 33 [Source:HGNC S |
| AL035410.1 | 1 | ribosomal protein L26 (RPL26) pseudogene |
| EIF2S2P5 | 1 | eukaryotic translation initiation factor 2 subur |
| Y_RNA | 1 | Y RNA [Source:RFAM;Acc:RF00019] |
| PKMP1 | 1 | pyruvate kinase M1/2 pseudogene 1 [Source:I |
| BCAS2 | 1 | BCAS2 pre-mRNA processing factor [Source:H |
| DENND2C | 1 | DENN domain containing 2C [Source:HGNC Sy |
| AMPD1 | 1 | adenosine monophosphate deaminase 1 [Sou |
| RN7SL432P | 1 | RNA, 7SL, cytoplasmic 432, pseudogene [Sour |
| NRAS | 1 | NRAS proto-oncogene, GTPase [Source:HGNC |
| CSDE1 | 1 | cold shock domain containing E1 [Source:HGN |
| RNY1P13 | 1 | RNY1 pseudogene 13 [Source:HGNC Symbol;A |
| SIKE1 | 1 | suppressor of IKBKE 1 [Source:HGNC Symbol;A |
| NR1H5P | 1 | nuclear receptor subfamily 1 group H membe |
| SYCP1 | 1 | synaptonemal complex protein 1 [Source:HGN |
| TSHB | 1 | thyroid stimulating hormone subunit beta [So |
| TSPAN2 | 1 | tetraspanin 2 [Source:HGNC Symbol;Acc:HGN |
| LINC01765 | 1 | long intergenic non-protein coding RNA 1765 |
| AL049825.1 | 1 | Novel transcript |
| NGF-AS1 | 1 | NGF antisense RNA 1 [Source:HGNC Symbol;A |
| NGF | 1 | nerve growth factor [Source:HGNC Symbol;Ac |
| AL512638.3 | 1 | novel transcript |
| AL512638.1 | 1 | renin (REN) pseudogene |
| AL512638.2 | 1 | ethanolamine kinase 2 (ETNK2) pseudogene |
| ELOCP20 | 1 | elongin C pseudogene 20 [Source:HGNC Symb |
| CNOT7P2 | 1 | CCR4-NOT transcription complex subunit 7 ps |
| AL592436.2 | 1 | tetraspanin 12 (TSPAN12) pseudogene |
| RN7SL420P | 1 | RNA, 7SL, cytoplasmic 420, pseudogene [Sour |
| AL592436.1 | 1 | NULL |
| VANGL1 | 1 | VANGL planar cell polarity protein 1 [Source:H |
| CASQ2 | 1 | calsequestrin 2 [Source:HGNC Symbol;Acc:HG |
| NHLH2 | 1 | nescient helix-loop-helix 2 [Source:HGNC Sym |
| HNRNPA1P43 | 1 | heterogeneous nuclear ribonucleoprotein A1 |

| | | |
|---|---|---|
| LINC01649 | 1 | long intergenic non-protein coding RNA 1649 |
| AL357137.1 | 1 | serine palmitoyltransferase, small subunit A (S |
| SLC22A15 | 1 | solute carrier family 22 member 15 [Source:H |
| AL365318.1 | 1 | novel transcript |
| MAB21L3 | 1 | mab-21 like 3 [Source:HGNC Symbol;Acc:HGN |
| LINC01779 | 1 | long intergenic non-protein coding RNA 1779 |
| U3 | 1 | Small nucleolar RNA U3 [Source:RFAM;Acc:RF |
| AL355538.1 | 1 | novel transcript |
| ATP1A1 | 1 | ATPase Na+/K+ transporting subunit alpha 1 [S |
| ATP1A1-AS1 | 1 | ATP1A1 antisense RNA 1 [Source:HGNC Symb |
| RNU6-817P | 1 | RNA, U6 small nuclear 817, pseudogene [Sour |
| LINC01762 | 1 | long intergenic non-protein coding RNA 1762 |
| AL136376.1 | 1 | novel transcript |
| Y_RNA | 1 | Y RNA [Source:RFAM;Acc:RF00019] |
| AL390066.1 | 1 | novel transcript |
| AL390066.2 | 1 | novel transcript |
| CD58 | 1 | CD58 molecule [Source:HGNC Symbol;Acc:HG |
| NAP1L4P1 | 1 | nucleosome assembly protein 1 like 4 pseudo |
| MIR548AC | 1 | microRNA 548ac [Source:HGNC Symbol;Acc:H |
| IGSF3 | 1 | immunoglobulin superfamily member 3 [Sour |
| MIR320B1 | 1 | microRNA 320b-1 [Source:HGNC Symbol;Acc:I |
| C1orf137 | 1 | chromosome 1 open reading frame 137 (puta |
| GAPDHP64 | 1 | glyceraldehyde-3-phosphate dehydrogenase p |
| NEFHP1 | 1 | neurofilament heavy pseudogene 1 [Source:H |
| CD2 | 1 | CD2 molecule [Source:HGNC Symbol;Acc:HGN |
| FTH1P22 | 1 | ferritin heavy chain 1 pseudogene 22 [Source: |
| AL157904.1 | 1 | novel transcript |
| PTGFRN | 1 | prostaglandin F2 receptor inhibitor [Source:H |
| RNA5SP55 | 1 | RNA, 5S ribosomal pseudogene 55 [Source:HG |
| CD101 | 1 | CD101 molecule [Source:HGNC Symbol;Acc:H |
| AL445231.1 | 1 | novel transcript |
| TTF2 | 1 | transcription termination factor 2 [Source:HG |
| MIR942 | 1 | microRNA 942 [Source:HGNC Symbol;Acc:HGI |
| TRIM45 | 1 | tripartite motif containing 45 [Source:HGNC S |
| RPS15AP9 | 1 | ribosomal protein S15a pseudogene 9 [Source |
| VTCN1 | 1 | V-set domain containing T cell activation inhib |
| Metazoa_SRP | 1 | Metazoan signal recognition particle RNA [Sou |
| LINC01525 | 1 | long intergenic non-protein coding RNA 1525 |
| AL358072.1 | 1 | novel transcript |
| MAN1A2 | 1 | mannosidase alpha class 1A member 2 [Sourc |
| AL157902.2 | 1 | TEC |
| VPS25P1 | 1 | VPS25 pseudogene 1 [Source:HGNC Symbol;A |
| AL157902.1 | 1 | novel transcript |
| TENT5C | 1 | terminal nucleotidyltransferase 5C [Source:H |
| VDAC2P3 | 1 | voltage dependent anion channel 2 pseudoge |
| AL390877.1 | 1 | NULL |
| AL390877.2 | 1 | anaphase promoting complex subunit 13 (AN |
| PNRC2P1 | 1 | proline rich nuclear receptor coactivator 2 pse |
| GDAP2 | 1 | ganglioside induced differentiation associated |
| WDR3 | 1 | WD repeat domain 3 [Source:HGNC Symbol;A |

| | | |
|---|---|---|
| SPAG17 | 1 | sperm associated antigen 17 [Source:HGNC Sy |
| AL391557.1 | 1 | novel transcript |
| RNA5SP56 | 1 | RNA, 5S ribosomal pseudogene 56 [Source:HG |
| PSMC1P12 | 1 | proteasome 26S subunit, ATPase 1 pseudoger |
| AL139148.1 | 1 | high density lipoprotein binding protein (vigili |
| TBX15 | 1 | T-box transcription factor 15 [Source:HGNC Sy |
| AL139420.1 | 1 | novel transcript |
| AL139420.2 | 1 | novel transcript |
| WARS2 | 1 | tryptophanyl tRNA synthetase 2, mitochondri |
| WARS2-IT1 | 1 | WARS2 intronic transcript 1 [Source:HGNC Syr |
| RBMX2P3 | 1 | RBMX2 pseudogene 3 [Source:HGNC Symbol;/ |
| RPS3AP12 | 1 | ribosomal protein S3a pseudogene 12 [Source |
| WARS2-AS1 | 1 | WARS2 antisense RNA 1 [Source:HGNC Symbc |
| RPL6P2 | 1 | ribosomal protein L6 pseudogene 2 [Source:H |
| AL359915.1 | 1 | novel transcript |
| LINC01780 | 1 | long intergenic non-protein coding RNA 1780 |
| RNU1-75P | 1 | RNA, U1 small nuclear 75, pseudogene [Sourc |
| HAO2 | 1 | hydroxyacid oxidase 2 [Source:HGNC Symbol;. |
| HAO2-IT1 | 1 | HAO2 intronic transcript 1 [Source:HGNC Sym |
| HSD3B2 | 1 | hydroxy-delta-5-steroid dehydrogenase, 3 bet |
| GAPDHP74 | 1 | glyceraldehyde-3-phosphate dehydrogenase p |
| HSD3BP2 | 1 | hydroxy-delta-5-steroid dehydrogenase, 3 bet |
| GAPDHP23 | 1 | glyceraldehyde 3 phosphate dehydrogenase p |
| HSD3BP1 | 1 | hydroxy-delta-5-steroid dehydrogenase, 3 bet |
| GAPDHP58 | 1 | glyceraldehyde 3 phosphate dehydrogenase p |
| HSD3B1 | 1 | hydroxy-delta-5-steroid dehydrogenase, 3 bet |
| GAPDHP32 | 1 | glyceraldehyde 3 phosphate dehydrogenase p |
| HSD3BP3 | 1 | hydroxy-delta-5-steroid dehydrogenase, 3 bet |
| GAPDHP27 | 1 | glyceraldehyde 3 phosphate dehydrogenase p |
| HSD3BP4 | 1 | hydroxy-delta-5-steroid dehydrogenase, 3 bet |
| GAPDHP33 | 1 | glyceraldehyde 3 phosphate dehydrogenase p |
| LINC00622 | 1 | long intergenic non-protein coding RNA 622 [S |
| HSD3BP5 | 1 | hydroxy-delta-5-steroid dehydrogenase, 3 bet |
| ZNF697 | 1 | zinc finger protein 697 [Source:HGNC Symbol; |
| PHGDH | 1 | phosphoglycerate dehydrogenase [Source:HG |
| HMGCS2 | 1 | 3-hydroxy-3-methylglutaryl-CoA synthase 2 [S |
| REG4 | 1 | regenerating family member 4 [Source:HGNC |
| NBPF7 | 1 | NBPF member 7 [Source:HGNC Symbol;Acc:H( |
| PFN1P9 | 1 | profilin 1 pseudogene 9 [Source:HGNC Symbo |
| NOTCH2P1 | 1 | notch 2 pseudogene 1 [Source:HGNC Symbol;. |
| ADAM30 | 1 | ADAM metallopeptidase domain 30 [Source:H |
| NOTCH2 | 1 | notch receptor 2 [Source:HGNC Symbol;Acc:H |
| AC245008.1 | 1 | novel transcript |
| RNU6-465P | 1 | RNA, U6 small nuclear 465, pseudogene [Sour |
| SEC22B | 1 | SEC22 homolog B, vesicle trafficking protein (g |
| AC244669.2 | 1 | phosphodiesterase 4D interacting protein (PD |
| 7SK | 1 | 7SK RNA [Source:RFAM;Acc:RF00100] |
| AC244669.1 | 1 | Poly [ADP-ribose] polymerase 8 (PARP8) pseu |
| U1 | 1 | U1 spliceosomal RNA [Source:RFAM;Acc:RF00 |
| NBPF8 | 1 | NBPF member 8 [Source:HGNC Symbol;Acc:H( |

| | | |
|---|---|---|
| PFN1P2 | 1 | profilin 1 pseudogene 2 [Source:HGNC Symbo |
| AC241952.1 | 1 | phosphodiesterase 4D interacting protein (PD |
| U2 | 1 | U2 spliceosomal RNA [Source:RFAM;Acc:RF00 |
| 7SK | 1 | 7SK RNA [Source:RFAM;Acc:RF00100] |
| AC253572.1 | 1 | notch 2 N-terminal like R |
| NBPF26 | 1 | NBPF member 26 [Source:HGNC Symbol;Acc:H |
| AC253572.2 | 1 | novel transcript |
| RNVU1-19 | 1 | RNA, variant U1 small nuclear 19 [Source:HGN |
| PPIAL4A | 1 | peptidylprolyl isomerase A like 4A [Source:HG |
| AC241377.3 | 1 | TEC |
| LINC00623 | 1 | long intergenic non-protein coding RNA 623 [S |
| RNVU1-4 | 1 | RNA, variant U1 small nuclear 4 [Source:HGNC |
| AC241377.4 | 1 | novel pseudogene, similar to phosphodiestera |
| AC241377.2 | 1 | phosphodiesterase 4D interacting protein (PD |
| U1 | 1 | U1 spliceosomal RNA [Source:RFAM;Acc:RF00 |
| AC241377.1 | 1 | ubiquitin-conjugating enzyme E2D 4 (putative |
| FCGR1B | 1 | Fc fragment of IgG receptor Ib [Source:HGNC S |
| AC244453.3 | 1 | novel transcript, antisense to FCGR1B |
| AC244453.2 | 1 | novel transcript |
| HIST2H2BA | 1 | histone cluster 2 H2B family member a (pseud |
| AC244453.1 | 1 | novel transcript |
| HIST2H3DP1 | 1 | histone cluster 2 H3 family member d pseudo |
| RPL22P6 | 1 | ribosomal protein L22 pseudogene 6 [Source: |
| FAM72B | 1 | family with sequence similarity 72 member B |
| SRGAP2C | 1 | SLIT-ROBO Rho GTPase activating protein 2C [ |
| AC243994.1 | 1 | RAP1, GTP-GDP dissociation stimulator 1 (RAP |
| SRGAP2-AS1 | 1 | SRGAP2 antisense RNA 1 [Source:HGNC Symb |
| LINC02798 | 1 | long intergenic non-protein coding RNA 2798 |
| AL592494.1 | 1 | novel transcript |
| MTIF2P1 | 1 | mitochondrial translational initiation factor 2 |
| AL592494.2 | 1 | novel transcript |
| EMBP1 | 1 | embigin pseudogene 1 [Source:HGNC Symbol; |
| LINC01691 | 1 | long intergenic non-protein coding RNA 1691 |
| AC242852.1 | 1 | transmembrane protein 128 (TMEM128) pseu |
| AC242852.2 | 1 | otopetrin 1 (OTOP1) pseudogene |
| AC239859.5 | 1 | lymphocyte-specific protein 1 (LSP1) pseudog |
| RNA5SP533 | 1 | RNA, 5S ribosomal pseudogene 533 [Source:H |
| AC239859.3 | 1 | dopamine receptor D5 (DRD5) pseudogene |
| AC239859.4 | 1 | lysine (K)-specific methyltransferase 2C (KMT2 |
| NKAIN1P1 | 1 | sodium/potassium transporting ATPase intera |
| AC239859.2 | 1 | kinase suppressor of ras 1 (KSR1) pseudogene |
| LINC02799 | 1 | long intergenic non-protein coding RNA 2799 |
| AC239859.1 | 1 | phosphodiesterase 4D interacting protein (my |
| NBPF17P | 1 | NBPF member 17, pseudogene [Source:HGNC |
| PFN1P12 | 1 | profilin 1 pseudogene 12 [Source:HGNC Symb |
| RNU1-114P | 1 | RNA, U1 small nuclear 114, pseudogene [Sour |
| RNVU1-17 | 1 | RNA, variant U1 small nuclear 17 [Source:HGN |
| RNU1-92P | 1 | RNA, U1 small nuclear 92, pseudogene [Sourc |
| RNVU1-18 | 1 | RNA, variant U1 small nuclear 18 [Source:HGN |
| AC239800.2 | 1 | novel transcript |

| | | |
|---|---|---|
| AC239800.1 | 1 | novel pseudogene, similar to phosphodiestera |
| FAM91A3P | 1 | family with sequence similarity 91 member A3 |
| AC239800.3 | 1 | NHP2 non-histone chromosome protein 2-like |
| LINC02591 | 1 | long intergenic non-protein coding RNA 2591 |
| RNU1-143P | 1 | RNA, U1 small nuclear 143, pseudogene [Sour |
| AC239798.1 | 1 | phosphodiesterase 4D interacting protein (PD |
| AC239798.3 | 1 | ubiquitin-conjugating enzyme E2D 4 (putative |
| FCGR1CP | 1 | Fc fragment of IgG receptor Ic, pseudogene [S |
| AC239798.2 | 1 | histone cluster 2, H2bb (pseudogene) |
| HIST2H3PS2 | 1 | histone cluster 2 H3 pseudogene 2 [Source:HC |
| AC246680.1 | 1 | novel transcript |
| RPL22P5 | 1 | ribosomal protein L22 pseudogene 5 [Source: |
| FAM72C | 1 | family with sequence similarity 72 member C |
| SRGAP2D | 1 | SLIT-ROBO Rho GTPase activating protein 2D ( |
| IGKV1OR1-1 | 1 | immunoglobulin kappa variable 1/OR1-1 (pseu |
| AC244015.1 | 1 | transmembrane protein 128 (TMEM128) pseu |
| AC244015.2 | 1 | otopetrin 1 (OTOP1) pseudogene |
| LINC02802 | 1 | long intergenic non-protein coding RNA 2802 |
| RNA5SP529 | 1 | RNA, 5S ribosomal pseudogene 529 [Source:H |
| DRD5P2 | 1 | dopamine receptor D5 pseudogene 2 [Source: |
| AC245595.1 | 1 | lysine (K)-specific methyltransferase 2C (KMT2 |
| PPIAL4E | 1 | peptidylprolyl isomerase A like 4E [Source:HG |
| AC246785.4 | 1 | HIG1 hypoxia inducible domain family, membe |
| RNVU1-15 | 1 | RNA, variant U1 small nuclear 15 [Source:HGN |
| AC246785.3 | 1 | novel transcript |
| AC246785.2 | 1 | novel pseudogene (FLJ39739) |
| NBPF15 | 1 | NBPF member 15 [Source:HGNC Symbol;Acc:H |
| PFN1P6 | 1 | profilin 1 pseudogene 6 [Source:HGNC Symbo |
| AC246785.1 | 1 | novel pseudogene |
| U1 | 1 | U1 spliceosomal RNA [Source:RFAM;Acc:RF00 |
| U1 | 1 | U1 spliceosomal RNA [Source:RFAM;Acc:RF00 |
| U1 | 1 | U1 spliceosomal RNA [Source:RFAM;Acc:RF00 |
| PPIAL4F | 1 | peptidylprolyl isomerase A like 4F [Source:HG |
| LINC01632 | 1 | long intergenic non-protein coding RNA 1632 |
| FP700111.1 | 1 | novel transcript |
| FP700107.1 | 1 | myeloid/lymphoid or mixed-lineage leukemia |
| RNA5SP59 | 1 | RNA, 5S ribosomal pseudogene 59 [Source:HG |
| FP700111.2 | 1 | otopetrin 1 (OTOP1) pseudogene |
| FP700111.3 | 1 | transmembrane protein 128 (TMEM128) pseu |
| SRGAP2B | 1 | SLIT-ROBO Rho GTPase activating protein 2B [ |
| AC242498.1 | 1 | RAP1, GTP-GDP dissociation stimulator 1 (RAP |
| FAM72D | 1 | family with sequence similarity 72 member D |
| AC241585.3 | 1 | phosphodiesterase 4D interacting protein (PD |
| LINC01145 | 1 | long intergenic non-protein coding RNA 1145 |
| U1 | 1 | U1 spliceosomal RNA [Source:RFAM;Acc:RF00 |
| AC241585.2 | 1 | TEC |
| AC241585.1 | 1 | EGFR-coamplified and overexpressed protein |
| PPIAL4D | 1 | peptidylprolyl isomerase A like 4D [Source:HG |
| AC245014.2 | 1 | HIG1 hypoxia inducible domain family, membe |
| RNVU1-14 | 1 | RNA, variant U1 small nuclear 14 [Source:HGN |

| | | |
|---|---|---|
| AC245014.3 | 1 | novel transcript |
| AC245014.1 | 1 | novel transcript |
| NBPF20 | 1 | NBPF member 20 [Source:HGNC Symbol;Acc:H |
| PFN1P3 | 1 | profilin 1 pseudogene 3 [Source:HGNC Symbo |
| U1 | 1 | U1 spliceosomal RNA [Source:RFAM;Acc:RF00 |
| AC239860.3 | 1 | TEC |
| U1 | 1 | U1 spliceosomal RNA [Source:RFAM;Acc:RF00 |
| AC239860.4 | 1 | novel transcript |
| AC239860.2 | 1 | phosphodiesterase 4D interacting protein (PD |
| AC239860.1 | 1 | phosphodiesterase 4D interacting protein (PD |
| NBPF25P | 1 | NBPF member 25, pseudogene [Source:HGNC |
| GPR89A | 1 | G protein-coupled receptor 89A [Source:HGN |
| PDZK1 | 1 | PDZ domain containing 1 [Source:HGNC Symb |
| CD160 | 1 | CD160 molecule [Source:HGNC Symbol;Acc:H |
| RNF115 | 1 | ring finger protein 115 [Source:HGNC Symbol; |
| POLR3C | 1 | RNA polymerase III subunit C [Source:HGNC Sy |
| NUDT17 | 1 | nudix hydrolase 17 [Source:HGNC Symbol;Acc |
| PIAS3 | 1 | protein inhibitor of activated STAT 3 [Source:H |
| MIR6736 | 1 | microRNA 6736 [Source:HGNC Symbol;Acc:HG |
| ANKRD35 | 1 | ankyrin repeat domain 35 [Source:HGNC Syml |
| ITGA10 | 1 | integrin subunit alpha 10 [Source:HGNC Symb |
| AC243547.1 | 1 | novel transcript, antisense to ITGA10 |
| PEX11B | 1 | peroxisomal biogenesis factor 11 beta [Source |
| RBM8A | 1 | RNA binding motif protein 8A [Source:HGNC S |
| GNRHR2 | 1 | gonadotropin releasing hormone receptor 2 (p |
| LIX1L-AS1 | 1 | LIX1L antisense RNA 1 [Source:HGNC Symbol;A |
| AC243547.3 | 1 | novel protein, lncRNA-POLR3GL readthrough |
| LIX1L | 1 | limb and CNS expressed 1 like [Source:HGNC S |
| ANKRD34A | 1 | ankyrin repeat domain 34A [Source:HGNC Syn |
| AC243547.2 | 1 | novel transcript, antisense to ANKRD34A |
| POLR3GL | 1 | RNA polymerase III subunit G like [Source:HGN |
| TXNIP | 1 | thioredoxin interacting protein [Source:HGNC |
| HJV | 1 | hemojuvelin BMP co-receptor [Source:HGNC S |
| AC239799.2 | 1 | novel transcript |
| RNVU1-6 | 1 | RNA, variant U1 small nuclear 6 [Source:HGN |
| LINC01719 | 1 | long intergenic non-protein coding RNA 1719 |
| NBPF10 | 1 | NBPF member 10 [Source:HGNC Symbol;Acc:H |
| NOTCH2NLA | 1 | notch 2 N-terminal like A [Source:HGNC Symb |
| AC239799.1 | 1 | novel transcript |
| RNU6-1071P | 1 | RNA, U6 small nuclear 1071, pseudogene [Sou |
| NUDT4P2 | 1 | nudix hydrolase 4 pseudogene 2 [Source:HGN |
| SEC22B4P | 1 | SEC22 homolog B4, pseudogene [Source:HGN |
| PPIAL4H | 1 | peptidylprolyl isomerase A like 4H [Source:HG |
| U1 | 1 | U1 spliceosomal RNA [Source:RFAM;Acc:RF00 |
| AC245407.2 | 1 | novel transcript |
| U1 | 1 | U1 spliceosomal RNA [Source:RFAM;Acc:RF00 |
| AC245407.1 | 1 | neuroblastoma breakpoint family, member 6 |
| HYDIN2 | 1 | HYDIN axonemal central pair apparatus protei |
| RNA5SP536 | 1 | RNA, 5S ribosomal pseudogene 536 [Source:H |
| AC241929.1 | 1 | NULL |

| | | |
|---|---|---|
| NBPF12 | 1 | NBPF member 12 [Source:HGNC Symbol;Acc:I |
| PFN1P8 | 1 | profilin 1 pseudogene 8 [Source:HGNC Symbo |
| AC244394.2 | 1 | novel transcript |
| NBPF13P | 1 | NBPF member 13, pseudogene [Source:HGNC |
| AC244394.1 | 1 | insulin-like growth factor 2 mRNA binding pro |
| RNU1-151P | 1 | RNA, U1 small nuclear 151, pseudogene [Sour |
| AC241584.1 | 1 | Sjogren syndrome antigen B (autoantigen La) |
| RNVU1-8 | 1 | RNA, variant U1 small nuclear 8 [Source:HGN |
| AC241584.2 | 1 | neuroblastoma breakpoint family, member 1 |
| PRKAB2 | 1 | protein kinase AMP-activated non-catalytic su |
| PDIA3P1 | 1 | protein disulfide isomerase family A member |
| AC242426.2 | 1 | novel transcript, antisense to FMO5 |
| FMO5 | 1 | flavin containing dimethylaniline monoxygena |
| CCT8P1 | 1 | chaperonin containing TCP1 subunit 8 pseudo |
| RPL7AP15 | 1 | ribosomal protein L7a pseudogene 15 [Source |
| CHD1L | 1 | chromodomain helicase DNA binding protein |
| LINC00624 | 1 | long intergenic non-protein coding RNA 624 [ |
| AC242426.1 | 1 | prostaglandin reductase 1 (PTGR1) pseudoger |
| OR13Z1P | 1 | olfactory receptor family 13 subfamily Z mem |
| Y_RNA | 1 | Y RNA [Source:RFAM;Acc:RF00019] |
| OR13Z2P | 1 | olfactory receptor family 13 subfamily Z mem |
| OR13Z3P | 1 | olfactory receptor family 13 subfamily Z mem |
| BCL9 | 1 | BCL9 transcription coactivator [Source:HGNC |
| ACP6 | 1 | acid phosphatase 6, lysophosphatidic [Source |
| RN7SL261P | 1 | RNA, 7SL, cytoplasmic 261, pseudogene [Sour |
| AC241644.1 | 1 | novel transcript |
| GJA5 | 1 | gap junction protein alpha 5 [Source:HGNC Sy |
| AC241644.3 | 1 | novel transcript, antisense to GJA5 |
| AC241644.2 | 1 | novel transcript |
| AC239801.1 | 1 | PRAME family member 17 (PRAMEF17) pseud |
| GJA8 | 1 | gap junction protein alpha 8 [Source:HGNC Sy |
| GPR89B | 1 | G protein-coupled receptor 89B [Source:HGN |
| PDZK1P1 | 1 | PDZ domain containing 1 pseudogene 1 [Sour |
| AC239803.1 | 1 | novel transcript |
| LINC02804 | 1 | long intergenic non-protein coding RNA 2804 |
| RNU1-129P | 1 | RNA, U1 small nuclear 129, pseudogene [Sour |
| RNVU1-7 | 1 | RNA, variant U1 small nuclear 7 [Source:HGN |
| PDE4DIPP1 | 1 | phosphodiesterase 4D interacting protein pse |
| NBPF11 | 1 | NBPF member 11 [Source:HGNC Symbol;Acc:I |
| PFN1P4 | 1 | profilin 1 pseudogene 4 [Source:HGNC Symbo |
| ABHD17AP1 | 1 | abhydrolase domain containing 17A pseudoge |
| LINC02805 | 1 | long intergenic non-protein coding RNA 2805 |
| AC239809.3 | 1 | novel transcript |
| RNA5SP57 | 1 | RNA, 5S ribosomal pseudogene 57 [Source:HG |
| AC239809.1 | 1 | neuroblastoma breakpoint family, member 4 |
| AC239809.2 | 1 | novel pseudogene |
| RNU1-120P | 1 | RNA, U1 small nuclear 120, pseudogene [Sour |
| LINC01731 | 1 | long intergenic non-protein coding RNA 1731 |
| AC245100.3 | 1 | novel pseudogene |
| LINC01138 | 1 | long intergenic non-protein coding RNA 1138 |

| | | |
|---|---|---|
| LINC02806 | 1 | long intergenic non-protein coding RNA 2806 |
| AC245100.1 | 1 | novel transcript |
| AC245100.5 | 1 | novel pseudogene |
| Y_RNA | 1 | Y RNA [Source:RFAM;Acc:RF00019] |
| MIR5087 | 1 | microRNA 5087 [Source:HGNC Symbol;Acc:HG |
| RNU1-122P | 1 | RNA, U1 small nuclear 122, pseudogene [Sour |
| AC245100.6 | 1 | novel transcript |
| RNVU1-1 | 1 | RNA, variant U1 small nuclear 1 [Source:HGN( |
| RNVU1-2 | 1 | RNA, variant U1 small nuclear 2 [Source:HGN( |
| MIR6077 | 1 | microRNA 6077 [Source:HGNC Symbol;Acc:HG |
| RNU1-13P | 1 | RNA, U1 small nuclear 13, pseudogene [Sourc |
| AC245100.4 | 1 | phosphodiesterase 4D interacting protein (PD |
| RNVU1-3 | 1 | RNA, variant U1 small nuclear 3 [Source:HGN( |
| AC245100.2 | 1 | family with sequence similarity 91, member A |
| AC245100.8 | 1 | poly(ADP-ribose) polymerase family member |
| AC245100.7 | 1 | TEC |
| PPIAL4G | 1 | peptidylprolyl isomerase A like 4G [Source:HG |
| U1 | 1 | U1 spliceosomal RNA [Source:RFAM;Acc:RF00 |
| NBPF14 | 1 | NBPF member 14 [Source:HGNC Symbol;Acc:H |
| NOTCH2NLB | 1 | notch 2 N-terminal like B [Source:HGNC Symb |
| RNU6-1171P | 1 | RNA, U6 small nuclear 1171, pseudogene [Sou |
| NUDT4B | 1 | nudix hydrolase 4B [Source:HGNC Symbol;Acc |
| SEC22B3 | 1 | SEC22 homolog B3 [Source:HGNC Symbol;Acc |
| AC245389.1 | 1 | CKLF-like MARVEL transmembrane domain co |
| PDE4DIP | 1 | phosphodiesterase 4D interacting protein [So |
| RN7SKP88 | 1 | RN7SK pseudogene 88 [Source:HGNC Symbol; |
| RNU2-38P | 1 | RNA, U2 small nuclear 38, pseudogene [Sourc |
| AC239802.1 | 1 | novel transcript |
| AC239802.2 | 1 | novel transcript |
| AC239804.1 | 1 | novel transcript |
| NBPF9 | 1 | NBPF member 9 [Source:HGNC Symbol;Acc:H( |
| AC239804.2 | 1 | Neuroblastoma breakpoint family (NBPF) pseu |
| RNU1-59P | 1 | RNA, U1 small nuclear 59, pseudogene [Sourc |
| AC245297.3 | 1 | novel transcript |
| AC245297.1 | 1 | phosphodiesterase 4D interacting protein (my |
| AC245297.2 | 1 | novel transcript |
| 7SK | 1 | 7SK RNA [Source:RFAM;Acc:RF00100] |
| SEC22B2 | 1 | SEC22 homolog B2 [Source:HGNC Symbol;Acc |
| NOTCH2NLC | 1 | notch 2 N-terminal like C [Source:HGNC Symb |
| AC242842.3 | 1 | novel protein, identical to neuroblastoma bre |
| NBPF19 | 1 | NBPF member 19 [Source:HGNC Symbol;Acc:H |
| PPIAL4C | 1 | peptidylprolyl isomerase A like 4C [Source:HG |
| LINC00869 | 1 | long intergenic non-protein coding RNA 869 [S |
| AC242842.2 | 1 | poly(ADP-ribose) polymerase family member |
| AC242842.1 | 1 | novel transcript |
| U1 | 1 | U1 spliceosomal RNA [Source:RFAM;Acc:RF00 |
| FAM91A2P | 1 | family with sequence similarity 91 member A2 |
| AC243772.3 | 1 | phosphodiesterase 4D interacting protein (PD |
| RNU1-68P | 1 | RNA, U1 small nuclear 68, pseudogene [Sourc |
| AC243772.1 | 1 | ubiquitin-conjugating enzyme E2D 1 (UBC4/5 |

| | | |
|---|---|---|
| FCGR1A | 1 | Fc fragment of IgG receptor Ia [Source:HGNC S |
| HIST2H2BF | 1 | histone cluster 2 H2B family member f [Sourc |
| AC243772.2 | 1 | novel transcript, antisense to FCGR1A |
| HIST2H3D | 1 | histone cluster 2 H3 family member d [Source |
| HIST2H4A | 1 | histone cluster 2 H4 family member a [Source |
| HIST2H3C | 1 | histone cluster 2 H3 family member c [Source |
| HIST2H2AA3 | 1 | histone cluster 2 H2A family member a3 [Sour |
| HIST2H2BD | 1 | histone cluster 2 H2B family member d (pseuc |
| HIST2H2BC | 1 | histone cluster 2 H2B family member c (pseuc |
| HIST2H2AA4 | 1 | histone cluster 2 H2A family member a4 [Sour |
| HIST2H3A | 1 | histone cluster 2 H3 family member a [Source |
| HIST2H4B | 1 | histone cluster 2 H4 family member b [Source |
| AC239868.1 | 1 | novel transcript, antisense HIST2H4A |
| HIST2H2BE | 1 | histone cluster 2 H2B family member e [Sourc |
| HIST2H2AC | 1 | histone cluster 2 H2A family member c [Sourc |
| HIST2H2AB | 1 | histone cluster 2 H2A family member b [Sourc |
| BOLA1 | 1 | bolA family member 1 [Source:HGNC Symbol; |
| SV2A | 1 | synaptic vesicle glycoprotein 2A [Source:HGN |
| SF3B4 | 1 | splicing factor 3b subunit 4 [Source:HGNC Sym |
| MTMR11 | 1 | myotubularin related protein 11 [Source:HGN |
| OTUD7B | 1 | OTU deubiquitinase 7B [Source:HGNC Symbol |
| AC244033.2 | 1 | novel transcript |
| AC244033.1 | 1 | ribosomal protein L6 (RPL6) pseudogene |
| VPS45 | 1 | vacuolar protein sorting 45 homolog [Source:I |
| PLEKHO1 | 1 | pleckstrin homology domain containing O1 [Sc |
| AC242988.2 | 1 | Novel transcript |
| RN7SL480P | 1 | RNA, 7SL, cytoplasmic 480, pseudogene [Sour |
| ANP32E | 1 | acidic nuclear phosphoprotein 32 family mem |
| RNU2-17P | 1 | RNA, U2 small nuclear 17, pseudogene [Sourc |
| AC242988.1 | 1 | novel transcript, antisense to CA14 |
| CA14 | 1 | carbonic anhydrase 14 [Source:HGNC Symbol; |
| APH1A | 1 | aph-1 homolog A, gamma-secretase subunit [S |
| C1orf54 | 1 | chromosome 1 open reading frame 54 [Sourc |
| CIART | 1 | circadian associated repressor of transcriptior |
| MRPS21 | 1 | mitochondrial ribosomal protein S21 [Source: |
| PRPF3 | 1 | pre-mRNA processing factor 3 [Source:HGNC S |
| RPRD2 | 1 | regulation of nuclear pre-mRNA domain conta |
| TARS2 | 1 | threonyl-tRNA synthetase 2, mitochondrial [Sc |
| MIR6878 | 1 | microRNA 6878 [Source:HGNC Symbol;Acc:HG |
| ECM1 | 1 | extracellular matrix protein 1 [Source:HGNC S |
| FALEC | 1 | focally amplified long non-coding RNA in epith |
| AL356356.1 | 1 | novel transcript, antisense to ADAMTSL4 |
| ADAMTSL4 | 1 | ADAMTS like 4 [Source:HGNC Symbol;Acc:HGI |
| MIR4257 | 1 | microRNA 4257 [Source:HGNC Symbol;Acc:HG |
| ADAMTSL4-AS1 | 1 | ADAMTSL4 antisense RNA 1 [Source:HGNC Syr |
| RN7SL473P | 1 | RNA, 7SL, cytoplasmic 473, pseudogene [Sour |
| RN7SL600P | 1 | RNA, 7SL, cytoplasmic 600, pseudogene [Sour |
| MCL1 | 1 | MCL1 apoptosis regulator, BCL2 family memb |
| AL356356.2 | 1 | NULL |
| ENSA | 1 | endosulfine alpha [Source:HGNC Symbol;Acc:I |

| | | |
|---|---|---|
| U4 | 1 | U4 spliceosomal RNA [Source:RFAM;Acc:RF00 |
| GOLPH3L | 1 | golgi phosphoprotein 3 like [Source:HGNC Syr |
| HORMAD1 | 1 | HORMA domain containing 1 [Source:HGNC S |
| RNU6-1042P | 1 | RNA, U6 small nuclear 1042, pseudogene [Sou |
| CTSS | 1 | cathepsin S [Source:HGNC Symbol;Acc:HGNC: |
| AL356292.1 | 1 | pseudogene similar to part of short coiled-coi |
| CTSK | 1 | cathepsin K [Source:HGNC Symbol;Acc:HGNC: |
| UBE2D3P3 | 1 | ubiquitin conjugating enzyme E2 D3 pseudoge |
| ARNT | 1 | aryl hydrocarbon receptor nuclear translocato |
| RNU6-1309P | 1 | RNA, U6 small nuclear 1309, pseudogene [Sou |
| RPS27AP6 | 1 | ribosomal protein S27a pseudogene 6 [Source |
| CTXND2 | 1 | cortexin domain containing 2 [Source:HGNC S |
| CYCSP51 | 1 | CYCS pseudogene 51 [Source:HGNC Symbol;A |
| SETDB1 | 1 | SET domain bifurcated histone lysine methyltr |
| CERS2 | 1 | ceramide synthase 2 [Source:HGNC Symbol;A |
| AL590133.2 | 1 | novel transcript, antisense to CERS2 |
| AL590133.1 | 1 | novel transcript, antisense to CERS2 |
| ANXA9 | 1 | annexin A9 [Source:HGNC Symbol;Acc:HGNC: |
| MINDY1 | 1 | MINDY lysine 48 deubiquitinase 1 [Source:HG |
| PRUNE1 | 1 | prune exopolyphosphatase 1 [Source:HGNC S |
| RNU6-884P | 1 | RNA, U6 small nuclear 884, pseudogene [Sour |
| BNIPL | 1 | BCL2 interacting protein like [Source:HGNC Sy |
| C1orf56 | 1 | chromosome 1 open reading frame 56 [Sourc |
| CDC42SE1 | 1 | CDC42 small effector 1 [Source:HGNC Symbol |
| MLLT11 | 1 | MLLT11 transcription factor 7 cofactor [Sourc |
| GABPB2 | 1 | GA binding protein transcription factor subuni |
| RPS29P29 | 1 | ribosomal protein S29 pseudogene 29 [Source |
| AL592424.1 | 1 | novel transcript, sense overlapping SEMA6C |
| SEMA6C | 1 | semaphorin 6C [Source:HGNC Symbol;Acc:HG |
| TNFAIP8L2 | 1 | TNF alpha induced protein 8 like 2 [Source:HG |
| SCNM1 | 1 | sodium channel modifier 1 [Source:HGNC Sym |
| LYSMD1 | 1 | LysM domain containing 1 [Source:HGNC Sym |
| TMOD4 | 1 | tropomodulin 4 [Source:HGNC Symbol;Acc:HG |
| VPS72 | 1 | vacuolar protein sorting 72 homolog [Source: |
| PIP5K1A | 1 | phosphatidylinositol-4-phosphate 5-kinase typ |
| PSMD4 | 1 | proteasome 26S subunit, non-ATPase 4 [Sourc |
| ZNF687-AS1 | 1 | ZNF687 antisense RNA 1 [Source:HGNC Symb |
| ZNF687 | 1 | zinc finger protein 687 [Source:HGNC Symbol; |
| PI4KB | 1 | phosphatidylinositol 4-kinase beta [Source:HG |
| RN7SL444P | 1 | RNA, 7SL, cytoplasmic 444, pseudogene [Sour |
| AL391069.3 | 1 | novel transcript |
| RFX5 | 1 | regulatory factor X5 [Source:HGNC Symbol;Ac |
| AL391069.1 | 1 | novel transcript |
| AL391069.2 | 1 | novel transcript |
| SELENBP1 | 1 | selenium binding protein 1 [Source:HGNC Sym |
| PSMB4 | 1 | proteasome subunit beta 4 [Source:HGNC Syn |
| POGZ | 1 | pogo transposable element derived with ZNF |
| RNY4P25 | 1 | RNY4 pseudogene 25 [Source:HGNC Symbol;A |
| CGN | 1 | cingulin [Source:HGNC Symbol;Acc:HGNC:174 |
| AL365436.3 | 1 | NULL |

| | | |
|---|---|---|
| TUFT1 | 1 | tuftelin 1 [Source:HGNC Symbol;Acc:HGNC:12 |
| AL365436.2 | 1 | novel transcript |
| MIR554 | 1 | microRNA 554 [Source:HGNC Symbol;Acc:HGI |
| AL365436.1 | 1 | ribosomal protein S10 (RPS10) pseudogene |
| SNX27 | 1 | sorting nexin 27 [Source:HGNC Symbol;Acc:H( |
| AL391335.1 | 1 | novel transcript, antisense to SNX27 |
| RNU6-1062P | 1 | RNA, U6 small nuclear 1062, pseudogene [Sou |
| CELF3 | 1 | CUGBP Elav-like family member 3 [Source:HGI |
| AL589765.1 | 1 | novel transcript |
| RIIAD1 | 1 | regulatory subunit of type II PKA R-subunit (RI |
| RNU6-662P | 1 | RNA, U6 small nuclear 662, pseudogene [Sour |
| AL589765.7 | 1 | novel transcript, antisense to MRPL9 |
| AL589765.3 | 1 | ribosomal protein S11 (RPS11) pseudogene |
| MRPL9 | 1 | mitochondrial ribosomal protein L9 [Source:H |
| OAZ3 | 1 | ornithine decarboxylase antizyme 3 [Source:H |
| AL589765.4 | 1 | novel transcript |
| AL589765.2 | 1 | novel transcript |
| AL589765.5 | 1 | novel transcript, antisense to OAZ3 |
| TDRKH | 1 | tudor and KH domain containing [Source:HGN |
| TDRKH-AS1 | 1 | TDRKH antisense RNA 1 [Source:HGNC Symbo |
| AL589765.6 | 1 | novel transcript |
| LINGO4 | 1 | leucine rich repeat and Ig domain containing 4 |
| RORC | 1 | RAR related orphan receptor C [Source:HGNC |
| C2CD4D | 1 | C2 calcium dependent domain containing 4D [ |
| Y_RNA | 1 | Y RNA [Source:RFAM;Acc:RF00019] |
| C2CD4D-AS1 | 1 | C2CD4D and THEM5 antisense RNA 1 [Source: |
| THEM5 | 1 | thioesterase superfamily member 5 [Source:H |
| THEM4 | 1 | thioesterase superfamily member 4 [Source:H |
| AL450992.2 | 1 | novel transcript, antisense to THEM4 |
| AL450992.3 | 1 | novel transcript |
| KRT8P28 | 1 | keratin 8 pseudogene 28 [Source:HGNC Symb |
| S100A10 | 1 | S100 calcium binding protein A10 [Source:HGI |
| AL450992.1 | 1 | novel transcript |
| NBPF18P | 1 | NBPF member 18, pseudogene [Source:HGNC |
| S100A11 | 1 | S100 calcium binding protein A11 [Source:HGI |
| SPTLC1P4 | 1 | serine palmitoyltransferase long chain base su |
| AL591893.1 | 1 | putative UPF0607 protein ENSP00000383783 |
| TCHHL1 | 1 | trichohyalin like 1 [Source:HGNC Symbol;Acc:I |
| TCHH | 1 | trichohyalin [Source:HGNC Symbol;Acc:HGNC: |
| AL589986.1 | 1 | novel transcript |
| PUDPP2 | 1 | pseudouridine 5'-phosphatase pseudogene 2 |
| RPTN | 1 | repetin [Source:HGNC Symbol;Acc:HGNC:268( |
| FLG-AS1 | 1 | FLG antisense RNA 1 [Source:HGNC Symbol;A( |
| AL589986.2 | 1 | novel transcript |
| HRNR | 1 | hornerin [Source:HGNC Symbol;Acc:HGNC:20 |
| FLG | 1 | filaggrin [Source:HGNC Symbol;Acc:HGNC:374 |
| FLG2 | 1 | filaggrin family member 2 [Source:HGNC Symb |
| HMGN3P1 | 1 | high mobility group nucleosomal binding dom |
| CRNN | 1 | cornulin [Source:HGNC Symbol;Acc:HGNC:123 |
| LCE5A | 1 | late cornified envelope 5A [Source:HGNC Sym |

| | | |
|---|---|---|
| CRCT1 | 1 | cysteine rich C-terminal 1 [Source:HGNC Symb |
| LCE3E | 1 | late cornified envelope 3E [Source:HGNC Sym |
| LCE3D | 1 | late cornified envelope 3D [Source:HGNC Sym |
| LCE3C | 1 | late cornified envelope 3C [Source:HGNC Sym |
| LCE3B | 1 | late cornified envelope 3B [Source:HGNC Sym |
| LCE3A | 1 | late cornified envelope 3A [Source:HGNC Sym |
| LCEP4 | 1 | late cornified envelope pseudogene 4 [Source |
| AL139247.1 | 1 | late cornified envelope family pseudogene |
| LCE2D | 1 | late cornified envelope 2D [Source:HGNC Sym |
| LCE2C | 1 | late cornified envelope 2C [Source:HGNC Sym |
| LCE2B | 1 | late cornified envelope 2B [Source:HGNC Sym |
| LCE2A | 1 | late cornified envelope 2A [Source:HGNC Sym |
| LCE4A | 1 | late cornified envelope 4A [Source:HGNC Sym |
| C1orf68 | 1 | chromosome 1 open reading frame 68 [Sourc |
| LCEP2 | 1 | late cornified envelope pseudogene 2 [Source |
| LCEP1 | 1 | late cornified envelope pseudogene 1 [Source |
| KPRP | 1 | keratinocyte proline rich protein [Source:HGN |
| LCE1F | 1 | late cornified envelope 1F [Source:HGNC Sym |
| LCE1E | 1 | late cornified envelope 1E [Source:HGNC Sym |
| LCE1D | 1 | late cornified envelope 1D [Source:HGNC Sym |
| LCE1C | 1 | late cornified envelope 1C [Source:HGNC Sym |
| LCE1B | 1 | late cornified envelope 1B [Source:HGNC Sym |
| LCE1A | 1 | late cornified envelope 1A [Source:HGNC Sym |
| LCE6A | 1 | late cornified envelope 6A [Source:HGNC Sym |
| AL162596.1 | 1 | novel late cornified envelope family protein |
| SMCP | 1 | sperm mitochondria associated cysteine rich p |
| IVL | 1 | involucrin [Source:HGNC Symbol;Acc:HGNC:6 |
| LINC01527 | 1 | long intergenic non-protein coding RNA 1527 |
| SPRR5 | 1 | small proline rich protein 5 [Source:HGNC Sym |
| SPRR4 | 1 | small proline rich protein 4 [Source:HGNC Sym |
| SPRR1A | 1 | small proline rich protein 1A [Source:HGNC Sy |
| SPRR3 | 1 | small proline rich protein 3 [Source:HGNC Sym |
| AL356867.2 | 1 | NULL |
| AL356867.1 | 1 | small proline-rich protein 3 (SPRR3) pseudoge |
| SPRR1B | 1 | small proline rich protein 1B [Source:HGNC Sy |
| SPRR2D | 1 | small proline rich protein 2D [Source:HGNC Sy |
| SPRR2A | 1 | small proline rich protein 2A [Source:HGNC Sy |
| SPRR2B | 1 | small proline rich protein 2B [Source:HGNC Sy |
| SPRR2E | 1 | small proline rich protein 2E [Source:HGNC Sy |
| SPRR2F | 1 | small proline rich protein 2F [Source:HGNC Sy |
| SPRR2C | 1 | small proline rich protein 2C (pseudogene) [Sc |
| SPRR2G | 1 | small proline rich protein 2G [Source:HGNC Sy |
| AL161636.1 | 1 | novel transcript |
| LELP1 | 1 | late cornified envelope like proline rich 1 [Sou |
| PRR9 | 1 | proline rich 9 [Source:HGNC Symbol;Acc:HGN |
| AL161636.2 | 1 | ribosomal protein, large, P0 (RPLP0) pseudoge |
| LOR | 1 | loricrin [Source:HGNC Symbol;Acc:HGNC:666 |
| PGLYRP3 | 1 | peptidoglycan recognition protein 3 [Source:H |
| PGLYRP4 | 1 | peptidoglycan recognition protein 4 [Source:H |
| RNU6-160P | 1 | RNA, U6 small nuclear 160, pseudogene [Sour |

| | | |
|---|---|---|
| S100A9 | 1 | S100 calcium binding protein A9 [Source:HGN |
| S100A12 | 1 | S100 calcium binding protein A12 [Source:HGI |
| LAPTM4BP1 | 1 | lysosomal protein transmembrane 4 beta pseu |
| S100A8 | 1 | S100 calcium binding protein A8 [Source:HGN |
| S100A15A | 1 | S100 calcium binding protein A15A (pseudoge |
| S100A7A | 1 | S100 calcium binding protein A7A [Source:HG |
| S100A7P1 | 1 | S100 calcium binding protein A7 pseudogene |
| S100A7L2 | 1 | S100 calcium binding protein A7 like 2 [Source |
| S100A7 | 1 | S100 calcium binding protein A7 [Source:HGN |
| RN7SL44P | 1 | RNA, 7SL, cytoplasmic 44, pseudogene [Source |
| BX470102.1 | 1 | novel transcript |
| S100A6 | 1 | S100 calcium binding protein A6 [Source:HGN |
| S100A5 | 1 | S100 calcium binding protein A5 [Source:HGN |
| S100A4 | 1 | S100 calcium binding protein A4 [Source:HGN |
| S100A3 | 1 | S100 calcium binding protein A3 [Source:HGN |
| S100A2 | 1 | S100 calcium binding protein A2 [Source:HGN |
| BX470102.2 | 1 | Novel transcript |
| S100A16 | 1 | S100 calcium binding protein A16 [Source:HGI |
| S100A14 | 1 | S100 calcium binding protein A14 [Source:HGI |
| S100A13 | 1 | S100 calcium binding protein A13 [Source:HGI |
| AL162258.1 | 1 | novel transcript |
| S100A1 | 1 | S100 calcium binding protein A1 [Source:HGN |
| AL162258.2 | 1 | novel transcript, antisense to S100A1 and CHT |
| CHTOP | 1 | chromatin target of PRMT1 [Source:HGNC Syr |
| SNAPIN | 1 | SNAP associated protein [Source:HGNC Symbo |
| ILF2 | 1 | interleukin enhancer binding factor 2 [Source: |
| NPR1 | 1 | natriuretic peptide receptor 1 [Source:HGNC S |
| MIR8083 | 1 | microRNA 8083 [Source:HGNC Symbol;Acc:HG |
| RN7SL372P | 1 | RNA, 7SL, cytoplasmic 372, pseudogene [Sour |
| GEMIN2P1 | 1 | gem nuclear organelle associated protein 2 ps |
| Y_RNA | 1 | Y RNA [Source:RFAM;Acc:RF00019] |
| INTS3 | 1 | integrator complex subunit 3 [Source:HGNC S |
| AL513523.4 | 1 | novel transcript |
| AL513523.3 | 1 | novel transcript |
| SLC27A3 | 1 | solute carrier family 27 member 3 [Source:HG |
| Y_RNA | 1 | Y RNA [Source:RFAM;Acc:RF00019] |
| GATAD2B | 1 | GATA zinc finger domain containing 2B [Sourc |
| AL513523.2 | 1 | immunoglobulin superfamily, DCC subclass, m |
| AL513523.1 | 1 | adipose differentiation-related protein (ADFP) |
| AL831737.1 | 1 | ribosomal protein L30 (RPL30) pseudogene |
| AL358472.5 | 1 | novel transcript, antisense to DENND4B |
| DENND4B | 1 | DENN domain containing 4B [Source:HGNC Sy |
| CRTC2 | 1 | CREB regulated transcription coactivator 2 [Sc |
| SLC39A1 | 1 | solute carrier family 39 member 1 [Source:HG |
| AL358472.7 | 1 | novel protein |
| MIR6737 | 1 | microRNA 6737 [Source:HGNC Symbol;Acc:HG |
| AL358472.4 | 1 | novel transcript, antisense to SLC39A1 |
| AL358472.3 | 1 | novel transcript, antisense to SLC39A1 |
| CREB3L4 | 1 | cAMP responsive element binding protein 3 li |
| JTB | 1 | jumping translocation breakpoint [Source:HGI |

| | | |
|---|---|---|
| AL358472.6 | 1 | novel protein |
| AL358472.2 | 1 | novel transcript |
| RAB13 | 1 | RAB13, member RAS oncogene family [Source |
| RPS27 | 1 | ribosomal protein S27 [Source:HGNC Symbol; |
| NUP210L | 1 | nucleoporin 210 like [Source:HGNC Symbol;Ac |
| AL358472.1 | 1 | ribosomal protein L34 (RPL34) pseudogene |
| RNU6-179P | 1 | RNA, U6 small nuclear 179, pseudogene [Sour |
| RPS7P2 | 1 | ribosomal protein S7 pseudogene 2 [Source:H |
| MIR5698 | 1 | microRNA 5698 [Source:HGNC Symbol;Acc:HG |
| TPM3 | 1 | tropomyosin 3 [Source:HGNC Symbol;Acc:HGI |
| RN7SL431P | 1 | RNA, 7SL, cytoplasmic 431, pseudogene [Sour |
| MIR190B | 1 | microRNA 190b [Source:HGNC Symbol;Acc:HG |
| C1orf189 | 1 | chromosome 1 open reading frame 189 [Sour |
| C1orf43 | 1 | chromosome 1 open reading frame 43 [Sourc |
| UBAP2L | 1 | ubiquitin associated protein 2 like [Source:HG |
| SNORA58B | 1 | small nucleolar RNA, H/ACA box 58B [Source:I |
| HAX1 | 1 | HCLS1 associated protein X-1 [Source:HGNC S |
| AL354980.1 | 1 | NULL |
| RNU6-239P | 1 | RNA, U6 small nuclear 239, pseudogene [Sour |
| RNU6-121P | 1 | RNA, U6 small nuclear 121, pseudogene [Sour |
| AL354980.2 | 1 | aquaporin 10 (AQP10) pseudogene |
| AQP10 | 1 | aquaporin 10 [Source:HGNC Symbol;Acc:HGN |
| ATP8B2 | 1 | ATPase phospholipid transporting 8B2 [Source |
| RNU7-57P | 1 | RNA, U7 small nuclear 57 pseudogene [Source |
| AL162591.1 | 1 | mitochondrial ribosomal protein S33 (MRPS33 |
| RPSAP17 | 1 | ribosomal protein SA pseudogene 17 [Source: |
| IL6R-AS1 | 1 | IL6R antisense RNA 1 [Source:HGNC Symbol;A |
| IL6R | 1 | interleukin 6 receptor [Source:HGNC Symbol; |
| PSMD8P1 | 1 | proteasome 26S subunit, non-ATPase, 8 pseu |
| SHE | 1 | Src homology 2 domain containing E [Source:I |
| AL162591.2 | 1 | novel transcript, antisense to SHE |
| TDRD10 | 1 | tudor domain containing 10 [Source:HGNC Sy |
| UBE2Q1 | 1 | ubiquitin conjugating enzyme E2 Q1 [Source:H |
| UBE2Q1-AS1 | 1 | UBE2Q1 antisense RNA 1 [Source:HGNC Symb |
| AL592078.2 | 1 | novel transcript |
| CHRNB2 | 1 | cholinergic receptor nicotinic beta 2 subunit [ |
| AL592078.1 | 1 | novel transcript |
| ADAR | 1 | adenosine deaminase RNA specific [Source:HG |
| AL606500.1 | 1 | novel transcript |
| KCNN3 | 1 | potassium calcium-activated channel subfami |
| PMVK | 1 | phosphomevalonate kinase [Source:HGNC Syr |
| AL451085.1 | 1 | novel transcript |
| PBXIP1 | 1 | PBX homeobox interacting protein 1 [Source:I |
| Metazoa_SRP | 1 | Metazoan signal recognition particle RNA [Sou |
| PYGO2 | 1 | pygopus family PHD finger 2 [Source:HGNC Sy |
| AL451085.2 | 1 | uncharacterized LOC101928120 [Source:NCBI |
| SHC1 | 1 | SHC adaptor protein 1 [Source:HGNC Symbol; |
| CKS1B | 1 | CDC28 protein kinase regulatory subunit 1B [S |
| MIR4258 | 1 | microRNA 4258 [Source:HGNC Symbol;Acc:HG |
| FLAD1 | 1 | flavin adenine dinucleotide synthetase 1 [Sour |

| | | |
|---|---|---|
| LENEP | 1 | lens epithelial protein [Source:HGNC Symbol;/ |
| ZBTB7B | 1 | zinc finger and BTB domain containing 7B [Sou |
| DCST2 | 1 | DC-STAMP domain containing 2 [Source:HGN( |
| DCST1 | 1 | DC-STAMP domain containing 1 [Source:HGN( |
| DCST1-AS1 | 1 | DCST1 antisense RNA 1 [Source:HGNC Symbol |
| ADAM15 | 1 | ADAM metallopeptidase domain 15 [Source:H |
| EFNA4 | 1 | ephrin A4 [Source:HGNC Symbol;Acc:HGNC:3Z |
| AL691442.1 | 1 | novel ephrin-A4 (EFNA4) and ephrin-A3 (EFNA |
| EFNA3 | 1 | ephrin A3 [Source:HGNC Symbol;Acc:HGNC:3Z |
| Y_RNA | 1 | Y RNA [Source:RFAM;Acc:RF00019] |
| EFNA1 | 1 | ephrin A1 [Source:HGNC Symbol;Acc:HGNC:3Z |
| SLC50A1 | 1 | solute carrier family 50 member 1 [Source:HG |
| DPM3 | 1 | dolichyl-phosphate mannosyltransferase subu |
| HMGN2P18 | 1 | high mobility group nucleosomal binding dom |
| KRTCAP2 | 1 | keratinocyte associated protein 2 [Source:HGI |
| AL713999.1 | 1 | novel protein |
| TRIM46 | 1 | tripartite motif containing 46 [Source:HGNC S' |
| MUC1 | 1 | mucin 1, cell surface associated [Source:HGN( |
| AC234582.1 | 1 | novel transcript, antisense to THBS3 |
| MIR92B | 1 | microRNA 92b [Source:HGNC Symbol;Acc:HGI |
| THBS3 | 1 | thrombospondin 3 [Source:HGNC Symbol;Acc |
| MTX1 | 1 | metaxin 1 [Source:HGNC Symbol;Acc:HGNC:7! |
| AC234582.2 | 1 | novel transcript |
| GBAP1 | 1 | glucosylceramidase beta pseudogene 1 [Sourc |
| MTX1P1 | 1 | metaxin 1 pseudogene 1 [Source:HGNC Symb |
| GBA | 1 | glucosylceramidase beta [Source:HGNC Symb |
| FAM189B | 1 | family with sequence similarity 189 member E |
| SCAMP3 | 1 | secretory carrier membrane protein 3 [Source |
| CLK2 | 1 | CDC like kinase 2 [Source:HGNC Symbol;Acc:H |
| HCN3 | 1 | hyperpolarization activated cyclic nucleotide ք |
| PKLR | 1 | pyruvate kinase L/R [Source:HGNC Symbol;Ac |
| FDPS | 1 | farnesyl diphosphate synthase [Source:HGNC |
| RUSC1-AS1 | 1 | RUSC1 antisense RNA 1 [Source:HGNC Symbo |
| RUSC1 | 1 | RUN and SH3 domain containing 1 [Source:HG |
| ASH1L | 1 | ASH1 like histone lysine methyltransferase [Sc |
| MIR555 | 1 | microRNA 555 [Source:HGNC Symbol;Acc:HGI |
| RNU6-106P | 1 | RNA, U6 small nuclear 106, pseudogene [Sour |
| ASH1L-IT1 | 1 | ASH1L intronic transcript 1 [Source:HGNC Sym |
| RNU6-1297P | 1 | RNA, U6 small nuclear 1297, pseudogene [Sou |
| POU5F1P4 | 1 | POU class 5 homeobox 1 pseudogene 4 [Sourc |
| ASH1L-AS1 | 1 | ASH1L antisense RNA 1 [Source:HGNC Symbol |
| AL353807.4 | 1 | ATP synthase, H+ transporting, mitochondrial |
| DAP3P1 | 1 | death associated protein 3 pseudogene 1 [Sou |
| AL353807.1 | 1 | claudin 6 (CLDN6) pseudogene |
| AL353807.2 | 1 | novel transcript, antisense to MSTO1 |
| MSTO1 | 1 | misato mitochondrial distribution and morpho |
| AL353807.3 | 1 | novel pseudogene |
| AL353807.5 | 1 | novel transcript |
| YY1AP1 | 1 | YY1 associated protein 1 [Source:HGNC Symb |
| DAP3 | 1 | death associated protein 3 [Source:HGNC Sym |

| | | |
|---|---|---|
| AL162734.1 | 1 | novel transcript, antisense to DAP3 |
| MSTO2P | 1 | misato family member 2, pseudogene [Source |
| GON4L | 1 | gon-4 like [Source:HGNC Symbol;Acc:HGNC:2[illegible] |
| AL139128.1 | 1 | proteasome (prosome, macropain) 26S subun |
| SYT11 | 1 | synaptotagmin 11 [Source:HGNC Symbol;Acc: |
| U4 | 1 | U4 spliceosomal RNA [Source:RFAM;Acc:RF00 |
| RIT1 | 1 | Ras like without CAAX 1 [Source:HGNC Symbo |
| KHDC4 | 1 | KH domain containing 4, pre-mRNA splicing fa |
| SNORA80E | 1 | small nucleolar RNA, H/ACA box 80E [Source:[illegible] |
| SCARNA4 | 1 | small Cajal body-specific RNA 4 [Source:HGNC |
| SCARNA4 | 1 | small Cajal body-specific RNA 4 [Source:HGNC |
| RXFP4 | 1 | relaxin family peptide/INSL5 receptor 4 [Sourc |
| ARHGEF2 | 1 | Rho/Rac guanine nucleotide exchange factor [illegible] |
| MIR6738 | 1 | microRNA 6738 [Source:HGNC Symbol;Acc:H[illegible] |
| AL355388.2 | 1 | novel transcript, antisense to ARHGEF2 |
| AL355388.1 | 1 | novel transcript |
| AL355388.3 | 1 | Novel transcript, antisense to SSR2 |
| SSR2 | 1 | signal sequence receptor subunit 2 [Source:H[illegible] |
| UBQLN4 | 1 | ubiquilin 4 [Source:HGNC Symbol;Acc:HGNC:1 |
| LAMTOR2 | 1 | late endosomal/lysosomal adaptor, MAPK an[illegible] |
| RAB25 | 1 | RAB25, member RAS oncogene family [Source |
| MEX3A | 1 | mex-3 RNA binding family member A [Source: |
| LMNA | 1 | lamin A/C [Source:HGNC Symbol;Acc:HGNC:6[illegible] |
| SEMA4A | 1 | semaphorin 4A [Source:HGNC Symbol;Acc:HG |
| AL135927.1 | 1 | NULL |
| SLC25A44 | 1 | solute carrier family 25 member 44 [Source:H |
| PMF1-BGLAP | 1 | PMF1-BGLAP readthrough [Source:HGNC Sym |
| PMF1 | 1 | polyamine modulated factor 1 [Source:HGNC |
| BGLAP | 1 | bone gamma-carboxyglutamate protein [Sour |
| PAQR6 | 1 | progestin and adipoQ receptor family membe |
| SMG5 | 1 | SMG5 nonsense mediated mRNA decay facto[illegible] |
| TMEM79 | 1 | transmembrane protein 79 [Source:HGNC Syn |
| GLMP | 1 | glycosylated lysosomal membrane protein [Sc |
| VHLL | 1 | VHL like [Source:HGNC Symbol;Acc:HGNC:306 |
| CCT3 | 1 | chaperonin containing TCP1 subunit 3 [Source |
| TSACC | 1 | TSSK6 activating cochaperone [Source:HGNC [illegible] |
| RHBG | 1 | Rh family B glycoprotein (gene/pseudogene) [ |
| AL139130.1 | 1 | novel transcript |
| C1orf61 | 1 | chromosome 1 open reading frame 61 [Sourc[illegible] |
| MIR9-1 | 1 | microRNA 9-1 [Source:HGNC Symbol;Acc:HG[illegible] |
| MEF2D | 1 | myocyte enhancer factor 2D [Source:HGNC Sy |
| Y_RNA | 1 | Y RNA [Source:RFAM;Acc:RF00019] |
| AL365181.1 | 1 | novel transcript |
| IQGAP3 | 1 | IQ motif containing GTPase activating protein |
| TTC24 | 1 | tetratricopeptide repeat domain 24 [Source:H |
| NAXE | 1 | NAD(P)HX epimerase [Source:HGNC Symbol;A |
| GPATCH4 | 1 | G-patch domain containing 4 [Source:HGNC S[illegible] |
| AL365181.4 | 1 | novel transcript |
| HAPLN2 | 1 | hyaluronan and proteoglycan link protein 2 [S |
| AL365181.2 | 1 | novel transcript |

| | | |
|---|---|---|
| BCAN | 1 | brevican [Source:HGNC Symbol;Acc:HGNC:23( |
| AL365181.3 | 1 | novel transcript, antisense to BCAN |
| AL590666.2 | 1 | novel transcript |
| NES | 1 | nestin [Source:HGNC Symbol;Acc:HGNC:7756] |
| AL590666.3 | 1 | novel transcript |
| AL590666.4 | 1 | Novel transcript, antisense to CRABP2 |
| CRABP2 | 1 | cellular retinoic acid binding protein 2 [Source |
| AL590666.1 | 1 | novel transcript |
| ISG20L2 | 1 | interferon stimulated exonuclease gene 20 lik |
| RRNAD1 | 1 | ribosomal RNA adenine dimethylase domain c |
| MRPL24 | 1 | mitochondrial ribosomal protein L24 [Source:I |
| HDGF | 1 | heparin binding growth factor [Source:HGNC S |
| PRCC | 1 | proline rich mitotic checkpoint control factor |
| AL590666.5 | 1 | novel transcript, antisense to PRCC |
| SH2D2A | 1 | SH2 domain containing 2A [Source:HGNC Sym |
| NTRK1 | 1 | neurotrophic receptor tyrosine kinase 1 [Sour |
| INSRR | 1 | insulin receptor related receptor [Source:HGN |
| PEAR1 | 1 | platelet endothelial aggregation receptor 1 [S |
| LRRC71 | 1 | leucine rich repeat containing 71 [Source:HGN |
| ARHGEF11 | 1 | Rho guanine nucleotide exchange factor 11 [S |
| MIR765 | 1 | microRNA 765 [Source:HGNC Symbol;Acc:HGI |
| RN7SL612P | 1 | RNA, 7SL, cytoplasmic 612, pseudogene [Sour |
| AL157713.1 | 1 | smu-1 suppressor of mec-8 and unc-52 homol |
| KRT8P45 | 1 | keratin 8 pseudogene 45 [Source:HGNC Symb |
| ETV3L | 1 | ETS variant 3 like [Source:HGNC Symbol;Acc:H |
| ETV3 | 1 | ETS variant 3 [Source:HGNC Symbol;Acc:HGN( |
| CYCSP52 | 1 | CYCS pseudogene 52 [Source:HGNC Symbol;A |
| AL357143.1 | 1 | novel SLAM family member pseudogene |
| AL357143.2 | 1 | novel SLAM family member pseudogene |
| AL138900.1 | 1 | novel transcript |
| LINC02772 | 1 | long intergenic non-protein coding RNA 2772 |
| AL138900.2 | 1 | SLAM family member 5-like pseudogene |
| AL732406.1 | 1 | SLAM family member 5-like pseudogene |
| AL353899.1 | 1 | SLAM family member 5-like pseudogene |
| FCRL5 | 1 | Fc receptor like 5 [Source:HGNC Symbol;Acc:H |
| FCRL4 | 1 | Fc receptor like 4 [Source:HGNC Symbol;Acc:H |
| AL135929.2 | 1 | pseudogene similar to part of Fc receptor-like |
| AL135929.1 | 1 | pseudogene similar to part of Fc receptor-like |
| FCRL3 | 1 | Fc receptor like 3 [Source:HGNC Symbol;Acc:H |
| AL356276.1 | 1 | novel transcript |
| AL356276.2 | 1 | pseudogene similar to part of SON DNA bindir |
| VDAC1P9 | 1 | voltage dependent anion channel 1 pseudoge |
| FCRL2 | 1 | Fc receptor like 2 [Source:HGNC Symbol;Acc:H |
| FCRL1 | 1 | Fc receptor like 1 [Source:HGNC Symbol;Acc:H |
| CD5L | 1 | CD5 molecule like [Source:HGNC Symbol;Acc:I |
| MRPS21P2 | 1 | mitochondrial ribosomal protein S21 pseudog |
| AL139010.1 | 1 | pseudogene similar to part of a immunoglobu |
| KIRREL1 | 1 | kirre like nephrin family adhesion molecule 1 |
| KIRREL1-IT1 | 1 | KIRREL1 intronic transcript 1 [Source:HGNC Sy |
| AL138899.3 | 1 | novel protein |

| | | |
|---|---|---|
| LINC01704 | 1 | long intergenic non-protein coding RNA 1704 |
| ELL2P1 | 1 | elongation factor for RNA polymerase II 2 pse |
| CD1D | 1 | CD1d molecule [Source:HGNC Symbol;Acc:HG |
| AL138899.2 | 1 | ribosomal protein S10 (RPS10) pseudogene |
| AL138899.1 | 1 | novel transcript |
| CD1A | 1 | CD1a molecule [Source:HGNC Symbol;Acc:HG |
| HMGN1P5 | 1 | high mobility group nucleosome binding dom |
| CD1C | 1 | CD1c molecule [Source:HGNC Symbol;Acc:HG |
| CD1B | 1 | CD1b molecule [Source:HGNC Symbol;Acc:HG |
| CD1E | 1 | CD1e molecule [Source:HGNC Symbol;Acc:HG |
| OR10T2 | 1 | olfactory receptor family 10 subfamily T mem |
| OR10K2 | 1 | olfactory receptor family 10 subfamily K mem |
| OR10T1P | 1 | olfactory receptor family 10 subfamily T mem |
| EI24P2 | 1 | EI24 pseudogene 2 [Source:HGNC Symbol;Acc |
| OR10K1 | 1 | olfactory receptor family 10 subfamily K mem |
| OR10R2 | 1 | olfactory receptor family 10 subfamily R mem |
| AL365440.2 | 1 | novel transcript |
| OR10R3P | 1 | olfactory receptor family 10 subfamily R mem |
| OR10R1P | 1 | olfactory receptor family 10 subfamily R mem |
| AL365440.1 | 1 | heat shock 90kDa protein 1, alpha (HSPCA) ps |
| OR6Y1 | 1 | olfactory receptor family 6 subfamily Y memb |
| OR6P1 | 1 | olfactory receptor family 6 subfamily P memb |
| OR10X1 | 1 | olfactory receptor family 10 subfamily X mem |
| OR10Z1 | 1 | olfactory receptor family 10 subfamily Z mem |
| SPTA1 | 1 | spectrin alpha, erythrocytic 1 [Source:HGNC S |
| OR6K1P | 1 | olfactory receptor family 6 subfamily K memb |
| OR6K2 | 1 | olfactory receptor family 6 subfamily K memb |
| OR6K3 | 1 | olfactory receptor family 6 subfamily K memb |
| OR6K4P | 1 | olfactory receptor family 6 subfamily K memb |
| OR6K5P | 1 | olfactory receptor family 6 subfamily K memb |
| OR6N1 | 1 | olfactory receptor family 6 subfamily N memb |
| OR6K6 | 1 | olfactory receptor family 6 subfamily K memb |
| OR6N2 | 1 | olfactory receptor family 6 subfamily N memb |
| OR2AQ1P | 1 | olfactory receptor family 2 subfamily AQ mem |
| OR10AA1P | 1 | olfactory receptor family 10 subfamily AA me |
| MNDA | 1 | myeloid cell nuclear differentiation antigen [S |
| PYHIN5P | 1 | pyrin and HIN domain family member 5, pseu |
| PYHIN1 | 1 | pyrin and HIN domain family member 1 [Sour |
| IFI16 | 1 | interferon gamma inducible protein 16 [Sourc |
| AIM2 | 1 | absent in melanoma 2 [Source:HGNC Symbol; |
| RAD1P2 | 1 | RAD1 pseudogene 2 [Source:HGNC Symbol;Ac |
| CADM3 | 1 | cell adhesion molecule 3 [Source:HGNC Symb |
| RNA5SP60 | 1 | RNA, 5S ribosomal pseudogene 60 [Source:HG |
| CADM3-AS1 | 1 | CADM3 antisense RNA 1 [Source:HGNC Symb |
| ACKR1 | 1 | atypical chemokine receptor 1 (Duffy blood gr |
| MPTX1 | 1 | mucosal pentraxin 1 (pseudogene) [Source:H |
| OR10J2P | 1 | olfactory receptor family 10 subfamily J mem |
| FCER1A | 1 | Fc fragment of IgE receptor Ia [Source:HGNC S |
| OR10J3 | 1 | olfactory receptor family 10 subfamily J mem |
| AL513323.1 | 1 | novel transcript |

| | | |
|---|---|---|
| OR10J7P | 1 | olfactory receptor family 10 subfamily J meml |
| OR10J8P | 1 | olfactory receptor family 10 subfamily J meml |
| OR10J9P | 1 | olfactory receptor family 10 subfamily J meml |
| OR10J4 | 1 | olfactory receptor family 10 subfamily J meml |
| OR10J1 | 1 | olfactory receptor family 10 subfamily J meml |
| LINC02819 | 1 | long intergenic non-protein coding RNA 2819 |
| OR10J5 | 1 | olfactory receptor family 10 subfamily J meml |
| AL606752.1 | 1 | amyloid P component, serum (APCS) pseudog |
| OR10AE1P | 1 | olfactory receptor family 10 subfamily AE mer |
| APCS | 1 | amyloid P component, serum [Source:HGNC S |
| OR10J6P | 1 | olfactory receptor family 10 subfamily J meml |
| AL445528.2 | 1 | elastin microfibril interfacer 2 (EMILIN2) pseu |
| CRPP1 | 1 | C-reactive protein pseudogene 1 [Source:HGN |
| CRP | 1 | C-reactive protein [Source:HGNC Symbol;Acc: |
| AL445528.1 | 1 | ribosomal protein L27 (RPL27) pseudogene |
| DUSP23 | 1 | dual specificity phosphatase 23 [Source:HGNC |
| FCRL6 | 1 | Fc receptor like 6 [Source:HGNC Symbol;Acc:H |
| SLAMF8 | 1 | SLAM family member 8 [Source:HGNC Symbo |
| AL590560.2 | 1 | novel protein |
| SNHG28 | 1 | small nucleolar RNA host gene 28 [Source:HGI |
| AL590560.1 | 1 | NULL |
| VSIG8 | 1 | V-set and immunoglobulin domain containing |
| AL590560.3 | 1 | novel transcript, antisense to VSIG8 |
| CFAP45 | 1 | cilia and flagella associated protein 45 [Source |
| MIR4259 | 1 | microRNA 4259 [Source:HGNC Symbol;Acc:HG |
| AL590560.5 | 1 | novel transcript |
| AL590560.4 | 1 | novel transcript |
| TAGLN2 | 1 | transgelin 2 [Source:HGNC Symbol;Acc:HGNC: |
| IGSF9 | 1 | immunoglobulin superfamily member 9 [Sour |
| SLAMF9 | 1 | SLAM family member 9 [Source:HGNC Symbo |
| LINC01133 | 1 | long intergenic non-protein coding RNA 1133 |
| AL513485.1 | 1 | Fc receptor-like 6 (FCRL6) pseudogene |
| KCNJ10 | 1 | potassium voltage-gated channel subfamily J r |
| AL513302.1 | 1 | ribosomal protein L27a (RPL27A) pseudogene |
| PIGM | 1 | phosphatidylinositol glycan anchor biosynthes |
| AL121987.1 | 1 | novel transcript |
| KCNJ9 | 1 | potassium voltage-gated channel subfamily J r |
| IGSF8 | 1 | immunoglobulin superfamily member 8 [Sour |
| ATP1A2 | 1 | ATPase Na+/K+ transporting subunit alpha 2 [ |
| ATP1A4 | 1 | ATPase Na+/K+ transporting subunit alpha 4 [ |
| CASQ1 | 1 | calsequestrin 1 [Source:HGNC Symbol;Acc:HG |
| AL121987.2 | 1 | novel transcript |
| PEA15 | 1 | proliferation and apoptosis adaptor protein 1 |
| DCAF8 | 1 | DDB1 and CUL4 associated factor 8 [Source:H |
| AL139011.2 | 1 | novel protein |
| AL139011.1 | 1 | novel transcript |
| RPSAP18 | 1 | ribosomal protein SA pseudogene 18 [Source: |
| PEX19 | 1 | peroxisomal biogenesis factor 19 [Source:HGN |
| COPA | 1 | coatomer protein complex subunit alpha [Sou |
| SUMO1P3 | 1 | SUMO1 pseudogene 3 [Source:HGNC Symbol; |

| | | |
|---|---|---|
| Y_RNA | 1 | Y RNA [Source:RFAM;Acc:RF00019] |
| NCSTN | 1 | nicastrin [Source:HGNC Symbol;Acc:HGNC:17( |
| NHLH1 | 1 | nescient helix-loop-helix 1 [Source:HGNC Sym |
| RNU4-42P | 1 | RNA, U4 small nuclear 42, pseudogene [Sourc |
| VANGL2 | 1 | VANGL planar cell polarity protein 2 [Source:H |
| SLAMF6 | 1 | SLAM family member 6 [Source:HGNC Symbo |
| AL138930.1 | 1 | novel transcript |
| CD84 | 1 | CD84 molecule [Source:HGNC Symbol;Acc:HG |
| SLAMF1 | 1 | signaling lymphocytic activation molecule fam |
| SETP9 | 1 | SET pseudogene 9 [Source:HGNC Symbol;Acc: |
| AL121985.1 | 1 | novel transcript, antisense to CD48 |
| CD48 | 1 | CD48 molecule [Source:HGNC Symbol;Acc:HG |
| SLAMF7 | 1 | SLAM family member 7 [Source:HGNC Symbo |
| AL121985.3 | 1 | SLAM family member 9 (SLAMF9) pseudogene |
| AL121985.2 | 1 | SLAM family member 9 (SLAMF9) pseudogene |
| LY9 | 1 | lymphocyte antigen 9 [Source:HGNC Symbol;A |
| CD244 | 1 | CD244 molecule [Source:HGNC Symbol;Acc:H |
| PPIAP37 | 1 | peptidylprolyl isomerase A pseudogene 37 [Sc |
| ITLN1 | 1 | intelectin 1 [Source:HGNC Symbol;Acc:HGNC: |
| AL354714.2 | 1 | spermine synthase (SMS) pseudogene |
| AL354714.3 | 1 | novel transcript |
| AL354714.1 | 1 | antisense to ITLN2 |
| AL354714.4 | 1 | pseudogene similar to part of DEAD (Asp-Glu- |
| ITLN2 | 1 | intelectin 2 [Source:HGNC Symbol;Acc:HGNC: |
| F11R | 1 | F11 receptor [Source:HGNC Symbol;Acc:HGN( |
| AL591806.4 | 1 | novel transcript, TSTD1 - F11R readthrough |
| AL591806.2 | 1 | glutaredoxin 5 (GLRX5) pseudogene |
| TSTD1 | 1 | thiosulfate sulfurtransferase like domain cont |
| USF1 | 1 | upstream transcription factor 1 [Source:HGN( |
| ARHGAP30 | 1 | Rho GTPase activating protein 30 [Source:HGI |
| NECTIN4 | 1 | nectin cell adhesion molecule 4 [Source:HGN( |
| AL591806.1 | 1 | novel transcript |
| KLHDC9 | 1 | kelch domain containing 9 [Source:HGNC Sym |
| PFDN2 | 1 | prefoldin subunit 2 [Source:HGNC Symbol;Acc |
| NIT1 | 1 | nitrilase 1 [Source:HGNC Symbol;Acc:HGNC:7 |
| DEDD | 1 | death effector domain containing [Source:HGI |
| AL591806.3 | 1 | NULL |
| UFC1 | 1 | ubiquitin-fold modifier conjugating enzyme 1 |
| AL590714.1 | 1 | novel transcript |
| USP21 | 1 | ubiquitin specific peptidase 21 [Source:HGNC |
| PPOX | 1 | protoporphyrinogen oxidase [Source:HGNC Sy |
| B4GALT3 | 1 | beta-1,4-galactosyltransferase 3 [Source:HGN |
| ADAMTS4 | 1 | ADAM metallopeptidase with thrombospondi |
| NDUFS2 | 1 | NADH:ubiquinone oxidoreductase core subun |
| FCER1G | 1 | Fc fragment of IgE receptor Ig [Source:HGNC S |
| APOA2 | 1 | apolipoprotein A2 [Source:HGNC Symbol;Acc: |
| TOMM40L | 1 | translocase of outer mitochondrial membrane |
| MIR5187 | 1 | microRNA 5187 [Source:HGNC Symbol;Acc:HG |
| NR1I3 | 1 | nuclear receptor subfamily 1 group I member |
| PCP4L1 | 1 | Purkinje cell protein 4 like 1 [Source:HGNC Syr |

| | | |
|---|---|---|
| MPZ | 1 | myelin protein zero [Source:HGNC Symbol;Ac |
| SDHC | 1 | succinate dehydrogenase complex subunit C [ |
| CFAP126 | 1 | cilia and flagella associated protein 126 [Sourc |
| RRM2P2 | 1 | ribonucleotide reductase M2 polypeptide pse |
| AL592295.4 | 1 | novel transcript |
| AL592295.5 | 1 | novel transcript |
| RNU6-481P | 1 | RNA, U6 small nuclear 481, pseudogene [Sour |
| AL592295.3 | 1 | novel transcript |
| AL592295.1 | 1 | ribosomal protein S2 (RPS2) pseudogene |
| AL592295.2 | 1 | COMM domain containing 6 (COMMD6) pseu |
| AL831711.1 | 1 | novel transcript |
| FCGR2A | 1 | Fc fragment of IgG receptor IIa [Source:HGNC |
| AL590385.2 | 1 | novel transcript, readthrough between FCGR2 |
| HSPA6 | 1 | heat shock protein family A (Hsp70) member |
| RPS23P10 | 1 | ribosomal protein S23 pseudogene 10 [Source |
| FCGR3A | 1 | Fc fragment of IgG receptor IIIa [Source:HGNC |
| AL590385.1 | 1 | novel transcript |
| FCGR2C | 1 | Fc fragment of IgG receptor IIc (gene/pseudog |
| HSPA7 | 1 | heat shock protein family A (Hsp70) member |
| RPS23P9 | 1 | ribosomal protein S23 pseudogene 9 [Source: |
| FCGR3B | 1 | Fc fragment of IgG receptor IIIb [Source:HGNC |
| FCGR2B | 1 | Fc fragment of IgG receptor IIb [Source:HGNC |
| AL451067.1 | 1 | novel transcript, antisense to FCGR2B |
| RPL31P11 | 1 | ribosomal protein L31 pseudogene 11 [Source |
| Y_RNA | 1 | Y RNA [Source:RFAM;Acc:RF00019] |
| FCRLA | 1 | Fc receptor like A [Source:HGNC Symbol;Acc:H |
| FCRLB | 1 | Fc receptor like B [Source:HGNC Symbol;Acc:H |
| RN7SL466P | 1 | RNA, 7SL, cytoplasmic 466, pseudogene [Sour |
| DUSP12 | 1 | dual specificity phosphatase 12 [Source:HGNC |
| AL359541.1 | 1 | novel transcript |
| ATF6 | 1 | activating transcription factor 6 [Source:HGNC |
| AL391825.1 | 1 | proteasome (prosome, macropain) activator s |
| OLFML2B | 1 | olfactomedin like 2B [Source:HGNC Symbol;Ac |
| AL590408.1 | 1 | novel transcript |
| NOS1AP | 1 | nitric oxide synthase 1 adaptor protein [Sourc |
| AL450163.1 | 1 | Novel transcript, antisense to NOS1AP |
| MIR4654 | 1 | microRNA 4654 [Source:HGNC Symbol;Acc:HG |
| AL512785.1 | 1 | novel transcript |
| RNA5SP61 | 1 | RNA, 5S ribosomal pseudogene 61 [Source:HG |
| MIR556 | 1 | microRNA 556 [Source:HGNC Symbol;Acc:HGN |
| AL512785.2 | 1 | novel protein |
| SPATA46 | 1 | spermatogenesis associated 46 [Source:HGNC |
| C1orf226 | 1 | chromosome 1 open reading frame 226 [Sour |
| SH2D1B | 1 | SH2 domain containing 1B [Source:HGNC Sym |
| SLAMF6P1 | 1 | SLAM family member 6 pseudogene 1 [Source |
| UHMK1 | 1 | U2AF homology motif kinase 1 [Source:HGNC |
| UQCRBP2 | 1 | ubiquinol-cytochrome c reductase binding pro |
| AL596325.1 | 1 | novel transcript, antisense to UAP1 |
| UAP1 | 1 | UDP-N-acetylglucosamine pyrophosphorylase |
| AL596325.2 | 1 | novel transcript, sense intronic to UAP1 |

| | | |
|---|---|---|
| DDR2 | 1 | discoidin domain receptor tyrosine kinase 2 [S |
| RN7SL861P | 1 | RNA, 7SL, cytoplasmic 861, pseudogene [Sour |
| HSD17B7 | 1 | hydroxysteroid 17-beta dehydrogenase 7 [Sou |
| CCDC190 | 1 | coiled-coil domain containing 190 [Source:HG |
| AL392003.2 | 1 | hydroxysteroid (17-beta) dehydrogenase 7 (H |
| AL392003.1 | 1 | cytochrome c oxidase subunit VIIb (COX7B) ps |
| RGS4 | 1 | regulator of G protein signaling 4 [Source:HGN |
| RGS5 | 1 | regulator of G protein signaling 5 [Source:HGN |
| AL499616.1 | 1 | novel transcript |
| AL451063.1 | 1 | tRNA nucleotidyl transferase, CCA-adding, 1 (T |
| AL592435.1 | 1 | novel transcript |
| AL592435.2 | 1 | novel transcript |
| NUF2 | 1 | NUF2 component of NDC80 kinetochore comp |
| AL603841.1 | 1 | eukaryotic translation initiation factor 3, subu |
| RNA5SP62 | 1 | RNA, 5S ribosomal pseudogene 62 [Source:HG |
| RNA5SP63 | 1 | RNA, 5S ribosomal pseudogene 63 [Source:HG |
| AL355862.1 | 1 | pseudogene similar to the likely ortholog of m |
| U3 | 1 | Small nucleolar RNA U3 [Source:RFAM;Acc:RF |
| NMNAT1P2 | 1 | NMNAT1 pseudogene 2 [Source:HGNC Symbo |
| RNU5F-6P | 1 | RNA, U5F small nuclear 6, pseudogene [Sourc |
| HMGB3P6 | 1 | high mobility group box 3 pseudogene 6 [Sour |
| PBX1 | 1 | PBX homeobox 1 [Source:HGNC Symbol;Acc:H |
| RNU6-171P | 1 | RNA, U6 small nuclear 171, pseudogene [Sour |
| AL391001.1 | 1 | novel transcript, sense intronic to PBX1 |
| Metazoa_SRP | 1 | Metazoan signal recognition particle RNA [Sou |
| AL357568.1 | 1 | novel transcript, antisense to PBX1 |
| AL357568.2 | 1 | novel transcript |
| Y_RNA | 1 | Y RNA [Source:RFAM;Acc:RF00019] |
| AL390119.1 | 1 | ribosomal protein L35a (RPL35A) pseudogene |
| RNU6-755P | 1 | RNA, U6 small nuclear 755, pseudogene [Sour |
| LMX1A | 1 | LIM homeobox transcription factor 1 alpha [So |
| AL390730.1 | 1 | novel transcript, antisense to LMX1A |
| AL390730.2 | 1 | novel transcript, antisense to LMX1A |
| RXRG | 1 | retinoid X receptor gamma [Source:HGNC Sym |
| LRRC52-AS1 | 1 | LRRC52 antisense RNA 1 [Source:HGNC Symbo |
| AL157714.1 | 1 | px19-like protein (PX19) pseudogene |
| LRRC52 | 1 | leucine rich repeat containing 52 [Source:HGN |
| AL356441.2 | 1 | ribosomal protein S2 (RPS2) pseudogene |
| AL356441.1 | 1 | novel transcript |
| MGST3 | 1 | microsomal glutathione S-transferase 3 [Sourc |
| ALDH9A1 | 1 | aldehyde dehydrogenase 9 family member A1 |
| Y_RNA | 1 | Y RNA [Source:RFAM;Acc:RF00019] |
| AL451074.3 | 1 | ribosomal protein S13 (RPS13) pseudogene |
| AL451074.2 | 1 | ribosomal protein L21 (RPL21) pseudogene |
| AL451074.1 | 1 | carbonic anhydrase XIV (CA14) pseudogene |
| AL451074.5 | 1 | novel transcript |
| TMCO1 | 1 | transmembrane and coiled-coil domains 1 [So |
| TMCO1-AS1 | 1 | TMCO1 antisense RNA 1 [Source:HGNC Symbo |
| AL451074.4 | 1 | ribosomal protein L26 (RPL26) pseudogene |
| UCK2 | 1 | uridine-cytidine kinase 2 [Source:HGNC Symbo |

| | | |
|---|---|---|
| AL358115.1 | 1 | novel transcript, antisense to UCK2 |
| MIR3658 | 1 | microRNA 3658 [Source:HGNC Symbol;Acc:HG |
| AL606495.2 | 1 | post-GPI attachment to proteins 2 (PGAP2) ps |
| AL606495.1 | 1 | ribosomal protein S3A (RPS3A) pseudogene |
| RNA5SP64 | 1 | RNA, 5S ribosomal pseudogene 64 [Source:HG |
| FAM78B | 1 | family with sequence similarity 78 member B |
| AL626787.1 | 1 | novel transcript |
| AL596087.3 | 1 | novel transcript |
| MIR921 | 1 | microRNA 921 [Source:HGNC Symbol;Acc:HGN |
| AL596087.2 | 1 | novel transcript |
| AL596087.1 | 1 | Ewing sarcoma breakpoint region 1 (EWSR1) p |
| AL583804.1 | 1 | novel transcript |
| FMO7P | 1 | flavin containing dimethylaniline monoxygena |
| LINC01675 | 1 | long intergenic non-protein coding RNA 1675 |
| FMO8P | 1 | flavin containing dimethylaniline monoxygena |
| FMO9P | 1 | flavin containing dimethylaniline monoxygena |
| FMO10P | 1 | flavin containing dimethylaniline monoxygena |
| RPL4P2 | 1 | ribosomal protein L4 pseudogene 2 [Source:H |
| FMO11P | 1 | flavin containing dimethylaniline monoxygena |
| CNN2P10 | 1 | calponin 2 pseudogene 10 [Source:HGNC Sym |
| POGK | 1 | pogo transposable element derived with KRAB |
| TADA1 | 1 | transcriptional adaptor 1 [Source:HGNC Symb |
| DUTP6 | 1 | deoxyuridine triphosphatase pseudogene 6 [S |
| ILDR2 | 1 | immunoglobulin like domain containing recep |
| MAEL | 1 | maelstrom spermatogenic transposon silence |
| RNA5SP65 | 1 | RNA, 5S ribosomal pseudogene 65 [Source:HG |
| AL158837.1 | 1 | novel transcript |
| GPA33 | 1 | glycoprotein A33 [Source:HGNC Symbol;Acc:H |
| DUSP27 | 1 | dual specificity phosphatase 27, atypical [Sour |
| AL451050.1 | 1 | ribosomal protein S17 (RPS17) pseudogene |
| LINC01363 | 1 | long intergenic non-protein coding RNA 1363 |
| AL451050.2 | 1 | novel transcript |
| POU2F1 | 1 | POU class 2 homeobox 1 [Source:HGNC Symb |
| AL136984.1 | 1 | novel transcript, sense intronic to POU2F1 |
| CD247 | 1 | CD247 molecule [Source:HGNC Symbol;Acc:H |
| AL359962.3 | 1 | novel transcript, antisense to CD247 |
| AL359962.1 | 1 | novel transcript |
| AL359962.2 | 1 | novel transcript |
| AKR1D1P1 | 1 | aldo-keto reductase family 1 member D1 pseu |
| CREG1 | 1 | cellular repressor of E1A stimulated genes 1 [S |
| AL031733.1 | 1 | ribosomal protein S18 (RPS18) pseudogene |
| AL031733.2 | 1 | novel transcript |
| RCSD1 | 1 | RCSD domain containing 1 [Source:HGNC Sym |
| MPZL1 | 1 | myelin protein zero like 1 [Source:HGNC Symb |
| ADCY10 | 1 | adenylate cyclase 10 [Source:HGNC Symbol;Ac |
| Z99943.2 | 1 | chromosome 14 open reading frame 24 (C14o |
| Z99943.1 | 1 | novel transcript |
| MPC2 | 1 | mitochondrial pyruvate carrier 2 [Source:HGN |
| DCAF6 | 1 | DDB1 and CUL4 associated factor 6 [Source:H |
| MIR1255B2 | 1 | microRNA 1255b-2 [Source:HGNC Symbol;Acc |

| | | |
|---|---|---|
| GCSHP5 | 1 | glycine cleavage system protein H pseudogene |
| GPR161 | 1 | G protein-coupled receptor 161 [Source:HGN( |
| TIPRL | 1 | TOR signaling pathway regulator [Source:HGN |
| RPL34P1 | 1 | ribosomal protein L34 pseudogene 1 [Source: |
| AL021397.1 | 1 | NADH dehydrogenase (ubiquinone) 1 alpha su |
| SFT2D2 | 1 | SFT2 domain containing 2 [Source:HGNC Syml |
| ANKRD36BP1 | 1 | ankyrin repeat domain 36B pseudogene 1 [So |
| RNU6-1310P | 1 | RNA, U6 small nuclear 1310, pseudogene [Sou |
| TBX19 | 1 | T-box transcription factor 19 [Source:HGNC Sy |
| AL022100.1 | 1 | dynactin 4 pseudogene |
| MIR557 | 1 | microRNA 557 [Source:HGNC Symbol;Acc:HGI |
| AL023755.1 | 1 | novel transcript |
| AL022100.2 | 1 | novel transcript |
| QRSL1P1 | 1 | QRSL1 pseudogene 1 [Source:HGNC Symbol;A |
| XCL2 | 1 | X-C motif chemokine ligand 2 [Source:HGNC S |
| AL031736.1 | 1 | ribosomal protein L7a (RPL7A) pseudogene |
| XCL1 | 1 | X-C motif chemokine ligand 1 [Source:HGNC S |
| AL031736.2 | 1 | ribosomal protein L7a (RPL7A) pseudogene |
| DPT | 1 | dermatopontin [Source:HGNC Symbol;Acc:HG |
| AL135926.1 | 1 | novel transcript |
| AL031275.1 | 1 | novel transcript |
| LINC00626 | 1 | long intergenic non-protein coding RNA 626 [S |
| SUMO1P2 | 1 | SUMO1 pseudogene 2 [Source:HGNC Symbol; |
| AL135926.2 | 1 | novel transcript |
| LINC00970 | 1 | long intergenic non-protein coding RNA 970 [S |
| RPL29P7 | 1 | ribosomal protein L29 pseudogene 7 [Source: |
| AL031726.2 | 1 | novel transcript |
| RNA5SP66 | 1 | RNA, 5S ribosomal pseudogene 66 [Source:HG |
| AL031726.1 | 1 | novel transcript |
| ATP1B1 | 1 | ATPase Na+/K+ transporting subunit beta 1 [S |
| NME7 | 1 | NME/NM23 family member 7 [Source:HGNC S |
| Z99758.1 | 1 | novel transcript |
| BLZF1 | 1 | basic leucine zipper nuclear factor 1 [Source:H |
| CCDC181 | 1 | coiled-coil domain containing 181 [Source:HG |
| SLC19A2 | 1 | solute carrier family 19 member 2 [Source:HG |
| AL021068.1 | 1 | ATP synthase 6 (MTATP6) pseudogene |
| Z99572.1 | 1 | novel transcript |
| F5 | 1 | coagulation factor V [Source:HGNC Symbol;Ac |
| SELP | 1 | selectin P [Source:HGNC Symbol;Acc:HGNC:10 |
| C1orf112 | 1 | chromosome 1 open reading frame 112 [Sour |
| SELL | 1 | selectin L [Source:HGNC Symbol;Acc:HGNC:10 |
| SELE | 1 | selectin E [Source:HGNC Symbol;Acc:HGNC:10 |
| AL021940.1 | 1 | novel transcript |
| METTL18 | 1 | methyltransferase like 18 [Source:HGNC Symb |
| SCYL3 | 1 | SCY1 like pseudokinase 3 [Source:HGNC Symb |
| RN7SL333P | 1 | RNA, 7SL, cytoplasmic 333, pseudogene [Sour |
| AL031297.1 | 1 | small lysine-rich protein 1 (SMKR1) pseudoger |
| KIFAP3 | 1 | kinesin associated protein 3 [Source:HGNC Syr |
| RN7SL269P | 1 | RNA, 7SL, cytoplasmic 269, pseudogene [Sour |
| MRPS10P1 | 1 | mitochondrial ribosomal protein S10 pseudog |

| | | |
|---|---|---|
| AL356475.1 | 1 | novel transcript |
| SIGLEC30P | 1 | sialic acid binding Ig like lectin 30, pseudogene |
| METTL11B | 1 | methyltransferase like 11B [Source:HGNC Sym |
| MIR3119-1 | 1 | microRNA 3119-1 [Source:HGNC Symbol;Acc:I |
| MIR3119-2 | 1 | microRNA 3119-2 [Source:HGNC Symbol;Acc:I |
| LINC01681 | 1 | long intergenic non-protein coding RNA 1681 |
| ISCUP1 | 1 | iron-sulfur cluster assembly enzyme pseudoge |
| LINC01142 | 1 | long intergenic non-protein coding RNA 1142 |
| HAUS4P1 | 1 | HAUS augmin like complex subunit 4 pseudog |
| GORAB-AS1 | 1 | GORAB antisense RNA 1 [Source:HGNC Symbo |
| GORAB | 1 | golgin, RAB6 interacting [Source:HGNC Symbo |
| AL162399.1 | 1 | novel transcript |
| AL023495.1 | 1 | novel transcript |
| PRRX1 | 1 | paired related homeobox 1 [Source:HGNC Syr |
| Z97200.1 | 1 | novel transcript |
| BX284613.2 | 1 | novel transcript, antisense to FMO1 |
| MROH9 | 1 | maestro heat like repeat family member 9 [Sc |
| BX284613.1 | 1 | novel pseudogene |
| FMO3 | 1 | flavin containing dimethylaniline monoxygena |
| MIR1295A | 1 | microRNA 1295a [Source:HGNC Symbol;Acc:H |
| MIR1295B | 1 | microRNA 1295b [Source:HGNC Symbol;Acc:H |
| FMO6P | 1 | flavin containing dimethylaniline monoxygena |
| FMO2 | 1 | flavin containing dimethylaniline monoxygena |
| AL021026.1 | 1 | novel transcript, antisense to FMO2 |
| FMO1 | 1 | flavin containing dimethylaniline monoxygena |
| Y_RNA | 1 | Y RNA [Source:RFAM;Acc:RF00019] |
| HMGB1P11 | 1 | high mobility group box 1 pseudogene 11 [Sou |
| FMO4 | 1 | flavin containing dimethylaniline monoxygena |
| AL031274.1 | 1 | DNA topoisomerase I pseudogene 1 |
| SRP14P4 | 1 | signal recognition particle 14 pseudogene 4 [S |
| GM2AP2 | 1 | GM2 ganglioside activator pseudogene 2 [Sou |
| CYCSP53 | 1 | CYCS pseudogene 53 [Source:HGNC Symbol;A |
| PRRC2C | 1 | proline rich coiled-coil 2C [Source:HGNC Symb |
| MYOCOS | 1 | myocilin opposite strand [Source:HGNC Symb |
| MYOC | 1 | myocilin [Source:HGNC Symbol;Acc:HGNC:76 |
| PFN1P1 | 1 | profilin 1 pseudogene 1 [Source:HGNC Symbo |
| RPL4P3 | 1 | ribosomal protein L4 pseudogene 3 [Source:H |
| VAMP4 | 1 | vesicle associated membrane protein 4 [Sourc |
| Z98751.1 | 1 | solute carrier family 5 (sodium/glucose cotrar |
| Z98751.3 | 1 | solute carrier family 5 (low affinity glucose co |
| Z98751.2 | 1 | RNA binding motif protein 3 (RBM3) pseudoge |
| AL135931.1 | 1 | NULL |
| EEF1AKNMT | 1 | eEF1A lysine and N-terminal methyltransferas |
| AL031864.1 | 1 | ribosomal protein S15 (RPS15) pseudogene |
| AL031864.2 | 1 | ribosomal protein, large, P1 (RPLP1) pseudoge |
| DNM3 | 1 | dynamin 3 [Source:HGNC Symbol;Acc:HGNC:2 |
| DNM3-IT1 | 1 | DNM3 intronic transcript 1 [Source:HGNC Sym |
| DNM3OS | 1 | DNM3 opposite strand/antisense RNA [Source |
| MIR214 | 1 | microRNA 214 [Source:HGNC Symbol;Acc:HGI |
| MIR3120 | 1 | microRNA 3120 [Source:HGNC Symbol;Acc:HG |

| | | |
|---|---|---|
| MIR199A2 | 1 | microRNA 199a-2 [Source:HGNC Symbol;Acc:I |
| AL137157.1 | 1 | novel transcript, antisense to DNM3 |
| RNU6-157P | 1 | RNA, U6 small nuclear 157, pseudogene [Sour |
| PIGC | 1 | phosphatidylinositol glycan anchor biosynthes |
| C1orf105 | 1 | chromosome 1 open reading frame 105 [Sour |
| SUCO | 1 | SUN domain containing ossification factor [So |
| RNU6-693P | 1 | RNA, U6 small nuclear 693, pseudogene [Sour |
| FASLG | 1 | Fas ligand [Source:HGNC Symbol;Acc:HGNC:1 |
| SLC25A38P1 | 1 | solute carrier family 25 member 38 pseudoge |
| Z98043.1 | 1 | TEC |
| AL031599.1 | 1 | novel transcript |
| AIMP1P2 | 1 | aminoacyl tRNA synthetase complex interacti |
| Z97198.1 | 1 | novel transcript |
| TNFSF18 | 1 | TNF superfamily member 18 [Source:HGNC Sy |
| GOT2P2 | 1 | glutamic-oxaloacetic transaminase 2 pseudog |
| AL022310.1 | 1 | novel transcript |
| TNFSF4 | 1 | TNF superfamily member 4 [Source:HGNC Syn |
| AL645568.3 | 1 | pseudogene similar to part of ribosomal prote |
| AL645568.2 | 1 | novel pseudogene |
| AL645568.1 | 1 | novel transcript |
| PRDX6 | 1 | peroxiredoxin 6 [Source:HGNC Symbol;Acc:HG |
| SLC9C2 | 1 | solute carrier family 9 member C2 (putative) [ |
| AL139142.1 | 1 | novel transcript |
| AL139142.2 | 1 | novel protein |
| ANKRD45 | 1 | ankyrin repeat domain 45 [Source:HGNC Syml |
| TEX50 | 1 | testis expressed 50 [Source:HGNC Symbol;Acc |
| KLHL20 | 1 | kelch like family member 20 [Source:HGNC Sy |
| BX248409.1 | 1 | ribosomal protein S27 (metallopanstimulin 1) |
| RN7SKP160 | 1 | RN7SK pseudogene 160 [Source:HGNC Symbo |
| CENPL | 1 | centromere protein L [Source:HGNC Symbol;A |
| Y_RNA | 1 | Y RNA [Source:RFAM;Acc:RF00019] |
| DARS2 | 1 | aspartyl-tRNA synthetase 2, mitochondrial [Sc |
| GAS5 | 1 | growth arrest specific 5 [Source:HGNC Symbo |
| GAS5-AS1 | 1 | GAS5 antisense RNA 1 [Source:HGNC Symbol; |
| ZBTB37 | 1 | zinc finger and BTB domain containing 37 [Sou |
| SERPINC1 | 1 | serpin family C member 1 [Source:HGNC Syml |
| RNA5SP67 | 1 | RNA, 5S ribosomal pseudogene 67 [Source:HG |
| RC3H1 | 1 | ring finger and CCCH-type domains 1 [Source: |
| RNA5SP68 | 1 | RNA, 5S ribosomal pseudogene 68 [Source:HG |
| RC3H1-IT1 | 1 | RC3H1 intronic transcript 1 [Source:HGNC Syn |
| LINC02776 | 1 | long intergenic non-protein coding RNA 2776 |
| RPL30P1 | 1 | ribosomal protein L30 pseudogene 1 [Source: |
| RABGAP1L-DT | 1 | RABGAP1L divergent transcript [Source:HGNC |
| RABGAP1L | 1 | RAB GTPase activating protein 1 like [Source:H |
| AL022400.1 | 1 | forty-two-three domain containing 1 (FYTTD1 |
| GPR52 | 1 | G protein-coupled receptor 52 [Source:HGNC |
| BANF1P4 | 1 | barrier to autointegration factor 1 pseudogen |
| NDUFAF4P4 | 1 | NADH:ubiquinone oxidoreductase complex as |
| Z99127.3 | 1 | transmembrane protein 56 (TMEM56) pseudc |
| RABGAP1L-IT1 | 1 | RABGAP1L intronic transcript 1 [Source:HGNC |

| | | |
|---|---|---|
| Z99127.2 | 1 | ribosomal protein S26 (RPS26) pseudogene |
| Z99127.1 | 1 | novel transcript |
| RNU6-307P | 1 | RNA, U6 small nuclear 307, pseudogene [Sour |
| Z99127.4 | 1 | novel transcript, antisense to CACYBP |
| CACYBP | 1 | calcyclin binding protein [Source:HGNC Symbo |
| MRPS14 | 1 | mitochondrial ribosomal protein S14 [Source: |
| Y_RNA | 1 | Y RNA [Source:RFAM;Acc:RF00019] |
| ENTR1P2 | 1 | ENTR1 pseudogene 2 [Source:HGNC Symbol;A |
| TNN | 1 | tenascin N [Source:HGNC Symbol;Acc:HGNC:2 |
| KIAA0040 | 1 | KIAA0040 [Source:HGNC Symbol;Acc:HGNC:2 |
| AL008626.1 | 1 | novel transcript |
| RPS29P4 | 1 | ribosomal protein S29 pseudogene 4 [Source: |
| Z94057.1 | 1 | novel transcript |
| TNR | 1 | tenascin R [Source:HGNC Symbol;Acc:HGNC:1 |
| TNR-IT1 | 1 | TNR intronic transcript 1 [Source:HGNC Symbo |
| LINC01657 | 1 | long intergenic non-protein coding RNA 1657 |
| LINC02803 | 1 | long intergenic non-protein coding RNA 2803 |
| RPS29P5 | 1 | ribosomal protein S29 pseudogene 5 [Source: |
| COP1 | 1 | COP1 E3 ubiquitin ligase [Source:HGNC Symbo |
| SCARNA3 | 1 | small Cajal body-specific RNA 3 [Source:HGNC |
| AL590723.1 | 1 | novel transcript |
| AL591043.2 | 1 | novel transcript |
| U7 | 1 | U7 small nuclear RNA [Source:RFAM;Acc:RF00 |
| AL359265.2 | 1 | ribosomal protein L13 (RPL13) pseudogene |
| RNU2-12P | 1 | RNA, U2 small nuclear 12, pseudogene [Sourc |
| AL359265.1 | 1 | novel pseudogene |
| AL591043.1 | 1 | proteolipid protein 2 (colonic epithelium-enri |
| MORF4L1P7 | 1 | mortality factor 4 like 1 pseudogene 7 [Source |
| PAPPA2 | 1 | pappalysin 2 [Source:HGNC Symbol;Acc:HGNC |
| PTP4A1P7 | 1 | PTP4A1 pseudogene 7 [Source:HGNC Symbol; |
| AL031290.1 | 1 | novel transcript, antisense to PAPPA2 |
| ASTN1 | 1 | astrotactin 1 [Source:HGNC Symbol;Acc:HGNC |
| MIR488 | 1 | microRNA 488 [Source:HGNC Symbol;Acc:HGN |
| BRINP2 | 1 | BMP/retinoic acid inducible neural specific 2 [ |
| LINC01645 | 1 | long intergenic non-protein coding RNA 1645 |
| AL136114.1 | 1 | novel transcript |
| LINC01741 | 1 | long intergenic non-protein coding RNA 1741 |
| SEC16B | 1 | SEC16 homolog B, endoplasmic reticulum exp |
| CRYZL2P-SEC16B | 1 | CRYZL2P-SEC16B readthrough [Source:HGNC S |
| CRYZL2P | 1 | crystallin zeta like 2, pseudogene [Source:HGN |
| AL359075.1 | 1 | ubiquitin-conjugating enzyme E2K (UBE2K) ps |
| RASAL2-AS1 | 1 | RASAL2 antisense RNA 1 [Source:HGNC Symbo |
| RASAL2 | 1 | RAS protein activator like 2 [Source:HGNC Syn |
| AL160281.1 | 1 | H3 histone, family 3C (H3F3C) pseudogene |
| AL365357.1 | 1 | ribosomal protein S14 (RPS14) pseudogene |
| CLEC20A | 1 | C-type lectin domain containing 20A [Source:I |
| Metazoa_SRP | 1 | Metazoan signal recognition particle RNA [Sou |
| AL513013.1 | 1 | novel transcript |
| TEX35 | 1 | testis expressed 35 [Source:HGNC Symbol;Acc |
| C1orf220 | 1 | chromosome 1 open reading frame 220 (puta |

| | | |
|---|---|---|
| RNA5SP69 | 1 | RNA, 5S ribosomal pseudogene 69 [Source:HG |
| AL137796.1 | 1 | novel transcript |
| MIR4424 | 1 | microRNA 4424 [Source:HGNC Symbol;Acc:HG |
| AL449106.1 | 1 | novel transcript, antisense to RALGPS2 |
| RALGPS2 | 1 | Ral GEF with PH domain and SH3 binding moti |
| PTPN2P1 | 1 | PTPN2 pseudogene 1 [Source:HGNC Symbol;A |
| SNORA63 | 1 | Small nucleolar RNA SNORA63 [Source:RFAM; |
| ANGPTL1 | 1 | angiopoietin like 1 [Source:HGNC Symbol;Acc: |
| FAM20B | 1 | FAM20B glycosaminoglycan xylosylkinase [Sou |
| AL139132.1 | 1 | ribosomal protein S26 (RPS26) pseudogene |
| TOR3A | 1 | torsin family 3 member A [Source:HGNC Symb |
| ABL2 | 1 | ABL proto-oncogene 2, non-receptor tyrosine |
| AL512326.5 | 1 | ubiquinol-cytochrome c reductase (6.4kD) sub |
| SETP10 | 1 | SET pseudogene 10 [Source:HGNC Symbol;Acc |
| AL512326.2 | 1 | NULL |
| AL512326.1 | 1 | NULL |
| EIF4A1P11 | 1 | eukaryotic translation initiation factor 4A1 pse |
| AL512326.3 | 1 | UDP-Gal:betaGlcNAc beta 1,3-galactosyltransf |
| COX5BP8 | 1 | cytochrome c oxidase subunit 5B pseudogene |
| AL512326.4 | 1 | ribosomal protein L39-like (RPL39L) pseudoge |
| SOAT1 | 1 | sterol O-acyltransferase 1 [Source:HGNC Syml |
| AXDND1 | 1 | axonemal dynein light chain domain containin |
| MEF2AP1 | 1 | myocyte enhancer factor 2A pseudogene 1 [S |
| HNRNPA1P54 | 1 | heterogeneous nuclear ribonucleoprotein A1 |
| AL160286.3 | 1 | novel transcript |
| NPHS2 | 1 | NPHS2 stomatin family member, podocin [Sou |
| RNU5F-2P | 1 | RNA, U5F small nuclear 2, pseudogene [Sourc |
| AL160286.1 | 1 | centaurin, gamma 2 (CENTG2) pseudogene |
| AL160286.2 | 1 | novel transcript, sense overlapping TDRD5 |
| TDRD5 | 1 | tudor domain containing 5 [Source:HGNC Sym |
| AL359853.2 | 1 | novel transcript |
| FAM163A | 1 | family with sequence similarity 163 member A |
| AL359853.1 | 1 | novel transcript |
| LINC02818 | 1 | long intergenic non-protein coding RNA 2818 |
| TOR1AIP2 | 1 | torsin 1A interacting protein 2 [Source:HGNC S |
| AL353708.3 | 1 | novel transcript, antisense to TOR1AIP1 |
| TOR1AIP1 | 1 | torsin 1A interacting protein 1 [Source:HGNC S |
| RN7SL230P | 1 | RNA, 7SL, cytoplasmic 230, pseudogene [Sour |
| AL353708.2 | 1 | novel transcript |
| AL353708.1 | 1 | novel transcript |
| CEP350 | 1 | centrosomal protein 350 [Source:HGNC Symb |
| RPSAP16 | 1 | ribosomal protein SA pseudogene 16 [Source: |
| AL590632.1 | 1 | ribosomal protein S24 (RPS24) pseudogene |
| AL390718.1 | 1 | novel transcript |
| QSOX1 | 1 | quiescin sulfhydryl oxidase 1 [Source:HGNC Sy |
| LHX4 | 1 | LIM homeobox 4 [Source:HGNC Symbol;Acc:H |
| ACBD6 | 1 | acyl-CoA binding domain containing 6 [Source |
| VDAC1P4 | 1 | voltage dependent anion channel 1 pseudoge |
| MIR3121 | 1 | microRNA 3121 [Source:HGNC Symbol;Acc:HG |
| Y_RNA | 1 | Y RNA [Source:RFAM;Acc:RF00019] |

| | | |
|---|---|---|
| OVAAL | 1 | ovarian adenocarcinoma amplified long non-c |
| XPR1 | 1 | xenotropic and polytropic retrovirus receptor |
| U6 | 1 | U6 spliceosomal RNA [Source:RFAM;Acc:RF00 |
| LINC02816 | 1 | long intergenic non-protein coding RNA 2816 |
| KIAA1614 | 1 | KIAA1614 [Source:HGNC Symbol;Acc:HGNC:2 |
| AL162431.1 | 1 | novel transcript |
| KIAA1614-AS1 | 1 | KIAA1614 antisense RNA 1 [Source:HGNC Sym |
| AL162431.2 | 1 | pseudogene similar to part of major histocom |
| AL162431.3 | 1 | cell division cycle associated 8 (CDCA8) pseud |
| STX6 | 1 | syntaxin 6 [Source:HGNC Symbol;Acc:HGNC:1 |
| MR1 | 1 | major histocompatibility complex, class I-relat |
| IER5 | 1 | immediate early response 5 [Source:HGNC Sy |
| LINC01732 | 1 | long intergenic non-protein coding RNA 1732 |
| AL358393.1 | 1 | novel transcript |
| LINC01699 | 1 | long intergenic non-protein coding RNA 1699 |
| CACNA1E | 1 | calcium voltage-gated channel subunit alpha1 |
| Metazoa_SRP | 1 | Metazoan signal recognition particle RNA [Sou |
| RNA5SP70 | 1 | RNA, 5S ribosomal pseudogene 70 [Source:HG |
| AL161734.2 | 1 | novel transcript |
| AL161734.1 | 1 | novel transcript |
| RN7SKP229 | 1 | RN7SK pseudogene 229 [Source:HGNC Symbo |
| ZNF648 | 1 | zinc finger protein 648 [Source:HGNC Symbol; |
| AL355482.1 | 1 | novel transcript (LOC149157) |
| AL355482.2 | 1 | novel transcript |
| LINC01344 | 1 | long intergenic non-protein coding RNA 1344 |
| AL390856.1 | 1 | novel transcript |
| YPEL5P1 | 1 | YPEL5 pseudogene 1 [Source:HGNC Symbol;A |
| RNU6-152P | 1 | RNA, U6 small nuclear 152, pseudogene [Sour |
| AL513480.1 | 1 | ribosomal protein L18 (RPL18) pseudogene |
| EIF1P3 | 1 | eukaryotic translation initiation factor 1 pseu |
| GLUL | 1 | glutamate-ammonia ligase [Source:HGNC Sym |
| TEDDM1 | 1 | transmembrane epididymal protein 1 [Source |
| LINC00272 | 1 | long intergenic non-protein coding RNA 272 [ |
| RGSL1 | 1 | regulator of G protein signaling like 1 [Source: |
| AL139344.1 | 1 | NOP10 ribonucleoprotein (NOP10) pseudoger |
| TEDDM2P | 1 | transmembrane epididymal protein 2, pseudo |
| RNASEL | 1 | ribonuclease L [Source:HGNC Symbol;Acc:HG |
| Metazoa_SRP | 1 | Metazoan signal recognition particle RNA [Sou |
| RGS16 | 1 | regulator of G protein signaling 16 [Source:HG |
| LINC01686 | 1 | long intergenic non-protein coding RNA 1686 |
| RGS8 | 1 | regulator of G protein signaling 8 [Source:HG |
| LINC01688 | 1 | long intergenic non-protein coding RNA 1688 |
| AL353778.1 | 1 | non-SMC element 4 homolog A (S. cerevisiae) |
| NPL | 1 | N-acetylneuraminate pyruvate lyase [Source: |
| AL355999.1 | 1 | novel transcript, antisense to DHX9 |
| DHX9 | 1 | DExH-box helicase 9 [Source:HGNC Symbol;A |
| SHCBP1L | 1 | SHC binding and spindle associated 1 like [Sou |
| HMGN1P4 | 1 | high mobility group nucleosome binding dom |
| AL450304.2 | 1 | NULL |
| RNA5SP71 | 1 | RNA, 5S ribosomal pseudogene 71 [Source:HG |

| | | |
|---|---|---|
| AL450304.1 | 1 | natural cytotoxicity triggering receptor 1 (NCR |
| KRT18P28 | 1 | keratin 18 pseudogene 28 [Source:HGNC Sym |
| RNU6-41P | 1 | RNA, U6 small nuclear 41, pseudogene [Sourc |
| LAMC1 | 1 | laminin subunit gamma 1 [Source:HGNC Symb |
| LAMC1-AS1 | 1 | LAMC1 antisense RNA 1 [Source:HGNC Symbo |
| LAMC2 | 1 | laminin subunit gamma 2 [Source:HGNC Symb |
| NMNAT2 | 1 | nicotinamide nucleotide adenylyltransferase 2 |
| AL354953.1 | 1 | novel transcript, antisense to NMNAT2 |
| AL356981.1 | 1 | pseudogene similar to ribosomal protein S3A |
| AL449223.1 | 1 | novel transcript, antisense to NMNAT2 |
| SMG7-AS1 | 1 | SMG7 antisense RNA 1 [Source:HGNC Symbol |
| SMG7 | 1 | SMG7 nonsense mediated mRNA decay factor |
| NCF2 | 1 | neutrophil cytosolic factor 2 [Source:HGNC Sy |
| AL137800.1 | 1 | novel transcript |
| ARPC5 | 1 | actin related protein 2/3 complex subunit 5 [S |
| RGL1 | 1 | ral guanine nucleotide dissociation stimulator |
| APOBEC4 | 1 | apolipoprotein B mRNA editing enzyme cataly |
| AL157899.1 | 1 | T, brachyury homolog (mouse) (T) pseudogene |
| AL590422.1 | 1 | novel transcript |
| COLGALT2 | 1 | collagen beta(1-O)galactosyltransferase 2 [Sou |
| TSEN15 | 1 | tRNA splicing endonuclease subunit 15 [Sourc |
| AL158011.1 | 1 | novel transcript, antisense to TSEN15 |
| Y_RNA | 1 | Y RNA [Source:RFAM;Acc:RF00019] |
| Y_RNA | 1 | Y RNA [Source:RFAM;Acc:RF00019] |
| AL445228.1 | 1 | novel transcript |
| RN7SL654P | 1 | RNA, 7SL, cytoplasmic 654, pseudogene [Sour |
| AL445228.2 | 1 | novel transcript |
| C1orf21 | 1 | chromosome 1 open reading frame 21 [Source |
| AL078645.1 | 1 | novel transcript |
| AL078645.2 | 1 | novel pseudogene |
| AL713852.1 | 1 | novel transcript |
| AL713852.2 | 1 | novel transcript |
| EDEM3 | 1 | ER degradation enhancing alpha-mannosidase |
| NIBAN1 | 1 | niban apoptosis regulator 1 [Source:HGNC Syr |
| RNU7-13P | 1 | RNA, U7 small nuclear 13 pseudogene [Source |
| LINC01633 | 1 | long intergenic non-protein coding RNA 1633 |
| RNA5SP72 | 1 | RNA, 5S ribosomal pseudogene 72 [Source:HG |
| RNF2 | 1 | ring finger protein 2 [Source:HGNC Symbol;Ac |
| FTH1P25 | 1 | ferritin heavy chain 1 pseudogene 25 [Source: |
| TRMT1L | 1 | tRNA methyltransferase 1 like [Source:HGNC S |
| SWT1 | 1 | SWT1 RNA endoribonuclease homolog [Source |
| RPL22P24 | 1 | ribosomal protein L22 pseudogene 24 [Source |
| RPL5P5 | 1 | ribosomal protein L5 pseudogene 5 [Source:H |
| Y_RNA | 1 | Y RNA [Source:RFAM;Acc:RF00019] |
| Y_RNA | 1 | Y RNA [Source:RFAM;Acc:RF00019] |
| AL356273.1 | 1 | tyrosine aminotransferase (TAT) pseudogene |
| Y_RNA | 1 | Y RNA [Source:RFAM;Acc:RF00019] |
| AL356273.2 | 1 | proteasome (prosome, macropain) 26S subun |
| AL356273.3 | 1 | TEC |
| IVNS1ABP | 1 | influenza virus NS1A binding protein [Source:H |

| | | |
|---|---|---|
| AL078644.2 | 1 | novel transcript |
| AL078644.1 | 1 | microtubule associated protein 1 light chain 3 |
| RNU7-183P | 1 | RNA, U7 small nuclear 183 pseudogene [Sourc |
| MCRIP2P2 | 1 | MAPK regulated corepressor interacting prote |
| AL133383.2 | 1 | TEC |
| LINC01350 | 1 | long intergenic non-protein coding RNA 1350 |
| Y_RNA | 1 | Y RNA [Source:RFAM;Acc:RF00019] |
| Y_RNA | 1 | Y RNA [Source:RFAM;Acc:RF00019] |
| AL133383.1 | 1 | novel transcript |
| HMCN1 | 1 | hemicentin 1 [Source:HGNC Symbol;Acc:HGN |
| AL133553.1 | 1 | novel transcript |
| AL133553.2 | 1 | novel transcript |
| PRG4 | 1 | proteoglycan 4 [Source:HGNC Symbol;Acc:HG |
| TPR | 1 | translocated promoter region, nuclear basket |
| RNU6-1240P | 1 | RNA, U6 small nuclear 1240, pseudogene [Sou |
| ODR4 | 1 | odr-4 GPCR localization factor homolog [Sourc |
| OCLM | 1 | oculomedin [Source:HGNC Symbol;Acc:HGNC: |
| AL096803.2 | 1 | novel transcript, antisense to PDC |
| PDC | 1 | phosducin [Source:HGNC Symbol;Acc:HGNC:8 |
| AL096803.4 | 1 | novel transcript |
| AL096803.1 | 1 | zinc-finger protein pseudogene |
| AL096803.3 | 1 | novel pseudogene |
| PTGS2 | 1 | prostaglandin-endoperoxide synthase 2 [Sour |
| PACERR | 1 | PTGS2 antisense NFKB1 complex-mediated ex |
| PLA2G4A | 1 | phospholipase A2 group IVA [Source:HGNC Sy |
| ERVMER61-1 | 1 | endogenous retrovirus group MER61 member |
| AL391813.1 | 1 | uridine-cytidine kinase 1 (UCK1) pseudogene |
| FDPSP1 | 1 | farnesyl diphosphate synthase pseudogene 1 |
| AL357559.1 | 1 | neuroepithelial cell transforming 1 (NET1) pse |
| SLC4A1APP2 | 1 | solute carrier family 4 member 1 adaptor prot |
| AL645474.1 | 1 | NIF3 NGG1 interacting factor 3-like 1 (S. pomb |
| AL136372.2 | 1 | novel transcript |
| AL136372.1 | 1 | novel zinc finger protein pseudogene |
| RN7SKP156 | 1 | RN7SK pseudogene 156 [Source:HGNC Symbo |
| AL596211.1 | 1 | novel transcript |
| AL592447.1 | 1 | DEAD (Asp-Glu-Ala-As) box polypeptide 19 (DI |
| AL929288.1 | 1 | novel transcript |
| AL929288.2 | 1 | apolipoprotein L, 3 (APOL3) pseudogene |
| RPS3AP9 | 1 | ribosomal protein S3a pseudogene 9 [Source: |
| AL691515.1 | 1 | novel transcript |
| AL691515.2 | 1 | novel transcript |
| LINC01035 | 1 | long intergenic non-protein coding RNA 1035 |
| CLPTM1LP1 | 1 | CLPTM1L pseudogene 1 [Source:HGNC Symbo |
| GAPDHP75 | 1 | glyceraldehyde-3-phosphate dehydrogenase p |
| RNA5SP73 | 1 | RNA, 5S ribosomal pseudogene 73 [Source:HG |
| LINC01701 | 1 | long intergenic non-protein coding RNA 1701 |
| AL591504.1 | 1 | novel transcript |
| AL591504.2 | 1 | novel transcript |
| AL359976.1 | 1 | DEAH (Asp-Glu-Ala-His) box polypeptide 29 (D |
| BRINP3 | 1 | BMP/retinoic acid inducible neural specific 3 [ |

| | | |
|---|---|---|
| AL354771.1 | 1 | novel transcript |
| LINC01351 | 1 | long intergenic non-protein coding RNA 1351 |
| AL391645.1 | 1 | novel transcript |
| LINC01720 | 1 | long intergenic non-protein coding RNA 1720 |
| AL139135.1 | 1 | coronin, actin binding protein, 1C (CORO1C) p |
| AL139135.2 | 1 | glycine cleavage system protein H (aminomet |
| AL138927.1 | 1 | novel transcript |
| HNRNPA1P46 | 1 | heterogeneous nuclear ribonucleoprotein A1 |
| AL713866.1 | 1 | novel transcript |
| AL713866.2 | 1 | small integral membrane protein 15 (SMIM15 |
| LINC01680 | 1 | long intergenic non-protein coding RNA 1680 |
| LINC02770 | 1 | long intergenic non-protein coding RNA 2770 |
| AL359081.1 | 1 | novel transcript |
| RGS18 | 1 | regulator of G protein signaling 18 [Source:HG |
| AL390957.1 | 1 | novel transcript |
| AL513175.1 | 1 | pseudogene similar to ATP-binding cassette, s |
| AL513175.2 | 1 | pseudogene similar to ATP-binding cassette, s |
| RGS21 | 1 | regulator of G protein signaling 21 [Source:HG |
| AL136987.1 | 1 | novel transcript |
| RGS1 | 1 | regulator of G protein signaling 1 [Source:HGN |
| RGS13 | 1 | regulator of G protein signaling 13 [Source:HG |
| AL136454.1 | 1 | novel protein similar to ribosomal protein S27 |
| MIR4426 | 1 | microRNA 4426 [Source:HGNC Symbol;Acc:HG |
| AL035407.2 | 1 | pseudogene similar to a hypothetical protein ( |
| AL035407.1 | 1 | pseudogene similar to integrin beta 1 binding |
| RGS2 | 1 | regulator of G protein signaling 2 [Source:HGN |
| RN7SKP126 | 1 | RN7SK pseudogene 126 [Source:HGNC Symbo |
| ZNF101P2 | 1 | zinc finger protein 101 pseudogene 2 [Source: |
| UCHL5 | 1 | ubiquitin C-terminal hydrolase L5 [Source:HGI |
| SCARNA18B | 1 | small Cajal body-specific RNA 18B [Source:HG |
| RO60 | 1 | Ro60, Y RNA binding protein [Source:HGNC Sy |
| GLRX2 | 1 | glutaredoxin 2 [Source:HGNC Symbol;Acc:HGI |
| CDC73 | 1 | cell division cycle 73 [Source:HGNC Symbol;Ac |
| MIR1278 | 1 | microRNA 1278 [Source:HGNC Symbol;Acc:HG |
| B3GALT2 | 1 | beta-1,3-galactosyltransferase 2 [Source:HGN |
| Metazoa_SRP | 1 | Metazoan signal recognition particle RNA [Sou |
| LINC01031 | 1 | long intergenic non-protein coding RNA 1031 |
| AL138778.1 | 1 | novel transcript |
| AL136456.1 | 1 | novel transcript |
| AL357793.1 | 1 | novel transcript |
| AL357793.2 | 1 | novel transcript |
| RPL23AP22 | 1 | ribosomal protein L23a pseudogene 22 [Sourc |
| EEF1A1P14 | 1 | eukaryotic translation elongation factor 1 alpl |
| AL513348.1 | 1 | novel transcript |
| RNU6-983P | 1 | RNA, U6 small nuclear 983, pseudogene [Sour |
| AL353072.1 | 1 | translocase of inner mitochondrial membrane |
| AL353072.2 | 1 | novel transcript |
| AL353709.1 | 1 | novel transcript |
| LINC01724 | 1 | long intergenic non-protein coding RNA 1724 |
| KCNT2 | 1 | potassium sodium-activated channel subfamil |

| | | |
|---|---|---|
| MIR4735 | 1 | microRNA 4735 [Source:HGNC Symbol;Acc:HG |
| CFH | 1 | complement factor H [Source:HGNC Symbol;A |
| CFHR3 | 1 | complement factor H related 3 [Source:HGNC |
| CFHR1 | 1 | complement factor H related 1 [Source:HGNC |
| BX248415.1 | 1 | complement factor H related 3 (CFHR3) pseuc |
| CFHR4 | 1 | complement factor H related 4 [Source:HGNC |
| CFHR2 | 1 | complement factor H related 2 [Source:HGNC |
| AL139418.1 | 1 | complement factor H-related famlily member |
| CFHR5 | 1 | complement factor H related 5 [Source:HGNC |
| F13B | 1 | coagulation factor XIII B chain [Source:HGNC S |
| ASPM | 1 | abnormal spindle microtubule assembly [Sour |
| SEPTIN14P12 | 1 | septin 14 pseudogene 12 [Source:HGNC Symb |
| ZBTB41 | 1 | zinc finger and BTB domain containing 41 [Sou |
| AL513325.1 | 1 | ATPase, H+ transporting, lysosomal accessory |
| CRB1 | 1 | crumbs cell polarity complex component 1 [Sc |
| MRPS21P3 | 1 | mitochondrial ribosomal protein S21 pseudog |
| AL136322.1 | 1 | novel transcript |
| DENND1B | 1 | DENN domain containing 1B [Source:HGNC Sy |
| Y_RNA | 1 | Y RNA [Source:RFAM;Acc:RF00019] |
| EEF1A1P32 | 1 | eukaryotic translation elongation factor 1 alpł |
| AL365258.2 | 1 | ribosomal protein L24 (RPL24) pseudogene |
| FAM204BP | 1 | family with sequence similarity 204 member E |
| AL365258.1 | 1 | novel transcript |
| C1orf53 | 1 | chromosome 1 open reading frame 53 [Sourc |
| LHX9 | 1 | LIM homeobox 9 [Source:HGNC Symbol;Acc:H |
| NEK7 | 1 | NIMA related kinase 7 [Source:HGNC Symbol; |
| PRR13P1 | 1 | proline rich 13 pseudogene 1 [Source:HGNC S |
| AL450352.1 | 1 | novel transcript |
| ATP6V1G3 | 1 | ATPase H+ transporting V1 subunit G3 [Source |
| AL157402.1 | 1 | novel transcript |
| PTPRC | 1 | protein tyrosine phosphatase receptor type C |
| AL157402.2 | 1 | novel transcript |
| PEBP1P3 | 1 | phosphatidylethanolamine binding protein 1 p |
| MIR181A1HG | 1 | MIR181A1 host gene [Source:HGNC Symbol;A |
| MIR181B1 | 1 | microRNA 181b-1 [Source:HGNC Symbol;Acc:I |
| MIR181A1 | 1 | microRNA 181a-1 [Source:HGNC Symbol;Acc:I |
| AC096631.2 | 1 | ribosomal protein S2 (RPS2) pseudogene |
| AC096631.1 | 1 | novel transcript |
| LINC01222 | 1 | long intergenic non-protein coding RNA 1222 |
| LINC01221 | 1 | long intergenic non-protein coding RNA 1221 |
| AC105941.1 | 1 | novel transcript |
| LINC02789 | 1 | long intergenic non-protein coding RNA 2789 |
| RPL23AP16 | 1 | ribosomal protein L23a pseudogene 16 [Sourc |
| AC099335.1 | 1 | eukaryotic translation elongation factor 1 alpł |
| AL596266.1 | 1 | translational inhibitor protein p14.5 (UK114) p |
| AL445687.1 | 1 | NADH dehydrogenase (ubiquinone) 1 alpha su |
| RNU6-778P | 1 | RNA, U6 small nuclear 778, pseudogene [Sour |
| AL445687.2 | 1 | phosphatidylinositol transfer protein, cytoplas |
| RNU6-716P | 1 | RNA, U6 small nuclear 716, pseudogene [Sour |
| RNU6-609P | 1 | RNA, U6 small nuclear 609, pseudogene [Sour |

| | | |
|---|---|---|
| NR5A2 | 1 | nuclear receptor subfamily 5 group A member |
| RNU6-570P | 1 | RNA, U6 small nuclear 570, pseudogene [Sour |
| AC096633.1 | 1 | collagen, type XXII, alpha 1 (COL22A1) pseudo |
| CCNQP1 | 1 | CCNQ pseudogene 1 [Source:HGNC Symbol;A |
| LINC00862 | 1 | long intergenic non-protein coding RNA 862 [ |
| AC097065.2 | 1 | novel transcript |
| AC097065.1 | 1 | cofilin 1 (non-muscle) (CFL1) pseudogene |
| AC104461.1 | 1 | novel transcript, antisense to ZNF281 |
| ZNF281 | 1 | zinc finger protein 281 [Source:HGNC Symbol; |
| AL359834.1 | 1 | novel transcript |
| KIF14 | 1 | kinesin family member 14 [Source:HGNC Syml |
| DDX59 | 1 | DEAD-box helicase 59 [Source:HGNC Symbol; |
| DDX59-AS1 | 1 | DDX59 antisense RNA 1 [Source:HGNC Symbo |
| CAMSAP2 | 1 | calmodulin regulated spectrin associated prot |
| RPL34P6 | 1 | ribosomal protein L34 pseudogene 6 [Source: |
| GPR25 | 1 | G protein-coupled receptor 25 [Source:HGNC |
| INAVA | 1 | innate immunity activator [Source:HGNC Sym |
| MROH3P | 1 | maestro heat like repeat family member 3, ps |
| RNU6-704P | 1 | RNA, U6 small nuclear 704, pseudogene [Sour |
| KIF21B | 1 | kinesin family member 21B [Source:HGNC Syr |
| AL358473.1 | 1 | novel transcript, antisense KIF21B |
| AL358473.2 | 1 | novel transcript, antisense CACNA1S |
| CACNA1S | 1 | calcium voltage-gated channel subunit alpha1 |
| ASCL5 | 1 | achaete-scute family bHLH transcription facto |
| TMEM9 | 1 | transmembrane protein 9 [Source:HGNC Syml |
| IGFN1 | 1 | immunoglobulin like and fibronectin type III d |
| AC103925.1 | 1 | novel transcript |
| PKP1 | 1 | plakophilin 1 [Source:HGNC Symbol;Acc:HGN |
| TNNT2 | 1 | troponin T2, cardiac type [Source:HGNC Symb |
| AC119427.2 | 1 | novel transcript |
| LAD1 | 1 | ladinin 1 [Source:HGNC Symbol;Acc:HGNC:64 |
| AC119427.1 | 1 | novel transcript, antisense to LAD1 |
| TNNI1 | 1 | troponin I1, slow skeletal type [Source:HGNC |
| AC096677.3 | 1 | protein (peptidyl-prolyl cis/trans isomerase) N |
| PHLDA3 | 1 | pleckstrin homology like domain family A mer |
| AC096677.2 | 1 | novel transcript |
| CSRP1 | 1 | cysteine and glycine rich protein 1 [Source:HG |
| AC096677.1 | 1 | novel transcript |
| RPS10P7 | 1 | ribosomal protein S10 pseudogene 7 [Source: |
| NAV1 | 1 | neuron navigator 1 [Source:HGNC Symbol;Acc |
| AC092800.1 | 1 | novel transcript |
| IPO9-AS1 | 1 | IPO9 antisense RNA 1 [Source:HGNC Symbol; |
| MIR5191 | 1 | microRNA 5191 [Source:HGNC Symbol;Acc:HG |
| AL645504.1 | 1 | novel transcript |
| RNU6-501P | 1 | RNA, U6 small nuclear 501, pseudogene [Sour |
| MIR1231 | 1 | microRNA 1231 [Source:HGNC Symbol;Acc:HG |
| IPO9 | 1 | importin 9 [Source:HGNC Symbol;Acc:HGNC:1 |
| MIR6739 | 1 | microRNA 6739 [Source:HGNC Symbol;Acc:HG |
| SHISA4 | 1 | shisa family member 4 [Source:HGNC Symbol; |
| AL513217.1 | 1 | novel transcript |

| | | |
|---|---|---|
| LMOD1 | 1 | leiomodin 1 [Source:HGNC Symbol;Acc:HGNC: |
| TIMM17A | 1 | translocase of inner mitochondrial membrane |
| SNORA70H | 1 | small nucleolar RNA, H/ACA box 70H [Source: |
| RPL10P4 | 1 | ribosomal protein L10 pseudogene 4 [Source: |
| RNPEP | 1 | arginyl aminopeptidase [Source:HGNC Symbo |
| ELF3-AS1 | 1 | ELF3 antisense RNA 1 [Source:HGNC Symbol;A |
| MIR6740 | 1 | microRNA 6740 [Source:HGNC Symbol;Acc:HG |
| ELF3 | 1 | E74 like ETS transcription factor 3 [Source:HG |
| AL691482.3 | 1 | novel transcript |
| AL691482.1 | 1 | LSM3 homolog, U6 small nuclear RNA associa |
| AL691482.2 | 1 | ATPase, H+ transporting, lysosomal 9kDa, V0 s |
| Y_RNA | 1 | Y RNA [Source:RFAM;Acc:RF00019] |
| CRIP1P3 | 1 | cysteine rich protein 1 pseudogene 3 [Source: |
| GPR37L1 | 1 | G protein-coupled receptor 37 like 1 [Source: |
| ARL8A | 1 | ADP ribosylation factor like GTPase 8A [Sourc |
| PTPN7 | 1 | protein tyrosine phosphatase non-receptor ty |
| PTPRVP | 1 | protein tyrosine phosphatase receptor type V, |
| LGR6 | 1 | leucine rich repeat containing G protein-coupl |
| UBE2T | 1 | ubiquitin conjugating enzyme E2 T [Source:HG |
| PPP1R12B | 1 | protein phosphatase 1 regulatory subunit 12B |
| CYCSP4 | 1 | CYCS pseudogene 4 [Source:HGNC Symbol;Ac |
| RNU6-89P | 1 | RNA, U6 small nuclear 89, pseudogene [Sourc |
| AC099336.1 | 1 | FK506 binding protein 8, 38kDa (FKBP8) pseuc |
| AC099336.2 | 1 | ribosomal protein S27 (metallopanstimulin 1) |
| SNORA70 | 1 | Small nucleolar RNA SNORA70 [Source:RFAM; |
| SYT2 | 1 | synaptotagmin 2 [Source:HGNC Symbol;Acc:H |
| AC104463.2 | 1 | novel transcript |
| AC104463.1 | 1 | novel transcript |
| KDM5B | 1 | lysine demethylase 5B [Source:HGNC Symbol; |
| AC098934.3 | 1 | cytochrome c oxidase subunit VIIc (COX7C) ps |
| SLC25A39P1 | 1 | solute carrier family 25 member 39 pseudoge |
| AC098934.4 | 1 | novel transcript, antisense to a novel transcri |
| PCAT6 | 1 | prostate cancer associated transcript 6 [Sourc |
| MGAT4EP | 1 | MGAT4 family member E, pseudogene [Sourc |
| AC098934.1 | 1 | tubulin, alpha (TUBA) pseudogene |
| AC098934.2 | 1 | actin, gamma 1 (ACTG1) pseudogene |
| RABIF | 1 | RAB interacting factor [Source:HGNC Symbol; |
| KLHL12 | 1 | kelch like family member 12 [Source:HGNC Sy |
| HNRNPA1P59 | 1 | heterogeneous nuclear ribonucleoprotein A1 |
| Y_RNA | 1 | Y RNA [Source:RFAM;Acc:RF00019] |
| ADIPOR1 | 1 | adiponectin receptor 1 [Source:HGNC Symbol |
| CYB5R1 | 1 | cytochrome b5 reductase 1 [Source:HGNC Syr |
| AC096632.2 | 1 | novel pseudogene |
| AC096632.1 | 1 | ribosomal protein S20 (RPS20) pseudogene |
| TMEM183A | 1 | transmembrane protein 183A [Source:HGNC S |
| PPFIA4 | 1 | PTPRF interacting protein alpha 4 [Source:HGI |
| MYOG | 1 | myogenin [Source:HGNC Symbol;Acc:HGNC:7 |
| ADORA1 | 1 | adenosine A1 receptor [Source:HGNC Symbol, |
| AC105940.2 | 1 | novel transcript |
| AC105940.1 | 1 | novel transcript |

| | | |
|---|---|---|
| MYBPH | 1 | myosin binding protein H [Source:HGNC Symb |
| CHI3L1 | 1 | chitinase 3 like 1 [Source:HGNC Symbol;Acc:H |
| CHIT1 | 1 | chitinase 1 [Source:HGNC Symbol;Acc:HGNC:1 |
| NPM1P40 | 1 | nucleophosmin 1 pseudogene 40 [Source:HGN |
| LINC01353 | 1 | long intergenic non-protein coding RNA 1353 |
| LINC01136 | 1 | long intergenic non-protein coding RNA 1136 |
| BTG2 | 1 | BTG anti-proliferation factor 2 [Source:HGNC |
| RNU6-487P | 1 | RNA, U6 small nuclear 487, pseudogene [Sour |
| FMOD | 1 | fibromodulin [Source:HGNC Symbol;Acc:HGN |
| AL359837.1 | 1 | small EDRK-rich factor 2 (SERF2) pseudogene |
| LARP7P1 | 1 | La ribonucleoprotein domain family member |
| PRELP | 1 | proline and arginine rich end leucine rich repe |
| OPTC | 1 | opticin [Source:HGNC Symbol;Acc:HGNC:815 |
| ATP2B4 | 1 | ATPase plasma membrane Ca2+ transporting |
| AL513343.1 | 1 | TGF beta-inducible nuclear protein 1 (TINP1) |
| SNORA77 | 1 | small nucleolar RNA, H/ACA box 77 [Source:H |
| LAX1 | 1 | lymphocyte transmembrane adaptor 1 [Sourc |
| ZC3H11A | 1 | zinc finger CCCH-type containing 11A [Source: |
| ZBED6 | 1 | zinc finger BED-type containing 6 [Source:HGN |
| AC114402.2 | 1 | SNF7 domain containing 2 (SNF7DC20) pseud |
| AC114402.1 | 1 | ribosomal protein L35a (RPL35A) pseudogene |
| SNRPE | 1 | small nuclear ribonucleoprotein polypeptide |
| KRT8P29 | 1 | keratin 8 pseudogene 29 [Source:HGNC Symb |
| HSPE1P6 | 1 | heat shock protein family E (Hsp10) member |
| CBX1P3 | 1 | chromobox 1 pseudogene 3 [Source:HGNC Sy |
| AC096645.1 | 1 | novel transcript |
| LINC00303 | 1 | long intergenic non-protein coding RNA 303 [ |
| AL592146.2 | 1 | novel transcript, antisense to SOX13 |
| SOX13 | 1 | SRY-box transcription factor 13 [Source:HGNC |
| ETNK2 | 1 | ethanolamine kinase 2 [Source:HGNC Symbol; |
| AL592146.1 | 1 | novel transcript |
| ERLNC1 | 1 | estrogen receptor responsive lncRNA 1 [Sourc |
| REN | 1 | renin [Source:HGNC Symbol;Acc:HGNC:9958] |
| AL592114.2 | 1 | novel pseudogene |
| KISS1 | 1 | KiSS-1 metastasis suppressor [Source:HGNC S |
| GOLT1A | 1 | golgi transport 1A [Source:HGNC Symbol;Acc: |
| PLEKHA6 | 1 | pleckstrin homology domain containing A6 [S |
| AL592114.3 | 1 | novel transcript |
| AL592114.1 | 1 | ribosomal protein L21 (RPL21) pseudogene |
| LINC00628 | 1 | long intergenic non-protein coding RNA 628 [ |
| AL606489.1 | 1 | novel transcript |
| AL606489.2 | 1 | small nuclear ribonucleoprotein polypeptide |
| PPP1R15B | 1 | protein phosphatase 1 regulatory subunit 15B |
| PIK3C2B | 1 | phosphatidylinositol-4-phosphate 3-kinase ca |
| MDM4 | 1 | MDM4 regulator of p53 [Source:HGNC Symbo |
| AL512306.1 | 1 | PEF protein with a long N-terminal hydropho |
| RNA5SP74 | 1 | RNA, 5S ribosomal pseudogene 74 [Source:H |
| AL512306.3 | 1 | novel transcript |
| LRRN2 | 1 | leucine rich repeat neuronal 2 [Source:HGNC |
| AL512306.2 | 1 | novel transcript |

| | | |
|---|---|---|
| AL161793.1 | 1 | novel transcript |
| RNA5SP75 | 1 | RNA, 5S ribosomal pseudogene 75 [Source:HG |
| AL161793.2 | 1 | NULL |
| AC096675.1 | 1 | novel transcript, antisense to NFASC |
| NFASC | 1 | neurofascin [Source:HGNC Symbol;Acc:HGNC: |
| AL391822.1 | 1 | ribosomal protein L13a (RPL13A) pseudogene |
| CNTN2 | 1 | contactin 2 [Source:HGNC Symbol;Acc:HGNC: |
| TMEM81 | 1 | transmembrane protein 81 [Source:HGNC Syn |
| RBBP5 | 1 | RB binding protein 5, histone lysine methyltra |
| AL583832.1 | 1 | glycogenin 2 (GYG2) pseudogene |
| AC093422.3 | 1 | signal peptidase 12kDa (SPC12) pseudogene |
| DSTYK | 1 | dual serine/threonine and tyrosine protein kir |
| AC093422.1 | 1 | ribosomal protein L17 (RPL17) pseudogene |
| TMCC2 | 1 | transmembrane and coiled-coil domain family |
| AC093422.2 | 1 | novel transcript |
| NUAK2 | 1 | NUAK family kinase 2 [Source:HGNC Symbol;A |
| KLHDC8A | 1 | kelch domain containing 8A [Source:HGNC Syr |
| SNRPGP10 | 1 | small nuclear ribonucleoprotein polypeptide ( |
| LEMD1-AS1 | 1 | LEMD1 antisense RNA 1 [Source:HGNC Symbo |
| LEMD1 | 1 | LEM domain containing 1 [Source:HGNC Symb |
| BLACAT1 | 1 | bladder cancer associated transcript 1 [Source |
| MIR135B | 1 | microRNA 135b [Source:HGNC Symbol;Acc:HG |
| LEMD1-DT | 1 | LEMD1 divergent transcript [Source:HGNC Syr |
| CDK18 | 1 | cyclin dependent kinase 18 [Source:HGNC Syn |
| RNU2-19P | 1 | RNA, U2 small nuclear 19, pseudogene [Sourc |
| MFSD4A | 1 | major facilitator superfamily domain containir |
| RNU6-418P | 1 | RNA, U6 small nuclear 418, pseudogene [Sour |
| ELK4 | 1 | ETS transcription factor ELK4 [Source:HGNC Sy |
| AC096533.1 | 1 | GABA(A) receptor-associated protein (GABAR |
| SLC45A3 | 1 | solute carrier family 45 member 3 [Source:HG |
| Metazoa_SRP | 1 | Metazoan signal recognition particle RNA [Sou |
| NUCKS1 | 1 | nuclear casein kinase and cyclin dependent kir |
| SNORA72 | 1 | Small nucleolar RNA SNORA72 [Source:RFAM; |
| RAB29 | 1 | RAB29, member RAS oncogene family [Source |
| AC119673.1 | 1 | novel transcript |
| SLC41A1 | 1 | solute carrier family 41 member 1 [Source:HG |
| AC119673.2 | 1 | novel transcript, antisense to PM20D1 |
| PM20D1 | 1 | peptidase M20 domain containing 1 [Source:H |
| SLC26A9 | 1 | solute carrier family 26 member 9 [Source:HG |
| AL713965.1 | 1 | novel transcript |
| RAB7B | 1 | RAB7B, member RAS oncogene family [Source |
| CTSE | 1 | cathepsin E [Source:HGNC Symbol;Acc:HGNC: |
| BX571818.1 | 1 | novel pseudogene |
| RHEX | 1 | regulator of hemoglobinization and erythroid |
| AC244035.3 | 1 | NULL |
| AVPR1B | 1 | arginine vasopressin receptor 1B [Source:HGN |
| AC244035.1 | 1 | novel transcript |
| AC244035.4 | 1 | novel transcript |
| RPL22P4 | 1 | ribosomal protein L22 pseudogene 4 [Source: |
| AC244035.2 | 1 | novel transcript |

| | | |
|---|---|---|
| FAM72A | 1 | family with sequence similarity 72 member A |
| SRGAP2 | 1 | SLIT-ROBO Rho GTPase activating protein 2 [S |
| AC244023.1 | 1 | RAP1, GTP-GDP dissociation stimulator 1 (RAF |
| IKBKE | 1 | inhibitor of nuclear factor kappa B kinase subu |
| MIR6769B | 1 | microRNA 6769b [Source:HGNC Symbol;Acc:H |
| C1orf147 | 1 | chromosome 1 open reading frame 147 [Sour |
| AC244034.2 | 1 | novel transcript |
| RASSF5 | 1 | Ras association domain family member 5 [Sou |
| AC244034.1 | 1 | ribosomal protein L7a (RPL7A) pseudogene |
| AC244034.3 | 1 | novel transcript, sense intronic to RASSF5 |
| EIF2D | 1 | eukaryotic translation initiation factor 2D [Sou |
| AL591846.2 | 1 | novel transcript |
| DYRK3 | 1 | dual specificity tyrosine phosphorylation regu |
| MAPKAPK2 | 1 | MAPK activated protein kinase 2 [Source:HGN |
| AL591846.1 | 1 | ribosomal protein S14 (RPS14) pseudogene |
| Y_RNA | 1 | Y RNA [Source:RFAM;Acc:RF00019] |
| IL10 | 1 | interleukin 10 [Source:HGNC Symbol;Acc:HGN |
| IL19 | 1 | interleukin 19 [Source:HGNC Symbol;Acc:HGN |
| IL20 | 1 | interleukin 20 [Source:HGNC Symbol;Acc:HGN |
| IL24 | 1 | interleukin 24 [Source:HGNC Symbol;Acc:HGN |
| FCMR | 1 | Fc fragment of IgM receptor [Source:HGNC Sy |
| AC098935.2 | 1 | La ribonucleoprotein domain family, member |
| AC098935.1 | 1 | ribosomal protein L13a (RPL13A) pseudogene |
| PIGR | 1 | polymeric immunoglobulin receptor [Source:H |
| FCAMR | 1 | Fc fragment of IgA and IgM receptor [Source:H |
| C1orf116 | 1 | chromosome 1 open reading frame 116 [Sour |
| PFKFB2 | 1 | 6-phosphofructo-2-kinase/fructose-2,6-bipho: |
| YOD1 | 1 | YOD1 deubiquitinase [Source:HGNC Symbol;A |
| C4BPB | 1 | complement component 4 binding protein be |
| C4BPA | 1 | complement component 4 binding protein alp |
| AL445493.3 | 1 | novel transcript, antisense to C4BPA |
| AL445493.1 | 1 | signal recognition particle 72kDa (SRP72) pseu |
| C4BPAP1 | 1 | C4BPA pseudogene 1 [Source:HGNC Symbol;A |
| AL445493.2 | 1 | novel transcript |
| C4BPAP2 | 1 | C4BPA pseudogene 2 [Source:HGNC Symbol;A |
| AL596218.1 | 1 | novel transcript |
| CD55 | 1 | CD55 molecule (Cromer blood group) [Source |
| AL391597.1 | 1 | novel transcript |
| CR2 | 1 | complement C3d receptor 2 [Source:HGNC Sy |
| CR1 | 1 | complement C3b/C4b receptor 1 (Knops bloo |
| AL137789.1 | 1 | novel transcript, antisense to CR1 |
| CR1L | 1 | complement C3b/C4b receptor 1 like [Source: |
| CD46P1 | 1 | CD46 molecule pseudogene 1 [Source:HGNC S |
| CDCA4P3 | 1 | cell division cycle associated 4 pseudogene 3 [ |
| RNA5SP534 | 1 | RNA, 5S ribosomal pseudogene 534 [Source:H |
| AL137789.2 | 1 | novel transcript |
| CD46 | 1 | CD46 molecule [Source:HGNC Symbol;Acc:HG |
| CDCA4P4 | 1 | cell division cycle associated 4 pseudogene 4 [ |
| MIR29B2CHG | 1 | MIR29B2 and MIR29C host gene [Source:HGN |
| MIR29C | 1 | microRNA 29c [Source:HGNC Symbol;Acc:HGN |

| | | |
|---|---|---|
| MIR29B2 | 1 | microRNA 29b-2 [Source:HGNC Symbol;Acc:H |
| CD34 | 1 | CD34 molecule [Source:HGNC Symbol;Acc:HG |
| AL356275.1 | 1 | novel transcript, antisense to CD34 |
| LINC02767 | 1 | long intergenic non-protein coding RNA 2767 |
| PLXNA2 | 1 | plexin A2 [Source:HGNC Symbol;Acc:HGNC:91 |
| AL590138.1 | 1 | novel transcript, antisense to PLXNA2 |
| AL606753.1 | 1 | heterogeneous nuclear ribonucleoprotein D-li |
| AL606753.2 | 1 | novel transcript |
| LINC01735 | 1 | long intergenic non-protein coding RNA 1735 |
| LINC02769 | 1 | long intergenic non-protein coding RNA 2769 |
| RPS26P13 | 1 | ribosomal protein S26 pseudogene 13 [Source |
| LINC01717 | 1 | long intergenic non-protein coding RNA 1717 |
| LINC01774 | 1 | long intergenic non-protein coding RNA 1774 |
| AC104465.1 | 1 | vacuole membrane protein 1 (VMP1) pseudog |
| AC092810.4 | 1 | novel transcript |
| AC092810.3 | 1 | novel transcript |
| AC092810.2 | 1 | GRB2-related adaptor protein (GRAP) pseudog |
| AC092810.1 | 1 | transcription factor Dp-1 (TFDP1) pseudogene |
| ATP5MC2P1 | 1 | ATP synthase membrane subunit c locus 2 pse |
| LINC01696 | 1 | long intergenic non-protein coding RNA 1696 |
| LINC01698 | 1 | long intergenic non-protein coding RNA 1698 |
| MIR205HG | 1 | MIR205 host gene [Source:HGNC Symbol;Acc: |
| MIR205 | 1 | microRNA 205 [Source:HGNC Symbol;Acc:HGN |
| AL023754.1 | 1 | novel transcript |
| AL023754.2 | 1 | novel transcript |
| CAMK1G | 1 | calcium/calmodulin dependent protein kinase |
| LAMB3 | 1 | laminin subunit beta 3 [Source:HGNC Symbol; |
| MIR4260 | 1 | microRNA 4260 [Source:HGNC Symbol;Acc:HG |
| HSD11B1-AS1 | 1 | HSD11B1 antisense RNA 1 [Source:HGNC Sym |
| G0S2 | 1 | G0/G1 switch 2 [Source:HGNC Symbol;Acc:HG |
| HSD11B1 | 1 | hydroxysteroid 11-beta dehydrogenase 1 [Sou |
| ADORA2BP1 | 1 | adenosine A2b receptor pseudogene 1 [Sourc |
| TRAF3IP3 | 1 | TRAF3 interacting protein 3 [Source:HGNC Syr |
| C1orf74 | 1 | chromosome 1 open reading frame 74 [Sourc |
| IRF6 | 1 | interferon regulatory factor 6 [Source:HGNC S |
| UTP25 | 1 | UTP25 small subunit processor component [S |
| SYT14 | 1 | synaptotagmin 14 [Source:HGNC Symbol;Acc: |
| AL022397.1 | 1 | novel transcript, antisense to SYT14 |
| SERTAD4-AS1 | 1 | SERTAD4 antisense RNA 1 [Source:HGNC Syml |
| SERTAD4 | 1 | SERTA domain containing 4 [Source:HGNC Syr |
| ST13P19 | 1 | ST13, Hsp70 interacting protein pseudogene 1 |
| AL035414.1 | 1 | ribonuclease H1 (RNASEH1) pseudogene |
| HHAT | 1 | hedgehog acyltransferase [Source:HGNC Syml |
| AL034351.4 | 1 | novel transcript, antisense to HHAT |
| RNU5A-8P | 1 | RNA, U5A small nuclear 8, pseudogene [Sourc |
| AL034351.3 | 1 | novel transcript, antisense to HHAT |
| AL034351.1 | 1 | inositol monophosphatase domain containing |
| KCNH1 | 1 | potassium voltage-gated channel subfamily H |
| AC096636.1 | 1 | novel transcript |
| AL590132.1 | 1 | novel protein |

| | | |
|---|---|---|
| IPO8P1 | 1 | importin 8 pseudogene 1 [Source:HGNC Symb |
| AC092017.3 | 1 | novel protein |
| AC092017.4 | 1 | novel transcript |
| KCNH1-IT1 | 1 | KCNH1 intronic transcript 1 [Source:HGNC Syr |
| AC092017.2 | 1 | pseudogene similar to ribosomal protein S25 ( |
| AC092017.1 | 1 | pseudogene similar to px19-like protein (PX19 |
| RCOR3 | 1 | REST corepressor 3 [Source:HGNC Symbol;Acc |
| TRAF5 | 1 | TNF receptor associated factor 5 [Source:HGN |
| AL590101.1 | 1 | novel transcript |
| LINC00467 | 1 | long intergenic non-protein coding RNA 467 [S |
| SNX25P1 | 1 | sorting nexin 25 pseudogene 1 [Source:HGNC |
| ARPC3P2 | 1 | actin related protein 2/3 complex subunit 3 ps |
| RD3 | 1 | retinal degeneration 3, GUCY2D regulator [Sou |
| AC105275.1 | 1 | novel transcript |
| SLC30A1 | 1 | solute carrier family 30 member 1 [Source:HG |
| AC105275.2 | 1 | novel transcript, sense overlapping SLC30A1 |
| AL356310.1 | 1 | novel transcript |
| LINC01693 | 1 | long intergenic non-protein coding RNA 1693 |
| NEK2 | 1 | NIMA related kinase 2 [Source:HGNC Symbol; |
| FDPSP8 | 1 | farnesyl diphosphate synthase pseudogene 8 |
| AC096637.2 | 1 | novel transcript |
| AC096637.1 | 1 | novel transcript |
| LPGAT1 | 1 | lysophosphatidylglycerol acyltransferase 1 [Sc |
| RN7SL344P | 1 | RNA, 7SL, cytoplasmic 344, pseudogene [Sour |
| Y_RNA | 1 | Y RNA [Source:RFAM;Acc:RF00019] |
| LPGAT1-AS1 | 1 | LPGAT1 antisense RNA 1 [Source:HGNC Symbo |
| AC092814.2 | 1 | protein (peptidyl-prolyl cis\\/trans isomerase) |
| INTS7 | 1 | integrator complex subunit 7 [Source:HGNC Sy |
| AC092814.1 | 1 | NULL |
| DTL | 1 | denticleless E3 ubiquitin protein ligase homolo |
| RPL21P28 | 1 | ribosomal protein L21 pseudogene 28 [Source |
| MIR3122 | 1 | microRNA 3122 [Source:HGNC Symbol;Acc:HG |
| RN7SKP98 | 1 | RN7SK pseudogene 98 [Source:HGNC Symbol; |
| AL606468.1 | 1 | novel transcript |
| LINC02608 | 1 | long intergenic non-protein coding RNA 2608 |
| PPP2R5A | 1 | protein phosphatase 2 regulatory subunit B'al |
| AL360091.2 | 1 | novel transcript, sense intronic to PPP2R5A |
| AL360091.3 | 1 | novel transcript, antisense to PPP2R5A |
| RPL23AP18 | 1 | ribosomal protein L23a pseudogene 18 [Sourc |
| SNORA16B | 1 | small nucleolar RNA, H/ACA box 16B [Source:I |
| AL360091.1 | 1 | novel transcript, sense intronic to PPP2R5A |
| PACC1 | 1 | proton activated chloride channel 1 [Source:H |
| AC092803.3 | 1 | novel transcript |
| NENF | 1 | neudesin neurotrophic factor [Source:HGNC S |
| LINC02771 | 1 | long intergenic non-protein coding RNA 2771 |
| AC092803.2 | 1 | novel transcript |
| LINC01740 | 1 | long intergenic non-protein coding RNA 1740 |
| AC092803.1 | 1 | novel transcript |
| AC092803.4 | 1 | novel transcript |
| ATF3 | 1 | activating transcription factor 3 [Source:HGN( |

| | | |
|---|---|---|
| AL590648.2 | 1 | novel transcript |
| FAM71A | 1 | family with sequence similarity 71 member A |
| AL590648.1 | 1 | riboflavin kinase pseudogene |
| LINC02773 | 1 | long intergenic non-protein coding RNA 2773 |
| BATF3 | 1 | basic leucine zipper ATF-like transcription fact |
| NSL1 | 1 | NSL1 component of MIS12 kinetochore compl |
| TATDN3 | 1 | TatD DNase domain containing 3 [Source:HGN |
| AC104333.3 | 1 | ribosomal protein S5 (RPS5) pseudogene |
| SPATA45 | 1 | spermatogenesis associated 45 [Source:HGNC |
| FLVCR1-DT | 1 | FLVCR1 divergent transcript [Source:HGNC Sy |
| AC104333.1 | 1 | pseudogene similar to part of ribosomal prote |
| AC104333.2 | 1 | ribosomal protein S28 (RPS28) pseudogene |
| FLVCR1 | 1 | feline leukemia virus subgroup C cellular rece |
| AC104333.4 | 1 | novel transcript |
| VASH2 | 1 | vasohibin 2 [Source:HGNC Symbol;Acc:HGNC: |
| ANGEL2 | 1 | angel homolog 2 [Source:HGNC Symbol;Acc:H |
| RPS6KC1 | 1 | ribosomal protein S6 kinase C1 [Source:HGNC |
| RPL31P13 | 1 | ribosomal protein L31 pseudogene 13 [Source |
| AL592402.1 | 1 | novel transcript |
| AC096639.1 | 1 | novel transcript |
| PROX1-AS1 | 1 | PROX1 antisense RNA 1 [Source:HGNC Symbo |
| LINC00538 | 1 | long intergenic non-protein coding RNA 538 [ |
| PROX1 | 1 | prospero homeobox 1 [Source:HGNC Symbol; |
| AC011700.1 | 1 | novel transcript, antisense to PROX1 |
| AL606537.1 | 1 | PROX1 antisense RNA 1 |
| LINC02775 | 1 | long intergenic non-protein coding RNA 2775 |
| SMYD2 | 1 | SET and MYND domain containing 2 [Source:H |
| AL929236.1 | 1 | novel transcript |
| PTPN14 | 1 | protein tyrosine phosphatase non-receptor ty |
| AL445305.1 | 1 | adaptor-related protein complex 3 pseudoger |
| KRT18P12 | 1 | keratin 18 pseudogene 12 [Source:HGNC Sym |
| CENPF | 1 | centromere protein F [Source:HGNC Symbol;A |
| ABHD17AP3 | 1 | abhydrolase domain containing 17A pseudoge |
| UBE2V1P13 | 1 | ubiquitin conjugating enzyme E2 V1 pseudoge |
| GAPDHP24 | 1 | glyceraldehyde 3 phosphate dehydrogenase p |
| AC099563.1 | 1 | translocase of inner mitochondrial membrane |
| AC099563.2 | 1 | novel transcript |
| KCNK2 | 1 | potassium two pore domain channel subfamil |
| VDAC1P10 | 1 | voltage dependent anion channel 1 pseudoge |
| AL450990.1 | 1 | novel transcript |
| KCTD3 | 1 | potassium channel tetramerization domain cc |
| USH2A | 1 | usherin [Source:HGNC Symbol;Acc:HGNC:126 |
| SNORD116 | 1 | Small nucleolar RNA SNORD116 [Source:RFAN |
| AL358452.1 | 1 | novel transcript |
| AC093581.1 | 1 | novel transcript |
| AC138024.1 | 1 | novel transcript |
| MRPS18BP1 | 1 | mitochondrial ribosomal protein S18B pseudo |
| ESRRG | 1 | estrogen related receptor gamma [Source:HG |
| GPATCH2 | 1 | G-patch domain containing 2 [Source:HGNC S |
| SPATA17 | 1 | spermatogenesis associated 17 [Source:HGNC |

| | | |
|---|---|---|
| SPATA17-AS1 | 1 | SPATA17 antisense RNA 1 [Source:HGNC Syml |
| UBBP2 | 1 | ubiquitin B pseudogene 2 [Source:HGNC Symb |
| LINC00210 | 1 | long intergenic non-protein coding RNA 210 [S |
| AL355526.1 | 1 | novel transcript |
| LINC01653 | 1 | long intergenic non-protein coding RNA 1653 |
| AL355526.2 | 1 | novel transcript |
| RNU1-141P | 1 | RNA, U1 small nuclear 141, pseudogene [Sour |
| RRP15 | 1 | ribosomal RNA processing 15 homolog [Sourc |
| AC096638.2 | 1 | novel pseudogene |
| AC096638.1 | 1 | ribosomal protein S26 (RPS26) pseudogenes |
| TGFB2-AS1 | 1 | TGFB2 antisense RNA 1 (head to head) [Sourc |
| TGFB2 | 1 | transforming growth factor beta 2 [Source:HG |
| TGFB2-OT1 | 1 | TGFB2 overlapping transcript 1 [Source:HGNC |
| C1orf143 | 1 | chromosome 1 open reading frame 143 (puta |
| U3 | 1 | Small nucleolar RNA U3 [Source:RFAM;Acc:RF |
| NXNP1 | 1 | nucleoredoxin pseudogene 1 [Source:HGNC S |
| LINC01710 | 1 | long intergenic non-protein coding RNA 1710 |
| LYPLAL1-DT | 1 | LYPLAL1 divergent transcript [Source:HGNC Sy |
| LYPLAL1 | 1 | lysophospholipase like 1 [Source:HGNC Symbo |
| RIMKLBP2 | 1 | ribosomal modification protein rimK like famil |
| AL360093.1 | 1 | novel transcript |
| AC096642.1 | 1 | novel transcript |
| AC096642.2 | 1 | novel transcript |
| LYPLAL1-AS1 | 1 | LYPLAL1 antisense RNA 1 [Source:HGNC Symb |
| AL356364.1 | 1 | novel transcript |
| ZC3H11B | 1 | zinc finger CCCH-type containing 11B [Source: |
| SLC30A10 | 1 | solute carrier family 30 member 10 [Source:H |
| RNA5SP76 | 1 | RNA, 5S ribosomal pseudogene 76 [Source:HG |
| U3 | 1 | Small nucleolar RNA U3 [Source:RFAM;Acc:RF |
| EPRS | 1 | glutamyl-prolyl-tRNA synthetase [Source:HGN |
| BPNT1 | 1 | 3'(2'), 5'-bisphosphate nucleotidase 1 [Source |
| IARS2 | 1 | isoleucyl-tRNA synthetase 2, mitochondrial [S |
| MIR215 | 1 | microRNA 215 [Source:HGNC Symbol;Acc:HGN |
| MIR194-1 | 1 | microRNA 194-1 [Source:HGNC Symbol;Acc:H |
| RPS15AP12 | 1 | ribosomal protein S15a pseudogene 12 [Sourc |
| RAB3GAP2 | 1 | RAB3 GTPase activating non-catalytic protein |
| MIR664A | 1 | microRNA 664a [Source:HGNC Symbol;Acc:HG |
| SNORA36B | 1 | small nucleolar RNA, H/ACA box 36B [Source:I |
| AL445435.1 | 1 | sorting nexin 2 (SNX2) pseudogene |
| KF455155.1 | 1 | GABA(A) receptor-associated protein-like 2 (G |
| MORF4L1P1 | 1 | mortality factor 4 like 1 pseudogene 1 [Source |
| AURKAP1 | 1 | aurora kinase A pseudogene 1 [Source:HGNC : |
| XRCC6P3 | 1 | X-ray repair cross complementing 6 pseudoge |
| AL451081.2 | 1 | ribosomal protein, large, P0 (RPLP0) pseudoge |
| AL451081.1 | 1 | novel transcript |
| AC096644.3 | 1 | novel transcript |
| AC096644.2 | 1 | ribosomal protein L7a (RPL7A) pseudogene |
| AC096644.1 | 1 | part of a cytochrome c oxidase subunit III (CO |
| PRELID3BP1 | 1 | PRELI domain containing 3B pseudogene 1 [Sc |
| LINC02779 | 1 | long intergenic non-protein coding RNA 2779 |

| | | |
|---|---|---|
| MARK1 | 1 | microtubule affinity regulating kinase 1 [Sourc |
| RN7SL464P | 1 | RNA, 7SL, cytoplasmic 464, pseudogene [Sour |
| HDAC1P2 | 1 | histone deacetylase 1 pseudogene 2 [Source:I |
| C1orf115 | 1 | chromosome 1 open reading frame 115 [Sour |
| MARC2 | 1 | mitochondrial amidoxime reducing componer |
| MARC1 | 1 | mitochondrial amidoxime reducing componer |
| AL445423.3 | 1 | novel protein |
| RNU6ATAC35P | 1 | RNA, U6atac small nuclear 35, pseudogene [S |
| AL445423.2 | 1 | NULL |
| AL445423.1 | 1 | novel transcript |
| LINC01352 | 1 | long intergenic non-protein coding RNA 1352 |
| HLX-AS1 | 1 | HLX antisense RNA 1 [Source:HGNC Symbol;A |
| HLX | 1 | H2.0 like homeobox [Source:HGNC Symbol;Ac |
| AL445466.1 | 1 | novel pseudogene |
| LINC02817 | 1 | long intergenic non-protein coding RNA 2817 |
| AL360013.2 | 1 | novel transcript |
| AL360013.3 | 1 | novel transcript |
| AL360013.1 | 1 | solute carrier family 35, member E3 (SLC35E3 |
| AL360013.4 | 1 | novel transcript |
| DUSP10 | 1 | dual specificity phosphatase 10 [Source:HGNC |
| LINC01655 | 1 | long intergenic non-protein coding RNA 1655 |
| RNU6-403P | 1 | RNA, U6 small nuclear 403, pseudogene [Sour |
| LINC02257 | 1 | long intergenic non-protein coding RNA 2257 |
| LINC02474 | 1 | long intergenic non-protein coding RNA 2474 |
| LINC01705 | 1 | long intergenic non-protein coding RNA 1705 |
| AL356108.1 | 1 | novel transcript |
| QRSL1P2 | 1 | QRSL1 pseudogene 2 [Source:HGNC Symbol;A |
| AL513314.1 | 1 | novel transcript |
| CICP13 | 1 | capicua transcriptional repressor pseudogene |
| AL513314.2 | 1 | novel transcript |
| RNU6-791P | 1 | RNA, U6 small nuclear 791, pseudogene [Sour |
| AL592148.2 | 1 | general transcription factor IIi (GTF2I) pseudo |
| HHIPL2 | 1 | HHIP like 2 [Source:HGNC Symbol;Acc:HGNC:2 |
| TAF1A | 1 | TATA-box binding protein associated factor, R |
| TAF1A-AS1 | 1 | TAF1A antisense RNA 1 [Source:HGNC Symbol |
| MIA3 | 1 | MIA SH3 domain ER export factor 3 [Source:H |
| AL592148.1 | 1 | prefoldin 5 (PFDN5) pseudogene |
| AL592148.3 | 1 | novel transcript, antisense to MIA3 |
| AIDA | 1 | axin interactor, dorsalization associated [Sour |
| BROX | 1 | BRO1 domain and CAAX motif containing [Sou |
| FAM177B | 1 | family with sequence similarity 177 member E |
| AL392172.2 | 1 | novel transcript, antisense to FAM177B |
| AL392172.1 | 1 | novel transcript |
| AC093152.1 | 1 | novel transcript, lincRNA-DISP1 readthrough |
| DISP1 | 1 | dispatched RND transporter family member 1 |
| NDUFB1P2 | 1 | NADH:ubiquinone oxidoreductase subunit B1 |
| AL929091.1 | 1 | novel transcript |
| TLR5 | 1 | toll like receptor 5 [Source:HGNC Symbol;Acc: |
| AL359979.2 | 1 | novel transcript |
| AL359979.1 | 1 | novel transcript |

| | | |
|---|---|---|
| SUSD4 | 1 | sushi domain containing 4 [Source:HGNC Sym |
| RNU4-57P | 1 | RNA, U4 small nuclear 57, pseudogene [Sourc |
| CCDC185 | 1 | coiled-coil domain containing 185 [Source:HG |
| CAPN8 | 1 | calpain 8 [Source:HGNC Symbol;Acc:HGNC:14 |
| SNRPEP10 | 1 | small nuclear ribonucleoprotein polypeptide E |
| RNU6-1248P | 1 | RNA, U6 small nuclear 1248, pseudogene [Sou |
| CAPN2 | 1 | calpain 2 [Source:HGNC Symbol;Acc:HGNC:14 |
| TP53BP2 | 1 | tumor protein p53 binding protein 2 [Source:H |
| PHBP11 | 1 | prohibitin pseudogene 11 [Source:HGNC Symb |
| ACTBP11 | 1 | ACTB pseudogene 11 [Source:HGNC Symbol;A |
| CICP5 | 1 | capicua transcriptional repressor pseudogene |
| GTF2IP20 | 1 | general transcription factor IIi pseudogene 20 |
| RNU6-1319P | 1 | RNA, U6 small nuclear 1319, pseudogene [Sou |
| AC138393.1 | 1 | septin 7 (SEPT7) pseudogene |
| AC138393.3 | 1 | novel transcript |
| AC138393.2 | 1 | F-box only protein 25 (FBXO25) pseudogene |
| RN7SKP49 | 1 | RN7SK pseudogene 49 [Source:HGNC Symbol; |
| FBXO28 | 1 | F-box protein 28 [Source:HGNC Symbol;Acc:H |
| AC092809.3 | 1 | novel pseudogene |
| DEGS1 | 1 | delta 4-desaturase, sphingolipid 1 [Source:HG |
| SNORA72 | 1 | Small nucleolar RNA SNORA72 [Source:RFAM; |
| AC092809.2 | 1 | novel transcript |
| AC092809.4 | 1 | novel transcript |
| NVL | 1 | nuclear VCP like [Source:HGNC Symbol;Acc:HG |
| MIR320B2 | 1 | microRNA 320b-2 [Source:HGNC Symbol;Acc:I |
| AC092809.1 | 1 | ribosomal protein L21 (RPL21) pseudogene |
| RNU6-1008P | 1 | RNA, U6 small nuclear 1008, pseudogene [Sou |
| CNIH4 | 1 | cornichon family AMPA receptor auxiliary pro |
| WDR26 | 1 | WD repeat domain 26 [Source:HGNC Symbol;A |
| MIR4742 | 1 | microRNA 4742 [Source:HGNC Symbol;Acc:HG |
| CNIH3 | 1 | cornichon family AMPA receptor auxiliary pro |
| AKR1B1P1 | 1 | aldehyde reductase family 1 member B1 pseu |
| AC096537.1 | 1 | novel transcript |
| AC096537.2 | 1 | DnaJ (Hsp40) homolog, subfamily B, member |
| AL596330.2 | 1 | novel transcript, antisense to CNIH3 |
| AL596330.1 | 1 | novel transcript |
| LINC02813 | 1 | long intergenic non-protein coding RNA 2813 |
| AL391811.1 | 1 | novel transcript |
| DNAH14 | 1 | dynein axonemal heavy chain 14 [Source:HGN |
| LBR | 1 | lamin B receptor [Source:HGNC Symbol;Acc:H |
| LINC02765 | 1 | long intergenic non-protein coding RNA 2765 |
| AC092811.1 | 1 | novel transcript |
| ENAH | 1 | ENAH actin regulator [Source:HGNC Symbol;A |
| AC099066.2 | 1 | novel transcript |
| AC099066.1 | 1 | novel transcript |
| RNU6-1304P | 1 | RNA, U6 small nuclear 1304, pseudogene [Sou |
| SRP9 | 1 | signal recognition particle 9 [Source:HGNC Syr |
| AC099066.3 | 1 | endometrial bleeding associated factor (left-ri |
| EPHX1 | 1 | epoxide hydrolase 1 [Source:HGNC Symbol;Ac |
| AL591895.1 | 1 | novel transcript |

| | | |
|---|---|---|
| TMEM63A | 1 | transmembrane protein 63A [Source:HGNC Sy |
| LEFTY1 | 1 | left-right determination factor 1 [Source:HGN |
| AL117348.2 | 1 | novel protein |
| PYCR2 | 1 | pyrroline-5-carboxylate reductase 2 [Source:H |
| MIR6741 | 1 | microRNA 6741 [Source:HGNC Symbol;Acc:HG |
| AL117348.1 | 1 | novel transcript |
| LEFTY2 | 1 | left-right determination factor 2 [Source:HGN |
| NDUFA3P3 | 1 | NADH:ubiquinone oxidoreductase subunit A3 |
| SDE2 | 1 | SDE2 telomere maintenance homolog [Source |
| AL512343.2 | 1 | novel transcript, antisense to H3F3A |
| H3F3A | 1 | H3 histone family member 3A [Source:HGNC S |
| LINC01703 | 1 | long intergenic non-protein coding RNA 1703 |
| AL512343.1 | 1 | chromosome 17 open reading frame 89 (C17c |
| ACBD3 | 1 | acyl-CoA binding domain containing 3 [Source |
| ACBD3-AS1 | 1 | ACBD3 antisense RNA 1 [Source:HGNC Symbo |
| AL592045.1 | 1 | ribosomal protein L34 (RPL34) pseudogene |
| MIXL1 | 1 | Mix paired-like homeobox [Source:HGNC Sym |
| LIN9 | 1 | lin-9 DREAM MuvB core complex component |
| YBX1P9 | 1 | Y-box binding protein 1 pseudogene 9 [Source |
| AL359742.1 | 1 | signal peptidase complex subunit 3 homolog ( |
| PARP1 | 1 | poly(ADP-ribose) polymerase 1 [Source:HGNC |
| AL359704.2 | 1 | SP100 nuclear antigen (SP100) pseudogene |
| AL359704.1 | 1 | ribosomal protein S3A (RPS3A) pseudogene |
| RN7SKP165 | 1 | RN7SK pseudogene 165 [Source:HGNC Symbo |
| CDKN2AIPNLP1 | 1 | CDKN2A interacting protein N-terminal like ps |
| AL359704.3 | 1 | novel transcript |
| STUM | 1 | stum, mechanosensory transduction mediator |
| ITPKB | 1 | inositol-trisphosphate 3-kinase B [Source:HGN |
| ITPKB-IT1 | 1 | ITPKB intronic transcript 1 [Source:HGNC Sym |
| ITPKB-AS1 | 1 | ITPKB antisense RNA 1 [Source:HGNC Symbol; |
| RPS27P5 | 1 | ribosomal protein S27 pseudogene 5 [Source: |
| AL359732.1 | 1 | novel transcript |
| PSEN2 | 1 | presenilin 2 [Source:HGNC Symbol;Acc:HGNC: |
| COQ8A | 1 | coenzyme Q8A [Source:HGNC Symbol;Acc:HG |
| AL353689.1 | 1 | pseudogene similar to part of NADH dehydrog |
| CDC42BPA | 1 | CDC42 binding protein kinase alpha [Source:H |
| AL353689.2 | 1 | novel transcript |
| AL353689.3 | 1 | novel transcript, antisense to CDC42BPA |
| AL451047.1 | 1 | novel transcript |
| AL627308.1 | 1 | cathepsin B (CTSB) pseudogene |
| AL627308.2 | 1 | novel pseudogene |
| AL627308.3 | 1 | novel transcript, antisense to CDC42BPA |
| LINC01641 | 1 | long intergenic non-protein coding RNA 1641 |
| NUCKS1P1 | 1 | nuclear casein kinase and cyclin dependent ki |
| AL606462.1 | 1 | ribosomal protein S18 (RPS18) pseudogene |
| BTF3P9 | 1 | basic transcription factor 3 pseudogene 9 [Sou |
| AL451054.2 | 1 | novel pseudogene |
| AL451054.3 | 1 | FAM166A, pseudogene 5 |
| TUBB8P10 | 1 | tubulin beta 8 class VIII pseudogene 10 [Sourc |
| TUBB8P9 | 1 | tubulin beta 8 class VIII pseudogene 9 [Source |

| | | |
|---|---|---|
| AL451054.1 | 1 | FAM166A, pseudogene 4 |
| AL451054.4 | 1 | novel transcript |
| RNA5SP77 | 1 | RNA, 5S ribosomal pseudogene 77 [Source:HG |
| ZNF678 | 1 | zinc finger protein 678 [Source:HGNC Symbol; |
| AL592310.1 | 1 | novel pseudogene |
| ZNF847P | 1 | zinc finger protein 847, pseudogene [Source:H |
| SNAP47 | 1 | synaptosome associated protein 47 [Source:H |
| JMJD4 | 1 | jumonji domain containing 4 [Source:HGNC Sy |
| SNAP47-AS1 | 1 | SNAP47 antisense RNA 1 [Source:HGNC Symb |
| AL731702.1 | 1 | novel transcript |
| PRSS38 | 1 | serine protease 38 [Source:HGNC Symbol;Acc |
| WNT9A | 1 | Wnt family member 9A [Source:HGNC Symbo |
| MIR5008 | 1 | microRNA 5008 [Source:HGNC Symbol;Acc:HG |
| CICP26 | 1 | capicua transcriptional repressor pseudogene |
| SEPTIN14P17 | 1 | septin 14 pseudogene 17 [Source:HGNC Symb |
| WNT3A | 1 | Wnt family member 3A [Source:HGNC Symbo |
| LINC02809 | 1 | long intergenic non-protein coding RNA 2809 |
| ARF1 | 1 | ADP ribosylation factor 1 [Source:HGNC Symb |
| MIR3620 | 1 | microRNA 3620 [Source:HGNC Symbol;Acc:HG |
| C1orf35 | 1 | chromosome 1 open reading frame 35 [Source |
| MRPL55 | 1 | mitochondrial ribosomal protein L55 [Source:I |
| FAM96AP2 | 1 | family with sequence similarity 96 member A |
| AL136379.1 | 1 | novel transcript |
| AL359510.2 | 1 | TEC |
| AL359510.1 | 1 | ribosomal protein S28 (RPS28) pseudogene |
| GUK1 | 1 | guanylate kinase 1 [Source:HGNC Symbol;Acc: |
| GJC2 | 1 | gap junction protein gamma 2 [Source:HGNC S |
| IBA57-DT | 1 | IBA57 divergent transcript [Source:HGNC Sym |
| IBA57 | 1 | iron-sulfur cluster assembly factor IBA57 [Sou |
| OBSCN-AS1 | 1 | OBSCN antisense RNA 1 [Source:HGNC Symbo |
| OBSCN | 1 | obscurin, cytoskeletal calmodulin and titin-int |
| AL353593.1 | 1 | novel transcript, antisense to OBSCN |
| AL353593.2 | 1 | novel transcript, antisense to OBSCN |
| AL353593.3 | 1 | novel transcript, sense intronic to OBSCN |
| AL670729.3 | 1 | novel transcript, antisense to OBSCN |
| AL670729.2 | 1 | novel transcript |
| TRIM11 | 1 | tripartite motif containing 11 [Source:HGNC S |
| AL670729.1 | 1 | novel transcript, antisense to TRIM11 |
| MIR6742 | 1 | microRNA 6742 [Source:HGNC Symbol;Acc:HG |
| AL139288.1 | 1 | novel transcript |
| TRIM17 | 1 | tripartite motif containing 17 [Source:HGNC S |
| HIST3H3 | 1 | histone cluster 3 H3 [Source:HGNC Symbol;Ac |
| RPL23AP15 | 1 | ribosomal protein L23a pseudogene 15 [Sourc |
| HIST3H2A | 1 | histone cluster 3 H2A [Source:HGNC Symbol;A |
| HIST3H2BB | 1 | histone cluster 3 H2B family member b [Sourc |
| MIR4666A | 1 | microRNA 4666a [Source:HGNC Symbol;Acc:H |
| HIST3H2BA | 1 | histone cluster 3 H2B family member a (pseud |
| AL139288.2 | 1 | novel transcript |
| RNF187 | 1 | ring finger protein 187 [Source:HGNC Symbol; |
| BTNL10 | 1 | butyrophilin like 10 [Source:HGNC Symbol;Acc |

| | | |
|---|---|---|
| RNA5SP19 | 1 | RNA, 5S ribosomal pseudogene 19 [Source:HG |
| RNA5SP162 | 1 | RNA, 5S ribosomal pseudogene 162 [Source:H |
| RNA5S1 | 1 | RNA, 5S ribosomal 1 [Source:HGNC Symbol;Ac |
| RNA5S2 | 1 | RNA, 5S ribosomal 2 [Source:HGNC Symbol;Ac |
| RNA5S3 | 1 | RNA, 5S ribosomal 3 [Source:HGNC Symbol;Ac |
| RNA5S4 | 1 | RNA, 5S ribosomal 4 [Source:HGNC Symbol;Ac |
| RNA5S5 | 1 | RNA, 5S ribosomal 5 [Source:HGNC Symbol;Ac |
| RNA5S6 | 1 | RNA, 5S ribosomal 6 [Source:HGNC Symbol;Ac |
| RNA5S7 | 1 | RNA, 5S ribosomal 7 [Source:HGNC Symbol;Ac |
| RNA5S8 | 1 | RNA, 5S ribosomal 8 [Source:HGNC Symbol;Ac |
| RNA5S9 | 1 | RNA, 5S ribosomal 9 [Source:HGNC Symbol;Ac |
| RNA5S10 | 1 | RNA, 5S ribosomal 10 [Source:HGNC Symbol;A |
| RNA5S11 | 1 | RNA, 5S ribosomal 11 [Source:HGNC Symbol;A |
| RNA5S12 | 1 | RNA, 5S ribosomal 12 [Source:HGNC Symbol;A |
| RNA5S13 | 1 | RNA, 5S ribosomal 13 [Source:HGNC Symbol;A |
| RNA5S14 | 1 | RNA, 5S ribosomal 14 [Source:HGNC Symbol;A |
| RNA5S15 | 1 | RNA, 5S ribosomal 15 [Source:HGNC Symbol;A |
| RNA5S16 | 1 | RNA, 5S ribosomal 16 [Source:HGNC Symbol;A |
| RNA5S17 | 1 | RNA, 5S ribosomal 17 [Source:HGNC Symbol;A |
| RNA5SP18 | 1 | RNA, 5S ribosomal pseudogene 18 [Source:HG |
| DUSP5P1 | 1 | dual specificity phosphatase 5 pseudogene 1 [ |
| AL713899.1 | 1 | NULL |
| FTH1P2 | 1 | ferritin heavy chain 1 pseudogene 2 [Source:H |
| RHOU | 1 | ras homolog family member U [Source:HGNC |
| AL078624.2 | 1 | eukaryotic translation elongation factor 1 alph |
| AL078624.1 | 1 | MAP/microtubule affinity-regulating kinase 2 |
| LINC02815 | 1 | long intergenic non-protein coding RNA 2815 |
| ISCA1P2 | 1 | iron-sulfur cluster assembly 1 pseudogene 2 [S |
| LINC02814 | 1 | long intergenic non-protein coding RNA 2814 |
| AL162595.2 | 1 | novel transcript |
| TMEM78 | 1 | transmembrane protein 78 [Source:HGNC Syn |
| AL162595.1 | 1 | novel transcript |
| RAB4A | 1 | RAB4A, member RAS oncogene family [Source |
| SPHAR | 1 | S-phase response (cyclin related) [Source:HGN |
| AL117350.1 | 1 | novel transcript |
| CCSAP | 1 | centriole, cilia and spindle associated protein |
| RNU6-180P | 1 | RNA, U6 small nuclear 180, pseudogene [Sour |
| RN7SKP276 | 1 | RN7SK pseudogene 276 [Source:HGNC Symbo |
| AL160004.2 | 1 | ribosomal protein L21 (RPL21) pseudogene |
| ACTA1 | 1 | actin alpha 1, skeletal muscle [Source:HGNC S |
| NUP133 | 1 | nucleoporin 133 [Source:HGNC Symbol;Acc:H |
| AL160004.1 | 1 | novel transcript |
| AL121990.1 | 1 | novel transcript |
| ABCB10 | 1 | ATP binding cassette subfamily B member 10 |
| RNU4-21P | 1 | RNA, U4 small nuclear 21, pseudogene [Sourc |
| RNA5SP78 | 1 | RNA, 5S ribosomal pseudogene 78 [Source:HG |
| HMGN2P19 | 1 | high mobility group nucleosomal binding dom |
| TAF5L | 1 | TATA-box binding protein associated factor 5 |
| URB2 | 1 | URB2 ribosome biogenesis homolog [Source:H |
| AL354983.1 | 1 | novel pseudogene |

| | | |
|---|---|---|
| HMGB1P26 | 1 | high mobility group box 1 pseudogene 26 [Sou |
| LINC01682 | 1 | long intergenic non-protein coding RNA 1682 |
| LINC01736 | 1 | long intergenic non-protein coding RNA 1736 |
| GALNT2 | 1 | polypeptide N-acetylgalactosaminyltransferas |
| AL136988.2 | 1 | novel transcript |
| AL136988.1 | 1 | novel transcript |
| PGBD5 | 1 | piggyBac transposable element derived 5 [Sou |
| AL133516.1 | 1 | novel transcript |
| LINC01737 | 1 | long intergenic non-protein coding RNA 1737 |
| AL158214.1 | 1 | ribosomal protein S24 (RPS24) pseudogene |
| COG2 | 1 | component of oligomeric golgi complex 2 [Sou |
| AL158214.2 | 1 | novel transcript, antisense to COG2 |
| AGT | 1 | angiotensinogen [Source:HGNC Symbol;Acc:H |
| AL512328.1 | 1 | novel transcript, antisense to CAPN9 |
| CAPN9 | 1 | calpain 9 [Source:HGNC Symbol;Acc:HGNC:14 |
| RNA5SP79 | 1 | RNA, 5S ribosomal pseudogene 79 [Source:HG |
| AL118511.2 | 1 | novel transcript |
| C1orf198 | 1 | chromosome 1 open reading frame 198 [Sour |
| AL118511.1 | 1 | novel transcript |
| AL118511.4 | 1 | novel transcript |
| AL118511.3 | 1 | novel transcript |
| RN7SL837P | 1 | RNA, 7SL, cytoplasmic 837, pseudogene [Sour |
| TTC13 | 1 | tetratricopeptide repeat domain 13 [Source:H |
| ARV1 | 1 | ARV1 homolog, fatty acid homeostasis modul |
| FAM89A | 1 | family with sequence similarity 89 member A |
| MIR1182 | 1 | microRNA 1182 [Source:HGNC Symbol;Acc:HG |
| AL732414.1 | 1 | ring finger protein 4 (RNF4) pseudogene |
| AL109810.1 | 1 | peptidylprolyl isomerase A (cyclophilin A) pseu |
| TRIM67 | 1 | tripartite motif containing 67 [Source:HGNC S |
| AL109810.2 | 1 | uncharacterized LOC149373 [Source:NCBI ger |
| C1orf131 | 1 | chromosome 1 open reading frame 131 [Sour |
| GNPAT | 1 | glyceronephosphate O-acyltransferase [Sourc |
| RNA5SP80 | 1 | RNA, 5S ribosomal pseudogene 80 [Source:HG |
| EXOC8 | 1 | exocyst complex component 8 [Source:HGNC |
| SPRTN | 1 | SprT-like N-terminal domain [Source:HGNC Sy |
| AL117352.1 | 1 | exocyst complex component 8 (EXOC8) pseud |
| EGLN1 | 1 | egl-9 family hypoxia inducible factor 1 [Source |
| AL445524.2 | 1 | novel protein |
| SNRPD2P2 | 1 | small nuclear ribonucleoprotein D2 pseudoge |
| AL445524.1 | 1 | novel transcript |
| TSNAX | 1 | translin associated factor X [Source:HGNC Syn |
| TSNAX-DISC1 | 1 | TSNAX-DISC1 readthrough (NMD candidate) [S |
| LINC00582 | 1 | long intergenic non-protein coding RNA 582 [S |
| DISC1 | 1 | DISC1 scaffold protein [Source:HGNC Symbol; |
| RNU5A-5P | 1 | RNA, U5A small nuclear 5, pseudogene [Sourc |
| AL136171.2 | 1 | disrupted in schizophrenia 2 |
| AL136171.1 | 1 | NULL |
| DISC1-IT1 | 1 | DISC1 intronic transcript 1 [Source:HGNC Sym |
| AL353052.1 | 1 | novel transcript |
| AL353052.2 | 1 | novel transcript |

| | | |
|---|---|---|
| BX323014.1 | 1 | DEAD (Asp-Glu-Ala-Asp) box polypeptide 60 (D |
| RN7SL299P | 1 | RNA, 7SL, cytoplasmic 299, pseudogene [Sour |
| SIPA1L2 | 1 | signal induced proliferation associated 1 like 2 |
| RNU6-1211P | 1 | RNA, U6 small nuclear 1211, pseudogene [Sou |
| LINC01745 | 1 | long intergenic non-protein coding RNA 1745 |
| LINC01744 | 1 | long intergenic non-protein coding RNA 1744 |
| MAP10 | 1 | microtubule associated protein 10 [Source:HG |
| RNU1-74P | 1 | RNA, U1 small nuclear 74, pseudogene [Sourc |
| AL122003.2 | 1 | novel transcript |
| AL122003.1 | 1 | chromosome 9 open reading frame 78 (C9orf |
| NTPCR | 1 | nucleoside-triphosphatase, cancer-related [Sc |
| PCNX2 | 1 | pecanex 2 [Source:HGNC Symbol;Acc:HGNC:8 |
| RPS7P3 | 1 | ribosomal protein S7 pseudogene 3 [Source:H |
| AL133380.1 | 1 | pecanex homolog (Drosophila) (PCNX) pseudo |
| MAP3K21 | 1 | mitogen-activated protein kinase kinase kinas |
| RNU4-77P | 1 | RNA, U4 small nuclear 77, pseudogene [Sourc |
| AL360006.1 | 1 | novel transcript |
| KCNK1 | 1 | potassium two pore domain channel subfamil |
| MIR4427 | 1 | microRNA 4427 [Source:HGNC Symbol;Acc:HG |
| AL357972.1 | 1 | novel transcript |
| AL663058.1 | 1 | acid phosphatase 1, soluble (ACP1) pseudoger |
| AL663058.2 | 1 | novel transcript |
| AL713868.1 | 1 | novel transcript, antisense to SLC35F3 |
| SLC35F3 | 1 | solute carrier family 35 member F3 [Source:H |
| RAC1P7 | 1 | Rac family small GTPase 1 pseudogene 7 [Sou |
| AL122008.3 | 1 | novel transcript |
| AL122008.1 | 1 | novel transcript |
| AL122008.4 | 1 | novel transcript |
| AL122008.2 | 1 | ribosomal protein L9 (RPL9) pseudogene |
| MIR4671 | 1 | microRNA 4671 [Source:HGNC Symbol;Acc:HG |
| AL355472.1 | 1 | ribosomal protein S15 (RPS15) pseudogene |
| AL355472.2 | 1 | novel transcript |
| AL355472.3 | 1 | novel transcript |
| COA6-AS1 | 1 | COA6 antisense RNA 1 [Source:HGNC Symbol; |
| COA6 | 1 | cytochrome c oxidase assembly factor 6 [Sour |
| TARBP1 | 1 | TAR (HIV-1) RNA binding protein 1 [Source:HG |
| LINC01354 | 1 | long intergenic non-protein coding RNA 1354 |
| AL161640.2 | 1 | novel transcript |
| AL161640.3 | 1 | novel transcript |
| AL161640.1 | 1 | novel transcript |
| U8 | 1 | U8 small nucleolar RNA [Source:RFAM;Acc:RF |
| IRF2BP2 | 1 | interferon regulatory factor 2 binding protein |
| AL160408.2 | 1 | novel transcript |
| LINC00184 | 1 | long intergenic non-protein coding RNA 184 [S |
| AL160408.5 | 1 | novel transcript |
| AL160408.1 | 1 | novel transcript |
| AL160408.4 | 1 | novel transcript |
| AL160408.3 | 1 | novel transcript |
| AL160408.6 | 1 | novel transcript |
| LINC01132 | 1 | long intergenic non-protein coding RNA 1132 |

| | | |
|---|---|---|
| AL360294.1 | 1 | novel transcript |
| AL391832.4 | 1 | novel transcript |
| RNY4P16 | 1 | RNY4 pseudogene 16 [Source:HGNC Symbol;A |
| RN7SL668P | 1 | RNA, 7SL, cytoplasmic 668, pseudogene [Sour |
| AL391832.1 | 1 | novel transcript |
| AL391832.2 | 1 | novel transcript |
| AL391832.3 | 1 | novel transcript |
| LINC01348 | 1 | long intergenic non-protein coding RNA 1348 |
| AL732292.1 | 1 | pseudogene similar to part of ribosomal prote |
| AL732292.2 | 1 | novel transcript |
| TOMM20 | 1 | translocase of outer mitochondrial membrane |
| SNORA14B | 1 | small nucleolar RNA, H/ACA box 14B [Source:I |
| RBM34 | 1 | RNA binding motif protein 34 [Source:HGNC S |
| ARID4B | 1 | AT-rich interaction domain 4B [Source:HGNC S |
| MIR4753 | 1 | microRNA 4753 [Source:HGNC Symbol;Acc:HG |
| RPL23AP23 | 1 | ribosomal protein L23a pseudogene 23 [Sourc |
| GGPS1 | 1 | geranylgeranyl diphosphate synthase 1 [Sourc |
| TBCE | 1 | tubulin folding cofactor E [Source:HGNC Symb |
| AL391994.1 | 1 | ubiquitin-conjugating enzyme E2G 1 (UBC7 ho |
| AL357556.1 | 1 | dentatorubral-pallidoluysian atrophy (atrophi |
| AL357556.4 | 1 | novel transcript, antisense to TBCE |
| TBCE | 1 | tubulin folding cofactor E [Source:HGNC Symb |
| RPS21P1 | 1 | ribosomal protein S21 pseudogene 1 [Source: |
| B3GALNT2 | 1 | beta-1,3-N-acetylgalactosaminyltransferase 2 |
| MTCYBP14 | 1 | MT-CYB pseudogene 14 [Source:HGNC Symbo |
| MTND6P14 | 1 | MT-ND6 pseudogene 14 [Source:HGNC Symbo |
| MTND5P19 | 1 | MT-ND5 pseudogene 19 [Source:HGNC Symbo |
| MTND4P10 | 1 | MT-ND4 pseudogene 10 [Source:HGNC Symbo |
| MTND4LP21 | 1 | MT-ND4L pseudogene 21 [Source:HGNC Symb |
| MTND3P8 | 1 | MT-ND3 pseudogene 8 [Source:HGNC Symbol |
| MTCO3P46 | 1 | MT-CO3 pseudogene 46 [Source:HGNC Symbo |
| GNG4 | 1 | G protein subunit gamma 4 [Source:HGNC Syr |
| FO393422.1 | 1 | methylenetetrahydrofolate dehydrogenase (N |
| AL583844.1 | 1 | solute carrier family 39 (zinc transporter), mer |
| LYST | 1 | lysosomal trafficking regulator [Source:HGNC |
| LDHAP2 | 1 | lactate dehydrogenase A pseudogene 2 [Sourc |
| LYST-AS1 | 1 | LYST antisense RNA 1 [Source:HGNC Symbol;A |
| MIR1537 | 1 | microRNA 1537 [Source:HGNC Symbol;Acc:HG |
| RNU6-968P | 1 | RNA, U6 small nuclear 968, pseudogene [Sour |
| AL139161.1 | 1 | novel transcript |
| LINC02768 | 1 | long intergenic non-protein coding RNA 2768 |
| NID1 | 1 | nidogen 1 [Source:HGNC Symbol;Acc:HGNC:78 |
| Y_RNA | 1 | Y RNA [Source:RFAM;Acc:RF00019] |
| AL122018.1 | 1 | aconitase 2, mitochondrial (ACO2) pseudogen |
| AL122018.2 | 1 | novel transcript |
| GPR137B | 1 | G protein-coupled receptor 137B [Source:HGN |
| ERO1B | 1 | endoplasmic reticulum oxidoreductase 1 beta |
| RNU2-70P | 1 | RNA, U2 small nuclear 70, pseudogene [Sourc |
| AL450309.1 | 1 | FK506 binding protein 3, 25kDa (FKBP3) pseuc |
| EDARADD | 1 | EDAR associated death domain [Source:HGNC |

| | | |
|---|---|---|
| ENO1P1 | 1 | enolase 1 pseudogene 1 [Source:HGNC Symbo |
| LGALS8 | 1 | galectin 8 [Source:HGNC Symbol;Acc:HGNC:65 |
| LGALS8-AS1 | 1 | LGALS8 antisense RNA 1 [Source:HGNC Symbo |
| AL359921.2 | 1 | novel transcript |
| AL359921.1 | 1 | novel transcript |
| HEATR1 | 1 | HEAT repeat containing 1 [Source:HGNC Symb |
| ACTN2 | 1 | actinin alpha 2 [Source:HGNC Symbol;Acc:HG |
| MTR | 1 | 5-methyltetrahydrofolate-homocysteine meth |
| RPSAP21 | 1 | ribosomal protein SA pseudogene 21 [Source: |
| RPL35P1 | 1 | ribosomal protein L35 pseudogene 1 [Source: |
| MT1HL1 | 1 | metallothionein 1H like 1 [Source:HGNC Symb |
| AL359259.1 | 1 | novel transcript |
| RYR2 | 1 | ryanodine receptor 2 [Source:HGNC Symbol;A |
| RN7SKP195 | 1 | RN7SK pseudogene 195 [Source:HGNC Symbo |
| MIR4428 | 1 | microRNA 4428 [Source:HGNC Symbol;Acc:HG |
| AL442065.1 | 1 | NULL |
| AL359924.1 | 1 | novel transcript |
| ZP4 | 1 | zona pellucida glycoprotein 4 [Source:HGNC S |
| AL590396.1 | 1 | actin, beta (ACTB) pseudogene |
| AL590396.3 | 1 | mitochondrially encoded cytochrome c oxidas |
| AL590396.2 | 1 | mitochondrially encoded NADH dehydrogenas |
| AL590396.4 | 1 | mitochondrially encoded NADH dehydrogenas |
| MTRNR2L11 | 1 | MT-RNR2 like 11 [Source:HGNC Symbol;Acc:H |
| MTCYBP15 | 1 | MT-CYB pseudogene 15 [Source:HGNC Symbo |
| MTND6P15 | 1 | MT-ND6 pseudogene 15 [Source:HGNC Symbo |
| MTND5P18 | 1 | MT-ND5 pseudogene 18 [Source:HGNC Symbo |
| YWHAQP9 | 1 | YWHAQ pseudogene 9 [Source:HGNC Symbol; |
| AL356010.2 | 1 | novel transcript |
| AL356010.1 | 1 | POM (POM121 homolog, rat) and ZP3 fusion ( |
| RNU6-725P | 1 | RNA, U6 small nuclear 725, pseudogene [Sour |
| LINC01139 | 1 | long intergenic non-protein coding RNA 1139 |
| AL359551.1 | 1 | novel transcript |
| KRT18P32 | 1 | keratin 18 pseudogene 32 [Source:HGNC Sym |
| MIPEPP2 | 1 | mitochondrial intermediate peptidase pseudo |
| AL583825.1 | 1 | novel transcript |
| AL590135.1 | 1 | ribosomal protein L39 (RPL39) pseudogene |
| AC093426.2 | 1 | novel transcript |
| AC093426.1 | 1 | novel transcript |
| CHRM3 | 1 | cholinergic receptor muscarinic 3 [Source:HG |
| CHRM3-AS2 | 1 | CHRM3 antisense RNA 2 [Source:HGNC Symbo |
| CHRM3-AS1 | 1 | CHRM3 antisense RNA 1 [Source:HGNC Symbo |
| AL356361.2 | 1 | novel transcript |
| AL356361.1 | 1 | 3-hydroxy-3-methylglutaryl-Coenzyme A synt |
| AL356361.3 | 1 | novel transcript |
| RPS7P5 | 1 | ribosomal protein S7 pseudogene 5 [Source:H |
| FMN2 | 1 | formin 2 [Source:HGNC Symbol;Acc:HGNC:14 |
| AL359918.1 | 1 | proteasome 26S non-ATPase subunit 2 (PSMD |
| Y_RNA | 1 | Y RNA [Source:RFAM;Acc:RF00019] |
| ADH5P3 | 1 | ADH5 pseudogene 3 [Source:HGNC Symbol;Ac |
| AL359918.2 | 1 | novel transcript |

| | | |
|---|---|---|
| Y_RNA | 1 | Y RNA [Source:RFAM;Acc:RF00019] |
| AL590490.1 | 1 | novel transcript, antisense to FMN2 |
| GREM2 | 1 | gremlin 2, DAN family BMP antagonist [Source |
| AL358176.1 | 1 | novel transcript, antisense to GREM2 |
| AL358176.2 | 1 | cytochrome c oxidase subunit VIb polypeptide |
| AL358176.4 | 1 | novel transcript |
| AL358176.3 | 1 | ribosomal protein S11 (RPS11) pseudogene |
| RNU5F-8P | 1 | RNA, U5F small nuclear 8, pseudogene [Sourc |
| AL358176.5 | 1 | ferredoxin pseudogene |
| Y_RNA | 1 | Y RNA [Source:RFAM;Acc:RF00019] |
| AL365184.2 | 1 | novel transcript |
| AL365184.1 | 1 | novel transcript, antisense to RGS7 |
| THAP12P8 | 1 | THAP domain containing 12 pseudogene 8 [Sc |
| RGS7 | 1 | regulator of G protein signaling 7 [Source:HGN |
| RFKP1 | 1 | riboflavin kinase pseudogene 1 [Source:HGNC |
| HNRNPA1P42 | 1 | heterogeneous nuclear ribonucleoprotein A1 |
| AL590682.1 | 1 | ribosomal protein, large, P1 (RPLP1) pseudoge |
| MIR3123 | 1 | microRNA 3123 [Source:HGNC Symbol;Acc:HG |
| AL592076.1 | 1 | ribosomal protein L36 (RPL36) pseudogene |
| AL359764.1 | 1 | novel transcript |
| AL359764.2 | 1 | novel transcript |
| AL359764.3 | 1 | novel transcript |
| FH | 1 | fumarate hydratase [Source:HGNC Symbol;Ac |
| KMO | 1 | kynurenine 3-monooxygenase [Source:HGNC |
| OPN3 | 1 | opsin 3 [Source:HGNC Symbol;Acc:HGNC:140 |
| CHML | 1 | CHM like Rab escort protein [Source:HGNC Sy |
| AL133390.1 | 1 | novel transcript |
| WDR64 | 1 | WD repeat domain 64 [Source:HGNC Symbol; |
| AL365366.1 | 1 | ribosomal protein L6 (RPL6) pseudogene |
| EXO1 | 1 | exonuclease 1 [Source:HGNC Symbol;Acc:HGN |
| RPL23AP20 | 1 | ribosomal protein L23a pseudogene 20 [Sourc |
| BECN2 | 1 | beclin 2 [Source:HGNC Symbol;Acc:HGNC:386 |
| CFL1P4 | 1 | cofilin 1 pseudogene 4 [Source:HGNC Symbol; |
| MAP1LC3C | 1 | microtubule associated protein 1 light chain 3 |
| TUBB8P6 | 1 | tubulin beta 8 class VIII pseudogene 6 [Source |
| BX571673.1 | 1 | FAM166A, pseudogene |
| PLD5 | 1 | phospholipase D family member 5 [Source:HG |
| AL591686.2 | 1 | novel transcript, antisense to PLD5 |
| RN7SKP12 | 1 | RN7SK pseudogene 12 [Source:HGNC Symbol; |
| AL591686.1 | 1 | novel transcript, antisense to PLD5 |
| AL360271.2 | 1 | pseudogene similar to part of excision repair c |
| RPL10AP5 | 1 | ribosomal protein L10a pseudogene 5 [Source |
| AL360271.1 | 1 | novel zinc finger protein pseudogene |
| AC099785.1 | 1 | Josephin domain containing 1 (JOSD1) pseudo |
| RSL24D1P4 | 1 | ribosomal L24 domain containing 1 pseudoger |
| AL445675.1 | 1 | regulator of chromosome condensation 2 (RC |
| AL445675.2 | 1 | chromosome 2 open reading frame 27B (C2or |
| AL603825.1 | 1 | ribosomal protein L41 (RPL41) pseudogene |
| AL606534.3 | 1 | novel transcript |
| SEPTIN14P21 | 1 | septin 14 pseudogene 21 [Source:HGNC Symb |

| | | |
|---|---|---|
| AL606534.5 | 1 | capicua transcriptional repressor (CIC) pseudc |
| AL606534.4 | 1 | novel pseudogene |
| LINC01347 | 1 | long intergenic non-protein coding RNA 1347 |
| RNU6-747P | 1 | RNA, U6 small nuclear 747, pseudogene [Sour |
| CEP170 | 1 | centrosomal protein 170 [Source:HGNC Symb |
| AL606534.1 | 1 | novel transcript |
| AL606534.2 | 1 | novel transcript, antisense to CEP170 |
| SDCCAG8 | 1 | serologically defined colon cancer antigen 8 [S |
| FCF1P7 | 1 | FCF1 pseudogene 7 [Source:HGNC Symbol;Acc |
| MIR4677 | 1 | microRNA 4677 [Source:HGNC Symbol;Acc:HG |
| AKT3 | 1 | AKT serine/threonine kinase 3 [Source:HGNC S |
| AL591721.1 | 1 | novel transcript |
| FABP7P1 | 1 | fatty acid binding protein 7 pseudogene 1 [Sou |
| AL662889.1 | 1 | novel transcript |
| AKT3-IT1 | 1 | AKT3 intronic transcript 1 [Source:HGNC Symb |
| LINC02774 | 1 | long intergenic non-protein coding RNA 2774 |
| ZBTB18 | 1 | zinc finger and BTB domain containing 18 [Sou |
| AL590483.3 | 1 | novel transcript |
| AL590483.1 | 1 | novel transcript |
| AL590483.4 | 1 | novel transcript |
| RN7SL148P | 1 | RNA, 7SL, cytoplasmic 148, pseudogene [Sour |
| AL590483.2 | 1 | novel transcript |
| AL358177.1 | 1 | novel transcript |
| C1orf100 | 1 | chromosome 1 open reading frame 100 [Sour |
| AL645465.1 | 1 | novel transcript, antisense to C1orf100 |
| TGIF2P1 | 1 | TGFB induced factor homeobox 2 pseudogene |
| ADSS | 1 | adenylosuccinate synthase [Source:HGNC Sym |
| CATSPERE | 1 | catsper channel auxiliary subunit epsilon [Sou |
| CYCSP5 | 1 | CYCS pseudogene 5 [Source:HGNC Symbol;Acc |
| DESI2 | 1 | desumoylating isopeptidase 2 [Source:HGNC S |
| AL451007.1 | 1 | ribosomal protein L37 (RPL37) pseudogene |
| AL451007.3 | 1 | novel protein |
| AL451007.2 | 1 | novel transcript |
| BX323046.2 | 1 | novel transcript |
| COX20 | 1 | cytochrome c oxidase assembly factor COX20 |
| HNRNPU | 1 | heterogeneous nuclear ribonucleoprotein U [S |
| BX323046.1 | 1 | novel transcript |
| RN7SKP55 | 1 | RN7SK pseudogene 55 [Source:HGNC Symbol; |
| RNU6-947P | 1 | RNA, U6 small nuclear 947, pseudogene [Sour |
| AL356512.1 | 1 | novel transcript, antisense to EFCAB2 |
| EFCAB2 | 1 | EF-hand calcium binding domain 2 [Source:HG |
| RNU6-1089P | 1 | RNA, U6 small nuclear 1089, pseudogene [Sou |
| RNU6-999P | 1 | RNA, U6 small nuclear 999, pseudogene [Sour |
| AL589763.1 | 1 | aldo-keto reductase family 1, member B1 (ald |
| RNU1-132P | 1 | RNA, U1 small nuclear 132, pseudogene [Sour |
| KIF26B | 1 | kinesin family member 26B [Source:HGNC Syr |
| KIF26B-AS1 | 1 | KIF26B antisense RNA 1 [Source:HGNC Symbo |
| DNAJC19P8 | 1 | DnaJ heat shock protein family (Hsp40) memb |
| AC104462.2 | 1 | novel transcript |
| AC104462.1 | 1 | novel transcript |

| | | |
|---|---|---|
| SMYD3 | 1 | SET and MYND domain containing 3 [Source:H |
| AC118555.1 | 1 | novel transcript, antisense to SMYD3 |
| AC092801.1 | 1 | novel transcript, antisense to SMYD3 |
| CHCHD4P5 | 1 | coiled-coil-helix-coiled-coil-helix domain conta |
| RNU6-1283P | 1 | RNA, U6 small nuclear 1283, pseudogene [Sou |
| SMYD3-IT1 | 1 | SMYD3 intronic transcript 1 [Source:HGNC Syr |
| Y_RNA | 1 | Y RNA [Source:RFAM;Acc:RF00019] |
| AL356583.1 | 1 | ubiquitin specific protease 6 (Tre-2 oncogene) |
| LINC01743 | 1 | long intergenic non-protein coding RNA 1743 |
| AL356583.3 | 1 | pseudogene similar to part of tumor necrosis |
| AL356583.2 | 1 | pseudogene similar to part of carboxypeptida |
| TFB2M | 1 | transcription factor B2, mitochondrial [Source |
| CNST | 1 | consortin, connexin sorting protein [Source:H |
| AL591623.1 | 1 | novel transcript |
| AL591623.2 | 1 | ribosomal protein L7a (RPL7A) pseudogene |
| AL591848.3 | 1 | novel transcript |
| AL591848.2 | 1 | novel transcript |
| SCCPDH | 1 | saccharopine dehydrogenase (putative) [Sour |
| AL591848.1 | 1 | ribosomal protein L35a (RPL35A) pseudogene |
| KIF28P | 1 | kinesin family member 28, pseudogene [Sour |
| AL591848.4 | 1 | novel transcript |
| LINC01341 | 1 | long intergenic non-protein coding RNA 1341 |
| AC113174.1 | 1 | pseudogene similar to part of microfibrillar-as |
| AC113174.2 | 1 | NULL |
| AHCTF1 | 1 | AT-hook containing transcription factor 1 [Sou |
| ZNF670-ZNF695 | 1 | ZNF670-ZNF695 readthrough (NMD candidate |
| ZNF695 | 1 | zinc finger protein 695 [Source:HGNC Symbol; |
| AL512637.2 | 1 | pseudogene similar to part of zinc finger prote |
| ZNF670 | 1 | zinc finger protein 670 [Source:HGNC Symbol; |
| AL512637.1 | 1 | primase polypeptide 1 49kDa (PRIM1) pseudo |
| AL627095.1 | 1 | pseudogene similar to part of fibroblast grow |
| ZNF669 | 1 | zinc finger protein 669 [Source:HGNC Symbol; |
| AL627095.2 | 1 | novel transcript |
| C1orf229 | 1 | chromosome 1 open reading frame 229 [Sour |
| FGFR3P6 | 1 | fibroblast growth factor receptor 3 pseudoger |
| ZNF124 | 1 | zinc finger protein 124 [Source:HGNC Symbol; |
| AL390728.3 | 1 | SMT3 suppressor of mif two 3 homolog 3 (yea |
| AL390728.2 | 1 | peroxisomal biogenesis factor 5 (PEX5) pseud |
| AL390728.6 | 1 | novel transcript |
| AL390728.4 | 1 | zinc finger protein pseudogene |
| MIR3916 | 1 | microRNA 3916 [Source:HGNC Symbol;Acc:HG |
| RNA5SP82 | 1 | RNA, 5S ribosomal pseudogene 82 [Source:HG |
| AL390728.5 | 1 | novel transcript |
| AL390728.1 | 1 | heat shock 70kDa protein 8 (HSPA8) pseudoge |
| VN1R17P | 1 | vomeronasal 1 receptor 17 pseudogene [Sour |
| VN1R5 | 1 | vomeronasal 1 receptor 5 (gene/pseudogene) |
| Y_RNA | 1 | Y RNA [Source:RFAM;Acc:RF00019] |
| ZNF496 | 1 | zinc finger protein 496 [Source:HGNC Symbol; |
| AC104335.1 | 1 | novel transcript |
| AC104335.2 | 1 | novel transcript |

| | | |
|---|---|---|
| NLRP3 | 1 | NLR family pyrin domain containing 3 [Source |
| OR2B11 | 1 | olfactory receptor family 2 subfamily B memb |
| OR2W5 | 1 | olfactory receptor family 2 subfamily W meml |
| GCSAML | 1 | germinal center associated signaling and moti |
| OR2C3 | 1 | olfactory receptor family 2 subfamily C memb |
| GCSAML-AS1 | 1 | GCSAML antisense RNA 1 [Source:HGNC Symb |
| AL606804.1 | 1 | novel transcript |
| OR2G2 | 1 | olfactory receptor family 2 subfamily G memb |
| OR2G3 | 1 | olfactory receptor family 2 subfamily G memb |
| U6 | 1 | U6 spliceosomal RNA [Source:RFAM;Acc:RF00 |
| OR14L1P | 1 | olfactory receptor family 14 subfamily L mem |
| AL390860.1 | 1 | novel transcript, antisense to OR13G1, OR6F1 |
| OR3D1P | 1 | olfactory receptor family 3 subfamily D memb |
| OR13G1 | 1 | olfactory receptor family 13 subfamily G mem |
| OR6F1 | 1 | olfactory receptor family 6 subfamily F memb |
| AC118470.1 | 1 | novel protein |
| OR14A2 | 1 | olfactory receptor family 14 subfamily A mem |
| OR14K1 | 1 | olfactory receptor family 14 subfamily K mem |
| OR1C1 | 1 | olfactory receptor family 1 subfamily C memb |
| OR9H1P | 1 | olfactory receptor family 9 subfamily H memb |
| OR14A16 | 1 | olfactory receptor family 14 subfamily A mem |
| HSD17B7P1 | 1 | hydroxysteroid 17-beta dehydrogenase 7 pseu |
| OR6R1P | 1 | olfactory receptor family 6 subfamily R memb |
| OR11L1 | 1 | olfactory receptor family 11 subfamily L mem |
| TRIM58 | 1 | tripartite motif containing 58 [Source:HGNC S |
| OR2W3 | 1 | olfactory receptor family 2 subfamily W meml |
| OR2T8 | 1 | olfactory receptor family 2 subfamily T memb |
| OR2AJ1 | 1 | olfactory receptor family 2 subfamily AJ meml |
| CLK3P2 | 1 | CDC like kinase 3 pseudogene 2 [Source:HGN( |
| OR2X1P | 1 | olfactory receptor family 2 subfamily X memb |
| OR2L8 | 1 | olfactory receptor family 2 subfamily L memb |
| OR2AK2 | 1 | olfactory receptor family 2 subfamily AK mem |
| OR2L9P | 1 | olfactory receptor family 2 subfamily L memb |
| OR2L1P | 1 | olfactory receptor family 2 subfamily L memb |
| Y_RNA | 1 | Y RNA [Source:RFAM;Acc:RF00019] |
| OR2L6P | 1 | olfactory receptor family 2 subfamily L memb |
| OR2L5 | 1 | olfactory receptor family 2 subfamily L memb |
| Y_RNA | 1 | Y RNA [Source:RFAM;Acc:RF00019] |
| OR2L2 | 1 | olfactory receptor family 2 subfamily L memb |
| Y_RNA | 1 | Y RNA [Source:RFAM;Acc:RF00019] |
| OR2L3 | 1 | olfactory receptor family 2 subfamily L memb |
| OR2T32P | 1 | olfactory receptor family 2 subfamily T memb |
| OR2L13 | 1 | olfactory receptor family 2 subfamily L memb |
| OR2M1P | 1 | olfactory receptor family 2 subfamily M meml |
| OR2M5 | 1 | olfactory receptor family 2 subfamily M meml |
| OR2M2 | 1 | olfactory receptor family 2 subfamily M meml |
| OR2M3 | 1 | olfactory receptor family 2 subfamily M meml |
| OR2M4 | 1 | olfactory receptor family 2 subfamily M meml |
| OR2T33 | 1 | olfactory receptor family 2 subfamily T memb |
| OR2T12 | 1 | olfactory receptor family 2 subfamily T memb |

| | | |
|---|---|---|
| OR2M7 | 1 | olfactory receptor family 2 subfamily M meml |
| OR14C36 | 1 | olfactory receptor family 14 subfamily C mem |
| OR2T4 | 1 | olfactory receptor family 2 subfamily T memb |
| OR2T6 | 1 | olfactory receptor family 2 subfamily T memb |
| OR2T1 | 1 | olfactory receptor family 2 subfamily T memb |
| OR2T7 | 1 | olfactory receptor family 2 subfamily T memb |
| OR2T2 | 1 | olfactory receptor family 2 subfamily T memb |
| OR2T3 | 1 | olfactory receptor family 2 subfamily T memb |
| AC138089.1 | 1 | novel transcript |
| OR2T5 | 1 | olfactory receptor family 2 subfamily T memb |
| OR2AS2P | 1 | olfactory receptor family 2 subfamily AS mem |
| OR2G6 | 1 | olfactory receptor family 2 subfamily G memb |
| AC098483.1 | 1 | novel transcript |
| OR2AS1P | 1 | olfactory receptor family 2 subfamily AS mem |
| OR2T29 | 1 | olfactory receptor family 2 subfamily T memb |
| OR2T34 | 1 | olfactory receptor family 2 subfamily T memb |
| OR2T10 | 1 | olfactory receptor family 2 subfamily T memb |
| Y_RNA | 1 | Y RNA [Source:RFAM;Acc:RF00019] |
| AC098483.2 | 1 | epithelial cell transforming sequence 2 oncog |
| OR2T11 | 1 | olfactory receptor family 2 subfamily T memb |
| OR2T35 | 1 | olfactory receptor family 2 subfamily T memb |
| OR2T27 | 1 | olfactory receptor family 2 subfamily T memb |
| CR589904.2 | 1 | novel protein |
| OR14I1 | 1 | olfactory receptor family 14 subfamily I meml |
| LYPD9P | 1 | LY6/PLAUR domain containing 9, pseudogene |
| CR589904.1 | 1 | TGF-beta activated kinase 1/MAP3K7 binding |
| AHCYP8 | 1 | adenosylhomocysteinase pseudogene 8 [Sour |
| LYPD8 | 1 | LY6/PLAUR domain containing 8 [Source:HGN |
| DPY19L4P1 | 1 | DPY19L4 pseudogene 1 [Source:HGNC Symbo |
| SH3BP5L | 1 | SH3 binding domain protein 5 like [Source:HG |
| MIR3124 | 1 | microRNA 3124 [Source:HGNC Symbol;Acc:HG |
| ZNF672 | 1 | zinc finger protein 672 [Source:HGNC Symbol; |
| ZNF692 | 1 | zinc finger protein 692 [Source:HGNC Symbol; |
| AL672291.1 | 1 | novel transcript |
| PGBD2 | 1 | piggyBac transposable element derived 2 [Sou |
| RNU6-1205P | 1 | RNA, U6 small nuclear 1205, pseudogene [Sou |
| RPL23AP25 | 1 | ribosomal protein L23a pseudogene 25 [Sourc |
| FAM110C | 2 | family with sequence similarity 110 member C |
| AC079779.1 | 2 | novel transcript |
| SH3YL1 | 2 | SH3 and SYLF domain containing 1 [Source:HG |
| ACP1 | 2 | acid phosphatase 1 [Source:HGNC Symbol;Acc |
| ALKAL2 | 2 | ALK and LTK ligand 2 [Source:HGNC Symbol;Ac |
| AC079779.2 | 2 | novel transcript |
| AC079779.3 | 2 | novel transcript |
| AC079779.4 | 2 | novel transcript |
| LINC01865 | 2 | long intergenic non-protein coding RNA 1865 |
| AC105393.2 | 2 | novel transcript |
| AC105393.1 | 2 | novel transcript |
| LINC01874 | 2 | long intergenic non-protein coding RNA 1874 |
| AC093326.2 | 2 | novel transcript |

| | | |
|---|---|---|
| LINC01875 | 2 | long intergenic non-protein coding RNA 1875 |
| AC093326.1 | 2 | novel transcript |
| TMEM18 | 2 | transmembrane protein 18 [Source:HGNC Syn |
| AC092159.2 | 2 | novel transcript |
| AC092159.4 | 2 | TEC |
| AC092159.3 | 2 | novel transcript |
| AC092159.1 | 2 | novel transcript |
| AC116609.3 | 2 | novel transcript |
| AC116609.2 | 2 | novel transcript |
| AC116609.1 | 2 | novel transcript |
| LINC01115 | 2 | long intergenic non-protein coding RNA 1115 |
| LINC01939 | 2 | long intergenic non-protein coding RNA 1939 |
| AC113607.1 | 2 | novel transcript |
| SNTG2-AS1 | 2 | SNTG2 antisense RNA 1 [Source:HGNC Symbo |
| SNTG2 | 2 | syntrophin gamma 2 [Source:HGNC Symbol;A |
| AC114808.1 | 2 | novel transcript |
| AC114808.2 | 2 | novel transcript |
| AC108462.1 | 2 | novel transcript |
| AC105450.1 | 2 | novel transcript |
| TPO | 2 | thyroid peroxidase [Source:HGNC Symbol;Acc |
| AC141930.1 | 2 | novel transcript |
| AC141930.2 | 2 | novel transcript |
| AC144450.1 | 2 | novel transcript |
| PXDN | 2 | peroxidasin [Source:HGNC Symbol;Acc:HGNC: |
| MYT1L | 2 | myelin transcription factor 1 like [Source:HGN |
| AC093390.2 | 2 | novel transcript, antisense to MYT1L |
| AC093390.1 | 2 | novel transcript |
| MYT1L-AS1 | 2 | MYT1L antisense RNA 1 [Source:HGNC Symbo |
| AC018685.2 | 2 | novel transcript |
| AC018685.3 | 2 | novel transcript |
| AC011995.2 | 2 | novel transcript |
| AC018685.1 | 2 | novel transcript |
| AC011995.1 | 2 | novel transcript |
| LINC01250 | 2 | long intergenic non-protein coding RNA 1250 |
| AC019118.2 | 2 | novel transcript |
| AC019118.1 | 2 | novel transcript |
| EIPR1 | 2 | EARP complex and GARP complex interacting |
| EIPR1-IT1 | 2 | EIPR1 intronic transcript 1 [Source:HGNC Sym |
| TRAPPC12 | 2 | trafficking protein particle complex 12 [Source |
| TRAPPC12-AS1 | 2 | TRAPPC12 antisense RNA 1 [Source:HGNC Syn |
| AC114810.1 | 2 | novel transcript |
| ADI1 | 2 | acireductone dioxygenase 1 [Source:HGNC Sy |
| AC231981.1 | 2 | novel transcript, antisense to ADI1 |
| AC108488.1 | 2 | novel transcript |
| AC108488.2 | 2 | novel protein |
| RNASEH1 | 2 | ribonuclease H1 [Source:HGNC Symbol;Acc:H |
| RNASEH1-AS1 | 2 | RNASEH1 antisense RNA 1 [Source:HGNC Sym |
| AC108488.3 | 2 | novel transcript |
| RPS7 | 2 | ribosomal protein S7 [Source:HGNC Symbol;A |
| COLEC11 | 2 | collectin subfamily member 11 [Source:HGNC |

| | | |
|---|---|---|
| AC010907.2 | 2 | novel transcript |
| TMSB4XP2 | 2 | TMSB4X pseudogene 2 [Source:HGNC Symbol |
| ALLC | 2 | allantoicase [Source:HGNC Symbol;Acc:HGNC: |
| GAPDHP48 | 2 | glyceraldehyde 3 phosphate dehydrogenase p |
| AC010907.1 | 2 | novel transcript |
| DCDC2C | 2 | doublecortin domain containing 2C [Source:H |
| LINC01304 | 2 | long intergenic non-protein coding RNA 1304 |
| AC012445.1 | 2 | novel transcript |
| NPM1P48 | 2 | nucleophosmin 1 pseudogene 48 [Source:HGN |
| LINC01249 | 2 | long intergenic non-protein coding RNA 1249 |
| AC092169.1 | 2 | NULL |
| RNU6-649P | 2 | RNA, U6 small nuclear 649, pseudogene [Sour |
| AC073143.1 | 2 | muscle RAS oncogene homolog (MRAS) pseud |
| AC107057.1 | 2 | novel transcript |
| LINC01248 | 2 | long intergenic non-protein coding RNA 1248 |
| AC108025.1 | 2 | novel transcript |
| SOX11 | 2 | SRY-box transcription factor 11 [Source:HGNC |
| AC010729.2 | 2 | novel transcript |
| AC010729.1 | 2 | novel transcript |
| LINC01810 | 2 | long intergenic non-protein coding RNA 1810 |
| SILC1 | 2 | sciatic injury induced lincRNA upregulator of S |
| MIR7158 | 2 | microRNA 7158 [Source:HGNC Symbol;Acc:HG |
| AC017053.1 | 2 | novel transcript |
| LINC01247 | 2 | long intergenic non-protein coding RNA 1247 |
| LINC01824 | 2 | long intergenic non-protein coding RNA 1824 |
| PIK3CDP1 | 2 | phosphatidylinositol-4,5-bisphosphate 3-kinas |
| MIR7515HG | 2 | MIR7515 host gene [Source:HGNC Symbol;Acc |
| AC097517.1 | 2 | novel transcript |
| MIR7515 | 2 | microRNA 7515 [Source:HGNC Symbol;Acc:HG |
| LINC00487 | 2 | long intergenic non-protein coding RNA 487 [S |
| NRIR | 2 | negative regulator of interferon response [Sou |
| CMPK2 | 2 | cytidine/uridine monophosphate kinase 2 [Sou |
| RSAD2 | 2 | radical S-adenosyl methionine domain contain |
| AC017076.1 | 2 | novel transcript |
| RNF144A-AS1 | 2 | RNF144A antisense RNA 1 [Source:HGNC Sym |
| RNF144A | 2 | ring finger protein 144A [Source:HGNC Symbo |
| AC068481.1 | 2 | novel transcript, antisense to RNF144A |
| RN7SKP112 | 2 | RN7SK pseudogene 112 [Source:HGNC Symbo |
| AC010904.2 | 2 | novel transcript |
| AC010904.1 | 2 | CDK2-associated protein 1 (CDK2AP1) pseudo |
| AC013460.1 | 2 | novel transcript |
| RNU6ATAC37P | 2 | RNA, U6atac small nuclear 37, pseudogene [So |
| AC092580.4 | 2 | novel transcript |
| AC092580.2 | 2 | novel transcript |
| LINC01871 | 2 | long intergenic non-protein coding RNA 1871 |
| AC092580.3 | 2 | pseudogene similar to part of proteasome (pr |
| AC092580.1 | 2 | proteasome (prosome, macropain) subunit, b |
| AC007463.1 | 2 | novel transcript |
| LINC00298 | 2 | long intergenic non-protein coding RNA 298 [S |
| AC007463.2 | 2 | novel transcript |

| | | |
|---|---|---|
| LINC00299 | 2 | long intergenic non-protein coding RNA 299 [S |
| U91324.1 | 2 | novel transcript |
| LINC01814 | 2 | long intergenic non-protein coding RNA 1814 |
| AC092617.1 | 2 | novel transcript |
| AC011747.1 | 2 | novel transcript |
| SNRPEP5 | 2 | small nuclear ribonucleoprotein polypeptide E |
| ID2-AS1 | 2 | ID2 antisense RNA 1 [Source:HGNC Symbol;Ac |
| ID2 | 2 | inhibitor of DNA binding 2 [Source:HGNC Sym |
| KIDINS220 | 2 | kinase D interacting substrate 220 [Source:HG |
| MBOAT2 | 2 | membrane bound O-acyltransferase domain c |
| HMGB1P25 | 2 | high mobility group box 1 pseudogene 25 [Sou |
| RPL30P3 | 2 | ribosomal protein L30 pseudogene 3 [Source: |
| AC093904.2 | 2 | novel transcript |
| AC093904.4 | 2 | novel transcript |
| AC093904.3 | 2 | novel transcript |
| AC093904.1 | 2 | novel transcript |
| ASAP2 | 2 | ArfGAP with SH3 domain, ankyrin repeat and |
| EIF1P7 | 2 | eukaryotic translation initiation factor 1 pseuc |
| ITGB1BP1 | 2 | integrin subunit beta 1 binding protein 1 [Sour |
| CPSF3 | 2 | cleavage and polyadenylation specific factor 3 |
| IAH1 | 2 | isoamyl acetate hydrolyzing esterase 1 (putati |
| ADAM17 | 2 | ADAM metallopeptidase domain 17 [Source:H |
| AC080162.1 | 2 | novel transcript |
| AC073195.1 | 2 | novel transcript |
| YWHAQ | 2 | tyrosine 3-monooxygenase/tryptophan 5-mor |
| Y_RNA | 2 | Y RNA [Source:RFAM;Acc:RF00019] |
| AC082651.1 | 2 | novel transcript |
| AC082651.4 | 2 | novel transcript |
| RNU4-73P | 2 | RNA, U4 small nuclear 73, pseudogene [Sourc |
| AC082651.2 | 2 | NADH dehydrogenase subunit 3 (ND3) pseudc |
| AC082651.3 | 2 | novel transcript |
| TAF1B | 2 | TATA-box binding protein associated factor, R |
| AC010969.2 | 2 | novel transcript |
| AC010969.3 | 2 | novel transcript |
| GRHL1 | 2 | grainyhead like transcription factor 1 [Source: |
| AC010969.1 | 2 | novel transcript |
| AC104794.4 | 2 | novel transcript |
| AC104794.3 | 2 | novel transcript |
| KLF11 | 2 | Kruppel like factor 11 [Source:HGNC Symbol;A |
| AC104794.5 | 2 | novel transcript, antisense to KLF11 |
| CYS1 | 2 | cystin 1 [Source:HGNC Symbol;Acc:HGNC:185 |
| AC104794.2 | 2 | novel transcript |
| AC104794.1 | 2 | NULL |
| RRM2 | 2 | ribonucleotide reductase regulatory subunit M |
| AC007240.1 | 2 | NULL |
| MIR4261 | 2 | microRNA 4261 [Source:HGNC Symbol;Acc:HG |
| RN7SL66P | 2 | RNA, 7SL, cytoplasmic 66, pseudogene [Sourc |
| AC007240.3 | 2 | novel transcript |
| AC007240.2 | 2 | NULL |
| HPCAL1 | 2 | hippocalcin like 1 [Source:HGNC Symbol;Acc:H |

| | | |
|---|---|---|
| ODC1 | 2 | ornithine decarboxylase 1 [Source:HGNC Syml |
| SNORA80B | 2 | small nucleolar RNA, H/ACA box 80B [Source:I |
| ODC1-DT | 2 | ODC1 divergent transcript [Source:HGNC Sym |
| NOL10 | 2 | nucleolar protein 10 [Source:HGNC Symbol;Ac |
| AC007314.1 | 2 | novel transcript |
| RN7SL832P | 2 | RNA, 7SL, cytoplasmic 832, pseudogene [Sour |
| Metazoa_SRP | 2 | Metazoan signal recognition particle RNA [Sou |
| AC092687.2 | 2 | novel transcript |
| ATP6V1C2 | 2 | ATPase H+ transporting V1 subunit C2 [Source |
| RNU7-138P | 2 | RNA, U7 small nuclear 138 pseudogene [Sourc |
| AC092687.3 | 2 | novel transcript |
| PDIA6 | 2 | protein disulfide isomerase family A member |
| RNU7-176P | 2 | RNA, U7 small nuclear 176 pseudogene [Sourc |
| LINC01954 | 2 | long intergenic non-protein coding RNA 1954 |
| AC092687.1 | 2 | novel transcript |
| AC079587.1 | 2 | ribosomal protein L32 (RPL32) pseudogene |
| KCNF1 | 2 | potassium voltage-gated channel modifier sub |
| AC062028.2 | 2 | ribosomal protein L6 (RPL6) pseudogene |
| AC062028.1 | 2 | novel transcript |
| C2orf50 | 2 | chromosome 2 open reading frame 50 [Sourc |
| SLC66A3 | 2 | solute carrier family 66 member 3 [Source:HG |
| ROCK2 | 2 | Rho associated coiled-coil containing protein l |
| RNU6-1081P | 2 | RNA, U6 small nuclear 1081, pseudogene [Sou |
| AC018463.1 | 2 | chromosome 1 open reading frame 80 (C1orf8 |
| PPIAP60 | 2 | peptidylprolyl isomerase A pseudogene 60 [Sc |
| AC099344.1 | 2 | novel transcript |
| AC099344.2 | 2 | novel transcript |
| LINC00570 | 2 | long intergenic non-protein coding RNA 570 [S |
| AC099344.3 | 2 | novel transcript |
| E2F6 | 2 | E2F transcription factor 6 [Source:HGNC Symb |
| GREB1 | 2 | growth regulating estrogen receptor binding 1 |
| RNA5SP84 | 2 | RNA, 5S ribosomal pseudogene 84 [Source:HG |
| MIR4429 | 2 | microRNA 4429 [Source:HGNC Symbol;Acc:HG |
| RNU2-13P | 2 | RNA, U2 small nuclear 13, pseudogene [Sourc |
| AC011994.1 | 2 | NULL |
| RNA5SP85 | 2 | RNA, 5S ribosomal pseudogene 85 [Source:HG |
| RN7SL674P | 2 | RNA, 7SL, cytoplasmic 674, pseudogene [Sour |
| NTSR2 | 2 | neurotensin receptor 2 [Source:HGNC Symbol |
| AC110754.1 | 2 | cyclin-dependent kinase 8 (CDK8) pseudogene |
| AC106875.2 | 2 | novel transcript |
| LPIN1 | 2 | lipin 1 [Source:HGNC Symbol;Acc:HGNC:1334! |
| AC106875.1 | 2 | uncharacterized LOC100506405 [Source:NCBI |
| AC012456.1 | 2 | novel transcript |
| AC012456.2 | 2 | novel transcript |
| MIR548S | 2 | microRNA 548s [Source:HGNC Symbol;Acc:HG |
| MIR3681HG | 2 | MIR3681 host gene [Source:HGNC Symbol;Acc |
| MIR4262 | 2 | microRNA 4262 [Source:HGNC Symbol;Acc:HG |
| SNORD18 | 2 | Small nucleolar RNA SNORD18 [Source:RFAM; |
| MIR3681 | 2 | microRNA 3681 [Source:HGNC Symbol;Acc:HG |
| RNU6-843P | 2 | RNA, U6 small nuclear 843, pseudogene [Sour |

| | | |
|---|---|---|
| AC096559.1 | 2 | novel transcript |
| AC009486.2 | 2 | novel transcript |
| AC009486.1 | 2 | novel transcript |
| TRIB2 | 2 | tribbles pseudokinase 2 [Source:HGNC Symbo |
| MIR3125 | 2 | microRNA 3125 [Source:HGNC Symbol;Acc:H( |
| AC064875.1 | 2 | novel transcript |
| AC093912.1 | 2 | novel transcript |
| AC073062.1 | 2 | novel transcript |
| LINC00276 | 2 | long intergenic non-protein coding RNA 276 [S |
| AC016730.1 | 2 | novel transcript |
| RNU6-1288P | 2 | RNA, U6 small nuclear 1288, pseudogene [Sou |
| Metazoa_SRP | 2 | Metazoan signal recognition particle RNA [Sou |
| AC011897.1 | 2 | novel transcript |
| LRATD1 | 2 | LRAT domain containing 1 [Source:HGNC Sym |
| AC068286.2 | 2 | novel transcript |
| AC068286.1 | 2 | novel transcript |
| NBAS | 2 | neuroblastoma amplified sequence [Source:H |
| AC007738.1 | 2 | ribosomal protein S26 (RPS26) pseudogene |
| AC008278.2 | 2 | novel transcript |
| DDX1 | 2 | DEAD-box helicase 1 [Source:HGNC Symbol;A( |
| AC008278.1 | 2 | novel transcript |
| LINC01804 | 2 | long intergenic non-protein coding RNA 1804 |
| AC113608.2 | 2 | novel transcript |
| RNU5E-7P | 2 | RNA, U5E small nuclear 7, pseudogene [Sourc |
| AC113608.1 | 2 | ribosomal protein, large, P1 (RPLP1) pseudoge |
| AC010145.2 | 2 | NULL |
| MYCNOS | 2 | MYCN opposite strand [Source:HGNC Symbol; |
| MYCNUT | 2 | MYCN upstream transcript [Source:HGNC Sym |
| MYCN | 2 | MYCN proto-oncogene, bHLH transcription fa( |
| RN7SL104P | 2 | RNA, 7SL, cytoplasmic 104, pseudogene [Sour |
| AC010145.1 | 2 | novel transcript |
| GACAT3 | 2 | gastric cancer associated transcript 3 [Source: |
| AC130710.1 | 2 | novel transcript |
| AC174048.1 | 2 | polymerase (RNA) II (DNA directed) polypepti( |
| AC142283.1 | 2 | NULL |
| AC010745.1 | 2 | novel transcript |
| AC010745.2 | 2 | novel transcript |
| AC010745.3 | 2 | novel transcript |
| AC010745.5 | 2 | novel transcript |
| AC010745.4 | 2 | novel transcript |
| AC010880.1 | 2 | novel transcript |
| AC104623.1 | 2 | novel transcript, antisense to FAM49A |
| AC104623.2 | 2 | TEC |
| FAM49A | 2 | family with sequence similarity 49 member A |
| AC008164.1 | 2 | novel transcript |
| AC008069.1 | 2 | tubulin, beta 2C (TUBB2C) pseudogene |
| LINC01866 | 2 | long intergenic non-protein coding RNA 1866 |
| RN7SKP168 | 2 | RN7SK pseudogene 168 [Source:HGNC Symbo |
| ZFYVE9P2 | 2 | zinc finger FYVE-type containing 9 pseudogen( |
| PSMC1P10 | 2 | proteasome 26S subunit, ATPase 1 pseudoger |

| | | |
|---|---|---|
| RAD51AP2 | 2 | RAD51 associated protein 2 [Source:HGNC Syr |
| VSNL1 | 2 | visinin like 1 [Source:HGNC Symbol;Acc:HGNC |
| SMC6 | 2 | structural maintenance of chromosomes 6 [Sc |
| GEN1 | 2 | GEN1 Holliday junction 5' flap endonuclease [! |
| MSGN1 | 2 | mesogenin 1 [Source:HGNC Symbol;Acc:HGN( |
| KCNS3 | 2 | potassium voltage-gated channel modifier sub |
| AC079802.1 | 2 | NULL |
| AC079148.3 | 2 | novel transcript |
| AC079148.2 | 2 | novel transcript |
| AC079148.1 | 2 | novel transcript |
| RDH14 | 2 | retinol dehydrogenase 14 [Source:HGNC Symb |
| NT5C1B-RDH14 | 2 | NT5C1B-RDH14 readthrough [Source:HGNC Sy |
| NT5C1B | 2 | 5'-nucleotidase, cytosolic IB [Source:HGNC Syr |
| RNU6-1215P | 2 | RNA, U6 small nuclear 1215, pseudogene [Sou |
| AC106053.1 | 2 | novel transcript |
| AC130814.1 | 2 | novel transcript |
| LINC01376 | 2 | long intergenic non-protein coding RNA 1376 |
| MIR4757 | 2 | microRNA 4757 [Source:HGNC Symbol;Acc:HG |
| OSR1 | 2 | odd-skipped related transcription factor 1 [So |
| AC010096.1 | 2 | novel transcript |
| LINC01808 | 2 | long intergenic non-protein coding RNA 1808 |
| AC019055.1 | 2 | novel transcript |
| CISD1P1 | 2 | CDGSH iron sulfur domain 1 pseudogene 1 [Sc |
| LINC00954 | 2 | long intergenic non-protein coding RNA 954 [S |
| TTC32 | 2 | tetratricopeptide repeat domain 32 [Source:H |
| AC013400.1 | 2 | novel transcript |
| WDR35 | 2 | WD repeat domain 35 [Source:HGNC Symbol;. |
| AC079145.1 | 2 | novel transcript |
| MATN3 | 2 | matrilin 3 [Source:HGNC Symbol;Acc:HGNC:69 |
| LAPTM4A | 2 | lysosomal protein transmembrane 4 alpha [Sc |
| LAPTM4A-DT | 2 | LAPTM4A divergent transcript [Source:HGNC ! |
| AC098828.2 | 2 | novel transcript |
| AC098828.1 | 2 | ribosomal protein S16 (RPS16) pseudogene |
| RN7SL140P | 2 | RNA, 7SL, cytoplasmic 140, pseudogene [Sour |
| RNU6-961P | 2 | RNA, U6 small nuclear 961, pseudogene [Sour |
| SDC1 | 2 | syndecan 1 [Source:HGNC Symbol;Acc:HGNC:: |
| AC007041.1 | 2 | developmentally regulated GTP binding prote |
| RNU7-113P | 2 | RNA, U7 small nuclear 113 pseudogene [Sourc |
| PUM2 | 2 | pumilio RNA binding family member 2 [Source |
| SLC7A15P | 2 | solute carrier family 7 member 15, pseudoger |
| RNA5SP86 | 2 | RNA, 5S ribosomal pseudogene 86 [Source:HG |
| RHOB | 2 | ras homolog family member B [Source:HGNC : |
| AC023137.1 | 2 | novel transcript |
| Y_RNA | 2 | Y RNA [Source:RFAM;Acc:RF00019] |
| AC012065.1 | 2 | novel transcript |
| AC012065.5 | 2 | NADH dehydrogenase (ubiquinone) complex I |
| HS1BP3 | 2 | HCLS1 binding protein 3 [Source:HGNC Symbc |
| AC012065.3 | 2 | novel transcript |
| HS1BP3-IT1 | 2 | HS1BP3 intronic transcript 1 [Source:HGNC Sy |
| AC012065.2 | 2 | ribosomal protein S25 (RPS25) pseudogene |

| | | |
|---|---|---|
| GDF7 | 2 | growth differentiation factor 7 [Source:HGNC |
| AC012065.4 | 2 | novel transcript |
| LDAH | 2 | lipid droplet associated hydrolase [Source:HG |
| LINC02850 | 2 | long intergenic non-protein coding RNA 2850 |
| AC115619.1 | 2 | novel transcript |
| APOB | 2 | apolipoprotein B [Source:HGNC Symbol;Acc:H |
| AC010872.1 | 2 | novel transcript, antisense to APOB |
| AC010872.2 | 2 | novel transcript |
| TDRD15 | 2 | tudor domain containing 15 [Source:HGNC Sy |
| AC067959.1 | 2 | tumor protein, translationally-controlled 1 (TF |
| AC018742.1 | 2 | novel transcript |
| AC011752.1 | 2 | novel transcript |
| NUTF2P8 | 2 | nuclear transport factor 2 pseudogene 8 [Sou |
| AC009411.2 | 2 | novel transcript |
| AC009411.1 | 2 | novel transcript |
| LINC01822 | 2 | long intergenic non-protein coding RNA 1822 |
| RN7SL117P | 2 | RNA, 7SL, cytoplasmic 117, pseudogene [Sour |
| AC096570.1 | 2 | long intergenic non-protein coding RNA 1830 |
| AC068044.1 | 2 | Mdm2, transformed 3T3 cell double minute 2 |
| AC099799.1 | 2 | mitochondrially encoded cytochrome b (MT-C |
| RNA5SP87 | 2 | RNA, 5S ribosomal pseudogene 87 [Source:HG |
| AC096570.2 | 2 | novel transcript |
| LINC01884 | 2 | long intergenic non-protein coding RNA 1884 |
| RN7SKP27 | 2 | RN7SK pseudogene 27 [Source:HGNC Symbol; |
| AC018467.1 | 2 | novel transcript |
| AC012506.1 | 2 | novel transcript |
| AC012506.3 | 2 | novel transcript |
| AC012506.2 | 2 | novel transcript |
| AC012506.4 | 2 | novel transcript |
| KLHL29 | 2 | kelch like family member 29 [Source:HGNC Sy |
| AC011239.1 | 2 | novel transcript, antisense to KLHL29 |
| AC011239.2 | 2 | TEC |
| AC009242.1 | 2 | novel transcript |
| ATAD2B | 2 | ATPase family AAA domain containing 2B [Sou |
| PGAM1P6 | 2 | phosphoglycerate mutase 1 pseudogene 6 [Sc |
| AC066692.1 | 2 | ribosomal protein S13 (RPS13) pseudogene |
| UBXN2A | 2 | UBX domain protein 2A [Source:HGNC Symbo |
| SDHCP3 | 2 | succinate dehydrogenase complex subunit C p |
| RN7SL610P | 2 | RNA, 7SL, cytoplasmic 610, pseudogene [Sour |
| MFSD2B | 2 | major facilitator superfamily domain containi |
| WDCP | 2 | WD repeat and coiled coil containing [Source: |
| RNU6-370P | 2 | RNA, U6 small nuclear 370, pseudogene [Sour |
| FKBP1B | 2 | FKBP prolyl isomerase 1B [Source:HGNC Symb |
| SF3B6 | 2 | splicing factor 3b subunit 6 [Source:HGNC Sym |
| FAM228B | 2 | family with sequence similarity 228 member B |
| TP53I3 | 2 | tumor protein p53 inducible protein 3 [Source |
| PFN4 | 2 | profilin family member 4 [Source:HGNC Symb |
| AC008073.3 | 2 | novel transcript |
| AC008073.2 | 2 | novel transcript |
| FAM228A | 2 | family with sequence similarity 228 member A |

| | | |
|---|---|---|
| AC008073.1 | 2 | novel transcript |
| ITSN2 | 2 | intersectin 2 [Source:HGNC Symbol;Acc:HGNC |
| AC009228.1 | 2 | novel transcript |
| AC009228.2 | 2 | NULL |
| HMGN2P20 | 2 | high mobility group nucleosomal binding dom |
| RPL36AP13 | 2 | ribosomal protein L36a pseudogene 13 [Sourc |
| AC093798.1 | 2 | novel transcript |
| NCOA1 | 2 | nuclear receptor coactivator 1 [Source:HGNC |
| RNA5SP88 | 2 | RNA, 5S ribosomal pseudogene 88 [Source:HG |
| RNU6-936P | 2 | RNA, U6 small nuclear 936, pseudogene [Sour |
| PTRHD1 | 2 | peptidyl-tRNA hydrolase domain containing 1 |
| CENPO | 2 | centromere protein O [Source:HGNC Symbol;A |
| ADCY3 | 2 | adenylate cyclase 3 [Source:HGNC Symbol;Acc |
| AC012073.1 | 2 | novel transcript, antisense to ADCY3 |
| DNAJC27 | 2 | DnaJ heat shock protein family (Hsp40) memb |
| AC013267.1 | 2 | ribosomal protein S13 (RPS13) pseudogene |
| SNORD14 | 2 | Small nucleolar RNA SNORD14 [Source:RFAM; |
| DNAJC27-AS1 | 2 | DNAJC27 antisense RNA 1 [Source:HGNC Syml |
| AC013267.2 | 2 | novel transcript |
| Y_RNA | 2 | Y RNA [Source:RFAM;Acc:RF00019] |
| EFR3B | 2 | EFR3 homolog B [Source:HGNC Symbol;Acc:H( |
| RN7SL856P | 2 | RNA, 7SL, cytoplasmic 856, pseudogene [Sour |
| SUCLA2P3 | 2 | succinate-CoA ligase ADP-forming beta subun |
| POMC | 2 | proopiomelanocortin [Source:HGNC Symbol;A |
| LINC01381 | 2 | long intergenic non-protein coding RNA 1381 |
| DNMT3A | 2 | DNA methyltransferase 3 alpha [Source:HGNC |
| MIR1301 | 2 | microRNA 1301 [Source:HGNC Symbol;Acc:HG |
| AC012074.1 | 2 | novel transcript |
| DTNB | 2 | dystrobrevin beta [Source:HGNC Symbol;Acc:I |
| AC104699.1 | 2 | novel transcript |
| Y_RNA | 2 | Y RNA [Source:RFAM;Acc:RF00019] |
| ASXL2 | 2 | ASXL transcriptional regulator 2 [Source:HGN( |
| TPM3P7 | 2 | tropomyosin 3 pseudogene 7 [Source:HGNC S |
| PTGES3P2 | 2 | prostaglandin E synthase 3 pseudogene 2 [Sou |
| AC064847.1 | 2 | enolase 2 (gamma, neuronal) (ENO2) pseudog |
| NDUFB4P4 | 2 | NADH:ubiquinone oxidoreductase subunit B4 |
| KIF3C | 2 | kinesin family member 3C [Source:HGNC Sym |
| UQCRHP2 | 2 | ubiquinol-cytochrome c reductase hinge prote |
| TRMT112P6 | 2 | tRNA methyltransferase subunit 11-2 pseudog |
| RAB10 | 2 | RAB10, member RAS oncogene family [Source |
| RNU6-942P | 2 | RNA, U6 small nuclear 942, pseudogene [Sour |
| AC011742.3 | 2 | ribosomal protein S2 (RPS2) pseudogene |
| AC011742.2 | 2 | epithelial membrane protein 2 (EMP2) pseudo |
| SMARCE1P6 | 2 | SWI/SNF related, matrix associated, actin dep |
| AC011742.1 | 2 | chromosome 11 open reading frame 73 (C11o |
| PPIL1P1 | 2 | peptidylprolyl isomerase like 1 pseudogene 1 |
| GAREM2 | 2 | GRB2 associated regulator of MAPK1 subtype |
| HADHA | 2 | hydroxyacyl-CoA dehydrogenase trifunctional |
| HADHB | 2 | hydroxyacyl-CoA dehydrogenase trifunctional |
| AC010896.1 | 2 | novel transcript |

| | | |
|---|---|---|
| ADGRF3 | 2 | adhesion G protein-coupled receptor F3 [Sour |
| SELENOI | 2 | selenoprotein I [Source:HGNC Symbol;Acc:HG |
| DRC1 | 2 | dynein regulatory complex subunit 1 [Source:I |
| OTOF | 2 | otoferlin [Source:HGNC Symbol;Acc:HGNC:85: |
| FAM166C | 2 | family with sequence similarity 166 member ( |
| CIB4 | 2 | calcium and integrin binding family member 4 |
| AC015977.2 | 2 | novel transcript |
| AC015977.1 | 2 | ribosomal protein L37 (RPL37) pseudogene |
| KCNK3 | 2 | potassium two pore domain channel subfamil |
| SLC35F6 | 2 | solute carrier family 35 member F6 [Source:H |
| CENPA | 2 | centromere protein A [Source:HGNC Symbol;/ |
| CDKN2AIPNLP2 | 2 | CDKN2A interacting protein N-terminal like ps |
| DPYSL5 | 2 | dihydropyrimidinase like 5 [Source:HGNC Sym |
| AC013472.2 | 2 | novel transcript |
| MAPRE3 | 2 | microtubule associated protein RP/EB family r |
| AC013472.1 | 2 | novel transcript |
| TMEM214 | 2 | transmembrane protein 214 [Source:HGNC Sy |
| AGBL5 | 2 | ATP/GTP binding protein like 5 [Source:HGNC |
| AGBL5-AS1 | 2 | AGBL5 antisense RNA 1 [Source:HGNC Symbo |
| AC013472.3 | 2 | novel transcript, antisense to AGBL5 |
| AGBL5-IT1 | 2 | AGBL5 intronic transcript 1 [Source:HGNC Syn |
| AC013403.2 | 2 | novel transcript, antisense to AGBL5 |
| OST4 | 2 | oligosaccharyltransferase complex subunit 4, |
| EMILIN1 | 2 | elastin microfibril interfacer 1 [Source:HGNC S |
| KHK | 2 | ketohexokinase [Source:HGNC Symbol;Acc:HG |
| CGREF1 | 2 | cell growth regulator with EF-hand domain 1 [ |
| ABHD1 | 2 | abhydrolase domain containing 1 [Source:HGI |
| PREB | 2 | prolactin regulatory element binding [Source: |
| PRR30 | 2 | proline rich 30 [Source:HGNC Symbol;Acc:HGI |
| TCF23 | 2 | transcription factor 23 [Source:HGNC Symbol; |
| AC013403.1 | 2 | pseudogene similar to part of solute carrier fa |
| SLC5A6 | 2 | solute carrier family 5 member 6 [Source:HGN |
| ATRAID | 2 | all-trans retinoic acid induced differentiation f |
| CAD | 2 | carbamoyl-phosphate synthetase 2, aspartate |
| SLC30A3 | 2 | solute carrier family 30 member 3 [Source:HG |
| DNAJC5G | 2 | DnaJ heat shock protein family (Hsp40) memb |
| TRIM54 | 2 | tripartite motif containing 54 [Source:HGNC S |
| UCN | 2 | urocortin [Source:HGNC Symbol;Acc:HGNC:12 |
| MPV17 | 2 | mitochondrial inner membrane protein MPV1 |
| GTF3C2 | 2 | general transcription factor IIIC subunit 2 [Sou |
| GTF3C2-AS1 | 2 | GTF3C2 antisense RNA 1 [Source:HGNC Symbo |
| AC074117.1 | 2 | novel transcript, antisense to GTF3C2 and EIF: |
| EIF2B4 | 2 | eukaryotic translation initiation factor 2B subu |
| SNX17 | 2 | sorting nexin 17 [Source:HGNC Symbol;Acc:H( |
| ZNF513 | 2 | zinc finger protein 513 [Source:HGNC Symbol; |
| PPM1G | 2 | protein phosphatase, Mg2+/Mn2+ dependent |
| FTH1P3 | 2 | ferritin heavy chain 1 pseudogene 3 [Source:H |
| NRBP1 | 2 | nuclear receptor binding protein 1 [Source:HG |
| KRTCAP3 | 2 | keratinocyte associated protein 3 [Source:HGI |
| IFT172 | 2 | intraflagellar transport 172 [Source:HGNC Syn |

| | | |
|---|---|---|
| AC074117.2 | 2 | ribosomal protein S15a (RPS15A) pseudogene |
| RNU6-986P | 2 | RNA, U6 small nuclear 986, pseudogene [Sour |
| FNDC4 | 2 | fibronectin type III domain containing 4 [Sour |
| GCKR | 2 | glucokinase regulator [Source:HGNC Symbol;A |
| C2orf16 | 2 | chromosome 2 open reading frame 16 [Source |
| ZNF512 | 2 | zinc finger protein 512 [Source:HGNC Symbol; |
| AC074091.2 | 2 | novel transcript |
| CCDC121 | 2 | coiled-coil domain containing 121 [Source:HG |
| GPN1 | 2 | GPN-loop GTPase 1 [Source:HGNC Symbol;Acc |
| AC074091.1 | 2 | NULL |
| SUPT7L | 2 | SPT7 like, STAGA complex gamma subunit [So |
| SLC4A1AP | 2 | solute carrier family 4 member 1 adaptor prot |
| LINC01460 | 2 | long intergenic non-protein coding RNA 1460 |
| MRPL33 | 2 | mitochondrial ribosomal protein L33 [Source: |
| RBKS | 2 | ribokinase [Source:HGNC Symbol;Acc:HGNC:3 |
| BABAM2 | 2 | BRISC and BRCA1 A complex member 2 [Sourc |
| AC021171.2 | 2 | chromosome 12 open reading frame 10 (C12c |
| MIR4263 | 2 | microRNA 4263 [Source:HGNC Symbol;Acc:HG |
| AC021171.1 | 2 | family with sequence similarity 133, member |
| AC096552.1 | 2 | NULL |
| AC093690.1 | 2 | novel transcript |
| RPL23AP34 | 2 | ribosomal protein L23a pseudogene 34 [Sourc |
| AC104695.2 | 2 | novel transcript, antisense to FOSL2 |
| FOSL2 | 2 | FOS like 2, AP-1 transcription factor subunit [S |
| AC104695.4 | 2 | novel transcript, sense intronic to FOSL2 |
| AC104695.3 | 2 | novel transcript |
| AC104695.1 | 2 | novel transcript |
| PLB1 | 2 | phospholipase B1 [Source:HGNC Symbol;Acc:I |
| SNRPGP7 | 2 | small nuclear ribonucleoprotein polypeptide C |
| AC074011.1 | 2 | novel transcript |
| RNA5SP89 | 2 | RNA, 5S ribosomal pseudogene 89 [Source:HG |
| AC092164.1 | 2 | novel transcript |
| PPP1CB | 2 | protein phosphatase 1 catalytic subunit beta [ |
| SPDYA | 2 | speedy/RINGO cell cycle regulator family mem |
| AC097724.1 | 2 | novel transcript |
| TRMT61B | 2 | tRNA methyltransferase 61B [Source:HGNC Sy |
| WDR43 | 2 | WD repeat domain 43 [Source:HGNC Symbol; |
| SNORD92 | 2 | small nucleolar RNA, C/D box 92 [Source:HGN |
| SNORD53 | 2 | small nucleolar RNA, C/D box 53 [Source:HGN |
| Y_RNA | 2 | Y RNA [Source:RFAM;Acc:RF00019] |
| SNORD53B | 2 | small nucleolar RNA, C/D box 53B [Source:HG |
| TOGARAM2 | 2 | TOG array regulator of axonemal microtubule |
| Y_RNA | 2 | Y RNA [Source:RFAM;Acc:RF00019] |
| PCARE | 2 | photoreceptor cilium actin regulator [Source: |
| AC105398.1 | 2 | novel transcript |
| CLIP4 | 2 | CAP-Gly domain containing linker protein fam |
| ALK | 2 | ALK receptor tyrosine kinase [Source:HGNC Sy |
| AC074096.1 | 2 | novel transcript, antisense to ALK |
| RN7SL516P | 2 | RNA, 7SL, cytoplasmic 516, pseudogene [Sour |
| AC016907.2 | 2 | novel transcript |

| | | |
|---|---|---|
| AC106870.2 | 2 | novel transcript |
| AC106870.1 | 2 | novel transcript |
| AC016907.1 | 2 | ubiquitin-conjugating enzyme E2I (UBC9 homo |
| YPEL5 | 2 | yippee like 5 [Source:HGNC Symbol;Acc:HGNC |
| SNORA10B | 2 | small nucleolar RNA, H/ACA box 10B [Source:I |
| AC104698.1 | 2 | H3 histone, family 3A (H3F3A) pseudogene |
| LBH | 2 | LBH regulator of WNT signaling pathway [Sour |
| AC109642.1 | 2 | novel transcript |
| LINC01936 | 2 | long intergenic non-protein coding RNA 1936 |
| AC073255.1 | 2 | TEC |
| LCLAT1 | 2 | lysocardiolipin acyltransferase 1 [Source:HGN |
| AC132154.1 | 2 | TEC |
| CAPN13 | 2 | calpain 13 [Source:HGNC Symbol;Acc:HGNC:1 |
| AC092569.1 | 2 | ribosomal protein L15 (RPL15) pseudogene |
| AC009301.1 | 2 | novel transcript |
| GALNT14 | 2 | polypeptide N-acetylgalactosaminyltransferas |
| AC009305.1 | 2 | novel transcript |
| CAPN14 | 2 | calpain 14 [Source:HGNC Symbol;Acc:HGNC:1 |
| RNA5SP90 | 2 | RNA, 5S ribosomal pseudogene 90 [Source:HG |
| EHD3 | 2 | EH domain containing 3 [Source:HGNC Symbo |
| AL121657.1 | 2 | ribosomal protein L21 (RPL21) pseudogene |
| XDH | 2 | xanthine dehydrogenase [Source:HGNC Symb |
| SRD5A2 | 2 | steroid 5 alpha-reductase 2 [Source:HGNC Syr |
| AL133247.1 | 2 | novel transcript |
| AL133247.2 | 2 | catenin (cadherin-associated protein), alpha 1 |
| LINC01946 | 2 | long intergenic non-protein coding RNA 1946 |
| KRT18P52 | 2 | keratin 18 pseudogene 52 [Source:HGNC Sym |
| AK2P2 | 2 | adenylate kinase 2 pseudogene 2 [Source:HGI |
| AL121652.1 | 2 | novel transcript |
| MEMO1 | 2 | mediator of cell motility 1 [Source:HGNC Sym |
| DPY30 | 2 | dpy-30 histone methyltransferase complex re |
| AL121655.1 | 2 | novel transcript, antisense to DPY30 |
| SPAST | 2 | spastin [Source:HGNC Symbol;Acc:HGNC:1123 |
| AL121658.1 | 2 | novel transcript |
| SLC30A6 | 2 | solute carrier family 30 member 6 [Source:HG |
| DDX50P1 | 2 | DEAD-box helicase 50 pseudogene 1 [Source:I |
| RNU6-647P | 2 | RNA, U6 small nuclear 647, pseudogene [Sour |
| NLRC4 | 2 | NLR family CARD domain containing 4 [Source |
| YIPF4 | 2 | Yip1 domain family member 4 [Source:HGNC |
| AL133245.1 | 2 | novel transcript |
| BIRC6 | 2 | baculoviral IAP repeat containing 6 [Source:H |
| BIRC6-AS1 | 2 | BIRC6 antisense RNA 1 [Source:HGNC Symbol; |
| AL133243.2 | 2 | novel transcript, sense intronic to BIRC6 |
| AL133243.3 | 2 | novel transcript, sense intronic to BIRC6 |
| MIR558 | 2 | microRNA 558 [Source:HGNC Symbol;Acc:HGI |
| AL133243.1 | 2 | novel transcript, sense intronic to BIRC6 |
| BIRC6-AS2 | 2 | BIRC6 antisense RNA 2 [Source:HGNC Symbol; |
| AL133243.4 | 2 | TEC |
| TTC27 | 2 | tetratricopeptide repeat domain 27 [Source:H |
| MIR4765 | 2 | microRNA 4765 [Source:HGNC Symbol;Acc:HG |

| | | |
|---|---|---|
| LINC00486 | 2 | long intergenic non-protein coding RNA 486 [S |
| AL133244.2 | 2 | novel transcript |
| Y_RNA | 2 | Y RNA [Source:RFAM;Acc:RF00019] |
| LTBP1 | 2 | latent transforming growth factor beta bindin |
| AL133244.1 | 2 | histone cluster 2, H2ac (HIST2H2AC) pseudoge |
| AC019127.1 | 2 | novel transcript, antisense to LTBP1 |
| RNA5SP91 | 2 | RNA, 5S ribosomal pseudogene 91 [Source:HG |
| RNA5SP92 | 2 | RNA, 5S ribosomal pseudogene 92 [Source:HG |
| MIR4430 | 2 | microRNA 4430 [Source:HGNC Symbol;Acc:HG |
| RASGRP3 | 2 | RAS guanyl releasing protein 3 [Source:HGNC |
| AC020594.1 | 2 | novel transcript |
| FAM98A | 2 | family with sequence similarity 98 member A |
| AC017050.1 | 2 | novel transcript |
| ATP6V0E1P3 | 2 | ATPase H+ transporting V0 subunit e1 pseudo |
| HNRNPA1P61 | 2 | heterogeneous nuclear ribonucleoprotein A1 |
| LINC01320 | 2 | long intergenic non-protein coding RNA 1320 |
| MYADML | 2 | myeloid associated differentiation marker like |
| SLC25A5P2 | 2 | solute carrier family 25 member 5 pseudogen |
| LINC01318 | 2 | long intergenic non-protein coding RNA 1318 |
| AC008170.1 | 2 | novel transcript |
| AC073218.1 | 2 | novel transcript |
| AC012593.1 | 2 | novel transcript |
| AC079352.1 | 2 | novel transcript |
| RN7SL602P | 2 | RNA, 7SL, cytoplasmic 602, pseudogene [Sour |
| AC019064.1 | 2 | novel transcript |
| AC013442.1 | 2 | chromosome 19 open reading frame 42 (C19o |
| MIR548AD | 2 | microRNA 548ad [Source:HGNC Symbol;Acc:H |
| RNU6-1117P | 2 | RNA, U6 small nuclear 1117, pseudogene [Sou |
| AC083939.1 | 2 | GTPase activating protein (SH3 domain) bindi |
| MRPL50P1 | 2 | mitochondrial ribosomal protein L50 pseudog |
| AC009414.1 | 2 | ribosomal protein L21 (RPL21) pseudogene |
| CRIM1-DT | 2 | CRIM1 divergent transcript [Source:HGNC Syn |
| CRIM1 | 2 | cysteine rich transmembrane BMP regulator 1 |
| AC007378.1 | 2 | novel transcript |
| FEZ2 | 2 | fasciculation and elongation protein zeta 2 [So |
| AC007363.1 | 2 | guanine nucleotide binding protein (G protein |
| VIT | 2 | vitrin [Source:HGNC Symbol;Acc:HGNC:12697 |
| STRN | 2 | striatin [Source:HGNC Symbol;Acc:HGNC:1142 |
| AC007382.1 | 2 | TEC |
| RNU6-577P | 2 | RNA, U6 small nuclear 577, pseudogene [Sour |
| HEATR5B | 2 | HEAT repeat containing 5B [Source:HGNC Sym |
| GPATCH11 | 2 | G-patch domain containing 11 [Source:HGNC |
| EIF2AK2 | 2 | eukaryotic translation initiation factor 2 alpha |
| AC007899.1 | 2 | chromosome 11 open reading frame 46 (C11o |
| SULT6B1 | 2 | sulfotransferase family 6B member 1 [Source: |
| AC007390.1 | 2 | ribosomal protein L31 (RPL31) pseudogene |
| CEBPZOS | 2 | CEBPZ opposite strand [Source:HGNC Symbol; |
| CEBPZ | 2 | CCAAT enhancer binding protein zeta [Source: |
| AC007390.2 | 2 | novel transcript |
| NDUFAF7 | 2 | NADH:ubiquinone oxidoreductase complex as |

| | | |
|---|---|---|
| PRKD3 | 2 | protein kinase D3 [Source:HGNC Symbol;Acc:[illegible] |
| AC007391.1 | 2 | novel transcript |
| RNU6-939P | 2 | RNA, U6 small nuclear 939, pseudogene [Sour |
| AC007391.2 | 2 | novel transcript, antisense to QPCT |
| QPCT | 2 | glutaminyl-peptide cyclotransferase [Source:[illegible] |
| RNU6-1116P | 2 | RNA, U6 small nuclear 1116, pseudogene [Sou |
| AC007391.3 | 2 | novel transcript |
| AC006369.1 | 2 | novel transcript |
| CDC42EP3 | 2 | CDC42 effector protein 3 [Source:HGNC Symb |
| AC006369.2 | 2 | novel transcript |
| AC010878.1 | 2 | serine/threonine kinase receptor associated [illegible] |
| LINC00211 | 2 | long intergenic non-protein coding RNA 211 [[illegible] |
| RMDN2 | 2 | regulator of microtubule dynamics 2 [Source:[illegible] |
| RMDN2-AS1 | 2 | RMDN2 antisense RNA 1 [Source:HGNC Symb[illegible] |
| CYP1B1 | 2 | cytochrome P450 family 1 subfamily B memb[illegible] |
| CYP1B1-AS1 | 2 | CYP1B1 antisense RNA 1 [Source:HGNC Symb[illegible] |
| AC009229.4 | 2 | novel transcript |
| AC009229.2 | 2 | novel transcript |
| RNU6-951P | 2 | RNA, U6 small nuclear 951, pseudogene [Sour |
| AC009229.3 | 2 | novel transcript |
| AC009229.1 | 2 | novel transcript |
| RPL7P12 | 2 | ribosomal protein L7 pseudogene 12 [Source:[illegible] |
| GAPDHP25 | 2 | glyceraldehyde 3 phosphate dehydrogenase [illegible] |
| ATL2 | 2 | atlastin GTPase 2 [Source:HGNC Symbol;Acc:[illegible] |
| LINC02613 | 2 | long intergenic non-protein coding RNA 2613 |
| AC016995.1 | 2 | novel transcript |
| LINC01883 | 2 | long intergenic non-protein coding RNA 1883 |
| RPLP0P6 | 2 | ribosomal protein lateral stalk subunit P0 pse[illegible] |
| AC011247.2 | 2 | cell death-inducing DFFA-like effector c (CIDE[illegible] |
| HNRNPLL | 2 | heterogeneous nuclear ribonucleoprotein L li[illegible] |
| AC011247.1 | 2 | novel transcript |
| GALM | 2 | galactose mutarotase [Source:HGNC Symbol;[illegible] |
| AC074366.1 | 2 | novel transcript |
| SRSF7 | 2 | serine and arginine rich splicing factor 7 [Sour |
| GEMIN6 | 2 | gem nuclear organelle associated protein 6 [S[illegible] |
| TTC39DP | 2 | tetratricopeptide repeat domain 39D, pseudo |
| AC018693.2 | 2 | N-acetylneuraminate pyruvate lyase (dihydro[illegible] |
| DHX57 | 2 | DExH-box helicase 57 [Source:HGNC Symbol;[illegible] |
| ASS1P2 | 2 | argininosuccinate synthetase 1 pseudogene 2 |
| AC018693.1 | 2 | novel transcript |
| MORN2 | 2 | MORN repeat containing 2 [Source:HGNC Sym |
| RNU6-851P | 2 | RNA, U6 small nuclear 851, pseudogene [Sour |
| ARHGEF33 | 2 | Rho guanine nucleotide exchange factor 33 [S |
| RN7SL96P | 2 | RNA, 7SL, cytoplasmic 96, pseudogene [Sourc[illegible] |
| AC019171.1 | 2 | novel transcript, antisense to ARHGEF33 |
| SOS1 | 2 | SOS Ras/Rac guanine nucleotide exchange fac |
| SOS1-IT1 | 2 | SOS1 intronic transcript 1 [Source:HGNC Symb[illegible] |
| RNU6-198P | 2 | RNA, U6 small nuclear 198, pseudogene [Sour |
| HSPE1P13 | 2 | heat shock protein family E (Hsp10) member [illegible] |
| Y_RNA | 2 | Y RNA [Source:RFAM;Acc:RF00019] |

| | | |
|---|---|---|
| CDKL4 | 2 | cyclin dependent kinase like 4 [Source:HGNC S |
| MAP4K3 | 2 | mitogen-activated protein kinase kinase kinas |
| AC007684.1 | 2 | NULL |
| AC007684.2 | 2 | novel transcript, sense intronic to MAP4K3 |
| MAP4K3-DT | 2 | MAP4K3 divergent transcript [Source:HGNC Sy |
| TMEM178A | 2 | transmembrane protein 178A [Source:HGNC S |
| THUMPD2 | 2 | THUMP domain containing 2 [Source:HGNC Sy |
| SLC8A1-AS1 | 2 | SLC8A1 antisense RNA 1 [Source:HGNC Symbo |
| AC007253.1 | 2 | zinc finger protein 19 (ZNF19) pseudogene |
| SLC8A1 | 2 | solute carrier family 8 member A1 [Source:HG |
| AC007877.1 | 2 | novel transcript, antisense to SLC8A1 |
| AC007317.1 | 2 | Novel transcript |
| AC007317.2 | 2 | novel transcript |
| LINC01794 | 2 | long intergenic non-protein coding RNA 1794 |
| HNRNPA1P57 | 2 | heterogeneous nuclear ribonucleoprotein A1 |
| AC009413.2 | 2 | novel transcript |
| LDHAP3 | 2 | lactate dehydrogenase A pseudogene 3 [Sourc |
| AC009413.1 | 2 | ribosomal protein S12 (RPS12) pseudogene |
| Y_RNA | 2 | Y RNA [Source:RFAM;Acc:RF00019] |
| LINC01913 | 2 | long intergenic non-protein coding RNA 1913 |
| RNU4-63P | 2 | RNA, U4 small nuclear 63, pseudogene [Sourc |
| LINC01914 | 2 | long intergenic non-protein coding RNA 1914 |
| C2orf91 | 2 | chromosome 2 open reading frame 91 [Sourc |
| AC013480.1 | 2 | novel transcript |
| PKDCC | 2 | protein kinase domain containing, cytoplasmi |
| AC083949.1 | 2 | novel transcript |
| EML4 | 2 | EMAP like 4 [Source:HGNC Symbol;Acc:HGNC: |
| COX7A2L | 2 | cytochrome c oxidase subunit 7A2 like [Sourc |
| AC025750.2 | 2 | NULL |
| KCNG3 | 2 | potassium voltage-gated channel modifier sub |
| VDAC1P13 | 2 | voltage dependent anion channel 1 pseudoge |
| AC025750.3 | 2 | ribosomal protein S13 (RPS13) pseudogene |
| MTA3 | 2 | metastasis associated 1 family member 3 [Sou |
| Metazoa_SRP | 2 | Metazoan signal recognition particle RNA [Sou |
| AC025750.1 | 2 | ribosomal protein L21 (RPL21) pseudogene |
| AC074375.1 | 2 | c-myc binding protein (MYCBP) pseudogene |
| RNU6-137P | 2 | RNA, U6 small nuclear 137, pseudogene [Sour |
| OXER1 | 2 | oxoeicosanoid receptor 1 [Source:HGNC Symb |
| HAAO | 2 | 3-hydroxyanthranilate 3,4-dioxygenase [Sourc |
| FTOP1 | 2 | FTO pseudogene 1 [Source:HGNC Symbol;Acc |
| AC098824.1 | 2 | istidine-rich domain (CHORD)-containing 1 (CH |
| AC016735.1 | 2 | novel transcript |
| LINC01819 | 2 | long intergenic non-protein coding RNA 1819 |
| LINC02590 | 2 | long intergenic non-protein coding RNA 2590 |
| RNU6-242P | 2 | RNA, U6 small nuclear 242, pseudogene [Sour |
| LINC02580 | 2 | long intergenic non-protein coding RNA 2580 |
| AC010883.2 | 2 | novel transcript |
| AC010883.3 | 2 | novel transcript, antisense to ZFP36L2 |
| ZFP36L2 | 2 | ZFP36 ring finger protein like 2 [Source:HGNC |
| LINC01126 | 2 | long intergenic non-protein coding RNA 1126 |

| | | |
|---|---|---|
| AC010883.1 | 2 | novel transcript |
| THADA | 2 | THADA armadillo repeat containing [Source:H |
| RNU6-958P | 2 | RNA, U6 small nuclear 958, pseudogene [Sour |
| RN7SL531P | 2 | RNA, 7SL, cytoplasmic 531, pseudogene [Sour |
| Y_RNA | 2 | Y RNA [Source:RFAM;Acc:RF00019] |
| PLEKHH2 | 2 | pleckstrin homology, MyTH4 and FERM doma |
| C1GALT1C1L | 2 | C1GALT1 specific chaperone 1 like [Source:HG |
| AC011242.1 | 2 | zinc finger protein 106 homolog (mouse) (ZFP |
| RN7SKP66 | 2 | RN7SK pseudogene 66 [Source:HGNC Symbol; |
| DYNC2LI1 | 2 | dynein cytoplasmic 2 light intermediate chain |
| ABCG5 | 2 | ATP binding cassette subfamily G member 5 [S |
| ABCG8 | 2 | ATP binding cassette subfamily G member 8 [S |
| LRPPRC | 2 | leucine rich pentatricopeptide repeat containi |
| RNU6-1048P | 2 | RNA, U6 small nuclear 1048, pseudogene [Sou |
| RN7SL455P | 2 | RNA, 7SL, cytoplasmic 455, pseudogene [Sour |
| AC019129.1 | 2 | poly(A) binding protein, cytoplasmic 1 (PABPC |
| RNU6-566P | 2 | RNA, U6 small nuclear 566, pseudogene [Sour |
| PDSS1P2 | 2 | decaprenyl diphosphate synthase subunit 1 ps |
| AC019129.2 | 2 | novel transcript |
| PPM1B | 2 | protein phosphatase, Mg2+/Mn2+ dependent |
| AC013717.1 | 2 | novel protein, PPM1B-SLC3A1 readthrough |
| RPL12P19 | 2 | ribosomal protein L12 pseudogene 19 [Source |
| SLC3A1 | 2 | solute carrier family 3 member 1 [Source:HGN |
| PREPL | 2 | prolyl endopeptidase like [Source:HGNC Symb |
| CAMKMT | 2 | calmodulin-lysine N-methyltransferase [Sourc |
| LINC01833 | 2 | long intergenic non-protein coding RNA 1833 |
| AC012354.10 | 2 | TEC |
| AC012354.3 | 2 | NULL |
| AC012354.8 | 2 | NULL |
| AC012354.6 | 2 | NULL |
| AC012354.9 | 2 | NULL |
| AC012354.5 | 2 | NULL |
| AC012354.4 | 2 | NULL |
| AC012354.7 | 2 | NULL |
| SIX3-AS1 | 2 | SIX3 antisense RNA 1 [Source:HGNC Symbol;A |
| SIX3 | 2 | SIX homeobox 3 [Source:HGNC Symbol;Acc:H |
| AC012354.1 | 2 | novel transcript |
| AC012354.2 | 2 | keratinocyte associated protein 2 (KRTCAP2) p |
| SIX2 | 2 | SIX homeobox 2 [Source:HGNC Symbol;Acc:H |
| AC093702.1 | 2 | novel transcript |
| LINC01121 | 2 | long intergenic non-protein coding RNA 1121 |
| AC093833.1 | 2 | novel transcript |
| AC009236.2 | 2 | novel transcript |
| AC009236.1 | 2 | novel transcript |
| SRBD1 | 2 | S1 RNA binding domain 1 [Source:HGNC Symb |
| RN7SL414P | 2 | RNA, 7SL, cytoplasmic 414, pseudogene [Sour |
| PRKCE | 2 | protein kinase C epsilon [Source:HGNC Symbo |
| U51244.1 | 2 | novel transcript |
| AC017078.1 | 2 | ribosomal protein L26 (RPL26) pseudogene |
| AC017006.2 | 2 | novel transcript |

| | | |
|---|---|---|
| AC017006.1 | 2 | novel transcript |
| AC016696.1 | 2 | ribosomal protein L36a (RPL36A) pseudogene |
| Metazoa_SRP | 2 | Metazoan signal recognition particle RNA [Sou |
| EPAS1 | 2 | endothelial PAS domain protein 1 [Source:HG |
| LINC01820 | 2 | long intergenic non-protein coding RNA 1820 |
| LINC02583 | 2 | long intergenic non-protein coding RNA 2583 |
| AC016912.1 | 2 | novel transcript |
| RN7SL817P | 2 | RNA, 7SL, cytoplasmic 817, pseudogene [Sour |
| TMEM247 | 2 | transmembrane protein 247 [Source:HGNC Sy |
| ATP6V1E2 | 2 | ATPase H+ transporting V1 subunit E2 [Source |
| AC018682.2 | 2 | novel transcript |
| RHOQ | 2 | ras homolog family member Q [Source:HGNC |
| AC018682.1 | 2 | novel transcript |
| PIGF | 2 | phosphatidylinositol glycan anchor biosynthes |
| CRIPT | 2 | CXXC repeat containing interactor of PDZ3 do |
| AC020604.1 | 2 | TEC |
| LINC01118 | 2 | long intergenic non-protein coding RNA 1118 |
| SOCS5 | 2 | suppressor of cytokine signaling 5 [Source:HG |
| LINC01119 | 2 | long intergenic non-protein coding RNA 1119 |
| AC016722.1 | 2 | novel transcript |
| AC016722.2 | 2 | novel transcript, antisense to MCFD2 |
| MCFD2 | 2 | multiple coagulation factor deficiency 2 [Sour |
| TTC7A | 2 | tetratricopeptide repeat domain 7A [Source:H |
| AC093732.2 | 2 | novel transcript |
| AC093732.1 | 2 | novel transcript |
| STPG4 | 2 | sperm-tail PG-rich repeat containing 4 [Source |
| AC073283.3 | 2 | novel transcript |
| AC073283.1 | 2 | novel transcript |
| CALM2 | 2 | calmodulin 2 [Source:HGNC Symbol;Acc:HGN |
| EPCAM-DT | 2 | EPCAM divergent transcript [Source:HGNC Sy |
| AC073283.2 | 2 | novel transcript |
| BCYRN1 | 2 | brain cytoplasmic RNA 1 [Source:HGNC Symb |
| EPCAM | 2 | epithelial cell adhesion molecule [Source:HGN |
| RN7SKP119 | 2 | RN7SK pseudogene 119 [Source:HGNC Symbo |
| MIR559 | 2 | microRNA 559 [Source:HGNC Symbol;Acc:HGN |
| Metazoa_SRP | 2 | Metazoan signal recognition particle RNA [Sou |
| MSH2 | 2 | mutS homolog 2 [Source:HGNC Symbol;Acc:H |
| KCNK12 | 2 | potassium two pore domain channel subfamil |
| AC138655.1 | 2 | novel transcript, antisense to KCNK12 |
| AC079250.1 | 2 | ribosomal protein L18a (RPL18A) pseudogene |
| MSH6 | 2 | mutS homolog 6 [Source:HGNC Symbol;Acc:H |
| AC006509.2 | 2 | non-metastatic cells 2, protein (NM23B) expre |
| AC006509.1 | 2 | novel pseudogene |
| U6 | 2 | U6 spliceosomal RNA [Source:RFAM;Acc:RF00 |
| FBXO11 | 2 | F-box protein 11 [Source:HGNC Symbol;Acc:H |
| RPL36AP15 | 2 | ribosomal protein L36a pseudogene 15 [Sourc |
| RPS27AP7 | 2 | ribosomal protein S27a pseudogene 7 [Source |
| AC079807.1 | 2 | novel transcript |
| AC092650.1 | 2 | novel transcript |
| PPIAP62 | 2 | peptidylprolyl isomerase A pseudogene 62 [Sc |

| | | |
|---|---|---|
| AC092650.2 | 2 | vomeronasal 1 receptor 4 (VN1R4) pseudoger |
| RN7SKP224 | 2 | RN7SK pseudogene 224 [Source:HGNC Symbo |
| FOXN2 | 2 | forkhead box N2 [Source:HGNC Symbol;Acc:H |
| RNU4-49P | 2 | RNA, U4 small nuclear 49, pseudogene [Sourc |
| AC093635.1 | 2 | novel transcript |
| PPP1R21 | 2 | protein phosphatase 1 regulatory subunit 21 [ |
| RNU6-282P | 2 | RNA, U6 small nuclear 282, pseudogene [Sour |
| STON1 | 2 | stonin 1 [Source:HGNC Symbol;Acc:HGNC:170 |
| STON1-GTF2A1L | 2 | STON1-GTF2A1L readthrough [Source:HGNC S |
| GTF2A1L | 2 | general transcription factor IIA subunit 1 like [ |
| AC073082.1 | 2 | novel transcript |
| TPT1P11 | 2 | tumor protein, translationally-controlled 1 pse |
| LHCGR | 2 | luteinizing hormone/choriogonadotropin rece |
| ELOBP3 | 2 | elongin B pseudogene 3 [Source:HGNC Symbo |
| AC009975.1 | 2 | novel transcript |
| CTBP2P5 | 2 | C-terminal binding protein 2 pseudogene 5 [So |
| FSHR | 2 | follicle stimulating hormone receptor [Source |
| MIR548BA | 2 | microRNA 548ba [Source:HGNC Symbol;Acc:H |
| AC009975.2 | 2 | novel transcript |
| RNU6-439P | 2 | RNA, U6 small nuclear 439, pseudogene [Sour |
| AC009971.1 | 2 | novel transcript |
| RPL7P13 | 2 | ribosomal protein L7 pseudogene 13 [Source: |
| SNORA75 | 2 | Small nucleolar RNA SNORA75 [Source:RFAM; |
| NRXN1 | 2 | neurexin 1 [Source:HGNC Symbol;Acc:HGNC:8 |
| AC068725.1 | 2 | Novel transcript, antisense to NRXN1 |
| MTCO1P42 | 2 | MT-CO1 pseudogene 42 [Source:HGNC Symbo |
| AC009234.1 | 2 | novel transcript, antisense to NRXN1 |
| MIR8485 | 2 | microRNA 8485 [Source:HGNC Symbol;Acc:HG |
| AC007560.1 | 2 | chromosome 20 open reading frame 30 (C20o |
| AC007682.1 | 2 | novel transcript, antisense to NRXN1 |
| AC007402.1 | 2 | novel transcript |
| AC007402.2 | 2 | novel transcript |
| KNOP1P3 | 2 | lysine rich nucleolar protein 1 pseudogene 3 [ |
| CRYGGP | 2 | crystallin gamma G, pseudogene [Source:HGN |
| AC097463.2 | 2 | biliverdin reductase B (flavin reductase (NADP |
| AC097463.1 | 2 | coiled-coil domain containing 12 (CCDC12) pse |
| AC079304.1 | 2 | novel transcript |
| ZNF863P | 2 | zinc finger protein 863, pseudogene [Source:H |
| Y_RNA | 2 | Y RNA [Source:RFAM;Acc:RF00019] |
| LINC01867 | 2 | long intergenic non-protein coding RNA 1867 |
| AC139712.2 | 2 | chromosome 7 open reading frame 24 (C7orf2 |
| AC139712.3 | 2 | novel transcript |
| AC139712.1 | 2 | CREB regulated transcription coactivator 1 (CF |
| FTH1P6 | 2 | ferritin heavy chain 1 pseudogene 6 [Source:H |
| MIR4431 | 2 | microRNA 4431 [Source:HGNC Symbol;Acc:HG |
| AC010967.1 | 2 | novel transcript |
| AC010967.2 | 2 | novel transcript |
| AC010967.3 | 2 | keratin associated protein 5-11 (KRTAP5-11) p |
| AC069157.2 | 2 | NULL |
| AC069157.1 | 2 | neuropilin (NRP) and tolloid (TLL)-like 2 (NETO |

| | | |
|---|---|---|
| ASB3 | 2 | ankyrin repeat and SOCS box containing 3 [Sou |
| RNU6-997P | 2 | RNA, U6 small nuclear 997, pseudogene [Sour |
| CHAC2 | 2 | ChaC cation transport regulator homolog 2 [S |
| ERLEC1 | 2 | endoplasmic reticulum lectin 1 [Source:HGNC |
| MIR3682 | 2 | microRNA 3682 [Source:HGNC Symbol;Acc:HG |
| GPR75 | 2 | G protein-coupled receptor 75 [Source:HGNC |
| PSME4 | 2 | proteasome activator subunit 4 [Source:HGNC |
| ACYP2 | 2 | acylphosphatase 2 [Source:HGNC Symbol;Acc: |
| AC008280.1 | 2 | ribosomal protein L21 (RPL21) pseudogene |
| HMGB1P31 | 2 | high mobility group box 1 pseudogene 31 [Sou |
| AC008280.2 | 2 | ribosomal protein S26 (RPS26) pseudogene |
| AC008280.3 | 2 | novel transcript |
| RNU7-172P | 2 | RNA, U7 small nuclear 172 pseudogene [Sourc |
| TSPYL6 | 2 | TSPY like 6 [Source:HGNC Symbol;Acc:HGNC:1 |
| C2orf73 | 2 | chromosome 2 open reading frame 73 [Sourc |
| SPTBN1 | 2 | spectrin beta, non-erythrocytic 1 [Source:HGN |
| AC092839.1 | 2 | novel transcript |
| RPL23AP32 | 2 | ribosomal protein L23a pseudogene 32 [Sourc |
| AC092839.2 | 2 | novel transcript |
| AC093110.1 | 2 | novel transcript, antisense to SPTBN1 |
| EML6 | 2 | EMAP like 6 [Source:HGNC Symbol;Acc:HGNC: |
| AC104781.1 | 2 | novel transcript |
| AC104781.2 | 2 | Novel transcript, antisense to EML6 |
| RNU7-81P | 2 | RNA, U7 small nuclear 81 pseudogene [Source |
| RTN4 | 2 | reticulon 4 [Source:HGNC Symbol;Acc:HGNC:1 |
| RNU6-433P | 2 | RNA, U6 small nuclear 433, pseudogene [Sour |
| CLHC1 | 2 | clathrin heavy chain linker domain containing |
| AC012358.1 | 2 | novel transcript |
| AC012358.2 | 2 | pseudogene similar to RIKEN cDNA 2210021J2 |
| RPS27A | 2 | ribosomal protein S27a [Source:HGNC Symbol |
| MTIF2 | 2 | mitochondrial translational initiation factor 2 |
| PRORSD1P | 2 | prolyl-tRNA synthetase associated domain cor |
| AC012358.3 | 2 | novel transcript, antisense to CCDC88A |
| Y_RNA | 2 | Y RNA [Source:RFAM;Acc:RF00019] |
| CCDC88A | 2 | coiled-coil domain containing 88A [Source:HG |
| BTF3P5 | 2 | basic transcription factor 3 pseudogene 5 [Sou |
| RNU6-775P | 2 | RNA, U6 small nuclear 775, pseudogene [Sour |
| RNU6-221P | 2 | RNA, U6 small nuclear 221, pseudogene [Sour |
| RNU6-634P | 2 | RNA, U6 small nuclear 634, pseudogene [Sour |
| CFAP36 | 2 | cilia and flagella associated protein 36 [Source |
| PPP4R3B | 2 | protein phosphatase 4 regulatory subunit 3B [ |
| AC015982.1 | 2 | NULL |
| AC015982.2 | 2 | novel transcript |
| PNPT1 | 2 | polyribonucleotide nucleotidyltransferase 1 [S |
| EFEMP1 | 2 | EGF containing fibulin extracellular matrix pro |
| RN7SKP208 | 2 | RN7SK pseudogene 208 [Source:HGNC Symbo |
| AC011306.1 | 2 | novel transcript |
| MIR217HG | 2 | MIR217 host gene [Source:HGNC Symbol;Acc: |
| MIR217 | 2 | microRNA 217 [Source:HGNC Symbol;Acc:HGN |
| MIR216A | 2 | microRNA 216a [Source:HGNC Symbol;Acc:HG |

| | | |
|---|---|---|
| MIR216B | 2 | microRNA 216b [Source:HGNC Symbol;Acc:HG |
| LINC01813 | 2 | long intergenic non-protein coding RNA 1813 |
| AC007744.1 | 2 | novel transcript |
| AC007743.1 | 2 | novel transcript, antisense to CCDC85A |
| CCDC85A | 2 | coiled-coil domain containing 85A [Source:HG |
| RNA5SP93 | 2 | RNA, 5S ribosomal pseudogene 93 [Source:HG |
| PPIAP63 | 2 | peptidylprolyl isomerase A pseudogene 63 [Sc |
| AC010738.1 | 2 | eukaryotic translation initiation factor 2, subu |
| AC132153.1 | 2 | Novel transcript |
| AC068276.1 | 2 | proteasome maturation protein (POMP) pseu |
| AC073875.1 | 2 | NULL |
| ACTG1P22 | 2 | actin gamma 1 pseudogene 22 [Source:HGNC |
| VRK2 | 2 | VRK serine/threonine kinase 2 [Source:HGNC |
| AC068193.1 | 2 | novel transcript, antisense to VRK2 |
| FANCL | 2 | FA complementation group L [Source:HGNC Sy |
| AC007250.1 | 2 | novel transcript |
| EIF3FP3 | 2 | eukaryotic translation initiation factor 3 subur |
| LINC01795 | 2 | long intergenic non-protein coding RNA 1795 |
| LINC01122 | 2 | long intergenic non-protein coding RNA 1122 |
| AC007238.1 | 2 | IK cytokine, down-regulator of HLA II (IK) pseu |
| AC007092.1 | 2 | novel transcript |
| LINC01793 | 2 | long intergenic non-protein coding RNA 1793 |
| AC007100.1 | 2 | novel transcript |
| AC007179.2 | 2 | novel transcript |
| AC007179.1 | 2 | novel transcript |
| AC007179.3 | 2 | LSM6 homolog, U6 small nuclear RNA associa |
| RNU6-508P | 2 | RNA, U6 small nuclear 508, pseudogene [Sour |
| RNA5SP94 | 2 | RNA, 5S ribosomal pseudogene 94 [Source:HG |
| AC007100.2 | 2 | novel transcript |
| AC007132.1 | 2 | Novel transcript |
| AC007381.1 | 2 | novel transcript |
| AC007381.2 | 2 | novel transcript |
| RNU1-32P | 2 | RNA, U1 small nuclear 32, pseudogene [Sourc |
| MIR4432 | 2 | microRNA 4432 [Source:HGNC Symbol;Acc:HG |
| BCL11A | 2 | BAF chromatin remodeling complex subunit B |
| AC009970.1 | 2 | novel transcript |
| RN7SL361P | 2 | RNA, 7SL, cytoplasmic 361, pseudogene [Sour |
| AC012498.2 | 2 | interferon induced transmembrane protein 3 |
| AC012498.1 | 2 | ribosomal protein L26 (RPL26) pseudogene |
| RNU6-612P | 2 | RNA, U6 small nuclear 612, pseudogene [Sour |
| ATP1B3P1 | 2 | ATPase Na+/K+ transporting subunit beta 3 ps |
| PAPOLG | 2 | poly(A) polymerase gamma [Source:HGNC Syr |
| LINC01185 | 2 | long intergenic non-protein coding RNA 1185 |
| RN7SL632P | 2 | RNA, 7SL, cytoplasmic 632, pseudogene [Sour |
| RPL21P33 | 2 | ribosomal protein L21 pseudogene 33 [Source |
| REL | 2 | REL proto-oncogene, NF-kB subunit [Source:H |
| RNU4-51P | 2 | RNA, U4 small nuclear 51, pseudogene [Sourc |
| AC010733.2 | 2 | novel transcript |
| NONOP2 | 2 | non-POU domain containing, octamer-binding |
| AC010733.1 | 2 | ribosomal protein S12 (RPS12) pseudogene |

| | | |
|---|---|---|
| PUS10 | 2 | pseudouridine synthase 10 [Source:HGNC Sym |
| RNA5SP95 | 2 | RNA, 5S ribosomal pseudogene 95 [Source:HG |
| PEX13 | 2 | peroxisomal biogenesis factor 13 [Source:HGN |
| KIAA1841 | 2 | KIAA1841 [Source:HGNC Symbol;Acc:HGNC:29 |
| AC016747.4 | 2 | novel transcript |
| AC016747.1 | 2 | novel transcript |
| C2orf74 | 2 | chromosome 2 open reading frame 74 [Source |
| AC016747.2 | 2 | novel transcript |
| AHSA2P | 2 | activator of HSP90 ATPase homolog 2, pseudo |
| USP34 | 2 | ubiquitin specific peptidase 34 [Source:HGNC |
| AC016747.3 | 2 | novel transcript |
| AC018889.1 | 2 | BCL2/adenovirus E1B 19kDa interacting prote |
| AC016894.1 | 2 | ribosomal protein L10-like (RPL10L) pseudoge |
| SNORA70B | 2 | small nucleolar RNA, H/ACA box 70B [Source:I |
| AC016727.1 | 2 | novel transcript, antisense to XPO1 |
| XPO1 | 2 | exportin 1 [Source:HGNC Symbol;Acc:HGNC:1 |
| AC016727.3 | 2 | novel transcript, antisense to XPO1 |
| AC016727.2 | 2 | proteasome (prosome, macropain) 26S subun |
| RPS29P10 | 2 | ribosomal protein S29 pseudogene 10 [Source |
| RNU6-1145P | 2 | RNA, U6 small nuclear 1145, pseudogene [Sou |
| AC108039.1 | 2 | ribosomal protein L14 (RPL14) pseudogene |
| RPS24P7 | 2 | ribosomal protein S24 pseudogene 7 [Source: |
| AC108039.2 | 2 | heterogeneous nuclear ribonucleoprotein A1 |
| FAM161A | 2 | FAM161 centrosomal protein A [Source:HGNC |
| AC107081.1 | 2 | ribosomal protein L31 (RPL31) pseudogene |
| AC107081.3 | 2 | serine/arginine repetitive matrix 1 (SRRM1) ps |
| CCT4 | 2 | chaperonin containing TCP1 subunit 4 [Source |
| AC107081.2 | 2 | novel transcript |
| AC107081.4 | 2 | novel transcript, antisense to CCT4 |
| COMMD1 | 2 | copper metabolism domain containing 1 [Sour |
| Y_RNA | 2 | Y RNA [Source:RFAM;Acc:RF00019] |
| AC018462.1 | 2 | novel transcript |
| RPSAP26 | 2 | ribosomal protein SA pseudogene 26 [Source: |
| AC018462.2 | 2 | phosphatidylinositol glycan anchor biosynthes |
| B3GNT2 | 2 | UDP-GlcNAc:betaGal beta-1,3-N-acetylglucosa |
| MIR5192 | 2 | microRNA 5192 [Source:HGNC Symbol;Acc:HG |
| RN7SL51P | 2 | RNA, 7SL, cytoplasmic 51, pseudogene [Source |
| AC093159.2 | 2 | eukaryotic translation initiation factor 3, subu |
| AC093159.1 | 2 | novel transcript |
| RN7SL18P | 2 | RNA, 7SL, cytoplasmic 18, pseudogene [Source |
| TMEM17 | 2 | transmembrane protein 17 [Source:HGNC Sym |
| AC107083.1 | 2 | ribosomal protein L37 pseudogene |
| AC092155.2 | 2 | ribosomal protein L21 (RPL21) pseudogene |
| AC092155.1 | 2 | novel transcript |
| PSAT1P2 | 2 | phosphoserine aminotransferase 1 pseudoger |
| RSL24D1P2 | 2 | ribosomal L24 domain containing 1 pseudoger |
| EHBP1 | 2 | EH domain binding protein 1 [Source:HGNC Sy |
| Y_RNA | 2 | Y RNA [Source:RFAM;Acc:RF00019] |
| AC092567.2 | 2 | novel transcript, antisense to EHBP1 |
| AC092567.1 | 2 | novel transcript |

| | | |
|---|---|---|
| AC007098.2 | 2 | ribosomal protein S20 (RPS20) pseudogene |
| AC007098.1 | 2 | novel transcript, antisense to EHBP1 |
| OTX1 | 2 | orthodenticle homeobox 1 [Source:HGNC Sym |
| AC009501.2 | 2 | BAI1-associated protein 2-like 1 (BAIAP2L1) ps |
| AC009501.3 | 2 | novel transcript, antisense to WDPCP |
| AC009501.1 | 2 | ribosomal protein L27 (RPL27) pseudogene |
| DBIL5P2 | 2 | diazepam binding inhibitor-like 5 pseudogene |
| WDPCP | 2 | WD repeat containing planar cell polarity effe |
| Metazoa_SRP | 2 | Metazoan signal recognition particle RNA [Sou |
| AC074367.1 | 2 | family with sequence similarity 54, member A |
| AC016734.2 | 2 | TEC |
| MDH1 | 2 | malate dehydrogenase 1 [Source:HGNC Symb |
| AC016734.1 | 2 | PRELI domain containing 1 (PRELID1) pseudog |
| RPS4XP5 | 2 | ribosomal protein S4X pseudogene 5 [Source: |
| AC096664.2 | 2 | ribosomal protein L27 (RPL27) pseudogene |
| AC096664.1 | 2 | pseudogene similar to part of citrate synthase |
| HNRNPA1P66 | 2 | heterogeneous nuclear ribonucleoprotein A1 |
| UGP2 | 2 | UDP-glucose pyrophosphorylase 2 [Source:HG |
| AC114748.1 | 2 | NULL |
| VPS54 | 2 | VPS54 subunit of GARP complex [Source:HGN |
| AC012368.2 | 2 | novel transcript |
| PELI1 | 2 | pellino E3 ubiquitin protein ligase 1 [Source:H |
| AC012368.1 | 2 | novel transcript, antisense to PELI1 |
| LINC00309 | 2 | long intergenic non-protein coding RNA 309 [S |
| AC012368.3 | 2 | family with sequence similarity 136, member |
| AC114752.1 | 2 | novel transcript |
| AC114752.2 | 2 | novel transcript |
| MIR4433B | 2 | microRNA 4433b [Source:HGNC Symbol;Acc:H |
| MIR4433A | 2 | microRNA 4433a [Source:HGNC Symbol;Acc:H |
| RPL23AP37 | 2 | ribosomal protein L23a pseudogene 37 [Sourc |
| LGALSL-DT | 2 | LGALSL divergent transcript [Source:HGNC Syr |
| AC008074.1 | 2 | novel pseudogene |
| LGALSL | 2 | galectin like [Source:HGNC Symbol;Acc:HGNC: |
| LINC01805 | 2 | long intergenic non-protein coding RNA 1805 |
| AC008074.2 | 2 | novel transcript |
| AFTPH | 2 | aftiphilin [Source:HGNC Symbol;Acc:HGNC:25 |
| MIR4434 | 2 | microRNA 4434 [Source:HGNC Symbol;Acc:HG |
| RNU6-100P | 2 | RNA, U6 small nuclear 100, pseudogene [Sour |
| LINC02579 | 2 | long intergenic non-protein coding RNA 2579 |
| SERTAD2 | 2 | SERTA domain containing 2 [Source:HGNC Syr |
| AC007365.1 | 2 | novel transcript |
| AC007365.2 | 2 | ribosomal protein S10 (RPS10) pseudogene |
| RN7SL341P | 2 | RNA, 7SL, cytoplasmic 341, pseudogene [Sour |
| Y_RNA | 2 | Y RNA [Source:RFAM;Acc:RF00019] |
| AC007880.1 | 2 | novel transcript |
| LINC01800 | 2 | long intergenic non-protein coding RNA 1800 |
| LINC02245 | 2 | long intergenic non-protein coding RNA 2245 |
| RN7SL211P | 2 | RNA, 7SL, cytoplasmic 211, pseudogene [Sour |
| AC007386.1 | 2 | ribosomal protein L11 (RPL11) pseudogene |
| SLC1A4 | 2 | solute carrier family 1 member 4 [Source:HGN |

| | | |
|---|---|---|
| RNU6-548P | 2 | RNA, U6 small nuclear 548, pseudogene [Sour |
| LINC02576 | 2 | long intergenic non-protein coding RNA 2576 |
| CEP68 | 2 | centrosomal protein 68 [Source:HGNC Symbo |
| RAB1A | 2 | RAB1A, member RAS oncogene family [Source |
| SNORA74 | 2 | Small nucleolar RNA SNORA74 [Source:RFAM; |
| AC007318.2 | 2 | marker of proliferation Ki-67 (MKI67) pseudog |
| AC007318.1 | 2 | voltage-dependent anion channel 2 (VDAC2) p |
| ACTR2 | 2 | actin related protein 2 [Source:HGNC Symbol; |
| SPRED2 | 2 | sprouty related EVH1 domain containing 2 [Sc |
| AC012370.1 | 2 | novel transcript, sense intronic to SPRED2 |
| AC007389.1 | 2 | novel transcript |
| AC012370.2 | 2 | novel transcript |
| AC007389.4 | 2 | DnaJ (Hsp40) homolog, subfamily B, member |
| AC007389.2 | 2 | ribosomal protein S15a (RPS15A) pseudogene |
| RN7SL635P | 2 | RNA, 7SL, cytoplasmic 635, pseudogene [Sour |
| VTRNA2-2P | 2 | vault RNA 2-2, pseudogene [Source:HGNC Syn |
| AC007389.3 | 2 | novel transcript |
| AC007389.5 | 2 | novel transcript |
| KRT18P33 | 2 | keratin 18 pseudogene 33 [Source:HGNC Sym |
| AC007741.1 | 2 | novel transcript |
| AC118345.1 | 2 | novel transcript |
| AC092669.1 | 2 | novel transcript |
| MIR4778 | 2 | microRNA 4778 [Source:HGNC Symbol;Acc:HG |
| LINC01873 | 2 | long intergenic non-protein coding RNA 1873 |
| MEIS1-AS3 | 2 | MEIS1 antisense RNA 3 [Source:HGNC Symbol |
| MEIS1 | 2 | Meis homeobox 1 [Source:HGNC Symbol;Acc: |
| MEIS1-AS2 | 2 | MEIS1 antisense RNA 2 [Source:HGNC Symbol |
| LINC01798 | 2 | long intergenic non-protein coding RNA 1798 |
| LINC01797 | 2 | long intergenic non-protein coding RNA 1797 |
| AC009474.1 | 2 | DNA (cytosine-5-)-methyltransferase 3 alpha ( |
| AC009474.2 | 2 | novel pseudogene |
| LINC01799 | 2 | long intergenic non-protein coding RNA 1799 |
| LINC01628 | 2 | long intergenic non-protein coding RNA 1628 |
| AC007403.1 | 2 | novel transcript |
| LINC01828 | 2 | long intergenic non-protein coding RNA 1828 |
| LINC01829 | 2 | long intergenic non-protein coding RNA 1829 |
| AC023115.1 | 2 | novel transcript |
| ETAA1 | 2 | ETAA1 activator of ATR kinase [Source:HGNC S |
| LINC02831 | 2 | long intergenic non-protein coding RNA 2831 |
| AC007422.1 | 2 | novel transcript |
| AC010987.1 | 2 | novel transcript |
| LINC01812 | 2 | long intergenic non-protein coding RNA 1812 |
| AC079112.1 | 2 | F-box and leucine-rich repeat protein 12 (FBXL |
| C1D | 2 | C1D nuclear receptor corepressor [Source:HG |
| AC017083.4 | 2 | novel pseudogene |
| WDR92 | 2 | WD repeat domain 92 [Source:HGNC Symbol;A |
| PPIAP64 | 2 | peptidylprolyl isomerase A pseudogene 64 [Sc |
| AC017083.3 | 2 | novel protein |
| PNO1 | 2 | partner of NOB1 homolog [Source:HGNC Sym |
| PPP3R1 | 2 | protein phosphatase 3 regulatory subunit B, a |

| | | |
|---|---|---|
| AC017083.2 | 2 | novel transcript |
| AC017083.1 | 2 | novel transcript |
| CNRIP1 | 2 | cannabinoid receptor interacting protein 1 [Sc |
| AC015969.1 | 2 | novel transcript, antisense to PLEK |
| PLEK | 2 | pleckstrin [Source:HGNC Symbol;Acc:HGNC:9( |
| AC127383.1 | 2 | WD repeat domain 4 (WDR4) pseudogene |
| FBXO48 | 2 | F-box protein 48 [Source:HGNC Symbol;Acc:H |
| APLF | 2 | aprataxin and PNKP like factor [Source:HGNC |
| AC130709.1 | 2 | chromosome 15 open reading frame 2 (C15or |
| PROKR1 | 2 | prokineticin receptor 1 [Source:HGNC Symbol |
| ARHGAP25 | 2 | Rho GTPase activating protein 25 [Source:HGI |
| LINC01890 | 2 | long intergenic non-protein coding RNA 1890 |
| LINC01888 | 2 | long intergenic non-protein coding RNA 1888 |
| BMP10 | 2 | bone morphogenetic protein 10 [Source:HGN( |
| GKN3P | 2 | gastrokine 3, pseudogene [Source:HGNC Syml |
| GKN2 | 2 | gastrokine 2 [Source:HGNC Symbol;Acc:HGNC |
| GKN1 | 2 | gastrokine 1 [Source:HGNC Symbol;Acc:HGNC |
| ANTXR1 | 2 | ANTXR cell adhesion molecule 1 [Source:HGN( |
| AC114802.1 | 2 | novel transcript, antisense to ANTXR1 |
| MIR3126 | 2 | microRNA 3126 [Source:HGNC Symbol;Acc:HG |
| RNA5SP96 | 2 | RNA, 5S ribosomal pseudogene 96 [Source:HG |
| RNU6-1216P | 2 | RNA, U6 small nuclear 1216, pseudogene [Sou |
| AC112230.1 | 2 | golgi reassembly stacking protein 2, 55kDa (G( |
| GFPT1 | 2 | glutamine--fructose-6-phosphate transaminas |
| Y_RNA | 2 | Y RNA [Source:RFAM;Acc:RF00019] |
| NFU1 | 2 | NFU1 iron-sulfur cluster scaffold [Source:HGN |
| AAK1 | 2 | AP2 associated kinase 1 [Source:HGNC Symbo |
| RN7SL604P | 2 | RNA, 7SL, cytoplasmic 604, pseudogene [Sour |
| SNORA36C | 2 | small nucleolar RNA, H/ACA box 36C [Source:I |
| AC092431.2 | 2 | ribosomal protein L36a (RPL36A) pseudogene |
| B3GALNT1P1 | 2 | beta-1,3-N-acetylgalactosaminyltransferase 1 |
| ANXA4 | 2 | annexin A4 [Source:HGNC Symbol;Acc:HGNC:! |
| AC092431.1 | 2 | novel transcript |
| GMCL1 | 2 | germ cell-less 1, spermatogenesis associated [ |
| AC019206.1 | 2 | Zinc finger protein 607 (ZNF607) pseudogene |
| RPL23AP92 | 2 | ribosomal protein L23a pseudogene 92 [Sourc |
| SNRNP27 | 2 | small nuclear ribonucleoprotein U4/U6.U5 sul |
| MXD1 | 2 | MAX dimerization protein 1 [Source:HGNC Syı |
| ASPRV1 | 2 | aspartic peptidase retroviral like 1 [Source:HG |
| PCBP1-AS1 | 2 | PCBP1 antisense RNA 1 [Source:HGNC Symbol |
| RN7SL470P | 2 | RNA, 7SL, cytoplasmic 470, pseudogene [Sour |
| PCBP1 | 2 | poly(rC) binding protein 1 [Source:HGNC Symt |
| AC016700.3 | 2 | novel transcript |
| AC016700.1 | 2 | mitochondrial ribosomal protein L36 (MRPL3( |
| LINC01816 | 2 | long intergenic non-protein coding RNA 1816 |
| C2orf42 | 2 | chromosome 2 open reading frame 42 [Sourc( |
| TIA1 | 2 | TIA1 cytotoxic granule associated RNA bindinį |
| MIR1285-2 | 2 | microRNA 1285-2 [Source:HGNC Symbol;Acc:I |
| AC016700.2 | 2 | ribosomal protein L39 (RPL39) pseudogene |
| PCYOX1 | 2 | prenylcysteine oxidase 1 [Source:HGNC Symb( |

| | | |
|---|---|---|
| SNRPG | 2 | small nuclear ribonucleoprotein polypeptide ( |
| FAM136A | 2 | family with sequence similarity 136 member / |
| AC022201.2 | 2 | novel transcript |
| BRD7P6 | 2 | bromodomain containing 7 pseudogene 6 [So |
| AC022201.1 | 2 | novel transcript |
| TGFA | 2 | transforming growth factor alpha [Source:HGI |
| TGFA-IT1 | 2 | TGFA intronic transcript 1 [Source:HGNC Syml |
| ADD2 | 2 | adducin 2 [Source:HGNC Symbol;Acc:HGNC:2 |
| AC005234.2 | 2 | novel transcript, antisense to ADD2 |
| AC005234.1 | 2 | novel transcript |
| FIGLA | 2 | folliculogenesis specific bHLH transcription fac |
| HMGN2P21 | 2 | high mobility group nucleosomal binding dom |
| CLEC4F | 2 | C-type lectin domain family 4 member F [Sour |
| AC007395.1 | 2 | pseudogene similar to part of MOB1, Mps On |
| CD207 | 2 | CD207 molecule [Source:HGNC Symbol;Acc:H |
| LINC01143 | 2 | long intergenic non-protein coding RNA 1143 |
| AC007040.3 | 2 | NULL |
| VAX2 | 2 | ventral anterior homeobox 2 [Source:HGNC S |
| ATP6V1B1 | 2 | ATPase H+ transporting V1 subunit B1 [Source |
| AC007040.2 | 2 | novel protein |
| ATP6V1B1-AS1 | 2 | ATP6V1B1 antisense RNA 1 [Source:HGNC Syr |
| ELOCP21 | 2 | elongin C pseudogene 21 [Source:HGNC Symb |
| RN7SL160P | 2 | RNA, 7SL, cytoplasmic 160, pseudogene [Sour |
| ANKRD53 | 2 | ankyrin repeat domain 53 [Source:HGNC Syml |
| TEX261 | 2 | testis expressed 261 [Source:HGNC Symbol;A |
| AC007040.1 | 2 | novel transcript |
| AC007881.2 | 2 | novel transcript, antisense to NAGK |
| OR7E91P | 2 | olfactory receptor family 7 subfamily E memb |
| OR7E46P | 2 | olfactory receptor family 7 subfamily E memb |
| OR7E62P | 2 | olfactory receptor family 7 subfamily E memb |
| NAGK | 2 | N-acetylglucosamine kinase [Source:HGNC Syı |
| AC007881.3 | 2 | novel transcript |
| MCEE | 2 | methylmalonyl-CoA epimerase [Source:HGNC |
| AC007881.4 | 2 | novel transcript, antisense to MCEE |
| MPHOSPH10 | 2 | M-phase phosphoprotein 10 [Source:HGNC Sy |
| AC007881.1 | 2 | pseudogene similar to part of N-acetylglucosa |
| PAIP2B | 2 | poly(A) binding protein interacting protein 2B |
| ZNF638 | 2 | zinc finger protein 638 [Source:HGNC Symbol; |
| AC007878.1 | 2 | novel transcript |
| RNU6-105P | 2 | RNA, U6 small nuclear 105, pseudogene [Sour |
| DYSF | 2 | dysferlin [Source:HGNC Symbol;Acc:HGNC:30 |
| RPS20P10 | 2 | ribosomal protein S20 pseudogene 10 [Source |
| CYP26B1 | 2 | cytochrome P450 family 26 subfamily B meml |
| EXOC6B | 2 | exocyst complex component 6B [Source:HGN( |
| AC006461.1 | 2 | ribosomal protein S15a (RPS15A) pseudogene |
| RNU2-39P | 2 | RNA, U2 small nuclear 39, pseudogene [Sourc |
| AC092630.1 | 2 | NULL |
| SPR | 2 | sepiapterin reductase [Source:HGNC Symbol;/ |
| EMX1 | 2 | empty spiracles homeobox 1 [Source:HGNC Sy |
| AC012366.1 | 2 | novel transcript, antisense to EMX1 |

| | | |
|---|---|---|
| SFXN5 | 2 | sideroflexin 5 [Source:HGNC Symbol;Acc:HGN |
| RAB11FIP5 | 2 | RAB11 family interacting protein 5 [Source:HG |
| AC010913.1 | 2 | novel transcript |
| NOTO | 2 | notochord homeobox [Source:HGNC Symbol;A |
| SMYD5 | 2 | SMYD family member 5 [Source:HGNC Symbo |
| PRADC1 | 2 | protease associated domain containing 1 [Sou |
| CCT7 | 2 | chaperonin containing TCP1 subunit 7 [Source |
| FBXO41 | 2 | F-box protein 41 [Source:HGNC Symbol;Acc:H |
| EGR4 | 2 | early growth response 4 [Source:HGNC Symbo |
| RNU6-111P | 2 | RNA, U6 small nuclear 111, pseudogene [Sour |
| AC069404.1 | 2 | novel transcript |
| AC074008.2 | 2 | novel transcript |
| RPSAP28 | 2 | ribosomal protein SA pseudogene 28 [Source: |
| ALMS1 | 2 | ALMS1 centrosome and basal body associated |
| ALMS1-IT1 | 2 | ALMS1 intronic transcript 1 [Source:HGNC Syr |
| AC074008.1 | 2 | novel transcript, sense intronic to ALMS1 |
| NAT8 | 2 | N-acetyltransferase 8 (putative) [Source:HGN |
| ALMS1P1 | 2 | ALMS1 pseudogene 1 [Source:HGNC Symbol;A |
| NAT8B | 2 | N-acetyltransferase 8B (putative, gene/pseud |
| TPRKB | 2 | TP53RK binding protein [Source:HGNC Symbo |
| AC092653.1 | 2 | novel transcript |
| AC092653.2 | 2 | TEC |
| DUSP11 | 2 | dual specificity phosphatase 11 [Source:HGNC |
| C2orf78 | 2 | chromosome 2 open reading frame 78 [Source |
| AC073046.4 | 2 | novel transcript |
| STAMBP | 2 | STAM binding protein [Source:HGNC Symbol;A |
| AC073046.3 | 2 | novel transcript, antisense to STAMBP |
| AC073046.2 | 2 | zinc finger, DHHC-type containing 6 (ZDHHC6) |
| ACTG2 | 2 | actin gamma 2, smooth muscle [Source:HGNC |
| DGUOK | 2 | deoxyguanosine kinase [Source:HGNC Symbol |
| DGUOK-AS1 | 2 | DGUOK antisense RNA 1 [Source:HGNC Symbo |
| RNA5SP97 | 2 | RNA, 5S ribosomal pseudogene 97 [Source:HG |
| AC073046.1 | 2 | novel transcript (LOC100132318) |
| TET3 | 2 | tet methylcytosine dioxygenase 3 [Source:HG |
| AC110801.1 | 2 | TEC |
| FNBP1P1 | 2 | formin binding protein 1 pseudogene 1 [Sourc |
| AC073263.1 | 2 | novel transcript, antisense to BOLA3 |
| BOLA3 | 2 | bolA family member 3 [Source:HGNC Symbol; |
| BOLA3-AS1 | 2 | BOLA3 divergent transcript [Source:HGNC Syn |
| MOB1A | 2 | MOB kinase activator 1A [Source:HGNC Symb |
| AC073263.3 | 2 | novel transcript |
| AC073263.2 | 2 | novel transcript |
| MTHFD2 | 2 | methylenetetrahydrofolate dehydrogenase (N |
| AC006030.1 | 2 | novel protein |
| SLC4A5 | 2 | solute carrier family 4 member 5 [Source:HGN |
| KRT18P26 | 2 | keratin 18 pseudogene 26 [Source:HGNC Sym |
| RNU6-542P | 2 | RNA, U6 small nuclear 542, pseudogene [Sour |
| NECAP1P2 | 2 | NECAP endocytosis associated 1 pseudogene |
| TAF13P2 | 2 | TATA-box binding protein associated factor 13 |
| DCTN1 | 2 | dynactin subunit 1 [Source:HGNC Symbol;Acc: |

| | | |
|---|---|---|
| DCTN1-AS1 | 2 | DCTN1 antisense RNA 1 [Source:HGNC Symbo |
| AC005041.1 | 2 | novel protein |
| AC005041.2 | 2 | novel retinol dehydrogenase pseudogene |
| C2orf81 | 2 | chromosome 2 open reading frame 81 [Sourc |
| AC005041.4 | 2 | novel transcript, antisense to C2orf81 |
| HMGA1P8 | 2 | high mobility group AT-hook 1 pseudogene 8 |
| WDR54 | 2 | WD repeat domain 54 [Source:HGNC Symbol; |
| RTKN | 2 | rhotekin [Source:HGNC Symbol;Acc:HGNC:10 |
| INO80B | 2 | INO80 complex subunit B [Source:HGNC Symb |
| INO80B-WBP1 | 2 | INO80B-WBP1 readthrough (NMD candidate) |
| WBP1 | 2 | WW domain binding protein 1 [Source:HGNC |
| MOGS | 2 | mannosyl-oligosaccharide glucosidase [Source |
| AC005041.5 | 2 | novel transcript, antisense to CCDC142and MI |
| MRPL53 | 2 | mitochondrial ribosomal protein L53 [Source: |
| CCDC142 | 2 | coiled-coil domain containing 142 [Source:HG |
| TTC31 | 2 | tetratricopeptide repeat domain 31 [Source:H |
| LBX2 | 2 | ladybird homeobox 2 [Source:HGNC Symbol;A |
| AC005041.3 | 2 | novel transcript, antisense to LBX2 |
| LBX2-AS1 | 2 | LBX2 antisense RNA 1 [Source:HGNC Symbol; |
| PCGF1 | 2 | polycomb group ring finger 1 [Source:HGNC S |
| TLX2 | 2 | T cell leukemia homeobox 2 [Source:HGNC Sy |
| DQX1 | 2 | DEAQ-box RNA dependent ATPase 1 [Source:I |
| AUP1 | 2 | AUP1 lipid droplet regulating VLDL assembly f |
| HTRA2 | 2 | HtrA serine peptidase 2 [Source:HGNC Symbo |
| LOXL3 | 2 | lysyl oxidase like 3 [Source:HGNC Symbol;Acc: |
| DOK1 | 2 | docking protein 1 [Source:HGNC Symbol;Acc:I |
| M1AP | 2 | meiosis 1 associated protein [Source:HGNC Sy |
| AC005033.1 | 2 | torsin family 1, member A (torsin A) (TOR1A) |
| TVP23BP2 | 2 | TVP23B pseudogene 2 [Source:HGNC Symbol; |
| SEMA4F | 2 | ssemaphorin 4F [Source:HGNC Symbol;Acc:H |
| AC007387.3 | 2 | novel transcript |
| AC007387.1 | 2 | ribosomal protein S28 (RPS28) pseudogene |
| AC007387.2 | 2 | novel transcript |
| AC019069.1 | 2 | novel transcript |
| HK2 | 2 | hexokinase 2 [Source:HGNC Symbol;Acc:HGN |
| LINC01291 | 2 | long intergenic non-protein coding RNA 1291 |
| AC104135.1 | 2 | novel transcript |
| LINC01293 | 2 | long intergenic non-protein coding RNA 1293 |
| POLE4 | 2 | DNA polymerase epsilon 4, accessory subunit |
| TACR1 | 2 | tachykinin receptor 1 [Source:HGNC Symbol;A |
| MIR5000 | 2 | microRNA 5000 [Source:HGNC Symbol;Acc:HG |
| AC007681.1 | 2 | novel transcript, antisense to TACR1 |
| GAPDHP57 | 2 | glyceraldehyde 3 phosphate dehydrogenase p |
| EVA1A | 2 | eva-1 homolog A, regulator of programmed c |
| AC007099.2 | 2 | novel transcript |
| AC007099.3 | 2 | NULL |
| AC007099.1 | 2 | novel transcript, antisense to EVA1A |
| AC005034.1 | 2 | homeobox B3 (HOXB3) pseudogene |
| U3 | 2 | Small nucleolar RNA U3 [Source:RFAM;Acc:RF |
| MRPL19 | 2 | mitochondrial ribosomal protein L19 [Source: |

| | | |
|---|---|---|
| SUPT4H1P1 | 2 | SPT4 homolog, DSIF elongation factor subunit |
| GCFC2 | 2 | GC-rich sequence DNA-binding factor 2 [Sourc |
| AC005034.3 | 2 | novel transcript, antisense to C2orf3 |
| AC005034.5 | 2 | novel transcript, sense intronic to MRPL19 |
| AC005034.2 | 2 | novel transcript, antisense to C2orf3 |
| AC005034.6 | 2 | novel transcript, antisense to GCFC2 |
| AC005034.4 | 2 | novel transcript |
| AC110614.1 | 2 | novel transcript |
| SUCLA2P2 | 2 | succinate-CoA ligase ADP-forming beta subun |
| AC073091.3 | 2 | novel transcript |
| AC073091.4 | 2 | novel transcript |
| AC073091.1 | 2 | nucleoside phosphorylase (NP) pseudogene |
| AC073091.2 | 2 | ubiquitin specific peptidase 21 (USP21) pseud |
| RN7SKP203 | 2 | RN7SK pseudogene 203 [Source:HGNC Symbo |
| RN7SKP164 | 2 | RN7SK pseudogene 164 [Source:HGNC Symbo |
| AC068616.2 | 2 | novel transcript |
| AC068616.3 | 2 | novel transcript |
| AC068616.1 | 2 | novel transcript |
| LRRTM4 | 2 | leucine rich repeat transmembrane neuronal |
| RNA5SP98 | 2 | RNA, 5S ribosomal pseudogene 98 [Source:HG |
| AC079117.2 | 2 | novel transcript, sense intronic to LRRTM4 |
| AC079117.1 | 2 | novel transcript |
| AC013716.1 | 2 | novel transcript, sense intronic to LRRTM4 |
| AC012494.1 | 2 | novel transcript |
| AC084149.1 | 2 | novel transcript |
| AC105399.1 | 2 | ribosomal protein L38 (PRL38) pseudogene |
| LINC01851 | 2 | long intergenic non-protein coding RNA 1851 |
| AC012494.2 | 2 | novel transcript |
| CYCSP6 | 2 | CYCS pseudogene 6 [Source:HGNC Symbol;Ac |
| AC064872.1 | 2 | ClpX caseinolytic peptidase X homolog (E. coli |
| AC092660.1 | 2 | novel transcript |
| AC092604.1 | 2 | novel transcript |
| RNU6-827P | 2 | RNA, U6 small nuclear 827, pseudogene [Sour |
| RNU6-812P | 2 | RNA, U6 small nuclear 812, pseudogene [Sour |
| REG3G | 2 | regenerating family member 3 gamma [Sourc |
| REG1B | 2 | regenerating family member 1 beta [Source:H |
| REG1A | 2 | regenerating family member 1 alpha [Source: |
| REG1CP | 2 | regenerating family member 1 gamma, pseud |
| REG3A | 2 | regenerating family member 3 alpha [Source: |
| AC011754.1 | 2 | novel transcript |
| CTNNA2 | 2 | catenin alpha 2 [Source:HGNC Symbol;Acc:HG |
| AC011754.2 | 2 | novel transcript, antisense to CTNNA2 |
| AC010975.1 | 2 | novel transcript |
| AC010975.3 | 2 | novel zinc finger protein pseudogene |
| AC010975.2 | 2 | guanine nucleotide binding protein (G protein |
| RNU6-561P | 2 | RNA, U6 small nuclear 561, pseudogene [Sour |
| MIR4264 | 2 | microRNA 4264 [Source:HGNC Symbol;Acc:HG |
| MIR8080 | 2 | microRNA 8080 [Source:HGNC Symbol;Acc:HG |
| AC016716.2 | 2 | novel transcript |
| AC016716.1 | 2 | RNA binding motif protein 7 (RBM7) pseudoge |

| | | |
|---|---|---|
| LRRTM1 | 2 | leucine rich repeat transmembrane neuronal |
| AC008067.1 | 2 | novel transcript |
| AC012355.1 | 2 | novel transcript |
| AC105342.1 | 2 | ankyrin repeat domain 11 (ANKRD11) pseudo |
| AC012075.1 | 2 | chromatin modifying protein 4A (CHMP4A) ps |
| LINC01815 | 2 | long intergenic non-protein coding RNA 1815 |
| RNA5SP99 | 2 | RNA, 5S ribosomal pseudogene 99 [Source:HG |
| AC013262.1 | 2 | mitochondrially encoded cytochrome c oxidas |
| RN7SL201P | 2 | RNA, 7SL, cytoplasmic 201, pseudogene [Sour |
| AC079896.1 | 2 | novel transcript |
| LYARP1 | 2 | Ly1 antibody reactive pseudogene 1 [Source:H |
| RNU6-685P | 2 | RNA, U6 small nuclear 685, pseudogene [Sour |
| Y_RNA | 2 | Y RNA [Source:RFAM;Acc:RF00019] |
| AC010105.1 | 2 | novel transcript |
| AC109638.1 | 2 | ring-box 1 (RBX1) pseudogene |
| MTND4P25 | 2 | MT-ND4 pseudogene 25 [Source:HGNC Symbo |
| MTND5P27 | 2 | MT-ND5 pseudogene 27 [Source:HGNC Symbo |
| MTND6P7 | 2 | MT-ND6 pseudogene 7 [Source:HGNC Symbol |
| MTCYBP7 | 2 | MT-CYB pseudogene 7 [Source:HGNC Symbol; |
| AC098817.1 | 2 | novel transcript |
| DHFRP3 | 2 | dihydrofolate reductase pseudogene 3 [Sourc |
| AC138623.1 | 2 | zinc finger protein 141 (ZNF141) pseudogene |
| LINC01809 | 2 | long intergenic non-protein coding RNA 1809 |
| RPL37P10 | 2 | ribosomal protein L37 pseudogene 10 [Source |
| RNU6-1312P | 2 | RNA, U6 small nuclear 1312, pseudogene [Sou |
| AC016908.1 | 2 | cytokine receptor-like factor 3 (CRLF3) pseudo |
| ST6GALNAC2P1 | 2 | ST6 N-acetylgalactosaminide alpha-2,6-sialyltr |
| FUNDC2P2 | 2 | FUN14 domain containing 2 pseudogene 2 [So |
| AC106874.1 | 2 | novel transcript |
| SUCLG1 | 2 | succinate-CoA ligase alpha subunit [Source:HG |
| DNAH6 | 2 | dynein axonemal heavy chain 6 [Source:HGNC |
| DUXAP1 | 2 | double homeobox A pseudogene 1 [Source:HG |
| LDHAP7 | 2 | lactate dehydrogenase A pseudogene 7 [Sourc |
| TRABD2A | 2 | TraB domain containing 2A [Source:HGNC Syn |
| AC010087.1 | 2 | family with sequence similarity 71, member B |
| RPL12P18 | 2 | ribosomal protein L12 pseudogene 18 [Source |
| TMSB10 | 2 | thymosin beta 10 [Source:HGNC Symbol;Acc:H |
| AC022210.1 | 2 | ribosomal protein S2 (RPS2) pseudogene |
| AC022210.2 | 2 | novel transcript |
| KCMF1 | 2 | potassium channel modulatory factor 1 [Sourc |
| LINC01964 | 2 | long intergenic non-protein coding RNA 1964 |
| AC078974.1 | 2 | Cytochrome c oxidase subunit 1 (COX1) pseud |
| LSM3P3 | 2 | LSM3 homolog, U6 small nuclear RNA and mR |
| TCF7L1 | 2 | transcription factor 7 like 1 [Source:HGNC Syn |
| TCF7L1-IT1 | 2 | TCF7L1 intronic transcript 1 [Source:HGNC Syr |
| RNU6-674P | 2 | RNA, U6 small nuclear 674, pseudogene [Sour |
| AC093162.1 | 2 | small nuclear ribonucleoprotein polypeptide E |
| AC093162.2 | 2 | retinol saturase (all-trans-retinol 13,14-reduct |
| TGOLN2 | 2 | trans-golgi network protein 2 [Source:HGNC S |
| Y_RNA | 2 | Y RNA [Source:RFAM;Acc:RF00019] |

| | | |
|---|---|---|
| PEBP1P2 | 2 | phosphatidylethanolamine binding protein 1 p |
| RETSAT | 2 | retinol saturase [Source:HGNC Symbol;Acc:HG |
| ELMOD3 | 2 | ELMO domain containing 3 [Source:HGNC Sym |
| AC062037.1 | 2 | NADH dehydrogenase (ubiquinone) 1 beta sub |
| Y_RNA | 2 | Y RNA [Source:RFAM;Acc:RF00019] |
| RN7SL113P | 2 | RNA, 7SL, cytoplasmic 113, pseudogene [Sour |
| AC062037.2 | 2 | novel transcript, antisense to ELMOD3 |
| CAPG | 2 | capping actin protein, gelsolin like [Source:HG |
| AC062037.3 | 2 | novel transcript |
| SH2D6 | 2 | SH2 domain containing 6 [Source:HGNC Symb |
| Y_RNA | 2 | Y RNA [Source:RFAM;Acc:RF00019] |
| RN7SL251P | 2 | RNA, 7SL, cytoplasmic 251, pseudogene [Sour |
| Y_RNA | 2 | Y RNA [Source:RFAM;Acc:RF00019] |
| RPSAP22 | 2 | ribosomal protein SA pseudogene 22 [Source: |
| PARTICL | 2 | promoter of MAT2A antisense radiation-induc |
| MAT2A | 2 | methionine adenosyltransferase 2A [Source:H |
| GGCX | 2 | gamma-glutamyl carboxylase [Source:HGNC S |
| VAMP8 | 2 | vesicle associated membrane protein 8 [Sourc |
| RN7SL126P | 2 | RNA, 7SL, cytoplasmic 126, pseudogene [Sour |
| VAMP5 | 2 | vesicle associated membrane protein 5 [Sourc |
| RNF181 | 2 | ring finger protein 181 [Source:HGNC Symbol; |
| TMEM150A | 2 | transmembrane protein 150A [Source:HGNC S |
| USP39 | 2 | ubiquitin specific peptidase 39 [Source:HGNC |
| C2orf68 | 2 | chromosome 2 open reading frame 68 [Source |
| SFTPB | 2 | surfactant protein B [Source:HGNC Symbol;Ac |
| GNLY | 2 | granulysin [Source:HGNC Symbol;Acc:HGNC:4 |
| AC012454.1 | 2 | novel pseudogene |
| RNU1-38P | 2 | RNA, U1 small nuclear 38, pseudogene [Sourc |
| ATOH8 | 2 | atonal bHLH transcription factor 8 [Source:HG |
| MIR6071 | 2 | microRNA 6071 [Source:HGNC Symbol;Acc:HG |
| AC105053.1 | 2 | novel transcript |
| RN7SKP83 | 2 | RN7SK pseudogene 83 [Source:HGNC Symbol; |
| ST3GAL5 | 2 | ST3 beta-galactoside alpha-2,3-sialyltransfera |
| ST3GAL5-AS1 | 2 | ST3GAL5 antisense RNA 1 (head to head) [Sou |
| AC012511.1 | 2 | novel transcript |
| AC012511.2 | 2 | novel transcript |
| POLR1A | 2 | RNA polymerase I subunit A [Source:HGNC Sy |
| PTCD3 | 2 | pentatricopeptide repeat domain 3 [Source:H |
| SNORD94 | 2 | small nucleolar RNA, C/D box 94 [Source:HGN |
| IMMT | 2 | inner membrane mitochondrial protein [Sourc |
| Y_RNA | 2 | Y RNA [Source:RFAM;Acc:RF00019] |
| MIR4779 | 2 | microRNA 4779 [Source:HGNC Symbol;Acc:HG |
| AC009309.1 | 2 | novel transcript, antisense to IMMT |
| MRPL35 | 2 | mitochondrial ribosomal protein L35 [Source: |
| REEP1 | 2 | receptor accessory protein 1 [Source:HGNC Sy |
| U8 | 2 | U8 small nucleolar RNA [Source:RFAM;Acc:RF |
| AC009408.1 | 2 | NULL |
| KDM3A | 2 | lysine demethylase 3A [Source:HGNC Symbol; |
| CHMP3 | 2 | charged multivesicular body protein 3 [Source |
| RNF103-CHMP3 | 2 | RNF103-CHMP3 readthrough [Source:HGNC S |

| | | |
|---|---|---|
| RNU6-640P | 2 | RNA, U6 small nuclear 640, pseudogene [Sour |
| AC015971.1 | 2 | novel transcript |
| RNF103 | 2 | ring finger protein 103 [Source:HGNC Symbol; |
| RMND5A | 2 | required for meiotic nuclear division 5 homolo |
| CD8A | 2 | CD8a molecule [Source:HGNC Symbol;Acc:HG |
| CD8B | 2 | CD8b molecule [Source:HGNC Symbol;Acc:HG |
| ANAPC1P1 | 2 | anaphase promoting complex subunit 1 pseuc |
| AC111200.1 | 2 | ribosomal protein S26 (RPS26) pseudogene |
| RGPD1 | 2 | RANBP2 like and GRIP domain containing 1 [S |
| AC111200.2 | 2 | WW domain binding protein 1 (WBP1) pseudo |
| NDUFB4P5 | 2 | NADH:ubiquinone oxidoreductase subunit B4 |
| PLGLB1 | 2 | plasminogen like B1 [Source:HGNC Symbol;Ac |
| AC125232.2 | 2 | novel transcript |
| AC125232.1 | 2 | anaphase promoting complex subunit 1 (ANAI |
| AC092651.2 | 2 | required for meiotic nuclear division 5 homolo |
| AC092651.3 | 2 | novel transcript |
| AC092651.1 | 2 | anaphase promoting complex subunit 1 (ANAI |
| AC083899.1 | 2 | anaphase promoting complex subunit 1 (ANAI |
| MIR4771-1 | 2 | microRNA 4771-1 [Source:HGNC Symbol;Acc:I |
| AC083899.2 | 2 | centromere protein N (CENPN) pseudogene |
| LINC01955 | 2 | long intergenic non-protein coding RNA 1955 |
| AC083899.3 | 2 | DBF4 homolog (S. cerevisiae) (DBF4) pseudog |
| AC068279.2 | 2 | novel transcript |
| IGKV3OR2-268 | 2 | immunoglobulin kappa variable 3/OR2-268 (n |
| AC068279.1 | 2 | novel pseudogene |
| LINC01943 | 2 | long intergenic non-protein coding RNA 1943 |
| CYTOR | 2 | cytoskeleton regulator RNA [Source:HGNC Syr |
| AC133644.3 | 2 | novel transcript |
| AC133644.2 | 2 | novel transcript |
| AC133644.1 | 2 | platelet-activating factor acetylhydrolase, isof |
| MIR4435-1 | 2 | microRNA 4435-1 [Source:HGNC Symbol;Acc:I |
| AC133965.1 | 2 | ribosomal protein S14 (RPS14) pseudogene |
| AC093616.1 | 2 | anaphase promoting complex subunit 1 (ANAI |
| PLGLB2 | 2 | plasminogen like B2 [Source:HGNC Symbol;Ac |
| RGPD2 | 2 | RANBP2 like and GRIP domain containing 2 [S |
| MTATP8P2 | 2 | MT-ATP8 pseudogene 2 [Source:HGNC Symbo |
| NDUFB4P7 | 2 | NADH:ubiquinone oxidoreductase subunit B4 |
| AC108479.1 | 2 | WW domain binding protein 1 (WBP1) pseudo |
| AC108479.2 | 2 | anaphase promoting complex subunit 1 (ANAI |
| AC108479.4 | 2 | lysine-rich coiled-coil 1 pseudogene (KRCC1) |
| RNU2-63P | 2 | RNA, U2 small nuclear 63, pseudogene [Sourc |
| AC108479.3 | 2 | novel transcript |
| KRCC1 | 2 | lysine rich coiled-coil 1 [Source:HGNC Symbol; |
| SMYD1 | 2 | SET and MYND domain containing 1 [Source:H |
| MIR4780 | 2 | microRNA 4780 [Source:HGNC Symbol;Acc:HG |
| FABP1 | 2 | fatty acid binding protein 1 [Source:HGNC Syn |
| THNSL2 | 2 | threonine synthase like 2 [Source:HGNC Symb |
| MRPL45P1 | 2 | mitochondrial ribosomal protein L45 pseudog |
| U6 | 2 | U6 spliceosomal RNA [Source:RFAM;Acc:RF00 |
| RNU6-1168P | 2 | RNA, U6 small nuclear 1168, pseudogene [Sou |

| | | |
|---|---|---|
| RNU6-1007P | 2 | RNA, U6 small nuclear 1007, pseudogene [Sou |
| AC012671.1 | 2 | ribosomal protein L38 (RPL38) pseudogene |
| FOXI3 | 2 | forkhead box I3 [Source:HGNC Symbol;Acc:HG |
| TEX37 | 2 | testis expressed 37 [Source:HGNC Symbol;Acc |
| AC104134.1 | 2 | novel transcript |
| EIF2AK3 | 2 | eukaryotic translation initiation factor 2 alpha |
| EIF2AK3-DT | 2 | EIF2AK3 divergent transcript [Source:HGNC Sy |
| AC062029.1 | 2 | novel transcript |
| RPIA | 2 | ribose 5-phosphate isomerase A [Source:HGN |
| ANKRD36BP2 | 2 | ankyrin repeat domain 36B pseudogene 2 [Sou |
| AC244205.1 | 2 | novel transcript |
| AC244205.2 | 2 | mal, T-cell differentiation protein-like (MALL) |
| MIR4436A | 2 | microRNA 4436a [Source:HGNC Symbol;Acc:H |
| IGKC | 2 | immunoglobulin kappa constant [Source:HGN |
| IGKJ5 | 2 | immunoglobulin kappa joining 5 [Source:HGN |
| IGKJ4 | 2 | immunoglobulin kappa joining 4 [Source:HGN |
| IGKJ3 | 2 | immunoglobulin kappa joining 3 [Source:HGN |
| IGKJ2 | 2 | immunoglobulin kappa joining 2 [Source:HGN |
| IGKJ1 | 2 | immunoglobulin kappa joining 1 [Source:HGN |
| IGKV4-1 | 2 | immunoglobulin kappa variable 4-1 [Source:H |
| IGKV5-2 | 2 | immunoglobulin kappa variable 5-2 [Source:H |
| IGKV7-3 | 2 | immunoglobulin kappa variable 7-3 (pseudoge |
| PGBD4P5 | 2 | piggyBac transposable element derived 4 pseu |
| IGKV2-4 | 2 | immunoglobulin kappa variable 2-4 (pseudoge |
| IGKV1-5 | 2 | immunoglobulin kappa variable 1-5 [Source:H |
| IGKV1-6 | 2 | immunoglobulin kappa variable 1-6 [Source:H |
| IGKV3-7 | 2 | immunoglobulin kappa variable 3-7 (non-func |
| IGKV1-8 | 2 | immunoglobulin kappa variable 1-8 [Source:H |
| IGKV1-9 | 2 | immunoglobulin kappa variable 1-9 [Source:H |
| IGKV2-10 | 2 | immunoglobulin kappa variable 2-10 (pseudog |
| IGKV3-11 | 2 | immunoglobulin kappa variable 3-11 [Source:I |
| IGKV1-12 | 2 | immunoglobulin kappa variable 1-12 [Source:I |
| IGKV1-13 | 2 | immunoglobulin kappa variable 1-13 (gene/ps |
| IGKV2-14 | 2 | immunoglobulin kappa variable 2-14 (pseudog |
| IGKV3-15 | 2 | immunoglobulin kappa variable 3-15 [Source:I |
| IGKV1-16 | 2 | immunoglobulin kappa variable 1-16 [Source:I |
| IGKV1-17 | 2 | immunoglobulin kappa variable 1-17 [Source:I |
| IGKV2-18 | 2 | immunoglobulin kappa variable 2-18 (pseudog |
| IGKV2-19 | 2 | immunoglobulin kappa variable 2-19 (pseudog |
| IGKV3-20 | 2 | immunoglobulin kappa variable 3-20 [Source:I |
| IGKV6-21 | 2 | immunoglobulin kappa variable 6-21 (non-fun |
| IGKV1-22 | 2 | immunoglobulin kappa variable 1-22 (pseudog |
| IGKV2-23 | 2 | immunoglobulin kappa variable 2-23 (pseudog |
| IGKV2-24 | 2 | immunoglobulin kappa variable 2-24 [Source:I |
| IGKV3-25 | 2 | immunoglobulin kappa variable 3-25 (pseudog |
| IGKV2-26 | 2 | immunoglobulin kappa variable 2-26 (pseudog |
| AC244255.1 | 2 | Ig kappa chain V-I region Lay pseudogene |
| IGKV1-27 | 2 | immunoglobulin kappa variable 1-27 [Source:I |
| IGKV2-28 | 2 | immunoglobulin kappa variable 2-28 [Source:I |
| IGKV2-29 | 2 | immunoglobulin kappa variable 2-29 (gene/ps |

| | | |
|---|---|---|
| IGKV2-30 | 2 | immunoglobulin kappa variable 2-30 [Source:I |
| IGKV3-31 | 2 | immunoglobulin kappa variable 3-31 (pseudog |
| IGKV1-32 | 2 | immunoglobulin kappa variable 1-32 (pseudog |
| IGKV1-33 | 2 | immunoglobulin kappa variable 1-33 [Source:I |
| IGKV3-34 | 2 | immunoglobulin kappa variable 3-34 (pseudog |
| IGKV1-35 | 2 | immunoglobulin kappa variable 1-35 (pseudog |
| IGKV2-36 | 2 | immunoglobulin kappa variable 2-36 (pseudog |
| IGKV1-37 | 2 | immunoglobulin kappa variable 1-37 (non-fun |
| IGKV2-38 | 2 | immunoglobulin kappa variable 2-38 (pseudog |
| IGKV1-39 | 2 | immunoglobulin kappa variable 1-39 (gene/ps |
| IGKV2-40 | 2 | immunoglobulin kappa variable 2-40 [Source:I |
| AC006453.4 | 2 | otopetrin 1 (OTPO1) pseudogene |
| AC006453.2 | 2 | lymphocyte-specific protein 1 (LSP1) pseudog |
| 5S_rRNA | 2 | 5S ribosomal RNA [Source:RFAM;Acc:RF00001 |
| AC006453.1 | 2 | lysine (K)-specific methyltransferase 2C (KMT2 |
| AC006453.3 | 2 | immunoglobulin superfamily member 3 (IGSF3 |
| AC006453.5 | 2 | novel solute carrier family 9, subfamily B (NH |
| IGKV2D-40 | 2 | immunoglobulin kappa variable 2D-40 [Source |
| IGKV1D-39 | 2 | immunoglobulin kappa variable 1D-39 [Source |
| IGKV2D-38 | 2 | immunoglobulin kappa variable 2D-38 (pseud |
| IGKV1D-37 | 2 | immunoglobulin kappa variable 1D-37 (non-fu |
| IGKV2D-36 | 2 | immunoglobulin kappa variable 2D-36 (pseud |
| IGKV1D-35 | 2 | immunoglobulin kappa variable 1D-35 (pseud |
| IGKV3D-34 | 2 | immunoglobulin kappa variable 3D-34 (pseud |
| IGKV1D-33 | 2 | immunoglobulin kappa variable 1D-33 [Source |
| IGKV1D-32 | 2 | immunoglobulin kappa variable 1D-32 (pseud |
| IGKV3D-31 | 2 | immunoglobulin kappa variable 3D-31 (pseud |
| IGKV2D-30 | 2 | immunoglobulin kappa variable 2D-30 [Source |
| IGKV2D-29 | 2 | immunoglobulin kappa variable 2D-29 [Source |
| IGKV2D-28 | 2 | immunoglobulin kappa variable 2D-28 [Source |
| IGKV1D-27 | 2 | immunoglobulin kappa variable 1D-27 (pseud |
| IGKV2D-26 | 2 | immunoglobulin kappa variable 2D-26 [Source |
| IGKV3D-25 | 2 | immunoglobulin kappa variable 3D-25 (pseud |
| IGKV2D-24 | 2 | immunoglobulin kappa variable 2D-24 (non-fu |
| IGKV2D-23 | 2 | immunoglobulin kappa variable 2D-23 (pseud |
| IGKV1D-22 | 2 | immunoglobulin kappa variable 1D-22 (pseud |
| IGKV6D-21 | 2 | immunoglobulin kappa variable 6D-21 (non-fu |
| IGKV3D-20 | 2 | immunoglobulin kappa variable 3D-20 [Source |
| IGKV2D-19 | 2 | immunoglobulin kappa variable 2D-19 (pseud |
| IGKV2D-18 | 2 | immunoglobulin kappa variable 2D-18 (pseud |
| IGKV6D-41 | 2 | immunoglobulin kappa variable 6D-41 (non-fu |
| IGKV1D-17 | 2 | immunoglobulin kappa variable 1D-17 [Source |
| IGKV1D-16 | 2 | immunoglobulin kappa variable 1D-16 [Source |
| IGKV3D-15 | 2 | immunoglobulin kappa variable 3D-15 (gene/p |
| IGKV2D-14 | 2 | immunoglobulin kappa variable 2D-14 (pseud |
| IGKV1D-13 | 2 | immunoglobulin kappa variable 1D-13 [Source |
| IGKV1D-12 | 2 | immunoglobulin kappa variable 1D-12 [Source |
| IGKV3D-11 | 2 | immunoglobulin kappa variable 3D-11 [Source |
| IGKV2D-10 | 2 | immunoglobulin kappa variable 2D-10 (pseud |
| IGKV1D-42 | 2 | immunoglobulin kappa variable 1D-42 (non-fu |

| | | |
|---|---|---|
| IGKV1D-43 | 2 | immunoglobulin kappa variable 1D-43 [Source |
| IGKV1D-8 | 2 | immunoglobulin kappa variable 1D-8 [Source: |
| IGKV3D-7 | 2 | immunoglobulin kappa variable 3D-7 [Source: |
| AC233263.2 | 2 | striatin, calmodulin binding protein (STRN) ps |
| IGKV1OR2-118 | 2 | immunoglobulin kappa variable 1/OR2-118 (p |
| AC233263.1 | 2 | pseudogene similar to part of kinase suppress |
| AC233263.6 | 2 | novel transcript |
| AC233266.1 | 2 | pseudogene similar to part of transmembrane |
| AC233266.2 | 2 | novel transcript |
| AC116050.1 | 2 | otopetrin 1 (OTOP1) pseudogene |
| LSP1P4 | 2 | LSP1 pseudogene 4 [Source:HGNC Symbol;Acc |
| RNA5SP100 | 2 | RNA, 5S ribosomal pseudogene 100 [Source:H |
| DRD5P1 | 2 | dopamine receptor D5 pseudogene 1 [Source: |
| AC027612.2 | 2 | pseudogene similar to part of myeloid/lymph |
| AC027612.4 | 2 | Na+/K+ transporting ATPase interacting 1 (NK |
| AC027612.1 | 2 | pseudogene similar to part of immunoglobulir |
| AC027612.3 | 2 | SET domain containing (lysine methyltransfer |
| AC027612.5 | 2 | novel transcript |
| GGT8P | 2 | gamma-glutamyltransferase 8 pseudogene [S |
| IGKV1OR2-1 | 2 | immunoglobulin kappa variable 1/OR2-1 (pse |
| ABCD1P5 | 2 | ATP binding cassette subfamily D member 1 p |
| AC127391.1 | 2 | pseudogene similar to part of CDGSH iron sulf |
| PABPC1P6 | 2 | poly(A) binding protein cytoplasmic 1 pseudog |
| SLC9B1P2 | 2 | solute carrier family 9 member B1 pseudogen |
| ACTR3BP2 | 2 | ACTR3B pseudogene 2 [Source:HGNC Symbol; |
| CHEK2P3 | 2 | checkpoint kinase 2 pseudogene 3 [Source:HG |
| AC128677.1 | 2 | pseudogene similar to part of Tripartite motif- |
| IGKV1OR2-2 | 2 | immunoglobulin kappa variable 1/OR2-2 (pse |
| AC026273.1 | 2 | BMS1, ribosome biogenesis factor (BMS1) pse |
| AL845331.3 | 2 | methylenetetrahydrofolate dehydrogenase (N |
| AL845331.2 | 2 | putative aquaporin-7-like protein 3 [Source:N |
| CNN2P11 | 2 | calponin 2 pseudogene 11 [Source:HGNC Sym |
| AL845331.1 | 2 | glycosyltransferase 8 domain containing 3 (GL |
| CNN2P8 | 2 | calponin 2 pseudogene 8 [Source:HGNC Symb |
| AC073464.2 | 2 | novel zinc finger protein pseudogene |
| CYP4F32P | 2 | cytochrome P450 family 4 subfamily F membe |
| AC073464.1 | 2 | novel pseudogene |
| SNX18P14 | 2 | sorting nexin 18 pseudogene 14 [Source:HGN |
| ANKRD20A8P | 2 | ankyrin repeat domain 20 family member A8, |
| RNU6-1320P | 2 | RNA, U6 small nuclear 1320, pseudogene [Sou |
| AC097374.2 | 2 | ankyrin repeat domain 57 pseudogene |
| AC097374.1 | 2 | pseudogene similar to part of asparagine synt |
| TEKT4 | 2 | tektin 4 [Source:HGNC Symbol;Acc:HGNC:310 |
| MTCO1P48 | 2 | MT-CO1 pseudogene 48 [Source:HGNC Symbo |
| MTCO3P45 | 2 | MT-CO3 pseudogene 45 [Source:HGNC Symbo |
| RPS24P6 | 2 | ribosomal protein S24 pseudogene 6 [Source: |
| AC103563.3 | 2 | immunoglobulin lambda variable group pseud |
| AC103563.4 | 2 | immunoglobulin heavy variable group pseudo |
| AC103563.5 | 2 | immunoglobulin heavy variable group pseudo |
| AC103563.1 | 2 | immunoglobulin heavy variable group pseudo |

| | | |
|---|---|---|
| AC103563.6 | 2 | pseudogene similar to part of V-set and immu |
| AC103563.8 | 2 | pseudogene similar to part of mal, T-cell differ |
| RN7SL575P | 2 | RNA, 7SL, cytoplasmic 575, pseudogene [Sour |
| AC103563.7 | 2 | novel transcript |
| MAL | 2 | mal, T cell differentiation protein [Source:HGN |
| AC103563.2 | 2 | putative novel transcript |
| AC103563.9 | 2 | novel transcript |
| MRPS5 | 2 | mitochondrial ribosomal protein S5 [Source:H |
| ZNF514 | 2 | zinc finger protein 514 [Source:HGNC Symbol; |
| ZNF2 | 2 | zinc finger protein 2 [Source:HGNC Symbol;Ac |
| SLC2AXP1 | 2 | solute carrier family 2 pseudogene 1 [Source:I |
| AC092835.1 | 2 | novel C2H2 type zinc finger protein |
| PROM2 | 2 | prominin 2 [Source:HGNC Symbol;Acc:HGNC:2 |
| KCNIP3 | 2 | potassium voltage-gated channel interacting p |
| FABP7P2 | 2 | fatty acid binding protein 7 pseudogene 2 [Sou |
| FAHD2A | 2 | fumarylacetoacetate hydrolase domain conta |
| AC009238.3 | 2 | novel pseudogene |
| AC009238.2 | 2 | pseudogene similar to part of Cdon homolog ( |
| AC009238.1 | 2 | pseudogene similar to part of Cdon homolog ( |
| UBTFL5 | 2 | UBTF like 5 (pseudogene) [Source:HGNC Symb |
| AC133104.2 | 2 | family with sequence similarity 178, member |
| AC133104.1 | 2 | pseudogene similar to part of a tripartite moti |
| TRIM43B | 2 | tripartite motif containing 43B [Source:HGNC |
| AC009237.11 | 2 | pseudogene similar to part of tripartite motif- |
| AC009237.1 | 2 | pseudogene similar to part of tripartite motif- |
| AC009237.10 | 2 | pseudogene similar to part of tripartite motif- |
| AC009237.2 | 2 | pseudogene similar to part of folate hydrolase |
| TRIM64FP | 2 | tripartite motif containing 64F, pseudogene [S |
| AC009237.14 | 2 | novel transcript |
| AC009237.3 | 2 | Ankyrin repeat domain-containing protein pse |
| AC009237.8 | 2 | pseudogene similar to part of tripartite motif- |
| AC009237.15 | 2 | novel transcript |
| AC009237.5 | 2 | pseudogene similar to part of tripartite motif- |
| OR7E102P | 2 | olfactory receptor family 7 subfamily E memb |
| TRIM51JP | 2 | tripartite motif-containing 51J, pseudogene [S |
| AC009237.16 | 2 | tripartite motif-containing 51B, pseudogene |
| AC009237.4 | 2 | pseudogene similar to part of tripartite motif- |
| TRIM43 | 2 | tripartite motif containing 43 [Source:HGNC S |
| AC009237.7 | 2 | pseudogene similar to part of tripartite motif- |
| AC009237.13 | 2 | family with sequence similarity 178, member |
| AC009237.12 | 2 | cAMP responsive element binding protein 1 (C |
| UBTFL3 | 2 | UBTF like 3 (pseudogene) [Source:HGNC Symb |
| AC009237.17 | 2 | tripartite motif containing 64C (TRIM64C) pse |
| AC009237.6 | 2 | pseudogene similar to part of Cdon homolog ( |
| AC009237.9 | 2 | pseudogene similar to part of Cdon homolog ( |
| AC008268.1 | 2 | novel transcript |
| GPAT2P1 | 2 | glycerol-3-phosphate acyltransferase 2 pseud |
| LINC00342 | 2 | long intergenic non-protein coding RNA 342 [S |
| ANKRD36C | 2 | ankyrin repeat domain 36C [Source:HGNC Syn |
| RN7SL210P | 2 | RNA, 7SL, cytoplasmic 210, pseudogene [Sour |

| | | |
|---|---|---|
| FAHD2CP | 2 | fumarylacetoacetate hydrolase domain conta |
| GPAT2 | 2 | glycerol-3-phosphate acyltransferase 2, mitoc |
| ADRA2B | 2 | adrenoceptor alpha 2B [Source:HGNC Symbol |
| ASTL | 2 | astacin like metalloendopeptidase [Source:HG |
| DUSP2 | 2 | dual specificity phosphatase 2 [Source:HGNC S |
| AC012307.1 | 2 | novel transcript |
| STARD7 | 2 | StAR related lipid transfer domain containing |
| STARD7-AS1 | 2 | STARD7 antisense RNA 1 [Source:HGNC Symb |
| TMEM127 | 2 | transmembrane protein 127 [Source:HGNC Sy |
| CIAO1 | 2 | cytosolic iron-sulfur assembly component 1 [S |
| SNRNP200 | 2 | small nuclear ribonucleoprotein U5 subunit 20 |
| AC021188.1 | 2 | novel transcript |
| ITPRIPL1 | 2 | ITPRIP like 1 [Source:HGNC Symbol;Acc:HGNC |
| NCAPH | 2 | non-SMC condensin I complex subunit H [Sour |
| NEURL3 | 2 | neuralized E3 ubiquitin protein ligase 3 [Sourc |
| AC013270.1 | 2 | novel transcript |
| ARID5A | 2 | AT-rich interaction domain 5A [Source:HGNC S |
| KANSL3 | 2 | KAT8 regulatory NSL complex subunit 3 [Sourc |
| FER1L5 | 2 | fer-1 like family member 5 [Source:HGNC Sym |
| LMAN2L | 2 | lectin, mannose binding 2 like [Source:HGNC S |
| CNNM4 | 2 | cyclin and CBS domain divalent metal cation t |
| MIR3127 | 2 | microRNA 3127 [Source:HGNC Symbol;Acc:HG |
| CNNM3-DT | 2 | CNNM3 divergent transcript [Source:HGNC Sy |
| CNNM3 | 2 | cyclin and CBS domain divalent metal cation t |
| ANKRD23 | 2 | ankyrin repeat domain 23 [Source:HGNC Syml |
| ANKRD39 | 2 | ankyrin repeat domain 39 [Source:HGNC Syml |
| SEMA4C | 2 | semaphorin 4C [Source:HGNC Symbol;Acc:HG |
| FAM178B | 2 | family with sequence similarity 178 member B |
| AC092636.1 | 2 | novel transcript, antisense to FAM178B |
| Metazoa_SRP | 2 | Metazoan signal recognition particle RNA [Sou |
| RNA5SP101 | 2 | RNA, 5S ribosomal pseudogene 101 [Source:H |
| AC079395.3 | 2 | ribosomal L24 domain containing 1 (RSL24D1) |
| AC079395.2 | 2 | cAMP responsive element binding protein 1 ( |
| AC079395.1 | 2 | pseudogene similar to part of a tripartite moti |
| TRIM43CP | 2 | tripartite motif containing 43C, pseudogene [S |
| AC018892.1 | 2 | pseudogene similar to part of a tripartite moti |
| AC018892.2 | 2 | pseudogene similar to part of a tripartite moti |
| IGKV2OR2-1 | 2 | immunoglobulin kappa variable 2/OR2-1 (pseu |
| IGKV2OR2-2 | 2 | immunoglobulin kappa variable 2/OR2-2 (pseu |
| IGKV1OR2-3 | 2 | immunoglobulin kappa variable 1/OR2-3 (pseu |
| GPAT2P2 | 2 | glycerol-3-phosphate acyltransferase 2 pseud |
| FAHD2B | 2 | fumarylacetoacetate hydrolase domain conta |
| AC018892.3 | 2 | TEC |
| RN7SL313P | 2 | RNA, 7SL, cytoplasmic 313, pseudogene [Sour |
| ANKRD36 | 2 | ankyrin repeat domain 36 [Source:HGNC Syml |
| AC159540.2 | 2 | novel transcript |
| AC159540.1 | 2 | novel transcript |
| IGKV1OR2-9 | 2 | immunoglobulin kappa variable 1/OR2-9 (pseu |
| IGKV2OR2-10 | 2 | immunoglobulin kappa variable 2/OR2-10 (ps |
| IGKV2OR2-7D | 2 | immunoglobulin kappa variable 2/OR2-7D (ps |

| | | |
|---|---|---|
| IGKV3OR2-5 | 2 | immunoglobulin kappa variable 3/OR2-5 (pseu |
| IGKV1OR2-6 | 2 | immunoglobulin kappa variable 1/OR2-6 (pseu |
| IGKV2OR2-7 | 2 | immunoglobulin kappa variable 2/OR2-7 (pseu |
| IGKV2OR2-8 | 2 | immunoglobulin kappa variable 2/OR2-8 (pseu |
| IGKV1OR2-11 | 2 | immunoglobulin kappa variable 1/OR2-11 (ps |
| AC092683.1 | 2 | novel transcript |
| AC092683.2 | 2 | novel transcript |
| ANKRD36B | 2 | ankyrin repeat domain 36B [Source:HGNC Syn |
| AC017099.1 | 2 | coiled-coil domain containing 144C (CCDC144 |
| AC017099.2 | 2 | novel transcript |
| UBTFL6 | 2 | UBTF like 6 (pseudogene) [Source:HGNC Symb |
| COX5B | 2 | cytochrome c oxidase subunit 5B [Source:HGN |
| ACTR1B | 2 | actin related protein 1B [Source:HGNC Symbo |
| C2orf92 | 2 | chromosome 2 open reading frame 92 [Sourc |
| RNU4-8P | 2 | RNA, U4 small nuclear 8, pseudogene [Source |
| ZAP70 | 2 | zeta chain of T cell receptor associated protei |
| TMEM131 | 2 | transmembrane protein 131 [Source:HGNC Sy |
| HMGN1P36 | 2 | high mobility group nucleosome binding doma |
| RNU7-96P | 2 | RNA, U7 small nuclear 96 pseudogene [Source |
| VWA3B | 2 | von Willebrand factor A domain containing 3E |
| ATP5F1BP1 | 2 | ATP synthase F1 subunit beta pseudogene 1 [S |
| AC092675.1 | 2 | novel transcript, antisense to CNGA3 |
| CNGA3 | 2 | cyclic nucleotide gated channel subunit alpha |
| AC092675.2 | 2 | novel transcript |
| INPP4A | 2 | inositol polyphosphate-4-phosphatase type I A |
| COA5 | 2 | cytochrome c oxidase assembly factor 5 [Sour |
| UNC50 | 2 | unc-50 inner nuclear membrane RNA binding |
| MGAT4A | 2 | alpha-1,3-mannosyl-glycoprotein 4-beta-N-ac |
| YWHAQP5 | 2 | YWHAQ pseudogene 5 [Source:HGNC Symbol; |
| LINC02611 | 2 | long intergenic non-protein coding RNA 2611 |
| RNU4-84P | 2 | RNA, U4 small nuclear 84, pseudogene [Sourc |
| KIAA1211L | 2 | KIAA1211 like [Source:HGNC Symbol;Acc:HGN |
| RNU7-46P | 2 | RNA, U7 small nuclear 46 pseudogene [Source |
| TSGA10 | 2 | testis specific 10 [Source:HGNC Symbol;Acc:H |
| AC019097.1 | 2 | structural maintenance of chromosomes 3 (SM |
| C2orf15 | 2 | chromosome 2 open reading frame 15 [Sourc |
| AC079447.1 | 2 | novel protein, C2orf15-MRPL30-novel lincRNA |
| LIPT1 | 2 | lipoyltransferase 1 [Source:HGNC Symbol;Acc: |
| AC092587.1 | 2 | novel LIPT1-MRPL30 readthrough |
| MITD1 | 2 | microtubule interacting and trafficking domai |
| MRPL30 | 2 | mitochondrial ribosomal protein L30 [Source: |
| LYG2 | 2 | lysozyme g2 [Source:HGNC Symbol;Acc:HGNC |
| LYG1 | 2 | lysozyme g1 [Source:HGNC Symbol;Acc:HGNC |
| TXNDC9 | 2 | thioredoxin domain containing 9 [Source:HGN |
| EIF5B | 2 | eukaryotic translation initiation factor 5B [Sou |
| REV1 | 2 | REV1 DNA directed polymerase [Source:HGNC |
| AC018690.1 | 2 | novel transcript |
| AFF3 | 2 | AF4/FMR2 family member 3 [Source:HGNC Sy |
| AC092667.1 | 2 | novel transcript |
| LINC01104 | 2 | long intergenic non-protein coding RNA 1104 |

| | | |
|---|---|---|
| LONRF2 | 2 | LON peptidase N-terminal domain and ring fir |
| AC012493.2 | 2 | cytochrome c, somatic (CYCS) pseudogene |
| AC012493.3 | 2 | ribosomal L24 domain containing 1 (RSL24D1) |
| CHST10 | 2 | carbohydrate sulfotransferase 10 [Source:HGI |
| AC012493.1 | 2 | ralA binding protein 1 (RALBP1) pseudogene |
| NMS | 2 | neuromedin S [Source:HGNC Symbol;Acc:HGN |
| AC068538.1 | 2 | cyclic AMP phosphoprotein, 19 kD (ARPP-19) |
| PDCL3 | 2 | phosducin like 3 [Source:HGNC Symbol;Acc:H( |
| HMGN2P22 | 2 | high mobility group nucleosomal binding dom |
| LINC01849 | 2 | long intergenic non-protein coding RNA 1849 |
| AC092845.1 | 2 | novel transcript |
| NANOGNBP1 | 2 | NANOGNB pseudogene 1 [Source:HGNC Symb |
| LINC01868 | 2 | long intergenic non-protein coding RNA 1868 |
| AC092168.1 | 2 | novel transcript |
| AC092168.3 | 2 | novel transcript |
| NPAS2 | 2 | neuronal PAS domain protein 2 [Source:HGNC |
| AC092168.2 | 2 | novel transcript |
| AC016738.2 | 2 | novel transcript, antisense to NPAS2 |
| AC016738.1 | 2 | novel transcript |
| RPL31 | 2 | ribosomal protein L31 [Source:HGNC Symbol;/ |
| TBC1D8 | 2 | TBC1 domain family member 8 [Source:HGNC |
| BBIP1P1 | 2 | BBSome interacting protein 1 pseudogene 1 [! |
| TBC1D8-AS1 | 2 | TBC1 domain family member 8 antisense RNA |
| CNOT11 | 2 | CCR4-NOT transcription complex subunit 11 [! |
| RNF149 | 2 | ring finger protein 149 [Source:HGNC Symbol; |
| SNORD89 | 2 | small nucleolar RNA, C/D box 89 [Source:HGN |
| MIR5696 | 2 | microRNA 5696 [Source:HGNC Symbol;Acc:HG |
| CREG2 | 2 | cellular repressor of E1A stimulated genes 2 [! |
| AC092570.1 | 2 | succinate dehydrogenase complex, subunit D, |
| RFX8 | 2 | RFX family member 8, lacking RFX DNA bindin |
| LINC01870 | 2 | long intergenic non-protein coding RNA 1870 |
| AC092570.3 | 2 | ribosomal protein S6 (RPS6) pseudogene |
| AC092570.2 | 2 | prolylcarboxypeptidase (angiotensinase C) (PF |
| AC093894.1 | 2 | novel transcript |
| AC093894.2 | 2 | novel transcript |
| MAP4K4 | 2 | mitogen-activated protein kinase kinase kinas |
| LINC01127 | 2 | long intergenic non-protein coding RNA 1127 |
| IL1R2 | 2 | interleukin 1 receptor type 2 [Source:HGNC Sy |
| AC007271.1 | 2 | novel transcript |
| IL1R1 | 2 | interleukin 1 receptor type 1 [Source:HGNC Sy |
| IL1R1-AS1 | 2 | IL1R1 antisense RNA 1 [Source:HGNC Symbol; |
| IL1RL2 | 2 | interleukin 1 receptor like 2 [Source:HGNC Syı |
| FAM183DP | 2 | family with sequence similarity 183 member [ |
| IL1RL1 | 2 | interleukin 1 receptor like 1 [Source:HGNC Syı |
| IL18R1 | 2 | interleukin 18 receptor 1 [Source:HGNC Symb |
| SDR42E1P5 | 2 | short chain dehydrogenase/reductase family 4 |
| IL18RAP | 2 | interleukin 18 receptor accessory protein [Sou |
| MIR4772 | 2 | microRNA 4772 [Source:HGNC Symbol;Acc:HG |
| AC007278.2 | 2 | novel transcript |
| AC007278.1 | 2 | novel transcript |

| | | |
|---|---|---|
| SLC9A4 | 2 | solute carrier family 9 member A4 [Source:HG |
| SLC9A2 | 2 | solute carrier family 9 member A2 [Source:HG |
| MFSD9 | 2 | major facilitator superfamily domain containir |
| TMEM182 | 2 | transmembrane protein 182 [Source:HGNC Sy |
| LINC01796 | 2 | long intergenic non-protein coding RNA 1796 |
| LINC01935 | 2 | long intergenic non-protein coding RNA 1935 |
| AC108051.1 | 2 | novel transcript |
| AC073987.1 | 2 | novel transcript |
| AC010881.1 | 2 | pseudogene similar to part of zinc finger, CCH |
| AC011593.1 | 2 | novel transcript |
| AC092568.1 | 2 | cytokine receptor-like factor 3 (CRLF3) pseudo |
| AC018880.1 | 2 | zinc finger protein 547 (ZNF547) pseudogene |
| AC018880.2 | 2 | capping protein (actin filament) muscle Z-line, |
| AC013727.1 | 2 | novel transcript |
| AC013727.2 | 2 | novel transcript |
| LINC01965 | 2 | long intergenic non-protein coding RNA 1965 |
| AC013727.3 | 2 | NULL |
| AC096554.1 | 2 | novel transcript |
| AHCYP3 | 2 | adenosylhomocysteinase pseudogene 3 [Sour |
| AC068535.1 | 2 | novel transcript |
| LINC01831 | 2 | long intergenic non-protein coding RNA 1831 |
| AC068535.2 | 2 | novel transcript |
| LINC01102 | 2 | long intergenic non-protein coding RNA 1102 |
| LINC01103 | 2 | long intergenic non-protein coding RNA 1103 |
| AC013402.2 | 2 | novel transcript |
| AC013402.3 | 2 | novel transcript |
| PANTR1 | 2 | POU3F3 adjacent non-coding transcript 1 [Sou |
| AC013402.1 | 2 | novel transcript |
| RPL23AP27 | 2 | ribosomal protein L23a pseudogene 27 [Sourc |
| AC068057.1 | 2 | novel transcript |
| AC068057.2 | 2 | TEC |
| SNORA72 | 2 | Small nucleolar RNA SNORA72 [Source:RFAM; |
| LINC01114 | 2 | long intergenic non-protein coding RNA 1114 |
| HMGB3P11 | 2 | high mobility group box 3 pseudogene 11 [Sou |
| AC018730.1 | 2 | novel transcript, sense overlapping POU3F3 |
| POU3F3 | 2 | POU class 3 homeobox 3 [Source:HGNC Symb |
| LINC01159 | 2 | long intergenic non-protein coding RNA 1159 |
| AC010884.2 | 2 | novel transcript |
| AC010884.1 | 2 | novel transcript |
| MRPS9 | 2 | mitochondrial ribosomal protein S9 [Source:H |
| AC104655.1 | 2 | novel transcript |
| LINC01918 | 2 | long intergenic non-protein coding RNA 1918 |
| GPR45 | 2 | G protein-coupled receptor 45 [Source:HGNC |
| AC012360.2 | 2 | novel transcript |
| TGFBRAP1 | 2 | transforming growth factor beta receptor asso |
| AC012360.1 | 2 | novel transcript |
| AC012360.3 | 2 | novel transcript |
| C2orf49 | 2 | chromosome 2 open reading frame 49 [Source |
| FHL2 | 2 | four and a half LIM domains 2 [Source:HGNC S |
| AC108058.1 | 2 | novel transcript, antisense to FHL2 |

| | | |
|---|---|---|
| AC018802.1 | 2 | novel transcript |
| AC018802.2 | 2 | novel transcript |
| NCK2 | 2 | NCK adaptor protein 2 [Source:HGNC Symbol; |
| AC010978.1 | 2 | novel transcript |
| AC009505.1 | 2 | novel transcript, antisense to NCK2 |
| ECRG4 | 2 | ECRG4 augurin precursor [Source:HGNC Symb |
| UXS1 | 2 | UDP-glucuronate decarboxylase 1 [Source:HG |
| AC018878.1 | 2 | novel transcript |
| AC092106.2 | 2 | novel pseudogene |
| RPL22P10 | 2 | ribosomal protein L22 pseudogene 10 [Source |
| AC092106.1 | 2 | splicing factor, arginine/serine-rich 3 (SFRS3) p |
| AC114755.3 | 2 | ribosomal protein S21 (RPS21) pseudogene |
| AC114755.4 | 2 | ribosomal protein L27a (RPL27A) pseudogene |
| AC114755.1 | 2 | chromosome 6 open reading frame 106 (C6or |
| AC114755.2 | 2 | ribosomal protein S26 (RPS26) pseudogene |
| AC114755.5 | 2 | pseudogene similar to part of anaphase prom |
| PLGLA | 2 | plasminogen like A (pseudogene) [Source:HGN |
| RGPD3 | 2 | RANBP2 like and GRIP domain containing 3 [So |
| AC097527.1 | 2 | pseudogene similar to part of anaphase prom |
| CD8B2 | 2 | CD8b2 molecule [Source:HGNC Symbol;Acc:H |
| AC108868.2 | 2 | novel transcript |
| AC108868.1 | 2 | novel transcript |
| EEF1A1P12 | 2 | eukaryotic translation elongation factor 1 alph |
| ST6GAL2 | 2 | ST6 beta-galactoside alpha-2,6-sialyltransfera |
| ST6GAL2-IT1 | 2 | ST6GAL2 intronic transcript 1 [Source:HGNC S |
| AC016994.1 | 2 | novel transcript |
| AC005040.2 | 2 | novel transcript |
| PPP1R2P5 | 2 | protein phosphatase 1 regulatory inhibitor sul |
| AC005040.1 | 2 | tripartite motif-containing 43 (TRIM43) pseud |
| LINC01789 | 2 | long intergenic non-protein coding RNA 1789 |
| AC096669.1 | 2 | novel transcript |
| LINC01885 | 2 | long intergenic non-protein coding RNA 1885 |
| LINC01886 | 2 | long intergenic non-protein coding RNA 1886 |
| AC064869.1 | 2 | novel transcript |
| GACAT1 | 2 | gastric cancer associated transcript 1 [Source: |
| RGPD4-AS1 | 2 | RGPD4 antisense RNA 1 (head to head) [Sourc |
| RGPD4 | 2 | RANBP2 like and GRIP domain containing 4 [So |
| AC009963.2 | 2 | pseudogene similar to part of plasminogen (Pl |
| AC009963.1 | 2 | pseudogene similar to part of anaphase prom |
| RPL22P8 | 2 | ribosomal protein L22 pseudogene 8 [Source: |
| AC009963.3 | 2 | splicing factor, arginine/serine-rich 3 (SFRS3) p |
| SLC5A7 | 2 | solute carrier family 5 member 7 [Source:HGN |
| LINC01593 | 2 | long intergenic non-protein coding RNA 1593 |
| ACTP1 | 2 | actin family pseudogene 1 [Source:HGNC Sym |
| LINC01594 | 2 | long intergenic non-protein coding RNA 1594 |
| SETD6P1 | 2 | SET domain containing 6 pseudogene 1 [Sourc |
| SULT1C3 | 2 | sulfotransferase family 1C member 3 [Source: |
| WASF1P1 | 2 | WASF1 pseudogene 1 [Source:HGNC Symbol;A |
| SULT1C2 | 2 | sulfotransferase family 1C member 2 [Source: |
| SULT1C2P2 | 2 | sulfotransferase family, cytosolic, 1C, member |

| | | |
|---|---|---|
| SULT1C2P1 | 2 | sulfotransferase family 1C member 2 pseudog |
| GMCL1P2 | 2 | GMCL1 pseudogene 2 [Source:HGNC Symbol;/ |
| SULT1C4 | 2 | sulfotransferase family 1C member 4 [Source: |
| SMIM12P1 | 2 | small integral membrane protein 12 pseudoge |
| GCC2 | 2 | GRIP and coiled-coil domain containing 2 [Sou |
| GCC2-AS1 | 2 | GCC2 antisense RNA 1 [Source:HGNC Symbol; |
| LIMS1 | 2 | LIM zinc finger domain containing 1 [Source:H |
| LIMS1-AS1 | 2 | LIMS1 antisense RNA 1 [Source:HGNC Symbol |
| AC010095.1 | 2 | cytochrome c oxidase subunit VIIb (COX7B) ps |
| RPL10P5 | 2 | ribosomal protein L10 pseudogene 5 [Source: |
| RANBP2 | 2 | RAN binding protein 2 [Source:HGNC Symbol;/ |
| CCDC138 | 2 | coiled-coil domain containing 138 [Source:HG |
| AC073415.1 | 2 | ribosomal protein L39 (RPL39) pseudogene |
| EDAR | 2 | ectodysplasin A receptor [Source:HGNC Symb |
| Metazoa_SRP | 2 | Metazoan signal recognition particle RNA [Sou |
| SH3RF3-AS1 | 2 | SH3RF3 antisense RNA 1 [Source:HGNC Symbo |
| SH3RF3 | 2 | SH3 domain containing ring finger 3 [Source:H |
| MIR4265 | 2 | microRNA 4265 [Source:HGNC Symbol;Acc:HG |
| SNRPGP9 | 2 | small nuclear ribonucleoprotein polypeptide G |
| MIR4266 | 2 | microRNA 4266 [Source:HGNC Symbol;Acc:HG |
| SEPTIN10 | 2 | septin 10 [Source:HGNC Symbol;Acc:HGNC:14 |
| AC011753.4 | 2 | ribosomal protein L37 (RPL37) pseudogene |
| AC011753.2 | 2 | novel transcript, sense intronic to SEPT10 |
| SOWAHC | 2 | sosondowah ankyrin repeat domain family me |
| AC011753.1 | 2 | ribosomal protein S21 (RPS21) pseudogene |
| BMS1P19 | 2 | BMS1 pseudogene 19 [Source:HGNC Symbol;/ |
| AC011753.3 | 2 | splicing factor, arginine/serine-rich 3 (SFRS3) p |
| RPL22P11 | 2 | ribosomal protein L22 pseudogene 11 [Source |
| AC074387.1 | 2 | novel transcript |
| RGPD5 | 2 | RANBP2 like and GRIP domain containing 5 [S |
| LIMS3 | 2 | LIM zinc finger domain containing 3 [Source:H |
| AC013271.1 | 2 | novel transcript |
| AC013268.1 | 2 | two-pore calcium channel 3 (TPCN3) pseudog |
| AC013268.5 | 2 | novel transcript |
| GPAA1P1 | 2 | glycosylphosphatidylinositol anchor attachme |
| ZBTB45P1 | 2 | zinc finger and BTB domain containing 45 pse |
| LINC01123 | 2 | long intergenic non-protein coding RNA 1123 |
| AC013268.3 | 2 | pseudogene similar to part of zinc finger prote |
| AC013268.2 | 2 | zinc finger protein 595 (ZNF595) pseudogene |
| AC013268.4 | 2 | novel pseudogene, similar to zonadhesin (Zan |
| MIR4267 | 2 | microRNA 4267 [Source:HGNC Symbol;Acc:HG |
| MALL | 2 | mal, T cell differentiation protein like [Source: |
| MIR4436B1 | 2 | microRNA 4436b-1 [Source:HGNC Symbol;Acc |
| NPHP1 | 2 | nephrocystin 1 [Source:HGNC Symbol;Acc:HG |
| AC140479.3 | 2 | ARP1 actin-related protein 1 homolog A, centr |
| MTLN | 2 | mitoregulin [Source:HGNC Symbol;Acc:HGNC: |
| AC140479.4 | 2 | novel ion transport protein domain-containing |
| AC140479.5 | 2 | novel transcript |
| AC140479.6 | 2 | mal, T-cell differentiation protein-like (MALL) |
| MIR4436B2 | 2 | microRNA 4436b-2 [Source:HGNC Symbol;Acc |

| | | |
|---|---|---|
| AC140479.7 | 2 | novel pseudogene, similar to zonadhesin (Zan |
| AC140479.1 | 2 | zinc finger protein pseudogene |
| AC140479.2 | 2 | zinc finger protein 532 (ZNF532) pseudogene |
| LINC01106 | 2 | long intergenic non-protein coding RNA 1106 |
| ZBTB45P2 | 2 | zinc finger and BTB domain containing 45 pseu |
| GPAA1P2 | 2 | glycosylphosphatidylinositol anchor attachme |
| AC112229.4 | 2 | novel transcript |
| AC112229.1 | 2 | two-pore calcium channel 3 (TPCN3) pseudog |
| AC112229.3 | 2 | novel transcript |
| LIMS4 | 2 | LIM zinc finger domain containing 4 [Source:H |
| AC112229.2 | 2 | novel transcript |
| AC108938.1 | 2 | pseudogene similar to part of tubulin tyrosine |
| RGPD6 | 2 | RANBP2 like and GRIP domain containing 6 [S |
| AC226101.1 | 2 | novel transcript |
| RPL22P12 | 2 | ribosomal protein L22 pseudogene 12 [Source |
| BUB1 | 2 | BUB1 mitotic checkpoint serine/threonine kin |
| AC114776.1 | 2 | novel transcript |
| ACOXL | 2 | acyl-CoA oxidase like [Source:HGNC Symbol;A |
| RPL5P9 | 2 | ribosomal protein L5 pseudogene 9 [Source:H |
| MIR4435-2HG | 2 | MIR4435-2 host gene [Source:HGNC Symbol;A |
| ACOXL-AS1 | 2 | ACOXL antisense RNA 1 [Source:HGNC Symbo |
| BCL2L11 | 2 | BCL2 like 11 [Source:HGNC Symbol;Acc:HGNC |
| AC108463.2 | 2 | novel transcript |
| AC108463.1 | 2 | pseudogene similar to part of ubiquitin carbox |
| AC108463.3 | 2 | novel transcript |
| AC068491.2 | 2 | novel transcript |
| AC068491.3 | 2 | novel transcript |
| RPS14P4 | 2 | ribosomal protein S14 pseudogene 4 [Source: |
| MIR4435-2 | 2 | microRNA 4435-2 [Source:HGNC Symbol;Acc:I |
| AC068491.4 | 2 | TEC |
| AC068491.1 | 2 | platelet-activating factor acetylhydrolase, isof |
| AC017002.6 | 2 | novel transcript |
| AC017002.5 | 2 | novel transcript |
| AC017002.3 | 2 | novel transcript |
| AC017002.1 | 2 | novel transcript |
| AC017002.4 | 2 | DBF4 homolog (S. cerevisiae) (DBF4) pseudog |
| AC017002.2 | 2 | novel transcript |
| ANAPC1 | 2 | anaphase promoting complex subunit 1 [Sour |
| AC093166.4 | 2 | ribosomal protein L34 (RPL34) pseudogene |
| AC093166.1 | 2 | EP300 interacting inhibitor of differentiation 2 |
| AC093166.2 | 2 | centromere protein N (CENPN) pseudogene |
| MIR4771-2 | 2 | microRNA 4771-2 [Source:HGNC Symbol;Acc:I |
| AC093166.3 | 2 | ribosomal protein S26 (RPS26) pseudogene |
| EEF1E1P1 | 2 | eukaryotic translation elongation factor 1 eps |
| MERTK | 2 | MER proto-oncogene, tyrosine kinase [Source |
| RN7SL297P | 2 | RNA, 7SL, cytoplasmic 297, pseudogene [Sour |
| AC104651.1 | 2 | chromosome 14 open reading frame 166 (C14 |
| AC104651.2 | 2 | solute carrier family 30 (zinc transporter), me |
| AC093675.1 | 2 | novel transcript, antisense to TMEM87B |
| TMEM87B | 2 | transmembrane protein 87B [Source:HGNC Sy |

| | | |
|---|---|---|
| AC093675.2 | 2 | novel transcript, antisense to TMEM87B |
| FBLN7 | 2 | fibulin 7 [Source:HGNC Symbol;Acc:HGNC:267 |
| AC092645.1 | 2 | TEC |
| ZC3H8 | 2 | zinc finger CCCH-type containing 8 [Source:HG |
| AC092645.2 | 2 | novel transcript, antisense to ZC3H8 |
| ZC3H6 | 2 | zinc finger CCCH-type containing 6 [Source:HG |
| NDUFB4P6 | 2 | NADH:ubiquinone oxidoreductase subunit B4 |
| AC115115.1 | 2 | axin 2 (conductin, axil) (AXIN2) pseudogene |
| RGPD8 | 2 | RANBP2 like and GRIP domain containing 8 [S |
| VINAC1P | 2 | vinculin/alpha-catenin family member 1, pseu |
| TTL | 2 | tubulin tyrosine ligase [Source:HGNC Symbol; |
| POLR1B | 2 | RNA polymerase I subunit B [Source:HGNC Sy |
| Y_RNA | 2 | Y RNA [Source:RFAM;Acc:RF00019] |
| CHCHD5 | 2 | coiled-coil-helix-coiled-coil-helix domain cont |
| AC012442.2 | 2 | novel transcript |
| AC012442.1 | 2 | novel transcript |
| AC079922.1 | 2 | ribosomal protein S20 (RPS20) pseudogene |
| AC079922.2 | 2 | novel transcript (FLJ42351) |
| SLC20A1 | 2 | solute carrier family 20 member 1 [Source:HG |
| NT5DC4 | 2 | 5'-nucleotidase domain containing 4 [Source: |
| CKAP2L | 2 | cytoskeleton associated protein 2 like [Source |
| IL1A | 2 | interleukin 1 alpha [Source:HGNC Symbol;Acc |
| AC079753.2 | 2 | novel transcript |
| IL1B | 2 | interleukin 1 beta [Source:HGNC Symbol;Acc: |
| AC079753.1 | 2 | TEC |
| XIAPP3 | 2 | X-linked inhibitor of apoptosis pseudogene 3 [ |
| IL37 | 2 | interleukin 37 [Source:HGNC Symbol;Acc:HGN |
| CDK8P2 | 2 | cyclin dependent kinase 8 pseudogene 2 [Sou |
| IL36G | 2 | interleukin 36 gamma [Source:HGNC Symbol; |
| HMGN2P23 | 2 | high mobility group nucleosomal binding dom |
| AC016724.1 | 2 | polymerase (RNA) II (DNA directed) polypepti |
| IL36A | 2 | interleukin 36 alpha [Source:HGNC Symbol;Ac |
| IL36B | 2 | interleukin 36 beta [Source:HGNC Symbol;Acc |
| IL36RN | 2 | interleukin 36 receptor antagonist [Source:HG |
| IL1F10 | 2 | interleukin 1 family member 10 [Source:HGN |
| RNU6-1180P | 2 | RNA, U6 small nuclear 1180, pseudogene [Sou |
| IL1RN | 2 | interleukin 1 receptor antagonist [Source:HGN |
| PSD4 | 2 | pleckstrin and Sec7 domain containing 4 [Sour |
| AC016683.1 | 2 | novel transcript, antisense to PSD4 |
| PAX8-AS1 | 2 | PAX8 antisense RNA 1 [Source:HGNC Symbol; |
| PAX8 | 2 | paired box 8 [Source:HGNC Symbol;Acc:HGNC |
| AC016745.1 | 2 | novel transcript |
| IGKV1OR2-108 | 2 | immunoglobulin kappa variable 1/OR2-108 (n |
| AC016745.2 | 2 | novel transcript |
| CBWD2 | 2 | COBW domain containing 2 [Source:HGNC Sy |
| FOXD4L1 | 2 | forkhead box D4 like 1 [Source:HGNC Symbol; |
| LINC01961 | 2 | long intergenic non-protein coding RNA 1961 |
| PGM5P4-AS1 | 2 | PGM5P4 antisense RNA 1 [Source:HGNC Symb |
| PGM5P4 | 2 | phosphoglucomutase 5 pseudogene 4 [Source |
| FAM138B | 2 | family with sequence similarity 138 member |

| | | |
|---|---|---|
| AL078621.4 | 2 | novel transcript |
| MIR1302-3 | 2 | microRNA 1302-3 [Source:HGNC Symbol;Acc:I |
| WASH2P | 2 | WASP family homolog 2, pseudogene [Source |
| DDX11L2 | 2 | DEAD/H-box helicase 11 like 2 [Source:HGNC S |
| AL078621.3 | 2 | TEC |
| RPL23AP7 | 2 | ribosomal protein L23a pseudogene 7 [Source |
| RABL2A | 2 | RAB, member of RAS oncogene family like 2A |
| AL078621.1 | 2 | small nuclear ribonucleoprotein polypeptide A |
| AL078621.2 | 2 | acrosin (ACR) pseudogene |
| AC017074.2 | 2 | novel transcript |
| AC017074.1 | 2 | novel transcript |
| RNU6-744P | 2 | RNA, U6 small nuclear 744, pseudogene [Sour |
| SLC35F5 | 2 | solute carrier family 35 member F5 [Source:H |
| MIR4782 | 2 | microRNA 4782 [Source:HGNC Symbol;Acc:HG |
| AC104653.2 | 2 | novel transcript |
| AC104653.1 | 2 | novel transcript |
| AC110769.2 | 2 | novel transcript |
| ACTR3 | 2 | actin related protein 3 [Source:HGNC Symbol; |
| AC110769.3 | 2 | TEC |
| LINC01191 | 2 | long intergenic non-protein coding RNA 1191 |
| AC110769.1 | 2 | novel transcript |
| U3 | 2 | Small nucleolar RNA U3 [Source:RFAM;Acc:RF |
| AC010982.2 | 2 | novel transcript |
| AC010982.1 | 2 | novel transcript |
| SEPHS1P7 | 2 | selenophosphate synthetase 1 pseudogene 7 |
| RNU2-41P | 2 | RNA, U2 small nuclear 41, pseudogene [Sourc |
| DPP10 | 2 | dipeptidyl peptidase like 10 [Source:HGNC Syr |
| DPP10-AS3 | 2 | DPP10 antisense RNA 3 [Source:HGNC Symbo |
| DPP10-AS2 | 2 | DPP10 antisense RNA 2 [Source:HGNC Symbo |
| AC093610.1 | 2 | novel transcript, antisense to DPP10 |
| DPP10-AS1 | 2 | DPP10 antisense RNA 1 [Source:HGNC Symbo |
| RPSAP23 | 2 | ribosomal protein SA pseudogene 23 [Source: |
| AC016721.1 | 2 | novel transcript, antisense to DPP10 |
| AC104408.1 | 2 | sideroflexin 5 (SFXN5) pseudogene |
| AC062016.1 | 2 | pseudogene similar to part of fibronectin type |
| MTCYBP39 | 2 | MT-CYB pseudogene 39 [Source:HGNC Symbo |
| AC062016.2 | 2 | pseudogene similar to part of piggyBac transp |
| MTND1P28 | 2 | MT-ND1 pseudogene 28 [Source:HGNC Symbo |
| MTND2P21 | 2 | MT-ND2 pseudogene 21 [Source:HGNC Symbo |
| MTCO1P43 | 2 | MT-CO1 pseudogene 43 [Source:HGNC Symbo |
| RNU7-190P | 2 | RNA, U7 small nuclear 190 pseudogene [Sourc |
| AC092170.1 | 2 | pseudogene similar to part of golgi autoantige |
| AC064856.1 | 2 | novel transcript |
| AC009312.1 | 2 | novel transcript |
| DDX18 | 2 | DEAD-box helicase 18 [Source:HGNC Symbol;A |
| AC009404.1 | 2 | novel transcript |
| HTR5BP | 2 | 5-hydroxytryptamine receptor 5B, pseudogen |
| CCDC93 | 2 | coiled-coil domain containing 93 [Source:HGN |
| AC009303.1 | 2 | nuclear factor (erythroid-derived 2)-like 2 (NF |
| AC009303.2 | 2 | novel transcript, antisense to CCDC93 |

| | | |
|---|---|---|
| AC009303.3 | 2 | novel transcript, antisense to CCDC93 |
| AC009303.4 | 2 | novel transcript |
| RN7SL111P | 2 | RNA, 7SL, cytoplasmic 111, pseudogene [Sour |
| INSIG2 | 2 | insulin induced gene 2 [Source:HGNC Symbol; |
| THORLNC | 2 | testis associated oncogenic lncRNA [Source:H |
| LINC01956 | 2 | long intergenic non-protein coding RNA 1956 |
| EN1 | 2 | engrailed homeobox 1 [Source:HGNC Symbol; |
| MARCO | 2 | macrophage receptor with collagenous struct |
| AC013457.1 | 2 | novel transcript, antisense to MARCO |
| C1QL2 | 2 | complement C1q like 2 [Source:HGNC Symbol |
| RN7SL468P | 2 | RNA, 7SL, cytoplasmic 468, pseudogene [Sour |
| STEAP3 | 2 | STEAP3 metalloreductase [Source:HGNC Symb |
| STEAP3-AS1 | 2 | STEAP3 antisense RNA 1 [Source:HGNC Symbo |
| C2orf76 | 2 | chromosome 2 open reading frame 76 [Sourc |
| DBI | 2 | diazepam binding inhibitor, acyl-CoA binding p |
| TMEM37 | 2 | transmembrane protein 37 [Source:HGNC Sym |
| SCTR | 2 | secretin receptor [Source:HGNC Symbol;Acc:H |
| AC013275.1 | 2 | novel transcript, antisense to SCTR |
| CFAP221 | 2 | cilia and flagella associated protein 221 [Sourc |
| TMEM177 | 2 | transmembrane protein 177 [Source:HGNC Sy |
| AC069154.1 | 2 | ribosomal protein L17 (RPL17) pseudogene |
| PTPN4 | 2 | protein tyrosine phosphatase non-receptor ty |
| AC016691.1 | 2 | ribosomal protein L27 (RPL27) pseudogene |
| EPB41L5 | 2 | erythrocyte membrane protein band 4.1 like 5 |
| AC012363.1 | 2 | novel transcript |
| Y_RNA | 2 | Y RNA [Source:RFAM;Acc:RF00019] |
| MTATP6P26 | 2 | MT-ATP6 pseudogene 26 [Source:HGNC Symb |
| MTCO3P43 | 2 | MT-CO3 pseudogene 43 [Source:HGNC Symbo |
| MTND3P10 | 2 | MT-ND3 pseudogene 10 [Source:HGNC Symbo |
| MTND4LP14 | 2 | MT-ND4L pseudogene 14 [Source:HGNC Symb |
| MTND4P26 | 2 | MT-ND4 pseudogene 26 [Source:HGNC Symbo |
| MTND5P28 | 2 | MT-ND5 pseudogene 28 [Source:HGNC Symbo |
| TMEM185B | 2 | transmembrane protein 185B [Source:HGNC S |
| RALB | 2 | RAS like proto-oncogene B [Source:HGNC Sym |
| AC012363.2 | 2 | novel transcript |
| INHBB | 2 | inhibin subunit beta B [Source:HGNC Symbol; |
| LINC01101 | 2 | long intergenic non-protein coding RNA 1101 |
| AC073257.2 | 2 | novel transcript |
| AC073257.1 | 2 | novel transcript |
| Y_RNA | 2 | Y RNA [Source:RFAM;Acc:RF00019] |
| AC018866.1 | 2 | novel transcript |
| GLI2 | 2 | GLI family zinc finger 2 [Source:HGNC Symbol; |
| AC017033.1 | 2 | novel transcript |
| Y_RNA | 2 | Y RNA [Source:RFAM;Acc:RF00019] |
| AC067960.1 | 2 | novel transcript |
| AC079988.1 | 2 | novel transcript |
| TFCP2L1 | 2 | transcription factor CP2 like 1 [Source:HGNC S |
| AC013399.1 | 2 | ribosomal protein S17 (RPS17) pseudogene |
| CLASP1 | 2 | cytoplasmic linker associated protein 1 [Sourc |
| AC012447.1 | 2 | novel transcript, antisense to CLASP1 |

| | | |
|---|---|---|
| RNU4ATAC | 2 | RNA, U4atac small nuclear (U12-dependent sp |
| Y_RNA | 2 | Y RNA [Source:RFAM;Acc:RF00019] |
| NIFK-AS1 | 2 | NIFK antisense RNA 1 [Source:HGNC Symbol;A |
| RPL12P15 | 2 | ribosomal protein L12 pseudogene 15 [Source |
| AC018737.2 | 2 | TEC |
| NPM1P32 | 2 | nucleophosmin 1 pseudogene 32 [Source:HGN |
| NIFK | 2 | nucleolar protein interacting with the FHA do |
| TSN | 2 | translin [Source:HGNC Symbol;Acc:HGNC:123 |
| AC018737.1 | 2 | ribosomal protein S18 (RPS18) pseudogene |
| LINC01823 | 2 | long intergenic non-protein coding RNA 1823 |
| AC011246.1 | 2 | novel transcript |
| AC073632.1 | 2 | novel transcript |
| AC062020.1 | 2 | novel transcript |
| LINC01826 | 2 | long intergenic non-protein coding RNA 1826 |
| AC073409.2 | 2 | novel transcript |
| AC073409.1 | 2 | novel transcript |
| AC092646.2 | 2 | transcription elongation factor B (SIII), polype |
| AC092646.1 | 2 | proteasome (prosome, macropain) 26S subun |
| AC064859.1 | 2 | TEC |
| RN7SKP102 | 2 | RN7SK pseudogene 102 [Source:HGNC Symbo |
| AC079154.1 | 2 | novel transcript |
| CNTNAP5 | 2 | contactin associated protein like 5 [Source:HG |
| MTND5P22 | 2 | MT-ND5 pseudogene 22 [Source:HGNC Symbo |
| AC019159.2 | 2 | novel transcript, antisense to CNTNAP5 |
| AC019159.1 | 2 | heat shock protein 90kDa alpha (cytosolic), cla |
| RNA5SP102 | 2 | RNA, 5S ribosomal pseudogene 102 [Source:H |
| RNU6-259P | 2 | RNA, U6 small nuclear 259, pseudogene [Sour |
| AC097499.2 | 2 | novel transcript |
| LINC01889 | 2 | long intergenic non-protein coding RNA 1889 |
| AC097499.1 | 2 | Wolf-Hirschhorn syndrome candidate 1 (WHS |
| LINC01941 | 2 | long intergenic non-protein coding RNA 1941 |
| AC142116.2 | 2 | novel transcript |
| AC023347.1 | 2 | novel transcript |
| AC023347.2 | 2 | solute carrier family 6 (amino acid transporte |
| YWHAZP2 | 2 | tyrosine 3-monooxygenase/tryptophan 5-mo |
| GYPC | 2 | glycophorin C (Gerbich blood group) [Source:I |
| AC013474.1 | 2 | novel pseudogene |
| RNU6-675P | 2 | RNA, U6 small nuclear 675, pseudogene [Sour |
| TEX51 | 2 | testis expressed 51 [Source:HGNC Symbol;Acc |
| RNU7-182P | 2 | RNA, U7 small nuclear 182 pseudogene [Sour |
| AC012508.2 | 2 | novel transcript |
| AC012508.3 | 2 | novel transcript |
| AC012508.1 | 2 | novel transcript |
| BIN1 | 2 | bridging integrator 1 [Source:HGNC Symbol;A |
| AC110926.2 | 2 | novel transcript |
| NIFKP9 | 2 | nucleolar protein interacting with the FHA do |
| CYP27C1 | 2 | cytochrome P450 family 27 subfamily C memb |
| AC110926.1 | 2 | WW domain binding protein 11 (WBP11) pseu |
| ERCC3 | 2 | ERCC excision repair 3, TFIIH core complex he |
| AC068282.1 | 2 | novel transcript, antisense to MAP3K2 |

| | | |
|---|---|---|
| MAP3K2 | 2 | mitogen-activated protein kinase kinase kinas |
| RNU6-1147P | 2 | RNA, U6 small nuclear 1147, pseudogene [Sou |
| MAP3K2-DT | 2 | MAP3K2 divergent transcript [Source:HGNC S |
| AC068282.2 | 2 | novel transcript |
| PROC | 2 | protein C, inactivator of coagulation factors Va |
| MIR4783 | 2 | microRNA 4783 [Source:HGNC Symbol;Acc:HG |
| IWS1 | 2 | interacts with SUPT6H, CTD assembly factor 1 |
| AC010976.1 | 2 | novel transcript, antisense to IWS1 |
| RNU4-48P | 2 | RNA, U4 small nuclear 48, pseudogene [Sourc |
| MYO7B | 2 | myosin VIIB [Source:HGNC Symbol;Acc:HGNC: |
| AC010976.2 | 2 | novel transcript, antisense to MYO7B |
| LIMS2 | 2 | LIM zinc finger domain containing 2 [Source:H |
| GPR17 | 2 | G protein-coupled receptor 17 [Source:HGNC |
| WDR33 | 2 | WD repeat domain 33 [Source:HGNC Symbol; |
| SFT2D3 | 2 | SFT2 domain containing 3 [Source:HGNC Syml |
| RNY4P7 | 2 | RNY4 pseudogene 7 [Source:HGNC Symbol;Ac |
| Y_RNA | 2 | Y RNA [Source:RFAM;Acc:RF00019] |
| AC006011.2 | 2 | ZFP91 zinc finger protein (ZFP91) pseudogene |
| POLR2D | 2 | RNA polymerase II subunit D [Source:HGNC Sy |
| RNU6-395P | 2 | RNA, U6 small nuclear 395, pseudogene [Sour |
| AC006011.1 | 2 | ribosomal protein S26 (RPS26) pseudogene |
| Metazoa_SRP | 2 | Metazoan signal recognition particle RNA [Sou |
| AMMECR1L | 2 | AMMECR1 like [Source:HGNC Symbol;Acc:HGI |
| AC012306.2 | 2 | novel transcript |
| AC012306.3 | 2 | novel transcript |
| AC012306.1 | 2 | archaelysin family metallopeptidase 2 (AMZ2) |
| SAP130 | 2 | Sin3A associated protein 130 [Source:HGNC S |
| AC108059.3 | 2 | enhancer of yellow 2 homolog (ENY2) pseudo |
| Y_RNA | 2 | Y RNA [Source:RFAM;Acc:RF00019] |
| UGGT1 | 2 | UDP-glucose glycoprotein glucosyltransferase |
| AC108059.1 | 2 | spermidine synthase (SRM) pseudogene |
| AC108059.2 | 2 | coiled-coil domain containing 72 (CCDC72) ps |
| DYNLT3P2 | 2 | dynein light chain Tctex-type 3 pseudogene 2 |
| AC017079.1 | 2 | ribosomal protein L14 (RPL14) pseudogene |
| AC017079.2 | 2 | ribosomal protein L21 (RPL21) pseudogene |
| HS6ST1 | 2 | heparan sulfate 6-O-sulfotransferase 1 [Sourc |
| Y_RNA | 2 | Y RNA [Source:RFAM;Acc:RF00019] |
| RNA5SP103 | 2 | RNA, 5S ribosomal pseudogene 103 [Source:H |
| ISCA1P6 | 2 | iron-sulfur cluster assembly 1 pseudogene 6 [ |
| AC012451.1 | 2 | novel transcript |
| AC068483.1 | 2 | novel transcript |
| LINC01854 | 2 | long intergenic non-protein coding RNA 1854 |
| PPIAP65 | 2 | peptidylprolyl isomerase A pseudogene 65 [Sc |
| LINC02572 | 2 | long intergenic non-protein coding RNA 2572 |
| AC079776.5 | 2 | novel transcript |
| PLAC9P1 | 2 | placenta associated 9 pseudogene 1 [Source:H |
| LINC01856 | 2 | long intergenic non-protein coding RNA 1856 |
| RPL22P7 | 2 | ribosomal protein L22 pseudogene 7 [Source: |
| RAB6C-AS1 | 2 | RAB6C antisense RNA 1 [Source:HGNC Symbo |
| RAB6C | 2 | RAB6C, member RAS oncogene family [Source |

| | | |
|---|---|---|
| AC079776.3 | 2 | pseudogene similar to part of fatty acyl CoA re |
| AC079776.4 | 2 | novel transcript |
| AC079776.2 | 2 | pseudogene similar to part of pleckstrin homo |
| AC079776.1 | 2 | single stranded DNA binding protein 3 (SSBP3) |
| ARHGAP42P2 | 2 | Rho GTPase activating protein 42 pseudogene |
| FAR2P1 | 2 | fatty acyl-CoA reductase 2 pseudogene 1 [Sou |
| AC018865.3 | 2 | novel transcript |
| KLF2P1 | 2 | Kruppel like factor 2 pseudogene 1 [Source:H( |
| CYP4F27P | 2 | cytochrome P450 family 4 subfamily F membe |
| POTEF | 2 | POTE ankyrin domain family member F [Sourc |
| AC018865.2 | 2 | pseudogene similar to part of nucleolar protei |
| AC018865.1 | 2 | novel transcript |
| RNU6-1049P | 2 | RNA, U6 small nuclear 1049, pseudogene [Sou |
| MED15P9 | 2 | mediator complex subunit 15 pseudogene 9 [S |
| CCDC74B | 2 | coiled-coil domain containing 74B [Source:HG |
| SMPD4 | 2 | sphingomyelin phosphodiesterase 4 [Source:H |
| MZT2B | 2 | mitotic spindle organizing protein 2B [Source: |
| TUBA3E | 2 | tubulin alpha 3e [Source:HGNC Symbol;Acc:H( |
| AC018804.1 | 2 | pseudogene similar to part of CDC42 binding p |
| RHOQP3 | 2 | ras homolog family member Q pseudogene 3 |
| NOC2LP1 | 2 | NOC2 like nucleolar associated transcriptional |
| AC134915.1 | 2 | pseudogene similar to part of ATPase, H+/K+ t |
| MED15P5 | 2 | mediator complex subunit 15 pseudogene 5 [S |
| MTND1P29 | 2 | MT-ND1 pseudogene 29 [Source:HGNC Symbo |
| MTND2P22 | 2 | MT-ND2 pseudogene 22 [Source:HGNC Symbo |
| MTCO1P7 | 2 | MT-CO1 pseudogene 7 [Source:HGNC Symbol |
| MTCO2P7 | 2 | MT-CO2 pseudogene 7 [Source:HGNC Symbol |
| MTATP6P7 | 2 | MT-ATP6 pseudogene 7 [Source:HGNC Symbo |
| MTCO3P7 | 2 | MT-CO3 pseudogene 7 [Source:HGNC Symbol |
| AC068137.3 | 2 | mitochondrially encoded NADH dehydrogenas |
| AC068137.4 | 2 | mitochondrially encoded NADH 4L dehydroge |
| MTND4P27 | 2 | MT-ND4 pseudogene 27 [Source:HGNC Symbo |
| MTND5P29 | 2 | MT-ND5 pseudogene 29 [Source:HGNC Symbo |
| MTND6P8 | 2 | MT-ND6 pseudogene 8 [Source:HGNC Symbol |
| MTCYBP8 | 2 | MT-CYB pseudogene 8 [Source:HGNC Symbol; |
| AC068137.1 | 2 | ribosomal protein L19 (RPL19) pseudogene |
| AC068137.2 | 2 | tektin 4 (TEKT4) pseudogene |
| AC132479.1 | 2 | pseudogene similar to part of YME1-like 1 (S. ( |
| CCDC115 | 2 | coiled-coil domain containing 115 [Source:HG |
| IMP4 | 2 | IMP U3 small nucleolar ribonucleoprotein 4 [S |
| PTPN18 | 2 | protein tyrosine phosphatase non-receptor ty |
| FAR2P2 | 2 | fatty acyl-CoA reductase 2 pseudogene 2 [Sou |
| KLF2P2 | 2 | Kruppel like factor 2 pseudogene 2 [Source:H( |
| CYP4F62P | 2 | cytochrome P450 family 4 subfamily F membe |
| POTEI | 2 | POTE ankyrin domain family member I [Sourc |
| RNU6-473P | 2 | RNA, U6 small nuclear 473, pseudogene [Sour |
| AC013269.2 | 2 | novel transcript |
| Metazoa_SRP | 2 | Metazoan signal recognition particle RNA [Sou |
| CFC1B | 2 | cripto, FRL-1, cryptic family 1B [Source:HGNC |
| AC013269.1 | 2 | novel transcript |

| | | |
|---|---|---|
| PRSS40B | 2 | serine protease 40B [Source:HGNC Symbol;Ac |
| PRSS40A | 2 | serine protease 40A [Source:HGNC Symbol;Ac |
| AC140481.2 | 2 | novel transcript |
| CFC1 | 2 | cripto, FRL-1, cryptic family 1 [Source:HGNC Sy |
| Metazoa_SRP | 2 | Metazoan signal recognition particle RNA [Sou |
| POTEJ | 2 | POTE ankyrin domain family member J [Source |
| RNU6-848P | 2 | RNA, U6 small nuclear 848, pseudogene [Sour |
| CYP4F30P | 2 | cytochrome P450 family 4 subfamily F membe |
| AC140481.3 | 2 | novel transcript |
| FAR2P3 | 2 | fatty acyl-CoA reductase 2 pseudogene 3 [Sou |
| KLF2P3 | 2 | Kruppel like factor 2 pseudogene 3 [Source:HG |
| GPR148 | 2 | G protein-coupled receptor 148 [Source:HGNC |
| AC140481.1 | 2 | novel transcript |
| AMER3 | 2 | APC membrane recruitment protein 3 [Source |
| CYCSP8 | 2 | CYCS pseudogene 8 [Source:HGNC Symbol;Acc |
| AC133785.1 | 2 | novel transcript, antisense to ARHGEF4 |
| ARHGEF4 | 2 | Rho guanine nucleotide exchange factor 4 [So |
| AC112786.1 | 2 | NULL |
| AC112786.2 | 2 | NULL |
| SMIM39 | 2 | small integral membrane protein 39 [Source:H |
| FAM168B | 2 | family with sequence similarity 168 member B |
| PLEKHB2 | 2 | pleckstrin homology domain containing B2 [So |
| NEK2P4 | 2 | NEK2 pseudogene 4 [Source:HGNC Symbol;Ac |
| AC009477.1 | 2 | nitric oxide synthase 2A (NOS2) pseudogene |
| NF1P8 | 2 | neurofibromin 1 pseudogene 8 [Source:HGNC |
| AC131180.1 | 2 | novel transcript |
| MED15P8 | 2 | mediator complex subunit 15 pseudogene 8 [S |
| POTEE | 2 | POTE ankyrin domain family member E [Sourc |
| RNU6-127P | 2 | RNA, U6 small nuclear 127, pseudogene [Sour |
| FAR2P4 | 2 | fatty acyl-CoA reductase 2 pseudogene 4 [Sou |
| KLF2P4 | 2 | Kruppel like factor 2 pseudogene 4 [Source:HG |
| AC073869.7 | 2 | novel transcript |
| ARHGAP42P1 | 2 | Rho GTPase activating protein 42 pseudogene |
| AC073869.9 | 2 | fatty acyl CoA reductase 2 (FAR2) pseudogene |
| AC073869.2 | 2 | single-stranded DNA binding protein 3 (SSBP3 |
| RAB6D | 2 | RAB6D, member RAS oncogene family [Source |
| AC073869.5 | 2 | novel transcript |
| MTCYBP10 | 2 | MT-CYB pseudogene 10 [Source:HGNC Symbo |
| MTND6P10 | 2 | MT-ND6 pseudogene 10 [Source:HGNC Symbo |
| MTND5P23 | 2 | MT-ND5 pseudogene 23 [Source:HGNC Symbo |
| MTND4P21 | 2 | MT-ND4 pseudogene 21 [Source:HGNC Symbo |
| AC073869.6 | 2 | mitochondrially encoded NADH 4L dehydroge |
| MTND3P18 | 2 | MT-ND3 pseudogene 18 [Source:HGNC Symbo |
| MTCO3P18 | 2 | MT-CO3 pseudogene 18 [Source:HGNC Symbo |
| MTATP6P4 | 2 | MT-ATP6 pseudogene 4 [Source:HGNC Symbo |
| MTCO2P18 | 2 | MT-CO2 pseudogene 18 [Source:HGNC Symbo |
| MTCO1P18 | 2 | MT-CO1 pseudogene 18 [Source:HGNC Symbo |
| MTND2P18 | 2 | MT-ND2 pseudogene 18 [Source:HGNC Symbo |
| MTND1P26 | 2 | MT-ND1 pseudogene 26 [Source:HGNC Symbo |
| MED15P3 | 2 | mediator complex subunit 15 pseudogene 3 [S |

| | | |
|---|---|---|
| AC073869.4 | 2 | ATPase, H+/K+ transporting, nongastric, alpha |
| LINC01120 | 2 | long intergenic non-protein coding RNA 1120 |
| AC073869.8 | 2 | secreted frizzled related protein 4 (SFRP4) pse |
| GNAQP1 | 2 | G protein subunit alpha q pseudogene 1 [Sour |
| NOC2LP2 | 2 | NOC2 like nucleolar associated transcriptional |
| RHOQP2 | 2 | ras homolog family member Q pseudogene 2 |
| AC073869.3 | 2 | novel transcript |
| MZT2A | 2 | mitotic spindle organizing protein 2A [Source: |
| TUBA3D | 2 | tubulin alpha 3d [Source:HGNC Symbol;Acc:H |
| MIR4784 | 2 | microRNA 4784 [Source:HGNC Symbol;Acc:HG |
| AC073869.1 | 2 | novel pseudogene |
| CCDC74A | 2 | coiled-coil domain containing 74A [Source:HG |
| MED15P4 | 2 | mediator complex subunit 15 pseudogene 4 [ |
| AC093838.1 | 2 | novel transcript |
| POTEKP | 2 | POTE ankyrin domain family member K, pseuc |
| RNU6-617P | 2 | RNA, U6 small nuclear 617, pseudogene [Sour |
| LINC01087 | 2 | long intergenic non-protein coding RNA 1087 |
| GRAMD4P8 | 2 | GRAM domain containing 4 pseudogene 8 [So |
| C2orf27A | 2 | chromosome 2 open reading frame 27A [Sour |
| AC093838.2 | 2 | novel pseudogene |
| C2orf27A | 2 | chromosome 2 open reading frame 27A [Sour |
| AC093724.1 | 2 | translocase of outer mitochondrial membrane |
| AC093724.3 | 2 | CMT1A duplicated region transcript 15-like 2 ( |
| C2orf27B | 2 | chromosome 2 open reading frame 27B [Sour |
| AC103564.3 | 2 | TBC1 domain family, member 26 (TBC1D26) p |
| AC103564.1 | 2 | novel transcript |
| AC103564.2 | 2 | novel pseudogene |
| AJ239322.2 | 2 | novel transcript |
| AJ239322.1 | 2 | novel transcript |
| LINC01945 | 2 | long intergenic non-protein coding RNA 1945 |
| AC093787.2 | 2 | novel transcript |
| AC093787.1 | 2 | pseudogene similar to part of kinase suppress |
| AC098826.3 | 2 | BCL2/adenovirus E1B 19kDa interacting prote |
| AC098826.1 | 2 | pseudogene similar to putative zinc finger pro |
| AC098826.2 | 2 | novel pseudogene similar to ankyrin repeat do |
| ANKRD30BL | 2 | ankyrin repeat domain 30B like [Source:HGNC |
| RNU6-1132P | 2 | RNA, U6 small nuclear 1132, pseudogene [Sou |
| RNA5-8SP5 | 2 | RNA, 5.8S ribosomal pseudogene 5 [Source:H |
| MIR663B | 2 | microRNA 663b [Source:HGNC Symbol;Acc:HG |
| CDC27P1 | 2 | cell division cycle 27 pseudogene 1 [Source:H |
| AC097532.1 | 2 | novel transcript |
| ZNF806 | 2 | zinc finger protein 806 [Source:HGNC Symbol; |
| AC097532.2 | 2 | novel transcript |
| AC097532.3 | 2 | novel transcript |
| FAM201B | 2 | family with sequence similarity 201 member E |
| RNU6-175P | 2 | RNA, U6 small nuclear 175, pseudogene [Sour |
| GPR39 | 2 | G protein-coupled receptor 39 [Source:HGNC |
| YBX1P7 | 2 | Y-box binding protein 1 pseudogene 7 [Source |
| RN7SKP103 | 2 | RN7SK pseudogene 103 [Source:HGNC Symbo |
| LYPD1 | 2 | LY6/PLAUR domain containing 1 [Source:HGN |

| | | |
|---|---|---|
| AC010974.1 | 2 | novel transcript |
| NCKAP5 | 2 | NCK associated protein 5 [Source:HGNC Symb |
| AC010974.2 | 2 | novel transcript, antisense to NCKAP5 |
| AC016909.1 | 2 | pseudogene similar to part of single-stranded |
| NCKAP5-AS1 | 2 | NCKAP5 antisense RNA 1 [Source:HGNC Symb |
| AC011755.1 | 2 | novel transcript |
| NCKAP5-AS2 | 2 | NCKAP5 antisense RNA 2 [Source:HGNC Symb |
| RNU6-579P | 2 | RNA, U6 small nuclear 579, pseudogene [Sour |
| RN7SKP154 | 2 | RN7SK pseudogene 154 [Source:HGNC Symbo |
| NCKAP5-IT1 | 2 | NCKAP5 intronic transcript 1 [Source:HGNC Sy |
| AC011243.1 | 2 | novel transcript |
| RN7SKP93 | 2 | RN7SK pseudogene 93 [Source:HGNC Symbol; |
| AC092623.1 | 2 | novel transcript |
| MGAT5 | 2 | alpha-1,6-mannosylglycoprotein 6-beta-N-ace |
| MIR3679 | 2 | microRNA 3679 [Source:HGNC Symbol;Acc:HG |
| RNA5SP104 | 2 | RNA, 5S ribosomal pseudogene 104 [Source:H |
| EDDM3CP | 2 | epididymal protein 3C, pseudogene [Source:H |
| TMEM163 | 2 | transmembrane protein 163 [Source:HGNC Sy |
| AC110620.2 | 2 | novel transcript, antisense to TMEM163 |
| CCNT2-AS1 | 2 | CCNT2 antisense RNA 1 [Source:HGNC Symbo |
| VDAC2P4 | 2 | voltage dependent anion channel 2 pseudoge |
| AC110620.1 | 2 | ribosomal protein L17 pseudogene |
| ACMSD | 2 | aminocarboxymuconate semialdehyde decarb |
| MIR5590 | 2 | microRNA 5590 [Source:HGNC Symbol;Acc:HG |
| CCNT2 | 2 | cyclin T2 [Source:HGNC Symbol;Acc:HGNC:16 |
| MAP3K19 | 2 | mitogen-activated protein kinase kinase kinas |
| RAB3GAP1 | 2 | RAB3 GTPase activating protein catalytic subu |
| SNORA40B | 2 | small nucleolar RNA, H/ACA box 40B [Source: |
| ZRANB3 | 2 | zinc finger RANBP2-type containing 3 [Source: |
| AC064850.1 | 2 | GTPase activating protein (SH3 domain) bindi |
| R3HDM1 | 2 | R3H domain containing 1 [Source:HGNC Symb |
| RNU6-512P | 2 | RNA, U6 small nuclear 512, pseudogene [Sour |
| MIR128-1 | 2 | microRNA 128-1 [Source:HGNC Symbol;Acc:H |
| UBXN4 | 2 | UBX domain protein 4 [Source:HGNC Symbol; |
| LCT | 2 | lactase [Source:HGNC Symbol;Acc:HGNC:653 |
| Y_RNA | 2 | Y RNA [Source:RFAM;Acc:RF00019] |
| AC011893.1 | 2 | novel transcript |
| MCM6 | 2 | minichromosome maintenance complex comp |
| AC011999.1 | 2 | mannosidase, endo-alpha-like (MANEAL) pseu |
| DARS | 2 | aspartyl-tRNA synthetase [Source:HGNC Symb |
| DARS-AS1 | 2 | DARS antisense RNA 1 [Source:HGNC Symbol; |
| AC068492.1 | 2 | novel transcript |
| CXCR4 | 2 | C-X-C motif chemokine receptor 4 [Source:HG |
| HNRNPKP2 | 2 | heterogeneous nuclear ribonucleoprotein K p |
| AC112255.1 | 2 | TEC |
| UBBP1 | 2 | ubiquitin B pseudogene 1 [Source:HGNC Symb |
| RN7SKP141 | 2 | RN7SK pseudogene 141 [Source:HGNC Symbo |
| SMC4P1 | 2 | structural maintenance of chromosomes 4 ps |
| THSD7B | 2 | thrombospondin type 1 domain containing 7B |
| RNA5SP105 | 2 | RNA, 5S ribosomal pseudogene 105 [Source:H |

| | | |
|---|---|---|
| Y_RNA | 2 | Y RNA [Source:RFAM;Acc:RF00019] |
| HNMT | 2 | histamine N-methyltransferase [Source:HGNC |
| LINC01832 | 2 | long intergenic non-protein coding RNA 1832 |
| AC097523.1 | 2 | ribosomal protein L15 (RPL15) pseudogene |
| YWHAEP5 | 2 | tyrosine 3-monooxygenase/tryptophan 5-mor |
| AC097523.2 | 2 | isopentenyl-diphosphate delta isomerase 1 (II |
| AC097721.1 | 2 | ring finger protein 14 (RNF14) pseudogene |
| AC114763.1 | 2 | novel transcript |
| AC114763.2 | 2 | novel transcript |
| SPOPL | 2 | speckle type BTB/POZ protein like [Source:HG |
| AC092620.1 | 2 | novel transcript |
| AC092620.2 | 2 | novel transcript |
| LINC02631 | 2 | long intergenic non-protein coding RNA 2631 |
| NXPH2 | 2 | neurexophilin 2 [Source:HGNC Symbol;Acc:HG |
| RN7SKP286 | 2 | RN7SK pseudogene 286 [Source:HGNC Symbo |
| YY1P2 | 2 | YY1 transcription factor pseudogene 2 [Source |
| AHCYP4 | 2 | adenosylhomocysteinase pseudogene 4 [Sour |
| AC023468.1 | 2 | novel transcript |
| AC013265.1 | 2 | novel transcript |
| AC062021.1 | 2 | novel transcript |
| AC016710.1 | 2 | novel transcript |
| SNORA72 | 2 | Small nucleolar RNA SNORA72 [Source:RFAM; |
| MRPS18BP2 | 2 | mitochondrial ribosomal protein S18B pseudo |
| AC012497.1 | 2 | ribosomal protein L9 (RPL9) pseudogene |
| AC078851.1 | 2 | novel transcript |
| RN7SL283P | 2 | RNA, 7SL, cytoplasmic 283, pseudogene [Sour |
| LINC01853 | 2 | long intergenic non-protein coding RNA 1853 |
| MTCO1P44 | 2 | MT-CO1 pseudogene 44 [Source:HGNC Symbo |
| MTND2P19 | 2 | MT-ND2 pseudogene 19 [Source:HGNC Symbo |
| MTND1P27 | 2 | MT-ND1 pseudogene 27 [Source:HGNC Symbo |
| LRP1B | 2 | LDL receptor related protein 1B [Source:HGN( |
| AC092156.1 | 2 | NULL |
| MIR7157 | 2 | microRNA 7157 [Source:HGNC Symbol;Acc:HG |
| AC117947.1 | 2 | chromosome 17 open reading frame 79 (C17c |
| AC010740.1 | 2 | leucine rich repeat containing 57 (LRRC57) pse |
| Y_RNA | 2 | Y RNA [Source:RFAM;Acc:RF00019] |
| RNU6-904P | 2 | RNA, U6 small nuclear 904, pseudogene [Sour |
| AC068287.1 | 2 | ribosomal protein S16 (RPS16) pseudogene |
| AC078882.1 | 2 | novel transcript, sense overlapping to LRP1B |
| UBE2V1P14 | 2 | ubiquitin conjugating enzyme E2 V1 pseudoge |
| AC013444.1 | 2 | RRN3 RNA polymerase I transcription factor h |
| KYNU | 2 | kynureninase [Source:HGNC Symbol;Acc:HGN |
| AC013444.2 | 2 | sideroflexin 4 (SFXN4) pseudogene |
| ARHGAP15 | 2 | Rho GTPase activating protein 15 [Source:HGI |
| MTCYBP11 | 2 | MT-CYB pseudogene 11 [Source:HGNC Symbo |
| MTND6P11 | 2 | MT-ND6 pseudogene 11 [Source:HGNC Symbo |
| MTND5P24 | 2 | MT-ND5 pseudogene 24 [Source:HGNC Symbo |
| MTND4P22 | 2 | MT-ND4 pseudogene 22 [Source:HGNC Symbo |
| MTND4LP12 | 2 | MT-ND4L pseudogene 12 [Source:HGNC Symb |
| MTND3P9 | 2 | MT-ND3 pseudogene 9 [Source:HGNC Symbol |

| | | |
|---|---|---|
| MTCO3P5 | 2 | MT-CO3 pseudogene 5 [Source:HGNC Symbol |
| AC013437.2 | 2 | mitochondrially encoded ATP synthase 6 (MT- |
| MTCO2P5 | 2 | MT-CO2 pseudogene 5 [Source:HGNC Symbol |
| AC013437.1 | 2 | novel transcript, antisense to ARHGAP15 |
| AC079793.1 | 2 | novel transcript, antisense to ARHGAP15 |
| AC092652.3 | 2 | novel transcript, sense intronic to ARHGAP15 |
| AC092652.1 | 2 | novel transcript, antisense to ARHGAP15 |
| AC092652.2 | 2 | novel transcript, antisense to ARHGAP15 |
| AC079584.2 | 2 | novel transcript, antisense to ARHGAP15 |
| AC079584.1 | 2 | novel transcript |
| Y_RNA | 2 | Y RNA [Source:RFAM;Acc:RF00019] |
| AC016910.1 | 2 | novel transcript |
| GTDC1 | 2 | glycosyltransferase like domain containing 1 [ |
| ZEB2 | 2 | zinc finger E-box binding homeobox 2 [Source |
| AC009951.5 | 2 | novel transcript, antisense to ZEB2 |
| AC009951.4 | 2 | TEC |
| ZEB2-AS1 | 2 | ZEB2 antisense RNA 1 [Source:HGNC Symbol; |
| AC009951.2 | 2 | NULL |
| AC009951.1 | 2 | NULL |
| AC009951.3 | 2 | NULL |
| AC009951.6 | 2 | novel transcript |
| LINC01412 | 2 | long intergenic non-protein coding RNA 1412 |
| TEX41 | 2 | testis expressed 41 [Source:HGNC Symbol;Acc |
| AC023128.1 | 2 | novel transcript |
| AC023128.3 | 2 | NULL |
| AC023128.2 | 2 | sarcoglycan, epsilon (SGCE) pseudogene |
| LINC01966 | 2 | long intergenic non-protein coding RNA 1966 |
| AC096666.1 | 2 | novel transcript |
| RPL6P5 | 2 | ribosomal protein L6 pseudogene 5 [Source:H |
| AC092484.1 | 2 | novel transcript |
| AC079163.2 | 2 | novel transcript |
| AC079163.1 | 2 | novel transcript |
| AC079248.2 | 2 | methionyl aminopeptidase 2 (METAP2) pseud |
| AC079248.1 | 2 | novel transcript |
| AC079248.3 | 2 | TEC |
| RNU7-2P | 2 | RNA, U7 small nuclear 2 pseudogene [Source: |
| RPL17P12 | 2 | ribosomal protein L17 pseudogene 12 [Source |
| AC093084.1 | 2 | orthodenticle homeobox 2 (OTX2) pseudogen |
| PABPC1P2 | 2 | poly(A) binding protein cytoplasmic 1 pseudo |
| AC103881.1 | 2 | glioblastoma amplified sequence (GBAS) pseu |
| LINC01911 | 2 | long intergenic non-protein coding RNA 1911 |
| AC062032.1 | 2 | novel transcript |
| AC018678.1 | 2 | novel transcript |
| RNU6-692P | 2 | RNA, U6 small nuclear 692, pseudogene [Sour |
| Y_RNA | 2 | Y RNA [Source:RFAM;Acc:RF00019] |
| RNU6-715P | 2 | RNA, U6 small nuclear 715, pseudogene [Sour |
| RNA5SP106 | 2 | RNA, 5S ribosomal pseudogene 106 [Source:H |
| AC009480.2 | 2 | ribosomal protein L26 (RPL26) pseudogene |
| ACVR2A | 2 | activin A receptor type 2A [Source:HGNC Sym |
| RNU6-1275P | 2 | RNA, U6 small nuclear 1275, pseudogene [Sou |

| | | |
|---|---|---|
| AC009480.1 | 2 | novel transcript |
| ORC4 | 2 | origin recognition complex subunit 4 [Source:I |
| MBD5 | 2 | methyl-CpG binding domain protein 5 [Source |
| AC019226.1 | 2 | TEC |
| AC019226.2 | 2 | TEC |
| AC019070.1 | 2 | NULL |
| AC019070.2 | 2 | Ubiquitin carboxyl-terminal hydrolase 12 (USF |
| RPS29P8 | 2 | ribosomal protein S29 pseudogene 8 [Source: |
| EPC2 | 2 | enhancer of polycomb homolog 2 [Source:HG |
| RNU2-9P | 2 | RNA, U2 small nuclear 9, pseudogene [Source |
| UBBP3 | 2 | ubiquitin B pseudogene 3 [Source:HGNC Symb |
| AC105402.3 | 2 | novel transcript |
| AC105402.2 | 2 | ribosomal protein S20 (RPS20) pseudogene |
| AC105402.1 | 2 | ubiquitin specific peptidase 8 (USP8) pseudog |
| KIF5C | 2 | kinesin family member 5C [Source:HGNC Sym |
| AC105402.4 | 2 | novel transcript, antisense to KIF5C |
| LYPD6B | 2 | LY6/PLAUR domain containing 6B [Source:HGI |
| TXNP5 | 2 | thioredoxin pseudogene 5 [Source:HGNC Sym |
| AC009230.1 | 2 | novel transcript, antisense to LYPD6B |
| FAM8A3P | 2 | family with sequence similarity 8 member A3, |
| LYPD6 | 2 | LY6/PLAUR domain containing 6 [Source:HGN |
| AC007394.1 | 2 | ribosomal protein pseudogene |
| MMADHC | 2 | metabolism of cobalamin associated D [Sourc |
| AC007364.1 | 2 | novel protein (LOC642340) |
| RNU6-601P | 2 | RNA, U6 small nuclear 601, pseudogene [Sour |
| LINC01931 | 2 | long intergenic non-protein coding RNA 1931 |
| AC016682.1 | 2 | novel transcript |
| LINC01818 | 2 | long intergenic non-protein coding RNA 1818 |
| LINC01817 | 2 | long intergenic non-protein coding RNA 1817 |
| RND3 | 2 | Rho family GTPase 3 [Source:HGNC Symbol;Ac |
| LINC01920 | 2 | long intergenic non-protein coding RNA 1920 |
| AC104777.2 | 2 | novel transcript |
| AC104777.1 | 2 | novel transcript |
| LINC02612 | 2 | long intergenic non-protein coding RNA 2612 |
| AC023469.2 | 2 | novel transcript |
| AC023469.1 | 2 | novel transcript |
| AC018731.1 | 2 | novel transcript |
| FABP5P10 | 2 | fatty acid binding protein 5 pseudogene 10 [S |
| RBM43 | 2 | RNA binding motif protein 43 [Source:HGNC S |
| NMI | 2 | N-myc and STAT interactor [Source:HGNC Sym |
| TNFAIP6 | 2 | TNF alpha induced protein 6 [Source:HGNC Sy |
| MIR4773-2 | 2 | microRNA 4773-2 [Source:HGNC Symbol;Acc:I |
| MIR4773-1 | 2 | microRNA 4773-1 [Source:HGNC Symbol;Acc:I |
| RN7SL124P | 2 | RNA, 7SL, cytoplasmic 124, pseudogene [Sour |
| RIF1 | 2 | replication timing regulatory factor 1 [Source: |
| NEB | 2 | nebulin [Source:HGNC Symbol;Acc:HGNC:772 |
| ARL5A | 2 | ADP ribosylation factor like GTPase 5A [Sourc |
| AC097448.1 | 2 | novel transcript, antisense to ARL5A |
| AC068547.1 | 2 | novel transcript |
| CACNB4 | 2 | calcium voltage-gated channel auxiliary subur |

| | | |
|---|---|---|
| AC079790.1 | 2 | novel transcript (LOC100127920) |
| STAM2 | 2 | signal transducing adaptor molecule 2 [Source |
| RPL30P2 | 2 | ribosomal protein L30 pseudogene 2 [Source: |
| AC079790.2 | 2 | novel transcript |
| Y_RNA | 2 | Y RNA [Source:RFAM;Acc:RF00019] |
| FMNL2 | 2 | formin like 2 [Source:HGNC Symbol;Acc:HGNC |
| AC012066.1 | 2 | nuclear distribution gene C (NUDC) pseudoger |
| AC012066.2 | 2 | novel transcript, antisense to FMNL2 |
| AC012443.1 | 2 | ribosomal protein L22 (RPL22) pseudogene |
| AC012443.2 | 2 | novel transcript, antisense to FMNL2 |
| PRPF40A | 2 | pre-mRNA processing factor 40 homolog A [So |
| ARL6IP6 | 2 | ADP ribosylation factor like GTPase 6 interacti |
| AC009969.1 | 2 | makorin, ring finger protein 1 (MKRN1) pseud |
| UBQLN4P2 | 2 | ubiquilin 4 pseudogene 2 [Source:HGNC Symb |
| ATP5PBP4 | 2 | ATP synthase peripheral stalk-membrane subu |
| AC011901.1 | 2 | novel transcript |
| AC079150.1 | 2 | pseudogene similar to part of phospholipase C |
| LINC01850 | 2 | long intergenic non-protein coding RNA 1850 |
| RPL23AP29 | 2 | ribosomal protein L23a pseudogene 29 [Sourc |
| AC012501.3 | 2 | tubulin, alpha 1b (TUBA1B) pseudogene |
| AC012501.2 | 2 | novel transcript |
| AC012501.1 | 2 | NULL |
| RPRM | 2 | reprimo, TP53 dependent G2 arrest mediator |
| DNAJA1P2 | 2 | DnaJ heat shock protein family (Hsp40) memb |
| GALNT13 | 2 | polypeptide N-acetylgalactosaminyltransferas |
| PHBP4 | 2 | prohibitin pseudogene 4 [Source:HGNC Symbo |
| AC009227.1 | 2 | novel transcript |
| AC009227.2 | 2 | novel transcript |
| RNA5SP107 | 2 | RNA, 5S ribosomal pseudogene 107 [Source:H |
| AC061961.1 | 2 | novel transcript |
| KCNJ3 | 2 | potassium voltage-gated channel subfamily J r |
| CBX3P6 | 2 | chromobox 3 pseudogene 6 [Source:HGNC Sy |
| AC092625.1 | 2 | novel transcript |
| RNU6-1001P | 2 | RNA, U6 small nuclear 1001, pseudogene [Sou |
| MTCO1P45 | 2 | MT-CO1 pseudogene 45 [Source:HGNC Symbo |
| MTND2P20 | 2 | MT-ND2 pseudogene 20 [Source:HGNC Symbo |
| ATP5F1AP2 | 2 | ATP synthase F1 subunit alpha pseudogene 2 |
| MTND4P28 | 2 | MT-ND4 pseudogene 28 [Source:HGNC Symbo |
| MTND5P30 | 2 | MT-ND5 pseudogene 30 [Source:HGNC Symbo |
| MTND6P9 | 2 | MT-ND6 pseudogene 9 [Source:HGNC Symbol |
| MTCYBP9 | 2 | MT-CYB pseudogene 9 [Source:HGNC Symbol; |
| RNU6-546P | 2 | RNA, U6 small nuclear 546, pseudogene [Sour |
| AC073225.1 | 2 | novel transcript |
| LINC01876 | 2 | long intergenic non-protein coding RNA 1876 |
| AC073551.1 | 2 | heme binding protein 2 (HEBP2) pseudogene |
| AC074099.1 | 2 | novel transcript |
| NR4A2 | 2 | nuclear receptor subfamily 4 group A member |
| GPD2 | 2 | glycerol-3-phosphate dehydrogenase 2 [Sourc |
| LINC01958 | 2 | long intergenic non-protein coding RNA 1958 |
| RPLP0P7 | 2 | ribosomal protein lateral stalk subunit P0 pseu |

| | | |
|---|---|---|
| AC096589.2 | 2 | novel transcript |
| AC096589.1 | 2 | novel transcript |
| AC108057.1 | 2 | novel transcript |
| CDK7P1 | 2 | cyclin dependent kinase 7 pseudogene 1 [Sour |
| GALNT5 | 2 | polypeptide N-acetylgalactosaminyltransferas |
| RN7SKP281 | 2 | RN7SK pseudogene 281 [Source:HGNC Symbo |
| ERMN | 2 | ermin [Source:HGNC Symbol;Acc:HGNC:2920 |
| FAM133DP | 2 | family with sequence similarity 133 member [ |
| CYTIP | 2 | cytohesin 1 interacting protein [Source:HGNC |
| ACVR1C | 2 | activin A receptor type 1C [Source:HGNC Sym |
| AC019186.1 | 2 | novel transcript |
| ACVR1 | 2 | activin A receptor type 1 [Source:HGNC Symb |
| UPP2 | 2 | uridine phosphorylase 2 [Source:HGNC Symbo |
| AC013731.1 | 2 | novel transcript |
| RNU6-436P | 2 | RNA, U6 small nuclear 436, pseudogene [Sour |
| AC091488.1 | 2 | novel transcript |
| AC005539.1 | 2 | metastasis associated 1 family, member 3 (M |
| RNU6-932P | 2 | RNA, U6 small nuclear 932, pseudogene [Sour |
| PTP4A1P1 | 2 | PTP4A1 pseudogene 1 [Source:HGNC Symbol; |
| UPP2-IT1 | 2 | UPP2 intronic transcript 1 [Source:HGNC Sym |
| CCDC148-AS1 | 2 | CCDC148 antisense RNA 1 [Source:HGNC Sym |
| CCDC148 | 2 | coiled-coil domain containing 148 [Source:HG |
| HNRNPDLP2 | 2 | HNRNPDL pseudogene 2 [Source:HGNC Symb |
| RN7SL393P | 2 | RNA, 7SL, cytoplasmic 393, pseudogene [Sour |
| PKP4 | 2 | plakophilin 4 [Source:HGNC Symbol;Acc:HGN |
| AC005042.1 | 2 | ribosomal protein L7a (RPL7A) pseudogene |
| PKP4-AS1 | 2 | PKP4 antisense RNA 1 [Source:HGNC Symbol; |
| AC005042.2 | 2 | novel transcript |
| AC005042.3 | 2 | nuclear receptor coactivator 6 (NCOA6) pseud |
| DAPL1 | 2 | death associated protein like 1 [Source:HGNC |
| AC064843.1 | 2 | novel transcript, antisense to DAPL1 |
| OR7E89P | 2 | olfactory receptor family 7 subfamily E memb |
| OR7E28P | 2 | olfactory receptor family 7 subfamily E memb |
| OR7E90P | 2 | olfactory receptor family 7 subfamily E memb |
| RNU2-21P | 2 | RNA, U2 small nuclear 21, pseudogene [Sourc |
| TANC1 | 2 | tetratricopeptide repeat, ankyrin repeat and c |
| BTF3L4P2 | 2 | basic transcription factor 3 like 4 pseudogene |
| RNU6-580P | 2 | RNA, U6 small nuclear 580, pseudogene [Sour |
| GSTM3P2 | 2 | glutathione S-transferase mu 3 pseudogene 2 |
| MIR6888 | 2 | microRNA 6888 [Source:HGNC Symbol;Acc:HG |
| WDSUB1 | 2 | WD repeat, sterile alpha motif and U-box dom |
| BAZ2B | 2 | bromodomain adjacent to zinc finger domain |
| AC008277.1 | 2 | novel transcript, antisense to BAZ2B |
| AC008277.2 | 2 | elongation factor Tu GTP binding domain cont |
| AC009506.2 | 2 | capping protein (actin filament) muscle Z-line, |
| Y_RNA | 2 | Y RNA [Source:RFAM;Acc:RF00019] |
| AC009506.1 | 2 | novel transcript, antisense to BAZ2B |
| AC009961.1 | 2 | novel transcript |
| AC009961.3 | 2 | ribosomal protein S3A (RPS3A) pseudogene |
| MARCH7 | 2 | membrane associated ring-CH-type finger 7 [S |

| | | |
|---|---|---|
| CD302 | 2 | CD302 molecule [Source:HGNC Symbol;Acc:H |
| LY75-CD302 | 2 | LY75-CD302 readthrough [Source:HGNC Symb |
| AC009961.4 | 2 | novel transcript, sense intronic to CD302and L |
| LY75 | 2 | lymphocyte antigen 75 [Source:HGNC Symbol |
| AC009961.2 | 2 | centrosomal protein 57kDa (CEP57) pseudoge |
| PLA2R1 | 2 | phospholipase A2 receptor 1 [Source:HGNC Sy |
| ITGB6 | 2 | integrin subunit beta 6 [Source:HGNC Symbol; |
| AC092153.1 | 2 | novel transcript |
| LINC02478 | 2 | long intergenic non-protein coding RNA 2478 |
| RBMS1 | 2 | RNA binding motif single stranded interacting |
| MIR4785 | 2 | microRNA 4785 [Source:HGNC Symbol;Acc:HG |
| AC096656.1 | 2 | TEC |
| RN7SL423P | 2 | RNA, 7SL, cytoplasmic 423, pseudogene [Sour |
| AC009313.2 | 2 | DEAD (Asp-Glu-Ala-Asp) box polypeptide 18 (D |
| AC009313.1 | 2 | novel transcript |
| TANK | 2 | TRAF family member associated NFKB activato |
| AC009299.3 | 2 | novel transcript, antisense to TANK |
| LINC01806 | 2 | long intergenic non-protein coding RNA 1806 |
| AC009299.1 | 2 | novel pseudogene |
| PSMD14 | 2 | proteasome 26S subunit, non-ATPase 14 [Sou |
| AC009299.2 | 2 | matrix-remodelling associated 7 (MXRA7) pse |
| RNA5SP108 | 2 | RNA, 5S ribosomal pseudogene 108 [Source:H |
| TBR1 | 2 | T-box brain transcription factor 1 [Source:HGN |
| AC009487.1 | 2 | novel transcript |
| AC009487.3 | 2 | novel transcript |
| SLC4A10 | 2 | solute carrier family 4 member 10 [Source:HG |
| AC009487.4 | 2 | TEC |
| AC009487.2 | 2 | AT hook containing transcription factor 1 (AH |
| AC092654.1 | 2 | NULL |
| AC062022.1 | 2 | chromosome 17 open reading frame 80 (C17o |
| AC062022.2 | 2 | novel transcript, antisense to SLC4A10 |
| KRT18P46 | 2 | keratin 18 pseudogene 46 [Source:HGNC Sym |
| RPEP5 | 2 | ribulose-5-phosphate-3-epimerase pseudoger |
| DPP4 | 2 | dipeptidyl peptidase 4 [Source:HGNC Symbol; |
| AC008063.1 | 2 | novel transcript |
| TIMM8AP1 | 2 | translocase of inner mitochondrial membrane |
| EIF3EP2 | 2 | eukaryotic translation initiation factor 3 subur |
| AC008063.2 | 2 | novel transcript |
| GCG | 2 | glucagon [Source:HGNC Symbol;Acc:HGNC:41 |
| AC007750.1 | 2 | novel transcript |
| FAP | 2 | fibroblast activation protein alpha [Source:HG |
| IFIH1 | 2 | interferon induced with helicase C domain 1 [ |
| GCA | 2 | grancalcin [Source:HGNC Symbol;Acc:HGNC:1 |
| KCNH7 | 2 | potassium voltage-gated channel subfamily H |
| RNA5SP109 | 2 | RNA, 5S ribosomal pseudogene 109 [Source:H |
| AC011900.1 | 2 | novel transcript |
| RPL7P61 | 2 | ribosomal protein L7 pseudogene 61 [Source: |
| RNU6-627P | 2 | RNA, U6 small nuclear 627, pseudogene [Sour |
| FIGN | 2 | fidgetin, microtubule severing factor [Source: |
| AC016766.1 | 2 | novel transcript |

| | | |
|---|---|---|
| PRPS1P1 | 2 | phosphoribosyl pyrophosphate synthetase 1 p |
| CYP2C56P | 2 | cytochrome P450 family 2 subfamily C membe |
| GRB14 | 2 | growth factor receptor bound protein 14 [Sou |
| AC107075.1 | 2 | eukaryotic translation initiation factor 3, subu |
| COBLL1 | 2 | cordon-bleu WH2 repeat protein like 1 [Sourc |
| AC019181.1 | 2 | pseudogene similar to part of ribosomal prote |
| SNORA70F | 2 | small nucleolar RNA, H/ACA box 70F [Source:H |
| AC019197.1 | 2 | novel transcript, antisense to SLC38A11 and S |
| RNA5SP110 | 2 | RNA, 5S ribosomal pseudogene 110 [Source:H |
| RNA5SP111 | 2 | RNA, 5S ribosomal pseudogene 111 [Source:H |
| SLC38A11 | 2 | solute carrier family 38 member 11 [Source:H |
| SCN3A | 2 | sodium voltage-gated channel alpha subunit 3 |
| SCN2A | 2 | sodium voltage-gated channel alpha subunit 2 |
| MAPRE1P3 | 2 | MAPRE1 pseudogene 3 [Source:HGNC Symbol |
| AC011303.1 | 2 | novel transcript |
| CSRNP3 | 2 | cysteine and serine rich nuclear protein 3 [Sou |
| GALNT3 | 2 | polypeptide N-acetylgalactosaminyltransferas |
| AC009495.3 | 2 | novel transcript |
| AC009495.1 | 2 | novel transcript |
| AC009495.2 | 2 | novel transcript |
| TTC21B | 2 | tetratricopeptide repeat domain 21B [Source: |
| TTC21B-AS1 | 2 | TTC21B antisense RNA 1 [Source:HGNC Symbo |
| SCN1A-AS1 | 2 | SCN1A and SCN9A antisense RNA 1 [Source:H |
| SCN1A | 2 | sodium voltage-gated channel alpha subunit 1 |
| Y_RNA | 2 | Y RNA [Source:RFAM;Acc:RF00019] |
| RN7SKP152 | 2 | RN7SK pseudogene 152 [Source:HGNC Symbo |
| SCN9A | 2 | sodium voltage-gated channel alpha subunit 9 |
| SCN7A | 2 | sodium voltage-gated channel alpha subunit 7 |
| AC074101.1 | 2 | mitochondrially encoded cytochrome c oxidas |
| XIRP2 | 2 | xin actin binding repeat containing 2 [Source:H |
| AC092601.1 | 2 | PDGFA associated protein 1 (PDAP1) pseudog |
| XIRP2-AS1 | 2 | XIRP2 antisense RNA 1 [Source:HGNC Symbol; |
| AC073050.1 | 2 | novel transcript (hCG1660379) |
| RNU7-148P | 2 | RNA, U7 small nuclear 148 pseudogene [Sourc |
| CTAGE14P | 2 | CTAGE family member 14, pseudogene [Sourc |
| AC016723.1 | 2 | novel transcript |
| B3GALT1 | 2 | beta-1,3-galactosyltransferase 1 [Source:HGN |
| STK39 | 2 | serine/threonine kinase 39 [Source:HGNC Sym |
| AC067940.1 | 2 | PHD finger protein 5A (PHF5A) pseudogene |
| RN7SL813P | 2 | RNA, 7SL, cytoplasmic 813, pseudogene [Sour |
| CERS6 | 2 | ceramide synthase 6 [Source:HGNC Symbol;Ac |
| MIR4774 | 2 | microRNA 4774 [Source:HGNC Symbol;Acc:HG |
| RNU6-766P | 2 | RNA, U6 small nuclear 766, pseudogene [Sour |
| CERS6-AS1 | 2 | CERS6 antisense RNA 1 [Source:HGNC Symbol |
| NOSTRIN | 2 | nitric oxide synthase trafficking [Source:HGNC |
| SPC25 | 2 | SPC25 component of NDC80 kinetochore com |
| G6PC2 | 2 | glucose-6-phosphatase catalytic subunit 2 [So |
| ABCB11 | 2 | ATP binding cassette subfamily B member 11 |
| DHRS9 | 2 | dehydrogenase/reductase 9 [Source:HGNC Sy |
| AC007556.1 | 2 | novel transcript |

| | | |
|---|---|---|
| UBE2V1P6 | 2 | ubiquitin conjugating enzyme E2 V1 pseudoge |
| LRP2 | 2 | LDL receptor related protein 2 [Source:HGNC |
| AC093899.2 | 2 | novel protein |
| BBS5 | 2 | Bardet-Biedl syndrome 5 [Source:HGNC Symb |
| KLHL41 | 2 | kelch like family member 41 [Source:HGNC Sy |
| FASTKD1 | 2 | FAST kinase domains 1 [Source:HGNC Symbol; |
| AC093899.1 | 2 | pseudogene similar to part of inositol polypho |
| PPIG | 2 | peptidylprolyl isomerase G [Source:HGNC Sym |
| CCDC173 | 2 | coiled-coil domain containing 173 [Source:HG |
| AC016772.1 | 2 | family with sequence similarity 49, member B |
| PHOSPHO2 | 2 | phosphatase, orphan 2 [Source:HGNC Symbol |
| KLHL23 | 2 | kelch like family member 23 [Source:HGNC Sy |
| AC009967.1 | 2 | NULL |
| PTCHD3P2 | 2 | patched domain containing 3 pseudogene 2 [S |
| AC009967.2 | 2 | cytochrome b5 type A (microsomal) (CYB5A) p |
| SSB | 2 | small RNA binding exonuclease protection fac |
| METTL5 | 2 | methyltransferase like 5 [Source:HGNC Symbo |
| UBR3 | 2 | ubiquitin protein ligase E3 component n-recog |
| AC092641.1 | 2 | TGF beta-inducible nuclear protein 1 (TINP1) p |
| RNU6-1006P | 2 | RNA, U6 small nuclear 1006, pseudogene [Sou |
| MYO3B | 2 | myosin IIIB [Source:HGNC Symbol;Acc:HGNC:1 |
| MYO3B-AS1 | 2 | MYO3B antisense RNA 1 [Source:HGNC Symbo |
| HMGB1P4 | 2 | high mobility group box 1 pseudogene 4 [Sour |
| AC007277.1 | 2 | novel transcript, antisense to MYO3B |
| AC007277.2 | 2 | NULL |
| AC007405.1 | 2 | novel transcript (FLJ13453) |
| LINC01124 | 2 | long intergenic non-protein coding RNA 1124 |
| SP5 | 2 | Sp5 transcription factor [Source:HGNC Symbo |
| EIF2S2P4 | 2 | eukaryotic translation initiation factor 2 subur |
| AC007405.3 | 2 | glutamate rich 2 [Source:NCBI gene;Acc:28514 |
| Y_RNA | 2 | Y RNA [Source:RFAM;Acc:RF00019] |
| ERICH2 | 2 | glutamate rich 2 [Source:HGNC Symbol;Acc:H |
| GAD1 | 2 | glutamate decarboxylase 1 [Source:HGNC Sym |
| AC007405.2 | 2 | novel transcript |
| AC007405.4 | 2 | novel transcript, antisense to GAD1 |
| AC010092.1 | 2 | novel transcript |
| GORASP2 | 2 | golgi reassembly stacking protein 2 [Source:H |
| TLK1 | 2 | tousled like kinase 1 [Source:HGNC Symbol;Ac |
| METTL8 | 2 | methyltransferase like 8 [Source:HGNC Symbo |
| AC008065.1 | 2 | ribosomal protein S26 (RPS26) pseudogene |
| DCAF17 | 2 | DDB1 and CUL4 associated factor 17 [Source:H |
| DAP3P2 | 2 | death associated protein 3 pseudogene 2 [Sou |
| AC007969.2 | 2 | ubiquitin-fold modifier 1 (UFM1) pseudogene |
| AC007969.1 | 2 | ribosomal protein S15 (RPS15) pseudogene |
| CYBRD1 | 2 | cytochrome b reductase 1 [Source:HGNC Sym |
| RPL21P38 | 2 | ribosomal protein L21 pseudogene 38 [Source |
| DYNC1I2 | 2 | dynein cytoplasmic 1 intermediate chain 2 [Sc |
| SLC25A12 | 2 | solute carrier family 25 member 12 [Source:H |
| RNU6-182P | 2 | RNA, U6 small nuclear 182, pseudogene [Sour |
| HAT1 | 2 | histone acetyltransferase 1 [Source:HGNC Syn |

| | | |
|---|---|---|
| METAP1D | 2 | methionyl aminopeptidase type 1D, mitochon |
| DLX1 | 2 | distal-less homeobox 1 [Source:HGNC Symbol |
| DLX2 | 2 | distal-less homeobox 2 [Source:HGNC Symbol |
| DLX2-DT | 2 | DLX2 divergent transcript [Source:HGNC Symb |
| AC104088.1 | 2 | novel transcript |
| AC104088.3 | 2 | novel transcript |
| AC104088.2 | 2 | novel transcript |
| AC078883.3 | 2 | TEC |
| ITGA6 | 2 | integrin subunit alpha 6 [Source:HGNC Symbo |
| AC078883.2 | 2 | novel transcript |
| ITGA6-AS1 | 2 | ITGA6 antisense RNA 1 [Source:HGNC Symbol |
| AC078883.1 | 2 | novel transcript |
| PDK1 | 2 | pyruvate dehydrogenase kinase 1 [Source:HG |
| Y_RNA | 2 | Y RNA [Source:RFAM;Acc:RF00019] |
| AC018712.1 | 2 | stem-loop binding protein (SLBP) pseudogene |
| RAPGEF4-AS1 | 2 | RAPGEF4 antisense RNA 1 [Source:HGNC Sym |
| AC018712.2 | 2 | NULL |
| RAPGEF4 | 2 | Rap guanine nucleotide exchange factor 4 [So |
| ALDH7A1P2 | 2 | aldehyde dehydrogenase 7 family member A1 |
| MAP3K20 | 2 | mitogen-activated protein kinase kinase kinas |
| MAP3K20-AS1 | 2 | MAP3K20 antisense RNA 1 [Source:HGNC Sym |
| AC092573.1 | 2 | ribosomal protein S2 (RPS2) pseudogene |
| AC092573.2 | 2 | novel pseudogene |
| CDCA7 | 2 | cell division cycle associated 7 [Source:HGNC |
| AC073465.1 | 2 | hematological and neurological expressed 1 ( |
| AC073465.2 | 2 | chibby homolog 1 (Drosophila) (CBY1) pseudo |
| PPIAP66 | 2 | peptidylprolyl isomerase A pseudogene 66 [Sc |
| AC013410.1 | 2 | heat shock protein 90kDa alpha (cytosolic), cl |
| AC013410.2 | 2 | dynein, axonemal, heavy chain 11 (DNAH11) p |
| AC106900.1 | 2 | novel transcript |
| RPL5P7 | 2 | ribosomal protein L5 pseudogene 7 [Source:H |
| SP3 | 2 | Sp3 transcription factor [Source:HGNC Symbo |
| AC016737.1 | 2 | novel transcript |
| AC016737.2 | 2 | novel transcript |
| LINC01960 | 2 | long intergenic non-protein coding RNA 1960 |
| RPSAP24 | 2 | ribosomal protein SA pseudogene 24 [Source: |
| OLA1 | 2 | Obg like ATPase 1 [Source:HGNC Symbol;Acc: |
| AC013467.2 | 2 | Y box binding protein 1 (YBX1) pseudogene |
| RN7SL65P | 2 | RNA, 7SL, cytoplasmic 65, pseudogene [Sourc |
| Y_RNA | 2 | Y RNA [Source:RFAM;Acc:RF00019] |
| AC013467.1 | 2 | leucine rich repeat containing 2 (LRRC2) pseu |
| HNRNPA1P39 | 2 | heterogeneous nuclear ribonucleoprotein A1 |
| LINC01305 | 2 | long intergenic non-protein coding RNA 1305 |
| AC018470.1 | 2 | novel transcript |
| SP9 | 2 | Sp9 transcription factor [Source:HGNC Symbo |
| CIR1 | 2 | corepressor interacting with RBPJ, 1 [Source: |
| SCRN3 | 2 | secernin 3 [Source:HGNC Symbol;Acc:HGNC:3 |
| GPR155 | 2 | G protein-coupled receptor 155 [Source:HGN |
| AC010894.2 | 2 | novel transcript |
| AC010894.1 | 2 | replication protein A3, 14kDa (RPA3) pseudog |

| | | |
|---|---|---|
| AC010894.5 | 2 | TEC |
| AC010894.3 | 2 | novel transcript, antisense to WIPF1 and CHRI |
| RNU6-5P | 2 | RNA, U6 small nuclear 5, pseudogene [Source |
| WIPF1 | 2 | WAS/WASL interacting protein family membe |
| AC010894.4 | 2 | novel transcript, antisense WIPF1 |
| H3F3AP4 | 2 | H3 histone, family 3A, pseudogene 4 [Source: |
| CHRNA1 | 2 | cholinergic receptor nicotinic alpha 1 subunit |
| CHN1 | 2 | chimerin 1 [Source:HGNC Symbol;Acc:HGNC:1 |
| RNU6-763P | 2 | RNA, U6 small nuclear 763, pseudogene [Sour |
| RNU6-1290P | 2 | RNA, U6 small nuclear 1290, pseudogene [Sou |
| AC007435.1 | 2 | ribosomal protein L21 (RPL21) pseudogene |
| ATF2 | 2 | activating transcription factor 2 [Source:HGN( |
| AC007435.2 | 2 | NULL |
| MIR933 | 2 | microRNA 933 [Source:HGNC Symbol;Acc:HGI |
| AC096649.1 | 2 | novel transcript |
| ATP5MC3 | 2 | ATP synthase membrane subunit c locus 3 [So |
| Y_RNA | 2 | Y RNA [Source:RFAM;Acc:RF00019] |
| AC096649.2 | 2 | ribosomal protein S15a (RPS15A) pseudogene |
| AC093459.1 | 2 | novel transcript |
| EXTL2P1 | 2 | exostosin like glycosyltransferase 2 pseudoge |
| AC016751.2 | 2 | novel transcript |
| AC016751.1 | 2 | novel transcript |
| LNPK | 2 | lunapark, ER junction formation factor [Sourc |
| EVX2 | 2 | even-skipped homeobox 2 [Source:HGNC Sym |
| HOXD13 | 2 | homeobox D13 [Source:HGNC Symbol;Acc:HG |
| HOXD12 | 2 | homeobox D12 [Source:HGNC Symbol;Acc:HG |
| HOXD11 | 2 | homeobox D11 [Source:HGNC Symbol;Acc:HG |
| HOXD10 | 2 | homeobox D10 [Source:HGNC Symbol;Acc:HG |
| HOXD-AS2 | 2 | HOXD cluster antisense RNA 2 [Source:HGNC : |
| HOXD9 | 2 | homeobox D9 [Source:HGNC Symbol;Acc:HGN |
| HOXD8 | 2 | homeobox D8 [Source:HGNC Symbol;Acc:HGN |
| HOXD3 | 2 | homeobox D3 [Source:HGNC Symbol;Acc:HGN |
| MIR10B | 2 | microRNA 10b [Source:HGNC Symbol;Acc:HGI |
| HOXD4 | 2 | homeobox D4 [Source:HGNC Symbol;Acc:HGN |
| HAGLR | 2 | HOXD antisense growth-associated long non-( |
| AC009336.1 | 2 | novel transcript, antisense to HOXD3 |
| HAGLROS | 2 | HAGLR opposite strand lncRNA [Source:HGNC |
| HOXD1 | 2 | homeobox D1 [Source:HGNC Symbol;Acc:HGN |
| MIR7704 | 2 | microRNA 7704 [Source:HGNC Symbol;Acc:HG |
| AC009336.2 | 2 | TEC |
| AC016739.1 | 2 | ribosomal protein, large, P1 (RPLP1) pseudoge |
| RPSAP25 | 2 | ribosomal protein SA pseudogene 25 [Source: |
| MTX2 | 2 | metaxin 2 [Source:HGNC Symbol;Acc:HGNC:7! |
| PPIAP67 | 2 | peptidylprolyl isomerase A pseudogene 67 [Sc |
| LINC01117 | 2 | long intergenic non-protein coding RNA 1117 |
| AC017048.1 | 2 | novel transcript |
| AC017048.2 | 2 | novel transcript |
| MIR1246 | 2 | microRNA 1246 [Source:HGNC Symbol;Acc:HG |
| AC017048.3 | 2 | novel transcript |
| LINC01116 | 2 | long intergenic non-protein coding RNA 1116 |

| | | |
|---|---|---|
| RNU6ATAC14P | 2 | RNA, U6atac small nuclear 14, pseudogene [So |
| AC092162.2 | 2 | novel transcript |
| FUCA1P1 | 2 | alpha-L-fucosidase 1 pseudogene 1 [Source:H |
| AC092162.1 | 2 | ribosomal protein L29 (RPL29) pseudogene |
| AC092162.3 | 2 | TEC |
| AC073636.1 | 2 | novel transcript |
| RNU6-187P | 2 | RNA, U6 small nuclear 187, pseudogene [Sour |
| AC074286.1 | 2 | novel transcript |
| AC079305.3 | 2 | novel transcript |
| RNA5SP112 | 2 | RNA, 5S ribosomal pseudogene 112 [Source:H |
| AC079305.2 | 2 | STIP1 homology and U-box containing protein |
| Y_RNA | 2 | Y RNA [Source:RFAM;Acc:RF00019] |
| KRT8P40 | 2 | keratin 8 pseudogene 40 [Source:HGNC Symb |
| HNRNPA3 | 2 | heterogeneous nuclear ribonucleoprotein A3 |
| MIR4444-1 | 2 | microRNA 4444-1 [Source:HGNC Symbol;Acc:I |
| NFE2L2 | 2 | nuclear factor, erythroid 2 like 2 [Source:HGN |
| DNAJC19P5 | 2 | DnaJ heat shock protein family (Hsp40) memb |
| MIR3128 | 2 | microRNA 3128 [Source:HGNC Symbol;Acc:HG |
| AC079305.1 | 2 | novel transcript |
| AC019080.1 | 2 | novel transcript |
| AC019080.4 | 2 | novel transcript, antisense to NFE2L2 |
| AC019080.5 | 2 | novel transcript, antisense to NFE2L2 |
| MIR6512 | 2 | microRNA 6512 [Source:HGNC Symbol;Acc:HG |
| AC019080.6 | 2 | novel transcript |
| AC019080.2 | 2 | Histone H3 pseudogene |
| AC019080.3 | 2 | NULL |
| AGPS | 2 | alkylglycerone phosphate synthase [Source:H |
| TTC30B | 2 | tetratricopeptide repeat domain 30B [Source: |
| AC073834.1 | 2 | novel transcript, antisense to TTC30A |
| TTC30A | 2 | tetratricopeptide repeat domain 30A [Source: |
| PDE11A | 2 | phosphodiesterase 11A [Source:HGNC Symbo |
| PDE11A | 2 | phosphodiesterase 11A [Source:NCBI gene;Ac |
| AC012499.1 | 2 | novel transcript |
| AC011998.3 | 2 | novel transcript, sense intronic to PDE11A |
| API5P2 | 2 | apoptosis inhibitor 5 pseudogene 2 [Source:H |
| AC011998.2 | 2 | succinate dehydrogenase complex, subunit D, |
| RNU6-629P | 2 | RNA, U6 small nuclear 629, pseudogene [Sour |
| CYCTP | 2 | cytochrome c, testis, pseudogene [Source:HG |
| AC011998.1 | 2 | peroxiredoxin 5 (PRDX5) pseudogene |
| RBM45 | 2 | RNA binding motif protein 45 [Source:HGNC S |
| RNU5E-9P | 2 | RNA, U5E small nuclear 9, pseudogene [Sourc |
| OSBPL6 | 2 | oxysterol binding protein like 6 [Source:HGNC |
| CHROMR | 2 | cholesterol induced regulator of metabolism I |
| PRKRA | 2 | protein activator of interferon induced protei |
| PJVK | 2 | pejvakin [Source:HGNC Symbol;Acc:HGNC:29! |
| AC009948.1 | 2 | nuclear distribution gene C homolog (A. nidula |
| FKBP7 | 2 | FKBP prolyl isomerase 7 [Source:HGNC Symbo |
| PLEKHA3 | 2 | pleckstrin homology domain containing A3 [So |
| TTN-AS1 | 2 | TTN antisense RNA 1 [Source:HGNC Symbol;A |
| TTN | 2 | titin [Source:HGNC Symbol;Acc:HGNC:12403] |

| | | |
|---|---|---|
| AC009948.3 | 2 | novel transcript, antisense to TTN |
| AC009948.2 | 2 | novel transcript, antisense to titin |
| AC009948.4 | 2 | novel transcript, antisense to titin |
| AC010680.3 | 2 | novel transcript, antisense to TTN |
| AC010680.2 | 2 | novel transcript, antisense to titin |
| AC010680.4 | 2 | novel transcript |
| AC010680.5 | 2 | novel transcript |
| AC010680.1 | 2 | novel transcript, antisense to titin |
| AC092640.1 | 2 | novel transcript, antisense to CCDC141and TT |
| CCDC141 | 2 | coiled-coil domain containing 141 [Source:HG |
| RNU7-104P | 2 | RNA, U7 small nuclear 104 pseudogene [Sourc |
| RPS6P2 | 2 | ribosomal protein S6 pseudogene 2 [Source:H |
| SESTD1 | 2 | SEC14 and spectrin domain containing 1 [Sour |
| AC093911.1 | 2 | novel transcript |
| AC068042.1 | 2 | RAD52 homolog (S. cerevisiae) (RAD52) pseud |
| ZNF385B | 2 | zinc finger protein 385B [Source:HGNC Symbo |
| AC092642.1 | 2 | TXNL4A thioredoxin-like 4A (TXNL4A) pseudog |
| MIR1258 | 2 | microRNA 1258 [Source:HGNC Symbol;Acc:HG |
| AC096587.2 | 2 | NULL |
| AC096587.1 | 2 | NULL |
| CWC22 | 2 | CWC22 spliceosome associated protein homo |
| AC012669.1 | 2 | ribosomal protein L17 (RPL17) pseudogene |
| AC009478.1 | 2 | novel transcript |
| SCHLAP1 | 2 | SWI/SNF complex antagonist associated with |
| AC104820.1 | 2 | TEC |
| FTH1P20 | 2 | ferritin heavy chain 1 pseudogene 20 [Source: |
| AC104076.2 | 2 | ribosomal protein L27a (RPL27A) pseudogene |
| UBE2E3 | 2 | ubiquitin conjugating enzyme E2 E3 [Source:H |
| AC104076.1 | 2 | novel transcript |
| AC068196.1 | 2 | novel transcript |
| LINC01934 | 2 | long intergenic non-protein coding RNA 1934 |
| MIR4437 | 2 | microRNA 4437 [Source:HGNC Symbol;Acc:HG |
| AC020595.1 | 2 | novel transcript |
| ITGA4 | 2 | integrin subunit alpha 4 [Source:HGNC Symbo |
| CERKL | 2 | ceramide kinase like [Source:HGNC Symbol;Ac |
| NEUROD1 | 2 | neuronal differentiation 1 [Source:HGNC Syml |
| AC013733.2 | 2 | novel transcript |
| AC013733.1 | 2 | novel protein (FLJ40163) |
| SAP18P2 | 2 | SAP18 pseudogene 2 [Source:HGNC Symbol;A |
| RNU6ATAC19P | 2 | RNA, U6atac small nuclear 19, pseudogene [S |
| AC009962.1 | 2 | novel transcript, antisense to SSFA2 |
| ITPRID2 | 2 | ITPR interacting domain containing 2 [Source: |
| PPP1R1C | 2 | protein phosphatase 1 regulatory inhibitor sul |
| KRT18P29 | 2 | keratin 18 pseudogene 29 [Source:HGNC Sym |
| RNA5SP113 | 2 | RNA, 5S ribosomal pseudogene 113 [Source:H |
| PDE1A | 2 | phosphodiesterase 1A [Source:HGNC Symbol; |
| AC012500.1 | 2 | novel transcript, antisense to PDE1A |
| RN7SL267P | 2 | RNA, 7SL, cytoplasmic 267, pseudogene [Sour |
| DNAJC10 | 2 | DnaJ heat shock protein family (Hsp40) memb |
| RPL31P15 | 2 | ribosomal protein L31 pseudogene 15 [Source |

| | | |
|---|---|---|
| Y_RNA | 2 | Y RNA [Source:RFAM;Acc:RF00019] |
| FRZB | 2 | frizzled related protein [Source:HGNC Symbol, |
| RNU6-1122P | 2 | RNA, U6 small nuclear 1122, pseudogene [Sou |
| NCKAP1 | 2 | NCK associated protein 1 [Source:HGNC Symb |
| Y_RNA | 2 | Y RNA [Source:RFAM;Acc:RF00019] |
| AC064871.1 | 2 | NULL |
| KRT8P10 | 2 | keratin 8 pseudogene 10 [Source:HGNC Symb |
| DUSP19 | 2 | dual specificity phosphatase 19 [Source:HGNC |
| AC064871.2 | 2 | novel transcript |
| NUP35 | 2 | nucleoporin 35 [Source:HGNC Symbol;Acc:HG |
| AC021851.1 | 2 | novel transcript |
| AC021851.2 | 2 | novel transcript |
| LIN28AP1 | 2 | LIN28A pseudogene 1 [Source:HGNC Symbol;A |
| CACYBPP2 | 2 | calcyclin binding protein pseudogene 2 [Sourc |
| AC093639.3 | 2 | novel transcript |
| AC093639.1 | 2 | novel transcript |
| AC093639.2 | 2 | novel transcript |
| AC096555.1 | 2 | novel transcript |
| AC020584.1 | 2 | novel transcript |
| MIR548AE1 | 2 | microRNA 548ae-1 [Source:HGNC Symbol;Acc |
| AC096667.1 | 2 | proline-rich protein 18-like [Source:NCBI gene |
| ZNF804A | 2 | zinc finger protein 804A [Source:HGNC Symbo |
| RPL23AP33 | 2 | ribosomal protein L23a pseudogene 33 [Sourc |
| AC009315.1 | 2 | novel transcript |
| AC080125.1 | 2 | pseudogene similar to part of E74-like factor 2 |
| FSIP2-AS1 | 2 | FSIP2 antisense RNA 1 [Source:HGNC Symbol; |
| U8 | 2 | U8 small nucleolar RNA [Source:RFAM;Acc:RF |
| FSIP2-AS2 | 2 | FSIP2 antisense RNA 2 [Source:HGNC Symbol; |
| FSIP2 | 2 | fibrous sheath interacting protein 2 [Source:H |
| AC097500.1 | 2 | novel transcript |
| RPL21P32 | 2 | ribosomal protein L21 pseudogene 32 [Source |
| LINC01473 | 2 | long intergenic non-protein coding RNA 1473 |
| AC104058.1 | 2 | ribosomal protein L29 (RPL29) pseudogene |
| RPL23AP35 | 2 | ribosomal protein L23a pseudogene 35 [Sourc |
| AC017071.1 | 2 | novel transcript |
| MED28P3 | 2 | mediator complex subunit 28 pseudogene 3 [S |
| ZC3H15 | 2 | zinc finger CCCH-type containing 15 [Source:H |
| DPRXP1 | 2 | divergent-paired related homeobox pseudoge |
| ITGAV | 2 | integrin subunit alpha V [Source:HGNC Symbo |
| AC017101.1 | 2 | novel transcript |
| FAM171B | 2 | family with sequence similarity 171 member B |
| ZSWIM2 | 2 | zinc finger SWIM-type containing 2 [Source:H( |
| AC018735.1 | 2 | IMP (inosine monophosphate) dehydrogenase |
| AC007319.1 | 2 | novel transcript, antisense to TFPI and CALCRL |
| RN7SKP42 | 2 | RN7SK pseudogene 42 [Source:HGNC Symbol; |
| RNU6-989P | 2 | RNA, U6 small nuclear 989, pseudogene [Sour |
| CALCRL | 2 | calcitonin receptor like receptor [Source:HGN |
| GAPDHP59 | 2 | glyceraldehyde-3-phosphate dehydrogenase p |
| TFPI | 2 | tissue factor pathway inhibitor [Source:HGNC |
| LINC01090 | 2 | long intergenic non-protein coding RNA 1090 |

| | | |
|---|---|---|
| ST13P2 | 2 | ST13, Hsp70 interacting protein pseudogene 2 |
| AC104131.1 | 2 | chromosome 18 open reading frame 25 (C18c |
| RNA5SP114 | 2 | RNA, 5S ribosomal pseudogene 114 [Source:H |
| GULP1 | 2 | GULP PTB domain containing engulfment adaj |
| MIR561 | 2 | microRNA 561 [Source:HGNC Symbol;Acc:HGI |
| AC092598.1 | 2 | novel transcript, overlapping DIRC1 |
| DIRC1 | 2 | disrupted in renal carcinoma 1 [Source:HGNC |
| COL3A1 | 2 | collagen type III alpha 1 chain [Source:HGNC S |
| MIR1245A | 2 | microRNA 1245a [Source:HGNC Symbol;Acc:H |
| MIR1245B | 2 | microRNA 1245b [Source:HGNC Symbol;Acc:H |
| MIR3606 | 2 | microRNA 3606 [Source:HGNC Symbol;Acc:HG |
| COL5A2 | 2 | collagen type V alpha 2 chain [Source:HGNC S |
| AC133106.1 | 2 | novel transcript |
| MIR3129 | 2 | microRNA 3129 [Source:HGNC Symbol;Acc:HG |
| KRT18P19 | 2 | keratin 18 pseudogene 19 [Source:HGNC Sym |
| WDR75 | 2 | WD repeat domain 75 [Source:HGNC Symbol; |
| AC013439.1 | 2 | jumonji domain containing 1A (JMJD1A) pseu |
| AC013439.2 | 2 | novel transcript |
| SLC40A1 | 2 | solute carrier family 40 member 1 [Source:HG |
| AC012488.1 | 2 | DnaJ (Hsp40) homolog, subfamily B, member |
| ASNSD1 | 2 | asparagine synthetase domain containing 1 [S |
| ASDURF | 2 | ASNSD1 upstream reading frame [Source:HGN |
| AC012488.2 | 2 | novel protein |
| ANKAR | 2 | ankyrin and armadillo repeat containing [Sour |
| OSGEPL1 | 2 | O-sialoglycoprotein endopeptidase like 1 [Sou |
| OSGEPL1-AS1 | 2 | OSGEPL1 antisense RNA 1 [Source:HGNC Sym |
| AC013468.1 | 2 | novel transcript |
| ORMDL1 | 2 | ORMDL sphingolipid biosynthesis regulator 1 [ |
| PMS1 | 2 | PMS1 homolog 1, mismatch repair system cor |
| C2orf88 | 2 | chromosome 2 open reading frame 88 [Sourc |
| AC019178.2 | 2 | pseudogene similar to part of telomeric repea |
| HNRNPCP2 | 2 | heterogeneous nuclear ribonucleoprotein C p |
| AC019178.1 | 2 | ring finger protein 11 (RNF11) pseudogene |
| MSTN | 2 | myostatin [Source:HGNC Symbol;Acc:HGNC:4 |
| HIBCH | 2 | 3-hydroxyisobutyryl-CoA hydrolase [Source:H |
| INPP1 | 2 | inositol polyphosphate-1-phosphatase [Sourc |
| MFSD6 | 2 | major facilitator superfamily domain containi |
| AC093388.1 | 2 | novel transcript, antisense to MFSD6 |
| NEMP2 | 2 | nuclear envelope integral membrane protein |
| AC108047.1 | 2 | novel transcript |
| RN7SKP179 | 2 | RN7SK pseudogene 179 [Source:HGNC Symbo |
| AC006460.2 | 2 | novel transcript |
| NAB1 | 2 | NGFI-A binding protein 1 [Source:HGNC Symb |
| AC006460.1 | 2 | novel transcript |
| AC005540.1 | 2 | novel transcript |
| GLS | 2 | glutaminase [Source:HGNC Symbol;Acc:HGNC |
| STAT1 | 2 | signal transducer and activator of transcriptio |
| AC067945.1 | 2 | RAB1A, member RAS oncogene family (RAB1A |
| AC067945.2 | 2 | novel transcript |
| AC067945.3 | 2 | novel transcript |

| | | |
|---|---|---|
| STAT4 | 2 | signal transducer and activator of transcriptio |
| AC067945.4 | 2 | novel transcript, antisense to STAT4 |
| RNU6-959P | 2 | RNA, U6 small nuclear 959, pseudogene [Sour |
| AC079777.1 | 2 | TEC |
| HMGB1P27 | 2 | high mobility group box 1 pseudogene 27 [Sou |
| AC092614.1 | 2 | novel transcript |
| MYO1B | 2 | myosin IB [Source:HGNC Symbol;Acc:HGNC:75 |
| RNU6-1045P | 2 | RNA, U6 small nuclear 1045, pseudogene [Sou |
| NABP1 | 2 | nucleic acid binding protein 1 [Source:HGNC S |
| AC114778.1 | 2 | NULL |
| AC098872.1 | 2 | novel transcript (hCG2045843) |
| CAVIN2 | 2 | caveolae associated protein 2 [Source:HGNC S |
| AC098617.1 | 2 | novel transcript, antisense to TMEFF2 |
| DNAJB1P1 | 2 | DnaJ heat shock protein family (Hsp40) memb |
| TMEFF2 | 2 | transmembrane protein with EGF like and two |
| AC062039.1 | 2 | novel transcript |
| AC013401.1 | 2 | novel transcript |
| PCGEM1 | 2 | PCGEM1 prostate-specific transcript [Source:H |
| RPS17P8 | 2 | ribosomal protein S17 pseudogene 8 [Source: |
| AC013401.2 | 2 | NULL |
| AC096647.1 | 2 | novel transcript |
| AC092638.1 | 2 | solute carrier family 44, member 3 (SLC44A3) |
| AC092638.2 | 2 | DnaJ (Hsp40) homolog, subfamily C, member |
| AC074290.1 | 2 | EndoMitotic Oocytes (EMO-1) pseudogene |
| AC074290.2 | 2 | novel transcript |
| AC068135.2 | 2 | novel transcript |
| AC068135.1 | 2 | S-phase kinase-associated protein 1 (SKP1) ps |
| GLULP6 | 2 | glutamate-ammonia ligase pseudogene 6 [Sou |
| HNRNPA1P47 | 2 | heterogeneous nuclear ribonucleoprotein A1 |
| LINC01821 | 2 | long intergenic non-protein coding RNA 1821 |
| AC106883.1 | 2 | COBW domain containing (CBWD) pseudogen |
| AC073973.1 | 2 | novel transcript |
| Metazoa_SRP | 2 | Metazoan signal recognition particle RNA [Sou |
| AC006196.1 | 2 | NULL |
| LINC01790 | 2 | long intergenic non-protein coding RNA 1790 |
| AC010983.1 | 2 | novel transcript |
| AC104823.1 | 2 | novel transcript |
| AC064834.1 | 2 | novel transcript |
| RNU6-169P | 2 | RNA, U6 small nuclear 169, pseudogene [Sour |
| LINC01825 | 2 | long intergenic non-protein coding RNA 1825 |
| LINC01827 | 2 | long intergenic non-protein coding RNA 1827 |
| SLC39A10 | 2 | solute carrier family 39 member 10 [Source:H |
| AHCYP5 | 2 | adenosylhomocysteinase pseudogene 5 [Sour |
| RNU6-915P | 2 | RNA, U6 small nuclear 915, pseudogene [Sour |
| DNAH7 | 2 | dynein axonemal heavy chain 7 [Source:HGNC |
| E2F3P2 | 2 | E2F transcription factor 3 pseudogene 2 [Sour |
| AC114760.1 | 2 | glutathione S-transferase pseudogene |
| STK17B | 2 | serine/threonine kinase 17b [Source:HGNC Sy |
| AC114760.2 | 2 | novel transcript, antisense to STK17B |
| HECW2 | 2 | HECT, C2 and WW domain containing E3 ubiq |

| | | |
|---|---|---|
| HECW2-AS1 | 2 | HECW2 antisense RNA 1 [Source:HGNC Symbo |
| RN7SL820P | 2 | RNA, 7SL, cytoplasmic 820, pseudogene [Sour |
| AC068544.2 | 2 | novel transcript, antisense to CCDC150 |
| CCDC150 | 2 | coiled-coil domain containing 150 [Source:HG |
| AC068544.1 | 2 | NULL |
| GTF3C3 | 2 | general transcription factor IIIC subunit 3 [Sou |
| C2orf66 | 2 | chromosome 2 open reading frame 66 [Sourc |
| AC012486.1 | 2 | TEC |
| PGAP1 | 2 | post-GPI attachment to proteins 1 [Source:HG |
| ANKRD44 | 2 | ankyrin repeat domain 44 [Source:HGNC Syml |
| HNRNPA3P15 | 2 | heterogeneous nuclear ribonucleoprotein A3 |
| AC017035.1 | 2 | ribosomal protein L4 (RPL4) pseudogene |
| AC013264.1 | 2 | novel transcript |
| ANKRD44-IT1 | 2 | ANKRD44 intronic transcript 1 [Source:HGNC S |
| ATP5MC2P3 | 2 | ATP synthase membrane subunit c locus 2 pse |
| AC010746.2 | 2 | novel transcript |
| NPM1P46 | 2 | nucleophosmin 1 pseudogene 46 [Source:HGN |
| SF3B1 | 2 | splicing factor 3b subunit 1 [Source:HGNC Sym |
| AC010746.1 | 2 | NULL |
| RNU6-1029P | 2 | RNA, U6 small nuclear 1029, pseudogene [Sou |
| COQ10B | 2 | coenzyme Q10B [Source:HGNC Symbol;Acc:H( |
| HSPD1 | 2 | heat shock protein family D (Hsp60) member |
| HSPE1 | 2 | heat shock protein family E (Hsp10) member : |
| HSPE1-MOB4 | 2 | HSPE1-MOB4 readthrough [Source:HGNC Sym |
| MOB4 | 2 | MOB family member 4, phocein [Source:HGN( |
| AC020550.1 | 2 | ribosomal protein L39 (RPL39) pseudogene |
| RFTN2 | 2 | raftlin family member 2 [Source:HGNC Symbo |
| AC020550.2 | 2 | novel transcript, antisense to RFTN2 |
| AC011997.1 | 2 | novel transcript, antisense to BOLL |
| MARS2 | 2 | methionyl-tRNA synthetase 2, mitochondrial [ |
| BOLL | 2 | boule homolog, RNA binding protein [Source:I |
| PLCL1 | 2 | phospholipase C like 1 (inactive) [Source:HGN |
| AC011997.2 | 2 | novel transcript, antisense to PLCL1 |
| AC020719.1 | 2 | novel transcript |
| LINC01923 | 2 | long intergenic non-protein coding RNA 1923 |
| AC019330.1 | 2 | novel transcript |
| AC020718.1 | 2 | novel transcript |
| AC018717.1 | 2 | pseudogene similar to part of CTAGE family, n |
| RNU7-147P | 2 | RNA, U7 small nuclear 147 pseudogene [Sourc |
| SATB2 | 2 | SATB homeobox 2 [Source:HGNC Symbol;Acc: |
| AC016746.1 | 2 | novel transcript, antisense to SATB homeobox |
| SATB2-AS1 | 2 | SATB2 antisense RNA 1 [Source:HGNC Symbol |
| LINC01877 | 2 | long intergenic non-protein coding RNA 1877 |
| SEPHS1P6 | 2 | selenophosphate synthetase 1 pseudogene 6 |
| FTCDNL1 | 2 | formiminotransferase cyclodeaminase N-term |
| AC097717.1 | 2 | novel transcript |
| AC108032.1 | 2 | cytochrome c oxidase, subunit VIc (COX6C) ps |
| RN7SL717P | 2 | RNA, 7SL, cytoplasmic 717, pseudogene [Sour |
| C2orf69 | 2 | chromosome 2 open reading frame 69 [Sourc |
| TYW5 | 2 | tRNA-yW synthesizing protein 5 [Source:HGN( |

| | | |
|---|---|---|
| MAIP1 | 2 | matrix AAA peptidase interacting protein 1 [S |
| AC004464.1 | 2 | novel transcript |
| SPATS2L | 2 | spermatogenesis associated serine rich 2 like |
| RNY4P34 | 2 | RNY4 pseudogene 34 [Source:HGNC Symbol;A |
| AC012459.1 | 2 | novel transcript, antisense to SPATS2L |
| KCTD18 | 2 | potassium channel tetramerization domain co |
| SGO2 | 2 | shugoshin 2 [Source:HGNC Symbol;Acc:HGNC |
| AOX1 | 2 | aldehyde oxidase 1 [Source:HGNC Symbol;Acc |
| AOX3P | 2 | aldehyde oxidase 3, pseudogene [Source:HGN |
| AOX3P-AOX2P | 2 | AOX3P-AOX2P readthrough, transcribed pseu |
| LINC01792 | 2 | long intergenic non-protein coding RNA 1792 |
| AOX2P | 2 | aldehyde oxidase 2 pseudogene [Source:HGN |
| RNU1-133P | 2 | RNA, U1 small nuclear 133, pseudogene [Sour |
| AC007163.1 | 2 | novel transcript |
| BZW1 | 2 | basic leucine zipper and W2 domains 1 [Sourc |
| RNU6-31P | 2 | RNA, U6 small nuclear 31, pseudogene [Sourc |
| BICD1P1 | 2 | BICD cargo adaptor 1 pseudogene 1 [Source:H |
| RNA5SP115 | 2 | RNA, 5S ribosomal pseudogene 115 [Source:H |
| CLK1 | 2 | CDC like kinase 1 [Source:HGNC Symbol;Acc:H |
| Y_RNA | 2 | Y RNA [Source:RFAM;Acc:RF00019] |
| PPIL3 | 2 | peptidylprolyl isomerase like 3 [Source:HGNC |
| RNU6-312P | 2 | RNA, U6 small nuclear 312, pseudogene [Sour |
| NIF3L1 | 2 | NGG1 interacting factor 3 like 1 [Source:HGN |
| RNU6-762P | 2 | RNA, U6 small nuclear 762, pseudogene [Sour |
| ORC2 | 2 | origin recognition complex subunit 2 [Source: |
| AC005037.1 | 2 | novel transcript |
| FAM126B | 2 | family with sequence similarity 126 member B |
| HNRNPA1P35 | 2 | heterogeneous nuclear ribonucleoprotein A1 |
| RPL23AP30 | 2 | ribosomal protein L23a pseudogene 30 [Sourc |
| NDUFB3 | 2 | NADH:ubiquinone oxidoreductase subunit B3 |
| RNU6-1206P | 2 | RNA, U6 small nuclear 1206, pseudogene [Sou |
| Metazoa_SRP | 2 | Metazoan signal recognition particle RNA [Sou |
| AC007272.1 | 2 | novel transcript |
| RPL17P10 | 2 | ribosomal protein L17 pseudogene 10 [Source |
| CFLAR | 2 | CASP8 and FADD like apoptosis regulator [Sou |
| IMPDH1P10 | 2 | inosine monophosphate dehydrogenase 1 pse |
| CFLAR-AS1 | 2 | CFLAR antisense RNA 1 [Source:HGNC Symbol |
| RNU7-45P | 2 | RNA, U7 small nuclear 45 pseudogene [Source |
| AC007283.2 | 2 | ribosomal protein L38 (RPL38) pseudogene |
| AC007283.1 | 2 | novel transcript |
| CASP10 | 2 | caspase 10 [Source:HGNC Symbol;Acc:HGNC: |
| MTND5P25 | 2 | MT-ND5 pseudogene 25 [Source:HGNC Symbo |
| MTND4P23 | 2 | MT-ND4 pseudogene 23 [Source:HGNC Symbo |
| MTND4LP13 | 2 | MT-ND4L pseudogene 13 [Source:HGNC Symb |
| CASP8 | 2 | caspase 8 [Source:HGNC Symbol;Acc:HGNC:1 |
| FLACC1 | 2 | flagellum associated containing coiled-coil do |
| TRAK2 | 2 | trafficking kinesin protein 2 [Source:HGNC Sy |
| STRADB | 2 | STE20 related adaptor beta [Source:HGNC Sy |
| AC007256.1 | 2 | SCY1-like 2 (S. cerevisiae) (SCYL2) pseudogene |
| Y_RNA | 2 | Y RNA [Source:RFAM;Acc:RF00019] |

| | | |
|---|---|---|
| C2CD6 | 2 | C2 calcium dependent domain containing 6 [S |
| MTCO2P16 | 2 | MT-CO2 pseudogene 16 [Source:HGNC Symbo |
| MTATP6P16 | 2 | MT-ATP6 pseudogene 16 [Source:HGNC Symb |
| MTCO3P16 | 2 | MT-CO3 pseudogene 16 [Source:HGNC Symbo |
| MTND3P16 | 2 | MT-ND3 pseudogene 16 [Source:HGNC Symbo |
| MTND4LP16 | 2 | MT-ND4L pseudogene 16 [Source:HGNC Symb |
| MTND4P29 | 2 | MT-ND4 pseudogene 29 [Source:HGNC Symbo |
| MTND5P31 | 2 | MT-ND5 pseudogene 31 [Source:HGNC Symbo |
| TMEM237 | 2 | transmembrane protein 237 [Source:HGNC Sy |
| ENO1P4 | 2 | enolase 1 pseudogene 4 [Source:HGNC Symbo |
| AC007279.2 | 2 | novel transcript, antisense to MPP4and TMEN |
| MPP4 | 2 | membrane palmitoylated protein 4 [Source:H |
| RNU6-651P | 2 | RNA, U6 small nuclear 651, pseudogene [Sour |
| ALS2 | 2 | alsin Rho guanine nucleotide exchange factor |
| AC007279.1 | 2 | ribosomal protein pseudogene |
| CDK15 | 2 | cyclin dependent kinase 15 [Source:HGNC Syn |
| Y_RNA | 2 | Y RNA [Source:RFAM;Acc:RF00019] |
| UBE2V1P11 | 2 | ubiquitin conjugating enzyme E2 V1 pseudoge |
| RNU6-440P | 2 | RNA, U6 small nuclear 440, pseudogene [Sour |
| AC069148.1 | 2 | novel transcript |
| FZD7 | 2 | frizzled class receptor 7 [Source:HGNC Symbo |
| KIAA2012 | 2 | KIAA2012 [Source:HGNC Symbol;Acc:HGNC:5: |
| KIAA2012-AS1 | 2 | KIAA2012 antisense RNA 1 [Source:HGNC Sym |
| AC079354.3 | 2 | novel transcript, antisense to KIAA2012 |
| AC079354.2 | 2 | novel transcript |
| PSMA2P3 | 2 | proteasome subunit alpha 2 pseudogene 3 [So |
| DAZAP2P1 | 2 | DAZ associated protein 2 pseudogene 1 [Sour |
| SUMO1 | 2 | small ubiquitin like modifier 1 [Source:HGNC S |
| AC079354.1 | 2 | ribosomal protein L39 (RPL39) pseudogene |
| NOP58 | 2 | NOP58 ribonucleoprotein [Source:HGNC Symb |
| SNORD70 | 2 | small nucleolar RNA, C/D box 70 [Source:HGN |
| SNORD70B | 2 | small nucleolar RNA, C/D box 70B [Source:HG |
| SNORD11B | 2 | small nucleolar RNA, C/D box 11B [Source:HG |
| SNORD11 | 2 | small nucleolar RNA, C/D box 11 [Source:HGN |
| AC064836.4 | 2 | novel transcript, antisense to NOP58 |
| RN7SL753P | 2 | RNA, 7SL, cytoplasmic 753, pseudogene [Sour |
| AC064836.1 | 2 | family with sequence similarity 64, member A |
| AC064836.2 | 2 | novel transcript |
| RN7SL40P | 2 | RNA, 7SL, cytoplasmic 40, pseudogene [Sourc |
| AC064836.3 | 2 | novel transcript |
| BMPR2 | 2 | bone morphogenetic protein receptor type 2 |
| AC073410.2 | 2 | H3 histone, family 3B (H3.3B) (H3F3B) pseudo |
| AC073410.1 | 2 | ribosomal protein L13a (RPL13A) pseudogene |
| MTCO1P17 | 2 | MT-CO1 pseudogene 17 [Source:HGNC Symbo |
| MTCO2P17 | 2 | MT-CO2 pseudogene 17 [Source:HGNC Symbo |
| MTATP6P17 | 2 | MT-ATP6 pseudogene 17 [Source:HGNC Symb |
| MTCO3P17 | 2 | MT-CO3 pseudogene 17 [Source:HGNC Symbo |
| MTND3P17 | 2 | MT-ND3 pseudogene 17 [Source:HGNC Symbo |
| MTND4LP17 | 2 | MT-ND4L pseudogene 17 [Source:HGNC Symb |
| MTND4P30 | 2 | MT-ND4 pseudogene 30 [Source:HGNC Symbo |

| | | |
|---|---|---|
| MTCO1P54 | 2 | MT-CO1 pseudogene 54 [Source:HGNC Symbo |
| FAM117B | 2 | family with sequence similarity 117 member E |
| ICA1L | 2 | islet cell autoantigen 1 like [Source:HGNC Sym |
| AC098831.1 | 2 | phosphatidylinositol glycan anchor biosynthes |
| KRT8P15 | 2 | keratin 8 pseudogene 15 [Source:HGNC Symb |
| WDR12 | 2 | WD repeat domain 12 [Source:HGNC Symbol;/ |
| CARF | 2 | calcium responsive transcription factor [Sourc |
| AC010900.1 | 2 | keratin 8 (KRT8) pseudogene |
| AC010900.2 | 2 | chromosome 18 open reading frame 32 (C18c |
| NBEAL1 | 2 | neurobeachin like 1 [Source:HGNC Symbol;Ac |
| RPL7P14 | 2 | ribosomal protein L7 pseudogene 14 [Source: |
| AC023271.1 | 2 | NULL |
| AC023271.2 | 2 | zinc finger protein 14 (ZNF14) pseudogene |
| RPL23AP36 | 2 | ribosomal protein L23a pseudogene 36 [Sourc |
| RPL12P16 | 2 | ribosomal protein L12 pseudogene 16 [Source |
| CYP20A1 | 2 | cytochrome P450 family 20 subfamily A meml |
| RN7SL670P | 2 | RNA, 7SL, cytoplasmic 670, pseudogene [Sour |
| MRPL50P2 | 2 | mitochondrial ribosomal protein L50 pseudog |
| ABI2 | 2 | abl interactor 2 [Source:HGNC Symbol;Acc:HG |
| AC080075.1 | 2 | novel transcript, antisense to abl interactor 2 |
| RAPH1 | 2 | Ras association (RalGDS/AF-6) and pleckstrin I |
| AC125238.2 | 2 | zinc finger protein 207 (ZNF207) pseudogene |
| AC125238.1 | 2 | NADH dehydrogenase (ubiquinone) 1 alpha su |
| CD28 | 2 | CD28 molecule [Source:HGNC Symbol;Acc:HG |
| KRT18P39 | 2 | keratin 18 pseudogene 39 [Source:HGNC Sym |
| NPM1P33 | 2 | nucleophosmin 1 pseudogene 33 [Source:HGN |
| RNU6-474P | 2 | RNA, U6 small nuclear 474, pseudogene [Sour |
| CTLA4 | 2 | cytotoxic T-lymphocyte associated protein 4 [ |
| ICOS | 2 | inducible T cell costimulator [Source:HGNC Sy |
| AC009965.2 | 2 | novel pseudogene similar to part of a zinc fing |
| AC009965.1 | 2 | novel transcript |
| AC009498.1 | 2 | novel transcript |
| DSTNP5 | 2 | DSTN pseudogene 5 [Source:HGNC Symbol;Ac |
| AC106901.1 | 2 | carboxymethylenebutenolidase homolog (Pse |
| AC016903.1 | 2 | novel transcript |
| AC016903.2 | 2 | novel transcript |
| PARD3B | 2 | par-3 family cell polarity regulator beta [Sourc |
| AC007736.1 | 2 | novel transcript, antisense to PARD3B |
| AC008171.1 | 2 | pseudogene similar to part of CAP-GLY domai |
| NRP2 | 2 | neuropilin 2 [Source:HGNC Symbol;Acc:HGNC |
| AC007362.1 | 2 | novel transcript |
| RN7SKP178 | 2 | RN7SK pseudogene 178 [Source:HGNC Symbo |
| PPIAP68 | 2 | peptidylprolyl isomerase A pseudogene 68 [Sc |
| AC007679.1 | 2 | novel transcript, antisense to INO80D |
| INO80D | 2 | INO80 complex subunit D [Source:HGNC Symb |
| Vault | 2 | Vault RNA [Source:RFAM;Acc:RF00006] |
| Y_RNA | 2 | Y RNA [Source:RFAM;Acc:RF00019] |
| AC007679.2 | 2 | ribosomal protein L27 (RPL27) pseudogene |
| AC007383.2 | 2 | novel transcript |
| NDUFS1 | 2 | NADH:ubiquinone oxidoreductase core subun |

| | | |
|---|---|---|
| AC007383.3 | 2 | novel transcript |
| GCSHP3 | 2 | glycine cleavage system protein H pseudogene |
| EEF1B2 | 2 | eukaryotic translation elongation factor 1 beta |
| AC007383.4 | 2 | NULL |
| snoZ196 | 2 | Small nucleolar RNA Z196/R39/R59 family [So |
| SNORD51 | 2 | small nucleolar RNA, C/D box 51 [Source:HGN |
| SNORA41 | 2 | small nucleolar RNA, H/ACA box 41 [Source:H |
| GPR1 | 2 | G protein-coupled receptor 1 [Source:HGNC S |
| GPR1-AS | 2 | GPR1 antisense RNA [Source:HGNC Symbol;Ac |
| RN7SKP200 | 2 | RN7SK pseudogene 200 [Source:HGNC Symbo |
| AC007383.1 | 2 | ATP synthase, H+ transporting, mitochondrial |
| HMGN1P6 | 2 | high mobility group nucleosome binding doma |
| RN7SKP260 | 2 | RN7SK pseudogene 260 [Source:HGNC Symbo |
| ZDBF2 | 2 | zinc finger DBF-type containing 2 [Source:HGN |
| AC017081.1 | 2 | N-acylsphingosine amidohydrolase 3-like (ASA |
| HNRNPA1P51 | 2 | heterogeneous nuclear ribonucleoprotein A1 |
| ADAM23 | 2 | ADAM metallopeptidase domain 23 [Source:H |
| AC010731.2 | 2 | novel transcript |
| AC010731.3 | 2 | novel transcript |
| FAM237A | 2 | family with sequence similarity 237 member A |
| DYTN | 2 | dystrotelin [Source:HGNC Symbol;Acc:HGNC:2 |
| AC010731.1 | 2 | Down syndrome critical region gene 3 (DSCR3 |
| Y_RNA | 2 | Y RNA [Source:RFAM;Acc:RF00019] |
| MDH1B | 2 | malate dehydrogenase 1B [Source:HGNC Sym |
| FASTKD2 | 2 | FAST kinase domains 2 [Source:HGNC Symbol; |
| MIR3130-1 | 2 | microRNA 3130-1 [Source:HGNC Symbol;Acc:I |
| MIR3130-2 | 2 | microRNA 3130-2 [Source:HGNC Symbol;Acc:I |
| AC008269.1 | 2 | novel transcript |
| AC008269.2 | 2 | novel transcript |
| CPO | 2 | carboxypeptidase O [Source:HGNC Symbol;Ac |
| KLF7 | 2 | Kruppel like factor 7 [Source:HGNC Symbol;Ac |
| MIR2355 | 2 | microRNA 2355 [Source:HGNC Symbol;Acc:HG |
| KLF7-IT1 | 2 | KLF7 intronic transcript 1 [Source:HGNC Symb |
| MYOSLID | 2 | myocardin-induced smooth muscle lncRNA, in |
| MIR7845 | 2 | microRNA 7845 [Source:HGNC Symbol;Acc:HG |
| AC007879.1 | 2 | novel transcript |
| AC007879.3 | 2 | novel transcript |
| AC007879.5 | 2 | novel transcript |
| AC007879.2 | 2 | novel transcript |
| AC007879.4 | 2 | novel transcript |
| AC009226.1 | 2 | novel transcript |
| LINC01802 | 2 | long intergenic non-protein coding RNA 1802 |
| MIR1302-4 | 2 | microRNA 1302-4 [Source:HGNC Symbol;Acc:I |
| CREB1 | 2 | cAMP responsive element binding protein 1 [S |
| METTL21A | 2 | methyltransferase like 21A [Source:HGNC Sym |
| RPS29P9 | 2 | ribosomal protein S29 pseudogene 9 [Source: |
| LINC01857 | 2 | long intergenic non-protein coding RNA 1857 |
| RNU6-664P | 2 | RNA, U6 small nuclear 664, pseudogene [Sour |
| PPP1R14BP2 | 2 | protein phosphatase 1 regulatory inhibitor sul |
| Y_RNA | 2 | Y RNA [Source:RFAM;Acc:RF00019] |

| | | |
|---|---|---|
| CCNYL1 | 2 | cyclin Y like 1 [Source:HGNC Symbol;Acc:HGN |
| AC096772.1 | 2 | novel transcript |
| MIR4775 | 2 | microRNA 4775 [Source:HGNC Symbol;Acc:HG |
| FZD5 | 2 | frizzled class receptor 5 [Source:HGNC Symbo |
| PLEKHM3 | 2 | pleckstrin homology domain containing M3 [S |
| AC083900.1 | 2 | novel transcript |
| RNU6-360P | 2 | RNA, U6 small nuclear 360, pseudogene [Sour |
| AC093698.1 | 2 | ribosomal protein L9 (RPL9) pseudogene |
| RNA5SP116 | 2 | RNA, 5S ribosomal pseudogene 116 [Source:H |
| RPL12P17 | 2 | ribosomal protein L12 pseudogene 17 [Source |
| CRYGEP | 2 | crystallin gamma E, pseudogene [Source:HGN |
| CRYGD | 2 | crystallin gamma D [Source:HGNC Symbol;Acc |
| CRYGC | 2 | crystallin gamma C [Source:HGNC Symbol;Acc |
| AC093698.2 | 2 | replication factor C (activator 1) 2, 40kDa (RF( |
| CRYGB | 2 | crystallin gamma B [Source:HGNC Symbol;Acc |
| CRYGA | 2 | crystallin gamma A [Source:HGNC Symbol;Acc |
| C2orf80 | 2 | chromosome 2 open reading frame 80 [Sourc |
| RPSAP27 | 2 | ribosomal protein SA pseudogene 27 [Source: |
| TPT1P2 | 2 | tumor protein, translationally-controlled 1 ps |
| IDH1 | 2 | isocitrate dehydrogenase (NADP(+)) 1, cytosol |
| IDH1-AS1 | 2 | IDH1 antisense RNA 1 [Source:HGNC Symbol;A |
| PIKFYVE | 2 | phosphoinositide kinase, FYVE-type zinc finge |
| MYL6BP1 | 2 | myosin light chain 6B pseudogene 1 [Source:H |
| PTH2R | 2 | parathyroid hormone 2 receptor [Source:HGN |
| AC012362.1 | 2 | actin related protein 2/3 complex, subunit 1A |
| AC019185.1 | 2 | H3 histone, family 3B (H3.3B) (H3F3B) pseudo |
| AC019185.2 | 2 | novel transcript |
| AC019185.3 | 2 | pseudogene similar to part of WD repeat dom |
| AC109824.1 | 2 | novel transcript |
| RNA5SP117 | 2 | RNA, 5S ribosomal pseudogene 117 [Source:H |
| HSPA8P6 | 2 | heat shock protein family A (Hsp70) member |
| CRYGFP | 2 | crystallin gamma F, pseudogene [Source:HGN |
| MEAF6P1 | 2 | MYST/Esa1-associated factor 6 pseudogene 1 |
| PKP4P1 | 2 | plakophilin 4 pseudogene 1 [Source:HGNC Syr |
| AC016743.1 | 2 | novel transcript |
| MAP2 | 2 | microtubule associated protein 2 [Source:HGN |
| RNA5SP118 | 2 | RNA, 5S ribosomal pseudogene 118 [Source:H |
| UNC80 | 2 | unc-80 homolog, NALCN channel complex sub |
| SNAI1P1 | 2 | snail family zinc finger 1 pseudogene 1 [Sourc |
| RPE | 2 | ribulose-5-phosphate-3-epimerase [Source:H( |
| KANSL1L | 2 | KAT8 regulatory NSL complex subunit 1 like [S |
| AC007038.2 | 2 | novel transcript, antisense to KANSL1L |
| AC007038.1 | 2 | novel transcript |
| AC006994.1 | 2 | ribosomal protein L6 (RPL6) pseudogene |
| AC006994.2 | 2 | novel transcript |
| ACADL | 2 | acyl-CoA dehydrogenase long chain [Source:H |
| Y_RNA | 2 | Y RNA [Source:RFAM;Acc:RF00019] |
| MYL1 | 2 | myosin light chain 1 [Source:HGNC Symbol;Ac |
| LANCL1-AS1 | 2 | LANCL1 antisense RNA 1 [Source:HGNC Symbo |
| LANCL1 | 2 | LanC like 1 [Source:HGNC Symbol;Acc:HGNC:6 |

| | | |
|---|---|---|
| CPS1 | 2 | carbamoyl-phosphate synthase 1 [Source:HGI |
| CPS1-IT1 | 2 | CPS1 intronic transcript 1 [Source:HGNC Symb |
| AC012491.1 | 2 | 5'-nucleotidase domain containing 2 (NT5DC2 |
| AC013404.1 | 2 | ribosomal protein S27 (RPS27) pseudogene |
| ERBB4 | 2 | erb-b2 receptor tyrosine kinase 4 [Source:HGI |
| RNA5SP119 | 2 | RNA, 5S ribosomal pseudogene 119 [Source:H |
| MTND2P23 | 2 | MT-ND2 pseudogene 23 [Source:HGNC Symbo |
| MTCO1P46 | 2 | MT-CO1 pseudogene 46 [Source:HGNC Symbo |
| MIR548F2 | 2 | microRNA 548f-2 [Source:HGNC Symbol;Acc:H |
| AC093865.1 | 2 | novel transcript, sense overlapping IKZF2 |
| LINC01878 | 2 | long intergenic non-protein coding RNA 1878 |
| PCED1CP | 2 | PC-esterase domain containing 1C, pseudoger |
| AC108066.1 | 2 | novel transcript |
| MIR4776-2 | 2 | microRNA 4776-2 [Source:HGNC Symbol;Acc:I |
| MIR4776-1 | 2 | microRNA 4776-1 [Source:HGNC Symbol;Acc:I |
| AC108066.2 | 2 | novel transcript |
| IKZF2 | 2 | IKAROS family zinc finger 2 [Source:HGNC Sym |
| AC079610.1 | 2 | novel transcript |
| LINC01953 | 2 | long intergenic non-protein coding RNA 1953 |
| AC079610.2 | 2 | novel transcript |
| SPAG16-DT | 2 | SPAG16 divergent transcript [Source:HGNC Sy |
| SPAG16 | 2 | sperm associated antigen 16 [Source:HGNC Sy |
| MIR4438 | 2 | microRNA 4438 [Source:HGNC Symbol;Acc:HG |
| AC068051.1 | 2 | novel transcript, antisense to VWC2L |
| RPL5P8 | 2 | ribosomal protein L5 pseudogene 8 [Source:H |
| VWC2L | 2 | von Willebrand factor C domain containing 2 l |
| VWC2L-IT1 | 2 | VWC2L intronic transcript 1 [Source:HGNC Syr |
| ENSAP3 | 2 | endosulfine alpha pseudogene 3 [Source:HGN |
| BARD1 | 2 | BRCA1 associated RING domain 1 [Source:HGI |
| SNHG31 | 2 | small nucleolar RNA host gene 31 [Source:HGI |
| SNORA70I | 2 | small nucleolar RNA, H/ACA box 70I [Source:H |
| RPL10P6 | 2 | ribosomal protein L10 pseudogene 6 [Source: |
| ABCA12 | 2 | ATP binding cassette subfamily A member 12 |
| AC072062.1 | 2 | novel transcript, antisense to ABCA12 |
| AC092844.1 | 2 | novel transcript |
| AC073284.1 | 2 | novel transcript |
| ATIC | 2 | 5-aminoimidazole-4-carboxamide ribonucleot |
| FN1 | 2 | fibronectin 1 [Source:HGNC Symbol;Acc:HGN( |
| AC012462.3 | 2 | novel transcript |
| AC012462.1 | 2 | novel transcript |
| AC012462.2 | 2 | novel transcript |
| AC012668.2 | 2 | novel transcript |
| AC012668.3 | 2 | novel transcript |
| AC012668.1 | 2 | novel transcript |
| LINC00607 | 2 | long intergenic non-protein coding RNA 607 [S |
| LINC01614 | 2 | long intergenic non-protein coding RNA 1614 |
| AC093850.1 | 2 | NADH dehydrogenase (ubiquinone) 1 alpha su |
| AC122136.1 | 2 | novel transcript |
| AC093382.1 | 2 | novel transcript |
| MREG | 2 | melanoregulin [Source:HGNC Symbol;Acc:HGI |

| | | |
|---|---|---|
| PECR | 2 | peroxisomal trans-2-enoyl-CoA reductase [Sou |
| TMEM169 | 2 | transmembrane protein 169 [Source:HGNC Sy |
| XRCC5 | 2 | X-ray repair cross complementing 5 [Source:H |
| POLHP1 | 2 | DNA polymerase eta pseudogene 1 [Source:H |
| AC012513.2 | 2 | heterogeneous nuclear ribonucleoprotein A3 |
| AC012513.3 | 2 | TEC |
| LINC01963 | 2 | long intergenic non-protein coding RNA 1963 |
| MARCH4 | 2 | membrane associated ring-CH-type finger 4 [S |
| AC012513.1 | 2 | novel transcript |
| AC069155.1 | 2 | novel transcript |
| Y_RNA | 2 | Y RNA [Source:RFAM;Acc:RF00019] |
| AC098820.2 | 2 | novel transcript |
| SMARCAL1 | 2 | SWI/SNF related, matrix associated, actin dep |
| AC098820.1 | 2 | novel transcript |
| AC098820.3 | 2 | novel transcript |
| RPL37A | 2 | ribosomal protein L37a [Source:HGNC Symbol |
| LINC01280 | 2 | long intergenic non-protein coding RNA 1280 |
| AC073321.1 | 2 | novel transcript |
| PSMB3P2 | 2 | proteasome subunit beta 3 pseudogene 2 [Sou |
| IGFBP2 | 2 | insulin like growth factor binding protein 2 [Sc |
| IGFBP5 | 2 | insulin like growth factor binding protein 5 [Sc |
| AC007563.2 | 2 | novel transcript, antisense to IGFBP5 and TNP |
| AC007563.3 | 2 | ribosomal protein L31 (RPL31) pseudogene |
| RNA5SP120 | 2 | RNA, 5S ribosomal pseudogene 120 [Source:H |
| AC007563.1 | 2 | novel transcript |
| FABP5P14 | 2 | fatty acid binding protein 5 pseudogene 14 [S |
| AC007563.4 | 2 | novel transcript |
| AC007557.2 | 2 | novel transcript |
| TNP1 | 2 | transition protein 1 [Source:HGNC Symbol;Acc |
| LINC01921 | 2 | long intergenic non-protein coding RNA 1921 |
| AC007557.4 | 2 | novel transcript |
| AC007557.1 | 2 | novel transcript |
| AC007557.3 | 2 | novel transcript |
| AC007749.1 | 2 | novel transcript |
| AC007749.2 | 2 | novel transcript |
| RN7SKP43 | 2 | RN7SK pseudogene 43 [Source:HGNC Symbol; |
| DIRC3-AS1 | 2 | DIRC3 antisense RNA 1 [Source:HGNC Symbol |
| AC009492.1 | 2 | NULL |
| DIRC3 | 2 | disrupted in renal carcinoma 3 [Source:HGNC |
| AC010886.1 | 2 | ATPase, H+/K+ transporting, nongastric, alpha |
| RPL7L1P9 | 2 | ribosomal protein L7 like 1 pseudogene 9 [Sou |
| TNS1 | 2 | tensin 1 [Source:HGNC Symbol;Acc:HGNC:119 |
| MIR6809 | 2 | microRNA 6809 [Source:HGNC Symbol;Acc:HG |
| AC010136.1 | 2 | novel transcript |
| RUFY4 | 2 | RUN and FYVE domain containing 4 [Source:H |
| CXCR2P1 | 2 | C-X-C motif chemokine receptor 2 pseudogen |
| CXCR2 | 2 | C-X-C motif chemokine receptor 2 [Source:HG |
| CXCR1 | 2 | C-X-C motif chemokine receptor 1 [Source:HG |
| HMGB1P9 | 2 | high mobility group box 1 pseudogene 9 [Sour |
| ARPC2 | 2 | actin related protein 2/3 complex subunit 2 [S |

| | | |
|---|---|---|
| AC021016.2 | 2 | novel transcript, overlapping to ARPC2 and GI |
| GPBAR1 | 2 | G protein-coupled bile acid receptor 1 [Source |
| AAMP | 2 | angio associated migratory cell protein [Sourc |
| PNKD | 2 | PNKD metallo-beta-lactamase domain contair |
| TMBIM1 | 2 | transmembrane BAX inhibitor motif containin |
| MIR6513 | 2 | microRNA 6513 [Source:HGNC Symbol;Acc:HG |
| CATIP-AS2 | 2 | CATIP antisense RNA 2 [Source:HGNC Symbol; |
| MIR6810 | 2 | microRNA 6810 [Source:HGNC Symbol;Acc:HG |
| AC021016.1 | 2 | ribosomal protein L19 (RPL19) pseudogene |
| CATIP | 2 | ciliogenesis associated TTC17 interacting prot |
| CATIP-AS1 | 2 | CATIP antisense RNA 1 [Source:HGNC Symbol; |
| SLC11A1 | 2 | solute carrier family 11 member 1 [Source:HG |
| CTDSP1 | 2 | CTD small phosphatase 1 [Source:HGNC Symb |
| AC021016.3 | 2 | novel transcript, antisense to CTDSP1 |
| MIR26B | 2 | microRNA 26b [Source:HGNC Symbol;Acc:HGI |
| VIL1 | 2 | villin 1 [Source:HGNC Symbol;Acc:HGNC:1269 |
| USP37 | 2 | ubiquitin specific peptidase 37 [Source:HGNC |
| RN7SKP38 | 2 | RN7SK pseudogene 38 [Source:HGNC Symbol; |
| CNOT9 | 2 | CCR4-NOT transcription complex subunit 9 [Sc |
| RNU6-136P | 2 | RNA, U6 small nuclear 136, pseudogene [Sour |
| PLCD4 | 2 | phospholipase C delta 4 [Source:HGNC Symbo |
| AC012510.1 | 2 | novel transcript, antisense to PLCD4 |
| ZNF142 | 2 | zinc finger protein 142 [Source:HGNC Symbol; |
| BCS1L | 2 | BCS1 homolog, ubiquinol-cytochrome c reduc |
| RNF25 | 2 | ring finger protein 25 [Source:HGNC Symbol;A |
| STK36 | 2 | serine/threonine kinase 36 [Source:HGNC Syn |
| TTLL4 | 2 | tubulin tyrosine ligase like 4 [Source:HGNC Sy |
| CYP27A1 | 2 | cytochrome P450 family 27 subfamily A meml |
| AC009974.2 | 2 | novel transcript, antisense to CYP27A1 |
| AC009974.1 | 2 | novel transcript |
| PRKAG3 | 2 | protein kinase AMP-activated non-catalytic su |
| MIR9500 | 2 | microRNA 9500 [Source:HGNC Symbol;Acc:HG |
| RPL23AP31 | 2 | ribosomal protein L23a pseudogene 31 [Sourc |
| WNT6 | 2 | Wnt family member 6 [Source:HGNC Symbol;/ |
| WNT10A | 2 | Wnt family member 10A [Source:HGNC Symb |
| LINC01494 | 2 | long intergenic non-protein coding RNA 1494 |
| RNU6-642P | 2 | RNA, U6 small nuclear 642, pseudogene [Sour |
| AC097468.2 | 2 | novel transcript, antisense to CDK5R2 |
| KRT8P30 | 2 | keratin 8 pseudogene 30 [Source:HGNC Symb |
| CDK5R2 | 2 | cyclin dependent kinase 5 regulatory subunit |
| LINC00608 | 2 | long intergenic non-protein coding RNA 608 [S |
| FEV | 2 | FEV transcription factor, ETS family member [ |
| CRYBA2 | 2 | crystallin beta A2 [Source:HGNC Symbol;Acc:H |
| MIR375 | 2 | microRNA 375 [Source:HGNC Symbol;Acc:HGI |
| AC097468.1 | 2 | novel transcript, antisense to CCDC108 |
| CFAP65 | 2 | cilia and flagella associated protein 65 [Source |
| IHH | 2 | Indian hedgehog signaling molecule [Source:H |
| MIR3131 | 2 | microRNA 3131 [Source:HGNC Symbol;Acc:HG |
| AC097468.4 | 2 | novel transcript |
| AC097468.3 | 2 | novel transcript |

| | | |
|---|---|---|
| NHEJ1 | 2 | non-homologous end joining factor 1 [Source: |
| AC068946.1 | 2 | novel protein, SLC23A3-NHEJ1 readthrough |
| RN7SL764P | 2 | RNA, 7SL, cytoplasmic 764, pseudogene [Sour |
| SLC23A3 | 2 | solute carrier family 23 member 3 [Source:HG |
| CNPPD1 | 2 | cyclin Pas1/PHO80 domain containing 1 [Sour |
| RETREG2 | 2 | reticulophagy regulator family member 2 [Sou |
| ZFAND2B | 2 | zinc finger AN1-type containing 2B [Source:HG |
| AC068946.3 | 2 | novel transcript, antisense to ZFAND2B |
| AC068946.2 | 2 | novel protein |
| ABCB6 | 2 | ATP binding cassette subfamily B member 6 (L |
| ATG9A | 2 | autophagy related 9A [Source:HGNC Symbol;A |
| ANKZF1 | 2 | ankyrin repeat and zinc finger peptidyl tRNA h |
| GLB1L | 2 | galactosidase beta 1 like [Source:HGNC Symbo |
| STK16 | 2 | serine/threonine kinase 16 [Source:HGNC Sym |
| TUBA4A | 2 | tubulin alpha 4a [Source:HGNC Symbol;Acc:H |
| TUBA4B | 2 | tubulin alpha 4b [Source:HGNC Symbol;Acc:H |
| DNAJB2 | 2 | DnaJ heat shock protein family (Hsp40) memb |
| PTPRN | 2 | protein tyrosine phosphatase receptor type N |
| MIR153-1 | 2 | microRNA 153-1 [Source:HGNC Symbol;Acc:H |
| AC114803.1 | 2 | novel transcript |
| RESP18 | 2 | regulated endocrine specific protein 18 [Sourc |
| AC053503.3 | 2 | pseudogene similar to part of protein tyrosine |
| DNPEP | 2 | aspartyl aminopeptidase [Source:HGNC Symb |
| AC053503.2 | 2 | novel transcript, antisense to DNPEP |
| DES | 2 | desmin [Source:HGNC Symbol;Acc:HGNC:277 |
| AC053503.4 | 2 | novel transcript |
| AC053503.6 | 2 | TEC |
| SPEG | 2 | striated muscle enriched protein kinase [Sour |
| AC053503.1 | 2 | novel transcript |
| AC053503.7 | 2 | novel protein |
| SPEGNB | 2 | SPEG neighbor [Source:HGNC Symbol;Acc:HG |
| AC053503.5 | 2 | novel transcript, antisense to SPEG |
| GMPPA | 2 | GDP-mannose pyrophosphorylase A [Source:H |
| ASIC4 | 2 | acid sensing ion channel subunit family memb |
| CHPF | 2 | chondroitin polymerizing factor [Source:HGN |
| TMEM198 | 2 | transmembrane protein 198 [Source:HGNC Sy |
| AC009955.3 | 2 | novel transcript, antisense to TMEM198 |
| MIR3132 | 2 | microRNA 3132 [Source:HGNC Symbol;Acc:HG |
| OBSL1 | 2 | obscurin like 1 [Source:HGNC Symbol;Acc:HGN |
| AC009955.4 | 2 | novel transcript, antisense to OBSL1 |
| INHA | 2 | inhibin subunit alpha [Source:HGNC Symbol;A |
| RN7SKP213 | 2 | RN7SK pseudogene 213 [Source:HGNC Symbo |
| STK11IP | 2 | serine/threonine kinase 11 interacting protein |
| AC009955.1 | 2 | novel transcript |
| SLC4A3 | 2 | solute carrier family 4 member 3 [Source:HGN |
| AC009955.2 | 2 | novel transcript |
| AC009502.1 | 2 | DnaJ (Hsp40) homolog, subfamily B, member |
| LINC02832 | 2 | long intergenic non-protein coding RNA 2832 |
| RPL23P4 | 2 | ribosomal protein L23 pseudogene 4 [Source: |
| LINC01803 | 2 | long intergenic non-protein coding RNA 1803 |

| | | |
|---|---|---|
| AC008281.1 | 2 | novel transcript |
| MIR4268 | 2 | microRNA 4268 [Source:HGNC Symbol;Acc:HG |
| AC009310.1 | 2 | pseudogene similar to part of mitochondrially |
| AC019211.1 | 2 | novel transcript |
| AC093083.1 | 2 | novel transcript |
| AC067956.1 | 2 | novel transcript |
| AC093843.1 | 2 | novel transcript |
| AC093843.2 | 2 | novel transcript |
| AC011233.1 | 2 | SMT3 suppressor of mif two 3 homolog 3 (S. c |
| AC011233.2 | 2 | novel transcript |
| EPHA4 | 2 | EPH receptor A4 [Source:HGNC Symbol;Acc:H |
| TMEM256P2 | 2 | transmembrane protein 256 pseudogene 2 [S |
| AC079834.2 | 2 | novel transcript |
| AC079834.1 | 2 | ribosomal protein L23 (RPL23) pseudogene |
| AC068489.1 | 2 | novel transcript |
| AC009302.1 | 2 | chromosome 12 open reading frame 31 (C12c |
| HSPA9P1 | 2 | heat shock protein family A (Hsp70) member |
| RPL23AP28 | 2 | ribosomal protein L23a pseudogene 28 [Sourc |
| PAX3 | 2 | paired box 3 [Source:HGNC Symbol;Acc:HGNC |
| CCDC140 | 2 | coiled-coil domain containing 140 [Source:HG |
| AC010980.1 | 2 | novel transcript |
| CT75 | 2 | cancer/testis associated transcript 75 [Source |
| Y_RNA | 2 | Y RNA [Source:RFAM;Acc:RF00019] |
| SGPP2 | 2 | sphingosine-1-phosphate phosphatase 2 [Sou |
| NANOGP2 | 2 | Nanog homeobox pseudogene 2 [Source:HGN |
| RNU6-619P | 2 | RNA, U6 small nuclear 619, pseudogene [Sour |
| GAPDHP49 | 2 | glyceraldehyde 3 phosphate dehydrogenase p |
| FARSB | 2 | phenylalanyl-tRNA synthetase subunit beta [S |
| MOGAT1 | 2 | monoacylglycerol O-acyltransferase 1 [Source |
| AC104772.2 | 2 | related RAS viral (r-ras) oncogene homolog 2 ( |
| AC104772.1 | 2 | C-type lectin domain family 1, member A (CLE |
| AC016712.1 | 2 | ribosomal protein L31 (RPL31) pseudogene |
| ACSL3 | 2 | acyl-CoA synthetase long chain family membe |
| ATG12P2 | 2 | autophagy related 12 pseudogene 2 [Source:H |
| AC013476.1 | 2 | novel transcript |
| AC013476.2 | 2 | zinc finger protein 354B (ZNF354B) pseudoger |
| KCNE4 | 2 | potassium voltage-gated channel subfamily E |
| RN7SL807P | 2 | RNA, 7SL, cytoplasmic 807, pseudogene [Sour |
| HIGD1AP4 | 2 | HIG1 hypoxia inducible domain family membe |
| AC013448.1 | 2 | novel transcript |
| AC013448.2 | 2 | novel transcript |
| SCG2 | 2 | secretogranin II [Source:HGNC Symbol;Acc:HG |
| AC012512.1 | 2 | MAX-like protein X (MLX) pseudogene |
| AP1S3 | 2 | adaptor related protein complex 1 subunit sig |
| AC093884.1 | 2 | novel protein, WDFY1-AP1S3 readthrough |
| WDFY1 | 2 | WD repeat and FYVE domain containing 1 [So |
| MRPL44 | 2 | mitochondrial ribosomal protein L44 [Source: |
| AC073641.1 | 2 | novel transcript |
| SERPINE2 | 2 | serpin family E member 2 [Source:HGNC Symb |
| AC019109.1 | 2 | ubiquitin specific peptidase 21 (USP21) pseud |

| | | |
|---|---|---|
| AC008072.1 | 2 | ribosomal protein S27a (RPS27A) pseudogene |
| FAM124B | 2 | family with sequence similarity 124 member E |
| AC073052.2 | 2 | novel transcript, antisense to CUL3 |
| CUL3 | 2 | cullin 3 [Source:HGNC Symbol;Acc:HGNC:2553 |
| AC073052.1 | 2 | ankyrin repeat domain 49 (ANKRD49) pseudo |
| CCDC195 | 2 | coiled-coil domain containing 195 [Source:HG |
| DOCK10 | 2 | dedicator of cytokinesis 10 [Source:HGNC Sym |
| MIR4439 | 2 | microRNA 4439 [Source:HGNC Symbol;Acc:HG |
| NYAP2 | 2 | neuronal tyrosine-phosphorylated phosphoin |
| AC016717.1 | 2 | similar to part of chromosome 7 open reading |
| AC016717.2 | 2 | novel transcript |
| AC016717.3 | 2 | TEC |
| AC068138.1 | 2 | similar to part of the heterogeneous nuclear r |
| AC062015.1 | 2 | novel transcript |
| MIR5702 | 2 | microRNA 5702 [Source:HGNC Symbol;Acc:HG |
| IRS1 | 2 | insulin receptor substrate 1 [Source:HGNC Syr |
| AC010735.2 | 2 | novel transcript |
| AC010735.1 | 2 | novel transcript |
| RHBDD1 | 2 | rhomboid domain containing 1 [Source:HGNC |
| AC073149.1 | 2 | NULL |
| COL4A4 | 2 | collagen type IV alpha 4 chain [Source:HGNC S |
| COL4A3 | 2 | collagen type IV alpha 3 chain [Source:HGNC S |
| AC097662.1 | 2 | novel transcript, antisense to COL4A3 |
| MFF | 2 | mitochondrial fission factor [Source:HGNC Syr |
| TM4SF20 | 2 | transmembrane 4 L six family member 20 [Sou |
| SCYGR1 | 2 | small cysteine and glycine repeat containing 1 |
| MIR5703 | 2 | microRNA 5703 [Source:HGNC Symbol;Acc:HG |
| AGFG1 | 2 | ArfGAP with FG repeats 1 [Source:HGNC Symb |
| AC064853.2 | 2 | novel keratin associated protein |
| SCYGR2 | 2 | small cysteine and glycine repeat containing 2 |
| SCYGR10 | 2 | small cysteine and glycine repeat containing 1 |
| C2orf83 | 2 | chromosome 2 open reading frame 83 [Sourc |
| SCYGR3 | 2 | small cysteine and glycine repeat containing 3 |
| SCYGR4 | 2 | small cysteine and glycine repeat containing 4 |
| AC064853.1 | 2 | novel transcript |
| SCYGR5 | 2 | small cysteine and glycine repeat containing 5 |
| SLC19A3 | 2 | solute carrier family 19 member 3 [Source:HG |
| SCYGR6 | 2 | small cysteine and glycine repeat containing 6 |
| SCYGR7 | 2 | small cysteine and glycine repeat containing 7 |
| SCYGR8 | 2 | small cysteine and glycine repeat containing 8 |
| RNA5SP121 | 2 | RNA, 5S ribosomal pseudogene 121 [Source:H |
| SNRPGP8 | 2 | small nuclear ribonucleoprotein polypeptide ( |
| AC073065.1 | 2 | ATP-binding cassette, sub-family G (WHITE), m |
| CCL20 | 2 | C-C motif chemokine ligand 20 [Source:HGNC |
| TDGF1P2 | 2 | teratocarcinoma-derived growth factor 1 pseu |
| DAW1 | 2 | dynein assembly factor with WD repeats 1 [Sc |
| AC073065.2 | 2 | NULL |
| SPHKAP | 2 | SPHK1 interactor, AKAP domain containing [Sc |
| RNU6-624P | 2 | RNA, U6 small nuclear 624, pseudogene [Sour |
| AC074019.1 | 2 | SNF8, ESCRT-II complex subunit, homolog (S. c |

| | | |
|---|---|---|
| AC009410.1 | 2 | novel transcript |
| LINC01807 | 2 | long intergenic non-protein coding RNA 1807 |
| AC012070.1 | 2 | novel transcript |
| RPL7L1P10 | 2 | ribosomal protein L7 like 1 pseudogene 10 [Sc |
| PID1 | 2 | phosphotyrosine interaction domain containir |
| AC007677.1 | 2 | ribosomal protein L17 (RPL17) pseudogene |
| RN7SKP283 | 2 | RN7SK pseudogene 283 [Source:HGNC Symbo |
| DNER | 2 | delta/notch like EGF repeat containing [Sourc |
| AC008273.1 | 2 | novel transcript, antisense to DNER |
| RNU7-9P | 2 | RNA, U7 small nuclear 9 pseudogene [Source: |
| AC007559.1 | 2 | thioredoxin domain containing 17 (TXNDC17) |
| TRIP12 | 2 | thyroid hormone receptor interactor 12 [Sour |
| RNU6-613P | 2 | RNA, U6 small nuclear 613, pseudogene [Sour |
| FBXO36 | 2 | F-box protein 36 [Source:HGNC Symbol;Acc:H |
| RNU6-964P | 2 | RNA, U6 small nuclear 964, pseudogene [Sour |
| FBXO36-IT1 | 2 | FBXO36 intronic transcript 1 [Source:HGNC Sy |
| RNU6-1027P | 2 | RNA, U6 small nuclear 1027, pseudogene [Sou |
| SLC16A14 | 2 | solute carrier family 16 member 14 [Source:H |
| RNY4P19 | 2 | RNY4 pseudogene 19 [Source:HGNC Symbol;A |
| AC009950.1 | 2 | novel transcript, antisense to SP110 |
| AC009950.2 | 2 | component of Sp100-rs (Csprs) pseudogene |
| SP110 | 2 | SP110 nuclear body protein [Source:HGNC Syr |
| SP140 | 2 | SP140 nuclear body protein [Source:HGNC Syr |
| SP140L | 2 | SP140 nuclear body protein like [Source:HGN( |
| AC009949.1 | 2 | TEC |
| SP100 | 2 | SP100 nuclear antigen [Source:HGNC Symbol; |
| RN7SL834P | 2 | RNA, 7SL, cytoplasmic 834, pseudogene [Sour |
| HMGB1P3 | 2 | high mobility group box 1 pseudogene 3 [Sour |
| AC010149.1 | 2 | novel transcript, antisense to SP100 |
| TPM3P8 | 2 | tropomyosin 3 pseudogene 8 [Source:HGNC S |
| AC010149.2 | 2 | Novel Sp100-like protein pseudogene |
| RNU6-451P | 2 | RNA, U6 small nuclear 451, pseudogene [Sour |
| LINC01907 | 2 | long intergenic non-protein coding RNA 1907 |
| CAB39 | 2 | calcium binding protein 39 [Source:HGNC Sym |
| BANF1P3 | 2 | barrier to autointegration factor 1 pseudogen |
| RNU6-268P | 2 | RNA, U6 small nuclear 268, pseudogene [Sour |
| AC012507.2 | 2 | coiled-coil domain containing 93 (CCDC93) ps |
| ITM2C | 2 | integral membrane protein 2C [Source:HGNC |
| GCSIR | 2 | GPR55 cis regulatory suppressor of immune re |
| GPR55 | 2 | G protein-coupled receptor 55 [Source:HGNC |
| AC012507.1 | 2 | novel transcript |
| AC012507.3 | 2 | novel transcript, antisense to GPR55 |
| COX20P2 | 2 | COX20 pseudogene 2 [Source:HGNC Symbol;A |
| SPATA3-AS1 | 2 | SPATA3 antisense RNA 1 (head to head) [Sour |
| SPATA3 | 2 | spermatogenesis associated 3 [Source:HGNC S |
| C2orf72 | 2 | chromosome 2 open reading frame 72 [Sourc |
| AC009407.1 | 2 | novel transcript |
| PSMD1 | 2 | proteasome 26S subunit, non-ATPase 1 [Sourc |
| HTR2B | 2 | 5-hydroxytryptamine receptor 2B [Source:HG |
| RNU1-93P | 2 | RNA, U1 small nuclear 93, pseudogene [Sourc |

| | | |
|---|---|---|
| ARMC9 | 2 | armadillo repeat containing 9 [Source:HGNC S |
| AC018738.1 | 2 | ribosomal protein S28 (RPS28) pseudogene |
| MIR4777 | 2 | microRNA 4777 [Source:HGNC Symbol;Acc:HG |
| AC017104.1 | 2 | novel transcript |
| B3GNT7 | 2 | UDP-GlcNAc:betaGal beta-1,3-N-acetylglucosa |
| ZBTB8OSP2 | 2 | zinc finger and BTB domain containing 8 oppo |
| AC017104.2 | 2 | HIG1 domain family, member 2A (HIGD2A) ps |
| AC017104.3 | 2 | novel transcript |
| NCL | 2 | nucleolin [Source:HGNC Symbol;Acc:HGNC:76 |
| SNORA75 | 2 | small nucleolar RNA, H/ACA box 75 [Source:H |
| SNORD20 | 2 | small nucleolar RNA, C/D box 20 [Source:HGN |
| SNORD82 | 2 | small nucleolar RNA, C/D box 82 [Source:HGN |
| AC017104.4 | 2 | GTF2I repeat domain containing 2(GTF2IRD2) |
| AC017104.5 | 2 | nucleolin (NCL) pseudogene |
| RNU2-22P | 2 | RNA, U2 small nuclear 22, pseudogene [Sourc |
| LINC00471 | 2 | long intergenic non-protein coding RNA 471 [S |
| AC017104.6 | 2 | novel transcript |
| NMUR1 | 2 | neuromedin U receptor 1 [Source:HGNC Symb |
| AC104634.1 | 2 | ribosomal protein L21 (RPL21) pseudogene |
| RPL23AP26 | 2 | ribosomal protein L23a pseudogene 26 [Sourc |
| AC104634.2 | 2 | novel transcript |
| TEX44 | 2 | testis expressed 44 [Source:HGNC Symbol;Acc |
| RN7SL499P | 2 | RNA, 7SL, cytoplasmic 499, pseudogene [Sour |
| PTMA | 2 | prothymosin alpha [Source:HGNC Symbol;Acc |
| U4 | 2 | U4 spliceosomal RNA [Source:RFAM;Acc:RF00 |
| MIR1244-1 | 2 | microRNA 1244-1 [Source:HGNC Symbol;Acc:H |
| PDE6D | 2 | phosphodiesterase 6D [Source:HGNC Symbol; |
| AC073476.4 | 2 | NULL |
| COPS7B | 2 | COP9 signalosome subunit 7B [Source:HGNC S |
| AC073476.1 | 2 | ribosomal protein L28 (RPL28) pseudogene |
| AC073476.3 | 2 | novel transcript |
| AC073476.2 | 2 | guanidinoacetate N-methyltransferase (GAMT |
| MIR1471 | 2 | microRNA 1471 [Source:HGNC Symbol;Acc:HG |
| NPPC | 2 | natriuretic peptide C [Source:HGNC Symbol;A |
| DIS3L2 | 2 | DIS3 like 3'-5' exoribonuclease 2 [Source:HGN |
| AC019130.1 | 2 | novel transcript |
| MIR562 | 2 | microRNA 562 [Source:HGNC Symbol;Acc:HGN |
| AC068134.3 | 2 | nuclear receptor binding factor 2 (NRBF2) pse |
| ECEL1P3 | 2 | endothelin converting enzyme like 1 pseudog |
| AC068134.4 | 2 | DIS3 mitotic control homolog (S. cerevisiae)-li |
| ALPP | 2 | alkaline phosphatase, placental [Source:HGNC |
| AC068134.1 | 2 | novel transcript |
| ECEL1P2 | 2 | endothelin converting enzyme like 1 pseudog |
| ALPG | 2 | alkaline phosphatase, germ cell [Source:HGNC |
| ECEL1P1 | 2 | endothelin converting enzyme like 1 pseudog |
| AC068134.2 | 2 | novel transcript |
| DIS3L2P1 | 2 | DIS3 like 3'-5' exoribonuclease 2 pseudogene |
| ALPI | 2 | alkaline phosphatase, intestinal [Source:HGNC |
| ECEL1 | 2 | endothelin converting enzyme like 1 [Source:H |
| PRSS56 | 2 | serine protease 56 [Source:HGNC Symbol;Acc |

| | | |
|---|---|---|
| CHRND | 2 | cholinergic receptor nicotinic delta subunit [S |
| CHRNG | 2 | cholinergic receptor nicotinic gamma subunit |
| TIGD1 | 2 | tigger transposable element derived 1 [Source |
| MIR5001 | 2 | microRNA 5001 [Source:HGNC Symbol;Acc:H |
| EIF4E2 | 2 | eukaryotic translation initiation factor 4E fam |
| AC073254.1 | 2 | novel transcript |
| EFHD1 | 2 | EF-hand domain family member D1 [Source:H |
| RN7SL359P | 2 | RNA, 7SL, cytoplasmic 359, pseudogene [Sour |
| GIGYF2 | 2 | GRB10 interacting GYF protein 2 [Source:HGN |
| EEF1B2P7 | 2 | eukaryotic translation elongation factor 1 bet |
| AC064852.1 | 2 | novel transcript |
| KCNJ13 | 2 | potassium voltage-gated channel subfamily J |
| RNU6-107P | 2 | RNA, U6 small nuclear 107, pseudogene [Sour |
| Y_RNA | 2 | Y RNA [Source:RFAM;Acc:RF00019] |
| SNORC | 2 | secondary ossification center associated regul |
| NGEF | 2 | neuronal guanine nucleotide exchange factor |
| AC106876.1 | 2 | novel transcript, antisense to NGEF |
| NEU2 | 2 | neuraminidase 2 [Source:HGNC Symbol;Acc:H |
| INPP5D | 2 | inositol polyphosphate-5-phosphatase D [Sou |
| RN7SL32P | 2 | RNA, 7SL, cytoplasmic 32, pseudogene [Sourc |
| ATG16L1 | 2 | autophagy related 16 like 1 [Source:HGNC Syr |
| SCARNA5 | 2 | small Cajal body-specific RNA 5 [Source:HGNC |
| SCARNA6 | 2 | small Cajal body-specific RNA 6 [Source:HGNC |
| SAG | 2 | S-antigen visual arrestin [Source:HGNC Symbo |
| AC013726.1 | 2 | novel transcript, antisense to DGKD |
| DGKD | 2 | diacylglycerol kinase delta [Source:HGNC Sym |
| Y_RNA | 2 | Y RNA [Source:RFAM;Acc:RF00019] |
| USP40 | 2 | ubiquitin specific peptidase 40 [Source:HGNC |
| AC019072.1 | 2 | protein tyrosine phosphatase, receptor type, |
| UGT1A12P | 2 | UDP glucuronosyltransferase family 1 membe |
| UGT1A11P | 2 | UDP glucuronosyltransferase family 1 membe |
| UGT1A8 | 2 | UDP glucuronosyltransferase family 1 membe |
| UGT1A10 | 2 | UDP glucuronosyltransferase family 1 membe |
| UGT1A13P | 2 | UDP glucuronosyltransferase family 1 membe |
| UGT1A9 | 2 | UDP glucuronosyltransferase family 1 membe |
| UGT1A7 | 2 | UDP glucuronosyltransferase family 1 membe |
| UGT1A6 | 2 | UDP glucuronosyltransferase family 1 membe |
| AC114812.2 | 2 | novel transcript |
| UGT1A5 | 2 | UDP glucuronosyltransferase family 1 membe |
| UGT1A4 | 2 | UDP glucuronosyltransferase family 1 membe |
| RPL17P11 | 2 | ribosomal protein L17 pseudogene 11 [Source |
| UGT1A3 | 2 | UDP glucuronosyltransferase family 1 membe |
| AC114812.3 | 2 | UDP glycosyltransferase 1 family, polypeptide |
| DNAJB3 | 2 | DnaJ heat shock protein family (Hsp40) memb |
| UGT1A2P | 2 | UDP glucuronosyltransferase family 1 membe |
| AC114812.4 | 2 | UDP glycosyltransferase 1 family, polypeptide |
| AC114812.1 | 2 | DnaJ (Hsp40) homolog, subfamily B, member |
| UGT1A1 | 2 | UDP glucuronosyltransferase family 1 membe |
| MROH2A | 2 | maestro heat like repeat family member 2A [S |
| HJURP | 2 | Holliday junction recognition protein [Source: |

| | | |
|---|---|---|
| MSL3P1 | 2 | MSL complex subunit 3 pseudogene 1 [Source |
| AC005538.2 | 2 | novel transcript |
| TRPM8 | 2 | transient receptor potential cation channel su |
| AC005538.1 | 2 | novel transcript |
| AC005538.3 | 2 | novel transcript, antisense to TRPM8 |
| SPP2 | 2 | secreted phosphoprotein 2 [Source:HGNC Syn |
| AC006037.2 | 2 | TEC |
| AC006037.1 | 2 | novel transcript |
| AC122134.1 | 2 | novel transcript |
| AC097713.2 | 2 | ribosomal protein S20 (RPS20) pseudogene |
| AC097713.3 | 2 | novel transcript |
| AC097713.1 | 2 | novel transcript |
| LINC01891 | 2 | long intergenic non-protein coding RNA 1891 |
| HSPE1P9 | 2 | heat shock protein family E (Hsp10) member 1 |
| ARL4C | 2 | ADP ribosylation factor like GTPase 4C [Source |
| LINC01173 | 2 | long intergenic non-protein coding RNA 1173 |
| AC104394.1 | 2 | heat shock 70kDa protein 8 (HSPA8) pseudoge |
| AC010148.1 | 2 | novel transcript |
| SH3BP4 | 2 | SH3 domain binding protein 4 [Source:HGNC S |
| AC114814.1 | 2 | novel transcript |
| AC114814.2 | 2 | chromosome 3 open reading frame 34 (C3orf3 |
| AC114814.3 | 2 | novel transcript |
| AGAP1 | 2 | ArfGAP with GTPase domain, ankyrin repeat a |
| AGAP1-IT1 | 2 | AGAP1 intronic transcript 1 [Source:HGNC Syr |
| AC064874.1 | 2 | novel transcript, antisense to AGAP1 |
| TMSB10P1 | 2 | thymosin beta 10 pseudogene 1 [Source:HGN |
| RNU7-127P | 2 | RNA, U7 small nuclear 127 pseudogene [Sourc |
| RN7SL204P | 2 | RNA, 7SL, cytoplasmic 204, pseudogene [Sour |
| Y_RNA | 2 | Y RNA [Source:RFAM;Acc:RF00019] |
| GBX2 | 2 | gastrulation brain homeobox 2 [Source:HGNC |
| AC019068.1 | 2 | novel transcript, antisense to ASB18 and GBX2 |
| ASB18 | 2 | ankyrin repeat and SOCS box containing 18 [So |
| RNU1-31P | 2 | RNA, U1 small nuclear 31, pseudogene [Sourc |
| IQCA1 | 2 | IQ motif containing with AAA domain 1 [Sourc |
| AC093915.1 | 2 | novel transcript, antisense to IQCA1 |
| IQCA1-AS1 | 2 | IQCA1 antisense RNA 1 [Source:HGNC Symbol |
| RPL3P5 | 2 | ribosomal protein L3 pseudogene 5 [Source:H |
| ACKR3 | 2 | atypical chemokine receptor 3 [Source:HGNC |
| AC084030.1 | 2 | novel transcript |
| AC012063.1 | 2 | novel transcript, antisense to COPS8 |
| RNU6-1051P | 2 | RNA, U6 small nuclear 1051, pseudogene [Sou |
| AC105760.1 | 2 | novel transcript |
| COPS8 | 2 | COP9 signalosome subunit 8 [Source:HGNC Sy |
| AC107079.1 | 2 | novel transcript |
| AC112715.1 | 2 | novel transcript |
| COL6A3 | 2 | collagen type VI alpha 3 chain [Source:HGNC S |
| AC112721.1 | 2 | novel transcript |
| AC112721.2 | 2 | novel protein (LOC728009) |
| MLPH | 2 | melanophilin [Source:HGNC Symbol;Acc:HGN( |
| MIR6811 | 2 | microRNA 6811 [Source:HGNC Symbol;Acc:HG |

| | | |
|---|---|---|
| RNU6-1140P | 2 | RNA, U6 small nuclear 1140, pseudogene [Sou |
| PRLH | 2 | prolactin releasing hormone [Source:HGNC Sy |
| RAB17 | 2 | RAB17, member RAS oncogene family [Source |
| AC104667.2 | 2 | novel transcript, antisense to RAB17 |
| AC104667.1 | 2 | novel transcript |
| LRRFIP1 | 2 | LRR binding FLII interacting protein 1 [Source: |
| AC012076.1 | 2 | novel transcript, antisense to LRRFIP1 |
| RBM44 | 2 | RNA binding motif protein 44 [Source:HGNC S |
| RAMP1 | 2 | receptor activity modifying protein 1 [Source: |
| SNORD39 | 2 | Small nucleolar RNA SNORD55/SNORD39 [Sou |
| UBE2F | 2 | ubiquitin conjugating enzyme E2 F (putative) [ |
| UBE2F-SCLY | 2 | UBE2F-SCLY readthrough (NMD candidate) [So |
| RNU6-1333P | 2 | RNA, U6 small nuclear 1333, pseudogene [Sou |
| SCLY | 2 | selenocysteine lyase [Source:HGNC Symbol;Ac |
| ESPNL | 2 | espin like [Source:HGNC Symbol;Acc:HGNC:27 |
| KLHL30 | 2 | kelch like family member 30 [Source:HGNC Sy |
| KLHL30-AS1 | 2 | KLHL30 antisense RNA 1 [Source:HGNC Symbo |
| ERFE | 2 | erythroferrone [Source:HGNC Symbol;Acc:HG |
| ILKAP | 2 | ILK associated serine/threonine phosphatase |
| LINC02610 | 2 | long intergenic non-protein coding RNA 2610 |
| AC012485.1 | 2 | novel transcript |
| HES6 | 2 | hes family bHLH transcription factor 6 [Source |
| PER2 | 2 | period circadian regulator 2 [Source:HGNC Syı |
| AC012485.3 | 2 | novel transcript, antisense to PER2 |
| AC012485.2 | 2 | novel transcript |
| TRAF3IP1 | 2 | TRAF3 interacting protein 1 [Source:HGNC Syr |
| RNU6-234P | 2 | RNA, U6 small nuclear 234, pseudogene [Sour |
| ASB1 | 2 | ankyrin repeat and SOCS box containing 1 [So |
| AC016999.1 | 2 | novel transcript |
| AC016999.2 | 2 | novel zinc finger protein pseudogene |
| LINC01107 | 2 | long intergenic non-protein coding RNA 1107 |
| LINC01937 | 2 | long intergenic non-protein coding RNA 1937 |
| AC113618.2 | 2 | novel transcript |
| AC113618.1 | 2 | ribosomal protein L5 (RPL5) pseudogene |
| AC145625.2 | 2 | novel transcript |
| AC145625.1 | 2 | novel transcript |
| TWIST2 | 2 | twist family bHLH transcription factor 2 [Sourc |
| LINC01940 | 2 | long intergenic non-protein coding RNA 1940 |
| HDAC4 | 2 | histone deacetylase 4 [Source:HGNC Symbol;A |
| MIR4440 | 2 | microRNA 4440 [Source:HGNC Symbol;Acc:HG |
| MIR4441 | 2 | microRNA 4441 [Source:HGNC Symbol;Acc:HG |
| AC017028.1 | 2 | novel transcript, antisense to HDAC4 |
| AC017028.2 | 2 | novel transcript, antisense to HDAC4 |
| MIR4269 | 2 | microRNA 4269 [Source:HGNC Symbol;Acc:HG |
| MIR2467 | 2 | microRNA 2467 [Source:HGNC Symbol;Acc:HG |
| HDAC4-AS1 | 2 | HDAC4 antisense RNA 1 [Source:HGNC Symbo |
| AC023787.2 | 2 | novel transcript |
| AC023787.3 | 2 | novel transcript |
| AC023787.1 | 2 | novel transcript |
| AC079612.1 | 2 | uncharacterized LOC401040 [Source:NCBI ger |

| | | |
|---|---|---|
| AC079612.2 | 2 | novel transcript |
| AC093802.2 | 2 | novel transcript |
| AC093802.1 | 2 | novel transcript |
| NDUFA10 | 2 | NADH:ubiquinone oxidoreductase subunit A1( |
| MIR4786 | 2 | microRNA 4786 [Source:HGNC Symbol;Acc:HG |
| OR6B2 | 2 | olfactory receptor family 6 subfamily B memb |
| OR6B3 | 2 | olfactory receptor family 6 subfamily B memb |
| OR9S24P | 2 | olfactory receptor family 9 subfamily S memb |
| OR5S1P | 2 | olfactory receptor family 5 subfamily S memb |
| COPS9 | 2 | COP9 signalosome subunit 9 [Source:HGNC Sy |
| OTOS | 2 | otospiralin [Source:HGNC Symbol;Acc:HGNC:2 |
| AC124861.3 | 2 | novel transcript |
| AC124861.2 | 2 | novel transcript |
| AC124861.1 | 2 | novel transcript |
| U3 | 2 | Small nucleolar RNA U3 [Source:RFAM;Acc:RF |
| GPC1 | 2 | glypican 1 [Source:HGNC Symbol;Acc:HGNC:4 |
| AC110619.1 | 2 | novel transcript, antisense to GPC1 |
| MIR149 | 2 | microRNA 149 [Source:HGNC Symbol;Acc:HGI |
| ANKMY1 | 2 | ankyrin repeat and MYND domain containing |
| AC124862.1 | 2 | novel transcript, antisense to ANKMY1 |
| DUSP28 | 2 | dual specificity phosphatase 28 [Source:HGNC |
| RNPEPL1 | 2 | arginyl aminopeptidase like 1 [Source:HGNC S |
| CAPN10-DT | 2 | CAPN10 divergent transcript [Source:HGNC Sy |
| CAPN10 | 2 | calpain 10 [Source:HGNC Symbol;Acc:HGNC:1 |
| GPR35 | 2 | G protein-coupled receptor 35 [Source:HGNC |
| AC011298.3 | 2 | galactose-3-O-sulfotransferase 1 (GAL3ST1) ps |
| AQP12B | 2 | aquaporin 12B [Source:HGNC Symbol;Acc:HGI |
| AC011298.1 | 2 | novel transcript |
| AQP12A | 2 | aquaporin 12A [Source:HGNC Symbol;Acc:HGI |
| AC011298.2 | 2 | galactose-3-O-sulfotransferase 4 (GAL3ST4) ps |
| KIF1A | 2 | kinesin family member 1A [Source:HGNC Sym |
| AGXT | 2 | alanine--glyoxylate and serine--pyruvate amin |
| MAB21L4 | 2 | mab-21 like 4 [Source:HGNC Symbol;Acc:HGN |
| CROCC2 | 2 | ciliary rootlet coiled-coil, rootletin family men |
| AC104809.2 | 2 | novel transcript |
| AC104809.1 | 2 | novel transcript |
| SNED1 | 2 | sushi, nidogen and EGF like domains 1 [Source |
| AC093585.1 | 2 | novel transcript |
| AC005237.1 | 2 | novel transcript |
| MTERF4 | 2 | mitochondrial transcription termination facto |
| PASK | 2 | PAS domain containing serine/threonine kinas |
| AC005237.2 | 2 | zinc finger CCCH-type containing 12A (ZC3H12 |
| PPP1R7 | 2 | protein phosphatase 1 regulatory subunit 7 [S |
| ANO7 | 2 | anoctamin 7 [Source:HGNC Symbol;Acc:HGNC |
| HDLBP | 2 | high density lipoprotein binding protein [Sour |
| SEPTIN2 | 2 | septin 2 [Source:HGNC Symbol;Acc:HGNC:772 |
| AC104841.1 | 2 | novel transcript, antisense to SEPT2 |
| AC005104.1 | 2 | novel transcript |
| FARP2 | 2 | FERM, ARH/RhoGEF and pleckstrin domain pr |
| AC005104.3 | 2 | novel transcript, antisense to FARP2 |

| | | |
|---|---|---|
| MIR3133 | 2 | microRNA 3133 [Source:HGNC Symbol;Acc:HG |
| AC005104.2 | 2 | potassium channel tetramerisation domain co |
| STK25 | 2 | serine/threonine kinase 25 [Source:HGNC Sym |
| BOK-AS1 | 2 | BOK antisense RNA 1 [Source:HGNC Symbol;A |
| BOK | 2 | BCL2 family apoptosis regulator BOK [Source:I |
| AC133528.1 | 2 | novel transcript |
| THAP4 | 2 | THAP domain containing 4 [Source:HGNC Sym |
| RNA5SP122 | 2 | RNA, 5S ribosomal pseudogene 122 [Source:H |
| ATG4B | 2 | autophagy related 4B cysteine peptidase [Sou |
| DTYMK | 2 | deoxythymidylate kinase [Source:HGNC Symb |
| ING5 | 2 | inhibitor of growth family member 5 [Source:I |
| AC114730.3 | 2 | novel transcript |
| D2HGDH | 2 | D-2-hydroxyglutarate dehydrogenase [Source |
| AC114730.1 | 2 | novel transcript |
| AC114730.2 | 2 | novel transcript |
| GAL3ST2 | 2 | galactose-3-O-sulfotransferase 2 [Source:HGN |
| AC131097.1 | 2 | novel transcript |
| AC131097.2 | 2 | novel transcript |
| NEU4 | 2 | neuraminidase 4 [Source:HGNC Symbol;Acc:H |
| AC131097.3 | 2 | novel transcript |
| PDCD1 | 2 | programmed cell death 1 [Source:HGNC Symb |
| RTP5 | 2 | receptor transporter protein 5 (putative) [Sou |
| LINC01237 | 2 | long intergenic non-protein coding RNA 1237 |
| FAM240C | 2 | family with sequence similarity 240 member C |
| LINC01238 | 2 | long intergenic non-protein coding RNA 1238 |
| AC093642.2 | 2 | TEC |
| AC093642.1 | 2 | novel transcript |
| AC093642.7 | 2 | novel transcript |
| LINC01880 | 2 | long intergenic non-protein coding RNA 1880 |
| LINC01238 | 2 | long intergenic non-protein coding RNA 1238 |
| LINC01881 | 2 | long intergenic non-protein coding RNA 1881 |
| CICP10 | 2 | capicua transcriptional repressor pseudogene |
| SEPTIN14P2 | 2 | septin 14 pseudogene 2 [Source:HGNC Symbo |
| RPL23AP88 | 2 | ribosomal protein L23a pseudogene 88 [Sourc |
| LINC01986 | 3 | long intergenic non-protein coding RNA 1986 |
| AC066595.1 | 3 | novel transcript |
| CHL1-AS2 | 3 | CHL1 antisense RNA 2 [Source:HGNC Symbol;/ |
| AC066595.2 | 3 | novel transcript |
| CHL1 | 3 | cell adhesion molecule L1 like [Source:HGNC S |
| RNU6-1194P | 3 | RNA, U6 small nuclear 1194, pseudogene [Sou |
| RPS8P6 | 3 | ribosomal protein S8 pseudogene 6 [Source:H |
| CHL1-AS1 | 3 | CHL1 antisense RNA 1 [Source:HGNC Symbol;/ |
| AC087428.1 | 3 | novel transcript |
| LINC01266 | 3 | long intergenic non-protein coding RNA 1266 |
| RPSAP32 | 3 | ribosomal protein SA pseudogene 32 [Source: |
| AC090044.1 | 3 | novel transcript |
| AC087430.1 | 3 | novel transcript |
| RN7SL120P | 3 | RNA, 7SL, cytoplasmic 120, pseudogene [Sour |
| CNTN6 | 3 | contactin 6 [Source:HGNC Symbol;Acc:HGNC:: |
| CRB3P1 | 3 | crumbs cell polarity complex component 3 ps |

| | | |
|---|---|---|
| RPL23AP38 | 3 | ribosomal protein L23a pseudogene 38 [Sourc |
| RPL23AP39 | 3 | ribosomal protein L23a pseudogene 39 [Sourc |
| RPL21P17 | 3 | ribosomal protein L21 pseudogene 17 [Source |
| AC018814.1 | 3 | novel transcript |
| RN7SKP144 | 3 | RN7SK pseudogene 144 [Source:HGNC Symbo |
| CNTN4 | 3 | contactin 4 [Source:HGNC Symbol;Acc:HGNC:: |
| CNTN4-AS2 | 3 | CNTN4 antisense RNA 2 [Source:HGNC Symbo |
| AC087427.1 | 3 | novel transcript, sense intronic to CNTN4 |
| HINT2P1 | 3 | histidine triad nucleotide binding protein 2 ps |
| DNAJC19P4 | 3 | DnaJ heat shock protein family (Hsp40) memb |
| CNTN4-AS1 | 3 | CNTN4 antisense RNA 1 [Source:HGNC Symbo |
| IL5RA | 3 | interleukin 5 receptor subunit alpha [Source:H |
| AC024060.1 | 3 | NULL |
| TRNT1 | 3 | tRNA nucleotidyl transferase 1 [Source:HGNC |
| CRBN | 3 | cereblon [Source:HGNC Symbol;Acc:HGNC:30 |
| AC024060.2 | 3 | novel transcript, antisense to CRBN |
| AC034195.1 | 3 | novel transcript |
| SUMF1 | 3 | sulfatase modifying factor 1 [Source:HGNC Syr |
| LRRN1 | 3 | leucine rich repeat neuronal 1 [Source:HGNC S |
| AC023480.1 | 3 | novel transcript, antisense to SUMF1 |
| PNPT1P1 | 3 | polyribonucleotide nucleotidyltransferase 1 ps |
| AC023483.1 | 3 | novel transcript, antisense to SETMAR |
| SETMAR | 3 | SET domain and mariner transposase fusion g |
| MRPS10P2 | 3 | mitochondrial ribosomal protein S10 pseudog |
| ITPR1-DT | 3 | ITPR1 divergent transcript [Source:HGNC Sym |
| ITPR1 | 3 | inositol 1,4,5-trisphosphate receptor type 1 [S |
| EGOT | 3 | eosinophil granule ontogeny transcript [Sourc |
| AC018816.2 | 3 | NULL |
| AC018816.1 | 3 | novel transcript, antisense to ITPR1 |
| BHLHE40-AS1 | 3 | BHLHE40 antisense RNA 1 [Source:HGNC Sym |
| BHLHE40 | 3 | basic helix-loop-helix family member e40 [Sou |
| RNF10P1 | 3 | ring finger protein 10 pseudogene 1 [Source:H |
| UBTFL8 | 3 | UBTF like 8 (pseudogene) [Source:HGNC Symb |
| Y_RNA | 3 | Y RNA [Source:RFAM;Acc:RF00019] |
| AC090955.1 | 3 | CD24 pseudogene |
| ARL8B | 3 | ADP ribosylation factor like GTPase 8B [Source |
| AC026202.2 | 3 | novel transcript, antisense to ARL8B |
| AC026202.3 | 3 | novel transcript, antisense to EDEM1 |
| EDEM1 | 3 | ER degradation enhancing alpha-mannosidase |
| MIR4790 | 3 | microRNA 4790 [Source:HGNC Symbol;Acc:HG |
| RN7SL553P | 3 | RNA, 7SL, cytoplasmic 553, pseudogene [Sour |
| AC026202.1 | 3 | novel transcript |
| AC087857.1 | 3 | novel transcript |
| AC026167.1 | 3 | novel transcript |
| AC027119.1 | 3 | pseudogene similar to part of MOCO sulphura |
| AC069277.1 | 3 | novel transcript |
| GRM7-AS3 | 3 | GRM7 antisense RNA 3 [Source:HGNC Symbol |
| AC069277.2 | 3 | tyrosine 3-monooxygenase/tryptophan 5-mor |
| GRM7 | 3 | glutamate metabotropic receptor 7 [Source:H |
| MRPS36P1 | 3 | mitochondrial ribosomal protein S36 pseudog |

| | | |
|---|---|---|
| GRM7-AS2 | 3 | GRM7 antisense RNA 2 [Source:HGNC Symbol |
| AC066585.1 | 3 | novel transcript, sense intronic to GRM7 |
| GRM7-AS1 | 3 | GRM7 antisense RNA 1 [Source:HGNC Symbol |
| AC077690.1 | 3 | novel transcript, antisense to GRM7 |
| LMCD1-AS1 | 3 | LMCD1 antisense RNA 1 [Source:HGNC Symbo |
| AC018832.1 | 3 | novel transcript |
| AC023481.1 | 3 | novel transcript |
| RNU4ATAC17P | 3 | RNA, U4atac small nuclear 17, pseudogene [S |
| LMCD1 | 3 | LIM and cysteine rich domains 1 [Source:HGN |
| AC034187.1 | 3 | novel transcript |
| SSUH2 | 3 | ssu-2 homolog [Source:HGNC Symbol;Acc:HG |
| OR7E122P | 3 | olfactory receptor family 7 subfamily E memb |
| CAV3 | 3 | caveolin 3 [Source:HGNC Symbol;Acc:HGNC:1 |
| OXTR | 3 | oxytocin receptor [Source:HGNC Symbol;Acc: |
| RAD18 | 3 | RAD18 E3 ubiquitin protein ligase [Source:HG |
| AC034186.2 | 3 | NULL |
| AC034186.1 | 3 | NULL |
| SRGAP3 | 3 | SLIT-ROBO Rho GTPase activating protein 3 [S |
| SRGAP3-AS1 | 3 | SRGAP3 antisense RNA 1 [Source:HGNC Symb |
| AC037193.1 | 3 | novel transcript, antisense to SRGAP3 |
| SRGAP3-AS2 | 3 | SRGAP3 antisense RNA 2 [Source:HGNC Symb |
| SRGAP3-AS3 | 3 | SRGAP3 antisense RNA 3 [Source:HGNC Symb |
| SRGAP3-AS4 | 3 | SRGAP3 antisense RNA 4 [Source:HGNC Symb |
| AC026191.1 | 3 | novel transcript, antisense to THUMPD3 |
| PGAM1P4 | 3 | phosphoglycerate mutase 1 pseudogene 4 [Sc |
| THUMPD3-AS1 | 3 | THUMPD3 antisense RNA 1 [Source:HGNC Syr |
| THUMPD3 | 3 | THUMP domain containing 3 [Source:HGNC Sy |
| SETD5 | 3 | SET domain containing 5 [Source:HGNC Symb |
| LHFPL4 | 3 | LHFPL tetraspan subfamily member 4 [Source |
| DUSP5P2 | 3 | dual specificity phosphatase 5 pseudogene 2 [ |
| MTMR14 | 3 | myotubularin related protein 14 [Source:HGN |
| AC022382.2 | 3 | novel transcript, antisense to CPNE9and MTM |
| CPNE9 | 3 | copine family member 9 [Source:HGNC Symb |
| BRPF1 | 3 | bromodomain and PHD finger containing 1 [S |
| OGG1 | 3 | 8-oxoguanine DNA glycosylase [Source:HGNC |
| CAMK1 | 3 | calcium/calmodulin dependent protein kinase |
| TADA3 | 3 | transcriptional adaptor 3 [Source:HGNC Symb |
| ARPC4 | 3 | actin related protein 2/3 complex subunit 4 [S |
| ARPC4-TTLL3 | 3 | ARPC4-TTLL3 readthrough [Source:HGNC Sym |
| TTLL3 | 3 | tubulin tyrosine ligase like 3 [Source:HGNC Sy |
| AC022382.1 | 3 | novel transcript, antisense to TTLL3 |
| RPUSD3 | 3 | RNA pseudouridine synthase D3 [Source:HGN |
| CIDEC | 3 | cell death inducing DFFA like effector c [Sourc |
| JAGN1 | 3 | jagunal homolog 1 [Source:HGNC Symbol;Acc: |
| IL17RE | 3 | interleukin 17 receptor E [Source:HGNC Symb |
| IL17RC | 3 | interleukin 17 receptor C [Source:HGNC Symb |
| RNU6-882P | 3 | RNA, U6 small nuclear 882, pseudogene [Sour |
| CRELD1 | 3 | cysteine rich with EGF like domains 1 [Source: |
| AC018809.1 | 3 | novel transcript, sense intronic to CRELD1 |
| PRRT3 | 3 | proline rich transmembrane protein 3 [Source |

| | | |
|---|---|---|
| PRRT3-AS1 | 3 | PRRT3 antisense RNA 1 [Source:HGNC Symbol |
| AC018809.2 | 3 | novel transcript, antisense to TMEM111 |
| EMC3 | 3 | ER membrane protein complex subunit 3 [Sou |
| EMC3-AS1 | 3 | EMC3 antisense RNA 1 [Source:HGNC Symbol; |
| CYCSP10 | 3 | CYCS pseudogene 10 [Source:HGNC Symbol;A |
| AC022007.1 | 3 | novel transcript |
| CIDECP1 | 3 | cell death inducing DFFA like effector c pseud |
| FANCD2 | 3 | FA complementation group D2 [Source:HGNC |
| RNU6-670P | 3 | RNA, U6 small nuclear 670, pseudogene [Sour |
| CYCSP11 | 3 | CYCS pseudogene 11 [Source:HGNC Symbol;A |
| FANCD2OS | 3 | FANCD2 opposite strand [Source:HGNC Symb |
| BRK1 | 3 | BRICK1 subunit of SCAR/WAVE actin nucleatin |
| VHL | 3 | von Hippel-Lindau tumor suppressor [Source: |
| AC034193.1 | 3 | novel transcript, antisense to VHL |
| IRAK2 | 3 | interleukin 1 receptor associated kinase 2 [So |
| TATDN2 | 3 | TatD DNase domain containing 2 [Source:HGN |
| AC022384.1 | 3 | novel protein |
| LINC00852 | 3 | long intergenic non-protein coding RNA 852 [S |
| GHRL | 3 | ghrelin and obestatin prepropeptide [Source: |
| GHRLOS | 3 | ghrelin opposite strand/antisense RNA [Sourc |
| AC022384.2 | 3 | NULL |
| SEC13 | 3 | SEC13 homolog, nuclear pore and COPII coat |
| ATP2B2 | 3 | ATPase plasma membrane Ca2+ transporting |
| MIR378B | 3 | microRNA 378b [Source:HGNC Symbol;Acc:H |
| MIR885 | 3 | microRNA 885 [Source:HGNC Symbol;Acc:HG |
| ATP2B2-IT1 | 3 | ATP2B2 intronic transcript 1 [Source:HGNC Sy |
| ATP2B2-IT2 | 3 | ATP2B2 intronic transcript 2 [Source:HGNC Sy |
| LINC00606 | 3 | long intergenic non-protein coding RNA 606 [S |
| AC018495.1 | 3 | novel transcript |
| SLC6A11 | 3 | solute carrier family 6 member 11 [Source:HG |
| AC027128.1 | 3 | novel transcript, antisense to SLC6A11 |
| SLC6A1 | 3 | solute carrier family 6 member 1 [Source:HGN |
| SLC6A1-AS1 | 3 | SLC6A1 antisense RNA 1 [Source:HGNC Symb |
| AC066580.1 | 3 | novel transcript |
| HRH1 | 3 | histamine receptor H1 [Source:HGNC Symbol; |
| AC083855.2 | 3 | Novel transcript, antisense to HRH1 |
| CHCHD4P4 | 3 | coiled-coil-helix-coiled-coil-helix domain cont |
| ATG7 | 3 | autophagy related 7 [Source:HGNC Symbol;Ac |
| AC083855.1 | 3 | ribosomal protein L13a (RPL13A) pseudogene |
| AC026185.1 | 3 | microtubule associated serine/threonine kina |
| AC022001.1 | 3 | cystatin B (stefin B), pseudogene |
| VGLL4 | 3 | vestigial like family member 4 [Source:HGNC |
| AC022001.2 | 3 | novel transcript, antisense to VGLL4 |
| AC022001.3 | 3 | novel transcript, antisense to VGLL4 |
| TAMM41 | 3 | TAM41 mitochondrial translocator assembly |
| FANCD2P2 | 3 | FANCD2 pseudogene 2 [Source:HGNC Symbol |
| CYCSP12 | 3 | CYCS pseudogene 12 [Source:HGNC Symbol;A |
| NUP210P2 | 3 | nucleoporin 210 pseudogene 2 [Source:HGNC |
| MARK2P14 | 3 | microtubule affinity regulating kinase 2 pseud |
| AC090958.1 | 3 | novel pseudogene |

| | | |
|---|---|---|
| RN7SL147P | 3 | RNA, 7SL, cytoplasmic 147, pseudogene [Sour |
| SYN2 | 3 | synapsin II [Source:HGNC Symbol;Acc:HGNC:1 |
| ACTG1P12 | 3 | actin gamma 1 pseudogene 12 [Source:HGNC |
| TIMP4 | 3 | TIMP metallopeptidase inhibitor 4 [Source:HG |
| MTCO1P5 | 3 | MT-CO1 pseudogene 5 [Source:HGNC Symbol |
| GSTM5P1 | 3 | glutathione S-transferase mu 5 pseudogene 1 |
| PPARG | 3 | peroxisome proliferator activated receptor ga |
| AC091492.1 | 3 | Prefoldin subunit 1 (PFD1) pseudogene |
| TSEN2 | 3 | tRNA splicing endonuclease subunit 2 [Source |
| RNA5SP123 | 3 | RNA, 5S ribosomal pseudogene 123 [Source:H |
| RNU6-377P | 3 | RNA, U6 small nuclear 377, pseudogene [Sour |
| MKRN2OS | 3 | MKRN2 opposite strand [Source:HGNC Symbo |
| MKRN2 | 3 | makorin ring finger protein 2 [Source:HGNC Sy |
| Metazoa_SRP | 3 | Metazoan signal recognition particle RNA [Sou |
| RAF1 | 3 | Raf-1 proto-oncogene, serine/threonine kinas |
| CRIP1P1 | 3 | cysteine rich protein 1 pseudogene 1 [Source: |
| TMEM40 | 3 | transmembrane protein 40 [Source:HGNC Syn |
| KRT18P17 | 3 | keratin 18 pseudogene 17 [Source:HGNC Sym |
| CAND2 | 3 | cullin associated and neddylation dissociated |
| AC034198.2 | 3 | novel transcript, antisense to CAND2 |
| RPL32 | 3 | ribosomal protein L32 [Source:HGNC Symbol; |
| SNORA7A | 3 | small nucleolar RNA, H/ACA box 7A [Source:H |
| AC034198.1 | 3 | intraflagellar transport 122 (IFT122) pseudoge |
| LINC02022 | 3 | long intergenic non-protein coding RNA 2022 |
| IQSEC1 | 3 | IQ motif and Sec7 domain 1 [Source:HGNC Sy |
| AC069271.1 | 3 | ribosomal protein L39 (RPL39) pseudogene |
| NUP210 | 3 | nucleoporin 210 [Source:HGNC Symbol;Acc:H |
| AC027124.2 | 3 | protein phosphatase 1, regulatory (inhibitor) s |
| AC027124.1 | 3 | proline rich 3 (PPR3) pseudogene |
| HDAC11-AS1 | 3 | HDAC11 antisense RNA 1 [Source:HGNC Symb |
| HDAC11 | 3 | histone deacetylase 11 [Source:HGNC Symbol |
| FBLN2 | 3 | fibulin 2 [Source:HGNC Symbol;Acc:HGNC:360 |
| LINC00620 | 3 | long intergenic non-protein coding RNA 620 [S |
| WNT7A | 3 | Wnt family member 7A [Source:HGNC Symbo |
| VN1R20P | 3 | vomeronasal 1 receptor 20 pseudogene [Sour |
| FGD5P1 | 3 | FYVE, RhoGEF and PH domain containing 5 ps |
| TPRXL | 3 | tetrapeptide repeat homeobox like [Source:H |
| AC090004.2 | 3 | novel transcript |
| VN1R21P | 3 | vomeronasal 1 receptor 21 pseudogene [Sour |
| CHCHD4 | 3 | coiled-coil-helix-coiled-coil-helix domain conta |
| TMEM43 | 3 | transmembrane protein 43 [Source:HGNC Syn |
| AC090004.1 | 3 | novel transcript |
| AC093495.1 | 3 | novel transcript, antisense to XPC |
| XPC | 3 | XPC complex subunit, DNA damage recognitio |
| LSM3 | 3 | LSM3 homolog, U6 small nuclear RNA and mR |
| AC093496.1 | 3 | novel transcript |
| LINC01267 | 3 | long intergenic non-protein coding RNA 1267 |
| RNA5SP124 | 3 | RNA, 5S ribosomal pseudogene 124 [Source:H |
| SLC6A6 | 3 | solute carrier family 6 member 6 [Source:HGN |
| GRIP2 | 3 | glutamate receptor interacting protein 2 [Sou |

| | | |
|---|---|---|
| RNU6-905P | 3 | RNA, U6 small nuclear 905, pseudogene [Sour |
| RHBDF1P1 | 3 | RHBDF1 pseudogene 1 [Source:HGNC Symbol, |
| AC090952.2 | 3 | novel transcript |
| AC090952.1 | 3 | novel transcript |
| CCDC174 | 3 | coiled-coil domain containing 174 [Source:HG |
| C3orf20 | 3 | chromosome 3 open reading frame 20 [Source |
| AC090957.1 | 3 | novel transcript |
| AC087591.1 | 3 | novel transcript |
| LINC02011 | 3 | long intergenic non-protein coding RNA 2011 |
| FGD5 | 3 | FYVE, RhoGEF and PH domain containing 5 [So |
| FGD5-AS1 | 3 | FGD5 antisense RNA 1 [Source:HGNC Symbol; |
| NR2C2 | 3 | nuclear receptor subfamily 2 group C member |
| MRPS25 | 3 | mitochondrial ribosomal protein S25 [Source: |
| RBSN | 3 | rabenosyn, RAB effector [Source:HGNC Symbo |
| AC090954.1 | 3 | ribosomal protein S24 (RPS24) pseudogene |
| RPS3AP53 | 3 | ribosomal protein S3a pseudogene 53 [Source |
| COL6A4P1 | 3 | collagen type VI alpha 4 pseudogene 1 [Source |
| CAPN7 | 3 | calpain 7 [Source:HGNC Symbol;Acc:HGNC:14 |
| SH3BP5-AS1 | 3 | SH3BP5 antisense RNA 1 [Source:HGNC Symbo |
| SH3BP5 | 3 | SH3 domain binding protein 5 [Source:HGNC S |
| RNU6-454P | 3 | RNA, U6 small nuclear 454, pseudogene [Sour |
| HMGN2P7 | 3 | high mobility group nucleosomal binding dom |
| METTL6 | 3 | methyltransferase like 6 [Source:HGNC Symbo |
| EAF1 | 3 | ELL associated factor 1 [Source:HGNC Symbol, |
| RNU6-1024P | 3 | RNA, U6 small nuclear 1024, pseudogene [Sou |
| EAF1-AS1 | 3 | EAF1 antisense RNA 1 [Source:HGNC Symbol;A |
| COLQ | 3 | collagen like tail subunit of asymmetric acetyl |
| MIR4270 | 3 | microRNA 4270 [Source:HGNC Symbol;Acc:HG |
| AC027129.1 | 3 | novel transcript, antisense to COLQ |
| RN7SL110P | 3 | RNA, 7SL, cytoplasmic 110, pseudogene [Sour |
| HACL1 | 3 | 2-hydroxyacyl-CoA lyase 1 [Source:HGNC Sym |
| BTD | 3 | biotinidase [Source:HGNC Symbol;Acc:HGNC:1 |
| ANKRD28 | 3 | ankyrin repeat domain 28 [Source:HGNC Symb |
| MIR3134 | 3 | microRNA 3134 [Source:HGNC Symbol;Acc:HG |
| AC090950.2 | 3 | TEC |
| AC090950.1 | 3 | chromosome 9 open reading frame 64 (C9orf6 |
| RN7SL4P | 3 | RNA, 7SL, cytoplasmic 4, pseudogene [Source: |
| AC090945.1 | 3 | novel transcript |
| MIR563 | 3 | microRNA 563 [Source:HGNC Symbol;Acc:HGN |
| IMPDH1P8 | 3 | inosine monophosphate dehydrogenase 1 pse |
| AC090946.1 | 3 | novel transcript |
| GALNT15 | 3 | polypeptide N-acetylgalactosaminyltransferas |
| AC090953.2 | 3 | Novel transcript |
| DPH3 | 3 | diphthamide biosynthesis 3 [Source:HGNC Syr |
| OXNAD1 | 3 | oxidoreductase NAD binding domain containi |
| AC090953.1 | 3 | THAP domain containing 5. pseudogene |
| RFTN1 | 3 | raftlin, lipid raft linker 1 [Source:HGNC Symbo |
| AC090948.1 | 3 | novel transcript, antisense to RFTN1 |
| AC090948.2 | 3 | novel transcript, antisense to RFTN1 |
| AC090948.3 | 3 | novel transcript, antisense to RFTN1 |

| | | |
|---|---|---|
| AC090948.4 | 3 | novel transcript, antisense to RFTN1 |
| AC010727.1 | 3 | novel transcript, antisense to RFTN1 |
| LINC00690 | 3 | long intergenic non-protein coding RNA 690 [S |
| AC010139.1 | 3 | novel transcript |
| DAZL | 3 | deleted in azoospermia like [Source:HGNC Syr |
| AC091493.1 | 3 | novel transcript |
| CDYLP1 | 3 | CDYL pseudogene 1 [Source:HGNC Symbol;Ac |
| PLCL2 | 3 | phospholipase C like 2 [Source:HGNC Symbol; |
| MIR3714 | 3 | microRNA 3714 [Source:HGNC Symbol;Acc:HG |
| PLCL2-AS1 | 3 | PLCL2 antisense RNA 1 [Source:HGNC Symbol, |
| AC090644.1 | 3 | Novel transcript, antisense to TBC1D5 |
| TBC1D5 | 3 | TBC1 domain family member 5 [Source:HGNC |
| AC090960.1 | 3 | chromosome 8 open reading frame 59 (C8orf! |
| AC140076.1 | 3 | ribosomal protein L31 (RPL31) pseudogene |
| RNU7-10P | 3 | RNA, U7 small nuclear 10 pseudogene [Source |
| AC104451.1 | 3 | pseudogene similar to part of aminopeptidase |
| AC104451.2 | 3 | novel transcript |
| AC104297.1 | 3 | pyrophosphatase (inorganic) 1 (PPA1) pseudo |
| PDCL3P3 | 3 | PDCL3 pseudogene 3 [Source:HGNC Symbol;A |
| AC132807.1 | 3 | novel transcript |
| AC132807.2 | 3 | novel transcript |
| SATB1 | 3 | SATB homeobox 1 [Source:HGNC Symbol;Acc: |
| AC144521.1 | 3 | novel transcript, sense intronic to SATB1 |
| SATB1-AS1 | 3 | SATB1 antisense RNA 1 [Source:HGNC Symbol |
| RAD23BP1 | 3 | RAD23B pseudogene 1 [Source:HGNC Symbol, |
| RNU6-138P | 3 | RNA, U6 small nuclear 138, pseudogene [Sour |
| KCNH8 | 3 | potassium voltage-gated channel subfamily H |
| MIR4791 | 3 | microRNA 4791 [Source:HGNC Symbol;Acc:HG |
| AC061958.1 | 3 | novel transcript, antisense to KCNH8 |
| EFHB | 3 | EF-hand domain family member B [Source:HG |
| HSPA8P18 | 3 | heat shock protein family A (Hsp70) member |
| RAB5A | 3 | RAB5A, member RAS oncogene family [Source |
| PP2D1 | 3 | protein phosphatase 2C like domain containin |
| RNU4-85P | 3 | RNA, U4 small nuclear 85, pseudogene [Sourc |
| SAP18P3 | 3 | SAP18 pseudogene 3 [Source:HGNC Symbol;A |
| RPL39P18 | 3 | ribosomal protein L39 pseudogene 18 [Source |
| KAT2B | 3 | lysine acetyltransferase 2B [Source:HGNC Sym |
| MIR3135A | 3 | microRNA 3135a [Source:HGNC Symbol;Acc:H |
| SGO1 | 3 | shugoshin 1 [Source:HGNC Symbol;Acc:HGNC: |
| SGO1-AS1 | 3 | SGO1 antisense RNA 1 [Source:HGNC Symbol; |
| RNU6-822P | 3 | RNA, U6 small nuclear 822, pseudogene [Sour |
| AC116096.1 | 3 | novel transcript |
| RNU6-815P | 3 | RNA, U6 small nuclear 815, pseudogene [Sour |
| AC099753.1 | 3 | novel transcript |
| VENTXP7 | 3 | VENT homeobox pseudogene 7 [Source:HGNC |
| ZNF385D | 3 | zinc finger protein 385D [Source:HGNC Symbo |
| ZNF385D-AS1 | 3 | ZNF385D antisense RNA 1 [Source:HGNC Sym |
| ZNF385D-AS2 | 3 | ZNF385D antisense RNA 2 [Source:HGNC Sym |
| HMGB1P5 | 3 | high mobility group box 1 pseudogene 5 [Sour |
| AC092421.1 | 3 | pseudogene similar to part of Fc receptor-like |

| | | |
|---|---|---|
| AC114477.1 | 3 | ribosome binding protein 1, pseudogene |
| RANP7 | 3 | RAN pseudogene 7 [Source:HGNC Symbol;Acc |
| SALL4P5 | 3 | spalt like transcription factor 4 pseudogene 5 |
| RPL24P7 | 3 | ribosomal protein L24 pseudogene 7 [Source: |
| UBE2E2-AS1 | 3 | UBE2E2 antisense RNA 1 (head to head) [Sour |
| UBE2E2 | 3 | ubiquitin conjugating enzyme E2 E2 [Source:H |
| Y_RNA | 3 | Y RNA [Source:RFAM;Acc:RF00019] |
| AC121251.1 | 3 | pseudogene similar to part of chromosome 1 |
| RNU6-922P | 3 | RNA, U6 small nuclear 922, pseudogene [Sour |
| RNU6-788P | 3 | RNA, U6 small nuclear 788, pseudogene [Sour |
| UBE2E1-AS1 | 3 | UBE2E1 antisense RNA 1 [Source:HGNC Symb |
| UBE2E1 | 3 | ubiquitin conjugating enzyme E2 E1 [Source:H |
| ARL4AP4 | 3 | ADP ribosylation factor like GTPase 4A pseudo |
| NKIRAS1 | 3 | NFKB inhibitor interacting Ras like 1 [Source:H |
| RPL15 | 3 | ribosomal protein L15 [Source:HGNC Symbol; |
| NR1D2 | 3 | nuclear receptor subfamily 1 group D membe |
| LINC00691 | 3 | long intergenic non-protein coding RNA 691 [ |
| NPM1P23 | 3 | nucleophosmin 1 pseudogene 23 [Source:HGN |
| THRB | 3 | thyroid hormone receptor beta [Source:HGNC |
| RPL31P20 | 3 | ribosomal protein L31 pseudogene 20 [Source |
| THRB-IT1 | 3 | THRB intronic transcript 1 [Source:HGNC Sym |
| THRB-AS1 | 3 | THRB antisense RNA 1 [Source:HGNC Symbol; |
| AC012087.2 | 3 | novel transcript |
| AC012087.1 | 3 | NULL |
| EIF3KP2 | 3 | eukaryotic translation initiation factor 3 subu |
| AC092422.1 | 3 | novel transcript |
| RN7SL216P | 3 | RNA, 7SL, cytoplasmic 216, pseudogene [Sour |
| RNA5SP125 | 3 | RNA, 5S ribosomal pseudogene 125 [Source:H |
| RARB | 3 | retinoic acid receptor beta [Source:HGNC Sym |
| CFL1P7 | 3 | cofilin 1 pseudogene 7 [Source:HGNC Symbol; |
| RNA5SP126 | 3 | RNA, 5S ribosomal pseudogene 126 [Source:H |
| AC133141.1 | 3 | H3 histone, family 3B (H3F3B) pseudogene |
| AC093416.1 | 3 | NULL |
| TOP2B | 3 | DNA topoisomerase II beta [Source:HGNC Sym |
| AC093416.2 | 3 | NULL |
| MIR4442 | 3 | microRNA 4442 [Source:HGNC Symbol;Acc:HG |
| CRIP1P2 | 3 | cysteine rich protein 1 pseudogene 2 [Source: |
| NGLY1 | 3 | N-glycanase 1 [Source:HGNC Symbol;Acc:HGN |
| AC092798.1 | 3 | ribosomal protein L32 (RPL32) pseudogene |
| TAF9BP1 | 3 | TATA-box binding protein associated factor 9b |
| OXSM | 3 | 3-oxoacyl-ACP synthase, mitochondrial [Sourc |
| LINC00692 | 3 | long intergenic non-protein coding RNA 692 [ |
| RPEP2 | 3 | ribulose-5-phosphate-3-epimerase pseudoger |
| HMGB3P12 | 3 | high mobility group box 3 pseudogene 12 [Sou |
| VENTXP4 | 3 | VENT homeobox pseudogene 4 [Source:HGNC |
| AC099754.1 | 3 | novel transcript |
| LRRC3B | 3 | leucine rich repeat containing 3B [Source:HGN |
| NEK10 | 3 | NIMA related kinase 10 [Source:HGNC Symbo |
| MICOS10P3 | 3 | MICOS10 pseudogene 3 [Source:HGNC Symbo |
| RNU6-342P | 3 | RNA, U6 small nuclear 342, pseudogene [Sour |

| | | |
|---|---|---|
| AC099535.2 | 3 | novel transcript, antisense to SLC4A7 |
| SLC4A7 | 3 | solute carrier family 4 member 7 [Source:HGN |
| RPS20P15 | 3 | ribosomal protein S20 pseudogene 15 [Source |
| AC099535.1 | 3 | ubiquitin pseudogene |
| RNU1-96P | 3 | RNA, U1 small nuclear 96, pseudogene [Sourc |
| AC137675.1 | 3 | frataxin (FXN) pseudogene |
| Y_RNA | 3 | Y RNA [Source:RFAM;Acc:RF00019] |
| AC098614.1 | 3 | ribosomal protein S27 (RPS27) pseudogene |
| AC098614.3 | 3 | tropomyosin 4 (TPM4) pseudogene |
| AC098614.2 | 3 | KIAA1143 (KIAA1143) pseudogene |
| LINC02084 | 3 | long intergenic non-protein coding RNA 2084 |
| EOMES | 3 | eomesodermin [Source:HGNC Symbol;Acc:HG |
| LINC01980 | 3 | long intergenic non-protein coding RNA 1980 |
| LINC01981 | 3 | long intergenic non-protein coding RNA 1981 |
| LINC01967 | 3 | long intergenic non-protein coding RNA 1967 |
| CMC1 | 3 | C-X9-C motif containing 1 [Source:HGNC Symb |
| AZI2 | 3 | 5-azacytidine induced 2 [Source:HGNC Symbo |
| ZCWPW2 | 3 | zinc finger CW-type and PWWP domain conta |
| AC098650.1 | 3 | novel protein |
| FAM96AP1 | 3 | family with sequence similarity 96 member A |
| AC093142.1 | 3 | ribosomal protein L34 (RPL34) pseudogene |
| RBMS3 | 3 | RNA binding motif single stranded interacting |
| RBMS3-AS3 | 3 | RBMS3 antisense RNA 3 [Source:HGNC Symbo |
| MESTP4 | 3 | mesoderm specific transcript pseudogene 4 [S |
| RPS12P5 | 3 | ribosomal protein S12 pseudogene 5 [Source: |
| RBMS3-AS2 | 3 | RBMS3 antisense RNA 2 [Source:HGNC Symbo |
| MTND4LP9 | 3 | MT-ND4L pseudogene 9 [Source:HGNC Symbo |
| RBMS3-AS1 | 3 | RBMS3 antisense RNA 1 [Source:HGNC Symbo |
| AC025614.2 | 3 | novel transcript |
| AC025614.1 | 3 | NULL |
| U3 | 3 | Small nucleolar RNA U3 [Source:RFAM;Acc:RF |
| LINC01985 | 3 | long intergenic non-protein coding RNA 1985 |
| AC116035.1 | 3 | novel transcript |
| TGFBR2 | 3 | transforming growth factor beta receptor 2 [S |
| AC096921.1 | 3 | ectodermal-neural cortex (ENC1) pseudogene |
| AC096921.2 | 3 | novel transcript |
| GADL1 | 3 | glutamate decarboxylase like 1 [Source:HGNC |
| MIR466 | 3 | microRNA 466 [Source:HGNC Symbol;Acc:HGN |
| CNN2P6 | 3 | calponin 2 pseudogene 6 [Source:HGNC Symb |
| RNA5SP127 | 3 | RNA, 5S ribosomal pseudogene 127 [Source:H |
| AC115283.1 | 3 | centromere protein A (CENPA) pseudogene |
| THRAP3P1 | 3 | thyroid hormone receptor associated protein |
| STT3B | 3 | STT3 oligosaccharyltransferase complex cataly |
| OSBPL10 | 3 | oxysterol binding protein like 10 [Source:HGN |
| OSBPL10-AS1 | 3 | OSBPL10 antisense RNA 1 [Source:HGNC Symb |
| ZNF587P1 | 3 | zinc finger protein 587 pseudogene 1 [Source: |
| ZNF860 | 3 | zinc finger protein 860 [Source:HGNC Symbol; |
| RPL21P40 | 3 | ribosomal protein L21 pseudogene 40 [Source |
| AC094019.1 | 3 | NULL |
| Y_RNA | 3 | Y RNA [Source:RFAM;Acc:RF00019] |

| | | |
|---|---|---|
| NIFKP7 | 3 | nucleolar protein interacting with the FHA do[illegible] |
| GPD1L | 3 | glycerol-3-phosphate dehydrogenase 1 like [S[illegible] |
| AC094019.2 | 3 | ribosomal protein L21 (RPL21) pseudogene |
| RPSAP11 | 3 | ribosomal protein SA pseudogene 11 [Source: |
| AC097639.1 | 3 | novel transcript, intronic to CMTM8 |
| CMTM8 | 3 | CKLF like MARVEL transmembrane domain co |
| AC097639.2 | 3 | small integral membrane protein 11, pseudog |
| KRT18P15 | 3 | keratin 18 pseudogene 15 [Source:HGNC Sym |
| CMTM7 | 3 | CKLF like MARVEL transmembrane domain co |
| CMTM6 | 3 | CKLF like MARVEL transmembrane domain co |
| MIR548AY | 3 | microRNA 548ay [Source:HGNC Symbol;Acc:H |
| AC104306.1 | 3 | ribosomal protein L31 (RPL31) pseudogene |
| DYNC1LI1 | 3 | dynein cytoplasmic 1 light intermediate chain |
| AC104306.2 | 3 | polymerase (DNA-directed), epsilon 4 (p12 su[illegible] |
| IGBP1P3 | 3 | immunoglobulin (CD79A) binding protein 1 ps |
| RPL30P4 | 3 | ribosomal protein L30 pseudogene 4 [Source:[illegible] |
| CNOT10 | 3 | CCR4-NOT transcription complex subunit 10 [[illegible] |
| CNOT10-AS1 | 3 | CNOT10 antisense RNA 1 [Source:HGNC Symb |
| SUGT1P2 | 3 | SUGT1 pseudogene 2 [Source:HGNC Symbol;[illegible] |
| RPL23AP43 | 3 | ribosomal protein L23a pseudogene 43 [Sourc |
| TRIM71 | 3 | tripartite motif containing 71 [Source:HGNC S[illegible] |
| CCR4 | 3 | C-C motif chemokine receptor 4 [Source:HGN[illegible] |
| GLB1 | 3 | galactosidase beta 1 [Source:HGNC Symbol;A[illegible] |
| SEC13P1 | 3 | SEC13 homolog, nuclear pore and COPII coat [illegible] |
| SUMO2P10 | 3 | SUMO2 pseudogene 10 [Source:HGNC Symbo |
| TMPPE | 3 | transmembrane protein with metallophospho |
| RN7SL296P | 3 | RNA, 7SL, cytoplasmic 296, pseudogene [Sour |
| CRTAP | 3 | cartilage associated protein [Source:HGNC Sy[illegible] |
| AC112211.1 | 3 | novel transcript |
| SUSD5 | 3 | sushi domain containing 5 [Source:HGNC Sym |
| AC123900.1 | 3 | pseudogene similar to part of cytochrome c, s |
| FBXL2 | 3 | F-box and leucine rich repeat protein 2 [Sourc |
| UBP1 | 3 | upstream binding protein 1 [Source:HGNC Sy[illegible] |
| RNU7-110P | 3 | RNA, U7 small nuclear 110 pseudogene [Sourc |
| RNA5SP128 | 3 | RNA, 5S ribosomal pseudogene 128 [Source:H |
| CLASP2 | 3 | cytoplasmic linker associated protein 2 [Sourc |
| COX6CP10 | 3 | cytochrome c oxidase subunit 6C pseudogene |
| AC112220.1 | 3 | interferon induced transmembrane protein 1 |
| SDAD1P3 | 3 | SDA1 domain containing 1 pseudogene 3 [Sou |
| AC112220.2 | 3 | novel transcript |
| PDCD6IP | 3 | programmed cell death 6 interacting protein [ |
| LINC01811 | 3 | long intergenic non-protein coding RNA 1811 |
| AC123023.2 | 3 | novel transcript |
| AC123023.1 | 3 | novel transcript |
| AC018359.1 | 3 | novel transcript |
| AC007483.1 | 3 | novel transcript |
| FECHP1 | 3 | ferrochelatase pseudogene 1 [Source:HGNC S[illegible] |
| KRT8P18 | 3 | keratin 8 pseudogene 18 [Source:HGNC Symb |
| RNU6-243P | 3 | RNA, U6 small nuclear 243, pseudogene [Sour |
| ARPP21 | 3 | cAMP regulated phosphoprotein 21 [Source:H |

| | | |
|---|---|---|
| ARPP21-AS1 | 3 | ARPP21 antisense RNA 1 [Source:HGNC Symb |
| MIR128-2 | 3 | microRNA 128-2 [Source:HGNC Symbol;Acc:H |
| AC104308.1 | 3 | ribosomal protein L36a (RPL36A) pseudogene |
| RFC3P1 | 3 | replication factor C 3 pseudogene 1 [Source:H |
| STAC | 3 | SH3 and cysteine rich domain [Source:HGNC S |
| NBPF21P | 3 | NBPF member 21, pseudogene [Source:HGNC |
| RN7SKP227 | 3 | RN7SK pseudogene 227 [Source:HGNC Symbo |
| DCLK3 | 3 | doublecortin like kinase 3 [Source:HGNC Symb |
| HSPD1P6 | 3 | heat shock protein family D (Hsp60) member |
| LINC02033 | 3 | long intergenic non-protein coding RNA 2033 |
| AC105749.1 | 3 | TEC |
| TRANK1 | 3 | tetratricopeptide repeat and ankyrin repeat c |
| AC011816.1 | 3 | chromobox homolog 3 (CBX3) pseudogene |
| RNU6ATAC4P | 3 | RNA, U6atac small nuclear 4, pseudogene [So |
| AC011816.2 | 3 | novel transcript |
| PRADC1P1 | 3 | protease associated domain containing 1 pseu |
| EPM2AIP1 | 3 | EPM2A interacting protein 1 [Source:HGNC Sy |
| MLH1 | 3 | mutL homolog 1 [Source:HGNC Symbol;Acc:H |
| RPL29P11 | 3 | ribosomal protein L29 pseudogene 11 [Source |
| LRRFIP2 | 3 | LRR binding FLII interacting protein 2 [Source: |
| Y_RNA | 3 | Y RNA [Source:RFAM;Acc:RF00019] |
| UBE2D3P2 | 3 | ubiquitin conjugating enzyme E2 D3 pseudoge |
| RNU6-1301P | 3 | RNA, U6 small nuclear 1301, pseudogene [Sou |
| UBE2FP1 | 3 | UBE2F pseudogene 1 [Source:HGNC Symbol;A |
| AC126118.1 | 3 | novel transcript, antisense to LRRFIP2 |
| AC097359.1 | 3 | ribosomal protein S16 (RPS16) pseudogene |
| AC097359.3 | 3 | novel transcript |
| AC097359.2 | 3 | novel transcript, antisense to GOLGA4 |
| GOLGA4 | 3 | golgin A4 [Source:HGNC Symbol;Acc:HGNC:44 |
| TCEA1P2 | 3 | transcription elongation factor A1 pseudogen |
| RNA5SP129 | 3 | RNA, 5S ribosomal pseudogene 129 [Source:H |
| C3orf35 | 3 | chromosome 3 open reading frame 35 [Sourc |
| ITGA9 | 3 | integrin subunit alpha 9 [Source:HGNC Symbo |
| RNU7-73P | 3 | RNA, U7 small nuclear 73 pseudogene [Source |
| ITGA9-AS1 | 3 | ITGA9 antisense RNA 1 [Source:HGNC Symbol |
| RPL21P135 | 3 | ribosomal protein L21 pseudogene 135 [Sourc |
| NDUFAF4P3 | 3 | NADH:ubiquinone oxidoreductase complex as |
| CTDSPL | 3 | CTD small phosphatase like [Source:HGNC Syr |
| MIR26A1 | 3 | microRNA 26a-1 [Source:HGNC Symbol;Acc:H |
| VILL | 3 | villin like [Source:HGNC Symbol;Acc:HGNC:30 |
| PLCD1 | 3 | phospholipase C delta 1 [Source:HGNC Symbo |
| Y_RNA | 3 | Y RNA [Source:RFAM;Acc:RF00019] |
| DLEC1 | 3 | DLEC1 cilia and flagella associated protein [So |
| PPP2R2DP1 | 3 | protein phosphatase 2 regulatory subunit Bde |
| ACAA1 | 3 | acetyl-CoA acyltransferase 1 [Source:HGNC Sy |
| Y_RNA | 3 | Y RNA [Source:RFAM;Acc:RF00019] |
| MYD88 | 3 | MYD88 innate immune signal transduction ad |
| OXSR1 | 3 | oxidative stress responsive kinase 1 [Source:H |
| SLC22A13 | 3 | solute carrier family 22 member 13 [Source:H |
| AP006193.1 | 3 | cell division cycle 42 (CDC42) pseudogene |

| | | |
|---|---|---|
| SLC22A14 | 3 | solute carrier family 22 member 14 [Source:H |
| RNU6-235P | 3 | RNA, U6 small nuclear 235, pseudogene [Sour |
| DLEC1P1 | 3 | DLEC1 pseudogene 1 [Source:HGNC Symbol;A |
| XYLB | 3 | xylulokinase [Source:HGNC Symbol;Acc:HGNC |
| ACVR2B-AS1 | 3 | ACVR2B antisense RNA 1 [Source:HGNC Symb |
| ACVR2B | 3 | activin A receptor type 2B [Source:HGNC Sym |
| EXOG | 3 | exo/endonuclease G [Source:HGNC Symbol;A |
| Y_RNA | 3 | Y RNA [Source:RFAM;Acc:RF00019] |
| DDTP1 | 3 | D-dopachrome tautomerase pseudogene 1 [S |
| RPL18AP7 | 3 | ribosomal protein L18a pseudogene 7 [Source |
| SCN5A | 3 | sodium voltage-gated channel alpha subunit 5 |
| SCN10A | 3 | sodium voltage-gated channel alpha subunit 1 |
| AC116038.1 | 3 | novel transcript |
| SCN11A | 3 | sodium voltage-gated channel alpha subunit 1 |
| RNU6-1227P | 3 | RNA, U6 small nuclear 1227, pseudogene [Sou |
| WDR48 | 3 | WD repeat domain 48 [Source:HGNC Symbol; |
| GORASP1 | 3 | golgi reassembly stacking protein 1 [Source:H |
| TTC21A | 3 | tetratricopeptide repeat domain 21A [Source: |
| MIR6822 | 3 | microRNA 6822 [Source:HGNC Symbol;Acc:HG |
| CSRNP1 | 3 | cysteine and serine rich nuclear protein 1 [Sou |
| AC092053.3 | 3 | novel transcript, antisense CSRNP1 |
| AC092053.2 | 3 | novel transcript, antisense CSRNP1 |
| Metazoa_SRP | 3 | Metazoan signal recognition particle RNA [Sou |
| AC092053.4 | 3 | novel transcript |
| XIRP1 | 3 | xin actin binding repeat containing 1 [Source:I |
| AC092053.6 | 3 | novel transcript |
| DSTNP4 | 3 | DSTN pseudogene 4 [Source:HGNC Symbol;Ac |
| AC092053.5 | 3 | novel transcript |
| CX3CR1 | 3 | C-X3-C motif chemokine receptor 1 [Source:H |
| AC092053.1 | 3 | Adenosine deaminase domain-containing prot |
| AC104850.2 | 3 | novel transcript, antisense to SLC25A38and C( |
| CCR8 | 3 | C-C motif chemokine receptor 8 [Source:HGN |
| HNRNPA1P21 | 3 | heterogeneous nuclear ribonucleoprotein A1 |
| AC104850.1 | 3 | centromere protein P (CENPP) pseudogene |
| EEF1A1P24 | 3 | eukaryotic translation elongation factor 1 alph |
| SLC25A38 | 3 | solute carrier family 25 member 38 [Source:H |
| RPSA | 3 | ribosomal protein SA [Source:HGNC Symbol;A |
| SNORA6 | 3 | small nucleolar RNA, H/ACA box 6 [Source:HG |
| SNORA62 | 3 | small nucleolar RNA, H/ACA box 62 [Source:H |
| AC099332.2 | 3 | novel transcript |
| AC099332.1 | 3 | novel transcript |
| MOBP | 3 | myelin associated oligodendrocyte basic prote |
| AC092058.1 | 3 | Novel transcript, antisense to MOBP |
| NFU1P1 | 3 | NFU1 iron-sulfur cluster scaffold pseudogene |
| MYRIP | 3 | myosin VIIA and Rab interacting protein [Sour |
| RNU4-56P | 3 | RNA, U4 small nuclear 56, pseudogene [Sourc |
| EIF1B-AS1 | 3 | EIF1B antisense RNA 1 [Source:HGNC Symbol; |
| RN7SL411P | 3 | RNA, 7SL, cytoplasmic 411, pseudogene [Sour |
| AC099331.1 | 3 | NULL |
| EIF1B | 3 | eukaryotic translation initiation factor 1B [Sou |

| | | |
|---|---|---|
| ENTPD3-AS1 | 3 | ENTPD3 antisense RNA 1 [Source:HGNC Symb |
| PGAM1P3 | 3 | phosphoglycerate mutase 1 pseudogene 3 [Sc |
| ENTPD3 | 3 | ectonucleoside triphosphate diphosphohydro |
| RPL14 | 3 | ribosomal protein L14 [Source:HGNC Symbol;/ |
| ZNF619 | 3 | zinc finger protein 619 [Source:HGNC Symbol; |
| ZNF620 | 3 | zinc finger protein 620 [Source:HGNC Symbol; |
| RNU5B-2P | 3 | RNA, U5B small nuclear 2, pseudogene [Sourc |
| ZNF621 | 3 | zinc finger protein 621 [Source:HGNC Symbol; |
| RPL5P10 | 3 | ribosomal protein L5 pseudogene 10 [Source: |
| EEF1GP3 | 3 | eukaryotic translation elongation factor 1 gam |
| AC122683.1 | 3 | novel transcript |
| HMGN2P24 | 3 | high mobility group nucleosomal binding dom |
| AC099560.1 | 3 | novel transcript |
| AC099560.2 | 3 | ribosomal protein S27 (RPS27) pseudogene |
| AC099541.2 | 3 | novel transcript |
| AC099541.1 | 3 | novel transcript |
| AC009743.1 | 3 | novel transcript |
| MRPS31P1 | 3 | mitochondrial ribosomal protein S31 pseudog |
| CTNNB1 | 3 | catenin beta 1 [Source:HGNC Symbol;Acc:HGI |
| ULK4 | 3 | unc-51 like kinase 4 [Source:HGNC Symbol;Ac |
| AC099537.1 | 3 | NULL |
| RN7SKP58 | 3 | RN7SK pseudogene 58 [Source:HGNC Symbol; |
| ATP6V0E1P2 | 3 | ATPase H+ transporting V0 subunit e1 pseudo |
| Y_RNA | 3 | Y RNA [Source:RFAM;Acc:RF00019] |
| U8 | 3 | U8 small nucleolar RNA [Source:RFAM;Acc:RF |
| RPL36P20 | 3 | ribosomal protein L36 pseudogene 20 [Source |
| GEMIN2P2 | 3 | gem nuclear organelle associated protein 2 ps |
| TRAK1 | 3 | trafficking kinesin protein 1 [Source:HGNC Syr |
| AC093414.1 | 3 | ribosomal protein L7a (RPL7A) pseudogene |
| TPMTP2 | 3 | thiopurine S-methyltransferase pseudogene 2 |
| RNU4-78P | 3 | RNA, U4 small nuclear 78, pseudogene [Sourc |
| CCK | 3 | cholecystokinin [Source:HGNC Symbol;Acc:HG |
| AC018358.1 | 3 | novel transcript, antisense to CCK |
| SALL4P6 | 3 | spalt like transcription factor 4 pseudogene 6 |
| EIF4BP4 | 3 | eukaryotic translation initiation factor 4B pseu |
| LYZL4 | 3 | lysozyme like 4 [Source:HGNC Symbol;Acc:HG |
| AC092047.1 | 3 | ribosomal protein L35a (RPL35A) pseudogene |
| VIPR1 | 3 | vasoactive intestinal peptide receptor 1 [Sour |
| VIPR1-AS1 | 3 | VIPR1 antisense RNA 1 [Source:HGNC Symbol; |
| SEC22C | 3 | SEC22 homolog C, vesicle trafficking protein [S |
| SS18L2 | 3 | SS18 like 2 [Source:HGNC Symbol;Acc:HGNC:1 |
| NKTR | 3 | natural killer cell triggering receptor [Source:H |
| AC006059.1 | 3 | novel transcript |
| ZBTB47 | 3 | zinc finger and BTB domain containing 47 [Sou |
| AC006059.5 | 3 | novel transcript, antisense to ZBTB47 |
| RN7SL567P | 3 | RNA, 7SL, cytoplasmic 567, pseudogene [Sour |
| KLHL40 | 3 | kelch like family member 40 [Source:HGNC Sy |
| AC006059.3 | 3 | novel protein |
| HHATL | 3 | hedgehog acyltransferase like [Source:HGNC S |
| HHATL-AS1 | 3 | HHATL antisense RNA 1 [Source:HGNC Symbo |

| | | |
|---|---|---|
| CCDC13 | 3 | coiled-coil domain containing 13 [Source:HGN |
| AC006059.2 | 3 | novel protein |
| AC006059.4 | 3 | novel transcript, antisense to CCDC13 |
| CCDC13-AS1 | 3 | CCDC13 antisense RNA 1 [Source:HGNC Symb |
| LINC02158 | 3 | long intergenic non-protein coding RNA 2158 |
| HIGD1A | 3 | HIG1 hypoxia inducible domain family membe |
| AC099329.1 | 3 | novel transcript, antisense to CCBP2 |
| ACKR2 | 3 | atypical chemokine receptor 2 [Source:HGNC |
| AC099329.2 | 3 | novel transcript |
| AC092042.3 | 3 | novel protein |
| KRBOX1 | 3 | KRAB box domain containing 1 [Source:HGNC |
| CYP8B1 | 3 | cytochrome P450 family 8 subfamily B membe |
| HNRNPA1P22 | 3 | heterogeneous nuclear ribonucleoprotein A1 |
| ZNF662 | 3 | zinc finger protein 662 [Source:HGNC Symbol; |
| KRBOX1-AS1 | 3 | KRBOX1 antisense RNA 1 [Source:HGNC Symb |
| GASK1A | 3 | golgi associated kinase 1A [Source:HGNC Sym |
| AC092042.2 | 3 | pseudogene similar to part of family with seq |
| POMGNT2 | 3 | protein O-linked mannose N-acetylglucosamir |
| AC092042.1 | 3 | novel transcript |
| AC092042.4 | 3 | Novel trranscript, antisense to POMGNT2 |
| SNRK | 3 | SNF related kinase [Source:HGNC Symbol;Acc: |
| RN7SL517P | 3 | RNA, 7SL, cytoplasmic 517, pseudogene [Sour |
| SNRK-AS1 | 3 | SNRK antisense RNA 1 [Source:HGNC Symbol; |
| ANO10 | 3 | anoctamin 10 [Source:HGNC Symbol;Acc:HGN |
| Y_RNA | 3 | Y RNA [Source:RFAM;Acc:RF00019] |
| AC097638.1 | 3 | ribosomal protein L18a (RPL18A) pseudogene |
| ABHD5 | 3 | abhydrolase domain containing 5 [Source:HGI |
| AC006055.1 | 3 | novel transcript |
| AC006058.2 | 3 | novel transcript |
| AC006058.3 | 3 | novel transcript |
| RNU6-367P | 3 | RNA, U6 small nuclear 367, pseudogene [Sour |
| MIR138-1 | 3 | microRNA 138-1 [Source:HGNC Symbol;Acc:H |
| AC006058.4 | 3 | TEC |
| AC006058.1 | 3 | novel transcript |
| TOPAZ1 | 3 | testis and ovary specific PAZ domain containir |
| AC104187.1 | 3 | novel transcript, antisense to TCAIM |
| TCAIM | 3 | T cell activation inhibitor, mitochondrial [Sour |
| C3orf86 | 3 | chromosome 3 open reading frame 86 [Sourc |
| LINC01988 | 3 | long intergenic non-protein coding RNA 1988 |
| ZNF445 | 3 | zinc finger protein 445 [Source:HGNC Symbol; |
| ZNF852 | 3 | zinc finger protein 852 [Source:HGNC Symbol; |
| EI24P3 | 3 | EI24 pseudogene 3 [Source:HGNC Symbol;Acc |
| AC099669.1 | 3 | zinc finger protein 347 (ZNF347) pseudogene |
| ZKSCAN7 | 3 | zinc finger with KRAB and SCAN domains 7 [Sc |
| ZKSCAN7-AS1 | 3 | ZKSCAN7 ZNF cluster antisense RNA 1 [Source |
| ZNF660 | 3 | zinc finger protein 660 [Source:HGNC Symbol; |
| MPRIPP1 | 3 | myosin phosphatase Rho interacting protein p |
| ZNF197 | 3 | zinc finger protein 197 [Source:HGNC Symbol; |
| ZNF197-AS1 | 3 | ZNF197 antisense RNA 1 [Source:HGNC Symbo |
| ZNF35 | 3 | zinc finger protein 35 [Source:HGNC Symbol;A |

| | | |
|---|---|---|
| AC124045.1 | 3 | novel transcript |
| SOCS5P3 | 3 | suppressor of cytokine signaling 5 pseudogene |
| ZNF502 | 3 | zinc finger protein 502 [Source:HGNC Symbol; |
| ZNF501 | 3 | zinc finger protein 501 [Source:HGNC Symbol; |
| KIAA1143 | 3 | KIAA1143 [Source:HGNC Symbol;Acc:HGNC:2 |
| KIF15 | 3 | kinesin family member 15 [Source:HGNC Sym |
| MIR564 | 3 | microRNA 564 [Source:HGNC Symbol;Acc:HG |
| TMEM42 | 3 | transmembrane protein 42 [Source:HGNC Syn |
| TGM4 | 3 | transglutaminase 4 [Source:HGNC Symbol;Acc |
| AC098649.1 | 3 | novel transcript |
| ZDHHC3 | 3 | zinc finger DHHC-type containing 3 [Source:H |
| RPL12P44 | 3 | ribosomal protein L12 pseudogene 44 [Source |
| EXOSC7 | 3 | exosome component 7 [Source:HGNC Symbol |
| CLEC3B | 3 | C-type lectin domain family 3 member B [Sour |
| RNU5B-3P | 3 | RNA, U5B small nuclear 3, pseudogene [Sourc |
| CDCP1 | 3 | CUB domain containing protein 1 [Source:HG |
| RPS24P8 | 3 | ribosomal protein S24 pseudogene 8 [Source: |
| TMEM158 | 3 | transmembrane protein 158 (gene/pseudoger |
| U3 | 3 | Small nucleolar RNA U3 [Source:RFAM;Acc:RF |
| LARS2 | 3 | leucyl-tRNA synthetase 2, mitochondrial [Sour |
| LARS2-AS1 | 3 | LARS2 antisense RNA 1 [Source:HGNC Symbol |
| LIMD1 | 3 | LIM domains containing 1 [Source:HGNC Sym |
| LIMD1-AS1 | 3 | LIMD1 antisense RNA 1 [Source:HGNC Symbo |
| SACM1L | 3 | SAC1 like phosphatidylinositide phosphatase [ |
| RN7SL145P | 3 | RNA, 7SL, cytoplasmic 145, pseudogene [Sour |
| AC098476.1 | 3 | TEC |
| SLC6A20 | 3 | solute carrier family 6 member 20 [Source:HG |
| LZTFL1 | 3 | leucine zipper transcription factor like 1 [Sour |
| AC099782.2 | 3 | Novel transcript |
| AC099782.1 | 3 | succinate dehydrogenase complex, subunit D, |
| CCR9 | 3 | C-C motif chemokine receptor 9 [Source:HGN |
| Y_RNA | 3 | Y RNA [Source:RFAM;Acc:RF00019] |
| FYCO1 | 3 | FYVE and coiled-coil domain containing 1 [Sou |
| CXCR6 | 3 | C-X-C motif chemokine receptor 6 [Source:HG |
| XCR1 | 3 | X-C motif chemokine receptor 1 [Source:HGN |
| NRBF2P2 | 3 | nuclear receptor binding factor 2 pseudogene |
| FLT1P1 | 3 | fms related tyrosine kinase 1 pseudogene 1 [S |
| CCR3 | 3 | C-C motif chemokine receptor 3 [Source:HGN |
| CCR1 | 3 | C-C motif chemokine receptor 1 [Source:HGN |
| UQCRC2P1 | 3 | ubiquinol-cytochrome c reductase core protei |
| CCR2 | 3 | C-C motif chemokine receptor 2 [Source:HGN |
| CCR5AS | 3 | CCR5 antisense RNA [Source:HGNC Symbol;Ac |
| CCR5 | 3 | C-C motif chemokine receptor 5 (gene/pseudo |
| CCRL2 | 3 | C-C motif chemokine receptor like 2 [Source:H |
| LINC02009 | 3 | long intergenic non-protein coding RNA 2009 |
| LTF | 3 | lactotransferrin [Source:HGNC Symbol;Acc:HG |
| RTP3 | 3 | receptor transporter protein 3 [Source:HGNC |
| LRRC2 | 3 | leucine rich repeat containing 2 [Source:HGN |
| LRRC2-AS1 | 3 | LRRC2 antisense RNA 1 [Source:HGNC Symbol |
| TDGF1 | 3 | teratocarcinoma-derived growth factor 1 [Sou |

| | | |
|---|---|---|
| AC104304.2 | 3 | novel transcript |
| FAM240A | 3 | family with sequence similarity 240 member A |
| AC104304.3 | 3 | novel transcript |
| AC104304.1 | 3 | NULL |
| ALS2CL | 3 | ALS2 C-terminal like [Source:HGNC Symbol;Ac |
| TMIE | 3 | transmembrane inner ear [Source:HGNC Symb |
| PRSS50 | 3 | serine protease 50 [Source:HGNC Symbol;Acc |
| PRSS50 | 3 | serine protease 50 [Source:NCBI gene;Acc:29 |
| PRSS46P | 3 | serine protease 46, pseudogene [Source:HGN |
| PRSS45P | 3 | serine protease 45, pseudogene [Source:HGN |
| AC109583.3 | 3 | novel transcript |
| AC134504.1 | 3 | serine protease 43, pseudogene |
| AC109583.1 | 3 | ATP synthase, H+ transporting, mitochondrial |
| PRSS44P | 3 | serine protease 44, pseudogene [Source:HGN |
| PRSS42P | 3 | serine protease 42, pseudogene [Source:HGN |
| MYL3 | 3 | myosin light chain 3 [Source:HGNC Symbol;Ac |
| PTH1R | 3 | parathyroid hormone 1 receptor [Source:HGN |
| AC109583.2 | 3 | biliverdin reductase B (flavin reductase (NADP |
| CCDC12 | 3 | coiled-coil domain containing 12 [Source:HGN |
| NBEAL2 | 3 | neurobeachin like 2 [Source:HGNC Symbol;Ac |
| NRADDP | 3 | neurotrophin receptor associated death doma |
| SETD2 | 3 | SET domain containing 2, histone lysine meth |
| MRPL57P3 | 3 | mitochondrial ribosomal protein L57 pseudog |
| KIF9-AS1 | 3 | KIF9 antisense RNA 1 [Source:HGNC Symbol;A |
| KIF9 | 3 | kinesin family member 9 [Source:HGNC Symb |
| SNORD13P3 | 3 | small nucleolar RNA, C/D box 13 pseudogene |
| KLHL18 | 3 | kelch like family member 18 [Source:HGNC Sy |
| AC104447.1 | 3 | TEC |
| AC099778.1 | 3 | novel transcript, antisense to PTPN23 |
| PTPN23 | 3 | protein tyrosine phosphatase non-receptor ty |
| SCAP | 3 | SREBF chaperone [Source:HGNC Symbol;Acc:H |
| AC099778.2 | 3 | novel pseudogene |
| ELP6 | 3 | elongator acetyltransferase complex subunit 6 |
| BOLA2P2 | 3 | bolA family member 2 pseudogene 2 [Source: |
| RN7SL870P | 3 | RNA, 7SL, cytoplasmic 870, pseudogene [Sour |
| CSPG5 | 3 | chondroitin sulfate proteoglycan 5 [Source:HG |
| SMARCC1 | 3 | SWI/SNF related, matrix associated, actin dep |
| AC112512.1 | 3 | novel transcript, sense intronic SMARCC1 |
| AC130472.1 | 3 | HIG1 domain family, member 2A (HIGD2A) ps |
| DHX30 | 3 | DExH-box helicase 30 [Source:HGNC Symbol;A |
| AC026318.1 | 3 | NULL |
| MIR1226 | 3 | microRNA 1226 [Source:HGNC Symbol;Acc:HG |
| MAP4 | 3 | microtubule associated protein 4 [Source:HGN |
| Y_RNA | 3 | Y RNA [Source:RFAM;Acc:RF00019] |
| VPS26BP1 | 3 | VPS26B pseudogene 1 [Source:HGNC Symbol; |
| AC124916.2 | 3 | ribosomal protein S27-like, pseudogene |
| RN7SL664P | 3 | RNA, 7SL, cytoplasmic 664, pseudogene [Sour |
| AC124916.1 | 3 | ribosomal protein L17 (RPL17) pseudogene |
| CDC25A | 3 | cell division cycle 25A [Source:HGNC Symbol;A |
| SNRPFP4 | 3 | small nuclear ribonucleoprotein polypeptide F |

| | | |
|---|---|---|
| RNU7-128P | 3 | RNA, U7 small nuclear 128 pseudogene [Sourc |
| NDUFB1P1 | 3 | NADH:ubiquinone oxidoreductase subunit B1 |
| MIR4443 | 3 | microRNA 4443 [Source:HGNC Symbol;Acc:HG |
| CAMP | 3 | cathelicidin antimicrobial peptide [Source:HGI |
| ZNF589 | 3 | zinc finger protein 589 [Source:HGNC Symbol; |
| MRPS18AP1 | 3 | mitochondrial ribosomal protein S18A pseudo |
| Y_RNA | 3 | Y RNA [Source:RFAM;Acc:RF00019] |
| NME6 | 3 | NME/NM23 nucleoside diphosphate kinase 6 |
| FCF1P2 | 3 | FCF1 pseudogene 2 [Source:HGNC Symbol;Acc |
| SPINK8 | 3 | serine peptidase inhibitor, Kazal type 8 (putat |
| MIR2115 | 3 | microRNA 2115 [Source:HGNC Symbol;Acc:HG |
| FBXW12 | 3 | F-box and WD repeat domain containing 12 [S |
| RN7SL321P | 3 | RNA, 7SL, cytoplasmic 321, pseudogene [Sour |
| PLXNB1 | 3 | plexin B1 [Source:HGNC Symbol;Acc:HGNC:91 |
| CCDC51 | 3 | coiled-coil domain containing 51 [Source:HGN |
| TMA7 | 3 | translation machinery associated 7 homolog [ |
| AC134772.1 | 3 | novel transcript, antisense to TMA7 |
| ATRIP | 3 | ATR interacting protein [Source:HGNC Symbol |
| TREX1 | 3 | three prime repair exonuclease 1 [Source:HGN |
| SHISA5 | 3 | shisa family member 5 [Source:HGNC Symbol; |
| PFKFB4 | 3 | 6-phosphofructo-2-kinase/fructose-2,6-bipho |
| MIR6823 | 3 | microRNA 6823 [Source:HGNC Symbol;Acc:HG |
| UCN2 | 3 | urocortin 2 [Source:HGNC Symbol;Acc:HGNC:1 |
| COL7A1 | 3 | collagen type VII alpha 1 chain [Source:HGNC |
| MIR711 | 3 | microRNA 711 [Source:HGNC Symbol;Acc:HGN |
| UQCRC1 | 3 | ubiquinol-cytochrome c reductase core protei |
| TMEM89 | 3 | transmembrane protein 89 [Source:HGNC Syn |
| SLC26A6 | 3 | solute carrier family 26 member 6 [Source:HG |
| MIR6824 | 3 | microRNA 6824 [Source:HGNC Symbol;Acc:HG |
| CELSR3 | 3 | cadherin EGF LAG seven-pass G-type receptor |
| MIR4793 | 3 | microRNA 4793 [Source:HGNC Symbol;Acc:HG |
| LINC02585 | 3 | long intergenic non-protein coding RNA 2585 |
| AC141002.1 | 3 | novel transcript |
| NCKIPSD | 3 | NCK interacting protein with SH3 domain [Sou |
| IP6K2 | 3 | inositol hexakisphosphate kinase 2 [Source:HG |
| PRKAR2A | 3 | protein kinase cAMP-dependent type II regula |
| PRKAR2A-AS1 | 3 | PRKAR2A antisense RNA 1 [Source:HGNC Sym |
| SLC25A20 | 3 | solute carrier family 25 member 20 [Source:H |
| ARIH2OS | 3 | ariadne RBR E3 ubiquitin protein ligase 2 oppo |
| ARIH2 | 3 | ariadne RBR E3 ubiquitin protein ligase 2 [Sou |
| AC137630.2 | 3 | novel transcript |
| AC137630.1 | 3 | novel transcript |
| AC137630.4 | 3 | novel transcript, antisense to ARIH2 |
| P4HTM | 3 | prolyl 4-hydroxylase, transmembrane [Source |
| AC137630.5 | 3 | novel pseudogene, ortholog of avian secreted |
| WDR6 | 3 | WD repeat domain 6 [Source:HGNC Symbol;A |
| DALRD3 | 3 | DALR anticodon binding domain containing 3 |
| MIR425 | 3 | microRNA 425 [Source:HGNC Symbol;Acc:HGN |
| NDUFAF3 | 3 | NADH:ubiquinone oxidoreductase complex as |
| MIR191 | 3 | microRNA 191 [Source:HGNC Symbol;Acc:HGN |

| | | |
|---|---|---|
| IMPDH2 | 3 | inosine monophosphate dehydrogenase 2 [So |
| AC137630.3 | 3 | novel transcript, antisense to IMPDH2 and QR |
| QRICH1 | 3 | glutamine rich 1 [Source:HGNC Symbol;Acc:H( |
| RN7SL182P | 3 | RNA, 7SL, cytoplasmic 182, pseudogene [Sour |
| QARS | 3 | glutaminyl-tRNA synthetase [Source:HGNC Sy |
| MIR6890 | 3 | microRNA 6890 [Source:HGNC Symbol;Acc:HG |
| USP19 | 3 | ubiquitin specific peptidase 19 [Source:HGNC |
| LAMB2 | 3 | laminin subunit beta 2 [Source:HGNC Symbol; |
| AC135506.1 | 3 | laminin, beta 1 (LAMB1) pseudogene |
| CCDC71 | 3 | coiled-coil domain containing 71 [Source:HGN |
| KLHDC8B | 3 | kelch domain containing 8B [Source:HGNC Syr |
| C3orf84 | 3 | chromosome 3 open reading frame 84 [Sourc |
| CCDC36 | 3 | coiled-coil domain containing 36 [Source:HGN |
| AC121247.2 | 3 | glycine cleavage system protein H (aminomet |
| AC121247.1 | 3 | novel transcript |
| C3orf62 | 3 | chromosome 3 open reading frame 62 [Sourc |
| Y_RNA | 3 | Y RNA [Source:RFAM;Acc:RF00019] |
| MIR4271 | 3 | microRNA 4271 [Source:HGNC Symbol;Acc:HG |
| USP4 | 3 | ubiquitin specific peptidase 4 [Source:HGNC S |
| GPX1 | 3 | glutathione peroxidase 1 [Source:HGNC Symb |
| RHOA | 3 | ras homolog family member A [Source:HGNC |
| RHOA-IT1 | 3 | RHOA intronic transcript 1 [Source:HGNC Sym |
| TCTA | 3 | T cell leukemia translocation altered [Source:I |
| AMT | 3 | aminomethyltransferase [Source:HGNC Symb |
| AC104452.1 | 3 | novel protein |
| NICN1 | 3 | nicolin 1 [Source:HGNC Symbol;Acc:HGNC:18 |
| RNA5SP130 | 3 | RNA, 5S ribosomal pseudogene 130 [Source:H |
| DAG1 | 3 | dystroglycan 1 [Source:HGNC Symbol;Acc:HGI |
| BSN-DT | 3 | BSN divergent transcript [Source:HGNC Symbo |
| BSN | 3 | bassoon presynaptic cytomatrix protein [Sour |
| BSN-AS1 | 3 | BSN antisense RNA 1 [Source:HGNC Symbol;A |
| APEH | 3 | acylaminoacyl-peptide hydrolase [Source:HGN |
| MST1 | 3 | macrophage stimulating 1 [Source:HGNC Sym |
| AC099668.1 | 3 | novel transcript |
| RNF123 | 3 | ring finger protein 123 [Source:HGNC Symbol; |
| AMIGO3 | 3 | adhesion molecule with Ig like domain 3 [Sour |
| GMPPB | 3 | GDP-mannose pyrophosphorylase B [Source:H |
| IP6K1 | 3 | inositol hexakisphosphate kinase 1 [Source:H( |
| COX6CP14 | 3 | cytochrome c oxidase subunit 6C pseudogene |
| AC139451.1 | 3 | PHD finger protein 5A (PHF5A) pseudogene |
| CDHR4 | 3 | cadherin related family member 4 [Source:HG |
| INKA1 | 3 | inka box actin regulator 1 [Source:HGNC Symb |
| UBA7 | 3 | ubiquitin like modifier activating enzyme 7 [So |
| MIR5193 | 3 | microRNA 5193 [Source:HGNC Symbol;Acc:HG |
| TRAIP | 3 | TRAF interacting protein [Source:HGNC Symbo |
| CAMKV | 3 | CaM kinase like vesicle associated [Source:HG |
| RN7SL217P | 3 | RNA, 7SL, cytoplasmic 217, pseudogene [Sour |
| ACTBP13 | 3 | ACTB pseudogene 13 [Source:HGNC Symbol;A |
| MST1R | 3 | macrophage stimulating 1 receptor [Source:H |
| AC105935.2 | 3 | novel transcript |

| | | |
|---|---|---|
| AC105935.1 | 3 | novel transcript, antisense to MON1A |
| MON1A | 3 | MON1 homolog A, secretory trafficking associ |
| RBM6 | 3 | RNA binding motif protein 6 [Source:HGNC Sy |
| RBM5 | 3 | RNA binding motif protein 5 [Source:HGNC Sy |
| RBM5-AS1 | 3 | RBM5 antisense RNA 1 [Source:HGNC Symbol |
| SEMA3F-AS1 | 3 | SEMA3F antisense RNA 1 [Source:HGNC Symb |
| SEMA3F | 3 | semaphorin 3F [Source:HGNC Symbol;Acc:HG |
| AC104450.1 | 3 | NULL |
| GNAT1 | 3 | G protein subunit alpha transducin 1 [Source: |
| SLC38A3 | 3 | solute carrier family 38 member 3 [Source:HG |
| GNAI2 | 3 | G protein subunit alpha i2 [Source:HGNC Sym |
| MIR5787 | 3 | microRNA 5787 [Source:HGNC Symbol;Acc:HG |
| U73169.1 | 3 | novel transcript, sense intronic to GNAI2 |
| U73166.1 | 3 | novel transcript |
| SEMA3B-AS1 | 3 | SEMA3B antisense RNA 1 (head to head) [Sou |
| SEMA3B | 3 | semaphorin 3B [Source:HGNC Symbol;Acc:HG |
| MIR6872 | 3 | microRNA 6872 [Source:HGNC Symbol;Acc:HG |
| LSMEM2 | 3 | leucine rich single-pass membrane protein 2 [ |
| IFRD2 | 3 | interferon related developmental regulator 2 |
| HYAL3 | 3 | hyaluronidase 3 [Source:HGNC Symbol;Acc:HG |
| NAA80 | 3 | N(alpha)-acetyltransferase 80, NatH catalytic |
| HYAL1 | 3 | hyaluronidase 1 [Source:HGNC Symbol;Acc:HG |
| HYAL2 | 3 | hyaluronidase 2 [Source:HGNC Symbol;Acc:HG |
| TUSC2 | 3 | tumor suppressor 2, mitochondrial calcium re |
| RASSF1 | 3 | Ras association domain family member 1 [Sou |
| RASSF1-AS1 | 3 | RASSF1 antisense RNA 1 [Source:HGNC Symbo |
| ZMYND10-AS1 | 3 | ZMYND10 antisense RNA 1 [Source:HGNC Sym |
| ZMYND10 | 3 | zinc finger MYND-type containing 10 [Source: |
| NPRL2 | 3 | NPR2 like, GATOR1 complex subunit [Source:H |
| CYB561D2 | 3 | cytochrome b561 family member D2 [Source: |
| Z84492.1 | 3 | novel protein |
| TMEM115 | 3 | transmembrane protein 115 [Source:HGNC Sy |
| CACNA2D2 | 3 | calcium voltage-gated channel auxiliary subun |
| CYB561D2 | 3 | cytochrome b561 family member D2 [Source: |
| RNA5SP131 | 3 | RNA, 5S ribosomal pseudogene 131 [Source:H |
| C3orf18 | 3 | chromosome 3 open reading frame 18 [Source |
| HEMK1 | 3 | HemK methyltransferase family member 1 [So |
| AC096920.1 | 3 | novel transcript |
| CISH | 3 | cytokine inducible SH2 containing protein [Sou |
| MAPKAPK3 | 3 | MAPK activated protein kinase 3 [Source:HGN |
| LINC02019 | 3 | long intergenic non-protein coding RNA 2019 |
| DOCK3 | 3 | dedicator of cytokinesis 3 [Source:HGNC Symb |
| MIR4787 | 3 | microRNA 4787 [Source:HGNC Symbol;Acc:HG |
| PPIAP69 | 3 | peptidylprolyl isomerase A pseudogene 69 [So |
| AC126120.1 | 3 | peptidylprolyl isomerase A (cyclophilin A) (PPI |
| ZNF652P1 | 3 | zinc finger protein 652 pseudogene 1 [Source: |
| ST13P14 | 3 | ST13, Hsp70 interacting protein pseudogene 1 |
| MANF | 3 | mesencephalic astrocyte derived neurotrophi |
| RBM15B | 3 | RNA binding motif protein 15B [Source:HGNC |
| DCAF1 | 3 | DDB1 and CUL4 associated factor 1 [Source:H |

| | | |
|---|---|---|
| RAD54L2 | 3 | RAD54 like 2 [Source:HGNC Symbol;Acc:HGNC |
| RNU6ATAC29P | 3 | RNA, U6atac small nuclear 29, pseudogene [So |
| TEX264 | 3 | testis expressed 264, ER-phagy receptor [Sour |
| AC099050.1 | 3 | novel transcript, antisense to RAD54L2and TE |
| RNA5SP132 | 3 | RNA, 5S ribosomal pseudogene 132 [Source:H |
| GRM2 | 3 | glutamate metabotropic receptor 2 [Source:H |
| IQCF6 | 3 | IQ motif containing F6 [Source:HGNC Symbol; |
| AC097636.2 | 3 | novel protein |
| AC097636.1 | 3 | IQ motif containing F4 (IQCF4) pseudogene |
| IQCF3 | 3 | IQ motif containing F3 [Source:HGNC Symbol; |
| IQCF2 | 3 | IQ motif containing F2 [Source:HGNC Symbol; |
| RN7SL504P | 3 | RNA, 7SL, cytoplasmic 504, pseudogene [Sour |
| IQCF5-AS1 | 3 | IQCF5 antisense RNA 1 [Source:HGNC Symbol |
| IQCF5 | 3 | IQ motif containing F5 [Source:HGNC Symbol; |
| IQCF1 | 3 | IQ motif containing F1 [Source:HGNC Symbol; |
| AC115284.4 | 3 | novel transcript |
| RRP9 | 3 | ribosomal RNA processing 9, U3 small nucleol |
| PARP3 | 3 | poly(ADP-ribose) polymerase family member |
| AC115284.2 | 3 | TEC |
| GPR62 | 3 | G protein-coupled receptor 62 [Source:HGNC |
| PCBP4 | 3 | poly(rC) binding protein 4 [Source:HGNC Symb |
| AC115284.1 | 3 | novel transcript |
| ABHD14B | 3 | abhydrolase domain containing 14B [Source:H |
| ABHD14A | 3 | abhydrolase domain containing 14A [Source:H |
| ABHD14A-ACY1 | 3 | ABHD14A-ACY1 readthrough [Source:HGNC Sy |
| ACY1 | 3 | aminoacylase 1 [Source:HGNC Symbol;Acc:HG |
| RPL29 | 3 | ribosomal protein L29 [Source:HGNC Symbol; |
| AC115284.3 | 3 | Novel trancript |
| DUSP7 | 3 | dual specificity phosphatase 7 [Source:HGNC S |
| POC1A | 3 | POC1 centriolar protein A [Source:HGNC Symb |
| AC097637.3 | 3 | novel transcript |
| ALDOAP1 | 3 | ALDOA pseudogene 1 [Source:HGNC Symbol;A |
| ALAS1 | 3 | 5'-aminolevulinate synthase 1 [Source:HGNC S |
| AC097637.2 | 3 | TEC |
| TLR9 | 3 | toll like receptor 9 [Source:HGNC Symbol;Acc: |
| AC097637.1 | 3 | novel twinfilin, actin-binding protein, homolo |
| TWF2 | 3 | twinfilin actin binding protein 2 [Source:HGNC |
| AC006252.1 | 3 | novel transcript, antisense to TWF2 |
| PPM1M | 3 | protein phosphatase, Mg2+/Mn2+ dependent |
| WDR82 | 3 | WD repeat domain 82 [Source:HGNC Symbol; |
| MIRLET7G | 3 | microRNA let-7g [Source:HGNC Symbol;Acc:H |
| GLYCTK | 3 | glycerate kinase [Source:HGNC Symbol;Acc:H |
| GLYCTK-AS1 | 3 | GLYCTK antisense RNA 1 [Source:HGNC Symbo |
| MIR135A1 | 3 | microRNA 135a-1 [Source:HGNC Symbol;Acc:H |
| DNAH1 | 3 | dynein axonemal heavy chain 1 [Source:HGNC |
| AC092045.1 | 3 | protein phosphatase 2, regulatory subunit B', |
| BAP1 | 3 | BRCA1 associated protein 1 [Source:HGNC Syr |
| PHF7 | 3 | PHD finger protein 7 [Source:HGNC Symbol;Ac |
| SEMA3G | 3 | semaphorin 3G [Source:HGNC Symbol;Acc:HG |
| TNNC1 | 3 | troponin C1, slow skeletal and cardiac type [So |

| | | |
|---|---|---|
| NISCH | 3 | nischarin [Source:HGNC Symbol;Acc:HGNC:18 |
| STAB1 | 3 | stabilin 1 [Source:HGNC Symbol;Acc:HGNC:18 |
| NT5DC2 | 3 | 5'-nucleotidase domain containing 2 [Source:I |
| SMIM4 | 3 | small integral membrane protein 4 [Source:H( |
| PBRM1 | 3 | polybromo 1 [Source:HGNC Symbol;Acc:HGN( |
| RNU6-856P | 3 | RNA, U6 small nuclear 856, pseudogene [Sour |
| RNU6ATAC16P | 3 | RNA, U6atac small nuclear 16, pseudogene [S( |
| GNL3 | 3 | G protein nucleolar 3 [Source:HGNC Symbol;A |
| AC104446.3 | 3 | NULL |
| AC104446.1 | 3 | NULL |
| SNORD19 | 3 | small nucleolar RNA, C/D box 19 [Source:HGN |
| SNORD19B | 3 | small nucleolar RNA, C/D box 19B [Source:HG |
| SNORD19C | 3 | small nucleolar RNA, C/D box 19C [Source:HG |
| SNORD69 | 3 | small nucleolar RNA, C/D box 69 [Source:HGN |
| GLT8D1 | 3 | glycosyltransferase 8 domain containing 1 [So |
| SPCS1 | 3 | signal peptidase complex subunit 1 [Source:H( |
| NEK4 | 3 | NIMA related kinase 4 [Source:HGNC Symbol;. |
| AC104446.2 | 3 | zinc finger protein 610 (ZNF610) pseudogene |
| AC006254.2 | 3 | Adenylate kinase isoenzyme 4, mitochondrial, |
| AC006254.3 | 3 | AKT interacting protein, pseudogene |
| ITIH1 | 3 | inter-alpha-trypsin inhibitor heavy chain 1 [So |
| ITIH3 | 3 | inter-alpha-trypsin inhibitor heavy chain 3 [So |
| ITIH4 | 3 | inter-alpha-trypsin inhibitor heavy chain 4 [So |
| AC006254.1 | 3 | novel MUSTN1-ITIH4 readthrough |
| ITIH4-AS1 | 3 | ITIH4 antisense RNA 1 [Source:HGNC Symbol;. |
| MUSTN1 | 3 | musculoskeletal, embryonic nuclear protein 1 |
| STIMATE-MUSTN1 | 3 | STIMATE-MUSTN1 readthrough [Source:HGN( |
| STIMATE | 3 | STIM activating enhancer [Source:HGNC Symb |
| MIR8064 | 3 | microRNA 8064 [Source:HGNC Symbol;Acc:H( |
| AC099667.1 | 3 | TEC |
| SFMBT1 | 3 | Scm like with four mbt domains 1 [Source:HGI |
| AC096887.1 | 3 | novel transcript |
| AC096887.2 | 3 | TEC |
| SERBP1P3 | 3 | SERPINE1 mRNA binding protein 1 pseudogen |
| RFT1 | 3 | RFT1 homolog [Source:HGNC Symbol;Acc:HGI |
| PRKCD | 3 | protein kinase C delta [Source:HGNC Symbol;/ |
| AC097015.1 | 3 | novel transcript |
| TKT | 3 | transketolase [Source:HGNC Symbol;Acc:HGN |
| DCP1A | 3 | decapping mRNA 1A [Source:HGNC Symbol;A( |
| Y_RNA | 3 | Y RNA [Source:RFAM;Acc:RF00019] |
| CACNA1D | 3 | calcium voltage-gated channel subunit alpha1 |
| SNORD38C | 3 | small nucleolar RNA, C/D box 38C [Source:HG |
| AC112218.1 | 3 | NULL |
| AC092035.1 | 3 | ribosomal protein S25 (RPS25) pseudogene |
| SNORD63 | 3 | Small nucleolar RNA SNORD63 [Source:RFAM; |
| AC012467.1 | 3 | novel transcript, antisense to CACNA1D |
| CHDH | 3 | choline dehydrogenase [Source:HGNC Symbol |
| IL17RB | 3 | interleukin 17 receptor B [Source:HGNC Symb |
| AC012467.2 | 3 | novel transcript, antisense to IL17RB |
| ACTR8 | 3 | actin related protein 8 [Source:HGNC Symbol; |

| | | |
|---|---|---|
| SELENOK | 3 | selenoprotein K [Source:HGNC Symbol;Acc:HG |
| AC115282.2 | 3 | novel transcript |
| CABYRP1 | 3 | calcium binding tyrosine phosphorylation regu |
| AC115282.1 | 3 | novel transcript, antisense to CACNA2D3 |
| CACNA2D3 | 3 | calcium voltage-gated channel auxiliary subun |
| RPS15P5 | 3 | ribosomal protein S15 pseudogene 5 [Source: |
| ESRG | 3 | embryonic stem cell related [Source:HGNC Sy |
| CACNA2D3-AS1 | 3 | CACNA2D3 antisense RNA 1 [Source:HGNC Syı |
| LRTM1 | 3 | leucine rich repeats and transmembrane dom |
| AC092059.1 | 3 | novel transcript |
| LINC02017 | 3 | long intergenic non-protein coding RNA 2017 |
| LINC02030 | 3 | long intergenic non-protein coding RNA 2030 |
| AC121764.2 | 3 | novel transcript |
| AC121764.3 | 3 | novel transcript |
| WNT5A | 3 | Wnt family member 5A [Source:HGNC Symbo |
| WNT5A-AS1 | 3 | WNT5A antisense RNA 1 [Source:HGNC Symb |
| AC121764.1 | 3 | novel transcript |
| ERC2 | 3 | ELKS/RAB6-interacting/CAST family member 2 |
| AC025572.1 | 3 | novel transcript |
| ERC2-IT1 | 3 | ERC2 intronic transcript 1 [Source:HGNC Symb |
| MIR3938 | 3 | microRNA 3938 [Source:HGNC Symbol;Acc:HG |
| RN7SKP45 | 3 | RN7SK pseudogene 45 [Source:HGNC Symbol; |
| RNA5SP133 | 3 | RNA, 5S ribosomal pseudogene 133 [Source:H |
| CCDC66 | 3 | coiled-coil domain containing 66 [Source:HGN |
| TASOR | 3 | transcription activation suppressor [Source:HG |
| ARHGEF3 | 3 | Rho guanine nucleotide exchange factor 3 [So |
| ARHGEF3-AS1 | 3 | ARHGEF3 antisense RNA 1 [Source:HGNC Sym |
| SPATA12 | 3 | spermatogenesis associated 12 [Source:HGNC |
| AC097358.2 | 3 | novel transcript, antisense to ARHGEF3 |
| IL17RD | 3 | interleukin 17 receptor D [Source:HGNC Symb |
| AC097358.1 | 3 | ribosomal protein L19 (RPL19) pseudogene |
| HESX1 | 3 | HESX homeobox 1 [Source:HGNC Symbol;Acc: |
| APPL1 | 3 | adaptor protein, phosphotyrosine interacting |
| ASB14 | 3 | ankyrin repeat and SOCS box containing 14 [S |
| AC093928.1 | 3 | novel transcript, antisense to DNAH12 |
| DNAH12 | 3 | dynein axonemal heavy chain 12 [Source:HGN |
| Y_RNA | 3 | Y RNA [Source:RFAM;Acc:RF00019] |
| RNU6-1181P | 3 | RNA, U6 small nuclear 1181, pseudogene [Sou |
| RNF7P1 | 3 | ring finger protein 7 pseudogene 1 [Source:HG |
| AC092418.1 | 3 | ribosomal protein S8 (RPS8) pseudogene |
| RNU6-108P | 3 | RNA, U6 small nuclear 108, pseudogene [Sour |
| RNU6-483P | 3 | RNA, U6 small nuclear 483, pseudogene [Sour |
| AC092418.2 | 3 | novel transcript |
| PDE12 | 3 | phosphodiesterase 12 [Source:HGNC Symbol;A |
| ARF4 | 3 | ADP ribosylation factor 4 [Source:HGNC Symb |
| ARF4-AS1 | 3 | ARF4 antisense RNA 1 [Source:HGNC Symbol;A |
| RNU6ATAC26P | 3 | RNA, U6atac small nuclear 26, pseudogene [S |
| DENND6A | 3 | DENN domain containing 6A [Source:HGNC Sy |
| DENND6A-AS1 | 3 | DENND6A antisense RNA 1 [Source:HGNC Sym |
| DENND6A-DT | 3 | DENND6A divergent transcript [Source:HGNC |

| | | |
|---|---|---|
| SLMAP | 3 | sarcolemma associated protein [Source:HGNC |
| PDHA1P1 | 3 | pyruvate dehydrogenase alpha 1 pseudogene |
| PPIAP16 | 3 | peptidylprolyl isomerase A pseudogene 16 [Sc |
| FLNB | 3 | filamin B [Source:HGNC Symbol;Acc:HGNC:37. |
| FLNB-AS1 | 3 | FLNB antisense RNA 1 [Source:HGNC Symbol;/ |
| AC137936.1 | 3 | novel transcript |
| AC137936.2 | 3 | novel pseudogene |
| DNASE1L3 | 3 | deoxyribonuclease 1 like 3 [Source:HGNC Sym |
| ABHD6 | 3 | abhydrolase domain containing 6 [Source:HGI |
| RPP14 | 3 | ribonuclease P/MRP subunit p14 [Source:HGN |
| HTD2 | 3 | hydroxyacyl-thioester dehydratase type 2 [Sou |
| AC098479.1 | 3 | novel transcript |
| PXK | 3 | PX domain containing serine/threonine kinase |
| AC135507.2 | 3 | WD repeat domain 53 (WDR53) pseudogene |
| PDHB | 3 | pyruvate dehydrogenase E1 beta subunit [Sou |
| AC135507.1 | 3 | novel transcript, antisense to PDHB |
| AC116036.2 | 3 | novel transcript |
| KCTD6 | 3 | potassium channel tetramerization domain cc |
| ACOX2 | 3 | acyl-CoA oxidase 2 [Source:HGNC Symbol;Acc |
| AC116036.1 | 3 | ribosomal protein L27 (RPL27) pseudogene |
| FAM107A | 3 | family with sequence similarity 107 member / |
| AC119424.1 | 3 | novel transcript |
| FAM3D-AS1 | 3 | FAM3D antisense RNA 1 [Source:HGNC Symbo |
| FAM3D | 3 | family with sequence similarity 3 member D [! |
| C3orf67 | 3 | chromosome 3 open reading frame 67 [Sourc |
| C3orf67-AS1 | 3 | C3orf67 antisense RNA 1 [Source:HGNC Symb |
| AC126121.3 | 3 | novel transcript |
| AC138057.1 | 3 | novel transcript |
| AC126121.2 | 3 | novel transcript |
| AC126121.1 | 3 | small nuclear ribonucleoprotein polypeptide E |
| Metazoa_SRP | 3 | Metazoan signal recognition particle RNA [Sou |
| FHIT | 3 | fragile histidine triad diadenosine triphosphat |
| AC093418.1 | 3 | novel transcript, antisense to FHIT |
| AC104164.2 | 3 | mitochondrial transcription termination facto |
| MIR548BB | 3 | microRNA 548bb [Source:HGNC Symbol;Acc:H |
| PPIAP70 | 3 | peptidylprolyl isomerase A pseudogene 70 [Sc |
| AC104164.1 | 3 | transmembrane emp24 domain trafficking prc |
| PPIAP71 | 3 | peptidylprolyl isomerase A pseudogene 71 [Sc |
| U3 | 3 | Small nucleolar RNA U3 [Source:RFAM;Acc:RF |
| PTPRG | 3 | protein tyrosine phosphatase receptor type G |
| Y_RNA | 3 | Y RNA [Source:RFAM;Acc:RF00019] |
| RPL10AP6 | 3 | ribosomal protein L10a pseudogene 6 [Source |
| RNU2-10P | 3 | RNA, U2 small nuclear 10, pseudogene [Sourc |
| AC104849.1 | 3 | inhibitor of DNA binding 2, dominant negative |
| PTPRG-AS1 | 3 | PTPRG antisense RNA 1 [Source:HGNC Symbo |
| C3orf14 | 3 | chromosome 3 open reading frame 14 [Sourc |
| FEZF2 | 3 | FEZ family zinc finger 2 [Source:HGNC Symbol |
| CADPS | 3 | calcium dependent secretion activator [Sourc |
| AC012519.1 | 3 | ribosomal protein L14 (RPL14) pseudogene |
| RN7SL863P | 3 | RNA, 7SL, cytoplasmic 863, pseudogene [Sour |

| | | |
|---|---|---|
| RNU6-139P | 3 | RNA, U6 small nuclear 139, pseudogene [Sour |
| AC096915.1 | 3 | ribosomal protein S10 (RPS10) pseudogene |
| LINC00698 | 3 | long intergenic non-protein coding RNA 698 [S |
| KRT8P35 | 3 | keratin 8 pseudogene 35 [Source:HGNC Symb |
| UBL5P3 | 3 | ubiquitin like 5 pseudogene 3 [Source:HGNC S |
| SYNPR | 3 | synaptoporin [Source:HGNC Symbol;Acc:HGN |
| SYNPR-AS1 | 3 | SYNPR antisense RNA 1 [Source:HGNC Symbol |
| RNA5SP134 | 3 | RNA, 5S ribosomal pseudogene 134 [Source:H |
| SNTN | 3 | sentan, cilia apical structure protein [Source:H |
| AC104162.1 | 3 | novel cadherin domain containing pseudogen |
| C3orf49 | 3 | chromosome 3 open reading frame 49 [Source |
| THOC7 | 3 | THO complex 7 [Source:HGNC Symbol;Acc:HG |
| THOC7-AS1 | 3 | THOC7 antisense RNA 1 [Source:HGNC Symbo |
| AC104162.2 | 3 | novel transcript |
| ATXN7 | 3 | ataxin 7 [Source:HGNC Symbol;Acc:HGNC:105 |
| ATXN7 | 3 | ataxin 7 [Source:HGNC Symbol;Acc:HGNC:105 |
| SCAANT1 | 3 | SCA7/ATXN7 antisense RNA 1 [Source:HGNC S |
| PSMD6-AS2 | 3 | PSMD6 antisense RNA 2 [Source:HGNC Symbo |
| AC012557.1 | 3 | novel transcript |
| PSMD6 | 3 | proteasome 26S subunit, non-ATPase 6 [Sourc |
| PSMD6-AS1 | 3 | PSMD6 antisense RNA 1 [Source:HGNC Symbo |
| AC012557.2 | 3 | novel transcript, antisense to PSMD6 |
| AC092040.1 | 3 | novel transcript |
| U3 | 3 | Small nucleolar RNA U3 [Source:RFAM;Acc:RF |
| LINC00994 | 3 | long intergenic non-protein coding RNA 994 [S |
| PRICKLE2 | 3 | prickle planar cell polarity protein 2 [Source:H |
| PRICKLE2-AS1 | 3 | PRICKLE2 antisense RNA 1 [Source:HGNC Sym |
| PRICKLE2-AS2 | 3 | PRICKLE2 antisense RNA 2 [Source:HGNC Sym |
| PRDX3P4 | 3 | peroxiredoxin 3 pseudogene 4 [Source:HGNC |
| PRICKLE2-AS3 | 3 | PRICKLE2 antisense RNA 3 [Source:HGNC Sym |
| PRICKLE2-DT | 3 | PRICKLE2 divergent transcript [Source:HGNC S |
| RNU6-739P | 3 | RNA, U6 small nuclear 739, pseudogene [Sour |
| ADAMTS9 | 3 | ADAM metallopeptidase with thrombospondi |
| ADAMTS9-AS1 | 3 | ADAMTS9 antisense RNA 1 [Source:HGNC Sym |
| ADAMTS9-AS2 | 3 | ADAMTS9 antisense RNA 2 [Source:HGNC Sym |
| LINC02040 | 3 | long intergenic non-protein coding RNA 2040 |
| MAGI1 | 3 | membrane associated guanylate kinase, WW |
| AC121493.1 | 3 | novel transcript, intronic to MAGI1 |
| ILF2P1 | 3 | interleukin enhancer binding factor 2 pseudog |
| RPL17P17 | 3 | ribosomal protein L17 pseudogene 17 [Source |
| MAGI1-IT1 | 3 | MAGI1 intronic transcript 1 [Source:HGNC Syr |
| MAGI1-AS1 | 3 | MAGI1 antisense RNA 1 [Source:HGNC Symbo |
| Y_RNA | 3 | Y RNA [Source:RFAM;Acc:RF00019] |
| AC106827.1 | 3 | novel pseudogene |
| SLC25A26 | 3 | solute carrier family 25 member 26 [Source:H |
| RNU6-787P | 3 | RNA, U6 small nuclear 787, pseudogene [Sour |
| RN7SL482P | 3 | RNA, 7SL, cytoplasmic 482, pseudogene [Sour |
| LRIG1 | 3 | leucine rich repeats and immunoglobulin like |
| AC104440.1 | 3 | transcription elongation factor A (SII)-like 8, p |
| RPL21P41 | 3 | ribosomal protein L21 pseudogene 41 [Source |

| | | |
|---|---|---|
| AC098969.2 | 3 | novel transcript |
| AC098969.1 | 3 | Novel transcript |
| AC020655.1 | 3 | novel transcript |
| KBTBD8 | 3 | kelch repeat and BTB domain containing 8 [So |
| MIR4272 | 3 | microRNA 4272 [Source:HGNC Symbol;Acc:HG |
| AC114401.1 | 3 | novel transcript |
| SUCLG2 | 3 | succinate-CoA ligase GDP-forming beta subun |
| NDUFB4P1 | 3 | NADH:ubiquinone oxidoreductase subunit B4 |
| SUCLG2-AS1 | 3 | SUCLG2 antisense RNA 1 (head to head) [Sour |
| TAFA1 | 3 | TAFA chemokine like family member 1 [Sourc |
| COPS8P2 | 3 | COP9 signalosome subunit 8 pseudogene 2 [S |
| AC107626.1 | 3 | pseudogene similar to part of ligand depende |
| AC104167.1 | 3 | novel transcript, sense intronic to FAM19A1 |
| AC096922.1 | 3 | ribosomal protein, large, P0 (RPLP0) pseudog |
| PSMC1P1 | 3 | proteasome 26S subunit, ATPase 1 pseudoger |
| TAFA4 | 3 | TAFA chemokine like family member 4 [Sourc |
| RNA5SP135 | 3 | RNA, 5S ribosomal pseudogene 135 [Source:H |
| EOGT | 3 | EGF domain specific O-linked N-acetylglucosa |
| AC109587.1 | 3 | novel transcript |
| TMF1 | 3 | TATA element modulatory factor 1 [Source:H |
| MIR3136 | 3 | microRNA 3136 [Source:HGNC Symbol;Acc:HG |
| UBA3 | 3 | ubiquitin like modifier activating enzyme 3 [S |
| ARL6IP5 | 3 | ADP ribosylation factor like GTPase 6 interacti |
| LMOD3 | 3 | leiomodin 3 [Source:HGNC Symbol;Acc:HGNC: |
| FRMD4B | 3 | FERM domain containing 4B [Source:HGNC Sy |
| RBM43P1 | 3 | RNA binding motif protein 43 pseudogene 1 [S |
| AC099328.1 | 3 | H1 histone family, member O, oocyte-specific |
| AC099328.2 | 3 | novel transcript |
| AC104445.1 | 3 | novel transcript |
| MITF | 3 | melanocyte inducing transcription factor [Sou |
| RN7SL418P | 3 | RNA, 7SL, cytoplasmic 418, pseudogene [Sour |
| SAMMSON | 3 | survival associated mitochondrial melanoma s |
| AC139700.1 | 3 | RNA-binding region (RNP1, RRM) containing 3 |
| UQCRHP4 | 3 | ubiquinol-cytochrome c reductase hinge prote |
| MDFIC2 | 3 | MyoD family inhibitor domain containing 2 [S |
| AC104633.1 | 3 | novel transcript |
| AC104633.2 | 3 | novel transcript |
| COX6CP6 | 3 | cytochrome c oxidase subunit 6C pseudogene |
| HMGB1P36 | 3 | high mobility group box 1 pseudogene 36 [Sou |
| RNU6-281P | 3 | RNA, U6 small nuclear 281, pseudogene [Sour |
| FOXP1 | 3 | forkhead box P1 [Source:HGNC Symbol;Acc:H |
| AC097634.4 | 3 | novel protein |
| FOXP1-AS1 | 3 | FOXP1 antisense RNA 1 [Source:HGNC Symbo |
| MIR1284 | 3 | microRNA 1284 [Source:HGNC Symbol;Acc:HG |
| AC097634.2 | 3 | novel transcript, antisense to FOXP1 |
| FOXP1-IT1 | 3 | FOXP1 intronic transcript 1 [Source:HGNC Sym |
| AC097634.3 | 3 | novel transcript, antisense to FOXP1 |
| AC097634.1 | 3 | novel transcript |
| EIF4E3 | 3 | eukaryotic translation initiation factor 4E fami |
| GPR27 | 3 | G protein-coupled receptor 27 [Source:HGNC |

| | | |
|---|---|---|
| PROK2 | 3 | prokineticin 2 [Source:HGNC Symbol;Acc:HGN |
| AC096970.1 | 3 | novel transcript |
| RN7SL271P | 3 | RNA, 7SL, cytoplasmic 271, pseudogene [Sour |
| LINC00877 | 3 | long intergenic non-protein coding RNA 877 [S |
| UBE2Q2P9 | 3 | ubiquitin conjugating enzyme E2 Q2 pseudoge |
| AC105265.3 | 3 | novel transcript |
| AC105265.4 | 3 | novel transcript |
| AC105265.1 | 3 | pseudogene similar to part of coiled-coil doma |
| AC105265.2 | 3 | coiled-coil domain containing 137 (CCDC137) |
| LINC00870 | 3 | long intergenic non-protein coding RNA 870 [S |
| AC112219.2 | 3 | novel transcript |
| AC112219.1 | 3 | novel transcript |
| AC134508.2 | 3 | novel transcript |
| AC134508.1 | 3 | novel transcript |
| RYBP | 3 | RING1 and YY1 binding protein [Source:HGNC |
| AC104435.2 | 3 | novel transcript |
| RNU1-62P | 3 | RNA, U1 small nuclear 62, pseudogene [Sourc |
| AC104435.1 | 3 | pseudogene similar to part of mitochondrially |
| RNA5SP136 | 3 | RNA, 5S ribosomal pseudogene 136 [Source:H |
| AC097369.2 | 3 | novel transcript |
| SHQ1 | 3 | SHQ1, H/ACA ribonucleoprotein assembly fact |
| AC134050.1 | 3 | U2-associated SR140 protein (SR140) pseudog |
| PSMD12P | 3 | proteasome 26S subunit, non-ATPase, 12 pseu |
| LAPTM4BP2 | 3 | lysosomal protein transmembrane 4 beta pse |
| GXYLT2 | 3 | glucoside xylosyltransferase 2 [Source:HGNC S |
| Metazoa_SRP | 3 | Metazoan signal recognition particle RNA [Sou |
| FTH1P23 | 3 | ferritin heavy chain 1 pseudogene 23 [Source: |
| PPP4R2 | 3 | protein phosphatase 4 regulatory subunit 2 [S |
| RNU7-19P | 3 | RNA, U7 small nuclear 19 pseudogene [Source |
| EBLN2 | 3 | endogenous Bornavirus like nucleoprotein 2 [S |
| RNU6-557P | 3 | RNA, U6 small nuclear 557, pseudogene [Sour |
| RNU2-64P | 3 | RNA, U2 small nuclear 64, pseudogene [Sourc |
| CCDC75P1 | 3 | coiled-coil domain containing 75 pseudogene |
| RNU6-1270P | 3 | RNA, U6 small nuclear 1270, pseudogene [Sou |
| PDZRN3 | 3 | PDZ domain containing ring finger 3 [Source:H |
| RNU7-119P | 3 | RNA, U7 small nuclear 119 pseudogene [Sourc |
| PDZRN3-AS1 | 3 | PDZRN3 antisense RNA 1 [Source:HGNC Symb |
| AC117489.1 | 3 | TEC |
| AC108725.1 | 3 | peroxiredoxin 5 (PRDX5) pseudogene |
| LINC02005 | 3 | long intergenic non-protein coding RNA 2005 |
| Metazoa_SRP | 3 | Metazoan signal recognition particle RNA [Sou |
| LINC02047 | 3 | long intergenic non-protein coding RNA 2047 |
| AC016930.1 | 3 | heat shock protein 90kDa alpha (cytosolic), cla |
| AKR1B1P2 | 3 | aldo-keto reductase family 1 member B1 pseu |
| CNTN3 | 3 | contactin 3 [Source:HGNC Symbol;Acc:HGNC:1 |
| NIPA2P2 | 3 | NIPA magnesium transporter 2 pseudogene 2 |
| RN7SL294P | 3 | RNA, 7SL, cytoplasmic 294, pseudogene [Sour |
| MIR4444-2 | 3 | microRNA 4444-2 [Source:HGNC Symbol;Acc:I |
| AC117481.1 | 3 | novel transcript |
| HNRNPA3P6 | 3 | heterogeneous nuclear ribonucleoprotein A3 |

| | | |
|---|---|---|
| MYLKP1 | 3 | myosin light chain kinase pseudogene 1 [Sour |
| OR7E66P | 3 | olfactory receptor family 7 subfamily E memb |
| OR7E22P | 3 | olfactory receptor family 7 subfamily E memb |
| OR7E55P | 3 | olfactory receptor family 7 subfamily E memb |
| AC139453.2 | 3 | chromosome 21 open reading frame 63 (C21c |
| LSP1P2 | 3 | LSP1 pseudogene 2 [Source:HGNC Symbol;Acc |
| AC139453.1 | 3 | pseudogene similar to part of ankyrin repeat |
| ALG1L6P | 3 | ALG1 like 6, pseudogene [Source:HGNC Symb |
| FAM86DP | 3 | family with sequence similarity 86 member D, |
| LINC02018 | 3 | long intergenic non-protein coding RNA 2018 |
| ENPP7P2 | 3 | ectonucleotide pyrophosphatase/phosphodie |
| SNRPCP10 | 3 | small nuclear ribonucleoprotein polypeptide |
| AC133041.1 | 3 | RNA binding protein, fox-1 homolog (C. elegar |
| RPS3AP15 | 3 | ribosomal protein S3a pseudogene 15 [Source |
| AC108724.1 | 3 | pseudogene similar to part of chromosome 1 |
| OR7E121P | 3 | olfactory receptor family 7 subfamily E memb |
| UNC93B3 | 3 | unc-93 homolog B3, pseudogene [Source:HGN |
| RPL23AP49 | 3 | ribosomal protein L23a pseudogene 49 [Sourc |
| AC108724.2 | 3 | major histocompatibility complex, class I, F (H |
| CLUHP10 | 3 | clustered mitochondria homolog pseudogene |
| MIR1324 | 3 | microRNA 1324 [Source:HGNC Symbol;Acc:HG |
| AGGF1P3 | 3 | angiogenic factor with G-patch and FHA doma |
| RARRES2P1 | 3 | retinoic acid receptor responder 2 pseudogen |
| FRG2C | 3 | FSHD region gene 2 family member C [Source: |
| DUX4L26 | 3 | double homeobox 4 like 26 (pseudogene) [Sou |
| LINC00960 | 3 | long intergenic non-protein coding RNA 960 [ |
| ZNF717 | 3 | zinc finger protein 717 [Source:HGNC Symbol; |
| MIR4273 | 3 | microRNA 4273 [Source:HGNC Symbol;Acc:HG |
| ROBO2 | 3 | roundabout guidance receptor 2 [Source:HGN |
| Y_RNA | 3 | Y RNA [Source:RFAM;Acc:RF00019] |
| AC026877.1 | 3 | CAP, adenylate cyclase-associated protein 1 ( |
| RNU6-386P | 3 | RNA, U6 small nuclear 386, pseudogene [Sour |
| VDAC1P7 | 3 | voltage dependent anion channel 1 pseudoge |
| AC133040.1 | 3 | novel transcript, antisense to ROBO2 |
| RNU6-217P | 3 | RNA, U6 small nuclear 217, pseudogene [Sour |
| LINC02077 | 3 | long intergenic non-protein coding RNA 2077 |
| AC117462.1 | 3 | novel transcript |
| RN7SL647P | 3 | RNA, 7SL, cytoplasmic 647, pseudogene [Sour |
| RN7SKP61 | 3 | RN7SK pseudogene 61 [Source:HGNC Symbol; |
| AC108752.1 | 3 | novel transcript |
| MRPS17P3 | 3 | mitochondrial ribosomal protein S17 pseudog |
| ROBO1 | 3 | roundabout guidance receptor 1 [Source:HGN |
| AC055731.1 | 3 | ribosomal protein S12 (RPS12) pseudogene |
| RN7SL751P | 3 | RNA, 7SL, cytoplasmic 751, pseudogene [Sour |
| AC117461.1 | 3 | novel transcript, antisense to ROBO1 |
| MIR3923 | 3 | microRNA 3923 [Source:HGNC Symbol;Acc:HG |
| HMGB1P38 | 3 | high mobility group box 1 pseudogene 38 [Sou |
| OSBPL9P1 | 3 | oxysterol binding protein like 9 pseudogene 1 |
| HNRNPA3P8 | 3 | heterogeneous nuclear ribonucleoprotein A3 |
| AC108740.1 | 3 | hydrocephalus inducing homolog (mouse) (HY |

| | | |
|---|---|---|
| AC108690.1 | 3 | novel transcript |
| AC068756.1 | 3 | novel transcript |
| LINC02050 | 3 | long intergenic non-protein coding RNA 2050 |
| LINC02027 | 3 | long intergenic non-protein coding RNA 2027 |
| AC099542.2 | 3 | novel transcript |
| AC099542.1 | 3 | novel transcript |
| AC107302.2 | 3 | novel transcript |
| AC107302.1 | 3 | ret finger protein-like pseudogene |
| GBE1 | 3 | 1,4-alpha-glucan branching enzyme 1 [Source: |
| RNU2-28P | 3 | RNA, U2 small nuclear 28, pseudogene [Sourc |
| SETP6 | 3 | SET pseudogene 6 [Source:HGNC Symbol;Acc: |
| AC017015.1 | 3 | SET nuclear oncogene (SET) pseudogene |
| AC017015.2 | 3 | novel transcript |
| AC129807.1 | 3 | novel transcript |
| LINC02008 | 3 | long intergenic non-protein coding RNA 2008 |
| RPL7AP23 | 3 | ribosomal protein L7a pseudogene 23 [Source |
| CYP51A1P1 | 3 | cytochrome P450 family 51 subfamily A meml |
| AC023503.1 | 3 | novel transcript |
| AC092825.1 | 3 | novel transcript |
| SRRM1P2 | 3 | serine/arginine repetitive matrix 1 pseudogen |
| AC117464.1 | 3 | novel transcript |
| LINC00971 | 3 | long intergenic non-protein coding RNA 971 [S |
| AC117482.1 | 3 | transmembrane protein 183B (TMEM183B) ps |
| LINC02025 | 3 | long intergenic non-protein coding RNA 2025 |
| AC132660.2 | 3 | novel transcript |
| CADM2 | 3 | cell adhesion molecule 2 [Source:HGNC Symb |
| MIR5688 | 3 | microRNA 5688 [Source:HGNC Symbol;Acc:HG |
| CADM2-AS2 | 3 | CADM2 antisense RNA 2 [Source:HGNC Symb |
| CADM2-AS1 | 3 | CADM2 antisense RNA 1 [Source:HGNC Symb |
| THAP12P2 | 3 | THAP domain containing 12 pseudogene 2 [Sc |
| AC107024.1 | 3 | NULL |
| RNU6-1129P | 3 | RNA, U6 small nuclear 1129, pseudogene [Sou |
| AC108733.1 | 3 | novel transcript |
| RN7SKP284 | 3 | RN7SK pseudogene 284 [Source:HGNC Symbo |
| LINC02070 | 3 | long intergenic non-protein coding RNA 2070 |
| AC108706.1 | 3 | zinc finger protein 418 (ZNF418) pseudogene |
| VGLL3 | 3 | vestigial like family member 3 [Source:HGNC S |
| PPATP1 | 3 | phosphoribosyl pyrophosphate amidotransfer |
| AC107204.1 | 3 | long intergenic non-protein coding RNA 506 |
| MIR4795 | 3 | microRNA 4795 [Source:HGNC Symbol;Acc:HG |
| CHMP2B | 3 | charged multivesicular body protein 2B [Sourc |
| POU1F1 | 3 | POU class 1 homeobox 1 [Source:HGNC Symb |
| KRT8P25 | 3 | keratin 8 pseudogene 25 [Source:HGNC Symb |
| APOOP2 | 3 | apolipoprotein O pseudogene 2 [Source:HGN( |
| PSMC1P6 | 3 | proteasome 26S subunit, ATPase 1 pseudoger |
| AC108749.1 | 3 | novel transcript |
| RNU6-873P | 3 | RNA, U6 small nuclear 873, pseudogene [Sour |
| RNU6ATAC6P | 3 | RNA, U6atac small nuclear 6, pseudogene [Sou |
| HTR1F | 3 | 5-hydroxytryptamine receptor 1F [Source:HGI |
| CGGBP1 | 3 | CGG triplet repeat binding protein 1 [Source:H |

| | | |
|---|---|---|
| ZNF654 | 3 | zinc finger protein 654 [Source:HGNC Symbol; |
| CBX5P1 | 3 | chromobox 5 pseudogene 1 [Source:HGNC Sy |
| C3orf38 | 3 | chromosome 3 open reading frame 38 [Sourc |
| ABCF2P1 | 3 | ATP binding cassette subfamily F member 2 p |
| CSNKA2IP | 3 | casein kinase 2 subunit alpha' interacting prot |
| Y_RNA | 3 | Y RNA [Source:RFAM;Acc:RF00019] |
| NDUFA5P5 | 3 | NADH:ubiquinone oxidoreductase subunit A5 |
| ICE2P2 | 3 | interactor of little elongation complex ELL sub |
| GAPDHP50 | 3 | glyceraldehyde 3 phosphate dehydrogenase p |
| EPHA3 | 3 | EPH receptor A3 [Source:HGNC Symbol;Acc:H |
| AC109129.1 | 3 | Novel transcript, antisense to EPHA3 |
| MTCO2P6 | 3 | MT-CO2 pseudogene 6 [Source:HGNC Symbol |
| MTCO1P6 | 3 | MT-CO1 pseudogene 6 [Source:HGNC Symbol |
| U3 | 3 | Small nucleolar RNA U3 [Source:RFAM;Acc:RF |
| RNU6-712P | 3 | RNA, U6 small nuclear 712, pseudogene [Sour |
| PROS2P | 3 | protein S (beta) pseudogene [Source:HGNC Sy |
| HSPE1P19 | 3 | heat shock protein family E (Hsp10) member : |
| ABBA01000933.1 | 3 | NULL |
| ABBA01000935.2 | 3 | primase, DNA, polypeptide 2 (58kDa) (PRIM2) |
| RNU6-488P | 3 | RNA, U6 small nuclear 488, pseudogene [Sour |
| PROS1 | 3 | protein S [Source:HGNC Symbol;Acc:HGNC:94 |
| Y_RNA | 3 | Y RNA [Source:RFAM;Acc:RF00019] |
| RNU6-511P | 3 | RNA, U6 small nuclear 511, pseudogene [Sour |
| ARL13B | 3 | ADP ribosylation factor like GTPase 13B [Sour |
| HMGN1P7 | 3 | high mobility group nucleosome binding dom |
| STX19 | 3 | syntaxin 19 [Source:HGNC Symbol;Acc:HGNC: |
| DHFR2 | 3 | dihydrofolate reductase 2 [Source:HGNC Sym |
| NSUN3 | 3 | NOP2/Sun RNA methyltransferase 3 [Source:H |
| RBBP4P2 | 3 | RBBP4 pseudogene 2 [Source:HGNC Symbol;A |
| ARMC10P1 | 3 | armadillo repeat containing 10 pseudogene 1 |
| AC140059.1 | 3 | WD repeat domain 82 (WDR82) pseudogene |
| WDR82P1 | 3 | WD repeat domain 82 pseudogene 1 [Source: |
| LINC00879 | 3 | long intergenic non-protein coding RNA 879 [S |
| RPS18P6 | 3 | ribosomal protein S18 pseudogene 6 [Source: |
| MTHFD2P1 | 3 | methylenetetrahydrofolate dehydrogenase (N |
| AC106719.1 | 3 | novel transcript |
| AC107304.1 | 3 | SYF2 homolog, RNA splicing factor (S. cerevisi |
| HNRNPKP4 | 3 | heterogeneous nuclear ribonucleoprotein K p |
| MIR8060 | 3 | microRNA 8060 [Source:HGNC Symbol;Acc:HG |
| AC107015.1 | 3 | retinoblastoma binding protein 4 (RBBP4) pse |
| RNU6-1094P | 3 | RNA, U6 small nuclear 1094, pseudogene [Sou |
| MTRNR2L12 | 3 | MT-RNR2 like 12 [Source:HGNC Symbol;Acc:H |
| RPL18AP8 | 3 | ribosomal protein L18a pseudogene 8 [Source |
| RCC2P5 | 3 | regulator of chromosome condensation 2 pse |
| CDV3P1 | 3 | CDV3 pseudogene 1 [Source:HGNC Symbol;Ac |
| AC109782.1 | 3 | novel transcript |
| EPHA6 | 3 | EPH receptor A6 [Source:HGNC Symbol;Acc:H |
| AC117470.1 | 3 | pseudogene similar to part of transmembrane |
| AC110491.1 | 3 | novel transcript |
| ARL6 | 3 | ADP ribosylation factor like GTPase 6 [Source: |

| | | |
|---|---|---|
| AC110491.3 | 3 | novel transcript, antisense to ARL6 |
| CRYBG3 | 3 | crystallin beta-gamma domain containing 3 [S |
| AC110491.2 | 3 | novel transcript, antisense to CRYBG3 |
| RIOX2 | 3 | ribosomal oxygenase 2 [Source:HGNC Symbol |
| AC026100.1 | 3 | Novel transcript |
| GABRR3 | 3 | gamma-aminobutyric acid type A receptor rho |
| OR5BM1P | 3 | olfactory receptor family 5 subfamily BM mem |
| OR5AC1 | 3 | olfactory receptor family 5 subfamily AC mem |
| OR5AC2 | 3 | olfactory receptor family 5 subfamily AC mem |
| AC117460.1 | 3 | novel transcript, antisense to OR5H1, OR5H14 |
| OR5AC4P | 3 | olfactory receptor family 5 subfamily AC mem |
| POU5F1P7 | 3 | POU class 5 homeobox 1 pseudogene 7 [Sour |
| OR5H1 | 3 | olfactory receptor family 5 subfamily H memb |
| OR5H14 | 3 | olfactory receptor family 5 subfamily H memb |
| OR5H15 | 3 | olfactory receptor family 5 subfamily H memb |
| OR5H5P | 3 | olfactory receptor family 5 subfamily H memb |
| OR5H3P | 3 | olfactory receptor family 5 subfamily H memb |
| OR5H4P | 3 | olfactory receptor family 5 subfamily H memb |
| AC117473.1 | 3 | novel transcript |
| OR5H7P | 3 | olfactory receptor family 5 subfamily H memb |
| OR5H6 | 3 | olfactory receptor family 5 subfamily H memb |
| OR5H2 | 3 | olfactory receptor family 5 subfamily H memb |
| OR5H8 | 3 | olfactory receptor family 5 subfamily H memb |
| OR5K4 | 3 | olfactory receptor family 5 subfamily K memb |
| OR5K3 | 3 | olfactory receptor family 5 subfamily K memb |
| UBFD1P1 | 3 | ubiquitin family domain containing 1 pseudog |
| OR5K1 | 3 | olfactory receptor family 5 subfamily K memb |
| OR5K2 | 3 | olfactory receptor family 5 subfamily K memb |
| CLDND1 | 3 | claudin domain containing 1 [Source:HGNC Sy |
| AC021660.3 | 3 | novel transcript |
| AC021660.2 | 3 | novel transcript |
| RPL38P4 | 3 | ribosomal protein L38 pseudogene 4 [Source: |
| GPR15 | 3 | G protein-coupled receptor 15 [Source:HGNC |
| AC021660.4 | 3 | novel transcript |
| CPOX | 3 | coproporphyrinogen oxidase [Source:HGNC Sy |
| AC021660.1 | 3 | ribosomal protein L19 (RPL19) pseudogene |
| WWP1P1 | 3 | WW domain containing E3 ubiquitin protein li |
| ST3GAL6-AS1 | 3 | ST3GAL6 antisense RNA 1 [Source:HGNC Symb |
| ST3GAL6 | 3 | ST3 beta-galactoside alpha-2,3-sialyltransfera |
| PDLIM1P4 | 3 | PDZ and LIM domain 1 pseudogene 4 [Source: |
| DCBLD2 | 3 | discoidin, CUB and LCCL domain containing 2 [ |
| RNU6-26P | 3 | RNA, U6 small nuclear 26, pseudogene [Sourc |
| AC091212.1 | 3 | novel transcript |
| RNU6-1263P | 3 | RNA, U6 small nuclear 1263, pseudogene [Sou |
| LINC00973 | 3 | long intergenic non-protein coding RNA 973 [S |
| U7 | 3 | U7 small nuclear RNA [Source:RFAM;Acc:RF00 |
| ACTG1P13 | 3 | actin gamma 1 pseudogene 13 [Source:HGNC |
| AC107297.1 | 3 | pseudogene similar to Rho-guanine nucleotide |
| AC078828.1 | 3 | novel transcript |
| AC107029.1 | 3 | novel transcript |

| | | |
|---|---|---|
| AC107029.2 | 3 | novel transcript |
| COL8A1 | 3 | collagen type VIII alpha 1 chain [Source:HGNC |
| AC069222.1 | 3 | novel transcript |
| CMSS1 | 3 | cms1 ribosomal small subunit homolog [Sourc |
| FILIP1L | 3 | filamin A interacting protein 1 like [Source:HG |
| MIR3921 | 3 | microRNA 3921 [Source:HGNC Symbol;Acc:HG |
| AC129803.1 | 3 | novel transcript, antisense to CMSS1 |
| TMEM30CP | 3 | transmembrane protein 30C, pseudogene [So |
| DUSP12P1 | 3 | dual specificity phosphatase 12 pseudogene 1 |
| VTI1BP1 | 3 | vesicle transport through interaction with t-SN |
| TBC1D23 | 3 | TBC1 domain family member 23 [Source:HGN |
| NIT2 | 3 | nitrilase family member 2 [Source:HGNC Symb |
| TOMM70 | 3 | translocase of outer mitochondrial membrane |
| LNP1 | 3 | leukemia NUP98 fusion partner 1 [Source:HGN |
| TMEM45A | 3 | transmembrane protein 45A [Source:HGNC Sy |
| ADGRG7 | 3 | adhesion G protein-coupled receptor G7 [Sour |
| AC069223.1 | 3 | NULL |
| GMFBP1 | 3 | glia maturation factor beta pseudogene 1 [Sou |
| TFG | 3 | trafficking from ER to golgi regulator [Source:H |
| ABI3BP | 3 | ABI family member 3 binding protein [Source: |
| AC080014.1 | 3 | family with sequence similarity 175, member |
| RNU6-865P | 3 | RNA, U6 small nuclear 865, pseudogene [Sour |
| ACTR3P3 | 3 | ACTR3 pseudogene 3 [Source:HGNC Symbol;A |
| IMPG2 | 3 | interphotoreceptor matrix proteoglycan 2 [So |
| SENP7 | 3 | SUMO specific peptidase 7 [Source:HGNC Sym |
| ZNF90P1 | 3 | zinc finger protein 90 pseudogene 1 [Source:H |
| AC110994.1 | 3 | family with sequence similarity 136, member |
| AC110994.2 | 3 | basic transcription factor 3 (BTF3) pseudogene |
| Y_RNA | 3 | Y RNA [Source:RFAM;Acc:RF00019] |
| FAM172BP | 3 | family with sequence similarity 172 member B |
| TRMT10C | 3 | tRNA methyltransferase 10C, mitochondrial R |
| PCNP | 3 | PEST proteolytic signal containing nuclear pro |
| AC073861.1 | 3 | ribosomal protein S18 (RPS18) pseudogene |
| AC084198.1 | 3 | ribosomal protein L32 (RPL32) pseudogene |
| RNU6-1256P | 3 | RNA, U6 small nuclear 1256, pseudogene [Sou |
| Y_RNA | 3 | Y RNA [Source:RFAM;Acc:RF00019] |
| Y_RNA | 3 | Y RNA [Source:RFAM;Acc:RF00019] |
| ZBTB11 | 3 | zinc finger and BTB domain containing 11 [Sou |
| RNY1P12 | 3 | RNY1 pseudogene 12 [Source:HGNC Symbol;A |
| ZBTB11-AS1 | 3 | ZBTB11 antisense RNA 1 [Source:HGNC Symbo |
| RPL24 | 3 | ribosomal protein L24 [Source:HGNC Symbol; |
| AC084198.2 | 3 | novel transcript |
| PDCL3P4 | 3 | PDCL3 pseudogene 4 [Source:HGNC Symbol;A |
| CEP97 | 3 | centrosomal protein 97 [Source:HGNC Symbo |
| NXPE3 | 3 | neurexophilin and PC-esterase domain family |
| AC020651.1 | 3 | novel transcript |
| NFKBIZ | 3 | NFKB inhibitor zeta [Source:HGNC Symbol;Acc |
| AC020651.2 | 3 | novel transcript |
| LINC02085 | 3 | long intergenic non-protein coding RNA 2085 |
| Y_RNA | 3 | Y RNA [Source:RFAM;Acc:RF00019] |

| | | |
|---|---|---|
| AC106712.1 | 3 | novel transcript |
| ZPLD1 | 3 | zona pellucida like domain containing 1 [Sourc |
| AC063938.1 | 3 | novel transcript |
| RNU6-461P | 3 | RNA, U6 small nuclear 461, pseudogene [Sour |
| RNU1-43P | 3 | RNA, U1 small nuclear 43, pseudogene [Sourc |
| NDUFA4P2 | 3 | NDUFA4 pseudogene 2 [Source:HGNC Symbol |
| MIR548AB | 3 | microRNA 548ab [Source:HGNC Symbol;Acc:H |
| AC046144.1 | 3 | synovial sarcoma, X breakpoint 2 interacting p |
| TUBBP11 | 3 | tubulin beta class I pseudogene 11 [Source:HG |
| RAP1BP2 | 3 | RAP1B, member of RAS oncogene family pseu |
| AC091804.1 | 3 | ezrin (EZR) pseudogene |
| ALCAM | 3 | activated leukocyte cell adhesion molecule [S |
| CBLB | 3 | Cbl proto-oncogene B [Source:HGNC Symbol;A |
| AC131237.1 | 3 | novel transcript |
| AC079382.1 | 3 | TM2 domain containing 1 (TM2D1) pseudogen |
| AC079382.2 | 3 | novel transcript |
| LINC00882 | 3 | long intergenic non-protein coding RNA 882 [S |
| Y_RNA | 3 | Y RNA [Source:RFAM;Acc:RF00019] |
| Y_RNA | 3 | Y RNA [Source:RFAM;Acc:RF00019] |
| FCF1P3 | 3 | FCF1 pseudogene 3 [Source:HGNC Symbol;Acc |
| MTND4P16 | 3 | MT-ND4 pseudogene 16 [Source:HGNC Symbo |
| MTND4LP3 | 3 | MT-ND4L pseudogene 3 [Source:HGNC Symbo |
| MTND3P6 | 3 | MT-ND3 pseudogene 6 [Source:HGNC Symbol |
| MTCO3P35 | 3 | MT-CO3 pseudogene 35 [Source:HGNC Symbo |
| MTATP6P22 | 3 | MT-ATP6 pseudogene 22 [Source:HGNC Symb |
| MTND5P16 | 3 | MT-ND5 pseudogene 16 [Source:HGNC Symbo |
| MTND6P6 | 3 | MT-ND6 pseudogene 6 [Source:HGNC Symbol |
| MTCO1P35 | 3 | MT-CO1 pseudogene 35 [Source:HGNC Symbo |
| MTND1P16 | 3 | MT-ND1 pseudogene 16 [Source:HGNC Symbo |
| MTND2P14 | 3 | MT-ND2 pseudogene 14 [Source:HGNC Symbo |
| RNU6-1308P | 3 | RNA, U6 small nuclear 1308, pseudogene [Sou |
| AC117435.1 | 3 | novel transcript |
| AC074043.1 | 3 | protein tyrosine phosphatase, non-receptor ty |
| DUBR | 3 | DPPA2 upstream binding RNA [Source:HGNC S |
| AC063944.1 | 3 | novel transcript |
| AC063944.2 | 3 | citrate synthase (CS) pseudogene |
| AC063944.3 | 3 | novel transcript |
| CCDC54 | 3 | coiled-coil domain containing 54 [Source:HGN |
| LINC01990 | 3 | long intergenic non-protein coding RNA 1990 |
| BBX | 3 | BBX high mobility group box domain containir |
| LINC00636 | 3 | long intergenic non-protein coding RNA 636 [S |
| LINC00635 | 3 | long intergenic non-protein coding RNA 635 [S |
| AC012020.1 | 3 | novel transcript |
| CD47 | 3 | CD47 molecule [Source:HGNC Symbol;Acc:HG |
| LINC01215 | 3 | long intergenic non-protein coding RNA 1215 |
| IFT57 | 3 | intraflagellar transport 57 [Source:HGNC Symb |
| HHLA2 | 3 | HERV-H LTR-associating 2 [Source:HGNC Symb |
| HNRNPA1P17 | 3 | heterogeneous nuclear ribonucleoprotein A1 |
| MYH15 | 3 | myosin heavy chain 15 [Source:HGNC Symbol; |
| AC069499.1 | 3 | ribosomal protein L13 (RPL13) pseudogene |

| | | |
|---|---|---|
| CIP2A | 3 | cell proliferation regulating inhibitor of protei |
| SNORA70 | 3 | Small nucleolar RNA SNORA70 [Source:RFAM; |
| AC069499.2 | 3 | ribosomal protein L10 (RPL10) pseudogene |
| DZIP3 | 3 | DAZ interacting zinc finger protein 3 [Source:H |
| AC092185.1 | 3 | resistin like beta (RETNLB) pseudogene |
| RETNLB | 3 | resistin like beta [Source:HGNC Symbol;Acc:H |
| TRAT1 | 3 | T cell receptor associated transmembrane ada |
| GUCA1C | 3 | guanylate cyclase activator 1C [Source:HGNC S |
| MORC1 | 3 | MORC family CW-type zinc finger 1 [Source:H |
| U3 | 3 | Small nucleolar RNA U3 [Source:RFAM;Acc:RF |
| MORC1-AS1 | 3 | MORC1 antisense RNA 1 [Source:HGNC Symbo |
| C3orf85 | 3 | chromosome 3 open reading frame 85 [Source |
| AC063923.2 | 3 | novel transcript |
| LINC00488 | 3 | long intergenic non-protein coding RNA 488 [S |
| RNU6-1236P | 3 | RNA, U6 small nuclear 1236, pseudogene [Sou |
| AC063923.1 | 3 | bromodomain containing 7 (BRD7) pseudogen |
| DPPA2 | 3 | developmental pluripotency associated 2 [Sou |
| Metazoa_SRP | 3 | Metazoan signal recognition particle RNA [Sou |
| DPPA4 | 3 | developmental pluripotency associated 4 [Sou |
| AC124945.1 | 3 | ribosomal protein L36 (RPL36) pseudogene |
| AC092905.1 | 3 | chromosome 12 open reading frame 29 (C12o |
| H3F3AP3 | 3 | H3 histone family member 3A pseudogene 3 [ |
| LINC01205 | 3 | long intergenic non-protein coding RNA 1205 |
| PPIAP15 | 3 | peptidylprolyl isomerase A pseudogene 15 [So |
| MIR4445 | 3 | microRNA 4445 [Source:HGNC Symbol;Acc:HG |
| AC078980.1 | 3 | novel transcript |
| DIMT1P1 | 3 | DIM1 dimethyladenosine transferase 1 homol |
| AC068781.1 | 3 | nuclear transcription factor Y, beta (NFYB) pse |
| AC078918.1 | 3 | pseudogene similar to part of zinc finger, AN1 |
| AC117430.1 | 3 | novel transcript |
| RNU6ATAC15P | 3 | RNA, U6atac small nuclear 15, pseudogene [So |
| RPSAP29 | 3 | ribosomal protein SA pseudogene 29 [Source: |
| Y_RNA | 3 | Y RNA [Source:RFAM;Acc:RF00019] |
| NECTIN3-AS1 | 3 | NECTIN3 antisense RNA 1 [Source:HGNC Symb |
| NECTIN3 | 3 | nectin cell adhesion molecule 3 [Source:HGNC |
| CD96 | 3 | CD96 molecule [Source:HGNC Symbol;Acc:HG |
| AC092916.1 | 3 | novel transcript |
| ATP6V0CP2 | 3 | ATPase H+ transporting V0 subunit c pseudog |
| AC092916.2 | 3 | pseudogene similar to part of poliovirus recep |
| ZBED2 | 3 | zinc finger BED-type containing 2 [Source:HGN |
| AC055748.1 | 3 | 5'-nucleotidase, cytosolic III (NT5C3) pseudog |
| PLCXD2 | 3 | phosphatidylinositol specific phospholipase C |
| PLCXD2-AS1 | 3 | PLCXD2 antisense RNA 1 [Source:HGNC Symbo |
| PHLDB2 | 3 | pleckstrin homology like domain family B mer |
| AC117509.1 | 3 | novel transcript, antisense to PHLDB2and PLC |
| ABHD10 | 3 | abhydrolase domain containing 10 [Source:HG |
| TAGLN3 | 3 | transgelin 3 [Source:HGNC Symbol;Acc:HGNC: |
| RFKP2 | 3 | riboflavin kinase pseudogene 2 [Source:HGNC |
| TMPRSS7 | 3 | transmembrane serine protease 7 [Source:HG |
| C3orf52 | 3 | chromosome 3 open reading frame 52 [Source |

| | | |
|---|---|---|
| MIR567 | 3 | microRNA 567 [Source:HGNC Symbol;Acc:HGI |
| GCSAM | 3 | germinal center associated signaling and moti |
| TBILA | 3 | TGF-beta induced lncRNA [Source:HGNC Symb |
| SLC9C1 | 3 | solute carrier family 9 member C1 [Source:HG |
| AC128688.1 | 3 | chromosome 1 open reading frame 106 (C1or |
| AC112487.1 | 3 | novel transcript |
| AC112487.2 | 3 | novel pseudogene |
| CD200 | 3 | CD200 molecule [Source:HGNC Symbol;Acc:H |
| AC092894.1 | 3 | novel transcript |
| BTLA | 3 | B and T lymphocyte associated [Source:HGNC |
| OR7E100P | 3 | olfactory receptor family 7 subfamily E memb |
| AC092692.1 | 3 | TEC |
| ATG3 | 3 | autophagy related 3 [Source:HGNC Symbol;Ac |
| SLC35A5 | 3 | solute carrier family 35 member A5 [Source:H |
| CCDC80 | 3 | coiled-coil domain containing 80 [Source:HGN |
| AC048334.1 | 3 | pseudogene similar to methionine adenosyltr |
| LINC02042 | 3 | long intergenic non-protein coding RNA 2042 |
| CD200R1L-AS1 | 3 | CD200R1L antisense RNA 1 [Source:HGNC Syn |
| CD200R1L | 3 | CD200 receptor 1 like [Source:HGNC Symbol; |
| CD200R1 | 3 | CD200 receptor 1 [Source:HGNC Symbol;Acc: |
| AC074044.1 | 3 | novel transcript, antisense to GTPBP8 |
| GTPBP8 | 3 | GTP binding protein 8 (putative) [Source:HGN |
| NEPRO | 3 | nucleolus and neural progenitor protein [Sour |
| AC078785.1 | 3 | uncharacterized LOC107986114 [Source:NCBI |
| AC078785.3 | 3 | DnaJ (Hsp40) homolog, subfamily B, member |
| AC078785.2 | 3 | novel transcript |
| LINC02044 | 3 | long intergenic non-protein coding RNA 2044 |
| BOC | 3 | BOC cell adhesion associated, oncogene regul |
| AC026329.1 | 3 | novel transcript, antisense to BOC |
| CFAP44 | 3 | cilia and flagella associated protein 44 [Source |
| AC026348.1 | 3 | Rab geranylgeranyltransferase, beta subunit ( |
| AC112128.1 | 3 | novel protein |
| CFAP44-AS1 | 3 | CFAP44 antisense RNA 1 [Source:HGNC Symb |
| MIR8076 | 3 | microRNA 8076 [Source:HGNC Symbol;Acc:HG |
| SPICE1 | 3 | spindle and centriole associated protein 1 [So |
| SIDT1 | 3 | SID1 transmembrane family member 1 [Sourc |
| SIDT1-AS1 | 3 | SIDT1 antisense RNA 1 [Source:HGNC Symbol; |
| MIR4446 | 3 | microRNA 4446 [Source:HGNC Symbol;Acc:HG |
| RN7SL767P | 3 | RNA, 7SL, cytoplasmic 767, pseudogene [Sour |
| USF3 | 3 | upstream transcription factor family member |
| NAA50 | 3 | N(alpha)-acetyltransferase 50, NatE catalytic s |
| AC108693.1 | 3 | pseudogene similar to part of dihydrolipoamic |
| AC108693.2 | 3 | novel transcript |
| ATP6V1A | 3 | ATPase H+ transporting V1 subunit A [Source: |
| GRAMD1C | 3 | GRAM domain containing 1C [Source:HGNC Sy |
| AC079944.1 | 3 | ribosomal protein S10 (RPS10) pseudogene |
| AC079944.2 | 3 | dullard homolog (Xenopus laevis) (DULLARD) |
| VPS26AP1 | 3 | VPS26A pseudogene 1 [Source:HGNC Symbol; |
| AC128687.2 | 3 | novel transcript, antisense to GRAMD1C |
| ZDHHC23 | 3 | zinc finger DHHC-type containing 23 [Source:H |

| | | |
|---|---|---|
| CCDC191 | 3 | coiled-coil domain containing 191 [Source:HG |
| AC128687.1 | 3 | ATP5F1 ATP synthase, H+ transporting, mitocl |
| AC128687.3 | 3 | novel transcript, antisense to CCDC191 |
| AC092896.2 | 3 | Novel trancript, antisense to CCDC191 |
| QTRT2 | 3 | queuine tRNA-ribosyltransferase accessory su |
| AC092896.1 | 3 | histone cluster 1 pseudogene |
| DRD3 | 3 | dopamine receptor D3 [Source:HGNC Symbol; |
| AC093010.2 | 3 | novel transcript, antisense to ZNF80 |
| AC093010.1 | 3 | novel pseudogene |
| ZNF80 | 3 | zinc finger protein 80 [Source:HGNC Symbol;A |
| TIGIT | 3 | T cell immunoreceptor with Ig and ITIM doma |
| ZBTB20 | 3 | zinc finger and BTB domain containing 20 [Sou |
| AC093010.3 | 3 | novel transcript |
| MIR568 | 3 | microRNA 568 [Source:HGNC Symbol;Acc:HGI |
| ZBTB20-AS1 | 3 | ZBTB20 antisense RNA 1 [Source:HGNC Symbo |
| ZBTB20-AS5 | 3 | ZBTB20 antisense RNA 5 [Source:HGNC Symbo |
| ZBTB20-AS2 | 3 | ZBTB20 antisense RNA 2 [Source:HGNC Symbo |
| MIR4796 | 3 | microRNA 4796 [Source:HGNC Symbol;Acc:HG |
| ZBTB20-AS3 | 3 | ZBTB20 antisense RNA 3 [Source:HGNC Symbo |
| YBX1P3 | 3 | Y-box binding protein 1 pseudogene 3 [Source |
| ZBTB20-AS4 | 3 | ZBTB20 antisense RNA 4 [Source:HGNC Symbo |
| AC026341.1 | 3 | novel transcript |
| EIF4E2P2 | 3 | eukaryotic translation initiation factor 4E fami |
| AC026341.2 | 3 | novel transcript |
| AC026341.3 | 3 | novel transcript |
| GAP43 | 3 | growth associated protein 43 [Source:HGNC S |
| AC092468.1 | 3 | novel transcript |
| LSAMP | 3 | limbic system associated membrane protein [ |
| RN7SL815P | 3 | RNA, 7SL, cytoplasmic 815, pseudogene [Sour |
| LSAMP-AS1 | 3 | LSAMP antisense RNA 1 [Source:HGNC Symbo |
| LINC00903 | 3 | long intergenic non-protein coding RNA 903 [S |
| RN7SL582P | 3 | RNA, 7SL, cytoplasmic 582, pseudogene [Sour |
| BZW1P2 | 3 | basic leucine zipper and W2 domains 1 pseud |
| TUSC7 | 3 | tumor suppressor candidate 7 [Source:HGNC S |
| AC108713.1 | 3 | NULL |
| MIR4447 | 3 | microRNA 4447 [Source:HGNC Symbol;Acc:HG |
| AC069504.1 | 3 | NULL |
| LINC00901 | 3 | long intergenic non-protein coding RNA 901 [S |
| RNU5E-8P | 3 | RNA, U5E small nuclear 8, pseudogene [Sourc |
| PTMAP8 | 3 | prothymosin alpha pseudogene 8 [Source:HGI |
| RNU6-1200P | 3 | RNA, U6 small nuclear 1200, pseudogene [Sou |
| AC092691.1 | 3 | novel transcript |
| AC092691.2 | 3 | developmentally regulated GTP binding prote |
| LINC02024 | 3 | long intergenic non-protein coding RNA 2024 |
| AC092691.3 | 3 | novel transcript |
| AC068633.1 | 3 | novel transcript |
| AC068633.2 | 3 | novel transcript |
| IGSF11 | 3 | immunoglobulin superfamily member 11 [Sou |
| IGSF11-AS1 | 3 | IGSF11 antisense RNA 1 [Source:HGNC Symbo |
| TEX55 | 3 | testis expressed 55 [Source:HGNC Symbol;Acc |

| | | |
|---|---|---|
| AC083800.1 | 3 | novel chromosome 3 open reading frame 30 ( |
| UPK1B | 3 | uroplakin 1B [Source:HGNC Symbol;Acc:HGNC |
| B4GALT4 | 3 | beta-1,4-galactosyltransferase 4 [Source:HGN |
| B4GALT4-AS1 | 3 | B4GALT4 antisense RNA 1 [Source:HGNC Syml |
| ARHGAP31 | 3 | Rho GTPase activating protein 31 [Source:HGN |
| RPS26P21 | 3 | ribosomal protein S26 pseudogene 21 [Source |
| ARHGAP31-AS1 | 3 | ARHGAP31 antisense RNA 1 [Source:HGNC Syı |
| RNU6-1127P | 3 | RNA, U6 small nuclear 1127, pseudogene [Sou |
| TMEM39A | 3 | transmembrane protein 39A [Source:HGNC Sy |
| POGLUT1 | 3 | protein O-glucosyltransferase 1 [Source:HGNC |
| AC073352.1 | 3 | novel transcript |
| TIMMDC1 | 3 | translocase of inner mitochondrial membrane |
| CD80 | 3 | CD80 molecule [Source:HGNC Symbol;Acc:HG |
| AC073352.2 | 3 | novel transcript, antisense to ADPRH |
| ADPRH | 3 | ADP-ribosylarginine hydrolase [Source:HGNC S |
| PLA1A | 3 | phospholipase A1 member A [Source:HGNC Sy |
| RPL10P7 | 3 | ribosomal protein L10 pseudogene 7 [Source: |
| POPDC2 | 3 | popeye domain containing 2 [Source:HGNC Sy |
| COX17 | 3 | cytochrome c oxidase copper chaperone COX: |
| AC023494.1 | 3 | novel transcript, antisense to COX17 |
| MAATS1 | 3 | MYCBP associated and testis expressed 1 [Sou |
| AC069444.2 | 3 | novel protein |
| AC069444.1 | 3 | novel transcript |
| NR1I2 | 3 | nuclear receptor subfamily 1 group I member |
| PHBP8 | 3 | prohibitin pseudogene 8 [Source:HGNC Symbo |
| GSK3B | 3 | glycogen synthase kinase 3 beta [Source:HGN |
| AC092910.1 | 3 | presenilin associated, rhomboid-like (PARL) ps |
| AC092910.2 | 3 | riboflavin kinase (RFK) pseudogene |
| AC092910.3 | 3 | novel transcript |
| RN7SL762P | 3 | RNA, 7SL, cytoplasmic 762, pseudogene [Sour |
| RN7SL397P | 3 | RNA, 7SL, cytoplasmic 397, pseudogene [Sour |
| GPR156 | 3 | G protein-coupled receptor 156 [Source:HGN |
| Y_RNA | 3 | Y RNA [Source:RFAM;Acc:RF00019] |
| AC063952.2 | 3 | prostate tumor overexpressed 1 (PTOV1) pseu |
| LRRC58 | 3 | leucine rich repeat containing 58 [Source:HGN |
| AC063952.1 | 3 | novel V-set domain-containing pseudogene |
| AC063952.4 | 3 | novel transcript |
| AC063952.3 | 3 | chromosome 20 open reading frame 149 (C20 |
| FSTL1 | 3 | follistatin like 1 [Source:HGNC Symbol;Acc:HG |
| MIR198 | 3 | microRNA 198 [Source:HGNC Symbol;Acc:HGI |
| AC126182.3 | 3 | novel transcript, antisense to FSTL1 |
| AC126182.1 | 3 | cyclin-dependent kinase 2 associated protein |
| AC126182.2 | 3 | ribosomal protein L34 (RPL34) pseudogene |
| NDUFB4 | 3 | NADH:ubiquinone oxidoreductase subunit B4 |
| HGD | 3 | homogentisate 1,2-dioxygenase [Source:HGN |
| RABL3 | 3 | RAB, member of RAS oncogene family like 3 [S |
| MTCO1P29 | 3 | MT-CO1 pseudogene 29 [Source:HGNC Symbo |
| MTCO2P29 | 3 | MT-CO2 pseudogene 29 [Source:HGNC Symbo |
| GTF2E1 | 3 | general transcription factor IIE subunit 1 [Sour |
| NAP1L1P3 | 3 | nucleosome assembly protein 1 like 1 pseudo |

| | | |
|---|---|---|
| AC072026.2 | 3 | novel transcript, antisense to STXBP5L |
| LINC02049 | 3 | long intergenic non-protein coding RNA 2049 |
| AC072026.3 | 3 | novel transcript |
| STXBP5L | 3 | syntaxin binding protein 5 like [Source:HGNC S |
| AC072026.1 | 3 | C18orf32 chromosome 18 open reading frame |
| MIR5682 | 3 | microRNA 5682 [Source:HGNC Symbol;Acc:HG |
| AC078857.1 | 3 | zinc finger protein pseudogene |
| AC079841.1 | 3 | actin related protein 2/3 complex, subunit 1A, |
| AC079841.2 | 3 | novel transcript, antisense to STXBP5L |
| POLQ | 3 | DNA polymerase theta [Source:HGNC Symbol; |
| RPL7AP11 | 3 | ribosomal protein L7a pseudogene 11 [Source |
| ARGFX | 3 | arginine-fifty homeobox [Source:HGNC Symbo |
| FBXO40 | 3 | F-box protein 40 [Source:HGNC Symbol;Acc:H |
| HCLS1 | 3 | hematopoietic cell-specific Lyn substrate 1 [So |
| RN7SL172P | 3 | RNA, 7SL, cytoplasmic 172, pseudogene [Sour |
| RNU4-62P | 3 | RNA, U4 small nuclear 62, pseudogene [Sourc |
| GOLGB1 | 3 | golgin B1 [Source:HGNC Symbol;Acc:HGNC:44 |
| IQCB1 | 3 | IQ motif containing B1 [Source:HGNC Symbol; |
| EAF2 | 3 | ELL associated factor 2 [Source:HGNC Symbol, |
| SLC15A2 | 3 | solute carrier family 15 member 2 [Source:HG |
| ILDR1 | 3 | immunoglobulin like domain containing recep |
| Y_RNA | 3 | Y RNA [Source:RFAM;Acc:RF00019] |
| CD86 | 3 | CD86 molecule [Source:HGNC Symbol;Acc:HG |
| AC068630.1 | 3 | tigger transposable element derived 1 (TIGD1 |
| CASR | 3 | calcium sensing receptor [Source:HGNC Symb |
| HNRNPA1P23 | 3 | heterogeneous nuclear ribonucleoprotein A1 |
| CSTA | 3 | cystatin A [Source:HGNC Symbol;Acc:HGNC:2 |
| CCDC58 | 3 | coiled-coil domain containing 58 [Source:HGN |
| FAM162A | 3 | family with sequence similarity 162 member A |
| WDR5B | 3 | WD repeat domain 5B [Source:HGNC Symbol; |
| AC083798.2 | 3 | novel transcript, antisense to KPNA1 |
| AC083798.1 | 3 | pseudogene containing WD repeat domains |
| KPNA1 | 3 | karyopherin subunit alpha 1 [Source:HGNC Sy |
| AC096861.1 | 3 | high mobility group nucleosome binding doma |
| AC096861.2 | 3 | novel transcript |
| PARP9 | 3 | poly(ADP-ribose) polymerase family member |
| DTX3L | 3 | deltex E3 ubiquitin ligase 3L [Source:HGNC Sy |
| PARP15 | 3 | poly(ADP-ribose) polymerase family member |
| EIF4BP8 | 3 | eukaryotic translation initiation factor 4B pseu |
| PARP14 | 3 | poly(ADP-ribose) polymerase family member |
| HSPBAP1 | 3 | HSPB1 associated protein 1 [Source:HGNC Syr |
| SLC49A4 | 3 | solute carrier family 49 member 4 [Source:HG |
| LINC02035 | 3 | long intergenic non-protein coding RNA 2035 |
| SEMA5B | 3 | semaphorin 5B [Source:HGNC Symbol;Acc:HG |
| PDIA5 | 3 | protein disulfide isomerase family A member |
| MIR7110 | 3 | microRNA 7110 [Source:HGNC Symbol;Acc:HG |
| SEC22A | 3 | SEC22 homolog A, vesicle trafficking protein [S |
| AC112503.2 | 3 | novel transcript |
| ADCY5 | 3 | adenylate cyclase 5 [Source:HGNC Symbol;Acc |
| AC112503.1 | 3 | novel transcript, antisense to ADCY5 |

| | | |
|---|---|---|
| HACD2 | 3 | 3-hydroxyacyl-CoA dehydratase 2 [Source:HG |
| MYLK-AS1 | 3 | MYLK antisense RNA 1 [Source:HGNC Symbol; |
| MYLK | 3 | myosin light chain kinase [Source:HGNC Symb |
| MYLK-AS2 | 3 | MYLK antisense RNA 2 [Source:HGNC Symbol; |
| AC020634.2 | 3 | novel transcript, antisense to MYLK |
| AC020634.1 | 3 | NULL |
| CCDC14 | 3 | coiled-coil domain containing 14 [Source:HGN |
| ROPN1 | 3 | rhophilin associated tail protein 1 [Source:HG |
| KALRN | 3 | kalirin RhoGEF kinase [Source:HGNC Symbol;A |
| MIR5002 | 3 | microRNA 5002 [Source:HGNC Symbol;Acc:HG |
| RPL7P15 | 3 | ribosomal protein L7 pseudogene 15 [Source: |
| MIR6083 | 3 | microRNA 6083 [Source:HGNC Symbol;Acc:HG |
| RNU6-143P | 3 | RNA, U6 small nuclear 143, pseudogene [Sour |
| AC022336.2 | 3 | novel transcript, overlapping to KALRN |
| UMPS | 3 | uridine monophosphate synthetase [Source:H |
| MIR544B | 3 | microRNA 544b [Source:HGNC Symbol;Acc:HG |
| AC022336.1 | 3 | ribosomal protein S26 (RPS26) pseudogene |
| ITGB5 | 3 | integrin subunit beta 5 [Source:HGNC Symbol; |
| ITGB5-AS1 | 3 | ITGB5 antisense RNA 1 [Source:HGNC Symbol; |
| AC022336.3 | 3 | TEC |
| 7SK | 3 | 7SK RNA [Source:RFAM;Acc:RF00100] |
| ENO1P3 | 3 | enolase 1 pseudogene 3 [Source:HGNC Symbo |
| MUC13 | 3 | mucin 13, cell surface associated [Source:HGN |
| HEG1 | 3 | heart development protein with EGF like dom |
| RNA5SP137 | 3 | RNA, 5S ribosomal pseudogene 137 [Source:H |
| AC117488.1 | 3 | novel transcript |
| SLC12A8 | 3 | solute carrier family 12 member 8 [Source:HG |
| RNU6-230P | 3 | RNA, U6 small nuclear 230, pseudogene [Sour |
| MIR5092 | 3 | microRNA 5092 [Source:HGNC Symbol;Acc:HG |
| ZNF148 | 3 | zinc finger protein 148 [Source:HGNC Symbol; |
| DUTP1 | 3 | deoxyuridine triphosphatase pseudogene 1 [S |
| AC108688.1 | 3 | DnaJ (Hsp40) homolog, subfamily B, member |
| RNU6-232P | 3 | RNA, U6 small nuclear 232, pseudogene [Sour |
| SNX4 | 3 | sorting nexin 4 [Source:HGNC Symbol;Acc:HG |
| Y_RNA | 3 | Y RNA [Source:RFAM;Acc:RF00019] |
| Y_RNA | 3 | Y RNA [Source:RFAM;Acc:RF00019] |
| OSBPL11 | 3 | oxysterol binding protein like 11 [Source:HGN |
| AF186996.3 | 3 | pseudogene similar to part of YTH domain fan |
| AF186996.2 | 3 | pseudogene similar to part of myosin light cha |
| OR7E130P | 3 | olfactory receptor family 7 subfamily E memb |
| OR7E29P | 3 | olfactory receptor family 7 subfamily E memb |
| OR7E93P | 3 | olfactory receptor family 7 subfamily E memb |
| OR7E53P | 3 | olfactory receptor family 7 subfamily E memb |
| OR7E97P | 3 | olfactory receptor family 7 subfamily E memb |
| AF186996.1 | 3 | pseudogene similar to part of chromosome 1: |
| AC092902.4 | 3 | novel transcript |
| AC092902.5 | 3 | RNA binding protein, fox-1 homolog (C. elegar |
| AC092902.2 | 3 | novel transcript |
| AC092902.1 | 3 | ribosomal protein S24 (RPS24) pseudogene |
| MIR548I1 | 3 | microRNA 548i-1 [Source:HGNC Symbol;Acc:H |

| | | |
|---|---|---|
| RPS3AP14 | 3 | ribosomal protein S3a pseudogene 14 [Source |
| AC092902.6 | 3 | novel transcript |
| SNRPCP11 | 3 | small nuclear ribonucleoprotein polypeptide C |
| LINC02614 | 3 | long intergenic non-protein coding RNA 2614 |
| ENPP7P4 | 3 | ectonucleotide pyrophosphatase/phosphodie |
| AC092903.2 | 3 | novel transcript |
| FAM86JP | 3 | family with sequence similarity 86 member J, |
| ALG1L | 3 | ALG1 chitobiosyldiphosphodolichol beta-man |
| AC092903.1 | 3 | pseudogene similar to part of fibrosin-like 1 (F |
| ROPN1B | 3 | rhophilin associated tail protein 1B [Source:H |
| SLC41A3 | 3 | solute carrier family 41 member 3 [Source:HG |
| AC117422.1 | 3 | novel transcript |
| AC079848.1 | 3 | novel transcript |
| ALDH1L1 | 3 | aldehyde dehydrogenase 1 family member L1 |
| ALDH1L1-AS1 | 3 | ALDH1L1 antisense RNA 1 [Source:HGNC Sym |
| RNU1-30P | 3 | RNA, U1 small nuclear 30, pseudogene [Sourc |
| ALDH1L1-AS2 | 3 | ALDH1L1 antisense RNA 2 [Source:HGNC Sym |
| AC079848.2 | 3 | novel transcript |
| AC016924.1 | 3 | novel transcript |
| AC063919.1 | 3 | novel transcript |
| AC063919.2 | 3 | novel transcript |
| KLF15 | 3 | Kruppel like factor 15 [Source:HGNC Symbol;A |
| CCDC37-DT | 3 | CCDC37 divergent transcript [Source:HGNC Sy |
| CFAP100 | 3 | cilia and flagella associated protein 100 [Sourc |
| AC073439.1 | 3 | TEC |
| ZXDC | 3 | ZXD family zinc finger C [Source:HGNC Symbo |
| UROC1 | 3 | urocanate hydratase 1 [Source:HGNC Symbol; |
| CHST13 | 3 | carbohydrate sulfotransferase 13 [Source:HGI |
| C3orf22 | 3 | chromosome 3 open reading frame 22 [Sourc |
| RNA5SP138 | 3 | RNA, 5S ribosomal pseudogene 138 [Source:H |
| AC024558.2 | 3 | Protein TXNRD3NB [Source:UniProtKB/Swiss- |
| TXNRD3 | 3 | thioredoxin reductase 3 [Source:HGNC Symbo |
| AC024558.1 | 3 | methyltransferase like 5 (METTL5) pseudogen |
| NUP210P1 | 3 | nucleoporin 210 pseudogene 1 [Source:HGNC |
| AC078867.1 | 3 | splicing factor 3a, subunit 2, 66kDa pseudoger |
| CHCHD6 | 3 | coiled-coil-helix-coiled-coil-helix domain conta |
| RCC2P4 | 3 | regulator of chromosome condensation 2 pse |
| AC011199.1 | 3 | TEC |
| PLXNA1 | 3 | plexin A1 [Source:HGNC Symbol;Acc:HGNC:90 |
| AC112482.2 | 3 | novel transcript, antisense to C3orf56 |
| C3orf56 | 3 | chromosome 3 open reading frame 56 [Sourc |
| Y_RNA | 3 | Y RNA [Source:RFAM;Acc:RF00019] |
| AC112482.1 | 3 | chromosome 3 open reading frame 56 (C3orf! |
| RNU6-1047P | 3 | RNA, U6 small nuclear 1047, pseudogene [Sou |
| AC133681.1 | 3 | novel transcript |
| PRR20G | 3 | proline rich 20G [Source:HGNC Symbol;Acc:H( |
| LINC02016 | 3 | long intergenic non-protein coding RNA 2016 |
| LINC01471 | 3 | long intergenic non-protein coding RNA 1471 |
| AC016968.1 | 3 | novel transcript |
| LINC02034 | 3 | long intergenic non-protein coding RNA 2034 |

| | | |
|---|---|---|
| TPRA1 | 3 | transmembrane protein adipocyte associated |
| AC084035.1 | 3 | novel transcript |
| MIR6825 | 3 | microRNA 6825 [Source:HGNC Symbol;Acc:HG |
| MIR7976 | 3 | microRNA 7976 [Source:HGNC Symbol;Acc:HG |
| MCM2 | 3 | minichromosome maintenance complex comp |
| AC023593.1 | 3 | Novel transcript, antisense to MCM2 |
| PODXL2 | 3 | podocalyxin like 2 [Source:HGNC Symbol;Acc: |
| ABTB1 | 3 | ankyrin repeat and BTB domain containing 1 [ |
| MGLL | 3 | monoglyceride lipase [Source:HGNC Symbol;A |
| AC117480.1 | 3 | novel transcript, antisense to MGLL |
| KBTBD12 | 3 | kelch repeat and BTB domain containing 12 [S |
| RNA5SP139 | 3 | RNA, 5S ribosomal pseudogene 139 [Source:H |
| SEC61A1 | 3 | SEC61 translocon alpha 1 subunit [Source:HGN |
| RUVBL1 | 3 | RuvB like AAA ATPase 1 [Source:HGNC Symbo |
| RNU2-37P | 3 | RNA, U2 small nuclear 37, pseudogene [Sourc |
| RUVBL1-AS1 | 3 | RUVBL1 antisense RNA 1 [Source:HGNC Symb |
| RNU6-823P | 3 | RNA, U6 small nuclear 823, pseudogene [Sour |
| EEFSEC | 3 | eukaryotic elongation factor, selenocysteine-t |
| AL449214.1 | 3 | Novel transcript, antisense to EEFSEC |
| DNAJB8 | 3 | DnaJ heat shock protein family (Hsp40) memb |
| DNAJB8-AS1 | 3 | DNAJB8 antisense RNA 1 [Source:HGNC Symb |
| GATA2 | 3 | GATA binding protein 2 [Source:HGNC Symbol |
| GATA2-AS1 | 3 | GATA2 antisense RNA 1 [Source:HGNC Symbo |
| TMED10P2 | 3 | transmembrane p24 trafficking protein 10 pse |
| AC080005.1 | 3 | novel transcript |
| LINC01565 | 3 | long intergenic non-protein coding RNA 1565 |
| RPN1 | 3 | ribophorin I [Source:HGNC Symbol;Acc:HGNC: |
| Metazoa_SRP | 3 | Metazoan signal recognition particle RNA [Sou |
| POU5F1P6 | 3 | POU class 5 homeobox 1 pseudogene 6 [Sourc |
| AC079945.1 | 3 | NULL |
| RAB7A | 3 | RAB7A, member RAS oncogene family [Source |
| MARK2P8 | 3 | microtubule affinity regulating kinase 2 pseud |
| FTH1P4 | 3 | ferritin heavy chain 1 pseudogene 4 [Source:H |
| RN7SL698P | 3 | RNA, 7SL, cytoplasmic 698, pseudogene [Sour |
| RPS15AP16 | 3 | ribosomal protein S15a pseudogene 16 [Sourc |
| MARK2P17 | 3 | microtubule affinity regulating kinase 2 pseud |
| MARK3P3 | 3 | microtubule affinity regulating kinase 3 pseud |
| MARK2P6 | 3 | microtubule affinity regulating kinase 2 pseud |
| AC112484.3 | 3 | novel transcript |
| AC112484.1 | 3 | novel transcript |
| AC112484.2 | 3 | cell death-inducing DFFA-like effector c (CIDE( |
| AC112484.5 | 3 | novel transcript |
| ACAD9 | 3 | acyl-CoA dehydrogenase family member 9 [Sc |
| KIAA1257 | 3 | KIAA1257 [Source:HGNC Symbol;Acc:HGNC:2( |
| AC112484.4 | 3 | TEC |
| EFCC1 | 3 | EF-hand and coiled-coil domain containing 1 [ |
| AC108673.1 | 3 | jumonji domain containing 4 (JMJD4) pseudog |
| GP9 | 3 | glycoprotein IX platelet [Source:HGNC Symbol |
| RAB43 | 3 | RAB43, member RAS oncogene family [Source |
| ISY1-RAB43 | 3 | ISY1-RAB43 readthrough [Source:HGNC Symb |

| | | |
|---|---|---|
| AC108673.2 | 3 | novel transcript, antisense to RAB43 |
| ISY1 | 3 | ISY1 splicing factor homolog [Source:HGNC Sy |
| AC108673.3 | 3 | novel transcript |
| CNBP | 3 | CCHC-type zinc finger nucleic acid binding pro |
| RPS27P12 | 3 | ribosomal protein S27 pseudogene 12 [Source |
| COPG1 | 3 | coatomer protein complex subunit gamma 1 [ |
| MIR6826 | 3 | microRNA 6826 [Source:HGNC Symbol;Acc:HG |
| AC137695.3 | 3 | novel transcript, antisense to COPG1 |
| HMCES | 3 | 5-hydroxymethylcytosine binding, ES cell spec |
| H1FX | 3 | H1 histone family member X [Source:HGNC Sy |
| H1FX-AS1 | 3 | H1FX antisense RNA 1 [Source:HGNC Symbol;/ |
| NUP210P3 | 3 | nucleoporin 210 pseudogene 3 [Source:HGNC |
| AC137695.1 | 3 | MAP/microtubule affinity-regulating kinase 2, |
| AC137695.2 | 3 | MAP/microtubule affinity-regulating kinase 2, |
| AL449212.1 | 3 | pseudogene similar to intraflagellar transport |
| RPL32P3 | 3 | ribosomal protein L32 pseudogene 3 [Source: |
| SNORA7B | 3 | small nucleolar RNA, H/ACA box 7B [Source:H |
| EFCAB12 | 3 | EF-hand calcium binding domain 12 [Source:H |
| MBD4 | 3 | methyl-CpG binding domain 4, DNA glycosylas |
| IFT122 | 3 | intraflagellar transport 122 [Source:HGNC Syn |
| RHO | 3 | rhodopsin [Source:HGNC Symbol;Acc:HGNC:1 |
| H1FOO | 3 | H1 histone family member O oocyte specific [ |
| PLXND1 | 3 | plexin D1 [Source:HGNC Symbol;Acc:HGNC:91 |
| RN7SL752P | 3 | RNA, 7SL, cytoplasmic 752, pseudogene [Sour |
| AC023162.1 | 3 | pseudogene similar to part of RGD motif, leuc |
| TMCC1 | 3 | transmembrane and coiled-coil domain family |
| RNY3P13 | 3 | RNY3 pseudogene 13 [Source:HGNC Symbol;A |
| U6 | 3 | U6 spliceosomal RNA [Source:RFAM;Acc:RF00 |
| AC083799.1 | 3 | novel transcript, sense intronic to TMCC1 |
| TMCC1-AS1 | 3 | TMCC1 antisense RNA 1 (head to head) [Sourc |
| AC083906.3 | 3 | novel transcript |
| TRH | 3 | thyrotropin releasing hormone [Source:HGNC |
| AC083906.1 | 3 | ribosomal protein S17 (RPS17) pseudogene |
| AC083906.5 | 3 | RNA binding protein, fox-1 homolog (C. elegar |
| AC083906.2 | 3 | pseudogene similar to part of chromosome 1 |
| OR7E129P | 3 | olfactory receptor family 7 subfamily E memb |
| OR7E21P | 3 | olfactory receptor family 7 subfamily E memb |
| AC083906.4 | 3 | eva-1 homolog C (EVA1C) pseudogene |
| ALG1L2 | 3 | ALG1 chitobiosyldiphosphodolichol beta-manı |
| LINC02014 | 3 | long intergenic non-protein coding RNA 2014 |
| FAM86HP | 3 | family with sequence similarity 86 member H, |
| LINC02021 | 3 | long intergenic non-protein coding RNA 2021 |
| ENPP7P3 | 3 | ectonucleotide pyrophosphatase/phosphodie |
| AC130888.1 | 3 | novel transcript |
| SNRPCP8 | 3 | small nuclear ribonucleoprotein polypeptide ( |
| COL6A4P2 | 3 | collagen type VI alpha 4 pseudogene 2 [Sourc |
| AC093004.1 | 3 | ATP synthase, H+ transporting, mitochondrial |
| COL6A5 | 3 | collagen type VI alpha 5 chain [Source:HGNC S |
| COL6A6 | 3 | collagen type VI alpha 6 chain [Source:HGNC S |
| PIK3R4 | 3 | phosphoinositide-3-kinase regulatory subunit |

| | | |
|---|---|---|
| RN7SKP212 | 3 | RN7SK pseudogene 212 [Source:HGNC Symbo |
| AC097105.1 | 3 | Novel transcript |
| GSTO3P | 3 | glutathione S-transferase omega 3, pseudoge |
| ATP2C1 | 3 | ATPase secretory pathway Ca2+ transporting |
| AC055733.4 | 3 | TEC |
| AC055733.2 | 3 | novel transcript, antisense to ATP2C1 |
| Y_RNA | 3 | Y RNA [Source:RFAM;Acc:RF00019] |
| AC055733.1 | 3 | cysteine and glycine-rich protein 2 pseudogen |
| AC055733.3 | 3 | pseudogene similar to part of makorin ring fin |
| ASTE1 | 3 | asteroid homolog 1 [Source:HGNC Symbol;Ac |
| NEK11 | 3 | NIMA related kinase 11 [Source:HGNC Symbo |
| AC116424.1 | 3 | novel transcript |
| RNU6-726P | 3 | RNA, U6 small nuclear 726, pseudogene [Sour |
| AC010210.1 | 3 | novel transcript |
| NUDT16P1 | 3 | nudix hydrolase 16 pseudogene 1 [Source:HG |
| NUDT16 | 3 | nudix hydrolase 16 [Source:HGNC Symbol;Acc |
| AC107027.3 | 3 | novel transcript, overlapping to MRPL3 |
| MRPL3 | 3 | mitochondrial ribosomal protein L3 [Source:H |
| SNORA58 | 3 | small nucleolar RNA, H/ACA box 58 [Source:H |
| AC107027.1 | 3 | novel transcript |
| BCL2L12P1 | 3 | BCL2 like 12 pseudogene 1 [Source:HGNC Sym |
| CPNE4 | 3 | copine 4 [Source:HGNC Symbol;Acc:HGNC:23 |
| AC107027.2 | 3 | pseudogene similar to part of copine IV |
| MIR5704 | 3 | microRNA 5704 [Source:HGNC Symbol;Acc:HG |
| AC117409.1 | 3 | 26S protease regulatory subunit 7 (PSMC2) ps |
| RPL7P16 | 3 | ribosomal protein L7 pseudogene 16 [Source: |
| ACPP | 3 | acid phosphatase, prostate [Source:HGNC Syn |
| AC020633.1 | 3 | G2/M-phase specific E3 ubiquitin ligase (G2E3 |
| NIP7P2 | 3 | NIP7 pseudogene 2 [Source:HGNC Symbol;Ac |
| DNAJC13 | 3 | DnaJ heat shock protein family (Hsp40) memb |
| ACAD11 | 3 | acyl-CoA dehydrogenase family member 11 [S |
| NPHP3-ACAD11 | 3 | NPHP3-ACAD11 readthrough (NMD candidate |
| ACKR4 | 3 | atypical chemokine receptor 4 [Source:HGNC |
| HSPA8P19 | 3 | heat shock protein family A (Hsp70) member |
| UBA5 | 3 | ubiquitin like modifier activating enzyme 5 [S |
| NPHP3 | 3 | nephrocystin 3 [Source:HGNC Symbol;Acc:HG |
| NPHP3-AS1 | 3 | NPHP3 antisense RNA 1 [Source:HGNC Symbo |
| AC079942.1 | 3 | novel transcript |
| TMEM108 | 3 | transmembrane protein 108 [Source:HGNC Sy |
| TMEM108-AS1 | 3 | TMEM108 antisense RNA 1 [Source:HGNC Syn |
| BFSP2 | 3 | beaded filament structural protein 2 [Source: |
| AC055753.1 | 3 | pseudogene similar to part of AF4/FMR2 fami |
| BFSP2-AS1 | 3 | BFSP2 antisense RNA 1 [Source:HGNC Symbol |
| AC022296.2 | 3 | pseudogene similar to part of MAF1 |
| AC022296.4 | 3 | novel transcript |
| AC022296.3 | 3 | pseudogene similar to part of FAM76A, family |
| AC022296.1 | 3 | NULL |
| CDV3 | 3 | CDV3 homolog [Source:HGNC Symbol;Acc:HG |
| TOPBP1 | 3 | DNA topoisomerase II binding protein 1 [Sour |
| INHCAP | 3 | inhibitor of carbonic anhydrase pseudogene [ |

| | | |
|---|---|---|
| RNU6-678P | 3 | RNA, U6 small nuclear 678, pseudogene [Sour |
| RNA5SP140 | 3 | RNA, 5S ribosomal pseudogene 140 [Source:H |
| TF | 3 | transferrin [Source:HGNC Symbol;Acc:HGNC:1 |
| AC080128.1 | 3 | acyl-CoA synthetase long-chain family membe |
| SRPRB | 3 | SRP receptor subunit beta [Source:HGNC Sym |
| AC080128.2 | 3 | novel transcript, antisense to SRPRB |
| RAB6B | 3 | RAB6B, member RAS oncogene family [Source |
| SLCO2A1 | 3 | solute carrier organic anion transporter family |
| LINC02000 | 3 | long intergenic non-protein coding RNA 2000 |
| RYK | 3 | receptor like tyrosine kinase [Source:HGNC Sy |
| HMGB3P14 | 3 | high mobility group box 3 pseudogene 14 [Sou |
| LINC02004 | 3 | long intergenic non-protein coding RNA 2004 |
| AC010207.1 | 3 | novel transcript |
| RPL39P5 | 3 | ribosomal protein L39 pseudogene 5 [Source: |
| AMOTL2 | 3 | angiomotin like 2 [Source:HGNC Symbol;Acc:H |
| MIR6827 | 3 | microRNA 6827 [Source:HGNC Symbol;Acc:HG |
| HMGN1P9 | 3 | high mobility group nucleosome binding doma |
| HMGB3P13 | 3 | high mobility group box 3 pseudogene 13 [Sou |
| MIR4788 | 3 | microRNA 4788 [Source:HGNC Symbol;Acc:HG |
| ANAPC13 | 3 | anaphase promoting complex subunit 13 [Sou |
| CEP63 | 3 | centrosomal protein 63 [Source:HGNC Symbo |
| AC026117.1 | 3 | developmental pluripotency associated 4 (DPI |
| EPHB1 | 3 | EPH receptor B1 [Source:HGNC Symbol;Acc:H |
| KY | 3 | kyphoscoliosis peptidase [Source:HGNC Symb |
| AC016931.1 | 3 | novel transcript, antisense to EPHB1 |
| RNU6-1174P | 3 | RNA, U6 small nuclear 1174, pseudogene [Sou |
| RNA5SP141 | 3 | RNA, 5S ribosomal pseudogene 141 [Source:H |
| AC092969.1 | 3 | novel transcript, antisense to EPHB1 |
| AC092989.1 | 3 | succinate dehydrogenase complex, subunit B, |
| NDUFS6P1 | 3 | NADH:ubiquinone oxidoreductase subunit S6 |
| PPP2R3A | 3 | protein phosphatase 2 regulatory subunit B''a |
| AC072039.1 | 3 | downstream neighbor of SON (DONSON) pseu |
| AC072039.2 | 3 | novel transcript, antisense to PPP2R3A |
| MSL2 | 3 | MSL complex subunit 2 [Source:HGNC Symbol |
| TDGF1P6 | 3 | teratocarcinoma-derived growth factor 1 pseu |
| AC092991.1 | 3 | ribosomal protein L31 (RPL31) pseudogene |
| PCCB | 3 | propionyl-CoA carboxylase subunit beta [Sour |
| STAG1 | 3 | stromal antigen 1 [Source:HGNC Symbol;Acc:H |
| RNU6-1284P | 3 | RNA, U6 small nuclear 1284, pseudogene [Sou |
| HMGN1P10 | 3 | high mobility group nucleosome binding doma |
| RNU6-789P | 3 | RNA, U6 small nuclear 789, pseudogene [Sour |
| AC117382.1 | 3 | DnaJ (Hsp40) homolog, subfamily C, member |
| AC117382.2 | 3 | novel transcript, antisense to STAG1 |
| AC096992.3 | 3 | novel transcript |
| AC096992.1 | 3 | sulfotransferase family, cytosolic, 1C, member |
| SLC35G2 | 3 | solute carrier family 35 member G2 [Source:H |
| NCK1-DT | 3 | NCK1 divergent transcript [Source:HGNC Syml |
| AC096992.2 | 3 | novel transcript, antisense to SLC35G2 |
| NCK1 | 3 | NCK adaptor protein 1 [Source:HGNC Symbol; |
| RAD51AP1P1 | 3 | RAD51AP1 pseudogene 1 [Source:HGNC Symb |

| | | |
|---|---|---|
| IL20RB | 3 | interleukin 20 receptor subunit beta [Source:H |
| IL20RB-AS1 | 3 | IL20RB antisense RNA 1 [Source:HGNC Symbo |
| RNA5SP142 | 3 | RNA, 5S ribosomal pseudogene 142 [Source:H |
| AC117440.1 | 3 | mitochondrial carrier homolog 2 (C. elegans) ( |
| NPM1P17 | 3 | nucleophosmin 1 pseudogene 17 [Source:HGN |
| SOX14 | 3 | SRY-box transcription factor 14 [Source:HGNC |
| LINC01210 | 3 | long intergenic non-protein coding RNA 1210 |
| AC007159.1 | 3 | novel transcript |
| HSPA8P9 | 3 | heat shock protein family A (Hsp70) member 8 |
| CLDN18 | 3 | claudin 18 [Source:HGNC Symbol;Acc:HGNC:2 |
| AC016252.1 | 3 | novel transcript, antisense to CLDN18 |
| DZIP1L | 3 | DAZ interacting zinc finger protein 1 like [Sour |
| KRT8P36 | 3 | keratin 8 pseudogene 36 [Source:HGNC Symb |
| A4GNT | 3 | alpha-1,4-N-acetylglucosaminyltransferase [So |
| AC023049.1 | 3 | alpha-1,4-N-acetylglucosaminyltransferase (A4 |
| DBR1 | 3 | debranching RNA lariats 1 [Source:HGNC Syml |
| ARMC8 | 3 | armadillo repeat containing 8 [Source:HGNC S |
| NME9 | 3 | NME/NM23 family member 9 [Source:HGNC S |
| MRAS | 3 | muscle RAS oncogene homolog [Source:HGNC |
| ESYT3 | 3 | extended synaptotagmin 3 [Source:HGNC Sym |
| AC022497.1 | 3 | TEC |
| CEP70 | 3 | centrosomal protein 70 [Source:HGNC Symbo |
| FAIM | 3 | Fas apoptotic inhibitory molecule [Source:HGI |
| PPIAP72 | 3 | peptidylprolyl isomerase A pseudogene 72 [So |
| PIK3CB | 3 | phosphatidylinositol-4,5-bisphosphate 3-kinas |
| GAPDHP39 | 3 | glyceraldehyde 3 phosphate dehydrogenase p |
| RPL23AP40 | 3 | ribosomal protein L23a pseudogene 40 [Sourc |
| EEF1A1P25 | 3 | eukaryotic translation elongation factor 1 alph |
| ATP5MC1P3 | 3 | ATP synthase membrane subunit c locus 1 pse |
| LINC01391 | 3 | long intergenic non-protein coding RNA 1391 |
| FOXL2 | 3 | forkhead box L2 [Source:HGNC Symbol;Acc:HC |
| FOXL2NB | 3 | FOXL2 neighbor [Source:HGNC Symbol;Acc:HC |
| PRR23A | 3 | proline rich 23A [Source:HGNC Symbol;Acc:HC |
| MRPS22 | 3 | mitochondrial ribosomal protein S22 [Source: |
| PRR23B | 3 | proline rich 23B [Source:HGNC Symbol;Acc:HC |
| PRR23C | 3 | proline rich 23C [Source:HGNC Symbol;Acc:HC |
| RPL7L1P7 | 3 | ribosomal protein L7 like 1 pseudogene 7 [Sou |
| BPESC1 | 3 | blepharophimosis, epicanthus inversus and pt |
| PISRT1 | 3 | PISRT1 lncRNA [Source:HGNC Symbol;Acc:HGI |
| AC119740.1 | 3 | NULL |
| AC024933.2 | 3 | novel transcript, antisense to MRPS22 |
| AC024933.1 | 3 | novel transcript, antisense to MRPS22 |
| COPB2 | 3 | coatomer protein complex subunit beta 2 [Sou |
| U6 | 3 | U6 spliceosomal RNA [Source:RFAM;Acc:RF00 |
| AC046134.2 | 3 | novel transcript, antisense to RBP2, RBP1 & N |
| RBP2 | 3 | retinol binding protein 2 [Source:HGNC Symbo |
| AC097103.1 | 3 | novel transcript |
| ACTG1P1 | 3 | actin gamma 1 pseudogene 1 [Source:HGNC S |
| RBP1 | 3 | retinol binding protein 1 [Source:HGNC Symbo |
| NMNAT3 | 3 | nicotinamide nucleotide adenylyltransferase 3 |

| | | |
|---|---|---|
| AC046134.1 | 3 | DTW domain containing 1 (DTWD1) pseudoge |
| RN7SKP124 | 3 | RN7SK pseudogene 124 [Source:HGNC Symbo |
| RN7SL724P | 3 | RNA, 7SL, cytoplasmic 724, pseudogene [Sour |
| AC110716.2 | 3 | novel transcript |
| AC110716.1 | 3 | novel transcript |
| AC016933.1 | 3 | novel transcript |
| TRMT112P5 | 3 | tRNA methyltransferase subunit 11-2 pseudog |
| CLSTN2 | 3 | calsyntenin 2 [Source:HGNC Symbol;Acc:HGN |
| AC010181.1 | 3 | novel transcript |
| AC010181.2 | 3 | novel transcript |
| CLSTN2-AS1 | 3 | CLSTN2 antisense RNA 1 [Source:HGNC Symbo |
| AC048346.1 | 3 | pseudogene similar to killer cell lectin-like rec |
| TRIM42 | 3 | tripartite motif containing 42 [Source:HGNC S |
| AC121333.1 | 3 | novel transcript |
| RPL23AP41 | 3 | ribosomal protein L23a pseudogene 41 [Sourc |
| SLC25A36 | 3 | solute carrier family 25 member 36 [Source:H |
| AC108727.1 | 3 | novel transcript |
| SPSB4 | 3 | splA/ryanodine receptor domain and SOCS bo |
| AC108727.2 | 3 | novel transcript |
| PXYLP1 | 3 | 2-phosphoxylose phosphatase 1 [Source:HGN |
| AC022215.2 | 3 | novel transcript, antisense to PXYLP1 |
| AC022215.1 | 3 | ribosomal protein L31 (RPL31) pseudogene |
| AC117383.1 | 3 | novel transcript, antisense to ZBTB38 & ACPL |
| AC117383.2 | 3 | PHD finger protein 5A (PHF5A) pseudogene |
| ZBTB38 | 3 | zinc finger and BTB domain containing 38 [Sou |
| KRT18P35 | 3 | keratin 18 pseudogene 35 [Source:HGNC Sym |
| RASA2 | 3 | RAS p21 protein activator 2 [Source:HGNC Sy |
| RASA2-IT1 | 3 | RASA2 intronic transcript 1 [Source:HGNC Syn |
| YWHAQP6 | 3 | YWHAQ pseudogene 6 [Source:HGNC Symbol; |
| LINC02618 | 3 | long intergenic non-protein coding RNA 2618 |
| TPT1P3 | 3 | tumor protein, translationally-controlled 1 ps |
| AC112771.1 | 3 | pseudogene similar to part of glyceraldehyde- |
| RNF7 | 3 | ring finger protein 7 [Source:HGNC Symbol;Ac |
| AC112504.2 | 3 | novel protein |
| GRK7 | 3 | G protein-coupled receptor kinase 7 [Source:H |
| AC112504.3 | 3 | novel transcript |
| HMGN2P25 | 3 | high mobility group nucleosomal binding dom |
| ATP1B3 | 3 | ATPase Na+/K+ transporting subunit beta 3 [S |
| RNU6-509P | 3 | RNA, U6 small nuclear 509, pseudogene [Sour |
| ATP1B3-AS1 | 3 | ATP1B3 antisense RNA 1 [Source:HGNC Symb |
| AC112504.1 | 3 | TEC |
| TFDP2 | 3 | transcription factor Dp-2 [Source:HGNC Symb |
| AC133435.1 | 3 | ribosomal protein L19 (RPL19) pseudogene |
| RNU6-425P | 3 | RNA, U6 small nuclear 425, pseudogene [Sour |
| GK5 | 3 | glycerol kinase 5 [Source:HGNC Symbol;Acc:H |
| XRN1 | 3 | 5'-3' exoribonuclease 1 [Source:HGNC Symbol |
| RNU6-1294P | 3 | RNA, U6 small nuclear 1294, pseudogene [Sou |
| RNU1-100P | 3 | RNA, U1 small nuclear 100, pseudogene [Sour |
| ATR | 3 | ATR serine/threonine kinase [Source:HGNC Sy |
| AC109992.1 | 3 | eukaryotic translation initiation factor 2-alpha |

| | | |
|---|---|---|
| AC109992.2 | 3 | novel transcript, sense overlapping to ATR |
| RPL6P9 | 3 | ribosomal protein L6 pseudogene 9 [Source:H |
| RNA5SP143 | 3 | RNA, 5S ribosomal pseudogene 143 [Source:H |
| PLS1 | 3 | plastin 1 [Source:HGNC Symbol;Acc:HGNC:90 |
| PLS1-AS1 | 3 | PLS1 antisense RNA 1 [Source:HGNC Symbol;A |
| RN7SKP25 | 3 | RN7SK pseudogene 25 [Source:HGNC Symbol; |
| AC072028.1 | 3 | novel transcript, antisense to PLS1 |
| TRPC1 | 3 | transient receptor potential cation channel su |
| RNU7-47P | 3 | RNA, U7 small nuclear 47 pseudogene [Source |
| PCOLCE2 | 3 | procollagen C-endopeptidase enhancer 2 [Sou |
| AC021074.1 | 3 | ribosomal protein L8 (RPL8) pseudogene |
| AC021074.3 | 3 | novel transcript |
| PAQR9 | 3 | progestin and adipoQ receptor family membe |
| AC021074.2 | 3 | pseudogene similar to part of ribosomal prote |
| PAQR9-AS1 | 3 | PAQR9 antisense RNA 1 [Source:HGNC Symbo |
| U2SURP | 3 | U2 snRNP associated SURP domain containing |
| AC026304.1 | 3 | novel transcript |
| AC018450.2 | 3 | novel transcript |
| CHST2 | 3 | carbohydrate sulfotransferase 2 [Source:HGN |
| AC018450.1 | 3 | novel transcript |
| PBX2P1 | 3 | PBX homeobox 2 pseudogene 1 [Source:HGN |
| SLC9A9 | 3 | solute carrier family 9 member A9 [Source:HG |
| AC131210.1 | 3 | novel transcript |
| SLC9A9-AS1 | 3 | SLC9A9 antisense RNA 1 [Source:HGNC Symbo |
| SLC9A9-AS2 | 3 | SLC9A9 antisense RNA 2 [Source:HGNC Symbo |
| GAPDHP47 | 3 | glyceraldehyde 3 phosphate dehydrogenase p |
| ST13P15 | 3 | ST13, Hsp70 interacting protein pseudogene 1 |
| AC073358.1 | 3 | ubiquitin domain containing 2 (UBTD2) pseud |
| AC107421.1 | 3 | ribosomal protein S17 (RPS17) pseudogene |
| DIPK2A | 3 | divergent protein kinase domain 2A [Source:H |
| RNA5SP144 | 3 | RNA, 5S ribosomal pseudogene 144 [Source:H |
| AC011592.1 | 3 | TEC |
| LARP7P4 | 3 | La ribonucleoprotein domain family member |
| GM2AP1 | 3 | GM2 ganglioside activator pseudogene 1 [Sou |
| RPL21P39 | 3 | ribosomal protein L21 pseudogene 39 [Source |
| AC055758.2 | 3 | novel transcript |
| AC055758.1 | 3 | novel transcript |
| AC107021.2 | 3 | novel transcript, overlapping to PLOD2 |
| AC107021.1 | 3 | novel transcript, antisense to PLOD2 |
| LNCSRLR | 3 | lncRNA sorafenib resistance in renal cell carci |
| PLOD2 | 3 | procollagen-lysine,2-oxoglutarate 5-dioxygen |
| PLSCR4 | 3 | phospholipid scramblase 4 [Source:HGNC Sym |
| AC130509.1 | 3 | pseudogene similar to part of phospholipid sc |
| AC069528.1 | 3 | phospholipid scramblase 2 (PLSCR2) pseudoge |
| PLSCR2 | 3 | phospholipid scramblase 2 [Source:HGNC Sym |
| AC069528.2 | 3 | TEC |
| PLSCR1 | 3 | phospholipid scramblase 1 [Source:HGNC Sym |
| RNU6-428P | 3 | RNA, U6 small nuclear 428, pseudogene [Sour |
| PLSCR5 | 3 | phospholipid scramblase family member 5 [Sc |
| PLSCR5-AS1 | 3 | PLSCR5 antisense RNA 1 [Source:HGNC Symbo |

| | | |
|---|---|---|
| AC092957.1 | 3 | novel transcript |
| LINC02010 | 3 | long intergenic non-protein coding RNA 2010 |
| RNU6-505P | 3 | RNA, U6 small nuclear 505, pseudogene [Sour |
| RPL21P71 | 3 | ribosomal protein L21 pseudogene 71 [Source |
| ZIC4 | 3 | Zic family member 4 [Source:HGNC Symbol;Ac |
| ZIC4-AS1 | 3 | ZIC4 antisense RNA 1 [Source:HGNC Symbol;A |
| ZIC1 | 3 | Zic family member 1 [Source:HGNC Symbol;Ac |
| NPM1P28 | 3 | nucleophosmin 1 pseudogene 28 [Source:HGN |
| AC092925.1 | 3 | novel transcript |
| AC092958.1 | 3 | novel transcript |
| AC092958.4 | 3 | novel transcript |
| LINC02032 | 3 | long intergenic non-protein coding RNA 2032 |
| AC092958.3 | 3 | novel transcript |
| AC092958.2 | 3 | novel transcript |
| AC025566.1 | 3 | novel transcript |
| HNRNPA1P20 | 3 | heterogeneous nuclear ribonucleoprotein A1 |
| LINC02045 | 3 | long intergenic non-protein coding RNA 2045 |
| LINC02046 | 3 | long intergenic non-protein coding RNA 2046 |
| AC069410.1 | 3 | novel transcript |
| RPL38P1 | 3 | ribosomal protein L38 pseudogene 1 [Source: |
| AGTR1 | 3 | angiotensin II receptor type 1 [Source:HGNC S |
| CPB1 | 3 | carboxypeptidase B1 [Source:HGNC Symbol;A |
| HMGB1P30 | 3 | high mobility group box 1 pseudogene 30 [Sou |
| AC092979.1 | 3 | novel transcript |
| CPA3 | 3 | carboxypeptidase A3 [Source:HGNC Symbol;A |
| AC092979.2 | 3 | novel transcript |
| UBQLN4P1 | 3 | ubiquilin 4 pseudogene 1 [Source:HGNC Symb |
| GYG1 | 3 | glycogenin 1 [Source:HGNC Symbol;Acc:HGNC |
| HLTF | 3 | helicase like transcription factor [Source:HGN |
| HLTF-AS1 | 3 | HLTF antisense RNA 1 [Source:HGNC Symbol;A |
| Y_RNA | 3 | Y RNA [Source:RFAM;Acc:RF00019] |
| MED28P2 | 3 | mediator complex subunit 28 pseudogene 2 [S |
| HPS3 | 3 | HPS3 biogenesis of lysosomal organelles comp |
| CP | 3 | ceruloplasmin [Source:HGNC Symbol;Acc:HGN |
| AC093001.2 | 3 | novel transcript |
| CPHL1P | 3 | ceruloplasmin and hephaestin like 1, pseudog |
| AC093001.1 | 3 | novel transcript |
| TM4SF18 | 3 | transmembrane 4 L six family member 18 [Sou |
| AC073522.1 | 3 | ribosomal protein S25 (RPS25) pseudogene |
| TM4SF1 | 3 | transmembrane 4 L six family member 1 [Sour |
| TM4SF1-AS1 | 3 | TM4SF1 antisense RNA 1 [Source:HGNC Symb |
| AC108751.4 | 3 | novel transcript |
| AC108751.2 | 3 | ribosomal protein L32 (RPL32) pseudogene |
| AC108751.3 | 3 | WD repeat domain 70 (WDR70) pseudogene |
| FKBP1AP4 | 3 | FKBP prolyl isomerase 1A pseudogene 4 [Sour |
| AC108751.1 | 3 | pseudogene similar to part of chromosome 10 |
| TM4SF4 | 3 | transmembrane 4 L six family member 4 [Sour |
| AC108751.5 | 3 | eukaryotic translation elongation factor 1 alph |
| WWTR1 | 3 | WW domain containing transcription regulato |
| RNU6-1098P | 3 | RNA, U6 small nuclear 1098, pseudogene [Sou |

| | | |
|---|---|---|
| WWTR1-IT1 | 3 | WWTR1 intronic transcript 1 [Source:HGNC Sy |
| WWTR1-AS1 | 3 | WWTR1 antisense RNA 1 [Source:HGNC Symb |
| COMMD2 | 3 | COMM domain containing 2 [Source:HGNC Sy |
| ANKUB1 | 3 | ankyrin repeat and ubiquitin domain containi |
| RNU6-507P | 3 | RNA, U6 small nuclear 507, pseudogene [Sour |
| RNF13 | 3 | ring finger protein 13 [Source:HGNC Symbol;A |
| RNU6-720P | 3 | RNA, U6 small nuclear 720, pseudogene [Sour |
| NPM1P29 | 3 | nucleophosmin 1 pseudogene 29 [Source:HGN |
| PFN2 | 3 | profilin 2 [Source:HGNC Symbol;Acc:HGNC:88 |
| AC117395.1 | 3 | novel transcript |
| TMEM183B | 3 | transmembrane protein 183B [Source:HGNC S |
| PPIAP73 | 3 | peptidylprolyl isomerase A pseudogene 73 [So |
| HNRNPA1P24 | 3 | heterogeneous nuclear ribonucleoprotein A1 |
| AC117386.2 | 3 | novel transcript, antisense to PFN2 |
| AC022494.1 | 3 | eukaryotic translation initiation factor 3, subu |
| LINC01998 | 3 | long intergenic non-protein coding RNA 1998 |
| RPL32P9 | 3 | ribosomal protein L32 pseudogene 9 [Source: |
| AC117386.1 | 3 | novel transcript |
| LINC01213 | 3 | long intergenic non-protein coding RNA 1213 |
| U2 | 3 | U2 spliceosomal RNA [Source:RFAM;Acc:RF00 |
| LINC01214 | 3 | long intergenic non-protein coding RNA 1214 |
| AC018545.1 | 3 | novel transcript |
| TSC22D2 | 3 | TSC22 domain family member 2 [Source:HGN |
| SERP1 | 3 | stress associated endoplasmic reticulum prote |
| EIF2A | 3 | eukaryotic translation initiation factor 2A [Sou |
| SELENOT | 3 | selenoprotein T [Source:HGNC Symbol;Acc:HG |
| AC069236.1 | 3 | Sin3A-associated protein, 30kDa, pseudogene |
| ERICH6 | 3 | glutamate rich 6 [Source:HGNC Symbol;Acc:H |
| ERICH6-AS1 | 3 | ERICH6 antisense RNA 1 [Source:HGNC Symbo |
| SNRPCP3 | 3 | small nuclear ribonucleoprotein polypeptide C |
| AC011317.1 | 3 | novel transcript |
| SIAH2 | 3 | siah E3 ubiquitin protein ligase 2 [Source:HGN |
| SIAH2-AS1 | 3 | SIAH2 antisense RNA 1 [Source:HGNC Symbol; |
| CLRN1-AS1 | 3 | CLRN1 antisense RNA 1 [Source:HGNC Symbol |
| MINDY4B | 3 | MINDY family member 4B [Source:HGNC Sym |
| AC020636.2 | 3 | CLRN1 and FAM188B2 readthrough transcript |
| AC020636.1 | 3 | novel transcript |
| CLRN1 | 3 | clarin 1 [Source:HGNC Symbol;Acc:HGNC:126 |
| AC108726.1 | 3 | NULL |
| MED12L | 3 | mediator complex subunit 12L [Source:HGNC |
| RNA5SP145 | 3 | RNA, 5S ribosomal pseudogene 145 [Source:H |
| GPR171 | 3 | G protein-coupled receptor 171 [Source:HGN |
| P2RY14 | 3 | purinergic receptor P2Y14 [Source:HGNC Sym |
| SETP11 | 3 | SET pseudogene 11 [Source:HGNC Symbol;Acc |
| AC078816.1 | 3 | novel transcript, antisense to GPR87 |
| GPR87 | 3 | G protein-coupled receptor 87 [Source:HGNC |
| P2RY13 | 3 | purinergic receptor P2Y13 [Source:HGNC Sym |
| P2RY12 | 3 | purinergic receptor P2Y12 [Source:HGNC Sym |
| IGSF10 | 3 | immunoglobulin superfamily member 10 [Sou |
| MRPL42P6 | 3 | mitochondrial ribosomal protein L42 pseudog |

| | | |
|---|---|---|
| AC117454.1 | 3 | pseudogene similar to part of zinc and ring fin |
| MIR5186 | 3 | microRNA 5186 [Source:HGNC Symbol;Acc:HG |
| LINC02066 | 3 | long intergenic non-protein coding RNA 2066 |
| AADACL2 | 3 | arylacetamide deacetylase like 2 [Source:HGN |
| AADACL2-AS1 | 3 | AADACL2 antisense RNA 1 [Source:HGNC Sym |
| AC068647.1 | 3 | UDP-N-acetyl-alpha-D-galactosamine:polypep |
| AADACP1 | 3 | arylacetamide deacetylase pseudogene 1 [Sou |
| AC068647.2 | 3 | arylacetamide deacetylase (esterase) (AADAC |
| AADAC | 3 | arylacetamide deacetylase [Source:HGNC Sym |
| SUCNR1 | 3 | succinate receptor 1 [Source:HGNC Symbol;Ac |
| AC108718.1 | 3 | novel transcript |
| MBNL1 | 3 | muscleblind like splicing regulator 1 [Source:H |
| MBNL1-AS1 | 3 | MBNL1 antisense RNA 1 [Source:HGNC Symbo |
| TMEM14EP | 3 | transmembrane protein 14E, pseudogene [Sou |
| Y_RNA | 3 | Y RNA [Source:RFAM;Acc:RF00019] |
| AC026347.1 | 3 | novel transcript |
| AC092924.1 | 3 | novel transcript |
| AC092924.2 | 3 | novel transcript |
| ATP5MGP5 | 3 | ATP synthase membrane subunit g pseudogen |
| P2RY1 | 3 | purinergic receptor P2Y1 [Source:HGNC Symb |
| HMGN2P13 | 3 | high mobility group nucleosomal binding dom |
| AC117394.1 | 3 | pseudogene similar to KIAA1328 |
| AC117394.2 | 3 | novel transcript |
| RAP2B | 3 | RAP2B, member of RAS oncogene family [Sou |
| RN7SL300P | 3 | RNA, 7SL, cytoplasmic 300, pseudogene [Sour |
| AC078788.1 | 3 | novel transcript |
| AC078788.2 | 3 | novel transcript |
| LINC02006 | 3 | long intergenic non-protein coding RNA 2006 |
| C3orf79 | 3 | chromosome 3 open reading frame 79 [Sourc |
| RNU6-901P | 3 | RNA, U6 small nuclear 901, pseudogene [Sour |
| Y_RNA | 3 | Y RNA [Source:RFAM;Acc:RF00019] |
| AC068985.1 | 3 | novel transcript |
| U8 | 3 | U8 small nucleolar RNA [Source:RFAM;Acc:RF |
| RPL21P42 | 3 | ribosomal protein L21 pseudogene 42 [Source |
| ARHGEF26-AS1 | 3 | ARHGEF26 antisense RNA 1 [Source:HGNC Syr |
| ARHGEF26 | 3 | Rho guanine nucleotide exchange factor 26 [S |
| DHX36 | 3 | DEAH-box helicase 36 [Source:HGNC Symbol; |
| AC134026.1 | 3 | NULL |
| AC134026.2 | 3 | TEC |
| GPR149 | 3 | G protein-coupled receptor 149 [Source:HGN |
| DDX50P2 | 3 | DEAD-box helicase 50 pseudogene 2 [Source: |
| AC092994.1 | 3 | novel transcript |
| SYPL1P1 | 3 | synaptophysin like 1 pseudogene 1 [Source:H |
| DYNLL1P5 | 3 | dynein light chain LC8-type 1 pseudogene 5 [S |
| RPL9P15 | 3 | ribosomal protein L9 pseudogene 15 [Source: |
| AC073359.2 | 3 | novel transcript |
| AC073359.1 | 3 | novel transcript |
| MME | 3 | membrane metalloendopeptidase [Source:HG |
| MME-AS1 | 3 | MME antisense RNA 1 [Source:HGNC Symbol; |
| LINC01487 | 3 | long intergenic non-protein coding RNA 1487 |

| | | |
|---|---|---|
| STRIT1 | 3 | small transmembrane regulator of ion transpo |
| AC104831.1 | 3 | novel transcript |
| PABPC1P10 | 3 | poly(A) binding protein cytoplasmic 1 pseudog |
| PLCH1 | 3 | phospholipase C eta 1 [Source:HGNC Symbol; |
| AC108729.3 | 3 | phospholipase C, eta 1 (PLCH1) pseudogene |
| AC108729.2 | 3 | DTW domain containing 1 (DTWD1) pseudoge |
| AC108729.1 | 3 | mitochondrially encoded cytochrome c oxidas |
| RN7SL715P | 3 | RNA, 7SL, cytoplasmic 715, pseudogene [Sour |
| PLCH1-AS1 | 3 | PLCH1 antisense RNA 1 [Source:HGNC Symbol |
| PLCH1-AS2 | 3 | PLCH1 antisense RNA 2 [Source:HGNC Symbol |
| RPL6P7 | 3 | ribosomal protein L6 pseudogene 7 [Source:H |
| AC104472.1 | 3 | novel transcript, sense intronic to PLCH1 |
| AC104472.2 | 3 | ribosomal protein L7a (RPL7A) pseudogene |
| C3orf33 | 3 | chromosome 3 open reading frame 33 [Sourc |
| AC104472.3 | 3 | novel transcript |
| SLC33A1 | 3 | solute carrier family 33 member 1 [Source:HG |
| AC104472.4 | 3 | novel transcript |
| GMPS | 3 | guanine monophosphate synthase [Source:HG |
| AC104472.5 | 3 | novel transcript, sense intronic to GMPS |
| SETP14 | 3 | SET pseudogene 14 [Source:HGNC Symbol;Acc |
| VN2R1P | 3 | vomeronasal 2 receptor 1 pseudogene [Sourc |
| KCNAB1 | 3 | potassium voltage-gated channel subfamily A |
| ALG1L15P | 3 | ALG1 like 15, pseudogene [Source:HGNC Syml |
| MRE11P1 | 3 | MRE11 homolog, double strand break repair r |
| AC091607.2 | 3 | novel transcript, antisense to SSR3 |
| AC091607.1 | 3 | small nuclear ribonucleoprotein 40kDa (U5) (S |
| KCNAB1-AS2 | 3 | KCNAB1 antisense RNA 2 [Source:HGNC Symb |
| KCNAB1-AS1 | 3 | KCNAB1 antisense RNA 1 [Source:HGNC Symb |
| AC084036.1 | 3 | novel transcript, antisense to KCNAB1 |
| SSR3 | 3 | signal sequence receptor subunit 3 [Source:H |
| TIPARP-AS1 | 3 | TIPARP antisense RNA 1 [Source:HGNC Symbo |
| TIPARP | 3 | TCDD inducible poly(ADP-ribose) polymerase |
| METTL15P1 | 3 | methyltransferase like 15 pseudogene 1 [Sour |
| LINC00886 | 3 | long intergenic non-protein coding RNA 886 [S |
| PA2G4P4 | 3 | proliferation-associated 2G4 pseudogene 4 [S |
| LEKR1 | 3 | leucine, glutamate and lysine rich 1 [Source:H |
| KLF3P2 | 3 | Kruppel like factor 3 pseudogene 2 [Source:H |
| AC117392.1 | 3 | CASP2 and RIPK1 domain containing adaptor v |
| RN7SKP177 | 3 | RN7SK pseudogene 177 [Source:HGNC Symbo |
| LINC00880 | 3 | long intergenic non-protein coding RNA 880 [S |
| LINC02029 | 3 | long intergenic non-protein coding RNA 2029 |
| LINC00881 | 3 | long intergenic non-protein coding RNA 881 [S |
| CCNL1 | 3 | cyclin L1 [Source:HGNC Symbol;Acc:HGNC:20 |
| Y_RNA | 3 | Y RNA [Source:RFAM;Acc:RF00019] |
| KRT18P34 | 3 | keratin 18 pseudogene 34 [Source:HGNC Sym |
| AC104411.1 | 3 | novel transcript |
| RNA5SP146 | 3 | RNA, 5S ribosomal pseudogene 146 [Source:H |
| AC092944.1 | 3 | novel transcript |
| AC092944.2 | 3 | poly(rC) binding protein 1, pseudogene |
| AC092944.3 | 3 | novel transcript |

| | | |
|---|---|---|
| VEPH1 | 3 | ventricular zone expressed PH domain contair |
| PTX3 | 3 | pentraxin 3 [Source:HGNC Symbol;Acc:HGNC: |
| SLC66A1L | 3 | solute carrier family 66 member 1 like [Source |
| AC084212.1 | 3 | novel transcript |
| AC079943.2 | 3 | novel transcript |
| RN7SKP46 | 3 | RN7SK pseudogene 46 [Source:HGNC Symbol; |
| AC079943.1 | 3 | novel transcript |
| SHOX2 | 3 | short stature homeobox 2 [Source:HGNC Sym |
| RSRC1 | 3 | arginine and serine rich coiled-coil 1 [Source:H |
| AC074276.1 | 3 | ribosomal protein L15 (RPL15) pseudogene |
| AC074276.2 | 3 | leukocyte immunoglobulin-like receptor, subf |
| AC106707.2 | 3 | ribosomal protein S12 (RPS12) pseudogene |
| AC106707.1 | 3 | novel transcript, antisense to RSRC1 |
| MLF1 | 3 | myeloid leukemia factor 1 [Source:HGNC Sym |
| MTAPP1 | 3 | methylthioadenosine phosphorylase pseudog |
| AC025033.1 | 3 | riboflavin kinase (RFK) pseudogene |
| GFM1 | 3 | G elongation factor mitochondrial 1 [Source:H |
| LXN | 3 | latexin [Source:HGNC Symbol;Acc:HGNC:1334 |
| AC080013.4 | 3 | novel transcript |
| AC080013.6 | 3 | novel transcript |
| RARRES1 | 3 | retinoic acid receptor responder 1 [Source:HG |
| MFSD1 | 3 | major facilitator superfamily domain containir |
| AC080013.1 | 3 | novel transcript, antisense to RARRES1 |
| AC080013.2 | 3 | ribosomal protein L35a (RPL35A) pseudogene |
| AC080013.3 | 3 | novel transcript, sense intronic to MFSD1 |
| AC080013.5 | 3 | novel transcript, antisense to MFSD1 |
| GPR79 | 3 | G protein-coupled receptor 79, pseudogene [S |
| AC092999.1 | 3 | TEC |
| IQCJ-SCHIP1 | 3 | IQCJ-SCHIP1 readthrough [Source:HGNC Symb |
| IQCJ | 3 | IQ motif containing J [Source:HGNC Symbol;A |
| MIR3919 | 3 | microRNA 3919 [Source:HGNC Symbol;Acc:HG |
| AC063955.1 | 3 | ribosomal protein S2 (RPS2) pseudogene |
| AC092943.2 | 3 | novel transcript, antisense to IQCJ-SCHIP1 |
| AC092943.1 | 3 | chromosome 9 open reading frame 78 (C9orf |
| IQCJ-SCHIP1-AS1 | 3 | IQCJ-SCHIP1 readthrough antisense RNA 1 [So |
| RNU2-31P | 3 | RNA, U2 small nuclear 31, pseudogene [Sourc |
| SCHIP1 | 3 | schwannomin interacting protein 1 [Source:H |
| IL12A-AS1 | 3 | IL12A antisense RNA 1 [Source:HGNC Symbol; |
| IL12A | 3 | interleukin 12A [Source:HGNC Symbol;Acc:HG |
| LINC01100 | 3 | long intergenic non-protein coding RNA 1100 |
| BRD7P2 | 3 | bromodomain containing 7 pseudogene 2 [So |
| C3orf80 | 3 | chromosome 3 open reading frame 80 [Sourc |
| AC079594.2 | 3 | TRIM59 and iFT80 readthrough |
| IFT80 | 3 | intraflagellar transport 80 [Source:HGNC Sym |
| AC079594.1 | 3 | ribosomal protein L35a (RPL35A) pseudogene |
| SMC4 | 3 | structural maintenance of chromosomes 4 [Sc |
| MIR15B | 3 | microRNA 15b [Source:HGNC Symbol;Acc:HGI |
| MIR16-2 | 3 | microRNA 16-2 [Source:HGNC Symbol;Acc:HG |
| TRIM59 | 3 | tripartite motif containing 59 [Source:HGNC S |
| B3GAT3P1 | 3 | beta-1,3-glucuronyltransferase 3 pseudogene |

| | | |
|---|---|---|
| RNU7-136P | 3 | RNA, U7 small nuclear 136 pseudogene [Sourc |
| KPNA4 | 3 | karyopherin subunit alpha 4 [Source:HGNC Sy |
| SCARNA7 | 3 | small Cajal body-specific RNA 7 [Source:HGNC |
| KRT8P12 | 3 | keratin 8 pseudogene 12 [Source:HGNC Symb |
| RPL6P8 | 3 | ribosomal protein L6 pseudogene 8 [Source:H |
| ARL14 | 3 | ADP ribosylation factor like GTPase 14 [Source |
| SNORA72 | 3 | Small nucleolar RNA SNORA72 [Source:RFAM; |
| AC069224.1 | 3 | novel transcript, antisense to PPM1L |
| PPM1L | 3 | protein phosphatase, Mg2+/Mn2+ dependent |
| B3GALNT1 | 3 | beta-1,3-N-acetylgalactosaminyltransferase 1 |
| NMD3 | 3 | NMD3 ribosome export adaptor [Source:HGN |
| AC108738.1 | 3 | signal recognition particle 14kDa (homologous |
| EEF1GP4 | 3 | eukaryotic translation elongation factor 1 gam |
| PSMC1P7 | 3 | proteasome 26S subunit, ATPase 1 pseudoger |
| SPTSSB | 3 | serine palmitoyltransferase small subunit B [S |
| LINC02067 | 3 | long intergenic non-protein coding RNA 2067 |
| AC112491.1 | 3 | novel transcript |
| RPL23AP42 | 3 | ribosomal protein L23a pseudogene 42 [Sourc |
| OTOL1 | 3 | otolin 1 [Source:HGNC Symbol;Acc:HGNC:340 |
| AC112770.1 | 3 | novel transcript |
| AC131211.1 | 3 | novel transcript |
| TOMM22P6 | 3 | TOMM22 pseudogene 6 [Source:HGNC Symbo |
| AC128716.1 | 3 | zinc finger protein (ZNF) pseudogene |
| AC128685.1 | 3 | novel transcript |
| LINC01192 | 3 | long intergenic non-protein coding RNA 1192 |
| RPS6P4 | 3 | ribosomal protein S6 pseudogene 4 [Source:H |
| AC079910.1 | 3 | novel transcript |
| RNU7-82P | 3 | RNA, U7 small nuclear 82 pseudogene [Source |
| AC084017.1 | 3 | neugrin, neurite outgrowth associated (NGRN |
| MIR1263 | 3 | microRNA 1263 [Source:HGNC Symbol;Acc:HG |
| LINC01323 | 3 | long intergenic non-protein coding RNA 1323 |
| LINC01324 | 3 | long intergenic non-protein coding RNA 1324 |
| SI | 3 | sucrase-isomaltase [Source:HGNC Symbol;Acc |
| Y_RNA | 3 | Y RNA [Source:RFAM;Acc:RF00019] |
| LINC02023 | 3 | long intergenic non-protein coding RNA 2023 |
| SLITRK3 | 3 | SLIT and NTRK like family member 3 [Source:H |
| LINC01322 | 3 | long intergenic non-protein coding RNA 1322 |
| BCHE | 3 | butyrylcholinesterase [Source:HGNC Symbol;A |
| MTND4P17 | 3 | MT-ND4 pseudogene 17 [Source:HGNC Symbo |
| MTND4LP10 | 3 | MT-ND4L pseudogene 10 [Source:HGNC Symb |
| MTND3P7 | 3 | MT-ND3 pseudogene 7 [Source:HGNC Symbol |
| MTCO3P38 | 3 | MT-CO3 pseudogene 38 [Source:HGNC Symbo |
| MCUR1P2 | 3 | MCUR1 pseudogene 2 [Source:HGNC Symbol; |
| AC104629.1 | 3 | Sjogren syndrome antigen B (autoantigen La) |
| LINC01326 | 3 | long intergenic non-protein coding RNA 1326 |
| AC072046.2 | 3 | protein kinase, X-linked (PRKX) pseudogene |
| RN7SKP298 | 3 | RN7SK pseudogene 298 [Source:HGNC Symbo |
| PSAT1P4 | 3 | phosphoserine aminotransferase 1 pseudoger |
| AC072046.1 | 3 | NADH dehydrogenase (ubiquinone) 1 alpha su |
| CBX1P5 | 3 | chromobox 1 pseudogene 5 [Source:HGNC Sy |

| | | |
|---|---|---|
| AC069439.2 | 3 | novel transcript |
| AC069439.1 | 3 | pseudogene similar to part of S-phase kinase-a |
| PPIAP74 | 3 | peptidylprolyl isomerase A pseudogene 74 [So |
| ZBBX | 3 | zinc finger B-box domain containing [Source:H |
| LINC01327 | 3 | long intergenic non-protein coding RNA 1327 |
| SERPINI2 | 3 | serpin family I member 2 [Source:HGNC Symb |
| WDR49 | 3 | WD repeat domain 49 [Source:HGNC Symbol;A |
| PDCD10 | 3 | programmed cell death 10 [Source:HGNC Sym |
| HMGN1P8 | 3 | high mobility group nucleosome binding doma |
| SERPINI1 | 3 | serpin family I member 1 [Source:HGNC Symb |
| AC079822.1 | 3 | novel transcript, antisense to SERPINI1 |
| LRRC77P | 3 | leucine rich repeat containing 77, pseudogene |
| AC026353.1 | 3 | novel transcript |
| MEMO1P3 | 3 | mediator of cell motility 1 pseudogene 3 [Sou |
| AC128713.1 | 3 | cytochrome c, somatic (CYCS) pseudogene |
| HMGN2P26 | 3 | high mobility group nucleosomal binding dom |
| GOLIM4 | 3 | golgi integral membrane protein 4 [Source:HG |
| AC069243.1 | 3 | novel transcript, sense intronic to GOLIM4 |
| EGFEM1P | 3 | EGF like and EMI domain containing 1, pseudo |
| AC124893.1 | 3 | novel transcript |
| RNU2-20P | 3 | RNA, U2 small nuclear 20, pseudogene [Sourc |
| MIR551B | 3 | microRNA 551b [Source:HGNC Symbol;Acc:HG |
| RPSAP33 | 3 | ribosomal protein SA pseudogene 33 [Source: |
| RPL21P43 | 3 | ribosomal protein L21 pseudogene 43 [Source |
| LINC02082 | 3 | long intergenic non-protein coding RNA 2082 |
| AC092954.1 | 3 | novel transcript |
| AC092954.2 | 3 | novel transcript |
| LINC01997 | 3 | long intergenic non-protein coding RNA 1997 |
| MECOM | 3 | MDS1 and EVI1 complex locus [Source:HGNC S |
| AC074033.1 | 3 | novel transcript |
| RPL22P1 | 3 | ribosomal protein L22 pseudogene 1 [Source: |
| AC007849.1 | 3 | novel transcript |
| AC007849.2 | 3 | succinate dehydrogenase complex, subunit D, |
| RNU6-637P | 3 | RNA, U6 small nuclear 637, pseudogene [Sour |
| TERC | 3 | telomerase RNA component [Source:HGNC Sy |
| Telomerase-vert | 3 | Vertebrate telomerase RNA [Source:RFAM;Ac |
| ACTRT3 | 3 | actin related protein T3 [Source:HGNC Symbo |
| AC078802.1 | 3 | novel transcript, antisense to ACTRT3 |
| MYNN | 3 | myoneurin [Source:HGNC Symbol;Acc:HGNC:1 |
| AC078795.1 | 3 | novel transcript, antisense to MYNN |
| LRRC34 | 3 | leucine rich repeat containing 34 [Source:HGN |
| AC078795.3 | 3 | novel transcript, antisense to LRRC34 |
| AC078795.2 | 3 | novel transcript, antisense to LRRC34 |
| LRRIQ4 | 3 | leucine rich repeats and IQ motif containing 4 |
| LRRC31 | 3 | leucine rich repeat containing 31 [Source:HGN |
| KRT18P43 | 3 | keratin 18 pseudogene 43 [Source:HGNC Sym |
| SAMD7 | 3 | sterile alpha motif domain containing 7 [Sourc |
| AC008040.3 | 3 | family with sequence similarity 20, member B |
| AC008040.1 | 3 | novel transcript, antisense to SEC62 |
| AC008040.4 | 3 | four and a half LIM domains 1 (FHL1) pseudog |

| | | |
|---|---|---|
| SEC62 | 3 | SEC62 homolog, preprotein translocation fact |
| SEC62-AS1 | 3 | SEC62 antisense RNA 1 [Source:HGNC Symbol |
| GPR160 | 3 | G protein-coupled receptor 160 [Source:HGN( |
| RNU4-38P | 3 | RNA, U4 small nuclear 38, pseudogene [Sourc |
| AC008040.2 | 3 | SET domain containing (lysine methyltransfera |
| PHC3 | 3 | polyhomeotic homolog 3 [Source:HGNC Symb |
| RNU6-315P | 3 | RNA, U6 small nuclear 315, pseudogene [Sour |
| PRKCI | 3 | protein kinase C iota [Source:HGNC Symbol;A |
| Y_RNA | 3 | Y RNA [Source:RFAM;Acc:RF00019] |
| AC073288.1 | 3 | novel transcript |
| SKIL | 3 | SKI like proto-oncogene [Source:HGNC Symbo |
| AC073288.2 | 3 | novel transcript |
| CLDN11 | 3 | claudin 11 [Source:HGNC Symbol;Acc:HGNC:8 |
| AC026316.4 | 3 | novel protein |
| MIR6828 | 3 | microRNA 6828 [Source:HGNC Symbol;Acc:HG |
| SLC7A14-AS1 | 3 | SLC7A14 antisense RNA 1 [Source:HGNC Symb |
| SLC7A14 | 3 | solute carrier family 7 member 14 [Source:HG |
| KRT8P13 | 3 | keratin 8 pseudogene 13 [Source:HGNC Symb |
| AC026316.1 | 3 | ribosomal protein L28 (RPL28) pseudogene |
| AC026316.3 | 3 | novel transcript |
| AC026316.5 | 3 | novel transcript |
| AC026316.2 | 3 | novel transcript |
| RPL22L1 | 3 | ribosomal protein L22 like 1 [Source:HGNC Sy |
| EIF5A2 | 3 | eukaryotic translation initiation factor 5A2 [Sc |
| AC061708.1 | 3 | novel transcript, antisense to SLC2A2 |
| KLF7P1 | 3 | kruppel like factor 7 pseudogene 1 [Source:H( |
| RNU1-70P | 3 | RNA, U1 small nuclear 70, pseudogene [Sourc |
| SLC2A2 | 3 | solute carrier family 2 member 2 [Source:HGN |
| TNIK | 3 | TRAF2 and NCK interacting kinase [Source:HG |
| MIR569 | 3 | microRNA 569 [Source:HGNC Symbol;Acc:HGI |
| Y_RNA | 3 | Y RNA [Source:RFAM;Acc:RF00019] |
| AC092919.1 | 3 | novel transcript |
| AC092919.2 | 3 | TEC |
| RNU6-348P | 3 | RNA, U6 small nuclear 348, pseudogene [Sour |
| AC008134.1 | 3 | pseudogene similar to part of mitochondrially |
| PLD1 | 3 | phospholipase D1 [Source:HGNC Symbol;Acc:I |
| TMEM212-AS1 | 3 | TMEM212 antisense RNA 1 [Source:HGNC Syn |
| TMEM212 | 3 | transmembrane protein 212 [Source:HGNC Sy |
| TMEM212-IT1 | 3 | TMEM212 intronic transcript 1 [Source:HGNC |
| AC055714.1 | 3 | novel transcript |
| FNDC3B | 3 | fibronectin type III domain containing 3B [Sou |
| RPS27AP8 | 3 | ribosomal protein S27a pseudogene 8 [Source |
| RN7SL141P | 3 | RNA, 7SL, cytoplasmic 141, pseudogene [Sour |
| AC092964.1 | 3 | novel transcript |
| BZW1P1 | 3 | basic leucine zipper and W2 domains 1 pseud |
| GHSR | 3 | growth hormone secretagogue receptor [Sour |
| TNFSF10 | 3 | TNF superfamily member 10 [Source:HGNC Sy |
| AC007919.1 | 3 | pseudogene similar to part of TBP-like 1 TBPL: |
| LINC02068 | 3 | long intergenic non-protein coding RNA 2068 |
| SLC31A1P1 | 3 | solute carrier family 31 member 1 pseudogen |

| | | |
|---|---|---|
| NCEH1 | 3 | neutral cholesterol ester hydrolase 1 [Source: |
| AC007919.2 | 3 | nucleosome assembly protein 1-like 5 (NAP1L |
| RNU6-547P | 3 | RNA, U6 small nuclear 547, pseudogene [Sour |
| AC108667.2 | 3 | novel transcript |
| ECT2 | 3 | epithelial cell transforming 2 [Source:HGNC Sy |
| SNORA72 | 3 | Small nucleolar RNA SNORA72 [Source:RFAM; |
| RNU4-4P | 3 | RNA, U4 small nuclear 4, pseudogene [Source |
| ATP5MC1P4 | 3 | ATP synthase membrane subunit c locus 1 pse |
| AC108667.1 | 3 | pseudogene similar to part of methylmalonic |
| SPATA16 | 3 | spermatogenesis associated 16 [Source:HGNC |
| AC068759.1 | 3 | novel transcript |
| NLGN1 | 3 | neuroligin 1 [Source:HGNC Symbol;Acc:HGNC |
| AC092967.1 | 3 | chromosome 14 open reading frame 83 (C14o |
| NLGN1-AS1 | 3 | NLGN1 antisense RNA 1 [Source:HGNC Symbo |
| RN7SKP234 | 3 | RN7SK pseudogene 234 [Source:HGNC Symbo |
| AC069218.2 | 3 | chromosome 11 open reading frame 51 (C11o |
| AC069218.1 | 3 | ribosomal protein L8 (RPL8) pseudogene |
| NAALADL2 | 3 | N-acetylated alpha-linked acidic dipeptidase li |
| RN7SKP40 | 3 | RN7SK pseudogene 40 [Source:HGNC Symbol; |
| EEF1B2P8 | 3 | eukaryotic translation elongation factor 1 beta |
| NAALADL2-AS3 | 3 | NAALADL2 antisense RNA 3 [Source:HGNC Syn |
| NAALADL2-AS2 | 3 | NAALADL2 antisense RNA 2 [Source:HGNC Syn |
| MIR4789 | 3 | microRNA 4789 [Source:HGNC Symbol;Acc:HG |
| RNU6-1233P | 3 | RNA, U6 small nuclear 1233, pseudogene [Sou |
| AC008180.2 | 3 | novel transcript, antisense to NAALADL2 |
| RNU4-91P | 3 | RNA, U4 small nuclear 91, pseudogene [Sourc |
| AC119744.1 | 3 | DEAD (Asp-Glu-Ala-Asp) box polypeptide 5 (DI |
| UBE2V1P2 | 3 | ubiquitin conjugating enzyme E2 V1 pseudoge |
| RNU6-1317P | 3 | RNA, U6 small nuclear 1317, pseudogene [Sou |
| NAALADL2-AS1 | 3 | NAALADL2 antisense RNA 1 [Source:HGNC Syn |
| AC104640.1 | 3 | novel transcript |
| ACTG1P23 | 3 | actin gamma 1 pseudogene 23 [Source:HGNC |
| EI24P1 | 3 | EI24 pseudogene 1 [Source:HGNC Symbol;Acc |
| AC104640.2 | 3 | etoposide induced 2.4, pseudogene |
| AC117434.1 | 3 | transmembrane protein 38B (TMEM38B) pseu |
| LINC01208 | 3 | long intergenic non-protein coding RNA 1208 |
| MIR7977 | 3 | microRNA 7977 [Source:HGNC Symbol;Acc:HG |
| RNA5SP147 | 3 | RNA, 5S ribosomal pseudogene 147 [Source:H |
| LINC01209 | 3 | long intergenic non-protein coding RNA 1209 |
| MTND5P15 | 3 | MT-ND5 pseudogene 15 [Source:HGNC Symbo |
| TBL1XR1 | 3 | transducin beta like 1 X-linked receptor 1 [Sou |
| TBL1XR1-AS1 | 3 | TBL1XR1 antisense RNA 1 [Source:HGNC Symb |
| Y_RNA | 3 | Y RNA [Source:RFAM;Acc:RF00019] |
| RNU6-681P | 3 | RNA, U6 small nuclear 681, pseudogene [Sour |
| LINC00501 | 3 | long intergenic non-protein coding RNA 501 [S |
| ASS1P7 | 3 | argininosuccinate synthetase 1 pseudogene 7 |
| AC117465.1 | 3 | novel transcript |
| LINC00578 | 3 | long intergenic non-protein coding RNA 578 [S |
| RN7SKP52 | 3 | RN7SK pseudogene 52 [Source:HGNC Symbol; |
| AC026355.3 | 3 | novel transcript |

| | | |
|---|---|---|
| AC026355.1 | 3 | NULL |
| AC026355.2 | 3 | novel transcript |
| AC026355.4 | 3 | fibroblast growth factor receptor 3 (FGFR3) ps |
| LINC02015 | 3 | long intergenic non-protein coding RNA 2015 |
| AC007953.1 | 3 | novel transcript |
| AC007953.2 | 3 | novel transcript |
| RNU6-1120P | 3 | RNA, U6 small nuclear 1120, pseudogene [Sou |
| AC110992.1 | 3 | novel transcript |
| AC117453.1 | 3 | novel transcript, antisense to KCNMB2 |
| KCNMB2 | 3 | potassium calcium-activated channel subfami |
| AC117457.1 | 3 | potassium large conductance calcium-activate |
| LINC01014 | 3 | long intergenic non-protein coding RNA 1014 |
| RNA5SP148 | 3 | RNA, 5S ribosomal pseudogene 148 [Source:H |
| PPIAP75 | 3 | peptidylprolyl isomerase A pseudogene 75 [Sc |
| KCNMB2-AS1 | 3 | KCNMB2 antisense RNA 1 [Source:HGNC Symb |
| ZMAT3 | 3 | zinc finger matrin-type 3 [Source:HGNC Symb |
| Y_RNA | 3 | Y RNA [Source:RFAM;Acc:RF00019] |
| AC076966.2 | 3 | novel transcript |
| PIK3CA | 3 | phosphatidylinositol-4,5-bisphosphate 3-kinas |
| AC076966.1 | 3 | NULL |
| KCNMB3 | 3 | potassium calcium-activated channel subfami |
| LRRFIP1P1 | 3 | LRR binding FLII interacting protein 1 pseudog |
| ZNF639 | 3 | zinc finger protein 639 [Source:HGNC Symbol; |
| AC007823.1 | 3 | novel transcript |
| MFN1 | 3 | mitofusin 1 [Source:HGNC Symbol;Acc:HGNC: |
| GNB4 | 3 | G protein subunit beta 4 [Source:HGNC Symb |
| AC007620.2 | 3 | novel transcript |
| AC007620.3 | 3 | novel transcript, antisense to GNB4 |
| AC007620.1 | 3 | pseudogene similar to part of guanine nucleot |
| MTHFD2P7 | 3 | methylenetetrahydrofolate dehydrogenase (N |
| ACTL6A | 3 | actin like 6A [Source:HGNC Symbol;Acc:HGNC |
| AC090425.3 | 3 | novel transcript, antisense to ACTL6A |
| AC090425.2 | 3 | novel transcript, antisense to ACTL6A |
| MRPL47 | 3 | mitochondrial ribosomal protein L47 [Source: |
| NDUFB5 | 3 | NADH:ubiquinone oxidoreductase subunit B5 |
| AC090425.1 | 3 | NULL |
| AC125604.1 | 3 | histone H3-like pseudogene |
| USP13 | 3 | ubiquitin specific peptidase 13 [Source:HGNC |
| PEX5L | 3 | peroxisomal biogenesis factor 5 like [Source:H |
| AC007687.1 | 3 | novel transcript |
| PEX5L-AS1 | 3 | PEX5L antisense RNA 1 [Source:HGNC Symbol |
| PEX5L-AS2 | 3 | PEX5L antisense RNA 2 [Source:HGNC Symbol |
| AC092939.1 | 3 | novel transcript, antisense to PEX5L |
| RNA5SP149 | 3 | RNA, 5S ribosomal pseudogene 149 [Source:H |
| AC134503.1 | 3 | pseudogene similar to part of glyceraldehyde- |
| GAPDHP36 | 3 | glyceraldehyde 3 phosphate dehydrogenase p |
| AC068760.1 | 3 | ralA binding protein 1 (RALBP1) pseudogene |
| LINC02053 | 3 | long intergenic non-protein coding RNA 2053 |
| U8 | 3 | U8 small nucleolar RNA [Source:RFAM;Acc:RF |
| AC125618.1 | 3 | novel transcript |

| | | |
|---|---|---|
| RNU6-486P | 3 | RNA, U6 small nuclear 486, pseudogene [Sour |
| AC068298.1 | 3 | pseudogene similar to part of proteasome (pr |
| TTC14 | 3 | tetratricopeptide repeat domain 14 [Source:H |
| CCDC39 | 3 | coiled-coil domain containing 39 [Source:HGN |
| CCDC39 | 3 | coiled-coil domain containing 39 [Source:HGN |
| CCDC39-AS1 | 3 | CCDC39 antisense RNA 1 [Source:HGNC Symb |
| AC108734.4 | 3 | novel transcript |
| RN7SL229P | 3 | RNA, 7SL, cytoplasmic 229, pseudogene [Sour |
| AC108734.3 | 3 | ring finger protein 13 (RNF13) pseudogene |
| AC108734.2 | 3 | actin, beta (ACTB) pseudogene |
| FLYWCH1P1 | 3 | FLYWCH-type zinc finger 1 pseudogene 1 [Sou |
| FXR1 | 3 | FMR1 autosomal homolog 1 [Source:HGNC Sy |
| AC108734.1 | 3 | ribosomal protein L32 (RPL32) pseudogene |
| DNAJC19 | 3 | DnaJ heat shock protein family (Hsp40) memb |
| SOX2-OT | 3 | SOX2 overlapping transcript [Source:HGNC Sy |
| AC083904.1 | 3 | matrin 3 (MATR3) pseudogene |
| RNU6-4P | 3 | RNA, U6 small nuclear 4, pseudogene [Source |
| FAUP2 | 3 | FAU pseudogene 2 [Source:HGNC Symbol;Acc |
| AC068308.1 | 3 | novel transcript |
| RPL7AP25 | 3 | ribosomal protein L7a pseudogene 25 [Source |
| AC125613.1 | 3 | novel transcript |
| AC125613.2 | 3 | NULL |
| AC117415.1 | 3 | NULL |
| SOX2 | 3 | SRY-box transcription factor 2 [Source:HGNC S |
| AC117415.2 | 3 | NULL |
| RN7SL703P | 3 | RNA, 7SL, cytoplasmic 703, pseudogene [Sour |
| AC007547.3 | 3 | NULL |
| RNA5SP150 | 3 | RNA, 5S ribosomal pseudogene 150 [Source:H |
| AC007547.2 | 3 | proteasome (prosome, macropain) assembly |
| AC007547.1 | 3 | zinc finger, CCHC domain containing 10 (ZCCH |
| LINC01206 | 3 | long intergenic non-protein coding RNA 1206 |
| RN7SKP265 | 3 | RN7SK pseudogene 265 [Source:HGNC Symbo |
| AC012081.2 | 3 | novel transcript |
| AC012081.1 | 3 | novel transcript |
| AC109131.1 | 3 | novel transcript |
| AC084211.1 | 3 | novel transcript |
| LINC01994 | 3 | long intergenic non-protein coding RNA 1994 |
| EIF4EP3 | 3 | eukaryotic translation initiation factor 4E pseu |
| LINC01995 | 3 | long intergenic non-protein coding RNA 1995 |
| AC021055.1 | 3 | novel transcript |
| RPL7L1P8 | 3 | ribosomal protein L7 like 1 pseudogene 8 [Sou |
| LINC02031 | 3 | long intergenic non-protein coding RNA 2031 |
| AC083801.1 | 3 | phospholipase A2, group X (PLA2G10) pseudo |
| AC083801.2 | 3 | novel transcript |
| AC069431.1 | 3 | novel transcript |
| ATP11B | 3 | ATPase phospholipid transporting 11B (putati |
| AC069431.2 | 3 | eukaryotic translation initiation factor 3, subu |
| AC092953.2 | 3 | novel transcript, antisense to ATP11B |
| DCUN1D1 | 3 | defective in cullin neddylation 1 domain conta |
| AC092953.1 | 3 | chromosome 9 open reading frame 85 (C9orf8 |

| | | |
|---|---|---|
| MCCC1 | 3 | methylcrotonoyl-CoA carboxylase 1 [Source:H |
| MCCC1-AS1 | 3 | MCCC1 antisense RNA 1 [Source:HGNC Symbo |
| LAMP3 | 3 | lysosomal associated membrane protein 3 [Sc |
| MCF2L2 | 3 | MCF.2 cell line derived transforming sequence |
| B3GNT5 | 3 | UDP-GlcNAc:betaGal beta-1,3-N-acetylglucosa |
| RNA5SP151 | 3 | RNA, 5S ribosomal pseudogene 151 [Source:H |
| LINC00888 | 3 | long intergenic non-protein coding RNA 888 [S |
| SNORA63D | 3 | small nucleolar RNA, H/ACA box 63D [Source: |
| AC092960.2 | 3 | NULL |
| SNORA63E | 3 | small nucleolar RNA, H/ACA box 63E [Source:I |
| AC092960.1 | 3 | NULL |
| AC092960.3 | 3 | TVP23B trans-golgi network vesicle protein 23 |
| KLHL6 | 3 | kelch like family member 6 [Source:HGNC Sym |
| KLHL6-AS1 | 3 | KLHL6 antisense RNA 1 [Source:HGNC Symbol |
| AC068769.1 | 3 | prickle homolog 1 (Drosophila) (PRICKLE1) pse |
| KLHL24 | 3 | kelch like family member 24 [Source:HGNC Sy |
| YEATS2 | 3 | YEATS domain containing 2 [Source:HGNC Syn |
| YEATS2-AS1 | 3 | YEATS2 antisense RNA 1 [Source:HGNC Symbo |
| AC131160.1 | 3 | presenilin associated rhomboid like [Source:N |
| MAP6D1 | 3 | MAP6 domain containing 1 [Source:HGNC Syn |
| PARL | 3 | presenilin associated rhomboid like [Source:H |
| RPSAP31 | 3 | ribosomal protein SA pseudogene 31 [Source: |
| MIR4448 | 3 | microRNA 4448 [Source:HGNC Symbol;Acc:HG |
| CYP2AB1P | 3 | cytochrome P450 family 2 subfamily AB meml |
| ABCC5 | 3 | ATP binding cassette subfamily C member 5 [S |
| ABCC5-AS1 | 3 | ABCC5 antisense RNA 1 [Source:HGNC Symbo |
| EEF1A1P8 | 3 | eukaryotic translation elongation factor 1 alpl |
| HTR3D | 3 | 5-hydroxytryptamine receptor 3D [Source:HG |
| HTR3C | 3 | 5-hydroxytryptamine receptor 3C [Source:HGI |
| HTR3C2P | 3 | 5-hydroxytryptamine receptor 3C2, pseudoge |
| Y_RNA | 3 | Y RNA [Source:RFAM;Acc:RF00019] |
| HTR3E-AS1 | 3 | HTR3E antisense RNA 1 [Source:HGNC Symbol |
| HTR3E | 3 | 5-hydroxytryptamine receptor 3E [Source:HGI |
| HSP90AA5P | 3 | heat shock protein 90 alpha family class A me |
| AC131235.3 | 3 | novel transcript |
| AC131235.2 | 3 | novel transcript, antisense to EIF2B5 |
| EIF2B5 | 3 | eukaryotic translation initiation factor 2B subu |
| AC131235.1 | 3 | ribosomal protein S3 (RPS3) pseudogene |
| DVL3 | 3 | dishevelled segment polarity protein 3 [Sourc |
| AP2M1 | 3 | adaptor related protein complex 2 subunit mu |
| ABCF3 | 3 | ATP binding cassette subfamily F member 3 [S |
| VWA5B2 | 3 | von Willebrand factor A domain containing 5E |
| MIR1224 | 3 | microRNA 1224 [Source:HGNC Symbol;Acc:HG |
| ALG3 | 3 | ALG3 alpha-1,3- mannosyltransferase [Source |
| EEF1AKMT4 | 3 | EEF1A lysine methyltransferase 4 [Source:HGI |
| EEF1AKMT4-ECE2 | 3 | EEF1AKMT4-ECE2 readthrough [Source:HGNC |
| CAMK2N2 | 3 | calcium/calmodulin dependent protein kinase |
| ECE2 | 3 | endothelin converting enzyme 2 [Source:HGN |
| PSMD2 | 3 | proteasome 26S subunit, non-ATPase 2 [Sourc |
| EIF4G1 | 3 | eukaryotic translation initiation factor 4 gamn |

| | | |
|---|---|---|
| SNORD66 | 3 | small nucleolar RNA, C/D box 66 [Source:HGN |
| FAM131A | 3 | family with sequence similarity 131 member A |
| CLCN2 | 3 | chloride voltage-gated channel 2 [Source:HGN |
| POLR2H | 3 | RNA polymerase II subunit H [Source:HGNC Sy |
| THPO | 3 | thrombopoietin [Source:HGNC Symbol;Acc:HG |
| CHRD | 3 | chordin [Source:HGNC Symbol;Acc:HGNC:194 |
| LINC02054 | 3 | long intergenic non-protein coding RNA 2054 |
| AC128714.1 | 3 | transmembrane epididymal protein 1 (TEDDM |
| LINC01839 | 3 | long intergenic non-protein coding RNA 1839 |
| LINC01840 | 3 | long intergenic non-protein coding RNA 1840 |
| EPHB3 | 3 | EPH receptor B3 [Source:HGNC Symbol;Acc:H |
| MAGEF1 | 3 | MAGE family member F1 [Source:HGNC Symb |
| AC107294.2 | 3 | uncharacterized LOC107986163 [Source:NCBI |
| LINC02069 | 3 | long intergenic non-protein coding RNA 2069 |
| AC107294.3 | 3 | novel transcript |
| AC107294.1 | 3 | novel transcript |
| VPS8 | 3 | VPS8 subunit of CORVET complex [Source:HGI |
| C3orf70 | 3 | chromosome 3 open reading frame 70 [Source |
| EHHADH-AS1 | 3 | EHHADH antisense RNA 1 [Source:HGNC Symb |
| EHHADH | 3 | enoyl-CoA hydratase and 3-hydroxyacyl CoA d |
| EIF2S2P2 | 3 | eukaryotic translation initiation factor 2 subur |
| MIR5588 | 3 | microRNA 5588 [Source:HGNC Symbol;Acc:HG |
| MAP3K13 | 3 | mitogen-activated protein kinase kinase kinas |
| RPL4P4 | 3 | ribosomal protein L4 pseudogene 4 [Source:H |
| TMEM41A | 3 | transmembrane protein 41A [Source:HGNC Sy |
| AC099661.1 | 3 | novel transcript |
| LIPH | 3 | lipase H [Source:HGNC Symbol;Acc:HGNC:184 |
| SENP2 | 3 | SUMO specific peptidase 2 [Source:HGNC Sym |
| AC133473.1 | 3 | ribosomal protein L34 (RPL34) pseudogene |
| IGF2BP2 | 3 | insulin like growth factor 2 mRNA binding pro |
| IGF2BP2-AS1 | 3 | IGF2BP2 antisense RNA 1 [Source:HGNC Symb |
| MIR548AQ | 3 | microRNA 548aq [Source:HGNC Symbol;Acc:H |
| TRA2B | 3 | transformer 2 beta homolog [Source:HGNC Sy |
| NMRAL2P | 3 | NmrA like redox sensor 2, pseudogene [Source |
| ETV5 | 3 | ETS variant 5 [Source:HGNC Symbol;Acc:HGN( |
| Metazoa_SRP | 3 | Metazoan signal recognition particle RNA [Sou |
| ETV5-AS1 | 3 | ETV5 antisense RNA 1 [Source:HGNC Symbol;A |
| DGKG | 3 | diacylglycerol kinase gamma [Source:HGNC Sy |
| LINC02020 | 3 | long intergenic non-protein coding RNA 2020 |
| LINC02052 | 3 | long intergenic non-protein coding RNA 2052 |
| LINC02051 | 3 | long intergenic non-protein coding RNA 2051 |
| CRYGS | 3 | crystallin gamma S [Source:HGNC Symbol;Acc |
| TBCCD1 | 3 | TBCC domain containing 1 [Source:HGNC Sym |
| DNAJB11 | 3 | DnaJ heat shock protein family (Hsp40) memb |
| AC068631.1 | 3 | novel transcript, antisense to KNG1, HRG, FET |
| AC068631.2 | 3 | novel protein |
| AC068631.3 | 3 | PET100 homolog (PET100) pseudogene |
| AHSG | 3 | alpha 2-HS glycoprotein [Source:HGNC Symbo |
| Metazoa_SRP | 3 | Metazoan signal recognition particle RNA [Sou |
| FETUB | 3 | fetuin B [Source:HGNC Symbol;Acc:HGNC:365 |

| | | |
|---|---|---|
| HRG | 3 | histidine rich glycoprotein [Source:HGNC Sym |
| AC109780.1 | 3 | histidine-rich glycoprotein (HRG) pseudogene |
| AC109780.2 | 3 | histidine-rich glycoprotein (HRG) pseudogene |
| KNG1 | 3 | kininogen 1 [Source:HGNC Symbol;Acc:HGNC: |
| PSMD10P2 | 3 | proteasome 26S subunit, non-ATPase, 10 pseu |
| RNU6-1105P | 3 | RNA, U6 small nuclear 1105, pseudogene [Sou |
| GPS2P2 | 3 | G protein pathway suppressor 2 pseudogene |
| AC112907.3 | 3 | novel transcript, antisense to eukaryotic trans |
| EIF4A2 | 3 | eukaryotic translation initiation factor 4A2 [Sc |
| SNORD2 | 3 | small nucleolar RNA, C/D box 2 [Source:HGNC |
| SNORA63B | 3 | small nucleolar RNA, H/ACA box 63B [Source: |
| MIR1248 | 3 | microRNA 1248 [Source:HGNC Symbol;Acc:HG |
| SNORA81 | 3 | small nucleolar RNA, H/ACA box 81 [Source:H |
| SNORA63 | 3 | small nucleolar RNA, H/ACA box 63 [Source:H |
| SNORA4 | 3 | small nucleolar RNA, H/ACA box 4 [Source:HG |
| RFC4 | 3 | replication factor C subunit 4 [Source:HGNC S |
| AC112907.1 | 3 | novel transcript |
| LINC02043 | 3 | long intergenic non-protein coding RNA 2043 |
| AC112907.2 | 3 | MCF.2 cell line derived transforming sequence |
| ADIPOQ | 3 | adiponectin, C1Q and collagen domain contai |
| ADIPOQ-AS1 | 3 | ADIPOQ antisense RNA 1 [Source:HGNC Symb |
| RPS20P14 | 3 | ribosomal protein S20 pseudogene 14 [Source |
| AC007690.1 | 3 | ribosomal protein L29 (RPL29) pseudogene |
| ST6GAL1 | 3 | ST6 beta-galactoside alpha-2,6-sialyltransfera |
| RPL39L | 3 | ribosomal protein L39 like [Source:HGNC Sym |
| AC007920.1 | 3 | novel transcript |
| RTP1 | 3 | receptor transporter protein 1 [Source:HGNC |
| MASP1 | 3 | mannan binding lectin serine peptidase 1 [Sou |
| AC007920.2 | 3 | novel transcript, antisense to MASP1 |
| RTP4 | 3 | receptor transporter protein 4 [Source:HGNC |
| AC068299.2 | 3 | TEC |
| AC068299.1 | 3 | NULL |
| LINC02041 | 3 | long intergenic non-protein coding RNA 2041 |
| SST | 3 | somatostatin [Source:HGNC Symbol;Acc:HGN |
| RTP2 | 3 | receptor transporter protein 2 [Source:HGNC |
| AC072022.1 | 3 | novel transcript, antisense to RTP2 and BCL6 |
| BCL6 | 3 | BCL6 transcription repressor [Source:HGNC Sy |
| AC072022.2 | 3 | BCL6 antisense 1 |
| AC108681.1 | 3 | novel transcript |
| AC068295.1 | 3 | novel transcript |
| LINC01991 | 3 | long intergenic non-protein coding RNA 1991 |
| AC092941.1 | 3 | novel transcript |
| AC092941.2 | 3 | novel transcript |
| AC022498.1 | 3 | novel transcript |
| LPP-AS2 | 3 | LPP antisense RNA 2 [Source:HGNC Symbol;Ac |
| LPP | 3 | LIM domain containing preferred translocatio |
| AC022498.2 | 3 | novel transcript |
| LPP-AS1 | 3 | LPP antisense RNA 1 [Source:HGNC Symbol;Ac |
| MIR28 | 3 | microRNA 28 [Source:HGNC Symbol;Acc:HGN |
| TPRG1-AS1 | 3 | TPRG1 antisense RNA 1 [Source:HGNC Symbo |

| | | |
|---|---|---|
| TPRG1 | 3 | tumor protein p63 regulated 1 [Source:HGNC |
| AC009319.1 | 3 | enolase-phosphatase 1 (ENOPH1) pseudogen |
| TPRG1-AS2 | 3 | TPRG1 antisense RNA 2 [Source:HGNC Symbo |
| TP63 | 3 | tumor protein p63 [Source:HGNC Symbol;Acc |
| AC078809.1 | 3 | serine/arginine-rich splicing factor 10 (SRSF10 |
| MIR944 | 3 | microRNA 944 [Source:HGNC Symbol;Acc:HGI |
| P3H2 | 3 | prolyl 3-hydroxylase 2 [Source:HGNC Symbol; |
| MTAPP2 | 3 | methylthioadenosine phosphorylase pseudog |
| P3H2-AS1 | 3 | P3H2 antisense RNA 1 [Source:HGNC Symbol; |
| RNU6-1109P | 3 | RNA, U6 small nuclear 1109, pseudogene [Sou |
| NMNAT1P3 | 3 | NMNAT1 pseudogene 3 [Source:HGNC Symbo |
| CLDN1 | 3 | claudin 1 [Source:HGNC Symbol;Acc:HGNC:20 |
| CLDN16 | 3 | claudin 16 [Source:HGNC Symbol;Acc:HGNC:2 |
| TMEM207 | 3 | transmembrane protein 207 [Source:HGNC Sy |
| IL1RAP | 3 | interleukin 1 receptor accessory protein [Sour |
| GCNT1P3 | 3 | glucosaminyl (N-acetyl) transferase 1 pseudog |
| RN7SKP296 | 3 | RN7SK pseudogene 296 [Source:HGNC Symbo |
| AC108747.1 | 3 | novel transcript |
| LINC02013 | 3 | long intergenic non-protein coding RNA 2013 |
| CCT6P4 | 3 | chaperonin containing TCP1 subunit 6 pseudo |
| AC092952.1 | 3 | enabled homolog (Drosophila) (ENAH) pseudc |
| GMNC | 3 | geminin coiled-coil domain containing [Source |
| OSTN | 3 | osteocrin [Source:HGNC Symbol;Acc:HGNC:29 |
| OSTN-AS1 | 3 | OSTN antisense RNA 1 [Source:HGNC Symbol; |
| UTS2B | 3 | urotensin 2B [Source:HGNC Symbol;Acc:HGN |
| CCDC50 | 3 | coiled-coil domain containing 50 [Source:HGN |
| AC073365.1 | 3 | novel transcript |
| PYDC2 | 3 | pyrin domain containing 2 [Source:HGNC Sym |
| Y_RNA | 3 | Y RNA [Source:RFAM;Acc:RF00019] |
| AC026320.2 | 3 | PERP, TP53 apoptosis effector (PERP) pseudo |
| AC026320.1 | 3 | novel transcript |
| AC026320.3 | 3 | novel transcript |
| RN7SKP222 | 3 | RN7SK pseudogene 222 [Source:HGNC Symbo |
| FGF12 | 3 | fibroblast growth factor 12 [Source:HGNC Syn |
| FGF12-AS1 | 3 | FGF12 antisense RNA 1 [Source:HGNC Symbol |
| FGF12-AS2 | 3 | FGF12 antisense RNA 2 [Source:HGNC Symbol |
| FGF12-AS3 | 3 | FGF12 antisense RNA 3 [Source:HGNC Symbol |
| AC026671.1 | 3 | NULL |
| RNU1-20P | 3 | RNA, U1 small nuclear 20, pseudogene [Sourc |
| MB21D2 | 3 | Mab-21 domain containing 2 [Source:HGNC S |
| AC092966.1 | 3 | novel transcript |
| VEZF1P1 | 3 | vascular endothelial zinc finger 1 pseudogene |
| PLAAT1 | 3 | phospholipase A and acyltransferase 1 [Sourc |
| ATP13A5 | 3 | ATPase 13A5 [Source:HGNC Symbol;Acc:HGN |
| ATP13A5-AS1 | 3 | ATP13A5 antisense RNA 1 [Source:HGNC Sym |
| ATP13A4 | 3 | ATPase 13A4 [Source:HGNC Symbol;Acc:HGN |
| ATP13A4-AS1 | 3 | ATP13A4 antisense RNA 1 [Source:HGNC Sym |
| AC048351.1 | 3 | eukaryotic translation elongation factor 1 alp |
| OPA1 | 3 | OPA1 mitochondrial dynamin like GTPase [Sou |
| OPA1-AS1 | 3 | OPA1 antisense RNA 1 [Source:HGNC Symbol; |

| | | |
|---|---|---|
| Y_RNA | 3 | Y RNA [Source:RFAM;Acc:RF00019] |
| RN7SL447P | 3 | RNA, 7SL, cytoplasmic 447, pseudogene [Sour |
| AC069421.1 | 3 | cytochrome c oxidase subunit VIa polypeptide |
| AC069421.2 | 3 | novel transcript |
| AC069421.3 | 3 | novel transcript |
| LINC02038 | 3 | long intergenic non-protein coding RNA 2038 |
| LINC02026 | 3 | long intergenic non-protein coding RNA 2026 |
| AC024559.1 | 3 | novel transcript |
| DPPA2P3 | 3 | developmental pluripotency associated 2 pseu |
| LINC02028 | 3 | long intergenic non-protein coding RNA 2028 |
| AC024559.2 | 3 | novel transcript |
| AC080129.1 | 3 | novel transcript |
| AC080129.2 | 3 | novel transcript |
| HES1 | 3 | hes family bHLH transcription factor 1 [Source |
| RN7SL215P | 3 | RNA, 7SL, cytoplasmic 215, pseudogene [Sour |
| LINC02036 | 3 | long intergenic non-protein coding RNA 2036 |
| LINC02037 | 3 | long intergenic non-protein coding RNA 2037 |
| LINC02048 | 3 | long intergenic non-protein coding RNA 2048 |
| LINC00887 | 3 | long intergenic non-protein coding RNA 887 [S |
| AC117469.1 | 3 | novel transcript |
| CPN2 | 3 | carboxypeptidase N subunit 2 [Source:HGNC S |
| LRRC15 | 3 | leucine rich repeat containing 15 [Source:HGN |
| GP5 | 3 | glycoprotein V platelet [Source:HGNC Symbol; |
| ATP13A3 | 3 | ATPase 13A3 [Source:HGNC Symbol;Acc:HGN( |
| AC108676.2 | 3 | UBC ubiquitin C, pseudogene |
| LINC00884 | 3 | long intergenic non-protein coding RNA 884 [S |
| AC108676.1 | 3 | novel transcript, sense overlapping to ATP13A |
| RNU6-1101P | 3 | RNA, U6 small nuclear 1101, pseudogene [Sou |
| RPL23AP93 | 3 | ribosomal protein L23a pseudogene 93 [Sourc |
| TMEM44-AS1 | 3 | TMEM44 antisense RNA 1 [Source:HGNC Sym |
| TMEM44 | 3 | transmembrane protein 44 [Source:HGNC Syn |
| AC046143.1 | 3 | novel transcript |
| AC046143.2 | 3 | novel transcript |
| LSG1 | 3 | large 60S subunit nuclear export GTPase 1 [So |
| FAM43A | 3 | family with sequence similarity 43 member A |
| AC106706.1 | 3 | novel transcript |
| AC106706.2 | 3 | novel transcript |
| LINC01968 | 3 | long intergenic non-protein coding RNA 1968 |
| AC090505.1 | 3 | family with sequence similarity 151, member |
| LINC01972 | 3 | long intergenic non-protein coding RNA 1972 |
| AC090505.2 | 3 | novel transcript |
| XXYLT1 | 3 | xyloside xylosyltransferase 1 [Source:HGNC Sy |
| XXYLT1-AS1 | 3 | XXYLT1 antisense RNA 1 [Source:HGNC Symbo |
| MIR3137 | 3 | microRNA 3137 [Source:HGNC Symbol;Acc:HG |
| RN7SL36P | 3 | RNA, 7SL, cytoplasmic 36, pseudogene [Sourc |
| XXYLT1-AS2 | 3 | XXYLT1 antisense RNA 2 [Source:HGNC Symbo |
| RNU6-25P | 3 | RNA, U6 small nuclear 25, pseudogene [Sourc |
| AC090018.2 | 3 | novel transcript, antisense to XXYLT1 |
| ACAP2 | 3 | ArfGAP with coiled-coil, ankyrin repeat and PH |
| ACAP2-IT1 | 3 | ACAP2 intronic transcript 1 [Source:HGNC Syn |

| | | |
|---|---|---|
| AC090018.1 | 3 | ribosomal protein L31 (RPL31) pseudogene |
| Y_RNA | 3 | Y RNA [Source:RFAM;Acc:RF00019] |
| PPP1R2 | 3 | protein phosphatase 1 regulatory inhibitor sul |
| RNU6ATAC24P | 3 | RNA, U6atac small nuclear 24, pseudogene [S |
| AC069213.1 | 3 | novel transcript |
| RN7SL73P | 3 | RNA, 7SL, cytoplasmic 73, pseudogene [Sourc |
| AC069213.3 | 3 | ribosomal protein L24 (RPL24) pseudogene |
| AC069213.2 | 3 | ribosomal protein L31 (RPL31) pseudogene |
| APOD | 3 | apolipoprotein D [Source:HGNC Symbol;Acc:H |
| MUC20P1 | 3 | mucin 20, cell surface associated pseudogene |
| AC233280.19 | 3 | programmed cell death 6 (PDCD6) pseudogen |
| AC233280.2 | 3 | protein phosphatase 4, regulatory subunit 2, p |
| AC233280.1 | 3 | novel transcript |
| MUC20-OT1 | 3 | MUC20 overlapping transcript [Source:HGNC S |
| SDHAP2 | 3 | succinate dehydrogenase complex flavoprotei |
| MIR570 | 3 | microRNA 570 [Source:HGNC Symbol;Acc:HGN |
| SMBD1P | 3 | somatomedin B domain containing 1, pseudog |
| MUC20 | 3 | mucin 20, cell surface associated [Source:HGN |
| MUC4 | 3 | mucin 4, cell surface associated [Source:HGNC |
| LINC01983 | 3 | long intergenic non-protein coding RNA 1983 |
| KIF3AP1 | 3 | kinesin family member 3A pseudogene 1 [Sou |
| TNK2 | 3 | tyrosine kinase non receptor 2 [Source:HGNC |
| MIR6829 | 3 | microRNA 6829 [Source:HGNC Symbol;Acc:HG |
| AC124944.3 | 3 | novel transcript, antisense to TNK2 |
| TNK2-AS1 | 3 | TNK2 antisense RNA 1 [Source:HGNC Symbol;A |
| AC124944.2 | 3 | programmed cell death 6 (PDCD6) pseudogen |
| RNU2-11P | 3 | RNA, U2 small nuclear 11, pseudogene [Sourc |
| AC124944.1 | 3 | protein phosphatase 4, regulatory subunit 2 (I |
| SDHAP1 | 3 | succinate dehydrogenase complex flavoprotei |
| AC024937.3 | 3 | novel transcript, antisense to TFRC |
| AC024937.2 | 3 | protein phosphatase 4, regulatory subunit 2 (I |
| TFRC | 3 | transferrin receptor [Source:HGNC Symbol;Ac |
| AC024937.1 | 3 | ADP-ribosylation factor-like 8B (ARL8B) pseud |
| RNU7-18P | 3 | RNA, U7 small nuclear 18 pseudogene [Source |
| LINC00885 | 3 | long intergenic non-protein coding RNA 885 [S |
| ZDHHC19 | 3 | zinc finger DHHC-type containing 19 [Source:H |
| Y_RNA | 3 | Y RNA [Source:RFAM;Acc:RF00019] |
| SLC51A | 3 | solute carrier family 51 alpha subunit [Source: |
| PCYT1A | 3 | phosphate cytidylyltransferase 1, choline, alph |
| AC069257.3 | 3 | novel protein |
| AC069257.1 | 3 | novel transcript |
| TCTEX1D2 | 3 | Tctex1 domain containing 2 [Source:HGNC Syn |
| AC069257.2 | 3 | ribosomal protein L39 (RPL39) pseudogene |
| TM4SF19-TCTEX1D2 | 3 | TM4SF19-TCTEX1D2 readthrough (NMD candi |
| TM4SF19-AS1 | 3 | TM4SF19 antisense RNA 1 [Source:HGNC Sym |
| TM4SF19 | 3 | transmembrane 4 L six family member 19 [Sou |
| RNU6-910P | 3 | RNA, U6 small nuclear 910, pseudogene [Sour |
| UBXN7 | 3 | UBX domain protein 7 [Source:HGNC Symbol;A |
| RNU6-1279P | 3 | RNA, U6 small nuclear 1279, pseudogene [Sou |
| RN7SL434P | 3 | RNA, 7SL, cytoplasmic 434, pseudogene [Sour |

| | | |
|---|---|---|
| RN7SL738P | 3 | RNA, 7SL, cytoplasmic 738, pseudogene [Sour |
| AC083822.1 | 3 | zinc finger, DHHC-type containing 1 (ZDHHC1) |
| UBXN7-AS1 | 3 | UBXN7 antisense RNA 1 [Source:HGNC Symbo |
| AC117490.1 | 3 | ring finger protein 168 (RNF168) pseudogene |
| RNF168 | 3 | ring finger protein 168 [Source:HGNC Symbol; |
| AC117490.2 | 3 | novel transcript, antisense to RNF168 |
| SMCO1 | 3 | single-pass membrane protein with coiled-coi |
| AC092933.1 | 3 | N-acetyltransferase 13 (GCN5-related) (NAT13 |
| RPS29P3 | 3 | ribosomal protein S29 pseudogene 3 [Source: |
| WDR53 | 3 | WD repeat domain 53 [Source:HGNC Symbol; |
| FBXO45 | 3 | F-box protein 45 [Source:HGNC Symbol;Acc:H |
| AC092933.2 | 3 | novel transcript |
| LINC01063 | 3 | long intergenic non-protein coding RNA 1063 |
| NRROS | 3 | negative regulator of reactive oxygen species |
| PIGX | 3 | phosphatidylinositol glycan anchor biosynthes |
| CEP19 | 3 | centrosomal protein 19 [Source:HGNC Symbo |
| RNU6-646P | 3 | RNA, U6 small nuclear 646, pseudogene [Sour |
| PAK2 | 3 | p21 (RAC1) activated kinase 2 [Source:HGNC S |
| RNU4-89P | 3 | RNA, U4 small nuclear 89, pseudogene [Sourc |
| RNU6-42P | 3 | RNA, U6 small nuclear 42, pseudogene [Sourc |
| SENP5 | 3 | SUMO specific peptidase 5 [Source:HGNC Sym |
| AC016949.1 | 3 | novel transcript |
| NCBP2 | 3 | nuclear cap binding protein subunit 2 [Source: |
| NCBP2-AS1 | 3 | NCBP2 antisense RNA 1 [Source:HGNC Symbo |
| NCBP2AS2 | 3 | NCBP2 antisense 2 (head to head) [Source:HG |
| PIGZ | 3 | phosphatidylinositol glycan anchor biosynthes |
| AC011322.1 | 3 | novel transcript |
| RPSAP69 | 3 | ribosomal protein SA pseudogene 69 [Source: |
| MELTF | 3 | melanotransferrin [Source:HGNC Symbol;Acc: |
| MELTF-AS1 | 3 | MELTF antisense RNA 1 [Source:HGNC Symbo |
| DLG1 | 3 | discs large MAGUK scaffold protein 1 [Source: |
| MIR4797 | 3 | microRNA 4797 [Source:HGNC Symbol;Acc:HG |
| DLG1-AS1 | 3 | DLG1 antisense RNA 1 [Source:HGNC Symbol; |
| AL121981.1 | 3 | novel transcript |
| AC128709.2 | 3 | novel transcript |
| AC128709.1 | 3 | novel transcript |
| AC128709.3 | 3 | novel transcript |
| LINC02012 | 3 | long intergenic non-protein coding RNA 2012 |
| BDH1 | 3 | 3-hydroxybutyrate dehydrogenase 1 [Source:I |
| AC132008.2 | 3 | succinate dehydrogenase complex, subunit A, |
| AC132008.1 | 3 | ribosomal protein L36 (RPL36) pseudogene |
| AC024560.2 | 3 | protein phosphatase 4, regulatory subunit 2 (I |
| AC024560.3 | 3 | novel transcript |
| AC024560.4 | 3 | novel transcript |
| AC024560.1 | 3 | novel transcript |
| AC024560.5 | 3 | novel transcript, antisense to RUBCN |
| RUBCN | 3 | rubicon autophagy regulator [Source:HGNC Sy |
| MIR922 | 3 | microRNA 922 [Source:HGNC Symbol;Acc:HGN |
| FYTTD1 | 3 | forty-two-three domain containing 1 [Source: |
| AC055764.2 | 3 | novel transcript |

| | | |
|---|---|---|
| LRCH3 | 3 | leucine rich repeats and calponin homology d |
| AC055764.1 | 3 | novel transcript |
| AC144530.1 | 3 | ribosomal protein L17 (RPL17) pseudogene |
| IQCG | 3 | IQ motif containing G [Source:HGNC Symbol; |
| RNU6-858P | 3 | RNA, U6 small nuclear 858, pseudogene [Sour |
| RPL35A | 3 | ribosomal protein L35a [Source:HGNC Symbol |
| LMLN | 3 | leishmanolysin like peptidase [Source:HGNC S |
| RNU6-621P | 3 | RNA, U6 small nuclear 621, pseudogene [Sour |
| AC135893.1 | 3 | novel transcript, antisense to LMLN |
| LMLN-AS1 | 3 | LMLN antisense RNA 1 [Source:HGNC Symbol; |
| ANKRD18DP | 3 | ankyrin repeat domain 18D, pseudogene [Sou |
| RNU6-821P | 3 | RNA, U6 small nuclear 821, pseudogene [Sour |
| FRG2FP | 3 | FSHD region gene 2 family member F, pseudo |
| TUBB8P8 | 3 | tubulin beta 8 class VIII pseudogene 8 [Source |
| AC073135.1 | 3 | FAM166A, pseudogene 10 |
| FAM157A | 3 | family with sequence similarity 157 member |
| RNU6-860P | 3 | RNA, U6 small nuclear 860, pseudogene [Sour |
| CICP6 | 3 | capicua transcriptional repressor pseudogene |
| SEPTIN14P3 | 3 | septin 14 pseudogene 3 [Source:HGNC Symbo |
| BNIP3P41 | 4 | BCL2 interacting protein 3 pseudogene 41 [So |
| ZNF595 | 4 | zinc finger protein 595 [Source:HGNC Symbol; |
| AC253576.1 | 4 | pseudogene similar to part of WEE1 homolog |
| ZNF718 | 4 | zinc finger protein 718 [Source:HGNC Symbol; |
| AC253576.2 | 4 | novel transcript |
| AC108475.1 | 4 | pseudogene similar to part of WEE1 homolog |
| ZNF876P | 4 | zinc finger protein 876, pseudogene [Source:H |
| AC079140.3 | 4 | novel zinc finger protein pseudogene |
| AC079140.5 | 4 | novel zinc finger protein pseudogene |
| ZNF732 | 4 | zinc finger protein 732 [Source:HGNC Symbol; |
| AC079140.6 | 4 | novel transcript |
| AC079140.4 | 4 | pseudogene similar to part of zinc finger prot |
| ZNF141 | 4 | zinc finger protein 141 [Source:HGNC Symbol; |
| AC079140.1 | 4 | novel pseudogene |
| AC079140.2 | 4 | pseudogene similar to part of ribosomal prote |
| MIR571 | 4 | microRNA 571 [Source:HGNC Symbol;Acc:HGI |
| AC092574.2 | 4 | novel transcript, antisense to ZNF721 |
| ZNF519P4 | 4 | zinc finger protein 519 pseudogene 4 [Source: |
| AC092574.1 | 4 | novel transcript |
| ABCA11P | 4 | ATP binding cassette subfamily A member 11, |
| ZNF721 | 4 | zinc finger protein 721 [Source:HGNC Symbol; |
| PIGG | 4 | phosphatidylinositol glycan anchor biosynthes |
| TMEM271 | 4 | transmembrane protein 271 [Source:HGNC Sy |
| AC116565.1 | 4 | novel transcript |
| PDE6B | 4 | phosphodiesterase 6B [Source:HGNC Symbol; |
| AC107464.1 | 4 | novel transcript, antisense to PDE6B |
| AC107464.3 | 4 | novel transcript, antisense to PDE6B |
| ATP5ME | 4 | ATP synthase membrane subunit e [Source:H( |
| MYL5 | 4 | myosin light chain 5 [Source:HGNC Symbol;Ac |
| SLC49A3 | 4 | solute carrier family 49 member 3 [Source:HG |
| PCGF3 | 4 | polycomb group ring finger 3 [Source:HGNC S |

| | | |
|---|---|---|
| AC107464.2 | 4 | novel transcript |
| AC139887.4 | 4 | novel transcript, antisense to PCGF3 |
| AC139887.2 | 4 | novel transcript, antisense to PCGF3 |
| AC139887.1 | 4 | novel transcript, antisense to PCGF3 |
| CPLX1 | 4 | complexin 1 [Source:HGNC Symbol;Acc:HGNC |
| AC139887.3 | 4 | novel transcript |
| AC139887.5 | 4 | novel transcript |
| GAK | 4 | cyclin G associated kinase [Source:HGNC Syml |
| TMEM175 | 4 | transmembrane protein 175 [Source:HGNC Sy |
| DGKQ | 4 | diacylglycerol kinase theta [Source:HGNC Sym |
| SLC26A1 | 4 | solute carrier family 26 member 1 [Source:HG |
| IDUA | 4 | iduronidase alpha-L- [Source:HGNC Symbol;Ac |
| FGFRL1 | 4 | fibroblast growth factor receptor like 1 [Sourc |
| AC019103.1 | 4 | novel transcript |
| RNF212 | 4 | ring finger protein 212 [Source:HGNC Symbol; |
| AC092535.3 | 4 | protein phosphatase 2, regulatory subunit B, ( |
| AC092535.4 | 4 | novel transcript |
| TMED11P | 4 | transmembrane p24 trafficking protein 11, ps |
| AC092535.1 | 4 | NULL |
| AC092535.2 | 4 | novel transcript |
| SPON2 | 4 | spondin 2 [Source:HGNC Symbol;Acc:HGNC:1 |
| AC092535.5 | 4 | novel transcript, antisense to SPON2 |
| CTBP1-AS | 4 | CTBP1 antisense RNA [Source:HGNC Symbol;A |
| CTBP1 | 4 | C-terminal binding protein 1 [Source:HGNC Sy |
| CTBP1-DT | 4 | CTBP1 divergent transcript [Source:HGNC Sym |
| MAEA | 4 | macrophage erythroblast attacher [Source:HG |
| UVSSA | 4 | UV stimulated scaffold protein A [Source:HGN |
| AC078852.2 | 4 | novel transcript |
| AC078852.1 | 4 | novel transcript |
| CRIPAK | 4 | cysteine rich PAK1 inhibitor [Source:HGNC Syr |
| NKX1-1 | 4 | NK1 homeobox 1 [Source:HGNC Symbol;Acc:H |
| AC147067.2 | 4 | novel transcript |
| AC147067.1 | 4 | novel transcript |
| FAM53A | 4 | family with sequence similarity 53 member A |
| Y_RNA | 4 | Y RNA [Source:RFAM;Acc:RF00019] |
| SLBP | 4 | stem-loop binding protein [Source:HGNC Sym |
| AC016773.1 | 4 | novel transcript |
| TACC3 | 4 | transforming acidic coiled-coil containing prot |
| TMEM129 | 4 | transmembrane protein 129 [Source:HGNC Sy |
| FGFR3 | 4 | fibroblast growth factor receptor 3 [Source:H( |
| LETM1 | 4 | leucine zipper and EF-hand containing transm |
| NSD2 | 4 | nuclear receptor binding SET domain protein : |
| SCARNA22 | 4 | small Cajal body-specific RNA 22 [Source:HGN |
| NELFA | 4 | negative elongation factor complex member / |
| MIR943 | 4 | microRNA 943 [Source:HGNC Symbol;Acc:HGI |
| C4orf48 | 4 | chromosome 4 open reading frame 48 [Sourc |
| NAT8L | 4 | N-acetyltransferase 8 like [Source:HGNC Symt |
| POLN | 4 | DNA polymerase nu [Source:HGNC Symbol;Ac |
| HAUS3 | 4 | HAUS augmin like complex subunit 3 [Source:I |
| AL136360.1 | 4 | pseudogene similar to part of HBS1-like |

| | | |
|---|---|---|
| AL136360.2 | 4 | novel transcript |
| COX6B1P5 | 4 | cytochrome c oxidase subunit 6B1 pseudogen |
| MXD4 | 4 | MAX dimerization protein 4 [Source:HGNC Sy |
| MIR4800 | 4 | microRNA 4800 [Source:HGNC Symbol;Acc:HG |
| ZFYVE28 | 4 | zinc finger FYVE-type containing 28 [Source:H |
| AL158068.2 | 4 | novel transcript |
| RN7SL589P | 4 | RNA, 7SL, cytoplasmic 589, pseudogene [Sour |
| AL158068.1 | 4 | pseudogene similar to Spi-C transcription fact |
| CFAP99 | 4 | cilia and flagella associated protein 99 [Source |
| RNF4 | 4 | ring finger protein 4 [Source:HGNC Symbol;Ac |
| AL645924.1 | 4 | novel transcript |
| AL645924.2 | 4 | ribosomal protein S3A (RPS3A) pseudogene |
| FAM193A | 4 | family with sequence similarity 193 member A |
| CR545473.1 | 4 | novel pseudogene |
| Y_RNA | 4 | Y RNA [Source:RFAM;Acc:RF00019] |
| TNIP2 | 4 | TNFAIP3 interacting protein 2 [Source:HGNC S |
| SH3BP2 | 4 | SH3 domain binding protein 2 [Source:HGNC S |
| ADD1 | 4 | adducin 1 [Source:HGNC Symbol;Acc:HGNC:24 |
| AL121750.1 | 4 | novel transcript, antisense to ADD1 |
| MFSD10 | 4 | major facilitator superfamily domain containi |
| NOP14-AS1 | 4 | NOP14 antisense RNA 1 [Source:HGNC Symbo |
| NOP14 | 4 | NOP14 nucleolar protein [Source:HGNC Symb |
| GRK4 | 4 | G protein-coupled receptor kinase 4 [Source:H |
| HTT | 4 | huntingtin [Source:HGNC Symbol;Acc:HGNC:4 |
| RNU6-204P | 4 | RNA, U6 small nuclear 204, pseudogene [Sour |
| HTT-AS | 4 | HTT antisense RNA [Source:HGNC Symbol;Acc |
| AL390065.2 | 4 | NULL |
| AL390065.1 | 4 | NULL |
| MSANTD1 | 4 | Myb/SANT DNA binding domain containing 1 [ |
| RGS12 | 4 | regulator of G protein signaling 12 [Source:HG |
| AL645949.1 | 4 | X antigen family, member 5 (XAGE5) pseudog |
| AL645949.2 | 4 | ribosomal protein L7a (RPL7A) pseudogene |
| HGFAC | 4 | HGF activator [Source:HGNC Symbol;Acc:HGN |
| AL590235.2 | 4 | novel transcript |
| DOK7 | 4 | docking protein 7 [Source:HGNC Symbol;Acc:H |
| LRPAP1 | 4 | LDL receptor related protein associated prote |
| AL590235.1 | 4 | novel transcript |
| LINC00955 | 4 | long intergenic non-protein coding RNA 955 [S |
| AL121796.1 | 4 | novel transcript |
| LINC02171 | 4 | long intergenic non-protein coding RNA 2171 |
| LINC02600 | 4 | long intergenic non-protein coding RNA 2600 |
| ADRA2C | 4 | adrenoceptor alpha 2C [Source:HGNC Symbol |
| OR7E163P | 4 | olfactory receptor family 7 subfamily E memb |
| OR7E162P | 4 | olfactory receptor family 7 subfamily E memb |
| AC147876.1 | 4 | novel pseudogene |
| AC226119.1 | 4 | novel transcript |
| AC226119.2 | 4 | novel pseudogene |
| ALG1L7P | 4 | ALG1 like 7, pseudogene [Source:HGNC Symb |
| FAM86EP | 4 | family with sequence similarity 86 member E, |
| ENPP7P9 | 4 | ectonucleotide pyrophosphatase/phosphodie |

| | | |
|---|---|---|
| AC116562.4 | 4 | RNA binding protein, fox-1 homolog (C. elegar |
| AC116562.1 | 4 | ribosomal protein S3A (RPS3A) pseudogene |
| AC116562.3 | 4 | ribosomal protein S24 (RPS24) pseudogene |
| AC116562.2 | 4 | chromosome 11 open reading frame 2 (C11or |
| OR4D12P | 4 | olfactory receptor family 4 subfamily D memb |
| UNC93B4 | 4 | unc-93 homolog B4, pseudogene [Source:HGN |
| OR7E99P | 4 | olfactory receptor family 7 subfamily E memb |
| OR7E43P | 4 | olfactory receptor family 7 subfamily E memb |
| OTOP1 | 4 | otopetrin 1 [Source:HGNC Symbol;Acc:HGNC: |
| TMEM128 | 4 | transmembrane protein 128 [Source:HGNC Sy |
| LYAR | 4 | Ly1 antibody reactive [Source:HGNC Symbol;A |
| ZBTB49 | 4 | zinc finger and BTB domain containing 49 [Sou |
| AC105415.1 | 4 | novel transcript |
| NSG1 | 4 | neuronal vesicle trafficking associated 1 [Sour |
| STX18 | 4 | syntaxin 18 [Source:HGNC Symbol;Acc:HGNC: |
| STX18-IT1 | 4 | STX18 intronic transcript 1 [Source:HGNC Sym |
| RPS7P15 | 4 | ribosomal protein S7 pseudogene 15 [Source: |
| STX18-AS1 | 4 | STX18 antisense RNA 1 (head to head) [Source |
| AC092437.1 | 4 | tec |
| LINC01396 | 4 | long intergenic non-protein coding RNA 1396 |
| MSX1 | 4 | msh homeobox 1 [Source:HGNC Symbol;Acc:H |
| LDHAP1 | 4 | lactate dehydrogenase A pseudogene 1 [Sourc |
| RN7SKP113 | 4 | RN7SK pseudogene 113 [Source:HGNC Symbo |
| CYTL1 | 4 | cytokine like 1 [Source:HGNC Symbol;Acc:HGN |
| STK32B | 4 | serine/threonine kinase 32B [Source:HGNC Sy |
| Y_RNA | 4 | Y RNA [Source:RFAM;Acc:RF00019] |
| RN7SKP275 | 4 | RN7SK pseudogene 275 [Source:HGNC Symbo |
| LINC01587 | 4 | long intergenic non-protein coding RNA 1587 |
| EVC2 | 4 | EvC ciliary complex subunit 2 [Source:HGNC S |
| EVC | 4 | EvC ciliary complex subunit 1 [Source:HGNC S |
| CRMP1 | 4 | collapsin response mediator protein 1 [Source |
| C4orf50 | 4 | chromosome 4 open reading frame 50 [Sourc |
| MIR378D1 | 4 | microRNA 378d-1 [Source:HGNC Symbol;Acc:I |
| JAKMIP1 | 4 | janus kinase and microtubule interacting prot |
| AC092442.1 | 4 | novel protein |
| AC113615.2 | 4 | novel transcript, antisense to C4orf50and JAK |
| LINC02495 | 4 | long intergenic non-protein coding RNA 2495 |
| AC113615.1 | 4 | novel transcript |
| WFS1 | 4 | wolframin ER transmembrane glycoprotein [S |
| AC116317.1 | 4 | novel transcript |
| PPP2R2C | 4 | protein phosphatase 2 regulatory subunit Bga |
| MAN2B2 | 4 | mannosidase alpha class 2B member 2 [Sourc |
| MRFAP1 | 4 | Morf4 family associated protein 1 [Source:HG |
| LINC02482 | 4 | long intergenic non-protein coding RNA 2482 |
| AC093323.4 | 4 | Morf4 family associated protein 1 (MRFAP1) p |
| AC093323.1 | 4 | Putative MORF4 family-associated protein 1-li |
| LINC002481 | 4 | long intergenic non-protein coding RNA 2481 |
| AC093323.2 | 4 | TEC |
| S100P | 4 | S100 calcium binding protein P [Source:HGNC |
| AC093323.3 | 4 | TEC |

| | | |
|---|---|---|
| MRFAP1L1 | 4 | Morf4 family associated protein 1 like 1 [Sour |
| BLOC1S4 | 4 | biogenesis of lysosomal organelles complex 1 |
| AC106045.1 | 4 | novel transcript |
| KIAA0232 | 4 | KIAA0232 [Source:HGNC Symbol;Acc:HGNC:2 |
| TBC1D14 | 4 | TBC1 domain family member 14 [Source:HGN |
| AC097382.3 | 4 | novel transcript, antisense to TBC1D14 |
| AC097382.1 | 4 | novel transcript, sense overlapping TBC1D14 |
| RN7SKP292 | 4 | RN7SK pseudogene 292 [Source:HGNC Symbo |
| AC097382.2 | 4 | novel transcript, antisense to TBC1D14 and T |
| CCDC96 | 4 | coiled-coil domain containing 96 [Source:HGN |
| TADA2B | 4 | transcriptional adaptor 2B [Source:HGNC Sym |
| GRPEL1 | 4 | GrpE like 1, mitochondrial [Source:HGNC Sym |
| LINC02447 | 4 | long intergenic non-protein coding RNA 2447 |
| RN7SKP36 | 4 | RN7SK pseudogene 36 [Source:HGNC Symbol; |
| SORCS2 | 4 | sortilin related VPS10 domain containing rece |
| MIR4798 | 4 | microRNA 4798 [Source:HGNC Symbol;Acc:H |
| PSAPL1 | 4 | prosaposin like 1 (gene/pseudogene) [Source: |
| MIR4274 | 4 | microRNA 4274 [Source:HGNC Symbol;Acc:H |
| AFAP1-AS1 | 4 | AFAP1 antisense RNA 1 [Source:HGNC Symbol |
| AFAP1 | 4 | actin filament associated protein 1 [Source:H |
| AC112254.1 | 4 | novel transcript, antisense to AFAP1 |
| AC097381.1 | 4 | uncharacterized LOC389199 [Source:NCBI ger |
| ABLIM2 | 4 | actin binding LIM protein family member 2 [S |
| MIR95 | 4 | microRNA 95 [Source:HGNC Symbol;Acc:HGN |
| AC097381.3 | 4 | novel transcript, antisense to ABLIM2 |
| AC097381.2 | 4 | novel transcript |
| GMPSP1 | 4 | guanine monophosphate synthase pseudogen |
| SH3TC1 | 4 | SH3 domain and tetratricopeptide repeats 1 [ |
| HTRA3 | 4 | HtrA serine peptidase 3 [Source:HGNC Symbo |
| LINC02517 | 4 | long intergenic non-protein coding RNA 2517 |
| AC104825.1 | 4 | novel transcript |
| ACOX3 | 4 | acyl-CoA oxidase 3, pristanoyl [Source:HGNC |
| RNA5SP152 | 4 | RNA, 5S ribosomal pseudogene 152 [Source:H |
| TRMT44 | 4 | tRNA methyltransferase 44 homolog [Source: |
| AC105345.2 | 4 | novel transcript |
| AC105345.1 | 4 | novel transcript |
| GPR78 | 4 | G protein-coupled receptor 78 [Source:HGNC |
| CPZ | 4 | carboxypeptidase Z [Source:HGNC Symbol;Ac |
| AC209005.1 | 4 | novel transcript |
| HMX1 | 4 | H6 family homeobox 1 [Source:HGNC Symbol; |
| AC116612.1 | 4 | novel transcript, sense intronic to HMX1 |
| AC073648.1 | 4 | olfactory receptor, family 7 pseudogene |
| UNC93B8 | 4 | unc-93 homolog B8, pseudogene [Source:HGN |
| AC073648.3 | 4 | olfactory receptor, family 7 pseudogene |
| AC073648.2 | 4 | pseudogene similar to part of chromosome 1 |
| AC073648.6 | 4 | RNA binding protein, fox-1 homolog (C. elegar |
| AC073648.5 | 4 | ribosomal protein S24 (RPS24) pseudogene |
| AC073648.4 | 4 | ribosomal protein S3A (RPS3A) pseudogene |
| AC108519.1 | 4 | novel transcript |
| SNRPCP16 | 4 | small nuclear ribonucleoprotein polypeptide |

| | | |
|---|---|---|
| AC073648.7 | 4 | novel transcript |
| ENPP7P10 | 4 | ectonucleotide pyrophosphatase/phosphodie |
| FAM86KP | 4 | family with sequence similarity 86 member K, |
| ALG1L14P | 4 | ALG1 like 14, pseudogene [Source:HGNC Syml |
| FAM90A26 | 4 | family with sequence similarity 90 member A2 |
| USP17L10 | 4 | ubiquitin specific peptidase 17 like family mer |
| USP17L11 | 4 | ubiquitin specific peptidase 17 like family mer |
| USP17L12 | 4 | ubiquitin specific peptidase 17 like family mer |
| USP17L13 | 4 | ubiquitin specific peptidase 17 like family mer |
| USP17L14P | 4 | ubiquitin specific peptidase 17 like family mer |
| USP17L15 | 4 | ubiquitin specific peptidase 17 like family mer |
| USP17L16P | 4 | ubiquitin specific peptidase 17 like family mer |
| USP17L17 | 4 | ubiquitin specific peptidase 17 like family mer |
| USP17L18 | 4 | ubiquitin specific peptidase 17 like family mer |
| USP17L19 | 4 | ubiquitin specific peptidase 17 like family mer |
| USP17L20 | 4 | ubiquitin specific peptidase 17 like family mer |
| USP17L21 | 4 | ubiquitin specific peptidase 17 like family mer |
| USP17L22 | 4 | ubiquitin specific peptidase 17 like family mer |
| USP17L23 | 4 | ubiquitin specific peptidase 17 like family mer |
| USP17L24 | 4 | ubiquitin specific peptidase 17 like family mer |
| USP17L25 | 4 | ubiquitin specific peptidase 17 like family mer |
| USP17L26 | 4 | ubiquitin specific peptidase 17 like family mer |
| USP17L5 | 4 | ubiquitin specific peptidase 17 like family mer |
| USP17L27 | 4 | ubiquitin specific peptidase 17 like family mer |
| USP17L28 | 4 | ubiquitin specific peptidase 17 like family mer |
| USP17L29 | 4 | ubiquitin specific peptidase 17 like family mer |
| USP17L9P | 4 | ubiquitin specific peptidase 17 like family mer |
| USP17L30 | 4 | ubiquitin specific peptidase 17 like family mer |
| USP17L6P | 4 | ubiquitin specific peptidase 17 like family mer |
| AC116655.1 | 4 | C2H2 zinc finger protein pseudogene |
| DEFB108F | 4 | defensin beta 108F (pseudogene) [Source:HGI |
| RNA5SP153 | 4 | RNA, 5S ribosomal pseudogene 153 [Source:H |
| DEFB130D | 4 | defensin beta 130D (pseudogene) [Source:HG |
| DEFB131A | 4 | defensin beta 131A [Source:HGNC Symbol;Acc |
| OR7E86P | 4 | olfactory receptor family 7 subfamily E memb |
| OR7E84P | 4 | olfactory receptor family 7 subfamily E memb |
| OR7E85P | 4 | olfactory receptor family 7 subfamily E memb |
| UNC93B7 | 4 | unc-93 homolog B7, pseudogene [Source:HGN |
| OR7E83P | 4 | olfactory receptor family 7 subfamily E memb |
| AC097493.3 | 4 | pseudogene similar to part of chromosome 1 |
| AC097493.4 | 4 | RNA binding protein, fox-1 homolog (C. elegar |
| AC097493.1 | 4 | ribosomal protein S24 (RPS24) pseudogene |
| MIR548I2 | 4 | microRNA 548i-2 [Source:HGNC Symbol;Acc:H |
| AC097493.2 | 4 | ribosomal protein S24 (RPS24) pseudogene |
| AC105916.1 | 4 | novel transcript |
| SNRPCP13 | 4 | small nuclear ribonucleoprotein polypeptide ( |
| ENPP7P11 | 4 | ectonucleotide pyrophosphatase/phosphodie |
| FAM86MP | 4 | family with sequence similarity 86 member M |
| ALG1L3P | 4 | ALG1 like 3, pseudogene [Source:HGNC Symb |
| Metazoa_SRP | 4 | Metazoan signal recognition particle RNA [Sou |

| | | |
|---|---|---|
| AC098976.1 | 4 | chromosome 21 open reading frame 63 (C21c |
| OR7E35P | 4 | olfactory receptor family 7 subfamily E memb |
| SLC2A9 | 4 | solute carrier family 2 member 9 [Source:HGN |
| DRD5 | 4 | dopamine receptor D5 [Source:HGNC Symbol; |
| RNA5SP154 | 4 | RNA, 5S ribosomal pseudogene 154 [Source:H |
| AC108199.1 | 4 | novel transcript |
| AC005674.1 | 4 | novel transcript, antisense to SLC2A9 |
| AC005674.2 | 4 | novel transcript, overlapping WDR1 |
| WDR1 | 4 | WD repeat domain 1 [Source:HGNC Symbol;A |
| MIR3138 | 4 | microRNA 3138 [Source:HGNC Symbol;Acc:HG |
| RNA5SP155 | 4 | RNA, 5S ribosomal pseudogene 155 [Source:H |
| AC093664.1 | 4 | G antigen pseudogene |
| AC006499.6 | 4 | novel developmental pluripotency associated |
| AC006499.1 | 4 | developmental pluripotency associated 4 (DPI |
| AC006499.8 | 4 | myxovirus (influenza virus) resistance 1, interf |
| RAF1P1 | 4 | RAF1 pseudogene 1 [Source:HGNC Symbol;Ac |
| AC006499.4 | 4 | pseudogene similar to part of myxovirus (influ |
| AC006499.2 | 4 | novel beta-tubulin pseudogene |
| AC006499.3 | 4 | novel myxovirus (influenza virus) resistance pı |
| AC006499.5 | 4 | pseudogene similar to part of myxovirus (influ |
| AC006499.7 | 4 | pseudogene similar to part of myxovirus (influ |
| AC110768.2 | 4 | dynein, light chain, Tctex-type 1 (DYNLT1) pse |
| ZNF518B | 4 | zinc finger protein 518B [Source:HGNC Symbo |
| AC005599.1 | 4 | novel transcript, antisense to ZNF518Band CLI |
| AC110768.1 | 4 | heat-responsive protein 12 (HRSP12) pseudog |
| CLNK | 4 | cytokine dependent hematopoietic cell linker |
| AC084048.1 | 4 | novel transcript |
| LINC02498 | 4 | long intergenic non-protein coding RNA 2498 |
| MIR572 | 4 | microRNA 572 [Source:HGNC Symbol;Acc:HGI |
| RNPS1P1 | 4 | RNA binding protein with serine rich domain 1 |
| HS3ST1 | 4 | heparan sulfate-glucosamine 3-sulfotransfera: |
| RNA5SP156 | 4 | RNA, 5S ribosomal pseudogene 156 [Source:H |
| AC006230.1 | 4 | novel transcript, antisense to HS3ST1 |
| AC025539.1 | 4 | novel transcript |
| AC005699.1 | 4 | novel transcript |
| LINC02360 | 4 | long intergenic non-protein coding RNA 2360 |
| AC108037.1 | 4 | novel transcript |
| LINC02270 | 4 | long intergenic non-protein coding RNA 2270 |
| RNU6-578P | 4 | RNA, U6 small nuclear 578, pseudogene [Sour |
| ECM1P2 | 4 | extracellular matrix protein 1 pseudogene 2 [S |
| AC093809.1 | 4 | pseudogene similar to part of mitochondrially |
| AC007370.2 | 4 | novel transcript |
| AC007370.1 | 4 | novel transcript |
| U6 | 4 | U6 spliceosomal RNA [Source:RFAM;Acc:RF00 |
| HSP90AB2P | 4 | heat shock protein 90 alpha family class B mei |
| RAB28 | 4 | RAB28, member RAS oncogene family [Source |
| AC006445.3 | 4 | novel transcript |
| LINC01097 | 4 | long intergenic non-protein coding RNA 1097 |
| NKX3-2 | 4 | NK3 homeobox 2 [Source:HGNC Symbol;Acc:H |
| LINC01096 | 4 | long intergenic non-protein coding RNA 1096 |

| | | |
|---|---|---|
| BOD1L1 | 4 | biorientation of chromosomes in cell division |
| MIR5091 | 4 | microRNA 5091 [Source:HGNC Symbol;Acc:HG |
| AC006445.2 | 4 | ribosomal protein L21 (RPL21) pseudogene |
| AC006445.1 | 4 | ribosomal protein L32 (RPL32) pseudogene |
| LINC01182 | 4 | long intergenic non-protein coding RNA 1182 |
| AC007126.1 | 4 | novel transcript |
| AC095052.1 | 4 | chromosome 9 open reading frame 78 (C9orf7 |
| LINC01085 | 4 | long intergenic non-protein coding RNA 1085 |
| AC073848.1 | 4 | novel transcript |
| AC092546.1 | 4 | novel transcript |
| AC006296.3 | 4 | novel transcript |
| AC006296.2 | 4 | novel transcript |
| AC006296.1 | 4 | novel transcript |
| LINC00504 | 4 | long intergenic non-protein coding RNA 504 [S |
| MTND2P31 | 4 | MT-ND2 pseudogene 31 [Source:HGNC Symbo |
| SNORA63 | 4 | Small nucleolar RNA SNORA63 [Source:RFAM; |
| CPEB2-DT | 4 | CPEB2 divergent transcript [Source:HGNC Sym |
| CPEB2 | 4 | cytoplasmic polyadenylation element binding |
| AC098829.1 | 4 | novel transcript |
| RN7SKP170 | 4 | RN7SK pseudogene 170 [Source:HGNC Symbo |
| C1QTNF7 | 4 | C1q and TNF related 7 [Source:HGNC Symbol; |
| AC099550.1 | 4 | novel transcript |
| CC2D2A | 4 | coiled-coil and C2 domain containing 2A [Sour |
| AC007016.1 | 4 | pseudogene similar to part of glyceraldehyde- |
| AC116651.1 | 4 | novel transcript, antisense to CC2D2A |
| FBXL5 | 4 | F-box and leucine rich repeat protein 5 [Sourc |
| FAM200B | 4 | family with sequence similarity 200 member E |
| BST1 | 4 | bone marrow stromal cell antigen 1 [Source:H |
| AC114744.1 | 4 | ribosomal protein L10a (RPL10A) pseudogene |
| AC005798.1 | 4 | prefoldin subunit 1 (PFDN1) pseudogene |
| CD38 | 4 | CD38 molecule [Source:HGNC Symbol;Acc:HG |
| HPRT1P1 | 4 | hypoxanthine phosphoribosyltransferase 1 ps |
| FGFBP1 | 4 | fibroblast growth factor binding protein 1 [Sou |
| FGFBP2 | 4 | fibroblast growth factor binding protein 2 [Sou |
| PROM1 | 4 | prominin 1 [Source:HGNC Symbol;Acc:HGNC:S |
| AC108063.2 | 4 | novel transcript |
| TAPT1 | 4 | transmembrane anterior posterior transforma |
| AC108063.1 | 4 | novel transcript |
| TAPT1-AS1 | 4 | TAPT1 antisense RNA 1 (head to head) [Sourc |
| RPS21P4 | 4 | ribosomal protein S21 pseudogene 4 [Source: |
| AC006427.1 | 4 | fibroblast growth factor binding protein 1 (FG |
| AC006427.2 | 4 | novel transcript |
| ZEB2P1 | 4 | zinc finger E-box binding homeobox 2 pseudo |
| AC097515.1 | 4 | novel transcript |
| LDB2 | 4 | LIM domain binding 2 [Source:HGNC Symbol;A |
| AC104656.1 | 4 | novel transcript, antisense to LDB2 |
| AC106894.1 | 4 | novel transcript |
| MTND5P4 | 4 | MT-ND5 pseudogene 4 [Source:HGNC Symbol |
| LINC02493 | 4 | long intergenic non-protein coding RNA 2493 |
| SNORA75B | 4 | small nucleolar RNA, H/ACA box 75B [Source:I |

| | | |
|---|---|---|
| RPS7P6 | 4 | ribosomal protein S7 pseudogene 6 [Source:H |
| QDPR | 4 | quinoid dihydropteridine reductase [Source:H |
| CLRN2 | 4 | clarin 2 [Source:HGNC Symbol;Acc:HGNC:3393 |
| NACAP5 | 4 | NACA pseudogene 5 [Source:HGNC Symbol;Ac |
| AC006160.2 | 4 | mitochondrially encoded cytochrome b (MT-C |
| LAP3 | 4 | leucine aminopeptidase 3 [Source:HGNC Syml |
| AC006160.1 | 4 | novel transcript, antisense to LAP3 |
| MED28 | 4 | mediator complex subunit 28 [Source:HGNC S |
| FAM184B | 4 | family with sequence similarity 184 member E |
| DCAF16 | 4 | DDB1 and CUL4 associated factor 16 [Source:I |
| NCAPG | 4 | non-SMC condensin I complex subunit G [Sour |
| LCORL | 4 | ligand dependent nuclear receptor corepresso |
| KRT18P63 | 4 | keratin 18 pseudogene 63 [Source:HGNC Sym |
| AC093898.1 | 4 | novel transcript |
| AC093871.1 | 4 | novel transcript |
| AC006840.1 | 4 | TEC |
| LINC02438 | 4 | long intergenic non-protein coding RNA 2438 |
| RNA5SP157 | 4 | RNA, 5S ribosomal pseudogene 157 [Source:H |
| AC024595.1 | 4 | TEC |
| AC024230.1 | 4 | novel transcript |
| AC110767.1 | 4 | novel transcript |
| AC114728.1 | 4 | ribosomal protein L21 (RPL21) pseudogene |
| AC098862.1 | 4 | high-mobility group nucleosome binding dom |
| SLIT2 | 4 | slit guidance ligand 2 [Source:HGNC Symbol;A |
| SLIT2-IT1 | 4 | SLIT2 intronic transcript 1 [Source:HGNC Symb |
| MIR218-1 | 4 | microRNA 218-1 [Source:HGNC Symbol;Acc:H |
| PACRGL | 4 | parkin coregulated like [Source:HGNC Symbol |
| KCNIP4 | 4 | potassium voltage-gated channel interacting p |
| AC097505.1 | 4 | novel transcript, antisense to KCNIP4 |
| AC110296.1 | 4 | novel transcript |
| MIR7978 | 4 | microRNA 7978 [Source:HGNC Symbol;Acc:HG |
| AC096576.6 | 4 | novel transcript, sense intronic to KCNIP4 |
| AC096576.3 | 4 | novel transcript |
| AC096576.1 | 4 | ribosomal protein L31 (RPL31) pseudogene |
| AC096576.2 | 4 | novel transcript |
| KCNIP4-IT1 | 4 | KCNIP4 intronic transcript 1 [Source:HGNC Syr |
| AC096719.1 | 4 | novel transcript |
| RNU6-420P | 4 | RNA, U6 small nuclear 420, pseudogene [Sour |
| ADGRA3 | 4 | adhesion G protein-coupled receptor A3 [Sour |
| AC098583.1 | 4 | ribosomal protein S27 (RPS27) pseudogene |
| AC098583.2 | 4 | chromosome 18 open reading frame 16 (C18c |
| GBA3 | 4 | glucosylceramidase beta 3 (gene/pseudogene |
| CDC42P6 | 4 | cell division cycle 42 pseudogene 6 [Source:H( |
| AC097512.1 | 4 | novel transcript |
| AC092440.1 | 4 | novel transcript |
| RFPL4AP3 | 4 | ret finger protein like 4A pseudogene 3 [Sourc |
| AC093607.1 | 4 | novel transcript, antisense to PPARGC1A |
| ERVH-1 | 4 | endogenous retrovirus group H member 1 [So |
| PPARGC1A | 4 | PPARG coactivator 1 alpha [Source:HGNC Sym |
| AC092834.1 | 4 | novel transcript |

| | | |
|---|---|---|
| DHX15 | 4 | DEAH-box helicase 15 [Source:HGNC Symbol;/ |
| MIR573 | 4 | microRNA 573 [Source:HGNC Symbol;Acc:HGI |
| RN7SL16P | 4 | RNA, 7SL, cytoplasmic 16, pseudogene [Sourc |
| ATP5MGP3 | 4 | ATP synthase membrane subunit g pseudoger |
| LINC02473 | 4 | long intergenic non-protein coding RNA 2473 |
| AC006390.1 | 4 | novel pseudogene |
| HNRNPA1P65 | 4 | heterogeneous nuclear ribonucleoprotein A1 |
| SOD3 | 4 | superoxide dismutase 3 [Source:HGNC Symbo |
| CCDC149 | 4 | coiled-coil domain containing 149 [Source:HG |
| LGI2 | 4 | leucine rich repeat LGI family member 2 [Sour |
| SEPSECS | 4 | Sep (O-phosphoserine) tRNA:Sec (selenocyste |
| SEPSECS-AS1 | 4 | SEPSECS antisense RNA 1 (head to head) [Sou |
| PI4K2B | 4 | phosphatidylinositol 4-kinase type 2 beta [Sou |
| AC104662.1 | 4 | novel transcript |
| U2 | 4 | U2 spliceosomal RNA [Source:RFAM;Acc:RF00 |
| ZCCHC4 | 4 | zinc finger CCHC-type containing 4 [Source:HG |
| ANAPC4 | 4 | anaphase promoting complex subunit 4 [Sour |
| AC133963.2 | 4 | ring finger protein 168 (RNF168) pseudogene |
| AC133963.1 | 4 | inositol-3-phosphate synthase 1 (ISYNA1) pse |
| AC105290.1 | 4 | novel transcript |
| RNU7-126P | 4 | RNA, U7 small nuclear 126 pseudogene [Sourc |
| AC092436.1 | 4 | S-phase kinase-associated protein 1 (SKP1) ps |
| SLC34A2 | 4 | solute carrier family 34 member 2 [Source:HG |
| RPS29P11 | 4 | ribosomal protein S29 pseudogene 11 [Source |
| AC092436.2 | 4 | chromosome 14 open reading frame 156 (C14 |
| MTND4P9 | 4 | MT-ND4 pseudogene 9 [Source:HGNC Symbol |
| MTND4LP22 | 4 | MT-ND4L pseudogene 22 [Source:HGNC Symb |
| MTND3P5 | 4 | MT-ND3 pseudogene 5 [Source:HGNC Symbol |
| MTCO3P44 | 4 | MT-CO3 pseudogene 44 [Source:HGNC Symbo |
| SEL1L3 | 4 | SEL1L family member 3 [Source:HGNC Symbol |
| AC092436.3 | 4 | novel transcript |
| SMIM20 | 4 | small integral membrane protein 20 [Source:H |
| AC133961.1 | 4 | novel transcript |
| LINC02357 | 4 | long intergenic non-protein coding RNA 2357 |
| AC097714.1 | 4 | zinc finger protein 248 (ZNF248) pseudogene |
| RBPJ | 4 | recombination signal binding protein for imm |
| CCKAR | 4 | cholecystokinin A receptor [Source:HGNC Sym |
| TBC1D19 | 4 | TBC1 domain family member 19 [Source:HGN |
| AC107390.1 | 4 | NULL |
| STIM2 | 4 | stromal interaction molecule 2 [Source:HGNC |
| STIM2-AS1 | 4 | STIM2 antisense RNA 1 [Source:HGNC Symbol |
| AC106047.1 | 4 | family with sequence similarity 64, member A |
| AC024132.2 | 4 | novel transcript |
| AC024132.1 | 4 | novel transcript |
| LINC02261 | 4 | long intergenic non-protein coding RNA 2261 |
| Y_RNA | 4 | Y RNA [Source:RFAM;Acc:RF00019] |
| AC024132.3 | 4 | novel transcript |
| IGBP1P5 | 4 | immunoglobulin (CD79A) binding protein 1 ps |
| AC007106.2 | 4 | novel transcript |
| AC007106.1 | 4 | novel transcript |

| | | |
|---|---|---|
| AC093791.2 | 4 | novel transcript |
| AC093791.1 | 4 | novel transcript |
| AC097480.1 | 4 | novel transcript |
| AC097480.2 | 4 | novel transcript |
| RN7SL101P | 4 | RNA, 7SL, cytoplasmic 101, pseudogene [Sour |
| MIR4275 | 4 | microRNA 4275 [Source:HGNC Symbol;Acc:HG |
| MESTP3 | 4 | mesoderm specific transcript pseudogene 3 [S |
| LINC02364 | 4 | long intergenic non-protein coding RNA 2364 |
| AC068944.2 | 4 | transmembrane protein 56 (TMEM56) pseudo |
| AC068944.1 | 4 | novel transcript |
| AC109349.1 | 4 | novel transcript |
| LINC02472 | 4 | long intergenic non-protein coding RNA 2472 |
| AC092593.2 | 4 | pseudogene similar to part of plectin 1, intern |
| AC092593.1 | 4 | protein inhibitor of activated STAT, 2 (PIAS2) p |
| U6 | 4 | U6 spliceosomal RNA [Source:RFAM;Acc:RF00 |
| EEF1A1P21 | 4 | eukaryotic translation elongation factor 1 alph |
| AC097510.1 | 4 | novel pseudogene |
| RPS3AP17 | 4 | ribosomal protein S3a pseudogene 17 [Source |
| AC106868.1 | 4 | cathepsin B (CTSB) pseudogene |
| AC098595.1 | 4 | novel transcript, antisense to PCDH7 |
| PCDH7 | 4 | protocadherin 7 [Source:HGNC Symbol;Acc:H( |
| AC097716.1 | 4 | novel transcript |
| MTCYBP43 | 4 | MT-CYB pseudogene 43 [Source:HGNC Symbo |
| LINC02497 | 4 | long intergenic non-protein coding RNA 2497 |
| AC104071.1 | 4 | novel transcript |
| LINC02501 | 4 | long intergenic non-protein coding RNA 2501 |
| LINC02506 | 4 | long intergenic non-protein coding RNA 2506 |
| AC097654.1 | 4 | novel transcript |
| LINC02353 | 4 | long intergenic non-protein coding RNA 2353 |
| AC116611.1 | 4 | novel transcript |
| AC017013.1 | 4 | microtubule-associated protein, RP/EB family, |
| AC093606.1 | 4 | novel transcript |
| AC093878.1 | 4 | pseudogene similar to part of calmodulin regu |
| AC097535.2 | 4 | keratin 8, type II (KRT8) pseudogene |
| AC097535.1 | 4 | novel transcript |
| AC079772.1 | 4 | novel transcript |
| AC016687.3 | 4 | novel transcript |
| AC016687.2 | 4 | novel transcript |
| AC016687.1 | 4 | ribosomal protein S3A (RPS3A) pseudogene |
| LINC02484 | 4 | long intergenic non-protein coding RNA 2484 |
| AC110790.1 | 4 | novel transcript |
| AC093689.1 | 4 | novel transcript |
| AC093913.1 | 4 | zinc finger protein 709 (ZNF709) pseudogene |
| AC020589.1 | 4 | NULL |
| SEC63P2 | 4 | SEC63 homolog, protein translocation regulat |
| RNU6-573P | 4 | RNA, U6 small nuclear 573, pseudogene [Sour |
| ARAP2 | 4 | ArfGAP with RhoGAP domain, ankyrin repeat a |
| AC098827.1 | 4 | pseudogene similar to part of LIM and senesc |
| AC104078.1 | 4 | novel transcript, antisense to CENTD1 |
| DTHD1 | 4 | death domain containing 1 [Source:HGNC Sym |

| | | |
|---|---|---|
| AC104078.2 | 4 | novel transcript |
| MIR1255B1 | 4 | microRNA 1255b-1 [Source:HGNC Symbol;Acc |
| LINC02505 | 4 | long intergenic non-protein coding RNA 2505 |
| AC125336.1 | 4 | F-box protein 38 (FBXO38) pseudogene |
| AC093746.1 | 4 | novel transcript |
| LINC02616 | 4 | long intergenic non-protein coding RNA 2616 |
| AL136537.1 | 4 | novel transcript |
| MIR4801 | 4 | microRNA 4801 [Source:HGNC Symbol;Acc:H |
| NWD2 | 4 | NACHT and WD repeat domain containing 2 [S |
| C4orf19 | 4 | chromosome 4 open reading frame 19 [Sourc |
| AC022463.1 | 4 | novel transcript, antisense to C4orf19 |
| AC027607.1 | 4 | putatitive novel transcript |
| RELL1 | 4 | RELT like 1 [Source:HGNC Symbol;Acc:HGNC:2 |
| Y_RNA | 4 | Y RNA [Source:RFAM;Acc:RF00019] |
| AC108022.1 | 4 | ribosomal protein L21 (RPL21) pseudogene |
| PGM2 | 4 | phosphoglucomutase 2 [Source:HGNC Symbo |
| AC021106.3 | 4 | TEC |
| AC021106.1 | 4 | synaptotagmin binding, cytoplasmic RNA inter |
| TBC1D1 | 4 | TBC1 domain family member 1 [Source:HGNC |
| PTTG2 | 4 | pituitary tumor-transforming 2 [Source:HGNC |
| AC021106.2 | 4 | proteasome (prosome, macropain) activator s |
| MRPS33P2 | 4 | mitochondrial ribosomal protein S33 pseudog |
| AC098680.2 | 4 | novel transcript |
| AC098680.1 | 4 | novel transcript |
| LINC02513 | 4 | long intergenic non-protein coding RNA 2513 |
| AC024023.1 | 4 | tec |
| LINC01258 | 4 | long intergenic non-protein coding RNA 1258 |
| LINC01259 | 4 | long intergenic non-protein coding RNA 1259 |
| LINC02278 | 4 | long intergenic non-protein coding RNA 2278 |
| KLF3-AS1 | 4 | KLF3 antisense RNA 1 [Source:HGNC Symbol;A |
| AC021860.1 | 4 | novel transcript |
| KLF3 | 4 | Kruppel like factor 3 [Source:HGNC Symbol;Ac |
| RNA5SP158 | 4 | RNA, 5S ribosomal pseudogene 158 [Source:H |
| TLR10 | 4 | toll like receptor 10 [Source:HGNC Symbol;Ac |
| TLR1 | 4 | toll like receptor 1 [Source:HGNC Symbol;Acc: |
| TLR6 | 4 | toll like receptor 6 [Source:HGNC Symbol;Acc: |
| FAM114A1 | 4 | family with sequence similarity 114 member A |
| MIR574 | 4 | microRNA 574 [Source:HGNC Symbol;Acc:HGN |
| TMEM156 | 4 | transmembrane protein 156 [Source:HGNC Sy |
| KLHL5 | 4 | kelch like family member 5 [Source:HGNC Sym |
| AC079921.1 | 4 | novel transcript |
| AC079921.2 | 4 | novel transcript |
| WDR19 | 4 | WD repeat domain 19 [Source:HGNC Symbol;A |
| RFC1 | 4 | replication factor C subunit 1 [Source:HGNC Sy |
| RNU6-32P | 4 | RNA, U6 small nuclear 32, pseudogene [Sourc |
| RNU6-887P | 4 | RNA, U6 small nuclear 887, pseudogene [Sour |
| KLB | 4 | klotho beta [Source:HGNC Symbol;Acc:HGNC: |
| MIR5591 | 4 | microRNA 5591 [Source:HGNC Symbol;Acc:HG |
| Y_RNA | 4 | Y RNA [Source:RFAM;Acc:RF00019] |
| RPL9 | 4 | ribosomal protein L9 [Source:HGNC Symbol;A |

| | | |
|---|---|---|
| LIAS | 4 | lipoic acid synthetase [Source:HGNC Symbol;A |
| AC021148.1 | 4 | WD repeat domain 5 (WDR5) pseudogene |
| UGDH | 4 | UDP-glucose 6-dehydrogenase [Source:HGNC |
| UGDH-AS1 | 4 | UGDH antisense RNA 1 [Source:HGNC Symbol |
| AC021148.2 | 4 | novel transcript |
| SMIM14 | 4 | small integral membrane protein 14 [Source:H |
| AC108471.3 | 4 | novel transcript, antisense to SMIM14 |
| RNU7-11P | 4 | RNA, U7 small nuclear 11 pseudogene [Source |
| AC108471.2 | 4 | novel transcript |
| UBE2K | 4 | ubiquitin conjugating enzyme E2 K [Source:HG |
| Y_RNA | 4 | Y RNA [Source:RFAM;Acc:RF00019] |
| AC108471.1 | 4 | ribosomal protein S7 (RPS7) pseudogene |
| RN7SL558P | 4 | RNA, 7SL, cytoplasmic 558, pseudogene [Sour |
| ZBTB12BP | 4 | zinc finger and BTB domain containing 12B, ps |
| PDS5A | 4 | PDS5 cohesin associated factor A [Source:HGN |
| ELOCP33 | 4 | elongin C pseudogene 33 [Source:HGNC Symb |
| RNA5SP159 | 4 | RNA, 5S ribosomal pseudogene 159 [Source:H |
| AC098591.3 | 4 | poly(A) binding protein, cytoplasmic 1 (PABPC |
| PABPC1P1 | 4 | poly(A) binding protein cytoplasmic 1 pseudog |
| KRT18P25 | 4 | keratin 18 pseudogene 25 [Source:HGNC Sym |
| AC098591.2 | 4 | acyl-CoA thioesterase 7 (ACOT7) pseudogene |
| N4BP2 | 4 | NEDD4 binding protein 2 [Source:HGNC Symb |
| RNU6-1112P | 4 | RNA, U6 small nuclear 1112, pseudogene [Sou |
| AC098591.1 | 4 | NULL |
| AC095057.1 | 4 | heterogeneous nuclear ribonucleoprotein A3 |
| AC095057.3 | 4 | novel transcript, overlapping N4BP2 |
| AC095057.4 | 4 | novel transcript |
| RHOH | 4 | ras homolog family member H [Source:HGNC |
| AC095057.2 | 4 | novel transcript |
| LINC02265 | 4 | long intergenic non-protein coding RNA 2265 |
| CHRNA9 | 4 | cholinergic receptor nicotinic alpha 9 subunit |
| RNU7-74P | 4 | RNA, U7 small nuclear 74 pseudogene [Source |
| RBM47 | 4 | RNA binding motif protein 47 [Source:HGNC S |
| AC098869.2 | 4 | novel transcript |
| AC098869.1 | 4 | ribosomal protein L37 (RPL37) pseudogene |
| MIR4802 | 4 | microRNA 4802 [Source:HGNC Symbol;Acc:HG |
| AC131953.2 | 4 | novel transcript, antisense to NSUN7 |
| NSUN7 | 4 | NOP2/Sun RNA methyltransferase family men |
| ARL4AP2 | 4 | ADP ribosylation factor like GTPase 4A pseudo |
| APBB2 | 4 | amyloid beta precursor protein binding family |
| AC131953.1 | 4 | novel transcript |
| Y_RNA | 4 | Y RNA [Source:RFAM;Acc:RF00019] |
| RNA5SP160 | 4 | RNA, 5S ribosomal pseudogene 160 [Source:H |
| RNU6-836P | 4 | RNA, U6 small nuclear 836, pseudogene [Sour |
| RNU6-1195P | 4 | RNA, U6 small nuclear 1195, pseudogene [Sou |
| UCHL1-AS1 | 4 | UCHL1 antisense RNA 1 (head to head) [Sourc |
| UCHL1 | 4 | ubiquitin C-terminal hydrolase L1 [Source:HGI |
| Y_RNA | 4 | Y RNA [Source:RFAM;Acc:RF00019] |
| LIMCH1 | 4 | LIM and calponin homology domains 1 [Sourc |
| RPL12P20 | 4 | ribosomal protein L12 pseudogene 20 [Source |

| | | |
|---|---|---|
| AC108050.1 | 4 | novel transcript, antisense to LIMCH1 |
| AC105389.2 | 4 | novel transcript |
| OR5M14P | 4 | olfactory receptor family 5 subfamily M meml |
| PHOX2B | 4 | paired like homeobox 2B [Source:HGNC Symb |
| AC105389.3 | 4 | novel transcript, antisense to PHOX2B |
| AC105389.1 | 4 | novel transcript |
| RNU1-49P | 4 | RNA, U1 small nuclear 49, pseudogene [Sourc |
| HMGB1P28 | 4 | high mobility group box 1 pseudogene 28 [Sou |
| LINC00682 | 4 | long intergenic non-protein coding RNA 682 [S |
| AC108210.1 | 4 | novel transcript |
| AC108210.2 | 4 | tetraspanin 17 (TSPAN17) pseudogene |
| TMEM33 | 4 | transmembrane protein 33 [Source:HGNC Syn |
| DCAF4L1 | 4 | DDB1 and CUL4 associated factor 4 like 1 [Sou |
| AC106052.1 | 4 | novel transcript |
| SLC30A9 | 4 | solute carrier family 30 member 9 [Source:HG |
| ATP1B1P1 | 4 | ATPase Na+/K+ transporting subunit beta 1 ps |
| BEND4 | 4 | BEN domain containing 4 [Source:HGNC Symb |
| AC111006.1 | 4 | novel transcript |
| AC024022.1 | 4 | novel transcript |
| SHISA3 | 4 | shisa family member 3 [Source:HGNC Symbol; |
| ATP8A1 | 4 | ATPase phospholipid transporting 8A1 [Source |
| RPS7P7 | 4 | ribosomal protein S7 pseudogene 7 [Source:H |
| CCNL2P1 | 4 | cyclin L2 pseudogene 1 [Source:HGNC Symbol |
| AC096734.2 | 4 | novel transcript, antisense to ATP8A1 |
| AC096734.1 | 4 | novel transcript |
| RN7SKP82 | 4 | RN7SK pseudogene 82 [Source:HGNC Symbol; |
| GRXCR1 | 4 | glutaredoxin and cysteine rich domain contair |
| AC111198.1 | 4 | novel transcript |
| AC097451.1 | 4 | novel transcript |
| AC098590.2 | 4 | major histocompatibility complex, class II, DP |
| AC098590.1 | 4 | ribosomal protein S2 (RPS2) pseudogene |
| LINC02383 | 4 | long intergenic non-protein coding RNA 2383 |
| AC114774.1 | 4 | piggyBac transposable element derived 3 (PGI |
| RN7SL691P | 4 | RNA, 7SL, cytoplasmic 691, pseudogene [Sour |
| AC080132.1 | 4 | novel transcript |
| RN7SL193P | 4 | RNA, 7SL, cytoplasmic 193, pseudogene [Sour |
| NDUFB4P12 | 4 | NADH:ubiquinone oxidoreductase subunit B4 |
| AC114757.1 | 4 | novel transcript |
| LINC02475 | 4 | long intergenic non-protein coding RNA 2475 |
| KCTD8 | 4 | potassium channel tetramerization domain cc |
| AC093852.1 | 4 | endoplasmic reticulum protein 44 (ERP44) pse |
| YIPF7 | 4 | Yip1 domain family member 7 [Source:HGNC S |
| GUF1 | 4 | GUF1 homolog, GTPase [Source:HGNC Symbo |
| GNPDA2 | 4 | glucosamine-6-phosphate deaminase 2 [Sourc |
| AC096586.2 | 4 | novel transcript, antisense to GUF1 |
| AC096586.1 | 4 | novel transcript, antisense to GNPDA2 |
| AC093755.1 | 4 | tubulin, alpha 1b (TUBA1B) pseudogene |
| PRDX4P1 | 4 | peroxiredoxin 4 pseudogene 1 [Source:HGNC |
| AC108467.2 | 4 | endogenous retrovirus group FRD, member 1 |
| AC108467.1 | 4 | novel transcript |

| | | |
|---|---|---|
| THAP12P9 | 4 | THAP domain containing 12 pseudogene 9 [So |
| AC108043.1 | 4 | ribosomal protein L4 (RPL4) pseudogene |
| RNU6-931P | 4 | RNA, U6 small nuclear 931, pseudogene [Sour |
| RN7SKP199 | 4 | RN7SK pseudogene 199 [Source:HGNC Symbo |
| GABRG1 | 4 | gamma-aminobutyric acid type A receptor gar |
| AC104072.1 | 4 | novel transcript, overlapping GABRA2 |
| GABRA2 | 4 | gamma-aminobutyric acid type A receptor alp |
| AC095060.1 | 4 | novel transcript |
| RNU6-412P | 4 | RNA, U6 small nuclear 412, pseudogene [Sour |
| RAC1P2 | 4 | Rac family small GTPase 1 pseudogene 2 [Sou |
| COX7B2 | 4 | cytochrome c oxidase subunit 7B2 [Source:HG |
| GABRA4 | 4 | gamma-aminobutyric acid type A receptor alp |
| GABRB1 | 4 | gamma-aminobutyric acid type A receptor bet |
| AC107398.3 | 4 | novel transcript, overlapping GABRB1 |
| COMMD8 | 4 | COMM domain containing 8 [Source:HGNC Sy |
| AC107398.4 | 4 | novel transcript |
| AC107398.5 | 4 | novel transcript |
| ATP10D | 4 | ATPase phospholipid transporting 10D (putati |
| AC107398.1 | 4 | ribosomal protein L21 (RPL21) pseudogene |
| AC107398.2 | 4 | novel transcript |
| CORIN | 4 | corin, serine peptidase [Source:HGNC Symbol |
| MIR8053 | 4 | microRNA 8053 [Source:HGNC Symbol;Acc:HG |
| AC092597.1 | 4 | ribosomal protein L15 (RPL15) pseudogene |
| AC107068.2 | 4 | novel transcript, antisense to CORIN and NFXL |
| AC107068.1 | 4 | novel transcript |
| NFXL1 | 4 | nuclear transcription factor, X-box binding like |
| NIPAL1 | 4 | NIPA like domain containing 1 [Source:HGNC S |
| CNGA1 | 4 | cyclic nucleotide gated channel subunit alpha |
| TXK | 4 | TXK tyrosine kinase [Source:HGNC Symbol;Acc |
| RNU6-838P | 4 | RNA, U6 small nuclear 838, pseudogene [Sour |
| U6 | 4 | U6 spliceosomal RNA [Source:RFAM;Acc:RF00 |
| TEC | 4 | tec protein tyrosine kinase [Source:HGNC Sym |
| Y_RNA | 4 | Y RNA [Source:RFAM;Acc:RF00019] |
| SLAIN2 | 4 | SLAIN motif family member 2 [Source:HGNC S |
| SLC10A4 | 4 | solute carrier family 10 member 4 [Source:HG |
| ZAR1 | 4 | zygote arrest 1 [Source:HGNC Symbol;Acc:HG |
| FRYL | 4 | FRY like transcription coactivator [Source:HGN |
| RNU5E-3P | 4 | RNA, U5E small nuclear 3, pseudogene [Sourc |
| OCIAD1 | 4 | OCIA domain containing 1 [Source:HGNC Sym |
| OCIAD1-AS1 | 4 | OCIAD1 antisense RNA 1 [Source:HGNC Symbo |
| OCIAD2 | 4 | OCIA domain containing 2 [Source:HGNC Sym |
| RNU6-158P | 4 | RNA, U6 small nuclear 158, pseudogene [Sour |
| AC020593.1 | 4 | novel pseudogene |
| CWH43 | 4 | cell wall biogenesis 43 C-terminal homolog [So |
| TPI1P4 | 4 | triosephosphate isomerase 1 pseudogene 4 [S |
| AC118282.3 | 4 | sorting nexin 18 (SNX18) pseudogene |
| AC118282.2 | 4 | ankyrin repeat domain 26 (ANKRD26) pseudog |
| AC118282.4 | 4 | tec |
| SNX18P23 | 4 | sorting nexin 18 pseudogene 23 [Source:HGN( |
| AC118282.1 | 4 | ankyrin repeat domain 18B (ANKRD18B) pseu |

| | | |
|---|---|---|
| MTCO3P39 | 4 | MT-CO3 pseudogene 39 [Source:HGNC Symbo |
| MTND3P22 | 4 | MT-ND3 pseudogene 22 [Source:HGNC Symbo |
| AC119751.3 | 4 | ankyrin repeat domain family (ANKRD) pseudo |
| ANKRD20A17P | 4 | ankyrin repeat domain 20 family member A17 |
| AC119751.5 | 4 | coiled-coil domain containing 144 family (CCD |
| AC119751.2 | 4 | family with sequence similarity 201, member |
| AC119751.8 | 4 | family with sequence similarity 201, member |
| AC119751.7 | 4 | NADH dehydrogenase, subunit 3 (complex I) ( |
| MTCO3P42 | 4 | MT-CO3 pseudogene 42 [Source:HGNC Symbo |
| AC119751.1 | 4 | ankyrin repeat domain family (ANKRD) pseudo |
| AC119751.6 | 4 | sorting nexin 18 (SNX18) pseudogene |
| SNX18P24 | 4 | sorting nexin 18 pseudogene 24 [Source:HGN |
| AC119751.4 | 4 | coiled-coil domain containing 144A (CCDC144 |
| SNX18P25 | 4 | sorting nexin 18 pseudogene 25 [Source:HGN |
| DCUN1D4 | 4 | defective in cullin neddylation 1 domain conta |
| DUTP7 | 4 | deoxyuridine triphosphatase pseudogene 7 [S |
| AC027271.1 | 4 | novel transcript |
| AC104784.1 | 4 | ribosomal protein L37a (RPL37A) pseudogene |
| LRRC66 | 4 | leucine rich repeat containing 66 [Source:HGN |
| SGCB | 4 | sarcoglycan beta [Source:HGNC Symbol;Acc:H |
| LINC02480 | 4 | long intergenic non-protein coding RNA 2480 |
| SPATA18 | 4 | spermatogenesis associated 18 [Source:HGNC |
| AC097522.2 | 4 | novel transcript |
| AC097522.1 | 4 | metallophosphoesterase 1 (MPPE1) pseudoge |
| RNU6-1252P | 4 | RNA, U6 small nuclear 1252, pseudogene [Sou |
| USP46 | 4 | ubiquitin specific peptidase 46 [Source:HGNC |
| USP46-AS1 | 4 | USP46 antisense RNA 1 [Source:HGNC Symbol |
| AC104066.3 | 4 | novel transcript |
| LINC01618 | 4 | long intergenic non-protein coding RNA 1618 |
| DANCR | 4 | differentiation antagonizing non-protein codir |
| MIR4449 | 4 | microRNA 4449 [Source:HGNC Symbol;Acc:HG |
| AC104066.4 | 4 | novel transcript |
| SNORA26 | 4 | small nucleolar RNA, H/ACA box 26 [Source:H |
| AC104066.2 | 4 | novel pseudogene |
| ERVMER34-1 | 4 | endogenous retrovirus group MER34 member |
| AC104066.1 | 4 | NULL |
| Y_RNA | 4 | Y RNA [Source:RFAM;Acc:RF00019] |
| RASL11B | 4 | RAS like family 11 member B [Source:HGNC Sy |
| SCFD2 | 4 | sec1 family domain containing 2 [Source:HGN |
| AC023154.1 | 4 | novel transcript |
| RNU6-310P | 4 | RNA, U6 small nuclear 310, pseudogene [Sour |
| FIP1L1 | 4 | factor interacting with PAPOLA and CPSF1 [So |
| AC058822.1 | 4 | novel FIP1L1-PDGFRA fusion protein |
| LNX1 | 4 | ligand of numb-protein X 1 [Source:HGNC Sym |
| LNX1-AS1 | 4 | LNX1 antisense RNA 1 [Source:HGNC Symbol; |
| AC095040.1 | 4 | COMM domain containing 5 (COMMD5) pseu |
| LNX1-AS2 | 4 | LNX1 antisense RNA 2 [Source:HGNC Symbol; |
| AC124017.1 | 4 | novel transcript |
| AC105384.1 | 4 | novel transcript |
| AC105384.2 | 4 | nucleic acid binding protein 2 (NABP2) pseudo |

| | | |
|---|---|---|
| RPL21P44 | 4 | ribosomal protein L21 pseudogene 44 [Source |
| AC110792.3 | 4 | novel transcript |
| CHIC2 | 4 | cysteine rich hydrophobic domain 2 [Source:H |
| AC110792.2 | 4 | novel transcript, antisense to CHIC2 |
| AC110792.4 | 4 | novel transcript, sense intronic to RP11-231C1 |
| AC110792.1 | 4 | cytochrome c oxidase subunit VIc (COX6C) pse |
| MORF4L2P1 | 4 | mortality factor 4 like 2 pseudogene 1 [Source |
| GSX2 | 4 | GS homeobox 2 [Source:HGNC Symbol;Acc:HG |
| RPL22P13 | 4 | ribosomal protein L22 pseudogene 13 [Source |
| PDGFRA | 4 | platelet derived growth factor receptor alpha |
| LINC02283 | 4 | long intergenic non-protein coding RNA 2283 |
| AC098587.1 | 4 | novel transcript |
| AC098868.1 | 4 | novel transcript |
| LINC02260 | 4 | long intergenic non-protein coding RNA 2260 |
| KIT | 4 | KIT proto-oncogene, receptor tyrosine kinase |
| AC097494.1 | 4 | novel transcript |
| LINC02358 | 4 | long intergenic non-protein coding RNA 2358 |
| AC111194.1 | 4 | novel transcript |
| RPL38P3 | 4 | ribosomal protein L38 pseudogene 3 [Source: |
| RNU6-410P | 4 | RNA, U6 small nuclear 410, pseudogene [Sour |
| AC111194.2 | 4 | novel transcript, antisense to KDR |
| RN7SL424P | 4 | RNA, 7SL, cytoplasmic 424, pseudogene [Sour |
| KDR | 4 | kinase insert domain receptor [Source:HGNC S |
| AC021220.2 | 4 | novel transcript |
| RN7SL822P | 4 | RNA, 7SL, cytoplasmic 822, pseudogene [Sour |
| AC021220.1 | 4 | methyltransferase like 5 (METTL5) pseudogen |
| RNU6-746P | 4 | RNA, U6 small nuclear 746, pseudogene [Sour |
| LRRC34P2 | 4 | leucine rich repeat containing 34 pseudogene |
| SRD5A3 | 4 | steroid 5 alpha-reductase 3 [Source:HGNC Syr |
| FCF1P8 | 4 | FCF1 pseudogene 8 [Source:HGNC Symbol;Acc |
| SRD5A3-AS1 | 4 | SRD5A3 antisense RNA 1 [Source:HGNC Symb |
| AC069200.1 | 4 | novel transcript |
| TMEM165 | 4 | transmembrane protein 165 [Source:HGNC Sy |
| Y_RNA | 4 | Y RNA [Source:RFAM;Acc:RF00019] |
| CLOCK | 4 | clock circadian regulator [Source:HGNC Symbo |
| Y_RNA | 4 | Y RNA [Source:RFAM;Acc:RF00019] |
| RN7SKP30 | 4 | RN7SK pseudogene 30 [Source:HGNC Symbol; |
| AC024243.1 | 4 | novel transcript, antisense to clock |
| PDCL2 | 4 | phosducin like 2 [Source:HGNC Symbol;Acc:H( |
| NMU | 4 | neuromedin U [Source:HGNC Symbol;Acc:HGN |
| RNU6-276P | 4 | RNA, U6 small nuclear 276, pseudogene [Sour |
| EXOC1L | 4 | exocyst complex component 1 like [Source:HC |
| EXOC1 | 4 | exocyst complex component 1 [Source:HGNC |
| RNU6-652P | 4 | RNA, U6 small nuclear 652, pseudogene [Sour |
| AC110611.1 | 4 | novel transcript, antisense to EXOC1 |
| AC110611.2 | 4 | novel transcript |
| CEP135 | 4 | centrosomal protein 135 [Source:HGNC Symb |
| KIAA1211 | 4 | KIAA1211 [Source:HGNC Symbol;Acc:HGNC:2 |
| RNA5SP161 | 4 | RNA, 5S ribosomal pseudogene 161 [Source:H |
| RNU6-197P | 4 | RNA, U6 small nuclear 197, pseudogene [Sour |

| | | |
|---|---|---|
| AC022267.1 | 4 | novel transcript |
| MRPL22P1 | 4 | mitochondrial ribosomal protein L22 pseudog |
| AASDH | 4 | aminoadipate-semialdehyde dehydrogenase [ |
| RPL7AP31 | 4 | ribosomal protein L7a pseudogene 31 [Source |
| AC068620.1 | 4 | novel transcript |
| PPAT | 4 | phosphoribosyl pyrophosphate amidotransfer |
| AC068620.2 | 4 | novel transcript |
| AC068620.3 | 4 | novel Myb/SANT-like DNA-binding domain co |
| PAICS | 4 | phosphoribosylaminoimidazole carboxylase a |
| SRP72 | 4 | signal recognition particle 72 [Source:HGNC S |
| ARL9 | 4 | ADP ribosylation factor like GTPase 9 [Source: |
| THEGL | 4 | theg spermatid protein like [Source:HGNC Syn |
| GLDCP1 | 4 | glycine decarboxylase pseudogene 1 [Source: |
| HOPX | 4 | HOP homeobox [Source:HGNC Symbol;Acc:H |
| AC108215.1 | 4 | TEC |
| RPL17P20 | 4 | ribosomal protein L17 pseudogene 20 [Source |
| AC022483.1 | 4 | thioredoxin domain containing 9 (TXNDC9) ps |
| RN7SL492P | 4 | RNA, 7SL, cytoplasmic 492, pseudogene [Sour |
| RN7SL357P | 4 | RNA, 7SL, cytoplasmic 357, pseudogene [Sour |
| SPINK2 | 4 | serine peptidase inhibitor, Kazal type 2 [Sourc |
| Metazoa_SRP | 4 | Metazoan signal recognition particle RNA [Sou |
| Y_RNA | 4 | Y RNA [Source:RFAM;Acc:RF00019] |
| REST | 4 | RE1 silencing transcription factor [Source:HGN |
| AC069307.1 | 4 | novel transcript |
| NOA1 | 4 | nitric oxide associated 1 [Source:HGNC Symbo |
| POLR2B | 4 | RNA polymerase II subunit B [Source:HGNC Sy |
| RNU6-998P | 4 | RNA, U6 small nuclear 998, pseudogene [Sour |
| IGFBP7 | 4 | insulin like growth factor binding protein 7 [So |
| UBE2CP3 | 4 | ubiquitin conjugating enzyme E2 C pseudogen |
| IGFBP7-AS1 | 4 | IGFBP7 antisense RNA 1 [Source:HGNC Symbo |
| AC111197.1 | 4 | novel transcript |
| AC105390.1 | 4 | novel transcript |
| RPS26P24 | 4 | ribosomal protein S26 pseudogene 24 [Source |
| LINC02380 | 4 | long intergenic non-protein coding RNA 2380 |
| AC013724.1 | 4 | novel transcript |
| AC107396.1 | 4 | novel transcript |
| AC093725.2 | 4 | novel transcript |
| AC093725.1 | 4 | novel transcript |
| AC104790.1 | 4 | novel transcript |
| SRIP1 | 4 | sorcin pseudogene 1 [Source:HGNC Symbol;A |
| AC096725.1 | 4 | echinoderm microtubule associated protein li |
| LINC02494 | 4 | long intergenic non-protein coding RNA 2494 |
| AC019133.1 | 4 | novel transcript |
| AC017091.1 | 4 | novel transcript |
| LINC02619 | 4 | long intergenic non-protein coding RNA 2619 |
| LINC02429 | 4 | long intergenic non-protein coding RNA 2429 |
| AC097501.1 | 4 | novel transcript |
| AC108517.2 | 4 | novel transcript |
| AC108517.1 | 4 | novel transcript |
| Y_RNA | 4 | Y RNA [Source:RFAM;Acc:RF00019] |

| | | |
|---|---|---|
| AC096588.1 | 4 | novel transcript |
| AC093857.1 | 4 | novel transcript |
| Y_RNA | 4 | Y RNA [Source:RFAM;Acc:RF00019] |
| RNU6-1325P | 4 | RNA, U6 small nuclear 1325, pseudogene [Sou |
| AC092596.1 | 4 | novel transcript |
| AC097655.1 | 4 | ribosomal protein, large, P1 (RPLP1) pseudoge |
| LINC02496 | 4 | long intergenic non-protein coding RNA 2496 |
| AC105420.1 | 4 | protein phosphatase 2 (formerly 2A), regulato |
| MIR548AG1 | 4 | microRNA 548ag-1 [Source:HGNC Symbol;Acc |
| LINC02271 | 4 | long intergenic non-protein coding RNA 2271 |
| ADGRL3 | 4 | adhesion G protein-coupled receptor L3 [Sour |
| RPL17P19 | 4 | ribosomal protein L17 pseudogene 19 [Source |
| AC020741.1 | 4 | novel transcript |
| AC108161.1 | 4 | ribosomal protein S12 (RPS12) pseudogene |
| Y_RNA | 4 | Y RNA [Source:RFAM;Acc:RF00019] |
| ADGRL3-AS1 | 4 | adhesion G protein-coupled receptor L3 antise |
| RPS15AP17 | 4 | ribosomal protein S15a pseudogene 17 [Sourc |
| RPL21P47 | 4 | ribosomal protein L21 pseudogene 47 [Source |
| AC105313.1 | 4 | ubiquitin-conjugating enzyme E2L 3 (UBE2L3) |
| AC074087.1 | 4 | coiled-coil domain containing 65 (CCDC65) pse |
| HMGN1P11 | 4 | high mobility group nucleosome binding doma |
| EXOC5P1 | 4 | exocyst complex component 5 pseudogene 1 |
| AC097491.1 | 4 | novel transcript |
| LARP1BP1 | 4 | La ribonucleoprotein domain family member 1 |
| AC093730.1 | 4 | novel transcript |
| TECRL | 4 | trans-2,3-enoyl-CoA reductase like [Source:HG |
| RNU6-191P | 4 | RNA, U6 small nuclear 191, pseudogene [Sour |
| DPP3P1 | 4 | dipeptidyl peptidase 3 pseudogene 1 [Source: |
| MTCO3P27 | 4 | MT-CO3 pseudogene 27 [Source:HGNC Symbo |
| MTCYBP16 | 4 | MT-CYB pseudogene 16 [Source:HGNC Symbo |
| MTND6P16 | 4 | MT-ND6 pseudogene 16 [Source:HGNC Symbo |
| MTND5P13 | 4 | MT-ND5 pseudogene 13 [Source:HGNC Symbo |
| MTND4LP31 | 4 | MT-ND4L pseudogene 31 [Source:HGNC Symb |
| MTND3P24 | 4 | MT-ND3 pseudogene 24 [Source:HGNC Symbo |
| MTCO3P28 | 4 | MT-CO3 pseudogene 28 [Source:HGNC Symbo |
| AC104689.1 | 4 | ribosomal protein S6 (RPS6) pseudogene |
| AC104689.2 | 4 | novel transcript |
| LINC02232 | 4 | long intergenic non-protein coding RNA 2232 |
| AC107058.1 | 4 | tripartite motif-containing 75 (TRIM75) pseud |
| EFL1P2 | 4 | elongation factor like GTPase 1 pseudogene 2 |
| AC096754.1 | 4 | ArfGAP with GTPase domain, ankyrin repeat a |
| LINC02835 | 4 | long intergenic non-protein coding RNA 2835 |
| EPHA5 | 4 | EPH receptor A5 [Source:HGNC Symbol;Acc:H |
| AC115223.1 | 4 | ribosomal protein L6 (RPL6) pseudogene |
| EPHA5-AS1 | 4 | EPHA5 antisense RNA 1 [Source:HGNC Symbo |
| AC104137.1 | 4 | novel transcript |
| RNU2-40P | 4 | RNA, U2 small nuclear 40, pseudogene [Sourc |
| AC097110.1 | 4 | novel transcript |
| AC116049.2 | 4 | novel transcript |
| AC096721.1 | 4 | novel transcript |

| | | |
|---|---|---|
| AC116049.1 | 4 | interferon induced transmembrane protein 3 |
| AC110809.1 | 4 | novel transcript |
| MIR1269A | 4 | microRNA 1269a [Source:HGNC Symbol;Acc:H |
| RPS23P3 | 4 | ribosomal protein S23 pseudogene 3 [Source: |
| RNU6-699P | 4 | RNA, U6 small nuclear 699, pseudogene [Sour |
| AC104806.2 | 4 | novel transcript |
| RNU1-63P | 4 | RNA, U1 small nuclear 63, pseudogene [Sourc |
| RNA5SP527 | 4 | RNA, 5S ribosomal pseudogene 527 [Source:H |
| AC104806.1 | 4 | CCR4-NOT transcription complex, subunit 7 (C |
| CENPC | 4 | centromere protein C [Source:HGNC Symbol;A |
| STAP1 | 4 | signal transducing adaptor family member 1 [S |
| AC096720.2 | 4 | tec |
| UBA6 | 4 | ubiquitin like modifier activating enzyme 6 [Sc |
| AC096720.1 | 4 | small EDRK-rich factor 2 (SERF2) pseudogene |
| UBA6-AS1 | 4 | UBA6 antisense RNA 1 (head to head) [Source |
| ST3GAL1P1 | 4 | ST3 beta-galactoside alpha-2,3-sialyltransfera: |
| AC079880.1 | 4 | HEAT repeat containing 2 (HEATR2) pseudoge |
| AC079880.2 | 4 | tubulin, alpha 4a (TUBA4A) pseudogene |
| GNRHR | 4 | gonadotropin releasing hormone receptor [So |
| SNORA62 | 4 | Small nucleolar RNA SNORA62/SNORA6 family |
| TMPRSS11CP | 4 | transmembrane serine protease 11C, pseudog |
| TMPRSS11D | 4 | transmembrane serine protease 11D [Source: |
| TMPRSS11A | 4 | transmembrane serine protease 11A [Source: |
| TMPRSS11GP | 4 | transmembrane serine protease 11G, pseudog |
| RNU6-95P | 4 | RNA, U6 small nuclear 95, pseudogene [Sourc |
| GCOM2 | 4 | GRINL1B complex locus 2 (pseudogene) [Sour |
| MTND2P41 | 4 | MT-ND2 pseudogene 41 [Source:HGNC Symbo |
| TMPRSS11F | 4 | transmembrane serine protease 11F [Source:I |
| SYT14P1 | 4 | synaptotagmin 14 pseudogene 1 [Source:HGN |
| FTLP10 | 4 | ferritin light chain pseudogene 10 [Source:HG |
| TMPRSS11BNL | 4 | TMPRSS11B N-terminal like, pseudogene [Sou |
| TMPRSS11B | 4 | transmembrane serine protease 11B [Source: |
| AC098799.3 | 4 | UDP glucuronosyltransferase 2 family, polype |
| AC098799.1 | 4 | UDP glucuronos...yltransferase 2 family, polyp |
| AC098799.2 | 4 | UDP glucuronosyltransferase 2 family, polype |
| APOOP4 | 4 | apolipoprotein O pseudogene 4 [Source:HGN( |
| YTHDC1 | 4 | YTH domain containing 1 [Source:HGNC Symb |
| MT2P1 | 4 | metallothionein 2 pseudogene 1 [Source:HGN |
| TMPRSS11E | 4 | transmembrane serine protease 11E [Source:I |
| UGT2B29P | 4 | UDP glucuronosyltransferase family 2 membe |
| UGT2B17 | 4 | UDP glucuronosyltransferase family 2 membe |
| AC147055.2 | 4 | pseudogene similar to part of RAS guanyl rele |
| AC147055.4 | 4 | UBX domain protein 2A (UBXN2A) pseudogen |
| AC147055.3 | 4 | UDP glucuronosyltransferase 2 family pseudog |
| UGT2B15 | 4 | UDP glucuronosyltransferase family 2 membe |
| AC147055.1 | 4 | pseudogene similar to part of UDP glucuronos |
| AC093720.1 | 4 | pseudogene similar to part of UDP glucuronos |
| AC021146.8 | 4 | pseudogene similar to part of UDP glucuronos |
| UGT2B10 | 4 | UDP glucuronosyltransferase family 2 membe |
| AC021146.9 | 4 | pseudogene similar to part of UDP glucuronos |

| | | |
|---|---|---|
| AC021146.4 | 4 | pseudogene similar to part of UDP glucuronos |
| AC021146.11 | 4 | pseudogene similar to part of UDP glucuronos |
| AC021146.2 | 4 | speckle-type POZ protein-like (SPOPL) pseudo |
| AC021146.3 | 4 | pseudogene similar to part of UDP glucuronos |
| AC021146.12 | 4 | novel transcript |
| AC021146.6 | 4 | pseudogene similar to part of UDP glucuronos |
| AC021146.1 | 4 | pseudogene similar to part of UDP glucuronos |
| UGT2A3 | 4 | UDP glucuronosyltransferase family 2 membe |
| AC021146.10 | 4 | pseudogene similar to part of UDP glucuronos |
| AC021146.5 | 4 | UDP glucuronosyltransferase 2 family pseudo |
| AC021146.7 | 4 | speckle-type POZ protein-like (SPOPL) pseudo |
| UGT2B27P | 4 | UDP glucuronosyltransferase family 2 membe |
| AC107401.1 | 4 | pseudogene similar to part of UDP glucuronos |
| UGT2B26P | 4 | UDP glucuronosyltransferase family 2 membe |
| UGT2B7 | 4 | UDP glucuronosyltransferase family 2 membe |
| AC111000.2 | 4 | UDP glucuronosyltransferase 2 family, polype |
| AC111000.5 | 4 | speckle-type POZ protein-like (SPOPL) pseudo |
| AC111000.3 | 4 | UDP glucuronosyltransferase 2 family pseudo |
| AC111000.6 | 4 | UDP glucuronosyltransferase 2 family pseudo |
| AC111000.1 | 4 | UDP glucuronosyltransferase 2 family, polype |
| AC111000.4 | 4 | novel transcript, antisense to UGT2B11 |
| UGT2B11 | 4 | UDP glucuronosyltransferase family 2 membe |
| AC114797.1 | 4 | UDP glucuronosyltransferase 2 family pseudo |
| AC114786.1 | 4 | UDP glucuronosyltransferase 2 family pseudo |
| UGT2B28 | 4 | UDP glucuronosyltransferase family 2 membe |
| AC114786.3 | 4 | UDP glucuronosyltransferase 2 family, polype |
| AC114786.2 | 4 | UDP glucuronosyltransferase 2 family pseudo |
| AC114786.4 | 4 | UDP glucuronosyltransferase 2 family pseudo |
| UGT2B25P | 4 | UDP glucuronosyltransferase family 2 membe |
| AC108078.1 | 4 | UDP glucuronosyltransferase 2 family pseudo |
| UGT2B24P | 4 | UDP glucuronosyltransferase family 2 membe |
| AC108078.3 | 4 | UDP glucuronosyltransferase 2 family pseudo |
| AC108078.2 | 4 | UDP glucuronosyltransferase 2 family pseudo |
| UGT2B4 | 4 | UDP glucuronosyltransferase family 2 membe |
| UGT2A3P7 | 4 | UDP glucuronosyltransferase family 2 membe |
| AC093829.1 | 4 | UDP glucuronosyltransferase 2 family pseudo |
| UGT2A1 | 4 | UDP glucuronosyltransferase family 2 membe |
| UGT2A2 | 4 | UDP glucuronosyltransferase family 2 membe |
| SULT1B1 | 4 | sulfotransferase family 1B member 1 [Source: |
| SULT1D1P | 4 | sulfotransferase family 1D member 1, pseudo |
| AC108941.2 | 4 | novel protein |
| SULT1E1 | 4 | sulfotransferase family 1E member 1 [Source: |
| AC108941.1 | 4 | solute carrier family 25 (mitochondrial carrier |
| CSN1S1 | 4 | casein alpha s1 [Source:HGNC Symbol;Acc:HG |
| CSN2 | 4 | casein beta [Source:HGNC Symbol;Acc:HGNC: |
| STATH | 4 | statherin [Source:HGNC Symbol;Acc:HGNC:11 |
| HTN3 | 4 | histatin 3 [Source:HGNC Symbol;Acc:HGNC:52 |
| HTN1 | 4 | histatin 1 [Source:HGNC Symbol;Acc:HGNC:52 |
| CSN1S2AP | 4 | casein alpha s2 like A, pseudogene [Source:HG |
| PRR27 | 4 | proline rich 27 [Source:HGNC Symbol;Acc:HGN |

| | | |
|---|---|---|
| ODAM | 4 | odontogenic, ameleoblast associated [Source:H |
| FDCSP | 4 | follicular dendritic cell secreted protein [Sourc |
| CSN3 | 4 | casein kappa [Source:HGNC Symbol;Acc:HGN( |
| CABS1 | 4 | calcium binding protein, spermatid associated |
| SMR3A | 4 | submaxillary gland androgen regulated protei |
| SMR3B | 4 | submaxillary gland androgen regulated protei |
| OPRPN | 4 | opiorphin prepropeptide [Source:HGNC Symb |
| MUC7 | 4 | mucin 7, secreted [Source:HGNC Symbol;Acc:I |
| AMTN | 4 | amelotin [Source:HGNC Symbol;Acc:HGNC:33 |
| AMBN | 4 | ameloblastin [Source:HGNC Symbol;Acc:HGN( |
| ENAM | 4 | enamelin [Source:HGNC Symbol;Acc:HGNC:33 |
| AC009570.2 | 4 | novel transcript |
| JCHAIN | 4 | joining chain of multimeric IgA and IgM [Sourc |
| UTP3 | 4 | UTP3 small subunit processome component [S |
| RNU6-520P | 4 | RNA, U6 small nuclear 520, pseudogene [Sour |
| RNU6-784P | 4 | RNA, U6 small nuclear 784, pseudogene [Sour |
| AC009570.1 | 4 | novel transcript, antisense to RUFY3 |
| RUFY3 | 4 | RUN and FYVE domain containing 3 [Source:H |
| GRSF1 | 4 | G-rich RNA sequence binding factor 1 [Source |
| U6 | 4 | U6 spliceosomal RNA [Source:RFAM;Acc:RF00 |
| RNU6-891P | 4 | RNA, U6 small nuclear 891, pseudogene [Sour |
| MOB1B | 4 | MOB kinase activator 1B [Source:HGNC Symb |
| DCK | 4 | deoxycytidine kinase [Source:HGNC Symbol;A |
| SLC4A4 | 4 | solute carrier family 4 member 4 [Source:HGN |
| AC096713.1 | 4 | solute carrier family 35, member A4 (SLC35A4 |
| LDHAL6EP | 4 | lactate dehydrogenase A like 6E, pseudogene |
| GC | 4 | GC vitamin D binding protein [Source:HGNC Sy |
| RNA5SP163 | 4 | RNA, 5S ribosomal pseudogene 163 [Source:H |
| AC068721.1 | 4 | novel transcript |
| NPFFR2 | 4 | neuropeptide FF receptor 2 [Source:HGNC Syr |
| ADAMTS3 | 4 | ADAM metallopeptidase with thrombospondi |
| AC095056.1 | 4 | novel transcript |
| HNRNPA1P67 | 4 | heterogeneous nuclear ribonucleoprotein A1 |
| RNU4ATAC9P | 4 | RNA, U4atac small nuclear 9, pseudogene [Sou |
| RNU6ATAC5P | 4 | RNA, U6atac small nuclear 5, pseudogene [Sou |
| COX18 | 4 | cytochrome c oxidase assembly factor COX18 |
| ANKRD17 | 4 | ankyrin repeat domain 17 [Source:HGNC Syml |
| HMGA1P2 | 4 | high mobility group AT-hook 1 pseudogene 2 |
| AC053527.2 | 4 | novel transcript |
| AC053527.1 | 4 | NULL |
| AC108157.1 | 4 | zinc finger, AN1-type domain 1 (ZFAND1) pseu |
| ALB | 4 | albumin [Source:HGNC Symbol;Acc:HGNC:399 |
| AFP | 4 | alpha fetoprotein [Source:HGNC Symbol;Acc:H |
| AFM | 4 | afamin [Source:HGNC Symbol;Acc:HGNC:316] |
| LINC02499 | 4 | long intergenic non-protein coding RNA 2499 |
| AC074250.1 | 4 | ribosomal protein L13a (RPL13A) pseudogene |
| RASSF6 | 4 | Ras association domain family member 6 [Sou |
| UMLILO | 4 | upstream master lncRNA of the inflammatory |
| CXCL8 | 4 | C-X-C motif chemokine ligand 8 [Source:HGN( |
| AC112518.1 | 4 | chromosome 11 open reading frame 73 (C11c |

| | | |
|---|---|---|
| CXCL6 | 4 | C-X-C motif chemokine ligand 6 [Source:HGN( |
| PPBPP1 | 4 | pro-platelet basic protein pseudogene 1 [Sour |
| PF4V1 | 4 | platelet factor 4 variant 1 [Source:HGNC Symb |
| CXCL1 | 4 | C-X-C motif chemokine ligand 1 [Source:HGN( |
| HNRNPA1P55 | 4 | heterogeneous nuclear ribonucleoprotein A1 |
| CXCL1P1 | 4 | C-X-C motif chemokine ligand 1 pseudogene 1 |
| PF4 | 4 | platelet factor 4 [Source:HGNC Symbol;Acc:H( |
| PPBP | 4 | pro-platelet basic protein [Source:HGNC Symb |
| CXCL5 | 4 | C-X-C motif chemokine ligand 5 [Source:HGN( |
| AC097709.1 | 4 | novel transcript, antisense to CXCL5 |
| RN7SL218P | 4 | RNA, 7SL, cytoplasmic 218, pseudogene [Sour |
| CXCL3 | 4 | C-X-C motif chemokine ligand 3 [Source:HGN( |
| PPBPP2 | 4 | pro-platelet basic protein pseudogene 2 [Sour |
| AC093677.3 | 4 | novel transcript |
| AC093677.1 | 4 | coiled-coil domain containing 90B (CCDC90B) |
| CXCL2 | 4 | C-X-C motif chemokine ligand 2 [Source:HGN( |
| MTHFD2L | 4 | methylenetetrahydrofolate dehydrogenase (N |
| AC093677.2 | 4 | novel transcript, antisense to MTHFD2L |
| EPGN | 4 | epithelial mitogen [Source:HGNC Symbol;Acc: |
| EREG | 4 | epiregulin [Source:HGNC Symbol;Acc:HGNC:3 |
| AC021180.1 | 4 | novel transcript |
| AREG | 4 | amphiregulin [Source:HGNC Symbol;Acc:HGN |
| AC239584.1 | 4 | novel transcript |
| BTC | 4 | betacellulin [Source:HGNC Symbol;Acc:HGNC: |
| AC098825.1 | 4 | novel transcript |
| HSPE1P23 | 4 | heat shock protein family E (Hsp10) member : |
| PARM1 | 4 | prostate androgen-regulated mucin-like prote |
| AC110760.2 | 4 | novel transcript |
| AC110760.1 | 4 | novel transcript |
| LINC02562 | 4 | long intergenic non-protein coding RNA 2562 |
| AC110760.3 | 4 | ubiquitin-conjugating enzyme E2D 4 (putative |
| AC093870.1 | 4 | zinc finger protein 99 (ZNF99) pseudogene |
| AC025244.1 | 4 | novel transcript |
| AC025244.2 | 4 | novel transcript |
| LINC02483 | 4 | long intergenic non-protein coding RNA 2483 |
| AC096759.1 | 4 | novel transcript |
| AC096759.2 | 4 | novel transcript |
| RCHY1 | 4 | ring finger and CHY zinc finger domain contair |
| THAP6 | 4 | THAP domain containing 6 [Source:HGNC Sym |
| ODAPH | 4 | odontogenesis associated phosphoprotein [Sc |
| CDKL2 | 4 | cyclin dependent kinase like 2 [Source:HGNC S |
| G3BP2 | 4 | G3BP stress granule assembly factor 2 [Source |
| Y_RNA | 4 | Y RNA [Source:RFAM;Acc:RF00019] |
| USO1 | 4 | USO1 vesicle transport factor [Source:HGNC S |
| AC110615.1 | 4 | novel transcript |
| RNU2-16P | 4 | RNA, U2 small nuclear 16, pseudogene [Sourc |
| PPEF2 | 4 | protein phosphatase with EF-hand domain 2 [ |
| NAAA | 4 | N-acylethanolamine acid amidase [Source:HG |
| SDAD1 | 4 | SDA1 domain containing 1 [Source:HGNC Sym |
| AC112719.2 | 4 | novel transcript |

| | | |
|---|---|---|
| CXCL9 | 4 | C-X-C motif chemokine ligand 9 [Source:HGNC |
| ART3 | 4 | ADP-ribosyltransferase 3 [Source:HGNC Symb |
| CXCL10 | 4 | C-X-C motif chemokine ligand 10 [Source:HGN |
| CXCL11 | 4 | C-X-C motif chemokine ligand 11 [Source:HGN |
| AC112719.1 | 4 | ribosomal protein L36 (RPL36) pseudogene |
| NUP54 | 4 | nucleoporin 54 [Source:HGNC Symbol;Acc:HG |
| AC110795.1 | 4 | novel transcript |
| SCARB2 | 4 | scavenger receptor class B member 2 [Source |
| FAM47E | 4 | family with sequence similarity 47 member E |
| AC034139.1 | 4 | novel transcript, antisense to STBD1and FAM4 |
| FAM47E-STBD1 | 4 | FAM47E-STBD1 readthrough [Source:HGNC Sy |
| STBD1 | 4 | starch binding domain 1 [Source:HGNC Symbo |
| CCDC158 | 4 | coiled-coil domain containing 158 [Source:HG |
| SNX5P1 | 4 | sorting nexin 5 pseudogene 1 [Source:HGNC S |
| RNU6-1000P | 4 | RNA, U6 small nuclear 1000, pseudogene [Sou |
| AC096743.2 | 4 | ribosomal protein L36a pseudogene 18 |
| AC096743.1 | 4 | NULL |
| SHROOM3 | 4 | shroom family member 3 [Source:HGNC Symb |
| AC112249.1 | 4 | ribosomal protein L18 (RPL18) pseudogene |
| RNU6-145P | 4 | RNA, U6 small nuclear 145, pseudogene [Sour |
| MIR4450 | 4 | microRNA 4450 [Source:HGNC Symbol;Acc:HG |
| MIR548AH | 4 | microRNA 548ah [Source:HGNC Symbol;Acc:H |
| AC107072.2 | 4 | novel transcript |
| AC107072.1 | 4 | novel transcript |
| AC104687.2 | 4 | NUAK family, SNF1-like kinase, 2 (NUAK2) pse |
| AC104687.1 | 4 | ribosomal protein L26 (RPL26) pseudogene |
| SOWAHB | 4 | sosondowah ankyrin repeat domain family me |
| SEPTIN11 | 4 | septin 11 [Source:HGNC Symbol;Acc:HGNC:25 |
| TXNP6 | 4 | thioredoxin pseudogene 6 [Source:HGNC Sym |
| CCNI | 4 | cyclin I [Source:HGNC Symbol;Acc:HGNC:1595 |
| RPL7P17 | 4 | ribosomal protein L7 pseudogene 17 [Source: |
| AC104771.1 | 4 | ubiquitin family domain containing 1 (UBFD1) |
| RNU6-1187P | 4 | RNA, U6 small nuclear 1187, pseudogene [Sou |
| CCNG2 | 4 | cyclin G2 [Source:HGNC Symbol;Acc:HGNC:15 |
| AC008638.3 | 4 | ferritin, light polypeptide (FTL) pseudogene |
| AC008638.2 | 4 | developmental pluripotency associated 2 (DP |
| AC008638.1 | 4 | developmental pluripotency associated 2 (DP |
| AC092674.1 | 4 | novel transcript |
| CXCL13 | 4 | C-X-C motif chemokine ligand 13 [Source:HGN |
| AC104701.1 | 4 | NULL |
| CNOT6L | 4 | CCR4-NOT transcription complex subunit 6 lik |
| MRPL1 | 4 | mitochondrial ribosomal protein L1 [Source:H |
| AC112225.1 | 4 | cytochrome c oxidase subunit VIc (COX6C) pse |
| AC114801.2 | 4 | FUS interacting protein (serine/arginine-rich) |
| HMGB1P44 | 4 | high mobility group box 1 pseudogene 44 [Sou |
| HNRNPA1P56 | 4 | heterogeneous nuclear ribonucleoprotein A1 |
| AC114801.1 | 4 | transmembrane protein 128 (TMEM128) pseu |
| AC114801.3 | 4 | novel pseudogene |
| FRAS1 | 4 | Fraser extracellular matrix complex subunit 1 |
| SERBP1P5 | 4 | SERPINE1 mRNA binding protein 1 pseudogen |

| | | |
|---|---|---|
| MICOS10P4 | 4 | MICOS10 pseudogene 4 [Source:HGNC Symbo |
| ANXA3 | 4 | annexin A3 [Source:HGNC Symbol;Acc:HGNC:! |
| Y_RNA | 4 | Y RNA [Source:RFAM;Acc:RF00019] |
| LINC01094 | 4 | long intergenic non-protein coding RNA 1094 |
| HIGD1AP13 | 4 | HIG1 hypoxia inducible domain family membe |
| AC112253.1 | 4 | novel transcript |
| AC098818.1 | 4 | ribosomal protein S21 (RPS21) pseudogene |
| AC098818.2 | 4 | novel transcript, antisense to BMP2K |
| BMP2K | 4 | BMP2 inducible kinase [Source:HGNC Symbol; |
| AC098818.3 | 4 | NULL |
| PAQR3 | 4 | progestin and adipoQ receptor family membe |
| RN7SL127P | 4 | RNA, 7SL, cytoplasmic 127, pseudogene [Sour |
| LINC01088 | 4 | long intergenic non-protein coding RNA 1088 |
| NAA11 | 4 | N(alpha)-acetyltransferase 11, NatA catalytic : |
| GK2 | 4 | glycerol kinase 2 [Source:HGNC Symbol;Acc:H |
| LINC00989 | 4 | long intergenic non-protein coding RNA 989 [S |
| OR7E94P | 4 | olfactory receptor family 7 subfamily E memb |
| AC092542.1 | 4 | novel transcript |
| LINC02469 | 4 | long intergenic non-protein coding RNA 2469 |
| PCAT4 | 4 | prostate cancer associated transcript 4 [Sourc |
| SNORA75 | 4 | Small nucleolar RNA SNORA75 [Source:RFAM; |
| ANTXR2 | 4 | ANTXR cell adhesion molecule 2 [Source:HGN( |
| KPNA2P1 | 4 | karyopherin subunit alpha 2 pseudogene 1 [Sc |
| RPSAP39 | 4 | ribosomal protein SA pseudogene 39 [Source: |
| AC021127.1 | 4 | putative novel transript |
| PRDM8 | 4 | PR/SET domain 8 [Source:HGNC Symbol;Acc:H |
| FGF5 | 4 | fibroblast growth factor 5 [Source:HGNC Symt |
| CFAP299 | 4 | cilia and flagella associated protein 299 [Sourc |
| AC105917.1 | 4 | zinc finger protein 341 (ZNF341) pseudogene |
| BMP3 | 4 | bone morphogenetic protein 3 [Source:HGNC |
| PRKG2 | 4 | protein kinase cGMP-dependent 2 [Source:HG |
| AC139722.1 | 4 | novel transcript |
| RNU5A-2P | 4 | RNA, U5A small nuclear 2, pseudogene [Sourc |
| RASGEF1B | 4 | RasGEF domain family member 1B [Source:HG |
| MTCYBP44 | 4 | MT-CYB pseudogene 44 [Source:HGNC Symbo |
| COX5BP1 | 4 | cytochrome c oxidase subunit 5B pseudogene |
| AC021863.1 | 4 | NULL |
| NPM1P41 | 4 | nucleophosmin 1 pseudogene 41 [Source:HGN |
| HNRNPA3P13 | 4 | heterogeneous nuclear ribonucleoprotein A3 |
| RNU6-499P | 4 | RNA, U6 small nuclear 499, pseudogene [Sour |
| BIN2P1 | 4 | bridging integrator 2 pseudogene 1 [Source:H |
| VAMP9P | 4 | vesicle associated membrane protein 9, pseuc |
| AC124016.2 | 4 | novel transcript |
| HNRNPD | 4 | heterogeneous nuclear ribonucleoprotein D [S |
| AC124016.1 | 4 | novel transcript, antisense to HNRNPD |
| IGBP1P4 | 4 | immunoglobulin (CD79A) binding protein 1 ps |
| SNORD42 | 4 | Small nucleolar RNA SNORD42 [Source:RFAM; |
| HNRNPDL | 4 | heterogeneous nuclear ribonucleoprotein D li |
| ENOPH1 | 4 | enolase-phosphatase 1 [Source:HGNC Symbol |
| TMEM150C | 4 | transmembrane protein 150C [Source:HGNC S |

| | | |
|---|---|---|
| RPL7AP26 | 4 | ribosomal protein L7a pseudogene 26 [Source |
| AC067942.1 | 4 | ribosomal protein S6 (RPS6) pseudogene |
| AC067942.3 | 4 | embryonic ectoderm development (EED) pseu |
| AC067942.2 | 4 | Tyrosine-protein phosphatase non-receptor ty |
| LINC00575 | 4 | long intergenic non-protein coding RNA 575 [S |
| AC067942.4 | 4 | novel transcript |
| SCD5 | 4 | stearoyl-CoA desaturase 5 [Source:HGNC Sym |
| AC073413.1 | 4 | PQ loop repeat containing 1 (PQLC1) pseudog |
| MIR575 | 4 | microRNA 575 [Source:HGNC Symbol;Acc:HGN |
| AC073413.2 | 4 | ribosomal protein S26 (RPS26) pseudogene |
| SEC31A | 4 | SEC31 homolog A, COPII coat complex compo |
| THAP9-AS1 | 4 | THAP9 antisense RNA 1 [Source:HGNC Symbo |
| THAP9 | 4 | THAP domain containing 9 [Source:HGNC Sym |
| LIN54 | 4 | lin-54 DREAM MuvB core complex componen |
| Y_RNA | 4 | Y RNA [Source:RFAM;Acc:RF00019] |
| RNU6-615P | 4 | RNA, U6 small nuclear 615, pseudogene [Sour |
| COPS4 | 4 | COP9 signalosome subunit 4 [Source:HGNC Sy |
| AC073840.1 | 4 | novel transcript |
| PLAC8 | 4 | placenta associated 8 [Source:HGNC Symbol;A |
| AC114781.2 | 4 | novel transcript |
| AC114781.3 | 4 | tumor protein, translationally-controlled 1 (TF |
| AC114781.1 | 4 | aminomethyltransferase (AMT) pseudogene |
| COQ2 | 4 | coenzyme Q2, polyprenyltransferase [Source: |
| HPSE | 4 | heparanase [Source:HGNC Symbol;Acc:HGNC: |
| AC114781.5 | 4 | ribosomal protein S23 (RPS23) pseudogene |
| AC114781.4 | 4 | chromosome 18 open reading frame 21 (C18c |
| HELQ | 4 | helicase, POLQ like [Source:HGNC Symbol;Acc |
| MRPS18C | 4 | mitochondrial ribosomal protein S18C [Source |
| ABRAXAS1 | 4 | abraxas 1, BRCA1 A complex subunit [Source:I |
| SLC25A14P1 | 4 | solute carrier family 25 member 14 pseudoge |
| RPL30P5 | 4 | ribosomal protein L30 pseudogene 5 [Source: |
| GPAT3 | 4 | glycerol-3-phosphate acyltransferase 3 [Sourc |
| Y_RNA | 4 | Y RNA [Source:RFAM;Acc:RF00019] |
| AC021192.1 | 4 | novel transcript |
| AC079160.1 | 4 | novel transcript |
| RNU6-774P | 4 | RNA, U6 small nuclear 774, pseudogene [Sour |
| AC025538.1 | 4 | pleiomorphic adenoma gene-like 2 (PLAGL2) p |
| AC104063.1 | 4 | novel transcript |
| NKX6-1 | 4 | NK6 homeobox 1 [Source:HGNC Symbol;Acc:H |
| RPL3P13 | 4 | ribosomal protein L3 pseudogene 13 [Source: |
| CDS1 | 4 | CDP-diacylglycerol synthase 1 [Source:HGNC S |
| WDFY3 | 4 | WD repeat and FYVE domain containing 3 [So |
| RN7SL552P | 4 | RNA, 7SL, cytoplasmic 552, pseudogene [Sour |
| WDFY3-AS1 | 4 | WDFY3 antisense RNA 1 [Source:HGNC Symbo |
| RNU6-469P | 4 | RNA, U6 small nuclear 469, pseudogene [Sour |
| WDFY3-AS2 | 4 | WDFY3 antisense RNA 2 [Source:HGNC Symbo |
| AC108021.1 | 4 | novel transcript |
| RN7SKP48 | 4 | RN7SK pseudogene 48 [Source:HGNC Symbol; |
| AC097488.1 | 4 | novel pseudogene |
| ARHGAP24 | 4 | Rho GTPase activating protein 24 [Source:HGI |

| | | |
|---|---|---|
| MIR4451 | 4 | microRNA 4451 [Source:HGNC Symbol;Acc:HG |
| MAPK10 | 4 | mitogen-activated protein kinase 10 [Source:H |
| AC104827.1 | 4 | novel transcript, antisense to MAPK10 |
| RN7SKP96 | 4 | RN7SK pseudogene 96 [Source:HGNC Symbol; |
| MIR4452 | 4 | microRNA 4452 [Source:HGNC Symbol;Acc:HG |
| PTPN13 | 4 | protein tyrosine phosphatase non-receptor ty |
| SLC10A6 | 4 | solute carrier family 10 member 6 [Source:HG |
| AC093827.1 | 4 | ribosomal protein L6 (RPL6) pseudogene |
| C4orf36 | 4 | chromosome 4 open reading frame 36 [Source |
| AC093827.2 | 4 | CCZ1 vacuolar protein trafficking and biogene |
| AC093827.5 | 4 | novel protein |
| AC093827.3 | 4 | actin related protein 2/3 complex, subunit 1A, |
| AC093827.4 | 4 | Novel transcript, antisense to AFF1 |
| AFF1 | 4 | AF4/FMR2 family member 1 [Source:HGNC Sy |
| TECRP1 | 4 | trans-2,3-enoyl-CoA reductase pseudogene 1 |
| KLHL8 | 4 | kelch like family member 8 [Source:HGNC Syn |
| AC092658.1 | 4 | uracil-DNA glycosylase (UNG) pseudogene |
| GAPDHP60 | 4 | glyceraldehyde-3-phosphate dehydrogenase p |
| AC108516.1 | 4 | novel transcript |
| MIR5705 | 4 | microRNA 5705 [Source:HGNC Symbol;Acc:HG |
| HSD17B13 | 4 | hydroxysteroid 17-beta dehydrogenase 13 [So |
| AC108516.2 | 4 | novel transcript |
| HSD17B11 | 4 | hydroxysteroid 17-beta dehydrogenase 11 [So |
| RN7SL681P | 4 | RNA, 7SL, cytoplasmic 681, pseudogene [Sour |
| AC112250.1 | 4 | mitochondrial ribosomal protein L19 (MRPL19 |
| NUDT9 | 4 | nudix hydrolase 9 [Source:HGNC Symbol;Acc:H |
| AC112250.2 | 4 | novel transcript |
| SPARCL1 | 4 | SPARC like 1 [Source:HGNC Symbol;Acc:HGNC |
| AC093895.1 | 4 | novel transcript |
| DSPP | 4 | dentin sialophosphoprotein [Source:HGNC Syn |
| DMP1 | 4 | dentin matrix acidic phosphoprotein 1 [Source |
| Y_RNA | 4 | Y RNA [Source:RFAM;Acc:RF00019] |
| CHCHD2P7 | 4 | coiled-coil-helix-coiled-coil-helix domain conta |
| IBSP | 4 | integrin binding sialoprotein [Source:HGNC Sy |
| MEPE | 4 | matrix extracellular phosphoglycoprotein [Sou |
| HSP90AB3P | 4 | heat shock protein 90 alpha family class B mem |
| AC131944.1 | 4 | novel transcript, antisense to PKD2and SPP1 |
| SPP1 | 4 | secreted phosphoprotein 1 [Source:HGNC Syn |
| RNU1-36P | 4 | RNA, U1 small nuclear 36, pseudogene [Sourc |
| PKD2 | 4 | polycystin 2, transient receptor potential catic |
| ABCG2 | 4 | ATP binding cassette subfamily G member 2 (J |
| AC097484.1 | 4 | ribosomal protein L31 (RPL31) pseudogene |
| RNU6-818P | 4 | RNA, U6 small nuclear 818, pseudogene [Sour |
| RNU6ATAC31P | 4 | RNA, U6atac small nuclear 31, pseudogene [So |
| AC097484.2 | 4 | ribosomal protein S27 (RPS27) pseudogene |
| RNU6-1298P | 4 | RNA, U6 small nuclear 1298, pseudogene [Sou |
| PPM1K | 4 | protein phosphatase, Mg2+/Mn2+ dependent |
| RNU6-112P | 4 | RNA, U6 small nuclear 112, pseudogene [Sour |
| PPM1K-DT | 4 | PPM1K divergent transcript [Source:HGNC Syn |
| Y_RNA | 4 | Y RNA [Source:RFAM;Acc:RF00019] |

| | | |
|---|---|---|
| AC107067.2 | 4 | novel transcript |
| AC107067.1 | 4 | novel transcript |
| HERC6 | 4 | HECT and RLD domain containing E3 ubiquitin |
| HERC5 | 4 | HECT and RLD domain containing E3 ubiquitin |
| NCOA4P2 | 4 | nuclear receptor coactivator 4 pseudogene 2 |
| PYURF | 4 | PIGY upstream reading frame [Source:HGNC S |
| PIGY | 4 | phosphatidylinositol glycan anchor biosynthes |
| AC098582.1 | 4 | HECT and RLD domain containing E3 ubiquitin |
| PIGY-DT | 4 | PIGY divergent transcript [Source:HGNC Symb |
| AC083829.1 | 4 | CD53 molecule (CD53) pseudogene |
| RN7SKP244 | 4 | RN7SK pseudogene 244 [Source:HGNC Symbo |
| HERC3 | 4 | HECT and RLD domain containing E3 ubiquitin |
| RNU6-33P | 4 | RNA, U6 small nuclear 33, pseudogene [Sourc |
| NAP1L5 | 4 | nucleosome assembly protein 1 like 5 [Source |
| FAM13A-AS1 | 4 | FAM13A antisense RNA 1 [Source:HGNC Symb |
| AC108065.2 | 4 | NULL |
| AC108065.1 | 4 | NULL |
| FAM13A | 4 | family with sequence similarity 13 member A |
| TIGD2 | 4 | tigger transposable element derived 2 [Source |
| AC104785.1 | 4 | novel transcript |
| RNU6-907P | 4 | RNA, U6 small nuclear 907, pseudogene [Sour |
| GPRIN3 | 4 | GPRIN family member 3 [Source:HGNC Symbo |
| AC093866.1 | 4 | novel transcript |
| SNCA | 4 | synuclein alpha [Source:HGNC Symbol;Acc:HG |
| AC097478.1 | 4 | novel transcript, antisense to SNCA |
| AC097478.3 | 4 | novel transcript, antisense to SNCA |
| AC097478.2 | 4 | novel transcript, antisense to SNCA |
| SNCA-AS1 | 4 | SNCA antisense RNA 1 [Source:HGNC Symbol; |
| MMRN1 | 4 | multimerin 1 [Source:HGNC Symbol;Acc:HGN( |
| AC105445.1 | 4 | novel transcript |
| CCSER1 | 4 | coiled-coil serine rich protein 1 [Source:HGNC |
| RN7SKP248 | 4 | RN7SK pseudogene 248 [Source:HGNC Symbo |
| AC093729.1 | 4 | Interleukin-9 (IL9) pseudogene |
| TMSB4XP8 | 4 | TMSB4X pseudogene 8 [Source:HGNC Symbol |
| AC004054.1 | 4 | novel transcript |
| AC074124.1 | 4 | novel transcript, antisense to CCSER1 |
| AC110774.1 | 4 | tec |
| KRT19P6 | 4 | keratin 19 pseudogene 6 [Source:HGNC Symb |
| AC093810.1 | 4 | novel transcript |
| PMPCAP1 | 4 | peptidase, mitochondrial processing alpha sul |
| AC097520.2 | 4 | novel transcript |
| AC097520.1 | 4 | tec |
| LNCPRESS2 | 4 | lncRNA p53 regulated and ESC associated 2 [S |
| AC112695.1 | 4 | novel transcript |
| GRID2 | 4 | glutamate ionotropic receptor delta type subu |
| MTND1P19 | 4 | MT-ND1 pseudogene 19 [Source:HGNC Symbo |
| AC095059.2 | 4 | Guanine nucleotide-binding protein subunit b |
| AC095059.1 | 4 | novel transcript |
| AC020699.1 | 4 | Tudor domain-containing protein 1 (TDRD1) p |
| AC110800.1 | 4 | novel transcript |

| | | |
|---|---|---|
| RNA5SP164 | 4 | RNA, 5S ribosomal pseudogene 164 [Source:H |
| ATOH1 | 4 | atonal bHLH transcription factor 1 [Source:HG |
| AC096746.1 | 4 | novel transcript |
| HMGB3P15 | 4 | high mobility group box 3 pseudogene 15 [Sou |
| SMARCAD1 | 4 | SWI/SNF-related, matrix-associated actin-dep |
| HPGDS | 4 | hematopoietic prostaglandin D synthase [Sour |
| AC109925.2 | 4 | novel transcript, antisense to HPGDS |
| AC109925.1 | 4 | ribosomal protein L35a (RPL35A) pseudogene |
| PDLIM5 | 4 | PDZ and LIM domain 5 [Source:HGNC Symbol; |
| AC108067.1 | 4 | novel transcript |
| BMPR1B-DT | 4 | BMPR1B divergent transcript [Source:HGNC Sy |
| BMPR1B | 4 | bone morphogenetic protein receptor type 1E |
| UNC5C | 4 | unc-5 netrin receptor C [Source:HGNC Symbol |
| AC106881.1 | 4 | novel transcript, antisense to UNC5C |
| RPL30P6 | 4 | ribosomal protein L30 pseudogene 6 [Source: |
| PDHA2 | 4 | pyruvate dehydrogenase E1 alpha 2 subunit [S |
| RNU6-1059P | 4 | RNA, U6 small nuclear 1059, pseudogene [Sou |
| RNU6-34P | 4 | RNA, U6 small nuclear 34, pseudogene [Sourc |
| AC096661.1 | 4 | talin 2 (TLN2) pseudogene |
| LINC02267 | 4 | long intergenic non-protein coding RNA 2267 |
| RN7SKP28 | 4 | RN7SK pseudogene 28 [Source:HGNC Symbol; |
| AC096585.1 | 4 | novel transcript |
| COX7A2P2 | 4 | cytochrome c oxidase subunit 7A2 pseudogen |
| AC108515.1 | 4 | novel transcript |
| STPG2 | 4 | sperm tail PG-rich repeat containing 2 [Source |
| AC034154.1 | 4 | novel transcript, sense overlapping to STPG2 |
| STPG2-AS1 | 4 | STPG2 antisense RNA 1 [Source:HGNC Symbol |
| CRYZP2 | 4 | crystallin zeta pseudogene 2 [Source:HGNC Sy |
| RPL5P12 | 4 | ribosomal protein L5 pseudogene 12 [Source: |
| DUTP8 | 4 | deoxyuridine triphosphatase pseudogene 8 [S |
| AC019077.1 | 4 | novel transcript |
| RAP1GDS1 | 4 | Rap1 GTPase-GDP dissociation stimulator 1 [S |
| AC058823.1 | 4 | ribosomal protein L21 (RPL21) pseudogene |
| TSPAN5 | 4 | tetraspanin 5 [Source:HGNC Symbol;Acc:HGN |
| AC108159.1 | 4 | novel transcript |
| AC114811.2 | 4 | novel transcript, antisense TSPAN5 |
| AC114811.1 | 4 | translocase of outer mitochondrial membrane |
| BTF3P13 | 4 | basic transcription factor 3 pseudogene 13 [Sc |
| EIF4E | 4 | eukaryotic translation initiation factor 4E [Sou |
| TBCAP3 | 4 | tubulin folding cofactor A pseudogene 3 [Sour |
| AC019131.1 | 4 | novel transcript, antisense to EIF4E |
| FAM177A1P1 | 4 | family with sequence similarity 177 member A |
| AC019131.3 | 4 | novel transcript |
| RNU7-149P | 4 | RNA, U7 small nuclear 149 pseudogene [Sourc |
| NDUFS5P4 | 4 | NADH:ubiquinone oxidoreductase subunit S5 |
| METAP1 | 4 | methionyl aminopeptidase 1 [Source:HGNC Sy |
| MIR3684 | 4 | microRNA 3684 [Source:HGNC Symbol;Acc:HG |
| ABT1P1 | 4 | activator of basal transcription 1 pseudogene |
| AC019131.2 | 4 | novel transcript |
| ADH5 | 4 | alcohol dehydrogenase 5 (class III), chi polype |

| | | |
|---|---|---|
| AP002026.1 | 4 | novel transcript, antisense to ADH4, ADH6 & / |
| ADH4 | 4 | alcohol dehydrogenase 4 (class II), pi polypept |
| PCNAP1 | 4 | proliferating cell nuclear antigen pseudogene |
| ADH6 | 4 | alcohol dehydrogenase 6 (class V) [Source:HG |
| ADH1A | 4 | alcohol dehydrogenase 1A (class I), alpha poly |
| ADH1B | 4 | alcohol dehydrogenase 1B (class I), beta polyp |
| ADH1C | 4 | alcohol dehydrogenase 1C (class I), gamma pc |
| ADH7 | 4 | alcohol dehydrogenase 7 (class IV), mu or sign |
| AP001960.1 | 4 | alcohol dehydrogenase (ADH) family pseudog |
| C4orf17 | 4 | chromosome 4 open reading frame 17 [Sourc |
| TRMT10A | 4 | tRNA methyltransferase 10A [Source:HGNC Sy |
| MTTP | 4 | microsomal triglyceride transfer protein [Sour |
| AC083902.1 | 4 | novel transcript |
| C4orf54 | 4 | chromosome 4 open reading frame 54 [Sourc |
| DAPP1 | 4 | dual adaptor of phosphotyrosine and 3-phosp |
| AP001962.1 | 4 | TEC |
| LAMTOR3 | 4 | late endosomal/lysosomal adaptor, MAPK anc |
| DNAJB14 | 4 | DnaJ heat shock protein family (Hsp40) memb |
| AC097460.3 | 4 | TEC |
| H2AFZ | 4 | H2A histone family member Z [Source:HGNC S |
| AC097460.1 | 4 | novel transcript |
| AC097460.2 | 4 | ribosomal protein L7-like 1 (RPL7L1) pseudoge |
| DYNLL1P6 | 4 | dynein light chain LC8-type 1 pseudogene 6 [S |
| DDIT4L | 4 | DNA damage inducible transcript 4 like [Sourc |
| AP001961.1 | 4 | novel transcript |
| EMCN | 4 | endomucin [Source:HGNC Symbol;Acc:HGNC: |
| AC097459.1 | 4 | novel transcript, antisense to EMCN |
| LINC01216 | 4 | long intergenic non-protein coding RNA 1216 |
| LINC01217 | 4 | long intergenic non-protein coding RNA 1217 |
| LINC01218 | 4 | long intergenic non-protein coding RNA 1218 |
| PPP3CA | 4 | protein phosphatase 3 catalytic subunit alpha |
| MIR8066 | 4 | microRNA 8066 [Source:HGNC Symbol;Acc:HG |
| MIR1255A | 4 | microRNA 1255a [Source:HGNC Symbol;Acc:H |
| AP001816.1 | 4 | novel transcript |
| BANK1 | 4 | B cell scaffold protein with ankyrin repeats 1 [ |
| AC084277.1 | 4 | novel transcript |
| RNU6-462P | 4 | RNA, U6 small nuclear 462, pseudogene [Sour |
| MTND5P5 | 4 | MT-ND5 pseudogene 5 [Source:HGNC Symbol |
| AP002075.1 | 4 | novel transcript |
| SLC39A8 | 4 | solute carrier family 39 member 8 [Source:HG |
| RN7SL728P | 4 | RNA, 7SL, cytoplasmic 728, pseudogene [Sour |
| AC098487.1 | 4 | novel transcript |
| AF213884.2 | 4 | zinc finger protein 705A (ZNF705A) pseudoge |
| AF213884.3 | 4 | novel transcript, antisense NFKB1 |
| NFKB1 | 4 | nuclear factor kappa B subunit 1 [Source:HGN |
| MANBA | 4 | mannosidase beta [Source:HGNC Symbol;Acc: |
| AF213884.1 | 4 | ribosomal protein L21 (RPL21) pseudogene |
| LRRC37A15P | 4 | leucine rich repeat containing 37 member A1! |
| KRT8P46 | 4 | keratin 8 pseudogene 46 [Source:HGNC Symb |
| AC018797.1 | 4 | mannosidase, beta A, lysosomal-like (MANBAI |

| | | |
|---|---|---|
| AC018797.2 | 4 | WD repeat domain 12 (WDR12) pseudogene |
| UBE2D3 | 4 | ubiquitin conjugating enzyme E2 D3 [Source:H |
| AC018797.3 | 4 | novel transcript, antisense to UBE2D3 |
| UBE2D3-AS1 | 4 | UBE2D3 antisense RNA 1 [Source:HGNC Symb |
| RNU7-151P | 4 | RNA, U7 small nuclear 151 pseudogene [Sourc |
| CISD2 | 4 | CDGSH iron sulfur domain 2 [Source:HGNC Sy |
| SLC9B1 | 4 | solute carrier family 9 member B1 [Source:HG |
| PABPC1P7 | 4 | poly(A) binding protein cytoplasmic 1 pseudog |
| ACTR3BP4 | 4 | ACTR3B pseudogene 4 [Source:HGNC Symbol; |
| SLC9B2 | 4 | solute carrier family 9 member B2 [Source:HG |
| BDH2 | 4 | 3-hydroxybutyrate dehydrogenase 2 [Source:I |
| CENPE | 4 | centromere protein E [Source:HGNC Symbol;A |
| LINC02428 | 4 | long intergenic non-protein coding RNA 2428 |
| AC105460.2 | 4 | novel transcript, antisense to TACR3 |
| AC105460.1 | 4 | novel transcript |
| DDX3P3 | 4 | DEAD-box helicase 3 pseudogene 3 [Source:H |
| TACR3 | 4 | tachykinin receptor 3 [Source:HGNC Symbol;A |
| AC004047.1 | 4 | novel transcript |
| RNU6-635P | 4 | RNA, U6 small nuclear 635, pseudogene [Sour |
| LINC02503 | 4 | long intergenic non-protein coding RNA 2503 |
| AC004052.1 | 4 | novel transcript |
| CXXC4 | 4 | CXXC finger protein 4 [Source:HGNC Symbol;A |
| CXXC4-AS1 | 4 | CXXC4 antisense RNA 1 [Source:HGNC Symbol |
| AC004063.1 | 4 | novel transcript |
| AC004053.1 | 4 | novel transcript |
| AC105918.1 | 4 | ribosomal protein L6 (RPL6) pseudogene |
| AC096577.1 | 4 | novel transcript |
| RNU6-351P | 4 | RNA, U6 small nuclear 351, pseudogene [Sour |
| AC096577.2 | 4 | nucleolar and coiled-body phosphoprotein 1 ( |
| AC004069.2 | 4 | nucleoporin 43kDa (NUP43) pseudogene |
| AC004069.3 | 4 | NULL |
| AC004069.1 | 4 | novel transcript |
| TET2 | 4 | tet methylcytosine dioxygenase 2 [Source:HG |
| TET2-AS1 | 4 | TET2 antisense RNA 1 [Source:HGNC Symbol;A |
| RN7SL89P | 4 | RNA, 7SL, cytoplasmic 89, pseudogene [Sourc |
| PPA2 | 4 | pyrophosphatase (inorganic) 2 [Source:HGNC |
| RNU6-553P | 4 | RNA, U6 small nuclear 553, pseudogene [Sour |
| EEF1A1P9 | 4 | eukaryotic translation elongation factor 1 alph |
| AC004066.1 | 4 | family with sequence similarity 64, member A |
| ATP5F1EP1 | 4 | ATP synthase F1 subunit epsilon pseudogene |
| AC004066.2 | 4 | novel transcript |
| ARHGEF38 | 4 | Rho guanine nucleotide exchange factor 38 [S |
| ARHGEF38-IT1 | 4 | ARHGEF38 intronic transcript 1 [Source:HGNC |
| AC105391.1 | 4 | DEAD (Asp-Glu-Ala-Asp) box polypeptide 50 (D |
| INTS12 | 4 | integrator complex subunit 12 [Source:HGNC |
| GSTCD | 4 | glutathione S-transferase C-terminal domain c |
| AC008243.1 | 4 | novel transcript |
| NPNT | 4 | nephronectin [Source:HGNC Symbol;Acc:HGN |
| AC109361.2 | 4 | novel transcript |
| AC109361.1 | 4 | novel transcript |

| | | |
|---|---|---|
| TBCK | 4 | TBC1 domain containing kinase [Source:HGNC |
| AIMP1 | 4 | aminoacyl tRNA synthetase complex interacti |
| GIMD1 | 4 | GIMAP family P-loop NTPase domain containi |
| LINC02173 | 4 | long intergenic non-protein coding RNA 2173 |
| AC092445.1 | 4 | novel transcript, antisense to DKK2 |
| ACTR6P1 | 4 | ACTR6 pseudogene 1 [Source:HGNC Symbol;A |
| DKK2 | 4 | dickkopf WNT signaling pathway inhibitor 2 [S |
| RAC1P5 | 4 | Rac family small GTPase 1 pseudogene 5 [Sou |
| AC104663.1 | 4 | novel transcript |
| RNU6-551P | 4 | RNA, U6 small nuclear 551, pseudogene [Sour |
| PAPSS1 | 4 | 3'-phosphoadenosine 5'-phosphosulfate synth |
| SGMS2 | 4 | sphingomyelin synthase 2 [Source:HGNC Syml |
| AC096564.1 | 4 | novel transcript |
| RNU6-733P | 4 | RNA, U6 small nuclear 733, pseudogene [Sour |
| CYP2U1 | 4 | cytochrome P450 family 2 subfamily U memb |
| AC096564.2 | 4 | novel transcript |
| HADH | 4 | hydroxyacyl-CoA dehydrogenase [Source:HGN |
| LEF1 | 4 | lymphoid enhancer binding factor 1 [Source:H |
| LEF1-AS1 | 4 | LEF1 antisense RNA 1 [Source:HGNC Symbol;A |
| RPSAP34 | 4 | ribosomal protein SA pseudogene 34 [Source: |
| ZACNP1 | 4 | zinc activated ion channel pseudogene 1 [Sou |
| EXOC7P1 | 4 | exocyst complex component 7 pseudogene 1 |
| RPL34-AS1 | 4 | RPL34 antisense RNA 1 (head to head) [Source |
| RPL34 | 4 | ribosomal protein L34 [Source:HGNC Symbol;A |
| OSTC | 4 | oligosaccharyltransferase complex non-cataly |
| RNU6-431P | 4 | RNA, U6 small nuclear 431, pseudogene [Sour |
| AC107071.1 | 4 | novel transcript |
| ETNPPL | 4 | ethanolamine-phosphate phospho-lyase [Sour |
| AC097473.1 | 4 | protein phosphatase 1, regulatory subunit 36 |
| RCC2P8 | 4 | regulator of chromosome condensation 2 pse |
| COL25A1 | 4 | collagen type XXV alpha 1 chain [Source:HGN( |
| ZCCHC23 | 4 | zinc finger CCHC-type containing 23 [Source:H |
| RBMXP4 | 4 | RBMX pseudogene 4 [Source:HGNC Symbol;A |
| SEC24B-AS1 | 4 | SEC24B antisense RNA 1 [Source:HGNC Symbo |
| SEC24B | 4 | SEC24 homolog B, COPII coat complex compo |
| RN7SL55P | 4 | RNA, 7SL, cytoplasmic 55, pseudogene [Sourc |
| MIR576 | 4 | microRNA 576 [Source:HGNC Symbol;Acc:HGN |
| SETP20 | 4 | SET pseudogene 20 [Source:HGNC Symbol;Acc |
| CDC42P4 | 4 | cell division cycle 42 pseudogene 4 [Source:H( |
| MCUB | 4 | mitochondrial calcium uniporter dominant ne |
| HIGD1AP14 | 4 | HIG1 hypoxia inducible domain family membe |
| CASP6 | 4 | caspase 6 [Source:HGNC Symbol;Acc:HGNC:1! |
| AC004067.1 | 4 | novel transcript, antisense to CASP6 |
| PLA2G12A | 4 | phospholipase A2 group XIIA [Source:HGNC Sy |
| AC126283.2 | 4 | novel protein |
| CFI | 4 | complement factor I [Source:HGNC Symbol;A( |
| AC126283.1 | 4 | novel transcript |
| GAR1 | 4 | GAR1 ribonucleoprotein [Source:HGNC Symbo |
| RRH | 4 | retinal pigment epithelium-derived rhodopsin |
| LRIT3 | 4 | leucine rich repeat, Ig-like and transmembran |

| | | |
|---|---|---|
| KRT19P3 | 4 | keratin 19 pseudogene 3 [Source:HGNC Symb |
| EGF | 4 | epidermal growth factor [Source:HGNC Symbo |
| RNU6-35P | 4 | RNA, U6 small nuclear 35, pseudogene [Sourc |
| ELOVL6 | 4 | ELOVL fatty acid elongase 6 [Source:HGNC Syr |
| RN7SL275P | 4 | RNA, 7SL, cytoplasmic 275, pseudogene [Sour |
| AC093770.1 | 4 | ribosomal protein L17 (RPL17) pseudogene |
| HSBP1P2 | 4 | heat shock factor binding protein 1 pseudoge |
| AC108145.1 | 4 | ring finger protein 14 (RNF14) pseudogene |
| RNU6-205P | 4 | RNA, U6 small nuclear 205, pseudogene [Sour |
| ZBED1P1 | 4 | zinc finger BED-type containing 1 pseudogene |
| ENPEP | 4 | glutamyl aminopeptidase [Source:HGNC Symb |
| RPL7L1P13 | 4 | ribosomal protein L7 like 1 pseudogene 13 [So |
| ZNF969P | 4 | zinc finger protein 969, pseudogene [Source:H |
| AC093872.1 | 4 | NULL |
| AC098798.1 | 4 | novel transcript, antisense to ENPEP |
| PANCR | 4 | PITX2 adjacent non-coding RNA [Source:HGNC |
| PITX2 | 4 | paired like homeodomain 2 [Source:HGNC Syr |
| LINC01438 | 4 | long intergenic non-protein coding RNA 1438 |
| MIR297 | 4 | microRNA 297 [Source:HGNC Symbol;Acc:HGN |
| LYPLA1P2 | 4 | LYPLA1 pseudogene 2 [Source:HGNC Symbol;A |
| AC004062.1 | 4 | eukaryotic translation elongation factor 1 beta |
| RNU6-289P | 4 | RNA, U6 small nuclear 289, pseudogene [Sour |
| AC083795.1 | 4 | ribosomal protein L36a (RPL36A) pseudogene |
| AC083795.2 | 4 | novel transcript |
| AC139718.2 | 4 | novel transcript |
| AC139718.1 | 4 | novel transcript |
| AC004704.1 | 4 | novel transcript |
| CCDC34P1 | 4 | coiled-coil domain containing 34 pseudogene |
| TUBB8P3 | 4 | tubulin beta 8 class VIII pseudogene 3 [Source |
| AC093663.1 | 4 | ribosomal protein S12 (RPS12) pseudogene |
| FAM241A | 4 | family with sequence similarity 241 member A |
| AC093663.2 | 4 | ribosomal protein L36a (RPL36A) pseudogene |
| AC109347.1 | 4 | novel transcript, antisense to AP1AR |
| AP1AR | 4 | adaptor related protein complex 1 associated |
| AC109347.2 | 4 | novel transcript |
| TIFA | 4 | TRAF interacting protein with forkhead associa |
| ALPK1 | 4 | alpha kinase 1 [Source:HGNC Symbol;Acc:HGN |
| RTEL1P1 | 4 | regulator of telomere elongation helicase 1 ps |
| TOX4P1 | 4 | TOX high mobility group box family member 4 |
| NEUROG2 | 4 | neurogenin 2 [Source:HGNC Symbol;Acc:HGN |
| AC023886.1 | 4 | novel transcript |
| RPL23AP94 | 4 | ribosomal protein L23a pseudogene 94 [Sourc |
| ZGRF1 | 4 | zinc finger GRF-type containing 1 [Source:HGN |
| AC023886.2 | 4 | H3 histone, family 3B (H3.3B) (H3F3B) pseudo |
| LARP7 | 4 | La ribonucleoprotein domain family member |
| MIR302CHG | 4 | miR-302/367 cluster host gene [Source:HGNC |
| MIR367 | 4 | microRNA 367 [Source:HGNC Symbol;Acc:HGN |
| MIR302D | 4 | microRNA 302d [Source:HGNC Symbol;Acc:HG |
| MIR302A | 4 | microRNA 302a [Source:HGNC Symbol;Acc:HG |
| MIR302C | 4 | microRNA 302c [Source:HGNC Symbol;Acc:HG |

| | | |
|---|---|---|
| MIR302B | 4 | microRNA 302b [Source:HGNC Symbol;Acc:HG |
| OSTCP4 | 4 | oligosaccharyltransferase complex subunit ps |
| AC106864.1 | 4 | novel transcript |
| WRBP1 | 4 | tryptophan rich basic protein pseudogene 1 [S |
| AC017007.3 | 4 | ribosomal protein L32 (RPL32) pseudogene |
| Y_RNA | 4 | Y RNA [Source:RFAM;Acc:RF00019] |
| RPL7AP30 | 4 | ribosomal protein L7a pseudogene 30 [Source |
| AC017007.4 | 4 | eukaryotic translation initiation factor 2, subu |
| ANK2 | 4 | ankyrin 2 [Source:HGNC Symbol;Acc:HGNC:49 |
| AC017007.2 | 4 | TAF4b RNA polymerase II, TATA box binding p |
| AC017007.5 | 4 | novel transcript, antisense to ANK2 |
| AC017007.1 | 4 | heterogeneous nuclear ribonucleoprotein A1 |
| AC093879.1 | 4 | novel transcript |
| AC093879.2 | 4 | novel transcript |
| MIR1243 | 4 | microRNA 1243 [Source:HGNC Symbol;Acc:HG |
| MIR8082 | 4 | microRNA 8082 [Source:HGNC Symbol;Acc:HG |
| AC004057.1 | 4 | ribosomal protein S26 (RPS26) pseudogene |
| RN7SL184P | 4 | RNA, 7SL, cytoplasmic 184, pseudogene [Sour |
| RNU1-138P | 4 | RNA, U1 small nuclear 138, pseudogene [Sour |
| CAMK2D | 4 | calcium/calmodulin dependent protein kinase |
| AC111193.1 | 4 | pseudogene similar to part of sterol-C4-methy |
| ARSJ | 4 | arylsulfatase family member J [Source:HGNC S |
| AC104779.1 | 4 | novel transcript |
| AC093815.1 | 4 | ribosomal protein L21 (RPL21) pseudogene |
| UGT8 | 4 | UDP glycosyltransferase 8 [Source:HGNC Syml |
| MIR577 | 4 | microRNA 577 [Source:HGNC Symbol;Acc:HGN |
| CIR1P2 | 4 | corepressor interacting with RBPJ, 1 pseudoge |
| RN7SL808P | 4 | RNA, 7SL, cytoplasmic 808, pseudogene [Sour |
| NDST4 | 4 | N-deacetylase and N-sulfotransferase 4 [Sourc |
| MRPS33P3 | 4 | mitochondrial ribosomal protein S33 pseudog |
| RPF2P2 | 4 | ribosome production factor 2 homolog pseud |
| PGAM4P2 | 4 | phosphoglycerate mutase family member 4 ps |
| KRT18P21 | 4 | keratin 18 pseudogene 21 [Source:HGNC Sym |
| AC027613.1 | 4 | rearranged L-myc fusion (RLF) pseudogene |
| EIF3KP3 | 4 | eukaryotic translation initiation factor 3 subur |
| TTC39CP1 | 4 | tetratricopeptide repeat domain 39C pseudog |
| MTRNR2L13 | 4 | MT-RNR2 like 13 [Source:HGNC Symbol;Acc:H |
| MIR1973 | 4 | microRNA 1973 [Source:HGNC Symbol;Acc:HG |
| TRMT112P1 | 4 | tRNA methyltransferase subunit 11-2 pseudog |
| CUL4AP1 | 4 | cullin 4A pseudogene 1 [Source:HGNC Symbol |
| AC106892.1 | 4 | novel transcript |
| ACTN4P1 | 4 | actinin alpha 4 pseudogene 1 [Source:HGNC S |
| AC093765.5 | 4 | novel transcript |
| AC093765.4 | 4 | novel transcript |
| AC093765.3 | 4 | novel transcript |
| AC093765.2 | 4 | novel transcript |
| RNU6-119P | 4 | RNA, U6 small nuclear 119, pseudogene [Sour |
| AC093765.1 | 4 | hepatitis A virus cellular receptor 1 (HAVCR1) |
| TRAM1L1 | 4 | translocation associated membrane protein 1 |
| AC107399.1 | 4 | ninein (GSK3B interacting protein) (NIN) pseu |

| | | |
|---|---|---|
| LINC02262 | 4 | long intergenic non-protein coding RNA 2262 |
| LINC02263 | 4 | long intergenic non-protein coding RNA 2263 |
| LINC01378 | 4 | long intergenic non-protein coding RNA 1378 |
| RPSAP35 | 4 | ribosomal protein SA pseudogene 35 [Source: |
| NT5C3AP1 | 4 | 5'-nucleotidase, cytosolic IIIA pseudogene 1 [S |
| AC092661.1 | 4 | novel transcript |
| LINC02264 | 4 | long intergenic non-protein coding RNA 2264 |
| NDST3 | 4 | N-deacetylase and N-sulfotransferase 3 [Sourc |
| FKBP4P1 | 4 | FKBP prolyl isomerase 4 pseudogene 1 [Sourc |
| SNHG8 | 4 | small nucleolar RNA host gene 8 [Source:HGN |
| SNORA24 | 4 | small nucleolar RNA, H/ACA box 24 [Source:H |
| PRSS12 | 4 | serine protease 12 [Source:HGNC Symbol;Acc |
| NDUFS5P5 | 4 | NADH:ubiquinone oxidoreductase subunit S5 |
| AC092670.1 | 4 | ribosomal protein L41 (RPL41) pseudogene |
| CEP170P1 | 4 | centrosomal protein 170 pseudogene 1 [Sourc |
| AC110079.1 | 4 | general transcription factor IIi (GTF2I) pseudo |
| RNU6-1054P | 4 | RNA, U6 small nuclear 1054, pseudogene [Sou |
| CICP16 | 4 | capicua transcriptional repressor pseudogene |
| SEPTIN14P4 | 4 | septin 14 pseudogene 4 [Source:HGNC Symbo |
| METTL14-DT | 4 | METTL14 divergent transcript [Source:HGNC S |
| METTL14 | 4 | methyltransferase like 14 [Source:HGNC Symb |
| SEC24D | 4 | SEC24 homolog D, COPII coat complex compo |
| SYNPO2 | 4 | synaptopodin 2 [Source:HGNC Symbol;Acc:HG |
| AC096745.1 | 4 | iron-sulfur cluster assembly 1 homolog (S. cer |
| MYOZ2 | 4 | myozenin 2 [Source:HGNC Symbol;Acc:HGNC: |
| AC092656.1 | 4 | mitochondrial ribosomal protein L42 (MRPL42 |
| USP53 | 4 | ubiquitin specific peptidase 53 [Source:HGNC |
| C4orf3 | 4 | chromosome 4 open reading frame 3 [Source: |
| FABP2 | 4 | fatty acid binding protein 2 [Source:HGNC Syn |
| KLHL2P1 | 4 | kelch like family member 2 pseudogene 1 [Sou |
| RNU4-33P | 4 | RNA, U4 small nuclear 33, pseudogene [Sourc |
| AC110373.1 | 4 | glycerol kinase 2 pseudogene |
| GTF2IP12 | 4 | general transcription factor IIi pseudogene 12 |
| AC093752.3 | 4 | novel transcript |
| RNU6-1217P | 4 | RNA, U6 small nuclear 1217, pseudogene [Sou |
| AC093752.1 | 4 | septin 4 (sept4) pseudogene |
| AC093752.2 | 4 | novel transcript |
| PDE5A | 4 | phosphodiesterase 5A [Source:HGNC Symbol; |
| AC080089.1 | 4 | NULL |
| AC080089.2 | 4 | novel transcript |
| LINC01365 | 4 | long intergenic non-protein coding RNA 1365 |
| AC097173.1 | 4 | Rho GTPase activating protein 20 (ARHGAP20 |
| LINC02502 | 4 | long intergenic non-protein coding RNA 2502 |
| Y_RNA | 4 | Y RNA [Source:RFAM;Acc:RF00019] |
| AC097173.2 | 4 | novel transcript |
| MAD2L1 | 4 | mitotic arrest deficient 2 like 1 [Source:HGNC |
| AC073475.1 | 4 | novel transcript, antisense to MAD2L1 |
| LTV1P1 | 4 | LTV1 ribosome biogenesis factor pseudogene |
| SAR1AP3 | 4 | secretion associated Ras related GTPase 1A ps |
| AC025741.1 | 4 | novel transcript |

| | | |
|---|---|---|
| PRDM5 | 4 | PR/SET domain 5 [Source:HGNC Symbol;Acc:H |
| RNU6-550P | 4 | RNA, U6 small nuclear 550, pseudogene [Sour |
| SETP12 | 4 | SET pseudogene 12 [Source:HGNC Symbol;Acc |
| NDNF | 4 | neuron derived neurotrophic factor [Source:H |
| RN7SKP137 | 4 | RN7SK pseudogene 137 [Source:HGNC Symbo |
| AC105254.1 | 4 | novel transcript |
| AC105254.2 | 4 | serine threonine kinase 39 (STK39) pseudoger |
| TNIP3 | 4 | TNFAIP3 interacting protein 3 [Source:HGNC S |
| RNU6-948P | 4 | RNA, U6 small nuclear 948, pseudogene [Sour |
| QRFPR | 4 | pyroglutamylated RFamide peptide receptor [ |
| Y_RNA | 4 | Y RNA [Source:RFAM;Acc:RF00019] |
| AC093816.1 | 4 | novel pseudogene |
| TUBB4BP5 | 4 | tubulin beta 4B class IVb pseudogene 5 [Sourc |
| ANXA5 | 4 | annexin A5 [Source:HGNC Symbol;Acc:HGNC:! |
| TMEM155 | 4 | transmembrane protein 155 [Source:HGNC Sy |
| AC079341.1 | 4 | novel transcript, antisense to TMEM155 |
| EXOSC9 | 4 | exosome component 9 [Source:HGNC Symbol |
| CCNA2 | 4 | cyclin A2 [Source:HGNC Symbol;Acc:HGNC:15 |
| BBS7 | 4 | Bardet-Biedl syndrome 7 [Source:HGNC Symb |
| AC079341.2 | 4 | novel transcript |
| TRPC3 | 4 | transient receptor potential cation channel su |
| AC097533.1 | 4 | pseudogene similar to part of ring finger prote |
| RN7SL335P | 4 | RNA, 7SL, cytoplasmic 335, pseudogene [Sour |
| KIAA1109 | 4 | KIAA1109 [Source:HGNC Symbol;Acc:HGNC:2( |
| ADAD1 | 4 | adenosine deaminase domain containing 1 [S( |
| IL2 | 4 | interleukin 2 [Source:HGNC Symbol;Acc:HGNC |
| IL21 | 4 | interleukin 21 [Source:HGNC Symbol;Acc:HGN |
| IL21-AS1 | 4 | IL21 antisense RNA 1 [Source:HGNC Symbol;A |
| AC053545.1 | 4 | novel transcript |
| CETN4P | 4 | centrin 4, pseudogene [Source:HGNC Symbol; |
| BBS12 | 4 | Bardet-Biedl syndrome 12 [Source:HGNC Sym |
| AC021205.2 | 4 | ribosomal protein L34 (RPL34) pseudogene |
| FGF2 | 4 | fibroblast growth factor 2 [Source:HGNC Symb |
| AC021205.1 | 4 | ribosomal protein S26 (RPS26) pseudogene |
| AC021205.3 | 4 | novel transcript, antisense to FGF2 |
| NUDT6 | 4 | nudix hydrolase 6 [Source:HGNC Symbol;Acc:I |
| SPATA5 | 4 | spermatogenesis associated 5 [Source:HGNC S |
| AC026402.1 | 4 | 40S ribosomal protein S13 (RPS13) pseudoger |
| AC026402.2 | 4 | novel transcript |
| SPRY1 | 4 | sprouty RTK signaling antagonist 1 [Source:HG |
| AC093821.1 | 4 | novel transcript |
| LINC02435 | 4 | long intergenic non-protein coding RNA 2435 |
| LINC01091 | 4 | long intergenic non-protein coding RNA 1091 |
| AC096732.1 | 4 | novel transcript |
| RPL21P50 | 4 | ribosomal protein L21 pseudogene 50 [Source |
| AC093767.1 | 4 | voltage-dependent anion channel 2 (VDAC2) p |
| AC108075.1 | 4 | novel transcript |
| PPIAP76 | 4 | peptidylprolyl isomerase A pseudogene 76 [So |
| AC121154.1 | 4 | novel transcript |
| AC133530.1 | 4 | novel transcript |

| | | |
|---|---|---|
| AC097474.1 | 4 | Tubulin alpha chain pseudogene |
| TECRP2 | 4 | trans-2,3-enoyl-CoA reductase pseudogene 2 |
| LINC02516 | 4 | long intergenic non-protein coding RNA 2516 |
| ANKRD50 | 4 | ankyrin repeat domain 50 [Source:HGNC Syml |
| FAT4 | 4 | FAT atypical cadherin 4 [Source:HGNC Symbol |
| Y_RNA | 4 | Y RNA [Source:RFAM;Acc:RF00019] |
| MIR2054 | 4 | microRNA 2054 [Source:HGNC Symbol;Acc:HG |
| AC104664.1 | 4 | novel transcript |
| NUP58P1 | 4 | nucleoporin 58 pseudogene 1 [Source:HGNC S |
| TMEM248P1 | 4 | transmembrane protein 248 pseudogene 1 [So |
| LINC02379 | 4 | long intergenic non-protein coding RNA 2379 |
| AC105758.1 | 4 | histone cluster 3, H3 (HIST3H3) pseudogene |
| AC097528.1 | 4 | novel transcript |
| RBM48P1 | 4 | RNA binding motif protein 48 pseudogene 1 [S |
| Y_RNA | 4 | Y RNA [Source:RFAM;Acc:RF00019] |
| AC096773.1 | 4 | novel transcript |
| AC093772.1 | 4 | novel transcript |
| AC093772.2 | 4 | novel transcript |
| AC097462.1 | 4 | TAF10 RNA polymerase II, TATA box binding p |
| AC097462.2 | 4 | spermidine/spermine N1-acetyltransferase 1 ( |
| AC097462.3 | 4 | novel transcript, antisense to INTU |
| INTU | 4 | inturned planar cell polarity protein [Source:H |
| SLC25A31 | 4 | solute carrier family 25 member 31 [Source:H |
| HSPA4L | 4 | heat shock protein family A (Hsp70) member 4 |
| AC093591.3 | 4 | INTS3 and NABP interacting protein (INIP) pse |
| AC093591.1 | 4 | ribosomal protein L21 (RPL21) pseudogene |
| AC093591.2 | 4 | novel transcript, antisense HSPA4L |
| PLK4 | 4 | polo like kinase 4 [Source:HGNC Symbol;Acc:H |
| RNU6-583P | 4 | RNA, U6 small nuclear 583, pseudogene [Sour |
| MFSD8 | 4 | major facilitator superfamily domain containir |
| ABHD18 | 4 | abhydrolase domain containing 18 [Source:HG |
| LARP1B | 4 | La ribonucleoprotein domain family member 1 |
| AC099340.1 | 4 | ribosomal protein L15 (RPL15) pseudogene |
| FOSL1P1 | 4 | FOSL1 pseudogene 1 [Source:HGNC Symbol;A |
| PGRMC2 | 4 | progesterone receptor membrane componen |
| LINC02615 | 4 | long intergenic non-protein coding RNA 2615 |
| AC110609.1 | 4 | novel transcript |
| AC078850.1 | 4 | novel transcript |
| AC078850.2 | 4 | novel transcript |
| JADRR | 4 | JADE1 adjacent regulatory RNA [Source:HGNC |
| JADE1 | 4 | jade family PHD finger 1 [Source:HGNC Symbo |
| SCLT1 | 4 | sodium channel and clathrin linker 1 [Source:H |
| AC093783.1 | 4 | zinc finger protein 93 (ZNF93) pseudogene |
| AC093826.2 | 4 | NULL |
| C4orf33 | 4 | chromosome 4 open reading frame 33 [Source |
| AC093826.1 | 4 | CMT1A duplicated region transcript 15-like 2 ( |
| ZSWIM5P3 | 4 | zinc finger SWIM-type containing 5 pseudoger |
| AC105417.1 | 4 | chromosome 4 open reading frame 33 (C4orf3 |
| AC082650.1 | 4 | novel transcript |
| AC082650.2 | 4 | novel transcript |

| | | |
|---|---|---|
| LINC02466 | 4 | long intergenic non-protein coding RNA 2466 |
| LINC02465 | 4 | long intergenic non-protein coding RNA 2465 |
| EEF1GP8 | 4 | eukaryotic translation elongation factor 1 gam |
| LINC02479 | 4 | long intergenic non-protein coding RNA 2479 |
| PGBD4P4 | 4 | piggyBac transposable element derived 4 pseu |
| GAPDHP56 | 4 | glyceraldehyde 3 phosphate dehydrogenase p |
| AC093831.2 | 4 | tec |
| AC093831.1 | 4 | novel transcript |
| AC093902.1 | 4 | tigger transposable element derived 1 (TIGD1 |
| AC025554.1 | 4 | COX17 cytochrome c oxidase assembly homol |
| RNU6-224P | 4 | RNA, U6 small nuclear 224, pseudogene [Sour |
| LINC02377 | 4 | long intergenic non-protein coding RNA 2377 |
| AC105424.1 | 4 | neuroblastoma amplified sequence (NBAS) ps |
| RN7SL205P | 4 | RNA, 7SL, cytoplasmic 205, pseudogene [Sour |
| SNHG27 | 4 | small nucleolar RNA host gene 27 [Source:HGI |
| SNORA70 | 4 | Small nucleolar RNA SNORA70 [Source:RFAM; |
| AC107393.1 | 4 | ribosomal protein L10-like (RPL10L) pseudoge |
| RPL7AP28 | 4 | ribosomal protein L7a pseudogene 28 [Source |
| AC096711.2 | 4 | novel transcript |
| AC093916.1 | 4 | novel transcript |
| AC096711.1 | 4 | novel transcript |
| ELL2P2 | 4 | elongation factor for RNA polymerase II 2 pse |
| AC114741.1 | 4 | novel transcript |
| LINC01256 | 4 | long intergenic non-protein coding RNA 1256 |
| AARSP1 | 4 | alanyl-tRNA synthetase pseudogene 1 [Source |
| AC115622.1 | 4 | novel transcript |
| AC110751.1 | 4 | novel transcript |
| AC105383.1 | 4 | novel transcript |
| R3HDM2P1 | 4 | R3H domain containing 2 pseudogene 1 [Sour |
| PCDH10 | 4 | protocadherin 10 [Source:HGNC Symbol;Acc:H |
| AC108052.1 | 4 | novel transcript |
| AC105252.1 | 4 | Yip1 domain family, member 3 (YIPF3) pseudo |
| AC079380.1 | 4 | novel transcript |
| AC015631.1 | 4 | novel transcript |
| AC096763.1 | 4 | novel transcript |
| PABPC4L | 4 | poly(A) binding protein cytoplasmic 4 like [Sou |
| PES1P1 | 4 | pescadillo ribosomal biogenesis factor 1 pseu |
| LINC02462 | 4 | long intergenic non-protein coding RNA 2462 |
| AC093722.1 | 4 | novel transcript |
| AC104619.2 | 4 | novel transcript |
| AC104619.4 | 4 | ubiquinol-cytochrome c reductase, complex II |
| AC104619.1 | 4 | ribosomal protein S2 (RPS2) pseudogene |
| AC104619.3 | 4 | eukaryotic translation elongation factor 1 alph |
| AC107463.1 | 4 | novel transcript |
| LINC02485 | 4 | long intergenic non-protein coding RNA 2485 |
| U6 | 4 | U6 spliceosomal RNA [Source:RFAM;Acc:RF00 |
| KRT18P54 | 4 | keratin 18 pseudogene 54 [Source:HGNC Sym |
| TARS2P1 | 4 | TARS2 pseudogene 1 [Source:HGNC Symbol;A |
| AC105362.1 | 4 | novel transcript |
| LINC00613 | 4 | long intergenic non-protein coding RNA 613 [S |

| | | |
|---|---|---|
| RNU1-89P | 4 | RNA, U1 small nuclear 89, pseudogene [Sourc |
| AC104136.1 | 4 | eukaryotic translation termination factor 1 (E |
| AC073429.2 | 4 | novel transcript |
| AC018680.1 | 4 | novel transcript |
| AC073429.1 | 4 | novel transcript |
| TERF1P3 | 4 | telomeric repeat binding factor 1 pseudogene |
| AC093875.1 | 4 | TEC |
| LINC02511 | 4 | long intergenic non-protein coding RNA 2511 |
| LINC02510 | 4 | long intergenic non-protein coding RNA 2510 |
| SERF1AP1 | 4 | small EDRK-rich factor 1A pseudogene 1 [Sour |
| STMN1P2 | 4 | stathmin 1 pseudogene 2 [Source:HGNC Symb |
| RPS23P2 | 4 | ribosomal protein S23 pseudogene 2 [Source: |
| PCDH18 | 4 | protocadherin 18 [Source:HGNC Symbol;Acc:H |
| LINC02172 | 4 | long intergenic non-protein coding RNA 2172 |
| AC116563.2 | 4 | TEC |
| AC116563.1 | 4 | novel transcript |
| AC131956.2 | 4 | novel transcript |
| AC131956.3 | 4 | novel transcript |
| AC131956.1 | 4 | novel transcript |
| AC097511.1 | 4 | novel transcript |
| LINC00616 | 4 | long intergenic non-protein coding RNA 616 [S |
| SLC7A11-AS1 | 4 | SLC7A11 antisense RNA 1 [Source:HGNC Symb |
| SLC7A11 | 4 | solute carrier family 7 member 11 [Source:HG |
| AC093903.1 | 4 | novel transcript |
| LINC00498 | 4 | long intergenic non-protein coding RNA 498 [S |
| LINC00499 | 4 | long intergenic non-protein coding RNA 499 [S |
| LINC00500 | 4 | long intergenic non-protein coding RNA 500 [S |
| AC105416.1 | 4 | armadillo repeat containing 1 (ARMC1) pseud |
| AC098859.1 | 4 | NIN1/RPN12 binding protein 1 homolog (S. ce |
| AC098859.2 | 4 | novel transcript |
| AC093766.1 | 4 | novel transcript |
| AC109927.1 | 4 | novel transcript |
| AC109927.2 | 4 | novel transcript |
| NOCT | 4 | nocturnin [Source:HGNC Symbol;Acc:HGNC:14 |
| ELF2 | 4 | E74 like ETS transcription factor 2 [Source:HG |
| RNU6-531P | 4 | RNA, U6 small nuclear 531, pseudogene [Sour |
| RN7SL382P | 4 | RNA, 7SL, cytoplasmic 382, pseudogene [Sour |
| PPP1R14BP3 | 4 | protein phosphatase 1 regulatory inhibitor sul |
| Metazoa_SRP | 4 | Metazoan signal recognition particle RNA [Sou |
| RN7SL311P | 4 | RNA, 7SL, cytoplasmic 311, pseudogene [Sour |
| Y_RNA | 4 | Y RNA [Source:RFAM;Acc:RF00019] |
| RNU6-506P | 4 | RNA, U6 small nuclear 506, pseudogene [Sour |
| RNU6-1074P | 4 | RNA, U6 small nuclear 1074, pseudogene [Sou |
| MGARP | 4 | mitochondria localized glutamic acid rich prot |
| NDUFC1 | 4 | NADH:ubiquinone oxidoreductase subunit C1 |
| NAA15 | 4 | N(alpha)-acetyltransferase 15, NatA auxiliary s |
| RNU6-1214P | 4 | RNA, U6 small nuclear 1214, pseudogene [Sou |
| AC097376.3 | 4 | novel transcript, antisense to RAB33B |
| AC097376.2 | 4 | glucose-fructose oxidoreductase domain cont |
| AC097376.1 | 4 | NULL |

| | | |
|---|---|---|
| RAB33B | 4 | RAB33B, member RAS oncogene family [Sourc |
| SETD7 | 4 | SET domain containing 7, histone lysine methy |
| FTH1P24 | 4 | ferritin heavy chain 1 pseudogene 24 [Source: |
| AC112236.2 | 4 | novel transcript, antisense to SETD7 |
| AC112236.3 | 4 | novel transcript, antisense to SETD7 |
| AC112236.1 | 4 | novel transcript |
| MGST2 | 4 | microsomal glutathione S-transferase 2 [Sourc |
| RN7SKP237 | 4 | RN7SK pseudogene 237 [Source:HGNC Symbo |
| H3F3AP6 | 4 | H3 histone, family 3A, pseudogene 6 [Source: |
| MAML3 | 4 | mastermind like transcriptional coactivator 3 |
| AC108053.1 | 4 | novel transcript, antisense to MAML3 |
| RN7SKP253 | 4 | RN7SK pseudogene 253 [Source:HGNC Symbo |
| AC131182.1 | 4 | novel transcript |
| SCOC | 4 | short coiled-coil protein [Source:HGNC Symbo |
| SCOC-AS1 | 4 | SCOC antisense RNA 1 [Source:HGNC Symbol; |
| NDUFB4P9 | 4 | NADH:ubiquinone oxidoreductase subunit B4 |
| RPL14P3 | 4 | ribosomal protein L14 pseudogene 3 [Source: |
| CLGN | 4 | calmegin [Source:HGNC Symbol;Acc:HGNC:20 |
| MGAT4D | 4 | MGAT4 family member D [Source:HGNC Symb |
| AC093671.1 | 4 | tec |
| ELMOD2 | 4 | ELMO domain containing 2 [Source:HGNC Syn |
| UCP1 | 4 | uncoupling protein 1 [Source:HGNC Symbol;A |
| AC108019.2 | 4 | novel transcript |
| RN7SL152P | 4 | RNA, 7SL, cytoplasmic 152, pseudogene [Sour |
| AC108019.1 | 4 | riboflavin kinase (RFK) pseudogene |
| TBC1D9 | 4 | TBC1 domain family member 9 [Source:HGNC |
| TNRC18P1 | 4 | trinucleotide repeat containing 18 pseudogen |
| RNY1P14 | 4 | RNY1 pseudogene 14 [Source:HGNC Symbol;A |
| AC096733.1 | 4 | novel transcript |
| AC096733.2 | 4 | novel transcript |
| Y_RNA | 4 | Y RNA [Source:RFAM;Acc:RF00019] |
| RNF150 | 4 | ring finger protein 150 [Source:HGNC Symbol; |
| ZNF330 | 4 | zinc finger protein 330 [Source:HGNC Symbol; |
| AC107220.1 | 4 | novel transcript |
| LINC02432 | 4 | long intergenic non-protein coding RNA 2432 |
| AC097504.1 | 4 | F-box and leucine-rich repeat protein 5 (FBXL5 |
| AC097504.2 | 4 | novel transcript |
| LINC02276 | 4 | long intergenic non-protein coding RNA 2276 |
| IL15 | 4 | interleukin 15 [Source:HGNC Symbol;Acc:HGN |
| INPP4B | 4 | inositol polyphosphate-4-phosphatase type II |
| RPL5P13 | 4 | ribosomal protein L5 pseudogene 13 [Source: |
| LSM3P4 | 4 | LSM3 homolog, U6 small nuclear RNA and mR |
| AC139720.2 | 4 | novel transcript |
| AC139720.1 | 4 | novel transcript |
| AC104596.1 | 4 | novel transcript |
| AC104596.2 | 4 | LysM, putative peptidoglycan-binding, domain |
| USP38 | 4 | ubiquitin specific peptidase 38 [Source:HGNC |
| AC097658.2 | 4 | novel transcript |
| Y_RNA | 4 | Y RNA [Source:RFAM;Acc:RF00019] |
| GAB1 | 4 | GRB2 associated binding protein 1 [Source:HG |

| | | |
|---|---|---|
| MIR3139 | 4 | microRNA 3139 [Source:HGNC Symbol;Acc:HG |
| AC097658.1 | 4 | ribosomal protein S2 (RPS2) pseudogene |
| AC097658.3 | 4 | novel transcript, antisense to GAB1 |
| RPSAP36 | 4 | ribosomal protein SA pseudogene 36 [Source: |
| AC104685.1 | 4 | torsin A interacting protein 2 (TOR1AIP2) pseu |
| SMARCA5-AS1 | 4 | SMARCA5 antisense RNA 1 [Source:HGNC Sym |
| SMARCA5 | 4 | SWI/SNF related, matrix associated, actin dep |
| AC139713.1 | 4 | heterogeneous nuclear ribonucleoprotein A1 |
| AC107223.1 | 4 | novel transcript, antisense to GYPA |
| GUSBP5 | 4 | GUSB pseudogene 5 [Source:HGNC Symbol;Ac |
| FREM3 | 4 | FRAS1 related extracellular matrix 3 [Source:H |
| AC104090.1 | 4 | ethanolamine kinase 1 (ETNK1) pseudogene |
| GYPE | 4 | glycophorin E (MNS blood group) [Source:HGN |
| AC093890.1 | 4 | salvador homolog 1 (Drosophila) (SAV1) pseud |
| GYPB | 4 | glycophorin B (MNS blood group) [Source:HGI |
| AC093890.2 | 4 | novel transcript, antisense to GYPB |
| GYPA | 4 | glycophorin A (MNS blood group) [Source:HGI |
| AC098588.3 | 4 | novel protein |
| AC098588.2 | 4 | novel transcript |
| AC106871.1 | 4 | novel transcript |
| KRT18P51 | 4 | keratin 18 pseudogene 51 [Source:HGNC Sym |
| HHIP-AS1 | 4 | HHIP antisense RNA 1 [Source:HGNC Symbol;A |
| HHIP | 4 | hedgehog interacting protein [Source:HGNC S |
| AC098588.1 | 4 | NULL |
| ANAPC10 | 4 | anaphase promoting complex subunit 10 [Sou |
| HSPD1P5 | 4 | heat shock protein family D (Hsp60) member |
| AC109811.1 | 4 | novel transcript |
| AC109811.2 | 4 | adrenocortical dysplasia homolog (mouse) (A( |
| RN7SKP235 | 4 | RN7SK pseudogene 235 [Source:HGNC Symbo |
| ABCE1 | 4 | ATP binding cassette subfamily E member 1 [S |
| OTUD4 | 4 | OTU deubiquitinase 4 [Source:HGNC Symbol;A |
| Y_RNA | 4 | Y RNA [Source:RFAM;Acc:RF00019] |
| RPS23P4 | 4 | ribosomal protein S23 pseudogene 4 [Source: |
| LINC02266 | 4 | long intergenic non-protein coding RNA 2266 |
| RTN3P1 | 4 | reticulon 3 pseudogene 1 [Source:HGNC Symb |
| AC079228.1 | 4 | tec |
| AC093796.1 | 4 | novel transcript |
| NMNAT1P4 | 4 | NMNAT1 pseudogene 4 [Source:HGNC Symbo |
| SMAD1 | 4 | SMAD family member 1 [Source:HGNC Symbo |
| SMAD1-AS2 | 4 | SMAD1 antisense RNA 2 [Source:HGNC Symbo |
| SMAD1-AS1 | 4 | SMAD1 antisense RNA 1 [Source:HGNC Symbo |
| MMAA | 4 | metabolism of cobalamin associated A [Sourc |
| NCOA4P3 | 4 | nuclear receptor coactivator 4 pseudogene 3 |
| AC093864.1 | 4 | novel transcript |
| C4orf51 | 4 | chromosome 4 open reading frame 51 [Sourc |
| ZNF827 | 4 | zinc finger protein 827 [Source:HGNC Symbol; |
| AC104791.2 | 4 | TEC |
| AC104791.1 | 4 | novel transcript |
| Y_RNA | 4 | Y RNA [Source:RFAM;Acc:RF00019] |
| AC108206.1 | 4 | novel transcript |

| | | |
|---|---|---|
| LINC01095 | 4 | long intergenic non-protein coding RNA 1095 |
| LSM6 | 4 | LSM6 homolog, U6 small nuclear RNA and mR |
| AC097372.2 | 4 | TEC |
| REELD1 | 4 | reeler domain containing 1 [Source:HGNC Syn |
| AC097372.3 | 4 | TEC |
| AC097372.1 | 4 | novel transcript |
| SLC10A7 | 4 | solute carrier family 10 member 7 [Source:HG |
| MIR7849 | 4 | microRNA 7849 [Source:HGNC Symbol;Acc:HG |
| RNU1-44P | 4 | RNA, U1 small nuclear 44, pseudogene [Sourc |
| POU4F2 | 4 | POU class 4 homeobox 2 [Source:HGNC Symb |
| TTC29 | 4 | tetratricopeptide repeat domain 29 [Source:H |
| AC092435.2 | 4 | novel transcript |
| AC092435.1 | 4 | novel transcript |
| AC092435.3 | 4 | novel transcript |
| AC092435.4 | 4 | tec |
| AC097450.1 | 4 | novel transcript |
| MIR548G | 4 | microRNA 548g [Source:HGNC Symbol;Acc:HG |
| AC010683.1 | 4 | ribosomal protein L31 (RPL31) pseudogene |
| PRMT5P1 | 4 | protein arginine methyltransferase 5 pseudog |
| EDNRA | 4 | endothelin receptor type A [Source:HGNC Syn |
| AC093908.1 | 4 | TEC |
| GTF2F2P1 | 4 | general transcription factor IIF subunit 2 pseu |
| LINC02507 | 4 | long intergenic non-protein coding RNA 2507 |
| AC093835.1 | 4 | novel transcript |
| TMEM184C | 4 | transmembrane protein 184C [Source:HGNC S |
| PRMT9 | 4 | protein arginine methyltransferase 9 [Source: |
| ARHGAP10 | 4 | Rho GTPase activating protein 10 [Source:HGN |
| RNA5SP165 | 4 | RNA, 5S ribosomal pseudogene 165 [Source:H |
| MIR4799 | 4 | microRNA 4799 [Source:HGNC Symbol;Acc:HG |
| RN7SL254P | 4 | RNA, 7SL, cytoplasmic 254, pseudogene [Sour |
| AC115621.1 | 4 | ret finger protein-like 4A (RFPL4A) pseudogen |
| NR3C2 | 4 | nuclear receptor subfamily 3 group C member |
| AC069272.1 | 4 | novel transcript |
| AC002460.2 | 4 | novel transcript |
| ASS1P8 | 4 | argininosuccinate synthetase 1 pseudogene 8 |
| RNA5SP166 | 4 | RNA, 5S ribosomal pseudogene 166 [Source:H |
| AC108935.1 | 4 | novel transcript |
| ATP5MGP4 | 4 | ATP synthase membrane subunit g pseudogen |
| AC093648.1 | 4 | TEC |
| AC002460.1 | 4 | novel transcript |
| RNU7-197P | 4 | RNA, U7 small nuclear 197 pseudogene [Sourc |
| AC108156.1 | 4 | novel transcript |
| LINC02430 | 4 | long intergenic non-protein coding RNA 2430 |
| LINC02355 | 4 | long intergenic non-protein coding RNA 2355 |
| IQCM | 4 | IQ motif containing M [Source:HGNC Symbol;A |
| AC108168.1 | 4 | novel transcript, antisense to RP11-526A4.1 |
| AC093893.1 | 4 | novel transcript |
| AKIRIN2P1 | 4 | akirin 2 pseudogene 1 [Source:HGNC Symbol;A |
| RNA5SP167 | 4 | RNA, 5S ribosomal pseudogene 167 [Source:H |
| RNU6-1230P | 4 | RNA, U6 small nuclear 1230, pseudogene [Sou |

| | | |
|---|---|---|
| AC105343.1 | 4 | novel transcript |
| AC105343.2 | 4 | novel transcript |
| DCLK2 | 4 | doublecortin like kinase 2 [Source:HGNC Symb |
| RNU7-194P | 4 | RNA, U7 small nuclear 194 pseudogene [Sourc |
| LRBA | 4 | LPS responsive beige-like anchor protein [Sou |
| AC092612.1 | 4 | nei like 2 (E. coli) (NEIL2) pseudogene |
| AC110813.1 | 4 | novel transcript, antisense to LRBA |
| MAB21L2 | 4 | mab-21 like 2 [Source:HGNC Symbol;Acc:HGN |
| ZBTB8OSP1 | 4 | zinc finger and BTB domain containing 8 oppo |
| RNA5SP168 | 4 | RNA, 5S ribosomal pseudogene 168 [Source:H |
| AK4P6 | 4 | adenylate kinase 4 pseudogene 6 [Source:HGI |
| RPS3A | 4 | ribosomal protein S3A [Source:HGNC Symbol; |
| SNORD73B | 4 | small nucleolar RNA, C/D box U73B [Source:H |
| SH3D19 | 4 | SH3 domain containing 19 [Source:HGNC Sym |
| SNORD73A | 4 | small nucleolar RNA, C/D box 73A [Source:HG |
| AC095055.1 | 4 | novel transcript, antisense to SH3D19 |
| AC104819.1 | 4 | trypsin domain contaiing protein pseudogene |
| PRSS48 | 4 | serine protease 48 [Source:HGNC Symbol;Acc |
| AC104819.2 | 4 | epidermis-specific serine protease-like proteir |
| RNU6-1282P | 4 | RNA, U6 small nuclear 1282, pseudogene [Sou |
| AC104819.3 | 4 | novel transcript |
| FAM160A1-DT | 4 | FAM160A1 divergent transcript [Source:HGNC |
| FAM160A1 | 4 | family with sequence similarity 160 member A |
| RN7SKP35 | 4 | RN7SK pseudogene 35 [Source:HGNC Symbol; |
| GATB | 4 | glutamyl-tRNA amidotransferase subunit B [So |
| AC092611.1 | 4 | novel transcript, antisense to PET112L |
| AC107074.1 | 4 | novel transcript |
| AC112242.2 | 4 | novel transcript |
| AC112242.1 | 4 | novel transcript |
| AC097375.2 | 4 | novel transcript |
| AC097375.1 | 4 | novel transcript |
| AC097375.4 | 4 | novel transcript |
| AC097375.3 | 4 | novel transcript |
| RNA5SP169 | 4 | RNA, 5S ribosomal pseudogene 169 [Source:H |
| AC097375.5 | 4 | novel transcript |
| LINC02273 | 4 | long intergenic non-protein coding RNA 2273 |
| AC079340.3 | 4 | novel transcript |
| AC079340.1 | 4 | novel transcript |
| AC079340.2 | 4 | novel transcript |
| RN7SL446P | 4 | RNA, 7SL, cytoplasmic 446, pseudogene [Sour |
| AC080078.1 | 4 | novel transcript |
| AC080078.2 | 4 | novel transcript |
| FBXW7 | 4 | F-box and WD repeat domain containing 7 [Sc |
| FBXW7-AS1 | 4 | FBXW7 antisense RNA 1 [Source:HGNC Symbo |
| AC023424.3 | 4 | novel transcript, antisense to FBXW7 |
| MIR3140 | 4 | microRNA 3140 [Source:HGNC Symbol;Acc:HG |
| MIR4453HG | 4 | MIR4453 host gene [Source:HGNC Symbol;Acc |
| MIR4453 | 4 | microRNA 4453 [Source:HGNC Symbol;Acc:HG |
| AC023424.2 | 4 | ribosomal protein S3A (RPS3A) pseudogene |
| AC023424.1 | 4 | ribosomal protein S14 (RPS14) pseudogene |

| | | |
|---|---|---|
| TMEM154 | 4 | transmembrane protein 154 [Source:HGNC Sy |
| AC106882.1 | 4 | novel transcript |
| TIGD4 | 4 | tigger transposable element derived 4 [Source |
| ARFIP1 | 4 | ADP ribosylation factor interacting protein 1 [ |
| AC113155.1 | 4 | NSA2 ribosome biogenesis homolog (S. cerevi |
| FAM192BP | 4 | family with sequence similarity 192 member E |
| AC093599.2 | 4 | novel transcript, antisense to FHDC1 |
| FHDC1 | 4 | FH2 domain containing 1 [Source:HGNC Symb |
| AC093599.1 | 4 | novel transcript |
| TRIM2 | 4 | tripartite motif containing 2 [Source:HGNC Sy |
| RNU6-1196P | 4 | RNA, U6 small nuclear 1196, pseudogene [Sou |
| Y_RNA | 4 | Y RNA [Source:RFAM;Acc:RF00019] |
| ANXA2P1 | 4 | annexin A2 pseudogene 1 [Source:HGNC Syml |
| MND1 | 4 | meiotic nuclear divisions 1 [Source:HGNC Sym |
| RN7SL419P | 4 | RNA, 7SL, cytoplasmic 419, pseudogene [Sour |
| TMEM131L | 4 | transmembrane 131 like [Source:HGNC Symbo |
| AC106865.2 | 4 | novel transcript, antisense to KIAA0922 |
| AC106865.1 | 4 | toll-like receptor 2 pseudogene |
| WDR45P1 | 4 | WD repeat domain 45 pseudogene 1 [Source: |
| TLR2 | 4 | toll like receptor 2 [Source:HGNC Symbol;Acc: |
| RNF175 | 4 | ring finger protein 175 [Source:HGNC Symbol; |
| AC020703.1 | 4 | novel transcript |
| SFRP2 | 4 | secreted frizzled related protein 2 [Source:HG |
| AC079298.3 | 4 | novel transcript, antisense to DCHS2 |
| DCHS2 | 4 | dachsous cadherin-related 2 [Source:HGNC Sy |
| AC079298.1 | 4 | novel transcript, antisense to DCHS2 |
| AC079298.2 | 4 | novel transcript, antisense to DCHS2 |
| AC110775.1 | 4 | kelch domain containing 4 pseudogene |
| AC110753.1 | 4 | WD repeat domain 77 (WDR77) pseudogene |
| PLRG1 | 4 | pleiotropic regulator 1 [Source:HGNC Symbol; |
| RNU6-1285P | 4 | RNA, U6 small nuclear 1285, pseudogene [Sou |
| AC107385.1 | 4 | pseudogene similar to part of 5-aminoimidazo |
| FGB | 4 | fibrinogen beta chain [Source:HGNC Symbol;A |
| FGA | 4 | fibrinogen alpha chain [Source:HGNC Symbol; |
| FGG | 4 | fibrinogen gamma chain [Source:HGNC Symbo |
| AC107385.2 | 4 | fibrinogen gamma chain (FGG) pseudogene |
| LRAT | 4 | lecithin retinol acyltransferase [Source:HGNC |
| NDUFB2P1 | 4 | NADH:ubiquinone oxidoreductase subunit B2 |
| AC009567.1 | 4 | novel transcript |
| AC009567.2 | 4 | pseudogene similar to part of ribosomal prote |
| RBM46 | 4 | RNA binding motif protein 46 [Source:HGNC S |
| RNU2-66P | 4 | RNA, U2 small nuclear 66, pseudogene [Sourc |
| RNU2-44P | 4 | RNA, U2 small nuclear 44, pseudogene [Sourc |
| AC097467.3 | 4 | novel transcript, antisense to MAP9 |
| AC104407.1 | 4 | novel transcript |
| NPY2R | 4 | neuropeptide Y receptor Y2 [Source:HGNC Syr |
| AC097467.1 | 4 | novel transcript |
| MAP9 | 4 | microtubule associated protein 9 [Source:HGN |
| YWHAEP4 | 4 | tyrosine 3-monooxygenase/tryptophan 5-mor |
| AC097467.2 | 4 | tumor necrosis factor receptor superfamily, m |

| | | |
|---|---|---|
| AC097467.4 | 4 | TBC1 domain family, member 3 (TBC1D3) pseu |
| AC097526.1 | 4 | novel transcript |
| MTCYBP17 | 4 | MT-CYB pseudogene 17 [Source:HGNC Symbo |
| MTND6P17 | 4 | MT-ND6 pseudogene 17 [Source:HGNC Symbo |
| MTND5P9 | 4 | MT-ND5 pseudogene 9 [Source:HGNC Symbol |
| MTND4P8 | 4 | MT-ND4 pseudogene 8 [Source:HGNC Symbol |
| MTND3P3 | 4 | MT-ND3 pseudogene 3 [Source:HGNC Symbol |
| MTCO3P9 | 4 | MT-CO3 pseudogene 9 [Source:HGNC Symbol |
| MTATP6P9 | 4 | MT-ATP6 pseudogene 9 [Source:HGNC Symbo |
| MTCO2P9 | 4 | MT-CO2 pseudogene 9 [Source:HGNC Symbol |
| MTCO1P9 | 4 | MT-CO1 pseudogene 9 [Source:HGNC Symbol |
| MTND2P33 | 4 | MT-ND2 pseudogene 33 [Source:HGNC Symbo |
| MTND1P22 | 4 | MT-ND1 pseudogene 22 [Source:HGNC Symbo |
| AC107208.1 | 4 | novel transcript |
| GUCY1A1 | 4 | guanylate cyclase 1 soluble subunit alpha 1 [S |
| AC104083.1 | 4 | novel transcript, overlapping GUCY1A3 |
| GUCY1B1 | 4 | guanylate cyclase 1 soluble subunit beta 1 [So |
| ASIC5 | 4 | acid sensing ion channel subunit family memb |
| TDO2 | 4 | tryptophan 2,3-dioxygenase [Source:HGNC Sy |
| CTSO | 4 | cathepsin O [Source:HGNC Symbol;Acc:HGNC: |
| FTH1P21 | 4 | ferritin heavy chain 1 pseudogene 21 [Source: |
| AC096736.3 | 4 | novel transcript |
| LINC02272 | 4 | long intergenic non-protein coding RNA 2272 |
| AC096736.2 | 4 | novel transcript |
| AC096736.1 | 4 | novel transcript |
| PDGFC | 4 | platelet derived growth factor C [Source:HGN |
| AC092608.1 | 4 | novel transcript |
| AC092608.3 | 4 | TEC |
| AC092608.2 | 4 | TEC |
| AC093325.1 | 4 | ribosomal protein L34 (RPL34) pseudogene |
| RPPH1-3P | 4 | ribonuclease P RNA component H1, 3 pseudo |
| Y_RNA | 4 | Y RNA [Source:RFAM;Acc:RF00019] |
| GLRB | 4 | glycine receptor beta [Source:HGNC Symbol;A |
| RNU6-582P | 4 | RNA, U6 small nuclear 582, pseudogene [Sour |
| GRIA2 | 4 | glutamate ionotropic receptor AMPA type sub |
| AC112240.1 | 4 | chromatin accessibility complex 1 (CHRAC1) p |
| AC093817.2 | 4 | novel transcript |
| LINC02433 | 4 | long intergenic non-protein coding RNA 2433 |
| AC093817.1 | 4 | novel transcript |
| Y_RNA | 4 | Y RNA [Source:RFAM;Acc:RF00019] |
| AC017037.1 | 4 | ATP-binding cassette, sub-family B (MDR/TAP |
| AC017037.3 | 4 | TEC |
| AC017037.5 | 4 | novel transcript |
| AC084740.1 | 4 | novel transcript |
| AC017037.4 | 4 | TEC |
| AC017037.2 | 4 | ribosomal protein L6 (RPL6) pseudogene |
| AC098679.3 | 4 | TEC |
| AC098679.4 | 4 | TEC |
| GASK1B | 4 | golgi associated kinase 1B [Source:HGNC Syml |
| FAM198B-AS1 | 4 | FAM198B antisense RNA 1 [Source:HGNC Sym |

| | | |
|---|---|---|
| AC098679.6 | 4 | novel transcript |
| NUDT19P5 | 4 | nudix hydrolase 19 pseudogene 5 [Source:HG |
| AC098679.1 | 4 | novel transcript |
| TMEM144 | 4 | transmembrane protein 144 [Source:HGNC Sy |
| AC098679.5 | 4 | novel transcript |
| AC098679.2 | 4 | axin interactor, dorsalization associated (AIDA |
| RXFP1 | 4 | relaxin family peptide receptor 1 [Source:HGN |
| RNU4-79P | 4 | RNA, U4 small nuclear 79, pseudogene [Sourc |
| AC121161.1 | 4 | GC-rich promoter binding protein 1 (GPBP1) p |
| AC121161.2 | 4 | novel transcript |
| C4orf46 | 4 | chromosome 4 open reading frame 46 [Sourc |
| ETFDH | 4 | electron transfer flavoprotein dehydrogenase |
| U3 | 4 | Small nucleolar RNA U3 [Source:RFAM;Acc:RF |
| PPID | 4 | peptidylprolyl isomerase D [Source:HGNC Sym |
| FNIP2 | 4 | folliculin interacting protein 2 [Source:HGNC S |
| RNU6-128P | 4 | RNA, U6 small nuclear 128, pseudogene [Sour |
| C4orf45 | 4 | chromosome 4 open reading frame 45 [Sourc |
| SNORD39 | 4 | Small nucleolar RNA SNORD55/SNORD39 [Sou |
| AC093700.1 | 4 | proteasome (prosome, macropain) activator s |
| FABP5P12 | 4 | fatty acid binding protein 5 pseudogene 12 [S |
| Y_RNA | 4 | Y RNA [Source:RFAM;Acc:RF00019] |
| RAPGEF2 | 4 | Rap guanine nucleotide exchange factor 2 [So |
| MIR3688-1 | 4 | microRNA 3688-1 [Source:HGNC Symbol;Acc: |
| MIR3688-2 | 4 | microRNA 3688-2 [Source:HGNC Symbol;Acc: |
| AC074344.1 | 4 | kinase non-catalytic C-lobe domain (KIND) cor |
| AC074344.2 | 4 | novel transcript |
| LINC02233 | 4 | long intergenic non-protein coding RNA 2233 |
| AC093853.1 | 4 | v-myb myeloblastosis viral oncogene homolog |
| LINC02477 | 4 | long intergenic non-protein coding RNA 2477 |
| RPS14P7 | 4 | ribosomal protein S14 pseudogene 7 [Source: |
| AC104793.1 | 4 | novel transcript |
| FSTL5 | 4 | follistatin like 5 [Source:HGNC Symbol;Acc:HG |
| AC023136.1 | 4 | novel transcript |
| MTHFD2P4 | 4 | methylenetetrahydrofolate dehydrogenase (N |
| TOMM22P4 | 4 | TOMM22 pseudogene 4 [Source:HGNC Symbo |
| AC021134.1 | 4 | novel transcript |
| AC021134.2 | 4 | novel transcript |
| AC084752.1 | 4 | chromosome 6 open reading frame 221 (C6or |
| MIR4454 | 4 | microRNA 4454 [Source:HGNC Symbol;Acc:H |
| AC022272.1 | 4 | novel transcript |
| NAF1 | 4 | nuclear assembly factor 1 ribonucleoprotein [ |
| AC079238.1 | 4 | mitochondrial ribosomal protein S5 (MRPS5) p |
| NPY1R | 4 | neuropeptide Y receptor Y1 [Source:HGNC Sy |
| NPY5R | 4 | neuropeptide Y receptor Y5 [Source:HGNC Sy |
| RPL35AP12 | 4 | ribosomal protein L35a pseudogene 12 [Sourc |
| TKTL2 | 4 | transketolase like 2 [Source:HGNC Symbol;Ac |
| TMA16 | 4 | translation machinery associated 16 homolog |
| MARCH1 | 4 | membrane associated ring-CH-type finger 1 [S |
| AC093788.1 | 4 | novel transcript |
| RN7SKP105 | 4 | RN7SK pseudogene 105 [Source:HGNC Symbo |

| | | |
|---|---|---|
| AC112247.1 | 4 | tyrosine 3-monooxygenase/tryptophan 5-mor |
| AC105250.1 | 4 | heat shock 70kDa protein 5 (glucose-regulate |
| ANP32C | 4 | acidic nuclear phosphoprotein 32 family mem |
| AC074198.2 | 4 | novel pseudogene |
| RNU6-284P | 4 | RNA, U6 small nuclear 284, pseudogene [Sour |
| AC074198.1 | 4 | cysteine and histidine-rich domain (CHORD)-c |
| SMIM31 | 4 | small integral membrane protein 31 [Source:H |
| RNU6-668P | 4 | RNA, U6 small nuclear 668, pseudogene [Sour |
| APELA | 4 | apelin receptor early endogenous ligand [Sou |
| TRIM60P14 | 4 | tripartite motif containing 60 pseudogene 14 |
| AC106872.3 | 4 | ribosomal protein L26 (RPL26) pseudogene |
| AC106872.12 | 4 | novel transcript |
| FAM218BP | 4 | family with sequence similarity 218 member B |
| AC106872.10 | 4 | immediate early response 3 (IER3) pseudogen |
| AC106872.5 | 4 | novel pseudogene |
| NACA3P | 4 | NACA family member 3, pseudogene [Source: |
| AC106872.1 | 4 | novel pseudogene containing zinc finger doma |
| TRIM61 | 4 | tripartite motif containing 61 [Source:HGNC S |
| FAM218A | 4 | family with sequence similarity 218 member A |
| AC106872.4 | 4 | novel pseudogene containing a SPRY domain |
| AC106872.9 | 4 | novel transcript |
| AC106872.8 | 4 | novel pseudogene containing zinc finger doma |
| AC106872.2 | 4 | ribosomal protein L21 (RPL21) pseudogene |
| TRIM60 | 4 | tripartite motif containing 60 [Source:HGNC S |
| AC106872.11 | 4 | tripartite motif containing 60 (TRIM60) pseud |
| AC106872.7 | 4 | novel pseudogene containing a zinc finger dor |
| AC106872.6 | 4 | ribosomal protein S10 (RPS10) pseudogene |
| TRIM75P | 4 | tripartite motif containing 75, pseudogene [Sc |
| TMEM192 | 4 | transmembrane protein 192 [Source:HGNC Sy |
| KLHL2 | 4 | kelch like family member 2 [Source:HGNC Sym |
| RNU4-87P | 4 | RNA, U4 small nuclear 87, pseudogene [Sourc |
| GK3P | 4 | glycerol kinase 3 pseudogene [Source:HGNC S |
| MSMO1 | 4 | methylsterol monooxygenase 1 [Source:HGNC |
| CPE | 4 | carboxypeptidase E [Source:HGNC Symbol;Acc |
| MIR578 | 4 | microRNA 578 [Source:HGNC Symbol;Acc:HGN |
| HADHAP1 | 4 | HADHA pseudogene 1 [Source:HGNC Symbol;A |
| SCGB1D5P | 4 | secretoglobin family 1D member 5, pseudoge |
| NOL8P1 | 4 | nucleolar protein 8 pseudogene 1 [Source:HG |
| AC080079.2 | 4 | novel transcript |
| AC080079.1 | 4 | tripartite motif-containing 38 (TRIM38) pseud |
| LINC01179 | 4 | long intergenic non-protein coding RNA 1179 |
| TLL1 | 4 | tolloid like 1 [Source:HGNC Symbol;Acc:HGNC |
| RNA5SP170 | 4 | RNA, 5S ribosomal pseudogene 170 [Source:H |
| Y_RNA | 4 | Y RNA [Source:RFAM;Acc:RF00019] |
| AC097487.1 | 4 | novel transcript |
| SPOCK3 | 4 | SPARC (osteonectin), cwcv and kazal like dom |
| RNA5SP171 | 4 | RNA, 5S ribosomal pseudogene 171 [Source:H |
| AC079349.1 | 4 | novel transcript |
| RN7SL776P | 4 | RNA, 7SL, cytoplasmic 776, pseudogene [Sour |
| PHBP14 | 4 | prohibitin pseudogene 14 [Source:HGNC Symb |

| | | |
|---|---|---|
| RN7SKP188 | 4 | RN7SK pseudogene 188 [Source:HGNC Symbo |
| AC105148.1 | 4 | protein phosphatase 2, catalytic subunit, beta |
| AC116634.1 | 4 | novel transcript |
| ANXA10 | 4 | annexin A10 [Source:HGNC Symbol;Acc:HGNC |
| AC068989.1 | 4 | novel transcript, antisense to ANXA10 |
| DDX60 | 4 | DExD/H-box helicase 60 [Source:HGNC Symbo |
| DDX60L | 4 | DExD/H-box 60 like [Source:HGNC Symbol;Acc |
| AC079926.1 | 4 | NULL |
| PALLD | 4 | palladin, cytoskeletal associated protein [Sour |
| BTF3L4P4 | 4 | basic transcription factor 3 like 4 pseudogene |
| AC079858.1 | 4 | novel transcript |
| RNU6-1336P | 4 | RNA, U6 small nuclear 1336, pseudogene [Sou |
| RPL9P16 | 4 | ribosomal protein L9 pseudogene 16 [Source: |
| AC080188.1 | 4 | novel transcript |
| AC080188.2 | 4 | TEC |
| CBR4 | 4 | carbonyl reductase 4 [Source:HGNC Symbol;A |
| RNU6-853P | 4 | RNA, U6 small nuclear 853, pseudogene [Sour |
| RNY4P17 | 4 | RNY4 pseudogene 17 [Source:HGNC Symbol;A |
| AC021151.1 | 4 | novel transcript |
| SH3RF1 | 4 | SH3 domain containing ring finger 1 [Source:H |
| RPL6P12 | 4 | ribosomal protein L6 pseudogene 12 [Source: |
| AC096741.1 | 4 | novel transcript |
| NEK1 | 4 | NIMA related kinase 1 [Source:HGNC Symbol; |
| AC084724.1 | 4 | novel transcript, antisense to NEK1 |
| CLCN3 | 4 | chloride voltage-gated channel 3 [Source:HGN |
| HPF1 | 4 | histone PARylation factor 1 [Source:HGNC Syr |
| PTGES3P3 | 4 | prostaglandin E synthase 3 pseudogene 3 [Sou |
| AC079768.1 | 4 | chloride channel, nucleotide-sensitive, 1A (CLI |
| AC079768.3 | 4 | novel transcript |
| AC079768.4 | 4 | novel transcript |
| LINC02275 | 4 | long intergenic non-protein coding RNA 2275 |
| AC079768.2 | 4 | novel transcript |
| AC084866.2 | 4 | novel transcript |
| AC084866.1 | 4 | novel transcript |
| MFAP3L | 4 | microfibril associated protein 3 like [Source:H |
| AADAT | 4 | aminoadipate aminotransferase [Source:HGN |
| APOBEC3AP1 | 4 | apolipoprotein B mRNA editing enzyme cataly |
| AC069306.1 | 4 | phosphoribosyl pyrophosphate synthetase 1 ( |
| LINC01612 | 4 | long intergenic non-protein coding RNA 1612 |
| LINC02512 | 4 | long intergenic non-protein coding RNA 2512 |
| AC074255.1 | 4 | DENN/MADD domain containing 3 (DENND3) |
| HSP90AA6P | 4 | heat shock protein 90 alpha family class A me |
| LINC02382 | 4 | long intergenic non-protein coding RNA 2382 |
| AC097486.1 | 4 | myeloid cell leukemia sequence 1 (BCL2-relate |
| RNU6ATAC13P | 4 | RNA, U6atac small nuclear 13, pseudogene [So |
| AC034159.2 | 4 | novel transcript |
| LINC02431 | 4 | long intergenic non-protein coding RNA 2431 |
| AC034159.1 | 4 | RWD domain containing 2B (RWDD2B) pseudo |
| AC092639.1 | 4 | unconventional SNARE in the ER 1 homolog (S |
| MIR6082 | 4 | microRNA 6082 [Source:HGNC Symbol;Acc:HG |

| | | |
|---|---|---|
| LINC02504 | 4 | long intergenic non-protein coding RNA 2504 |
| AC074254.2 | 4 | novel transcript |
| LINC02174 | 4 | long intergenic non-protein coding RNA 2174 |
| AC074254.1 | 4 | novel transcript |
| GALNTL6 | 4 | polypeptide N-acetylgalactosaminyltransferas |
| AC108064.1 | 4 | ubiquitin-conjugating enzyme E2N (UBC13 ho |
| AC105285.1 | 4 | novel transcript, antisense to GALNT7 |
| RN7SL253P | 4 | RNA, 7SL, cytoplasmic 253, pseudogene [Sour |
| GALNT7 | 4 | polypeptide N-acetylgalactosaminyltransferas |
| MIR548T | 4 | microRNA 548t [Source:HGNC Symbol;Acc:HG |
| AC097534.1 | 4 | novel transcript |
| HMGB2 | 4 | high mobility group box 2 [Source:HGNC Symb |
| AC097534.2 | 4 | novel transcript, antisense to SAP30 |
| SAP30 | 4 | Sin3A associated protein 30 [Source:HGNC Sy |
| SCRG1 | 4 | stimulator of chondrogenesis 1 [Source:HGNC |
| AC093849.4 | 4 | novel transcript, antisense to SCRG1 |
| RPL5P11 | 4 | ribosomal protein L5 pseudogene 11 [Source: |
| AC093849.3 | 4 | novel transcript |
| RNU6-1096P | 4 | RNA, U6 small nuclear 1096, pseudogene [Sou |
| AC093849.1 | 4 | novel transcript |
| AC093849.2 | 4 | novel transcript |
| HAND2 | 4 | heart and neural crest derivatives expressed 2 |
| HAND2-AS1 | 4 | HAND2 antisense RNA 1 [Source:HGNC Symbo |
| MORF4 | 4 | mortality factor 4 (pseudogene) [Source:HGN |
| RANP6 | 4 | RAN pseudogene 6 [Source:HGNC Symbol;Acc |
| LINC02269 | 4 | long intergenic non-protein coding RNA 2269 |
| AC106895.1 | 4 | novel transcript |
| AC106895.2 | 4 | novel transcript |
| LINC02268 | 4 | long intergenic non-protein coding RNA 2268 |
| AC012055.1 | 4 | novel transcript |
| AC012055.2 | 4 | novel transcript |
| FBXO8 | 4 | F-box protein 8 [Source:HGNC Symbol;Acc:HG |
| CEP44 | 4 | centrosomal protein 44 [Source:HGNC Symbo |
| AC116616.1 | 4 | novel transcript |
| MIR4276 | 4 | microRNA 4276 [Source:HGNC Symbol;Acc:HC |
| HPGD | 4 | 15-hydroxyprostaglandin dehydrogenase [Sou |
| AC096751.2 | 4 | novel transcript |
| AC096751.1 | 4 | Zinc finger protein pseudogene |
| GLRA3 | 4 | glycine receptor alpha 3 [Source:HGNC Symbo |
| ADAM29 | 4 | ADAM metallopeptidase domain 29 [Source:H |
| AC105914.1 | 4 | GC-rich promoter binding protein 1-like 1 (GP |
| AC105914.2 | 4 | novel transcript |
| AC022325.1 | 4 | ADAM metallopeptidase domain 29 (ADAM29 |
| AC022325.2 | 4 | novel transcript |
| AC095050.2 | 4 | PC4 and SFRS1 interacting protein 1 (PSIP1) ps |
| AC095050.1 | 4 | novel transcript |
| AC131094.1 | 4 | novel transcript |
| TSEN2P1 | 4 | tRNA splicing endonuclease subunit 2 pseudo |
| ADAM20P2 | 4 | ADAM metallopeptidase domain 20 pseudoge |
| GPM6A | 4 | glycoprotein M6A [Source:HGNC Symbol;Acc: |

| | | |
|---|---|---|
| AC097537.1 | 4 | novel transcript, antisense to GPM6A |
| AC110794.1 | 4 | Rac GTPase activating protein 1 (RACGAP1) ps |
| MARK2P4 | 4 | microtubule affinity regulating kinase 2 pseud |
| WDR17 | 4 | WD repeat domain 17 [Source:HGNC Symbol;/ |
| AC093605.1 | 4 | NULL |
| SPATA4 | 4 | spermatogenesis associated 4 [Source:HGNC S |
| ASB5 | 4 | ankyrin repeat and SOCS box containing 5 [So |
| AC019163.1 | 4 | novel transcript |
| SPCS3 | 4 | signal peptidase complex subunit 3 [Source:H( |
| AC093801.2 | 4 | CTAGE family, member 5 (CTAGE5) pseudoger |
| AC093801.1 | 4 | novel transcript |
| VEGFC | 4 | vascular endothelial growth factor C [Source:H |
| AC097518.2 | 4 | novel transcript |
| AC092673.1 | 4 | novel transcript |
| LINC02509 | 4 | long intergenic non-protein coding RNA 2509 |
| RN7SKP136 | 4 | RN7SK pseudogene 136 [Source:HGNC Symbo |
| AC097518.1 | 4 | novel transcript |
| NEIL3 | 4 | nei like DNA glycosylase 3 [Source:HGNC Syml |
| AC027627.1 | 4 | novel transcript |
| AGA | 4 | aspartylglucosaminidase [Source:HGNC Symb( |
| AC078881.1 | 4 | novel transcript |
| AC078881.2 | 4 | NULL |
| RNA5SP172 | 4 | RNA, 5S ribosomal pseudogene 172 [Source:H |
| AC103879.1 | 4 | novel transcript |
| LINC01098 | 4 | long intergenic non-protein coding RNA 1098 |
| LINC01099 | 4 | long intergenic non-protein coding RNA 1099 |
| RNU1-45P | 4 | RNA, U1 small nuclear 45, pseudogene [Sourc |
| AC095041.1 | 4 | ribosomal protein L19 (RPL19) pseudogene |
| RNA5SP173 | 4 | RNA, 5S ribosomal pseudogene 173 [Source:H |
| AC017087.1 | 4 | NULL |
| AC018710.1 | 4 | mitochondrially encoded cytochrome c oxidas |
| AC020551.1 | 4 | novel transcript |
| AC021193.1 | 4 | ankyrin repeat, SAM and basic leucine zipper ( |
| AC108169.1 | 4 | novel transcript |
| AC096747.1 | 4 | novel transcript |
| NDUFB5P1 | 4 | NADH:ubiquinone oxidoreductase subunit B5 |
| AC104803.1 | 4 | novel transcript |
| LINC00290 | 4 | long intergenic non-protein coding RNA 290 [S |
| AC019235.1 | 4 | mitochondrially encoded cytochrome c oxidas |
| LINC02500 | 4 | long intergenic non-protein coding RNA 2500 |
| AC093840.1 | 4 | epithelial cell adhesion molecule (EPCAM) pse |
| AC093840.2 | 4 | novel transcript |
| AC084741.1 | 4 | novel transcript |
| TEMN3-AS1 | 4 | TENM3 antisense transcript 1 [Source:HGNC S |
| RN7SKP13 | 4 | RN7SK pseudogene 13 [Source:HGNC Symbol; |
| TENM3-AS1 | 4 | TENM3 antisense RNA 1 [Source:HGNC Symbo |
| CCNHP1 | 4 | cyclin H pseudogene 1 [Source:HGNC Symbol; |
| TENM3 | 4 | teneurin transmembrane protein 3 [Source:H( |
| AC108142.1 | 4 | novel transcript |
| MIR1305 | 4 | microRNA 1305 [Source:HGNC Symbol;Acc:HG |

| | | |
|---|---|---|
| RNU2-34P | 4 | RNA, U2 small nuclear 34, pseudogene [Sourc |
| RN7SKP67 | 4 | RN7SK pseudogene 67 [Source:HGNC Symbol; |
| AC079226.2 | 4 | novel transcript |
| AC079226.1 | 4 | novel transcript |
| AC114798.1 | 4 | novel trasncript |
| AC079766.1 | 4 | novel transcript |
| DCTD | 4 | dCMP deaminase [Source:HGNC Symbol;Acc:H |
| AC019193.4 | 4 | novel transcript |
| AC019193.1 | 4 | survival of motor neuron 1, telomeric (SMN1) |
| FAM92A1P2 | 4 | family with sequence similarity 92 member A1 |
| AC019193.3 | 4 | novel transcript |
| AC019193.2 | 4 | novel transcript |
| WWC2-AS2 | 4 | WWC2 antisense RNA 2 [Source:HGNC Symbo |
| WWC2 | 4 | WW and C2 domain containing 2 [Source:HGN |
| WWC2-AS1 | 4 | WWC2 antisense RNA 1 [Source:HGNC Symbo |
| CLDN22 | 4 | claudin 22 [Source:HGNC Symbol;Acc:HGNC:2 |
| CLDN24 | 4 | claudin 24 [Source:HGNC Symbol;Acc:HGNC:3 |
| AC093844.1 | 4 | chimerin (chimaerin) 1 (CHN1) pseudogene |
| AC112698.1 | 4 | fibrillarin (FBL) pseudogene |
| CDKN2AIP | 4 | CDKN2A interacting protein [Source:HGNC Syr |
| VTI1BP2 | 4 | vesicle transport through interaction with t-SN |
| AC107214.1 | 4 | novel transcript |
| ING2 | 4 | inhibitor of growth family member 2 [Source:I |
| AC107214.2 | 4 | novel transcript |
| RWDD4 | 4 | RWD domain containing 4 [Source:HGNC Sym |
| RNU6-479P | 4 | RNA, U6 small nuclear 479, pseudogene [Sour |
| TRAPPC11 | 4 | trafficking protein particle complex 11 [Source |
| RNU6-335P | 4 | RNA, U6 small nuclear 335, pseudogene [Sour |
| RNU6-1053P | 4 | RNA, U6 small nuclear 1053, pseudogene [Sou |
| AC108477.2 | 4 | novel transcript |
| AC108477.1 | 4 | novel transcript |
| STOX2 | 4 | storkhead box 2 [Source:HGNC Symbol;Acc:H( |
| AC074194.1 | 4 | novel transcript |
| AC074194.2 | 4 | TEC |
| AC107222.1 | 4 | low density lipoprotein receptor-related prote |
| ENPP6 | 4 | ectonucleotide pyrophosphatase/phosphodie |
| AC107222.2 | 4 | novel transcript, antisense to ENPP6 |
| AC079080.1 | 4 | ribosomal protein L6 (RPL6) pseudogene |
| RN7SL28P | 4 | RNA, 7SL, cytoplasmic 28, pseudogene [Sourc |
| MYL12BP2 | 4 | myosin light chain 12B pseudogene 2 [Source: |
| LINC02363 | 4 | long intergenic non-protein coding RNA 2363 |
| LINC02362 | 4 | long intergenic non-protein coding RNA 2362 |
| IRF2 | 4 | interferon regulatory factor 2 [Source:HGNC S |
| SNORD79 | 4 | Small nucleolar RNA SNORD79 [Source:RFAM; |
| AC099343.2 | 4 | novel transcript, sense intronic to IRF2 |
| AC099343.3 | 4 | novel transcript |
| AC099343.4 | 4 | novel transcript |
| LINC02427 | 4 | long intergenic non-protein coding RNA 2427 |
| AC099343.1 | 4 | GAR1 ribonucleoprotein homolog (yeast) (GAI |
| LINC02365 | 4 | long intergenic non-protein coding RNA 2365 |

| | | |
|---|---|---|
| CASP3 | 4 | caspase 3 [Source:HGNC Symbol;Acc:HGNC:15 |
| PRIMPOL | 4 | primase and DNA directed polymerase [Sourc |
| CENPU | 4 | centromere protein U [Source:HGNC Symbol;/ |
| ACSL1 | 4 | acyl-CoA synthetase long chain family membe |
| SLED1 | 4 | proteoglycan 3, pro eosinophil major basic pr |
| AC084871.1 | 4 | novel transcript, antisense to ACSL1 |
| AC084871.3 | 4 | novel transcript, antisense to ACSL1 |
| MIR3945HG | 4 | MIR3945 host gene [Source:HGNC Symbol;Acc |
| MIR3945 | 4 | microRNA 3945 [Source:HGNC Symbol;Acc:HG |
| LINC01093 | 4 | long intergenic non-protein coding RNA 1093 |
| AC084871.2 | 4 | transient receptor potential cation channel, su |
| MIR4455 | 4 | microRNA 4455 [Source:HGNC Symbol;Acc:HG |
| AC112243.1 | 4 | TEC |
| LINC02437 | 4 | long intergenic non-protein coding RNA 2437 |
| HELT | 4 | helt bHLH transcription factor [Source:HGNC S |
| LINC02436 | 4 | long intergenic non-protein coding RNA 2436 |
| AC093824.1 | 4 | ribosomal protein L22-like 1 (RPL22L1) pseudo |
| AC093824.2 | 4 | novel transcript |
| SLC25A4 | 4 | solute carrier family 25 member 4 [Source:HG |
| CFAP97 | 4 | cilia and flagella associated protein 97 [Source |
| SNX25 | 4 | sorting nexin 25 [Source:HGNC Symbol;Acc:H( |
| LRP2BP | 4 | LRP2 binding protein [Source:HGNC Symbol;A |
| AC112722.1 | 4 | novel transcript |
| ANKRD37 | 4 | ankyrin repeat domain 37 [Source:HGNC Syml |
| UFSP2 | 4 | UFM1 specific peptidase 2 [Source:HGNC Sym |
| Y_RNA | 4 | Y RNA [Source:RFAM;Acc:RF00019] |
| C4orf47 | 4 | chromosome 4 open reading frame 47 [Sourc |
| CCDC110 | 4 | coiled-coil domain containing 110 [Source:HG |
| AC106897.1 | 4 | novel transcript, antisense to CCDC110 |
| PDLIM3 | 4 | PDZ and LIM domain 3 [Source:HGNC Symbol; |
| SORBS2 | 4 | sorbin and SH3 domain containing 2 [Source:H |
| AC093797.1 | 4 | novel transcript, antisense to SORBS2 |
| AC108472.1 | 4 | novel transcript |
| Y_RNA | 4 | Y RNA [Source:RFAM;Acc:RF00019] |
| AC104805.1 | 4 | general transcription factor IIH subunit 5 (GTF |
| AC096659.1 | 4 | novel transcript |
| RNU4-64P | 4 | RNA, U4 small nuclear 64, pseudogene [Sourc |
| TLR3 | 4 | toll like receptor 3 [Source:HGNC Symbol;Acc: |
| FAM149A | 4 | family with sequence similarity 149 member / |
| AC104070.1 | 4 | NULL |
| RPSAP70 | 4 | ribosomal protein SA pseudogene 70 [Source: |
| ORAOV1P1 | 4 | oral cancer overexpressed 1 pseudogene 1 [So |
| AC110771.1 | 4 | novel transcript |
| CYP4V2 | 4 | cytochrome P450 family 4 subfamily V memb |
| KLKB1 | 4 | kallikrein B1 [Source:HGNC Symbol;Acc:HGNC |
| F11 | 4 | coagulation factor XI [Source:HGNC Symbol;A |
| F11-AS1 | 4 | F11 antisense RNA 1 [Source:HGNC Symbol;Ac |
| AC109517.1 | 4 | novel transcript |
| SLC25A5P6 | 4 | solute carrier family 25 member 5 pseudogen |
| AC018709.1 | 4 | novel protein |

| | | |
|---|---|---|
| RNU6-1055P | 4 | RNA, U6 small nuclear 1055, pseudogene [Sou |
| MTNR1A | 4 | melatonin receptor 1A [Source:HGNC Symbol; |
| FAT1 | 4 | FAT atypical cadherin 1 [Source:HGNC Symbol |
| AC107050.1 | 4 | novel transcript, antisense to FAT1 |
| AC108865.1 | 4 | novel transcript |
| AC108865.2 | 4 | novel transcript |
| MRPS36P2 | 4 | mitochondrial ribosomal protein S36 pseudog |
| AC110772.1 | 4 | novel transcript |
| AC110772.2 | 4 | novel transcript |
| AC108046.1 | 4 | novel transcript |
| LINC02374 | 4 | long intergenic non-protein coding RNA 2374 |
| AC097652.1 | 4 | novel transcript |
| LINC02514 | 4 | long intergenic non-protein coding RNA 2514 |
| LINC02515 | 4 | long intergenic non-protein coding RNA 2515 |
| AC093763.2 | 4 | COP9 constitutive photomorphogenic homolo |
| AC093763.1 | 4 | novel transcript |
| LINC02492 | 4 | long intergenic non-protein coding RNA 2492 |
| AC097521.1 | 4 | novel transcript |
| ADAM20P3 | 4 | ADAM metallopeptidase domain 20 pseudoge |
| AC108073.3 | 4 | novel transcript |
| AC108073.1 | 4 | peroxiredoxin 6 (PRDX6) pseudogene |
| ZFP42 | 4 | ZFP42 zinc finger protein [Source:HGNC Symb |
| TRIML2 | 4 | tripartite motif family like 2 [Source:HGNC Syr |
| AC108073.2 | 4 | Finkel-Biskis-Reilly murine sarcoma virus (FBR- |
| RNU6-173P | 4 | RNA, U6 small nuclear 173, pseudogene [Sour |
| TRIML1 | 4 | tripartite motif family like 1 [Source:HGNC Syr |
| AC138781.1 | 4 | novel transcript |
| LINC02434 | 4 | long intergenic non-protein coding RNA 2434 |
| AC093789.1 | 4 | ribosomal protein L7a (RPL7A) pseudogene |
| ICE2P1 | 4 | interactor of little elongation complex ELL sub |
| LINC01060 | 4 | long intergenic non-protein coding RNA 1060 |
| AC093909.3 | 4 | pseudogene similar to part of cleavage and po |
| AC093909.5 | 4 | heat shock 70kDa protein 8 (HSPA8) pseudoge |
| RNU7-192P | 4 | RNA, U7 small nuclear 192 pseudogene [Sourc |
| AC093909.1 | 4 | mitochondrial translational release factor 1-lik |
| AC093909.6 | 4 | tec |
| AC093909.2 | 4 | novel transcript |
| LINC02508 | 4 | long intergenic non-protein coding RNA 2508 |
| AC093909.4 | 4 | novel transcript |
| AC122138.1 | 4 | novel transcript |
| AC107391.1 | 4 | novel transcript |
| AC105924.1 | 4 | novel transcript |
| HSP90AA4P | 4 | heat shock protein 90 alpha family class A me |
| LINC01262 | 4 | long intergenic non-protein coding RNA 1262 |
| AF250324.1 | 4 | novel transcript |
| RNU1-51P | 4 | RNA, U1 small nuclear 51, pseudogene [Sourc |
| FRG1-DT | 4 | FRG1 divergent transcript [Source:HGNC Symb |
| LINC01596 | 4 | long intergenic non-protein coding RNA 1596 |
| FRG1 | 4 | FSHD region gene 1 [Source:HGNC Symbol;Ac |
| MLLT10P2 | 4 | MLLT10 pseudogene 2 [Source:HGNC Symbol; |

| | | |
|---|---|---|
| AF146191.1 | 4 | family with sequence similarity 166, member |
| TUBB7P | 4 | tubulin beta 7 pseudogene [Source:HGNC Sym |
| RNA5SP174 | 4 | RNA, 5S ribosomal pseudogene 174 [Source:H |
| RNA5SP175 | 4 | RNA, 5S ribosomal pseudogene 175 [Source:H |
| DUX4L9 | 4 | double homeobox 4 like 9 (pseudogene) [Sour |
| FRG2 | 4 | FSHD region gene 2 [Source:HGNC Symbol;Ac |
| RARRES2P4 | 4 | retinoic acid receptor responder 2 pseudogen |
| AGGF1P1 | 4 | angiogenic factor with G-patch and FHA doma |
| CLUHP4 | 4 | clustered mitochondria homolog pseudogene |
| DBET | 4 | D4Z4 binding element transcript [Source:HGN |
| U85056.1 | 4 | double homeobox 4 like 30 |
| DUX4L8 | 4 | double homeobox 4 like 8 (pseudogene) [Sour |
| DUX4L7 | 4 | double homeobox 4 like 7 (pseudogene) [Sour |
| DUX4L6 | 4 | double homeobox 4 like 6 (pseudogene) [Sour |
| DUX4L5 | 4 | double homeobox 4 like 5 (pseudogene) [Sour |
| DUX4L4 | 4 | double homeobox 4 like 4 (pseudogene) [Sour |
| DUX4L1 | 4 | double homeobox 4 like 1 (pseudogene) [Sour |
| DUX4L3 | 4 | double homeobox 4 like 3 (pseudogene) [Sour |
| DUX4L2 | 4 | double homeobox 4 like 2 (pseudogene) [Sour |
| DUX4 | 4 | double homeobox 4 [Source:HGNC Symbol;Ac |
| RPL23AP84 | 4 | ribosomal protein L23a pseudogene 84 [Sourc |
| AC215524.1 | 4 | major histocompatibility complex, class I, F (H |
| AC113430.1 | 5 | Spi-C transcription factor (Spi-1/PU.1 related) |
| PLEKHG4B | 5 | pleckstrin homology and RhoGEF domain cont |
| Y_RNA | 5 | Y RNA [Source:RFAM;Acc:RF00019] |
| LRRC14B | 5 | leucine rich repeat containing 14B [Source:HG |
| AC021087.4 | 5 | novel transcript, overlapping LRRC14B |
| CCDC127 | 5 | coiled-coil domain containing 127 [Source:HG |
| AC021087.3 | 5 | novel transcript |
| SDHA | 5 | succinate dehydrogenase complex flavoprotei |
| AC021087.5 | 5 | novel protein |
| AC021087.1 | 5 | novel transcript |
| PDCD6 | 5 | programmed cell death 6 [Source:HGNC Symb |
| AHRR | 5 | aryl-hydrocarbon receptor repressor [Source: |
| AC021087.2 | 5 | protein phosphatase 4, regulatory subunit 2 ( |
| AHRR | 5 | aryl-hydrocarbon receptor repressor [Source: |
| AC010442.2 | 5 | mitochondrial translational initiation factor 3 |
| AC010442.3 | 5 | family with sequence similarity 136, member |
| EXOC3-AS1 | 5 | EXOC3 antisense RNA 1 [Source:HGNC Symbo |
| EXOC3 | 5 | exocyst complex component 3 [Source:HGNC |
| AC010442.1 | 5 | novel transcript, antisense to EXOC3 |
| SLC9A3 | 5 | solute carrier family 9 member A3 [Source:HG |
| SLC9A3-AS1 | 5 | SLC9A3 antisense RNA 1 [Source:HGNC Symbo |
| AC106772.2 | 5 | novel transcript, antisense to SLC9A3 |
| MIR4456 | 5 | microRNA 4456 [Source:HGNC Symbol;Acc:HG |
| AC106772.1 | 5 | novel transcript |
| CEP72 | 5 | centrosomal protein 72 [Source:HGNC Symbo |
| TPPP | 5 | tubulin polymerization promoting protein [So |
| AC026740.1 | 5 | novel transcript, antisense to TPPP |
| ZDHHC11B | 5 | zinc finger DHHC-type containing 11B [Source |

| | | |
|---|---|---|
| AC026740.3 | 5 | zinc finger DHHC-type containing 11B [Source |
| BRD9P2 | 5 | bromodomain containing 9 pseudogene 2 [So |
| AC026740.2 | 5 | tubulin polymerization promoting protein (TP |
| ZDHHC11 | 5 | zinc finger DHHC-type containing 11 [Source:H |
| SPCS2P3 | 5 | signal peptidase complex subunit 2 pseudoger |
| BRD9 | 5 | bromodomain containing 9 [Source:HGNC Syr |
| TRIP13 | 5 | thyroid hormone receptor interactor 13 [Sour |
| AC122719.3 | 5 | novel transcript, antisense to TRIP13 |
| AC122719.1 | 5 | novel transcript |
| AC122719.2 | 5 | novel transcript |
| AC116351.1 | 5 | novel transcript |
| AC116351.2 | 5 | novel transcript |
| NKD2 | 5 | NKD inhibitor of WNT signaling pathway 2 [So |
| SLC12A7 | 5 | solute carrier family 12 member 7 [Source:HG |
| MIR4635 | 5 | microRNA 4635 [Source:HGNC Symbol;Acc:HG |
| AC114291.1 | 5 | novel transcript |
| SLC6A19 | 5 | solute carrier family 6 member 19 [Source:HG |
| SLC6A18 | 5 | solute carrier family 6 member 18 [Source:HG |
| AC114291.2 | 5 | novel transcript |
| TERT | 5 | telomerase reverse transcriptase [Source:HGN |
| MIR4457 | 5 | microRNA 4457 [Source:HGNC Symbol;Acc:HG |
| CLPTM1L | 5 | CLPTM1 like [Source:HGNC Symbol;Acc:HGNC |
| AC026748.3 | 5 | novel transcript |
| LINC01511 | 5 | long intergenic non-protein coding RNA 1511 |
| MTCO2P32 | 5 | MT-CO2 pseudogene 32 [Source:HGNC Symbo |
| AC026748.1 | 5 | tec |
| SLC6A3 | 5 | solute carrier family 6 member 3 [Source:HGN |
| LPCAT1 | 5 | lysophosphatidylcholine acyltransferase 1 [So |
| MIR6075 | 5 | microRNA 6075 [Source:HGNC Symbol;Acc:HG |
| AC091849.2 | 5 | novel transcript |
| SDHAP3 | 5 | succinate dehydrogenase complex flavoprotei |
| AC026412.3 | 5 | novel transcript |
| AC091849.1 | 5 | coiled-coil domain containing 127 (CCDC127) |
| AC026412.1 | 5 | programmed cell death 6 (PDCD6) pseudogen |
| AC026412.2 | 5 | protein phosphatase 4, regulatory subunit 2 (I |
| MIR4277 | 5 | microRNA 4277 [Source:HGNC Symbol;Acc:HG |
| AC112176.1 | 5 | novel transcript |
| MRPL36 | 5 | mitochondrial ribosomal protein L36 [Source:I |
| NDUFS6 | 5 | NADH:ubiquinone oxidoreductase subunit S6 |
| AC025183.3 | 5 | novel transcript |
| LINC02116 | 5 | long intergenic non-protein coding RNA 2116 |
| AC025183.4 | 5 | tec |
| IRX4 | 5 | iroquois homeobox 4 [Source:HGNC Symbol;A |
| AC025183.1 | 5 | novel transcript, antisense to IRX4 |
| AC025183.2 | 5 | novel transcript, antisense to IRX4 |
| AC126768.3 | 5 | novel transcript |
| AC124852.1 | 5 | novel transcript |
| AC126768.1 | 5 | novel transcript |
| AC126768.2 | 5 | novel transcript |
| Y_RNA | 5 | Y RNA [Source:RFAM;Acc:RF00019] |

| | | |
|---|---|---|
| AC092266.1 | 5 | ornithine decarboxylase 1 (ODC1) pseudogen |
| LSINCT5 | 5 | long stress-induced non-coding transcript 5 [S |
| AC091891.1 | 5 | novel transcript |
| IRX2 | 5 | iroquois homeobox 2 [Source:HGNC Symbol;A |
| C5orf38 | 5 | chromosome 5 open reading frame 38 [Sourc |
| AC116359.1 | 5 | 3-hydroxyisobutyryl-Coenzyme A hydrolase (H |
| AC094105.1 | 5 | novel transcript |
| AC094105.2 | 5 | novel transcript |
| AC091832.1 | 5 | long intergenic non-protein coding RNA 1377 |
| LINC01019 | 5 | long intergenic non-protein coding RNA 1019 |
| LINC02162 | 5 | long intergenic non-protein coding RNA 2162 |
| LINC01017 | 5 | long intergenic non-protein coding RNA 1017 |
| IRX1 | 5 | iroquois homeobox 1 [Source:HGNC Symbol;A |
| AC016595.1 | 5 | novel transcript, antisense to IRX1 |
| AC025773.1 | 5 | novel transcript |
| AC025187.1 | 5 | novel transcript |
| LINC02063 | 5 | long intergenic non-protein coding RNA 2063 |
| AC106799.3 | 5 | novel transcript |
| AC106799.2 | 5 | novel transcript |
| AC106799.1 | 5 | novel transcript |
| AC010634.1 | 5 | novel transcript |
| LINC02114 | 5 | long intergenic non-protein coding RNA 2114 |
| AC026719.1 | 5 | novel transcript |
| AC026415.1 | 5 | novel transcript |
| AC010451.3 | 5 | novel transcript |
| AC010451.1 | 5 | novel transcript |
| LINC01020 | 5 | long intergenic non-protein coding RNA 1020 |
| AC010451.2 | 5 | novel transcript |
| LINC02121 | 5 | long intergenic non-protein coding RNA 2121 |
| RN7SKP73 | 5 | RN7SK pseudogene 73 [Source:HGNC Symbol; |
| AC022424.1 | 5 | novel transcript |
| ADAMTS16 | 5 | ADAM metallopeptidase with thrombospondi |
| AC022424.2 | 5 | novel transcript |
| ALG3P1 | 5 | ALG3 pseudogene 1 [Source:HGNC Symbol;Ac |
| AC091978.1 | 5 | novel transcript, antisense to ICE1 |
| MTCO1P30 | 5 | MT-CO1 pseudogene 30 [Source:HGNC Symbo |
| MTCO2P30 | 5 | MT-CO2 pseudogene 30 [Source:HGNC Symbo |
| MTCO1P31 | 5 | MT-CO1 pseudogene 31 [Source:HGNC Symbo |
| ICE1 | 5 | interactor of little elongation complex ELL sub |
| AC026736.1 | 5 | novel transcript |
| AC010266.2 | 5 | novel transcript |
| AC010266.1 | 5 | novel transcript |
| AC026797.1 | 5 | tec |
| HMGB3P3 | 5 | high mobility group box 3 pseudogene 3 [Sour |
| LINC02145 | 5 | long intergenic non-protein coding RNA 2145 |
| MED10 | 5 | mediator complex subunit 10 [Source:HGNC S |
| AC093307.1 | 5 | novel transcript |
| UBE2QL1 | 5 | ubiquitin conjugating enzyme E2 Q family like |
| AC027334.1 | 5 | histone cluster 3, H3 pseudogene (HIST3H3) p |
| LINC01018 | 5 | long intergenic non-protein coding RNA 1018 |

| | | |
|---|---|---|
| NSUN2 | 5 | NOP2/Sun RNA methyltransferase 2 [Source:H |
| SRD5A1 | 5 | steroid 5 alpha-reductase 1 [Source:HGNC Syr |
| LINC02102 | 5 | long intergenic non-protein coding RNA 2102 |
| TENT4A | 5 | terminal nucleotidyltransferase 4A [Source:HC |
| LINC02236 | 5 | long intergenic non-protein coding RNA 2236 |
| AC122710.2 | 5 | novel transcript |
| MIR4278 | 5 | microRNA 4278 [Source:HGNC Symbol;Acc:HG |
| LINC02196 | 5 | long intergenic non-protein coding RNA 2196 |
| RN7SKP79 | 5 | RN7SK pseudogene 79 [Source:HGNC Symbol; |
| AC122710.1 | 5 | novel transcript |
| RNA5SP176 | 5 | RNA, 5S ribosomal pseudogene 176 [Source:H |
| AC025756.1 | 5 | novel transcript |
| AC091889.1 | 5 | novel transcript |
| Metazoa_SRP | 5 | Metazoan signal recognition particle RNA [Sou |
| AC091951.4 | 5 | novel transcript |
| AC091951.2 | 5 | chromosome 16 open reading frame 61 (C16c |
| AC091951.3 | 5 | pseudogene similar to part of hect domain an |
| AC091951.1 | 5 | golgi autoantigen, golgin subfamily a, 6-like 2 |
| AC027343.3 | 5 | novel transcript |
| LINC02123 | 5 | long intergenic non-protein coding RNA 2123 |
| AC027343.1 | 5 | novel transcript |
| AC027343.2 | 5 | novel transcript |
| LINC02142 | 5 | long intergenic non-protein coding RNA 2142 |
| ADCY2 | 5 | adenylate cyclase 2 [Source:HGNC Symbol;Acc |
| AC093305.1 | 5 | novel transcript |
| C5orf49 | 5 | chromosome 5 open reading frame 49 [Source |
| MTRR | 5 | 5-methyltetrahydrofolate-homocysteine meth |
| FASTKD3 | 5 | FAST kinase domains 3 [Source:HGNC Symbol; |
| AC025174.1 | 5 | novel transcript |
| RNU1-76P | 5 | RNA, U1 small nuclear 76, pseudogene [Sourc |
| AC116361.1 | 5 | novel transcript |
| AC091941.1 | 5 | tec |
| AC091912.2 | 5 | tec |
| AC091912.1 | 5 | MYC binding protein 2 (MYCBP2) pseudogene |
| AC091912.3 | 5 | novel transcript |
| LINC02226 | 5 | long intergenic non-protein coding RNA 2226 |
| AC091965.1 | 5 | novel transcript |
| AC091965.3 | 5 | sema domain, seven thrombospondin repeats |
| AC091965.4 | 5 | novel transcript |
| MIR4458HG | 5 | MIR4458 host gene [Source:HGNC Symbol;Acc |
| AC091965.2 | 5 | FAST kinase domains 3 (FASTKD3) pseudogene |
| AC091965.5 | 5 | TEC |
| MIR4458 | 5 | microRNA 4458 [Source:HGNC Symbol;Acc:HG |
| AC091953.1 | 5 | novel transcript |
| AC091953.3 | 5 | tec |
| AC091953.5 | 5 | tec |
| MTND6P2 | 5 | MT-ND6 pseudogene 2 [Source:HGNC Symbol |
| MTCYBP37 | 5 | MT-CYB pseudogene 37 [Source:HGNC Symbo |
| AC091953.2 | 5 | tec |
| AC091953.6 | 5 | tec |

| | | |
|---|---|---|
| AC091953.4 | 5 | TEC |
| AC091932.4 | 5 | tec |
| AC091932.2 | 5 | tec |
| AC091932.1 | 5 | novel transcript |
| AC091932.3 | 5 | tec |
| LINC02199 | 5 | long intergenic non-protein coding RNA 2199 |
| AC091895.1 | 5 | tec |
| AC021088.1 | 5 | novel transcript |
| SEMA5A | 5 | semaphorin 5A [Source:HGNC Symbol;Acc:HG |
| MIR4636 | 5 | microRNA 4636 [Source:HGNC Symbol;Acc:HG |
| AC091906.1 | 5 | novel transcript |
| AC027335.2 | 5 | TEC |
| SEMA5A-AS1 | 5 | SEMA5A antisense RNA 1 [Source:HGNC Symb |
| AC027335.1 | 5 | novel transcript |
| SNHG18 | 5 | small nucleolar RNA host gene 18 [Source:HG |
| SNORD123 | 5 | small nucleolar RNA, C/D box 123 [Source:HG |
| AC026787.1 | 5 | novel transcript, overlapping TAS2R1 |
| TAS2R1 | 5 | taste 2 receptor member 1 [Source:HGNC Sym |
| LINC02112 | 5 | long intergenic non-protein coding RNA 2112 |
| RNA5SP177 | 5 | RNA, 5S ribosomal pseudogene 177 [Source:H |
| AC091838.1 | 5 | novel transcript |
| LINC02221 | 5 | long intergenic non-protein coding RNA 2221 |
| AC091905.2 | 5 | tec |
| AC091905.3 | 5 | tec |
| AC091905.1 | 5 | tec |
| AC091905.4 | 5 | tec |
| AC034229.1 | 5 | novel transcript |
| AC034229.6 | 5 | tec |
| AC034229.5 | 5 | tec |
| AC034229.2 | 5 | novel transcript |
| AC034229.3 | 5 | novel transcript |
| AC034229.4 | 5 | novel transcript |
| ATPSCKMT | 5 | ATP synthase c subunit lysine N-methyltransfe |
| AC012640.1 | 5 | novel transcript |
| CCT5 | 5 | chaperonin containing TCP1 subunit 5 [Source |
| AC012640.4 | 5 | novel transcript, antisense to CCT5 |
| AC012640.3 | 5 | novel transcript |
| CMBL | 5 | carboxymethylenebutenolidase homolog [Sou |
| Y_RNA | 5 | Y RNA [Source:RFAM;Acc:RF00019] |
| Y_RNA | 5 | Y RNA [Source:RFAM;Acc:RF00019] |
| AC012640.5 | 5 | tec |
| AC012640.2 | 5 | novel transcript, antisense to MARCH6 |
| MARCH6 | 5 | membrane associated ring-CH-type finger 6 [S |
| ROPN1L-AS1 | 5 | ROPN1L antisense RNA 1 [Source:HGNC Symb |
| ROPN1L | 5 | rhophilin associated tail protein 1 like [Source |
| MIR6131 | 5 | microRNA 6131 [Source:HGNC Symbol;Acc:HG |
| LINC01513 | 5 | long intergenic non-protein coding RNA 1513 |
| RPL30P7 | 5 | ribosomal protein L30 pseudogene 7 [Source: |
| LINC02212 | 5 | long intergenic non-protein coding RNA 2212 |
| AC092336.1 | 5 | novel transcript |

| | | |
|---|---|---|
| LINC02213 | 5 | long intergenic non-protein coding RNA 2213 |
| AC112200.2 | 5 | TEC |
| AC112200.1 | 5 | TEC |
| ANKRD33B | 5 | ankyrin repeat domain 33B [Source:HGNC Syn |
| ANKRD33B-AS1 | 5 | ANKRD33B antisense RNA 1 [Source:HGNC Syı |
| AC106760.1 | 5 | novel pseudogene |
| AC106760.2 | 5 | Intersectin-1 (ITSN1) pseudogene |
| AC012629.4 | 5 | tec |
| AC012629.3 | 5 | tec |
| DAP | 5 | death associated protein [Source:HGNC Symb |
| AC012629.2 | 5 | novel transcript, antisense to DAP |
| AC012629.1 | 5 | novel transcript |
| CTNND2 | 5 | catenin delta 2 [Source:HGNC Symbol;Acc:HG |
| RNU6-429P | 5 | RNA, U6 small nuclear 429, pseudogene [Sour |
| RNU6-679P | 5 | RNA, U6 small nuclear 679, pseudogene [Sour |
| AC106771.1 | 5 | novel transcript |
| LINC01194 | 5 | long intergenic non-protein coding RNA 1194 |
| AC106794.2 | 5 | chromosome X open reading frame 67 (CXorf( |
| AC106794.1 | 5 | chromosome X open reading frame 67 (CXorf( |
| LINC02220 | 5 | long intergenic non-protein coding RNA 2220 |
| AC016553.1 | 5 | novel transcript |
| RPS23P5 | 5 | ribosomal protein S23 pseudogene 5 [Source: |
| NENFP3 | 5 | neudesin neurotrophic factor pseudogene 3 [! |
| AC016546.1 | 5 | ribosomal protein L29 (RPL29) pseudogene |
| DNAH5 | 5 | dynein axonemal heavy chain 5 [Source:HGNC |
| AC016576.1 | 5 | novel transcript |
| TRIO | 5 | trio Rho guanine nucleotide exchange factor [ |
| AC016549.1 | 5 | novel transcript, antisense to TRIO |
| OTULINL | 5 | OTU deubiquitinase with linear linkage specifi |
| AC010627.1 | 5 | pleckstrin homology domain containing, famil |
| CCT6P2 | 5 | chaperonin containing TCP1 subunit 6 pseudo |
| EEF1A1P13 | 5 | eukaryotic translation elongation factor 1 alpł |
| AC010491.1 | 5 | novel transcript, antisense to FAM105B |
| OTULIN | 5 | OTU deubiquitinase with linear linkage specifi |
| ANKH | 5 | ANKH inorganic pyrophosphate transport regu |
| AC010491.2 | 5 | novel transcript, antisense to ANKH |
| RBBP4P1 | 5 | RBBP4 pseudogene 1 [Source:HGNC Symbol;A |
| MIR4637 | 5 | microRNA 4637 [Source:HGNC Symbol;Acc:HG |
| AC016575.1 | 5 | novel transcript |
| UQCRBP3 | 5 | ubiquinol-cytochrome c reductase binding prc |
| HNRNPKP5 | 5 | heterogeneous nuclear ribonucleoprotein K p: |
| SEPHS2P1 | 5 | selenophosphate synthetase 2 pseudogene 1 |
| AC016575.2 | 5 | novel transcript |
| U8 | 5 | U8 small nucleolar RNA [Source:RFAM;Acc:RF( |
| LINC02149 | 5 | long intergenic non-protein coding RNA 2149 |
| MARK2P5 | 5 | microtubule affinity regulating kinase 2 pseud |
| AC114964.2 | 5 | novel transcript |
| AC114964.1 | 5 | actin, gamma 1 (ACTG1) pseudogene |
| FBXL7 | 5 | F-box and leucine rich repeat protein 7 [Sourc |
| AC010638.1 | 5 | novel transcript |

| | | |
|---|---|---|
| MIR887 | 5 | microRNA 887 [Source:HGNC Symbol;Acc:HGI |
| RNA5SP178 | 5 | RNA, 5S ribosomal pseudogene 178 [Source:H |
| MARCH11 | 5 | membrane associated ring-CH-type finger 11 [ |
| AC016650.1 | 5 | novel transcript |
| AC092335.1 | 5 | novel transcript, antisense to MARCH11 |
| NACAP6 | 5 | NACA pseudogene 6 [Source:HGNC Symbol;Ac |
| LINC02150 | 5 | long intergenic non-protein coding RNA 2150 |
| AC020980.1 | 5 | novel transcript |
| ZNF622 | 5 | zinc finger protein 622 [Source:HGNC Symbol; |
| RETREG1 | 5 | reticulophagy regulator 1 [Source:HGNC Symb |
| AC024588.1 | 5 | novel transcript, antisense to FAM134B |
| AC022113.1 | 5 | novel transcript |
| Y_RNA | 5 | Y RNA [Source:RFAM;Acc:RF00019] |
| MYO10 | 5 | myosin X [Source:HGNC Symbol;Acc:HGNC:75 |
| RNA5SP179 | 5 | RNA, 5S ribosomal pseudogene 179 [Source:H |
| RPS26P28 | 5 | ribosomal protein S26 pseudogene 28 [Source |
| RNU6-660P | 5 | RNA, U6 small nuclear 660, pseudogene [Sour |
| BASP1 | 5 | brain abundant membrane attached signal pro |
| BASP1-AS1 | 5 | BASP1 antisense RNA 1 [Source:HGNC Symbol |
| AC026790.1 | 5 | novel transcript |
| Y_RNA | 5 | Y RNA [Source:RFAM;Acc:RF00019] |
| RNA5SP180 | 5 | RNA, 5S ribosomal pseudogene 180 [Source:H |
| DCAF13P2 | 5 | DDB1 and CUL4 associated factor 13 pseudog |
| RNU6-1003P | 5 | RNA, U6 small nuclear 1003, pseudogene [Sou |
| AC026785.1 | 5 | aurora kinase A interacting protein 1 (AURKAI |
| RN7SKP133 | 5 | RN7SK pseudogene 133 [Source:HGNC Symbo |
| FTH1P10 | 5 | ferritin heavy chain 1 pseudogene 10 [Source: |
| AC026785.2 | 5 | novel transcript |
| AC026785.3 | 5 | novel transcript |
| LINC02111 | 5 | long intergenic non-protein coding RNA 2111 |
| LINC02217 | 5 | long intergenic non-protein coding RNA 2217 |
| LINC02218 | 5 | long intergenic non-protein coding RNA 2218 |
| AC106774.4 | 5 | Putative histone H3.X [Source:UniProtKB/Swi |
| AC106774.3 | 5 | TAF11-like protein pseudogene |
| TAF11L2 | 5 | TATA-box binding protein associated factor 11 |
| AC106774.1 | 5 | H3 histone family 3 pseudogene |
| AC106774.2 | 5 | H3 histone family 3 pseudogene |
| TAF11L3 | 5 | TATA-box binding protein associated factor 11 |
| TAF11L4 | 5 | TATA-box binding protein associated factor 11 |
| TAF11L5 | 5 | TATA-box binding protein associated factor 11 |
| TAF11L6 | 5 | TATA-box binding protein associated factor 11 |
| TAF11L7 | 5 | TATA-box binding protein associated factor 11 |
| AC233724.7 | 5 | TAF11-like protein pseudogene |
| TAF11L8 | 5 | TATA-box binding protein associated factor 11 |
| TAF11L9 | 5 | TATA-box binding protein associated factor 11 |
| TAF11L10 | 5 | TATA-box binding protein associated factor 11 |
| AC233724.3 | 5 | TAF11-like protein pseudogene |
| AC233724.6 | 5 | TAF11-like protein pseudogene |
| TAF11L11 | 5 | TATA-box binding protein associated factor 11 |
| TAF11L12 | 5 | TATA-box binding protein associated factor 11 |

| | | |
|---|---|---|
| AC233724.11 | 5 | H3 histone family 3 pseudogene |
| AC233724.8 | 5 | H3 histone family 3 pseudogene |
| AC233724.9 | 5 | H3 histone family 3 pseudogene |
| TAF11L13 | 5 | TATA-box binding protein associated factor 11 |
| TAF11L14 | 5 | TATA-box binding protein associated factor 11 |
| AC233724.4 | 5 | TAF11-like protein pseudogene |
| AC233724.1 | 5 | TAF11-like protein pseudogene |
| AC233724.5 | 5 | TAF11-like protein pseudogene |
| AC233724.2 | 5 | TAF11-like protein pseudogene |
| H3.Y | 5 | histone cluster 2, H3c pseudogene [Source:NC |
| AC233724.12 | 5 | TAF11-like protein pseudogene |
| AC233724.10 | 5 | H3 histone family 3 pseudogene |
| LINC02223 | 5 | long intergenic non-protein coding RNA 2223 |
| RPL36AP21 | 5 | ribosomal protein L36a pseudogene 21 [Sourc |
| SNORD81 | 5 | Small nucleolar RNA SNORD81 [Source:RFAM; |
| RN7SL58P | 5 | RNA, 7SL, cytoplasmic 58, pseudogene [Sourc |
| LINC02100 | 5 | long intergenic non-protein coding RNA 2100 |
| UBE2V1P12 | 5 | ubiquitin conjugating enzyme E2 V1 pseudoge |
| AC025768.1 | 5 | novel pseudogene |
| AC114981.1 | 5 | novel pseudogene |
| Metazoa_SRP | 5 | Metazoan signal recognition particle RNA [Sou |
| AC106744.1 | 5 | novel transcript |
| AC106744.2 | 5 | novel transcript |
| RPL32P14 | 5 | ribosomal protein L32 pseudogene 14 [Source |
| HSPD1P15 | 5 | heat shock protein family D (Hsp60) member |
| CDH18 | 5 | cadherin 18 [Source:HGNC Symbol;Acc:HGNC: |
| AC093303.1 | 5 | novel pseudogene |
| AC094103.1 | 5 | nucleoporin 50kDa (NUP50) pseudogene |
| CDH18-AS1 | 5 | CDH18 antisense RNA 1 [Source:HGNC Symbo |
| LINC02146 | 5 | long intergenic non-protein coding RNA 2146 |
| LINC02241 | 5 | long intergenic non-protein coding RNA 2241 |
| AC138938.1 | 5 | UBX domain protein 2A (UBXN2A) pseudogen |
| AC140168.1 | 5 | novel transcript |
| AC140172.1 | 5 | stearoyl-CoA desaturase (delta-9-desaturase) |
| AC093274.1 | 5 | novel transcript |
| GUSBP1 | 5 | GUSB pseudogene 1 [Source:HGNC Symbol;Ac |
| AC138951.1 | 5 | POM121 membrane glycoprotein (POM121) p |
| AC138951.2 | 5 | TEC |
| AC091946.2 | 5 | novel transcript |
| AC091946.1 | 5 | novel transcript |
| CDH12 | 5 | cadherin 12 [Source:HGNC Symbol;Acc:HGNC: |
| HSPD1P1 | 5 | heat shock protein family D (Hsp60) member |
| AC139497.1 | 5 | novel transcript |
| PMCHL1 | 5 | pro-melanin concentrating hormone like 1 (ps |
| AC138854.1 | 5 | novel transcript, antisense to CDH12 |
| GCNT1P2 | 5 | glucosaminyl (N-acetyl) transferase 1, core 2 p |
| AC091938.1 | 5 | novel transcript, antisense to CDH12 |
| AC010445.1 | 5 | novel transcript |
| AC010460.2 | 5 | zinc finger CCCH-type containing 14 (ZC3H14) |
| Y_RNA | 5 | Y RNA [Source:RFAM;Acc:RF00019] |

| | | |
|---|---|---|
| AC010460.3 | 5 | protein tyrosine phosphatase, non-receptor ty |
| AC010460.1 | 5 | non-SMC condensin I complex, subunit G (NCA |
| PRDM9 | 5 | PR/SET domain 9 [Source:HGNC Symbol;Acc:H |
| AC109445.1 | 5 | cerebellar degeneration-related protein 2, 62l |
| AC108103.1 | 5 | transient receptor potential cation channel, su |
| C5orf17 | 5 | chromosome 5 open reading frame 17 (putati |
| AC026784.2 | 5 | NULL |
| AC026784.1 | 5 | alcohol dehydrogenase 5 (class III), chi polype |
| Y_RNA | 5 | Y RNA [Source:RFAM;Acc:RF00019] |
| AC114930.1 | 5 | novel transcript |
| AC010342.1 | 5 | AKT interacting protein (AKTIP) pseudogene |
| AC010387.1 | 5 | novel transcript |
| Metazoa_SRP | 5 | Metazoan signal recognition particle RNA [Sou |
| AC010387.2 | 5 | novel transcript |
| CDH10 | 5 | cadherin 10 [Source:HGNC Symbol;Acc:HGNC: |
| AC091885.2 | 5 | novel transcript |
| AC091885.1 | 5 | FERM domain containing 7 (FRMD7) pseudoge |
| AC091893.2 | 5 | novel transcript |
| AC091893.1 | 5 | transient receptor potential cation channel su |
| AC106821.2 | 5 | novel transcript |
| BTG4P1 | 5 | BTG anti-proliferation factor 4 pseudogene 1 [ |
| SNORD29 | 5 | Small nucleolar RNA SNORD29 [Source:RFAM; |
| LINC02239 | 5 | long intergenic non-protein coding RNA 2239 |
| AC106821.1 | 5 | novel transcript |
| Y_RNA | 5 | Y RNA [Source:RFAM;Acc:RF00019] |
| LINC02228 | 5 | long intergenic non-protein coding RNA 2228 |
| LINC02211 | 5 | long intergenic non-protein coding RNA 2211 |
| AC099499.2 | 5 | novel transcript |
| AC099499.1 | 5 | novel transcript |
| AC106754.1 | 5 | novel transcript |
| AC106754.2 | 5 | TEC |
| AC022140.1 | 5 | novel transcript |
| AC022140.2 | 5 | TEC |
| RNU6-374P | 5 | RNA, U6 small nuclear 374, pseudogene [Sour |
| AC022418.1 | 5 | TEC |
| MSNP1 | 5 | moesin pseudogene 1 [Source:HGNC Symbol;A |
| AC113370.1 | 5 | novel transcript |
| RNU4-43P | 5 | RNA, U4 small nuclear 43, pseudogene [Sourc |
| AC025475.1 | 5 | novel transcript |
| AC025475.2 | 5 | novel transcript |
| AC113347.1 | 5 | novel pseudogene |
| AC113347.2 | 5 | ankyrin repeat domain 11 (ANKRD11) pseudog |
| AC113347.3 | 5 | glycine cleavage system protein H (aminometh |
| AC113347.4 | 5 | novel transcript |
| CCNB3P1 | 5 | cyclin B3 pseudogene 1 [Source:HGNC Symbo |
| CDH9 | 5 | cadherin 9 [Source:HGNC Symbol;Acc:HGNC:1 |
| RNU6-738P | 5 | RNA, U6 small nuclear 738, pseudogene [Sour |
| PURPL | 5 | p53 upregulated regulator of p53 levels [Sour |
| AC113355.1 | 5 | novel transcript |
| AC093300.1 | 5 | T, brachyury homolog (mouse) (T) pseudogen |

| | | |
|---|---|---|
| LINC02103 | 5 | long intergenic non-protein coding RNA 2103 |
| AC008825.1 | 5 | novel transcript |
| AC109472.1 | 5 | novel transcript |
| RNU6-909P | 5 | RNA, U6 small nuclear 909, pseudogene [Sour |
| AC008825.2 | 5 | PQ loop repeat containing 1 (PQLC1) pseudog |
| LINC02109 | 5 | long intergenic non-protein coding RNA 2109 |
| AC010385.2 | 5 | developmental pluripotency associated 3 (DPI |
| LSP1P3 | 5 | LSP1 pseudogene 3 [Source:HGNC Symbol;Acc |
| AC010385.1 | 5 | striatin, calmodulin binding protein (STRN) ps |
| AC010385.3 | 5 | ankyrin repeat domain 36C (ANKRD36C) pseu |
| SUCLG2P4 | 5 | succinate-CoA ligase GDP-forming beta subun |
| AC024589.2 | 5 | NULL |
| AC024589.1 | 5 | aldehyde dehydrogenase 6 family, member A |
| AC024589.3 | 5 | coiled-coil domain containing 176 (CCDC176) |
| AC112172.2 | 5 | novel transcript |
| LINC02064 | 5 | long intergenic non-protein coding RNA 2064 |
| AC027345.1 | 5 | aurora kinase A interacting protein 1 (AURKAI |
| UBL5P1 | 5 | ubiquitin like 5 pseudogene 1 [Source:HGNC S |
| AC108082.1 | 5 | cytochrome c oxidase subunit VIa polypeptide |
| AC010374.2 | 5 | zinc finger, CCHC domain containing 10 (ZCCH |
| AC010374.1 | 5 | piggyBac transposable element derived 3 (PGI |
| RN7SKP207 | 5 | RN7SK pseudogene 207 [Source:HGNC Symbo |
| AC018765.1 | 5 | tec |
| HPRT1P2 | 5 | hypoxanthine phosphoribosyltransferase 1 ps |
| AC114300.1 | 5 | dynein, cytoplasmic 1, light intermediate chai |
| AC099517.1 | 5 | novel transcript |
| AC026433.1 | 5 | novel transcript |
| RPL19P11 | 5 | ribosomal protein L19 pseudogene 11 [Source |
| AC026421.1 | 5 | mitochondrially encoded cytochrome b (MT- |
| AC113386.1 | 5 | novel transcript |
| CDH6 | 5 | cadherin 6 [Source:HGNC Symbol;Acc:HGNC:1 |
| DUX4L51 | 5 | double homeobox 4 like 51 (pseudogene) [Sou |
| DROSHA | 5 | drosha ribonuclease III [Source:HGNC Symbol; |
| Y_RNA | 5 | Y RNA [Source:RFAM;Acc:RF00019] |
| C5orf22 | 5 | chromosome 5 open reading frame 22 [Sourc |
| AC022417.1 | 5 | tec |
| RNU6-363P | 5 | RNA, U6 small nuclear 363, pseudogene [Sour |
| PDZD2 | 5 | PDZ domain containing 2 [Source:HGNC Symb |
| AC022447.2 | 5 | novel transcript |
| AC022447.4 | 5 | FUS interacting protein (serine/arginine-rich) |
| AC022447.5 | 5 | novel transcript |
| AC022447.3 | 5 | novel transcript |
| AC022447.1 | 5 | novel transcript |
| AC022447.7 | 5 | novel transcript |
| AC022447.6 | 5 | novel transcript |
| RNU6-358P | 5 | RNA, U6 small nuclear 358, pseudogene [Sour |
| AC010469.1 | 5 | ribosomal protein L9 (RPL9) pseudogene |
| RNU6-760P | 5 | RNA, U6 small nuclear 760, pseudogene [Sour |
| AC010469.2 | 5 | ribosomal protein L21 (RPL21) pseudogene |
| RPL5P14 | 5 | ribosomal protein L5 pseudogene 14 [Source: |

| | | |
|---|---|---|
| TPT1P5 | 5 | tumor protein, translationally-controlled 1 ps |
| AC025458.1 | 5 | translocase of outer mitochondrial membrane |
| MIR4279 | 5 | microRNA 4279 [Source:HGNC Symbol;Acc:H |
| AC025178.1 | 5 | novel transcript |
| GOLPH3 | 5 | golgi phosphoprotein 3 [Source:HGNC Symbol |
| Y_RNA | 5 | Y RNA [Source:RFAM;Acc:RF00019] |
| AC025181.2 | 5 | novel transcript |
| AC025181.1 | 5 | ribosomal protein L27 (RPL27) pseudogene |
| MTMR12 | 5 | myotubularin related protein 12 [Source:HGN |
| RNU6-1079P | 5 | RNA, U6 small nuclear 1079, pseudogene [Sou |
| RNU6-378P | 5 | RNA, U6 small nuclear 378, pseudogene [Sour |
| ZFR | 5 | zinc finger RNA binding protein [Source:HGNC |
| MIR579 | 5 | microRNA 579 [Source:HGNC Symbol;Acc:HG |
| AC074134.1 | 5 | transmembrane protein 183A (TMEM183A) p |
| SUB1 | 5 | SUB1 regulator of transcription [Source:HGNC |
| LINC02061 | 5 | long intergenic non-protein coding RNA 2061 |
| AC008766.1 | 5 | novel transcript |
| NPR3 | 5 | natriuretic peptide receptor 3 [Source:HGNC S |
| AC025459.1 | 5 | ribosomal protein L17 (RPL17) pseudogene |
| AC010343.3 | 5 | novel transcript |
| LINC02120 | 5 | long intergenic non-protein coding RNA 2120 |
| AC034223.1 | 5 | novel transcript |
| AC034223.2 | 5 | novel transcript |
| AC010343.2 | 5 | tripartite motif-containing 28 (TRIM28) pseud |
| AC010343.1 | 5 | ribosomal protein S8 (RPS8) pseudogene |
| LINC02160 | 5 | long intergenic non-protein coding RNA 2160 |
| AC034231.1 | 5 | novel transcript |
| TARS | 5 | threonyl-tRNA synthetase [Source:HGNC Syml |
| AC034232.1 | 5 | translocase of outer mitochondrial membrane |
| AC034232.2 | 5 | novel transcript |
| ADAMTS12 | 5 | ADAM metallopeptidase with thrombospondi |
| RNU6-923P | 5 | RNA, U6 small nuclear 923, pseudogene [Sour |
| RXFP3 | 5 | relaxin family peptide receptor 3 [Source:HGN |
| SLC45A2 | 5 | solute carrier family 45 member 2 [Source:HG |
| AC139783.2 | 5 | proteasome (prosome, macropain) 26S subun |
| AMACR | 5 | alpha-methylacyl-CoA racemase [Source:HGN |
| C1QTNF3-AMACR | 5 | C1QTNF3-AMACR readthrough (NMD candida |
| AC139783.1 | 5 | fumarate hydratase (FH) pseudogene |
| C1QTNF3 | 5 | C1q and TNF related 3 [Source:HGNC Symbol; |
| AC139792.2 | 5 | TEC |
| AC139792.3 | 5 | TEC |
| AC139792.1 | 5 | novel transcript |
| AC138409.2 | 5 | Putative beta-glucuronidase-like protein FLJ7 |
| AC138409.1 | 5 | POM121-like protein 1 pseudogene |
| AC138409.3 | 5 | novel transcript |
| AC138853.1 | 5 | UBX domain protein 2A (UBXN2A) pseudogen |
| AC025754.2 | 5 | novel transcript |
| RAI14 | 5 | retinoic acid induced 14 [Source:HGNC Symbo |
| AC025754.1 | 5 | novel transcript |
| AC026801.2 | 5 | novel transcript, antisense to TTC23L |

| | | |
|---|---|---|
| TTC23L | 5 | tetratricopeptide repeat domain 23 like [Sour |
| AC026801.1 | 5 | ribosomal protein L26-like 1 (RPL26L1) pseud |
| RPL21P54 | 5 | ribosomal protein L21 pseudogene 54 [Source |
| RAD1 | 5 | RAD1 checkpoint DNA exonuclease [Source:H |
| BRIX1 | 5 | biogenesis of ribosomes BRX1 [Source:HGNC S |
| DNAJC21 | 5 | DnaJ heat shock protein family (Hsp40) memb |
| AGXT2 | 5 | alanine--glyoxylate aminotransferase 2 [Sourc |
| PRLR | 5 | prolactin receptor [Source:HGNC Symbol;Acc: |
| AC010368.1 | 5 | glyceraldehyde-3-phosphate dehydrogenase ( |
| U3 | 5 | Small nucleolar RNA U3 [Source:RFAM;Acc:RF |
| SPEF2 | 5 | sperm flagellar 2 [Source:HGNC Symbol;Acc:H |
| RNU7-130P | 5 | RNA, U7 small nuclear 130 pseudogene [Sourc |
| AC137810.1 | 5 | novel transcript, antisense to SPEF2 |
| IL7R | 5 | interleukin 7 receptor [Source:HGNC Symbol;A |
| AC112204.3 | 5 | TEC |
| AC112204.1 | 5 | calcyphosine-like (CAPSL) pseudogene |
| CAPSL | 5 | calcyphosine like [Source:HGNC Symbol;Acc:H |
| AC112204.2 | 5 | novel transcript |
| UGT3A1 | 5 | UDP glycosyltransferase family 3 member A1 |
| UGT3A2 | 5 | UDP glycosyltransferase family 3 member A2 |
| AC016612.1 | 5 | UDP glycosyltransferase 3 family, polypeptide |
| LMBRD2 | 5 | LMBR1 domain containing 2 [Source:HGNC Sy |
| MIR580 | 5 | microRNA 580 [Source:HGNC Symbol;Acc:HGN |
| SKP2 | 5 | S-phase kinase associated protein 2 [Source:H |
| RNU6-1305P | 5 | RNA, U6 small nuclear 1305, pseudogene [Sou |
| NADK2 | 5 | NAD kinase 2, mitochondrial [Source:HGNC Sy |
| NADK2-AS1 | 5 | NADK2 antisense RNA 1 [Source:HGNC Symbo |
| RANBP3L | 5 | RAN binding protein 3 like [Source:HGNC Sym |
| AC114277.1 | 5 | novel transcript |
| RNA5SP181 | 5 | RNA, 5S ribosomal pseudogene 181 [Source:H |
| AC010631.1 | 5 | ribosomal protein L39 (RPL39) pseudogene |
| SLC1A3 | 5 | solute carrier family 1 member 3 [Source:HGN |
| AC008957.2 | 5 | novel transcript, antisense SLC1A3 |
| AC008957.1 | 5 | novel transcript |
| AC008957.3 | 5 | TEC |
| AC026741.1 | 5 | novel transcript |
| NIPBL-DT | 5 | NIPBL divergent transcript [Source:HGNC Sym |
| NIPBL | 5 | NIPBL cohesin loading factor [Source:HGNC Sy |
| KRT18P31 | 5 | keratin 18 pseudogene 31 [Source:HGNC Sym |
| RPS4XP6 | 5 | ribosomal protein S4X pseudogene 6 [Source: |
| CPLANE1 | 5 | ciliogenesis and planar polarity effector 1 [Sou |
| RBISP2 | 5 | RBIS pseudogene 2 [Source:HGNC Symbol;Acc |
| OFD1P17 | 5 | OFD1 pseudogene 17 [Source:HGNC Symbol;A |
| RN7SL37P | 5 | RNA, 7SL, cytoplasmic 37, pseudogene [Sourc |
| AC025449.2 | 5 | novel transcript, antisense to CPLANE1 |
| AC025449.1 | 5 | chromobox homolog 3 (CBX3) pseudogene |
| NUP155 | 5 | nucleoporin 155 [Source:HGNC Symbol;Acc:H |
| RNU7-75P | 5 | RNA, U7 small nuclear 75 pseudogene [Source |
| AC117532.1 | 5 | novel pseudogene |
| WDR70 | 5 | WD repeat domain 70 [Source:HGNC Symbol; |

| | | |
|---|---|---|
| KCTD9P5 | 5 | potassium channel tetramerization domain cc |
| RNU6-1190P | 5 | RNA, U6 small nuclear 1190, pseudogene [Sou |
| RNU6-484P | 5 | RNA, U6 small nuclear 484, pseudogene [Sour |
| GDNF-AS1 | 5 | GDNF antisense RNA 1 [Source:HGNC Symbol; |
| GDNF | 5 | glial cell derived neurotrophic factor [Source:I |
| LINC02117 | 5 | long intergenic non-protein coding RNA 2117 |
| AC008869.1 | 5 | novel transcript |
| LINC02110 | 5 | long intergenic non-protein coding RNA 2110 |
| AC034226.1 | 5 | novel transcript |
| LINC02119 | 5 | long intergenic non-protein coding RNA 2119 |
| AC010338.1 | 5 | peroxiredoxin 4 (PRDX4) pseudogene |
| EGFLAM | 5 | EGF like, fibronectin type III and laminin G dor |
| EGFLAM-AS4 | 5 | EGFLAM antisense RNA 4 [Source:HGNC Symb |
| EGFLAM-AS3 | 5 | EGFLAM antisense RNA 3 [Source:HGNC Symb |
| EGFLAM-AS2 | 5 | EGFLAM antisense RNA 2 [Source:HGNC Symb |
| EGFLAM-AS1 | 5 | EGFLAM antisense RNA 1 [Source:HGNC Symb |
| AC091839.1 | 5 | ribosomal protein L27 (RPL27) pseudogene |
| AC010457.1 | 5 | novel transcript |
| LIFR | 5 | LIF receptor subunit alpha [Source:HGNC Sym |
| LIFR-AS1 | 5 | LIFR antisense RNA 1 [Source:HGNC Symbol;A |
| MIR3650 | 5 | microRNA 3650 [Source:HGNC Symbol;Acc:HG |
| AC010425.1 | 5 | novel transcript |
| OSMR-AS1 | 5 | OSMR antisense RNA 1 (head to head) [Source |
| AC091435.1 | 5 | transmembrane protein 126A (TMEM126A) ps |
| AC091435.2 | 5 | novel transcript |
| OSMR | 5 | oncostatin M receptor [Source:HGNC Symbol; |
| RICTOR | 5 | RPTOR independent companion of MTOR com |
| AIG1P1 | 5 | androgen induced 1 pseudogene 1 [Source:HG |
| AC008964.1 | 5 | novel transcript |
| FYB1 | 5 | FYN binding protein 1 [Source:HGNC Symbol;A |
| GOLGA5P1 | 5 | golgin A5 pseudogene 1 [Source:HGNC Symbo |
| C9 | 5 | complement C9 [Source:HGNC Symbol;Acc:HG |
| DAB2 | 5 | DAB adaptor protein 2 [Source:HGNC Symbol; |
| LINC02104 | 5 | long intergenic non-protein coding RNA 2104 |
| CCDC11P1 | 5 | coiled-coil domain containing 11 pseudogene |
| INTS6P1 | 5 | integrator complex subunit 6 pseudogene 1 [S |
| GCSHP1 | 5 | glycine cleavage system protein H pseudogene |
| LINC00603 | 5 | long intergenic non-protein coding RNA 603 [S |
| KRT18P56 | 5 | keratin 18 pseudogene 56 [Source:HGNC Sym |
| LINC00604 | 5 | long intergenic non-protein coding RNA 604 [S |
| RNU1-150P | 5 | RNA, U1 small nuclear 150, pseudogene [Sour |
| AC108105.1 | 5 | novel transcript |
| AC093277.1 | 5 | novel transcript |
| AC114977.1 | 5 | novel transcript |
| AC114977.2 | 5 | novel transcript |
| TTC33 | 5 | tetratricopeptide repeat domain 33 [Source:H |
| SNORA63 | 5 | Small nucleolar RNA SNORA63 [Source:RFAM; |
| PTGER4 | 5 | prostaglandin E receptor 4 [Source:HGNC Sym |
| PRKAA1 | 5 | protein kinase AMP-activated catalytic subuni |
| AC008810.1 | 5 | NULL |

| | | |
|---|---|---|
| RPL37 | 5 | ribosomal protein L37 [Source:HGNC Symbol;/ |
| SNORD72 | 5 | small nucleolar RNA, C/D box 72 [Source:HGN |
| CARD6 | 5 | caspase recruitment domain family member 6 |
| C7 | 5 | complement C7 [Source:HGNC Symbol;Acc:HC |
| RNU7-161P | 5 | RNA, U7 small nuclear 161 pseudogene [Sourc |
| AC114967.1 | 5 | ribosomal protein L5 pseudogene |
| MROH2B | 5 | maestro heat like repeat family member 2B [S |
| C6 | 5 | complement C6 [Source:HGNC Symbol;Acc:HC |
| PLCXD3 | 5 | phosphatidylinositol specific phospholipase C |
| AC008817.1 | 5 | t-complex 1 (TCP1) pseudogene |
| OXCT1 | 5 | 3-oxoacid CoA-transferase 1 [Source:HGNC Sy |
| AC034222.2 | 5 | novel transcript, antisense to OXCT1 |
| OXCT1-AS1 | 5 | OXCT1 antisense RNA 1 [Source:HGNC Symbo |
| AC034222.1 | 5 | ribosomal protein S2 (RPS2) pseudogene |
| C5orf51 | 5 | chromosome 5 open reading frame 51 [Sourc |
| FBXO4 | 5 | F-box protein 4 [Source:HGNC Symbol;Acc:HG |
| RPSAP38 | 5 | ribosomal protein SA pseudogene 38 [Source: |
| Y_RNA | 5 | Y RNA [Source:RFAM;Acc:RF00019] |
| MTHFD2P6 | 5 | methylenetetrahydrofolate dehydrogenase (N |
| AC108099.1 | 5 | novel transcript |
| GHR | 5 | growth hormone receptor [Source:HGNC Sym |
| SERBP1P6 | 5 | SERPINE1 mRNA binding protein 1 pseudogen |
| AC093225.1 | 5 | novel zinc finger protein pseudogene |
| AC113368.1 | 5 | NADH dehydrogenase (ubiquinone) 1 beta sub |
| CCDC152 | 5 | coiled-coil domain containing 152 [Source:HG |
| SELENOP | 5 | selenoprotein P [Source:HGNC Symbol;Acc:HC |
| AC008945.1 | 5 | novel transcript, antisense to SEPP1 |
| AC008945.2 | 5 | novel transcript, antisense to SEPP1 |
| PPIAP77 | 5 | peptidylprolyl isomerase A pseudogene 77 [Sc |
| PRELID3BP4 | 5 | PRELI domain containing 3B pseudogene 4 [Sc |
| PRELID3BP5 | 5 | PRELI domain containing 3B pseudogene 5 [Sc |
| PRELID3BP6 | 5 | PRELI domain containing 3B pseudogene 6 [Sc |
| PRELID3BP7 | 5 | PRELI domain containing 3B pseudogene 7 [Sc |
| AC008875.3 | 5 | novel transcript |
| AC008875.2 | 5 | novel transcript |
| AC008875.1 | 5 | novel transcript |
| AC025171.1 | 5 | novel transcript, sense overlapping to ANXA2I |
| AC025171.3 | 5 | novel transcript |
| AC025171.5 | 5 | novel transcript |
| ANXA2R | 5 | annexin A2 receptor [Source:HGNC Symbol;Ac |
| AC025171.2 | 5 | novel transcript, antisense to ANXA2R |
| AC025171.4 | 5 | novel transcript |
| ZNF131 | 5 | zinc finger protein 131 [Source:HGNC Symbol; |
| AC106800.4 | 5 | eukaryotic translation elongation factor 1 beta |
| NIM1K | 5 | NIM1 serine/threonine protein kinase [Source |
| AC106800.2 | 5 | spermine synthase (SMS) pseudogene |
| AC106800.3 | 5 | NADH dehydrogenase (ubiquinone) 1 alpha su |
| AC106800.1 | 5 | tubulin, alpha 1b (TUBA1B) pseudogene |
| HMGCS1 | 5 | 3-hydroxy-3-methylglutaryl-CoA synthase 1 [S |
| AC114947.2 | 5 | novel transcript, antisense to HMGCS1 |

| | | |
|---|---|---|
| AC114947.1 | 5 | ribosomal protein, large, P1 (RPLP1) pseudoge |
| CCL28 | 5 | C-C motif chemokine ligand 28 [Source:HGNC |
| TMEM267 | 5 | transmembrane protein 267 [Source:HGNC Sy |
| AC114956.3 | 5 | novel transcript |
| C5orf34 | 5 | chromosome 5 open reading frame 34 [Sourc |
| EEF1A1P19 | 5 | eukaryotic translation elongation factor 1 alpl |
| AC114956.2 | 5 | novel transcript |
| AC114956.1 | 5 | novel transcript |
| PAIP1 | 5 | poly(A) binding protein interacting protein 1 [ |
| NNT-AS1 | 5 | NNT antisense RNA 1 [Source:HGNC Symbol;A |
| AC010435.1 | 5 | TEC |
| AMD1P3 | 5 | adenosylmethionine decarboxylase 1 pseudog |
| NNT | 5 | nicotinamide nucleotide transhydrogenase [S |
| RPL29P12 | 5 | ribosomal protein L29 pseudogene 12 [Source |
| AC104119.1 | 5 | novel transcript |
| RNU6-381P | 5 | RNA, U6 small nuclear 381, pseudogene [Sour |
| FGF10 | 5 | fibroblast growth factor 10 [Source:HGNC Syn |
| FGF10-AS1 | 5 | FGF10 antisense RNA 1 [Source:HGNC Symbol |
| LINC02224 | 5 | long intergenic non-protein coding RNA 2224 |
| AC093292.1 | 5 | novel transcript |
| RN7SL383P | 5 | RNA, 7SL, cytoplasmic 383, pseudogene [Sour |
| MRPS30-DT | 5 | MRPS30 divergent transcript [Source:HGNC Sy |
| AC093297.1 | 5 | novel transcript |
| MRPS30 | 5 | mitochondrial ribosomal protein S30 [Source: |
| AC093297.2 | 5 | novel transcript |
| AC114954.1 | 5 | novel transcript |
| AC116350.1 | 5 | novel transcript |
| HCN1 | 5 | hyperpolarization activated cyclic nucleotide g |
| AC117529.2 | 5 | tigger transposable element derived 1 (TIGD1 |
| AC117529.1 | 5 | tigger transposable element derived 1 (TIGD1 |
| AC122694.1 | 5 | novel transcript |
| EMB | 5 | embigin [Source:HGNC Symbol;Acc:HGNC:304 |
| AC112187.2 | 5 | HIG1 hypoxia inducible domain family, memb |
| AC112187.1 | 5 | PPIAL4A peptidylprolyl isomerase A (cyclophil |
| AC112187.3 | 5 | novel transcript |
| PARP8 | 5 | poly(ADP-ribose) polymerase family member |
| AC022441.1 | 5 | novel transcript |
| AC022441.3 | 5 | ribosomal protein, large, P1 (RPLP1) pseudoge |
| AC022441.2 | 5 | novel transcript |
| AC008808.2 | 5 | novel transcript |
| LINC02106 | 5 | long intergenic non-protein coding RNA 2106 |
| AC008808.1 | 5 | novel transcript |
| RNU6-480P | 5 | RNA, U6 small nuclear 480, pseudogene [Sour |
| AC091834.1 | 5 | transmembrane protein 69 (TMEM69) pseudo |
| RNU6-1296P | 5 | RNA, U6 small nuclear 1296, pseudogene [Sou |
| AC026798.1 | 5 | ribosomal protein S10 (RPS10) pseudogene |
| AC010478.1 | 5 | novel transcript, antisense to ISL1 |
| ISL1 | 5 | ISL LIM homeobox 1 [Source:HGNC Symbol;Ac |
| HMGB1P47 | 5 | high mobility group box 1 pseudogene 47 [Sou |
| AC091860.1 | 5 | mago-nashi homolog B (Drosophila) (MAGOH |

| | | |
|---|---|---|
| AC022433.1 | 5 | novel transcript |
| AC091860.2 | 5 | novel transcript |
| RNA5SP182 | 5 | RNA, 5S ribosomal pseudogene 182 [Source:H |
| KATNBL1P4 | 5 | katanin regulatory subunit B1 like 1 pseudoge |
| AC091903.1 | 5 | TEC |
| LINC02118 | 5 | long intergenic non-protein coding RNA 2118 |
| RPS17P11 | 5 | ribosomal protein S17 pseudogene 11 [Source |
| MFSD4BP1 | 5 | major facilitator superfamily domain containi |
| AC022126.1 | 5 | novel transcript |
| ITGA1 | 5 | integrin subunit alpha 1 [Source:HGNC Symbo |
| PELO | 5 | pelota mRNA surveillance and ribosome rescu |
| B3GNTL1P1 | 5 | B3GNTL1 pseudogene 1 [Source:HGNC Symbo |
| AC022133.1 | 5 | ribosomal protein L17 (RPL17) pseudogene |
| AC025180.1 | 5 | novel transcript |
| ITGA2 | 5 | integrin subunit alpha 2 [Source:HGNC Symbo |
| AC008966.3 | 5 | novel transcript, antisense to ITGA2 |
| AC008966.2 | 5 | novel transcript, antisense to ITGA2 |
| MOCS2 | 5 | molybdenum cofactor synthesis 2 [Source:HG |
| AC008966.1 | 5 | novel transcript, antisense to MOCS2 |
| AC026477.1 | 5 | ribosomal protein L13a (RPL13A) pseudogene |
| AC027329.1 | 5 | novel transcript |
| AC108114.1 | 5 | ribosomal protein S19 (RPS19) pseudogene |
| FST | 5 | follistatin [Source:HGNC Symbol;Acc:HGNC:3 |
| NDUFS4 | 5 | NADH:ubiquinone oxidoreductase subunit S4 |
| LINC02105 | 5 | long intergenic non-protein coding RNA 2105 |
| ASS1P9 | 5 | argininosuccinate synthetase 1 pseudogene 9 |
| AC025175.1 | 5 | novel transcript |
| ARL15 | 5 | ADP ribosylation factor like GTPase 15 [Source |
| AC025175.2 | 5 | novel transcript, antisense to ARL15 |
| RNU6-272P | 5 | RNA, U6 small nuclear 272, pseudogene [Sour |
| MIR581 | 5 | microRNA 581 [Source:HGNC Symbol;Acc:HGI |
| RN7SL801P | 5 | RNA, 7SL, cytoplasmic 801, pseudogene [Sour |
| AC016601.1 | 5 | NULL |
| AC008873.1 | 5 | TEC |
| LINC01033 | 5 | long intergenic non-protein coding RNA 1033 |
| RPL37P25 | 5 | ribosomal protein L37 pseudogene 25 [Source |
| HSPB3 | 5 | heat shock protein family B (small) member 3 |
| SNX18 | 5 | sorting nexin 18 [Source:HGNC Symbol;Acc:H |
| AC016583.1 | 5 | tec |
| AC016583.2 | 5 | novel transcript |
| AC112198.2 | 5 | novel pseudogene |
| AC112198.1 | 5 | novel zinc finger protein pseudogene |
| AC112198.4 | 5 | thiosulfate sulfurtransferase (rhodanese)-like |
| AC112198.3 | 5 | novel transcript |
| ESM1 | 5 | endothelial cell specific molecule 1 [Source:H |
| AC034238.1 | 5 | novel transcript, antisense to GZMK |
| GZMK | 5 | granzyme K [Source:HGNC Symbol;Acc:HGNC: |
| Y_RNA | 5 | Y RNA [Source:RFAM;Acc:RF00019] |
| AC034238.3 | 5 | novel transcript |
| AC034238.2 | 5 | novel transcript |

| | | |
|---|---|---|
| GZMAP1 | 5 | granzyme A pseudogene 1 [Source:HGNC Sym |
| GZMA | 5 | granzyme A [Source:HGNC Symbol;Acc:HGNC: |
| CDC20B | 5 | cell division cycle 20B [Source:HGNC Symbol;A |
| GPX8 | 5 | glutathione peroxidase 8 (putative) [Source:H |
| MIR449A | 5 | microRNA 449a [Source:HGNC Symbol;Acc:HG |
| MIR449B | 5 | microRNA 449b [Source:HGNC Symbol;Acc:HG |
| MIR449C | 5 | microRNA 449c [Source:HGNC Symbol;Acc:HG |
| MCIDAS | 5 | multiciliate differentiation and DNA synthesis |
| CCNO | 5 | cyclin O [Source:HGNC Symbol;Acc:HGNC:185 |
| AC026704.1 | 5 | novel transcript |
| DHX29 | 5 | DExH-box helicase 29 [Source:HGNC Symbol;A |
| MTREX | 5 | Mtr4 exosome RNA helicase [Source:HGNC Sy |
| AC020728.1 | 5 | protein serine kinase H1 (PSKH1) pseudogene |
| PLPP1 | 5 | phospholipid phosphatase 1 [Source:HGNC Sy |
| AC010480.1 | 5 | TEC |
| MIR5687 | 5 | microRNA 5687 [Source:HGNC Symbol;Acc:HG |
| RNF138P1 | 5 | ring finger protein 138 pseudogene 1 [Source: |
| AK4P2 | 5 | adenylate kinase 4 pseudogene 2 [Source:HGI |
| SLC38A9 | 5 | solute carrier family 38 member 9 [Source:HG |
| DDX4 | 5 | DEAD-box helicase 4 [Source:HGNC Symbol;A |
| AC016632.1 | 5 | S-phase kinase-associated protein 1 (SKP1) ps |
| RNA5SP183 | 5 | RNA, 5S ribosomal pseudogene 183 [Source:H |
| HNRNPH1P3 | 5 | heterogeneous nuclear ribonucleoprotein H1 |
| IL31RA | 5 | interleukin 31 receptor A [Source:HGNC Symb |
| IL6ST | 5 | interleukin 6 signal transducer [Source:HGNC |
| AC008914.1 | 5 | novel transcript, antisense to IL6ST |
| AC016596.3 | 5 | ribosomal protein L41 (RPL41) pseudogene |
| AC008892.1 | 5 | novel transcript |
| AC016596.1 | 5 | novel transcript, antisense to IL6ST |
| AC016596.2 | 5 | NULL |
| ANKRD55 | 5 | ankyrin repeat domain 55 [Source:HGNC Syml |
| RNU6-299P | 5 | RNA, U6 small nuclear 299, pseudogene [Sour |
| RPL17P22 | 5 | ribosomal protein L17 pseudogene 22 [Source |
| RNA5SP184 | 5 | RNA, 5S ribosomal pseudogene 184 [Source:H |
| C1GALT1P2 | 5 | C1GALT1 pseudogene 2 [Source:HGNC Symbo |
| RNA5SP185 | 5 | RNA, 5S ribosomal pseudogene 185 [Source:H |
| PSMC1P4 | 5 | proteasome 26S subunit, ATPase 1 pseudoger |
| RNU6ATAC2P | 5 | RNA, U6atac small nuclear 2, pseudogene [Sou |
| AC034245.1 | 5 | novel transcript |
| HMGN1P17 | 5 | high mobility group nucleosome binding doma |
| LINC01948 | 5 | long intergenic non-protein coding RNA 1948 |
| AC008391.1 | 5 | novel transcript |
| RPL26P19 | 5 | ribosomal protein L26 pseudogene 19 [Source |
| C5orf67 | 5 | chromosome 5 open reading frame 67 [Sourc |
| AC022431.1 | 5 | novel transcript |
| AC008940.1 | 5 | novel transcript |
| MAP3K1 | 5 | mitogen-activated protein kinase kinase kinas |
| AC008937.2 | 5 | novel transcript |
| AC008937.1 | 5 | novel transcript |
| SETD9 | 5 | SET domain containing 9 [Source:HGNC Symb |

| | | |
|---|---|---|
| MIER3 | 5 | MIER family member 3 [Source:HGNC Symbol |
| AC008937.3 | 5 | novel transcript, antisense to MIER3 |
| AC016644.1 | 5 | novel transcript |
| AC114973.1 | 5 | small nuclear ribonucleoprotein pseudogene |
| AC034244.1 | 5 | cytochrome c oxidase subunit VIc (COX6C) pse |
| AC034244.2 | 5 | chromosome 12 open reading frame 29 (C12c |
| Y_RNA | 5 | Y RNA [Source:RFAM;Acc:RF00019] |
| GPBP1 | 5 | GC-rich promoter binding protein 1 [Source:H |
| AC025470.1 | 5 | TSR1, 20S rRNA accumulation, homolog (S. ce |
| AC025470.2 | 5 | novel transcript, antisense to ACTBL2 |
| SALL4P1 | 5 | spalt like transcription factor 4 pseudogene 1 |
| ACTBL2 | 5 | actin beta like 2 [Source:HGNC Symbol;Acc:HG |
| AC008780.1 | 5 | novel transcript |
| AC008780.2 | 5 | novel transcript |
| AC106822.1 | 5 | novel transcript |
| LINC02225 | 5 | long intergenic non-protein coding RNA 2225 |
| LINC02101 | 5 | long intergenic non-protein coding RNA 2101 |
| AC008786.1 | 5 | novel transcript |
| PGAM1P1 | 5 | phosphoglycerate mutase 1 pseudogene 1 [Sc |
| PLK2 | 5 | polo like kinase 2 [Source:HGNC Symbol;Acc:H |
| GAPT | 5 | GRB2 binding adaptor protein, transmembran |
| AC008814.1 | 5 | prenyl (decaprenyl) diphosphate synthase, sul |
| MIR548AE2 | 5 | microRNA 548ae-2 [Source:HGNC Symbol;Acc |
| LINC02108 | 5 | long intergenic non-protein coding RNA 2108 |
| RAB3C | 5 | RAB3C, member RAS oncogene family [Source |
| AC016642.1 | 5 | novel transcript |
| RPL5P15 | 5 | ribosomal protein L5 pseudogene 15 [Source: |
| AC008852.1 | 5 | novel transcript |
| PDE4D | 5 | phosphodiesterase 4D [Source:HGNC Symbol; |
| AC092343.1 | 5 | novel transcript |
| AC008833.1 | 5 | endogenous retrovirus group V, member pse |
| NDUFB4P2 | 5 | NADH:ubiquinone oxidoreductase subunit B4 |
| MIR582 | 5 | microRNA 582 [Source:HGNC Symbol;Acc:HGN |
| AC034234.1 | 5 | novel transcript |
| RNU6-806P | 5 | RNA, U6 small nuclear 806, pseudogene [Sour |
| AC109486.1 | 5 | ribosomal protein L31 (RPL31) pseudogene |
| SETP21 | 5 | SET pseudogene 21 [Source:HGNC Symbol;Acc |
| AC109486.2 | 5 | NULL |
| PART1 | 5 | prostate androgen-regulated transcript 1 [Sou |
| AC109486.3 | 5 | NULL |
| AC016591.1 | 5 | NULL |
| DEPDC1B | 5 | DEP domain containing 1B [Source:HGNC Sym |
| CAB39P1 | 5 | calcium binding protein 39 pseudogene 1 [Sou |
| AC109133.1 | 5 | novel transcript, antisense to DEPDC1B |
| KRT8P31 | 5 | keratin 8 pseudogene 31 [Source:HGNC Symb |
| ELOVL7 | 5 | ELOVL fatty acid elongase 7 [Source:HGNC Syr |
| AC104113.1 | 5 | novel transcript |
| ERCC8 | 5 | ERCC excision repair 8, CSA ubiquitin ligase co |
| GNL3LP1 | 5 | G protein nucleolar 3 like pseudogene 1 [Sour |
| AC022445.1 | 5 | novel transcript |

| | | |
|---|---|---|
| NDUFAF2 | 5 | NADH:ubiquinone oxidoreductase complex as |
| AC008498.2 | 5 | novel transcript |
| AC008498.1 | 5 | protocadherin 19 (PCDH19) pseudogene |
| SMIM15 | 5 | small integral membrane protein 15 [Source:H |
| SMIM15-AS1 | 5 | SMIM15 antisense RNA 1 [Source:HGNC Symb |
| LINC02057 | 5 | long intergenic non-protein coding RNA 2057 |
| ZSWIM6 | 5 | zinc finger SWIM-type containing 6 [Source:H |
| AC122718.2 | 5 | DEAD (Asp-Glu-Ala-Asp) box polypeptide 25 (D |
| AC122718.1 | 5 | family with sequence similarity 133, member |
| RPL3P6 | 5 | ribosomal protein L3 pseudogene 6 [Source:H |
| C5orf64 | 5 | chromosome 5 open reading frame 64 [Source |
| AC026746.1 | 5 | novel transcript |
| RNU6-913P | 5 | RNA, U6 small nuclear 913, pseudogene [Sour |
| C5orf64-AS1 | 5 | C5orf64 antisense RNA 1 [Source:HGNC Symb |
| AC026434.1 | 5 | actin, gamma 1 (ACTG1) pseudogene |
| AC008877.1 | 5 | novel transcript |
| RN7SKP157 | 5 | RN7SK pseudogene 157 [Source:HGNC Symbo |
| KIF2A | 5 | kinesin family member 2A [Source:HGNC Sym |
| DIMT1 | 5 | DIMT1 rRNA methyltransferase and ribosome |
| IPO11 | 5 | importin 11 [Source:HGNC Symbol;Acc:HGNC: |
| RNU6-661P | 5 | RNA, U6 small nuclear 661, pseudogene [Sour |
| CKS1BP3 | 5 | CDC28 protein kinase regulatory subunit 1B ps |
| AC114982.2 | 5 | ribosomal protein L31 (RPL31) pseudogene |
| AC114982.1 | 5 | ribosomal protein L35a (RPL35A) pseudogene |
| LRRC70 | 5 | leucine rich repeat containing 70 [Source:HGN |
| ISCA1P1 | 5 | iron-sulfur cluster assembly 1 pseudogene 1 [S |
| AC113420.1 | 5 | secretory carrier membrane protein 1 (SCAMP |
| AC025445.1 | 5 | dedicator of cytokinesis family pseudogene |
| AC008558.1 | 5 | novel transcript |
| AC122707.1 | 5 | novel transcript, sense overlapping HTR1A |
| HTR1A | 5 | 5-hydroxytryptamine receptor 1A [Source:HG |
| RNF180 | 5 | ring finger protein 180 [Source:HGNC Symbol; |
| AC091862.1 | 5 | pseudogene similar to 4933425L06Rik |
| RGS7BP | 5 | regulator of G protein signaling 7 binding prot |
| RN7SL169P | 5 | RNA, 7SL, cytoplasmic 169, pseudogene [Sour |
| RNU6-294P | 5 | RNA, U6 small nuclear 294, pseudogene [Sour |
| MRPL49P1 | 5 | mitochondrial ribosomal protein L49 pseudog |
| SHISAL2B | 5 | shisa like 2B [Source:HGNC Symbol;Acc:HGNC |
| SREK1IP1 | 5 | SREK1 interacting protein 1 [Source:HGNC Syr |
| CWC27 | 5 | CWC27 spliceosome associated cyclophilin [So |
| AC092354.2 | 5 | novel transcript |
| AC092354.1 | 5 | novel transcript |
| Y_RNA | 5 | Y RNA [Source:RFAM;Acc:RF00019] |
| ADAMTS6 | 5 | ADAM metallopeptidase with thrombospondi |
| AC008868.1 | 5 | ribosomal protein L5 (RPL5) pseudogene |
| RPEP1 | 5 | ribulose-5-phosphate-3-epimerase pseudoger |
| AC025186.1 | 5 | ribosomal protein S2 (RPS2) pseudogene |
| AC025176.1 | 5 | novel transcript |
| CENPK | 5 | centromere protein K [Source:HGNC Symbol;A |
| PPWD1 | 5 | peptidylprolyl isomerase domain and WD repe |

| | | |
|---|---|---|
| TRIM23 | 5 | tripartite motif containing 23 [Source:HGNC S |
| RNU6-540P | 5 | RNA, U6 small nuclear 540, pseudogene [Sour |
| SHLD3 | 5 | shieldin complex subunit 3 [Source:HGNC Sym |
| TRAPPC13 | 5 | trafficking protein particle complex 13 [Source |
| SGTB | 5 | small glutamine rich tetratricopeptide repeat |
| NLN | 5 | neurolysin [Source:HGNC Symbol;Acc:HGNC:1 |
| AC008958.1 | 5 | PRAME family pseudogene |
| AC010359.2 | 5 | novel transcript |
| ERBIN | 5 | erbb2 interacting protein [Source:HGNC Symb |
| AC010359.1 | 5 | NULL |
| AC010359.3 | 5 | novel transcript, sense intronic to ERBIN |
| AC025442.2 | 5 | novel transcipt |
| SREK1 | 5 | splicing regulatory glutamic acid and lysine ric |
| AC025442.1 | 5 | novel transcript |
| LINC02065 | 5 | long intergenic non-protein coding RNA 2065 |
| AC092353.1 | 5 | novel transcript |
| AC092353.2 | 5 | novel transcript |
| LINC02229 | 5 | long intergenic non-protein coding RNA 2229 |
| PPIAP78 | 5 | peptidylprolyl isomerase A pseudogene 78 [So |
| MAST4 | 5 | microtubule associated serine/threonine kina |
| AC092349.1 | 5 | ribosomal protein L15 pseudogene |
| MAST4-IT1 | 5 | MAST4 intronic transcript 1 [Source:HGNC Syr |
| MAST4-AS1 | 5 | MAST4 antisense RNA 1 [Source:HGNC Symbo |
| CD180 | 5 | CD180 molecule [Source:HGNC Symbol;Acc:H |
| AC010420.1 | 5 | novel transcript |
| AC008459.1 | 5 | novel transcript |
| AC112206.2 | 5 | novel transcript |
| AC079467.1 | 5 | novel transcript |
| AC112206.1 | 5 | ribosomal protein L21 (RPL21) pseudogene |
| LINC02242 | 5 | long intergenic non-protein coding RNA 2242 |
| BCL9P1 | 5 | BCL9 pseudogene 1 [Source:HGNC Symbol;Ac |
| AC112206.4 | 5 | mitochondrial ribosomal protein L39 (MRPL39 |
| AC112206.3 | 5 | ribosomal protein L23a (RPL23A) pseudogene |
| AC106798.1 | 5 | novel transcript |
| AC024581.1 | 5 | novel transcript |
| AC024581.2 | 5 | dicer 1, ribonuclease type III (DICER1) pseudo |
| AC010376.1 | 5 | DnaJ (Hsp40) homolog, subfamily C, member |
| RNU6-1232P | 5 | RNA, U6 small nuclear 1232, pseudogene [Sou |
| EEF1B2P2 | 5 | eukaryotic translation elongation factor 1 bet |
| LINC02219 | 5 | long intergenic non-protein coding RNA 2219 |
| PIK3R1 | 5 | phosphoinositide-3-kinase regulatory subunit |
| AC024579.1 | 5 | novel transcript |
| AC010280.2 | 5 | novel transcript |
| AC010280.1 | 5 | novel transcript |
| AC010280.3 | 5 | novel transcript |
| AC093523.1 | 5 | novel transcript |
| VWA8P1 | 5 | von Willebrand factor A domain containing 8 |
| U8 | 5 | U8 small nucleolar RNA [Source:RFAM;Acc:RF |
| AC093248.1 | 5 | novel transcript |
| RN7SKP251 | 5 | RN7SK pseudogene 251 [Source:HGNC Symbo |

| | | |
|---|---|---|
| LINC02198 | 5 | long intergenic non-protein coding RNA 2198 |
| AC010273.3 | 5 | novel transcript |
| SUMO2P4 | 5 | SUMO2 pseudogene 4 [Source:HGNC Symbol; |
| SLC30A5 | 5 | solute carrier family 30 member 5 [Source:HG |
| AC010273.2 | 5 | novel transcript |
| AC010273.1 | 5 | NULL |
| SNORA50D | 5 | small nucleolar RNA, H/ACA box 50D [Source: |
| CCNB1 | 5 | cyclin B1 [Source:HGNC Symbol;Acc:HGNC:15 |
| AC022107.1 | 5 | TEC |
| CENPH | 5 | centromere protein H [Source:HGNC Symbol; |
| MRPS36 | 5 | mitochondrial ribosomal protein S36 [Source: |
| CDK7 | 5 | cyclin dependent kinase 7 [Source:HGNC Sym |
| CCDC125 | 5 | coiled-coil domain containing 125 [Source:HG |
| CFL1P5 | 5 | cofilin 1 pseudogene 5 [Source:HGNC Symbol; |
| CHCHD2P2 | 5 | coiled-coil-helix-coiled-coil-helix domain cont |
| NDUFB9P1 | 5 | NADH:ubiquinone oxidoreductase subunit B9 |
| AK6 | 5 | adenylate kinase 6 [Source:HGNC Symbol;Acc |
| TAF9 | 5 | TATA-box binding protein associated factor 9 |
| RAD17 | 5 | RAD17 checkpoint clamp loader component [ |
| MARVELD2 | 5 | MARVEL domain containing 2 [Source:HGNC S |
| RPS27P14 | 5 | ribosomal protein S27 pseudogene 14 [Source |
| AC145146.1 | 5 | novel transcript, antisense to OCLN |
| RN7SL616P | 5 | RNA, 7SL, cytoplasmic 616, pseudogene [Sour |
| OCLN | 5 | occludin [Source:HGNC Symbol;Acc:HGNC:81 |
| RNU6-724P | 5 | RNA, U6 small nuclear 724, pseudogene [Sour |
| GTF2H2C | 5 | GTF2H2 family member C [Source:HGNC Sym |
| AC139495.1 | 5 | NLR family, apoptosis inhibitory protein (NAIP |
| AC139495.3 | 5 | POM121 membrane glycoprotein-like 1 (POM |
| GUSBP3 | 5 | GUSB pseudogene 3 [Source:HGNC Symbol;A |
| AC139495.2 | 5 | POM121 membrane glycoprotein-like 1 (POM |
| AC131392.1 | 5 | glucuronidase, beta (GUSB) pseudogene |
| AC131392.2 | 5 | POM121 membrane glycoprotein-like (POM1 |
| CDH12P2 | 5 | cadherin 12 pseudogene 2 [Source:HGNC Sym |
| SERF1B | 5 | small EDRK-rich factor 1B [Source:HGNC Sym |
| SMN2 | 5 | survival of motor neuron 2, centromeric [Sour |
| AC140134.2 | 5 | novel transcript, antisense to SMN2 |
| AC140134.1 | 5 | NLR family, apoptosis inhibitory protein (NAIP |
| CDH12P3 | 5 | cadherin 12 pseudogene 3 [Source:HGNC Sym |
| AC138866.2 | 5 | glucuronidase, beta (GUSB) pseudogene |
| AC138866.1 | 5 | glucuronidase, beta (GUSB) pseudogene |
| GTF2H2B | 5 | general transcription factor IIH subunit 2B (ps |
| AC146944.4 | 5 | novel transcript |
| AC146944.2 | 5 | NLR family, apoptosis inhibitory protein (NAIP |
| AC146944.3 | 5 | glucuronidase, beta (GUSB) pseudogene |
| AC146944.1 | 5 | glucuronidase, beta (GUSB) pseudogene |
| AC139272.1 | 5 | glucuronidase, beta (GUSB) pseudogene |
| AC139272.2 | 5 | POM121 transmembrane nucleoporin C (POM |
| CDH12P1 | 5 | cadherin 12 pseudogene 1 [Source:HGNC Sym |
| SERF1A | 5 | small EDRK-rich factor 1A [Source:HGNC Sym |
| SMN1 | 5 | survival of motor neuron 1, telomeric [Source |

| | | |
|---|---|---|
| AC139834.1 | 5 | novel transcript, antisense to SMN1 |
| NAIP | 5 | NLR family apoptosis inhibitory protein [Sourc |
| GTF2H2 | 5 | general transcription factor IIH subunit 2 [Sou |
| OCLNP1 | 5 | OCLN pseudogene 1 [Source:HGNC Symbol;Ac |
| AC145138.1 | 5 | NLR family, apoptosis inhibitory protein (NAIP |
| CDH12P4 | 5 | cadherin 12 pseudogene 4 [Source:HGNC Sym |
| GUSBP9 | 5 | GUSB pseudogene 9 [Source:HGNC Symbol;Ac |
| AC146949.1 | 5 | glucuronidase, beta (GUSB) pseudogene |
| LINC02197 | 5 | long intergenic non-protein coding RNA 2197 |
| AC145141.2 | 5 | novel transcript |
| PMCHL2 | 5 | pro-melanin concentrating hormone like 2 (ps |
| AC145141.1 | 5 | novel transcript |
| BDP1 | 5 | B double prime 1, subunit of RNA polymerase |
| AC138832.1 | 5 | sterol carrier protein (SCP2) pseudogene |
| HMGN1P12 | 5 | high mobility group nucleosome binding doma |
| MCCC2 | 5 | methylcrotonoyl-CoA carboxylase 2 [Source:H |
| CARTPT | 5 | CART prepropeptide [Source:HGNC Symbol;Ac |
| AC143336.1 | 5 | catenin (cadherin-associated protein), beta 1, |
| AC025774.1 | 5 | novel transcript |
| MAP1B | 5 | microtubule associated protein 1B [Source:HG |
| MIR4803 | 5 | microRNA 4803 [Source:HGNC Symbol;Acc:HG |
| MRPS27 | 5 | mitochondrial ribosomal protein S27 [Source: |
| RN7SL153P | 5 | RNA, 7SL, cytoplasmic 153, pseudogene [Sour |
| PTCD2 | 5 | pentatricopeptide repeat domain 2 [Source:H |
| AC093278.1 | 5 | ribosomal protein L10a (RPL10A) pseudogene |
| YBX1P5 | 5 | Y-box binding protein 1 pseudogene 5 [Source |
| AC093278.2 | 5 | novel transcript, overlapping ZNF366 |
| ZNF366 | 5 | zinc finger protein 366 [Source:HGNC Symbol; |
| LINC02056 | 5 | long intergenic non-protein coding RNA 2056 |
| AC063979.2 | 5 | novel transcript |
| RPL7P22 | 5 | ribosomal protein L7 pseudogene 22 [Source: |
| AC063979.1 | 5 | novel pseudogene |
| Y_RNA | 5 | Y RNA [Source:RFAM;Acc:RF00019] |
| AC035140.1 | 5 | novel transcript |
| TNPO1 | 5 | transportin 1 [Source:HGNC Symbol;Acc:HGN |
| MIR4804 | 5 | microRNA 4804 [Source:HGNC Symbol;Acc:HG |
| AC008953.1 | 5 | ribosomal protein L35a (RPL35A) pseudogene |
| AC008972.1 | 5 | novel transcript |
| AC008972.2 | 5 | novel transcript |
| FCHO2 | 5 | FCH domain only 2 [Source:HGNC Symbol;Acc |
| AC020893.2 | 5 | dullard homolog (Xenopus laevis) (DULLARD) |
| AC020893.1 | 5 | calcium binding protein P22 (CHP) pseudogen |
| TMEM171 | 5 | transmembrane protein 171 [Source:HGNC Sy |
| AC116345.3 | 5 | novel transcript |
| AC116345.2 | 5 | ribosomal protein S3 (RPS3 ) pseudogene |
| TMEM174 | 5 | transmembrane protein 174 [Source:HGNC Sy |
| AC116345.4 | 5 | novel transcript |
| AC116345.1 | 5 | novel transcript |
| LINC02230 | 5 | long intergenic non-protein coding RNA 2230 |
| AC099522.1 | 5 | High mobility group nucleosome-binding dom |

| | | |
|---|---|---|
| FOXD1 | 5 | forkhead box D1 [Source:HGNC Symbol;Acc:H |
| FOXD1-AS1 | 5 | FOXD1 antisense RNA 1 [Source:HGNC Symbo |
| LINC01385 | 5 | long intergenic non-protein coding RNA 1385 |
| LINC01386 | 5 | long intergenic non-protein coding RNA 1386 |
| AC099522.2 | 5 | novel transcript |
| BTF3 | 5 | basic transcription factor 3 [Source:HGNC Sym |
| FUNDC2P1 | 5 | FUN14 domain containing 2 pseudogene 1 [So |
| ANKRA2 | 5 | ankyrin repeat family A member 2 [Source:HG |
| UTP15 | 5 | UTP15 small subunit processome component |
| ARHGEF28 | 5 | Rho guanine nucleotide exchange factor 28 [S |
| AC091868.2 | 5 | novel transcript |
| AC091868.1 | 5 | ATP synthase, H+ transporting, mitochondrial |
| RNU7-196P | 5 | RNA, U7 small nuclear 196 pseudogene [Sourc |
| AC093283.1 | 5 | novel transcript |
| LINC02122 | 5 | long intergenic non-protein coding RNA 2122 |
| LINC01331 | 5 | long intergenic non-protein coding RNA 1331 |
| LINC01335 | 5 | long intergenic non-protein coding RNA 1335 |
| RN7SL814P | 5 | RNA, 7SL, cytoplasmic 814, pseudogene [Sour |
| LINC01333 | 5 | long intergenic non-protein coding RNA 1333 |
| LINC01332 | 5 | long intergenic non-protein coding RNA 1332 |
| ENC1 | 5 | ectodermal-neural cortex 1 [Source:HGNC Syr |
| HEXB | 5 | hexosaminidase subunit beta [Source:HGNC S |
| GFM2 | 5 | G elongation factor mitochondrial 2 [Source:H |
| RNU6-658P | 5 | RNA, U6 small nuclear 658, pseudogene [Sour |
| NSA2 | 5 | NSA2 ribosome biogenesis factor [Source:HGN |
| FAM169A | 5 | family with sequence similarity 169 member A |
| RNU6-1330P | 5 | RNA, U6 small nuclear 1330, pseudogene [Sou |
| AC093214.1 | 5 | actin related protein 2/3 complex, subunit 1A, |
| AC010501.2 | 5 | novel transcript, antisense to FAM169A |
| AC010501.1 | 5 | NULL |
| AC116337.4 | 5 | novel transcript |
| AC116337.3 | 5 | novel transcript |
| AC116337.1 | 5 | zinc finger protein pseudogene |
| AC116337.2 | 5 | ribosomal protein L39 (RPL39) pseudogene |
| RPL27AP5 | 5 | ribosomal protein L27a pseudogene 5 [Source |
| AC093259.1 | 5 | novel pseudogene |
| GCNT4 | 5 | glucosaminyl (N-acetyl) transferase 4 [Source: |
| ANKRD31 | 5 | ankyrin repeat domain 31 [Source:HGNC Syml |
| SUMO2P5 | 5 | SUMO2 pseudogene 5 [Source:HGNC Symbol; |
| AC008897.2 | 5 | novel transcript |
| HMGCR | 5 | 3-hydroxy-3-methylglutaryl-CoA reductase [So |
| CERT1 | 5 | ceramide transporter 1 [Source:HGNC Symbol |
| AC008897.3 | 5 | novel transcript |
| AC008897.1 | 5 | ribosomal protein S26 (RPS26) pseudogene |
| POLK | 5 | DNA polymerase kappa [Source:HGNC Symbo |
| RNU7-175P | 5 | RNA, U7 small nuclear 175 pseudogene [Sourc |
| AC010245.1 | 5 | novel transcript |
| AC010245.2 | 5 | novel transcript |
| ANKDD1B | 5 | ankyrin repeat and death domain containing 1 |
| POC5 | 5 | POC5 centriolar protein [Source:HGNC Symbo |

| | | |
|---|---|---|
| RNU6-680P | 5 | RNA, U6 small nuclear 680, pseudogene [Sour |
| SLC25A5P9 | 5 | solute carrier family 25 member 5 pseudogen |
| AC108120.3 | 5 | karyopherin alpha pseudogene |
| AC108120.2 | 5 | bridging integrator 2 (BIN2) pseudogene |
| BIN2P2 | 5 | bridging integrator 2 pseudogene 2 [Source:H |
| AC108120.1 | 5 | aconitase 2, mitochondrial (ACO2) pseudogen |
| SAP18P1 | 5 | SAP18 pseudogene 1 [Source:HGNC Symbol;A |
| AC026774.2 | 5 | spermine synthase (SMS) pseudogene |
| AC026774.1 | 5 | novel transcript |
| SV2C | 5 | synaptic vesicle glycoprotein 2C [Source:HGN( |
| RNA5SP186 | 5 | RNA, 5S ribosomal pseudogene 186 [Source:H |
| AC113404.2 | 5 | pseudogene similar to part of microtubule ass |
| AC113404.3 | 5 | RAP1B, member of RAS oncogene family pseu |
| HMGN2P4 | 5 | high mobility group nucleosomal binding dom |
| PDCD5P2 | 5 | programmed cell death 5 pseudogene 2 [Sour |
| AC113404.1 | 5 | novel transcript |
| SNRPCP2 | 5 | small nuclear ribonucleoprotein polypeptide ( |
| IQGAP2 | 5 | IQ motif containing GTPase activating protein |
| AC112173.1 | 5 | sulfiredoxin 1 homolog (S. cerevisiae) (SRXN1) |
| AC026725.1 | 5 | novel transcript |
| F2RL2 | 5 | coagulation factor II thrombin receptor like 2 |
| AC025188.1 | 5 | novel transcript, antisense to F2R |
| AC025188.2 | 5 | NULL |
| AC025188.3 | 5 | NULL |
| F2R | 5 | coagulation factor II thrombin receptor [Sourc |
| F2RL1 | 5 | F2R like trypsin receptor 1 [Source:HGNC Sym |
| RN7SL208P | 5 | RNA, 7SL, cytoplasmic 208, pseudogene [Sour |
| S100Z | 5 | S100 calcium binding protein Z [Source:HGNC |
| RNU6ATAC36P | 5 | RNA, U6atac small nuclear 36, pseudogene [S( |
| CRHBP | 5 | corticotropin releasing hormone binding prot( |
| AGGF1 | 5 | angiogenic factor with G-patch and FHA doma |
| AC008581.2 | 5 | novel protein, AGGF1-ZBED3-AS1 readthrough |
| ZBED3 | 5 | zinc finger BED-type containing 3 [Source:HGN |
| AC008581.1 | 5 | novel transcript |
| SNORA47 | 5 | small nucleolar RNA, H/ACA box 47 [Source:H( |
| ZBED3-AS1 | 5 | ZBED3 antisense RNA 1 [Source:HGNC Symbol |
| AC022414.1 | 5 | phosphodiesterase 8B [Source:NCBI gene;Acc |
| HMGB1P35 | 5 | high mobility group box 1 pseudogene 35 [Sou |
| PDE8B | 5 | phosphodiesterase 8B [Source:HGNC Symbol; |
| ALDH7A1P1 | 5 | aldehyde dehydrogenase 7 family member A1 |
| WDR41 | 5 | WD repeat domain 41 [Source:HGNC Symbol; |
| RPS2P24 | 5 | ribosomal protein S2 pseudogene 24 [Source: |
| RPL7P23 | 5 | ribosomal protein L7 pseudogene 23 [Source: |
| OTP | 5 | orthopedia homeobox [Source:HGNC Symbol; |
| TBCA | 5 | tubulin folding cofactor A [Source:HGNC Symb |
| ACTBP2 | 5 | ACTB pseudogene 2 [Source:HGNC Symbol;Ac |
| AC108482.1 | 5 | novel transcript |
| AC108482.2 | 5 | novel transcript |
| AP3B1 | 5 | adaptor related protein complex 3 subunit be |
| AC024568.1 | 5 | heat shock protein 90kDa alpha (cytosolic), cl |

| | | |
|---|---|---|
| ATP6V1G1P6 | 5 | ATPase H+ transporting V1 subunit G1 pseudc |
| AC104108.1 | 5 | NADH dehydrogenase [ubiquinone] 1 alpha su |
| RNU6-183P | 5 | RNA, U6 small nuclear 183, pseudogene [Sour |
| Y_RNA | 5 | Y RNA [Source:RFAM;Acc:RF00019] |
| SCAMP1-AS1 | 5 | SCAMP1 antisense RNA 1 [Source:HGNC Symb |
| SCAMP1 | 5 | secretory carrier membrane protein 1 [Source |
| LHFPL2 | 5 | LHFPL tetraspan subfamily member 2 [Source |
| HMGB1P21 | 5 | high mobility group box 1 pseudogene 21 [Sou |
| AC008802.1 | 5 | ribosomal protein S3A (RPS3A) pseudogene |
| AC025755.1 | 5 | SMT3 suppressor of mif two 3 homolog 2 (S. c |
| ARSB | 5 | arylsulfatase B [Source:HGNC Symbol;Acc:HGI |
| DMGDH | 5 | dimethylglycine dehydrogenase [Source:HGN( |
| AC020937.1 | 5 | Impact homolog (mouse) (IMPACT) pseudoge |
| BHMT2 | 5 | betaine--homocysteine S-methyltransferase 2 |
| BHMT | 5 | betaine--homocysteine S-methyltransferase [ |
| RNY3P1 | 5 | RNY3 pseudogene 1 [Source:HGNC Symbol;Ac |
| AC016559.2 | 5 | ubiquinol-cytochrome c reductase hinge prote |
| AC016559.1 | 5 | NULL |
| AC016559.3 | 5 | GABA(A) receptor-associated protein-like 2 (G |
| JMY | 5 | junction mediating and regulatory protein, p5 |
| AC020898.1 | 5 | ribosomal protein S3A (RPS3A) pseudogene |
| HOMER1 | 5 | homer scaffold protein 1 [Source:HGNC Symb |
| AC093270.1 | 5 | ribosomal protein L29 (RPL29) pseudogene |
| AC093270.2 | 5 | ribosomal protein L7a (RPL7A) pseudogene |
| Y_RNA | 5 | Y RNA [Source:RFAM;Acc:RF00019] |
| Y_RNA | 5 | Y RNA [Source:RFAM;Acc:RF00019] |
| TENT2 | 5 | terminal nucleotidyltransferase 2 [Source:HGI |
| CMYA5 | 5 | cardiomyopathy associated 5 [Source:HGNC S |
| AC008496.2 | 5 | novel transcript, antisense to CMYA5 |
| AC008496.3 | 5 | novel transcript |
| AC008496.1 | 5 | ribosomal protein L10 (RPL10) pseudogene |
| LINC01455 | 5 | long intergenic non-protein coding RNA 1455 |
| MTX3 | 5 | metaxin 3 [Source:HGNC Symbol;Acc:HGNC:2 |
| THBS4 | 5 | thrombospondin 4 [Source:HGNC Symbol;Acc |
| AC112184.1 | 5 | ring-box 1 (RBX1) pseudogene |
| TRMT112P2 | 5 | tRNA methyltransferase subunit 11-2 pseudog |
| AC012636.1 | 5 | novel transcript |
| SERINC5 | 5 | serine incorporator 5 [Source:HGNC Symbol;A |
| AC010260.1 | 5 | novel transcript |
| KRT18P45 | 5 | keratin 18 pseudogene 45 [Source:HGNC Sym |
| AC026410.1 | 5 | actin, beta (ACTB) pseudogene |
| AC026410.4 | 5 | NULL |
| AC026410.2 | 5 | ribosomal protein L39 (RPL39) pseudogene |
| SPZ1 | 5 | spermatogenic leucine zipper 1 [Source:HGNC |
| RBMX2P5 | 5 | RBMX2 pseudogene 5 [Source:HGNC Symbol; |
| AC026410.3 | 5 | mediator complex subunit 27 (MED27) pseud |
| HNRNPA1P12 | 5 | heterogeneous nuclear ribonucleoprotein A1 |
| RNU6-211P | 5 | RNA, U6 small nuclear 211, pseudogene [Sour |
| ZFYVE16 | 5 | zinc finger FYVE-type containing 16 [Source:H |
| AC008771.1 | 5 | novel transcript |

| | | |
|---|---|---|
| FAM151B | 5 | family with sequence similarity 151 member B |
| RPS27AP9 | 5 | ribosomal protein S27a pseudogene 9 [Source |
| RPL7P24 | 5 | ribosomal protein L7 pseudogene 24 [Source: |
| ANKRD34B | 5 | ankyrin repeat domain 34B [Source:HGNC Syn |
| DBIP2 | 5 | DBI pseudogene 2 [Source:HGNC Symbol;Acc: |
| LINC01337 | 5 | long intergenic non-protein coding RNA 1337 |
| DHFR | 5 | dihydrofolate reductase [Source:HGNC Symbo |
| AC008434.1 | 5 | novel transcript |
| MTRNR2L2 | 5 | MT-RNR2 like 2 [Source:HGNC Symbol;Acc:HG |
| MSH3 | 5 | mutS homolog 3 [Source:HGNC Symbol;Acc:H |
| AC022493.2 | 5 | S-phase kinase-associated protein 1 (SKP1) ps |
| AC022493.1 | 5 | ribosomal protein S26 (RPS26) pseudogene |
| RASGRF2-AS1 | 5 | RASGRF2 antisense RNA 1 [Source:HGNC Sym |
| RASGRF2 | 5 | Ras protein specific guanine nucleotide releas |
| AC026427.2 | 5 | peroxiredoxin 3 (PRDX3) pseudogene |
| AC026427.1 | 5 | novel transcript |
| CKMT2-AS1 | 5 | CKMT2 antisense RNA 1 [Source:HGNC Symbo |
| CKMT2 | 5 | creatine kinase, mitochondrial 2 [Source:HGN |
| AC026436.1 | 5 | ribosomal protein S12 (RPS12) pseudogene |
| ZCCHC9 | 5 | zinc finger CCHC-type containing 9 [Source:HG |
| ACOT12 | 5 | acyl-CoA thioesterase 12 [Source:HGNC Symb |
| AC010623.1 | 5 | novel transcript |
| SSBP2 | 5 | single stranded DNA binding protein 2 [Source |
| AC026726.2 | 5 | novel transcript |
| AC026726.1 | 5 | novel transcript |
| SEM1P1 | 5 | SEM1 pseudogene 1 [Source:HGNC Symbol;Ac |
| ATG10 | 5 | autophagy related 10 [Source:HGNC Symbol;A |
| ATG10-IT1 | 5 | ATG10 intronic transcript 1 [Source:HGNC Syn |
| PPIAP11 | 5 | peptidylprolyl isomerase A pseudogene 11 [Sc |
| ATG10-AS1 | 5 | ATG10 antisense RNA 1 [Source:HGNC Symbo |
| RN7SL378P | 5 | RNA, 7SL, cytoplasmic 378, pseudogene [Sour |
| AC010598.1 | 5 | ribosomal protein S23 pseudogene 5 |
| RPS23 | 5 | ribosomal protein S23 [Source:HGNC Symbol; |
| ATP6AP1L | 5 | ATPase H+ transporting accessory protein 1 lil |
| AC026782.1 | 5 | novel transcript |
| AC026782.2 | 5 | novel transcript |
| AC008885.2 | 5 | novel transcript |
| RPL5P16 | 5 | ribosomal protein L5 pseudogene 16 [Source: |
| AC008885.1 | 5 | pancreatic progenitor cell differentiation and |
| MIR3977 | 5 | microRNA 3977 [Source:HGNC Symbol;Acc:HG |
| LINC01338 | 5 | long intergenic non-protein coding RNA 1338 |
| AC108174.1 | 5 | novel transcript |
| AC027338.2 | 5 | novel transcript |
| ST13P12 | 5 | ST13, Hsp70 interacting protein pseudogene 1 |
| AC027338.1 | 5 | novel transcript |
| AC104118.1 | 5 | novel transcript |
| TMEM167A | 5 | transmembrane protein 167A [Source:HGNC S |
| SCARNA18 | 5 | small Cajal body-specific RNA 18 [Source:HGN |
| XRCC4 | 5 | X-ray repair cross complementing 4 [Source:H |
| AC094085.1 | 5 | ribosomal protein L13 (RPL13) pseudogene |

| | | |
|---|---|---|
| COQ10BP2 | 5 | coenzyme Q10B pseudogene 2 [Source:HGNC |
| FTH1P9 | 5 | ferritin heavy chain 1 pseudogene 9 [Source:H |
| VCAN | 5 | versican [Source:HGNC Symbol;Acc:HGNC:246 |
| VCAN-AS1 | 5 | VCAN antisense RNA 1 [Source:HGNC Symbol; |
| HAPLN1 | 5 | hyaluronan and proteoglycan link protein 1 [S |
| RN7SKP295 | 5 | RN7SK pseudogene 295 [Source:HGNC Symbo |
| RNU6-620P | 5 | RNA, U6 small nuclear 620, pseudogene [Sour |
| AC020899.1 | 5 | ribosomal protein L13a (RPL13A) pseudogene |
| RNU4-11P | 5 | RNA, U4 small nuclear 11, pseudogene [Sourc |
| AC093286.1 | 5 | zona pellucida glycoprotein 3 (sperm receptor |
| EDIL3 | 5 | EGF like repeats and discoidin domains 3 [Sou |
| RNU6-448P | 5 | RNA, U6 small nuclear 448, pseudogene [Sour |
| AC113383.1 | 5 | novel transcript |
| PPIAP79 | 5 | peptidylprolyl isomerase A pseudogene 79 [Sc |
| AC114928.1 | 5 | novel transcript |
| RBBP4P6 | 5 | RBBP4 pseudogene 6 [Source:HGNC Symbol;A |
| AC117522.3 | 5 | ribosomal protein L14 pseudogene |
| AC117522.2 | 5 | novel transcript |
| AC117522.1 | 5 | cytochrome b pseudogene |
| AC010486.1 | 5 | novel transcript |
| AC010486.2 | 5 | novel transcript |
| AC010486.3 | 5 | novel transcript |
| RPL5P17 | 5 | ribosomal protein L5 pseudogene 17 [Source: |
| AC026700.1 | 5 | PDGFA associated protein 1 pseudogene |
| AC010595.1 | 5 | novel transcript |
| RPS2P25 | 5 | ribosomal protein S2 pseudogene 25 [Source: |
| AC026414.1 | 5 | novel transcript |
| PTP4A1P4 | 5 | PTP4A1 pseudogene 4 [Source:HGNC Symbol; |
| NBPF22P | 5 | NBPF member 22, pseudogene [Source:HGNC |
| AC020923.1 | 5 | novel transcript |
| AC016550.1 | 5 | novel transcript |
| AC016550.2 | 5 | novel transcript |
| AC016550.3 | 5 | novel transcript |
| AC026444.1 | 5 | ash2 (absent, small, or homeotic)-like (Drosop |
| COX7C | 5 | cytochrome c oxidase subunit 7C [Source:HGN |
| SNORD138 | 5 | small nucleolar RNA, C/D box 138 [Source:HGI |
| RNU6-804P | 5 | RNA, U6 small nuclear 804, pseudogene [Sour |
| LINC02059 | 5 | long intergenic non-protein coding RNA 2059 |
| AC008539.1 | 5 | novel transcript |
| AC091429.1 | 5 | ribosomal protein L10a (RPL10A) pseudogene |
| AC109492.1 | 5 | novel transcript |
| LINC01949 | 5 | long intergenic non-protein coding RNA 1949 |
| RN7SKP34 | 5 | RN7SK pseudogene 34 [Source:HGNC Symbol; |
| MIR4280 | 5 | microRNA 4280 [Source:HGNC Symbol;Acc:HG |
| RASA1 | 5 | RAS p21 protein activator 1 [Source:HGNC Syr |
| RN7SL629P | 5 | RNA, 7SL, cytoplasmic 629, pseudogene [Sour |
| RNU6-606P | 5 | RNA, U6 small nuclear 606, pseudogene [Sour |
| CCNH | 5 | cyclin H [Source:HGNC Symbol;Acc:HGNC:159 |
| AC018754.1 | 5 | novel transcript, antisense to CCNH |
| RNU6-727P | 5 | RNA, U6 small nuclear 727, pseudogene [Sour |

| | | |
|---|---|---|
| LINC02488 | 5 | long intergenic non-protein coding RNA 2488 |
| LINC02144 | 5 | long intergenic non-protein coding RNA 2144 |
| AC020930.1 | 5 | novel transcript |
| AC114971.1 | 5 | novel transcript |
| TMEM161B | 5 | transmembrane protein 161B [Source:HGNC S |
| TMEM161B-AS1 | 5 | TMEM161B antisense RNA 1 [Source:HGNC Sy |
| RNA5SP187 | 5 | RNA, 5S ribosomal pseudogene 187 [Source:H |
| RPS3AP22 | 5 | ribosomal protein S3a pseudogene 22 [Source |
| SNORA70 | 5 | Small nucleolar RNA SNORA70 [Source:RFAM; |
| AC091826.1 | 5 | 60S ribosomal protein L10 (RPL10) pseudogen |
| LINC02060 | 5 | long intergenic non-protein coding RNA 2060 |
| AC091826.2 | 5 | novel transcript |
| LINC00461 | 5 | long intergenic non-protein coding RNA 461 [S |
| AC008526.1 | 5 | H3 histone family 3 pseudogene |
| MIR9-2 | 5 | microRNA 9-2 [Source:HGNC Symbol;Acc:HGN |
| MEF2C-AS2 | 5 | MEF2C antisense RNA 2 [Source:HGNC Symbo |
| AC008525.1 | 5 | novel transcript |
| AC008525.2 | 5 | novel transcript |
| MEF2C | 5 | myocyte enhancer factor 2C [Source:HGNC Sy |
| MEF2C-AS1 | 5 | MEF2C antisense RNA 1 [Source:HGNC Symbo |
| AC074131.1 | 5 | novel transcript |
| LINC02161 | 5 | long intergenic non-protein coding RNA 2161 |
| AC113167.1 | 5 | novel transcript |
| MIR3660 | 5 | microRNA 3660 [Source:HGNC Symbol;Acc:HG |
| LINC01339 | 5 | long intergenic non-protein coding RNA 1339 |
| AC099554.1 | 5 | novel transcript |
| AC093510.2 | 5 | novel transcript, overlapping CETN3 |
| CETN3 | 5 | centrin 3 [Source:HGNC Symbol;Acc:HGNC:18 |
| AC093510.1 | 5 | novel transcript |
| MBLAC2 | 5 | metallo-beta-lactamase domain containing 2 [ |
| POLR3G | 5 | RNA polymerase III subunit G [Source:HGNC S |
| LYSMD3 | 5 | LysM domain containing 3 [Source:HGNC Sym |
| ADGRV1 | 5 | adhesion G protein-coupled receptor V1 [Sour |
| TMEM251P1 | 5 | transmembrane protein 251 pseudogene 1 [So |
| LUCAT1 | 5 | lung cancer associated transcript 1 [Source:HG |
| AC074132.1 | 5 | novel transcript, antisense to ADGRV1 |
| AC093281.1 | 5 | novel transcript |
| AC093281.2 | 5 | novel transcript |
| RNU4-90P | 5 | RNA, U4 small nuclear 90, pseudogene [Sourc |
| AC123595.1 | 5 | novel transcript |
| AC123595.2 | 5 | novel transcript |
| Metazoa_SRP | 5 | Metazoan signal recognition particle RNA [Sou |
| AC008799.1 | 5 | ribosomal protein S18 (RPS18) pseudogene |
| AC008799.2 | 5 | novel transcript |
| ARRDC3 | 5 | arrestin domain containing 3 [Source:HGNC Sy |
| ARRDC3-AS1 | 5 | ARRDC3 antisense RNA 1 [Source:HGNC Symb |
| RAB5CP2 | 5 | RAB5C, member RAS oncogene family pseudo |
| AC093298.2 | 5 | novel transcript |
| PCBP2P3 | 5 | poly(rC) binding protein 2 pseudogene 3 [Sour |
| AC093298.1 | 5 | NULL |

| | | |
|---|---|---|
| AC114316.1 | 5 | novel transcript |
| AC124854.1 | 5 | novel transcript |
| AC026780.1 | 5 | novel transcript |
| AC026780.2 | 5 | novel transcript |
| AC008517.1 | 5 | novel transcript |
| LDHBP3 | 5 | lactate dehydrogenase B pseudogene 3 [Sourc |
| CCT7P2 | 5 | chaperonin containing TCP1 subunit 7 pseudo |
| LINC02058 | 5 | long intergenic non-protein coding RNA 2058 |
| AC012625.1 | 5 | novel transcript |
| AC026408.1 | 5 | novel transcript |
| POLD2P1 | 5 | DNA polymerase delta 2, accessory subunit ps |
| NR2F1-AS1 | 5 | NR2F1 antisense RNA 1 [Source:HGNC Symbo |
| NR2F1 | 5 | nuclear receptor subfamily 2 group F member |
| AC106818.1 | 5 | novel transcript |
| FAM172A | 5 | family with sequence similarity 172 member A |
| MIR2277 | 5 | microRNA 2277 [Source:HGNC Symbol;Acc:HG |
| AC106818.2 | 5 | novel transcript, antisense to FAM172A |
| NPM1P27 | 5 | nucleophosmin 1 pseudogene 27 [Source:HGN |
| POU5F2 | 5 | POU domain class 5, transcription factor 2 [So |
| AC108102.1 | 5 | novel transcript |
| AC099501.1 | 5 | novel transcript, antisense to FAM172A |
| AC114980.1 | 5 | novel transcript |
| AC117528.1 | 5 | novel transcript, antisense to KIAA0825 |
| KIAA0825 | 5 | KIAA0825 [Source:HGNC Symbol;Acc:HGNC:28 |
| MTND5P12 | 5 | MT-ND5 pseudogene 12 [Source:HGNC Symbo |
| MTND6P3 | 5 | MT-ND6 pseudogene 3 [Source:HGNC Symbol |
| MTCYBP35 | 5 | MT-CYB pseudogene 35 [Source:HGNC Symbo |
| AC093311.1 | 5 | FAST kinase domains 1 (FASTKD1) pseudogene |
| AC025766.1 | 5 | novel transcript, sense intronic to C5orf36 |
| SLF1 | 5 | SMC5-SMC6 complex localization factor 1 [Sou |
| MCTP1 | 5 | multiple C2 and transmembrane domain cont |
| AC008534.1 | 5 | novel transcript |
| RPL7P18 | 5 | ribosomal protein L7 pseudogene 18 [Source: |
| AC008534.2 | 5 | NULL |
| MCTP1-AS1 | 5 | MCTP1 antisense RNA 1 [Source:HGNC Symbo |
| FAM81B | 5 | family with sequence similarity 81 member B |
| AC090071.1 | 5 | chromosome 14 open reading frame 166 (C14 |
| TTC37 | 5 | tetratricopeptide repeat domain 37 [Source:H |
| RNU6-308P | 5 | RNA, U6 small nuclear 308, pseudogene [Sour |
| ARSK | 5 | arylsulfatase family member K [Source:HGNC |
| GPR150 | 5 | G protein-coupled receptor 150 [Source:HGN |
| RFESD | 5 | Rieske Fe-S domain containing [Source:HGNC |
| SPATA9 | 5 | spermatogenesis associated 9 [Source:HGNC S |
| AC008840.1 | 5 | novel transcript |
| RHOBTB3 | 5 | Rho related BTB domain containing 3 [Source: |
| GLRX | 5 | glutaredoxin [Source:HGNC Symbol;Acc:HGNC |
| HSPD1P11 | 5 | heat shock protein family D (Hsp60) member |
| AC008592.3 | 5 | gamma-glutamyl cyclotransferase (GGCT) pse |
| AC008592.4 | 5 | novel transcript |
| LINC01554 | 5 | long intergenic non-protein coding RNA 1554 |

| | | |
|---|---|---|
| AC008592.5 | 5 | novel transcript |
| AC008592.1 | 5 | novel transcript |
| ELL2 | 5 | elongation factor for RNA polymerase II 2 [Sou |
| AC104123.1 | 5 | novel transcript, antisense to PCSK1 |
| AC008592.2 | 5 | novel transcript |
| FABP5P5 | 5 | fatty acid binding protein 5 pseudogene 5 [Sou |
| MIR583HG | 5 | MIR583 host gene [Source:HGNC Symbol;Acc: |
| MIR583 | 5 | microRNA 583 [Source:HGNC Symbol;Acc:HGN |
| RNU6-524P | 5 | RNA, U6 small nuclear 524, pseudogene [Sour |
| AC099509.1 | 5 | novel transcript |
| AC099509.2 | 5 | novel transcript |
| PCSK1 | 5 | proprotein convertase subtilisin/kexin type 1 [ |
| CAST | 5 | calpastatin [Source:HGNC Symbol;Acc:HGNC:1 |
| AC020900.1 | 5 | novel transcript |
| ERAP1 | 5 | endoplasmic reticulum aminopeptidase 1 [Sou |
| AC008906.1 | 5 | novel transcript |
| AC008906.2 | 5 | novel transcript, antisense to ERAP1 |
| AC009126.1 | 5 | novel transcript, antisense to LNPEP and ERAP |
| ERAP2 | 5 | endoplasmic reticulum aminopeptidase 2 [Sou |
| LNPEP | 5 | leucyl and cystinyl aminopeptidase [Source:H( |
| AC008850.1 | 5 | ribosomal protein S20 (RPS20) pseudogene |
| Y_RNA | 5 | Y RNA [Source:RFAM;Acc:RF00019] |
| SETP22 | 5 | SET pseudogene 22 [Source:HGNC Symbol;Acc |
| LIX1-AS1 | 5 | LIX1 and RIOK2 antisense RNA 1 [Source:HGN |
| LIX1 | 5 | limb and CNS expressed 1 [Source:HGNC Syml |
| RIOK2 | 5 | RIO kinase 2 [Source:HGNC Symbol;Acc:HGNC |
| RNU1-73P | 5 | RNA, U1 small nuclear 73, pseudogene [Sourc |
| AC008883.3 | 5 | novel transcript |
| AC008883.1 | 5 | novel transcript |
| AC008883.2 | 5 | novel transcript |
| AC074133.1 | 5 | ubiquitin-conjugating enzyme E2 variant 1 (UE |
| YTHDF1P1 | 5 | YTH domain family member 1 pseudogene 1 [ |
| AC106748.1 | 5 | NHP2 non-histone chromosome protein 2-like |
| LINC01340 | 5 | long intergenic non-protein coding RNA 1340 |
| MTCYBP40 | 5 | MT-CYB pseudogene 40 [Source:HGNC Symbo |
| AC112203.1 | 5 | novel DHHC-type zinc finger pseudogene |
| AC122697.1 | 5 | novel transcript |
| LINC02234 | 5 | long intergenic non-protein coding RNA 2234 |
| AC008834.1 | 5 | selenophosphate synthetase 1 (SEPHS1) pseu |
| LINC01846 | 5 | long intergenic non-protein coding RNA 1846 |
| AC116347.2 | 5 | RB1-inducible coiled-coil 1 (RB1CC1) pseudog |
| AC116347.1 | 5 | proteasome (prosome, macropain) activator s |
| AC233964.1 | 5 | actin, beta (ACTB) pseudogene |
| KRT8P32 | 5 | keratin 8 pseudogene 32 [Source:HGNC Symb |
| MRPS35P2 | 5 | mitochondrial ribosomal protein S35 pseudog |
| MTCO2P24 | 5 | MT-CO2 pseudogene 24 [Source:HGNC Symbo |
| MTCO1P24 | 5 | MT-CO1 pseudogene 24 [Source:HGNC Symbo |
| CTBP2P4 | 5 | C-terminal binding protein 2 pseudogene 4 [S |
| DDX18P4 | 5 | DEAD-box helicase 18 pseudogene 4 [Source:I |
| RGMB | 5 | repulsive guidance molecule BMP co-receptor |

| | | |
|---|---|---|
| RGMB-AS1 | 5 | RGMB antisense RNA 1 [Source:HGNC Symbol |
| AC008522.1 | 5 | novel transcript, antisense to RGMB |
| CSNK1A1P3 | 5 | casein kinase 1 alpha 1 pseudogene 3 [Source |
| CHD1 | 5 | chromodomain helicase DNA binding protein |
| RNU6-402P | 5 | RNA, U6 small nuclear 402, pseudogene [Sour |
| LINC02062 | 5 | long intergenic non-protein coding RNA 2062 |
| Y_RNA | 5 | Y RNA [Source:RFAM;Acc:RF00019] |
| AC022121.1 | 5 | ribosomal protein S9 (RPS9) pseudogene |
| AC114324.2 | 5 | WD repeat domain 70 (WDR70) pseudogene |
| AC114324.1 | 5 | putative POM121-like protein 1-like [Source:N |
| GUSBP8 | 5 | GUSB pseudogene 8 [Source:HGNC Symbol;Ac |
| LINC02113 | 5 | long intergenic non-protein coding RNA 2113 |
| AC008837.1 | 5 | Centromere protein O (CENP-O) pseudogene |
| AC008837.2 | 5 | Centromere protein O (CENP-O) pseudogene |
| AC092284.1 | 5 | cytokine receptor-like factor 3 (CRLF3) pseudo |
| EEF1A1P20 | 5 | eukaryotic translation elongation factor 1 alph |
| Y_RNA | 5 | Y RNA [Source:RFAM;Acc:RF00019] |
| MTCYBP22 | 5 | MT-CYB pseudogene 22 [Source:HGNC Symbo |
| MTND6P22 | 5 | MT-ND6 pseudogene 22 [Source:HGNC Symbo |
| MTND5P10 | 5 | MT-ND5 pseudogene 10 [Source:HGNC Symbo |
| MTND4P35 | 5 | MT-ND4 pseudogene 35 [Source:HGNC Symbo |
| MTND4LP5 | 5 | MT-ND4L pseudogene 5 [Source:HGNC Symbo |
| MTND3P19 | 5 | MT-ND3 pseudogene 19 [Source:HGNC Symbo |
| MTCO3P22 | 5 | MT-CO3 pseudogene 22 [Source:HGNC Symbo |
| MTATP6P2 | 5 | MT-ATP6 pseudogene 2 [Source:HGNC Symbo |
| MTCO2P22 | 5 | MT-CO2 pseudogene 22 [Source:HGNC Symbo |
| MTCO1P22 | 5 | MT-CO1 pseudogene 22 [Source:HGNC Symbo |
| AC092278.1 | 5 | glucuronidase, beta (GUSB) pseudogene |
| RNU6-1119P | 5 | RNA, U6 small nuclear 1119, pseudogene [Sou |
| AC113385.3 | 5 | glucuronidase, beta (GUSB) pseudogene |
| AC113385.2 | 5 | POM121 transmembrane nucleoporin (POM1 |
| AC113385.1 | 5 | novel transcript |
| AC021086.1 | 5 | novel transcript |
| FAM174A | 5 | family with sequence similarity 174 member A |
| AC027315.1 | 5 | novel transcript |
| RN7SKP62 | 5 | RN7SK pseudogene 62 [Source:HGNC Symbol; |
| ST8SIA4 | 5 | ST8 alpha-N-acetyl-neuraminide alpha-2,8-sia |
| MIR548P | 5 | microRNA 548p [Source:HGNC Symbol;Acc:HG |
| RN7SL802P | 5 | RNA, 7SL, cytoplasmic 802, pseudogene [Sour |
| OR7H2P | 5 | olfactory receptor family 7 subfamily H memb |
| AC117525.1 | 5 | novel ankyrin repeat-containing pseudogene |
| RNA5SP188 | 5 | RNA, 5S ribosomal pseudogene 188 [Source:H |
| AC008948.1 | 5 | novel transcript |
| SLCO4C1 | 5 | solute carrier organic anion transporter family |
| RN7SKP68 | 5 | RN7SK pseudogene 68 [Source:HGNC Symbol; |
| SLCO6A1 | 5 | solute carrier organic anion transporter family |
| AC094108.1 | 5 | novel transcript |
| LINC00492 | 5 | long intergenic non-protein coding RNA 492 [S |
| LINC00491 | 5 | long intergenic non-protein coding RNA 491 [S |
| AC099487.1 | 5 | novel transcript |

| | | |
|---|---|---|
| AC099487.2 | 5 | novel transcript |
| PAM | 5 | peptidylglycine alpha-amidating monooxygen |
| EIF3KP1 | 5 | eukaryotic translation initiation factor 3 subur |
| GIN1 | 5 | gypsy retrotransposon integrase 1 [Source:HG |
| PPIP5K2 | 5 | diphosphoinositol pentakisphosphate kinase 2 |
| AC011362.1 | 5 | novel transcript |
| C5orf30 | 5 | chromosome 5 open reading frame 30 [Source |
| AC010406.1 | 5 | novel transcript |
| PDZPH1P | 5 | PDZ and pleckstrin homology domains 1, pseu |
| LINC02115 | 5 | long intergenic non-protein coding RNA 2115 |
| NUDT12 | 5 | nudix hydrolase 12 [Source:HGNC Symbol;Acc |
| AC008505.1 | 5 | novel transcript |
| LINC02163 | 5 | long intergenic non-protein coding RNA 2163 |
| RNU1-140P | 5 | RNA, U1 small nuclear 140, pseudogene [Sour |
| RN7SL255P | 5 | RNA, 7SL, cytoplasmic 255, pseudogene [Sour |
| AC099520.1 | 5 | novel transcript |
| AC091931.1 | 5 | novel transcript |
| AC099520.2 | 5 | novel transcript |
| RNU6-334P | 5 | RNA, U6 small nuclear 334, pseudogene [Sour |
| AC091987.1 | 5 | novel transcript |
| RAB9BP1 | 5 | RAB9B, member RAS oncogene family pseudo |
| RNA5SP189 | 5 | RNA, 5S ribosomal pseudogene 189 [Source:H |
| AC027313.1 | 5 | RUN and FYVE domain containing 2 (RUFY2) p |
| AC114940.1 | 5 | zinc finger protein family pseudogene |
| AC114940.2 | 5 | NULL |
| LINC01950 | 5 | long intergenic non-protein coding RNA 1950 |
| PSMC1P5 | 5 | proteasome 26S subunit, ATPase 1 pseudoger |
| EFNA5 | 5 | ephrin A5 [Source:HGNC Symbol;Acc:HGNC:3 |
| AC024587.2 | 5 | novel transcript |
| AC024587.1 | 5 | ubiquitin specific peptidase 7 (herpes virus-as |
| RN7SL782P | 5 | RNA, 7SL, cytoplasmic 782, pseudogene [Sour |
| RN7SKP122 | 5 | RN7SK pseudogene 122 [Source:HGNC Symbo |
| FBXL17 | 5 | F-box and leucine rich repeat protein 17 [Sour |
| AC008462.1 | 5 | novel transcript |
| AC133134.1 | 5 | ribosomal protein S20 (RPS20) pseudogene |
| LINC01023 | 5 | long intergenic non-protein coding RNA 1023 |
| FER | 5 | FER tyrosine kinase [Source:HGNC Symbol;Acc |
| AC034207.1 | 5 | guanine nucleotide binding protein (G protein |
| AC008871.1 | 5 | novel transcript |
| Y_RNA | 5 | Y RNA [Source:RFAM;Acc:RF00019] |
| AC109481.1 | 5 | LSM2 homolog, U6 small nuclear RNA associa |
| RNU6-47P | 5 | RNA, U6 small nuclear 47, pseudogene [Sourc |
| GJA1P1 | 5 | gap junction protein alpha 1 pseudogene 1 [So |
| AC116428.1 | 5 | novel transcript, antisense to FER |
| AC008467.1 | 5 | novel transcript |
| PJA2 | 5 | praja ring finger ubiquitin ligase 2 [Source:HGI |
| AC010625.1 | 5 | DnaJ (Hsp40) homolog family pseudogene |
| AC091917.3 | 5 | novel transcript |
| AC091917.2 | 5 | novel transcript |
| AC091917.1 | 5 | novel transcript |

| | | |
|---|---|---|
| KRT18P42 | 5 | keratin 18 pseudogene 42 [Source:HGNC Sym |
| AC012603.1 | 5 | novel transcript |
| MAN2A1 | 5 | mannosidase alpha class 2A member 1 [Sourc |
| RN7SKP230 | 5 | RN7SK pseudogene 230 [Source:HGNC Symbo |
| AC008417.1 | 5 | cancer/testis antigen family 45, member A5 p |
| LINC01848 | 5 | long intergenic non-protein coding RNA 1848 |
| PGAM5P1 | 5 | PGAM family member 5, mitochondrial serine |
| TMEM232 | 5 | transmembrane protein 232 [Source:HGNC Sy |
| MIR548F3 | 5 | microRNA 548f-3 [Source:HGNC Symbol;Acc:H |
| AC008650.1 | 5 | NULL |
| SLC25A46 | 5 | solute carrier family 25 member 46 [Source:H |
| AC008782.1 | 5 | DEAD (Asp-Glu-Ala-Asp) box polypeptide 4 (DI |
| BCLAF1P1 | 5 | BCL2 associated transcription factor 1 pseudo |
| AC010395.1 | 5 | novel transcript |
| AC010395.2 | 5 | integrin alpha FG-GAP repeat containing 2 (ITI |
| TSLP | 5 | thymic stromal lymphopoietin [Source:HGNC |
| AC008572.1 | 5 | novel transcript |
| WDR36 | 5 | WD repeat domain 36 [Source:HGNC Symbol; |
| AC010468.3 | 5 | proline-rich nuclear receptor coactivator 2 (PI |
| AC010468.1 | 5 | ribosomal protein S3A (RPS3A) pseudogene |
| CAMK4 | 5 | calcium/calmodulin dependent protein kinase |
| AC010468.2 | 5 | novel transcript |
| AC010275.1 | 5 | novel transcript |
| STARD4 | 5 | StAR related lipid transfer domain containing |
| STARD4-AS1 | 5 | STARD4 antisense RNA 1 [Source:HGNC Symb |
| HMGN1P13 | 5 | high mobility group nucleosome binding dom |
| AC008967.1 | 5 | high-mobility group nucleosome binding dom |
| NREP | 5 | neuronal regeneration related protein [Sourc |
| RN7SKP57 | 5 | RN7SK pseudogene 57 [Source:HGNC Symbol; |
| HMGN1P14 | 5 | high mobility group nucleosome binding dom |
| NREP-AS1 | 5 | NREP antisense RNA 1 [Source:HGNC Symbol; |
| EPB41L4A | 5 | erythrocyte membrane protein band 4.1 like |
| EPB41L4A-AS1 | 5 | EPB41L4A antisense RNA 1 [Source:HGNC Sym |
| SNORA13 | 5 | small nucleolar RNA, H/ACA box 13 [Source:H |
| AC010261.1 | 5 | novel transcript |
| AC010261.2 | 5 | novel transcript |
| AC104126.1 | 5 | novel transcript |
| AC010265.1 | 5 | heat shock factor binding protein 1 (HSBP1) p |
| EPB41L4A-DT | 5 | EPB41L4A divergent transcript [Source:HGNC |
| HMGB3P16 | 5 | high mobility group box 3 pseudogene 16 [Sou |
| AC137549.1 | 5 | transmembrane protein 183 (TMEM183) pseu |
| LINC02200 | 5 | long intergenic non-protein coding RNA 2200 |
| APC | 5 | APC regulator of WNT signaling pathway [Sour |
| CBX3P3 | 5 | chromobox 3 pseudogene 3 [Source:HGNC Sy |
| RNU6-482P | 5 | RNA, U6 small nuclear 482, pseudogene [Sour |
| AC008575.1 | 5 | novel protein |
| SRP19 | 5 | signal recognition particle 19 [Source:HGNC S |
| REEP5 | 5 | receptor accessory protein 5 [Source:HGNC Sy |
| XBP1P1 | 5 | X-box binding protein 1 pseudogene 1 [Sourc |
| ZRSR2P1 | 5 | ZRSR2 pseudogene 1 [Source:HGNC Symbol;A |

| | | |
|---|---|---|
| DCP2 | 5 | decapping mRNA 2 [Source:HGNC Symbol;Acc |
| MCC | 5 | MCC regulator of WNT signaling pathway [Sou |
| AC079465.1 | 5 | novel transcript |
| TSSK1B | 5 | testis specific serine kinase 1B [Source:HGNC : |
| RNU4ATAC13P | 5 | RNA, U4atac small nuclear 13, pseudogene [S( |
| YTHDC2 | 5 | YTH domain containing 2 [Source:HGNC Symb |
| AC093240.1 | 5 | novel transcript |
| KCNN2 | 5 | potassium calcium-activated channel subfami |
| RN7SKP89 | 5 | RN7SK pseudogene 89 [Source:HGNC Symbol; |
| AC010230.1 | 5 | artifact |
| LINC01957 | 5 | long intergenic non-protein coding RNA 1957 |
| AC094104.2 | 5 | novel transcript |
| AC094104.1 | 5 | novel transcript |
| TRIM36 | 5 | tripartite motif containing 36 [Source:HGNC S |
| TRIM36-IT1 | 5 | TRIM36 intronic transcript 1 [Source:HGNC Sy |
| AC008494.2 | 5 | novel transcript |
| PGGT1B | 5 | protein geranylgeranyltransferase type I subu |
| AC008494.3 | 5 | novel transcript, antisense to PGGT1B |
| AC008494.1 | 5 | ribosomal protein S2 (RPS2) pseudogene |
| CCDC112 | 5 | coiled-coil domain containing 112 [Source:HG |
| HMGN1P15 | 5 | high mobility group nucleosome binding doma |
| CTNNA1P1 | 5 | catenin alpha 1 pseudogene 1 [Source:HGNC ! |
| AK3P4 | 5 | adenylate kinase 3 pseudogene 4 [Source:HGI |
| AC008628.2 | 5 | COX assembly mitochondrial protein homolog |
| AC008628.1 | 5 | high-mobility group box 1-like 1 pseudogene |
| CCT5P1 | 5 | chaperonin containing TCP1 subunit 5 pseudo |
| FEM1C | 5 | fem-1 homolog C [Source:HGNC Symbol;Acc:H |
| TMED7-TICAM2 | 5 | TMED7-TICAM2 readthrough [Source:HGNC S |
| TICAM2 | 5 | toll like receptor adaptor molecule 2 [Source:I |
| AC010226.1 | 5 | novel transcript |
| TMED7 | 5 | transmembrane p24 trafficking protein 7 [Sou |
| AC008549.2 | 5 | novel transcript |
| AC008549.1 | 5 | novel transcript |
| AC008549.3 | 5 | H3 histone, family 3A pseudogene |
| RNU2-49P | 5 | RNA, U2 small nuclear 49, pseudogene [Sourc |
| CDO1 | 5 | cysteine dioxygenase type 1 [Source:HGNC Sy |
| ATG12 | 5 | autophagy related 12 [Source:HGNC Symbol;A |
| AP3S1 | 5 | adaptor related protein complex 3 subunit sig |
| LINCADL | 5 | lincRNA adipogenesis and lipogenesis associat |
| LVRN | 5 | laeverin [Source:HGNC Symbol;Acc:HGNC:269 |
| DDX43P1 | 5 | DEAD-box helicase 43 pseudogene 1 [Source:I |
| ARL14EPL | 5 | ADP ribosylation factor like GTPase 14 effecto |
| AC034236.1 | 5 | ribosomal protein S25 (RPS25) pseudogene |
| AC034236.3 | 5 | novel transcript |
| AC034236.2 | 5 | novel transcript, antisense to COMMD10 |
| COMMD10 | 5 | COMM domain containing 10 [Source:HGNC S |
| RNU6-644P | 5 | RNA, U6 small nuclear 644, pseudogene [Sour |
| HMGN2P27 | 5 | high mobility group nucleosomal binding dom |
| AC018752.1 | 5 | novel transcript |
| SEMA6A | 5 | semaphorin 6A [Source:HGNC Symbol;Acc:HG |

| | | |
|---|---|---|
| SEMA6A-AS1 | 5 | SEMA6A antisense RNA 1 [Source:HGNC Symb |
| RPS14P8 | 5 | ribosomal protein S14 pseudogene 8 [Source: |
| SEMA6A-AS2 | 5 | SEMA6A antisense RNA 2 [Source:HGNC Symb |
| RPS17P2 | 5 | ribosomal protein S17 pseudogene 2 [Source: |
| LINC02214 | 5 | long intergenic non-protein coding RNA 2214 |
| AC010267.1 | 5 | novel transcript |
| AC093534.1 | 5 | thymosin beta 15a (TMSB15A) pseudogene |
| AC093534.2 | 5 | novel transcript |
| RPL35AP15 | 5 | ribosomal protein L35a pseudogene 15 [Sourc |
| AC093295.1 | 5 | Novel transcript |
| LINC00992 | 5 | long intergenic non-protein coding RNA 992 [S |
| AC114322.1 | 5 | protein tyrosine phosphatase, receptor type f |
| LINC02147 | 5 | long intergenic non-protein coding RNA 2147 |
| RPL7L1P4 | 5 | ribosomal protein L7 like 1 pseudogene 4 [Sou |
| LINC02208 | 5 | long intergenic non-protein coding RNA 2208 |
| LINC02148 | 5 | long intergenic non-protein coding RNA 2148 |
| LINC02216 | 5 | long intergenic non-protein coding RNA 2216 |
| AC114311.1 | 5 | novel transcript |
| LINC02215 | 5 | long intergenic non-protein coding RNA 2215 |
| RNU7-34P | 5 | RNA, U7 small nuclear 34 pseudogene [Source |
| AC093206.1 | 5 | novel transcript |
| DTWD2 | 5 | DTW domain containing 2 [Source:HGNC Sym |
| PTMAP2 | 5 | prothymosin alpha pseudogene 2 [Source:HGI |
| MIR1244-2 | 5 | microRNA 1244-2 [Source:HGNC Symbol;Acc:I |
| AC008629.1 | 5 | novel transcript |
| RNU6-373P | 5 | RNA, U6 small nuclear 373, pseudogene [Sour |
| AC008629.2 | 5 | N-acetylneuraminic acid phosphatase (NANP) |
| AC008629.3 | 5 | cation channel, sperm associated 2 (CATSPER |
| DMXL1 | 5 | Dmx like 1 [Source:HGNC Symbol;Acc:HGNC:2 |
| AC118465.1 | 5 | septin family pseudogene |
| LAMTOR3P2 | 5 | late endosomal/lysosomal adaptor, MAPK and |
| RNU6-701P | 5 | RNA, U6 small nuclear 701, pseudogene [Sour |
| MIR5706 | 5 | microRNA 5706 [Source:HGNC Symbol;Acc:HG |
| TNFAIP8 | 5 | TNF alpha induced protein 8 [Source:HGNC Sy |
| RN7SL174P | 5 | RNA, 7SL, cytoplasmic 174, pseudogene [Sour |
| RNA5SP190 | 5 | RNA, 5S ribosomal pseudogene 190 [Source:H |
| HSD17B4 | 5 | hydroxysteroid 17-beta dehydrogenase 4 [Sou |
| AC010409.1 | 5 | ribosomal protein L21 (RPL21) pseudogene |
| FABP5P6 | 5 | fatty acid binding protein 5 pseudogene 6 [So |
| FAM170A | 5 | family with sequence similarity 170 member A |
| AC010409.2 | 5 | tubulin, alpha 1c pseudogene |
| AC008550.1 | 5 | mitochondrial ribosomal protein L10 pseudog |
| AC008574.1 | 5 | novel transcript |
| RNU6-718P | 5 | RNA, U6 small nuclear 718, pseudogene [Sour |
| AC113418.1 | 5 | novel transcript |
| PRR16 | 5 | proline rich 16 [Source:HGNC Symbol;Acc:HGI |
| PRELID3BP8 | 5 | PRELI domain containing 3B pseudogene 8 [So |
| RNU4-69P | 5 | RNA, U4 small nuclear 69, pseudogene [Sourc |
| AC114284.1 | 5 | novel transcript |
| AC008565.1 | 5 | serum response factor binding protein 1 (SRFE |

| | | |
|---|---|---|
| AC113352.1 | 5 | novel zinc finger pseudogene |
| AC008568.1 | 5 | novel transcript |
| AC010350.1 | 5 | Rho GTPase activating protein 5 (ARHGAP5) p |
| AC008568.2 | 5 | tec |
| RPL23AP44 | 5 | ribosomal protein L23a pseudogene 44 [Sourc |
| AC106806.1 | 5 | ribosomal protein L18 (RPL18) pseudogene |
| AC106806.2 | 5 | ribosomal L24 domain containing 1 (RSL24D1) |
| FTMT | 5 | ferritin mitochondrial [Source:HGNC Symbol;A |
| SRFBP1 | 5 | serum response factor binding protein 1 [Sour |
| LOX | 5 | lysyl oxidase [Source:HGNC Symbol;Acc:HGNC |
| AC010255.1 | 5 | novel transcript |
| ZNF474 | 5 | zinc finger protein 474 [Source:HGNC Symbol; |
| AC010255.3 | 5 | zinc finger protein 474-like [Source:NCBI gene |
| AC010255.2 | 5 | novel transcript |
| AC113349.2 | 5 | novel transcript, antisense to SNCAIP |
| SNCAIP | 5 | synuclein alpha interacting protein [Source:HC |
| AC113349.1 | 5 | novel transcript |
| AC119150.1 | 5 | novel transcript |
| AC119150.2 | 5 | novel anceint vomeronasal 1 receptor pseudo |
| AC022101.1 | 5 | novel transcript |
| LINC02201 | 5 | long intergenic non-protein coding RNA 2201 |
| ARGFXP1 | 5 | arginine-fifty homeobox pseudogene 1 [Sourc |
| SNX2 | 5 | sorting nexin 2 [Source:HGNC Symbol;Acc:HGI |
| AC008669.1 | 5 | novel transcript, overlapping SNX2 |
| SNX24 | 5 | sorting nexin 24 [Source:HGNC Symbol;Acc:HC |
| RN7SL689P | 5 | RNA, 7SL, cytoplasmic 689, pseudogene [Sour |
| PPIC | 5 | peptidylprolyl isomerase C [Source:HGNC Sym |
| AC106786.2 | 5 | novel transcript |
| RN7SL711P | 5 | RNA, 7SL, cytoplasmic 711, pseudogene [Sour |
| AC106786.1 | 5 | novel transcript |
| PRDM6 | 5 | PR/SET domain 6 [Source:HGNC Symbol;Acc:H |
| AC010493.1 | 5 | SMT3 suppressor of mif two 3 homolog 1 (S. c |
| CEP120 | 5 | centrosomal protein 120 [Source:HGNC Symb |
| KRT8P33 | 5 | keratin 8 pseudogene 33 [Source:HGNC Symb |
| HMGB3P17 | 5 | high mobility group box 3 pseudogene 17 [Sou |
| AC026422.1 | 5 | tec |
| CSNK1G3 | 5 | casein kinase 1 gamma 3 [Source:HGNC Symb |
| KRT18P16 | 5 | keratin 18 pseudogene 16 [Source:HGNC Sym |
| LINC01170 | 5 | long intergenic non-protein coding RNA 1170 |
| HMGB1P29 | 5 | high mobility group box 1 pseudogene 29 [Sou |
| AC016556.1 | 5 | novel transcript |
| AC025465.2 | 5 | novel transcript |
| AC025465.4 | 5 | novel transcript |
| AC025465.3 | 5 | novel transcript |
| AC025465.1 | 5 | novel transcript |
| ZNF608 | 5 | zinc finger protein 608 [Source:HGNC Symbol; |
| AC112196.1 | 5 | novel transcript, antisense to ZNF608 |
| AC113398.2 | 5 | novel transcript |
| AC113398.1 | 5 | ribosomal protein L28 (RPL28) pseudogene |
| LINC02240 | 5 | long intergenic non-protein coding RNA 2240 |

| | | |
|---|---|---|
| AC109464.1 | 5 | novel transcript |
| AC109464.2 | 5 | novel transcript |
| AC109458.2 | 5 | novel transcript |
| AC109458.1 | 5 | novel pseudogene |
| HMGB1P22 | 5 | high mobility group box 1 pseudogene 22 [Sou |
| AC116362.1 | 5 | novel transcript |
| RN7SKP117 | 5 | RN7SK pseudogene 117 [Source:HGNC Symbo |
| AC109471.1 | 5 | novel transcript |
| AC116362.2 | 5 | Novel transcript |
| AC010587.2 | 5 | ribosomal protein S3a pseudogene |
| AC010587.1 | 5 | chromosome 1 open reading frame 146 pseuc |
| RPSAP37 | 5 | ribosomal protein SA pseudogene 37 [Source: |
| LINC02039 | 5 | long intergenic non-protein coding RNA 2039 |
| GRAMD2B | 5 | GRAM domain containing 2B [Source:HGNC Sy |
| AC093535.1 | 5 | novel transcript, antisense to GRAMD3 |
| AC093535.2 | 5 | tec |
| ALDH7A1 | 5 | aldehyde dehydrogenase 7 family member A1 |
| RNU6-290P | 5 | RNA, U6 small nuclear 290, pseudogene [Sour |
| PHAX | 5 | phosphorylated adaptor for RNA export [Sour |
| AC099513.1 | 5 | ribosomal protein, large, P1 (RPLP1) pseudoge |
| TEX43 | 5 | testis expressed 43 [Source:HGNC Symbol;Acc |
| BOLA3P3 | 5 | bolA family member 3 pseudogene 3 [Source: |
| HSPE1P10 | 5 | heat shock protein family E (Hsp10) member |
| LMNB1-DT | 5 | LMNB1 divergent transcript [Source:HGNC Syr |
| RNU6-752P | 5 | RNA, U6 small nuclear 752, pseudogene [Sour |
| LMNB1 | 5 | lamin B1 [Source:HGNC Symbol;Acc:HGNC:66 |
| MARCH3 | 5 | membrane associated ring-CH-type finger 3 [S |
| C5orf63 | 5 | chromosome 5 open reading frame 63 [Sourc |
| AC005915.1 | 5 | tec |
| MRPS5P3 | 5 | mitochondrial ribosomal protein S5 pseudoge |
| AC011416.1 | 5 | RNA binding motif protein 15B pseudogene |
| SELENOTP2 | 5 | selenoprotein T pseudogene 2 [Source:HGNC |
| AC011416.2 | 5 | eukaryotic translation elongation factor 1 alpl |
| AC011416.3 | 5 | novel transcript |
| AC011416.4 | 5 | novel transcript |
| MEGF10 | 5 | multiple EGF like domains 10 [Source:HGNC S |
| AC010424.1 | 5 | ARP3 actin-related protein 3 homolog (yeast) |
| AC010424.2 | 5 | novel transcript |
| HNRNPKP1 | 5 | heterogeneous nuclear ribonucleoprotein K p |
| PRRC1 | 5 | proline rich coiled-coil 1 [Source:HGNC Symbo |
| AC010424.3 | 5 | novel transcript |
| CTXN3 | 5 | cortexin 3 [Source:HGNC Symbol;Acc:HGNC:3 |
| AC022118.1 | 5 | novel transcript |
| CCDC192 | 5 | coiled-coil domain containing 192 [Source:HG |
| CUL1P1 | 5 | cullin 1 pseudogene 1 [Source:HGNC Symbol;/ |
| AC068658.1 | 5 | novel transcript |
| LINC01184 | 5 | long intergenic non-protein coding RNA 1184 |
| SLC12A2 | 5 | solute carrier family 12 member 2 [Source:HG |
| KDELC1P1 | 5 | KDEL motif containing 1 pseudogene 1 [Sourc |
| FBN2 | 5 | fibrillin 2 [Source:HGNC Symbol;Acc:HGNC:36 |

| | | |
|---|---|---|
| Y_RNA | 5 | Y RNA [Source:RFAM;Acc:RF00019] |
| SLC27A6 | 5 | solute carrier family 27 member 6 [Source:HG |
| AC008588.1 | 5 | novel transcript |
| ISOC1 | 5 | isochorismatase domain containing 1 [Source: |
| MIR4633 | 5 | microRNA 4633 [Source:HGNC Symbol;Acc:HG |
| AC008679.1 | 5 | novel transcript |
| MIR4460 | 5 | microRNA 4460 [Source:HGNC Symbol;Acc:HG |
| ADAMTS19-AS1 | 5 | ADAMTS19 antisense RNA 1 [Source:HGNC Sy |
| ADAMTS19 | 5 | ADAM metallopeptidase with thrombospondi |
| AC008591.1 | 5 | novel transcript |
| MINAR2 | 5 | membrane integral NOTCH2 associated recep |
| CHSY3 | 5 | chondroitin sulfate synthase 3 [Source:HGNC |
| RNU6ATAC10P | 5 | RNA, U6atac small nuclear 10, pseudogene [So |
| RNA5SP191 | 5 | RNA, 5S ribosomal pseudogene 191 [Source:H |
| HSPA8P4 | 5 | heat shock protein family A (Hsp70) member |
| RNU7-53P | 5 | RNA, U7 small nuclear 53 pseudogene [Source |
| ARL2BPP4 | 5 | ADP ribosylation factor like GTPase 2 binding |
| AC005741.1 | 5 | frataxin (FXN) pseudogene |
| AC006525.1 | 5 | novel transcript |
| AC113367.1 | 5 | ribosomal protein L11 (RPL11) pseudogene |
| AC113367.3 | 5 | splicing factor 4 (SF4) pseudogene |
| AC113367.2 | 5 | leucine rich repeat containing 59 (LRRC59) pse |
| HINT1 | 5 | histidine triad nucleotide binding protein 1 [So |
| LYRM7 | 5 | LYR motif containing 7 [Source:HGNC Symbol; |
| CDC42SE2 | 5 | CDC42 small effector 2 [Source:HGNC Symbol |
| AC004777.1 | 5 | TEC |
| RAPGEF6 | 5 | Rap guanine nucleotide exchange factor 6 [So |
| AC008695.1 | 5 | novel protein |
| AC008497.1 | 5 | NULL |
| FNIP1 | 5 | folliculin interacting protein 1 [Source:HGNC S |
| AC008695.2 | 5 | actin, beta pseudogene 4 |
| AC005593.1 | 5 | TEC |
| MEIKIN | 5 | meiotic kinetochore factor [Source:HGNC Sym |
| AC034228.4 | 5 | novel transcript |
| AC034228.3 | 5 | novel transcript |
| ACSL6 | 5 | acyl-CoA synthetase long chain family membe |
| AC034228.1 | 5 | novel transcript |
| AC034228.2 | 5 | novel transcript |
| IL3 | 5 | interleukin 3 [Source:HGNC Symbol;Acc:HGNC |
| CSF2 | 5 | colony stimulating factor 2 [Source:HGNC Sym |
| AC034216.1 | 5 | NULL |
| AC063976.3 | 5 | procollagen-proline, 2-oxoglutarate 4-dioxyge |
| AC063976.4 | 5 | Chediak-Higashi syndrome 1 (CHS10) pseudog |
| AC063976.1 | 5 | novel transcript |
| P4HA2-AS1 | 5 | P4HA2 antisense RNA 1 [Source:HGNC Symbo |
| P4HA2 | 5 | prolyl 4-hydroxylase subunit alpha 2 [Source:H |
| AC063976.2 | 5 | cytoplasmic linker associated protein 2 (CLASI |
| MIR6830 | 5 | microRNA 6830 [Source:HGNC Symbol;Acc:HG |
| PDLIM4 | 5 | PDZ and LIM domain 4 [Source:HGNC Symbol; |
| SLC22A4 | 5 | solute carrier family 22 member 4 [Source:HG |

| | | |
|---|---|---|
| MIR3936HG | 5 | MIR3936 host gene [Source:HGNC Symbol;Acc |
| MIR3936 | 5 | microRNA 3936 [Source:HGNC Symbol;Acc:HG |
| SLC22A5 | 5 | solute carrier family 22 member 5 [Source:HG |
| C5orf56 | 5 | chromosome 5 open reading frame 56 [Source |
| AC116366.3 | 5 | novel protein |
| AC116366.2 | 5 | novel transcript |
| Y_RNA | 5 | Y RNA [Source:RFAM;Acc:RF00019] |
| AC116366.1 | 5 | novel transcript |
| IRF1 | 5 | interferon regulatory factor 1 [Source:HGNC S |
| IL5 | 5 | interleukin 5 [Source:HGNC Symbol;Acc:HGNC |
| RAD50 | 5 | RAD50 double strand break repair protein [So |
| TH2LCRR | 5 | T helper type 2 locus control region associated |
| IL13 | 5 | interleukin 13 [Source:HGNC Symbol;Acc:HGN |
| IL4 | 5 | interleukin 4 [Source:HGNC Symbol;Acc:HGNC |
| AC004039.1 | 5 | novel transcript, antisense to KIF3A |
| KIF3A | 5 | kinesin family member 3A [Source:HGNC Sym |
| CCNI2 | 5 | cyclin I family member 2 [Source:HGNC Symbo |
| SEPTIN8 | 5 | septin 8 [Source:HGNC Symbol;Acc:HGNC:165 |
| SOWAHA | 5 | sosondowah ankyrin repeat domain family me |
| AC004775.1 | 5 | novel transcript |
| SHROOM1 | 5 | shroom family member 1 [Source:HGNC Symb |
| RNA5SP192 | 5 | RNA, 5S ribosomal pseudogene 192 [Source:H |
| GDF9 | 5 | growth differentiation factor 9 [Source:HGNC |
| UQCRQ | 5 | ubiquinol-cytochrome c reductase complex III |
| LEAP2 | 5 | liver enriched antimicrobial peptide 2 [Source |
| AFF4 | 5 | AF4/FMR2 family member 4 [Source:HGNC Sy |
| AC010240.2 | 5 | eukaryotic translation elongation factor 1 alph |
| Y_RNA | 5 | Y RNA [Source:RFAM;Acc:RF00019] |
| ATP6V0E1P1 | 5 | ATPase H+ transporting V0 subunit e1 pseudo |
| AC010240.3 | 5 | novel transcript |
| AC010240.1 | 5 | putative nucleic acid binding protein RY-1 (RY |
| Y_RNA | 5 | Y RNA [Source:RFAM;Acc:RF00019] |
| ZCCHC10 | 5 | zinc finger CCHC-type containing 10 [Source:H |
| AC113410.1 | 5 | NADH dehydrogenase (ubiquinone) 1 alpha su |
| AC113410.5 | 5 | novel transcript, antisense to ZCCHC10 |
| AC113410.4 | 5 | novel transcript |
| HSPA4 | 5 | heat shock protein family A (Hsp70) member |
| AC113410.3 | 5 | TEC |
| AC113410.2 | 5 | novel transcript |
| AC010307.2 | 5 | novel transcript |
| AC010307.1 | 5 | ribosomal protein L6 (RPL6) pseudogene |
| FSTL4 | 5 | follistatin like 4 [Source:HGNC Symbol;Acc:HG |
| AC010307.3 | 5 | novel transcript |
| AC010307.4 | 5 | novel transcript |
| AC010608.1 | 5 | novel transcript |
| MIR1289-2 | 5 | microRNA 1289-2 [Source:HGNC Symbol;Acc: |
| AC010608.3 | 5 | TEC |
| AC010608.2 | 5 | TEC |
| AC005178.1 | 5 | DnaJ (Hsp40) homolog, subfamily C, member |
| WSPAR | 5 | WNT signaling pathway activating non-coding |

| | | |
|---|---|---|
| C5orf15 | 5 | chromosome 5 open reading frame 15 [Sourc |
| VDAC1 | 5 | voltage dependent anion channel 1 [Source:H |
| AC008608.2 | 5 | novel transcript |
| AC008608.1 | 5 | chromosome 7 open reading frame 73 (C7orf |
| TCF7 | 5 | transcription factor 7 [Source:HGNC Symbol;A |
| SKP1 | 5 | S-phase kinase associated protein 1 [Source:H |
| AC104109.3 | 5 | novel protein |
| PPP2CA | 5 | protein phosphatase 2 catalytic subunit alpha |
| AC104109.4 | 5 | novel transcript |
| MIR3661 | 5 | microRNA 3661 [Source:HGNC Symbol;Acc:H |
| AC104109.2 | 5 | novel transcript |
| AC104109.1 | 5 | ribosomal protein S13 (RPS13) pseudogene |
| CDKL3 | 5 | cyclin dependent kinase like 3 [Source:HGNC |
| UBE2B | 5 | ubiquitin conjugating enzyme E2 B [Source:H |
| AC109454.4 | 5 | TEC |
| AC109454.2 | 5 | novel transcript |
| CDKN2AIPNL | 5 | CDKN2A interacting protein N-terminal like [S |
| AC109454.1 | 5 | ribosomal protein S10 (RPS10) pseudogene |
| AC109454.3 | 5 | novel transcript |
| AC005355.1 | 5 | novel transcript |
| RNU6-456P | 5 | RNA, U6 small nuclear 456, pseudogene [Sour |
| LINC01843 | 5 | long intergenic non-protein coding RNA 1843 |
| RN7SL541P | 5 | RNA, 7SL, cytoplasmic 541, pseudogene [Sour |
| AC005355.2 | 5 | TEC |
| JADE2 | 5 | jade family PHD finger 2 [Source:HGNC Symbo |
| SAR1B | 5 | secretion associated Ras related GTPase 1B [S |
| RNU6-1311P | 5 | RNA, U6 small nuclear 1311, pseudogene [Sou |
| SEC24A | 5 | SEC24 homolog A, COPII coat complex compo |
| RNU6-1164P | 5 | RNA, U6 small nuclear 1164, pseudogene [Sou |
| RNU6-757P | 5 | RNA, U6 small nuclear 757, pseudogene [Sour |
| CAMLG | 5 | calcium modulating ligand [Source:HGNC Sym |
| DDX46 | 5 | DEAD-box helicase 46 [Source:HGNC Symbol; |
| AC010301.1 | 5 | ribosomal protein L34 (RPL34) pseudogene |
| C5orf24 | 5 | chromosome 5 open reading frame 24 [Sourc |
| AC006077.1 | 5 | ribosomal protein L31 (RPL31) pseudogene |
| TXNDC15 | 5 | thioredoxin domain containing 15 [Source:HG |
| PCBD2 | 5 | pterin-4 alpha-carbinolamine dehydratase 2 [ |
| AC006077.2 | 5 | TEC |
| MTCYBP18 | 5 | MT-CYB pseudogene 18 [Source:HGNC Symbo |
| MTND6P4 | 5 | MT-ND6 pseudogene 4 [Source:HGNC Symbol |
| MTND5P11 | 5 | MT-ND5 pseudogene 11 [Source:HGNC Symbo |
| MTND4P12 | 5 | MT-ND4 pseudogene 12 [Source:HGNC Symbo |
| MTND4LP30 | 5 | MT-ND4L pseudogene 30 [Source:HGNC Symb |
| AC008670.1 | 5 | NULL |
| MTND3P25 | 5 | MT-ND3 pseudogene 25 [Source:HGNC Symbo |
| CATSPER3 | 5 | cation channel sperm associated 3 [Source:H |
| PITX1 | 5 | paired like homeodomain 1 [Source:HGNC Sy |
| C5orf66 | 5 | chromosome 5 open reading frame 66 [Sourc |
| AC008406.3 | 5 | novel transcript, antisense to PITX1 |
| C5orf66-AS1 | 5 | C5orf66 antisense RNA 1 [Source:HGNC Symb |

| | | |
|---|---|---|
| AC008406.2 | 5 | novel transcript |
| AC008406.1 | 5 | novel transcript |
| C5orf66-AS2 | 5 | C5orf66 antisense RNA 2 [Source:HGNC Symb |
| H2AFY | 5 | H2A histone family member Y [Source:HGNC S |
| AC026691.1 | 5 | novel transcript, antisense to H2AFY |
| DCANP1 | 5 | dendritic cell associated nuclear protein [Sour |
| TIFAB | 5 | TIFA inhibitor [Source:HGNC Symbol;Acc:HGN |
| AC022092.1 | 5 | novel transcript, antisense to TIFAB |
| NEUROG1 | 5 | neurogenin 1 [Source:HGNC Symbol;Acc:HGN |
| AC034206.1 | 5 | novel transcript |
| CXCL14 | 5 | C-X-C motif chemokine ligand 14 [Source:HGN |
| SLC25A48 | 5 | solute carrier family 25 member 48 [Source:H |
| SLC25A48-AS1 | 5 | SLC25A48 antisense RNA 1 [Source:HGNC Sym |
| AC011431.1 | 5 | nucleotide binding protein 1 (MinD homolog, |
| MIR5692C1 | 5 | microRNA 5692c-1 [Source:HGNC Symbol;Acc |
| AC114296.1 | 5 | uncharacterized LOC107986453 [Source:NCBI |
| IL9 | 5 | interleukin 9 [Source:HGNC Symbol;Acc:HGNC |
| AC002428.2 | 5 | novel transcript |
| AC002428.1 | 5 | novel transcript |
| LECT2 | 5 | leukocyte cell derived chemotaxin 2 [Source:H |
| FBXL21P | 5 | F-box and leucine rich repeat protein 21, pseu |
| TGFBI | 5 | transforming growth factor beta induced [Sou |
| VTRNA2-1 | 5 | vault RNA 2-1 [Source:HGNC Symbol;Acc:HGN |
| Vault | 5 | Vault RNA [Source:RFAM;Acc:RF00006] |
| SMAD5-AS1 | 5 | SMAD5 antisense RNA 1 [Source:HGNC Symbo |
| AC009014.4 | 5 | NULL |
| SMAD5 | 5 | SMAD family member 5 [Source:HGNC Symbo |
| AC009014.2 | 5 | NULL |
| AC009014.3 | 5 | NULL |
| AC009014.1 | 5 | NULL |
| SMIM32 | 5 | small integral membrane protein 32 [Source:H |
| TRPC7 | 5 | transient receptor potential cation channel su |
| TRPC7-AS1 | 5 | TRPC7 antisense RNA 1 [Source:HGNC Symbol |
| TRPC7-AS2 | 5 | TRPC7 antisense RNA 2 [Source:HGNC Symbol |
| AC063980.1 | 5 | novel transcript |
| HSPD1P18 | 5 | heat shock protein family D (Hsp60) member |
| HNRNPA1P13 | 5 | heterogeneous nuclear ribonucleoprotein A1 |
| AC112178.1 | 5 | novel transcript |
| AC109439.2 | 5 | novel transcript |
| AC109439.1 | 5 | ankyrin repeat domain 49 (ANKRD49) pseudog |
| RNA5SP193 | 5 | RNA, 5S ribosomal pseudogene 193 [Source:H |
| SPOCK1 | 5 | SPARC (osteonectin), cwcv and kazal like dom |
| KLHL3 | 5 | kelch like family member 3 [Source:HGNC Sym |
| MIR874 | 5 | microRNA 874 [Source:HGNC Symbol;Acc:HGN |
| HNRNPA0 | 5 | heterogeneous nuclear ribonucleoprotein A0 |
| AC106791.3 | 5 | zinc finger BED-type containing 9 (ZBED9) pse |
| AC106791.2 | 5 | novel transcript |
| NPY6R | 5 | neuropeptide Y receptor Y6 (pseudogene) [So |
| AC106791.1 | 5 | novel transcript |
| MYOT | 5 | myotilin [Source:HGNC Symbol;Acc:HGNC:123 |

| | | |
|---|---|---|
| PKD2L2 | 5 | polycystin 2 like 2, transient receptor potentia |
| FAM13B | 5 | family with sequence similarity 13 member B |
| AC113382.1 | 5 | novel transcript, antisense to FAM13B |
| AC113382.2 | 5 | novel transcript |
| RNU6-1148P | 5 | RNA, U6 small nuclear 1148, pseudogene [Sou |
| WNT8A | 5 | Wnt family member 8A [Source:HGNC Symbo |
| NME5 | 5 | NME/NM23 family member 5 [Source:HGNC S |
| RNU6-460P | 5 | RNA, U6 small nuclear 460, pseudogene [Sour |
| RNU6-888P | 5 | RNA, U6 small nuclear 888, pseudogene [Sour |
| BRD8 | 5 | bromodomain containing 8 [Source:HGNC Syr |
| KIF20A | 5 | kinesin family member 20A [Source:HGNC Syr |
| CDC23 | 5 | cell division cycle 23 [Source:HGNC Symbol;Ac |
| GFRA3 | 5 | GDNF family receptor alpha 3 [Source:HGNC S |
| RPS27AP18 | 5 | ribosomal protein S27a pseudogene 18 [Sourc |
| CDC25C | 5 | cell division cycle 25C [Source:HGNC Symbol;A |
| FAM53C | 5 | family with sequence similarity 53 member C |
| AC104116.1 | 5 | novel transcript |
| KDM3B | 5 | lysine demethylase 3B [Source:HGNC Symbol; |
| REEP2 | 5 | receptor accessory protein 2 [Source:HGNC Sy |
| AC113403.1 | 5 | tubulin tyrosine ligase-like family, member 12 |
| EGR1 | 5 | early growth response 1 [Source:HGNC Symbo |
| RPL7P19 | 5 | ribosomal protein L7 pseudogene 19 [Source: |
| ETF1 | 5 | eukaryotic translation termination factor 1 [Sc |
| HSPA9 | 5 | heat shock protein family A (Hsp70) member 9 |
| SNORD63B | 5 | small nucleolar RNA, C/D box 63B [Source:HG |
| SNORD63 | 5 | small nucleolar RNA, C/D box 63 [Source:HGN |
| CTNNA1 | 5 | catenin alpha 1 [Source:HGNC Symbol;Acc:HG |
| AC113340.1 | 5 | ribosomal protein L10a (RPL10A) pseudogene |
| AC113340.2 | 5 | NULL |
| AC034243.1 | 5 | novel transcript |
| LRRTM2 | 5 | leucine rich repeat transmembrane neuronal 2 |
| SIL1 | 5 | SIL1 nucleotide exchange factor [Source:HGN( |
| RNA5SP194 | 5 | RNA, 5S ribosomal pseudogene 194 [Source:H |
| AC011405.2 | 5 | NULL |
| AC011405.1 | 5 | novel transcript |
| RPL12P21 | 5 | ribosomal protein L12 pseudogene 21 [Source |
| RNU6-572P | 5 | RNA, U6 small nuclear 572, pseudogene [Sour |
| MATR3 | 5 | matrin 3 [Source:NCBI gene;Acc:9782] |
| SNHG4 | 5 | small nucleolar RNA host gene 4 [Source:HGN |
| SNORA74D | 5 | small nucleolar RNA, H/ACA box 74D [Source: |
| SNORA74A | 5 | small nucleolar RNA, H/ACA box 74A [Source: |
| MATR3 | 5 | matrin 3 [Source:HGNC Symbol;Acc:HGNC:69 |
| RNA5SP195 | 5 | RNA, 5S ribosomal pseudogene 195 [Source:H |
| RN7SKP64 | 5 | RN7SK pseudogene 64 [Source:HGNC Symbol; |
| PAIP2 | 5 | poly(A) binding protein interacting protein 2 [ |
| AC135457.1 | 5 | novel transcript, antisense to PAIP2 |
| SLC23A1 | 5 | solute carrier family 23 member 1 [Source:HG |
| MZB1 | 5 | marginal zone B and B1 cell specific protein [S |
| PROB1 | 5 | proline rich basic protein 1 [Source:HGNC Syn |
| SPATA24 | 5 | spermatogenesis associated 24 [Source:HGNC |

| | | |
|---|---|---|
| DNAJC18 | 5 | DnaJ heat shock protein family (Hsp40) memb |
| AC142391.1 | 5 | novel transcript |
| RNU5B-4P | 5 | RNA, U5B small nuclear 4, pseudogene [Sourc |
| ECSCR | 5 | endothelial cell surface expressed chemotaxis |
| SMIM33 | 5 | small integral membrane protein 33 [Source:H |
| TMEM173 | 5 | transmembrane protein 173 [Source:HGNC Sy |
| AC138517.1 | 5 | chromosome 1 open reading frame 57 pseudo |
| NCOA4P4 | 5 | nuclear receptor coactivator 4 pseudogene 4 |
| AC010378.2 | 5 | novel transcript |
| UBE2D2 | 5 | ubiquitin conjugating enzyme E2 D2 [Source:H |
| Y_RNA | 5 | Y RNA [Source:RFAM;Acc:RF00019] |
| AC010378.1 | 5 | histone cluster 3, H3 (HIST3H3) pseudogene |
| AC113361.1 | 5 | novel transcript |
| CXXC5 | 5 | CXXC finger protein 5 [Source:HGNC Symbol;A |
| CXXC5-AS1 | 5 | CXXC5 antisense RNA 1 [Source:HGNC Symbol |
| AC008667.1 | 5 | novel transcript |
| RNU6-236P | 5 | RNA, U6 small nuclear 236, pseudogene [Sour |
| PSD2-AS1 | 5 | PSD2 antisense RNA 1 [Source:HGNC Symbol; |
| AC008667.3 | 5 | novel transcript |
| AC008667.4 | 5 | novel transcript |
| PSD2 | 5 | pleckstrin and Sec7 domain containing 2 [Sour |
| NRG2 | 5 | neuregulin 2 [Source:HGNC Symbol;Acc:HGNC |
| AC008667.2 | 5 | novel transcript |
| Y_RNA | 5 | Y RNA [Source:RFAM;Acc:RF00019] |
| MALINC1 | 5 | mitosis associated long intergenic non-coding |
| PURA | 5 | purine rich element binding protein A [Source |
| IGIP | 5 | IgA inducing protein [Source:HGNC Symbol;Ac |
| AC011379.2 | 5 | novel transcript |
| CYSTM1 | 5 | cysteine rich transmembrane module contain |
| AC011379.1 | 5 | novel transcript |
| PFDN1 | 5 | prefoldin subunit 1 [Source:HGNC Symbol;Acc |
| HBEGF | 5 | heparin binding EGF like growth factor [Sourc |
| AC008438.2 | 5 | novel transcript |
| SLC4A9 | 5 | solute carrier family 4 member 9 [Source:HGN |
| AC008438.1 | 5 | novel transcript |
| RNU4-14P | 5 | RNA, U4 small nuclear 14, pseudogene [Sourc |
| ANKHD1 | 5 | ankyrin repeat and KH domain containing 1 [S |
| ANKHD1-EIF4EBP3 | 5 | ANKHD1-EIF4EBP3 readthrough [Source:HGN |
| AC011399.1 | 5 | NULL |
| SRA1 | 5 | steroid receptor RNA activator 1 [Source:HGN |
| EIF4EBP3 | 5 | eukaryotic translation initiation factor 4E bind |
| APBB3 | 5 | amyloid beta precursor protein binding family |
| AC116353.5 | 5 | novel transcript |
| MIR6831 | 5 | microRNA 6831 [Source:HGNC Symbol;Acc:HG |
| SLC35A4 | 5 | solute carrier family 35 member A4 [Source:H |
| AC116353.6 | 5 | novel pseudogene |
| AC116353.1 | 5 | NULL |
| HAUS1P1 | 5 | HAUS augmin like complex subunit 1 pseudog |
| AC116353.2 | 5 | ribosomal protein L36 (RPL36) pseudogene |
| CD14 | 5 | CD14 molecule [Source:HGNC Symbol;Acc:HG |

| | | |
|---|---|---|
| NDUFA2 | 5 | NADH:ubiquinone oxidoreductase subunit A2 |
| TMCO6 | 5 | transmembrane and coiled-coil domains 6 [So |
| IK | 5 | IK cytokine [Source:HGNC Symbol;Acc:HGNC:5 |
| MIR3655 | 5 | microRNA 3655 [Source:HGNC Symbol;Acc:HG |
| WDR55 | 5 | WD repeat domain 55 [Source:HGNC Symbol; |
| DND1 | 5 | DND microRNA-mediated repression inhibitor |
| HARS | 5 | histidyl-tRNA synthetase [Source:HGNC Symb |
| HARS2 | 5 | histidyl-tRNA synthetase 2, mitochondrial [Sou |
| ZMAT2 | 5 | zinc finger matrin-type 2 [Source:HGNC Symb |
| VTRNA1-1 | 5 | vault RNA 1-1 [Source:HGNC Symbol;Acc:HGN |
| VTRNA1-2 | 5 | vault RNA 1-2 [Source:HGNC Symbol;Acc:HGN |
| VTRNA1-3 | 5 | vault RNA 1-3 [Source:HGNC Symbol;Acc:HGN |
| AC116353.4 | 5 | rearranged L-myc fusion pseudogene |
| AC116353.3 | 5 | zinc finger protein 470 pseudogene |
| AC005609.5 | 5 | novel transcript, antisense to PCDHA5, PCDHA |
| PCDHA1 | 5 | protocadherin alpha 1 [Source:HGNC Symbol; |
| PCDHA2 | 5 | protocadherin alpha 2 [Source:HGNC Symbol; |
| PCDHA3 | 5 | protocadherin alpha 3 [Source:HGNC Symbol; |
| PCDHA4 | 5 | protocadherin alpha 4 [Source:HGNC Symbol; |
| PCDHA5 | 5 | protocadherin alpha 5 [Source:HGNC Symbol; |
| PCDHA6 | 5 | protocadherin alpha 6 [Source:HGNC Symbol; |
| PCDHA7 | 5 | protocadherin alpha 7 [Source:HGNC Symbol; |
| PCDHA8 | 5 | protocadherin alpha 8 [Source:HGNC Symbol; |
| PCDHA9 | 5 | protocadherin alpha 9 [Source:HGNC Symbol; |
| AC005609.2 | 5 | novel transcript, antisense to PCDHA9 |
| PCDHA10 | 5 | protocadherin alpha 10 [Source:HGNC Symbo |
| PCDHA14 | 5 | protocadherin alpha 14 pseudogene [Source:H |
| AC005609.4 | 5 | novel transcript, antisense to PCDHA10 |
| PCDHA11 | 5 | protocadherin alpha 11 [Source:HGNC Symbo |
| PCDHA12 | 5 | protocadherin alpha 12 [Source:HGNC Symbo |
| AC005609.3 | 5 | novel transcript, antisense to PCDHA12 |
| AC005609.1 | 5 | novel transcript, antisense to PCDHA13 |
| PCDHA13 | 5 | protocadherin alpha 13 [Source:HGNC Symbo |
| PCDHAC1 | 5 | protocadherin alpha subfamily C, 1 [Source:H |
| AC010223.1 | 5 | novel transcript, antisense to PCDHAC2 |
| PCDHAC2 | 5 | protocadherin alpha subfamily C, 2 [Source:H |
| AC244517.5 | 5 | novel transcript, antisense to PCDHB1 and PC |
| PCDHB1 | 5 | protocadherin beta 1 [Source:HGNC Symbol;A |
| AC244517.9 | 5 | novel transcript |
| PCDHB2 | 5 | protocadherin beta 2 [Source:HGNC Symbol;A |
| AC244517.2 | 5 | novel transcript, antisense to PCDHB3, PCDHE |
| PCDHB3 | 5 | protocadherin beta 3 [Source:HGNC Symbol;A |
| AC244517.1 | 5 | novel transcript |
| PCDHB4 | 5 | protocadherin beta 4 [Source:HGNC Symbol;A |
| PCDHB5 | 5 | protocadherin beta 5 [Source:HGNC Symbol;A |
| AC244517.11 | 5 | novel transcript, antisense to PCDHB5, PCDHE |
| PCDHB6 | 5 | protocadherin beta 6 [Source:HGNC Symbol;A |
| PCDHB17P | 5 | protocadherin beta 17 pseudogene [Source:H |
| AC244517.12 | 5 | novel transcript |
| PCDHB7 | 5 | protocadherin beta 7 [Source:HGNC Symbol;A |

| | | |
|---|---|---|
| PCDHB8 | 5 | protocadherin beta 8 [Source:HGNC Symbol;A |
| AC244517.8 | 5 | novel transcript |
| PCDHB16 | 5 | protocadherin beta 16 [Source:HGNC Symbol; |
| AC244517.6 | 5 | novel transcript |
| AC244517.4 | 5 | novel transcript, antisense to PCDHB16, PCDH |
| PCDHB9 | 5 | protocadherin beta 9 [Source:HGNC Symbol;A |
| AC244517.7 | 5 | novel transcript, antisense to PCDHB10 |
| PCDHB10 | 5 | protocadherin beta 10 [Source:HGNC Symbol; |
| PCDHB11 | 5 | protocadherin beta 11 [Source:HGNC Symbol; |
| PCDHB12 | 5 | protocadherin beta 12 [Source:HGNC Symbol; |
| PCDHB13 | 5 | protocadherin beta 13 [Source:HGNC Symbol; |
| PCDHB14 | 5 | protocadherin beta 14 [Source:HGNC Symbol; |
| AC244517.10 | 5 | novel protein |
| PCDHB18P | 5 | protocadherin beta 18 pseudogene [Source:H |
| PCDHB19P | 5 | protocadherin beta 19 pseudogene [Source:H |
| PCDHB15 | 5 | protocadherin beta 15 [Source:HGNC Symbol; |
| AC244517.3 | 5 | novel zinc finger pseudogene |
| TAF7 | 5 | TATA-box binding protein associated factor 7 |
| SLC25A2 | 5 | solute carrier family 25 member 2 [Source:HG |
| AC005618.4 | 5 | novel transcript |
| AC005618.1 | 5 | novel transcript |
| PCDHGA1 | 5 | protocadherin gamma subfamily A, 1 [Source: |
| PCDHGA2 | 5 | protocadherin gamma subfamily A, 2 [Source: |
| PCDHGA3 | 5 | protocadherin gamma subfamily A, 3 [Source: |
| PCDHGB1 | 5 | protocadherin gamma subfamily B, 1 [Source: |
| AC005618.3 | 5 | novel transcript, antisense to PCDHGB1 |
| PCDHGA4 | 5 | protocadherin gamma subfamily A, 4 [Source: |
| PCDHGB2 | 5 | protocadherin gamma subfamily B, 2 [Source: |
| PCDHGA5 | 5 | protocadherin gamma subfamily A, 5 [Source: |
| PCDHGB3 | 5 | protocadherin gamma subfamily B, 3 [Source: |
| PCDHGA6 | 5 | protocadherin gamma subfamily A, 6 [Source: |
| PCDHGA7 | 5 | protocadherin gamma subfamily A, 7 [Source: |
| PCDHGB4 | 5 | protocadherin gamma subfamily B, 4 [Source: |
| PCDHGA8 | 5 | protocadherin gamma subfamily A, 8 [Source: |
| PCDHGB5 | 5 | protocadherin gamma subfamily B, 5 [Source: |
| PCDHGA9 | 5 | protocadherin gamma subfamily A, 9 [Source: |
| PCDHGB6 | 5 | protocadherin gamma subfamily B, 6 [Source: |
| PCDHGA10 | 5 | protocadherin gamma subfamily A, 10 [Source |
| PCDHGB7 | 5 | protocadherin gamma subfamily B, 7 [Source: |
| PCDHGA11 | 5 | protocadherin gamma subfamily A, 11 [Source |
| PCDHGB8P | 5 | protocadherin gamma subfamily B, 8 pseudog |
| AC005618.2 | 5 | novel transcript, antisense to protocadherin g |
| PCDHGA12 | 5 | protocadherin gamma subfamily A, 12 [Source |
| PCDHGB9P | 5 | protocadherin gamma subfamily B, 9 pseudog |
| AC008781.3 | 5 | novel transcript, sense intronic to PCDHG clus |
| PCDHGC3 | 5 | protocadherin gamma subfamily C, 3 [Source: |
| RN7SL68P | 5 | RNA, 7SL, cytoplasmic 68, pseudogene [Sourc |
| PCDHGC4 | 5 | protocadherin gamma subfamily C, 4 [Source: |
| PCDHGC5 | 5 | protocadherin gamma subfamily C, 5 [Source: |
| DIAPH1 | 5 | diaphanous related formin 1 [Source:HGNC Sy |

| | | |
|---|---|---|
| AC008781.2 | 5 | novel transcript |
| RPS27AP10 | 5 | ribosomal protein S27a pseudogene 10 [Sourc |
| AC008781.1 | 5 | novel transcript |
| HDAC3 | 5 | histone deacetylase 3 [Source:HGNC Symbol;A |
| RELL2 | 5 | RELT like 2 [Source:HGNC Symbol;Acc:HGNC:2 |
| FCHSD1 | 5 | FCH and double SH3 domains 1 [Source:HGNC |
| ARAP3 | 5 | ArfGAP with RhoGAP domain, ankyrin repeat a |
| AC005753.2 | 5 | novel transcript |
| AC005753.3 | 5 | novel transcript |
| AC005753.1 | 5 | novel transcript |
| PCDH1 | 5 | protocadherin 1 [Source:HGNC Symbol;Acc:HG |
| AC005740.5 | 5 | novel transcript |
| AC005740.1 | 5 | glycine cleavage system protein H (aminomet |
| DELE1 | 5 | DAP3 binding cell death enhancer 1 [Source:H |
| PCDH12 | 5 | protocadherin 12 [Source:HGNC Symbol;Acc:H |
| AC005740.4 | 5 | novel transcript, antisense to PCDH12 |
| RNF14 | 5 | ring finger protein 14 [Source:HGNC Symbol;A |
| AC005740.3 | 5 | novel transcript |
| GNPDA1 | 5 | glucosamine-6-phosphate deaminase 1 [Sourc |
| AC005740.2 | 5 | mitochondrial ribosomal protein L11 (MRPL11 |
| NDFIP1 | 5 | Nedd4 family interacting protein 1 [Source:HG |
| AC091825.3 | 5 | TEC |
| SPRY4 | 5 | sprouty RTK signaling antagonist 4 [Source:HG |
| AC091825.2 | 5 | NULL |
| AC091825.1 | 5 | NULL |
| SPRY4-AS1 | 5 | SPRY4 antisense RNA 1 [Source:HGNC Symbol |
| RPS12P10 | 5 | ribosomal protein S12 pseudogene 10 [Source |
| FGF1 | 5 | fibroblast growth factor 1 [Source:HGNC Symb |
| AC005592.1 | 5 | novel transcript |
| LINC01844 | 5 | long intergenic non-protein coding RNA 1844 |
| AC005592.2 | 5 | novel transcript |
| ARHGAP26 | 5 | Rho GTPase activating protein 26 [Source:HGN |
| ARHGAP26-AS1 | 5 | ARHGAP26 antisense RNA 1 [Source:HGNC Syr |
| AC008533.1 | 5 | TEC |
| ARHGAP26-IT1 | 5 | ARHGAP26 intronic transcript 1 [Source:HGNC |
| NR3C1 | 5 | nuclear receptor subfamily 3 group C member |
| RPL7P21 | 5 | ribosomal protein L7 pseudogene 21 [Source: |
| AC091925.2 | 5 | TEC |
| AC091925.1 | 5 | TEC |
| AC016598.1 | 5 | novel transcript |
| AC016598.2 | 5 | novel transcript |
| RNU7-156P | 5 | RNA, U7 small nuclear 156 pseudogene [Sourc |
| AC008696.2 | 5 | novel transcript, sense overlapping HMHB1 |
| MIR5197 | 5 | microRNA 5197 [Source:HGNC Symbol;Acc:HG |
| AC008696.1 | 5 | TNF receptor-associated factor (TNF) pseudog |
| HMHB1 | 5 | histocompatibility minor HB-1 [Source:HGNC S |
| RN7SL87P | 5 | RNA, 7SL, cytoplasmic 87, pseudogene [Sourc |
| YIPF5 | 5 | Yip1 domain family member 5 [Source:HGNC S |
| KCTD16 | 5 | potassium channel tetramerization domain co |
| AC008652.1 | 5 | novel transcript |

| | | |
|---|---|---|
| AC109441.1 | 5 | Novel transcript, antisense to KCTD16 |
| CKS1BP5 | 5 | CDC28 protein kinase regulatory subunit 1B p[illegible] |
| AC008716.1 | 5 | ribosomal protein L9 (RPL9) pseudogene |
| RN7SKP246 | 5 | RN7SK pseudogene 246 [Source:HGNC Symbo |
| AC132803.1 | 5 | novel transcript |
| NAMPTP2 | 5 | nicotinamide phosphoribosyltransferase pseu |
| ASS1P10 | 5 | argininosuccinate synthetase 1 pseudogene 1[illegible] |
| AC137770.1 | 5 | novel transcript |
| AC008700.1 | 5 | novel transcript |
| PRELID2 | 5 | PRELI domain containing 2 [Source:HGNC Sym |
| AC011411.1 | 5 | transient receptor potential cation channel fa |
| GRXCR2 | 5 | glutaredoxin and cysteine rich domain contair |
| SH3RF2 | 5 | SH3 domain containing ring finger 2 [Source:H |
| AC005351.1 | 5 | apolipoprotein O-like (APOOL) pseudogene |
| PLAC8L1 | 5 | PLAC8 like 1 [Source:HGNC Symbol;Acc:HGNC |
| AC091887.1 | 5 | novel transcript |
| LARS | 5 | leucyl-tRNA synthetase [Source:HGNC Symbol |
| AC091959.2 | 5 | ribosomal protein L35a (RPL35A) pseudogene |
| AC091959.1 | 5 | ribosomal protein L35a (RPL35A) pseudogene |
| AC091959.3 | 5 | novel readthrough transcript |
| RBM27 | 5 | RNA binding motif protein 27 [Source:HGNC S |
| RNU7-180P | 5 | RNA, U7 small nuclear 180 pseudogene [Sourc |
| AC011396.2 | 5 | novel transcript, antisense to POU4F3 |
| POU4F3 | 5 | POU class 4 homeobox 3 [Source:HGNC Symb |
| AC011396.3 | 5 | novel transcript |
| AC011396.1 | 5 | novel transcript |
| TCERG1 | 5 | transcription elongation regulator 1 [Source:H |
| AC011375.1 | 5 | NULL |
| GPR151 | 5 | G protein-coupled receptor 151 [Source:HGN[illegible] |
| AC008728.1 | 5 | novel transcript, antisense to PPP2R2B |
| PPP2R2B | 5 | protein phosphatase 2 regulatory subunit Bbe |
| RNA5SP196 | 5 | RNA, 5S ribosomal pseudogene 196 [Source:H |
| RN7SL791P | 5 | RNA, 7SL, cytoplasmic 791, pseudogene [Sour |
| KRT8P48 | 5 | keratin 8 pseudogene 48 [Source:HGNC Symb |
| PPP2R2B-IT1 | 5 | PPP2R2B intronic transcript 1 [Source:HGNC S |
| AC010251.1 | 5 | novel transcript |
| STK32A-AS1 | 5 | STK32A antisense RNA 1 [Source:HGNC Symbo |
| STK32A | 5 | serine/threonine kinase 32A [Source:HGNC Sy |
| DPYSL3 | 5 | dihydropyrimidinase like 3 [Source:HGNC Sym |
| AC011373.1 | 5 | novel transcript, antisense to DPYSL3 |
| JAKMIP2-AS1 | 5 | JAKMIP2 antisense RNA 1 [Source:HGNC Symb |
| JAKMIP2 | 5 | janus kinase and microtubule interacting prot |
| AC011370.1 | 5 | novel transcript, antisense to JAKMIP2 |
| SPINK1 | 5 | serine peptidase inhibitor, Kazal type 1 [Sourc |
| AC011352.2 | 5 | formyl peptide receptor 3 pseudogene |
| SCGB3A2 | 5 | secretoglobin family 3A member 2 [Source:HG |
| C5orf46 | 5 | chromosome 5 open reading frame 46 [Sourc |
| AC011352.1 | 5 | novel transcript |
| AC011352.3 | 5 | novel transcript |
| EEF1GP2 | 5 | eukaryotic translation elongation factor 1 gam |

| | | |
|---|---|---|
| SPINK5 | 5 | serine peptidase inhibitor, Kazal type 5 [Sourc |
| AC011346.1 | 5 | novel transcript, antisense to SPINK9, SPINK7 |
| SPINK14 | 5 | serine peptidase inhibitor, Kazal type 14 (puta |
| SPINK6 | 5 | serine peptidase inhibitor, Kazal type 6 [Sourc |
| HMGN1P16 | 5 | high mobility group nucleosome binding dom |
| MARCOL | 5 | MARCO like [Source:HGNC Symbol;Acc:HGNC: |
| SPINK13 | 5 | serine peptidase inhibitor, Kazal type 13 (puta |
| PGBD4P3 | 5 | piggyBac transposable element derived 4 pseu |
| SPINK7 | 5 | serine peptidase inhibitor, Kazal type 7 (putat |
| SPINK9 | 5 | serine peptidase inhibitor, Kazal type 9 [Sourc |
| AC091948.1 | 5 | novel ovomucoid-like pseudogene |
| FBXO38 | 5 | F-box protein 38 [Source:HGNC Symbol;Acc:H |
| Metazoa_SRP | 5 | Metazoan signal recognition particle RNA [Sou |
| AC114939.1 | 5 | novel transcript, antisense to FBXO38 |
| HTR4 | 5 | 5-hydroxytryptamine receptor 4 [Source:HGN |
| AC008627.1 | 5 | novel transcript |
| ADRB2 | 5 | adrenoceptor beta 2 [Source:HGNC Symbol;A |
| SH3TC2 | 5 | SH3 domain and tetratricopeptide repeats 2 [ |
| AC116312.1 | 5 | novel transcript, antisense to SH3TC2 |
| RNU6-732P | 5 | RNA, U6 small nuclear 732, pseudogene [Sour |
| MIR584 | 5 | microRNA 584 [Source:HGNC Symbol;Acc:HGI |
| SH3TC2-DT | 5 | SH3TC2 divergent transcript [Source:HGNC Sy |
| RN7SKP145 | 5 | RN7SK pseudogene 145 [Source:HGNC Symbo |
| ABLIM3 | 5 | actin binding LIM protein family member 3 [S |
| AC012613.1 | 5 | novel transcript |
| AC012613.2 | 5 | novel transcript, antisense to ABLIM3 |
| AFAP1L1 | 5 | actin filament associated protein 1 like 1 [Sou |
| AC131025.3 | 5 | Novel transcript |
| GRPEL2 | 5 | GrpE like 2, mitochondrial [Source:HGNC Sym |
| GRPEL2-AS1 | 5 | GRPEL2 antisense RNA 1 [Source:HGNC Symb |
| PCYOX1L | 5 | prenylcysteine oxidase 1 like [Source:HGNC Sy |
| IL17B | 5 | interleukin 17B [Source:HGNC Symbol;Acc:HG |
| AC131025.1 | 5 | novel transcript |
| CARMN | 5 | cardiac mesoderm enhancer-associated non-c |
| AC131025.2 | 5 | novel transcript |
| MIR143 | 5 | microRNA 143 [Source:HGNC Symbol;Acc:HGI |
| MIR145 | 5 | microRNA 145 [Source:HGNC Symbol;Acc:HGI |
| CSNK1A1 | 5 | casein kinase 1 alpha 1 [Source:HGNC Symbol |
| AC021078.1 | 5 | novel transcript |
| RPL29P14 | 5 | ribosomal protein L29 pseudogene 14 [Source |
| ARHGEF37 | 5 | Rho guanine nucleotide exchange factor 37 [S |
| RNU6-588P | 5 | RNA, U6 small nuclear 588, pseudogene [Sour |
| AC022100.1 | 5 | novel transcript |
| U3 | 5 | Small nucleolar RNA U3 [Source:RFAM;Acc:RF |
| RN7SL868P | 5 | RNA, 7SL, cytoplasmic 868, pseudogene [Sour |
| PPARGC1B | 5 | PPARG coactivator 1 beta [Source:HGNC Symb |
| MIR378A | 5 | microRNA 378a [Source:HGNC Symbol;Acc:HG |
| PDE6A | 5 | phosphodiesterase 6A [Source:HGNC Symbol; |
| AC008427.1 | 5 | mitochondrial fission factor (MFF) pseudogen |
| Y_RNA | 5 | Y RNA [Source:RFAM;Acc:RF00019] |

| | | |
|---|---|---|
| SLC26A2 | 5 | solute carrier family 26 member 2 [Source:HG |
| TIGD6 | 5 | tigger transposable element derived 6 [Source |
| HMGXB3 | 5 | HMG-box containing 3 [Source:HGNC Symbol; |
| AC011406.1 | 5 | Transmembrane protein pseudogene |
| RPS20P4 | 5 | ribosomal protein S20 pseudogene 4 [Source: |
| CSF1R | 5 | colony stimulating factor 1 receptor [Source:H |
| RPL7P1 | 5 | ribosomal protein L7 pseudogene 1 [Source:H |
| Y_RNA | 5 | Y RNA [Source:RFAM;Acc:RF00019] |
| PDGFRB | 5 | platelet derived growth factor receptor beta [ |
| CDX1 | 5 | caudal type homeobox 1 [Source:HGNC Symb |
| SLC6A7 | 5 | solute carrier family 6 member 7 [Source:HGN |
| CAMK2A | 5 | calcium/calmodulin dependent protein kinase |
| ARSI | 5 | arylsulfatase family member I [Source:HGNC S |
| TCOF1 | 5 | treacle ribosome biogenesis factor 1 [Source:I |
| CD74 | 5 | CD74 molecule [Source:HGNC Symbol;Acc:HG |
| AC011388.1 | 5 | novel transcript |
| RPS14 | 5 | ribosomal protein S14 [Source:HGNC Symbol;. |
| NDST1-AS1 | 5 | NDST1 antisense RNA 1 [Source:HGNC Symbo |
| NDST1 | 5 | N-deacetylase and N-sulfotransferase 1 [Sourc |
| SYNPO | 5 | synaptopodin [Source:HGNC Symbol;Acc:HGN |
| AC011383.1 | 5 | novel transcript |
| AC008453.2 | 5 | novel transcript, antisense to SYNPO |
| MYOZ3 | 5 | myozenin 3 [Source:HGNC Symbol;Acc:HGNC: |
| AC008453.1 | 5 | novel transcript |
| RBM22 | 5 | RNA binding motif protein 22 [Source:HGNC S |
| DCTN4 | 5 | dynactin subunit 4 [Source:HGNC Symbol;Acc: |
| SMIM3 | 5 | small integral membrane protein 3 [Source:HG |
| IRGM | 5 | immunity related GTPase M [Source:HGNC Sy |
| ZNF300 | 5 | zinc finger protein 300 [Source:HGNC Symbol; |
| ZNF300P1 | 5 | zinc finger protein 300 pseudogene 1 [Source: |
| AC022106.1 | 5 | novel zinc finger protein pseudogene |
| AC022106.2 | 5 | novel zinc finger protein pseudogene |
| GPX3 | 5 | glutathione peroxidase 3 [Source:HGNC Symb |
| TNIP1 | 5 | TNFAIP3 interacting protein 1 [Source:HGNC S |
| ANXA6 | 5 | annexin A6 [Source:HGNC Symbol;Acc:HGNC: |
| AC008641.1 | 5 | novel transcript |
| AC011342.1 | 5 | novel platelet-derived growth factor receptor |
| CCDC69 | 5 | coiled-coil domain containing 69 [Source:HGN |
| GM2A | 5 | GM2 ganglioside activator [Source:HGNC Sym |
| SLC36A3 | 5 | solute carrier family 36 member 3 [Source:HG |
| SLC36A2 | 5 | solute carrier family 36 member 2 [Source:HG |
| AC008385.1 | 5 | ATPase, H+ transporting, lysosomal 13kDa, V1 |
| AC034205.2 | 5 | solute carrier family 36 (proton/amino acid sy |
| AC034205.1 | 5 | mediator complex subunit 13 (MED13) pseud |
| AC034205.3 | 5 | RUN domain containing 1 (RUNDC1) pseudog |
| SLC36A1 | 5 | solute carrier family 36 member 1 [Source:HG |
| RNA5SP197 | 5 | RNA, 5S ribosomal pseudogene 197 [Source:H |
| FAT2 | 5 | FAT atypical cadherin 2 [Source:HGNC Symbol |
| AC011337.1 | 5 | novel transcript, antisense to FAT2 |
| MIR6499 | 5 | microRNA 6499 [Source:HGNC Symbol;Acc:HG |

| | | |
|---|---|---|
| AC011374.3 | 5 | novel transcript |
| AC011374.1 | 5 | novel transcript, antisense to SPARC |
| SPARC | 5 | secreted protein acidic and cysteine rich [Sour |
| CLMAT3 | 5 | colorectal liver metastasis associated transcri |
| RN7SKP232 | 5 | RN7SK pseudogene 232 [Source:HGNC Symbo |
| AC011374.2 | 5 | novel transcript |
| ATOX1 | 5 | antioxidant 1 copper chaperone [Source:HGN |
| AC091982.1 | 5 | novel transcript |
| RPLP1P6 | 5 | ribosomal protein lateral stalk subunit P1 pse |
| AC091982.3 | 5 | novel transcript |
| G3BP1 | 5 | G3BP stress granule assembly factor 1 [Source |
| GLRA1 | 5 | glycine receptor alpha 1 [Source:HGNC Symbo |
| AC091982.2 | 5 | glutamate-ammonia ligase (GLUL) pseudogen |
| RNA5SP198 | 5 | RNA, 5S ribosomal pseudogene 198 [Source:H |
| LINC01933 | 5 | long intergenic non-protein coding RNA 1933 |
| AC008571.2 | 5 | novel transcript, antisense to NMUR2 |
| NMUR2 | 5 | neuromedin U receptor 2 [Source:HGNC Symb |
| AC008571.1 | 5 | ribosomal protein L36a (RPL36A) pseudogene |
| LINC01470 | 5 | long intergenic non-protein coding RNA 1470 |
| GRIA1 | 5 | glutamate ionotropic receptor AMPA type sub |
| RN7SL177P | 5 | RNA, 7SL, cytoplasmic 177, pseudogene [Sour |
| LINC01861 | 5 | long intergenic non-protein coding RNA 1861 |
| AC091962.1 | 5 | novel transcript |
| FAM114A2 | 5 | family with sequence similarity 114 member |
| MFAP3 | 5 | microfibril associated protein 3 [Source:HGNC |
| GALNT10 | 5 | polypeptide N-acetylgalactosaminyltransferas |
| SAP30L-AS1 | 5 | SAP30L antisense RNA 1 (head to head) [Sour |
| HNRNPA3P7 | 5 | heterogeneous nuclear ribonucleoprotein A3 |
| AC008625.1 | 5 | novel transcript, sense intronic to GALNT10 |
| MIR1294 | 5 | microRNA 1294 [Source:HGNC Symbol;Acc:H |
| RN7SL655P | 5 | RNA, 7SL, cytoplasmic 655, pseudogene [Sour |
| SAP30L | 5 | SAP30 like [Source:HGNC Symbol;Acc:HGNC:2 |
| HAND1 | 5 | heart and neural crest derivatives expressed 1 |
| AC026688.2 | 5 | novel transcript |
| AC026688.1 | 5 | Vacuolar protein sorting-associated protein 3 |
| CIR1P1 | 5 | corepressor interacting with RBPJ, 1 pseudoge |
| MIR3141 | 5 | microRNA 3141 [Source:HGNC Symbol;Acc:H |
| MIR1303 | 5 | microRNA 1303 [Source:HGNC Symbol;Acc:H |
| Y_RNA | 5 | Y RNA [Source:RFAM;Acc:RF00019] |
| LARP1 | 5 | La ribonucleoprotein domain family member |
| RN7SL803P | 5 | RNA, 7SL, cytoplasmic 803, pseudogene [Sour |
| AC008379.1 | 5 | ribosomal protein L21 (RPL21) pseudogene |
| FAXDC2 | 5 | fatty acid hydroxylase domain containing 2 [S |
| Metazoa_SRP | 5 | Metazoan signal recognition particle RNA [Sou |
| MIR378H | 5 | microRNA 378h [Source:HGNC Symbol;Acc:H |
| CNOT8 | 5 | CCR4-NOT transcription complex subunit 8 [S |
| GEMIN5 | 5 | gem nuclear organelle associated protein 5 [S |
| MRPL22 | 5 | mitochondrial ribosomal protein L22 [Source: |
| AC008410.1 | 5 | cytochrome c oxidase subunit VIIa polypeptid |
| KIF4B | 5 | kinesin family member 4B [Source:HGNC Sym |

| | | |
|---|---|---|
| AC010476.2 | 5 | novel transcript |
| AC010476.1 | 5 | ribosomal protein L6 (RPL6) pseudogene |
| AC010591.1 | 5 | oligonucleotide/oligosaccharide-binding fold c |
| AC008725.1 | 5 | peptidylprolyl isomerase G (cyclophilin G) (PPI |
| RNA5SP199 | 5 | RNA, 5S ribosomal pseudogene 199 [Source:H |
| SGCD | 5 | sarcoglycan delta [Source:HGNC Symbol;Acc:H |
| AC011351.1 | 5 | novel transcript, antisense to SGCD |
| AC027308.1 | 5 | pseudogene similar to part of WW and C2 dor |
| AC025434.1 | 5 | novel transcript |
| RNU6-556P | 5 | RNA, U6 small nuclear 556, pseudogene [Sour |
| PPP1R2B | 5 | PPP1R2 family member B [Source:HGNC Symb |
| TIMD4 | 5 | T cell immunoglobulin and mucin domain con |
| APOOP1 | 5 | apolipoprotein O pseudogene 1 [Source:HGN( |
| HAVCR1 | 5 | hepatitis A virus cellular receptor 1 [Source:H( |
| HAVCR2 | 5 | hepatitis A virus cellular receptor 2 [Source:H( |
| AC011377.1 | 5 | novel transcript |
| MED7 | 5 | mediator complex subunit 7 [Source:HGNC Sy |
| ITK | 5 | IL2 inducible T cell kinase [Source:HGNC Symb |
| FAM71B | 5 | family with sequence similarity 71 member B |
| AC010609.1 | 5 | novel transcript |
| AC009185.1 | 5 | novel transcript |
| CYFIP2 | 5 | cytoplasmic FMR1 interacting protein 2 [Sourc |
| AC008676.3 | 5 | Novel protein |
| FNDC9 | 5 | fibronectin type III domain containing 9 [Sourc |
| AC016571.1 | 5 | novel transcript, sense overlapping ADAM19 |
| AC008676.1 | 5 | novel transcript, antisense to CYFIP2 |
| ADAM19 | 5 | ADAM metallopeptidase domain 19 [Source:H |
| AC008676.2 | 5 | ribosomal protein L26 (RPL26) pseudogene |
| NIPAL4 | 5 | NIPA like domain containing 4 [Source:HGNC S |
| AC106801.1 | 5 | novel transcript |
| RNU6-390P | 5 | RNA, U6 small nuclear 390, pseudogene [Sour |
| AC008694.2 | 5 | pseudogene similar to part of ribosomal prote |
| SOX30 | 5 | SRY-box transcription factor 30 [Source:HGNC |
| C5orf52 | 5 | chromosome 5 open reading frame 52 [Sourc |
| AC008694.1 | 5 | SRY (sex determining region Y)-box 30 (Sox30) |
| THG1L | 5 | tRNA-histidine guanylyltransferase 1 like [Sour |
| LSM11 | 5 | LSM11, U7 small nuclear RNA associated [Sou |
| AC026407.1 | 5 | pseudogene similar to part of chromosome 7 |
| CLINT1 | 5 | clathrin interactor 1 [Source:HGNC Symbol;Ac |
| RNU6-260P | 5 | RNA, U6 small nuclear 260, pseudogene [Sour |
| AC008677.2 | 5 | ribosomal protein, large, P2 (RPLP2) pseudoge |
| RNU2-48P | 5 | RNA, U2 small nuclear 48, pseudogene [Sourc |
| MARK2P11 | 5 | microtubule affinity regulating kinase 2 pseud |
| AC008677.1 | 5 | cell division cycle 37 homolog (S. cerevisiae) (( |
| AC008677.3 | 5 | latexin (LXN) pseudogene |
| AC025437.5 | 5 | novel transcript |
| AC025437.4 | 5 | novel transcript |
| AC025437.6 | 5 | uracil-DNA glycosylase (UNG) pseudogene |
| AC025437.3 | 5 | novel transcript |
| AC025437.2 | 5 | novel transcript |

| | | |
|---|---|---|
| AC025437.1 | 5 | novel transcript |
| LINC02227 | 5 | long intergenic non-protein coding RNA 2227 |
| AC091979.1 | 5 | novel transcript |
| AC091979.2 | 5 | novel transcript |
| AC091939.1 | 5 | novel transcript |
| EBF1 | 5 | EBF transcription factor 1 [Source:HGNC Symb |
| AC136424.2 | 5 | novel transcript |
| AC136424.1 | 5 | novel transcript |
| LINC02202 | 5 | long intergenic non-protein coding RNA 2202 |
| AC134043.3 | 5 | TEC |
| RNF145 | 5 | ring finger protein 145 [Source:HGNC Symbol; |
| AC134043.2 | 5 | novel transcript |
| LINC01932 | 5 | long intergenic non-protein coding RNA 1932 |
| AC134043.1 | 5 | NULL |
| UBLCP1 | 5 | ubiquitin like domain containing CTD phospha |
| AC008691.1 | 5 | novel transcript |
| IL12B | 5 | interleukin 12B [Source:HGNC Symbol;Acc:HG |
| RNU4ATAC2P | 5 | RNA, U4atac small nuclear 2, pseudogene [So |
| LINC01845 | 5 | long intergenic non-protein coding RNA 1845 |
| AC008703.1 | 5 | novel transcript |
| LINC01847 | 5 | long intergenic non-protein coding RNA 1847 |
| ADRA1B | 5 | adrenoceptor alpha 1B [Source:HGNC Symbol |
| GAPDHP40 | 5 | glyceraldehyde 3 phosphate dehydrogenase p |
| Y_RNA | 5 | Y RNA [Source:RFAM;Acc:RF00019] |
| TTC1 | 5 | tetratricopeptide repeat domain 1 [Source:HG |
| AC068657.1 | 5 | CCHC domain zinc finger protein pseudogene |
| PWWP2A | 5 | PWWP domain containing 2A [Source:HGNC S |
| FABP6 | 5 | fatty acid binding protein 6 [Source:HGNC Syn |
| AC008609.1 | 5 | novel transcript |
| AC112191.1 | 5 | acid phosphatase 1, soluble (ACP1) pseudoger |
| CCNJL | 5 | cyclin J like [Source:HGNC Symbol;Acc:HGNC:2 |
| AC112191.2 | 5 | ubiquinol-cytochrome c reductase binding pro |
| AC112191.3 | 5 | small nuclear ribonucleoprotein polypeptide E |
| C1QTNF2 | 5 | C1q and TNF related 2 [Source:HGNC Symbol; |
| ZBED8 | 5 | zinc finger BED-type containing 8 [Source:HGN |
| SLU7 | 5 | SLU7 homolog, splicing factor [Source:HGNC S |
| PTTG1 | 5 | PTTG1 regulator of sister chromatid separatio |
| MIR3142HG | 5 | MIR3142 host gene [Source:HGNC Symbol;Acc |
| MIR3142 | 5 | microRNA 3142 [Source:HGNC Symbol;Acc:HG |
| MIR146A | 5 | microRNA 146a [Source:HGNC Symbol;Acc:HG |
| ATP10B | 5 | ATPase phospholipid transporting 10B (putativ |
| AC008456.1 | 5 | novel transcript |
| AC011363.1 | 5 | novel transcript |
| LINC02159 | 5 | long intergenic non-protein coding RNA 2159 |
| GABRB2 | 5 | gamma-aminobutyric acid type A receptor bet |
| GABRA6 | 5 | gamma-aminobutyric acid type A receptor alp |
| AC091944.1 | 5 | novel transcript |
| GLRXP3 | 5 | glutaredoxin pseudogene 3 [Source:HGNC Syr |
| GABRA1 | 5 | gamma-aminobutyric acid type A receptor alp |
| LINC01202 | 5 | long intergenic non-protein coding RNA 1202 |

| | | |
|---|---|---|
| GABRG2 | 5 | gamma-aminobutyric acid type A receptor gar |
| AC113414.1 | 5 | novel transcript |
| RNU6-164P | 5 | RNA, U6 small nuclear 164, pseudogene [Sour |
| ARL2BPP5 | 5 | ADP ribosylation factor like GTPase 2 binding |
| MRPL57P6 | 5 | mitochondrial ribosomal protein L57 pseudog |
| AC008723.1 | 5 | lin-28 homolog B (C. elegans) (LIN28B) pseudc |
| CCNG1 | 5 | cyclin G1 [Source:HGNC Symbol;Acc:HGNC:15 |
| NUDCD2 | 5 | NudC domain containing 2 [Source:HGNC Sym |
| HMMR | 5 | hyaluronan mediated motility receptor [Sourc |
| HMMR-AS1 | 5 | HMMR antisense RNA 1 [Source:HGNC Symbo |
| MAT2B | 5 | methionine adenosyltransferase 2B [Source:H |
| AC010291.1 | 5 | Novel transcript |
| AC116917.1 | 5 | pseudogene similar to part of tropomyosin 2 ( |
| RNU6-168P | 5 | RNA, U6 small nuclear 168, pseudogene [Sour |
| AC008432.1 | 5 | novel zinc finger protein pseudogene |
| LSM1P2 | 5 | LSM1 homolog, mRNA degradation associatec |
| AC008662.1 | 5 | novel transcript |
| AC008662.2 | 5 | novel transcript |
| AC109466.1 | 5 | novel transcript |
| LINC02143 | 5 | long intergenic non-protein coding RNA 2143 |
| AC008446.1 | 5 | ribosomal protein S15 (RPS15) pseudogene |
| RNU6-209P | 5 | RNA, U6 small nuclear 209, pseudogene [Sour |
| AC091973.1 | 5 | Novel transcript |
| LINC01938 | 5 | long intergenic non-protein coding RNA 1938 |
| AC008415.1 | 5 | novel transcript |
| RN7SKP60 | 5 | RN7SK pseudogene 60 [Source:HGNC Symbol; |
| AC008562.1 | 5 | CDGSH iron sulfur domain 1 (CISD1) pseudoge |
| AC008489.1 | 5 | ribosomal protein L21 (RPL21) pseudogene |
| RPL7P20 | 5 | ribosomal protein L7 pseudogene 20 [Source: |
| AC114321.1 | 5 | novel transcript |
| AC026403.1 | 5 | 60S acidic ribosomal protein (RPLP0) pseudog |
| LINC01947 | 5 | long intergenic non-protein coding RNA 1947 |
| AC091819.2 | 5 | novel transcript |
| AC091819.1 | 5 | novel transcript |
| TENM2 | 5 | teneurin transmembrane protein 2 [Source:H |
| AC091820.2 | 5 | novel transcript |
| AC091820.1 | 5 | novel transcript |
| AC008601.1 | 5 | novel transcript |
| AC008708.2 | 5 | novel transcript |
| AC008708.1 | 5 | novel transcript |
| AC008464.1 | 5 | novel transcript |
| AC008705.2 | 5 | novel transcript |
| AC008705.1 | 5 | NULL |
| AC011369.1 | 5 | novel transcript |
| AC011369.2 | 5 | TEC |
| AC026689.1 | 5 | novel transcript |
| WWC1 | 5 | WW and C2 domain containing 1 [Source:HGN |
| RARS | 5 | arginyl-tRNA synthetase [Source:HGNC Symbo |
| FBLL1 | 5 | fibrillarin like 1 [Source:HGNC Symbol;Acc:HG |
| PANK3 | 5 | pantothenate kinase 3 [Source:HGNC Symbol; |

| | | |
|---|---|---|
| SLC2A3P1 | 5 | solute carrier family 2 member 3 pseudogene |
| MIR103A1 | 5 | microRNA 103a-1 [Source:HGNC Symbol;Acc:H |
| MIR103B1 | 5 | microRNA 103b-1 [Source:HGNC Symbol;Acc:H |
| RPL10P9 | 5 | ribosomal protein L10 pseudogene 9 [Source: |
| AC011365.2 | 5 | novel transcript |
| SLIT3 | 5 | slit guidance ligand 3 [Source:HGNC Symbol;A |
| AC011365.1 | 5 | novel transcript |
| MIR218-2 | 5 | microRNA 218-2 [Source:HGNC Symbol;Acc:H |
| Y_RNA | 5 | Y RNA [Source:RFAM;Acc:RF00019] |
| AC011389.2 | 5 | novel transcript |
| AC011389.3 | 5 | TEC |
| AC011389.1 | 5 | novel transcript |
| MIR585 | 5 | microRNA 585 [Source:HGNC Symbol;Acc:HGN |
| RNU6-477P | 5 | RNA, U6 small nuclear 477, pseudogene [Sour |
| SPDL1 | 5 | spindle apparatus coiled-coil protein 1 [Source |
| DOCK2 | 5 | dedicator of cytokinesis 2 [Source:HGNC Symb |
| AC008680.1 | 5 | novel transcript |
| INSYN2B | 5 | inhibitory synaptic factor family member 2B [S |
| MIR378E | 5 | microRNA 378e [Source:HGNC Symbol;Acc:HG |
| FOXI1 | 5 | forkhead box I1 [Source:HGNC Symbol;Acc:HG |
| KRT18P41 | 5 | keratin 18 pseudogene 41 [Source:HGNC Sym |
| LINC01187 | 5 | long intergenic non-protein coding RNA 1187 |
| C5orf58 | 5 | chromosome 5 open reading frame 58 [Source |
| LCP2 | 5 | lymphocyte cytosolic protein 2 [Source:HGNC |
| LINC01366 | 5 | long intergenic non-protein coding RNA 1366 |
| AC034199.1 | 5 | novel transcript |
| KCNIP1 | 5 | potassium voltage-gated channel interacting p |
| KCNMB1 | 5 | potassium calcium-activated channel subfami |
| AC027306.1 | 5 | novel transcript |
| AC008619.1 | 5 | novel transcript |
| AC027312.1 | 5 | novel transcript |
| AC008514.1 | 5 | novel transcript |
| GABRP | 5 | gamma-aminobutyric acid type A receptor pi s |
| RANBP17 | 5 | RAN binding protein 17 [Source:HGNC Symbol |
| RN7SL623P | 5 | RNA, 7SL, cytoplasmic 623, pseudogene [Sour |
| AC008514.2 | 5 | tubulin, epsilon 1 (TUBE1) pseudogene |
| AC010306.1 | 5 | ubiquitin specific peptidase 46 (USP46) pseud |
| AC091980.2 | 5 | novel transcript, antisense to TLX3 |
| TLX3 | 5 | T cell leukemia homeobox 3 [Source:HGNC Sy |
| AC091980.1 | 5 | pseudogene similar to part of ribosomal prote |
| RN7SL339P | 5 | RNA, 7SL, cytoplasmic 339, pseudogene [Sour |
| RPSAP71 | 5 | ribosomal protein SA pseudogene 71 [Source: |
| SNORA70J | 5 | small nucleolar RNA, H/ACA box 70J [Source:H |
| RPL10P8 | 5 | ribosomal protein L10 pseudogene 8 [Source: |
| MIR3912 | 5 | microRNA 3912 [Source:HGNC Symbol;Acc:HG |
| NPM1 | 5 | nucleophosmin 1 [Source:HGNC Symbol;Acc:H |
| FGF18 | 5 | fibroblast growth factor 18 [Source:HGNC Sym |
| AC022440.1 | 5 | Novel transcript |
| AC011410.1 | 5 | novel transcript |
| SMIM23 | 5 | small integral membrane protein 23 [Source:H |

| | | |
|---|---|---|
| FBXW11 | 5 | F-box and WD repeat domain containing 11 [S |
| STK10 | 5 | serine/threonine kinase 10 [Source:HGNC Sym |
| EFCAB9 | 5 | EF-hand calcium binding domain 9 [Source:HG |
| UBTD2 | 5 | ubiquitin domain containing 2 [Source:HGNC S |
| AC024561.1 | 5 | NULL |
| KLF3P1 | 5 | Kruppel like factor 3 pseudogene 1 [Source:H( |
| SH3PXD2B | 5 | SH3 and PX domains 2B [Source:HGNC Symbo |
| AC011407.1 | 5 | novel transcript |
| LINC01944 | 5 | long intergenic non-protein coding RNA 1944 |
| AC027309.3 | 5 | novel transcript |
| NEURL1B | 5 | neuralized E3 ubiquitin protein ligase 1B [Sour |
| AC027309.2 | 5 | coiled-coil domain containing 72 (CCDC72) ps |
| MIR5003 | 5 | microRNA 5003 [Source:HGNC Symbol;Acc:HG |
| AC027309.1 | 5 | novel transcript |
| AC022217.2 | 5 | novel transcript |
| AC022217.1 | 5 | family with sequence similarity 58, member A |
| AC022217.3 | 5 | novel transcript, antisense to DUSP1 |
| DUSP1 | 5 | dual specificity phosphatase 1 [Source:HGNC S |
| Y_RNA | 5 | Y RNA [Source:RFAM;Acc:RF00019] |
| AC022217.4 | 5 | phosphatidylinositol glycan anchor biosynthes |
| AC110011.1 | 5 | novel transcript |
| ERGIC1 | 5 | endoplasmic reticulum-golgi intermediate cor |
| AC008429.1 | 5 | novel transcript, antisense to RPL26L1 |
| AC008429.3 | 5 | novel transcript |
| RPL26L1 | 5 | ribosomal protein L26 like 1 [Source:HGNC Sy |
| AC008429.2 | 5 | pyrophosphatase (inorganic) 1 (PPA1) pseudo |
| ATP6V0E1 | 5 | ATPase H+ transporting V0 subunit e1 [Source |
| AC008429.4 | 5 | coiled-coil domain containing 72 (CCDC72) ps |
| SNORA74B | 5 | small nucleolar RNA, H/ACA box 74B [Source:I |
| Y_RNA | 5 | Y RNA [Source:RFAM;Acc:RF00019] |
| CREBRF | 5 | CREB3 regulatory factor [Source:HGNC Symbo |
| CDC42P5 | 5 | cell division cycle 42 pseudogene 5 [Source:H( |
| AC008378.1 | 5 | novel transcript |
| BNIP1 | 5 | BCL2 interacting protein 1 [Source:HGNC Sym |
| RPL7AP33 | 5 | ribosomal protein L7a pseudogene 33 [Source |
| NKX2-5 | 5 | NK2 homeobox 5 [Source:HGNC Symbol;Acc:H |
| Y_RNA | 5 | Y RNA [Source:RFAM;Acc:RF00019] |
| RNA5SP200 | 5 | RNA, 5S ribosomal pseudogene 200 [Source:H |
| STC2 | 5 | stanniocalcin 2 [Source:HGNC Symbol;Acc:HG |
| MIR8056 | 5 | microRNA 8056 [Source:HGNC Symbol;Acc:HG |
| RNU6-500P | 5 | RNA, U6 small nuclear 500, pseudogene [Sour |
| AC016573.1 | 5 | novel transcript |
| AC008632.1 | 5 | novel transcript |
| AC008663.1 | 5 | novel transcript |
| AC008663.3 | 5 | novel transcript |
| AC008663.2 | 5 | novel transcript |
| BOD1 | 5 | biorientation of chromosomes in cell division |
| LINC01863 | 5 | long intergenic non-protein coding RNA 1863 |
| LINC01942 | 5 | long intergenic non-protein coding RNA 1942 |
| LINC01484 | 5 | long intergenic non-protein coding RNA 1484 |

| | | |
|---|---|---|
| AC008674.1 | 5 | novel transcript |
| LINC01485 | 5 | long intergenic non-protein coding RNA 1485 |
| CPEB4 | 5 | cytoplasmic polyadenylation element binding |
| C5orf47 | 5 | chromosome 5 open reading frame 47 [Sourc |
| RPL12P22 | 5 | ribosomal protein L12 pseudogene 22 [Source |
| NSG2 | 5 | neuronal vesicle trafficking associated 2 [Sour |
| AC113423.1 | 5 | novel transcript |
| AC113423.2 | 5 | novel transcript |
| AC011333.1 | 5 | novel transcript |
| AC025752.1 | 5 | novel transcript |
| LINC01411 | 5 | long intergenic non-protein coding RNA 1411 |
| GAPDHP71 | 5 | glyceraldehyde-3-phosphate dehydrogenase p |
| SUMO2P6 | 5 | SUMO2 pseudogene 6 [Source:HGNC Symbol; |
| AC108112.1 | 5 | novel transcript |
| HIGD1AP3 | 5 | HIG1 hypoxia inducible domain family membe |
| MSX2 | 5 | msh homeobox 2 [Source:HGNC Symbol;Acc:H |
| AC113346.2 | 5 | novel transcript |
| MIR4634 | 5 | microRNA 4634 [Source:HGNC Symbol;Acc:HG |
| AC113346.1 | 5 | novel transcript |
| LINC01951 | 5 | long intergenic non-protein coding RNA 1951 |
| NIFKP2 | 5 | nucleolar protein interacting with the FHA dor |
| AC008413.1 | 5 | SIN3 homolog A, transcription regulator (yeas |
| AC008413.2 | 5 | novel transcript |
| ARL2BPP6 | 5 | ADP ribosylation factor like GTPase 2 binding |
| DRD1 | 5 | dopamine receptor D1 [Source:HGNC Symbol; |
| AC091393.2 | 5 | novel transcript |
| SFXN1 | 5 | sideroflexin 1 [Source:HGNC Symbol;Acc:HGN |
| AC091393.1 | 5 | nuclear transcription factor Y, beta (NFYB) pse |
| RN7SKP148 | 5 | RN7SK pseudogene 148 [Source:HGNC Symbo |
| HRH2 | 5 | histamine receptor H2 [Source:HGNC Symbol; |
| RNU6-226P | 5 | RNA, U6 small nuclear 226, pseudogene [Sour |
| CPLX2 | 5 | complexin 2 [Source:HGNC Symbol;Acc:HGNC |
| AC138965.3 | 5 | TEC |
| AC138965.2 | 5 | novel transcript, sense overlapping to THOC3 |
| THOC3 | 5 | THO complex 3 [Source:HGNC Symbol;Acc:HG |
| AC138965.1 | 5 | novel transcript |
| OR1X1P | 5 | olfactory receptor family 1 subfamily X memb |
| AC139491.3 | 5 | novel transcript |
| AC139491.1 | 5 | novel transcript, antisense to FAM153B |
| FAM153B | 5 | family with sequence similarity 153 member E |
| AC139491.7 | 5 | novel transcript |
| AC139491.5 | 5 | centrosomal protein 192 (CEP192), pseudoge |
| AC139491.4 | 5 | novel transcript |
| AC139491.6 | 5 | centrosomal protein 192kDa (CEP192) pseudo |
| AC139491.2 | 5 | novel transcript |
| AC139491.8 | 5 | novel transcript |
| AC139493.2 | 5 | novel transcript |
| AC139493.1 | 5 | selenophosphate synthetase 1 (SEPHS1) pseu |
| SIMC1 | 5 | SUMO interacting motifs containing 1 [Source |
| BRCC3P1 | 5 | BRCA1/BRCA2-containing complex subunit 3 p |

| | | |
|---|---|---|
| KIAA1191 | 5 | KIAA1191 [Source:HGNC Symbol;Acc:HGNC:2 |
| AC138956.1 | 5 | novel transcript |
| AC138956.2 | 5 | novel transcript |
| ARL10 | 5 | ADP ribosylation factor like GTPase 10 [Source |
| MIR1271 | 5 | microRNA 1271 [Source:HGNC Symbol;Acc:HG |
| NOP16 | 5 | NOP16 nucleolar protein [Source:HGNC Symb |
| HIGD2A | 5 | HIG1 hypoxia inducible domain family membe |
| CLTB | 5 | clathrin light chain B [Source:HGNC Symbol;Ac |
| AC010297.1 | 5 | tec |
| FAF2 | 5 | Fas associated factor family member 2 [Sourc |
| RNF44 | 5 | ring finger protein 44 [Source:HGNC Symbol;A |
| CDHR2 | 5 | cadherin related family member 2 [Source:HG |
| RN7SL684P | 5 | RNA, 7SL, cytoplasmic 684, pseudogene [Sour |
| Y_RNA | 5 | Y RNA [Source:RFAM;Acc:RF00019] |
| GPRIN1 | 5 | G protein regulated inducer of neurite outgro |
| SNCB | 5 | synuclein beta [Source:HGNC Symbol;Acc:HGN |
| MIR4281 | 5 | microRNA 4281 [Source:HGNC Symbol;Acc:HG |
| EIF4E1B | 5 | eukaryotic translation initiation factor 4E fami |
| TSPAN17 | 5 | tetraspanin 17 [Source:HGNC Symbol;Acc:HGI |
| AC113391.1 | 5 | novel transcript |
| AC113391.2 | 5 | novel transcript |
| LINC01574 | 5 | long intergenic non-protein coding RNA 1574 |
| UNC5A | 5 | unc-5 netrin receptor A [Source:HGNC Symbo |
| HK3 | 5 | hexokinase 3 [Source:HGNC Symbol;Acc:HGN |
| UIMC1 | 5 | ubiquitin interaction motif containing 1 [Sourc |
| ZNF346 | 5 | zinc finger protein 346 [Source:HGNC Symbol; |
| ZNF346-IT1 | 5 | ZNF346 intronic transcript 1 [Source:HGNC Sy |
| FGFR4 | 5 | fibroblast growth factor receptor 4 [Source:H |
| NSD1 | 5 | nuclear receptor binding SET domain protein |
| AC146507.3 | 5 | novel transcript, sense intronic to NSD1 |
| AC146507.2 | 5 | ribosomal protein L21 (RPL21) pseudogene |
| PRMT1P1 | 5 | protein arginine methyltransferase 1 pseudog |
| RAB24 | 5 | RAB24, member RAS oncogene family [Source |
| MXD3 | 5 | MAX dimerization protein 3 [Source:HGNC Sy |
| PRELID1 | 5 | PRELI domain containing 1 [Source:HGNC Sym |
| LMAN2 | 5 | lectin, mannose binding 2 [Source:HGNC Sym |
| AC146507.1 | 5 | ribosomal protein S20 (RPS20) pseudogene |
| RGS14 | 5 | regulator of G protein signaling 14 [Source:HG |
| SLC34A1 | 5 | solute carrier family 34 member 1 [Source:HG |
| PFN3 | 5 | profilin 3 [Source:HGNC Symbol;Acc:HGNC:18 |
| F12 | 5 | coagulation factor XII [Source:HGNC Symbol;A |
| GRK6 | 5 | G protein-coupled receptor kinase 6 [Source:H |
| PRR7-AS1 | 5 | PRR7 antisense RNA 1 [Source:HGNC Symbol; |
| PRR7 | 5 | proline rich 7, synaptic [Source:HGNC Symbol; |
| DBN1 | 5 | drebrin 1 [Source:HGNC Symbol;Acc:HGNC:26 |
| AC145098.2 | 5 | TEC |
| PDLIM7 | 5 | PDZ and LIM domain 7 [Source:HGNC Symbol; |
| AC145098.1 | 5 | novel transcript |
| DOK3 | 5 | docking protein 3 [Source:HGNC Symbol;Acc:H |
| DDX41 | 5 | DEAD-box helicase 41 [Source:HGNC Symbol; |

| | | |
|---|---|---|
| FAM193B | 5 | family with sequence similarity 193 member E |
| AC139795.3 | 5 | novel transcript |
| TMED9 | 5 | transmembrane p24 trafficking protein 9 [Sou |
| B4GALT7 | 5 | beta-1,4-galactosyltransferase 7 [Source:HGN |
| AC139795.2 | 5 | novel transcript |
| AC139795.1 | 5 | SUMO interacting motifs containing 1 (SIMC1) |
| AC138819.1 | 5 | novel transcript |
| FAM153A | 5 | family with sequence similarity 153 member A |
| AC140125.2 | 5 | novel transcript, antisense to FAM153A |
| AC140125.1 | 5 | novel transcript |
| AC106795.1 | 5 | THO complex 3 (THOC3) pseudogene |
| OR1X5P | 5 | olfactory receptor family 1 subfamily X memb |
| AC140125.3 | 5 | novel transcript |
| AC106795.3 | 5 | novel transcript |
| AC106795.2 | 5 | novel transcript, antisense to PROP1 |
| AC106795.5 | 5 | novel transcript |
| SUDS3P1 | 5 | SDS3 homolog, SIN3A corepressor complex cc |
| PROP1 | 5 | PROP paired-like homeobox 1 [Source:HGNC S |
| FAM153CP | 5 | family with sequence similarity 153 member C |
| AC136632.1 | 5 | ribosomal protein L19 (RPL19) pseudogene |
| N4BP3 | 5 | NEDD4 binding protein 3 [Source:HGNC Symb |
| RMND5B | 5 | required for meiotic nuclear division 5 homolo |
| NHP2 | 5 | NHP2 ribonucleoprotein [Source:HGNC Symbo |
| AC136632.2 | 5 | apolipoprotein A-I binding protein (APOA1BP) |
| GMCL2 | 5 | germ cell-less 2, spermatogenesis associated [ |
| HNRNPAB | 5 | heterogeneous nuclear ribonucleoprotein A/E |
| PHYKPL | 5 | 5-phosphohydroxy-L-lysine phospho-lyase [So |
| COL23A1 | 5 | collagen type XXIII alpha 1 chain [Source:HGN |
| AC136601.2 | 5 | mitochondrial ribosomal protein L50 (MRPL50 |
| AC136601.1 | 5 | novel transcript |
| RN7SL646P | 5 | RNA, 7SL, cytoplasmic 646, pseudogene [Sour |
| AC008659.1 | 5 | novel transcript |
| CLK4 | 5 | CDC like kinase 4 [Source:HGNC Symbol;Acc:H |
| RN7SKP70 | 5 | RN7SK pseudogene 70 [Source:HGNC Symbol; |
| AC113348.3 | 5 | novel transcript |
| AC113348.1 | 5 | uncharacterized LOC102724657 [Source:NCBI |
| AC113348.2 | 5 | novel transcript |
| ZNF354A | 5 | zinc finger protein 354A [Source:HGNC Symbo |
| AACSP1 | 5 | acetoacetyl-CoA synthetase pseudogene 1 [So |
| AC126915.2 | 5 | immunoglobulin lambda variable 5-48 (non-fu |
| AC126915.1 | 5 | coiled-coil domain containing 86 (CCDC86) pse |
| AC126915.3 | 5 | coiled-coil domain containing 86 (CCDC86) pse |
| ZNF354B | 5 | zinc finger protein 354B [Source:HGNC Symbo |
| RNU1-39P | 5 | RNA, U1 small nuclear 39, pseudogene [Sourc |
| ZFP2 | 5 | ZFP2 zinc finger protein [Source:HGNC Symbo |
| AC104117.1 | 5 | DCN1, defective in cullin neddylation 1, doma |
| PIGFP1 | 5 | phosphatidylinositol glycan anchor biosynthes |
| AC104117.5 | 5 | novel transcript |
| ZNF454 | 5 | zinc finger protein 454 [Source:HGNC Symbol; |
| AC104117.3 | 5 | novel transcript |

| | | |
|---|---|---|
| GRM6 | 5 | glutamate metabotropic receptor 6 [Source:H |
| AC104117.4 | 5 | F-box and WD repeat domain containing 2 (FB |
| ZNF879 | 5 | zinc finger protein 879 [Source:HGNC Symbol; |
| ZNF354C | 5 | zinc finger protein 354C [Source:HGNC Symbo |
| AC104117.2 | 5 | pseudogene similat to part of cold shock dom |
| ADAMTS2 | 5 | ADAM metallopeptidase with thrombospondi |
| AC109479.3 | 5 | novel transcript |
| AC109479.1 | 5 | novel transcript |
| AC109479.2 | 5 | pseudogene similar to part of 5'-nucleotidase |
| AC136628.1 | 5 | novel transcript |
| U4 | 5 | U4 spliceosomal RNA [Source:RFAM;Acc:RF00 |
| AC136628.4 | 5 | novel transcript |
| AC136628.2 | 5 | novel pseudogene |
| AC136628.3 | 5 | 6-pyruvoyltetrahydropterin synthase (PTS) ps |
| RUFY1 | 5 | RUN and FYVE domain containing 1 [Source:H |
| PRDX2P3 | 5 | peroxiredoxin 2 pseudogene 3 [Source:HGNC |
| AC136604.2 | 5 | novel transcript |
| HNRNPH1 | 5 | heterogeneous nuclear ribonucleoprotein H1 |
| Metazoa_SRP | 5 | Metazoan signal recognition particle RNA [Sou |
| C5orf60 | 5 | chromosome 5 open reading frame 60 [Sourc |
| AC136604.1 | 5 | chromosome 5 open reading frame 60 (C5orf |
| AC136604.3 | 5 | novel transcript |
| CBY3 | 5 | chibby family member 3 [Source:HGNC Symbo |
| CANX | 5 | calnexin [Source:HGNC Symbol;Acc:HGNC:147 |
| HMGB3P22 | 5 | high mobility group box 3 pseudogene 22 [Sou |
| MAML1 | 5 | mastermind like transcriptional coactivator 1 |
| LTC4S | 5 | leukotriene C4 synthase [Source:HGNC Symbo |
| MGAT4B | 5 | alpha-1,3-mannosyl-glycoprotein 4-beta-N-ac |
| MIR1229 | 5 | microRNA 1229 [Source:HGNC Symbol;Acc:HG |
| SQSTM1 | 5 | sequestosome 1 [Source:HGNC Symbol;Acc:H |
| MRNIP | 5 | MRN complex interacting protein [Source:HG |
| RN7SKP150 | 5 | RN7SK pseudogene 150 [Source:HGNC Symbo |
| Y_RNA | 5 | Y RNA [Source:RFAM;Acc:RF00019] |
| AC008393.1 | 5 | novel transcript |
| TBC1D9B | 5 | TBC1 domain family member 9B [Source:HGN |
| RNF130 | 5 | ring finger protein 130 [Source:HGNC Symbol; |
| RPS15AP18 | 5 | ribosomal protein S15a pseudogene 18 [Sourc |
| AC010285.2 | 5 | DnaJ (Hsp40) homolog, subfamily B, member |
| AC010285.1 | 5 | novel transcript |
| AC010285.3 | 5 | Novel transcript, antisense to RNF130 |
| MIR340 | 5 | microRNA 340 [Source:HGNC Symbol;Acc:HG |
| AC122713.2 | 5 | tec |
| AC122713.1 | 5 | chromosome 4 open reading frame 27 (C4orf |
| RASGEF1C | 5 | RasGEF domain family member 1C [Source:HG |
| AC104115.1 | 5 | ribosomal protein S8 (RPS8) pseudogene |
| MAPK9 | 5 | mitogen-activated protein kinase 9 [Source:H |
| AC104115.2 | 5 | zinc finger protein pseudogene |
| AC008610.1 | 5 | novel transcript |
| GFPT2 | 5 | glutamine-fructose-6-phosphate transaminas |
| AC034213.1 | 5 | novel transcript |

| | | |
|---|---|---|
| RNU6-525P | 5 | RNA, U6 small nuclear 525, pseudogene [Sour |
| CNOT6 | 5 | CCR4-NOT transcription complex subunit 6 [So |
| AC122714.1 | 5 | proteolipid protein 2 (colonic epithelium-enric |
| SCGB3A1 | 5 | secretoglobin family 3A member 1 [Source:HG |
| FLT4 | 5 | fms related tyrosine kinase 4 [Source:HGNC Sy |
| AC122714.2 | 5 | coiled-coil domain containing 86 (CCDC86) pse |
| LINC02222 | 5 | long intergenic non-protein coding RNA 2222 |
| OR2AI1P | 5 | olfactory receptor family 2 subfamily AI memb |
| RNU1-17P | 5 | RNA, U1 small nuclear 17, pseudogene [Sourc |
| OR2Y1 | 5 | olfactory receptor family 2 subfamily Y memb |
| MGAT1 | 5 | mannosyl (alpha-1,3-)-glycoprotein beta-1,2-N |
| AC022413.1 | 5 | TEC |
| HEIH | 5 | hepatocellular carcinoma up-regulated EZH2-a |
| LINC00847 | 5 | long intergenic non-protein coding RNA 847 [S |
| AC022413.2 | 5 | novel transcript |
| ZFP62 | 5 | ZFP62 zinc finger protein [Source:HGNC Symb |
| BTNL8 | 5 | butyrophilin like 8 [Source:HGNC Symbol;Acc: |
| Y_RNA | 5 | Y RNA [Source:RFAM;Acc:RF00019] |
| RPS29P12 | 5 | ribosomal protein S29 pseudogene 12 [Source |
| AC091874.3 | 5 | telomeric repeat binding factor 2, interacting |
| AC091874.2 | 5 | transmembrane protein 69 (TMEM69) pseudo |
| AC091874.1 | 5 | cAMP-regulated phosphoprotein, 19kDa (ARP |
| BTNL3 | 5 | butyrophilin like 3 [Source:HGNC Symbol;Acc: |
| RNU6-1036P | 5 | RNA, U6 small nuclear 1036, pseudogene [Sou |
| BTNL9 | 5 | butyrophilin like 9 [Source:HGNC Symbol;Acc: |
| MIR8089 | 5 | microRNA 8089 [Source:HGNC Symbol;Acc:HG |
| AC008620.1 | 5 | ribosomal protein L13 (RPL13) pseudogene |
| FOXO1B | 5 | forkhead box O1B (pseudogene) [Source:HGN |
| AC008620.2 | 5 | transmembrane and tetratricopeptide repeat |
| OR2V1 | 5 | olfactory receptor family 2 subfamily V memb |
| OR2V2 | 5 | olfactory receptor family 2 subfamily V memb |
| LINC01962 | 5 | long intergenic non-protein coding RNA 1962 |
| AC008443.5 | 5 | novel transcript |
| TRIM7 | 5 | tripartite motif containing 7 [Source:HGNC Sy |
| AC008443.6 | 5 | novel transcript |
| AC008443.4 | 5 | novel transcript |
| AC008443.9 | 5 | novel transcript |
| TRIM41 | 5 | tripartite motif containing 41 [Source:HGNC S |
| MIR4638 | 5 | microRNA 4638 [Source:HGNC Symbol;Acc:HG |
| AC008443.2 | 5 | novel transcript |
| RACK1 | 5 | receptor for activated C kinase 1 [Source:HGN |
| SNORD96A | 5 | small nucleolar RNA, C/D box 96A [Source:HG |
| SNORD95 | 5 | small nucleolar RNA, C/D box 95 [Source:HGN |
| AC008443.1 | 5 | novel transcript, antisense to TRIM52 |
| TRIM52 | 5 | tripartite motif containing 52 [Source:HGNC S |
| TRIM52-AS1 | 5 | TRIM52 antisense RNA 1 (head to head) [Sour |
| AC008443.8 | 5 | novel transcript |
| AC008443.3 | 5 | novel transcript |
| AC008443.7 | 5 | general transcription factor IIi (GTF2I) pseudo |
| RNU6-705P | 5 | RNA, U6 small nuclear 705, pseudogene [Sour |

| | | |
|---|---|---|
| AC138035.1 | 5 | novel transcript |
| AC138035.3 | 5 | capicua transcriptional repressor (CIC) pseudc |
| AC138035.2 | 5 | novel transcript |
| OR4F3 | 5 | olfactory receptor family 4 subfamily F memb |
| WBP1LP4 | 5 | WBP1L pseudogene 4 [Source:HGNC Symbol;/ |
| RPL23AP45 | 5 | ribosomal protein L23a pseudogene 45 [Sourc |
| WBP1LP12 | 6 | WBP1L pseudogene 12 [Source:HGNC Symbol |
| OR4F1P | 6 | olfactory receptor family 4 subfamily F memb |
| CICP18 | 6 | capicua transcriptional repressor pseudogene |
| AL035696.2 | 6 | novel Ankyrin repeat domain-containing prote |
| AL035696.3 | 6 | novel transcript |
| AL035696.4 | 6 | Novel transcript |
| AL035696.1 | 6 | novel transcript |
| AL365272.1 | 6 | novel transcript, antisense to DUSP22 |
| DUSP22 | 6 | dual specificity phosphatase 22 [Source:HGNC |
| IRF4 | 6 | interferon regulatory factor 4 [Source:HGNC S |
| AL512308.1 | 6 | novel transcript |
| EXOC2 | 6 | exocyst complex component 2 [Source:HGNC |
| AL031770.1 | 6 | novel transcript |
| HUS1B | 6 | HUS1 checkpoint clamp component B [Source |
| AL357054.1 | 6 | novel serine/threonine kinase pseudogene |
| AL357054.2 | 6 | novel transcript |
| AL357054.4 | 6 | novel transcript |
| AL357054.5 | 6 | novel transcript |
| AL357054.3 | 6 | novel transcript |
| AL392183.1 | 6 | novel transcript |
| AL356130.1 | 6 | novel transcript |
| LINC01622 | 6 | long intergenic non-protein coding RNA 1622 |
| AL033381.2 | 6 | novel transcript |
| AL033381.1 | 6 | novel transcript |
| AL033381.3 | 6 | novel transcript |
| FOXQ1 | 6 | forkhead box Q1 [Source:HGNC Symbol;Acc:H |
| LINC01394 | 6 | long intergenic non-protein coding RNA 1394 |
| AL034346.1 | 6 | novel transcript, antisense to FOXF2 |
| FOXF2 | 6 | forkhead box F2 [Source:HGNC Symbol;Acc:H( |
| MIR6720 | 6 | microRNA 6720 [Source:HGNC Symbol;Acc:HG |
| RN7SL352P | 6 | RNA, 7SL, cytoplasmic 352, pseudogene [Sour |
| AL512329.1 | 6 | E74-like factor 2 (ets domain transcription fac |
| AL512329.2 | 6 | novel transcript |
| FOXCUT | 6 | FOXC1 upstream transcript [Source:HGNC Syn |
| FOXC1 | 6 | forkhead box C1 [Source:HGNC Symbol;Acc:H( |
| GMDS | 6 | GDP-mannose 4,6-dehydratase [Source:HGNC |
| AL035693.1 | 6 | novel transcript, antisense to GMDS |
| GMDS-DT | 6 | GMDS divergent transcript [Source:HGNC Sym |
| HMGN2P28 | 6 | high mobility group nucleosomal binding dom |
| AL031768.2 | 6 | novel transcript |
| AL031768.1 | 6 | novel transcript |
| AL359852.2 | 6 | novel transcript |
| AL359852.1 | 6 | TEC |
| LINC02521 | 6 | long intergenic non-protein coding RNA 2521 |

| | | |
|---|---|---|
| LINC01600 | 6 | long intergenic non-protein coding RNA 1600 |
| MYLK4 | 6 | myosin light chain kinase family member 4 [Sc |
| WRNIP1 | 6 | WRN helicase interacting protein 1 [Source:H( |
| SERPINB1 | 6 | serpin family B member 1 [Source:HGNC Syml |
| SERPINB9P1 | 6 | serpin family B member 9 pseudogene 1 [Sour |
| MIR4645 | 6 | microRNA 4645 [Source:HGNC Symbol;Acc:H( |
| SERPINB9 | 6 | serpin family B member 9 [Source:HGNC Syml |
| AL133351.3 | 6 | pseudogene similar to part of serine (or cystei |
| AL133351.2 | 6 | novel transcript |
| SERPINB6 | 6 | serpin family B member 6 [Source:HGNC Syml |
| LINC01011 | 6 | long intergenic non-protein coding RNA 1011 |
| NQO2 | 6 | N-ribosyldihydronicotinamide:quinone reduct |
| AL133351.1 | 6 | novel transcript |
| HTATSF1P2 | 6 | HIV-1 Tat specific factor 1 pseudogene 2 [Sour |
| AL133351.4 | 6 | NAD(P)H dehydrogenase, quinone 2 (NQO2) p |
| FAM136BP | 6 | family with sequence similarity 136 member E |
| SERPINB8P1 | 6 | serpin family B member 8 pseudogene 1 [Sour |
| AL031963.2 | 6 | novel transcript |
| RIPK1 | 6 | receptor interacting serine/threonine kinase 1 |
| AL031963.3 | 6 | novel transcript, antisense to RIPK1 |
| RNA5SP201 | 6 | RNA, 5S ribosomal pseudogene 201 [Source:H |
| BPHL | 6 | biphenyl hydrolase like [Source:HGNC Symbol |
| AL031963.1 | 6 | novel transcript |
| TUBB2A | 6 | tubulin beta 2A class IIa [Source:HGNC Symbo |
| TUBB2BP1 | 6 | tubulin beta 2B class IIb pseudogene 1 [Source |
| LINC02525 | 6 | long intergenic non-protein coding RNA 2525 |
| TUBB2B | 6 | tubulin beta 2B class IIb [Source:HGNC Symbo |
| PSMG4 | 6 | proteasome assembly chaperone 4 [Source:H( |
| SLC22A23 | 6 | solute carrier family 22 member 23 [Source:H |
| AL445309.1 | 6 | novel transcript, antisense to SLC22A23 |
| AL033523.1 | 6 | novel transcript |
| AL033523.2 | 6 | translocase of outer mitochondrial membrane |
| PXDC1 | 6 | PX domain containing 1 [Source:HGNC Symbo |
| AL391422.4 | 6 | novel transcript, antisense to PXDC1 |
| AL391422.1 | 6 | FUN14 domain containing 1 (FUNDC1) pseudc |
| AL391422.3 | 6 | novel transcript |
| AL391422.2 | 6 | novel transcript |
| FAM50B | 6 | family with sequence similarity 50 member B |
| AL590004.3 | 6 | novel transcript |
| AL590004.2 | 6 | ribosomal protein S25 (RPS25) pseudogene |
| AL590004.1 | 6 | high mobility group nucleosome binding dom |
| TDGF1P4 | 6 | teratocarcinoma-derived growth factor 1 pseu |
| GLRX3P2 | 6 | glutaredoxin 3 pseudogene 2 [Source:HGNC S |
| AL138831.3 | 6 | TEC |
| AL138831.2 | 6 | novel transcript |
| AL138831.1 | 6 | novel transcript |
| PRPF4B | 6 | pre-mRNA processing factor 4B [Source:HGNC |
| FAM217A | 6 | family with sequence similarity 217 member / |
| C6orf201 | 6 | chromosome 6 open reading frame 201 [Sour |
| ECI2 | 6 | enoyl-CoA delta isomerase 2 [Source:HGNC Sy |

| | | |
|---|---|---|
| AL136309.4 | 6 | novel transcript, antisense to ECI2 |
| AL136309.2 | 6 | novel transcript |
| AL136309.3 | 6 | novel transcript |
| AL136309.1 | 6 | N(alpha)-acetyltransferase 40, NatD catalytic : |
| AL162718.1 | 6 | novel transcript |
| AL159166.1 | 6 | novel transcript |
| AL162718.3 | 6 | novel transcript |
| RNA5SP202 | 6 | RNA, 5S ribosomal pseudogene 202 [Source:H |
| AL162718.2 | 6 | novel transcript |
| LINC02533 | 6 | long intergenic non-protein coding RNA 2533 |
| AL133393.1 | 6 | SRSF protein kinase 2 (SRPK2) pseudogene |
| AL034376.2 | 6 | novel transcript |
| AL034376.1 | 6 | novel transcript |
| PSMC1P11 | 6 | proteasome 26S subunit, ATPase 1 pseudoger |
| CDYL | 6 | chromodomain Y like [Source:HGNC Symbol;A |
| AL356747.1 | 6 | novel transcript, antisense to CDYL |
| AL022725.1 | 6 | novel transcript |
| AL359643.1 | 6 | ribosomal protein S18 (RPS18) pseudogene |
| RPP40 | 6 | ribonuclease P/MRP subunit p40 [Source:HGN |
| AL359643.3 | 6 | novel transcript |
| AL359643.2 | 6 | novel transcript |
| AL139094.1 | 6 | transcription factor B2, mitochondrial (TFB2M |
| PPP1R3G | 6 | protein phosphatase 1 regulatory subunit 3G [ |
| LYRM4 | 6 | LYR motif containing 4 [Source:HGNC Symbol; |
| MIR3691 | 6 | microRNA 3691 [Source:HGNC Symbol;Acc:HG |
| FARS2 | 6 | phenylalanyl-tRNA synthetase 2, mitochondri |
| AL121978.1 | 6 | microtubule-associated protein 1 light chain 3 |
| AL021328.1 | 6 | novel transcript, antisense to FARS2 |
| HNRNPA1P37 | 6 | heterogeneous nuclear ribonucleoprotein A1 |
| AL022097.1 | 6 | novel transcript, antisense to FARS2 |
| AL136968.2 | 6 | ribosomal protein L34 (RPL34) pseudogene |
| AL136968.1 | 6 | Glyceraldehyde 3-phosphate dehydrogenase ( |
| AL136307.1 | 6 | novel transcript |
| RN7SL221P | 6 | RNA, 7SL, cytoplasmic 221, pseudogene [Sour |
| PKMP5 | 6 | pyruvate kinase M1/2 pseudogene 5 [Source:I |
| NRN1 | 6 | neuritin 1 [Source:HGNC Symbol;Acc:HGNC:17 |
| F13A1 | 6 | coagulation factor XIII A chain [Source:HGNC S |
| MIR7853 | 6 | microRNA 7853 [Source:HGNC Symbol;Acc:HG |
| MIR5683 | 6 | microRNA 5683 [Source:HGNC Symbol;Acc:HG |
| LY86-AS1 | 6 | LY86 antisense RNA 1 [Source:HGNC Symbol;A |
| SNAPC5P1 | 6 | small nuclear RNA activating complex polypep |
| AL162719.1 | 6 | novel transcript |
| CNN3P1 | 6 | calponin 3 pseudogene 1 [Source:HGNC Symb |
| LY86 | 6 | lymphocyte antigen 86 [Source:HGNC Symbol |
| AL031123.2 | 6 | novel transcript |
| AL031123.3 | 6 | TEC |
| AL031123.1 | 6 | novel transcript |
| AL031123.4 | 6 | novel transcript |
| AL031123.5 | 6 | novel transcript |
| AL136361.1 | 6 | novel transcript |

| | | |
|---|---|---|
| BTF3P7 | 6 | basic transcription factor 3 pseudogene 7 [Sou |
| AL158817.1 | 6 | novel transcript |
| RN7SL554P | 6 | RNA, 7SL, cytoplasmic 554, pseudogene [Sour |
| AL139390.1 | 6 | novel transcript |
| RREB1 | 6 | ras responsive element binding protein 1 [Sou |
| AL355336.1 | 6 | novel transcript |
| Y_RNA | 6 | Y RNA [Source:RFAM;Acc:RF00019] |
| AL589644.1 | 6 | ras responsive element binding protein 1 (RRE |
| AL139095.5 | 6 | novel transcript |
| SSR1 | 6 | signal sequence receptor subunit 1 [Source:H( |
| AL139095.4 | 6 | novel transcript |
| CAGE1 | 6 | cancer antigen 1 [Source:HGNC Symbol;Acc:H |
| AL139095.1 | 6 | ATP synthase, H+ transporting, mitochondrial |
| AL139095.2 | 6 | ribosomal protein S3 (RPS3) pseudogene |
| AL139095.3 | 6 | 1-aminocyclopropane-1-carboxylate synthase |
| RIOK1 | 6 | RIO kinase 1 [Source:HGNC Symbol;Acc:HGNC |
| HNRNPLP1 | 6 | heterogeneous nuclear ribonucleoprotein L ps |
| AL138878.1 | 6 | ribosomal protein S26 pseudogene |
| AL138878.2 | 6 | isocitrate dehydrogenase 1 (NADP+), soluble ( |
| AL031058.1 | 6 | novel transcript, antisense to DSP |
| DSP | 6 | desmoplakin [Source:HGNC Symbol;Acc:HGN( |
| SNRNP48 | 6 | small nuclear ribonucleoprotein U11/U12 sub |
| AL390026.1 | 6 | ribosomal protein L29 (RPL29) pseudogene |
| BMP6 | 6 | bone morphogenetic protein 6 [Source:HGNC |
| TXNDC5 | 6 | thioredoxin domain containing 5 [Source:HGN |
| BLOC1S5-TXNDC5 | 6 | BLOC1S5-TXNDC5 readthrough (NMD candida |
| AL096800.1 | 6 | coxsackie virus and adenovirus receptor (CXAI |
| PIP5K1P1 | 6 | phosphatidylinositol-4-phosphate 5-kinase typ |
| BLOC1S5 | 6 | biogenesis of lysosomal organelles complex 1 |
| EEF1E1-BLOC1S5 | 6 | EEF1E1-BLOC1S5 readthrough (NMD candidat |
| EEF1E1 | 6 | eukaryotic translation elongation factor 1 eps |
| AL355499.1 | 6 | novel transcript |
| AL355499.2 | 6 | novel transcript |
| AL359378.1 | 6 | novel transcript |
| SLC35B3 | 6 | solute carrier family 35 member B3 [Source:H |
| HULC | 6 | hepatocellular carcinoma up-regulated long n |
| AL161437.1 | 6 | NULL |
| AL137220.1 | 6 | novel transcript |
| OFCC1 | 6 | orofacial cleft 1 candidate 1 [Source:HGNC Sy |
| AL136226.1 | 6 | ribosomal protein L7A (RPL7A) pseudogene |
| AL354868.1 | 6 | ribosomal protein L21 (RPL21) pseudogene |
| RNU6ATAC21P | 6 | RNA, U6atac small nuclear 21, pseudogene [S |
| AL138885.1 | 6 | ribosomal protein L7 (RPL7) pseudogene |
| TFAP2A | 6 | transcription factor AP-2 alpha [Source:HGNC |
| TFAP2A-AS2 | 6 | TFAP2A antisense RNA 2 [Source:NCBI gene;A |
| TFAP2A-AS1 | 6 | TFAP2A antisense RNA 1 [Source:HGNC Symb |
| AL138885.2 | 6 | novel transcript |
| AL138885.3 | 6 | novel transcript |
| LINC00518 | 6 | long intergenic non-protein coding RNA 518 [S |
| MIR5689HG | 6 | MIR5689 host gene [Source:HGNC Symbol;Ac |

| | | |
|---|---|---|
| MIR5689 | 6 | microRNA 5689 [Source:HGNC Symbol;Acc:HG |
| MRPL48P1 | 6 | mitochondrial ribosomal protein L48 pseudog |
| LINC02522 | 6 | long intergenic non-protein coding RNA 2522 |
| GCNT2 | 6 | glucosaminyl (N-acetyl) transferase 2 (I blood |
| AL139039.3 | 6 | novel transcript, antisense to GCNT2 |
| AL139039.1 | 6 | glucosaminyl (N-acetyl) transferase 1, core 2 ( |
| AL139039.2 | 6 | ribosomal protein L21 (RPL21) pseudogene |
| GCNT2P1 | 6 | GCNT2 pseudogene 1 [Source:HGNC Symbol;A |
| AL358777.1 | 6 | novel transcript, antisense to C6orf52 |
| C6orf52 | 6 | chromosome 6 open reading frame 52 [Sourc |
| Y_RNA | 6 | Y RNA [Source:RFAM;Acc:RF00019] |
| PAK1IP1 | 6 | PAK1 interacting protein 1 [Source:HGNC Sym |
| TMEM14C | 6 | transmembrane protein 14C [Source:HGNC Sy |
| AL024498.1 | 6 | novel transcript |
| TMEM14B | 6 | transmembrane protein 14B [Source:HGNC Sy |
| AL024498.2 | 6 | novel transcript, TMEM14B-SYCP2L readthrou |
| RNA5SP203 | 6 | RNA, 5S ribosomal pseudogene 203 [Source:H |
| MAK | 6 | male germ cell associated kinase [Source:HGN |
| GCM2 | 6 | glial cells missing transcription factor 2 [Sourc |
| AL357497.1 | 6 | novel transcript, antisense to GCM2 |
| SYCP2L | 6 | synaptonemal complex protein 2 like [Source: |
| ELOVL2 | 6 | ELOVL fatty acid elongase 2 [Source:HGNC Syr |
| AL121955.1 | 6 | siah E3 ubiquitin protein ligase family membe |
| ELOVL2-AS1 | 6 | ELOVL2 antisense RNA 1 [Source:HGNC Symbo |
| SMIM13 | 6 | small integral membrane protein 13 [Source:H |
| ERVFRD-1 | 6 | endogenous retrovirus group FRD member 1, |
| AL136139.1 | 6 | P antigen family, member 4 (prostate associat |
| AL139807.1 | 6 | novel transcript |
| NEDD9 | 6 | neural precursor cell expressed, development |
| AL022098.1 | 6 | novel transcript, antisense to NEDD9 |
| AL445430.2 | 6 | novel transcript |
| AL445430.1 | 6 | novel transcript |
| RNU1-64P | 6 | RNA, U1 small nuclear 64, pseudogene [Sourc |
| THAP12P5 | 6 | THAP domain containing 12 pseudogene 5 [Sc |
| TMEM170B | 6 | transmembrane protein 170B [Source:HGNC S |
| AL357518.1 | 6 | novel transcript |
| Metazoa_SRP | 6 | Metazoan signal recognition particle RNA [Sou |
| AL022724.1 | 6 | NULL |
| ADTRP | 6 | androgen dependent TFPI regulating protein [ |
| AL022724.3 | 6 | novel transcript, antisense to ADTRP |
| AL022724.2 | 6 | novel transcript |
| AMD1P4 | 6 | adenosylmethionine decarboxylase 1 pseudog |
| AL157373.1 | 6 | novel transcript |
| AL157373.2 | 6 | novel transcript |
| HIVEP1 | 6 | human immunodeficiency virus type I enhanc |
| EDN1 | 6 | endothelin 1 [Source:HGNC Symbol;Acc:HGNC |
| SUMO2P12 | 6 | SUMO2 pseudogene 12 [Source:HGNC Symbo |
| RN7SKP293 | 6 | RN7SK pseudogene 293 [Source:HGNC Symbo |
| RPL15P3 | 6 | ribosomal protein L15 pseudogene 3 [Source: |
| LINC02530 | 6 | long intergenic non-protein coding RNA 2530 |

| | | |
|---|---|---|
| PHACTR1 | 6 | phosphatase and actin regulator 1 [Source:HG |
| RNU1-11P | 6 | RNA, U1 small nuclear 11, pseudogene [Sourc |
| AL008729.1 | 6 | novel transcript, antisense to PHACTR1 |
| TBC1D7 | 6 | TBC1 domain family member 7 [Source:HGNC |
| AL008729.2 | 6 | novel transcript |
| GFOD1 | 6 | glucose-fructose oxidoreductase domain cont |
| GFOD1-AS1 | 6 | GFOD1 antisense RNA 1 [Source:HGNC Symbo |
| RPS4XP7 | 6 | ribosomal protein S4X pseudogene 7 [Source: |
| RN7SKP204 | 6 | RN7SK pseudogene 204 [Source:HGNC Symbo |
| SIRT5 | 6 | sirtuin 5 [Source:HGNC Symbol;Acc:HGNC:14 |
| AL441883.1 | 6 | novel transcript, antisense to NOL7 |
| NOL7 | 6 | nucleolar protein 7 [Source:HGNC Symbol;Acc |
| RANBP9 | 6 | RAN binding protein 9 [Source:HGNC Symbol; |
| Z93020.1 | 6 | novel transcript, antisense to RANBP9 |
| MCUR1 | 6 | mitochondrial calcium uniporter regulator 1 [ |
| AL023583.1 | 6 | novel transcript |
| RNF182 | 6 | ring finger protein 182 [Source:HGNC Symbol; |
| MRPL35P1 | 6 | mitochondrial ribosomal protein L35 pseudog |
| AL022396.1 | 6 | novel transcript |
| CD83 | 6 | CD83 molecule [Source:HGNC Symbol;Acc:HG |
| AL133259.1 | 6 | novel transcript |
| RNU7-133P | 6 | RNA, U7 small nuclear 133 pseudogene [Sour |
| AL353152.2 | 6 | novel transcript |
| AL353152.1 | 6 | novel transcript |
| LINC01108 | 6 | long intergenic non-protein coding RNA 1108 |
| AL080313.2 | 6 | novel transcript |
| AL080313.1 | 6 | novel transcript |
| AL009177.1 | 6 | novel transcript |
| AL109914.1 | 6 | novel transcript |
| RNU6-793P | 6 | RNA, U6 small nuclear 793, pseudogene [Sour |
| AL138720.1 | 6 | novel transcript |
| AL050335.1 | 6 | ribosomal protein L6 (RPL6) pseudogene |
| RN7SL332P | 6 | RNA, 7SL, cytoplasmic 332, pseudogene [Sour |
| AL136162.1 | 6 | novel transcript |
| JARID2 | 6 | jumonji and AT-rich interaction domain conta |
| JARID2-AS1 | 6 | JARID2 antisense RNA 1 [Source:HGNC Symbo |
| RNU6-522P | 6 | RNA, U6 small nuclear 522, pseudogene [Sour |
| RNU6-645P | 6 | RNA, U6 small nuclear 645, pseudogene [Sour |
| 5S_rRNA | 6 | 5S ribosomal RNA [Source:RFAM;Acc:RF00001 |
| DTNBP1 | 6 | dystrobrevin binding protein 1 [Source:HGNC |
| AL021978.1 | 6 | novel transcript, antisense to DTNBP1 |
| ARPC3P5 | 6 | actin related protein 2/3 complex subunit 3 p |
| LINC02543 | 6 | long intergenic non-protein coding RNA 2543 |
| AL365265.1 | 6 | mitochondrial calcium uptake 2 (MICU2) pseu |
| AL021407.2 | 6 | malate dehydrogenase pseudogene |
| MYLIP | 6 | myosin regulatory light chain interacting prot |
| MIR4639 | 6 | microRNA 4639 [Source:HGNC Symbol;Acc:HG |
| AL021407.3 | 6 | PERP, TP53 apoptosis effector (PERP) pseudog |
| AL021407.1 | 6 | pseudogene similar to part of eukaryotic tran |
| MRPL42P2 | 6 | mitochondrial ribosomal protein L42 pseudog |

| | | |
|---|---|---|
| RNU6-1114P | 6 | RNA, U6 small nuclear 1114, pseudogene [Sou |
| Y_RNA | 6 | Y RNA [Source:RFAM;Acc:RF00019] |
| GMPR | 6 | guanosine monophosphate reductase [Source |
| AL009031.1 | 6 | novel transcript, antisense to GMPR |
| ATXN1 | 6 | ataxin 1 [Source:HGNC Symbol;Acc:HGNC:105 |
| AL137003.1 | 6 | novel transcript |
| AL137003.2 | 6 | novel transcript |
| AL133405.2 | 6 | novel transcript |
| AL133405.1 | 6 | novel transcript |
| AL138740.1 | 6 | novel transcript |
| STMND1 | 6 | stathmin domain containing 1 [Source:HGNC S |
| AL136305.1 | 6 | novel transcript, antisense to RBM24 |
| RBM24 | 6 | RNA binding motif protein 24 [Source:HGNC S |
| AL034372.1 | 6 | ESD (esterase D/formylglutathione hydrolase) |
| CAP2 | 6 | cyclase associated actin cytoskeleton regulato |
| RPL7P26 | 6 | ribosomal protein L7 pseudogene 26 [Source: |
| SUMO2P13 | 6 | SUMO2 pseudogene 13 [Source:HGNC Symbo |
| AL138824.1 | 6 | novel transcript |
| FAM8A1 | 6 | family with sequence similarity 8 member A1 |
| NUP153 | 6 | nucleoporin 153 [Source:HGNC Symbol;Acc:H |
| RNU6-190P | 6 | RNA, U6 small nuclear 190, pseudogene [Sour |
| AL138724.1 | 6 | novel transcript, antisense to NUP153 |
| RNA5SP204 | 6 | RNA, 5S ribosomal pseudogene 204 [Source:H |
| Y_RNA | 6 | Y RNA [Source:RFAM;Acc:RF00019] |
| KIF13A | 6 | kinesin family member 13A [Source:HGNC Syr |
| AL023807.1 | 6 | ribosomal protein S12 (RPS12) pseudogene |
| NHLRC1 | 6 | NHL repeat containing E3 ubiquitin protein lig |
| TPMT | 6 | thiopurine S-methyltransferase [Source:HGNC |
| KDM1B | 6 | lysine demethylase 1B [Source:HGNC Symbol; |
| DEK | 6 | DEK proto-oncogene [Source:HGNC Symbol;A |
| Y_RNA | 6 | Y RNA [Source:RFAM;Acc:RF00019] |
| AL138825.1 | 6 | chromosome 15 open reading frame 40 (C15o |
| RNU6-263P | 6 | RNA, U6 small nuclear 263, pseudogene [Sour |
| DDX18P3 | 6 | DEAD-box helicase 18 pseudogene 3 [Source: |
| AL138725.1 | 6 | IMP (inosine monophosphate) dehydrogenase |
| RNF144B | 6 | ring finger protein 144B [Source:HGNC Symbo |
| MIR548A1HG | 6 | MIR548A1 host gene [Source:HGNC Symbol;A |
| MIR548A1 | 6 | microRNA 548a-1 [Source:HGNC Symbol;Acc: |
| AL589647.1 | 6 | ribosomal protein L21 (RPL21) pseudogene |
| AL357052.1 | 6 | novel transcript |
| AL022068.1 | 6 | novel transcript, antisense to ID4 |
| RPL5P20 | 6 | ribosomal protein L5 pseudogene 20 [Source: |
| RNA5SP205 | 6 | RNA, 5S ribosomal pseudogene 205 [Source:H |
| KRT18P38 | 6 | keratin 18 pseudogene 38 [Source:HGNC Sym |
| UQCRFS1P3 | 6 | ubiquinol-cytochrome c reductase, Rieske iror |
| RNU6-801P | 6 | RNA, U6 small nuclear 801, pseudogene [Sour |
| ID4 | 6 | inhibitor of DNA binding 4, HLH protein [Sourc |
| AL022726.1 | 6 | novel transcript |
| AL008627.1 | 6 | ribosomal protein L29 (RPL29) pseudogene |
| MBOAT1 | 6 | membrane bound O-acyltransferase domain c |

| | | |
|---|---|---|
| AL158198.1 | 6 | novel transcript, antisense to MBOAT1 |
| AL158198.2 | 6 | novel transcript |
| AL132775.1 | 6 | novel transcript |
| AL132775.2 | 6 | novel transcript |
| E2F3 | 6 | E2F transcription factor 3 [Source:HGNC Symb |
| RN7SL128P | 6 | RNA, 7SL, cytoplasmic 128, pseudogene [Sour |
| E2F3-IT1 | 6 | E2F3 intronic transcript 1 [Source:HGNC Symb |
| RNU6-141P | 6 | RNA, U6 small nuclear 141, pseudogene [Sour |
| Y_RNA | 6 | Y RNA [Source:RFAM;Acc:RF00019] |
| CDKAL1 | 6 | CDK5 regulatory subunit associated protein 1 |
| AL035090.1 | 6 | ribosomal protein L36A (RPL36A) pseudogene |
| AL513188.1 | 6 | novel transcript |
| RNU6-150P | 6 | RNA, U6 small nuclear 150, pseudogene [Sour |
| AL451080.1 | 6 | novel transcript |
| AL031767.1 | 6 | novel transcript |
| AL031767.2 | 6 | novel transcript |
| LINC00581 | 6 | long intergenic non-protein coding RNA 581 [S |
| AL512380.1 | 6 | novel transcript |
| AL512380.2 | 6 | novel transcript, antisense to SOX4 |
| SOX4 | 6 | SRY-box transcription factor 4 [Source:HGNC S |
| BOLA2P3 | 6 | bolA family member 2 pseudogene 3 [Source: |
| CASC15 | 6 | cancer susceptibility 15 [Source:HGNC Symbo |
| AL136313.1 | 6 | TEC |
| RN7SKP240 | 6 | RN7SK pseudogene 240 [Source:HGNC Symbo |
| NBAT1 | 6 | neuroblastoma associated transcript 1 [Sourc |
| AL359694.1 | 6 | neugrin, neurite outgrowth associated (NGRN |
| PRL | 6 | prolactin [Source:HGNC Symbol;Acc:HGNC:94 |
| HDGFL1 | 6 | HDGF like 1 [Source:HGNC Symbol;Acc:HGNC: |
| AL033539.1 | 6 | novel transcript |
| AL033539.2 | 6 | novel transcript |
| AL035401.1 | 6 | novel transcript |
| AL591416.1 | 6 | ribosomal protein L6 (RPL6) pseudogene |
| RNU6-1060P | 6 | RNA, U6 small nuclear 1060, pseudogene [Sou |
| AL139231.1 | 6 | novel transcript |
| AL133270.1 | 6 | ferritin pseudogene |
| AL139093.1 | 6 | nuclear pore associated protein 1 (NPAP1) pse |
| SPTLC1P2 | 6 | serine palmitoyltransferase long chain base su |
| AL032822.1 | 6 | interferon stimulated exonuclease gene 20kD |
| HNRNPA1P58 | 6 | heterogeneous nuclear ribonucleoprotein A1 |
| NRSN1 | 6 | neurensin 1 [Source:HGNC Symbol;Acc:HGNC: |
| FO393410.1 | 6 | NULL |
| DCDC2 | 6 | doublecortin domain containing 2 [Source:HG |
| KAAG1 | 6 | kidney associated antigen 1 [Source:HGNC Syr |
| RNU6-391P | 6 | RNA, U6 small nuclear 391, pseudogene [Sour |
| MRS2 | 6 | magnesium transporter MRS2 [Source:HGNC S |
| GPLD1 | 6 | glycosylphosphatidylinositol specific phosphol |
| ALDH5A1 | 6 | aldehyde dehydrogenase 5 family member A1 |
| KIAA0319 | 6 | KIAA0319 [Source:HGNC Symbol;Acc:HGNC:2 |
| KRT8P43 | 6 | keratin 8 pseudogene 43 [Source:HGNC Symb |
| TDP2 | 6 | tyrosyl-DNA phosphodiesterase 2 [Source:HG |

| | | |
|---|---|---|
| ACOT13 | 6 | acyl-CoA thioesterase 13 [Source:HGNC Symb |
| AL031775.1 | 6 | novel transcript, antisense to ACOT13 |
| C6orf62 | 6 | chromosome 6 open reading frame 62 [Sourc |
| AL031775.2 | 6 | novel transcript, antisense to C6orf62 |
| LINC02828 | 6 | long intergenic non-protein coding RNA 2828 |
| AL133264.1 | 6 | adaptor-related protein complex 3, sigma 1 su |
| GMNN | 6 | geminin DNA replication inhibitor [Source:HG |
| ARMH2 | 6 | armadillo like helical domain containing 2 [Sou |
| AL512428.1 | 6 | novel protein |
| RIPOR2 | 6 | RHO family interacting cell polarization regula |
| MTCO2P33 | 6 | MT-CO2 pseudogene 33 [Source:HGNC Symbo |
| PPIAP29 | 6 | peptidylprolyl isomerase A pseudogene 29 [So |
| AL160400.1 | 6 | novel transcript, antisense to RIPOR2 |
| ASS1P1 | 6 | argininosuccinate synthetase 1 pseudogene 1 |
| RN7SL334P | 6 | RNA, 7SL, cytoplasmic 334, pseudogene [Sour |
| AL133268.3 | 6 | novel transcript |
| AL133268.2 | 6 | novel transcript |
| CMAHP | 6 | cytidine monophospho-N-acetylneuraminic ac |
| AL133268.4 | 6 | Novel transcript, antisense to CMAHP |
| AL133268.1 | 6 | nucleoporin 50kD (NUP50) pseudogene |
| AL590084.3 | 6 | calcium activated nucleotidase 1 (CANT1) pseu |
| AL590084.1 | 6 | ubiquitin-conjugating enzyme pseudogene |
| AL590084.2 | 6 | aryl hydrocarbon receptor (AHR) pseudogene |
| KATNBL1P5 | 6 | katanin regulatory subunit B1 like 1 pseudoge |
| AL024509.2 | 6 | 60S Ribosomal Protein L21 (RPL21) pseudoger |
| AL024509.1 | 6 | heterogeneous nuclear ribonucleoprotein pse |
| CARMIL1 | 6 | capping protein regulator and myosin 1 linker |
| Y_RNA | 6 | Y RNA [Source:RFAM;Acc:RF00019] |
| RNU6-987P | 6 | RNA, U6 small nuclear 987, pseudogene [Sour |
| AL022170.1 | 6 | novel transcript |
| SCGN | 6 | secretagogin, EF-hand calcium binding proteir |
| PRELID1P2 | 6 | PRELI domain containing 1 pseudogene 2 [Sou |
| HIST1H2AA | 6 | histone cluster 1 H2A family member a [Sourc |
| HIST1H2BA | 6 | histone cluster 1 H2B family member a [Sourc |
| HIST1H2BPS1 | 6 | histone cluster 1 H2B pseudogene 1 [Source:H |
| HIST1H2APS1 | 6 | histone cluster 1 H2A pseudogene 1 [Source:H |
| SLC17A4 | 6 | solute carrier family 17 member 4 [Source:HG |
| SLC17A1 | 6 | solute carrier family 17 member 1 [Source:HG |
| SLC17A3 | 6 | solute carrier family 17 member 3 [Source:HG |
| HIST1H2APS2 | 6 | histone cluster 1 H2A pseudogene 2 [Source:H |
| AL138726.1 | 6 | histone cluster 1, H2bo (HIST1H2BO) pseudog |
| SLC17A2 | 6 | solute carrier family 17 member 2 [Source:HG |
| TRIM38 | 6 | tripartite motif containing 38 [Source:HGNC S |
| U91328.2 | 6 | novel transcript, antisense to TRIM38 |
| U91328.1 | 6 | novel transcript |
| U91328.3 | 6 | novel transcript |
| HIST1H1PS2 | 6 | histone cluster 1 H1 pseudogene 2 [Source:HC |
| HIST1H1A | 6 | histone cluster 1 H1 family member a [Source |
| HIST1H3A | 6 | histone cluster 1 H3 family member a [Source |
| HIST1H4A | 6 | histone cluster 1 H4 family member a [Source |

| | | |
|---|---|---|
| HIST1H4B | 6 | histone cluster 1 H4 family member b [Source |
| HIST1H3B | 6 | histone cluster 1 H3 family member b [Source |
| HIST1H2AB | 6 | histone cluster 1 H2A family member b [Sourc |
| HIST1H2BB | 6 | histone cluster 1 H2B family member b [Sourc |
| HIST1H2APS5 | 6 | histone cluster 1 H2A pseudogene 5 [Source:H |
| HIST1H3C | 6 | histone cluster 1 H3 family member c [Source |
| HIST1H1C | 6 | histone cluster 1 H1 family member c [Source |
| HFE | 6 | homeostatic iron regulator [Source:HGNC Sym |
| HIST1H4C | 6 | histone cluster 1 H4 family member c [Source |
| HIST1H1T | 6 | histone cluster 1 H1 family member t [Source: |
| HIST1H2BC | 6 | histone cluster 1 H2B family member c [Sourc |
| HIST1H2AC | 6 | histone cluster 1 H2A family member c [Sourc |
| HIST1H1E | 6 | histone cluster 1 H1 family member e [Source |
| HIST1H2BD | 6 | histone cluster 1 H2B family member d [Sourc |
| AL353759.1 | 6 | novel transcript, antisense to HIST1H2BD |
| LARP1P1 | 6 | La ribonucleoprotein domain family member : |
| HIST1H2BE | 6 | histone cluster 1 H2B family member e [Sourc |
| HIST1H4D | 6 | histone cluster 1 H4 family member d [Source |
| HIST1H1PS1 | 6 | histone cluster 1 H1 pseudogene 1 [Source:HC |
| AL031777.3 | 6 | novel protein |
| HIST1H3D | 6 | histone cluster 1 H3 family member d [Source |
| HIST1H2AD | 6 | histone cluster 1 H2A family member d [Sourc |
| HIST1H2BF | 6 | histone cluster 1 H2B family member f [Sourc |
| AL031777.1 | 6 | ribosomal protein S10 (RPS10) pseudogene |
| HIST1H4E | 6 | histone cluster 1 H4 family member e [Source |
| HIST1H2BG | 6 | histone cluster 1 H2B family member g [Sourc |
| HIST1H2AE | 6 | histone cluster 1 H2A family member e [Sourc |
| AL031777.2 | 6 | histone cluster 1, H3b (HIST1H3B) pseudogen |
| HIST1H3E | 6 | histone cluster 1 H3 family member e [Source |
| HIST1H2APS3 | 6 | histone cluster 1 H2A pseudogene 3 [Source:H |
| HIST1H1D | 6 | histone cluster 1 H1 family member d [Source |
| HIST1H4F | 6 | histone cluster 1 H4 family member f [Source: |
| HIST1H4G | 6 | histone cluster 1 H4 family member g [Source |
| HIST1H3F | 6 | histone cluster 1 H3 family member f [Source: |
| HIST1H2BH | 6 | histone cluster 1 H2B family member h [Sourc |
| HIST1H3G | 6 | histone cluster 1 H3 family member g [Source |
| HIST1H2APS4 | 6 | histone cluster 1 H2A pseudogene 4 [Source:H |
| HIST1H2BI | 6 | histone cluster 1 H2B family member i [Sourc |
| HIST1H4H | 6 | histone cluster 1 H4 family member h [Source |
| HIST1H3PS1 | 6 | histone cluster 1 H3 pseudogene 1 [Source:HC |
| AL021917.1 | 6 | novel transcript |
| BTN3A2 | 6 | butyrophilin subfamily 3 member A2 [Source: |
| BTN2A2 | 6 | butyrophilin subfamily 2 member A2 [Source: |
| BTN3A1 | 6 | butyrophilin subfamily 3 member A1 [Source: |
| BTN2A3P | 6 | butyrophilin subfamily 2 member A3, pseudog |
| BTN3A3 | 6 | butyrophilin subfamily 3 member A3 [Source: |
| BTN2A1 | 6 | butyrophilin subfamily 2 member A1 [Source: |
| AL121936.2 | 6 | butyrophilin family pseudogene |
| BTN1A1 | 6 | butyrophilin subfamily 1 member A1 [Source: |
| RNU6-502P | 6 | RNA, U6 small nuclear 502, pseudogene [Sour |

| | | |
|---|---|---|
| HCG11 | 6 | HLA complex group 11 [Source:HGNC Symbol; |
| HMGN4 | 6 | high mobility group nucleosomal binding dom |
| AL121936.1 | 6 | novel transcript |
| ABT1 | 6 | activator of basal transcription 1 [Source:HGN |
| AL513548.3 | 6 | novel transcript |
| AL513548.2 | 6 | POM121 transmembrane nucleoporin C (POM |
| ZNF322 | 6 | zinc finger protein 322 [Source:HGNC Symbol; |
| AL513548.4 | 6 | novel transcript |
| AL513548.1 | 6 | novel transcript |
| GUSBP2 | 6 | GUSB pseudogene 2 [Source:HGNC Symbol;Ac |
| POM121L6P | 6 | POM121 transmembrane nucleoporin like 6, p |
| LINC00240 | 6 | long intergenic non-protein coding RNA 240 [S |
| AL133255.1 | 6 | novel transcript |
| AL590062.1 | 6 | novel transcript |
| VN1R12P | 6 | vomeronasal 1 receptor 12, pseudogene [Sou |
| VN1R13P | 6 | vomeronasal 1 receptor 13 pseudogene [Sour |
| VN1R11P | 6 | vomeronasal 1 receptor 11, pseudogene [Sou |
| RNU2-62P | 6 | RNA, U2 small nuclear 62, pseudogene [Sourc |
| AL021807.1 | 6 | novel transcript |
| HIST1H2BJ | 6 | histone cluster 1 H2B family member j [Source |
| HIST1H2AG | 6 | histone cluster 1 H2A family member g [Sourc |
| HIST1H4I | 6 | histone cluster 1 H4 family member i [Source: |
| HIST1H2BK | 6 | histone cluster 1 H2B family member k [Sourc |
| HIST1H2AH | 6 | histone cluster 1 H2A family member h [Sourc |
| MIR3143 | 6 | microRNA 3143 [Source:HGNC Symbol;Acc:HG |
| RPL10P2 | 6 | ribosomal protein L10 pseudogene 2 [Source: |
| PRSS16 | 6 | serine protease 16 [Source:HGNC Symbol;Acc |
| AL021808.1 | 6 | cell division cycle associated 7 (CDCA7) pseud |
| POM121L2 | 6 | POM121 transmembrane nucleoporin like 2 [S |
| VN1R10P | 6 | vomeronasal 1 receptor 10 pseudogene [Sour |
| ZNF204P | 6 | zinc finger protein 204, pseudogene [Source:H |
| ZNF391 | 6 | zinc finger protein 391 [Source:HGNC Symbol; |
| AL031118.1 | 6 | novel transcript |
| MCFD2P1 | 6 | multiple coagulation factor deficiency 2 pseuc |
| ZNF184 | 6 | zinc finger protein 184 [Source:HGNC Symbol; |
| AL021918.4 | 6 | novel transcript, antisense to ZNF184 |
| AL021918.5 | 6 | novel transcript |
| AL021918.3 | 6 | novel transcript |
| AL021918.2 | 6 | sterile alpha motif domain containing 12 (SAM |
| AL021918.1 | 6 | WD repeat domain 59 (WDR59) pseudogene |
| HNRNPA1P1 | 6 | heterogeneous nuclear ribonucleoprotein A1 |
| CD83P1 | 6 | CD83 molecule pseudogene 1 [Source:HGNC S |
| RNU6-471P | 6 | RNA, U6 small nuclear 471, pseudogene [Sour |
| AL031229.1 | 6 | 60S ribosomal protein L8 (RPL8) pseudogene |
| AL009179.3 | 6 | novel transcript |
| AL009179.1 | 6 | long intergenic non-protein coding RNA 1012 |
| GPR89P | 6 | G protein-coupled receptor 89 pseudogene [S |
| AL009179.2 | 6 | novel transcript |
| RSL24D1P1 | 6 | ribosomal L24 domain containing 1 pseudoger |
| HIST1H4PS1 | 6 | histone cluster 1 H4 pseudogene 1 [Source:HG |

| | | |
|---|---|---|
| HIST1H2BL | 6 | histone cluster 1 H2B family member l [Source |
| HIST1H2AI | 6 | histone cluster 1 H2A family member i [Source |
| HIST1H3H | 6 | histone cluster 1 H3 family member h [Source |
| HIST1H2AJ | 6 | histone cluster 1 H2A family member j [Source |
| HIST1H2BM | 6 | histone cluster 1 H2B family member m [Sour |
| HIST1H4J | 6 | histone cluster 1 H4 family member j [Source: |
| HIST1H4K | 6 | histone cluster 1 H4 family member k [Source |
| HIST1H2BN | 6 | histone cluster 1 H2B family member n [Sourc |
| HIST1H2AK | 6 | histone cluster 1 H2A family member k [Sourc |
| HIST1H2BPS2 | 6 | histone cluster 1 H2B pseudogene 2 [Source:H |
| HIST1H2AL | 6 | histone cluster 1 H2A family member l [Source |
| HIST1H1B | 6 | histone cluster 1 H1 family member b [Source |
| HIST1H3I | 6 | histone cluster 1 H3 family member i [Source: |
| HIST1H4L | 6 | histone cluster 1 H4 family member l [Source: |
| HIST1H3J | 6 | histone cluster 1 H3 family member j [Source: |
| HIST1H2AM | 6 | histone cluster 1 H2A family member m [Sour |
| HIST1H2BO | 6 | histone cluster 1 H2B family member o [Sourc |
| RNU7-26P | 6 | RNA, U7 small nuclear 26 pseudogene [Source |
| OR2B2 | 6 | olfactory receptor family 2 subfamily B memb |
| OR2W6P | 6 | olfactory receptor family 2 subfamily W meml |
| OR2B6 | 6 | olfactory receptor family 2 subfamily B memb |
| RPLP2P1 | 6 | ribosomal protein lateral stalk subunit P2 pseu |
| OR2W4P | 6 | olfactory receptor family 2 subfamily W meml |
| IQCB2P | 6 | IQ motif containing B2 pseudogene [Source:H |
| ZSCAN16-AS1 | 6 | ZSCAN16 antisense RNA 1 [Source:HGNC Syml |
| U3 | 6 | Small nucleolar RNA U3 [Source:RFAM;Acc:RF |
| OR2W2P | 6 | olfactory receptor family 2 subfamily W meml |
| OR2B7P | 6 | olfactory receptor family 2 subfamily B memb |
| OR2B8P | 6 | olfactory receptor family 2 subfamily B memb |
| OR1F12 | 6 | olfactory receptor family 1 subfamily F memb |
| AL121944.1 | 6 | novel transcript, antisense to ZNF165 |
| ZNF165 | 6 | zinc finger protein 165 [Source:HGNC Symbol; |
| ZSCAN12P1 | 6 | zinc finger and SCAN domain containing 12 ps |
| ZNF602P | 6 | zinc finger protein 602, pseudogene [Source:H |
| ZSCAN16 | 6 | zinc finger and SCAN domain containing 16 [So |
| AL358933.1 | 6 | novel transcript, overlapping to ZNF435 |
| ZKSCAN8 | 6 | zinc finger with KRAB and SCAN domains 8 [Sc |
| ZNF192P1 | 6 | zinc finger protein 192 pseudogene 1 [Source: |
| AL022393.1 | 6 | TEC |
| ZNF603P | 6 | zinc finger protein 603, pseudogene [Source:H |
| ZNF192P2 | 6 | zinc finger protein 192 pseudogene 2 [Source: |
| TOB2P1 | 6 | transducer of ERBB2, 2 pseudogene 1 [Source |
| ZSCAN9 | 6 | zinc finger and SCAN domain containing 9 [Sou |
| ZKSCAN4 | 6 | zinc finger with KRAB and SCAN domains 4 [Sc |
| NKAPL | 6 | NFKB activating protein like [Source:HGNC Syr |
| ZSCAN26 | 6 | zinc finger and SCAN domain containing 26 [So |
| AL021997.3 | 6 | novel protein |
| PGBD1 | 6 | piggyBac transposable element derived 1 [Sou |
| AL021997.2 | 6 | zinc finger and SCAN domain containing 16 (ZS |
| AL021997.1 | 6 | small integral membrane protein 15 (SMIM15 |

| | | |
|---|---|---|
| ZSCAN31 | 6 | zinc finger and SCAN domain containing 31 [Sc |
| ZKSCAN3 | 6 | zinc finger with KRAB and SCAN domains 3 [Sc |
| ZSCAN12 | 6 | zinc finger and SCAN domain containing 12 [Sc |
| RNU2-45P | 6 | RNA, U2 small nuclear 45, pseudogene [Sourc |
| ZSCAN23 | 6 | zinc finger and SCAN domain containing 23 [Sc |
| COX11P1 | 6 | COX11 pseudogene 1 [Source:HGNC Symbol;A |
| OR2E1P | 6 | olfactory receptor family 2 subfamily E memb |
| Z98745.2 | 6 | novel transcript |
| Z98745.1 | 6 | novel transcript |
| GPX6 | 6 | glutathione peroxidase 6 [Source:HGNC Symb |
| GPX5 | 6 | glutathione peroxidase 5 [Source:HGNC Symb |
| ZBED9 | 6 | zinc finger BED-type containing 9 [Source:HGN |
| AL049543.1 | 6 | novel transcript |
| AL121932.1 | 6 | cytochrome c oxidase subunit VIIIC (COX8C) p |
| AL121932.2 | 6 | novel transcript |
| LINC00533 | 6 | long intergenic non-protein coding RNA 533 [S |
| RPSAP2 | 6 | ribosomal protein SA pseudogene 2 [Source:H |
| NOP56P1 | 6 | NOP56 ribonucleoprotein pseudogene 1 [Sour |
| AL662890.1 | 6 | novel transcript |
| LINC01623 | 6 | long intergenic non-protein coding RNA 1623 |
| RPL13P | 6 | ribosomal protein L13 pseudogene [Source:H( |
| ZNF90P2 | 6 | zinc finger protein 90 pseudogene 2 [Source:H |
| HCG14 | 6 | HLA complex group 14 [Source:HGNC Symbol; |
| TRIM27 | 6 | tripartite motif containing 27 [Source:HGNC S |
| RNU6-930P | 6 | RNA, U6 small nuclear 930, pseudogene [Sour |
| LINC01556 | 6 | long intergenic non-protein coding RNA 1556 |
| KRT18P1 | 6 | keratin 18 pseudogene 1 [Source:HGNC Symb |
| RN7SL471P | 6 | RNA, 7SL, cytoplasmic 471, pseudogene [Sour |
| HCG15 | 6 | HLA complex group 15 [Source:HGNC Symbol; |
| AL662791.1 | 6 | HLA complex group 16 |
| ZNF311 | 6 | zinc finger protein 311 [Source:HGNC Symbol; |
| OR2AD1P | 6 | olfactory receptor family 2 subfamily AD mem |
| AL662791.2 | 6 | novel transcript, antisense to OR2W1 |
| OR2W1 | 6 | olfactory receptor family 2 subfamily W meml |
| OR2P1P | 6 | olfactory receptor family 2 subfamily P memb |
| SAR1AP1 | 6 | secretion associated Ras related GTPase 1A ps |
| OR2B3 | 6 | olfactory receptor family 2 subfamily B memb |
| OR2J1 | 6 | olfactory receptor family 2 subfamily J membe |
| OR2J3 | 6 | olfactory receptor family 2 subfamily J membe |
| AL645937.2 | 6 | novel transcript |
| AL645937.1 | 6 | olfactory receptor, family 2, subfamily N, men |
| AL645937.4 | 6 | novel transcript |
| OR2J2 | 6 | olfactory receptor family 2 subfamily J membe |
| OR2J4P | 6 | olfactory receptor family 2 subfamily J membe |
| AL645937.3 | 6 | olfactory receptor, family 2, subfamily H, men |
| AL672167.1 | 6 | olfactory receptor, family 2, subfamily B, mem |
| OR2G1P | 6 | olfactory receptor family 2 subfamily G memb |
| OR2U1P | 6 | olfactory receptor family 2 subfamily U memb |
| OR2U2P | 6 | olfactory receptor family 2 subfamily U memb |
| OR14J1 | 6 | olfactory receptor family 14 subfamily J meml |

| | | |
|---|---|---|
| DDX6P1 | 6 | DEAD-box helicase 6 pseudogene 1 [Source:H |
| OR5V1 | 6 | olfactory receptor family 5 subfamily V memb |
| OR12D3 | 6 | olfactory receptor family 12 subfamily D mem |
| OR12D2 | 6 | olfactory receptor family 12 subfamily D mem |
| OR12D1 | 6 | olfactory receptor family 12 subfamily D mem |
| OR11A1 | 6 | olfactory receptor family 11 subfamily A mem |
| OR10C1 | 6 | olfactory receptor family 10 subfamily C mem |
| AL645927.1 | 6 | YTH domain family, member 1 (YTHDF1) pseu |
| OR2H1 | 6 | olfactory receptor family 2 subfamily H memb |
| UBDP1 | 6 | ubiquitin D pseudogene 1 [Source:HGNC Symb |
| MAS1LP1 | 6 | MAS1L pseudogene 1 [Source:HGNC Symbol;A |
| MAS1L | 6 | MAS1 proto-oncogene like, G protein-couplec |
| RPS17P1 | 6 | ribosomal protein S17 pseudogene 1 [Source: |
| LINC02829 | 6 | long intergenic non-protein coding RNA 2829 |
| LINC01015 | 6 | long intergenic non-protein coding RNA 1015 |
| GPR53P | 6 | G protein-coupled receptor 53, pseudogene [S |
| OR2I1P | 6 | olfactory receptor family 2 subfamily I membe |
| UBD | 6 | ubiquitin D [Source:HGNC Symbol;Acc:HGNC:1 |
| GABBR1 | 6 | gamma-aminobutyric acid type B receptor sub |
| OR2H5P | 6 | olfactory receptor family 2 subfamily H memb |
| TMEM183AP1 | 6 | transmembrane protein 183A pseudogene 1 [ |
| SNORD32B | 6 | small nucleolar RNA, C/D box 32B [Source:HG |
| RPL13AP | 6 | ribosomal protein L13a pseudogene [Source:H |
| OR2H2 | 6 | olfactory receptor family 2 subfamily H memb |
| SUMO2P1 | 6 | SUMO2 pseudogene 1 [Source:HGNC Symbol; |
| MOG | 6 | myelin oligodendrocyte glycoprotein [Source: |
| ZFP57 | 6 | ZFP57 zinc finger protein [Source:HGNC Symb |
| ZDHHC20P1 | 6 | zinc finger DHHC-type containing 20 pseudoge |
| HLA-F | 6 | major histocompatibility complex, class I, F [S |
| AL645939.1 | 6 | HLA complex group 4 pseudogene 11 |
| HLA-F-AS1 | 6 | HLA-F antisense RNA 1 [Source:HGNC Symbol; |
| RPL23AP1 | 6 | ribosomal protein L23a pseudogene 1 [Source |
| MICE | 6 | MHC class I polypeptide-related sequence E (p |
| AL645939.2 | 6 | HLA complex group 9 pseudogene 5 |
| Y_RNA | 6 | Y RNA [Source:RFAM;Acc:RF00019] |
| IFITM4P | 6 | interferon induced transmembrane protein 4 |
| AL645939.4 | 6 | novel transcript |
| AL645939.5 | 6 | novel transcript |
| AL645939.3 | 6 | mitochondrial coiled-coil domain 1 (MCCD1) p |
| HLA-V | 6 | major histocompatibility complex, class I, V (p |
| HCG4 | 6 | HLA complex group 4 [Source:HGNC Symbol;A |
| HLA-P | 6 | major histocompatibility complex, class I, P (p |
| RPL7AP7 | 6 | ribosomal protein L7a pseudogene 7 [Source: |
| MICG | 6 | MHC class I polypeptide-related sequence G ( |
| HLA-G | 6 | major histocompatibility complex, class I, G [S |
| HCG4P8 | 6 | HLA complex group 4 pseudogene 8 [Source:H |
| AL645929.3 | 6 | MHC class I polypeptide-related sequence F ps |
| AL645929.2 | 6 | HLA complex P5B |
| AL645929.1 | 6 | HLA complex group 4 pseudogene 7 |
| HLA-H | 6 | major histocompatibility complex, class I, H (p |

| | | |
|---|---|---|
| HLA-T | 6 | major histocompatibility complex, class I, T (p |
| DDX39BP1 | 6 | DEAD-box helicase 39B pseudogene 1 [Source |
| MCCD1P1 | 6 | mitochondrial coiled-coil domain 1 pseudoger |
| HCG4B | 6 | HLA complex group 4B [Source:HGNC Symbol; |
| HLA-K | 6 | major histocompatibility complex, class I, K (p |
| HLA-U | 6 | major histocompatibility complex, class I, U (p |
| HLA-A | 6 | major histocompatibility complex, class I, A [S |
| AL671277.1 | 6 | HLA complex group 4 pseudogene 5 |
| HLA-W | 6 | major histocompatibility complex, class I, W (p |
| MICD | 6 | MHC class I polypeptide-related sequence D (p |
| HCG9 | 6 | HLA complex group 9 [Source:HGNC Symbol;A |
| DDX39BP2 | 6 | DEAD-box helicase 39B pseudogene 2 [Source |
| MCCD1P2 | 6 | mitochondrial coiled-coil domain 1 pseudoger |
| ZNRD1ASP | 6 | zinc ribbon domain containing 1 antisense, ps |
| HLA-J | 6 | major histocompatibility complex, class I, J (ps |
| AL671277.2 | 6 | HLA complex group 4 pseudogene 3 |
| ETF1P1 | 6 | eukaryotic translation termination factor 1 ps |
| AL669914.3 | 6 | NULL |
| AL669914.2 | 6 | NULL |
| ZNRD1 | 6 | zinc ribbon domain containing 1 [Source:HGN |
| AL669914.4 | 6 | NULL |
| PPP1R11 | 6 | protein phosphatase 1 regulatory inhibitor sul |
| RNF39 | 6 | ring finger protein 39 [Source:HGNC Symbol;A |
| TRIM31 | 6 | tripartite motif containing 31 [Source:HGNC S |
| TRIM31-AS1 | 6 | TRIM31 antisense RNA 1 [Source:HGNC Symb |
| AL669914.1 | 6 | NULL |
| TRIM40 | 6 | tripartite motif containing 40 [Source:HGNC S |
| TRIM10 | 6 | tripartite motif containing 10 [Source:HGNC S |
| TRIM15 | 6 | tripartite motif containing 15 [Source:HGNC S |
| TRIM26 | 6 | tripartite motif containing 26 [Source:HGNC S |
| PAIP1P1 | 6 | poly(A) binding protein interacting protein 1 p |
| HCG17 | 6 | HLA complex group 17 [Source:HGNC Symbol; |
| TRIM26BP | 6 | tripartite motif containing 26B, pseudogene [S |
| HLA-L | 6 | major histocompatibility complex, class I, L (ps |
| AL662795.2 | 6 | TEC |
| HCG18 | 6 | HLA complex group 18 [Source:HGNC Symbol; |
| TRIM39 | 6 | tripartite motif containing 39 [Source:HGNC S |
| TRIM39-RPP21 | 6 | TRIM39-RPP21 readthrough [Source:HGNC Sy |
| RPP21 | 6 | ribonuclease P/MRP subunit p21 [Source:HGN |
| HLA-N | 6 | major histocompatibility complex, class I, N (p |
| AL662795.1 | 6 | HLA complex group 19 pseudogene |
| UBQLN1P1 | 6 | ubiquilin 1 pseudogene 1 [Source:HGNC Symb |
| MICC | 6 | MHC class I polypeptide-related sequence C (p |
| AL662873.1 | 6 | novel transcript |
| TMPOP1 | 6 | thymopoietin pseudogene 1 [Source:HGNC Sy |
| SUCLA2P1 | 6 | succinate-CoA ligase ADP-forming beta subun |
| RANP1 | 6 | RAN pseudogene 1 [Source:HGNC Symbol;Acc |
| HLA-E | 6 | major histocompatibility complex, class I, E [S |
| LINC02569 | 6 | long intergenic non-protein coding RNA 2569 |
| GNL1 | 6 | G protein nucleolar 1 (putative) [Source:HGN |

| | | |
|---|---|---|
| PRR3 | 6 | proline rich 3 [Source:HGNC Symbol;Acc:HGN |
| ABCF1 | 6 | ATP binding cassette subfamily F member 1 [S |
| MIR877 | 6 | microRNA 877 [Source:HGNC Symbol;Acc:HGN |
| PPP1R10 | 6 | protein phosphatase 1 regulatory subunit 10 [ |
| MRPS18B | 6 | mitochondrial ribosomal protein S18B [Source |
| ATAT1 | 6 | alpha tubulin acetyltransferase 1 [Source:HGN |
| PTMAP1 | 6 | prothymosin alpha pseudogene 1 [Source:HGI |
| C6orf136 | 6 | chromosome 6 open reading frame 136 [Sour |
| DHX16 | 6 | DEAH-box helicase 16 [Source:HGNC Symbol;A |
| PPP1R18 | 6 | protein phosphatase 1 regulatory subunit 18 [ |
| NRM | 6 | nurim [Source:HGNC Symbol;Acc:HGNC:8003] |
| RPL7P4 | 6 | ribosomal protein L7 pseudogene 4 [Source:H |
| MDC1 | 6 | mediator of DNA damage checkpoint 1 [Sourc |
| MDC1-AS1 | 6 | MDC1 antisense RNA 1 [Source:HGNC Symbol |
| TUBB | 6 | tubulin beta class I [Source:HGNC Symbol;Acc |
| AL662797.1 | 6 | novel transcript, antisense to TUBB |
| FLOT1 | 6 | flotillin 1 [Source:HGNC Symbol;Acc:HGNC:37 |
| Y_RNA | 6 | Y RNA [Source:RFAM;Acc:RF00019] |
| IER3-AS1 | 6 | IER3 antisense RNA 1 [Source:HGNC Symbol;A |
| IER3 | 6 | immediate early response 3 [Source:HGNC Sy |
| HCG20 | 6 | HLA complex group 20 [Source:HGNC Symbol; |
| RN7SL353P | 6 | RNA, 7SL, cytoplasmic 353, pseudogene [Sour |
| LINC00243 | 6 | long intergenic non-protein coding RNA 243 [S |
| LINC02570 | 6 | long intergenic non-protein coding RNA 2570 |
| RN7SKP186 | 6 | RN7SK pseudogene 186 [Source:HGNC Symbo |
| DDR1-DT | 6 | DDR1 divergent transcript [Source:HGNC Sym |
| DDR1 | 6 | discoidin domain receptor tyrosine kinase 1 [S |
| MIR4640 | 6 | microRNA 4640 [Source:HGNC Symbol;Acc:HG |
| GTF2H4 | 6 | general transcription factor IIH subunit 4 [Sou |
| AL669830.1 | 6 | novel protein |
| VARS2 | 6 | valyl-tRNA synthetase 2, mitochondrial [Sourc |
| SFTA2 | 6 | surfactant associated 2 [Source:HGNC Symbol |
| Y_RNA | 6 | Y RNA [Source:RFAM;Acc:RF00019] |
| MUCL3 | 6 | mucin like 3 [Source:HGNC Symbol;Acc:HGNC |
| HCG21 | 6 | HLA complex group 21 [Source:HGNC Symbol; |
| NAPGP2 | 6 | N-ethylmaleimide-sensitive factor attachment |
| MUC21 | 6 | mucin 21, cell surface associated [Source:HGN |
| MUC22 | 6 | mucin 22 [Source:HGNC Symbol;Acc:HGNC:39 |
| HCG22 | 6 | HLA complex group 22 (gene/pseudogene) [So |
| RNU6-1133P | 6 | RNA, U6 small nuclear 1133, pseudogene [Sou |
| C6orf15 | 6 | chromosome 6 open reading frame 15 [Sourc |
| PSORS1C1 | 6 | psoriasis susceptibility 1 candidate 1 [Source:I |
| CDSN | 6 | corneodesmosin [Source:HGNC Symbol;Acc:H |
| PSORS1C2 | 6 | psoriasis susceptibility 1 candidate 2 [Source:I |
| POLR2LP1 | 6 | RNA polymerase II subunit L pseudogene 1 [So |
| CCHCR1 | 6 | coiled-coil alpha-helical rod protein 1 [Source: |
| TCF19 | 6 | transcription factor 19 [Source:HGNC Symbol; |
| POU5F1 | 6 | POU class 5 homeobox 1 [Source:HGNC Symb |
| PSORS1C3 | 6 | psoriasis susceptibility 1 candidate 3 [Source:I |
| AL662844.4 | 6 | novel transcript, antisense to HCG27 |

| | | |
|---|---|---|
| HCG27 | 6 | HLA complex group 27 [Source:HGNC Symbol; |
| AL662844.3 | 6 | novel transcript |
| AL662844.1 | 6 | mitochondrial coiled-coil domain 1 (MCCD1) p |
| AL662844.2 | 6 | HLA-B associated transcript 1 (BAT1) pseudog |
| HLA-C | 6 | major histocompatibility complex, class I, C [S |
| HLA-B | 6 | major histocompatibility complex, class I, B [S |
| USP8P1 | 6 | ubiquitin specific peptidase 8 pseudogene 1 [S |
| RPL3P2 | 6 | ribosomal protein L3 pseudogene 2 [Source:H |
| WASF5P | 6 | WASP family member 5, pseudogene [Source: |
| LINC02571 | 6 | long intergenic non-protein coding RNA 2571 |
| AL671883.1 | 6 | novel HLA class I pseudogene fragment |
| MIR6891 | 6 | microRNA 6891 [Source:HGNC Symbol;Acc:HG |
| AL671883.2 | 6 | HLA complex group 4 (HCG4) pseudogene |
| DHFRP2 | 6 | dihydrofolate reductase pseudogene 2 [Sourc |
| AL671883.3 | 6 | novel transcript |
| RNU6-283P | 6 | RNA, U6 small nuclear 283, pseudogene [Sour |
| FGFR3P1 | 6 | fibroblast growth factor receptor 3 pseudoger |
| ZDHHC20P2 | 6 | zinc finger DHHC-type containing 20 pseudoge |
| HLA-S | 6 | major histocompatibility complex, class I, S (p: |
| AL645933.2 | 6 | novel transcript |
| MICA | 6 | MHC class I polypeptide-related sequence A [S |
| HCP5 | 6 | HLA complex P5 [Source:HGNC Symbol;Acc:H( |
| Y_RNA | 6 | Y RNA [Source:RFAM;Acc:RF00019] |
| LINC01149 | 6 | long intergenic non-protein coding RNA 1149 |
| AL645933.1 | 6 | HLA complex group 4 (HCG4) pseudogene |
| AL645933.3 | 6 | novel transcript |
| MICB | 6 | MHC class I polypeptide-related sequence B [S |
| Y_RNA | 6 | Y RNA [Source:RFAM;Acc:RF00019] |
| AL663061.1 | 6 | HLA pseudogene |
| PPIAP9 | 6 | peptidylprolyl isomerase A pseudogene 9 [Sou |
| RPL15P4 | 6 | ribosomal protein L15 pseudogene 4 [Source: |
| MCCD1 | 6 | mitochondrial coiled-coil domain 1 [Source:H( |
| ATP6V1G2-DDX39B | 6 | ATP6V1G2-DDX39B readthrough (NMD candic |
| DDX39B | 6 | DExD-box helicase 39B [Source:HGNC Symbol, |
| SNORD117 | 6 | small nucleolar RNA, C/D box 117 [Source:HGI |
| SNORD84 | 6 | small nucleolar RNA, C/D box 84 [Source:HGN |
| DDX39B-AS1 | 6 | DDX39B antisense RNA 1 [Source:HGNC Symb |
| ATP6V1G2 | 6 | ATPase H+ transporting V1 subunit G2 [Source |
| NFKBIL1 | 6 | NFKB inhibitor like 1 [Source:HGNC Symbol;Ac |
| LTA | 6 | lymphotoxin alpha [Source:HGNC Symbol;Acc |
| TNF | 6 | tumor necrosis factor [Source:HGNC Symbol;A |
| LTB | 6 | lymphotoxin beta [Source:HGNC Symbol;Acc:I |
| LST1 | 6 | leukocyte specific transcript 1 [Source:HGNC S |
| NCR3 | 6 | natural cytotoxicity triggering receptor 3 [Sou |
| UQCRHP1 | 6 | ubiquinol-cytochrome c reductase hinge prote |
| AIF1 | 6 | allograft inflammatory factor 1 [Source:HGNC |
| PRRC2A | 6 | proline rich coiled-coil 2A [Source:HGNC Symb |
| SNORA38 | 6 | small nucleolar RNA, H/ACA box 38 [Source:H( |
| MIR6832 | 6 | microRNA 6832 [Source:HGNC Symbol;Acc:HG |
| BAG6 | 6 | BCL2 associated athanogene 6 [Source:HGNC |

| | | |
|---|---|---|
| APOM | 6 | apolipoprotein M [Source:HGNC Symbol;Acc:I |
| C6orf47 | 6 | chromosome 6 open reading frame 47 [Sourc |
| C6orf47-AS1 | 6 | C6orf47 antisense RNA 1 [Source:HGNC Symb |
| GPANK1 | 6 | G-patch domain and ankyrin repeats 1 [Sourc |
| Y_RNA | 6 | Y RNA [Source:RFAM;Acc:RF00019] |
| CSNK2B | 6 | casein kinase 2 beta [Source:HGNC Symbol;Ac |
| AL662899.2 | 6 | novel protein |
| LY6G5B | 6 | lymphocyte antigen 6 family member G5B [So |
| LY6G5C | 6 | lymphocyte antigen 6 family member G5C [So |
| ABHD16A | 6 | abhydrolase domain containing 16A [Source:I |
| AL662899.1 | 6 | novel transcript |
| MIR4646 | 6 | microRNA 4646 [Source:HGNC Symbol;Acc:H |
| LY6G6F | 6 | lymphocyte antigen 6 family member G6F [So |
| LY6G6F-LY6G6D | 6 | LY6G6F-LY6G6D readthrough [Source:NCBI ge |
| LY6G6E | 6 | lymphocyte antigen 6 family member G6E [So |
| LY6G6D | 6 | lymphocyte antigen 6 family member G6D [Sc |
| MPIG6B | 6 | megakaryocyte and platelet inhibitory recept |
| LY6G6C | 6 | lymphocyte antigen 6 family member G6C [So |
| DDAH2 | 6 | dimethylarginine dimethylaminohydrolase 2 [ |
| CLIC1 | 6 | chloride intracellular channel 1 [Source:HGNC |
| MSH5 | 6 | mutS homolog 5 [Source:HGNC Symbol;Acc:H |
| MSH5-SAPCD1 | 6 | MSH5-SAPCD1 readthrough (NMD candidate) |
| RNU6-850P | 6 | RNA, U6 small nuclear 850, pseudogene [Sour |
| SAPCD1 | 6 | suppressor APC domain containing 1 [Source: |
| SAPCD1-AS1 | 6 | SAPCD1 antisense RNA 1 [Source:HGNC Symb |
| VWA7 | 6 | von Willebrand factor A domain containing 7 |
| VARS | 6 | valyl-tRNA synthetase [Source:HGNC Symbol; |
| Y_RNA | 6 | Y RNA [Source:RFAM;Acc:RF00019] |
| LSM2 | 6 | LSM2 homolog, U6 small nuclear RNA and mR |
| HSPA1L | 6 | heat shock protein family A (Hsp70) member |
| HSPA1A | 6 | heat shock protein family A (Hsp70) member |
| HSPA1B | 6 | heat shock protein family A (Hsp70) member |
| AL671762.1 | 6 | novel transcript |
| SNHG32 | 6 | small nucleolar RNA host gene 32 [Source:HG |
| SNORD48 | 6 | small nucleolar RNA, C/D box 48 [Source:HGN |
| SNORD52 | 6 | small nucleolar RNA, C/D box 52 [Source:HGN |
| NEU1 | 6 | neuraminidase 1 [Source:HGNC Symbol;Acc:H |
| SLC44A4 | 6 | solute carrier family 44 member 4 [Source:HG |
| EHMT2-AS1 | 6 | EHMT2 and SLC44A4 antisense RNA 1 [Source |
| EHMT2 | 6 | euchromatic histone lysine methyltransferase |
| C2 | 6 | complement C2 [Source:HGNC Symbol;Acc:H |
| ZBTB12 | 6 | zinc finger and BTB domain containing 12 [Sou |
| AL645922.1 | 6 | novel complement component 2 (C2) and con |
| C2-AS1 | 6 | C2 antisense RNA 1 [Source:HGNC Symbol;Acc |
| CFB | 6 | complement factor B [Source:HGNC Symbol;A |
| NELFE | 6 | negative elongation factor complex member |
| MIR1236 | 6 | microRNA 1236 [Source:HGNC Symbol;Acc:H |
| SKIV2L | 6 | Ski2 like RNA helicase [Source:HGNC Symbol; |
| DXO | 6 | decapping exoribonuclease [Source:HGNC Sy |
| STK19 | 6 | serine/threonine kinase 19 [Source:HGNC Sym |

| | | |
|---|---|---|
| C4A | 6 | complement C4A (Rodgers blood group) [Sour |
| C4A-AS1 | 6 | C4A antisense RNA 1 [Source:HGNC Symbol;A |
| CYP21A1P | 6 | cytochrome P450 family 21 subfamily A meml |
| TNXA | 6 | tenascin XA (pseudogene) [Source:HGNC Sym |
| STK19B | 6 | serine/threonine kinase 19B (pseudogene) [Sc |
| C4B | 6 | complement C4B (Chido blood group) [Source |
| C4B-AS1 | 6 | C4B antisense RNA 1 [Source:HGNC Symbol;A |
| CYP21A2 | 6 | cytochrome P450 family 21 subfamily A meml |
| TNXB | 6 | tenascin XB [Source:HGNC Symbol;Acc:HGNC: |
| RNA5SP206 | 6 | RNA, 5S ribosomal pseudogene 206 [Source:H |
| AL662884.2 | 6 | novel transcript |
| AL662884.5 | 6 | novel transcript, antisense to TNXB |
| ATF6B | 6 | activating transcription factor 6 beta [Source:I |
| FKBPL | 6 | FKBP prolyl isomerase like [Source:HGNC Sym |
| PRRT1 | 6 | proline rich transmembrane protein 1 [Source |
| AL662884.4 | 6 | novel protein |
| AL662884.3 | 6 | novel transcript |
| PPT2 | 6 | palmitoyl-protein thioesterase 2 [Source:HGN |
| PPT2-EGFL8 | 6 | PPT2-EGFL8 readthrough (NMD candidate) [Sc |
| EGFL8 | 6 | EGF like domain multiple 8 [Source:HGNC Syn |
| AGPAT1 | 6 | 1-acylglycerol-3-phosphate O-acyltransferase |
| MIR6721 | 6 | microRNA 6721 [Source:HGNC Symbol;Acc:HG |
| RNF5 | 6 | ring finger protein 5 [Source:HGNC Symbol;Ac |
| MIR6833 | 6 | microRNA 6833 [Source:HGNC Symbol;Acc:HG |
| AGER | 6 | advanced glycosylation end-product specific r |
| PBX2 | 6 | PBX homeobox 2 [Source:HGNC Symbol;Acc:H |
| AL662884.1 | 6 | novel transcript |
| GPSM3 | 6 | G protein signaling modulator 3 [Source:HGN( |
| NOTCH4 | 6 | notch receptor 4 [Source:HGNC Symbol;Acc:H |
| TSBP1-AS1 | 6 | TSBP1 and BTNL2 antisense RNA 1 [Source:HG |
| TSBP1 | 6 | testis expressed basic protein 1 [Source:HGN( |
| HNRNPA1P2 | 6 | heterogeneous nuclear ribonucleoprotein A1 |
| RNU6-603P | 6 | RNA, U6 small nuclear 603, pseudogene [Sour |
| BTNL2 | 6 | butyrophilin like 2 [Source:HGNC Symbol;Acc: |
| HLA-DRA | 6 | major histocompatibility complex, class II, DR |
| HLA-DRB9 | 6 | major histocompatibility complex, class II, DR |
| HLA-DRB5 | 6 | major histocompatibility complex, class II, DR |
| RNU1-61P | 6 | RNA, U1 small nuclear 61, pseudogene [Sourc |
| HLA-DRB6 | 6 | major histocompatibility complex, class II, DR |
| HLA-DRB1 | 6 | major histocompatibility complex, class II, DR |
| HLA-DQA1 | 6 | major histocompatibility complex, class II, DQ |
| HLA-DQB1 | 6 | major histocompatibility complex, class II, DQ |
| HLA-DQB1-AS1 | 6 | HLA-DQB1 antisense RNA 1 [Source:HGNC Syr |
| MTCO3P1 | 6 | MT-CO3 pseudogene 1 [Source:HGNC Symbol |
| AL662789.1 | 6 | novel transcript |
| HLA-DQB3 | 6 | major histocompatibility complex, class II, DQ |
| HLA-DQA2 | 6 | major histocompatibility complex, class II, DQ |
| MIR3135B | 6 | microRNA 3135b [Source:HGNC Symbol;Acc:H |
| HLA-DQB2 | 6 | major histocompatibility complex, class II, DQ |
| HLA-DOB | 6 | major histocompatibility complex, class II, DO |

| | | |
|---|---|---|
| AL669918.1 | 6 | novel protein, TAP2-HLA-DOB readthrough |
| TAP2 | 6 | transporter 2, ATP binding cassette subfamily |
| PSMB8 | 6 | proteasome subunit beta 8 [Source:HGNC Syn |
| PSMB8-AS1 | 6 | PSMB8 antisense RNA 1 (head to head) [Sourc |
| PSMB9 | 6 | proteasome subunit beta 9 [Source:HGNC Syn |
| TAP1 | 6 | transporter 1, ATP binding cassette subfamily |
| PPP1R2P1 | 6 | protein phosphatase 1 regulatory inhibitor sul |
| HLA-Z | 6 | major histocompatibility complex, class I, Z (p |
| HLA-DMB | 6 | major histocompatibility complex, class II, DM |
| AL645941.2 | 6 | novel protein |
| HLA-DMA | 6 | major histocompatibility complex, class II, DM |
| BRD2 | 6 | bromodomain containing 2 [Source:HGNC Syn |
| AL645941.1 | 6 | novel transcript |
| AL645941.3 | 6 | novel transcript, antisense to BRD2 |
| HLA-DOA | 6 | major histocompatibility complex, class II, DO |
| HLA-DPA1 | 6 | major histocompatibility complex, class II, DP |
| HLA-DPB1 | 6 | major histocompatibility complex, class II, DP |
| RPL32P1 | 6 | ribosomal protein L32 pseudogene 1 [Source: |
| HLA-DPA2 | 6 | major histocompatibility complex, class II, DP |
| COL11A2P1 | 6 | collagen type XI alpha 2 pseudogene 1 [Sourc |
| HLA-DPB2 | 6 | major histocompatibility complex, class II, DP |
| HLA-DPA3 | 6 | major histocompatibility complex, class II, DP |
| HCG24 | 6 | HLA complex group 24 [Source:HGNC Symbol; |
| COL11A2 | 6 | collagen type XI alpha 2 chain [Source:HGNC S |
| RXRB | 6 | retinoid X receptor beta [Source:HGNC Symbo |
| RNY4P10 | 6 | RNY4 pseudogene 10 [Source:HGNC Symbol;A |
| SLC39A7 | 6 | solute carrier family 39 member 7 [Source:HG |
| HSD17B8 | 6 | hydroxysteroid 17-beta dehydrogenase 8 [Sou |
| MIR219A1 | 6 | microRNA 219a-1 [Source:HGNC Symbol;Acc:H |
| RING1 | 6 | ring finger protein 1 [Source:HGNC Symbol;Ac |
| ZNF70P1 | 6 | zinc finger protein 70 pseudogene 1 [Source:H |
| HTATSF1P1 | 6 | HIV-1 Tat specific factor 1 pseudogene 1 [Sour |
| AL645940.1 | 6 | novel transcript |
| HCG25 | 6 | HLA complex group 25 [Source:HGNC Symbol; |
| VPS52 | 6 | VPS52 subunit of GARP complex [Source:HGN |
| RPS18 | 6 | ribosomal protein S18 [Source:HGNC Symbol;. |
| B3GALT4 | 6 | beta-1,3-galactosyltransferase 4 [Source:HGN |
| WDR46 | 6 | WD repeat domain 46 [Source:HGNC Symbol;. |
| MIR6873 | 6 | microRNA 6873 [Source:HGNC Symbol;Acc:HG |
| PFDN6 | 6 | prefoldin subunit 6 [Source:HGNC Symbol;Acc |
| MIR6834 | 6 | microRNA 6834 [Source:HGNC Symbol;Acc:HG |
| RGL2 | 6 | ral guanine nucleotide dissociation stimulator |
| TAPBP | 6 | TAP binding protein [Source:HGNC Symbol;Ac |
| ZBTB22 | 6 | zinc finger and BTB domain containing 22 [Sou |
| DAXX | 6 | death domain associated protein [Source:HGN |
| SMIM40 | 6 | small integral membrane protein 40 [Source:H |
| SMIM40 | 6 | small integral membrane protein 40 [Source:N |
| MYL8P | 6 | myosin light chain 8, pseudogene [Source:HGI |
| LYPLA2P1 | 6 | LYPLA2 pseudogene 1 [Source:HGNC Symbol;A |
| RPL35AP4 | 6 | ribosomal protein L35a pseudogene 4 [Source |

| | | |
|---|---|---|
| KIFC1 | 6 | kinesin family member C1 [Source:HGNC Sym |
| RPL12P1 | 6 | ribosomal protein L12 pseudogene 1 [Source: |
| PHF1 | 6 | PHD finger protein 1 [Source:HGNC Symbol;Ac |
| CUTA | 6 | cutA divalent cation tolerance homolog [Sourc |
| SYNGAP1 | 6 | synaptic Ras GTPase activating protein 1 [Sour |
| SYNGAP1-AS1 | 6 | SYNGAP1 antisense RNA 1 [Source:HGNC Sym |
| MIR5004 | 6 | microRNA 5004 [Source:HGNC Symbol;Acc:HG |
| ZBTB9 | 6 | zinc finger and BTB domain containing 9 [Sour |
| GGNBP1 | 6 | gametogenetin binding protein 1 (pseudogene |
| RN7SL26P | 6 | RNA, 7SL, cytoplasmic 26, pseudogene [Sourc |
| Metazoa_SRP | 6 | Metazoan signal recognition particle RNA [Sou |
| BAK1 | 6 | BCL2 antagonist/killer 1 [Source:HGNC Symbo |
| LINC00336 | 6 | long intergenic non-protein coding RNA 336 [S |
| ITPR3 | 6 | inositol 1,4,5-trisphosphate receptor type 3 [S |
| UQCC2 | 6 | ubiquinol-cytochrome c reductase complex as |
| MIR3934 | 6 | microRNA 3934 [Source:HGNC Symbol;Acc:HG |
| IP6K3 | 6 | inositol hexakisphosphate kinase 3 [Source:HG |
| LEMD2 | 6 | LEM domain containing 2 [Source:HGNC Symb |
| MLN | 6 | motilin [Source:HGNC Symbol;Acc:HGNC:7141 |
| AL138889.3 | 6 | novel transcript, antisense to MLN |
| LINC01016 | 6 | long intergenic non-protein coding RNA 1016 |
| AL138889.2 | 6 | ankyrin repeat domain 36C (ANKRD36C) pseu |
| AL138889.1 | 6 | novel transcript |
| MIR7159 | 6 | microRNA 7159 [Source:HGNC Symbol;Acc:HG |
| MIR1275 | 6 | microRNA 1275 [Source:HGNC Symbol;Acc:HG |
| GRM4 | 6 | glutamate metabotropic receptor 4 [Source:H |
| KRT18P9 | 6 | keratin 18 pseudogene 9 [Source:HGNC Symb |
| CYCSP55 | 6 | CYCS pseudogene 55 [Source:HGNC Symbol;A |
| HMGA1 | 6 | high mobility group AT-hook 1 [Source:HGNC |
| MIR6835 | 6 | microRNA 6835 [Source:HGNC Symbol;Acc:HG |
| SMIM29 | 6 | small integral membrane protein 29 [Source:H |
| AL354740.1 | 6 | novel transcript, antisense to C6orf1 |
| RPL35P2 | 6 | ribosomal protein L35 pseudogene 2 [Source: |
| NUDT3 | 6 | nudix hydrolase 3 [Source:HGNC Symbol;Acc:H |
| RPS10-NUDT3 | 6 | RPS10-NUDT3 readthrough [Source:HGNC Syr |
| RPS10 | 6 | ribosomal protein S10 [Source:HGNC Symbol; |
| PACSIN1 | 6 | protein kinase C and casein kinase substrate ir |
| SPDEF | 6 | SAM pointed domain containing ETS transcrip |
| AL031577.1 | 6 | interferon induced transmembrane protein (II |
| ILRUN | 6 | inflammation and lipid regulator with UBA-like |
| RPL7P25 | 6 | ribosomal protein L7 pseudogene 25 [Source: |
| RN7SL200P | 6 | RNA, 7SL, cytoplasmic 200, pseudogene [Sour |
| AL031577.2 | 6 | ribosomal protein L44 (RPL44) pseudogene |
| AL451165.2 | 6 | novel transcript, antisense to C6orf106 |
| AL451165.1 | 6 | ATPase, H+ transporting, lysosomal (vacuolar |
| AL139100.1 | 6 | ribosomal protein S10 (RPS10) pseudogene |
| SNRPC | 6 | small nuclear ribonucleoprotein polypeptide C |
| UHRF1BP1 | 6 | UHRF1 binding protein 1 [Source:HGNC Symbo |
| Y_RNA | 6 | Y RNA [Source:RFAM;Acc:RF00019] |
| RNY3P15 | 6 | RNY3 pseudogene 15 [Source:HGNC Symbol;A |

| | | |
|---|---|---|
| TAF11 | 6 | TATA-box binding protein associated factor 1[illegible] |
| ANKS1A | 6 | ankyrin repeat and sterile alpha motif domain |
| HSPE1P11 | 6 | heat shock protein family E (Hsp10) member [illegible] |
| AL591002.1 | 6 | novel transcript, antisense to ANKS1A |
| AL138721.1 | 6 | ribosomal protein S12 (RPS12) pseudogene |
| TCP11 | 6 | t-complex 11 [Source:HGNC Symbol;Acc:HGN[illegible] |
| SCUBE3 | 6 | signal peptide, CUB domain and EGF like dom[illegible] |
| Z97832.2 | 6 | novel transcript, antisense to SCUBE3 |
| ZNF76 | 6 | zinc finger protein 76 [Source:HGNC Symbol;[illegible] |
| Z97832.1 | 6 | ribosomal protein L35A (RPL35A) pseudogene |
| DEF6 | 6 | DEF6 guanine nucleotide exchange factor [So[illegible] |
| PPARD | 6 | peroxisome proliferator activated receptor de |
| AL022721.1 | 6 | makorin, ring finger protein, pseudogene 2 |
| FANCE | 6 | FA complementation group E [Source:HGNC S |
| RPL10A | 6 | ribosomal protein L10a [Source:HGNC Symbol |
| MIR7111 | 6 | microRNA 7111 [Source:HGNC Symbol;Acc:H[illegible] |
| TEAD3 | 6 | TEA domain transcription factor 3 [Source:HG |
| TULP1 | 6 | TUB like protein 1 [Source:HGNC Symbol;Acc:[illegible] |
| AL033519.4 | 6 | novel transcript |
| AL033519.3 | 6 | ribosomal protein S15A (RPS15A) pseudogene |
| FKBP5 | 6 | FKBP prolyl isomerase 5 [Source:HGNC Symbo |
| AL033519.2 | 6 | ribosomal protein L36 (RPL36) pseudogene |
| AL033519.5 | 6 | novel transcript, antisense to FKBP5 |
| AL033519.1 | 6 | NULL |
| MIR5690 | 6 | microRNA 5690 [Source:HGNC Symbol;Acc:H[illegible] |
| AL157823.2 | 6 | novel transcript |
| ARMC12 | 6 | armadillo repeat containing 12 [Source:HGNC |
| AL157823.1 | 6 | cytidylate kinase (CMPK) pseudogene |
| CLPSL2 | 6 | colipase like 2 [Source:HGNC Symbol;Acc:HG[illegible] |
| CLPSL1 | 6 | colipase like 1 [Source:HGNC Symbol;Acc:HG[illegible] |
| CLPS | 6 | colipase [Source:HGNC Symbol;Acc:HGNC:20[illegible] |
| LHFPL5 | 6 | LHFPL tetraspan subfamily member 5 [Source |
| SRPK1 | 6 | SRSF protein kinase 1 [Source:HGNC Symbol;[illegible] |
| SLC26A8 | 6 | solute carrier family 26 member 8 [Source:HG |
| DPRXP2 | 6 | divergent-paired related homeobox pseudoge |
| MAPK14 | 6 | mitogen-activated protein kinase 14 [Source:[illegible] |
| Z95152.1 | 6 | nudix (nucleoside diphosphate linked moiety [illegible] |
| MAPK13 | 6 | mitogen-activated protein kinase 13 [Source:[illegible] |
| Z84485.1 | 6 | novel transcript, antisense to BRPF3 |
| BRPF3 | 6 | bromodomain and PHD finger containing 3 [S[illegible] |
| PNPLA1 | 6 | patatin like phospholipase domain containing |
| BNIP5 | 6 | BCL2 interacting protein 5 [Source:HGNC Sym |
| ETV7 | 6 | ETS variant 7 [Source:HGNC Symbol;Acc:HGN[illegible] |
| Z84484.1 | 6 | novel transcript |
| PXT1 | 6 | peroxisomal testis enriched protein 1 [Source |
| KCTD20 | 6 | potassium channel tetramerization domain co |
| RN7SL502P | 6 | RNA, 7SL, cytoplasmic 502, pseudogene [Sour |
| STK38 | 6 | serine/threonine kinase 38 [Source:HGNC Sym |
| RN7SL748P | 6 | RNA, 7SL, cytoplasmic 748, pseudogene [Sour |
| SRSF3 | 6 | serine and arginine rich splicing factor 3 [Sour |

| | | |
|---|---|---|
| MIR3925 | 6 | microRNA 3925 [Source:HGNC Symbol;Acc:HG |
| RNU1-88P | 6 | RNA, U1 small nuclear 88, pseudogene [Sourc |
| Y_RNA | 6 | Y RNA [Source:RFAM;Acc:RF00019] |
| PANDAR | 6 | promoter of CDKN1A antisense DNA damage a |
| LAP3P2 | 6 | leucine aminopeptidase 3 pseudogene 2 [Sour |
| CDKN1A | 6 | cyclin dependent kinase inhibitor 1A [Source:H |
| DINOL | 6 | damage induced long noncoding RNA [Source |
| RAB44 | 6 | RAB44, member RAS oncogene family [Source |
| Z85996.1 | 6 | galanin receptor pseudogene |
| CPNE5 | 6 | copine 5 [Source:HGNC Symbol;Acc:HGNC:23 |
| Z85996.2 | 6 | novel transcript |
| Z85996.3 | 6 | novel transcript |
| PPIL1 | 6 | peptidylprolyl isomerase like 1 [Source:HGNC |
| C6orf89 | 6 | chromosome 6 open reading frame 89 [Sourc |
| AL122034.1 | 6 | novel transcript |
| PI16 | 6 | peptidase inhibitor 16 [Source:HGNC Symbol;A |
| MTCH1 | 6 | mitochondrial carrier 1 [Source:HGNC Symbol |
| FGD2 | 6 | FYVE, RhoGEF and PH domain containing 2 [So |
| COX6A1P2 | 6 | cytochrome c oxidase subunit 6A1 pseudogen |
| RPL12P2 | 6 | ribosomal protein L12 pseudogene 2 [Source: |
| PIM1 | 6 | Pim-1 proto-oncogene, serine/threonine kinas |
| TMEM217 | 6 | transmembrane protein 217 [Source:HGNC Sy |
| AL353579.1 | 6 | transmembrane protein 217 [Source:NCBI gen |
| TBC1D22B | 6 | TBC1 domain family member 22B [Source:HG |
| AL096712.1 | 6 | immediate early response 3 interacting protei |
| AL096712.2 | 6 | novel transcript, antisense to TBC1D22B |
| RNF8 | 6 | ring finger protein 8 [Source:HGNC Symbol;Ac |
| RN7SL273P | 6 | RNA, 7SL, cytoplasmic 273, pseudogene [Sour |
| CMTR1 | 6 | cap methyltransferase 1 [Source:HGNC Symbo |
| CCDC167 | 6 | coiled-coil domain containing 167 [Source:HG |
| LINC02520 | 6 | long intergenic non-protein coding RNA 2520 |
| AL353597.2 | 6 | ribosomal protein S20 (RPS20) pseudogene |
| AL353597.1 | 6 | novel transcript |
| MIR4462 | 6 | microRNA 4462 [Source:HGNC Symbol;Acc:HG |
| AL353597.3 | 6 | TEC |
| MDGA1 | 6 | MAM domain containing glycosylphosphatidy |
| AL121574.1 | 6 | novel transcript |
| ZFAND3 | 6 | zinc finger AN1-type containing 3 [Source:HGN |
| RN7SL285P | 6 | RNA, 7SL, cytoplasmic 285, pseudogene [Sour |
| RNU1-87P | 6 | RNA, U1 small nuclear 87, pseudogene [Sourc |
| AL589655.1 | 6 | pseudogene similar to an integrin Alpha-E pre |
| BTBD9 | 6 | BTB domain containing 9 [Source:HGNC Symb |
| AL033518.1 | 6 | NULL |
| BTBD9-AS1 | 6 | BTBD9 antisense RNA 1 [Source:HGNC Symbo |
| Y_RNA | 6 | Y RNA [Source:RFAM;Acc:RF00019] |
| AL079341.1 | 6 | TRK-fused gene (TFG) pseudogene |
| GLO1 | 6 | glyoxalase I [Source:HGNC Symbol;Acc:HGNC: |
| AL391415.1 | 6 | novel transcript |
| DNAH8 | 6 | dynein axonemal heavy chain 8 [Source:HGNC |
| RN7SL465P | 6 | RNA, 7SL, cytoplasmic 465, pseudogene [Sour |

| | | |
|---|---|---|
| AL035555.2 | 6 | zuotin related factor-1 (ZRF1) pseudogene |
| AL035555.1 | 6 | NULL |
| AL034345.2 | 6 | novel transcript |
| AL034345.1 | 6 | pseudogene similar to part of 40S Ribosomal p |
| AL035690.1 | 6 | 40S ribosomal protein S2 (RPS2) pseudogene |
| GLP1R | 6 | glucagon like peptide 1 receptor [Source:HGN |
| SAYSD1 | 6 | SAYSVFN motif domain containing 1 [Source:H |
| ANKRD18EP | 6 | ankyrin repeat domain 18E, pseudogene [Sour |
| KCNK5 | 6 | potassium two pore domain channel subfamil |
| KCNK17 | 6 | potassium two pore domain channel subfamil |
| KCNK16 | 6 | potassium two pore domain channel subfamil |
| KIF6 | 6 | kinesin family member 6 [Source:HGNC Symb |
| AL136087.1 | 6 | novel pseudogene |
| E2F4P1 | 6 | E2F transcription factor 4 pseudogene 1 [Sour |
| RNU1-54P | 6 | RNA, U1 small nuclear 54, pseudogene [Sourc |
| DAAM2 | 6 | dishevelled associated activator of morphoger |
| AL592158.1 | 6 | TEC |
| AL590999.1 | 6 | novel transcript |
| MOCS1 | 6 | molybdenum cofactor synthesis 1 [Source:HG |
| TUBBP9 | 6 | tubulin beta class I pseudogene 9 [Source:HGI |
| FO393411.1 | 6 | pseudogene similar to part of ribosomal prote |
| AL139275.2 | 6 | novel transcript |
| TDRG1 | 6 | testis development related 1 [Source:HGNC Sy |
| LINC00951 | 6 | long intergenic non-protein coding RNA 951 [S |
| AL139275.1 | 6 | novel transcript |
| LRFN2 | 6 | leucine rich repeat and fibronectin type III dor |
| RNU6-250P | 6 | RNA, U6 small nuclear 250, pseudogene [Sour |
| AL359475.1 | 6 | novel transcript |
| AL033380.1 | 6 | novel transcript |
| AL583854.1 | 6 | novel transcript |
| UNC5CL | 6 | unc-5 family C-terminal like [Source:HGNC Syr |
| OARD1 | 6 | O-acyl-ADP-ribose deacylase 1 [Source:HGNC |
| TSPO2 | 6 | translocator protein 2 [Source:HGNC Symbol;/ |
| APOBEC2 | 6 | apolipoprotein B mRNA editing enzyme cataly |
| NFYA | 6 | nuclear transcription factor Y subunit alpha [S |
| AL031778.1 | 6 | novel transcript, antisense to NFYA |
| ADCY10P1 | 6 | ADCY10 pseudogene 1 [Source:HGNC Symbol; |
| TREML1 | 6 | triggering receptor expressed on myeloid cells |
| TREM2 | 6 | triggering receptor expressed on myeloid cells |
| TREML2 | 6 | triggering receptor expressed on myeloid cells |
| TREML3P | 6 | triggering receptor expressed on myeloid cells |
| TREML4 | 6 | triggering receptor expressed on myeloid cells |
| RNA5SP207 | 6 | RNA, 5S ribosomal pseudogene 207 [Source:H |
| TREML5P | 6 | triggering receptor expressed on myeloid cells |
| AL391903.2 | 6 | natural cytotoxicity triggering receptor 2 (NCF |
| AL391903.3 | 6 | triggering receptor expressed on myeloid cells |
| TREM1 | 6 | triggering receptor expressed on myeloid cells |
| AL391903.1 | 6 | triggering receptor expressed on myeloid cells |
| RNU6-643P | 6 | RNA, U6 small nuclear 643, pseudogene [Sour |
| AL672212.1 | 6 | ribosomal protein L32 (RPL32) pseudogene |

| | | |
|---|---|---|
| NCR2 | 6 | natural cytotoxicity triggering receptor 2 [Sou |
| AL136967.1 | 6 | TEC gene |
| AL136967.2 | 6 | TEC gene |
| FOXP4-AS1 | 6 | FOXP4 antisense RNA 1 [Source:HGNC Symbo |
| LINC01276 | 6 | long intergenic non-protein coding RNA 1276 |
| FOXP4 | 6 | forkhead box P4 [Source:HGNC Symbol;Acc:H |
| MIR4641 | 6 | microRNA 4641 [Source:HGNC Symbol;Acc:HG |
| MDFI | 6 | MyoD family inhibitor [Source:HGNC Symbol;/ |
| NPM1P51 | 6 | nucleophosmin 1 pseudogene 51 [Source:HGN |
| TFEB | 6 | transcription factor EB [Source:HGNC Symbol; |
| AL035588.1 | 6 | novel transcript |
| PGC | 6 | progastricsin [Source:HGNC Symbol;Acc:HGN( |
| AL365205.4 | 6 | novel transcript |
| FRS3 | 6 | fibroblast growth factor receptor substrate 3 [ |
| AL365205.1 | 6 | novel protein |
| PRICKLE4 | 6 | prickle planar cell polarity protein 4 [Source:H |
| TOMM6 | 6 | translocase of outer mitochondrial membrane |
| USP49 | 6 | ubiquitin specific peptidase 49 [Source:HGNC |
| AL365205.3 | 6 | novel transcript, antisense to USP49 |
| AL365205.2 | 6 | NULL |
| AL160163.1 | 6 | novel transcript |
| MED20 | 6 | mediator complex subunit 20 [Source:HGNC S |
| Y_RNA | 6 | Y RNA [Source:RFAM;Acc:RF00019] |
| BYSL | 6 | bystin like [Source:HGNC Symbol;Acc:HGNC:1 |
| CCND3 | 6 | cyclin D3 [Source:HGNC Symbol;Acc:HGNC:15 |
| RNU6-761P | 6 | RNA, U6 small nuclear 761, pseudogene [Sour |
| AL513008.1 | 6 | novel transcript |
| TAF8 | 6 | TATA-box binding protein associated factor 8 |
| AL512274.1 | 6 | novel transcript |
| C6orf132 | 6 | chromosome 6 open reading frame 132 [Sour |
| AL096814.2 | 6 | guanylate cyclase activator 1A [Source:NCBI g |
| GUCA1A | 6 | guanylate cyclase activator 1A [Source:HGNC |
| AL096814.1 | 6 | Uncharacterized protein 114841037 [Source:N |
| GUCA1B | 6 | guanylate cyclase activator 1B [Source:HGNC |
| MRPS10 | 6 | mitochondrial ribosomal protein S10 [Source: |
| TRERF1 | 6 | transcriptional regulating factor 1 [Source:HG |
| AL591473.1 | 6 | ribosomal protein L36a (RPL36A) pseudogene |
| UBR2 | 6 | ubiquitin protein ligase E3 component n-recog |
| RNU6-890P | 6 | RNA, U6 small nuclear 890, pseudogene [Sour |
| PRPH2 | 6 | peripherin 2 [Source:HGNC Symbol;Acc:HGNC |
| ATP6V0CP3 | 6 | ATPase H+ transporting V0 subunit c pseudog |
| TBCC | 6 | tubulin folding cofactor C [Source:HGNC Symb |
| BICRAL | 6 | BRD4 interacting chromatin remodeling comp |
| RPL7L1 | 6 | ribosomal protein L7 like 1 [Source:HGNC Sym |
| C6orf226 | 6 | chromosome 6 open reading frame 226 [Sour |
| AL035587.3 | 6 | novel transcript |
| PTCRA | 6 | pre T cell antigen receptor alpha [Source:HGN |
| AL035587.2 | 6 | novel transcript, antisense to CNPY3 |
| CNPY3 | 6 | canopy FGF signaling regulator 3 [Source:HGN |
| AL035587.1 | 6 | novel transcript |

| | | |
|---|---|---|
| RPL24P4 | 6 | ribosomal protein L24 pseudogene 4 [Source: |
| GNMT | 6 | glycine N-methyltransferase [Source:HGNC Sy |
| PEX6 | 6 | peroxisomal biogenesis factor 6 [Source:HGN( |
| PPP2R5D | 6 | protein phosphatase 2 regulatory subunit B'de |
| MEA1 | 6 | male-enhanced antigen 1 [Source:HGNC Symb |
| KLHDC3 | 6 | kelch domain containing 3 [Source:HGNC Sym |
| RRP36 | 6 | ribosomal RNA processing 36 [Source:HGNC S |
| AL136304.1 | 6 | novel transcript, antisense to RRP36 |
| RN7SL403P | 6 | RNA, 7SL, cytoplasmic 403, pseudogene [Sour |
| CUL7 | 6 | cullin 7 [Source:HGNC Symbol;Acc:HGNC:2102 |
| KLC4 | 6 | kinesin light chain 4 [Source:HGNC Symbol;Ac |
| MRPL2 | 6 | mitochondrial ribosomal protein L2 [Source:H |
| AL355385.1 | 6 | novel transcript, antisense to KLC4 |
| PTK7 | 6 | protein tyrosine kinase 7 (inactive) [Source:H( |
| SRF | 6 | serum response factor [Source:HGNC Symbol; |
| CUL9 | 6 | cullin 9 [Source:HGNC Symbol;Acc:HGNC:1598 |
| AL133375.1 | 6 | novel transcript |
| DNPH1 | 6 | 2'-deoxynucleoside 5'-phosphate N-hydrolase |
| TTBK1 | 6 | tau tubulin kinase 1 [Source:HGNC Symbol;Ac |
| SLC22A7 | 6 | solute carrier family 22 member 7 [Source:HG |
| CRIP3 | 6 | cysteine rich protein 3 [Source:HGNC Symbol; |
| ZNF318 | 6 | zinc finger protein 318 [Source:HGNC Symbol; |
| RPL12P47 | 6 | ribosomal protein L12 pseudogene 47 [Source |
| AL583834.1 | 6 | ribosomal protein L34 (RPL34) pseudogene |
| RPS2P28 | 6 | ribosomal protein S2 pseudogene 28 [Source: |
| AL590383.1 | 6 | ribosomal protein S2 (RPS2) pseudogene |
| AL359813.1 | 6 | novel transcript |
| AL359813.2 | 6 | novel transcript |
| ABCC10 | 6 | ATP binding cassette subfamily C member 10 |
| MIR6780B | 6 | microRNA 6780b [Source:HGNC Symbol;Acc:H |
| DLK2 | 6 | delta like non-canonical Notch ligand 2 [Sourc |
| RNU6-1113P | 6 | RNA, U6 small nuclear 1113, pseudogene [Sou |
| TJAP1 | 6 | tight junction associated protein 1 [Source:HG |
| LRRC73 | 6 | leucine rich repeat containing 73 [Source:HGN |
| POLR1C | 6 | RNA polymerase I and III subunit C [Source:HG |
| YIPF3 | 6 | Yip1 domain family member 3 [Source:HGNC S |
| AL355802.3 | 6 | novel transcript, antisense to POLR1C |
| XPO5 | 6 | exportin 5 [Source:HGNC Symbol;Acc:HGNC:1 |
| AL355802.1 | 6 | 40S ribosomal protein S2 (RPS2) pseudogene |
| AL355802.2 | 6 | NULL |
| POLH | 6 | DNA polymerase eta [Source:HGNC Symbol;A |
| POLH-AS1 | 6 | POLH antisense RNA 1 [Source:HGNC Symbol; |
| GTPBP2 | 6 | GTP binding protein 2 [Source:HGNC Symbol;A |
| MAD2L1BP | 6 | MAD2L1 binding protein [Source:HGNC Symb |
| RSPH9 | 6 | radial spoke head 9 homolog [Source:HGNC Sy |
| MRPS18A | 6 | mitochondrial ribosomal protein S18A [Source |
| AL136131.1 | 6 | OPA-interacting protein (OIP21) pseudogene |
| AL136131.2 | 6 | novel transcript |
| VEGFA | 6 | vascular endothelial growth factor A [Source:H |
| AL136131.3 | 6 | novel transcript, antisense to VEGFA |

| | | |
|---|---|---|
| AL157371.2 | 6 | novel transcript |
| LINC02537 | 6 | long intergenic non-protein coding RNA 2537 |
| AL157371.1 | 6 | novel transcript |
| AL109615.3 | 6 | novel transcript, antisense to C6orf223 |
| C6orf223 | 6 | chromosome 6 open reading frame 223 [Sour |
| AL109615.2 | 6 | novel transcript |
| AL109615.1 | 6 | ribosomal protein L29 (RPL29) pseudogene |
| AL109615.4 | 6 | novel transcript |
| MRPL14 | 6 | mitochondrial ribosomal protein L14 [Source: |
| TMEM63B | 6 | transmembrane protein 63B [Source:HGNC Sy |
| CAPN11 | 6 | calpain 11 [Source:HGNC Symbol;Acc:HGNC:1 |
| RN7SL811P | 6 | RNA, 7SL, cytoplasmic 811, pseudogene [Sour |
| MYMX | 6 | myomixer, myoblast fusion factor [Source:HG |
| SLC29A1 | 6 | solute carrier family 29 member 1 (Augustine |
| HSP90AB1 | 6 | heat shock protein 90 alpha family class B me |
| SLC35B2 | 6 | solute carrier family 35 member B2 [Source:H |
| MIR4647 | 6 | microRNA 4647 [Source:HGNC Symbol;Acc:HG |
| NFKBIE | 6 | NFKB inhibitor epsilon [Source:HGNC Symbol; |
| TMEM151B | 6 | transmembrane protein 151B [Source:HGNC S |
| AL353588.1 | 6 | novel protein, TMEM151B-SPATS1 readtrough |
| TCTE1 | 6 | t-complex-associated-testis-expressed 1 [Sour |
| AARS2 | 6 | alanyl-tRNA synthetase 2, mitochondrial [Sou |
| SPATS1 | 6 | spermatogenesis associated serine rich 1 [Sou |
| CDC5L | 6 | cell division cycle 5 like [Source:HGNC Symbol |
| MIR4642 | 6 | microRNA 4642 [Source:HGNC Symbol;Acc:HG |
| AL136140.1 | 6 | novel transcript |
| AL136140.2 | 6 | TEC |
| AL133334.1 | 6 | novel transcript, antisense to SUPT3H |
| SUPT3H | 6 | SPT3 homolog, SAGA and STAGA complex com |
| NUDT19P4 | 6 | nudix hydrolase 19 pseudogene 4 [Source:HG |
| AL138880.2 | 6 | ribonuclease P/MRP 40kDa subunit (RPP40) p |
| RBM22P4 | 6 | RNA binding motif protein 22 pseudogene 4 [S |
| AL138880.1 | 6 | pseudogene similar to part of poly(rC) binding |
| MIR586 | 6 | microRNA 586 [Source:HGNC Symbol;Acc:HGN |
| RUNX2 | 6 | RUNX family transcription factor 2 [Source:HG |
| AL096865.1 | 6 | novel transcript, antisense to RUNX2 |
| RUNX2-AS1 | 6 | RUNX2 antisense RNA 1 [Source:HGNC Symbo |
| CLIC5 | 6 | chloride intracellular channel 5 [Source:HGNC |
| RNU6-754P | 6 | RNA, U6 small nuclear 754, pseudogene [Sour |
| AL035701.1 | 6 | novel transcript |
| ENPP4 | 6 | ectonucleotide pyrophosphatase/phosphodie |
| ENPP5 | 6 | ectonucleotide pyrophosphatase/phosphodie |
| ACTG1P9 | 6 | actin gamma 1 pseudogene 9 [Source:HGNC S |
| RCAN2 | 6 | regulator of calcineurin 2 [Source:HGNC Symb |
| AL359633.1 | 6 | ribosomal protein L36 (RPL36) pseudogene |
| AL035670.1 | 6 | novel transcript |
| CYP39A1 | 6 | cytochrome P450 family 39 subfamily A meml |
| SLC25A27 | 6 | solute carrier family 25 member 27 [Source:H |
| AL591242.1 | 6 | novel transcript, antisense to SLC25A27 |
| TDRD6 | 6 | tudor domain containing 6 [Source:HGNC Sym |

| | | |
|---|---|---|
| PLA2G7 | 6 | phospholipase A2 group VII [Source:HGNC Syr |
| ANKRD66 | 6 | ankyrin repeat domain 66 [Source:HGNC Syml |
| AL161618.1 | 6 | novel transcript |
| MEP1A | 6 | meprin A subunit alpha [Source:HGNC Symbol |
| ADGRF5 | 6 | adhesion G protein-coupled receptor F5 [Sour |
| AL096772.1 | 6 | novel transcript |
| ADGRF1 | 6 | adhesion G protein-coupled receptor F1 [Sour |
| TNFRSF21 | 6 | TNF receptor superfamily member 21 [Source |
| B3GNTL1P2 | 6 | B3GNTL1 pseudogene 2 [Source:HGNC Symbc |
| AL451166.1 | 6 | novel transcript |
| AL355353.1 | 6 | novel transcript |
| CD2AP | 6 | CD2 associated protein [Source:HGNC Symbol |
| Y_RNA | 6 | Y RNA [Source:RFAM;Acc:RF00019] |
| AL358178.1 | 6 | 40S ribosomal protein S12 (RPS12) pseudoger |
| ADGRF2 | 6 | adhesion G protein-coupled receptor F2 [Sour |
| ADGRF4 | 6 | adhesion G protein-coupled receptor F4 [Sour |
| RN7SKP116 | 6 | RN7SK pseudogene 116 [Source:HGNC Symbo |
| AL356421.2 | 6 | novel transcript |
| AL356421.1 | 6 | ribosomal protein L27A (RPL27A) pseudogene |
| OPN5 | 6 | opsin 5 [Source:HGNC Symbol;Acc:HGNC:199 |
| RNU1-105P | 6 | RNA, U1 small nuclear 105, pseudogene [Sour |
| PTCHD4 | 6 | patched domain containing 4 [Source:HGNC S |
| AL353138.1 | 6 | novel transcript |
| HNRNPA3P4 | 6 | heterogeneous nuclear ribonucleoprotein A3 |
| RBMXP1 | 6 | RBMX pseudogene 1 [Source:HGNC Symbol;A |
| FO393412.1 | 6 | patched domain containing 4 (PTCHD4) pseud |
| AL391538.1 | 6 | novel transcript |
| AL022099.1 | 6 | proline rich 11 (PRR11) pseudogene |
| AL589994.1 | 6 | pseudogene similar to part of myotubularin re |
| AL589994.2 | 6 | piggyBac transposable element derived 4 (PGI |
| FO393413.1 | 6 | zinc finger pseudogene |
| RNU7-65P | 6 | RNA, U7 small nuclear 65 pseudogene [Source |
| EEF1A1P42 | 6 | eukaryotic translation elongation factor 1 alpl |
| MMUT | 6 | methylmalonyl-CoA mutase [Source:HGNC Syı |
| CENPQ | 6 | centromere protein Q [Source:HGNC Symbol;/ |
| GLYATL3 | 6 | glycine-N-acyltransferase like 3 [Source:HGNC |
| C6orf141 | 6 | chromosome 6 open reading frame 141 [Sour |
| CYP2AC1P | 6 | cytochrome P450 family 2 subfamily AC meml |
| RHAG | 6 | Rh associated glycoprotein [Source:HGNC Syn |
| CRISP2 | 6 | cysteine rich secretory protein 2 [Source:HGN |
| AL121974.1 | 6 | novel transcript |
| AL121950.1 | 6 | TEC |
| CRISP3 | 6 | cysteine rich secretory protein 3 [Source:HGN |
| PGK2 | 6 | phosphoglycerate kinase 2 [Source:HGNC Syn |
| AL359458.1 | 6 | novel transcript |
| CRISP1 | 6 | cysteine rich secretory protein 1 [Source:HGN |
| DEFB133 | 6 | defensin beta 133 [Source:HGNC Symbol;Acc: |
| DEFB114 | 6 | defensin beta 114 [Source:HGNC Symbol;Acc: |
| DEFB113 | 6 | defensin beta 113 [Source:HGNC Symbol;Acc: |
| DEFB110 | 6 | defensin beta 110 [Source:HGNC Symbol;Acc: |

| | | |
|---|---|---|
| DEFB112 | 6 | defensin beta 112 [Source:HGNC Symbol;Acc: |
| AL138826.1 | 6 | uncharacterized LOC100505985 [Source:NCBI |
| AL132799.1 | 6 | novel transcript |
| AL360175.1 | 6 | novel transcript |
| AL135908.1 | 6 | novel transcript |
| TFAP2D | 6 | transcription factor AP-2 delta [Source:HGNC |
| TFAP2B | 6 | transcription factor AP-2 beta [Source:HGNC S |
| RPS17P5 | 6 | ribosomal protein S17 pseudogene 5 [Source: |
| AL049693.1 | 6 | pseudogene similar to part of nuclear transpo |
| FTH1P5 | 6 | ferritin heavy chain 1 pseudogene 5 [Source:H |
| AL158050.2 | 6 | TIA1 cytotoxic granule-associated RNA binding |
| AL158050.1 | 6 | Wilms' tumour 1-associating protein (WTAP) p |
| AL365219.1 | 6 | NULL |
| AL445529.1 | 6 | ribosomal protein S15a (RPS15A) pseudogene |
| AL355997.1 | 6 | novel transcript |
| PKHD1 | 6 | PKHD1 ciliary IPT domain containing fibrocysti |
| RN7SL580P | 6 | RNA, 7SL, cytoplasmic 580, pseudogene [Sour |
| MIR206 | 6 | microRNA 206 [Source:HGNC Symbol;Acc:HGN |
| LINCMD1 | 6 | long intergenic non-protein coding RNA, musc |
| MIR133B | 6 | microRNA 133b [Source:HGNC Symbol;Acc:HG |
| IL17A | 6 | interleukin 17A [Source:HGNC Symbol;Acc:HG |
| IL17F | 6 | interleukin 17F [Source:HGNC Symbol;Acc:HG |
| SLC25A20P1 | 6 | solute carrier family 25 member 20 pseudoge |
| MCM3 | 6 | minichromosome maintenance complex comp |
| PAQR8 | 6 | progestin and adipoQ receptor family membe |
| EFHC1 | 6 | EF-hand domain containing 1 [Source:HGNC S |
| TRAM2 | 6 | translocation associated membrane protein 2 |
| TRAM2-AS1 | 6 | TRAM2 antisense RNA 1 (head to head) [Sourc |
| AL109918.2 | 6 | SH3 domain binding glutamic acid-rich protein |
| AL109918.1 | 6 | heterogeneous nuclear ribonucleoprotein A/B |
| TMEM14A | 6 | transmembrane protein 14A [Source:HGNC Sy |
| GSTA8P | 6 | glutathione S-transferase alpha 8, pseudogene |
| AL109918.3 | 6 | SREK1-interacting protein 1 (SREK1IP1) pseud |
| GSTA7P | 6 | glutathione S-transferase alpha 7, pseudogene |
| GSTA2 | 6 | glutathione S-transferase alpha 2 [Source:HGN |
| GSTA12P | 6 | glutathione S-transferase alpha 12, pseudogen |
| GSTA1 | 6 | glutathione S-transferase alpha 1 [Source:HGN |
| GSTA6P | 6 | glutathione S-transferase alpha 6, pseudogene |
| GSTA5 | 6 | glutathione S-transferase alpha 5 [Source:HGN |
| GSTA11P | 6 | glutathione S-transferase alpha 11, pseudogen |
| GSTA10P | 6 | glutathione S-transferase alpha 10, pseudogen |
| GSTA3 | 6 | glutathione S-transferase alpha 3 [Source:HGN |
| GSTA9P | 6 | glutathione S-transferase alpha 9, pseudogene |
| AL121969.1 | 6 | family with sequence similarity 110 member A |
| GSTA4 | 6 | glutathione S-transferase alpha 4 [Source:HGN |
| 7SK | 6 | 7SK RNA [Source:RFAM;Acc:RF00100] |
| RN7SK | 6 | RNA component of 7SK nuclear ribonucleopro |
| ICK | 6 | intestinal cell kinase [Source:HGNC Symbol;Ac |
| FBXO9 | 6 | F-box protein 9 [Source:HGNC Symbol;Acc:HG |
| RN7SL244P | 6 | RNA, 7SL, cytoplasmic 244, pseudogene [Sour |

| | | |
|---|---|---|
| AL512347.1 | 6 | novel transcript, overlapping to GCM1 |
| GCM1 | 6 | glial cells missing transcription factor 1 [Sourc |
| RNU6-464P | 6 | RNA, U6 small nuclear 464, pseudogene [Sour |
| SOD1P1 | 6 | superoxide dismutase 1 pseudogene 1 [Source |
| AL512378.1 | 6 | regulator of chromosome condensation 1 (RC |
| RNU1-136P | 6 | RNA, U1 small nuclear 136, pseudogene [Sour |
| HMGB1P20 | 6 | high mobility group box 1 pseudogene 20 [Sou |
| ELOVL5 | 6 | ELOVL fatty acid elongase 5 [Source:HGNC Syr |
| MIR5685 | 6 | microRNA 5685 [Source:HGNC Symbol;Acc:HG |
| RPS16P5 | 6 | ribosomal protein S16 pseudogene 5 [Source: |
| AL034374.1 | 6 | novel transcript |
| RPL31P28 | 6 | ribosomal protein L31 pseudogene 28 [Source |
| RPL31P33 | 6 | ribosomal protein L31 pseudogene 33 [Source |
| AL591034.1 | 6 | replication protein A3 (RPA3) pseudogene |
| AL591034.2 | 6 | WW domain binding protein 11 (WBP11) pseu |
| RN7SKP256 | 6 | RN7SK pseudogene 256 [Source:HGNC Symbo |
| NANOGP3 | 6 | Nanog homeobox pseudogene 3 [Source:HGN |
| GCLC | 6 | glutamate-cysteine ligase catalytic subunit [Sc |
| AL033397.2 | 6 | novel transcript |
| AL033397.1 | 6 | novel transcript |
| LINC01564 | 6 | long intergenic non-protein coding RNA 1564 |
| KLHL31 | 6 | kelch like family member 31 [Source:HGNC Sy |
| AL590652.1 | 6 | novel transcript |
| LRRC1 | 6 | leucine rich repeat containing 1 [Source:HGN( |
| AL033384.2 | 6 | novel transcript, antisense to LRRC1 |
| MLIP | 6 | muscular LMNA interacting protein [Source:H |
| AL033384.1 | 6 | novel transcript |
| MLIP-AS1 | 6 | MLIP antisense RNA 1 [Source:HGNC Symbol;/ |
| MLIP-IT1 | 6 | MLIP intronic transcript 1 [Source:HGNC Symb |
| ERHP2 | 6 | ERH pseudogene 2 [Source:HGNC Symbol;Acc |
| AL359380.1 | 6 | novel transcript, sense intronic to MLIP |
| TINAG | 6 | tubulointerstitial nephritis antigen [Source:HG |
| AL589946.1 | 6 | novel transcript |
| CLNS1AP1 | 6 | chloride nucleotide-sensitive channel 1A pseu |
| RPL10P10 | 6 | ribosomal protein L10 pseudogene 10 [Source |
| RPSAP44 | 6 | ribosomal protein SA pseudogene 44 [Source: |
| KRASP1 | 6 | KRAS proto-oncogene, GTPase pseudogene 1 |
| RNU6-1023P | 6 | RNA, U6 small nuclear 1023, pseudogene [Sou |
| AL512363.1 | 6 | novel transcript |
| FAM83B | 6 | family with sequence similarity 83 member B |
| AL049555.1 | 6 | novel transcript, overlapping to FAM83B |
| HCRTR2 | 6 | hypocretin receptor 2 [Source:HGNC Symbol;/ |
| GFRAL | 6 | GDNF family receptor alpha like [Source:HGN( |
| HMGCLL1 | 6 | 3-hydroxymethyl-3-methylglutaryl-CoA lyase l |
| AL590290.1 | 6 | NIFU-like protein pseudogene |
| BMP5 | 6 | bone morphogenetic protein 5 [Source:HGNC |
| NPM1P36 | 6 | nucleophosmin 1 pseudogene 36 [Source:HGI |
| COL21A1 | 6 | collagen type XXI alpha 1 chain [Source:HGNC |
| DHFRP6 | 6 | dihydrofolate reductase pseudogene 6 [Sourc |
| AL031779.1 | 6 | novel transcript |

| | | |
|---|---|---|
| RCC2P7 | 6 | regulator of chromosome condensation 2 pse |
| DST | 6 | dystonin [Source:HGNC Symbol;Acc:HGNC:10 |
| AL512422.1 | 6 | novel transcript |
| RPL17P26 | 6 | ribosomal protein L17 pseudogene 26 [Source |
| RNU6-626P | 6 | RNA, U6 small nuclear 626, pseudogene [Sour |
| BEND6 | 6 | BEN domain containing 6 [Source:HGNC Symb |
| OSTCP6 | 6 | oligosaccharyltransferase complex subunit ps |
| FTH1P15 | 6 | ferritin heavy chain 1 pseudogene 15 [Source: |
| MRPL30P1 | 6 | mitochondrial ribosomal protein L30 pseudog |
| KIAA1586 | 6 | KIAA1586 [Source:HGNC Symbol;Acc:HGNC:2 |
| ZNF451 | 6 | zinc finger protein 451 [Source:HGNC Symbol; |
| ZNF451-AS1 | 6 | ZNF451 regulatory antisense RNA 1 [Source:H |
| BAG2 | 6 | BCL2 associated athanogene 2 [Source:HGNC |
| RAB23 | 6 | RAB23, member RAS oncogene family [Source |
| PRIM2 | 6 | DNA primase subunit 2 [Source:HGNC Symbol |
| MIR548U | 6 | microRNA 548u [Source:HGNC Symbol;Acc:HG |
| FO680682.1 | 6 | novel transcript |
| AL021368.3 | 6 | novel transcript |
| AL021368.5 | 6 | novel transcript |
| AL021368.1 | 6 | novel transcript |
| AL021368.2 | 6 | novel transcript |
| GUSBP4 | 6 | GUSB pseudogene 4 [Source:HGNC Symbol;Ac |
| AL021368.4 | 6 | novel transcript |
| POM121L14P | 6 | POM121 transmembrane nucleoporin like 14, |
| LINC00680 | 6 | long intergenic non-protein coding RNA 680 [ |
| AL445250.1 | 6 | novel transcript |
| GAPDHP15 | 6 | glyceraldehyde-3-phosphate dehydrogenase |
| AL390237.1 | 6 | SUMO1/sentrin specific peptidase 6 (SENP6) |
| RBBP4P4 | 6 | RBBP4 pseudogene 4 [Source:HGNC Symbol;A |
| AL603755.1 | 6 | RAB5A, member RAS oncogene family (RAB5A |
| AC244258.1 | 6 | primase, DNA, polypeptide 2 (58kDa) (PRIM2) |
| GAPDHP41 | 6 | glyceraldehyde 3 phosphate dehydrogenase |
| AL590227.1 | 6 | novel transcript |
| AL512427.2 | 6 | novel transcript |
| AL512427.1 | 6 | SUMO1/sentrin specific peptidase 6 (SENP6) |
| RBBP4P3 | 6 | RBBP4 pseudogene 3 [Source:HGNC Symbol;A |
| AL356131.1 | 6 | proteasome subunit alpha type 4 (PROS29) ps |
| MTRNR2L9 | 6 | MT-RNR2 like 9 [Source:HGNC Symbol;Acc:HG |
| KHDRBS2-OT | 6 | KHDRBS2 3'UTR overlapping transcript 1 [Sou |
| KHDRBS2 | 6 | KH RNA binding domain containing, signal tra |
| AL355347.1 | 6 | novel transcript, antisense to KHDRBS2 |
| DHFRP5 | 6 | dihydrofolate reductase pseudogene 5 [Sourc |
| AL355375.1 | 6 | ribosomal protein S17 (RPS17) pseudogene |
| AL355375.2 | 6 | endogenous retrovirus group FRD, member 1 |
| AL450346.1 | 6 | keratin 8 (KRT8) pseudogene |
| FKBP1C | 6 | FKBP prolyl isomerase 1C [Source:HGNC Symb |
| SPTLC1P3 | 6 | serine palmitoyltransferase long chain base s |
| AL450346.2 | 6 | novel pseudogene |
| LGSN | 6 | lengsin, lens protein with glutamine syntheta |
| AL121949.3 | 6 | crumbs homolog 3 (Drosophila) (CRB3) pseud |

| | | |
|---|---|---|
| AL121949.2 | 6 | DENN/MADD domain containing 1C (DENND1 |
| AL121949.1 | 6 | leucine rich repeat containing 59 (LRRC59) pse |
| EEF1B2P5 | 6 | eukaryotic translation elongation factor 1 beta |
| PTP4A1 | 6 | protein tyrosine phosphatase 4A1 [Source:HG |
| RPL7AP34 | 6 | ribosomal protein L7a pseudogene 34 [Source |
| AL135905.1 | 6 | novel transcript, antisense to PTP4A1 |
| AL135905.2 | 6 | novel protein |
| AL513043.1 | 6 | ribosomal protein S7 (RPS7) pseudogene |
| RPL9P18 | 6 | ribosomal protein L9 pseudogene 18 [Source: |
| PHF3 | 6 | PHD finger protein 3 [Source:HGNC Symbol;Ac |
| EYS | 6 | eyes shut homolog [Source:HGNC Symbol;Acc |
| AL354719.1 | 6 | platelet-activating factor acetylhydrolase 1b, r |
| SCAT8 | 6 | S-phase cancer associated transcript 8 [Source |
| GCNT1P4 | 6 | glucosaminyl (N-acetyl) transferase 1 pseudog |
| AL357375.1 | 6 | novel transcript |
| HNRNPDP2 | 6 | heterogeneous nuclear ribonucleoprotein D p |
| AL355357.1 | 6 | transmembrane protein 98 (TMEM98) pseudo |
| AL365217.1 | 6 | zinc finger CCCH-type containing 11A (ZC3H11 |
| SLC25A51P1 | 6 | SLC25A51 pseudogene 1 [Source:HGNC Symb |
| ADH5P4 | 6 | ADH5 pseudogene 4 [Source:HGNC Symbol;Ac |
| NUFIP1P1 | 6 | NUFIP1 pseudogene 1 [Source:HGNC Symbol; |
| AL450336.1 | 6 | transcription elongation regulator 1 (TCERG1) |
| RNU7-66P | 6 | RNA, U7 small nuclear 66 pseudogene [Source |
| AL591004.1 | 6 | NULL |
| AL365503.1 | 6 | SH3-domain binding protein 4 (SH3BP4) pseud |
| RNA5SP208 | 6 | RNA, 5S ribosomal pseudogene 208 [Source:H |
| RNU6-280P | 6 | RNA, U6 small nuclear 280, pseudogene [Sour |
| Y_RNA | 6 | Y RNA [Source:RFAM;Acc:RF00019] |
| AL713998.1 | 6 | novel transcript |
| AL713998.3 | 6 | novel transcript |
| AL713998.4 | 6 | novel transcript |
| AL713998.2 | 6 | suppressor of variegation 3-9 homolog 2 (Dros |
| AL646090.2 | 6 | novel transcript |
| AL646090.1 | 6 | novel transcript |
| AL593844.1 | 6 | novel transcript |
| LINC02549 | 6 | long intergenic non-protein coding RNA 2549 |
| AL606923.2 | 6 | novel transcript |
| AL606923.1 | 6 | zinc finger protein 791 (ZNF791) pseudogene |
| AL391807.1 | 6 | novel transcript |
| ADGRB3 | 6 | adhesion G protein-coupled receptor B3 [Sour |
| LMBRD1 | 6 | LMBR1 domain containing 1 [Source:HGNC Sy |
| NPM1P37 | 6 | nucleophosmin 1 pseudogene 37 [Source:HGN |
| GAPDHP42 | 6 | glyceraldehyde 3 phosphate dehydrogenase p |
| COL19A1 | 6 | collagen type XIX alpha 1 chain [Source:HGNC |
| RPL37P15 | 6 | ribosomal protein L37 pseudogene 15 [Source |
| COL9A1 | 6 | collagen type IX alpha 1 chain [Source:HGNC S |
| AL160262.1 | 6 | novel transcript |
| AL080275.1 | 6 | novel transcript |
| AL365226.2 | 6 | novel transcript |
| AL365226.1 | 6 | novel transcript |

| | | |
|---|---|---|
| FAM135A | 6 | family with sequence similarity 135 member A |
| RNU7-48P | 6 | RNA, U7 small nuclear 48 pseudogene [Source |
| SDHAF4 | 6 | succinate dehydrogenase complex assembly f |
| AL583856.2 | 6 | novel transcript |
| AL583856.1 | 6 | spermine synthase (SMS) pseudogene |
| SLC25A6P6 | 6 | solute carrier family 25 member 6 pseudogen |
| SMAP1 | 6 | small ArfGAP 1 [Source:HGNC Symbol;Acc:HG |
| AL354943.1 | 6 | transmembrane protein 192 (TMEM192) pseu |
| NDUFAB1P1 | 6 | NADH:ubiquinone oxidoreductase subunit AB |
| B3GAT2 | 6 | beta-1,3-glucuronyltransferase 2 [Source:HGN |
| AL096709.1 | 6 | novel transcript |
| BECN1P2 | 6 | beclin 1 pseudogene 2 [Source:HGNC Symbol; |
| U3 | 6 | Small nucleolar RNA U3 [Source:RFAM;Acc:RF |
| LYPLA1P3 | 6 | LYPLA1 pseudogene 3 [Source:HGNC Symbol;A |
| RNU6-411P | 6 | RNA, U6 small nuclear 411, pseudogene [Sour |
| AL589935.1 | 6 | novel transcript, antisense OGFRL1 |
| AL589935.3 | 6 | novel transcript |
| AL589935.2 | 6 | coiled-coil domain containing 112 (CCDC112) |
| OGFRL1 | 6 | opioid growth factor receptor like 1 [Source:H |
| AL136164.2 | 6 | novel transcript |
| AL136164.1 | 6 | novel transcript |
| LINC00472 | 6 | long intergenic non-protein coding RNA 472 [S |
| MIR30C2 | 6 | microRNA 30c-2 [Source:HGNC Symbol;Acc:H |
| AL136164.3 | 6 | TEC |
| MIR30A | 6 | microRNA 30a [Source:HGNC Symbol;Acc:HGN |
| LINC01626 | 6 | long intergenic non-protein coding RNA 1626 |
| AL035467.2 | 6 | novel transcript |
| AL035467.1 | 6 | neugrin, neurite outgrowth associated (NGRN |
| KRT19P1 | 6 | keratin 19 pseudogene 1 [Source:HGNC Symb |
| RNU4-66P | 6 | RNA, U4 small nuclear 66, pseudogene [Sourc |
| RIMS1 | 6 | regulating synaptic membrane exocytosis 1 [S |
| AL035633.1 | 6 | forkhead box H1 (FOXH1) pseudogene |
| FO393414.2 | 6 | polymerase (DNA directed), delta 1, catalytic s |
| FO393414.1 | 6 | HORMA domain containing 1 (HORMAD1) pse |
| FO393414.3 | 6 | novel transcript |
| KCNQ5 | 6 | potassium voltage-gated channel subfamily Q |
| KCNQ5-IT1 | 6 | KCNQ5 intronic transcript 1 [Source:HGNC Syr |
| MIR4282 | 6 | microRNA 4282 [Source:HGNC Symbol;Acc:HG |
| KNOP1P4 | 6 | lysine rich nucleolar protein 1 pseudogene 4 [ |
| PGAM1P10 | 6 | phosphoglycerate mutase 1 pseudogene 10 [S |
| KCNQ5-AS1 | 6 | KCNQ5 antisense RNA 1 [Source:HGNC Symbo |
| RBPMS2P1 | 6 | RBPMS2 pseudogene 1 [Source:HGNC Symbol |
| KHDC1P1 | 6 | KH domain containing 1 pseudogene 1 [Sourc |
| AL365232.1 | 6 | novel protein |
| KHDC1L | 6 | KH domain containing 1 like [Source:HGNC Syr |
| KHDC1 | 6 | KH domain containing 1 [Source:HGNC Symbo |
| AC019205.1 | 6 | novel transcript, antisense to KHDC1 |
| RPSAP41 | 6 | ribosomal protein SA pseudogene 41 [Source: |
| EIF3EP1 | 6 | eukaryotic translation initiation factor 3 subur |
| SDCBP2P1 | 6 | syndecan binding protein 2 pseudogene 1 [Sou |

| | | |
|---|---|---|
| PAICSP3 | 6 | phosphoribosylaminoimidazole carboxylase, p |
| DPPA5 | 6 | developmental pluripotency associated 5 [Sou |
| KHDC3L | 6 | KH domain containing 3 like, subcortical mate |
| OOEP | 6 | oocyte expressed protein [Source:HGNC Symb |
| OOEP-AS1 | 6 | OOEP antisense RNA 1 [Source:HGNC Symbol; |
| RPL39P3 | 6 | ribosomal protein L39 pseudogene 3 [Source: |
| RPS6P8 | 6 | ribosomal protein S6 pseudogene 8 [Source:H |
| DDX43 | 6 | DEAD-box helicase 43 [Source:HGNC Symbol;A |
| CGAS | 6 | cyclic GMP-AMP synthase [Source:HGNC Symb |
| MTO1 | 6 | mitochondrial tRNA translation optimization 1 |
| RNU6-975P | 6 | RNA, U6 small nuclear 975, pseudogene [Sour |
| RN7SL827P | 6 | RNA, 7SL, cytoplasmic 827, pseudogene [Sour |
| AL603910.1 | 6 | novel transcript |
| EEF1A1 | 6 | eukaryotic translation elongation factor 1 alph |
| AL121972.1 | 6 | novel transcript |
| AL603910.2 | 6 | thymosin, beta (TMSNB) pseudogene |
| SLC17A5 | 6 | solute carrier family 17 member 5 [Source:HG |
| RPS27P15 | 6 | ribosomal protein S27 pseudogene 15 [Source |
| AL590428.1 | 6 | novel transcript |
| CD109 | 6 | CD109 molecule [Source:HGNC Symbol;Acc:H |
| AL590552.1 | 6 | TEC |
| TXNP7 | 6 | thioredoxin pseudogene 7 [Source:HGNC Sym |
| AL357507.1 | 6 | novel transcript |
| AL357563.1 | 6 | family with sequence similarity 136, member |
| AL356277.3 | 6 | novel transcript |
| AL356277.1 | 6 | ubiquinol-cytochrome c reductase core protei |
| AL356277.2 | 6 | novel transcript |
| AL356473.1 | 6 | nucleoporin 50kDa (NUP50) pseudogene |
| COL12A1 | 6 | collagen type XII alpha 1 chain [Source:HGNC |
| COX7A2 | 6 | cytochrome c oxidase subunit 7A2 [Source:HG |
| TMEM30A | 6 | transmembrane protein 30A [Source:HGNC Sy |
| AL080250.1 | 6 | novel transcript |
| FILIP1 | 6 | filamin A interacting protein 1 [Source:HGNC |
| HMGB1P39 | 6 | high mobility group box 1 pseudogene 39 [Sou |
| AL445465.1 | 6 | novel transcript |
| U3 | 6 | Small nucleolar RNA U3 [Source:RFAM;Acc:RF |
| MIR4463 | 6 | microRNA 4463 [Source:HGNC Symbol;Acc:HG |
| AL445465.2 | 6 | novel transcript |
| UBE2V1P15 | 6 | ubiquitin conjugating enzyme E2 V1 pseudoge |
| RNU1-34P | 6 | RNA, U1 small nuclear 34, pseudogene [Sourc |
| RPL26P20 | 6 | ribosomal protein L26 pseudogene 20 [Source |
| AL356057.2 | 6 | H3 histone, family 3C (H3F3C) pseudogene |
| RNU6-1338P | 6 | RNA, U6 small nuclear 1338, pseudogene [Sou |
| SENP6 | 6 | SUMO specific peptidase 6 [Source:HGNC Sym |
| AL356057.1 | 6 | LSM14A, SCD6 homolog A (S. cerevisiae) (LSM |
| RNU6-1016P | 6 | RNA, U6 small nuclear 1016, pseudogene [Sou |
| RN7SKP163 | 6 | RN7SK pseudogene 163 [Source:HGNC Symbo |
| AL109897.1 | 6 | novel oocyte-secreted protein family member |
| MYO6 | 6 | myosin VI [Source:HGNC Symbol;Acc:HGNC:76 |
| RNU6-155P | 6 | RNA, U6 small nuclear 155, pseudogene [Sour |

| | | |
|---|---|---|
| RNA5SP209 | 6 | RNA, 5S ribosomal pseudogene 209 [Source:H |
| IMPG1 | 6 | interphotoreceptor matrix proteoglycan 1 [So |
| Y_RNA | 6 | Y RNA [Source:RFAM;Acc:RF00019] |
| RNU6-248P | 6 | RNA, U6 small nuclear 248, pseudogene [Sour |
| RNU6-261P | 6 | RNA, U6 small nuclear 261, pseudogene [Sour |
| LINC02540 | 6 | long intergenic non-protein coding RNA 2540 |
| RNU6-84P | 6 | RNA, U6 small nuclear 84, pseudogene [Sourc |
| AL591030.1 | 6 | novel transcript |
| AL590426.1 | 6 | novel pseudogene |
| AL355612.1 | 6 | novel transcript |
| AL365222.1 | 6 | chromosome 12 open reading frame 76 (C12c |
| HTR1B | 6 | 5-hydroxytryptamine receptor 1B [Source:HG |
| RPS6P7 | 6 | ribosomal protein S6 pseudogene 7 [Source:H |
| MEI4 | 6 | meiotic double-stranded break formation pro |
| AL121718.1 | 6 | novel transcript |
| AL450327.1 | 6 | novel transcript |
| IRAK1BP1 | 6 | interleukin 1 receptor associated kinase 1 bin |
| PHIP | 6 | pleckstrin homology domain interacting prote |
| AL356776.1 | 6 | nucleoporin 155kDa pseudogene |
| AL356776.2 | 6 | novel transcript, antisense to PHIP |
| HMGN3 | 6 | high mobility group nucleosomal binding dom |
| HMGN3-AS1 | 6 | HMGN3 antisense RNA 1 [Source:HGNC Symb |
| AL355796.1 | 6 | ribosomal protein S18 (RPS18) pseudogene |
| Y_RNA | 6 | Y RNA [Source:RFAM;Acc:RF00019] |
| LCAL1 | 6 | lung cancer associated lncRNA 1 [Source:HGN |
| AL355379.1 | 6 | high mobility group nucleosomal binding dom |
| AL078601.1 | 6 | high mobility group nucleosomal binding dom |
| AL078601.2 | 6 | stomatin (STOM) pseudogene |
| AL391840.1 | 6 | novel transcript |
| DBIP1 | 6 | DBI pseudogene 1 [Source:HGNC Symbol;Acc: |
| LCA5 | 6 | lebercilin LCA5 [Source:HGNC Symbol;Acc:HG |
| AL391840.3 | 6 | novel transcript |
| AL391840.2 | 6 | solute carrier family 19, member 3 (SLC19A3) |
| AL451064.1 | 6 | novel transcript |
| AL451064.2 | 6 | TEC |
| SH3BGRL2 | 6 | SH3 domain binding glutamate rich protein lik |
| LINC01621 | 6 | long intergenic non-protein coding RNA 1621 |
| AL132875.1 | 6 | cell division cycle 73, Paf1/RNA polymerase II |
| AL132875.3 | 6 | novel transcript |
| AL132875.2 | 6 | novel transcript |
| ELOVL4 | 6 | ELOVL fatty acid elongase 4 [Source:HGNC Syr |
| AL133475.1 | 6 | novel transcript |
| GAPDHP63 | 6 | glyceraldehyde 3 phosphate dehydrogenase p |
| RPL35AP18 | 6 | ribosomal protein L35a pseudogene 18 [Sourc |
| TTK | 6 | TTK protein kinase [Source:HGNC Symbol;Acc: |
| AL591135.2 | 6 | novel transcript |
| AL591135.1 | 6 | Rho GTPase activating protein 21 (ARHGAP21 |
| AK4P5 | 6 | adenylate kinase 4 pseudogene 5 [Source:HGI |
| BCKDHB | 6 | branched chain keto acid dehydrogenase E1 s |
| AL359715.3 | 6 | novel transcript |

| | | |
|---|---|---|
| RPL17P25 | 6 | ribosomal protein L17 pseudogene 25 [Source |
| AL359715.4 | 6 | TEC |
| AL359715.1 | 6 | novel transcript |
| AL359715.2 | 6 | novel transcript |
| RPSAP72 | 6 | ribosomal protein SA pseudogene 72 [Source: |
| AL596028.1 | 6 | pituitary tumor-transforming 1 (PTTG1) pseud |
| AL590824.1 | 6 | citrate synthase (CS) pseudogene |
| TENT5A | 6 | terminal nucleotidyltransferase 5A [Source:H( |
| AL122017.1 | 6 | novel transcript, sense intronic to FAM46A |
| AL359693.1 | 6 | novel transcript |
| RNA5SP210 | 6 | RNA, 5S ribosomal pseudogene 210 [Source:H |
| AL078599.2 | 6 | novel transcript |
| SNORA70 | 6 | Small nucleolar RNA SNORA70 [Source:RFAM; |
| AL078599.1 | 6 | ribosomal protein L10 (RPL10) pseudogene |
| LINC01526 | 6 | long intergenic non-protein coding RNA 1526 |
| LINC02542 | 6 | long intergenic non-protein coding RNA 2542 |
| AL360157.1 | 6 | novel transcript |
| IBTK | 6 | inhibitor of Bruton tyrosine kinase [Source:HG |
| RNU6-130P | 6 | RNA, U6 small nuclear 130, pseudogene [Sour |
| AL121977.1 | 6 | single stranded DNA binding protein pseudog |
| AL121977.2 | 6 | novel transcript, antisense to TPBG |
| TPBG | 6 | trophoblast glycoprotein [Source:HGNC Symb |
| UBE3D | 6 | ubiquitin protein ligase E3D [Source:HGNC Syr |
| LAP3P1 | 6 | leucine aminopeptidase 3 pseudogene 1 [Sour |
| AL355613.1 | 6 | novel transcript |
| DOP1A | 6 | DOP1 leucine zipper like protein A [Source:HG |
| PGM3 | 6 | phosphoglucomutase 3 [Source:HGNC Symbo |
| RWDD2A | 6 | RWD domain containing 2A [Source:HGNC Syr |
| ME1 | 6 | malic enzyme 1 [Source:HGNC Symbol;Acc:HG |
| AL391416.1 | 6 | glyceraldehyde-3-phosphate dehydrogenase ( |
| PRSS35 | 6 | serine protease 35 [Source:HGNC Symbol;Acc |
| SNAP91 | 6 | synaptosome associated protein 91 [Source:H |
| AL136972.1 | 6 | novel transcript |
| AL139232.1 | 6 | novel transcript, antisense to RIPPLY2 |
| RIPPLY2 | 6 | ripply transcriptional repressor 2 [Source:HGN |
| CYB5R4 | 6 | cytochrome b5 reductase 4 [Source:HGNC Syr |
| AL034347.1 | 6 | novel transcript |
| LINC02857 | 6 | long intergenic non-protein coding RNA 2857 |
| MRAP2 | 6 | melanocortin 2 receptor accessory protein 2 [ |
| CEP162 | 6 | centrosomal protein 162 [Source:HGNC Symb |
| LINC01611 | 6 | long intergenic non-protein coding RNA 1611 |
| SMARCE1P2 | 6 | SWI/SNF related, matrix associated, actin dep |
| TBX18 | 6 | T-box transcription factor 18 [Source:HGNC Sy |
| TBX18-AS1 | 6 | TBX18 antisense RNA 1 [Source:HGNC Symbol |
| AL590143.1 | 6 | pseudogene similar to part of cytochrome c o: |
| AL035694.1 | 6 | novel transcript |
| AL109941.1 | 6 | novel pseudogene |
| AL139806.1 | 6 | pseudogene similar to part of hormonally upr |
| AL122016.1 | 6 | ribosomal protein L31 (RPL31) pseudogene |
| AL135903.1 | 6 | karyopherin alpha 5 (importin alpha 6) (KPNA! |

| | | |
|---|---|---|
| KRT18P64 | 6 | keratin 18 pseudogene 64 [Source:HGNC Sym |
| LINC02535 | 6 | long intergenic non-protein coding RNA 2535 |
| DUTP5 | 6 | deoxyuridine triphosphatase pseudogene 5 [S |
| TPT1P6 | 6 | tumor protein, translationally-controlled 1 ps |
| NT5E | 6 | 5'-nucleotidase ecto [Source:HGNC Symbol;Ac |
| AL589666.2 | 6 | TEC |
| SNX14 | 6 | sorting nexin 14 [Source:HGNC Symbol;Acc:H |
| AL589666.1 | 6 | novel protein, SYNCRIP-SNX14 readthrough |
| SYNCRIP | 6 | synaptotagmin binding cytoplasmic RNA inter |
| SNHG5 | 6 | small nucleolar RNA host gene 5 [Source:HGN |
| PKMP3 | 6 | pyruvate kinase M1/2 pseudogene 3 [Source: |
| SNORD50B | 6 | small nucleolar RNA, C/D box 50B [Source:HG |
| AL355615.1 | 6 | pseudogene similar to part of heat shock 90k |
| SMIM11P1 | 6 | small integral membrane protein 11 pseudoge |
| RNU4-72P | 6 | RNA, U4 small nuclear 72, pseudogene [Sourc |
| Y_RNA | 6 | Y RNA [Source:RFAM;Acc:RF00019] |
| AL450338.1 | 6 | ras-related C3 botulinum toxin substrate 3 (rh |
| AL450338.2 | 6 | novel transcript |
| RNU4-12P | 6 | RNA, U4 small nuclear 12, pseudogene [Sourc |
| RPL7P27 | 6 | ribosomal protein L7 pseudogene 27 [Source: |
| AL049545.1 | 6 | RAB1A, member RAS oncogene family (RAB1A |
| NDUFA5P9 | 6 | NADH:ubiquinone oxidoreductase subunit A5 |
| AL391417.1 | 6 | novel pseudogene |
| RN7SL643P | 6 | RNA, 7SL, cytoplasmic 643, pseudogene [Sour |
| AL353133.1 | 6 | ribosomal protein L29 (RPL29) pseudogene |
| AL353133.2 | 6 | novel pseudogene |
| AL157777.1 | 6 | suppression of tumorigenicity 13, Hsp70-inter |
| AL157777.2 | 6 | pseudogene similar to part of Tec protein-tyr |
| HTR1E | 6 | 5-hydroxytryptamine receptor 1E [Source:HG |
| RPL7P29 | 6 | ribosomal protein L7 pseudogene 29 [Source: |
| MTHFD2P2 | 6 | methylenetetrahydrofolate dehydrogenase (N |
| AL138827.1 | 6 | 60S ribosomal protein L21 (RPL21) pseudogen |
| RN7SKP209 | 6 | RN7SK pseudogene 209 [Source:HGNC Symbo |
| CGA | 6 | glycoprotein hormones, alpha polypeptide [So |
| RCN1P1 | 6 | reticulocalbin 1 pseudogene 1 [Source:HGNC S |
| AL139274.2 | 6 | novel transcript, antisense to ZNF292 |
| ZNF292 | 6 | zinc finger protein 292 [Source:HGNC Symbol; |
| AL139274.1 | 6 | novel TWIST neighbor (TWISTNB) pseudogene |
| GJB7 | 6 | gap junction protein beta 7 [Source:HGNC Syn |
| HSPD1P10 | 6 | heat shock protein family D (Hsp60) member |
| SMIM8 | 6 | small integral membrane protein 8 [Source:H |
| C6orf163 | 6 | chromosome 6 open reading frame 163 [Sour |
| AL096817.1 | 6 | novel transcript, sense intronic to C6orf163 |
| CFAP206 | 6 | cilia and flagella associated protein 206 [Sourc |
| AL049697.1 | 6 | novel transcript, C6orf165-SLC35A1 readthrou |
| AL049697.2 | 6 | TAF13 RNA polymerase II, TATA box binding p |
| AL049697.3 | 6 | S-adenosylmethionine decarboxylase 1 (AMD |
| ST13P16 | 6 | ST13, Hsp70 interacting protein pseudogene 1 |
| SLC35A1 | 6 | solute carrier family 35 member A1 [Source:H |
| RNU6-444P | 6 | RNA, U6 small nuclear 444, pseudogene [Sour |

| | | |
|---|---|---|
| RARS2 | 6 | arginyl-tRNA synthetase 2, mitochondrial [Sou |
| ORC3 | 6 | origin recognition complex subunit 3 [Source:I |
| AKIRIN2 | 6 | akirin 2 [Source:HGNC Symbol;Acc:HGNC:214( |
| AL133211.1 | 6 | NULL |
| Y_RNA | 6 | Y RNA [Source:RFAM;Acc:RF00019] |
| RN7SL183P | 6 | RNA, 7SL, cytoplasmic 183, pseudogene [Sour |
| AL590392.1 | 6 | proteasome (prosome, macropain) subunit, al |
| AL136096.1 | 6 | novel transcript |
| SPACA1 | 6 | sperm acrosome associated 1 [Source:HGNC S |
| CNR1 | 6 | cannabinoid receptor 1 [Source:HGNC Symbol |
| AL121835.2 | 6 | TEC |
| AL139042.1 | 6 | novel transcript |
| AL121835.1 | 6 | ribosomal protein S14 (RPS14) pseudogene |
| ACTBP8 | 6 | ACTB pseudogene 8 [Source:HGNC Symbol;Ac |
| AL139090.1 | 6 | novel transcript |
| RNGTT | 6 | RNA guanylyltransferase and 5'-phosphatase [ |
| SNORA73 | 6 | Small nucleolar RNA SNORA73 family [Source: |
| AL079342.3 | 6 | TEC |
| AL079342.1 | 6 | small EDRK-rich factor 2 (SERF2) pseudogene |
| AL079342.2 | 6 | adaptor-related (clathrin-associated) protein c |
| RNU2-61P | 6 | RNA, U2 small nuclear 61, pseudogene [Sourc |
| AL353135.1 | 6 | novel transcript, antisense to PNRC1 |
| PNRC1 | 6 | proline rich nuclear receptor coactivator 1 [So |
| RN7SL336P | 6 | RNA, 7SL, cytoplasmic 336, pseudogene [Sour |
| SRSF12 | 6 | serine and arginine rich splicing factor 12 [Sou |
| AL353135.2 | 6 | novel transcript |
| PM20D2 | 6 | peptidase M20 domain containing 2 [Source:H |
| GABRR1 | 6 | gamma-aminobutyric acid type A receptor rho |
| GABRR2 | 6 | gamma-aminobutyric acid type A receptor rho |
| TUBB3P1 | 6 | tubulin beta 3 class III pseudogene 1 [Source:H |
| UBE2J1 | 6 | ubiquitin conjugating enzyme E2 J1 [Source:H |
| RRAGD | 6 | Ras related GTP binding D [Source:HGNC Symb |
| NACAP7 | 6 | NACA pseudogene 7 [Source:HGNC Symbol;Ac |
| ANKRD6 | 6 | ankyrin repeat domain 6 [Source:HGNC Symbo |
| RN7SL11P | 6 | RNA, 7SL, cytoplasmic 11, pseudogene [Sourc |
| AL159174.1 | 6 | novel transcript |
| LYRM2 | 6 | LYR motif containing 2 [Source:HGNC Symbol; |
| MDN1 | 6 | midasin AAA ATPase 1 [Source:HGNC Symbol; |
| AL096678.1 | 6 | novel transcript |
| DNAJC19P6 | 6 | DnaJ heat shock protein family (Hsp40) memb |
| AL353692.1 | 6 | family with sequence similarity 64, member A |
| CASP8AP2 | 6 | caspase 8 associated protein 2 [Source:HGNC |
| Y_RNA | 6 | Y RNA [Source:RFAM;Acc:RF00019] |
| RPL22P14 | 6 | ribosomal protein L22 pseudogene 14 [Source |
| AL353692.2 | 6 | G protein pathway suppressor 1 (GPS1) pseud |
| GJA10 | 6 | gap junction protein alpha 10 [Source:HGNC S |
| Y_RNA | 6 | Y RNA [Source:RFAM;Acc:RF00019] |
| BACH2 | 6 | BTB domain and CNC homolog 2 [Source:HGN |
| AL121787.1 | 6 | novel transcript |
| RN7SKP110 | 6 | RN7SK pseudogene 110 [Source:HGNC Symbo |

| | | |
|---|---|---|
| AL158136.1 | 6 | novel transcript, antisense to BACH2 |
| AL117342.1 | 6 | A kinase (PRKA) interacting protein 1 (AKIP1) |
| AL132996.1 | 6 | novel transcript |
| MIR4464 | 6 | microRNA 4464 [Source:HGNC Symbol;Acc:HG |
| MAP3K7 | 6 | mitogen-activated protein kinase kinase kinas |
| AL080284.1 | 6 | novel transcript |
| MIR4643 | 6 | microRNA 4643 [Source:HGNC Symbol;Acc:HG |
| CASC6 | 6 | cancer susceptibility 6 [Source:HGNC Symbol; |
| MTATP6P25 | 6 | MT-ATP6 pseudogene 25 [Source:HGNC Symb |
| MTND4LP19 | 6 | MT-ND4L pseudogene 19 [Source:HGNC Symb |
| MTCO1P56 | 6 | MT-CO1 pseudogene 56 [Source:HGNC Symbo |
| RN7SL415P | 6 | RNA, 7SL, cytoplasmic 415, pseudogene [Sour |
| AL590635.1 | 6 | ras homologue gene family pseudogene |
| AL590814.1 | 6 | novel transcript |
| RPL5P19 | 6 | ribosomal protein L5 pseudogene 19 [Source: |
| U3 | 6 | Small nucleolar RNA U3 [Source:RFAM;Acc:RF |
| AL133457.1 | 6 | novel transcript |
| AL359987.1 | 6 | novel transcript |
| LINC02531 | 6 | long intergenic non-protein coding RNA 2531 |
| U3 | 6 | Small nucleolar RNA U3 [Source:RFAM;Acc:RF |
| ATF1P1 | 6 | activating transcription factor 1 pseudogene 1 |
| AL138731.1 | 6 | novel transcript |
| COPS5P1 | 6 | COP9 signalosome subunit 5 pseudogene 1 [S |
| EPHA7 | 6 | EPH receptor A7 [Source:HGNC Symbol;Acc:H |
| AL591519.1 | 6 | novel transcript, antisense to EPHA7 |
| AL356094.2 | 6 | novel transcript |
| AL356094.1 | 6 | TEC |
| AL391336.2 | 6 | NULL |
| AL391336.1 | 6 | novel transcript |
| AL157378.1 | 6 | novel transcript |
| AL078595.1 | 6 | ribosomal protein (RPL7A) pseudogene |
| AL078595.3 | 6 | TEC |
| AL078595.2 | 6 | zinc finger CCCH-type containing 14 (ZC3H14) |
| MTCYBP36 | 6 | MT-CYB pseudogene 36 [Source:HGNC Symbo |
| CYCSP17 | 6 | CYCS pseudogene 17 [Source:HGNC Symbol;A |
| MANEA-DT | 6 | MANEA divergent transcript [Source:HGNC Sy |
| MANEA | 6 | mannosidase endo-alpha [Source:HGNC Symb |
| AL607077.1 | 6 | pseudogene similar to part of achalsia, adrenc |
| AL034348.1 | 6 | novel transcript |
| KRT18P50 | 6 | keratin 18 pseudogene 50 [Source:HGNC Sym |
| FUT9 | 6 | fucosyltransferase 9 [Source:HGNC Symbol;Ac |
| UFL1-AS1 | 6 | UFL1 antisense RNA 1 [Source:HGNC Symbol; |
| RN7SL797P | 6 | RNA, 7SL, cytoplasmic 797, pseudogene [Sour |
| UFL1 | 6 | UFM1 specific ligase 1 [Source:HGNC Symbol; |
| FHL5 | 6 | four and a half LIM domains 5 [Source:HGNC S |
| RPS7P8 | 6 | ribosomal protein S7 pseudogene 8 [Source:H |
| RNU4-70P | 6 | RNA, U4 small nuclear 70, pseudogene [Sourc |
| TYMSP1 | 6 | thymidylate synthetase pseudogene 1 [Source |
| AL355143.1 | 6 | protein phosphatase 1, catalytic subunit (PPP1 |
| EEF1GP6 | 6 | eukaryotic translation elongation factor 1 gan |

| | | |
|---|---|---|
| AL033379.1 | 6 | novel transcript |
| GPR63 | 6 | G protein-coupled receptor 63 [Source:HGNC |
| NDUFAF4 | 6 | NADH:ubiquinone oxidoreductase complex as |
| RN7SL509P | 6 | RNA, 7SL, cytoplasmic 509, pseudogene [Sour |
| KLHL32 | 6 | kelch like family member 32 [Source:HGNC Sy |
| MMS22L | 6 | MMS22 like, DNA repair protein [Source:HGN |
| AL589740.1 | 6 | novel transcript |
| AL080316.1 | 6 | novel transcript |
| MIR2113 | 6 | microRNA 2113 [Source:HGNC Symbol;Acc:HG |
| EIF4EBP2P3 | 6 | eukaryotic translation initiation factor 4E bind |
| AL590727.1 | 6 | novel transcript |
| AL589826.1 | 6 | 60S ribosomal protein L6 (RPL6) pseudogene |
| AL589826.2 | 6 | novel transcript |
| POU3F2 | 6 | POU class 3 homeobox 2 [Source:HGNC Symb |
| FBXL4 | 6 | F-box and leucine rich repeat protein 4 [Sourc |
| AL078603.1 | 6 | diaphanous homolog 2 (Drosophila) (DIAPH2) |
| MIR548AI | 6 | microRNA 548ai [Source:HGNC Symbol;Acc:H |
| BDH2P1 | 6 | 3-hydroxybutyrate dehydrogenase 2, pseudog |
| FAXC | 6 | failed axon connections homolog, metaxin like |
| COQ3 | 6 | coenzyme Q3, methyltransferase [Source:HGN |
| PNISR | 6 | PNN interacting serine and arginine rich prote |
| AL513550.1 | 6 | novel transcript |
| USP45 | 6 | ubiquitin specific peptidase 45 [Source:HGNC |
| TSTD3 | 6 | thiosulfate sulfurtransferase like domain cont |
| Y_RNA | 6 | Y RNA [Source:RFAM;Acc:RF00019] |
| CCNC | 6 | cyclin C [Source:HGNC Symbol;Acc:HGNC:158 |
| AL137784.1 | 6 | 40S ribososmal protein 3 (RPS3) pseudogene |
| PRDM13 | 6 | PR/SET domain 13 [Source:HGNC Symbol;Acc: |
| Y_RNA | 6 | Y RNA [Source:RFAM;Acc:RF00019] |
| MCHR2 | 6 | melanin concentrating hormone receptor 2 [S |
| MCHR2-AS1 | 6 | MCHR2 antisense RNA 1 [Source:HGNC Symbo |
| NPM1P38 | 6 | nucleophosmin 1 pseudogene 38 [Source:HGN |
| AL080285.2 | 6 | tetratricopeptide repeat domain 22 (TTC22) p |
| PRDX2P4 | 6 | peroxiredoxin 2 pseudogene 4 [Source:HGNC |
| AL080285.1 | 6 | arginase, type II (ARG2) pseudogene |
| SIM1 | 6 | SIM bHLH transcription factor 1 [Source:HGN |
| Z86062.2 | 6 | novel transcript |
| ASCC3 | 6 | activating signal cointegrator 1 complex subu |
| Z86062.1 | 6 | RP42 homolog (RP42) pseudogene |
| AL133338.1 | 6 | novel transcript, antisense to ASCC3 |
| AL133338.2 | 6 | mitogen-activated protein kinase 1 interacting |
| GRIK2 | 6 | glutamate ionotropic receptor kainate type su |
| ACTG1P18 | 6 | actin gamma 1 pseudogene 18 [Source:HGNC |
| AP002528.1 | 6 | novel transcript, antisense to GRIK2 |
| AP002530.1 | 6 | laminin receptor 1 (67kD, ribosomal protein S |
| AL357139.2 | 6 | novel transcript |
| AL357139.1 | 6 | pseudogene similar to part of heterogeneous |
| Z98755.1 | 6 | ClpX caseinolytic protease X homolog (E. coli) |
| AL591387.1 | 6 | NULL |
| AL590608.1 | 6 | novel transcript |

| | | |
|---|---|---|
| R3HDM2P2 | 6 | R3H domain containing 2 pseudogene 2 [Sour |
| NPM1P10 | 6 | nucleophosmin 1 pseudogene 10 [Source:HGN |
| AL357522.1 | 6 | novel transcript |
| AL356967.1 | 6 | novel pseudogene |
| HACE1 | 6 | HECT domain and ankyrin repeat containing E |
| RNU6-897P | 6 | RNA, U6 small nuclear 897, pseudogene [Sour |
| AL357315.1 | 6 | novel transcript |
| AL357315.2 | 6 | poly(A) binding protein, cytoplasmic 4 (inducil |
| LIN28B-AS1 | 6 | LIN28B antisense RNA 1 [Source:HGNC Symbo |
| LIN28B | 6 | lin-28 homolog B [Source:HGNC Symbol;Acc:H |
| RNU6-1106P | 6 | RNA, U6 small nuclear 1106, pseudogene [Sou |
| BVES | 6 | blood vessel epicardial substance [Source:HGI |
| BVES-AS1 | 6 | BVES antisense RNA 1 [Source:HGNC Symbol;A |
| POPDC3 | 6 | popeye domain containing 3 [Source:HGNC Sy |
| PREP | 6 | prolyl endopeptidase [Source:HGNC Symbol;A |
| AL133406.2 | 6 | novel transcript |
| AL133406.1 | 6 | ribosomal protein L7 (RPL7) pseudogene |
| RPL35P3 | 6 | ribosomal protein L35 pseudogene 3 [Source: |
| AL096794.1 | 6 | novel transcript |
| LINC02836 | 6 | long intergenic non-protein coding RNA 2836 |
| Z97206.1 | 6 | chromatin assembly factor 1, p60 subunit B (C |
| AL591518.1 | 6 | novel transcript |
| RN7SKP211 | 6 | RN7SK pseudogene 211 [Source:HGNC Symbo |
| PRDM1 | 6 | PR/SET domain 1 [Source:HGNC Symbol;Acc:H |
| ATG5 | 6 | autophagy related 5 [Source:HGNC Symbol;Ac |
| AL022067.1 | 6 | novel transcript, sense intronic to PRDM1 |
| U4 | 6 | U4 spliceosomal RNA [Source:RFAM;Acc:RF00 |
| RN7SL47P | 6 | RNA, 7SL, cytoplasmic 47, pseudogene [Sourc |
| RNU6-344P | 6 | RNA, U6 small nuclear 344, pseudogene [Sour |
| AL356859.1 | 6 | novel transcript |
| CRYBG1 | 6 | crystallin beta-gamma domain containing 1 [S |
| Y_RNA | 6 | Y RNA [Source:RFAM;Acc:RF00019] |
| RNA5SP211 | 6 | RNA, 5S ribosomal pseudogene 211 [Source:H |
| AL109920.1 | 6 | novel transcript, antisense to AIM1 |
| Y_RNA | 6 | Y RNA [Source:RFAM;Acc:RF00019] |
| RTN4IP1 | 6 | reticulon 4 interacting protein 1 [Source:HGN |
| RNU6-527P | 6 | RNA, U6 small nuclear 527, pseudogene [Sour |
| QRSL1 | 6 | glutaminyl-tRNA amidotransferase subunit QF |
| RPL21P65 | 6 | ribosomal protein L21 pseudogene 65 [Source |
| LINC02526 | 6 | long intergenic non-protein coding RNA 2526 |
| LINC02532 | 6 | long intergenic non-protein coding RNA 2532 |
| RNU6-117P | 6 | RNA, U6 small nuclear 117, pseudogene [Sour |
| MIR587 | 6 | microRNA 587 [Source:HGNC Symbol;Acc:HGI |
| CD24 | 6 | CD24 molecule [Source:HGNC Symbol;Acc:HG |
| MTRES1 | 6 | mitochondrial transcription rescue factor 1 [S |
| BEND3 | 6 | BEN domain containing 3 [Source:HGNC Symb |
| RNU6-1299P | 6 | RNA, U6 small nuclear 1299, pseudogene [Sou |
| PDSS2 | 6 | decaprenyl diphosphate synthase subunit 2 [S |
| AL591516.1 | 6 | ribosomal L1 domain containing 1 (RSL1D1) ps |
| RPS24P12 | 6 | ribosomal protein S24 pseudogene 12 [Source |

| | | |
|---|---|---|
| SOBP | 6 | sine oculis binding protein homolog [Source:H |
| AL121957.1 | 6 | novel transcript |
| AL096816.1 | 6 | TEC |
| SCML4 | 6 | Scm polycomb group protein like 4 [Source:H( |
| SEC63 | 6 | SEC63 homolog, protein translocation regulat( |
| RNU6-437P | 6 | RNA, U6 small nuclear 437, pseudogene [Sour |
| RPL23AP50 | 6 | ribosomal protein L23a pseudogene 50 [Sourc |
| AL024507.2 | 6 | novel transcript, antisense to SEC63 |
| AL024507.1 | 6 | ribosomal protein L14 (RPL14) pseudogene |
| SNORA73 | 6 | Small nucleolar RNA SNORA73 family [Source: |
| MTHFD2P3 | 6 | methylenetetrahydrofolate dehydrogenase (N |
| RPL3P7 | 6 | ribosomal protein L3 pseudogene 7 [Source:H |
| Z98200.1 | 6 | novel transcript |
| OSTM1 | 6 | osteoclastogenesis associated transmembran( |
| Y_RNA | 6 | Y RNA [Source:RFAM;Acc:RF00019] |
| OSTM1-AS1 | 6 | OSTM1 antisense RNA 1 [Source:HGNC Symbo |
| NR2E1 | 6 | nuclear receptor subfamily 2 group E member |
| AL078596.1 | 6 | TEC |
| SNX3 | 6 | sorting nexin 3 [Source:HGNC Symbol;Acc:HGI |
| Z98742.2 | 6 | pseudogene similar to hypothetical protein HS |
| RNA5SP212 | 6 | RNA, 5S ribosomal pseudogene 212 [Source:H |
| Z98742.4 | 6 | novel transcript |
| Z98742.1 | 6 | novel transcript |
| RNU6-1144P | 6 | RNA, U6 small nuclear 1144, pseudogene [Sou |
| AFG1L | 6 | AFG1 like ATPase [Source:HGNC Symbol;Acc:H |
| Z98742.3 | 6 | ribosomal protein L36a (RPL36A) pseudogene |
| AL356121.2 | 6 | TEC |
| AL356121.1 | 6 | novel pseudogene |
| AL353706.1 | 6 | RAB22A, member RAS oncogene family (RAB2 |
| RNU6-770P | 6 | RNA, U6 small nuclear 770, pseudogene [Sour |
| AL139106.1 | 6 | novel transcript, antisense to LACE1 |
| AL391646.1 | 6 | MACRO domain containing 2 (MACROD2) pseu |
| FOXO3 | 6 | forkhead box O3 [Source:HGNC Symbol;Acc:H |
| SUMO2P8 | 6 | SUMO2 pseudogene 8 [Source:HGNC Symbol; |
| LINC00222 | 6 | long intergenic non-protein coding RNA 222 [S |
| ZNF259P1 | 6 | zinc finger protein 259 pseudogene 1 [Source: |
| Z95118.2 | 6 | novel transcript |
| Z95118.1 | 6 | ribosomal protein L37a (RPL37A) pseudogene |
| ARMC2 | 6 | armadillo repeat containing 2 [Source:HGNC S |
| AL445189.1 | 6 | ribosomal protein S12 (RPS12) pseudogene |
| AL445189.2 | 6 | ATP synthase, H+ transporting, mitochondrial |
| ARMC2-AS1 | 6 | ARMC2 antisense RNA 1 [Source:HGNC Symbo |
| SESN1 | 6 | sestrin 1 [Source:HGNC Symbol;Acc:HGNC:21! |
| AL390208.1 | 6 | novel transcript, antisense to SESN1 |
| RNU6-653P | 6 | RNA, U6 small nuclear 653, pseudogene [Sour |
| CEP57L1 | 6 | centrosomal protein 57 like 1 [Source:HGNC S |
| CCDC162P | 6 | coiled-coil domain containing 162, pseudogen |
| AL359711.1 | 6 | RAB18, member RAS oncogene family (RAB18 |
| PTCHD3P3 | 6 | patched domain containing 3 pseudogene 3 [S |
| RNY3P11 | 6 | RNY3 pseudogene 11 [Source:HGNC Symbol;A |

| | | |
|---|---|---|
| RPL7P28 | 6 | ribosomal protein L7 pseudogene 28 [Source: |
| CD164 | 6 | CD164 molecule [Source:HGNC Symbol;Acc:H |
| AL359711.2 | 6 | novel transcript, antisense to CD164 |
| PPIL6 | 6 | peptidylprolyl isomerase like 6 [Source:HGNC |
| SMPD2 | 6 | sphingomyelin phosphodiesterase 2 [Source:H |
| MICAL1 | 6 | microtubule associated monooxygenase, calp |
| ZBTB24 | 6 | zinc finger and BTB domain containing 24 [Sou |
| AL109947.1 | 6 | novel transcript |
| AK9 | 6 | adenylate kinase 9 [Source:HGNC Symbol;Acc |
| FIG4 | 6 | FIG4 phosphoinositide 5-phosphatase [Source |
| AL512303.1 | 6 | novel transcript |
| GPR6 | 6 | G protein-coupled receptor 6 [Source:HGNC S |
| AL590009.1 | 6 | novel transcript |
| WASF1 | 6 | WASP family member 1 [Source:HGNC Symbo |
| CDC40 | 6 | cell division cycle 40 [Source:HGNC Symbol;Ac |
| METTL24 | 6 | methyltransferase like 24 [Source:HGNC Symb |
| AL050350.1 | 6 | novel transcript, antisense to METTL24 |
| DDO | 6 | D-aspartate oxidase [Source:HGNC Symbol;Ac |
| SLC22A16 | 6 | solute carrier family 22 member 16 [Source:H |
| RN7SL617P | 6 | RNA, 7SL, cytoplasmic 617, pseudogene [Sour |
| AC002464.1 | 6 | novel transcript, antisense to SLC22A16 |
| AL512430.3 | 6 | ribosomal protein S19 (RPS19) pseudogene |
| AL512430.4 | 6 | abhydrolase domain containing 10 (ABHD10) |
| CDK19 | 6 | cyclin dependent kinase 19 [Source:HGNC Syn |
| AL512430.1 | 6 | novel pseudogene |
| AL512430.2 | 6 | ribosomal protein S20 (RPS20) pseudogene |
| AL603914.1 | 6 | ribosomal protein L21 (RPL21) pseudogene |
| RNU6-957P | 6 | RNA, U6 small nuclear 957, pseudogene [Sour |
| SNORA40C | 6 | small nucleolar RNA, H/ACA box 40C [Source: |
| RNU6-1115P | 6 | RNA, U6 small nuclear 1115, pseudogene [Sou |
| CNN2P9 | 6 | calponin 2 pseudogene 9 [Source:HGNC Symb |
| Z84480.1 | 6 | novel pseudogene |
| AMD1 | 6 | adenosylmethionine decarboxylase 1 [Source: |
| AL357515.1 | 6 | PDGFA associated protein 1 (PDGFA) pseudog |
| GTF3C6 | 6 | general transcription factor IIIC subunit 6 [Sou |
| RPF2 | 6 | ribosome production factor 2 homolog [Sourc |
| GSTM2P1 | 6 | glutathione S-transferase mu 2 pseudogene 1 |
| SLC16A10 | 6 | solute carrier family 16 member 10 [Source:H |
| RNU6-960P | 6 | RNA, U6 small nuclear 960, pseudogene [Sour |
| AL360227.1 | 6 | novel transcript |
| MFSD4B | 6 | major facilitator superfamily domain containi |
| AL080317.1 | 6 | novel transcript |
| REV3L | 6 | REV3 like, DNA directed polymerase zeta cata |
| AL080317.2 | 6 | novel transcript, antisense to REV3L |
| FCF1P5 | 6 | FCF1 pseudogene 5 [Source:HGNC Symbol;Ac |
| REV3L-IT1 | 6 | REV3L intronic transcript 1 [Source:HGNC Sym |
| BRD7P4 | 6 | bromodomain containing 7 pseudogene 4 [So |
| TRAF3IP2-AS1 | 6 | TRAF3IP2 antisense RNA 1 [Source:HGNC Sym |
| AL136310.1 | 6 | novel pseudogene |
| AL008730.1 | 6 | novel transcript |

| | | |
|---|---|---|
| TRAF3IP2 | 6 | TRAF3 interacting protein 2 [Source:HGNC Syr |
| Z97989.1 | 6 | chromosome 6 open reading frame 3 |
| FYN | 6 | FYN proto-oncogene, Src family tyrosine kinas |
| LINC02527 | 6 | long intergenic non-protein coding RNA 2527 |
| CCN6 | 6 | cellular communication network factor 6 [Sou |
| TUBE1 | 6 | tubulin epsilon 1 [Source:HGNC Symbol;Acc:H |
| FAM229B | 6 | family with sequence similarity 229 member E |
| LAMA4 | 6 | laminin subunit alpha 4 [Source:HGNC Symbol |
| Z99289.1 | 6 | novel transcript |
| RNU6-1226P | 6 | RNA, U6 small nuclear 1226, pseudogene [Sou |
| Z99289.2 | 6 | novel transcript, sense intronic LAMA4 |
| Z99289.3 | 6 | novel transcript, sense intronic LAMA4 |
| AL365214.2 | 6 | novel transcript |
| AL365214.3 | 6 | novel ret finger protein-like 4B |
| AL365214.1 | 6 | hepatocellular carcinoma-related putative tur |
| RFPL4B | 6 | ret finger protein like 4B [Source:HGNC Symbo |
| RPSAP45 | 6 | ribosomal protein SA pseudogene 45 [Source: |
| KRT18P65 | 6 | keratin 18 pseudogene 65 [Source:HGNC Sym |
| FEM1AP3 | 6 | fem-1 homolog A pseudogene 3 [Source:HGN |
| AL365214.4 | 6 | novel transcript |
| AL357514.1 | 6 | novel transcript |
| AL358942.1 | 6 | proliferation-associated 2G4, 38kD (PA2G4) ps |
| AL360085.1 | 6 | family with sequence similarity 98, member B |
| RNU6-1163P | 6 | RNA, U6 small nuclear 1163, pseudogene [Sou |
| AL360015.1 | 6 | novel transcript |
| FCF1P10 | 6 | FCF1 pseudogene 10 [Source:HGNC Symbol;A |
| SOCS5P5 | 6 | suppressor of cytokine signaling 5 pseudogen |
| AL589684.1 | 6 | novel transcript |
| LINC02518 | 6 | long intergenic non-protein coding RNA 2518 |
| AL513123.1 | 6 | novel transcript |
| RPS27AP11 | 6 | ribosomal protein S27a pseudogene 11 [Sourc |
| LINC02541 | 6 | long intergenic non-protein coding RNA 2541 |
| FO393415.3 | 6 | novel transcript |
| FO393415.2 | 6 | ribosomal protein L30 (RPL30) pseudogene |
| MARCKS | 6 | myristoylated alanine rich protein kinase C sul |
| LINC01268 | 6 | long intergenic non-protein coding RNA 1268 |
| FO393415.1 | 6 | novel transcript |
| HDAC2 | 6 | histone deacetylase 2 [Source:HGNC Symbol;A |
| HDAC2-AS2 | 6 | HDAC2 and HS3ST5 antisense RNA 2 [Source:H |
| NUDT19P3 | 6 | nudix hydrolase 19 pseudogene 3 [Source:HGI |
| HS3ST5 | 6 | heparan sulfate-glucosamine 3-sulfotransfera: |
| RPSAP43 | 6 | ribosomal protein SA pseudogene 43 [Source: |
| RNA5SP213 | 6 | RNA, 5S ribosomal pseudogene 213 [Source:H |
| DNAJA1P4 | 6 | DnaJ heat shock protein family (Hsp40) memb |
| AL138830.2 | 6 | novel transcript |
| AL138830.1 | 6 | novel transcript |
| AL590550.1 | 6 | uncharacterized LOC107986639 [Source:NCBI |
| RNU6-475P | 6 | RNA, U6 small nuclear 475, pseudogene [Sour |
| AL606845.1 | 6 | ATP synthase, H+ transporting, mitochondrial |
| LINC02534 | 6 | long intergenic non-protein coding RNA 2534 |

| | | |
|---|---|---|
| FRK | 6 | fyn related Src family tyrosine kinase [Source: |
| AL357141.1 | 6 | novel transcript, antisense to FRK |
| TPI1P3 | 6 | triosephosphate isomerase 1 pseudogene 3 [S |
| NT5DC1 | 6 | 5'-nucleotidase domain containing 1 [Source:H |
| COL10A1 | 6 | collagen type X alpha 1 chain [Source:HGNC S |
| NIP7P3 | 6 | NIP7 pseudogene 3 [Source:HGNC Symbol;Acc |
| AL050331.2 | 6 | novel transcript |
| TSPYL4 | 6 | TSPY like 4 [Source:HGNC Symbol;Acc:HGNC:2 |
| DSE | 6 | dermatan sulfate epimerase [Source:HGNC Sy |
| AL050331.1 | 6 | RPS5 (40S Ribosomal Protein S5) pseudogene |
| TSPYL1 | 6 | TSPY like 1 [Source:HGNC Symbol;Acc:HGNC:1 |
| Z84488.2 | 6 | novel protein |
| CBX3P9 | 6 | chromobox 3 pseudogene 9 [Source:HGNC Sy |
| KRT18P22 | 6 | keratin 18 pseudogene 22 [Source:HGNC Sym |
| Z84488.1 | 6 | novel transcript |
| CALHM6 | 6 | calcium homeostasis modulator family memb |
| AL445224.1 | 6 | novel transcript, antisense to BET3L |
| TRAPPC3L | 6 | trafficking protein particle complex 3 like [Sou |
| CALHM5 | 6 | calcium homeostasis modulator family memb |
| CALHM4 | 6 | calcium homeostasis modulator family memb |
| RWDD1 | 6 | RWD domain containing 1 [Source:HGNC Sym |
| RSPH4A | 6 | radial spoke head component 4A [Source:HGN |
| ZUP1 | 6 | zinc finger containing ubiquitin peptidase 1 [S |
| KPNA5 | 6 | karyopherin subunit alpha 5 [Source:HGNC Sy |
| FAM162B | 6 | family with sequence similarity 162 member B |
| GPRC6A | 6 | G protein-coupled receptor class C group 6 m |
| RFX6 | 6 | regulatory factor X6 [Source:HGNC Symbol;Ac |
| RPS29P13 | 6 | ribosomal protein S29 pseudogene 13 [Source |
| RNA5SP214 | 6 | RNA, 5S ribosomal pseudogene 214 [Source:H |
| Z98880.1 | 6 | novel transcript |
| VGLL2 | 6 | vestigial like family member 2 [Source:HGNC S |
| ROS1 | 6 | ROS proto-oncogene 1, receptor tyrosine kina |
| RN7SKP18 | 6 | RN7SK pseudogene 18 [Source:HGNC Symbol; |
| RN7SKP51 | 6 | RN7SK pseudogene 51 [Source:HGNC Symbol; |
| AL132671.2 | 6 | novel protein, GOPC-ROS1 readthrough |
| RAP1BP3 | 6 | RAP1B, member of RAS oncogene family pseu |
| AL132671.1 | 6 | novel transcript |
| DCBLD1 | 6 | discoidin, CUB and LCCL domain containing 1 [ |
| RNU6-253P | 6 | RNA, U6 small nuclear 253, pseudogene [Sour |
| GOPC | 6 | golgi associated PDZ and coiled-coil motif con |
| NEPNP | 6 | nephrocan, pseudogene [Source:HGNC Symbo |
| NUS1 | 6 | NUS1 dehydrodolichyl diphosphate synthase s |
| SLC35F1 | 6 | solute carrier family 35 member F1 [Source:H |
| AL450405.1 | 6 | ribosomal protein L29 (RPL29) pseudogene |
| AL589993.1 | 6 | DEAD/H (Asp-Glu-Ala-Asp/His) box polypeptid |
| CEP85L | 6 | centrosomal protein 85 like [Source:HGNC Syr |
| Z99496.1 | 6 | bromodomain-containing 7 (BRD7) pseudoger |
| PLN | 6 | phospholamban [Source:HGNC Symbol;Acc:H |
| Z99496.2 | 6 | novel transcript, antisense to CEP85L |
| SSXP10 | 6 | SSX family pseudogene 10 [Source:HGNC Sym |

| | | |
|---|---|---|
| SELENOKP3 | 6 | selenoprotein K pseudogene 3 [Source:HGNC |
| MCM9 | 6 | minichromosome maintenance 9 homologous |
| ASF1A | 6 | anti-silencing function 1A histone chaperone [ |
| AL137009.1 | 6 | novel transcript, antisense to FAM184A |
| FAM184A | 6 | family with sequence similarity 184 member A |
| Y_RNA | 6 | Y RNA [Source:RFAM;Acc:RF00019] |
| MIR548B | 6 | microRNA 548b [Source:HGNC Symbol;Acc:HG |
| AL022722.2 | 6 | cytochrome c oxidase polypeptide I (COX) pse |
| MAN1A1 | 6 | mannosidase alpha class 1A member 1 [Sourc |
| AL022722.1 | 6 | 60S Ribosomal Protein L13A (RPL13) pseudoge |
| RNU6-194P | 6 | RNA, U6 small nuclear 194, pseudogene [Sour |
| AL078600.1 | 6 | novel transcript |
| MIR3144 | 6 | microRNA 3144 [Source:HGNC Symbol;Acc:HG |
| AL096854.1 | 6 | novel transcript |
| RNU6-214P | 6 | RNA, U6 small nuclear 214, pseudogene [Sour |
| RNA5SP215 | 6 | RNA, 5S ribosomal pseudogene 215 [Source:H |
| COX6A1P3 | 6 | cytochrome c oxidase subunit 6A1 pseudogen |
| AL357513.1 | 6 | novel transcript |
| TBC1D32 | 6 | TBC1 domain family member 32 [Source:HGN |
| RNU6-1286P | 6 | RNA, U6 small nuclear 1286, pseudogene [Sou |
| Y_RNA | 6 | Y RNA [Source:RFAM;Acc:RF00019] |
| AL139098.1 | 6 | ribosomal protein S15a (RPS15A) pseudogene |
| AL139098.2 | 6 | sialic acid binding Ig-like lectin 7 (SIGLEC7) pse |
| GJA1 | 6 | gap junction protein alpha 1 [Source:HGNC Sy |
| RNU4-35P | 6 | RNA, U4 small nuclear 35, pseudogene [Sourc |
| AL138729.1 | 6 | transcription factor A, mitochondrial (TFAM) p |
| RNU4-76P | 6 | RNA, U4 small nuclear 76, pseudogene [Sourc |
| RNU2-8P | 6 | RNA, U2 small nuclear 8, pseudogene [Source |
| AL603865.2 | 6 | cytochrome c oxidase subunit VIc (COX6C) pse |
| SLC25A5P7 | 6 | solute carrier family 25 member 5 pseudogen |
| RPL23AP48 | 6 | ribosomal protein L23a pseudogene 48 [Sourc |
| AL603865.1 | 6 | LSM14A, SCD6 homolog A (S. cerevisiae) (LSM |
| HMGB3P18 | 6 | high mobility group box 3 pseudogene 18 [Sou |
| RNU1-18P | 6 | RNA, U1 small nuclear 18, pseudogene [Sourc |
| HSF2 | 6 | heat shock transcription factor 2 [Source:HGN |
| Z99129.4 | 6 | TEC |
| SERINC1 | 6 | serine incorporator 1 [Source:HGNC Symbol;A |
| Z99129.1 | 6 | mitogen-activated protein kinase 6 (MAPK6) p |
| PKIB | 6 | cAMP-dependent protein kinase inhibitor beta |
| Z99129.3 | 6 | TEC |
| Z99129.2 | 6 | RAB GTPase activating protein 1-like (RABGAP |
| AL512283.2 | 6 | novel transcript, antisense to PKIB |
| AL512283.1 | 6 | novel transcript |
| AL512283.3 | 6 | novel transcript, antisense to PKIB |
| RN7SL564P | 6 | RNA, 7SL, cytoplasmic 564, pseudogene [Sour |
| FABP7 | 6 | fatty acid binding protein 7 [Source:HGNC Syn |
| SMPDL3A | 6 | sphingomyelin phosphodiesterase acid like 3A |
| ATP5MGP2 | 6 | ATP synthase membrane subunit g pseudoger |
| AL591428.1 | 6 | novel transcript |
| CLVS2 | 6 | clavesin 2 [Source:HGNC Symbol;Acc:HGNC:23 |

| | | |
|---|---|---|
| TRDN | 6 | triadin [Source:HGNC Symbol;Acc:HGNC:1226 |
| TRDN-AS1 | 6 | TRDN antisense RNA 1 [Source:HGNC Symbol; |
| AL445259.1 | 6 | novel transcript, antisense to TRDN |
| AL133257.1 | 6 | novel transcript, antisense to TRDN |
| NKAIN2 | 6 | sodium/potassium transporting ATPase intera |
| AL354936.1 | 6 | novel transcript |
| RNF217-AS1 | 6 | RNF217 antisense RNA 1 (head to head) [Sour |
| RNF217 | 6 | ring finger protein 217 [Source:HGNC Symbol; |
| TPD52L1 | 6 | TPD52 like 1 [Source:HGNC Symbol;Acc:HGNC |
| HDDC2 | 6 | HD domain containing 2 [Source:HGNC Symbo |
| AL121938.1 | 6 | novel transcript, antisense to HDDC2 |
| AL450332.1 | 6 | novel transcript |
| AL365259.1 | 6 | novel transcript, antisense to HEY2 |
| LINC02523 | 6 | long intergenic non-protein coding RNA 2523 |
| HEY2 | 6 | hes related family bHLH transcription factor w |
| NCOA7 | 6 | nuclear receptor coactivator 7 [Source:HGNC |
| NCOA7-AS1 | 6 | NCOA7 antisense RNA 1 [Source:HGNC Symbo |
| RN7SKP56 | 6 | RN7SK pseudogene 56 [Source:HGNC Symbol; |
| HINT3 | 6 | histidine triad nucleotide binding protein 3 [So |
| RNA5SP216 | 6 | RNA, 5S ribosomal pseudogene 216 [Source:H |
| TRMT11 | 6 | tRNA methyltransferase 11 homolog [Source: |
| PPP1R14BP5 | 6 | protein phosphatase 1 regulatory inhibitor sul |
| AL356534.1 | 6 | vimentin 2, pseudogene |
| CENPW | 6 | centromere protein W [Source:HGNC Symbol; |
| MIR588 | 6 | microRNA 588 [Source:HGNC Symbol;Acc:HGN |
| RNU6-200P | 6 | RNA, U6 small nuclear 200, pseudogene [Sour |
| AL359535.1 | 6 | Yes-associated protein 1 (YAP1) pseudogene |
| PRELID1P1 | 6 | PRELI domain containing 1 pseudogene 1 [Sou |
| RPS4XP9 | 6 | ribosomal protein S4X pseudogene 9 [Source: |
| AL035603.1 | 6 | novel transcript |
| RSPO3 | 6 | R-spondin 3 [Source:HGNC Symbol;Acc:HGNC: |
| RNF146 | 6 | ring finger protein 146 [Source:HGNC Symbol; |
| ECHDC1 | 6 | ethylmalonyl-CoA decarboxylase 1 [Source:HG |
| RNA5SP217 | 6 | RNA, 5S ribosomal pseudogene 217 [Source:H |
| YWHAZP4 | 6 | tyrosine 3-monooxygenase/tryptophan 5-mor |
| RPL5P18 | 6 | ribosomal protein L5 pseudogene 18 [Source: |
| AL590002.1 | 6 | cornichon homolog 4 (Drosophila) (CNIH4) pse |
| AL096711.1 | 6 | ribosomal protein L17 (RPL17) pseudogene |
| AL096711.2 | 6 | novel protein |
| KIAA0408 | 6 | KIAA0408 [Source:HGNC Symbol;Acc:HGNC:2 |
| SOGA3 | 6 | SOGA family member 3 [Source:HGNC Symbo |
| C6orf58 | 6 | chromosome 6 open reading frame 58 [Sourc |
| AL592291.1 | 6 | novel pseudogene |
| AL356432.1 | 6 | transducin (beta)-like 1 X-linked receptor 1 (T |
| AL356432.3 | 6 | transducin (beta)-like 1 X-linked receptor 1 (T |
| AL356432.2 | 6 | transducin (beta)-like 1 X-linked receptor 1 (T |
| LINC02536 | 6 | long intergenic non-protein coding RNA 2536 |
| THEMIS | 6 | thymocyte selection associated [Source:HGN |
| MRPS17P5 | 6 | mitochondrial ribosomal protein S17 pseudog |
| PTPRK | 6 | protein tyrosine phosphatase receptor type K |

| | | |
|---|---|---|
| AL590006.1 | 6 | novel transcript, antisense to PTPRK |
| AL034349.1 | 6 | novel transcript, antisense to PTPRK |
| EEF1DP5 | 6 | eukaryotic translation elongation factor 1 delt |
| Y_RNA | 6 | Y RNA [Source:RFAM;Acc:RF00019] |
| AL080315.1 | 6 | ribosomal protein L21 (RPL21) pseudogene |
| Y_RNA | 6 | Y RNA [Source:RFAM;Acc:RF00019] |
| LAMA2 | 6 | laminin subunit alpha 2 [Source:HGNC Symbol |
| MESTP1 | 6 | mesoderm specific transcript pseudogene 1 [S |
| BMPR1AP1 | 6 | bone morphogenetic protein receptor type 1/ |
| RNU6-861P | 6 | RNA, U6 small nuclear 861, pseudogene [Sour |
| AL356124.1 | 6 | novel transcript |
| AL356124.2 | 6 | novel transcript |
| ARHGAP18 | 6 | Rho GTPase activating protein 18 [Source:HGN |
| RPL5P21 | 6 | ribosomal protein L5 pseudogene 21 [Source: |
| AL451046.2 | 6 | SMT3 suppressor of mif two 3 homolog 1 (S. c |
| B3GALNT2P1 | 6 | beta-1,3-N-acetylgalactosaminyltransferase 2 |
| AL451046.1 | 6 | pyridoxal (pyridoxine, vitamin B6) kinase (PDX |
| TMEM244 | 6 | transmembrane protein 244 [Source:HGNC Sy |
| L3MBTL3 | 6 | L3MBTL histone methyl-lysine binding protein |
| AL355581.1 | 6 | novel transcript, antisense to L3MBTL3 |
| SAMD3 | 6 | sterile alpha motif domain containing 3 [Sourc |
| TMEM200A | 6 | transmembrane protein 200A [Source:HGNC S |
| Y_RNA | 6 | Y RNA [Source:RFAM;Acc:RF00019] |
| AL583803.1 | 6 | novel transcript |
| SMLR1 | 6 | small leucine rich protein 1 [Source:HGNC Syn |
| EPB41L2 | 6 | erythrocyte membrane protein band 4.1 like 2 |
| AKAP7 | 6 | A-kinase anchoring protein 7 [Source:HGNC Sy |
| AL137222.1 | 6 | D4, zinc and double PHD fingers family 2 (DPF |
| RPL21P67 | 6 | ribosomal protein L21 pseudogene 67 [Source |
| ARG1 | 6 | arginase 1 [Source:HGNC Symbol;Acc:HGNC:6 |
| MED23 | 6 | mediator complex subunit 23 [Source:HGNC S |
| ENPP3 | 6 | ectonucleotide pyrophosphatase/phosphodie |
| RNU4-18P | 6 | RNA, U4 small nuclear 18, pseudogene [Sourc |
| OR2A4 | 6 | olfactory receptor family 2 subfamily A memb |
| CTAGE9 | 6 | CTAGE family member 9 [Source:HGNC Symbo |
| AC005587.1 | 6 | ribosomal protein L15 (RPL15) pseudogene |
| MIR548H5 | 6 | microRNA 548h-5 [Source:HGNC Symbol;Acc:I |
| ENPP1 | 6 | ectonucleotide pyrophosphatase/phosphodie |
| SELENOKP2 | 6 | selenoprotein K pseudogene 2 [Source:HGNC |
| RN7SKP245 | 6 | RN7SK pseudogene 245 [Source:HGNC Symbo |
| AL139805.1 | 6 | T-cell activation leucine repeat-rich protein (T |
| AL117378.1 | 6 | novel transcript |
| AL117378.2 | 6 | TEC |
| CCN2 | 6 | cellular communication network factor 2 [Sou |
| AL133346.1 | 6 | novel transcript |
| AL021408.1 | 6 | novel transcript |
| MIR548AJ1 | 6 | microRNA 548aj-1 [Source:HGNC Symbol;Acc: |
| AL021408.2 | 6 | TEC |
| LINC01013 | 6 | long intergenic non-protein coding RNA 1013 |
| EEF1A1P36 | 6 | eukaryotic translation elongation factor 1 alph |

| | | |
|---|---|---|
| MOXD1 | 6 | monooxygenase DBH like 1 [Source:HGNC Syr |
| STX7 | 6 | syntaxin 7 [Source:HGNC Symbol;Acc:HGNC:1 |
| RPL21P66 | 6 | ribosomal protein L21 pseudogene 66 [Source |
| TAAR9 | 6 | trace amine associated receptor 9 (gene/pseu |
| TAAR8 | 6 | trace amine associated receptor 8 [Source:HG |
| TAAR7P | 6 | trace amine associated receptor 7, pseudoger |
| TAAR6 | 6 | trace amine associated receptor 6 [Source:HG |
| TAAR5 | 6 | trace amine associated receptor 5 [Source:HG |
| TAAR4P | 6 | trace amine associated receptor 4, pseudoger |
| AL513524.1 | 6 | novel transcript |
| TAAR3P | 6 | trace amine associated receptor 3, pseudoger |
| TAAR2 | 6 | trace amine associated receptor 2 (gene/pseu |
| TAAR1 | 6 | trace amine associated receptor 1 [Source:HG |
| HLFP1 | 6 | HLF pseudogene 1 [Source:HGNC Symbol;Acc: |
| VNN1 | 6 | vanin 1 [Source:HGNC Symbol;Acc:HGNC:127( |
| CCNG1P1 | 6 | cyclin G1 pseudogene 1 [Source:HGNC Symbo |
| VNN3 | 6 | vanin 3 [Source:HGNC Symbol;Acc:HGNC:164 |
| VNN2 | 6 | vanin 2 [Source:HGNC Symbol;Acc:HGNC:127( |
| AL032821.1 | 6 | novel transcript |
| RBM11P1 | 6 | RNA binding motif protein 11 pseudogene 1 [ |
| SLC18B1 | 6 | solute carrier family 18 member B1 [Source:H |
| RPS12 | 6 | ribosomal protein S12 [Source:HGNC Symbol; |
| SNORD101 | 6 | small nucleolar RNA, C/D box 101 [Source:HG |
| SNORD100 | 6 | small nucleolar RNA, C/D box 100 [Source:HG |
| SNORA33 | 6 | small nucleolar RNA, H/ACA box 33 [Source:H |
| HMGB1P13 | 6 | high mobility group box 1 pseudogene 13 [Sou |
| AL137783.1 | 6 | TEC |
| LINC00326 | 6 | long intergenic non-protein coding RNA 326 [ |
| RPL23AP46 | 6 | ribosomal protein L23a pseudogene 46 [Sourc |
| AL591115.1 | 6 | novel transcript |
| MTCYBP4 | 6 | MT-CYB pseudogene 4 [Source:HGNC Symbol; |
| EYA4 | 6 | EYA transcriptional coactivator and phosphata |
| AL024497.1 | 6 | novel transcript, antisense to EYA4 |
| AL024497.2 | 6 | novel transcript |
| TARID | 6 | TCF21 antisense RNA inducing promoter dem |
| HSPE1P21 | 6 | heat shock protein family E (Hsp10) member |
| AL450270.1 | 6 | novel transcript |
| FTH1P26 | 6 | ferritin heavy chain 1 pseudogene 26 [Source: |
| LINC01312 | 6 | long intergenic non-protein coding RNA 1312 |
| TCF21 | 6 | transcription factor 21 [Source:HGNC Symbol; |
| TBPL1 | 6 | TATA-box binding protein like 1 [Source:HGN( |
| SLC2A12 | 6 | solute carrier family 2 member 12 [Source:HG |
| AL449363.1 | 6 | epithelial splicing regulatory protein 1 (ESRP1) |
| AL449363.2 | 6 | novel transcript |
| HMGA1P7 | 6 | high mobility group AT-hook 1 pseudogene 7 |
| RN7SL408P | 6 | RNA, 7SL, cytoplasmic 408, pseudogene [Sour |
| SGK1 | 6 | serum/glucocorticoid regulated kinase 1 [Sour |
| RPS29P32 | 6 | ribosomal protein S29 pseudogene 32 [Source |
| RNA5SP218 | 6 | RNA, 5S ribosomal pseudogene 218 [Source:H |
| Y_RNA | 6 | Y RNA [Source:RFAM;Acc:RF00019] |

| | | |
|---|---|---|
| AL355881.1 | 6 | novel transcript, antisense to SGK1 |
| KRT8P42 | 6 | keratin 8 pseudogene 42 [Source:HGNC Symb |
| LINC01010 | 6 | long intergenic non-protein coding RNA 1010 |
| Z84486.1 | 6 | novel transcript |
| CHCHD2P4 | 6 | coiled-coil-helix-coiled-coil-helix domain conta |
| CT69 | 6 | cancer/testis associated transcript 69 [Source |
| AL078590.2 | 6 | novel transcript |
| FAM8A6P | 6 | family with sequence similarity 8 member A6, |
| AL078590.1 | 6 | novel transcript |
| AL596188.1 | 6 | novel transcript |
| AL121970.1 | 6 | novel transcript |
| AL021939.1 | 6 | mediator of cell motility 1 (MEMO1) pseudoge |
| ALDH8A1 | 6 | aldehyde dehydrogenase 8 family member A1 |
| AL445190.1 | 6 | novel transcript |
| HBS1L | 6 | HBS1 like translational GTPase [Source:HGNC |
| MIR3662 | 6 | microRNA 3662 [Source:HGNC Symbol;Acc:HG |
| AL353596.1 | 6 | novel transcript |
| MYB | 6 | MYB proto-oncogene, transcription factor [So |
| MYB-AS1 | 6 | MYB antisense RNA 1 [Source:HGNC Symbol;A |
| MIR548A2 | 6 | microRNA 548a-2 [Source:HGNC Symbol;Acc:H |
| AL023693.1 | 6 | novel transcript |
| AHI1 | 6 | Abelson helper integration site 1 [Source:HGN |
| AL049552.1 | 6 | novel transcript |
| AL049552.2 | 6 | novel transcript, antisense to AHI1 |
| Y_RNA | 6 | Y RNA [Source:RFAM;Acc:RF00019] |
| LINC00271 | 6 | long intergenic non-protein coding RNA 271 [S |
| GAPDHP73 | 6 | glyceraldehyde-3-phosphate dehydrogenase p |
| AL035604.1 | 6 | novel transcript |
| HMGB1P17 | 6 | high mobility group box 1 pseudogene 17 [Sou |
| AL133319.1 | 6 | novel transcript |
| PDE7B | 6 | phosphodiesterase 7B [Source:HGNC Symbol; |
| AL360178.1 | 6 | novel transcript, antisense to PDE7B |
| AL360178.2 | 6 | novel transcript, antisense to PDE7B |
| AL138828.1 | 6 | novel transcript |
| COX5BP2 | 6 | cytochrome c oxidase subunit 5B pseudogene |
| AL138828.2 | 6 | family with sequence similarity 98, member A |
| MTFR2 | 6 | mitochondrial fission regulator 2 [Source:HGN |
| BCLAF1 | 6 | BCL2 associated transcription factor 1 [Source |
| AL023284.1 | 6 | eukaryotic translation elongation factor 1 alph |
| AL023284.4 | 6 | novel transcript |
| MAP7 | 6 | microtubule associated protein 7 [Source:HGN |
| AL023284.3 | 6 | ribosomal protein, large, P1 (RPLP1) pseudoge |
| AL023284.2 | 6 | phosphopantothenoylcysteine synthetase (PP |
| NDUFS5P1 | 6 | NADH:ubiquinone oxidoreductase subunit S5 |
| RN7SKP299 | 6 | RN7SK pseudogene 299 [Source:HGNC Symbo |
| AL024508.2 | 6 | novel transcript, antisense to MAP7 |
| MAP3K5 | 6 | mitogen-activated protein kinase kinase kinas |
| AL024508.1 | 6 | novel transcript |
| RNA5SP219 | 6 | RNA, 5S ribosomal pseudogene 219 [Source:H |
| AL121933.2 | 6 | novel transcript, antisense to MAP3K5 |

| | | |
|---|---|---|
| PEX7 | 6 | peroxisomal biogenesis factor 7 [Source:HGN( |
| AL121933.1 | 6 | NULL |
| AL365223.1 | 6 | ribosomal protein L7a (RPL7A) pseudogene |
| SLC35D3 | 6 | solute carrier family 35 member D3 [Source:H |
| Y_RNA | 6 | Y RNA [Source:RFAM;Acc:RF00019] |
| RPL35AP3 | 6 | ribosomal protein L35a pseudogene 3 [Source |
| AL135902.1 | 6 | novel transcript |
| AL135902.2 | 6 | novel transcript |
| IL20RA | 6 | interleukin 20 receptor subunit alpha [Source: |
| IL22RA2 | 6 | interleukin 22 receptor subunit alpha 2 [Sourc |
| IFNGR1 | 6 | interferon gamma receptor 1 [Source:HGNC S |
| OLIG3 | 6 | oligodendrocyte transcription factor 3 [Source |
| BTF3L4P3 | 6 | basic transcription factor 3 like 4 pseudogene |
| AL356234.2 | 6 | novel transcript |
| AL356234.3 | 6 | novel transcript |
| AL356234.1 | 6 | protein tyrosine phosphatase, non-receptor ty |
| LINC02539 | 6 | long intergenic non-protein coding RNA 2539 |
| Y_RNA | 6 | Y RNA [Source:RFAM;Acc:RF00019] |
| WAKMAR2 | 6 | wound and keratinocyte migration associated |
| TNFAIP3 | 6 | TNF alpha induced protein 3 [Source:HGNC Sy |
| AL591468.2 | 6 | novel transcript |
| AL591468.1 | 6 | novel transcript |
| LINC02528 | 6 | long intergenic non-protein coding RNA 2528 |
| RPSAP42 | 6 | ribosomal protein SA pseudogene 42 [Source: |
| PERP | 6 | p53 apoptosis effector related to PMP22 [Sou |
| ARFGEF3 | 6 | ARFGEF family member 3 [Source:HGNC Symb |
| PBOV1 | 6 | prostate and breast cancer overexpressed 1 [S |
| SMIM28 | 6 | small integral membrane protein 28 [Source:H |
| AL031003.1 | 6 | pseudogene similar to Mus musculus upregula |
| MARCKSL1P2 | 6 | MARCKS like 1 pseudogene 2 [Source:HGNC S |
| HEBP2 | 6 | heme binding protein 2 [Source:HGNC Symbo |
| NHSL1 | 6 | NHS like 1 [Source:HGNC Symbol;Acc:HGNC:2 |
| MIR3145 | 6 | microRNA 3145 [Source:HGNC Symbol;Acc:HG |
| AL391669.1 | 6 | transmembrane protein 80 (TMEM80) pseudo |
| AL590617.1 | 6 | mitochondrial carrier homolog 1 (MTCH1) pse |
| AL590617.2 | 6 | novel transcript |
| GVQW2 | 6 | GVQW motif containing 2 [Source:HGNC Symb |
| CCDC28A | 6 | coiled-coil domain containing 28A [Source:HG |
| ECT2L | 6 | epithelial cell transforming 2 like [Source:HGN |
| ACKR4P1 | 6 | ACKR4 pseudogene 1 [Source:HGNC Symbol;A |
| AL121834.1 | 6 | SMT3 suppressor of mif two 3 homolog 2 (S. c |
| REPS1 | 6 | RALBP1 associated Eps domain containing 1 [S |
| ABRACL | 6 | ABRA C-terminal like [Source:HGNC Symbol;A |
| HECA | 6 | hdc homolog, cell cycle regulator [Source:HGN |
| AL031772.1 | 6 | novel transcript |
| AL158850.1 | 6 | novel transcript |
| TXLNB | 6 | taxilin beta [Source:HGNC Symbol;Acc:HGNC:2 |
| AL592429.2 | 6 | novel transcript |
| AL592429.1 | 6 | DnaJ (Hsp40) homolog, subfamily C, member |
| CITED2 | 6 | Cbp/p300 interacting transactivator with Glu/ |

| | | |
|---|---|---|
| LINC01625 | 6 | long intergenic non-protein coding RNA 1625 |
| ATP5PBP6 | 6 | ATP synthase peripheral stalk-membrane subu |
| AL390205.1 | 6 | thyroid hormone receptor interactor 4 (TRIP4 |
| FILNC1 | 6 | FOXO induced long non-coding RNA 1 [Source |
| AL050338.1 | 6 | novel transcript |
| AL050338.2 | 6 | novel transcript |
| AL357146.1 | 6 | novel transcript |
| RNA5SP220 | 6 | RNA, 5S ribosomal pseudogene 220 [Source:H |
| MIR3668 | 6 | microRNA 3668 [Source:HGNC Symbol;Acc:HG |
| AL035446.2 | 6 | novel transcript |
| MIR4465 | 6 | microRNA 4465 [Source:HGNC Symbol;Acc:HG |
| RPS3AP24 | 6 | ribosomal protein S3a pseudogene 24 [Source |
| AL035446.1 | 6 | novel transcript |
| AL357084.1 | 6 | ribosomal protein S18 (RPS18) pseudogene |
| AL357080.1 | 6 | glioblastoma amplified sequence (GBAS) pseu |
| AL355596.2 | 6 | novel transcript |
| AL355596.1 | 6 | novel transcript |
| RN7SKP106 | 6 | RN7SK pseudogene 106 [Source:HGNC Symbo |
| RPS3AP23 | 6 | ribosomal protein S3a pseudogene 23 [Source |
| NMBR | 6 | neuromedin B receptor [Source:HGNC Symbo |
| AL589674.1 | 6 | ubiquitin-like modifier activating enzyme 2 (U |
| NMBR-AS1 | 6 | NMBR antisense RNA 1 [Source:HGNC Symbol |
| GJE1 | 6 | gap junction protein epsilon 1 [Source:HGNC S |
| VTA1 | 6 | vesicle trafficking 1 [Source:HGNC Symbol;Acc |
| AL360007.1 | 6 | novel transcript |
| ADGRG6 | 6 | adhesion G protein-coupled receptor G6 [Sour |
| AL359313.1 | 6 | novel transcript |
| AL355337.1 | 6 | novel transcript |
| AL023584.1 | 6 | novel transcript |
| HIVEP2 | 6 | human immunodeficiency virus type I enhanc |
| AL023584.2 | 6 | novel transcript |
| AL355304.1 | 6 | novel transcript |
| LINC01277 | 6 | long intergenic non-protein coding RNA 1277 |
| AL023581.2 | 6 | novel transcript |
| AIG1 | 6 | androgen induced 1 [Source:HGNC Symbol;Ac |
| AL023581.1 | 6 | novel transcript |
| AL136116.2 | 6 | actin protein pseudogene |
| AL136116.1 | 6 | ribosomal protein L31 (RPL31) pseudogene |
| AL136116.3 | 6 | pseudogene similar to presenilin associated, r |
| AL031320.1 | 6 | chromosome 4 open reading frame 46 (C4orf4 |
| ADAT2 | 6 | adenosine deaminase tRNA specific 2 [Source |
| TUBB8P2 | 6 | tubulin beta 8 class VIII pseudogene 2 [Source |
| RNA5SP221 | 6 | RNA, 5S ribosomal pseudogene 221 [Source:H |
| PEX3 | 6 | peroxisomal biogenesis factor 3 [Source:HGN( |
| AL031320.2 | 6 | novel transcript |
| VDAC1P8 | 6 | voltage dependent anion channel 1 pseudoge |
| FUCA2 | 6 | alpha-L-fucosidase 2 [Source:HGNC Symbol;Ac |
| PHACTR2 | 6 | phosphatase and actin regulator 2 [Source:HG |
| PHACTR2-AS1 | 6 | PHACTR2 antisense RNA 1 [Source:HGNC Sym |
| AL049844.1 | 6 | novel protein |

| | | |
|---|---|---|
| LTV1 | 6 | LTV1 ribosome biogenesis factor [Source:HGN |
| AL049844.3 | 6 | novel transcript |
| AL049844.2 | 6 | synaptophysin-like 1 (SYPL1) pseudogene |
| ZC2HC1B | 6 | zinc finger C2HC-type containing 1B [Source:H |
| PLAGL1 | 6 | PLAG1 like zinc finger 1 [Source:HGNC Symbol |
| HYMAI | 6 | hydatidiform mole associated and imprinted [ |
| AL109755.2 | 6 | NULL |
| AL109755.1 | 6 | FK506 binding protein 7 (FKBP7) pseudogene |
| SF3B5 | 6 | splicing factor 3b subunit 5 [Source:HGNC Sym |
| MRPL42P3 | 6 | mitochondrial ribosomal protein L42 pseudog |
| STX11 | 6 | syntaxin 11 [Source:HGNC Symbol;Acc:HGNC: |
| TPT1P4 | 6 | tumor protein, translationally-controlled 1 ps |
| AL024474.1 | 6 | single-stranded DNA binding protein 1 (SSBP1 |
| UTRN | 6 | utrophin [Source:HGNC Symbol;Acc:HGNC:12 |
| AL024474.2 | 6 | novel transcript, antisense to UTRN |
| AL590704.1 | 6 | rhomboid domain containing 1 (RHBDD1) pse |
| AL513475.1 | 6 | 5'-nucleotidase domain containing 1 (NT5DC1 |
| AL513475.2 | 6 | 5'-nucleotidase domain containing 1 (NT5DC1 |
| AL023283.1 | 6 | novel transcript |
| EPM2A | 6 | EPM2A glucan phosphatase, laforin [Source:H |
| RNU1-33P | 6 | RNA, U1 small nuclear 33, pseudogene [Sourc |
| AL031119.1 | 6 | BTB/POZ domain zinc finger protein pseudoge |
| AL023806.1 | 6 | novel transcript |
| AL023806.2 | 6 | novel transcript |
| AL356599.1 | 6 | novel transcript, antisense to SHPRH |
| AL023806.4 | 6 | novel transcript |
| AL023806.3 | 6 | TEC |
| FBXO30 | 6 | F-box protein 30 [Source:HGNC Symbol;Acc:H |
| SHPRH | 6 | SNF2 histone linker PHD RING helicase [Sourc |
| GRM1 | 6 | glutamate metabotropic receptor 1 [Source:H |
| FUNDC2P3 | 6 | FUN14 domain containing 2 pseudogene 3 [Sc |
| RNA5SP222 | 6 | RNA, 5S ribosomal pseudogene 222 [Source:H |
| AL035698.1 | 6 | phosphoribosyl pyrophosphate synthetase (P |
| RAB32 | 6 | RAB32, member RAS oncogene family [Source |
| AL359547.1 | 6 | novel transcript |
| ADGB | 6 | androglobin [Source:HGNC Symbol;Acc:HGNC |
| AL359547.2 | 6 | piggyBac transposable element derived 4 (PG |
| RNU6-906P | 6 | RNA, U6 small nuclear 906, pseudogene [Sour |
| KATNBL1P6 | 6 | katanin regulatory subunit B1 like 1 pseudoge |
| STXBP5-AS1 | 6 | STXBP5 antisense RNA 1 [Source:HGNC Symbo |
| AL138916.1 | 6 | novel transcript |
| AL138916.2 | 6 | novel transcript |
| LUADT1 | 6 | lung adenocarcinoma associated transcript 1 [ |
| STXBP5 | 6 | syntaxin binding protein 5 [Source:HGNC Sym |
| AL355365.1 | 6 | heat shock 70kDa protein 8 (HSPA8) pseudoge |
| YAP1P1 | 6 | Yes associated protein 1 pseudogene 1 [Sourc |
| SAMD5 | 6 | sterile alpha motif domain containing 5 [Sourc |
| AL033504.1 | 6 | novel transcript |
| AL365271.1 | 6 | novel transcript |
| AL445123.2 | 6 | novel transcript |

| | | |
|---|---|---|
| AL445123.1 | 6 | novel transcript |
| AL359382.1 | 6 | novel transcript |
| AL359382.2 | 6 | novel transcript |
| AL513164.1 | 6 | novel transcript |
| SASH1 | 6 | SAM and SH3 domain containing 1 [Source:HG |
| RNU6-1222P | 6 | RNA, U6 small nuclear 1222, pseudogene [Sou |
| CYP51A1P3 | 6 | cytochrome P450 family 51 subfamily A meml |
| SNRPEP6 | 6 | small nuclear ribonucleoprotein polypeptide E |
| RPSAP40 | 6 | ribosomal protein SA pseudogene 40 [Source: |
| UST | 6 | uronyl 2-sulfotransferase [Source:HGNC Symb |
| UST-AS1 | 6 | UST antisense RNA 1 [Source:HGNC Symbol;A |
| AL357992.1 | 6 | novel transcript |
| AL603766.1 | 6 | small nuclear ribonucleoprotein polypeptide E |
| AL031056.1 | 6 | novel transcript |
| TAB2 | 6 | TGF-beta activated kinase 1 (MAP3K7) binding |
| TAB2-AS1 | 6 | TAB2 antisense RNA 1 [Source:HGNC Symbol;A |
| AL031056.2 | 6 | novel transcript |
| RN7SL234P | 6 | RNA, 7SL, cytoplasmic 234, pseudogene [Sour |
| FABP12P1 | 6 | fatty acid binding protein 12 pseudogene 1 [S |
| SUMO4 | 6 | small ubiquitin like modifier 4 [Source:HGNC S |
| ZC3H12D | 6 | zinc finger CCCH-type containing 12D [Source: |
| AL031133.1 | 6 | coiled-coil domain containing 59 (CCDC59) ps |
| PPIL4 | 6 | peptidylprolyl isomerase like 4 [Source:HGNC |
| RNU7-3P | 6 | RNA, U7 small nuclear 3 pseudogene [Source: |
| AL078581.3 | 6 | novel transcript |
| GINM1 | 6 | glycoprotein integral membrane 1 [Source:HG |
| AL078581.4 | 6 | novel transcript |
| AL078581.2 | 6 | novel transcript |
| RPS18P9 | 6 | ribosomal protein S18 pseudogene 9 [Source: |
| AL078581.1 | 6 | NULL |
| KATNA1 | 6 | katanin catalytic subunit A1 [Source:HGNC Syr |
| Y_RNA | 6 | Y RNA [Source:RFAM;Acc:RF00019] |
| LATS1 | 6 | large tumor suppressor kinase 1 [Source:HGN |
| AL358852.1 | 6 | novel transcript, antisense to LATS1 |
| NUP43 | 6 | nucleoporin 43 [Source:HGNC Symbol;Acc:HG |
| PCMT1 | 6 | protein-L-isoaspartate (D-aspartate) O-methyl |
| BTBD10P2 | 6 | BTB domain containing 10 pseudogene 2 [Sou |
| AL355312.2 | 6 | novel transcript |
| AL355312.5 | 6 | novel transcript |
| AL355312.6 | 6 | novel transcript |
| LRP11 | 6 | LDL receptor related protein 11 [Source:HGN( |
| AL355312.4 | 6 | novel transcript, antisense to LRP11 |
| AL355312.3 | 6 | novel transcript, antisense to LRP11, RAET1E a |
| CCT7P1 | 6 | chaperonin containing TCP1 subunit 7 pseudo |
| RAET1E | 6 | retinoic acid early transcript 1E [Source:HGNC |
| RAET1E-AS1 | 6 | RAET1E antisense RNA 1 [Source:HGNC Symbo |
| AL355312.1 | 6 | multimerin 2 (MMRN2) pseudogene |
| RAET1G | 6 | retinoic acid early transcript 1G [Source:HGN( |
| AL583835.2 | 6 | pseudogene similar to small nuclear RNA activ |
| ULBP2 | 6 | UL16 binding protein 2 [Source:HGNC Symbol |

| | | |
|---|---|---|
| AL583835.1 | 6 | pseudogene similar to small nuclear RNA activ |
| ULBP1 | 6 | UL16 binding protein 1 [Source:HGNC Symbol; |
| AL355497.1 | 6 | pseudogene similar to UL16 binding protein 1 |
| BTF3P10 | 6 | basic transcription factor 3 pseudogene 10 [Sc |
| RAET1K | 6 | retinoic acid early transcript 1K pseudogene [S |
| RAET1L | 6 | retinoic acid early transcript 1L [Source:HGNC |
| RAET1M | 6 | retinoic acid early transcript 1M pseudogene [ |
| AL355497.2 | 6 | novel transcript |
| PHBP1 | 6 | prohibitin pseudogene 1 [Source:HGNC Symbo |
| ULBP3 | 6 | UL16 binding protein 3 [Source:HGNC Symbol; |
| PPP1R14C | 6 | protein phosphatase 1 regulatory inhibitor sul |
| AL355497.3 | 6 | NULL |
| RNU4-7P | 6 | RNA, U4 small nuclear 7, pseudogene [Source |
| IYD | 6 | iodotyrosine deiodinase [Source:HGNC Symbo |
| SSR1P1 | 6 | signal sequence receptor subunit 1 pseudoger |
| AL451061.1 | 6 | alpha-methylacyl-CoA racemase (AMACR) pse |
| PLEKHG1 | 6 | pleckstrin homology and RhoGEF domain cont |
| AL450344.3 | 6 | novel transcript |
| AL450344.1 | 6 | novel transcript |
| AL450344.2 | 6 | novel transcript |
| PDCL3P5 | 6 | PDCL3 pseudogene 5 [Source:HGNC Symbol;A |
| MTHFD1L | 6 | methylenetetrahydrofolate dehydrogenase (N |
| ARL4AP5 | 6 | ADP ribosylation factor like GTPase 4A pseudo |
| RNU6-302P | 6 | RNA, U6 small nuclear 302, pseudogene [Sour |
| AL133260.2 | 6 | novel transcript |
| AL138733.1 | 6 | novel transcript, antisense to MTHFD1L |
| AL133260.1 | 6 | 60S ribosomal protein L32 (RPL32) pseudogen |
| AL138733.2 | 6 | novel transcript |
| RNU6-300P | 6 | RNA, U6 small nuclear 300, pseudogene [Sour |
| AL356535.1 | 6 | ribosomal protein S12 (RPS12) pseudogene |
| AKAP12 | 6 | A-kinase anchoring protein 12 [Source:HGNC S |
| RNU6-1247P | 6 | RNA, U6 small nuclear 1247, pseudogene [Sou |
| RNY4P20 | 6 | RNY4 pseudogene 20 [Source:HGNC Symbol;A |
| RN7SKP268 | 6 | RN7SK pseudogene 268 [Source:HGNC Symbo |
| ZBTB2 | 6 | zinc finger and BTB domain containing 2 [Sour |
| Y_RNA | 6 | Y RNA [Source:RFAM;Acc:RF00019] |
| RMND1 | 6 | required for meiotic nuclear division 1 homolo |
| HSPA8P15 | 6 | heat shock protein family A (Hsp70) member 8 |
| ARMT1 | 6 | acidic residue methyltransferase 1 [Source:HG |
| CCDC170 | 6 | coiled-coil domain containing 170 [Source:HG |
| RNU6-813P | 6 | RNA, U6 small nuclear 813, pseudogene [Sour |
| ESR1 | 6 | estrogen receptor 1 [Source:HGNC Symbol;Ac |
| AL356311.1 | 6 | novel transcript |
| AL078582.2 | 6 | novel transcript, antisense to ESR1 |
| AL078582.1 | 6 | novel transcript, sense intronic to ESR1 |
| SYNE1 | 6 | spectrin repeat containing nuclear envelope p |
| SYNE1-AS1 | 6 | SYNE1 antisense RNA 1 [Source:HGNC Symbol |
| AL049548.1 | 6 | novel transcript |
| RNA5SP223 | 6 | RNA, 5S ribosomal pseudogene 223 [Source:H |
| NANOGP11 | 6 | Nanog homeobox pseudogene 11 [Source:HG |

| | | |
|---|---|---|
| MYCT1 | 6 | MYC target 1 [Source:HGNC Symbol;Acc:HGN( |
| HSPD1P16 | 6 | heat shock protein family D (Hsp60) member |
| VIP | 6 | vasoactive intestinal peptide [Source:HGNC Sy |
| LINC02840 | 6 | long intergenic non-protein coding RNA 2840 |
| TUBB4BP7 | 6 | tubulin beta 4B class IVb pseudogene 7 [Sourc |
| AL080276.1 | 6 | chromosome 11 open reading frame 48 (C11c |
| FBXO5 | 6 | F-box protein 5 [Source:HGNC Symbol;Acc:HG |
| AL080276.2 | 6 | novel transcript |
| MTRF1L | 6 | mitochondrial translational release factor 1 lik |
| RGS17 | 6 | regulator of G protein signaling 17 [Source:HG |
| RNA5SP224 | 6 | RNA, 5S ribosomal pseudogene 224 [Source:H |
| AL590867.1 | 6 | novel transcript |
| AL590867.2 | 6 | 60S ribosomal protein L27a (RPL27A) pseudog |
| AL358134.1 | 6 | novel transcript |
| RNA5SP225 | 6 | RNA, 5S ribosomal pseudogene 225 [Source:H |
| MTCO2P31 | 6 | MT-CO2 pseudogene 31 [Source:HGNC Symbo |
| AL603908.1 | 6 | mitochondrially encoded ATP synthase 8 (MT- |
| MTATP6P31 | 6 | MT-ATP6 pseudogene 31 [Source:HGNC Symb |
| MTCO3P31 | 6 | MT-CO3 pseudogene 31 [Source:HGNC Symbo |
| MTND3P20 | 6 | MT-ND3 pseudogene 20 [Source:HGNC Symbo |
| MTND4LP20 | 6 | MT-ND4L pseudogene 20 [Source:HGNC Symb |
| MTND4P13 | 6 | MT-ND4 pseudogene 13 [Source:HGNC Symbo |
| RNU6-896P | 6 | RNA, U6 small nuclear 896, pseudogene [Sour |
| HMGB3P19 | 6 | high mobility group box 3 pseudogene 19 [Sou |
| OPRM1 | 6 | opioid receptor mu 1 [Source:HGNC Symbol;A |
| IPCEF1 | 6 | interaction protein for cytohesin exchange fac |
| AL357075.5 | 6 | CNK3/IPCEF1 fusion protein [Source:UniProtK |
| AL445220.1 | 6 | ribosomal protein L17 (RPL17) pseudogene |
| CNKSR3 | 6 | CNKSR family member 3 [Source:HGNC Symbo |
| AL357075.3 | 6 | novel transcript, antisense to CNKSR3 |
| AL357075.4 | 6 | novel transcript |
| AL357075.2 | 6 | La ribonucleoprotein domain family, member |
| AL357075.1 | 6 | chromosome 6 open reading frame 203 (C6or |
| RPS4XP8 | 6 | ribosomal protein S4X pseudogene 8 [Source: |
| AL591419.1 | 6 | ribosomal protein L31 (RPL31) pseudogene |
| AL591499.1 | 6 | stathmin-like 2 (STMN2) pseudogene |
| SCAF8 | 6 | SR-related CTD associated factor 8 [Source:HG |
| RNU6-824P | 6 | RNA, U6 small nuclear 824, pseudogene [Sour |
| TIAM2 | 6 | T cell lymphoma invasion and metastasis 2 [Sc |
| AL121952.1 | 6 | 60S ribosomal protein L17 (RPL17) pseudogen |
| MIR1273C | 6 | microRNA 1273c [Source:HGNC Symbol;Acc:H |
| U8 | 6 | U8 small nucleolar RNA [Source:RFAM;Acc:RF( |
| AL136228.1 | 6 | lysophosphatidylcholine acyltransferase 3 (LP( |
| AL596202.1 | 6 | novel transcript |
| TFB1M | 6 | transcription factor B1, mitochondrial [Source |
| CLDN20 | 6 | claudin 20 [Source:HGNC Symbol;Acc:HGNC:2 |
| AL031773.1 | 6 | novel transcript |
| NOX3 | 6 | NADPH oxidase 3 [Source:HGNC Symbol;Acc:H |
| RNU7-152P | 6 | RNA, U7 small nuclear 152 pseudogene [Sourc |
| AL589693.1 | 6 | novel transcript |

| | | |
|---|---|---|
| MIR1202 | 6 | microRNA 1202 [Source:HGNC Symbol;Acc:HG |
| AL512658.2 | 6 | novel transcript |
| SNORD28B | 6 | small nucleolar RNA, C/D box 28B [Source:HG |
| AL512658.1 | 6 | novel transcript |
| AL355297.1 | 6 | H3 histone, family 3A (H3F3A) pseudogene |
| AL355297.3 | 6 | novel transcript |
| ARID1B | 6 | AT-rich interaction domain 1B [Source:HGNC S |
| AL355297.4 | 6 | novel transcript, antisense to ARID1B |
| MIR4466 | 6 | microRNA 4466 [Source:HGNC Symbol;Acc:HG |
| AL355297.2 | 6 | novel transcript, antisense to ARID1B |
| AL162578.1 | 6 | ribosomal protein S15a (RPS15A) pseudogene |
| AL049820.1 | 6 | novel transcript |
| TMEM242 | 6 | transmembrane protein 242 [Source:HGNC Sy |
| LDHAL6FP | 6 | lactate dehydrogenase A like 6F, pseudogene |
| AL390955.2 | 6 | novel transcript |
| AL390955.1 | 6 | steroid dehydrogenase homolog pseudogene |
| AL117344.1 | 6 | novel pseudogene |
| AL117344.2 | 6 | novel transcript |
| ZDHHC14 | 6 | zinc finger DHHC-type containing 14 [Source:H |
| MIR3692 | 6 | microRNA 3692 [Source:HGNC Symbol;Acc:HG |
| SNX9 | 6 | sorting nexin 9 [Source:HGNC Symbol;Acc:HGI |
| AL391863.2 | 6 | novel transcript |
| AL391863.1 | 6 | novel transcript |
| AL035634.1 | 6 | novel transcript |
| HSPE1P26 | 6 | heat shock protein family E (Hsp10) member 1 |
| RNU6-786P | 6 | RNA, U6 small nuclear 786, pseudogene [Sour |
| SYNJ2 | 6 | synaptojanin 2 [Source:HGNC Symbol;Acc:HGI |
| AL139330.1 | 6 | novel transcript |
| SYNJ2-IT1 | 6 | SYNJ2 intronic transcript 1 [Source:HGNC Sym |
| SERAC1 | 6 | serine active site containing 1 [Source:HGNC S |
| GTF2H5 | 6 | general transcription factor IIH subunit 5 [Sou |
| TULP4 | 6 | TUB like protein 4 [Source:HGNC Symbol;Acc:I |
| SRP72P2 | 6 | signal recognition particle 72 pseudogene 2 [S |
| AL360169.2 | 6 | novel transcript, sense overlapping TULP4 |
| AL360169.3 | 6 | novel transcript |
| AL360169.1 | 6 | novel transcript |
| RN7SL173P | 6 | RNA, 7SL, cytoplasmic 173, pseudogene [Sour |
| CACYBPP3 | 6 | calcyclin binding protein pseudogene 3 [Sourc |
| TMEM181 | 6 | transmembrane protein 181 [Source:HGNC Sy |
| TATDN2P2 | 6 | TatD DNase domain containing 2 pseudogene |
| MIR7161 | 6 | microRNA 7161 [Source:HGNC Symbol;Acc:HG |
| DYNLT1 | 6 | dynein light chain Tctex-type 1 [Source:HGNC |
| SYTL3 | 6 | synaptotagmin like 3 [Source:HGNC Symbol;A |
| AMZ2P2 | 6 | archaelysin family metallopeptidase 2 pseudo |
| MIR3918 | 6 | microRNA 3918 [Source:HGNC Symbol;Acc:HG |
| EZR | 6 | ezrin [Source:HGNC Symbol;Acc:HGNC:12691] |
| EZR-AS1 | 6 | EZR antisense RNA 1 [Source:HGNC Symbol;Ac |
| OSTCP1 | 6 | oligosaccharyltransferase complex subunit ps |
| AL627422.2 | 6 | novel transcript |
| C6orf99 | 6 | chromosome 6 open reading frame 99 [Sourc |

| | | |
|---|---|---|
| AL627422.1 | 6 | SNARE Vti1a-beta protein pseudogene |
| RNU6-293P | 6 | RNA, U6 small nuclear 293, pseudogene [Sour |
| RSPH3 | 6 | radial spoke head 3 [Source:HGNC Symbol;Acc |
| AL035530.2 | 6 | novel transcript, antisense to TAGAP |
| TAGAP | 6 | T cell activation RhoGTPase activating protein |
| AL035530.1 | 6 | novel transcript |
| AL356417.3 | 6 | uncharacterized LOC112267968 [Source:NCBI |
| AL356417.1 | 6 | novel transcript |
| AL356417.2 | 6 | novel transcript, antisense to FNDC1 |
| FNDC1 | 6 | fibronectin type III domain containing 1 [Sourc |
| FNDC1-IT1 | 6 | FNDC1 intronic transcript 1 [Source:HGNC Sym |
| AL357832.1 | 6 | novel transcript |
| LINC02529 | 6 | long intergenic non-protein coding RNA 2529 |
| AL078604.4 | 6 | novel transcript |
| AL078604.1 | 6 | 60S Ribosomal protein L21 (RPL21) pseudoger |
| AL078604.3 | 6 | cullin 2 (CUL2) pseudogene |
| AL078604.2 | 6 | novel transcript |
| SOD2 | 6 | superoxide dismutase 2 [Source:HGNC Symbo |
| SOD2 | 6 | superoxide dismutase 2 [Source:NCBI gene;Ac |
| HNRNPH1P1 | 6 | heterogeneous nuclear ribonucleoprotein H1 |
| RNU4ATAC18P | 6 | RNA, U4atac small nuclear 18, pseudogene [So |
| WTAP | 6 | WT1 associated protein [Source:HGNC Symbo |
| SOD2-OT1 | 6 | SOD2 overlapping transcript 1 [Source:HGNC S |
| ACAT2 | 6 | acetyl-CoA acetyltransferase 2 [Source:HGNC |
| TCP1 | 6 | t-complex 1 [Source:HGNC Symbol;Acc:HGNC: |
| SNORA20 | 6 | small nucleolar RNA, H/ACA box 20 [Source:H |
| SNORA29 | 6 | small nucleolar RNA, H/ACA box 29 [Source:H |
| MRPL18 | 6 | mitochondrial ribosomal protein L18 [Source: |
| PNLDC1 | 6 | PARN like, ribonuclease domain containing 1 [ |
| AL035691.1 | 6 | novel transcript |
| MAS1 | 6 | MAS1 proto-oncogene, G protein-coupled rec |
| IGF2R | 6 | insulin like growth factor 2 receptor [Source:H |
| AIRN | 6 | antisense of IGF2R non-protein coding RNA [S |
| AL353625.1 | 6 | calcium binding protein P22 (CHP) pseudogen |
| SLC22A1 | 6 | solute carrier family 22 member 1 [Source:HG |
| AL645733.1 | 6 | novel pseudogene |
| SLC22A2 | 6 | solute carrier family 22 member 2 [Source:HG |
| AL162582.1 | 6 | novel transcript |
| SLC22A3 | 6 | solute carrier family 22 member 3 [Source:HG |
| AL591069.1 | 6 | novel transcript, antisense to SLC22A3 |
| LPAL2 | 6 | lipoprotein(a) like 2, pseudogene [Source:HGN |
| LPA | 6 | lipoprotein(a) [Source:HGNC Symbol;Acc:HGN |
| AL109933.2 | 6 | pseudogene similar to part of plasminogen PL |
| AL109933.1 | 6 | novel transcript |
| PLG | 6 | plasminogen [Source:HGNC Symbol;Acc:HGNC |
| AL109933.5 | 6 | novel transcript, antisense to PLG |
| AL109933.4 | 6 | solute carrier family 22 (organic cation transp |
| AL109933.3 | 6 | plasminogen (PLG) pseudogene |
| AL391361.2 | 6 | novel transcript |
| AL391361.1 | 6 | pseudogene similar to lipoproteins (contains k |

| | | |
|---|---|---|
| AL139393.2 | 6 | novel transcript |
| AL139393.1 | 6 | novel transcript |
| AL139393.3 | 6 | novel transcript, antisense to MAP3K4 |
| MAP3K4 | 6 | mitogen-activated protein kinase kinase kinas |
| AGPAT4 | 6 | 1-acylglycerol-3-phosphate O-acyltransferase |
| PRKN | 6 | parkin RBR E3 ubiquitin protein ligase [Source |
| AL132982.1 | 6 | novel transcript, antisense to PARK2 |
| AL138716.1 | 6 | novel transcript, antisense to PARK2 |
| KRT8P44 | 6 | keratin 8 pseudogene 44 [Source:HGNC Symb |
| PACRG | 6 | parkin coregulated [Source:HGNC Symbol;Acc |
| AL590286.1 | 6 | ribosomal protein L34 (RPL34) pseudogene |
| AL590286.2 | 6 | novel transcript, antisense to PACRG |
| PACRG-AS2 | 6 | PACRG antisense RNA 2 [Source:HGNC Symbo |
| AL137182.1 | 6 | novel transcript |
| PACRG-AS3 | 6 | PACRG antisense RNA 3 [Source:HGNC Symbo |
| PACRG-AS1 | 6 | PACRG antisense RNA 1 [Source:HGNC Symbo |
| AL031121.2 | 6 | novel transcript |
| AL031121.3 | 6 | novel transcript |
| AL031121.1 | 6 | RPL37A (60S Ribosomal Protein L37A) pseudo |
| CAHM | 6 | colon adenocarcinoma hypermethylated [Sou |
| QKI | 6 | QKI, KH domain containing RNA binding [Sour |
| AL445307.1 | 6 | novel transcript |
| AL078602.1 | 6 | novel transcript |
| AL137005.1 | 6 | novel transcript |
| RN7SL366P | 6 | RNA, 7SL, cytoplasmic 366, pseudogene [Sour |
| AL136225.1 | 6 | novel transcript |
| Z97205.2 | 6 | novel transcript |
| Z97205.1 | 6 | novel transcript |
| AL358972.1 | 6 | novel transcript |
| AL450345.2 | 6 | novel transcript |
| AL450345.1 | 6 | novel transcript |
| AL133538.1 | 6 | novel transcript |
| AL136100.1 | 6 | pseudogene similar to zinc finger proteins |
| C6orf118 | 6 | chromosome 6 open reading frame 118 [Sour |
| PDE10A | 6 | phosphodiesterase 10A [Source:HGNC Symbo |
| RNA5SP226 | 6 | RNA, 5S ribosomal pseudogene 226 [Source:H |
| RNU6-730P | 6 | RNA, U6 small nuclear 730, pseudogene [Sour |
| AL590302.2 | 6 | novel transcript, antisense to PDE10A |
| AL590302.1 | 6 | novel transcript |
| LINC00602 | 6 | long intergenic non-protein coding RNA 602 [S |
| GAPDHP72 | 6 | glyceraldehyde-3-phosphate dehydrogenase p |
| AL627443.3 | 6 | novel transcript |
| AL627443.1 | 6 | novel transcript |
| AL627443.2 | 6 | NULL |
| TBXT | 6 | T-box transcription factor T [Source:HGNC Syr |
| RNU6-153P | 6 | RNA, U6 small nuclear 153, pseudogene [Sour |
| AL121956.5 | 6 | heterogeneous nuclear ribonucleoprotein D (/ |
| AL121956.1 | 6 | novel transcript |
| GNG5P1 | 6 | G protein subunit gamma 5 pseudogene 1 [So |
| AL121956.4 | 6 | novel transcript |

| | | |
|---|---|---|
| PRR18 | 6 | proline rich 18 [Source:HGNC Symbol;Acc:HGI |
| AL121956.6 | 6 | novel transcript |
| SFT2D1 | 6 | SFT2 domain containing 1 [Source:HGNC Syml |
| HNRNPA1P49 | 6 | heterogeneous nuclear ribonucleoprotein A1 |
| AL022069.3 | 6 | novel transcript |
| MPC1 | 6 | mitochondrial pyruvate carrier 1 [Source:HGN |
| AL022069.1 | 6 | novel transcript, antisense to BRP44L |
| AL022069.2 | 6 | TEC |
| RPS6KA2 | 6 | ribosomal protein S6 kinase A2 [Source:HGNC |
| RPS6KA2-IT1 | 6 | RPS6KA2 intronic transcript 1 [Source:HGNC S |
| MIR1913 | 6 | microRNA 1913 [Source:HGNC Symbol;Acc:H( |
| RAMACL | 6 | RNA guanine-7 methyltransferase activating s |
| AL023775.2 | 6 | novel transcript, antisense to RPS6KA2 |
| AL023775.1 | 6 | IGF-II mRNA-binding protein 3 (KOC1) pseudo; |
| AL159163.1 | 6 | novel protein |
| RPS6KA2-AS1 | 6 | RPS6KA2 antisense RNA 1 [Source:HGNC Syml |
| RNASET2 | 6 | ribonuclease T2 [Source:HGNC Symbol;Acc:H( |
| Z94721.1 | 6 | novel transcript |
| Z94721.3 | 6 | novel transcript |
| MIR3939 | 6 | microRNA 3939 [Source:HGNC Symbol;Acc:H( |
| FGFR1OP | 6 | FGFR1 oncogene partner [Source:HGNC Symb |
| Z94721.2 | 6 | novel transcript |
| CCR6 | 6 | C-C motif chemokine receptor 6 [Source:HGN( |
| AL121935.1 | 6 | novel transcript |
| AL121935.2 | 6 | novel transcript |
| TCP10L2 | 6 | t-complex 10 like 2 [Source:HGNC Symbol;Acc |
| GPR31 | 6 | G protein-coupled receptor 31 [Source:HGNC |
| AL121935.3 | 6 | novel transcript, antisense to TCP10L2 |
| AL353747.1 | 6 | novel pseudogene |
| AL353747.4 | 6 | novel transcript |
| AL353747.3 | 6 | novel transcript |
| AL353747.2 | 6 | novel transcript |
| UNC93A | 6 | unc-93 homolog A [Source:HGNC Symbol;Acc: |
| TTLL2 | 6 | tubulin tyrosine ligase like 2 [Source:HGNC Sy |
| AL021331.1 | 6 | MAD2 mitotic arrest deficient-like 1 (yeast) (N |
| TCP10 | 6 | t-complex 10 [Source:HGNC Symbol;Acc:HGN( |
| AL356123.1 | 6 | novel transcript |
| LINC02538 | 6 | long intergenic non-protein coding RNA 2538 |
| AL356123.2 | 6 | novel transcript |
| LINC02487 | 6 | long intergenic non-protein coding RNA 2487 |
| AL009178.1 | 6 | serine palmitoyltransferase, long chain base s |
| LINC01558 | 6 | long intergenic non-protein coding RNA 1558 |
| AL009178.2 | 6 | novel transcript |
| AL009178.3 | 6 | novel transcript |
| AFDN-DT | 6 | AFDN divergent transcript [Source:HGNC Sym |
| AFDN | 6 | afadin, adherens junction formation factor [S( |
| HGC6.3 | 6 | uncharacterized LOC100128124 [Source:NCBI |
| KIF25-AS1 | 6 | KIF25 antisense RNA 1 [Source:HGNC Symbol; |
| KIF25 | 6 | kinesin family member 25 [Source:HGNC Syml |
| FRMD1 | 6 | FERM domain containing 1 [Source:HGNC Sym |

| | | |
|---|---|---|
| AL611929.1 | 6 | novel transcript |
| AL606970.5 | 6 | novel transcript |
| AL606970.4 | 6 | novel transcript |
| AL606970.2 | 6 | novel transcript |
| AL606970.3 | 6 | novel transcript |
| CTAGE13P | 6 | CTAGE family member 13, pseudogene [Sourc |
| DACT2 | 6 | dishevelled binding antagonist of beta catenir |
| AL606970.1 | 6 | novel transcript |
| AL138918.1 | 6 | novel transcript |
| SMOC2 | 6 | SPARC related modular calcium binding 2 [Sou |
| AL136099.1 | 6 | novel transcript |
| AL513210.2 | 6 | TEC |
| AL513210.1 | 6 | novel transcript |
| AL109924.4 | 6 | novel transcript |
| AL109924.3 | 6 | TEC |
| AL109924.1 | 6 | novel transcript |
| AL109924.2 | 6 | novel transcript |
| AL136129.1 | 6 | novel transcript |
| LINC01615 | 6 | long intergenic non-protein coding RNA 1615 |
| LINC02544 | 6 | long intergenic non-protein coding RNA 2544 |
| BX322234.1 | 6 | novel transcript |
| THBS2 | 6 | thrombospondin 2 [Source:HGNC Symbol;Acc |
| BX322234.2 | 6 | novel transcript |
| VTA1P1 | 6 | vesicle trafficking 1 pseudogene 1 [Source:HG |
| LINC02519 | 6 | long intergenic non-protein coding RNA 2519 |
| AL009176.1 | 6 | novel transcript |
| AL031315.1 | 6 | novel protein |
| AL135910.1 | 6 | novel transcript |
| WDR27 | 6 | WD repeat domain 27 [Source:HGNC Symbol; |
| C6orf120 | 6 | chromosome 6 open reading frame 120 [Sour |
| PHF10 | 6 | PHD finger protein 10 [Source:HGNC Symbol; |
| AL354892.2 | 6 | novel transcript |
| AL354892.1 | 6 | novel transcript |
| TCTE3 | 6 | t-complex-associated-testis-expressed 3 [Sour |
| ERMARD | 6 | ER membrane associated RNA degradation [So |
| AL354892.3 | 6 | novel transcript |
| LINC00242 | 6 | long intergenic non-protein coding RNA 242 [S |
| LINC00574 | 6 | long intergenic non-protein coding RNA 574 [S |
| AL049612.1 | 6 | novel transcript |
| AL603783.1 | 6 | novel transcript |
| AL596442.4 | 6 | novel transcript |
| AL596442.3 | 6 | novel transcript |
| AL596442.1 | 6 | novel transcript |
| AL596442.2 | 6 | novel transcript |
| RPL12P23 | 6 | ribosomal protein L12 pseudogene 23 [Source |
| AL109910.1 | 6 | novel transcript |
| AL109910.2 | 6 | novel transcript |
| LINC01624 | 6 | long intergenic non-protein coding RNA 1624 |
| DLL1 | 6 | delta like canonical Notch ligand 1 [Source:HG |
| FAM120B | 6 | family with sequence similarity 120B [Source: |

| | | |
|---|---|---|
| AL078605.1 | 6 | novel transcript |
| MIR4644 | 6 | microRNA 4644 [Source:HGNC Symbol;Acc:HG |
| AL008628.1 | 6 | novel transcript, overlapping to FAM120B |
| PSMB1 | 6 | proteasome subunit beta 1 [Source:HGNC Syn |
| TBP | 6 | TATA-box binding protein [Source:HGNC Symb |
| PDCD2 | 6 | programmed cell death 2 [Source:HGNC Symb |
| AL031259.1 | 6 | novel transcript |
| AL672310.1 | 6 | novel transcript |
| OR4F7P | 6 | olfactory receptor family 4 subfamily F memb |
| WBP1LP8 | 6 | WBP1L pseudogene 8 [Source:HGNC Symbol;A |
| AL731661.1 | 6 | novel transcript |
| AL731684.1 | 6 | novel transcript |
| RPL23AP47 | 6 | ribosomal protein L23a pseudogene 47 [Sourc |
| AC215522.3 | 7 | novel transcript |
| AC215522.2 | 7 | novel transcript similar to family member witl |
| AC215522.1 | 7 | von Hippel-Lindau binding protein 1 (VBP1) ps |
| AC093627.1 | 7 | novel transcript |
| AC093627.6 | 7 | novel transcript |
| AC093627.22 | 7 | novel transcript |
| AC093627.5 | 7 | novel transcript |
| AC093627.4 | 7 | novel transcript |
| AC093627.7 | 7 | novel transcript |
| AC093627.2 | 7 | novel transcript |
| AC093627.3 | 7 | novel transcript |
| FAM20C | 7 | FAM20C golgi associated secretory pathway k |
| AC145676.1 | 7 | TEC |
| AC187653.1 | 7 | novel forkhead box family protein |
| AC188616.1 | 7 | novel transcript |
| AC226118.1 | 7 | novel transcript |
| AC188617.1 | 7 | novel transcript |
| AC188617.2 | 7 | novel transcript |
| AC147651.3 | 7 | novel transcript |
| PDGFA | 7 | platelet derived growth factor subunit A [Sour |
| AC147651.1 | 7 | novel transcript |
| PRKAR1B | 7 | protein kinase cAMP-dependent type I regulat |
| AC147651.2 | 7 | novel transcript |
| AC147651.4 | 7 | novel transcript |
| DNAAF5 | 7 | dynein axonemal assembly factor 5 [Source:H |
| SUN1 | 7 | Sad1 and UNC84 domain containing 1 [Source |
| GET4 | 7 | guided entry of tail-anchored proteins factor 4 |
| AC073957.3 | 7 | novel transcript, antisense to GET4 |
| ADAP1 | 7 | ArfGAP with dual PH domains 1 [Source:HGN( |
| COX19 | 7 | cytochrome c oxidase assembly factor COX19 |
| CYP2W1 | 7 | cytochrome P450 family 2 subfamily W memb |
| C7orf50 | 7 | chromosome 7 open reading frame 50 [Sourc |
| MIR339 | 7 | microRNA 339 [Source:HGNC Symbol;Acc:HGN |
| AC073957.1 | 7 | novel transcript |
| GPR146 | 7 | G protein-coupled receptor 146 [Source:HGN( |
| AC073957.2 | 7 | novel transcript, antisense to GPR146 |
| AC091729.1 | 7 | novel transcript |

| | | |
|---|---|---|
| AC091729.2 | 7 | novel transcript |
| GPER1 | 7 | G protein-coupled estrogen receptor 1 [Sourc |
| ZFAND2A | 7 | zinc finger AN1-type containing 2A [Source:H( |
| AC091729.3 | 7 | novel transcript |
| UNCX | 7 | UNC homeobox [Source:HGNC Symbol;Acc:H( |
| AC073094.1 | 7 | novel transcript |
| MICALL2 | 7 | MICAL like 2 [Source:HGNC Symbol;Acc:HGNC |
| AC102953.1 | 7 | novel transcript |
| AC102953.2 | 7 | novel transcript |
| INTS1 | 7 | integrator complex subunit 1 [Source:HGNC S |
| AC093734.1 | 7 | novel transcript |
| MAFK | 7 | MAF bZIP transcription factor K [Source:HGNC |
| TMEM184A | 7 | transmembrane protein 184A [Source:HGNC S |
| PSMG3 | 7 | proteasome assembly chaperone 3 [Source:H( |
| PSMG3-AS1 | 7 | PSMG3 antisense RNA 1 (head to head) [Sour( |
| TFAMP1 | 7 | transcription factor A, mitochondrial pseudog |
| AC074389.2 | 7 | novel transcript |
| ELFN1 | 7 | extracellular leucine rich repeat and fibronect |
| AC074389.3 | 7 | novel transcript |
| AC074389.1 | 7 | novel transcript |
| ELFN1-AS1 | 7 | ELFN1 antisense RNA 1 [Source:HGNC Symbol |
| MAD1L1 | 7 | mitotic arrest deficient 1 like 1 [Source:HGNC |
| AC069288.1 | 7 | Novel protein |
| AC104129.1 | 7 | novel transcript, antisense to MAD1L1 |
| MIR4655 | 7 | microRNA 4655 [Source:HGNC Symbol;Acc:H( |
| MRM2 | 7 | mitochondrial rRNA methyltransferase 2 [Sou |
| NUDT1 | 7 | nudix hydrolase 1 [Source:HGNC Symbol;Acc:I |
| SNX8 | 7 | sorting nexin 8 [Source:HGNC Symbol;Acc:HGI |
| MIR6836 | 7 | microRNA 6836 [Source:HGNC Symbol;Acc:H( |
| IMMP1LP3 | 7 | inner mitochondrial membrane peptidase sub |
| EIF3B | 7 | eukaryotic translation initiation factor 3 subur |
| AC004840.1 | 7 | kinesin family member pseudogene |
| AC004840.2 | 7 | neugrin, neurite outgrowth associated (NGRN |
| CHST12 | 7 | carbohydrate sulfotransferase 12 [Source:HGI |
| GRIFIN | 7 | galectin-related inter-fiber protein [Source:H( |
| LFNG | 7 | LFNG O-fucosylpeptide 3-beta-N-acetylglucos |
| MIR4648 | 7 | microRNA 4648 [Source:HGNC Symbol;Acc:H( |
| BRAT1 | 7 | BRCA1 associated ATM activator 1 [Source:H( |
| IQCE | 7 | IQ motif containing E [Source:HGNC Symbol;A |
| TTYH3 | 7 | tweety family member 3 [Source:HGNC Symb |
| AMZ1 | 7 | archaelysin family metallopeptidase 1 [Source |
| GNA12 | 7 | G protein subunit alpha 12 [Source:HGNC Sym |
| AC006028.1 | 7 | prefoldin subunit 4 (PFDN4) pseudogene |
| CARD11 | 7 | caspase recruitment domain family member 1 |
| AC004906.1 | 7 | novel transcript, antisense to CARD11 |
| RN7SKP130 | 7 | RN7SK pseudogene 130 [Source:HGNC Symbo |
| AC024028.1 | 7 | novel transcript |
| AC073316.1 | 7 | novel transcript |
| AC073316.2 | 7 | novel transcript |
| AC073316.3 | 7 | novel transcript |

| | | |
|---|---|---|
| SDK1 | 7 | sidekick cell adhesion molecule 1 [Source:HGN |
| AC092427.1 | 7 | ribosomal protein L21 (RPL21) pseudogene |
| AC011284.1 | 7 | novel transcript |
| AC017000.1 | 7 | novel transcript, antisense to SDK1 |
| CYP3A54P | 7 | cytochrome P450 family 3 subfamily A membe |
| AC072054.1 | 7 | novel transcript |
| FOXK1 | 7 | forkhead box K1 [Source:HGNC Symbol;Acc:H |
| AC092428.1 | 7 | novel transcript, sense intronic to FOXK1 |
| AP5Z1 | 7 | adaptor related protein complex 5 subunit zet |
| MIR4656 | 7 | microRNA 4656 [Source:HGNC Symbol;Acc:HG |
| AC092610.1 | 7 | Ras association and DIL domains (RADIL) pseu |
| RADIL | 7 | Rap associating with DIL domain [Source:HGN |
| Y_RNA | 7 | Y RNA [Source:RFAM;Acc:RF00019] |
| PAPOLB | 7 | poly(A) polymerase beta [Source:HGNC Symb |
| RPL22P16 | 7 | ribosomal protein L22 pseudogene 16 [Source |
| MMD2 | 7 | monocyte to macrophage differentiation asso |
| RNF216P1 | 7 | ring finger protein 216 pseudogene 1 [Source: |
| RBAK-RBAKDN | 7 | RBAK-RBAKDN readthrough [Source:HGNC Sy |
| AC092032.1 | 7 | solute carrier family 25 (mitochondrial carrier |
| SPDYE19P | 7 | speedy/RINGO cell cycle regulator family men |
| RBAK | 7 | RB associated KRAB zinc finger [Source:HGNC |
| RBAKDN | 7 | RBAK downstream neighbor [Source:HGNC Sy |
| OR10AH1P | 7 | olfactory receptor family 10 subfamily AH me |
| ZNF890P | 7 | zinc finger protein 890, pseudogene [Source:H |
| RNU6-215P | 7 | RNA, U6 small nuclear 215, pseudogene [Sour |
| WIPI2 | 7 | WD repeat domain, phosphoinositide interact |
| SLC29A4 | 7 | solute carrier family 29 member 4 [Source:HG |
| TNRC18 | 7 | trinucleotide repeat containing 18 [Source:HG |
| AC093620.1 | 7 | novel transcript |
| AC092171.1 | 7 | novel transcript |
| AC092171.4 | 7 | novel transcript |
| AC092171.3 | 7 | uncharacterized LOC100129484 [Source:NCBI |
| AC092171.5 | 7 | novel transcript |
| FBXL18 | 7 | F-box and leucine rich repeat protein 18 [Sour |
| AC092171.2 | 7 | novel transcript |
| MIR589 | 7 | microRNA 589 [Source:HGNC Symbol;Acc:HGN |
| ACTB | 7 | actin beta [Source:HGNC Symbol;Acc:HGNC:1 |
| AC006483.2 | 7 | novel transcript, antisense to ACTB |
| AC006483.1 | 7 | S100 calcium binding protein A11 (S100A11) p |
| FSCN1 | 7 | fascin actin-bundling protein 1 [Source:HGNC |
| RNF216 | 7 | ring finger protein 216 [Source:HGNC Symbol; |
| RNF216-IT1 | 7 | RNF216 intronic transcript 1 [Source:HGNC Sy |
| MIR6874 | 7 | microRNA 6874 [Source:HGNC Symbol;Acc:HG |
| ZNF815P | 7 | zinc finger protein 815, pseudogene [Source:H |
| RN7SL556P | 7 | RNA, 7SL, cytoplasmic 556, pseudogene [Sour |
| OCM | 7 | oncomodulin [Source:HGNC Symbol;Acc:HGN |
| CCZ1 | 7 | CCZ1 homolog, vacuolar protein trafficking an |
| RSPH10B | 7 | radial spoke head 10 homolog B [Source:HGN |
| PMS2 | 7 | PMS1 homolog 2, mismatch repair system co |
| Y_RNA | 7 | Y RNA [Source:RFAM;Acc:RF00019] |

| | | |
|---|---|---|
| AIMP2 | 7 | aminoacyl tRNA synthetase complex interacti |
| SNORA80D | 7 | small nucleolar RNA, H/ACA box 80D [Source: |
| EIF2AK1 | 7 | eukaryotic translation initiation factor 2 alpha |
| ANKRD61 | 7 | ankyrin repeat domain 61 [Source:HGNC Sym |
| RN7SL851P | 7 | RNA, 7SL, cytoplasmic 851, pseudogene [Sour |
| RNU6-218P | 7 | RNA, U6 small nuclear 218, pseudogene [Sour |
| AC004895.1 | 7 | novel transcript |
| USP42 | 7 | ubiquitin specific peptidase 42 [Source:HGNC |
| CYTH3 | 7 | cytohesin 3 [Source:HGNC Symbol;Acc:HGNC: |
| Y_RNA | 7 | Y RNA [Source:RFAM;Acc:RF00019] |
| FAM220A | 7 | family with sequence similarity 220 member |
| AC009412.1 | 7 | novel protein |
| RPSAP73 | 7 | ribosomal protein SA pseudogene 73 [Source: |
| RAC1 | 7 | Rac family small GTPase 1 [Source:HGNC Sym |
| DAGLB | 7 | diacylglycerol lipase beta [Source:HGNC Symb |
| KDELR2 | 7 | KDEL endoplasmic reticulum protein retentio |
| AC072052.1 | 7 | ribosomal protein L31 (RPL31) pseudogene |
| GRID2IP | 7 | Grid2 interacting protein [Source:HGNC Symb |
| ZDHHC4 | 7 | zinc finger DHHC-type containing 4 [Source:H |
| AC079742.1 | 7 | novel transcript |
| C7orf26 | 7 | chromosome 7 open reading frame 26 [Sourc |
| ZNF853 | 7 | zinc finger protein 853 [Source:HGNC Symbol; |
| ZNF316 | 7 | zinc finger protein 316 [Source:HGNC Symbol; |
| AC073343.2 | 7 | novel transcript, antisense to ZNF12 |
| AC073343.1 | 7 | zinc finger protein pseudogene |
| ZNF12 | 7 | zinc finger protein 12 [Source:HGNC Symbol; |
| PMS2CL | 7 | PMS2 C-terminal like pseudogene [Source:HG |
| SPDYE20P | 7 | speedy/RINGO cell cycle regulator family men |
| RSPH10B2 | 7 | radial spoke head 10 homolog B2 [Source:HG |
| CCZ1B | 7 | CCZ1 homolog B, vacuolar protein trafficking |
| OR7E39P | 7 | olfactory receptor family 7 subfamily E memb |
| UNC93B2 | 7 | unc-93 homolog B2, pseudogene [Source:HGN |
| OR7E136P | 7 | olfactory receptor family 7 subfamily E memb |
| OR7E59P | 7 | olfactory receptor family 7 subfamily E memb |
| AC079804.2 | 7 | novel pseudogene |
| ALG1L5P | 7 | ALG1 like 5, pseudogene [Source:HGNC Symb |
| AC079804.1 | 7 | novel pseudogene |
| FAM86LP | 7 | family with sequence similarity 86 member L, |
| AC079804.3 | 7 | novel transcript |
| MIR3683 | 7 | microRNA 3683 [Source:HGNC Symbol;Acc:H |
| AC092104.1 | 7 | zinc finger protein 655 (ZNF655) pseudogene |
| Y_RNA | 7 | Y RNA [Source:RFAM;Acc:RF00019] |
| C1GALT1 | 7 | core 1 synthase, glycoprotein-N-acetylgalacto |
| AC005532.2 | 7 | pseudogene similar to part of UTP6, small sub |
| AC005532.1 | 7 | novel transcript |
| AC079448.1 | 7 | microtubule-associated protein 1 light chain 3 |
| COL28A1 | 7 | collagen type XXVIII alpha 1 chain [Source:HG |
| AC004982.1 | 7 | novel transcript |
| AC004982.2 | 7 | novel transcript |
| MIOS | 7 | meiosis regulator for oocyte development [So |

| | | |
|---|---|---|
| AC004948.1 | 7 | novel transcript |
| RPA3 | 7 | replication protein A3 [Source:HGNC Symbol;/ |
| UMAD1 | 7 | UBAP1-MVB12-associated (UMA) domain con |
| AC007161.3 | 7 | novel transcript |
| AC007161.2 | 7 | 5-azacytidine induced 2 (AZI2) pseudogene |
| AC007161.1 | 7 | ribosomal protein L23a (RPL23A) pseudogene |
| RNU6-534P | 7 | RNA, U6 small nuclear 534, pseudogene [Sour |
| CCNB2P1 | 7 | cyclin B2 pseudogene 1 [Source:HGNC Symbo |
| AC006042.4 | 7 | novel transcript |
| AC006042.2 | 7 | novel transcript |
| GLCCI1 | 7 | glucocorticoid induced 1 [Source:HGNC Symb |
| AC006042.3 | 7 | peptidylprolyl isomerase (cyclophilin)-like 4 (P |
| ICA1 | 7 | islet cell autoantigen 1 [Source:HGNC Symbol; |
| AC006042.1 | 7 | novel transcript |
| AC007009.1 | 7 | novel transcript, antisense to ICA1 |
| AC007128.1 | 7 | novel transcript, antisense to ICA1 |
| AC007128.2 | 7 | novel transcript, antisense to ICA1 |
| NXPH1 | 7 | neurexophilin 1 [Source:HGNC Symbol;Acc:HG |
| AC009500.1 | 7 | ribosomal protein L9 (RPL9) pseudogene |
| AC004852.2 | 7 | novel transcript |
| AC004852.1 | 7 | LYR motif containing 9 (LYRM9) pseudogene |
| AC004852.3 | 7 | ret finger protein-like 4A (RFPL4A) pseudogen |
| GAPDHP68 | 7 | glyceraldehyde 3 phosphate dehydrogenase p |
| AC060834.2 | 7 | novel transcript |
| AC096553.1 | 7 | period circadian clock 3, pseudogene |
| AC060834.1 | 7 | C3 and PZP-like, alpha-2-macroglobulin domai |
| AC006363.1 | 7 | eukaryotic translation elongation factor 1 alph |
| AC010991.1 | 7 | novel transcript |
| AC004936.1 | 7 | novel transcript |
| AC004879.1 | 7 | novel transcript |
| U3 | 7 | Small nucleolar RNA U3 [Source:RFAM;Acc:RF |
| AC004949.1 | 7 | novel transcript |
| HSPA8P8 | 7 | heat shock protein family A (Hsp70) member 8 |
| AC009945.1 | 7 | CDK2-associated protein 2 (CDK2AP2) pseudo |
| AC004415.1 | 7 | novel transcript |
| Y_RNA | 7 | Y RNA [Source:RFAM;Acc:RF00019] |
| NDUFA4 | 7 | NDUFA4 mitochondrial complex associated [S |
| AC007029.1 | 7 | novel transcript |
| PHF14 | 7 | PHD finger protein 14 [Source:HGNC Symbol;/ |
| RPL23AP52 | 7 | ribosomal protein L23a pseudogene 52 [Sourc |
| AC004160.1 | 7 | novel transcript, antisense to THSD7A |
| NPM1P11 | 7 | nucleophosmin 1 pseudogene 11 [Source:HGN |
| THSD7A | 7 | thrombospondin type 1 domain containing 7A |
| AC004160.2 | 7 | novel transcript |
| AC073109.1 | 7 | BCL2-associated transcription factor 1 (BCLAF |
| TMEM106B | 7 | transmembrane protein 106B [Source:HGNC S |
| VWDE | 7 | von Willebrand factor D and EGF domains [So |
| AC013470.4 | 7 | CTAGE family, member 5 (CTAGE5) pseudoger |
| AC013470.2 | 7 | novel protein |
| AC013470.3 | 7 | Sjogren syndrome antigen B (autoantigen La) |

| | | |
|---|---|---|
| TAS2R2P | 7 | taste 2 receptor member 2, pseudogene [Sour |
| AC013470.1 | 7 | PC4 and SFRS1 interacting protein 1 (PSIP1) ps |
| AC005281.1 | 7 | novel transcript |
| SCIN | 7 | scinderin [Source:HGNC Symbol;Acc:HGNC:21 |
| AC011891.2 | 7 | novel transcript |
| AC011891.3 | 7 | apoptosis-inducing, TAF9-like domain 1, pseu |
| ARL4A | 7 | ADP ribosylation factor like GTPase 4A [Source |
| AC011891.1 | 7 | NULL |
| AC011287.1 | 7 | novel transcript |
| RN7SKP228 | 7 | RN7SK pseudogene 228 [Source:HGNC Symbo |
| RBMX2P4 | 7 | RBMX2 pseudogene 4 [Source:HGNC Symbol; |
| AC011287.2 | 7 | novel transcript |
| AC005019.2 | 7 | novel transcript |
| AC005019.1 | 7 | ribosomal protein L26 (RPL26) pseudogene |
| ETV1 | 7 | ETS variant 1 [Source:HGNC Symbol;Acc:HGN |
| Y_RNA | 7 | Y RNA [Source:RFAM;Acc:RF00019] |
| RPL6P21 | 7 | ribosomal protein L6 pseudogene 21 [Source: |
| DGKB | 7 | diacylglycerol kinase beta [Source:HGNC Symb |
| EEF1A1P26 | 7 | eukaryotic translation elongation factor 1 alp |
| AC006150.1 | 7 | novel transcript |
| GTF3AP5 | 7 | general transcription factor IIIA pseudogene 5 |
| AC006458.1 | 7 | novel transcript |
| AGMO | 7 | alkylglycerol monooxygenase [Source:HGNC S |
| MEOX2 | 7 | mesenchyme homeobox 2 [Source:HGNC Sym |
| AC005550.2 | 7 | novel transcript, antisense to MEOX2 |
| LINC02587 | 7 | long intergenic non-protein coding RNA 2587 |
| AC005550.1 | 7 | novel transcript |
| RPL36AP26 | 7 | ribosomal protein L36a pseudogene 26 [Sourc |
| AC006041.2 | 7 | novel transcript |
| AC006041.1 | 7 | novel transcript |
| CRPPA | 7 | CDP-L-ribitol pyrophosphorylase A [Source:H |
| RPL36AP29 | 7 | ribosomal protein L36a pseudogene 29 [Sourc |
| CRPPA-AS1 | 7 | CRPPA antisense RNA 1 [Source:HGNC Symbo |
| SOSTDC1 | 7 | sclerostin domain containing 1 [Source:HGNC |
| AC005014.3 | 7 | novel transcript |
| LRRC72 | 7 | leucine rich repeat containing 72 [Source:HGN |
| AC005014.1 | 7 | ataxin 2-like (ATXN2L) pseudogene |
| AC005014.2 | 7 | novel transcript |
| ANKMY2 | 7 | ankyrin repeat and MYND domain containing |
| AC005014.4 | 7 | novel transcript, antisense to ANKMY2 |
| BZW2 | 7 | basic leucine zipper and W2 domains 2 [Sourc |
| AC073333.1 | 7 | novel transcript |
| TSPAN13 | 7 | tetraspanin 13 [Source:HGNC Symbol;Acc:HG |
| AGR2 | 7 | anterior gradient 2, protein disulphide isomer |
| AC073333.2 | 7 | Rho GTPase activating protein 20 (ARHGAP20 |
| AGR3 | 7 | anterior gradient 3, protein disulphide isomer |
| RAD17P1 | 7 | RAD17 pseudogene 1 [Source:HGNC Symbol; |
| AC098592.2 | 7 | LSM5 homolog, U6 small nuclear RNA associa |
| AHR | 7 | aryl hydrocarbon receptor [Source:HGNC Sym |
| AC098592.1 | 7 | pseudogene similar to part of serine/threonin |

| | | |
|---|---|---|
| AC073332.1 | 7 | novel transcript |
| BRWD1P3 | 7 | bromodomain and WD repeat domain contair |
| Metazoa_SRP | 7 | Metazoan signal recognition particle RNA [Sou |
| AC019117.3 | 7 | novel protein |
| SNORA63 | 7 | Small nucleolar RNA SNORA63 [Source:RFAM; |
| AC019117.2 | 7 | renal clear cell carcinoma-associated transcrip |
| AC019117.1 | 7 | novel transcript |
| AC006482.1 | 7 | novel transcript |
| SNX13 | 7 | sorting nexin 13 [Source:HGNC Symbol;Acc:H( |
| AC080080.1 | 7 | tec |
| RNMTL1P2 | 7 | RNA methyltransferase like 1 pseudogene 2 [S |
| AC080080.2 | 7 | tec |
| PRPS1L1 | 7 | phosphoribosyl pyrophosphate synthetase 1 l |
| HDAC9 | 7 | histone deacetylase 9 [Source:HGNC Symbol;A |
| MIR1302-6 | 7 | microRNA 1302-6 [Source:HGNC Symbol;Acc:I |
| AC010082.1 | 7 | novel transcript |
| RN7SKP266 | 7 | RN7SK pseudogene 266 [Source:HGNC Symbo |
| AC004994.1 | 7 | novel transcript |
| NPM1P13 | 7 | nucleophosmin 1 pseudogene 13 [Source:HGI |
| TWIST1 | 7 | twist family bHLH transcription factor 1 [Sourc |
| AC003986.2 | 7 | novel transcript |
| AC003986.3 | 7 | novel transcript |
| AC003986.1 | 7 | novel transcript, antisense to FERD3L |
| FERD3L | 7 | Fer3 like bHLH transcription factor [Source:H( |
| AC007091.1 | 7 | novel transcript |
| TWISTNB | 7 | TWIST neighbor [Source:HGNC Symbol;Acc:H( |
| MIR3146 | 7 | microRNA 3146 [Source:HGNC Symbol;Acc:H( |
| TMEM196 | 7 | transmembrane protein 196 [Source:HGNC Sy |
| AC004543.1 | 7 | novel transcript |
| AC005062.1 | 7 | novel transcript, sense overlapping to MACC1 |
| RPL21P75 | 7 | ribosomal protein L21 pseudogene 75 [Source |
| AC007001.1 | 7 | novel transcript |
| MACC1 | 7 | MET transcriptional regulator MACC1 [Source |
| MACC1-AS1 | 7 | MACC1 antisense RNA 1 [Source:HGNC Symbo |
| AC005083.1 | 7 | novel transcript |
| AC099342.1 | 7 | novel transcript |
| AC004130.2 | 7 | novel transcript, antisense to ITGB8 |
| ITGB8 | 7 | integrin subunit beta 8 [Source:HGNC Symbol; |
| AC004130.1 | 7 | NULL |
| EEF1A1P27 | 7 | eukaryotic translation elongation factor 1 alpl |
| AC002486.2 | 7 | v-ral simian leukemia viral oncogene homolog |
| ABCB5 | 7 | ATP binding cassette subfamily B member 5 [S |
| AC002486.1 | 7 | pseudogene similar to part of pituitary tumor- |
| SP8 | 7 | Sp8 transcription factor [Source:HGNC Symbo |
| RPL23P8 | 7 | ribosomal protein L23 pseudogene 8 [Source: |
| LINC01162 | 7 | long intergenic non-protein coding RNA 1162 |
| RPS26P30 | 7 | ribosomal protein S26 pseudogene 30 [Source |
| AC080068.1 | 7 | novel transcript |
| RN7SL542P | 7 | RNA, 7SL, cytoplasmic 542, pseudogene [Sour |
| ASS1P11 | 7 | argininosuccinate synthetase 1 pseudogene 1 |

| | | |
|---|---|---|
| RNU1-15P | 7 | RNA, U1 small nuclear 15, pseudogene [Sourc |
| AC005094.1 | 7 | ribosomal protein L31 (RPL31) pseudogene |
| SP4 | 7 | Sp4 transcription factor [Source:HGNC Symbo |
| MIR1183 | 7 | microRNA 1183 [Source:HGNC Symbol;Acc:HG |
| DNAH11 | 7 | dynein axonemal heavy chain 11 [Source:HGN |
| CDCA7L | 7 | cell division cycle associated 7 like [Source:HG |
| RAPGEF5 | 7 | Rap guanine nucleotide exchange factor 5 [So |
| RNA5SP227 | 7 | RNA, 5S ribosomal pseudogene 227 [Source:H |
| STEAP1B | 7 | STEAP family member 1B [Source:HGNC Symb |
| EEF1A1P6 | 7 | eukaryotic translation elongation factor 1 alph |
| AC002480.1 | 7 | novel transcript, antisense to STEAP1B |
| AC002480.2 | 7 | novel transcript, antisense to STEAP1B |
| AC073072.1 | 7 | novel transcript |
| IL6 | 7 | interleukin 6 [Source:HGNC Symbol;Acc:HGNC |
| MTCYBP42 | 7 | MT-CYB pseudogene 42 [Source:HGNC Symbo |
| AC073072.2 | 7 | ribosomal protein S26 (RPS26) pseudogene |
| TOMM7 | 7 | translocase of outer mitochondrial membrane |
| AC005682.2 | 7 | novel transcript |
| SNHG26 | 7 | small nucleolar RNA host gene 26 [Source:HGI |
| SNORD93 | 7 | small nucleolar RNA, C/D box 93 [Source:HGN |
| AC005682.1 | 7 | solute carrier family 40 (iron-regulated transp |
| RPL12P10 | 7 | ribosomal protein L12 pseudogene 10 [Source |
| FAM126A | 7 | family with sequence similarity 126 member A |
| KLHL7-DT | 7 | KLHL7 divergent transcript [Source:HGNC Sym |
| KLHL7 | 7 | kelch like family member 7 [Source:HGNC Sym |
| AK3P3 | 7 | adenylate kinase 3 pseudogene 3 [Source:HGI |
| NUP42 | 7 | nucleoporin 42 [Source:HGNC Symbol;Acc:HG |
| AC005082.2 | 7 | Novel transcript, antisense to NUPL2 |
| AC005082.1 | 7 | novel transcript |
| GPNMB | 7 | glycoprotein nmb [Source:HGNC Symbol;Acc:I |
| MALSU1 | 7 | mitochondrial assembly of ribosomal large sul |
| IGF2BP3 | 7 | insulin like growth factor 2 mRNA binding pro |
| AC021876.1 | 7 | ribosomal protein L31 (RPL31) pseudogene |
| SNORD65C | 7 | small nucleolar RNA, C/D box 65C [Source:HG |
| RNU7-143P | 7 | RNA, U7 small nuclear 143 pseudogene [Sourc |
| AC079780.1 | 7 | chromosome 5 open reading frame 13 (C5orf1 |
| RPS2P32 | 7 | ribosomal protein S2 pseudogene 32 [Source: |
| Y_RNA | 7 | Y RNA [Source:RFAM;Acc:RF00019] |
| TRA2A | 7 | transformer 2 alpha homolog [Source:HGNC S |
| AC006026.1 | 7 | ATP-binding cassette, sub-family E (OABP), me |
| CLK2P1 | 7 | CDC like kinase 2, pseudogene 1 [Source:HGN |
| CCDC126 | 7 | coiled-coil domain containing 126 [Source:HG |
| FCF1P1 | 7 | FCF1 pseudogene 1 [Source:HGNC Symbol;Acc |
| AC006026.2 | 7 | riboflavin kinase (RFK) pseudogene |
| FAM221A | 7 | family with sequence similarity 221 member A |
| AC006026.3 | 7 | novel transcript |
| STK31 | 7 | serine/threonine kinase 31 [Source:HGNC Sym |
| PCMTD1P3 | 7 | protein-L-isoaspartate (D-aspartate) O-methyl |
| TPT1P7 | 7 | tumor protein, translationally-controlled 1 pse |
| AC003087.1 | 7 | novel transcript |

| | | |
|---|---|---|
| RNA5SP228 | 7 | RNA, 5S ribosomal pseudogene 228 [Source:H |
| AC004485.1 | 7 | novel transcript |
| NPY | 7 | neuropeptide Y [Source:HGNC Symbol;Acc:HG |
| AC003044.1 | 7 | novel transcript |
| RNU6-1103P | 7 | RNA, U6 small nuclear 1103, pseudogene [Sou |
| MPP6 | 7 | membrane palmitoylated protein 6 [Source:H |
| SUMO2P14 | 7 | SUMO2 pseudogene 14 [Source:HGNC Symbo |
| AC005084.1 | 7 | signal recognition particle 14kDa (homologous |
| GSDME | 7 | gasdermin E [Source:HGNC Symbol;Acc:HGNC |
| AC003093.1 | 7 | novel transcript |
| OSBPL3 | 7 | oxysterol binding protein like 3 [Source:HGNC |
| SNRPCP19 | 7 | small nuclear ribonucleoprotein polypeptide C |
| CYCS | 7 | cytochrome c, somatic [Source:HGNC Symbol; |
| C7orf31 | 7 | chromosome 7 open reading frame 31 [Source |
| AC004129.2 | 7 | ubiquitin specific peptidase 22 (USP22) pseud |
| AC004129.1 | 7 | ribosomal protein L7a (RPL7A) pseudogene |
| NPVF | 7 | neuropeptide VF precursor [Source:HGNC Syn |
| AC004129.3 | 7 | novel transcript |
| U3 | 7 | Small nucleolar RNA U3 [Source:RFAM;Acc:RF |
| TSEN15P3 | 7 | tRNA splicing endonuclease subunit 15 pseudo |
| AC003985.1 | 7 | novel transcript |
| AC005100.1 | 7 | novel transcript |
| AC005100.2 | 7 | novel transcript |
| AC091705.1 | 7 | novel transcript |
| AC005165.1 | 7 | novel transcript |
| AC005165.2 | 7 | ubiquitin A-52 residue ribosomal protein fusio |
| AC005165.3 | 7 | novel transcript |
| AC018706.1 | 7 | novel transcript |
| AC010719.1 | 7 | novel transcript |
| MIR148A | 7 | microRNA 148a [Source:HGNC Symbol;Acc:HG |
| NFE2L3 | 7 | nuclear factor, erythroid 2 like 3 [Source:HGN |
| HNRNPA2B1 | 7 | heterogeneous nuclear ribonucleoprotein A2/ |
| CBX3 | 7 | chromobox 3 [Source:HGNC Symbol;Acc:HGN |
| AC010677.2 | 7 | novel transcript |
| AC010677.1 | 7 | ribosomal protein L23 (RPL23) pseudogene |
| SNX10 | 7 | sorting nexin 10 [Source:HGNC Symbol;Acc:H |
| AC004540.2 | 7 | novel transcript |
| AC004540.1 | 7 | novel transcript |
| KIAA0087 | 7 | KIAA0087 lncRNA [Source:HGNC Symbol;Acc:H |
| AC004947.2 | 7 | tec |
| AC004947.1 | 7 | novel transcript |
| AC004947.3 | 7 | novel transcript |
| LINC02860 | 7 | long intergenic non-protein coding RNA 2860 |
| SKAP2 | 7 | src kinase associated phosphoprotein 2 [Sourc |
| RPL7AP38 | 7 | ribosomal protein L7a pseudogene 38 [Source |
| HMGB3P20 | 7 | high mobility group box 3 pseudogene 20 [Sou |
| TPM3P4 | 7 | tropomyosin 3 pseudogene 4 [Source:HGNC S |
| NHP2P2 | 7 | NHP2 ribonucleoprotein pseudogene 2 [Sourc |
| HOXA1 | 7 | homeobox A1 [Source:HGNC Symbol;Acc:HGN |
| HOTAIRM1 | 7 | HOXA transcript antisense RNA, myeloid-spec |

| | | |
|---|---|---|
| AC004079.3 | 7 | NULL |
| AC004079.4 | 7 | NULL |
| AC004079.2 | 7 | NULL |
| AC004079.1 | 7 | NULL |
| AC004079.5 | 7 | NULL |
| HOXA2 | 7 | homeobox A2 [Source:HGNC Symbol;Acc:HGN |
| HOXA3 | 7 | homeobox A3 [Source:HGNC Symbol;Acc:HGN |
| HOXA-AS2 | 7 | HOXA cluster antisense RNA 2 [Source:HGNC S |
| HOXA4 | 7 | homeobox A4 [Source:HGNC Symbol;Acc:HGN |
| AC004080.6 | 7 | novel transcript |
| HOXA-AS3 | 7 | HOXA cluster antisense RNA 3 [Source:HGNC S |
| HOXA5 | 7 | homeobox A5 [Source:HGNC Symbol;Acc:HGN |
| HOXA6 | 7 | homeobox A6 [Source:HGNC Symbol;Acc:HGN |
| AC004080.5 | 7 | novel transcript |
| HOXA7 | 7 | homeobox A7 [Source:HGNC Symbol;Acc:HGN |
| AC004080.4 | 7 | novel transcript |
| HOXA9 | 7 | homeobox A9 [Source:HGNC Symbol;Acc:HGN |
| AC004080.3 | 7 | HOXA10-HOXA9 readthrough |
| HOXA10-AS | 7 | HOXA10 antisense RNA [Source:HGNC Symbo |
| MIR196B | 7 | microRNA 196b [Source:HGNC Symbol;Acc:HG |
| HOXA10 | 7 | homeobox A10 [Source:HGNC Symbol;Acc:HG |
| HOXA11 | 7 | homeobox A11 [Source:HGNC Symbol;Acc:HG |
| HOXA11-AS | 7 | HOXA11 antisense RNA [Source:HGNC Symbo |
| AC004080.7 | 7 | NULL |
| AC004080.14 | 7 | NULL |
| AC004080.8 | 7 | NULL |
| AC004080.2 | 7 | novel transcript |
| AC004080.12 | 7 | NULL |
| AC004080.15 | 7 | NULL |
| AC004080.13 | 7 | NULL |
| HOXA13 | 7 | homeobox A13 [Source:HGNC Symbol;Acc:HG |
| HOTTIP | 7 | HOXA distal transcript antisense RNA [Source: |
| AC004080.9 | 7 | NULL |
| AC004080.16 | 7 | NULL |
| AC004080.10 | 7 | NULL |
| AC004080.11 | 7 | NULL |
| AC004080.17 | 7 | novel transcript |
| AC004080.1 | 7 | novel transcript |
| EVX1-AS | 7 | EVX1 antisense RNA [Source:HGNC Symbol;Ac |
| EVX1 | 7 | even-skipped homeobox 1 [Source:HGNC Sym |
| RPL35P4 | 7 | ribosomal protein L35 pseudogene 4 [Source: |
| HNRNPA1P73 | 7 | heterogeneous nuclear ribonucleoprotein A1 |
| AC004009.1 | 7 | novel transcript |
| AC004009.3 | 7 | TEC |
| AC004009.2 | 7 | TEC |
| EIF4HP1 | 7 | eukaryotic translation initiation factor 4H pse |
| PSMC1P2 | 7 | proteasome 26S subunit, ATPase 1 pseudoger |
| AC073150.1 | 7 | coiled-coil domain containing 86 (CCDC86) ps |
| HIBADH | 7 | 3-hydroxyisobutyrate dehydrogenase [Source |
| AC007130.1 | 7 | novel transcript, antisense to HIBADH |

| | | |
|---|---|---|
| AC005091.1 | 7 | novel transcript, antisense to TAX1BP1 |
| TAX1BP1 | 7 | Tax1 binding protein 1 [Source:HGNC Symbol; |
| JAZF1 | 7 | JAZF zinc finger 1 [Source:HGNC Symbol;Acc:H |
| AC004549.1 | 7 | novel transcript, antisense to JAZF1 |
| RNU6-979P | 7 | RNA, U6 small nuclear 979, pseudogene [Sour |
| JAZF1-AS1 | 7 | JAZF1 antisense RNA 1 [Source:HGNC Symbol; |
| PPIAP80 | 7 | peptidylprolyl isomerase A pseudogene 80 [Sc |
| CREB5 | 7 | cAMP responsive element binding protein 5 [S |
| AC005105.1 | 7 | ribosomal protein S2 (RPS2) pseudogene |
| TRIL | 7 | TLR4 interactor with leucine rich repeats [Sou |
| AC005013.1 | 7 | novel transcript |
| AC005162.3 | 7 | novel transcript, antisense to CPVL |
| AC005162.2 | 7 | novel transcript |
| CPVL | 7 | carboxypeptidase vitellogenic like [Source:HG |
| AC005162.1 | 7 | uncharacterized LOC107986703 [Source:NCBI |
| AC005232.1 | 7 | novel transcript, antisense to CPVL |
| AC004593.3 | 7 | chimerin 2 [Source:NCBI gene;Acc:1124] |
| CHN2 | 7 | chimerin 2 [Source:HGNC Symbol;Acc:HGNC:1 |
| AC004593.2 | 7 | Novel transcript, sense intronic to CPVL |
| AC004593.1 | 7 | novel transcript |
| NANOGP4 | 7 | Nanog homeobox pseudogene 4 [Source:HGN |
| AC007255.1 | 7 | novel transcript, antisense to CHN2 |
| AC007255.2 | 7 | general transcription factor IIIC, polypeptide 6 |
| PRR15 | 7 | proline rich 15 [Source:HGNC Symbol;Acc:HGI |
| ZNRF2P2 | 7 | zinc and ring finger 2 pseudogene 2 [Source:H |
| TMSB4XP3 | 7 | TMSB4X pseudogene 3 [Source:HGNC Symbol |
| DPY19L2P3 | 7 | DPY19L2 pseudogene 3 [Source:HGNC Symbo |
| MIR550A3 | 7 | microRNA 550a-3 [Source:HGNC Symbol;Acc:I |
| WIPF3 | 7 | WAS/WASL interacting protein family membe |
| AC004912.1 | 7 | NULL |
| SCRN1 | 7 | secernin 1 [Source:HGNC Symbol;Acc:HGNC:2 |
| AC007285.1 | 7 | novel transcript |
| FKBP14 | 7 | FKBP prolyl isomerase 14 [Source:HGNC Symb |
| PLEKHA8 | 7 | pleckstrin homology domain containing A8 [Sc |
| AC007285.2 | 7 | novel transcript |
| AC007036.4 | 7 | NULL |
| AC007036.5 | 7 | novel transcript, antisense to PLEKHA8 |
| MTURN | 7 | maturin, neural progenitor differentiation reg |
| AC007036.2 | 7 | novel transcript |
| AC007036.3 | 7 | novel transcript |
| RPS27P16 | 7 | ribosomal protein S27 pseudogene 16 [Source |
| AC007036.1 | 7 | novel transcript |
| ZNRF2 | 7 | zinc and ring finger 2 [Source:HGNC Symbol;A |
| MIR550B1 | 7 | microRNA 550b-1 [Source:HGNC Symbol;Acc:I |
| MIR550A1 | 7 | microRNA 550a-1 [Source:HGNC Symbol;Acc:I |
| AC006978.2 | 7 | novel protein, ZNRF2-LINC01176 readthrough |
| AC006978.1 | 7 | proteasome (prosome, macropain) 26S subun |
| LINC01176 | 7 | long intergenic non-protein coding RNA 1176 |
| NOD1 | 7 | nucleotide binding oligomerization domain co |
| AC006027.1 | 7 | novel transcript, antisense to NOD1 |

| | | |
|---|---|---|
| GGCT | 7 | gamma-glutamylcyclotransferase [Source:HGI |
| AC005154.5 | 7 | novel protein |
| GARS-DT | 7 | GARS divergent transcript [Source:HGNC Syml |
| AC005154.2 | 7 | pseudogene similar golgi autoantigen, golgin s |
| AC005154.4 | 7 | novel transcript |
| AC005154.1 | 7 | pseudogene similar to part of transient recept |
| AC005154.3 | 7 | novel transcript |
| GARS | 7 | glycyl-tRNA synthetase [Source:HGNC Symbol |
| CRHR2 | 7 | corticotropin releasing hormone receptor 2 [S |
| INMT | 7 | indolethylamine N-methyltransferase [Source |
| AC006022.1 | 7 | indolethylamine N-methyltransferase (INMT) |
| INMT-MINDY4 | 7 | INMT-MINDY4 readthrough (NMD candidate) |
| MINDY4 | 7 | MINDY lysine 48 deubiquitinase 4 [Source:HG |
| AC004691.1 | 7 | novel transcript |
| AC004691.2 | 7 | novel protein, MINDY4 and AQP1 readthrougł |
| AQP1 | 7 | aquaporin 1 (Colton blood group) [Source:HGI |
| GHRHR | 7 | growth hormone releasing hormone receptor |
| ADCYAP1R1 | 7 | ADCYAP receptor type I [Source:HGNC Symbo |
| AC006466.1 | 7 | novel pseudogene |
| AC006398.1 | 7 | novel transcript |
| AC006380.1 | 7 | guanine monphosphate synthetase, pseudoge |
| NEUROD6 | 7 | neuronal differentiation 6 [Source:HGNC Syml |
| AC005090.1 | 7 | novel transcript |
| ITPRID1 | 7 | ITPR interacting domain containing 1 [Source: |
| PPP1R17 | 7 | protein phosphatase 1 regulatory subunit 17 [ |
| PDE1C | 7 | phosphodiesterase 1C [Source:HGNC Symbol; |
| SNX2P2 | 7 | sorting nexin 2 pseudogene 2 [Source:HGNC S |
| AC018637.1 | 7 | novel transcript |
| AC018641.1 | 7 | phosphoribosyl pyrophosphate synthetase 2 ( |
| SLC25A5P5 | 7 | solute carrier family 25 member 5 pseudogen |
| LSM5 | 7 | LSM5 homolog, U6 small nuclear RNA and mR |
| AVL9 | 7 | AVL9 cell migration associated [Source:HGNC |
| DPY19L1P1 | 7 | DPY19L1 pseudogene 1 [Source:HGNC Symbo |
| ZNRF2P1 | 7 | zinc and ring finger 2 pseudogene 1 [Source:H |
| MIR550A2 | 7 | microRNA 550a-2 [Source:HGNC Symbol;Acc:H |
| MIR550B2 | 7 | microRNA 550b-2 [Source:HGNC Symbol;Acc:I |
| AC018645.3 | 7 | novel transcript |
| LINC00997 | 7 | long intergenic non-protein coding RNA 997 [S |
| AC018645.1 | 7 | neuropeptide S receptor 1 (NPSR1) pseudoge |
| AC018645.2 | 7 | NULL |
| DPY19L1P2 | 7 | DPY19L1 pseudogene 2 [Source:HGNC Symbo |
| AC018648.1 | 7 | novel transcript |
| KBTBD2 | 7 | kelch repeat and BTB domain containing 2 [So |
| RP9P | 7 | RP9 pseudogene [Source:HGNC Symbol;Acc:H |
| AC083863.1 | 7 | pseudogene similar to part of ribosomal prote |
| FKBP9 | 7 | FKBP prolyl isomerase 9 [Source:HGNC Symbo |
| RNU6-388P | 7 | RNA, U6 small nuclear 388, pseudogene [Sour |
| NT5C3A | 7 | 5'-nucleotidase, cytosolic IIIA [Source:HGNC S |
| RPS29P14 | 7 | ribosomal protein S29 pseudogene 14 [Source |
| RN7SL505P | 7 | RNA, 7SL, cytoplasmic 505, pseudogene [Sour |

| | | |
|---|---|---|
| RP9 | 7 | RP9 pre-mRNA splicing factor [Source:HGNC S |
| BBS9 | 7 | Bardet-Biedl syndrome 9 [Source:HGNC Symb |
| AC074338.1 | 7 | pseudogene similar to part of 5'-nucleotidase, |
| RNA5SP229 | 7 | RNA, 5S ribosomal pseudogene 229 [Source:H |
| AC007312.1 | 7 | pseudogene similar to part of ubiquitin specifi |
| AC006195.1 | 7 | RAB3A interacting protein (rabin3)-like 1 (RAB |
| AC008080.1 | 7 | novel transcript |
| AC008080.2 | 7 | novel transcript |
| AC008080.4 | 7 | novel transcript |
| AC008080.3 | 7 | novel transcript |
| BMPER | 7 | BMP binding endothelial regulator [Source:HG |
| AC007350.1 | 7 | novel transcript, antisense to BMPER |
| AC009262.1 | 7 | novel transcript |
| RNU6-438P | 7 | RNA, U6 small nuclear 438, pseudogene [Sour |
| NPSR1-AS1 | 7 | NPSR1 antisense RNA 1 [Source:HGNC Symbo |
| NPSR1 | 7 | neuropeptide S receptor 1 [Source:HGNC Sym |
| RPL7P31 | 7 | ribosomal protein L7 pseudogene 31 [Source: |
| NCAPD2P1 | 7 | non-SMC condensin I complex subunit D2 pse |
| RN7SL132P | 7 | RNA, 7SL, cytoplasmic 132, pseudogene [Sour |
| AC004788.1 | 7 | NULL |
| DPY19L1 | 7 | dpy-19 like C-mannosyltransferase 1 [Source: |
| MIR548N | 7 | microRNA 548n [Source:HGNC Symbol;Acc:HG |
| AC005400.1 | 7 | novel transcript, antisense to DPY19L1 |
| DPY19L2P1 | 7 | DPY19L2 pseudogene 1 [Source:HGNC Symbo |
| S100A11P2 | 7 | S100 calcium binding protein A11 pseudogene |
| TBX20 | 7 | T-box transcription factor 20 [Source:HGNC Sy |
| AC009531.1 | 7 | novel transcript |
| AC007652.3 | 7 | novel transcript |
| AC007652.2 | 7 | novel transcript |
| AC007652.1 | 7 | novel transcript |
| AC018647.3 | 7 | novel transcript |
| HERPUD2 | 7 | HERPUD family member 2 [Source:HGNC Sym |
| AC018647.2 | 7 | novel transcript |
| AC018647.1 | 7 | novel transcript |
| SEPTIN7-AS1 | 7 | SEPT7 antisense RNA 1 (head to head) [Source |
| AC007551.1 | 7 | glycine cleavage system protein H (aminomet |
| SEPTIN7 | 7 | septin 7 [Source:HGNC Symbol;Acc:HGNC:171 |
| AC007551.2 | 7 | gem (nuclear organelle) associated protein 7 ( |
| RNU6-1085P | 7 | RNA, U6 small nuclear 1085, pseudogene [Sou |
| AC087072.2 | 7 | zinc finger protein 383 (ZNF383) pseudogene |
| SEPTIN7P3 | 7 | septin 7 pseudogene 3 [Source:HGNC Symbol; |
| AC087072.1 | 7 | protein phosphatase 1, regulatory (inhibitor) s |
| AC083864.5 | 7 | novel transcript |
| AC083864.2 | 7 | novel transcript |
| AC083864.4 | 7 | novel transcript |
| AC083864.1 | 7 | novel transcript |
| AC083864.3 | 7 | novel transcript |
| MARK2P13 | 7 | microtubule affinity regulating kinase 2 pseud |
| AC078841.1 | 7 | MAP/microtubule affinity-regulating kinase 2. |
| EEPD1 | 7 | endonuclease/exonuclease/phosphatase fami |

| | | |
|---|---|---|
| AC007327.1 | 7 | pseudogene similar to part of mitochondrially |
| AC007327.2 | 7 | novel transcript |
| AC006960.1 | 7 | ribosomal protein L39 (RPL39) pseudogene |
| AC006960.3 | 7 | novel transcript, antisense to KIAA0895 |
| KIAA0895 | 7 | KIAA0895 [Source:HGNC Symbol;Acc:HGNC:2 |
| Y_RNA | 7 | Y RNA [Source:RFAM;Acc:RF00019] |
| ANLN | 7 | anillin actin binding protein [Source:HGNC Syr |
| AC006960.2 | 7 | novel transcript |
| AOAH | 7 | acyloxyacyl hydrolase [Source:HGNC Symbol;/ |
| AOAH-IT1 | 7 | AOAH intronic transcript 1 [Source:HGNC Sym |
| AC007349.4 | 7 | novel transcript |
| AC007349.1 | 7 | novel transcript |
| AC007349.3 | 7 | novel transcript |
| NPM1P18 | 7 | nucleophosmin 1 pseudogene 18 [Source:HGI |
| AC007349.2 | 7 | novel transcript |
| ELMO1 | 7 | engulfment and cell motility 1 [Source:HGNC |
| MIR1200 | 7 | microRNA 1200 [Source:HGNC Symbol;Acc:HC |
| ELMO1-AS1 | 7 | ELMO1 antisense RNA 1 [Source:HGNC Symbc |
| RPS17P13 | 7 | ribosomal protein S17 pseudogene 13 [Source |
| RNU6-565P | 7 | RNA, U6 small nuclear 565, pseudogene [Sour |
| RPS10P14 | 7 | ribosomal protein S10 pseudogene 14 [Source |
| AC078843.1 | 7 | NULL |
| Y_RNA | 7 | Y RNA [Source:RFAM;Acc:RF00019] |
| NECAP1P1 | 7 | NECAP endocytosis associated 1 pseudogene |
| GPR141 | 7 | G protein-coupled receptor 141 [Source:HGN( |
| EPDR1 | 7 | ependymin related 1 [Source:HGNC Symbol;A |
| GPR141BP | 7 | G protein-coupled receptor 141B, pseudogene |
| AC018634.1 | 7 | pseudogene similar to part of cell division cycl |
| NME8 | 7 | NME/NM23 family member 8 [Source:HGNC S |
| SFRP4 | 7 | secreted frizzled related protein 4 [Source:HG |
| STARD3NL | 7 | STARD3 N-terminal like [Source:HGNC Symbol |
| TRGC2 | 7 | T cell receptor gamma constant 2 [Source:HGI |
| TRGJ2 | 7 | T cell receptor gamma joining 2 [Source:HGN( |
| TRGJP2 | 7 | T cell receptor gamma joining P2 [Source:HGN |
| TRGC1 | 7 | T cell receptor gamma constant 1 [Source:HGI |
| TRGJ1 | 7 | T cell receptor gamma joining 1 [Source:HGN( |
| TRGJP | 7 | T cell receptor gamma joining P [Source:HGN( |
| TRGJP1 | 7 | T cell receptor gamma joining P1 [Source:HGN |
| TRGV11 | 7 | T cell receptor gamma variable 11 (non-functi |
| TRGVB | 7 | T cell receptor gamma variable B (pseudogene |
| TRGV10 | 7 | T cell receptor gamma variable 10 (non-functi |
| AC006033.1 | 7 | coiled-coil-helix-coiled-coil-helix domain conta |
| TRGV9 | 7 | T cell receptor gamma variable 9 [Source:HGN |
| TRGVA | 7 | T cell receptor gamma variable A (pseudogene |
| AC006033.2 | 7 | novel transcript |
| TRGV8 | 7 | T cell receptor gamma variable 8 [Source:HGN |
| TRGV7 | 7 | T cell receptor gamma variable 7 (pseudogene |
| TRGV6 | 7 | T cell receptor gamma variable 6 (pseudogene |
| TRG-AS1 | 7 | T cell receptor gamma locus antisense RNA 1 |
| TRGV5P | 7 | T cell receptor gamma variable 5P (pseudoger |

| | | |
|---|---|---|
| TRGV5 | 7 | T cell receptor gamma variable 5 [Source:HGN |
| TRGV4 | 7 | T cell receptor gamma variable 4 [Source:HGN |
| TRGV3 | 7 | T cell receptor gamma variable 3 [Source:HGN |
| TRGV2 | 7 | T cell receptor gamma variable 2 [Source:HGN |
| TRGV1 | 7 | T cell receptor gamma variable 1 (non-functio |
| AMPH | 7 | amphiphysin [Source:HGNC Symbol;Acc:HGN( |
| RN7SL83P | 7 | RNA, 7SL, cytoplasmic 83, pseudogene [Sourc |
| KRT8P20 | 7 | keratin 8 pseudogene 20 [Source:HGNC Symb |
| FAM183BP | 7 | family with sequence similarity 183 member E |
| VPS41 | 7 | VPS41 subunit of HOPS complex [Source:HGN |
| POU6F2 | 7 | POU class 6 homeobox 2 [Source:HGNC Symb |
| POU6F2-AS2 | 7 | POU6F2 antisense RNA 2 [Source:HGNC Symb |
| SNORA20B | 7 | small nucleolar RNA, H/ACA box 20B [Source:I |
| POU6F2-AS1 | 7 | POU6F2 antisense RNA 1 [Source:HGNC Symb |
| AC011290.1 | 7 | novel transcript, antisense to POU6F2 |
| YAE1 | 7 | YAE1 maturation factor of ABCE1 [Source:HGI |
| AC011290.2 | 7 | thioredoxin domain containing 1 (TXNDC1) ps |
| AC004837.3 | 7 | novel transcript |
| AC004837.1 | 7 | akirin 1 (AKIRIN1) pseudogene |
| AC004837.4 | 7 | novel transcript |
| RALA | 7 | RAS like proto-oncogene A [Source:HGNC Sym |
| AC004837.2 | 7 | novel transcript |
| LINC00265 | 7 | long intergenic non-protein coding RNA 265 [S |
| RNU6-719P | 7 | RNA, U6 small nuclear 719, pseudogene [Sour |
| AC004987.4 | 7 | calmodulin 1 (phosphorylase kinase, delta) (C/ |
| CICP22 | 7 | capicua transcriptional repressor pseudogene |
| AC004987.3 | 7 | pseudogene similar to part of a member of AI |
| AC004987.1 | 7 | pseudogene similar to part of pleckstrin homc |
| AC004987.2 | 7 | single stranded DNA binding protein 3 (SSBP3) |
| RWDD4P2 | 7 | RWD domain containing 4 pseudogene 2 [Sou |
| RN7SL496P | 7 | RNA, 7SL, cytoplasmic 496, pseudogene [Sour |
| AC072061.1 | 7 | novel transcript, antisense to CDC2L5 |
| CDK13 | 7 | cyclin dependent kinase 13 [Source:HGNC Syn |
| AC006023.2 | 7 | methionyl aminopeptidase 2 (METAP2) pseud |
| AC006023.1 | 7 | HIG1 hypoxia inducible domain family, membe |
| MPLKIP | 7 | M-phase specific PLK1 interacting protein [Sou |
| Y_RNA | 7 | Y RNA [Source:RFAM;Acc:RF00019] |
| SUGCT | 7 | succinyl-CoA:glutarate-CoA transferase [Sourc |
| THUMPD3P1 | 7 | THUMP domain containing 3 pseudogene 1 [S |
| AC004988.1 | 7 | novel transcript |
| AC005160.1 | 7 | novel transcript |
| LINC01450 | 7 | long intergenic non-protein coding RNA 1450 |
| LINC01449 | 7 | long intergenic non-protein coding RNA 1449 |
| INHBA | 7 | inhibin subunit beta A [Source:HGNC Symbol;/ |
| INHBA-AS1 | 7 | INHBA antisense RNA 1 [Source:HGNC Symbol |
| GLI3 | 7 | GLI family zinc finger 3 [Source:HGNC Symbol; |
| HMGN2P30 | 7 | high mobility group nucleosomal binding dom |
| LINC01448 | 7 | long intergenic non-protein coding RNA 1448 |
| TCP1P1 | 7 | t-complex 1 pseudogene 1 [Source:HGNC Sym |
| AC010132.1 | 7 | novel transcript |

| | | |
|---|---|---|
| AC010132.4 | 7 | novel transcript |
| C7orf25 | 7 | chromosome 7 open reading frame 25 [Sourc |
| AC010132.3 | 7 | novel PSMA2 and C7orf25 readthrough |
| PSMA2 | 7 | proteasome subunit alpha 2 [Source:HGNC Sy |
| AC010132.2 | 7 | serine/threonine kinase receptor associated p |
| MRPL32 | 7 | mitochondrial ribosomal protein L32 [Source: |
| AC005537.1 | 7 | novel transcript |
| Y_RNA | 7 | Y RNA [Source:RFAM;Acc:RF00019] |
| HECW1 | 7 | HECT, C2 and WW domain containing E3 ubiq |
| HECW1-IT1 | 7 | HECW1 intronic transcript 1 [Source:HGNC Sy |
| MIR3943 | 7 | microRNA 3943 [Source:HGNC Symbol;Acc:HG |
| RNU7-35P | 7 | RNA, U7 small nuclear 35 pseudogene [Source |
| RNU6-575P | 7 | RNA, U6 small nuclear 575, pseudogene [Sour |
| AC004692.2 | 7 | novel transcript |
| AC004692.1 | 7 | ribosomal protein L18a (RPL18A) pseudogene |
| LUARIS | 7 | lncRNA upregulator of antiviral response inter |
| STK17A | 7 | serine/threonine kinase 17a [Source:HGNC Sy |
| COA1 | 7 | cytochrome c oxidase assembly factor 1 homo |
| BLVRA | 7 | biliverdin reductase A [Source:HGNC Symbol; |
| AC005189.1 | 7 | novel pseudogene similar to LOC203547 |
| URGCP-MRPS24 | 7 | URGCP-MRPS24 readthrough [Source:HGNC S |
| MRPS24 | 7 | mitochondrial ribosomal protein S24 [Source: |
| URGCP | 7 | upregulator of cell proliferation [Source:HGN |
| TUBG1P | 7 | tubulin gamma 1 pseudogene [Source:HGNC S |
| UBE2D4 | 7 | ubiquitin conjugating enzyme E2 D4 (putative |
| POLR2J4 | 7 | RNA polymerase II subunit J4, pseudogene [So |
| AC004951.4 | 7 | novel transcript, POLR2J4 pseudogene-RASA4 |
| AC004951.3 | 7 | RAS p21 protein activator 4 (RASA4) pseudoge |
| AC004951.2 | 7 | uroplakin-like protein (UPLP) pseudogene |
| SPDYE1 | 7 | speedy/RINGO cell cycle regulator family mem |
| AC004951.1 | 7 | novel transcript |
| AC017116.2 | 7 | novel transcript |
| POLR2J4 | 7 | RNA polymerase II subunit J4, pseudogene [So |
| RASA4CP | 7 | RAS p21 protein activator 4C, pseudogene [So |
| LINC00957 | 7 | long intergenic non-protein coding RNA 957 [S |
| DBNL | 7 | drebrin like [Source:HGNC Symbol;Acc:HGNC: |
| MIR6837 | 7 | microRNA 6837 [Source:HGNC Symbol;Acc:HG |
| PGAM2 | 7 | phosphoglycerate mutase 2 [Source:HGNC Sym |
| AC017116.1 | 7 | novel transcript, sense overlapping DBNL |
| POLM | 7 | DNA polymerase mu [Source:HGNC Symbol;Ac |
| MIR6838 | 7 | microRNA 6838 [Source:HGNC Symbol;Acc:HG |
| AEBP1 | 7 | AE binding protein 1 [Source:HGNC Symbol;Ac |
| MIR4649 | 7 | microRNA 4649 [Source:HGNC Symbol;Acc:HG |
| POLD2 | 7 | DNA polymerase delta 2, accessory subunit [S |
| RNA5SP230 | 7 | RNA, 5S ribosomal pseudogene 230 [Source:H |
| MYL7 | 7 | myosin light chain 7 [Source:HGNC Symbol;Ac |
| GCK | 7 | glucokinase [Source:HGNC Symbol;Acc:HGNC: |
| YKT6 | 7 | YKT6 v-SNARE homolog [Source:HGNC Symbo |
| CAMK2B | 7 | calcium/calmodulin dependent protein kinase |
| AC004453.2 | 7 | tec |

| | | |
|---|---|---|
| NUDCD3 | 7 | NudC domain containing 3 [Source:HGNC Sym |
| AC004453.1 | 7 | ribosomal protein L32 (RPL32) pseudogene |
| RNU6-1097P | 7 | RNA, U6 small nuclear 1097, pseudogene [Sou |
| AC004938.1 | 7 | ribosomal protein L36a (RPL36a) pseudogene |
| NPC1L1 | 7 | NPC1 like intracellular cholesterol transporter |
| DDX56 | 7 | DEAD-box helicase 56 [Source:HGNC Symbol;/ |
| TMED4 | 7 | transmembrane p24 trafficking protein 4 [Sou |
| AC004938.2 | 7 | novel transcript |
| AC011894.1 | 7 | ribosomal protein L18a, pseudogene |
| OGDH | 7 | oxoglutarate dehydrogenase [Source:HGNC Sy |
| ZMIZ2 | 7 | zinc finger MIZ-type containing 2 [Source:HGN |
| AC013436.1 | 7 | novel transcript |
| PPIA | 7 | peptidylprolyl isomerase A [Source:HGNC Sym |
| H2AFV | 7 | H2A histone family member V [Source:HGNC S |
| LINC01952 | 7 | long intergenic non-protein coding RNA 1952 |
| PURB | 7 | purine rich element binding protein B [Source |
| MIR4657 | 7 | microRNA 4657 [Source:HGNC Symbol;Acc:HG |
| AC004854.2 | 7 | novel transcript, antisense to PURB |
| AC004854.1 | 7 | mitochondrial ribosomal protein S23 (MRPS23 |
| AC004847.1 | 7 | novel transcript, overlapping MYO1G |
| MYO1G | 7 | myosin IG [Source:HGNC Symbol;Acc:HGNC:1: |
| SNHG15 | 7 | small nucleolar RNA host gene 15 [Source:HGI |
| SNORA9 | 7 | small nucleolar RNA, H/ACA box 9 [Source:HG |
| CCM2 | 7 | CCM2 scaffold protein [Source:HGNC Symbol; |
| NACAD | 7 | NAC alpha domain containing [Source:HGNC S |
| TBRG4 | 7 | transforming growth factor beta regulator 4 [S |
| SNORA5A | 7 | small nucleolar RNA, H/ACA box 5A [Source:H |
| SNORA5C | 7 | small nucleolar RNA, H/ACA box 5C [Source:H |
| SNORA5B | 7 | small nucleolar RNA, H/ACA box 5B [Source:H |
| RAMP3 | 7 | receptor activity modifying protein 3 [Source: |
| AC073968.2 | 7 | novel transcript |
| AC073968.1 | 7 | zinc finger protein 729 (ZNF729) pseudogene |
| ELK1P1 | 7 | ELK1 pseudogene 1 [Source:HGNC Symbol;Acc |
| AC073325.1 | 7 | novel transcript |
| AC073325.2 | 7 | novel ubiquitin-like ribosomal protein pseudo |
| ADCY1 | 7 | adenylate cyclase 1 [Source:HGNC Symbol;Acc |
| SEPTIN7P2 | 7 | septin 7 pseudogene 2 [Source:HGNC Symbol; |
| GTF2IP13 | 7 | general transcription factor IIi pseudogene 13 |
| RNU6-241P | 7 | RNA, U6 small nuclear 241, pseudogene [Sour |
| CICP20 | 7 | capicua transcriptional repressor pseudogene |
| AC096582.1 | 7 | chromosome 20 open reading frame 69 (C20c |
| AC096582.2 | 7 | novel pseudogene |
| RNU6-326P | 7 | RNA, U6 small nuclear 326, pseudogene [Sour |
| CCDC201 | 7 | coiled-coil domain containing 201 [Source:HG |
| IGFBP1 | 7 | insulin like growth factor binding protein 1 [Sc |
| IGFBP3 | 7 | insulin like growth factor binding protein 3 [Sc |
| AC073115.2 | 7 | novel transcript |
| AC073115.1 | 7 | novel transcript |
| FTLP15 | 7 | ferritin light chain pseudogene 15 [Source:HG |
| TTC4P1 | 7 | tetratricopeptide repeat domain 4 pseudogen |

| | | |
|---|---|---|
| ZNF619P1 | 7 | zinc finger protein 619 pseudogene 1 [Source: |
| AC023669.1 | 7 | novel transcript |
| AC023669.2 | 7 | novel transcript |
| AC004869.2 | 7 | novel transcript |
| AC004869.1 | 7 | novel transcript |
| HMGN1P19 | 7 | high mobility group nucleosome binding doma |
| AC011294.1 | 7 | novel transcript |
| EPS15P1 | 7 | epidermal growth factor receptor pathway su |
| AC004870.4 | 7 | novel transcript |
| AC004870.2 | 7 | novel transcript |
| AC004870.3 | 7 | novel transcript |
| MRPL42P4 | 7 | mitochondrial ribosomal protein L42 pseudog |
| AC004870.1 | 7 | splicing factor, arginine/serine-rich 2B (SFRS2I |
| AC087175.1 | 7 | novel transcript |
| TNS3 | 7 | tensin 3 [Source:HGNC Symbol;Acc:HGNC:216 |
| LINC01447 | 7 | long intergenic non-protein coding RNA 1447 |
| C7orf65 | 7 | chromosome 7 open reading frame 65 [Sourc |
| PKD1L1 | 7 | polycystin 1 like 1, transient receptor potentia |
| LINC00525 | 7 | long intergenic non-protein coding RNA 525 [S |
| C7orf69 | 7 | chromosome 7 open reading frame 69 [Sourc |
| AC069282.1 | 7 | pseudogene similar to part of TTK protein kina |
| HUS1 | 7 | HUS1 checkpoint clamp component [Source:H |
| SUN3 | 7 | Sad1 and UNC84 domain containing 3 [Source |
| C7orf57 | 7 | chromosome 7 open reading frame 57 [Sourc |
| UPP1 | 7 | uridine phosphorylase 1 [Source:HGNC Symbo |
| ABCA13 | 7 | ATP binding cassette subfamily A member 13 |
| AC091770.1 | 7 | novel transcript |
| LINC02838 | 7 | long intergenic non-protein coding RNA 2838 |
| AC004899.1 | 7 | novel pseudogene similar to CDC14 cell divisic |
| AC004899.2 | 7 | TEC |
| GDI2P1 | 7 | GDP dissociation inhibitor 2 pseudogene 1 [Sc |
| CDC14C | 7 | cell division cycle 14C [Source:HGNC Symbol;A |
| AC091730.1 | 7 | novel transcript |
| DDX43P2 | 7 | DEAD-box helicase 43 pseudogene 2 [Source:I |
| AC092448.1 | 7 | novel transcript |
| AC093775.1 | 7 | tec |
| VWC2 | 7 | von Willebrand factor C domain containing 2 [ |
| ZPBP | 7 | zona pellucida binding protein [Source:HGNC S |
| GNL2P1 | 7 | G protein nucleolar 2 pseudogene 1 [Source:H |
| AC034148.1 | 7 | novel transcript, antisense to ZPBP |
| SPATA48 | 7 | spermatogenesis associated 48 [Source:HGNC |
| AC020743.1 | 7 | novel transcript |
| AC020743.2 | 7 | novel transcript |
| AC020743.3 | 7 | novel transcript |
| IKZF1 | 7 | IKAROS family zinc finger 1 [Source:HGNC Sym |
| AC124014.1 | 7 | novel transcript, antisense to IKZF1 |
| RNU6-1091P | 7 | RNA, U6 small nuclear 1091, pseudogene [Sou |
| FIGNL1 | 7 | fidgetin like 1 [Source:HGNC Symbol;Acc:HGN |
| DDC | 7 | dopa decarboxylase [Source:HGNC Symbol;Ac |
| DDC-AS1 | 7 | DDC antisense RNA 1 [Source:HGNC Symbol;A |

| | | |
|---|---|---|
| GRB10 | 7 | growth factor receptor bound protein 10 [Sou |
| AC005153.1 | 7 | novel transcript, antisense to GRB10 |
| AC004920.1 | 7 | novel transcript |
| AC009296.1 | 7 | ribosomal protein L39 (RPL39) pseudogene |
| AC004830.2 | 7 | novel transcript, antisense to COBL |
| AC004830.1 | 7 | NULL |
| COBL | 7 | cordon-bleu WH2 repeat protein [Source:HGN |
| RPL7L1P2 | 7 | ribosomal protein L7 like 1 pseudogene 2 [Sou |
| CICP17 | 7 | capicua transcriptional repressor pseudogene |
| AC012441.2 | 7 | novel pseudogene |
| AC012441.1 | 7 | novel pseudogene |
| AC005999.2 | 7 | novel transcript |
| ROBO2P1 | 7 | roundabout guidance receptor 2 pseudogene |
| AC005999.1 | 7 | novel transcript |
| RN7SL292P | 7 | RNA, 7SL, cytoplasmic 292, pseudogene [Sour |
| AC006478.1 | 7 | novel transcript |
| AC006478.2 | 7 | novel transcript |
| AC079763.1 | 7 | novel transcript |
| AC006320.1 | 7 | NULL |
| AC006459.1 | 7 | novel transcript |
| SGO1P2 | 7 | shugoshin 1 pseudogene 2 [Source:HGNC Sym |
| AC074397.1 | 7 | pseudogene similar to part of ribosome bindir |
| POM121L12 | 7 | POM121 transmembrane nucleoporin like 12 |
| HAUS6P1 | 7 | HAUS augmin like complex subunit 6 pseudog |
| RNF138P2 | 7 | ring finger protein 138 pseudogene 2 [Source: |
| RNU1-14P | 7 | RNA, U1 small nuclear 14, pseudogene [Sourc |
| RN7SKP218 | 7 | RN7SK pseudogene 218 [Source:HGNC Symbo |
| AC009468.2 | 7 | novel transcript |
| AC009468.1 | 7 | novel transcript |
| LINC01446 | 7 | long intergenic non-protein coding RNA 1446 |
| RNU2-29P | 7 | RNA, U2 small nuclear 29, pseudogene [Sourc |
| RAC1P9 | 7 | Rac family small GTPase 1 pseudogene 9 [Sou |
| AC074348.1 | 7 | pseudogene similar to part of ataxia telangiec |
| HAUS6P3 | 7 | HAUS augmin like complex subunit 6 pseudog |
| LINC02854 | 7 | long intergenic non-protein coding RNA 2854 |
| AC092848.2 | 7 | novel transcript |
| AC092848.1 | 7 | human papillomavirus (type 18) E5 central sec |
| LINC01445 | 7 | long intergenic non-protein coding RNA 1445 |
| SLC25A5P3 | 7 | solute carrier family 25 member 5 pseudogen |
| VSTM2A | 7 | V-set and transmembrane domain containing |
| VSTM2A-OT1 | 7 | VSTM2A overlapping transcript 1 [Source:HGN |
| AC011228.1 | 7 | novel transcript |
| RNU6-1125P | 7 | RNA, U6 small nuclear 1125, pseudogene [Sou |
| RPL31P35 | 7 | ribosomal protein L31 pseudogene 35 [Source |
| AC011228.2 | 7 | novel transcript |
| SEC61G | 7 | SEC61 translocon gamma subunit [Source:HGI |
| AC074351.1 | 7 | novel transcript |
| AC006971.1 | 7 | Rho GTPase activating protein 5 (ARHGAP5) p |
| EGFR | 7 | epidermal growth factor receptor [Source:HG |
| EGFR-AS1 | 7 | EGFR antisense RNA 1 [Source:HGNC Symbol;/ |

| | | |
|---|---|---|
| ELDR | 7 | EGFR long non-coding downstream RNA [Sour |
| AC073324.1 | 7 | solute carrier family 19, member 3 (SLC19A3) |
| CALM1P2 | 7 | calmodulin 1 pseudogene 2 [Source:HGNC Syr |
| AC073347.1 | 7 | REX1, RNA exonuclease 1 homolog (S. cerevisi |
| LANCL2 | 7 | LanC like 2 [Source:HGNC Symbol;Acc:HGNC:6 |
| VOPP1 | 7 | VOPP1 WW domain binding protein [Source:H |
| AC099681.1 | 7 | novel transcript |
| AC099681.3 | 7 | novel transcript |
| AC099681.2 | 7 | novel transcript |
| CDC42P2 | 7 | cell division cycle 42 pseudogene 2 [Source:H |
| TUBBP6 | 7 | tubulin beta class I pseudogene 6 [Source:HGI |
| AC091812.1 | 7 | coiled-coil domain containing 144C (CCDC144 |
| FKBP9P1 | 7 | FKBP prolyl isomerase 9 pseudogene 1 [Sourc |
| RNU6-389P | 7 | RNA, U6 small nuclear 389, pseudogene [Sour |
| Metazoa_SRP | 7 | Metazoan signal recognition particle RNA [Sou |
| SUMO2P3 | 7 | SUMO2 pseudogene 3 [Source:HGNC Symbol; |
| CICP11 | 7 | capicua transcriptional repressor pseudogene |
| AC091812.3 | 7 | capicua homolog (Drosophila) (CIC) pseudoge |
| AC091812.2 | 7 | capicua homolog (Drosophila) (CIC) pseudoge |
| PSPHP1 | 7 | phosphoserine phosphatase pseudogene 1 [S |
| RNU6-1126P | 7 | RNA, U6 small nuclear 1126, pseudogene [Sou |
| SEPTIN14 | 7 | septin 14 [Source:HGNC Symbol;Acc:HGNC:33 |
| AC092647.4 | 7 | novel pseudogene |
| CICP12 | 7 | capicua transcriptional repressor pseudogene |
| AC092647.3 | 7 | cell division cycle associated 8 (CDCA8) pseud |
| AC092647.1 | 7 | pseudogene similar to part of cell division cycl |
| AC092647.2 | 7 | pseudogene similar to part of cell division cycl |
| AC092647.5 | 7 | novel zinc finger protein 713 (ZNF713) and mi |
| ZNF713 | 7 | zinc finger protein 713 [Source:HGNC Symbol; |
| NIPSNAP2 | 7 | nipsnap homolog 2 [Source:HGNC Symbol;Acc |
| MRPS17 | 7 | mitochondrial ribosomal protein S17 [Source: |
| PSPH | 7 | phosphoserine phosphatase [Source:HGNC Sy |
| CCT6A | 7 | chaperonin containing TCP1 subunit 6A [Sourc |
| SNORA22B | 7 | small nucleolar RNA, H/ACA box 22B [Source:I |
| SNORA15 | 7 | small nucleolar RNA, H/ACA box 15 [Source:H |
| SUMF2 | 7 | sulfatase modifying factor 2 [Source:HGNC Syr |
| PHKG1 | 7 | phosphorylase kinase catalytic subunit gamma |
| CHCHD2 | 7 | coiled-coil-helix-coiled-coil-helix domain conta |
| NUPR2 | 7 | nuclear protein 2, transcriptional regulator [S |
| AC006970.1 | 7 | interferon induced transmembrane protein 3 |
| AC006970.2 | 7 | ribonuclease T2 (RNASET2) pseudogene |
| AC006970.3 | 7 | novel transcript |
| CCNJP1 | 7 | cyclin J pseudogene 1 [Source:HGNC Symbol;A |
| AC073136.1 | 7 | exportin, tRNA (nuclear export receptor for tR |
| AC073136.2 | 7 | protein phosphatase 2, regulatory subunit B', |
| AC093392.1 | 7 | pseudogene similar to part of septin 13 SEPT1 |
| RNU6-1335P | 7 | RNA, U6 small nuclear 1335, pseudogene [Sou |
| SEPTIN14P24 | 7 | septin 14 pseudogene 24 [Source:HGNC Symb |
| CICP8 | 7 | capicua transcriptional repressor pseudogene |
| AC093392.2 | 7 | calmodulin 3 (phosphorylase kinase, delta) (C/ |

| | | |
|---|---|---|
| RNU6-1052P | 7 | RNA, U6 small nuclear 1052, pseudogene [Sou |
| AC092423.1 | 7 | pseudogene similar to part of general transcri |
| AC092447.7 | 7 | pseudogene similar to part of coiled-coil doma |
| AC092447.10 | 7 | zinc finger protein 195 (ZNF195) pseudogene |
| AC092447.4 | 7 | novel transcript |
| RBM22P3 | 7 | RNA binding motif protein 22 pseudogene 3 [S |
| AC092447.8 | 7 | novel transcript |
| NMD3P2 | 7 | NMD3 ribosome export adaptor pseudogene |
| AC092447.3 | 7 | C2H2 zinc finger protein pseudogene |
| AC092447.5 | 7 | novel transcript |
| VN1R25P | 7 | vomeronasal 1 receptor 25 pseudogene [Sour |
| AC092447.2 | 7 | tyrosine 3-monooxygenase/tryptophan 5-mor |
| AC092447.6 | 7 | pseudogene similar to part of a C2H2 zinc fing |
| AC092447.1 | 7 | insulin-like growth factor 2 mRNA binding pro |
| AC092447.9 | 7 | chromosome 7 open reading frame 73 (C7orf7 |
| AC095038.5 | 7 | novel transcript |
| TRIM60P16 | 7 | tripartite motif containing 60 pseudogene 16 |
| AC095038.1 | 7 | C2H2 zinc finger protein pseudogene |
| AC095038.4 | 7 | novel transcript |
| AC095038.2 | 7 | zinc finger protein 730 (ZNF730) pseudogene |
| AC095038.3 | 7 | translation machinery associated 7 homolog ( |
| AC118758.2 | 7 | capicua transcriptional repressor (CIC) pseudo |
| AC118758.3 | 7 | novel piRNA host transcript |
| AC118758.1 | 7 | pseudogene similar to part of phosphorylase k |
| AC069152.1 | 7 | pseudogene similar to part of solute carrier fa |
| MIR4283-1 | 7 | microRNA 4283-1 [Source:HGNC Symbol;Acc:I |
| TNRC18P3 | 7 | trinucleotide repeat containing 18 pseudogen |
| SLC29A4P1 | 7 | solute carrier family 29 member 4 pseudogen |
| PHKG1P4 | 7 | phosphorylase kinase catalytic subunit gamma |
| AC122133.1 | 7 | septin 7 (SEPT7) pseudogene |
| ZNF479 | 7 | zinc finger protein 479 [Source:HGNC Symbol; |
| AC099654.15 | 7 | novel transcript |
| AC099654.2 | 7 | novel pseudogene |
| AC099654.6 | 7 | novel zinc finger protein pseudogene |
| AC099654.12 | 7 | mitochondrially encoded ATP synthase 8 (MT- |
| AC099654.7 | 7 | mitochondrially encoded ATP synthase 6 (MT- |
| MTCO3P10 | 7 | MT-CO3 pseudogene 10 [Source:HGNC Symbo |
| AC099654.5 | 7 | mitochondrially encoded NADH dehydrogenas |
| MTND4LP32 | 7 | MT-ND4L pseudogene 32 [Source:HGNC Symb |
| MTND4P4 | 7 | MT-ND4 pseudogene 4 [Source:HGNC Symbol |
| MTND5P6 | 7 | MT-ND5 pseudogene 6 [Source:HGNC Symbol |
| MTND6P29 | 7 | MT-ND6 pseudogene 29 [Source:HGNC Symbo |
| MTCYBP29 | 7 | MT-CYB pseudogene 29 [Source:HGNC Symbo |
| GUSBP10 | 7 | GUSB pseudogene 10 [Source:HGNC Symbol;A |
| MTND1P4 | 7 | MT-ND1 pseudogene 4 [Source:HGNC Symbol |
| MTND2P6 | 7 | MT-ND2 pseudogene 6 [Source:HGNC Symbol |
| AC099654.3 | 7 | mitochondrially encoded cytochrome c oxidas |
| AC099654.11 | 7 | mitochondrially encoded cytochrome c oxidas |
| AC099654.9 | 7 | mitochondrially encoded ATP synthase 8 (MT- |
| AC099654.8 | 7 | mitochondrially encoded ATP synthase 6 (MT- |

| | | |
|---|---|---|
| MTCO3P4 | 7 | MT-CO3 pseudogene 4 [Source:HGNC Symbol |
| AC099654.13 | 7 | mitochondrially encoded NADH dehydrogenas |
| AC099654.10 | 7 | mitochondrially encoded NADH 4L dehydroge |
| MTND4P5 | 7 | MT-ND4 pseudogene 5 [Source:HGNC Symbol |
| MTND5P7 | 7 | MT-ND5 pseudogene 7 [Source:HGNC Symbol |
| AC099654.14 | 7 | mitochondrially encoded NADH dehydrogenas |
| MTCYBP5 | 7 | MT-CYB pseudogene 5 [Source:HGNC Symbol; |
| GUSBP12 | 7 | GUSB pseudogene 12 [Source:HGNC Symbol;A |
| AC099654.1 | 7 | novel transcript |
| RNU7-157P | 7 | RNA, U7 small nuclear 157 pseudogene [Sourc |
| AC099654.16 | 7 | novel transcript |
| AC099654.4 | 7 | zinc finger protein 195 (ZNF195) pseudogene |
| AC237221.2 | 7 | TEC |
| AC237221.1 | 7 | novel transcript |
| MIR3147 | 7 | microRNA 3147 [Source:HGNC Symbol;Acc:HG |
| VN1R28P | 7 | vomeronasal 1 receptor 28 pseudogene [Sour |
| SAPCD2P2 | 7 | suppressor APC domain containing 2 pseudog |
| ZNF716 | 7 | zinc finger protein 716 [Source:HGNC Symbol; |
| AC092175.1 | 7 | family with sequence similarity 27, member A |
| NCOR1P3 | 7 | nuclear receptor corepressor 1 pseudogene 3 |
| AC064862.6 | 7 | pseudogene similar to part of family with sequ |
| AC064862.7 | 7 | abhydrolase domain containing 15, pseudoger |
| BSNDP4 | 7 | barttin CLCNK type accessory beta subunit pse |
| AC064862.1 | 7 | family with sequence similarity 27 (FAM27) ps |
| AC064862.5 | 7 | family with sequence similarity 27 (FAM27) ps |
| AC064862.2 | 7 | family with sequence similarity 27 (FAM27) ps |
| AC064862.4 | 7 | family with sequence similarity 27 (FAM27) ps |
| AC064862.3 | 7 | family with sequence similarity 27 (FAM27) ps |
| AC023141.4 | 7 | family with sequence similarity 27 (FAM27) ps |
| AC023141.8 | 7 | family with sequence similarity 27-like (FAM2 |
| AC023141.5 | 7 | family with sequence similarity 27 (FAM27) ps |
| AC023141.12 | 7 | family with sequence similarity 27-like (FAM2 |
| AC023141.2 | 7 | family with sequence similarity 27 (FAM27) ps |
| AC023141.9 | 7 | family with sequence similarity 27 (FAM27) ps |
| AC023141.11 | 7 | family with sequence similarity 27-like (FAM2 |
| AC023141.3 | 7 | family with sequence similarity 27 (FAM27) ps |
| AC023141.1 | 7 | family with sequence similarity 27 (FAM27) ps |
| AC023141.10 | 7 | family with sequence similarity 27 (FAM27) ps |
| AC023141.6 | 7 | family with sequence similarity 27 (FAM27) ps |
| AC023141.7 | 7 | family with sequence similarity 182, member |
| AC023141.13 | 7 | abhydrolase domain containing 15 (ABHD15) |
| AC128676.1 | 7 | kinase suppressor of ras 1 (KSR1) pseudogene |
| RNU6-417P | 7 | RNA, U6 small nuclear 417, pseudogene [Sour |
| AC069285.2 | 7 | novel transcript |
| AC069285.3 | 7 | phosphoserine phosphatase (PSPH) pseudoge |
| ZNF90P3 | 7 | zinc finger protein 90 pseudogene 3 [Source:H |
| SAPCD2P4 | 7 | suppressor APC domain containing 2 pseudog |
| SEPTIN14P1 | 7 | septin 14 pseudogene 1 [Source:HGNC Symbo |
| AC069285.1 | 7 | NULL |
| AC092001.1 | 7 | glutamyl aminopeptidase (aminopeptidase A) |

| | | |
|---|---|---|
| PHKG1P1 | 7 | phosphorylase kinase catalytic subunit gamma |
| AC006455.8 | 7 | novel transcript |
| SEPTIN7P4 | 7 | septin 7 pseudogene 4 [Source:HGNC Symbol; |
| AC006455.7 | 7 | gem (nuclear organelle) associated protein 7 ( |
| ZNF733P | 7 | zinc finger protein 733, pseudogene [Source:H |
| AC006455.1 | 7 | novel transcript |
| ARAFP3 | 7 | ARAF pseudogene 3 [Source:HGNC Symbol;Ac |
| AC006455.6 | 7 | vomeronasal 1 receptor family pseudogene |
| AC006455.5 | 7 | novel transcript |
| AC006455.2 | 7 | solute carrier family 25 (mitochondrial carrier |
| AC006455.3 | 7 | TP53 target 3 (TP53TG3) pseudogene |
| AC006455.4 | 7 | novel transcript |
| VN1R31P | 7 | vomeronasal 1 receptor 31 pseudogene [Sour |
| VN1R32P | 7 | vomeronasal 1 receptor 32 pseudogene [Sour |
| AC073188.3 | 7 | novel transcript |
| AC073188.4 | 7 | novel transcript |
| AC073188.2 | 7 | novel transcript |
| AC073188.1 | 7 | TP53 target 3 (TP53TG3) pseudogene |
| AC073188.5 | 7 | novel transcript |
| AC073188.6 | 7 | novel transcript |
| SLC25A1P2 | 7 | solute carrier family 25 member 1 pseudogen |
| VN1R33P | 7 | vomeronasal 1 receptor 33 pseudogene [Sour |
| ARAFP2 | 7 | ARAF pseudogene 2 [Source:HGNC Symbol;Ac |
| ZNF734P | 7 | zinc finger protein 734, pseudogene [Source:H |
| AC006457.1 | 7 | gem (nuclear organelle) associated protein 7 ( |
| SEPTIN7P5 | 7 | septin 7 pseudogene 5 [Source:HGNC Symbol; |
| PHKG1P2 | 7 | phosphorylase kinase catalytic subunit gamma |
| SLC29A4P2 | 7 | solute carrier family 29 member 4 pseudogen |
| TNRC18P2 | 7 | trinucleotide repeat containing 18 pseudogen |
| MIR4283-2 | 7 | microRNA 4283-2 [Source:HGNC Symbol;Acc:I |
| AC006015.1 | 7 | solute carrier family 29 (nucleoside transporte |
| AC079355.1 | 7 | phosphorylase kinase, gamma 1 (muscle) (PHI |
| CICP24 | 7 | capicua transcriptional repressor pseudogene |
| AC079355.2 | 7 | vitamin K epoxide reductase complex, subunit |
| AC092634.5 | 7 | coiled-coil domain containing 72 (CCDC72) ps |
| AC092634.2 | 7 | zinc finger protein (ZNF) family pseudogene |
| AC092634.4 | 7 | novel transcript |
| GABPAP | 7 | GABPA pseudogene [Source:HGNC Symbol;Ac |
| LINC02848 | 7 | long intergenic non-protein coding RNA 2848 |
| NMD3P1 | 7 | NMD3 ribosome export adaptor pseudogene |
| AC092634.3 | 7 | TP53 target 3 (TP53TG3) pseudogene |
| AC092634.8 | 7 | novel transcript |
| SLC25A1P3 | 7 | solute carrier family 25 member 1 pseudogen |
| VN1R34P | 7 | vomeronasal 1 receptor 34 pseudogene [Sour |
| ARAFP1 | 7 | ARAF pseudogene 1 [Source:HGNC Symbol;Ac |
| AC092634.7 | 7 | novel transcript |
| VN1R35P | 7 | vomeronasal 1 receptor 35 pseudogene [Sour |
| VN1R36P | 7 | vomeronasal 1 receptor 36 pseudogene [Sour |
| AC092634.1 | 7 | zinc finger protein family (ZNF) pseudogene |
| AC092634.6 | 7 | zinc finger protein 679 (ZNF679) pseudogene |

| | | |
|---|---|---|
| AC115220.1 | 7 | novel zinc finger protein |
| LINC01005 | 7 | long intergenic non-protein coding RNA 1005 |
| AC115220.3 | 7 | novel transcript |
| AC115220.2 | 7 | novel transcript |
| ZNF727 | 7 | zinc finger protein 727 [Source:HGNC Symbol; |
| TRIM60P17 | 7 | tripartite motif containing 60 pseudogene 17 |
| GUSBP6 | 7 | GUSB pseudogene 6 [Source:HGNC Symbol;Ac |
| MTND4P2 | 7 | MT-ND4 pseudogene 2 [Source:HGNC Symbol |
| MTND4LP2 | 7 | MT-ND4L pseudogene 2 [Source:HGNC Symbo |
| MTND3P2 | 7 | MT-ND3 pseudogene 2 [Source:HGNC Symbol |
| MTCO3P8 | 7 | MT-CO3 pseudogene 8 [Source:HGNC Symbol |
| MTATP6P18 | 7 | MT-ATP6 pseudogene 18 [Source:HGNC Symb |
| MTCO2P8 | 7 | MT-CO2 pseudogene 8 [Source:HGNC Symbol |
| MTCO1P8 | 7 | MT-CO1 pseudogene 8 [Source:HGNC Symbol |
| MTND2P4 | 7 | MT-ND2 pseudogene 4 [Source:HGNC Symbol |
| MTND1P2 | 7 | MT-ND1 pseudogene 2 [Source:HGNC Symbol |
| AC091685.1 | 7 | novel transcript |
| AC091685.2 | 7 | ribosomal protein L6 (RPL6) pseudogene |
| VN1R37P | 7 | vomeronasal 1 receptor 37 pseudogene [Sour |
| VN1R38P | 7 | vomeronasal 1 receptor 38 pseudogene [Sour |
| SAPCD2P1 | 7 | suppressor APC domain containing 2 pseudog |
| ZNF735 | 7 | zinc finger protein 735 [Source:HGNC Symbol; |
| ZNF679 | 7 | zinc finger protein 679 [Source:HGNC Symbol; |
| AC073270.1 | 7 | novel transcript |
| ZNF736 | 7 | zinc finger protein 736 [Source:HGNC Symbol; |
| TRIM60P18 | 7 | tripartite motif containing 60 pseudogene 18 |
| AC073270.2 | 7 | novel transcript |
| AC022202.1 | 7 | novel zinc finger protein pseudogene |
| YWHAEP1 | 7 | tyrosine 3-monooxygenase/tryptophan 5-mon |
| VN1R40P | 7 | vomeronasal 1 receptor 40 pseudogene [Sour |
| ZNF680P1 | 7 | ZNF680 pseudogene 1 [Source:HGNC Symbol; |
| HNRNPCP7 | 7 | heterogeneous nuclear ribonucleoprotein C p |
| ZNF680 | 7 | zinc finger protein 680 [Source:HGNC Symbol; |
| BNIP3P42 | 7 | BCL2 interacting protein 3 pseudogene 42 [So |
| AC016769.3 | 7 | P antigen family, member 4 (prostate associat |
| AC016769.1 | 7 | novel pseudogene |
| AC016769.2 | 7 | glycine cleavage system protein H (aminometh |
| AC016769.4 | 7 | BCL2/adenovirus E1B 19kDa interacting prote |
| AC016769.5 | 7 | novel transcript |
| AC016769.6 | 7 | novel transcript |
| ZNF107 | 7 | zinc finger protein 107 [Source:HGNC Symbol; |
| BNIP3P11 | 7 | BCL2 interacting protein 3 pseudogene 11 [So |
| MIR6839 | 7 | microRNA 6839 [Source:HGNC Symbol;Acc:HG |
| AC091799.1 | 7 | myelin protein zero-like 1 (MPZL1) pseudogen |
| AC073349.3 | 7 | novel zinc finger protein pseudogene |
| ZNF138 | 7 | zinc finger protein 138 [Source:HGNC Symbol; |
| AC073349.4 | 7 | novel transcript, antisense to ZNF138 |
| AC073349.2 | 7 | endogenous retroviral family W, env(C7), mer |
| SEPHS1P1 | 7 | selenophosphate synthetase 1 pseudogene 1 |
| EEF1DP4 | 7 | eukaryotic translation elongation factor 1 delt |

| | | |
|---|---|---|
| AC073349.5 | 7 | novel transcript |
| ZNF273 | 7 | zinc finger protein 273 [Source:HGNC Symbol; |
| AC073349.1 | 7 | uncharacterized LOC441239 [Source:NCBI ger |
| VN1R42P | 7 | vomeronasal 1 receptor 42 pseudogene [Sour |
| MTDHP1 | 7 | metadherin pseudogene 1 [Source:HGNC Sym |
| AC073210.3 | 7 | novel transcript |
| ZNF117 | 7 | zinc finger protein 117 [Source:HGNC Symbol; |
| ERV3-1 | 7 | endogenous retrovirus group 3 member 1, en |
| RNU6-1229P | 7 | RNA, U6 small nuclear 1229, pseudogene [Sou |
| CCT6P3 | 7 | chaperonin containing TCP1 subunit 6 pseudo |
| AC073210.2 | 7 | pseudogene similar to part of vitamin K epoxi |
| SNORA22C | 7 | small nucleolar RNA, H/ACA box 22C [Source:I |
| SNORA15B-1 | 7 | small nucleolar RNA, H/ACA box 15B-1 [Sourc |
| AC073210.1 | 7 | pseudogene similar to part of sulfatase modif |
| GTF2IP14 | 7 | general transcription factor IIi pseudogene 14 |
| INTS4P1 | 7 | integrator complex subunit 4 pseudogene 1 [S |
| AC104073.2 | 7 | pseudogene similar to part of fibroblast grow |
| AC104073.1 | 7 | pseudogene similar to part of 24-dehydrochol |
| AC104073.4 | 7 | novel transcript |
| AC104073.3 | 7 | novel zinc finger protein pseudogene |
| RSL24D1P3 | 7 | ribosomal L24 domain containing 1 pseudoger |
| AC092685.1 | 7 | novel pseudogene containing a Zinc finger, C2 |
| ZNF92 | 7 | zinc finger protein 92 [Source:HGNC Symbol;A |
| AC114501.3 | 7 | novel transcript |
| AC114501.1 | 7 | novel pseudogene |
| AC114501.2 | 7 | novel pseudogene similar to part of integrato |
| AC104057.1 | 7 | pseudogene similar to part of glutamate rece |
| AC073107.1 | 7 | nuclear protein 2, transcriptional regulator (N |
| INTS4P2 | 7 | integrator complex subunit 4 pseudogene 2 [S |
| CCT6P1 | 7 | chaperonin containing TCP1 subunit 6 pseudo |
| SNORA22 | 7 | small nucleolar RNA, H/ACA box 22 [Source:H |
| SNORA15B-2 | 7 | small nucleolar RNA, H/ACA box 15B-2 [Sourc |
| AC073107.2 | 7 | sulfatase modifying factor 2 (SUMF2) pseudog |
| GTF2IP5 | 7 | general transcription factor IIi pseudogene 5 [ |
| RNU6-912P | 7 | RNA, U6 small nuclear 912, pseudogene [Sour |
| AC093582.1 | 7 | novel transcript |
| RNU6-973P | 7 | RNA, U6 small nuclear 973, pseudogene [Sour |
| VKORC1L1 | 7 | vitamin K epoxide reductase complex subunit |
| GUSB | 7 | glucuronidase beta [Source:HGNC Symbol;Acc |
| AC073261.1 | 7 | vitamin K epoxide reductase complex, subunit |
| AC068533.2 | 7 | novel pseudogene similar to part of CGRP rec |
| AC068533.1 | 7 | golgi autoantigen, golgin subfamily a, 7 (GOLG |
| ASL | 7 | argininosuccinate lyase [Source:HGNC Symbol |
| AC068533.4 | 7 | novel protein |
| CRCP | 7 | CGRP receptor component [Source:HGNC Sym |
| AC068533.3 | 7 | novel transcript |
| TPST1 | 7 | tyrosylprotein sulfotransferase 1 [Source:HGN |
| RNU6-313P | 7 | RNA, U6 small nuclear 313, pseudogene [Sour |
| LINC00174 | 7 | long intergenic non-protein coding RNA 174 [S |
| GTF2IP9 | 7 | general transcription factor IIi pseudogene 9 [ |

| | | |
|---|---|---|
| SKP1P1 | 7 | S-phase kinase associated protein 1 pseudoge |
| AC008267.2 | 7 | RAB guanine nucleotide exchange factor 1 (RA |
| AC008267.4 | 7 | GrpE-like 1, mitochondrial (E. coli) (GRPEL1) p |
| AC008267.8 | 7 | TEC |
| AC008267.6 | 7 | general transcription factor II, i (GTF2I) pseud |
| AC008267.7 | 7 | general transcription factor II, i (GTF2I) pseud |
| AC008267.5 | 7 | novel transcript |
| AC008267.1 | 7 | guanine nucleotide binding protein (G protein |
| AC008267.3 | 7 | GTF2I repeat domain containing 1 (GTF2IRD1) |
| AC006001.3 | 7 | RAB guanine nucleotide exchange factor (GEF |
| AC006001.4 | 7 | translocase of outer mitochondrial membrane |
| SAPCD2P3 | 7 | suppressor APC domain containing 2 pseudog |
| AC006001.1 | 7 | novel pseudogene |
| RPL35P5 | 7 | ribosomal protein L35 pseudogene 5 [Source: |
| KCTD7 | 7 | potassium channel tetramerization domain cc |
| AC027644.4 | 7 | novel transcript |
| AC006001.2 | 7 | novel transcript |
| RABGEF1 | 7 | RAB guanine nucleotide exchange factor 1 [So |
| AC027644.3 | 7 | novel transcript, antisense to RABGEF1 |
| AC027644.2 | 7 | coiled-coil-helix-coiled-coil-helix domain cont |
| GTF2IRD1P1 | 7 | GTF2I repeat domain containing 1 pseudogen |
| AC027644.1 | 7 | pseudogene similar to part of general transcri |
| GTF2IP23 | 7 | general transcription factor IIi pseudogene 23 |
| RNU6-1254P | 7 | RNA, U6 small nuclear 1254, pseudogene [Sou |
| LINC02604 | 7 | long intergenic non-protein coding RNA 2604 |
| AC073335.1 | 7 | ribosomal protein L31 (Rpl31) pseudogene |
| Metazoa_SRP | 7 | Metazoan signal recognition particle RNA [Sou |
| TMEM248 | 7 | transmembrane protein 248 [Source:HGNC Sy |
| RN7SL43P | 7 | RNA, 7SL, cytoplasmic 43, pseudogene [Sourc |
| SBDS | 7 | SBDS ribosome maturation factor [Source:HG |
| TYW1 | 7 | tRNA-yW synthesizing protein 1 homolog [Sou |
| AC073089.1 | 7 | HLA-B associated transcript 2 (BAT2) pseudog |
| MIR4650-1 | 7 | microRNA 4650-1 [Source:HGNC Symbol;Acc: |
| SPDYE21P | 7 | speedy/RINGO cell cycle regulator family men |
| PMS2P4 | 7 | PMS1 homolog 2, mismatch repair system cor |
| Y_RNA | 7 | Y RNA [Source:RFAM;Acc:RF00019] |
| STAG3L4 | 7 | stromal antigen 3-like 4 (pseudogene) [Source |
| AC006480.1 | 7 | ribosomal protein L29 (RPL29) pseudogene |
| AC006480.2 | 7 | novel transcript |
| MTATP6P21 | 7 | MT-ATP6 pseudogene 21 [Source:HGNC Symb |
| MTCO3P41 | 7 | MT-CO3 pseudogene 41 [Source:HGNC Symbo |
| AC005482.1 | 7 | novel transcript |
| AC092648.1 | 7 | prefoldin subunit 4 (PFDN4) pseudogene |
| AC092637.1 | 7 | novel transcript |
| AC006013.1 | 7 | novel transcript |
| MTCO1P57 | 7 | MT-CO1 pseudogene 57 [Source:HGNC Symbo |
| AC093655.1 | 7 | novel transcript |
| AC093655.2 | 7 | septin 13 (SEPT13) pseudogene |
| MTND4P3 | 7 | MT-ND4 pseudogene 3 [Source:HGNC Symbol |
| AC004910.1 | 7 | ribosomal protein L21 (RPL21) pseudogene |

| | | |
|---|---|---|
| RNA5SP231 | 7 | RNA, 5S ribosomal pseudogene 231 [Source:H |
| AC104688.1 | 7 | novel transcript |
| RNU6-832P | 7 | RNA, U6 small nuclear 832, pseudogene [Sour |
| AC069280.2 | 7 | novel transcript |
| AC092100.1 | 7 | novel transcript |
| AC069280.1 | 7 | novel transcript |
| MTCO2P25 | 7 | MT-CO2 pseudogene 25 [Source:HGNC Symbo |
| MTCO1P25 | 7 | MT-CO1 pseudogene 25 [Source:HGNC Symbo |
| RNU6-229P | 7 | RNA, U6 small nuclear 229, pseudogene [Sour |
| Y_RNA | 7 | Y RNA [Source:RFAM;Acc:RF00019] |
| CT66 | 7 | cancer/testis associated transcript 66 [Source |
| AUTS2 | 7 | activator of transcription and developmental r |
| AC073873.1 | 7 | neuroepithelial cell transforming 1 (NET1) pse |
| GALNT17 | 7 | polypeptide N-acetylgalactosaminyltransferas |
| MIR3914-1 | 7 | microRNA 3914-1 [Source:HGNC Symbol;Acc:H |
| MIR3914-2 | 7 | microRNA 3914-2 [Source:HGNC Symbol;Acc:H |
| RN7SKP75 | 7 | RN7SK pseudogene 75 [Source:HGNC Symbol; |
| CALN1 | 7 | calneuron 1 [Source:HGNC Symbol;Acc:HGNC: |
| RNA5SP232 | 7 | RNA, 5S ribosomal pseudogene 232 [Source:H |
| AC005011.1 | 7 | ribosomal protein S28 (RPS28) pseudogene |
| ABCF2P2 | 7 | ATP binding cassette subfamily F member 2 ps |
| TYW1B | 7 | tRNA-yW synthesizing protein 1 homolog B [So |
| MIR4650-2 | 7 | microRNA 4650-2 [Source:HGNC Symbol;Acc:H |
| AC211469.2 | 7 | proline-rich coiled-coil 2A, pseudogene |
| AC211469.1 | 7 | transmembrane protein 68, pseudogene |
| RN7SL377P | 7 | RNA, 7SL, cytoplasmic 377, pseudogene [Sour |
| SBDSP1 | 7 | SBDS pseudogene 1 [Source:HGNC Symbol;Ac |
| RN7SL625P | 7 | RNA, 7SL, cytoplasmic 625, pseudogene [Sour |
| SPDYE7P | 7 | speedy/RINGO cell cycle regulator family men |
| POM121 | 7 | POM121 transmembrane nucleoporin [Source |
| AC211476.2 | 7 | novel transcript, antisense to POM121 |
| NSUN5P2 | 7 | NSUN5 pseudogene 2 [Source:HGNC Symbol;A |
| AC211476.6 | 7 | novel transcript |
| TRIM74 | 7 | tripartite motif containing 74 [Source:HGNC S |
| STAG3L3 | 7 | stromal antigen 3-like 3 (pseudogene) [Source |
| AC211476.1 | 7 | FK506 binding protein 6, 36kDa (FKBP6 ) pseu |
| Y_RNA | 7 | Y RNA [Source:RFAM;Acc:RF00019] |
| PMS2P7 | 7 | PMS1 homolog 2, mismatch repair system cor |
| Y_RNA | 7 | Y RNA [Source:RFAM;Acc:RF00019] |
| SPDYE8P | 7 | speedy/RINGO cell cycle regulator family men |
| AC211476.3 | 7 | PMS2 postmeiotic segregation increased 2 (S. |
| Y_RNA | 7 | Y RNA [Source:RFAM;Acc:RF00019] |
| AC211476.5 | 7 | novel transcript |
| SPDYE11 | 7 | speedy/RINGO cell cycle regulator family men |
| SPDYE17 | 7 | speedy/RINGO cell cycle regulator family men |
| AC211476.4 | 7 | PMS2 postmeiotic segregation increased 2 (S. |
| Y_RNA | 7 | Y RNA [Source:RFAM;Acc:RF00019] |
| SPDYE9P | 7 | speedy/RINGO cell cycle regulator family men |
| PMS2P6 | 7 | PMS1 homolog 2, mismatch repair system cor |
| Y_RNA | 7 | Y RNA [Source:RFAM;Acc:RF00019] |

| | | |
|---|---|---|
| SPDYE10P | 7 | speedy/RINGO cell cycle regulator family mem |
| GTF2IP4 | 7 | general transcription factor IIi pseudogene 4 [ |
| PHBP5 | 7 | prohibitin pseudogene 5 [Source:HGNC Symbo |
| NCF1B | 7 | neutrophil cytosolic factor 1B pseudogene [So |
| GTF2IRD2P1 | 7 | GTF2I repeat domain containing 2 pseudogene |
| POM121B | 7 | POM121 transmembrane nucleoporin B (pseu |
| NSUN5 | 7 | NOP2/Sun RNA methyltransferase 5 [Source:H |
| TRIM50 | 7 | tripartite motif containing 50 [Source:HGNC S |
| AC211485.1 | 7 | ribosomal protein L7a (RPL7A) pseudogene |
| FKBP6 | 7 | FKBP prolyl isomerase 6 [Source:HGNC Symbo |
| RNU6-1080P | 7 | RNA, U6 small nuclear 1080, pseudogene [Sou |
| Y_RNA | 7 | Y RNA [Source:RFAM;Acc:RF00019] |
| FZD9 | 7 | frizzled class receptor 9 [Source:HGNC Symbo |
| BAZ1B | 7 | bromodomain adjacent to zinc finger domain |
| RNU6-1198P | 7 | RNA, U6 small nuclear 1198, pseudogene [Sou |
| BCL7B | 7 | BAF chromatin remodeling complex subunit B |
| TBL2 | 7 | transducin beta like 2 [Source:HGNC Symbol;A |
| MLXIPL | 7 | MLX interacting protein like [Source:HGNC Sy |
| AC005089.1 | 7 | novel transcript |
| VPS37D | 7 | VPS37D subunit of ESCRT-I [Source:HGNC Sym |
| DNAJC30 | 7 | DnaJ heat shock protein family (Hsp40) memb |
| BUD23 | 7 | BUD23 rRNA methyltransferase and ribosome |
| STX1A | 7 | syntaxin 1A [Source:HGNC Symbol;Acc:HGNC: |
| MIR4284 | 7 | microRNA 4284 [Source:HGNC Symbol;Acc:HG |
| RN7SL265P | 7 | RNA, 7SL, cytoplasmic 265, pseudogene [Sour |
| ABHD11-AS1 | 7 | ABHD11 antisense RNA 1 (tail to tail) [Source: |
| ABHD11 | 7 | abhydrolase domain containing 11 [Source:HG |
| CLDN3 | 7 | claudin 3 [Source:HGNC Symbol;Acc:HGNC:20 |
| CLDN4 | 7 | claudin 4 [Source:HGNC Symbol;Acc:HGNC:20 |
| METTL27 | 7 | methyltransferase like 27 [Source:HGNC Symb |
| TMEM270 | 7 | transmembrane protein 270 [Source:HGNC Sy |
| AC099398.1 | 7 | novel transcript |
| ELN | 7 | elastin [Source:HGNC Symbol;Acc:HGNC:3327 |
| ELN-AS1 | 7 | ELN antisense RNA 1 [Source:HGNC Symbol;A |
| LIMK1 | 7 | LIM domain kinase 1 [Source:HGNC Symbol;A |
| EIF4H | 7 | eukaryotic translation initiation factor 4H [Sou |
| MIR590 | 7 | microRNA 590 [Source:HGNC Symbol;Acc:HGN |
| LAT2 | 7 | linker for activation of T cells family member |
| RFC2 | 7 | replication factor C subunit 2 [Source:HGNC S |
| AC005081.1 | 7 | novel transcript |
| CLIP2 | 7 | CAP-Gly domain containing linker protein 2 [S |
| GTF2IRD1 | 7 | GTF2I repeat domain containing 1 [Source:HG |
| RNA5SP233 | 7 | RNA, 5S ribosomal pseudogene 233 [Source:H |
| AC211433.2 | 7 | novel transcript |
| AC211433.3 | 7 | novel transcript |
| GTF2I | 7 | general transcription factor IIi [Source:HGNC S |
| AC211433.1 | 7 | novel transcript, antisense to GTF2I |
| PHBP15 | 7 | prohibitin pseudogene 15 [Source:HGNC Symb |
| NCF1 | 7 | neutrophil cytosolic factor 1 [Source:HGNC Sy |
| GTF2IRD2 | 7 | GTF2I repeat domain containing 2 [Source:HG |

| | | |
|---|---|---|
| STAG3L2 | 7 | stromal antigen 3-like 2 (pseudogene) [Source |
| PMS2P5 | 7 | PMS1 homolog 2, mismatch repair system cor |
| Y_RNA | 7 | Y RNA [Source:RFAM;Acc:RF00019] |
| SPDYE12P | 7 | speedy/RINGO cell cycle regulator family men |
| CASTOR2 | 7 | cytosolic arginine sensor for mTORC1 subunit |
| RCC1L | 7 | RCC1 like [Source:HGNC Symbol;Acc:HGNC:14 |
| Metazoa_SRP | 7 | Metazoan signal recognition particle RNA [Sou |
| GTF2IRD2B | 7 | GTF2I repeat domain containing 2B [Source:H |
| NCF1C | 7 | neutrophil cytosolic factor 1C pseudogene [So |
| GTF2IP1 | 7 | general transcription factor IIi pseudogene 1 [ |
| PHBP6 | 7 | prohibitin pseudogene 6 [Source:HGNC Symbo |
| AC211486.5 | 7 | novel transcript |
| AC211486.2 | 7 | novel transcript |
| AC211486.4 | 7 | GATS protein-like 2 (GATSL2) pseudogene |
| AC211486.7 | 7 | novel speedy/RINGO cell cycle regulator famil |
| PMS2P10 | 7 | PMS1 homolog 2, mismatch repair system cor |
| Y_RNA | 7 | Y RNA [Source:RFAM;Acc:RF00019] |
| AC211486.8 | 7 | novel speedy/RINGO cell cycle regulator famil |
| AC211486.3 | 7 | postmeiotic segregation increased 2 pseudoge |
| Y_RNA | 7 | Y RNA [Source:RFAM;Acc:RF00019] |
| AC211486.6 | 7 | novel speedy/RINGO cell cycle regulator famil |
| PMS2P2 | 7 | PMS1 homolog 2, mismatch repair system cor |
| Y_RNA | 7 | Y RNA [Source:RFAM;Acc:RF00019] |
| STAG3L1 | 7 | stromal antigen 3-like 1 (pseudogene) [Source |
| Y_RNA | 7 | Y RNA [Source:RFAM;Acc:RF00019] |
| AC211486.1 | 7 | FK506 binding protein 6, 36kDa (FKBP6 ) pseu |
| TRIM73 | 7 | tripartite motif containing 73 [Source:HGNC S |
| NSUN5P1 | 7 | NSUN5 pseudogene 1 [Source:HGNC Symbol;A |
| POM121C | 7 | POM121 transmembrane nucleoporin C [Sour |
| AC211429.1 | 7 | postmeiotic segregation increased 2-like 3 (PM |
| SPDYE5 | 7 | speedy/RINGO cell cycle regulator family men |
| PMS2P3 | 7 | PMS1 homolog 2, mismatch repair system cor |
| Y_RNA | 7 | Y RNA [Source:RFAM;Acc:RF00019] |
| HIP1 | 7 | huntingtin interacting protein 1 [Source:HGNC |
| AC004491.1 | 7 | TEC |
| CCL26 | 7 | C-C motif chemokine ligand 26 [Source:HGNC |
| CCL24 | 7 | C-C motif chemokine ligand 24 [Source:HGNC |
| AC005102.1 | 7 | THO complex 4 (THOC4) pseudogene |
| RHBDD2 | 7 | rhomboid domain containing 2 [Source:HGNC |
| POR | 7 | cytochrome p450 oxidoreductase [Source:HG |
| MIR4651 | 7 | microRNA 4651 [Source:HGNC Symbol;Acc:HG |
| RPL7L1P3 | 7 | ribosomal protein L7 like 1 pseudogene 3 [Sou |
| SNORA14A | 7 | small nucleolar RNA, H/ACA box 14A [Source:H |
| TMEM120A | 7 | transmembrane protein 120A [Source:HGNC S |
| STYXL1 | 7 | serine/threonine/tyrosine interacting like 1 [S |
| AC006330.1 | 7 | TEC |
| MDH2 | 7 | malate dehydrogenase 2 [Source:HGNC Symb |
| AC005077.4 | 7 | hypothetical protein LOC285908 (LOC285908) |
| RNU6-863P | 7 | RNA, U6 small nuclear 863, pseudogene [Sour |
| GTF2IP7 | 7 | general transcription factor IIi pseudogene 7 [ |

| | | |
|---|---|---|
| AC005077.2 | 7 | pseudogene similar to part of RAB guanine nu |
| AC005077.1 | 7 | GrpE-like 1, mitochondrial (E. coli) (GRPEL1) p |
| AC005077.3 | 7 | pseudogene similar to part of GTF2I repeat dc |
| SRRM3 | 7 | serine/arginine repetitive matrix 3 [Source:HG |
| HSPB1 | 7 | heat shock protein family B (small) member 1 |
| YWHAG | 7 | tyrosine 3-monooxygenase/tryptophan 5-mor |
| PPIAP81 | 7 | peptidylprolyl isomerase A pseudogene 81 [Sc |
| SSC4D | 7 | scavenger receptor cysteine rich family memb |
| ZP3 | 7 | zona pellucida glycoprotein 3 [Source:HGNC S |
| DTX2 | 7 | deltex E3 ubiquitin ligase 2 [Source:HGNC Syn |
| FDPSP2 | 7 | farnesyl diphosphate synthase pseudogene 2 |
| AC005522.1 | 7 | novel transcript, antisense to DTX2 |
| UPK3B | 7 | uroplakin 3B [Source:HGNC Symbol;Acc:HGN( |
| AC004980.4 | 7 | pseudogene similar to part of postmeiotic seg |
| Y_RNA | 7 | Y RNA [Source:RFAM;Acc:RF00019] |
| AC004980.3 | 7 | postmeiotic segregation increased 2-like 5 (PN |
| SPDYE16 | 7 | speedy/RINGO cell cycle regulator family men |
| AC004980.1 | 7 | pseudogene similar to part of postmeiotic seg |
| POMZP3 | 7 | POM121 and ZP3 fusion [Source:HGNC Symbc |
| AC004980.2 | 7 | pseudogene similar to part of RB-associated K |
| AC006972.1 | 7 | novel transcript |
| AC007003.1 | 7 | novel transcript |
| DTX2P1-UPK3BP1-PMS2P | 7 | DTX2P1-UPK3BP1-PMS2P11 readthrough, trai |
| AC114737.3 | 7 | novel transcript |
| FDPSP7 | 7 | farnesyl diphosphate synthase pseudogene 7 |
| AC114737.2 | 7 | pseudogene similar to part of chromosome 1 |
| AC114737.1 | 7 | novel transcript |
| DTX2P1 | 7 | DTX2 pseudogene 1 [Source:HGNC Symbol;Ac |
| UPK3BP1 | 7 | uroplakin 3B pseudogene 1 [Source:HGNC Syr |
| PMS2P11 | 7 | PMS1 homolog 2, mismatch repair system cor |
| Y_RNA | 7 | Y RNA [Source:RFAM;Acc:RF00019] |
| SPDYE17 | 7 | speedy/RINGO cell cycle regulator family men |
| PMS2P9 | 7 | PMS1 homolog 2, mismatch repair system cor |
| Y_RNA | 7 | Y RNA [Source:RFAM;Acc:RF00019] |
| AC007000.3 | 7 | novel transcript |
| SPDYE18 | 7 | speedy/RINGO cell cycle regulator family men |
| AC007000.2 | 7 | pseudogene similar to part of SWI/SNF relate( |
| FAM185BP | 7 | family with sequence similarity 185 member E |
| AC007000.1 | 7 | ribosomal protein L7a (RPL7A) pseudogene |
| CCDC146 | 7 | coiled-coil domain containing 146 [Source:HG |
| FGL2 | 7 | fibrinogen like 2 [Source:HGNC Symbol;Acc:H( |
| AC098851.1 | 7 | novel transcript |
| GSAP | 7 | gamma-secretase activating protein [Source:H |
| AC004921.1 | 7 | novel transcript |
| GCNT1P5 | 7 | glucosaminyl (N-acetyl) transferase 1 pseudog |
| AC004921.2 | 7 | novel transcript |
| PTPN12 | 7 | protein tyrosine phosphatase non-receptor ty |
| APTR | 7 | Alu-mediated CDKN1A/p21 transcriptional reg |
| RSBN1L | 7 | round spermatid basic protein 1 like [Source:H |
| TMEM60 | 7 | transmembrane protein 60 [Source:HGNC Syn |

| | | |
|---|---|---|
| PHTF2 | 7 | putative homeodomain transcription factor 2 |
| Y_RNA | 7 | Y RNA [Source:RFAM;Acc:RF00019] |
| AC004990.1 | 7 | novel transcript |
| MAGI2 | 7 | membrane associated guanylate kinase, WW |
| AC004808.2 | 7 | TEC |
| AC004808.1 | 7 | TEC |
| RPL13AP17 | 7 | ribosomal protein L13a pseudogene 17 [Sourc |
| AC006043.1 | 7 | novel transcript |
| AC007237.1 | 7 | ubiquitin-fold modifier conjugating enzyme 1 |
| MAGI2-AS1 | 7 | MAGI2 antisense RNA 1 [Source:HGNC Symbo |
| MAGI2-AS2 | 7 | MAGI2 antisense RNA 2 [Source:HGNC Symbo |
| RNU6-337P | 7 | RNA, U6 small nuclear 337, pseudogene [Sour |
| AC006355.1 | 7 | heterogeneous nuclear ribonucleoprotein A1- |
| AC006355.2 | 7 | novel transcript, antisense to MAGI2 |
| AC074024.1 | 7 | novel transcript |
| RNU6-530P | 7 | RNA, U6 small nuclear 530, pseudogene [Sour |
| AC004945.1 | 7 | pseudogene similar to part of transmembrane |
| MAGI2-AS3 | 7 | MAGI2 antisense RNA 3 [Source:HGNC Symbo |
| NUP35P2 | 7 | nucleoporin 35 pseudogene 2 [Source:HGNC S |
| RNA5SP234 | 7 | RNA, 5S ribosomal pseudogene 234 [Source:H |
| AC073048.1 | 7 | novel transcript |
| GNAI1 | 7 | G protein subunit alpha i1 [Source:HGNC Sym |
| RNU6-849P | 7 | RNA, U6 small nuclear 849, pseudogene [Sour |
| RPL10P11 | 7 | ribosomal protein L10 pseudogene 11 [Source |
| RN7SL869P | 7 | RNA, 7SL, cytoplasmic 869, pseudogene [Sour |
| AC003988.1 | 7 | novel transcript |
| AC004862.1 | 7 | novel transcript, antisense to CD36 |
| RN7SL35P | 7 | RNA, 7SL, cytoplasmic 35, pseudogene [Sourc |
| CD36 | 7 | CD36 molecule [Source:HGNC Symbol;Acc:HG |
| SNRPBP1 | 7 | small nuclear ribonucleoprotein polypeptides |
| GNAT3 | 7 | G protein subunit alpha transducin 3 [Source: |
| AC073850.1 | 7 | pseudogene similar to part of transcription fa |
| SEMA3C | 7 | semaphorin 3C [Source:HGNC Symbol;Acc:HG |
| AC004972.1 | 7 | tec |
| AC005008.2 | 7 | novel transcript |
| AC005008.1 | 7 | voltage-dependent anion channel 3 (Vdac3) p |
| AC004866.1 | 7 | pseudogene similar to part of mediator compl |
| AC004866.2 | 7 | pseudogene similar to part of mediator compl |
| EIF4EP4 | 7 | eukaryotic translation initiation factor 4E pseu |
| AC008163.1 | 7 | novel pseudogene containing cadherin repeat |
| DDX43P3 | 7 | DEAD-box helicase 43 pseudogene 3 [Source: |
| HGF | 7 | hepatocyte growth factor [Source:HGNC Symb |
| CACNA2D1 | 7 | calcium voltage-gated channel auxiliary subun |
| AC006145.1 | 7 | novel transcript |
| MTHFD2P5 | 7 | methylenetetrahydrofolate dehydrogenase (N |
| AC004006.1 | 7 | pseudogene similar to part of anterior pharyn |
| PCLO | 7 | piccolo presynaptic cytomatrix protein [Sourc |
| RNA5SP235 | 7 | RNA, 5S ribosomal pseudogene 235 [Source:H |
| AC079799.1 | 7 | novel transcript |
| SEMA3E | 7 | semaphorin 3E [Source:HGNC Symbol;Acc:HG |

| | | |
|---|---|---|
| AC079799.2 | 7 | TEC |
| AC079987.1 | 7 | pseudogene similar to part of proteasome (pr |
| RAD23BP2 | 7 | RAD23B pseudogene 2 [Source:HGNC Symbol; |
| SEMA3A | 7 | semaphorin 3A [Source:HGNC Symbol;Acc:HG |
| RPL7P30 | 7 | ribosomal protein L7 pseudogene 30 [Source: |
| AC003984.1 | 7 | novel transcript |
| AC093716.1 | 7 | pseudogene similar to part of selenoprotein K |
| HMGN2P11 | 7 | high mobility group nucleosomal binding dom |
| AC074183.2 | 7 | novel transcript similar to small nucleolar RNA |
| AC074183.1 | 7 | novel transcript similar to small nucleolar RNA |
| HNRNPA1P8 | 7 | heterogeneous nuclear ribonucleoprotein A1 |
| SEMA3D | 7 | semaphorin 3D [Source:HGNC Symbol;Acc:HG |
| HSPA8P16 | 7 | heat shock protein family A (Hsp70) member 8 |
| DYNLL1P7 | 7 | dynein light chain LC8-type 1 pseudogene 7 [S |
| LINC00972 | 7 | long intergenic non-protein coding RNA 972 [S |
| AC092013.1 | 7 | leukemia NUP98 fusion partner 1 (LNP1) pseu |
| SOCS5P1 | 7 | suppressor of cytokine signaling 5 pseudogene |
| GRM3 | 7 | glutamate metabotropic receptor 3 [Source:H |
| AC005009.1 | 7 | novel transcript |
| AC005009.2 | 7 | novel transcript |
| KIAA1324L | 7 | KIAA1324 like [Source:HGNC Symbol;Acc:HGN |
| AC004023.1 | 7 | novel transcript |
| AC005076.1 | 7 | novel transcript, antisense to DMTF1 |
| DMTF1 | 7 | cyclin D binding myb like transcription factor 1 |
| AC005076.2 | 7 | NULL |
| TMEM243 | 7 | transmembrane protein 243 [Source:HGNC Sy |
| Y_RNA | 7 | Y RNA [Source:RFAM;Acc:RF00019] |
| TP53TG1 | 7 | TP53 target 1 [Source:HGNC Symbol;Acc:HGN |
| AC005045.1 | 7 | NULL |
| AC005045.2 | 7 | NULL |
| CROT | 7 | carnitine O-octanoyltransferase [Source:HGNC |
| ABCB4 | 7 | ATP binding cassette subfamily B member 4 [S |
| ABCB1 | 7 | ATP binding cassette subfamily B member 1 [S |
| HNRNPA1P9 | 7 | heterogeneous nuclear ribonucleoprotein A1 |
| RUNDC3B | 7 | RUN domain containing 3B [Source:HGNC Sym |
| SLC25A40 | 7 | solute carrier family 25 member 40 [Source:H |
| DBF4 | 7 | DBF4 zinc finger [Source:HGNC Symbol;Acc:HG |
| ADAM22 | 7 | ADAM metallopeptidase domain 22 [Source:H |
| SRI | 7 | sorcin [Source:HGNC Symbol;Acc:HGNC:1129 |
| AC003991.2 | 7 | novel transcript |
| AC003991.1 | 7 | novel transcript, antisense to STEAP4 |
| STEAP4 | 7 | STEAP4 metalloreductase [Source:HGNC Symb |
| AC002069.2 | 7 | novel transcript |
| EIF4A1P13 | 7 | eukaryotic translation initiation factor 4A1 pse |
| AC002069.1 | 7 | NULL |
| AC002069.4 | 7 | polymerase (RNA) III (DNA directed) polypepti |
| AC002069.3 | 7 | CSRP2 binding protein (CSRP2BP) pseudogene |
| KPNA2P2 | 7 | karyopherin subunit alpha 2 pseudogene 2 [So |
| PQLC1P1 | 7 | PQ loop repeat containing 1 pseudogene 1 [So |
| EEF1A1P28 | 7 | eukaryotic translation elongation factor 1 alph |

| | | |
|---|---|---|
| AC002127.1 | 7 | cytochrome c oxidase subunit VIa polypeptide |
| ZNF804B | 7 | zinc finger protein 804B [Source:HGNC Symbo |
| TEX47 | 7 | testis expressed 47 [Source:HGNC Symbol;Acc |
| AC002383.1 | 7 | novel transcript |
| RNU6-274P | 7 | RNA, U6 small nuclear 274, pseudogene [Sour |
| STEAP2-AS1 | 7 | STEAP2 antisense RNA 1 [Source:HGNC Symbo |
| AC004969.1 | 7 | novel transcript |
| DPY19L2P4 | 7 | DPY19L2 pseudogene 4 [Source:HGNC Symbo |
| STEAP1 | 7 | STEAP family member 1 [Source:HGNC Symbo |
| STEAP2 | 7 | STEAP2 metalloreductase [Source:HGNC Symb |
| CFAP69 | 7 | cilia and flagella associated protein 69 [Source |
| Y_RNA | 7 | Y RNA [Source:RFAM;Acc:RF00019] |
| AC002064.2 | 7 | novel transcript |
| AC002064.1 | 7 | novel transcript |
| FAM237B | 7 | family with sequence similarity 237 member E |
| GTPBP10 | 7 | GTP binding protein 10 [Source:HGNC Symbol |
| AC002064.3 | 7 | TEC |
| CLDN12 | 7 | claudin 12 [Source:HGNC Symbol;Acc:HGNC:2 |
| AC006153.1 | 7 | novel transcript |
| CDK14 | 7 | cyclin dependent kinase 14 [Source:HGNC Syn |
| AC002456.2 | 7 | novel pseudogene |
| AC002456.1 | 7 | novel transcript |
| TVP23CP1 | 7 | TVP23C pseudogene 1 [Source:HGNC Symbol; |
| AC002458.1 | 7 | novel transcript, antisense to CDK14 |
| PTP4A1P3 | 7 | PTP4A1 pseudogene 3 [Source:HGNC Symbol; |
| FZD1 | 7 | frizzled class receptor 1 [Source:HGNC Symbo |
| AC079760.2 | 7 | novel transcript |
| NIPA2P1 | 7 | NIPA magnesium transporter 2 pseudogene 1 |
| AC079760.1 | 7 | novel transcript |
| AC000058.1 | 7 | novel transcript |
| MTERF1 | 7 | mitochondrial transcription termination facto |
| AC003086.1 | 7 | novel transcript |
| AKAP9 | 7 | A-kinase anchoring protein 9 [Source:HGNC Sy |
| CYP51A1 | 7 | cytochrome P450 family 51 subfamily A meml |
| AC000120.2 | 7 | novel protein |
| AC000120.3 | 7 | novel protein |
| CYP51A1-AS1 | 7 | CYP51A1 antisense RNA 1 [Source:HGNC Syml |
| LRRD1 | 7 | leucine rich repeats and death domain contair |
| KRIT1 | 7 | KRIT1 ankyrin repeat containing [Source:HGN( |
| AC000120.1 | 7 | novel transcript |
| Y_RNA | 7 | Y RNA [Source:RFAM;Acc:RF00019] |
| MIR1285-1 | 7 | microRNA 1285-1 [Source:HGNC Symbol;Acc:I |
| ANKIB1 | 7 | ankyrin repeat and IBR domain containing 1 [S |
| TMBIM7P | 7 | transmembrane BAX inhibitor motif containin |
| AC007566.2 | 7 | coiled-coil-helix-coiled-coil-helix domain conta |
| GATAD1 | 7 | GATA zinc finger domain containing 1 [Source |
| AC007566.1 | 7 | novel transcript, antisense to ERVW-1 and PE) |
| ERVW-1 | 7 | endogenous retrovirus group W member 1, er |
| PEX1 | 7 | peroxisomal biogenesis factor 1 [Source:HGN( |
| RBM48 | 7 | RNA binding motif protein 48 [Source:HGNC S |

| | | |
|---|---|---|
| AC005156.1 | 7 | EF-hand calcium binding domain 1 (EFCAB1) p |
| FAM133B | 7 | family with sequence similarity 133 member E |
| CDK6 | 7 | cyclin dependent kinase 6 [Source:HGNC Symb |
| AC000065.2 | 7 | novel transcript, sense intronic to CDK6 |
| AC000065.1 | 7 | novel transcript, antisense to CDK6 |
| RNU6-10P | 7 | RNA, U6 small nuclear 10, pseudogene [Sourc |
| AC002454.1 | 7 | novel transcript |
| RN7SL7P | 7 | RNA, 7SL, cytoplasmic 7, pseudogene [Source: |
| SAMD9 | 7 | sterile alpha motif domain containing 9 [Sourc |
| SAMD9L | 7 | sterile alpha motif domain containing 9 like [S |
| HEPACAM2 | 7 | HEPACAM family member 2 [Source:HGNC Sy |
| VPS50 | 7 | VPS50 subunit of EARP/GARPII complex [Sour |
| CALCR | 7 | calcitonin receptor [Source:HGNC Symbol;Acc |
| MIR653 | 7 | microRNA 653 [Source:HGNC Symbol;Acc:HGI |
| MIR489 | 7 | microRNA 489 [Source:HGNC Symbol;Acc:HGI |
| GNGT1 | 7 | G protein subunit gamma transducin 1 [Sourc |
| AC002075.2 | 7 | ribosomal protein S27 (RPS27) pseudogene |
| MIR4652 | 7 | microRNA 4652 [Source:HGNC Symbol;Acc:HC |
| AC002075.1 | 7 | translin (TSN) pseudogene |
| NDUFAF4P2 | 7 | NADH:ubiquinone oxidoreductase complex as |
| TFPI2 | 7 | tissue factor pathway inhibitor 2 [Source:HGN |
| AC002076.1 | 7 | novel transcript |
| AC002076.2 | 7 | guanine nucleotide binding protein (G protein |
| GNG11 | 7 | G protein subunit gamma 11 [Source:HGNC Sy |
| AC006378.2 | 7 | tec |
| BET1 | 7 | Bet1 golgi vesicular membrane trafficking pro |
| AC006378.1 | 7 | novel transcript |
| AC003092.1 | 7 | novel transcript |
| AC003092.2 | 7 | novel transcript |
| AC002074.1 | 7 | COL1A2 antisense RNA 1 |
| AC002074.2 | 7 | novel transcript |
| COL1A2 | 7 | collagen type I alpha 2 chain [Source:HGNC Sy |
| RNU6-1328P | 7 | RNA, U6 small nuclear 1328, pseudogene [Sou |
| CASD1 | 7 | CAS1 domain containing 1 [Source:HGNC Sym |
| SGCE | 7 | sarcoglycan epsilon [Source:HGNC Symbol;Acc |
| PEG10 | 7 | paternally expressed 10 [Source:HGNC Symbo |
| RPS3AP25 | 7 | ribosomal protein S3a pseudogene 25 [Source |
| RNU6-956P | 7 | RNA, U6 small nuclear 956, pseudogene [Sour |
| ATP5PBP2 | 7 | ATP synthase peripheral stalk-membrane subu |
| GRPEL2P3 | 7 | GRPEL2 pseudogene 3 [Source:HGNC Symbol; |
| RN7SKP129 | 7 | RN7SK pseudogene 129 [Source:HGNC Symbo |
| ARF1P1 | 7 | ADP ribosylation factor 1 pseudogene 1 [Sour |
| PPP1R9A | 7 | protein phosphatase 1 regulatory subunit 9A [ |
| HINT1P2 | 7 | histidine triad nucleotide binding protein 1 ps |
| AC002429.1 | 7 | histidine triad nucleotide binding protein 1 (H |
| AC002429.2 | 7 | novel transcript, antisense to PPP1R9A |
| RNU4-16P | 7 | RNA, U4 small nuclear 16, pseudogene [Sourc |
| PON1 | 7 | paraoxonase 1 [Source:HGNC Symbol;Acc:HGI |
| AC004022.2 | 7 | HERV-FRD provirus ancestral Env polyprotein( |
| AC004022.1 | 7 | HERV-FRD provirus ancestral Env polyprotein( |

| | | |
|---|---|---|
| PON3 | 7 | paraoxonase 3 [Source:HGNC Symbol;Acc:HGI |
| PON2 | 7 | paraoxonase 2 [Source:HGNC Symbol;Acc:HGI |
| AC005021.1 | 7 | novel transcript, antisense to PON2 |
| AC004012.1 | 7 | novel transcript |
| ASB4 | 7 | ankyrin repeat and SOCS box containing 4 [So |
| AC002451.2 | 7 | TEC |
| AC002451.1 | 7 | novel transcript |
| PDK4 | 7 | pyruvate dehydrogenase kinase 4 [Source:HG |
| DYNC1I1 | 7 | dynein cytoplasmic 1 intermediate chain 1 [Sc |
| AC002540.1 | 7 | tec |
| SLC25A13 | 7 | solute carrier family 25 member 13 [Source:H |
| MIR591 | 7 | microRNA 591 [Source:HGNC Symbol;Acc:HGI |
| AC096775.1 | 7 | ribosomal protein L2(RPL21) pseudogene |
| AC084368.1 | 7 | small EDRK-rich factor 2 (SERF2) pseudogene |
| RNU6-532P | 7 | RNA, U6 small nuclear 532, pseudogene [Sour |
| RNU6-364P | 7 | RNA, U6 small nuclear 364, pseudogene [Sour |
| RNU7-188P | 7 | RNA, U7 small nuclear 188 pseudogene [Sourc |
| SEM1 | 7 | SEM1 26S proteasome complex subunit [Sour |
| MARK2P10 | 7 | microtubule affinity regulating kinase 2 pseud |
| RN7SL252P | 7 | RNA, 7SL, cytoplasmic 252, pseudogene [Sour |
| DLX6-AS1 | 7 | DLX6 antisense RNA 1 [Source:HGNC Symbol;/ |
| AC004774.1 | 7 | NULL |
| AC004774.3 | 7 | NULL |
| DLX6 | 7 | distal-less homeobox 6 [Source:HGNC Symbol |
| AC004774.2 | 7 | NULL |
| DLX5 | 7 | distal-less homeobox 5 [Source:HGNC Symbol |
| SDHAF3 | 7 | succinate dehydrogenase complex assembly f |
| HMGB3P21 | 7 | high mobility group box 3 pseudogene 21 [Sou |
| AC073900.1 | 7 | ribosomal protein L7a (RPL7A) pseudogene |
| AP1S2P1 | 7 | AP1S2 pseudogene 1 [Source:HGNC Symbol;A |
| RN7SKP104 | 7 | RN7SK pseudogene 104 [Source:HGNC Symbo |
| TAC1 | 7 | tachykinin precursor 1 [Source:HGNC Symbol; |
| ASNS | 7 | asparagine synthetase (glutamine-hydrolyzing |
| AC079781.5 | 7 | novel transcript |
| AC005326.1 | 7 | pseudogene similar to asparagine synthetase |
| SNRPCP9 | 7 | small nuclear ribonucleoprotein polypeptide C |
| RPS3AP29 | 7 | ribosomal protein S3a pseudogene 29 [Source |
| AC079781.1 | 7 | novel pseudogene |
| AC079781.2 | 7 | novel pseudogene |
| AC079781.4 | 7 | RNA binding protein, fox-1 homolog 1 (RBFOX |
| AC079781.3 | 7 | novel pseudogene |
| OR7E7P | 7 | olfactory receptor family 7 subfamily E memb |
| MIR5692A1 | 7 | microRNA 5692a-1 [Source:HGNC Symbol;Acc |
| MIR5692C2 | 7 | microRNA 5692c-2 [Source:HGNC Symbol;Acc |
| OR7E38P | 7 | olfactory receptor family 7 subfamily E memb |
| AC004967.2 | 7 | novel transcript |
| AC004967.1 | 7 | CCZ1 homolog, vacuolar protein trafficking an |
| OCM2 | 7 | oncomodulin 2 [Source:HGNC Symbol;Acc:HG |
| RN7SL478P | 7 | RNA, 7SL, cytoplasmic 478, pseudogene [Sour |
| LMTK2 | 7 | lemur tyrosine kinase 2 [Source:HGNC Symbo |

| | | |
|---|---|---|
| BHLHA15 | 7 | basic helix-loop-helix family member a15 [Sou |
| TECPR1 | 7 | tectonin beta-propeller repeat containing 1 [S |
| BRI3 | 7 | brain protein I3 [Source:HGNC Symbol;Acc:HG |
| BAIAP2L1 | 7 | BAI1 associated protein 2 like 1 [Source:HGNC |
| AC093799.1 | 7 | novel transcript, antisense to BAIAP2L1 |
| RPS3AP26 | 7 | ribosomal protein S3a pseudogene 26 [Source |
| PPIAP82 | 7 | peptidylprolyl isomerase A pseudogene 82 [Sc |
| AC074121.1 | 7 | ribosomal protein S26 (RPS26) pseudogene |
| NPTX2 | 7 | neuronal pentraxin 2 [Source:HGNC Symbol;A |
| RNU6-393P | 7 | RNA, U6 small nuclear 393, pseudogene [Sour |
| TMEM130 | 7 | transmembrane protein 130 [Source:HGNC Sy |
| TRRAP | 7 | transformation/transcription domain associat |
| MIR3609 | 7 | microRNA 3609 [Source:HGNC Symbol;Acc:HG |
| AC004893.3 | 7 | novel transcript, antisense to TRRAP |
| AC004893.1 | 7 | ring finger protein 14(RNF14)pseudogene |
| SMURF1 | 7 | SMAD specific E3 ubiquitin protein ligase 1 [Sc |
| AC004893.2 | 7 | transformation/transcription domain-associat |
| KPNA7 | 7 | karyopherin subunit alpha 7 [Source:HGNC Sy |
| MYH16 | 7 | myosin heavy chain 16 pseudogene [Source:H |
| AC004834.1 | 7 | novel transcript |
| AC004922.1 | 7 | novel protein, ARPC1A and ARPC1B readthrou |
| ARPC1A | 7 | actin related protein 2/3 complex subunit 1A |
| ARPC1B | 7 | actin related protein 2/3 complex subunit 1B |
| PDAP1 | 7 | PDGFA associated protein 1 [Source:HGNC Syı |
| BUD31 | 7 | BUD31 homolog [Source:HGNC Symbol;Acc:H |
| PTCD1 | 7 | pentatricopeptide repeat domain 1 [Source:H |
| ATP5MF-PTCD1 | 7 | ATP5MF-PTCD1 readthrough [Source:HGNC Sy |
| CPSF4 | 7 | cleavage and polyadenylation specific factor 4 |
| AC073063.1 | 7 | cytokine induced protein 29 kDa (CIP29) pseu |
| ATP5MF | 7 | ATP synthase membrane subunit f [Source:HG |
| ZNF789 | 7 | zinc finger protein 789 [Source:HGNC Symbol; |
| ZNF394 | 7 | zinc finger protein 394 [Source:HGNC Symbol; |
| ZKSCAN5 | 7 | zinc finger with KRAB and SCAN domains 5 [Sc |
| FAM200A | 7 | family with sequence similarity 200 member A |
| ZNF655 | 7 | zinc finger protein 655 [Source:HGNC Symbol; |
| AC005020.2 | 7 | novel protein |
| TMEM225B | 7 | transmembrane protein 225B [Source:HGNC S |
| ZSCAN25 | 7 | zinc finger and SCAN domain containing 25 [Sc |
| AC005020.1 | 7 | ADP-ribosylation factor 6 (ARF6) pseudogene |
| CYP3A5 | 7 | cytochrome P450 family 3 subfamily A membe |
| CYP3A7 | 7 | cytochrome P450 family 3 subfamily A membe |
| CYP3A51P | 7 | cytochrome P450 family 3 subfamily A membe |
| CYP3A7-CYP3A51P | 7 | CYP3A7-CYP3A51P readthrough [Source:HGN( |
| CYP3A4 | 7 | cytochrome P450 family 3 subfamily A membe |
| AC069294.1 | 7 | novel transcript, antisense to CYP3A4 |
| CYP3A137P | 7 | cytochrome P450 family 3 subfamily A membe |
| CYP3A43 | 7 | cytochrome P450 family 3 subfamily A membe |
| AC011904.1 | 7 | cytochrome P450, family 3, subfamily A family |
| CYP3A52P | 7 | cytochrome P450 family 3 subfamily A membe |
| OR2AE1 | 7 | olfactory receptor family 2 subfamily AE mem |

| | | |
|---|---|---|
| TRIM4 | 7 | tripartite motif containing 4 [Source:HGNC Sy |
| GJC3 | 7 | gap junction protein gamma 3 [Source:HGNC ! |
| AC004522.3 | 7 | novel transcript |
| Y_RNA | 7 | Y RNA [Source:RFAM;Acc:RF00019] |
| AC004522.1 | 7 | NULL |
| AZGP1 | 7 | alpha-2-glycoprotein 1, zinc-binding [Source:H |
| AC004522.4 | 7 | novel transcript, antisense to AZGP1 |
| AZGP1P1 | 7 | AZGP1 pseudogene 1 [Source:HGNC Symbol;A |
| AC004522.2 | 7 | gap junction protein, gamma 3, 30.2kDa (GJC3 |
| AC004522.5 | 7 | novel transcript |
| ZKSCAN1 | 7 | zinc finger with KRAB and SCAN domains 1 [Sc |
| ZSCAN21 | 7 | zinc finger and SCAN domain containing 21 [S |
| ZNF3 | 7 | zinc finger protein 3 [Source:HGNC Symbol;Ac |
| COPS6 | 7 | COP9 signalosome subunit 6 [Source:HGNC Sy |
| MCM7 | 7 | minichromosome maintenance complex comp |
| MIR25 | 7 | microRNA 25 [Source:HGNC Symbol;Acc:HGN |
| MIR93 | 7 | microRNA 93 [Source:HGNC Symbol;Acc:HGN |
| MIR106B | 7 | microRNA 106b [Source:HGNC Symbol;Acc:HG |
| AP4M1 | 7 | adaptor related protein complex 4 subunit mu |
| TAF6 | 7 | TATA-box binding protein associated factor 6 |
| AC073842.2 | 7 | novel transcript |
| CNPY4 | 7 | canopy FGF signaling regulator 4 [Source:HGN |
| MBLAC1 | 7 | metallo-beta-lactamase domain containing 1 [ |
| AC073842.1 | 7 | novel transcript |
| RPL7P60 | 7 | ribosomal protein L7 pseudogene 60 [Source: |
| LAMTOR4 | 7 | late endosomal/lysosomal adaptor, MAPK and |
| MAP11 | 7 | microtubule associated protein 11 [Source:HG |
| MIR4658 | 7 | microRNA 4658 [Source:HGNC Symbol;Acc:HG |
| GAL3ST4 | 7 | galactose-3-O-sulfotransferase 4 [Source:HGN |
| GPC2 | 7 | glypican 2 [Source:HGNC Symbol;Acc:HGNC:4 |
| STAG3 | 7 | stromal antigen 3 [Source:HGNC Symbol;Acc:H |
| CASTOR3 | 7 | CASTOR family member 3 [Source:HGNC Symb |
| PVRIG | 7 | PVR related immunoglobulin domain containi |
| SPDYE3 | 7 | speedy/RINGO cell cycle regulator family mem |
| PMS2P1 | 7 | PMS1 homolog 2, mismatch repair system cor |
| Y_RNA | 7 | Y RNA [Source:RFAM;Acc:RF00019] |
| STAG3L5P | 7 | stromal antigen 3-like 5 pseudogene [Source:I |
| STAG3L5P-PVRIG2P-PILRE | 7 | STAG3L5P-PVRIG2P-PILRB readthrough [Sourc |
| PILRB | 7 | paired immunoglobin like type 2 receptor bet |
| PVRIG2P | 7 | poliovirus receptor related immunoglobulin d |
| MIR6840 | 7 | microRNA 6840 [Source:HGNC Symbol;Acc:HG |
| PILRA | 7 | paired immunoglobin like type 2 receptor alph |
| AC005071.1 | 7 | novel transcript, antisense to PILRA |
| ZCWPW1 | 7 | zinc finger CW-type and PWWP domain conta |
| MEPCE | 7 | methylphosphate capping enzyme [Source:HG |
| AC092849.2 | 7 | novel transcript |
| PPP1R35 | 7 | protein phosphatase 1 regulatory subunit 35 [ |
| AC092849.1 | 7 | novel transcript |
| C7orf61 | 7 | chromosome 7 open reading frame 61 [Sourc |
| TSC22D4 | 7 | TSC22 domain family member 4 [Source:HGN |

| | | |
|---|---|---|
| AC092849.3 | 7 | novel transcript |
| NYAP1 | 7 | neuronal tyrosine phosphorylated phosphoin |
| AC069281.1 | 7 | novel transcript |
| RN7SL416P | 7 | RNA, 7SL, cytoplasmic 416, pseudogene [Sour |
| AGFG2 | 7 | ArfGAP with FG repeats 2 [Source:HGNC Symb |
| IRS3P | 7 | insulin receptor substrate 3, pseudogene [Sou |
| SAP25 | 7 | Sin3A associated protein 25 [Source:HGNC Sy |
| AC069281.2 | 7 | novel transcript |
| LRCH4 | 7 | leucine rich repeats and calponin homology d |
| FBXO24 | 7 | F-box protein 24 [Source:HGNC Symbol;Acc:H |
| PCOLCE-AS1 | 7 | PCOLCE antisense RNA 1 [Source:HGNC Symb |
| PCOLCE | 7 | procollagen C-endopeptidase enhancer [Sour |
| MOSPD3 | 7 | motile sperm domain containing 3 [Source:HG |
| TFR2 | 7 | transferrin receptor 2 [Source:HGNC Symbol; |
| ACTL6B | 7 | actin like 6B [Source:HGNC Symbol;Acc:HGNC |
| GNB2 | 7 | G protein subunit beta 2 [Source:HGNC Symb |
| GIGYF1 | 7 | GRB10 interacting GYF protein 1 [Source:HGN |
| POP7 | 7 | POP7 homolog, ribonuclease P/MRP subunit [ |
| EPO | 7 | erythropoietin [Source:HGNC Symbol;Acc:HG |
| ZAN | 7 | zonadhesin (gene/pseudogene) [Source:HGN |
| EPHB4 | 7 | EPH receptor B4 [Source:HGNC Symbol;Acc:H |
| RN7SL750P | 7 | RNA, 7SL, cytoplasmic 750, pseudogene [Sour |
| SLC12A9 | 7 | solute carrier family 12 member 9 [Source:HG |
| SLC12A9-AS1 | 7 | SLC12A9 antisense RNA 1 [Source:HGNC Symb |
| TRIP6 | 7 | thyroid hormone receptor interactor 6 [Sourc |
| MIR6875 | 7 | microRNA 6875 [Source:HGNC Symbol;Acc:HG |
| SRRT | 7 | serrate, RNA effector molecule [Source:HGNC |
| UFSP1 | 7 | UFM1 specific peptidase 1 (inactive) [Source: |
| ACHE | 7 | acetylcholinesterase (Cartwright blood group) |
| RN7SL549P | 7 | RNA, 7SL, cytoplasmic 549, pseudogene [Sour |
| RPS29P15 | 7 | ribosomal protein S29 pseudogene 15 [Source |
| MUC3A | 7 | mucin 3A, cell surface associated [Source:HGN |
| AC254629.1 | 7 | novel transcript |
| MUC12 | 7 | mucin 12, cell surface associated [Source:HGN |
| AC105446.1 | 7 | novel transcript |
| MUC17 | 7 | mucin 17, cell surface associated [Source:HGN |
| RN7SKP54 | 7 | RN7SK pseudogene 54 [Source:HGNC Symbol; |
| TRIM56 | 7 | tripartite motif containing 56 [Source:HGNC S |
| SERPINE1 | 7 | serpin family E member 1 [Source:HGNC Symb |
| AP1S1 | 7 | adaptor related protein complex 1 subunit sig |
| MIR4653 | 7 | microRNA 4653 [Source:HGNC Symbol;Acc:HG |
| VGF | 7 | VGF nerve growth factor inducible [Source:HG |
| NAT16 | 7 | N-acetyltransferase 16 (putative) [Source:HGN |
| MOGAT3 | 7 | monoacylglycerol O-acyltransferase 3 [Source |
| DGAT2L7P | 7 | diacylglycerol O-acyltransferase 2 like 7, pseu |
| RPSAP46 | 7 | ribosomal protein SA pseudogene 46 [Source: |
| PLOD3 | 7 | procollagen-lysine,2-oxoglutarate 5-dioxygen |
| ZNHIT1 | 7 | zinc finger HIT-type containing 1 [Source:HGN |
| CLDN15 | 7 | claudin 15 [Source:HGNC Symbol;Acc:HGNC:2 |
| FIS1 | 7 | fission, mitochondrial 1 [Source:HGNC Symbo |

| | | |
|---|---|---|
| RNU6-1104P | 7 | RNA, U6 small nuclear 1104, pseudogene [Sou |
| AC006329.2 | 7 | zinc finger protein 397 (ZNF397) pseudogene |
| AZGP1P2 | 7 | AZGP1 pseudogene 2 [Source:HGNC Symbol;A |
| LNCPRESS1 | 7 | lncRNA p53 regulated and ESC associated 1 [S |
| AC006329.1 | 7 | novel transcript |
| IFT22 | 7 | intraflagellar transport 22 [Source:HGNC Syml |
| COL26A1 | 7 | collagen type XXVI alpha 1 chain [Source:HGN |
| AC004965.1 | 7 | pseudogene similar to part of ribosomal prote |
| LINC01007 | 7 | long intergenic non-protein coding RNA 1007 |
| MYL10 | 7 | myosin light chain 10 [Source:HGNC Symbol;A |
| CUX1 | 7 | cut like homeobox 1 [Source:HGNC Symbol;Ac |
| AC005096.1 | 7 | novel transcript, antisense to CUX1 |
| AC005072.1 | 7 | novel transcript, sense intronic to CUX1 |
| AC005086.2 | 7 | novel transcript, antisense to CUX1 |
| AC005086.5 | 7 | NULL |
| AC005086.4 | 7 | NULL |
| AC005086.3 | 7 | NULL |
| AC005086.1 | 7 | NULL |
| AC005088.1 | 7 | ribosomal protein L10 (RPL10) pseudogene |
| SH2B2 | 7 | SH2B adaptor protein 2 [Source:HGNC Symbo |
| MIR4285 | 7 | microRNA 4285 [Source:HGNC Symbol;Acc:HG |
| AC091390.1 | 7 | pseudogene similar to part of stromal antigen |
| Y_RNA | 7 | Y RNA [Source:RFAM;Acc:RF00019] |
| AC091390.2 | 7 | postmeiotic segregation increased 2-like prote |
| SPDYE6 | 7 | speedy/RINGO cell cycle regulator family mem |
| PRKRIP1 | 7 | PRKR interacting protein 1 [Source:HGNC Sym |
| AC091390.4 | 7 | uncharacterized LOC100289561 [Source:NCBI |
| AC091390.3 | 7 | pseudogene similar to part of stromal antigen |
| MIR548O | 7 | microRNA 548o [Source:HGNC Symbol;Acc:HG |
| AC073517.1 | 7 | novel transcript |
| ORAI2 | 7 | ORAI calcium release-activated calcium modu |
| ALKBH4 | 7 | alkB homolog 4, lysine demethylase [Source:H |
| LRWD1 | 7 | leucine rich repeats and WD repeat domain co |
| MIR5090 | 7 | microRNA 5090 [Source:HGNC Symbol;Acc:HG |
| MIR4467 | 7 | microRNA 4467 [Source:HGNC Symbol;Acc:HG |
| POLR2J | 7 | RNA polymerase II subunit J [Source:HGNC Syr |
| RASA4B | 7 | RAS p21 protein activator 4B [Source:HGNC Sy |
| POLR2J3 | 7 | RNA polymerase II subunit J3 [Source:HGNC S |
| UPK3BL2 | 7 | uroplakin 3B like 2 [Source:HGNC Symbol;Acc: |
| AC093668.1 | 7 | novel protein |
| SPDYE2 | 7 | speedy/RINGO cell cycle regulator family mem |
| POLR2J3 | 7 | RNA polymerase II subunit J3 [Source:HGNC S |
| RASA4 | 7 | RAS p21 protein activator 4 [Source:HGNC Syr |
| AC105052.4 | 7 | novel transcript, sense overlapping POLR2J2, U |
| AC105052.1 | 7 | novel protein, UPK3BL-RASA4 readthrough |
| UPK3BL1 | 7 | uroplakin 3B like 1 [Source:HGNC Symbol;Acc: |
| AC105052.3 | 7 | novel protein, POLR2J2-UPK3BL readthrough |
| SPDYE2B | 7 | speedy/RINGO cell cycle regulator family mem |
| POLR2J2 | 7 | RNA polymerase II subunit J2 [Source:HGNC S |
| RASA4DP | 7 | RAS p21 protein activator 4CD, pseudogene [S |

| | | |
|---|---|---|
| AC105052.5 | 7 | novel transcript |
| FAM185A | 7 | family with sequence similarity 185 member A |
| AC105052.2 | 7 | ribosomal protein L7a (RPL7A) pseudogene |
| FBXL13 | 7 | F-box and leucine rich repeat protein 13 [Sour |
| RNU6-1136P | 7 | RNA, U6 small nuclear 1136, pseudogene [Sou |
| RN7SKP198 | 7 | RN7SK pseudogene 198 [Source:HGNC Symbo |
| LRRC17 | 7 | leucine rich repeat containing 17 [Source:HGN |
| NFE4 | 7 | nuclear factor, erythroid 4 [Source:HGNC Sym |
| AC073127.1 | 7 | novel transcript |
| ARMC10 | 7 | armadillo repeat containing 10 [Source:HGNC |
| CRYZP1 | 7 | crystallin zeta pseudogene 1 [Source:HGNC Sy |
| NAPEPLD | 7 | N-acyl phosphatidylethanolamine phospholipa |
| AC007683.1 | 7 | ribosomal protein L19 (RPL19) pseudogene |
| RPL23AP95 | 7 | ribosomal protein L23a pseudogene 95 [Sourc |
| AC007683.2 | 7 | tec |
| DPY19L2P2 | 7 | DPY19L2 pseudogene 2 [Source:HGNC Symbo |
| S100A11P1 | 7 | S100 calcium binding protein A11 pseudogene |
| PMPCB | 7 | peptidase, mitochondrial processing beta subu |
| DNAJC2 | 7 | DnaJ heat shock protein family (Hsp40) memb |
| PSMC2 | 7 | proteasome 26S subunit, ATPase 2 [Source:HG |
| RPS29P16 | 7 | ribosomal protein S29 pseudogene 16 [Source |
| SLC26A5 | 7 | solute carrier family 26 member 5 [Source:HG |
| Y_RNA | 7 | Y RNA [Source:RFAM;Acc:RF00019] |
| Y_RNA | 7 | Y RNA [Source:RFAM;Acc:RF00019] |
| AC005064.1 | 7 | novel transcript, antisense to RELN and SLC26 |
| RELN | 7 | reelin [Source:HGNC Symbol;Acc:HGNC:9957] |
| RN7SKP86 | 7 | RN7SK pseudogene 86 [Source:HGNC Symbol; |
| ORC5 | 7 | origin recognition complex subunit 5 [Source:I |
| LHFPL3 | 7 | LHFPL tetraspan subfamily member 3 [Source |
| EIF4BP6 | 7 | eukaryotic translation initiation factor 4B pseu |
| LHFPL3-AS1 | 7 | LHFPL3 antisense RNA 1 [Source:HGNC Symbo |
| AC091286.1 | 7 | novel pseudogene KIAA1191 |
| LHFPL3-AS2 | 7 | LHFPL3 antisense RNA 2 [Source:HGNC Symbo |
| RN7SL8P | 7 | RNA, 7SL, cytoplasmic 8, pseudogene [Source: |
| AC007384.1 | 7 | novel transcript |
| LINC01004 | 7 | long intergenic non-protein coding RNA 1004 |
| KMT2E-AS1 | 7 | KMT2E antisense RNA 1 [Source:HGNC Symbo |
| KMT2E | 7 | lysine methyltransferase 2E [Source:HGNC Syr |
| AC005070.3 | 7 | novel transcript, antisense to MLL5 |
| SRPK2 | 7 | SRSF protein kinase 2 [Source:HGNC Symbol;A |
| AC005070.1 | 7 | ribosomal protein, large, P1, pseudogene |
| AC005070.2 | 7 | peroxiredoxin 3, pseudogene |
| AC004884.1 | 7 | ribosomal protein L36 (RPL36) pseudogene |
| RWDD4P1 | 7 | RWD domain containing 4 pseudogene 1 [Sou |
| AC004884.2 | 7 | novel transcript |
| RNU6-1322P | 7 | RNA, U6 small nuclear 1322, pseudogene [Sou |
| PUS7 | 7 | pseudouridine synthase 7 [Source:HGNC Symb |
| AC073073.1 | 7 | ring finger protein 181 (RNF181) pseudogene |
| RINT1 | 7 | RAD50 interactor 1 [Source:HGNC Symbol;Acc |
| EFCAB10 | 7 | EF-hand calcium binding domain 10 [Source:H |

| | | |
|---|---|---|
| AC073073.2 | 7 | novel transcript, antisense to EFCAB10 |
| YBX1P2 | 7 | Y-box binding protein 1 pseudogene 2 [Source |
| ATXN7L1 | 7 | ataxin 7 like 1 [Source:HGNC Symbol;Acc:HGN |
| AC005099.1 | 7 | ribosomal protein L13a (RPL13A) pseudogene |
| CDHR3 | 7 | cadherin related family member 3 [Source:HG |
| AC004836.1 | 7 | TEC |
| SYPL1 | 7 | synaptophysin like 1 [Source:HGNC Symbol;Ac |
| DCAF13P1 | 7 | DDB1 and CUL4 associated factor 13 pseudog |
| RNU6-392P | 7 | RNA, U6 small nuclear 392, pseudogene [Sour |
| NAMPT | 7 | nicotinamide phosphoribosyltransferase [Sou |
| AC007032.1 | 7 | novel transcript, antisense to NAMPT |
| LARP1BP2 | 7 | La ribonucleoprotein domain family member |
| AC004917.1 | 7 | novel transcript, sense overlapping CCDC71L |
| AC005050.3 | 7 | novel transcript |
| AC005050.1 | 7 | novel transcript |
| AC005050.2 | 7 | novel transcript |
| CCDC71L | 7 | coiled-coil domain containing 71 like [Source: |
| LINC02577 | 7 | long intergenic non-protein coding RNA 2577 |
| RNA5SP236 | 7 | RNA, 5S ribosomal pseudogene 236 [Source:H |
| PIK3CG | 7 | phosphatidylinositol-4,5-bisphosphate 3-kinas |
| PRKAR2B | 7 | protein kinase cAMP-dependent type II regula |
| AC006387.1 | 7 | novel transcript, antisense to PRKAR2B |
| HBP1 | 7 | HMG-box transcription factor 1 [Source:HGNC |
| AC004492.1 | 7 | novel transcript, antisense to HBP1 |
| COG5 | 7 | component of oligomeric golgi complex 5 [Sou |
| AC002381.1 | 7 | ribosomal protein L37a (RPL37A) pseudogene |
| GPR22 | 7 | G protein-coupled receptor 22 [Source:HGNC |
| DUS4L | 7 | dihydrouridine synthase 4 like [Source:HGNC S |
| AC004839.2 | 7 | novel transcript, antisense to BCAP29 |
| BCAP29 | 7 | B cell receptor associated protein 29 [Source: |
| AC004839.1 | 7 | NULL |
| WBP1LP2 | 7 | WBP1L pseudogene 2 [Source:HGNC Symbol;A |
| BANF1P5 | 7 | barrier to autointegration factor 1 pseudogen |
| SLC26A4-AS1 | 7 | SLC26A4 antisense RNA 1 [Source:HGNC Symb |
| SLC26A4 | 7 | solute carrier family 26 member 4 [Source:HG |
| AC002467.1 | 7 | novel transcript, antisense to CBLL1 |
| CBLL1 | 7 | Cbl proto-oncogene like 1 [Source:HGNC Syml |
| SLC26A3 | 7 | solute carrier family 26 member 3 [Source:HG |
| PIGCP2 | 7 | phosphatidylinositol glycan anchor biosynthes |
| DLD | 7 | dihydrolipoamide dehydrogenase [Source:HG |
| LAMB1 | 7 | laminin subunit beta 1 [Source:HGNC Symbol; |
| AC005046.1 | 7 | novel transcript, antisense to LAMB1 |
| U3 | 7 | Small nucleolar RNA U3 [Source:RFAM;Acc:RF |
| LAMB4 | 7 | laminin subunit beta 4 [Source:HGNC Symbol; |
| NRCAM | 7 | neuronal cell adhesion molecule [Source:HGN |
| PNPLA8 | 7 | patatin like phospholipase domain containing |
| RPL7P32 | 7 | ribosomal protein L7 pseudogene 32 [Source: |
| THAP5 | 7 | THAP domain containing 5 [Source:HGNC Sym |
| DNAJB9 | 7 | DnaJ heat shock protein family (Hsp40) memb |
| AC005487.1 | 7 | novel transcript |

| | | |
|---|---|---|
| AC002487.1 | 7 | sorbitol dehydrogenase, pseudogene |
| C7orf66 | 7 | chromosome 7 open reading frame 66 [Sourc |
| AC004014.2 | 7 | TEC |
| AC004014.1 | 7 | novel transcript |
| BUB3P1 | 7 | BUB3 mitotic checkpoint protein pseudogene |
| AC002386.1 | 7 | novel transcript |
| AC073071.1 | 7 | novel transcript |
| AC073387.1 | 7 | STAM binding protein-like 1 (STAMBPL1) pseu |
| AC073387.2 | 7 | immunoglobulin (CD79A) binding protein 1, p |
| EIF3IP1 | 7 | eukaryotic translation initiation factor 3 subu |
| RPL3P8 | 7 | ribosomal protein L3 pseudogene 8 [Source:H |
| AC073114.1 | 7 | novel transcript |
| AC073114.2 | 7 | novel zinc finger protein pseudogene |
| AC073326.2 | 7 | novel transcript |
| IMMP2L | 7 | inner mitochondrial membrane peptidase sub |
| AC073326.1 | 7 | novel transcript |
| LRRN3 | 7 | leucine rich repeat neuronal 3 [Source:HGNC |
| AC003989.2 | 7 | pseudogene similar to part of adenosine dean |
| AC003989.1 | 7 | pseudogene similar to part of argininosuccina |
| DOCK4 | 7 | dedicator of cytokinesis 4 [Source:HGNC Symb |
| AC003077.1 | 7 | novel transcript, antisense to DOCK4 |
| DOCK4-AS1 | 7 | DOCK4 antisense RNA 1 [Source:HGNC Symbo |
| RNA5SP237 | 7 | RNA, 5S ribosomal pseudogene 237 [Source:H |
| AC003080.1 | 7 | ribosomal protein L8 (RPL8) pseudogene |
| ZNF277 | 7 | zinc finger protein 277 [Source:HGNC Symbol; |
| RN7SKP187 | 7 | RN7SK pseudogene 187 [Source:HGNC Symbo |
| AC004112.1 | 7 | novel transcript, antisense to ZNF277 |
| MTND5P8 | 7 | MT-ND5 pseudogene 8 [Source:HGNC Symbol |
| MTND6P24 | 7 | MT-ND6 pseudogene 24 [Source:HGNC Symbo |
| MTCYBP24 | 7 | MT-CYB pseudogene 24 [Source:HGNC Symbo |
| IFRD1 | 7 | interferon related developmental regulator 1 |
| AC079741.1 | 7 | ribosomal protein S27a (RPS27A) pseudogene |
| LSMEM1 | 7 | leucine rich single-pass membrane protein 1 [ |
| NPM1P14 | 7 | nucleophosmin 1 pseudogene 14 [Source:HGI |
| AC002463.1 | 7 | novel transcript |
| MIPEPP1 | 7 | mitochondrial intermediate peptidase pseudo |
| TMEM168 | 7 | transmembrane protein 168 [Source:HGNC Sy |
| BMT2 | 7 | base methyltransferase of 25S rRNA 2 homolo |
| AC018464.1 | 7 | novel transcript |
| AC073346.2 | 7 | tec |
| GPR85 | 7 | G protein-coupled receptor 85 [Source:HGNC |
| AC073346.1 | 7 | novel transcript |
| SMIM30 | 7 | small integral membrane protein 30 [Source:H |
| AC073137.1 | 7 | VAMP (vesicle-associated membrane protein) |
| AC073348.1 | 7 | TEC |
| AC073348.2 | 7 | novel transcript |
| PPP1R3A | 7 | protein phosphatase 1 regulatory subunit 3A [ |
| FOXP2 | 7 | forkhead box P2 [Source:HGNC Symbol;Acc:H |
| AC073626.2 | 7 | ribosomal protein L36 (RPL36) pseudogene |
| AC073626.1 | 7 | novel transcript |

| | | |
|---|---|---|
| AC003992.1 | 7 | novel transcript, sense intronic to FOXP2 |
| RNA5SP238 | 7 | RNA, 5S ribosomal pseudogene 238 [Source:H |
| MIR3666 | 7 | microRNA 3666 [Source:HGNC Symbol;Acc:HG |
| MDFIC | 7 | MyoD family inhibitor domain containing [Sou |
| LINC01393 | 7 | long intergenic non-protein coding RNA 1393 |
| LINC01392 | 7 | long intergenic non-protein coding RNA 1392 |
| RAC1P6 | 7 | Rac family small GTPase 1 pseudogene 6 [Sou |
| Y_RNA | 7 | Y RNA [Source:RFAM;Acc:RF00019] |
| AC073901.1 | 7 | polymerase (RNA) II (DNA directed) polypepti |
| SNORA25B | 7 | small nucleolar RNA, H/ACA box 25B [Source:I |
| AC092590.1 | 7 | novel transcript |
| AC093714.1 | 7 | novel transcript |
| AC093714.2 | 7 | novel transcript |
| Y_RNA | 7 | Y RNA [Source:RFAM;Acc:RF00019] |
| TFEC | 7 | transcription factor EC [Source:HGNC Symbol; |
| AC073130.3 | 7 | TEC |
| TES | 7 | testin LIM domain protein [Source:HGNC Sym |
| AC073130.2 | 7 | novel transcript |
| AC002066.1 | 7 | novel transcript |
| AC073130.1 | 7 | uncharacterized LOC102724434 [Source:NCBI |
| CAV2 | 7 | caveolin 2 [Source:HGNC Symbol;Acc:HGNC:1 |
| Metazoa_SRP | 7 | Metazoan signal recognition particle RNA [Sou |
| CAV1 | 7 | caveolin 1 [Source:HGNC Symbol;Acc:HGNC:1 |
| AC006159.1 | 7 | novel transcript |
| LINC01510 | 7 | long intergenic non-protein coding RNA 1510 |
| AC006159.2 | 7 | novel transcript |
| MET | 7 | MET proto-oncogene, receptor tyrosine kinas |
| CAPZA2 | 7 | capping actin protein of muscle Z-line subunit |
| AC002543.1 | 7 | novel pseudogene |
| Y_RNA | 7 | Y RNA [Source:RFAM;Acc:RF00019] |
| RNA5SP239 | 7 | RNA, 5S ribosomal pseudogene 239 [Source:H |
| ST7-AS1 | 7 | ST7 antisense RNA 1 [Source:HGNC Symbol;Ac |
| AC106873.5 | 7 | NULL |
| AC106873.7 | 7 | NULL |
| ST7 | 7 | suppression of tumorigenicity 7 [Source:HGN( |
| ST7-OT4 | 7 | ST7 overlapping transcript 4 [Source:HGNC Sy |
| AC106873.3 | 7 | NULL |
| AC106873.2 | 7 | NULL |
| AC106873.6 | 7 | NULL |
| AC106873.4 | 7 | NULL |
| AC106873.1 | 7 | novel transcript, antisense to ST7 |
| TPM3P1 | 7 | tropomyosin 3 pseudogene 1 [Source:HGNC S |
| MIR6132 | 7 | microRNA 6132 [Source:HGNC Symbol;Acc:HG |
| ST7-AS2 | 7 | ST7 antisense RNA 2 [Source:HGNC Symbol;Ac |
| AC002542.6 | 7 | novel transcript, antisense to ST7 |
| AC002542.2 | 7 | NULL |
| AC002542.3 | 7 | NULL |
| AC002542.1 | 7 | SMT3 suppressor of mif two 3 homolog 2 (S. c |
| AC002542.5 | 7 | NULL |
| AC002542.4 | 7 | NULL |

| | | |
|---|---|---|
| AC006326.1 | 7 | heterogeneous nuclear ribonucleoprotein A1 |
| MTND4P6 | 7 | MT-ND4 pseudogene 6 [Source:HGNC Symbol |
| MTCYBP6 | 7 | MT-CYB pseudogene 6 [Source:HGNC Symbol; |
| WNT2 | 7 | Wnt family member 2 [Source:HGNC Symbol;/ |
| AC002465.1 | 7 | novel transcript |
| AC002465.2 | 7 | novel transcript |
| ASZ1 | 7 | ankyrin repeat, SAM and basic leucine zipper ( |
| AC003045.1 | 7 | fetal globin-inducing factor (FGIF) pseudogene |
| CFTR | 7 | CF transmembrane conductance regulator [Sc |
| AC000111.2 | 7 | ribosomal protein L32 (RPL32) pseudogene |
| CFTR-AS1 | 7 | CFTR antisense RNA 1 [Source:HGNC Symbol;/ |
| AC000111.1 | 7 | dynein, cytoplasmic, light polypeptide 1 (DNCl |
| AC000061.1 | 7 | novel transcript, antisense to CFTR |
| CTTNBP2 | 7 | cortactin binding protein 2 [Source:HGNC Sym |
| AC007568.1 | 7 | splicing factor, arginine/serine-rich 3 (SFRS3) p |
| AC003084.1 | 7 | novel transcript |
| LSM8 | 7 | LSM8 homolog, U6 small nuclear RNA associa |
| ANKRD7 | 7 | ankyrin repeat domain 7 [Source:HGNC Symb |
| AC002529.1 | 7 | novel transcript |
| GTF3AP6 | 7 | general transcription factor IIIA pseudogene 6 |
| AC092098.1 | 7 | eukaryotic translation initiation factor 3, subu |
| AC079791.1 | 7 | cereblon, pseudogene |
| LINC02476 | 7 | long intergenic non-protein coding RNA 2476 |
| RNU1-29P | 7 | RNA, U1 small nuclear 29, pseudogene [Sourc |
| AC004946.1 | 7 | novel transcript |
| AC004946.2 | 7 | novel transcript |
| KCND2 | 7 | potassium voltage-gated channel subfamily D |
| RNU6-581P | 7 | RNA, U6 small nuclear 581, pseudogene [Sour |
| AC004888.1 | 7 | novel transcript |
| TSPAN12 | 7 | tetraspanin 12 [Source:HGNC Symbol;Acc:HGI |
| ING3 | 7 | inhibitor of growth family member 3 [Source:l |
| RNA5SP240 | 7 | RNA, 5S ribosomal pseudogene 240 [Source:H |
| CPED1 | 7 | cadherin like and PC-esterase domain contain |
| HMGN1P18 | 7 | high mobility group nucleosome binding doma |
| RNA5SP241 | 7 | RNA, 5S ribosomal pseudogene 241 [Source:H |
| RNU6-517P | 7 | RNA, U6 small nuclear 517, pseudogene [Sour |
| AC006364.1 | 7 | novel transcript |
| WNT16 | 7 | Wnt family member 16 [Source:HGNC Symbol |
| FAM3C | 7 | family with sequence similarity 3 member C [S |
| CYCSP19 | 7 | CYCS pseudogene 19 [Source:HGNC Symbol;A |
| AC074085.1 | 7 | CKLF-like MARVEL transmembrane domain co |
| AC074085.2 | 7 | ribosomal protein L18 (RPL18) pseudogene |
| AC004875.1 | 7 | novel transcript |
| RN7SKP277 | 7 | RN7SK pseudogene 277 [Source:HGNC Symbo |
| PNPT1P2 | 7 | polyribonucleotide nucleotidyltransferase 1 ps |
| PTPRZ1 | 7 | protein tyrosine phosphatase receptor type Z1 |
| AASS | 7 | aminoadipate-semialdehyde synthase [Source |
| RNU7-154P | 7 | RNA, U7 small nuclear 154 pseudogene [Sourc |
| AC015983.2 | 7 | novel transcript |
| AC006020.1 | 7 | tetratricopeptide repeat domain 14 (TTC14) p |

| | | |
|---|---|---|
| AC015983.1 | 7 | ribosomal protein L31 (RPL31) pseudogene |
| FEZF1 | 7 | FEZ family zinc finger 1 [Source:HGNC Symbol; |
| FEZF1-AS1 | 7 | FEZF1 antisense RNA 1 [Source:HGNC Symbol; |
| CADPS2 | 7 | calcium dependent secretion activator 2 [Sour |
| AC004594.1 | 7 | novel transcript, antisense to CADPS2 |
| AC006463.1 | 7 | pseudogene similar to part of dedicator of cyt |
| RPS26P31 | 7 | ribosomal protein S26 pseudogene 31 [Source |
| RNF133 | 7 | ring finger protein 133 [Source:HGNC Symbol; |
| RNF148 | 7 | ring finger protein 148 [Source:HGNC Symbol; |
| AC004986.1 | 7 | v-rel reticuloendotheliosis viral oncogene hon |
| TAS2R16 | 7 | taste 2 receptor member 16 [Source:HGNC Sy |
| AC073311.1 | 7 | novel transcript |
| SLC13A1 | 7 | solute carrier family 13 member 1 [Source:HG |
| LYPLA1P1 | 7 | LYPLA1 pseudogene 1 [Source:HGNC Symbol; |
| IQUB | 7 | IQ motif and ubiquitin domain containing [Sou |
| AC073323.1 | 7 | novel transcript |
| RNU6-296P | 7 | RNA, U6 small nuclear 296, pseudogene [Sour |
| NDUFA5 | 7 | NADH:ubiquinone oxidoreductase subunit A5 |
| ASB15 | 7 | ankyrin repeat and SOCS box containing 15 [S |
| AC006333.1 | 7 | novel transcript |
| LMOD2 | 7 | leiomodin 2 [Source:HGNC Symbol;Acc:HGNC: |
| WASL | 7 | WASP like actin nucleation promoting factor [ |
| AC006333.2 | 7 | novel transcript, antisense to WASL |
| RNU6-11P | 7 | RNA, U6 small nuclear 11, pseudogene [Sourc |
| HYAL6P | 7 | hyaluronidase 6, pseudogene [Source:HGNC S |
| HYAL4 | 7 | hyaluronidase 4 [Source:HGNC Symbol;Acc:H( |
| SPAM1 | 7 | sperm adhesion molecule 1 [Source:HGNC Syr |
| AC004690.1 | 7 | cytochrome c oxidase subunit VIa polypeptide |
| AC004690.2 | 7 | novel transcript |
| TMEM229A | 7 | transmembrane protein 229A [Source:HGNC S |
| AC006148.1 | 7 | novel transcript |
| AC006148.2 | 7 | novel transcript |
| AC004930.1 | 7 | novel transcript |
| SSU72P8 | 7 | SSU72 pseudogene 8 [Source:HGNC Symbol;A |
| AC005521.1 | 7 | ribosomal protein S2 (RPS2) pseudogene |
| RNU6-102P | 7 | RNA, U6 small nuclear 102, pseudogene [Sour |
| AC004925.1 | 7 | tec |
| GPR37 | 7 | G protein-coupled receptor 37 [Source:HGNC |
| C7orf77 | 7 | chromosome 7 open reading frame 77 [Sourc |
| POT1 | 7 | protection of telomeres 1 [Source:HGNC Syml |
| POT1-AS1 | 7 | POT1 antisense RNA 1 [Source:HGNC Symbol; |
| EEF1GP1 | 7 | eukaryotic translation elongation factor 1 gan |
| LINC02830 | 7 | long intergenic non-protein coding RNA 2830 |
| AC019155.1 | 7 | chromosome 11 open reading frame 58 (C11c |
| AC019155.2 | 7 | novel transcript |
| AC010099.1 | 7 | ribosomal protein L31 (RPL31) pseudogene |
| AC010099.3 | 7 | metaxin 2 (MTX2) pseudogene |
| AC010099.2 | 7 | peptidylprolyl isomerase A (cyclophilin A) (PPI |
| AC003975.1 | 7 | novel transcript |
| AC000372.1 | 7 | novel transcript |

| | | |
|---|---|---|
| GRM8 | 7 | glutamate metabotropic receptor 8 [Source:H |
| AC002057.2 | 7 | novel transcript, antisense to GRM8 |
| AC000374.1 | 7 | poly(A) binding protein, cytoplasmic 1 (PABPC |
| AC002057.1 | 7 | novel transcript, antisense to GRM8 |
| AC000367.1 | 7 | tigger transposable element derived 1 (TIGD1 |
| AC000362.1 | 7 | ATP synthase, H+ transporting, mitochondrial |
| MIR592 | 7 | microRNA 592 [Source:HGNC Symbol;Acc:HGI |
| AC000099.1 | 7 | novel transcript |
| PRELID3BP10 | 7 | PRELI domain containing 3B pseudogene 10 [S |
| ZNF800 | 7 | zinc finger protein 800 [Source:HGNC Symbol; |
| AC000123.2 | 7 | TEC |
| AC000123.1 | 7 | novel transcript |
| AC000123.3 | 7 | TEC |
| AC000124.1 | 7 | novel transcript |
| GCC1 | 7 | GRIP and coiled-coil domain containing 1 [Sou |
| ARF5 | 7 | ADP ribosylation factor 5 [Source:HGNC Symb |
| FSCN3 | 7 | fascin actin-bundling protein 3 [Source:HGNC |
| PAX4 | 7 | paired box 4 [Source:HGNC Symbol;Acc:HGNC |
| AC073934.1 | 7 | novel transcript, antisense to SND1 |
| SND1 | 7 | staphylococcal nuclease and tudor domain co |
| AC006529.1 | 7 | pseudogene similar to part of CD2 (cytoplasm |
| SND1-IT1 | 7 | SND1 intronic transcript 1 [Source:HGNC Syml |
| LRRC4 | 7 | leucine rich repeat containing 4 [Source:HGN( |
| MIR593 | 7 | microRNA 593 [Source:HGNC Symbol;Acc:HGI |
| MIR129-1 | 7 | microRNA 129-1 [Source:HGNC Symbol;Acc:H( |
| LEP | 7 | leptin [Source:HGNC Symbol;Acc:HGNC:6553] |
| AC018635.2 | 7 | novel transcript |
| RBM28 | 7 | RNA binding motif protein 28 [Source:HGNC S |
| AC018635.1 | 7 | aurora kinase A (AURKA) pseudogene |
| RNU7-27P | 7 | RNA, U7 small nuclear 27 pseudogene [Source |
| PRRT4 | 7 | proline rich transmembrane protein 4 [Source |
| IMPDH1 | 7 | inosine monophosphate dehydrogenase 1 [So |
| AC010655.1 | 7 | ribosomal protein L37 (RPL37) pseudogene |
| RNU7-54P | 7 | RNA, U7 small nuclear 54 pseudogene [Source |
| AC010655.4 | 7 | novel transcript, readthrough between HILPD/ |
| HILPDA | 7 | hypoxia inducible lipid droplet associated [Sou |
| AC010655.2 | 7 | novel transcript |
| AC010655.3 | 7 | EF-hand calcium binding domain 3 (EFCAB3) p |
| METTL2B | 7 | methyltransferase like 2B [Source:HGNC Symb |
| Metazoa_SRP | 7 | Metazoan signal recognition particle RNA [Sou |
| AC090114.2 | 7 | novel transcript |
| AC090114.3 | 7 | pseudogene similar to part of tousled-like kin |
| AC090114.1 | 7 | ribosomal protein S10 (RPS10) pseudogene |
| AC108010.1 | 7 | novel transcript |
| AC018638.8 | 7 | pseudogene, similar to general transcription f |
| RNU6-177P | 7 | RNA, U6 small nuclear 177, pseudogene [Sour |
| AC018638.4 | 7 | novel pseudogene |
| AC018638.2 | 7 | novel pseudogene |
| AC018638.1 | 7 | novel pseudogene |
| AC018638.5 | 7 | novel pseudogene |

| | | |
|---|---|---|
| CICP14 | 7 | capicua transcriptional repressor pseudogene |
| AC018638.7 | 7 | novel transcript |
| AC018638.3 | 7 | pseudogene similar to part of EMG1 nucleolar |
| FAM71F2 | 7 | family with sequence similarity 71 member F2 |
| AC018638.6 | 7 | novel transcript |
| IMP3P2 | 7 | IMP3 pseudogene 2 [Source:HGNC Symbol;Ac |
| RNA5SP242 | 7 | RNA, 5S ribosomal pseudogene 242 [Source:H |
| RNA5SP243 | 7 | RNA, 5S ribosomal pseudogene 243 [Source:H |
| FAM71F1 | 7 | family with sequence similarity 71 member F1 |
| CALU | 7 | calumenin [Source:HGNC Symbol;Acc:HGNC:1 |
| RN7SL81P | 7 | RNA, 7SL, cytoplasmic 81, pseudogene [Sourc |
| OPN1SW | 7 | opsin 1, short wave sensitive [Source:HGNC Sy |
| CCDC136 | 7 | coiled-coil domain containing 136 [Source:HG |
| FLNC | 7 | filamin C [Source:HGNC Symbol;Acc:HGNC:37 |
| FLNC-AS1 | 7 | FLNC antisense RNA 1 [Source:HGNC Symbol; |
| KCP | 7 | kielin cysteine rich BMP regulator [Source:HG |
| ATP6V1F | 7 | ATPase H+ transporting V1 subunit F [Source: |
| ATP6V1FNB | 7 | ATP6V1F neighbor [Source:HGNC Symbol;Acc: |
| AC025594.1 | 7 | pseudogene similar to part of filamin C, gamm |
| IRF5 | 7 | interferon regulatory factor 5 [Source:HGNC S |
| AC025594.2 | 7 | novel transcript |
| TNPO3 | 7 | transportin 3 [Source:HGNC Symbol;Acc:HGN |
| RN7SL306P | 7 | RNA, 7SL, cytoplasmic 306, pseudogene [Sour |
| ODCP | 7 | ornithine decarboxylase pseudogene [Source: |
| TPI1P2 | 7 | triosephosphate isomerase 1 pseudogene 2 [S |
| AC018639.1 | 7 | pseudogene similar to part of Sec61 alpha 1 su |
| AC011005.2 | 7 | pseudogene similar to part of vomeronasal 1 r |
| AC011005.3 | 7 | zinc finger pseudogene |
| CYCSP20 | 7 | CYCS pseudogene 20 [Source:HGNC Symbol;A |
| AC011005.1 | 7 | novel protein similar to mitogen-activated pro |
| TSPAN33 | 7 | tetraspanin 33 [Source:HGNC Symbol;Acc:HGI |
| RNY1P11 | 7 | RNY1 pseudogene 11 [Source:HGNC Symbol;A |
| SMO | 7 | smoothened, frizzled class receptor [Source:H |
| AC011005.4 | 7 | novel transcript |
| AHCYL2 | 7 | adenosylhomocysteinase like 2 [Source:HGNC |
| AC009244.2 | 7 | novel transcript, antisense to AHCYL2 |
| AC009244.1 | 7 | cell division cycle 26 homolog (S. cerevisiae) ( |
| STRIP2 | 7 | striatin interacting protein 2 [Source:HGNC Sy |
| SNRPGP3 | 7 | small nuclear ribonucleoprotein polypeptide G |
| RNU1-72P | 7 | RNA, U1 small nuclear 72, pseudogene [Sourc |
| SMKR1 | 7 | small lysine rich protein 1 [Source:HGNC Symb |
| AC078846.1 | 7 | novel transcript |
| NRF1 | 7 | nuclear respiratory factor 1 [Source:HGNC Syr |
| RNA5SP244 | 7 | RNA, 5S ribosomal pseudogene 244 [Source:H |
| AC084864.2 | 7 | novel transcript |
| MIR182 | 7 | microRNA 182 [Source:HGNC Symbol;Acc:HGI |
| MIR96 | 7 | microRNA 96 [Source:HGNC Symbol;Acc:HGN |
| MIR183 | 7 | microRNA 183 [Source:HGNC Symbol;Acc:HGI |
| AC084864.1 | 7 | novel transcript |
| UBE2H | 7 | ubiquitin conjugating enzyme E2 H [Source:HG |

| | | |
|---|---|---|
| AC073320.1 | 7 | novel transcript, antisense to ZC3HC1 |
| Y_RNA | 7 | Y RNA [Source:RFAM;Acc:RF00019] |
| ZC3HC1 | 7 | zinc finger C3HC-type containing 1 [Source:HG |
| RNA5SP245 | 7 | RNA, 5S ribosomal pseudogene 245 [Source:H |
| KLHDC10 | 7 | kelch domain containing 10 [Source:HGNC Syr |
| AC087071.1 | 7 | novel transcript |
| TMEM209 | 7 | transmembrane protein 209 [Source:HGNC Sy |
| AC087071.2 | 7 | novel transcript |
| SSMEM1 | 7 | serine rich single-pass membrane protein 1 [S |
| AC024085.1 | 7 | ribosomal protein L29, pseudogene |
| CPA2 | 7 | carboxypeptidase A2 [Source:HGNC Symbol;A |
| CPA4 | 7 | carboxypeptidase A4 [Source:HGNC Symbol;A |
| CPA5 | 7 | carboxypeptidase A5 [Source:HGNC Symbol;A |
| CPA1 | 7 | carboxypeptidase A1 [Source:HGNC Symbol;A |
| CEP41 | 7 | centrosomal protein 41 [Source:HGNC Symbo |
| AC007938.1 | 7 | novel transcript |
| MESTIT1 | 7 | MEST intronic transcript 1, antisense RNA [Sou |
| MEST | 7 | mesoderm specific transcript [Source:HGNC S |
| AC007938.5 | 7 | NULL |
| AC007938.6 | 7 | NULL |
| AC007938.4 | 7 | NULL |
| AC007938.2 | 7 | novel transcript, antisense to MEST |
| MIR335 | 7 | microRNA 335 [Source:HGNC Symbol;Acc:HGN |
| COPG2 | 7 | coatomer protein complex subunit gamma 2 [ |
| AC007938.3 | 7 | novel transcript, antisense to COPG2 |
| AUXG01000058.1 | 7 | COPG2 imprinted transcript 1 |
| RNA5SP246 | 7 | RNA, 5S ribosomal pseudogene 246 [Source:H |
| AC234644.1 | 7 | glutamyl-tRNA(Gln) amidotransferase, subuni |
| TSGA13 | 7 | testis specific 13 [Source:HGNC Symbol;Acc:H |
| AC234644.2 | 7 | novel transcript, antisense to TSGA13 |
| KLF14 | 7 | Kruppel like factor 14 [Source:HGNC Symbol;A |
| AC016831.6 | 7 | novel transcript |
| AC016831.7 | 7 | novel transcript |
| AC016831.2 | 7 | armadillo repeat containing, X-linked 2 (ARMC |
| AC016831.4 | 7 | histone cluster H4 pseudogene |
| AC016831.3 | 7 | zinc finger protein 131 (ZNF131) pseudogene |
| LINC00513 | 7 | long intergenic non-protein coding RNA 513 [S |
| MIR29A | 7 | microRNA 29a [Source:HGNC Symbol;Acc:HGN |
| AC016831.1 | 7 | novel transcript |
| MIR29B1 | 7 | microRNA 29b-1 [Source:HGNC Symbol;Acc:H |
| AC016831.5 | 7 | novel transcript |
| AC058791.1 | 7 | novel transcript |
| LINC-PINT | 7 | long intergenic non-protein coding RNA, p53 i |
| RNU6-1010P | 7 | RNA, U6 small nuclear 1010, pseudogene [Sou |
| MKLN1 | 7 | muskelin 1 [Source:HGNC Symbol;Acc:HGNC:7 |
| AC009362.1 | 7 | ribosomal protein L27 (RPL27) pseudogene |
| MKLN1-AS | 7 | MKLN1 antisense RNA [Source:HGNC Symbol; |
| AC008264.2 | 7 | novel transcript, antisense to MKLN1 |
| PODXL | 7 | podocalyxin like [Source:HGNC Symbol;Acc:HG |
| AC008264.1 | 7 | ribosomal protein L31 (RPL31) pseudogene |

| | | |
|---|---|---|
| EEF1B2P6 | 7 | eukaryotic translation elongation factor 1 beta |
| AC093106.1 | 7 | ribosomal protein S14 (RPS14) pseudogene |
| AC093106.2 | 7 | ribosomal protein S15a (RPS15A) pseudogene |
| NDUFB9P2 | 7 | NADH:ubiquinone oxidoreductase subunit B9 |
| AC009518.2 | 7 | capping protein (actin filament) muscle Z-line, |
| AC009518.3 | 7 | capping protein (actin filament) muscle Z-line, |
| AC009518.1 | 7 | novel transcript |
| AC105443.1 | 7 | zinc finger protein pseudogene |
| PLXNA4 | 7 | plexin A4 [Source:HGNC Symbol;Acc:HGNC:91 |
| AC018643.1 | 7 | novel transcript |
| AC011625.1 | 7 | novel transcript |
| AC009365.1 | 7 | novel transcript |
| AC009365.2 | 7 | NULL |
| AC009365.3 | 7 | novel transcript |
| CHCHD3 | 7 | coiled-coil-helix-coiled-coil-helix domain conta |
| AC009365.4 | 7 | novel transcript |
| AC008038.1 | 7 | eukaryotic translation elongation factor 1 gam |
| MIR3654 | 7 | microRNA 3654 [Source:HGNC Symbol;Acc:HG |
| RNU6-92P | 7 | RNA, U6 small nuclear 92, pseudogene [Sourc |
| AC083873.1 | 7 | suppression of tumorigenicity 13 (colon carcir |
| EXOC4 | 7 | exocyst complex component 4 [Source:HGNC |
| MIR6133 | 7 | microRNA 6133 [Source:HGNC Symbol;Acc:HG |
| AC083875.1 | 7 | ribosomal protein S15a(RPS15A)pseudogene |
| COX5BP3 | 7 | cytochrome c oxidase subunit 5B pseudogene |
| AC007790.1 | 7 | ribosomal protein S3A (RPS3A) pseudogene |
| LRGUK | 7 | leucine rich repeats and guanylate kinase dom |
| AC008154.2 | 7 | novel transcript |
| SLC35B4 | 7 | solute carrier family 35 member B4 [Source:H |
| AC008154.1 | 7 | novel transcript |
| AC009275.1 | 7 | novel transcript |
| AKR1B1 | 7 | aldo-keto reductase family 1 member B [Sourc |
| AKR1B10 | 7 | aldo-keto reductase family 1 member B10 [So |
| AKR1B15 | 7 | aldo-keto reductase family 1 member B15 [So |
| BPGM | 7 | bisphosphoglycerate mutase [Source:HGNC Sy |
| AC009276.1 | 7 | novel transcript |
| TUBB3P2 | 7 | tubulin beta 3 class III pseudogene 2 [Source:H |
| CALD1 | 7 | caldesmon 1 [Source:HGNC Symbol;Acc:HGNC |
| AC083870.1 | 7 | novel transcript, antisense to AGBL3and CALD |
| AGBL3 | 7 | ATP/GTP binding protein like 3 [Source:HGNC |
| CYREN | 7 | cell cycle regulator of NHEJ [Source:HGNC Syn |
| AC083862.1 | 7 | ring finger protein 14 (RNF14) pseudogene |
| TMEM140 | 7 | transmembrane protein 140 [Source:HGNC Sy |
| AC083862.2 | 7 | novel transcript, antisense to C7orf49 |
| AC083862.3 | 7 | novel transcript |
| WDR91 | 7 | WD repeat domain 91 [Source:HGNC Symbol;A |
| AC009542.1 | 7 | novel transcript, antisense to WDR91 |
| MIR6509 | 7 | microRNA 6509 [Source:HGNC Symbol;Acc:HG |
| STRA8 | 7 | stimulated by retinoic acid 8 [Source:HGNC Sy |
| AC009542.2 | 7 | solute carrier family 23 (ascorbic acid transpo |
| CNOT4 | 7 | CCR4-NOT transcription complex subunit 4 [So |

| | | |
|---|---|---|
| SDHDP2 | 7 | succinate dehydrogenase complex subunit D p |
| NUP205 | 7 | nucleoporin 205 [Source:HGNC Symbol;Acc:H |
| AC093107.2 | 7 | MYC associated factor X, pseudogene |
| AC093107.1 | 7 | ribosomal protein L15 (RPL15) pseudogene |
| STMP1 | 7 | short transmembrane mitochondrial protein 1 |
| RNU6-1154P | 7 | RNA, U6 small nuclear 1154, pseudogene [Sou |
| SLC13A4 | 7 | solute carrier family 13 member 4 [Source:HG |
| AC091736.1 | 7 | novel transcript, antisense to SLC13A4 |
| FAM180A | 7 | family with sequence similarity 180 member A |
| AC015987.1 | 7 | novel transcript, antisense to MTPN |
| MTPN | 7 | myotrophin [Source:HGNC Symbol;Acc:HGNC: |
| LUZP6 | 7 | leucine zipper protein 6 [Source:HGNC Symbo |
| RNU6-223P | 7 | RNA, U6 small nuclear 223, pseudogene [Sour |
| AC024084.1 | 7 | novel transcript |
| AC078845.1 | 7 | novel transcript |
| Y_RNA | 7 | Y RNA [Source:RFAM;Acc:RF00019] |
| AC009784.1 | 7 | transient receptor potential cation channel, su |
| AC009541.1 | 7 | zona pellucida glycoprotein 3 (sperm receptor |
| AC009264.1 | 7 | novel transcript, antisense to CHRM2 |
| PSMC1P3 | 7 | proteasome 26S subunit, ATPase 1 pseudogen |
| CHRM2 | 7 | cholinergic receptor muscarinic 2 [Source:HGN |
| MIR490 | 7 | microRNA 490 [Source:HGNC Symbol;Acc:HGN |
| KRT8P51 | 7 | keratin 8 pseudogene 51 [Source:HGNC Symb |
| AC009517.1 | 7 | ribosomal protein L18 (RPL18) pseudogene |
| PTN | 7 | pleiotrophin [Source:HGNC Symbol;Acc:HGNC |
| SNORD81 | 7 | Small nucleolar RNA SNORD81 [Source:RFAM; |
| AC078842.2 | 7 | novel transcript |
| AC078842.1 | 7 | novel transcript |
| DGKI | 7 | diacylglycerol kinase iota [Source:HGNC Symb |
| AC090498.1 | 7 | ribosomal protein L41 (RPL41) pseudogene |
| AC009245.1 | 7 | ribosomal protein L6 (RPL6) pseudogene |
| CREB3L2 | 7 | cAMP responsive element binding protein 3 li |
| CREB3L2-AS1 | 7 | CREB3L2 antisense RNA 1 [Source:HGNC Symb |
| AKR1D1 | 7 | aldo-keto reductase family 1 member D1 [Sou |
| AC009263.1 | 7 | protein phosphatase 2, regulatory subunit B', |
| AC024082.1 | 7 | Der1-like domain family, member 2 (DERL2) p |
| AC024082.2 | 7 | pseudogene similiar to part of tumor protein, |
| RN7SKP223 | 7 | RN7SK pseudogene 223 [Source:HGNC Symbo |
| RCC2P3 | 7 | regulator of chromosome condensation 2 pse |
| MIR4468 | 7 | microRNA 4468 [Source:HGNC Symbol;Acc:HG |
| AC083867.2 | 7 | novel transcript |
| Y_RNA | 7 | Y RNA [Source:RFAM;Acc:RF00019] |
| AC083867.1 | 7 | NULL |
| AC008155.1 | 7 | ribosomal protein S17 (RPS17) pseudogene |
| PTMAP10 | 7 | prothymosin alpha pseudogene 10 [Source:HG |
| IMPDH1P3 | 7 | inosine monophosphate dehydrogenase 1 pse |
| TRIM24 | 7 | tripartite motif containing 24 [Source:HGNC S |
| AC013429.2 | 7 | ribosomal protein S3A (RPS3A) pseudogene |
| SVOPL | 7 | SVOP like [Source:HGNC Symbol;Acc:HGNC:27 |
| AC013429.3 | 7 | NULL |

| | | |
|---|---|---|
| AC013429.1 | 7 | ribosomal protein L21 (RPL21) pseudogene |
| AC020983.1 | 7 | ribosomal protein L17 (RPL17) pseudogene |
| UQCRFS1P2 | 7 | ubiquinol-cytochrome c reductase, Rieske iror |
| ATP6V0A4 | 7 | ATPase H+ transporting V0 subunit a4 [Source |
| TMEM213 | 7 | transmembrane protein 213 [Source:HGNC Sy |
| KIAA1549 | 7 | KIAA1549 [Source:HGNC Symbol;Acc:HGNC:2 |
| RNU6-1272P | 7 | RNA, U6 small nuclear 1272, pseudogene [Sou |
| ZC3HAV1L | 7 | zinc finger CCCH-type containing, antiviral 1 lil |
| ZC3HAV1 | 7 | zinc finger CCCH-type containing, antiviral 1 [S |
| AC018644.1 | 7 | ribosomal protein L17 (RPL17) pseudogene |
| TTC26 | 7 | tetratricopeptide repeat domain 26 [Source:H |
| AC009220.2 | 7 | Male-specific lethal-3 homolog 1 (Msl3l1) pse |
| AC009220.3 | 7 | FCF1 small subunit (SSU) processome compor |
| AC009220.1 | 7 | ribosomal protein L37a (RPL37A) pseudogene |
| MZT1P2 | 7 | mitotic spindle organizing protein 1 pseudoge |
| UBN2 | 7 | ubinuclein 2 [Source:HGNC Symbol;Acc:HGNC |
| RNU6-206P | 7 | RNA, U6 small nuclear 206, pseudogene [Sour |
| FMC1 | 7 | formation of mitochondrial complex V assemk |
| LUC7L2 | 7 | LUC7 like 2, pre-mRNA splicing factor [Source: |
| FMC1-LUC7L2 | 7 | FMC1-LUC7L2 readthrough [Source:HGNC Syn |
| AC083880.1 | 7 | novel transcript, antisense to LUC7L2 |
| RNU6-911P | 7 | RNA, U6 small nuclear 911, pseudogene [Sour |
| KLRG2 | 7 | killer cell lectin like receptor G2 [Source:HGN( |
| AC005531.1 | 7 | suppression of tumorigenicity 13 (colon carcir |
| CLEC2L | 7 | C-type lectin domain family 2 member L [Sour |
| ERHP1 | 7 | ERH pseudogene 1 [Source:HGNC Symbol;Acc |
| HIPK2 | 7 | homeodomain interacting protein kinase 2 [S( |
| TBXAS1 | 7 | thromboxane A synthase 1 [Source:HGNC Sym |
| PARP12 | 7 | poly(ADP-ribose) polymerase family member |
| KDM7A | 7 | lysine demethylase 7A [Source:HGNC Symbol; |
| Y_RNA | 7 | Y RNA [Source:RFAM;Acc:RF00019] |
| KDM7A-DT | 7 | KDM7A divergent transcript [Source:HGNC Sy |
| RNU6-797P | 7 | RNA, U6 small nuclear 797, pseudogene [Sour |
| RNU1-58P | 7 | RNA, U1 small nuclear 58, pseudogene [Sourc |
| AC005377.1 | 7 | ribosomal protein L9, pseudogene |
| PPP1R2P6 | 7 | protein phosphatase 1 regulatory inhibitor sul |
| SLC37A3 | 7 | solute carrier family 37 member 3 [Source:HG |
| RNA5SP247 | 7 | RNA, 5S ribosomal pseudogene 247 [Source:H |
| RNA5SP248 | 7 | RNA, 5S ribosomal pseudogene 248 [Source:H |
| RAB19 | 7 | RAB19, member RAS oncogene family [Source |
| AC069335.1 | 7 | non-metastatic cells 4, protein expressed in (N |
| MKRN1 | 7 | makorin ring finger protein 1 [Source:HGNC Sy |
| DENND2A | 7 | DENN domain containing 2A [Source:HGNC Sy |
| Y_RNA | 7 | Y RNA [Source:RFAM;Acc:RF00019] |
| AC006452.1 | 7 | novel transcript |
| RN7SL771P | 7 | RNA, 7SL, cytoplasmic 771, pseudogene [Sour |
| ADCK2 | 7 | aarF domain containing kinase 2 [Source:HGN |
| NDUFB2 | 7 | NADH:ubiquinone oxidoreductase subunit B2 |
| NDUFB2-AS1 | 7 | NDUFB2 antisense RNA 1 [Source:HGNC Symb |
| BRAF | 7 | B-Raf proto-oncogene, serine/threonine kinas |

| | | |
|---|---|---|
| RNU6-85P | 7 | RNA, U6 small nuclear 85, pseudogene [Sourc |
| AC006006.1 | 7 | mediator complex subunit 1, pseudogene |
| CCT4P1 | 7 | chaperonin containing TCP1 subunit 4 pseudo |
| MRPS33 | 7 | mitochondrial ribosomal protein S33 [Source: |
| RNU4-74P | 7 | RNA, U4 small nuclear 74, pseudogene [Sourc |
| TMEM178B | 7 | transmembrane protein 178B [Source:HGNC S |
| AC006362.1 | 7 | mitochondrially encoded NADH dehydrogenas |
| NDUFB10P2 | 7 | NADH:ubiquinone oxidoreductase subunit B1( |
| AC005692.3 | 7 | novel transcript |
| AC005692.1 | 7 | novel transcript |
| AC005692.2 | 7 | novel transcript, antisense to novel TMEM ge |
| AC073878.1 | 7 | novel transcript, antisense to AGK |
| AC073878.2 | 7 | novel transcript |
| AGK | 7 | acylglycerol kinase [Source:HGNC Symbol;Acc |
| AC004918.1 | 7 | novel transcript, antisense to AGK |
| DENND11 | 7 | DENN domain containing 11 [Source:HGNC Sy |
| AC004918.4 | 7 | novel transcript |
| WEE2-AS1 | 7 | WEE2 antisense RNA 1 [Source:HGNC Symbol; |
| WEE2 | 7 | WEE1 homolog 2 [Source:HGNC Symbol;Acc:H |
| RNU1-82P | 7 | RNA, U1 small nuclear 82, pseudogene [Sourc |
| SSBP1 | 7 | single stranded DNA binding protein 1 [Source |
| TAS2R3 | 7 | taste 2 receptor member 3 [Source:HGNC Syn |
| TAS2R4 | 7 | taste 2 receptor member 4 [Source:HGNC Syn |
| TAS2R6P | 7 | taste 2 receptor member 6 pseudogene [Sour |
| TAS2R5 | 7 | taste 2 receptor member 5 [Source:HGNC Syn |
| MTCO1P55 | 7 | MT-CO1 pseudogene 55 [Source:HGNC Symbo |
| MTND2P5 | 7 | MT-ND2 pseudogene 5 [Source:HGNC Symbol |
| MTND1P3 | 7 | MT-ND1 pseudogene 3 [Source:HGNC Symbol |
| MYL6P4 | 7 | myosin light chain 6 pseudogene 4 [Source:HG |
| PRSS37 | 7 | serine protease 37 [Source:HGNC Symbol;Acc |
| OR9A3P | 7 | olfactory receptor family 9 subfamily A memb |
| OR9A1P | 7 | olfactory receptor family 9 subfamily A memb |
| MGAM | 7 | maltase-glucoamylase [Source:HGNC Symbol; |
| OR9N1P | 7 | olfactory receptor family 9 subfamily N memb |
| OR9A4 | 7 | olfactory receptor family 9 subfamily A memb |
| CLEC5A | 7 | C-type lectin domain containing 5A [Source:H |
| TAS2R38 | 7 | taste 2 receptor member 38 [Source:HGNC Sy |
| MGAM2 | 7 | maltase-glucoamylase 2 (putative) [Source:HG |
| MOXD2P | 7 | monooxygenase DBH like 2, pseudogene [Sou |
| PRSS58 | 7 | serine protease 58 [Source:HGNC Symbol;Acc |
| AC245088.1 | 7 | trypsinogen-like pseudogene |
| AC245088.2 | 7 | novel transcript |
| PRSS3P3 | 7 | PRSS3 pseudogene 3 [Source:HGNC Symbol;A |
| TRBV1 | 7 | T cell receptor beta variable 1 (pseudogene) [ |
| TRBV2 | 7 | T cell receptor beta variable 2 [Source:HGNC S |
| TRBV3-1 | 7 | T cell receptor beta variable 3-1 [Source:HGN |
| TRBV4-1 | 7 | T cell receptor beta variable 4-1 [Source:HGN |
| TRBV5-1 | 7 | T cell receptor beta variable 5-1 [Source:HGN |
| TRBV6-1 | 7 | T cell receptor beta variable 6-1 [Source:HGN |
| TRBV7-1 | 7 | T cell receptor beta variable 7-1 (non-functior |

| | | |
|---|---|---|
| TRBV4-2 | 7 | T cell receptor beta variable 4-2 [Source:HGN( |
| TRBV6-2 | 7 | T cell receptor beta variable 6-2 (gene/pseudo |
| TRBV7-2 | 7 | T cell receptor beta variable 7-2 [Source:HGN( |
| TRBV8-1 | 7 | T cell receptor beta variable 8-1 (pseudogene) |
| TRBV5-2 | 7 | T cell receptor beta variable 5-2 (pseudogene) |
| TRBV6-4 | 7 | T cell receptor beta variable 6-4 [Source:HGN( |
| TRBV7-3 | 7 | T cell receptor beta variable 7-3 [Source:HGN( |
| TRBV8-2 | 7 | T cell receptor beta variable 8-2 (pseudogene) |
| TRBV5-3 | 7 | T cell receptor beta variable 5-3 (non-functior |
| TRBV9 | 7 | T cell receptor beta variable 9 [Source:HGNC S |
| TRBV10-1 | 7 | T cell receptor beta variable 10-1(gene/pseud |
| TRBV11-1 | 7 | T cell receptor beta variable 11-1 [Source:HGN |
| TRBV12-1 | 7 | T cell receptor beta variable 12-1 (pseudogen |
| TRBV10-2 | 7 | T cell receptor beta variable 10-2 [Source:HGN |
| TRBV11-2 | 7 | T cell receptor beta variable 11-2 [Source:HGN |
| TRBV12-2 | 7 | T cell receptor beta variable 12-2 (pseudogen |
| TRBV6-5 | 7 | T cell receptor beta variable 6-5 [Source:HGN( |
| TRBV7-4 | 7 | T cell receptor beta variable 7-4 (gene/pseudo |
| TRBV5-4 | 7 | T cell receptor beta variable 5-4 [Source:HGN( |
| TRBV6-6 | 7 | T cell receptor beta variable 6-6 [Source:HGN( |
| TRBV7-5 | 7 | T cell receptor beta variable 7-5 (pseudogene) |
| TRBV5-5 | 7 | T cell receptor beta variable 5-5 [Source:HGN( |
| TRBV6-7 | 7 | T cell receptor beta variable 6-7 (non-functior |
| TRBV7-6 | 7 | T cell receptor beta variable 7-6 [Source:HGN( |
| TRBV5-6 | 7 | T cell receptor beta variable 5-6 [Source:HGN( |
| TRBV6-8 | 7 | T cell receptor beta variable 6-8 [Source:HGN( |
| TRBV7-7 | 7 | T cell receptor beta variable 7-7 [Source:HGN( |
| TRBV5-7 | 7 | T cell receptor beta variable 5-7 (non-functior |
| TRBV7-9 | 7 | T cell receptor beta variable 7-9 [Source:HGN( |
| TRBV13 | 7 | T cell receptor beta variable 13 [Source:HGNC |
| TRBV10-3 | 7 | T cell receptor beta variable 10-3 [Source:HGN |
| TRBV11-3 | 7 | T cell receptor beta variable 11-3 [Source:HGN |
| TRBV12-3 | 7 | T cell receptor beta variable 12-3 [Source:HGN |
| TRBV12-4 | 7 | T cell receptor beta variable 12-4 [Source:HGN |
| TRBV12-5 | 7 | T cell receptor beta variable 12-5 [Source:HGN |
| TRBV14 | 7 | T cell receptor beta variable 14 [Source:HGNC |
| TRBV15 | 7 | T cell receptor beta variable 15 [Source:HGNC |
| TRBV16 | 7 | T cell receptor beta variable 16 (gene/pseudo |
| TRBV17 | 7 | T cell receptor beta variable 17 (non-functiona |
| TRBV18 | 7 | T cell receptor beta variable 18 [Source:HGNC |
| TRBV19 | 7 | T cell receptor beta variable 19 [Source:HGNC |
| TRBV20-1 | 7 | T cell receptor beta variable 20-1 [Source:HGN |
| TRBV21-1 | 7 | T cell receptor beta variable 21-1 (pseudogen |
| TRBV22-1 | 7 | T cell receptor beta variable 22-1 (pseudogen |
| TRBV23-1 | 7 | T cell receptor beta variable 23-1 (non-functic |
| TRBV24-1 | 7 | T cell receptor beta variable 24-1 [Source:HGN |
| MTRNR2L6 | 7 | MT-RNR2 like 6 [Source:HGNC Symbol;Acc:HG |
| TRBV25-1 | 7 | T cell receptor beta variable 25-1 [Source:HGN |
| TRBVA | 7 | T cell receptor beta variable A (pseudogene) [ |
| TRBV26 | 7 | T cell receptor beta variable 26 (pseudogene) |

| | | |
|---|---|---|
| TRBVB | 7 | T cell receptor beta variable B (pseudogene) [S |
| TRBV27 | 7 | T cell receptor beta variable 27 [Source:HGNC |
| TRBV28 | 7 | T cell receptor beta variable 28 [Source:HGNC |
| PGBD4P1 | 7 | piggyBac transposable element derived 4 pseu |
| AC244472.1 | 7 | novel transcript |
| TRBV29-1 | 7 | T cell receptor beta variable 29-1 [Source:HGN |
| PRSS1 | 7 | serine protease 1 [Source:HGNC Symbol;Acc:H |
| PRSS2 | 7 | serine protease 2 [Source:HGNC Symbol;Acc:H |
| PRSS3P1 | 7 | PRSS3 pseudogene 1 [Source:HGNC Symbol;A |
| WBP1LP1 | 7 | WBP1L pseudogene 1 [Source:HGNC Symbol;A |
| TRBD1 | 7 | T cell receptor beta diversity 1 [Source:HGNC |
| TRBJ1-1 | 7 | T cell receptor beta joining 1-1 [Source:HGNC |
| TRBJ1-2 | 7 | T cell receptor beta joining 1-2 [Source:HGNC |
| TRBJ1-3 | 7 | T cell receptor beta joining 1-3 [Source:HGNC |
| TRBJ1-4 | 7 | T cell receptor beta joining 1-4 [Source:HGNC |
| TRBJ1-5 | 7 | T cell receptor beta joining 1-5 [Source:HGNC |
| TRBJ1-6 | 7 | T cell receptor beta joining 1-6 [Source:HGNC |
| TRBC1 | 7 | T cell receptor beta constant 1 [Source:HGNC |
| TRBJ2-1 | 7 | T cell receptor beta joining 2-1 [Source:HGNC |
| TRBJ2-2 | 7 | T cell receptor beta joining 2-2 [Source:HGNC |
| TRBJ2-2P | 7 | T cell receptor beta joining 2-2P (non-function |
| TRBJ2-3 | 7 | T cell receptor beta joining 2-3 [Source:HGNC |
| TRBJ2-4 | 7 | T cell receptor beta joining 2-4 [Source:HGNC |
| TRBJ2-5 | 7 | T cell receptor beta joining 2-5 [Source:HGNC |
| TRBJ2-6 | 7 | T cell receptor beta joining 2-6 [Source:HGNC |
| TRBJ2-7 | 7 | T cell receptor beta joining 2-7 [Source:HGNC |
| TRBC2 | 7 | T cell receptor beta constant 2 [Source:HGNC |
| TRBV30 | 7 | T cell receptor beta variable 30 (gene/pseudo |
| EPHB6 | 7 | EPH receptor B6 [Source:HGNC Symbol;Acc:H |
| TRPV6 | 7 | transient receptor potential cation channel su |
| AC245427.1 | 7 | novel transcript, antisense to TRPV6 |
| AC245427.2 | 7 | novel transcript |
| TRPV5 | 7 | transient receptor potential cation channel su |
| LLCFC1 | 7 | LLLL and CFNLAS motif containing 1 [Source:H |
| KEL | 7 | Kell metallo-endopeptidase (Kell blood group) |
| OR9A2 | 7 | olfactory receptor family 9 subfamily A memb |
| OR9P1P | 7 | olfactory receptor family 9 subfamily P memb |
| OR6V1 | 7 | olfactory receptor family 6 subfamily V memb |
| OR6W1P | 7 | olfactory receptor family 6 subfamily W meml |
| PIP | 7 | prolactin induced protein [Source:HGNC Symb |
| TAS2R39 | 7 | taste 2 receptor member 39 [Source:HGNC Sy |
| AC073342.2 | 7 | novel transcript, antisense to TAS2R40 |
| TAS2R40 | 7 | taste 2 receptor member 40 [Source:HGNC Sy |
| AC073342.3 | 7 | tec |
| GSTK1 | 7 | glutathione S-transferase kappa 1 [Source:HG |
| AC073342.1 | 7 | novel transcript |
| TMEM139 | 7 | transmembrane protein 139 [Source:HGNC Sy |
| CASP2 | 7 | caspase 2 [Source:HGNC Symbol;Acc:HGNC:15 |
| RN7SL535P | 7 | RNA, 7SL, cytoplasmic 535, pseudogene [Sour |
| RN7SL481P | 7 | RNA, 7SL, cytoplasmic 481, pseudogene [Sour |

| | | |
|---|---|---|
| HINT1P1 | 7 | histidine triad nucleotide binding protein 1 ps |
| CLCN1 | 7 | chloride voltage-gated channel 1 [Source:HGN |
| FAM131B | 7 | family with sequence similarity 131 member E |
| AC093673.2 | 7 | novel transcript |
| AC093673.1 | 7 | novel transcript |
| ZYX | 7 | zyxin [Source:HGNC Symbol;Acc:HGNC:13200] |
| MIR6892 | 7 | microRNA 6892 [Source:HGNC Symbol;Acc:HG |
| EPHA1 | 7 | EPH receptor A1 [Source:HGNC Symbol;Acc:H |
| EPHA1-AS1 | 7 | EPHA1 antisense RNA 1 [Source:HGNC Symbo |
| TAS2R62P | 7 | taste 2 receptor member 62 pseudogene [Sou |
| TAS2R60 | 7 | taste 2 receptor member 60 [Source:HGNC Sy |
| TAS2R41 | 7 | taste 2 receptor member 41 [Source:HGNC Sy |
| OR2R1P | 7 | olfactory receptor family 2 subfamily R memb |
| OR10AC1 | 7 | olfactory receptor family 10 subfamily AC mer |
| AC073264.1 | 7 | ribosomal protein L26 (RPL26) pseudogene |
| PAICSP5 | 7 | phosphoribosylaminoimidazole carboxylase, p |
| CTAGE15 | 7 | CTAGE family member 15 [Source:HGNC Symb |
| RNU6-162P | 7 | RNA, U6 small nuclear 162, pseudogene [Sour |
| AC073264.2 | 7 | pseudogene similar to part of thioredoxin dor |
| TCAF1P1 | 7 | TRPM8 channel associated factor 1 pseudoger |
| AC073264.3 | 7 | novel TRPM8 channel-associated factor pseuc |
| TCAF2 | 7 | TRPM8 channel associated factor 2 [Source:H |
| TCAF2C | 7 | TRPM8 channel associated factor 2C [Source:I |
| RNU6-267P | 7 | RNA, U6 small nuclear 267, pseudogene [Sour |
| CTAGE6 | 7 | CTAGE family member 6 [Source:HGNC Symbo |
| AC099548.2 | 7 | family with sequence similarity 115, member |
| PAICSP6 | 7 | phosphoribosylaminoimidazole carboxylase, p |
| AC099548.1 | 7 | ribosomal protein L26 (RPL26) pseudogene |
| TCAF2P1 | 7 | TRPM8 channel associated factor 2 pseudoger |
| TCAF1 | 7 | TRPM8 channel associated factor 1 [Source:H |
| OR2F2 | 7 | olfactory receptor family 2 subfamily F memb |
| OR2F1 | 7 | olfactory receptor family 2 subfamily F memb |
| AC004853.1 | 7 | capping protein (actin filament) muscle Z-line, |
| OR2Q1P | 7 | olfactory receptor family 2 subfamily Q memb |
| SLC16A1P1 | 7 | SLC16A1 pseudogene 1 [Source:HGNC Symbol |
| OR6B1 | 7 | olfactory receptor family 6 subfamily B memb |
| OR2A5 | 7 | olfactory receptor family 2 subfamily A memb |
| OR2A25 | 7 | olfactory receptor family 2 subfamily A memb |
| OR2A41P | 7 | olfactory receptor family 2 subfamily A memb |
| OR2A12 | 7 | olfactory receptor family 2 subfamily A memb |
| OR2A2 | 7 | olfactory receptor family 2 subfamily A memb |
| OR2A15P | 7 | olfactory receptor family 2 subfamily A memb |
| OR2A14 | 7 | olfactory receptor family 2 subfamily A memb |
| OR2A13P | 7 | olfactory receptor family 2 subfamily A memb |
| OR2A3P | 7 | olfactory receptor family 2 subfamily A memb |
| OR2AO1P | 7 | olfactory receptor family 2 subfamily AO mer |
| CTAGE4 | 7 | CTAGE family member 4 [Source:HGNC Symbo |
| ARHGEF35 | 7 | Rho guanine nucleotide exchange factor 35 [S |
| AC004889.1 | 7 | novel transcript, antisense to ARHGEF35, OR2 |
| OR2A42 | 7 | olfactory receptor family 2 subfamily A memb |

| | | |
|---|---|---|
| OR2A20P | 7 | olfactory receptor family 2 subfamily A memb |
| AC074386.1 | 7 | novel transcript, antisense to OR2A1 and ARH |
| OR2A7 | 7 | olfactory receptor family 2 subfamily A memb |
| CTAGE8 | 7 | CTAGE family member 8 [Source:HGNC Symbo |
| ARHGEF34P | 7 | Rho guanine nucleotide exchange factor 34, p |
| OR2A9P | 7 | olfactory receptor family 2 subfamily A memb |
| OR2A1-AS1 | 7 | OR2A1 antisense RNA 1 [Source:HGNC Symbo |
| OR2A1 | 7 | olfactory receptor family 2 subfamily A memb |
| ARHGEF5 | 7 | Rho guanine nucleotide exchange factor 5 [So |
| NOBOX | 7 | NOBOX oogenesis homeobox [Source:HGNC S |
| PPIAP83 | 7 | peptidylprolyl isomerase A pseudogene 83 [Sc |
| RNU6ATAC40P | 7 | RNA, U6atac small nuclear 40, pseudogene [S |
| TPK1 | 7 | thiamin pyrophosphokinase 1 [Source:HGNC S |
| EEF1A1P10 | 7 | eukaryotic translation elongation factor 1 alpl |
| RN7SKP174 | 7 | RN7SK pseudogene 174 [Source:HGNC Symbo |
| EI24P4 | 7 | EI24 pseudogene 4 [Source:HGNC Symbol;Acc |
| AC073310.1 | 7 | karyopherin alpha 2 (RAG cohort 1, importin a |
| RPL7P59 | 7 | ribosomal protein L7 pseudogene 59 [Source: |
| AC004911.1 | 7 | cytoskeleton-associated protein 4 (CKAP4) pse |
| DPY19L4P2 | 7 | DPY19L4 pseudogene 2 [Source:HGNC Symbo |
| AC006007.1 | 7 | novel transcript |
| AC073308.1 | 7 | tec |
| CNTNAP2 | 7 | contactin associated protein like 2 [Source:HG |
| AC073418.1 | 7 | NULL |
| Y_RNA | 7 | Y RNA [Source:RFAM;Acc:RF00019] |
| AC006004.1 | 7 | novel transcript |
| DUTP3 | 7 | deoxyuridine triphosphatase pseudogene 3 [S |
| RANP2 | 7 | RAN pseudogene 2 [Source:HGNC Symbol;Acc |
| MIR548F4 | 7 | microRNA 548f-4 [Source:HGNC Symbol;Acc:H |
| AC005518.1 | 7 | novel transcript |
| AC005518.2 | 7 | phosphoribosylaminoimidazole carboxylase, p |
| RNU6-1184P | 7 | RNA, U6 small nuclear 1184, pseudogene [Sou |
| RNA5SP249 | 7 | RNA, 5S ribosomal pseudogene 249 [Source:H |
| RN7SL456P | 7 | RNA, 7SL, cytoplasmic 456, pseudogene [Sour |
| U3 | 7 | Small nucleolar RNA U3 [Source:RFAM;Acc:RF |
| RN7SL72P | 7 | RNA, 7SL, cytoplasmic 72, pseudogene [Sourc |
| AC083849.1 | 7 | exosome component 6 (EXOSC6) pseudogene |
| AC006974.1 | 7 | novel transcript |
| AC006974.2 | 7 | novel transcript |
| AC005229.2 | 7 | ribosomal protein L32 (RPL32) pseudogene |
| AC005229.5 | 7 | novel transcript, antisense to C7orf33 |
| C7orf33 | 7 | chromosome 7 open reading frame 33 [Sourc |
| AC005229.1 | 7 | Huntingtin interacting protein K (HYPK) pseud |
| AC005229.3 | 7 | ACOT13 acyl-CoA thioesterase 13, pseudogen |
| AC005229.4 | 7 | novel transcript, antisense to CUL1 |
| CUL1 | 7 | cullin 1 [Source:HGNC Symbol;Acc:HGNC:2551 |
| EZH2 | 7 | enhancer of zeste 2 polycomb repressive com |
| RNU7-20P | 7 | RNA, U7 small nuclear 20 pseudogene [Source |
| Metazoa_SRP | 7 | Metazoan signal recognition particle RNA [Sou |
| RN7SL569P | 7 | RNA, 7SL, cytoplasmic 569, pseudogene [Sour |

| | | |
|---|---|---|
| AC073140.2 | 7 | novel transcript |
| AC073140.1 | 7 | RNA, Ro-associated Y5 |
| RNY4 | 7 | RNA, Ro60-associated Y4 [Source:HGNC Symb |
| RNY3 | 7 | RNA, Ro60-associated Y3 [Source:HGNC Symb |
| RNY1 | 7 | RNA, Ro60-associated Y1 [Source:HGNC Symb |
| GHET1 | 7 | gastric carcinoma proliferation enhancing trar |
| PDIA4 | 7 | protein disulfide isomerase family A member |
| RNU6-650P | 7 | RNA, U6 small nuclear 650, pseudogene [Sour |
| COX6B1P1 | 7 | cytochrome c oxidase subunit 6B1 pseudogen |
| ZNF786 | 7 | zinc finger protein 786 [Source:HGNC Symbol; |
| ZNF425 | 7 | zinc finger protein 425 [Source:HGNC Symbol; |
| RN7SL521P | 7 | RNA, 7SL, cytoplasmic 521, pseudogene [Sour |
| ZNF398 | 7 | zinc finger protein 398 [Source:HGNC Symbol; |
| AC004890.1 | 7 | ribosomal protein L36a (RPL36A) pseudogene |
| ZNF282 | 7 | zinc finger protein 282 [Source:HGNC Symbol; |
| ZNF212 | 7 | zinc finger protein 212 [Source:HGNC Symbol; |
| ZNF783 | 7 | zinc finger family member 783 [Source:HGNC |
| AC004890.3 | 7 | SECIS binding protein 2, pseudogene |
| AC004890.2 | 7 | zinc finger protein pseudogene |
| AC004941.2 | 7 | karyopherin (importin) beta 1 (KPNB1) pseudo |
| NPM1P12 | 7 | nucleophosmin 1 pseudogene 12 [Source:HGI |
| AC004941.1 | 7 | uncharacterized LOC101928733 [Source:NCBI |
| AC073314.1 | 7 | novel transcript |
| ZNF777 | 7 | zinc finger protein 777 [Source:HGNC Symbol; |
| ZNF746 | 7 | zinc finger protein 746 [Source:HGNC Symbol; |
| ZNF767P | 7 | zinc finger family member 767, pseudogene [S |
| KRBA1 | 7 | KRAB-A domain containing 1 [Source:HGNC Sy |
| ZNF467 | 7 | zinc finger protein 467 [Source:HGNC Symbol; |
| SSPO | 7 | SCO-spondin [Source:HGNC Symbol;Acc:HGN( |
| ZNF862 | 7 | zinc finger protein 862 [Source:HGNC Symbol; |
| AC004877.2 | 7 | TEC |
| AC004877.1 | 7 | novel transcript, antisense to ZNF862 |
| ATP6V0E2-AS1 | 7 | ATP6V0E2 antisense RNA 1 [Source:HGNC Syn |
| ATP6V0E2 | 7 | ATPase H+ transporting V0 subunit e2 [Source |
| AC093458.1 | 7 | novel transcript |
| AC093458.2 | 7 | tec |
| AC092681.3 | 7 | novel transcript |
| AC092681.1 | 7 | pseudogene similar to part of ARP3 actin-relat |
| AC092681.2 | 7 | novel transcript |
| AC092666.1 | 7 | novel transcript |
| AC092666.2 | 7 | dipeptidyl-peptidase 6 (DPP6) pseudogene |
| AC006008.1 | 7 | novel transcript |
| ACTR3C | 7 | actin related protein 3C [Source:HGNC Symbo |
| ZBED6CL | 7 | ZBED6 C-terminal like [Source:HGNC Symbol;A |
| LRRC61 | 7 | leucine rich repeat containing 61 [Source:HGI |
| AC005586.2 | 7 | novel transcript, overlapping LRRC61 |
| RARRES2 | 7 | retinoic acid receptor responder 2 [Source:HG |
| AC005586.1 | 7 | novel transcript |
| REPIN1 | 7 | replication initiator 1 [Source:HGNC Symbol;A |
| ZNF775 | 7 | zinc finger protein 775 [Source:HGNC Symbol; |

| | | |
|---|---|---|
| AC073111.3 | 7 | novel transcript |
| AC073111.1 | 7 | novel transcript |
| AC073111.4 | 7 | novel zinc finger protein |
| LINC00996 | 7 | long intergenic non-protein coding RNA 996 [S |
| GIMAP8 | 7 | GTPase, IMAP family member 8 [Source:HGN( |
| AC073111.2 | 7 | EIF2A eukaryotic translation initiation factor 2 |
| STRADBP1 | 7 | STE20-related kinase adaptor beta pseudogen |
| GIMAP7 | 7 | GTPase, IMAP family member 7 [Source:HGN( |
| ALDH7A1P3 | 7 | aldehyde dehydrogenase 7 family member A1 |
| GIMAP4 | 7 | GTPase, IMAP family member 4 [Source:HGN( |
| AC069304.3 | 7 | EIF2A eukaryotic translation initiation factor 2 |
| AC069304.1 | 7 | pseudogene similar to part of transient recept |
| GIMAP6 | 7 | GTPase, IMAP family member 6 [Source:HGN( |
| AC069304.2 | 7 | eukaryotic translation initiation factor 2A, 65k |
| GIMAP2 | 7 | GTPase, IMAP family member 2 [Source:HGN( |
| GIMAP1 | 7 | GTPase, IMAP family member 1 [Source:HGN( |
| GIMAP1-GIMAP5 | 7 | GIMAP1-GIMAP5 readthrough [Source:HGNC |
| GIMAP5 | 7 | GTPase, IMAP family member 5 [Source:HGN( |
| GIMAP3P | 7 | GTPase, IMAP family member 3 pseudogene [ |
| BET1P1 | 7 | Bet1 golgi vesicular membrane trafficking pro |
| TMEM176B | 7 | transmembrane protein 176B [Source:HGNC S |
| TMEM176A | 7 | transmembrane protein 176A [Source:HGNC S |
| AOC1 | 7 | amine oxidase copper containing 1 [Source:H( |
| KCNH2 | 7 | potassium voltage-gated channel subfamily H |
| NOS3 | 7 | nitric oxide synthase 3 [Source:HGNC Symbol; |
| ATG9B | 7 | autophagy related 9B [Source:HGNC Symbol;A |
| ABCB8 | 7 | ATP binding cassette subfamily B member 8 [S |
| AC010973.1 | 7 | novel transcript |
| ASIC3 | 7 | acid sensing ion channel subunit 3 [Source:HG |
| CDK5 | 7 | cyclin dependent kinase 5 [Source:HGNC Syml |
| SLC4A2 | 7 | solute carrier family 4 member 2 [Source:HGN |
| AC010973.2 | 7 | novel transcript |
| FASTK | 7 | Fas activated serine/threonine kinase [Source |
| TMUB1 | 7 | transmembrane and ubiquitin like domain cor |
| AGAP3 | 7 | ArfGAP with GTPase domain, ankyrin repeat a |
| GBX1 | 7 | gastrulation brain homeobox 1 [Source:HGNC |
| ASB10 | 7 | ankyrin repeat and SOCS box containing 10 [S |
| IQCA1L | 7 | IQ motif containing with AAA domain 1 like [S |
| AC021097.2 | 7 | novel histone H2B-like protein [Source:NCBI g |
| ABCF2 | 7 | ATP binding cassette subfamily F member 2 [S |
| ABCF2 | 7 | ATP binding cassette subfamily F member 2 [S |
| CHPF2 | 7 | chondroitin polymerizing factor 2 [Source:HGI |
| MIR671 | 7 | microRNA 671 [Source:HGNC Symbol;Acc:HGI |
| SMARCD3 | 7 | SWI/SNF related, matrix associated, actin dep |
| AC021097.1 | 7 | novel transcript, antisense to SMARCD3 |
| AC005486.1 | 7 | novel transcript |
| NUB1 | 7 | negative regulator of ubiquitin like proteins 1 |
| WDR86 | 7 | WD repeat domain 86 [Source:HGNC Symbol;. |
| WDR86-AS1 | 7 | WDR86 antisense RNA 1 [Source:HGNC Symbo |
| AC005996.1 | 7 | novel transcript |

| | | |
|---|---|---|
| CRYGN | 7 | crystallin gamma N [Source:HGNC Symbol;Acc |
| MIR3907 | 7 | microRNA 3907 [Source:HGNC Symbol;Acc:HG |
| RN7SL76P | 7 | RNA, 7SL, cytoplasmic 76, pseudogene [Sourc |
| RHEB | 7 | Ras homolog, mTORC1 binding [Source:HGNC |
| ETF1P2 | 7 | eukaryotic translation termination factor 1 ps |
| PRKAG2 | 7 | protein kinase AMP-activated non-catalytic su |
| AC093583.1 | 7 | uncharacterized LOC644090 [Source:NCBI ger |
| PRKAG2-AS1 | 7 | PRKAG2 antisense RNA 1 [Source:HGNC Symb |
| RNU6-604P | 7 | RNA, U6 small nuclear 604, pseudogene [Sour |
| Y_RNA | 7 | Y RNA [Source:RFAM;Acc:RF00019] |
| AC074257.1 | 7 | novel transcript |
| GALNTL5 | 7 | polypeptide N-acetylgalactosaminyltransferas |
| Metazoa_SRP | 7 | Metazoan signal recognition particle RNA [Sou |
| GALNT11 | 7 | polypeptide N-acetylgalactosaminyltransferas |
| AC006017.1 | 7 | novel transcript |
| YBX1P4 | 7 | Y-box binding protein 1 pseudogene 4 [Source |
| KMT2C | 7 | lysine methyltransferase 2C [Source:HGNC Sy |
| AC104692.2 | 7 | nuclear protein, transcriptional regulator, 1-lil |
| AC104692.1 | 7 | pseudogene similar to phosphoserine phosph |
| SEPTIN7P6 | 7 | septin 7 pseudogene 6 [Source:HGNC Symbol; |
| Y_RNA | 7 | Y RNA [Source:RFAM;Acc:RF00019] |
| FABP5P3 | 7 | fatty acid binding protein 5 pseudogene 3 [Sou |
| CCT8L1P | 7 | chaperonin containing TCP1 subunit 8 like 1, p |
| LINC01003 | 7 | long intergenic non-protein coding RNA 1003 |
| AC104843.2 | 7 | ribosomal protein S20 (RPS20) pseudogene |
| AC104843.3 | 7 | NULL |
| RNA5SP250 | 7 | RNA, 5S ribosomal pseudogene 250 [Source:H |
| AC104843.1 | 7 | ribosomal protein L36a (RPL36A) pseudogene |
| AC003109.1 | 7 | NULL |
| XRCC2 | 7 | X-ray repair cross complementing 2 [Source:H |
| ATP5PBP3 | 7 | ATP synthase peripheral stalk-membrane subu |
| RN7SL845P | 7 | RNA, 7SL, cytoplasmic 845, pseudogene [Sour |
| ACTR3B | 7 | actin related protein 3B [Source:HGNC Symbo |
| AC006348.1 | 7 | novel transcript |
| AC079809.1 | 7 | SMT3 suppressor of mif two 3 homolog 2 (S. c |
| LINC01287 | 7 | long intergenic non-protein coding RNA 1287 |
| AC073236.1 | 7 | novel transcript |
| PAXBP1P1 | 7 | PAX3 and PAX7 binding protein 1 pseudogene |
| AC005998.1 | 7 | novel transcript |
| DPP6 | 7 | dipeptidyl peptidase like 6 [Source:HGNC Sym |
| AC005588.1 | 7 | NADH dehydrogenase subunit 6 (ND6) pseudo |
| AC006019.3 | 7 | novel transcript |
| AC006019.1 | 7 | novel transcript |
| AC006019.2 | 7 | novel transcript |
| AC024730.1 | 7 | novel transcript, antisense to DPP6 |
| AC073336.1 | 7 | novel transcript |
| PAXIP1-AS2 | 7 | PAXIP1 antisense RNA 2 [Source:HGNC Symbo |
| PAXIP1 | 7 | PAX interacting protein 1 [Source:HGNC Symb |
| AC093726.1 | 7 | novel transcript, antisense to PAXIP1 |
| AC093726.2 | 7 | novel transcript, antisense to PAXIP1 |

| | | |
|---|---|---|
| PAXIP1-AS1 | 7 | PAXIP1 antisense RNA 1 (head to head) [Sourc |
| HTR5A-AS1 | 7 | HTR5A antisense RNA 1 [Source:HGNC Symbo |
| HTR5A | 7 | 5-hydroxytryptamine receptor 5A [Source:HG |
| AC093726.3 | 7 | mitochondrially encoded cytochrome c oxidas |
| AC092628.2 | 7 | zinc finger protein 532 (ZNF532) pseudogene |
| AC092628.1 | 7 | zinc finger protein 532 (ZNF532) pseudogene |
| RN7SKP280 | 7 | RN7SK pseudogene 280 [Source:HGNC Symbo |
| AC099552.1 | 7 | novel transcript |
| AC099552.3 | 7 | novel transcript |
| AC099552.5 | 7 | novel transcript |
| AC099552.4 | 7 | tec |
| AC099552.2 | 7 | novel transcript |
| AC144652.1 | 7 | novel transcript |
| INSIG1 | 7 | insulin induced gene 1 [Source:HGNC Symbol; |
| BLACE | 7 | B cell acute lymphoblastic leukemia expressed |
| AC008060.1 | 7 | novel transcript |
| AC008060.4 | 7 | novel transcript, overlapping novel protein |
| AC008060.5 | 7 | novel transcript |
| AC008060.3 | 7 | novel transcript |
| EN2 | 7 | engrailed homeobox 2 [Source:HGNC Symbol; |
| CNPY1 | 7 | canopy FGF signaling regulator 1 [Source:HGN |
| AC009403.2 | 7 | novel transcript |
| AC008060.2 | 7 | novel transcript, antisense to CNPY1 |
| AC009403.3 | 7 | novel transcript, antisense to CNPY1 |
| AC009403.1 | 7 | novel transcript, antisense to RBM33 |
| RBM33 | 7 | RNA binding motif protein 33 [Source:HGNC S |
| SHH | 7 | sonic hedgehog signaling molecule [Source:H( |
| AC021218.1 | 7 | uncharacterized LOC389602 [Source:NCBI ger |
| Y_RNA | 7 | Y RNA [Source:RFAM;Acc:RF00019] |
| AC093813.1 | 7 | ribosomal protein S24, pseudogene |
| LINC01006 | 7 | long intergenic non-protein coding RNA 1006 |
| LINC00244 | 7 | long intergenic non-protein coding RNA 244 [S |
| AC005534.2 | 7 | ribosomal protein L26-like 1 (RPL26L1) pseudo |
| RNF32 | 7 | ring finger protein 32 [Source:HGNC Symbol;A |
| AC005534.1 | 7 | novel transcript |
| LMBR1 | 7 | limb development membrane protein 1 [Sour |
| RNU4-31P | 7 | RNA, U4 small nuclear 31, pseudogene [Sourc |
| AC006357.1 | 7 | novel transcript |
| NOM1 | 7 | nucleolar protein with MIF4G domain 1 [Sour |
| MNX1 | 7 | motor neuron and pancreas homeobox 1 [Sou |
| MNX1-AS2 | 7 | MNX1 antisense RNA 2 [Source:HGNC Symbol |
| MNX1-AS1 | 7 | MNX1 antisense RNA 1 (head to head) [Sourc |
| AC006967.2 | 7 | novel transcript |
| AC006967.1 | 7 | adenosylmethionine decarboxylase 1 (AMD1) |
| AC006967.3 | 7 | novel transcript |
| UBE3C | 7 | ubiquitin protein ligase E3C [Source:HGNC Syr |
| RPS27AP12 | 7 | ribosomal protein S27a pseudogene 12 [Sourc |
| AC004898.1 | 7 | UBX domain protein 2B (UBXN2B) pseudogen |
| AC004975.1 | 7 | ribosomal protein L36 (RPL36A) pseudogene |
| AC004975.2 | 7 | novel transcript |

| | | |
|---|---|---|
| DNAJB6 | 7 | DnaJ heat shock protein family (Hsp40) memb |
| AC006372.3 | 7 | novel transcript |
| AC006372.1 | 7 | novel transcript |
| AC006372.2 | 7 | novel transcript |
| AC006372.4 | 7 | TEC |
| PTPRN2 | 7 | protein tyrosine phosphatase receptor type N |
| MIR153-2 | 7 | microRNA 153-2 [Source:HGNC Symbol;Acc:H |
| AC005481.1 | 7 | novel transcript, antisense to PTPRN2 |
| AC006003.1 | 7 | novel transcript |
| AC011899.2 | 7 | novel transcript, antisense to PTPRN2 |
| AC011899.3 | 7 | novel transcript, antisense to PTPRN2 |
| AC011899.4 | 7 | novel transcript, antisense to PTPRN2 |
| AC011899.1 | 7 | novel transcript |
| MIR595 | 7 | microRNA 595 [Source:HGNC Symbol;Acc:HGI |
| AC078942.1 | 7 | novel transcript |
| LINC01022 | 7 | long intergenic non-protein coding RNA 1022 |
| MIR5707 | 7 | microRNA 5707 [Source:HGNC Symbol;Acc:HG |
| THAP5P1 | 7 | THAP domain containing 5 pseudogene 1 [Sou |
| NCAPG2 | 7 | non-SMC condensin II complex subunit G2 [So |
| AC019084.1 | 7 | ribosomal protein L21 (RPL21) pseudogene |
| ESYT2 | 7 | extended synaptotagmin 2 [Source:HGNC Sym |
| WDR60 | 7 | WD repeat domain 60 [Source:HGNC Symbol; |
| AC004863.1 | 7 | novel transcript |
| LINC00689 | 7 | long intergenic non-protein coding RNA 689 [S |
| VIPR2 | 7 | vasoactive intestinal peptide receptor 2 [Sour |
| AC007269.1 | 7 | novel transcript, antisense to VIPR2 |
| PIP5K1P2 | 7 | phosphatidylinositol-4-phosphate 5-kinase typ |
| AC144568.1 | 8 | family with sequence similarity 41, member C |
| AC144568.2 | 8 | novel transcript |
| AC131281.2 | 8 | microtubule associated serine/threonine kina |
| WBP1LP3 | 8 | WBP1L pseudogene 3 [Source:HGNC Symbol;A |
| OR4F21 | 8 | olfactory receptor family 4 subfamily F memb |
| SEPTIN14P8 | 8 | septin 14 pseudogene 8 [Source:HGNC Symbo |
| AC131281.1 | 8 | family with sequence similarity 41, member C |
| RPL23AP53 | 8 | ribosomal protein L23a pseudogene 53 [Sourc |
| ZNF596 | 8 | zinc finger protein 596 [Source:HGNC Symbol; |
| AC004908.2 | 8 | novel transcript, sense intronic to ZNF596 |
| AC004908.1 | 8 | novel transcript, antisense to ZNF596 |
| AC004908.3 | 8 | novel transcript, antisense to ZNF596 |
| AC136777.2 | 8 | ring finger protein 11 (RNF11) pseudogene |
| AC136777.1 | 8 | novel transcript |
| FAM87A | 8 | family with sequence similarity 87 member A |
| FBXO25 | 8 | F-box protein 25 [Source:HGNC Symbol;Acc:H |
| AC083964.1 | 8 | novel transcript, antisense to FBXO25 |
| AC083964.2 | 8 | TEC |
| TDRP | 8 | testis development related protein [Source:H |
| ERICH1 | 8 | glutamate rich 1 [Source:HGNC Symbol;Acc:H |
| AC100797.1 | 8 | novel transcript |
| DLGAP2 | 8 | DLG associated protein 2 [Source:HGNC Symb |
| AC100797.5 | 8 | novel transcript |

| | | |
|---|---|---|
| AC026950.2 | 8 | novel transcript |
| AC129915.3 | 8 | novel transcript |
| AC110288.1 | 8 | novel transcript |
| AF067845.2 | 8 | novel transcript |
| AF067845.1 | 8 | novel transcript |
| AF067845.5 | 8 | novel transcript |
| AF067845.4 | 8 | novel transcript |
| DLGAP2-AS1 | 8 | DLGAP2 antisense RNA 1 [Source:HGNC Symb |
| AC100810.6 | 8 | mitochondrially encoded cytochrome c oxidas |
| AC100810.2 | 8 | DNA fragmentation factor, 45kDa, alpha polyp |
| CLN8 | 8 | CLN8 transmembrane ER and ERGIC protein [S |
| AC100810.3 | 8 | novel transcript |
| AC100810.1 | 8 | novel transcript |
| AC019257.8 | 8 | novel transcript |
| MIR3674 | 8 | microRNA 3674 [Source:HGNC Symbol;Acc:HG |
| AC100810.7 | 8 | novel transcript |
| MIR596 | 8 | microRNA 596 [Source:HGNC Symbol;Acc:HGN |
| ARHGEF10 | 8 | Rho guanine nucleotide exchange factor 10 [S |
| AC019257.9 | 8 | novel transcript, antisense to ARHGEF10 |
| AC019257.1 | 8 | novel transcript |
| KBTBD11-OT1 | 8 | KBTBD11 overlapping transcript 1 [Source:HG |
| KBTBD11 | 8 | kelch repeat and BTB domain containing 11 [S |
| AC019257.2 | 8 | novel transcript |
| AC245164.1 | 8 | novel transcript |
| MYOM2 | 8 | myomesin 2 [Source:HGNC Symbol;Acc:HGNC |
| MIR7160 | 8 | microRNA 7160 [Source:HGNC Symbol;Acc:HG |
| AC245164.2 | 8 | novel transcript, antisense to MYOM2 |
| AC245519.1 | 8 | novel transcript |
| AC245519.2 | 8 | novel transcript |
| AC245187.2 | 8 | novel transcript |
| AC245187.1 | 8 | novel transcript |
| AC245123.1 | 8 | novel transcript |
| AC246817.2 | 8 | novel transcript |
| AC246817.1 | 8 | novel transcript |
| AC115485.1 | 8 | novel transcript |
| CSMD1 | 8 | CUB and Sushi multiple domains 1 [Source:HG |
| AC026991.1 | 8 | novel transcript |
| AC026991.2 | 8 | novel transcript, antisense to CSMD1 |
| RNA5SP251 | 8 | RNA, 5S ribosomal pseudogene 251 [Source:H |
| AC027251.1 | 8 | novel transcript, antisense to CSMD1 |
| AC010941.1 | 8 | novel transcript, antisense to CSMD1 |
| AC022068.1 | 8 | novel transcript, antisense to CSMD1 |
| PAICSP4 | 8 | phosphoribosylaminoimidazole carboxylase, p |
| AC019176.1 | 8 | ribosomal L24 domain containing 1 (RSL24D1) |
| AC019176.2 | 8 | ribosomal protein L10 (RPL10) pseudogene |
| SNORA70 | 8 | Small nucleolar RNA SNORA70 [Source:RFAM; |
| RN7SL318P | 8 | RNA, 7SL, cytoplasmic 318, pseudogene [Sour |
| AC007718.1 | 8 | peroxisomal membrane protein 2, 22k (PXMP |
| AC091193.1 | 8 | ribosomal protein L26 (RPL26) pseudogene |
| RPL23AP54 | 8 | ribosomal protein L23a pseudogene 54 [Sourc |

| | | |
|---|---|---|
| AC004944.1 | 8 | TEC |
| AC084768.1 | 8 | novel transcript |
| AC084768.2 | 8 | ring finger protein 175 (RNF175) pseudogene |
| AC087369.1 | 8 | transcriptional adaptor 1 (TADA1) pseudogen |
| AC087369.2 | 8 | novel transcript |
| RN7SKP159 | 8 | RN7SK pseudogene 159 [Source:HGNC Symbo |
| AC079054.1 | 8 | novel transcript |
| AC009435.1 | 8 | novel transcript |
| AC016065.1 | 8 | novel transcript |
| MCPH1 | 8 | microcephalin 1 [Source:HGNC Symbol;Acc:H |
| AC016065.2 | 8 | TAF9 RNA polymerase II (TAF9) pseudogene |
| ANGPT2 | 8 | angiopoietin 2 [Source:HGNC Symbol;Acc:HGI |
| AC018398.1 | 8 | novel transcript, antisense to MCPH1 |
| AF287957.1 | 8 | novel transcript |
| MCPH1-AS1 | 8 | MCPH1 antisense RNA 1 [Source:HGNC Symb |
| MIR8055 | 8 | microRNA 8055 [Source:HGNC Symbol;Acc:H |
| AGPAT5 | 8 | 1-acylglycerol-3-phosphate O-acyltransferase |
| MIR4659A | 8 | microRNA 4659a [Source:HGNC Symbol;Acc:H |
| MIR4659B | 8 | microRNA 4659b [Source:HGNC Symbol;Acc:H |
| AF233439.3 | 8 | novel transcript |
| XKR5 | 8 | XK related 5 [Source:HGNC Symbol;Acc:HGNC |
| AF233439.1 | 8 | novel transcript |
| DEFB1 | 8 | defensin beta 1 [Source:HGNC Symbol;Acc:HG |
| RPL23AP96 | 8 | ribosomal protein L23a pseudogene 96 [Sourc |
| DEFA6 | 8 | defensin alpha 6 [Source:HGNC Symbol;Acc:H |
| AF233439.2 | 8 | novel transcript |
| DEFA4 | 8 | defensin alpha 4 [Source:HGNC Symbol;Acc:H |
| DEFA8P | 8 | defensin alpha 8, pseudogene [Source:HGNC S |
| DEFA9P | 8 | defensin alpha 9, pseudogene [Source:HGNC S |
| DEFA10P | 8 | defensin alpha 10, pseudogene [Source:HGNC |
| DEFA1 | 8 | defensin alpha 1 [Source:HGNC Symbol;Acc:H |
| DEFT1P | 8 | defensin theta 1, pseudogene [Source:HGNC S |
| DEFA1B | 8 | defensin alpha 1B [Source:HGNC Symbol;Acc:I |
| DEFT1P2 | 8 | defensin theta 1, pseudogene 2 [Source:HGN |
| DEFA3 | 8 | defensin alpha 3 [Source:HGNC Symbol;Acc:H |
| DEFA11P | 8 | defensin alpha 11, pseudogene [Source:HGNC |
| DEFA7P | 8 | defensin alpha 7, pseudogene [Source:HGNC S |
| DEFA5 | 8 | defensin alpha 5 [Source:HGNC Symbol;Acc:H |
| AF238378.1 | 8 | chromosome 11 open reading frame 2 (C11or |
| AF228730.5 | 8 | RNA binding protein, fox-1 homolog (C. elegar |
| AF228730.2 | 8 | Ribosomal protein S24 pseudogene |
| RPS3AP30 | 8 | ribosomal protein S3a pseudogene 30 [Source |
| SNRPCP6 | 8 | small nuclear ribonucleoprotein polypeptide ( |
| SNRPCP15 | 8 | small nuclear ribonucleoprotein polypeptide ( |
| RPS3AP33 | 8 | ribosomal protein S3a pseudogene 33 [Source |
| AF228730.1 | 8 | ribosomal protein S24 (RPS24) pseudogene |
| AF228730.4 | 8 | RNA binding protein, fox-1 homolog (C. elegar |
| AF228730.6 | 8 | RNA binding protein, fox-1 homolog (C. elegar |
| AF228730.3 | 8 | chromosome 11 open reading frame 2 (C11or |
| OR7E125P | 8 | olfactory receptor family 7 subfamily E memb |

| | | |
|---|---|---|
| FAM90A15P | 8 | family with sequence similarity 90 member A1 |
| FAM90A3P | 8 | family with sequence similarity 90 member A3 |
| FAM90A4P | 8 | family with sequence similarity 90 member A4 |
| FAM90A13P | 8 | family with sequence similarity 90 member A1 |
| FAM90A5P | 8 | family with sequence similarity 90 member A5 |
| FAM90A20P | 8 | family with sequence similarity 90 member A2 |
| FAM66B | 8 | family with sequence similarity 66 member B |
| DEFB109B | 8 | defensin beta 109B [Source:HGNC Symbol;Acc |
| USP17L1 | 8 | ubiquitin specific peptidase 17 like family mer |
| USP17L4 | 8 | ubiquitin specific peptidase 17 like family mer |
| AC130360.1 | 8 | ubiquitin-specific protease (USP) pseudogene |
| ZNF705G | 8 | zinc finger protein 705G [Source:HGNC Symbo |
| DEFB108C | 8 | defensin beta 108C [Source:HGNC Symbol;Acc |
| DEFB4B | 8 | defensin beta 4B [Source:HGNC Symbol;Acc:H |
| HSPD1P3 | 8 | heat shock protein family D (Hsp60) member |
| DEFB103B | 8 | defensin beta 103B [Source:HGNC Symbol;Acc |
| SPAG11B | 8 | sperm associated antigen 11B [Source:HGNC S |
| DEFB104B | 8 | defensin beta 104B [Source:HGNC Symbol;Acc |
| DEFB106B | 8 | defensin beta 106B [Source:HGNC Symbol;Acc |
| DEFB105B | 8 | defensin beta 105B [Source:HGNC Symbol;Acc |
| DEFB107B | 8 | defensin beta 107B [Source:HGNC Symbol;Acc |
| AC134684.7 | 8 | proline rich 23 domain containing 1 (PRR23D1 |
| PRR23D1 | 8 | proline rich 23 domain containing 1 [Source:H |
| AC134684.10 | 8 | proline rich 23 domain containing 1 (PRR23D1 |
| FAM90A6P | 8 | family with sequence similarity 90 member A6 |
| AC134684.11 | 8 | Putative protein FAM90A7P [Source:UniProtK |
| FAM90A21P | 8 | family with sequence similarity 90 member A2 |
| AC134684.8 | 8 | Putative protein FAM90A22P [Source:UniProt |
| AC134684.9 | 8 | Putative protein FAM90A23P [Source:UniProt |
| OR7E157P | 8 | olfactory receptor family 7 subfamily E memb |
| AC134684.1 | 8 | chromosome 11 open reading frame 2 (C11or |
| AC134684.6 | 8 | RNA binding protein, fox-1 homolog (C. elegar |
| AC084121.3 | 8 | RNA binding protein, fox-1 homolog (C. elegar |
| AC084121.4 | 8 | RNA binding protein, fox-1 homolog (C. elegar |
| AC084121.2 | 8 | chromosome 11 open reading frame 2 (C11or |
| OR7E154P | 8 | olfactory receptor family 7 subfamily E memb |
| AC084121.9 | 8 | Putative protein FAM90A14P [Source:UniProt |
| AC084121.11 | 8 | Putative protein FAM90A18P/FAM90A19P [S |
| AC084121.6 | 8 | Putative protein FAM90A16P/FAM90A17P [S |
| AC084121.12 | 8 | Putative protein FAM90A8P [Source:UniProtK |
| AC084121.8 | 8 | Putative protein FAM90A16P/FAM90A17P [S |
| AC084121.7 | 8 | Putative protein FAM90A18P/FAM90A19P [S |
| AC084121.5 | 8 | FAM90A pseudogene [Source:NCBI gene;Acc:1 |
| AC084121.13 | 8 | Putative protein FAM90A10P [Source:UniProt |
| AC084121.10 | 8 | proline rich 23 domain containing 1 (PRR23D1 |
| PRR23D2 | 8 | proline rich 23 domain containing 2 [Source:H |
| AC084121.1 | 8 | novel pseudogene |
| DEFB107A | 8 | defensin beta 107A [Source:HGNC Symbol;Acc |
| DEFB105A | 8 | defensin beta 105A [Source:HGNC Symbol;Acc |
| DEFB106A | 8 | defensin beta 106A [Source:HGNC Symbol;Acc |

| | | |
|---|---|---|
| DEFB104A | 8 | defensin beta 104A [Source:HGNC Symbol;Acc |
| SPAG11A | 8 | sperm associated antigen 11A [Source:HGNC S |
| DEFB103A | 8 | defensin beta 103A [Source:HGNC Symbol;Acc |
| HSPD1P2 | 8 | heat shock protein family D (Hsp60) member |
| DEFB4A | 8 | defensin beta 4A [Source:HGNC Symbol;Acc:H |
| ZNF705B | 8 | zinc finger protein 705B [Source:HGNC Symbo |
| DEFB108A | 8 | defensin beta 108A [Source:HGNC Symbol;Acc |
| FAM66E | 8 | family with sequence similarity 66 member E |
| AC130365.1 | 8 | ubiquitin specific peptidase 17 (USP17) pseud |
| USP17L8 | 8 | ubiquitin specific peptidase 17 like family mer |
| USP17L3 | 8 | ubiquitin specific peptidase 17 like family mer |
| DEFB109C | 8 | defensin beta 109C [Source:HGNC Symbol;Acc |
| FAM90A11P | 8 | family with sequence similarity 90 member A1 |
| FAM90A24P | 8 | family with sequence similarity 90 member A2 |
| FAM90A12P | 8 | family with sequence similarity 90 member A1 |
| OR7E96P | 8 | olfactory receptor family 7 subfamily E memb |
| AC105233.2 | 8 | chromosome 11 open reading frame 2 (C11or |
| AC105233.4 | 8 | RNA binding protein, fox-1 homolog (C. elegar |
| AC105233.5 | 8 | novel transcript |
| AC105233.6 | 8 | RNA binding protein, fox-1 homolog (C. elegar |
| AC105233.3 | 8 | ribosomal protein S24 pseudogene |
| MIR548I3 | 8 | microRNA 548i-3 [Source:HGNC Symbol;Acc:H |
| AC105233.1 | 8 | ribosomal protein S3A (RPS3A) pseudogene |
| SNRPCP17 | 8 | small nuclear ribonucleoprotein polypeptide C |
| ENPP7P1 | 8 | ectonucleotide pyrophosphatase/phosphodie |
| FAM85B | 8 | family with sequence similarity 85 member B |
| AC068020.1 | 8 | novel transcript |
| FAM86B3P | 8 | family with sequence similarity 86 member B3 |
| ALG1L13P | 8 | ALG1 like 13, pseudogene [Source:HGNC Syml |
| PRAG1 | 8 | PEAK1 related, kinase-activating pseudokinase |
| AC103957.1 | 8 | novel transcript |
| AC103957.2 | 8 | novel transcript |
| AC114550.2 | 8 | novel transcript |
| AC114550.3 | 8 | novel transcript |
| AC114550.1 | 8 | novel transcript |
| RN7SL178P | 8 | RNA, 7SL, cytoplasmic 178, pseudogene [Sour |
| AC087269.3 | 8 | novel transcript |
| CLDN23 | 8 | claudin 23 [Source:HGNC Symbol;Acc:HGNC:1 |
| AC087269.2 | 8 | novel lipoprotein receptor-related protein pse |
| AC087269.1 | 8 | novel transcript |
| MFHAS1 | 8 | malignant fibrous histiocytoma amplified sequ |
| RNU6-682P | 8 | RNA, U6 small nuclear 682, pseudogene [Sour |
| AC087763.1 | 8 | mitochondrial ribosomal protein S18C (MRPS1 |
| RPL10P19 | 8 | ribosomal protein L10 pseudogene 19 [Source |
| SNORA70 | 8 | Small nucleolar RNA SNORA70 [Source:RFAM; |
| ERI1 | 8 | exoribonuclease 1 [Source:HGNC Symbol;Acc: |
| MIR4660 | 8 | microRNA 4660 [Source:HGNC Symbol;Acc:HG |
| RNU7-55P | 8 | RNA, U7 small nuclear 55 pseudogene [Source |
| Metazoa_SRP | 8 | Metazoan signal recognition particle RNA [Sou |
| PPP1R3B | 8 | protein phosphatase 1 regulatory subunit 3B [ |

| | | |
|---|---|---|
| AC022784.5 | 8 | novel transcript |
| AC022784.1 | 8 | novel transcript |
| AC022784.6 | 8 | novel transcript |
| AC022784.2 | 8 | novel transcript |
| AC022784.4 | 8 | ubiquitin-conjugating enzyme E2 variant 1 (UE |
| AC022784.3 | 8 | novel transcript |
| RNU6-1151P | 8 | RNA, U6 small nuclear 1151, pseudogene [Sou |
| AC021736.1 | 8 | novel transcript |
| AC021242.2 | 8 | novel transcript |
| RNU6-526P | 8 | RNA, U6 small nuclear 526, pseudogene [Sour |
| AC021242.1 | 8 | DENN/MADD domain containing 4A (DENND4 |
| AC021242.3 | 8 | novel transcript, antisense to TNKS |
| TNKS | 8 | tankyrase [Source:HGNC Symbol;Acc:HGNC:1 |
| AC103834.1 | 8 | microtubule associated serine/threonine kina |
| MIR597 | 8 | microRNA 597 [Source:HGNC Symbol;Acc:HGN |
| LINC00599 | 8 | long intergenic non-protein coding RNA 599 [S |
| MIR124-1 | 8 | microRNA 124-1 [Source:HGNC Symbol;Acc:H |
| AC034111.2 | 8 | novel transcript, antisense to MSRA |
| AC034111.1 | 8 | novel transcript, antisense to MSRA |
| MSRA | 8 | methionine sulfoxide reductase A [Source:HG |
| AC023385.1 | 8 | novel transcript |
| AC079200.1 | 8 | novel transcript |
| AC112673.1 | 8 | transmembrane protein 126A (TMEM126A) p |
| AC104964.3 | 8 | novel transcript, overlapping to MSRA |
| AC104964.1 | 8 | novel transcript |
| AC104964.5 | 8 | novel protease, serine (PRSS) gene family pse |
| RNU6-729P | 8 | RNA, U6 small nuclear 729, pseudogene [Sour |
| AC104964.2 | 8 | novel transcript |
| PRSS51 | 8 | serine protease 51 [Source:HGNC Symbol;Acc |
| AC104964.4 | 8 | novel transcript |
| PRSS55 | 8 | serine protease 55 [Source:HGNC Symbol;Acc |
| RP1L1 | 8 | RP1 like 1 [Source:HGNC Symbol;Acc:HGNC:1 |
| MIR4286 | 8 | microRNA 4286 [Source:HGNC Symbol;Acc:HG |
| C8orf74 | 8 | chromosome 8 open reading frame 74 [Sourc |
| RNA5SP252 | 8 | RNA, 5S ribosomal pseudogene 252 [Source:H |
| SOX7 | 8 | SRY-box transcription factor 7 [Source:HGNC S |
| PINX1 | 8 | PIN2 (TERF1) interacting telomerase inhibitor |
| AC105001.1 | 8 | novel transcript, antisense to SOX7 and PINX1 |
| PINX1 | 8 | PIN2 (TERF1) interacting telomerase inhibitor |
| MIR1322 | 8 | microRNA 1322 [Source:HGNC Symbol;Acc:HG |
| AC011008.1 | 8 | novel transcript |
| AC011008.2 | 8 | TEC |
| XKR6 | 8 | XK related 6 [Source:HGNC Symbol;Acc:HGNC |
| MIR598 | 8 | microRNA 598 [Source:HGNC Symbol;Acc:HGN |
| AF131215.3 | 8 | novel transcript |
| AF131215.6 | 8 | novel transcript |
| AF131215.5 | 8 | novel transcript |
| AF131215.4 | 8 | novel transcript |
| AF131215.2 | 8 | novel transcript |
| AF131215.1 | 8 | ribosomal protein L17 (RPL17) pseudogene |

| | | |
|---|---|---|
| AF131215.7 | 8 | novel transcript |
| LINC00529 | 8 | long intergenic non-protein coding RNA 529 [S |
| RPL19P13 | 8 | ribosomal protein L19 pseudogene 13 [Source |
| AF131216.2 | 8 | HERV-FRD provirus ancestral Env polyprotein |
| AF131216.4 | 8 | novel transcript, antisense to MTMR9 |
| MTMR9 | 8 | myotubularin related protein 9 [Source:HGNC |
| AF131216.1 | 8 | uncharacterized LOC101929290 [Source:NCBI |
| SLC35G5 | 8 | solute carrier family 35 member G5 [Source:H |
| TDH | 8 | L-threonine dehydrogenase (pseudogene) [So |
| AF131216.3 | 8 | novel transcript, antisense to TDH |
| FAM167A-AS1 | 8 | FAM167A antisense RNA 1 [Source:HGNC Sym |
| RN7SL293P | 8 | RNA, 7SL, cytoplasmic 293, pseudogene [Sour |
| RNU6-1084P | 8 | RNA, U6 small nuclear 1084, pseudogene [Sou |
| FAM167A | 8 | family with sequence similarity 167 member A |
| AF131216.5 | 8 | novel transcript |
| BLK | 8 | BLK proto-oncogene, Src family tyrosine kinas |
| AC022239.1 | 8 | novel transcript, antisense to BLK |
| LINC00208 | 8 | long intergenic non-protein coding RNA 208 [S |
| AC022239.2 | 8 | novel transcript |
| GATA4 | 8 | GATA binding protein 4 [Source:HGNC Symbol |
| C8orf49 | 8 | chromosome 8 open reading frame 49 [Source |
| NEIL2 | 8 | nei like DNA glycosylase 2 [Source:HGNC Syml |
| SUB1P1 | 8 | SUB1 pseudogene 1 [Source:HGNC Symbol;Ac |
| FDFT1 | 8 | farnesyl-diphosphate farnesyltransferase 1 [So |
| AC069185.1 | 8 | novel transcript |
| CTSB | 8 | cathepsin B [Source:HGNC Symbol;Acc:HGNC: |
| AC025857.2 | 8 | novel transcript, sense intronic to CTSB |
| AC025857.1 | 8 | transmembrane protein 126A (TMEM126A) p |
| OR7E158P | 8 | olfactory receptor family 7 subfamily E memb |
| OR7E161P | 8 | olfactory receptor family 7 subfamily E memb |
| AC107918.5 | 8 | novel transcript |
| AC107918.6 | 8 | novel transcript |
| DEFB136 | 8 | defensin beta 136 [Source:HGNC Symbol;Acc: |
| DEFB135 | 8 | defensin beta 135 [Source:HGNC Symbol;Acc: |
| DEFB134 | 8 | defensin beta 134 [Source:HGNC Symbol;Acc: |
| AC107918.3 | 8 | Wolf-Hirschhorn syndrome candidate 1-like 1 |
| DEFB131E | 8 | defensin beta 131E (pseudogene) [Source:HG |
| AC107918.4 | 8 | RNA binding protein, fox-1 homolog (C. elegar |
| AC107918.2 | 8 | chromosome 11 open reading frame 2 (C11or |
| AC107918.1 | 8 | olfactory receptor, family 7, subfamily E, mem |
| DEFB131C | 8 | defensin beta 131C (pseudogene) [Source:HG |
| DEFB130B | 8 | defensin beta 130B [Source:HGNC Symbol;Acc |
| RNA5SP253 | 8 | RNA, 5S ribosomal pseudogene 253 [Source:H |
| DEFB108D | 8 | defensin beta 108D (pseudogene) [Source:HG |
| ZNF705D | 8 | zinc finger protein 705D [Source:HGNC Symbo |
| FAM66D | 8 | family with sequence similarity 66 member D |
| AC130366.1 | 8 | deubiquitinating enzyme 3 pseudogene |
| USP17L7 | 8 | ubiquitin specific peptidase 17 like family mer |
| USP17L2 | 8 | ubiquitin specific peptidase 17 like family mer |
| DEFB109D | 8 | defensin beta 109D (pseudogene) [Source:HG |

| | | |
|---|---|---|
| FAM90A2P | 8 | family with sequence similarity 90 member A2 |
| ALG1L11P | 8 | ALG1 like 11, pseudogene [Source:HGNC Syml |
| FAM86B1 | 8 | family with sequence similarity 86 member B1 |
| AC145124.1 | 8 | novel transcript |
| ENPP7P12 | 8 | ectonucleotide pyrophosphatase/phosphodie |
| DEFB131D | 8 | defensin beta 131D (pseudogene) [Source:HG |
| DEFB130A | 8 | defensin beta 130A [Source:HGNC Symbol;Acc |
| RNA5SP254 | 8 | RNA, 5S ribosomal pseudogene 254 [Source:H |
| DEFB108E | 8 | defensin beta 108E (pseudogene) [Source:HGI |
| ZNF705CP | 8 | zinc finger protein 705C, pseudogene [Source: |
| FAM66A | 8 | family with sequence similarity 66 member A |
| AC087203.1 | 8 | ubiquitin specific peptidase 17-like 2 ( USP17L |
| AC087203.2 | 8 | ubiquitin specific peptidase 17-like 2 (USP17L2 |
| DEFB109A | 8 | defensin beta 109A (pseudogene) [Source:HG |
| AC087203.3 | 8 | novel transcript |
| FAM90A25P | 8 | family with sequence similarity 90 member A2 |
| ALG1L12P | 8 | ALG1 like 12, pseudogene [Source:HGNC Syml |
| FAM86B2 | 8 | family with sequence similarity 86 member B2 |
| ENPP7P6 | 8 | ectonucleotide pyrophosphatase/phosphodie |
| AC068587.4 | 8 | novel transcript |
| AC130352.1 | 8 | novel transcript |
| RPS3AP34 | 8 | ribosomal protein S3a pseudogene 34 [Source |
| AC068587.2 | 8 | 40S ribosomal protein S24 (Rps24) pseudogen |
| AC068587.8 | 8 | RNA binding protein, fox-1 homolog (C. elegar |
| AC068587.6 | 8 | RNA binding protein, fox-1 homolog (C. elegar |
| RPS3AP35 | 8 | ribosomal protein S3a pseudogene 35 [Source |
| AC068587.3 | 8 | ribosomal protein S24 (Rps24) pseudogene |
| AC068587.7 | 8 | RNA binding protein, fox-1 homolog (C. elegar |
| AC068587.5 | 8 | RNA binding protein, fox-1 homolog (C. elegar |
| AC068587.1 | 8 | novel pseudogene |
| OR7E8P | 8 | olfactory receptor family 7 subfamily E memb |
| OR7E15P | 8 | olfactory receptor family 7 subfamily E memb |
| OR7E10P | 8 | olfactory receptor family 7 subfamily E memb |
| MIR5692A2 | 8 | microRNA 5692a-2 [Source:HGNC Symbol;Acc |
| LONRF1 | 8 | LON peptidase N-terminal domain and ring fir |
| MIR3926-1 | 8 | microRNA 3926-1 [Source:HGNC Symbol;Acc:I |
| MIR3926-2 | 8 | microRNA 3926-2 [Source:HGNC Symbol;Acc:I |
| AC123777.1 | 8 | novel transcript |
| LINC00681 | 8 | long intergenic non-protein coding RNA 681 [S |
| TRMT9B | 8 | tRNA methyltransferase 9B (putative) [Source |
| AC135352.1 | 8 | regulatory solute carrier protein, family 1, me |
| RNU6-842P | 8 | RNA, U6 small nuclear 842, pseudogene [Sour |
| DLC1 | 8 | DLC1 Rho GTPase activating protein [Source:H |
| AC106845.1 | 8 | novel transcript, antisense to DLC1 |
| Y_RNA | 8 | Y RNA [Source:RFAM;Acc:RF00019] |
| AC019270.1 | 8 | novel transcript |
| MTND4P7 | 8 | MT-ND4 pseudogene 7 [Source:HGNC Symbol |
| RNA5SP255 | 8 | RNA, 5S ribosomal pseudogene 255 [Source:H |
| C8orf48 | 8 | chromosome 8 open reading frame 48 [Sourc |
| AC022832.1 | 8 | nuclear transcription factor Y, gamma (NFYC) |

| | | |
|---|---|---|
| AC022832.2 | 8 | novel transcript |
| AC022880.2 | 8 | septin 8 (SEPT8) pseudogene |
| AC022880.1 | 8 | zinc finger protein 654 (ZNF654) pseudogene |
| AC022690.2 | 8 | novel transcript |
| AC022690.1 | 8 | small nuclear ribonucleoprotein polypeptide E |
| AC023538.1 | 8 | structural maintenance of chromosomes 5 (SM |
| SGCZ | 8 | sarcoglycan zeta [Source:HGNC Symbol;Acc:H |
| AC022039.1 | 8 | novel transcript |
| EIF4EP5 | 8 | eukaryotic translation initiation factor 4E pseu |
| RNU7-153P | 8 | RNA, U7 small nuclear 153 pseudogene [Sourc |
| Y_RNA | 8 | Y RNA [Source:RFAM;Acc:RF00019] |
| RNU6-397P | 8 | RNA, U6 small nuclear 397, pseudogene [Sour |
| AC040926.1 | 8 | diazepam binding inhibitor (GABA receptor m |
| MIR383 | 8 | microRNA 383 [Source:HGNC Symbol;Acc:HGN |
| AC084838.1 | 8 | novel transcript |
| AC103851.1 | 8 | pseudogene similar to part of polymerase (DN |
| TUSC3 | 8 | tumor suppressor candidate 3 [Source:HGNC S |
| AC100850.1 | 8 | clathrin, light chain (Lca) (CLTA) pseudogene |
| PPM1AP1 | 8 | protein phosphatase, Mg2+/Mn2+ dependent |
| AC018437.3 | 8 | novel transcript |
| AC018437.1 | 8 | ribosomal protein L32 (RPL32) pseudogene |
| AC018437.2 | 8 | novel transcript |
| MSR1 | 8 | macrophage scavenger receptor 1 [Source:HG |
| AC087360.1 | 8 | chaperonin containing TCP1, subunit 3 (gamm |
| MRPL49P2 | 8 | mitochondrial ribosomal protein L49 pseudog |
| AC011586.2 | 8 | novel transcript |
| AC011586.1 | 8 | novel transcript |
| RN7SL474P | 8 | RNA, 7SL, cytoplasmic 474, pseudogene [Sour |
| FGF20 | 8 | fibroblast growth factor 20 [Source:HGNC Sym |
| MICU3 | 8 | mitochondrial calcium uptake family member |
| AC079193.2 | 8 | novel transcript |
| AC079193.1 | 8 | ribosomal protein L9 (RPL9) pseudogene |
| ZDHHC2 | 8 | zinc finger DHHC-type containing 2 [Source:HG |
| CNOT7 | 8 | CCR4-NOT transcription complex subunit 7 [Sc |
| VPS37A | 8 | VPS37A subunit of ESCRT-I [Source:HGNC Sym |
| MTMR7 | 8 | myotubularin related protein 7 [Source:HGNC |
| ADAM24P | 8 | ADAM metallopeptidase domain 24, pseudoge |
| SLC7A2 | 8 | solute carrier family 7 member 2 [Source:HGN |
| AP006248.2 | 8 | TEC |
| AP006248.5 | 8 | TEC |
| AP006248.3 | 8 | TEC |
| AP006248.4 | 8 | novel transcript, sense intronic SLC7A2 |
| PDGFRL | 8 | platelet derived growth factor receptor like [S |
| AP006248.1 | 8 | GRB2-related adaptor protein (GRAP) pseudog |
| AP006248.6 | 8 | novel transcript, antisense to PDGFRL |
| MTUS1 | 8 | microtubule associated scaffold protein 1 [Sou |
| MIR548V | 8 | microRNA 548v [Source:HGNC Symbol;Acc:HG |
| AC124069.1 | 8 | novel transcript |
| RNA5SP256 | 8 | RNA, 5S ribosomal pseudogene 256 [Source:H |
| AC027117.2 | 8 | novel transcript |

| | | |
|---|---|---|
| AC027117.1 | 8 | novel transcript |
| FGL1 | 8 | fibrinogen like 1 [Source:HGNC Symbol;Acc:H( |
| AC087273.1 | 8 | novel transcript |
| AC087273.2 | 8 | novel transcript |
| PCM1 | 8 | pericentriolar material 1 [Source:HGNC Symbo |
| ASAH1 | 8 | N-acylsphingosine amidohydrolase 1 [Source:I |
| AC124242.1 | 8 | novel transcript |
| AC124242.2 | 8 | family with sequence similarity 175, member |
| AC124242.3 | 8 | novel transcript |
| NAT1 | 8 | N-acetyltransferase 1 [Source:HGNC Symbol;A |
| MTND4LP26 | 8 | MT-ND4L pseudogene 26 [Source:HGNC Symb |
| NATP | 8 | N-acetyltransferase pseudogene [Source:HGN |
| AC025062.1 | 8 | ribosomal protein L10a (RPL10A) pseudogene |
| AC025062.3 | 8 | novel transcript |
| AC025062.2 | 8 | SURF6 (surfeit 6) pseudogene |
| NAT2 | 8 | N-acetyltransferase 2 [Source:HGNC Symbol;A |
| PSD3 | 8 | pleckstrin and Sec7 domain containing 3 [Sour |
| AC100800.1 | 8 | novel transcript, antisense to PSD3 |
| RPL35P6 | 8 | ribosomal protein L35 pseudogene 6 [Source: |
| AC087821.2 | 8 | mitochondrially encoded cytochrome b (MT-C |
| AC087821.1 | 8 | TEC |
| SNORA62 | 8 | Small nucleolar RNA SNORA62/SNORA6 family |
| AC009884.1 | 8 | novel transcript |
| AC009884.2 | 8 | chromosome 8 open reading frame 46 (C8orf4 |
| AC100849.1 | 8 | novel transcript |
| AC100849.2 | 8 | novel transcript |
| AC068880.1 | 8 | ATPase, Ca++ transporting, plasma membrane |
| AC068880.3 | 8 | novel transcript |
| AC068880.2 | 8 | novel transcript |
| AC068880.5 | 8 | TEC |
| SH2D4A | 8 | SH2 domain containing 4A [Source:HGNC Sym |
| AC068880.4 | 8 | TEC |
| CSGALNACT1 | 8 | chondroitin sulfate N-acetylgalactosaminyltra |
| AC090541.1 | 8 | novel transcript |
| INTS10 | 8 | integrator complex subunit 10 [Source:HGNC |
| LPL | 8 | lipoprotein lipase [Source:HGNC Symbol;Acc:H |
| AC100802.1 | 8 | novel transcript |
| RPL30P9 | 8 | ribosomal protein L30 pseudogene 9 [Source: |
| SLC18A1 | 8 | solute carrier family 18 member A1 [Source:H |
| ATP6V1B2 | 8 | ATPase H+ transporting V1 subunit B2 [Source |
| RNU6-892P | 8 | RNA, U6 small nuclear 892, pseudogene [Sour |
| LZTS1 | 8 | leucine zipper tumor suppressor 1 [Source:HG |
| LZTS1-AS1 | 8 | LZTS1 antisense RNA 1 [Source:HGNC Symbol; |
| RNA5SP257 | 8 | RNA, 5S ribosomal pseudogene 257 [Source:H |
| AC023403.1 | 8 | novel transcript |
| AC022559.1 | 8 | mitochondrially encoded ATP synthase 6 (MT- |
| AC018541.1 | 8 | novel transcript |
| TMEM97P2 | 8 | transmembrane protein 97 pseudogene 2 [Sou |
| AC015468.3 | 8 | novel transcript |
| AC021613.1 | 8 | novel transcript |

| | | |
|---|---|---|
| AC015468.1 | 8 | novel transcript |
| AC015468.2 | 8 | novel transcript |
| LINC02153 | 8 | long intergenic non-protein coding RNA 2153 |
| AC103719.1 | 8 | novel transcript |
| AC021613.2 | 8 | TEC |
| AC021613.3 | 8 | novel transcript |
| AC021355.1 | 8 | novel transcript |
| GFRA2 | 8 | GDNF family receptor alpha 2 [Source:HGNC S |
| OR6R2P | 8 | olfactory receptor family 6 subfamily R memb |
| DOK2 | 8 | docking protein 2 [Source:HGNC Symbol;Acc:H |
| XPO7 | 8 | exportin 7 [Source:HGNC Symbol;Acc:HGNC:1 |
| NPM2 | 8 | nucleophosmin/nucleoplasmin 2 [Source:HGN |
| FGF17 | 8 | fibroblast growth factor 17 [Source:HGNC Sym |
| DMTN | 8 | dematin actin binding protein [Source:HGNC S |
| FAM160B2 | 8 | family with sequence similarity 160 member B |
| NUDT18 | 8 | nudix hydrolase 18 [Source:HGNC Symbol;Acc |
| HR | 8 | HR lysine demethylase and nuclear receptor c |
| REEP4 | 8 | receptor accessory protein 4 [Source:HGNC Sy |
| LGI3 | 8 | leucine rich repeat LGI family member 3 [Sour |
| SFTPC | 8 | surfactant protein C [Source:HGNC Symbol;Ac |
| BMP1 | 8 | bone morphogenetic protein 1 [Source:HGNC |
| AC105206.1 | 8 | novel transcript, sense overlapping BMP1 |
| AC105206.3 | 8 | Novel transcript, antisense to BMP1 |
| PHYHIP | 8 | phytanoyl-CoA 2-hydroxylase interacting prot |
| MIR320A | 8 | microRNA 320a [Source:HGNC Symbol;Acc:HG |
| POLR3D | 8 | RNA polymerase III subunit D [Source:HGNC S |
| AC105206.2 | 8 | novel transcript |
| PIWIL2 | 8 | piwi like RNA-mediated gene silencing 2 [Sour |
| SLC39A14 | 8 | solute carrier family 39 member 14 [Source:H |
| AC105910.1 | 8 | ribosomal protein L21 (RPL21) pseudogene |
| RNU6-336P | 8 | RNA, U6 small nuclear 336, pseudogene [Sour |
| PPP3CC | 8 | protein phosphatase 3 catalytic subunit gamm |
| AC087854.1 | 8 | novel transcript |
| AC087854.2 | 8 | basic transcription factor 3-like (BTF3L) pseud |
| SORBS3 | 8 | sorbin and SH3 domain containing 3 [Source:H |
| AC037459.4 | 8 | novel transcript, sense intronic to SORBS3 |
| AC037459.3 | 8 | novel transcript |
| PDLIM2 | 8 | PDZ and LIM domain 2 [Source:HGNC Symbol; |
| AC037459.1 | 8 | novel protein |
| C8orf58 | 8 | chromosome 8 open reading frame 58 [Source |
| CCAR2 | 8 | cell cycle and apoptosis regulator 2 [Source:H |
| AC037459.2 | 8 | novel transcript, antisense to KIAA1967 |
| BIN3 | 8 | bridging integrator 3 [Source:HGNC Symbol;A |
| AC105046.1 | 8 | novel transcript, overlapping to EGR3 |
| EGR3 | 8 | early growth response 3 [Source:HGNC Symbo |
| AC055854.1 | 8 | novel transcript |
| AC105046.2 | 8 | novel transcript |
| PEBP4 | 8 | phosphatidylethanolamine binding protein 4 [ |
| AC037441.1 | 8 | novel transcript |
| AC037441.2 | 8 | novel transcript |

| | | |
|---|---|---|
| RN7SL303P | 8 | RNA, 7SL, cytoplasmic 303, pseudogene [Sour |
| AC107959.1 | 8 | novel transcript, antisense to RHOBTB2 |
| RHOBTB2 | 8 | Rho related BTB domain containing 2 [Source: |
| TNFRSF10B | 8 | TNF receptor superfamily member 10b [Sourc |
| AC107959.2 | 8 | novel transcript |
| AC107959.3 | 8 | novel transcript |
| AC107959.4 | 8 | novel transcript |
| AC107959.5 | 8 | novel protein |
| TNFRSF10C | 8 | TNF receptor superfamily member 10c [Sourc |
| TNFRSF10D | 8 | TNF receptor superfamily member 10d [Sourc |
| TNFRSF10A-AS1 | 8 | TNFRSF10A antisense RNA 1 [Source:HGNC Sy |
| TNFRSF10A | 8 | TNF receptor superfamily member 10a [Sourc |
| RPL23AP55 | 8 | ribosomal protein L23a pseudogene 55 [Sourc |
| AC100861.1 | 8 | novel transcript, antisense to TNFRSF10A |
| AC100861.2 | 8 | novel transcript |
| CHMP7 | 8 | charged multivesicular body protein 7 [Source |
| R3HCC1 | 8 | R3H domain and coiled-coil containing 1 [Sour |
| LOXL2 | 8 | lysyl oxidase like 2 [Source:HGNC Symbol;Acc: |
| AC090197.1 | 8 | novel transcript |
| ENTPD4 | 8 | ectonucleoside triphosphate diphosphohydro |
| AC104561.1 | 8 | novel transcript |
| AC104561.2 | 8 | DnaJ (Hsp40) homolog, subfamily C, member |
| AC104561.4 | 8 | novel transcript |
| AC104561.3 | 8 | novel transcript |
| SLC25A37 | 8 | solute carrier family 25 member 37 [Source:H |
| RNU4-71P | 8 | RNA, U4 small nuclear 71, pseudogene [Sourc |
| SINHCAFP3 | 8 | SINHCAF pseudogene 3 [Source:HGNC Symbol |
| NKX3-1 | 8 | NK3 homeobox 1 [Source:HGNC Symbol;Acc:H |
| NKX2-6 | 8 | NK2 homeobox 6 [Source:HGNC Symbol;Acc:H |
| AC012574.1 | 8 | novel transcript |
| AC012574.2 | 8 | novel transcript |
| AC012119.2 | 8 | eukaryotic translation initiation factor 4A1 (EI |
| AC012119.1 | 8 | NULL |
| STC1 | 8 | stanniocalcin 1 [Source:HGNC Symbol;Acc:HG |
| RNU1-148P | 8 | RNA, U1 small nuclear 148, pseudogene [Sour |
| AC023202.1 | 8 | novel transcript |
| ADAM28 | 8 | ADAM metallopeptidase domain 28 [Source:H |
| AC120193.1 | 8 | novel transcript |
| ADAMDEC1 | 8 | ADAM like decysin 1 [Source:HGNC Symbol;Ac |
| ADAM7 | 8 | ADAM metallopeptidase domain 7 [Source:HG |
| AC024958.1 | 8 | novel transcript |
| AF106564.1 | 8 | novel transcript, antisense to NEFM |
| NEFM | 8 | neurofilament medium [Source:HGNC Symbol |
| NEFL | 8 | neurofilament light [Source:HGNC Symbol;Acc |
| MIR6841 | 8 | microRNA 6841 [Source:HGNC Symbol;Acc:HG |
| AC107373.2 | 8 | novel transcript, antisense to NEFL |
| AC107373.1 | 8 | novel transcript |
| AC107373.3 | 8 | novel transcript |
| AC073581.1 | 8 | ependymin related protein 1 (zebrafish) (EPDI |
| AC041005.1 | 8 | novel transcript |

| | | |
|---|---|---|
| DOCK5 | 8 | dedicator of cytokinesis 5 [Source:HGNC Symt |
| AC091185.2 | 8 | novel oocyte-secreted protein family member |
| MIR6876 | 8 | microRNA 6876 [Source:HGNC Symbol;Acc:HG |
| GNRH1 | 8 | gonadotropin releasing hormone 1 [Source:H( |
| AC091185.1 | 8 | novel transcript |
| KCTD9 | 8 | potassium channel tetramerization domain cc |
| CDCA2 | 8 | cell division cycle associated 2 [Source:HGNC : |
| AC009623.1 | 8 | novel transcript |
| AC009623.2 | 8 | novel transcript |
| AC009623.3 | 8 | novel transcript |
| AC090103.1 | 8 | novel transcript |
| EBF2 | 8 | EBF transcription factor 2 [Source:HGNC Symt |
| RNA5SP258 | 8 | RNA, 5S ribosomal pseudogene 258 [Source:H |
| AC079097.1 | 8 | matrix metallopeptidase 12 (macrophage elas |
| PPP2R2A | 8 | protein phosphatase 2 regulatory subunit Balp |
| SDAD1P1 | 8 | SDA1 domain containing 1 pseudogene 1 [Sou |
| BNIP3L | 8 | BCL2 interacting protein 3 like [Source:HGNC : |
| AC011726.2 | 8 | novel transcript |
| AC011726.1 | 8 | DnaJ (Hsp40) homolog, subfamily B, member |
| AC011726.3 | 8 | novel transcript |
| PNMA2 | 8 | PNMA family member 2 [Source:HGNC Symbc |
| DPYSL2 | 8 | dihydropyrimidinase like 2 [Source:HGNC Sym |
| AC015743.1 | 8 | proteasome (prosome, macropain) activator s |
| ADRA1A | 8 | adrenoceptor alpha 1A [Source:HGNC Symbol |
| AC091675.1 | 8 | ribosomal protein, large, P1 (RPLP1) pseudoge |
| COX6B1P4 | 8 | cytochrome c oxidase subunit 6B1 pseudogen |
| AC067904.3 | 8 | serine palmitoyltransferase, long chain base s |
| AC067904.1 | 8 | chromosome 15 open reading frame 2 (C15or |
| MIR548H4 | 8 | microRNA 548h-4 [Source:HGNC Symbol;Acc:I |
| AC067904.2 | 8 | novel pseudogene |
| AC090150.2 | 8 | novel transcript |
| AC090150.1 | 8 | novel transcript |
| STMN4 | 8 | stathmin 4 [Source:HGNC Symbol;Acc:HGNC:1 |
| TRIM35 | 8 | tripartite motif containing 35 [Source:HGNC S |
| PTK2B | 8 | protein tyrosine kinase 2 beta [Source:HGNC : |
| MIR6842 | 8 | microRNA 6842 [Source:HGNC Symbol;Acc:HG |
| CHRNA2 | 8 | cholinergic receptor nicotinic alpha 2 subunit |
| EPHX2 | 8 | epoxide hydrolase 2 [Source:HGNC Symbol;Ac |
| GULOP | 8 | gulonolactone (L-) oxidase, pseudogene [Sour |
| CLU | 8 | clusterin [Source:HGNC Symbol;Acc:HGNC:20! |
| MIR6843 | 8 | microRNA 6843 [Source:HGNC Symbol;Acc:HG |
| SCARA3 | 8 | scavenger receptor class A member 3 [Source |
| RNU6-1086P | 8 | RNA, U6 small nuclear 1086, pseudogene [Sou |
| AC013643.1 | 8 | coenzyme Q6 homolog, monooxygenase (S. c |
| AC013643.3 | 8 | TEC |
| MIR3622B | 8 | microRNA 3622b [Source:HGNC Symbol;Acc:H |
| MIR3622A | 8 | microRNA 3622a [Source:HGNC Symbol;Acc:H |
| AC013643.2 | 8 | novel transcript |
| CCDC25 | 8 | coiled-coil domain containing 25 [Source:HGN |
| ESCO2 | 8 | establishment of sister chromatid cohesion N- |

| | | |
|---|---|---|
| RNU6-1276P | 8 | RNA, U6 small nuclear 1276, pseudogene [Sou |
| PBK | 8 | PDZ binding kinase [Source:HGNC Symbol;Acc |
| AC104997.1 | 8 | TAF9B RNA polymerase II, TATA box binding p |
| SCARA5 | 8 | scavenger receptor class A member 5 [Source |
| MIR4287 | 8 | microRNA 4287 [Source:HGNC Symbol;Acc:HG |
| AC069113.1 | 8 | mitochondrially encoded cytochrome b (MT-C |
| AC069113.3 | 8 | mitochondrially encoded NADH dehydrogenas |
| AC069113.2 | 8 | novel transcript |
| NUGGC | 8 | nuclear GTPase, germinal center associated [S |
| ELP3 | 8 | elongator acetyltransferase complex subunit 3 |
| AC021678.1 | 8 | degenerative spermatocyte homolog 1, lipid c |
| AC021678.3 | 8 | PDZ and LIM domain 7 (enigma) (PDLIM7) pse |
| AC021678.2 | 8 | novel transcript |
| RPL5P22 | 8 | ribosomal protein L5 pseudogene 22 [Source: |
| PNOC | 8 | prepronociceptin [Source:HGNC Symbol;Acc:H |
| ZNF395 | 8 | zinc finger protein 395 [Source:HGNC Symbol; |
| FBXO16 | 8 | F-box protein 16 [Source:HGNC Symbol;Acc:H |
| AC025871.3 | 8 | novel transcript, sense intronic to FBXO16 |
| AC025871.2 | 8 | novel transcript |
| RNU6-178P | 8 | RNA, U6 small nuclear 178, pseudogene [Sour |
| AC025871.1 | 8 | novel transcript |
| FZD3 | 8 | frizzled class receptor 3 [Source:HGNC Symbo |
| MIR4288 | 8 | microRNA 4288 [Source:HGNC Symbol;Acc:HG |
| RNA5SP259 | 8 | RNA, 5S ribosomal pseudogene 259 [Source:H |
| EXTL3 | 8 | exostosin like glycosyltransferase 3 [Source:H |
| EXTL3-AS1 | 8 | EXTL3 antisense RNA 1 [Source:HGNC Symbol; |
| INTS9 | 8 | integrator complex subunit 9 [Source:HGNC Sy |
| INTS9-AS1 | 8 | INTS9 antisense RNA 1 [Source:HGNC Symbol; |
| AC040975.1 | 8 | ribosomal protein L36a (RPL36A) pseudogene |
| HMBOX1 | 8 | homeobox containing 1 [Source:HGNC Symbo |
| Metazoa_SRP | 8 | Metazoan signal recognition particle RNA [Sou |
| HMBOX1-IT1 | 8 | HMBOX1 intronic transcript 1 [Source:HGNC S |
| RNA5SP260 | 8 | RNA, 5S ribosomal pseudogene 260 [Source:H |
| AC108449.2 | 8 | novel transcript, sense intronic to HMBOX1 |
| KIF13B | 8 | kinesin family member 13B [Source:HGNC Syr |
| AC108449.3 | 8 | novel transcript, antisense to KIF13B |
| AC108449.1 | 8 | novel transcript, antisense to KIF13B |
| HMGB1P23 | 8 | high mobility group box 1 pseudogene 23 [Sou |
| DUSP4 | 8 | dual specificity phosphatase 4 [Source:HGNC S |
| AC084262.2 | 8 | novel transcript |
| AC084262.1 | 8 | novel transcript, overlapping to DUSP4 |
| AC084026.2 | 8 | novel transcript |
| AC084026.1 | 8 | novel transcript |
| AC084026.3 | 8 | TEC |
| RPL17P33 | 8 | ribosomal protein L17 pseudogene 33 [Source |
| LINC00589 | 8 | long intergenic non-protein coding RNA 589 [S |
| LINC02099 | 8 | long intergenic non-protein coding RNA 2099 |
| AC131254.1 | 8 | novel transcript |
| AC131254.3 | 8 | novel transcript |
| AC131254.2 | 8 | novel transcript |

| | | |
|---|---|---|
| LINC02209 | 8 | long intergenic non-protein coding RNA 2209 |
| MIR3148 | 8 | microRNA 3148 [Source:HGNC Symbol;Acc:HG |
| MAP2K1P1 | 8 | mitogen-activated protein kinase kinase 1 pse |
| RNU6-1218P | 8 | RNA, U6 small nuclear 1218, pseudogene [Sou |
| SARAF | 8 | store-operated calcium entry associated regul |
| AC044849.1 | 8 | novel transcript, antisense to TMEM66 |
| LEPROTL1 | 8 | leptin receptor overlapping transcript like 1 [S |
| RPS15AP24 | 8 | ribosomal protein S15a pseudogene 24 [Sourc |
| MBOAT4 | 8 | membrane bound O-acyltransferase domain c |
| AC026979.2 | 8 | novel transcript |
| DCTN6 | 8 | dynactin subunit 6 [Source:HGNC Symbol;Acc: |
| AC026979.1 | 8 | novel transcript |
| AC026979.4 | 8 | novel transcript |
| AC026979.3 | 8 | novel transcript |
| HSPA8P11 | 8 | heat shock protein family A (Hsp70) member 8 |
| AC090820.1 | 8 | transgelin 2 (TAGLN2) pseudogene |
| AC090820.2 | 8 | novel transcript |
| AC109329.1 | 8 | general transcription factor IIA, 2, 12kDa (GTF |
| PPIAP84 | 8 | peptidylprolyl isomerase A pseudogene 84 [Sc |
| TUBBP1 | 8 | tubulin beta pseudogene 1 [Source:HGNC Sym |
| RBPMS-AS1 | 8 | RBPMS antisense RNA 1 [Source:HGNC Symbo |
| RBPMS | 8 | RNA binding protein, mRNA processing factor |
| AC102945.2 | 8 | TEC |
| GTF2E2 | 8 | general transcription factor IIE subunit 2 [Sour |
| AC102945.1 | 8 | novel transcript |
| SMIM18 | 8 | small integral membrane protein 18 [Source:H |
| RNU5A-3P | 8 | RNA, U5A small nuclear 3, pseudogene [Sourc |
| GSR | 8 | glutathione-disulfide reductase [Source:HGNC |
| UBXN8 | 8 | UBX domain protein 8 [Source:HGNC Symbol;A |
| AC009314.1 | 8 | chromosome 11 open reading frame 73 (C11o |
| PPP2CB | 8 | protein phosphatase 2 catalytic subunit beta [ |
| TEX15 | 8 | testis expressed 15, meiosis and synapsis asso |
| AC090281.1 | 8 | sorting nexin 5 (SNX5) pseudogene |
| AC008066.1 | 8 | lysophosphatidylglycerol acyltransferase 1 (LP |
| PURG | 8 | purine rich element binding protein G [Source |
| WRN | 8 | WRN RecQ like helicase [Source:HGNC Symbo |
| SUMO2P16 | 8 | SUMO2 pseudogene 16 [Source:HGNC Symbo |
| AC009563.1 | 8 | novel transcript |
| KCTD9P6 | 8 | potassium channel tetramerization domain co |
| AC068672.2 | 8 | novel transcript |
| AC068672.3 | 8 | novel transcript |
| AC068672.1 | 8 | novel transcript |
| AC090816.1 | 8 | novel transcript |
| RNA5SP261 | 8 | RNA, 5S ribosomal pseudogene 261 [Source:H |
| NRG1 | 8 | neuregulin 1 [Source:HGNC Symbol;Acc:HGNC |
| AC068931.1 | 8 | novel transcript, antisense to NRG1 |
| NRG1-IT1 | 8 | NRG1 intronic transcript 1 [Source:HGNC Sym |
| RNA5SP262 | 8 | RNA, 5S ribosomal pseudogene 262 [Source:H |
| RNA5SP263 | 8 | RNA, 5S ribosomal pseudogene 263 [Source:H |
| NRG1-IT3 | 8 | NRG1 intronic transcript 3 [Source:HGNC Sym |

| | | |
|---|---|---|
| AC083977.1 | 8 | novel protein |
| RNU6-663P | 8 | RNA, U6 small nuclear 663, pseudogene [Sour |
| AC090204.1 | 8 | novel transcript |
| MTND1P6 | 8 | MT-ND1 pseudogene 6 [Source:HGNC Symbol |
| MTND2P32 | 8 | MT-ND2 pseudogene 32 [Source:HGNC Symbo |
| RANP9 | 8 | RAN pseudogene 9 [Source:HGNC Symbol;Acc |
| AC104027.1 | 8 | endoplasmic reticulum protein 44 (ERP44) pse |
| RNU6-528P | 8 | RNA, U6 small nuclear 528, pseudogene [Sour |
| AC067838.1 | 8 | novel transcript |
| FUT10 | 8 | fucosyltransferase 10 [Source:HGNC Symbol;A |
| AC091144.2 | 8 | RNA binding motif protein 4 (RBM4) pseudoge |
| TTI2 | 8 | TELO2 interacting protein 2 [Source:HGNC Syr |
| MAK16 | 8 | MAK16 homolog [Source:HGNC Symbol;Acc:H |
| SNORD13 | 8 | small nucleolar RNA, C/D box 13 [Source:HGN |
| AC091144.1 | 8 | thymosin beta 4, X-linked (TMSB4X) pseudoge |
| SNORA70 | 8 | Small nucleolar RNA SNORA70 [Source:RFAM; |
| RPL10P18 | 8 | ribosomal protein L10 pseudogene 18 [Source |
| RNF122 | 8 | ring finger protein 122 [Source:HGNC Symbol; |
| RN7SL621P | 8 | RNA, 7SL, cytoplasmic 621, pseudogene [Sour |
| DUSP26 | 8 | dual specificity phosphatase 26 [Source:HGNC |
| AF279873.3 | 8 | novel transcript |
| BUD31P1 | 8 | BUD31 homolog pseudogene 1 [Source:HGNC |
| RN7SL457P | 8 | RNA, 7SL, cytoplasmic 457, pseudogene [Sour |
| VENTXP5 | 8 | VENT homeobox pseudogene 5 [Source:HGNC |
| AF279873.2 | 8 | ribosomal protein L6 (RPL6) pseudogene |
| AF279873.1 | 8 | cytochrome c, somatic (CYCS) pseudogene |
| AF279873.4 | 8 | novel transcript |
| AC087855.2 | 8 | novel transcript |
| AC090993.1 | 8 | novel transcript |
| AC087855.1 | 8 | novel transcript |
| RPL10AP3 | 8 | ribosomal protein L10a pseudogene 3 [Source |
| LINC01288 | 8 | long intergenic non-protein coding RNA 1288 |
| AC087343.1 | 8 | ribosomal protein L21 (RPl21) pseudogene |
| AC099685.1 | 8 | novel transcript |
| UNC5D | 8 | unc-5 netrin receptor D [Source:HGNC Symbo |
| AC090740.1 | 8 | TEC |
| LSM12P1 | 8 | LSM12 pseudogene 1 [Source:HGNC Symbol;A |
| AC105230.1 | 8 | novel transcript, antisense to UNC5D |
| AC124290.1 | 8 | novel transcript |
| AC124290.2 | 8 | novel transcript |
| AC124290.3 | 8 | zinc finger, DHHC-type containing 7 (ZDHHC7) |
| MTCYBP19 | 8 | MT-CYB pseudogene 19 [Source:HGNC Symbo |
| MTND6P19 | 8 | MT-ND6 pseudogene 19 [Source:HGNC Symbo |
| MTND5P41 | 8 | MT-ND5 pseudogene 41 [Source:HGNC Symbo |
| AP006245.1 | 8 | methylenetetrahydrofolate dehydrogenase (N |
| RNU6-533P | 8 | RNA, U6 small nuclear 533, pseudogene [Sour |
| AP006245.2 | 8 | novel pseudogene |
| AC090809.1 | 8 | novel transcript |
| RPL23P10 | 8 | ribosomal protein L23 pseudogene 10 [Source |
| RNA5SP264 | 8 | RNA, 5S ribosomal pseudogene 264 [Source:H |

| | | |
|---|---|---|
| KCNU1 | 8 | potassium calcium-activated channel subfami |
| AC124076.1 | 8 | mitochondrial ribosomal protein S7 (MRPS7) p |
| TPT1P8 | 8 | tumor protein, translationally-controlled 1 ps |
| AC090453.1 | 8 | ribosomal protein L26 (RPL26) pseudogene |
| AC092818.1 | 8 | novel transcript |
| SMARCE1P4 | 8 | SWI/SNF related, matrix associated, actin dep |
| AC091182.1 | 8 | novel transcript |
| AC091182.2 | 8 | novel transcript |
| LINC01605 | 8 | long intergenic non-protein coding RNA 1605 |
| AC124067.1 | 8 | novel transcript |
| AC124067.2 | 8 | novel transcript |
| AC124067.3 | 8 | novel transcript |
| AC124067.4 | 8 | novel transcript |
| AC137579.2 | 8 | novel transcript |
| AC137579.1 | 8 | novel transcript |
| RNU6-607P | 8 | RNA, U6 small nuclear 607, pseudogene [Sour |
| ZNF703 | 8 | zinc finger protein 703 [Source:HGNC Symbol; |
| AC138356.3 | 8 | novel transcript |
| AC138356.1 | 8 | novel transcript, antisense to ERLIN2 |
| ERLIN2 | 8 | ER lipid raft associated 2 [Source:HGNC Symb |
| AC138356.2 | 8 | chromosome X open reading frame 56 (CXorf |
| PLPBP | 8 | pyridoxal phosphate binding protein [Source: |
| ADGRA2 | 8 | adhesion G protein-coupled receptor A2 [Sour |
| BRF2 | 8 | BRF2 RNA polymerase III transcription initiati |
| RAB11FIP1 | 8 | RAB11 family interacting protein 1 [Source:H |
| AC130304.1 | 8 | novel transcript |
| RN7SL709P | 8 | RNA, 7SL, cytoplasmic 709, pseudogene [Sour |
| GOT1L1 | 8 | glutamic-oxaloacetic transaminase 1 like 1 [Sc |
| AC144573.1 | 8 | ADRB3-GOT1L1 readthrough |
| ADRB3 | 8 | adrenoceptor beta 3 [Source:HGNC Symbol;A |
| RPL12P48 | 8 | ribosomal protein L12 pseudogene 48 [Source |
| EIF4EBP1 | 8 | eukaryotic translation initiation factor 4E bind |
| AP006545.1 | 8 | novel transcript, overlapping to EIF4EBP1 |
| AP006545.3 | 8 | novel transcript |
| AP006545.2 | 8 | novel transcript |
| ASH2L | 8 | ASH2 like, histone lysine methyltransferase co |
| RNU6-988P | 8 | RNA, U6 small nuclear 988, pseudogene [Sour |
| AC084024.1 | 8 | novel transcript |
| STAR | 8 | steroidogenic acute regulatory protein [Sourc |
| AC084024.3 | 8 | novel transcript |
| LSM1 | 8 | LSM1 homolog, mRNA degradation associatec |
| RNU6-323P | 8 | RNA, U6 small nuclear 323, pseudogene [Sour |
| BAG4 | 8 | BCL2 associated athanogene 4 [Source:HGNC |
| AC084024.2 | 8 | origin recognition complex, subunit 3-like (yea |
| AC084024.4 | 8 | novel transcript |
| DDHD2 | 8 | DDHD domain containing 2 [Source:HGNC Syn |
| PLPP5 | 8 | phospholipid phosphatase 5 [Source:HGNC Sy |
| NSD3 | 8 | nuclear receptor binding SET domain protein : |
| AC087362.2 | 8 | novel transcript |
| AC087362.1 | 8 | novel transcript, antisense to NSD3 |

| | | |
|---|---|---|
| AC087623.2 | 8 | novel transcript |
| LETM2 | 8 | leucine zipper and EF-hand containing transm |
| FGFR1 | 8 | fibroblast growth factor receptor 1 [Source:H( |
| AC087623.3 | 8 | novel transcript, antisense to LETM2 |
| AC087623.1 | 8 | novel transcript |
| RPS20P22 | 8 | ribosomal protein S20 pseudogene 22 [Source |
| C8orf86 | 8 | chromosome 8 open reading frame 86 [Sourc |
| AC069120.1 | 8 | novel transcript |
| RNF5P1 | 8 | ring finger protein 5 pseudogene 1 [Source:H( |
| AC016813.1 | 8 | novel transcript |
| TACC1 | 8 | transforming acidic coiled-coil containing prot |
| Y_RNA | 8 | Y RNA [Source:RFAM;Acc:RF00019] |
| AC067817.1 | 8 | novel transcript |
| AC067817.2 | 8 | novel transcript |
| PLEKHA2 | 8 | pleckstrin homology domain containing A2 [S |
| AC108863.2 | 8 | novel transcript |
| HTRA4 | 8 | HtrA serine peptidase 4 [Source:HGNC Symbo |
| TM2D2 | 8 | TM2 domain containing 2 [Source:HGNC Symb |
| ADAM9 | 8 | ADAM metallopeptidase domain 9 [Source:HG |
| AC108863.1 | 8 | NULL |
| ADAM32 | 8 | ADAM metallopeptidase domain 32 [Source:H |
| RPL3P10 | 8 | ribosomal protein L3 pseudogene 10 [Source: |
| AC105185.1 | 8 | K(lysine) acetyltransferase 2B (KAT2B) pseudo |
| ADAM5 | 8 | ADAM metallopeptidase domain 5 (pseudoge |
| ADAM3A | 8 | ADAM metallopeptidase domain 3A (pseudog |
| AC123767.1 | 8 | ADAM metallopeptidase domain 2 (ADAM2) p |
| ADAM18 | 8 | ADAM metallopeptidase domain 18 [Source:H |
| ADAM2 | 8 | ADAM metallopeptidase domain 2 [Source:HG |
| AP005902.2 | 8 | ribosomal protein L7a (RPL7A) pseudogene |
| AP005902.1 | 8 | Nipped-B homolog (Drosophila) (NIPBL) pseuc |
| IDO1 | 8 | indoleamine 2,3-dioxygenase 1 [Source:HGNC |
| AC007991.3 | 8 | novel transcript, antisense to IDO1 and IDO2 |
| AC007991.2 | 8 | novel transcript |
| AC007991.4 | 8 | novel transcript |
| IDO2 | 8 | indoleamine 2,3-dioxygenase 2 [Source:HGNC |
| AC007991.1 | 8 | nuclear transcription factor Y, beta (NFYB) pse |
| AC087518.1 | 8 | novel transcript |
| AC022733.1 | 8 | novel transcript |
| TCIM | 8 | transcriptional and immune response regulat( |
| AC022733.2 | 8 | novel transcript |
| SIRLNT | 8 | SIRT1 regulating lncRNA tumor promoter [Sou |
| AC105999.1 | 8 | novel transcript |
| AC010857.1 | 8 | novel transcript |
| ZMAT4 | 8 | zinc finger matrin-type 4 [Source:HGNC Symb |
| AC048387.1 | 8 | novel transcript |
| RNU6-356P | 8 | RNA, U6 small nuclear 356, pseudogene [Sour |
| SFRP1 | 8 | secreted frizzled related protein 1 [Source:HG |
| MIR548AO | 8 | microRNA 548ao [Source:HGNC Symbol;Acc:H |
| RPS29P2 | 8 | ribosomal protein S29 pseudogene 2 [Source: |
| AC104393.1 | 8 | novel transcript |

| | | |
|---|---|---|
| RNU6-895P | 8 | RNA, U6 small nuclear 895, pseudogene [Sour |
| SNORD65B | 8 | small nucleolar RNA, C/D box 65B [Source:HG |
| AC016868.1 | 8 | uncharacterized LOC105379391 [Source:NCBI |
| GOLGA7 | 8 | golgin A7 [Source:HGNC Symbol;Acc:HGNC:24 |
| AC009630.1 | 8 | novel transcript, antisense to AGPAT6 and GIN |
| KRT18P37 | 8 | keratin 18 pseudogene 37 [Source:HGNC Sym |
| GINS4 | 8 | GINS complex subunit 4 [Source:HGNC Symbo |
| AC009630.2 | 8 | novel transcript, antisense to GINS4 |
| AC009630.4 | 8 | novel transcript |
| GPAT4 | 8 | glycerol-3-phosphate acyltransferase 4 [Sourc |
| AC009630.3 | 8 | novel transcript, antisense to AGPAT6 |
| NKX6-3 | 8 | NK6 homeobox 3 [Source:HGNC Symbol;Acc:H |
| ANK1 | 8 | ankyrin 1 [Source:HGNC Symbol;Acc:HGNC:49 |
| MIR486-1 | 8 | microRNA 486-1 [Source:HGNC Symbol;Acc:H |
| MIR486-2 | 8 | microRNA 486-2 [Source:HGNC Symbol;Acc:H |
| AC113133.1 | 8 | novel transcript, antisense to ANK1 |
| Y_RNA | 8 | Y RNA [Source:RFAM;Acc:RF00019] |
| AC027702.1 | 8 | novel transcript |
| RN7SL149P | 8 | RNA, 7SL, cytoplasmic 149, pseudogene [Sour |
| KAT6A | 8 | lysine acetyltransferase 6A [Source:HGNC Sym |
| AC090571.1 | 8 | NULL |
| AC103724.1 | 8 | ribosomal protein L17 (RPL17) pseudogene |
| AC103724.2 | 8 | myosin, light chain 9, regulatory (MYL9) pseuc |
| AC103724.4 | 8 | novel transcript |
| AC103724.3 | 8 | novel transcript, antisense to AP3M2 |
| AP3M2 | 8 | adaptor related protein complex 3 subunit mu |
| PLAT | 8 | plasminogen activator, tissue type [Source:HG |
| AC083973.1 | 8 | novel transcript |
| IKBKB | 8 | inhibitor of nuclear factor kappa B kinase subu |
| POLB | 8 | DNA polymerase beta [Source:HGNC Symbol;A |
| RPL5P23 | 8 | ribosomal protein L5 pseudogene 23 [Source: |
| DKK4 | 8 | dickkopf WNT signaling pathway inhibitor 4 [S |
| VDAC3 | 8 | voltage dependent anion channel 3 [Source:H |
| SLC20A2 | 8 | solute carrier family 20 member 2 [Source:HG |
| AC093367.1 | 8 | glucosidase, beta (bile acid) 2 (GBA2) pseudog |
| AC090739.1 | 8 | novel transcript |
| SMIM19 | 8 | small integral membrane protein 19 [Source:H |
| CHRNB3 | 8 | cholinergic receptor nicotinic beta 3 subunit [S |
| AC103843.1 | 8 | novel transcript |
| AC103843.2 | 8 | ribosomal protein L7-like 1 (RPL7L1) pseudoge |
| CHRNA6 | 8 | cholinergic receptor nicotinic alpha 6 subunit |
| THAP1 | 8 | THAP domain containing 1 [Source:HGNC Sym |
| AC087533.1 | 8 | novel transcript, antisense to THAP1 |
| RNF170 | 8 | ring finger protein 170 [Source:HGNC Symbol; |
| RN7SL806P | 8 | RNA, 7SL, cytoplasmic 806, pseudogene [Sour |
| MIR4469 | 8 | microRNA 4469 [Source:HGNC Symbol;Acc:HG |
| HOOK3 | 8 | hook microtubule tethering protein 3 [Source |
| Y_RNA | 8 | Y RNA [Source:RFAM;Acc:RF00019] |
| AC110275.1 | 8 | novel protein |
| FNTA | 8 | farnesyltransferase, CAAX box, alpha [Source: |

| | | |
|---|---|---|
| RNU1-124P | 8 | RNA, U1 small nuclear 124, pseudogene [Sour |
| POMK | 8 | protein O-mannose kinase [Source:HGNC Sym |
| AC113191.1 | 8 | Family with sequence similarity 9, member C ( |
| HGSNAT | 8 | heparan-alpha-glucosaminide N-acetyltransfe |
| VN1R46P | 8 | vomeronasal 1 receptor 46 pseudogene [Sour |
| AC022616.5 | 8 | chromosome 9 open reading frame 122 (C9or |
| AC022616.7 | 8 | Ankyrin repeat domain-containing protein (AN |
| AC022616.3 | 8 | ankyrin repeat domain (ANKRD) pseudogene |
| AC022616.2 | 8 | coiled-coil domain containing 144 (CCDC144) |
| AFG3L2P1 | 8 | AFG3 like AAA ATPase 2 pseudogene 1 [Sourc |
| AC022616.1 | 8 | spermatogenesis associated 5-like 1 (SPATA5L |
| AC022616.4 | 8 | S-phase response (cyclin related) (SPHAR) pse |
| POTEA | 8 | POTE ankyrin domain family member A [Sourc |
| RNU6-104P | 8 | RNA, U6 small nuclear 104, pseudogene [Sour |
| AC022616.8 | 8 | coiled-coil domain containing 144C (CCDC144 |
| AC022616.6 | 8 | POTE ankyrin domain family, member A (POTI |
| U3 | 8 | Small nucleolar RNA U3 [Source:RFAM;Acc:RF |
| AC134698.1 | 8 | Rho GTPase activating protein 20 (ARHGAP20 |
| AC134698.4 | 8 | ankyrin repeat domain 26 (ANKRD26) pseudo |
| AC134698.3 | 8 | novel pseudogene |
| SNX18P27 | 8 | sorting nexin 18 pseudogene 27 [Source:HGN |
| AC134698.5 | 8 | chromosome 2 open reading frame 14 (C2orf |
| CYP4F44P | 8 | cytochrome P450 family 4 subfamily F membe |
| AC134698.2 | 8 | cytochrome P450, family 4, subfamily F, polyp |
| AC139365.2 | 8 | dynein, axonemal, heavy chain 11 (DNAH11) p |
| AC139365.1 | 8 | Cell division cycle-associated 7-like protein (C |
| AC118650.1 | 8 | NULL |
| HSPA8P13 | 8 | heat shock protein family A (Hsp70) member |
| ASNSP1 | 8 | asparagine synthetase pseudogene 1 [Source: |
| AC104576.1 | 8 | heat shock 70kDa protein (HSPA) pseudogene |
| ASNSP4 | 8 | asparagine synthetase pseudogene 4 [Source: |
| AC021451.1 | 8 | similar to zinc finger protein 682 |
| TRIM60P15 | 8 | tripartite motif containing 60 pseudogene 15 |
| AC021451.2 | 8 | novel transcript |
| AC021451.3 | 8 | novel transcript |
| LINC00293 | 8 | long intergenic non-protein coding RNA 293 [S |
| MTND2P38 | 8 | MT-ND2 pseudogene 38 [Source:HGNC Symbo |
| MTND1P7 | 8 | MT-ND1 pseudogene 7 [Source:HGNC Symbol |
| RNU6-656P | 8 | RNA, U6 small nuclear 656, pseudogene [Sour |
| MTCYBP20 | 8 | MT-CYB pseudogene 20 [Source:HGNC Symbo |
| MTND6P20 | 8 | MT-ND6 pseudogene 20 [Source:HGNC Symbo |
| AC091163.3 | 8 | similar to zinc finger protein 682 (ZNF682) pse |
| AC091163.1 | 8 | novel transcript |
| AC091163.2 | 8 | uncharacterized LOC105375814 [Source:NCBI |
| MAPK6P4 | 8 | mitogen-activated protein kinase 6 pseudoge |
| RN7SKP32 | 8 | RN7SK pseudogene 32 [Source:HGNC Symbol; |
| AC120036.1 | 8 | POTE ankyrin domain family pseudogene |
| AC120036.5 | 8 | novel transcript |
| RNU6-819P | 8 | RNA, U6 small nuclear 819, pseudogene [Sour |
| RPL10AP2 | 8 | ribosomal protein L10a pseudogene 2 [Source |

| | | |
|---|---|---|
| NDUFA5P12 | 8 | NADH:ubiquinone oxidoreductase subunit A5 |
| AC120036.3 | 8 | patched homolog 1 (Drosophila) (PTCH1) pseu |
| AC120036.4 | 8 | novel transcript |
| ATP6V1G1P2 | 8 | ATPase H+ transporting V1 subunit G1 pseudo |
| AC120036.2 | 8 | pre-B lymphocyte 1 (VPREB1) pseudogene |
| IGLV8OR8-1 | 8 | immunoglobulin lambda variable 8/OR8-1 (ps |
| SPIDR | 8 | scaffold protein involved in DNA repair [Sourc |
| AC024451.2 | 8 | NSFL1 (p97) cofactor (p47) (NSFL1C) pseudog |
| AC024451.4 | 8 | novel transcript, sense intronic to KIAA0146 |
| AC024451.1 | 8 | ribosomal protein S4, X-linked (RPS4X) pseudo |
| AC024451.3 | 8 | ubiquitin-conjugating enzyme E2N (UBC13 ho |
| RNU6-665P | 8 | RNA, U6 small nuclear 665, pseudogene [Sour |
| CEBPD | 8 | CCAAT enhancer binding protein delta [Source |
| PRKDC | 8 | protein kinase, DNA-activated, catalytic subur |
| Y_RNA | 8 | Y RNA [Source:RFAM;Acc:RF00019] |
| AC021236.1 | 8 | novel transcript, antisense to PRKDC |
| MCM4 | 8 | minichromosome maintenance complex comp |
| RNU6-519P | 8 | RNA, U6 small nuclear 519, pseudogene [Sour |
| UBE2V2 | 8 | ubiquitin conjugating enzyme E2 V2 [Source:H |
| AC104989.1 | 8 | cell division cycle associated 8 (CDCA8) pseud |
| AC105029.1 | 8 | isopentenyl-diphosphate delta isomerase 1 (ID |
| RNU6-295P | 8 | RNA, U6 small nuclear 295, pseudogene [Sour |
| RNA5SP531 | 8 | RNA, 5S ribosomal pseudogene 531 [Source:H |
| RPL29P19 | 8 | ribosomal protein L29 pseudogene 19 [Source |
| AC041040.1 | 8 | TEC |
| AC026904.1 | 8 | novel transcript |
| AC026904.2 | 8 | novel transcript |
| AC022915.2 | 8 | novel transcript |
| LINC02599 | 8 | long intergenic non-protein coding RNA 2599 |
| LINC02847 | 8 | long intergenic non-protein coding RNA 2847 |
| AC022915.1 | 8 | novel transcript |
| AC022915.3 | 8 | novel transcript |
| EFCAB1 | 8 | EF-hand calcium binding domain 1 [Source:HG |
| AC100805.1 | 8 | novel transcript |
| AC013701.1 | 8 | erythrocyte membrane protein band 4.1 like 5 |
| SNAI2 | 8 | snail family transcriptional repressor 2 [Source |
| AC013701.2 | 8 | novel transcript |
| AC044893.2 | 8 | novel transcript |
| PPDPFL | 8 | pancreatic progenitor cell differentiation and |
| AC044893.1 | 8 | novel transcript |
| RN7SKP294 | 8 | RN7SK pseudogene 294 [Source:HGNC Symbo |
| RFPL4AP7 | 8 | ret finger protein like 4A pseudogene 7 [Sourc |
| AC090155.2 | 8 | novel transcript |
| AC090155.1 | 8 | novel transcript |
| PSAT1P1 | 8 | phosphoserine aminotransferase 1 pseudoger |
| AC113145.1 | 8 | novel transcript |
| AC023762.1 | 8 | novel transcript |
| SNTG1 | 8 | syntrophin gamma 1 [Source:HGNC Symbol;A |
| AC090539.1 | 8 | novel transcript |
| AC090539.2 | 8 | coagulation factor III (thromboplastin, tissue f |

| | | |
|---|---|---|
| CYCSP22 | 8 | CYCS pseudogene 22 [Source:HGNC Symbol;A |
| AC087272.1 | 8 | programmed cell death 10 (PDCD10) pseudog |
| AC090919.1 | 8 | NULL |
| AC012413.1 | 8 | novel transcript |
| PXDNL | 8 | peroxidasin like [Source:HGNC Symbol;Acc:HG |
| BRIX1P1 | 8 | BRIX1 pseudogene 1 [Source:HGNC Symbol;Ac |
| BTF3P1 | 8 | BTF3 pseudogene 1 [Source:HGNC Symbol;Acc |
| AC090186.1 | 8 | novel transcript |
| PCMTD1 | 8 | protein-L-isoaspartate (D-aspartate) O-methyl |
| AC103769.1 | 8 | novel transcript |
| AC064807.1 | 8 | novel transcript |
| AC064807.4 | 8 | novel transcript |
| AC064807.2 | 8 | novel transcript |
| AC064807.3 | 8 | novel transcript |
| ST18 | 8 | ST18 C2H2C-type zinc finger transcription fact |
| AC021915.2 | 8 | novel transcript |
| AC021915.1 | 8 | novel transcript |
| AC103831.1 | 8 | novel transcript |
| RPL34P17 | 8 | ribosomal protein L34 pseudogene 17 [Source |
| ALKAL1 | 8 | ALK and LTK ligand 1 [Source:HGNC Symbol;Ac |
| RB1CC1 | 8 | RB1 inducible coiled-coil 1 [Source:HGNC Sym |
| AC113139.1 | 8 | novel transcript, sense intronic KANK1 |
| NPBWR1 | 8 | neuropeptides B and W receptor 1 [Source:HG |
| AC009800.1 | 8 | ring finger protein 2 (RNF2) pseudogene |
| AC009646.1 | 8 | histone acetyltransferase 1 (HAT1) pseudogen |
| AC009646.2 | 8 | novel transcript |
| OPRK1 | 8 | opioid receptor kappa 1 [Source:HGNC Symbo |
| AC131902.1 | 8 | novel transcript |
| AC022034.3 | 8 | novel transcript |
| AC131902.2 | 8 | polymerase (DNA directed), epsilon 2, accesso |
| AC022034.1 | 8 | novel transcript |
| AC022034.4 | 8 | novel transcript |
| AC022034.2 | 8 | ribosomal protein L21 (RPL21) pseudogene |
| MAPK6P1 | 8 | mitogen-activated protein kinase 6 pseudogen |
| AC103778.1 | 8 | novel pseudogene |
| ATP6V1H | 8 | ATPase H+ transporting V1 subunit H [Source: |
| RGS20 | 8 | regulator of G protein signaling 20 [Source:HG |
| AC113194.1 | 8 | novel transcript, antisense to RGS20 |
| RPS27AP13 | 8 | ribosomal protein S27a pseudogene 13 [Sourc |
| RNU6-1331P | 8 | RNA, U6 small nuclear 1331, pseudogene [Sou |
| TCEA1 | 8 | transcription elongation factor A1 [Source:HG |
| AC100821.1 | 8 | ESF1, nucleolar pre-rRNA processing protein, l |
| AC100821.2 | 8 | novel transcript, overlapping to LYPLA1 |
| LYPLA1 | 8 | lysophospholipase 1 [Source:HGNC Symbol;Ac |
| TDGF1P5 | 8 | teratocarcinoma-derived growth factor 1 pseu |
| Metazoa_SRP | 8 | Metazoan signal recognition particle RNA [Sou |
| MRPL15 | 8 | mitochondrial ribosomal protein L15 [Source:I |
| RNU6ATAC32P | 8 | RNA, U6atac small nuclear 32, pseudogene [So |
| AC060764.1 | 8 | coiled-coil-helix-coiled-coil-helix domain conta |
| AC044836.1 | 8 | dolichyl-phosphate mannosyltransferase poly |

| | | |
|---|---|---|
| RNU105C | 8 | RNA, U105C small nucleolar [Source:HGNC Sy |
| AC027250.1 | 8 | complement component 7 (C7) pseudogene |
| AC027250.2 | 8 | novel transcript |
| RN7SL250P | 8 | RNA, 7SL, cytoplasmic 250, pseudogene [Sour |
| SOX17 | 8 | SRY-box transcription factor 17 [Source:HGNC |
| AC091076.1 | 8 | SRY (sex determining region Y)-box 17 (SOX17 |
| TRMT112P7 | 8 | tRNA methyltransferase subunit 11-2 pseudog |
| RP1 | 8 | RP1 axonemal microtubule associated [Source |
| SEC11B | 8 | SEC11 homolog B, signal peptidase complex su |
| AC090151.1 | 8 | ribosomal protein L7-like 1 (RPL7L1) pseudoge |
| AC084834.1 | 8 | novel transcript |
| XKR4 | 8 | XK related 4 [Source:HGNC Symbol;Acc:HGNC |
| AC022679.2 | 8 | novel transcript |
| AC022679.1 | 8 | novel transcript |
| SBF1P1 | 8 | SET binding factor 1 pseudogene 1 [Source:HG |
| AC090200.1 | 8 | novel transcript |
| TMEM68 | 8 | transmembrane protein 68 [Source:HGNC Syn |
| RNA5SP265 | 8 | RNA, 5S ribosomal pseudogene 265 [Source:H |
| TGS1 | 8 | trimethylguanosine synthase 1 [Source:HGNC |
| LYN | 8 | LYN proto-oncogene, Src family tyrosine kinas |
| AC018607.1 | 8 | novel transcript |
| SNORA1B | 8 | small nucleolar RNA, H/ACA box 1B [Source:H |
| RN7SL798P | 8 | RNA, 7SL, cytoplasmic 798, pseudogene [Sour |
| RN7SL323P | 8 | RNA, 7SL, cytoplasmic 323, pseudogene [Sour |
| AC046176.1 | 8 | proteasome (prosome, macropain) 26S subun |
| RPS20 | 8 | ribosomal protein S20 [Source:HGNC Symbol; |
| SNORD54 | 8 | small nucleolar RNA, C/D box 54 [Source:HGN |
| CERNA3 | 8 | competing endogenous lncRNA 3 for miR-645 |
| NPM1P21 | 8 | nucleophosmin 1 pseudogene 21 [Source:HGN |
| MOS | 8 | MOS proto-oncogene, serine/threonine kinas |
| AC107376.1 | 8 | NULL |
| PLAG1 | 8 | PLAG1 zinc finger [Source:HGNC Symbol;Acc:H |
| CHCHD7 | 8 | coiled-coil-helix-coiled-coil-helix domain cont |
| AC107952.2 | 8 | novel transcript |
| AC107952.1 | 8 | SPRYD4 SPRY domain containing 4 (SPRYD4) p |
| SDR16C5 | 8 | short chain dehydrogenase/reductase family 1 |
| SDR16C6P | 8 | short chain dehydrogenase/reductase family 1 |
| PENK | 8 | proenkephalin [Source:HGNC Symbol;Acc:HGN |
| AC012349.1 | 8 | novel transcript |
| SEPTIN10P1 | 8 | septin 10 pseudogene 1 [Source:HGNC Symbo |
| LINC00968 | 8 | long intergenic non-protein coding RNA 968 [S |
| AC013644.1 | 8 | novel transcript |
| RPL37P6 | 8 | ribosomal protein L37 pseudogene 6 [Source: |
| AC009597.1 | 8 | novel transcript |
| AC009597.2 | 8 | impact RWD domain protein (IMPACT) pseudo |
| RNU6-13P | 8 | RNA, U6 small nuclear 13, pseudogene [Sourc |
| IMPAD1 | 8 | inositol monophosphatase domain containing |
| RNA5SP266 | 8 | RNA, 5S ribosomal pseudogene 266 [Source:H |
| LINC01606 | 8 | long intergenic non-protein coding RNA 1606 |
| AC025674.1 | 8 | ankyrin repeat domain 26 (ANKRD26) pseudog |

| | | |
|---|---|---|
| AC025674.2 | 8 | novel transcript |
| LINC00588 | 8 | long intergenic non-protein coding RNA 588 [S |
| RNU6-596P | 8 | RNA, U6 small nuclear 596, pseudogene [Sour |
| AC068075.1 | 8 | novel transcript |
| RPL30P10 | 8 | ribosomal protein L30 pseudogene 10 [Source |
| AC068075.2 | 8 | novel transcript |
| AC090796.1 | 8 | novel transcript |
| AC104051.1 | 8 | novel transcript |
| AC104051.2 | 8 | novel transcript |
| LINC01602 | 8 | long intergenic non-protein coding RNA 1602 |
| AC012103.1 | 8 | novel transcript |
| FAM110B | 8 | family with sequence similarity 110 member B |
| AC104350.1 | 8 | novel transcript |
| AC027698.1 | 8 | novel transcript |
| RPL26P26 | 8 | ribosomal protein L26 pseudogene 26 [Source |
| AC092819.3 | 8 | novel transcript |
| AC092819.1 | 8 | novel transcript |
| AC092819.2 | 8 | novel transcript |
| UBXN2B | 8 | UBX domain protein 2B [Source:HGNC Symbo |
| AC009927.1 | 8 | protein tyrosine phosphatase, non-receptor ty |
| CYP7A1 | 8 | cytochrome P450 family 7 subfamily A membe |
| PPIAP85 | 8 | peptidylprolyl isomerase A pseudogene 85 [Sc |
| SDCBP | 8 | syndecan binding protein [Source:HGNC Symb |
| NSMAF | 8 | neutral sphingomyelinase activation associate |
| AC068522.1 | 8 | ribosomal protein S26 (RPS26) pseudogene |
| TOX | 8 | thymocyte selection associated high mobility |
| RNU4-50P | 8 | RNA, U4 small nuclear 50, pseudogene [Sourc |
| AC105150.1 | 8 | novel transcript |
| AC090152.1 | 8 | novel transcript |
| AC087698.1 | 8 | NULL |
| RNA5SP267 | 8 | RNA, 5S ribosomal pseudogene 267 [Source:H |
| AC087664.3 | 8 | nudix (nucleoside diphosphate linked moiety X |
| AC087664.1 | 8 | novel transcript |
| AC087664.2 | 8 | novel transcript |
| AC107934.1 | 8 | novel pseudogene |
| AC107934.2 | 8 | sirtuin (silent mating type information regulat |
| AC022709.1 | 8 | betaine--homocysteine S-methyltransferase 2 |
| AC021393.1 | 8 | uncharacterized LOC105375861 [Source:NCBI |
| SLC2A13P1 | 8 | SLC2A13 pseudogene 1 [Source:HGNC Symbol |
| CA8 | 8 | carbonic anhydrase 8 [Source:HGNC Symbol;A |
| LINC01301 | 8 | long intergenic non-protein coding RNA 1301 |
| PDCL3P1 | 8 | PDCL3 pseudogene 1 [Source:HGNC Symbol;A |
| RAB2A | 8 | RAB2A, member RAS oncogene family [Source |
| AC068389.3 | 8 | novel transcript |
| AC068389.1 | 8 | novel transcript |
| AC068389.2 | 8 | novel transcript |
| CHD7 | 8 | chromodomain helicase DNA binding protein |
| AC113143.1 | 8 | novel transcript |
| AC113143.3 | 8 | novel transcript |
| AC113143.2 | 8 | interferon induced transmembrane protein 3 |

| | | |
|---|---|---|
| AC022182.1 | 8 | novel transcript |
| AC022182.3 | 8 | nuclear autoantigenic sperm protein (histone- |
| AC022182.2 | 8 | novel transcript |
| CLVS1 | 8 | clavesin 1 [Source:HGNC Symbol;Acc:HGNC:2 |
| NPM1P6 | 8 | nucleophosmin 1 pseudogene 6 [Source:HGN |
| AC023866.2 | 8 | novel transcript |
| AC023866.1 | 8 | novel transcript |
| ASPH | 8 | aspartate beta-hydroxylase [Source:HGNC Syr |
| KRT8P3 | 8 | keratin 8 pseudogene 3 [Source:HGNC Symbo |
| RN7SKP97 | 8 | RN7SK pseudogene 97 [Source:HGNC Symbol; |
| MIR4470 | 8 | microRNA 4470 [Source:HGNC Symbol;Acc:HG |
| AC091173.1 | 8 | novel transcript |
| LINC02842 | 8 | long intergenic non-protein coding RNA 2842 |
| AC025524.1 | 8 | novel transcript |
| LINC02155 | 8 | long intergenic non-protein coding RNA 2155 |
| C1GALT1P3 | 8 | C1GALT1 pseudogene 3 [Source:HGNC Symbo |
| NARSP2 | 8 | asparaginyl-tRNA synthetase pseudogene 2 [S |
| AC104852.1 | 8 | polo-like kinase 1 (Drosophila) (PLK1) pseudog |
| AC023095.1 | 8 | novel transcript |
| NKAIN3 | 8 | sodium/potassium transporting ATPase intera |
| AC018861.2 | 8 | novel transcript |
| AC018861.1 | 8 | ubiquitin-conjugating enzyme E2L 3 (UBE2L3) |
| AC090577.1 | 8 | novel transcript, antisense to NKAIN3 |
| XRCC6P4 | 8 | X-ray repair cross complementing 6 pseudoge |
| SRPK2P | 8 | SRSF protein kinase 2 pseudogene [Source:HG |
| NKAIN3-IT1 | 8 | NKAIN3 intronic transcript [Source:HGNC Sym |
| AC120042.1 | 8 | novel transcript |
| GGH | 8 | gamma-glutamyl hydrolase [Source:HGNC Syr |
| AC120042.2 | 8 | novel transcript |
| TTPA | 8 | alpha tocopherol transfer protein [Source:HG |
| AC011978.1 | 8 | TAR DNA binding protein (TARDBP) pseudoge |
| YTHDF3-AS1 | 8 | YTHDF3 antisense RNA 1 (head to head) [Sour |
| YTHDF3 | 8 | YTH N6-methyladenosine RNA binding protei |
| AC011978.2 | 8 | novel transcript |
| RN7SL135P | 8 | RNA, 7SL, cytoplasmic 135, pseudogene [Sour |
| AC011124.1 | 8 | novel transcript |
| IFITM8P | 8 | interferon induced transmembrane protein 8 |
| AC011124.2 | 8 | novel transcript |
| AC018953.1 | 8 | novel transcript |
| RN7SKP135 | 8 | RN7SK pseudogene 135 [Source:HGNC Symbo |
| AC018953.2 | 8 | novel transcript |
| LINC01414 | 8 | long intergenic non-protein coding RNA 1414 |
| AC069133.1 | 8 | novel transcript |
| LINC01289 | 8 | long intergenic non-protein coding RNA 1289 |
| COX6CP8 | 8 | cytochrome c oxidase subunit 6C pseudogene |
| MIR124-2HG | 8 | MIR124-2 host gene [Source:HGNC Symbol;Ac |
| MIR124-2 | 8 | microRNA 124-2 [Source:HGNC Symbol;Acc:H |
| AC012535.1 | 8 | novel transcript |
| AC090136.1 | 8 | NADH dehydrogenase (ubiquinone) 1 alpha su |
| AC090136.2 | 8 | 60S ribosomal protein L23 (RPL23) pseudoger |

| | | |
|---|---|---|
| AC090136.3 | 8 | novel transcript |
| BHLHE22 | 8 | basic helix-loop-helix family member e22 [Sou |
| CYP7B1 | 8 | cytochrome P450 family 7 subfamily B membe |
| AC104232.2 | 8 | novel transcript |
| AC104232.3 | 8 | novel transcript |
| AC104232.1 | 8 | novel transcript |
| AC087808.1 | 8 | interferon induced transmembrane protein 3 |
| RPL31P41 | 8 | ribosomal protein L31 pseudogene 41 [Source |
| LINC00251 | 8 | long intergenic non-protein coding RNA 251 [S |
| PPIAP86 | 8 | peptidylprolyl isomerase A pseudogene 86 [So |
| LINC01299 | 8 | long intergenic non-protein coding RNA 1299 |
| AC100814.1 | 8 | novel transcript |
| Y_RNA | 8 | Y RNA [Source:RFAM;Acc:RF00019] |
| ARMC1 | 8 | armadillo repeat containing 1 [Source:HGNC S |
| MTFR1 | 8 | mitochondrial fission regulator 1 [Source:HGN |
| AC055822.1 | 8 | novel transcript, antisense to MTFR1 |
| PDE7A | 8 | phosphodiesterase 7A [Source:HGNC Symbol; |
| AC100812.1 | 8 | novel transcript |
| DNAJC5B | 8 | DnaJ heat shock protein family (Hsp40) memb |
| AC084082.1 | 8 | novel transcript |
| TRIM55 | 8 | tripartite motif containing 55 [Source:HGNC S |
| CRH | 8 | corticotropin releasing hormone [Source:HGN |
| LINC00967 | 8 | long intergenic non-protein coding RNA 967 [S |
| AC009879.4 | 8 | novel transcript, antisense to RRS1 |
| RRS1-AS1 | 8 | RRS1 antisense RNA 1 (head to head) [Source: |
| RRS1 | 8 | ribosome biogenesis regulator 1 homolog [Sou |
| AC009879.2 | 8 | novel protein |
| AC009879.3 | 8 | novel transcript |
| ADHFE1 | 8 | alcohol dehydrogenase iron containing 1 [Sou |
| AC009879.1 | 8 | TAF9 RNA polymerase II, TATA box binding pro |
| VXN | 8 | vexin [Source:HGNC Symbol;Acc:HGNC:28498 |
| RNU6-1324P | 8 | RNA, U6 small nuclear 1324, pseudogene [Sou |
| MYBL1 | 8 | MYB proto-oncogene like 1 [Source:HGNC Syr |
| VCPIP1 | 8 | valosin containing protein interacting protein |
| Y_RNA | 8 | Y RNA [Source:RFAM;Acc:RF00019] |
| SGK3 | 8 | serum/glucocorticoid regulated kinase family |
| PTTG3P | 8 | pituitary tumor-transforming 3, pseudogene [ |
| MCMDC2 | 8 | minichromosome maintenance domain conta |
| SNHG6 | 8 | small nucleolar RNA host gene 6 [Source:HGN |
| SNORD87 | 8 | small nucleolar RNA, C/D box 87 [Source:HGN |
| TCF24 | 8 | transcription factor 24 [Source:HGNC Symbol; |
| PPP1R42 | 8 | protein phosphatase 1 regulatory subunit 42 [ |
| AC110998.1 | 8 | PRP3 pre-mRNA processing factor 3 homolog |
| COPS5 | 8 | COP9 signalosome subunit 5 [Source:HGNC Sy |
| CSPP1 | 8 | centrosome and spindle pole associated prote |
| RNA5SP268 | 8 | RNA, 5S ribosomal pseudogene 268 [Source:H |
| AC087359.1 | 8 | tigger transposable element derived 1 (TIGD1 |
| ARFGEF1 | 8 | ADP ribosylation factor guanine nucleotide ex |
| AC021321.2 | 8 | novel transcript, sense intronic to ARFGEF1 |
| AC021321.1 | 8 | novel transcript |

| | | |
|---|---|---|
| AC011037.1 | 8 | ribosomal protein L17 (RPL17) pseudogene |
| CPA6 | 8 | carboxypeptidase A6 [Source:HGNC Symbol;A |
| NACAP10 | 8 | NACA pseudogene 10 [Source:HGNC Symbol;/ |
| AC022874.1 | 8 | novel transcript |
| NDUFS5P6 | 8 | NADH:ubiquinone oxidoreductase subunit S5 |
| PREX2 | 8 | phosphatidylinositol-3,4,5-trisphosphate depe |
| AC011853.2 | 8 | novel transcript |
| AC011853.1 | 8 | NADH dehydrogenase [ubiquinone] flavoprote |
| Y_RNA | 8 | Y RNA [Source:RFAM;Acc:RF00019] |
| RPL31P40 | 8 | ribosomal protein L31 pseudogene 40 [Source |
| C8orf34-AS1 | 8 | C8orf34 antisense RNA 1 [Source:HGNC Symb |
| C8orf34 | 8 | chromosome 8 open reading frame 34 [Sourc |
| RPS15AP25 | 8 | ribosomal protein S15a pseudogene 25 [Sourc |
| RNA5SP269 | 8 | RNA, 5S ribosomal pseudogene 269 [Source:H |
| AC083967.1 | 8 | novel transcript |
| LINC01592 | 8 | long intergenic non-protein coding RNA 1592 |
| RNU7-102P | 8 | RNA, U7 small nuclear 102 pseudogene [Sourc |
| AF201337.1 | 8 | tropomyosin 4 (TPM4) pseudogene |
| AC021785.1 | 8 | novel transcript |
| LINC01603 | 8 | long intergenic non-protein coding RNA 1603 |
| AC022790.1 | 8 | guanine nucleotide binding protein (G protein |
| SULF1 | 8 | sulfatase 1 [Source:HGNC Symbol;Acc:HGNC:2 |
| SLCO5A1 | 8 | solute carrier organic anion transporter family |
| RN7SKP29 | 8 | RN7SK pseudogene 29 [Source:HGNC Symbol; |
| AC091047.1 | 8 | novel transcript |
| AC079089.3 | 8 | thioredoxin-related transmembrane protein 1 |
| AC079089.1 | 8 | novel transcript, antisense to SLCO5A1 |
| Metazoa_SRP | 8 | Metazoan signal recognition particle RNA [Sou |
| AC079089.2 | 8 | BAG family molecular chaperone regulator 4 ( |
| RN7SL675P | 8 | RNA, 7SL, cytoplasmic 675, pseudogene [Sour |
| RNA5SP270 | 8 | RNA, 5S ribosomal pseudogene 270 [Source:H |
| AC090574.1 | 8 | TEC |
| SDCBPP2 | 8 | syndecan binding protein pseudogene 2 [Sour |
| SUMO2P20 | 8 | SUMO2 pseudogene 20 [Source:HGNC Symbo |
| PRDM14 | 8 | PR/SET domain 14 [Source:HGNC Symbol;Acc: |
| RNU1-101P | 8 | RNA, U1 small nuclear 101, pseudogene [Sour |
| H2AFZP2 | 8 | H2A histone family member Z pseudogene 2 [ |
| NCOA2 | 8 | nuclear receptor coactivator 2 [Source:HGNC |
| BTF3P12 | 8 | basic transcription factor 3 pseudogene 12 [Sc |
| RNY3P14 | 8 | RNY3 pseudogene 14 [Source:HGNC Symbol;A |
| AC022730.2 | 8 | ribosomal protein S18 (RPS18) pseudogene |
| AC022730.3 | 8 | novel transcript |
| AC022730.4 | 8 | novel transcript |
| RN7SL203P | 8 | RNA, 7SL, cytoplasmic 203, pseudogene [Sour |
| AC022730.1 | 8 | ribosomal protein L13 (RPL13) pseudogene |
| AC120194.1 | 8 | novel transcript |
| TRAM1 | 8 | translocation associated membrane protein 1 |
| LACTB2-AS1 | 8 | LACTB2 antisense RNA 1 [Source:HGNC Symbo |
| LACTB2 | 8 | lactamase beta 2 [Source:HGNC Symbol;Acc:H |
| RN7SL19P | 8 | RNA, 7SL, cytoplasmic 19, pseudogene [Sourc |

| | | |
|---|---|---|
| XKR9 | 8 | XK related 9 [Source:HGNC Symbol;Acc:HGNC |
| AC015687.1 | 8 | novel transcript |
| Y_RNA | 8 | Y RNA [Source:RFAM;Acc:RF00019] |
| AC022858.1 | 8 | novel transcript |
| EYA1 | 8 | EYA transcriptional coactivator and phosphata |
| TRAPPC2P2 | 8 | trafficking protein particle complex 2 pseudog |
| AC009446.1 | 8 | novel transcript |
| AC104012.1 | 8 | novel transcript |
| U8 | 8 | U8 small nucleolar RNA [Source:RFAM;Acc:RF( |
| MSC-AS1 | 8 | MSC antisense RNA 1 [Source:HGNC Symbol;A |
| MSC | 8 | musculin [Source:HGNC Symbol;Acc:HGNC:73 |
| RPS20P20 | 8 | ribosomal protein S20 pseudogene 20 [Source |
| TRPA1 | 8 | transient receptor potential cation channel su |
| AC078906.1 | 8 | novel transcript |
| AC022905.1 | 8 | transient receptor potential cation channel, su |
| RNA5SP271 | 8 | RNA, 5S ribosomal pseudogene 271 [Source:H |
| AC124293.1 | 8 | nucleolar pre-rRNA processing protein, homol |
| AC011131.1 | 8 | novel transcript |
| KCNB2 | 8 | potassium voltage-gated channel subfamily B |
| HAUS1P3 | 8 | HAUS augmin like complex subunit 1 pseudog |
| AC013562.2 | 8 | CTAGE family, member 5 (CTAGE5) pseudoger |
| AC013562.1 | 8 | novel transcript |
| AC090735.1 | 8 | novel transcript |
| AC022893.2 | 8 | novel transcript |
| TERF1 | 8 | telomeric repeat binding factor 1 [Source:HGI |
| RNU6-285P | 8 | RNA, U6 small nuclear 285, pseudogene [Sour |
| AC022893.3 | 8 | novel transcript, antisense to TERF1 |
| AC022893.1 | 8 | novel transcript |
| SBSPON | 8 | somatomedin B and thrombospondin type 1 c |
| AC100823.1 | 8 | novel transcript |
| AC100823.2 | 8 | heat shock protein 90kDa alpha (cytosolic), cla |
| C8orf89 | 8 | chromosome 8 open reading frame 89 [Sourc |
| PRXL2AP2 | 8 | peroxiredoxin like 2A pseudogene 2 [Source:H |
| RPL7 | 8 | ribosomal protein L7 [Source:HGNC Symbol;A |
| RDH10 | 8 | retinol dehydrogenase 10 [Source:HGNC Symb |
| RDH10-AS1 | 8 | RDH10 antisense RNA 1 [Source:HGNC Symbo |
| AC111149.1 | 8 | novel transcript |
| AC111149.2 | 8 | novel transcript |
| STAU2-AS1 | 8 | STAU2 antisense RNA 1 [Source:HGNC Symbo |
| STAU2 | 8 | staufen double-stranded RNA binding protein |
| AC018620.1 | 8 | chromatin accessibility complex protein 1 (CH |
| AC100784.1 | 8 | 60S ribosomal protein L36 (RPL36) pseudogen |
| VENTXP6 | 8 | VENT homeobox pseudogene 6 [Source:HGNC |
| AC027018.1 | 8 | novel transcript |
| AC022826.2 | 8 | novel protein, UBE2W-STAU2 readthrough |
| UBE2W | 8 | ubiquitin conjugating enzyme E2 W [Source:H |
| AC022826.1 | 8 | regulator of G-protein signaling 10 (RGS10) ps |
| RN7SL760P | 8 | RNA, 7SL, cytoplasmic 760, pseudogene [Sour |
| AC022868.2 | 8 | long intergenic non-protein coding RNA 1617 |
| GYG1P1 | 8 | glycogenin 1 pseudogene 1 [Source:HGNC Syn |

| | | |
|---|---|---|
| ELOC | 8 | elongin C [Source:HGNC Symbol;Acc:HGNC:11 |
| TMEM70 | 8 | transmembrane protein 70 [Source:HGNC Syn |
| RPS20P21 | 8 | ribosomal protein S20 pseudogene 21 [Source |
| AC022868.1 | 8 | destrin (actin depolymerizing factor) (DSTN) p |
| LY96 | 8 | lymphocyte antigen 96 [Source:HGNC Symbol |
| RNU6-1300P | 8 | RNA, U6 small nuclear 1300, pseudogene [Sou |
| AC087672.2 | 8 | novel transcript |
| RPS3AP32 | 8 | ribosomal protein S3a pseudogene 32 [Source |
| RNU6-1197P | 8 | RNA, U6 small nuclear 1197, pseudogene [Sou |
| AC087627.1 | 8 | novel transcript |
| AC087672.1 | 8 | novel transcript |
| JPH1 | 8 | junctophilin 1 [Source:HGNC Symbol;Acc:HGN |
| GDAP1 | 8 | ganglioside induced differentiation associated |
| AC103952.1 | 8 | novel transcript |
| Y_RNA | 8 | Y RNA [Source:RFAM;Acc:RF00019] |
| MIR5681A | 8 | microRNA 5681a [Source:HGNC Symbol;Acc:H |
| MIR5681B | 8 | microRNA 5681b [Source:HGNC Symbol;Acc:H |
| MIR2052HG | 8 | MIR2052 host gene [Source:HGNC Symbol;Acc |
| AC115837.1 | 8 | poly(rC) binding protein 2 (PCBP2) pseudogen |
| AC115837.2 | 8 | novel transcript |
| MIR2052 | 8 | microRNA 2052 [Source:HGNC Symbol;Acc:HG |
| AC011632.1 | 8 | novel transcript, antisense to PI15 |
| AC026616.1 | 8 | RAB9A, member RAS oncogene family (RAB9A |
| PI15 | 8 | peptidase inhibitor 15 [Source:HGNC Symbol; |
| AC011632.2 | 8 | novel transcript |
| CRISPLD1 | 8 | cysteine rich secretory protein LCCL domain c |
| AC100782.1 | 8 | novel transcript |
| AC022274.1 | 8 | novel transcript |
| CASC9 | 8 | cancer susceptibility 9 [Source:HGNC Symbol; |
| HIGD1AP6 | 8 | HIG1 hypoxia inducible domain family membe |
| PKMP4 | 8 | pyruvate kinase M1/2 pseudogene 4 [Source: |
| HNF4G | 8 | hepatocyte nuclear factor 4 gamma [Source:H |
| AC011029.1 | 8 | novel transcript |
| RNU2-54P | 8 | RNA, U2 small nuclear 54, pseudogene [Sourc |
| AC067773.1 | 8 | novel transcript |
| LINC01109 | 8 | long intergenic non-protein coding RNA 1109 |
| LINC01111 | 8 | long intergenic non-protein coding RNA 1111 |
| ZFHX4-AS1 | 8 | ZFHX4 antisense RNA 1 [Source:HGNC Symbol |
| AC013509.1 | 8 | novel transcript |
| MRPL9P1 | 8 | mitochondrial ribosomal protein L9 pseudoge |
| ZFHX4 | 8 | zinc finger homeobox 4 [Source:HGNC Symbo |
| AC011716.1 | 8 | NULL |
| AC023200.1 | 8 | novel transcript |
| MIR3149 | 8 | microRNA 3149 [Source:HGNC Symbol;Acc:HG |
| PEX2 | 8 | peroxisomal biogenesis factor 2 [Source:HGN |
| HIGD1AP18 | 8 | HIG1 hypoxia inducible domain family membe |
| AC062004.1 | 8 | novel transcript |
| AC084706.1 | 8 | novel transcript |
| PKIA-AS1 | 8 | PKIA antisense RNA 1 [Source:HGNC Symbol;A |
| RNU6-1220P | 8 | RNA, U6 small nuclear 1220, pseudogene [Sou |

| | | |
|---|---|---|
| PKIA | 8 | cAMP-dependent protein kinase inhibitor alph |
| AC068700.1 | 8 | novel transcript, overlapping to PKIA |
| AC068700.2 | 8 | novel transcript |
| ZC2HC1A | 8 | zinc finger C2HC-type containing 1A [Source:H |
| IL7 | 8 | interleukin 7 [Source:HGNC Symbol;Acc:HGNC |
| AC100854.1 | 8 | chromosome 4 open reading frame 46 (C4orf4 |
| THAP12P7 | 8 | THAP domain containing 12 pseudogene 7 [So |
| AC083837.1 | 8 | novel transcript |
| LINC02605 | 8 | long intergenic non-protein coding RNA 2605 |
| AC138646.1 | 8 | novel transcript |
| AC100870.1 | 8 | family with sequence similarity 71, member C |
| RPL3P9 | 8 | ribosomal protein L3 pseudogene 9 [Source:H |
| STMN2 | 8 | stathmin 2 [Source:HGNC Symbol;Acc:HGNC:1 |
| AC016240.1 | 8 | novel transcript |
| HEY1 | 8 | hes related family bHLH transcription factor w |
| LINC01607 | 8 | long intergenic non-protein coding RNA 1607 |
| AC036214.1 | 8 | novel transcript |
| RNU7-85P | 8 | RNA, U7 small nuclear 85 pseudogene [Source |
| AC036214.4 | 8 | novel transcript, antisense to MRPS28 |
| MRPS28 | 8 | mitochondrial ribosomal protein S28 [Source: |
| AC036214.3 | 8 | novel protein, TPD52-MRPS28 readthrough |
| AC036214.2 | 8 | novel transcript, sense intronic to MRPS28 |
| AC009686.2 | 8 | novel transcript |
| TPD52 | 8 | tumor protein D52 [Source:HGNC Symbol;Acc |
| AC009686.1 | 8 | novel transcript |
| RN7SL41P | 8 | RNA, 7SL, cytoplasmic 41, pseudogene [Sourc |
| MIR5708 | 8 | microRNA 5708 [Source:HGNC Symbol;Acc:HG |
| AC104212.3 | 8 | ribosomal protein L13a (RPL13A) pseudogene |
| AC034114.2 | 8 | novel transcript |
| AC104212.2 | 8 | ribosomal protein S5 (RPS5) pseudogene |
| AC104212.1 | 8 | NULL |
| AC034114.1 | 8 | ADP-ribosylation-like factor 6 interacting prot |
| RNU6-1213P | 8 | RNA, U6 small nuclear 1213, pseudogene [Sou |
| AC009812.1 | 8 | novel transcript, antisense to ZBTB10 |
| Y_RNA | 8 | Y RNA [Source:RFAM;Acc:RF00019] |
| ZBTB10 | 8 | zinc finger and BTB domain containing 10 [Sou |
| AC009812.3 | 8 | novel transcript |
| AC009812.4 | 8 | novel transcript |
| RPSAP47 | 8 | ribosomal protein SA pseudogene 47 [Source: |
| RNU7-174P | 8 | RNA, U7 small nuclear 174 pseudogene [Sourc |
| AC009812.2 | 8 | OCIA domain containing 2 (OCIAD2) pseudoge |
| RNU2-71P | 8 | RNA, U2 small nuclear 71, pseudogene [Sourc |
| SLC25A51P3 | 8 | SLC25A51 pseudogene 3 [Source:HGNC Symbo |
| RN7SL107P | 8 | RNA, 7SL, cytoplasmic 107, pseudogene [Sour |
| ZNF704 | 8 | zinc finger protein 704 [Source:HGNC Symbol; |
| CKS1BP7 | 8 | CDC28 protein kinase regulatory subunit 1B ps |
| RNU11-6P | 8 | RNA, U11 small nuclear 6, pseudogene [Sourc |
| RN7SL308P | 8 | RNA, 7SL, cytoplasmic 308, pseudogene [Sour |
| HMGB1P41 | 8 | high mobility group box 1 pseudogene 41 [Sou |
| AC012533.1 | 8 | novel transcript |

| | | |
|---|---|---|
| PAG1 | 8 | phosphoprotein membrane anchor with glyco |
| AC022778.1 | 8 | novel transcript |
| AC079209.2 | 8 | novel transcript |
| AC079209.1 | 8 | novel transcript |
| AC009902.1 | 8 | ribosomal protein L17 (RPL17) pseudogene |
| UBE2HP1 | 8 | ubiquitin conjugating enzyme E2 H pseudoger |
| AC009902.3 | 8 | novel transcript |
| AC009902.2 | 8 | novel transcript |
| FABP5 | 8 | fatty acid binding protein 5 [Source:HGNC Syn |
| AC018616.1 | 8 | novel transcript |
| PMP2 | 8 | peripheral myelin protein 2 [Source:HGNC Syr |
| FABP9 | 8 | fatty acid binding protein 9 [Source:HGNC Syn |
| FABP4 | 8 | fatty acid binding protein 4 [Source:HGNC Syn |
| AC023644.1 | 8 | novel transcript |
| FTH1P11 | 8 | ferritin heavy chain 1 pseudogene 11 [Source: |
| FABP12 | 8 | fatty acid binding protein 12 [Source:HGNC Sy |
| IMPA1P1 | 8 | inositol monophosphatase 1 pseudogene 1 [S |
| NIPA2P4 | 8 | NIPA magnesium transporter 2 pseudogene 4 |
| RPS26P34 | 8 | ribosomal protein S26 pseudogene 34 [Source |
| SLC10A5P1 | 8 | SLC10A5 pseudogene 1 [Source:HGNC Symbol |
| IMPA1 | 8 | inositol monophosphatase 1 [Source:HGNC Sy |
| SLC10A5 | 8 | solute carrier family 10 member 5 [Source:HG |
| AC132219.2 | 8 | TEC |
| ZFAND1 | 8 | zinc finger AN1-type containing 1 [Source:HGI |
| CHMP4C | 8 | charged multivesicular body protein 4C [Sourc |
| AC132219.1 | 8 | novel transcript, antisense to SNX16 |
| SNX16 | 8 | sorting nexin 16 [Source:HGNC Symbol;Acc:H |
| HNRNPA1P36 | 8 | heterogeneous nuclear ribonucleoprotein A1 |
| LINC02235 | 8 | long intergenic non-protein coding RNA 2235 |
| AC060765.2 | 8 | novel transcript |
| LINC02839 | 8 | long intergenic non-protein coding RNA 2839 |
| HNRNPA1P4 | 8 | heterogeneous nuclear ribonucleoprotein A1 |
| AC060765.1 | 8 | novel transcript |
| AC105031.2 | 8 | novel transcript |
| AC105031.1 | 8 | developmental pluripotency associated 3 (DPI |
| AC090095.1 | 8 | solute carrier family 25, member 43 (SLC25A4 |
| AC090132.1 | 8 | sulfite oxidase (SUOX) pseudogene |
| AC090132.2 | 8 | zinc finger protein 511 (ZNF511) pseudogene |
| LINC01419 | 8 | long intergenic non-protein coding RNA 1419 |
| AC015522.1 | 8 | novel transcript |
| TPM3P3 | 8 | tropomyosin 3 pseudogene 3 [Source:HGNC S |
| AC012400.1 | 8 | tropomyosin 3 (TPM3) pseudogene |
| RALYL | 8 | RALY RNA binding protein like [Source:HGNC S |
| RNU6-1040P | 8 | RNA, U6 small nuclear 1040, pseudogene [Sou |
| AC092709.1 | 8 | proteasome (prosome, macropain) 26S subun |
| AC009901.1 | 8 | glucosamine-phosphate N-acetyltransferase 1 |
| AC009901.2 | 8 | extra spindle pole bodies homolog 1 (S. cerevi |
| ACTBP6 | 8 | ACTB pseudogene 6 [Source:HGNC Symbol;Ac |
| JCHAINP1 | 8 | joining chain of multimeric IgA and IgM pseud |
| AC091175.1 | 8 | solute carrier family 7, (cationic amino acid tra |

| | | |
|---|---|---|
| LRRCC1 | 8 | leucine rich repeat and coiled-coil centrosoma |
| AC011773.4 | 8 | novel transcript, antisense to E2F5 |
| E2F5 | 8 | E2F transcription factor 5 [Source:HGNC Symb |
| AC011773.1 | 8 | novel transcript, antisense to E2F5 |
| RBIS | 8 | ribosomal biogenesis factor [Source:HGNC Syr |
| CA13 | 8 | carbonic anhydrase 13 [Source:HGNC Symbol; |
| AC011773.3 | 8 | novel transcript, antisense to CA13 |
| AC011773.2 | 8 | novel transcript, antisense to CA13 |
| CA1 | 8 | carbonic anhydrase 1 [Source:HGNC Symbol;A |
| CA3 | 8 | carbonic anhydrase 3 [Source:HGNC Symbol;A |
| CA3-AS1 | 8 | CA3 antisense RNA 1 [Source:HGNC Symbol;A |
| CA2 | 8 | carbonic anhydrase 2 [Source:HGNC Symbol;A |
| AC084734.1 | 8 | novel transcript |
| AC093331.1 | 8 | solute carrier family 33 (acetyl-CoA transporte |
| REXO1L8P | 8 | REXO1 like 8, pseudogene [Source:HGNC Sym |
| REXO1L3P | 8 | REXO1 like 3, pseudogene [Source:HGNC Sym |
| REXO1L1P | 8 | REXO1 like 1, pseudogene [Source:HGNC Sym |
| REXO1L12P | 8 | REXO1 like 12, pseudogene [Source:HGNC Syr |
| REXO1L11P | 8 | REXO1 like 11, pseudogene [Source:HGNC Syr |
| REXO1L10P | 8 | REXO1 like 10, pseudogene [Source:HGNC Syr |
| REXO1L9P | 8 | REXO1 like 9, pseudogene [Source:HGNC Sym |
| REXO1L2P | 8 | REXO1 like 2, pseudogene [Source:HGNC Sym |
| AC232323.1 | 8 | EX1, RNA exonuclease 1 homolog (S. cerevisia |
| REXO1L4P | 8 | REXO1 like 4, pseudogene [Source:HGNC Sym |
| REXO1L5P | 8 | REXO1 like 5, pseudogene [Source:HGNC Sym |
| REXO1L6P | 8 | REXO1 like 6, pseudogene [Source:HGNC Sym |
| AC100801.1 | 8 | novel transcript |
| LINC02849 | 8 | long intergenic non-protein coding RNA 2849 |
| AC100801.2 | 8 | uncharacterized LOC107986954 [Source:NCBI |
| ATP6V0D2 | 8 | ATPase H+ transporting V0 subunit d2 [Source |
| AC023194.1 | 8 | ribosomal protein L36a (RPL36A) pseudogene |
| AC023194.2 | 8 | ribosomal protein L32 (RPL32) pseudogene |
| PSKH2 | 8 | protein serine kinase H2 [Source:HGNC Symbo |
| AC023194.3 | 8 | novel transcript |
| AC084128.1 | 8 | novel transcript |
| SLC7A13 | 8 | solute carrier family 7 member 13 [Source:HG |
| AC103760.1 | 8 | novel transcript |
| WWP1 | 8 | WW domain containing E3 ubiquitin protein li |
| RMDN1 | 8 | regulator of microtubule dynamics 1 [Source:I |
| NTAN1P2 | 8 | N-terminal asparagine amidase pseudogene 2 |
| SLC2A3P4 | 8 | solute carrier family 2 member 3 pseudogene |
| CPNE3 | 8 | copine 3 [Source:HGNC Symbol;Acc:HGNC:23 |
| CNGB3 | 8 | cyclic nucleotide gated channel subunit beta 3 |
| MIOXP1 | 8 | myo-inositol oxygenase pseudogene 1 [Source |
| GOLGA2P1 | 8 | GOLGA2 pseudogene 1 [Source:HGNC Symbol |
| UBE2Q2P10 | 8 | ubiquitin conjugating enzyme E2 Q2 pseudoge |
| AC090572.3 | 8 | novel transcript |
| AC090572.2 | 8 | novel transcript |
| AC090572.1 | 8 | SET and MYND domain containing 3 (SMYD3) |
| CNBD1 | 8 | cyclic nucleotide binding domain containing 1 |

| | | |
|---|---|---|
| AF121898.1 | 8 | novel transcript, antisense to CNBD1 |
| VTA1P2 | 8 | vesicle trafficking 1 pseudogene 2 [Source:HG |
| IARS2P1 | 8 | IARS2 pseudogene 1 [Source:HGNC Symbol;Ac |
| SOX5P1 | 8 | SOX5 pseudogene 1 [Source:HGNC Symbol;Ac |
| AC005066.1 | 8 | zinc finger protein 773 (ZNF773) pseudogene |
| DCAF4L2 | 8 | DDB1 and CUL4 associated factor 4 like 2 [Sou |
| AC037450.1 | 8 | novel transcript |
| MMP16 | 8 | matrix metallopeptidase 16 [Source:HGNC Syr |
| AC090578.1 | 8 | novel transcript, antisense to MMP16 |
| RNA5SP272 | 8 | RNA, 5S ribosomal pseudogene 272 [Source:H |
| AC090568.1 | 8 | chromosome 12 open reading frame 29 (C12c |
| AC090578.2 | 8 | novel transcript |
| AC090578.3 | 8 | novel transcript |
| RPSAP74 | 8 | ribosomal protein SA pseudogene 74 [Source: |
| KRT8P4 | 8 | keratin 8 pseudogene 4 [Source:HGNC Symbo |
| AF117829.1 | 8 | novel transcript |
| RIPK2 | 8 | receptor interacting serine/threonine kinase 2 |
| COX6B1P6 | 8 | cytochrome c oxidase subunit 6B1 pseudogen |
| AF117829.2 | 8 | solute carrier family 7 (amino acid transporter |
| RNU6-925P | 8 | RNA, U6 small nuclear 925, pseudogene [Sour |
| OSGIN2 | 8 | oxidative stress induced growth inhibitor fami |
| NBN | 8 | nibrin [Source:HGNC Symbol;Acc:HGNC:7652] |
| DECR1 | 8 | 2,4-dienoyl-CoA reductase 1 [Source:HGNC Sy |
| CALB1 | 8 | calbindin 1 [Source:HGNC Symbol;Acc:HGNC:1 |
| AC004083.1 | 8 | novel transcript |
| LINC00534 | 8 | long intergenic non-protein coding RNA 534 [S |
| RNA5SP273 | 8 | RNA, 5S ribosomal pseudogene 273 [Source:H |
| LINC01030 | 8 | long intergenic non-protein coding RNA 1030 |
| TMEM64 | 8 | transmembrane protein 64 [Source:HGNC Syn |
| AC106038.1 | 8 | novel transcript |
| AC106038.2 | 8 | slowmo homolog 2 (Drosophila) (SLMO2) pseu |
| NECAB1 | 8 | N-terminal EF-hand calcium binding protein 1 |
| AC103770.1 | 8 | novel transcript |
| C8orf88 | 8 | chromosome 8 open reading frame 88 [Source |
| PIP4P2 | 8 | phosphatidylinositol-4,5-bisphosphate 4-phos |
| AC087439.1 | 8 | novel transcript |
| AC087439.2 | 8 | novel transcript |
| OTUD6B-AS1 | 8 | OTUD6B antisense RNA 1 (head to head) [Sou |
| OTUD6B | 8 | OTU domain containing 6B [Source:HGNC Sym |
| LRRC69 | 8 | leucine rich repeat containing 69 [Source:HGN |
| AC104966.1 | 8 | ceruloplasmin (ferroxidase) (CP) pseudogene |
| MIR4661 | 8 | microRNA 4661 [Source:HGNC Symbol;Acc:HG |
| SLC26A7 | 8 | solute carrier family 26 member 7 [Source:HG |
| AC103409.1 | 8 | novel transcript |
| RN7SKP231 | 8 | RN7SK pseudogene 231 [Source:HGNC Symbo |
| PRR13P7 | 8 | proline rich 13 pseudogene 7 [Source:HGNC S |
| MRPS16P1 | 8 | mitochondrial ribosomal protein S16 pseudog |
| RUNX1T1 | 8 | RUNX1 partner transcriptional co-repressor 1 |
| AF181450.1 | 8 | novel transcript |
| AC104339.1 | 8 | ribosomal protein S26 (RPS26) pseudogene |

| | | |
|---|---|---|
| AC022695.1 | 8 | NADH dehydrogenase (ubiquinone) 1 alpha su |
| AC022695.2 | 8 | novel transcript |
| AC022695.3 | 8 | novel transcript |
| AC091096.1 | 8 | novel transcript |
| AC104211.2 | 8 | novel transcript |
| AC104211.4 | 8 | novel transcript |
| AC104211.1 | 8 | novel transcript |
| AC104211.3 | 8 | novel transcript |
| AC117834.2 | 8 | novel transcript |
| AC117834.1 | 8 | novel transcript |
| TRIQK | 8 | triple QxxK/R motif containing [Source:HGNC |
| IRF5P1 | 8 | interferon regulatory factor 5 pseudogene 1 [S |
| MIR8084 | 8 | microRNA 8084 [Source:HGNC Symbol;Acc:HG |
| C8orf87 | 8 | chromosome 8 open reading frame 87 [Source |
| LINC00535 | 8 | long intergenic non-protein coding RNA 535 [S |
| AC016885.1 | 8 | novel transcript |
| AC016885.3 | 8 | variable charge X-linked protein pseudogene |
| AC016885.2 | 8 | novel transcript |
| RNA5SP274 | 8 | RNA, 5S ribosomal pseudogene 274 [Source:H |
| ZNF317P1 | 8 | zinc finger protein 317 pseudogene 1 [Source: |
| FAM92A | 8 | family with sequence similarity 92 member A |
| AC120053.1 | 8 | novel transcript, antisense to FAM92A1 |
| AC010834.3 | 8 | novel transcript |
| RBM12B | 8 | RNA binding motif protein 12B [Source:HGNC |
| AC010834.1 | 8 | novel transcript |
| RBM12B-AS1 | 8 | RBM12B antisense RNA 1 [Source:HGNC Symb |
| AC010834.2 | 8 | novel transcript |
| TMEM67 | 8 | transmembrane protein 67 [Source:HGNC Syn |
| AC084346.1 | 8 | novel transcript |
| MYL12AP1 | 8 | myosin light chain 12A pseudogene 1 [Source: |
| PDP1 | 8 | pyruvate dehyrogenase phosphatase catalytic |
| MIR378D2HG | 8 | MIR378D2 host gene [Source:HGNC Symbol;A |
| MIR378D2 | 8 | microRNA 378d-2 [Source:HGNC Symbol;Acc:I |
| PSMA2P2 | 8 | proteasome subunit alpha 2 pseudogene 2 [So |
| RPL34P18 | 8 | ribosomal protein L34 pseudogene 18 [Source |
| AC105081.1 | 8 | glutamic-oxaloacetic transaminase 1, soluble ( |
| AP003351.1 | 8 | novel transcript |
| CDH17 | 8 | cadherin 17 [Source:HGNC Symbol;Acc:HGNC: |
| AP003478.1 | 8 | ribosomal protein L6 (RPL6) pseudogene |
| GEM | 8 | GTP binding protein overexpressed in skeletal |
| RPS4XP10 | 8 | ribosomal protein S4X pseudogene 10 [Source |
| RAD54B | 8 | RAD54 homolog B [Source:HGNC Symbol;Acc: |
| FSBP | 8 | fibrinogen silencer binding protein [Source:HG |
| VIRMA | 8 | vir like m6A methyltransferase associated [So |
| AC023632.5 | 8 | novel transcript |
| AC023632.2 | 8 | novel transcript |
| AC023632.1 | 8 | ribosomal protein L31 (RPL31) pseudogene |
| AC023632.3 | 8 | microtubule associated serine/threonine kina |
| AC023632.4 | 8 | ATP synthase, H+ transporting, mitochondrial |
| AC108860.2 | 8 | novel transcript, antisense to ESRP1 |

| | | |
|---|---|---|
| ESRP1 | 8 | epithelial splicing regulatory protein 1 [Source |
| Metazoa_SRP | 8 | Metazoan signal recognition particle RNA [Sou |
| AC108860.1 | 8 | ribosomal protein S15a (RPS15A) pseudogene |
| AP005660.1 | 8 | ribosomal protein L21 (RPL21) pseudogene |
| Y_RNA | 8 | Y RNA [Source:RFAM;Acc:RF00019] |
| DPY19L4 | 8 | dpy-19 like 4 [Source:HGNC Symbol;Acc:HGN( |
| AP003692.1 | 8 | novel transcript |
| INTS8 | 8 | integrator complex subunit 8 [Source:HGNC Sy |
| CCNE2 | 8 | cyclin E2 [Source:HGNC Symbol;Acc:HGNC:15! |
| AC087752.4 | 8 | novel transcript, antisense to CCNE2 |
| AC087752.1 | 8 | ribosomal protein L17 (RPL17) pseudogene |
| NDUFAF6 | 8 | NADH:ubiquinone oxidoreductase complex as |
| TP53INP1 | 8 | tumor protein p53 inducible nuclear protein 1 |
| AC087752.3 | 8 | novel transcript |
| AC087752.2 | 8 | ORM1-like 2 (S. cerevisiae) (ORMDL2) pseudo |
| AC068189.2 | 8 | novel transcript, antisense to NDUFAF6 |
| RNU6-1209P | 8 | RNA, U6 small nuclear 1209, pseudogene [Sou |
| MIR3150BHG | 8 | MIR3150B host gene [Source:HGNC Symbol;A |
| AC068189.1 | 8 | novel transcript, antisense to NDUFAF6 |
| MIR3150B | 8 | microRNA 3150b [Source:HGNC Symbol;Acc:H |
| MIR3150A | 8 | microRNA 3150a [Source:HGNC Symbol;Acc:H |
| PLEKHF2 | 8 | pleckstrin homology and FYVE domain contair |
| C8orf37-AS1 | 8 | C8orf37 antisense RNA 1 [Source:HGNC Symb |
| LINC01298 | 8 | long intergenic non-protein coding RNA 1298 |
| C8orf37 | 8 | chromosome 8 open reading frame 37 [Sourc |
| AC024995.1 | 8 | ribosomal protein L36a-like (RPL36AL) pseudc |
| AC083836.1 | 8 | novel transcript |
| RNU6-690P | 8 | RNA, U6 small nuclear 690, pseudogene [Sour |
| AC012339.2 | 8 | splicing factor, arginine/serine-rich 3 (SFRS3) p |
| AC012339.1 | 8 | novel transcript |
| AC007992.1 | 8 | glyceraldehyde-3-phosphate dehydrogenase ( |
| AC007992.2 | 8 | tubulin, beta class I (TUBB) pseudogene |
| AP003465.2 | 8 | novel transcript |
| GDF6 | 8 | growth differentiation factor 6 [Source:HGNC |
| UQCRB | 8 | ubiquinol-cytochrome c reductase binding prc |
| AP003465.1 | 8 | novel transcript |
| MTERF3 | 8 | mitochondrial transcription termination facto |
| PTDSS1 | 8 | phosphatidylserine synthase 1 [Source:HGNC |
| Y_RNA | 8 | Y RNA [Source:RFAM;Acc:RF00019] |
| AP003548.1 | 8 | novel transcript |
| RNU6-1172P | 8 | RNA, U6 small nuclear 1172, pseudogene [Sou |
| SDC2 | 8 | syndecan 2 [Source:HGNC Symbol;Acc:HGNC:: |
| CPQ | 8 | carboxypeptidase Q [Source:HGNC Symbol;Ac |
| AP003117.2 | 8 | novel transcript, antisense to CPQ |
| AP003117.1 | 8 | novel transcript, sense intronic to PGCP |
| AP003115.1 | 8 | novel transcript |
| TSPYL5 | 8 | TSPY like 5 [Source:HGNC Symbol;Acc:HGNC:2 |
| SNORD3H | 8 | small nucleolar RNA, C/D box 3H [Source:HGN |
| AP002982.1 | 8 | ribosomal protein S2 (RPS2) pseudogene |
| AP002982.2 | 8 | novel transcript |

| | | |
|---|---|---|
| MTDH | 8 | metadherin [Source:HGNC Symbol;Acc:HGNC: |
| Y_RNA | 8 | Y RNA [Source:RFAM;Acc:RF00019] |
| LAPTM4B | 8 | lysosomal protein transmembrane 4 beta [Sou |
| AP003357.1 | 8 | ribosomal protein L39 (RPL39) pseudogene |
| AP002906.1 | 8 | transmembrane protein 69 (TMEM69) pseudo |
| SUMO2P18 | 8 | SUMO2 pseudogene 18 [Source:HGNC Symbo |
| RPS23P1 | 8 | ribosomal protein S23 pseudogene 1 [Source: |
| MATN2 | 8 | matrilin 2 [Source:HGNC Symbol;Acc:HGNC:69 |
| RPL30 | 8 | ribosomal protein L30 [Source:HGNC Symbol;A |
| AP003352.1 | 8 | novel transcript |
| SNORA72 | 8 | small nucleolar RNA, H/ACA box 72 [Source:H |
| RNU6-703P | 8 | RNA, U6 small nuclear 703, pseudogene [Sour |
| ERICH5 | 8 | glutamate rich 5 [Source:HGNC Symbol;Acc:H |
| RIDA | 8 | reactive intermediate imine deaminase A hom |
| POP1 | 8 | POP1 homolog, ribonuclease P/MRP subunit [ |
| NIPAL2 | 8 | NIPA like domain containing 2 [Source:HGNC S |
| RNU6-914P | 8 | RNA, U6 small nuclear 914, pseudogene [Sour |
| STK3 | 8 | serine/threonine kinase 3 [Source:HGNC Symb |
| AP003355.3 | 8 | TEC |
| AP003355.1 | 8 | coiled-coil domain containing 53 (CCDC53) ps |
| KCNS2 | 8 | potassium voltage-gated channel modifier sub |
| AP003355.2 | 8 | novel transcript, antisense to STK3 |
| RNU6-748P | 8 | RNA, U6 small nuclear 748, pseudogene [Sour |
| AP003467.1 | 8 | novel transcript |
| AP003467.2 | 8 | ATP synthase, H+ transporting, mitochondrial |
| RPL19P14 | 8 | ribosomal protein L19 pseudogene 14 [Source |
| AC016877.2 | 8 | RNA binding motif protein 4 family pseudoger |
| MRPL57P7 | 8 | mitochondrial ribosomal protein L57 pseudog |
| RN7SKP85 | 8 | RN7SK pseudogene 85 [Source:HGNC Symbol; |
| AC016877.3 | 8 | novel transcript |
| OSR2 | 8 | odd-skipped related transciption factor 2 [Sou |
| VPS13B-DT | 8 | VPS13B divergent transcript [Source:HGNC Sy |
| AC016877.1 | 8 | novel transcript |
| AC104986.1 | 8 | ubiquitin-conjugating enzyme E2 variant 1 (UE |
| VPS13B | 8 | vacuolar protein sorting 13 homolog B [Source |
| AC107909.2 | 8 | novel transcript |
| AC107909.1 | 8 | novel transcript |
| AP004290.1 | 8 | insulin-like growth factor 2 mRNA binding pro |
| AP004289.2 | 8 | male-specific lethal 3-like 1 (Drosophila) (MSL |
| AP004289.1 | 8 | aconitase 2, mitochondrial (ACO2) pseudogen |
| Y_RNA | 8 | Y RNA [Source:RFAM;Acc:RF00019] |
| MIR599 | 8 | microRNA 599 [Source:HGNC Symbol;Acc:HGN |
| MIR875 | 8 | microRNA 875 [Source:HGNC Symbol;Acc:HGN |
| RN7SL350P | 8 | RNA, 7SL, cytoplasmic 350, pseudogene [Sour |
| AC018442.1 | 8 | leucine zipper-EF-hand containing transmemb |
| AC018442.2 | 8 | novel transcript |
| AC018442.3 | 8 | ribosomal protein L18a (RPL18A) pseudogene |
| COX6C | 8 | cytochrome c oxidase subunit 6C [Source:HGN |
| AC105328.1 | 8 | novel transcript, antisense to COX6C |
| RGS22 | 8 | regulator of G protein signaling 22 [Source:HG |

| | | |
|---|---|---|
| SNORD77B | 8 | small nucleolar RNA, C/D box 77B [Source:HG |
| AC021590.1 | 8 | NULL |
| FBXO43 | 8 | F-box protein 43 [Source:HGNC Symbol;Acc:H |
| POLR2K | 8 | RNA polymerase II subunit K [Source:HGNC Sy |
| SPAG1 | 8 | sperm associated antigen 1 [Source:HGNC Syr |
| Y_RNA | 8 | Y RNA [Source:RFAM;Acc:RF00019] |
| AC025647.1 | 8 | ubiquitin-fold modifier 1 (UFM1) pseudogene |
| Y_RNA | 8 | Y RNA [Source:RFAM;Acc:RF00019] |
| RNF19A | 8 | ring finger protein 19A, RBR E3 ubiquitin prote |
| AP001574.1 | 8 | novel transcript |
| AP003472.1 | 8 | novel transcript |
| MIR4471 | 8 | microRNA 4471 [Source:HGNC Symbol;Acc:HG |
| AP003472.2 | 8 | novel transcript |
| AP000424.1 | 8 | novel transcript |
| AP000424.2 | 8 | novel transcript |
| ANKRD46 | 8 | ankyrin repeat domain 46 [Source:HGNC Syml |
| GAPDHP62 | 8 | glyceraldehyde 3 phosphate dehydrogenase p |
| SNX31 | 8 | sorting nexin 31 [Source:HGNC Symbol;Acc:H( |
| AP001205.1 | 8 | novel transcript |
| RNU6-1092P | 8 | RNA, U6 small nuclear 1092, pseudogene [Sou |
| PABPC1 | 8 | poly(A) binding protein cytoplasmic 1 [Source |
| MIR7705 | 8 | microRNA 7705 [Source:HGNC Symbol;Acc:HG |
| RNU6ATAC41P | 8 | RNA, U6atac small nuclear 41, pseudogene [S( |
| Y_RNA | 8 | Y RNA [Source:RFAM;Acc:RF00019] |
| RPS20P23 | 8 | ribosomal protein S20 pseudogene 23 [Source |
| RNU4-83P | 8 | RNA, U4 small nuclear 83, pseudogene [Sourc |
| RPS26P6 | 8 | ribosomal protein S26 pseudogene 6 [Source: |
| Metazoa_SRP | 8 | Metazoan signal recognition particle RNA [Sou |
| AC027373.1 | 8 | novel transcript, overlapping YWHAZ |
| YWHAZ | 8 | tyrosine 3-monooxygenase/tryptophan 5-mor |
| RN7SL685P | 8 | RNA, 7SL, cytoplasmic 685, pseudogene [Sour |
| AC018781.1 | 8 | novel transcript |
| AP003469.2 | 8 | novel transcript |
| AP003469.1 | 8 | novel transcript |
| RN7SKP249 | 8 | RN7SK pseudogene 249 [Source:HGNC Symbo |
| AP003469.3 | 8 | novel transcript |
| AP003469.4 | 8 | novel transcript |
| ZNF706 | 8 | zinc finger protein 706 [Source:HGNC Symbol; |
| AP001330.4 | 8 | novel transcript |
| AP001330.2 | 8 | myeloid leukemia factor 2 (MLF2) pseudogene |
| LINC02844 | 8 | long intergenic non-protein coding RNA 2844 |
| AP001330.5 | 8 | novel transcript |
| AP001330.1 | 8 | novel transcript |
| RNU7-67P | 8 | RNA, U7 small nuclear 67 pseudogene [Source |
| AP001330.3 | 8 | ribosomal protein L39 (RPL39) pseudogene |
| NACA4P | 8 | NACA family member 4, pseudogene [Source: |
| DUXAP2 | 8 | double homeobox A pseudogene 2 [Source:H( |
| LINC02845 | 8 | long intergenic non-protein coding RNA 2845 |
| RN7SL563P | 8 | RNA, 7SL, cytoplasmic 563, pseudogene [Sour |
| AP001208.1 | 8 | novel transcript |

| | | |
|---|---|---|
| AP001207.3 | 8 | novel transcript |
| GRHL2 | 8 | grainyhead like transcription factor 2 [Source: |
| AP001207.2 | 8 | SAP domain containing ribonucleoprotein (SA |
| AP001207.1 | 8 | ribosomal L24 domain containing 1 (RSL24D1) |
| NCALD | 8 | neurocalcin delta [Source:HGNC Symbol;Acc:H |
| AP000426.1 | 8 | novel transcript |
| AP001329.1 | 8 | NULL |
| PDCL3P2 | 8 | PDCL3 pseudogene 2 [Source:HGNC Symbol;A |
| MIR5680 | 8 | microRNA 5680 [Source:HGNC Symbol;Acc:HG |
| AP001328.1 | 8 | 6.8 kDa mitochondrial proteolipid (MP68) pse |
| RRM2B | 8 | ribonucleotide reductase regulatory TP53 indu |
| UBR5-AS1 | 8 | UBR5 antisense RNA 1 [Source:HGNC Symbol; |
| SUMO2P19 | 8 | SUMO2 pseudogene 19 [Source:HGNC Symbo |
| UBR5 | 8 | ubiquitin protein ligase E3 component n-recog |
| AP002907.1 | 8 | novel transcript, antisense to UBR5 |
| AP002981.1 | 8 | heat shock 10kDa protein 1 (chaperonin 10) (I |
| RNU6ATAC8P | 8 | RNA, U6atac small nuclear 8, pseudogene [Sou |
| HSPE1P14 | 8 | heat shock protein family E (Hsp10) member 1 |
| RPS12P15 | 8 | ribosomal protein S12 pseudogene 15 [Source |
| AP002852.1 | 8 | novel transcript |
| RNU6-1224P | 8 | RNA, U6 small nuclear 1224, pseudogene [Sou |
| ODF1 | 8 | outer dense fiber of sperm tails 1 [Source:HGI |
| POU5F1P2 | 8 | POU class 5 homeobox 1 pseudogene 2 [Sourc |
| KLF10 | 8 | Kruppel like factor 10 [Source:HGNC Symbol;A |
| AP002851.1 | 8 | novel transcript |
| ADI1P2 | 8 | acireductone dioxygenase 1 pseudogene 2 [Sc |
| GASAL1 | 8 | growth arrest associated lncRNA 1 [Source:HG |
| AZIN1 | 8 | antizyme inhibitor 1 [Source:HGNC Symbol;Ac |
| AP003696.1 | 8 | novel transcript |
| AZIN1-AS1 | 8 | AZIN1 antisense RNA 1 [Source:HGNC Symbol; |
| AP003354.1 | 8 | novel transcript |
| RPL5P24 | 8 | ribosomal protein L5 pseudogene 24 [Source: |
| Y_RNA | 8 | Y RNA [Source:RFAM;Acc:RF00019] |
| NPM1P52 | 8 | nucleophosmin 1 pseudogene 52 [Source:HGI |
| AP003550.1 | 8 | novel transcript |
| ATP6V1C1 | 8 | ATPase H+ transporting V1 subunit C1 [Source |
| MTND1P5 | 8 | MT-ND1 pseudogene 5 [Source:HGNC Symbol |
| MTCO1P4 | 8 | MT-CO1 pseudogene 4 [Source:HGNC Symbol |
| MTCO2P4 | 8 | MT-CO2 pseudogene 4 [Source:HGNC Symbol |
| LINC01181 | 8 | long intergenic non-protein coding RNA 1181 |
| BAALC-AS2 | 8 | BAALC antisense RNA 2 [Source:HGNC Symbol |
| BAALC | 8 | BAALC binder of MAP3K1 and KLF4 [Source:H |
| BAALC-AS1 | 8 | BAALC antisense RNA 1 [Source:HGNC Symbol |
| MIR3151 | 8 | microRNA 3151 [Source:HGNC Symbol;Acc:HG |
| AC025370.1 | 8 | novel transcript |
| FZD6 | 8 | frizzled class receptor 6 [Source:HGNC Symbo |
| AC025370.2 | 8 | novel transcript, antisense to FZD6 |
| CTHRC1 | 8 | collagen triple helix repeat containing 1 [Sourc |
| AC012213.3 | 8 | novel transcript |
| RNU6-1011P | 8 | RNA, U6 small nuclear 1011, pseudogene [Sou |

| | | |
|---|---|---|
| SLC25A32 | 8 | solute carrier family 25 member 32 [Source:H |
| AC012213.5 | 8 | novel protein |
| DCAF13 | 8 | DDB1 and CUL4 associated factor 13 [Source:I |
| AC012213.2 | 8 | novel transcript |
| AC012213.4 | 8 | novel transcript |
| AC012213.1 | 8 | novel transcript |
| RIMS2 | 8 | regulating synaptic membrane exocytosis 2 [S |
| PTMAP15 | 8 | prothymosin alpha pseudogene 15 [Source:HC |
| AC007751.1 | 8 | mediator complex subunit 21 (MED21) pseud |
| AC090686.1 | 8 | ribosomal protein S6 (RPS6) pseudogene |
| DPYS | 8 | dihydropyrimidinase [Source:HGNC Symbol;A |
| DCSTAMP | 8 | dendrocyte expressed seven transmembrane |
| AP003471.1 | 8 | novel transcript, sense intronic to DPYS |
| AP002847.1 | 8 | TEC |
| MIR548A3 | 8 | microRNA 548a-3 [Source:HGNC Symbol;Acc:I |
| LRP12 | 8 | LDL receptor related protein 12 [Source:HGN( |
| NDUFA5P2 | 8 | NADH:ubiquinone oxidoreductase subunit A5 |
| ZFPM2 | 8 | zinc finger protein, FOG family member 2 [Sou |
| AC012564.1 | 8 | novel transcript, antisense to ZFPM2 |
| AC090142.2 | 8 | ribosomal protein L23 (RPL23) pseudogene |
| AC090142.3 | 8 | chibby homolog 1 (Drosophila) (CBY1) pseudo |
| AC090142.1 | 8 | ribosomal protein L29 (RPL29) pseudogene |
| AC021546.1 | 8 | novel transcript |
| TMCC1P1 | 8 | transmembrane and coiled-coil domain family |
| AC041039.1 | 8 | ribosomal protein L17 (RPL17) pseudogene |
| ZFPM2-AS1 | 8 | ZFPM2 antisense RNA 1 [Source:HGNC Symbo |
| AC103853.1 | 8 | novel transcript |
| AC103853.2 | 8 | novel transcript |
| RPL12P24 | 8 | ribosomal protein L12 pseudogene 24 [Source |
| AC090802.1 | 8 | novel transcript |
| Y_RNA | 8 | Y RNA [Source:RFAM;Acc:RF00019] |
| SLC16A14P1 | 8 | solute carrier family 16 member 14 pseudoge |
| AC027031.1 | 8 | novel transcript |
| AC027031.2 | 8 | novel transcript |
| OXR1 | 8 | oxidation resistance 1 [Source:HGNC Symbol;/ |
| AC090579.1 | 8 | novel transcript |
| TAGLN2P1 | 8 | transgelin 2 pseudogene 1 [Source:HGNC Sym |
| RNU7-84P | 8 | RNA, U7 small nuclear 84 pseudogene [Source |
| ABRA | 8 | actin binding Rho activating protein [Source:H |
| AP003789.1 | 8 | Kruppel-like factor 8 (KLF8) pseudogene |
| AP000428.2 | 8 | novel transcript |
| HMGB1P46 | 8 | high mobility group box 1 pseudogene 46 [Sou |
| AP000428.1 | 8 | novel transcript |
| ANGPT1 | 8 | angiopoietin 1 [Source:HGNC Symbol;Acc:HGI |
| AC091010.1 | 8 | novel transcript |
| PGAM1P13 | 8 | phosphoglycerate mutase 1 pseudogene 13 [S |
| AC025508.1 | 8 | novel transcript, antisense to RSPO2 |
| RNA5SP275 | 8 | RNA, 5S ribosomal pseudogene 275 [Source:H |
| RSPO2 | 8 | R-spondin 2 [Source:HGNC Symbol;Acc:HGNC: |
| NRBF2P4 | 8 | nuclear receptor binding factor 2 pseudogene |

| | | |
|---|---|---|
| AP001331.1 | 8 | aurora kinase B (AURKB) pseudogene |
| EIF3E | 8 | eukaryotic translation initiation factor 3 subur |
| AC087620.1 | 8 | novel transcript |
| RPS17P14 | 8 | ribosomal protein S17 pseudogene 14 [Source |
| EMC2 | 8 | ER membrane protein complex subunit 2 [Sou |
| AC022634.1 | 8 | DnaJ (Hsp40) homolog, subfamily B, member |
| AC022634.2 | 8 | novel transcript |
| TMEM74 | 8 | transmembrane protein 74 [Source:HGNC Syn |
| TRMT10BP1 | 8 | tRNA methyltransferase 10B pseudogene 1 [S |
| AC104248.1 | 8 | novel transcript |
| TRHR | 8 | thyrotropin releasing hormone receptor [Sour |
| NUDCD1 | 8 | NudC domain containing 1 [Source:HGNC Sym |
| AC021237.1 | 8 | novel transcript |
| ENY2 | 8 | ENY2 transcription and export complex 2 subu |
| PKHD1L1 | 8 | PKHD1 like 1 [Source:HGNC Symbol;Acc:HGNC |
| MAPK6P5 | 8 | mitogen-activated protein kinase 6 pseudoger |
| EBAG9 | 8 | estrogen receptor binding site associated anti |
| SYBU | 8 | syntabulin [Source:HGNC Symbol;Acc:HGNC:2 |
| AC079061.1 | 8 | novel transcript, antisense to SYBU |
| AC023245.2 | 8 | Smg-5 homolog, nonsense mediated mRNA de |
| U2 | 8 | U2 spliceosomal RNA [Source:RFAM;Acc:RF00 |
| AC023245.1 | 8 | ribosomal protein L18 (RPL18) pseudogene |
| KCNV1 | 8 | potassium voltage-gated channel modifier sub |
| AC027451.1 | 8 | novel transcript |
| RPSAP48 | 8 | ribosomal protein SA pseudogene 48 [Source: |
| AC073023.1 | 8 | novel transcript |
| AC090819.1 | 8 | ataxin 3 (ATXN3) pseudogene |
| AC025366.1 | 8 | novel transcript |
| AP005357.1 | 8 | novel transcript |
| NDUFB9P3 | 8 | NADH:ubiquinone oxidoreductase subunit B9 |
| LINC01608 | 8 | long intergenic non-protein coding RNA 1608 |
| MTCO1P47 | 8 | MT-CO1 pseudogene 47 [Source:HGNC Symbo |
| LINC01609 | 8 | long intergenic non-protein coding RNA 1609 |
| AC009930.1 | 8 | novel transcript |
| EEF1A1P37 | 8 | eukaryotic translation elongation factor 1 alph |
| LINC02237 | 8 | long intergenic non-protein coding RNA 2237 |
| SERPINA15P | 8 | serpin family A member 15, pseudogene [Sou |
| AC022360.1 | 8 | H2A histone family, member Z (H2AFZ) pseud |
| AC022360.2 | 8 | novel transcript |
| RNU4-37P | 8 | RNA, U4 small nuclear 37, pseudogene [Sourc |
| CSMD3 | 8 | CUB and Sushi multiple domains 3 [Source:HG |
| AC024996.1 | 8 | novel transcript, antisense to CSMD3 |
| RPL30P16 | 8 | ribosomal protein L30 pseudogene 16 [Source |
| MIR2053 | 8 | microRNA 2053 [Source:HGNC Symbol;Acc:HG |
| AC107890.1 | 8 | ribosomal protein L18 (RPL18) pseudogene |
| AC055788.2 | 8 | novel transcript, antisense to CSMD3 |
| AC055788.1 | 8 | novel transcript |
| AC103993.1 | 8 | novel transcript |
| Y_RNA | 8 | Y RNA [Source:RFAM;Acc:RF00019] |
| AC064802.1 | 8 | novel transcript |

| | | |
|---|---|---|
| AC025881.1 | 8 | chromosome 2 open reading frame 56 (C2orf! |
| CARSP2 | 8 | cysteinyl-tRNA synthetase pseudogene 2 [Sou |
| TRPS1 | 8 | transcriptional repressor GATA binding 1 [Sou |
| AF178030.1 | 8 | novel transcript |
| LINC00536 | 8 | long intergenic non-protein coding RNA 536 [S |
| RNA5SP276 | 8 | RNA, 5S ribosomal pseudogene 276 [Source:H |
| AC090994.1 | 8 | novel transcript |
| AC105177.1 | 8 | novel transcript |
| EIF3H | 8 | eukaryotic translation initiation factor 3 subur |
| UTP23 | 8 | UTP23 small subunit processome component |
| RAD21 | 8 | RAD21 cohesin complex component [Source:H |
| RAD21-AS1 | 8 | RAD21 antisense RNA 1 [Source:HGNC Symbo |
| MIR3610 | 8 | microRNA 3610 [Source:HGNC Symbol;Acc:HG |
| AARD | 8 | alanine and arginine rich domain containing p |
| SLC30A8 | 8 | solute carrier family 30 member 8 [Source:HG |
| RN7SL228P | 8 | RNA, 7SL, cytoplasmic 228, pseudogene [Sour |
| AC027419.1 | 8 | Nuclear receptor coactivator 3 (NCoA-3) pseu |
| AC027419.2 | 8 | novel transcript |
| RN7SL826P | 8 | RNA, 7SL, cytoplasmic 826, pseudogene [Sour |
| AC084114.1 | 8 | novel transcript |
| AP002848.1 | 8 | NULL |
| MED30 | 8 | mediator complex subunit 30 [Source:HGNC S |
| RPS10P16 | 8 | ribosomal protein S10 pseudogene 16 [Source |
| EXT1 | 8 | exostosin glycosyltransferase 1 [Source:HGNC |
| SAMD12 | 8 | sterile alpha motif domain containing 12 [Sou |
| AC023590.1 | 8 | novel transcript, antisense to SAMD12 |
| SAMD12-AS1 | 8 | SAMD12 antisense RNA 1 [Source:HGNC Symb |
| RPS26P35 | 8 | ribosomal protein S26 pseudogene 35 [Source |
| TNFRSF11B | 8 | TNF receptor superfamily member 11b [Sourc |
| RNU6-12P | 8 | RNA, U6 small nuclear 12, pseudogene [Sourc |
| AC107953.1 | 8 | GABA(A) receptor-associated protein-like 2 (G |
| COLEC10 | 8 | collectin subfamily member 10 [Source:HGNC |
| AC107953.2 | 8 | novel transcript |
| MAL2 | 8 | mal, T cell differentiation protein 2 (gene/pse |
| MAL2-AS1 | 8 | MAL2 antisense RNA 1 [Source:HGNC Symbol; |
| MIR548AZ | 8 | microRNA 548az [Source:HGNC Symbol;Acc:H |
| AC021733.4 | 8 | novel transcript, antisense to NOV |
| AC021733.1 | 8 | NULL |
| CCN3 | 8 | cellular communication network factor 3 [Sou |
| AC021733.2 | 8 | novel transcript |
| AC021733.3 | 8 | zinc finger protein 532 (ZNF532) pseudogene |
| ENPP2 | 8 | ectonucleotide pyrophosphatase/phosphodie |
| CYCSP23 | 8 | CYCS pseudogene 23 [Source:HGNC Symbol;A |
| RN7SKP153 | 8 | RN7SK pseudogene 153 [Source:HGNC Symbo |
| TAF2 | 8 | TATA-box binding protein associated factor 2 |
| AP005717.2 | 8 | novel transcript, antisense to DSCC1 |
| DSCC1 | 8 | DNA replication and sister chromatid cohesior |
| AC021945.1 | 8 | TEC |
| RN7SL396P | 8 | RNA, 7SL, cytoplasmic 396, pseudogene [Sour |
| AP005717.1 | 8 | novel transcript |

| | | |
|---|---|---|
| DEPTOR | 8 | DEP domain containing MTOR interacting prot |
| RNA5SP277 | 8 | RNA, 5S ribosomal pseudogene 277 [Source:H |
| AC091563.1 | 8 | novel transcript |
| COL14A1 | 8 | collagen type XIV alpha 1 chain [Source:HGNC |
| MRPL13 | 8 | mitochondrial ribosomal protein L13 [Source: |
| MTBP | 8 | MDM2 binding protein [Source:HGNC Symbol |
| SNTB1 | 8 | syntrophin beta 1 [Source:HGNC Symbol;Acc: |
| NCAPGP1 | 8 | non-SMC condensin I complex subunit G pseu |
| AC104958.1 | 8 | novel transcript |
| AC104958.2 | 8 | novel transcript |
| AC068413.1 | 8 | novel transcript |
| AC011626.1 | 8 | glucosamine-phosphate N-acetyltransferase 1 |
| RPL35AP19 | 8 | ribosomal protein L35a pseudogene 19 [Sourc |
| HAS2 | 8 | hyaluronan synthase 2 [Source:HGNC Symbol; |
| HAS2-AS1 | 8 | HAS2 antisense RNA 1 [Source:HGNC Symbol; |
| LINC02855 | 8 | long intergenic non-protein coding RNA 2855 |
| AC037486.1 | 8 | novel transcript |
| MRPS36P3 | 8 | mitochondrial ribosomal protein S36 pseudog |
| SMILR | 8 | smooth muscle induced lncRNA, enhancer of p |
| LINC01151 | 8 | long intergenic non-protein coding RNA 1151 |
| AC108136.1 | 8 | novel transcript |
| AC100872.1 | 8 | NULL |
| AC100872.2 | 8 | novel transcript |
| CDK5P1 | 8 | cyclin dependent kinase 5 pseudogene 1 [Sou |
| AC016405.3 | 8 | novel transcript |
| AC016405.2 | 8 | novel transcript |
| ZHX2 | 8 | zinc fingers and homeoboxes 2 [Source:HGNC |
| AC016405.1 | 8 | novel transcript |
| AC104316.2 | 8 | novel transcript |
| AC104316.1 | 8 | novel transcript, antisense to DERL1 |
| DERL1 | 8 | derlin 1 [Source:HGNC Symbol;Acc:HGNC:284 |
| TBC1D31 | 8 | TBC1 domain family member 31 [Source:HGN |
| HMGB1P19 | 8 | high mobility group box 1 pseudogene 19 [Sou |
| AC068228.3 | 8 | novel transcript |
| FAM83A | 8 | family with sequence similarity 83 member A |
| U3 | 8 | Small nucleolar RNA U3 [Source:RFAM;Acc:RF |
| AC068228.1 | 8 | novel transcript, antisense to FAM83A |
| AC068228.2 | 8 | WD repeat domain 67 (WDR67) pseudogene |
| FAM83A-AS1 | 8 | FAM83A antisense RNA 1 [Source:HGNC Symb |
| MIR4663 | 8 | microRNA 4663 [Source:HGNC Symbol;Acc:HG |
| C8orf76 | 8 | chromosome 8 open reading frame 76 [Sourc |
| ZHX1-C8orf76 | 8 | ZHX1-C8orf76 readthrough [Source:HGNC Sym |
| UBA52P5 | 8 | ubiquitin A-52 residue ribosomal protein fusic |
| ZHX1 | 8 | zinc fingers and homeoboxes 1 [Source:HGNC |
| RNU6-628P | 8 | RNA, U6 small nuclear 628, pseudogene [Sour |
| AC018992.1 | 8 | hydroxysteroid (17-beta) dehydrogenase 4 (H |
| ATAD2 | 8 | ATPase family AAA domain containing 2 [Sour |
| RNU6-875P | 8 | RNA, U6 small nuclear 875, pseudogene [Sour |
| MIR548AA1 | 8 | microRNA 548aa-1 [Source:HGNC Symbol;Acc |
| MIR548D1 | 8 | microRNA 548d-1 [Source:HGNC Symbol;Acc: |

| | | |
|---|---|---|
| DUTP2 | 8 | deoxyuridine triphosphatase pseudogene 2 [S |
| IMPDH1P6 | 8 | inosine monophosphate dehydrogenase 1 pse |
| WDYHV1 | 8 | WDYHV motif containing 1 [Source:HGNC Sym |
| U4 | 8 | U4 spliceosomal RNA [Source:RFAM;Acc:RF00 |
| FBXO32 | 8 | F-box protein 32 [Source:HGNC Symbol;Acc:H |
| AC090193.1 | 8 | novel transcript |
| RN7SKP155 | 8 | RN7SK pseudogene 155 [Source:HGNC Symbo |
| AC135166.1 | 8 | novel transcript |
| KLHL38 | 8 | kelch like family member 38 [Source:HGNC Sy |
| ANXA13 | 8 | annexin A13 [Source:HGNC Symbol;Acc:HGNC |
| FAM91A1 | 8 | family with sequence similarity 91 member A |
| FER1L6 | 8 | fer-1 like family member 6 [Source:HGNC Sym |
| AC090753.1 | 8 | novel transcript |
| FER1L6-AS1 | 8 | FER1L6 antisense RNA 1 [Source:HGNC Symbo |
| AC100871.2 | 8 | novel transcript |
| FER1L6-AS2 | 8 | FER1L6 antisense RNA 2 [Source:HGNC Symbo |
| AC100871.1 | 8 | neuropilin (NRP) and tolloid (TLL)-like 2 (NETC |
| ARF1P3 | 8 | ADP ribosylation factor 1 pseudogene 3 [Sour |
| AC090921.1 | 8 | novel transcript |
| RNU6-756P | 8 | RNA, U6 small nuclear 756, pseudogene [Sour |
| AC090192.2 | 8 | novel transcript |
| AC090192.1 | 8 | SRY (sex determining region Y)-box 2 (SOX2) p |
| TMEM65 | 8 | transmembrane protein 65 [Source:HGNC Syn |
| TRMT12 | 8 | tRNA methyltransferase 12 homolog [Source: |
| RNF139-AS1 | 8 | RNF139 antisense RNA 1 (head to head) [Sour |
| RNF139 | 8 | ring finger protein 139 [Source:HGNC Symbol; |
| TATDN1 | 8 | TatD DNase domain containing 1 [Source:HGN |
| AC090198.1 | 8 | novel transcript, antisense to TATDN1 |
| MIR6844 | 8 | microRNA 6844 [Source:HGNC Symbol;Acc:HG |
| NDUFB9 | 8 | NADH:ubiquinone oxidoreductase subunit B9 |
| MTSS1 | 8 | MTSS I-BAR domain containing 1 [Source:HGN |
| AC100858.3 | 8 | uncharacterized LOC105375743 [Source:NCBI |
| MIR4662B | 8 | microRNA 4662b [Source:HGNC Symbol;Acc:H |
| MIR4662A | 8 | microRNA 4662a [Source:HGNC Symbol;Acc:H |
| LINC00964 | 8 | long intergenic non-protein coding RNA 964 [S |
| AC100858.1 | 8 | MRS2 magnesium homeostasis factor homolo |
| AC100858.2 | 8 | novel transcript |
| AC100858.4 | 8 | novel transcript |
| ZNF572 | 8 | zinc finger protein 572 [Source:HGNC Symbol; |
| AC009908.1 | 8 | novel transcript |
| SQLE | 8 | squalene epoxidase [Source:HGNC Symbol;Ac |
| WASHC5 | 8 | WASH complex subunit 5 [Source:HGNC Symb |
| WASHC5-AS1 | 8 | WASHC5 antisense RNA 1 [Source:HGNC Symb |
| NSMCE2 | 8 | NSE2 (MMS21) homolog, SMC5-SMC6 comple |
| RN7SL329P | 8 | RNA, 7SL, cytoplasmic 329, pseudogene [Sour |
| AC084083.1 | 8 | novel transcript |
| TRIB1 | 8 | tribbles pseudokinase 1 [Source:HGNC Symbo |
| AC091114.1 | 8 | novel transcript |
| AC016074.2 | 8 | novel transcript |
| AC016074.1 | 8 | novel transcript |

| | | |
|---|---|---|
| LINC00861 | 8 | long intergenic non-protein coding RNA 861 [S |
| RNU6-442P | 8 | RNA, U6 small nuclear 442, pseudogene [Sour |
| SOD1P3 | 8 | superoxide dismutase 1 pseudogene 3 [Source |
| AC024681.2 | 8 | novel transcript |
| KNOP1P5 | 8 | lysine rich nucleolar protein 1 pseudogene 5 [ |
| AC024681.1 | 8 | Transmembrane protein C12orf67 pseudogen |
| RFPL4AP5 | 8 | ret finger protein like 4A pseudogene 5 [Sourc |
| AC087667.1 | 8 | novel transcript |
| AC084116.2 | 8 | novel transcript |
| AC084116.1 | 8 | novel transcript |
| RNU6-869P | 8 | RNA, U6 small nuclear 869, pseudogene [Sour |
| AC084116.3 | 8 | novel transcript |
| LRATD2 | 8 | LRAT domain containing 2 [Source:HGNC Sym |
| PCAT1 | 8 | prostate cancer associated transcript 1 [Sourc |
| AC083923.1 | 8 | zinc finger protein pseudogene |
| RNU11-4P | 8 | RNA, U11 small nuclear 4, pseudogene [Sourc |
| AC024382.1 | 8 | novel transcript |
| CASC19 | 8 | cancer susceptibility 19 [Source:HGNC Symbo |
| PRNCR1 | 8 | prostate cancer associated non-coding RNA 1 |
| AC020688.1 | 8 | serine/arginine repetitive matrix 1 (SRRM1) p |
| AC018714.2 | 8 | novel transcript |
| CASC8 | 8 | cancer susceptibility 8 [Source:HGNC Symbol; |
| AC018714.1 | 8 | small nuclear ribonucleoprotein polypeptide ( |
| POU5F1B | 8 | POU class 5 homeobox 1B [Source:HGNC Sym |
| CCAT2 | 8 | colon cancer associated transcript 2 [Source:H |
| AC104370.1 | 8 | novel transcript |
| AC108925.1 | 8 | novel transcript |
| CASC11 | 8 | cancer susceptibility 11 [Source:HGNC Symbo |
| MYC | 8 | MYC proto-oncogene, bHLH transcription fact |
| PVT1 | 8 | Pvt1 oncogene [Source:HGNC Symbol;Acc:HG |
| MIR1204 | 8 | microRNA 1204 [Source:HGNC Symbol;Acc:HG |
| AC084123.1 | 8 | NULL |
| TMEM75 | 8 | transmembrane protein 75 [Source:HGNC Syn |
| MIR1205 | 8 | microRNA 1205 [Source:HGNC Symbol;Acc:HG |
| RNU1-106P | 8 | RNA, U1 small nuclear 106, pseudogene [Sour |
| MIR1206 | 8 | microRNA 1206 [Source:HGNC Symbol;Acc:HG |
| AC103705.1 | 8 | protein tyrosine phosphatase type IVA, memb |
| RNU4-25P | 8 | RNA, U4 small nuclear 25, pseudogene [Sourc |
| MIR1207 | 8 | microRNA 1207 [Source:HGNC Symbol;Acc:HG |
| MIR1208 | 8 | microRNA 1208 [Source:HGNC Symbol;Acc:HG |
| RN7SKP226 | 8 | RN7SK pseudogene 226 [Source:HGNC Symbo |
| AC100822.1 | 8 | novel transcript |
| LINC00824 | 8 | long intergenic non-protein coding RNA 824 [S |
| CCDC26 | 8 | CCDC26 long non-coding RNA [Source:HGNC S |
| AC103833.2 | 8 | novel transcript |
| AC103833.1 | 8 | minichromosome maintenance complex comp |
| RN7SKP206 | 8 | RN7SK pseudogene 206 [Source:HGNC Symbo |
| AC011257.1 | 8 | novel transcript |
| MIR3686 | 8 | microRNA 3686 [Source:HGNC Symbol;Acc:HG |
| AC103718.1 | 8 | novel transcript |

| | | |
|---|---|---|
| MTRF1LP2 | 8 | mitochondrial translational release factor 1 lik |
| GSDMC | 8 | gasdermin C [Source:HGNC Symbol;Acc:HGNC |
| AC022973.5 | 8 | novel transcript |
| FAM49B | 8 | family with sequence similarity 49 member B |
| AC022973.2 | 8 | ribosomal protein L15 (RPL15) pseudogene |
| AC022973.1 | 8 | NULL |
| AC022973.4 | 8 | novel transcript |
| AC022973.3 | 8 | novel transcript |
| AC131568.1 | 8 | novel transcript, antisense to FAM49B |
| RNU7-181P | 8 | RNA, U7 small nuclear 181 pseudogene [Sourc |
| MIR5194 | 8 | microRNA 5194 [Source:HGNC Symbol;Acc:HG |
| ASAP1 | 8 | ArfGAP with SH3 domain, ankyrin repeat and |
| RNU6-1255P | 8 | RNA, U6 small nuclear 1255, pseudogene [Sou |
| ASAP1-IT2 | 8 | ASAP1 intronic transcript 2 [Source:HGNC Sym |
| AC103725.1 | 8 | NULL |
| AC009682.1 | 8 | novel transcript, antisense to ASAP1 |
| AC139019.1 | 8 | novel transcript, antisense to ASAP1 |
| AC090987.1 | 8 | novel transcript |
| ADCY8 | 8 | adenylate cyclase 8 [Source:HGNC Symbol;Acc |
| AC103726.1 | 8 | novel transcript |
| AC103726.2 | 8 | novel transcript |
| AC087341.1 | 8 | novel transcript |
| AC104257.1 | 8 | novel transcript |
| SNORA72 | 8 | Small nucleolar RNA SNORA72 [Source:RFAM; |
| AC060788.1 | 8 | RNA binding motif protein 45 (RBM45) pseudo |
| EFR3A | 8 | EFR3 homolog A [Source:HGNC Symbol;Acc:H |
| OC90 | 8 | otoconin 90 [Source:HGNC Symbol;Acc:HGNC |
| AC100868.1 | 8 | novel protein |
| HHLA1 | 8 | HERV-H LTR-associating 1 [Source:HGNC Symb |
| KCNQ3 | 8 | potassium voltage-gated channel subfamily Q |
| HPYR1 | 8 | Helicobacter pylori responsive 1 [Source:HGN |
| LRRC6 | 8 | leucine rich repeat containing 6 [Source:HGN |
| TMEM71 | 8 | transmembrane protein 71 [Source:HGNC Syn |
| AF228727.1 | 8 | novel transcript, antisense to TMEM71 |
| PHF20L1 | 8 | PHD finger protein 20 like 1 [Source:HGNC Syr |
| AF230666.2 | 8 | novel transcript, intronic to PHF20L1 |
| AF230666.1 | 8 | novel transcript, antisense to PHF20L1 |
| TG | 8 | thyroglobulin [Source:HGNC Symbol;Acc:HGN |
| AF305872.1 | 8 | ribosomal protein L21 (RPL21) pseudogene |
| SLA | 8 | Src like adaptor [Source:HGNC Symbol;Acc:HG |
| MIR7848 | 8 | microRNA 7848 [Source:HGNC Symbol;Acc:HG |
| AF305872.2 | 8 | papillary thyroid carcinoma susceptibility canc |
| CCN4 | 8 | cellular communication network factor 4 [Sou |
| NDRG1 | 8 | N-myc downstream regulated 1 [Source:HGN |
| KC877373.1 | 8 | ribosomal protein L32 (RPL32) pseudogene |
| AF186190.1 | 8 | developmental pluripotency associated 3 pseu |
| ST13P6 | 8 | ST13, Hsp70 interacting protein pseudogene 6 |
| ST3GAL1 | 8 | ST3 beta-galactoside alpha-2,3-sialyltransfera |
| AC103706.2 | 8 | mitochondrially encoded NADH dehydrogenas |
| AC103706.3 | 8 | mitochondrially encoded NADH 4L dehydroge |

| | | |
|---|---|---|
| AC103706.1 | 8 | novel transcript |
| AC090821.1 | 8 | NULL |
| Metazoa_SRP | 8 | Metazoan signal recognition particle RNA [Sou |
| MTND2P7 | 8 | MT-ND2 pseudogene 7 [Source:HGNC Symbol |
| MTCO1P49 | 8 | MT-CO1 pseudogene 49 [Source:HGNC Symbo |
| AC133634.1 | 8 | novel transcript |
| AC090821.2 | 8 | novel transcript |
| AC090821.3 | 8 | non-metastatic cells 2, protein (NM23B) expre |
| AC110741.1 | 8 | novel transcript |
| AC110741.3 | 8 | CTAGE family, member 5 (CTAGE5) pseudoger |
| AC110741.2 | 8 | NIMA-related kinase 4 (NEK4) pseudogene |
| AC105180.2 | 8 | novel transcript |
| AC105180.1 | 8 | novel transcript |
| ZFAT | 8 | zinc finger and AT-hook domain containing [So |
| AC015599.1 | 8 | NULL |
| ZFAT-AS1 | 8 | ZFAT antisense RNA 1 [Source:HGNC Symbol;A |
| AC015599.2 | 8 | NULL |
| AC015599.3 | 8 | NULL |
| AC087045.1 | 8 | developmental pluripotency associated 3 (DPI |
| AC087045.2 | 8 | novel transcript |
| AC087045.3 | 8 | TEC |
| AC083843.4 | 8 | TEC |
| AC083843.3 | 8 | novel transcript |
| MIR30B | 8 | microRNA 30b [Source:HGNC Symbol;Acc:HGI |
| MIR30D | 8 | microRNA 30d [Source:HGNC Symbol;Acc:HGI |
| AC083843.2 | 8 | novel transcript |
| AC083843.1 | 8 | novel transcript |
| AC103764.1 | 8 | novel transcript |
| RPL23AP56 | 8 | ribosomal protein L23a pseudogene 56 [Sourc |
| LINC01591 | 8 | long intergenic non-protein coding RNA 1591 |
| AC040914.1 | 8 | novel transcript |
| KHDRBS3 | 8 | KH RNA binding domain containing, signal trar |
| MAPRE1P1 | 8 | MAPRE1 pseudogene 1 [Source:HGNC Symbol |
| RNU1-35P | 8 | RNA, U1 small nuclear 35, pseudogene [Sourc |
| LINC02055 | 8 | long intergenic non-protein coding RNA 2055 |
| AC103955.1 | 8 | novel transcript |
| AC023781.1 | 8 | Novel transcript |
| RNU6-144P | 8 | RNA, U6 small nuclear 144, pseudogene [Sour |
| ZYXP1 | 8 | zyxin pseudogene 1 [Source:HGNC Symbol;Ac |
| AC105213.1 | 8 | Novel transcript |
| AC110053.1 | 8 | nucleolar and coiled-body phosphoprotein 1 ( |
| AC046195.1 | 8 | novel transcript |
| AC046195.2 | 8 | novel transcript |
| AC079015.1 | 8 | novel transcript |
| FAM135B | 8 | family with sequence similarity 135 member E |
| COL22A1 | 8 | collagen type XXII alpha 1 chain [Source:HGN( |
| AC027541.1 | 8 | novel transcript |
| AC100807.1 | 8 | NULL |
| AC100807.2 | 8 | novel transcript |
| AC087354.1 | 8 | novel transcript |

| | | |
|---|---|---|
| AC090093.1 | 8 | MAD1 mitotic arrest deficient-like 1 (yeast) (M |
| KCNK9 | 8 | potassium two pore domain channel subfamil |
| TRAPPC9 | 8 | trafficking protein particle complex 9 [Source: |
| AC021744.1 | 8 | novel transcript |
| C8orf17 | 8 | chromosome 8 open reading frame 17 [Source |
| AC040978.1 | 8 | glyceraldehyde-3-phosphate dehydrogenase ( |
| PEG13 | 8 | paternally expressed 13 [Source:HGNC Symbo |
| AC107375.1 | 8 | novel transcript, antisense to CHRAC1 |
| CHRAC1 | 8 | chromatin accessibility complex subunit 1 [Sou |
| AGO2 | 8 | argonaute RISC catalytic component 2 [Source |
| AC067931.1 | 8 | TEC |
| AC067931.2 | 8 | TEC |
| ERICD | 8 | E2F1-regulated inhibitor of cell death [Source: |
| PTK2 | 8 | protein tyrosine kinase 2 [Source:HGNC Symb |
| MIR151A | 8 | microRNA 151a [Source:HGNC Symbol;Acc:HG |
| AC105235.1 | 8 | TEC |
| RNA5SP278 | 8 | RNA, 5S ribosomal pseudogene 278 [Source:H |
| AC100860.1 | 8 | novel transcript |
| DENND3 | 8 | DENN domain containing 3 [Source:HGNC Sym |
| AC040970.1 | 8 | novel transcript |
| SLC45A4 | 8 | solute carrier family 45 member 4 [Source:HG |
| AC011676.5 | 8 | TEC |
| AC011676.1 | 8 | novel transcript |
| AC011676.2 | 8 | novel transcript, antisense to SLC45A4 |
| AC011676.3 | 8 | novel transcript |
| AC011676.4 | 8 | novel transcript |
| LINC01300 | 8 | long intergenic non-protein coding RNA 1300 |
| AC100803.3 | 8 | novel transcript, overlapping to GPR20 |
| GPR20 | 8 | G protein-coupled receptor 20 [Source:HGNC |
| AC100803.2 | 8 | novel transcript, antisense to PTP4A3 |
| PTP4A3 | 8 | protein tyrosine phosphatase 4A3 [Source:HG |
| MROH5 | 8 | maestro heat like repeat family member 5 (ge |
| AC100803.4 | 8 | novel transcript, antisense to MROH5 |
| AC100803.1 | 8 | NULL |
| HNRNPA1P38 | 8 | heterogeneous nuclear ribonucleoprotein A1p |
| AC138647.2 | 8 | novel transcript |
| AC138647.1 | 8 | novel protein |
| AC025839.1 | 8 | novel transcript |
| MIR1302-7 | 8 | microRNA 1302-7 [Source:HGNC Symbol;Acc:H |
| AC104417.1 | 8 | DnaJ (Hsp40) homolog, subfamily C, member |
| AC104417.2 | 8 | novel transcript |
| AC103758.1 | 8 | novel transcript |
| MIR4472-1 | 8 | microRNA 4472-1 [Source:HGNC Symbol;Acc:H |
| LINC00051 | 8 | long intergenic non-protein coding RNA 51 [Sc |
| TSNARE1 | 8 | t-SNARE domain containing 1 [Source:HGNC S |
| RN7SL260P | 8 | RNA, 7SL, cytoplasmic 260, pseudogene [Sour |
| AC134682.1 | 8 | novel transcript, antisense to TSNARE1 |
| ADGRB1 | 8 | adhesion G protein-coupled receptor B1 [Sour |
| MROH4P | 8 | maestro heat like repeat family member 4, ps |
| ARC | 8 | activity regulated cytoskeleton associated pro |

| | | |
|---|---|---|
| AP006547.1 | 8 | novel transcript, antisense to ARC |
| AC108002.2 | 8 | novel transcript |
| JRK | 8 | Jrk helix-turn-helix protein [Source:HGNC Sym |
| PSCA | 8 | prostate stem cell antigen [Source:HGNC Sym |
| LY6K | 8 | lymphocyte antigen 6 family member K [Sourc |
| LNCOC1 | 8 | lncRNA associated with ovarian cancer 1 [Sour |
| THEM6 | 8 | thioesterase superfamily member 6 [Source:H |
| AC108002.1 | 8 | novel transcript |
| SLURP1 | 8 | secreted LY6/PLAUR domain containing 1 [Sou |
| LYPD2 | 8 | LY6/PLAUR domain containing 2 [Source:HGN |
| AC083841.1 | 8 | novel transcript |
| LYNX1-SLURP2 | 8 | LYNX1-SLURP2 readthrough [Source:HGNC Sy |
| SLURP2 | 8 | secreted LY6/PLAUR domain containing 2 [Sou |
| LYNX1 | 8 | Ly6/neurotoxin 1 [Source:HGNC Symbol;Acc:H |
| LY6D | 8 | lymphocyte antigen 6 family member D [Sour |
| AC083841.2 | 8 | novel transcript |
| AC083841.4 | 8 | lymphocyte antigen 6 complex, locus K (LY6K) |
| AC083841.3 | 8 | novel transcript |
| GML | 8 | glycosylphosphatidylinositol anchored molecu |
| ZNHIT1P1 | 8 | zinc finger HIT-type containing 1 pseudogene |
| CYP11B1 | 8 | cytochrome P450 family 11 subfamily B meml |
| CYP11B2 | 8 | cytochrome P450 family 11 subfamily B meml |
| LY6E-DT | 8 | LY6E divergent transcript [Source:HGNC Symb |
| CDC42P3 | 8 | cell division cycle 42 pseudogene 3 [Source:H |
| LY6E | 8 | lymphocyte antigen 6 family member E [Sourc |
| C8orf31 | 8 | chromosome 8 open reading frame 31 [Sourc |
| AK3P2 | 8 | adenylate kinase 3 pseudogene 2 [Source:HGI |
| AC083982.1 | 8 | ribosomal protein S26 (RPS26) pseudogene |
| LY6L | 8 | lymphocyte antigen 6 family member L [Sourc |
| LY6H | 8 | lymphocyte antigen 6 family member H [Sour |
| GPIHBP1 | 8 | glycosylphosphatidylinositol anchored high de |
| ZFP41 | 8 | ZFP41 zinc finger protein [Source:HGNC Symb |
| AC138696.1 | 8 | novel protein |
| GLI4 | 8 | GLI family zinc finger 4 [Source:HGNC Symbol; |
| MINCR | 8 | MYC-induced long non-coding RNA [Source:H |
| ZNF696 | 8 | zinc finger protein 696 [Source:HGNC Symbol; |
| AC138696.2 | 8 | novel transcript, antisense to ZNF696 |
| TOP1MT | 8 | DNA topoisomerase I mitochondrial [Source:H |
| RNU6-220P | 8 | RNA, U6 small nuclear 220, pseudogene [Sour |
| RHPN1-AS1 | 8 | RHPN1 antisense RNA 1 (head to head) [Sourc |
| RHPN1 | 8 | rhophilin Rho GTPase binding protein 1 [Sourc |
| AC105118.1 | 8 | novel transcript |
| MAFA-AS1 | 8 | MAFA antisense RNA 1 [Source:HGNC Symbol |
| MAFA | 8 | MAF bZIP transcription factor A [Source:HGN( |
| ZC3H3 | 8 | zinc finger CCCH-type containing 3 [Source:HG |
| AC067930.8 | 8 | novel transcript |
| AC067930.3 | 8 | novel transcript |
| RN7SKP175 | 8 | RN7SK pseudogene 175 [Source:HGNC Symbo |
| GSDMD | 8 | gasdermin D [Source:HGNC Symbol;Acc:HGNC |
| MROH6 | 8 | maestro heat like repeat family member 6 [Sc |

| | | |
|---|---|---|
| AC067930.4 | 8 | novel transcript |
| NAPRT | 8 | nicotinate phosphoribosyltransferase [Source |
| AC067930.5 | 8 | TEC |
| AC067930.1 | 8 | novel transcript |
| EEF1D | 8 | eukaryotic translation elongation factor 1 delt |
| TIGD5 | 8 | tigger transposable element derived 5 [Source |
| PYCR3 | 8 | pyrroline-5-carboxylate reductase 3 [Source:H |
| TSTA3 | 8 | tissue specific transplantation antigen P35B [S |
| AC067930.2 | 8 | novel transcript |
| ZNF623 | 8 | zinc finger protein 623 [Source:HGNC Symbol; |
| ZNF707 | 8 | zinc finger protein 707 [Source:HGNC Symbol; |
| AC105219.1 | 8 | breast cancer estrogen-induced apoptosis 2 (B |
| CCDC166 | 8 | coiled-coil domain containing 166 [Source:HG |
| AC105219.3 | 8 | novel transcript |
| MAPK15 | 8 | mitogen-activated protein kinase 15 [Source:H |
| AC105219.2 | 8 | novel transcript |
| FAM83H | 8 | family with sequence similarity 83 member H |
| MIR4664 | 8 | microRNA 4664 [Source:HGNC Symbol;Acc:HG |
| IQANK1 | 8 | IQ motif and ankyrin repeat containing 1 [Sou |
| AC105219.4 | 8 | novel transcript |
| SCRIB | 8 | scribble planar cell polarity protein [Source:H( |
| MIR937 | 8 | microRNA 937 [Source:HGNC Symbol;Acc:HGN |
| PUF60 | 8 | poly(U) binding splicing factor 60 [Source:HGN |
| AC234917.3 | 8 | novel transcript, antisense to PUF60 |
| AC234917.1 | 8 | novel transcript, sense overlapping NRBP2 3' U |
| NRBP2 | 8 | nuclear receptor binding protein 2 [Source:HG |
| MIR6845 | 8 | microRNA 6845 [Source:HGNC Symbol;Acc:HG |
| AC234917.2 | 8 | Novel transcript, antisense to NRBP2 |
| EPPK1 | 8 | epiplakin 1 [Source:HGNC Symbol;Acc:HGNC:1 |
| AC109322.2 | 8 | novel transcript |
| PLEC | 8 | plectin [Source:HGNC Symbol;Acc:HGNC:9069 |
| MIR661 | 8 | microRNA 661 [Source:HGNC Symbol;Acc:HGN |
| PARP10 | 8 | poly(ADP-ribose) polymerase family member |
| GRINA | 8 | glutamate ionotropic receptor NMDA type sul |
| SPATC1 | 8 | spermatogenesis and centriole associated 1 [S |
| SMPD5 | 8 | sphingomyelin phosphodiesterase 5 [Source:H |
| OPLAH | 8 | 5-oxoprolinase, ATP-hydrolysing [Source:HGN |
| MIR6846 | 8 | microRNA 6846 [Source:HGNC Symbol;Acc:HG |
| AC109322.1 | 8 | TEC |
| EXOSC4 | 8 | exosome component 4 [Source:HGNC Symbol |
| MIR6847 | 8 | microRNA 6847 [Source:HGNC Symbol;Acc:HG |
| GPAA1 | 8 | glycosylphosphatidylinositol anchor attachme |
| CYC1 | 8 | cytochrome c1 [Source:HGNC Symbol;Acc:HG |
| SHARPIN | 8 | SHANK associated RH domain interactor [Sour |
| MAF1 | 8 | MAF1 homolog, negative regulator of RNA po |
| WDR97 | 8 | WD repeat domain 97 [Source:HGNC Symbol;A |
| HGH1 | 8 | HGH1 homolog [Source:HGNC Symbol;Acc:HG |
| TSSK5P | 8 | testis specific serine kinase 5, pseudogene [So |
| MROH1 | 8 | maestro heat like repeat family member 1 [Sc |
| BOP1 | 8 | BOP1 ribosomal biogenesis factor [Source:HG |

| | | |
|---|---|---|
| MIR7112 | 8 | microRNA 7112 [Source:HGNC Symbol;Acc:HG |
| SCX | 8 | scleraxis bHLH transcription factor [Source:HG |
| HSF1 | 8 | heat shock transcription factor 1 [Source:HGN |
| DGAT1 | 8 | diacylglycerol O-acyltransferase 1 [Source:HG |
| AC233992.1 | 8 | novel transcript |
| MIR6848 | 8 | microRNA 6848 [Source:HGNC Symbol;Acc:HG |
| SCRT1 | 8 | scratch family transcriptional repressor 1 [Sou |
| TMEM249 | 8 | transmembrane protein 249 [Source:HGNC Sy |
| AC233992.2 | 8 | novel transcript |
| SLC52A2 | 8 | solute carrier family 52 member 2 [Source:HG |
| FBXL6 | 8 | F-box and leucine rich repeat protein 6 [Sourc |
| ADCK5 | 8 | aarF domain containing kinase 5 [Source:HGN |
| CPSF1 | 8 | cleavage and polyadenylation specific factor 1 |
| MIR939 | 8 | microRNA 939 [Source:HGNC Symbol;Acc:HGN |
| MIR1234 | 8 | microRNA 1234 [Source:HGNC Symbol;Acc:HG |
| MIR6849 | 8 | microRNA 6849 [Source:HGNC Symbol;Acc:HG |
| AC233992.3 | 8 | novel transcript, antisense to SLC39A4 |
| SLC39A4 | 8 | solute carrier family 39 member 4 [Source:HG |
| VPS28 | 8 | VPS28 subunit of ESCRT-I [Source:HGNC Symb |
| TONSL | 8 | tonsoku like, DNA repair protein [Source:HGN |
| MIR6893 | 8 | microRNA 6893 [Source:HGNC Symbol;Acc:HG |
| TONSL-AS1 | 8 | TONSL antisense RNA 1 [Source:HGNC Symbo |
| CYHR1 | 8 | cysteine and histidine rich 1 [Source:HGNC Sy |
| AC084125.1 | 8 | novel transcript, antisense to CYHR1 |
| KIFC2 | 8 | kinesin family member C2 [Source:HGNC Sym |
| FOXH1 | 8 | forkhead box H1 [Source:HGNC Symbol;Acc:H |
| PPP1R16A | 8 | protein phosphatase 1 regulatory subunit 16A |
| AC084125.2 | 8 | novel transcript, antisense to PPP1R16A |
| GPT | 8 | glutamic--pyruvic transaminase [Source:HGNC |
| MFSD3 | 8 | major facilitator superfamily domain containi |
| RECQL4 | 8 | RecQ like helicase 4 [Source:HGNC Symbol;Ac |
| AC084125.4 | 8 | novel transcript, antisense to RECQL4 |
| LRRC14 | 8 | leucine rich repeat containing 14 [Source:HGN |
| LRRC24 | 8 | leucine rich repeat containing 24 [Source:HGN |
| C8orf82 | 8 | chromosome 8 open reading frame 82 [Sourc |
| ARHGAP39 | 8 | Rho GTPase activating protein 39 [Source:HGN |
| AC084125.3 | 8 | novel transcript, antisense to ARHGAP39 |
| Metazoa_SRP | 8 | Metazoan signal recognition particle RNA [Sou |
| AF186192.1 | 8 | novel transcript |
| AF186192.2 | 8 | zinc finger protein 154 (ZNF154) pseudogene |
| AF186192.3 | 8 | novel transcript |
| ZNF251 | 8 | zinc finger protein 251 [Source:HGNC Symbol; |
| ZNF34 | 8 | zinc finger protein 34 [Source:HGNC Symbol;A |
| RN7SL395P | 8 | RNA, 7SL, cytoplasmic 395, pseudogene [Sour |
| RPL8 | 8 | ribosomal protein L8 [Source:HGNC Symbol;A |
| MIR6850 | 8 | microRNA 6850 [Source:HGNC Symbol;Acc:HG |
| ZNF517 | 8 | zinc finger protein 517 [Source:HGNC Symbol; |
| ZNF7 | 8 | zinc finger protein 7 [Source:HGNC Symbol;Ac |
| COMMD5 | 8 | COMM domain containing 5 [Source:HGNC Sy |
| AF235103.3 | 8 | novel transcript |

| | | |
|---|---|---|
| ZNF250 | 8 | zinc finger protein 250 [Source:HGNC Symbol; |
| ZNF16 | 8 | zinc finger protein 16 [Source:HGNC Symbol;A |
| ZNF252P | 8 | zinc finger protein 252, pseudogene [Source:H |
| AF235103.1 | 8 | novel transcript, antisense to ZNF252 |
| TMED10P1 | 8 | transmembrane p24 trafficking protein 10 pse |
| ZNF252P-AS1 | 8 | ZNF252P antisense RNA 1 [Source:HGNC Syml |
| AF235103.2 | 8 | zinc finger with KRAB and SCAN domains 1 (ZK |
| AC139103.1 | 8 | zinc finger protein pseudogene |
| C8orf33 | 8 | chromosome 8 open reading frame 33 [Sourc |
| DDX11L5 | 9 | DEAD/H-box helicase 11 like 5 [Source:HGNC S |
| WASHC1 | 9 | WASH complex subunit 1 [Source:HGNC Symb |
| AL928970.1 | 9 | novel transcript |
| MIR1302-9 | 9 | microRNA 1302-9 [Source:HGNC Symbol;Acc:H |
| FAM138C | 9 | family with sequence similarity 138 member C |
| AL449043.2 | 9 | phosphoglucomutase 5 (PGM5) pseudogene |
| PGM5P3-AS1 | 9 | PGM5P3 antisense RNA 1 [Source:HGNC Symb |
| AL449043.1 | 9 | novel transcript |
| LINC01388 | 9 | long intergenic non-protein coding RNA 1388 |
| FOXD4 | 9 | forkhead box D4 [Source:HGNC Symbol;Acc:H |
| CBWD1 | 9 | COBW domain containing 1 [Source:HGNC Syr |
| AL158832.2 | 9 | novel transcript |
| DOCK8-AS1 | 9 | DOCK8 antisense RNA 1 [Source:HGNC Symbo |
| DOCK8 | 9 | dedicator of cytokinesis 8 [Source:HGNC Symb |
| AL158832.1 | 9 | novel transcript |
| AL161725.1 | 9 | novel transcript |
| KANK1 | 9 | KN motif and ankyrin repeat domains 1 [Sourc |
| RPL12P25 | 9 | ribosomal protein L12 pseudogene 25 [Source |
| AL161725.2 | 9 | novel transcript |
| AL392089.1 | 9 | novel transcript |
| RNU6-1327P | 9 | RNA, U6 small nuclear 1327, pseudogene [Sou |
| EIF1P1 | 9 | eukaryotic translation initiation factor 1 pseu |
| AL136979.1 | 9 | novel transcript |
| DMRT1 | 9 | doublesex and mab-3 related transcription fa |
| DMRT3 | 9 | doublesex and mab-3 related transcription fa |
| RNU6-1073P | 9 | RNA, U6 small nuclear 1073, pseudogene [Sou |
| AL358976.1 | 9 | H3 histone pseudogene |
| AL358976.2 | 9 | novel transcript |
| LINC01230 | 9 | long intergenic non-protein coding RNA 1230 |
| DMRT2 | 9 | doublesex and mab-3 related transcription fa |
| AL356498.1 | 9 | DiGeorge syndrome-related protein pseudoge |
| RPS27AP14 | 9 | ribosomal protein S27a pseudogene 14 [Sourc |
| AL358934.1 | 9 | transcription factor IIA small 12 kDa subunit ( |
| RNA5SP279 | 9 | RNA, 5S ribosomal pseudogene 279 [Source:H |
| AL358053.1 | 9 | novel transcript |
| AL591644.1 | 9 | laminin receptor 1 (67kD, ribosomal protein S |
| SMARCA2 | 9 | SWI/SNF related, matrix associated, actin dep |
| AL359076.1 | 9 | novel transcript |
| RNU2-25P | 9 | RNA, U2 small nuclear 25, pseudogene [Sourc |
| RN7SL592P | 9 | RNA, 7SL, cytoplasmic 592, pseudogene [Sour |
| VLDLR-AS1 | 9 | VLDLR antisense RNA 1 [Source:HGNC Symbol |

| | | |
|---|---|---|
| AL353614.1 | 9 | novel transcript |
| VLDLR | 9 | very low density lipoprotein receptor [Source: |
| AL450467.1 | 9 | novel transcript, antisense to VLDLR |
| KCNV2 | 9 | potassium voltage-gated channel modifier sub |
| PUM3 | 9 | pumilio RNA binding family member 3 [Source |
| GPS2P1 | 9 | G protein pathway suppressor 2 pseudogene : |
| ATP5PDP2 | 9 | ATP synthase peripheral stalk subunit d pseud |
| CARM1P1 | 9 | coactivator associated arginine methyltransfe |
| LINC01231 | 9 | long intergenic non-protein coding RNA 1231 |
| RFX3 | 9 | regulatory factor X3 [Source:HGNC Symbol;Ac |
| AL354941.1 | 9 | ribosomal protein L7 (RPL7) pseudogene |
| RFX3-AS1 | 9 | RFX3 antisense RNA 1 [Source:HGNC Symbol;/ |
| AL354977.2 | 9 | novel transcript |
| AL354977.1 | 9 | novel transcript |
| GLIS3 | 9 | GLIS family zinc finger 3 [Source:HGNC Symbo |
| AL137071.1 | 9 | novel transcript |
| GLIS3-AS1 | 9 | GLIS3 antisense RNA 1 [Source:HGNC Symbol; |
| AL359095.1 | 9 | novel transcript |
| AL162419.1 | 9 | novel transcript |
| RNU6-694P | 9 | RNA, U6 small nuclear 694, pseudogene [Sour |
| SLC1A1 | 9 | solute carrier family 1 member 1 [Source:HGN |
| SPATA6L | 9 | spermatogenesis associated 6 like [Source:HG |
| AL136231.1 | 9 | ribosomal protein S6 (RPS6) pseudogene |
| PLPP6 | 9 | phospholipid phosphatase 6 [Source:HGNC Sy |
| CDC37L1-DT | 9 | CDC37L1 divergent transcript [Source:HGNC S |
| CDC37L1 | 9 | cell division cycle 37 like 1 [Source:HGNC Sym |
| AK3 | 9 | adenylate kinase 3 [Source:HGNC Symbol;Acc |
| AL353151.2 | 9 | novel transcript, antisense to AK3 |
| ECM1P1 | 9 | extracellular matrix protein 1 pseudogene 1 [S |
| AL353151.1 | 9 | ribosomal protein S5 (RPS5) pseudogene |
| RCL1 | 9 | RNA terminal phosphate cyclase like 1 [Source |
| KLF4P1 | 9 | Kruppel like factor 4 pseudogene 1 [Source:HG |
| MIR101-2 | 9 | microRNA 101-2 [Source:HGNC Symbol;Acc:H |
| AL158147.1 | 9 | miR-101-precursor-9 micro RNA |
| HNRNPA1P41 | 9 | heterogeneous nuclear ribonucleoprotein A1 |
| JAK2 | 9 | Janus kinase 2 [Source:HGNC Symbol;Acc:HGN |
| CSNK1G2P1 | 9 | casein kinase 1 gamma 2 pseudogene 1 [Sour |
| PDSS1P1 | 9 | decaprenyl diphosphate synthase subunit 1 ps |
| MTND6P5 | 9 | MT-ND6 pseudogene 5 [Source:HGNC Symbol |
| MTND1P11 | 9 | MT-ND1 pseudogene 11 [Source:HGNC Symbo |
| MTCO1P11 | 9 | MT-CO1 pseudogene 11 [Source:HGNC Symbo |
| MTCO2P11 | 9 | MT-CO2 pseudogene 11 [Source:HGNC Symbo |
| MTATP6P11 | 9 | MT-ATP6 pseudogene 11 [Source:HGNC Symb |
| MTCO3P11 | 9 | MT-CO3 pseudogene 11 [Source:HGNC Symbo |
| MTND4P14 | 9 | MT-ND4 pseudogene 14 [Source:HGNC Symbo |
| MTND5P14 | 9 | MT-ND5 pseudogene 14 [Source:HGNC Symbo |
| TCF3P1 | 9 | transcription factor 3 pseudogene 1 [Source:H |
| IGHEP2 | 9 | immunoglobulin heavy constant epsilon P2 (p |
| INSL6 | 9 | insulin like 6 [Source:HGNC Symbol;Acc:HGNC |
| AL133547.1 | 9 | insulin-like 4 (placenta) (INSL4) pseudogene |

| | | |
|---|---|---|
| INSL4 | 9 | insulin like 4 [Source:HGNC Symbol;Acc:HGNC |
| RLN2 | 9 | relaxin 2 [Source:HGNC Symbol;Acc:HGNC:10( |
| HMGN2P31 | 9 | high mobility group nucleosomal binding dom |
| RLN1 | 9 | relaxin 1 [Source:HGNC Symbol;Acc:HGNC:10( |
| AL135786.2 | 9 | novel transcript |
| AL135786.1 | 9 | novel pseudogene |
| PLGRKT | 9 | plasminogen receptor with a C-terminal lysine |
| RNF152P1 | 9 | ring finger protein 152 pseudogene 1 [Source: |
| CD274 | 9 | CD274 molecule [Source:HGNC Symbol;Acc:H |
| AL162253.2 | 9 | novel transcript |
| PDCD1LG2 | 9 | programmed cell death 1 ligand 2 [Source:HG |
| AL162253.1 | 9 | novel transcript |
| RIC1 | 9 | RIC1 homolog, RAB6A GEF complex partner 1 |
| AL136980.1 | 9 | novel transcript |
| ERMP1 | 9 | endoplasmic reticulum metallopeptidase 1 [So |
| AK4P4 | 9 | adenylate kinase 4 pseudogene 4 [Source:HGI |
| KIAA2026 | 9 | KIAA2026 [Source:HGNC Symbol;Acc:HGNC:23 |
| MLANA | 9 | melan-A [Source:HGNC Symbol;Acc:HGNC:712 |
| MIR4665 | 9 | microRNA 4665 [Source:HGNC Symbol;Acc:HG |
| RANBP6 | 9 | RAN binding protein 6 [Source:HGNC Symbol;A |
| AL162384.1 | 9 | novel transcript |
| GTF3AP1 | 9 | general transcription factor IIIA pseudogene 1 |
| IL33 | 9 | interleukin 33 [Source:HGNC Symbol;Acc:HGN |
| AL360221.1 | 9 | glycine cleavage system protein H (aminometl |
| SELENOTP1 | 9 | selenoprotein T pseudogene 1 [Source:HGNC |
| TPD52L3 | 9 | TPD52 like 3 [Source:HGNC Symbol;Acc:HGNC |
| UHRF2 | 9 | ubiquitin like with PHD and ring finger domair |
| AL133480.1 | 9 | novel transcript |
| GLDC | 9 | glycine decarboxylase [Source:HGNC Symbol;A |
| RN7SL25P | 9 | RNA, 7SL, cytoplasmic 25, pseudogene [Sourc |
| AL353718.1 | 9 | pseudogene similar to part of DCN1, defective |
| RPL23AP57 | 9 | ribosomal protein L23a pseudogene 57 [Sourc |
| AL162411.1 | 9 | novel transcript |
| RN7SL123P | 9 | RNA, 7SL, cytoplasmic 123, pseudogene [Sour |
| RPS3AP54 | 9 | ribosomal protein S3a pseudogene 54 [Source |
| RNF2P1 | 9 | ring finger protein 2 pseudogene 1 [Source:HG |
| AL354707.2 | 9 | 60S ribosomal protein L35a (RPL35A) pseudog |
| LINC02851 | 9 | long intergenic non-protein coding RNA 2851 |
| AL354707.1 | 9 | novel transcript |
| KDM4C | 9 | lysine demethylase 4C [Source:HGNC Symbol; |
| PRELID3BP11 | 9 | PRELI domain containing 3B pseudogene 11 [S |
| SNRPEP2 | 9 | small nuclear ribonucleoprotein polypeptide E |
| ACTG1P14 | 9 | actin gamma 1 pseudogene 14 [Source:HGNC |
| AL513412.1 | 9 | novel transcript |
| AL161443.1 | 9 | novel transcript |
| AL157703.1 | 9 | novel transcript |
| RPL4P5 | 9 | ribosomal protein L4 pseudogene 5 [Source:H |
| PPIAP33 | 9 | peptidylprolyl isomerase A pseudogene 33 [So |
| AL354694.1 | 9 | novel transcript |
| DMAC1 | 9 | distal membrane arm assembly complex 1 [So |

| | | |
|---|---|---|
| AL135923.2 | 9 | novel transcript |
| AL135923.1 | 9 | novel transcript |
| PTPRD | 9 | protein tyrosine phosphatase receptor type D |
| AL583805.2 | 9 | novel transcript |
| AL583805.1 | 9 | ribosomal protein 18 (RPL18) pseudogene |
| RNU7-185P | 9 | RNA, U7 small nuclear 185 pseudogene [Sourc |
| PTPRD-AS1 | 9 | PTPRD antisense RNA 1 [Source:HGNC Symbo |
| AL596451.1 | 9 | novel transcript |
| RPS26P3 | 9 | ribosomal protein S26 pseudogene 3 [Source: |
| RN7SL5P | 9 | RNA, 7SL, cytoplasmic 5, pseudogene [Source: |
| AL513422.1 | 9 | novel transcript |
| AL135790.2 | 9 | ribosomal protein S12 (RPS12) pseudogene |
| PTPRD-AS2 | 9 | PTPRD antisense RNA 2 (head to head) [Sourc |
| AL135790.1 | 9 | novel transcript |
| AL161788.1 | 9 | pseudogene similar to part of ribosomal prote |
| IMP3P1 | 9 | IMP3 pseudogene 1 [Source:HGNC Symbol;Ac |
| AL162413.1 | 9 | novel transcript |
| AKAP8P1 | 9 | A-kinase anchoring protein 8 pseudogene 1 [S |
| AL451129.1 | 9 | novel transcript |
| AL355672.1 | 9 | novel transcript |
| AL592227.1 | 9 | cytochrome c oxidase subunit VIIIC (COX8C) p |
| AL353595.1 | 9 | Novel transcript |
| AL589678.1 | 9 | novel transcript |
| JKAMPP1 | 9 | JNK1/MAPK8-associated membrane protein p |
| RNU2-47P | 9 | RNA, U2 small nuclear 47, pseudogene [Sourc |
| TYRP1 | 9 | tyrosinase related protein 1 [Source:HGNC Sy |
| LURAP1L-AS1 | 9 | LURAP1L antisense RNA 1 [Source:HGNC Syml |
| RN7SL849P | 9 | RNA, 7SL, cytoplasmic 849, pseudogene [Sour |
| LURAP1L | 9 | leucine rich adaptor protein 1 like [Source:HG |
| Y_RNA | 9 | Y RNA [Source:RFAM;Acc:RF00019] |
| AL161449.2 | 9 | Novel transcript |
| PRDX1P1 | 9 | peroxiredoxin 1 pseudogene 1 [Source:HGNC |
| AL161449.1 | 9 | Sjogren syndrome antigen B (autoantigen La) |
| MPDZ | 9 | multiple PDZ domain crumbs cell polarity com |
| AL162386.2 | 9 | novel transcript |
| AL162386.1 | 9 | inhibitor of growth family, member 1 (ING1) p |
| AL583785.1 | 9 | novel transcript |
| LINC01235 | 9 | long intergenic non-protein coding RNA 1235 |
| LINC00583 | 9 | long intergenic non-protein coding RNA 583 [S |
| PES1P2 | 9 | pescadillo ribosomal biogenesis factor 1 pseu |
| AL360089.1 | 9 | novel transcript |
| RPL3P11 | 9 | ribosomal protein L3 pseudogene 11 [Source: |
| ATP5PDP3 | 9 | ATP synthase peripheral stalk subunit d pseud |
| NFIB | 9 | nuclear factor I B [Source:HGNC Symbol;Acc:H |
| AL441963.1 | 9 | ribosomal protein L7a (RPL7A) pseudogene |
| AL136366.1 | 9 | novel transcript |
| AL137017.1 | 9 | novel transcript, antisense to NFIB |
| AL159169.1 | 9 | pseudogene similar to part of nuclear factor I/ |
| AL159169.3 | 9 | novel transcript |
| AL159169.2 | 9 | novel transcript |

| | | |
|---|---|---|
| CDCA4P1 | 9 | cell division cycle associated 4 pseudogene 1 [ |
| ZDHHC21 | 9 | zinc finger DHHC-type containing 21 [Source:H |
| CER1 | 9 | cerberus 1, DAN family BMP antagonist [Sourc |
| FREM1 | 9 | FRAS1 related extracellular matrix 1 [Source:H |
| AL512643.2 | 9 | novel transcript, antisense to FREM1 |
| AL512643.1 | 9 | novel transcript, antisense to FREM1 |
| RNU6-1260P | 9 | RNA, U6 small nuclear 1260, pseudogene [Sou |
| CLCN3P1 | 9 | chloride voltage-gated channel 3 pseudogene |
| LDHAP4 | 9 | lactate dehydrogenase A pseudogene 4 [Sourc |
| AL592293.1 | 9 | chromosome 4 open reading frame 27 (C4orf2 |
| AL592293.2 | 9 | PC4 and SFRS1 interacting protein 1 (PSIP1) ps |
| RNU6-559P | 9 | RNA, U6 small nuclear 559, pseudogene [Sour |
| TTC39B | 9 | tetratricopeptide repeat domain 39B [Source: |
| RPL7P33 | 9 | ribosomal protein L7 pseudogene 33 [Source: |
| SNAPC3 | 9 | small nuclear RNA activating complex polypep |
| RNU6-319P | 9 | RNA, U6 small nuclear 319, pseudogene [Sour |
| PSIP1 | 9 | PC4 and SFRS1 interacting protein 1 [Source:H |
| AL513423.1 | 9 | thymosin beta 10 (TMSB10) pseudogene |
| RN7SL98P | 9 | RNA, 7SL, cytoplasmic 98, pseudogene [Sourc |
| FTH1P12 | 9 | ferritin heavy chain 1 pseudogene 12 [Source: |
| RNU6-246P | 9 | RNA, U6 small nuclear 246, pseudogene [Sour |
| CCDC171 | 9 | coiled-coil domain containing 171 [Source:HG |
| HMGN2P16 | 9 | high mobility group nucleosomal binding dom |
| RNU6-14P | 9 | RNA, U6 small nuclear 14, pseudogene [Sourc |
| C9orf92 | 9 | chromosome 9 open reading frame 92 [Sourc |
| BNC2 | 9 | basonuclin 2 [Source:HGNC Symbol;Acc:HGNC |
| AL449983.1 | 9 | novel transcript |
| AL450003.1 | 9 | novel transcript |
| BNC2-AS1 | 9 | BNC2 antisense RNA 1 [Source:HGNC Symbol; |
| LSM1P1 | 9 | LSM1 homolog, mRNA degradation associatec |
| AL162725.2 | 9 | novel transcript |
| AL358117.1 | 9 | novel transcript |
| RN7SL720P | 9 | RNA, 7SL, cytoplasmic 720, pseudogene [Sour |
| RPS29P33 | 9 | ribosomal protein S29 pseudogene 33 [Source |
| AL162725.1 | 9 | ribosomal protein L31 (RPL31) pseudogene |
| CNTLN | 9 | centlein [Source:HGNC Symbol;Acc:HGNC:234 |
| SAMM50P1 | 9 | SAMM50 sorting and assembly machinery cor |
| SH3GL2 | 9 | SH3 domain containing GRB2 like 2, endophili |
| PABPC1P11 | 9 | poly(A) binding protein cytoplasmic 1 pseudog |
| ADAMTSL1 | 9 | ADAMTS like 1 [Source:HGNC Symbol;Acc:HGI |
| AL442638.1 | 9 | novel transcript |
| MIR3152 | 9 | microRNA 3152 [Source:HGNC Symbol;Acc:HG |
| RN7SKP258 | 9 | RN7SK pseudogene 258 [Source:HGNC Symbo |
| RAP1BP1 | 9 | RAP1B, member of RAS oncogene family pseu |
| SAXO1 | 9 | stabilizer of axonemal microtubules 1 [Source |
| AL356000.1 | 9 | proteasome (prosome, macropain) 26S subun |
| RRAGA | 9 | Ras related GTP binding A [Source:HGNC Syml |
| HAUS6 | 9 | HAUS augmin like complex subunit 6 [Source: |
| SCARNA8 | 9 | small Cajal body-specific RNA 8 [Source:HGNC |
| RNU6-264P | 9 | RNA, U6 small nuclear 264, pseudogene [Sour |

| | | |
|---|---|---|
| Y_RNA | 9 | Y RNA [Source:RFAM;Acc:RF00019] |
| PLIN2 | 9 | perilipin 2 [Source:HGNC Symbol;Acc:HGNC:2 |
| AL161909.1 | 9 | ribosomal protein S6 (RPS6) pseudogene |
| DENND4C | 9 | DENN domain containing 4C [Source:HGNC Sy |
| AL161909.2 | 9 | novel transcript |
| AL391834.1 | 9 | novel transcript, antisense to DENND4C |
| AL391834.2 | 9 | novel transcript, antisense to RPS6 |
| RPS6 | 9 | ribosomal protein S6 [Source:HGNC Symbol;A |
| NDUFA5P3 | 9 | NADH:ubiquinone oxidoreductase subunit A5 |
| AL391834.3 | 9 | Novel transcript |
| ACER2 | 9 | alkaline ceramidase 2 [Source:HGNC Symbol; |
| AL158206.1 | 9 | novel transcript, overlapping ACER2 |
| MAP1LC3BP1 | 9 | microtubule associated protein 1 light chain 3 |
| SLC24A2 | 9 | solute carrier family 24 member 2 [Source:HG |
| C11orf98P1 | 9 | chromosome 11 open reading frame 98 pseu |
| AL158077.2 | 9 | novel transcript |
| AL158077.1 | 9 | CDC-like kinase 3 (CLK3) pseudogene |
| AL591222.1 | 9 | novel transcript |
| AL591222.2 | 9 | KIAA1191 (KIAA1191) pseudogene |
| AL591222.3 | 9 | glutaredoxin 3 (GLRX3) pseudogene |
| AL512635.1 | 9 | survival of motor neuron 1, telomeric (SMN1) |
| MLLT3 | 9 | MLLT3 super elongation complex subunit [Sou |
| MIR4473 | 9 | microRNA 4473 [Source:HGNC Symbol;Acc:H |
| RNU4-26P | 9 | RNA, U4 small nuclear 26, pseudogene [Sourc |
| MIR4474 | 9 | microRNA 4474 [Source:HGNC Symbol;Acc:H |
| AL163193.1 | 9 | ribosomal protein L7a (RPL7A) pseudogene |
| FOCAD | 9 | focadhesin [Source:HGNC Symbol;Acc:HGNC:2 |
| FOCAD-AS1 | 9 | FOCAD antisense RNA 1 [Source:HGNC Symbo |
| MIR491 | 9 | microRNA 491 [Source:HGNC Symbol;Acc:HG |
| SNORA30B | 9 | small nucleolar RNA, H/ACA box 30B [Source: |
| HACD4 | 9 | 3-hydroxyacyl-CoA dehydratase 4 [Source:HG |
| IFNNP1 | 9 | interferon nu 1, pseudogene [Source:HGNC Sy |
| IFNB1 | 9 | interferon beta 1 [Source:HGNC Symbol;Acc:H |
| IFNWP4 | 9 | interferon omega 1 pseudogene 4 [Source:HG |
| AL390882.1 | 9 | DnaJ (Hsp40) homolog, subfamily B, member |
| IFNW1 | 9 | interferon omega 1 [Source:HGNC Symbol;Ac |
| Y_RNA | 9 | Y RNA [Source:RFAM;Acc:RF00019] |
| IFNA21 | 9 | interferon alpha 21 [Source:HGNC Symbol;Ac |
| IFNWP15 | 9 | interferon omega 1 pseudogene 15 [Source:H |
| IFNA4 | 9 | interferon alpha 4 [Source:HGNC Symbol;Acc: |
| IFNWP9 | 9 | interferon omega 1 pseudogene 9 [Source:HG |
| IFNA7 | 9 | interferon alpha 7 [Source:HGNC Symbol;Acc: |
| IFNA10 | 9 | interferon alpha 10 [Source:HGNC Symbol;Ac |
| IFNWP18 | 9 | interferon omega 1 pseudogene 18 [Source:H |
| IFNA16 | 9 | interferon alpha 16 [Source:HGNC Symbol;Ac |
| IFNA17 | 9 | interferon alpha 17 [Source:HGNC Symbol;Ac |
| IFNWP5 | 9 | interferon omega 1 pseudogene 5 [Source:HG |
| IFNA14 | 9 | interferon alpha 14 [Source:HGNC Symbol;Ac |
| IFNA22P | 9 | interferon alpha 22, pseudogene [Source:HG |
| IFNA5 | 9 | interferon alpha 5 [Source:HGNC Symbol;Acc: |

| | | |
|---|---|---|
| IFNA20P | 9 | interferon alpha 20, pseudogene [Source:HGN |
| KLHL9 | 9 | kelch like family member 9 [Source:HGNC Sym |
| IFNA6 | 9 | interferon alpha 6 [Source:HGNC Symbol;Acc: |
| IFNA13 | 9 | interferon alpha 13 [Source:HGNC Symbol;Acc |
| IFNA2 | 9 | interferon alpha 2 [Source:HGNC Symbol;Acc: |
| AL353732.1 | 9 | novel transcript |
| IFNA11P | 9 | interferon alpha 11, pseudogene [Source:HGN |
| IFNA12P | 9 | interferon alpha 12, pseudogene [Source:HGN |
| IFNA8 | 9 | interferon alpha 8 [Source:HGNC Symbol;Acc: |
| MIR31HG | 9 | MIR31 host gene [Source:HGNC Symbol;Acc:H |
| IFNWP2 | 9 | interferon omega 1 pseudogene 2 [Source:HG |
| IFNA1 | 9 | interferon alpha 1 [Source:HGNC Symbol;Acc: |
| IFNWP19 | 9 | interferon omega 1 pseudogene 19 [Source:H |
| IFNE | 9 | interferon epsilon [Source:HGNC Symbol;Acc: |
| MIR31 | 9 | microRNA 31 [Source:HGNC Symbol;Acc:HGN |
| SNORD39 | 9 | Small nucleolar RNA SNORD55/SNORD39 [Sou |
| AL137022.1 | 9 | histone H3 pseudogene |
| KHSRPP1 | 9 | KH-type splicing regulatory protein pseudoger |
| RN7SL151P | 9 | RNA, 7SL, cytoplasmic 151, pseudogene [Sour |
| MTAP | 9 | methylthioadenosine phosphorylase [Source:I |
| AL359922.1 | 9 | novel transcript |
| TUBB8P1 | 9 | tubulin beta 8 class VIII pseudogene 1 [Source |
| AL359922.2 | 9 | novel transcript, antisense MTAP |
| AL359922.3 | 9 | TEC |
| ERVFRD-3 | 9 | endogenous retrovirus group FRD member 3 [ |
| CDKN2A-DT | 9 | CDKN2A divergent transcript [Source:HGNC Sy |
| CDKN2A | 9 | cyclin dependent kinase inhibitor 2A [Source:I |
| CDKN2B-AS1 | 9 | CDKN2B antisense RNA 1 [Source:HGNC Symb |
| AL449423.1 | 9 | novel transcript, sense intronic CDKN2B-AS1 |
| CDKN2B | 9 | cyclin dependent kinase inhibitor 2B [Source:I |
| UBA52P6 | 9 | ubiquitin A-52 residue ribosomal protein fusic |
| AL354709.1 | 9 | NULL |
| AL354709.2 | 9 | NULL |
| AL157937.1 | 9 | novel transcript |
| DMRTA1 | 9 | DMRT like family A1 [Source:HGNC Symbol;Ac |
| LINC01239 | 9 | long intergenic non-protein coding RNA 1239 |
| AL391117.1 | 9 | novel transcript |
| CLIC4P1 | 9 | chloride intracellular channel 4 pseudogene 1 |
| AC017067.1 | 9 | novel transcript |
| AL357614.1 | 9 | novel transcript |
| AL445623.2 | 9 | novel transcript |
| SUMO2P2 | 9 | SUMO2 pseudogene 2 [Source:HGNC Symbol; |
| AL445623.1 | 9 | novel transcript |
| NOP56P2 | 9 | NOP56 ribonucleoprotein pseudogene 2 [Sour |
| ELAVL2 | 9 | ELAV like RNA binding protein 2 [Source:HGN( |
| AL161628.1 | 9 | novel transcript |
| AL365204.1 | 9 | novel transcript |
| AL365204.2 | 9 | novel transcript |
| AL365204.3 | 9 | novel zinc finger protein pseudogene |
| AL513317.1 | 9 | novel transcript |

| | | |
|---|---|---|
| IZUMO3 | 9 | IZUMO family member 3 [Source:HGNC Symb |
| AL353811.1 | 9 | novel transcript |
| RMRPP5 | 9 | RNA component of mitochondrial RNA proces |
| RN7SKP120 | 9 | RN7SK pseudogene 120 [Source:HGNC Symbo |
| AL353730.1 | 9 | novel transcript |
| TUSC1 | 9 | tumor suppressor candidate 1 [Source:HGNC S |
| LINC01241 | 9 | long intergenic non-protein coding RNA 1241 |
| FAM71BP1 | 9 | family with sequence similarity 71 member B |
| AL353753.1 | 9 | putative deoxyuridine 5'-triphosphate nucleot |
| AL442639.1 | 9 | tec |
| AL451137.2 | 9 | novel transcript |
| AL451137.1 | 9 | NULL |
| AL356133.2 | 9 | novel transcript |
| CAAP1 | 9 | caspase activity and apoptosis inhibitor 1 [Sou |
| AL356133.1 | 9 | H3 histone, family 3B (H3.3B) (H3F3B) pseudo |
| RN7SL100P | 9 | RNA, 7SL, cytoplasmic 100, pseudogene [Sour |
| PLAA | 9 | phospholipase A2 activating protein [Source:H |
| IFT74 | 9 | intraflagellar transport 74 [Source:HGNC Syml |
| IFT74-AS1 | 9 | IFT74 antisense RNA 1 [Source:HGNC Symbol; |
| LRRC19 | 9 | leucine rich repeat containing 19 [Source:HGN |
| AL355432.1 | 9 | novel transcript |
| TEK | 9 | TEK receptor tyrosine kinase [Source:HGNC Sy |
| RNA5SP280 | 9 | RNA, 5S ribosomal pseudogene 280 [Source:H |
| AL355433.1 | 9 | ribosomal protein L36a (RPL36A) pseudogene |
| LINC00032 | 9 | long intergenic non-protein coding RNA 32 [So |
| EQTN | 9 | equatorin [Source:HGNC Symbol;Acc:HGNC:1 |
| MOB3B | 9 | MOB kinase activator 3B [Source:HGNC Symb |
| AL163192.1 | 9 | novel transcript |
| AL451123.2 | 9 | novel transcript, sense intronic to MOB3B |
| IFNK | 9 | interferon kappa [Source:HGNC Symbol;Acc:H |
| AL451123.1 | 9 | novel transcript |
| C9orf72 | 9 | C9orf72-SMCR8 complex subunit [Source:HGN |
| CTAGE12P | 9 | CTAGE family member 12, pseudogene [Sourc |
| AL360014.1 | 9 | novel transcript |
| AL353746.1 | 9 | novel transcript, overlapping LINGO2 |
| LINGO2 | 9 | leucine rich repeat and Ig domain containing 2 |
| AL451124.1 | 9 | novel pseudogene |
| AL445526.1 | 9 | novel transcript, antisense to LINGO2 |
| KCTD10P1 | 9 | potassium channel tetramerization domain co |
| MIR876 | 9 | microRNA 876 [Source:HGNC Symbol;Acc:HGN |
| MIR873 | 9 | microRNA 873 [Source:HGNC Symbol;Acc:HGN |
| AL162388.2 | 9 | novel transcript |
| AL162388.1 | 9 | NHP2 non-histone chromosome protein 2-like |
| AL353684.1 | 9 | pyruvate dehydrogenase kinase, isozyme 1 (P |
| AL354676.1 | 9 | pseudogene similar to part of non-metastatic |
| ME2P1 | 9 | malic enzyme 2 pseudogene 1 [Source:HGNC S |
| LINC01242 | 9 | long intergenic non-protein coding RNA 1242 |
| SLC4A1APP1 | 9 | solute carrier family 4 member 1 adaptor prot |
| C2orf27AP2 | 9 | chromosome 2 open reading frame 27A pseud |
| RBMXP2 | 9 | RBMX pseudogene 2 [Source:HGNC Symbol;A |

| | | |
|---|---|---|
| KRT18P66 | 9 | keratin 18 pseudogene 66 [Source:HGNC Sym |
| KRT18P36 | 9 | keratin 18 pseudogene 36 [Source:HGNC Sym |
| RPS26P2 | 9 | ribosomal protein S26 pseudogene 2 [Source: |
| FTLP4 | 9 | ferritin light chain pseudogene 4 [Source:HGN |
| HSPA8P17 | 9 | heat shock protein family A (Hsp70) member |
| AL354854.1 | 9 | pseudogene similar to part of a novel protein |
| SLC25A6P2 | 9 | solute carrier family 25 member 6 pseudogen |
| LINC01243 | 9 | long intergenic non-protein coding RNA 1243 |
| MTATP6P30 | 9 | MT-ATP6 pseudogene 30 [Source:HGNC Symb |
| MTCO3P30 | 9 | MT-CO3 pseudogene 30 [Source:HGNC Symbo |
| HMGB3P23 | 9 | high mobility group box 3 pseudogene 23 [Sou |
| RNA5SP281 | 9 | RNA, 5S ribosomal pseudogene 281 [Source:H |
| SLC25A5P8 | 9 | solute carrier family 25 member 5 pseudogen |
| ACO1 | 9 | aconitase 1 [Source:HGNC Symbol;Acc:HGNC: |
| DDX58 | 9 | DExD/H-box helicase 58 [Source:HGNC Symbo |
| TOPORS | 9 | TOP1 binding arginine/serine rich protein [Sou |
| SMIM27 | 9 | small integral membrane protein 27 [Source:H |
| NDUFB6 | 9 | NADH:ubiquinone oxidoreductase subunit B6 |
| DFFBP1 | 9 | DNA fragmentation factor subunit beta pseud |
| TAF1L | 9 | TATA-box binding protein associated factor 1 |
| AL589642.1 | 9 | novel transcript |
| AL589642.2 | 9 | novel transcript |
| SNRPCP1 | 9 | small nuclear ribonucleoprotein polypeptide C |
| AL157884.1 | 9 | ribosomal protein S11 (RPS11) pseudogene |
| TMEM215 | 9 | transmembrane protein 215 [Source:HGNC Sy |
| Y_RNA | 9 | Y RNA [Source:RFAM;Acc:RF00019] |
| AL157884.3 | 9 | pseudogene similar to part of membrane guar |
| AL157884.2 | 9 | novel transcript |
| APTX | 9 | aprataxin [Source:HGNC Symbol;Acc:HGNC:15 |
| BOLA3P4 | 9 | bolA family member 3 pseudogene 4 [Source: |
| ASS1P12 | 9 | argininosuccinate synthetase 1 pseudogene 1 |
| TCEA1P4 | 9 | transcription elongation factor A1 pseudogene |
| LAGE3P1 | 9 | L antigen family member 3 pseudogene 1 [Sou |
| DNAJA1 | 9 | DnaJ heat shock protein family (Hsp40) memb |
| SMU1 | 9 | SMU1 DNA replication regulator and spliceoso |
| B4GALT1 | 9 | beta-1,4-galactosyltransferase 1 [Source:HGN |
| B4GALT1-AS1 | 9 | B4GALT1 antisense RNA 1 [Source:HGNC Symb |
| RNU4ATAC15P | 9 | RNA, U4atac small nuclear 15, pseudogene [So |
| SPINK4 | 9 | serine peptidase inhibitor, Kazal type 4 [Sourc |
| BAG1 | 9 | BCL2 associated athanogene 1 [Source:HGNC |
| CHMP5 | 9 | charged multivesicular body protein 5 [Source |
| NFX1 | 9 | nuclear transcription factor, X-box binding 1 [S |
| Y_RNA | 9 | Y RNA [Source:RFAM;Acc:RF00019] |
| AQP7 | 9 | aquaporin 7 [Source:HGNC Symbol;Acc:HGNC |
| AL356218.2 | 9 | novel transcript |
| AQP3 | 9 | aquaporin 3 (Gill blood group) [Source:HGNC S |
| AL356218.1 | 9 | heat-responsive protein 12 (HRSP12) pseudog |
| NOL6 | 9 | nucleolar protein 6 [Source:HGNC Symbol;Acc |
| MIR6851 | 9 | microRNA 6851 [Source:HGNC Symbol;Acc:HG |
| SUGT1P1 | 9 | SUGT1 pseudogene 1 [Source:HGNC Symbol;A |

| | | |
|---|---|---|
| AL139008.2 | 9 | novel transcript |
| ANKRD18B | 9 | ankyrin repeat domain 18B [Source:HGNC Syn |
| AL139008.1 | 9 | novel pseudogene |
| SNX18P7 | 9 | sorting nexin 18 pseudogene 7 [Source:HGNC |
| CYP4F26P | 9 | cytochrome P450 family 4 subfamily F membe |
| PRSS3P4 | 9 | PRSS3 pseudogene 4 [Source:HGNC Symbol;A |
| TRBV20OR9-2 | 9 | T cell receptor beta variable 20/OR9-2 (non-fu |
| ANXA2P2 | 9 | annexin A2 pseudogene 2 [Source:HGNC Syml |
| TRBV21OR9-2 | 9 | T cell receptor beta variable 21/OR9-2 (pseud |
| TRBV22OR9-2 | 9 | T cell receptor beta variable 22/OR9-2 (pseud |
| TRBV23OR9-2 | 9 | T cell receptor beta variable 23/OR9-2 (pseud |
| TRBV24OR9-2 | 9 | T cell receptor beta variable 24/OR9-2 (pseud |
| TRBV25OR9-2 | 9 | T cell receptor beta variable 25/OR9-2 (pseud |
| PTENP1 | 9 | phosphatase and tensin homolog pseudogene |
| PTENP1-AS | 9 | PTENP1 antisense RNA [Source:HGNC Symbol, |
| TRBV26OR9-2 | 9 | T cell receptor beta variable 26/OR9-2 (pseud |
| AL356489.2 | 9 | novel transcript |
| AL356489.3 | 9 | novel transcript |
| AL356489.1 | 9 | novel transcript |
| AL356489.4 | 9 | novel transcript |
| LINC01251 | 9 | long intergenic non-protein coding RNA 1251 |
| PRSS3 | 9 | serine protease 3 [Source:HGNC Symbol;Acc:H |
| UBE2R2-AS1 | 9 | UBE2R2 antisense RNA 1 [Source:HGNC Symb |
| TRBV29OR9-2 | 9 | T cell receptor beta variable 29/OR9-2 (non-fu |
| UBE2R2 | 9 | ubiquitin conjugating enzyme E2 R2 [Source:H |
| Y_RNA | 9 | Y RNA [Source:RFAM;Acc:RF00019] |
| RNU4ATAC11P | 9 | RNA, U4atac small nuclear 11, pseudogene [S |
| UBAP2 | 9 | ubiquitin associated protein 2 [Source:HGNC S |
| SNORD121B | 9 | small nucleolar RNA, C/D box 121B [Source:H |
| SNORD121A | 9 | small nucleolar RNA, C/D box 121A [Source:H |
| OSTCP8 | 9 | oligosaccharyltransferase complex subunit ps |
| RN7SKP114 | 9 | RN7SK pseudogene 114 [Source:HGNC Symbo |
| AL354989.1 | 9 | novel transcript |
| DCAF12 | 9 | DDB1 and CUL4 associated factor 12 [Source:I |
| TUBB4BP2 | 9 | tubulin beta 4B class IVb pseudogene 2 [Sourc |
| AL354989.2 | 9 | IMP (inosine monophosphate) dehydrogenase |
| UBAP1 | 9 | ubiquitin associated protein 1 [Source:HGNC S |
| AL353662.1 | 9 | ribosomal protein L35A (RPL35A) pseudogene |
| AL353662.3 | 9 | NULL |
| RNA5SP282 | 9 | RNA, 5S ribosomal pseudogene 282 [Source:H |
| AL353662.2 | 9 | ribosomal protein S8 (RPS8) pseudogene |
| KIF24 | 9 | kinesin family member 24 [Source:HGNC Syml |
| RNU2-50P | 9 | RNA, U2 small nuclear 50, pseudogene [Sourc |
| SERPINH1P1 | 9 | serpin family H member 1 pseudogene 1 [Sou |
| NUDT2 | 9 | nudix hydrolase 2 [Source:HGNC Symbol;Acc:I |
| MYORG | 9 | myogenesis regulating glycosidase (putative) [ |
| C9orf24 | 9 | chromosome 9 open reading frame 24 [Sourc |
| Y_RNA | 9 | Y RNA [Source:RFAM;Acc:RF00019] |
| FAM219A | 9 | family with sequence similarity 219 member A |
| DNAI1 | 9 | dynein axonemal intermediate chain 1 [Sourc |

| | | |
|---|---|---|
| RN7SKP24 | 9 | RN7SK pseudogene 24 [Source:HGNC Symbol; |
| ENHO | 9 | energy homeostasis associated [Source:HGNC |
| AL160270.1 | 9 | novel transcript |
| AL160270.2 | 9 | novel pseudogene |
| CNTFR | 9 | ciliary neurotrophic factor receptor [Source:H |
| CNTFR-AS1 | 9 | CNTFR antisense RNA 1 [Source:HGNC Symbo |
| RPP25L | 9 | ribonuclease P/MRP subunit p25 like [Source: |
| DCTN3 | 9 | dynactin subunit 3 [Source:HGNC Symbol;Acc: |
| ARID3C | 9 | AT-rich interaction domain 3C [Source:HGNC S |
| SIGMAR1 | 9 | sigma non-opioid intracellular receptor 1 [Sou |
| GALT | 9 | galactose-1-phosphate uridylyltransferase [So |
| AL162231.3 | 9 | novel protein (GALT-IL11RA readthrough) |
| IL11RA | 9 | interleukin 11 receptor subunit alpha [Source: |
| CCL27 | 9 | C-C motif chemokine ligand 27 [Source:HGNC |
| AL162231.4 | 9 | novel transcript (novel gene-CCL27 readthrou |
| AL162231.1 | 9 | uncharacterized LOC730098 [Source:NCBI ger |
| AL162231.2 | 9 | novel transcript |
| CCL19 | 9 | C-C motif chemokine ligand 19 [Source:HGNC |
| CCL21 | 9 | C-C motif chemokine ligand 21 [Source:HGNC |
| FAM205A | 9 | family with sequence similarity 205 member A |
| AL589645.1 | 9 | novel transcript, antisense to FAM205C |
| FAM205BP | 9 | family with sequence similarity 205 member E |
| FAM205C | 9 | family with sequence similarity 205 member C |
| GLULP4 | 9 | glutamate-ammonia ligase pseudogene 4 [Sou |
| YWHAZP6 | 9 | tyrosine 3-monooxygenase/tryptophan 5-mor |
| PHF24 | 9 | PHD finger protein 24 [Source:HGNC Symbol;A |
| DNAJB5-DT | 9 | DNAJB5 divergent transcript [Source:HGNC Sy |
| DNAJB5 | 9 | DnaJ heat shock protein family (Hsp40) memb |
| AL355377.1 | 9 | SPATA31 subfamily D, member 1 (SPATA31D1 |
| AL355377.2 | 9 | tec |
| SYF2P2 | 9 | SYF2 pre-mRNA splicing factor pseudogene 2 [ |
| AL355377.3 | 9 | novel transcript |
| AL355377.4 | 9 | novel transcript |
| AL353795.3 | 9 | TEC |
| C9orf131 | 9 | chromosome 9 open reading frame 131 [Sour |
| RN7SL338P | 9 | RNA, 7SL, cytoplasmic 338, pseudogene [Sour |
| VCP | 9 | valosin containing protein [Source:HGNC Sym |
| FANCG | 9 | FA complementation group G [Source:HGNC S |
| PIGO | 9 | phosphatidylinositol glycan anchor biosynthes |
| AL353795.2 | 9 | novel transcript |
| STOML2 | 9 | stomatin like 2 [Source:HGNC Symbol;Acc:HGI |
| FAM214B | 9 | family with sequence similarity 214 member E |
| AL353795.1 | 9 | zinc finger protein pseudogene |
| UNC13B | 9 | unc-13 homolog B [Source:HGNC Symbol;Acc: |
| AL160274.1 | 9 | TEC |
| ATP8B5P | 9 | ATPase phospholipid transporting 8B5, pseud |
| ZFAND6P1 | 9 | zinc finger AN1-type containing 6 pseudogene |
| RUSC2 | 9 | RUN and SH3 domain containing 2 [Source:HG |
| AL133476.1 | 9 | novel transcript, antisense to RUSC2 |
| RPL36AP33 | 9 | ribosomal protein L36a pseudogene 33 [Sourc |

| | | |
|---|---|---|
| FAM166B | 9 | family with sequence similarity 166 member |
| RPS29P17 | 9 | ribosomal protein S29 pseudogene 17 [Source |
| TESK1 | 9 | testis associated actin remodelling kinase 1 [S |
| MIR4667 | 9 | microRNA 4667 [Source:HGNC Symbol;Acc:HG |
| CD72 | 9 | CD72 molecule [Source:HGNC Symbol;Acc:HG |
| AL357874.2 | 9 | novel transcript |
| AL357874.1 | 9 | novel transcript |
| SIT1 | 9 | signaling threshold regulating transmembrane |
| RMRP | 9 | RNA component of mitochondrial RNA proces |
| RMRP | 9 | RNA component of mitochondrial RNA proces |
| CCDC107 | 9 | coiled-coil domain containing 107 [Source:HG |
| ARHGEF39 | 9 | Rho guanine nucleotide exchange factor 39 [S |
| RN7SL22P | 9 | RNA, 7SL, cytoplasmic 22, pseudogene [Sourc |
| CA9 | 9 | carbonic anhydrase 9 [Source:HGNC Symbol; |
| TPM2 | 9 | tropomyosin 2 [Source:HGNC Symbol;Acc:HG |
| TLN1 | 9 | talin 1 [Source:HGNC Symbol;Acc:HGNC:1184 |
| MIR6852 | 9 | microRNA 6852 [Source:HGNC Symbol;Acc:HG |
| CREB3 | 9 | cAMP responsive element binding protein 3 [ |
| MIR6853 | 9 | microRNA 6853 [Source:HGNC Symbol;Acc:HG |
| GBA2 | 9 | glucosylceramidase beta 2 [Source:HGNC Sym |
| RGP1 | 9 | RGP1 homolog, RAB6A GEF complex partner 1 |
| MSMP | 9 | microseminoprotein, prostate associated [Sou |
| AL133410.2 | 9 | novel transcript |
| AL133410.1 | 9 | novel transcript |
| NPR2 | 9 | natriuretic peptide receptor 2 [Source:HGNC |
| SPAG8 | 9 | sperm associated antigen 8 [Source:HGNC Sy |
| HINT2 | 9 | histidine triad nucleotide binding protein 2 [S |
| AL133410.3 | 9 | Novel protein |
| TMEM8B | 9 | transmembrane protein 8B [Source:HGNC Sy |
| FAM221B | 9 | family with sequence similarity 221 member |
| OR13E1P | 9 | olfactory receptor family 13 subfamily E mem |
| OR13J1 | 9 | olfactory receptor family 13 subfamily J mem |
| NDUFA5P4 | 9 | NADH:ubiquinone oxidoreductase subunit A5 |
| HRCT1 | 9 | histidine rich carboxyl terminus 1 [Source:HG |
| SPAAR | 9 | small regulatory polypeptide of amino acid re |
| AL135841.1 | 9 | novel transcript, antisense to SPAAR |
| PGAM1P2 | 9 | phosphoglycerate mutase 1 pseudogene 2 [S |
| OR2S2 | 9 | olfactory receptor family 2 subfamily S memb |
| OR2S1P | 9 | olfactory receptor family 2 subfamily S memb |
| YBX1P10 | 9 | Y-box binding protein 1 pseudogene 10 [Sour |
| OR13C6P | 9 | olfactory receptor family 13 subfamily C mem |
| OR13C7 | 9 | olfactory receptor family 13 subfamily C mem |
| OR2AM1P | 9 | olfactory receptor family 2 subfamily AM me |
| RECK | 9 | reversion inducing cysteine rich protein with |
| AL138834.1 | 9 | NULL |
| GLIPR2 | 9 | GLI pathogenesis related 2 [Source:HGNC Sym |
| CCIN | 9 | calicin [Source:HGNC Symbol;Acc:HGNC:1568 |
| CLTA | 9 | clathrin light chain A [Source:HGNC Symbol;A |
| GNE | 9 | glucosamine (UDP-N-acetyl)-2-epimerase/N-a |
| RNU4-53P | 9 | RNA, U4 small nuclear 53, pseudogene [Sourc |

| | | |
|---|---|---|
| HMGB3P24 | 9 | high mobility group box 3 pseudogene 24 [Sou |
| RNF38 | 9 | ring finger protein 38 [Source:HGNC Symbol;A |
| MELK | 9 | maternal embryonic leucine zipper kinase [So |
| AL450267.2 | 9 | novel transcript |
| MIR4475 | 9 | microRNA 4475 [Source:HGNC Symbol;Acc:HG |
| PAX5 | 9 | paired box 5 [Source:HGNC Symbol;Acc:HGNC |
| AL450267.1 | 9 | novel transcript, antisense to PAX5 |
| MIR4540 | 9 | microRNA 4540 [Source:HGNC Symbol;Acc:HG |
| MIR4476 | 9 | microRNA 4476 [Source:HGNC Symbol;Acc:HG |
| AL161781.2 | 9 | novel transcript, antisense to PAX5 |
| AL161781.1 | 9 | ribosomal protein L32 (RPL32) pseudogene |
| LINC01400 | 9 | long intergenic non-protein coding RNA 1400 |
| AL512604.3 | 9 | novel transcript |
| EBLN3P | 9 | endogenous Bornavirus like nucleoprotein 3, p |
| AL512604.2 | 9 | novel transcript |
| ZCCHC7 | 9 | zinc finger CCHC-type containing 7 [Source:HG |
| Y_RNA | 9 | Y RNA [Source:RFAM;Acc:RF00019] |
| AL512604.1 | 9 | ribosomal protein L21 (RPL21) pseudogene |
| Y_RNA | 9 | Y RNA [Source:RFAM;Acc:RF00019] |
| RNU6-677P | 9 | RNA, U6 small nuclear 677, pseudogene [Sour |
| LINC01627 | 9 | long intergenic non-protein coding RNA 1627 |
| GRHPR | 9 | glyoxylate and hydroxypyruvate reductase [Sc |
| CHCHD4P3 | 9 | coiled-coil-helix-coiled-coil-helix domain conta |
| ZBTB5 | 9 | zinc finger and BTB domain containing 5 [Sour |
| KCTD9P3 | 9 | potassium channel tetramerization domain cc |
| POLR1E | 9 | RNA polymerase I subunit E [Source:HGNC Syr |
| AL158156.1 | 9 | ribosomal protein L21 (RPL21) pseudogene |
| AL513165.1 | 9 | novel transcript |
| FBXO10 | 9 | F-box protein 10 [Source:HGNC Symbol;Acc:H |
| AL513165.2 | 9 | novel transcript |
| TOMM5 | 9 | translocase of outer mitochondrial membrane |
| AL138752.2 | 9 | novel transcript |
| RAB1C | 9 | RAB1C, member RAS oncogene family pseudo |
| FRMPD1 | 9 | FERM and PDZ domain containing 1 [Source:H |
| RN7SKP171 | 9 | RN7SK pseudogene 171 [Source:HGNC Symbo |
| TRMT10B | 9 | tRNA methyltransferase 10B [Source:HGNC Sy |
| EXOSC3 | 9 | exosome component 3 [Source:HGNC Symbol |
| DCAF10 | 9 | DDB1 and CUL4 associated factor 10 [Source:I |
| AL138752.1 | 9 | mitochondrial ribosomal protein S10 (MRPS1( |
| PAICSP1 | 9 | phosphoribosylaminoimidazole carboxylase, p |
| SLC25A51 | 9 | solute carrier family 25 member 51 [Source:H |
| TMX2P1 | 9 | thioredoxin related transmembrane protein 2 |
| SHB | 9 | SH2 domain containing adaptor protein B [Sou |
| RNU7-124P | 9 | RNA, U7 small nuclear 124 pseudogene [Sourc |
| U8 | 9 | U8 small nucleolar RNA [Source:RFAM;Acc:RF( |
| AL135785.1 | 9 | novel transcript |
| ALDH1B1 | 9 | aldehyde dehydrogenase 1 family member B1 |
| IGFBPL1 | 9 | insulin like growth factor binding protein like : |
| AL390726.5 | 9 | STRA6-like (Stra6l) pseudogene |
| ARMC8P1 | 9 | armadillo repeat containing 8 pseudogene 1 [! |

| | | |
|---|---|---|
| TCEA1P3 | 9 | transcription elongation factor A1 pseudogen |
| GAS2L1P1 | 9 | growth arrest specific 2 like 1 pseudogene 1 [S |
| VN1R48P | 9 | vomeronasal 1 receptor 48 pseudogene [Sour |
| AL390726.1 | 9 | family with sequence similarity 220, member |
| FAM220BP | 9 | family with sequence similarity 220 member E |
| AL390726.4 | 9 | cytochrome P450, family 4, subfamily F family |
| FAM95C | 9 | family with sequence similarity 95 member C |
| AL390726.3 | 9 | novel transcript |
| AL390726.2 | 9 | novel transcript |
| SNX18P3 | 9 | sorting nexin 18 pseudogene 3 [Source:HGNC |
| ANKRD18A | 9 | ankyrin repeat domain 18A [Source:HGNC Syr |
| FAM201A | 9 | family with sequence similarity 201 member A |
| YWHABP1 | 9 | tyrosine 3-monooxygenase/tryptophan 5-mor |
| AL591543.1 | 9 | novel transcript |
| RNU6-765P | 9 | RNA, U6 small nuclear 765, pseudogene [Sour |
| FAM240B | 9 | family with sequence similarity 240 member E |
| AL845311.1 | 9 | novel transcript |
| AL953883.1 | 9 | novel transcript |
| VN2R3P | 9 | vomeronasal 2 receptor 3, pseudogene [Sourc |
| CNTNAP3 | 9 | contactin associated protein like 3 [Source:HG |
| RBM17P1 | 9 | RNA binding motif protein 17 pseudogene 1 [S |
| AL353729.2 | 9 | CAS1 domain containing 1 (CASD1) pseudoger |
| AL353729.1 | 9 | TGF beta-inducible nuclear protein 1 (TINP1) p |
| RN7SL640P | 9 | RNA, 7SL, cytoplasmic 640, pseudogene [Sour |
| SPATA31A1 | 9 | SPATA31 subfamily A member 1 [Source:HGN |
| BX005214.3 | 9 | 6-pyruvoyltetrahydropterin synthase (PTS) ps |
| FAM74A1 | 9 | family with sequence similarity 74 member A1 |
| BX005214.1 | 9 | novel transcript |
| BX005214.2 | 9 | novel transcript |
| ZNF658B | 9 | zinc finger protein 658B (pseudogene) [Source |
| SKP1P3 | 9 | S-phase kinase associated protein 1 pseudoge |
| FKBP4P2 | 9 | FKBP prolyl isomerase 4 pseudogene 2 [Sourc |
| SDR42E1P1 | 9 | short chain dehydrogenase/reductase family 4 |
| ATP5F1AP8 | 9 | ATP synthase F1 subunit alpha pseudogene 8 |
| RAB28P1 | 9 | RAB28, member RAS oncogene family pseudo |
| RBPJP5 | 9 | RBPJ pseudogene 5 [Source:HGNC Symbol;Acc |
| GLIDR | 9 | glioblastoma down-regulated RNA [Source:HG |
| BX664615.1 | 9 | novel pseudogene |
| FGF7P3 | 9 | fibroblast growth factor 7 pseudogene 3 [Sour |
| BX664615.2 | 9 | Uncharacterized protein FLJ76381 [Source:Ur |
| RN7SL763P | 9 | RNA, 7SL, cytoplasmic 763, pseudogene [Sour |
| SNORA70 | 9 | Small nucleolar RNA SNORA70 [Source:RFAM; |
| CR769767.1 | 9 | pseudogene similar to part of ribosomal prote |
| AL772307.1 | 9 | novel transcript |
| RPL7AP49 | 9 | ribosomal protein L7a pseudogene 49 [Source |
| AL773545.3 | 9 | novel pseudogene |
| C2orf27AP3 | 9 | chromosome 2 open reading frame 27A pseud |
| AL773545.1 | 9 | novel transcript |
| AL773545.2 | 9 | novel pseudogene |
| AL773545.4 | 9 | adhesion G protein-coupled receptor F5 (ADG |

| | | |
|---|---|---|
| MEP1AP1 | 9 | meprin A subunit alpha pseudogene 1 [Source |
| ANKRD20A2 | 9 | ankyrin repeat domain 20 family member A2 [ |
| BX664727.3 | 9 | novel transcript |
| RNU6-1269P | 9 | RNA, U6 small nuclear 1269, pseudogene [Sou |
| BX664727.2 | 9 | melanoma antigen (LOC51152) pseudogene |
| SNX18P8 | 9 | sorting nexin 18 pseudogene 8 [Source:HGNC |
| FAM95B1 | 9 | family with sequence similarity 95 member B1 |
| BX664727.1 | 9 | zinc finger pseudogene |
| CNN2P2 | 9 | calponin 2 pseudogene 2 [Source:HGNC Symb |
| GXYLT1P3 | 9 | glucoside xylosyltransferase 1 pseudogene 3 [ |
| AL591471.1 | 9 | calponin 2 (CNN2) pseudogene |
| AQP7P5 | 9 | aquaporin 7 pseudogene 5 [Source:HGNC Sym |
| FP885919.1 | 9 | aquaporin 7 (AQP7) pseudogene |
| AMYH02020865.1 | 9 | methylenetetrahydrofolate dehydrogenase (N |
| BMS1P14 | 9 | BMS1 pseudogene 14 [Source:HGNC Symbol;A |
| FP325318.1 | 9 | novel transcript |
| IGKV1OR9-2 | 9 | immunoglobulin kappa variable 1/OR9-2 (pseu |
| AL353626.3 | 9 | ankyrin repeat domain 30B (ANKRD30B) pseu |
| RNU6-156P | 9 | RNA, U6 small nuclear 156, pseudogene [Sour |
| AL353626.1 | 9 | novel transcript |
| AL353626.2 | 9 | lymphocyte-specific protein 1 (LSP1) pseudog |
| MIR1299 | 9 | microRNA 1299 [Source:HGNC Symbol;Acc:HG |
| FRG1HP | 9 | FSHD region gene 1 family member H, pseudo |
| PGM5P2 | 9 | phosphoglucomutase 5 pseudogene 2 [Source |
| AL353763.2 | 9 | TEC |
| AL353763.1 | 9 | TEC |
| BX255923.1 | 9 | novel transcript |
| BX255923.2 | 9 | novel transcript |
| FOXD4L6 | 9 | forkhead box D4 like 6 [Source:HGNC Symbol; |
| CBWD6 | 9 | COBW domain containing 6 [Source:HGNC Syr |
| MIR4477A | 9 | microRNA 4477a [Source:HGNC Symbol;Acc:H |
| RNA5SP530 | 9 | RNA, 5S ribosomal pseudogene 530 [Source:H |
| AL845472.2 | 9 | TEC |
| AL845472.1 | 9 | novel transcript |
| CDK2AP2P1 | 9 | cyclin dependent kinase 2 associated protein : |
| MYO5BP1 | 9 | myosin VB pseudogene 1 [Source:HGNC Symb |
| AL354718.2 | 9 | novel transcript |
| AL354718.3 | 9 | CMT1A duplicated region transcript 15 (CDRT: |
| AL354718.1 | 9 | ribosomal protein L7a (RPL7A) pseudogene |
| AL591926.3 | 9 | ribosomal protein L10 (RPL10) pseudogene |
| SNORA70 | 9 | Small nucleolar RNA SNORA70 [Source:RFAM; |
| RN7SL565P | 9 | RNA, 7SL, cytoplasmic 565, pseudogene [Sour |
| AL591926.7 | 9 | family with sequence similarity 27 member E3 |
| AL591926.6 | 9 | zinc finger protein 829 (ZNF829) pseudogene |
| AL591926.5 | 9 | novel pseudogene |
| AL591926.2 | 9 | novel transcript |
| FAM242F | 9 | family with sequence similarity 242 member F |
| Y_RNA | 9 | Y RNA [Source:RFAM;Acc:RF00019] |
| AL591926.1 | 9 | family with sequence similarity 27-like (FAM2 |
| AL591926.4 | 9 | family with sequence similarity 27, member D |

| | | |
|---|---|---|
| AL162731.1 | 9 | novel transcript |
| CNTNAP3B | 9 | contactin associated protein like 3B [Source:H |
| RBM17P3 | 9 | RNA binding motif protein 17 pseudogene 3 [S |
| CR788268.1 | 9 | CAS1 domain containing 1 (CASD1) pseudoger |
| RN7SL343P | 9 | RNA, 7SL, cytoplasmic 343, pseudogene [Sour |
| SPATA31A6 | 9 | SPATA31 subfamily A member 6 [Source:HGN |
| AL445584.1 | 9 | 6-pyruvoyltetrahydropterin synthase (PTPS) (I |
| FAM74A7 | 9 | family with sequence similarity 74 member A7 |
| AL445584.3 | 9 | developmental pluripotency associated 3 (DPI |
| AL445584.2 | 9 | novel transcript |
| FKBP4P6 | 9 | FKBP prolyl isomerase 4 pseudogene 6 [Sourc |
| SDR42E1P2 | 9 | short chain dehydrogenase/reductase family 4 |
| ATP5F1AP1 | 9 | ATP synthase F1 subunit alpha pseudogene 1 |
| RAB28P3 | 9 | RAB28, member RAS oncogene family pseudo |
| RBPJP6 | 9 | RBPJ pseudogene 6 [Source:HGNC Symbol;Acc |
| BX088651.4 | 9 | novel transcript |
| BX088651.1 | 9 | novel pseudogene |
| BX088651.2 | 9 | novel transcript |
| FGF7P5 | 9 | fibroblast growth factor 7 pseudogene 5 [Sour |
| BX088651.3 | 9 | novel pseudogene |
| FGF7P4 | 9 | fibroblast growth factor 7 pseudogene 4 [Sour |
| BX664718.1 | 9 | novel transcript |
| BX664718.2 | 9 | contactin associated protein like 3B (CNTNAP3 |
| ADGRF5P2 | 9 | adhesion G protein-coupled receptor F5 pseu |
| MEP1AP4 | 9 | meprin A subunit alpha pseudogene 4 [Source |
| ANKRD20A7P | 9 | ankyrin repeat domain 20 family member A7, |
| CR848007.3 | 9 | poly(ADP-ribose) glycohydrolase (PARG) pseu |
| RNU6-599P | 9 | RNA, U6 small nuclear 599, pseudogene [Sour |
| CR848007.2 | 9 | novel pseudogene |
| SNX18P5 | 9 | sorting nexin 18 pseudogene 5 [Source:HGNC |
| CYP4F60P | 9 | cytochrome P450 family 4 subfamily F membe |
| CR848007.1 | 9 | novel zinc finger pseudogene |
| CNN2P4 | 9 | calponin 2 pseudogene 4 [Source:HGNC Symb |
| AL591441.1 | 9 | glucoside xylosyltransferase 1 pseudogene 5 |
| FP325317.1 | 9 | aquaporin 7 (AQP7) pseudogene |
| FP325317.2 | 9 | methylenetetrahydrofolate dehydrogenase (N |
| ABBA01045074.2 | 9 | mitochondrially encoded NADH dehydrogenas |
| ABBA01045074.1 | 9 | novel tripartite motif-containing protein famil |
| BX088702.1 | 9 | lymphocyte-specific protein 1 (LSP1) pseudoge |
| SPATA31A5 | 9 | SPATA31 subfamily A member 5 [Source:HGN |
| AL590491.1 | 9 | uncharacterized LOC107987066 [Source:NCBI |
| FAM74A6 | 9 | family with sequence similarity 74 member A6 |
| AL590491.2 | 9 | novel transcript |
| AL590491.3 | 9 | developmental pluripotency associated 3 (DPI |
| BX005040.4 | 9 | contactin associated protein-like 3B (CNTNAP3 |
| SPATA31A7 | 9 | SPATA31 subfamily A member 7 [Source:HGN |
| BX005040.3 | 9 | novel transcript, antisense to SPATA31A7 |
| BX005040.2 | 9 | 6-pyruvoyltetrahydropterin synthase (PTS) pse |
| FAM74A4 | 9 | family with sequence similarity 74 member A4 |
| BX005040.1 | 9 | novel transcript |

| | | |
|---|---|---|
| CNTNAP3C | 9 | contactin associated protein like 3C [Source:H |
| RN7SL462P | 9 | RNA, 7SL, cytoplasmic 462, pseudogene [Sour |
| AL935212.3 | 9 | novel zinc finger protein pseudogene |
| FAM242D | 9 | family with sequence similarity 242 member D |
| Y_RNA | 9 | Y RNA [Source:RFAM;Acc:RF00019] |
| AL935212.1 | 9 | novel transcript |
| AL935212.2 | 9 | novel pseudogene |
| RN7SL722P | 9 | RNA, 7SL, cytoplasmic 722, pseudogene [Sour |
| FAM27C | 9 | family with sequence similarity 27 member C |
| AL391987.2 | 9 | NULL |
| AL391987.1 | 9 | NULL |
| AL391987.4 | 9 | novel transcript |
| AL391987.3 | 9 | novel transcript |
| Y_RNA | 9 | Y RNA [Source:RFAM;Acc:RF00019] |
| FAM242E | 9 | family with sequence similarity 242 member E |
| CR769775.1 | 9 | zinc finger protein pseudogene |
| CR769775.2 | 9 | TEC |
| AL590399.4 | 9 | pseudogene similar to part of ribosomal prote |
| AL590399.3 | 9 | novel transcript |
| AL590399.6 | 9 | pseudogene similar to part of ribosomal prote |
| SNORA70 | 9 | Small nucleolar RNA SNORA70 [Source:RFAM; |
| RN7SL544P | 9 | RNA, 7SL, cytoplasmic 544, pseudogene [Sour |
| AL590399.1 | 9 | novel transcript |
| FGF7P6 | 9 | fibroblast growth factor 7 pseudogene 6 [Sour |
| AL590399.5 | 9 | novel transcript |
| AL590399.2 | 9 | novel pseudogene |
| BX664725.1 | 9 | novel pseudogene |
| LINC01189 | 9 | long intergenic non-protein coding RNA 1189 |
| FGF7P7 | 9 | fibroblast growth factor 7 pseudogene 7 [Sour |
| BX005266.2 | 9 | novel transcript |
| BX005266.1 | 9 | novel pseudogene |
| CNTNAP3P5 | 9 | CNTNAP3 pseudogene 5 [Source:HGNC Symbo |
| RBPJP7 | 9 | RBPJ pseudogene 7 [Source:HGNC Symbol;Acc |
| RAB28P2 | 9 | RAB28, member RAS oncogene family pseudo |
| BX005266.3 | 9 | G protein-coupled receptor, family C, group 5, |
| ATP5F1AP7 | 9 | ATP synthase F1 subunit alpha pseudogene 7 |
| SDR42E1P3 | 9 | short chain dehydrogenase/reductase family 4 |
| FKBP4P7 | 9 | FKBP prolyl isomerase 4 pseudogene 7 [Sourc |
| LINC01410 | 9 | long intergenic non-protein coding RNA 1410 |
| AL512625.3 | 9 | novel transcript |
| RNA5SP283 | 9 | RNA, 5S ribosomal pseudogene 283 [Source:H |
| AL512625.2 | 9 | novel transcript |
| PTGER4P2 | 9 | prostaglandin E receptor 4 pseudogene 2 [Sou |
| CDK2AP2P2 | 9 | cyclin dependent kinase 2 associated protein 2 |
| LERFS | 9 | lncRNA negative regulator of fibroblast-like sy |
| MYO5BP2 | 9 | myosin VB pseudogene 2 [Source:HGNC Symb |
| ADGRF5P1 | 9 | adhesion G protein-coupled receptor F5 pseu |
| FGF7P8 | 9 | fibroblast growth factor 7 pseudogene 8 [Sour |
| AL512625.1 | 9 | novel transcript |
| CNTNAP3P1 | 9 | CNTNAP3 pseudogene 1 [Source:HGNC Symbo |

| | | |
|---|---|---|
| AL591379.2 | 9 | pseudogene similar to part of tripartite motif- |
| BMS1P9 | 9 | BMS1 pseudogene 9 [Source:HGNC Symbol;Ac |
| AL591379.1 | 9 | methylenetetrahydrofolate dehydrogenase (N |
| AQP7P4 | 9 | aquaporin 7 pseudogene 4 [Source:HGNC Sym |
| BMS1P10 | 9 | BMS1 pseudogene 10 [Source:HGNC Symbol;A |
| AQP7P1 | 9 | aquaporin 7 pseudogene 1 [Source:HGNC Sym |
| AL845321.1 | 9 | methylenetetrahydrofolate dehydrogenase (N |
| AL591438.1 | 9 | ribosomal protein L7a (RPL7A) pseudogene |
| AL591438.2 | 9 | novel pseudogene |
| AL772155.3 | 9 | novel pseudogene |
| AL772155.1 | 9 | novel transcript |
| AL772155.2 | 9 | novel pseudogene |
| MYO5BP3 | 9 | myosin VB pseudogene 3 [Source:HGNC Symb |
| CDK2AP2P3 | 9 | cyclin dependent kinase 2 associated protein 2 |
| PTGER4P3 | 9 | prostaglandin E receptor 4 pseudogene 3 [Sou |
| CR786580.1 | 9 | novel transcript |
| RNA5SP284 | 9 | RNA, 5S ribosomal pseudogene 284 [Source:H |
| DUX4L50 | 9 | double homeobox 4 like 50 (pseudogene) [Sou |
| MIR4477B | 9 | microRNA 4477b [Source:HGNC Symbol;Acc:H |
| FRG1JP | 9 | FSHD region gene 1 family member J, pseudog |
| AL163540.1 | 9 | COBW domain containing protein pseudogene |
| U6 | 9 | U6 spliceosomal RNA [Source:RFAM;Acc:RF00 |
| CR769776.3 | 9 | Novel transcript |
| ANKRD20A4 | 9 | ankyrin repeat domain 20 family member A4 [ |
| RNU6-1193P | 9 | RNA, U6 small nuclear 1193, pseudogene [Sou |
| SNX18P9 | 9 | sorting nexin 18 pseudogene 9 [Source:HGNC |
| CR769776.2 | 9 | novel pseudogene |
| CYP4F25P | 9 | cytochrome P450 family 4 subfamily F membe |
| CR769776.1 | 9 | novel zinc finger pseudogene |
| CNN2P3 | 9 | calponin 2 pseudogene 3 [Source:HGNC Symb |
| GXYLT1P6 | 9 | glucoside xylosyltransferase 1 pseudogene 6 [ |
| AL773524.1 | 9 | calponin 2 (CNN2) pseudogene |
| AQP7P2 | 9 | aquaporin 7 pseudogene 2 [Source:HGNC Sym |
| AL445665.1 | 9 | methylenetetrahydrofolate dehydrogenase (N |
| BMS1P11 | 9 | BMS1 pseudogene 11 [Source:HGNC Symbol;A |
| AL359955.2 | 9 | pseudogene similar to tripartite motif-contain |
| IGKV1OR-2 | 9 | immunoglobulin kappa variable 1/OR-2 (pseu |
| BNIP3P4 | 9 | BCL2 interacting protein 3 pseudogene 4 [Sou |
| AL359955.1 | 9 | zinc finger pseudogene |
| AC125634.1 | 9 | novel ankyrin repeat domain protein pseudog |
| RNU6-1293P | 9 | RNA, U6 small nuclear 1293, pseudogene [Sou |
| FO082814.1 | 9 | methylenetetrahydrofolate dehydrogenase (N |
| BMS1P12 | 9 | BMS1 pseudogene 12 [Source:HGNC Symbol;A |
| AL512605.1 | 9 | tripartite motif-containing protein pseudogen |
| AL512605.2 | 9 | NADH dehydrogenase pseudogene |
| IGKV1OR9-1 | 9 | immunoglobulin kappa variable 1/OR9-1 (pseu |
| FOXD4L5 | 9 | forkhead box D4 like 5 [Source:HGNC Symbol; |
| CBWD4P | 9 | COBW domain containing 4 pseudogene [Sour |
| FP326651.1 | 9 | novel transcript |
| Y_RNA | 9 | Y RNA [Source:RFAM;Acc:RF00019] |

| | | |
|---|---|---|
| FRG1KP | 9 | FSHD region gene 1 family member K, pseudo |
| CBWD5 | 9 | COBW domain containing 5 [Source:HGNC Syr |
| FOXD4L4 | 9 | forkhead box D4 like 4 [Source:HGNC Symbol; |
| IGKV1OR-3 | 9 | immunoglobulin kappa variable 1/OR-3 (pseu |
| BX284632.2 | 9 | pseudogene similar to part of tripartite motif- |
| BX284632.1 | 9 | novel pseudogene |
| BMS1P13 | 9 | BMS1 pseudogene 13 [Source:HGNC Symbol;/ |
| AC129778.1 | 9 | methylenetetrahydrofolate dehydrogenase (N |
| AQP7P3 | 9 | aquaporin 7 pseudogene 3 [Source:HGNC Sym |
| AL136317.2 | 9 | novel transcript |
| AL136317.1 | 9 | calponin 2 (CNN2) pseudogene |
| GXYLT1P4 | 9 | glucoside xylosyltransferase 1 pseudogene 4 [ |
| AL136317.3 | 9 | calponin 2 (CNN2) pseudogene |
| AL669942.2 | 9 | zinc finger protein 114 (ZNF114) pseudogene |
| AL669942.1 | 9 | novel cytochrome P450 family pseudogene |
| AL513478.4 | 9 | novel transcript |
| SNX18P4 | 9 | sorting nexin 18 pseudogene 4 [Source:HGNC |
| AL513478.1 | 9 | novel pseudogene |
| ANKRD20A3 | 9 | ankyrin repeat domain 20 family member A3 [ |
| RNU6-538P | 9 | RNA, U6 small nuclear 538, pseudogene [Sour |
| AL513478.2 | 9 | meprin A subunit alpha (MEP1A) pseudogene |
| AL513478.3 | 9 | adhesion G protein-coupled receptor F5 (ADG |
| AL591479.1 | 9 | CDRT15 protein pseudogene |
| RBPJP2 | 9 | RBPJ pseudogene 2 [Source:HGNC Symbol;Acc |
| RAB28P4 | 9 | RAB28, member RAS oncogene family pseudo |
| ATP5F1AP10 | 9 | ATP synthase F1 subunit alpha pseudogene 10 |
| SDR42E1P4 | 9 | short chain dehydrogenase/reductase family 4 |
| FKBP4P8 | 9 | FKBP prolyl isomerase 4 pseudogene 8 [Sourc |
| ZNF658 | 9 | zinc finger protein 658 [Source:HGNC Symbol; |
| BX664608.1 | 9 | S-phase kinase-associated protein 1 (SKP1) ps |
| AL353770.4 | 9 | novel transcript |
| AL353770.2 | 9 | developmental pluripotency associated 3 (DPI |
| FAM74A3 | 9 | family with sequence similarity 74 member A3 |
| AL353770.3 | 9 | novel transcript |
| AL353770.1 | 9 | 6-pyruvoyltetrahydropterin synthase (PTS-P1) |
| SPATA31A3 | 9 | SPATA31 subfamily A member 3 [Source:HGN |
| CNTNAP3P2 | 9 | CNTNAP3 pseudogene 2 [Source:HGNC Symbo |
| RN7SL422P | 9 | RNA, 7SL, cytoplasmic 422, pseudogene [Sour |
| AL162233.1 | 9 | CAS1 domain containing 1 (CASD1) pseudoger |
| RBM17P2 | 9 | RNA binding motif protein 17 pseudogene 2 [S |
| BX005195.1 | 9 | calcium-sensing receptor (CASR) pseudogene |
| BX664730.1 | 9 | novel transcript |
| Y_RNA | 9 | Y RNA [Source:RFAM;Acc:RF00019] |
| FAM27E3 | 9 | family with sequence similarity 27 member E3 |
| AL627230.3 | 9 | NULL |
| FAM27B | 9 | family with sequence similarity 27 member B |
| AL627230.1 | 9 | NULL |
| RN7SL787P | 9 | RNA, 7SL, cytoplasmic 787, pseudogene [Sour |
| SNORA70 | 9 | Small nucleolar RNA SNORA70 [Source:RFAM; |
| AL627230.2 | 9 | ribosomal protein L10 (RPL10) pseudogene |

| | | |
|---|---|---|
| ANKRD20A1 | 9 | ankyrin repeat domain 20 family member A1 [ |
| RNU6-368P | 9 | RNA, U6 small nuclear 368, pseudogene [Sour |
| AL353608.2 | 9 | novel transcript |
| CBWD3 | 9 | COBW domain containing 3 [Source:HGNC Syr |
| AL353608.4 | 9 | TEC |
| FOXD4L3 | 9 | forkhead box D4 like 3 [Source:HGNC Symbol; |
| AL353608.3 | 9 | novel transcript |
| AL353608.1 | 9 | novel transcript |
| PGM5 | 9 | phosphoglucomutase 5 [Source:HGNC Symbo |
| PGM5-AS1 | 9 | PGM5 antisense RNA 1 [Source:HGNC Symbol |
| AL161457.2 | 9 | novel transcript, antisense to PGM5 |
| AL161457.1 | 9 | novel transcript |
| TMEM252 | 9 | transmembrane protein 252 [Source:HGNC Sy |
| TMEM252-DT | 9 | TMEM252 divergent transcript [Source:HGNC |
| LINC01506 | 9 | long intergenic non-protein coding RNA 1506 |
| AL353616.1 | 9 | dermatan sulfate epimerase (DSE) pseudogen |
| PIP5K1B | 9 | phosphatidylinositol-4-phosphate 5-kinase tyı |
| FAM122A | 9 | family with sequence similarity 122A [Source: |
| AL354794.2 | 9 | adaptor-related protein complex 3, mu 2 subu |
| AL354794.1 | 9 | novel transcript |
| RNU6-820P | 9 | RNA, U6 small nuclear 820, pseudogene [Sour |
| PRKACG | 9 | protein kinase cAMP-activated catalytic subur |
| FXN | 9 | frataxin [Source:HGNC Symbol;Acc:HGNC:395 |
| AL358113.1 | 9 | novel protein |
| AL162730.1 | 9 | adenylate kinase 3 (AK3) pseudogene |
| TJP2 | 9 | tight junction protein 2 [Source:HGNC Symbol |
| BANCR | 9 | BRAF-activated non-protein coding RNA [Sour |
| FAM189A2 | 9 | family with sequence similarity 189 member A |
| APBA1 | 9 | amyloid beta precursor protein binding family |
| AL355140.1 | 9 | novel transcript |
| AL353693.1 | 9 | novel transcript |
| AL162412.1 | 9 | novel transcript |
| PTAR1 | 9 | protein prenyltransferase alpha subunit repea |
| C9orf135-DT | 9 | C9orf135 divergent transcript [Source:HGNC S |
| C9orf135 | 9 | chromosome 9 open reading frame 135 [Sour |
| RN7SL570P | 9 | RNA, 7SL, cytoplasmic 570, pseudogene [Sour |
| MAMDC2-AS1 | 9 | MAMDC2 antisense RNA 1 [Source:HGNC Sym |
| MAMDC2 | 9 | MAM domain containing 2 [Source:HGNC Sym |
| RNU2-5P | 9 | RNA, U2 small nuclear 5, pseudogene [Source |
| SMC5-AS1 | 9 | SMC5 antisense RNA 1 (head to head) [Source |
| RPL24P8 | 9 | ribosomal protein L24 pseudogene 8 [Source: |
| SMC5 | 9 | structural maintenance of chromosomes 5 [Sc |
| AL162390.1 | 9 | novel transcript |
| Y_RNA | 9 | Y RNA [Source:RFAM;Acc:RF00019] |
| KLF9 | 9 | Kruppel like factor 9 [Source:HGNC Symbol;Ac |
| TRPM3 | 9 | transient receptor potential cation channel su |
| RN7SL726P | 9 | RNA, 7SL, cytoplasmic 726, pseudogene [Sour |
| AL159990.2 | 9 | novel transcript, antisense TRPM3 |
| AL159990.1 | 9 | novel transcript, antisense TRPM3 |
| MIR204 | 9 | microRNA 204 [Source:HGNC Symbol;Acc:HGI |

| | | |
|---|---|---|
| PIGUP1 | 9 | phosphatidylinositol glycan anchor biosynthes |
| RPL35AP21 | 9 | ribosomal protein L35a pseudogene 21 [Sourc |
| CEMIP2 | 9 | cell migration inducing hyaluronidase 2 [Sourc |
| AL671309.1 | 9 | glucosamine-6-phosphate deaminase 2 (GNPD |
| ABHD17B | 9 | abhydrolase domain containing 17B [Source:H |
| Y_RNA | 9 | Y RNA [Source:RFAM;Acc:RF00019] |
| C9orf85 | 9 | chromosome 9 open reading frame 85 [Sourc |
| HSPB1P1 | 9 | heat shock protein family B (small) member 1 |
| C9orf57 | 9 | chromosome 9 open reading frame 57 [Sourc |
| AL583829.1 | 9 | novel transcript, antisense to C9orf57 |
| BTF3P4 | 9 | basic transcription factor 3 pseudogene 4 [Sou |
| GDA | 9 | guanine deaminase [Source:HGNC Symbol;Acc |
| AL135924.2 | 9 | novel transcript |
| AL135924.1 | 9 | non-metastatic cells 2, protein (NM23B) expre |
| RNA5SP285 | 9 | RNA, 5S ribosomal pseudogene 285 [Source:H |
| LINC01504 | 9 | long intergenic non-protein coding RNA 1504 |
| ZFAND5 | 9 | zinc finger AN1-type containing 5 [Source:HGN |
| AL445646.1 | 9 | pseudogene similar to part of RAB7, member |
| AL596448.1 | 9 | ribosomal protein S12 (RPS12) pseudogene |
| TMC1 | 9 | transmembrane channel like 1 [Source:HGNC |
| RPS20P24 | 9 | ribosomal protein S20 pseudogene 24 [Source |
| RPS27AP15 | 9 | ribosomal protein S27a pseudogene 15 [Sourc |
| LINC01474 | 9 | long intergenic non-protein coding RNA 1474 |
| ALDH1A1 | 9 | aldehyde dehydrogenase 1 family member A1 |
| CYP1D1P | 9 | cytochrome P450 family 1 subfamily D memb |
| ANXA1 | 9 | annexin A1 [Source:HGNC Symbol;Acc:HGNC: |
| DPP3P2 | 9 | dipeptidyl peptidase 3 pseudogene 2 [Source: |
| AL451127.2 | 9 | novel transcript |
| AL451127.1 | 9 | novel transcript |
| AL512594.1 | 9 | pseudogene similar to part of a GTPase-activa |
| RNA5SP286 | 9 | RNA, 5S ribosomal pseudogene 286 [Source:H |
| AL513124.1 | 9 | novel pseudogene |
| AL355674.1 | 9 | novel transcript |
| RORB-AS1 | 9 | RORB antisense RNA 1 [Source:HGNC Symbol; |
| RORB | 9 | RAR related orphan receptor B [Source:HGNC |
| AL137018.1 | 9 | novel transcript, antisense to RORB |
| TRPM6 | 9 | transient receptor potential cation channel su |
| RN7SKP47 | 9 | RN7SK pseudogene 47 [Source:HGNC Symbol; |
| RPSAP75 | 9 | ribosomal protein SA pseudogene 75 [Source: |
| RNU6-445P | 9 | RNA, U6 small nuclear 445, pseudogene [Sour |
| AL158825.1 | 9 | novel pseudogene |
| C9orf40 | 9 | chromosome 9 open reading frame 40 [Sourc |
| CARNMT1-AS1 | 9 | CARNMT1 antisense RNA 1 [Source:HGNC Syn |
| CARNMT1 | 9 | carnosine N-methyltransferase 1 [Source:HGN |
| AL158825.2 | 9 | novel transcript |
| AL158825.3 | 9 | NULL |
| NMRK1 | 9 | nicotinamide riboside kinase 1 [Source:HGNC |
| OSTF1 | 9 | osteoclast stimulating factor 1 [Source:HGNC |
| Y_RNA | 9 | Y RNA [Source:RFAM;Acc:RF00019] |
| RNU6-1228P | 9 | RNA, U6 small nuclear 1228, pseudogene [Sou |

| | | |
|---|---|---|
| AL158073.1 | 9 | cell division cycle associated 7 (CDCA7) pseud |
| AL353780.1 | 9 | novel transcript |
| OTX2P1 | 9 | orthodenticle homeobox 2 pseudogene 1 [Sou |
| PCSK5 | 9 | proprotein convertase subtilisin/kexin type 5 [ |
| RBM22P5 | 9 | RNA binding motif protein 22 pseudogene 5 [S |
| RFK | 9 | riboflavin kinase [Source:HGNC Symbol;Acc:H |
| RPSAP9 | 9 | ribosomal protein SA pseudogene 9 [Source:H |
| GCNT1 | 9 | glucosaminyl (N-acetyl) transferase 1 [Source: |
| AL590664.1 | 9 | H3 histone, family 3A (H3F3A) pseudogene |
| PPIAP87 | 9 | peptidylprolyl isomerase A pseudogene 87 [So |
| PRUNE2 | 9 | prune homolog 2 with BCH domain [Source:H |
| PCA3 | 9 | prostate cancer associated 3 [Source:HGNC Sy |
| AL359314.1 | 9 | NULL |
| AL390239.1 | 9 | NULL |
| AL353637.2 | 9 | novel transcript |
| DNAJB5P1 | 9 | DnaJ heat shock protein family (Hsp40) memb |
| LYPLA2P3 | 9 | LYPLA2 pseudogene 3 [Source:HGNC Symbol; |
| AL353637.1 | 9 | eukaryotic translation initiation factor 3, subu |
| FOXB2 | 9 | forkhead box B2 [Source:HGNC Symbol;Acc:H |
| ATP5MFP3 | 9 | ATP synthase membrane subunit f pseudogen |
| RFC5P1 | 9 | replication factor C 5 pseudogene 1 [Source:H |
| VPS13A-AS1 | 9 | VPS13A antisense RNA 1 [Source:HGNC Symb |
| VPS13A | 9 | vacuolar protein sorting 13 homolog A [Sourc |
| GNA14 | 9 | G protein subunit alpha 14 [Source:HGNC Sym |
| GNA14-AS1 | 9 | GNA14 antisense RNA 1 [Source:HGNC Symbo |
| NUTF2P3 | 9 | nuclear transport factor 2 pseudogene 3 [Sou |
| RNU6-1303P | 9 | RNA, U6 small nuclear 1303, pseudogene [Sou |
| GNAQ | 9 | G protein subunit alpha q [Source:HGNC Symb |
| AL353705.2 | 9 | synaptogyrin 2 (SYNGR2) pseudogene |
| RN7SKP59 | 9 | RN7SK pseudogene 59 [Source:HGNC Symbol; |
| AL353705.3 | 9 | argininosuccinate synthetase (ASS) pseudoger |
| AL353705.1 | 9 | ribosomal protein L21 (RPL21) pseudogene |
| AL353705.4 | 9 | CDC28 protein kinase regulatory subunit 2 (CK |
| CEP78 | 9 | centrosomal protein 78 [Source:HGNC Symbo |
| PSAT1 | 9 | phosphoserine aminotransferase 1 [Source:H |
| AL592221.1 | 9 | CTAGE family, member 5 (CTAGE5) pseudoger |
| MTND2P8 | 9 | MT-ND2 pseudogene 8 [Source:HGNC Symbol |
| KRT18P24 | 9 | keratin 18 pseudogene 24 [Source:HGNC Sym |
| AL512634.1 | 9 | novel transcript |
| CHCHD2P9 | 9 | coiled-coil-helix-coiled-coil-helix domain conta |
| LNCARSR | 9 | lncRNA regulator of Akt signaling associated w |
| TLE4 | 9 | TLE family member 4, transcriptional corepres |
| AL161912.3 | 9 | ankyrin repeat domain 26 (ANKRD26) pseudo |
| NPAP1P6 | 9 | nuclear pore associated protein 1 pseudogene |
| AL161912.1 | 9 | novel transcript |
| AL161912.2 | 9 | novel transcript |
| AL161912.4 | 9 | tec |
| AL161782.1 | 9 | novel transcript |
| LINC01507 | 9 | long intergenic non-protein coding RNA 1507 |
| NPAP1P4 | 9 | nuclear pore associated protein 1 pseudogene |

| | | |
|---|---|---|
| MTCO1P51 | 9 | MT-CO1 pseudogene 51 [Source:HGNC Symbo |
| MTND2P9 | 9 | MT-ND2 pseudogene 9 [Source:HGNC Symbol |
| RPS19P6 | 9 | ribosomal protein S19 pseudogene 6 [Source: |
| AL138749.1 | 9 | putatitive novel transcript |
| RPS20P25 | 9 | ribosomal protein S20 pseudogene 25 [Source |
| AL161727.1 | 9 | novel pseudogene |
| TLE1 | 9 | TLE family member 1, transcriptional corepres |
| AL591368.1 | 9 | novel transcript |
| RNU6-1035P | 9 | RNA, U6 small nuclear 1035, pseudogene [Sou |
| RNA5SP287 | 9 | RNA, 5S ribosomal pseudogene 287 [Source:H |
| AL158154.1 | 9 | NULL |
| SPATA31D5P | 9 | SPATA31 subfamily D member 5, pseudogene |
| AL158154.2 | 9 | novel transcript |
| SPATA31D4 | 9 | SPATA31 subfamily D member 4 [Source:HGN |
| AL158154.3 | 9 | novel transcript |
| SPATA31D3 | 9 | SPATA31 subfamily D member 3 [Source:HGN |
| SPATA31D2P | 9 | SPATA31 subfamily D member 2, pseudogene |
| AL162726.3 | 9 | serine palmitoyltransferase long chain base su |
| SPATA31D1 | 9 | SPATA31 subfamily D member 1 [Source:HGN |
| SPATA31B1P | 9 | SPATA31 subfamily B member 1, pseudogene |
| DDX10P2 | 9 | DEAD-box helicase 10 pseudogene 2 [Source:I |
| AL158047.1 | 9 | ribosomal protein S2 (RPS2) pseudogene |
| AL162726.1 | 9 | novel transcript |
| MTCO3P40 | 9 | MT-CO3 pseudogene 40 [Source:HGNC Symbo |
| AL162726.2 | 9 | novel pseudogene |
| AL162726.4 | 9 | novel transcript |
| AL356490.1 | 9 | tec |
| AL353774.1 | 9 | novel transcript |
| RPS6P12 | 9 | ribosomal protein S6 pseudogene 12 [Source: |
| RASEF | 9 | RAS and EF-hand domain containing [Source:H |
| AL499602.1 | 9 | novel transcript |
| RN7SKP242 | 9 | RN7SK pseudogene 242 [Source:HGNC Symbo |
| AL137847.3 | 9 | ribosomal protein L10 (RPL10) pseudogene |
| AL137847.1 | 9 | novel transcript |
| FRMD3 | 9 | FERM domain containing 3 [Source:HGNC Sym |
| AL137847.2 | 9 | novel transcript |
| RNU4-29P | 9 | RNA, U4 small nuclear 29, pseudogene [Sourc |
| RNU4-15P | 9 | RNA, U4 small nuclear 15, pseudogene [Sourc |
| IDNK | 9 | IDNK gluconokinase [Source:HGNC Symbol;Ac |
| UBQLN1 | 9 | ubiquilin 1 [Source:HGNC Symbol;Acc:HGNC:1 |
| AL354920.1 | 9 | novel transcript, antisense to UBQLN1 |
| GKAP1 | 9 | G kinase anchoring protein 1 [Source:HGNC Sy |
| AL354733.1 | 9 | novel transcript, antisense to KIF27 |
| AL354733.2 | 9 | novel transcript, antisense to KIF27 |
| KIF27 | 9 | kinesin family member 27 [Source:HGNC Sym |
| C9orf64 | 9 | chromosome 9 open reading frame 64 [Sourc |
| HNRNPK | 9 | heterogeneous nuclear ribonucleoprotein K [S |
| MIR7-1 | 9 | microRNA 7-1 [Source:HGNC Symbol;Acc:HGN |
| AL354733.3 | 9 | novel transcript |
| RMI1 | 9 | RecQ mediated genome instability 1 [Source:H |

| | | |
|---|---|---|
| Y_RNA | 9 | Y RNA [Source:RFAM;Acc:RF00019] |
| AL390838.1 | 9 | novel transcript |
| AL390838.2 | 9 | high mobility group nucleosomal binding dom |
| SLC28A3 | 9 | solute carrier family 28 member 3 [Source:HG |
| AL356134.1 | 9 | novel transcript |
| AL157886.1 | 9 | Novel transcript, antisense to SLC28A3 |
| AL354692.1 | 9 | novel pseudogene |
| NTRK2 | 9 | neurotrophic receptor tyrosine kinase 2 [Sour |
| UBE2V1P10 | 9 | ubiquitin conjugating enzyme E2 V1 pseudoge |
| AL354897.1 | 9 | Novel transcript |
| AL354897.2 | 9 | novel transcript |
| AL583827.1 | 9 | Novel transcript |
| STK33P1 | 9 | serine/threonine kinase 33 pseudogene 1 [Sou |
| AGTPBP1 | 9 | ATP/GTP binding protein 1 [Source:HGNC Sym |
| AL353743.4 | 9 | kinesin family member 27 (KIF27) pseudogene |
| AL157882.1 | 9 | novel transcript |
| AL353743.2 | 9 | novel transcript |
| AL353743.1 | 9 | kinesin family member 27 (KIF27) pseudogene |
| PHBP7 | 9 | prohibitin pseudogene 7 [Source:HGNC Symbo |
| AL353743.3 | 9 | novel pseudogene |
| NAA35 | 9 | N(alpha)-acetyltransferase 35, NatC auxiliary s |
| GOLM1 | 9 | golgi membrane protein 1 [Source:HGNC Sym |
| AL161447.2 | 9 | novel transcript |
| AL161447.1 | 9 | novel pseudogene |
| C9orf153 | 9 | chromosome 9 open reading frame 153 [Sour |
| AL137849.1 | 9 | protein tyrosine phosphatase, receptor type, |
| RN7SKP264 | 9 | RN7SK pseudogene 264 [Source:HGNC Symbo |
| ISCA1 | 9 | iron-sulfur cluster assembly 1 [Source:HGNC S |
| TUT7 | 9 | terminal uridylyl transferase 7 [Source:HGNC |
| AL353678.1 | 9 | ribosomal protein S6 (RPS6) pseudogene |
| AL158828.1 | 9 | novel transcript |
| RNU2-36P | 9 | RNA, U2 small nuclear 36, pseudogene [Sourc |
| AL353613.1 | 9 | novel transcript |
| LINC02834 | 9 | long intergenic non-protein coding RNA 2834 |
| GAS1 | 9 | growth arrest specific 1 [Source:HGNC Symbo |
| GAS1RR | 9 | GAS1 adjacent regulatory RNA [Source:HGNC |
| AL513318.2 | 9 | novel transcript |
| CDC20P1 | 9 | cell division cycle 20 pseudogene 1 [Source:H |
| AL513318.1 | 9 | nuclear transcription factor Y, gamma (NFYC) |
| C9orf170 | 9 | chromosome 9 open reading frame 170 [Sour |
| AL136367.1 | 9 | NULL |
| AL353752.1 | 9 | NADH dehydrogenase (ubiquinone) 1 alpha (N |
| AL353752.2 | 9 | novel transcript |
| DAPK1 | 9 | death associated protein kinase 1 [Source:HG |
| AL161787.1 | 9 | ribosomal protein S29 (RPS29) pseudogene |
| DAPK1-IT1 | 9 | DAPK1 intronic transcript 1 [Source:HGNC Syn |
| CTSL | 9 | cathepsin L [Source:HGNC Symbol;Acc:HGNC: |
| AL160279.1 | 9 | eukaryotic translation initiation factor 3, subu |
| CTSL3P | 9 | cathepsin L family member 3, pseudogene [Sc |
| AL772337.4 | 9 | death-associated protein kinase 1 (DAPK1) ps |

| | | |
|---|---|---|
| FBP2P1 | 9 | fructose-bisphosphatase 2 pseudogene 1 [Sou |
| ELF2P3 | 9 | ELF2 pseudogene 3 [Source:HGNC Symbol;Acc |
| NPAP1P7 | 9 | nuclear pore associated protein 1 pseudogene |
| CTSLP8 | 9 | cathepsin L pseudogene 8 [Source:HGNC Syml |
| AL772337.2 | 9 | novel transcript |
| AL772337.1 | 9 | nuclear pore associated protein 1 (NPAP1) pse |
| AL772337.3 | 9 | cathepsin L (CTSL) pseudogene |
| SPATA31E1 | 9 | SPATA31 subfamily E member 1 [Source:HGN |
| SPATA31C1 | 9 | SPATA31 subfamily C member 1 [Source:HGN |
| AL353572.2 | 9 | novel pseudogene |
| AL353572.3 | 9 | novel protein |
| CDK20 | 9 | cyclin dependent kinase 20 [Source:HGNC Syn |
| RNU6-86P | 9 | RNA, U6 small nuclear 86, pseudogene [Sourc |
| RPS10P3 | 9 | ribosomal protein S10 pseudogene 3 [Source: |
| AL353572.4 | 9 | novel transcript |
| AL353572.1 | 9 | proline-rich nuclear receptor coactivator 2 (PN |
| SPATA31C2 | 9 | SPATA31 subfamily C member 2 [Source:HGN |
| AL353726.1 | 9 | arginine/serine-rich coiled-coil 1 (RSRC1) pseu |
| AL353726.2 | 9 | tec |
| AL451142.1 | 9 | potassium channel tetramerisation domain cc |
| AL451142.2 | 9 | mitofusin 1 (MFN1) pseudogene |
| RPSAP49 | 9 | ribosomal protein SA pseudogene 49 [Source: |
| AL353748.3 | 9 | novel transcript, antisense to SPIN1 |
| SPIN1 | 9 | spindlin 1 [Source:HGNC Symbol;Acc:HGNC:11 |
| AL353748.2 | 9 | developmental pluripotency associated 3 (DPI |
| HNRNPA1P14 | 9 | heterogeneous nuclear ribonucleoprotein A1 |
| AL353748.1 | 9 | ribosomal protein L21 (RPL21) pseudogene |
| NXNL2 | 9 | nucleoredoxin like 2 [Source:HGNC Symbol;Ac |
| LINC02843 | 9 | long intergenic non-protein coding RNA 2843 |
| AL390791.1 | 9 | novel transcript |
| MIR4289 | 9 | microRNA 4289 [Source:HGNC Symbol;Acc:HG |
| PCNPP2 | 9 | PEST containing nuclear protein pseudogene 2 |
| AL772202.1 | 9 | novel transcript |
| S1PR3 | 9 | sphingosine-1-phosphate receptor 3 [Source:H |
| SHC3 | 9 | SHC adaptor protein 3 [Source:HGNC Symbol;. |
| AL353150.1 | 9 | novel transcript |
| CKS2 | 9 | CDC28 protein kinase regulatory subunit 2 [So |
| MIR3153 | 9 | microRNA 3153 [Source:HGNC Symbol;Acc:HG |
| SECISBP2 | 9 | SECIS binding protein 2 [Source:HGNC Symbol |
| SEMA4D | 9 | semaphorin 4D [Source:HGNC Symbol;Acc:HG |
| PA2G4P6 | 9 | proliferation-associated 2G4 pseudogene 6 [S |
| AL590233.1 | 9 | ubiquitin-conjugating enzyme E2 pseudogene |
| AL161910.1 | 9 | novel transcript |
| GADD45G | 9 | growth arrest and DNA damage inducible gam |
| AL606807.1 | 9 | novel transcript |
| BX470209.2 | 9 | novel transcript |
| BX470209.1 | 9 | novel transcript |
| AL161629.1 | 9 | novel transcript |
| MIR4290HG | 9 | MIR4290 host gene [Source:HGNC Symbol;Acc |
| MIR4290 | 9 | microRNA 4290 [Source:HGNC Symbol;Acc:HG |

| | | |
|---|---|---|
| IL6RP1 | 9 | interleukin 6 receptor pseudogene 1 [Source:I |
| AL161629.2 | 9 | peptidylprolyl isomerase F (PPIF) pseudogene |
| AL353649.1 | 9 | tec |
| OR7E31P | 9 | olfactory receptor family 7 subfamily E memb |
| OR7E116P | 9 | olfactory receptor family 7 subfamily E memb |
| LINC01508 | 9 | long intergenic non-protein coding RNA 1508 |
| LINC01501 | 9 | long intergenic non-protein coding RNA 1501 |
| DIRAS2 | 9 | DIRAS family GTPase 2 [Source:HGNC Symbol; |
| OR7E109P | 9 | olfactory receptor family 7 subfamily E memb |
| OR7E108P | 9 | olfactory receptor family 7 subfamily E memb |
| SYK | 9 | spleen associated tyrosine kinase [Source:HGI |
| AL162585.1 | 9 | novel transcript |
| AL355607.1 | 9 | novel transcript |
| AL355607.2 | 9 | novel transcript |
| AL158071.4 | 9 | novel transcript |
| AL158071.3 | 9 | novel transcript |
| AL158071.1 | 9 | novel transcript |
| LINC00484 | 9 | long intergenic non-protein coding RNA 484 [S |
| AL158071.2 | 9 | peptidylpolyl isomerase A (cyclophilin A) (PPIA |
| AL158071.5 | 9 | novel transcript |
| AUH | 9 | AU RNA binding methylglutaconyl-CoA hydrat |
| NFIL3 | 9 | nuclear factor, interleukin 3 regulated [Source |
| AL353764.1 | 9 | novel transcript |
| ROR2 | 9 | receptor tyrosine kinase like orphan receptor |
| PAICSP2 | 9 | phosphoribosylaminoimidazole carboxylase, p |
| MIR3910-1 | 9 | microRNA 3910-1 [Source:HGNC Symbol;Acc:H |
| MIR3910-2 | 9 | microRNA 3910-2 [Source:HGNC Symbol;Acc:H |
| SPTLC1 | 9 | serine palmitoyltransferase long chain base su |
| AL391219.1 | 9 | ribosomal protein L21 (RPL21) pseudogene |
| HSPE1P22 | 9 | heat shock protein family E (Hsp10) member 1 |
| AL354751.3 | 9 | novel pseudogene |
| MTATP6P29 | 9 | MT-ATP6 pseudogene 29 [Source:HGNC Symb |
| MTCO3P29 | 9 | MT-CO3 pseudogene 29 [Source:HGNC Symbo |
| MTND3P23 | 9 | MT-ND3 pseudogene 23 [Source:HGNC Symbo |
| MTND4LP7 | 9 | MT-ND4L pseudogene 7 [Source:HGNC Symbo |
| MTND4P15 | 9 | MT-ND4 pseudogene 15 [Source:HGNC Symbo |
| AL354751.1 | 9 | protein tyrosine phosphatase (similar to roden |
| LINC00475 | 9 | long intergenic non-protein coding RNA 475 [S |
| AL354751.2 | 9 | transcription elongation regulator 1 like (TCER |
| BEND3P2 | 9 | BEN domain containing 3 pseudogene 2 [Sour |
| PRSS47 | 9 | serine protease 47 [Source:HGNC Symbol;Acc |
| AL136097.2 | 9 | novel transcript |
| IARS | 9 | isoleucyl-tRNA synthetase [Source:HGNC Sym |
| MIR3651 | 9 | microRNA 3651 [Source:HGNC Symbol;Acc:HG |
| SNORA84 | 9 | small nucleolar RNA, H/ACA box 84 [Source:H |
| NOL8 | 9 | nucleolar protein 8 [Source:HGNC Symbol;Acc |
| CENPP | 9 | centromere protein P [Source:HGNC Symbol;A |
| AL136097.1 | 9 | novel pseudogene |
| OGN | 9 | osteoglycin [Source:HGNC Symbol;Acc:HGNC: |
| OMD | 9 | osteomodulin [Source:HGNC Symbol;Acc:HGN |

| | | |
|---|---|---|
| ASPN | 9 | asporin [Source:HGNC Symbol;Acc:HGNC:148 |
| ECM2 | 9 | extracellular matrix protein 2 [Source:HGNC S |
| MIR4670 | 9 | microRNA 4670 [Source:HGNC Symbol;Acc:HG |
| AL157827.1 | 9 | pseudogene similar to part of cytochrome c o |
| AL157827.2 | 9 | Novel transcript, antisense to CENPP |
| IPPK | 9 | inositol-pentakisphosphate 2-kinase [Source:H |
| AL137074.1 | 9 | ribosomal protein L21 (RPL21) pseudogene |
| BICD2 | 9 | BICD cargo adaptor 2 [Source:HGNC Symbol;A |
| AL136981.3 | 9 | novel transcript |
| AL136981.4 | 9 | novel transcript |
| ANKRD19P | 9 | ankyrin repeat domain 19, pseudogene [Sourc |
| RNU6-714P | 9 | RNA, U6 small nuclear 714, pseudogene [Sour |
| EEF1DP2 | 9 | eukaryotic translation elongation factor 1 delt |
| AL136981.2 | 9 | novel zinc finger protein pseudogene |
| ZNF484 | 9 | zinc finger protein 484 [Source:HGNC Symbol; |
| AL136981.1 | 9 | pseudogene similar to family with sequence s |
| SNX18P2 | 9 | sorting nexin 18 pseudogene 2 [Source:HGNC |
| ALOX15P2 | 9 | arachidonate 15-lipoxygenase pseudogene 2 [ |
| FGD3 | 9 | FYVE, RhoGEF and PH domain containing 3 [So |
| SUSD3 | 9 | sushi domain containing 3 [Source:HGNC Sym |
| AL451065.1 | 9 | novel transcript |
| CARD19 | 9 | caspase recruitment domain family member 1 |
| NINJ1 | 9 | ninjurin 1 [Source:HGNC Symbol;Acc:HGNC:78 |
| AL390760.1 | 9 | novel transcript |
| WNK2 | 9 | WNK lysine deficient protein kinase 2 [Source |
| C9orf129 | 9 | chromosome 9 open reading frame 129 [Sour |
| RNU6-829P | 9 | RNA, U6 small nuclear 829, pseudogene [Sour |
| Y_RNA | 9 | Y RNA [Source:RFAM;Acc:RF00019] |
| AL583839.1 | 9 | novel transcript |
| FAM120AOS | 9 | family with sequence similarity 120A opposite |
| FAM120A | 9 | family with sequence similarity 120A [Source: |
| PHF2 | 9 | PHD finger protein 2 [Source:HGNC Symbol;Ac |
| MIR548AU | 9 | microRNA 548au [Source:HGNC Symbol;Acc:H |
| AL442224.1 | 9 | novel transcript |
| MIR4291 | 9 | microRNA 4291 [Source:HGNC Symbol;Acc:HG |
| BARX1 | 9 | BARX homeobox 1 [Source:HGNC Symbol;Acc: |
| BARX1-DT | 9 | BARX1 divergent transcript [Source:HGNC Sym |
| PTPDC1 | 9 | protein tyrosine phosphatase domain contain |
| CYCSP24 | 9 | CYCS pseudogene 24 [Source:HGNC Symbol;A |
| AL360020.1 | 9 | novel transcript |
| AL158152.2 | 9 | tripartite motif-containing protein 54 pseudog |
| AL158152.1 | 9 | novel transcript |
| MIRLET7A1 | 9 | microRNA let-7a-1 [Source:HGNC Symbol;Acc: |
| MIRLET7F1 | 9 | microRNA let-7f-1 [Source:HGNC Symbol;Acc: |
| LINC02603 | 9 | long intergenic non-protein coding RNA 2603 |
| MIRLET7D | 9 | microRNA let-7d [Source:HGNC Symbol;Acc:H |
| ZNF169 | 9 | zinc finger protein 169 [Source:HGNC Symbol; |
| VDAC1P11 | 9 | voltage dependent anion channel 1 pseudoge |
| NUTM2F | 9 | NUT family member 2F [Source:HGNC Symbol |
| AL691447.2 | 9 | novel transcript |

| | | |
|---|---|---|
| AL691447.1 | 9 | proline-rich nuclear receptor coactivator 2 (PN |
| PTMAP12 | 9 | prothymosin alpha pseudogene 12 [Source:HG |
| MFSD14B | 9 | major facilitator superfamily domain containi |
| YRDCP1 | 9 | yrdC N(6)-threonylcarbamoyltransferase dom |
| AL358232.2 | 9 | novel transcript |
| AL358232.1 | 9 | leishmanolysin-like (metallopeptidase M8 fam |
| RNU6-669P | 9 | RNA, U6 small nuclear 669, pseudogene [Sour |
| PCAT7 | 9 | prostate cancer associated transcript 7 [Sourc |
| FBP2 | 9 | fructose-bisphosphatase 2 [Source:HGNC Sym |
| FBP1 | 9 | fructose-bisphosphatase 1 [Source:HGNC Sym |
| AL157936.1 | 9 | novel transcript |
| AOPEP | 9 | aminopeptidase O (putative) [Source:HGNC Sy |
| MIR2278 | 9 | microRNA 2278 [Source:HGNC Symbol;Acc:HG |
| AL807757.1 | 9 | novel transcript |
| AL807757.2 | 9 | novel transcript |
| AL353768.1 | 9 | novel transcript |
| AL353768.2 | 9 | Novel transcript, antisense to C9orf3 |
| MIR6081 | 9 | microRNA 6081 [Source:HGNC Symbol;Acc:HG |
| MIR23B | 9 | microRNA 23b [Source:HGNC Symbol;Acc:HGN |
| MIR27B | 9 | microRNA 27b [Source:HGNC Symbol;Acc:HGN |
| MIR3074 | 9 | microRNA 3074 [Source:HGNC Symbol;Acc:HG |
| MIR24-1 | 9 | microRNA 24-1 [Source:HGNC Symbol;Acc:HG |
| FANCC | 9 | FA complementation group C [Source:HGNC S |
| AL354893.2 | 9 | novel transcript |
| AL354893.1 | 9 | ribosomal protein S26 (RPS26) pseudogene |
| RNA5SP288 | 9 | RNA, 5S ribosomal pseudogene 288 [Source:H |
| AL157384.1 | 9 | ATM interactor (ATMIN) pseudogene |
| AL161729.3 | 9 | novel transcript |
| MT1P1 | 9 | metallothionein 1 pseudogene 1 [Source:HGN |
| PTCH1 | 9 | patched 1 [Source:HGNC Symbol;Acc:HGNC:9 |
| AL161729.2 | 9 | novel transcript, antisense to PTCH1 |
| AL161729.1 | 9 | novel transcript, antisense to PTCH1 |
| AL161729.4 | 9 | novel transcript, antisense to PTCH1 |
| AL392185.1 | 9 | novel transcript |
| AL354861.2 | 9 | novel transcript |
| AL354861.1 | 9 | proteasome subunit, alpha type, 7 (PSMA7) ps |
| LINC00476 | 9 | long intergenic non-protein coding RNA 476 [S |
| AL354861.3 | 9 | novel transcript |
| RNU2-46P | 9 | RNA, U2 small nuclear 46, pseudogene [Sourc |
| ERCC6L2 | 9 | ERCC excision repair 6 like 2 [Source:HGNC Sy |
| RNA5SP289 | 9 | RNA, 5S ribosomal pseudogene 289 [Source:H |
| AL161454.1 | 9 | parkinson disease 7 (PARK7) pseudogene |
| LINC00092 | 9 | long intergenic non-protein coding RNA 92 [So |
| AL449403.1 | 9 | novel transcript |
| Y_RNA | 9 | Y RNA [Source:RFAM;Acc:RF00019] |
| AL449403.2 | 9 | novel transcript |
| AL449403.3 | 9 | novel transcript |
| EIF4BP3 | 9 | eukaryotic translation initiation factor 4B pseu |
| HSD17B3 | 9 | hydroxysteroid 17-beta dehydrogenase 3 [Sou |
| AL160269.1 | 9 | solute carrier family 35 member D2 [Source:N |

| | | |
|---|---|---|
| HSD17B3-AS1 | 9 | HSD17B3 antisense RNA 1 [Source:HGNC Sym |
| RNU6-1160P | 9 | RNA, U6 small nuclear 1160, pseudogene [Sou |
| SLC35D2 | 9 | solute carrier family 35 member D2 [Source:H |
| ZNF367 | 9 | zinc finger protein 367 [Source:HGNC Symbol; |
| AL133477.1 | 9 | NSA2 ribosome biogenesis homolog (S. cerevi |
| AL133477.2 | 9 | novel transcript, sense intronic to ZNF367 |
| AL133477.3 | 9 | novel transcript |
| HABP4 | 9 | hyaluronan binding protein 4 [Source:HGNC S |
| CDC14B | 9 | cell division cycle 14B [Source:HGNC Symbol;A |
| PRXL2C | 9 | peroxiredoxin like 2C [Source:HGNC Symbol;A |
| AL589843.1 | 9 | novel transcript |
| AL589843.2 | 9 | makorin, ring finger protein, 2 (MKRN2) pseuc |
| ZNF510 | 9 | zinc finger protein 510 [Source:HGNC Symbol; |
| ZNF782 | 9 | zinc finger protein 782 [Source:HGNC Symbol; |
| MFSD14C | 9 | major facilitator superfamily domain containir |
| PTMAP11 | 9 | prothymosin alpha pseudogene 11 [Source:HC |
| AL158827.1 | 9 | proline-rich nuclear receptor coactivator 2 (PI |
| NUTM2G | 9 | NUT family member 2G [Source:HGNC Symbo |
| YRDCP2 | 9 | yrdC N(6)-threonylcarbamoyltransferase dom |
| CTSV | 9 | cathepsin V [Source:HGNC Symbol;Acc:HGNC: |
| TCEA1P1 | 9 | transcription elongation factor A1 pseudogen |
| GAS2L1P2 | 9 | growth arrest specific 2 like 1 pseudogene 2 [S |
| VN1R51P | 9 | vomeronasal 1 receptor 51 pseudogene [Sour |
| FAM220CP | 9 | family with sequence similarity 220 member C |
| AL445670.1 | 9 | sorting nexin 18 (SNX18) pseudogene |
| ANKRD18CP | 9 | ankyrin repeat domain 18C, pseudogene [Sou |
| RNU6-798P | 9 | RNA, U6 small nuclear 798, pseudogene [Sour |
| AL590705.3 | 9 | novel transcript |
| ZNF322P1 | 9 | zinc finger protein 322 pseudogene 1 [Source: |
| AL590705.1 | 9 | novel transcript |
| AL590705.2 | 9 | novel transcript |
| SUGT1P4-STRA6LP | 9 | SUGT1P4-STRA6LP readthrough [Source:HGN( |
| STRA6LP | 9 | STRA6 like, pseudogene [Source:HGNC Symbo |
| SUGT1P4 | 9 | SUGT1 pseudogene 4 [Source:HGNC Symbol;A |
| AL512590.2 | 9 | novel pseudogene |
| CCDC180 | 9 | coiled-coil domain containing 180 [Source:HG |
| MIR1302-8 | 9 | microRNA 1302-8 [Source:HGNC Symbol;Acc:I |
| AL512590.1 | 9 | pseudogene similar to part of ral guanine nucl |
| AL512590.3 | 9 | novel transcript |
| TDRD7 | 9 | tudor domain containing 7 [Source:HGNC Sym |
| TMOD1 | 9 | tropomodulin 1 [Source:HGNC Symbol;Acc:HG |
| AL162385.1 | 9 | novel transcript |
| TSTD2 | 9 | thiosulfate sulfurtransferase like domain cont |
| NCBP1 | 9 | nuclear cap binding protein subunit 1 [Source: |
| AL162385.2 | 9 | novel transcript |
| XPA | 9 | XPA, DNA damage recognition and repair fact |
| KRT18P13 | 9 | keratin 18 pseudogene 13 [Source:HGNC Sym |
| PTCSC2 | 9 | papillary thyroid carcinoma susceptibility canc |
| AL445531.1 | 9 | novel transcript |
| FOXE1 | 9 | forkhead box E1 [Source:HGNC Symbol;Acc:H( |

| | | |
|---|---|---|
| TRMO | 9 | tRNA methyltransferase O [Source:HGNC Sym |
| Y_RNA | 9 | Y RNA [Source:RFAM;Acc:RF00019] |
| AL499604.1 | 9 | novel transcript, sense intronic to TRMO |
| HEMGN | 9 | hemogen [Source:HGNC Symbol;Acc:HGNC:17 |
| ANP32B | 9 | acidic nuclear phosphoprotein 32 family mem |
| AL354726.1 | 9 | novel transcript |
| RNU6-918P | 9 | RNA, U6 small nuclear 918, pseudogene [Sour |
| NANS | 9 | N-acetylneuraminate synthase [Source:HGNC |
| TRIM14 | 9 | tripartite motif containing 14 [Source:HGNC S |
| Metazoa_SRP | 9 | Metazoan signal recognition particle RNA [Sou |
| CORO2A | 9 | coronin 2A [Source:HGNC Symbol;Acc:HGNC:2 |
| TBC1D2 | 9 | TBC1 domain family member 2 [Source:HGNC |
| MIR6854 | 9 | microRNA 6854 [Source:HGNC Symbol;Acc:HG |
| AL360081.1 | 9 | ADP-ribosylation factor-like 8B (ARL8B) pseud |
| GABBR2 | 9 | gamma-aminobutyric acid type B receptor sub |
| SEPTIN7P7 | 9 | septin 7 pseudogene 7 [Source:HGNC Symbol; |
| ANKS6 | 9 | ankyrin repeat and sterile alpha motif domain |
| AL807776.1 | 9 | Novel transcript |
| GALNT12 | 9 | polypeptide N-acetylgalactosaminyltransferas |
| AL136084.2 | 9 | novel transcript, antisense to GALNT12 |
| AL136084.1 | 9 | non-metastatic cells 2, protein (NM23B) expre |
| COL15A1 | 9 | collagen type XV alpha 1 chain [Source:HGNC |
| AL136084.3 | 9 | novel transcript, antisense to COL15A1 |
| TGFBR1 | 9 | transforming growth factor beta receptor 1 [S |
| RNA5SP290 | 9 | RNA, 5S ribosomal pseudogene 290 [Source:H |
| AL162427.1 | 9 | novel transcript |
| RN7SL794P | 9 | RNA, 7SL, cytoplasmic 794, pseudogene [Sour |
| ALG2 | 9 | ALG2 alpha-1,3/1,6-mannosyltransferase [Sou |
| SEC61B | 9 | SEC61 translocon beta subunit [Source:HGNC |
| RN7SKP225 | 9 | RN7SK pseudogene 225 [Source:HGNC Symbo |
| KRT8P11 | 9 | keratin 8 pseudogene 11 [Source:HGNC Symb |
| NAMA | 9 | non-protein coding RNA, associated with MAF |
| AL137067.2 | 9 | NULL |
| AL137067.3 | 9 | novel transcript |
| AL137067.1 | 9 | NULL |
| AL359710.1 | 9 | novel transcript |
| STX17-AS1 | 9 | STX17 antisense RNA 1 [Source:HGNC Symbol |
| AL162394.1 | 9 | novel transcript |
| NR4A3 | 9 | nuclear receptor subfamily 4 group A member |
| STX17 | 9 | syntaxin 17 [Source:HGNC Symbol;Acc:HGNC: |
| AL358937.1 | 9 | novel transcript |
| ERP44 | 9 | endoplasmic reticulum protein 44 [Source:HG |
| UPF3AP3 | 9 | UPF3A pseudogene 3 [Source:HGNC Symbol;A |
| INVS | 9 | inversin [Source:HGNC Symbol;Acc:HGNC:178 |
| RN7SL75P | 9 | RNA, 7SL, cytoplasmic 75, pseudogene [Sourc |
| RPS2P35 | 9 | ribosomal protein S2 pseudogene 35 [Source: |
| AL137072.1 | 9 | developmental pluripotency associated 3 (DPI |
| NANOGP5 | 9 | Nanog homeobox pseudogene 5 [Source:HGN |
| TEX10 | 9 | testis expressed 10 [Source:HGNC Symbol;Acc |
| MSANTD3 | 9 | Myb/SANT DNA binding domain containing 3 [ |

| | | |
|---|---|---|
| MSANTD3-TMEFF1 | 9 | MSANTD3-TMEFF1 readthrough [Source:HGN |
| TMEFF1 | 9 | transmembrane protein with EGF like and two |
| CAVIN4 | 9 | caveolae associated protein 4 [Source:HGNC S |
| RN7SKP87 | 9 | RN7SK pseudogene 87 [Source:HGNC Symbol; |
| ACTG1P19 | 9 | actin gamma 1 pseudogene 19 [Source:HGNC |
| AL162395.1 | 9 | ribosomal protein L24 (RPL24) pseudogene |
| GAPDHP26 | 9 | glyceraldehyde 3 phosphate dehydrogenase p |
| PLPPR1 | 9 | phospholipid phosphatase related 1 [Source:H |
| AL161631.1 | 9 | pseudogene similar to part of the transient re |
| AL359893.2 | 9 | NULL |
| ACNATP | 9 | acyl-CoA:amino acid N-acyltransferase pseudo |
| TRMT112P4 | 9 | tRNA methyltransferase subunit 11-2 pseudog |
| AL359893.1 | 9 | zinc finger and BTB domain containing protein |
| BAAT | 9 | bile acid-CoA:amino acid N-acyltransferase [So |
| FYTTD1P1 | 9 | forty-two-three domain containing 1 pseudog |
| MRPL50 | 9 | mitochondrial ribosomal protein L50 [Source:I |
| ZNF189 | 9 | zinc finger protein 189 [Source:HGNC Symbol; |
| ALDOB | 9 | aldolase, fructose-bisphosphate B [Source:HG |
| AL353621.2 | 9 | zinc finger protein 551 (ZNF551) pseudogene |
| TMEM246-AS1 | 9 | TMEM246 antisense RNA 1 [Source:HGNC Syn |
| TMEM246 | 9 | transmembrane protein 246 [Source:HGNC Sy |
| AL353621.1 | 9 | NADH dehydrogenase (ubiquinone) 1 alpha su |
| RNF20 | 9 | ring finger protein 20 [Source:HGNC Symbol;A |
| GRIN3A | 9 | glutamate ionotropic receptor NMDA type sul |
| PPP3R2 | 9 | protein phosphatase 3 regulatory subunit B, b |
| MTND3P4 | 9 | MT-ND3 pseudogene 4 [Source:HGNC Symbol |
| ARL2BPP7 | 9 | ADP ribosylation factor like GTPase 2 binding |
| RNU6-329P | 9 | RNA, U6 small nuclear 329, pseudogene [Sour |
| AL442644.2 | 9 | CEBPZ opposite strand (CEBPZOS) pseudogene |
| AL442644.1 | 9 | family with sequence similarity 190, member |
| LINC00587 | 9 | long intergenic non-protein coding RNA 587 [S |
| ZYG11AP1 | 9 | zyg-11 family member A, cell cycle regulator p |
| AL365396.1 | 9 | phosphoribosyl pyrophosphate synthetase 2 ( |
| CYLC2 | 9 | cylicin 2 [Source:HGNC Symbol;Acc:HGNC:258 |
| AL589823.1 | 9 | pseudogene similar to part of NADH dehydrog |
| LINC01492 | 9 | long intergenic non-protein coding RNA 1492 |
| AL390962.1 | 9 | novel transcript |
| RNA5SP291 | 9 | RNA, 5S ribosomal pseudogene 291 [Source:H |
| AL590381.1 | 9 | novel transcript |
| SMC2-AS1 | 9 | SMC2 antisense RNA 1 (head to head) [Source |
| SMC2 | 9 | structural maintenance of chromosomes 2 [So |
| TOPORSLP | 9 | topoisomerase I binding, arginine and serine r |
| AL512646.1 | 9 | WDR45-like (WDR45L) pseudogene |
| OR13F1 | 9 | olfactory receptor family 13 subfamily F mem |
| AL450426.1 | 9 | novel transcript, antisense to OR13C3 |
| OR13C4 | 9 | olfactory receptor family 13 subfamily C mem |
| OR13C3 | 9 | olfactory receptor family 13 subfamily C mem |
| OR13C8 | 9 | olfactory receptor family 13 subfamily C mem |
| OR13D2P | 9 | olfactory receptor family 13 subfamily D mem |
| OR13C5 | 9 | olfactory receptor family 13 subfamily C mem |

| | | |
|---|---|---|
| OR13C2 | 9 | olfactory receptor family 13 subfamily C mem |
| OR13C9 | 9 | olfactory receptor family 13 subfamily C mem |
| OR13I1P | 9 | olfactory receptor family 13 subfamily I meml |
| OR13C1P | 9 | olfactory receptor family 13 subfamily C mem |
| OR13D1 | 9 | olfactory receptor family 13 subfamily D mem |
| OR13D3P | 9 | olfactory receptor family 13 subfamily D mem |
| NIPSNAP3A | 9 | nipsnap homolog 3A [Source:HGNC Symbol;A |
| NIPSNAP3B | 9 | nipsnap homolog 3B [Source:HGNC Symbol;A |
| ABCA1 | 9 | ATP binding cassette subfamily A member 1 [S |
| AL359182.1 | 9 | novel transcript |
| CT70 | 9 | cancer/testis associated transcript 70 [Source |
| AL591506.1 | 9 | novel transcript |
| RN7SKP191 | 9 | RN7SK pseudogene 191 [Source:HGNC Symbo |
| SLC44A1 | 9 | solute carrier family 44 member 1 [Source:HG |
| FSD1L | 9 | fibronectin type III and SPRY domain containir |
| RALGAPA1P1 | 9 | Ral GTPase activating protein catalytic alpha s |
| AL158070.1 | 9 | novel transcript |
| FKTN | 9 | fukutin [Source:HGNC Symbol;Acc:HGNC:362 |
| AL158070.2 | 9 | tec |
| TAL2 | 9 | TAL bHLH transcription factor 2 [Source:HGN |
| TMEM38B | 9 | transmembrane protein 38B [Source:HGNC Sy |
| AL592437.1 | 9 | FUS interacting protein (serine-arginine rich) 1 |
| AL592437.2 | 9 | DEP domain containing 1 (DEPDC1) pseudoge |
| SLC25A6P5 | 9 | solute carrier family 25 member 6 pseudogen |
| LINC01505 | 9 | long intergenic non-protein coding RNA 1505 |
| AL732323.1 | 9 | novel transcript |
| AL451140.1 | 9 | novel transcript |
| MIR8081 | 9 | microRNA 8081 [Source:HGNC Symbol;Acc:H |
| AL512649.2 | 9 | novel transcript |
| LINC01505 | 9 | long intergenic non-protein coding RNA 1505 |
| RN7SKP77 | 9 | RN7SK pseudogene 77 [Source:HGNC Symbol; |
| AL512649.1 | 9 | pseudogene similar to part of constitutive ph |
| RNA5SP292 | 9 | RNA, 5S ribosomal pseudogene 292 [Source:H |
| ZNF462 | 9 | zinc finger protein 462 [Source:HGNC Symbol; |
| AL807761.4 | 9 | novel transcript |
| AL512593.1 | 9 | ribosomal protein L7a (RPL7A) pseudogene |
| AL807761.1 | 9 | novel pseudogene |
| AL807761.2 | 9 | MMRP19 pseudogene |
| AL807761.3 | 9 | novel transcript |
| AL445487.1 | 9 | karyopherin alpha 2 (RAG cohort 1, importin a |
| RAD23B | 9 | RAD23 homolog B, nucleotide excision repair |
| LINC01509 | 9 | long intergenic non-protein coding RNA 1509 |
| AL359552.1 | 9 | novel transcript |
| HMGN2P32 | 9 | high mobility group nucleosomal binding dom |
| RNU6-492P | 9 | RNA, U6 small nuclear 492, pseudogene [Sour |
| KLF4 | 9 | Kruppel like factor 4 [Source:HGNC Symbol;A |
| AL389915.1 | 9 | ribosomal protein S16 (RPS16) pseudogene |
| PPIAP88 | 9 | peptidylprolyl isomerase A pseudogene 88 [S |
| RNU6-996P | 9 | RNA, U6 small nuclear 996, pseudogene [Sour |
| RNU6-1064P | 9 | RNA, U6 small nuclear 1064, pseudogene [So |

| | | |
|---|---|---|
| RN7SL659P | 9 | RNA, 7SL, cytoplasmic 659, pseudogene [Sour |
| RPL36AP6 | 9 | ribosomal protein L36a pseudogene 6 [Source |
| AL162389.1 | 9 | small EDRK-rich factor 2 (SERF2) pseudogene |
| RNA5SP293 | 9 | RNA, 5S ribosomal pseudogene 293 [Source:H |
| RPS15AP27 | 9 | ribosomal protein S15a pseudogene 27 [Sourc |
| AC068050.1 | 9 | ribosomal protein L31 (RPL31) pseudogene |
| CHCHD4P2 | 9 | coiled-coil-helix-coiled-coil-helix domain conta |
| AL353742.1 | 9 | novel transcript |
| AL669983.1 | 9 | ribosomal protein L36 (RPL36) pseudogene |
| AL358779.1 | 9 | novel transcript |
| RPL36AP35 | 9 | ribosomal protein L36a pseudogene 35 [Sourc |
| AL359692.1 | 9 | ATP synthase F chain (mitochondrial) pseudog |
| ACTL7B | 9 | actin like 7B [Source:HGNC Symbol;Acc:HGNC |
| ACTL7A | 9 | actin like 7A [Source:HGNC Symbol;Acc:HGNC |
| ELP1 | 9 | elongator complex protein 1 [Source:HGNC Sy |
| ABITRAM | 9 | actin binding transcription modulator [Source |
| CTNNAL1 | 9 | catenin alpha like 1 [Source:HGNC Symbol;Acc |
| TMEM245 | 9 | transmembrane protein 245 [Source:HGNC Sy |
| Y_RNA | 9 | Y RNA [Source:RFAM;Acc:RF00019] |
| MIR32 | 9 | microRNA 32 [Source:HGNC Symbol;Acc:HGN |
| FRRS1L | 9 | ferric chelate reductase 1 like [Source:HGNC S |
| EPB41L4B | 9 | erythrocyte membrane protein band 4.1 like 4 |
| RNU6-984P | 9 | RNA, U6 small nuclear 984, pseudogene [Sour |
| MTND2P11 | 9 | MT-ND2 pseudogene 11 [Source:HGNC Symbo |
| Y_RNA | 9 | Y RNA [Source:RFAM;Acc:RF00019] |
| PTPN3 | 9 | protein tyrosine phosphatase non-receptor ty |
| MIR3927 | 9 | microRNA 3927 [Source:HGNC Symbol;Acc:HG |
| YBX1P6 | 9 | Y-box binding protein 1 pseudogene 6 [Source |
| PALM2-AKAP2 | 9 | PALM2-AKAP2 fusion [Source:HGNC Symbol;A |
| AL158829.1 | 9 | novel transcript |
| AL158823.1 | 9 | ribosomal protein L21 (RPL21) pseudogene |
| C9orf152 | 9 | chromosome 9 open reading frame 152 [Sour |
| TXN | 9 | thioredoxin [Source:HGNC Symbol;Acc:HGNC: |
| TXNDC8 | 9 | thioredoxin domain containing 8 [Source:HGN |
| SVEP1 | 9 | sushi, von Willebrand factor type A, EGF and p |
| RNU6-1039P | 9 | RNA, U6 small nuclear 1039, pseudogene [Sou |
| AL592463.1 | 9 | novel transcript |
| MUSK | 9 | muscle associated receptor tyrosine kinase [S |
| AL513328.1 | 9 | ribosomal protein S21 (RPS21) pseudogene |
| LPAR1 | 9 | lysophosphatidic acid receptor 1 [Source:HGN |
| RNU6-432P | 9 | RNA, U6 small nuclear 432, pseudogene [Sour |
| Y_RNA | 9 | Y RNA [Source:RFAM;Acc:RF00019] |
| RNY4P18 | 9 | RNY4 pseudogene 18 [Source:HGNC Symbol;A |
| AL162414.1 | 9 | novel transcript |
| MIR7702 | 9 | microRNA 7702 [Source:HGNC Symbol;Acc:HG |
| OR2K2 | 9 | olfactory receptor family 2 subfamily K memb |
| ECPAS | 9 | Ecm29 proteasome adaptor and scaffold [Sou |
| RNA5SP294 | 9 | RNA, 5S ribosomal pseudogene 294 [Source:H |
| ZNF483 | 9 | zinc finger protein 483 [Source:HGNC Symbol; |
| PTGR1 | 9 | prostaglandin reductase 1 [Source:HGNC Sym |

| | | |
|---|---|---|
| AL135787.1 | 9 | novel transcript |
| LRRC37A5P | 9 | leucine rich repeat containing 37 member A5, |
| DNAJC25 | 9 | DnaJ heat shock protein family (Hsp40) memb |
| DNAJC25-GNG10 | 9 | DNAJC25-GNG10 readthrough [Source:HGNC : |
| GNG10 | 9 | G protein subunit gamma 10 [Source:HGNC Sy |
| SHOC1 | 9 | shortage in chiasmata 1 [Source:HGNC Symbo |
| AL354877.1 | 9 | pseudogene similar to part of HLA-B associate |
| RNU6-1013P | 9 | RNA, U6 small nuclear 1013, pseudogene [Sou |
| UGCG | 9 | UDP-glucose ceramide glucosyltransferase [Sc |
| MIR4668 | 9 | microRNA 4668 [Source:HGNC Symbol;Acc:HG |
| RNU6-710P | 9 | RNA, U6 small nuclear 710, pseudogene [Sour |
| AL138756.1 | 9 | novel transcript, overlapping SUSD1 |
| SUSD1 | 9 | sushi domain containing 1 [Source:HGNC Sym |
| RNU6-855P | 9 | RNA, U6 small nuclear 855, pseudogene [Sour |
| Y_RNA | 9 | Y RNA [Source:RFAM;Acc:RF00019] |
| AL158824.1 | 9 | ribosomal protein L29 (RPL29) pseudogene |
| PTBP3 | 9 | polypyrimidine tract binding protein 3 [Source |
| RN7SL57P | 9 | RNA, 7SL, cytoplasmic 57, pseudogene [Sourc |
| RNA5SP295 | 9 | RNA, 5S ribosomal pseudogene 295 [Source:H |
| RN7SL430P | 9 | RNA, 7SL, cytoplasmic 430, pseudogene [Sour |
| HSPE1P28 | 9 | heat shock protein family E (Hsp10) member : |
| AL359073.1 | 9 | ribosomal protein L32 (RPL32) pseudogene |
| HSDL2 | 9 | hydroxysteroid dehydrogenase like 2 [Source: |
| C9orf147 | 9 | chromosome 9 open reading frame 147 [Sour |
| KIAA1958 | 9 | KIAA1958 [Source:HGNC Symbol;Acc:HGNC:2: |
| AL445187.1 | 9 | vacuolar protein sorting 33B (VPS33B) pseudo |
| INIP | 9 | INTS3 and NABP interacting protein [Source:H |
| AL390067.1 | 9 | novel transcript |
| SNX30 | 9 | sorting nexin family member 30 [Source:HGN |
| SLC46A2 | 9 | solute carrier family 46 member 2 [Source:HG |
| AL139041.1 | 9 | novel transcript, antisense to SLC46A2 |
| ZNF883 | 9 | zinc finger protein 883 [Source:HGNC Symbol; |
| MUPP | 9 | major urinary protein, pseudogene [Source:H |
| ZNF883 | 9 | zinc finger protein 883 [Source:NCBI gene;Acc |
| ZFP37 | 9 | ZFP37 zinc finger protein [Source:HGNC Symb |
| AL162588.1 | 9 | novel pseudogene |
| AL449105.1 | 9 | novel pseudogene |
| AL449105.3 | 9 | novel pseudogene |
| AL449105.4 | 9 | novel pseudogene |
| AL449105.2 | 9 | novel pseudogene |
| AL449105.5 | 9 | novel pseudogene |
| FAM225B | 9 | family with sequence similarity 225 member E |
| FAM225A | 9 | family with sequence similarity 225 member / |
| SLC31A2 | 9 | solute carrier family 31 member 2 [Source:HG |
| FKBP15 | 9 | FKBP prolyl isomerase 15 [Source:HGNC Symb |
| SLC31A1 | 9 | solute carrier family 31 member 1 [Source:HG |
| CDC26 | 9 | cell division cycle 26 [Source:HGNC Symbol;Ac |
| PRPF4 | 9 | pre-mRNA processing factor 4 [Source:HGNC : |
| RNF183 | 9 | ring finger protein 183 [Source:HGNC Symbol; |
| WDR31 | 9 | WD repeat domain 31 [Source:HGNC Symbol;. |

| | | |
|---|---|---|
| BSPRY | 9 | B-box and SPRY domain containing [Source:H( |
| HDHD3 | 9 | haloacid dehalogenase like hydrolase domain |
| ALAD | 9 | aminolevulinate dehydratase [Source:HGNC S |
| POLE3 | 9 | DNA polymerase epsilon 3, accessory subunit |
| C9orf43 | 9 | chromosome 9 open reading frame 43 [Sourc |
| RGS3 | 9 | regulator of G protein signaling 3 [Source:HGN |
| AL162727.2 | 9 | novel transcript |
| AL162727.3 | 9 | TEC |
| AL162727.1 | 9 | novel transcript |
| AL157702.2 | 9 | novel transcript |
| AL157702.1 | 9 | novel pseudogene |
| AL162393.1 | 9 | programmed cell death 2 (PDCD2) pseudogen |
| ZNF618 | 9 | zinc finger protein 618 [Source:HGNC Symbol; |
| AMBP | 9 | alpha-1-microglobulin/bikunin precursor [Sou |
| KIF12 | 9 | kinesin family member 12 [Source:HGNC Syml |
| COL27A1 | 9 | collagen type XXVII alpha 1 chain [Source:HGN |
| MIR455 | 9 | microRNA 455 [Source:HGNC Symbol;Acc:HGI |
| ORM1 | 9 | orosomucoid 1 [Source:HGNC Symbol;Acc:HG |
| ORM2 | 9 | orosomucoid 2 [Source:HGNC Symbol;Acc:HG |
| AKNA | 9 | AT-hook transcription factor [Source:HGNC Sy |
| AL138895.1 | 9 | novel transcript |
| WHRN | 9 | whirlin [Source:HGNC Symbol;Acc:HGNC:1636 |
| AL138895.2 | 9 | small EDRK-rich factor 2 (SERF2) pseudogene |
| AL160275.2 | 9 | pseudogene similar to part of ARP3 actin-relat |
| ATP6V1G1 | 9 | ATPase H+ transporting V1 subunit G1 [Source |
| TMEM268 | 9 | transmembrane protein 268 [Source:HGNC Sy |
| Y_RNA | 9 | Y RNA [Source:RFAM;Acc:RF00019] |
| TEX53 | 9 | testis expressed 53 [Source:HGNC Symbol;Acc |
| TEX48 | 9 | testis expressed 48 [Source:HGNC Symbol;Acc |
| AL160275.1 | 9 | pseudogene similar to part of ribosomal prote |
| AL390240.1 | 9 | pumilio RNA-binding family member 2 (PUM2 |
| TNFSF15 | 9 | TNF superfamily member 15 [Source:HGNC Sy |
| AL133412.1 | 9 | PRP4 pre-mRNA processing factor 4 homolog |
| DEC1 | 9 | deleted in esophageal cancer 1 [Source:HGNC |
| TNFSF8 | 9 | TNF superfamily member 8 [Source:HGNC Syn |
| TNC | 9 | tenascin C [Source:HGNC Symbol;Acc:HGNC:5 |
| AL162425.1 | 9 | novel transcript |
| AL355601.1 | 9 | novel transcript |
| AL691420.1 | 9 | novel transcript, antisense to DEC1 |
| U2 | 9 | U2 spliceosomal RNA [Source:RFAM;Acc:RF00 |
| LINC00474 | 9 | long intergenic non-protein coding RNA 474 [S |
| AL691426.1 | 9 | novel transcript |
| PAPPA | 9 | pappalysin 1 [Source:HGNC Symbol;Acc:HGNC |
| PAPPA-AS2 | 9 | PAPPA antisense RNA 2 [Source:HGNC Symbo |
| AL137024.1 | 9 | novel transcript |
| PAPPA-AS1 | 9 | PAPPA antisense RNA 1 [Source:HGNC Symbo |
| ASTN2 | 9 | astrotactin 2 [Source:HGNC Symbol;Acc:HGNC |
| ASTN2-AS1 | 9 | ASTN2 antisense RNA 1 [Source:HGNC Symbo |
| AL133284.1 | 9 | novel transcript |
| TRIM32 | 9 | tripartite motif containing 32 [Source:HGNC S |

| | | |
|---|---|---|
| AL157829.1 | 9 | Novel transcript, antisense to ASTN2 |
| RPL10P3 | 9 | ribosomal protein L10 pseudogene 3 [Source: |
| SNORA70C | 9 | small nucleolar RNA, H/ACA box 70C [Source:I |
| RN7SKP128 | 9 | RN7SK pseudogene 128 [Source:HGNC Symbo |
| RN7SKP125 | 9 | RN7SK pseudogene 125 [Source:HGNC Symbo |
| AL445644.1 | 9 | novel transcript |
| AL161630.1 | 9 | novel transcript |
| RPL35AP22 | 9 | ribosomal protein L35a pseudogene 22 [Sourc |
| AL160272.2 | 9 | novel protein |
| TLR4 | 9 | toll like receptor 4 [Source:HGNC Symbol;Acc: |
| RNU6-1082P | 9 | RNA, U6 small nuclear 1082, pseudogene [Sou |
| AL160272.1 | 9 | novel transcript |
| AL354754.1 | 9 | novel transcript |
| TPT1P9 | 9 | tumor protein, translationally-controlled 1 ps |
| AL355592.1 | 9 | novel transcript |
| LINC02578 | 9 | long intergenic non-protein coding RNA 2578 |
| TUBB4BP6 | 9 | tubulin beta 4B class IVb pseudogene 6 [Sourc |
| BRINP1 | 9 | BMP/retinoic acid inducible neural specific 1 [ |
| AC006288.1 | 9 | novel transcript |
| LINC01613 | 9 | long intergenic non-protein coding RNA 1613 |
| AL441989.1 | 9 | novel transcript |
| MIR147A | 9 | microRNA 147a [Source:HGNC Symbol;Acc:HG |
| CDK5RAP2 | 9 | CDK5 regulatory subunit associated protein 2 |
| MEGF9 | 9 | multiple EGF like domains 9 [Source:HGNC Syı |
| AHCYP2 | 9 | adenosylhomocysteinase pseudogene 2 [Sour |
| FBXW2 | 9 | F-box and WD repeat domain containing 2 [Sc |
| B3GNT10 | 9 | UDP-GlcNAc:betaGal beta-1,3-N-acetylglucosa |
| AL161911.1 | 9 | chromatin assembly factor 1, subunit A (p150) |
| PSMD5 | 9 | proteasome 26S subunit, non-ATPase 5 [Sourc |
| CUTALP | 9 | cutA divalent cation tolerance homolog-like, p |
| PHF19 | 9 | PHD finger protein 19 [Source:HGNC Symbol;/ |
| TRAF1 | 9 | TNF receptor associated factor 1 [Source:HGN |
| C5 | 9 | complement C5 [Source:HGNC Symbol;Acc:HC |
| CNTRL | 9 | centriolin [Source:HGNC Symbol;Acc:HGNC:18 |
| RAB14 | 9 | RAB14, member RAS oncogene family [Source |
| RN7SL181P | 9 | RNA, 7SL, cytoplasmic 181, pseudogene [Sour |
| GSN | 9 | gelsolin [Source:HGNC Symbol;Acc:HGNC:462 |
| AL513122.1 | 9 | ATPase, H+ transporting, lysosomal 56/58kDa |
| AL513122.2 | 9 | novel transcript |
| GSN-AS1 | 9 | GSN antisense RNA 1 [Source:HGNC Symbol;A |
| STOM | 9 | stomatin [Source:HGNC Symbol;Acc:HGNC:33 |
| AL359644.1 | 9 | novel transcript, antisense STOM |
| GGTA1P | 9 | glycoprotein alpha-galactosyltransferase 1, ps |
| RN7SL187P | 9 | RNA, 7SL, cytoplasmic 187, pseudogene [Sour |
| HMGB1P37 | 9 | high mobility group box 1 pseudogene 37 [Sou |
| DAB2IP | 9 | DAB2 interacting protein [Source:HGNC Symb |
| AL357936.1 | 9 | novel transcript |
| AL596244.1 | 9 | novel transcript, overlapping TTLL11 |
| TTLL11 | 9 | tubulin tyrosine ligase like 11 [Source:HGNC S |
| TTLL11-IT1 | 9 | TTLL11 intronic transcript 1 [Source:HGNC Syr |

| | | |
|---|---|---|
| MIR4478 | 9 | microRNA 4478 [Source:HGNC Symbol;Acc:HG |
| NDUFA8 | 9 | NADH:ubiquinone oxidoreductase subunit A8 |
| MORN5 | 9 | MORN repeat containing 5 [Source:HGNC Sym |
| LHX6 | 9 | LIM homeobox 6 [Source:HGNC Symbol;Acc:H |
| RBM18 | 9 | RNA binding motif protein 18 [Source:HGNC S |
| MRRF | 9 | mitochondrial ribosome recycling factor [Sour |
| PTGS1 | 9 | prostaglandin-endoperoxide synthase 1 [Sour |
| AL162424.1 | 9 | novel transcript, sense intronic to PTGS1 |
| AL359636.1 | 9 | novel transcript |
| AL359636.2 | 9 | novel transcript |
| OR1J1 | 9 | olfactory receptor family 1 subfamily J membe |
| OR1J2 | 9 | olfactory receptor family 1 subfamily J membe |
| OR1J4 | 9 | olfactory receptor family 1 subfamily J membe |
| OR1N1 | 9 | olfactory receptor family 1 subfamily N memb |
| OR1N2 | 9 | olfactory receptor family 1 subfamily N memb |
| OR1L8 | 9 | olfactory receptor family 1 subfamily L memb |
| OR1H1P | 9 | olfactory receptor family 1 subfamily H memb |
| OR1Q1 | 9 | olfactory receptor family 1 subfamily Q memb |
| OR1B1 | 9 | olfactory receptor family 1 subfamily B memb |
| OR1L1 | 9 | olfactory receptor family 1 subfamily L memb |
| OR1L3 | 9 | olfactory receptor family 1 subfamily L memb |
| TLK1P1 | 9 | tousled like kinase 1 pseudogene 1 [Source:H |
| OR1L4 | 9 | olfactory receptor family 1 subfamily L memb |
| OR1L6 | 9 | olfactory receptor family 1 subfamily L memb |
| SKA2P1 | 9 | spindle and kinetochore associated complex s |
| OR5C1 | 9 | olfactory receptor family 5 subfamily C memb |
| PDCL | 9 | phosducin like [Source:HGNC Symbol;Acc:HGN |
| OR1K1 | 9 | olfactory receptor family 1 subfamily K memb |
| KRT18P67 | 9 | keratin 18 pseudogene 67 [Source:HGNC Sym |
| RC3H2 | 9 | ring finger and CCCH-type domains 2 [Source: |
| SNORD90 | 9 | small nucleolar RNA, C/D box 90 [Source:HGN |
| ZBTB6 | 9 | zinc finger and BTB domain containing 6 [Sour |
| ZBTB26 | 9 | zinc finger and BTB domain containing 26 [Sou |
| AC007066.2 | 9 | novel transcript, overlapping RABGAP1 |
| AC007066.3 | 9 | novel transcript, antisense to RABGAP1 |
| RABGAP1 | 9 | RAB GTPase activating protein 1 [Source:HGN |
| RNY1P15 | 9 | RNY1 pseudogene 15 [Source:HGNC Symbol;A |
| AC007066.1 | 9 | ribosomal protein S25 (RPS25) pseudogene |
| GPR21 | 9 | G protein-coupled receptor 21 [Source:HGNC |
| AL365338.1 | 9 | COP9 constitutive photomorphogenic homolo |
| MIR600HG | 9 | MIR600 host gene [Source:HGNC Symbol;Acc: |
| STRBP | 9 | spermatid perinuclear RNA binding protein [S |
| MIR600 | 9 | microRNA 600 [Source:HGNC Symbol;Acc:HGN |
| CRB2 | 9 | crumbs cell polarity complex component 2 [So |
| DENND1A | 9 | DENN domain containing 1A [Source:HGNC Sy |
| MIR601 | 9 | microRNA 601 [Source:HGNC Symbol;Acc:HGN |
| MIR7150 | 9 | microRNA 7150 [Source:HGNC Symbol;Acc:HG |
| AL445489.1 | 9 | novel transcript |
| AL390774.2 | 9 | novel transcript |
| AL390774.1 | 9 | LSM5 homolog, U6 small nuclear RNA associa |

| | | |
|---|---|---|
| PIGFP2 | 9 | phosphatidylinositol glycan anchor biosynthes |
| LHX2 | 9 | LIM homeobox 2 [Source:HGNC Symbol;Acc:H |
| AC006450.3 | 9 | novel transcript, antisense to LHX2 |
| AC006450.1 | 9 | novel transcript |
| AC006450.2 | 9 | novel transcript, antisense to LHX2 |
| NEK6 | 9 | NIMA related kinase 6 [Source:HGNC Symbol; |
| AL162724.2 | 9 | novel transcript |
| AL162724.1 | 9 | novel transcript |
| PSMB7 | 9 | proteasome subunit beta 7 [Source:HGNC Syn |
| ADGRD2 | 9 | adhesion G protein-coupled receptor D2 [Sou |
| NR5A1 | 9 | nuclear receptor subfamily 5 group A membe |
| AL354979.1 | 9 | novel transcript |
| NR6A1 | 9 | nuclear receptor subfamily 6 group A membe |
| AL669818.1 | 9 | RAN binding protein 1 (RANBP1) pseudogene |
| RN7SL302P | 9 | RNA, 7SL, cytoplasmic 302, pseudogene [Sour |
| MIR181A2HG | 9 | MIR181A2 host gene [Source:HGNC Symbol;A |
| MIR181A2 | 9 | microRNA 181a-2 [Source:HGNC Symbol;Acc: |
| MIR181B2 | 9 | microRNA 181b-2 [Source:HGNC Symbol;Acc: |
| AL354928.1 | 9 | novel transcript |
| OLFML2A | 9 | olfactomedin like 2A [Source:HGNC Symbol;A |
| WDR38 | 9 | WD repeat domain 38 [Source:HGNC Symbol; |
| RPL35 | 9 | ribosomal protein L35 [Source:HGNC Symbol; |
| ARPC5L | 9 | actin related protein 2/3 complex subunit 5 li |
| GOLGA1 | 9 | golgin A1 [Source:HGNC Symbol;Acc:HGNC:44 |
| RNU4-82P | 9 | RNA, U4 small nuclear 82, pseudogene [Sourc |
| SCAI | 9 | suppressor of cancer cell invasion [Source:HG |
| FXNP2 | 9 | frataxin pseuodgene 2 [Source:HGNC Symbol; |
| PPP6C | 9 | protein phosphatase 6 catalytic subunit [Sour |
| PRPS1P2 | 9 | phosphoribosyl pyrophosphate synthetase 1 p |
| RPSAP76 | 9 | ribosomal protein SA pseudogene 76 [Source: |
| RABEPK | 9 | Rab9 effector protein with kelch motifs [Sour |
| HSPA5 | 9 | heat shock protein family A (Hsp70) member |
| AL354710.2 | 9 | novel transcript |
| GAPVD1 | 9 | GTPase activating protein and VPS9 domains |
| AL354710.1 | 9 | ribosomal protein S10 (RPS10) pseudogene |
| RNU6-1020P | 9 | RNA, U6 small nuclear 1020, pseudogene [Sou |
| AL627223.1 | 9 | DnaJ (Hsp40) homolog, subfamily C, member |
| Metazoa_SRP | 9 | Metazoan signal recognition particle RNA [Sou |
| AL359632.1 | 9 | novel transcript, antisense to MAPKAP1 |
| MAPKAP1 | 9 | MAPK associated protein 1 [Source:HGNC Syn |
| RN7SL30P | 9 | RNA, 7SL, cytoplasmic 30, pseudogene [Sourc |
| AL162584.1 | 9 | novel transcript |
| HNRNPA1P15 | 9 | heterogeneous nuclear ribonucleoprotein A1 |
| AL358074.1 | 9 | novel transcript |
| PBX3 | 9 | PBX homeobox 3 [Source:HGNC Symbol;Acc:H |
| AL589923.1 | 9 | novel transcript |
| AL162391.1 | 9 | novel transcript |
| MVB12B | 9 | multivesicular body subunit 12B [Source:HGN |
| NRON | 9 | non-coding repressor of NFAT [Source:HGNC S |
| SNORD116 | 9 | Small nucleolar RNA SNORD116 [Source:RFAM |

| | | |
|---|---|---|
| AL356309.2 | 9 | novel transcript |
| AL356309.3 | 9 | tec |
| AL356309.1 | 9 | novel transcript |
| AL161908.1 | 9 | novel transcript, antisense to LIM1B |
| LMX1B | 9 | LIM homeobox transcription factor 1 beta [So |
| AL161908.2 | 9 | novel transcript, sense intronic to LMX1B |
| AL161731.1 | 9 | novel transcript |
| ZBTB43 | 9 | zinc finger and BTB domain containing 43 [Sou |
| ZBTB34 | 9 | zinc finger and BTB domain containing 34 [Sou |
| RALGPS1 | 9 | Ral GEF with PH domain and SH3 binding moti |
| ANGPTL2 | 9 | angiopoietin like 2 [Source:HGNC Symbol;Acc: |
| AL450263.1 | 9 | novel transcript, antisense to RALGPS1 |
| GARNL3 | 9 | GTPase activating Rap/RanGAP domain like 3 |
| AL450263.2 | 9 | density-regulated protein (DENR) pseudogene |
| AL445222.1 | 9 | novel transcript |
| SLC2A8 | 9 | solute carrier family 2 member 8 [Source:HGN |
| ZNF79 | 9 | zinc finger protein 79 [Source:HGNC Symbol;A |
| RPL12 | 9 | ribosomal protein L12 [Source:HGNC Symbol;A |
| SNORA65 | 9 | small nucleolar RNA, H/ACA box 65 [Source:H |
| LRSAM1 | 9 | leucine rich repeat and sterile alpha motif con |
| Y_RNA | 9 | Y RNA [Source:RFAM;Acc:RF00019] |
| NIBAN2 | 9 | niban apoptosis regulator 2 [Source:HGNC Syr |
| STXBP1 | 9 | syntaxin binding protein 1 [Source:HGNC Sym |
| AL162426.1 | 9 | novel transcript |
| PTRH1 | 9 | peptidyl-tRNA hydrolase 1 homolog [Source:H |
| MIR3911 | 9 | microRNA 3911 [Source:HGNC Symbol;Acc:HG |
| CFAP157 | 9 | cilia and flagella associated protein 157 [Sourc |
| TTC16 | 9 | tetratricopeptide repeat domain 16 [Source:H |
| TOR2A | 9 | torsin family 2 member A [Source:HGNC Symb |
| SH2D3C | 9 | SH2 domain containing 3C [Source:HGNC Sym |
| AL162586.2 | 9 | Novel transcript, antisense to SH2D3C |
| CDK9 | 9 | cyclin dependent kinase 9 [Source:HGNC Symb |
| MIR3960 | 9 | microRNA 3960 [Source:HGNC Symbol;Acc:HG |
| MIR2861 | 9 | microRNA 2861 [Source:HGNC Symbol;Acc:HG |
| FPGS | 9 | folylpolyglutamate synthase [Source:HGNC Sy |
| ENG | 9 | endoglin [Source:HGNC Symbol;Acc:HGNC:33 |
| AL162586.1 | 9 | novel transcript |
| Y_RNA | 9 | Y RNA [Source:RFAM;Acc:RF00019] |
| RNA5SP296 | 9 | RNA, 5S ribosomal pseudogene 296 [Source:H |
| AK1 | 9 | adenylate kinase 1 [Source:HGNC Symbol;Acc |
| AL157935.2 | 9 | novel protein |
| MIR4672 | 9 | microRNA 4672 [Source:HGNC Symbol;Acc:HG |
| ST6GALNAC6 | 9 | ST6 N-acetylgalactosaminide alpha-2,6-sialyltr |
| ST6GALNAC4 | 9 | ST6 N-acetylgalactosaminide alpha-2,6-sialyltr |
| PIP5KL1 | 9 | phosphatidylinositol-4-phosphate 5-kinase lik |
| AL157935.1 | 9 | novel transcript |
| DPM2 | 9 | dolichyl-phosphate mannosyltransferase subu |
| FAM102A | 9 | family with sequence similarity 102 member A |
| AL360268.2 | 9 | novel transcript |
| NAIF1 | 9 | nuclear apoptosis inducing factor 1 [Source:H |

| | | |
|---|---|---|
| SLC25A25 | 9 | solute carrier family 25 member 25 [Source:H |
| AL360268.1 | 9 | novel transcript |
| SLC25A25-AS1 | 9 | SLC25A25 antisense RNA 1 [Source:HGNC Syn |
| PTGES2 | 9 | prostaglandin E synthase 2 [Source:HGNC Syn |
| AL590708.1 | 9 | PTGES2 antisense RNA 1 |
| LCN2 | 9 | lipocalin 2 [Source:HGNC Symbol;Acc:HGNC:6 |
| C9orf16 | 9 | chromosome 9 open reading frame 16 [Sourc |
| CIZ1 | 9 | CDKN1A interacting zinc finger protein 1 [Sour |
| DNM1 | 9 | dynamin 1 [Source:HGNC Symbol;Acc:HGNC:2 |
| MIR199B | 9 | microRNA 199b [Source:HGNC Symbol;Acc:HG |
| MIR3154 | 9 | microRNA 3154 [Source:HGNC Symbol;Acc:HG |
| GOLGA2 | 9 | golgin A2 [Source:HGNC Symbol;Acc:HGNC:44 |
| SWI5 | 9 | SWI5 homologous recombination repair prote |
| TRUB2 | 9 | TruB pseudouridine synthase family member |
| AL359091.4 | 9 | novel transcript, antisense to TRUB2 |
| COQ4 | 9 | coenzyme Q4 [Source:HGNC Symbol;Acc:HGN |
| SLC27A4 | 9 | solute carrier family 27 member 4 [Source:HG |
| TMSB4XP4 | 9 | TMSB4X pseudogene 4 [Source:HGNC Symbol |
| URM1 | 9 | ubiquitin related modifier 1 [Source:HGNC Syr |
| AL359091.3 | 9 | novel transcript |
| AL359091.1 | 9 | novel transcript |
| MIR219A2 | 9 | microRNA 219a-2 [Source:HGNC Symbol;Acc:H |
| MIR219B | 9 | microRNA 219b [Source:HGNC Symbol;Acc:HG |
| CERCAM | 9 | cerebral endothelial cell adhesion molecule [S |
| AL359091.5 | 9 | novel transcript, sense intronic to CERCAM |
| AL359091.2 | 9 | novel pseudogene |
| ODF2 | 9 | outer dense fiber of sperm tails 2 [Source:HGI |
| ODF2-AS1 | 9 | ODF2 antisense RNA 1 [Source:HGNC Symbol; |
| GLE1 | 9 | GLE1 RNA export mediator [Source:HGNC Sym |
| AL356481.1 | 9 | novel transcript |
| RNU7-171P | 9 | RNA, U7 small nuclear 171 pseudogene [Sourc |
| SPTAN1 | 9 | spectrin alpha, non-erythrocytic 1 [Source:HG |
| AL356481.2 | 9 | novel transcript, sense intronic to SPTAN1 |
| AL356481.3 | 9 | novel transcript, antisense to SPTAN1 |
| WDR34 | 9 | WD repeat domain 34 [Source:HGNC Symbol; |
| VTI1BP4 | 9 | vesicle transport through interaction with t-SI |
| HMGA1P4 | 9 | high mobility group AT-hook 1 pseudogene 4 |
| SET | 9 | SET nuclear proto-oncogene [Source:HGNC Sy |
| PKN3 | 9 | protein kinase N3 [Source:HGNC Symbol;Acc:I |
| RN7SL560P | 9 | RNA, 7SL, cytoplasmic 560, pseudogene [Sour |
| ZDHHC12 | 9 | zinc finger DHHC-type containing 12 [Source:H |
| AL441992.1 | 9 | novel transcript |
| ZER1 | 9 | zyg-11 related cell cycle regulator [Source:HGI |
| Metazoa_SRP | 9 | Metazoan signal recognition particle RNA [Sou |
| TBC1D13 | 9 | TBC1 domain family member 13 [Source:HGN |
| ENDOG | 9 | endonuclease G [Source:HGNC Symbol;Acc:H( |
| SPOUT1 | 9 | SPOUT domain containing methyltransferase |
| AL441992.2 | 9 | kynurenine aminotransferase 1 [Source:NCBI |
| KYAT1 | 9 | kynurenine aminotransferase 1 [Source:HGNC |
| LRRC8A | 9 | leucine rich repeat containing 8 VRAC subunit |

| | | |
|---|---|---|
| PHYHD1 | 9 | phytanoyl-CoA dioxygenase domain containin |
| AL672142.1 | 9 | novel protein |
| DOLK | 9 | dolichol kinase [Source:HGNC Symbol;Acc:HG |
| NUP188 | 9 | nucleoporin 188 [Source:HGNC Symbol;Acc:H |
| AL592211.1 | 9 | novel transcript |
| SH3GLB2 | 9 | SH3 domain containing GRB2 like, endophilin |
| MIGA2 | 9 | mitoguardin 2 [Source:HGNC Symbol;Acc:HGN |
| AL592211.2 | 9 | novel transcript |
| DOLPP1 | 9 | dolichyldiphosphatase 1 [Source:HGNC Symbo |
| CRAT | 9 | carnitine O-acetyltransferase [Source:HGNC S |
| AL158151.2 | 9 | novel transcript |
| PTPA | 9 | protein phosphatase 2 phosphatase activator |
| AL158151.4 | 9 | novel transcript |
| IER5L | 9 | immediate early response 5 like [Source:HGN |
| AL158151.3 | 9 | novel transcript, antisense to IER5L |
| AL158151.1 | 9 | novel transcript |
| AL161785.2 | 9 | novel transcript |
| AL161785.3 | 9 | novel transcript |
| AL161785.1 | 9 | novel transcript |
| C9orf106 | 9 | chromosome 9 open reading frame 106 (puta |
| AL353803.5 | 9 | novel transcript |
| LINC01503 | 9 | long intergenic non-protein coding RNA 1503 |
| RN7SL159P | 9 | RNA, 7SL, cytoplasmic 159, pseudogene [Sour |
| AL353803.3 | 9 | novel pseudogene |
| AL353803.2 | 9 | novel transcript |
| AL353803.1 | 9 | novel transcript |
| LINC00963 | 9 | long intergenic non-protein coding RNA 963 [S |
| AL353803.4 | 9 | novel transcript |
| AL391056.2 | 9 | novel transcript |
| AL391056.1 | 9 | novel transcript |
| NTMT1 | 9 | N-terminal Xaa-Pro-Lys N-methyltransferase 1 |
| C9orf50 | 9 | chromosome 9 open reading frame 50 [Sourc |
| ASB6 | 9 | ankyrin repeat and SOCS box containing 6 [So |
| AL590369.1 | 9 | novel transcript |
| PRRX2 | 9 | paired related homeobox 2 [Source:HGNC Syr |
| PRRX2-AS1 | 9 | PRRX2 antisense RNA 1 [Source:HGNC Symbol |
| PTGES | 9 | prostaglandin E synthase [Source:HGNC Symb |
| UBE2V1P4 | 9 | ubiquitin conjugating enzyme E2 V1 pseudoge |
| TOR1B | 9 | torsin family 1 member B [Source:HGNC Symb |
| TOR1A | 9 | torsin family 1 member A [Source:HGNC Symb |
| C9orf78 | 9 | chromosome 9 open reading frame 78 [Sourc |
| USP20 | 9 | ubiquitin specific peptidase 20 [Source:HGNC |
| MIR6855 | 9 | microRNA 6855 [Source:HGNC Symbol;Acc:HG |
| AL158207.1 | 9 | 60S ribosomal protein L34 (RPL34) pseudogen |
| FNBP1 | 9 | formin binding protein 1 [Source:HGNC Symbo |
| AL158207.2 | 9 | novel transcript |
| AL136141.1 | 9 | novel transcript |
| GPR107 | 9 | G protein-coupled receptor 107 [Source:HGN |
| Y_RNA | 9 | Y RNA [Source:RFAM;Acc:RF00019] |
| GPRACR | 9 | GPR107 adjacent cis regulating lncRNA [Sourc |

| | | |
|---|---|---|
| NCS1 | 9 | neuronal calcium sensor 1 [Source:HGNC Sym |
| HMCN2 | 9 | hemicentin 2 [Source:HGNC Symbol;Acc:HGN |
| RN7SL665P | 9 | RNA, 7SL, cytoplasmic 665, pseudogene [Sour |
| ASS1 | 9 | argininosuccinate synthase 1 [Source:HGNC Sy |
| FUBP3 | 9 | far upstream element binding protein 3 [Sourc |
| MIR6856 | 9 | microRNA 6856 [Source:HGNC Symbol;Acc:HG |
| AL359092.2 | 9 | RPL19 (ribosomal protein L19) pseudogene |
| PRDM12 | 9 | PR/SET domain 12 [Source:HGNC Symbol;Acc: |
| EXOSC2 | 9 | exosome component 2 [Source:HGNC Symbol |
| ABL1 | 9 | ABL proto-oncogene 1, non-receptor tyrosine |
| AL359092.1 | 9 | ribosomal protein L37 (RPL37) pseudogene |
| QRFP | 9 | pyroglutamylated RFamide peptide [Source:H |
| FIBCD1 | 9 | fibrinogen C domain containing 1 [Source:HGN |
| AL161733.1 | 9 | novel transcript |
| LAMC3 | 9 | laminin subunit gamma 3 [Source:HGNC Symb |
| AL583807.1 | 9 | NULL |
| AIF1L | 9 | allograft inflammatory factor 1 like [Source:HG |
| NUP214 | 9 | nucleoporin 214 [Source:HGNC Symbol;Acc:HG |
| RNU6-881P | 9 | RNA, U6 small nuclear 881, pseudogene [Sour |
| AL157938.2 | 9 | novel transcript, antisense to NUP214 |
| AL157938.3 | 9 | novel transcript |
| AL157938.1 | 9 | ribosomal protein L13a (RPL13A) pseudogene |
| FAM78A | 9 | family with sequence similarity 78 member A |
| AL157938.4 | 9 | novel transcript |
| PLPP7 | 9 | phospholipid phosphatase 7 (inactive) [Source |
| AL354855.1 | 9 | novel pseudogene similar to avian transmemb |
| PRRC2B | 9 | proline rich coiled-coil 2B [Source:HGNC Symb |
| SNORD62A | 9 | small nucleolar RNA, C/D box 62A [Source:HG |
| SNORD62B | 9 | small nucleolar RNA, C/D box 62B [Source:HG |
| AL358781.1 | 9 | novel transcript, antisense to PRRC2B |
| POMT1 | 9 | protein O-mannosyltransferase 1 [Source:HGN |
| AL358781.2 | 9 | novel transcript |
| UCK1 | 9 | uridine-cytidine kinase 1 [Source:HGNC Symbo |
| PRRT1B | 9 | proline rich transmembrane protein 1B [Sourc |
| RAPGEF1 | 9 | Rap guanine nucleotide exchange factor 1 [So |
| RN7SL328P | 9 | RNA, 7SL, cytoplasmic 328, pseudogene [Sour |
| EIF4A1P3 | 9 | eukaryotic translation initiation factor 4A1 pse |
| AL160271.1 | 9 | NULL |
| AL160271.2 | 9 | novel transcript |
| MED27 | 9 | mediator complex subunit 27 [Source:HGNC S |
| NTNG2 | 9 | netrin G2 [Source:HGNC Symbol;Acc:HGNC:14 |
| SETX | 9 | senataxin [Source:HGNC Symbol;Acc:HGNC:44 |
| TTF1 | 9 | transcription termination factor 1 [Source:HG |
| CFAP77 | 9 | cilia and flagella associated protein 77 [Source |
| RNU5D-2P | 9 | RNA, U5D small nuclear 2, pseudogene [Sourc |
| BARHL1 | 9 | BarH like homeobox 1 [Source:HGNC Symbol; |
| DDX31 | 9 | DEAD-box helicase 31 [Source:HGNC Symbol;A |
| GTF3C4 | 9 | general transcription factor IIIC subunit 4 [Sou |
| AK8 | 9 | adenylate kinase 8 [Source:HGNC Symbol;Acc |
| AL445645.1 | 9 | novel transcript |

| | | |
|---|---|---|
| SPACA9 | 9 | sperm acrosome associated 9 [Source:HGNC S |
| TSC1 | 9 | TSC complex subunit 1 [Source:HGNC Symbol; |
| GFI1B | 9 | growth factor independent 1B transcriptional |
| MIR548AW | 9 | microRNA 548aw [Source:HGNC Symbol;Acc:H |
| AL593851.1 | 9 | ribosomal protein L39 (RPL39) pseudogene |
| RNU7-21P | 9 | RNA, U7 small nuclear 21 pseudogene [Source |
| EEF1A1P5 | 9 | eukaryotic translation elongation factor 1 alph |
| GTF3C5 | 9 | general transcription factor IIIC subunit 5 [Sou |
| MIR6877 | 9 | microRNA 6877 [Source:HGNC Symbol;Acc:HG |
| CEL | 9 | carboxyl ester lipase [Source:HGNC Symbol;Ac |
| CELP | 9 | carboxyl ester lipase pseudogene [Source:HGN |
| RALGDS | 9 | ral guanine nucleotide dissociation stimulator |
| AL162417.1 | 9 | novel protein |
| GBGT1 | 9 | globoside alpha-1,3-N-acetylgalactosaminyltra |
| OBP2B | 9 | odorant binding protein 2B [Source:HGNC Syn |
| LCN1P1 | 9 | lipocalin 1 pseudogene 1 [Source:HGNC Symb |
| ABO | 9 | ABO, alpha 1-3-N-acetylgalactosaminyltransfe |
| Y_RNA | 9 | Y RNA [Source:RFAM;Acc:RF00019] |
| LCN1P2 | 9 | lipocalin 1 pseudogene 2 [Source:HGNC Symb |
| SURF6 | 9 | surfeit 6 [Source:HGNC Symbol;Acc:HGNC:114 |
| Y_RNA | 9 | Y RNA [Source:RFAM;Acc:RF00019] |
| MED22 | 9 | mediator complex subunit 22 [Source:HGNC S |
| AL772161.1 | 9 | ribosomal protein L21 (RPL21) pseudogene |
| RPL7A | 9 | ribosomal protein L7a [Source:HGNC Symbol;A |
| SNORD24 | 9 | small nucleolar RNA, C/D box 24 [Source:HGN |
| SNORD36B | 9 | small nucleolar RNA, C/D box 36B [Source:HG |
| SNORD36A | 9 | small nucleolar RNA, C/D box 36A [Source:HG |
| SNORD36C | 9 | small nucleolar RNA, C/D box 36C [Source:HG |
| SURF1 | 9 | SURF1 cytochrome c oxidase assembly factor |
| SURF2 | 9 | surfeit 2 [Source:HGNC Symbol;Acc:HGNC:114 |
| SURF4 | 9 | surfeit 4 [Source:HGNC Symbol;Acc:HGNC:114 |
| STKLD1 | 9 | serine/threonine kinase like domain containin |
| REXO4 | 9 | REX4 homolog, 3'-5' exonuclease [Source:HGN |
| ADAMTS13 | 9 | ADAM metallopeptidase with thrombospondi |
| CACFD1 | 9 | calcium channel flower domain containing 1 [S |
| SLC2A6 | 9 | solute carrier family 2 member 6 [Source:HGN |
| MYMK | 9 | myomaker, myoblast fusion factor [Source:HG |
| ADAMTSL2 | 9 | ADAMTS like 2 [Source:HGNC Symbol;Acc:HGI |
| FAM163B | 9 | family with sequence similarity 163 member E |
| AC002101.1 | 9 | novel transcript, antisense to DBH |
| DBH | 9 | dopamine beta-hydroxylase [Source:HGNC Sy |
| DBH-AS1 | 9 | DBH antisense RNA 1 [Source:HGNC Symbol;A |
| SARDH | 9 | sarcosine dehydrogenase [Source:HGNC Symb |
| VAV2 | 9 | vav guanine nucleotide exchange factor 2 [Sou |
| BRD3OS | 9 | BRD3 opposite strand [Source:HGNC Symbol;A |
| BRD3 | 9 | bromodomain containing 3 [Source:HGNC Syn |
| AL445931.1 | 9 | novel transcript, antisense to BRD3 |
| ARF4P1 | 9 | ADP ribosylation factor 4 pseudogene 1 [Sour |
| BX649632.1 | 9 | novel transcript, antisense to WDR5 |
| WDR5 | 9 | WD repeat domain 5 [Source:HGNC Symbol;A |

| | | |
|---|---|---|
| RNU6ATAC | 9 | RNA, U6atac small nuclear (U12-dependent sp |
| BX649601.1 | 9 | novel transcript |
| LINC02247 | 9 | long intergenic non-protein coding RNA 2247 |
| AL354796.1 | 9 | tec |
| RXRA | 9 | retinoid X receptor alpha [Source:HGNC Symb |
| MIR4669 | 9 | microRNA 4669 [Source:HGNC Symbol;Acc:HG |
| AL669970.1 | 9 | novel transcript |
| AL669970.3 | 9 | novel transcript |
| AL669970.2 | 9 | novel transcript |
| COL5A1 | 9 | collagen type V alpha 1 chain [Source:HGNC S |
| COL5A1-AS1 | 9 | COL5A1 antisense RNA 1 [Source:HGNC Symb |
| AL645768.1 | 9 | novel transcript, antisense to COL5A1 |
| MIR3689C | 9 | microRNA 3689c [Source:HGNC Symbol;Acc:H |
| MIR3689A | 9 | microRNA 3689a [Source:HGNC Symbol;Acc:H |
| MIR3689D1 | 9 | microRNA 3689d-1 [Source:HGNC Symbol;Acc |
| MIR3689B | 9 | microRNA 3689b [Source:HGNC Symbol;Acc:H |
| MIR3689D2 | 9 | microRNA 3689d-2 [Source:HGNC Symbol;Acc |
| MIR3689E | 9 | microRNA 3689e [Source:HGNC Symbol;Acc:H |
| MIR3689F | 9 | microRNA 3689f [Source:HGNC Symbol;Acc:H |
| AL603650.1 | 9 | pseudogene similar to part of a ficolin family p |
| FCN2 | 9 | ficolin 2 [Source:HGNC Symbol;Acc:HGNC:362 |
| FCN1 | 9 | ficolin 1 [Source:HGNC Symbol;Acc:HGNC:362 |
| AL353611.1 | 9 | novel transcript |
| AL353611.2 | 9 | tec |
| OLFM1 | 9 | olfactomedin 1 [Source:HGNC Symbol;Acc:HG |
| AL390778.1 | 9 | novel transcript |
| AL390778.2 | 9 | novel transcript |
| C9orf62 | 9 | chromosome 9 open reading frame 62 [Sourc |
| AL353615.1 | 9 | novel transcript |
| SOCS5P2 | 9 | suppressor of cytokine signaling 5 pseudogen |
| PPP1R26-AS1 | 9 | PPP1R26 antisense RNA 1 [Source:HGNC Syml |
| PPP1R26 | 9 | protein phosphatase 1 regulatory subunit 26 [ |
| C9orf116 | 9 | chromosome 9 open reading frame 116 [Sour |
| MRPS2 | 9 | mitochondrial ribosomal protein S2 [Source:H |
| AL161452.1 | 9 | novel transcript |
| LCN1 | 9 | lipocalin 1 [Source:HGNC Symbol;Acc:HGNC:6 |
| OBP2A | 9 | odorant binding protein 2A [Source:HGNC Syn |
| PAEP | 9 | progestagen associated endometrial protein [ |
| LINC01502 | 9 | long intergenic non-protein coding RNA 1502 |
| PAEPP1 | 9 | progestagen associated endometrial protein p |
| AL354761.1 | 9 | lipocalin 1-like [Source:NCBI gene;Acc:102723 |
| GLT6D1 | 9 | glycosyltransferase 6 domain containing 1 [So |
| LCN9 | 9 | lipocalin 9 [Source:HGNC Symbol;Acc:HGNC:1 |
| SOHLH1 | 9 | spermatogenesis and oogenesis specific basic |
| KCNT1 | 9 | potassium sodium-activated channel subfamil |
| CAMSAP1 | 9 | calmodulin regulated spectrin associated prot |
| AL355574.1 | 9 | novel transcript |
| UBAC1 | 9 | UBA domain containing 1 [Source:HGNC Symb |
| NACC2 | 9 | NACC family member 2 [Source:HGNC Symbol |
| LINC02846 | 9 | long intergenic non-protein coding RNA 2846 |

| | | |
|---|---|---|
| TMEM250 | 9 | transmembrane protein 250 [Source:HGNC Sy |
| AL138781.1 | 9 | novel transcript |
| LHX3 | 9 | LIM homeobox 3 [Source:HGNC Symbol;Acc:H |
| QSOX2 | 9 | quiescin sulfhydryl oxidase 2 [Source:HGNC Sy |
| CCDC187 | 9 | coiled-coil domain containing 187 [Source:HG |
| CR392000.1 | 9 | TEC |
| CR392000.2 | 9 | novel transcript, antisense to a novel protein |
| AC174065.1 | 9 | novel transcript |
| GPSM1 | 9 | G protein signaling modulator 1 [Source:HGN( |
| DNLZ | 9 | DNL-type zinc finger [Source:HGNC Symbol;Ac |
| CARD9 | 9 | caspase recruitment domain family member 9 |
| SNAPC4 | 9 | small nuclear RNA activating complex polypep |
| ENTR1 | 9 | endosome associated trafficking regulator 1 [S |
| PMPCA | 9 | peptidase, mitochondrial processing alpha sub |
| INPP5E | 9 | inositol polyphosphate-5-phosphatase E [Sour |
| SEC16A | 9 | SEC16 homolog A, endoplasmic reticulum exp |
| C9orf163 | 9 | chromosome 9 open reading frame 163 [Sour |
| NOTCH1 | 9 | notch receptor 1 [Source:HGNC Symbol;Acc:H |
| MIR4673 | 9 | microRNA 4673 [Source:HGNC Symbol;Acc:HG |
| AL592301.1 | 9 | novel transcript |
| MIR4674 | 9 | microRNA 4674 [Source:HGNC Symbol;Acc:HG |
| NALT1 | 9 | NOTCH1 associated lncRNA in T cell acute lym |
| AL590226.2 | 9 | long intergenic non-protein coding RNA 1451 |
| AL590226.1 | 9 | novel transcript |
| EGFL7 | 9 | EGF like domain multiple 7 [Source:HGNC Sym |
| MIR126 | 9 | microRNA 126 [Source:HGNC Symbol;Acc:HGN |
| AGPAT2 | 9 | 1-acylglycerol-3-phosphate O-acyltransferase |
| DIPK1B | 9 | divergent protein kinase domain 1B [Source:H |
| SNHG7 | 9 | small nucleolar RNA host gene 7 [Source:HGN |
| SNORA17B | 9 | small nucleolar RNA, H/ACA box 17B [Source:I |
| SNORA17B | 9 | small nucleolar RNA, H/ACA box 17B [Source:I |
| SNORA17A | 9 | small nucleolar RNA, H/ACA box 17A [Source:I |
| AL355987.6 | 9 | NULL |
| AL355987.5 | 9 | ral guanine nucleotide dissociation stimulator |
| LCN10 | 9 | lipocalin 10 [Source:HGNC Symbol;Acc:HGNC: |
| AL355987.1 | 9 | novel protein |
| LCN6 | 9 | lipocalin 6 [Source:HGNC Symbol;Acc:HGNC:1 |
| MIR6722 | 9 | microRNA 6722 [Source:HGNC Symbol;Acc:HG |
| LCN8 | 9 | lipocalin 8 [Source:HGNC Symbol;Acc:HGNC:2 |
| LCN15 | 9 | lipocalin 15 [Source:HGNC Symbol;Acc:HGNC: |
| ATP6V1G1P3 | 9 | ATPase H+ transporting V1 subunit G1 pseudo |
| TMEM141 | 9 | transmembrane protein 141 [Source:HGNC Sy |
| AL355987.3 | 9 | novel protein |
| CCDC183 | 9 | coiled-coil domain containing 183 [Source:HG |
| AL355987.4 | 9 | novel transcript |
| AL355987.2 | 9 | novel transcript |
| CCDC183-AS1 | 9 | CCDC183 antisense RNA 1 [Source:HGNC Sym |
| RABL6 | 9 | RAB, member RAS oncogene family like 6 [Sou |
| NCLP1 | 9 | nucleolin pseudogene 1 [Source:HGNC Symbo |
| MIR4292 | 9 | microRNA 4292 [Source:HGNC Symbol;Acc:HG |

| | | |
|---|---|---|
| AJM1 | 9 | apical junction component 1 homolog [Source |
| PHPT1 | 9 | phosphohistidine phosphatase 1 [Source:HGN |
| MAMDC4 | 9 | MAM domain containing 4 [Source:HGNC Sym |
| EDF1 | 9 | endothelial differentiation related factor 1 [Sc |
| TRAF2 | 9 | TNF receptor associated factor 2 [Source:HGN |
| MIR4479 | 9 | microRNA 4479 [Source:HGNC Symbol;Acc:HG |
| AL807752.1 | 9 | ribosomal protein L30 (RPL30) pseudogene |
| AL807752.5 | 9 | novel transcript, overlapping FBXW5 |
| FBXW5 | 9 | F-box and WD repeat domain containing 5 [Sc |
| C8G | 9 | complement C8 gamma chain [Source:HGNC S |
| LCN12 | 9 | lipocalin 12 [Source:HGNC Symbol;Acc:HGNC: |
| LINC02692 | 9 | long intergenic non-protein coding RNA 2692 |
| PTGDS | 9 | prostaglandin D2 synthase [Source:HGNC Sym |
| AL807752.7 | 9 | novel transcript |
| LCNL1 | 9 | lipocalin like 1 [Source:HGNC Symbol;Acc:HGN |
| PAXX | 9 | PAXX non-homologous end joining factor [Sou |
| CLIC3 | 9 | chloride intracellular channel 3 [Source:HGNC |
| ABCA2 | 9 | ATP binding cassette subfamily A member 2 [S |
| C9orf139 | 9 | chromosome 9 open reading frame 139 [Sour |
| FUT7 | 9 | fucosyltransferase 7 [Source:HGNC Symbol;Ac |
| AL807752.6 | 9 | novel protein |
| NPDC1 | 9 | neural proliferation, differentiation and contr |
| ENTPD2 | 9 | ectonucleoside triphosphate diphosphohydro |
| AL807752.4 | 9 | novel transcript |
| AL807752.2 | 9 | novel transcript, antisense to SAPCD2 |
| SAPCD2 | 9 | suppressor APC domain containing 2 [Source:I |
| AL807752.3 | 9 | novel transcript |
| UAP1L1 | 9 | UDP-N-acetylglucosamine pyrophosphorylase |
| MAN1B1-DT | 9 | MAN1B1 divergent transcript [Source:HGNC S |
| MAN1B1 | 9 | mannosidase alpha class 1B member 1 [Sourc |
| SNORD62 | 9 | Small nucleolar RNA SNORD62 [Source:RFAM; |
| DPP7 | 9 | dipeptidyl peptidase 7 [Source:HGNC Symbol; |
| GRIN1 | 9 | glutamate ionotropic receptor NMDA type sul |
| LRRC26 | 9 | leucine rich repeat containing 26 [Source:HGN |
| AL929554.1 | 9 | novel transcript, TMEM210 and LRRC26 readt |
| MIR3621 | 9 | microRNA 3621 [Source:HGNC Symbol;Acc:HG |
| TMEM210 | 9 | transmembrane protein 210 [Source:HGNC Sy |
| ANAPC2 | 9 | anaphase promoting complex subunit 2 [Sour |
| SSNA1 | 9 | SS nuclear autoantigen 1 [Source:HGNC Symb |
| TPRN | 9 | taperin [Source:HGNC Symbol;Acc:HGNC:2689 |
| TMEM203 | 9 | transmembrane protein 203 [Source:HGNC Sy |
| NDOR1 | 9 | NADPH dependent diflavin oxidoreductase 1 [ |
| BX255925.3 | 9 | novel RING finger protein |
| RNF208 | 9 | ring finger protein 208 [Source:HGNC Symbol; |
| CYSRT1 | 9 | cysteine rich tail 1 [Source:HGNC Symbol;Acc: |
| RNF224 | 9 | ring finger protein 224 [Source:HGNC Symbol; |
| SLC34A3 | 9 | solute carrier family 34 member 3 [Source:HG |
| TUBB4B | 9 | tubulin beta 4B class IVb [Source:HGNC Symbo |
| FAM166A | 9 | family with sequence similarity 166 member A |
| STPG3-AS1 | 9 | STPG3 antisense RNA 1 [Source:HGNC Symbol |

| | | |
|---|---|---|
| STPG3 | 9 | sperm-tail PG-rich repeat containing 3 [Source |
| NELFB | 9 | negative elongation factor complex member B |
| TOR4A | 9 | torsin family 4 member A [Source:HGNC Symb |
| BX255925.2 | 9 | DSN1, MIND kinetochore complex component |
| BX255925.1 | 9 | novel transcript, overlapping NRARP |
| BX255925.4 | 9 | novel transcript, antisense to NRARP |
| NRARP | 9 | NOTCH regulated ankyrin repeat protein [Sou |
| EXD3 | 9 | exonuclease 3'-5' domain containing 3 [Source |
| NOXA1 | 9 | NADPH oxidase activator 1 [Source:HGNC Sym |
| ENTPD8 | 9 | ectonucleoside triphosphate diphosphohydro |
| NSMF | 9 | NMDA receptor synaptonuclear signaling and |
| MIR7114 | 9 | microRNA 7114 [Source:HGNC Symbol;Acc:HG |
| PNPLA7 | 9 | patatin like phospholipase domain containing |
| MRPL41 | 9 | mitochondrial ribosomal protein L41 [Source: |
| DPH7 | 9 | diphthamide biosynthesis 7 [Source:HGNC Syr |
| ZMYND19 | 9 | zinc finger MYND-type containing 19 [Source: |
| ARRDC1 | 9 | arrestin domain containing 1 [Source:HGNC Sy |
| AL590627.2 | 9 | novel transcript |
| ARRDC1-AS1 | 9 | ARRDC1 antisense RNA 1 [Source:HGNC Symb |
| EHMT1 | 9 | euchromatic histone lysine methyltransferase |
| Metazoa_SRP | 9 | Metazoan signal recognition particle RNA [Sou |
| SETP5 | 9 | SET pseudogene 5 [Source:HGNC Symbol;Acc: |
| AL590627.1 | 9 | forkhead box H1 (FOXH1) pseudogene |
| MIR602 | 9 | microRNA 602 [Source:HGNC Symbol;Acc:HGN |
| AL772363.1 | 9 | novel transcript, antisense to CACNA1B and El |
| CACNA1B | 9 | calcium voltage-gated channel subunit alpha1 |
| AL591424.2 | 9 | polymerase (RNA) III (DNA directed) polypepti |
| AL591424.1 | 9 | pseudogene similar to part of interleukin 9 re |
| TUBBP5 | 9 | tubulin beta pseudogene 5 [Source:HGNC Sym |
| AL591424.3 | 9 | family with sequence similarity 166, member |
| AL954642.1 | 9 | novel transcript |
| FAM157B | 9 | family with sequence similarity 157 member B |
| RNU6-785P | 9 | RNA, U6 small nuclear 785, pseudogene [Sour |
| FO082796.1 | 9 | septin 14 pseudogene 9 |
| FO082796.2 | 9 | novel olfactory receptor pseudogene |
| AC215217.1 | 10 | novel transcript |
| AL713922.1 | 10 | FAM166A, pseudogene 7 |
| TUBB8 | 10 | tubulin beta 8 class VIII [Source:HGNC Symbol |
| IL9RP2 | 10 | interleukin 9 receptor pseudogene 2 [Source: |
| ZMYND11 | 10 | zinc finger MYND-type containing 11 [Source: |
| AL713922.2 | 10 | DEAD (Asp-Glu-Ala-Asp) box polypeptide 20 (D |
| RNA5SP297 | 10 | RNA, 5S ribosomal pseudogene 297 [Source:H |
| DIP2C | 10 | disco interacting protein 2 homolog C [Source |
| RNA5SP298 | 10 | RNA, 5S ribosomal pseudogene 298 [Source:H |
| RN7SL754P | 10 | RNA, 7SL, cytoplasmic 754, pseudogene [Sour |
| AL669841.1 | 10 | novel transcript |
| AL358216.1 | 10 | novel transcript |
| MIR5699 | 10 | microRNA 5699 [Source:HGNC Symbol;Acc:HG |
| PRR26 | 10 | proline rich 26 [Source:HGNC Symbol;Acc:HGN |
| AL157709.1 | 10 | novel transcript |

| | | |
|---|---|---|
| LARP4B | 10 | La ribonucleoprotein domain family member 4 |
| AL359878.2 | 10 | novel transcript |
| AL359878.1 | 10 | novel transcript, antisense to GTPBP4 |
| GTPBP4 | 10 | GTP binding protein 4 [Source:HGNC Symbol;A |
| IDI2 | 10 | isopentenyl-diphosphate delta isomerase 2 [S |
| IDI2-AS1 | 10 | IDI2 antisense RNA 1 [Source:HGNC Symbol;A |
| IDI1 | 10 | isopentenyl-diphosphate delta isomerase 1 [S |
| WDR37 | 10 | WD repeat domain 37 [Source:HGNC Symbol;A |
| LINC00200 | 10 | long intergenic non-protein coding RNA 200 [S |
| ADARB2 | 10 | adenosine deaminase RNA specific B2 (inactiv |
| AL392083.2 | 10 | novel transcript, sense intronic to ADARB2 |
| AL392083.1 | 10 | novel transcript, sense intronic to ADARB2 |
| AL513304.1 | 10 | novel transcript |
| ADARB2-AS1 | 10 | ADARB2 antisense RNA 1 [Source:HGNC Symb |
| AL355597.1 | 10 | novel transcript |
| LINC00700 | 10 | long intergenic non-protein coding RNA 700 [S |
| AL441943.1 | 10 | novel transcript |
| MIR6072 | 10 | microRNA 6072 [Source:HGNC Symbol;Acc:HG |
| RNU6-889P | 10 | RNA, U6 small nuclear 889, pseudogene [Sour |
| AL441943.2 | 10 | novel transcript (FLJ31539) |
| LINC02662 | 10 | long intergenic non-protein coding RNA 2662 |
| RNU6-576P | 10 | RNA, U6 small nuclear 576, pseudogene [Sour |
| LINC00701 | 10 | long intergenic non-protein coding RNA 701 [S |
| LINC02645 | 10 | long intergenic non-protein coding RNA 2645 |
| AL713851.2 | 10 | novel transcript |
| AL713851.1 | 10 | novel transcript |
| AL355361.1 | 10 | novel transcript |
| AC026396.1 | 10 | novel pseudogene |
| AL731533.1 | 10 | novel transcript |
| AL731533.3 | 10 | novel transcript |
| AL731533.2 | 10 | novel transcript |
| PFKP | 10 | phosphofructokinase, platelet [Source:HGNC S |
| AL451164.2 | 10 | novel readthrough transcript, antisense to PIT |
| PITRM1 | 10 | pitrilysin metallopeptidase 1 [Source:HGNC Sy |
| PITRM1-AS1 | 10 | PITRM1 antisense RNA 1 [Source:HGNC Symbo |
| LINC02668 | 10 | long intergenic non-protein coding RNA 2668 |
| AL451164.1 | 10 | novel transcript |
| AL139280.1 | 10 | novel transcript |
| AL139280.2 | 10 | novel transcript |
| AL356433.1 | 10 | novel transcript |
| LINC02669 | 10 | long intergenic non-protein coding RNA 2669 |
| AL357833.1 | 10 | novel transcript |
| AL450322.2 | 10 | novel transcript |
| AL450322.1 | 10 | novel transcript |
| KLF6 | 10 | Kruppel like factor 6 [Source:HGNC Symbol;Ac |
| AL513303.1 | 10 | novel transcript |
| LINC02639 | 10 | long intergenic non-protein coding RNA 2639 |
| LINC02660 | 10 | long intergenic non-protein coding RNA 2660 |
| MIR6078 | 10 | microRNA 6078 [Source:HGNC Symbol;Acc:HG |
| AC025822.1 | 10 | novel transcript |

| | | |
|---|---|---|
| AC025822.2 | 10 | novel transcript (FLJ31241) |
| LINC00702 | 10 | long intergenic non-protein coding RNA 702 [S |
| LINC00703 | 10 | long intergenic non-protein coding RNA 703 [S |
| AC022535.1 | 10 | novel transcript |
| AC022535.3 | 10 | novel transcript |
| AC022535.2 | 10 | novel transcript |
| RNU6-163P | 10 | RNA, U6 small nuclear 163, pseudogene [Sour |
| MANCR | 10 | mitotically associated long non coding RNA [S |
| LINC00705 | 10 | long intergenic non-protein coding RNA 705 [S |
| AC018978.1 | 10 | novel transcript |
| AKR1E2 | 10 | aldo-keto reductase family 1 member E2 [Sou |
| AKR1C6P | 10 | aldo-keto reductase family 1 member C6, pse |
| U8 | 10 | U8 small nucleolar RNA [Source:RFAM;Acc:RF |
| AKR1C1 | 10 | aldo-keto reductase family 1 member C1 [Sou |
| AKR1C2 | 10 | aldo-keto reductase family 1 member C2 [Sou |
| AL391427.1 | 10 | novel transcript, antisense to AKR1C2 |
| U8 | 10 | U8 small nucleolar RNA [Source:RFAM;Acc:RF |
| AKR1C3 | 10 | aldo-keto reductase family 1 member C3 [Sou |
| U8 | 10 | U8 small nucleolar RNA [Source:RFAM;Acc:RF |
| AKR1C5P | 10 | aldo-keto reductase family 1 member C5, pse |
| AKR1C8P | 10 | aldo-keto reductase family 1 member C8, pse |
| AKR1C4 | 10 | aldo-keto reductase family 1 member C4 [Sou |
| ARL4AP3 | 10 | ADP ribosylation factor like GTPase 4A pseudo |
| AL355303.2 | 10 | novel transcript, antisense to AKR1C4 |
| AL355303.1 | 10 | novel transcript |
| LINC02561 | 10 | long intergenic non-protein coding RNA 2561 |
| AKR1C7P | 10 | aldo-keto reductase family 1 member C7, pse |
| RPL26P28 | 10 | ribosomal protein L26 pseudogene 28 [Source |
| AL683826.1 | 10 | TEC |
| UCN3 | 10 | urocortin 3 [Source:HGNC Symbol;Acc:HGNC: |
| TUBAL3 | 10 | tubulin alpha like 3 [Source:HGNC Symbol;Acc |
| NET1 | 10 | neuroepithelial cell transforming 1 [Source:H |
| CALML5 | 10 | calmodulin like 5 [Source:HGNC Symbol;Acc:H |
| CALML3-AS1 | 10 | CALML3 antisense RNA 1 [Source:HGNC Symb |
| CALML3 | 10 | calmodulin like 3 [Source:HGNC Symbol;Acc:H |
| AL732437.1 | 10 | novel transcript |
| AL732437.3 | 10 | novel transcript |
| AL732437.2 | 10 | novel transcript |
| LINC02657 | 10 | long intergenic non-protein coding RNA 2657 |
| LINC02678 | 10 | long intergenic non-protein coding RNA 2678 |
| LINC02677 | 10 | long intergenic non-protein coding RNA 2677 |
| AL365356.2 | 10 | ribosomal protein L23a (RPL23A) pseudogene |
| RN7SL445P | 10 | RNA, 7SL, cytoplasmic 445, pseudogene [Sour |
| ASB13 | 10 | ankyrin repeat and SOCS box containing 13 [S |
| TASOR2 | 10 | transcription activation suppressor family me |
| AL365356.1 | 10 | novel transcript |
| GDI2 | 10 | GDP dissociation inhibitor 2 [Source:HGNC Sy |
| NRBF2P5 | 10 | nuclear receptor binding factor 2 pseudogene |
| AL596094.1 | 10 | novel transcript |
| ANKRD16 | 10 | ankyrin repeat domain 16 [Source:HGNC Syml |

| | | |
|---|---|---|
| RPL12P28 | 10 | ribosomal protein L12 pseudogene 28 [Source |
| FBH1 | 10 | F-box DNA helicase 1 [Source:HGNC Symbol;A |
| AL137186.2 | 10 | novel transcript, antisense to FBXO18 |
| IL15RA | 10 | interleukin 15 receptor subunit alpha [Source: |
| IL2RA | 10 | interleukin 2 receptor subunit alpha [Source:H |
| AL137186.3 | 10 | NULL |
| AL137186.1 | 10 | novel transcript |
| RPL32P23 | 10 | ribosomal protein L32 pseudogene 23 [Source |
| RBM17 | 10 | RNA binding motif protein 17 [Source:HGNC S |
| PFKFB3 | 10 | 6-phosphofructo-2-kinase/fructose-2,6-bipho: |
| RN7SKP78 | 10 | RN7SK pseudogene 78 [Source:HGNC Symbol; |
| MIR3155A | 10 | microRNA 3155a [Source:HGNC Symbol;Acc:H |
| MIR3155B | 10 | microRNA 3155b [Source:HGNC Symbol;Acc:H |
| AL157395.1 | 10 | novel transcript |
| Y_RNA | 10 | Y RNA [Source:RFAM;Acc:RF00019] |
| LINC02649 | 10 | long intergenic non-protein coding RNA 2649 |
| SDCBPP1 | 10 | syndecan binding protein pseudogene 1 [Sour |
| LINC02656 | 10 | long intergenic non-protein coding RNA 2656 |
| PRKCQ | 10 | protein kinase C theta [Source:HGNC Symbol;. |
| PRKCQ-AS1 | 10 | PRKCQ antisense RNA 1 [Source:HGNC Symbo |
| LINC02648 | 10 | long intergenic non-protein coding RNA 2648 |
| AL158210.1 | 10 | novel transcript, LINP1-LINC00707 readthroug |
| LINP1 | 10 | lncRNA in non-homologous end joining pathw |
| AL392086.2 | 10 | novel transcript, LINP1-LINC00707 readthroug |
| LINC00706 | 10 | long intergenic non-protein coding RNA 706 [S |
| LINC00707 | 10 | long intergenic non-protein coding RNA 707 [S |
| AL392086.1 | 10 | novel transcript |
| AL392086.3 | 10 | novel transcript |
| LINC02665 | 10 | long intergenic non-protein coding RNA 2665 |
| AL590095.1 | 10 | ribosomal protein L21 (RPL21) pseudogene |
| SFMBT2 | 10 | Scm like with four mbt domains 2 [Source:HG |
| COX6CP17 | 10 | cytochrome c oxidase subunit 6C pseudogene |
| AL139125.1 | 10 | TEC |
| LINC02642 | 10 | long intergenic non-protein coding RNA 2642 |
| RNU6-535P | 10 | RNA, U6 small nuclear 535, pseudogene [Sour |
| AL445070.1 | 10 | novel transcript |
| ITIH5 | 10 | inter-alpha-trypsin inhibitor heavy chain 5 [So |
| ITIH2 | 10 | inter-alpha-trypsin inhibitor heavy chain 2 [So |
| KIN | 10 | Kin17 DNA and RNA binding protein [Source:H |
| ATP5F1C | 10 | ATP synthase F1 subunit gamma [Source:HGN |
| TAF3 | 10 | TATA-box binding protein associated factor 3 |
| AL353754.1 | 10 | mitochondrial ribosomal protein L13 (MRPL13 |
| GATA3 | 10 | GATA binding protein 3 [Source:HGNC Symbol |
| GATA3-AS1 | 10 | GATA3 antisense RNA 1 [Source:HGNC Symbo |
| AL390294.1 | 10 | novel transcript |
| PRPF38AP1 | 10 | PRP38 domain containing A pseudogene 1 [So |
| LINC00708 | 10 | long intergenic non-protein coding RNA 708 [S |
| AL390835.1 | 10 | novel transcript |
| AC025946.1 | 10 | novel transcript |
| AC025946.2 | 10 | SGT1, suppressor of G2 allele of SKP1 (S. cere\ |

| | | |
|---|---|---|
| KRT8P37 | 10 | keratin 8 pseudogene 37 [Source:HGNC Symb |
| CHCHD3P1 | 10 | coiled-coil-helix-coiled-coil-helix domain conta |
| RNA5SP299 | 10 | RNA, 5S ribosomal pseudogene 299 [Source:H |
| AL355333.1 | 10 | oocyte expressed protein (OOEP) pseudogene |
| LINC02676 | 10 | long intergenic non-protein coding RNA 2676 |
| AC044784.1 | 10 | novel transcript |
| LINC00709 | 10 | long intergenic non-protein coding RNA 709 [S |
| AL138773.1 | 10 | novel transcript |
| HSP90AB7P | 10 | heat shock protein 90 alpha family class B mer |
| LINC02663 | 10 | long intergenic non-protein coding RNA 2663 |
| LINC02670 | 10 | long intergenic non-protein coding RNA 2670 |
| ELOCP3 | 10 | elongin C pseudogene 3 [Source:HGNC Symbo |
| Y_RNA | 10 | Y RNA [Source:RFAM;Acc:RF00019] |
| CUX2P1 | 10 | cut like homeobox 2 pseudogene 1 [Source:H( |
| AL354916.1 | 10 | novel transcript |
| AL583859.1 | 10 | novel transcript |
| CELF2-DT | 10 | CELF2 divergent transript [Source:HGNC Symb |
| AL583859.2 | 10 | TEC |
| AL583859.3 | 10 | novel transcript |
| AL136452.1 | 10 | ORM1-like (S. cerevisiae) pseudogene |
| SFTA1P | 10 | surfactant associated 1, pseudogene [Source:I |
| CELF2 | 10 | CUGBP Elav-like family member 2 [Source:HGI |
| LINC00710 | 10 | long intergenic non-protein coding RNA 710 [S |
| AL136369.2 | 10 | novel transcript |
| AL136369.1 | 10 | novel transcript |
| AL162408.1 | 10 | novel transcript |
| CELF2-AS2 | 10 | CELF2 antisense RNA 2 [Source:HGNC Symbol, |
| AL136320.1 | 10 | novel transcript |
| CELF2-AS1 | 10 | CELF2 antisense RNA 1 [Source:HGNC Symbol, |
| AC026887.1 | 10 | TEC |
| USP6NL | 10 | USP6 N-terminal like [Source:HGNC Symbol;A( |
| AL512631.2 | 10 | novel transcript, antisense to USP6NL |
| AL512631.1 | 10 | novel transcript |
| ECHDC3 | 10 | enoyl-CoA hydratase domain containing 3 [So |
| PROSER2 | 10 | proline and serine rich 2 [Source:HGNC Symbo |
| PROSER2-AS1 | 10 | PROSER2 antisense RNA 1 [Source:HGNC Sym |
| UPF2 | 10 | UPF2 regulator of nonsense mediated mRNA ( |
| RNU6-1095P | 10 | RNA, U6 small nuclear 1095, pseudogene [Sou |
| DHTKD1 | 10 | dehydrogenase E1 and transketolase domain ( |
| RNU6-88P | 10 | RNA, U6 small nuclear 88, pseudogene [Sourc |
| SEC61A2 | 10 | SEC61 translocon alpha 2 subunit [Source:HGI |
| MIR548AK | 10 | microRNA 548ak [Source:HGNC Symbol;Acc:H |
| NUDT5 | 10 | nudix hydrolase 5 [Source:HGNC Symbol;Acc:I |
| CDC123 | 10 | cell division cycle 123 [Source:HGNC Symbol;A |
| AL512770.1 | 10 | novel transcript |
| RN7SL198P | 10 | RNA, 7SL, cytoplasmic 198, pseudogene [Sour |
| CAMK1D | 10 | calcium/calmodulin dependent protein kinase |
| AL391314.1 | 10 | NULL |
| RNU6ATAC39P | 10 | RNA, U6atac small nuclear 39, pseudogene [S( |
| AL731559.1 | 10 | novel transcript, sense intronic to CAMK1D |

| | | |
|---|---|---|
| MIR4480 | 10 | microRNA 4480 [Source:HGNC Symbol;Acc:HG |
| MIR4481 | 10 | microRNA 4481 [Source:HGNC Symbol;Acc:HG |
| MIR548Q | 10 | microRNA 548q [Source:HGNC Symbol;Acc:HG |
| AL353586.1 | 10 | novel transcript, antisense to CCDC3 |
| CCDC3 | 10 | coiled-coil domain containing 3 [Source:HGNC |
| RNA5SP300 | 10 | RNA, 5S ribosomal pseudogene 300 [Source:H |
| RPL5P25 | 10 | ribosomal protein L5 pseudogene 25 [Source: |
| OPTN | 10 | optineurin [Source:HGNC Symbol;Acc:HGNC:1 |
| SNRPGP5 | 10 | small nuclear ribonucleoprotein polypeptide ( |
| AL355355.2 | 10 | pseudogene similar to part of chromodomain |
| AL355355.1 | 10 | pseudogene similar to part of COX10 homolog |
| BTBD7P1 | 10 | BTB domain containing 7 pseudogene 1 [Sour |
| RPL36AP36 | 10 | ribosomal protein L36a pseudogene 36 [Sourc |
| MCM10 | 10 | minichromosome maintenance 10 replication |
| RNU6-6P | 10 | RNA, U6 small nuclear 6, pseudogene [Source |
| UCMA | 10 | upper zone of growth plate and cartilage matr |
| PHYH | 10 | phytanoyl-CoA 2-hydroxylase [Source:HGNC S |
| RBISP1 | 10 | RBIS pseudogene 1 [Source:HGNC Symbol;Acc |
| SEPHS1 | 10 | selenophosphate synthetase 1 [Source:HGNC |
| AL355870.2 | 10 | novel transcript |
| AL355870.1 | 10 | novel transcript |
| BEND7 | 10 | BEN domain containing 7 [Source:HGNC Symb |
| Y_RNA | 10 | Y RNA [Source:RFAM;Acc:RF00019] |
| AL590677.1 | 10 | novel transcript |
| PRPF18 | 10 | pre-mRNA processing factor 18 [Source:HGNC |
| RPL6P24 | 10 | ribosomal protein L6 pseudogene 24 [Source: |
| AL157392.5 | 10 | novel protein |
| AL157392.3 | 10 | novel transcript |
| FRMD4A | 10 | FERM domain containing 4A [Source:HGNC Sy |
| AL157392.4 | 10 | novel transcript, antisense to FRMD4A |
| AL157392.1 | 10 | novel transcript |
| RNA5SP301 | 10 | RNA, 5S ribosomal pseudogene 301 [Source:H |
| AL157392.2 | 10 | novel transcript |
| NUTF2P5 | 10 | nuclear transport factor 2 pseudogene 5 [Sour |
| AC044781.1 | 10 | novel transcript |
| AL157896.1 | 10 | novel transcript |
| AL139405.1 | 10 | novel transcript |
| MIR4293 | 10 | microRNA 4293 [Source:HGNC Symbol;Acc:HG |
| Metazoa_SRP | 10 | Metazoan signal recognition particle RNA [Sou |
| MIR1265 | 10 | microRNA 1265 [Source:HGNC Symbol;Acc:HG |
| FAM107B | 10 | family with sequence similarity 107 member B |
| AL158168.1 | 10 | novel transcript, antisense to FAM107B |
| RNA5SP302 | 10 | RNA, 5S ribosomal pseudogene 302 [Source:H |
| RPSAP7 | 10 | ribosomal protein SA pseudogene 7 [Source:H |
| CDNF | 10 | cerebral dopamine neurotrophic factor [Sourc |
| HSPA14 | 10 | heat shock protein family A (Hsp70) member 1 |
| HSPA14 | 10 | heat shock protein family A (Hsp70) member 1 |
| AC069544.1 | 10 | novel transcript |
| SUV39H2 | 10 | suppressor of variegation 3-9 homolog 2 [Sou |
| DCLRE1C | 10 | DNA cross-link repair 1C [Source:HGNC Symbo |

| | | |
|---|---|---|
| MEIG1 | 10 | meiosis/spermiogenesis associated 1 [Source: |
| OR7E110P | 10 | olfactory receptor family 7 subfamily E memb |
| OR7E26P | 10 | olfactory receptor family 7 subfamily E memb |
| OR7E115P | 10 | olfactory receptor family 7 subfamily E memb |
| DCLRE1CP1 | 10 | DNA cross-link repair 1C pseudogene 1 [Sourc |
| OLAH | 10 | oleoyl-ACP hydrolase [Source:HGNC Symbol;A |
| ACBD7 | 10 | acyl-CoA binding domain containing 7 [Source |
| GAPDHP45 | 10 | glyceraldehyde 3 phosphate dehydrogenase p |
| RPP38-DT | 10 | RPP38 divergent transcript [Source:HGNC Sym |
| RPP38 | 10 | ribonuclease P/MRP subunit p38 [Source:HGN |
| NMT2 | 10 | N-myristoyltransferase 2 [Source:HGNC Symb |
| PPIAP30 | 10 | peptidylprolyl isomerase A pseudogene 30 [So |
| AL590365.1 | 10 | NADH dehydrogenase (ubiquinone) 1 alpha su |
| FAM171A1 | 10 | family with sequence similarity 171 member A |
| AL607028.1 | 10 | novel transcript |
| ITGA8 | 10 | integrin subunit alpha 8 [Source:HGNC Symbo |
| MINDY3 | 10 | MINDY lysine 48 deubiquitinase 3 [Source:HG |
| AL358033.1 | 10 | NULL |
| FTLP19 | 10 | ferritin light chain pseudogene 19 [Source:HG |
| RNU6-1075P | 10 | RNA, U6 small nuclear 1075, pseudogene [Sou |
| LINC02654 | 10 | long intergenic non-protein coding RNA 2654 |
| PTER | 10 | phosphotriesterase related [Source:HGNC Syr |
| RNU2-18P | 10 | RNA, U2 small nuclear 18, pseudogene [Sourc |
| C1QL3 | 10 | complement C1q like 3 [Source:HGNC Symbol |
| AL353576.1 | 10 | novel transcript, antisense to C1QL3 |
| RSU1 | 10 | Ras suppressor protein 1 [Source:HGNC Symb |
| AC073367.1 | 10 | novel transcript, sense intronic to RSU1 |
| AL365215.1 | 10 | NULL |
| AL365215.2 | 10 | novel transcript |
| CUBN | 10 | cubilin [Source:HGNC Symbol;Acc:HGNC:2548 |
| AC067747.1 | 10 | novel transcript |
| TRDMT1 | 10 | tRNA aspartic acid methyltransferase 1 [Sourc |
| VIM-AS1 | 10 | VIM antisense RNA 1 [Source:HGNC Symbol;A |
| VIM | 10 | vimentin [Source:HGNC Symbol;Acc:HGNC:12 |
| AL133415.1 | 10 | novel transcript |
| ST8SIA6 | 10 | ST8 alpha-N-acetyl-neuraminide alpha-2,8-sia |
| ST8SIA6-AS1 | 10 | ST8SIA6 antisense RNA 1 [Source:HGNC Symb |
| Y_RNA | 10 | Y RNA [Source:RFAM;Acc:RF00019] |
| PRPF38AP2 | 10 | PRP38 domain containing A pseudogene 2 [So |
| HACD1 | 10 | 3-hydroxyacyl-CoA dehydratase 1 [Source:HGI |
| AC069542.2 | 10 | NULL |
| STAM-AS1 | 10 | STAM antisense RNA 1 (head to head) [Source |
| STAM | 10 | signal transducing adaptor molecule [Source:I |
| AC069542.1 | 10 | novel transcript |
| TMEM236 | 10 | transmembrane protein 236 [Source:HGNC Sy |
| MRC1 | 10 | mannose receptor C-type 1 [Source:HGNC Syr |
| MIR511 | 10 | microRNA 511 [Source:HGNC Symbol;Acc:HGI |
| AC069023.1 | 10 | novel transcript |
| SLC39A12 | 10 | solute carrier family 39 member 12 [Source:H |
| SLC39A12-AS1 | 10 | SLC39A12 antisense RNA 1 [Source:HGNC Sym |

| | | |
|---|---|---|
| CACNB2 | 10 | calcium voltage-gated channel auxiliary subun |
| AL390783.1 | 10 | novel transcript |
| AL450384.2 | 10 | novel transcript, antisense to CACNB2 |
| AL450384.1 | 10 | novel transcript |
| NSUN6 | 10 | NOP2/Sun RNA methyltransferase 6 [Source:H |
| ARL5B | 10 | ADP ribosylation factor like GTPase 5B [Source |
| AL512641.1 | 10 | novel transcript |
| AIFM1P1 | 10 | AIFM1 pseudogene 1 [Source:HGNC Symbol;A |
| AL357520.1 | 10 | pseudogene similar to part of glycogen syntha |
| MALRD1 | 10 | MAM and LDL receptor class A domain contai |
| UBE2V2P1 | 10 | ubiquitin conjugating enzyme E2 V2 pseudoge |
| AL590378.1 | 10 | novel transcript |
| HMGN1P20 | 10 | high mobility group nucleosome binding doma |
| AL354695.1 | 10 | pseudogene similar to part of ATPase, Ca++ tr |
| U3 | 10 | Small nucleolar RNA U3 [Source:RFAM;Acc:RF |
| RNA5SP303 | 10 | RNA, 5S ribosomal pseudogene 303 [Source:H |
| AL157895.1 | 10 | novel transcript, antisense to C10orf112 |
| MTND1P37 | 10 | MT-ND1 pseudogene 37 [Source:HGNC Symbo |
| MTND2P16 | 10 | MT-ND2 pseudogene 16 [Source:HGNC Symbo |
| RNU6-1212P | 10 | RNA, U6 small nuclear 1212, pseudogene [Sou |
| AL353147.1 | 10 | novel transcript |
| PLXDC2 | 10 | plexin domain containing 2 [Source:HGNC Syn |
| AC069549.1 | 10 | novel transcript |
| AC069549.2 | 10 | novel transcript, antisense to PLXDC2 |
| AMD1P1 | 10 | adenosylmethionine decarboxylase 1 pseudog |
| AL590032.1 | 10 | chromosome 21 open reading frame 59 (C21o |
| MIR4675 | 10 | microRNA 4675 [Source:HGNC Symbol;Acc:HG |
| NEBL | 10 | nebulette [Source:HGNC Symbol;Acc:HGNC:16 |
| MTND1P21 | 10 | MT-ND1 pseudogene 21 [Source:HGNC Symbo |
| AL157398.1 | 10 | pseudogene similar to part of MTCO3 (COIII)( |
| EIF4BP2 | 10 | eukaryotic translation initiation factor 4B pseu |
| NPM1P30 | 10 | nucleophosmin 1 pseudogene 30 [Source:HGN |
| C10orf113 | 10 | chromosome 10 open reading frame 113 [Sou |
| NEBL-AS1 | 10 | NEBL antisense RNA 1 [Source:HGNC Symbol;A |
| AL731547.1 | 10 | ribosomal protein L15 (RPL15) pseudogene |
| LUZP4P1 | 10 | leucine zipper protein 4 pseudogene 1 [Source |
| RNU6-15P | 10 | RNA, U6 small nuclear 15, pseudogene [Sourc |
| LINC02643 | 10 | long intergenic non-protein coding RNA 2643 |
| RNMTL1P1 | 10 | RNA methyltransferase like 1 pseudogene 1 [S |
| AL158209.1 | 10 | adenosine deaminase domain containing 1 (te |
| Metazoa_SRP | 10 | Metazoan signal recognition particle RNA [Sou |
| Y_RNA | 10 | Y RNA [Source:RFAM;Acc:RF00019] |
| MIR1915HG | 10 | MIR1915 host gene [Source:HGNC Symbol;Acc |
| MIR1915 | 10 | microRNA 1915 [Source:HGNC Symbol;Acc:HG |
| SKIDA1 | 10 | SKI/DACH domain containing 1 [Source:HGNC |
| MLLT10 | 10 | MLLT10 histone lysine methyltransferase DOT |
| AL358780.1 | 10 | novel transcript, sense intronic to MLLT10 |
| RNU6-306P | 10 | RNA, U6 small nuclear 306, pseudogene [Sour |
| HNRNPRP1 | 10 | heterogeneous nuclear ribonucleoprotein R p |
| RNU6-1141P | 10 | RNA, U6 small nuclear 1141, pseudogene [Sou |

| | | |
|---|---|---|
| DNAJC1 | 10 | DnaJ heat shock protein family (Hsp40) memb |
| RN7SKP219 | 10 | RN7SK pseudogene 219 [Source:HGNC Symbo |
| AL359697.1 | 10 | novel transcript, sense intronic to DNAJC1 |
| RN7SKP37 | 10 | RN7SK pseudogene 37 [Source:HGNC Symbol; |
| ADIPOR1P1 | 10 | adiponectin receptor 1 pseudogene 1 [Source |
| AL157831.1 | 10 | ribosomal protein L29 (RPL29) pseudogene |
| EBLN1 | 10 | endogenous Bornavirus like nucleoprotein 1 [ |
| AL157831.2 | 10 | novel transcript |
| PSME2P6 | 10 | proteasome activator subunit 2 pseudogene 6 |
| AL157831.3 | 10 | novel transcript |
| AL158211.5 | 10 | TEC |
| AL158211.1 | 10 | novel transcript |
| RPL31P45 | 10 | ribosomal protein L31 pseudogene 45 [Source |
| COMMD3 | 10 | COMM domain containing 3 [Source:HGNC Sy |
| COMMD3-BMI1 | 10 | COMMD3-BMI1 readthrough [Source:HGNC S |
| BMI1 | 10 | BMI1 proto-oncogene, polycomb ring finger [S |
| AL158211.4 | 10 | novel transcript |
| AL158211.2 | 10 | novel transcript |
| AL158211.3 | 10 | novel transcript |
| SPAG6 | 10 | sperm associated antigen 6 [Source:HGNC Syr |
| AL513128.1 | 10 | novel transcript, sense overlapping SPAG6 |
| AL513128.3 | 10 | novel transcript |
| AL513128.2 | 10 | novel transcript |
| PIP4K2A | 10 | phosphatidylinositol-5-phosphate 4-kinase typ |
| AL390318.1 | 10 | TEC |
| RNU6-413P | 10 | RNA, U6 small nuclear 413, pseudogene [Sour |
| ARMC3 | 10 | armadillo repeat containing 3 [Source:HGNC S |
| AL139815.1 | 10 | novel transcript, antisense to ARMC3and MSR |
| RNA5SP304 | 10 | RNA, 5S ribosomal pseudogene 304 [Source:H |
| MSRB2 | 10 | methionine sulfoxide reductase B2 [Source:HC |
| YWHAZP3 | 10 | tyrosine 3-monooxygenase/tryptophan 5-mor |
| AL139281.2 | 10 | NULL |
| PTF1A | 10 | pancreas associated transcription factor 1a [S |
| C10orf67 | 10 | chromosome 10 open reading frame 67 [Sour |
| AL139281.1 | 10 | novel pseudogene |
| AL606469.1 | 10 | novel transcript |
| AL606469.2 | 10 | NULL |
| OTUD1 | 10 | OTU deubiquitinase 1 [Source:HGNC Symbol;A |
| AL512603.2 | 10 | novel transcript |
| AL512603.1 | 10 | novel transcript |
| KIAA1217 | 10 | KIAA1217 [Source:HGNC Symbol;Acc:HGNC:2 |
| NUP35P1 | 10 | nucleoporin 35 pseudogene 1 [Source:HGNC S |
| AC063961.1 | 10 | NULL |
| MIR603 | 10 | microRNA 603 [Source:HGNC Symbol;Acc:HGI |
| AL356113.1 | 10 | novel transcript, antisense to KIAA1217 |
| AL353583.1 | 10 | TEC |
| ARHGAP21 | 10 | Rho GTPase activating protein 21 [Source:HGI |
| RNA5SP305 | 10 | RNA, 5S ribosomal pseudogene 305 [Source:H |
| AL157385.2 | 10 | novel transcript |
| PRTFDC1 | 10 | phosphoribosyl transferase domain containin |

| | | |
|---|---|---|
| AL157385.1 | 10 | pseudogene similar to part of NADH dehydrog |
| RN7SKP241 | 10 | RN7SK pseudogene 241 [Source:HGNC Symbo |
| AL512598.2 | 10 | novel transcript |
| AL512598.1 | 10 | novel transcript |
| ENKUR | 10 | enkurin, TRPC channel interacting protein [Sou |
| THNSL1 | 10 | threonine synthase like 1 [Source:HGNC Symb |
| LINC01516 | 10 | long intergenic non-protein coding RNA 1516 |
| GPR158-AS1 | 10 | GPR158 antisense RNA 1 [Source:HGNC Symb |
| GPR158 | 10 | G protein-coupled receptor 158 [Source:HGN( |
| AL354976.1 | 10 | PR domain containing 9 (PRDM9) pseudogene |
| RN7SKP220 | 10 | RN7SK pseudogene 220 [Source:HGNC Symbo |
| GPN3P1 | 10 | GPN-loop GTPase 3 pseudogene 1 [Source:HG |
| LINC00836 | 10 | long intergenic non-protein coding RNA 836 [S |
| HIRAP1 | 10 | histone cell cycle regulator pseudogene 1 [Sou |
| RNA5SP306 | 10 | RNA, 5S ribosomal pseudogene 306 [Source:H |
| RNU6-632P | 10 | RNA, U6 small nuclear 632, pseudogene [Sour |
| AL358612.1 | 10 | novel transcript |
| MYO3A | 10 | myosin IIIA [Source:HGNC Symbol;Acc:HGNC:7 |
| GAD2 | 10 | glutamate decarboxylase 2 [Source:HGNC Sym |
| AL355798.1 | 10 | novel pseudogene, ortholog of LOC10309363! |
| APBB1IP | 10 | amyloid beta precursor protein binding family |
| RNA5SP307 | 10 | RNA, 5S ribosomal pseudogene 307 [Source:H |
| AL160287.1 | 10 | novel pseudogene |
| FAM238A | 10 | family with sequence similarity 238 member A |
| AL390961.1 | 10 | novel transcript |
| FAM238B | 10 | family with sequence similarity 238 member E |
| SELENOOLP | 10 | selenoprotein O like, pseudogene [Source:HG |
| AL390961.3 | 10 | TEC |
| PDSS1 | 10 | decaprenyl diphosphate synthase subunit 1 [S |
| HSPA8P3 | 10 | heat shock protein family A (Hsp70) member 8 |
| AL390961.2 | 10 | novel transcript |
| ABI1 | 10 | abl interactor 1 [Source:HGNC Symbol;Acc:HG |
| AL139404.1 | 10 | NULL |
| RNU6-946P | 10 | RNA, U6 small nuclear 946, pseudogene [Sour |
| FAM238C | 10 | family with sequence similarity 238 member C |
| ANKRD26 | 10 | ankyrin repeat domain 26 [Source:HGNC Syml |
| RNU6-490P | 10 | RNA, U6 small nuclear 490, pseudogene [Sour |
| YME1L1 | 10 | YME1 like 1 ATPase [Source:HGNC Symbol;Acc |
| MASTL | 10 | microtubule associated serine/threonine kina: |
| RNU2-24P | 10 | RNA, U2 small nuclear 24, pseudogene [Sourc |
| ACBD5 | 10 | acyl-CoA binding domain containing 5 [Source |
| RNU7-12P | 10 | RNA, U7 small nuclear 12 pseudogene [Source |
| AL160291.1 | 10 | novel transcript |
| LRRC37A6P | 10 | leucine rich repeat containing 37 member A6, |
| ARMC4P1 | 10 | armadillo repeat containing 4 pseudogene 1 [S |
| AL355493.3 | 10 | ankyrin repeat domain 30B-like (ANKRD30BL) |
| RNU6-666P | 10 | RNA, U6 small nuclear 666, pseudogene [Sour |
| AL355493.1 | 10 | pseudogene similar to part of breast cancer ar |
| RNU6-452P | 10 | RNA, U6 small nuclear 452, pseudogene [Sour |
| LINC02673 | 10 | long intergenic non-protein coding RNA 2673 |

| | | |
|---|---|---|
| FAM210CP | 10 | family with sequence similarity 210 member ( |
| TRIAP1P1 | 10 | TP53 regulated inhibitor of apoptosis 1 pseud |
| PTCHD3 | 10 | patched domain containing 3 [Source:HGNC S |
| RAB18 | 10 | RAB18, member RAS oncogene family [Source |
| LINC02680 | 10 | long intergenic non-protein coding RNA 2680 |
| MKX | 10 | mohawk homeobox [Source:HGNC Symbol;Ac |
| MKX-AS1 | 10 | MKX antisense RNA 1 [Source:HGNC Symbol;/ |
| ARMC4 | 10 | armadillo repeat containing 4 [Source:HGNC S |
| RN7SKP132 | 10 | RN7SK pseudogene 132 [Source:HGNC Symbo |
| RPL36AP55 | 10 | ribosomal protein L36a pseudogene 55 [Sourc |
| AL390866.1 | 10 | novel transcript |
| MRPS21P5 | 10 | mitochondrial ribosomal protein S21 pseudog |
| MPP7 | 10 | membrane palmitoylated protein 7 [Source:H |
| AL355501.1 | 10 | novel transcript, antisense to MPP7 |
| MIR8086 | 10 | microRNA 8086 [Source:HGNC Symbol;Acc:HG |
| MPP7-DT | 10 | MPP7 divergent transcript [Source:HGNC Sym |
| ZNF101P1 | 10 | zinc finger protein 101 pseudogene 1 [Source: |
| U3 | 10 | Small nucleolar RNA U3 [Source:RFAM;Acc:RF |
| LINC02652 | 10 | long intergenic non-protein coding RNA 2652 |
| RPSAP10 | 10 | ribosomal protein SA pseudogene 10 [Source: |
| RN7SKP39 | 10 | RN7SK pseudogene 39 [Source:HGNC Symbol; |
| WAC-AS1 | 10 | WAC antisense RNA 1 (head to head) [Source: |
| WAC | 10 | WW domain containing adaptor with coiled-c |
| RNU4ATAC6P | 10 | RNA, U4atac small nuclear 6, pseudogene [So |
| TPRKBP1 | 10 | TP53RK binding protein pseudogene 1 [Source |
| RNU6-1067P | 10 | RNA, U6 small nuclear 1067, pseudogene [Sou |
| BAMBI | 10 | BMP and activin membrane bound inhibitor [S |
| LINC01517 | 10 | long intergenic non-protein coding RNA 1517 |
| AL355376.1 | 10 | novel transcript |
| RNU6-270P | 10 | RNA, U6 small nuclear 270, pseudogene [Sour |
| LINC00837 | 10 | long intergenic non-protein coding RNA 837 [S |
| AL355376.2 | 10 | neural proliferation, differentiation and contr |
| RNA5SP308 | 10 | RNA, 5S ribosomal pseudogene 308 [Source:H |
| RPL21P93 | 10 | ribosomal protein L21 pseudogene 93 [Source |
| LYZL1 | 10 | lysozyme like 1 [Source:HGNC Symbol;Acc:HG |
| AL158167.1 | 10 | novel transcript |
| SVIL-AS1 | 10 | SVIL antisense RNA 1 [Source:HGNC Symbol;A |
| PTCHD3P1 | 10 | patched domain containing 3 pseudogene 1 [S |
| SVIL | 10 | supervillin [Source:HGNC Symbol;Acc:HGNC:1 |
| MIR604 | 10 | microRNA 604 [Source:HGNC Symbol;Acc:HGN |
| SNORD115 | 10 | Small nucleolar RNA SNORD115 [Source:RFAN |
| MIR938 | 10 | microRNA 938 [Source:HGNC Symbol;Acc:HGN |
| CKS1BP2 | 10 | CDC28 protein kinase regulatory subunit 1B p: |
| RNU6-908P | 10 | RNA, U6 small nuclear 908, pseudogene [Sour |
| AL353093.1 | 10 | single-stranded DNA binding protein (SSBP1)(S |
| JCAD | 10 | junctional cadherin 5 associated [Source:HGN |
| RNU6-598P | 10 | RNA, U6 small nuclear 598, pseudogene [Sour |
| EEF1A1P39 | 10 | eukaryotic translation elongation factor 1 alpl |
| AL353796.1 | 10 | novel transcript, overlapping to MTPAP |
| MTPAP | 10 | mitochondrial poly(A) polymerase [Source:HG |

| | | |
|---|---|---|
| AL353796.2 | 10 | novel transcript, antisense to MTPAP |
| DNM1P17 | 10 | dynamin 1 pseudogene 17 [Source:HGNC Sym |
| GOLGA2P6 | 10 | GOLGA2 pseudogene 6 [Source:HGNC Symbol |
| RN7SL241P | 10 | RNA, 7SL, cytoplasmic 241, pseudogene [Sour |
| MIR7162 | 10 | microRNA 7162 [Source:HGNC Symbol;Acc:HG |
| NIFKP1 | 10 | nucleolar protein interacting with the FHA dor |
| AL161651.1 | 10 | pseudogene similar to part of SIN3 homolog A |
| CCND3P1 | 10 | cyclin D3 pseudogene 1 [Source:HGNC Symbo |
| MAP3K8 | 10 | mitogen-activated protein kinase kinase kinas |
| HNRNPA1P32 | 10 | heterogeneous nuclear ribonucleoprotein A1 |
| AL590068.4 | 10 | novel transcript |
| AL590068.1 | 10 | novel transcript |
| RN7SL63P | 10 | RNA, 7SL, cytoplasmic 63, pseudogene [Sourc |
| AL590068.2 | 10 | novel transcript |
| AL590068.3 | 10 | novel transcript |
| LYZL2 | 10 | lysozyme like 2 [Source:HGNC Symbol;Acc:HG |
| SVIL2P | 10 | supervillin family member 2, pseudogene [Sou |
| LINC02644 | 10 | long intergenic non-protein coding RNA 2644 |
| ZNF438 | 10 | zinc finger protein 438 [Source:HGNC Symbol; |
| AL359532.1 | 10 | novel transcript, sense intronic to ZNF438 |
| DDX10P1 | 10 | DEAD-box helicase 10 pseudogene 1 [Source:I |
| AL359546.1 | 10 | novel transcript |
| LINC02664 | 10 | long intergenic non-protein coding RNA 2664 |
| ZEB1-AS1 | 10 | ZEB1 antisense RNA 1 [Source:HGNC Symbol;A |
| RNA5SP309 | 10 | RNA, 5S ribosomal pseudogene 309 [Source:H |
| ZEB1 | 10 | zinc finger E-box binding homeobox 1 [Source |
| SPTLC1P1 | 10 | serine palmitoyltransferase long chain base su |
| AL117340.1 | 10 | novel pseudogene similar to nuclear pore asso |
| AL161935.1 | 10 | novel transcript |
| AL161935.3 | 10 | novel transcript |
| AL161935.2 | 10 | glutamate dehydrogenase pseudogene 5 |
| MACORIS | 10 | macrophage enriched lincRNA repressor of IFI |
| Y_RNA | 10 | Y RNA [Source:RFAM;Acc:RF00019] |
| ARHGAP12 | 10 | Rho GTPase activating protein 12 [Source:HGI |
| HMGB1P7 | 10 | high mobility group box 1 pseudogene 7 [Sour |
| RPL34P19 | 10 | ribosomal protein L34 pseudogene 19 [Source |
| RN7SL825P | 10 | RNA, 7SL, cytoplasmic 825, pseudogene [Sour |
| KIF5B | 10 | kinesin family member 5B [Source:HGNC Sym |
| Y_RNA | 10 | Y RNA [Source:RFAM;Acc:RF00019] |
| RPS4XP11 | 10 | ribosomal protein S4X pseudogene 11 [Source |
| AL161932.2 | 10 | part of a novel pseudogene |
| AL161932.1 | 10 | pseudogene similar to part of RAB18 (RAB18, |
| RNU7-22P | 10 | RNA, U7 small nuclear 22 pseudogene [Source |
| PPIAP31 | 10 | peptidylprolyl isomerase A pseudogene 31 [So |
| RPS24P13 | 10 | ribosomal protein S24 pseudogene 13 [Source |
| AL158834.2 | 10 | novel transcript |
| EPC1 | 10 | enhancer of polycomb homolog 1 [Source:HG |
| AL158834.1 | 10 | novel transcript, antisense to EPC1 |
| AL391839.2 | 10 | novel transcript, antisense to EPC1 |
| AL391839.1 | 10 | novel transcript |

| | | |
|---|---|---|
| RNU6-1244P | 10 | RNA, U6 small nuclear 1244, pseudogene [Sou |
| AL391839.3 | 10 | novel transcript |
| CCDC7 | 10 | coiled-coil domain containing 7 [Source:HGNC |
| C1DP1 | 10 | C1D nuclear receptor corepressor pseudogen |
| AL365203.2 | 10 | novel transcript |
| ITGB1 | 10 | integrin subunit beta 1 [Source:HGNC Symbol; |
| AK3P5 | 10 | adenylate kinase 3 pseudogene 5 [Source:HGI |
| ITGB1-DT | 10 | ITGB1 divergent transcript [Source:HGNC Sym |
| MTND4LP11 | 10 | MT-ND4L pseudogene 11 [Source:HGNC Symb |
| RN7SL847P | 10 | RNA, 7SL, cytoplasmic 847, pseudogene [Sour |
| AL365203.1 | 10 | pseudogene similar to part of integrin, beta 2 |
| RPL7AP53 | 10 | ribosomal protein L7a pseudogene 53 [Source |
| IATPR | 10 | ITGB1 adjacent tumor promoting lncRNA [Sou |
| NRP1 | 10 | neuropilin 1 [Source:HGNC Symbol;Acc:HGNC |
| AL121748.1 | 10 | novel transcript, antisense to NRP1 |
| RN7SL398P | 10 | RNA, 7SL, cytoplasmic 398, pseudogene [Sour |
| AL353600.1 | 10 | novel transcript |
| AL353600.2 | 10 | novel transcript |
| LINC02628 | 10 | long intergenic non-protein coding RNA 2628 |
| AL450313.1 | 10 | novel transcript |
| LINC00838 | 10 | long intergenic non-protein coding RNA 838 [S |
| RPL23P11 | 10 | ribosomal protein L23 pseudogene 11 [Source |
| LINC02629 | 10 | long intergenic non-protein coding RNA 2629 |
| PARD3 | 10 | par-3 family cell polarity regulator [Source:HG |
| Y_RNA | 10 | Y RNA [Source:RFAM;Acc:RF00019] |
| ELOBP4 | 10 | elongin B pseudogene 4 [Source:HGNC Symbo |
| RPL37P18 | 10 | ribosomal protein L37 pseudogene 18 [Source |
| RPS12P16 | 10 | ribosomal protein S12 pseudogene 16 [Source |
| PARD3-AS1 | 10 | PARD3 antisense RNA 1 [Source:HGNC Symbo |
| SS18L2P1 | 10 | SS18 like 2 pseudogene 1 [Source:HGNC Symb |
| RNU6-847P | 10 | RNA, U6 small nuclear 847, pseudogene [Sour |
| RNU6-193P | 10 | RNA, U6 small nuclear 193, pseudogene [Sour |
| PRDX2P2 | 10 | peroxiredoxin 2 pseudogene 2 [Source:HGNC |
| LINC02635 | 10 | long intergenic non-protein coding RNA 2635 |
| CUL2 | 10 | cullin 2 [Source:HGNC Symbol;Acc:HGNC:2552 |
| MIR3611 | 10 | microRNA 3611 [Source:HGNC Symbol;Acc:HG |
| AL392046.1 | 10 | novel transcript |
| CREM | 10 | cAMP responsive element modulator [Source: |
| RNU7-77P | 10 | RNA, U7 small nuclear 77 pseudogene [Source |
| ATP6V1G1P4 | 10 | ATPase H+ transporting V1 subunit G1 pseudo |
| AL117336.1 | 10 | novel transcript, sense intronic to CREM |
| LINC02634 | 10 | long intergenic non-protein coding RNA 2634 |
| RNU6-794P | 10 | RNA, U6 small nuclear 794, pseudogene [Sour |
| CCNY | 10 | cyclin Y [Source:HGNC Symbol;Acc:HGNC:233! |
| RNU6-1167P | 10 | RNA, U6 small nuclear 1167, pseudogene [Sou |
| AL117336.2 | 10 | novel transcript, antisense to CCNY |
| AL603824.1 | 10 | novel transcript, antisense to CCNY |
| AL121749.1 | 10 | novel transcript |
| GJD4 | 10 | gap junction protein delta 4 [Source:HGNC Sy |
| FZD8 | 10 | frizzled class receptor 8 [Source:HGNC Symbo |

| | | |
|---|---|---|
| AL121749.2 | 10 | novel transcript, antisense to FZD8 |
| MIR4683 | 10 | microRNA 4683 [Source:HGNC Symbol;Acc:HG |
| RPL7P37 | 10 | ribosomal protein L7 pseudogene 37 [Source: |
| PCAT5 | 10 | prostate cancer associated transcript 5 [Sourc |
| LINC02630 | 10 | long intergenic non-protein coding RNA 2630 |
| AL355300.1 | 10 | novel transcript |
| MTND5P17 | 10 | MT-ND5 pseudogene 17 [Source:HGNC Symbo |
| MTND4P18 | 10 | MT-ND4 pseudogene 18 [Source:HGNC Symbo |
| AL590730.1 | 10 | novel transcript |
| AL590730.2 | 10 | NULL |
| NAMPTP1 | 10 | nicotinamide phosphoribosyltransferase pseu |
| Y_RNA | 10 | Y RNA [Source:RFAM;Acc:RF00019] |
| AL390061.1 | 10 | novel transcript |
| MKNK2P1 | 10 | MAP kinase interacting serine/threonine kina |
| ARL6IP1P2 | 10 | ADP ribosylation factor like GTPase 6 interacti |
| ANKRD30A | 10 | ankyrin repeat domain 30A [Source:HGNC Syr |
| RNU6-811P | 10 | RNA, U6 small nuclear 811, pseudogene [Sour |
| AL157387.1 | 10 | novel transcript |
| LINC00993 | 10 | long intergenic non-protein coding RNA 993 [S |
| RN7SL314P | 10 | RNA, 7SL, cytoplasmic 314, pseudogene [Sour |
| Y_RNA | 10 | Y RNA [Source:RFAM;Acc:RF00019] |
| TMEM161BP1 | 10 | transmembrane protein 161B pseudogene 1 [ |
| VN1R53P | 10 | vomeronasal 1 receptor 53 pseudogene [Sour |
| TACC1P1 | 10 | transforming acidic coiled-coil containing prot |
| MTND1P18 | 10 | MT-ND1 pseudogene 18 [Source:HGNC Symbo |
| MTRNR2L7 | 10 | MT-RNR2 like 7 [Source:HGNC Symbol;Acc:HG |
| AL132657.1 | 10 | novel transcript |
| ZNF248 | 10 | zinc finger protein 248 [Source:HGNC Symbol; |
| AL132657.2 | 10 | novel transcript |
| ZNF33BP1 | 10 | zinc finger protein 33B pseudogene 1 [Source: |
| TLK2P2 | 10 | tousled like kinase 2 pseudogene 2 [Source:H( |
| AL135791.1 | 10 | novel transcript |
| ZNF37CP | 10 | zinc finger protein 37C, pseudogene [Source:H |
| ZNF33CP | 10 | zinc finger protein 33C, pseudogene [Source:H |
| ZNF25 | 10 | zinc finger protein 25 [Source:HGNC Symbol;A |
| ZNF25-DT | 10 | ZNF25 divergent transcript [Source:HGNC Sym |
| Y_RNA | 10 | Y RNA [Source:RFAM;Acc:RF00019] |
| ZNF33A | 10 | zinc finger protein 33A [Source:HGNC Symbol; |
| RNU6-795P | 10 | RNA, U6 small nuclear 795, pseudogene [Sour |
| AL161931.1 | 10 | immunoglobulin lambda variable 4-3 (IGLV4-3 |
| EIF3LP3 | 10 | eukaryotic translation initiation factor 3 subur |
| FXYD6P2 | 10 | FXYD domain containing ion transport regulat |
| ZNF37A | 10 | zinc finger protein 37A [Source:HGNC Symbol; |
| AL117339.5 | 10 | novel transcript |
| AL117339.4 | 10 | novel transcript |
| AL117339.2 | 10 | par-3 partitioning defective 3 homolog (C.eleg |
| AL117339.1 | 10 | novel cyclin Y-like pseudogene |
| AL133217.1 | 10 | phospholipase D family member 5 (PLD5) pseu |
| AL117339.3 | 10 | pseudogene similar to part of excision repair c |
| HSD17B7P2 | 10 | hydroxysteroid 17-beta dehydrogenase 7 pseu |

| | | |
|---|---|---|
| SEPTIN7P9 | 10 | septin 7 pseudogene 9 [Source:HGNC Symbol; |
| AL133216.2 | 10 | general transcription factor IIi (GTF2I) pseudo |
| RNU6-1118P | 10 | RNA, U6 small nuclear 1118, pseudogene [Sou |
| CICP9 | 10 | capicua transcriptional repressor pseudogene |
| AL133216.1 | 10 | novel transcript |
| ABCD1P2 | 10 | ATP binding cassette subfamily D member 1 p |
| AL133173.2 | 10 | CDGSH iron sulfur domain 2 (CISD2) pseudoge |
| PABPC1P12 | 10 | poly(A) binding protein cytoplasmic 1 pseudo |
| SLC9B1P3 | 10 | solute carrier family 9 member B1 pseudogen |
| ACTR3BP5 | 10 | ACTR3B pseudogene 5 [Source:HGNC Symbol; |
| CHEK2P5 | 10 | checkpoint kinase 2 pseudogene 5 [Source:HG |
| AL133173.1 | 10 | ring finger protein 18 (RNF18) pseudogene |
| AL590623.1 | 10 | N-acetylated alpha-linked acidic dipeptidase 2 |
| KSR1P1 | 10 | kinase suppressor of ras 1 pseudogene 1 [Sou |
| IGKV1OR10-1 | 10 | immunoglobulin kappa variable 1/OR10-1 (ps |
| AL031601.1 | 10 | tripartite motif containing 43 (TRIM43) pseud |
| AL031601.2 | 10 | solute carrier family 9, subfamily B (NHA2, cat |
| PABPC1P8 | 10 | poly(A) binding protein cytoplasmic 1 pseudo |
| BX322639.1 | 10 | zinc finger protein pseudogene |
| CCNYL2 | 10 | cyclin Y like 2 (pseudogene) [Source:HGNC Sy |
| AL391099.1 | 10 | par-3 family cell polarity regulator (PARD3) ps |
| LINC00839 | 10 | long intergenic non-protein coding RNA 839 [ |
| ZNF37BP | 10 | zinc finger protein 37B, pseudogene [Source:H |
| FXYD6P1 | 10 | FXYD domain containing ion transport regulat |
| EIF3LP2 | 10 | eukaryotic translation initiation factor 3 subu |
| ZNF33B | 10 | zinc finger protein 33B [Source:HGNC Symbol; |
| AL022345.3 | 10 | mmunoglobulin lambda variable pseudogene |
| AL022345.1 | 10 | double homeobox 4 (DUX4) pseudogene |
| AL022345.2 | 10 | double homeobox 4 (DUX4) pseudogene |
| RNU6-1170P | 10 | RNA, U6 small nuclear 1170, pseudogene [Sou |
| AL022345.4 | 10 | novel transcript |
| LINC01518 | 10 | long intergenic non-protein coding RNA 1518 |
| VN1R54P | 10 | vomeronasal 1 receptor 54 pseudogene [Sour |
| CUBNP1 | 10 | cubilin pseudogene 1 [Source:HGNC Symbol;A |
| LINC02632 | 10 | long intergenic non-protein coding RNA 2632 |
| RSU1P1 | 10 | Ras suppressor protein 1 pseudogene 1 [Sour |
| DUXAP3 | 10 | double homeobox A pseudogene 3 [Source:H |
| AL022344.1 | 10 | novel transcript |
| BMS1 | 10 | BMS1 ribosome biogenesis factor [Source:HG |
| RNU6-885P | 10 | RNA, U6 small nuclear 885, pseudogene [Sour |
| LINC02623 | 10 | long intergenic non-protein coding RNA 2623 |
| LINC01264 | 10 | long intergenic non-protein coding RNA 1264 |
| MIR5100 | 10 | microRNA 5100 [Source:HGNC Symbol;Acc:HG |
| RET | 10 | ret proto-oncogene [Source:HGNC Symbol;Ac |
| AC010864.1 | 10 | novel transcript |
| CSGALNACT2 | 10 | chondroitin sulfate N-acetylgalactosaminyltra |
| RASGEF1A | 10 | RasGEF domain family member 1A [Source:HG |
| LINC02633 | 10 | long intergenic non-protein coding RNA 2633 |
| AC068707.1 | 10 | novel transcript |
| RNU6ATAC11P | 10 | RNA, U6atac small nuclear 11, pseudogene [S |

| | | |
|---|---|---|
| FXYD4 | 10 | FXYD domain containing ion transport regulat |
| HNRNPF | 10 | heterogeneous nuclear ribonucleoprotein F [S |
| SNORD3J | 10 | small nucleolar RNA, C/D box 3J [Source:HGN( |
| AL450326.1 | 10 | novel transcript |
| ZNF487 | 10 | zinc finger protein 487 [Source:HGNC Symbol; |
| RNU7-193P | 10 | RNA, U7 small nuclear 193 pseudogene [Sourc |
| AL450326.2 | 10 | zinc finger protein pseudogene |
| ZNF239 | 10 | zinc finger protein 239 [Source:HGNC Symbol; |
| AL450326.3 | 10 | novel transcript |
| CAP1P2 | 10 | CAP1 pseudogene 2 [Source:HGNC Symbol;Ac |
| ZNF485 | 10 | zinc finger protein 485 [Source:HGNC Symbol; |
| ZNF32-AS3 | 10 | ZNF32 antisense RNA 3 [Source:HGNC Symbol |
| AL645634.2 | 10 | novel transcript |
| RPL21P88 | 10 | ribosomal protein L21 pseudogene 88 [Source |
| ZNF32 | 10 | zinc finger protein 32 [Source:HGNC Symbol;A |
| ZNF32-AS1 | 10 | ZNF32 antisense RNA 1 [Source:HGNC Symbol |
| ZNF32-AS2 | 10 | ZNF32 antisense RNA 2 [Source:HGNC Symbol |
| AL645634.1 | 10 | novel transcript |
| AL355989.1 | 10 | chromosome 12 open reading frame 49 (C12o |
| UQCRHP3 | 10 | ubiquinol-cytochrome c reductase hinge prote |
| ELOCP30 | 10 | elongin C pseudogene 30 [Source:HGNC Symb |
| LINC02658 | 10 | long intergenic non-protein coding RNA 2658 |
| LINC00840 | 10 | long intergenic non-protein coding RNA 840 [S |
| HNRNPA3P1 | 10 | heterogeneous nuclear ribonucleoprotein A3 |
| LINC02659 | 10 | long intergenic non-protein coding RNA 2659 |
| LINC00841 | 10 | long intergenic non-protein coding RNA 841 [S |
| AL139237.1 | 10 | novel transcript |
| AL137026.2 | 10 | novel transcript |
| AL137026.1 | 10 | novel transcript |
| C10orf142 | 10 | chromosome 10 open reading frame 142 [Sou |
| AL137026.3 | 10 | novel transcript |
| CXCL12 | 10 | C-X-C motif chemokine ligand 12 [Source:HGN |
| RPL9P21 | 10 | ribosomal protein L9 pseudogene 21 [Source: |
| AL356157.2 | 10 | RAP1, GTP-GDP dissociation stimulator 1 (RAF |
| AL356157.3 | 10 | novel transcript |
| AL356157.1 | 10 | novel transcript |
| TMEM72-AS1 | 10 | TMEM72 antisense RNA 1 [Source:HGNC Sym |
| EIF2AP4 | 10 | eukaryotic translation initiation factor 2A pseu |
| AL353801.2 | 10 | novel transcript |
| TMEM72 | 10 | transmembrane protein 72 [Source:HGNC Syn |
| AL353801.1 | 10 | novel transcript |
| AL353801.3 | 10 | novel transcript |
| RASSF4 | 10 | Ras association domain family member 4 [Sou |
| DEPP1 | 10 | DEPP1 autophagy regulator [Source:HGNC Syr |
| C10orf25 | 10 | chromosome 10 open reading frame 25 [Sour |
| ZNF22 | 10 | zinc finger protein 22 [Source:HGNC Symbol;A |
| CEP164P1 | 10 | centrosomal protein 164 pseudogene 1 [Sourc |
| AL358394.3 | 10 | novel transcript |
| DUXAP4 | 10 | double homeobox A pseudogene 4 [Source:H( |
| RPS19P7 | 10 | ribosomal protein S19 pseudogene 7 [Source: |

| | | |
|---|---|---|
| RSU1P2 | 10 | Ras suppressor protein 1 pseudogene 2 [Sourc |
| CUBNP3 | 10 | cubilin pseudogene 3 [Source:HGNC Symbol;A |
| AL358394.2 | 10 | novel transcript |
| ANKRD54P1 | 10 | ankyrin repeat domain 54 pseudogene 1 [Sour |
| ANKRD30BP3 | 10 | ankyrin repeat domain 30B pseudogene 3 [Sou |
| MIR3156-1 | 10 | microRNA 3156-1 [Source:HGNC Symbol;Acc:I |
| AL358394.1 | 10 | novel transcript |
| AL512324.1 | 10 | ankyrin repeat domain 30B-like (ANKRD30BL) |
| AL512324.3 | 10 | ankyrin repeat domain 30B-like (ANKRD30BL) |
| RNU6-1207P | 10 | RNA, U6 small nuclear 1207, pseudogene [Sou |
| CUBNP2 | 10 | cubilin pseudogene 2 [Source:HGNC Symbol;A |
| OR6D1P | 10 | olfactory receptor family 6 subfamily D memb |
| OR13A1 | 10 | olfactory receptor family 13 subfamily A mem |
| ALOX5 | 10 | arachidonate 5-lipoxygenase [Source:HGNC Sy |
| AL731567.1 | 10 | novel transcript |
| MARCH8 | 10 | membrane associated ring-CH-type finger 8 [S |
| ZFAND4 | 10 | zinc finger AN1-type containing 4 [Source:HGN |
| AGAP10P | 10 | ArfGAP with GTPase domain, ankyrin repeat a |
| FAM21FP | 10 | family with sequence similarity 21 member F, |
| AL645998.1 | 10 | novel pseudogene |
| WASHC2C | 10 | WASH complex subunit 2C [Source:HGNC Sym |
| AC012044.1 | 10 | ataxin 1-like (ATXN1L) pseudogene |
| FAM25E | 10 | family with sequence similarity 25 member E |
| AGAP4 | 10 | ArfGAP with GTPase domain, ankyrin repeat a |
| PARGP1 | 10 | poly(ADP-ribose) glycohydrolase pseudogene |
| RNA5SP310 | 10 | RNA, 5S ribosomal pseudogene 310 [Source:H |
| RPL35AP23 | 10 | ribosomal protein L35a pseudogene 23 [Sourc |
| TIMM23 | 10 | translocase of inner mitochondrial membrane |
| SNORA74C-1 | 10 | small nucleolar RNA, H/ACA box 74C-1 [Sourc |
| NCOA4 | 10 | nuclear receptor coactivator 4 [Source:HGNC |
| MSMB | 10 | microseminoprotein beta [Source:HGNC Symb |
| RPL23AP61 | 10 | ribosomal protein L23a pseudogene 61 [Sourc |
| AGAP7P | 10 | ArfGAP with GTPase domain, ankyrin repeat a |
| ANTXRLP1 | 10 | ANTXR like pseudogene 1 [Source:HGNC Symb |
| ANTXRL | 10 | ANTXR like [Source:HGNC Symbol;Acc:HGNC:2 |
| AGAP14P | 10 | ArfGAP with GTPase domain, ankyrin repeat a |
| FAM25BP | 10 | family with sequence similarity 25 member B, |
| AC244230.1 | 10 | novel transcript |
| ANXA8L1 | 10 | annexin A8 like 1 [Source:HGNC Symbol;Acc:H |
| AC244230.2 | 10 | novel transcript |
| LINC00842 | 10 | long intergenic non-protein coding RNA 842 [S |
| HNRNPA1P33 | 10 | heterogeneous nuclear ribonucleoprotein A1 |
| NPY4R | 10 | neuropeptide Y receptor Y4 [Source:HGNC Syr |
| GPRIN2 | 10 | G protein regulated inducer of neurite outgrov |
| SYT15 | 10 | synaptotagmin 15 [Source:HGNC Symbol;Acc: |
| AL356056.2 | 10 | novel transcript |
| AL356056.1 | 10 | novel transcript |
| SHLD2P1 | 10 | shieldin complex subunit 2 pseudogene 1 [Sou |
| LINC02637 | 10 | long intergenic non-protein coding RNA 2637 |
| RHEBP1 | 10 | RHEB pseudogene 1 [Source:HGNC Symbol;Ac |

| | | |
|---|---|---|
| RN7SL248P | 10 | RNA, 7SL, cytoplasmic 248, pseudogene [Sour |
| FAM245B | 10 | family with sequence similarity 245 member E |
| CTSLP3 | 10 | cathepsin L pseudogene 3 [Source:HGNC Syml |
| GLUD1P2 | 10 | glutamate dehydrogenase 1 pseudogene 2 [Sc |
| AL731733.1 | 10 | dual specificity phosphatase 8 pseudogene 2 |
| BMS1P1 | 10 | BMS1 pseudogene 1 [Source:HGNC Symbol;Ac |
| RNA5SP311 | 10 | RNA, 5S ribosomal pseudogene 311 [Source:H |
| AGAP13P | 10 | ArfGAP with GTPase domain, ankyrin repeat a |
| FRMPD2B | 10 | FERM and PDZ domain containing 2B, pseudo |
| PTPN20 | 10 | protein tyrosine phosphatase non-receptor ty |
| GDF10 | 10 | growth differentiation factor 10 [Source:HGN |
| GDF2 | 10 | growth differentiation factor 2 [Source:HGNC |
| RBP3 | 10 | retinol binding protein 3 [Source:HGNC Symbo |
| ZNF488 | 10 | zinc finger protein 488 [Source:HGNC Symbol; |
| ANXA8 | 10 | annexin A8 [Source:HGNC Symbol;Acc:HGNC:! |
| AL591684.3 | 10 | ataxin 1-like (ATXN1L) pseudogene |
| FAM25G | 10 | family with sequence similarity 25 member G |
| AL591684.1 | 10 | novel transcript |
| AGAP9 | 10 | ArfGAP with GTPase domain, ankyrin repeat a |
| AL591684.2 | 10 | novel transcript |
| RNA5SP312 | 10 | RNA, 5S ribosomal pseudogene 312 [Source:H |
| BMS1P2 | 10 | BMS1 pseudogene 2 [Source:HGNC Symbol;Ac |
| BX547991.2 | 10 | glutamate dehydrogenase pseudogene 8 |
| BX547991.1 | 10 | dual specificity phosphatase 8 (DUSP8) pseud |
| CTSLP2 | 10 | cathepsin L pseudogene 2 [Source:HGNC Syml |
| LINC02675 | 10 | long intergenic non-protein coding RNA 2675 |
| SHLD2P3 | 10 | shieldin complex subunit 2 pseudogene 3 [Sou |
| RN7SL453P | 10 | RNA, 7SL, cytoplasmic 453, pseudogene [Sour |
| RHEBP2 | 10 | RHEB pseudogene 2 [Source:HGNC Symbol;Ac |
| FO681492.1 | 10 | synaptotagmin-15 [Source:NCBI gene;Acc:102 |
| AC245041.2 | 10 | novel transcript |
| NPY4R2 | 10 | neuropeptide Y receptor Y4-2 [Source:HGNC S |
| AC245041.1 | 10 | novel transcript |
| FAM25C | 10 | family with sequence similarity 25 member C |
| AGAP12P | 10 | ArfGAP with GTPase domain, ankyrin repeat a |
| RNA5SP315 | 10 | RNA, 5S ribosomal pseudogene 315 [Source:H |
| BMS1P7 | 10 | BMS1 pseudogene 7 [Source:HGNC Symbol;Ac |
| PTPN20CP | 10 | protein tyrosine phosphatase non-receptor ty |
| FRMPD2 | 10 | FERM and PDZ domain containing 2 [Source:H |
| AC074325.1 | 10 | novel transcript, antisense to FRMPD2 |
| RPS6P14 | 10 | ribosomal protein S6 pseudogene 14 [Source: |
| MAPK8 | 10 | mitogen-activated protein kinase 8 [Source:H |
| AC016397.2 | 10 | TEC |
| ARHGAP22 | 10 | Rho GTPase activating protein 22 [Source:HGI |
| ARHGAP22-IT1 | 10 | ARHGAP22 intronic transcript 1 [Source:HGNC |
| AC016397.1 | 10 | cytochrome c oxidase subunit VIIb (COX7B) ps |
| AC068898.2 | 10 | novel transcript |
| AC068898.1 | 10 | novel transcript |
| WDFY4 | 10 | WDFY family member 4 [Source:HGNC Symbo |
| RPL13AP19 | 10 | ribosomal protein L13a pseudogene 19 [Sourc |

| | | |
|---|---|---|
| AC035139.2 | 10 | novel transcript, antisense to WDFY4 |
| AC035139.1 | 10 | novel transcript |
| AC060234.3 | 10 | novel transcript, sense intronic to LRRC18 |
| LRRC18 | 10 | leucine rich repeat containing 18 [Source:HGN |
| AC060234.2 | 10 | novel transcript |
| AC060234.1 | 10 | novel transcript |
| MIR4294 | 10 | microRNA 4294 [Source:HGNC Symbol;Acc:HG |
| VSTM4 | 10 | V-set and transmembrane domain containing |
| FAM170B-AS1 | 10 | FAM170B antisense RNA 1 [Source:HGNC Sym |
| FAM170B | 10 | family with sequence similarity 170 member B |
| TMEM273 | 10 | transmembrane protein 273 [Source:HGNC Sy |
| C10orf71-AS1 | 10 | C10orf71 antisense RNA 1 [Source:HGNC Sym |
| C10orf71 | 10 | chromosome 10 open reading frame 71 [Sour |
| DRGX | 10 | dorsal root ganglia homeobox [Source:HGNC S |
| AL138760.1 | 10 | novel transcript |
| ERCC6 | 10 | ERCC excision repair 6, chromatin remodeling |
| HMGB1P50 | 10 | high mobility group box 1 pseudogene 50 [Sou |
| CHAT | 10 | choline O-acetyltransferase [Source:HGNC Syr |
| SLC18A3 | 10 | solute carrier family 18 member A3 [Source:H |
| C10orf53 | 10 | chromosome 10 open reading frame 53 [Sour |
| OGDHL | 10 | oxoglutarate dehydrogenase like [Source:HGN |
| MAPK6P6 | 10 | mitogen-activated protein kinase 6 pseudoger |
| RPL21P89 | 10 | ribosomal protein L21 pseudogene 89 [Source |
| PARG | 10 | poly(ADP-ribose) glycohydrolase [Source:HGN |
| RPL35AP24 | 10 | ribosomal protein L35a pseudogene 24 [Sourc |
| TIMM23B | 10 | translocase of inner mitochondrial membrane |
| TIMM23B-AGAP6 | 10 | TIMM23B-AGAP6 readthrough [Source:HGNC |
| RNA5SP317 | 10 | RNA, 5S ribosomal pseudogene 317 [Source:H |
| AL442003.1 | 10 | novel transcript, antisense to AGAP6 |
| AGAP6 | 10 | ArfGAP with GTPase domain, ankyrin repeat a |
| FAM21EP | 10 | family with sequence similarity 21 member E, |
| SLC9A3P3 | 10 | solute carrier family 9 member 3 pseudogene |
| WASHC2A | 10 | WASH complex subunit 2A [Source:HGNC Sym |
| SLC9A3P1 | 10 | solute carrier family 9 member 3 pseudogene |
| ASAH2 | 10 | N-acylsphingosine amidohydrolase 2 [Source:I |
| DYNC1I2P1 | 10 | dynein cytoplasmic 1 intermediate chain 2 pse |
| SGMS1 | 10 | sphingomyelin synthase 1 [Source:HGNC Syml |
| AC069547.2 | 10 | novel transcript, antisense to SGMS1 |
| AC069547.1 | 10 | TEC |
| AL117341.1 | 10 | novel transcript |
| RNU7-107P | 10 | RNA, U7 small nuclear 107 pseudogene [Sourc |
| SGMS1-AS1 | 10 | SGMS1 antisense RNA 1 [Source:HGNC Symbo |
| SHQ1P1 | 10 | SHQ1, H/ACA ribonucleoprotein assembly fact |
| BEND3P1 | 10 | BEN domain containing 3 pseudogene 1 [Sour |
| NUTM2HP | 10 | NUT family member 2H, pseudogene [Source: |
| AL589794.1 | 10 | novel pseudogene |
| CTSLP4 | 10 | cathepsin L pseudogene 4 [Source:HGNC Syml |
| PGGT1BP1 | 10 | protein geranylgeranyltransferase type I subu |
| AL589794.2 | 10 | novel transcript, antisense to ASAH2B |
| ASAH2B | 10 | N-acylsphingosine amidohydrolase 2B [Source |

| | | |
|---|---|---|
| A1CF | 10 | APOBEC1 complementation factor [Source:HG |
| AL512366.1 | 10 | novel transcript |
| CCDC58P2 | 10 | coiled-coil domain containing 58 pseudogene |
| AL731537.2 | 10 | novel transcript |
| PRKG1 | 10 | protein kinase cGMP-dependent 1 [Source:HG |
| AL731537.1 | 10 | novel transcript |
| AC022537.1 | 10 | novel transcript |
| MIR605 | 10 | microRNA 605 [Source:HGNC Symbol;Acc:HGN |
| AC069079.1 | 10 | novel transcript |
| RSU1P3 | 10 | Ras suppressor protein 1 pseudogene 3 [Sourc |
| CSTF2T | 10 | cleavage stimulation factor subunit 2 tau varia |
| PRKG1-AS1 | 10 | PRKG1 antisense RNA 1 [Source:HGNC Symbo |
| DKK1 | 10 | dickkopf WNT signaling pathway inhibitor 1 [S |
| RPL31P44 | 10 | ribosomal protein L31 pseudogene 44 [Source |
| THAP12P3 | 10 | THAP domain containing 12 pseudogene 3 [Sc |
| LNCAROD | 10 | lncRNA activating regulator of DKK1 [Source:H |
| MBL2 | 10 | mannose binding lectin 2 [Source:HGNC Symb |
| Y_RNA | 10 | Y RNA [Source:RFAM;Acc:RF00019] |
| AC073174.1 | 10 | novel transcript |
| LINC02672 | 10 | long intergenic non-protein coding RNA 2672 |
| SNRPEP8 | 10 | small nuclear ribonucleoprotein polypeptide E |
| AC036101.1 | 10 | novel transcript |
| RNA5SP318 | 10 | RNA, 5S ribosomal pseudogene 318 [Source:H |
| AL365496.1 | 10 | pseudogene similar to part of pleckstrin homo |
| PCDH15 | 10 | protocadherin related 15 [Source:HGNC Symb |
| RNU6-687P | 10 | RNA, U6 small nuclear 687, pseudogene [Sour |
| AL353784.1 | 10 | novel transcript, antisense to PCDH15 |
| NEFMP1 | 10 | neurofilament medium pseudogene 1 [Source |
| MIR548F1 | 10 | microRNA 548f-1 [Source:HGNC Symbol;Acc:H |
| AC051618.1 | 10 | novel transcript |
| AC016822.1 | 10 | novel transcript |
| AL355314.1 | 10 | novel transcript |
| AL355314.2 | 10 | novel transcript |
| MTRNR2L5 | 10 | MT-RNR2 like 5 [Source:HGNC Symbol;Acc:HG |
| GAPDHP21 | 10 | glyceraldehyde 3 phosphate dehydrogenase p |
| AC069545.1 | 10 | zinc finger, MYM-type 4 (ZMYM4) pseudogene |
| ZWINT | 10 | ZW10 interacting kinetochore protein [Source |
| AC010996.1 | 10 | novel transcript |
| AC025039.1 | 10 | NULL |
| MIR3924 | 10 | microRNA 3924 [Source:HGNC Symbol;Acc:HG |
| AL731534.1 | 10 | pseudogene similar to part of tubulin, beta, 2 |
| AC006484.1 | 10 | novel pseudogene (KIAA0733) |
| MRPS35P3 | 10 | mitochondrial ribosomal protein S35 pseudog |
| AC026884.1 | 10 | dual specificity phosphatase 18 (DUSP18) pseu |
| IPMK | 10 | inositol polyphosphate multikinase [Source:H( |
| TPT1P10 | 10 | tumor protein, translationally-controlled 1 pse |
| CISD1 | 10 | CDGSH iron sulfur domain 1 [Source:HGNC Syr |
| AC016396.1 | 10 | novel transcript |
| AC016396.2 | 10 | novel transcript |
| UBE2D1 | 10 | ubiquitin conjugating enzyme E2 D1 [Source:H |

| | | |
|---|---|---|
| TFAM | 10 | transcription factor A, mitochondrial [Source:I |
| BICC1 | 10 | BicC family RNA binding protein 1 [Source:HGI |
| FAM133CP | 10 | family with sequence similarity 133 member C |
| RN7SKP196 | 10 | RN7SK pseudogene 196 [Source:HGNC Symbo |
| LINC00844 | 10 | long intergenic non-protein coding RNA 844 [S |
| RPLP1P10 | 10 | ribosomal protein lateral stalk subunit P1 pseu |
| TRAF6P1 | 10 | TNF receptor associated factor 6 pseudogene |
| PHYHIPL | 10 | phytanoyl-CoA 2-hydroxylase interacting prot |
| FAM13C | 10 | family with sequence similarity 13 member C |
| AC025038.1 | 10 | novel transcript |
| AL355474.1 | 10 | novel transcript, antisense to FAM13C |
| MRPL50P4 | 10 | mitochondrial ribosomal protein L50 pseudog |
| AC026391.1 | 10 | novel transcript |
| SLC16A9 | 10 | solute carrier family 16 member 9 [Source:HG |
| MRLN | 10 | myoregulin [Source:HGNC Symbol;Acc:HGNC: |
| CCDC6 | 10 | coiled-coil domain containing 6 [Source:HGNC |
| LINC01553 | 10 | long intergenic non-protein coding RNA 1553 |
| ANK3 | 10 | ankyrin 3 [Source:HGNC Symbol;Acc:HGNC:49 |
| AL592430.1 | 10 | novel transcript, antisense to ANK3 |
| Y_RNA | 10 | Y RNA [Source:RFAM;Acc:RF00019] |
| AL592430.2 | 10 | novel transcript, antisense to ANK3 |
| ARL4AP1 | 10 | ADP ribosylation factor like GTPase 4A pseudo |
| ANK3-DT | 10 | ANK3 divergent transcript [Source:HGNC Syml |
| CDK1 | 10 | cyclin dependent kinase 1 [Source:HGNC Syml |
| RHOBTB1 | 10 | Rho related BTB domain containing 1 [Source: |
| RNU2-72P | 10 | RNA, U2 small nuclear 72, pseudogene [Sourc |
| LINC00845 | 10 | long intergenic non-protein coding RNA 845 [S |
| TMEM26 | 10 | transmembrane protein 26 [Source:HGNC Syn |
| TMEM26-AS1 | 10 | TMEM26 antisense RNA 1 [Source:HGNC Sym |
| AL356952.1 | 10 | novel transcript |
| CABCOCO1 | 10 | ciliary associated calcium binding coiled-coil 1 |
| AL451049.1 | 10 | novel transcript, antisense to C10orf107 |
| LINC02625 | 10 | long intergenic non-protein coding RNA 2625 |
| ARID5B | 10 | AT-rich interaction domain 5B [Source:HGNC S |
| RTKN2 | 10 | rhotekin 2 [Source:HGNC Symbol;Acc:HGNC:1 |
| LINC02621 | 10 | long intergenic non-protein coding RNA 2621 |
| AC024597.1 | 10 | novel transcript, antisense to ZNF365 |
| RN7SL591P | 10 | RNA, 7SL, cytoplasmic 591, pseudogene [Sour |
| ZNF365 | 10 | zinc finger protein 365 [Source:HGNC Symbol; |
| AC024598.1 | 10 | novel transcript |
| AC067752.1 | 10 | zinc finger protein 365 [Source:NCBI gene;Acc |
| AC067751.1 | 10 | novel transcript |
| ALDH7A1P4 | 10 | aldehyde dehydrogenase 7 family member A1 |
| ADO | 10 | 2-aminoethanethiol dioxygenase [Source:HGN |
| EGR2 | 10 | early growth response 2 [Source:HGNC Symbo |
| RNU6-543P | 10 | RNA, U6 small nuclear 543, pseudogene [Sour |
| AL590502.1 | 10 | novel transcript |
| NRBF2 | 10 | nuclear receptor binding factor 2 [Source:HGN |
| JMJD1C | 10 | jumonji domain containing 1C [Source:HGNC S |
| TATDN1P1 | 10 | TatD DNase domain containing 1 pseudogene |

| | | |
|---|---|---|
| AC022022.2 | 10 | leucine rich repeat containing 40 (LRRC40) pse |
| MIR1296 | 10 | microRNA 1296 [Source:HGNC Symbol;Acc:HG |
| PRELID1P3 | 10 | PRELI domain containing 1 pseudogene 3 [Sou |
| JMJD1C-AS1 | 10 | JMJD1C antisense RNA 1 [Source:HGNC Symb |
| AC022022.1 | 10 | pseudogene similar to part of heat shock 10kD |
| REEP3 | 10 | receptor accessory protein 3 [Source:HGNC Sy |
| AC022387.2 | 10 | novel transcript |
| MRPL35P2 | 10 | mitochondrial ribosomal protein L35 pseudog |
| AC022387.1 | 10 | StAR-related lipid transfer (START) domain co |
| AC013287.1 | 10 | novel transcript |
| RPL7AP50 | 10 | ribosomal protein L7a pseudogene 50 [Source |
| AC012558.1 | 10 | novel transcript |
| DBF4P1 | 10 | DBF4 zinc finger pseudogene 1 [Source:HGNC |
| RPL17P35 | 10 | ribosomal protein L17 pseudogene 35 [Source |
| CYP2C61P | 10 | cytochrome P450 family 2 subfamily C membe |
| ANXA2P3 | 10 | annexin A2 pseudogene 3 [Source:HGNC Syml |
| LINC02671 | 10 | long intergenic non-protein coding RNA 2671 |
| NEK4P3 | 10 | NIMA-related kinase 4 pseudogene 3 [Source: |
| AL513321.2 | 10 | novel transcript |
| MYL6P3 | 10 | myosin light chain 6 pseudogene 3 [Source:HG |
| AL513321.1 | 10 | pseudogene similar to part of itchy homolog E |
| AL592466.1 | 10 | novel transcript |
| LINC01515 | 10 | long intergenic non-protein coding RNA 1515 |
| CTNNA3 | 10 | catenin alpha 3 [Source:HGNC Symbol;Acc:HG |
| AC022017.1 | 10 | novel transcript, antisense to CTNNA3 |
| AL607023.1 | 10 | leucine rich repeat protein 1 (LRR1) pseudoge |
| LRRTM3 | 10 | leucine rich repeat transmembrane neuronal |
| AL139240.1 | 10 | novel transcript |
| RPL7AP51 | 10 | ribosomal protein L7a pseudogene 51 [Source |
| MIR7151 | 10 | microRNA 7151 [Source:HGNC Symbol;Acc:HG |
| AKR1B10P1 | 10 | aldo-keto reductase family 1 member B10 pse |
| RPL21P92 | 10 | ribosomal protein L21 pseudogene 92 [Source |
| DNAJC12 | 10 | DnaJ heat shock protein family (Hsp40) memb |
| RNU6-1250P | 10 | RNA, U6 small nuclear 1250, pseudogene [Sou |
| RN7SL394P | 10 | RNA, 7SL, cytoplasmic 394, pseudogene [Sour |
| DNAJC19P1 | 10 | DnaJ heat shock protein family (Hsp40) memb |
| AL133551.1 | 10 | novel transcript |
| RNU6-523P | 10 | RNA, U6 small nuclear 523, pseudogene [Sour |
| RPL12P8 | 10 | ribosomal protein L12 pseudogene 8 [Source: |
| SIRT1 | 10 | sirtuin 1 [Source:HGNC Symbol;Acc:HGNC:149 |
| HERC4 | 10 | HECT and RLD domain containing E3 ubiquitin |
| RPS3AP38 | 10 | ribosomal protein S3a pseudogene 38 [Source |
| RN7SL220P | 10 | RNA, 7SL, cytoplasmic 220, pseudogene [Sour |
| POU5F1P5 | 10 | POU class 5 homeobox 1 pseudogene 5 [Sourc |
| MYPN | 10 | myopalladin [Source:HGNC Symbol;Acc:HGNC |
| RN7SKP202 | 10 | RN7SK pseudogene 202 [Source:HGNC Symbo |
| ATOH7 | 10 | atonal bHLH transcription factor 7 [Source:HG |
| LINC02640 | 10 | long intergenic non-protein coding RNA 2640 |
| KRT19P4 | 10 | keratin 19 pseudogene 4 [Source:HGNC Symb |
| PBLD | 10 | phenazine biosynthesis like protein domain co |

| | | |
|---|---|---|
| HNRNPH3 | 10 | heterogeneous nuclear ribonucleoprotein H3 |
| RUFY2 | 10 | RUN and FYVE domain containing 2 [Source:H |
| DNA2 | 10 | DNA replication helicase/nuclease 2 [Source:H |
| RPL26P29 | 10 | ribosomal protein L26 pseudogene 29 [Source |
| RNA5SP319 | 10 | RNA, 5S ribosomal pseudogene 319 [Source:H |
| SLC25A16 | 10 | solute carrier family 25 member 16 [Source:H |
| RPL26P27 | 10 | ribosomal protein L26 pseudogene 27 [Source |
| Y_RNA | 10 | Y RNA [Source:RFAM;Acc:RF00019] |
| TMEM14DP | 10 | transmembrane protein 14D, pseudogene [So |
| TET1 | 10 | tet methylcytosine dioxygenase 1 [Source:HG |
| COX20P1 | 10 | COX20 pseudogene 1 [Source:HGNC Symbol;A |
| RPS3AP37 | 10 | ribosomal protein S3a pseudogene 37 [Source |
| AL513534.2 | 10 | novel transcript, overlapping to TET1 |
| AL513534.3 | 10 | heterogeneous nuclear ribonucleoprotein A1 |
| CCAR1 | 10 | cell division cycle and apoptosis regulator 1 [S |
| Y_RNA | 10 | Y RNA [Source:RFAM;Acc:RF00019] |
| SNORD98 | 10 | small nucleolar RNA, C/D box 98 [Source:HGN |
| AL513534.1 | 10 | NULL |
| STOX1 | 10 | storkhead box 1 [Source:HGNC Symbol;Acc:H |
| RNU6-697P | 10 | RNA, U6 small nuclear 697, pseudogene [Sour |
| AL359844.1 | 10 | novel transcript |
| DDX50 | 10 | DExD-box helicase 50 [Source:HGNC Symbol;A |
| RNU6-571P | 10 | RNA, U6 small nuclear 571, pseudogene [Sour |
| DDX21 | 10 | DExD-box helicase 21 [Source:HGNC Symbol;A |
| KIF1BP | 10 | KIF1 binding protein [Source:HGNC Symbol;Ac |
| MED28P1 | 10 | mediator complex subunit 28 pseudogene 1 [S |
| ACTBP14 | 10 | ACTB pseudogene 14 [Source:HGNC Symbol;A |
| SRGN | 10 | serglycin [Source:HGNC Symbol;Acc:HGNC:93 |
| Metazoa_SRP | 10 | Metazoan signal recognition particle RNA [Sou |
| VPS26A | 10 | VPS26, retromer complex component A [Sour |
| RPS12P17 | 10 | ribosomal protein S12 pseudogene 17 [Source |
| SUPV3L1 | 10 | Suv3 like RNA helicase [Source:HGNC Symbol; |
| AL596223.1 | 10 | novel transcript, antisense to HKDC1 |
| HKDC1 | 10 | hexokinase domain containing 1 [Source:HGN |
| AL596223.2 | 10 | novel transcript |
| HK1 | 10 | hexokinase 1 [Source:HGNC Symbol;Acc:HGN |
| RPS15AP28 | 10 | ribosomal protein S15a pseudogene 28 [Sourc |
| TACR2 | 10 | tachykinin receptor 2 [Source:HGNC Symbol;A |
| ATP5MC1P7 | 10 | ATP synthase membrane subunit c locus 1 pse |
| TSPAN15 | 10 | tetraspanin 15 [Source:HGNC Symbol;Acc:HG |
| AC016821.1 | 10 | novel transcript |
| TMEM256P1 | 10 | transmembrane protein 256 pseudogene 1 [S |
| NEUROG3 | 10 | neurogenin 3 [Source:HGNC Symbol;Acc:HGN |
| AL450311.1 | 10 | novel transcript |
| MTATP6P23 | 10 | MT-ATP6 pseudogene 23 [Source:HGNC Symb |
| MTCO2P23 | 10 | MT-CO2 pseudogene 23 [Source:HGNC Symbo |
| MTCO1P23 | 10 | MT-CO1 pseudogene 23 [Source:HGNC Symbo |
| MTND2P15 | 10 | MT-ND2 pseudogene 15 [Source:HGNC Symbo |
| MTND1P20 | 10 | MT-ND1 pseudogene 20 [Source:HGNC Symbo |
| FAM241B | 10 | family with sequence similarity 241 member B |

| | | |
|---|---|---|
| LINC02651 | 10 | long intergenic non-protein coding RNA 2651 |
| RPL5P26 | 10 | ribosomal protein L5 pseudogene 26 [Source: |
| COL13A1 | 10 | collagen type XIII alpha 1 chain [Source:HGNC |
| LINC02636 | 10 | long intergenic non-protein coding RNA 2636 |
| H2AFY2 | 10 | H2A histone family member Y2 [Source:HGNC |
| AIFM2 | 10 | apoptosis inducing factor mitochondria associ |
| TYSND1 | 10 | trypsin domain containing 1 [Source:HGNC Sy |
| SAR1A | 10 | secretion associated Ras related GTPase 1A [S |
| CALM2P2 | 10 | calmodulin 2 pseudogene 2 [Source:HGNC Syr |
| RPS25P9 | 10 | ribosomal protein S25 pseudogene 9 [Source: |
| PPA1 | 10 | pyrophosphatase (inorganic) 1 [Source:HGNC |
| NPFFR1 | 10 | neuropeptide FF receptor 1 [Source:HGNC Syr |
| LRRC20 | 10 | leucine rich repeat containing 20 [Source:HGN |
| CEP57L1P1 | 10 | centrosomal protein 57 like 1 pseudogene 1 [S |
| EIF4EBP2 | 10 | eukaryotic translation initiation factor 4E bind |
| NODAL | 10 | nodal growth differentiation factor [Source:H |
| AC022532.1 | 10 | TEC |
| PALD1 | 10 | phosphatase domain containing paladin 1 [Sou |
| YY1P1 | 10 | YY1 transcription factor pseudogene 1 [Source |
| PRF1 | 10 | perforin 1 [Source:HGNC Symbol;Acc:HGNC:9 |
| ADAMTS14 | 10 | ADAM metallopeptidase with thrombospondi |
| TBATA | 10 | thymus, brain and testes associated [Source:H |
| RPS26P40 | 10 | ribosomal protein S26 pseudogene 40 [Source |
| SGPL1 | 10 | sphingosine-1-phosphate lyase 1 [Source:HGN |
| PCBD1 | 10 | pterin-4 alpha-carbinolamine dehydratase 1 [S |
| AC073176.2 | 10 | novel transcript |
| AC073176.1 | 10 | novel transcript |
| LINC02622 | 10 | long intergenic non-protein coding RNA 2622 |
| UNC5B | 10 | unc-5 netrin receptor B [Source:HGNC Symbol |
| UNC5B-AS1 | 10 | UNC5B antisense RNA 1 [Source:HGNC Symbo |
| SLC29A3 | 10 | solute carrier family 29 member 3 [Source:HG |
| AL359183.1 | 10 | TEC |
| CDH23 | 10 | cadherin related 23 [Source:HGNC Symbol;Acc |
| CDH23-AS1 | 10 | CDH23 antisense RNA 1 [Source:HGNC Symbo |
| C10orf105 | 10 | chromosome 10 open reading frame 105 [Sou |
| VSIR | 10 | V-set immunoregulatory receptor [Source:HG |
| MIR7152 | 10 | microRNA 7152 [Source:HGNC Symbol;Acc:HG |
| PSAP | 10 | prosaposin [Source:HGNC Symbol;Acc:HGNC:9 |
| RNU7-38P | 10 | RNA, U7 small nuclear 38 pseudogene [Source |
| AC073370.1 | 10 | novel transcript |
| CHST3 | 10 | carbohydrate sulfotransferase 3 [Source:HGN |
| AC022392.1 | 10 | novel transcript |
| SPOCK2 | 10 | SPARC (osteonectin), cwcv and kazal like dom |
| ASCC1 | 10 | activating signal cointegrator 1 complex subu |
| AL607035.1 | 10 | NULL |
| RPL15P14 | 10 | ribosomal protein L15 pseudogene 14 [Source |
| ANAPC16 | 10 | anaphase promoting complex subunit 16 [Sou |
| Y_RNA | 10 | Y RNA [Source:RFAM;Acc:RF00019] |
| DDIT4 | 10 | DNA damage inducible transcript 4 [Source:H |
| DDIT4-AS1 | 10 | DDIT4 antisense RNA 1 [Source:HGNC Symbol |

| | | |
|---|---|---|
| DNAJB12 | 10 | DnaJ heat shock protein family (Hsp40) memb |
| MICU1 | 10 | mitochondrial calcium uptake 1 [Source:HGN( |
| RNU6-805P | 10 | RNA, U6 small nuclear 805, pseudogene [Sour |
| AC091769.2 | 10 | novel transcript, sense-intronic to MICU1 |
| AC091769.1 | 10 | sorting nexin 19 (SNX19) pseudogene |
| AL513185.1 | 10 | novel transcript |
| AL513185.2 | 10 | cytochrome c oxidase subunit VIIc (COX7C) ps |
| RN7SL840P | 10 | RNA, 7SL, cytoplasmic 840, pseudogene [Sour |
| HMGN2P34 | 10 | high mobility group nucleosomal binding dom |
| AL513185.3 | 10 | novel pseudogene |
| Y_RNA | 10 | Y RNA [Source:RFAM;Acc:RF00019] |
| MCU | 10 | mitochondrial calcium uniporter [Source:HGN |
| MIR4676 | 10 | microRNA 4676 [Source:HGNC Symbol;Acc:HG |
| AC016542.3 | 10 | TEC |
| AC016542.2 | 10 | TEC |
| AC016542.1 | 10 | novel transcript, sense intronic with MCU |
| OIT3 | 10 | oncoprotein induced transcript 3 [Source:HGN |
| NPM1P24 | 10 | nucleophosmin 1 pseudogene 24 [Source:HGN |
| PLA2G12B | 10 | phospholipase A2 group XIIB [Source:HGNC Sy |
| RPL17P50 | 10 | ribosomal protein L17 pseudogene 50 [Source |
| P4HA1 | 10 | prolyl 4-hydroxylase subunit alpha 1 [Source:H |
| AL731563.1 | 10 | novel pseudogene |
| AL731563.2 | 10 | novel transcript, antisense to P4HA1 |
| Y_RNA | 10 | Y RNA [Source:RFAM;Acc:RF00019] |
| AL731563.3 | 10 | novel transcript |
| NUDT13 | 10 | nudix hydrolase 13 [Source:HGNC Symbol;Acc |
| AC016394.2 | 10 | novel transcript to NUDT13 |
| SNORA11F | 10 | small nucleolar RNA, H/ACA box 11F [Source:H |
| ECD | 10 | ecdysoneless cell cycle regulator [Source:HGN |
| FAM149B1 | 10 | family with sequence similarity 149 member B |
| DNAJC9 | 10 | DnaJ heat shock protein family (Hsp40) memb |
| Y_RNA | 10 | Y RNA [Source:RFAM;Acc:RF00019] |
| EIF4A2P2 | 10 | eukaryotic translation initiation factor 4A2 pse |
| DNAJC9-AS1 | 10 | DNAJC9 antisense RNA 1 [Source:HGNC Symb |
| MRPS16 | 10 | mitochondrial ribosomal protein S16 [Source: |
| AC016394.1 | 10 | novel transcript, antisense to TTC18 |
| CFAP70 | 10 | cilia and flagella associated protein 70 [Source |
| RNU6-833P | 10 | RNA, U6 small nuclear 833, pseudogene [Sour |
| Y_RNA | 10 | Y RNA [Source:RFAM;Acc:RF00019] |
| ANXA7 | 10 | annexin A7 [Source:HGNC Symbol;Acc:HGNC:! |
| AL512656.1 | 10 | novel transcript |
| RPL26P6 | 10 | ribosomal protein L26 pseudogene 6 [Source: |
| MSS51 | 10 | MSS51 mitochondrial translational activator [! |
| PPP3CB | 10 | protein phosphatase 3 catalytic subunit beta [ |
| PPP3CB-AS1 | 10 | PPP3CB antisense RNA 1 (head to head) [Sour |
| USP54 | 10 | ubiquitin specific peptidase 54 [Source:HGNC |
| RNU6-883P | 10 | RNA, U6 small nuclear 883, pseudogene [Sour |
| RPS26P41 | 10 | ribosomal protein S26 pseudogene 41 [Source |
| AC073389.1 | 10 | novel transcript, antisense to USP54 |
| AC073389.3 | 10 | novel transcript |

| | | |
|---|---|---|
| MYOZ1 | 10 | myozenin 1 [Source:HGNC Symbol;Acc:HGNC: |
| SYNPO2L | 10 | synaptopodin 2 like [Source:HGNC Symbol;Ac |
| AC073389.2 | 10 | novel transcript, antisense to SYNPO2L and A( |
| AGAP5 | 10 | ArfGAP with GTPase domain, ankyrin repeat a |
| BMS1P4-AGAP5 | 10 | BMS1P4-AGAP5 readthough [Source:NCBI ger |
| BMS1P4 | 10 | BMS1 pseudogene 4 [Source:NCBI gene;Acc:7 |
| AC022400.4 | 10 | novel transcript |
| RNA5SP320 | 10 | RNA, 5S ribosomal pseudogene 320 [Source:H |
| BMS1P4 | 10 | BMS1 pseudogene 4 [Source:HGNC Symbol;A |
| GLUD1P3 | 10 | glutamate dehydrogenase 1 pseudogene 3 [S |
| DUSP8P5 | 10 | dual specificity phosphatase 8 pseudogene 5 [ |
| AC022400.3 | 10 | ataxin 1-like (ATXN1L) pseudogene |
| AC022400.6 | 10 | TEC |
| SEC24C | 10 | SEC24 homolog C, COPII coat complex compo |
| AC022400.7 | 10 | TEC |
| FUT11 | 10 | fucosyltransferase 11 [Source:HGNC Symbol;A |
| CHCHD1 | 10 | coiled-coil-helix-coiled-coil-helix domain conta |
| ZSWIM8 | 10 | zinc finger SWIM-type containing 8 [Source:H |
| AC022400.8 | 10 | TEC |
| AC022400.5 | 10 | novel transcript |
| ZSWIM8-AS1 | 10 | ZSWIM8 antisense RNA 1 [Source:HGNC Symb |
| NDST2 | 10 | N-deacetylase and N-sulfotransferase 2 [Sour |
| CAMK2G | 10 | calcium/calmodulin dependent protein kinase |
| AC022400.1 | 10 | novel transcript |
| AC022400.2 | 10 | novel transcript, antisense to CAMK2G |
| PLAU | 10 | plasminogen activator, urokinase [Source:HGI |
| C10orf55 | 10 | chromosome 10 open reading frame 55 [Sour |
| VCL | 10 | vinculin [Source:HGNC Symbol;Acc:HGNC:126 |
| AL596247.1 | 10 | novel transcript |
| AP3M1 | 10 | adaptor related protein complex 3 subunit mu |
| ADK | 10 | adenosine kinase [Source:HGNC Symbol;Acc:H |
| TIMM9P1 | 10 | TIMM9 pseudogene 1 [Source:HGNC Symbol;A |
| RPSAP6 | 10 | ribosomal protein SA pseudogene 6 [Source:H |
| RAB5CP1 | 10 | RAB5C, member RAS oncogene family pseudo |
| AC022540.1 | 10 | novel transcript |
| MRPL35P3 | 10 | mitochondrial ribosomal protein L35 pseudog |
| Y_RNA | 10 | Y RNA [Source:RFAM;Acc:RF00019] |
| NDUFA8P1 | 10 | NADH:ubiquinone oxidoreductase subunit A8 |
| POLR3DP1 | 10 | RNA polymerase III subunit D pseudogene 1 [S |
| FAM32CP | 10 | family with sequence similarity 32 member C, |
| AC063962.1 | 10 | novel transcript |
| KAT6B | 10 | lysine acetyltransferase 6B [Source:HGNC Sym |
| AC018511.2 | 10 | novel transcript |
| AC018511.1 | 10 | novel transcript |
| DUPD1 | 10 | dual specificity phosphatase and pro isomeras |
| AC018511.5 | 10 | novel transcript, antisense to DUPD1 |
| PPIAP13 | 10 | peptidylprolyl isomerase A pseudogene 13 [S |
| DUSP13 | 10 | dual specificity phosphatase 13 [Source:HGNC |
| SAMD8 | 10 | sterile alpha motif domain containing 8 [Sour |
| RPS26P42 | 10 | ribosomal protein S26 pseudogene 42 [Source |

| | | |
|---|---|---|
| VDAC2 | 10 | voltage dependent anion channel 2 [Source:H |
| COMTD1 | 10 | catechol-O-methyltransferase domain contair |
| ZNF503-AS1 | 10 | ZNF503 antisense RNA 1 [Source:HGNC Symb |
| RPL39P25 | 10 | ribosomal protein L39 pseudogene 25 [Source |
| HMGA1P5 | 10 | high mobility group AT-hook 1 pseudogene 5 |
| AC010997.3 | 10 | novel transcript |
| AC010997.1 | 10 | peroxiredoxin 5 (PRDX5) pseudogene |
| SPA17P1 | 10 | sperm autoantigenic protein 17 pseudogene 1 |
| ZNF503 | 10 | zinc finger protein 503 [Source:HGNC Symbol; |
| ZNF503-AS2 | 10 | ZNF503 antisense RNA 2 [Source:HGNC Symb |
| AC010997.5 | 10 | novel transcript |
| AC010997.4 | 10 | novel transcript |
| AC010997.2 | 10 | novel transcript, antisense to C10orf11 |
| LRMDA | 10 | leucine rich melanocyte differentiation associ |
| AC010997.6 | 10 | TEC |
| MIR606 | 10 | microRNA 606 [Source:HGNC Symbol;Acc:HGI |
| AL589863.2 | 10 | novel transcript |
| AL589863.1 | 10 | novel transcript, sense intronic to C10orf11 |
| AL731568.1 | 10 | novel transcript, sense intronic to C10orf11 |
| RN7SL518P | 10 | RNA, 7SL, cytoplasmic 518, pseudogene [Sour |
| RNU6-673P | 10 | RNA, U6 small nuclear 673, pseudogene [Sour |
| AC012048.1 | 10 | TEC |
| AC013286.1 | 10 | novel transcript, antisense to C10orf11 |
| AC024603.1 | 10 | novel transcript, antisense to C10orf11 |
| AC013738.1 | 10 | NULL |
| ATP5MC1P8 | 10 | ATP synthase membrane subunit c locus 1 pse |
| KCNMA1 | 10 | potassium calcium-activated channel subfami |
| KCNMA1-AS1 | 10 | KCNMA1 antisense RNA 1 [Source:HGNC Syml |
| KCNMA1-AS2 | 10 | KCNMA1 antisense RNA 2 [Source:HGNC Syml |
| AL731575.1 | 10 | novel transcript, antisense to KCNMA1 |
| KCNMA1-AS3 | 10 | KCNMA1 antisense RNA 3 [Source:HGNC Syml |
| COX6CP15 | 10 | cytochrome c oxidase subunit 6C pseudogene |
| RNA5SP321 | 10 | RNA, 5S ribosomal pseudogene 321 [Source:H |
| AL731556.1 | 10 | novel zinc finger pseudogene (KIAA1629) |
| AL731556.2 | 10 | retinoblastoma binding protein 6 (RBBP6) pse |
| RNU6-1266P | 10 | RNA, U6 small nuclear 1266, pseudogene [Sou |
| IMPDH1P5 | 10 | inosine monophosphate dehydrogenase 1 pse |
| AL450306.1 | 10 | novel transcript |
| DLG5 | 10 | discs large MAGUK scaffold protein 5 [Source: |
| AL391421.1 | 10 | novel transcript, antisense to DLG5 |
| RN7SL284P | 10 | RNA, 7SL, cytoplasmic 284, pseudogene [Sour |
| DLG5-AS1 | 10 | DLG5 antisense RNA 1 [Source:HGNC Symbol; |
| H2AFZP5 | 10 | H2A histone family member Z pseudogene 5 [ |
| Metazoa_SRP | 10 | Metazoan signal recognition particle RNA [Sou |
| POLR3A | 10 | RNA polymerase III subunit A [Source:HGNC S |
| RPS24 | 10 | ribosomal protein S24 [Source:HGNC Symbol; |
| GNAI2P2 | 10 | G protein subunit alpha i2 pseudogene 2 [Sou |
| AL731555.1 | 10 | novel transcript |
| AC012560.1 | 10 | novel transcript |
| LINC00595 | 10 | long intergenic non-protein coding RNA 595 [S |

| | | |
|---|---|---|
| AC010163.3 | 10 | novel transcript |
| AC010163.1 | 10 | novel transcript |
| AC010163.2 | 10 | novel transcript |
| SNORA71 | 10 | Small nucleolar RNA SNORA71 [Source:RFAM; |
| AL583852.1 | 10 | novel transcript |
| AC016820.1 | 10 | novel transcript |
| ZMIZ1-AS1 | 10 | ZMIZ1 antisense RNA 1 [Source:HGNC Symbol |
| AL356753.1 | 10 | novel transcript |
| ZMIZ1 | 10 | zinc finger MIZ-type containing 1 [Source:HGN |
| PPIF | 10 | peptidylprolyl isomerase F [Source:HGNC Sym |
| ZCCHC24 | 10 | zinc finger CCHC-type containing 24 [Source:H |
| AL133481.1 | 10 | novel transcript |
| TPRX1P1 | 10 | tetrapeptide repeat homeobox 1 pseudogene |
| AL133481.3 | 10 | novel transcript |
| AL133481.2 | 10 | nuclear pore complex interacting protein (NPI |
| EIF5AL1 | 10 | eukaryotic translation initiation factor 5A like |
| BX248123.1 | 10 | acyl-CoA synthetase medium-chain family me |
| RPS12P18 | 10 | ribosomal protein S12 pseudogene 18 [Source |
| SFTPA2 | 10 | surfactant protein A2 [Source:HGNC Symbol;A |
| MBL3P | 10 | mannose-binding lectin family member 3, pse |
| SFTPA3P | 10 | surfactant protein A3, pseudogene [Source:H( |
| SFTPA1 | 10 | surfactant protein A1 [Source:HGNC Symbol;A |
| LINC02679 | 10 | long intergenic non-protein coding RNA 2679 |
| AL132656.2 | 10 | novel transcript |
| NUTM2B-AS1 | 10 | NUTM2B antisense RNA 1 [Source:HGNC Syml |
| AL132656.3 | 10 | novel transcript |
| AL132656.4 | 10 | novel transcript |
| BEND3P3 | 10 | BEN domain containing 3 pseudogene 3 [Sour |
| AL132656.1 | 10 | novel pseudogene |
| NUTM2B | 10 | NUT family member 2B [Source:HGNC Symbo |
| AL135925.1 | 10 | novel transcript |
| NUTM2E | 10 | NUT family member 2E [Source:HGNC Symbol |
| NPAP1P2 | 10 | nuclear pore associated protein 1 pseudogene |
| CTSLP6 | 10 | cathepsin L pseudogene 6 [Source:HGNC Syml |
| PGGT1BP2 | 10 | protein geranylgeranyltransferase type I subu |
| AL512662.2 | 10 | novel transcript |
| BMS1P21 | 10 | BMS1 pseudogene 21 [Source:HGNC Symbol;A |
| MBL1P | 10 | mannose binding lectin 1, pseudogene [Sourc |
| SFTPD | 10 | surfactant protein D [Source:HGNC Symbol;Ac |
| SFTPD-AS1 | 10 | SFTPD antisense RNA 1 [Source:HGNC Symbol |
| AL512662.1 | 10 | novel transcript |
| ZNRF2P3 | 10 | zinc and ring finger 2 pseudogene 3 [Source:H |
| DPY19L2P5 | 10 | DPY19L2 pseudogene 5 [Source:HGNC Symbo |
| AL356095.1 | 10 | double homeobox (DUX) pseudogene |
| C1DP3 | 10 | C1D nuclear receptor corepressor pseudogene |
| C1DP2 | 10 | C1D nuclear receptor corepressor pseudogene |
| C1DP4 | 10 | C1D nuclear receptor corepressor pseudogene |
| TMEM254-AS1 | 10 | TMEM254 antisense RNA 1 [Source:HGNC Syr |
| AL356095.3 | 10 | double homeobox (DUX) pseudogene |
| TMEM254 | 10 | transmembrane protein 254 [Source:HGNC Sy |

| | | |
|---|---|---|
| AL356095.2 | 10 | novel transcript |
| RPL22P18 | 10 | ribosomal protein L22 pseudogene 18 [Source |
| PLAC9 | 10 | placenta associated 9 [Source:HGNC Symbol;A |
| ANXA11 | 10 | annexin A11 [Source:HGNC Symbol;Acc:HGNC |
| LINC00857 | 10 | long intergenic non-protein coding RNA 857 [S |
| RPS12P2 | 10 | ribosomal protein S12 pseudogene 2 [Source: |
| AL513174.1 | 10 | acyl-CoA synthetase medium-chain family me |
| EIF5AP4 | 10 | eukaryotic translation initiation factor 5A pseu |
| AL359195.2 | 10 | TEC |
| MAT1A | 10 | methionine adenosyltransferase 1A [Source:H |
| ZNF519P1 | 10 | zinc finger protein 519 pseudogene 1 [Source: |
| AL359195.1 | 10 | novel transcript |
| DYDC1 | 10 | DPY30 domain containing 1 [Source:HGNC Syr |
| DYDC2 | 10 | DPY30 domain containing 2 [Source:HGNC Syr |
| PRXL2A | 10 | peroxiredoxin like 2A [Source:HGNC Symbol;A |
| TSPAN14 | 10 | tetraspanin 14 [Source:HGNC Symbol;Acc:HGI |
| AC021028.1 | 10 | novel transcript |
| SH2D4B | 10 | SH2 domain containing 4B [Source:HGNC Sym |
| LINC02655 | 10 | long intergenic non-protein coding RNA 2655 |
| RPS7P9 | 10 | ribosomal protein S7 pseudogene 9 [Source:H |
| AC027673.1 | 10 | eukaryotic translation initiation factor 5A2 pse |
| FARSBP1 | 10 | FARSB pseudogene 1 [Source:HGNC Symbol;A |
| WARS2P1 | 10 | tryptophanyl tRNA synthetase 2 pseudogene : |
| RPA2P2 | 10 | replication protein A2 pseudogene 2 [Source:I |
| AL096706.1 | 10 | novel transcript |
| NRG3 | 10 | neuregulin 3 [Source:HGNC Symbol;Acc:HGNC |
| AC010157.2 | 10 | novel transcript, antisense to NRG3 |
| AC010157.1 | 10 | novel transcript |
| NRG3-AS1 | 10 | NRG3 antisense RNA 1 [Source:HGNC Symbol; |
| RNU6-441P | 10 | RNA, U6 small nuclear 441, pseudogene [Sour |
| RNU6-478P | 10 | RNA, U6 small nuclear 478, pseudogene [Sour |
| MARK2P15 | 10 | microtubule affinity regulating kinase 2 pseud |
| LINC02650 | 10 | long intergenic non-protein coding RNA 2650 |
| AC069540.2 | 10 | novel transcript |
| AC069540.1 | 10 | T cell receptor associated transmembrane ada |
| RNU6-129P | 10 | RNA, U6 small nuclear 129, pseudogene [Sour |
| AL390786.1 | 10 | novel transcript |
| RNU1-65P | 10 | RNA, U1 small nuclear 65, pseudogene [Sourc |
| HMGN2P8 | 10 | high mobility group nucleosomal binding dom |
| AL603756.1 | 10 | novel transcript, antisense to GHITM |
| GHITM | 10 | growth hormone inducible transmembrane pr |
| CERNA2 | 10 | competing endogenous lncRNA 2 for microRN |
| C10orf99 | 10 | chromosome 10 open reading frame 99 [Sour |
| CDHR1 | 10 | cadherin related family member 1 [Source:HG |
| LRIT2 | 10 | leucine rich repeat, Ig-like and transmembran |
| RGR | 10 | retinal G protein coupled receptor [Source:HG |
| LRIT1 | 10 | leucine rich repeat, Ig-like and transmembran |
| LINC00858 | 10 | long intergenic non-protein coding RNA 858 [S |
| CCSER2 | 10 | coiled-coil serine rich protein 2 [Source:HGNC |
| RPL12P29 | 10 | ribosomal protein L12 pseudogene 29 [Source |

| | | |
|---|---|---|
| TNPO1P1 | 10 | transportin 1 pseudogene 1 [Source:HGNC Syı |
| CACYBPP1 | 10 | calcyclin binding protein pseudogene 1 [Sourc |
| RPS3AP5 | 10 | ribosomal protein S3A pseudogene 5 [Source: |
| AL358787.2 | 10 | SET domain and mariner transposase fusion g |
| LINC01519 | 10 | long intergenic non-protein coding RNA 1519 |
| AL358787.1 | 10 | novel transcript |
| LINC02647 | 10 | long intergenic non-protein coding RNA 2647 |
| AC025428.1 | 10 | novel transcript |
| LINC01520 | 10 | long intergenic non-protein coding RNA 1520 |
| AL731532.1 | 10 | TEC |
| RNU6-325P | 10 | RNA, U6 small nuclear 325, pseudogene [Sour |
| AL731532.2 | 10 | TEC |
| GRID1-AS1 | 10 | GRID1 antisense RNA 1 [Source:HGNC Symbol |
| GRID1 | 10 | glutamate ionotropic receptor delta type subu |
| AC022028.3 | 10 | NULL |
| AC022028.2 | 10 | novel transcript, antisense to GRID1 |
| RN7SKP238 | 10 | RN7SK pseudogene 238 [Source:HGNC Symbo |
| AC022028.1 | 10 | acid phosphatase 1, soluble (ACP1) pseudoger |
| RNA5SP322 | 10 | RNA, 5S ribosomal pseudogene 322 [Source:H |
| MIR346 | 10 | microRNA 346 [Source:HGNC Symbol;Acc:HGI |
| AL844892.2 | 10 | novel transcript |
| AL844892.1 | 10 | novel transcript |
| WAPL | 10 | WAPL cohesin release factor [Source:HGNC Sy |
| RNU6-780P | 10 | RNA, U6 small nuclear 780, pseudogene [Sour |
| AL731569.1 | 10 | novel transcript |
| RPL7AP8 | 10 | ribosomal protein L7a pseudogene 8 [Source:I |
| OPN4 | 10 | opsin 4 [Source:HGNC Symbol;Acc:HGNC:144 |
| LDB3 | 10 | LIM domain binding 3 [Source:HGNC Symbol;/ |
| AC067750.1 | 10 | novel transcript, antisense to BMPR1A |
| BMPR1A | 10 | bone morphogenetic protein receptor type 1/ |
| RPAP2P1 | 10 | RNA polymerase II associated protein 2 pseud |
| RNU1-19P | 10 | RNA, U1 small nuclear 19, pseudogene [Sourc |
| MMRN2 | 10 | multimerin 2 [Source:HGNC Symbol;Acc:HGN( |
| AC025268.1 | 10 | TEC |
| SNCG | 10 | synuclein gamma [Source:HGNC Symbol;Acc:I |
| ADIRF-AS1 | 10 | ADIRF antisense RNA 1 [Source:HGNC Symbol |
| ADIRF | 10 | adipogenesis regulatory factor [Source:HGNC |
| AL136982.7 | 10 | novel transcript, antisense to ADIRF |
| AGAP11 | 10 | ArfGAP with GTPase domain, ankyrin repeat a |
| BMS1P3 | 10 | BMS1 pseudogene 3 [Source:HGNC Symbol;A( |
| AL136982.1 | 10 | novel transcript |
| AL136982.6 | 10 | novel transcript |
| AL136982.4 | 10 | ArfGAP with GTPase domain, ankyrin repeat a |
| AL136982.2 | 10 | novel transcript |
| FAM25A | 10 | family with sequence similarity 25 member A |
| AL136982.5 | 10 | ataxin 1-like (ATXN1L) pseudogene |
| RNU6-529P | 10 | RNA, U6 small nuclear 529, pseudogene [Sour |
| GLUD1 | 10 | glutamate dehydrogenase 1 [Source:HGNC Sy |
| SHLD2 | 10 | shieldin complex subunit 2 [Source:HGNC Sym |
| AL136982.3 | 10 | novel transcript |

| | | |
|---|---|---|
| RN7SL733P | 10 | RNA, 7SL, cytoplasmic 733, pseudogene [Sour |
| NUTM2A-AS1 | 10 | NUTM2A antisense RNA 1 [Source:HGNC Sym |
| AL157893.2 | 10 | novel transcript |
| AL157893.1 | 10 | synaptotagmin 15 (SYT15) pseudogene |
| NUTM2A | 10 | NUT family member 2A [Source:HGNC Symbo |
| LINC00863 | 10 | long intergenic non-protein coding RNA 863 [S |
| NUTM2D | 10 | NUT family member 2D [Source:HGNC Symbo |
| NPAP1P3 | 10 | nuclear pore associated protein 1 pseudogene |
| CTSLP1 | 10 | cathepsin L pseudogene 1 [Source:HGNC Syml |
| FAM245A | 10 | family with sequence similarity 245 member A |
| AL355334.1 | 10 | pseudogene similar to part of ring finger prote |
| AL355334.2 | 10 | novel transcript |
| MIR4678 | 10 | microRNA 4678 [Source:HGNC Symbol;Acc:HG |
| MINPP1 | 10 | multiple inositol-polyphosphate phosphatase |
| AL138767.2 | 10 | novel transcript |
| AL138767.3 | 10 | novel transcript |
| AL138767.1 | 10 | novel transcript |
| RPS26P38 | 10 | ribosomal protein S26 pseudogene 38 [Source |
| PAPSS2 | 10 | 3'-phosphoadenosine 5'-phosphosulfate synth |
| ATAD1 | 10 | ATPase family AAA domain containing 1 [Sour |
| CFL1P1 | 10 | cofilin 1 pseudogene 1 [Source:HGNC Symbol; |
| RN7SL78P | 10 | RNA, 7SL, cytoplasmic 78, pseudogene [Source |
| KLLN | 10 | killin, p53 regulated DNA replication inhibitor |
| PTEN | 10 | phosphatase and tensin homolog [Source:HGI |
| AC063965.2 | 10 | novel transcript, sense intronic to PTEN |
| RPL11P3 | 10 | ribosomal protein L11 pseudogene 3 [Source: |
| AC063965.1 | 10 | NULL |
| MED6P1 | 10 | mediator complex subunit 6 pseudogene 1 [So |
| AL353149.1 | 10 | vomeronasal 1 receptor 1 (VN1R1) pseudoger |
| RNLS | 10 | renalase, FAD dependent amine oxidase [Sour |
| Y_RNA | 10 | Y RNA [Source:RFAM;Acc:RF00019] |
| LIPJ | 10 | lipase family member J [Source:HGNC Symbol |
| RPL7P34 | 10 | ribosomal protein L7 pseudogene 34 [Source: |
| LIPF | 10 | lipase F, gastric type [Source:HGNC Symbol;Ac |
| NAPGP1 | 10 | N-ethylmaleimide-sensitive factor attachment |
| LIPK | 10 | lipase family member K [Source:HGNC Symbo |
| KRT8P38 | 10 | keratin 8 pseudogene 38 [Source:HGNC Symb |
| LIPN | 10 | lipase family member N [Source:HGNC Symbo |
| RCBTB2P1 | 10 | RCC1 and BTB domain containing protein 2 ps |
| LIPM | 10 | lipase family member M [Source:HGNC Symbo |
| ANKRD22 | 10 | ankyrin repeat domain 22 [Source:HGNC Syml |
| PTCD2P2 | 10 | pentatricopeptide repeat domain 2 pseudoge |
| STAMBPL1 | 10 | STAM binding protein like 1 [Source:HGNC Syr |
| ACTA2-AS1 | 10 | ACTA2 antisense RNA 1 [Source:HGNC Symbo |
| ACTA2 | 10 | actin alpha 2, smooth muscle [Source:HGNC S |
| AL157394.3 | 10 | FAS antisense RNA 1 |
| FAS | 10 | Fas cell surface death receptor [Source:HGNC |
| AL157394.2 | 10 | NULL |
| AL157394.1 | 10 | novel transcript, overlapping FAS |
| MIR4679-2 | 10 | microRNA 4679-2 [Source:HGNC Symbol;Acc:I |

| | | |
|---|---|---|
| MIR4679-1 | 10 | microRNA 4679-1 [Source:HGNC Symbol;Acc:I |
| AL513533.1 | 10 | novel zinc finger pseudogene |
| CH25H | 10 | cholesterol 25-hydroxylase [Source:HGNC Syn |
| LIPA | 10 | lipase A, lysosomal acid type [Source:HGNC Sy |
| IFIT2 | 10 | interferon induced protein with tetratricopep |
| AL353751.1 | 10 | novel transcript, antisense to LIPA |
| IFIT3 | 10 | interferon induced protein with tetratricopep |
| IFIT6P | 10 | interferon induced protein with tetratricopep |
| IFIT1B | 10 | interferon induced protein with tetratricopep |
| IFIT1 | 10 | interferon induced protein with tetratricopep |
| IFIT5 | 10 | interferon induced protein with tetratricopep |
| SLC16A12 | 10 | solute carrier family 16 member 12 [Source:H |
| SLC16A12-AS1 | 10 | SLC16A12 antisense RNA 1 [Source:HGNC Sym |
| PANK1 | 10 | pantothenate kinase 1 [Source:HGNC Symbol; |
| MIR107 | 10 | microRNA 107 [Source:HGNC Symbol;Acc:HGI |
| AL157400.5 | 10 | pantothenate kinase 1 (PANK1) pseudogene |
| AL157400.2 | 10 | novel transcript |
| AL157400.1 | 10 | novel transcript |
| AL157400.3 | 10 | novel transcript |
| AL157400.4 | 10 | novel transcript |
| KIF20B | 10 | kinesin family member 20B [Source:HGNC Syr |
| MTND5P42 | 10 | MT-ND5 pseudogene 42 [Source:HGNC Symbo |
| LINC00865 | 10 | long intergenic non-protein coding RNA 865 [S |
| MARK2P16 | 10 | microtubule affinity regulating kinase 2 pseud |
| LINC01374 | 10 | long intergenic non-protein coding RNA 1374 |
| LINC01375 | 10 | long intergenic non-protein coding RNA 1375 |
| SNRPD2P1 | 10 | small nuclear ribonucleoprotein D2 pseudoge |
| AL139340.1 | 10 | novel transcript |
| RN7SKP143 | 10 | RN7SK pseudogene 143 [Source:HGNC Symbo |
| LINC02653 | 10 | long intergenic non-protein coding RNA 2653 |
| AL391704.1 | 10 | novel transcript |
| AL391704.2 | 10 | novel transcript |
| AL390862.1 | 10 | novel transcript |
| HTR7 | 10 | 5-hydroxytryptamine receptor 7 [Source:HGN |
| RPP30 | 10 | ribonuclease P/MRP subunit p30 [Source:HGN |
| Y_RNA | 10 | Y RNA [Source:RFAM;Acc:RF00019] |
| ANKRD1 | 10 | ankyrin repeat domain 1 [Source:HGNC Symbo |
| RNU6-740P | 10 | RNA, U6 small nuclear 740, pseudogene [Sour |
| AL365434.1 | 10 | novel transcript |
| AL365434.2 | 10 | novel transcript |
| LINC00502 | 10 | long intergenic non-protein coding RNA 502 [S |
| DDX18P6 | 10 | DEAD-box helicase 18 pseudogene 6 [Source:I |
| NUDT9P1 | 10 | nudix hydrolase 9 pseudogene 1 [Source:HGN |
| AL731553.1 | 10 | novel transcript |
| PCGF5 | 10 | polycomb group ring finger 5 [Source:HGNC S |
| HECTD2 | 10 | HECT domain E3 ubiquitin protein ligase 2 [So |
| AL161798.1 | 10 | transmembrane protein 230 (TMEM230) pseu |
| RPS27P1 | 10 | ribosomal protein S27 pseudogene 1 [Source: |
| PPP1R3C | 10 | protein phosphatase 1 regulatory subunit 3C [ |
| GAPDHP28 | 10 | glyceraldehyde 3 phosphate dehydrogenase p |

| | | |
|---|---|---|
| FAF2P1 | 10 | Fas associated factor family member 2 pseudo |
| TNKS2-AS1 | 10 | TNKS2 antisense RNA 1 (head to head) [Sourc |
| TNKS2 | 10 | tankyrase 2 [Source:HGNC Symbol;Acc:HGNC: |
| SRP9P1 | 10 | signal recognition particle 9 pseudogene 1 [So |
| AL359198.2 | 10 | DnaJ (Hsp40) homolog, subfamily C, member |
| FGFBP3 | 10 | fibroblast growth factor binding protein 3 [Sou |
| AL359198.1 | 10 | novel transcript, antisense to FGFBP3 |
| BTAF1 | 10 | B-TFIID TATA-box binding protein associated f |
| CPEB3 | 10 | cytoplasmic polyadenylation element binding |
| AL365398.1 | 10 | NULL |
| SDHCP2 | 10 | succinate dehydrogenase complex subunit C p |
| EIF4A1P8 | 10 | eukaryotic translation initiation factor 4A1 pse |
| AL158040.1 | 10 | NULL |
| NHP2P1 | 10 | NHP2 ribonucleoprotein pseudogene 1 [Sourc |
| Y_RNA | 10 | Y RNA [Source:RFAM;Acc:RF00019] |
| MARCH5 | 10 | membrane associated ring-CH-type finger 5 [S |
| RNY3P12 | 10 | RNY3 pseudogene 12 [Source:HGNC Symbol;A |
| MARK2P9 | 10 | microtubule affinity regulating kinase 2 pseud |
| AL161652.1 | 10 | novel transcript |
| IDE | 10 | insulin degrading enzyme [Source:HGNC Symb |
| KIF11 | 10 | kinesin family member 11 [Source:HGNC Sym |
| RPL11P4 | 10 | ribosomal protein L11 pseudogene 4 [Source: |
| RN7SL644P | 10 | RNA, 7SL, cytoplasmic 644, pseudogene [Sour |
| EIF2S2P3 | 10 | eukaryotic translation initiation factor 2 subur |
| HHEX | 10 | hematopoietically expressed homeobox [Sour |
| Y_RNA | 10 | Y RNA [Source:RFAM;Acc:RF00019] |
| EXOC6 | 10 | exocyst complex component 6 [Source:HGNC |
| AL392103.1 | 10 | pseudogene similar to part of nuclease sensiti |
| AL358613.1 | 10 | novel transcript |
| CYP26C1 | 10 | cytochrome P450 family 26 subfamily C meml |
| AL358613.3 | 10 | novel transcript |
| AL358613.2 | 10 | novel transcript |
| CYP26A1 | 10 | cytochrome P450 family 26 subfamily A meml |
| NIP7P1 | 10 | NIP7 pseudogene 1 [Source:HGNC Symbol;Acc |
| XRCC6P1 | 10 | X-ray repair cross complementing 6 pseudoge |
| RPL17P34 | 10 | ribosomal protein L17 pseudogene 34 [Source |
| MYOF | 10 | myoferlin [Source:HGNC Symbol;Acc:HGNC:3( |
| CEP55 | 10 | centrosomal protein 55 [Source:HGNC Symbo |
| RNA5SP323 | 10 | RNA, 5S ribosomal pseudogene 323 [Source:H |
| FFAR4 | 10 | free fatty acid receptor 4 [Source:HGNC Symb |
| RBP4 | 10 | retinol binding protein 4 [Source:HGNC Symbo |
| PDE6C | 10 | phosphodiesterase 6C [Source:HGNC Symbol; |
| FRA10AC1 | 10 | FRA10A associated CGG repeat 1 [Source:HGN |
| LGI1 | 10 | leucine rich glioma inactivated 1 [Source:HGN |
| AL157396.1 | 10 | novel transcript, sense intronic to LGI1 |
| AL358154.1 | 10 | novel transcript, antisense to LGI1 |
| RAB11AP1 | 10 | RAB11A, member RAS oncogene family pseud |
| SLC35G1 | 10 | solute carrier family 35 member G1 [Source:H |
| PIPSL | 10 | PIP5K1A and PSMD4 like (pseudogene) [Sourc |
| PLCE1 | 10 | phospholipase C epsilon 1 [Source:HGNC Sym |

| | | |
|---|---|---|
| PLCE1-AS2 | 10 | PLCE1 antisense RNA 2 [Source:HGNC Symbol |
| AL139118.1 | 10 | novel transcript |
| HDAC1P1 | 10 | histone deacetylase 1 pseudogene 1 [Source:H |
| RNU6-657P | 10 | RNA, U6 small nuclear 657, pseudogene [Sour |
| AL389885.1 | 10 | novel transcript |
| PLCE1-AS1 | 10 | PLCE1 antisense RNA 1 [Source:HGNC Symbol |
| AL139124.1 | 10 | novel transcript, antisense to PLCE1 |
| NOC3L | 10 | NOC3 like DNA replication regulator [Source:H |
| TBC1D12 | 10 | TBC1 domain family member 12 [Source:HGN |
| HELLS | 10 | helicase, lymphoid specific [Source:HGNC Sym |
| AL138759.1 | 10 | novel transcript |
| CTBP2P2 | 10 | C-terminal binding protein 2 pseudogene 2 [S |
| CYP2C18 | 10 | cytochrome P450 family 2 subfamily C membe |
| AL583836.1 | 10 | novel transcript |
| CYP2C19 | 10 | cytochrome P450 family 2 subfamily C membe |
| MTND4P19 | 10 | MT-ND4 pseudogene 19 [Source:HGNC Symbo |
| CYP2C58P | 10 | cytochrome P450 family 2 subfamily C membe |
| RPL7AP52 | 10 | ribosomal protein L7a pseudogene 52 [Source |
| CYP2C9 | 10 | cytochrome P450 family 2 subfamily C membe |
| MTND4P20 | 10 | MT-ND4 pseudogene 20 [Source:HGNC Symbo |
| CYP2C59P | 10 | cytochrome P450 family 2 subfamily C membe |
| CYP2C60P | 10 | cytochrome P450 family 2 subfamily C membe |
| AL359672.1 | 10 | novel pseudogene |
| CYP2C8 | 10 | cytochrome P450 family 2 subfamily C membe |
| AL157834.1 | 10 | novel transcript |
| AL157834.3 | 10 | nuclear pore complex interacting protein fami |
| PAWRP1 | 10 | pro-apoptotic WT1 regulator pseudogene 1 [S |
| ACSM6 | 10 | acyl-CoA synthetase medium chain family mei |
| AL157834.2 | 10 | novel transcript, antisense to C10orf129 |
| PDLIM1 | 10 | PDZ and LIM domain 1 [Source:HGNC Symbol; |
| SORBS1 | 10 | sorbin and SH3 domain containing 1 [Source:H |
| RPS3AP36 | 10 | ribosomal protein S3a pseudogene 36 [Source |
| ALDH18A1 | 10 | aldehyde dehydrogenase 18 family member A |
| TCTN3 | 10 | tectonic family member 3 [Source:HGNC Syml |
| ENTPD1 | 10 | ectonucleoside triphosphate diphosphohydro |
| ENTPD1-AS1 | 10 | ENTPD1 antisense RNA 1 [Source:HGNC Symb |
| AL365273.1 | 10 | novel transcript, sense intronic to ENTPD1 |
| AL365273.2 | 10 | novel transcript |
| CC2D2B | 10 | coiled-coil and C2 domain containing 2B [Sour |
| RPL21P90 | 10 | ribosomal protein L21 pseudogene 90 [Source |
| CCNJ | 10 | cyclin J [Source:HGNC Symbol;Acc:HGNC:2343 |
| MIR3157 | 10 | microRNA 3157 [Source:HGNC Symbol;Acc:HG |
| RNU6-271P | 10 | RNA, U6 small nuclear 271, pseudogene [Sour |
| AC021037.1 | 10 | novel transcript, antisense to ZNF518A |
| ZNF518A | 10 | zinc finger protein 518A [Source:HGNC Symbo |
| NPM1P25 | 10 | nucleophosmin 1 pseudogene 25 [Source:HGN |
| BLNK | 10 | B cell linker [Source:HGNC Symbol;Acc:HGNC: |
| AL136181.1 | 10 | novel transcript, antisense to DNTT |
| DNTT | 10 | DNA nucleotidylexotransferase [Source:HGNC |
| OPALIN | 10 | oligodendrocytic myelin paranodal and inner I |

| | | |
|---|---|---|
| TLL2 | 10 | tolloid like 2 [Source:HGNC Symbol;Acc:HGNC |
| TM9SF3 | 10 | transmembrane 9 superfamily member 3 [Sou |
| NPM1P26 | 10 | nucleophosmin 1 pseudogene 26 [Source:HGN |
| PIK3AP1 | 10 | phosphoinositide-3-kinase adaptor protein 1 [ |
| RNA5SP324 | 10 | RNA, 5S ribosomal pseudogene 324 [Source:H |
| RPS2P36 | 10 | ribosomal protein S2 pseudogene 36 [Source: |
| RNU6-1274P | 10 | RNA, U6 small nuclear 1274, pseudogene [Sou |
| RPL13AP5 | 10 | ribosomal protein L13a pseudogene 5 [Source |
| MIR607 | 10 | microRNA 607 [Source:HGNC Symbol;Acc:HGN |
| LCOR | 10 | ligand dependent nuclear receptor corepresso |
| HMGN2P35 | 10 | high mobility group nucleosomal binding dom |
| SLIT1 | 10 | slit guidance ligand 1 [Source:HGNC Symbol;A |
| SLIT1-AS1 | 10 | SLIT1 antisense RNA 1 [Source:HGNC Symbol;A |
| ARHGAP19-SLIT1 | 10 | ARHGAP19-SLIT1 readthrough (NMD candidat |
| AL512424.1 | 10 | novel transcript |
| ARHGAP19 | 10 | Rho GTPase activating protein 19 [Source:HGN |
| AL359385.1 | 10 | CD164 antigen, sialomucin (CD164) pseudoge |
| Metazoa_SRP | 10 | Metazoan signal recognition particle RNA [Sou |
| RPL12P27 | 10 | ribosomal protein L12 pseudogene 27 [Source |
| FRAT1 | 10 | FRAT regulator of WNT signaling pathway 1 [S |
| FRAT2 | 10 | FRAT regulator of WNT signaling pathway 2 [S |
| AL355490.1 | 10 | novel transcript |
| RRP12 | 10 | ribosomal RNA processing 12 homolog [Sourc |
| RPL34P20 | 10 | ribosomal protein L34 pseudogene 20 [Source |
| AL355490.2 | 10 | novel transcript |
| Y_RNA | 10 | Y RNA [Source:RFAM;Acc:RF00019] |
| PGAM1 | 10 | phosphoglycerate mutase 1 [Source:HGNC Sy |
| EXOSC1 | 10 | exosome component 1 [Source:HGNC Symbol |
| ZDHHC16 | 10 | zinc finger DHHC-type containing 16 [Source:H |
| MMS19 | 10 | MMS19 homolog, cytosolic iron-sulfur assemb |
| UBTD1 | 10 | ubiquitin domain containing 1 [Source:HGNC S |
| ANKRD2 | 10 | ankyrin repeat domain 2 [Source:HGNC Symbo |
| HOGA1 | 10 | 4-hydroxy-2-oxoglutarate aldolase 1 [Source:H |
| AL355315.1 | 10 | novel protein |
| C10orf62 | 10 | chromosome 10 open reading frame 62 [Sour |
| MORN4 | 10 | MORN repeat containing 4 [Source:HGNC Sym |
| PI4K2A | 10 | phosphatidylinositol 4-kinase type 2 alpha [So |
| AVPI1 | 10 | arginine vasopressin induced 1 [Source:HGNC |
| MARVELD1 | 10 | MARVEL domain containing 1 [Source:HGNC S |
| ZFYVE27 | 10 | zinc finger FYVE-type containing 27 [Source:H |
| SFRP5 | 10 | secreted frizzled related protein 5 [Source:HG |
| LINC00866 | 10 | long intergenic non-protein coding RNA 866 [S |
| GOLGA7B | 10 | golgin A7 family member B [Source:HGNC Syn |
| CRTAC1 | 10 | cartilage acidic protein 1 [Source:HGNC Symb |
| AL355301.1 | 10 | NULL |
| R3HCC1L | 10 | R3H domain and coiled-coil containing 1 like [ |
| LOXL4 | 10 | lysyl oxidase like 4 [Source:HGNC Symbol;Acc: |
| AL139241.1 | 10 | novel transcript |
| PYROXD2 | 10 | pyridine nucleotide-disulphide oxidoreductase |
| MIR1287 | 10 | microRNA 1287 [Source:HGNC Symbol;Acc:HG |

| | | |
|---|---|---|
| HPS1 | 10 | HPS1 biogenesis of lysosomal organelles comp |
| MIR4685 | 10 | microRNA 4685 [Source:HGNC Symbol;Acc:HG |
| AL139243.1 | 10 | novel transcript, antisense to HPS1 |
| HPSE2 | 10 | heparanase 2 (inactive) [Source:HGNC Symbo |
| MIR6507 | 10 | microRNA 6507 [Source:HGNC Symbol;Acc:HG |
| ARL5AP2 | 10 | ADP ribosylation factor like GTPase 5A pseudo |
| RPL7P36 | 10 | ribosomal protein L7 pseudogene 36 [Source: |
| CNNM1 | 10 | cyclin and CBS domain divalent metal cation t |
| GOT1 | 10 | glutamic-oxaloacetic transaminase 1 [Source: |
| AL391684.1 | 10 | novel transcript |
| LINC01475 | 10 | long intergenic non-protein coding RNA 1475 |
| AL513542.1 | 10 | novel transcript |
| NKX2-3 | 10 | NK2 homeobox 3 [Source:HGNC Symbol;Acc:H |
| SLC25A28 | 10 | solute carrier family 25 member 28 [Source:H |
| AL353719.1 | 10 | novel transcript |
| AL133353.2 | 10 | novel transcript |
| AL133353.1 | 10 | novel transcript |
| ENTPD7 | 10 | ectonucleoside triphosphate diphosphohydro |
| EBAG9P1 | 10 | EBAG9 pseudogene 1 [Source:HGNC Symbol;A |
| CUTC | 10 | cutC copper transporter [Source:HGNC Symbo |
| COX15 | 10 | cytochrome c oxidase assembly homolog COX |
| ABCC2 | 10 | ATP binding cassette subfamily C member 2 [S |
| NANOGP6 | 10 | Nanog homeobox pseudogene 6 [Source:HGN |
| DNMBP | 10 | dynamin binding protein [Source:HGNC Symb |
| DNMBP-AS1 | 10 | DNMBP antisense RNA 1 [Source:HGNC Symb |
| CPN1 | 10 | carboxypeptidase N subunit 1 [Source:HGNC S |
| CYP2C23P | 10 | cytochrome P450 family 2 subfamily C membe |
| TPM4P1 | 10 | tropomyosin 4 pseudogene 1 [Source:HGNC S |
| SPCS2P2 | 10 | signal peptidase complex subunit 2 pseudoge |
| ERLIN1 | 10 | ER lipid raft associated 1 [Source:HGNC Symb |
| CHUK | 10 | component of inhibitor of nuclear factor kapp |
| AL138921.2 | 10 | novel transcript, antisense to CHUK |
| AL138921.1 | 10 | novel transcript |
| CWF19L1 | 10 | CWF19 like cell cycle control factor 1 [Source: |
| SNORA12 | 10 | small nucleolar RNA, H/ACA box 12 [Source:H |
| PHBP9 | 10 | prohibitin pseudogene 9 [Source:HGNC Symb |
| RNU6-422P | 10 | RNA, U6 small nuclear 422, pseudogene [Sour |
| BLOC1S2 | 10 | biogenesis of lysosomal organelles complex 1 |
| PKD2L1 | 10 | polycystin 2 like 1, transient receptor potentia |
| AL139819.1 | 10 | novel transcript |
| SCD | 10 | stearoyl-CoA desaturase [Source:HGNC Symb |
| OLMALINC | 10 | oligodendrocyte maturation-associated long i |
| Y_RNA | 10 | Y RNA [Source:RFAM;Acc:RF00019] |
| WNT8B | 10 | Wnt family member 8B [Source:HGNC Symbol |
| SEC31B | 10 | SEC31 homolog B, COPII coat complex compo |
| AL133352.1 | 10 | NADH dehydrogenase (ubiquinone) 1 beta sub |
| NDUFB8 | 10 | NADH:ubiquinone oxidoreductase subunit B8 |
| HIF1AN | 10 | hypoxia inducible factor 1 subunit alpha inhibi |
| Metazoa_SRP | 10 | Metazoan signal recognition particle RNA [Sou |
| PAX2 | 10 | paired box 2 [Source:HGNC Symbol;Acc:HGNC |

| | | |
|---|---|---|
| Y_RNA | 10 | Y RNA [Source:RFAM;Acc:RF00019] |
| AL138762.1 | 10 | novel transcript, antisense to FAM178A |
| SLF2 | 10 | SMC5-SMC6 complex localization factor 2 [Sou |
| AL133215.3 | 10 | novel transcript |
| MRPL43 | 10 | mitochondrial ribosomal protein L43 [Source: |
| SEMA4G | 10 | semaphorin 4G [Source:HGNC Symbol;Acc:HG |
| MIR608 | 10 | microRNA 608 [Source:HGNC Symbol;Acc:HGN |
| AL133215.1 | 10 | novel transcript, antisense to SEMA4G |
| TWNK | 10 | twinkle mtDNA helicase [Source:HGNC Symbo |
| LZTS2 | 10 | leucine zipper tumor suppressor 2 [Source:HG |
| PDZD7 | 10 | PDZ domain containing 7 [Source:HGNC Symb |
| SFXN3 | 10 | sideroflexin 3 [Source:HGNC Symbol;Acc:HGN |
| AL133215.2 | 10 | novel transcript |
| KAZALD1 | 10 | Kazal type serine peptidase inhibitor domain 1 |
| TLX1NB | 10 | TLX1 neighbor [Source:HGNC Symbol;Acc:HGN |
| TLX1 | 10 | T cell leukemia homeobox 1 [Source:HGNC Sy |
| LINC01514 | 10 | long intergenic non-protein coding RNA 1514 |
| SMARCE1P7 | 10 | SWI/SNF related, matrix associated, actin dep |
| LBX1 | 10 | ladybird homeobox 1 [Source:HGNC Symbol;A |
| LBX1-AS1 | 10 | LBX1 antisense RNA 1 [Source:HGNC Symbol;A |
| LINC02681 | 10 | long intergenic non-protein coding RNA 2681 |
| RNU2-43P | 10 | RNA, U2 small nuclear 43, pseudogene [Sourc |
| AL133387.1 | 10 | novel transcript |
| AL133387.2 | 10 | novel transcript |
| BTRC | 10 | beta-transducin repeat containing E3 ubiquitin |
| RNU2-59P | 10 | RNA, U2 small nuclear 59, pseudogene [Sourc |
| DPCD | 10 | deleted in primary ciliary dyskinesia homolog |
| POLL | 10 | DNA polymerase lambda [Source:HGNC Symb |
| MIR3158-2 | 10 | microRNA 3158-2 [Source:HGNC Symbol;Acc:H |
| MIR3158-1 | 10 | microRNA 3158-1 [Source:HGNC Symbol;Acc:H |
| FBXW4 | 10 | F-box and WD repeat domain containing 4 [Sc |
| RNU6-1165P | 10 | RNA, U6 small nuclear 1165, pseudogene [Sou |
| AC010789.2 | 10 | novel transcript, antisense to FBXW4 |
| AC010789.1 | 10 | novel transcript |
| FGF8 | 10 | fibroblast growth factor 8 [Source:HGNC Symb |
| NPM3 | 10 | nucleophosmin/nucleoplasmin 3 [Source:HGN |
| OGA | 10 | O-GlcNAcase [Source:HGNC Symbol;Acc:HGN( |
| KCNIP2-AS1 | 10 | KCNIP2 antisense RNA 1 [Source:HGNC Symbo |
| KCNIP2 | 10 | potassium voltage-gated channel interacting p |
| ARMH3 | 10 | armadillo like helical domain containing 3 [Sou |
| Metazoa_SRP | 10 | Metazoan signal recognition particle RNA [Sou |
| Y_RNA | 10 | Y RNA [Source:RFAM;Acc:RF00019] |
| HPS6 | 10 | HPS6 biogenesis of lysosomal organelles comp |
| AL500527.1 | 10 | transcription elongation factor B (SIII), polype |
| AL500527.2 | 10 | transcription elongation factor B (SIII) pseudog |
| LDB1 | 10 | LIM domain binding 1 [Source:HGNC Symbol;A |
| PPRC1 | 10 | PPARG related coactivator 1 [Source:HGNC Sy |
| NOLC1 | 10 | nucleolar and coiled-body phosphoprotein 1 [ |
| ELOVL3 | 10 | ELOVL fatty acid elongase 3 [Source:HGNC Syr |
| PITX3 | 10 | paired like homeodomain 3 [Source:HGNC Syr |

| | | |
|---|---|---|
| GBF1 | 10 | golgi brefeldin A resistant guanine nucleotide |
| NFKB2 | 10 | nuclear factor kappa B subunit 2 [Source:HGN |
| PSD | 10 | pleckstrin and Sec7 domain containing [Sourc |
| FBXL15 | 10 | F-box and leucine rich repeat protein 15 [Sour |
| CUEDC2 | 10 | CUE domain containing 2 [Source:HGNC Symb |
| MIR146B | 10 | microRNA 146b [Source:HGNC Symbol;Acc:HG |
| RPARP-AS1 | 10 | RPARP antisense RNA 1 [Source:HGNC Symbo |
| C10orf95 | 10 | chromosome 10 open reading frame 95 [Sour |
| MFSD13A | 10 | major facilitator superfamily domain containi |
| ACTR1A | 10 | actin related protein 1A [Source:HGNC Symbo |
| AL121928.1 | 10 | novel transcript, antisense to ACTR1A |
| SUFU | 10 | SUFU negative regulator of hedgehog signalin |
| RPL23AP58 | 10 | ribosomal protein L23a pseudogene 58 [Sourc |
| RNU6-43P | 10 | RNA, U6 small nuclear 43, pseudogene [Sourc |
| TRIM8 | 10 | tripartite motif containing 8 [Source:HGNC Sy |
| AL391121.1 | 10 | novel transcript |
| ARL3 | 10 | ADP ribosylation factor like GTPase 3 [Source: |
| SFXN2 | 10 | sideroflexin 2 [Source:HGNC Symbol;Acc:HGN |
| WBP1L | 10 | WW domain binding protein 1 like [Source:HG |
| RNU6-1231P | 10 | RNA, U6 small nuclear 1231, pseudogene [Sou |
| CYP17A1 | 10 | cytochrome P450 family 17 subfamily A meml |
| PFN1P11 | 10 | profilin 1 pseudogene 11 [Source:HGNC Symb |
| PTGES3P4 | 10 | prostaglandin E synthase 3 pseudogene 4 [Sou |
| AL358790.1 | 10 | novel transcript, antisense to C10orf32 |
| BORCS7 | 10 | BLOC-1 related complex subunit 7 [Source:HG |
| BORCS7-ASMT | 10 | BORCS7-ASMT readthrough (NMD candidate) |
| U6 | 10 | U6 spliceosomal RNA [Source:RFAM;Acc:RF00 |
| AS3MT | 10 | arsenite methyltransferase [Source:HGNC Syn |
| RPL22P17 | 10 | ribosomal protein L22 pseudogene 17 [Source |
| AL356608.3 | 10 | novel transcript, antisense to CNNM2and AS3 |
| AL356608.1 | 10 | novel transcript |
| CNNM2 | 10 | cyclin and CBS domain divalent metal cation t |
| AL356608.2 | 10 | NULL |
| NT5C2 | 10 | 5'-nucleotidase, cytosolic II [Source:HGNC Syn |
| RPS15AP29 | 10 | ribosomal protein S15a pseudogene 29 [Sourc |
| MARCKSL1P1 | 10 | MARCKS like 1 pseudogene 1 [Source:HGNC S |
| ST13P13 | 10 | ST13, Hsp70 interacting protein pseudogene 1 |
| RPEL1 | 10 | ribulose-5-phosphate-3-epimerase like 1 [Sou |
| INA | 10 | internexin neuronal intermediate filament pro |
| PCGF6 | 10 | polycomb group ring finger 6 [Source:HGNC S |
| RNU11-3P | 10 | RNA, U11 small nuclear 3, pseudogene [Sourc |
| TAF5 | 10 | TATA-box binding protein associated factor 5 |
| ATP5MD | 10 | ATP synthase membrane subunit DAPIT [Sourc |
| MIR1307 | 10 | microRNA 1307 [Source:HGNC Symbol;Acc:HG |
| PDCD11 | 10 | programmed cell death 11 [Source:HGNC Sym |
| CALHM2 | 10 | calcium homeostasis modulator family memb |
| AL139339.2 | 10 | novel transcript, antisense to CALHM2 |
| AL139339.1 | 10 | novel transcript |
| CALHM1 | 10 | calcium homeostasis modulator 1 [Source:HG |
| CALHM3 | 10 | calcium homeostasis modulator 3 [Source:HG |

| | | |
|---|---|---|
| NEURL1-AS1 | 10 | NEURL1 antisense RNA 1 [Source:HGNC Symb |
| NEURL1 | 10 | neuralized E3 ubiquitin protein ligase 1 [Sourc |
| AL121929.1 | 10 | novel transcript |
| AL121929.3 | 10 | novel transcript, antisense to NEURL1 |
| SH3PXD2A | 10 | SH3 and PX domains 2A [Source:HGNC Symbo |
| AL121929.2 | 10 | novel transcript, antisense to SH3PXD2A |
| Y_RNA | 10 | Y RNA [Source:RFAM;Acc:RF00019] |
| SH3PXD2A-AS1 | 10 | SH3PXD2A antisense RNA 1 [Source:HGNC Syr |
| AL133355.1 | 10 | novel transcript, overlapping to OBFC1 |
| STN1 | 10 | STN1 subunit of CST complex [Source:HGNC S |
| SLK | 10 | STE20 like kinase [Source:HGNC Symbol;Acc:H |
| RN7SL524P | 10 | RNA, 7SL, cytoplasmic 524, pseudogene [Sour |
| COL17A1 | 10 | collagen type XVII alpha 1 chain [Source:HGN( |
| MIR936 | 10 | microRNA 936 [Source:HGNC Symbol;Acc:HGI |
| SFR1 | 10 | SWI5 dependent homologous recombination |
| CFAP43 | 10 | cilia and flagella associated protein 43 [Source |
| MIR609 | 10 | microRNA 609 [Source:HGNC Symbol;Acc:HGI |
| GSTO1 | 10 | glutathione S-transferase omega 1 [Source:HG |
| MIR4482 | 10 | microRNA 4482 [Source:HGNC Symbol;Acc:HG |
| GSTO2 | 10 | glutathione S-transferase omega 2 [Source:HG |
| ITPRIP | 10 | inositol 1,4,5-trisphosphate receptor interacti |
| AL162742.1 | 10 | novel transcript, antisense to ITPRIP |
| AL162742.2 | 10 | inositol 1,4,5-triphosphate receptor interactin |
| ITPRIP-AS1 | 10 | ITPRIP antisense RNA 1 [Source:HGNC Symbol |
| CFAP58-DT | 10 | CFAP58 divergent transcript [Source:HGNC Sy |
| CFAP58 | 10 | cilia and flagella associated protein 58 [Source |
| LINC02620 | 10 | long intergenic non-protein coding RNA 2620 |
| AL161646.1 | 10 | novel transcript |
| SORCS3 | 10 | sortilin related VPS10 domain containing rece |
| SORCS3-AS1 | 10 | SORCS3 antisense RNA 1 [Source:HGNC Symb |
| RNU6-463P | 10 | RNA, U6 small nuclear 463, pseudogene [Sour |
| PPIAP38 | 10 | peptidylprolyl isomerase A pseudogene 38 [Sc |
| YWHAZP5 | 10 | tyrosine 3-monooxygenase/tryptophan 5-mor |
| LINC02627 | 10 | long intergenic non-protein coding RNA 2627 |
| AL731574.1 | 10 | novel transcript |
| AL589166.1 | 10 | novel transcript |
| LINC02624 | 10 | long intergenic non-protein coding RNA 2624 |
| RPL23AP59 | 10 | ribosomal protein L23a pseudogene 59 [Sourc |
| SORCS1 | 10 | sortilin related VPS10 domain containing rece |
| AL133395.1 | 10 | novel transcript, antisense to SORCS1 |
| RNA5SP325 | 10 | RNA, 5S ribosomal pseudogene 325 [Source:H |
| RNA5SP326 | 10 | RNA, 5S ribosomal pseudogene 326 [Source:H |
| LINC01435 | 10 | long intergenic non-protein coding RNA 1435 |
| PTGES3P5 | 10 | prostaglandin E synthase 3 pseudogene 5 [Sou |
| AL353740.1 | 10 | novel transcript |
| LINC02661 | 10 | long intergenic non-protein coding RNA 2661 |
| MAPKAPK5P1 | 10 | MAPKAPK5 pseudogene 1 [Source:HGNC Sym |
| RN7SKP278 | 10 | RN7SK pseudogene 278 [Source:HGNC Symbo |
| RNU5B-6P | 10 | RNA, U5B small nuclear 6, pseudogene [Sourc |
| RNU6-839P | 10 | RNA, U6 small nuclear 839, pseudogene [Sour |

| | | |
|---|---|---|
| PHB2P1 | 10 | prohibitin 2 pseudogene 1 [Source:HGNC Sym |
| BTF3P15 | 10 | basic transcription factor 3 pseudogene 15 [So |
| AL390123.1 | 10 | dpy-19-like 3 (C. elegans) (DPY19L3) pseudoge |
| XIAPP1 | 10 | X-linked inhibitor of apoptosis pseudogene 1 [ |
| RPL21P91 | 10 | ribosomal protein L21 pseudogene 91 [Source |
| XPNPEP1 | 10 | X-prolyl aminopeptidase 1 [Source:HGNC Sym |
| RNU4-5P | 10 | RNA, U4 small nuclear 5, pseudogene [Source |
| ADD3-AS1 | 10 | ADD3 antisense RNA 1 [Source:HGNC Symbol; |
| ADD3 | 10 | adducin 3 [Source:HGNC Symbol;Acc:HGNC:24 |
| SNRPGP12 | 10 | small nuclear ribonucleoprotein polypeptide C |
| MXI1 | 10 | MAX interactor 1, dimerization protein [Sourc |
| AL360182.2 | 10 | novel transcript |
| AL360182.1 | 10 | ribosomal protein L21 (RPL21) pseudogene |
| SMNDC1 | 10 | survival motor neuron domain containing 1 [S |
| AL355512.1 | 10 | novel transcript |
| HMGB3P5 | 10 | high mobility group box 3 pseudogene 5 [Sour |
| DUSP5 | 10 | dual specificity phosphatase 5 [Source:HGNC S |
| RPL7P35 | 10 | ribosomal protein L7 pseudogene 35 [Source: |
| SMC3 | 10 | structural maintenance of chromosomes 3 [So |
| Y_RNA | 10 | Y RNA [Source:RFAM;Acc:RF00019] |
| RBM20 | 10 | RNA binding motif protein 20 [Source:HGNC S |
| RN7SKP288 | 10 | RN7SK pseudogene 288 [Source:HGNC Symbo |
| RNA5SP327 | 10 | RNA, 5S ribosomal pseudogene 327 [Source:H |
| AL136368.1 | 10 | novel transcript |
| PDCD4-AS1 | 10 | PDCD4 antisense RNA 1 [Source:HGNC Symbo |
| PDCD4 | 10 | programmed cell death 4 [Source:HGNC Symb |
| MIR4680 | 10 | microRNA 4680 [Source:HGNC Symbol;Acc:HG |
| BBIP1 | 10 | BBSome interacting protein 1 [Source:HGNC S |
| AL158163.2 | 10 | novel transcript, antisense to BBIP1 |
| AL158163.1 | 10 | novel transcript, antisense to BBIP1 |
| SHOC2 | 10 | SHOC2 leucine rich repeat scaffold protein [So |
| RPL13AP6 | 10 | ribosomal protein L13a pseudogene 6 [Source |
| MIR548E | 10 | microRNA 548e [Source:HGNC Symbol;Acc:HG |
| ADRA2A | 10 | adrenoceptor alpha 2A [Source:HGNC Symbol |
| AL355863.1 | 10 | HSPC023 protein (HSPC023) pseudogene |
| BTBD7P2 | 10 | BTB domain containing 7 pseudogene 2 [Sour |
| AC021035.1 | 10 | novel transcript |
| RPS6P15 | 10 | ribosomal protein S6 pseudogene 15 [Source: |
| AL136119.1 | 10 | novel transcript |
| GPAM | 10 | glycerol-3-phosphate acyltransferase, mitocho |
| TECTB | 10 | tectorin beta [Source:HGNC Symbol;Acc:HGNC |
| MIR6715A | 10 | microRNA 6715a [Source:HGNC Symbol;Acc:H |
| MIR6715B | 10 | microRNA 6715b [Source:HGNC Symbol;Acc:H |
| GUCY2GP | 10 | guanylate cyclase 2G, pseudogene [Source:HG |
| ACSL5 | 10 | acyl-CoA synthetase long chain family membe |
| AL157786.1 | 10 | novel transcript, antisense to ACSL5 |
| ZDHHC6 | 10 | zinc finger DHHC-type containing 6 [Source:HG |
| VTI1A | 10 | vesicle transport through interaction with t-SN |
| AC022018.1 | 10 | pseudogene similar to part of eukaryotic trans |
| MIR4295 | 10 | microRNA 4295 [Source:HGNC Symbol;Acc:HG |

| | | |
|---|---|---|
| AL139120.1 | 10 | novel transcript |
| AL158212.5 | 10 | novel transcript, antisense to VTI1A |
| AL158212.3 | 10 | novel transcript, overlapping to VTI1A |
| AL158212.4 | 10 | TEC |
| AL158212.1 | 10 | novel transcript |
| TCF7L2 | 10 | transcription factor 7 like 2 [Source:HGNC Syn |
| AL158212.2 | 10 | novel transcript |
| RPS15AP30 | 10 | ribosomal protein S15a pseudogene 30 [Sourc |
| RNU7-165P | 10 | RNA, U7 small nuclear 165 pseudogene [Sourc |
| AL133482.1 | 10 | novel transcript |
| PPIAP39 | 10 | peptidylprolyl isomerase A pseudogene 39 [Sc |
| HABP2 | 10 | hyaluronan binding protein 2 [Source:HGNC S |
| NRAP | 10 | nebulin related anchoring protein [Source:HG |
| CASP7 | 10 | caspase 7 [Source:HGNC Symbol;Acc:HGNC:15 |
| AL592546.2 | 10 | novel transcript |
| PLEKHS1 | 10 | pleckstrin homology domain containing S1 [Sc |
| MIR4483 | 10 | microRNA 4483 [Source:HGNC Symbol;Acc:HG |
| AL592546.1 | 10 | NULL |
| DCLRE1A | 10 | DNA cross-link repair 1A [Source:HGNC Symbo |
| NHLRC2 | 10 | NHL repeat containing 2 [Source:HGNC Symbo |
| AL592546.3 | 10 | novel transcript, antisense to NHLRC2 |
| ADRB1 | 10 | adrenoceptor beta 1 [Source:HGNC Symbol;Ac |
| AC022023.2 | 10 | novel transcript |
| RNU6-709P | 10 | RNA, U6 small nuclear 709, pseudogene [Sour |
| UBE2V1P5 | 10 | ubiquitin conjugating enzyme E2 V1 pseudoge |
| CCDC186 | 10 | coiled-coil domain containing 186 [Source:HG |
| AC022023.1 | 10 | NULL |
| MIR2110 | 10 | microRNA 2110 [Source:HGNC Symbol;Acc:HG |
| TDRD1 | 10 | tudor domain containing 1 [Source:HGNC Sym |
| VWA2 | 10 | von Willebrand factor A domain containing 2 [ |
| AURKAP2 | 10 | aurora kinase A pseudogene 2 [Source:HGNC S |
| AC005383.1 | 10 | novel transcript, antisense to VWA2 |
| AFAP1L2 | 10 | actin filament associated protein 1 like 2 [Sou |
| RN7SL384P | 10 | RNA, 7SL, cytoplasmic 384, pseudogene [Sour |
| AL133384.2 | 10 | TEC |
| ABLIM1 | 10 | actin binding LIM protein 1 [Source:HGNC Sym |
| AL133384.1 | 10 | pseudogene similar to part of small nuclear ril |
| PPIAP19 | 10 | peptidylprolyl isomerase A pseudogene 19 [Sc |
| AL137025.1 | 10 | novel transcript |
| TAF9BP2 | 10 | TATA-box binding protein associated factor 9b |
| FAM160B1 | 10 | family with sequence similarity 160 member E |
| RPL15P13 | 10 | ribosomal protein L15 pseudogene 13 [Source |
| TRUB1 | 10 | TruB pseudouridine synthase family member 1 |
| RNU6-1121P | 10 | RNA, U6 small nuclear 1121, pseudogene [Sou |
| LINC02626 | 10 | long intergenic non-protein coding RNA 2626 |
| ATRNL1 | 10 | attractin like 1 [Source:HGNC Symbol;Acc:HGN |
| AC016042.1 | 10 | novel transcript, antisense to ATRNL1 |
| NTAN1P1 | 10 | N-terminal asparagine amidase pseudogene 1 |
| GFRA1 | 10 | GDNF family receptor alpha 1 [Source:HGNC S |
| AC012470.1 | 10 | novel transcript |

| | | |
|---|---|---|
| CCDC172 | 10 | coiled-coil domain containing 172 [Source:HG |
| SNRPGP6 | 10 | small nuclear ribonucleoprotein polypeptide ( |
| PNLIPRP3 | 10 | pancreatic lipase related protein 3 [Source:HG |
| HMGB3P8 | 10 | high mobility group box 3 pseudogene 8 [Sour |
| RNU6-1090P | 10 | RNA, U6 small nuclear 1090, pseudogene [Sou |
| PNLIP | 10 | pancreatic lipase [Source:HGNC Symbol;Acc:H |
| PNLIPP1 | 10 | pancreatic lipase pseudogene 1 [Source:HGN( |
| PNLIPRP1 | 10 | pancreatic lipase related protein 1 [Source:HG |
| PNLIPRP2 | 10 | pancreatic lipase related protein 2 (gene/pseu |
| C10orf82 | 10 | chromosome 10 open reading frame 82 [Sour |
| AC016825.1 | 10 | novel transcript, antisense to HSPA12A |
| HSPA12A | 10 | heat shock protein family A (Hsp70) member : |
| RPL5P27 | 10 | ribosomal protein L5 pseudogene 27 [Source: |
| ENO4 | 10 | enolase 4 [Source:HGNC Symbol;Acc:HGNC:31 |
| SHTN1 | 10 | shootin 1 [Source:HGNC Symbol;Acc:HGNC:29 |
| AC023283.1 | 10 | novel transcript, antisense to SHTN1 |
| AC012308.1 | 10 | novel pseudogene |
| VAX1 | 10 | ventral anterior homeobox 1 [Source:HGNC S |
| MIR3663HG | 10 | MIR3663 host gene [Source:HGNC Symbol;Acc |
| MIR3663 | 10 | microRNA 3663 [Source:HGNC Symbol;Acc:HG |
| RPL12P26 | 10 | ribosomal protein L12 pseudogene 26 [Source |
| KCNK18 | 10 | potassium two pore domain channel subfamil |
| AL731557.1 | 10 | novel transcript |
| SLC18A2 | 10 | solute carrier family 18 member A2 [Source:H |
| AL391988.1 | 10 | novel transcript, antisense to SLC18A2 |
| PDZD8 | 10 | PDZ domain containing 8 [Source:HGNC Symb |
| AC005871.2 | 10 | novel transcript |
| EMX2OS | 10 | EMX2 opposite strand/antisense RNA [Source |
| EMX2 | 10 | empty spiracles homeobox 2 [Source:HGNC S |
| AC005871.1 | 10 | novel transcript |
| AL356419.1 | 10 | novel transcript |
| LINC02674 | 10 | long intergenic non-protein coding RNA 2674 |
| AL513324.1 | 10 | novel transcript |
| RAB11FIP2 | 10 | RAB11 family interacting protein 2 [Source:HG |
| AC022395.1 | 10 | novel transcript, antisense to RAB11FIP2 |
| CASC2 | 10 | cancer susceptibility 2 [Source:HGNC Symbol;. |
| AL354863.1 | 10 | novel transcript |
| FAM204A | 10 | family with sequence similarity 204 member / |
| LINC00867 | 10 | long intergenic non-protein coding RNA 867 [S |
| SLC25A18P1 | 10 | solute carrier family 25 member 18 pseudoge |
| PRLHR | 10 | prolactin releasing hormone receptor [Source |
| TOMM22P5 | 10 | TOMM22 pseudogene 5 [Source:HGNC Symbo |
| CACUL1 | 10 | CDK2 associated cullin domain 1 [Source:HGN |
| AL139407.1 | 10 | novel transcript, sense intronic to CACUL1 |
| AL157388.1 | 10 | novel transcript |
| RPL17P36 | 10 | ribosomal protein L17 pseudogene 36 [Source |
| LDHAP5 | 10 | lactate dehydrogenase A pseudogene 5 [Sourc |
| NANOS1 | 10 | nanos C2HC-type zinc finger 1 [Source:HGNC S |
| EIF3A | 10 | eukaryotic translation initiation factor 3 subur |
| SNORA19 | 10 | small nucleolar RNA, H/ACA box 19 [Source:H( |

| | | |
|---|---|---|
| AL355598.1 | 10 | NULL |
| AL355598.2 | 10 | ribosomal protein L37 (RPL37) pseudogene |
| DENND10 | 10 | DENN domain containing 10 [Source:HGNC Sy |
| SFXN4 | 10 | sideroflexin 4 [Source:HGNC Symbol;Acc:HGN |
| PRDX3 | 10 | peroxiredoxin 3 [Source:HGNC Symbol;Acc:HG |
| GRK5 | 10 | G protein-coupled receptor kinase 5 [Source:H |
| GRK5-IT1 | 10 | GRK5 intronic transcript 1 [Source:HGNC Syml |
| RN7SL749P | 10 | RNA, 7SL, cytoplasmic 749, pseudogene [Sour |
| AL583824.1 | 10 | novel transcript |
| MIR4681 | 10 | microRNA 4681 [Source:HGNC Symbol;Acc:HG |
| RGS10 | 10 | regulator of G protein signaling 10 [Source:HG |
| AC012468.1 | 10 | developmental pluripotency associated 3 (DPI |
| TIAL1 | 10 | TIA1 cytotoxic granule associated RNA binding |
| RAD1P1 | 10 | RAD1 pseudogene 1 [Source:HGNC Symbol;Ac |
| RPS8P4 | 10 | ribosomal protein S8 pseudogene 4 [Source:H |
| BAG3 | 10 | BCL2 associated athanogene 3 [Source:HGNC |
| TXNP1 | 10 | thioredoxin pseudogene 1 [Source:HGNC Sym |
| INPP5F | 10 | inositol polyphosphate-5-phosphatase F [Sour |
| RN7SL846P | 10 | RNA, 7SL, cytoplasmic 846, pseudogene [Sour |
| PHACTR2P1 | 10 | phosphatase and actin regulator 2 pseudogen |
| AL133461.1 | 10 | VMA21 vacuolar H+-ATPase homolog (S. cerev |
| MCMBP | 10 | minichromosome maintenance complex bindi |
| SEC23IP | 10 | SEC23 interacting protein [Source:HGNC Symb |
| MIR4682 | 10 | microRNA 4682 [Source:HGNC Symbol;Acc:HG |
| NACAP2 | 10 | NACA pseudogene 2 [Source:HGNC Symbol;Ac |
| AL355298.1 | 10 | ribosomal protein L3 (RPL3) pseudogene |
| RPL21P16 | 10 | ribosomal protein L21 pseudogene 16 [Source |
| AC073587.1 | 10 | spermatogenesis associated 7 (SPATA7) pseuc |
| PLPP4 | 10 | phospholipid phosphatase 4 [Source:HGNC Sy |
| LINC01561 | 10 | long intergenic non-protein coding RNA 1561 |
| AC023282.1 | 10 | novel transcript |
| WDR11-AS1 | 10 | WDR11 antisense RNA 1 [Source:HGNC Symbo |
| AL391425.1 | 10 | novel transcript |
| WDR11 | 10 | WD repeat domain 11 [Source:HGNC Symbol; |
| AC010998.2 | 10 | novel transcript, antisense to WDR11 |
| AC010998.1 | 10 | novel transcript |
| AC010998.3 | 10 | novel transcript |
| RPL19P16 | 10 | ribosomal protein L19 pseudogene 16 [Source |
| LINC01153 | 10 | long intergenic non-protein coding RNA 1153 |
| RN7SKP167 | 10 | RN7SK pseudogene 167 [Source:HGNC Symbo |
| FGFR2 | 10 | fibroblast growth factor receptor 2 [Source:H( |
| AC009988.1 | 10 | novel transcript |
| RPS15AP5 | 10 | ribosomal protein S15a pseudogene 5 [Source |
| AC025947.1 | 10 | novel transcript |
| ATE1 | 10 | arginyltransferase 1 [Source:HGNC Symbol;Ac |
| Y_RNA | 10 | Y RNA [Source:RFAM;Acc:RF00019] |
| ATE1-AS1 | 10 | ATE1 antisense RNA 1 [Source:HGNC Symbol;/ |
| AL731566.1 | 10 | NULL |
| AL731566.3 | 10 | novel transcript, antisense to NSMCE4A |
| NSMCE4A | 10 | NSE4 homolog A, SMC5-SMC6 complex compo |

| | | |
|---|---|---|
| AL731566.2 | 10 | novel transcript, antisense to NSMCE4A |
| TACC2 | 10 | transforming acidic coiled-coil containing prot |
| AC063960.1 | 10 | novel transcript, antisense to TACC2 |
| AC063960.2 | 10 | novel transcript, antisense to TACC2 |
| BTBD16 | 10 | BTB domain containing 16 [Source:HGNC Sym |
| RNU6-728P | 10 | RNA, U6 small nuclear 728, pseudogene [Sour |
| PLEKHA1 | 10 | pleckstrin homology domain containing A1 [S |
| MIR3941 | 10 | microRNA 3941 [Source:HGNC Symbol;Acc:H |
| BX842242.1 | 10 | novel transcript |
| ARMS2 | 10 | age-related maculopathy susceptibility 2 [Sou |
| HTRA1 | 10 | HtrA serine peptidase 1 [Source:HGNC Symbo |
| DMBT1 | 10 | deleted in malignant brain tumors 1 [Source:H |
| AL603764.1 | 10 | novel pseudogene similar to AQN-1 protein A |
| AL603764.2 | 10 | carbohydrate-binding protein AQN-1-like [Sou |
| C10orf120 | 10 | chromosome 10 open reading frame 120 [Sou |
| DMBT1P1 | 10 | deleted in malignant brain tumors 1 pseudoge |
| CUZD1 | 10 | CUB and zona pellucida like domains 1 [Source |
| AC073585.2 | 10 | novel protein |
| FAM24B | 10 | family with sequence similarity 24 member B |
| AC073585.1 | 10 | chromosome 10 open reading frame 88 (C10c |
| FAM24A | 10 | family with sequence similarity 24 member A |
| C10orf88 | 10 | chromosome 10 open reading frame 88 [Sour |
| PSTK | 10 | phosphoseryl-tRNA kinase [Source:HGNC Sym |
| IKZF5 | 10 | IKAROS family zinc finger 5 [Source:HGNC Sym |
| ACADSB | 10 | acyl-CoA dehydrogenase short/branched chai |
| HMX3 | 10 | H6 family homeobox 3 [Source:HGNC Symbol; |
| HMX2 | 10 | H6 family homeobox 2 [Source:HGNC Symbol; |
| BUB3 | 10 | BUB3 mitotic checkpoint protein [Source:HGN |
| RPS26P39 | 10 | ribosomal protein S26 pseudogene 39 [Source |
| AL160290.2 | 10 | cytochrome c oxidase subunit VIIc (COX7C) ps |
| LINC02641 | 10 | long intergenic non-protein coding RNA 2641 |
| AL160290.1 | 10 | novel transcript |
| AL357127.2 | 10 | novel transcript |
| AL357127.1 | 10 | novel transcript |
| GPR26 | 10 | G protein-coupled receptor 26 [Source:HGNC |
| CPXM2 | 10 | carboxypeptidase X, M14 family member 2 [S |
| AC009987.1 | 10 | novel transcript |
| AC068058.1 | 10 | novel transcript |
| BX469938.1 | 10 | novel transcript |
| YBX2P1 | 10 | Y-box binding protein 2 pseudogene 1 [Source |
| AL683842.1 | 10 | novel pseudogene |
| CHST15 | 10 | carbohydrate sulfotransferase 15 [Source:HG |
| OAT | 10 | ornithine aminotransferase [Source:HGNC Syr |
| NKX1-2 | 10 | NK1 homeobox 2 [Source:HGNC Symbol;Acc:H |
| AL445237.1 | 10 | novel transcript, antisense to NKX1-2 |
| LHPP | 10 | phospholysine phosphohistidine inorganic pyr |
| RPS10P18 | 10 | ribosomal protein S10 pseudogene 18 [Source |
| AC068896.1 | 10 | novel transcript, METTL10-FAM53B readthrou |
| FAM53B | 10 | family with sequence similarity 53 member B |
| AL513190.1 | 10 | novel transcript, antisense to FAM53B |

| | | |
|---|---|---|
| FAM53B-AS1 | 10 | FAM53B antisense RNA 1 [Source:HGNC Symb |
| EEF1AKMT2 | 10 | EEF1A lysine methyltransferase 2 [Source:HGI |
| Y_RNA | 10 | Y RNA [Source:RFAM;Acc:RF00019] |
| ABRAXAS2 | 10 | abraxas 2, BRISC complex subunit [Source:HGI |
| Y_RNA | 10 | Y RNA [Source:RFAM;Acc:RF00019] |
| NPM1P31 | 10 | nucleophosmin 1 pseudogene 31 [Source:HGI |
| AL731577.2 | 10 | novel transcript, sense overlapping ZRANB1 |
| ZRANB1 | 10 | zinc finger RANBP2-type containing 1 [Source: |
| AL731577.1 | 10 | novel transcript |
| CTBP2 | 10 | C-terminal binding protein 2 [Source:HGNC Sy |
| AL731571.1 | 10 | novel transcript, antisense to CTBP2 |
| MIR4296 | 10 | microRNA 4296 [Source:HGNC Symbol;Acc:HG |
| AL157888.1 | 10 | novel transcript |
| MRPS21P6 | 10 | mitochondrial ribosomal protein S21 pseudog |
| RPS27P18 | 10 | ribosomal protein S27 pseudogene 18 [Source |
| TEX36-AS1 | 10 | TEX36 antisense RNA 1 [Source:HGNC Symbol |
| TEX36 | 10 | testis expressed 36 [Source:HGNC Symbol;Acc |
| ALDOAP2 | 10 | ALDOA pseudogene 2 [Source:HGNC Symbol;A |
| AL158835.1 | 10 | novel transcript |
| EDRF1-DT | 10 | EDRF1 divergent transcript [Source:HGNC Sym |
| AL158835.3 | 10 | novel transcript, antisense to C10orf137 |
| AL158835.2 | 10 | novel transcript |
| EDRF1 | 10 | erythroid differentiation regulatory factor 1 [S |
| EDRF1-AS1 | 10 | EDRF1 antisense RNA 1 [Source:HGNC Symbol |
| MMP21 | 10 | matrix metallopeptidase 21 [Source:HGNC Syr |
| UROS | 10 | uroporphyrinogen III synthase [Source:HGNC S |
| MIR4484 | 10 | microRNA 4484 [Source:HGNC Symbol;Acc:HG |
| BCCIP | 10 | BRCA2 and CDKN1A interacting protein [Sourc |
| DHX32 | 10 | DEAH-box helicase 32 (putative) [Source:HGN |
| Metazoa_SRP | 10 | Metazoan signal recognition particle RNA [Sou |
| RNU2-42P | 10 | RNA, U2 small nuclear 42, pseudogene [Sourc |
| FANK1 | 10 | fibronectin type III and ankyrin repeat domain |
| GNG10P1 | 10 | G protein subunit gamma 10 pseudogene 1 [S |
| FANK1-AS1 | 10 | FANK1 antisense RNA 1 [Source:HGNC Symbo |
| ADAM12 | 10 | ADAM metallopeptidase domain 12 [Source:H |
| RNA5SP328 | 10 | RNA, 5S ribosomal pseudogene 328 [Source:H |
| SAR1AP2 | 10 | secretion associated Ras related GTPase 1A ps |
| AL589787.2 | 10 | novel transcript |
| LINC00601 | 10 | long intergenic non-protein coding RNA 601 [S |
| C10orf90 | 10 | chromosome 10 open reading frame 90 [Sour |
| AL589787.1 | 10 | Novel transcript, antisense to C10orf90 |
| AL512272.1 | 10 | NULL |
| DOCK1 | 10 | dedicator of cytokinesis 1 [Source:HGNC Symb |
| AL359094.1 | 10 | novel transcript |
| AL359094.2 | 10 | novel transcript, antisense to DOCK1 |
| INSYN2A | 10 | inhibitory synaptic factor 2A [Source:HGNC Sy |
| NPS | 10 | neuropeptide S [Source:HGNC Symbol;Acc:HG |
| AL391005.1 | 10 | TEC |
| FOXI2 | 10 | forkhead box I2 [Source:HGNC Symbol;Acc:HG |
| BUB1P1 | 10 | BUB1 mitotic checkpoint serine/threonine kin |

| | | |
|---|---|---|
| CLRN3 | 10 | clarin 3 [Source:HGNC Symbol;Acc:HGNC:207[illegible] |
| PTPRE | 10 | protein tyrosine phosphatase receptor type E |
| AL158166.2 | 10 | novel transcript |
| AL158166.1 | 10 | novel transcript |
| AL390236.1 | 10 | novel transcript, antisense to PTPRE |
| MKI67 | 10 | marker of proliferation Ki-67 [Source:HGNC S[illegible] |
| LINC01163 | 10 | long intergenic non-protein coding RNA 1163 |
| AL390763.1 | 10 | novel transcript |
| LINC02667 | 10 | long intergenic non-protein coding RNA 2667 |
| AL355537.1 | 10 | novel transcript |
| AL355537.2 | 10 | novel transcript |
| AL391869.1 | 10 | novel transcript |
| AL359508.1 | 10 | novel transcript |
| MGMT | 10 | O-6-methylguanine-DNA methyltransferase [S[illegible] |
| AL355531.1 | 10 | novel transcript, antisense to MGMT |
| AL157832.2 | 10 | novel transcript |
| AL157832.1 | 10 | novel transcript |
| LINC02666 | 10 | long intergenic non-protein coding RNA 2666 |
| EBF3 | 10 | EBF transcription factor 3 [Source:HGNC Sym[illegible] |
| AL354950.2 | 10 | novel transcript, antisense to EBF3 |
| MIR4297 | 10 | microRNA 4297 [Source:HGNC Symbol;Acc:H[illegible] |
| AL354950.1 | 10 | novel transcript, antisense to EBF3 |
| AL354950.3 | 10 | novel transcript, antisense to EBF3 |
| C10orf143 | 10 | chromosome 10 open reading frame 143 [Sou[illegible] |
| AL139123.1 | 10 | novel transcript |
| CTAGE7P | 10 | CTAGE family member 7, pseudogene [Source[illegible] |
| PPIAP32 | 10 | peptidylprolyl isomerase A pseudogene 32 [S[illegible] |
| GLRX3 | 10 | glutaredoxin 3 [Source:HGNC Symbol;Acc:HG[illegible] |
| LINC02646 | 10 | long intergenic non-protein coding RNA 2646 |
| AL157712.1 | 10 | zinc finger protein pseudogene |
| Y_RNA | 10 | Y RNA [Source:RFAM;Acc:RF00019] |
| AC016816.1 | 10 | novel transcript |
| MIR378C | 10 | microRNA 378c [Source:HGNC Symbol;Acc:H[illegible] |
| TCERG1L | 10 | transcription elongation regulator 1 like [Sour[illegible] |
| TCERG1L-AS1 | 10 | TCERG1L antisense RNA 1 [Source:HGNC Sym[illegible] |
| LINC01164 | 10 | long intergenic non-protein coding RNA 1164 |
| AL450307.1 | 10 | novel transcript |
| AL607033.1 | 10 | TEC |
| FP565171.1 | 10 | novel transcript |
| PPP2R2D | 10 | protein phosphatase 2 regulatory subunit Bde[illegible] |
| BNIP3 | 10 | BCL2 interacting protein 3 [Source:HGNC Sym[illegible] |
| AL162274.2 | 10 | novel transcript, antisense to BNIP3 |
| AL162274.1 | 10 | novel transcript, antisense to BNIP3 |
| AL162274.3 | 10 | TEC |
| JAKMIP3 | 10 | Janus kinase and microtubule interacting prot[illegible] |
| AL512622.1 | 10 | novel transcript |
| DPYSL4 | 10 | dihydropyrimidinase like 4 [Source:HGNC Sym[illegible] |
| STK32C | 10 | serine/threonine kinase 32C [Source:HGNC Sy[illegible] |
| LRRC27 | 10 | leucine rich repeat containing 27 [Source:HG[illegible] |
| PWWP2B | 10 | PWWP domain containing 2B [Source:HGNC S[illegible] |

| | | |
|---|---|---|
| AL451069.2 | 10 | novel transcript |
| AL451069.3 | 10 | novel transcript |
| C10orf91 | 10 | chromosome 10 open reading frame 91 (puta |
| AL451069.1 | 10 | novel transcript |
| LINC01165 | 10 | long intergenic non-protein coding RNA 1165 |
| INPP5A | 10 | inositol polyphosphate-5-phosphatase A [Sour |
| NKX6-2 | 10 | NK6 homeobox 2 [Source:HGNC Symbol;Acc:H |
| CFAP46 | 10 | cilia and flagella associated protein 46 [Source |
| LINC01166 | 10 | long intergenic non-protein coding RNA 1166 |
| LINC01167 | 10 | long intergenic non-protein coding RNA 1167 |
| LINC01168 | 10 | long intergenic non-protein coding RNA 1168 |
| ADGRA1 | 10 | adhesion G protein-coupled receptor A1 [Sour |
| ADGRA1-AS1 | 10 | adhesion G protein-coupled receptor A1 antis |
| RPL5P28 | 10 | ribosomal protein L5 pseudogene 28 [Source: |
| KNDC1 | 10 | kinase non-catalytic C-lobe domain containing |
| UTF1 | 10 | undifferentiated embryonic cell transcription |
| VENTX | 10 | VENT homeobox [Source:HGNC Symbol;Acc:H |
| MIR202HG | 10 | MIR202 host gene [Source:HGNC Symbol;Acc: |
| MIR202 | 10 | microRNA 202 [Source:HGNC Symbol;Acc:HGN |
| AL592071.1 | 10 | novel transcript |
| ADAM8 | 10 | ADAM metallopeptidase domain 8 [Source:HG |
| TUBGCP2 | 10 | tubulin gamma complex associated protein 2 |
| AL360181.2 | 10 | novel transcript |
| ZNF511 | 10 | zinc finger protein 511 [Source:HGNC Symbol; |
| ZNF511-PRAP1 | 10 | ZNF511-PRAP1 readthrough [Source:HGNC Sy |
| CALY | 10 | calcyon neuron specific vesicular protein [Sou |
| BANF1P2 | 10 | barrier to autointegration factor 1 pseudogen |
| AL360181.1 | 10 | novel transcript, antisense to PRAP1 |
| PRAP1 | 10 | proline rich acidic protein 1 [Source:HGNC Syr |
| FUOM | 10 | fucose mutarotase [Source:HGNC Symbol;Acc |
| ECHS1 | 10 | enoyl-CoA hydratase, short chain 1 [Source:H( |
| MIR3944 | 10 | microRNA 3944 [Source:HGNC Symbol;Acc:HG |
| AL360181.4 | 10 | novel transcript |
| PAOX | 10 | polyamine oxidase [Source:HGNC Symbol;Acc |
| AL360181.3 | 10 | novel protein |
| MTG1 | 10 | mitochondrial ribosome associated GTPase 1 |
| SPRN | 10 | shadow of prion protein [Source:HGNC Symbo |
| OR6L2P | 10 | olfactory receptor family 6 subfamily L memb |
| SCART1 | 10 | scavenger receptor family member expressed |
| OR7M1P | 10 | olfactory receptor family 7 subfamily M meml |
| CYP2E1 | 10 | cytochrome P450 family 2 subfamily E membe |
| AL161645.1 | 10 | novel transcript, antisense to CYP2E1 |
| AL161645.2 | 10 | novel transcript, sense intronic to CYP2E1 |
| SYCE1 | 10 | synaptonemal complex central element prote |
| AL731769.2 | 10 | novel transcript, antisense to FRG2Band SYCE |
| OR6L1P | 10 | olfactory receptor family 6 subfamily L memb |
| AL731769.1 | 10 | novel transcript |
| FRG2B | 10 | FSHD region gene 2 family member B [Source: |
| RARRES2P2 | 10 | retinoic acid receptor responder 2 pseudogen |
| AGGF1P2 | 10 | angiogenic factor with G-patch and FHA doma |

| | | |
|---|---|---|
| CLUHP5 | 10 | clustered mitochondria homolog pseudogene |
| DUX4L28 | 10 | double homeobox 4 like 28 (pseudogene) [Sou |
| DUX4L25 | 10 | double homeobox 4 like 25 (pseudogene) [Sou |
| DUX4L24 | 10 | double homeobox 4 like 24 (pseudogene) [Sou |
| DUX4L23 | 10 | double homeobox 4 like 23 (pseudogene) [Sou |
| DUX4L22 | 10 | double homeobox 4 like 22 (pseudogene) [Sou |
| DUX4L21 | 10 | double homeobox 4 like 21 (pseudogene) [Sou |
| DUX4L20 | 10 | double homeobox 4 like 20 (pseudogene) [Sou |
| DUX4L29 | 10 | double homeobox 4 like 29 (pseudogene) [Sou |
| DUX4L10 | 10 | double homeobox 4 like 10 (pseudogene) [Sou |
| DUX4L11 | 10 | double homeobox 4 like 11 (pseudogene) [Sou |
| DUX4L12 | 10 | double homeobox 4 like 12 (pseudogene) [Sou |
| DUX4L13 | 10 | double homeobox 4 like 13 (pseudogene) [Sou |
| DUX4L14 | 10 | double homeobox 4 like 14 (pseudogene) [Sou |
| DUX4L15 | 10 | double homeobox 4 like 15 (pseudogene) [Sou |
| RPL23AP60 | 10 | ribosomal protein L23a pseudogene 60 [Sourc |
| BX322534.1 | 10 | major histocompatibility complex, class I, F (H |
| WBP1LP10 | 11 | WBP1L pseudogene 10 [Source:HGNC Symbol |
| OR4F2P | 11 | olfactory receptor family 4 subfamily F memb |
| AC069287.1 | 11 | novel transcript |
| CICP23 | 11 | capicua transcriptional repressor pseudogene |
| LINC01001 | 11 | long intergenic non-protein coding RNA 1001 |
| AC069287.3 | 11 | novel transcript, antisense to BET1L |
| RNU6-447P | 11 | RNA, U6 small nuclear 447, pseudogene [Sour |
| BET1L | 11 | Bet1 golgi vesicular membrane trafficking pro |
| SCGB1C1 | 11 | secretoglobin family 1C member 1 [Source:HG |
| ODF3 | 11 | outer dense fiber of sperm tails 3 [Source:HGI |
| AC069287.2 | 11 | novel transcript |
| RIC8A | 11 | RIC8 guanine nucleotide exchange factor A [So |
| MIR6743 | 11 | microRNA 6743 [Source:HGNC Symbol;Acc:HG |
| SIRT3 | 11 | sirtuin 3 [Source:HGNC Symbol;Acc:HGNC:149 |
| PSMD13 | 11 | proteasome 26S subunit, non-ATPase 13 [Sou |
| AC136475.10 | 11 | signal recognition particle 19kDa (SRP19) pseu |
| NLRP6 | 11 | NLR family pyrin domain containing 6 [Source |
| AC136475.3 | 11 | novel transcript |
| PGGHG | 11 | protein-glucosylgalactosylhydroxylysine gluco |
| IFITM5 | 11 | interferon induced transmembrane protein 5 |
| AC136475.6 | 11 | mitochondrial ribosomal protein S24 (MRPS24 |
| IFITM2 | 11 | interferon induced transmembrane protein 2 |
| AC136475.2 | 11 | novel transcript |
| IFITM1 | 11 | interferon induced transmembrane protein 1 |
| AC136475.1 | 11 | novel transcript, antisense to IFITM3 |
| IFITM3 | 11 | interferon induced transmembrane protein 3 |
| AC136475.4 | 11 | novel transcript, antisense to IFITM3 |
| AC136475.8 | 11 | novel transcript |
| AC136475.7 | 11 | novel transcript |
| AC136475.5 | 11 | novel transcript |
| AC136475.9 | 11 | novel transcript |
| B4GALNT4 | 11 | beta-1,4-N-acetyl-galactosaminyltransferase 4 |
| PKP3 | 11 | plakophilin 3 [Source:HGNC Symbol;Acc:HGN( |

| | | |
|---|---|---|
| SIGIRR | 11 | single Ig and TIR domain containing [Source:H |
| ANO9 | 11 | anoctamin 9 [Source:HGNC Symbol;Acc:HGNC |
| RN7SL838P | 11 | RNA, 7SL, cytoplasmic 838, pseudogene [Sour |
| PTDSS2 | 11 | phosphatidylserine synthase 2 [Source:HGNC |
| AC138230.1 | 11 | novel transcript |
| RNH1 | 11 | ribonuclease/angiogenin inhibitor 1 [Source:H |
| AC137894.3 | 11 | novel transcript |
| AC137894.1 | 11 | novel transcript |
| HRAS | 11 | HRas proto-oncogene, GTPase [Source:HGNC |
| LRRC56 | 11 | leucine rich repeat containing 56 [Source:HGN |
| LMNTD2 | 11 | lamin tail domain containing 2 [Source:HGNC |
| AP006284.1 | 11 | novel transcript |
| RASSF7 | 11 | Ras association domain family member 7 [Sou |
| MIR210HG | 11 | MIR210 host gene [Source:HGNC Symbol;Acc: |
| MIR210 | 11 | microRNA 210 [Source:HGNC Symbol;Acc:HGN |
| PHRF1 | 11 | PHD and ring finger domains 1 [Source:HGNC |
| IRF7 | 11 | interferon regulatory factor 7 [Source:HGNC S |
| CDHR5 | 11 | cadherin related family member 5 [Source:HG |
| SCT | 11 | secretin [Source:HGNC Symbol;Acc:HGNC:106 |
| DRD4 | 11 | dopamine receptor D4 [Source:HGNC Symbol; |
| DEAF1 | 11 | DEAF1 transcription factor [Source:HGNC Sym |
| AC131934.1 | 11 | novel transcript |
| EPS8L2 | 11 | EPS8 like 2 [Source:HGNC Symbol;Acc:HGNC:2 |
| TMEM80 | 11 | transmembrane protein 80 [Source:HGNC Syn |
| AP006621.4 | 11 | novel transcript, antisense to EPS8L2 |
| TALDO1 | 11 | transaldolase 1 [Source:HGNC Symbol;Acc:HG |
| GATD1 | 11 | glutamine amidotransferase like class 1 doma |
| AP006621.3 | 11 | novel transcript |
| AP006621.5 | 11 | TEC |
| AP006621.2 | 11 | novel transcript |
| CEND1 | 11 | cell cycle exit and neuronal differentiation 1 [S |
| SLC25A22 | 11 | solute carrier family 25 member 22 [Source:H |
| PANO1 | 11 | proapoptotic nucleolar protein 1 [Source:HGN |
| PIDD1 | 11 | p53-induced death domain protein 1 [Source: |
| RPLP2 | 11 | ribosomal protein lateral stalk subunit P2 [Sou |
| SNORA52 | 11 | small nucleolar RNA, H/ACA box 52 [Source:H |
| PNPLA2 | 11 | patatin like phospholipase domain containing |
| AP006621.1 | 11 | novel transcript |
| CRACR2B | 11 | calcium release activated channel regulator 2I |
| CD151 | 11 | CD151 molecule (Raph blood group) [Source:I |
| POLR2L | 11 | RNA polymerase II subunit L [Source:HGNC Sy |
| TSPAN4 | 11 | tetraspanin 4 [Source:HGNC Symbol;Acc:HGN |
| AP006623.1 | 11 | novel transcript |
| CHID1 | 11 | chitinase domain containing 1 [Source:HGNC S |
| AP2A2 | 11 | adaptor related protein complex 2 subunit alp |
| RNU6-1025P | 11 | RNA, U6 small nuclear 1025, pseudogene [Sou |
| MUC6 | 11 | mucin 6, oligomeric mucus/gel-forming [Sourc |
| LINC02688 | 11 | long intergenic non-protein coding RNA 2688 |
| MUC2 | 11 | mucin 2, oligomeric mucus/gel-forming [Sourc |
| MUC5AC | 11 | mucin 5AC, oligomeric mucus/gel-forming [So |

| | | |
|---|---|---|
| AC061979.1 | 11 | novel transcript |
| MUC5B | 11 | mucin 5B, oligomeric mucus/gel-forming [Sou |
| MUC5B-AS1 | 11 | MUC5B antisense RNA 1 [Source:HGNC Symbo |
| MIR6744 | 11 | microRNA 6744 [Source:HGNC Symbol;Acc:HG |
| TOLLIP | 11 | toll interacting protein [Source:HGNC Symbol; |
| TOLLIP-AS1 | 11 | TOLLIP antisense RNA 1 (head to head) [Sourc |
| AC136297.1 | 11 | novel transcript |
| LINC02689 | 11 | long intergenic non-protein coding RNA 2689 |
| BRSK2 | 11 | BR serine/threonine kinase 2 [Source:HGNC S |
| AC091196.1 | 11 | novel transcript |
| MOB2 | 11 | MOB kinase activator 2 [Source:HGNC Symbol |
| DUSP8 | 11 | dual specificity phosphatase 8 [Source:HGNC S |
| KRTAP5-AS1 | 11 | KRTAP5-1/KRTAP5-2 antisense RNA 1 [Source |
| KRTAP5-1 | 11 | keratin associated protein 5-1 [Source:HGNC S |
| KRTAP5-2 | 11 | keratin associated protein 5-2 [Source:HGNC S |
| KRTAP5-3 | 11 | keratin associated protein 5-3 [Source:HGNC S |
| KRTAP5-4 | 11 | keratin associated protein 5-4 [Source:HGNC S |
| KRTAP5-5 | 11 | keratin associated protein 5-5 [Source:HGNC S |
| AP006285.1 | 11 | novel transcript |
| FAM99A | 11 | family with sequence similarity 99 member A |
| FAM99B | 11 | family with sequence similarity 99 member B |
| LINC02708 | 11 | long intergenic non-protein coding RNA 2708 |
| KRTAP5-6 | 11 | keratin associated protein 5-6 [Source:HGNC S |
| IFITM10 | 11 | interferon induced transmembrane protein 1( |
| AC068580.4 | 11 | novel protein |
| CTSD | 11 | cathepsin D [Source:HGNC Symbol;Acc:HGNC: |
| AC068580.3 | 11 | novel transcript |
| AC068580.1 | 11 | novel transcript |
| AC068580.5 | 11 | novel transcript |
| AC068580.2 | 11 | novel transcript |
| RPL36AP39 | 11 | ribosomal protein L36a pseudogene 39 [Sourc |
| AC139143.1 | 11 | actin pseudogene |
| SYT8 | 11 | synaptotagmin 8 [Source:HGNC Symbol;Acc:H |
| TNNI2 | 11 | troponin I2, fast skeletal type [Source:HGNC S |
| LSP1 | 11 | lymphocyte specific protein 1 [Source:HGNC S |
| MIR4298 | 11 | microRNA 4298 [Source:HGNC Symbol;Acc:HG |
| AC051649.1 | 11 | novel transcript, sense overlapping LSP1 |
| MIR7847 | 11 | microRNA 7847 [Source:HGNC Symbol;Acc:HG |
| PRR33 | 11 | proline rich 33 [Source:HGNC Symbol;Acc:HGN |
| LINC01150 | 11 | long intergenic non-protein coding RNA 1150 |
| TNNT3 | 11 | troponin T3, fast skeletal type [Source:HGNC S |
| MRPL23 | 11 | mitochondrial ribosomal protein L23 [Source: |
| MRPL23-AS1 | 11 | MRPL23 antisense RNA 1 [Source:HGNC Symb |
| LINC01219 | 11 | long intergenic non-protein coding RNA 1219 |
| H19 | 11 | H19 imprinted maternally expressed transcrip |
| AC051649.3 | 11 | NULL |
| MIR675 | 11 | microRNA 675 [Source:HGNC Symbol;Acc:HGN |
| AC051649.2 | 11 | NULL |
| IGF2 | 11 | insulin like growth factor 2 [Source:HGNC Sym |
| AC132217.2 | 11 | novel protein |

| | | |
|---|---|---|
| AC132217.1 | 11 | novel transcript, 3' overlapping ncRNA IGF2 |
| INS-IGF2 | 11 | INS-IGF2 readthrough [Source:HGNC Symbol;/ |
| MIR483 | 11 | microRNA 483 [Source:HGNC Symbol;Acc:HGI |
| IGF2-AS | 11 | IGF2 antisense RNA [Source:HGNC Symbol;Ac |
| INS | 11 | insulin [Source:HGNC Symbol;Acc:HGNC:6081 |
| TH | 11 | tyrosine hydroxylase [Source:HGNC Symbol;A |
| MIR4686 | 11 | microRNA 4686 [Source:HGNC Symbol;Acc:HG |
| ASCL2 | 11 | achaete-scute family bHLH transcription facto |
| C11orf21 | 11 | chromosome 11 open reading frame 21 [Sour |
| TSPAN32 | 11 | tetraspanin 32 [Source:HGNC Symbol;Acc:HGI |
| RNU6-878P | 11 | RNA, U6 small nuclear 878, pseudogene [Sour |
| CD81-AS1 | 11 | CD81 antisense RNA 1 [Source:HGNC Symbol; |
| RPL26P30 | 11 | ribosomal protein L26 pseudogene 30 [Source |
| Y_RNA | 11 | Y RNA [Source:RFAM;Acc:RF00019] |
| CD81 | 11 | CD81 molecule [Source:HGNC Symbol;Acc:HG |
| TSSC4 | 11 | tumor suppressing subtransferable candidate |
| AC124057.1 | 11 | novel transcript |
| TRPM5 | 11 | transient receptor potential cation channel su |
| KCNQ1 | 11 | potassium voltage-gated channel subfamily Q |
| KCNQ1OT1 | 11 | KCNQ1 opposite strand/antisense transcript 1 |
| AC021424.1 | 11 | NULL |
| AC021424.3 | 11 | NULL |
| AC021424.4 | 11 | NULL |
| AC021424.2 | 11 | NULL |
| COX6CP18 | 11 | cytochrome c oxidase subunit 6C pseudogene |
| KCNQ1-AS1 | 11 | KCNQ1 antisense RNA 1 [Source:HGNC Symbo |
| KCNQ1DN | 11 | KCNQ1 downstream neighbor [Source:HGNC S |
| AC013791.1 | 11 | NULL |
| CDKN1C | 11 | cyclin dependent kinase inhibitor 1C [Source:I |
| SLC22A18AS | 11 | solute carrier family 22 member 18 antisense |
| SLC22A18 | 11 | solute carrier family 22 member 18 [Source:H |
| PHLDA2 | 11 | pleckstrin homology like domain family A mer |
| NAP1L4 | 11 | nucleosome assembly protein 1 like 4 [Source |
| SNORA54 | 11 | small nucleolar RNA, H/ACA box 54 [Source:H |
| AC131971.1 | 11 | novel transcript, antisense to NAP1L4 |
| CARS | 11 | cysteinyl-tRNA synthetase [Source:HGNC Sym |
| CARS-AS1 | 11 | CARS antisense RNA 1 [Source:HGNC Symbol; |
| RNU1-91P | 11 | RNA, U1 small nuclear 91, pseudogene [Sourc |
| AC108448.3 | 11 | novel transcript, antisense to CARS |
| AC108448.2 | 11 | proteasome (prosome, macropain) subunit, al |
| OSBPL5 | 11 | oxysterol binding protein like 5 [Source:HGNC |
| AC108448.1 | 11 | novel transcript |
| MRGPRG | 11 | MAS related GPR family member G [Source:H |
| MRGPRG-AS1 | 11 | MRGPRG antisense RNA 1 [Source:HGNC Sym |
| MRGPRE | 11 | MAS related GPR family member E [Source:H |
| AC109309.1 | 11 | novel transcript |
| NDUFA5P8 | 11 | NADH:ubiquinone oxidoreductase subunit A5 |
| AC109309.2 | 11 | mastermind-like 1 (Drosophila) (MAML1) pseu |
| NDUFA5P1 | 11 | NADH:ubiquinone oxidoreductase subunit A5 |
| AC123788.2 | 11 | mastermind-like 1 (Drosophila) (MAML1) pseu |

| | | |
|---|---|---|
| AC123788.1 | 11 | novel transcript |
| ZNF195 | 11 | zinc finger protein 195 [Source:HGNC Symbol; |
| TSSC2 | 11 | tumor suppressing subtransferable candidate |
| OR7E12P | 11 | olfactory receptor family 7 subfamily E memb |
| FAM86GP | 11 | family with sequence similarity 86 member G, |
| AC127526.3 | 11 | ectonucleotide pyrophosphatase/phosphodie |
| AC127526.1 | 11 | novel transcript |
| AC127526.5 | 11 | novel transcript |
| AC127526.2 | 11 | novel transcript |
| AC127526.4 | 11 | novel transcript |
| SNRPCP5 | 11 | small nuclear ribonucleoprotein polypeptide ( |
| RPS3AP39 | 11 | ribosomal protein S3a pseudogene 39 [Source |
| RPS24P14 | 11 | ribosomal protein S24 pseudogene 14 [Source |
| AP006294.2 | 11 | RNA binding protein, fox-1 homolog (C. elegar |
| AP006294.1 | 11 | novel pseudogene (C11orf2) |
| OR7E117P | 11 | olfactory receptor family 7 subfamily E memb |
| TRPC2 | 11 | transient receptor potential cation channel su |
| ART5 | 11 | ADP-ribosyltransferase 5 [Source:HGNC Symb |
| ART1 | 11 | ADP-ribosyltransferase 1 [Source:HGNC Symb |
| Y_RNA | 11 | Y RNA [Source:RFAM;Acc:RF00019] |
| Y_RNA | 11 | Y RNA [Source:RFAM;Acc:RF00019] |
| CHRNA10 | 11 | cholinergic receptor nicotinic alpha 10 subuni |
| NUP98 | 11 | nucleoporin 98 [Source:HGNC Symbol;Acc:HG |
| RNU6-1143P | 11 | RNA, U6 small nuclear 1143, pseudogene [Sou |
| RNU7-50P | 11 | RNA, U7 small nuclear 50 pseudogene [Source |
| PGAP2 | 11 | post-GPI attachment to proteins 2 [Source:HG |
| RHOG | 11 | ras homolog family member G [Source:HGNC |
| AC090587.1 | 11 | novel transcript |
| STIM1 | 11 | stromal interaction molecule 1 [Source:HGNC |
| AC090587.2 | 11 | novel transcript, sense overlapping STIM1 |
| MIR4687 | 11 | microRNA 4687 [Source:HGNC Symbol;Acc:HG |
| PPIAP40 | 11 | peptidylprolyl isomerase A pseudogene 40 [Sc |
| AC087441.1 | 11 | NULL |
| AC087441.3 | 11 | Acyl-CoA-binding domain-containing protein 7 |
| HNRNPA1P76 | 11 | heterogeneous nuclear ribonucleoprotein A1 |
| AC087441.2 | 11 | Non-histone chromosomal protein HMG-14 (H |
| RPS29P20 | 11 | ribosomal protein S29 pseudogene 20 [Source |
| RRM1 | 11 | ribonucleotide reductase catalytic subunit M1 |
| RRM1-AS1 | 11 | RRM1 antisense RNA 1 [Source:HGNC Symbol |
| OR55B1P | 11 | olfactory receptor family 55 subfamily B mem |
| LINC02749 | 11 | long intergenic non-protein coding RNA 2749 |
| AC015689.1 | 11 | radixin (RDX) pseudogene |
| SSU72P5 | 11 | SSU72 pseudogene 5 [Source:HGNC Symbol;A |
| SSU72P2 | 11 | SSU72 pseudogene 2 [Source:HGNC Symbol;A |
| SSU72P6 | 11 | SSU72 pseudogene 6 [Source:HGNC Symbol;A |
| SSU72P4 | 11 | SSU72 pseudogene 4 [Source:HGNC Symbol;A |
| SSU72P3 | 11 | SSU72 pseudogene 3 [Source:HGNC Symbol;A |
| SSU72P7 | 11 | SSU72 pseudogene 7 [Source:HGNC Symbol;A |
| OR52B4 | 11 | olfactory receptor family 52 subfamily B mem |
| OR52B3P | 11 | olfactory receptor family 52 subfamily B mem |

| | | |
|---|---|---|
| TRIM21 | 11 | tripartite motif containing 21 [Source:HGNC S |
| OR52K3P | 11 | olfactory receptor family 52 subfamily K mem |
| OR51R1P | 11 | olfactory receptor family 51 subfamily R mem |
| OR52P2P | 11 | olfactory receptor family 52 subfamily P mem |
| OR52K2 | 11 | olfactory receptor family 52 subfamily K mem |
| OR52K1 | 11 | olfactory receptor family 52 subfamily K mem |
| OR52M2P | 11 | olfactory receptor family 52 subfamily M mem |
| OR52M1 | 11 | olfactory receptor family 52 subfamily M mem |
| C11orf40 | 11 | chromosome 11 open reading frame 40 (puta |
| OR52I2 | 11 | olfactory receptor family 52 subfamily I memb |
| OR52I1 | 11 | olfactory receptor family 52 subfamily I memb |
| TRIM68 | 11 | tripartite motif containing 68 [Source:HGNC S |
| OR51D1 | 11 | olfactory receptor family 51 subfamily D mem |
| OR51E1 | 11 | olfactory receptor family 51 subfamily E mem |
| OR51A9P | 11 | olfactory receptor family 51 subfamily A mem |
| OR51E2 | 11 | olfactory receptor family 51 subfamily E mem |
| OR51C1P | 11 | olfactory receptor family 51 subfamily C mem |
| MMP26 | 11 | matrix metallopeptidase 26 [Source:HGNC Syr |
| OR51F5P | 11 | olfactory receptor family 51 subfamily F mem |
| OR51C4P | 11 | olfactory receptor family 51 subfamily C mem |
| KRT8P49 | 11 | keratin 8 pseudogene 49 [Source:HGNC Symb |
| OR51F3P | 11 | olfactory receptor family 51 subfamily F mem |
| OR51F4P | 11 | olfactory receptor family 51 subfamily F mem |
| SNORA62 | 11 | Small nucleolar RNA SNORA62/SNORA6 family |
| OR51F1 | 11 | olfactory receptor family 51 subfamily F mem |
| OR51N1P | 11 | olfactory receptor family 51 subfamily N mem |
| OR52Y1P | 11 | olfactory receptor family 52 subfamily Y mem |
| OR52R1 | 11 | olfactory receptor family 52 subfamily R mem |
| OR51F2 | 11 | olfactory receptor family 51 subfamily F mem |
| OR51A8P | 11 | olfactory receptor family 51 subfamily A mem |
| OR51S1 | 11 | olfactory receptor family 51 subfamily S mem |
| OR51H1 | 11 | olfactory receptor family 51 subfamily H mem |
| OR51H2P | 11 | olfactory receptor family 51 subfamily H mem |
| OR51T1 | 11 | olfactory receptor family 51 subfamily T mem |
| OR51A6P | 11 | olfactory receptor family 51 subfamily A mem |
| OR51A7 | 11 | olfactory receptor family 51 subfamily A mem |
| OR51G2 | 11 | olfactory receptor family 51 subfamily G mem |
| OR51G1 | 11 | olfactory receptor family 51 subfamily G mem |
| OR51A3P | 11 | olfactory receptor family 51 subfamily A mem |
| OR51A4 | 11 | olfactory receptor family 51 subfamily A mem |
| OR51A2 | 11 | olfactory receptor family 51 subfamily A mem |
| OR51A5P | 11 | olfactory receptor family 51 subfamily A mem |
| OR51L1 | 11 | olfactory receptor family 51 subfamily L mem |
| OR51P1P | 11 | olfactory receptor family 51 subfamily P mem |
| OR52J2P | 11 | olfactory receptor family 52 subfamily J memb |
| OR52J3 | 11 | olfactory receptor family 52 subfamily J memb |
| OR52E2 | 11 | olfactory receptor family 52 subfamily E mem |
| OR52E1 | 11 | olfactory receptor family 52 subfamily E mem |
| OR52S1P | 11 | olfactory receptor family 52 subfamily S mem |
| OR52E3P | 11 | olfactory receptor family 52 subfamily E mem |

| | | |
|---|---|---|
| OR52J1P | 11 | olfactory receptor family 52 subfamily J meml |
| AC113331.1 | 11 | novel transcript |
| OR52A4P | 11 | olfactory receptor family 52 subfamily A mem |
| OR52A5 | 11 | olfactory receptor family 52 subfamily A mem |
| OR52A1 | 11 | olfactory receptor family 52 subfamily A mem |
| OR51A1P | 11 | olfactory receptor family 51 subfamily A mem |
| OR52Z1 | 11 | olfactory receptor family 52 subfamily Z mem |
| OR51V1 | 11 | olfactory receptor family 51 subfamily V mem |
| AC104389.2 | 11 | novel transcript |
| AC104389.7 | 11 | TEC |
| AC104389.1 | 11 | NULL |
| HBB | 11 | hemoglobin subunit beta [Source:HGNC Symb |
| HBD | 11 | hemoglobin subunit delta [Source:HGNC Symb |
| HBBP1 | 11 | hemoglobin subunit beta pseudogene 1 [Sour |
| BGLT3 | 11 | beta globin locus transcript 3 [Source:HGNC Sy |
| HBG1 | 11 | hemoglobin subunit gamma 1 [Source:HGNC S |
| AC104389.6 | 11 | novel protein |
| HBG2 | 11 | hemoglobin subunit gamma 2 [Source:HGNC S |
| AC104389.5 | 11 | novel transcript |
| HBE1 | 11 | hemoglobin subunit epsilon 1 [Source:HGNC S |
| OR51AB1P | 11 | olfactory receptor family 51 subfamily AB mer |
| OR51B4 | 11 | olfactory receptor family 51 subfamily B mem |
| OR51B5 | 11 | olfactory receptor family 51 subfamily B mem |
| OR51B3P | 11 | olfactory receptor family 51 subfamily B mem |
| OR51B2 | 11 | olfactory receptor family 51 subfamily B mem |
| OR51B8P | 11 | olfactory receptor family 51 subfamily B mem |
| OR51B6 | 11 | olfactory receptor family 51 subfamily B mem |
| AC104389.3 | 11 | amphiphysin (Stiff-Man syndrome with breast |
| AC104389.4 | 11 | chromosome 11 open reading frame 10 (C11c |
| OR51M1 | 11 | olfactory receptor family 51 subfamily M mer |
| OR51J1 | 11 | olfactory receptor family 51 subfamily J meml |
| OR51Q1 | 11 | olfactory receptor family 51 subfamily Q mer |
| OR51K1P | 11 | olfactory receptor family 51 subfamily K mem |
| OR51I1 | 11 | olfactory receptor family 51 subfamily I meml |
| OR51I2 | 11 | olfactory receptor family 51 subfamily I meml |
| OR51A10P | 11 | olfactory receptor family 51 subfamily A mem |
| OR52D1 | 11 | olfactory receptor family 52 subfamily D mem |
| UBQLN3 | 11 | ubiquilin 3 [Source:HGNC Symbol;Acc:HGNC:1 |
| UBQLNL | 11 | ubiquilin like [Source:HGNC Symbol;Acc:HGNC |
| OLFM5P | 11 | olfactomedin family member 5, pseudogene [ |
| OR52V1P | 11 | olfactory receptor family 52 subfamily V mem |
| OR52H1 | 11 | olfactory receptor family 52 subfamily H mem |
| OR52H2P | 11 | olfactory receptor family 52 subfamily H mem |
| OR52B5P | 11 | olfactory receptor family 52 subfamily B mem |
| OR52T1P | 11 | olfactory receptor family 52 subfamily T mem |
| HNRNPA1P53 | 11 | heterogeneous nuclear ribonucleoprotein A1 |
| OR52B6 | 11 | olfactory receptor family 52 subfamily B mem |
| TRIM6 | 11 | tripartite motif containing 6 [Source:HGNC Syr |
| TRIM6-TRIM34 | 11 | TRIM6-TRIM34 readthrough [Source:HGNC Sy |
| TRIM34 | 11 | tripartite motif containing 34 [Source:HGNC S |

| | | |
|---|---|---|
| TRIM5 | 11 | tripartite motif containing 5 [Source:HGNC Sy |
| TRIM22 | 11 | tripartite motif containing 22 [Source:HGNC S |
| OR52U1P | 11 | olfactory receptor family 52 subfamily U mem |
| OR52P1P | 11 | olfactory receptor family 52 subfamily P mem |
| OR56B1 | 11 | olfactory receptor family 56 subfamily B mem |
| OR52N4 | 11 | olfactory receptor family 52 subfamily N mem |
| OR56B2P | 11 | olfactory receptor family 56 subfamily B mem |
| OR52N5 | 11 | olfactory receptor family 52 subfamily N mem |
| OR52N1 | 11 | olfactory receptor family 52 subfamily N mem |
| OR52N3P | 11 | olfactory receptor family 52 subfamily N mem |
| OR52N2 | 11 | olfactory receptor family 52 subfamily N mem |
| OR52E6 | 11 | olfactory receptor family 52 subfamily E mem |
| OR52E8 | 11 | olfactory receptor family 52 subfamily E mem |
| OR52E7P | 11 | olfactory receptor family 52 subfamily E mem |
| OR52E4 | 11 | olfactory receptor family 52 subfamily E mem |
| OR52E5 | 11 | olfactory receptor family 52 subfamily E mem |
| OR52Q1P | 11 | olfactory receptor family 52 subfamily Q mem |
| OR56A3 | 11 | olfactory receptor family 56 subfamily A mem |
| OR56A5 | 11 | olfactory receptor family 56 subfamily A mem |
| OR52L1 | 11 | olfactory receptor family 52 subfamily L mem |
| KRT18P58 | 11 | keratin 18 pseudogene 58 [Source:HGNC Sym |
| OR56A4 | 11 | olfactory receptor family 56 subfamily A mem |
| RNA5SP329 | 11 | RNA, 5S ribosomal pseudogene 329 [Source:H |
| OR56A1 | 11 | olfactory receptor family 56 subfamily A mem |
| OR56A7P | 11 | olfactory receptor family 56 subfamily A mem |
| OR52L2P | 11 | olfactory receptor family 52 subfamily L mem |
| OR52X1P | 11 | olfactory receptor family 52 subfamily X mem |
| OR56B4 | 11 | olfactory receptor family 56 subfamily B mem |
| AC022762.1 | 11 | novel transcript, sense overlapping OR52B2 |
| OR56B3P | 11 | olfactory receptor family 56 subfamily B mem |
| OR52B1P | 11 | olfactory receptor family 52 subfamily B mem |
| OR52B2 | 11 | olfactory receptor family 52 subfamily B mem |
| OR52W1 | 11 | olfactory receptor family 52 subfamily W men |
| AC022762.2 | 11 | novel transcript |
| C11orf42 | 11 | chromosome 11 open reading frame 42 [Sour |
| FAM160A2 | 11 | family with sequence similarity 160 member A |
| CNGA4 | 11 | cyclic nucleotide gated channel subunit alpha |
| CCKBR | 11 | cholecystokinin B receptor [Source:HGNC Sym |
| AC068733.2 | 11 | TBC1 domain family, member 12, pseudogene |
| CAVIN3 | 11 | caveolae associated protein 3 [Source:HGNC S |
| AC068733.3 | 11 | novel transcript, antisense to PRKCDBP |
| AC068733.1 | 11 | novel transcript |
| SMPD1 | 11 | sphingomyelin phosphodiesterase 1 [Source:H |
| APBB1 | 11 | amyloid beta precursor protein binding family |
| HPX | 11 | hemopexin [Source:HGNC Symbol;Acc:HGNC:! |
| TRIM3 | 11 | tripartite motif containing 3 [Source:HGNC Sy |
| ARFIP2 | 11 | ADP ribosylation factor interacting protein 2 [ |
| AC084337.2 | 11 | novel transcript |
| TIMM10B | 11 | translocase of inner mitochondrial membrane |
| AC084337.1 | 11 | voltage-dependent anion channel 1(VDAC1) p |

| | | |
|---|---|---|
| DNHD1 | 11 | dynein heavy chain domain 1 [Source:HGNC S |
| RRP8 | 11 | ribosomal RNA processing 8 [Source:HGNC Sy |
| AC091564.2 | 11 | novel transcript |
| ILK | 11 | integrin linked kinase [Source:HGNC Symbol;A |
| TAF10 | 11 | TATA-box binding protein associated factor 1( |
| AC091564.3 | 11 | novel transcript, sense overlapping TAF10 |
| AC091564.7 | 11 | novel transcript |
| TPP1 | 11 | tripeptidyl peptidase 1 [Source:HGNC Symbol; |
| AC091564.6 | 11 | novel transcript |
| DCHS1 | 11 | dachsous cadherin-related 1 [Source:HGNC Sy |
| AC091564.4 | 11 | novel transcript |
| AC091564.5 | 11 | novel transcript |
| MRPL17 | 11 | mitochondrial ribosomal protein L17 [Source:I |
| AC091564.1 | 11 | ribosomal protein L21 (RPL21) pseudogene |
| GVINP1 | 11 | GTPase, very large interferon inducible pseud |
| GVINP2 | 11 | GTPase, very large interferon inducible pseud |
| OR2AG2 | 11 | olfactory receptor family 2 subfamily AG mem |
| OR2AG1 | 11 | olfactory receptor family 2 subfamily AG mem |
| OR6A2 | 11 | olfactory receptor family 6 subfamily A memb |
| AC087280.2 | 11 | novel transcript |
| AC087280.1 | 11 | phosphorylated adaptor for RNA export (PHA) |
| OR10A5 | 11 | olfactory receptor family 10 subfamily A mem |
| OR10A2 | 11 | olfactory receptor family 10 subfamily A mem |
| OR10A4 | 11 | olfactory receptor family 10 subfamily A mem |
| OR2D2 | 11 | olfactory receptor family 2 subfamily D memb |
| OR2D3 | 11 | olfactory receptor family 2 subfamily D memb |
| ZNF215 | 11 | zinc finger protein 215 [Source:HGNC Symbol; |
| ZNF214 | 11 | zinc finger protein 214 [Source:HGNC Symbol; |
| NLRP14 | 11 | NLR family pyrin domain containing 14 [Sourc |
| RBMXL2 | 11 | RBMX like 2 [Source:HGNC Symbol;Acc:HGNC |
| AC027804.1 | 11 | novel transcript |
| MIR302E | 11 | microRNA 302e [Source:HGNC Symbol;Acc:HG |
| SYT9 | 11 | synaptotagmin 9 [Source:HGNC Symbol;Acc:H |
| AC107884.1 | 11 | novel transcript, antisense to OLFML1 |
| OLFML1 | 11 | olfactomedin like 1 [Source:HGNC Symbol;Acc |
| PPFIBP2 | 11 | PPFIA binding protein 2 [Source:HGNC Symbo |
| AC107884.2 | 11 | novel transcript |
| CYB5R2 | 11 | cytochrome b5 reductase 2 [Source:HGNC Syr |
| OVCH2 | 11 | ovochymase 2 (gene/pseudogene) [Source:HG |
| AC104237.2 | 11 | novel transcript, sense intronic to OVCH2 |
| AC104237.3 | 11 | novel transcript, sense intronic to OVCH2 |
| AC044810.3 | 11 | novel transcript, sense overlapping OR5P2, OI |
| AC104237.1 | 11 | novel transcript |
| OR10AB1P | 11 | olfactory receptor family 10 subfamily AB me |
| OR5P4P | 11 | olfactory receptor family 5 subfamily P memb |
| AC044810.2 | 11 | novel transcript, sense overlapping OR5P2, OI |
| OR5P1P | 11 | olfactory receptor family 5 subfamily P memb |
| AC044810.1 | 11 | ribosomal protein L7a (RPL7A) pseudogene |
| OR5P2 | 11 | olfactory receptor family 5 subfamily P memb |
| OR5P3 | 11 | olfactory receptor family 5 subfamily P memb |

| | | |
|---|---|---|
| OR5E1P | 11 | olfactory receptor family 5 subfamily E memb |
| RNU6-943P | 11 | RNA, U6 small nuclear 943, pseudogene [Sour |
| OR10A6 | 11 | olfactory receptor family 10 subfamily A mem |
| OR10A3 | 11 | olfactory receptor family 10 subfamily A mem |
| NLRP10 | 11 | NLR family pyrin domain containing 10 [Sourc |
| EIF3F | 11 | eukaryotic translation initiation factor 3 subur |
| COX6CP5 | 11 | cytochrome c oxidase subunit 6C pseudogene |
| CASC23 | 11 | cancer susceptibility 23 [Source:HGNC Symbo |
| TUB | 11 | TUB bipartite transcription factor [Source:HGI |
| AC116456.1 | 11 | novel transcript |
| TUB-AS1 | 11 | TUB antisense RNA 1 [Source:HGNC Symbol;A |
| AC116456.2 | 11 | TEC |
| RIC3 | 11 | RIC3 acetylcholine receptor chaperone [Sourc |
| RIC3-DT | 11 | RIC3 divergent transcript [Source:HGNC Symb |
| LMO1 | 11 | LIM domain only 1 [Source:HGNC Symbol;Acc |
| STK33 | 11 | serine/threonine kinase 33 [Source:HGNC Sym |
| AP006442.1 | 11 | NULL |
| TRIM66 | 11 | tripartite motif containing 66 [Source:HGNC S |
| AC091053.2 | 11 | novel transcript |
| RPL27A | 11 | ribosomal protein L27a [Source:HGNC Symbol |
| SNORA3A | 11 | small nucleolar RNA, H/ACA box 3A [Source:H |
| SNORA3B | 11 | small nucleolar RNA, H/ACA box 3B [Source:H |
| ST5 | 11 | suppression of tumorigenicity 5 [Source:HGN( |
| AC091053.1 | 11 | novel transcript |
| AC026894.1 | 11 | novel transcript |
| RNA5SP330 | 11 | RNA, 5S ribosomal pseudogene 330 [Source:H |
| Y_RNA | 11 | Y RNA [Source:RFAM;Acc:RF00019] |
| AKIP1 | 11 | A-kinase interacting protein 1 [Source:HGNC S |
| C11orf16 | 11 | chromosome 11 open reading frame 16 [Sour |
| ASCL3 | 11 | achaete-scute family bHLH transcription facto |
| Y_RNA | 11 | Y RNA [Source:RFAM;Acc:RF00019] |
| TMEM9B | 11 | TMEM9 domain family member B [Source:HG |
| TMEM9B-AS1 | 11 | TMEM9B antisense RNA 1 [Source:HGNC Sym |
| NRIP3 | 11 | nuclear receptor interacting protein 3 [Source |
| AC079296.1 | 11 | novel transcript |
| SCUBE2 | 11 | signal peptide, CUB domain and EGF like dom; |
| AC079296.2 | 11 | novel transcript, antisense to SCUBE2 |
| MIR5691 | 11 | microRNA 5691 [Source:HGNC Symbol;Acc:HG |
| KRT8P41 | 11 | keratin 8 pseudogene 41 [Source:HGNC Symb |
| DENND5A | 11 | DENN domain containing 5A [Source:HGNC Sy |
| AP006259.1 | 11 | novel transcript |
| Metazoa_SRP | 11 | Metazoan signal recognition particle RNA [Sou |
| TMEM41B | 11 | transmembrane protein 41B [Source:HGNC Sy |
| PRR13P2 | 11 | proline rich 13 pseudogene 2 [Source:HGNC S |
| IPO7 | 11 | importin 7 [Source:HGNC Symbol;Acc:HGNC:9 |
| SNORA23 | 11 | small nucleolar RNA, H/ACA box 23 [Source:H |
| AC132192.1 | 11 | novel transcript |
| AC132192.2 | 11 | novel transcript, antisense to ZNF143 |
| ZNF143 | 11 | zinc finger protein 143 [Source:HGNC Symbol; |
| WEE1 | 11 | WEE1 G2 checkpoint kinase [Source:HGNC Sy |

| | | |
|---|---|---|
| AC122179.1 | 11 | NULL |
| RPL23AP65 | 11 | ribosomal protein L23a pseudogene 65 [Sourc |
| AC011979.2 | 11 | chondroitin sulfate synthase 1 (CHSY1) pseud |
| AC011979.1 | 11 | ribosomal protein L21 (RPL21) pseudogene |
| SWAP70 | 11 | switching B cell complex subunit SWAP70 [Sou |
| RN7SKP50 | 11 | RN7SK pseudogene 50 [Source:HGNC Symbol; |
| LINC02709 | 11 | long intergenic non-protein coding RNA 2709 |
| SBF2-AS1 | 11 | SBF2 antisense RNA 1 [Source:HGNC Symbol;A |
| SBF2 | 11 | SET binding factor 2 [Source:HGNC Symbol;Ac |
| AC011092.2 | 11 | novel transcript |
| RNU7-28P | 11 | RNA, U7 small nuclear 28 pseudogene [Source |
| AC011092.1 | 11 | glutamate-ammonia ligase (GLUL) pseudogen |
| AC011092.3 | 11 | novel transcript, antisense to SBF2 |
| AC100763.1 | 11 | novel transcript |
| AC080023.2 | 11 | guanine nucleotide binding protein (G protein |
| AC080023.1 | 11 | novel transcript |
| ADM | 11 | adrenomedullin [Source:HGNC Symbol;Acc:H( |
| AMPD3 | 11 | adenosine monophosphate deaminase 3 [Sou |
| RNU6ATAC33P | 11 | RNA, U6atac small nuclear 33, pseudogene [S |
| MTRNR2L8 | 11 | MT-RNR2 like 8 [Source:HGNC Symbol;Acc:HG |
| MIR4485 | 11 | microRNA 4485 [Source:HGNC Symbol;Acc:H( |
| RNF141 | 11 | ring finger protein 141 [Source:HGNC Symbol; |
| MRVI1-AS1 | 11 | MRVI1 antisense RNA 1 [Source:HGNC Symbo |
| LYVE1 | 11 | lymphatic vessel endothelial hyaluronan recep |
| MRVI1 | 11 | murine retrovirus integration site 1 homolog [ |
| Y_RNA | 11 | Y RNA [Source:RFAM;Acc:RF00019] |
| CTR9 | 11 | CTR9 homolog, Paf1/RNA polymerase II comp |
| EIF4G2 | 11 | eukaryotic translation initiation factor 4 gamn |
| SNORD97 | 11 | small nucleolar RNA, C/D box 97 [Source:HGN |
| AC116535.1 | 11 | novel transcript, antisense to ZBED5 |
| ZBED5 | 11 | zinc finger BED-type containing 5 [Source:HGN |
| ZBED5-AS1 | 11 | ZBED5 antisense RNA 1 [Source:HGNC Symbol |
| AC069360.1 | 11 | novel transcript |
| LINC02752 | 11 | long intergenic non-protein coding RNA 2752 |
| AC111188.1 | 11 | activating transcription factor 7 interacting pr |
| AC111188.2 | 11 | high mobility group nucleosome binding doma |
| MTND5P21 | 11 | MT-ND5 pseudogene 21 [Source:HGNC Symbo |
| KC877392.1 | 11 | novel transcript |
| GALNT18 | 11 | polypeptide N-acetylgalactosaminyltransferas |
| CSNK2A3 | 11 | casein kinase 2 alpha 3 [Source:HGNC Symbol |
| AC023946.1 | 11 | novel transcript |
| AC104031.1 | 11 | novel transcript |
| MIR4299 | 11 | microRNA 4299 [Source:HGNC Symbol;Acc:H( |
| AC131935.2 | 11 | TEC |
| AC131935.1 | 11 | ubiquitin-conjugating enzyme E2D 2 (UBC4/5 |
| AC104383.2 | 11 | novel transcript |
| MIR8070 | 11 | microRNA 8070 [Source:HGNC Symbol;Acc:H( |
| USP47 | 11 | ubiquitin specific peptidase 47 [Source:HGNC |
| AC104383.1 | 11 | H3 histone, family 3A (H3F3A) pseudogene |
| DKK3 | 11 | dickkopf WNT signaling pathway inhibitor 3 [S |

| | | |
|---|---|---|
| LINC02547 | 11 | long intergenic non-protein coding RNA 2547 |
| AC124276.1 | 11 | novel transcript |
| AC124276.2 | 11 | novel transcript |
| MICAL2 | 11 | microtubule associated monooxygenase, calp |
| MIR6124 | 11 | microRNA 6124 [Source:HGNC Symbol;Acc:HG |
| AC079329.1 | 11 | novel transcript |
| AC025300.1 | 11 | novel transcript |
| PARVA | 11 | parvin alpha [Source:HGNC Symbol;Acc:HGNC |
| AC009806.1 | 11 | novel transcript |
| AC107881.1 | 11 | novel transcript |
| TEAD1 | 11 | TEA domain transcription factor 1 [Source:HG |
| AC013549.3 | 11 | novel transcript |
| AC013549.4 | 11 | novel transcript |
| AC013549.2 | 11 | NULL |
| AC013549.1 | 11 | novel transcript |
| LINC00958 | 11 | long intergenic non-protein coding RNA 958 [S |
| RASSF10-DT | 11 | RASSF10 divergent transcript [Source:HGNC Sy |
| RASSF10 | 11 | Ras association domain family member 10 [So |
| AC013762.1 | 11 | novel transcript |
| AC022878.1 | 11 | aldo-keto reductase family 1, member A1 (ald |
| ARNTL | 11 | aryl hydrocarbon receptor nuclear translocato |
| RN7SKP151 | 11 | RN7SK pseudogene 151 [Source:HGNC Symbo |
| BTBD10 | 11 | BTB domain containing 10 [Source:HGNC Sym |
| AC021269.3 | 11 | novel transcript |
| AC021269.1 | 11 | tripartite motif-containing 4 (TRIM4) pseudog |
| AC021269.2 | 11 | peptidase (mitochondrial processing) alpha (P |
| PTH | 11 | parathyroid hormone [Source:HGNC Symbol;A |
| HMGN2P36 | 11 | high mobility group nucleosomal binding dom |
| FAR1 | 11 | fatty acyl-CoA reductase 1 [Source:HGNC Sym |
| FAR1-IT1 | 11 | FAR1 intronic transcript 1 [Source:HGNC Symb |
| RPL39P26 | 11 | ribosomal protein L39 pseudogene 26 [Source |
| Y_RNA | 11 | Y RNA [Source:RFAM;Acc:RF00019] |
| CENPUP1 | 11 | centromere protein U pseudogene 1 [Source:I |
| AC013828.1 | 11 | coiled-coil domain containing 111 (CCDC111) |
| LINC02548 | 11 | long intergenic non-protein coding RNA 2548 |
| AC027779.1 | 11 | novel transcript |
| LINC02545 | 11 | long intergenic non-protein coding RNA 2545 |
| RNA5SP331 | 11 | RNA, 5S ribosomal pseudogene 331 [Source:H |
| LINC02683 | 11 | long intergenic non-protein coding RNA 2683 |
| SPON1 | 11 | spondin 1 [Source:HGNC Symbol;Acc:HGNC:1 |
| RNA5SP332 | 11 | RNA, 5S ribosomal pseudogene 332 [Source:H |
| SPON1-AS1 | 11 | SPON1 antisense RNA 1 [Source:HGNC Symbo |
| RRAS2 | 11 | RAS related 2 [Source:HGNC Symbol;Acc:HGN |
| AC018523.1 | 11 | ribosomal protein L13a pseudogene |
| COPB1 | 11 | coatomer protein complex subunit beta 1 [So |
| RNU7-49P | 11 | RNA, U7 small nuclear 49 pseudogene [Source |
| AC018523.2 | 11 | novel protein |
| PSMA1 | 11 | proteasome subunit alpha 1 [Source:HGNC Sy |
| PDE3B | 11 | phosphodiesterase 3B [Source:HGNC Symbol; |
| MORF4L1P3 | 11 | mortality factor 4 like 1 pseudogene 3 [Source |

| | | |
|---|---|---|
| CYP2R1 | 11 | cytochrome P450 family 2 subfamily R membe |
| CALCB | 11 | calcitonin related polypeptide beta [Source:H |
| CALCP | 11 | calcitonin pseudogene [Source:HGNC Symbol; |
| CALCA | 11 | calcitonin related polypeptide alpha [Source:H |
| OR7E41P | 11 | olfactory receptor family 7 subfamily E memb |
| INSC | 11 | INSC spindle orientation adaptor protein [Sou |
| AC104361.1 | 11 | NULL |
| LINC02751 | 11 | long intergenic non-protein coding RNA 2751 |
| AC073172.1 | 11 | novel transcript |
| AC087379.2 | 11 | uncharacterized LOC102724957 [Source:NCBI |
| AC087379.1 | 11 | novel transcript |
| AC009869.1 | 11 | novel transcript |
| LINC02682 | 11 | long intergenic non-protein coding RNA 2682 |
| SOX6 | 11 | SRY-box transcription factor 6 [Source:HGNC S |
| MIR6073 | 11 | microRNA 6073 [Source:HGNC Symbol;Acc:HG |
| AC103794.1 | 11 | novel transcript |
| AKR1B1P3 | 11 | aldo-keto reductase family 1 member B1 pseu |
| RN7SL188P | 11 | RNA, 7SL, cytoplasmic 188, pseudogene [Sour |
| C11orf58 | 11 | chromosome 11 open reading frame 58 [Sour |
| PLEKHA7 | 11 | pleckstrin homology domain containing A7 [So |
| RN7SKP90 | 11 | RN7SK pseudogene 90 [Source:HGNC Symbol; |
| AC116533.1 | 11 | ribosomal protein L36a (RPL36A) pseudogene |
| OR7E14P | 11 | olfactory receptor family 7 subfamily E memb |
| RPS13 | 11 | ribosomal protein S13 [Source:HGNC Symbol;. |
| SNORD14A | 11 | small nucleolar RNA, C/D box 14A [Source:HG |
| SNORD14B | 11 | small nucleolar RNA, C/D box 14B [Source:HG |
| PIK3C2A | 11 | phosphatidylinositol-4-phosphate 3-kinase ca |
| RNU6-593P | 11 | RNA, U6 small nuclear 593, pseudogene [Sour |
| AC126389.1 | 11 | ribosomal protein L34 (RPL34) pseudogene |
| AC107956.2 | 11 | ribosomal protein L29 (RPL29) pseudogene |
| NUCB2 | 11 | nucleobindin 2 [Source:HGNC Symbol;Acc:HG |
| AC107956.1 | 11 | ribosomal protein S2 (RPS2) pseudogene |
| AC124798.2 | 11 | novel transcript, antisense to NUCB2 |
| NCR3LG1 | 11 | natural killer cell cytotoxicity receptor 3 ligan |
| AC124798.1 | 11 | antisense to KCNJ11 and overlapping to a nov |
| KCNJ11 | 11 | potassium voltage-gated channel subfamily J |
| ABCC8 | 11 | ATP binding cassette subfamily C member 8 [S |
| SDHCP4 | 11 | succinate dehydrogenase complex subunit C p |
| AC124798.3 | 11 | novel transcript |
| USH1C | 11 | USH1 protein network component harmonin |
| OTOG | 11 | otogelin [Source:HGNC Symbol;Acc:HGNC:851 |
| AC124301.1 | 11 | novel transcript |
| LINC02729 | 11 | long intergenic non-protein coding RNA 2729 |
| MYOD1 | 11 | myogenic differentiation 1 [Source:HGNC Sym |
| KCNC1 | 11 | potassium voltage-gated channel subfamily C |
| SERGEF | 11 | secretion regulating guanine nucleotide excha |
| AC055860.1 | 11 | novel transcript |
| TPH1 | 11 | tryptophan hydroxylase 1 [Source:HGNC Symb |
| SAAL1 | 11 | serum amyloid A like 1 [Source:HGNC Symbol; |
| HIGD1AP5 | 11 | HIG1 hypoxia inducible domain family membe |

| | | |
|---|---|---|
| SAA3P | 11 | serum amyloid A3, pseudogene [Source:HGN( |
| MRGPRX3 | 11 | MAS related GPR family member X3 [Source:H |
| AC090099.1 | 11 | novel transcript, antisense to MRGPRX4 |
| AC090099.2 | 11 | novel transcript |
| MRGPRX12P | 11 | MAS related GPR family member X12, pseudo |
| MRGPRX4 | 11 | MAS related GPR family member X4 [Source:H |
| GLTPP1 | 11 | glycolipid transfer protein pseudogene 1 [Sou |
| MRGPRX13P | 11 | MAS related GPR family member X13, pseudo |
| SLC25A51P4 | 11 | SLC25A51 pseudogene 4 [Source:HGNC Symb |
| SAA4 | 11 | serum amyloid A4, constitutive [Source:HGNC |
| SAA2-SAA4 | 11 | SAA2-SAA4 readthrough [Source:HGNC Symb |
| SAA2 | 11 | serum amyloid A2 [Source:HGNC Symbol;Acc: |
| RNA5SP333 | 11 | RNA, 5S ribosomal pseudogene 333 [Source:H |
| ST13P5 | 11 | ST13, Hsp70 interacting protein pseudogene 5 |
| SAA1 | 11 | serum amyloid A1 [Source:HGNC Symbol;Acc: |
| RNA5SP334 | 11 | RNA, 5S ribosomal pseudogene 334 [Source:H |
| HPS5 | 11 | HPS5 biogenesis of lysosomal organelles comp |
| GTF2H1 | 11 | general transcription factor IIH subunit 1 [Sou |
| MIR3159 | 11 | microRNA 3159 [Source:HGNC Symbol;Acc:HG |
| LDHA | 11 | lactate dehydrogenase A [Source:HGNC Symb |
| AC084117.1 | 11 | novel transcript |
| LDHC | 11 | lactate dehydrogenase C [Source:HGNC Symb |
| AC027544.3 | 11 | WW domain binding protein 2 (WBP2) pseudo |
| LDHAL6A | 11 | lactate dehydrogenase A like 6A [Source:HGN |
| TSG101 | 11 | tumor susceptibility 101 [Source:HGNC Symb |
| YWHABP2 | 11 | tyrosine 3-monooxygenase/tryptophan 5-mor |
| AC027544.2 | 11 | novel transcript |
| AC027544.1 | 11 | glycine cleavage system protein H (aminomet |
| UEVLD | 11 | UEV and lactate/malate dehyrogenase domai |
| Metazoa_SRP | 11 | Metazoan signal recognition particle RNA [Sou |
| SPTY2D1OS | 11 | SPTY2D1 opposite strand [Source:HGNC Symb |
| AC112694.1 | 11 | mitochondrial carrier homolog 1 (C. elegans) ( |
| AC112694.2 | 11 | novel transcript |
| SPTY2D1 | 11 | SPT2 chromatin protein domain containing 1 [ |
| AP006296.1 | 11 | serine/arginine-rich splicing factor 3 (SRSF3) p |
| TMEM86A | 11 | transmembrane protein 86A [Source:HGNC Sy |
| IGSF22 | 11 | immunoglobulin superfamily member 22 [Sou |
| AC103974.1 | 11 | novel transcript |
| PTPN5 | 11 | protein tyrosine phosphatase non-receptor ty |
| AC103974.2 | 11 | novel transcript, antisense to PTPN5 |
| MRGPRX8P | 11 | MAS related GPR family member X8, pseudog |
| MRGPRX7P | 11 | MAS related GPR family member X7, pseudog |
| AC023078.2 | 11 | methylenetetrahydrofolate dehydrogenase (N |
| MRGPRX6P | 11 | MAS related GPR family member X6, pseudog |
| MRGPRX5P | 11 | MAS related GPR family member X5, pseudog |
| MRGPRX1 | 11 | MAS related GPR family member X1 [Source:H |
| AC023078.1 | 11 | novel transcript |
| MRGPRX11P | 11 | MAS related GPR family member X11, pseudo |
| MRGPRX10P | 11 | MAS related GPR family member X10, pseudo |
| MRGPRX9P | 11 | MAS related GPR family member X9, pseudog |

| | | |
|---|---|---|
| MRGPRX2 | 11 | MAS related GPR family member X2 [Source:H |
| ZDHHC13 | 11 | zinc finger DHHC-type containing 13 [Source:H |
| CSRP3 | 11 | cysteine and glycine rich protein 3 [Source:HG |
| CSRP3-AS1 | 11 | CSRP3 and E2F8 antisense RNA 1 [Source:HGN |
| E2F8 | 11 | E2F transcription factor 8 [Source:HGNC Symb |
| PCNAP4 | 11 | proliferating cell nuclear antigen pseudogene |
| AC009652.1 | 11 | novel transcript |
| NAV2 | 11 | neuron navigator 2 [Source:HGNC Symbol;Acc |
| NAV2-IT1 | 11 | NAV2 intronic transcript 1 [Source:HGNC Sym |
| RNA5SP335 | 11 | RNA, 5S ribosomal pseudogene 335 [Source:H |
| NAV2-AS5 | 11 | NAV2 antisense RNA 5 [Source:HGNC Symbol; |
| NAV2-AS4 | 11 | NAV2 antisense RNA 4 [Source:HGNC Symbol; |
| MIR4486 | 11 | microRNA 4486 [Source:HGNC Symbol;Acc:HG |
| AC023950.1 | 11 | NULL |
| AC009549.1 | 11 | novel transcript |
| MIR4694 | 11 | microRNA 4694 [Source:HGNC Symbol;Acc:HG |
| NAV2-AS3 | 11 | NAV2 antisense RNA 3 [Source:HGNC Symbol; |
| NAV2-AS2 | 11 | NAV2 antisense RNA 2 [Source:HGNC Symbol; |
| NAV2-AS1 | 11 | NAV2 antisense RNA 1 [Source:HGNC Symbol; |
| AC068860.1 | 11 | TEC |
| DBX1 | 11 | developing brain homeobox 1 [Source:HGNC S |
| HTATIP2 | 11 | HIV-1 Tat interactive protein 2 [Source:HGNC |
| PRMT3 | 11 | protein arginine methyltransferase 3 [Source: |
| HMGB1P40 | 11 | high mobility group box 1 pseudogene 40 [Sou |
| AC090707.2 | 11 | novel transcript |
| SLC6A5 | 11 | solute carrier family 6 member 5 [Source:HGN |
| NELL1 | 11 | neural EGFL like 1 [Source:HGNC Symbol;Acc:I |
| AC090707.1 | 11 | novel transcript, antisense to NELL1 |
| RNA5SP336 | 11 | RNA, 5S ribosomal pseudogene 336 [Source:H |
| AC090857.1 | 11 | novel pseudogene |
| AC090857.2 | 11 | novel transcript |
| AC099730.1 | 11 | isocitrate dehydrogenase 3 (NAD+) gamma (ID |
| RNA5SP337 | 11 | RNA, 5S ribosomal pseudogene 337 [Source:H |
| AC130307.1 | 11 | novel transcript |
| ANO5 | 11 | anoctamin 5 [Source:HGNC Symbol;Acc:HGNC |
| AC116534.1 | 11 | novel transcript |
| AC107882.1 | 11 | novel transcript |
| AC107886.1 | 11 | novel transcript |
| AC104009.1 | 11 | novel transcript |
| SLC17A6 | 11 | solute carrier family 17 member 6 [Source:HG |
| AC040936.1 | 11 | novel transcript |
| LINC01495 | 11 | long intergenic non-protein coding RNA 1495 |
| AC055878.1 | 11 | novel transcript |
| FANCF | 11 | FA complementation group F [Source:HGNC S |
| GAS2 | 11 | growth arrest specific 2 [Source:HGNC Symbo |
| RNA5SP338 | 11 | RNA, 5S ribosomal pseudogene 338 [Source:H |
| SVIP | 11 | small VCP interacting protein [Source:HGNC S |
| AC006299.1 | 11 | novel transcript, antisense to SVIP |
| CCDC179 | 11 | coiled-coil domain containing 179 [Source:HG |
| LINC02718 | 11 | long intergenic non-protein coding RNA 2718 |

| | | |
|---|---|---|
| AC100767.2 | 11 | programmed cell death 2-like, pseudogene |
| WIZP1 | 11 | WIZ pseudogene 1 [Source:HGNC Symbol;Acc |
| MIR8054 | 11 | microRNA 8054 [Source:HGNC Symbol;Acc:HG |
| AC100767.1 | 11 | retinitis pigmentosa GTPase regulator (RPGR) |
| THAP12P4 | 11 | THAP domain containing 12 pseudogene 4 [Sc |
| AC068472.1 | 11 | ribosomal protein S2 (RPS2) pseudogene |
| LINC02726 | 11 | long intergenic non-protein coding RNA 2726 |
| AC100768.1 | 11 | novel transcript |
| RNU6-783P | 11 | RNA, U6 small nuclear 783, pseudogene [Sour |
| AC099842.1 | 11 | protein-kinase, interferon-inducible double st |
| LINC02686 | 11 | long intergenic non-protein coding RNA 2686 |
| Y_RNA | 11 | Y RNA [Source:RFAM;Acc:RF00019] |
| LUZP2 | 11 | leucine zipper protein 2 [Source:HGNC Symbo |
| AC115990.1 | 11 | high-mobility group nucleosome binding dom |
| AC087373.1 | 11 | N-acetylneuraminic acid phosphatase (NANP) |
| RPL36AP40 | 11 | ribosomal protein L36a pseudogene 40 [Sourc |
| AC100770.1 | 11 | ATPase, H+ transporting, lysosomal 13kDa, V1 |
| LINC02699 | 11 | long intergenic non-protein coding RNA 2699 |
| AC013799.1 | 11 | novel transcript |
| AC024341.1 | 11 | NEDD8 activating enzyme E1 subunit 1 (NAE1) |
| ANO3 | 11 | anoctamin 3 [Source:HGNC Symbol;Acc:HGNC |
| AC021698.1 | 11 | novel transcript |
| ANO3-AS1 | 11 | ANO3 antisense RNA 1 [Source:HGNC Symbol; |
| AC079064.1 | 11 | sex comb on midleg homolog 1 (Drosophila) (S |
| MUC15 | 11 | mucin 15, cell surface associated [Source:HGN |
| SLC5A12 | 11 | solute carrier family 5 member 12 [Source:HG |
| FIBIN | 11 | fin bud initiation factor homolog [Source:HGN |
| BBOX1 | 11 | gamma-butyrobetaine hydroxylase 1 [Source: |
| BBOX1-AS1 | 11 | BBOX1 antisense RNA 1 [Source:HGNC Symbo |
| CCDC34 | 11 | coiled-coil domain containing 34 [Source:HGN |
| LGR4 | 11 | leucine rich repeat containing G protein-coupl |
| AC100771.2 | 11 | novel transcript |
| AC100771.1 | 11 | ribosomal protein L37a (RPL37A) pseudogene |
| LIN7C | 11 | lin-7 homolog C, crumbs cell polarity complex |
| BDNF-AS | 11 | BDNF antisense RNA [Source:HGNC Symbol;Ac |
| MIR8087 | 11 | microRNA 8087 [Source:HGNC Symbol;Acc:HG |
| RNA5SP339 | 11 | RNA, 5S ribosomal pseudogene 339 [Source:H |
| AC104563.1 | 11 | novel pseudogene |
| LINC00678 | 11 | long intergenic non-protein coding RNA 678 [S |
| BDNF | 11 | brain derived neurotrophic factor [Source:HGI |
| AC103796.1 | 11 | novel transcript, sense overlapping BDNF |
| CBX3P1 | 11 | chromobox 3 pseudogene 1 [Source:HGNC Syr |
| HSP90AA2P | 11 | heat shock protein 90 alpha family class A me |
| AC090159.1 | 11 | novel transcript |
| KIF18A | 11 | kinesin family member 18A [Source:HGNC Syr |
| MIR610 | 11 | microRNA 610 [Source:HGNC Symbol;Acc:HGI |
| METTL15 | 11 | methyltransferase like 15 [Source:HGNC Symb |
| AC023206.1 | 11 | ATP synthase, H+ transporting, mitochondrial |
| RN7SKP158 | 11 | RN7SK pseudogene 158 [Source:HGNC Symbo |
| AC104978.1 | 11 | ribosomal protein S15a (RPS15A) pseudogene |

| | | |
|---|---|---|
| MIR8068 | 11 | microRNA 8068 [Source:HGNC Symbol;Acc:HG |
| AC013714.1 | 11 | novel transcript |
| LINC02758 | 11 | long intergenic non-protein coding RNA 2758 |
| LINC02742 | 11 | long intergenic non-protein coding RNA 2742 |
| OR2BH1P | 11 | olfactory receptor family 2 subfamily BH mem |
| AC090791.1 | 11 | novel transcript |
| AC110056.1 | 11 | eukaryotic translation elongation factor 1 alph |
| LINC02755 | 11 | long intergenic non-protein coding RNA 2755 |
| AC090124.2 | 11 | serine/threonine kinase family pseudogene |
| AC090124.1 | 11 | novel transcript |
| AC110058.1 | 11 | novel transcript |
| LINC02546 | 11 | long intergenic non-protein coding RNA 2546 |
| HNRNPRP2 | 11 | heterogeneous nuclear ribonucleoprotein R p |
| AC107973.1 | 11 | novel transcript |
| RN7SL240P | 11 | RNA, 7SL, cytoplasmic 240, pseudogene [Sour |
| RPL7AP58 | 11 | ribosomal protein L7a pseudogene 58 [Source |
| LINC01616 | 11 | long intergenic non-protein coding RNA 1616 |
| KCNA4 | 11 | potassium voltage-gated channel subfamily A |
| AL358944.1 | 11 | novel transcript |
| AL358944.2 | 11 | follicle stimulating hormone, beta polypeptide |
| FSHB | 11 | follicle stimulating hormone subunit beta [Sou |
| ARL14EP | 11 | ADP ribosylation factor like GTPase 14 effecto |
| RPL12P30 | 11 | ribosomal protein L12 pseudogene 30 [Source |
| MPPED2 | 11 | metallophosphoesterase domain containing 2 |
| AL353699.1 | 11 | novel transcript |
| MPPED2-AS1 | 11 | MPPED2 antisense RNA 1 [Source:HGNC Symb |
| AL122014.1 | 11 | novel transcript, antisense to DCDC1 |
| DCDC1 | 11 | doublecortin domain containing 1 [Source:HG |
| CYCSP25 | 11 | CYCS pseudogene 25 [Source:HGNC Symbol;A |
| AL137804.1 | 11 | novel transcript |
| DNAJC24 | 11 | DnaJ heat shock protein family (Hsp40) memb |
| IMMP1L | 11 | inner mitochondrial membrane peptidase sub |
| ELP4 | 11 | elongator acetyltransferase complex subunit 4 |
| AC131571.1 | 11 | novel transcript, antisense to ELP4 |
| PAX6 | 11 | paired box 6 [Source:HGNC Symbol;Acc:HGNC |
| PAUPAR | 11 | PAX6 upstream antisense RNA [Source:HGNC |
| AL035078.4 | 11 | novel protein |
| AL035078.1 | 11 | novel transcript |
| AL035078.3 | 11 | novel transcript |
| AL035078.2 | 11 | novel transcript |
| U3 | 11 | Small nucleolar RNA U3 [Source:RFAM;Acc:RF |
| EIF4A2P5 | 11 | eukaryotic translation initiation factor 4A2 pse |
| RCN1 | 11 | reticulocalbin 1 [Source:HGNC Symbol;Acc:HG |
| AL078612.1 | 11 | novel transcript |
| THEM7P | 11 | thioesterase superfamily member 7, pseudoge |
| AL078612.2 | 11 | novel transcript |
| WT1 | 11 | WT1 transcription factor [Source:HGNC Symbo |
| WT1-AS | 11 | WT1 antisense RNA [Source:HGNC Symbol;Acc |
| AL049692.6 | 11 | NULL |
| AL049692.1 | 11 | NULL |

| | | |
|---|---|---|
| AL049692.5 | 11 | NULL |
| AL049692.3 | 11 | NULL |
| AL049692.2 | 11 | NULL |
| AL049692.4 | 11 | NULL |
| EIF3M | 11 | eukaryotic translation initiation factor 3 subur |
| HNRNPA3P9 | 11 | heterogeneous nuclear ribonucleoprotein A3 |
| CCDC73 | 11 | coiled-coil domain containing 73 [Source:HGN |
| AL049714.1 | 11 | ribosomal protein L34 (RPL34) pseudogene |
| PRRG4 | 11 | proline rich and Gla domain 4 [Source:HGNC S |
| QSER1 | 11 | glutamine and serine rich 1 [Source:HGNC Syn |
| Y_RNA | 11 | Y RNA [Source:RFAM;Acc:RF00019] |
| DEPDC7 | 11 | DEP domain containing 7 [Source:HGNC Symb |
| TCP11L1 | 11 | t-complex 11 like 1 [Source:HGNC Symbol;Acc |
| PIGCP1 | 11 | phosphatidylinositol glycan anchor biosynthes |
| LINC00294 | 11 | long intergenic non-protein coding RNA 294 [S |
| CSTF3 | 11 | cleavage stimulation factor subunit 3 [Source: |
| CSTF3-DT | 11 | CSTF3 divergent transcript [Source:HGNC Sym |
| Y_RNA | 11 | Y RNA [Source:RFAM;Acc:RF00019] |
| RPL29P23 | 11 | ribosomal protein L29 pseudogene 23 [Source |
| Y_RNA | 11 | Y RNA [Source:RFAM;Acc:RF00019] |
| AL136126.1 | 11 | ribosomal protein S24 (RPS24) pseudogene |
| HIPK3 | 11 | homeodomain interacting protein kinase 3 [Sc |
| KIAA1549L | 11 | KIAA1549 like [Source:HGNC Symbol;Acc:HGN |
| AL137161.1 | 11 | novel pseudogene |
| AL133399.1 | 11 | S100 calcium binding protein A11 (S100A11) p |
| AL049629.1 | 11 | novel transcript |
| AL049629.2 | 11 | novel protein |
| C11orf91 | 11 | chromosome 11 open reading frame 91 [Sour |
| CD59 | 11 | CD59 molecule (CD59 blood group) [Source:H |
| FBXO3 | 11 | F-box protein 3 [Source:HGNC Symbol;Acc:HG |
| FBXO3-DT | 11 | FBXO3 divergent transcript [Source:HGNC Syn |
| AC113192.3 | 11 | novel transcript |
| AC113192.2 | 11 | Spi transcription factor (Spi-1/PU.1 related) ps |
| LINC02722 | 11 | long intergenic non-protein coding RNA 2722 |
| AC113192.1 | 11 | novel transcript |
| LINC02721 | 11 | long intergenic non-protein coding RNA 2721 |
| LMO2 | 11 | LIM domain only 2 [Source:HGNC Symbol;Acc |
| AC132216.1 | 11 | novel transcript, antisense to LMO2 |
| AC090469.1 | 11 | novel transcript, antisense to CAPRIN1 |
| CAPRIN1 | 11 | cell cycle associated protein 1 [Source:HGNC S |
| NAT10 | 11 | N-acetyltransferase 10 [Source:HGNC Symbol; |
| ABTB2 | 11 | ankyrin repeat and BTB domain containing 2 [ |
| Y_RNA | 11 | Y RNA [Source:RFAM;Acc:RF00019] |
| AL139174.1 | 11 | methylmalonic aciduria (cobalamin deficiency |
| AL035079.1 | 11 | pseudogene similar to part of DnaJ (Hsp40) hc |
| CIR1P3 | 11 | corepressor interacting with RBPJ, 1 pseudoge |
| CAT | 11 | catalase [Source:HGNC Symbol;Acc:HGNC:151 |
| ELF5 | 11 | E74 like ETS transcription factor 5 [Source:HG |
| AL137224.1 | 11 | novel transcript |
| LINC02707 | 11 | long intergenic non-protein coding RNA 2707 |

| | | |
|---|---|---|
| EHF | 11 | ETS homologous factor [Source:HGNC Symbol |
| AC087783.1 | 11 | NADH dehydrogenase (ubiquinone) 1 beta sul |
| AL359999.1 | 11 | Rho GTPase activating protein 26, pseudogen |
| APIP | 11 | APAF1 interacting protein [Source:HGNC Syml |
| PDHX | 11 | pyruvate dehydrogenase complex component |
| MIR1343 | 11 | microRNA 1343 [Source:HGNC Symbol;Acc:HG |
| AL356215.1 | 11 | novel transcript |
| CD44 | 11 | CD44 molecule (Indian blood group) [Source:H |
| CD44-AS1 | 11 | CD44 antisense RNA 1 [Source:HGNC Symbol; |
| AL133330.1 | 11 | novel transcript |
| SLC1A2 | 11 | solute carrier family 1 member 2 [Source:HGN |
| AL133330.2 | 11 | novel transcript |
| AC090625.1 | 11 | novel pseudogene |
| AC090625.2 | 11 | novel transcript |
| PAMR1 | 11 | peptidase domain containing associated with |
| AL135934.1 | 11 | novel transcript |
| FJX1 | 11 | four-jointed box kinase 1 [Source:HGNC Symb |
| AL138812.1 | 11 | novel transcript |
| TRIM44 | 11 | tripartite motif containing 44 [Source:HGNC S |
| AC090692.2 | 11 | novel transcript |
| KRT18P14 | 11 | keratin 18 pseudogene 14 [Source:HGNC Sym |
| AC090692.3 | 11 | novel transcript |
| AC090692.1 | 11 | novel transcript |
| LDLRAD3 | 11 | low density lipoprotein receptor class A doma |
| AL136146.1 | 11 | WEE1 homolog (S. pombe) (WEE1) pseudoger |
| AL136146.2 | 11 | novel transcript |
| RPL12P31 | 11 | ribosomal protein L12 pseudogene 31 [Source |
| MIR3973 | 11 | microRNA 3973 [Source:HGNC Symbol;Acc:HG |
| AC129502.1 | 11 | TEC |
| COMMD9 | 11 | COMM domain containing 9 [Source:HGNC Sy |
| PRR5L | 11 | proline rich 5 like [Source:HGNC Symbol;Acc:H |
| AC087277.1 | 11 | novel transcript |
| AC087277.2 | 11 | novel transcript |
| AC009656.1 | 11 | novel transcript |
| TRAF6 | 11 | TNF receptor associated factor 6 [Source:HGN |
| AC061999.1 | 11 | TEC |
| RAG1 | 11 | recombination activating 1 [Source:HGNC Sym |
| RAG2 | 11 | recombination activating 2 [Source:HGNC Sym |
| C11orf74 | 11 | chromosome 11 open reading frame 74 [Sour |
| AC104042.1 | 11 | transforming, acidic coiled-coil containing pro |
| AC061997.1 | 11 | NULL |
| RPL7AP56 | 11 | ribosomal protein L7a pseudogene 56 [Source |
| LINC02760 | 11 | long intergenic non-protein coding RNA 2760 |
| AC103798.1 | 11 | TEC |
| LRRC6P1 | 11 | leucine rich repeat containing 6 pseudogene 1 |
| AC021713.1 | 11 | novel transcript |
| LINC02759 | 11 | long intergenic non-protein coding RNA 2759 |
| LINC01493 | 11 | long intergenic non-protein coding RNA 1493 |
| AC021723.2 | 11 | novel transcript |
| AC021723.1 | 11 | ribosomal protein L18 (RPL18) pseudogene |

| | | |
|---|---|---|
| RNU6-99P | 11 | RNA, U6 small nuclear 99, pseudogene [Sourc |
| AC021749.1 | 11 | adaptor-related protein complex 5, mu 1 subu |
| AC021820.1 | 11 | zinc finger, CCHC domain containing 9 (ZCCHC |
| AC080100.1 | 11 | TEC |
| LRRC4C | 11 | leucine rich repeat containing 4C [Source:HGN |
| Y_RNA | 11 | Y RNA [Source:RFAM;Acc:RF00019] |
| AC090720.1 | 11 | zinc finger protein pseudogene |
| RNU6-365P | 11 | RNA, U6 small nuclear 365, pseudogene [Sour |
| AC090138.1 | 11 | novel transcript |
| LINC02741 | 11 | long intergenic non-protein coding RNA 2741 |
| AC021006.1 | 11 | ribosomal protein L9 (RPL9) pseudogene |
| AC021006.2 | 11 | damage-specific DNA binding protein 2, 48kDa |
| LINC01499 | 11 | long intergenic non-protein coding RNA 1499 |
| AC023442.1 | 11 | zinc finger and BTB domain containing 5 (ZBTE |
| AC023442.3 | 11 | novel transcript |
| AC023442.2 | 11 | novel transcript |
| LINC02745 | 11 | long intergenic non-protein coding RNA 2745 |
| LINC02740 | 11 | long intergenic non-protein coding RNA 2740 |
| AC109810.1 | 11 | cytochrome c oxidase subunit VIc (COX6C) pse |
| AC009643.2 | 11 | MAN1B1 mannosidase, alpha, class 1B, memb |
| AC009643.1 | 11 | mannosidase, alpha, class 1B, member 1 (MAI |
| HNRNPKP3 | 11 | heterogeneous nuclear ribonucleoprotein K ps |
| API5 | 11 | apoptosis inhibitor 5 [Source:HGNC Symbol;Ac |
| AC087276.2 | 11 | novel transcript, antisense to API5 and TTC17 |
| Y_RNA | 11 | Y RNA [Source:RFAM;Acc:RF00019] |
| TTC17 | 11 | tetratricopeptide repeat domain 17 [Source:H |
| AC087276.3 | 11 | novel transcript |
| AC087276.1 | 11 | novel transcript, sense overlapping TTC17 |
| RN7SKP287 | 11 | RN7SK pseudogene 287 [Source:HGNC Symbo |
| PPIAP41 | 11 | peptidylprolyl isomerase A pseudogene 41 [Sc |
| CTBP2P6 | 11 | C-terminal binding protein 2 pseudogene 6 [S |
| AC068205.1 | 11 | novel transcript |
| MIR670 | 11 | microRNA 670 [Source:HGNC Symbol;Acc:HGN |
| MIR670HG | 11 | MIR670 host gene [Source:HGNC Symbol;Acc: |
| AC068205.2 | 11 | novel transcript |
| MIR129-2 | 11 | microRNA 129-2 [Source:HGNC Symbol;Acc:H |
| HSD17B12 | 11 | hydroxysteroid 17-beta dehydrogenase 12 [Sc |
| AC068205.3 | 11 | ribosomal protein L23a (RPL23A) pseudogene |
| RPL23AP63 | 11 | ribosomal protein L23a pseudogene 63 [Sourc |
| PHBP2 | 11 | prohibitin pseudogene 2 [Source:HGNC Symbo |
| AC068205.4 | 11 | prohibitin (PHB) pseudogene |
| AC087521.2 | 11 | novel transcript |
| AC087521.4 | 11 | novel transcript, antisense HSD17B12 |
| ALKBH3 | 11 | alkB homolog 3, alpha-ketoglutaratedepender |
| SEC14L1P1 | 11 | SEC14 like 1 pseudogene 1 [Source:HGNC Sym |
| ALKBH3-AS1 | 11 | ALKBH3 antisense RNA 1 [Source:HGNC Symb |
| AC087521.3 | 11 | novel transcript, sense overlapping C11orf96 |
| C11orf96 | 11 | chromosome 11 open reading frame 96 [Sour |
| AC087521.1 | 11 | novel transcript |
| ACCSL | 11 | 1-aminocyclopropane-1-carboxylate synthase |

| | | |
|---|---|---|
| ACCS | 11 | 1-aminocyclopropane-1-carboxylate synthase |
| AC134775.1 | 11 | novel transcript |
| EXT2 | 11 | exostosin glycosyltransferase 2 [Source:HGNC |
| ALX4 | 11 | ALX homeobox 4 [Source:HGNC Symbol;Acc:H |
| AC010768.4 | 11 | novel transcript |
| AC010768.3 | 11 | ribosomal protein L7a (RPL7A) pseudogene |
| CD82 | 11 | CD82 molecule [Source:HGNC Symbol;Acc:HG |
| AC010768.1 | 11 | novel transcript |
| AC010768.2 | 11 | novel transcript |
| RPL34P22 | 11 | ribosomal protein L34 pseudogene 22 [Source |
| AC104010.1 | 11 | LYR motif containing 2 (LYRM2) pseudogene |
| LINC02704 | 11 | long intergenic non-protein coding RNA 2704 |
| TSPAN18-AS1 | 11 | TSPAN18 antisense RNA 1 [Source:HGNC Syml |
| TSPAN18 | 11 | tetraspanin 18 [Source:HGNC Symbol;Acc:HGI |
| TP53I11 | 11 | tumor protein p53 inducible protein 11 [Sourc |
| LINC02685 | 11 | long intergenic non-protein coding RNA 2685 |
| AC068858.1 | 11 | novel transcript |
| PRDM11 | 11 | PR/SET domain 11 [Source:HGNC Symbol;Acc: |
| AC103681.2 | 11 | novel transcript |
| SYT13 | 11 | synaptotagmin 13 [Source:HGNC Symbol;Acc: |
| AC103681.1 | 11 | novel transcript |
| LINC02696 | 11 | long intergenic non-protein coding RNA 2696 |
| LINC02687 | 11 | long intergenic non-protein coding RNA 2687 |
| AC103855.3 | 11 | novel transcript |
| AC018716.2 | 11 | novel transcript |
| AC018716.1 | 11 | novel transcript |
| AC103855.2 | 11 | novel transcript |
| AC103855.4 | 11 | high-mobility group nucleosomal binding dom |
| AC103855.1 | 11 | novel transcript |
| CHST1 | 11 | carbohydrate sulfotransferase 1 [Source:HGN |
| AC087442.1 | 11 | novel transcript |
| MIR7154 | 11 | microRNA 7154 [Source:HGNC Symbol;Acc:HG |
| AC044839.1 | 11 | novel transcript |
| AC044839.2 | 11 | novel transcript |
| LINC02690 | 11 | long intergenic non-protein coding RNA 2690 |
| LINC02716 | 11 | long intergenic non-protein coding RNA 2716 |
| SLC35C1 | 11 | solute carrier family 35 member C1 [Source:H |
| AC044839.3 | 11 | novel transcript |
| CRY2 | 11 | cryptochrome circadian regulator 2 [Source:H |
| MAPK8IP1 | 11 | mitogen-activated protein kinase 8 interacting |
| AC068385.1 | 11 | novel transcript |
| C11orf94 | 11 | chromosome 11 open reading frame 94 [Sour |
| PEX16 | 11 | peroxisomal biogenesis factor 16 [Source:HGI |
| LARGE2 | 11 | LARGE xylosyl- and glucuronyltransferase 2 [S |
| PHF21A | 11 | PHD finger protein 21A [Source:HGNC Symbol |
| AC024475.1 | 11 | novel transcript |
| AC024475.3 | 11 | novel transcript |
| AC024475.2 | 11 | filamin binding LIM protein 1 (FBLIM1) pseudo |
| LINC02710 | 11 | long intergenic non-protein coding RNA 2710 |
| AC116021.1 | 11 | novel transcript |

| | | |
|---|---|---|
| LINC02489 | 11 | long intergenic non-protein coding RNA 2489 |
| CREB3L1 | 11 | cAMP responsive element binding protein 3 li |
| DGKZ | 11 | diacylglycerol kinase zeta [Source:HGNC Symb |
| MIR4688 | 11 | microRNA 4688 [Source:HGNC Symbol;Acc:HG |
| MDK | 11 | midkine [Source:HGNC Symbol;Acc:HGNC:697 |
| CHRM4 | 11 | cholinergic receptor muscarinic 4 [Source:HGI |
| AMBRA1 | 11 | autophagy and beclin 1 regulator 1 [Source:H( |
| AC024293.1 | 11 | ribosomal protein S10 (RPS10) pseudogene |
| MIR3160-1 | 11 | microRNA 3160-1 [Source:HGNC Symbol;Acc:I |
| MIR3160-2 | 11 | microRNA 3160-2 [Source:HGNC Symbol;Acc:I |
| AC127035.1 | 11 | novel transcript, antisense to AMBRA1 |
| HARBI1 | 11 | harbinger transposase derived 1 [Source:HGN |
| ATG13 | 11 | autophagy related 13 [Source:HGNC Symbol;A |
| ARHGAP1 | 11 | Rho GTPase activating protein 1 [Source:HGN( |
| ZNF408 | 11 | zinc finger protein 408 [Source:HGNC Symbol; |
| F2 | 11 | coagulation factor II, thrombin [Source:HGNC |
| CKAP5 | 11 | cytoskeleton associated protein 5 [Source:HG |
| MIR5582 | 11 | microRNA 5582 [Source:HGNC Symbol;Acc:HG |
| AC115088.1 | 11 | NULL |
| SNORD67 | 11 | small nucleolar RNA, C/D box 67 [Source:HGN |
| LRP4-AS1 | 11 | LRP4 antisense RNA 1 [Source:HGNC Symbol;A |
| LRP4 | 11 | LDL receptor related protein 4 [Source:HGNC |
| AC021573.1 | 11 | novel transcript |
| C11orf49 | 11 | chromosome 11 open reading frame 49 [Sour |
| AC103792.1 | 11 | arsA arsenite transporter, ATP-binding, homol |
| AC090589.2 | 11 | novel transcript |
| ARFGAP2 | 11 | ADP ribosylation factor GTPase activating prot |
| AC090589.3 | 11 | novel transcript |
| PACSIN3 | 11 | protein kinase C and casein kinase substrate ir |
| MIR6745 | 11 | microRNA 6745 [Source:HGNC Symbol;Acc:HG |
| AC090589.1 | 11 | ribosomal protein S20 (RPS20) pseudogene |
| DDB2 | 11 | damage specific DNA binding protein 2 [Sourc |
| AC018410.2 | 11 | novel transcript |
| ACP2 | 11 | acid phosphatase 2, lysosomal [Source:HGNC |
| NR1H3 | 11 | nuclear receptor subfamily 1 group H member |
| MADD | 11 | MAP kinase activating death domain [Source:I |
| AC018410.1 | 11 | novel transcript |
| AC090582.1 | 11 | mitochondrially encoded NADH dehydrogenas |
| MYBPC3 | 11 | myosin binding protein C, cardiac [Source:HGI |
| SPI1 | 11 | Spi-1 proto-oncogene [Source:HGNC Symbol;A |
| AC090559.1 | 11 | novel transcript |
| MIR4487 | 11 | microRNA 4487 [Source:HGNC Symbol;Acc:HG |
| SLC39A13 | 11 | solute carrier family 39 member 13 [Source:H |
| PSMC3 | 11 | proteasome 26S subunit, ATPase 3 [Source:HG |
| RAPSN | 11 | receptor associated protein of the synapse [Sc |
| CELF1 | 11 | CUGBP Elav-like family member 1 [Source:HGI |
| AC090559.2 | 11 | novel transcript |
| RN7SL652P | 11 | RNA, 7SL, cytoplasmic 652, pseudogene [Sour |
| NDUFS3 | 11 | NADH:ubiquinone oxidoreductase core subun |
| PTPMT1 | 11 | protein tyrosine phosphatase mitochondrial 1 |

| | | |
|---|---|---|
| KBTBD4 | 11 | kelch repeat and BTB domain containing 4 [So |
| RNU5E-10P | 11 | RNA, U5E small nuclear 10, pseudogene [Sour |
| KF459542.1 | 11 | novel transcript, antisense to KBTBD4 |
| FAM180B | 11 | family with sequence similarity 180 member E |
| C1QTNF4 | 11 | C1q and TNF related 4 [Source:HGNC Symbol; |
| MTCH2 | 11 | mitochondrial carrier 2 [Source:HGNC Symbol |
| AGBL2 | 11 | ATP/GTP binding protein like 2 [Source:HGNC |
| FNBP4 | 11 | formin binding protein 4 [Source:HGNC Symbo |
| Y_RNA | 11 | Y RNA [Source:RFAM;Acc:RF00019] |
| NUP160 | 11 | nucleoporin 160 [Source:HGNC Symbol;Acc:H |
| AC021443.1 | 11 | ribosomal protein L6 (RPL6) pseudogene |
| RNA5SP340 | 11 | RNA, 5S ribosomal pseudogene 340 [Source:H |
| Y_RNA | 11 | Y RNA [Source:RFAM;Acc:RF00019] |
| YPEL5P2 | 11 | YPEL5 pseudogene 2 [Source:HGNC Symbol;A |
| AC023232.1 | 11 | Sec61 beta subunit (SEC61B) pseudogene |
| PTPRJ | 11 | protein tyrosine phosphatase receptor type J |
| PTPRJ-AS1 | 11 | PTPRJ antisense RNA 1 [Source:HGNC Symbol; |
| MIR3161 | 11 | microRNA 3161 [Source:HGNC Symbol;Acc:HG |
| OR4B1 | 11 | olfactory receptor family 4 subfamily B memb |
| OR4B2P | 11 | olfactory receptor family 4 subfamily B memb |
| OR4X2 | 11 | olfactory receptor family 4 subfamily X memb |
| OR4X1 | 11 | olfactory receptor family 4 subfamily X memb |
| OR4S1 | 11 | olfactory receptor family 4 subfamily S memb |
| OR4C3 | 11 | olfactory receptor family 4 subfamily C memb |
| OR4C4P | 11 | olfactory receptor family 4 subfamily C memb |
| OR4C5 | 11 | olfactory receptor family 4 subfamily C memb |
| OR4C2P | 11 | olfactory receptor family 4 subfamily C memb |
| OR4C10P | 11 | olfactory receptor family 4 subfamily C memb |
| OR4C9P | 11 | olfactory receptor family 4 subfamily C memb |
| OR4R1P | 11 | olfactory receptor family 4 subfamily R memb |
| OR4A47 | 11 | olfactory receptor family 4 subfamily A memb |
| OR4A48P | 11 | olfactory receptor family 4 subfamily A memb |
| OR4A46P | 11 | olfactory receptor family 4 subfamily A memb |
| OR4A40P | 11 | olfactory receptor family 4 subfamily A memb |
| OR4A43P | 11 | olfactory receptor family 4 subfamily A memb |
| OR4A45P | 11 | olfactory receptor family 4 subfamily A memb |
| OR4A41P | 11 | olfactory receptor family 4 subfamily A memb |
| OR4A42P | 11 | olfactory receptor family 4 subfamily A memb |
| OR4A44P | 11 | olfactory receptor family 4 subfamily A memb |
| AC027369.6 | 11 | novel pseudogene |
| AC027369.3 | 11 | ankyrin repeat domain 33B (ANKRD33B) pseu |
| AC027369.4 | 11 | tripartite motif-containing pseudogene |
| AC027369.5 | 11 | tripartite motif-containing 43 (TRIM43) pseud |
| TRIM51CP | 11 | tripartite motif-containing 51C, pseudogene [ |
| AC027369.2 | 11 | tripartite motif-containing pseudogene |
| TRIM51GP | 11 | tripartite motif-containing 51G, pseudogene [ |
| TRIM53CP | 11 | tripartite motif containing 53C, pseudogene [ |
| AC027369.1 | 11 | tripartite motif-containing pseudogene |
| TRIM49B | 11 | tripartite motif containing 49B [Source:HGNC |
| AC084851.4 | 11 | ankyrin repeat domain 33B (ANKRD33B) pseu |

| | | |
|---|---|---|
| TRIM64C | 11 | tripartite motif containing 64C [Source:HGNC |
| AC084851.3 | 11 | tripartite motif containing 49C (TRIM49C) pse |
| AC084851.1 | 11 | establishment of cohesion 1 homolog 1 (S. cer |
| UBTFL7 | 11 | UBTF like 7 (pseudogene) [Source:HGNC Symb |
| AC084851.2 | 11 | tripartite motif-containing pseudogene |
| AC118273.1 | 11 | Cdon homolog (mouse) (CDON) pseudogene |
| AC118273.2 | 11 | tripartite motif-containing pseudogene |
| TRIM77BP | 11 | tripartite motif containing 77B, pseudogene [S |
| UBTFL9 | 11 | UBTF like 9 (pseudogene) [Source:HGNC Symb |
| FOLH1 | 11 | folate hydrolase 1 [Source:HGNC Symbol;Acc: |
| AC118942.1 | 11 | NADPH oxidase 4 pseudogene |
| TYRL | 11 | tyrosinase like (pseudogene) [Source:HGNC Sy |
| CBX3P8 | 11 | chromobox 3 pseudogene 8 [Source:HGNC Syr |
| AC135977.1 | 11 | solute carrier family 25, member 33 (SLC25A3 |
| AC136759.1 | 11 | glutamate receptor, metabotropic 5 (GRM5) p |
| AC130364.1 | 11 | novel pseudogene |
| AC130364.2 | 11 | tripartite motif-containing protein pseudogen |
| TRIM51FP | 11 | tripartite motif-containing 51F, pseudogene [S |
| AP006587.5 | 11 | tripartite motif-containing pseudogene |
| AP006587.6 | 11 | novel transcript |
| AP006587.4 | 11 | novel pseudogene |
| AP006587.3 | 11 | tripartite motif-containing pseudogene |
| TRIM51DP | 11 | tripartite motif-containing 51D, pseudogene [ |
| AP006587.1 | 11 | tripartite motif-containing pseudogene |
| AP006587.2 | 11 | Ankyrin repeat domain-containing protein 33 |
| OR4A1P | 11 | olfactory receptor family 4 subfamily A memb |
| OR4A49P | 11 | olfactory receptor family 4 subfamily A memb |
| OR4A18P | 11 | olfactory receptor family 4 subfamily A memb |
| OR4A19P | 11 | olfactory receptor family 4 subfamily A memb |
| OR4R3P | 11 | olfactory receptor family 4 subfamily R memb |
| OR4C13 | 11 | olfactory receptor family 4 subfamily C memb |
| OR4C12 | 11 | olfactory receptor family 4 subfamily C memb |
| OR4C48P | 11 | olfactory receptor family 4 subfamily C memb |
| OR4C45 | 11 | olfactory receptor family 4 subfamily C memb |
| OR4C49P | 11 | olfactory receptor family 4 subfamily C memb |
| GTF2IP11 | 11 | general transcription factor IIi pseudogene 11 |
| AC109635.4 | 11 | glutamyl aminopeptidase (aminopeptidase A) |
| AC109635.7 | 11 | novel transcript |
| AC109635.6 | 11 | novel transcript |
| AC109635.1 | 11 | Phosphorylase b kinase gamma catalytic chair |
| AC109635.5 | 11 | pantothenate kinase 3 (PANK3) pseudogene |
| AC109635.3 | 11 | septin 7 (SEPT7) pseudogene |
| AC109635.2 | 11 | novel pseudogene |
| LINC02750 | 11 | long intergenic non-protein coding RNA 2750 |
| AC024405.2 | 11 | General transcription factor II-I (GTF2I) pseud |
| AC024405.1 | 11 | DEAH (Asp-Glu-Ala-His) box polypeptide 9 (DH |
| OR4C50P | 11 | olfactory receptor family 4 subfamily C memb |
| OR4C46 | 11 | olfactory receptor family 4 subfamily C memb |
| OR4C7P | 11 | olfactory receptor family 4 subfamily C memb |
| OR4R2P | 11 | olfactory receptor family 4 subfamily R memb |

| | | |
|---|---|---|
| OR4A4P | 11 | olfactory receptor family 4 subfamily A memb |
| OR4A3P | 11 | olfactory receptor family 4 subfamily A memb |
| OR4A2P | 11 | olfactory receptor family 4 subfamily A memb |
| OR4A8 | 11 | olfactory receptor family 4 subfamily A memb |
| OR4A7P | 11 | olfactory receptor family 4 subfamily A memb |
| OR4A5 | 11 | olfactory receptor family 4 subfamily A memb |
| OR4A6P | 11 | olfactory receptor family 4 subfamily A memb |
| TRIM48 | 11 | tripartite motif containing 48 [Source:HGNC S |
| AP005597.4 | 11 | Tripartite motif-containing protein pseudoger |
| AP005597.3 | 11 | novel pseudogene |
| AP005597.2 | 11 | novel pseudogene |
| TRIM51HP | 11 | tripartite motif-containing 51H, pseudogene [ |
| AP005597.1 | 11 | tripartite motif-containing protein pseudogen |
| OR4A11P | 11 | olfactory receptor family 4 subfamily A memb |
| OR4A12P | 11 | olfactory receptor family 4 subfamily A memb |
| OR4A16 | 11 | olfactory receptor family 4 subfamily A memb |
| OR4A15 | 11 | olfactory receptor family 4 subfamily A memb |
| OR4A9P | 11 | olfactory receptor family 4 subfamily A memb |
| OR4X7P | 11 | olfactory receptor family 4 subfamily X memb |
| OR4A10P | 11 | olfactory receptor family 4 subfamily A memb |
| OR4A17P | 11 | olfactory receptor family 4 subfamily A memb |
| OR4A13P | 11 | olfactory receptor family 4 subfamily A memb |
| OR4A50P | 11 | olfactory receptor family 4 subfamily A memb |
| OR4A21P | 11 | olfactory receptor family 4 subfamily A memb |
| OR4C1P | 11 | olfactory receptor family 4 subfamily C memb |
| OR4C14P | 11 | olfactory receptor family 4 subfamily C memb |
| OR4C15 | 11 | olfactory receptor family 4 subfamily C memb |
| OR4C16 | 11 | olfactory receptor family 4 subfamily C memb |
| OR4C11 | 11 | olfactory receptor family 4 subfamily C memb |
| OR4P4 | 11 | olfactory receptor family 4 subfamily P memb |
| OR4S2 | 11 | olfactory receptor family 4 subfamily S memb |
| OR4C6 | 11 | olfactory receptor family 4 subfamily C memb |
| OR4V1P | 11 | olfactory receptor family 4 subfamily V memb |
| OR4P1P | 11 | olfactory receptor family 4 subfamily P memb |
| AP006437.1 | 11 | novel transcript |
| OR5D2P | 11 | olfactory receptor family 5 subfamily D memb |
| OR5D3P | 11 | olfactory receptor family 5 subfamily D memb |
| OR5D17P | 11 | olfactory receptor family 5 subfamily D memb |
| OR5D13 | 11 | olfactory receptor family 5 subfamily D memb |
| OR5D15P | 11 | olfactory receptor family 5 subfamily D memb |
| OR5D14 | 11 | olfactory receptor family 5 subfamily D memb |
| OR5L1 | 11 | olfactory receptor family 5 subfamily L memb |
| OR5D18 | 11 | olfactory receptor family 5 subfamily D memb |
| OR5L2 | 11 | olfactory receptor family 5 subfamily L memb |
| OR5D16 | 11 | olfactory receptor family 5 subfamily D memb |
| OR9M1P | 11 | olfactory receptor family 9 subfamily M meml |
| AC036111.2 | 11 | tripartite motif-containing pseudogene |
| AC036111.3 | 11 | Tripartite motif-containing pseudogene |
| AC036111.1 | 11 | Ankyrin repeat domain-containing pseudogen |
| TRIM51 | 11 | tripartite motif-containing 51 [Source:HGNC S |

| | | |
|---|---|---|
| OR5W1P | 11 | olfactory receptor family 5 subfamily W meml |
| OR5W2 | 11 | olfactory receptor family 5 subfamily W meml |
| OR5I1 | 11 | olfactory receptor family 5 subfamily I membe |
| OR10AF1P | 11 | olfactory receptor family 10 subfamily AF mer |
| OR10AK1P | 11 | olfactory receptor family 10 subfamily AK mer |
| OR10AG1 | 11 | olfactory receptor family 10 subfamily AG mer |
| OR7E5P | 11 | olfactory receptor family 7 subfamily E memb |
| OR5F1 | 11 | olfactory receptor family 5 subfamily F memb |
| OR5F2P | 11 | olfactory receptor family 5 subfamily F memb |
| OR5AS1 | 11 | olfactory receptor family 5 subfamily AS mem |
| OR5AQ1P | 11 | olfactory receptor family 5 subfamily AQ mem |
| OR5J1P | 11 | olfactory receptor family 5 subfamily J membe |
| OR5BE1P | 11 | olfactory receptor family 5 subfamily BE mem |
| OR8I2 | 11 | olfactory receptor family 8 subfamily I membe |
| OR8I4P | 11 | olfactory receptor family 8 subfamily I membe |
| OR8H2 | 11 | olfactory receptor family 8 subfamily H memb |
| OR5BN2P | 11 | olfactory receptor family 5 subfamily BN mem |
| OR8H3 | 11 | olfactory receptor family 8 subfamily H memb |
| OR5BN1P | 11 | olfactory receptor family 5 subfamily BN mem |
| OR8J3 | 11 | olfactory receptor family 8 subfamily J membe |
| OR8K4P | 11 | olfactory receptor family 8 subfamily K memb |
| OR8K5 | 11 | olfactory receptor family 8 subfamily K memb |
| OR5J7P | 11 | olfactory receptor family 5 subfamily J membe |
| OR5J2 | 11 | olfactory receptor family 5 subfamily J membe |
| OR8V1P | 11 | olfactory receptor family 8 subfamily V memb |
| AC022882.1 | 11 | N(alpha)-acetyltransferase 50, NatE catalytic s |
| OR8J2 | 11 | olfactory receptor family 8 subfamily J membe |
| OR5T2 | 11 | olfactory receptor family 5 subfamily T memb |
| OR5T3 | 11 | olfactory receptor family 5 subfamily T memb |
| OR5T1 | 11 | olfactory receptor family 5 subfamily T memb |
| OR8H1 | 11 | olfactory receptor family 8 subfamily H memb |
| OR8I1P | 11 | olfactory receptor family 8 subfamily I membe |
| OR8K3 | 11 | olfactory receptor family 8 subfamily K memb |
| FAM8A2P | 11 | family with sequence similarity 8 member A2, |
| OR8K2P | 11 | olfactory receptor family 8 subfamily K memb |
| OR8K1 | 11 | olfactory receptor family 8 subfamily K memb |
| OR8J1 | 11 | olfactory receptor family 8 subfamily J membe |
| RPL5P29 | 11 | ribosomal protein L5 pseudogene 29 [Source: |
| OR8U1 | 11 | olfactory receptor family 8 subfamily U memb |
| OR8L1P | 11 | olfactory receptor family 8 subfamily L membe |
| OR5AL2P | 11 | olfactory receptor family 5 subfamily AL mem |
| OR5AL1 | 11 | olfactory receptor family 5 subfamily AL mem |
| OR5R1 | 11 | olfactory receptor family 5 subfamily R memb |
| OR5M4P | 11 | olfactory receptor family 5 subfamily M meml |
| AP002512.3 | 11 | novel olfactory receptor |
| OR5M9 | 11 | olfactory receptor family 5 subfamily M meml |
| OR5M3 | 11 | olfactory receptor family 5 subfamily M meml |
| OR5M2P | 11 | olfactory receptor family 5 subfamily M meml |
| OR5M8 | 11 | olfactory receptor family 5 subfamily M meml |
| AP002512.1 | 11 | ubiquitin carboxyl-terminal esterase L1 (ubiqu |

| | | |
|---|---|---|
| OR5M7P | 11 | olfactory receptor family 5 subfamily M meml |
| AP002512.2 | 11 | ubiquitin carboxyl-terminal esterase L1 (ubiqu |
| OR5M6P | 11 | olfactory receptor family 5 subfamily M meml |
| OR5M5P | 11 | olfactory receptor family 5 subfamily M meml |
| OR5M11 | 11 | olfactory receptor family 5 subfamily M meml |
| OR5M10 | 11 | olfactory receptor family 5 subfamily M meml |
| OR5M13P | 11 | olfactory receptor family 5 subfamily M meml |
| OR5M1 | 11 | olfactory receptor family 5 subfamily M meml |
| OR5AM1P | 11 | olfactory receptor family 5 subfamily AM mer |
| OR5M12P | 11 | olfactory receptor family 5 subfamily M meml |
| OR5AP1P | 11 | olfactory receptor family 5 subfamily AP mem |
| OR5AP2 | 11 | olfactory receptor family 5 subfamily AP mem |
| OR5AR1 | 11 | olfactory receptor family 5 subfamily AR mem |
| OR2AH1P | 11 | olfactory receptor family 2 subfamily AH mer |
| AP004245.1 | 11 | chromosome 20 open reading frame 30 (C20c |
| OR9G1 | 11 | olfactory receptor family 9 subfamily G memb |
| AP001803.1 | 11 | GNAS complex locus (GNAS) pseudogene |
| OR9G3P | 11 | olfactory receptor family 9 subfamily G memb |
| OR9G4 | 11 | olfactory receptor family 9 subfamily G memb |
| MIR6128 | 11 | microRNA 6128 [Source:HGNC Symbol;Acc:HG |
| AP001803.3 | 11 | novel olfactory protein pseudogene |
| OR9G2P | 11 | olfactory receptor family 9 subfamily G memb |
| OR5G1P | 11 | olfactory receptor family 5 subfamily G memb |
| OR5G4P | 11 | olfactory receptor family 5 subfamily G memb |
| OR5G5P | 11 | olfactory receptor family 5 subfamily G memb |
| OR5G3 | 11 | olfactory receptor family 5 subfamily G memb |
| LINC02735 | 11 | long intergenic non-protein coding RNA 2735 |
| FADS2P1 | 11 | fatty acid desaturase 2 pseudogene 1 [Source |
| OR5AK3P | 11 | olfactory receptor family 5 subfamily AK mem |
| OR5AK2 | 11 | olfactory receptor family 5 subfamily AK mem |
| OR5AK1P | 11 | olfactory receptor family 5 subfamily AK mem |
| OR5BQ1P | 11 | olfactory receptor family 5 subfamily BQ mer |
| OR5AK4P | 11 | olfactory receptor family 5 subfamily AK mem |
| OR5AO1P | 11 | olfactory receptor family 5 subfamily AO mer |
| OR5BP1P | 11 | olfactory receptor family 5 subfamily BP mem |
| LRRC55 | 11 | leucine rich repeat containing 55 [Source:HGN |
| APLNR | 11 | apelin receptor [Source:HGNC Symbol;Acc:HG |
| TNKS1BP1 | 11 | tankyrase 1 binding protein 1 [Source:HGNC S |
| AP000781.1 | 11 | novel transcript |
| SSRP1 | 11 | structure specific recognition protein 1 [Sourc |
| P2RX3 | 11 | purinergic receptor P2X 3 [Source:HGNC Symb |
| PRG3 | 11 | proteoglycan 3, pro eosinophil major basic pro |
| PRG2 | 11 | proteoglycan 2, pro eosinophil major basic pro |
| AP000781.2 | 11 | novel protein |
| SLC43A3 | 11 | solute carrier family 43 member 3 [Source:HG |
| RNA5SP341 | 11 | RNA, 5S ribosomal pseudogene 341 [Source:H |
| RN7SKP259 | 11 | RN7SK pseudogene 259 [Source:HGNC Symbo |
| RTN4RL2 | 11 | reticulon 4 receptor like 2 [Source:HGNC Syml |
| AP002893.1 | 11 | novel transcript |
| SLC43A1 | 11 | solute carrier family 43 member 1 [Source:HG |

| | | |
|---|---|---|
| RN7SL605P | 11 | RNA, 7SL, cytoplasmic 605, pseudogene [Sour |
| TIMM10 | 11 | translocase of inner mitochondrial membrane |
| SMTNL1 | 11 | smoothelin like 1 [Source:HGNC Symbol;Acc:H |
| UBE2L6 | 11 | ubiquitin conjugating enzyme E2 L6 [Source:H |
| RPS4XP13 | 11 | ribosomal protein S4X pseudogene 13 [Source |
| SERPING1 | 11 | serpin family G member 1 [Source:HGNC Sym |
| AP000662.1 | 11 | novel transcript, sense overlapping CLP1 |
| MIR130A | 11 | microRNA 130a [Source:HGNC Symbol;Acc:HG |
| YPEL4 | 11 | yippee like 4 [Source:HGNC Symbol;Acc:HGNC |
| CLP1 | 11 | cleavage and polyadenylation factor I subunit |
| ZDHHC5 | 11 | zinc finger DHHC-type containing 5 [Source:HG |
| MED19 | 11 | mediator complex subunit 19 [Source:HGNC S |
| TMX2 | 11 | thioredoxin related transmembrane protein 2 |
| TMX2-CTNND1 | 11 | TMX2-CTNND1 readthrough (NMD candidate) |
| PPIAP42 | 11 | peptidylprolyl isomerase A pseudogene 42 [So |
| SELENOH | 11 | selenoprotein H [Source:HGNC Symbol;Acc:HG |
| AP001931.1 | 11 | novel protein, C11orf31-CTNND1 readthrough |
| BTBD18 | 11 | BTB domain containing 18 [Source:HGNC Sym |
| CTNND1 | 11 | catenin delta 1 [Source:HGNC Symbol;Acc:HG |
| OR5BA1P | 11 | olfactory receptor family 5 subfamily BA mem |
| AP003484.1 | 11 | NADH dehydrogenase (ubiquinone) 1 alpha su |
| OR5AZ1P | 11 | olfactory receptor family 5 subfamily AZ mem |
| OR5BD1P | 11 | olfactory receptor family 5 subfamily BD mem |
| CYCSP26 | 11 | CYCS pseudogene 26 [Source:HGNC Symbol;A |
| OR9Q1 | 11 | olfactory receptor family 9 subfamily Q memb |
| RNU6-899P | 11 | RNA, U6 small nuclear 899, pseudogene [Sour |
| OR6Q1 | 11 | olfactory receptor family 6 subfamily Q memb |
| VN2R9P | 11 | vomeronasal 2 receptor 9, pseudogene [Sourc |
| AP004247.2 | 11 | novel transcript |
| AP004247.1 | 11 | tRNA methyltransferase 11 homolog (S. cerev |
| OR9L1P | 11 | olfactory receptor family 9 subfamily L membe |
| OR9I3P | 11 | olfactory receptor family 9 subfamily I membe |
| OR9I1 | 11 | olfactory receptor family 9 subfamily I membe |
| OR9I2P | 11 | olfactory receptor family 9 subfamily I membe |
| OR5BL1P | 11 | olfactory receptor family 5 subfamily BL mem |
| OR9Q2 | 11 | olfactory receptor family 9 subfamily Q memb |
| OR1S2 | 11 | olfactory receptor family 1 subfamily S memb |
| OR1S1 | 11 | olfactory receptor family 1 subfamily S memb |
| OR10Q1 | 11 | olfactory receptor family 10 subfamily Q mem |
| EIF4A2P3 | 11 | eukaryotic translation initiation factor 4A2 pse |
| OR10W1 | 11 | olfactory receptor family 10 subfamily W men |
| OR10Q2P | 11 | olfactory receptor family 10 subfamily Q mem |
| OR5BC1P | 11 | olfactory receptor family 5 subfamily BC mem |
| OR5B19P | 11 | olfactory receptor family 5 subfamily B memb |
| OR5B10P | 11 | olfactory receptor family 5 subfamily B memb |
| AP000435.1 | 11 | dihydropyrimidinase-like 5 pseudogene |
| OR5B17 | 11 | olfactory receptor family 5 subfamily B memb |
| OR5B1P | 11 | olfactory receptor family 5 subfamily B memb |
| OR5B15P | 11 | olfactory receptor family 5 subfamily B memb |
| OR5B3 | 11 | olfactory receptor family 5 subfamily B memb |

| | | |
|---|---|---|
| OR5B2 | 11 | olfactory receptor family 5 subfamily B memb |
| OR5B12 | 11 | olfactory receptor family 5 subfamily B memb |
| AP003557.2 | 11 | novel transcript |
| AP003557.1 | 11 | novel transcript |
| OR5B21 | 11 | olfactory receptor family 5 subfamily B memb |
| LPXN | 11 | leupaxin [Source:HGNC Symbol;Acc:HGNC:14( |
| ZFP91 | 11 | ZFP91 zinc finger protein [Source:HGNC Symb |
| ZFP91-CNTF | 11 | ZFP91-CNTF readthrough (NMD candidate) [S( |
| AP001350.1 | 11 | novel transcript, antisense to ZFP91 |
| CNTF | 11 | ciliary neurotrophic factor [Source:HGNC Sym |
| AP001350.2 | 11 | TEC |
| GLYAT | 11 | glycine-N-acyltransferase [Source:HGNC Symb |
| AP000445.1 | 11 | glycine-N-acyltransferase (GLYAT) pseudogen( |
| AP000445.2 | 11 | glycine-N-acyltransferase (GLYAT) pseudogen( |
| TMA16P1 | 11 | translation machinery associated 16 homolog |
| GLYATL2 | 11 | glycine-N-acyltransferase like 2 [Source:HGNC |
| GLYATL1P2 | 11 | glycine-N-acyltransferase like 1 pseudogene 2 |
| GLYATL1 | 11 | glycine-N-acyltransferase like 1 [Source:HGNC |
| AP001652.1 | 11 | novel transcript |
| AP001636.3 | 11 | novel transcript |
| AP001636.2 | 11 | cAMP-dependent protein kinase type II-alpha |
| GLYATL1P1 | 11 | glycine-N-acyltransferase like 1 pseudogene 1 |
| AP001636.1 | 11 | Glycine N-acyltransferase-like protein 1 (GLYA |
| GLYATL1P4 | 11 | glycine-N-acyltransferase like 1 pseudogene 4 |
| GLYATL1B | 11 | glycine-N-acyltransferase like 1B [Source:HGN |
| FAM111B | 11 | family with sequence similarity 111 member E |
| FAM111A-DT | 11 | FAM111A divergent transcript [Source:HGNC : |
| AP001258.1 | 11 | protein kinase, cAMP-dependent, regulatory, |
| FAM111A | 11 | family with sequence similarity 111 member / |
| DTX4 | 11 | deltex E3 ubiquitin ligase 4 [Source:HGNC Sym |
| MPEG1 | 11 | macrophage expressed 1 [Source:HGNC Symb |
| RN7SL42P | 11 | RNA, 7SL, cytoplasmic 42, pseudogene [Sourc( |
| WARSP1 | 11 | tryptophanyl-tRNA synthetase pseudogene 1 |
| SLC25A47P1 | 11 | solute carrier family 25 member 47 pseudoge |
| AP002358.1 | 11 | novel transcript |
| AP002358.2 | 11 | macrophage expressed 1 (MPEG1) pseudogen |
| RN7SL435P | 11 | RNA, 7SL, cytoplasmic 435, pseudogene [Sour |
| OR5AN2P | 11 | olfactory receptor family 5 subfamily AN mem |
| OR5BR1P | 11 | olfactory receptor family 5 subfamily BR mem |
| OR5AN1 | 11 | olfactory receptor family 5 subfamily AN mem |
| OR5BB1P | 11 | olfactory receptor family 5 subfamily BB mem |
| OR5A2 | 11 | olfactory receptor family 5 subfamily A memb |
| OR5A1 | 11 | olfactory receptor family 5 subfamily A memb |
| OR4D6 | 11 | olfactory receptor family 4 subfamily D memb |
| OR4D10 | 11 | olfactory receptor family 4 subfamily D memb |
| OR4D8P | 11 | olfactory receptor family 4 subfamily D memb |
| OR4D11 | 11 | olfactory receptor family 4 subfamily D memb |
| OR4D9 | 11 | olfactory receptor family 4 subfamily D memb |
| OR4D7P | 11 | olfactory receptor family 4 subfamily D memb |
| AP003778.1 | 11 | novel transcript |

| | | |
|---|---|---|
| LINC02739 | 11 | long intergenic non-protein coding RNA 2739 |
| AP000442.1 | 11 | novel transcript |
| OSBP | 11 | oxysterol binding protein [Source:HGNC Symb |
| MIR3162 | 11 | microRNA 3162 [Source:HGNC Symbol;Acc:HG |
| PATL1 | 11 | PAT1 homolog 1, processing body mRNA deca |
| AP000640.2 | 11 | novel transcript |
| RN7SKP192 | 11 | RN7SK pseudogene 192 [Source:HGNC Symbo |
| OR10V1 | 11 | olfactory receptor family 10 subfamily V mem |
| STX3 | 11 | syntaxin 3 [Source:HGNC Symbol;Acc:HGNC:1 |
| OR10Y1P | 11 | olfactory receptor family 10 subfamily Y mem |
| OR10V3P | 11 | olfactory receptor family 10 subfamily V mem |
| OR10V2P | 11 | olfactory receptor family 10 subfamily V mem |
| AP000640.1 | 11 | novel transcript |
| FABP5P7 | 11 | fatty acid binding protein 5 pseudogene 7 [Sou |
| MRPL16 | 11 | mitochondrial ribosomal protein L16 [Source: |
| CBLIF | 11 | cobalamin binding intrinsic factor [Source:HGI |
| TCN1 | 11 | transcobalamin 1 [Source:HGNC Symbol;Acc:H |
| OOSP3 | 11 | oocyte secreted protein family member 3 [Sou |
| SRD5A3P1 | 11 | steroid 5 alpha-reductase 3 pseudogene 1 [So |
| OOSP1 | 11 | oocyte secreted protein 1 [Source:HGNC Syml |
| AP000790.1 | 11 | novel transcript |
| OOSP4A | 11 | oocyte secreted protein family member 4A [S |
| OOSP4B | 11 | oocyte secreted protein family member 4B [S |
| OOSP2 | 11 | oocyte secreted protein 2 [Source:HGNC Syml |
| MS4A3 | 11 | membrane spanning 4-domains A3 [Source:H |
| AP000790.2 | 11 | tyrosine 3-monooxygenase/tryptophan 5-mor |
| MS4A2 | 11 | membrane spanning 4-domains A2 [Source:H |
| LINC02705 | 11 | long intergenic non-protein coding RNA 2705 |
| MS4A6A | 11 | membrane spanning 4-domains A6A [Source:H |
| MS4A4A | 11 | membrane spanning 4-domains A4A [Source:H |
| MS4A4E | 11 | membrane spanning 4-domains A4E [Source:H |
| MIR6503 | 11 | microRNA 6503 [Source:HGNC Symbol;Acc:HG |
| MS4A6E | 11 | membrane spanning 4-domains A6E [Source:H |
| MS4A7 | 11 | membrane spanning 4-domains A7 [Source:H |
| MS4A14 | 11 | membrane spanning 4-domains A14 [Source:H |
| MS4A5 | 11 | membrane spanning 4-domains A5 [Source:H |
| MS4A1 | 11 | membrane spanning 4-domains A1 [Source:H |
| MS4A12 | 11 | membrane spanning 4-domains A12 [Source:H |
| MS4A13 | 11 | membrane spanning 4-domains A13 [Source:H |
| AP003399.1 | 11 | membrane-spanning 4-domains subfamily A n |
| LINC00301 | 11 | long intergenic non-protein coding RNA 301 [S |
| MS4A8 | 11 | membrane spanning 4-domains A8 [Source:H |
| MS4A18 | 11 | membrane spanning 4-domains A18 [Source:H |
| MS4A15 | 11 | membrane spanning 4-domains A15 [Source:H |
| MS4A10 | 11 | membrane spanning 4-domains A10 [Source:H |
| AP000777.1 | 11 | novel transcript |
| AP000777.2 | 11 | novel transcript, antisense to CCDC86 |
| AP000777.3 | 11 | novel transcript |
| CCDC86 | 11 | coiled-coil domain containing 86 [Source:HGN |
| PTGDR2 | 11 | prostaglandin D2 receptor 2 [Source:HGNC Sy |

| | | |
|---|---|---|
| ZP1 | 11 | zona pellucida glycoprotein 1 [Source:HGNC S |
| PRPF19 | 11 | pre-mRNA processing factor 19 [Source:HGNC |
| AP003721.4 | 11 | novel transcript |
| AP003721.3 | 11 | novel transcript |
| TMEM109 | 11 | transmembrane protein 109 [Source:HGNC Sy |
| AP003721.1 | 11 | novel transcript |
| TMEM132A | 11 | transmembrane protein 132A [Source:HGNC S |
| SLC15A3 | 11 | solute carrier family 15 member 3 [Source:HG |
| CD6 | 11 | CD6 molecule [Source:HGNC Symbol;Acc:HGN |
| RNU6-933P | 11 | RNA, U6 small nuclear 933, pseudogene [Sour |
| AP003721.2 | 11 | novel transcript |
| CD5 | 11 | CD5 molecule [Source:HGNC Symbol;Acc:HGN |
| VPS37C | 11 | VPS37C subunit of ESCRT-I [Source:HGNC Sym |
| AP000437.1 | 11 | novel transcript |
| PGA3 | 11 | pepsinogen 3, group I (pepsinogen A) [Source: |
| PGA4 | 11 | pepsinogen 4, group I (pepsinogen A) [Source: |
| AP003037.1 | 11 | novel transcript |
| PGA5 | 11 | pepsinogen 5, group I (pepsinogen A) [Source: |
| VWCE | 11 | von Willebrand factor C and EGF domains [Sou |
| DDB1 | 11 | damage specific DNA binding protein 1 [Sourc |
| TKFC | 11 | triokinase and FMN cyclase [Source:HGNC Syr |
| CYB561A3 | 11 | cytochrome b561 family member A3 [Source: |
| AP003108.3 | 11 | novel transcript |
| TMEM138 | 11 | transmembrane protein 138 [Source:HGNC Sy |
| TMEM216 | 11 | transmembrane protein 216 [Source:HGNC Sy |
| CPSF7 | 11 | cleavage and polyadenylation specific factor 7 |
| AP003108.4 | 11 | novel transcript |
| AP003108.2 | 11 | novel transcript |
| SDHAF2 | 11 | succinate dehydrogenase complex assembly f |
| RN7SL23P | 11 | RNA, 7SL, cytoplasmic 23, pseudogene [Sourc |
| PPP1R32 | 11 | protein phosphatase 1 regulatory subunit 32 [ |
| AP003108.1 | 11 | novel transcript |
| MIR4488 | 11 | microRNA 4488 [Source:HGNC Symbol;Acc:HG |
| LRRC10B | 11 | leucine rich repeat containing 10B [Source:HG |
| SYT7 | 11 | synaptotagmin 7 [Source:HGNC Symbol;Acc:H |
| AP003108.5 | 11 | novel transcript |
| AP003559.1 | 11 | novel transcript |
| AP002754.1 | 11 | novel transcript |
| RPLP0P2 | 11 | ribosomal protein lateral stalk subunit P0 pseu |
| AP002754.2 | 11 | novel transcript |
| DAGLA | 11 | diacylglycerol lipase alpha [Source:HGNC Sym |
| MYRF-AS1 | 11 | MYRF antisense RNA 1 [Source:HGNC Symbol; |
| MYRF | 11 | myelin regulatory factor [Source:HGNC Symbo |
| TMEM258 | 11 | transmembrane protein 258 [Source:HGNC Sy |
| MIR611 | 11 | microRNA 611 [Source:HGNC Symbol;Acc:HGN |
| FEN1 | 11 | flap structure-specific endonuclease 1 [Source |
| FADS2 | 11 | fatty acid desaturase 2 [Source:HGNC Symbol; |
| FADS1 | 11 | fatty acid desaturase 1 [Source:HGNC Symbol; |
| MIR1908 | 11 | microRNA 1908 [Source:HGNC Symbol;Acc:HG |
| FADS3 | 11 | fatty acid desaturase 3 [Source:HGNC Symbol; |

| | | |
|---|---|---|
| MIR6746 | 11 | microRNA 6746 [Source:HGNC Symbol;Acc:HG |
| RAB3IL1 | 11 | RAB3A interacting protein like 1 [Source:HGN |
| RNU6-1243P | 11 | RNA, U6 small nuclear 1243, pseudogene [Sou |
| BEST1 | 11 | bestrophin 1 [Source:HGNC Symbol;Acc:HGNC |
| FTH1 | 11 | ferritin heavy chain 1 [Source:HGNC Symbol;A |
| AP003733.3 | 11 | TEC |
| AP003733.1 | 11 | ribosomal protein S2 (RPS2) pseudogene |
| LINC02733 | 11 | long intergenic non-protein coding RNA 2733 |
| AP003733.2 | 11 | Ribosome biogenesis protein NSA2 homolog ( |
| AP002793.1 | 11 | novel transcript |
| INCENP | 11 | inner centromere protein [Source:HGNC Symb |
| EEF1DP8 | 11 | eukaryotic translation elongation factor 1 delt |
| SCGB1D1 | 11 | secretoglobin family 1D member 1 [Source:HG |
| SCGB2A1 | 11 | secretoglobin family 2A member 1 [Source:HG |
| AP003306.1 | 11 | novel transcript |
| SCGB1D2 | 11 | secretoglobin family 1D member 2 [Source:HG |
| SCGB2A2 | 11 | secretoglobin family 2A member 2 [Source:HG |
| SCGB1D4 | 11 | secretoglobin family 1D member 4 [Source:HG |
| NPM1P35 | 11 | nucleophosmin 1 pseudogene 35 [Source:HGN |
| AP003306.2 | 11 | novel transcript |
| ASRGL1 | 11 | asparaginase and isoaspartyl peptidase 1 [Sou |
| RCC2P6 | 11 | regulator of chromosome condensation 2 pse |
| AP003064.1 | 11 | novel transcript |
| SCGB1A1 | 11 | secretoglobin family 1A member 1 [Source:HG |
| AP003064.2 | 11 | novel transcript |
| AHNAK | 11 | AHNAK nucleoprotein [Source:HGNC Symbol;A |
| AP001363.1 | 11 | novel transcript |
| AP001363.2 | 11 | novel transcript |
| EEF1G | 11 | eukaryotic translation elongation factor 1 gam |
| AP002990.1 | 11 | novel protein |
| MIR6747 | 11 | microRNA 6747 [Source:HGNC Symbol;Acc:HG |
| TUT1 | 11 | terminal uridylyl transferase 1, U6 snRNA-spe |
| MTA2 | 11 | metastasis associated 1 family member 2 [Sou |
| EML3 | 11 | EMAP like 3 [Source:HGNC Symbol;Acc:HGNC: |
| AP001458.1 | 11 | novel transcript |
| ROM1 | 11 | retinal outer segment membrane protein 1 [S |
| B3GAT3 | 11 | beta-1,3-glucuronyltransferase 3 [Source:HGN |
| GANAB | 11 | glucosidase II alpha subunit [Source:HGNC Syr |
| INTS5 | 11 | integrator complex subunit 5 [Source:HGNC S |
| AP001458.2 | 11 | novel protein |
| C11orf98 | 11 | chromosome 11 open reading frame 98 [Sour |
| LBHD1 | 11 | LBH domain containing 1 [Source:HGNC Symb |
| CSKMT | 11 | citrate synthase lysine methyltransferase [Sou |
| SNORA57 | 11 | small nucleolar RNA, H/ACA box 57 [Source:H |
| UQCC3 | 11 | ubiquinol-cytochrome c reductase complex as |
| UBXN1 | 11 | UBX domain protein 1 [Source:HGNC Symbol; |
| LRRN4CL | 11 | LRRN4 C-terminal like [Source:HGNC Symbol;A |
| BSCL2 | 11 | BSCL2 lipid droplet biogenesis associated, seip |
| HNRNPUL2-BSCL2 | 11 | HNRNPUL2-BSCL2 readthrough (NMD candida |
| GNG3 | 11 | G protein subunit gamma 3 [Source:HGNC Syr |

| | | |
|---|---|---|
| HNRNPUL2 | 11 | heterogeneous nuclear ribonucleoprotein U li |
| TTC9C | 11 | tetratricopeptide repeat domain 9C [Source:H |
| ZBTB3 | 11 | zinc finger and BTB domain containing 3 [Sour |
| POLR2G | 11 | RNA polymerase II subunit G [Source:HGNC Sy |
| AP001160.2 | 11 | novel transcript, antisense to TAF6L |
| TAF6L | 11 | TATA-box binding protein associated factor 6 |
| TMEM223 | 11 | transmembrane protein 223 [Source:HGNC Sy |
| AP001160.3 | 11 | novel transcript, antisense to TAF6L |
| TMEM179B | 11 | transmembrane protein 179B [Source:HGNC S |
| MIR6748 | 11 | microRNA 6748 [Source:HGNC Symbol;Acc:HG |
| NXF1 | 11 | nuclear RNA export factor 1 [Source:HGNC Sy |
| MIR6514 | 11 | microRNA 6514 [Source:HGNC Symbol;Acc:HG |
| STX5 | 11 | syntaxin 5 [Source:HGNC Symbol;Acc:HGNC:1 |
| AP001160.4 | 11 | novel transcript, sense intronic to STX5 |
| RNU6-118P | 11 | RNA, U6 small nuclear 118, pseudogene [Sour |
| AP001160.1 | 11 | novel transcript |
| TEX54 | 11 | testis expressed 54 [Source:HGNC Symbol;Acc |
| WDR74 | 11 | WD repeat domain 74 [Source:HGNC Symbol; |
| RNU2-2P | 11 | RNA, U2 small nuclear 2, pseudogene [Source |
| SNHG1 | 11 | small nucleolar RNA host gene 1 [Source:HGN |
| SNORD22 | 11 | small nucleolar RNA, C/D box 22 [Source:HGN |
| SNORD30 | 11 | small nucleolar RNA, C/D box 30 [Source:HGN |
| SNORD22 | 11 | Small nucleolar RNA SNORD22 [Source:RFAM; |
| SNORD28 | 11 | small nucleolar RNA, C/D box 28 [Source:HGN |
| SNORD27 | 11 | small nucleolar RNA, C/D box 27 [Source:HGN |
| SNORD26 | 11 | small nucleolar RNA, C/D box 26 [Source:HGN |
| SNORD25 | 11 | small nucleolar RNA, C/D box 25 [Source:HGN |
| SLC3A2 | 11 | solute carrier family 3 member 2 [Source:HGN |
| CHRM1 | 11 | cholinergic receptor muscarinic 1 [Source:HGI |
| AP000438.1 | 11 | novel transcript |
| RN7SL259P | 11 | RNA, 7SL, cytoplasmic 259, pseudogene [Sour |
| SLC22A6 | 11 | solute carrier family 22 member 6 [Source:HG |
| AP000438.2 | 11 | solute carrier family pseudogene |
| SLC22A8 | 11 | solute carrier family 22 member 8 [Source:HG |
| AP001858.2 | 11 | pituitary tumor-transforming pseudogene |
| AP001858.1 | 11 | tubulin alpha chain pseudogene |
| SLC22A24 | 11 | solute carrier family 22 member 24 [Source:H |
| RPL29P22 | 11 | ribosomal protein L29 pseudogene 22 [Source |
| SLC22A10 | 11 | solute carrier family 22 member 10 [Source:H |
| SLC22A25 | 11 | solute carrier family 22 member 25 [Source:H |
| AP001880.2 | 11 | solute carrier (SLC) family 22 (organic anion tr |
| CCND2P1 | 11 | cyclin D2 pseudogene 1 [Source:HGNC Symbo |
| AP001880.1 | 11 | solute carrier (SLC) family 22 (organic anion tr |
| SLC22A9 | 11 | solute carrier family 22 member 9 [Source:HG |
| PLAAT5 | 11 | phospholipase A and acyltransferase 5 [Source |
| AP000484.1 | 11 | zinc finger pseudogene |
| AP001591.1 | 11 | novel transcript |
| LGALS12 | 11 | galectin 12 [Source:HGNC Symbol;Acc:HGNC:1 |
| PLAAT4 | 11 | phospholipase A and acyltransferase 4 [Source |
| PLAAT2 | 11 | phospholipase A and acyltransferase 2 [Source |

| | | |
|---|---|---|
| PLAAT3 | 11 | phospholipase A and acyltransferase 3 [Sourc |
| AP000753.1 | 11 | novel transcript |
| ATL3 | 11 | atlastin GTPase 3 [Source:HGNC Symbol;Acc:H |
| IMMP1LP1 | 11 | inner mitochondrial membrane peptidase sub |
| AP000753.2 | 11 | novel transcript |
| RTN3 | 11 | reticulon 3 [Source:HGNC Symbol;Acc:HGNC:1 |
| AP000753.3 | 11 | ATPase, H+ transporting, lysosomal 56/58kDa |
| C11orf95 | 11 | chromosome 11 open reading frame 95 [Sour |
| RN7SL596P | 11 | RNA, 7SL, cytoplasmic 596, pseudogene [Sour |
| SPINDOC | 11 | spindlin interactor and repressor of chromatir |
| AP006289.1 | 11 | ATP synthase subunit g, mitochondrial (ATP5L |
| MARK2 | 11 | microtubule affinity regulating kinase 2 [Sourc |
| RNU6-1306P | 11 | RNA, U6 small nuclear 1306, pseudogene [Sou |
| RCOR2 | 11 | REST corepressor 2 [Source:HGNC Symbol;Acc |
| NAA40 | 11 | N(alpha)-acetyltransferase 40, NatD catalytic s |
| RNU6-45P | 11 | RNA, U6 small nuclear 45, pseudogene [Sourc |
| AP000721.1 | 11 | novel transcript |
| COX8A | 11 | cytochrome c oxidase subunit 8A [Source:HGN |
| OTUB1 | 11 | OTU deubiquitinase, ubiquitin aldehyde bindir |
| MACROD1 | 11 | mono-ADP ribosylhydrolase 1 [Source:HGNC S |
| AP000721.2 | 11 | novel transcript, antisense to MACROD1 |
| AP006333.2 | 11 | novel transcript, antisense to MACROD1 |
| FLRT1 | 11 | fibronectin leucine rich transmembrane prote |
| AP006333.1 | 11 | novel transcript |
| STIP1 | 11 | stress induced phosphoprotein 1 [Source:HGN |
| FERMT3 | 11 | fermitin family member 3 [Source:HGNC Symb |
| TRPT1 | 11 | tRNA phosphotransferase 1 [Source:HGNC Syr |
| NUDT22 | 11 | nudix hydrolase 22 [Source:HGNC Symbol;Acc |
| AP001453.1 | 11 | novel transcript |
| DNAJC4 | 11 | DnaJ heat shock protein family (Hsp40) memb |
| VEGFB | 11 | vascular endothelial growth factor B [Source:H |
| FKBP2 | 11 | FKBP prolyl isomerase 2 [Source:HGNC Symbo |
| AP001453.3 | 11 | Uncharacterized protein LOC114841035 [Sour |
| PPP1R14B | 11 | protein phosphatase 1 regulatory inhibitor sul |
| PPP1R14B-AS1 | 11 | PPP1R14B antisense RNA 1 [Source:HGNC Syn |
| AP001453.2 | 11 | novel transcript |
| PLCB3 | 11 | phospholipase C beta 3 [Source:HGNC Symbol |
| BAD | 11 | BCL2 associated agonist of cell death [Source: |
| GPR137 | 11 | G protein-coupled receptor 137 [Source:HGN( |
| KCNK4 | 11 | potassium two pore domain channel subfamil |
| KCNK4-TEX40 | 11 | KCNK4-TEX40 readthrough [Source:NCBI gene |
| Y_RNA | 11 | Y RNA [Source:RFAM;Acc:RF00019] |
| CATSPERZ | 11 | catsper channel auxiliary subunit zeta [Source |
| ESRRA | 11 | estrogen related receptor alpha [Source:HGN( |
| TRMT112 | 11 | tRNA methyltransferase subunit 11-2 [Source: |
| PRDX5 | 11 | peroxiredoxin 5 [Source:HGNC Symbol;Acc:HG |
| AP003774.2 | 11 | novel transcript |
| CCDC88B | 11 | coiled-coil domain containing 88B [Source:HG |
| MIR7155 | 11 | microRNA 7155 [Source:HGNC Symbol;Acc:HG |
| RPS6KA4 | 11 | ribosomal protein S6 kinase A4 [Source:HGNC |

| | | |
|---|---|---|
| MIR1237 | 11 | microRNA 1237 [Source:HGNC Symbol;Acc:HG |
| LINC02723 | 11 | long intergenic non-protein coding RNA 2723 |
| AP003774.1 | 11 | novel transcript |
| LINC02724 | 11 | long intergenic non-protein coding RNA 2724 |
| AP005273.2 | 11 | novel transcript |
| AP005273.1 | 11 | novel transcript |
| AP006288.1 | 11 | novel pseudogene |
| SLC22A11 | 11 | solute carrier family 22 member 11 [Source:H |
| SLC22A12 | 11 | solute carrier family 22 member 12 [Source:H |
| NRXN2 | 11 | neurexin 2 [Source:HGNC Symbol;Acc:HGNC:8 |
| AP001092.1 | 11 | novel transcript |
| RASGRP2 | 11 | RAS guanyl releasing protein 2 [Source:HGNC |
| PYGM | 11 | glycogen phosphorylase, muscle associated [S |
| SF1 | 11 | splicing factor 1 [Source:HGNC Symbol;Acc:HG |
| AP001462.1 | 11 | novel transcript |
| MAP4K2 | 11 | mitogen-activated protein kinase kinase kinas |
| MEN1 | 11 | menin 1 [Source:HGNC Symbol;Acc:HGNC:701 |
| CDC42BPG | 11 | CDC42 binding protein kinase gamma [Source |
| EHD1 | 11 | EH domain containing 1 [Source:HGNC Symbo |
| AP001187.1 | 11 | ribosomal protein S16 (RPS16) pseudogene |
| MIR194-2HG | 11 | MIR194-2 host gene [Source:HGNC Symbol;Ac |
| MIR192 | 11 | microRNA 192 [Source:HGNC Symbol;Acc:HGN |
| MIR194-2 | 11 | microRNA 194-2 [Source:HGNC Symbol;Acc:H |
| ATG2A | 11 | autophagy related 2A [Source:HGNC Symbol;A |
| MIR6750 | 11 | microRNA 6750 [Source:HGNC Symbol;Acc:HG |
| MIR6749 | 11 | microRNA 6749 [Source:HGNC Symbol;Acc:HG |
| PPP2R5B | 11 | protein phosphatase 2 regulatory subunit B'be |
| GPHA2 | 11 | glycoprotein hormone subunit alpha 2 [Source |
| MAJIN | 11 | membrane anchored junction protein [Source |
| BATF2 | 11 | basic leucine zipper ATF-like transcription fact |
| ARL2 | 11 | ADP ribosylation factor like GTPase 2 [Source: |
| ARL2-SNX15 | 11 | ARL2-SNX15 readthrough (NMD candidate) [S |
| MIR6879 | 11 | microRNA 6879 [Source:HGNC Symbol;Acc:HG |
| AP000436.1 | 11 | eukaryotic translation elongation factor 1 alph |
| SNX15 | 11 | sorting nexin 15 [Source:HGNC Symbol;Acc:H |
| SAC3D1 | 11 | SAC3 domain containing 1 [Source:HGNC Sym |
| NAALADL1 | 11 | N-acetylated alpha-linked acidic dipeptidase li |
| CDCA5 | 11 | cell division cycle associated 5 [Source:HGNC S |
| ZFPL1 | 11 | zinc finger protein like 1 [Source:HGNC Symbo |
| TMEM262 | 11 | transmembrane protein 262 [Source:HGNC Sy |
| VPS51 | 11 | VPS51 subunit of GARP complex [Source:HGN |
| AP003068.1 | 11 | novel transcript |
| TM7SF2 | 11 | transmembrane 7 superfamily member 2 [Sou |
| ZNHIT2 | 11 | zinc finger HIT-type containing 2 [Source:HGN |
| AP003068.3 | 11 | novel transcript |
| AP003068.4 | 11 | TEC |
| FAU | 11 | FAU ubiquitin like and ribosomal protein S30 f |
| SYVN1 | 11 | synoviolin 1 [Source:HGNC Symbol;Acc:HGNC: |
| MRPL49 | 11 | mitochondrial ribosomal protein L49 [Source: |
| MIR6751 | 11 | microRNA 6751 [Source:HGNC Symbol;Acc:HG |

| | | |
|---|---|---|
| HIGD1AP10 | 11 | HIG1 hypoxia inducible domain family membe |
| RNU2-23P | 11 | RNA, U2 small nuclear 23, pseudogene [Sourc |
| PDCL2P2 | 11 | PDCL2 pseudogene 2 [Source:HGNC Symbol;A |
| SPDYC | 11 | speedy/RINGO cell cycle regulator family men |
| PGAM1P8 | 11 | phosphoglycerate mutase 1 pseudogene 8 [Sc |
| AP003068.2 | 11 | novel transcript, antisense to CAPN1 |
| CAPN1 | 11 | calpain 1 [Source:HGNC Symbol;Acc:HGNC:14 |
| SLC22A20P | 11 | solute carrier family 22 member 20, pseudoge |
| AP000944.6 | 11 | novel transcript |
| POLA2 | 11 | DNA polymerase alpha 2, accessory subunit [S |
| AP000944.5 | 11 | novel protein, POLA2-CDC42EP2 readthrough |
| AP000944.7 | 11 | novel transcript, antisense to POLA2 |
| CDC42EP2 | 11 | CDC42 effector protein 2 [Source:HGNC Symb |
| DPF2 | 11 | double PHD fingers 2 [Source:HGNC Symbol;A |
| TIGD3 | 11 | tigger transposable element derived 3 [Source |
| AP000944.1 | 11 | novel transcript |
| SLC25A45 | 11 | solute carrier family 25 member 45 [Source:H |
| FRMD8 | 11 | FERM domain containing 8 [Source:HGNC Sym |
| NEAT1 | 11 | nuclear paraspeckle assembly transcript 1 [So |
| AP000944.4 | 11 | NULL |
| AP000944.3 | 11 | NULL |
| AP000944.2 | 11 | NULL |
| MIR612 | 11 | microRNA 612 [Source:HGNC Symbol;Acc:HGI |
| AP000769.4 | 11 | NULL |
| AP000769.1 | 11 | Finkel-Biskis-Reilly murine sarcoma virus (FBR |
| LINC02736 | 11 | long intergenic non-protein coding RNA 2736 |
| AP000769.6 | 11 | novel transcript |
| MALAT1 | 11 | metastasis associated lung adenocarcinoma tı |
| AP000769.2 | 11 | novel transcript |
| AP000769.5 | 11 | NULL |
| AP000769.3 | 11 | NULL |
| SNRPGP19 | 11 | small nuclear ribonucleoprotein polypeptide ( |
| SCYL1 | 11 | SCY1 like pseudokinase 1 [Source:HGNC Symb |
| LTBP3 | 11 | latent transforming growth factor beta bindin |
| AP001362.1 | 11 | TEC |
| SSSCA1-AS1 | 11 | SSSCA1 antisense RNA 1 (head to head) [Sour |
| ZNRD2 | 11 | zinc ribbon domain containing 2 [Source:HGN |
| FAM89B | 11 | family with sequence similarity 89 member B |
| EHBP1L1 | 11 | EH domain binding protein 1 like 1 [Source:HG |
| KCNK7 | 11 | potassium two pore domain channel subfamil |
| MAP3K11 | 11 | mitogen-activated protein kinase kinase kinas |
| PCNX3 | 11 | pecanex 3 [Source:HGNC Symbol;Acc:HGNC:1 |
| MIR4690 | 11 | microRNA 4690 [Source:HGNC Symbol;Acc:HG |
| SIPA1 | 11 | signal-induced proliferation-associated 1 [Sou |
| MIR4489 | 11 | microRNA 4489 [Source:HGNC Symbol;Acc:HG |
| RELA | 11 | RELA proto-oncogene, NF-kB subunit [Source: |
| RELA-DT | 11 | RELA divergent transcript [Source:HGNC Symb |
| RN7SL309P | 11 | RNA, 7SL, cytoplasmic 309, pseudogene [Sour |
| KAT5 | 11 | lysine acetyltransferase 5 [Source:HGNC Symb |
| RNASEH2C | 11 | ribonuclease H2 subunit C [Source:HGNC Sym |

| | | |
|---|---|---|
| KRT8P26 | 11 | keratin 8 pseudogene 26 [Source:HGNC Symb |
| AP001266.2 | 11 | novel transcript |
| AP5B1 | 11 | adaptor related protein complex 5 subunit be |
| OVOL1 | 11 | ovo like transcriptional repressor 1 [Source:H( |
| OVOL1-AS1 | 11 | OVOL1 antisense RNA 1 [Source:HGNC Symbo |
| AP001266.1 | 11 | novel transcript |
| CFL1 | 11 | cofilin 1 [Source:HGNC Symbol;Acc:HGNC:187 |
| SNX32 | 11 | sorting nexin 32 [Source:HGNC Symbol;Acc:H( |
| MUS81 | 11 | MUS81 structure-specific endonuclease subur |
| EFEMP2 | 11 | EGF containing fibulin extracellular matrix pro |
| CTSW | 11 | cathepsin W [Source:HGNC Symbol;Acc:HGNC |
| FIBP | 11 | FGF1 intracellular binding protein [Source:HGI |
| CCDC85B | 11 | coiled-coil domain containing 85B [Source:HG |
| FOSL1 | 11 | FOS like 1, AP-1 transcription factor subunit [S |
| C11orf68 | 11 | chromosome 11 open reading frame 68 [Sour |
| DRAP1 | 11 | DR1 associated protein 1 [Source:HGNC Symb |
| TSGA10IP | 11 | testis specific 10 interacting protein [Source:H |
| SART1 | 11 | spliceosome associated factor 1, recruiter of U |
| AP006287.1 | 11 | transmembrane protein 55B (TMEM55B) pseu |
| EIF1AD | 11 | eukaryotic translation initiation factor 1A dom |
| BANF1 | 11 | barrier to autointegration factor 1 [Source:HG |
| CST6 | 11 | cystatin E/M [Source:HGNC Symbol;Acc:HGNC |
| CATSPER1 | 11 | cation channel sperm associated 1 [Source:HG |
| GAL3ST3 | 11 | galactose-3-O-sulfotransferase 3 [Source:HGN |
| AP006287.3 | 11 | novel transcript, antisense to GAL3ST3 |
| SF3B2 | 11 | splicing factor 3b subunit 2 [Source:HGNC Sym |
| AP006287.2 | 11 | novel transcript, antisense to SF3B2 |
| PACS1 | 11 | phosphofurin acidic cluster sorting protein 1 [ |
| AP000759.1 | 11 | novel transcript |
| KLC2 | 11 | kinesin light chain 2 [Source:HGNC Symbol;Ac |
| AP001107.4 | 11 | novel transcript |
| AP001107.8 | 11 | novel transcript |
| AP001107.7 | 11 | novel transcript |
| RAB1B | 11 | RAB1B, member RAS oncogene family [Source |
| AP001107.1 | 11 | novel transcript |
| AP001107.2 | 11 | novel transcript |
| CNIH2 | 11 | cornichon family AMPA receptor auxiliary pro |
| YIF1A | 11 | Yip1 interacting factor homolog A, membrane |
| TMEM151A | 11 | transmembrane protein 151A [Source:HGNC S |
| AP001107.3 | 11 | novel transcript |
| CD248 | 11 | CD248 molecule [Source:HGNC Symbol;Acc:H( |
| RIN1 | 11 | Ras and Rab interactor 1 [Source:HGNC Symb |
| AP001107.6 | 11 | novel transcript |
| BRMS1 | 11 | BRMS1 transcriptional repressor and anoikis r |
| B4GAT1 | 11 | beta-1,4-glucuronyltransferase 1 [Source:HGN |
| AP001107.9 | 11 | novel transcript, antisense to SLC29A2 |
| SLC29A2 | 11 | solute carrier family 29 member 2 [Source:HG |
| Metazoa_SRP | 11 | Metazoan signal recognition particle RNA [Sou |
| BRD9P1 | 11 | bromodomain containing 9 pseudogene 1 [So |
| RNU1-84P | 11 | RNA, U1 small nuclear 84, pseudogene [Sourc |

| | | |
|---|---|---|
| AP001107.5 | 11 | novel transcript |
| NPAS4 | 11 | neuronal PAS domain protein 4 [Source:HGNC |
| AP002748.1 | 11 | NULL |
| MRPL11 | 11 | mitochondrial ribosomal protein L11 [Source:I |
| AP002748.3 | 11 | glutaredoxin-like protein YDR286C homolog p |
| PELI3 | 11 | pellino E3 ubiquitin protein ligase family mem |
| AP002748.4 | 11 | novel transcript |
| DPP3 | 11 | dipeptidyl peptidase 3 [Source:HGNC Symbol; |
| AP002748.5 | 11 | novel protein |
| BBS1 | 11 | Bardet-Biedl syndrome 1 [Source:HGNC Symb |
| AP002748.6 | 11 | novel transcript, antisense to BBS1 |
| ZDHHC24 | 11 | zinc finger DHHC-type containing 24 [Source:H |
| ACTN3 | 11 | actinin alpha 3 (gene/pseudogene) [Source:H( |
| AP002748.2 | 11 | novel transcript |
| CTSF | 11 | cathepsin F [Source:HGNC Symbol;Acc:HGNC: |
| CCDC87 | 11 | coiled-coil domain containing 87 [Source:HGN |
| CCS | 11 | copper chaperone for superoxide dismutase [: |
| RNU4-39P | 11 | RNA, U4 small nuclear 39, pseudogene [Sourc |
| RBM14 | 11 | RNA binding motif protein 14 [Source:HGNC S |
| RBM14-RBM4 | 11 | RBM14-RBM4 readthrough [Source:HGNC Syr |
| RBM4 | 11 | RNA binding motif protein 4 [Source:HGNC Sy |
| RBM4B | 11 | RNA binding motif protein 4B [Source:HGNC S |
| SPTBN2 | 11 | spectrin beta, non-erythrocytic 2 [Source:HGN |
| RN7SL12P | 11 | RNA, 7SL, cytoplasmic 12, pseudogene [Sourc |
| C11orf80 | 11 | chromosome 11 open reading frame 80 [Sour |
| C1QBPP2 | 11 | complement C1q binding protein pseudogene |
| FTLP6 | 11 | ferritin light chain pseudogene 6 [Source:HGN |
| RCE1 | 11 | Ras converting CAAX endopeptidase 1 [Source |
| PC | 11 | pyruvate carboxylase [Source:HGNC Symbol;A |
| LRFN4 | 11 | leucine rich repeat and fibronectin type III dor |
| RNU7-23P | 11 | RNA, U7 small nuclear 23 pseudogene [Source |
| MIR3163 | 11 | microRNA 3163 [Source:HGNC Symbol;Acc:HG |
| C11orf86 | 11 | chromosome 11 open reading frame 86 [Sour |
| SYT12 | 11 | synaptotagmin 12 [Source:HGNC Symbol;Acc: |
| MIR6860 | 11 | microRNA 6860 [Source:HGNC Symbol;Acc:HG |
| RHOD | 11 | ras homolog family member D [Source:HGNC |
| KDM2A | 11 | lysine demethylase 2A [Source:HGNC Symbol; |
| AP001885.1 | 11 | tetratricopeptide repeat domain 3 (TTC3) pseu |
| AP001885.3 | 11 | novel transcript, antisense to KDM2A |
| GRK2 | 11 | G protein-coupled receptor kinase 2 [Source:H |
| ANKRD13D | 11 | ankyrin repeat domain 13D [Source:HGNC Syr |
| SSH3 | 11 | slingshot protein phosphatase 3 [Source:HGN |
| AP001885.2 | 11 | novel transcript |
| RAD9A | 11 | RAD9 checkpoint clamp component A [Source |
| POLD4 | 11 | DNA polymerase delta 4, accessory subunit [S |
| AP003419.1 | 11 | novel protein |
| AP003419.2 | 11 | novel transcript |
| RN7SKP239 | 11 | RN7SK pseudogene 239 [Source:HGNC Symbo |
| CLCF1 | 11 | cardiotrophin like cytokine factor 1 [Source:H( |
| AP003419.3 | 11 | novel transcript |

| | | |
|---|---|---|
| RNU6-1238P | 11 | RNA, U6 small nuclear 1238, pseudogene [Sou |
| PPP1CA | 11 | protein phosphatase 1 catalytic subunit alpha |
| TBC1D10C | 11 | TBC1 domain family member 10C [Source:HGI |
| CARNS1 | 11 | carnosine synthase 1 [Source:HGNC Symbol;A |
| RPS6KB2 | 11 | ribosomal protein S6 kinase B2 [Source:HGNC |
| RPS6KB2-AS1 | 11 | RPS6KB2 antisense RNA 1 [Source:HGNC Symb |
| CORO1B | 11 | coronin 1B [Source:HGNC Symbol;Acc:HGNC:2 |
| PTPRCAP | 11 | protein tyrosine phosphatase receptor type C |
| GPR152 | 11 | G protein-coupled receptor 152 [Source:HGN( |
| CABP4 | 11 | calcium binding protein 4 [Source:HGNC Symb |
| TMEM134 | 11 | transmembrane protein 134 [Source:HGNC Sy |
| AIP | 11 | aryl hydrocarbon receptor interacting protein |
| MIR6752 | 11 | microRNA 6752 [Source:HGNC Symbol;Acc:HG |
| PITPNM1 | 11 | phosphatidylinositol transfer protein membra |
| CDK2AP2 | 11 | cyclin dependent kinase 2 associated protein : |
| CABP2 | 11 | calcium binding protein 2 [Source:HGNC Symb |
| AP001184.1 | 11 | glutathione S-transferase pi 1 (GSTP1) pseudo |
| GSTP1 | 11 | glutathione S-transferase pi 1 [Source:HGNC S |
| C11orf72 | 11 | chromosome 11 open reading frame 72 (puta |
| AP003385.3 | 11 | novel transcript |
| NDUFV1 | 11 | NADH:ubiquinone oxidoreductase core subun |
| DOC2GP | 11 | double C2 domain gamma, pseudogene [Sour |
| NUDT8 | 11 | nudix hydrolase 8 [Source:HGNC Symbol;Acc:I |
| TBX10 | 11 | T-box transcription factor 10 [Source:HGNC Sy |
| ACY3 | 11 | aminoacylase 3 [Source:HGNC Symbol;Acc:HG |
| AP003385.1 | 11 | novel pseudogene (similar to 2700097O09Rik) |
| ALDH3B2 | 11 | aldehyde dehydrogenase 3 family member B2 |
| RPL37P2 | 11 | ribosomal protein L37 pseudogene 2 [Source: |
| UNC93B5 | 11 | unc-93 homolog B5, pseudogene [Source:HGN |
| OR7E145P | 11 | olfactory receptor family 7 subfamily E memb |
| OR7E11P | 11 | olfactory receptor family 7 subfamily E memb |
| AP003385.5 | 11 | olfactory receptor, family 7, subfamily E, mem |
| AP003385.2 | 11 | novel pseudogene |
| AP003385.4 | 11 | eva-1 homolog Cv ( EVA1C) pseudogene |
| ALG1L8P | 11 | ALG1 like 8, pseudogene [Source:HGNC Symbo |
| FAM86C2P | 11 | family with sequence similarity 86 member C2 |
| ENPP7P7 | 11 | ectonucleotide pyrophosphatase/phosphodie |
| AP003716.1 | 11 | novel pseudogene |
| LINC02754 | 11 | long intergenic non-protein coding RNA 2754 |
| RNU6-46P | 11 | RNA, U6 small nuclear 46, pseudogene [Sourc |
| RPS3AP40 | 11 | ribosomal protein S3a pseudogene 40 [Source |
| AC004923.1 | 11 | RNA binding protein, fox-1 homolog (C. elegar |
| AC004923.2 | 11 | ribosomal protein S24 (RPS24) pseudogene |
| AC004923.3 | 11 | novel pseudogene (C11orf2) |
| OR7E1P | 11 | olfactory receptor family 7 subfamily E memb |
| UNC93B1 | 11 | unc-93 homolog B1, TLR signaling regulator [S |
| ALDH3B1 | 11 | aldehyde dehydrogenase 3 family member B1 |
| AC004923.4 | 11 | novel transcript |
| NDUFS8 | 11 | NADH:ubiquinone oxidoreductase core subun |
| MIR7113 | 11 | microRNA 7113 [Source:HGNC Symbol;Acc:HG |

| | | |
|---|---|---|
| MIR4691 | 11 | microRNA 4691 [Source:HGNC Symbol;Acc:HG |
| TCIRG1 | 11 | T cell immune regulator 1, ATPase H+ transpo |
| MIR6753 | 11 | microRNA 6753 [Source:HGNC Symbol;Acc:HG |
| AP002807.1 | 11 | novel transcript, antisense to CHKA |
| CHKA | 11 | choline kinase alpha [Source:HGNC Symbol;Ac |
| AP002992.1 | 11 | novel transcript |
| KMT5B | 11 | lysine methyltransferase 5B [Source:HGNC Sy |
| C11orf24 | 11 | chromosome 11 open reading frame 24 [Sour |
| AP002813.1 | 11 | novel transcript |
| LRP5 | 11 | LDL receptor related protein 5 [Source:HGNC |
| PPP6R3 | 11 | protein phosphatase 6 regulatory subunit 3 [S |
| NDUFA3P2 | 11 | NADH:ubiquinone oxidoreductase subunit A3 |
| AP003096.1 | 11 | novel transcript |
| GAL | 11 | galanin and GMAP prepropeptide [Source:HG |
| TESMIN | 11 | testis expressed metallothionein like protein [ |
| CPT1A | 11 | carnitine palmitoyltransferase 1A [Source:HG |
| LINC02701 | 11 | long intergenic non-protein coding RNA 2701 |
| MRPL21 | 11 | mitochondrial ribosomal protein L21 [Source: |
| IGHMBP2 | 11 | immunoglobulin mu DNA binding protein 2 [S |
| AP000808.1 | 11 | novel transcript |
| MRGPRD | 11 | MAS related GPR family member D [Source:H |
| AP003071.4 | 11 | novel transcript, overlapping to MRGPRF |
| AP003071.3 | 11 | novel transcript, antisense to MRGPRF |
| MRGPRF | 11 | MAS related GPR family member F [Source:H |
| MRGPRF-AS1 | 11 | MRGPRF antisense RNA 1 [Source:HGNC Symb |
| TPCN2 | 11 | two pore segment channel 2 [Source:HGNC Sy |
| AP003071.5 | 11 | novel protein, TPCN2 - SMIM38 readthrough |
| MIR3164 | 11 | microRNA 3164 [Source:HGNC Symbol;Acc:HG |
| AP003071.1 | 11 | novel transcript |
| AP003071.2 | 11 | novel transcript |
| SMIM38 | 11 | small integral membrane protein 38 [Source:H |
| MYEOV | 11 | myeloma overexpressed [Source:HGNC Symbo |
| IFITM9P | 11 | interferon induced transmembrane protein 9 |
| AP005233.2 | 11 | novel transcript |
| AP005233.1 | 11 | novel transcript |
| AP000439.2 | 11 | uncharacterized LOC102724265 [Source:NCBI |
| AP000439.1 | 11 | novel transcript |
| AP000439.3 | 11 | novel transcript |
| LINC02747 | 11 | long intergenic non-protein coding RNA 2747 |
| LINC01488 | 11 | long intergenic non-protein coding RNA 1488 |
| CCND1 | 11 | cyclin D1 [Source:HGNC Symbol;Acc:HGNC:15 |
| LTO1 | 11 | LTO1 maturation factor of ABCE1 [Source:HG |
| FGF19 | 11 | fibroblast growth factor 19 [Source:HGNC Sym |
| AP001888.1 | 11 | DnaJ (Hsp40) homolog, subfamily B, member |
| FGF4 | 11 | fibroblast growth factor 4 [Source:HGNC Symb |
| FGF3 | 11 | fibroblast growth factor 3 [Source:HGNC Symb |
| AP007216.1 | 11 | novel transcript |
| AP007216.2 | 11 | novel transcript |
| AP003555.2 | 11 | novel transcript |
| AP003555.1 | 11 | novel transcript |

| | | |
|---|---|---|
| LINC02753 | 11 | long intergenic non-protein coding RNA 2753 |
| LINC02584 | 11 | long intergenic non-protein coding RNA 2584 |
| RNU6-1175P | 11 | RNA, U6 small nuclear 1175, pseudogene [Sou |
| ANO1 | 11 | anoctamin 1 [Source:HGNC Symbol;Acc:HGNC |
| AP000879.2 | 11 | novel transcript, sense overlapping ANO1 |
| ANO1-AS1 | 11 | ANO1 antisense RNA 1 [Source:HGNC Symbol; |
| FADD | 11 | Fas associated via death domain [Source:HGN |
| AP000879.1 | 11 | novel transcript |
| PPFIA1 | 11 | PTPRF interacting protein alpha 1 [Source:HGI |
| H2AFZP4 | 11 | H2A histone family member Z pseudogene 4 [ |
| AP002336.2 | 11 | novel transcript |
| MIR548K | 11 | microRNA 548k [Source:HGNC Symbol;Acc:HG |
| AP002336.1 | 11 | novel transcript |
| AP002336.3 | 11 | novel transcript |
| AP000487.2 | 11 | novel transcript |
| AP000487.1 | 11 | novel transcript |
| CTTN | 11 | cortactin [Source:HGNC Symbol;Acc:HGNC:33 |
| SHANK2 | 11 | SH3 and multiple ankyrin repeat domains 2 [S |
| AP001271.1 | 11 | novel transcript |
| AP001271.2 | 11 | TEC |
| SHANK2-AS1 | 11 | SHANK2 antisense RNA 1 [Source:HGNC Symb |
| SHANK2-AS2 | 11 | SHANK2 antisense RNA 2 [Source:HGNC Symb |
| Y_RNA | 11 | Y RNA [Source:RFAM;Acc:RF00019] |
| AP000590.1 | 11 | TEC |
| SHANK2-AS3 | 11 | SHANK2 antisense RNA 3 [Source:HGNC Symb |
| MIR3664 | 11 | microRNA 3664 [Source:HGNC Symbol;Acc:HG |
| AP003783.1 | 11 | novel transcript, antisense to SHANK2 |
| AP002387.1 | 11 | novel actin domain containing pseudogene |
| DHCR7 | 11 | 7-dehydrocholesterol reductase [Source:HGN |
| AP002387.2 | 11 | novel transcript |
| NADSYN1 | 11 | NAD synthetase 1 [Source:HGNC Symbol;Acc:I |
| MIR6754 | 11 | microRNA 6754 [Source:HGNC Symbol;Acc:HG |
| AP000867.2 | 11 | S100 calcium binding protein A11 (S100A11) p |
| KRTAP5-7 | 11 | keratin associated protein 5-7 [Source:HGNC S |
| KRTAP5-8 | 11 | keratin associated protein 5-8 [Source:HGNC S |
| KRTAP5-9 | 11 | keratin associated protein 5-9 [Source:HGNC S |
| AP000867.5 | 11 | novel transcript, antisense to KRTAP5-10 |
| KRTAP5-10 | 11 | keratin associated protein 5-10 [Source:HGNC |
| AP000867.1 | 11 | novel keratin associated protein (KRTAP) pseu |
| KRTAP5-11 | 11 | keratin associated protein 5-11 [Source:HGNC |
| KRTAP5-14P | 11 | keratin associated protein 5-14, pseudogene [ |
| OR7E87P | 11 | olfactory receptor family 7 subfamily E memb |
| UNC93B6 | 11 | unc-93 homolog B6, pseudogene [Source:HGN |
| OR7E4P | 11 | olfactory receptor family 7 subfamily E memb |
| AP000867.3 | 11 | chromosome 11 open reading frame 2 (C11or |
| AP000867.4 | 11 | RNA binding protein, fox-1 homolog (C. elegar |
| RPS3AP41 | 11 | ribosomal protein S3a pseudogene 41 [Source |
| ALG1L9P | 11 | ALG1 like 9, pseudogene [Source:NCBI gene;A |
| SNRPCP14 | 11 | small nuclear ribonucleoprotein polypeptide C |
| AP003498.1 | 11 | novel transcript |

| | | |
|---|---|---|
| ENPP7P8 | 11 | ectonucleotide pyrophosphatase/phosphodie |
| AP003498.2 | 11 | novel transcript |
| FAM86C1 | 11 | family with sequence similarity 86 member C1 |
| ALG1L9P | 11 | ALG1 like 9, pseudogene [Source:HGNC Symb |
| ZNF705E | 11 | zinc finger protein 705E [Source:HGNC Symbo |
| DEFB108B | 11 | defensin beta 108B [Source:HGNC Symbol;Acc |
| RNA5SP342 | 11 | RNA, 5S ribosomal pseudogene 342 [Source:H |
| DEFB130C | 11 | defensin beta 130C (pseudogene) [Source:HG |
| AP002495.1 | 11 | Putative short transient receptor potential ch |
| DEFB131B | 11 | defensin beta 131B [Source:HGNC Symbol;Acc |
| OR7E128P | 11 | olfactory receptor family 7 subfamily E memb |
| OR7E126P | 11 | olfactory receptor family 7 subfamily E memb |
| RNF121 | 11 | ring finger protein 121 [Source:HGNC Symbol; |
| AP002490.2 | 11 | ribosomal protein L17 (RPL17) pseudogene |
| IL18BP | 11 | interleukin 18 binding protein [Source:HGNC S |
| NUMA1 | 11 | nuclear mitotic apparatus protein 1 [Source:H |
| AP002490.1 | 11 | novel transcript |
| RNU6-72P | 11 | RNA, U6 small nuclear 72, pseudogene [Sourc |
| MIR3165 | 11 | microRNA 3165 [Source:HGNC Symbol;Acc:HG |
| LRTOMT | 11 | leucine rich transmembrane and O-methyltra |
| AP000812.5 | 11 | novel protein |
| LAMTOR1 | 11 | late endosomal/lysosomal adaptor, MAPK and |
| AP000812.4 | 11 | novel protein, ortholog of mouse Tomt |
| ANAPC15 | 11 | anaphase promoting complex subunit 15 [Sou |
| FOLR3 | 11 | folate receptor gamma [Source:HGNC Symbol |
| RPEP6 | 11 | ribulose-5-phosphate-3-epimerase pseudoger |
| AP000812.3 | 11 | folate receptor 1 (adult) (FOLR1) pseudogene |
| AP000812.1 | 11 | novel transcript |
| AP000812.2 | 11 | folate receptor 3 (gamma) (FOLR3) pseudoger |
| FOLR1 | 11 | folate receptor alpha [Source:HGNC Symbol;A |
| FOLR2 | 11 | folate receptor beta [Source:HGNC Symbol;Ac |
| INPPL1 | 11 | inositol polyphosphate phosphatase like 1 [So |
| PHOX2A | 11 | paired like homeobox 2A [Source:HGNC Symb |
| AP000593.2 | 11 | docking protein 1, 62kDa (downstream of tyro |
| AP000593.4 | 11 | novel transcript |
| AP000593.1 | 11 | U2 small nuclear RNA auxiliary factor 1 (U2AF |
| CLPB | 11 | ClpB homolog, mitochondrial AAA ATPase cha |
| AP000593.3 | 11 | novel transcript |
| AP002892.2 | 11 | novel transcript |
| AP002892.1 | 11 | novel transcript |
| AP003785.1 | 11 | novel transcript |
| ART2BP | 11 | ADP-ribosyltransferase 2B, pseudogene [Sour |
| ART2P | 11 | ADP-ribosyltransferase 2, pseudogene [Source |
| AP005019.3 | 11 | cyclic nucleotide gated channel alpha 1 (CNGA |
| AP005019.2 | 11 | ADP-ribosyltransferase 5 (ART5) pseudogene |
| LINC01537 | 11 | long intergenic non-protein coding RNA 1537 |
| PDE2A | 11 | phosphodiesterase 2A [Source:HGNC Symbol; |
| AP005019.1 | 11 | novel transcript |
| MIR139 | 11 | microRNA 139 [Source:HGNC Symbol;Acc:HGN |
| RNU7-105P | 11 | RNA, U7 small nuclear 105 pseudogene [Sourc |

| | | |
|---|---|---|
| AP003065.1 | 11 | novel transcript |
| ARAP1 | 11 | ArfGAP with RhoGAP domain, ankyrin repeat a |
| ARAP1-AS1 | 11 | ARAP1 antisense RNA 1 [Source:HGNC Symbo |
| AP003065.2 | 11 | novel transcript, antisense to ARAP1 |
| ARAP1-AS2 | 11 | ARAP1 antisense RNA 2 [Source:HGNC Symbo |
| RPS12P20 | 11 | ribosomal protein S12 pseudogene 20 [Source |
| STARD10 | 11 | StAR related lipid transfer domain containing |
| Y_RNA | 11 | Y RNA [Source:RFAM;Acc:RF00019] |
| MIR4692 | 11 | microRNA 4692 [Source:HGNC Symbol;Acc:HG |
| AP002381.2 | 11 | Novel transcript |
| ATG16L2 | 11 | autophagy related 16 like 2 [Source:HGNC Syr |
| FCHSD2 | 11 | FCH and double SH3 domains 2 [Source:HGNC |
| RNU6-672P | 11 | RNA, U6 small nuclear 672, pseudogene [Sour |
| AP002381.1 | 11 | ribosomal protein L15 (RPL15) pseudogene |
| AP002761.2 | 11 | novel transcript |
| AP002761.1 | 11 | novel transcript |
| P2RY2 | 11 | purinergic receptor P2Y2 [Source:HGNC Symb |
| AP002761.4 | 11 | novel transcript, overlapping to P2RY2 |
| OR8R1P | 11 | olfactory receptor family 8 subfamily R memb |
| P2RY6 | 11 | pyrimidinergic receptor P2Y6 [Source:HGNC S |
| AP002761.3 | 11 | novel transcript |
| ARHGEF17 | 11 | Rho guanine nucleotide exchange factor 17 [S |
| RELT | 11 | RELT TNF receptor [Source:HGNC Symbol;Acc: |
| AP000763.4 | 11 | novel transcript |
| FAM168A | 11 | family with sequence similarity 168 member A |
| AP000763.3 | 11 | novel transcript |
| AP000763.1 | 11 | nucleophosmin/nucleoplasmin 3 (NPM3) pseu |
| AP000763.2 | 11 | mitochondrially encoded cytochrome c oxidas |
| HMGN2P38 | 11 | high mobility group nucleosomal binding dom |
| AP000860.1 | 11 | CTAGE family, member 5 (CTAGE5) pseudoger |
| PLEKHB1 | 11 | pleckstrin homology domain containing B1 [Sc |
| RAB6A | 11 | RAB6A, member RAS oncogene family [Source |
| AP002993.1 | 11 | novel transcript |
| AP002770.1 | 11 | novel transcript |
| MRPL48 | 11 | mitochondrial ribosomal protein L48 [Source:I |
| RN7SKP243 | 11 | RN7SK pseudogene 243 [Source:HGNC Symbo |
| CCDC58P5 | 11 | coiled-coil domain containing 58 pseudogene |
| COA4 | 11 | cytochrome c oxidase assembly factor 4 homc |
| PAAF1 | 11 | proteasomal ATPase associated factor 1 [Sour |
| AP002770.2 | 11 | actin related protein 2/3 complex, subunit 3, 2 |
| DNAJB13 | 11 | DnaJ heat shock protein family (Hsp40) memb |
| AP003717.1 | 11 | novel transcript |
| UCP2 | 11 | uncoupling protein 2 [Source:HGNC Symbol;A |
| AP003717.4 | 11 | novel transcript |
| AP003717.2 | 11 | developmental pluripotency associated 4 (DPI |
| AP003717.3 | 11 | family with sequence similarity 91, member A |
| UCP3 | 11 | uncoupling protein 3 [Source:HGNC Symbol;A |
| C2CD3 | 11 | C2 domain containing 3 centriole elongation r |
| AP002392.1 | 11 | cytochrome c oxidase subunit VIc (COX6C) pse |
| PPME1 | 11 | protein phosphatase methylesterase 1 [Sourc |

| | | |
|---|---|---|
| RNA5SP343 | 11 | RNA, 5S ribosomal pseudogene 343 [Source:H |
| AP000577.2 | 11 | TEC |
| P4HA3 | 11 | prolyl 4-hydroxylase subunit alpha 3 [Source:H |
| AP000577.1 | 11 | NULL |
| P4HA3-AS1 | 11 | P4HA3 antisense RNA 1 [Source:HGNC Symbo |
| PGM2L1 | 11 | phosphoglucomutase 2 like 1 [Source:HGNC S |
| HNRNPA1P40 | 11 | heterogeneous nuclear ribonucleoprotein A1 |
| AP001372.1 | 11 | novel transcript, antisense to PGM2L1 |
| AP001085.1 | 11 | novel transcript |
| MIR548AL | 11 | microRNA 548al [Source:HGNC Symbol;Acc:H |
| KCNE3 | 11 | potassium voltage-gated channel subfamily E |
| AP001372.3 | 11 | novel transcript |
| CYCSP27 | 11 | CYCS pseudogene 27 [Source:HGNC Symbol;A |
| AP001372.4 | 11 | ribosomal L24 domain containing 1 (RSL24D1) |
| LIPT2 | 11 | lipoyl(octanoyl) transferase 2 [Source:HGNC S |
| AP001372.2 | 11 | uncharacterized LOC100287896 [Source:NCBI |
| POLD3 | 11 | DNA polymerase delta 3, accessory subunit [S |
| RANP3 | 11 | RAN pseudogene 3 [Source:HGNC Symbol;Acc |
| RN7SKP297 | 11 | RN7SK pseudogene 297 [Source:HGNC Symbo |
| CHRDL2 | 11 | chordin like 2 [Source:HGNC Symbol;Acc:HGN |
| AP001324.3 | 11 | TEC |
| MIR4696 | 11 | microRNA 4696 [Source:HGNC Symbol;Acc:H |
| AP001324.2 | 11 | ribosomal protein L36a (RPL36A) pseudogene |
| AP001324.1 | 11 | ribosomal protein S12 (RPS12) pseudogene |
| RNF169 | 11 | ring finger protein 169 [Source:HGNC Symbol; |
| XRRA1 | 11 | X-ray radiation resistance associated 1 [Sourc |
| AP000560.1 | 11 | TEC |
| RN7SL239P | 11 | RNA, 7SL, cytoplasmic 239, pseudogene [Sour |
| AP001992.1 | 11 | ribosomal protein L31 (RPL31) pseudogene |
| AP001992.2 | 11 | novel transcript, antisense to XRRA1 |
| SPCS2 | 11 | signal peptidase complex subunit 2 [Source:H |
| RNU6-216P | 11 | RNA, U6 small nuclear 216, pseudogene [Sour |
| NEU3 | 11 | neuraminidase 3 [Source:HGNC Symbol;Acc:H |
| AP003175.1 | 11 | novel transcript |
| OR2AT2P | 11 | olfactory receptor family 2 subfamily AT mem |
| NPM1P50 | 11 | nucleophosmin 1 pseudogene 50 [Source:HGN |
| OR2AT4 | 11 | olfactory receptor family 2 subfamily AT mem |
| AP001972.4 | 11 | novel transcript, antisense to SLCO2B1 |
| SLCO2B1 | 11 | solute carrier organic anion transporter family |
| OR2AT1P | 11 | olfactory receptor family 2 subfamily AT mem |
| AP001972.3 | 11 | novel transcript |
| ZDHHC20P3 | 11 | zinc finger DHHC-type containing 20 pseudoge |
| TPBGL | 11 | trophoblast glycoprotein like [Source:HGNC Sy |
| ARRB1 | 11 | arrestin beta 1 [Source:HGNC Symbol;Acc:HG |
| AP001972.1 | 11 | novel transcript |
| AP001972.5 | 11 | TEC |
| AP001972.2 | 11 | novel transcript |
| MIR326 | 11 | microRNA 326 [Source:HGNC Symbol;Acc:HGN |
| RPS3 | 11 | ribosomal protein S3 [Source:HGNC Symbol;A |
| SNORD15A | 11 | small nucleolar RNA, C/D box 15A [Source:HG |

| | | |
|---|---|---|
| SNORD15B | 11 | small nucleolar RNA, C/D box 15B [Source:HG |
| KLHL35 | 11 | kelch like family member 35 [Source:HGNC Sy |
| GDPD5 | 11 | glycerophosphodiester phosphodiesterase do |
| AP002815.1 | 11 | novel transcript |
| SERPINH1 | 11 | serpin family H member 1 [Source:HGNC Syml |
| AP001922.5 | 11 | novel transcript |
| MAP6 | 11 | microtubule associated protein 6 [Source:HGN |
| AP001922.6 | 11 | novel transcript |
| AP001922.2 | 11 | novel transcript |
| AP001922.4 | 11 | ribosomal protein L37 (RPL37) pseudogene |
| MOGAT2 | 11 | monoacylglycerol O-acyltransferase 2 [Source |
| RN7SL786P | 11 | RNA, 7SL, cytoplasmic 786, pseudogene [Sour |
| AP001922.1 | 11 | novel transcript, antisense to DGAT2 |
| DGAT2 | 11 | diacylglycerol O-acyltransferase 2 [Source:HG |
| AP001922.3 | 11 | novel transcript |
| AP003031.1 | 11 | novel transcript |
| UVRAG-DT | 11 | UVRAG divergent transcript [Source:HGNC Syr |
| UVRAG | 11 | UV radiation resistance associated [Source:HG |
| Y_RNA | 11 | Y RNA [Source:RFAM;Acc:RF00019] |
| Y_RNA | 11 | Y RNA [Source:RFAM;Acc:RF00019] |
| PPP1R1AP1 | 11 | protein phosphatase 1 regulatory inhibitor sul |
| AP003168.2 | 11 | novel transcript |
| RNA5SP344 | 11 | RNA, 5S ribosomal pseudogene 344 [Source:H |
| AP003168.1 | 11 | SMT3 suppressor of mif two 3 homolog 1 (S. c |
| AP002340.1 | 11 | novel transcript |
| WNT11 | 11 | Wnt family member 11 [Source:HGNC Symbol |
| AP000785.1 | 11 | novel transcript |
| LINC02761 | 11 | long intergenic non-protein coding RNA 2761 |
| THAP12 | 11 | THAP domain containing 12 [Source:HGNC Syr |
| GVQW3 | 11 | GVQW motif containing 3 [Source:HGNC Symb |
| Y_RNA | 11 | Y RNA [Source:RFAM;Acc:RF00019] |
| AP002360.1 | 11 | novel transcript |
| EMSY | 11 | EMSY transcriptional repressor, BRCA2 intera |
| AP001189.2 | 11 | nucleolin (NCL) pseudogene |
| LINC02757 | 11 | long intergenic non-protein coding RNA 2757 |
| AP001189.5 | 11 | novel transcript |
| AP001189.3 | 11 | novel transcript |
| AP001189.1 | 11 | novel transcript |
| LRRC32 | 11 | leucine rich repeat containing 32 [Source:HGN |
| AP001189.4 | 11 | novel transcript |
| GUCY2EP | 11 | guanylate cyclase 2E, pseudogene [Source:HG |
| AP003119.1 | 11 | novel transcript |
| TSKU | 11 | tsukushi, small leucine rich proteoglycan [Sou |
| AP003119.2 | 11 | novel transcript |
| AP003119.3 | 11 | novel transcript, overlapping to TSKU |
| ACER3 | 11 | alkaline ceramidase 3 [Source:HGNC Symbol;A |
| AP002498.1 | 11 | novel transcript |
| B3GNT6 | 11 | UDP-GlcNAc:betaGal beta-1,3-N-acetylglucosa |
| CAPN5 | 11 | calpain 5 [Source:HGNC Symbol;Acc:HGNC:14 |
| OMP | 11 | olfactory marker protein [Source:HGNC Symb |

| | | |
|---|---|---|
| MYO7A | 11 | myosin VIIA [Source:HGNC Symbol;Acc:HGNC: |
| GDPD4 | 11 | glycerophosphodiester phosphodiesterase do |
| TOMM20P1 | 11 | TOMM20 pseudogene 1 [Source:HGNC Symbo |
| PAK1 | 11 | p21 (RAC1) activated kinase 1 [Source:HGNC S |
| AP003680.1 | 11 | novel transcript, antisense to PAK1 |
| CLNS1A | 11 | chloride nucleotide-sensitive channel 1A [Sou |
| RNU7-59P | 11 | RNA, U7 small nuclear 59 pseudogene [Source |
| AQP11 | 11 | aquaporin 11 [Source:HGNC Symbol;Acc:HGN |
| RSF1 | 11 | remodeling and spacing factor 1 [Source:HGN |
| Y_RNA | 11 | Y RNA [Source:RFAM;Acc:RF00019] |
| RSF1-IT2 | 11 | RSF1 intronic transcript 2 [Source:HGNC Symb |
| FTH1P16 | 11 | ferritin heavy chain 1 pseudogene 16 [Source: |
| RSF1-IT1 | 11 | RSF1 intronic transcript 1 [Source:HGNC Symb |
| AP000580.1 | 11 | ribosomal protein S20 (RPS20) pseudogene |
| AAMDC | 11 | adipogenesis associated Mth938 domain cont |
| AP002812.2 | 11 | novel transcript, antisense to AAMDC |
| AP002812.3 | 11 | novel transcript |
| AP002812.5 | 11 | novel transcript, antisense to AAMDC |
| AP002812.1 | 11 | ribosomal protein L21 (RPL21) pseudogene |
| INTS4 | 11 | integrator complex subunit 4 [Source:HGNC S |
| AP002812.4 | 11 | family with sequence similarity 162, member |
| AP003032.2 | 11 | novel transcript |
| KCTD14 | 11 | potassium channel tetramerization domain co |
| NDUFC2-KCTD14 | 11 | NDUFC2-KCTD14 readthrough [Source:HGNC S |
| AP003032.1 | 11 | novel transcript |
| THRSP | 11 | thyroid hormone responsive [Source:HGNC Sy |
| NDUFC2 | 11 | NADH:ubiquinone oxidoreductase subunit C2 |
| ALG8 | 11 | ALG8 alpha-1,3-glucosyltransferase [Source:H |
| RNU6-126P | 11 | RNA, U6 small nuclear 126, pseudogene [Sour |
| KCTD21-AS1 | 11 | KCTD21 antisense RNA 1 [Source:HGNC Symb |
| KCTD21 | 11 | potassium channel tetramerization domain co |
| USP35 | 11 | ubiquitin specific peptidase 35 [Source:HGNC |
| GAB2 | 11 | GRB2 associated binding protein 2 [Source:HG |
| AP003086.1 | 11 | novel transcript |
| AP003086.3 | 11 | zinc finger protein 75C, pseudogene |
| AP003086.2 | 11 | novel transcript |
| LINC02728 | 11 | long intergenic non-protein coding RNA 2728 |
| NARS2 | 11 | asparaginyl-tRNA synthetase 2, mitochondrial |
| AP003110.1 | 11 | novel transcript |
| RNU6-311P | 11 | RNA, U6 small nuclear 311, pseudogene [Sour |
| COPS8P3 | 11 | COP9 signalosome subunit 8 pseudogene 3 [S |
| TENM4 | 11 | teneurin transmembrane protein 4 [Source:H |
| AP002768.1 | 11 | novel transcript |
| AP002957.1 | 11 | novel transcript |
| AP001547.1 | 11 | novel transcript |
| MIR708 | 11 | microRNA 708 [Source:HGNC Symbol;Acc:HGN |
| MIR5579 | 11 | microRNA 5579 [Source:HGNC Symbol;Acc:HG |
| AP001978.1 | 11 | novel transcript |
| AP001541.1 | 11 | TEC |
| AP001284.1 | 11 | origin recognition complex, subunit 3-like (yea |

| | | |
|---|---|---|
| AP006295.1 | 11 | novel transcript |
| RNU6-544P | 11 | RNA, U6 small nuclear 544, pseudogene [Sour |
| ARL6IP1P3 | 11 | ADP ribosylation factor like GTPase 6 interacti |
| LINC02720 | 11 | long intergenic non-protein coding RNA 2720 |
| AP003398.1 | 11 | cytochrome c oxidase subunit VIa polypeptide |
| AP003398.2 | 11 | novel transcript |
| AP003464.1 | 11 | novel transcript |
| MTND4LP18 | 11 | MT-ND4L pseudogene 18 [Source:HGNC Symb |
| MTND6P25 | 11 | MT-ND6 pseudogene 25 [Source:HGNC Symbo |
| MIR4300HG | 11 | MIR4300 host gene [Source:HGNC Symbol;Acc |
| AP002802.2 | 11 | ribosomal protein S12, pseudogene |
| MIR4300 | 11 | microRNA 4300 [Source:HGNC Symbol;Acc:HG |
| AP002802.1 | 11 | novel transcript |
| AP000793.1 | 11 | novel transcript |
| RPS28P7 | 11 | ribosomal protein S28 pseudogene 7 [Source: |
| FAM181B | 11 | family with sequence similarity 181 member B |
| LINC02734 | 11 | long intergenic non-protein coding RNA 2734 |
| AP001646.2 | 11 | RNA binding motif protein, X-linked (RBMX) ps |
| PRCP | 11 | prolylcarboxypeptidase [Source:HGNC Symbo |
| AP001646.1 | 11 | eukaryotic translation initiation factor 2, subu |
| DDIAS | 11 | DNA damage induced apoptosis suppressor [S |
| AP000893.1 | 11 | ribosomal protein L7a (RPL7A) pseudogene |
| AP000893.2 | 11 | novel transcript, sense overlapping RAB30 |
| RAB30 | 11 | RAB30, member RAS oncogene family [Source |
| AP001767.1 | 11 | ribosomal protein L10 (RPL10) pseudogene |
| SNORA70E | 11 | small nucleolar RNA, H/ACA box 70E [Source:H |
| RAB30-DT | 11 | RAB30 divergent transcript [Source:HGNC Sym |
| AP001767.3 | 11 | novel transcript |
| AP001767.4 | 11 | TEC |
| AP001767.2 | 11 | cytochrome c oxidase subunit Vb (COX5B) pse |
| C1DP5 | 11 | C1D nuclear receptor corepressor pseudogene |
| BCAS2P1 | 11 | BCAS2 pseudogene 1 [Source:HGNC Symbol;A |
| PCF11 | 11 | PCF11 cleavage and polyadenylation factor su |
| AP000873.4 | 11 | novel transcript, sense overlapping PCF11 |
| AP000873.2 | 11 | novel transcript |
| PCF11-AS1 | 11 | PCF11 antisense RNA 1 [Source:HGNC Symbol |
| ANKRD42 | 11 | ankyrin repeat domain 42 [Source:HGNC Symb |
| AP000873.5 | 11 | CDC28 protein kinase regulatory subunit 1B (C |
| AP000873.3 | 11 | novel transcript |
| AP000873.1 | 11 | ribosomal protein L32 (RPL32) pseudogene |
| CCDC90B | 11 | coiled-coil domain containing 90B [Source:HG |
| AP000446.1 | 11 | novel transcript, antisense to CCDC90B |
| CYCSP28 | 11 | CYCS pseudogene 28 [Source:HGNC Symbol;A |
| AP000446.2 | 11 | NULL |
| DLG2 | 11 | discs large MAGUK scaffold protein 2 [Source: |
| AP002370.2 | 11 | novel transcript |
| AP002370.1 | 11 | suppressor of cytokine signaling 6 (SOCS6) pse |
| LDHAL6DP | 11 | lactate dehydrogenase A like 6D, pseudogene |
| HNRNPA1P72 | 11 | heterogeneous nuclear ribonucleoprotein A1 |
| AP000852.1 | 11 | ribosomal protein L9 (RPL9) pseudogene |

| | | |
|---|---|---|
| AP001825.1 | 11 | novel transcript |
| AP001825.2 | 11 | NULL |
| AP001825.3 | 11 | NULL |
| AP000857.2 | 11 | novel transcript |
| AP000857.1 | 11 | ras homolog gene family, member B (RHOB) p |
| HNRNPCP6 | 11 | heterogeneous nuclear ribonucleoprotein C p |
| AP002803.1 | 11 | REST corepressor 1 (RCOR1) pseudogene |
| AP003035.1 | 11 | DEAH (Asp-Glu-Ala-His) box polypeptide 9 (DH |
| RNU6-1292P | 11 | RNA, U6 small nuclear 1292, pseudogene [Sou |
| TMEM126B | 11 | transmembrane protein 126B [Source:HGNC S |
| TMEM126A | 11 | transmembrane protein 126A [Source:HGNC S |
| CREBZF | 11 | CREB/ATF bZIP transcription factor [Source:HG |
| CCDC89 | 11 | coiled-coil domain containing 89 [Source:HGN |
| SYTL2 | 11 | synaptotagmin like 2 [Source:HGNC Symbol;A |
| AP000974.1 | 11 | TEC |
| CCDC83 | 11 | coiled-coil domain containing 83 [Source:HGN |
| AP003128.1 | 11 | novel transcript |
| SLC25A1P1 | 11 | solute carrier family 25 member 1 pseudogen |
| PICALM | 11 | phosphatidylinositol binding clathrin assembly |
| RNU6-560P | 11 | RNA, U6 small nuclear 560, pseudogene [Sour |
| LINC02695 | 11 | long intergenic non-protein coding RNA 2695 |
| AP003097.1 | 11 | farnesyltransferase, CAAX box, alpha-like 1 |
| Metazoa_SRP | 11 | Metazoan signal recognition particle RNA [Sou |
| EED | 11 | embryonic ectoderm development [Source:HG |
| MIR6755 | 11 | microRNA 6755 [Source:HGNC Symbol;Acc:HG |
| AP003084.1 | 11 | immunoglobin superfamily, member 21 (IGSF2 |
| SETP17 | 11 | SET pseudogene 17 [Source:HGNC Symbol;Acc |
| HIKESHI | 11 | heat shock protein nuclear import factor hike |
| RN7SL225P | 11 | RNA, 7SL, cytoplasmic 225, pseudogene [Sour |
| CCDC81 | 11 | coiled-coil domain containing 81 [Source:HGN |
| AP001831.1 | 11 | novel transcript, antisense to ME3 |
| PTP4A1P6 | 11 | PTP4A1 pseudogene 6 [Source:HGNC Symbol; |
| AP001148.1 | 11 | novel transcript |
| ME3 | 11 | malic enzyme 3 [Source:HGNC Symbol;Acc:HG |
| AP003059.1 | 11 | novel transcript |
| AP003059.2 | 11 | novel transcript |
| PRSS23 | 11 | serine protease 23 [Source:HGNC Symbol;Acc |
| AP000654.2 | 11 | MOB family member 4, phocein, pseudogene |
| OR7E13P | 11 | olfactory receptor family 7 subfamily E memb |
| AP000654.1 | 11 | novel transcript |
| OR7E2P | 11 | olfactory receptor family 7 subfamily E memb |
| AP001528.1 | 11 | novel transcript |
| AP001528.2 | 11 | novel transcript |
| FZD4 | 11 | frizzled class receptor 4 [Source:HGNC Symbo |
| FZD4-DT | 11 | FZD4 divergent transcript [Source:HGNC Symb |
| HNRNPCP8 | 11 | heterogeneous nuclear ribonucleoprotein C p |
| TMEM135 | 11 | transmembrane protein 135 [Source:HGNC Sy |
| XIAPP2 | 11 | X-linked inhibitor of apoptosis pseudogene 2 [ |
| AP002967.1 | 11 | novel transcript |
| AP001102.1 | 11 | translocase of outer mitochondrial membrane |

| | | |
|---|---|---|
| AP000811.1 | 11 | coiled-coil domain containing 90B (CCDC90B) |
| PSMA2P1 | 11 | proteasome subunit alpha 2 pseudogene 1 [So |
| AP003497.1 | 11 | high-mobility group box 1 (HMGB1) pseudoge |
| AP003497.2 | 11 | novel transcript |
| AP001784.1 | 11 | novel transcript |
| RNU6-1135P | 11 | RNA, U6 small nuclear 1135, pseudogene [Sou |
| U6 | 11 | U6 spliceosomal RNA [Source:RFAM;Acc:RF00 |
| LINC02711 | 11 | long intergenic non-protein coding RNA 2711 |
| AP000756.1 | 11 | novel transcript |
| MTCYBP41 | 11 | MT-CYB pseudogene 41 [Source:HGNC Symbo |
| AP000676.2 | 11 | mitochondrially encoded NADH dehydrogenas |
| AP000676.3 | 11 | novel transcript |
| AP000676.5 | 11 | novel transcript |
| AP000676.4 | 11 | novel transcript |
| AP000676.1 | 11 | mitochondrially encoded cytochrome b (MT-C |
| AP005436.3 | 11 | novel transcript |
| AP005436.1 | 11 | novel transcript |
| AP005436.2 | 11 | novel transcript |
| Y_RNA | 11 | Y RNA [Source:RFAM;Acc:RF00019] |
| RAB38 | 11 | RAB38, member RAS oncogene family [Source |
| AP001642.1 | 11 | novel transcript |
| MIR3166 | 11 | microRNA 3166 [Source:HGNC Symbol;Acc:HG |
| CTSC | 11 | cathepsin C [Source:HGNC Symbol;Acc:HGNC: |
| GAPDHP70 | 11 | glyceraldehyde 3 phosphate dehydrogenase p |
| AP003120.1 | 11 | novel transcript |
| GRM5-AS1 | 11 | GRM5 antisense RNA 1 [Source:HGNC Symbol |
| GRM5 | 11 | glutamate metabotropic receptor 5 [Source:H |
| RNU6-16P | 11 | RNA, U6 small nuclear 16, pseudogene [Sourc |
| TYR | 11 | tyrosinase [Source:HGNC Symbol;Acc:HGNC:1 |
| CBX3P7 | 11 | chromobox 3 pseudogene 7 [Source:HGNC Sy |
| AP000720.1 | 11 | novel transcript |
| AP000720.2 | 11 | novel transcript |
| NOX4 | 11 | NADPH oxidase 4 [Source:HGNC Symbol;Acc:H |
| AP002404.1 | 11 | histone cluster 3, H3 (HIST3H3) pseudogene |
| AP003400.4 | 11 | novel transcript |
| AP003400.5 | 11 | novel transcript |
| AP003400.1 | 11 | novel transcript |
| AP003400.2 | 11 | novel transcript |
| AP003400.3 | 11 | novel transcript |
| AP003400.6 | 11 | novel transcript |
| FOLH1B | 11 | folate hydrolase 1B [Source:HGNC Symbol;Acc |
| UBTFL10 | 11 | UBTF like 10 (pseudogene) [Source:HGNC Sym |
| AP003122.6 | 11 | novel transcript |
| TRIM77 | 11 | tripartite motif containing 77 [Source:HGNC S |
| AP003122.4 | 11 | tripartite motif-containing protein (TRIM) pse |
| AP003122.5 | 11 | Cdon homolog (mouse) (CDON) pseudogene |
| UBTFL2 | 11 | UBTF like 2 (pseudogene) [Source:HGNC Symb |
| AP003122.2 | 11 | novel transcript |
| AP003122.3 | 11 | establishment of cohesion 1 homolog 2 (S. ce |
| AP003122.1 | 11 | tripartite motif-containing protein 43 (TRIM43 |

| | | |
|---|---|---|
| TRIM64DP | 11 | tripartite motif containing 64D, pseudogene [S |
| AP004833.2 | 11 | ankyrin repeat domain 33B (ANKRD33B ) pseu |
| AP004833.3 | 11 | novel pseudogene |
| TRIM49 | 11 | tripartite motif containing 49 [Source:HGNC S |
| AP004833.1 | 11 | tripartite motif-containing 43 (TRIM43) pseud |
| TRIM53BP | 11 | tripartite motif containing 53B, pseudogene [S |
| TRIM51BP | 11 | tripartite motif-containing 51B, pseudogene [S |
| AP005435.1 | 11 | ankyrin repeat domain-containing protein 33B |
| TRIM64B | 11 | tripartite motif containing 64B [Source:HGNC |
| AP005435.4 | 11 | folate hydrolase (prostate-specific membrane |
| AP005435.2 | 11 | tripartite motif-containing (TRIM) pseudogene |
| AP005435.3 | 11 | Cdon homolog (mouse) (CDON) pseudogene |
| MTND1P35 | 11 | MT-ND1 pseudogene 35 [Source:HGNC Symbo |
| TRIM49D1 | 11 | tripartite motif containing 49D1 [Source:HGN |
| TRIM49D2 | 11 | tripartite motif containing 49D2 [Source:HGN |
| AP004607.3 | 11 | Cdon homolog (mouse) (CDON) pseudogene |
| AP004607.9 | 11 | tripartite motif-containing 51 (TRIM51) pseud |
| AP004607.8 | 11 | tripartite motif-containing protein (TRIM) pse |
| AP004607.1 | 11 | N-acetylated alpha-linked acidic dipeptidase-li |
| TRIM64 | 11 | tripartite motif containing 64 [Source:HGNC S |
| AP004607.7 | 11 | Ankyrin repeat domain-containing protein 33B |
| TRIM51EP | 11 | tripartite motif-containing 51E, pseudogene [S |
| TRIM53AP | 11 | tripartite motif containing 53A, pseudogene [S |
| AP004607.6 | 11 | tripartite motif-containing protein 43 (TRIM43 |
| TRIM49C | 11 | tripartite motif containing 49C [Source:HGNC |
| AP004607.2 | 11 | novel pseudogene |
| AP004607.4 | 11 | ankyrin repeat domain 33B (ANKRD33B) pseu |
| TRIM64EP | 11 | tripartite motif containing 64E, pseudogene [S |
| AP004607.5 | 11 | tripartite motif-containg 49-like (TRIM49L) ps |
| AP000827.1 | 11 | establishment of cohesion 1 homolog 1 (S. ce |
| UBTFL1 | 11 | upstream binding transcription factor like 1 [S |
| AP000648.2 | 11 | tripartite motif-containing protein 43 (TRIM43 |
| AP000648.9 | 11 | novel transcript |
| AP000648.1 | 11 | NULL |
| AP000648.3 | 11 | novel transcript |
| NAALAD2 | 11 | N-acetylated alpha-linked acidic dipeptidase 2 |
| AP000648.4 | 11 | novel transcript |
| CHORDC1 | 11 | cysteine and histidine rich domain containing |
| AP002364.1 | 11 | novel transcript |
| DISC1FP1 | 11 | DISC1 fusion partner 1 [Source:HGNC Symbol; |
| TUBAP2 | 11 | tubulin alpha pseudogene 2 [Source:HGNC Sy |
| MIR4490 | 11 | microRNA 4490 [Source:HGNC Symbol;Acc:HG |
| AP002782.1 | 11 | novel transcript |
| MIR1261 | 11 | microRNA 1261 [Source:HGNC Symbol;Acc:HG |
| AP001002.3 | 11 | novel transcript |
| AP001002.1 | 11 | novel transcript |
| AP001002.2 | 11 | novel transcript |
| OSBPL9P2 | 11 | oxysterol binding protein like 9 pseudogene 2 |
| OSBPL9P3 | 11 | oxysterol binding protein like 9 pseudogene 3 |
| LINC02748 | 11 | long intergenic non-protein coding RNA 2748 |

| | | |
|---|---|---|
| AP003485.2 | 11 | novel transcript |
| AP003485.1 | 11 | novel transcript |
| AP002791.1 | 11 | novel transcript |
| LINC02756 | 11 | long intergenic non-protein coding RNA 2756 |
| TUBB4BP4 | 11 | tubulin beta 4B class IVb pseudogene 4 [Sourc |
| AP002799.1 | 11 | novel transcript |
| RPL7AP57 | 11 | ribosomal protein L7a pseudogene 57 [Source |
| NDUFB11P1 | 11 | NADH:ubiquinone oxidoreductase subunit B1 |
| FAT3 | 11 | FAT atypical cadherin 3 [Source:HGNC Symbol |
| PGAM1P9 | 11 | phosphoglycerate mutase 1 pseudogene 9 [Sc |
| AP000722.1 | 11 | novel transcript |
| RPS3AP42 | 11 | ribosomal protein S3a pseudogene 42 [Source |
| AP003718.1 | 11 | novel transcript |
| LINC02746 | 11 | long intergenic non-protein coding RNA 2746 |
| AP003171.2 | 11 | eukaryotic translation elongation factor 1 alpl |
| SNRPGP16 | 11 | small nuclear ribonucleoprotein polypeptide ( |
| AP003171.1 | 11 | novel transcript |
| MTNR1B | 11 | melatonin receptor 1B [Source:HGNC Symbol; |
| RPL26P31 | 11 | ribosomal protein L26 pseudogene 31 [Source |
| AP002371.1 | 11 | novel transcript |
| AP003072.4 | 11 | novel transcript |
| SLC36A4 | 11 | solute carrier family 36 member 4 [Source:HG |
| AP003072.2 | 11 | aminoacyl tRNA synthetase complex-interacti |
| AP003072.5 | 11 | novel transcript |
| AP003072.3 | 11 | novel transcript |
| AP003072.1 | 11 | chromosome 14 open reading frame 2 (C14or |
| AP003969.1 | 11 | novel transcript |
| AP003969.2 | 11 | novel transcript |
| DEUP1 | 11 | deuterosome assembly protein 1 [Source:HGI |
| SMCO4 | 11 | single-pass membrane protein with coiled-coi |
| SRP14P2 | 11 | signal recognition particle 14 pseudogene 2 [S |
| RN7SL223P | 11 | RNA, 7SL, cytoplasmic 223, pseudogene [Sour |
| AP003499.5 | 11 | novel transcript |
| AP003499.4 | 11 | novel transcript |
| AP003499.1 | 11 | N-6 adenine-specific DNA methyltransferase 1 |
| CEP295 | 11 | centrosomal protein 295 [Source:HGNC Symb |
| Y_RNA | 11 | Y RNA [Source:RFAM;Acc:RF00019] |
| SCARNA9 | 11 | small Cajal body-specific RNA 9 [Source:HGNC |
| AP003499.3 | 11 | NULL |
| AP003499.2 | 11 | NULL |
| AP001273.1 | 11 | novel transcript |
| TAF1D | 11 | TATA-box binding protein associated factor, R |
| SNORA25 | 11 | small nucleolar RNA, H/ACA box 25 [Source:H |
| SNORA32 | 11 | small nucleolar RNA, H/ACA box 32 [Source:H |
| SNORD6 | 11 | small nucleolar RNA, C/D box 6 [Source:HGNC |
| SNORA1 | 11 | small nucleolar RNA, H/ACA box 1 [Source:HG |
| SNORA8 | 11 | small nucleolar RNA, H/ACA box 8 [Source:HG |
| SNORD5 | 11 | small nucleolar RNA, C/D box 5 [Source:HGNC |
| SNORA18 | 11 | small nucleolar RNA, H/ACA box 18 [Source:H |
| MIR1304 | 11 | microRNA 1304 [Source:HGNC Symbol;Acc:HG |

| | | |
|---|---|---|
| SNORA40 | 11 | small nucleolar RNA, H/ACA box 40 [Source:H |
| C11orf54 | 11 | chromosome 11 open reading frame 54 [Sour |
| AP001273.2 | 11 | novel protein, C11orf54-MED17 readthrough |
| MED17 | 11 | mediator complex subunit 17 [Source:HGNC S |
| VSTM5 | 11 | V-set and transmembrane domain containing |
| Y_RNA | 11 | Y RNA [Source:RFAM;Acc:RF00019] |
| AP002795.1 | 11 | hypoxanthine phosphoribosyltransferase pseu |
| HPRT1P3 | 11 | hypoxanthine phosphoribosyltransferase 1 ps |
| HEPHL1 | 11 | hephaestin like 1 [Source:HGNC Symbol;Acc:H |
| PHBP16 | 11 | prohibitin pseudogene 16 [Source:HGNC Symb |
| PANX1 | 11 | pannexin 1 [Source:HGNC Symbol;Acc:HGNC: |
| AP002784.1 | 11 | novel transcript |
| AP002784.2 | 11 | actin related protein 2/3 complex, subunit 3, |
| IZUMO1R | 11 | IZUMO1 receptor, JUNO [Source:HGNC Symbo |
| GPR83 | 11 | G protein-coupled receptor 83 [Source:HGNC |
| MRE11 | 11 | MRE11 homolog, double strand break repair r |
| MIR548L | 11 | microRNA 548l [Source:HGNC Symbol;Acc:HG |
| AP000786.1 | 11 | novel transcript |
| ANKRD49 | 11 | ankyrin repeat domain 49 [Source:HGNC Syml |
| C11orf97 | 11 | chromosome 11 open reading frame 97 [Sour |
| PIWIL4 | 11 | piwi like RNA-mediated gene silencing 4 [Sour |
| FUT4 | 11 | fucosyltransferase 4 [Source:HGNC Symbol;Ac |
| AP000943.2 | 11 | novel transcript |
| AP000943.3 | 11 | TEC |
| AP000943.1 | 11 | novel transcript |
| LINC02700 | 11 | long intergenic non-protein coding RNA 2700 |
| AMOTL1 | 11 | angiomotin like 1 [Source:HGNC Symbol;Acc:H |
| AP002383.2 | 11 | novel transcript |
| ST13P11 | 11 | ST13, Hsp70 interacting protein pseudogene 1 |
| CWC15 | 11 | CWC15 spliceosome associated protein homo |
| KDM4D | 11 | lysine demethylase 4D [Source:HGNC Symbol; |
| AP002383.1 | 11 | lysine (K)-specific demethylase 4D (KDM4D) p |
| KDM4E | 11 | lysine demethylase 4E [Source:HGNC Symbol; |
| AP002383.5 | 11 | lysine (K)-specific demethylase 4D (KDM4D) p |
| AP002383.3 | 11 | serine/arginine-rich splicing factor 8 (SRSF8) p |
| AP002383.4 | 11 | lysine (K)-specific demethylase 4D (KDM4D) p |
| KDM4F | 11 | lysine demethylase 4F [Source:HGNC Symbol; |
| AP001264.1 | 11 | lysine (K)-specific demethylase 4D (KDM4D) p |
| SRSF8 | 11 | serine and arginine rich splicing factor 8 [Sour |
| ENDOD1 | 11 | endonuclease domain containing 1 [Source:H |
| BUD13P1 | 11 | BUD13 homolog pseudogene 1 [Source:HGNC |
| AP000787.2 | 11 | splicing factor 3a, subunit 2, 66kDa, pseudoge |
| AP000787.1 | 11 | novel transcript, antisense to SESN3 |
| SESN3 | 11 | sestrin 3 [Source:HGNC Symbol;Acc:HGNC:23 |
| AP000787.3 | 11 | ribosomal protein S16 (RPS16) pseudogene |
| AP001790.1 | 11 | novel transcript |
| AP000820.2 | 11 | novel transcript |
| AP000820.1 | 11 | novel transcript |
| AP001877.1 | 11 | fibroblast growth factor receptor 3 (FGFR3) ps |
| FAM76B | 11 | family with sequence similarity 76 member B |

| | | |
|---|---|---|
| CEP57 | 11 | centrosomal protein 57 [Source:HGNC Symbo |
| MTMR2 | 11 | myotubularin related protein 2 [Source:HGNC |
| RNA5SP345 | 11 | RNA, 5S ribosomal pseudogene 345 [Source:H |
| AP000870.1 | 11 | ribosomal protein L32 (RPL32) pseudogene |
| MAML2 | 11 | mastermind like transcriptional coactivator 2 |
| AP000779.1 | 11 | novel transcript |
| MIR1260B | 11 | microRNA 1260b [Source:HGNC Symbol;Acc:H |
| CCDC82 | 11 | coiled-coil domain containing 82 [Source:HGN |
| JRKL | 11 | JRK like [Source:HGNC Symbol;Acc:HGNC:620 |
| JRKL-AS1 | 11 | JRKL antisense RNA 1 [Source:HGNC Symbol;A |
| RNA5SP346 | 11 | RNA, 5S ribosomal pseudogene 346 [Source:H |
| LINC02737 | 11 | long intergenic non-protein coding RNA 2737 |
| AP003066.1 | 11 | novel transcript |
| AP003066.2 | 11 | AKT interacting protein (AKTIP) pseudogene |
| MED28P5 | 11 | mediator complex subunit 28 pseudogene 5 [ |
| AP001836.1 | 11 | phosphoribosyl pyrophosphate synthetase 1-l |
| LINC02553 | 11 | long intergenic non-protein coding RNA 2553 |
| RNA5SP347 | 11 | RNA, 5S ribosomal pseudogene 347 [Source:H |
| LINC02713 | 11 | long intergenic non-protein coding RNA 2713 |
| AP003730.1 | 11 | high-mobility group nucleosome binding dom |
| AP001317.1 | 11 | novel pseudogene containing a Zinc finger, C2 |
| AP003038.1 | 11 | zinc finger protein 614 (ZNF614) pseudogene |
| AP003715.1 | 11 | novel transcript |
| AP002428.1 | 11 | novel transcript |
| CNTN5 | 11 | contactin 5 [Source:HGNC Symbol;Acc:HGNC: |
| RN7SKP53 | 11 | RN7SK pseudogene 53 [Source:HGNC Symbol; |
| RPA2P3 | 11 | replication protein A2 pseudogene 3 [Source: |
| RN7SL222P | 11 | RNA, 7SL, cytoplasmic 222, pseudogene [Sour |
| PPIAP43 | 11 | peptidylprolyl isomerase A pseudogene 43 [S |
| AP001351.1 | 11 | novel transcript, antisense to ARHGAP42 |
| ARHGAP42 | 11 | Rho GTPase activating protein 42 [Source:HG |
| RN7SKP115 | 11 | RN7SK pseudogene 115 [Source:HGNC Symbo |
| PGR | 11 | progesterone receptor [Source:HGNC Symbol |
| PGR-AS1 | 11 | PGR antisense RNA 1 [Source:HGNC Symbol;A |
| AP001533.1 | 11 | myotubularin related protein 12 (MTMR12) p |
| TRPC6 | 11 | transient receptor potential cation channel su |
| MIR3920 | 11 | microRNA 3920 [Source:HGNC Symbol;Acc:H |
| AP003080.1 | 11 | novel pseudogene |
| AP000776.1 | 11 | plastin 1 (PLS1) pseudogene |
| ANGPTL5 | 11 | angiopoietin like 5 [Source:HGNC Symbol;Acc: |
| CEP126 | 11 | centrosomal protein 126 [Source:HGNC Symb |
| CFAP300 | 11 | cilia and flagella associated protein 300 [Sour |
| RNA5SP535 | 11 | RNA, 5S ribosomal pseudogene 535 [Source:H |
| AP001527.1 | 11 | novel transcript |
| AP001527.2 | 11 | novel transcript |
| YAP1 | 11 | Yes associated protein 1 [Source:HGNC Symb |
| AP000942.2 | 11 | novel transcript |
| AP000942.1 | 11 | ribosomal protein S6 (RPS6) pseudogene |
| AP000942.3 | 11 | cytochrome c oxidase subunit VIIb, pseudoge |
| AP000942.4 | 11 | cytochrome c oxidase subunit VIIb, pseudoge |

| | | |
|---|---|---|
| RNU6-952P | 11 | RNA, U6 small nuclear 952, pseudogene [Sour |
| BIRC3 | 11 | baculoviral IAP repeat containing 3 [Source:H( |
| BIRC2 | 11 | baculoviral IAP repeat containing 2 [Source:H( |
| TMEM123 | 11 | transmembrane protein 123 [Source:HGNC Sy |
| AP001830.1 | 11 | novel transcript |
| AP001830.2 | 11 | novel transcript, antisense to TMEM123 |
| MMP7 | 11 | matrix metallopeptidase 7 [Source:HGNC Sym |
| MMP20 | 11 | matrix metallopeptidase 20 [Source:HGNC Syr |
| AP000851.1 | 11 | novel transcript |
| AP000851.2 | 11 | novel transcript |
| MMP27 | 11 | matrix metallopeptidase 27 [Source:HGNC Syr |
| MMP8 | 11 | matrix metallopeptidase 8 [Source:HGNC Sym |
| WTAPP1 | 11 | Wilms tumor 1 associated protein pseudogen |
| AP000619.1 | 11 | matrix metallopeptidase 1 (MMP1) pseudoger |
| AP000619.3 | 11 | spastic ataxia of Charlevoix-Saguenay (SACS) p |
| MMP10 | 11 | matrix metallopeptidase 10 [Source:HGNC Syr |
| MMP1 | 11 | matrix metallopeptidase 1 [Source:HGNC Sym |
| AP000619.2 | 11 | casein kinase 1, alpha 1 (CSNK1A1) pseudoger |
| MMP3 | 11 | matrix metallopeptidase 3 [Source:HGNC Sym |
| MMP12 | 11 | matrix metallopeptidase 12 [Source:HGNC Syr |
| BOLA3P1 | 11 | bolA family member 3 pseudogene 1 [Source: |
| RNU7-159P | 11 | RNA, U7 small nuclear 159 pseudogene [Sourc |
| MMP13 | 11 | matrix metallopeptidase 13 [Source:HGNC Syr |
| AP001486.3 | 11 | matrix metallopeptidase 8 (neutrophil collage |
| AP001486.1 | 11 | ribosomal protein L21 (RPL21) pseudogene |
| DCUN1D5 | 11 | defective in cullin neddylation 1 domain conta |
| AP001486.2 | 11 | novel transcript, overlapping to DCUN1D5 |
| DYNC2H1 | 11 | dynein cytoplasmic 2 heavy chain 1 [Source:H |
| AP002961.1 | 11 | novel transcript, antisense to DYNC2H1 |
| MTCO3P15 | 11 | MT-CO3 pseudogene 15 [Source:HGNC Symbo |
| MTATP6P15 | 11 | MT-ATP6 pseudogene 15 [Source:HGNC Symb |
| MTCO2P15 | 11 | MT-CO2 pseudogene 15 [Source:HGNC Symbo |
| MTCO1P15 | 11 | MT-CO1 pseudogene 15 [Source:HGNC Symbo |
| MTND2P26 | 11 | MT-ND2 pseudogene 26 [Source:HGNC Symbo |
| MTND1P36 | 11 | MT-ND1 pseudogene 36 [Source:HGNC Symbo |
| AP003461.1 | 11 | mitochondrially encoded NADH dehydrogenas |
| AP002989.1 | 11 | novel transcript |
| MIR4693 | 11 | microRNA 4693 [Source:HGNC Symbol;Acc:HG |
| PDGFD | 11 | platelet derived growth factor D [Source:HGN |
| AP003043.1 | 11 | novel transcript |
| DDI1 | 11 | DNA damage inducible 1 homolog 1 [Source:H |
| AP003304.1 | 11 | NULL |
| LINC02552 | 11 | long intergenic non-protein coding RNA 2552 |
| AP001485.1 | 11 | nucleophosmin (nucleolar phosphoprotein B2 |
| AP002004.1 | 11 | dynein, axonemal, heavy chain 2 (DNAH2) pse |
| CASP12 | 11 | caspase 12 (gene/pseudogene) [Source:HGNC |
| CASP17P | 11 | caspase 17, pseudogene [Source:HGNC Symbo |
| CASP4 | 11 | caspase 4 [Source:HGNC Symbol;Acc:HGNC:15 |
| AP001153.1 | 11 | novel caspase family pseudogene |
| CASP5 | 11 | caspase 5 [Source:HGNC Symbol;Acc:HGNC:15 |

| | | |
|---|---|---|
| CASP1 | 11 | caspase 1 [Source:HGNC Symbol;Acc:HGNC:1 |
| CARD16 | 11 | caspase recruitment domain family member 1 |
| AP001153.2 | 11 | caspase 1, apoptosis-related cysteine peptida |
| CASP1P2 | 11 | caspase 1 pseudogene 2 [Source:HGNC Symb |
| CARD17 | 11 | caspase recruitment domain family member 1 |
| CASP1P1 | 11 | caspase 1 pseudogene 1 [Source:HGNC Symb |
| CARD18 | 11 | caspase recruitment domain family member 1 |
| OR2AL1P | 11 | olfactory receptor family 2 subfamily AL mem |
| AP003181.1 | 11 | olfactory receptor pseudogene |
| Metazoa_SRP | 11 | Metazoan signal recognition particle RNA [Sou |
| GRIA4 | 11 | glutamate ionotropic receptor AMPA type sub |
| AP000641.1 | 11 | heat shock 60kDa protein 1 (chaperonin) (HSP |
| RNU4-55P | 11 | RNA, U4 small nuclear 55, pseudogene [Sourc |
| RNU6-277P | 11 | RNA, U6 small nuclear 277, pseudogene [Sour |
| AP000813.1 | 11 | novel transcript, antisense to MSANTD4 |
| MSANTD4 | 11 | Myb/SANT DNA binding domain containing 4 |
| KBTBD3 | 11 | kelch repeat and BTB domain containing 3 [So |
| AASDHPPT | 11 | aminoadipate-semialdehyde dehydrogenase-p |
| AP001001.1 | 11 | novel transcript |
| AP001001.2 | 11 | SET domain and mariner transposase fusion g |
| LINC02719 | 11 | long intergenic non-protein coding RNA 2719 |
| AP002001.3 | 11 | novel transcript |
| AP002001.1 | 11 | zinc finger protein family pseudogene |
| AP002001.2 | 11 | G1 to S phase transition 1 (GSPT1) pseudogen |
| AP003173.1 | 11 | novel transcript |
| GUCY1A2 | 11 | guanylate cyclase 1 soluble subunit alpha 2 [S |
| AP001282.2 | 11 | RecQ protein-like (DNA helicase Q1-like) (REC |
| AP001282.1 | 11 | pyridine nucleotide-disulphide oxidoreductase |
| AP003057.1 | 11 | argininosuccinate synthase 1 (ASS1) pseudoge |
| AP000766.1 | 11 | novel transcript |
| CWF19L2 | 11 | CWF19 like cell cycle control factor 2 [Source: |
| SMARCE1P1 | 11 | SWI/SNF related, matrix associated, actin dep |
| ALKBH8 | 11 | alkB homolog 8, tRNA methyltransferase [Sou |
| AP001823.1 | 11 | succinate-CoA ligase, GDP-forming, beta subu |
| ELMOD1 | 11 | ELMO domain containing 1 [Source:HGNC Syn |
| AP000889.2 | 11 | methyltransferase like 5 (METTL5) pseudogen |
| AP000889.1 | 11 | MOB1, Mps One Binder kinase activator-like 3 |
| SLN | 11 | sarcolipin [Source:HGNC Symbol;Acc:HGNC:1 |
| AP002353.1 | 11 | sorting nexin 7 (SNX7) pseudogene |
| SLC35F2 | 11 | solute carrier family 35 member F2 [Source:H |
| AP001024.1 | 11 | ribosomal protein, large, P2 (RPLP2) pseudoge |
| RAB39A | 11 | RAB39A, member RAS oncogene family [Sourc |
| AP003307.1 | 11 | novel transcript, antisense to CUL5 |
| CUL5 | 11 | cullin 5 [Source:HGNC Symbol;Acc:HGNC:255 |
| Y_RNA | 11 | Y RNA [Source:RFAM;Acc:RF00019] |
| AP002433.1 | 11 | novel transcript |
| ACAT1 | 11 | acetyl-CoA acetyltransferase 1 [Source:HGNC |
| AP002433.2 | 11 | novel transcript, antisense to ACAT1 |
| NPAT | 11 | nuclear protein, coactivator of histone transcr |
| ATM | 11 | ATM serine/threonine kinase [Source:HGNC S |

| | | |
|---|---|---|
| Y_RNA | 11 | Y RNA [Source:RFAM;Acc:RF00019] |
| C11orf65 | 11 | chromosome 11 open reading frame 65 [Sour |
| POGLUT3 | 11 | protein O-glucosyltransferase 3 [Source:HGN( |
| EXPH5 | 11 | exophilin 5 [Source:HGNC Symbol;Acc:HGNC:3 |
| DDX10 | 11 | DEAD-box helicase 10 [Source:HGNC Symbol;/ |
| RPS2P39 | 11 | ribosomal protein S2 pseudogene 39 [Source: |
| AP002453.1 | 11 | adenosine deaminase-like (ADAL) pseudogene |
| CYCSP29 | 11 | CYCS pseudogene 29 [Source:HGNC Symbol;A |
| AP003027.1 | 11 | novel transcript |
| AP003123.1 | 11 | novel transcript |
| RNU6-654P | 11 | RNA, U6 small nuclear 654, pseudogene [Sour |
| RNA5SP349 | 11 | RNA, 5S ribosomal pseudogene 349 [Source:H |
| SNORD39 | 11 | Small nucleolar RNA SNORD55/SNORD39 [Sou |
| AP003049.2 | 11 | novel transcript, antisense to C11orf87 |
| C11orf87 | 11 | chromosome 11 open reading frame 87 [Sour |
| AP003049.1 | 11 | CD47 molecule (CD47) pseudogene |
| AP003102.1 | 11 | chaperonin containing TCP1, subunit 6A (zeta |
| LINC02715 | 11 | long intergenic non-protein coding RNA 2715 |
| RDX | 11 | radixin [Source:HGNC Symbol;Acc:HGNC:9944 |
| TFAMP2 | 11 | transcription factor A, mitochondrial pseudog |
| AP001981.2 | 11 | novel transcript, antisense to RDX |
| RPSAP50 | 11 | ribosomal protein SA pseudogene 50 [Source: |
| AP001981.1 | 11 | retinoblastoma binding protein 8, pseudogene |
| AP001889.1 | 11 | polymerase (RNA) II (DNA directed) polypeptio |
| ZC3H12C | 11 | zinc finger CCCH-type containing 12C [Source: |
| LINC02732 | 11 | long intergenic non-protein coding RNA 2732 |
| FDX1 | 11 | ferredoxin 1 [Source:HGNC Symbol;Acc:HGNC |
| ARHGAP20 | 11 | Rho GTPase activating protein 20 [Source:HGI |
| HNRNPA1P60 | 11 | heterogeneous nuclear ribonucleoprotein A1 |
| AP002963.1 | 11 | mitochondrially encoded cytochrome b (MT-C |
| AP003973.4 | 11 | novel transcript |
| RNA5SP350 | 11 | RNA, 5S ribosomal pseudogene 350 [Source:H |
| AP003973.2 | 11 | novel transcript |
| LINC02550 | 11 | long intergenic non-protein coding RNA 2550 |
| AP003973.3 | 11 | novel transcript |
| RPS17P15 | 11 | ribosomal protein S17 pseudogene 15 [Source |
| AP003973.1 | 11 | PEST proteolytic signal containing nuclear pro |
| C11orf53 | 11 | chromosome 11 open reading frame 53 [Sour |
| COLCA1 | 11 | colorectal cancer associated 1 [Source:HGNC : |
| COLCA2 | 11 | colorectal cancer associated 2 [Source:HGNC : |
| AP002448.1 | 11 | novel transcript |
| MIR4491 | 11 | microRNA 4491 [Source:HGNC Symbol;Acc:HG |
| POU2AF1 | 11 | POU class 2 homeobox associating factor 1 [S |
| RNU2-60P | 11 | RNA, U2 small nuclear 60, pseudogene [Sourc |
| AP002008.3 | 11 | novel transcript, sense intronic to POU2AF1 |
| AP002008.2 | 11 | novel pseudogene |
| BTG4 | 11 | BTG anti-proliferation factor 4 [Source:HGNC : |
| AP002008.4 | 11 | novel transcript, antisense to BTG4 |
| MIR34B | 11 | microRNA 34b [Source:HGNC Symbol;Acc:HGI |
| MIR34C | 11 | microRNA 34c [Source:HGNC Symbol;Acc:HGI |

| | | |
|---|---|---|
| AP002008.1 | 11 | novel transcript |
| C11orf88 | 11 | chromosome 11 open reading frame 88 [Sour |
| LAYN | 11 | layilin [Source:HGNC Symbol;Acc:HGNC:29471 |
| SIK2 | 11 | salt inducible kinase 2 [Source:HGNC Symbol;/ |
| AP000925.1 | 11 | keratin 8 (KRT8) pseudogene |
| RN7SKP273 | 11 | RN7SK pseudogene 273 [Source:HGNC Symbo |
| PPP2R1B | 11 | protein phosphatase 2 scaffold subunit Abeta |
| AP001781.1 | 11 | novel transcript |
| ALG9 | 11 | ALG9 alpha-1,2-mannosyltransferase [Source: |
| AP001781.2 | 11 | novel protein |
| ALG9-IT1 | 11 | ALG9 intronic transcript 1 [Source:HGNC Symb |
| GNG5P3 | 11 | G protein subunit gamma 5 pseudogene 3 [So |
| FDXACB1 | 11 | ferredoxin-fold anticodon binding domain con |
| C11orf1 | 11 | chromosome 11 open reading frame 1 [Source |
| RPL37AP8 | 11 | ribosomal protein L37a pseudogene 8 [Source |
| CRYAB | 11 | crystallin alpha B [Source:HGNC Symbol;Acc:H |
| HSPB2 | 11 | heat shock protein family B (small) member 2 |
| HSPB2-C11orf52 | 11 | HSPB2-C11orf52 readthrough (NMD candidate |
| C11orf52 | 11 | chromosome 11 open reading frame 52 [Sour |
| DIXDC1 | 11 | DIX domain containing 1 [Source:HGNC Symbo |
| RNA5SP351 | 11 | RNA, 5S ribosomal pseudogene 351 [Source:H |
| AP000907.1 | 11 | ribosomal protein L34 (RPL34) pseudogene |
| AP000907.2 | 11 | novel transcript |
| DLAT | 11 | dihydrolipoamide S-acetyltransferase [Source |
| PPIHP1 | 11 | peptidylprolyl isomerase H pseudogene 1 [Sou |
| RNU6-893P | 11 | RNA, U6 small nuclear 893, pseudogene [Sour |
| AP000907.3 | 11 | ATP synthase, H+ transporting, mitochondrial |
| PIH1D2 | 11 | PIH1 domain containing 2 [Source:HGNC Symb |
| NKAPD1 | 11 | NKAP domain containing 1 [Source:HGNC Sym |
| TIMM8B | 11 | translocase of inner mitochondrial membrane |
| SDHD | 11 | succinate dehydrogenase complex subunit D [ |
| AP002884.2 | 11 | novel transcript |
| IL18 | 11 | interleukin 18 [Source:HGNC Symbol;Acc:HGN |
| AP002884.1 | 11 | novel transcript |
| TEX12 | 11 | testis expressed 12 [Source:HGNC Symbol;Acc |
| BCO2 | 11 | beta-carotene oxygenase 2 [Source:HGNC Syn |
| KCTD9P4 | 11 | potassium channel tetramerization domain cc |
| MRPS36P4 | 11 | mitochondrial ribosomal protein S36 pseudog |
| RPS12P21 | 11 | ribosomal protein S12 pseudogene 21 [Source |
| PTS | 11 | 6-pyruvoyltetrahydropterin synthase [Source: |
| RPS6P16 | 11 | ribosomal protein S6 pseudogene 16 [Source: |
| PLET1 | 11 | placenta expressed transcript 1 [Source:HGNC |
| AP002884.3 | 11 | novel transcript, antisense to C11orf34 |
| ST13P10 | 11 | ST13, Hsp70 interacting protein pseudogene 1 |
| LINC02762 | 11 | long intergenic non-protein coding RNA 2762 |
| RNU6-44P | 11 | RNA, U6 small nuclear 44, pseudogene [Sourc |
| LINC02763 | 11 | long intergenic non-protein coding RNA 2763 |
| RPL23AP62 | 11 | ribosomal protein L23a pseudogene 62 [Sourc |
| LINC02764 | 11 | long intergenic non-protein coding RNA 2764 |
| AP003063.1 | 11 | NULL |

| | | |
|---|---|---|
| AP003100.2 | 11 | novel transcript |
| AP003100.3 | 11 | novel transcript |
| AP003100.1 | 11 | novel transcript |
| AC016902.1 | 11 | novel transcript |
| AP000802.1 | 11 | novel transcript, antisense to NCAM1 |
| NCAM1 | 11 | neural cell adhesion molecule 1 [Source:HGN( |
| RNU7-187P | 11 | RNA, U7 small nuclear 187 pseudogene [Sourc |
| NCAM1-AS1 | 11 | NCAM1 antisense RNA1 [Source:HGNC Symbo |
| AP000880.1 | 11 | novel transcript |
| TTC12 | 11 | tetratricopeptide repeat domain 12 [Source:H |
| AP002840.2 | 11 | novel transcript |
| ANKK1 | 11 | ankyrin repeat and kinase domain containing |
| AP002840.1 | 11 | novel transcript |
| DRD2 | 11 | dopamine receptor D2 [Source:HGNC Symbol; |
| MIR4301 | 11 | microRNA 4301 [Source:HGNC Symbol;Acc:HG |
| TMPRSS5 | 11 | transmembrane serine protease 5 [Source:HG |
| MTRF1LP1 | 11 | mitochondrial translational release factor 1 lik |
| ZW10 | 11 | zw10 kinetochore protein [Source:HGNC Syml |
| RPS29P19 | 11 | ribosomal protein S29 pseudogene 19 [Source |
| AP003170.5 | 11 | cytochrome c oxidase subunit VIIa polypeptid |
| AP003170.4 | 11 | novel transcript |
| CLDN25 | 11 | claudin 25 [Source:HGNC Symbol;Acc:HGNC:3 |
| ATF4P4 | 11 | activating transcription factor 4 pseudogene 4 |
| AP003170.2 | 11 | SH3-domain GRB2-like 1 (SH3GL1) pseudogen |
| LRRC37A13P | 11 | leucine rich repeat containing 37 member A1: |
| USP28 | 11 | ubiquitin specific peptidase 28 [Source:HGNC |
| AP003170.3 | 11 | novel transcript |
| AP003170.1 | 11 | interferon stimulated exonuclease gene 20kD. |
| RNU6-1107P | 11 | RNA, U6 small nuclear 1107, pseudogene [Sou |
| HTR3B | 11 | 5-hydroxytryptamine receptor 3B [Source:HG |
| HTR3A | 11 | 5-hydroxytryptamine receptor 3A [Source:HG |
| ZBTB16 | 11 | zinc finger and BTB domain containing 16 [Sou |
| AP002755.1 | 11 | novel transcript, antisense to ZBTB16 |
| AP002518.2 | 11 | novel transcript |
| NNMT | 11 | nicotinamide N-methyltransferase [Source:HC |
| AP002518.1 | 11 | novel transcript |
| C11orf71 | 11 | chromosome 11 open reading frame 71 [Sour |
| RBM7 | 11 | RNA binding motif protein 7 [Source:HGNC Sy |
| AP002373.1 | 11 | novel transcript, RBM7-REXO2 readthrough |
| REXO2 | 11 | RNA exonuclease 2 [Source:HGNC Symbol;Acc |
| AP002373.2 | 11 | CCR4 carbon catabolite repression 4-like (S. ce |
| NXPE2P1 | 11 | neurexophilin and PC-esterase domain family |
| AC020549.3 | 11 | phospholipase A2, group XIIA, pseudogene |
| NXPE1 | 11 | neurexophilin and PC-esterase domain family |
| AC020549.1 | 11 | NULL |
| NXPE4 | 11 | neurexophilin and PC-esterase domain family |
| AC020549.2 | 11 | histone deacetylase 1 (HDAC1) pseudogene |
| NXPE2 | 11 | neurexophilin and PC-esterase domain family |
| AP003179.1 | 11 | NULL |
| CADM1 | 11 | cell adhesion molecule 1 [Source:HGNC Symb |

| | | |
|---|---|---|
| AP000465.1 | 11 | novel transcript, sense intronic to CADM1 |
| AP000462.3 | 11 | novel transcript |
| AP000462.1 | 11 | novel transcript |
| AP000462.2 | 11 | novel transcript |
| AP003174.2 | 11 | novel transcript |
| AP003174.3 | 11 | ferritin, light polypeptide (FTL) pseudogene |
| RPL12P46 | 11 | ribosomal protein L12 pseudogene 46 [Source |
| AP003174.1 | 11 | novel transcript |
| AP000997.1 | 11 | NULL |
| AP000997.2 | 11 | novel transcript |
| AP000997.3 | 11 | novel transcript |
| AP000997.4 | 11 | novel transcript |
| LINC02698 | 11 | long intergenic non-protein coding RNA 2698 |
| AP002991.1 | 11 | novel transcript |
| AP002991.2 | 11 | novel transcript |
| LINC00900 | 11 | long intergenic non-protein coding RNA 900 [S |
| LINC02703 | 11 | long intergenic non-protein coding RNA 2703 |
| AP000797.2 | 11 | ribosomal protein L15 (RPL15) pseudogene |
| AP000797.1 | 11 | ribosomal protein L5 (RPL5) pseudogene |
| LINC02151 | 11 | long intergenic non-protein coding RNA 2151 |
| AP000770.1 | 11 | TEC |
| LINC02702 | 11 | long intergenic non-protein coding RNA 2702 |
| BUD13 | 11 | BUD13 homolog [Source:HGNC Symbol;Acc:H |
| AP006216.1 | 11 | novel transcript |
| ZPR1 | 11 | ZPR1 zinc finger [Source:HGNC Symbol;Acc:HC |
| APOA5 | 11 | apolipoprotein A5 [Source:HGNC Symbol;Acc: |
| AP006216.2 | 11 | novel transcript |
| AP006216.3 | 11 | novel transcript, antisense to APOA4 |
| APOA4 | 11 | apolipoprotein A4 [Source:HGNC Symbol;Acc: |
| APOC3 | 11 | apolipoprotein C3 [Source:HGNC Symbol;Acc: |
| APOA1 | 11 | apolipoprotein A1 [Source:HGNC Symbol;Acc: |
| APOA1-AS | 11 | APOA1 antisense RNA [Source:HGNC Symbol;A |
| SIK3 | 11 | SIK family kinase 3 [Source:HGNC Symbol;Acc: |
| RNY4P6 | 11 | RNY4 pseudogene 6 [Source:HGNC Symbol;Ac |
| AP000936.3 | 11 | ribosomal protein S27 (metallopanstimulin 1) |
| AP000936.1 | 11 | novel transcript |
| AP000936.2 | 11 | cleavage and polyadenylation specific factor 5 |
| AP005018.2 | 11 | proprotein convertase subtilisin/kexin type 7 ( |
| AP005018.1 | 11 | chromosome 18 open reading frame 12 (C18o |
| PAFAH1B2 | 11 | platelet activating factor acetylhydrolase 1b ca |
| SIDT2 | 11 | SID1 transmembrane family member 2 [Sourc |
| TAGLN | 11 | transgelin [Source:HGNC Symbol;Acc:HGNC:1 |
| PCSK7 | 11 | proprotein convertase subtilisin/kexin type 7 [ |
| AP000892.4 | 11 | novel transcript |
| RNF214 | 11 | ring finger protein 214 [Source:HGNC Symbol; |
| AP000892.2 | 11 | NULL |
| BACE1 | 11 | beta-secretase 1 [Source:HGNC Symbol;Acc:H |
| BACE1-AS | 11 | BACE1 antisense RNA [Source:HGNC Symbol;A |
| AP000892.3 | 11 | novel transcript, sense intronic to BACE1 |
| CEP164 | 11 | centrosomal protein 164 [Source:HGNC Symb |

| | | |
|---|---|---|
| AP000892.1 | 11 | novel transcript, antisense to CEP164 |
| PRR13P3 | 11 | proline rich 13 pseudogene 3 [Source:HGNC S |
| DSCAML1 | 11 | DS cell adhesion molecule like 1 [Source:HGN |
| AP000711.1 | 11 | novel transcript |
| AP001554.1 | 11 | novel transcript |
| FXYD2 | 11 | FXYD domain containing ion transport regulat |
| AP000757.2 | 11 | novel transcript, antisense to FXYD2 |
| FXYD6-FXYD2 | 11 | FXYD6-FXYD2 readthrough [Source:HGNC Sym |
| AP000757.1 | 11 | novel transcript, antisense to FXYD6 |
| FXYD6 | 11 | FXYD domain containing ion transport regulat |
| TMPRSS13 | 11 | transmembrane serine protease 13 [Source:H |
| IL10RA | 11 | interleukin 10 receptor subunit alpha [Source: |
| SMIM35 | 11 | small integral membrane protein 35 [Source:H |
| RN7SL828P | 11 | RNA, 7SL, cytoplasmic 828, pseudogene [Sour |
| TMPRSS4 | 11 | transmembrane serine protease 4 [Source:HG |
| SCN4B | 11 | sodium voltage-gated channel beta subunit 4 |
| SCN2B | 11 | sodium voltage-gated channel beta subunit 2 |
| JAML | 11 | junction adhesion molecule like [Source:HGN |
| HSPE1P18 | 11 | heat shock protein family E (Hsp10) member |
| MPZL3 | 11 | myelin protein zero like 3 [Source:HGNC Symb |
| MPZL2 | 11 | myelin protein zero like 2 [Source:HGNC Symb |
| AP002800.1 | 11 | TEC |
| CD3E | 11 | CD3e molecule [Source:HGNC Symbol;Acc:HG |
| AP001582.1 | 11 | carbohydrate (N-acetylglucosamine 6-O) sulfo |
| CD3D | 11 | CD3d molecule [Source:HGNC Symbol;Acc:HG |
| CD3G | 11 | CD3g molecule [Source:HGNC Symbol;Acc:HG |
| UBE4A | 11 | ubiquitination factor E4A [Source:HGNC Symb |
| AP001267.1 | 11 | novel transcript, antisense to ATP5L and UBE4 |
| AP001267.5 | 11 | novel protein |
| ATP5MG | 11 | ATP synthase membrane subunit g [Source:H |
| AP001267.4 | 11 | novel transcript |
| AP001267.2 | 11 | novel transcript |
| KMT2A | 11 | lysine methyltransferase 2A [Source:HGNC Sy |
| AP001267.3 | 11 | novel transcript, antisense to MLL and TTC36 |
| TTC36 | 11 | tetratricopeptide repeat domain 36 [Source:H |
| TMEM25 | 11 | transmembrane protein 25 [Source:HGNC Syn |
| IFT46 | 11 | intraflagellar transport 46 [Source:HGNC Sym |
| RPL5P30 | 11 | ribosomal protein L5 pseudogene 30 [Source: |
| ARCN1 | 11 | archain 1 [Source:HGNC Symbol;Acc:HGNC:64 |
| RNU6-1157P | 11 | RNA, U6 small nuclear 1157, pseudogene [Sou |
| PHLDB1 | 11 | pleckstrin homology like domain family B mer |
| AP000941.1 | 11 | novel transcript |
| MIR6716 | 11 | microRNA 6716 [Source:HGNC Symbol;Acc:HG |
| TREH | 11 | trehalase [Source:HGNC Symbol;Acc:HGNC:12 |
| TREHP1 | 11 | trehalase pseudogene 1 [Source:HGNC Symbo |
| AP002954.1 | 11 | novel transcript |
| RNU6-376P | 11 | RNA, U6 small nuclear 376, pseudogene [Sour |
| AP002954.2 | 11 | novel transcript |
| DDX6 | 11 | DEAD-box helicase 6 [Source:HGNC Symbol;A |
| AP004609.3 | 11 | novel transcript |

| | | |
|---|---|---|
| AP004609.2 | 11 | cytochrome c oxidase subunit VIIc (COX7C) ps |
| SETP16 | 11 | SET pseudogene 16 [Source:HGNC Symbol;Acc |
| Y_RNA | 11 | Y RNA [Source:RFAM;Acc:RF00019] |
| CXCR5 | 11 | C-X-C motif chemokine receptor 5 [Source:HG |
| AP004609.1 | 11 | novel transcript, antisense to CXCR5 |
| BCL9L | 11 | BCL9 like [Source:HGNC Symbol;Acc:HGNC:23 |
| MIR4492 | 11 | microRNA 4492 [Source:HGNC Symbol;Acc:HG |
| UPK2 | 11 | uroplakin 2 [Source:HGNC Symbol;Acc:HGNC: |
| RN7SL688P | 11 | RNA, 7SL, cytoplasmic 688, pseudogene [Sour |
| Y_RNA | 11 | Y RNA [Source:RFAM;Acc:RF00019] |
| FOXR1 | 11 | forkhead box R1 [Source:HGNC Symbol;Acc:H |
| Metazoa_SRP | 11 | Metazoan signal recognition particle RNA [Sou |
| Y_RNA | 11 | Y RNA [Source:RFAM;Acc:RF00019] |
| CCDC84-DT | 11 | CCDC84 divergent transcript [Source:HGNC Sy |
| CCDC84 | 11 | coiled-coil domain containing 84 [Source:HGN |
| RPL23AP64 | 11 | ribosomal protein L23a pseudogene 64 [Sourc |
| AP003392.1 | 11 | novel transcript, antisense to CCDC84 |
| AP003392.2 | 11 | potassium channel tetramerisation domain co |
| RPS25 | 11 | ribosomal protein S25 [Source:HGNC Symbol; |
| TRAPPC4 | 11 | trafficking protein particle complex 4 [Source: |
| AP003392.11 | 11 | NULL |
| SLC37A4 | 11 | solute carrier family 37 member 4 [Source:HG |
| HYOU1 | 11 | hypoxia up-regulated 1 [Source:HGNC Symbol |
| AP003392.3 | 11 | novel transcript, antisense to HYOU1 |
| AP003392.4 | 11 | novel transcript |
| AP003392.5 | 11 | novel transcript, antisense to VPS11 |
| VPS11 | 11 | VPS11 core subunit of CORVET and HOPS com |
| HMBS | 11 | hydroxymethylbilane synthase [Source:HGNC |
| H2AFX | 11 | H2A histone family member X [Source:HGNC S |
| DPAGT1 | 11 | dolichyl-phosphate N-acetylglucosaminephos |
| C2CD2L | 11 | C2CD2 like [Source:HGNC Symbol;Acc:HGNC:2 |
| HINFP | 11 | histone H4 transcription factor [Source:HGNC |
| ABCG4 | 11 | ATP binding cassette subfamily G member 4 [S |
| NLRX1 | 11 | NLR family member X1 [Source:HGNC Symbol |
| PDZD3 | 11 | PDZ domain containing 3 [Source:HGNC Symb |
| CCDC153 | 11 | coiled-coil domain containing 153 [Source:HG |
| CBL | 11 | Cbl proto-oncogene [Source:HGNC Symbol;Ac |
| RNU6-262P | 11 | RNA, U6 small nuclear 262, pseudogene [Sour |
| MCAM | 11 | melanoma cell adhesion molecule [Source:HG |
| MIR6756 | 11 | microRNA 6756 [Source:HGNC Symbol;Acc:HG |
| AP002956.1 | 11 | NULL |
| RNF26 | 11 | ring finger protein 26 [Source:HGNC Symbol;A |
| AP003396.1 | 11 | novel transcript, antisense to RNF26 |
| MFRP | 11 | membrane frizzled-related protein [Source:HG |
| C1QTNF5 | 11 | C1q and TNF related 5 [Source:HGNC Symbol; |
| USP2 | 11 | ubiquitin specific peptidase 2 [Source:HGNC S |
| USP2-AS1 | 11 | USP2 antisense RNA 1 (head to head) [Source |
| AP003396.3 | 11 | novel transcript, antisense to USP2 |
| AP003396.2 | 11 | hippocampus abundant transcript 1 (HIAT1) p |
| AP003396.4 | 11 | novel pseudogene |

| | | |
|---|---|---|
| THY1 | 11 | Thy-1 cell surface antigen [Source:HGNC Symt |
| AP003396.5 | 11 | novel transcript |
| DUXAP5 | 11 | double homeobox A pseudogene 5 [Source:H( |
| KRT8P7 | 11 | keratin 8 pseudogene 7 [Source:HGNC Symbo |
| AP003393.1 | 11 | novel transcript, antisense to PVRL1 |
| NECTIN1 | 11 | nectin cell adhesion molecule 1 [Source:HGN( |
| RNU6-1123P | 11 | RNA, U6 small nuclear 1123, pseudogene [Sou |
| NECTIN1-AS1 | 11 | NECTIN1 antisense RNA 1 [Source:HGNC Symt |
| AP003390.1 | 11 | novel transcript |
| AP001994.1 | 11 | HLA-B associated transcript 1 (BAT1) pseudog |
| AP001994.2 | 11 | novel transcript |
| AP001994.3 | 11 | novel transcript |
| AP001360.3 | 11 | novel transcript |
| LINC02744 | 11 | long intergenic non-protein coding RNA 2744 |
| AP001360.1 | 11 | novel transcript |
| AP001360.2 | 11 | novel transcript |
| TRIM29 | 11 | tripartite motif containing 29 [Source:HGNC S |
| AP000679.1 | 11 | novel transcript, antisense to TRIM29 |
| OAF | 11 | out at first homolog [Source:HGNC Symbol;Ac |
| POU2F3 | 11 | POU class 2 homeobox 3 [Source:HGNC Symb |
| AP001150.1 | 11 | novel transcript, antisense to POU2F3 |
| TLCD5 | 11 | TLC domain containing 5 [Source:HGNC Symb( |
| ARHGEF12 | 11 | Rho guanine nucleotide exchange factor 12 [S |
| 7SK | 11 | 7SK RNA [Source:RFAM;Acc:RF00100] |
| GRIK4 | 11 | glutamate ionotropic receptor kainate type su |
| ELOCP22 | 11 | elongin C pseudogene 22 [Source:HGNC Symb |
| AP004147.1 | 11 | novel transcript, antisense to GRIK4 |
| HMGB1P42 | 11 | high mobility group box 1 pseudogene 42 [Sou |
| TBCEL | 11 | tubulin folding cofactor E like [Source:HGNC S |
| AP000646.1 | 11 | novel protein |
| TECTA | 11 | tectorin alpha [Source:HGNC Symbol;Acc:HGN |
| AP001124.1 | 11 | ransient receptor potential cation channel, su |
| RPS4XP12 | 11 | ribosomal protein S4X pseudogene 12 [Source |
| SC5D | 11 | sterol-C5-desaturase [Source:HGNC Symbol;A |
| BMPR1AP2 | 11 | bone morphogenetic protein receptor type 1/ |
| AP002365.1 | 11 | programmed cell death 2, pseudogene |
| AP000977.1 | 11 | novel transcript, antisense to SORL1 |
| SORL1 | 11 | sortilin related receptor 1 [Source:HGNC Symt |
| AP001977.1 | 11 | novel transcript |
| AP001924.1 | 11 | novel transcript |
| RNU6-256P | 11 | RNA, U6 small nuclear 256, pseudogene [Sour |
| AP001924.2 | 11 | novel transcript |
| MIR100HG | 11 | mir-100-let-7a-2-mir-125b-1 cluster host gene |
| MIR125B1 | 11 | microRNA 125b-1 [Source:HGNC Symbol;Acc:I |
| BLID | 11 | BH3-like motif containing, cell death inducer [ |
| MIRLET7A2 | 11 | microRNA let-7a-2 [Source:HGNC Symbol;Acc: |
| MIR100 | 11 | microRNA 100 [Source:HGNC Symbol;Acc:HGI |
| AP000755.1 | 11 | novel transcript |
| AP000755.2 | 11 | novel transcript |
| RNU4ATAC10P | 11 | RNA, U4atac small nuclear 10, pseudogene [S( |

| | | |
|---|---|---|
| RNU4ATAC5P | 11 | RNA, U4atac small nuclear 5, pseudogene [Sou |
| RNU6-592P | 11 | RNA, U6 small nuclear 592, pseudogene [Sour |
| GLULP3 | 11 | glutamate-ammonia ligase pseudogene 3 [Sou |
| AP002469.1 | 11 | scaffold attachment factor B (SAFB) pseudoge |
| UBASH3B | 11 | ubiquitin associated and SH3 domain containi |
| AP002762.3 | 11 | novel transcript, antisense to UBASH3B |
| AP002762.1 | 11 | NULL |
| AP002762.2 | 11 | novel transcript, antisense to CRTAM |
| CRTAM | 11 | cytotoxic and regulatory T cell molecule [Sour |
| JHY | 11 | junctional cadherin complex regulator [Source |
| RNU4-23P | 11 | RNA, U4 small nuclear 23, pseudogene [Sourc |
| ATP5PBP5 | 11 | ATP synthase peripheral stalk-membrane subu |
| BSX | 11 | brain specific homeobox [Source:HGNC Symbo |
| AP003040.1 | 11 | SUMO-1 activating enzyme subunit 1 (SAE1) p |
| RPL34P23 | 11 | ribosomal protein L34 pseudogene 23 [Source |
| RPS26P43 | 11 | ribosomal protein S26 pseudogene 43 [Source |
| RPL31P47 | 11 | ribosomal protein L31 pseudogene 47 [Source |
| HSPA8 | 11 | heat shock protein family A (Hsp70) member |
| SNORD14E | 11 | small nucleolar RNA, C/D box 14E [Source:HG |
| SNORD14D | 11 | small nucleolar RNA, C/D box 14D [Source:HG |
| SNORD14C | 11 | small nucleolar RNA, C/D box 14C [Source:HG |
| AP000926.2 | 11 | novel transcript, antisense to HSPA8and CLMI |
| CLMP | 11 | CXADR like membrane protein [Source:HGNC |
| AP000926.1 | 11 | FUN14 domain containing 1 (FUNDC1) pseudo |
| AP001970.1 | 11 | novel transcript |
| U8 | 11 | U8 small nucleolar RNA [Source:RFAM;Acc:RF |
| LINC02727 | 11 | long intergenic non-protein coding RNA 2727 |
| PHBP17 | 11 | prohibitin pseudogene 17 [Source:HGNC Syml |
| GRAMD1B | 11 | GRAM domain containing 1B [Source:HGNC Sy |
| MIR4493 | 11 | microRNA 4493 [Source:HGNC Symbol;Acc:HG |
| SF3A3P2 | 11 | splicing factor 3a, subunit 3 pseudogene 2 [So |
| SCN3B | 11 | sodium voltage-gated channel beta subunit 3 |
| AP002765.1 | 11 | novel transcript |
| ZNF202 | 11 | zinc finger protein 202 [Source:HGNC Symbol; |
| OR6X1 | 11 | olfactory receptor family 6 subfamily X memb |
| RNU1-21P | 11 | RNA, U1 small nuclear 21, pseudogene [Sourc |
| OR6M1 | 11 | olfactory receptor family 6 subfamily M meml |
| OR6M2P | 11 | olfactory receptor family 6 subfamily M meml |
| OR6M3P | 11 | olfactory receptor family 6 subfamily M meml |
| TMEM225 | 11 | transmembrane protein 225 [Source:HGNC Sy |
| OR8D4 | 11 | olfactory receptor family 8 subfamily D memb |
| OR4D5 | 11 | olfactory receptor family 4 subfamily D memb |
| OR6T1 | 11 | olfactory receptor family 6 subfamily T memb |
| OR10S1 | 11 | olfactory receptor family 10 subfamily S mem |
| OR10G6 | 11 | olfactory receptor family 10 subfamily G mem |
| OR10G5P | 11 | olfactory receptor family 10 subfamily G mem |
| OR10G4 | 11 | olfactory receptor family 10 subfamily G mem |
| OR10G9 | 11 | olfactory receptor family 10 subfamily G mem |
| OR10G8 | 11 | olfactory receptor family 10 subfamily G mem |
| OR10G7 | 11 | olfactory receptor family 10 subfamily G mem |

| | | |
|---|---|---|
| OR10D5P | 11 | olfactory receptor family 10 subfamily D mem |
| OR10D4P | 11 | olfactory receptor family 10 subfamily D mem |
| OR10N1P | 11 | olfactory receptor family 10 subfamily N mem |
| VWA5A | 11 | von Willebrand factor A domain containing 5A |
| OR10D1P | 11 | olfactory receptor family 10 subfamily D mem |
| OR10D3 | 11 | olfactory receptor family 10 subfamily D mem |
| OR8F1P | 11 | olfactory receptor family 8 subfamily F memb |
| OR8G3P | 11 | olfactory receptor family 8 subfamily G memb |
| OR8G2P | 11 | olfactory receptor family 8 subfamily G memb |
| OR8G7P | 11 | olfactory receptor family 8 subfamily G memb |
| OR8G1 | 11 | olfactory receptor family 8 subfamily G memb |
| OR8G5 | 11 | olfactory receptor family 8 subfamily G memb |
| SLC5A4P1 | 11 | solute carrier family 5 member 4 pseudogene |
| OR8D1 | 11 | olfactory receptor family 8 subfamily D memb |
| OR8D2 | 11 | olfactory receptor family 8 subfamily D memb |
| OR8B7P | 11 | olfactory receptor family 8 subfamily B memb |
| OR8B6P | 11 | olfactory receptor family 8 subfamily B memb |
| OR8B5P | 11 | olfactory receptor family 8 subfamily B memb |
| OR8B1P | 11 | olfactory receptor family 8 subfamily B memb |
| OR8C1P | 11 | olfactory receptor family 8 subfamily C memb |
| OR8B2 | 11 | olfactory receptor family 8 subfamily B memb |
| OR8B3 | 11 | olfactory receptor family 8 subfamily B memb |
| OR8X1P | 11 | olfactory receptor family 8 subfamily X memb |
| AP001804.1 | 11 | novel transcript |
| OR8B4 | 11 | olfactory receptor family 8 subfamily B memb |
| OR8B8 | 11 | olfactory receptor family 8 subfamily B memb |
| OR8A2P | 11 | olfactory receptor family 8 subfamily A memb |
| OR8B9P | 11 | olfactory receptor family 8 subfamily B memb |
| OR8B10P | 11 | olfactory receptor family 8 subfamily B memb |
| OR8A3P | 11 | olfactory receptor family 8 subfamily A memb |
| AP000916.1 | 11 | olfactory receptor, family 8, subfamily B, mem |
| OR8B12 | 11 | olfactory receptor family 8 subfamily B memb |
| OR8A1 | 11 | olfactory receptor family 8 subfamily A memb |
| OR8Q1P | 11 | olfactory receptor family 8 subfamily Q memb |
| PANX3 | 11 | pannexin 3 [Source:HGNC Symbol;Acc:HGNC:2 |
| TBRG1 | 11 | transforming growth factor beta regulator 1 [S |
| SIAE | 11 | sialic acid acetylesterase [Source:HGNC Symbo |
| RNA5SP352 | 11 | RNA, 5S ribosomal pseudogene 352 [Source:H |
| SPA17 | 11 | sperm autoantigenic protein 17 [Source:HGNC |
| NRGN | 11 | neurogranin [Source:HGNC Symbol;Acc:HGNC |
| AP000866.5 | 11 | novel transcript |
| VSIG2 | 11 | V-set and immunoglobulin domain containing |
| ESAM | 11 | endothelial cell adhesion molecule [Source:HG |
| AP000866.2 | 11 | novel transcript, antisense to ESAM |
| MSANTD2 | 11 | Myb/SANT DNA binding domain containing 2 [ |
| AP000866.6 | 11 | novel transcript, sense intronic to MSANTD2 |
| AP000866.3 | 11 | endothelial cell adhesion molecule (ESAM) pse |
| AP000866.1 | 11 | novel transcript, antisense to C11orf61 |
| AP000866.4 | 11 | ring finger protein 181 (RNF181) pseudogene |
| ROBO3 | 11 | roundabout guidance receptor 3 [Source:HGN |

| | | |
|---|---|---|
| ROBO4 | 11 | roundabout guidance receptor 4 [Source:HGN |
| AP003501.1 | 11 | novel transcript |
| AP003501.2 | 11 | novel transcript, antisense to ROBO4 |
| HEPACAM | 11 | hepatic and glial cell adhesion molecule [Sour |
| HEPN1 | 11 | hepatocellular carcinoma, down-regulated 1 [ |
| CCDC15-DT | 11 | CCDC15 divergent transcript [Source:HGNC Sy |
| CCDC15 | 11 | coiled-coil domain containing 15 [Source:HGN |
| SLC37A2 | 11 | solute carrier family 37 member 2 [Source:HG |
| TMEM218 | 11 | transmembrane protein 218 [Source:HGNC Sy |
| KRT18P59 | 11 | keratin 18 pseudogene 59 [Source:HGNC Sym |
| AP001007.2 | 11 | novel transcript |
| PKNOX2-AS1 | 11 | PKNOX2 antisense RNA 1 (head to head) [Sour |
| PKNOX2 | 11 | PBX/knotted 1 homeobox 2 [Source:HGNC Syr |
| AP001007.3 | 11 | novel transcript, sense intronic to PKNOX2 |
| AP001007.1 | 11 | novel transcript, antisense to PKNOX2 |
| RNU6-321P | 11 | RNA, U6 small nuclear 321, pseudogene [Sour |
| FEZ1 | 11 | fasciculation and elongation protein zeta 1 [Sc |
| AP000708.1 | 11 | novel transcript, antisense to FEZ1 |
| EI24 | 11 | EI24 autophagy associated transmembrane pr |
| RNU6-1156P | 11 | RNA, U6 small nuclear 1156, pseudogene [Sou |
| STT3A | 11 | STT3 oligosaccharyltransferase complex cataly |
| CHEK1 | 11 | checkpoint kinase 1 [Source:HGNC Symbol;Ac |
| RNU2-35P | 11 | RNA, U2 small nuclear 35, pseudogene [Sourc |
| ACRV1 | 11 | acrosomal vesicle protein 1 [Source:HGNC Syr |
| PATE1 | 11 | prostate and testis expressed 1 [Source:HGNC |
| PATE2 | 11 | prostate and testis expressed 2 [Source:HGNC |
| PATE3 | 11 | prostate and testis expressed 3 [Source:HGNC |
| PATE4 | 11 | prostate and testis expressed 4 [Source:HGNC |
| AP000842.3 | 11 | novel transcript |
| HYLS1 | 11 | HYLS1 centriolar and ciliogenesis associated [S |
| PUS3 | 11 | pseudouridine synthase 3 [Source:HGNC Symb |
| AP000842.2 | 11 | novel transcript, antisense to DDX25 |
| DDX25 | 11 | DEAD-box helicase 25 [Source:HGNC Symbol;A |
| AP000842.1 | 11 | novel transcript |
| VSIG10L2 | 11 | V-set and immunoglobulin domain containing |
| CDON | 11 | cell adhesion associated, oncogene regulated |
| NAP1L1P1 | 11 | nucleosome assembly protein 1 like 1 pseudo |
| AP000821.1 | 11 | novel transcript |
| AP000821.2 | 11 | novel transcript |
| AP001893.3 | 11 | novel transcript |
| AP001893.2 | 11 | EA domain family member 1 (SV40 transcripti |
| RPUSD4 | 11 | RNA pseudouridine synthase D4 [Source:HGN |
| AP001893.1 | 11 | novel transcript |
| FAM118B | 11 | family with sequence similarity 118 member E |
| RNU4-86P | 11 | RNA, U4 small nuclear 86, pseudogene [Sourc |
| RN7SL351P | 11 | RNA, 7SL, cytoplasmic 351, pseudogene [Sour |
| SRPRA | 11 | SRP receptor subunit alpha [Source:HGNC Syn |
| FOXRED1 | 11 | FAD dependent oxidoreductase domain conta |
| RPL35AP26 | 11 | ribosomal protein L35a pseudogene 26 [Sourc |
| TIRAP | 11 | TIR domain containing adaptor protein [Sourc |

| | | |
|---|---|---|
| AP001318.1 | 11 | novel transcript, antisense to TIRAP |
| AP001318.2 | 11 | novel transcript, antisense to DCPS and TIRAP |
| DCPS | 11 | decapping enzyme, scavenger [Source:HGNC S |
| GSEC | 11 | G-quadruplex forming sequence containing ln |
| ST3GAL4 | 11 | ST3 beta-galactoside alpha-2,3-sialyltransfera: |
| AP001318.3 | 11 | Fc fragment of IgG, low affinity IIa, receptor (( |
| KIRREL3 | 11 | kirre like nephrin family adhesion molecule 3 |
| KIRREL3-AS1 | 11 | KIRREL3 antisense RNA 1 [Source:HGNC Symb |
| AP001783.1 | 11 | novel transcript, antisense to KIRREL3 |
| AP002833.1 | 11 | novel transcript, sense intronic to KIRREL3 |
| AP002833.3 | 11 | TEC |
| KIRREL3-AS2 | 11 | KIRREL3 antisense RNA 2 [Source:HGNC Symb |
| MIR3167 | 11 | microRNA 3167 [Source:HGNC Symbol;Acc:HC |
| AP002833.2 | 11 | novel transcript, antisense to KIRREL3 |
| KIRREL3-AS3 | 11 | KIRREL3 antisense RNA 3 [Source:HGNC Symb |
| AP001993.1 | 11 | novel transcript |
| AP003121.1 | 11 | novel transcript |
| LINC02712 | 11 | long intergenic non-protein coding RNA 2712 |
| RN7SKP121 | 11 | RN7SK pseudogene 121 [Source:HGNC Symbo |
| RN7SKP279 | 11 | RN7SK pseudogene 279 [Source:HGNC Symbo |
| AP003532.1 | 11 | HIG1 hypoxia inducible domain family (HIG1) |
| AP001000.1 | 11 | similar to DnaJ (Hsp40) homolog, subfamily B, |
| LINC02725 | 11 | long intergenic non-protein coding RNA 2725 |
| LINC02098 | 11 | long intergenic non-protein coding RNA 2098 |
| ETS1 | 11 | ETS proto-oncogene 1, transcription factor [Sc |
| MIR6090 | 11 | microRNA 6090 [Source:HGNC Symbol;Acc:HC |
| ETS1-AS1 | 11 | ETS1 antisense RNA 1 [Source:HGNC Symbol;/ |
| AP001122.1 | 11 | novel transcript, antisense to FLI1 |
| FLI1 | 11 | Fli-1 proto-oncogene, ETS transcription factor |
| SENCR | 11 | smooth muscle and endothelial cell enriched r |
| KCNJ1 | 11 | potassium voltage-gated channel subfamily J r |
| AP000920.1 | 11 | SRY (sex determining region Y)-box 15 (SOX15 |
| KCNJ5 | 11 | potassium voltage-gated channel subfamily J r |
| C11orf45 | 11 | chromosome 11 open reading frame 45 [Sour |
| TP53AIP1 | 11 | tumor protein p53 regulated apoptosis induci |
| ARHGAP32 | 11 | Rho GTPase activating protein 32 [Source:HGI |
| RNU6-876P | 11 | RNA, U6 small nuclear 876, pseudogene [Sour |
| Y_RNA | 11 | Y RNA [Source:RFAM;Acc:RF00019] |
| RNU6-874P | 11 | RNA, U6 small nuclear 874, pseudogene [Sour |
| ZNF123P | 11 | zinc finger protein 123, pseudogene [Source:H |
| BARX2 | 11 | BARX homeobox 2 [Source:HGNC Symbol;Acc: |
| RPS27P20 | 11 | ribosomal protein S27 pseudogene 20 [Source |
| AP003500.1 | 11 | novel transcript |
| LINC01395 | 11 | long intergenic non-protein coding RNA 1395 |
| AP003327.2 | 11 | novel transcript |
| AP003327.1 | 11 | zinc finger and BTB domain containing 24 (ZBT |
| TMEM45B | 11 | transmembrane protein 45B [Source:HGNC Sy |
| NFRKB | 11 | nuclear factor related to kappaB binding prote |
| PRDM10 | 11 | PR/SET domain 10 [Source:HGNC Symbol;Acc: |
| LINC00167 | 11 | long intergenic non-protein coding RNA 167 [S |

| | | |
|---|---|---|
| AP003041.1 | 11 | zinc finger protein 260 (ZNF260) pseudogene |
| ELOBP2 | 11 | elongin B pseudogene 2 [Source:HGNC Symbo |
| AP003041.2 | 11 | similar to ribosomal protein L10 (RPL10) pseu |
| APLP2 | 11 | amyloid beta precursor like protein 2 [Source: |
| RPL34P21 | 11 | ribosomal protein L34 pseudogene 21 [Source |
| U4 | 11 | U4 spliceosomal RNA [Source:RFAM;Acc:RF00 |
| ST14 | 11 | suppression of tumorigenicity 14 [Source:HGN |
| ZBTB44 | 11 | zinc finger and BTB domain containing 44 [Sou |
| AC019227.1 | 11 | NADH dehydrogenase (ubiquinone) 1 alpha su |
| DDX18P5 | 11 | DEAD-box helicase 18 pseudogene 5 [Source:H |
| RN7SL778P | 11 | RNA, 7SL, cytoplasmic 778, pseudogene [Sour |
| ZBTB44-DT | 11 | ZBTB44 divergent transcript [Source:HGNC Sy |
| ADAMTS8 | 11 | ADAM metallopeptidase with thrombospondi |
| ADAMTS15 | 11 | ADAM metallopeptidase with thrombospondi |
| AP004371.1 | 11 | translocase of outer mitochondrial membrane |
| BAK1P2 | 11 | BCL2 antagonist/killer 1 pseudogene 2 [Source |
| MIR8052 | 11 | microRNA 8052 [Source:HGNC Symbol;Acc:HG |
| C11orf44 | 11 | chromosome 11 open reading frame 44 [Sour |
| AP003486.2 | 11 | novel transcript |
| PPP1R10P1 | 11 | protein phosphatase 1 regulatory subunit 10 p |
| LINC02551 | 11 | long intergenic non-protein coding RNA 2551 |
| AP003486.1 | 11 | novel transcript |
| SNX19 | 11 | sorting nexin 19 [Source:HGNC Symbol;Acc:HG |
| RN7SL167P | 11 | RNA, 7SL, cytoplasmic 167, pseudogene [Sour |
| AP002856.3 | 11 | novel transcript |
| AP002856.1 | 11 | novel transcript |
| AP002856.4 | 11 | novel transcript |
| AP002856.2 | 11 | novel transcript |
| NTM | 11 | neurotrimin [Source:HGNC Symbol;Acc:HGNC |
| RNU6ATAC12P | 11 | RNA, U6atac small nuclear 12, pseudogene [So |
| AP003025.1 | 11 | novel transcript |
| AP003025.2 | 11 | novel transcript |
| NTM-AS1 | 11 | NTM antisense RNA 1 [Source:HGNC Symbol;A |
| AP004372.1 | 11 | GPN-loop GTPase 1, pseudogene |
| AP000844.2 | 11 | novel transcript |
| AP000844.1 | 11 | novel transcript |
| NTM-IT | 11 | NTM intronic transcript [Source:HGNC Symbo |
| AP000843.1 | 11 | TEC |
| OPCML | 11 | opioid binding protein/cell adhesion molecule |
| U6 | 11 | U6 spliceosomal RNA [Source:RFAM;Acc:RF00 |
| AP003784.1 | 11 | novel transcript |
| OPCML-IT2 | 11 | OPCML intronic transcript 2 [Source:HGNC Sy |
| AP004782.1 | 11 | novel transcript |
| AP003972.1 | 11 | novel transcript |
| OPCML-IT1 | 11 | OPCML intronic transcript 1 [Source:HGNC Sy |
| AP004606.1 | 11 | novel transcript, antisense to OPCML |
| LINC02743 | 11 | long intergenic non-protein coding RNA 2743 |
| SPATA19 | 11 | spermatogenesis associated 19 [Source:HGNC |
| AP001979.1 | 11 | calponin 3, acidic (CNN3) pseudogene |
| IGSF9B | 11 | immunoglobulin superfamily member 9B [Sou |

| | | |
|---|---|---|
| MIR4697 | 11 | microRNA 4697 [Source:HGNC Symbol;Acc:HG |
| LINC02730 | 11 | long intergenic non-protein coding RNA 2730 |
| LINC02731 | 11 | long intergenic non-protein coding RNA 2731 |
| AP000911.1 | 11 | novel transcript |
| JAM3 | 11 | junctional adhesion molecule 3 [Source:HGNC |
| PTP4A2P2 | 11 | PTP4A2 pseudogene 2 [Source:HGNC Symbol; |
| NCAPD3 | 11 | non-SMC condensin II complex subunit D3 [So |
| AP001775.1 | 11 | Zinc finger protein 628 (ZNF628) pseudogene |
| AP001775.2 | 11 | novel transcript |
| VPS26B | 11 | VPS26, retromer complex component B [Sour |
| THYN1 | 11 | thymocyte nuclear protein 1 [Source:HGNC Sy |
| ACAD8 | 11 | acyl-CoA dehydrogenase family member 8 [Sc |
| GLB1L3 | 11 | galactosidase beta 1 like 3 [Source:HGNC Sym |
| AP000859.1 | 11 | ATPase, H+/K+ exchanging, beta polypeptide ( |
| GLB1L2 | 11 | galactosidase beta 1 like 2 [Source:HGNC Sym |
| B3GAT1 | 11 | beta-1,3-glucuronyltransferase 1 [Source:HGN |
| AP004608.1 | 11 | novel transcript |
| AP004550.1 | 11 | novel transcript |
| AP001999.3 | 11 | novel transcript |
| AP001999.1 | 11 | novel transcript |
| AP001999.2 | 11 | novel transcript |
| LINC02706 | 11 | long intergenic non-protein coding RNA 2706 |
| LINC02714 | 11 | long intergenic non-protein coding RNA 2714 |
| AP003062.2 | 11 | novel transcript |
| LINC02697 | 11 | long intergenic non-protein coding RNA 2697 |
| AP003062.1 | 11 | novel transcript |
| LINC02717 | 11 | long intergenic non-protein coding RNA 2717 |
| LINC02684 | 11 | long intergenic non-protein coding RNA 2684 |
| DDX11L8 | 12 | DEAD/H-box helicase 11 like 8 [Source:HGNC S |
| WASH8P | 12 | WAS protein family homolog 8, pseudogene [S |
| FAM138D | 12 | family with sequence similarity 138 member D |
| IQSEC3 | 12 | IQ motif and Sec7 domain 3 [Source:HGNC Syr |
| AC026369.3 | 12 | novel transcript |
| AC026369.1 | 12 | novel transcript, antisense to IQSEC3 |
| AC026369.2 | 12 | novel transcript |
| AC007406.3 | 12 | novel transcript |
| SLC6A12 | 12 | solute carrier family 6 member 12 [Source:HG |
| AC007406.1 | 12 | novel transcript |
| AC007406.4 | 12 | novel transcript |
| SLC6A13 | 12 | solute carrier family 6 member 13 [Source:HG |
| AC007406.2 | 12 | novel transcript |
| AC007406.5 | 12 | novel transcript |
| KDM5A | 12 | lysine demethylase 5A [Source:HGNC Symbol; |
| CCDC77 | 12 | coiled-coil domain containing 77 [Source:HGN |
| B4GALNT3 | 12 | beta-1,4-N-acetyl-galactosaminyltransferase 3 |
| NINJ2 | 12 | ninjurin 2 [Source:HGNC Symbol;Acc:HGNC:78 |
| AC006205.2 | 12 | novel transcript |
| RNU7-103P | 12 | RNA, U7 small nuclear 103 pseudogene [Sourc |
| AC006205.1 | 12 | novel transcript |
| AC021054.1 | 12 | novel transcript |

| | | |
|---|---|---|
| LINC02455 | 12 | long intergenic non-protein coding RNA 2455 |
| WNK1 | 12 | WNK lysine deficient protein kinase 1 [Source |
| RNU4ATAC16P | 12 | RNA, U4atac small nuclear 16, pseudogene [S |
| AC004765.1 | 12 | novel transcript, antisense to WNK1 |
| RAD52 | 12 | RAD52 homolog, DNA repair protein [Source:I |
| AC004803.1 | 12 | novel transcript |
| ERC1 | 12 | ELKS/RAB6-interacting/CAST family member 1 |
| HTR1DP1 | 12 | 5-hydroxytryptamine receptor 1D pseudogen |
| AC004672.1 | 12 | novel transcript |
| AC004672.2 | 12 | novel transcript, sense intronic to ERC1 |
| LINC00942 | 12 | long intergenic non-protein coding RNA 942 [S |
| WNT5B | 12 | Wnt family member 5B [Source:HGNC Symbol |
| FBXL14 | 12 | F-box and leucine rich repeat protein 14 [Sour |
| MIR3649 | 12 | microRNA 3649 [Source:HGNC Symbol;Acc:HG |
| ADIPOR2 | 12 | adiponectin receptor 2 [Source:HGNC Symbol |
| AC005183.1 | 12 | SDA1 domain containing 1 (SDAD1) pseudoge |
| AC005343.1 | 12 | novel transcript, antisense to ADIPOR2 |
| RPS4XP14 | 12 | ribosomal protein S4X pseudogene 14 [Source |
| CACNA2D4 | 12 | calcium voltage-gated channel auxiliary subun |
| AC005343.4 | 12 | novel transcript, antisense to CACNA2D4 |
| LRTM2 | 12 | leucine rich repeats and transmembrane dom |
| AC005342.2 | 12 | novel transcript |
| LINC00940 | 12 | long intergenic non-protein coding RNA 940 [S |
| DCP1B | 12 | decapping mRNA 1B [Source:HGNC Symbol;Ac |
| CACNA1C | 12 | calcium voltage-gated channel subunit alpha1 |
| AC005342.1 | 12 | novel transcript |
| CACNA1C-IT1 | 12 | CACNA1C intronic transcript 1 [Source:HGNC S |
| CACNA1C-IT2 | 12 | CACNA1C intronic transcript 2 [Source:HGNC S |
| AC005344.1 | 12 | novel transcript, antisense to CACNA1C |
| CACNA1C-AS4 | 12 | CACNA1C antisense RNA 4 [Source:HGNC Sym |
| CACNA1C-IT3 | 12 | CACNA1C intronic transcript 3 [Source:HGNC S |
| AC005293.1 | 12 | novel transcript, antisense to CACNA1C |
| AC005414.1 | 12 | novel transcript, antisense to CACNA1C |
| CACNA1C-AS3 | 12 | CACNA1C antisense RNA 3 [Source:HGNC Sym |
| AC007618.1 | 12 | ribosomal protein S6 (RPS6) pseudogene |
| CACNA1C-AS2 | 12 | CACNA1C antisense RNA 2 [Source:HGNC Sym |
| CACNA1C-AS1 | 12 | CACNA1C antisense RNA 1 [Source:HGNC Sym |
| ITFG2-AS1 | 12 | ITFG2 antisense RNA 1 [Source:HGNC Symbol; |
| AC092471.2 | 12 | gap junction beta family pseudogene |
| LINC02371 | 12 | long intergenic non-protein coding RNA 2371 |
| IQSEC3P1 | 12 | IQ motif and Sec7 domain 3 pseudogene 1 [Sc |
| AC092471.1 | 12 | maestro heat-like repeat-containing protein fa |
| RPL23AP14 | 12 | ribosomal protein L23a pseudogene 14 [Sourc |
| CBX3P4 | 12 | chromobox 3 pseudogene 4 [Source:HGNC Sy |
| FKBP4 | 12 | FKBP prolyl isomerase 4 [Source:HGNC Symbo |
| ITFG2-AS1 | 12 | ITFG2 antisense RNA 1 [Source:HGNC Symbol; |
| AC005841.1 | 12 | novel transcript, antisense to FKBP4 |
| ITFG2 | 12 | integrin alpha FG-GAP repeat containing 2 [So |
| NRIP2 | 12 | nuclear receptor interacting protein 2 [Source |
| TEX52 | 12 | testis expressed 52 [Source:HGNC Symbol;Acc |

| | | |
|---|---|---|
| FOXM1 | 12 | forkhead box M1 [Source:HGNC Symbol;Acc:H |
| RHNO1 | 12 | RAD9-HUS1-RAD1 interacting nuclear orphan |
| TULP3 | 12 | TUB like protein 3 [Source:HGNC Symbol;Acc: |
| AC005911.1 | 12 | novel transcript |
| RNU6-1315P | 12 | RNA, U6 small nuclear 1315, pseudogene [Sou |
| TEAD4 | 12 | TEA domain transcription factor 4 [Source:HG |
| AC125807.2 | 12 | novel transcript |
| AC125807.1 | 12 | ribosomal protein L13a (RPL13A) pseudogene |
| TSPAN9 | 12 | tetraspanin 9 [Source:HGNC Symbol;Acc:HGN |
| Metazoa_SRP | 12 | Metazoan signal recognition particle RNA [Sou |
| TSPAN9-IT1 | 12 | TSPAN9 intronic transcript 1 [Source:HGNC Sy |
| AC005912.2 | 12 | novel transcript |
| AC005912.1 | 12 | ribosomal protein S27 (RPS27) pseudogene |
| AC005865.1 | 12 | tetraspanin 11 (TSPAN11) pseudogene |
| AC005865.2 | 12 | novel transcript |
| LINC02827 | 12 | long intergenic non-protein coding RNA 2827 |
| LINC02417 | 12 | long intergenic non-protein coding RNA 2417 |
| PRMT8 | 12 | protein arginine methyltransferase 8 [Source: |
| AC005908.2 | 12 | novel transcript |
| AC005908.1 | 12 | ribosomal protein S26 (RPS26) pseudogene |
| AC005908.3 | 12 | novel transcript, antisense to PRMT8 |
| CRACR2A | 12 | calcium release activated channel regulator 2/ |
| AC005831.1 | 12 | TEC |
| RNU6-174P | 12 | RNA, U6 small nuclear 174, pseudogene [Sour |
| AC006207.1 | 12 | novel transcript, antisense to EFCAB4B |
| PARP11 | 12 | poly(ADP-ribose) polymerase family member |
| OTUD4P1 | 12 | OTUD4 pseudogene 1 [Source:HGNC Symbol;/ |
| AC005842.1 | 12 | novel transcript, antisense to PARP11 |
| AC084375.1 | 12 | novel transcript |
| HSPA8P5 | 12 | heat shock protein family A (Hsp70) member |
| AC007207.1 | 12 | ribosomal protein L18 (RPL18) pseudogene |
| CCND2-AS1 | 12 | CCND2 antisense RNA 1 [Source:HGNC Symbo |
| CCND2 | 12 | cyclin D2 [Source:HGNC Symbol;Acc:HGNC:15 |
| AC008012.1 | 12 | novel transcript |
| TIGAR | 12 | TP53 induced glycolysis regulatory phosphata |
| AC006122.1 | 12 | ribosomal protein S15 (RPS15) pseudogene |
| FGF23 | 12 | fibroblast growth factor 23 [Source:HGNC Syn |
| FGF6 | 12 | fibroblast growth factor 6 [Source:HGNC Symb |
| C12orf4 | 12 | chromosome 12 open reading frame 4 [Sourc |
| RAD51AP1 | 12 | RAD51 associated protein 1 [Source:HGNC Syr |
| DYRK4 | 12 | dual specificity tyrosine phosphorylation regu |
| AC005832.2 | 12 | chromosome 1 open reading frame 114 (C1or |
| AC005832.4 | 12 | novel protein |
| AKAP3 | 12 | A-kinase anchoring protein 3 [Source:HGNC S |
| AC005832.3 | 12 | RAB5A, member RAS oncogene family (RAB5A |
| AC005832.1 | 12 | novel transcript, sense intronic to RP11-500M |
| NDUFA9 | 12 | NADH:ubiquinone oxidoreductase subunit A9 |
| AC005833.1 | 12 | novel protein |
| GAU1 | 12 | GALNT8 antisense upstream 1 [Source:HGNC |
| GALNT8 | 12 | polypeptide N-acetylgalactosaminyltransferas |

| | | |
|---|---|---|
| AC005833.2 | 12 | new variant |
| KCNA6 | 12 | potassium voltage-gated channel subfamily A |
| AC006063.1 | 12 | novel transcript |
| AC006063.3 | 12 | novel transcript |
| AC006063.4 | 12 | novel transcript |
| AC005906.2 | 12 | novel transcript, sense overlapping KCNA1 |
| KCNA1 | 12 | potassium voltage-gated channel subfamily A |
| AC006063.2 | 12 | novel transcript |
| AC005906.1 | 12 | eukaryotic translation initiation factor 3, subu |
| KCNA5 | 12 | potassium voltage-gated channel subfamily A |
| AC005906.3 | 12 | novel transcript |
| LINC02443 | 12 | long intergenic non-protein coding RNA 2443 |
| AC006206.2 | 12 | novel transcript |
| AC006206.1 | 12 | novel transcript |
| AC007848.2 | 12 | novel transcript |
| AC007848.1 | 12 | novel transcript |
| NTF3 | 12 | neurotrophin 3 [Source:HGNC Symbol;Acc:HG |
| ANO2 | 12 | anoctamin 2 [Source:HGNC Symbol;Acc:HGNC |
| AC137627.1 | 12 | dynein, light chain, roadblock-type 1 (DYNLRB |
| VWF | 12 | von Willebrand factor [Source:HGNC Symbol;/ |
| RN7SL69P | 12 | RNA, 7SL, cytoplasmic 69, pseudogene [Sourc |
| AC005845.1 | 12 | novel transcript |
| CD9 | 12 | CD9 molecule [Source:HGNC Symbol;Acc:HGN |
| Y_RNA | 12 | Y RNA [Source:RFAM;Acc:RF00019] |
| ATP5MFP5 | 12 | ATP synthase membrane subunit f pseudogen |
| PLEKHG6 | 12 | pleckstrin homology and RhoGEF domain cont |
| TNFRSF1A | 12 | TNF receptor superfamily member 1A [Source |
| RN7SL391P | 12 | RNA, 7SL, cytoplasmic 391, pseudogene [Sour |
| SCNN1A | 12 | sodium channel epithelial 1 alpha subunit [Sou |
| LTBR | 12 | lymphotoxin beta receptor [Source:HGNC Sym |
| AC005840.2 | 12 | novel transcript |
| AC005840.1 | 12 | ribosomal protein L31 (RPL31) pseudogene |
| SRP14P1 | 12 | signal recognition particle 14 pseudogene 1 [S |
| CD27-AS1 | 12 | CD27 antisense RNA 1 [Source:HGNC Symbol;. |
| CD27 | 12 | CD27 molecule [Source:HGNC Symbol;Acc:HG |
| TAPBPL | 12 | TAP binding protein like [Source:HGNC Symbo |
| VAMP1 | 12 | vesicle associated membrane protein 1 [Sourc |
| AC005840.4 | 12 | novel transcript, antisense to VAMP1 |
| AC005840.3 | 12 | plakophilin 2 (PKP2) pseudogene |
| MRPL51 | 12 | mitochondrial ribosomal protein L51 [Source:I |
| NCAPD2 | 12 | non-SMC condensin I complex subunit D2 [Sou |
| SCARNA10 | 12 | small Cajal body-specific RNA 10 [Source:HGN |
| AC006064.5 | 12 | small Cajal body-specific RNA 10 |
| AC006064.3 | 12 | novel transcript |
| GAPDH | 12 | glyceraldehyde-3-phosphate dehydrogenase [ |
| AC006064.4 | 12 | novel transcript, antisense to GAPDH |
| IFFO1 | 12 | intermediate filament family orphan 1 [Source |
| AC006064.1 | 12 | novel transcript |
| NOP2 | 12 | NOP2 nucleolar protein [Source:HGNC Symbo |
| AC006064.6 | 12 | novel transcript |

| | | |
|---|---|---|
| CHD4 | 12 | chromodomain helicase DNA binding protein 4 |
| AC006064.2 | 12 | novel transcript, antisense to CHD4 |
| SCARNA11 | 12 | small Cajal body-specific RNA 11 [Source:HGN |
| LPAR5 | 12 | lysophosphatidic acid receptor 5 [Source:HGN |
| ACRBP | 12 | acrosin binding protein [Source:HGNC Symbol |
| ING4 | 12 | inhibitor of growth family member 4 [Source:I |
| AC125494.1 | 12 | novel transcript |
| ZNF384 | 12 | zinc finger protein 384 [Source:HGNC Symbol; |
| PIANP | 12 | PILR alpha associated neural protein [Source:I |
| RNU6-781P | 12 | RNA, U6 small nuclear 781, pseudogene [Sour |
| AC125494.3 | 12 | novel transcript, antisense to ZNF384 |
| COPS7A | 12 | COP9 signalosome subunit 7A [Source:HGNC S |
| AC125494.2 | 12 | novel transcript |
| MLF2 | 12 | myeloid leukemia factor 2 [Source:HGNC Sym |
| PTMS | 12 | parathymosin [Source:HGNC Symbol;Acc:HGN |
| LAG3 | 12 | lymphocyte activating 3 [Source:HGNC Symbo |
| RN7SL380P | 12 | RNA, 7SL, cytoplasmic 380, pseudogene [Sour |
| CD4 | 12 | CD4 molecule [Source:HGNC Symbol;Acc:HGN |
| GPR162 | 12 | G protein-coupled receptor 162 [Source:HGN |
| P3H3 | 12 | prolyl 3-hydroxylase 3 [Source:HGNC Symbol;A |
| GNB3 | 12 | G protein subunit beta 3 [Source:HGNC Symbo |
| CDCA3 | 12 | cell division cycle associated 3 [Source:HGNC S |
| USP5 | 12 | ubiquitin specific peptidase 5 [Source:HGNC S |
| TPI1 | 12 | triosephosphate isomerase 1 [Source:HGNC Sy |
| SPSB2 | 12 | splA/ryanodine receptor domain and SOCS bo |
| RPL13P5 | 12 | ribosomal protein L13 pseudogene 5 [Source:I |
| LRRC23 | 12 | leucine rich repeat containing 23 [Source:HGN |
| DSTNP2 | 12 | DSTN pseudogene 2 [Source:HGNC Symbol;Ac |
| U47924.3 | 12 | novel transcript |
| ENO2 | 12 | enolase 2 [Source:HGNC Symbol;Acc:HGNC:33 |
| ATN1 | 12 | atrophin 1 [Source:HGNC Symbol;Acc:HGNC:3 |
| C12orf57 | 12 | chromosome 12 open reading frame 57 [Sour |
| U47924.2 | 12 | novel transcript, antisense to C12orf57 |
| RNU7-1 | 12 | RNA, U7 small nuclear 1 [Source:HGNC Symbo |
| PTPN6 | 12 | protein tyrosine phosphatase non-receptor ty |
| MIR200CHG | 12 | MIR200C and MIR141 host gene [Source:HGN |
| MIR200C | 12 | microRNA 200c [Source:HGNC Symbol;Acc:HG |
| MIR141 | 12 | microRNA 141 [Source:HGNC Symbol;Acc:HGN |
| U47924.1 | 12 | novel transcript, antisense to PHB2 |
| PHB2 | 12 | prohibitin 2 [Source:HGNC Symbol;Acc:HGNC: |
| SCARNA12 | 12 | small Cajal body-specific RNA 12 [Source:HGN |
| EMG1 | 12 | EMG1 N1-specific pseudouridine methyltransf |
| LPCAT3 | 12 | lysophosphatidylcholine acyltransferase 3 [So |
| C1S | 12 | complement C1s [Source:HGNC Symbol;Acc:H |
| AC006512.1 | 12 | ribosomal protein L37 (RPL37) pseudogene |
| AC006512.2 | 12 | zinc finger protein 655 (ZNF655) pseudogene |
| C1R | 12 | complement C1r [Source:HGNC Symbol;Acc:H |
| C1RL | 12 | complement C1r subcomponent like [Source:I |
| C1RL-AS1 | 12 | C1RL antisense RNA 1 [Source:HGNC Symbol;A |
| RBP5 | 12 | retinol binding protein 5 [Source:HGNC Symbo |

| | | |
|---|---|---|
| RNU6-485P | 12 | RNA, U6 small nuclear 485, pseudogene [Sour |
| AC018653.3 | 12 | novel transcript |
| CLSTN3 | 12 | calsyntenin 3 [Source:HGNC Symbol;Acc:HGN |
| AC018653.4 | 12 | novel transcript, antisense to CLSTN3 |
| AC018653.1 | 12 | novel transcript |
| PEX5 | 12 | peroxisomal biogenesis factor 5 [Source:HGN |
| AC018653.2 | 12 | chromosome 10 open reading frame 129 (C1C |
| ACSM4 | 12 | acyl-CoA synthetase medium chain family mer |
| CD163L1 | 12 | CD163 molecule like 1 [Source:HGNC Symbol; |
| AC131206.1 | 12 | M-phase phosphoprotein 8 (MPHOSPH8) pseu |
| CD163 | 12 | CD163 molecule [Source:HGNC Symbol;Acc:H |
| GAPDHP31 | 12 | glyceraldehyde 3 phosphate dehydrogenase p |
| NIFKP3 | 12 | nucleolar protein interacting with the FHA dor |
| AC006927.1 | 12 | clathrin, light chain A (CLTA) pseudogene |
| AC006927.2 | 12 | mitochondrially encoded cytochrome c oxidas |
| AC006927.4 | 12 | mitochondrially encoded cytochrome c oxidas |
| AC006927.3 | 12 | mitochondrially encoded NADH dehydrogenas |
| AC006927.5 | 12 | mitochondrially encoded cytochrome c oxidas |
| APOBEC1 | 12 | apolipoprotein B mRNA editing enzyme cataly |
| GDF3 | 12 | growth differentiation factor 3 [Source:HGNC |
| DPPA3 | 12 | developmental pluripotency associated 3 [Sou |
| CLEC4C | 12 | C-type lectin domain family 4 member C [Sour |
| NANOGNB | 12 | NANOG neighbor homeobox [Source:HGNC Sy |
| Y_RNA | 12 | Y RNA [Source:RFAM;Acc:RF00019] |
| NANOG | 12 | Nanog homeobox [Source:HGNC Symbol;Acc: |
| Y_RNA | 12 | Y RNA [Source:RFAM;Acc:RF00019] |
| SLC2A14 | 12 | solute carrier family 2 member 14 [Source:HG |
| AC006517.2 | 12 | novel transcript, antisense to SLC2A14 |
| Y_RNA | 12 | Y RNA [Source:RFAM;Acc:RF00019] |
| Metazoa_SRP | 12 | Metazoan signal recognition particle RNA [Sou |
| AC006517.1 | 12 | ribosomal protein S20 (RPS20) pseudogene |
| AC124891.1 | 12 | myosin, light chain 9, regulatory (MYL9) pseuc |
| NANOGP1 | 12 | Nanog homeobox pseudogene 1 [Source:HGN |
| AC007536.1 | 12 | Nanog homeobox (NANOG) pseudogene |
| SLC2A3 | 12 | solute carrier family 2 member 3 [Source:HGN |
| AC006511.6 | 12 | novel transcript |
| Y_RNA | 12 | Y RNA [Source:RFAM;Acc:RF00019] |
| AC006511.1 | 12 | ribosomal protein S20 (RPS20) pseudogene |
| AC006511.4 | 12 | novel transcript |
| HSPD1P12 | 12 | heat shock protein family D (Hsp60) member |
| AC006511.2 | 12 | heat shock 60kDa protein 1 (chaperonin) (HSP |
| FOXJ2 | 12 | forkhead box J2 [Source:HGNC Symbol;Acc:H |
| C3AR1 | 12 | complement C3a receptor 1 [Source:HGNC Sy |
| NECAP1 | 12 | NECAP endocytosis associated 1 [Source:HGN |
| AC006511.3 | 12 | TEC |
| AC006511.5 | 12 | novel transcript |
| CLEC4A | 12 | C-type lectin domain family 4 member A [Sour |
| POU5F1P3 | 12 | POU class 5 homeobox 1 pseudogene 3 [Sourc |
| AC092111.3 | 12 | novel protein |
| GCSHP4 | 12 | glycine cleavage system protein H pseudogen |

| | | |
|---|---|---|
| ZNF705A | 12 | zinc finger protein 705A [Source:HGNC Symbc |
| FAM66C | 12 | family with sequence similarity 66 member C |
| DEFB109F | 12 | defensin beta 109F (pseudogene) [Source:HGI |
| AC092111.2 | 12 | novel transcript |
| AC092111.1 | 12 | novel transcript |
| FAM90A1 | 12 | family with sequence similarity 90 member A: |
| ALG1L10P | 12 | ALG1 like 10, pseudogene [Source:HGNC Syml |
| FAM86FP | 12 | family with sequence similarity 86 member F, |
| LINC02449 | 12 | long intergenic non-protein coding RNA 2449 |
| AC092745.5 | 12 | novel transcript |
| ENPP7P5 | 12 | ectonucleotide pyrophosphatase/phosphodie |
| AC092745.4 | 12 | novel transcript |
| LINC00937 | 12 | long intergenic non-protein coding RNA 937 [S |
| AC092745.2 | 12 | ribosomal protein S24 (RPS24) pseudogene |
| AC092745.1 | 12 | novel transcript |
| AC092745.3 | 12 | small nuclear ribonucleoprotein polypeptide ( |
| RPS3AP43 | 12 | ribosomal protein S3a pseudogene 43 [Source |
| AC092865.4 | 12 | RNA binding protein, fox-1 homolog (C. elegar |
| AC092865.3 | 12 | RNA binding protein, fox-1 homolog (C. elegar |
| AC092865.2 | 12 | chromosome 11 open reading frame 2 (C11or |
| OR7E140P | 12 | olfactory receptor family 7 subfamily E memb |
| OR7E148P | 12 | olfactory receptor family 7 subfamily E memb |
| OR7E149P | 12 | olfactory receptor family 7 subfamily E memb |
| CLEC6A | 12 | C-type lectin domain containing 6A [Source:H( |
| AC092865.1 | 12 | ribosomal protein L15 (RPL15) pseudogene |
| RNU6-275P | 12 | RNA, U6 small nuclear 275, pseudogene [Sour |
| CLEC4D | 12 | C-type lectin domain family 4 member D [Sou |
| SUPT4H1P2 | 12 | SPT4 homolog, DSIF elongation factor subunit |
| CLEC4E | 12 | C-type lectin domain family 4 member E [Sour |
| AC092746.1 | 12 | novel transcript |
| Y_RNA | 12 | Y RNA [Source:RFAM;Acc:RF00019] |
| AC092184.1 | 12 | NULL |
| AICDA | 12 | activation induced cytidine deaminase [Sourc |
| AC092184.2 | 12 | TEC |
| HADHAP2 | 12 | HADHA pseudogene 2 [Source:HGNC Symbol;/ |
| MFAP5 | 12 | microfibril associated protein 5 [Source:HGNC |
| AC092490.2 | 12 | novel transcript |
| AC092490.3 | 12 | chromosome 19 open reading frame 40 (C19c |
| RIMKLB | 12 | ribosomal modification protein rimK like famil |
| RPSAP51 | 12 | ribosomal protein SA pseudogene 51 [Source: |
| A2ML1-AS1 | 12 | A2ML1 antisense RNA 1 [Source:HGNC Symbc |
| AC092490.1 | 12 | novel transcript |
| A2ML1-AS2 | 12 | A2ML1 antisense RNA 2 [Source:HGNC Symbc |
| A2ML1 | 12 | alpha-2-macroglobulin like 1 [Source:HGNC Sy |
| AC006581.2 | 12 | novel transcript, antisense to PHC1 and A2ML |
| PHC1 | 12 | polyhomeotic homolog 1 [Source:HGNC Symb |
| M6PR | 12 | mannose-6-phosphate receptor, cation deper |
| KLRG1 | 12 | killer cell lectin like receptor G1 [Source:HGN( |
| AC006581.1 | 12 | novel transcript |
| HNRNPA1P34 | 12 | heterogeneous nuclear ribonucleoprotein A1 |

| | | |
|---|---|---|
| VDAC2P2 | 12 | voltage dependent anion channel 2 pseudoge |
| LINC00612 | 12 | long intergenic non-protein coding RNA 612 [S |
| A2M-AS1 | 12 | A2M antisense RNA 1 [Source:HGNC Symbol;A |
| A2M | 12 | alpha-2-macroglobulin [Source:HGNC Symbol; |
| KRT17P8 | 12 | keratin 17 pseudogene 8 [Source:HGNC Symb |
| BTG1P1 | 12 | BTG anti-proliferation factor 1 pseudogene 1 [ |
| PZP | 12 | PZP alpha-2-macroglobulin like [Source:HGNC |
| TPT1P12 | 12 | tumor protein, translationally-controlled 1 ps |
| A2MP1 | 12 | alpha-2-macroglobulin pseudogene 1 [Source: |
| MIR1244-3 | 12 | microRNA 1244-3 [Source:HGNC Symbol;Acc:I |
| PTMAP4 | 12 | prothymosin alpha pseudogene 4 [Source:HGI |
| LINC00987 | 12 | long intergenic non-protein coding RNA 987 [S |
| AC010175.1 | 12 | novel transcript |
| AC009533.1 | 12 | DEAD/H (Asp-Glu-Ala-Asp/His) box polypeptid |
| SNORA75 | 12 | Small nucleolar RNA SNORA75 [Source:RFAM; |
| AC009533.2 | 12 | WAS protein family homolog pseudogene |
| AC009533.3 | 12 | novel transcript |
| LINC02367 | 12 | long intergenic non-protein coding RNA 2367 |
| AC009533.4 | 12 | novel transcript |
| AC141557.1 | 12 | ovostatin (OVOS) pseudogene |
| AC141557.2 | 12 | WAS protein family homolog 1 (WASH1) pseu |
| DDX12P | 12 | DEAD/H-box helicase 12, pseudogene [Source |
| SNORA75 | 12 | Small nucleolar RNA SNORA75 [Source:RFAM; |
| AC092821.3 | 12 | novel transcript |
| AC092821.1 | 12 | alpha-2-macroglobulin like 1 pseudogene |
| AC092821.2 | 12 | RAN binding protein (RANBP) pseudogene |
| KLRB1 | 12 | killer cell lectin like receptor B1 [Source:HGNC |
| RNU6-700P | 12 | RNA, U6 small nuclear 700, pseudogene [Sour |
| GOT2P3 | 12 | glutamic-oxaloacetic transaminase 2 pseudog |
| AC010186.4 | 12 | novel transcript |
| AC010186.1 | 12 | novel transcript |
| AC010186.2 | 12 | C-type lectin domain family 2 (CLEC2) pseudo |
| AC010186.3 | 12 | novel transcript |
| CLEC2D | 12 | C-type lectin domain family 2 member D [Sou |
| AC007068.1 | 12 | nucleophosmin (nucleolar phosphoprotein B2 |
| LINC02390 | 12 | long intergenic non-protein coding RNA 2390 |
| CLECL1 | 12 | C-type lectin like 1 [Source:HGNC Symbol;Acc: |
| CD69 | 12 | CD69 molecule [Source:HGNC Symbol;Acc:HG |
| AC007068.2 | 12 | acidic repeat containing (ACRC) pseudogene |
| KLRF1 | 12 | killer cell lectin like receptor F1 [Source:HGNC |
| CLEC2B | 12 | C-type lectin domain family 2 member B [Sour |
| AC091814.1 | 12 | novel transcript |
| KLRF2 | 12 | killer cell lectin like receptor F2 [Source:HGNC |
| CLEC2A | 12 | C-type lectin domain family 2 member A [Sour |
| LINC02470 | 12 | long intergenic non-protein coding RNA 2470 |
| CLEC12A-AS1 | 12 | CLEC12A antisense RNA 1 [Source:HGNC Symb |
| CLEC12A | 12 | C-type lectin domain family 12 member A [Sou |
| CLEC1B | 12 | C-type lectin domain family 1 member B [Sour |
| CLEC12B | 12 | C-type lectin domain family 12 member B [Sou |
| AC024224.2 | 12 | novel transcript |

| | | |
|---|---|---|
| CLEC9A | 12 | C-type lectin domain containing 9A [Source:H( |
| AC024224.1 | 12 | C-type lectin domain family 1, member B pseu |
| CLEC1A | 12 | C-type lectin domain family 1 member A [Sour |
| RN7SKP161 | 12 | RN7SK pseudogene 161 [Source:HGNC Symbo |
| HNRNPABP1 | 12 | heterogeneous nuclear ribonucleoprotein A/E |
| CLEC7A | 12 | C-type lectin domain containing 7A [Source:H( |
| OLR1 | 12 | oxidized low density lipoprotein receptor 1 [S( |
| TMEM52B | 12 | transmembrane protein 52B [Source:HGNC Sy |
| GABARAPL1 | 12 | GABA type A receptor associated protein like |
| AC115676.1 | 12 | novel transcript |
| KLRD1 | 12 | killer cell lectin like receptor D1 [Source:HGN( |
| LINC02617 | 12 | long intergenic non-protein coding RNA 2617 |
| LINC02598 | 12 | long intergenic non-protein coding RNA 2598 |
| AC022075.1 | 12 | novel transcript, antisense to KLRK1 |
| KLRK1 | 12 | killer cell lectin like receptor K1 [Source:HGN( |
| KLRC4-KLRK1 | 12 | KLRC4-KLRK1 readthrough [Source:HGNC Sym |
| KLRC4 | 12 | killer cell lectin like receptor C4 [Source:HGN( |
| KLRC3 | 12 | killer cell lectin like receptor C3 [Source:HGN( |
| AC068775.1 | 12 | novel protein |
| KLRC2 | 12 | killer cell lectin like receptor C2 [Source:HGN( |
| KLRC1 | 12 | killer cell lectin like receptor C1 [Source:HGN( |
| EIF2S3B | 12 | eukaryotic translation initiation factor 2 subur |
| SLC25A39P2 | 12 | SLC25A39 pseudogene 2 [Source:HGNC Symb( |
| LINC02446 | 12 | long intergenic non-protein coding RNA 2446 |
| KLRA1P | 12 | killer cell lectin like receptor A1, pseudogene [ |
| MAGOHB | 12 | mago homolog B, exon junction complex subu |
| STYK1 | 12 | serine/threonine/tyrosine kinase 1 [Source:H( |
| YBX3 | 12 | Y-box binding protein 3 [Source:HGNC Symbol |
| LINC02366 | 12 | long intergenic non-protein coding RNA 2366 |
| HSPE1P12 | 12 | heat shock protein family E (Hsp10) member 1 |
| TAS2R7 | 12 | taste 2 receptor member 7 [Source:HGNC Syn |
| TAS2R8 | 12 | taste 2 receptor member 8 [Source:HGNC Syn |
| TAS2R9 | 12 | taste 2 receptor member 9 [Source:HGNC Syn |
| PRH1 | 12 | proline rich protein HaeIII subfamily 1 [Source |
| TAS2R10 | 12 | taste 2 receptor member 10 [Source:HGNC Sy |
| AC006518.1 | 12 | taste receptor, type 2 pseudogene |
| PRR4 | 12 | proline rich 4 [Source:HGNC Symbol;Acc:HGN( |
| AC018630.2 | 12 | PRH1-PRR4 readthrough |
| AC006518.2 | 12 | taste receptor, type 2, member 12 pseudoger |
| TAS2R13 | 12 | taste 2 receptor member 13 [Source:HGNC Sy |
| PRH2 | 12 | proline rich protein HaeIII subfamily 2 [Source |
| TAS2R14 | 12 | taste 2 receptor member 14 [Source:HGNC Sy |
| TAS2R15P | 12 | taste 2 receptor member 15 pseudogene [Sou |
| TAS2R50 | 12 | taste 2 receptor member 50 [Source:HGNC Sy |
| TAS2R20 | 12 | taste 2 receptor member 20 [Source:HGNC Sy |
| TAS2R19 | 12 | taste 2 receptor member 19 [Source:HGNC Sy |
| TAS2R31 | 12 | taste 2 receptor member 31 [Source:HGNC Sy |
| TAS2R63P | 12 | taste 2 receptor member 63 pseudogene [Sou |
| TAS2R46 | 12 | taste 2 receptor member 46 [Source:HGNC Sy |
| TAS2R64P | 12 | taste 2 receptor member 64 pseudogene [Sou |

| | | |
|---|---|---|
| TAS2R43 | 12 | taste 2 receptor member 43 [Source:HGNC Sy |
| AC018630.1 | 12 | iron-responsive element binding protein 2 (IR |
| TAS2R30 | 12 | taste 2 receptor member 30 [Source:HGNC Sy |
| AC134349.2 | 12 | taste receptor, type 2, member 18 pseudoger |
| AC134349.1 | 12 | novel transcript |
| SMIM10L1 | 12 | small integral membrane protein 10 like 1 [So |
| TAS2R42 | 12 | taste 2 receptor member 42 [Source:HGNC Sy |
| AC244131.1 | 12 | taste receptor, type 2, member 7 pseudogene |
| AC244131.2 | 12 | novel transcript |
| PRB3 | 12 | proline rich protein BstNI subfamily 3 [Source: |
| PRB4 | 12 | proline rich protein BstNI subfamily 4 [Source: |
| PRB1 | 12 | proline rich protein BstNI subfamily 1 (gene/p |
| PRB2 | 12 | proline rich protein BstNI subfamily 2 [Source: |
| AC078950.1 | 12 | novel transcript |
| HIGD1AP8 | 12 | HIG1 hypoxia inducible domain family membe |
| AC007450.4 | 12 | IQ motif and Sec7 domain 3 (IQSEC3) pseudog |
| AC007450.3 | 12 | ovostatin (OVOS) pseudogene |
| DDX55P1 | 12 | DEAD-box helicase 55 pseudogene 1 [Source:I |
| AC007450.2 | 12 | tolloid-like 1 (TLL1) pseudogene |
| AC007450.1 | 12 | novel transcript |
| RNU7-60P | 12 | RNA, U7 small nuclear 60 pseudogene [Source |
| LINC01252 | 12 | long intergenic non-protein coding RNA 1252 |
| ETV6 | 12 | ETS variant 6 [Source:HGNC Symbol;Acc:HGN( |
| BCL2L14 | 12 | BCL2 like 14 [Source:HGNC Symbol;Acc:HGNC |
| PTMAP9 | 12 | prothymosin alpha pseudogene 9 [Source:HGI |
| MIR1244-4 | 12 | microRNA 1244-4 [Source:HGNC Symbol;Acc:I |
| LRP6 | 12 | LDL receptor related protein 6 [Source:HGNC |
| AC007537.1 | 12 | ribosomal protein L21 (RPL21) pseudogene |
| RNU6-545P | 12 | RNA, U6 small nuclear 545, pseudogene [Sour |
| AC007621.1 | 12 | ubiquitin-conjugating enzyme E2 variant 2 (UE |
| RNU6-318P | 12 | RNA, U6 small nuclear 318, pseudogene [Sour |
| MANSC1 | 12 | MANSC domain containing 1 [Source:HGNC Sy |
| RPL23AP66 | 12 | ribosomal protein L23a pseudogene 66 [Sourc |
| LOH12CR2 | 12 | loss of heterozygosity, 12, chromosomal regic |
| BORCS5 | 12 | BLOC-1 related complex subunit 5 [Source:HG |
| AC007619.2 | 12 | adaptor-related protein complex 3, sigma 1 su |
| DUSP16 | 12 | dual specificity phosphatase 16 [Source:HGNC |
| AC007619.1 | 12 | novel transcript |
| AC092824.1 | 12 | ribosomal protein L19 (RPL19) pseudogene |
| CREBL2 | 12 | cAMP responsive element binding protein like |
| RPL21P136 | 12 | ribosomal protein L21 pseudogene 136 [Sourc |
| AC008115.4 | 12 | novel transcript |
| GPR19 | 12 | G protein-coupled receptor 19 [Source:HGNC |
| AC008115.2 | 12 | novel transcript |
| CDKN1B | 12 | cyclin dependent kinase inhibitor 1B [Source:I |
| AC008115.3 | 12 | novel transcript |
| AC008115.1 | 12 | novel transcript |
| APOLD1 | 12 | apolipoprotein L domain containing 1 [Source |
| MIR613 | 12 | microRNA 613 [Source:HGNC Symbol;Acc:HGI |
| STX8P1 | 12 | syntaxin 8 pseudogene 1 [Source:HGNC Symb |

| | | |
|---|---|---|
| DDX47 | 12 | DEAD-box helicase 47 [Source:HGNC Symbol;/ |
| AC007215.1 | 12 | ribosomal protein L37a (RPL37A) pseudogene |
| AC007688.2 | 12 | ribosomal protein S6 (RPS6) pseudogene |
| RPL13AP20 | 12 | ribosomal protein L13a pseudogene 20 [Sourc |
| GPRC5A | 12 | G protein-coupled receptor class C group 5 m |
| MIR614 | 12 | microRNA 614 [Source:HGNC Symbol;Acc:HGI |
| GPRC5D-AS1 | 12 | GPRC5D and HEBP1 antisense RNA 1 [Source:I |
| GPRC5D | 12 | G protein-coupled receptor class C group 5 m |
| AC007688.3 | 12 | novel transcript |
| AC007688.1 | 12 | NULL |
| HEBP1 | 12 | heme binding protein 1 [Source:HGNC Symbo |
| HTR7P1 | 12 | 5-hydroxytryptamine receptor 7 pseudogene |
| AC023790.2 | 12 | novel transcript |
| AC023790.1 | 12 | ribosomal protein S26 (RPS26) pseudogene |
| FAM234B | 12 | family with sequence similarity 234 member E |
| GSG1 | 12 | germ cell associated 1 [Source:HGNC Symbol;/ |
| EMP1 | 12 | epithelial membrane protein 1 [Source:HGNC |
| AC022276.1 | 12 | novel transcript |
| LINC01559 | 12 | long intergenic non-protein coding RNA 1559 |
| GRIN2B | 12 | glutamate ionotropic receptor NMDA type sul |
| RNA5SP353 | 12 | RNA, 5S ribosomal pseudogene 353 [Source:H |
| RNU6-590P | 12 | RNA, U6 small nuclear 590, pseudogene [Sour |
| AC007527.1 | 12 | novel transcript, antisense to GRIN2B |
| AC007527.2 | 12 | novel transcript |
| AC007535.1 | 12 | novel transcript, antisense to GRIN2B |
| RN7SKP162 | 12 | RN7SK pseudogene 162 [Source:HGNC Symbo |
| AC092470.1 | 12 | Nucleoside diphosphate kinase B (NME2) pseu |
| AC135001.1 | 12 | Non-histone chromosomal protein HMG-14 (I |
| AC092112.1 | 12 | novel transcript |
| MRPS18CP4 | 12 | mitochondrial ribosomal protein S18C pseudo |
| RNU6-491P | 12 | RNA, U6 small nuclear 491, pseudogene [Sour |
| GNAI2P1 | 12 | G protein subunit alpha i2 pseudogene 1 [Sou |
| RPL30P11 | 12 | ribosomal protein L30 pseudogene 11 [Source |
| ATF7IP | 12 | activating transcription factor 7 interacting pr |
| PLBD1 | 12 | phospholipase B domain containing 1 [Source |
| AC008114.1 | 12 | novel transcript, antisense to PLBD1 |
| RN7SKP134 | 12 | RN7SK pseudogene 134 [Source:HGNC Symbo |
| PLBD1-AS1 | 12 | PLBD1 antisense RNA 1 [Source:HGNC Symbol |
| GUCY2C | 12 | guanylate cyclase 2C [Source:HGNC Symbol;A |
| AC010168.1 | 12 | novel transcript |
| AC010168.2 | 12 | novel transcript, overlapping HIST4H4 |
| HIST4H4 | 12 | histone cluster 4 H4 [Source:HGNC Symbol;Ac |
| H2AFJ | 12 | H2A histone family member J [Source:HGNC S |
| WBP11 | 12 | WW domain binding protein 11 [Source:HGN( |
| C12orf60 | 12 | chromosome 12 open reading frame 60 [Sour |
| SMCO3 | 12 | single-pass membrane protein with coiled-coi |
| AC007655.2 | 12 | novel transcript |
| ART4 | 12 | ADP-ribosyltransferase 4 (Dombrock blood gr |
| AC007655.1 | 12 | novel transcript |
| MGP | 12 | matrix Gla protein [Source:HGNC Symbol;Acc: |

| | | |
|---|---|---|
| ERP27 | 12 | endoplasmic reticulum protein 27 [Source:HG |
| ARHGDIB | 12 | Rho GDP dissociation inhibitor beta [Source:H |
| PDE6H | 12 | phosphodiesterase 6H [Source:HGNC Symbol; |
| LINC01489 | 12 | long intergenic non-protein coding RNA 1489 |
| RERG | 12 | RAS like estrogen regulated growth inhibitor [ |
| RERG-IT1 | 12 | RERG intronic transcript 1 [Source:HGNC Syml |
| RERG-AS1 | 12 | RERG antisense RNA 1 [Source:HGNC Symbol; |
| METTL8P1 | 12 | methyltransferase like 8 pseudogene 1 [Sourc |
| PTPRO | 12 | protein tyrosine phosphatase receptor type O |
| AC092183.1 | 12 | ribosomal protein S2 (RPS2) pseudogene |
| EPS8 | 12 | epidermal growth factor receptor pathway su |
| RNU6-251P | 12 | RNA, U6 small nuclear 251, pseudogene [Sour |
| AC073651.1 | 12 | novel transcript |
| AC073651.2 | 12 | mitchondrial ribosomal protein S7 (MRPS7) ps |
| STRAP | 12 | serine/threonine kinase receptor associated p |
| DERA | 12 | deoxyribose-phosphate aldolase [Source:HGN |
| EGLN3P1 | 12 | egl-9 family hypoxia-inducible factor 3 pseudo |
| SLC15A5 | 12 | solute carrier family 15 member 5 [Source:HG |
| MGST1 | 12 | microsomal glutathione S-transferase 1 [Sourc |
| SUPT16HP1 | 12 | SPT16 homolog, facilitates chromatin remode |
| AC007528.1 | 12 | novel transcript |
| AC007552.1 | 12 | glutamic-oxaloacetic transaminase 2, mitocho |
| LMO3 | 12 | LIM domain only 3 [Source:HGNC Symbol;Acc |
| AC007552.2 | 12 | novel transcript |
| AC007529.2 | 12 | novel transcript |
| AC007529.1 | 12 | novel transcript |
| AC023512.1 | 12 | S-phase kinase-associated protein 1 (SKP1) ps |
| EEF1A1P16 | 12 | eukaryotic translation elongation factor 1 alph |
| AC092793.1 | 12 | novel transcript |
| RNU6-837P | 12 | RNA, U6 small nuclear 837, pseudogene [Sour |
| AC092110.1 | 12 | novel transcript |
| RPL7P40 | 12 | ribosomal protein L7 pseudogene 40 [Source: |
| PSMC1P8 | 12 | proteasome 26S subunit, ATPase 1 pseudoger |
| TIMM17BP1 | 12 | translocase of inner mitochondrial membrane |
| AC048352.1 | 12 | novel transcript |
| LINC02378 | 12 | long intergenic non-protein coding RNA 2378 |
| MIR3974 | 12 | microRNA 3974 [Source:HGNC Symbol;Acc:HG |
| Y_RNA | 12 | Y RNA [Source:RFAM;Acc:RF00019] |
| RERGL | 12 | RERG like [Source:HGNC Symbol;Acc:HGNC:26 |
| PIK3C2G | 12 | phosphatidylinositol-4-phosphate 3-kinase ca |
| NDFIP1P1 | 12 | Nedd4 family interacting protein 1 pseudogen |
| AC092851.1 | 12 | F-box only protein 3 (FBXO3) pseudogene |
| ZKSCAN7P1 | 12 | zinc finger with KRAB and SCAN domains 7 ps |
| PLCZ1 | 12 | phospholipase C zeta 1 [Source:HGNC Symbol |
| PSMC1P9 | 12 | proteasome 26S subunit, ATPase 1 pseudoger |
| AC087242.1 | 12 | novel transcript, antisense to PLCZ1 |
| CAPZA3 | 12 | capping actin protein of muscle Z-line subunit |
| RPL7P6 | 12 | ribosomal protein L7 pseudogene 6 [Source:H |
| MEF2BNBP1 | 12 | MEF2B neighbor pseudogene 1 [Source:HGNC |
| PLEKHA5 | 12 | pleckstrin homology domain containing A5 [So |

| | | |
|---|---|---|
| AC092828.1 | 12 | novel transcript |
| RN7SL459P | 12 | RNA, 7SL, cytoplasmic 459, pseudogene [Sour |
| RN7SL67P | 12 | RNA, 7SL, cytoplasmic 67, pseudogene [Sourc |
| AEBP2 | 12 | AE binding protein 2 [Source:HGNC Symbol;Ac |
| PDCD5P1 | 12 | programmed cell death 5 pseudogene 1 [Sour |
| AC091805.2 | 12 | phosphoprotein enriched in astrocytes 15 (PE |
| RNU6-254P | 12 | RNA, U6 small nuclear 254, pseudogene [Sour |
| EEF1A1P4 | 12 | eukaryotic translation elongation factor 1 alpl |
| AC091805.1 | 12 | ribosomal protein L34 (RPL34) pseudogene |
| AC137561.1 | 12 | novel transcript, sense intronic to AEBP2 |
| RNU1-146P | 12 | RNA, U1 small nuclear 146, pseudogene [Sour |
| AC024901.1 | 12 | novel transcript |
| TCP1P3 | 12 | t-complex 1 pseudogene 3 [Source:HGNC Sym |
| LINC02398 | 12 | long intergenic non-protein coding RNA 2398 |
| AC126468.1 | 12 | novel transcript |
| LINC02468 | 12 | long intergenic non-protein coding RNA 2468 |
| AC129102.1 | 12 | novel transcript |
| PDE3A | 12 | phosphodiesterase 3A [Source:HGNC Symbol; |
| UBE2L2 | 12 | ubiquitin conjugating enzyme E2 L2 (pseudoge |
| AC112777.1 | 12 | ubiquitin-like with PHD and ring finger domair |
| SLCO1C1 | 12 | solute carrier organic anion transporter family |
| SLCO1B3 | 12 | solute carrier organic anion transporter family |
| AC011604.2 | 12 | novel protein |
| AC011604.1 | 12 | proteasomal ATPase-associated factor 1 (PAA |
| SLCO1B7 | 12 | solute carrier organic anion transporter family |
| AC022335.1 | 12 | novel transcript |
| SLCO1B1 | 12 | solute carrier organic anion transporter family |
| SLCO1A2 | 12 | solute carrier organic anion transporter family |
| IAPP | 12 | islet amyloid polypeptide [Source:HGNC Symb |
| PYROXD1 | 12 | pyridine nucleotide-disulphide oxidoreductase |
| ELOCP31 | 12 | elongin C pseudogene 31 [Source:HGNC Symb |
| RECQL | 12 | RecQ like helicase [Source:HGNC Symbol;Acc: |
| GOLT1B | 12 | golgi transport 1B [Source:HGNC Symbol;Acc:I |
| SPX | 12 | spexin hormone [Source:HGNC Symbol;Acc:H |
| GYS2 | 12 | glycogen synthase 2 [Source:HGNC Symbol;Ac |
| AC010197.2 | 12 | novel transcript |
| LDHB | 12 | lactate dehydrogenase B [Source:HGNC Symb |
| AC010197.1 | 12 | novel transcript |
| AC010185.1 | 12 | novel transcript |
| KCNJ8 | 12 | potassium voltage-gated channel subfamily J r |
| ABCC9 | 12 | ATP binding cassette subfamily C member 9 [S |
| AC008250.1 | 12 | novel transcript |
| AC008250.2 | 12 | novel transcript, sense intronic to ABCC9 |
| AC092862.1 | 12 | thioesterase superfamily member 4 (THEM4) |
| CMAS | 12 | cytidine monophosphate N-acetylneuraminic |
| ST8SIA1 | 12 | ST8 alpha-N-acetyl-neuraminide alpha-2,8-sia |
| AC007671.1 | 12 | novel transcript, sense intronic to ST8SIA1 |
| SULT6B2P | 12 | sulfotransferase family 6B member 2, pseudo |
| RNU1-149P | 12 | RNA, U1 small nuclear 149, pseudogene [Sour |
| AC087318.1 | 12 | novel transcript |

| | | |
|---|---|---|
| AC053513.1 | 12 | novel transcript |
| C2CD5 | 12 | C2 calcium dependent domain containing 5 [S |
| AC053513.2 | 12 | novel transcript |
| LRRC34P1 | 12 | leucine rich repeat containing 34 pseudogene |
| AC087241.1 | 12 | novel pseudogene |
| AC087241.2 | 12 | novel transcript, antisense to ETNK1 |
| ETNK1 | 12 | ethanolamine kinase 1 [Source:HGNC Symbol; |
| AC084816.1 | 12 | novel transcript |
| RPS27P22 | 12 | ribosomal protein S27 pseudogene 22 [Source |
| AC084819.1 | 12 | novel transcript |
| AC087258.1 | 12 | N(alpha)-acetyltransferase 20, NatB catalytic s |
| AC087235.2 | 12 | novel transcript |
| AC087235.1 | 12 | novel transcript |
| SOX5 | 12 | SRY-box transcription factor 5 [Source:HGNC S |
| AC087260.1 | 12 | novel transcript, antisense to SOX5 |
| MIR920 | 12 | microRNA 920 [Source:HGNC Symbol;Acc:HGN |
| SOX5-AS1 | 12 | SOX5 antisense RNA 1 [Source:HGNC Symbol;A |
| LINC00477 | 12 | long intergenic non-protein coding RNA 477 [S |
| KNOP1P1 | 12 | lysine rich nucleolar protein 1 pseudogene 1 [ |
| AC087312.1 | 12 | ribosomal protein L21 (RPL21) pseudogene |
| AC023796.1 | 12 | novel transcript |
| RN7SL38P | 12 | RNA, 7SL, cytoplasmic 38, pseudogene [Source |
| BCAT1 | 12 | branched chain amino acid transaminase 1 [So |
| AC023796.2 | 12 | novel transcript, antisense to BCAT1 |
| AC026310.3 | 12 | novel transcript |
| AC026310.2 | 12 | novel transcript, antisense to BCAT1 |
| AC026310.1 | 12 | brain protein I3 (BR13) pseudogene |
| C12orf77 | 12 | chromosome 12 open reading frame 77 [Sour |
| LRMP | 12 | lymphoid restricted membrane protein [Sourc |
| CENPUP2 | 12 | centromere protein U pseudogene 2 [Source:I |
| AC023510.1 | 12 | novel transcript, antisense to LRMP |
| AC023510.2 | 12 | novel transcript, antisense to LRMP |
| CASC1 | 12 | cancer susceptibility 1 [Source:HGNC Symbol;A |
| ETFRF1 | 12 | electron transfer flavoprotein regulatory facto |
| KRAS | 12 | KRAS proto-oncogene, GTPase [Source:HGNC |
| AC092794.1 | 12 | novel transcript, antisense to KRAS |
| AC092794.2 | 12 | novel transcript, antisense to KRAS |
| AC087239.1 | 12 | novel transcript |
| RNU4-67P | 12 | RNA, U4 small nuclear 67, pseudogene [Sourc |
| AC092451.1 | 12 | ribosomal protein L39 (RPL39) pseudogene |
| LMNTD1 | 12 | lamin tail domain containing 1 [Source:HGNC |
| AC092451.3 | 12 | novel transcript |
| AC092451.2 | 12 | family with sequence similarity 133, member |
| TUBB4BP1 | 12 | tubulin beta 4B class IVb pseudogene 1 [Sourc |
| AC022367.1 | 12 | novel transcript |
| RN7SKP262 | 12 | RN7SK pseudogene 262 [Source:HGNC Symbo |
| AC019209.3 | 12 | novel transcript |
| TDGP1 | 12 | thymine-DNA glycosylase pseudogene 1 [Sour |
| AC019209.2 | 12 | novel transcript |
| MIR4302 | 12 | microRNA 4302 [Source:HGNC Symbol;Acc:HG |

| | | |
|---|---|---|
| AC019209.1 | 12 | novel transcript |
| RASSF8-AS1 | 12 | RASSF8 antisense RNA 1 [Source:HGNC Symbo |
| RASSF8 | 12 | Ras association domain family member 8 [Sou |
| BHLHE41 | 12 | basic helix-loop-helix family member e41 [Sou |
| SSPN | 12 | sarcospan [Source:HGNC Symbol;Acc:HGNC:1 |
| AC022509.3 | 12 | novel transcript, antisense to SSPN |
| AC022509.5 | 12 | novel transcript, antisense to SSPN |
| AC022509.2 | 12 | novel transcript, antisense to SSPN and ITPR2 |
| AC022509.1 | 12 | novel transcript, antisense to SSPN |
| AC022509.4 | 12 | novel transcript, antisense to SSPN |
| AC055720.1 | 12 | novel transcript |
| AC024145.1 | 12 | novel transcript to ITPR2 |
| ITPR2 | 12 | inositol 1,4,5-trisphosphate receptor type 2 [S |
| AC055720.2 | 12 | novel transcript |
| RNA5SP354 | 12 | RNA, 5S ribosomal pseudogene 354 [Source:H |
| AC023051.1 | 12 | novel transcript, antisense to ITPR2 |
| AC023051.2 | 12 | Enoyl-CoA hydratase domain-containing prote |
| INTS13 | 12 | integrator complex subunit 13 [Source:HGNC |
| FGFR1OP2 | 12 | FGFR1 oncogene partner 2 [Source:HGNC Sym |
| TM7SF3 | 12 | transmembrane 7 superfamily member 3 [Sou |
| AC024896.1 | 12 | novel transcript, antisense to TM7SF3 |
| MED21 | 12 | mediator complex subunit 21 [Source:HGNC S |
| AC092747.4 | 12 | novel transcript |
| C12orf71 | 12 | chromosome 12 open reading frame 71 [Sour |
| AC092747.3 | 12 | high mobility group nucleosome binding doma |
| AC092747.1 | 12 | novel transcript |
| AC092747.2 | 12 | liprin-beta-1-like (PPFIBP1) pseudogene |
| STK38L | 12 | serine/threonine kinase 38 like [Source:HGNC |
| ARNTL2 | 12 | aryl hydrocarbon receptor nuclear translocato |
| ARNTL2-AS1 | 12 | ARNTL2 antisense RNA 1 [Source:HGNC Symb |
| SMCO2 | 12 | single-pass membrane protein with coiled-coi |
| RARSP1 | 12 | arginyl-tRNA synthetase pseudogene 1 [Sourc |
| PPFIBP1 | 12 | PPFIA binding protein 1 [Source:HGNC Symbo |
| AC087257.1 | 12 | novel transcript |
| AC087257.2 | 12 | chromosome 12 open reading frame 71 (C12o |
| AC009509.3 | 12 | high-mobility group nucleosome binding dom |
| AC009509.1 | 12 | novel transcript |
| REP15 | 12 | RAB15 effector protein [Source:HGNC Symbol |
| AC009509.4 | 12 | novel transcript, antisense to a lncRNA |
| HMGB1P49 | 12 | high mobility group box 1 pseudogene 49 [Sou |
| AC009509.2 | 12 | novel transcript |
| MRPS35 | 12 | mitochondrial ribosomal protein S35 [Source: |
| Y_RNA | 12 | Y RNA [Source:RFAM;Acc:RF00019] |
| MANSC4 | 12 | MANSC domain containing 4 [Source:HGNC Sy |
| AC009511.2 | 12 | novel transcript |
| KLHL42 | 12 | kelch like family member 42 [Source:HGNC Sy |
| AC009511.1 | 12 | novel transcript |
| RN7SKP15 | 12 | RN7SK pseudogene 15 [Source:HGNC Symbol; |
| AC009511.3 | 12 | NULL |
| AC008011.1 | 12 | pseudogene similar to part of aldehyde dehyd |

| | | |
|---|---|---|
| PTHLH | 12 | parathyroid hormone like hormone [Source:H |
| AC008011.2 | 12 | novel transcript, antisense to PTHLH |
| CCDC91 | 12 | coiled-coil domain containing 91 [Source:HGN |
| AC022364.1 | 12 | novel transcript, antisense to CCDC91 |
| AC022364.2 | 12 | intraflagellar transport 57 homolog (Chlamydo |
| AC022079.2 | 12 | novel transcript, antisense to CCDC91 |
| AC022079.1 | 12 | novel transcript, sense intronic to CCDC91 |
| RNU4-54P | 12 | RNA, U4 small nuclear 54, pseudogene [Sourc |
| RNA5SP355 | 12 | RNA, 5S ribosomal pseudogene 355 [Source:H |
| AC084754.1 | 12 | ribosomal protein L29 (RPL29) pseudogene |
| AC022081.1 | 12 | novel transcript |
| AC024255.1 | 12 | actin-like 8 (ACTL8) pseudogene |
| AC012150.1 | 12 | novel transcript |
| FAR2 | 12 | fatty acyl-CoA reductase 2 [Source:HGNC Sym |
| AC012150.2 | 12 | novel transcript, antisense to FAR2 |
| AC009318.4 | 12 | novel transcript, antisense to FAR2 |
| AC009318.1 | 12 | novel transcript, antisense to FAR2 |
| AC009318.3 | 12 | novel transcript, antisense to FAR2 |
| AC009318.2 | 12 | novel transcript, antisense to FAR2 |
| ERGIC2 | 12 | ERGIC and golgi 2 [Source:HGNC Symbol;Acc:H |
| OVCH1-AS1 | 12 | OVCH1 antisense RNA 1 [Source:HGNC Symbo |
| OVCH1 | 12 | ovochymase 1 [Source:HGNC Symbol;Acc:HGN |
| TMTC1 | 12 | transmembrane O-mannosyltransferase targe |
| AC009320.1 | 12 | novel transcript |
| RPL21P99 | 12 | ribosomal protein L21 pseudogene 99 [Source |
| AC023511.3 | 12 | novel transcript |
| AC023511.2 | 12 | transmembrane and tetratricopeptide repeat |
| AC023511.1 | 12 | novel transcript |
| LINC02386 | 12 | long intergenic non-protein coding RNA 2386 |
| AC078776.2 | 12 | maestro heat-like repeat-containing protein fa |
| RNA5SP356 | 12 | RNA, 5S ribosomal pseudogene 356 [Source:H |
| AC078776.1 | 12 | novel transcript |
| AC026371.1 | 12 | pseudogene similar to part of transmembrane |
| IPO8 | 12 | importin 8 [Source:HGNC Symbol;Acc:HGNC:9 |
| AC012673.1 | 12 | novel transcript |
| CAPRIN2 | 12 | caprin family member 2 [Source:HGNC Symbo |
| AC010198.1 | 12 | novel transcript |
| LINC00941 | 12 | long intergenic non-protein coding RNA 941 [S |
| AC010198.2 | 12 | long intergenic non-protein coding RNA 941 |
| PPIAP44 | 12 | peptidylprolyl isomerase A pseudogene 44 [So |
| TSPAN11 | 12 | tetraspanin 11 [Source:HGNC Symbol;Acc:HGN |
| AC008013.1 | 12 | tetraspanin family pseudogene |
| AC008013.2 | 12 | mitochondrially encoded cytochrome c oxidas |
| DDX11-AS1 | 12 | DDX11 antisense RNA 1 [Source:HGNC Symbo |
| DDX11 | 12 | DEAD/H-box helicase 11 [Source:HGNC Symbo |
| SNORA75 | 12 | Small nucleolar RNA SNORA75 [Source:RFAM; |
| AC008013.3 | 12 | WAS protein family homolog 4 pseudogene |
| AC024940.2 | 12 | ovostatin 2 |
| AC024940.5 | 12 | NADH dehydrogenase (ubiquinone) 1 alpha su |
| AC024940.3 | 12 | ribosomal protein L13a (RPL13A) pseudogene |

| | | |
|---|---|---|
| MREGP1 | 12 | melanoregulin pseudogene 1 [Source:HGNC S |
| AC024940.6 | 12 | novel transcript, overlapping 3'UTR of FAM60. |
| SINHCAF | 12 | SIN3-HDAC complex associated factor [Source |
| AC024940.4 | 12 | solute carrier family 25, member 13 (citrin) (S |
| AC024940.1 | 12 | novel transcript, antisense to FAM60A |
| LINC02387 | 12 | long intergenic non-protein coding RNA 2387 |
| U6 | 12 | U6 spliceosomal RNA [Source:RFAM;Acc:RF00 |
| DENND5B | 12 | DENN domain containing 5B [Source:HGNC Sy |
| RNU6-618P | 12 | RNA, U6 small nuclear 618, pseudogene [Sour |
| AC068792.1 | 12 | novel transcript, sense intronic to DENND5B |
| AC022080.1 | 12 | TAF10 RNA polymerase II, TATA box binding p |
| AC022080.3 | 12 | mitochondrial ribosomal protein L30 (MRPL3C |
| Y_RNA | 12 | Y RNA [Source:RFAM;Acc:RF00019] |
| AC022080.2 | 12 | ankyrin repeat domain 49 (ANKRD49) pseudo |
| DENND5B-AS1 | 12 | DENND5B antisense RNA 1 [Source:HGNC Syn |
| RNU5F-4P | 12 | RNA, U5F small nuclear 4, pseudogene [Sourc |
| RPL31P50 | 12 | ribosomal protein L31 pseudogene 50 [Source |
| AK4P3 | 12 | adenylate kinase 4 pseudogene 3 [Source:HGI |
| AC068774.1 | 12 | novel transcript, antisense to C12orf72 |
| ETFBKMT | 12 | electron transfer flavoprotein subunit beta lys |
| AMN1 | 12 | antagonist of mitotic exit network 1 homolog |
| STMN1P1 | 12 | stathmin 1 pseudogene 1 [Source:HGNC Symb |
| AC023157.3 | 12 | novel transcript, antisense to AMN1 |
| AC023157.2 | 12 | dpy-30 homolog (C. elegans) (DPY30) pseudog |
| AC023157.1 | 12 | interferon induced transmembrane protein 3 |
| H3F3C | 12 | H3 histone family member 3C [Source:HGNC S |
| RNU6-1069P | 12 | RNA, U6 small nuclear 1069, pseudogene [Sou |
| AC023050.2 | 12 | NULL |
| AC023050.3 | 12 | basic transcription factor 3-like 4 (BTF3L4) pse |
| LINC02422 | 12 | long intergenic non-protein coding RNA 2422 |
| AC023050.4 | 12 | Impact homolog (mouse) (IMPACT) pseudoge |
| AC023050.1 | 12 | ribosomal protein, large, P2 (RPLP2) pseudog |
| RPL12P32 | 12 | ribosomal protein L12 pseudogene 32 [Source |
| AC023050.5 | 12 | family with sequence similarity 98, member B |
| RESF1 | 12 | retroelement silencing factor 1 [Source:HGNC |
| AC016957.2 | 12 | novel transcript |
| AC016957.1 | 12 | COX assembly mitochondrial protein homolog |
| AC048344.2 | 12 | gamma-aminobutyric acid (GABA) A receptor, |
| AC048344.3 | 12 | aurora kinase B (AURKB) pseudogene |
| AC048344.1 | 12 | novel transcript |
| BICD1 | 12 | BICD cargo adaptor 1 [Source:HGNC Symbol;A |
| AC048344.4 | 12 | novel transcript, sense intronic to BICD1 |
| AC016954.1 | 12 | WW domain binding protein 2 (WBP2) pseudo |
| AC026356.1 | 12 | novel transcript, sense intronic to BICD1 |
| AC026356.2 | 12 | novel transcript, sense intronic to BICD1 |
| Y_RNA | 12 | Y RNA [Source:RFAM;Acc:RF00019] |
| FGD4 | 12 | FYVE, RhoGEF and PH domain containing 4 [So |
| RNU6-494P | 12 | RNA, U6 small nuclear 494, pseudogene [Sour |
| DNM1L | 12 | dynamin 1 like [Source:HGNC Symbol;Acc:HGI |
| AC084824.2 | 12 | mitochondrial assembly of ribosomal large sul |

| | | |
|---|---|---|
| Y_RNA | 12 | Y RNA [Source:RFAM;Acc:RF00019] |
| AC084824.5 | 12 | novel transcript, antisense to DNM1L |
| AC084824.1 | 12 | nucleosome assembly protein 1-like 1 (NAP1L |
| YARS2 | 12 | tyrosyl-tRNA synthetase 2 [Source:HGNC Sym |
| AC084824.3 | 12 | novel transcript, sense intronic to YARS2 |
| AC084824.4 | 12 | novel transcript, sense intronic to YARS2 |
| AC087588.3 | 12 | novel transcript, antisense to YARS2 |
| PKP2 | 12 | plakophilin 2 [Source:HGNC Symbol;Acc:HGN( |
| AC087588.2 | 12 | novel transcript, sense intronic to PKP2 |
| AC087588.1 | 12 | ribosomal protein L35a (RPL35A) pseudogene |
| AC087311.1 | 12 | argininosuccinate synthase 1 (ASS1) pseudoge |
| AC087311.2 | 12 | novel transcript |
| SYT10 | 12 | synaptotagmin 10 [Source:HGNC Symbol;Acc: |
| AC023158.1 | 12 | novel transcript, sense intronic to SYT10 |
| AC023158.2 | 12 | novel transcript, sense intronic to SYT10 |
| AC016956.1 | 12 | suppression of tumorigenicity 13 (colon carcir |
| RNU6-400P | 12 | RNA, U6 small nuclear 400, pseudogene [Sour |
| AC024153.1 | 12 | R3H domain and coiled-coil containing 1-like ( |
| RNU6-472P | 12 | RNA, U6 small nuclear 472, pseudogene [Sour |
| Y_RNA | 12 | Y RNA [Source:RFAM;Acc:RF00019] |
| AC046130.1 | 12 | novel transcript, antisense to ALG10 |
| ALG10 | 12 | ALG10 alpha-1,2-glucosyltransferase [Source:I |
| AC046130.2 | 12 | novel transcript |
| AC140847.1 | 12 | novel transcript |
| AC140847.3 | 12 | FAM166A, pseudogene 2 |
| TUBB8P4 | 12 | tubulin beta 8 class VIII pseudogene 4 [Source |
| AC140847.2 | 12 | novel transcript |
| RNA5SP357 | 12 | RNA, 5S ribosomal pseudogene 357 [Source:H |
| DUX4L27 | 12 | double homeobox 4 like 27 (pseudogene) [Sou |
| AK6P1 | 12 | adenylate kinase 6 pseudogene 1 [Source:HGI |
| ZNF970P | 12 | zinc finger protein 970, pseudogene [Source:H |
| AK6P2 | 12 | adenylate kinase 6 pseudogene 2 [Source:HGI |
| CLUHP8 | 12 | clustered mitochondria homolog pseudogene |
| AC079460.1 | 12 | NULL |
| AC079460.2 | 12 | novel transcript |
| RNA5SP358 | 12 | RNA, 5S ribosomal pseudogene 358 [Source:H |
| RNA5SP359 | 12 | RNA, 5S ribosomal pseudogene 359 [Source:H |
| TUBB8P5 | 12 | tubulin beta 8 class VIII pseudogene 5 [Source |
| AC117372.1 | 12 | FAM166A, pseudogene 9 |
| NF1P12 | 12 | neurofibromin 1 pseudogene 12 [Source:HGN |
| Y_RNA | 12 | Y RNA [Source:RFAM;Acc:RF00019] |
| ALG10B | 12 | ALG10 alpha-1,2-glucosyltransferase B [Sourc |
| AC087897.1 | 12 | ring finger protein 141 (RNF141) pseudogene |
| AC087897.2 | 12 | novel transcript |
| CPNE8 | 12 | copine 8 [Source:HGNC Symbol;Acc:HGNC:23 |
| CPNE8-AS1 | 12 | CPNE8 antisense RNA 1 [Source:HGNC Symbo |
| AC018448.2 | 12 | maltase-glucoamylase (alpha-glucosidase) (M |
| AC018448.1 | 12 | maltase-glucoamylase (alpha-glucosidase) (M |
| LINC02406 | 12 | long intergenic non-protein coding RNA 2406 |
| AC084373.1 | 12 | cytochrome c oxidase subunit VIIb (COX7B) ps |

| | | |
|---|---|---|
| KIF21A | 12 | kinesin family member 21A [Source:HGNC Syr |
| AC121334.4 | 12 | novel transcript |
| AC121334.1 | 12 | ribosomal protein L7a (RPL7A) pseudogene |
| AC121334.2 | 12 | transmembrane protein 230 (TMEM230) pseu |
| ABCD2 | 12 | ATP binding cassette subfamily D member 2 [S |
| AC121334.3 | 12 | novel transcript, antisense to ABCD2 |
| C12orf40 | 12 | chromosome 12 open reading frame 40 [Sour |
| SLC2A13 | 12 | solute carrier family 2 member 13 [Source:HG |
| AC121336.1 | 12 | NULL |
| RPL30P13 | 12 | ribosomal protein L30 pseudogene 13 [Source |
| LINC02555 | 12 | long intergenic non-protein coding RNA 2555 |
| LINC02471 | 12 | long intergenic non-protein coding RNA 2471 |
| AC079630.1 | 12 | novel transcript, antisense to LRRK2 |
| LRRK2 | 12 | leucine rich repeat kinase 2 [Source:HGNC Syr |
| AC084290.1 | 12 | pseudogene similar to part of the cytochrome |
| MUC19 | 12 | mucin 19, oligomeric [Source:HGNC Symbol;A |
| AC107023.1 | 12 | novel transcript |
| RNU6-713P | 12 | RNA, U6 small nuclear 713, pseudogene [Sour |
| CNTN1 | 12 | contactin 1 [Source:HGNC Symbol;Acc:HGNC:: |
| AC016144.1 | 12 | methionyl aminopeptidase 1 (METAP1) pseud |
| AC015540.1 | 12 | novel transcript, sense intronic to CNTN1 |
| PDZRN4 | 12 | PDZ domain containing ring finger 4 [Source:H |
| AC090531.1 | 12 | novel transcript |
| RNA5SP360 | 12 | RNA, 5S ribosomal pseudogene 360 [Source:H |
| MTND2P17 | 12 | MT-ND2 pseudogene 17 [Source:HGNC Symbo |
| MTND1P24 | 12 | MT-ND1 pseudogene 24 [Source:HGNC Symbo |
| LINC02400 | 12 | long intergenic non-protein coding RNA 2400 |
| AC090630.1 | 12 | novel transcript |
| AC006197.1 | 12 | tetratricopeptide repeat domain 5 (TTC5) pse |
| AC006197.2 | 12 | novel transcript |
| AC023513.1 | 12 | family with sequence similarity 173, member |
| GXYLT1 | 12 | glucoside xylosyltransferase 1 [Source:HGNC S |
| YAF2 | 12 | YY1 associated factor 2 [Source:HGNC Symbol |
| PPHLN1 | 12 | periphilin 1 [Source:HGNC Symbol;Acc:HGNC: |
| AC020629.1 | 12 | insulin-like growth factor 2 mRNA binding pro |
| ZCRB1 | 12 | zinc finger CCHC-type and RNA binding motif c |
| MIR7851 | 12 | microRNA 7851 [Source:HGNC Symbol;Acc:HG |
| SNORA74 | 12 | Small nucleolar RNA SNORA74 [Source:RFAM; |
| AC079684.1 | 12 | novel transcript, sense intronic to PPHLN1 |
| RNU6-249P | 12 | RNA, U6 small nuclear 249, pseudogene [Sour |
| AC079601.2 | 12 | fragile X mental retardation, autosomal homo |
| Y_RNA | 12 | Y RNA [Source:RFAM;Acc:RF00019] |
| PRICKLE1 | 12 | prickle planar cell polarity protein 1 [Source:H |
| AC079601.1 | 12 | novel transcript, antisense to PRICKLE1 |
| AC079600.3 | 12 | novel transcript, sense intronic to PRICKLE1 |
| LINC02402 | 12 | long intergenic non-protein coding RNA 2402 |
| AC079600.2 | 12 | DNA binding protein A pseudogene |
| AC079600.1 | 12 | CDGSH iron sulfur domain pseudogene |
| LINC02451 | 12 | long intergenic non-protein coding RNA 2451 |
| RPS27P21 | 12 | ribosomal protein S27 pseudogene 21 [Source |

| | | |
|---|---|---|
| LINC02450 | 12 | long intergenic non-protein coding RNA 2450 |
| AC068802.1 | 12 | aminolevulinate, delta-, synthase 1 (ALAS1) ps |
| AC012038.2 | 12 | novel transcript |
| AC012038.1 | 12 | novel transcript |
| MRPS36P5 | 12 | mitochondrial ribosomal protein S36 pseudog |
| LINC02461 | 12 | long intergenic non-protein coding RNA 2461 |
| ADAMTS20 | 12 | ADAM metallopeptidase with thrombospondi |
| AC090525.1 | 12 | ribosomal protein L21 (RPL21) pseudogene |
| AC090525.2 | 12 | protein-kinase, interferon-inducible double st |
| AC090525.3 | 12 | ribosomal protein S2 (RPS2) pseudogene |
| EEF1A1P17 | 12 | eukaryotic translation elongation factor 1 alph |
| PUS7L | 12 | pseudouridine synthase 7 like [Source:HGNC S |
| AC093012.1 | 12 | La ribonucleoprotein domain family, member |
| IRAK4 | 12 | interleukin 1 receptor associated kinase 4 [Sou |
| TWF1 | 12 | twinfilin actin binding protein 1 [Source:HGNC |
| TMEM117 | 12 | transmembrane protein 117 [Source:HGNC Sy |
| ZNF75BP | 12 | zinc finger protein 75B, pseudogene [Source:H |
| AC025030.2 | 12 | novel transcript, antisense to TMEM117 |
| AC025030.1 | 12 | IQ motif containing F1 (IQCF1) pseudogene |
| AC025253.1 | 12 | novel transcript |
| AC025253.2 | 12 | novel transcript |
| NELL2 | 12 | neural EGFL like 2 [Source:HGNC Symbol;Acc:I |
| Y_RNA | 12 | Y RNA [Source:RFAM;Acc:RF00019] |
| DBX2 | 12 | developing brain homeobox 2 [Source:HGNC S |
| AC008127.1 | 12 | novel transcript, antisense to DBX2 |
| RACGAP1P | 12 | Rac GTPase activating protein 1 pseudogene [ |
| AC008127.2 | 12 | Sjogren syndrome antigen B (autoantigen La) |
| RNA5SP361 | 12 | RNA, 5S ribosomal pseudogene 361 [Source:H |
| AC009248.3 | 12 | novel transcript |
| PLEKHA8P1 | 12 | pleckstrin homology domain containing A8 ps |
| AC009248.1 | 12 | carbonyl reductase 3 (CBR3) pseudogene |
| ANO6 | 12 | anoctamin 6 [Source:HGNC Symbol;Acc:HGNC |
| AC009248.2 | 12 | novel transcript, sense intronic to ANO6 |
| AC063924.1 | 12 | mesoderm development candidate 2 (MESDC |
| AC079950.1 | 12 | novel transcript |
| U6 | 12 | U6 spliceosomal RNA [Source:RFAM;Acc:RF00 |
| AC008124.1 | 12 | novel transcript |
| ARID2 | 12 | AT-rich interaction domain 2 [Source:HGNC Sy |
| AC009464.1 | 12 | novel transcript, sense intronic to ARID2 |
| RN7SL246P | 12 | RNA, 7SL, cytoplasmic 246, pseudogene [Sour |
| KNOP1P2 | 12 | lysine rich nucleolar protein 1 pseudogene 2 [ |
| SCAF11 | 12 | SR-related CTD associated factor 11 [Source:H |
| AC084878.1 | 12 | ribosomal protein L13a (RPL13A) pseudogene |
| SLC38A1 | 12 | solute carrier family 38 member 1 [Source:HG |
| AC025031.3 | 12 | novel transcript, sense intronic to SLC38A1 |
| SLC38A2 | 12 | solute carrier family 38 member 2 [Source:HG |
| AC025031.2 | 12 | novel transcript, antisense to SLC38A2 |
| AC008014.1 | 12 | novel transcript |
| AC025031.1 | 12 | novel transcript |
| AC025031.4 | 12 | novel transcript |

| | | |
|---|---|---|
| AC025031.5 | 12 | TEC |
| AC008035.1 | 12 | novel transcript |
| OR7A19P | 12 | olfactory receptor family 7 subfamily A memb |
| MARK3P1 | 12 | microtubule affinity regulating kinase 3 pseud |
| SLC38A4 | 12 | solute carrier family 38 member 4 [Source:HG |
| AC079906.1 | 12 | novel transcript |
| AMIGO2 | 12 | adhesion molecule with Ig like domain 2 [Sour |
| PCED1B | 12 | PC-esterase domain containing 1B [Source:HG |
| AC119044.1 | 12 | interferon induced transmembrane protein 3 |
| MIR4698 | 12 | microRNA 4698 [Source:HGNC Symbol;Acc:HG |
| PCED1B-AS1 | 12 | PCED1B antisense RNA 1 [Source:HGNC Symb |
| AC008083.2 | 12 | novel transcript, antisense to FAM113B |
| AC008083.3 | 12 | novel transcript |
| AC008083.4 | 12 | novel transcript |
| PPIAP45 | 12 | peptidylprolyl isomerase A pseudogene 45 [Sc |
| AC008083.1 | 12 | NULL |
| LINC02416 | 12 | long intergenic non-protein coding RNA 2416 |
| MIR4494 | 12 | microRNA 4494 [Source:HGNC Symbol;Acc:HG |
| LINC02156 | 12 | long intergenic non-protein coding RNA 2156 |
| ADI1P3 | 12 | acireductone dioxygenase 1 pseudogene 3 [Sc |
| AC003686.1 | 12 | starch binding domain 1 (STBD1) pseudogene |
| AC004486.1 | 12 | ubiquitin family Domain containing pseudoger |
| RPAP3 | 12 | RNA polymerase II associated protein 3 [Sourc |
| AC004241.3 | 12 | novel transcript, sense intronic to RPAP3 |
| AC004241.1 | 12 | novel transcript, antisense to ENDOU and RAF |
| ENDOU | 12 | endonuclease, poly(U) specific [Source:HGNC |
| AC004241.5 | 12 | TEC |
| AC004241.2 | 12 | novel transcript |
| RAPGEF3 | 12 | Rap guanine nucleotide exchange factor 3 [So |
| SLC48A1 | 12 | solute carrier family 48 member 1 [Source:HG |
| AC004241.4 | 12 | novel transcript, sense intronic to SLC48A1 |
| HDAC7 | 12 | histone deacetylase 7 [Source:HGNC Symbol;/ |
| AC004466.1 | 12 | novel transcript, antisense to HDAC7 |
| AC004466.3 | 12 | novel transcript, antisense to HDAC7 |
| AC004466.2 | 12 | novel transcript, sense intronic to HDAC7 |
| LINC02354 | 12 | long intergenic non-protein coding RNA 2354 |
| VDR | 12 | vitamin D receptor [Source:HGNC Symbol;Acc |
| AC121338.1 | 12 | novel transcript, antisense to VDR |
| AC121338.2 | 12 | novel transcript |
| AC004801.7 | 12 | TEC |
| TMEM106C | 12 | transmembrane protein 106C [Source:HGNC S |
| COL2A1 | 12 | collagen type II alpha 1 chain [Source:HGNC Sy |
| AC004801.5 | 12 | novel transcript |
| AC004801.4 | 12 | novel transcript, antisense to a novel protein |
| AC004801.3 | 12 | novel transcript |
| AC004801.6 | 12 | novel transcript |
| SENP1 | 12 | SUMO specific peptidase 1 [Source:HGNC Sym |
| AC004801.2 | 12 | ribosomal protein L37 (RPL37) pseudogene |
| RNU6-1203P | 12 | RNA, U6 small nuclear 1203, pseudogene [Sou |
| AC004801.1 | 12 | ATPase, H+ transporting, lysosomal 14kDa, V1 |

| | | |
|---|---|---|
| PFKM | 12 | phosphofructokinase, muscle [Source:HGNC S |
| MIR6505 | 12 | microRNA 6505 [Source:HGNC Symbol;Acc:HG |
| ASB8 | 12 | ankyrin repeat and SOCS box containing 8 [Sou |
| AC074029.3 | 12 | novel transcript, antisense to ASB8 |
| AC074029.2 | 12 | SCO cytochrome oxidase deficient homolog 1 |
| PHBP18 | 12 | prohibitin pseudogene 18 [Source:HGNC Symb |
| AC074029.1 | 12 | ERO1-like (S. cerevisiae) (ERO1L) pseudogene |
| CCDC184 | 12 | coiled-coil domain containing 184 [Source:HG |
| AC074029.4 | 12 | TEC |
| AC024257.3 | 12 | novel transcript, antisense to OR10AD1 |
| OR10AD1 | 12 | olfactory receptor family 10 subfamily AD me |
| AC024257.1 | 12 | novel transcript |
| AC024257.5 | 12 | novel transcript |
| H1FNT | 12 | H1 histone family member N, testis specific [S |
| AC024257.2 | 12 | pseudogene similar to part of microtubule ass |
| ZNF641 | 12 | zinc finger protein 641 [Source:HGNC Symbol; |
| AC090115.1 | 12 | novel transcript, antisense to ZNF641 |
| OR5BK1P | 12 | olfactory receptor family 5 subfamily BK mem |
| AC024257.4 | 12 | novel transcript |
| OR5BT1P | 12 | olfactory receptor family 5 subfamily BT mem |
| OR5BJ1P | 12 | olfactory receptor family 5 subfamily BJ meml |
| OR8S21P | 12 | olfactory receptor family 8 subfamily S memb |
| OR8T1P | 12 | olfactory receptor family 8 subfamily T memb |
| ANP32D | 12 | acidic nuclear phosphoprotein 32 family mem |
| C12orf54 | 12 | chromosome 12 open reading frame 54 [Sour |
| AC089987.2 | 12 | novel transcript, antisense to C12orf54 |
| RPS10P20 | 12 | ribosomal protein S10 pseudogene 20 [Source |
| AC089987.1 | 12 | biorientation of chromosomes in cell division |
| OR8S1 | 12 | olfactory receptor family 8 subfamily S memb |
| OR5BS1P | 12 | olfactory receptor family 5 subfamily BS mem |
| LALBA | 12 | lactalbumin alpha [Source:HGNC Symbol;Acc:I |
| OR11M1P | 12 | olfactory receptor family 11 subfamily M mem |
| KANSL2 | 12 | KAT8 regulatory NSL complex subunit 2 [Sourc |
| SNORA2C | 12 | small nucleolar RNA, H/ACA box 2C [Source:H |
| MIR1291 | 12 | microRNA 1291 [Source:HGNC Symbol;Acc:HG |
| SNORA2A | 12 | small nucleolar RNA, H/ACA box 2A [Source:H |
| SNORA2B | 12 | small nucleolar RNA, H/ACA box 2B [Source:H |
| CCNT1 | 12 | cyclin T1 [Source:HGNC Symbol;Acc:HGNC:15 |
| TEX49 | 12 | testis expressed 49 [Source:HGNC Symbol;Acc |
| ADCY6 | 12 | adenylate cyclase 6 [Source:HGNC Symbol;Acc |
| AC117498.1 | 12 | novel transcript, antisense to ADCY6 |
| MIR4701 | 12 | microRNA 4701 [Source:HGNC Symbol;Acc:HG |
| ADCY6-DT | 12 | ADCY6 divergent transcript [Source:HGNC Syn |
| AC117498.2 | 12 | chromosome 20 open reading frame 27 (C20o |
| Y_RNA | 12 | Y RNA [Source:RFAM;Acc:RF00019] |
| CACNB3 | 12 | calcium voltage-gated channel auxiliary subun |
| DDX23 | 12 | DEAD-box helicase 23 [Source:HGNC Symbol;A |
| RND1 | 12 | Rho family GTPase 1 [Source:HGNC Symbol;Ac |
| RNU6-600P | 12 | RNA, U6 small nuclear 600, pseudogene [Sour |
| AC073610.1 | 12 | ribosomal protein L32 (RPL32) pseudogene |

| | | |
|---|---|---|
| AC073610.2 | 12 | novel protein |
| CCDC65 | 12 | coiled-coil domain containing 65 [Source:HGN |
| FKBP11 | 12 | FKBP prolyl isomerase 11 [Source:HGNC Symb |
| ARF3 | 12 | ADP ribosylation factor 3 [Source:HGNC Symb |
| WNT10B | 12 | Wnt family member 10B [Source:HGNC Symb |
| RNU6-940P | 12 | RNA, U6 small nuclear 940, pseudogene [Sour |
| WNT1 | 12 | Wnt family member 1 [Source:HGNC Symbol; |
| DDN | 12 | dendrin [Source:HGNC Symbol;Acc:HGNC:244 |
| AC011603.3 | 12 | novel transcript, antisense to DDN |
| DDN-AS1 | 12 | DDN and PRKAG1 antisense RNA 1 [Source:H |
| PRKAG1 | 12 | protein kinase AMP-activated non-catalytic su |
| KMT2D | 12 | lysine methyltransferase 2D [Source:HGNC Sy |
| RHEBL1 | 12 | RHEB like 1 [Source:HGNC Symbol;Acc:HGNC: |
| AC011603.4 | 12 | tubulin, alpha 4a (TUBA4A) pseudogene |
| DHH | 12 | desert hedgehog signaling molecule [Source:H |
| AC011603.1 | 12 | novel transcript, antisense to DHH |
| LMBR1L | 12 | limb development membrane protein 1 like [S |
| AC011603.2 | 12 | novel transcript, antisense to TUBA1B |
| TUBA1B | 12 | tubulin alpha 1b [Source:HGNC Symbol;Acc:H |
| Y_RNA | 12 | Y RNA [Source:RFAM;Acc:RF00019] |
| TUBA1A | 12 | tubulin alpha 1a [Source:HGNC Symbol;Acc:H |
| TUBA1C | 12 | tubulin alpha 1c [Source:HGNC Symbol;Acc:H |
| Metazoa_SRP | 12 | Metazoan signal recognition particle RNA [Sou |
| AC010173.1 | 12 | novel transcript, antisense to TUBA1C |
| AC125611.3 | 12 | novel transcript, antisense to TUBA1C |
| AC125611.4 | 12 | novel transcript, antisense to PRPH |
| PRPH | 12 | peripherin [Source:HGNC Symbol;Acc:HGNC:9 |
| TROAP | 12 | trophinin associated protein [Source:HGNC Sy |
| C1QL4 | 12 | complement C1q like 4 [Source:HGNC Symbol |
| DNAJC22 | 12 | DnaJ heat shock protein family (Hsp40) memb |
| SPATS2 | 12 | spermatogenesis associated serine rich 2 [Sou |
| AC125611.2 | 12 | FGFR1 oncogene partner 2 (FGFR1OP2) pseud |
| AC125611.1 | 12 | Non-histone chromosomal protein HMG-14 (H |
| AC020612.4 | 12 | novel transcript |
| KCNH3 | 12 | potassium voltage-gated channel subfamily H |
| MCRS1 | 12 | microspherule protein 1 [Source:HGNC Symbo |
| PRPF40B | 12 | pre-mRNA processing factor 40 homolog B [So |
| AC020612.3 | 12 | novel transcript, antisense to PRPF40B |
| FAM186B | 12 | family with sequence similarity 186 member B |
| RNU6-834P | 12 | RNA, U6 small nuclear 834, pseudogene [Sour |
| POLR2KP1 | 12 | RNA polymerase II subunit K pseudogene 1 [So |
| AC020612.1 | 12 | Protein DJ-1 (PARK7) pseudogene |
| HIGD1AP9 | 12 | HIG1 hypoxia inducible domain family membe |
| FMNL3 | 12 | formin like 3 [Source:HGNC Symbol;Acc:HGNC |
| TMBIM6 | 12 | transmembrane BAX inhibitor motif containin |
| AC020612.2 | 12 | Cytochrome c oxidase subunit 5B, mitochondr |
| LSM6P2 | 12 | LSM6 homolog, U6 small nuclear RNA and mR |
| NCKAP5L | 12 | NCK associated protein 5 like [Source:HGNC S |
| BCDIN3D-AS1 | 12 | BCDIN3D antisense RNA 1 [Source:HGNC Sym |
| BCDIN3D | 12 | BCDIN3 domain containing RNA methyltransfe |

| | | |
|---|---|---|
| AC131157.1 | 12 | novel transcript |
| RPL35AP28 | 12 | ribosomal protein L35a pseudogene 28 [Sourc |
| FAIM2 | 12 | Fas apoptotic inhibitory molecule 2 [Source:H |
| LINC02395 | 12 | long intergenic non-protein coding RNA 2395 |
| LINC02396 | 12 | long intergenic non-protein coding RNA 2396 |
| AQP2 | 12 | aquaporin 2 [Source:HGNC Symbol;Acc:HGNC |
| AC025154.2 | 12 | novel transcript, antisense to AQP5 |
| AC025154.1 | 12 | novel transcript, antisense to AQP5 and AQP2 |
| AQP5 | 12 | aquaporin 5 [Source:HGNC Symbol;Acc:HGNC |
| AQP6 | 12 | aquaporin 6 [Source:HGNC Symbol;Acc:HGNC |
| RACGAP1 | 12 | Rac GTPase activating protein 1 [Source:HGN( |
| ASIC1 | 12 | acid sensing ion channel subunit 1 [Source:HG |
| SMARCD1 | 12 | SWI/SNF related, matrix associated, actin dep |
| GPD1 | 12 | glycerol-3-phosphate dehydrogenase 1 [Sourc |
| COX14 | 12 | cytochrome c oxidase assembly factor COX14 |
| AC074032.1 | 12 | novel transcript, antisense to CERS5 |
| CERS5 | 12 | ceramide synthase 5 [Source:HGNC Symbol;A( |
| LIMA1 | 12 | LIM domain and actin binding 1 [Source:HGN( |
| AC008147.2 | 12 | novel transcript, sense intronic to LIMA1 |
| AC008147.3 | 12 | 40S ribosomal protein S26 (RPS26) pseudoger |
| AC008147.1 | 12 | novel transcript, antisense to LIMA1 |
| MIR1293 | 12 | microRNA 1293 [Source:HGNC Symbol;Acc:HG |
| RNU6-1093P | 12 | RNA, U6 small nuclear 1093, pseudogene [Sou |
| AC008147.4 | 12 | novel pseudogene |
| FAM186A | 12 | family with sequence similarity 186 member / |
| AC090058.1 | 12 | ribosomal protein L26 (RPL26) pseudogene |
| LARP4 | 12 | La ribonucleoprotein domain family member ‹ |
| DIP2B | 12 | disco interacting protein 2 homolog B [Source |
| RNU6-769P | 12 | RNA, U6 small nuclear 769, pseudogene [Sour |
| RNU6-238P | 12 | RNA, U6 small nuclear 238, pseudogene [Sour |
| Y_RNA | 12 | Y RNA [Source:RFAM;Acc:RF00019] |
| AC013244.1 | 12 | mitochondrial inner membrane organizing sys |
| ATF1 | 12 | activating transcription factor 1 [Source:HGN( |
| AC013244.2 | 12 | chromosome 12 open reading frame 76 (C12c |
| RN7SL519P | 12 | RNA, 7SL, cytoplasmic 519, pseudogene [Sour |
| TMPRSS12 | 12 | transmembrane serine protease 12 [Source:H |
| AC013244.3 | 12 | ubiquitin-conjugating enzyme E2L 3 (UBE2L3) |
| METTL7A | 12 | methyltransferase like 7A [Source:HGNC Symb |
| AC008121.2 | 12 | novel transcript |
| Y_RNA | 12 | Y RNA [Source:RFAM;Acc:RF00019] |
| AC008121.1 | 12 | family with sequence similarity 32, member A |
| HIGD1C | 12 | HIG1 hypoxia inducible domain family membe |
| AC008121.3 | 12 | novel transcript, antisense to HIGD1C |
| SLC11A2 | 12 | solute carrier family 11 member 2 [Source:HG |
| RNU6-87P | 12 | RNA, U6 small nuclear 87, pseudogene [Sourc |
| RNU6-1273P | 12 | RNA, U6 small nuclear 1273, pseudogene [Sou |
| LETMD1 | 12 | LETM1 domain containing 1 [Source:HGNC Sy |
| CSRNP2 | 12 | cysteine and serine rich nuclear protein 2 [Sou |
| TFCP2 | 12 | transcription factor CP2 [Source:HGNC Symbo |
| AC087884.1 | 12 | ribosomal protein L35a (RPL35A) pseudogene |

| | | |
|---|---|---|
| PHBP19 | 12 | prohibitin pseudogene 19 [Source:HGNC Symb |
| Y_RNA | 12 | Y RNA [Source:RFAM;Acc:RF00019] |
| RNU6-199P | 12 | RNA, U6 small nuclear 199, pseudogene [Sour |
| POU6F1 | 12 | POU class 6 homeobox 1 [Source:HGNC Symb |
| AC139768.1 | 12 | novel transcript |
| DAZAP2 | 12 | DAZ associated protein 2 [Source:HGNC Symb |
| SMAGP | 12 | small cell adhesion glycoprotein [Source:HGN |
| BIN2 | 12 | bridging integrator 2 [Source:HGNC Symbol;A |
| CELA1 | 12 | chymotrypsin like elastase 1 [Source:HGNC Sy |
| GALNT6 | 12 | polypeptide N-acetylgalactosaminyltransferas |
| SLC4A8 | 12 | solute carrier family 4 member 8 [Source:HGN |
| AC107031.1 | 12 | novel transcript, antisense to SLC4A8 |
| SCN8A | 12 | sodium voltage-gated channel alpha subunit 8 |
| HNRNPA3P10 | 12 | heterogeneous nuclear ribonucleoprotein A3 |
| AC068987.1 | 12 | novel transcript |
| C12orf81 | 12 | chromosome 12 open reading frame 81 [Sour |
| AC068987.2 | 12 | novel transcript |
| FIGNL2 | 12 | fidgetin like 2 [Source:HGNC Symbol;Acc:HGN |
| AC068987.3 | 12 | novel transcript |
| FIGNL2-DT | 12 | FIGNL2 divergent transcript [Source:HGNC Syr |
| ANKRD33 | 12 | ankyrin repeat domain 33 [Source:HGNC Symb |
| AC025259.2 | 12 | filamin B, beta (FLNB) pseudogene |
| ACVRL1 | 12 | activin A receptor like type 1 [Source:HGNC Sy |
| ACVR1B | 12 | activin A receptor type 1B [Source:HGNC Sym |
| RNU6-574P | 12 | RNA, U6 small nuclear 574, pseudogene [Sour |
| GRASP | 12 | general receptor for phosphoinositides 1 asso |
| NR4A1 | 12 | nuclear receptor subfamily 4 group A member |
| AC025259.3 | 12 | novel transcript |
| ATG101 | 12 | autophagy related 101 [Source:HGNC Symbol; |
| AC025259.1 | 12 | novel transcript, antisense to C12orf44 |
| SMIM41 | 12 | small integral membrane protein 41 [Source:H |
| OR7E47P | 12 | olfactory receptor family 7 subfamily E memb |
| AC078864.1 | 12 | novel transcript |
| LINC00592 | 12 | long intergenic non-protein coding RNA 592 [S |
| KRT80 | 12 | keratin 80 [Source:HGNC Symbol;Acc:HGNC:2 |
| C12orf80 | 12 | chromosome 12 open reading frame 80 [Sour |
| METTL7AP1 | 12 | methyltransferase like 7A pseudogene 1 [Sour |
| KRT7 | 12 | keratin 7 [Source:HGNC Symbol;Acc:HGNC:64 |
| KRT7-AS | 12 | KRT7 antisense RNA 1 [Source:HGNC Symbol;A |
| KRT86 | 12 | keratin 86 [Source:HGNC Symbol;Acc:HGNC:6 |
| KRT87P | 12 | keratin 87 pseudogene [Source:HGNC Symbol |
| AC021066.2 | 12 | keratin protein pseudogene |
| AC021066.1 | 12 | novel transcript, antisense to KRT86 |
| AC021066.3 | 12 | keratin protein pseudogene |
| KRT81 | 12 | keratin 81 [Source:HGNC Symbol;Acc:HGNC:6 |
| AC121757.2 | 12 | novel transcript, antisense to KRT86 |
| AC121757.1 | 12 | novel transcript, antisense to KRT86 |
| KRT83 | 12 | keratin 83 [Source:HGNC Symbol;Acc:HGNC:6 |
| KRT89P | 12 | keratin 89 pseudogene [Source:HGNC Symbol |
| KRT85 | 12 | keratin 85 [Source:HGNC Symbol;Acc:HGNC:6 |

| | | |
|---|---|---|
| KRT84 | 12 | keratin 84 [Source:HGNC Symbol;Acc:HGNC:6 |
| AC078865.1 | 12 | novel transcript, antisense to KRT84 and KRT8 |
| KRT82 | 12 | keratin 82 [Source:HGNC Symbol;Acc:HGNC:6 |
| AC055736.1 | 12 | novel transcript, antisense to KRT75 and KRT8 |
| KRT90P | 12 | keratin 90 pseudogene [Source:HGNC Symbol |
| KRT75 | 12 | keratin 75 [Source:HGNC Symbol;Acc:HGNC:2 |
| KRT6B | 12 | keratin 6B [Source:HGNC Symbol;Acc:HGNC:6 |
| KRT6C | 12 | keratin 6C [Source:HGNC Symbol;Acc:HGNC:2 |
| KRT6A | 12 | keratin 6A [Source:HGNC Symbol;Acc:HGNC:6 |
| KRT5 | 12 | keratin 5 [Source:HGNC Symbol;Acc:HGNC:64 |
| KRT71 | 12 | keratin 71 [Source:HGNC Symbol;Acc:HGNC:2 |
| KRT74 | 12 | keratin 74 [Source:HGNC Symbol;Acc:HGNC:2 |
| KRT72 | 12 | keratin 72 [Source:HGNC Symbol;Acc:HGNC:2 |
| KRT73-AS1 | 12 | KRT73 antisense RNA 1 [Source:HGNC Symbol |
| KRT73 | 12 | keratin 73 [Source:HGNC Symbol;Acc:HGNC:2 |
| KRT128P | 12 | keratin 128 pseudogene [Source:HGNC Symbo |
| KRT2 | 12 | keratin 2 [Source:HGNC Symbol;Acc:HGNC:64 |
| KRT1 | 12 | keratin 1 [Source:HGNC Symbol;Acc:HGNC:64 |
| KRT77 | 12 | keratin 77 [Source:HGNC Symbol;Acc:HGNC:2 |
| AC055716.3 | 12 | novel transcript, antisense to KRT77 |
| KRT126P | 12 | keratin 126 pseudogene [Source:HGNC Symbo |
| KRT125P | 12 | keratin 125 pseudogene [Source:HGNC Symbo |
| AC055716.2 | 12 | DEAD (Asp-Glu-Ala-Asp) box polypeptide 23 (D |
| AC055716.1 | 12 | keratin pseudogene |
| BTBD10P1 | 12 | BTB domain containing 10 pseudogene 1 [Sou |
| KRT127P | 12 | keratin 127 pseudogene [Source:HGNC Symbo |
| ARL2BPP2 | 12 | ADP ribosylation factor like GTPase 2 binding |
| KRT76 | 12 | keratin 76 [Source:HGNC Symbol;Acc:HGNC:2 |
| AC107016.2 | 12 | ribosomal protein S28 (RPS28) pseudogene |
| KRT3 | 12 | keratin 3 [Source:HGNC Symbol;Acc:HGNC:64 |
| KRT4 | 12 | keratin 4 [Source:HGNC Symbol;Acc:HGNC:64 |
| KRT79 | 12 | keratin 79 [Source:HGNC Symbol;Acc:HGNC:2 |
| AC107016.1 | 12 | ribosomal protein S26 (RPS26) pseudogene |
| KRT78 | 12 | keratin 78 [Source:HGNC Symbol;Acc:HGNC:2 |
| RPL7P41 | 12 | ribosomal protein L7 pseudogene 41 [Source: |
| KRT8 | 12 | keratin 8 [Source:HGNC Symbol;Acc:HGNC:64 |
| KRT18 | 12 | keratin 18 [Source:HGNC Symbol;Acc:HGNC:6 |
| EIF4B | 12 | eukaryotic translation initiation factor 4B [Sou |
| AC068888.2 | 12 | novel transcript |
| AC068888.1 | 12 | novel transcript, antisense to TENC1 & EIF4B |
| TNS2 | 12 | tensin 2 [Source:HGNC Symbol;Acc:HGNC:197 |
| MIR6757 | 12 | microRNA 6757 [Source:HGNC Symbol;Acc:HG |
| SPRYD3 | 12 | SPRY domain containing 3 [Source:HGNC Sym |
| IGFBP6 | 12 | insulin like growth factor binding protein 6 [So |
| SOAT2 | 12 | sterol O-acyltransferase 2 [Source:HGNC Symb |
| VTI1BP3 | 12 | vesicle transport through interaction with t-SN |
| RNU6-333P | 12 | RNA, U6 small nuclear 333, pseudogene [Sour |
| HIGD1AP1 | 12 | HIG1 hypoxia inducible domain family membe |
| EIF4A1P4 | 12 | eukaryotic translation initiation factor 4A1 ps |
| CSAD | 12 | cysteine sulfinic acid decarboxylase [Source:H |

| | | |
|---|---|---|
| AC073573.1 | 12 | novel transcript |
| ZNF740 | 12 | zinc finger protein 740 [Source:HGNC Symbol; |
| ITGB7 | 12 | integrin subunit beta 7 [Source:HGNC Symbol; |
| RARG | 12 | retinoic acid receptor gamma [Source:HGNC S |
| AC021072.1 | 12 | novel protein |
| MFSD5 | 12 | major facilitator superfamily domain containin |
| ESPL1 | 12 | extra spindle pole bodies like 1, separase [Sou |
| PFDN5 | 12 | prefoldin subunit 5 [Source:HGNC Symbol;Acc |
| AC073611.1 | 12 | novel transcript, antisense to PFDN5 and C12c |
| C12orf10 | 12 | chromosome 12 open reading frame 10 [Sour |
| AAAS | 12 | aladin WD repeat nucleoporin [Source:HGNC S |
| SP7 | 12 | Sp7 transcription factor [Source:HGNC Symbo |
| SP1 | 12 | Sp1 transcription factor [Source:HGNC Symbo |
| AMHR2 | 12 | anti-Mullerian hormone receptor type 2 [Sour |
| PRR13 | 12 | proline rich 13 [Source:HGNC Symbol;Acc:HGI |
| AC023509.1 | 12 | novel transcript |
| PCBP2 | 12 | poly(rC) binding protein 2 [Source:HGNC Symb |
| PCBP2-OT1 | 12 | PCBP2 overlapping transcript 1 [Source:HGNC |
| AC023509.4 | 12 | NULL |
| MAP3K12 | 12 | mitogen-activated protein kinase kinase kinas |
| AC023509.3 | 12 | novel transcript, antisense to TARBP2 |
| TARBP2 | 12 | TARBP2 subunit of RISC loading complex [Sour |
| ATF7-NPFF | 12 | ATF7-NPFF readthrough [Source:NCBI gene;Ac |
| NPFF | 12 | neuropeptide FF-amide peptide precursor [So |
| ATF7 | 12 | activating transcription factor 7 [Source:HGN( |
| AC023509.2 | 12 | novel transcript, antisense to ATF7 |
| AC023509.6 | 12 | Novel transcript, antisense to ATF7 |
| AC023509.5 | 12 | zinc finger protein 337 (ZNF337) pseudogene |
| ATP5MC2 | 12 | ATP synthase membrane subunit c locus 2 [So |
| AC073594.1 | 12 | ribosomal protein L31 (RPL31) pseudogene |
| CALCOCO1 | 12 | calcium binding and coiled-coil domain 1 [Sou |
| AC076968.2 | 12 | novel transcript |
| CISTR | 12 | chondrogenesis-associated transcript [Source: |
| AC076968.1 | 12 | novel transcript |
| RN7SKP289 | 12 | RN7SK pseudogene 289 [Source:HGNC Symbo |
| HOXC13-AS | 12 | HOXC13 antisense RNA [Source:HGNC Symbol |
| HOXC13 | 12 | homeobox C13 [Source:HGNC Symbol;Acc:HG |
| HOXC12 | 12 | homeobox C12 [Source:HGNC Symbol;Acc:HG |
| HOTAIR | 12 | HOX transcript antisense RNA [Source:HGNC S |
| AC012531.5 | 12 | NULL |
| AC012531.7 | 12 | NULL |
| AC012531.4 | 12 | NULL |
| AC012531.6 | 12 | NULL |
| HOXC11 | 12 | homeobox C11 [Source:HGNC Symbol;Acc:HG |
| HOXC-AS3 | 12 | HOXC cluster antisense RNA 3 [Source:HGNC S |
| HOXC10 | 12 | homeobox C10 [Source:HGNC Symbol;Acc:HG |
| AC012531.3 | 12 | novel protein, readthrough between HOXC10 |
| HOXC6 | 12 | homeobox C6 [Source:HGNC Symbol;Acc:HGN |
| MIR196A2 | 12 | microRNA 196a-2 [Source:HGNC Symbol;Acc:H |
| HOXC-AS2 | 12 | HOXC cluster antisense RNA 2 [Source:HGNC S |

| | | |
|---|---|---|
| HOXC9 | 12 | homeobox C9 [Source:HGNC Symbol;Acc:HGN |
| HOXC-AS1 | 12 | HOXC cluster antisense RNA 1 [Source:HGNC S |
| HOXC8 | 12 | homeobox C8 [Source:HGNC Symbol;Acc:HGN |
| HOXC4 | 12 | homeobox C4 [Source:HGNC Symbol;Acc:HGN |
| AC012531.2 | 12 | novel transcript |
| AC012531.1 | 12 | novel transcript |
| HOXC5 | 12 | homeobox C5 [Source:HGNC Symbol;Acc:HGN |
| MIR615 | 12 | microRNA 615 [Source:HGNC Symbol;Acc:HGN |
| AC023794.1 | 12 | novel transcript |
| AC023794.4 | 12 | novel pseudogene |
| AC023794.3 | 12 | novel transcript |
| FAM242C | 12 | family with sequence similarity 242 member C |
| Y_RNA | 12 | Y RNA [Source:RFAM;Acc:RF00019] |
| SMUG1 | 12 | single-strand-selective monofunctional uracil- |
| LINC02381 | 12 | long intergenic non-protein coding RNA 2381 |
| AC023794.6 | 12 | novel transcript |
| AC023794.5 | 12 | novel transcript |
| AC023794.2 | 12 | novel transcript, antisense to SMUG1 |
| CBX5 | 12 | chromobox 5 [Source:HGNC Symbol;Acc:HGN |
| MIR3198-2 | 12 | microRNA 3198-2 [Source:HGNC Symbol;Acc:H |
| RN7SL390P | 12 | RNA, 7SL, cytoplasmic 390, pseudogene [Sour |
| SCAT2 | 12 | S-phase cancer associated transcript 2 [Source |
| AC078778.1 | 12 | novel transcript, sense overlapping to COPZ1 a |
| HNRNPA1 | 12 | heterogeneous nuclear ribonucleoprotein A1 |
| NFE2 | 12 | nuclear factor, erythroid 2 [Source:HGNC Sym |
| COPZ1 | 12 | coatomer protein complex subunit zeta 1 [Sou |
| RNU6-950P | 12 | RNA, U6 small nuclear 950, pseudogene [Sour |
| MIR148B | 12 | microRNA 148b [Source:HGNC Symbol;Acc:HG |
| RN7SL744P | 12 | RNA, 7SL, cytoplasmic 744, pseudogene [Sour |
| AC079313.2 | 12 | novel transcript, antisense to ZNF385A, GPR84 |
| AC079313.1 | 12 | novel transcript, antisense to ZNF385A, GPR84 |
| GPR84 | 12 | G protein-coupled receptor 84 [Source:HGNC |
| ZNF385A | 12 | zinc finger protein 385A [Source:HGNC Symbo |
| ITGA5 | 12 | integrin subunit alpha 5 [Source:HGNC Symbo |
| LINC01154 | 12 | long intergenic non-protein coding RNA 1154 |
| GTSF1 | 12 | gametocyte specific factor 1 [Source:HGNC Sy |
| NCKAP1L | 12 | NCK associated protein 1 like [Source:HGNC Sy |
| AC068789.1 | 12 | novel transcript, antisense to NCKAP1L |
| PDE1B | 12 | phosphodiesterase 1B [Source:HGNC Symbol; |
| PPP1R1A | 12 | protein phosphatase 1 regulatory inhibitor sub |
| GLYCAM1 | 12 | glycosylation dependent cell adhesion molecu |
| LACRT | 12 | lacritin [Source:HGNC Symbol;Acc:HGNC:1643 |
| DCD | 12 | dermcidin [Source:HGNC Symbol;Acc:HGNC:14 |
| AC079310.1 | 12 | novel transcript |
| VDAC1P5 | 12 | voltage dependent anion channel 1 pseudoge |
| MUCL1 | 12 | mucin like 1 [Source:HGNC Symbol;Acc:HGNC |
| TESPA1 | 12 | thymocyte expressed, positive selection assoc |
| AC027287.2 | 12 | novel transcript |
| NEUROD4 | 12 | neuronal differentiation 4 [Source:HGNC Syml |
| OR9K1P | 12 | olfactory receptor family 9 subfamily K memb |

| | | |
|---|---|---|
| AC027287.1 | 12 | heat shock transcription factor 2 (HSF2) pseuc |
| OR9K2 | 12 | olfactory receptor family 9 subfamily K memb |
| OR9R1P | 12 | olfactory receptor family 9 subfamily R memb |
| OR10U1P | 12 | olfactory receptor family 10 subfamily U mem |
| OR10A7 | 12 | olfactory receptor family 10 subfamily A mem |
| OR6C74 | 12 | olfactory receptor family 6 subfamily C memb |
| OR6C69P | 12 | olfactory receptor family 6 subfamily C memb |
| OR6C72P | 12 | olfactory receptor family 6 subfamily C memb |
| OR6C6 | 12 | olfactory receptor family 6 subfamily C memb |
| OR6C5P | 12 | olfactory receptor family 6 subfamily C memb |
| OR6C1 | 12 | olfactory receptor family 6 subfamily C memb |
| OR6C3 | 12 | olfactory receptor family 6 subfamily C memb |
| OR6C7P | 12 | olfactory receptor family 6 subfamily C memb |
| OR6C75 | 12 | olfactory receptor family 6 subfamily C memb |
| OR6C71P | 12 | olfactory receptor family 6 subfamily C memb |
| OR6C66P | 12 | olfactory receptor family 6 subfamily C memb |
| OR6C73P | 12 | olfactory receptor family 6 subfamily C memb |
| OR6C65 | 12 | olfactory receptor family 6 subfamily C memb |
| PHC1P1 | 12 | polyhomeotic homolog 1 pseudogene 1 [Sour |
| OR6C76 | 12 | olfactory receptor family 6 subfamily C memb |
| AC122685.1 | 12 | novel transcript, antisense to OR2AP1, OR6C4 |
| OR6C2 | 12 | olfactory receptor family 6 subfamily C memb |
| OR6C70 | 12 | olfactory receptor family 6 subfamily C memb |
| OR6C68 | 12 | olfactory receptor family 6 subfamily C memb |
| OR6C64P | 12 | olfactory receptor family 6 subfamily C memb |
| OR6C4 | 12 | olfactory receptor family 6 subfamily C memb |
| OR2AP1 | 12 | olfactory receptor family 2 subfamily AP mem |
| OR6U2P | 12 | olfactory receptor family 6 subfamily U memb |
| OR10P1 | 12 | olfactory receptor family 10 subfamily P mem |
| AC009779.5 | 12 | novel transcript, antisense to ITGA7 |
| OR10AE3P | 12 | olfactory receptor family 10 subfamily AE mer |
| AC009779.4 | 12 | proteasome (prosome, macropain) subunit, be |
| AC009779.6 | 12 | ribosomal protein S4, X-linked (RPS4X) pseudo |
| METTL7B | 12 | methyltransferase like 7B [Source:HGNC Symb |
| ITGA7 | 12 | integrin subunit alpha 7 [Source:HGNC Symbo |
| AC009779.3 | 12 | novel protein |
| BLOC1S1 | 12 | biogenesis of lysosomal organelles complex 1 |
| RDH5 | 12 | retinol dehydrogenase 5 [Source:HGNC Symbo |
| CD63 | 12 | CD63 molecule [Source:HGNC Symbol;Acc:HG |
| AC009779.2 | 12 | novel transcript, antisense to CD63 |
| Metazoa_SRP | 12 | Metazoan signal recognition particle RNA [Sou |
| AC009779.1 | 12 | inositol polyphosphate-1-phosphatase (INPP1 |
| GDF11 | 12 | growth differentiation factor 11 [Source:HGN( |
| SARNP | 12 | SAP domain containing ribonucleoprotein [Sou |
| AC023055.1 | 12 | novel protein |
| AC073487.1 | 12 | novel transcript, sense intronic to SARNP |
| ORMDL2 | 12 | ORMDL sphingolipid biosynthesis regulator 2 [ |
| DNAJC14 | 12 | DnaJ heat shock protein family (Hsp40) memb |
| TMEM198B | 12 | transmembrane protein 198B (pseudogene) [S |
| MMP19 | 12 | matrix metallopeptidase 19 [Source:HGNC Syr |

| | | |
|---|---|---|
| OLA1P3 | 12 | OLA1 pseudogene 3 [Source:HGNC Symbol;Ac |
| GSTP1P1 | 12 | glutathione S-transferase pi 1 pseudogene 1 [S |
| PYM1 | 12 | PYM homolog 1, exon junction complex assoc |
| DGKA | 12 | diacylglycerol kinase alpha [Source:HGNC Sym |
| AC025162.2 | 12 | novel transcript, sense intronic to DGKA |
| PMEL | 12 | premelanosome protein [Source:HGNC Symbo |
| CDK2 | 12 | cyclin dependent kinase 2 [Source:HGNC Syml |
| AC025162.1 | 12 | novel transcript, antisense to CDK2 |
| RAB5B | 12 | RAB5B, member RAS oncogene family [Source |
| AC034102.1 | 12 | RAB13 member RAS oncogene family pseudog |
| SUOX | 12 | sulfite oxidase [Source:HGNC Symbol;Acc:HGN |
| IKZF4 | 12 | IKAROS family zinc finger 4 [Source:HGNC Sym |
| AC034102.8 | 12 | novel transcript, antisence to IKZF4 |
| AC034102.3 | 12 | novel transcript , antisense to IKZF4 |
| RPS26 | 12 | ribosomal protein S26 [Source:HGNC Symbol; |
| ERBB3 | 12 | erb-b2 receptor tyrosine kinase 3 [Source:HGI |
| AC034102.2 | 12 | novel protein |
| PA2G4 | 12 | proliferation-associated 2G4 [Source:HGNC Sy |
| AC034102.4 | 12 | novel transcript, antisense to PA2G4 |
| RPL41 | 12 | ribosomal protein L41 [Source:HGNC Symbol; |
| ESYT1 | 12 | extended synaptotagmin 1 [Source:HGNC Sym |
| ZC3H10 | 12 | zinc finger CCCH-type containing 10 [Source:H |
| AC034102.7 | 12 | novel transcript, antisense to ZC3H10 and ES |
| AC034102.6 | 12 | novel transcript, antisense to ZC3H10 and ESY |
| AC034102.5 | 12 | novel transcript, antisense to MYL6B |
| MYL6B | 12 | myosin light chain 6B [Source:HGNC Symbol;A |
| MYL6 | 12 | myosin light chain 6 [Source:HGNC Symbol;Ac |
| SMARCC2 | 12 | SWI/SNF related, matrix associated, actin dep |
| AC073896.4 | 12 | novel transcript, sense overlapping to SMARC |
| RNF41 | 12 | ring finger protein 41 [Source:HGNC Symbol;A |
| NABP2 | 12 | nucleic acid binding protein 2 [Source:HGNC S |
| SLC39A5 | 12 | solute carrier family 39 member 5 [Source:HG |
| ANKRD52 | 12 | ankyrin repeat domain 52 [Source:HGNC Syml |
| COQ10A | 12 | coenzyme Q10A [Source:HGNC Symbol;Acc:H |
| AC073896.5 | 12 | novel transcript, antisense to COQ10A |
| CS | 12 | citrate synthase [Source:HGNC Symbol;Acc:H |
| AC073896.1 | 12 | novel protein |
| AC073896.2 | 12 | novel transcript, antisense to CNPY2 and CS |
| AC073896.3 | 12 | novel transcript, antisense to CNPY2 |
| CNPY2 | 12 | canopy FGF signaling regulator 2 [Source:HGN |
| PAN2 | 12 | poly(A) specific ribonuclease subunit PAN2 [S |
| IL23A | 12 | interleukin 23 subunit alpha [Source:HGNC Sy |
| STAT2 | 12 | signal transducer and activator of transcriptio |
| RNU7-40P | 12 | RNA, U7 small nuclear 40 pseudogene [Source |
| APOF | 12 | apolipoprotein F [Source:HGNC Symbol;Acc:H |
| APONP | 12 | apolipoprotein N, pseudogene [Source:HGNC |
| AC025574.1 | 12 | TBCC domain containing 1 (TBCCD1) pseudoge |
| AC024884.2 | 12 | novel transcript |
| AC024884.1 | 12 | apolipoprotein F (APOF) pseudogene |
| TIMELESS | 12 | timeless circadian regulator [Source:HGNC Sy |

| | | |
|---|---|---|
| MIP | 12 | major intrinsic protein of lens fiber [Source:H( |
| AC097104.1 | 12 | novel protein |
| SPRYD4 | 12 | SPRY domain containing 4 [Source:HGNC Sym |
| GLS2 | 12 | glutaminase 2 [Source:HGNC Symbol;Acc:HGN |
| HSPD1P4 | 12 | heat shock protein family D (Hsp60) member |
| RBMS2 | 12 | RNA binding motif single stranded interacting |
| RNU6-343P | 12 | RNA, U6 small nuclear 343, pseudogene [Sour |
| BAZ2A | 12 | bromodomain adjacent to zinc finger domain |
| ATP5F1B | 12 | ATP synthase F1 subunit beta [Source:HGNC S |
| SNORD59A | 12 | small nucleolar RNA, C/D box 59A [Source:HG |
| PTGES3 | 12 | prostaglandin E synthase 3 [Source:HGNC Sym |
| RN7SL809P | 12 | RNA, 7SL, cytoplasmic 809, pseudogene [Sour |
| NACA | 12 | nascent polypeptide associated complex subu |
| AC117378.1 | 12 | novel protein |
| PRIM1 | 12 | DNA primase subunit 1 [Source:HGNC Symbol |
| HSD17B6 | 12 | hydroxysteroid 17-beta dehydrogenase 6 [Sou |
| AC121758.2 | 12 | novel YWHA pseudogene |
| AC121758.4 | 12 | hydroxysteroid (17-beta) dehydrogenase 6 (H: |
| AC121758.3 | 12 | novel retinol dehydrogenase pseudogene |
| AC121758.1 | 12 | NULL |
| SDR9C7 | 12 | short chain dehydrogenase/reductase family ! |
| AC026120.2 | 12 | hydroxysteroid (17-beta) dehydrogenase 6 (H: |
| RDH16 | 12 | retinol dehydrogenase 16 [Source:HGNC Symb |
| AC026120.1 | 12 | hydroxysteroid (17-beta) dehydrogenase 6 (H: |
| AC026120.3 | 12 | novel transcript |
| GPR182 | 12 | G protein-coupled receptor 182 [Source:HGN( |
| ZBTB39 | 12 | zinc finger and BTB domain containing 39 [Sou |
| TAC3 | 12 | tachykinin 3 [Source:HGNC Symbol;Acc:HGNC |
| MYO1A | 12 | myosin IA [Source:HGNC Symbol;Acc:HGNC:7! |
| NEMP1 | 12 | nuclear envelope integral membrane protein |
| NAB2 | 12 | NGFI-A binding protein 2 [Source:HGNC Symb |
| STAT6 | 12 | signal transducer and activator of transcriptio |
| LRP1 | 12 | LDL receptor related protein 1 [Source:HGNC |
| LRP1-AS | 12 | LRP1 antisense RNA [Source:HGNC Symbol;Ac |
| MIR1228 | 12 | microRNA 1228 [Source:HGNC Symbol;Acc:H( |
| NXPH4 | 12 | neurexophilin 4 [Source:HGNC Symbol;Acc:H( |
| AC137834.2 | 12 | novel transcript, antisense to SHMT2 |
| SHMT2 | 12 | serine hydroxymethyltransferase 2 [Source:H( |
| NDUFA4L2 | 12 | NDUFA4 mitochondrial complex associated lik |
| STAC3 | 12 | SH3 and cysteine rich domain 3 [Source:HGN( |
| AC137834.1 | 12 | novel protein |
| R3HDM2 | 12 | R3H domain containing 2 [Source:HGNC Symb |
| RNU6-879P | 12 | RNA, U6 small nuclear 879, pseudogene [Sour |
| AC126614.1 | 12 | novel transcript |
| INHBC | 12 | inhibin subunit beta C [Source:HGNC Symbol;/ |
| INHBE | 12 | inhibin subunit beta E [Source:HGNC Symbol;/ |
| AC022506.2 | 12 | novel transcript, antisense to INHBEand GLI1 |
| GLI1 | 12 | GLI family zinc finger 1 [Source:HGNC Symbol; |
| ARHGAP9 | 12 | Rho GTPase activating protein 9 [Source:HGN( |
| MARS | 12 | methionyl-tRNA synthetase [Source:HGNC Syr |

| | | |
|---|---|---|
| MIR6758 | 12 | microRNA 6758 [Source:HGNC Symbol;Acc:HG |
| DDIT3 | 12 | DNA damage inducible transcript 3 [Source:H( |
| AC022506.1 | 12 | DDIT3 upstream open reading frame protein |
| MIR616 | 12 | microRNA 616 [Source:HGNC Symbol;Acc:HGI |
| MBD6 | 12 | methyl-CpG binding domain protein 6 [Source |
| DCTN2 | 12 | dynactin subunit 2 [Source:HGNC Symbol;Acc: |
| KIF5A | 12 | kinesin family member 5A [Source:HGNC Sym |
| PIP4K2C | 12 | phosphatidylinositol-5-phosphate 4-kinase typ |
| DTX3 | 12 | deltex E3 ubiquitin ligase 3 [Source:HGNC Sym |
| ARHGEF25 | 12 | Rho guanine nucleotide exchange factor 25 [S |
| AC025165.1 | 12 | novel transcript, antisense to ARHGEF25 |
| AC025165.6 | 12 | novel protein |
| SLC26A10 | 12 | solute carrier family 26 member 10 [Source:H |
| B4GALNT1 | 12 | beta-1,4-N-acetyl-galactosaminyltransferase 1 |
| RPL13AP23 | 12 | ribosomal protein L13a pseudogene 23 [Sourc |
| OS9 | 12 | OS9 endoplasmic reticulum lectin [Source:HGI |
| AC025165.2 | 12 | novel transcript, antisense to OS9 |
| AGAP2 | 12 | ArfGAP with GTPase domain, ankyrin repeat a |
| AGAP2-AS1 | 12 | AGAP2 antisense RNA 1 [Source:HGNC Symbo |
| TSPAN31 | 12 | tetraspanin 31 [Source:HGNC Symbol;Acc:HGI |
| CDK4 | 12 | cyclin dependent kinase 4 [Source:HGNC Syml |
| MIR6759 | 12 | microRNA 6759 [Source:HGNC Symbol;Acc:HG |
| MARCH9 | 12 | membrane associated ring-CH-type finger 9 [S |
| CYP27B1 | 12 | cytochrome P450 family 27 subfamily B meml |
| METTL1 | 12 | methyltransferase like 1 [Source:HGNC Symbo |
| EEF1AKMT3 | 12 | EEF1A lysine methyltransferase 3 [Source:HGI |
| AC025165.3 | 12 | novel protein |
| TSFM | 12 | Ts translation elongation factor, mitochondria |
| AVIL | 12 | advillin [Source:HGNC Symbol;Acc:HGNC:141[illegible] |
| AC025165.5 | 12 | novel transcript |
| RNU6-1083P | 12 | RNA, U6 small nuclear 1083, pseudogene [Sou |
| AC025165.4 | 12 | novel transcript |
| CTDSP2 | 12 | CTD small phosphatase 2 [Source:HGNC Symb |
| MIR26A2 | 12 | microRNA 26a-2 [Source:HGNC Symbol;Acc:H |
| AC083805.2 | 12 | novel transcript, antisense to CTDSP2 |
| AC083805.1 | 12 | novel transcript |
| AC083805.3 | 12 | novel transcript, antisense to Linc |
| ATP23 | 12 | ATP23 metallopeptidase and ATP synthase as |
| GIHCG | 12 | GIHCG inhibitor of miR-200b/200a/429 expres |
| AC084033.1 | 12 | zinc finger protein pseudogene |
| RN7SKP65 | 12 | RN7SK pseudogene 65 [Source:HGNC Symbol; |
| AC084033.2 | 12 | suppression of tumorigenicity 13 (colon carcir |
| LINC02403 | 12 | long intergenic non-protein coding RNA 2403 |
| AC090643.1 | 12 | TEC |
| AC090643.2 | 12 | TEC |
| RPL21P103 | 12 | ribosomal protein L21 pseudogene 103 [Sourc |
| AC025578.1 | 12 | ADP-ribosylation factor 1 (ARF1) pseudogene |
| AC020637.1 | 12 | novel transcript |
| LINC02388 | 12 | long intergenic non-protein coding RNA 2388 |
| LRIG3 | 12 | leucine rich repeats and immunoglobulin like |

| | | |
|---|---|---|
| AC068305.2 | 12 | novel transcript, antisense to LRIG3 |
| RPS6P22 | 12 | ribosomal protein S6 pseudogene 22 [Source: |
| AC068305.1 | 12 | methyltransferase 5 domain containing 1 (ME |
| AC046129.1 | 12 | novel transcript |
| AC108721.2 | 12 | novel transcript |
| AC108721.1 | 12 | novel transcript |
| RNU6-279P | 12 | RNA, U6 small nuclear 279, pseudogene [Sour |
| RNU6-871P | 12 | RNA, U6 small nuclear 871, pseudogene [Sour |
| LINC02448 | 12 | long intergenic non-protein coding RNA 2448 |
| RNU4-20P | 12 | RNA, U4 small nuclear 20, pseudogene [Sourc |
| SLC16A7 | 12 | solute carrier family 16 member 7 [Source:HG |
| AC079905.2 | 12 | esterase D (ESD) pseudogene |
| AC079905.1 | 12 | poly(A) binding protein, cytoplasmic 4 (inducil |
| AC080011.1 | 12 | novel transcript |
| AC087883.2 | 12 | novel transcript |
| AC087883.1 | 12 | RAB2A, member RAS oncogene family (RAB2A |
| Y_RNA | 12 | Y RNA [Source:RFAM;Acc:RF00019] |
| AC090022.1 | 12 | NULL |
| AC090022.2 | 12 | X antigen family, member 5 (XAGE5) pseudoge |
| AC090017.1 | 12 | piggyBac transposable element derived 3 (PGI |
| DUX4L52 | 12 | double homeobox 4 like 52 (pseudogene) [Sou |
| AC090629.1 | 12 | novel transcript |
| TAFA2 | 12 | TAFA chemokine like family member 2 [Sourc |
| RPL21P104 | 12 | ribosomal protein L21 pseudogene 104 [Sourc |
| KRT8P19 | 12 | keratin 8 pseudogene 19 [Source:HGNC Symb |
| RPS3P6 | 12 | ribosomal protein S3 pseudogene 6 [Source:H |
| AC078789.1 | 12 | novel transcript, sense intronic to FAM19A2 |
| KLF17P1 | 12 | Kruppel like factor 17 pseudogene 1 [Source:H |
| USP15 | 12 | ubiquitin specific peptidase 15 [Source:HGNC |
| MIR6125 | 12 | microRNA 6125 [Source:HGNC Symbol;Acc:HG |
| RNU6-595P | 12 | RNA, U6 small nuclear 595, pseudogene [Sour |
| Metazoa_SRP | 12 | Metazoan signal recognition particle RNA [Sou |
| MON2 | 12 | MON2 homolog, regulator of endosome-to-Go |
| AC079035.1 | 12 | novel transcript, antisense to MON2 |
| RNU6-399P | 12 | RNA, U6 small nuclear 399, pseudogene [Sour |
| LINC01465 | 12 | long intergenic non-protein coding RNA 1465 |
| AC048341.1 | 12 | novel transcript |
| MIRLET7I | 12 | microRNA let-7i [Source:HGNC Symbol;Acc:HC |
| AC048341.2 | 12 | novel transcript |
| PPM1H | 12 | protein phosphatase, Mg2+/Mn2+ dependent |
| RPL32P26 | 12 | ribosomal protein L32 pseudogene 26 [Source |
| GAPDHP44 | 12 | glyceraldehyde 3 phosphate dehydrogenase p |
| RNU1-83P | 12 | RNA, U1 small nuclear 83, pseudogene [Sourc |
| Y_RNA | 12 | Y RNA [Source:RFAM;Acc:RF00019] |
| RPL14P1 | 12 | ribosomal protein L14 pseudogene 1 [Source: |
| AC078814.2 | 12 | chromosome 6 open reading frame 35 (C6orf3 |
| LDHAL6CP | 12 | lactate dehydrogenase A like 6C, pseudogene |
| AC078814.1 | 12 | novel transcript |
| RSL24D1P5 | 12 | ribosomal L24 domain containing 1 pseudoger |
| AVPR1A | 12 | arginine vasopressin receptor 1A [Source:HGN |

| | | |
|---|---|---|
| AC135584.1 | 12 | TEC |
| HNRNPA1P69 | 12 | heterogeneous nuclear ribonucleoprotein A1 |
| AC026116.1 | 12 | novel transcript |
| DPY19L2 | 12 | dpy-19 like 2 [Source:HGNC Symbol;Acc:HGN( |
| AC084357.2 | 12 | antisense to TMEM5 |
| AC027667.1 | 12 | S100 calcium binding protein A11 (S100A11) p |
| AC084357.3 | 12 | T-box 20 (TBX20) pseudogene |
| AC084357.1 | 12 | ribosomal protein S27 (RPS27) pseudogene |
| RXYLT1 | 12 | ribitol xylosyltransferase 1 [Source:HGNC Sym |
| RXYLT1-AS1 | 12 | RXYLT1 antisense RNA 1 [Source:HGNC Symbo |
| PABPC1P4 | 12 | poly(A) binding protein cytoplasmic 1 pseudo |
| SRGAP1 | 12 | SLIT-ROBO Rho GTPase activating protein 1 [S |
| RPL36AP41 | 12 | ribosomal protein L36a pseudogene 41 [Sourc |
| AC079866.2 | 12 | novel transcript |
| AC079866.1 | 12 | CTAGE family, member 5 (CTAGE5) pseudoger |
| AC020611.1 | 12 | chromosome 3 open reading frame 19 (C3orf1 |
| AC020611.2 | 12 | novel transcript |
| AC025576.3 | 12 | novel transcript, antisense to SRGAP1 |
| AC025576.2 | 12 | novel transcript |
| AC025576.1 | 12 | novel transcript |
| C12orf66 | 12 | chromosome 12 open reading frame 66 [Sour |
| RNU6-1009P | 12 | RNA, U6 small nuclear 1009, pseudogene [Sou |
| AC012158.1 | 12 | ribosomal protein S11 pseudogene 6 |
| RNU5A-7P | 12 | RNA, U5A small nuclear 7, pseudogene [Sourc |
| C12orf56 | 12 | chromosome 12 open reading frame 56 [Sour |
| AC012158.2 | 12 | oocyte expressed protein homolog (dog) (OOI |
| AC135279.3 | 12 | ATPase, H+ transporting, lysosomal 31kDa, V1 |
| AC135279.1 | 12 | NULL |
| XPOT | 12 | exportin for tRNA [Source:HGNC Symbol;Acc:H |
| AC135279.2 | 12 | mitochondrial ribosomal protein S25 (MRPS25 |
| AC135279.4 | 12 | novel transcript, antisense to TBK1 |
| TBK1 | 12 | TANK binding kinase 1 [Source:HGNC Symbol; |
| RASSF3 | 12 | Ras association domain family member 3 [Sou |
| AC078962.2 | 12 | novel transcript, antisense to RASSF3 |
| SNORD83 | 12 | Small nucleolar RNA SNORD83 [Source:RFAM; |
| AC078962.4 | 12 | novel transcript |
| MIR548C | 12 | microRNA 548c [Source:HGNC Symbol;Acc:HG |
| MIR548Z | 12 | microRNA 548z [Source:HGNC Symbol;Acc:HG |
| AC078962.1 | 12 | novel transcript |
| AC078962.3 | 12 | novel transcript |
| AC025262.3 | 12 | novel transcript |
| AC025262.2 | 12 | TEC |
| GNS | 12 | glucosamine (N-acetyl)-6-sulfatase [Source:HC |
| AC025262.1 | 12 | novel transcript |
| TBC1D30 | 12 | TBC1 domain family member 30 [Source:HGN |
| AC025262.4 | 12 | novel transcript, antisense to TBC1D30 |
| RPL7P39 | 12 | ribosomal protein L7 pseudogene 39 [Source: |
| LINC02389 | 12 | long intergenic non-protein coding RNA 2389 |
| LINC02231 | 12 | long intergenic non-protein coding RNA 2231 |
| RNU6ATAC42P | 12 | RNA, U6atac small nuclear 42, pseudogene [S |

| | | |
|---|---|---|
| AC135895.1 | 12 | novel transcript, antisense to WIF1 |
| WIF1 | 12 | WNT inhibitory factor 1 [Source:HGNC Symbo |
| RNU6-166P | 12 | RNA, U6 small nuclear 166, pseudogene [Sour |
| APOOP3 | 12 | apolipoprotein O pseudogene 3 [Source:HGN( |
| LEMD3 | 12 | LEM domain containing 3 [Source:HGNC Symb |
| AC026124.2 | 12 | novel transcript, sense intronic to LEMD3 |
| MSRB3 | 12 | methionine sulfoxide reductase B3 [Source:H( |
| AC026124.1 | 12 | novel transcript |
| KRT18P60 | 12 | keratin 18 pseudogene 60 [Source:HGNC Sym |
| AC025419.1 | 12 | novel transcript |
| AC090023.1 | 12 | novel transcript |
| LINC02454 | 12 | long intergenic non-protein coding RNA 2454 |
| AC090023.2 | 12 | novel transcript |
| PCNPP3 | 12 | PEST containing nuclear protein pseudogene 3 |
| RPSAP52 | 12 | ribosomal protein SA pseudogene 52 [Source: |
| HMGA2 | 12 | high mobility group AT-hook 2 [Source:HGNC |
| AC107308.1 | 12 | novel transcript, sense intronic to HMGA2 |
| HMGA2-AS1 | 12 | HMGA2 antisense RNA 1 [Source:HGNC Symb |
| AC090673.1 | 12 | novel transcript, antisense to HMGA2 |
| MIR6074 | 12 | microRNA 6074 [Source:HGNC Symbol;Acc:HG |
| LINC02425 | 12 | long intergenic non-protein coding RNA 2425 |
| RPL21P18 | 12 | ribosomal protein L21 pseudogene 18 [Source |
| RNA5SP362 | 12 | RNA, 5S ribosomal pseudogene 362 [Source:H |
| LLPH | 12 | LLP homolog, long-term synaptic facilitation fa |
| AC078927.1 | 12 | novel protein |
| LLPH-DT | 12 | LLPH divergent transcript [Source:HGNC Symb |
| TMBIM4 | 12 | transmembrane BAX inhibitor motif containin |
| IRAK3 | 12 | interleukin 1 receptor associated kinase 3 [Sou |
| RBMS1P1 | 12 | RNA binding motif single stranded interacting |
| MIR6502 | 12 | microRNA 6502 [Source:HGNC Symbol;Acc:HG |
| AC078889.1 | 12 | novel transcript |
| PDCL3P7 | 12 | PDCL3 pseudogene 7 [Source:HGNC Symbol;A |
| RN7SKP166 | 12 | RN7SK pseudogene 166 [Source:HGNC Symbo |
| HELB | 12 | DNA helicase B [Source:HGNC Symbol;Acc:HG |
| GRIP1 | 12 | glutamate receptor interacting protein 1 [Sou |
| OSBPL9P5 | 12 | oxysterol binding protein like 9 pseudogene 5 |
| OSBPL9P4 | 12 | oxysterol binding protein like 9 pseudogene 4 |
| AC073530.1 | 12 | novel transcript |
| AC073530.2 | 12 | TEC |
| RAB11AP2 | 12 | RAB11A, member RAS oncogene family pseud |
| GGTA2P | 12 | glycoprotein alpha-galactosyltransferase 2, ps |
| CAND1 | 12 | cullin associated and neddylation dissociated |
| AC078983.1 | 12 | novel transcript |
| MRPL40P1 | 12 | mitochondrial ribosomal protein L40 pseudog |
| NTAN1P3 | 12 | N-terminal asparagine amidase pseudogene 3 |
| LINC02420 | 12 | long intergenic non-protein coding RNA 2420 |
| LINC02408 | 12 | long intergenic non-protein coding RNA 2408 |
| LINC02442 | 12 | long intergenic non-protein coding RNA 2442 |
| DYRK2 | 12 | dual specificity tyrosine phosphorylation regu |
| AC078777.1 | 12 | novel transcript |

| | | |
|---|---|---|
| LINC02421 | 12 | long intergenic non-protein coding RNA 2421 |
| AC022513.1 | 12 | TWC |
| LINC01479 | 12 | long intergenic non-protein coding RNA 1479 |
| AC005294.1 | 12 | zinc finger protein 766 (ZNF766) pseudogene |
| IFNG-AS1 | 12 | IFNG antisense RNA 1 [Source:HGNC Symbol;/ |
| RPL39P28 | 12 | ribosomal protein L39 pseudogene 28 [Source |
| HNRNPA1P70 | 12 | heterogeneous nuclear ribonucleoprotein A1 |
| AC007458.1 | 12 | mitochondrial ribosomal protein L21 (MRPL21 |
| IFNG | 12 | interferon gamma [Source:HGNC Symbol;Acc: |
| IL26 | 12 | interleukin 26 [Source:HGNC Symbol;Acc:HGN |
| IL22 | 12 | interleukin 22 [Source:HGNC Symbol;Acc:HGN |
| MDM1 | 12 | Mdm1 nuclear protein [Source:HGNC Symbol; |
| LINC02384 | 12 | long intergenic non-protein coding RNA 2384 |
| AC022511.1 | 12 | novel transcript |
| AC008033.3 | 12 | novel transcript |
| AC008033.1 | 12 | proteasome (prosome, macropain) 26S subun |
| AC008033.2 | 12 | chromosome 5 open reading frame 62 (C5orf( |
| RPSAP12 | 12 | ribosomal protein SA pseudogene 12 [Source: |
| Metazoa_SRP | 12 | Metazoan signal recognition particle RNA [Sou |
| RAP1B | 12 | RAP1B, member of RAS oncogene family [Sou |
| RPL10P12 | 12 | ribosomal protein L10 pseudogene 12 [Source |
| SNORA70G | 12 | small nucleolar RNA, H/ACA box 70G [Source: |
| ATP5PDP4 | 12 | ATP synthase peripheral stalk subunit d pseud |
| AC090061.1 | 12 | novel transcript |
| RPL7P42 | 12 | ribosomal protein L7 pseudogene 42 [Source: |
| NUP107 | 12 | nucleoporin 107 [Source:HGNC Symbol;Acc:H |
| KRT8P39 | 12 | keratin 8 pseudogene 39 [Source:HGNC Symb |
| SLC35E3 | 12 | solute carrier family 35 member E3 [Source:H |
| AC025423.3 | 12 | chromosome 1 open reading frame 144 (C1or |
| MDM2 | 12 | MDM2 proto-oncogene [Source:HGNC Symbo |
| AC025423.1 | 12 | novel transcript |
| AC025423.4 | 12 | novel transcript |
| AC025423.2 | 12 | ribosomal L24 domain containing 1 (RSL24D1) |
| CPM | 12 | carboxypeptidase M [Source:HGNC Symbol;A( |
| RNU7-4P | 12 | RNA, U7 small nuclear 4 pseudogene [Source: |
| AC025423.5 | 12 | guanine nucleotide binding protein, alpha trar |
| PRELID2P1 | 12 | PRELI domain containing 2 pseudogene 1 [Sou |
| AC127894.1 | 12 | novel transcript |
| CPSF6 | 12 | cleavage and polyadenylation specific factor 6 |
| MIR1279 | 12 | microRNA 1279 [Source:HGNC Symbol;Acc:HG |
| C1GALT1P1 | 12 | C1GALT1 pseudogene 1 [Source:HGNC Symbo |
| AC020656.2 | 12 | novel transcript |
| LYZ | 12 | lysozyme [Source:HGNC Symbol;Acc:HGNC:67 |
| AC020656.1 | 12 | novel transcript, antisense to LYZ |
| YEATS4 | 12 | YEATS domain containing 4 [Source:HGNC Syn |
| RN7SL804P | 12 | RNA, 7SL, cytoplasmic 804, pseudogene [Sour |
| RPS26P45 | 12 | ribosomal protein S26 pseudogene 45 [Source |
| LINC02373 | 12 | long intergenic non-protein coding RNA 2373 |
| FRS2 | 12 | fibroblast growth factor receptor substrate 2 [ |
| MIR3913-1 | 12 | microRNA 3913-1 [Source:HGNC Symbol;Acc:I |

| | | |
|---|---|---|
| MIR3913-2 | 12 | microRNA 3913-2 [Source:HGNC Symbol;Acc:I |
| CCT2 | 12 | chaperonin containing TCP1 subunit 2 [Source |
| LRRC10 | 12 | leucine rich repeat containing 10 [Source:HGN |
| BEST3 | 12 | bestrophin 3 [Source:HGNC Symbol;Acc:HGN( |
| AC025263.3 | 12 | TEC |
| AC025263.1 | 12 | novel transcript, antisense to RAB3IP |
| RAB3IP | 12 | RAB3A interacting protein [Source:HGNC Sym |
| AC025263.2 | 12 | novel protein |
| MYRFL | 12 | myelin regulatory factor like [Source:HGNC Sy |
| AC025159.1 | 12 | novel transcript |
| AC078922.1 | 12 | novel transcript, antisense to MYRFL |
| LINC02821 | 12 | long intergenic non-protein coding RNA 2821 |
| LINC01481 | 12 | long intergenic non-protein coding RNA 1481 |
| AC092881.2 | 12 | novel transcript |
| CNOT2 | 12 | CCR4-NOT transcription complex subunit 2 [S |
| AC092881.1 | 12 | TEC |
| KCNMB4 | 12 | potassium calcium-activated channel subfami |
| RNU4-65P | 12 | RNA, U4 small nuclear 65, pseudogene [Sourc |
| AC025569.1 | 12 | novel transcript, antisense to PTPRB |
| PTPRB | 12 | protein tyrosine phosphatase receptor type B |
| AC083809.1 | 12 | novel transcript, antisense to PTPRB |
| PTPRR | 12 | protein tyrosine phosphatase receptor type R |
| FAHD2P1 | 12 | fumarylacetoacetate hydrolase domain conta |
| Y_RNA | 12 | Y RNA [Source:RFAM;Acc:RF00019] |
| AC123905.1 | 12 | novel transcript |
| AC025575.1 | 12 | novel transcript |
| AC025575.2 | 12 | novel transcript |
| TSPAN8 | 12 | tetraspanin 8 [Source:HGNC Symbol;Acc:HGN |
| LGR5 | 12 | leucine rich repeat containing G protein-coupl |
| AC090116.1 | 12 | novel transcript, sense intronic to LGR5 |
| AC078860.2 | 12 | novel transcript, antisense to LGR5 |
| ZFC3H1 | 12 | zinc finger C3H1-type containing [Source:HGN |
| AC078860.1 | 12 | NULL |
| THAP2 | 12 | THAP domain containing 2 [Source:HGNC Sym |
| AC073612.1 | 12 | novel protein |
| TMEM19 | 12 | transmembrane protein 19 [Source:HGNC Syn |
| AC011601.1 | 12 | novel transcript |
| RAB21 | 12 | RAB21, member RAS oncogene family [Source |
| AC089984.1 | 12 | novel transcript |
| AC089984.3 | 12 | vacuolar protein sorting 24 homolog (S. cerev |
| TBC1D15 | 12 | TBC1 domain family member 15 [Source:HGN |
| AC089984.2 | 12 | MRS2 magnesium homeostasis factor homolo |
| TPH2 | 12 | tryptophan hydroxylase 2 [Source:HGNC Symb |
| AC090109.1 | 12 | novel transcript, antisense to TPH2 |
| TRHDE | 12 | thyrotropin releasing hormone degrading enz |
| TRHDE-AS1 | 12 | TRHDE antisense RNA 1 [Source:HGNC Symbo |
| AC016256.1 | 12 | histone cluster 3, H3 (HIST3H3) pseudogene |
| CHCHD3P2 | 12 | coiled-coil-helix-coiled-coil-helix domain conta |
| AC133480.1 | 12 | novel transcript |
| AC090503.2 | 12 | novel transcript |

| | | |
|---|---|---|
| LINC02444 | 12 | long intergenic non-protein coding RNA 2444 |
| AC090503.1 | 12 | novel transcript |
| AC087879.1 | 12 | ribosomal protein L31 (RPL31) pseudogene |
| RNU6-1012P | 12 | RNA, U6 small nuclear 1012, pseudogene [Sou |
| LINC02445 | 12 | long intergenic non-protein coding RNA 2445 |
| U8 | 12 | U8 small nucleolar RNA [Source:RFAM;Acc:RF |
| AC090015.1 | 12 | novel transcript |
| AC136188.1 | 12 | novel transcript |
| LINC02394 | 12 | long intergenic non-protein coding RNA 2394 |
| AC090502.3 | 12 | SURP and G patch domain containing 1 (SUGP |
| AC090502.2 | 12 | novel transcript |
| AC090502.1 | 12 | DnaJ (Hsp40) homolog, subfamily C, member |
| VENTXP3 | 12 | VENT homeobox pseudogene 3 [Source:HGNC |
| ATXN7L3B | 12 | ataxin 7 like 3B [Source:HGNC Symbol;Acc:HG |
| AC025257.1 | 12 | novel transcript, antisense to ATXN7L3B |
| AC123904.2 | 12 | cysteine-rich PDZ-binding protein (CRIPT) pseu |
| AC123904.1 | 12 | microtubule-associated protein 1 light chain 3 |
| AC123904.3 | 12 | TAF9 RNA polymerase II, TATA box binding pro |
| AC073525.1 | 12 | novel transcript, antisense to KCNC2 |
| KCNC2 | 12 | potassium voltage-gated channel subfamily C |
| AC091534.1 | 12 | novel transcript |
| CCNG2P1 | 12 | cyclin G2 pseudogene 1 [Source:HGNC Symbo |
| CAPS2 | 12 | calcyphosine 2 [Source:HGNC Symbol;Acc:HGI |
| AC092552.1 | 12 | necdin homolog (mouse) (NDN) pseudogene |
| AC121761.2 | 12 | novel transcript, sense intronic to CAPS2 |
| GLIPR1L1 | 12 | GLIPR1 like 1 [Source:HGNC Symbol;Acc:HGN |
| GLIPR1L2 | 12 | GLIPR1 like 2 [Source:HGNC Symbol;Acc:HGN |
| GLIPR1 | 12 | GLI pathogenesis related 1 [Source:HGNC Sym |
| AC121761.1 | 12 | novel transcript, antisense to GLIPR1 |
| KRR1 | 12 | KRR1 small subunit processome component h |
| AC078923.1 | 12 | novel transcript |
| AC022507.1 | 12 | phosphogluconate dehydrogenase (PGD) pseu |
| AC078820.2 | 12 | novel transcript |
| RPL10P13 | 12 | ribosomal protein L10 pseudogene 13 [Source |
| SNORA70 | 12 | Small nucleolar RNA SNORA70 [Source:RFAM; |
| AC078820.1 | 12 | novel transcript |
| AC011611.6 | 12 | transmembrane protein 141 (TMEM141) pseu |
| RN7SL734P | 12 | RNA, 7SL, cytoplasmic 734, pseudogene [Sour |
| AC011611.2 | 12 | novel transcript |
| AC011611.1 | 12 | NOP10 ribonucleoprotein homolog (yeast) (N |
| PHLDA1 | 12 | pleckstrin homology like domain family A mer |
| AC011611.3 | 12 | novel transcript, antisense to PHLDA1 |
| AC011611.4 | 12 | novel transcript, antisense to PHLDA1 |
| NAP1L1 | 12 | nucleosome assembly protein 1 like 1 [Source |
| AC011611.5 | 12 | density-regulated protein (DENR) pseudogene |
| AC233290.2 | 12 | polycomb group ring finger 1 (PCGF1) pseudo |
| AC233290.1 | 12 | reticulon 4 interacting protein 1 (RTN4IP1) pse |
| RNU6-1271P | 12 | RNA, U6 small nuclear 1271, pseudogene [Sou |
| LNCOG | 12 | lncRNA osteogenesis associated [Source:HGN |
| RN7SKP172 | 12 | RN7SK pseudogene 172 [Source:HGNC Symbo |

| | | |
|---|---|---|
| BBS10 | 12 | Bardet-Biedl syndrome 10 [Source:HGNC Sym |
| OSBPL8 | 12 | oxysterol binding protein like 8 [Source:HGNC |
| AC107032.2 | 12 | novel transcript |
| AC107032.3 | 12 | novel transcript |
| RPL7AP9 | 12 | ribosomal protein L7a pseudogene 9 [Source:I |
| YWHAQP7 | 12 | YWHAQ pseudogene 7 [Source:HGNC Symbol; |
| AC107032.1 | 12 | ribosomal protein L21 (RPL21) pseudogene |
| RPL7P43 | 12 | ribosomal protein L7 pseudogene 43 [Source:I |
| AC093014.1 | 12 | RWD domain containing 1 (RWDD1) pseudoge |
| ZDHHC17 | 12 | zinc finger DHHC-type containing 17 [Source:H |
| CSRP2 | 12 | cysteine and glycine rich protein 2 [Source:HG |
| AC124784.1 | 12 | novel transcript, antisense to CSRP2 |
| AC025161.1 | 12 | RAN binding protein 1 (RANBP1) pseudogene |
| E2F7 | 12 | E2F transcription factor 7 [Source:HGNC Symb |
| AC079030.1 | 12 | novel transcript |
| LINC02464 | 12 | long intergenic non-protein coding RNA 2464 |
| NAV3 | 12 | neuron navigator 3 [Source:HGNC Symbol;Acc |
| AC073591.1 | 12 | novel transcript |
| AC138331.1 | 12 | novel transcript |
| AC073571.1 | 12 | novel transcript, antisense to NAV3 |
| PRXL2AP1 | 12 | peroxiredoxin like 2A pseudogene 1 [Source:H |
| LINC02424 | 12 | long intergenic non-protein coding RNA 2424 |
| AC128707.1 | 12 | novel transcript |
| AC130415.1 | 12 | novel transcript |
| AC079362.1 | 12 | novel transcript |
| AC068993.1 | 12 | v-ral simian leukemia viral oncogene homolog |
| AC068993.2 | 12 | novel transcript |
| SYT1 | 12 | synaptotagmin 1 [Source:HGNC Symbol;Acc:H |
| AC090709.1 | 12 | novel transcript, antisense to SYT1 |
| AC027288.3 | 12 | novel transcript |
| MIR1252 | 12 | microRNA 1252 [Source:HGNC Symbol;Acc:HG |
| AC027288.2 | 12 | zinc finger protein 254 (ZNF254) pseudogene |
| AC027288.1 | 12 | novel transcript |
| NOP56P3 | 12 | NOP56 ribonucleoprotein pseudogene 3 [Sour |
| PAWR | 12 | pro-apoptotic WT1 regulator [Source:HGNC Sy |
| AC073569.2 | 12 | novel transcript |
| PPP1R12A | 12 | protein phosphatase 1 regulatory subunit 12A |
| AC073569.3 | 12 | novel transcript, antisense to PPP1R12A |
| AC073569.1 | 12 | novel transcript, antisense to PPP1R12A |
| RNA5SP363 | 12 | RNA, 5S ribosomal pseudogene 363 [Source:H |
| PPP1R12A-AS1 | 12 | PPP1R12A antisense RNA 1 [Source:HGNC Syn |
| SNRPGP20 | 12 | small nuclear ribonucleoprotein polypeptide ( |
| RPL7P38 | 12 | ribosomal protein L7 pseudogene 38 [Source:I |
| RNU7-106P | 12 | RNA, U7 small nuclear 106 pseudogene [Sourc |
| OTOGL | 12 | otogelin like [Source:HGNC Symbol;Acc:HGNC |
| AC078817.1 | 12 | ribosomal protein L26 (RPL26) pseudogene |
| RN7SKP261 | 12 | RN7SK pseudogene 261 [Source:HGNC Symbo |
| PTPRQ | 12 | protein tyrosine phosphatase receptor type Q |
| AKIRIN1P1 | 12 | akirin 1 pseudogene 1 [Source:HGNC Symbol;/ |
| AC074031.1 | 12 | novel transcript, antisense to PTPRQ |

| | | |
|---|---|---|
| MYF6 | 12 | myogenic factor 6 [Source:HGNC Symbol;Acc: |
| MYF5 | 12 | myogenic factor 5 [Source:HGNC Symbol;Acc: |
| LINC01490 | 12 | long intergenic non-protein coding RNA 1490 |
| LIN7A | 12 | lin-7 homolog A, crumbs cell polarity complex |
| MIR617 | 12 | microRNA 617 [Source:HGNC Symbol;Acc:HGI |
| AC078886.1 | 12 | tripartite motif containing 61 (TRIM61) pseud |
| MIR618 | 12 | microRNA 618 [Source:HGNC Symbol;Acc:HGI |
| ACSS3 | 12 | acyl-CoA synthetase short chain family memb |
| AC078955.1 | 12 | novel transcript, antisense to ACSS3 |
| MIR4699 | 12 | microRNA 4699 [Source:HGNC Symbol;Acc:HG |
| PPFIA2 | 12 | PTPRF interacting protein alpha 2 [Source:HGI |
| PPFIA2-AS1 | 12 | PPFIA2 antisense RNA 1 [Source:HGNC Symbo |
| AC069228.1 | 12 | novel transcript, antisense to PPFIA2 |
| AC079363.1 | 12 | transmembrane protein 66 (TMEM66) pseudo |
| AC091515.1 | 12 | vomeronasal 1 receptor 3 (gene/pseudogene) |
| LINC02426 | 12 | long intergenic non-protein coding RNA 2426 |
| AC011602.1 | 12 | peroxisomal biogenesis factor 2 (PEX2) pseud |
| CCDC59 | 12 | coiled-coil domain containing 59 [Source:HGN |
| METTL25 | 12 | methyltransferase like 25 [Source:HGNC Symb |
| AC089998.3 | 12 | F-box and WD repeat domain containing 12 (F |
| AC089998.4 | 12 | novel transcript |
| AC089998.2 | 12 | novel transcript |
| AC089998.1 | 12 | novel transcript |
| AC091214.1 | 12 | novel transcript |
| TMTC2 | 12 | transmembrane O-mannosyltransferase targe |
| RNU6-977P | 12 | RNA, U6 small nuclear 977, pseudogene [Sour |
| RPL6P25 | 12 | ribosomal protein L6 pseudogene 25 [Source: |
| AC090680.1 | 12 | novel transcript |
| AC093025.1 | 12 | novel transcript |
| AC090679.3 | 12 | novel transcript |
| AC090679.2 | 12 | novel transcript |
| AC090679.1 | 12 | NULL |
| AC087888.1 | 12 | WD repeat domain 12 (WDR12) pseudogene |
| AC093027.1 | 12 | cytochrome c oxidase subunit Va (COX5A) pse |
| SLC6A15 | 12 | solute carrier family 6 member 15 [Source:HG |
| AC128657.1 | 12 | small nuclear ribonucleoprotein D2 polypeptic |
| TSPAN19 | 12 | tetraspanin 19 [Source:HGNC Symbol;Acc:HGI |
| LRRIQ1 | 12 | leucine rich repeats and IQ motif containing 1 |
| ALX1 | 12 | ALX homeobox 1 [Source:HGNC Symbol;Acc:H |
| LINC02820 | 12 | long intergenic non-protein coding RNA 2820 |
| AC126471.1 | 12 | p53 and DNA-damage regulated 1 (PDRG1) ps |
| AC137768.1 | 12 | TM2 domain containing 3 (TM2D3) pseudoge |
| RASSF9 | 12 | Ras association domain family member 9 [Sou |
| NTS | 12 | neurotensin [Source:HGNC Symbol;Acc:HGNC |
| MGAT4C | 12 | MGAT4 family member C [Source:HGNC Symb |
| AC016993.1 | 12 | novel transcript |
| AC139697.1 | 12 | SH3-domain binding protein 4 (SH3BP4) pseu |
| AC010196.1 | 12 | novel transcript |
| AC079597.1 | 12 | novel transcript |
| RPL23AP68 | 12 | ribosomal protein L23a pseudogene 68 [Sourc |

| | | |
|---|---|---|
| AC079598.4 | 12 | cytochrome c, somatic (CYCS) pseudogene |
| CYCSP30 | 12 | CYCS pseudogene 30 [Source:HGNC Symbol;A |
| AC079598.2 | 12 | makorin ring finger protein (MKRN) pseudoge |
| MKRN9P | 12 | makorin ring finger protein 9, pseudogene [Sc |
| LINC02258 | 12 | long intergenic non-protein coding RNA 2258 |
| AC079598.1 | 12 | novel transcript |
| AC079598.3 | 12 | small nuclear ribonucleoprotein D1 polypeptic |
| RPS4XP15 | 12 | ribosomal protein S4X pseudogene 15 [Source |
| C12orf50 | 12 | chromosome 12 open reading frame 50 [Sour |
| C12orf29 | 12 | chromosome 12 open reading frame 29 [Sour |
| CEP290 | 12 | centrosomal protein 290 [Source:HGNC Symb |
| RNA5SP364 | 12 | RNA, 5S ribosomal pseudogene 364 [Source:H |
| AC091516.1 | 12 | transmembrane protein 38A (TMEM38A) |
| TMTC3 | 12 | transmembrane O-mannosyltransferase targe |
| Y_RNA | 12 | Y RNA [Source:RFAM;Acc:RF00019] |
| KITLG | 12 | KIT ligand [Source:HGNC Symbol;Acc:HGNC:63 |
| AC024941.2 | 12 | novel transcript, antisense to KITLG |
| RNU1-117P | 12 | RNA, U1 small nuclear 117, pseudogene [Sour |
| AC024941.1 | 12 | novel transcript |
| LINC02458 | 12 | long intergenic non-protein coding RNA 2458 |
| AC006199.1 | 12 | microtubule-associated protein 1 light chain 3 |
| RNU7-120P | 12 | RNA, U7 small nuclear 120 pseudogene [Sourc |
| MRPS6P4 | 12 | mitochondrial ribosomal protein S6 pseudoge |
| DUSP6 | 12 | dual specificity phosphatase 6 [Source:HGNC S |
| AC024909.1 | 12 | novel transcript, antisense to DUSP6 |
| AC010201.3 | 12 | novel transcript |
| AC010201.2 | 12 | novel transcript |
| AC010201.1 | 12 | novel transcript |
| POC1B | 12 | POC1 centriolar protein B [Source:HGNC Symb |
| CENPCP1 | 12 | centromere protein C pseudogene 1 [Source:H |
| POC1B-GALNT4 | 12 | POC1B-GALNT4 readthrough [Source:HGNC Sy |
| GALNT4 | 12 | polypeptide N-acetylgalactosaminyltransferas |
| POC1B-AS1 | 12 | POC1B antisense RNA 1 [Source:HGNC Symbo |
| AC025034.2 | 12 | TEC |
| AC025034.1 | 12 | novel transcript, antisense to ATP2B1 |
| ATP2B1 | 12 | ATPase plasma membrane Ca2+ transporting |
| ATP2B1-AS1 | 12 | ATP2B1 antisense RNA 1 [Source:HGNC Symbo |
| AC009522.1 | 12 | novel transcript |
| RNA5SP365 | 12 | RNA, 5S ribosomal pseudogene 365 [Source:H |
| MRPL2P1 | 12 | mitochondrial ribosomal protein L2 pseudoge |
| RNU6-148P | 12 | RNA, U6 small nuclear 148, pseudogene [Sour |
| BRWD1P2 | 12 | bromodomain and WD repeat domain contair |
| LINC02399 | 12 | long intergenic non-protein coding RNA 2399 |
| AC126178.1 | 12 | novel transcript |
| LINC02392 | 12 | long intergenic non-protein coding RNA 2392 |
| LINC02822 | 12 | long intergenic non-protein coding RNA 2822 |
| AC112481.2 | 12 | novel transcript |
| AC084365.1 | 12 | novel transcript |
| AC112481.1 | 12 | novel transcript |
| CCER1 | 12 | coiled-coil glutamate rich protein 1 [Source:H |

| | | |
|---|---|---|
| LINC00615 | 12 | long intergenic non-protein coding RNA 615 [S |
| EPYC | 12 | epiphycan [Source:HGNC Symbol;Acc:HGNC:3 |
| KERA | 12 | keratocan [Source:HGNC Symbol;Acc:HGNC:6 |
| LUM | 12 | lumican [Source:HGNC Symbol;Acc:HGNC:672 |
| DCN | 12 | decorin [Source:HGNC Symbol;Acc:HGNC:270 |
| AC007115.1 | 12 | GABA(A) receptor-associated protein (GABAR |
| LINC02823 | 12 | long intergenic non-protein coding RNA 2823 |
| AC090016.1 | 12 | novel transcript |
| AC126175.1 | 12 | NULL |
| AC126175.2 | 12 | novel transcript |
| AC025254.1 | 12 | novel transcript |
| AC090049.2 | 12 | alcohol dehydrogenase 5 (class III), chi polype |
| LINC02404 | 12 | long intergenic non-protein coding RNA 2404 |
| AC090049.1 | 12 | novel transcript |
| LINC01619 | 12 | long intergenic non-protein coding RNA 1619 |
| AC123512.1 | 12 | novel transcript |
| AC025164.2 | 12 | novel transcript |
| BTG1 | 12 | BTG anti-proliferation factor 1 [Source:HGNC |
| AC025164.1 | 12 | novel transcript, antisense to BTG1 |
| RPL21P106 | 12 | ribosomal protein L21 pseudogene 106 [Sour |
| Y_RNA | 12 | Y RNA [Source:RFAM;Acc:RF00019] |
| LINC02391 | 12 | long intergenic non-protein coding RNA 2391 |
| CLLU1OS | 12 | chronic lymphocytic leukemia up-regulated 1 |
| AC063949.2 | 12 | novel transcript, antisense to CLLU1OS |
| CLLU1 | 12 | chronic lymphocytic leukemia up-regulated 1 |
| LINC02397 | 12 | long intergenic non-protein coding RNA 2397 |
| AC063949.1 | 12 | mitochondrial carrier homolog 2 (MTCH2) pse |
| C12orf74 | 12 | chromosome 12 open reading frame 74 [Sour |
| EEA1 | 12 | early endosome antigen 1 [Source:HGNC Sym |
| AC026111.1 | 12 | methionine adenosyltransferase II, alpha (MA |
| HNRNPA1P50 | 12 | heterogeneous nuclear ribonucleoprotein A1 |
| RNU6-1329P | 12 | RNA, U6 small nuclear 1329, pseudogene [Sou |
| LINC02413 | 12 | long intergenic non-protein coding RNA 2413 |
| AC138123.1 | 12 | novel transcript, antisense to novel protein |
| Y_RNA | 12 | Y RNA [Source:RFAM;Acc:RF00019] |
| AC021646.1 | 12 | developmental pluripotency associated 3 (DP |
| RPL41P5 | 12 | ribosomal protein L41 pseudogene 5 [Source: |
| AC138123.2 | 12 | novel transcript |
| NACAP8 | 12 | NACA pseudogene 8 [Source:HGNC Symbol;A |
| LINC02412 | 12 | long intergenic non-protein coding RNA 2412 |
| AC126172.1 | 12 | NULL |
| AC124947.2 | 12 | anaphase promoting complex subunit 10 (AN |
| AC124947.1 | 12 | novel transcript, antisense to NUDT4 |
| NUDT4 | 12 | nudix hydrolase 4 [Source:HGNC Symbol;Acc: |
| UBE2N | 12 | ubiquitin conjugating enzyme E2 N [Source:H |
| Y_RNA | 12 | Y RNA [Source:RFAM;Acc:RF00019] |
| MRPL42 | 12 | mitochondrial ribosomal protein L42 [Source: |
| RN7SL737P | 12 | RNA, 7SL, cytoplasmic 737, pseudogene [Sour |
| AC025260.1 | 12 | RNA binding motif protein, X-linked (RBMX) p |
| SOCS2-AS1 | 12 | SOCS2 antisense RNA 1 [Source:HGNC Symbol |

| | | |
|---|---|---|
| SOCS2 | 12 | suppressor of cytokine signaling 2 [Source:HG |
| AC012085.1 | 12 | phosphoglycerate mutase 1 (brain) (PGAM1) p |
| CRADD | 12 | CASP2 and RIPK1 domain containing adaptor v |
| AC012085.2 | 12 | novel transcript |
| RN7SL630P | 12 | RNA, 7SL, cytoplasmic 630, pseudogene [Sour |
| AC012464.2 | 12 | novel transcript |
| AC012464.1 | 12 | novel transcript, antisense to RRP12 |
| AC012464.3 | 12 | novel transcript |
| RN7SKP263 | 12 | RN7SK pseudogene 263 [Source:HGNC Symbo |
| AC123567.1 | 12 | REST corepressor 2 (RCOR2) pseudogene |
| PLXNC1 | 12 | plexin C1 [Source:HGNC Symbol;Acc:HGNC:91 |
| AC123567.2 | 12 | novel transcript, antisense to PLXNC1 |
| AC073655.3 | 12 | novel transcript, antisense to PLXNC1 |
| AC073655.1 | 12 | novel transcript, antisense to PLXNC1 |
| AC073655.2 | 12 | novel transcript, antisense to PLXNC1 |
| CEP83 | 12 | centrosomal protein 83 [Source:HGNC Symbo |
| RN7SL330P | 12 | RNA, 7SL, cytoplasmic 330, pseudogene [Sour |
| RBMS2P1 | 12 | RNA binding motif single stranded interacting |
| CEP83-DT | 12 | CEP83 divergent transcript [Source:HGNC Syn |
| AC023161.1 | 12 | novel transcript |
| RN7SL483P | 12 | RNA, 7SL, cytoplasmic 483, pseudogene [Sour |
| AC023161.2 | 12 | NSA2 ribosome biogenesis homolog (S. cerevi |
| SUCLG2P2 | 12 | succinate-CoA ligase GDP-forming beta subun |
| MIR5700 | 12 | microRNA 5700 [Source:HGNC Symbol;Acc:HG |
| TMCC3 | 12 | transmembrane and coiled-coil domain family |
| MIR7844 | 12 | microRNA 7844 [Source:HGNC Symbol;Acc:HG |
| LSM3P2 | 12 | LSM3 homolog, U6 small nuclear RNA and mR |
| KRT19P2 | 12 | keratin 19 pseudogene 2 [Source:HGNC Symb |
| MIR492 | 12 | microRNA 492 [Source:HGNC Symbol;Acc:HGN |
| NDUFA12 | 12 | NADH:ubiquinone oxidoreductase subunit A1 |
| NR2C1 | 12 | nuclear receptor subfamily 2 group C member |
| FGD6 | 12 | FYVE, RhoGEF and PH domain containing 6 [So |
| AC126615.2 | 12 | peroxiredoxin 6 (PRDX6) pseudogene |
| AC126615.1 | 12 | ribosomal protein L14 (RPL14) pseudogene |
| VEZT | 12 | vezatin, adherens junctions transmembrane p |
| Y_RNA | 12 | Y RNA [Source:RFAM;Acc:RF00019] |
| CBX3P5 | 12 | chromobox 3 pseudogene 5 [Source:HGNC Sy |
| RNU6-808P | 12 | RNA, U6 small nuclear 808, pseudogene [Sour |
| MIR331 | 12 | microRNA 331 [Source:HGNC Symbol;Acc:HGN |
| MIR3685 | 12 | microRNA 3685 [Source:HGNC Symbol;Acc:HG |
| AC084879.1 | 12 | novel pseudogene |
| AC084879.2 | 12 | novel transcript |
| RNU6-735P | 12 | RNA, U6 small nuclear 735, pseudogene [Sour |
| AC018475.1 | 12 | ribosomal protein L29 (RPL29) pseudogene |
| METAP2 | 12 | methionyl aminopeptidase 2 [Source:HGNC Sy |
| USP44 | 12 | ubiquitin specific peptidase 44 [Source:HGNC |
| PGAM1P5 | 12 | phosphoglycerate mutase 1 pseudogene 5 [So |
| Y_RNA | 12 | Y RNA [Source:RFAM;Acc:RF00019] |
| NTN4 | 12 | netrin 4 [Source:HGNC Symbol;Acc:HGNC:136 |
| AC090001.1 | 12 | novel transcript |

| | | |
|---|---|---|
| RNU6-247P | 12 | RNA, U6 small nuclear 247, pseudogene [Sour |
| LINC02410 | 12 | long intergenic non-protein coding RNA 2410 |
| SNRPF | 12 | small nuclear ribonucleoprotein polypeptide F |
| CCDC38 | 12 | coiled-coil domain containing 38 [Source:HGN |
| AC090001.2 | 12 | novel oocyte-secreted protein family member |
| AMDHD1 | 12 | amidohydrolase domain containing 1 [Source: |
| HAL | 12 | histidine ammonia-lyase [Source:HGNC Symbo |
| AC007298.2 | 12 | novel transcript, antisense to LTA4H |
| LTA4H | 12 | leukotriene A4 hydrolase [Source:HGNC Symb |
| AC007298.1 | 12 | novel transcript, antisense to LTA4H |
| YPEL5P3 | 12 | YPEL5 pseudogene 3 [Source:HGNC Symbol;A |
| RN7SL88P | 12 | RNA, 7SL, cytoplasmic 88, pseudogene [Sourc |
| LINC02452 | 12 | long intergenic non-protein coding RNA 2452 |
| ELK3 | 12 | ETS transcription factor ELK3 [Source:HGNC S |
| AC008149.1 | 12 | novel transcript, antisense to ELK3 |
| AC008149.2 | 12 | novel transcript, antisense to ELK3 |
| CDK17 | 12 | cyclin dependent kinase 17 [Source:HGNC Syn |
| RNU4-24P | 12 | RNA, U4 small nuclear 24, pseudogene [Sourc |
| AC125612.1 | 12 | small nuclear ribonucleoprotein D1 polypeptic |
| AC007513.1 | 12 | novel transcript |
| RN7SKP11 | 12 | RN7SK pseudogene 11 [Source:HGNC Symbol; |
| CFAP54 | 12 | cilia and flagella associated protein 54 [Source |
| EEF1A1P33 | 12 | eukaryotic translation elongation factor 1 alph |
| NEDD1 | 12 | NEDD1 gamma-tubulin ring complex targeting |
| Y_RNA | 12 | Y RNA [Source:RFAM;Acc:RF00019] |
| AC007656.2 | 12 | novel transcript |
| AC007656.1 | 12 | novel transcript |
| AC007656.3 | 12 | novel transcript |
| AC007656.4 | 12 | novel transcript |
| LINC02409 | 12 | long intergenic non-protein coding RNA 2409 |
| RMST | 12 | rhabdomyosarcoma 2 associated transcript [S |
| MIR1251 | 12 | microRNA 1251 [Source:HGNC Symbol;Acc:H |
| AC007351.2 | 12 | NULL |
| AC007351.3 | 12 | NULL |
| AC007351.4 | 12 | NULL |
| AC007351.1 | 12 | NULL |
| AC018659.4 | 12 | NULL |
| AC018659.3 | 12 | NULL |
| AC018659.1 | 12 | NULL |
| AC018659.7 | 12 | NULL |
| AC018659.5 | 12 | NULL |
| AC018659.8 | 12 | TEC |
| MIR135A2 | 12 | microRNA 135a-2 [Source:HGNC Symbol;Acc:I |
| AC018659.6 | 12 | NULL |
| AC018659.2 | 12 | protein phosphatase, EF-hand calcium binding |
| AC007424.1 | 12 | platelet-activating factor acetylhydrolase 1b, |
| RNU6-36P | 12 | RNA, U6 small nuclear 36, pseudogene [Sourc |
| MIR4495 | 12 | microRNA 4495 [Source:HGNC Symbol;Acc:H |
| MIR4303 | 12 | microRNA 4303 [Source:HGNC Symbol;Acc:H |
| AC016152.1 | 12 | novel transcript |

| | | |
|---|---|---|
| AC008055.1 | 12 | family with sequence similarity 60, member A |
| RNU4-41P | 12 | RNA, U4 small nuclear 41, pseudogene [Sourc |
| AC008055.2 | 12 | novel transcript |
| SLC9A7P1 | 12 | solute carrier family 9 member 7 pseudogene |
| LINC02453 | 12 | long intergenic non-protein coding RNA 2453 |
| TMPO-AS1 | 12 | TMPO antisense RNA 1 [Source:HGNC Symbol |
| TMPO | 12 | thymopoietin [Source:HGNC Symbol;Acc:HGN |
| PPIAP8 | 12 | peptidylprolyl isomerase A pseudogene 8 [Sou |
| SLC25A3 | 12 | solute carrier family 25 member 3 [Source:HG |
| SNORA53 | 12 | small nucleolar RNA, H/ACA box 53 [Source:H |
| IKBIP | 12 | IKBKB interacting protein [Source:HGNC Symb |
| APAF1 | 12 | apoptotic peptidase activating factor 1 [Sourc |
| ANKS1B | 12 | ankyrin repeat and sterile alpha motif domain |
| AC008126.1 | 12 | Mdm2 p53 binding protein homolog (mouse) |
| AC069437.1 | 12 | novel transcript, antisense to ANKS1B |
| RNA5SP366 | 12 | RNA, 5S ribosomal pseudogene 366 [Source:H |
| AC117377.1 | 12 | novel transcript, antisense & intronic to ANKS |
| FAM71C | 12 | family with sequence similarity 71 member C |
| AC078916.1 | 12 | novel transcript |
| RPS4XP1 | 12 | ribosomal protein S4X pseudogene 1 [Source: |
| AC126474.1 | 12 | TEC |
| UHRF1BP1L | 12 | UHRF1 binding protein 1 like [Source:HGNC Sy |
| AC126474.2 | 12 | TEC |
| AC010203.2 | 12 | novel transcript, antisense to UHRF1BP1L |
| GOLGA2P5 | 12 | GOLGA2 pseudogene 5 [Source:HGNC Symbol |
| RN7SL176P | 12 | RNA, 7SL, cytoplasmic 176, pseudogene [Sour |
| AC010203.1 | 12 | TBC1 domain family, member 2B (TBC1D2B) p |
| MIR1827 | 12 | microRNA 1827 [Source:HGNC Symbol;Acc:HG |
| ACTR6 | 12 | actin related protein 6 [Source:HGNC Symbol; |
| DEPDC4 | 12 | DEP domain containing 4 [Source:HGNC Symb |
| Y_RNA | 12 | Y RNA [Source:RFAM;Acc:RF00019] |
| SCYL2 | 12 | SCY1 like pseudokinase 2 [Source:HGNC Symb |
| SLC17A8 | 12 | solute carrier family 17 member 8 [Source:HG |
| NR1H4 | 12 | nuclear receptor subfamily 1 group H membe |
| AC010200.2 | 12 | MAP/microtubule affinity-regulating kinase 2 |
| GAS2L3 | 12 | growth arrest specific 2 like 3 [Source:HGNC S |
| AC010200.1 | 12 | phosphatidylinositol glycan anchor biosynthes |
| ANO4 | 12 | anoctamin 4 [Source:HGNC Symbol;Acc:HGNC |
| AC138360.1 | 12 | novel transcript, antisense to ANO4 |
| AC063947.1 | 12 | novel transcript, antisense to ANO4 |
| AC063947.2 | 12 | sorting nexin 5 (SNX5) pseudogene |
| SLC5A8 | 12 | solute carrier family 5 member 8 [Source:HGN |
| RNU6-768P | 12 | RNA, U6 small nuclear 768, pseudogene [Sour |
| AC079953.1 | 12 | ribosomal protein S6 (RPS6) pseudogene |
| UTP20 | 12 | UTP20 small subunit processome component |
| ARL1 | 12 | ADP ribosylation factor like GTPase 1 [Source: |
| AC063948.1 | 12 | novel transcript |
| RNU6-1068P | 12 | RNA, U6 small nuclear 1068, pseudogene [Sou |
| RPS27P23 | 12 | ribosomal protein S27 pseudogene 23 [Source |
| Y_RNA | 12 | Y RNA [Source:RFAM;Acc:RF00019] |

| | | |
|---|---|---|
| RNU5E-5P | 12 | RNA, U5E small nuclear 5, pseudogene [Sourc |
| RNA5SP367 | 12 | RNA, 5S ribosomal pseudogene 367 [Source:H |
| SPIC | 12 | Spi-C transcription factor [Source:HGNC Symb |
| MYBPC1 | 12 | myosin binding protein C, slow type [Source:H |
| AC117505.1 | 12 | novel transcript, antisense to MYBPC1 |
| AC010205.1 | 12 | novel transcript |
| CHPT1 | 12 | choline phosphotransferase 1 [Source:HGNC S |
| RNY1P16 | 12 | RNY1 pseudogene 16 [Source:HGNC Symbol;A |
| SYCP3 | 12 | synaptonemal complex protein 3 [Source:HGN |
| GNPTAB | 12 | N-acetylglucosamine-1-phosphate transferase |
| RNU6-101P | 12 | RNA, U6 small nuclear 101, pseudogene [Sour |
| AC063950.1 | 12 | suppression of tumorigenicity 13 (colon carcir |
| RNA5SP368 | 12 | RNA, 5S ribosomal pseudogene 368 [Source:H |
| RNU6-172P | 12 | RNA, U6 small nuclear 172, pseudogene [Sour |
| RNA5SP369 | 12 | RNA, 5S ribosomal pseudogene 369 [Source:H |
| RNU6-1183P | 12 | RNA, U6 small nuclear 1183, pseudogene [Sou |
| HSPE1P4 | 12 | heat shock protein family E (Hsp10) member 1 |
| DRAM1 | 12 | DNA damage regulated autophagy modulator |
| AC084398.2 | 12 | novel transcript, antisense to DRAM1 |
| AC084398.1 | 12 | ribosomal protein L9 (RPL9) pseudogene |
| AC079907.2 | 12 | novel transcript |
| NENFP2 | 12 | neudesin neurotrophic factor pseudogene 2 [S |
| WASHC3 | 12 | WASH complex subunit 3 [Source:HGNC Symb |
| AC079907.1 | 12 | novel transcript, antisense to WASHC3 and NU |
| NUP37 | 12 | nucleoporin 37 [Source:HGNC Symbol;Acc:HG |
| PARPBP | 12 | PARP1 binding protein [Source:HGNC Symbol; |
| PMCH | 12 | pro-melanin concentrating hormone [Source:I |
| HELLPAR | 12 | HELLP associated long non-coding RNA [Sourc |
| RN7SL793P | 12 | RNA, 7SL, cytoplasmic 793, pseudogene [Sour |
| AC093023.1 | 12 | reticulon 3 (RTN3) pseudogene |
| LINC02456 | 12 | long intergenic non-protein coding RNA 2456 |
| IGF1 | 12 | insulin like growth factor 1 [Source:HGNC Sym |
| AC010202.1 | 12 | NADH dehydrogenase (ubiquinone) 1 alpha su |
| LINC00485 | 12 | long intergenic non-protein coding RNA 485 [S |
| PAH | 12 | phenylalanine hydroxylase [Source:HGNC Sym |
| AC026108.1 | 12 | novel transcript, sense intronic to PAH |
| ASCL1 | 12 | achaete-scute family bHLH transcription facto |
| AC026108.2 | 12 | novel transcript |
| AC068643.1 | 12 | novel transcript |
| AC068643.2 | 12 | novel transcript |
| C12orf42 | 12 | chromosome 12 open reading frame 42 [Sour |
| AC084364.1 | 12 | glutaryl-CoA dehydrogenase (GCDH) pseudoge |
| AC084364.2 | 12 | synaptonemal complex central element prote |
| AC084364.3 | 12 | novel transcript |
| LINC02401 | 12 | long intergenic non-protein coding RNA 2401 |
| STAB2 | 12 | stabilin 2 [Source:HGNC Symbol;Acc:HGNC:18 |
| U8 | 12 | U8 small nucleolar RNA [Source:RFAM;Acc:RF( |
| AC025265.2 | 12 | novel transcript, antisense to STAB2 |
| AC025265.3 | 12 | novel transcript, antisense to STAB2 |
| AC025265.1 | 12 | novel transcript, antisense to STAB2 |

| | | |
|---|---|---|
| NT5DC3 | 12 | 5'-nucleotidase domain containing 3 [Source:H |
| AC012555.2 | 12 | novel transcript, antisense to NT5DC3 |
| AC012555.1 | 12 | novel transcript |
| TTC41P | 12 | tetratricopeptide repeat domain 41, pseudoge |
| HSP90B1 | 12 | heat shock protein 90 beta family member 1 [ |
| MIR3652 | 12 | microRNA 3652 [Source:HGNC Symbol;Acc:HG |
| C12orf73 | 12 | chromosome 12 open reading frame 73 [Sour |
| TDG | 12 | thymine DNA glycosylase [Source:HGNC Symb |
| GLT8D2 | 12 | glycosyltransferase 8 domain containing 2 [So |
| AC078819.1 | 12 | phosphoglycerate mutase 1 (brain) (PGAM1) p |
| HCFC2 | 12 | host cell factor C2 [Source:HGNC Symbol;Acc: |
| NFYB | 12 | nuclear transcription factor Y subunit beta [So |
| RNA5SP370 | 12 | RNA, 5S ribosomal pseudogene 370 [Source:H |
| LINC02385 | 12 | long intergenic non-protein coding RNA 2385 |
| AC089983.2 | 12 | novel transcript |
| TXNRD1 | 12 | thioredoxin reductase 1 [Source:HGNC Symbo |
| AC089983.1 | 12 | novel transcript, antisense to TXNRD1 |
| RPL18AP3 | 12 | ribosomal protein L18a pseudogene 3 [Source |
| EID3 | 12 | EP300 interacting inhibitor of differentiation 3 |
| CHST11 | 12 | carbohydrate sulfotransferase 11 [Source:HGN |
| AC079316.1 | 12 | ATP synthase, H+ transporting, mitochondrial |
| AC079316.2 | 12 | TEC |
| MIR3922 | 12 | microRNA 3922 [Source:HGNC Symbol;Acc:HG |
| SLC41A2 | 12 | solute carrier family 41 member 2 [Source:HG |
| Y_RNA | 12 | Y RNA [Source:RFAM;Acc:RF00019] |
| AC089985.1 | 12 | novel transcript, antisense to SLC41A2 |
| KRT18P20 | 12 | keratin 18 pseudogene 20 [Source:HGNC Sym |
| C12orf45 | 12 | chromosome 12 open reading frame 45 [Sour |
| ALDH1L2 | 12 | aldehyde dehydrogenase 1 family member L2 |
| WASHC4 | 12 | WASH complex subunit 4 [Source:HGNC Symb |
| APPL2 | 12 | adaptor protein, phosphotyrosine interacting |
| C12orf75 | 12 | chromosome 12 open reading frame 75 [Sour |
| AC078874.1 | 12 | novel transcript, sense intronic to C12orf75 |
| AC011313.1 | 12 | novel transcript, antisense to C12orf75 |
| AC079851.1 | 12 | novel transcript |
| CASC18 | 12 | cancer susceptibility 18 [Source:HGNC Symbo |
| ST13P3 | 12 | ST13, Hsp70 interacting protein pseudogene 3 |
| AC011595.2 | 12 | novel transcript |
| NUAK1 | 12 | NUAK family kinase 1 [Source:HGNC Symbol;A |
| AC011595.1 | 12 | novel transcript, antisense to NUAK1 |
| AC010182.1 | 12 | thyroid hormone receptor interactor 11 (TRIP |
| CKAP4 | 12 | cytoskeleton associated protein 4 [Source:HG |
| AC079174.1 | 12 | novel transcript, sense intronic to CKAP4 |
| AC079174.2 | 12 | novel transcript, antisense to CKAP4 |
| RNU7-94P | 12 | RNA, U7 small nuclear 94 pseudogene [Source |
| TCP11L2 | 12 | t-complex 11 like 2 [Source:HGNC Symbol;Acc |
| POLR3B | 12 | RNA polymerase III subunit B [Source:HGNC Sy |
| AC079385.1 | 12 | novel transcript, antisense to POLR3B, RFX4 a |
| RFX4 | 12 | regulatory factor X4 [Source:HGNC Symbol;Ac |
| AC079385.2 | 12 | novel transcript, antisense to RFX4 |

| | | |
|---|---|---|
| AC079385.3 | 12 | novel transcript, antisense to RFX4 |
| RIC8B | 12 | RIC8 guanine nucleotide exchange factor B [So |
| AC007695.1 | 12 | novel transcript, antisense to RIC8B |
| EEF1B2P4 | 12 | eukaryotic translation elongation factor 1 beta |
| RPL30P12 | 12 | ribosomal protein L30 pseudogene 12 [Source |
| AC007541.1 | 12 | novel transcript, antisense to C12orf23 |
| TMEM263 | 12 | transmembrane protein 263 [Source:HGNC Sy |
| MTERF2 | 12 | mitochondrial transcription termination facto |
| CRY1 | 12 | cryptochrome circadian regulator 1 [Source:H |
| AC078929.1 | 12 | novel transcript, antisense to CRY1 |
| AC007649.1 | 12 | novel transcript |
| SETP7 | 12 | SET pseudogene 7 [Source:HGNC Symbol;Acc: |
| BTBD11 | 12 | BTB domain containing 11 [Source:HGNC Sym |
| AC009774.1 | 12 | NULL |
| RNA5SP371 | 12 | RNA, 5S ribosomal pseudogene 371 [Source:H |
| AC007540.1 | 12 | novel transcript, antisense to BTBD11 |
| Y_RNA | 12 | Y RNA [Source:RFAM;Acc:RF00019] |
| PWP1 | 12 | PWP1 homolog, endonuclein [Source:HGNC S |
| AC007622.1 | 12 | ribosomal protein L17 (RPL17) pseudogene |
| PRDM4 | 12 | PR/SET domain 4 [Source:HGNC Symbol;Acc:H |
| AC007622.2 | 12 | novel transcript, antisense to PRDM4 |
| ASCL4 | 12 | achaete-scute family bHLH transcription facto |
| AC126177.3 | 12 | novel transcript |
| AC126177.5 | 12 | ret finger protein-like 2 (RFPL2) pseudogene |
| AC126177.2 | 12 | ret finger protein-like 2 (RFPL2) pseudogene |
| AC126177.4 | 12 | coronin, actin binding protein, 1C (CORO1C) p |
| AC126177.6 | 12 | programmed cell death 2 (PDCD2) pseudogen |
| AC126177.1 | 12 | POM121 membrane glycoprotein (POM121) p |
| WSCD2 | 12 | WSC domain containing 2 [Source:HGNC Symb |
| AC009729.1 | 12 | apolipoprotein B mRNA editing enzyme, cataly |
| CMKLR1 | 12 | chemerin chemokine-like receptor 1 [Source:H |
| LINC01498 | 12 | long intergenic non-protein coding RNA 1498 |
| FICD | 12 | FIC domain containing [Source:HGNC Symbol; |
| SART3 | 12 | spliceosome associated factor 3, U4/U6 recyc |
| AC008119.1 | 12 | novel transcript, antisense to SART3 |
| ISCU | 12 | iron-sulfur cluster assembly enzyme [Source:H |
| TMEM119 | 12 | transmembrane protein 119 [Source:HGNC Sy |
| SELPLG | 12 | selectin P ligand [Source:HGNC Symbol;Acc:HG |
| AC007569.1 | 12 | novel transcript, antisense to SELPLG |
| MIR4496 | 12 | microRNA 4496 [Source:HGNC Symbol;Acc:HG |
| CORO1C | 12 | coronin 1C [Source:HGNC Symbol;Acc:HGNC:2 |
| RNU7-169P | 12 | RNA, U7 small nuclear 169 pseudogene [Sourc |
| Y_RNA | 12 | Y RNA [Source:RFAM;Acc:RF00019] |
| AC087893.1 | 12 | NULL |
| SSH1 | 12 | slingshot protein phosphatase 1 [Source:HGN |
| AC087893.3 | 12 | TEC |
| AC087893.2 | 12 | novel transcript, sense intronic to SSH1 |
| MIR619 | 12 | microRNA 619 [Source:HGNC Symbol;Acc:HGN |
| DAO | 12 | D-amino acid oxidase [Source:HGNC Symbol;A |
| SVOP | 12 | SV2 related protein [Source:HGNC Symbol;Acc |

| | | |
|---|---|---|
| RNU6-361P | 12 | RNA, U6 small nuclear 361, pseudogene [Sour |
| AC190387.1 | 12 | novel transcript |
| USP30 | 12 | ubiquitin specific peptidase 30 [Source:HGNC |
| USP30-AS1 | 12 | USP30 antisense RNA 1 [Source:HGNC Symbol |
| RNA5SP372 | 12 | RNA, 5S ribosomal pseudogene 372 [Source:H |
| ALKBH2 | 12 | alkB homolog 2, alpha-ketoglutarate depende |
| UNG | 12 | uracil DNA glycosylase [Source:HGNC Symbol; |
| AC007637.1 | 12 | novel transcript |
| ACACB | 12 | acetyl-CoA carboxylase beta [Source:HGNC Sy |
| FOXN4 | 12 | forkhead box N4 [Source:HGNC Symbol;Acc:H |
| MYO1H | 12 | myosin IH [Source:HGNC Symbol;Acc:HGNC:1 |
| LINC01486 | 12 | long intergenic non-protein coding RNA 1486 |
| AC007570.1 | 12 | novel transcript, antisense to MYO1H |
| KCTD10 | 12 | potassium channel tetramerization domain cc |
| UBE3B | 12 | ubiquitin protein ligase E3B [Source:HGNC Syr |
| MMAB | 12 | metabolism of cobalamin associated B [Sourc |
| RNU4-32P | 12 | RNA, U4 small nuclear 32, pseudogene [Sourc |
| MVK | 12 | mevalonate kinase [Source:HGNC Symbol;Acc |
| RN7SKP250 | 12 | RN7SK pseudogene 250 [Source:HGNC Symbo |
| FAM222A | 12 | family with sequence similarity 222 member A |
| FAM222A-AS1 | 12 | FAM222A antisense RNA 1 [Source:HGNC Syn |
| TRPV4 | 12 | transient receptor potential cation channel su |
| MIR4497 | 12 | microRNA 4497 [Source:HGNC Symbol;Acc:HG |
| GLTP | 12 | glycolipid transfer protein [Source:HGNC Syml |
| AC007834.2 | 12 | chromosome 8 open reading frame 59 (C8orf5 |
| RN7SL441P | 12 | RNA, 7SL, cytoplasmic 441, pseudogene [Sour |
| AC007834.1 | 12 | novel transcript |
| TCHP | 12 | trichoplein keratin filament binding [Source:H |
| GIT2 | 12 | GIT ArfGAP 2 [Source:HGNC Symbol;Acc:HGN( |
| AC084876.1 | 12 | novel transcript, antisense to GIT2 |
| AC084876.2 | 12 | TEC |
| ANKRD13A | 12 | ankyrin repeat domain 13A [Source:HGNC Syr |
| C12orf76 | 12 | chromosome 12 open reading frame 76 [Sour |
| AC007546.1 | 12 | novel transcript, sense intronic to ANKRD13A |
| AC007546.3 | 12 | purinergic receptor P2X, ligand-gated ion char |
| AC007546.2 | 12 | novel transcript |
| IFT81 | 12 | intraflagellar transport 81 [Source:HGNC Syml |
| ATP2A2 | 12 | ATPase sarcoplasmic/endoplasmic reticulum ( |
| RN7SL769P | 12 | RNA, 7SL, cytoplasmic 769, pseudogene [Sour |
| ANAPC7 | 12 | anaphase promoting complex subunit 7 [Sour |
| AC144548.1 | 12 | novel transcript |
| HMGA1P3 | 12 | high mobility group AT-hook 1 pseudogene 3 |
| ARPC3 | 12 | actin related protein 2/3 complex subunit 3 [S |
| GPN3 | 12 | GPN-loop GTPase 3 [Source:HGNC Symbol;Acc |
| RPL31P49 | 12 | ribosomal protein L31 pseudogene 49 [Source |
| FAM216A | 12 | family with sequence similarity 216 member A |
| VPS29 | 12 | VPS29 retromer complex component [Source: |
| AC002350.1 | 12 | NULL |
| AC002350.2 | 12 | novel transcript |
| RAD9B | 12 | RAD9 checkpoint clamp component B [Source |

| | | |
|---|---|---|
| PPTC7 | 12 | PTC7 protein phosphatase homolog [Source:H |
| TCTN1 | 12 | tectonic family member 1 [Source:HGNC Symb |
| RN7SL387P | 12 | RNA, 7SL, cytoplasmic 387, pseudogene [Sour |
| HVCN1 | 12 | hydrogen voltage gated channel 1 [Source:HG |
| PPP1CC | 12 | protein phosphatase 1 catalytic subunit gamm |
| AC144522.1 | 12 | novel transcript |
| AC002375.1 | 12 | novel transcript |
| RPL29P25 | 12 | ribosomal protein L29 pseudogene 25 [Source |
| CCDC63 | 12 | coiled-coil domain containing 63 [Source:HGN |
| MYL2 | 12 | myosin light chain 2 [Source:HGNC Symbol;Ac |
| LINC01405 | 12 | long intergenic non-protein coding RNA 1405 |
| AC002351.1 | 12 | novel transcript |
| CUX2 | 12 | cut like homeobox 2 [Source:HGNC Symbol;Ac |
| MIR6760 | 12 | microRNA 6760 [Source:HGNC Symbol;Acc:HG |
| RNA5SP373 | 12 | RNA, 5S ribosomal pseudogene 373 [Source:H |
| PHETA1 | 12 | PH domain containing endocytic trafficking ad |
| LINC02356 | 12 | long intergenic non-protein coding RNA 2356 |
| HSPA8P14 | 12 | heat shock protein family A (Hsp70) member 8 |
| SH2B3 | 12 | SH2B adaptor protein 3 [Source:HGNC Symbo |
| ATXN2 | 12 | ataxin 2 [Source:HGNC Symbol;Acc:HGNC:105 |
| U7 | 12 | U7 small nuclear RNA [Source:RFAM;Acc:RF00 |
| AC137055.1 | 12 | interferon induced transmembrane protein 3 |
| ATXN2-AS | 12 | ATXN2 antisense RNA [Source:HGNC Symbol;A |
| BRAP | 12 | BRCA1 associated protein [Source:HGNC Symb |
| PCNPP1 | 12 | PEST containing nuclear protein pseudogene 1 |
| ACAD10 | 12 | acyl-CoA dehydrogenase family member 10 [S |
| AC002996.1 | 12 | novel protein |
| ALDH2 | 12 | aldehyde dehydrogenase 2 family member [So |
| MIR6761 | 12 | microRNA 6761 [Source:HGNC Symbol;Acc:HG |
| MAPKAPK5-AS1 | 12 | MAPKAPK5 antisense RNA 1 [Source:HGNC Sy |
| MAPKAPK5 | 12 | MAPK activated protein kinase 5 [Source:HGN |
| RPS2P41 | 12 | ribosomal protein S2 pseudogene 41 [Source: |
| ADAM1A | 12 | ADAM metallopeptidase domain 1A (pseudog |
| ADAM1B | 12 | ADAM metallopeptidase domain 1B (pseudog |
| TMEM116 | 12 | transmembrane protein 116 [Source:HGNC Sy |
| SLC25A3P2 | 12 | solute carrier family 25 member 3 pseudogen |
| AC004024.1 | 12 | immediate early response 3 interacting protei |
| ERP29 | 12 | endoplasmic reticulum protein 29 [Source:HG |
| AC073575.2 | 12 | novel transcript, antisense to ERP29 |
| NAA25 | 12 | N(alpha)-acetyltransferase 25, NatB auxiliary s |
| MIR3657 | 12 | microRNA 3657 [Source:HGNC Symbol;Acc:HG |
| AC073575.1 | 12 | novel transcript, antisense to NAA25 |
| Y_RNA | 12 | Y RNA [Source:RFAM;Acc:RF00019] |
| TRAFD1 | 12 | TRAF-type zinc finger domain containing 1 [So |
| Y_RNA | 12 | Y RNA [Source:RFAM;Acc:RF00019] |
| HECTD4 | 12 | HECT domain E3 ubiquitin protein ligase 4 [So |
| MIR6861 | 12 | microRNA 6861 [Source:HGNC Symbol;Acc:HG |
| AC004217.1 | 12 | novel transcript |
| RN7SKP71 | 12 | RN7SK pseudogene 71 [Source:HGNC Symbol; |
| RPL7AP60 | 12 | ribosomal protein L7a pseudogene 60 [Source |

| | | |
|---|---|---|
| AC004217.2 | 12 | DnaJ (Hsp40) homolog, subfamily C, member |
| RPL6 | 12 | ribosomal protein L6 [Source:HGNC Symbol;A |
| PTPN11 | 12 | protein tyrosine phosphatase non-receptor ty |
| RPH3A | 12 | rabphilin 3A [Source:HGNC Symbol;Acc:HGNC |
| MIR1302-1 | 12 | microRNA 1302-1 [Source:HGNC Symbol;Acc:I |
| OAS1 | 12 | 2'-5'-oligoadenylate synthetase 1 [Source:HGI |
| AC004551.1 | 12 | novel transcript, antisense to OAS1, OAS2 and |
| OAS3 | 12 | 2'-5'-oligoadenylate synthetase 3 [Source:HGI |
| OAS2 | 12 | 2'-5'-oligoadenylate synthetase 2 [Source:HGI |
| IMMP1LP2 | 12 | inner mitochondrial membrane peptidase sub |
| RPS15AP32 | 12 | ribosomal protein S15a pseudogene 32 [Sourc |
| DTX1 | 12 | deltex E3 ubiquitin ligase 1 [Source:HGNC Sym |
| RASAL1 | 12 | RAS protein activator like 1 [Source:HGNC Syn |
| CFAP73 | 12 | cilia and flagella associated protein 73 [Source |
| Y_RNA | 12 | Y RNA [Source:RFAM;Acc:RF00019] |
| DDX54 | 12 | DEAD-box helicase 54 [Source:HGNC Symbol;/ |
| MIR7106 | 12 | microRNA 7106 [Source:HGNC Symbol;Acc:HG |
| Y_RNA | 12 | Y RNA [Source:RFAM;Acc:RF00019] |
| RITA1 | 12 | RBPJ interacting and tubulin associated 1 [Sou |
| AC089999.1 | 12 | novel transcript, antisense to C12orf52 |
| IQCD | 12 | IQ motif containing D [Source:HGNC Symbol;A |
| TPCN1 | 12 | two pore segment channel 1 [Source:HGNC Sy |
| AC089999.2 | 12 | novel transcript, sense intronic to TPCN1 |
| MIR6762 | 12 | microRNA 6762 [Source:HGNC Symbol;Acc:HG |
| SLC8B1 | 12 | solute carrier family 8 member B1 [Source:HG |
| PLBD2 | 12 | phospholipase B domain containing 2 [Source |
| SDS | 12 | serine dehydratase [Source:HGNC Symbol;Acc |
| SDSL | 12 | serine dehydratase like [Source:HGNC Symbol |
| LHX5 | 12 | LIM homeobox 5 [Source:HGNC Symbol;Acc:H |
| LHX5-AS1 | 12 | LHX5 antisense RNA 1 [Source:HGNC Symbol;/ |
| LINC01234 | 12 | long intergenic non-protein coding RNA 1234 |
| DYNLL1P4 | 12 | dynein light chain LC8-type 1 pseudogene 4 [S |
| RBM19 | 12 | RNA binding motif protein 19 [Source:HGNC S |
| AC009731.1 | 12 | novel transcript, antisense to RBM19 |
| AC073863.1 | 12 | novel transcript |
| AC010183.1 | 12 | novel transcript |
| AC010183.2 | 12 | novel transcript |
| HAUS8P1 | 12 | HAUS augmin like complex subunit 8 pseudog |
| GLULP5 | 12 | glutamate-ammonia ligase pseudogene 5 [Sou |
| LINC02459 | 12 | long intergenic non-protein coding RNA 2459 |
| TBX5 | 12 | T-box transcription factor 5 [Source:HGNC Syr |
| TBX5-AS1 | 12 | TBX5 antisense RNA 1 [Source:HGNC Symbol;/ |
| RN7SKP216 | 12 | RN7SK pseudogene 216 [Source:HGNC Symbo |
| AC069240.1 | 12 | microtubule-associated protein 1 light chain 3 |
| OSTF1P1 | 12 | osteoclast stimulating factor 1 pseudogene 1 |
| AC010177.1 | 12 | novel transcript |
| Y_RNA | 12 | Y RNA [Source:RFAM;Acc:RF00019] |
| TBX3 | 12 | T-box transcription factor 3 [Source:HGNC Syr |
| AC026765.3 | 12 | novel transcript |
| AC026765.4 | 12 | novel transcript |

| | | |
|---|---|---|
| AC026765.1 | 12 | NULL |
| AC026765.2 | 12 | novel transcript |
| AC009804.2 | 12 | novel transcript |
| AC009804.1 | 12 | zinc finger protein 717 (ZNF717) pseudogene |
| AC008125.1 | 12 | novel transcript |
| AC078880.1 | 12 | novel transcript |
| AC078880.3 | 12 | novel transcript |
| AC078880.4 | 12 | novel transcript |
| AC078880.5 | 12 | novel transcript |
| AC078880.2 | 12 | novel transcript |
| AC009803.2 | 12 | novel transcript |
| AC009803.1 | 12 | novel transcript |
| UBA52P7 | 12 | ubiquitin A-52 residue ribosomal protein fusic |
| RN7SL865P | 12 | RNA, 7SL, cytoplasmic 865, pseudogene [Sour |
| AC009387.1 | 12 | novel transcript |
| LINC02463 | 12 | long intergenic non-protein coding RNA 2463 |
| AC012157.1 | 12 | NULL |
| AC012157.3 | 12 | novel transcript |
| MED13L | 12 | mediator complex subunit 13L [Source:HGNC |
| AC012157.2 | 12 | novel transcript, antisense to MED13L |
| RNU6-1188P | 12 | RNA, U6 small nuclear 1188, pseudogene [Sou |
| MIR620 | 12 | microRNA 620 [Source:HGNC Symbol;Acc:HGI |
| SNRPGP18 | 12 | small nuclear ribonucleoprotein polypeptide ( |
| AC130895.1 | 12 | novel transcript, sense intronic to MED13L |
| AC079384.2 | 12 | novel transcript |
| AC079384.1 | 12 | novel transcript |
| MIR4472-2 | 12 | microRNA 4472-2 [Source:HGNC Symbol;Acc:I |
| LINC02457 | 12 | long intergenic non-protein coding RNA 2457 |
| LINC00173 | 12 | long intergenic non-protein coding RNA 173 [S |
| MAP1LC3B2 | 12 | microtubule associated protein 1 light chain 3 |
| AC125603.4 | 12 | novel transcript |
| AC125603.3 | 12 | novel transcript |
| AC125603.2 | 12 | novel transcript |
| AC125603.1 | 12 | novel transcript |
| C12orf49 | 12 | chromosome 12 open reading frame 49 [Sour |
| RNFT2 | 12 | ring finger protein, transmembrane 2 [Source: |
| AC090013.1 | 12 | ribosomal protein L21 (RPL21) pseudogene |
| RNU6-558P | 12 | RNA, U6 small nuclear 558, pseudogene [Sour |
| AC083806.3 | 12 | novel transcript |
| HRK | 12 | harakiri, BCL2 interacting protein [Source:HGI |
| AC083806.1 | 12 | ribosomal protein L36 (RPL36) pseudogene |
| FBXW8 | 12 | F-box and WD repeat domain containing 8 [Sc |
| AC083806.2 | 12 | novel transcript, antisense to FBXW8 |
| AC127164.1 | 12 | novel transcript |
| AC026368.1 | 12 | novel transcript, sense intronic to FBXW8 |
| TESC | 12 | tescalcin [Source:HGNC Symbol;Acc:HGNC:26( |
| TESC-AS1 | 12 | TESC antisense RNA 1 (head to head) [Source: |
| FBXO21 | 12 | F-box protein 21 [Source:HGNC Symbol;Acc:H |
| NOS1 | 12 | nitric oxide synthase 1 [Source:HGNC Symbol; |
| ELOCP32 | 12 | elongin C pseudogene 32 [Source:HGNC Symb |

| | | |
|---|---|---|
| AC073864.1 | 12 | novel transcript |
| KSR2 | 12 | kinase suppressor of ras 2 [Source:HGNC Syml |
| AC084291.1 | 12 | novel transcript |
| RFC5 | 12 | replication factor C subunit 5 [Source:HGNC S |
| AC131159.2 | 12 | novel transcript, antisense to RFC5 |
| WSB2 | 12 | WD repeat and SOCS box containing 2 [Source |
| AC131159.1 | 12 | novel transcript, antisense to WSB2 |
| VSIG10 | 12 | V-set and immunoglobulin domain containing |
| AC131238.1 | 12 | novel transcript, antisense to VSIG10 |
| PEBP1 | 12 | phosphatidylethanolamine binding protein 1 [ |
| TAOK3 | 12 | TAO kinase 3 [Source:HGNC Symbol;Acc:HGN |
| RPS2P5 | 12 | ribosomal protein S2 pseudogene 5 [Source:H |
| AC026367.1 | 12 | novel transcript |
| SUDS3 | 12 | SDS3 homolog, SIN3A corepressor complex cc |
| AC026367.3 | 12 | novel transcript |
| AC026367.2 | 12 | novel transcript |
| LINC02460 | 12 | long intergenic non-protein coding RNA 2460 |
| RPL17P37 | 12 | ribosomal protein L17 pseudogene 37 [Source |
| LINC02423 | 12 | long intergenic non-protein coding RNA 2423 |
| LINC02440 | 12 | long intergenic non-protein coding RNA 2440 |
| LINC02439 | 12 | long intergenic non-protein coding RNA 2439 |
| RNA5SP374 | 12 | RNA, 5S ribosomal pseudogene 374 [Source:H |
| AC087863.2 | 12 | novel transcript |
| AC087863.1 | 12 | NULL |
| SRRM4 | 12 | serine/arginine repetitive matrix 4 [Source:HG |
| AC087885.1 | 12 | novel transcript, antisense to SRRM4 |
| RN7SL508P | 12 | RNA, 7SL, cytoplasmic 508, pseudogene [Sour |
| AC084361.1 | 12 | novel transcript, antisense to SRRM4 |
| AC084880.4 | 12 | novel transcript |
| HSPB8 | 12 | heat shock protein family B (small) member 8 |
| AC084880.3 | 12 | novel transcript |
| AC084880.1 | 12 | PDGFA associated protein 1 (PDAP1) pseudog |
| AC084880.2 | 12 | novel transcript |
| LINC00934 | 12 | long intergenic non-protein coding RNA 934 [S |
| CCDC60 | 12 | coiled-coil domain containing 60 [Source:HGN |
| AC002070.1 | 12 | novel transcript |
| TMEM233 | 12 | transmembrane protein 233 [Source:HGNC Sy |
| RN7SKP197 | 12 | RN7SK pseudogene 197 [Source:HGNC Symbo |
| PRKAB1 | 12 | protein kinase AMP-activated non-catalytic su |
| CIT | 12 | citron rho-interacting serine/threonine kinase |
| AC002563.1 | 12 | novel transcript, antisense to CIT |
| MIR1178 | 12 | microRNA 1178 [Source:HGNC Symbol;Acc:HG |
| RPL35AP30 | 12 | ribosomal protein L35a pseudogene 30 [Sourc |
| BICDL1 | 12 | BICD family like cargo adaptor 1 [Source:HGN |
| Metazoa_SRP | 12 | Metazoan signal recognition particle RNA [Sou |
| AC004812.1 | 12 | ADP-ribosylation factor-like 2 binding protein |
| RAB35 | 12 | RAB35, member RAS oncogene family [Source |
| AC004812.2 | 12 | novel transcript, antisense to RAB35 |
| GCN1 | 12 | GCN1 activator of EIF2AK4 [Source:HGNC Sym |
| MIR4498 | 12 | microRNA 4498 [Source:HGNC Symbol;Acc:HG |

| | | |
|---|---|---|
| RPLP0 | 12 | ribosomal protein lateral stalk subunit P0 [Sou |
| PXN-AS1 | 12 | PXN antisense RNA 1 [Source:HGNC Symbol;A |
| PXN | 12 | paxillin [Source:HGNC Symbol;Acc:HGNC:9718 |
| AC004263.2 | 12 | novel transcript, antisense to PXN |
| AC004263.1 | 12 | novel transcript, antisense to PXN |
| RPS20P31 | 12 | ribosomal protein S20 pseudogene 31 [Source |
| NME2P1 | 12 | NME/NM23 nucleoside diphosphate kinase 2 |
| RNU4-2 | 12 | RNA, U4 small nuclear 2 [Source:HGNC Symbo |
| RNU4-1 | 12 | RNA, U4 small nuclear 1 [Source:HGNC Symbo |
| SIRT4 | 12 | sirtuin 4 [Source:HGNC Symbol;Acc:HGNC:149 |
| RNU6-1088P | 12 | RNA, U6 small nuclear 1088, pseudogene [Sou |
| PLA2G1B | 12 | phospholipase A2 group IB [Source:HGNC Sym |
| MSI1 | 12 | musashi RNA binding protein 1 [Source:HGNC |
| RPS27P25 | 12 | ribosomal protein S27 pseudogene 25 [Source |
| AC003982.1 | 12 | novel transcript |
| COX6A1 | 12 | cytochrome c oxidase subunit 6A1 [Source:HG |
| AL021546.1 | 12 | novel protein |
| TRIAP1 | 12 | TP53 regulated inhibitor of apoptosis 1 [Sourc |
| GATC | 12 | glutamyl-tRNA amidotransferase subunit C [So |
| RPL31P52 | 12 | ribosomal protein L31 pseudogene 52 [Source |
| SRSF9 | 12 | serine and arginine rich splicing factor 9 [Sour |
| DYNLL1 | 12 | dynein light chain LC8-type 1 [Source:HGNC Sy |
| NRAV | 12 | negative regulator of antiviral response [Sourc |
| AC063943.1 | 12 | novel transcript |
| COQ5 | 12 | coenzyme Q5, methyltransferase [Source:HGN |
| Y_RNA | 12 | Y RNA [Source:RFAM;Acc:RF00019] |
| RPL29P24 | 12 | ribosomal protein L29 pseudogene 24 [Source |
| RNF10 | 12 | ring finger protein 10 [Source:HGNC Symbol;A |
| POP5 | 12 | POP5 homolog, ribonuclease P/MRP subunit [ |
| RPL11P5 | 12 | ribosomal protein L11 pseudogene 5 [Source: |
| AC125616.1 | 12 | novel transcript, antisense to CABP1 |
| CABP1 | 12 | calcium binding protein 1 [Source:HGNC Symb |
| MLEC | 12 | malectin [Source:HGNC Symbol;Acc:HGNC:289 |
| AC069234.2 | 12 | novel transcript, antisense to MLEC |
| AC069234.5 | 12 | novel transcript |
| AC069234.4 | 12 | novel transcript |
| UNC119B | 12 | unc-119 lipid binding chaperone B [Source:HG |
| AC069234.3 | 12 | novel transcript, antisense to UNC119B |
| MIR4700 | 12 | microRNA 4700 [Source:HGNC Symbol;Acc:HG |
| ACADS | 12 | acyl-CoA dehydrogenase short chain [Source:H |
| AC069234.1 | 12 | novel transcript |
| SPPL3 | 12 | signal peptide peptidase like 3 [Source:HGNC |
| ARF1P2 | 12 | ADP ribosylation factor 1 pseudogene 2 [Sour |
| AC069214.1 | 12 | novel transcript |
| CLIC1P1 | 12 | chloride intracellular channel 1 pseudogene 1 |
| RPL12P33 | 12 | ribosomal protein L12 pseudogene 33 [Source |
| HNF1A-AS1 | 12 | HNF1A antisense RNA 1 [Source:HGNC Symbo |
| HNF1A | 12 | HNF1 homeobox A [Source:HGNC Symbol;Acc |
| C12orf43 | 12 | chromosome 12 open reading frame 43 [Sour |
| OASL | 12 | 2'-5'-oligoadenylate synthetase like [Source:H |

| | | |
|---|---|---|
| OASL2P | 12 | 2'-5' oligoadenylate synthetase like 2, pseudog |
| AC079602.2 | 12 | pancreatic progenitor cell differentiation and |
| AC079602.3 | 12 | novel transcript |
| SNORA70 | 12 | Small nucleolar RNA SNORA70 [Source:RFAM; |
| AC079602.1 | 12 | ribosomal protein L10 (RPL10) |
| P2RX7 | 12 | purinergic receptor P2X 7 [Source:HGNC Symb |
| AC069209.2 | 12 | novel transcript, antisense to P2RX7 |
| AC069209.1 | 12 | novel transcript |
| P2RX4 | 12 | purinergic receptor P2X 4 [Source:HGNC Symb |
| CAMKK2 | 12 | calcium/calmodulin dependent protein kinase |
| ANAPC5 | 12 | anaphase promoting complex subunit 5 [Sour |
| AC048337.1 | 12 | novel transcript |
| RNF34 | 12 | ring finger protein 34 [Source:HGNC Symbol;A |
| KDM2B | 12 | lysine demethylase 2B [Source:HGNC Symbol; |
| MIR7107 | 12 | microRNA 7107 [Source:HGNC Symbol;Acc:HG |
| KDM2B-DT | 12 | KDM2B divergent transcript [Source:HGNC Sy |
| RNU6-1004P | 12 | RNA, U6 small nuclear 1004, pseudogene [Sou |
| ORAI1 | 12 | ORAI calcium release-activated calcium modu |
| MORN3 | 12 | MORN repeat containing 3 [Source:HGNC Sym |
| AC084018.4 | 12 | novel transcript |
| TMEM120B | 12 | transmembrane protein 120B [Source:HGNC S |
| RHOF | 12 | ras homolog family member F, filopodia assoc |
| LINC01089 | 12 | long intergenic non-protein coding RNA 1089 |
| AC084018.1 | 12 | novel transcript |
| AC084018.2 | 12 | novel transcript |
| SETD1B | 12 | SET domain containing 1B, histone lysine metl |
| HPD | 12 | 4-hydroxyphenylpyruvate dioxygenase [Sourc |
| AC079360.1 | 12 | novel transcript |
| AC069503.3 | 12 | novel transcript |
| AC069503.4 | 12 | novel transcript |
| PSMD9 | 12 | proteasome 26S subunit, non-ATPase 9 [Sourc |
| AC069503.2 | 12 | novel transcript |
| Metazoa_SRP | 12 | Metazoan signal recognition particle RNA [Sou |
| RNU7-170P | 12 | RNA, U7 small nuclear 170 pseudogene [Sourc |
| WDR66 | 12 | WD repeat domain 66 [Source:HGNC Symbol;. |
| AC069503.1 | 12 | novel transcript |
| BCL7A | 12 | BAF chromatin remodeling complex subunit B |
| AC156455.1 | 12 | novel transcript |
| MLXIP | 12 | MLX interacting protein [Source:HGNC Symbo |
| LRRC43 | 12 | leucine rich repeat containing 43 [Source:HGN |
| IL31 | 12 | interleukin 31 [Source:HGNC Symbol;Acc:HGN |
| B3GNT4 | 12 | UDP-GlcNAc:betaGal beta-1,3-N-acetylglucosa |
| AC048338.2 | 12 | Diablo homolog, mitochondrial [Source:UniPr |
| DIABLO | 12 | diablo IAP-binding mitochondrial protein [Sou |
| AC048338.1 | 12 | novel protein |
| VPS33A | 12 | VPS33A core subunit of CORVET and HOPS co |
| CLIP1 | 12 | CAP-Gly domain containing linker protein 1 [S |
| RPL21P1 | 12 | ribosomal protein L21 pseudogene 1 [Source: |
| CLIP1-AS1 | 12 | CLIP1 antisense RNA 1 [Source:HGNC Symbol; |
| ZCCHC8 | 12 | zinc finger CCHC-type containing 8 [Source:HG |

| | | |
|---|---|---|
| AC127002.1 | 12 | NULL |
| AC127002.2 | 12 | novel transcript |
| RSRC2 | 12 | arginine and serine rich coiled-coil 2 [Source:H |
| KNTC1 | 12 | kinetochore associated 1 [Source:HGNC Symb |
| AC026333.1 | 12 | chemokine (C-C motif) receptor-like 2 (CCRL2) |
| AC026333.4 | 12 | novel transcript |
| AC026333.3 | 12 | novel transcript |
| HCAR2 | 12 | hydroxycarboxylic acid receptor 2 [Source:HG |
| HCAR3 | 12 | hydroxycarboxylic acid receptor 3 [Source:HG |
| HCAR1 | 12 | hydroxycarboxylic acid receptor 1 [Source:HG |
| AC026333.2 | 12 | novel transcript |
| DENR | 12 | density regulated re-initiation and release fac |
| Y_RNA | 12 | Y RNA [Source:RFAM;Acc:RF00019] |
| CCDC62 | 12 | coiled-coil domain containing 62 [Source:HGN |
| HIP1R | 12 | huntingtin interacting protein 1 related [Sourc |
| VPS37B | 12 | VPS37B subunit of ESCRT-I [Source:HGNC Sym |
| AC027290.1 | 12 | novel transcript |
| AC027290.2 | 12 | novel transcript |
| ABCB9 | 12 | ATP binding cassette subfamily B member 9 [S |
| OGFOD2 | 12 | 2-oxoglutarate and iron dependent oxygenase |
| AC026362.1 | 12 | novel transcript |
| ARL6IP4 | 12 | ADP ribosylation factor like GTPase 6 interacti |
| PITPNM2 | 12 | phosphatidylinositol transfer protein membra |
| MIR4304 | 12 | microRNA 4304 [Source:HGNC Symbol;Acc:HG |
| PITPNM2-AS1 | 12 | PITPNM2 antisense RNA 1 [Source:HGNC Sym |
| AC026362.2 | 12 | novel transcript |
| MPHOSPH9 | 12 | M-phase phosphoprotein 9 [Source:HGNC Syr |
| AC073857.1 | 12 | novel transcript |
| C12orf65 | 12 | chromosome 12 open reading frame 65 [Sour |
| CDK2AP1 | 12 | cyclin dependent kinase 2 associated protein : |
| AC068768.1 | 12 | novel transcript, antisense to CDK2AP1 |
| RNA5SP375 | 12 | RNA, 5S ribosomal pseudogene 375 [Source:H |
| SBNO1 | 12 | strawberry notch homolog 1 [Source:HGNC Sy |
| Y_RNA | 12 | Y RNA [Source:RFAM;Acc:RF00019] |
| SBNO1-AS1 | 12 | SBNO1 antisense RNA 1 [Source:HGNC Symbo |
| MIR8072 | 12 | microRNA 8072 [Source:HGNC Symbol;Acc:HG |
| KMT5A | 12 | lysine methyltransferase 5A [Source:HGNC Syr |
| RILPL2 | 12 | Rab interacting lysosomal protein like 2 [Sourc |
| AC145423.1 | 12 | COP9 constitutive photomorphogenic homolo |
| SNRNP35 | 12 | small nuclear ribonucleoprotein U11/U12 sub |
| RILPL1 | 12 | Rab interacting lysosomal protein like 1 [Sourc |
| AC145423.2 | 12 | novel transcript, sense intronic to RILPL1 |
| AC145423.3 | 12 | novel transcript, sense intronic to RILPL1 |
| MIR3908 | 12 | microRNA 3908 [Source:HGNC Symbol;Acc:HG |
| TMED2-DT | 12 | TMED2 divergent transcript [Source:HGNC Syr |
| TMED2 | 12 | transmembrane p24 trafficking protein 2 [Sou |
| DDX55 | 12 | DEAD-box helicase 55 [Source:HGNC Symbol;A |
| SNORA9B | 12 | small nucleolar RNA, H/ACA box 9B [Source:H |
| EIF2B1 | 12 | eukaryotic translation initiation factor 2B subu |
| GTF2H3 | 12 | general transcription factor IIH subunit 3 [Sou |

| | | |
|---|---|---|
| AC117503.3 | 12 | novel transcript |
| AC117503.2 | 12 | novel transcript |
| TCTN2 | 12 | tectonic family member 2 [Source:HGNC Syml |
| AC117503.1 | 12 | novel transcript |
| ATP6V0A2 | 12 | ATPase H+ transporting V0 subunit a2 [Source |
| AC117503.5 | 12 | novel transcript, antisense to ATP6V0A2 |
| RPL27P12 | 12 | ribosomal protein L27 pseudogene 12 [Source |
| AC117503.4 | 12 | novel transcript |
| DNAH10 | 12 | dynein axonemal heavy chain 10 [Source:HGN |
| CCDC92 | 12 | coiled-coil domain containing 92 [Source:HGN |
| DNAH10OS | 12 | dynein axonemal heavy chain 10 opposite stra |
| AC068790.7 | 12 | novel transcript, sense intronic to CCDC92 |
| AC068790.4 | 12 | novel transcript, sense intronic to CCDC92 |
| AC068790.3 | 12 | novel transcript, sense intronic to CCDC92 |
| AC068790.2 | 12 | novel transcript, sense intronic to CCDC92 |
| AC068790.5 | 12 | novel transcript, sense intronic to CCDC92 |
| AC068790.6 | 12 | novel transcript, sense intronic to CCDC92 |
| ZNF664 | 12 | zinc finger protein 664 [Source:HGNC Symbol; |
| AC068790.8 | 12 | novel protein |
| RFLNA | 12 | refilin A [Source:HGNC Symbol;Acc:HGNC:270 |
| AC068790.1 | 12 | spindle and kinetochore associated complex s |
| AC068790.9 | 12 | novel transcript |
| AC026358.1 | 12 | novel transcript |
| NCOR2 | 12 | nuclear receptor corepressor 2 [Source:HGNC |
| MIR6880 | 12 | microRNA 6880 [Source:HGNC Symbol;Acc:HG |
| AC073916.1 | 12 | chromosome 12 open reading frame 10 (C12c |
| AC073592.1 | 12 | novel transcript |
| AC073592.7 | 12 | novel transcript |
| AC073592.3 | 12 | novel transcript |
| AC073592.9 | 12 | novel transcript |
| AC073592.5 | 12 | novel transcript |
| AC073592.6 | 12 | novel transcript |
| AC073592.8 | 12 | novel transcript |
| AC073592.2 | 12 | novel transcript |
| AC073592.10 | 12 | novel transcript |
| AC073592.4 | 12 | novel transcript |
| AC073593.1 | 12 | novel transcript |
| SCARB1 | 12 | scavenger receptor class B member 1 [Source |
| AC073593.2 | 12 | novel transcript, antisense to SCARB1 |
| RNU6-927P | 12 | RNA, U6 small nuclear 927, pseudogene [Sour |
| UBC | 12 | ubiquitin C [Source:HGNC Symbol;Acc:HGNC:1 |
| MIR5188 | 12 | microRNA 5188 [Source:HGNC Symbol;Acc:HG |
| RPL22P19 | 12 | ribosomal protein L22 pseudogene 19 [Source |
| DHX37 | 12 | DEAH-box helicase 37 [Source:HGNC Symbol;A |
| BRI3BP | 12 | BRI3 binding protein [Source:HGNC Symbol;Ac |
| THRIL | 12 | TNF and HNRNPL related immunoregulatory lo |
| AC122688.1 | 12 | translocase of outer mitochondrial membrane |
| AACS | 12 | acetoacetyl-CoA synthetase [Source:HGNC Syr |
| AC122688.3 | 12 | novel transcript |
| AC122688.2 | 12 | novel transcript |

| | | |
|---|---|---|
| AC122688.5 | 12 | novel transcript |
| AC122688.4 | 12 | novel transcript |
| TMEM132B | 12 | transmembrane protein 132B [Source:HGNC [illegible] |
| AC093028.1 | 12 | novel transcript |
| AC005252.2 | 12 | novel transcript |
| AC005252.4 | 12 | novel transcript |
| AC005252.3 | 12 | novel transcript |
| AC005252.1 | 12 | CTAGE family, member 9 (CTAGE9) pseudoge[illegible] |
| LINC00939 | 12 | long intergenic non-protein coding RNA 939 [[illegible] |
| LINC02826 | 12 | long intergenic non-protein coding RNA 2826 |
| LINC02359 | 12 | long intergenic non-protein coding RNA 2359 |
| AC005858.2 | 12 | novel transcript |
| AC005858.3 | 12 | novel transcript |
| AC005858.1 | 12 | novel transcript |
| AC005888.1 | 12 | novel transcript |
| AC007368.1 | 12 | uncharacterized LOC101927531 [Source:NCBI |
| LINC02825 | 12 | long intergenic non-protein coding RNA 2825 |
| LINC02347 | 12 | long intergenic non-protein coding RNA 2347 |
| AC005868.1 | 12 | family with sequence similarity 32, member A |
| AC006065.1 | 12 | novel pseudogene |
| NDUFA5P6 | 12 | NADH:ubiquinone oxidoreductase subunit A5 |
| AC006065.4 | 12 | novel transcript |
| AC006065.2 | 12 | riboflavin kinase (RFK) pseudogene |
| AC006065.3 | 12 | novel transcript |
| AC006065.5 | 12 | novel transcript |
| AC006065.6 | 12 | novel transcript |
| LINC02824 | 12 | long intergenic non-protein coding RNA 2824 |
| LINC00943 | 12 | long intergenic non-protein coding RNA 943 [[illegible] |
| LINC00944 | 12 | long intergenic non-protein coding RNA 944 [[illegible] |
| AC078878.1 | 12 | transmembrane protein 132B (TMEM132B) p[illegible] |
| LINC02372 | 12 | long intergenic non-protein coding RNA 2372 |
| AC078878.2 | 12 | novel transcript |
| HSPE1P20 | 12 | heat shock protein family E (Hsp10) member [illegible] |
| LINC02405 | 12 | long intergenic non-protein coding RNA 2405 |
| AC079949.3 | 12 | novel transcript |
| AC079949.1 | 12 | novel transcript |
| AC079949.5 | 12 | novel transcript |
| AC079949.2 | 12 | novel transcript |
| AC079949.6 | 12 | novel transcript |
| LINC02376 | 12 | long intergenic non-protein coding RNA 2376 |
| AC073913.1 | 12 | transmembrane protein 132B (TMEM132B) p[illegible] |
| RNU1-104P | 12 | RNA, U1 small nuclear 104, pseudogene [Sour |
| LINC02375 | 12 | long intergenic non-protein coding RNA 2375 |
| AC073913.2 | 12 | transmembrane protein 132B (TMEM 132B) p |
| AC048347.1 | 12 | novel transcript |
| AC068787.4 | 12 | novel transcript |
| AC068787.1 | 12 | novel transcript |
| AC068787.3 | 12 | novel transcript |
| AC068787.2 | 12 | novel transcript |
| AC068787.5 | 12 | novel transcript |

| | | |
|---|---|---|
| AC025252.1 | 12 | novel transcript |
| Y_RNA | 12 | Y RNA [Source:RFAM;Acc:RF00019] |
| AC025252.2 | 12 | Transmembrane protein 132B (TMEM132B) p |
| LINC02411 | 12 | long intergenic non-protein coding RNA 2411 |
| AC025160.1 | 12 | novel transcript |
| AC087894.2 | 12 | novel transcript |
| AC087894.1 | 12 | fibroblast growth factor receptor 1 (FGFR1) |
| LINC02393 | 12 | long intergenic non-protein coding RNA 2393 |
| LINC00507 | 12 | long intergenic non-protein coding RNA 507 [S |
| LINC00508 | 12 | long intergenic non-protein coding RNA 508 [S |
| LINC02441 | 12 | long intergenic non-protein coding RNA 2441 |
| AC140121.1 | 12 | novel transcript |
| LINC02369 | 12 | long intergenic non-protein coding RNA 2369 |
| LINC02368 | 12 | long intergenic non-protein coding RNA 2368 |
| AC061709.2 | 12 | novel transcript |
| AC061709.1 | 12 | NULL |
| TMEM132C | 12 | transmembrane protein 132C [Source:HGNC S |
| AC061709.3 | 12 | novel transcript |
| MIR3612 | 12 | microRNA 3612 [Source:HGNC Symbol;Acc:HG |
| AC023595.1 | 12 | novel transcript |
| AC090424.1 | 12 | ribosomal protein S6 (RPS6) pseudogene |
| SLC15A4 | 12 | solute carrier family 15 member 4 [Source:HG |
| AC108704.2 | 12 | novel transcript |
| AC108704.1 | 12 | novel transcript |
| GLT1D1 | 12 | glycosyltransferase 1 domain containing 1 [So |
| AC069262.1 | 12 | adenylate kinase 3 (AK3) pseudogene |
| NLRP9P1 | 12 | NLR family pyrin domain containing 9 pseudog |
| TMEM132D | 12 | transmembrane protein 132D [Source:HGNC S |
| TMEM132D-AS1 | 12 | TMEM132D antisense RNA 1 [Source:HGNC Sy |
| AC107033.1 | 12 | NULL |
| TMEM132D-AS2 | 12 | TMEM132D antisense RNA 2 [Source:HGNC Sy |
| AC079456.1 | 12 | novel transcript |
| AC130404.1 | 12 | novel transcript |
| AC130404.2 | 12 | novel transcript, antisense to TMEM132D |
| AC117373.1 | 12 | novel transcript |
| AC117373.2 | 12 | novel transcript, antisense to TMEM132D |
| AC055717.3 | 12 | novel transcript |
| AC055717.1 | 12 | novel transcript |
| AC055717.2 | 12 | novel transcript |
| LINC02418 | 12 | long intergenic non-protein coding RNA 2418 |
| AC135388.1 | 12 | novel transcript |
| LINC02419 | 12 | long intergenic non-protein coding RNA 2419 |
| AC026336.4 | 12 | novel transcript |
| AC026336.3 | 12 | novel transcript |
| FZD10-AS1 | 12 | FZD10 antisense divergent transcript [Source: |
| AC026336.5 | 12 | novel transcript |
| FZD10 | 12 | frizzled class receptor 10 [Source:HGNC Symb |
| AC026336.1 | 12 | ribosomal protein S20 (RPS20) pseudogene |
| AC026336.2 | 12 | novel transcript |
| PIWIL1 | 12 | piwi like RNA-mediated gene silencing 1 [Sour |

| | | |
|---|---|---|
| AC127071.1 | 12 | novel transcript |
| AC127071.2 | 12 | novel transcript |
| RIMBP2 | 12 | RIMS binding protein 2 [Source:HGNC Symbol |
| AC063926.1 | 12 | novel transcript |
| AC063926.3 | 12 | novel transcript |
| AC063926.2 | 12 | novel transcript |
| AC095350.1 | 12 | novel transcript |
| AC073912.3 | 12 | novel transcript |
| STX2 | 12 | syntaxin 2 [Source:HGNC Symbol;Acc:HGNC:3 |
| AC073912.1 | 12 | novel transcript |
| AC073912.2 | 12 | novel transcript |
| RNU6-1077P | 12 | RNA, U6 small nuclear 1077, pseudogene [Sou |
| RAN | 12 | RAN, member RAS oncogene family [Source:H |
| Y_RNA | 12 | Y RNA [Source:RFAM;Acc:RF00019] |
| ADGRD1 | 12 | adhesion G protein-coupled receptor D1 [Sou |
| AC073862.3 | 12 | novel transcript |
| AC073862.4 | 12 | novel transcript |
| AC073862.5 | 12 | novel transcript |
| AC073862.1 | 12 | novel transcript |
| AC073862.2 | 12 | novel transcript |
| AC078925.4 | 12 | novel transcript |
| ADGRD1-AS1 | 12 | ADGRD1 antisense RNA 1 [Source:HGNC Symb |
| AC078925.2 | 12 | novel transcript |
| AC078925.3 | 12 | novel transcript |
| AC078925.1 | 12 | novel transcript |
| LINC01257 | 12 | long intergenic non-protein coding RNA 1257 |
| AC126564.1 | 12 | novel transcript |
| RPS6P20 | 12 | ribosomal protein S6 pseudogene 20 [Source: |
| AC092850.1 | 12 | novel transcript |
| LINC02415 | 12 | long intergenic non-protein coding RNA 2415 |
| RNA5SP376 | 12 | RNA, 5S ribosomal pseudogene 376 [Source:H |
| AC092850.2 | 12 | novel transcript |
| LINC02370 | 12 | long intergenic non-protein coding RNA 2370 |
| AC140118.2 | 12 | novel transcript |
| AC140118.1 | 12 | novel transcript |
| AC073578.4 | 12 | novel transcript |
| AC073578.2 | 12 | novel transcript |
| AC073578.5 | 12 | novel transcript |
| AC073578.1 | 12 | novel transcript |
| AC073578.3 | 12 | novel transcript |
| AC140063.1 | 12 | G protein-coupled receptor 133 (GPR133) pse |
| AC117500.5 | 12 | novel transcript |
| AC117500.3 | 12 | novel transcript |
| LINC02414 | 12 | long intergenic non-protein coding RNA 2414 |
| AC117500.6 | 12 | novel transcript |
| AC117500.4 | 12 | novel transcript |
| RNA5SP377 | 12 | RNA, 5S ribosomal pseudogene 377 [Source:H |
| AC117500.1 | 12 | novel transcript |
| RPS6P21 | 12 | ribosomal protein S6 pseudogene 21 [Source: |
| SFSWAP | 12 | splicing factor SWAP [Source:HGNC Symbol;A |

| | | |
|---|---|---|
| AC117500.2 | 12 | novel transcript |
| RNA5SP378 | 12 | RNA, 5S ribosomal pseudogene 378 [Source:H |
| RNU6-1017P | 12 | RNA, U6 small nuclear 1017, pseudogene [Sou |
| MMP17 | 12 | matrix metallopeptidase 17 [Source:HGNC Syr |
| AC131009.2 | 12 | novel transcript |
| ULK1 | 12 | unc-51 like autophagy activating kinase 1 [Sou |
| AC131009.4 | 12 | novel transcript |
| AC131009.1 | 12 | novel transcript |
| PUS1 | 12 | pseudouridine synthase 1 [Source:HGNC Symb |
| AC131009.3 | 12 | novel transcript, sense intronic to PUS1 |
| EP400 | 12 | E1A binding protein p400 [Source:HGNC Symb |
| SNORA49 | 12 | small nucleolar RNA, H/ACA box 49 [Source:H |
| AC137590.1 | 12 | novel transcript |
| AC137590.2 | 12 | novel transcript |
| EP400P1 | 12 | EP400 pseudogene 1 [Source:HGNC Symbol;A |
| AC138466.5 | 12 | novel transcript |
| AC138466.2 | 12 | placenta-specific 8 (PLAC8) pseudogene |
| DDX51 | 12 | DEAD-box helicase 51 [Source:HGNC Symbol;A |
| NOC4L | 12 | nucleolar complex associated 4 homolog [Sou |
| AC138466.3 | 12 | novel transcript |
| LINC02361 | 12 | long intergenic non-protein coding RNA 2361 |
| AC138466.1 | 12 | novel transcript |
| GALNT9 | 12 | polypeptide N-acetylgalactosaminyltransferas |
| AC138466.4 | 12 | novel transcript |
| AC148477.3 | 12 | novel transcript |
| AC148477.2 | 12 | novel transcript |
| AC148477.5 | 12 | novel transcript |
| AC148477.4 | 12 | novel transcript |
| AC148477.1 | 12 | novel transcript |
| AC148477.7 | 12 | novel transcript |
| AC148477.10 | 12 | novel transcript |
| AC148477.8 | 12 | novel transcript |
| AC148477.6 | 12 | novel transcript |
| AC148477.9 | 12 | novel transcript |
| AC148476.1 | 12 | novel transcript |
| AC079031.1 | 12 | novel transcript |
| AC079031.2 | 12 | novel transcript |
| FBRSL1 | 12 | fibrosin like 1 [Source:HGNC Symbol;Acc:HGN |
| AC131212.3 | 12 | novel transcript |
| AC131212.2 | 12 | novel transcript |
| MIR6763 | 12 | microRNA 6763 [Source:HGNC Symbol;Acc:HG |
| AC131212.1 | 12 | novel transcript |
| LRCOL1 | 12 | leucine rich colipase like 1 [Source:HGNC Sym |
| P2RX2 | 12 | purinergic receptor P2X 2 [Source:HGNC Symb |
| AC131212.4 | 12 | novel transcript |
| POLE | 12 | DNA polymerase epsilon, catalytic subunit [So |
| PXMP2 | 12 | peroxisomal membrane protein 2 [Source:HG |
| AC135586.2 | 12 | novel transcript |
| PGAM5 | 12 | PGAM family member 5, mitochondrial serine |
| RNA5SP379 | 12 | RNA, 5S ribosomal pseudogene 379 [Source:H |

| | | |
|---|---|---|
| ANKLE2 | 12 | ankyrin repeat and LEM domain containing 2 [ |
| AC135586.1 | 12 | tumor protein D52-like 2 (TPD52L2) pseudoge |
| GOLGA3 | 12 | golgin A3 [Source:HGNC Symbol;Acc:HGNC:44 |
| CHFR | 12 | checkpoint with forkhead and ring finger dom |
| RPS11P5 | 12 | ribosomal protein S11 pseudogene 5 [Source: |
| RNU6-327P | 12 | RNA, U6 small nuclear 327, pseudogene [Sour |
| AC127070.4 | 12 | novel transcript |
| AC127070.1 | 12 | novel transcript |
| AC127070.3 | 12 | NHP2 non-histone chromosome protein 2-like |
| AC127070.2 | 12 | novel transcript |
| ZNF605 | 12 | zinc finger protein 605 [Source:HGNC Symbol; |
| NANOGNBP2 | 12 | NANOGNB pseudogene 2 [Source:HGNC Symb |
| ZNF26 | 12 | zinc finger protein 26 [Source:HGNC Symbol;A |
| RNU4ATAC12P | 12 | RNA, U4atac small nuclear 12, pseudogene [So |
| ZNF84-DT | 12 | ZNF84 divergent transcript [Source:HGNC Sym |
| PTP4A1P2 | 12 | PTP4A1 pseudogene 2 [Source:HGNC Symbol; |
| ZNF84 | 12 | zinc finger protein 84 [Source:HGNC Symbol;A |
| AC073911.1 | 12 | novel transcript |
| AC073911.2 | 12 | novel transcript |
| ZNF140 | 12 | zinc finger protein 140 [Source:HGNC Symbol; |
| ZNF891 | 12 | zinc finger protein 891 [Source:HGNC Symbol; |
| AC026786.2 | 12 | novel transcript, antisense to ZNF891 |
| RPL23AP67 | 12 | ribosomal protein L23a pseudogene 67 [Sourc |
| ZNF10 | 12 | zinc finger protein 10 [Source:HGNC Symbol;A |
| AC026786.1 | 12 | novel protein |
| ZNF268 | 12 | zinc finger protein 268 [Source:HGNC Symbol; |
| AC226150.1 | 12 | interleukin 9 receptor (IL9R) pseudogene |
| ANHX | 12 | anomalous homeobox [Source:HGNC Symbol; |
| RNU6-717P | 12 | RNA, U6 small nuclear 717, pseudogene [Sour |
| Y_RNA | 12 | Y RNA [Source:RFAM;Acc:RF00019] |
| AL356585.4 | 13 | novel ankyrin repeat domain-containing prote |
| FAM230C | 13 | family with sequence similarity 230 member C |
| AL356585.1 | 13 | eukaryotic translation initiation factor 3 subur |
| AL356585.10 | 13 | gamma-glutamyltransferase 5 (GGT5) pseudo |
| GGT4P | 13 | gamma-glutamyltransferase 4 pseudogene [So |
| AL356585.2 | 13 | immunoglobulin superfamily, member 3 (IGSF |
| KMT5AP1 | 13 | KMT5A pseudogene 1 [Source:HGNC Symbol;A |
| AL356585.3 | 13 | lysine (K)-specific methyltransferase 2C (KMT2 |
| FO681491.1 | 13 | ankyrin repeat domain 30B (ANKRD30B) pseu |
| U6 | 13 | U6 spliceosomal RNA [Source:RFAM;Acc:RF00 |
| ZNF962P | 13 | zinc finger protein 962, pseudogene [Source:H |
| BNIP3P7 | 13 | BCL2 interacting protein 3 pseudogene 7 [Sou |
| LINC00349 | 13 | long intergenic non-protein coding RNA 349 [S |
| LONRF2P2 | 13 | LON peptidase N-terminal domain and ring fir |
| LINC00388 | 13 | long intergenic non-protein coding RNA 388 [S |
| FEM1AP4 | 13 | fem-1 homolog A pseudogene 4 [Source:HGN |
| LINC00387 | 13 | long intergenic non-protein coding RNA 387 [S |
| TERF1P5 | 13 | telomeric repeat binding factor 1 pseudogene |
| FAM207BP | 13 | family with sequence similarity 207 member B |
| GXYLT1P1 | 13 | glucoside xylosyltransferase 1 pseudogene 1 [ |

| | | |
|---|---|---|
| CNN2P12 | 13 | calponin 2 pseudogene 12 [Source:HGNC Sym |
| ZNF965P | 13 | zinc finger protein 965, pseudogene [Source:H |
| CYP4F34P | 13 | cytochrome P450 family 4 subfamily F membe |
| SNX18P26 | 13 | sorting nexin 18 pseudogene 26 [Source:HGN |
| LINC00328-2P | 13 | long intergenic non-protein coding RNA 328-2 |
| ANKRD20A9P | 13 | ankyrin repeat domain 20 family member A9, |
| RHOT1P3 | 13 | ras homolog family member T1 pseudogene 3 |
| RNU6-55P | 13 | RNA, U6 small nuclear 55, pseudogene [Sourc |
| RNU6-76P | 13 | RNA, U6 small nuclear 76, pseudogene [Sourc |
| SNX19P2 | 13 | sorting nexin 19 pseudogene 2 [Source:HGNC |
| LINC00408 | 13 | long intergenic non-protein coding RNA 408 [S |
| AL355516.1 | 13 | novel transcript |
| PHF2P2 | 13 | PHD finger protein 2 pseudogene 2 [Source:H |
| LINC00442 | 13 | long intergenic non-protein coding RNA 442 [S |
| USP24P1 | 13 | ubiquitin specific peptidase 24 pseudogene 1 |
| GTF2IP3 | 13 | general transcription factor IIi pseudogene 3 [ |
| AL137001.1 | 13 | novel transcript |
| AL137001.2 | 13 | novel transcript |
| RNA5SP24 | 13 | RNA, 5S ribosomal pseudogene 24 [Source:HG |
| CENPIP1 | 13 | centromere protein I pseudogene 1 [Source:H |
| AL139327.3 | 13 | novel pseudogene |
| SMPD4P2 | 13 | sphingomyelin phosphodiesterase 4 pseudoge |
| AL139327.4 | 13 | mitotic spindle organizing protein 2B (MZT2B) |
| TUBA3C | 13 | tubulin alpha 3c [Source:HGNC Symbol;Acc:H |
| AL139327.1 | 13 | CDC42 binding protein kinase beta (DMPK-like |
| PSPC1P1 | 13 | paraspeckle component 1 pseudogene 1 [Sou |
| AL139327.2 | 13 | novel transcript |
| ANKRD26P3 | 13 | ankyrin repeat domain 26 pseudogene 3 [Sou |
| MRPL3P1 | 13 | mitochondrial ribosomal protein L3 pseudoge |
| LINC00421 | 13 | long intergenic non-protein coding RNA 421 [S |
| PARP4P2 | 13 | poly(ADP-ribose) polymerase family member |
| CCNQP3 | 13 | CCNQ pseudogene 3 [Source:HGNC Symbol;Ac |
| AL356259.1 | 13 | novel transcript |
| SLC25A15P2 | 13 | solute carrier family 25 member 15 pseudoge |
| TPTE2 | 13 | transmembrane phosphoinositide 3-phosphat |
| CYCSP32 | 13 | CYCS pseudogene 32 [Source:HGNC Symbol;A |
| MRPS31P2 | 13 | mitochondrial ribosomal protein S31 pseudog |
| ESRRAP1 | 13 | estrogen-related receptor alpha pseudogene |
| LINC00350 | 13 | long intergenic non-protein coding RNA 350 [S |
| CASC4P1 | 13 | cancer susceptibility 4 pseudogene 1 [Source: |
| MPHOSPH8 | 13 | M-phase phosphoprotein 8 [Source:HGNC Syr |
| PSPC1-AS2 | 13 | PSPC1 antisense RNA 2 [Source:HGNC Symbol |
| PSPC1 | 13 | paraspeckle component 1 [Source:HGNC Syml |
| RN7SL166P | 13 | RNA, 7SL, cytoplasmic 166, pseudogene [Sour |
| ST6GALNAC4P1 | 13 | ST6 N-acetylgalactosaminide alpha-2,6-sialyltr |
| ZMYM5 | 13 | zinc finger MYM-type containing 5 [Source:HG |
| AL355001.1 | 13 | novel transcript |
| AL355001.2 | 13 | novel transcript |
| ZMYM2 | 13 | zinc finger MYM-type containing 2 [Source:HG |
| KRR1P1 | 13 | KRR1 pseudogene 1 [Source:HGNC Symbol;Ac |

| | | |
|---|---|---|
| AL138688.2 | 13 | TEC |
| LINC01072 | 13 | long intergenic non-protein coding RNA 1072 |
| GJA3 | 13 | gap junction protein alpha 3 [Source:HGNC Sy |
| PPIAP28 | 13 | peptidylprolyl isomerase A pseudogene 28 [Sc |
| LINC00556 | 13 | long intergenic non-protein coding RNA 556 [S |
| GJB2 | 13 | gap junction protein beta 2 [Source:HGNC Syn |
| AL138688.1 | 13 | TEC |
| GJB6 | 13 | gap junction protein beta 6 [Source:HGNC Syn |
| CRYL1 | 13 | crystallin lambda 1 [Source:HGNC Symbol;Acc |
| MIR4499 | 13 | microRNA 4499 [Source:HGNC Symbol;Acc:HG |
| AL590096.1 | 13 | novel transcript, antisense to IFT88 |
| IFT88 | 13 | intraflagellar transport 88 [Source:HGNC Syml |
| SLC35E1P1 | 13 | solute carrier family 35 member E1 pseudoge |
| RNU2-7P | 13 | RNA, U2 small nuclear 7, pseudogene [Source |
| AL161772.1 | 13 | novel transcript, antisense to IL17D |
| IL17D | 13 | interleukin 17D [Source:HGNC Symbol;Acc:HG |
| AL512652.2 | 13 | TEC |
| EEF1AKMT1 | 13 | EEF1A lysine methyltransferase 1 [Source:HGI |
| RANP8 | 13 | RAN pseudogene 8 [Source:HGNC Symbol;Acc |
| AL512652.1 | 13 | novel transcript, sense intronic to N6AMT2 |
| XPO4 | 13 | exportin 4 [Source:HGNC Symbol;Acc:HGNC:1 |
| CNOT4P1 | 13 | CCR4-NOT transcription complex subunit 4 ps |
| HNRNPA1P30 | 13 | heterogeneous nuclear ribonucleoprotein A1 |
| PPIAP27 | 13 | peptidylprolyl isomerase A pseudogene 27 [Sc |
| RPSAP54 | 13 | ribosomal protein SA pseudogene 54 [Source: |
| LATS2 | 13 | large tumor suppressor kinase 2 [Source:HGN |
| LATS2-AS1 | 13 | LATS2 antisense RNA 1 [Source:HGNC Symbol |
| RNU4-9P | 13 | RNA, U4 small nuclear 9, pseudogene [Source |
| RPS12P23 | 13 | ribosomal protein S12 pseudogene 23 [Source |
| IPPKP1 | 13 | inositol 1,3,4,5,6-pentakisphosphate 2-kinase |
| AL161613.1 | 13 | NULL |
| SNORD27 | 13 | Small nucleolar RNA SNORD27 [Source:RFAM; |
| SAP18 | 13 | Sin3A associated protein 18 [Source:HGNC Syı |
| SKA3 | 13 | spindle and kinetochore associated complex s |
| MRPL57 | 13 | mitochondrial ribosomal protein L57 [Source:I |
| AL158032.1 | 13 | ribosomal protein L34 (RPL34) pseudogene |
| LINC01046 | 13 | long intergenic non-protein coding RNA 1046 |
| ESRRAP2 | 13 | estrogen-related receptor alpha pseudogene : |
| AL158032.2 | 13 | POTE ankyrin domain family, member C (POTI |
| MIPEPP3 | 13 | mitochondrial intermediate peptidase pseudo |
| LINC00539 | 13 | long intergenic non-protein coding RNA 539 [S |
| GRK6P1 | 13 | G protein-coupled receptor kinase 6 pseudoge |
| GAPDHP52 | 13 | glyceraldehyde 3 phosphate dehydrogenase p |
| RNA5SP25 | 13 | RNA, 5S ribosomal pseudogene 25 [Source:HG |
| ZDHHC20 | 13 | zinc finger DHHC-type containing 20 [Source:H |
| ZDHHC20-IT1 | 13 | ZDHHC20 intronic transcript 1 [Source:HGNC S |
| HIST1H2BPS3 | 13 | HIST1H2B pseudogene 3 [Source:HGNC Symbo |
| MICU2 | 13 | mitochondrial calcium uptake 2 [Source:HGN( |
| FNTAP2 | 13 | farnesyltransferase, CAAX box, alpha pseudog |
| RNU6-59P | 13 | RNA, U6 small nuclear 59, pseudogene [Sourc |

| | | |
|---|---|---|
| RPS7P10 | 13 | ribosomal protein S7 pseudogene 10 [Source: |
| FGF9 | 13 | fibroblast growth factor 9 [Source:HGNC Symb |
| RN7SL766P | 13 | RNA, 7SL, cytoplasmic 766, pseudogene [Sour |
| LINC00424 | 13 | long intergenic non-protein coding RNA 424 [S |
| AL136962.1 | 13 | long intergenic non-protein coding RNA 540 |
| NME1P1 | 13 | NME/NM23 nucleoside diphosphate kinase 1 |
| MTND3P1 | 13 | MT-ND3 pseudogene 1 [Source:HGNC Symbol |
| AL512484.1 | 13 | novel transcript |
| FTH1P7 | 13 | ferritin heavy chain 1 pseudogene 7 [Source:H |
| DDX39AP1 | 13 | DEAD-box helicase 39A pseudogene 1 [Source |
| SNORD36 | 13 | Small nucleolar RNA SNORD36 [Source:RFAM; |
| RPL7AP73 | 13 | ribosomal protein L7a pseudogene 73 [Source |
| IPMKP1 | 13 | inositol polyphosphate multikinase pseudoger |
| RFESDP1 | 13 | Rieske (Fe-S) domain containing pseudogene 1 |
| LINC00621 | 13 | long intergenic non-protein coding RNA 621 [S |
| BASP1P1 | 13 | BASP1 pseudogene 1 [Source:HGNC Symbol;A |
| NUS1P2 | 13 | NUS1 pseudogene 2 [Source:HGNC Symbol;Ac |
| AL157931.1 | 13 | novel transcript |
| AL157931.2 | 13 | novel transcript |
| HMGA1P6 | 13 | high mobility group AT-hook 1 pseudogene 6 |
| RNY3P4 | 13 | RNY3 pseudogene 4 [Source:HGNC Symbol;Ac |
| LINC00362 | 13 | long intergenic non-protein coding RNA 362 [S |
| SGCG | 13 | sarcoglycan gamma [Source:HGNC Symbol;Ac |
| RNU6-58P | 13 | RNA, U6 small nuclear 58, pseudogene [Sourc |
| TATDN2P3 | 13 | TatD DNase domain containing 2 pseudogene |
| SDAD1P4 | 13 | SDA1 domain containing 1 pseudogene 4 [Sou |
| SACS | 13 | sacsin molecular chaperone [Source:HGNC Sy |
| RPLP1P13 | 13 | ribosomal protein lateral stalk subunit P1 pseu |
| SACS-AS1 | 13 | SACS antisense RNA 1 [Source:HGNC Symbol;A |
| LINC00327 | 13 | long intergenic non-protein coding RNA 327 [S |
| LINC00352 | 13 | long intergenic non-protein coding RNA 352 [S |
| TNFRSF19 | 13 | TNF receptor superfamily member 19 [Source |
| MIPEP | 13 | mitochondrial intermediate peptidase [Source |
| MTCO3P2 | 13 | MT-CO3 pseudogene 2 [Source:HGNC Symbol |
| PCOTH | 13 | Pro-X-Gly collagen triple helix like repeat cont |
| C1QTNF9B | 13 | C1q and TNF related 9B [Source:HGNC Symbo |
| ANKRD20A19P | 13 | ankyrin repeat domain 20 family member A19 |
| AL445985.2 | 13 | novel transcript |
| AL359736.1 | 13 | novel transcript |
| SPATA13 | 13 | spermatogenesis associated 13 [Source:HGNC |
| AL445985.1 | 13 | novel transcript |
| IPO7P2 | 13 | importin 7 pseudogene 2 [Source:HGNC Symb |
| MIR2276 | 13 | microRNA 2276 [Source:HGNC Symbol;Acc:HG |
| SPATA13-AS1 | 13 | SPATA13 antisense RNA 1 [Source:HGNC Symb |
| C1QTNF9 | 13 | C1q and TNF related 9 [Source:HGNC Symbol; |
| C1QTNF9-AS1 | 13 | C1QTNF9 antisense RNA 1 [Source:HGNC Sym |
| AL359736.2 | 13 | cell cycle exit and neuronal differentiation 1 (( |
| NUS1P3 | 13 | NUS1 pseudogene 3 [Source:HGNC Symbol;Ac |
| LINC00566 | 13 | long intergenic non-protein coding RNA 566 [S |
| CYCSP33 | 13 | CYCS pseudogene 33 [Source:HGNC Symbol;A |

| | | |
|---|---|---|
| TPTE2P6 | 13 | transmembrane phosphoinositide 3-phosphat |
| PARP4 | 13 | poly(ADP-ribose) polymerase family member |
| AL359538.4 | 13 | novel transcript |
| AL359538.3 | 13 | novel transcript |
| PSPC1P2 | 13 | paraspeckle component 1 pseudogene 2 [Sou |
| AL359538.1 | 13 | novel transcript |
| AL359538.2 | 13 | novel atrophin-1 superfamily protein pseudog |
| ATP12A | 13 | ATPase H+/K+ transporting non-gastric alpha2 |
| RNY1P7 | 13 | RNY1 pseudogene 7 [Source:HGNC Symbol;Ac |
| RPL26P34 | 13 | ribosomal protein L26 pseudogene 34 [Source |
| AL391560.1 | 13 | novel transcript |
| IRX1P1 | 13 | iroquois homeobox 1 pseudogene 1 [Source:H |
| AL391560.2 | 13 | novel transcript |
| ANKRD20A10P | 13 | ankyrin repeat domain 20 family member A10 |
| RNF17 | 13 | ring finger protein 17 [Source:HGNC Symbol;A |
| CENPJ | 13 | centromere protein J [Source:HGNC Symbol;A |
| TPTE2P1 | 13 | transmembrane phosphoinositide 3-phosphat |
| AL354798.1 | 13 | info tpoly (ADP-ribose) polymerase family, me |
| SLC25A15P3 | 13 | solute carrier family 25 member 15 pseudoge |
| RPL34P27 | 13 | ribosomal protein L34 pseudogene 27 [Source |
| LSP1P1 | 13 | LSP1 pseudogene 1 [Source:HGNC Symbol;Acc |
| AL590099.1 | 13 | ankyrin repeat domain 36C (ANKRD36C) pseu |
| PABPC3 | 13 | poly(A) binding protein cytoplasmic 3 [Source |
| AL359757.2 | 13 | TEC |
| AMER2 | 13 | APC membrane recruitment protein 2 [Source |
| LINC01053 | 13 | long intergenic non-protein coding RNA 1053 |
| LINC00463 | 13 | long intergenic non-protein coding RNA 463 [S |
| AL359757.1 | 13 | TEC |
| LINC01076 | 13 | long intergenic non-protein coding RNA 1076 |
| RPL23AP69 | 13 | ribosomal protein L23a pseudogene 69 [Sourc |
| MTMR6 | 13 | myotubularin related protein 6 [Source:HGNC |
| AL590787.1 | 13 | novel transcript, antisense to NUPL1 |
| NUP58 | 13 | nucleoporin 58 [Source:HGNC Symbol;Acc:HG |
| ELOBP1 | 13 | elongin B pseudogene 1 [Source:HGNC Symbo |
| ATP8A2 | 13 | ATPase phospholipid transporting 8A2 [Source |
| RNU6-78P | 13 | RNA, U6 small nuclear 78, pseudogene [Sourc |
| AL157366.1 | 13 | novel pseudogene |
| RN7SL741P | 13 | RNA, 7SL, cytoplasmic 741, pseudogene [Sour |
| RNY1P3 | 13 | RNY1 pseudogene 3 [Source:HGNC Symbol;Ac |
| SHISA2 | 13 | shisa family member 2 [Source:HGNC Symbol; |
| LINC00415 | 13 | long intergenic non-protein coding RNA 415 [S |
| ATP8A2P3 | 13 | ATPase phospholipid transporting 8A2 pseudo |
| PRUNEP1 | 13 | prune exopolyphosphatase pseudogene 1 [So |
| RNF6 | 13 | ring finger protein 6 [Source:HGNC Symbol;Ac |
| AL138966.2 | 13 | novel transcript, antisense to ring finger prote |
| AL138966.1 | 13 | synovial sarcoma translocation gene on chron |
| THAP12P6 | 13 | THAP domain containing 12 pseudogene 6 [Sc |
| CDK8 | 13 | cyclin dependent kinase 8 [Source:HGNC Syml |
| AL159159.1 | 13 | NULL |
| AL353789.1 | 13 | novel transcript |

| | | |
|---|---|---|
| WASF3 | 13 | WASP family member 3 [Source:HGNC Symbo |
| RPS3AP44 | 13 | ribosomal protein S3a pseudogene 44 [Source |
| WASF3-AS1 | 13 | WASF3 antisense RNA 1 [Source:HGNC Symbo |
| AL159978.1 | 13 | novel transcript, antisense to WASF3 |
| GPR12 | 13 | G protein-coupled receptor 12 [Source:HGNC |
| FGFR1OP2P1 | 13 | FGFR1 oncogene partner 2 pseudogene 1 [Sou |
| AL160035.1 | 13 | novel transcript |
| RPS21P8 | 13 | ribosomal protein S21 pseudogene 8 [Source: |
| RPS20P32 | 13 | ribosomal protein S20 pseudogene 32 [Source |
| USP12 | 13 | ubiquitin specific peptidase 12 [Source:HGNC |
| AL158062.1 | 13 | retinoblastoma binding protein 8 (RBBP8) pse |
| USP12-AS1 | 13 | USP12 antisense RNA 1 [Source:HGNC Symbol |
| USP12-AS2 | 13 | USP12 antisense RNA 2 (head to head) [Sourc |
| LINC02340 | 13 | long intergenic non-protein coding RNA 2340 |
| LINC00412 | 13 | long intergenic non-protein coding RNA 412 [S |
| RPL21 | 13 | ribosomal protein L21 [Source:HGNC Symbol; |
| SNORD102 | 13 | small nucleolar RNA, C/D box 102 [Source:HG |
| SNORA27 | 13 | small nucleolar RNA, H/ACA box 27 [Source:H |
| RASL11A | 13 | RAS like family 11 member A [Source:HGNC Sy |
| RNU6-70P | 13 | RNA, U6 small nuclear 70, pseudogene [Sourc |
| LINC01079 | 13 | long intergenic non-protein coding RNA 1079 |
| RNY1P1 | 13 | RNY1 pseudogene 1 [Source:HGNC Symbol;Ac |
| GTF3A | 13 | general transcription factor IIIA [Source:HGNC |
| MTIF3 | 13 | mitochondrial translational initiation factor 3 |
| RNU6-63P | 13 | RNA, U6 small nuclear 63, pseudogene [Sourc |
| LNX2 | 13 | ligand of numb-protein X 2 [Source:HGNC Syn |
| POLR1D | 13 | RNA polymerase I and III subunit D [Source:HC |
| AL136439.1 | 13 | novel transcript |
| NPM1P4 | 13 | nucleophosmin 1 pseudogene 4 [Source:HGN( |
| GSX1 | 13 | GS homeobox 1 [Source:HGNC Symbol;Acc:HC |
| PLUT | 13 | PDX1 associated lncRNA, upregulator of trans |
| RNU6-73P | 13 | RNA, U6 small nuclear 73, pseudogene [Sourc |
| PDX1 | 13 | pancreatic and duodenal homeobox 1 [Source |
| ATP5F1EP2 | 13 | ATP synthase F1 subunit epsilon pseudogene |
| LINC00543 | 13 | long intergenic non-protein coding RNA 543 [S |
| CDX2 | 13 | caudal type homeobox 2 [Source:HGNC Symb |
| URAD | 13 | ureidoimidazoline (2-oxo-4-hydroxy-4-carbox |
| RN7SL272P | 13 | RNA, 7SL, cytoplasmic 272, pseudogene [Sour |
| FLT3 | 13 | fms related tyrosine kinase 3 [Source:HGNC S |
| KATNBL1P1 | 13 | katanin regulatory subunit B1 like 1 pseudoge |
| CHCHD2P8 | 13 | coiled-coil-helix-coiled-coil-helix domain conta |
| PAN3-AS1 | 13 | PAN3 antisense RNA 1 [Source:HGNC Symbol; |
| PAN3 | 13 | poly(A) specific ribonuclease subunit PAN3 [Sc |
| EEF1A1P3 | 13 | eukaryotic translation elongation factor 1 alpl |
| RNU6-82P | 13 | RNA, U6 small nuclear 82, pseudogene [Sourc |
| FLT1 | 13 | fms related tyrosine kinase 1 [Source:HGNC S |
| AL139005.1 | 13 | TEC |
| EIF4A1P7 | 13 | eukaryotic translation initiation factor 4A1 ps |
| Y_RNA | 13 | Y RNA [Source:RFAM;Acc:RF00019] |
| POMP | 13 | proteasome maturation protein [Source:HGN( |

| | | |
|---|---|---|
| SLC46A3 | 13 | solute carrier family 46 member 3 [Source:HG |
| RNU6-53P | 13 | RNA, U6 small nuclear 53, pseudogene [Sourc |
| CYP51A1P2 | 13 | cytochrome P450 family 51 subfamily A meml |
| AL359741.1 | 13 | POM121 transmembrane nucleoporin-like 2 (I |
| POM121L13P | 13 | POM121 transmembrane nucleoporin like 13, |
| MTUS2 | 13 | microtubule associated scaffold protein 2 [Sou |
| MTUS2-AS2 | 13 | MTUS2 antisense RNA 2 [Source:HGNC Symbo |
| GAPDHP69 | 13 | glyceraldehyde 3 phosphate dehydrogenase p |
| AL596092.1 | 13 | carbonic anhydrase VB pseudogene 1 (CA5BP: |
| MTUS2-AS1 | 13 | MTUS2 antisense RNA 1 [Source:HGNC Symbo |
| SLC7A1 | 13 | solute carrier family 7 member 1 [Source:HGN |
| UBL3 | 13 | ubiquitin like 3 [Source:HGNC Symbol;Acc:HGI |
| Metazoa_SRP | 13 | Metazoan signal recognition particle RNA [Sou |
| TIMM8BP1 | 13 | translocase of inner mitochondrial membrane |
| LINC00297 | 13 | long intergenic non-protein coding RNA 297 [S |
| LINC00572 | 13 | long intergenic non-protein coding RNA 572 [S |
| LINC00544 | 13 | long intergenic non-protein coding RNA 544 [S |
| AL354674.1 | 13 | novel transcript |
| LINC00365 | 13 | long intergenic non-protein coding RNA 365 [S |
| LINC00384 | 13 | long intergenic non-protein coding RNA 384 [S |
| LINC00385 | 13 | long intergenic non-protein coding RNA 385 [S |
| AL157770.1 | 13 | TEC |
| KATNAL1 | 13 | katanin catalytic subunit A1 like 1 [Source:HGI |
| RNU6-64P | 13 | RNA, U6 small nuclear 64, pseudogene [Sourc |
| PRDX2P1 | 13 | peroxiredoxin 2 pseudogene 1 [Source:HGNC |
| LINC00427 | 13 | long intergenic non-protein coding RNA 427 [S |
| LINC00426 | 13 | long intergenic non-protein coding RNA 426 [S |
| AL356750.1 | 13 | TEC |
| AL161893.1 | 13 | novel transcript |
| LINC01058 | 13 | long intergenic non-protein coding RNA 1058 |
| UBE2L5 | 13 | ubiquitin conjugating enzyme E2 L5 [Source:H |
| HMGB1 | 13 | high mobility group box 1 [Source:HGNC Symb |
| AL353648.1 | 13 | novel transcript, antisense to HMGB1 |
| RBM22P2 | 13 | RNA binding motif protein 22 pseudogene 2 [S |
| MFAP1P1 | 13 | microfibril associated protein 1 pseudogene 1 |
| PTPN2P2 | 13 | PTPN2 pseudogene 2 [Source:HGNC Symbol;A |
| USPL1 | 13 | ubiquitin specific peptidase like 1 [Source:HGI |
| Metazoa_SRP | 13 | Metazoan signal recognition particle RNA [Sou |
| ALOX5AP | 13 | arachidonate 5-lipoxygenase activating protei |
| LINC00398 | 13 | long intergenic non-protein coding RNA 398 [S |
| LINC00545 | 13 | long intergenic non-protein coding RNA 545 [S |
| TEX26-AS1 | 13 | TEX26 antisense RNA 1 [Source:HGNC Symbol |
| MEDAG | 13 | mesenteric estrogen dependent adipogenesis |
| TEX26 | 13 | testis expressed 26 [Source:HGNC Symbol;Acc |
| AL353680.1 | 13 | novel transcript |
| LINC01066 | 13 | long intergenic non-protein coding RNA 1066 |
| AL158065.1 | 13 | novel transcript |
| WDR95P | 13 | WD repeat domain 95, pseudogene [Source:H |
| HSPH1 | 13 | heat shock protein family H (Hsp110) member |
| B3GLCT | 13 | beta 3-glucosyltransferase [Source:HGNC Sym |

| | | |
|---|---|---|
| ANKRD26P4 | 13 | ankyrin repeat domain 26 pseudogene 4 [Sour |
| AL161616.2 | 13 | novel transcript |
| AL161616.1 | 13 | ubiquitin specific peptidase 34 (USP34) pseud |
| RXFP2 | 13 | relaxin family peptide receptor 2 [Source:HGN |
| AL138708.1 | 13 | novel transcript |
| FRY | 13 | FRY microtubule binding protein [Source:HGN |
| EEF1DP3 | 13 | eukaryotic translation elongation factor 1 delt |
| AC002525.1 | 13 | TEC |
| Metazoa_SRP | 13 | Metazoan signal recognition particle RNA [Sou |
| FRY-AS1 | 13 | FRY antisense RNA 1 [Source:HGNC Symbol;A |
| ZAR1L | 13 | zygote arrest 1 like [Source:HGNC Symbol;Acc |
| BRCA2 | 13 | BRCA2 DNA repair associated [Source:HGNC S |
| IFIT1P1 | 13 | interferon induced protein with tetratricopep |
| N4BP2L1 | 13 | NEDD4 binding protein 2 like 1 [Source:HGNC |
| AL137247.1 | 13 | NULL |
| AL137247.2 | 13 | novel transcript |
| N4BP2L2 | 13 | NEDD4 binding protein 2 like 2 [Source:HGNC |
| ATP8A2P2 | 13 | ATPase phospholipid transporting 8A2 pseudo |
| N4BP2L2-IT2 | 13 | N4BPL2 intronic transcript 2 [Source:HGNC Sy |
| PDS5B | 13 | PDS5 cohesin associated factor B [Source:HGN |
| RNY1P4 | 13 | RNY1 pseudogene 4 [Source:HGNC Symbol;Ac |
| AL138820.1 | 13 | novel transcript |
| LINC00423 | 13 | long intergenic non-protein coding RNA 423 [S |
| TOMM22P3 | 13 | TOMM22 pseudogene 3 [Source:HGNC Symbo |
| KL | 13 | klotho [Source:HGNC Symbol;Acc:HGNC:6344 |
| STARD13 | 13 | StAR related lipid transfer domain containing |
| STARD13-IT1 | 13 | STARD13 intronic transcript 1 [Source:HGNC S |
| STARD13-AS | 13 | STARD13 antisense RNA [Source:HGNC Symbo |
| AL627232.1 | 13 | novel transcript, antisense to STARD13 |
| LINC02344 | 13 | long intergenic non-protein coding RNA 2344 |
| AL138999.2 | 13 | novel transcript, antisense to STARD13 |
| AL138999.1 | 13 | novel transcript, sense intronic to STARD13 |
| AL139383.1 | 13 | novel transcript |
| AL161719.1 | 13 | novel transcript |
| AL139081.1 | 13 | novel transcript |
| AL139081.2 | 13 | novel transcript |
| RFC3 | 13 | replication factor C subunit 3 [Source:HGNC S |
| RNU5A-4P | 13 | RNA, U5A small nuclear 4, pseudogene [Sourc |
| AL161891.1 | 13 | novel transcript, sense intronic to RFC3 |
| AL160394.2 | 13 | TEC |
| VDAC1P12 | 13 | voltage dependent anion channel 1 pseudoge |
| AL160394.1 | 13 | NULL |
| AL356321.1 | 13 | KIAA1551 (KIAA1551) pseudogene |
| LINC02343 | 13 | long intergenic non-protein coding RNA 2343 |
| LINC00457 | 13 | long intergenic non-protein coding RNA 457 [S |
| AL161716.1 | 13 | novel transcript |
| GAMTP2 | 13 | guanidinoacetate N-methyltransferase pseudo |
| Metazoa_SRP | 13 | Metazoan signal recognition particle RNA [Sou |
| AL138690.1 | 13 | novel transcript |
| NBEA | 13 | neurobeachin [Source:HGNC Symbol;Acc:HGN |

| | | |
|---|---|---|
| SCAND3P1 | 13 | SCAN domain containing 3 pseudogene 1 [Sou |
| PHBP13 | 13 | prohibitin pseudogene 13 [Source:HGNC Syml |
| MAB21L1 | 13 | mab-21 like 1 [Source:HGNC Symbol;Acc:HGN |
| AL390071.1 | 13 | novel transcript, antisense to NBEA |
| LINC00445 | 13 | long intergenic non-protein coding RNA 445 [S |
| DCLK1 | 13 | doublecortin like kinase 1 [Source:HGNC Symb |
| SOHLH2 | 13 | spermatogenesis and oogenesis specific basic |
| CCDC169-SOHLH2 | 13 | CCDC169-SOHLH2 readthrough [Source:HGNC |
| CCDC169 | 13 | coiled-coil domain containing 169 [Source:HG |
| RNU6-71P | 13 | RNA, U6 small nuclear 71, pseudogene [Sourc |
| SPART | 13 | spartin [Source:HGNC Symbol;Acc:HGNC:1851 |
| SPART-AS1 | 13 | SPART antisense RNA 1 [Source:HGNC Symbol |
| CCNA1 | 13 | cyclin A1 [Source:HGNC Symbol;Acc:HGNC:15 |
| HIST1H2APS6 | 13 | HIST1H2A pseudogene 6 [Source:HGNC Symb |
| SERTM1 | 13 | serine rich and transmembrane domain conta |
| GAPDHP34 | 13 | glyceraldehyde 3 phosphate dehydrogenase p |
| TCEAL4P1 | 13 | transcription elongation factor A like 4 pseud |
| ARL2BPP3 | 13 | ADP ribosylation factor like GTPase 2 binding |
| NDE1P2 | 13 | nudE neurodevelopment protein 1 pseudoger |
| RFXAP | 13 | regulatory factor X associated protein [Source |
| SMAD9 | 13 | SMAD family member 9 [Source:HGNC Symbo |
| SMAD9-IT1 | 13 | SMAD9 intronic transcript 1 [Source:HGNC Sy |
| LAMTOR3P1 | 13 | late endosomal/lysosomal adaptor, MAPK and |
| RPL29P28 | 13 | ribosomal protein L29 pseudogene 28 [Source |
| EIF4A1P5 | 13 | eukaryotic translation initiation factor 4A1 ps |
| ALG5 | 13 | ALG5 dolichyl-phosphate beta-glucosyltransfe |
| EXOSC8 | 13 | exosome component 8 [Source:HGNC Symbol |
| SUPT20H | 13 | SPT20 homolog, SAGA complex component [S |
| CSNK1A1L | 13 | casein kinase 1 alpha 1 like [Source:HGNC Sym |
| AL356274.1 | 13 | cutaneous T-cell lymphoma-associated antige |
| AL356274.2 | 13 | TEC |
| RN7SKP1 | 13 | RN7SK pseudogene 1 [Source:HGNC Symbol;A |
| RPS12P24 | 13 | ribosomal protein S12 pseudogene 24 [Source |
| LINC01048 | 13 | long intergenic non-protein coding RNA 1048 |
| LINC00547 | 13 | long intergenic non-protein coding RNA 547 [S |
| POSTN | 13 | periostin [Source:HGNC Symbol;Acc:HGNC:16 |
| TRPC4 | 13 | transient receptor potential cation channel su |
| RNA5SP26 | 13 | RNA, 5S ribosomal pseudogene 26 [Source:HG |
| AL356495.1 | 13 | novel transcript, antisense to TRPC4 |
| LINC02334 | 13 | long intergenic non-protein coding RNA 2334 |
| HSPD1P9 | 13 | heat shock protein family D (Hsp60) member |
| LINC00571 | 13 | long intergenic non-protein coding RNA 571 [S |
| UFM1 | 13 | ubiquitin fold modifier 1 [Source:HGNC Symbo |
| AL356863.1 | 13 | TEC |
| LINC00437 | 13 | long intergenic non-protein coding RNA 437 [S |
| LINC00366 | 13 | long intergenic non-protein coding RNA 366 [S |
| PRDX3P3 | 13 | peroxiredoxin 3 pseudogene 3 [Source:HGNC |
| FREM2 | 13 | FRAS1 related extracellular matrix 2 [Source:H |
| RNU6-56P | 13 | RNA, U6 small nuclear 56, pseudogene [Sourc |
| FREM2-AS1 | 13 | FREM2 antisense RNA 1 [Source:HGNC Symbo |

| | | |
|---|---|---|
| PLA2G12AP2 | 13 | phospholipase A2 group XIIA pseudogene 2 [S |
| ANKRD26P2 | 13 | ankyrin repeat domain 26 pseudogene 2 [Sour |
| STOML3 | 13 | stomatin like 3 [Source:HGNC Symbol;Acc:HG |
| PROSER1 | 13 | proline and serine rich 1 [Source:HGNC Symbo |
| NHLRC3 | 13 | NHL repeat containing 3 [Source:HGNC Symbo |
| AL354809.1 | 13 | novel transcript |
| NXT1P1 | 13 | nuclear transport factor 2 like export factor 1 |
| LHFPL6 | 13 | LHFPL tetraspan subfamily member 6 [Source |
| COG6 | 13 | component of oligomeric golgi complex 6 [Sou |
| MIR4305 | 13 | microRNA 4305 [Source:HGNC Symbol;Acc:HG |
| RNY4P14 | 13 | RNY4 pseudogene 14 [Source:HGNC Symbol;A |
| CDKN2AIPNLP3 | 13 | CDKN2A interacting protein N-terminal like ps |
| AZU1P1 | 13 | azurocidin 1 pseudogene 1 [Source:HGNC Sym |
| SNORD116 | 13 | Small nucleolar RNA SNORD116 [Source:RFAM |
| LINC00598 | 13 | long intergenic non-protein coding RNA 598 [S |
| LINC00332 | 13 | long intergenic non-protein coding RNA 332 [S |
| RPL17P51 | 13 | ribosomal protein L17 pseudogene 51 [Source |
| RNY3P9 | 13 | RNY3 pseudogene 9 [Source:HGNC Symbol;Ac |
| RN7SKP2 | 13 | RN7SK pseudogene 2 [Source:HGNC Symbol;A |
| FOXO1 | 13 | forkhead box O1 [Source:HGNC Symbol;Acc:H |
| RLIMP1 | 13 | ring finger protein, LIM domain interacting ps |
| MIR320D1 | 13 | microRNA 320d-1 [Source:HGNC Symbol;Acc:H |
| MRPS31 | 13 | mitochondrial ribosomal protein S31 [Source: |
| Metazoa_SRP | 13 | Metazoan signal recognition particle RNA [Sou |
| SLC25A15 | 13 | solute carrier family 25 member 15 [Source:H |
| MIR621 | 13 | microRNA 621 [Source:HGNC Symbol;Acc:HGN |
| TPTE2P5 | 13 | transmembrane phosphoinositide 3-phosphat |
| CYCSP34 | 13 | CYCS pseudogene 34 [Source:HGNC Symbol;A |
| AL590064.1 | 13 | novel transcript |
| SUGT1P3 | 13 | SUGT1 pseudogene 3 [Source:HGNC Symbol;A |
| ELF1 | 13 | E74 like ETS transcription factor 1 [Source:HG |
| RGS17P1 | 13 | regulator of G protein signaling 17 pseudogen |
| WBP4 | 13 | WW domain binding protein 4 [Source:HGNC |
| MIR3168 | 13 | microRNA 3168 [Source:HGNC Symbol;Acc:HG |
| RN7SL597P | 13 | RNA, 7SL, cytoplasmic 597, pseudogene [Sour |
| KBTBD6 | 13 | kelch repeat and BTB domain containing 6 [So |
| AL354696.2 | 13 | novel transcript, antisense to KBTBD7 and MT |
| CALM2P3 | 13 | calmodulin 2 pseudogene 3 [Source:HGNC Syr |
| KBTBD7 | 13 | kelch repeat and BTB domain containing 7 [So |
| MORF4L1P4 | 13 | mortality factor 4 like 1 pseudogene 4 [Source |
| MTRF1 | 13 | mitochondrial translation release factor 1 [Sou |
| AL354696.1 | 13 | novel transcript, sense intronic to MTRF1 |
| RAC1P3 | 13 | Rac family small GTPase 1 pseudogene 3 [Sou |
| NAA16 | 13 | N(alpha)-acetyltransferase 16, NatA auxiliary s |
| RNU6-57P | 13 | RNA, U6 small nuclear 57, pseudogene [Sourc |
| TUBBP2 | 13 | tubulin beta pseudogene 2 [Source:HGNC Sym |
| OR7E36P | 13 | olfactory receptor family 7 subfamily E memb |
| OR7E155P | 13 | olfactory receptor family 7 subfamily E memb |
| OR7E37P | 13 | olfactory receptor family 7 subfamily E memb |
| RGCC | 13 | regulator of cell cycle [Source:HGNC Symbol;A |

| | | |
|---|---|---|
| VWA8 | 13 | von Willebrand factor A domain containing 8 [ |
| MIR5006 | 13 | microRNA 5006 [Source:HGNC Symbol;Acc:HG |
| RN7SL515P | 13 | RNA, 7SL, cytoplasmic 515, pseudogene [Sour |
| RNU6-74P | 13 | RNA, U6 small nuclear 74, pseudogene [Sourc |
| KARSP1 | 13 | lysyl-tRNA synthetase pseudogene 1 [Source:H |
| VWA8-AS1 | 13 | VWA8 antisense RNA 1 (head to head) [Source |
| RPS28P8 | 13 | ribosomal protein S28 pseudogene 8 [Source: |
| DGKH | 13 | diacylglycerol kinase eta [Source:HGNC Symbo |
| AL157932.1 | 13 | novel transcript |
| MAPK6P3 | 13 | mitogen-activated protein kinase 6 pseudogen |
| CHCHD2P11 | 13 | coiled-coil-helix-coiled-coil-helix domain conta |
| FHP1 | 13 | fumarate hydratase pseudogene 1 [Source:HG |
| AKAP11 | 13 | A-kinase anchoring protein 11 [Source:HGNC S |
| LINC02341 | 13 | long intergenic non-protein coding RNA 2341 |
| FABP3P2 | 13 | fatty acid binding protein 3 pseudogene 2 [Sou |
| TNFSF11 | 13 | TNF superfamily member 11 [Source:HGNC Sy |
| FAM216B | 13 | family with sequence similarity 216 member B |
| LINC01050 | 13 | long intergenic non-protein coding RNA 1050 |
| LINC00428 | 13 | long intergenic non-protein coding RNA 428 [S |
| EPSTI1 | 13 | epithelial stromal interaction 1 [Source:HGNC |
| ZDHHC4P1 | 13 | zinc finger DHHC-type containing 4 pseudogen |
| DNAJC15 | 13 | DnaJ heat shock protein family (Hsp40) memb |
| LINC00400 | 13 | long intergenic non-protein coding RNA 400 [S |
| AL138709.1 | 13 | molybdenum cofactor synthesis 3 (MOCS3) ps |
| ENOX1 | 13 | ecto-NOX disulfide-thiol exchanger 1 [Source: |
| RPL36P19 | 13 | ribosomal protein L36 pseudogene 19 [Source |
| ENOX1-AS2 | 13 | ENOX1 antisense RNA 2 [Source:HGNC Symbo |
| ENOX1-AS1 | 13 | ENOX1 antisense RNA 1 [Source:HGNC Symbo |
| AL161714.1 | 13 | novel transcript, sense intronic to ENOX1 |
| AL162713.2 | 13 | novel transcript, antisense to ENOX1 |
| AL162713.1 | 13 | novel transcript |
| CCDC122 | 13 | coiled-coil domain containing 122 [Source:HG |
| AL512506.1 | 13 | novel transcript, sense intronic to CCDC122 |
| LACC1 | 13 | laccase domain containing 1 [Source:HGNC Sy |
| AL512506.3 | 13 | novel transcript |
| AL512506.2 | 13 | TEC |
| NRAD1 | 13 | non-coding RNA in the aldehyde dehydrogena |
| DGKZP1 | 13 | diacylglycerol kinase zeta pseudogene 1 [Sour |
| LINC00390 | 13 | long intergenic non-protein coding RNA 390 [S |
| SMIM2-AS1 | 13 | SMIM2 antisense RNA 1 [Source:HGNC Symbo |
| SMIM2 | 13 | small integral membrane protein 2 [Source:HG |
| SMIM2-IT1 | 13 | SMIM2 intronic transcript 1 [Source:HGNC Syn |
| AL589745.2 | 13 | novel transcript |
| MIR8079 | 13 | microRNA 8079 [Source:HGNC Symbol;Acc:HG |
| AL589745.1 | 13 | novel transcript |
| AL138960.1 | 13 | novel transcript |
| SERP2 | 13 | stress associated endoplasmic reticulum prote |
| TUSC8 | 13 | tumor suppressor candidate 8 [Source:HGNC S |
| TSC22D1 | 13 | TSC22 domain family member 1 [Source:HGN |
| AL138960.2 | 13 | TEC |

| | | |
|---|---|---|
| AL139184.1 | 13 | filamin A interacting protein 1-like (FILIP1L) ps |
| TSC22D1-AS1 | 13 | TSC22D1 antisense RNA 1 [Source:HGNC Symb |
| SMARCE1P5 | 13 | SWI/SNF related, matrix associated, actin dep |
| LINC00407 | 13 | long intergenic non-protein coding RNA 407 [S |
| AL356515.1 | 13 | novel transcript |
| LINC00330 | 13 | long intergenic non-protein coding RNA 330 [S |
| NUFIP1 | 13 | nuclear FMR1 interacting protein 1 [Source:H |
| AL359706.1 | 13 | novel transcript |
| GPALPP1 | 13 | GPALPP motifs containing 1 [Source:HGNC Syr |
| RN7SL49P | 13 | RNA, 7SL, cytoplasmic 49, pseudogene [Sourc |
| RN7SKP3 | 13 | RN7SK pseudogene 3 [Source:HGNC Symbol;A |
| GTF2F2 | 13 | general transcription factor IIF subunit 2 [Sour |
| RN7SKP4 | 13 | RN7SK pseudogene 4 [Source:HGNC Symbol;A |
| AL138693.1 | 13 | glyceraldehyde-3-phosphate dehydrogenase ( |
| KCTD4 | 13 | potassium channel tetramerization domain cc |
| AL138963.3 | 13 | cytochrome c oxidase subunit VIc (COX6C) pse |
| TPT1 | 13 | tumor protein, translationally-controlled 1 [Sc |
| AL138963.2 | 13 | NULL |
| SNORA31 | 13 | small nucleolar RNA, H/ACA box 31 [Source:H |
| AL138963.4 | 13 | novel transcript, antisense to TPT1 |
| TPT1-AS1 | 13 | TPT1 antisense RNA 1 [Source:HGNC Symbol;A |
| AL138963.1 | 13 | novel transcript |
| RCN1P2 | 13 | reticulocalbin 1 pseudogene 2 [Source:HGNC S |
| SLC25A30 | 13 | solute carrier family 25 member 30 [Source:H |
| SLC25A30-AS1 | 13 | SLC25A30 antisense RNA 1 [Source:HGNC Sym |
| PPIAP25 | 13 | peptidylprolyl isomerase A pseudogene 25 [Sc |
| COG3 | 13 | component of oligomeric golgi complex 3 [Sou |
| ERICH6B | 13 | glutamate rich 6B [Source:HGNC Symbol;Acc:I |
| RNA5SP27 | 13 | RNA, 5S ribosomal pseudogene 27 [Source:HG |
| TIMM9P3 | 13 | TIMM9 pseudogene 3 [Source:HGNC Symbol;A |
| COX4I1P2 | 13 | cytochrome c oxidase subunit 4I1 pseudogene |
| LINC01055 | 13 | long intergenic non-protein coding RNA 1055 |
| AKR1B1P4 | 13 | aldo-keto reductase family 1 member B1 pseu |
| SPERT | 13 | spermatid associated [Source:HGNC Symbol;A |
| Metazoa_SRP | 13 | Metazoan signal recognition particle RNA [Sou |
| SIAH3 | 13 | siah E3 ubiquitin protein ligase family membe |
| ZC3H13 | 13 | zinc finger CCCH-type containing 13 [Source:H |
| CPB2-AS1 | 13 | CPB2 antisense RNA 1 [Source:HGNC Symbol;A |
| CPB2 | 13 | carboxypeptidase B2 [Source:HGNC Symbol;A |
| LCP1 | 13 | lymphocyte cytosolic protein 1 [Source:HGNC |
| RN7SL288P | 13 | RNA, 7SL, cytoplasmic 288, pseudogene [Sour |
| LRRC63 | 13 | leucine rich repeat containing 63 [Source:HGN |
| RN7SKP5 | 13 | RN7SK pseudogene 5 [Source:HGNC Symbol;A |
| ABITRAMP1 | 13 | ABITRAM pseudogene 1 [Source:HGNC Symbo |
| AL139801.1 | 13 | novel transcript |
| LINC00563 | 13 | long intergenic non-protein coding RNA 563 [S |
| RUBCNL | 13 | rubicon like autophagy enhancer [Source:HGN |
| RNU2-6P | 13 | RNA, U2 small nuclear 6, pseudogene [Source |
| RNU6-68P | 13 | RNA, U6 small nuclear 68, pseudogene [Sourc |
| PPP1R2P4 | 13 | protein phosphatase 1 regulatory inhibitor sul |

| | | |
|---|---|---|
| LINC01198 | 13 | long intergenic non-protein coding RNA 1198 |
| AL138686.2 | 13 | novel keratin associated protein pseudogene |
| OR7E101P | 13 | olfactory receptor family 7 subfamily E memb |
| AL138686.1 | 13 | novel transcript |
| COX17P1 | 13 | COX17 pseudogene 1 [Source:HGNC Symbol;A |
| FKBP1AP3 | 13 | FKBP prolyl isomerase 1A pseudogene 3 [Sour |
| LRCH1 | 13 | leucine rich repeats and calponin homology d |
| AL359880.1 | 13 | novel transcript, sense intronic to LRCH1 |
| ESD | 13 | esterase D [Source:HGNC Symbol;Acc:HGNC:3 |
| HTR2A | 13 | 5-hydroxytryptamine receptor 2A [Source:HG |
| HTR2A-AS1 | 13 | HTR2A antisense RNA 1 [Source:HGNC Symbo |
| GNG5P5 | 13 | G protein subunit gamma 5 pseudogene 5 [So |
| RN7SL700P | 13 | RNA, 7SL, cytoplasmic 700, pseudogene [Sour |
| SUCLA2 | 13 | succinate-CoA ligase ADP-forming beta subun |
| NAP1L4P3 | 13 | nucleosome assembly protein 1 like 4 pseudo |
| RPL27AP8 | 13 | ribosomal protein L27a pseudogene 8 [Source |
| AL138962.1 | 13 | novel transcript |
| LINC00562 | 13 | long intergenic non-protein coding RNA 562 [S |
| AL157369.1 | 13 | novel transcript, antisense to SUCLA2 |
| SUCLA2-AS1 | 13 | SUCLA2 antisense RNA 1 [Source:HGNC Symb |
| NUDT15 | 13 | nudix hydrolase 15 [Source:HGNC Symbol;Acc |
| AL158196.1 | 13 | novel transcript, antisense to NUDT15 |
| MED4 | 13 | mediator complex subunit 4 [Source:HGNC Sy |
| MED4-AS1 | 13 | MED4 antisense RNA 1 [Source:HGNC Symbol |
| POLR2KP2 | 13 | RNA polymerase II subunit K pseudogene 2 [S |
| ITM2B | 13 | integral membrane protein 2B [Source:HGNC |
| RB1-DT | 13 | RB1 divergent transcript [Source:HGNC Symb |
| RB1 | 13 | RB transcriptional corepressor 1 [Source:HGN |
| PPP1R26P1 | 13 | protein phosphatase 1 regulatory subunit 26 p |
| PCNPP5 | 13 | PEST containing nuclear protein pseudogene 5 |
| AL392048.1 | 13 | novel transcript, antisense to RB1 |
| LPAR6 | 13 | lysophosphatidic acid receptor 6 [Source:HGN |
| Metazoa_SRP | 13 | Metazoan signal recognition particle RNA [Sou |
| RCBTB2 | 13 | RCC1 and BTB domain containing protein 2 [S |
| AL157813.2 | 13 | novel transcript, antisense to RCBTB2 |
| AL157813.1 | 13 | novel transcript, sense intronic to RCBTB2 |
| LINC01077 | 13 | long intergenic non-protein coding RNA 1077 |
| LINC00462 | 13 | long intergenic non-protein coding RNA 462 [S |
| CYSLTR2 | 13 | cysteinyl leukotriene receptor 2 [Source:HGN |
| PSME2P2 | 13 | proteasome activator subunit 2 pseudogene 2 |
| AL161421.1 | 13 | novel transcript, antisense to FNDC3A |
| FNDC3A | 13 | fibronectin type III domain containing 3A [Sou |
| RNU6-60P | 13 | RNA, U6 small nuclear 60, pseudogene [Sourc |
| RAD17P2 | 13 | RAD17 pseudogene 2 [Source:HGNC Symbol;A |
| RNY3P2 | 13 | RNY3 pseudogene 2 [Source:HGNC Symbol;Ac |
| COX7CP1 | 13 | cytochrome c oxidase subunit 7C pseudogene |
| OGFOD1P1 | 13 | 2-oxoglutarate and iron dependent oxygenase |
| MLNR | 13 | motilin receptor [Source:HGNC Symbol;Acc:H |
| AL137000.1 | 13 | protein-kinase, interferon-inducible double st |
| CDADC1 | 13 | cytidine and dCMP deaminase domain contair |

| | | |
|---|---|---|
| CAB39L | 13 | calcium binding protein 39 like [Source:HGNC |
| SETDB2 | 13 | SET domain bifurcated histone lysine methyltı |
| SNRPGP14 | 13 | small nuclear ribonucleoprotein polypeptide ( |
| PHF11 | 13 | PHD finger protein 11 [Source:HGNC Symbol;/ |
| RCBTB1 | 13 | RCC1 and BTB domain containing protein 1 [S( |
| AL135901.1 | 13 | novel pseudogene |
| ARL11 | 13 | ADP ribosylation factor like GTPase 11 [Source |
| EBPL | 13 | EBP like [Source:HGNC Symbol;Acc:HGNC:180 |
| KPNA3 | 13 | karyopherin subunit alpha 3 [Source:HGNC Sy |
| AL136301.1 | 13 | novel transcript, antisense to KPNA3 |
| RNY4P30 | 13 | RNY4 pseudogene 30 [Source:HGNC Symbol;A |
| CTAGE10P | 13 | CTAGE family member 10, pseudogene [Sourc |
| RNY4P9 | 13 | RNY4 pseudogene 9 [Source:HGNC Symbol;Ac |
| SPRYD7 | 13 | SPRY domain containing 7 [Source:HGNC Sym |
| DLEU2 | 13 | deleted in lymphocytic leukemia 2 [Source:HG |
| TRIM13 | 13 | tripartite motif containing 13 [Source:HGNC S |
| MIR3613 | 13 | microRNA 3613 [Source:HGNC Symbol;Acc:HG |
| KCNRG | 13 | potassium channel regulator [Source:HGNC Sy |
| AL137060.9 | 13 | NULL |
| AL137060.7 | 13 | NULL |
| MIR16-1 | 13 | microRNA 16-1 [Source:HGNC Symbol;Acc:HG |
| MIR15A | 13 | microRNA 15a [Source:HGNC Symbol;Acc:HGN |
| AL137060.4 | 13 | NULL |
| AL137060.2 | 13 | NULL |
| AL137060.8 | 13 | NULL |
| AL137060.5 | 13 | NULL |
| DLEU1 | 13 | deleted in lymphocytic leukemia 1 [Source:HG |
| AL137060.6 | 13 | NULL |
| RPL18P10 | 13 | ribosomal protein L18 pseudogene 10 [Source |
| AL137060.1 | 13 | NULL |
| AL137060.3 | 13 | novel transcript, sense intronic to DLEU1 |
| ST13P4 | 13 | ST13, Hsp70 interacting protein pseudogene 4 |
| RPL34P26 | 13 | ribosomal protein L34 pseudogene 26 [Source |
| AL158195.1 | 13 | novel transcript |
| DLEU7 | 13 | deleted in lymphocytic leukemia 7 [Source:HG |
| RNA5SP28 | 13 | RNA, 5S ribosomal pseudogene 28 [Source:HG |
| RNASEH2B-AS1 | 13 | RNASEH2B antisense RNA 1 [Source:HGNC Syı |
| RNASEH2B | 13 | ribonuclease H2 subunit B [Source:HGNC Sym |
| GUCY1B2 | 13 | guanylate cyclase 1 soluble subunit beta 2 (ps |
| RNA5SP29 | 13 | RNA, 5S ribosomal pseudogene 29 [Source:HG |
| C13orf42 | 13 | chromosome 13 open reading frame 42 [Sour |
| RPL5P31 | 13 | ribosomal protein L5 pseudogene 31 [Source: |
| AL157817.1 | 13 | novel transcript |
| PRELID3BP2 | 13 | PRELI domain containing 3B pseudogene 2 [Sc |
| FAM124A | 13 | family with sequence similarity 124 member / |
| SERPINE3 | 13 | serpin family E member 3 [Source:HGNC Symb |
| MIR5693 | 13 | microRNA 5693 [Source:HGNC Symbol;Acc:HG |
| INTS6 | 13 | integrator complex subunit 6 [Source:HGNC S |
| INTS6-AS1 | 13 | INTS6 antisense RNA 1 [Source:HGNC Symbol; |
| RPS4XP16 | 13 | ribosomal protein S4X pseudogene 16 [Source |

| | | |
|---|---|---|
| RN7SL320P | 13 | RNA, 7SL, cytoplasmic 320, pseudogene [Sour |
| SNRPGP11 | 13 | small nuclear ribonucleoprotein polypeptide ( |
| MIR4703 | 13 | microRNA 4703 [Source:HGNC Symbol;Acc:HG |
| RNU6-65P | 13 | RNA, U6 small nuclear 65, pseudogene [Sourc |
| WDFY2 | 13 | WD repeat and FYVE domain containing 2 [Sou |
| RNY1P6 | 13 | RNY1 pseudogene 6 [Source:HGNC Symbol;Ac |
| RN7SL413P | 13 | RNA, 7SL, cytoplasmic 413, pseudogene [Sour |
| ATP5PBP1 | 13 | ATP synthase peripheral stalk-membrane subu |
| AL162377.3 | 13 | novel transcript |
| DHRS12 | 13 | dehydrogenase/reductase 12 [Source:HGNC S |
| AL162377.1 | 13 | novel transcript |
| TMEM272 | 13 | transmembrane protein 272 [Source:HGNC Sy |
| CCDC70 | 13 | coiled-coil domain containing 70 [Source:HGN |
| CTAGE3P | 13 | CTAGE family member 3, pseudogene [Source |
| AL162377.2 | 13 | TEC |
| ATP7B | 13 | ATPase copper transporting beta [Source:HGN |
| FABP5P2 | 13 | fatty acid binding protein 5 pseudogene 2 [Sou |
| ALG11 | 13 | ALG11 alpha-1,2-mannosyltransferase [Source |
| UTP14C | 13 | UTP14C small subunit processome componen |
| NEK5 | 13 | NIMA related kinase 5 [Source:HGNC Symbol; |
| AL139082.1 | 13 | novel transcript, antisense to NEK3 & NEK5 |
| NEK3 | 13 | NIMA related kinase 3 [Source:HGNC Symbol; |
| AL158066.1 | 13 | novel transcript |
| MRPS31P5 | 13 | mitochondrial ribosomal protein S31 pseudog |
| AL139082.2 | 13 | thrombospondin type 1 domain containing 1 ( |
| AL158066.2 | 13 | SGT1 homolog, MIS12 kinetochore complex as |
| TPTE2P2 | 13 | transmembrane phosphoinositide 3-phosphat |
| LINC02333 | 13 | long intergenic non-protein coding RNA 2333 |
| THSD1 | 13 | thrombospondin type 1 domain containing 1 [ |
| RNY4P24 | 13 | RNY4 pseudogene 24 [Source:HGNC Symbol;A |
| VPS36 | 13 | vacuolar protein sorting 36 homolog [Source:I |
| AL359513.1 | 13 | novel transcript |
| CKAP2 | 13 | cytoskeleton associated protein 2 [Source:HG |
| LINC00345 | 13 | long intergenic non-protein coding RNA 345 [S |
| AL137058.1 | 13 | transmembrane phosphatase with tensin hom |
| AL137058.3 | 13 | TEC |
| AL137058.2 | 13 | novel transcript |
| MRPS31P4 | 13 | mitochondrial ribosomal protein S31 pseudog |
| HNRNPA1L2 | 13 | heterogeneous nuclear ribonucleoprotein A1 |
| AL139089.1 | 13 | novel transcript |
| SUGT1 | 13 | SGT1 homolog, MIS12 kinetochore complex as |
| CNMD | 13 | chondromodulin [Source:HGNC Symbol;Acc:H |
| MIR759 | 13 | microRNA 759 [Source:HGNC Symbol;Acc:HGN |
| PPIAP26 | 13 | peptidylprolyl isomerase A pseudogene 26 [Sc |
| PCDH8 | 13 | protocadherin 8 [Source:HGNC Symbol;Acc:HG |
| OLFM4 | 13 | olfactomedin 4 [Source:HGNC Symbol;Acc:HG |
| LINC01065 | 13 | long intergenic non-protein coding RNA 1065 |
| AL136359.1 | 13 | novel transcript |
| PCDH8P1 | 13 | protocadherin 8 pseudogene 1 [Source:HGNC |
| AL356295.1 | 13 | novel transcript |

| | | |
|---|---|---|
| RN7SL618P | 13 | RNA, 7SL, cytoplasmic 618, pseudogene [Sour |
| AL450423.2 | 13 | novel transcript |
| AL450423.1 | 13 | novel transcript |
| ZNF646P1 | 13 | zinc finger protein 646 pseudogene 1 [Source: |
| LINC00558 | 13 | long intergenic non-protein coding RNA 558 [S |
| LINC00458 | 13 | long intergenic non-protein coding RNA 458 [S |
| AL356052.1 | 13 | chromosome X open reading frame 40A (CXor |
| MIR1297 | 13 | microRNA 1297 [Source:HGNC Symbol;Acc:HG |
| RPL13AP25 | 13 | ribosomal protein L13a pseudogene 25 [Sourc |
| AL442636.1 | 13 | novel transcript |
| AL512655.1 | 13 | novel transcript |
| AL390964.1 | 13 | novel transcript |
| LINC02335 | 13 | long intergenic non-protein coding RNA 2335 |
| AL139038.1 | 13 | NULL |
| MIR5007 | 13 | microRNA 5007 [Source:HGNC Symbol;Acc:HG |
| AL354821.1 | 13 | novel transcript |
| HNF4GP1 | 13 | hepatocyte nuclear factor 4 gamma pseudoge |
| AL138997.1 | 13 | novel transcript |
| SPATA2P1 | 13 | spermatogenesis associated 2 pseudogene 1 [ |
| RN7SKP6 | 13 | RN7SK pseudogene 6 [Source:HGNC Symbol;A |
| PRR20A | 13 | proline rich 20A [Source:HGNC Symbol;Acc:HG |
| PRR20C | 13 | proline rich 20C [Source:HGNC Symbol;Acc:HG |
| PRR20B | 13 | proline rich 20B [Source:HGNC Symbol;Acc:HG |
| PRR20D | 13 | proline rich 20D [Source:HGNC Symbol;Acc:HG |
| PRR20E | 13 | proline rich 20E [Source:HGNC Symbol;Acc:HG |
| PRR20FP | 13 | proline rich 20F, pseudogene [Source:HGNC Sy |
| MTCO2P3 | 13 | MT-CO2 pseudogene 3 [Source:HGNC Symbol |
| SLC25A5P4 | 13 | solute carrier family 25 member 5 pseudogen |
| RPL31P53 | 13 | ribosomal protein L31 pseudogene 53 [Source |
| PCDH17 | 13 | protocadherin 17 [Source:HGNC Symbol;Acc:H |
| AL445288.1 | 13 | novel transcript, antisense to PCDH17 |
| RNA5SP30 | 13 | RNA, 5S ribosomal pseudogene 30 [Source:HG |
| LINC02338 | 13 | long intergenic non-protein coding RNA 2338 |
| LINC00374 | 13 | long intergenic non-protein coding RNA 374 [S |
| AL161423.1 | 13 | exosome component 3 (EXOSC3) |
| RNY4P29 | 13 | RNY4 pseudogene 29 [Source:HGNC Symbol;A |
| CTAGE16P | 13 | CTAGE family member 16, pseudogene [Sourc |
| DNAJA1P1 | 13 | DnaJ heat shock protein family (Hsp40) memb |
| HMGN2P39 | 13 | high mobility group nucleosomal binding dom |
| POLR3KP1 | 13 | RNA polymerase III subunit K pseudogene 1 [S |
| AL159156.1 | 13 | uncharacterized LOC105370224 [Source:NCBI |
| RPP40P2 | 13 | ribonuclease P/MRP subunit p40 pseudogene |
| RNU7-88P | 13 | RNA, U7 small nuclear 88 pseudogene [Source |
| DIAPH3 | 13 | diaphanous related formin 3 [Source:HGNC Sy |
| DIAPH3-AS1 | 13 | DIAPH3 antisense RNA 1 [Source:HGNC Symbo |
| RN7SL375P | 13 | RNA, 7SL, cytoplasmic 375, pseudogene [Sour |
| DIAPH3-AS2 | 13 | DIAPH3 antisense RNA 2 [Source:HGNC Symbo |
| RNY4P28 | 13 | RNY4 pseudogene 28 [Source:HGNC Symbol;A |
| LINC00434 | 13 | long intergenic non-protein coding RNA 434 [S |
| TARDBPP2 | 13 | TAR DNA binding protein pseudogene 2 [Sourc |

| | | |
|---|---|---|
| AL359920.1 | 13 | phosphoglycerate mutase 1 (brain) (PGAM1) p |
| TDRD3 | 13 | tudor domain containing 3 [Source:HGNC Sym |
| RNA5SP31 | 13 | RNA, 5S ribosomal pseudogene 31 [Source:HG |
| LINC00378 | 13 | long intergenic non-protein coding RNA 378 [S |
| EIF4A1P6 | 13 | eukaryotic translation initiation factor 4A1 pse |
| AL161901.1 | 13 | TEC |
| RNY3P5 | 13 | RNY3 pseudogene 5 [Source:HGNC Symbol;Ac |
| LINC01442 | 13 | long intergenic non-protein coding RNA 1442 |
| MIR3169 | 13 | microRNA 3169 [Source:HGNC Symbol;Acc:HG |
| PCDH20 | 13 | protocadherin 20 [Source:HGNC Symbol;Acc:H |
| AL592490.1 | 13 | novel protein |
| LINC02339 | 13 | long intergenic non-protein coding RNA 2339 |
| RAC1P8 | 13 | Rac family small GTPase 1 pseudogene 8 [Sou |
| AL353765.1 | 13 | cyclin-dependent kinase 2 associated protein |
| LINC00358 | 13 | long intergenic non-protein coding RNA 358 [S |
| LINC01075 | 13 | long intergenic non-protein coding RNA 1075 |
| LINC01074 | 13 | long intergenic non-protein coding RNA 1074 |
| LINC00459 | 13 | long intergenic non-protein coding RNA 459 [S |
| RPL32P28 | 13 | ribosomal protein L32 pseudogene 28 [Source |
| AL356102.1 | 13 | sequestosome 1 (SQSTM1) pseudogene |
| SQSTM1P1 | 13 | sequestosome 1 pseudogene 1 [Source:HGNC |
| AL445668.1 | 13 | novel transcript |
| LINC00448 | 13 | long intergenic non-protein coding RNA 448 [S |
| AL354810.1 | 13 | novel transcript |
| LINC00376 | 13 | long intergenic non-protein coding RNA 376 [S |
| AL359208.1 | 13 | novel transcript |
| LINC00395 | 13 | long intergenic non-protein coding RNA 395 [S |
| OR7E156P | 13 | olfactory receptor family 7 subfamily E memb |
| AL445989.2 | 13 | novel transcript |
| AL445989.1 | 13 | keratin-associated protein 21-1-like [Source:N |
| RNU6-81P | 13 | RNA, U6 small nuclear 81, pseudogene [Sourc |
| PPP1R2P10 | 13 | protein phosphatase 1 regulatory inhibitor sul |
| AL445238.2 | 13 | novel transcript |
| OR7E104P | 13 | olfactory receptor family 7 subfamily E memb |
| AL445238.1 | 13 | keratin-associated protein 21-1 [Source:NCBI |
| LINC00355 | 13 | long intergenic non-protein coding RNA 355 [S |
| NFYAP1 | 13 | nuclear transcription factor Y subunit alpha ps |
| LGMNP1 | 13 | legumain pseudogene 1 [Source:HGNC Symbo |
| STARP1 | 13 | steroidogenic acute regulatory protein pseud |
| HNRNPA3P5 | 13 | heterogeneous nuclear ribonucleoprotein A3 |
| LINC01052 | 13 | long intergenic non-protein coding RNA 1052 |
| AL158038.1 | 13 | novel transcript |
| MIR548X2 | 13 | microRNA 548x-2 [Source:HGNC Symbol;Acc:H |
| MIR4704 | 13 | microRNA 4704 [Source:HGNC Symbol;Acc:HG |
| TRIM60P19 | 13 | tripartite motif containing 60 pseudogene 19 |
| PCDH9 | 13 | protocadherin 9 [Source:HGNC Symbol;Acc:H |
| PCDH9-AS1 | 13 | PCDH9 antisense RNA 1 [Source:HGNC Symbo |
| RNU7-87P | 13 | RNA, U7 small nuclear 87 pseudogene [Source |
| PCDH9-AS2 | 13 | PCDH9 antisense RNA 2 [Source:HGNC Symbo |
| PCDH9-AS3 | 13 | PCDH9 antisense RNA 3 [Source:HGNC Symbo |

| | | |
|---|---|---|
| PCDH9-AS4 | 13 | PCDH9 antisense RNA 4 [Source:HGNC Symbo |
| AL160254.1 | 13 | novel transcript, antisense to PCDH9 |
| RPSAP53 | 13 | ribosomal protein SA pseudogene 53 [Source: |
| LINC00364 | 13 | long intergenic non-protein coding RNA 364 [S |
| AL512452.1 | 13 | novel transcript |
| BCRP9 | 13 | BCR pseudogene 9 [Source:HGNC Symbol;Acc |
| NPM1P22 | 13 | nucleophosmin 1 pseudogene 22 [Source:HGN |
| OR7E111P | 13 | olfactory receptor family 7 subfamily E memb |
| OR7E33P | 13 | olfactory receptor family 7 subfamily E memb |
| AL590027.1 | 13 | novel transcript |
| ELL2P3 | 13 | elongation factor for RNA polymerase II 2 pse |
| HNRNPA1P18 | 13 | heterogeneous nuclear ribonucleoprotein A1 |
| RPL37P21 | 13 | ribosomal protein L37 pseudogene 21 [Source |
| RPS3AP52 | 13 | ribosomal protein S3a pseudogene 52 [Source |
| RPL12P34 | 13 | ribosomal protein L12 pseudogene 34 [Source |
| RN7SL761P | 13 | RNA, 7SL, cytoplasmic 761, pseudogene [Sour |
| LINC00550 | 13 | long intergenic non-protein coding RNA 550 [S |
| LINC02342 | 13 | long intergenic non-protein coding RNA 2342 |
| ZDHHC20P4 | 13 | zinc finger DHHC-type containing 20 pseudoge |
| SNRPFP3 | 13 | small nuclear ribonucleoprotein polypeptide F |
| LINC00383 | 13 | long intergenic non-protein coding RNA 383 [S |
| SRSF1P1 | 13 | serine and arginine rich splicing factor 1 pseu |
| KLHL1 | 13 | kelch like family member 1 [Source:HGNC Sym |
| RNY3P10 | 13 | RNY3 pseudogene 10 [Source:HGNC Symbol;A |
| PSMC1P13 | 13 | proteasome 26S subunit, ATPase 1 pseudoger |
| ATXN8OS | 13 | ATXN8 opposite strand lncRNA [Source:HGNC |
| AL160391.1 | 13 | ataxin 8 |
| RNU6-54P | 13 | RNA, U6 small nuclear 54, pseudogene [Sourc |
| AL445264.1 | 13 | SYF2 pre-mRNA-splicing factor (SYF2) pseudog |
| MTCL1P1 | 13 | MTCL1 pseudogene 1 [Source:HGNC Symbol;A |
| LINC00348 | 13 | long intergenic non-protein coding RNA 348 [S |
| RABEPKP1 | 13 | Rab9 effector protein with kelch motifs pseud |
| DACH1 | 13 | dachshund family transcription factor 1 [Sourc |
| H3F3BP1 | 13 | H3 histone, family 3B pseudogene 1 [Source:H |
| RPL21P109 | 13 | ribosomal protein L21 pseudogene 109 [Sourc |
| RPL35AP31 | 13 | ribosomal protein L35a pseudogene 31 [Sourc |
| RPS10P21 | 13 | ribosomal protein S10 pseudogene 21 [Source |
| AL139003.1 | 13 | NULL |
| RPL18AP17 | 13 | ribosomal protein L18a pseudogene 17 [Sourc |
| AL139003.2 | 13 | NULL |
| AL356754.2 | 13 | NULL |
| AL356754.1 | 13 | NULL |
| RPL21P110 | 13 | ribosomal protein L21 pseudogene 110 [Sourc |
| RNU6-80P | 13 | RNA, U6 small nuclear 80, pseudogene [Sourc |
| MZT1 | 13 | mitotic spindle organizing protein 1 [Source:H |
| BORA | 13 | BORA aurora kinase A activator [Source:HGNC |
| DIS3 | 13 | DIS3 homolog, exosome endoribonuclease an |
| PIBF1 | 13 | progesterone immunomodulatory binding fac |
| RNU6-79P | 13 | RNA, U6 small nuclear 79, pseudogene [Sourc |
| PSMD10P3 | 13 | proteasome 26S subunit, non-ATPase, 10 pseu |

| | | |
|---|---|---|
| KLF5 | 13 | Kruppel like factor 5 [Source:HGNC Symbol;Ac |
| FABP5P1 | 13 | fatty acid binding protein 5 pseudogene 1 [So |
| RNU6-66P | 13 | RNA, U6 small nuclear 66, pseudogene [Sourc |
| RNU4-10P | 13 | RNA, U4 small nuclear 10, pseudogene [Sourc |
| RNY1P8 | 13 | RNY1 pseudogene 8 [Source:HGNC Symbol;Ac |
| AL162376.1 | 13 | TEC |
| MARK2P12 | 13 | microtubule affinity regulating kinase 2 pseud |
| LINC00393 | 13 | long intergenic non-protein coding RNA 393 [S |
| LINC00392 | 13 | long intergenic non-protein coding RNA 392 [S |
| KLF12 | 13 | Kruppel like factor 12 [Source:HGNC Symbol;A |
| AL138713.1 | 13 | TEC |
| AL159972.1 | 13 | novel transcript, sense intronic to KLF12 |
| LINC00402 | 13 | long intergenic non-protein coding RNA 402 [S |
| AL353660.1 | 13 | novel transcript |
| RNY1P5 | 13 | RNY1 pseudogene 5 [Source:HGNC Symbol;Ac |
| RPL21P108 | 13 | ribosomal protein L21 pseudogene 108 [Sourc |
| AL355390.1 | 13 | novel transcript |
| LINC00381 | 13 | long intergenic non-protein coding RNA 381 [S |
| AL355390.2 | 13 | novel transcript |
| LINC00347 | 13 | long intergenic non-protein coding RNA 347 [S |
| RIOK3P1 | 13 | RIO kinase 3 pseudogene 1 [Source:HGNC Sym |
| RNU6-38P | 13 | RNA, U6 small nuclear 38, pseudogene [Sourc |
| SSR1P2 | 13 | signal sequence receptor subunit 1 pseudoger |
| CTAGE11P | 13 | CTAGE family member 11, pseudogene [Sourc |
| LINC01078 | 13 | long intergenic non-protein coding RNA 1078 |
| TBC1D4 | 13 | TBC1 domain family member 4 [Source:HGNC |
| AL139230.1 | 13 | novel transcript |
| COMMD6 | 13 | COMM domain containing 6 [Source:HGNC Sy |
| UCHL3 | 13 | ubiquitin C-terminal hydrolase L3 [Source:HGI |
| AL137782.1 | 13 | novel transcript |
| LMO7-AS1 | 13 | LMO7 antisense RNA 1 [Source:HGNC Symbol |
| LMO7 | 13 | LIM domain 7 [Source:HGNC Symbol;Acc:HGN |
| AL137244.1 | 13 | novel transcript, sense intronic to LMO7 |
| FAM204CP | 13 | family with sequence similarity 204 member C |
| LMO7DN | 13 | LMO7 downstream neighbor [Source:HGNC Sy |
| LMO7DN-IT1 | 13 | LMO7DN intronic transcript 1 [Source:HGNC S |
| LINC00561 | 13 | long intergenic non-protein coding RNA 561 [S |
| LINC01034 | 13 | long intergenic non-protein coding RNA 1034 |
| RN7SL571P | 13 | RNA, 7SL, cytoplasmic 571, pseudogene [Sour |
| AL136441.1 | 13 | novel transcript |
| AL161717.1 | 13 | novel transcript |
| AL365394.1 | 13 | novel transcript |
| KCTD12 | 13 | potassium channel tetramerization domain cc |
| AC000403.1 | 13 | novel transcript |
| BTF3P11 | 13 | basic transcription factor 3 pseudogene 11 [Sc |
| ACOD1 | 13 | aconitate decarboxylase 1 [Source:HGNC Sym |
| RPL7P44 | 13 | ribosomal protein L7 pseudogene 44 [Source: |
| DHX9P1 | 13 | DEAH-box helicase 9 pseudogene 1 [Source:H |
| CLN5 | 13 | CLN5 intracellular trafficking protein [Source:I |
| AC001226.2 | 13 | novel protein |

| | | |
|---|---|---|
| FBXL3 | 13 | F-box and leucine rich repeat protein 3 [Sourc |
| MYCBP2-AS1 | 13 | MYCBP2 antisense RNA 1 [Source:HGNC Symb |
| AC001226.1 | 13 | novel transcript |
| MYCBP2 | 13 | MYC binding protein 2 [Source:HGNC Symbol; |
| MYCBP2-AS2 | 13 | MYCBP2 antisense RNA 2 [Source:HGNC Symb |
| AL159158.1 | 13 | novel transcript |
| SCEL | 13 | sciellin [Source:HGNC Symbol;Acc:HGNC:1057 |
| AL137140.1 | 13 | novel transcript, antisense to SCEL |
| SCEL-AS1 | 13 | SCEL antisense RNA 1 [Source:HGNC Symbol;/ |
| RNY3P7 | 13 | RNY3 pseudogene 7 [Source:HGNC Symbol;Ac |
| SPTLC1P5 | 13 | serine palmitoyltransferase long chain base su |
| AL450447.1 | 13 | putative UPF0607 protein ENSP00000383144 |
| SLAIN1 | 13 | SLAIN motif family member 1 [Source:HGNC S |
| MIR3665 | 13 | microRNA 3665 [Source:HGNC Symbol;Acc:HG |
| EDNRB-AS1 | 13 | EDNRB antisense RNA 1 [Source:HGNC Symbo |
| EDNRB | 13 | endothelin receptor type B [Source:HGNC Syn |
| OBI1-AS1 | 13 | OBI1 antisense RNA 1 [Source:HGNC Symbol;/ |
| AL139002.1 | 13 | novel transcript |
| RN7SL810P | 13 | RNA, 7SL, cytoplasmic 810, pseudogene [Sour |
| LINC01069 | 13 | long intergenic non-protein coding RNA 1069 |
| LINC00446 | 13 | long intergenic non-protein coding RNA 446 [S |
| AL138957.1 | 13 | novel transcript |
| SRGNP1 | 13 | serglycin pseudogene 1 [Source:HGNC Symbo |
| RNY3P3 | 13 | RNY3 pseudogene 3 [Source:HGNC Symbol;Ac |
| RPL31P54 | 13 | ribosomal protein L31 pseudogene 54 [Source |
| ELOCP23 | 13 | elongin C pseudogene 23 [Source:HGNC Symb |
| AL445209.1 | 13 | novel transcript, antisense to POU4F1 |
| POU4F1 | 13 | POU class 4 homeobox 1 [Source:HGNC Symb |
| OBI1 | 13 | ORC ubiquitin ligase 1 [Source:HGNC Symbol;/ |
| RPL21P111 | 13 | ribosomal protein L21 pseudogene 111 [Sourc |
| LINC00331 | 13 | long intergenic non-protein coding RNA 331 [S |
| HSPD1P8 | 13 | heat shock protein family D (Hsp60) member |
| CCT5P2 | 13 | chaperonin containing TCP1 subunit 5 pseudo |
| NIPA2P5 | 13 | NIPA magnesium transporter 2 pseudogene 5 |
| BCAS2P3 | 13 | BCAS2 pseudogene 3 [Source:HGNC Symbol;A |
| RBM26 | 13 | RNA binding motif protein 26 [Source:HGNC S |
| RNA5SP33 | 13 | RNA, 5S ribosomal pseudogene 33 [Source:HG |
| RBM26-AS1 | 13 | RBM26 antisense RNA 1 [Source:HGNC Symbo |
| NDFIP2-AS1 | 13 | NDFIP2 antisense RNA 1 [Source:HGNC Symbo |
| NDFIP2 | 13 | Nedd4 family interacting protein 2 [Source:HG |
| LINC01068 | 13 | long intergenic non-protein coding RNA 1068 |
| AL136442.1 | 13 | novel transcript |
| LINC01038 | 13 | long intergenic non-protein coding RNA 1038 |
| LINC00382 | 13 | long intergenic non-protein coding RNA 382 [S |
| AL158064.1 | 13 | novel transcript |
| LINC01080 | 13 | long intergenic non-protein coding RNA 1080 |
| AL137781.1 | 13 | novel transcript |
| SPRY2 | 13 | sprouty RTK signaling antagonist 2 [Source:HG |
| RNU6-61P | 13 | RNA, U6 small nuclear 61, pseudogene [Sourc |
| HNRNPA1P31 | 13 | heterogeneous nuclear ribonucleoprotein A1 |

| | | |
|---|---|---|
| PWWP2AP1 | 13 | PWWP domain containing 2A pseudogene 1 [S |
| AL590807.1 | 13 | novel transcript |
| ARF4P4 | 13 | ADP ribosylation factor 4 pseudogene 4 [Sour |
| LINC00377 | 13 | long intergenic non-protein coding RNA 377 [S |
| DPPA3P3 | 13 | developmental pluripotency associated 3 pseu |
| LINC00564 | 13 | long intergenic non-protein coding RNA 564 [S |
| AL353633.1 | 13 | novel transcript |
| RNU6-77P | 13 | RNA, U6 small nuclear 77, pseudogene [Sourc |
| HIGD1AP2 | 13 | HIG1 hypoxia inducible domain family membe |
| PTMAP5 | 13 | prothymosin alpha pseudogene 5 [Source:HGI |
| AL445255.1 | 13 | novel transcript |
| GYG1P2 | 13 | glycogenin 1 pseudogene 2 [Source:HGNC Syn |
| AL512782.1 | 13 | ubiquitin specific peptidase 7 (herpes virus-as |
| RNU6-67P | 13 | RNA, U6 small nuclear 67, pseudogene [Sourc |
| AL353688.1 | 13 | novel transcript |
| SLITRK1 | 13 | SLIT and NTRK like family member 1 [Source:H |
| AL355481.1 | 13 | novel transcript, antisense to SLITRK1 |
| VENTXP2 | 13 | VENT homeobox pseudogene 2 [Source:HGNC |
| AL590681.1 | 13 | novel transcript |
| UBE2D3P4 | 13 | ubiquitin conjugating enzyme E2 D3 pseudoge |
| AL445604.1 | 13 | novel transcript |
| AL162493.1 | 13 | novel transcript |
| MTND4P1 | 13 | MT-ND4 pseudogene 1 [Source:HGNC Symbol |
| MTND5P3 | 13 | MT-ND5 pseudogene 3 [Source:HGNC Symbol |
| LINC00333 | 13 | long intergenic non-protein coding RNA 333 [S |
| AL445588.1 | 13 | novel transcript |
| AL356313.1 | 13 | novel transcript |
| LINC00375 | 13 | long intergenic non-protein coding RNA 375 [S |
| AL160154.1 | 13 | novel transcript |
| LINC00351 | 13 | long intergenic non-protein coding RNA 351 [S |
| AL356413.1 | 13 | novel transcript |
| SLITRK6 | 13 | SLIT and NTRK like family member 6 [Source:H |
| AL162373.1 | 13 | piggyBac transposable element derived 4 (PGI |
| MOB1AP1 | 13 | MOB kinase activator 1A pseudogene 1 [Sourc |
| AL354994.1 | 13 | novel transcript |
| FO624990.1 | 13 | IQ motif containing K (IQCK) pseudogene |
| DDX6P2 | 13 | DEAD-box helicase 6 pseudogene 2 [Source:H |
| AL445207.1 | 13 | novel transcript |
| TXNL1P1 | 13 | thioredoxin like 1 pseudogene 1 [Source:HGN |
| LINC00430 | 13 | long intergenic non-protein coding RNA 430 [S |
| UBBP5 | 13 | ubiquitin B pseudogene 5 [Source:HGNC Symb |
| LIN28AP2 | 13 | LIN28A pseudogene 2 [Source:HGNC Symbol;A |
| AL360267.2 | 13 | novel transcript |
| AL360267.1 | 13 | novel transcript |
| MIR4500HG | 13 | MIR4500 host gene [Source:HGNC Symbol;Acc |
| AL355578.1 | 13 | novel transcript |
| MIR4500 | 13 | microRNA 4500 [Source:HGNC Symbol;Acc:HG |
| SLITRK5 | 13 | SLIT and NTRK like family member 5 [Source:H |
| AL445647.2 | 13 | nudix (nucleoside diphosphate linked moiety X |
| LINC00397 | 13 | long intergenic non-protein coding RNA 397 [S |

| | | |
|---|---|---|
| AL445647.1 | 13 | novel transcript |
| TET1P1 | 13 | tet methylcytosine dioxygenase 1 pseudogene |
| LINC00373 | 13 | long intergenic non-protein coding RNA 373 [S |
| RPL29P29 | 13 | ribosomal protein L29 pseudogene 29 [Source |
| AL355677.1 | 13 | novel transcript |
| LINC00433 | 13 | long intergenic non-protein coding RNA 433 [S |
| AL445242.1 | 13 | novel transcript |
| AL445984.1 | 13 | mitochondrially encoded NADH dehydrogenas |
| LINC00560 | 13 | long intergenic non-protein coding RNA 560 [S |
| GRPEL2P1 | 13 | GRPEL2 pseudogene 1 [Source:HGNC Symbol; |
| LINC00440 | 13 | long intergenic non-protein coding RNA 440 [S |
| LINC01047 | 13 | long intergenic non-protein coding RNA 1047 |
| TRIM60P13 | 13 | tripartite motif containing 60 pseudogene 13 |
| SP3P | 13 | Sp3 transcription factor pseudogene [Source:H |
| LINC02336 | 13 | long intergenic non-protein coding RNA 2336 |
| LINC01040 | 13 | long intergenic non-protein coding RNA 1040 |
| LINC00353 | 13 | long intergenic non-protein coding RNA 353 [S |
| RPL7L1P1 | 13 | ribosomal protein L7 like 1 pseudogene 1 [Sou |
| AL355821.1 | 13 | NULL |
| PEX12P1 | 13 | peroxisomal biogenesis factor 12 pseudogene |
| LINC00559 | 13 | long intergenic non-protein coding RNA 559 [S |
| RNA5SP34 | 13 | RNA, 5S ribosomal pseudogene 34 [Source:HG |
| FAR1P1 | 13 | fatty acyl-CoA reductase 1 pseudogene 1 [Sou |
| KRT18P27 | 13 | keratin 18 pseudogene 27 [Source:HGNC Sym |
| MIR622 | 13 | microRNA 622 [Source:HGNC Symbol;Acc:HGN |
| LINC01049 | 13 | long intergenic non-protein coding RNA 1049 |
| RNU6-75P | 13 | RNA, U6 small nuclear 75, pseudogene [Sourc |
| LINC00410 | 13 | long intergenic non-protein coding RNA 410 [S |
| BRK1P2 | 13 | BRK1 pseudogene 2 [Source:HGNC Symbol;Ac |
| LINC00380 | 13 | long intergenic non-protein coding RNA 380 [S |
| LINC00379 | 13 | long intergenic non-protein coding RNA 379 [S |
| PPIAP23 | 13 | peptidylprolyl isomerase A pseudogene 23 [So |
| MIR17HG | 13 | miR-17-92a-1 cluster host gene [Source:HGNC |
| MIR17 | 13 | microRNA 17 [Source:HGNC Symbol;Acc:HGN |
| MIR18A | 13 | microRNA 18a [Source:HGNC Symbol;Acc:HGN |
| MIR19A | 13 | microRNA 19a [Source:HGNC Symbol;Acc:HGN |
| MIR20A | 13 | microRNA 20a [Source:HGNC Symbol;Acc:HGN |
| MIR19B1 | 13 | microRNA 19b-1 [Source:HGNC Symbol;Acc:H |
| MIR92A1 | 13 | microRNA 92a-1 [Source:HGNC Symbol;Acc:H |
| GPC5 | 13 | glypican 5 [Source:HGNC Symbol;Acc:HGNC:4 |
| AL356118.1 | 13 | TEC |
| AL162456.1 | 13 | novel transcript |
| RNU4ATAC3P | 13 | RNA, U4atac small nuclear 3, pseudogene [So |
| FABP5P4 | 13 | fatty acid binding protein 5 pseudogene 4 [So |
| AL157815.1 | 13 | TEC |
| GPC5-AS2 | 13 | GPC5 antisense RNA 2 [Source:HGNC Symbol; |
| AL159152.1 | 13 | TEC |
| GPC5-IT1 | 13 | GPC5 intronic transcript 1 [Source:HGNC Symb |
| MIR548AS | 13 | microRNA 548as [Source:HGNC Symbol;Acc:H |
| AL356737.2 | 13 | novel transcript, antisense to GPC5 |

| | | |
|---|---|---|
| AL356737.1 | 13 | TEC |
| GPC5-AS1 | 13 | GPC5 antisense RNA 1 [Source:HGNC Symbol; |
| LINC00363 | 13 | long intergenic non-protein coding RNA 363 [S |
| AL354811.2 | 13 | novel transcript |
| AL354811.1 | 13 | novel transcript, antisense to GPC6 |
| GPC6 | 13 | glypican 6 [Source:HGNC Symbol;Acc:HGNC:4 |
| AL160159.1 | 13 | novel transcript, antisense to GPC6 |
| HNRNPA1P29 | 13 | heterogeneous nuclear ribonucleoprotein A1 |
| GPC6-AS2 | 13 | GPC6 antisense RNA 2 [Source:HGNC Symbol; |
| RNA5SP35 | 13 | RNA, 5S ribosomal pseudogene 35 [Source:HG |
| GPC6-AS1 | 13 | GPC6 antisense RNA 1 [Source:HGNC Symbol; |
| DCT | 13 | dopachrome tautomerase [Source:HGNC Sym |
| TGDS | 13 | TDP-glucose 4,6-dehydratase [Source:HGNC S |
| GPR180 | 13 | G protein-coupled receptor 180 [Source:HGN |
| RNA5SP36 | 13 | RNA, 5S ribosomal pseudogene 36 [Source:HG |
| LINC00391 | 13 | long intergenic non-protein coding RNA 391 [S |
| RN7SL585P | 13 | RNA, 7SL, cytoplasmic 585, pseudogene [Sour |
| SOX21-AS1 | 13 | SOX21 antisense divergent transcript 1 [Sourc |
| SOX21 | 13 | SRY-box transcription factor 21 [Source:HGNC |
| BRD7P5 | 13 | bromodomain containing 7 pseudogene 5 [So |
| RPL21P112 | 13 | ribosomal protein L21 pseudogene 112 [Sourc |
| LINC00557 | 13 | long intergenic non-protein coding RNA 557 [S |
| AL139381.1 | 13 | novel transcript |
| RNU6-62P | 13 | RNA, U6 small nuclear 62, pseudogene [Sourc |
| ABCC4 | 13 | ATP binding cassette subfamily C member 4 [S |
| RNY3P8 | 13 | RNY3 pseudogene 8 [Source:HGNC Symbol;Ac |
| RNY4P27 | 13 | RNY4 pseudogene 27 [Source:HGNC Symbol;A |
| MEMO1P5 | 13 | mediator of cell motility 1 pseudogene 5 [Sou |
| CLDN10 | 13 | claudin 10 [Source:HGNC Symbol;Acc:HGNC:2 |
| CLDN10-AS1 | 13 | CLDN10 antisense RNA 1 [Source:HGNC Symb |
| DZIP1 | 13 | DAZ interacting zinc finger protein 1 [Source:H |
| DNAJC3-DT | 13 | DNAJC3 divergent transcript [Source:HGNC Sy |
| DNAJC3 | 13 | DnaJ heat shock protein family (Hsp40) memb |
| MTND5P2 | 13 | MT-ND5 pseudogene 2 [Source:HGNC Symbol |
| MTND6P18 | 13 | MT-ND6 pseudogene 18 [Source:HGNC Symbo |
| MTCYBP3 | 13 | MT-CYB pseudogene 3 [Source:HGNC Symbol; |
| AL138955.1 | 13 | novel transcript, sense intronic to DNAJC3 |
| UGGT2 | 13 | UDP-glucose glycoprotein glucosyltransferase |
| HMGN1P24 | 13 | high mobility group nucleosome binding doma |
| HS6ST3 | 13 | heparan sulfate 6-O-sulfotransferase 3 [Sourc |
| RN7SL164P | 13 | RNA, 7SL, cytoplasmic 164, pseudogene [Sour |
| MIR4501 | 13 | microRNA 4501 [Source:HGNC Symbol;Acc:HG |
| AMMECR1LP1 | 13 | AMMECR1 like pseudogene 1 [Source:HGNC S |
| HSP90AB6P | 13 | heat shock protein 90 alpha family class B me |
| TULP3P1 | 13 | TUB like protein 3 pseudogene 1 [Source:HGN |
| AL353581.1 | 13 | novel transcript |
| RN7SKP7 | 13 | RN7SK pseudogene 7 [Source:HGNC Symbol;A |
| OXGR1 | 13 | oxoglutarate receptor 1 [Source:HGNC Symbo |
| LINC00456 | 13 | long intergenic non-protein coding RNA 456 [S |
| MBNL2 | 13 | muscleblind like splicing regulator 2 [Source:H |

| | | |
|---|---|---|
| AL161430.1 | 13 | novel transcript, antisense to MBNL2 |
| RNA5SP37 | 13 | RNA, 5S ribosomal pseudogene 37 [Source:HG |
| AL442067.3 | 13 | novel transcript |
| RAP2A | 13 | RAP2A, member of RAS oncogene family [Sou |
| AL442067.1 | 13 | novel transcript, sense intronic to RAP2A |
| AL442067.2 | 13 | novel transcript, sense intronic to RAP2A |
| PSMA6P4 | 13 | proteasome subunit alpha 6 pseudogene 4 [Sc |
| AL359502.1 | 13 | novel transcript |
| AL359502.2 | 13 | novel transcript |
| RPL7AP61 | 13 | ribosomal protein L7a pseudogene 61 [Source |
| AL356580.1 | 13 | novel transcript |
| IPO5 | 13 | importin 5 [Source:HGNC Symbol;Acc:HGNC:6 |
| FTLP8 | 13 | ferritin light chain pseudogene 8 [Source:HGN |
| FARP1 | 13 | FERM, ARH/RhoGEF and pleckstrin domain pr |
| RNF113B | 13 | ring finger protein 113B [Source:HGNC Symbo |
| RN7SKP8 | 13 | RN7SK pseudogene 8 [Source:HGNC Symbol;A |
| MIR3170 | 13 | microRNA 3170 [Source:HGNC Symbol;Acc:HG |
| AL445223.1 | 13 | novel transcript, sense intronic to FARP1 |
| AL161896.1 | 13 | novel transcript, antisense to FARP1 |
| FARP1-AS1 | 13 | FARP1 antisense RNA 1 [Source:HGNC Symbol |
| STK24 | 13 | serine/threonine kinase 24 [Source:HGNC Syn |
| AL356423.1 | 13 | high mobility group box 1 (HMGB1) pseudoge |
| STK24-AS1 | 13 | STK24 antisense RNA 1 [Source:HGNC Symbol |
| NUS1P4 | 13 | NUS1 pseudogene 4 [Source:HGNC Symbol;Ac |
| CYCSP35 | 13 | CYCS pseudogene 35 [Source:HGNC Symbol;A |
| CALM2P4 | 13 | calmodulin 2 pseudogene 4 [Source:HGNC Syr |
| RN7SL60P | 13 | RNA, 7SL, cytoplasmic 60, pseudogene [Sourc |
| SLC15A1 | 13 | solute carrier family 15 member 1 [Source:HG |
| DOCK9 | 13 | dedicator of cytokinesis 9 [Source:HGNC Symb |
| DOCK9-AS1 | 13 | DOCK9 antisense RNA 1 [Source:HGNC Symbo |
| AL161420.1 | 13 | keratin associated protein 3-1 (KRTAP3-1) pse |
| RPL7L1P12 | 13 | ribosomal protein L7 like 1 pseudogene 12 [Sc |
| RNU6-83P | 13 | RNA, U6 small nuclear 83, pseudogene [Sourc |
| DOCK9-DT | 13 | DOCK9 divergent transcript [Source:HGNC Syr |
| RPS6P23 | 13 | ribosomal protein S6 pseudogene 23 [Source: |
| UBAC2-AS1 | 13 | UBAC2 antisense RNA 1 [Source:HGNC Symbo |
| GAPDHP22 | 13 | glyceraldehyde 3 phosphate dehydrogenase p |
| UBAC2 | 13 | UBA domain containing 2 [Source:HGNC Symb |
| RN7SKP9 | 13 | RN7SK pseudogene 9 [Source:HGNC Symbol;A |
| H2AFZP3 | 13 | H2A histone family member Z pseudogene 3 [ |
| GPR18 | 13 | G protein-coupled receptor 18 [Source:HGNC |
| GPR183 | 13 | G protein-coupled receptor 183 [Source:HGN |
| MIR623 | 13 | microRNA 623 [Source:HGNC Symbol;Acc:HGI |
| HMGB3P4 | 13 | high mobility group box 3 pseudogene 4 [Sour |
| CCR12P | 13 | C-C motif chemokine receptor 12, pseudogen |
| AL136961.1 | 13 | novel transcript |
| TM9SF2 | 13 | transmembrane 9 superfamily member 2 [Sou |
| LINC01232 | 13 | long intergenic non-protein coding RNA 1232 |
| AL139035.1 | 13 | novel transcript |
| LINC00449 | 13 | long intergenic non-protein coding RNA 449 [S |

| | | |
|---|---|---|
| LINC01039 | 13 | long intergenic non-protein coding RNA 1039 |
| CFL1P8 | 13 | cofilin 1 pseudogene 8 [Source:HGNC Symbol; |
| AL139035.2 | 13 | novel transcript |
| CLYBL | 13 | citrate lyase beta like [Source:HGNC Symbol;A |
| MIR4306 | 13 | microRNA 4306 [Source:HGNC Symbol;Acc:HG |
| CLYBL-AS2 | 13 | CLYBL antisense RNA 2 [Source:HGNC Symbol, |
| CLYBL-AS1 | 13 | CLYBL antisense RNA 1 [Source:HGNC Symbol, |
| AL137139.1 | 13 | NULL |
| AL137139.2 | 13 | novel transcript, antisense to CLYBL |
| ZIC5 | 13 | Zic family member 5 [Source:HGNC Symbol;Ac |
| AL355338.1 | 13 | novel transcript |
| ZIC2 | 13 | Zic family member 2 [Source:HGNC Symbol;Ac |
| LINC00554 | 13 | long intergenic non-protein coding RNA 554 [S |
| NDUFA12P1 | 13 | NADH:ubiquinone oxidoreductase subunit A1 |
| PCCA-DT | 13 | PCCA divergent transcript [Source:HGNC Symb |
| ASNSP3 | 13 | asparagine synthetase pseudogene 3 [Source: |
| PCCA | 13 | propionyl-CoA carboxylase subunit alpha [Sou |
| RPL15P18 | 13 | ribosomal protein L15 pseudogene 18 [Source |
| AL356575.1 | 13 | novel pseudogene |
| AL353697.1 | 13 | novel transcript, antisense to PCCA |
| PCCA-AS1 | 13 | PCCA antisense RNA 1 [Source:HGNC Symbol; |
| GGACT | 13 | gamma-glutamylamine cyclotransferase [Sour |
| AL356966.1 | 13 | novel transcript |
| AL136526.1 | 13 | novel transcript, antisense to GGACT |
| RPS26P47 | 13 | ribosomal protein S26 pseudogene 47 [Source |
| TMTC4 | 13 | transmembrane O-mannosyltransferase targe |
| COX5BP6 | 13 | cytochrome c oxidase subunit 5B pseudogene |
| NALCN-AS1 | 13 | NALCN antisense RNA 1 [Source:HGNC Symbo |
| ARF4P3 | 13 | ADP ribosylation factor 4 pseudogene 3 [Sour |
| LINC00411 | 13 | long intergenic non-protein coding RNA 411 [S |
| NALCN | 13 | sodium leak channel, non-selective [Source:H |
| ITGBL1 | 13 | integrin subunit beta like 1 [Source:HGNC Sym |
| FGF14 | 13 | fibroblast growth factor 14 [Source:HGNC Sym |
| AL160153.1 | 13 | novel transcript, antisense to FGF14 |
| RNU1-24P | 13 | RNA, U1 small nuclear 24, pseudogene [Sourc |
| HMGB3P7 | 13 | high mobility group box 3 pseudogene 7 [Sour |
| MIR2681 | 13 | microRNA 2681 [Source:HGNC Symbol;Acc:HG |
| LIPT1P1 | 13 | lipoyltransferase 1 pseudogene 1 [Source:HGN |
| RNY1P2 | 13 | RNY1 pseudogene 2 [Source:HGNC Symbol;Ac |
| MIR4705 | 13 | microRNA 4705 [Source:HGNC Symbol;Acc:HG |
| RPL39P29 | 13 | ribosomal protein L39 pseudogene 29 [Source |
| FGF14-IT1 | 13 | FGF14 intronic transcript 1 [Source:HGNC Sym |
| AL356266.1 | 13 | TEC |
| FGF14-AS1 | 13 | FGF14 antisense RNA 1 [Source:HGNC Symbol |
| FGF14-AS2 | 13 | FGF14 antisense RNA 2 [Source:HGNC Symbol |
| LINC00555 | 13 | long intergenic non-protein coding RNA 555 [S |
| AL158063.1 | 13 | novel transcript |
| TPP2 | 13 | tripeptidyl peptidase 2 [Source:HGNC Symbol; |
| METTL21C | 13 | methyltransferase like 21C [Source:HGNC Sym |
| CCDC168 | 13 | coiled-coil domain containing 168 [Source:HG |

| | | |
|---|---|---|
| LINC00283 | 13 | long intergenic non-protein coding RNA 283 [S |
| AL157769.1 | 13 | novel transcript, antisense to TEX30 |
| TEX30 | 13 | testis expressed 30 [Source:HGNC Symbol;Acc |
| POGLUT2 | 13 | protein O-glucosyltransferase 2 [Source:HGNC |
| BIVM-ERCC5 | 13 | BIVM-ERCC5 readthrough [Source:HGNC Symb |
| BIVM | 13 | basic, immunoglobulin-like variable motif con |
| ERCC5 | 13 | ERCC excision repair 5, endonuclease [Source: |
| METTL21EP | 13 | methyltransferase like 21E, pseudogene [Sour |
| AL137246.1 | 13 | novel transcript |
| AL137246.2 | 13 | novel transcript |
| SLC10A2 | 13 | solute carrier family 10 member 2 [Source:HG |
| AL162717.1 | 13 | novel transcript |
| LINC01309 | 13 | long intergenic non-protein coding RNA 1309 |
| ATP6V1G1P7 | 13 | ATPase H+ transporting V1 subunit G1 pseudo |
| AL136524.1 | 13 | novel transcript |
| RPL7P45 | 13 | ribosomal protein L7 pseudogene 45 [Source: |
| AL138954.1 | 13 | novel transcript |
| DAOA-AS1 | 13 | DAOA antisense RNA 1 [Source:HGNC Symbol |
| AL359751.2 | 13 | NULL |
| AL359751.1 | 13 | NULL |
| DAOA | 13 | D-amino acid oxidase activator [Source:HGNC |
| AL359745.1 | 13 | novel transcript |
| LINC00343 | 13 | long intergenic non-protein coding RNA 343 [S |
| AL138701.1 | 13 | NULL |
| AL138701.2 | 13 | novel transcript |
| RNA5SP38 | 13 | RNA, 5S ribosomal pseudogene 38 [Source:HG |
| AL139379.1 | 13 | novel transcript |
| LINC00460 | 13 | long intergenic non-protein coding RNA 460 [S |
| RPL35P9 | 13 | ribosomal protein L35 pseudogene 9 [Source: |
| EFNB2 | 13 | ephrin B2 [Source:HGNC Symbol;Acc:HGNC:32 |
| AL138689.2 | 13 | novel transcript, antisense to EFNB2 |
| AL138689.1 | 13 | novel transcript, sense intronic to EFNB2 |
| ARGLU1 | 13 | arginine and glutamate rich 1 [Source:HGNC S |
| LINC00551 | 13 | long intergenic non-protein coding RNA 551 [S |
| LINC00443 | 13 | long intergenic non-protein coding RNA 443 [S |
| ATP5MC1P5 | 13 | ATP synthase membrane subunit c locus 1 pse |
| AL163534.1 | 13 | novel transcript |
| AL162574.2 | 13 | novel transcript |
| PPIAP24 | 13 | peptidylprolyl isomerase A pseudogene 24 [So |
| AL162574.1 | 13 | novel transcript |
| AL162574.3 | 13 | TEC |
| AL354741.1 | 13 | novel transcript |
| FAM155A | 13 | family with sequence similarity 155 member A |
| SNORD31B | 13 | small nucleolar RNA, C/D box 31B [Source:HG |
| AL445649.1 | 13 | novel transcript, sense intronic to FAM155A |
| MIR1267 | 13 | microRNA 1267 [Source:HGNC Symbol;Acc:HG |
| FAM155A-IT1 | 13 | FAM155A intronic transcript 1 [Source:HGNC S |
| AL359436.1 | 13 | novel transcript |
| AL136964.1 | 13 | novel transcript |
| LIG4 | 13 | DNA ligase 4 [Source:HGNC Symbol;Acc:HGNC |

| | | |
|---|---|---|
| ABHD13 | 13 | abhydrolase domain containing 13 [Source:HG |
| TNFSF13B | 13 | TNF superfamily member 13b [Source:HGNC S |
| AL157762.1 | 13 | novel transcript, sense intronic to TNFSF13B |
| RNA5SP39 | 13 | RNA, 5S ribosomal pseudogene 39 [Source:HG |
| AL138694.1 | 13 | novel transcript |
| HCFC2P1 | 13 | host cell factor C2 pseudogene 1 [Source:HGN |
| MYO16 | 13 | myosin XVI [Source:HGNC Symbol;Acc:HGNC:2 |
| MYO16-AS2 | 13 | MYO16 antisense RNA 2 [Source:HGNC Symbo |
| MYO16-AS1 | 13 | MYO16 antisense RNA 1 [Source:HGNC Symbo |
| AL161431.1 | 13 | novel transcript |
| LINC00370 | 13 | long intergenic non-protein coding RNA 370 [S |
| LINC01067 | 13 | long intergenic non-protein coding RNA 1067 |
| LINC00399 | 13 | long intergenic non-protein coding RNA 399 [S |
| AL163541.1 | 13 | novel transcript |
| LINC00676 | 13 | long intergenic non-protein coding RNA 676 [S |
| IRS2 | 13 | insulin receptor substrate 2 [Source:HGNC Syr |
| AL162497.1 | 13 | novel transcript, antisense to IRS2 |
| AL355810.1 | 13 | novel transcript |
| RN7SKP10 | 13 | RN7SK pseudogene 10 [Source:HGNC Symbol; |
| AL355974.1 | 13 | novel transcript |
| AL390755.1 | 13 | novel transcript |
| AL355974.2 | 13 | novel transcript |
| LINC00396 | 13 | long intergenic non-protein coding RNA 396 [S |
| RN7SL783P | 13 | RNA, 7SL, cytoplasmic 783, pseudogene [Sour |
| AL390755.2 | 13 | TEC |
| AL390755.3 | 13 | novel transcript |
| COL4A1 | 13 | collagen type IV alpha 1 chain [Source:HGNC S |
| AL161773.1 | 13 | TEC |
| COL4A2 | 13 | collagen type IV alpha 2 chain [Source:HGNC S |
| MIR8073 | 13 | microRNA 8073 [Source:HGNC Symbol;Acc:HG |
| COL4A2-AS2 | 13 | COL4A2 antisense 2 [Source:HGNC Symbol;Ac |
| COL4A2-AS1 | 13 | COL4A2 antisense RNA 1 [Source:HGNC Symb |
| RAB20 | 13 | RAB20, member RAS oncogene family [Source |
| AL139385.1 | 13 | novel transcript, antisense to CARKD |
| NAXD | 13 | NAD(P)HX dehydratase [Source:HGNC Symbol |
| CARS2 | 13 | cysteinyl-tRNA synthetase 2, mitochondrial [S |
| ING1 | 13 | inhibitor of growth family member 1 [Source:I |
| RPL21P107 | 13 | ribosomal protein L21 pseudogene 107 [Sourc |
| LINC00567 | 13 | long intergenic non-protein coding RNA 567 [S |
| LINC00346 | 13 | long intergenic non-protein coding RNA 346 [S |
| ANKRD10 | 13 | ankyrin repeat domain 10 [Source:HGNC Syml |
| ANKRD10-IT1 | 13 | ANKRD10 intronic transcript 1 [Source:HGNC S |
| AL442128.2 | 13 | novel transcript |
| PARP1P1 | 13 | poly(ADP-ribose) polymerase 1 pseudogene 1 |
| AL442128.1 | 13 | sperm acrosome associated 7 (SPACA7) pseuc |
| LINC00431 | 13 | long intergenic non-protein coding RNA 431 [S |
| LINC00368 | 13 | long intergenic non-protein coding RNA 368 [S |
| AL353704.2 | 13 | novel transcript |
| ARHGEF7-AS2 | 13 | ARHGEF7 antisense RNA 2 [Source:HGNC Sym |
| ARHGEF7 | 13 | Rho guanine nucleotide exchange factor 7 [So |

| | | |
|---|---|---|
| ARHGEF7-IT1 | 13 | ARHGEF7 intronic transcript 1 [Source:HGNC S |
| ARHGEF7-AS1 | 13 | ARHGEF7 antisense RNA 1 [Source:HGNC Sym |
| AL353704.1 | 13 | novel transcript, antisense to ARHGEF7 |
| AL390754.1 | 13 | spalt-like transcription factor 4 (SALL4) pseudo |
| TEX29 | 13 | testis expressed 29 [Source:HGNC Symbol;Acc |
| AL359649.1 | 13 | novel transcript |
| LINC02337 | 13 | long intergenic non-protein coding RNA 2337 |
| LINC00354 | 13 | long intergenic non-protein coding RNA 354 [S |
| AL138691.1 | 13 | NULL |
| SOX1-OT | 13 | SOX1 overlapping transcript [Source:HGNC Sy |
| SOX1 | 13 | SRY-box transcription factor 1 [Source:HGNC S |
| AL356961.1 | 13 | novel transcript |
| LINC00404 | 13 | long intergenic non-protein coding RNA 404 [S |
| LINC01070 | 13 | long intergenic non-protein coding RNA 1070 |
| LINC01043 | 13 | long intergenic non-protein coding RNA 1043 |
| LINC01044 | 13 | long intergenic non-protein coding RNA 1044 |
| SPACA7 | 13 | sperm acrosome associated 7 [Source:HGNC S |
| TUBGCP3 | 13 | tubulin gamma complex associated protein 3 |
| AL139384.2 | 13 | novel transcript |
| ATP11AUN | 13 | ATP11A upstream neighbor [Source:HGNC Syr |
| AL139384.1 | 13 | novel transcript |
| ATP11A | 13 | ATPase phospholipid transporting 11A [Source |
| ATP11A-AS1 | 13 | ATP11A antisense RNA 1 [Source:HGNC Symb |
| AL356752.1 | 13 | novel transcript, sense intronic to ATP11A |
| MCF2L | 13 | MCF.2 cell line derived transforming sequence |
| AL356740.1 | 13 | novel transcript |
| MCF2L-AS1 | 13 | MCF2L antisense RNA 1 [Source:HGNC Symbo |
| AL356740.2 | 13 | novel transcript |
| AL356740.3 | 13 | novel transcript |
| F7 | 13 | coagulation factor VII [Source:HGNC Symbol;A |
| F10 | 13 | coagulation factor X [Source:HGNC Symbol;Ac |
| F10-AS1 | 13 | F10 antisense RNA 1 [Source:HGNC Symbol;Ac |
| KARSP2 | 13 | lysyl-tRNA synthetase pseudogene 2 [Source:H |
| AL137002.2 | 13 | WAS/WASL-interacting protein family membe |
| PROZ | 13 | protein Z, vitamin K dependent plasma glycop |
| AL137002.1 | 13 | novel transcript, antisense to PROZ |
| PCID2 | 13 | PCI domain containing 2 [Source:HGNC Symbo |
| CUL4A | 13 | cullin 4A [Source:HGNC Symbol;Acc:HGNC:25 |
| MIR8075 | 13 | microRNA 8075 [Source:HGNC Symbol;Acc:HG |
| LDHBP1 | 13 | lactate dehydrogenase B pseudogene 1 [Sourc |
| LAMP1 | 13 | lysosomal associated membrane protein 1 [Sc |
| GRTP1 | 13 | growth hormone regulated TBC protein 1 [Sou |
| AL136221.1 | 13 | novel transcript, sense intronic to GRTP1 |
| GRTP1-AS1 | 13 | GRTP1 antisense RNA 1 [Source:HGNC Symbo |
| ADPRHL1 | 13 | ADP-ribosylhydrolase like 1 [Source:HGNC Syr |
| DCUN1D2 | 13 | defective in cullin neddylation 1 domain conta |
| DCUN1D2-AS | 13 | DCUN1D2 antisense RNA [Source:HGNC Symb |
| RNU1-16P | 13 | RNA, U1 small nuclear 16, pseudogene [Sourc |
| TMCO3 | 13 | transmembrane and coiled-coil domains 3 [So |
| AL442125.2 | 13 | novel transcript, sense intronic |

| | | |
|---|---|---|
| AL442125.1 | 13 | novel transcript, sense intronic to TMCO3 |
| TFDP1 | 13 | transcription factor Dp-1 [Source:HGNC Symb |
| ATP4B | 13 | ATPase H+/K+ transporting subunit beta [Sour |
| GRK1 | 13 | G protein-coupled receptor kinase 1 [Source:H |
| LINC00552 | 13 | long intergenic non-protein coding RNA 552 [S |
| TMEM255B | 13 | transmembrane protein 255B [Source:HGNC S |
| GAS6-AS1 | 13 | GAS6 antisense RNA 1 [Source:HGNC Symbol; |
| GAS6 | 13 | growth arrest specific 6 [Source:HGNC Symbo |
| GAS6-DT | 13 | GAS6 divergent transcript [Source:HGNC Symb |
| BX072579.1 | 13 | novel transcript |
| LINC00454 | 13 | long intergenic non-protein coding RNA 454 [S |
| BX072579.2 | 13 | long intergenic non-protein coding RNA 453 |
| LINC00452 | 13 | long intergenic non-protein coding RNA 452 [S |
| LINC00565 | 13 | long intergenic non-protein coding RNA 565 [S |
| C13orf46 | 13 | chromosome 13 open reading frame 46 [Sour |
| RASA3 | 13 | RAS p21 protein activator 3 [Source:HGNC Syr |
| RASA3-IT1 | 13 | RASA3 intronic transcript 1 [Source:HGNC Syn |
| CFAP97D2 | 13 | CFAP97 domain containing 2 [Source:HGNC Sy |
| AL160396.1 | 13 | novel transcript, antisense to novel protein |
| CDC16 | 13 | cell division cycle 16 [Source:HGNC Symbol;Ac |
| MIR548AR | 13 | microRNA 548ar [Source:HGNC Symbol;Acc:H |
| MIR4502 | 13 | microRNA 4502 [Source:HGNC Symbol;Acc:HG |
| AL160396.2 | 13 | novel transcript |
| UPF3A | 13 | UPF3A regulator of nonsense mediated mRNA |
| CLCP2 | 13 | Charcot-Leyden crystal protein pseudogene 2 |
| CHAMP1 | 13 | chromosome alignment maintaining phospho |
| LINC01054 | 13 | long intergenic non-protein coding RNA 1054 |
| RPL23AP97 | 13 | ribosomal protein L23a pseudogene 97 [Sourc |
| 5_8S_rRNA | 14 | 5.8S ribosomal RNA [Source:RFAM;Acc:RF000 |
| CR383658.1 | 14 | ankyrin repeat domain 7 (ANKRD7) pseudoger |
| RNU6-458P | 14 | RNA, U6 small nuclear 458, pseudogene [Sour |
| CR383658.2 | 14 | zinc finger protein 253 (ZNF253) pseudogene |
| BNIP3P6 | 14 | BCL2 interacting protein 3 pseudogene 6 [Sou |
| CR383656.10 | 14 | novel transcript, antisense to OR11H12 |
| CR383656.7 | 14 | membrane protein, palmitoylated 7 (MAGUK |
| CR383656.13 | 14 | YME1-like 1 (S. cerevisiae) (YME1L1) pseudog |
| CR383656.9 | 14 | cyclin-dependent kinase 2 (CDK2) pseudogene |
| OR11H12 | 14 | olfactory receptor family 11 subfamily H mem |
| CR383656.6 | 14 | fatty acyl CoA reductase 2 (FAR2) pseudogene |
| ARHGAP42P5 | 14 | Rho GTPase activating protein 42 pseudogene |
| LINC02297 | 14 | long intergenic non-protein coding RNA 2297 |
| CR383656.5 | 14 | fatty acyl CoA reductase 2 (FAR2) pseudogene |
| CR383656.1 | 14 | ubiquitin specific peptidase 10 (USP10) pseud |
| CR383656.12 | 14 | novel transcript |
| CR383656.8 | 14 | single stranded DNA binding protein 3 (SSBP3) |
| CR383656.11 | 14 | pleckstrin homology domain containing, famil |
| CR383656.3 | 14 | chromosome 10open reading frame 57 (C10o |
| RPL22P20 | 14 | ribosomal protein L22 pseudogene 20 [Source |
| CR383656.4 | 14 | NIMA (never in mitosis gene a)-related kinase |
| CR383656.2 | 14 | nitric oxide synthase 2, inducible (NOS2) pseu |

| | | |
|---|---|---|
| NF1P10 | 14 | neurofibromin 1 pseudogene 10 [Source:HGN |
| AL929601.4 | 14 | NIMA-related kinase 2 (NEK2) pseudogene |
| NF1P7 | 14 | neurofibromin 1 pseudogene 7 [Source:HGNC |
| MED15P1 | 14 | mediator complex subunit 15 pseudogene 1 [S |
| AL929601.1 | 14 | novel transcript |
| AL929601.2 | 14 | novel transcript |
| POTEM | 14 | POTE ankyrin domain family member M [Sour |
| AL929601.3 | 14 | novel transcript, antisense to POTEG |
| RNU6-1239P | 14 | RNA, U6 small nuclear 1239, pseudogene [Sou |
| AL589182.1 | 14 | novel transcript |
| GRAMD4P4 | 14 | GRAM domain containing 4 pseudogene 4 [So |
| DUXAP9 | 14 | double homeobox A pseudogene 9 [Source:H( |
| AL589182.3 | 14 | novel pseudogene |
| AL589743.4 | 14 | novel transcript |
| NBEAP5 | 14 | neurobeachin pseudogene 5 [Source:HGNC Sy |
| AL589182.2 | 14 | translocase of outer mitochondrial membrane |
| OR11H13P | 14 | olfactory receptor family 11 subfamily H mem |
| AL589743.3 | 14 | tetratricopeptide repeat domain 34 (TTC34) p |
| AL589743.6 | 14 | TBC1 domain family, member 26 (TBC1D26) p |
| AL589743.2 | 14 | CMT1A duplicated region transcript 15 like 2 ( |
| AL589743.7 | 14 | neurobeachin (NBEA) pseudogene |
| AL589743.1 | 14 | novel transcript |
| AL589743.5 | 14 | translocase of outer mitochondrial membrane |
| DUXAP10 | 14 | double homeobox A pseudogene 10 [Source:N |
| BMS1P17 | 14 | BMS1 pseudogene 17 [Source:HGNC Symbol;A |
| AL929602.1 | 14 | novel pseudogene |
| LINC01297 | 14 | long intergenic non-protein coding RNA 1297 |
| GRAMD4P3 | 14 | GRAM domain containing 4 pseudogene 3 [So |
| POTEG | 14 | POTE ankyrin domain family member G [Sour( |
| AL512624.2 | 14 | novel transcript, antisense to POTEM |
| RNU6-1268P | 14 | RNA, U6 small nuclear 1268, pseudogene [Sou |
| MED15P6 | 14 | mediator complex subunit 15 pseudogene 6 [S |
| NF1P4 | 14 | neurofibromin 1 pseudogene 4 [Source:HGNC |
| AL512310.11 | 14 | novel transcript |
| AL512624.1 | 14 | NIMA-related kinase 2 (NEK2) pseudogene |
| NF1P11 | 14 | neurofibromin 1 pseudogene 11 [Source:HGN |
| AL512310.3 | 14 | NIMA-related kinase 2 (NEK2) pseudogene |
| AL512310.10 | 14 | novel transcript |
| AL512310.8 | 14 | chromosome 10 open reading frame 57 (C10c |
| AL512310.4 | 14 | fatty acyl CoA reductase 2 (FAR2) pseudogene |
| AL512310.5 | 14 | pleckstrin homology domain containing, famil |
| AL512310.6 | 14 | single stranded DNA binding protein 3 (SSBP3) |
| AL512310.9 | 14 | novel transcript |
| AL512310.7 | 14 | ubiquitin specific peptidase 10 (USP10) pseud |
| AL512310.2 | 14 | novel transcript |
| ARHGAP42P4 | 14 | Rho GTPase activating protein 42 pseudogene |
| AL512310.1 | 14 | fatty acyl CoA reductase 2 (FAR2) pseudogene |
| OR11H2 | 14 | olfactory receptor family 11 subfamily H mem |
| OR4N2 | 14 | olfactory receptor family 4 subfamily N memb |
| OR11K2P | 14 | olfactory receptor family 11 subfamily K mem |

| | | |
|---|---|---|
| OR4Q3 | 14 | olfactory receptor family 4 subfamily Q memb |
| OR4H12P | 14 | olfactory receptor family 4 subfamily H memb |
| OR4M1 | 14 | olfactory receptor family 4 subfamily M meml |
| OR4N1P | 14 | olfactory receptor family 4 subfamily N memb |
| OR4K6P | 14 | olfactory receptor family 4 subfamily K memb |
| OR4K3 | 14 | olfactory receptor family 4 subfamily K memb |
| OR4K2 | 14 | olfactory receptor family 4 subfamily K memb |
| OR4K4P | 14 | olfactory receptor family 4 subfamily K memb |
| AL391156.1 | 14 | ATPase, Na+/K+ transporting, beta 3 polypept |
| OR4K5 | 14 | olfactory receptor family 4 subfamily K memb |
| OR4K1 | 14 | olfactory receptor family 4 subfamily K memb |
| OR4K16P | 14 | olfactory receptor family 4 subfamily K memb |
| OR4K15 | 14 | olfactory receptor family 4 subfamily K memb |
| OR4Q2 | 14 | olfactory receptor family 4 subfamily Q memb |
| OR4K14 | 14 | olfactory receptor family 4 subfamily K memb |
| OR4K13 | 14 | olfactory receptor family 4 subfamily K memb |
| OR4U1P | 14 | olfactory receptor family 4 subfamily U memb |
| OR4L1 | 14 | olfactory receptor family 4 subfamily L memb |
| RNA5SP380 | 14 | RNA, 5S ribosomal pseudogene 380 [Source:H |
| OR4T1P | 14 | olfactory receptor family 4 subfamily T memb |
| OR4K17 | 14 | olfactory receptor family 4 subfamily K memb |
| OR4N5 | 14 | olfactory receptor family 4 subfamily N memb |
| PSMB7P1 | 14 | PSMB7 pseudogene 1 [Source:HGNC Symbol;/ |
| GTF2IP22 | 14 | general transcription factor IIi pseudogene 22 |
| OR11G1P | 14 | olfactory receptor family 11 subfamily G mem |
| OR11P1P | 14 | olfactory receptor family 11 subfamily P mem |
| OR11G2 | 14 | olfactory receptor family 11 subfamily G mem |
| OR11H5P | 14 | olfactory receptor family 11 subfamily H mem |
| OR11H6 | 14 | olfactory receptor family 11 subfamily H mem |
| OR11H7 | 14 | olfactory receptor family 11 subfamily H mem |
| OR11H4 | 14 | olfactory receptor family 11 subfamily H mem |
| TTC5 | 14 | tetratricopeptide repeat domain 5 [Source:HG |
| AL356019.2 | 14 | novel transcript |
| AL356019.1 | 14 | novel transcript, antisense to CCNB1IP1 |
| CCNB1IP1 | 14 | cyclin B1 interacting protein 1 [Source:HGNC S |
| AL355075.6 | 14 | novel transcript, antisense to CCNB1IP1 |
| SNORA79B | 14 | small nucleolar RNA, H/ACA box 79B [Source:I |
| SNORD126 | 14 | small nucleolar RNA, C/D box 126 [Source:HGI |
| AL355075.4 | 14 | ribonuclease P RNA component H1 |
| RPPH1 | 14 | ribonuclease P RNA component H1 [Source:H( |
| PARP2 | 14 | poly(ADP-ribose) polymerase 2 [Source:HGNC |
| AL355075.1 | 14 | novel transcript |
| TEP1 | 14 | telomerase associated protein 1 [Source:HGN |
| RNA5SP382 | 14 | RNA, 5S ribosomal pseudogene 382 [Source:H |
| KLHL33 | 14 | kelch like family member 33 [Source:HGNC Sy |
| AL355075.5 | 14 | solute carrier family 12 (sodium/chloride tran |
| OSGEP | 14 | O-sialoglycoprotein endopeptidase [Source:H( |
| AL355075.2 | 14 | novel transcript, antisense to OSGEP |
| APEX1 | 14 | apurinic/apyrimidinic endodeoxyribonuclease |
| PIP4P1 | 14 | phosphatidylinositol-4,5-bisphosphate 4-phos |

| | | |
|---|---|---|
| PNP | 14 | purine nucleoside phosphorylase [Source:HGN |
| AL355075.3 | 14 | novel transcript |
| RNASE10 | 14 | ribonuclease A family member 10 (inactive) [S |
| PTCD2P1 | 14 | pentatricopeptide repeat domain 2 pseudoge |
| SETP1 | 14 | SET pseudogene 1 [Source:HGNC Symbol;Acc: |
| RNASE9 | 14 | ribonuclease A family member 9 (inactive) [Sc |
| AL163195.3 | 14 | novel protein |
| RNASE11 | 14 | ribonuclease A family member 11 (inactive) [S |
| AL163195.2 | 14 | novel transcript, antisense to RNASE11 & RNA |
| RNASE12 | 14 | ribonuclease A family member 12 (inactive) [S |
| AL163195.1 | 14 | ribosomal protein S26 (RPS26) pseudogene |
| OR6S1 | 14 | olfactory receptor family 6 subfamily S memb |
| Y_RNA | 14 | Y RNA [Source:RFAM;Acc:RF00019] |
| ANG | 14 | angiogenin [Source:HGNC Symbol;Acc:HGNC:4 |
| RNASE4 | 14 | ribonuclease A family member 4 [Source:HGN |
| AL163636.2 | 14 | novel protein, ANG-RNASE4 readthrough |
| AL163636.1 | 14 | novel transcript, antisense to RNASE4 and AN |
| RANBP20P | 14 | RAN binding protein 20 pseudogene [Source:H |
| EDDM3DP | 14 | epididymal protein 3D, pseudogene [Source:H |
| EDDM3A | 14 | epididymal protein 3A [Source:HGNC Symbol; |
| EDDM3B | 14 | epididymal protein 3B [Source:HGNC Symbol; |
| RNASE6 | 14 | ribonuclease A family member k6 [Source:HG |
| AL133371.2 | 14 | novel transcript, antisense to RNASE1 |
| RNASE1 | 14 | ribonuclease A family member 1, pancreatic [ |
| AL133371.3 | 14 | novel transcript |
| AL133371.1 | 14 | novel transcript, antisense to RNASE3 |
| RNASE3 | 14 | ribonuclease A family member 3 [Source:HGN |
| AL355922.4 | 14 | novel transcript |
| AL355922.3 | 14 | novel transcript |
| AL355922.1 | 14 | novel transcript |
| RNASE2 | 14 | ribonuclease A family member 2 [Source:HGN |
| RN7SL189P | 14 | RNA, 7SL, cytoplasmic 189, pseudogene [Sour |
| AL355922.2 | 14 | coiled-coil domain containing 107 (CCDC107) |
| METTL17 | 14 | methyltransferase like 17 [Source:HGNC Symb |
| AL161668.3 | 14 | novel transcript, antisense to SLC39A2 |
| SLC39A2 | 14 | solute carrier family 39 member 2 [Source:HG |
| NDRG2 | 14 | NDRG family member 2 [Source:HGNC Symbo |
| MIR6717 | 14 | microRNA 6717 [Source:HGNC Symbol;Acc:HG |
| TPPP2 | 14 | tubulin polymerization promoting protein fam |
| AL161668.4 | 14 | novel transcript |
| RNASE13 | 14 | ribonuclease A family member 13 (inactive) [S |
| AL161668.2 | 14 | novel transcript, antisense to RNASE13 |
| RNASE7 | 14 | ribonuclease A family member 7 [Source:HGN |
| RNASE8 | 14 | ribonuclease A family member 8 [Source:HGN |
| ARHGEF40 | 14 | Rho guanine nucleotide exchange factor 40 [S |
| ZNF219 | 14 | zinc finger protein 219 [Source:HGNC Symbol; |
| TMEM253 | 14 | transmembrane protein 253 [Source:HGNC Sy |
| AL161668.1 | 14 | novel transcript |
| CKAP2P1 | 14 | cytoskeleton associated protein 2 pseudogen |
| RNU6-252P | 14 | RNA, U6 small nuclear 252, pseudogene [Sour |

| | | |
|---|---|---|
| OR5AU1 | 14 | olfactory receptor family 5 subfamily AU mem |
| SMARCE1P3 | 14 | SWI/SNF related, matrix associated, actin dep |
| AL157687.1 | 14 | myoneurin (MYNN) pseudogene |
| LINC00641 | 14 | long intergenic non-protein coding RNA 641 [S |
| HNRNPC | 14 | heterogeneous nuclear ribonucleoprotein C (C |
| RPGRIP1 | 14 | RPGR interacting protein 1 [Source:HGNC Sym |
| SUPT16H | 14 | SPT16 homolog, facilitates chromatin remode |
| AL135744.1 | 14 | novel transcript, antisense to CHD8 and SUPT1 |
| CHD8 | 14 | chromodomain helicase DNA binding protein 8 |
| SNORD9 | 14 | small nucleolar RNA, C/D box 9 [Source:HGNC |
| SNORD8 | 14 | small nucleolar RNA, C/D box 8 [Source:HGNC |
| AL161747.1 | 14 | ubiquitin A-52 residue ribosomal protein fusic |
| EIF4EBP1P1 | 14 | eukaryotic translation initiation factor 4E bind |
| RN7SL650P | 14 | RNA, 7SL, cytoplasmic 650, pseudogene [Sour |
| RAB2B | 14 | RAB2B, member RAS oncogene family [Source |
| TOX4 | 14 | TOX high mobility group box family member 4 |
| METTL3 | 14 | methyltransferase like 3 [Source:HGNC Symbo |
| SALL2 | 14 | spalt like transcription factor 2 [Source:HGNC |
| AL161747.2 | 14 | novel transcript |
| RBBP4P5 | 14 | RBBP4 pseudogene 5 [Source:HGNC Symbol;A |
| OR10G3 | 14 | olfactory receptor family 10 subfamily G mem |
| AC243972.2 | 14 | ARP3 actin-related protein 3 homolog (yeast) |
| OR10G1P | 14 | olfactory receptor family 10 subfamily G mem |
| UBE2NP1 | 14 | ubiquitin conjugating enzyme E2 N pseudoger |
| TRAV1-1 | 14 | T cell receptor alpha variable 1-1 [Source:HGN |
| OR10G2 | 14 | olfactory receptor family 10 subfamily G mem |
| TRAV1-2 | 14 | T cell receptor alpha variable 1-2 [Source:HGN |
| ARL6IP1P1 | 14 | ADP ribosylation factor like GTPase 6 interacti |
| OR4E2 | 14 | olfactory receptor family 4 subfamily E memb |
| OR4E1 | 14 | olfactory receptor family 4 subfamily E memb |
| AC243972.3 | 14 | pseudogene similar to part of anoctamin 10 (A |
| TRAV2 | 14 | T cell receptor alpha variable 2 [Source:HGNC |
| AC243972.1 | 14 | novel transcript |
| TRAV3 | 14 | T cell receptor alpha variable 3 (gene/pseudog |
| TRAV4 | 14 | T cell receptor alpha variable 4 [Source:HGNC |
| TRAV5 | 14 | T cell receptor alpha variable 5 [Source:HGNC |
| RPL4P1 | 14 | ribosomal protein L4 pseudogene 1 [Source:H |
| TRAV6 | 14 | T cell receptor alpha variable 6 [Source:HGNC |
| TRAV7 | 14 | T cell receptor alpha variable 7 [Source:HGNC |
| TRAV8-1 | 14 | T cell receptor alpha variable 8-1 [Source:HGN |
| TRAV9-1 | 14 | T cell receptor alpha variable 9-1 [Source:HGN |
| TRAV10 | 14 | T cell receptor alpha variable 10 [Source:HGN |
| TRAV11 | 14 | T cell receptor alpha variable 11 (pseudogene |
| TRAV12-1 | 14 | T cell receptor alpha variable 12-1 [Source:HG |
| TRAV8-2 | 14 | T cell receptor alpha variable 8-2 [Source:HGN |
| TRAV8-3 | 14 | T cell receptor alpha variable 8-3 [Source:HGN |
| TRAV13-1 | 14 | T cell receptor alpha variable 13-1 [Source:HG |
| TRAV12-2 | 14 | T cell receptor alpha variable 12-2 [Source:HG |
| TRAV8-4 | 14 | T cell receptor alpha variable 8-4 [Source:HGN |
| TRAV8-5 | 14 | T cell receptor alpha variable 8-5 (pseudogene |

| | | |
|---|---|---|
| TRAV13-2 | 14 | T cell receptor alpha variable 13-2 [Source:HG |
| TRAV14DV4 | 14 | T cell receptor alpha variable 14/delta variabl |
| TRAV9-2 | 14 | T cell receptor alpha variable 9-2 [Source:HGN |
| TRAV15 | 14 | T cell receptor alpha variable 15 (pseudogene |
| AC245505.1 | 14 | ubiquitin-conjugating enzyme E2 variant 1 (UE |
| TRAV12-3 | 14 | T cell receptor alpha variable 12-3 [Source:HG |
| TRAV8-6 | 14 | T cell receptor alpha variable 8-6 [Source:HGN |
| TRAV16 | 14 | T cell receptor alpha variable 16 [Source:HGN |
| TRAV17 | 14 | T cell receptor alpha variable 17 [Source:HGN |
| TRAV18 | 14 | T cell receptor alpha variable 18 [Source:HGN |
| TRAV19 | 14 | T cell receptor alpha variable 19 [Source:HGN |
| AC245505.2 | 14 | sal-like 4 (Drosophila) (SALL4) pseudogene |
| TRAV20 | 14 | T cell receptor alpha variable 20 [Source:HGN |
| TRAV21 | 14 | T cell receptor alpha variable 21 [Source:HGN |
| AC245505.3 | 14 | T cell receptor pseudogene |
| TRAV22 | 14 | T cell receptor alpha variable 22 [Source:HGN |
| TRAV23DV6 | 14 | T cell receptor alpha variable 23/delta variabl |
| TRDV1 | 14 | T cell receptor delta variable 1 [Source:HGNC |
| TRAV24 | 14 | T cell receptor alpha variable 24 [Source:HGN |
| TRAV25 | 14 | T cell receptor alpha variable 25 [Source:HGN |
| TRAV26-1 | 14 | T cell receptor alpha variable 26-1 [Source:HG |
| TRAV8-7 | 14 | T cell receptor alpha variable 8-7 (non-functio |
| TRAV27 | 14 | T cell receptor alpha variable 27 [Source:HGN |
| TRAV28 | 14 | T cell receptor alpha variable 28 (pseudogene |
| TRAV29DV5 | 14 | T cell receptor alpha variable 29/delta variabl |
| TRAV30 | 14 | T cell receptor alpha variable 30 [Source:HGN |
| TRAV31 | 14 | T cell receptor alpha variable 31 (pseudogene |
| TRAV32 | 14 | T cell receptor alpha variable 32 (pseudogene |
| TRAV33 | 14 | T cell receptor alpha variable 33 (pseudogene |
| AC244502.1 | 14 | novel transcript, antisense to IG_gene_cluster |
| TRAV26-2 | 14 | T cell receptor alpha variable 26-2 [Source:HG |
| TRAV34 | 14 | T cell receptor alpha variable 34 [Source:HGN |
| TRAV35 | 14 | T cell receptor alpha variable 35 [Source:HGN |
| TRAV36DV7 | 14 | T cell receptor alpha variable 36/delta variabl |
| TRAV37 | 14 | T cell receptor alpha variable 37 (pseudogene |
| TRAV38-1 | 14 | T cell receptor alpha variable 38-1 [Source:HG |
| TRAV38-2DV8 | 14 | T cell receptor alpha variable 38-2/delta varia |
| TRAV39 | 14 | T cell receptor alpha variable 39 [Source:HGN |
| TRAV40 | 14 | T cell receptor alpha variable 40 [Source:HGN |
| TRAV41 | 14 | T cell receptor alpha variable 41 [Source:HGN |
| AC244502.2 | 14 | translin-associated factor X (TSNAX) pseudoge |
| AC244502.3 | 14 | novel transcript |
| TRDV2 | 14 | T cell receptor delta variable 2 [Source:HGNC |
| TRDD1 | 14 | T cell receptor delta diversity 1 [Source:HGNC |
| TRDD2 | 14 | T cell receptor delta diversity 2 [Source:HGNC |
| TRDD3 | 14 | T cell receptor delta diversity 3 [Source:HGNC |
| TRDJ1 | 14 | T cell receptor delta joining 1 [Source:HGNC S |
| TRDJ4 | 14 | T cell receptor delta joining 4 [Source:HGNC S |
| TRDJ2 | 14 | T cell receptor delta joining 2 [Source:HGNC S |
| TRDJ3 | 14 | T cell receptor delta joining 3 [Source:HGNC S |

| | | |
|---|---|---|
| TRDC | 14 | T cell receptor delta constant [Source:HGNC S |
| TRDV3 | 14 | T cell receptor delta variable 3 [Source:HGNC |
| TRAJ61 | 14 | T cell receptor alpha joining 61 (non-functiona |
| TRAJ60 | 14 | T cell receptor alpha joining 60 (pseudogene) |
| TRAJ59 | 14 | T cell receptor alpha joining 59 (non-functiona |
| TRAJ58 | 14 | T cell receptor alpha joining 58 (non-functiona |
| TRAJ57 | 14 | T cell receptor alpha joining 57 [Source:HGNC |
| TRAJ56 | 14 | T cell receptor alpha joining 56 [Source:HGNC |
| TRAJ55 | 14 | T cell receptor alpha joining 55 (pseudogene) |
| TRAJ54 | 14 | T cell receptor alpha joining 54 [Source:HGNC |
| TRAJ53 | 14 | T cell receptor alpha joining 53 [Source:HGNC |
| TRAJ52 | 14 | T cell receptor alpha joining 52 [Source:HGNC |
| TRAJ51 | 14 | T cell receptor alpha joining 51 (pseudogene) |
| TRAJ50 | 14 | T cell receptor alpha joining 50 [Source:HGNC |
| TRAJ49 | 14 | T cell receptor alpha joining 49 [Source:HGNC |
| TRAJ48 | 14 | T cell receptor alpha joining 48 [Source:HGNC |
| TRAJ47 | 14 | T cell receptor alpha joining 47 [Source:HGNC |
| TRAJ46 | 14 | T cell receptor alpha joining 46 [Source:HGNC |
| TRAJ45 | 14 | T cell receptor alpha joining 45 [Source:HGNC |
| TRAJ44 | 14 | T cell receptor alpha joining 44 [Source:HGNC |
| TRAJ43 | 14 | T cell receptor alpha joining 43 [Source:HGNC |
| TRAJ42 | 14 | T cell receptor alpha joining 42 [Source:HGNC |
| TRAJ41 | 14 | T cell receptor alpha joining 41 [Source:HGNC |
| TRAJ40 | 14 | T cell receptor alpha joining 40 [Source:HGNC |
| TRAJ39 | 14 | T cell receptor alpha joining 39 [Source:HGNC |
| TRAJ38 | 14 | T cell receptor alpha joining 38 [Source:HGNC |
| TRAJ37 | 14 | T cell receptor alpha joining 37 [Source:HGNC |
| TRAJ36 | 14 | T cell receptor alpha joining 36 [Source:HGNC |
| TRAJ35 | 14 | T cell receptor alpha joining 35 (non-functiona |
| TRAJ34 | 14 | T cell receptor alpha joining 34 [Source:HGNC |
| TRAJ33 | 14 | T cell receptor alpha joining 33 [Source:HGNC |
| TRAJ32 | 14 | T cell receptor alpha joining 32 [Source:HGNC |
| TRAJ31 | 14 | T cell receptor alpha joining 31 [Source:HGNC |
| TRAJ30 | 14 | T cell receptor alpha joining 30 [Source:HGNC |
| TRAJ29 | 14 | T cell receptor alpha joining 29 [Source:HGNC |
| TRAJ28 | 14 | T cell receptor alpha joining 28 [Source:HGNC |
| TRAJ27 | 14 | T cell receptor alpha joining 27 [Source:HGNC |
| TRAJ26 | 14 | T cell receptor alpha joining 26 [Source:HGNC |
| TRAJ25 | 14 | T cell receptor alpha joining 25 (non-functiona |
| TRAJ24 | 14 | T cell receptor alpha joining 24 [Source:HGNC |
| TRAJ23 | 14 | T cell receptor alpha joining 23 [Source:HGNC |
| TRAJ22 | 14 | T cell receptor alpha joining 22 [Source:HGNC |
| TRAJ21 | 14 | T cell receptor alpha joining 21 [Source:HGNC |
| TRAJ20 | 14 | T cell receptor alpha joining 20 [Source:HGNC |
| TRAJ19 | 14 | T cell receptor alpha joining 19 (non-functiona |
| TRAJ18 | 14 | T cell receptor alpha joining 18 [Source:HGNC |
| TRAJ17 | 14 | T cell receptor alpha joining 17 [Source:HGNC |
| TRAJ16 | 14 | T cell receptor alpha joining 16 [Source:HGNC |
| TRAJ14 | 14 | T cell receptor alpha joining 14 [Source:HGNC |
| TRAJ13 | 14 | T cell receptor alpha joining 13 [Source:HGNC |

| | | |
|---|---|---|
| TRAJ12 | 14 | T cell receptor alpha joining 12 [Source:HGNC |
| TRAJ11 | 14 | T cell receptor alpha joining 11 [Source:HGNC |
| TRAJ10 | 14 | T cell receptor alpha joining 10 [Source:HGNC |
| TRAJ9 | 14 | T cell receptor alpha joining 9 [Source:HGNC S |
| TRAJ8 | 14 | T cell receptor alpha joining 8 [Source:HGNC S |
| TRAJ7 | 14 | T cell receptor alpha joining 7 [Source:HGNC S |
| TRAJ6 | 14 | T cell receptor alpha joining 6 [Source:HGNC S |
| TRAJ5 | 14 | T cell receptor alpha joining 5 [Source:HGNC S |
| TRAJ4 | 14 | T cell receptor alpha joining 4 [Source:HGNC S |
| TRAJ3 | 14 | T cell receptor alpha joining 3 [Source:HGNC S |
| TRAJ2 | 14 | T cell receptor alpha joining 2 (non-functional |
| TRAJ1 | 14 | T cell receptor alpha joining 1 (non-functional |
| TRAC | 14 | T cell receptor alpha constant [Source:HGNC S |
| LINC02332 | 14 | long intergenic non-protein coding RNA 2332 |
| AC243965.1 | 14 | novel transcript |
| DAD1 | 14 | defender against cell death 1 [Source:HGNC Sy |
| AC243965.2 | 14 | novel transcript, antisense ABHD4 |
| ABHD4 | 14 | abhydrolase domain containing 4 [Source:HGN |
| OR6J1 | 14 | olfactory receptor family 6 subfamily J membe |
| AL160314.1 | 14 | ribosomal protein L26 (RPL26) pseudogene |
| AL160314.2 | 14 | novel transcript, antisense to OXA1L |
| OR6E1P | 14 | olfactory receptor family 6 subfamily E memb |
| AL160314.3 | 14 | von Willebrand factor A domain containing 5E |
| AL135998.1 | 14 | NULL |
| OXA1L | 14 | OXA1L mitochondrial inner membrane protein |
| SLC7A7 | 14 | solute carrier family 7 member 7 [Source:HGN |
| MRPL52 | 14 | mitochondrial ribosomal protein L52 [Source:I |
| MMP14 | 14 | matrix metallopeptidase 14 [Source:HGNC Syr |
| Y_RNA | 14 | Y RNA [Source:RFAM;Acc:RF00019] |
| Y_RNA | 14 | Y RNA [Source:RFAM;Acc:RF00019] |
| LRP10 | 14 | LDL receptor related protein 10 [Source:HGNC |
| REM2 | 14 | RRAD and GEM like GTPase 2 [Source:HGNC S |
| RBM23 | 14 | RNA binding motif protein 23 [Source:HGNC S |
| PRMT5-AS1 | 14 | PRMT5 antisense RNA 1 [Source:HGNC Symbo |
| PRMT5 | 14 | protein arginine methyltransferase 5 [Source: |
| AL132780.1 | 14 | novel transcript, antisense to HAUS4 |
| HAUS4 | 14 | HAUS augmin like complex subunit 4 [Source:I |
| AL132780.3 | 14 | novel protein |
| MIR4707 | 14 | microRNA 4707 [Source:HGNC Symbol;Acc:HG |
| AJUBA | 14 | ajuba LIM protein [Source:HGNC Symbol;Acc:I |
| AL132780.2 | 14 | novel transcript, antisense to C14orf93 |
| C14orf93 | 14 | chromosome 14 open reading frame 93 [Sour |
| AL132780.5 | 14 | TEC |
| PSMB5 | 14 | proteasome subunit beta 5 [Source:HGNC Syn |
| AL132780.4 | 14 | TEC |
| PSMB11 | 14 | proteasome subunit beta 11 [Source:HGNC Sy |
| CDH24 | 14 | cadherin 24 [Source:HGNC Symbol;Acc:HGNC: |
| ACIN1 | 14 | apoptotic chromatin condensation inducer 1 [ |
| C14orf119 | 14 | chromosome 14 open reading frame 119 [Sou |
| LMLN2 | 14 | leishmanolysin like peptidase 2 [Source:HGNC |

| | | |
|---|---|---|
| CEBPE | 14 | CCAAT enhancer binding protein epsilon [Sour |
| SLC7A8 | 14 | solute carrier family 7 member 8 [Source:HGN |
| RNU6-1138P | 14 | RNA, U6 small nuclear 1138, pseudogene [Sou |
| RNF212B | 14 | ring finger protein 212B [Source:HGNC Symbo |
| AL049829.1 | 14 | ribosomal protein L39 (RPL39) pseudogene |
| RNU6-1046P | 14 | RNA, U6 small nuclear 1046, pseudogene [Sou |
| HOMEZ | 14 | homeobox and leucine zipper encoding [Sourc |
| H3F3AP2 | 14 | H3 histone family member 3A pseudogene 2 [ |
| PPP1R3E | 14 | protein phosphatase 1 regulatory subunit 3E [ |
| BCL2L2 | 14 | BCL2 like 2 [Source:HGNC Symbol;Acc:HGNC:9 |
| BCL2L2-PABPN1 | 14 | BCL2L2-PABPN1 readthrough [Source:HGNC S |
| PABPN1 | 14 | poly(A) binding protein nuclear 1 [Source:HGN |
| SLC22A17 | 14 | solute carrier family 22 member 17 [Source:H |
| EFS | 14 | embryonal Fyn-associated substrate [Source:H |
| AL049829.2 | 14 | novel transcript, antisense to EFS |
| IL25 | 14 | interleukin 25 [Source:HGNC Symbol;Acc:HGN |
| CMTM5 | 14 | CKLF like MARVEL transmembrane domain co |
| MYH6 | 14 | myosin heavy chain 6 [Source:HGNC Symbol;A |
| MIR208A | 14 | microRNA 208a [Source:HGNC Symbol;Acc:HG |
| MYH7 | 14 | myosin heavy chain 7 [Source:HGNC Symbol;A |
| AL132855.1 | 14 | novel transcript, antisense to MYH7 |
| MIR208B | 14 | microRNA 208b [Source:HGNC Symbol;Acc:HG |
| NGDN | 14 | neuroguidin [Source:HGNC Symbol;Acc:HGNC |
| ZFHX2-AS1 | 14 | ZFHX2 antisense RNA 1 [Source:HGNC Symbol |
| ZFHX2 | 14 | zinc finger homeobox 2 [Source:HGNC Symbo |
| AL135999.2 | 14 | TEC |
| THTPA | 14 | thiamine triphosphatase [Source:HGNC Symbo |
| AP1G2 | 14 | adaptor related protein complex 1 subunit ga |
| AL135999.1 | 14 | novel transcript, antisense to AP1G2 |
| JPH4 | 14 | junctophilin 4 [Source:HGNC Symbol;Acc:HGN |
| AL135999.3 | 14 | novel transcript |
| DHRS2 | 14 | dehydrogenase/reductase 2 [Source:HGNC Sy |
| AL160237.1 | 14 | novel transcript |
| BRD7P1 | 14 | bromodomain containing 7 pseudogene 1 [So |
| AL160237.3 | 14 | suppressor of zeste 12 homolog (Drosophila) ( |
| RN7SKP205 | 14 | RN7SK pseudogene 205 [Source:HGNC Symbo |
| AL160237.2 | 14 | dehydrogenase/reductase (SDR family) memb |
| AL136419.2 | 14 | novel transcript |
| LINC00596 | 14 | long intergenic non-protein coding RNA 596 [S |
| DHRS4-AS1 | 14 | DHRS4 antisense RNA 1 [Source:HGNC Symbo |
| DHRS4 | 14 | dehydrogenase/reductase 4 [Source:HGNC Sy |
| DHRS4L2 | 14 | dehydrogenase/reductase 4 like 2 [Source:HG |
| AL136419.1 | 14 | dehydrogenase/reductase 4 like 1 [Source:NC |
| AL136419.3 | 14 | novel transcript |
| DHRS4L1 | 14 | dehydrogenase/reductase 4 like 1 [Source:HG |
| CARMIL3 | 14 | capping protein regulator and myosin 1 linker |
| CPNE6 | 14 | copine 6 [Source:HGNC Symbol;Acc:HGNC:23 |
| NRL | 14 | neural retina leucine zipper [Source:HGNC Syr |
| PCK2 | 14 | phosphoenolpyruvate carboxykinase 2, mitoc |
| DCAF11 | 14 | DDB1 and CUL4 associated factor 11 [Source:H |

| | | |
|---|---|---|
| AL136295.3 | 14 | novel protein |
| FITM1 | 14 | fat storage inducing transmembrane protein 1 |
| PSME1 | 14 | proteasome activator subunit 1 [Source:HGN( |
| EMC9 | 14 | ER membrane protein complex subunit 9 [Sou |
| AL136295.2 | 14 | novel transcript |
| PSME2 | 14 | proteasome activator subunit 2 [Source:HGN( |
| MIR7703 | 14 | microRNA 7703 [Source:HGNC Symbol;Acc:HG |
| RNF31 | 14 | ring finger protein 31 [Source:HGNC Symbol;A |
| AL136295.5 | 14 | novel transcript |
| RNA5SP383 | 14 | RNA, 5S ribosomal pseudogene 383 [Source:H |
| IRF9 | 14 | interferon regulatory factor 9 [Source:HGNC S |
| REC8 | 14 | REC8 meiotic recombination protein [Source:I |
| IPO4 | 14 | importin 4 [Source:HGNC Symbol;Acc:HGNC:1 |
| AL136295.4 | 14 | novel protein |
| AL136295.17 | 14 | novel transcript, antisense to TM9SF1 |
| TM9SF1 | 14 | transmembrane 9 superfamily member 1 [Sou |
| AL136295.1 | 14 | novel protein |
| AL136295.6 | 14 | novel transcript |
| AL136295.7 | 14 | novel transcript |
| TSSK4 | 14 | testis specific serine kinase 4 [Source:HGNC Sy |
| CHMP4A | 14 | charged multivesicular body protein 4A [Sourc |
| AL096870.2 | 14 | novel transcript |
| MDP1 | 14 | magnesium dependent phosphatase 1 [Source |
| NEDD8-MDP1 | 14 | NEDD8-MDP1 readthrough [Source:HGNC Sym |
| NEDD8 | 14 | NEDD8 ubiquitin like modifier [Source:HGNC S |
| GMPR2 | 14 | guanosine monophosphate reductase 2 [Sour |
| TINF2 | 14 | TERF1 interacting nuclear factor 2 [Source:HG |
| TGM1 | 14 | transglutaminase 1 [Source:HGNC Symbol;Acc |
| RABGGTA | 14 | Rab geranylgeranyltransferase subunit alpha [ |
| AL096870.8 | 14 | novel transcript, antisense to RABGGTAand DI |
| AL096870.6 | 14 | novel transcript, antisense to RABGGTA |
| DHRS1 | 14 | dehydrogenase/reductase 1 [Source:HGNC Sy |
| NOP9 | 14 | NOP9 nucleolar protein [Source:HGNC Symbo |
| CIDEB | 14 | cell death inducing DFFA like effector b [Sourc |
| LTB4R2 | 14 | leukotriene B4 receptor 2 [Source:HGNC Symb |
| LTB4R | 14 | leukotriene B4 receptor [Source:HGNC Symbo |
| ADCY4 | 14 | adenylate cyclase 4 [Source:HGNC Symbol;Acc |
| AL096870.1 | 14 | novel transcript |
| RIPK3 | 14 | receptor interacting serine/threonine kinase 3 |
| U6 | 14 | U6 spliceosomal RNA [Source:RFAM;Acc:RF00 |
| NFATC4 | 14 | nuclear factor of activated T cells 4 [Source:H( |
| NYNRIN | 14 | NYN domain and retroviral integrase containir |
| CBLN3 | 14 | cerebellin 3 precursor [Source:HGNC Symbol;A |
| KHNYN | 14 | KH and NYN domain containing [Source:HGNC |
| SDR39U1 | 14 | short chain dehydrogenase/reductase family 3 |
| AL132800.1 | 14 | novel transcript, antisense to CMA1 |
| CMA1 | 14 | chymase 1 [Source:HGNC Symbol;Acc:HGNC:2 |
| CTSG | 14 | cathepsin G [Source:HGNC Symbol;Acc:HGNC: |
| AL136018.1 | 14 | novel transcript, antisense to GZMB |
| GZMH | 14 | granzyme H [Source:HGNC Symbol;Acc:HGNC: |

| | | |
|---|---|---|
| GZMB | 14 | granzyme B [Source:HGNC Symbol;Acc:HGNC: |
| STXBP6 | 14 | syntaxin binding protein 6 [Source:HGNC Sym |
| AL161663.2 | 14 | novel transcript, antisense to STXBP6 |
| AL161663.1 | 14 | novel transcript, antisense to STXBP6 |
| LINC02286 | 14 | long intergenic non-protein coding RNA 2286 |
| HMGN2P6 | 14 | high mobility group nucleosomal binding dom |
| OR7K1P | 14 | olfactory receptor family 7 subfamily K memb |
| AL079352.1 | 14 | 1-acylglycerol-3-phosphate O-acyltransferase |
| LINC02306 | 14 | long intergenic non-protein coding RNA 2306 |
| AL359396.1 | 14 | NULL |
| AL359396.2 | 14 | TEC |
| AL132633.1 | 14 | novel transcript |
| CYB5AP5 | 14 | cytochrome b5 type A pseudogene 5 [Source: |
| NOVA1 | 14 | NOVA alternative splicing regulator 1 [Source: |
| AL079343.1 | 14 | novel transcript, sense intronic to NOVA1 |
| LINC02588 | 14 | long intergenic non-protein coding RNA 2588 |
| AL110292.1 | 14 | novel transcript |
| LINC02294 | 14 | long intergenic non-protein coding RNA 2294 |
| LINC02293 | 14 | long intergenic non-protein coding RNA 2293 |
| AL132718.1 | 14 | novel transcript |
| MIR4307HG | 14 | MIR4307 host gene [Source:HGNC Symbol;Acc |
| MIR4307 | 14 | microRNA 4307 [Source:HGNC Symbol;Acc:HG |
| AL390334.1 | 14 | ribosomal protein S27a (RPS27A) pseudogene |
| LINC00645 | 14 | long intergenic non-protein coding RNA 645 [S |
| MIR3171 | 14 | microRNA 3171 [Source:HGNC Symbol;Acc:HG |
| BNIP3P1 | 14 | BCL2 interacting protein 3 pseudogene 1 [Sou |
| AL139023.1 | 14 | novel transcript |
| RPL26P3 | 14 | ribosomal protein L26 pseudogene 3 [Source: |
| LINC02300 | 14 | long intergenic non-protein coding RNA 2300 |
| EIF4A1P12 | 14 | eukaryotic translation initiation factor 4A1 pse |
| BTF3P2 | 14 | BTF3 pseudogene 2 [Source:HGNC Symbol;Acc |
| FOXG1-AS1 | 14 | FOXG1 antisense RNA 1 [Source:HGNC Symbo |
| FOXG1 | 14 | forkhead box G1 [Source:HGNC Symbol;Acc:H |
| LINC01551 | 14 | long intergenic non-protein coding RNA 1551 |
| LINC02282 | 14 | long intergenic non-protein coding RNA 2282 |
| LINC02281 | 14 | long intergenic non-protein coding RNA 2281 |
| LINC02327 | 14 | long intergenic non-protein coding RNA 2327 |
| AL122126.1 | 14 | Vac14 homolog (S. cerevisiae) (VAC14) pseudo |
| LINC02326 | 14 | long intergenic non-protein coding RNA 2326 |
| Y_RNA | 14 | Y RNA [Source:RFAM;Acc:RF00019] |
| AL135878.1 | 14 | novel transcript |
| RNU6-864P | 14 | RNA, U6 small nuclear 864, pseudogene [Sour |
| AL133166.1 | 14 | novel transcript |
| AL158058.1 | 14 | novel transcript |
| AL158058.2 | 14 | novel transcript |
| RNU11-5P | 14 | RNA, U11 small nuclear 5, pseudogene [Sourc |
| PRKD1 | 14 | protein kinase D1 [Source:HGNC Symbol;Acc:H |
| AL356756.1 | 14 | novel transcript, antisense to PRKD1 |
| RNU6-1234P | 14 | RNA, U6 small nuclear 1234, pseudogene [Sou |
| AL355053.1 | 14 | novel transcript, antisense to PRKD1 |

| | | |
|---|---|---|
| AL133372.3 | 14 | novel transcript, antisense to PRKD1 |
| AL133372.2 | 14 | novel transcript |
| AL133372.1 | 14 | ribosomal protein S6 (RPS6) pseudogene |
| AL079305.1 | 14 | TEC |
| G2E3-AS1 | 14 | G2E3 antisense RNA 1 [Source:HGNC Symbol;A |
| AL117355.1 | 14 | NULL |
| SYF2P1 | 14 | SYF2 pre-mRNA splicing factor pseudogene 1 [ |
| G2E3 | 14 | G2/M-phase specific E3 ubiquitin protein ligas |
| SCFD1 | 14 | sec1 family domain containing 1 [Source:HGN |
| UBE2CP1 | 14 | ubiquitin conjugating enzyme E2 C pseudogen |
| AL121852.1 | 14 | novel transcript, antisense to SCFD1 |
| RPL12P5 | 14 | ribosomal protein L12 pseudogene 5 [Source: |
| AL049830.2 | 14 | ribosomal protein L27 (RPL27) pseudogene |
| AL049830.1 | 14 | ribosomal protein L27 (RPL27) pseudogene |
| COCH | 14 | cochlin [Source:HGNC Symbol;Acc:HGNC:218( |
| AL049830.3 | 14 | novel transcript, antisense to COCH |
| STRN3 | 14 | striatin 3 [Source:HGNC Symbol;Acc:HGNC:15 |
| HIGD1AP17 | 14 | HIG1 hypoxia inducible domain family membe |
| MIR624 | 14 | microRNA 624 [Source:HGNC Symbol;Acc:HGN |
| AL049830.4 | 14 | intraflagellar transport 20 homolog (Chlamydo |
| AP4S1 | 14 | adaptor related protein complex 4 subunit sig |
| HECTD1 | 14 | HECT domain E3 ubiquitin protein ligase 1 [So |
| RPL21P5 | 14 | ribosomal protein L21 pseudogene 5 [Source: |
| RNU6-541P | 14 | RNA, U6 small nuclear 541, pseudogene [Sour |
| Y_RNA | 14 | Y RNA [Source:RFAM;Acc:RF00019] |
| AL136418.1 | 14 | novel transcript, antisense to HEATR5A |
| Y_RNA | 14 | Y RNA [Source:RFAM;Acc:RF00019] |
| HEATR5A | 14 | HEAT repeat containing 5A [Source:HGNC Sym |
| AL139353.3 | 14 | ATP synthase, H+ transporting, mitochondrial |
| AL139353.1 | 14 | novel protein |
| AL139353.2 | 14 | novel transcript, antisense to C14orf126 |
| DTD2 | 14 | D-aminoacyl-tRNA deacylase 2 [Source:HGNC |
| AL163973.1 | 14 | ribosomal protein L9 (RPL9) pseudogene |
| GPR33 | 14 | G protein-coupled receptor 33 (gene/pseudog |
| NUBPL | 14 | nucleotide binding protein like [Source:HGNC |
| AL163973.3 | 14 | novel transcript, antisense to NUBPL |
| AL163973.2 | 14 | novel transcript, antisense to NUBPL |
| RNU6-602P | 14 | RNA, U6 small nuclear 602, pseudogene [Sour |
| RNU6-455P | 14 | RNA, U6 small nuclear 455, pseudogene [Sour |
| AL355112.1 | 14 | novel transcript |
| LINC02313 | 14 | long intergenic non-protein coding RNA 2313 |
| AL352984.2 | 14 | zinc finger, AN1-type domain 2A (ZFAND2A) p |
| AL352984.1 | 14 | novel transcript |
| ARHGAP5-AS1 | 14 | ARHGAP5 antisense RNA 1 (head to head) [So |
| ARHGAP5 | 14 | Rho GTPase activating protein 5 [Source:HGN( |
| AL161665.1 | 14 | novel transcript, antisense to ARHGAP5 |
| AL161665.2 | 14 | novel transcript |
| RNU6-7 | 14 | RNA, U6 small nuclear 7 [Source:HGNC Symbo |
| RNU6-8 | 14 | RNA, U6 small nuclear 8 [Source:HGNC Symbo |
| AL136298.2 | 14 | HERV-FRD provirus ancestral Env polyprotein |

| | | |
|---|---|---|
| AL136298.3 | 14 | KIAA1143 (KIAA1143) pseudogene |
| AKAP6 | 14 | A-kinase anchoring protein 6 [Source:HGNC Sy |
| AL136298.1 | 14 | novel transcript, antisense to AKAP6 |
| RN7SL660P | 14 | RNA, 7SL, cytoplasmic 660, pseudogene [Sour |
| MTCO1P2 | 14 | MT-CO1 pseudogene 2 [Source:HGNC Symbol |
| AL049781.1 | 14 | TEC |
| NPAS3 | 14 | neuronal PAS domain protein 3 [Source:HGNC |
| AL161851.1 | 14 | novel transcript, antisense to NPAS3 |
| AL161851.2 | 14 | novel transcript, antisense to NPAS3 |
| EGLN3 | 14 | egl-9 family hypoxia inducible factor 3 [Source |
| EGLN3-AS1 | 14 | EGLN3 antisense RNA 1 [Source:HGNC Symbo |
| AL356806.1 | 14 | sulfotransferase family 3A, member 1 (Sult3a1 |
| SPTSSA | 14 | serine palmitoyltransferase small subunit A [S |
| RPL23AP71 | 14 | ribosomal protein L23a pseudogene 71 [Sourc |
| EAPP | 14 | E2F associated phosphoprotein [Source:HGNC |
| AL445363.3 | 14 | TEC |
| AL445363.1 | 14 | novel transcript |
| RNU1-27P | 14 | RNA, U1 small nuclear 27, pseudogene [Sourc |
| RNU1-28P | 14 | RNA, U1 small nuclear 28, pseudogene [Sourc |
| AL445363.2 | 14 | novel transcript, antisense to SNX6 |
| SNX6 | 14 | sorting nexin 6 [Source:HGNC Symbol;Acc:HGI |
| RPS19P3 | 14 | ribosomal protein S19 pseudogene 3 [Source: |
| Metazoa_SRP | 14 | Metazoan signal recognition particle RNA [Sou |
| RNU6-1261P | 14 | RNA, U6 small nuclear 1261, pseudogene [Sou |
| RPL23AP8 | 14 | ribosomal protein L23a pseudogene 8 [Source |
| CFL2 | 14 | cofilin 2 [Source:HGNC Symbol;Acc:HGNC:187 |
| RPL12P6 | 14 | ribosomal protein L12 pseudogene 6 [Source: |
| BAZ1A | 14 | bromodomain adjacent to zinc finger domain |
| RNU7-41P | 14 | RNA, U7 small nuclear 41 pseudogene [Source |
| AL121603.2 | 14 | novel transcript, antisense to BAZ1A |
| SRP54-AS1 | 14 | SRP54 antisense RNA 1 (head to head) [Source |
| AL121603.1 | 14 | SMT3 suppressor of mif two 3 homolog 4 (S. c |
| IGBP1P1 | 14 | immunoglobulin (CD79A) binding protein 1 ps |
| SRP54 | 14 | signal recognition particle 54 [Source:HGNC Sy |
| FAM177A1 | 14 | family with sequence similarity 177 member A |
| PPP2R3C | 14 | protein phosphatase 2 regulatory subunit B''g |
| AL049776.1 | 14 | TEC |
| PRORP | 14 | protein only RNase P catalytic subunit [Source |
| AL121594.1 | 14 | novel transcript |
| RPL23AP70 | 14 | ribosomal protein L23a pseudogene 70 [Sourc |
| SEPTIN7P1 | 14 | septin 7 pseudogene 1 [Source:HGNC Symbol; |
| MRPL57P8 | 14 | mitochondrial ribosomal protein L57 pseudog |
| DPRXP3 | 14 | divergent-paired related homeobox pseudoge |
| RPL7AP3 | 14 | ribosomal protein L7a pseudogene 3 [Source:I |
| RPL9P3 | 14 | ribosomal protein L9 pseudogene 3 [Source:H |
| PSMA6 | 14 | proteasome subunit alpha 6 [Source:HGNC Sy |
| AL133163.2 | 14 | novel transcript |
| AL133163.1 | 14 | ribosomal protein, large, P0 (RPLP0) pseudoge |
| NFKBIA | 14 | NFKB inhibitor alpha [Source:HGNC Symbol;Ac |
| DNAJC8P1 | 14 | DnaJ heat shock protein family (Hsp40) memb |

| | | |
|---|---|---|
| AL133163.3 | 14 | novel transcript |
| RPS3AP4 | 14 | ribosomal protein S3A pseudogene 4 [Source: |
| KRT18P6 | 14 | keratin 18 pseudogene 6 [Source:HGNC Symb |
| INSM2 | 14 | INSM transcriptional repressor 2 [Source:HGN |
| RALGAPA1 | 14 | Ral GTPase activating protein catalytic alpha s |
| AL137818.1 | 14 | NULL |
| QRSL1P3 | 14 | QRSL1 pseudogene 3 [Source:HGNC Symbol;A |
| AL162311.3 | 14 | novel transcript, antisense to BRMS1L |
| NUTF2P2 | 14 | nuclear transport factor 2 pseudogene 2 [Sou |
| BRMS1L | 14 | BRMS1 like transcriptional repressor [Source:I |
| AL162311.1 | 14 | novel transcript |
| AL162311.2 | 14 | RNA-binding protein EWS (EWSR1) pseudoger |
| AL133304.3 | 14 | novel transcript, sense overlapping BRMS1L |
| AL133304.1 | 14 | novel transcript, antisense to BRMS1L |
| ILF2P2 | 14 | interleukin enhancer binding factor 2 pseudog |
| AL133304.2 | 14 | novel transcript |
| LINC00609 | 14 | long intergenic non-protein coding RNA 609 [S |
| PTCSC3 | 14 | papillary thyroid carcinoma susceptibility canc |
| AL162511.1 | 14 | novel transcript |
| RN7SKP21 | 14 | RN7SK pseudogene 21 [Source:HGNC Symbol; |
| MBIP | 14 | MAP3K12 binding inhibitory protein 1 [Source |
| AL132857.1 | 14 | novel transcript |
| DPPA3P2 | 14 | developmental pluripotency associated 3 pseu |
| RNU7-93P | 14 | RNA, U7 small nuclear 93 pseudogene [Source |
| SFTA3 | 14 | surfactant associated 3 [Source:NCBI gene;Acc |
| SFTA3 | 14 | surfactant associated 3 [Source:HGNC Symbol |
| NKX2-1 | 14 | NK2 homeobox 1 [Source:HGNC Symbol;Acc:H |
| NKX2-1-AS1 | 14 | NKX2-1 antisense RNA 1 [Source:HGNC Symbo |
| PHKBP2 | 14 | phosphorylase kinase regulatory subunit beta |
| RPL29P3 | 14 | ribosomal protein L29 pseudogene 3 [Source:I |
| NKX2-8 | 14 | NK2 homeobox 8 [Source:HGNC Symbol;Acc:H |
| RN7SKP257 | 14 | RN7SK pseudogene 257 [Source:HGNC Symbo |
| AL079303.1 | 14 | novel transcript, antisense to PAX9 |
| PAX9 | 14 | paired box 9 [Source:HGNC Symbol;Acc:HGNC |
| SLC25A21 | 14 | solute carrier family 25 member 21 [Source:H |
| AL162464.2 | 14 | novel transcript, antisense to SLC25A21 |
| AL162464.1 | 14 | novel transcript, antisense to SLC25A21 |
| MIR4503 | 14 | microRNA 4503 [Source:HGNC Symbol;Acc:HG |
| AL079304.1 | 14 | novel transcript, sense intronic to SLC25A21 |
| SLC25A21-AS1 | 14 | SLC25A21 antisense RNA 1 [Source:HGNC Sym |
| RNU6-273P | 14 | RNA, U6 small nuclear 273, pseudogene [Sour |
| MIPOL1 | 14 | mirror-image polydactyly 1 [Source:HGNC Syn |
| RNU6-886P | 14 | RNA, U6 small nuclear 886, pseudogene [Sour |
| AL121790.2 | 14 | novel transcript |
| AL121790.1 | 14 | novel transcript |
| FOXA1 | 14 | forkhead box A1 [Source:HGNC Symbol;Acc:H |
| TTC6 | 14 | tetratricopeptide repeat domain 6 [Source:HG |
| AL136296.1 | 14 | NULL |
| RNU6-1277P | 14 | RNA, U6 small nuclear 1277, pseudogene [Sou |
| LINC00517 | 14 | long intergenic non-protein coding RNA 517 [S |

| | | |
|---|---|---|
| AL359233.1 | 14 | novel transcript, antisense to TTC6 |
| AL392023.2 | 14 | novel transcript, antisense to TTC6 |
| AL392023.1 | 14 | novel transcript |
| SSTR1 | 14 | somatostatin receptor 1 [Source:HGNC Symbo |
| CLEC14A | 14 | C-type lectin domain containing 14A [Source:I |
| AL355835.1 | 14 | novel transcript |
| AL357094.1 | 14 | general transcription factor IIH, polypeptide 3 |
| KRT8P1 | 14 | keratin 8 pseudogene 1 [Source:HGNC Symbo |
| LINC00639 | 14 | long intergenic non-protein coding RNA 639 [S |
| AL132994.1 | 14 | zinc finger protein 41 homolog (mouse) (ZFP4 |
| AL132994.2 | 14 | Down syndrome critical region gene 6 (DSCR6 |
| Y_RNA | 14 | Y RNA [Source:RFAM;Acc:RF00019] |
| SEC23A | 14 | Sec23 homolog A, coat complex II component |
| AL109628.1 | 14 | ribosomal protein, large, P1 (RPLP1) pseudoge |
| SEC23A-AS1 | 14 | SEC23A antisense RNA 1 [Source:HGNC Symbo |
| PPIAP4 | 14 | peptidylprolyl isomerase A pseudogene 4 [Sou |
| GEMIN2 | 14 | gem nuclear organelle associated protein 2 [S |
| TRAPPC6B | 14 | trafficking protein particle complex 6B [Source |
| AL109628.2 | 14 | novel transcript, antisense to TRAPPC6B |
| RPL7AP2 | 14 | ribosomal protein L7a pseudogene 2 [Source:I |
| AL132639.3 | 14 | novel transcript, antisense to PNN |
| PNN | 14 | pinin, desmosome associated protein [Source |
| AL132639.1 | 14 | transmembrane protein 92 (TMEM92) pseudo |
| YTHDF2P1 | 14 | YTH domain family member 2 pseudogene 1 [ |
| MIA2 | 14 | MIA SH3 domain ER export factor 2 [Source:H |
| AL132639.2 | 14 | novel transcript, antisense to CTAGE5 |
| AL157791.2 | 14 | glutaredoxin 5 (GLRX5) pseudogene |
| AL157791.1 | 14 | coilin (COIL) pseudogene |
| FBXO33 | 14 | F-box protein 33 [Source:HGNC Symbol;Acc:H |
| AL049828.1 | 14 | novel transcript |
| AL049828.2 | 14 | novel transcript |
| AL390800.1 | 14 | novel transcript |
| LINC02315 | 14 | long intergenic non-protein coding RNA 2315 |
| AL391516.1 | 14 | WD repeat-containing protein 75 (WDR75) ps |
| AL356596.1 | 14 | TEC |
| AL121821.3 | 14 | ubiquitin-conjugating enzyme E2D 3 (UBE2D3 |
| AL121821.2 | 14 | novel transcript |
| AL121821.1 | 14 | NULL |
| LRFN5 | 14 | leucine rich repeat and fibronectin type III dor |
| FKBP1BP1 | 14 | FKBP prolyl isomerase 1B pseudogene 1 [Sour |
| AL445074.1 | 14 | novel transcript |
| AL442163.1 | 14 | novel transcript, sense overlapping to a novel |
| AL450442.1 | 14 | novel transcript |
| AL442163.2 | 14 | leucine-rich pentatricopeptide repeat contain |
| AL442163.3 | 14 | endogenous retrovirus group FRD, member 1 |
| AL356800.1 | 14 | 14-3-3 protein theta (YWHAQ) pseudogene |
| AL163153.1 | 14 | UDP glucuronosyltransferase 2 family, polype |
| AL445383.1 | 14 | Tubulin alpha-1B chain (TUB1A) pseudogene |
| HNRNPUP1 | 14 | heterogeneous nuclear ribonucleoprotein U p |
| AL583809.1 | 14 | novel transcript |

| | | |
|---|---|---|
| KRT8P2 | 14 | keratin 8 pseudogene 2 [Source:HGNC Symbo |
| AL358913.1 | 14 | Rho GTPase activating protein 17 (ARHGAP17 |
| AL358913.2 | 14 | Rho GTPase activating protein 17 (ARHGAP17 |
| LINC02307 | 14 | long intergenic non-protein coding RNA 2307 |
| EIF4BP1 | 14 | eukaryotic translation initiation factor 4B pseu |
| AL109766.1 | 14 | chromosome 11 open reading frame 58 (C11c |
| YWHAZP1 | 14 | tyrosine 3-monooxygenase/tryptophan 5-mor |
| AL161752.1 | 14 | novel transcript |
| LINC02277 | 14 | long intergenic non-protein coding RNA 2277 |
| FSCB | 14 | fibrous sheath CABYR binding protein [Source |
| AL356022.1 | 14 | novel transcript, antisense to FSCB |
| LINC02302 | 14 | long intergenic non-protein coding RNA 2302 |
| DOCK11P1 | 14 | dedicator of cytokinesis 11 pseudogene 1 [Sou |
| AL049870.2 | 14 | novel transcript, antisense to C14orf28 |
| AL049870.1 | 14 | DET1 and DDB1 associated 1 (DDA1) pseudog |
| AL049870.4 | 14 | Ras-related GTP binding A (RRAGA) pseudogen |
| C14orf28 | 14 | chromosome 14 open reading frame 28 [Sour |
| AL049870.3 | 14 | novel transcript, antisense to C14orf28 |
| KLHL28 | 14 | kelch like family member 28 [Source:HGNC Sy |
| TOGARAM1 | 14 | TOG array regulator of axonemal microtubule |
| AL121809.2 | 14 | novel transcript |
| AL121809.1 | 14 | novel transcript, antisense to PRPF39 |
| PRPF39 | 14 | pre-mRNA processing factor 39 [Source:HGNC |
| SNORD58 | 14 | Small nucleolar RNA SNORD58 [Source:RFAM; |
| SNORD127 | 14 | small nucleolar RNA, C/D box 127 [Source:HG |
| FKBP3 | 14 | FKBP prolyl isomerase 3 [Source:HGNC Symbo |
| FANCM | 14 | FA complementation group M [Source:HGNC S |
| MIS18BP1 | 14 | MIS18 binding protein 1 [Source:HGNC Symbo |
| RNU6-552P | 14 | RNA, U6 small nuclear 552, pseudogene [Sour |
| DNAJC19P9 | 14 | DnaJ heat shock protein family (Hsp40) memb |
| AL162632.3 | 14 | novel transcript |
| AL162632.1 | 14 | novel transcript |
| AL162632.2 | 14 | Rac GTPase activating protein 1 (RACGAP1) ps |
| AL139354.1 | 14 | novel transcript |
| LINC02303 | 14 | long intergenic non-protein coding RNA 2303 |
| LINC00871 | 14 | long intergenic non-protein coding RNA 871 [S |
| RPL10L | 14 | ribosomal protein L10 like [Source:HGNC Sym |
| MDGA2 | 14 | MAM domain containing glycosylphosphatidy |
| AL359951.1 | 14 | NULL |
| RPA2P1 | 14 | replication protein A2 pseudogene 1 [Source:I |
| RPL13AP2 | 14 | ribosomal protein L13a pseudogene 2 [Source |
| RPS15AP3 | 14 | ribosomal protein S15a pseudogene 3 [Source |
| MIR548Y | 14 | microRNA 548y [Source:HGNC Symbol;Acc:HG |
| LINC00648 | 14 | long intergenic non-protein coding RNA 648 [S |
| AL121576.1 | 14 | WD repeat and SOCS box containing 2 (WSB2) |
| AL359212.1 | 14 | ataxin 3 (ATXN3) pseudogene |
| AL512358.1 | 14 | novel transcript |
| AL512358.2 | 14 | novel transcript |
| AL358335.3 | 14 | mitochondrial ribosomal protein L54 (MRPL54 |
| RNU6-297P | 14 | RNA, U6 small nuclear 297, pseudogene [Sour |

| | | |
|---|---|---|
| AL358335.2 | 14 | novel transcript |
| AL358335.1 | 14 | STT3, subunit of the oligosaccharyltransferase |
| RPL18P1 | 14 | ribosomal protein L18 pseudogene 1 [Source: |
| AL512360.1 | 14 | nuclear factor (erythroid-derived 2)-like 3 (NF |
| AL110505.1 | 14 | novel transcript |
| AL512361.1 | 14 | ATP synthase, H+ transporting, mitochondrial |
| RNA5SP384 | 14 | RNA, 5S ribosomal pseudogene 384 [Source:H |
| RPS29 | 14 | ribosomal protein S29 [Source:HGNC Symbol; |
| RPL32P29 | 14 | ribosomal protein L32 pseudogene 29 [Source |
| RN7SL1 | 14 | RNA component of signal recognition particle |
| Y_RNA | 14 | Y RNA [Source:RFAM;Acc:RF00019] |
| LRR1 | 14 | leucine rich repeat protein 1 [Source:HGNC Sy |
| RHOQP1 | 14 | ras homolog family member Q pseudogene 1 |
| AL139099.3 | 14 | phosphatidylinositol glycan anchor biosynthes |
| RPL36AL | 14 | ribosomal protein L36a like [Source:HGNC Syr |
| MGAT2 | 14 | mannosyl (alpha-1,6-)-glycoprotein beta-1,2-N |
| AL139099.1 | 14 | novel transcript |
| DNAAF2 | 14 | dynein axonemal assembly factor 2 [Source:H |
| AL139099.2 | 14 | novel transcript |
| POLE2 | 14 | DNA polymerase epsilon 2, accessory subunit |
| AL591767.3 | 14 | serine/threonine kinase 16 (STK16) pseudoge |
| AL591767.2 | 14 | actin-like 6 (ACTL6) pseudogene |
| RNU6ATAC30P | 14 | RNA, U6atac small nuclear 30, pseudogene [S |
| KLHDC1 | 14 | kelch domain containing 1 [Source:HGNC Sym |
| AL591767.1 | 14 | novel transcript |
| KLHDC2 | 14 | kelch domain containing 2 [Source:HGNC Sym |
| NEMF | 14 | nuclear export mediator factor [Source:HGNC |
| Y_RNA | 14 | Y RNA [Source:RFAM;Acc:RF00019] |
| RNU6-539P | 14 | RNA, U6 small nuclear 539, pseudogene [Sour |
| RN7SL3 | 14 | RNA component of signal recognition particle |
| AL627171.2 | 14 | novel transcript |
| RN7SL2 | 14 | RNA component of signal recognition particle |
| AL627171.1 | 14 | novel transcript |
| ARF6 | 14 | ADP ribosylation factor 6 [Source:HGNC Symb |
| RNU6-189P | 14 | RNA, U6 small nuclear 189, pseudogene [Sour |
| Metazoa_SRP | 14 | Metazoan signal recognition particle RNA [Sou |
| LINC01588 | 14 | long intergenic non-protein coding RNA 1588 |
| MIR6076 | 14 | microRNA 6076 [Source:HGNC Symbol;Acc:HG |
| PDLIM1P1 | 14 | PDZ and LIM domain 1 pseudogene 1 [Source: |
| AL117692.1 | 14 | ribosomal protein S15a (RPS15A) pseudogene |
| RN7SKP193 | 14 | RN7SK pseudogene 193 [Source:HGNC Symbo |
| Y_RNA | 14 | Y RNA [Source:RFAM;Acc:RF00019] |
| VCPKMT | 14 | valosin containing protein lysine methyltransf |
| SOS2 | 14 | SOS Ras/Rho guanine nucleotide exchange fac |
| L2HGDH | 14 | L-2-hydroxyglutarate dehydrogenase [Source: |
| MIR4504 | 14 | microRNA 4504 [Source:HGNC Symbol;Acc:HG |
| DMAC2L | 14 | distal membrane arm assembly complex 2 like |
| AL359397.2 | 14 | novel transcript |
| CDKL1 | 14 | cyclin dependent kinase like 1 [Source:HGNC S |
| AL359397.1 | 14 | novel transcript |

| | | |
|---|---|---|
| MAP4K5 | 14 | mitogen-activated protein kinase kinase kinas |
| AL118556.1 | 14 | novel transcript |
| Y_RNA | 14 | Y RNA [Source:RFAM;Acc:RF00019] |
| AL118556.2 | 14 | zinc finger protein 124 (ZNF124) pseudogene |
| ATL1 | 14 | atlastin GTPase 1 [Source:HGNC Symbol;Acc:H |
| SNRPGP1 | 14 | small nuclear ribonucleoprotein polypeptide ( |
| SAV1 | 14 | salvador family WW domain containing protei |
| RN7SL452P | 14 | RNA, 7SL, cytoplasmic 452, pseudogene [Sour |
| AL606834.1 | 14 | novel transcript, sense intronic to SAV1 |
| ZFP64P1 | 14 | ZFP64 zinc finger protein pseudogene 1 [Sour |
| NIN | 14 | ninein [Source:HGNC Symbol;Acc:HGNC:1490 |
| AL606834.2 | 14 | novel transcript, sense intronic to NIN |
| AL133485.2 | 14 | novel transcript, antisense to NIN |
| AL133485.3 | 14 | novel transcript |
| AL133485.1 | 14 | NULL |
| AL358334.1 | 14 | novel transcript, antisense to PYGL |
| PYGL | 14 | glycogen phosphorylase L [Source:HGNC Symb |
| ABHD12B | 14 | abhydrolase domain containing 12B [Source:H |
| AL358334.3 | 14 | novel transcript, antisense to PYGL |
| AL358334.2 | 14 | novel transcript |
| TRIM9 | 14 | tripartite motif containing 9 [Source:HGNC Sy |
| AL358334.4 | 14 | novel transcript, antisense to TRIM9 |
| AL591770.1 | 14 | novel transcript, antisense to TRIM9 |
| TMX1 | 14 | thioredoxin related transmembrane protein 1 |
| SNORA70 | 14 | Small nucleolar RNA SNORA70 [Source:RFAM; |
| Y_RNA | 14 | Y RNA [Source:RFAM;Acc:RF00019] |
| LINC00519 | 14 | long intergenic non-protein coding RNA 519 [S |
| LINC00640 | 14 | long intergenic non-protein coding RNA 640 [S |
| AL358332.1 | 14 | novel transcript |
| LINC02310 | 14 | long intergenic non-protein coding RNA 2310 |
| SETP2 | 14 | SET pseudogene 2 [Source:HGNC Symbol;Acc: |
| FRMD6-AS2 | 14 | FRMD6 antisense RNA 2 [Source:HGNC Symbo |
| FRMD6 | 14 | FERM domain containing 6 [Source:HGNC Sym |
| RNU6-1291P | 14 | RNA, U6 small nuclear 1291, pseudogene [Sou |
| AL122125.1 | 14 | novel transcript, antisense to FRMD6 |
| FRMD6-AS1 | 14 | FRMD6 antisense RNA 1 [Source:HGNC Symbo |
| RNA5SP385 | 14 | RNA, 5S ribosomal pseudogene 385 [Source:H |
| RNU6-301P | 14 | RNA, U6 small nuclear 301, pseudogene [Sour |
| AL079307.2 | 14 | novel transcript |
| OR7E105P | 14 | olfactory receptor family 7 subfamily E memb |
| OR7E106P | 14 | olfactory receptor family 7 subfamily E memb |
| AL079307.1 | 14 | novel transcript |
| OR7E159P | 14 | olfactory receptor family 7 subfamily E memb |
| AL079307.3 | 14 | protein kinase-like protein SgK196 (SGK196) p |
| GNG2 | 14 | G protein subunit gamma 2 [Source:HGNC Syr |
| AL358333.2 | 14 | novel transcript, antisense to GNG2 |
| AL358333.3 | 14 | novel transcript, antisense to GNG2 |
| AL358333.1 | 14 | novel transcript, antisense to GNG2 and C14o |
| RTRAF | 14 | RNA transcription, translation and transport fa |
| NID2 | 14 | nidogen 2 [Source:HGNC Symbol;Acc:HGNC:1 |

| | | |
|---|---|---|
| LINC02319 | 14 | long intergenic non-protein coding RNA 2319 |
| COX5AP2 | 14 | cytochrome c oxidase subunit 5A pseudogene |
| AL118557.1 | 14 | integrin, alpha 3 (antigen CD49C, alpha 3 subu |
| PTGDR | 14 | prostaglandin D2 receptor [Source:HGNC Sym |
| AL365475.1 | 14 | novel Pumilio domain-containing protein KIAA |
| PTGER2 | 14 | prostaglandin E receptor 2 [Source:HGNC Sym |
| TXNDC16 | 14 | thioredoxin domain containing 16 [Source:HG |
| GPR137C | 14 | G protein-coupled receptor 137C [Source:HGN |
| ERO1A | 14 | endoplasmic reticulum oxidoreductase 1 alpha |
| AL133453.1 | 14 | novel transcript, antisense to ERO1L |
| PSMC6 | 14 | proteasome 26S subunit, ATPase 6 [Source:HG |
| STYX | 14 | serine/threonine/tyrosine interacting protein |
| GNPNAT1 | 14 | glucosamine-phosphate N-acetyltransferase 1 |
| AL139317.3 | 14 | novel transcript, antisense to GNPNAT1 |
| AL139317.5 | 14 | novel transcript, antisense to FERMT2 |
| AL139317.1 | 14 | novel transcript |
| AL139317.2 | 14 | myeloma overexpressed 2 (MYEOV2) pseudog |
| FERMT2 | 14 | fermitin family member 2 [Source:HGNC Symb |
| AL139317.4 | 14 | golgi membrane protein 1 (GOLM1) pseudoge |
| AL352979.4 | 14 | novel transcript, antisense to FERMT2 |
| AL352979.1 | 14 | NADH dehydrogenase (ubiquinone) 1 beta sub |
| AL352979.3 | 14 | centromere protein W (CENPW) pseudogene |
| DDHD1 | 14 | DDHD domain containing 1 [Source:HGNC Sym |
| AL352979.2 | 14 | novel transcript, antisense to DDHD1 |
| AL356020.1 | 14 | novel transcript, antisense to DDHD1 |
| AL365295.1 | 14 | novel transcript |
| AL365295.2 | 14 | novel transcript |
| AL163953.1 | 14 | novel transcript |
| RPS3AP46 | 14 | ribosomal protein S3a pseudogene 46 [Source |
| LINC02331 | 14 | long intergenic non-protein coding RNA 2331 |
| AL138479.1 | 14 | ribosomal protein L23a (RPL23A) pseudogene |
| MIR5580 | 14 | microRNA 5580 [Source:HGNC Symbol;Acc:HG |
| BMP4 | 14 | bone morphogenetic protein 4 [Source:HGNC |
| AL138479.2 | 14 | novel transcript, antisense to BMP4 |
| ATP5F1CP1 | 14 | ATP synthase F1 subunit gamma pseudogene |
| CDKN3 | 14 | cyclin dependent kinase inhibitor 3 [Source:HG |
| CNIH1 | 14 | cornichon family AMPA receptor auxiliary pro |
| AL359792.1 | 14 | novel transcript |
| GMFB | 14 | glia maturation factor beta [Source:HGNC Sym |
| CGRRF1 | 14 | cell growth regulator with ring finger domain |
| SAMD4A | 14 | sterile alpha motif domain containing 4A [Sou |
| AL133444.1 | 14 | novel transcript |
| GCH1 | 14 | GTP cyclohydrolase 1 [Source:HGNC Symbol;A |
| RNU6ATAC9P | 14 | RNA, U6atac small nuclear 9, pseudogene [Sou |
| MIR4308 | 14 | microRNA 4308 [Source:HGNC Symbol;Acc:HG |
| FDPSP3 | 14 | farnesyl diphosphate synthase pseudogene 3 |
| WDHD1 | 14 | WD repeat and HMG-box DNA binding protein |
| RPSAP13 | 14 | ribosomal protein SA pseudogene 13 [Source: |
| SOCS4 | 14 | suppressor of cytokine signaling 4 [Source:HG |
| MAPK1IP1L | 14 | mitogen-activated protein kinase 1 interacting |

| | | |
|---|---|---|
| LGALS3 | 14 | galectin 3 [Source:HGNC Symbol;Acc:HGNC:6[illegible] |
| DLGAP5 | 14 | DLG associated protein 5 [Source:HGNC Symb |
| AL139316.1 | 14 | novel transcript |
| AL158801.5 | 14 | ubiquitin-conjugating enzyme E2L 3 (UBE2L3) |
| AL158801.4 | 14 | cytochrome c oxidase subunit Va (COX5A) pse |
| AL158801.2 | 14 | novel transcript |
| FBXO34 | 14 | F-box protein 34 [Source:HGNC Symbol;Acc:H |
| AL158801.1 | 14 | ribosomal protein L34 (RPL34) pseudogene |
| Metazoa_SRP | 14 | Metazoan signal recognition particle RNA [Sou |
| CHMP4BP1 | 14 | charged multivesicular body protein 4B pseud |
| AL158801.3 | 14 | novel transcript |
| ATG14 | 14 | autophagy related 14 [Source:HGNC Symbol;[illegible] |
| AL356801.1 | 14 | high mobility group nucleosome binding pseu |
| TBPL2 | 14 | TATA-box binding protein like 2 [Source:HGN[illegible] |
| RPL21P6 | 14 | ribosomal protein L21 pseudogene 6 [Source:[illegible] |
| KTN1-AS1 | 14 | KTN1 antisense RNA 1 [Source:HGNC Symbol;[illegible] |
| RPL7AP4 | 14 | ribosomal protein L7a pseudogene 4 [Source:[illegible] |
| ABI1P1 | 14 | abl interactor 1 pseudogene 1 [Source:HGNC [illegible] |
| KTN1 | 14 | kinectin 1 [Source:HGNC Symbol;Acc:HGNC:6[illegible] |
| Y_RNA | 14 | Y RNA [Source:RFAM;Acc:RF00019] |
| AL355773.1 | 14 | novel transcript |
| RPL13AP3 | 14 | ribosomal protein L13a pseudogene 3 [Source |
| LINC00520 | 14 | long intergenic non-protein coding RNA 520 [[illegible] |
| AL163952.1 | 14 | novel transcript |
| AL138995.1 | 14 | novel transcript |
| PELI2 | 14 | pellino E3 ubiquitin protein ligase family mem |
| AL355073.1 | 14 | novel transcript |
| AL355073.2 | 14 | novel transcript |
| LINC02284 | 14 | long intergenic non-protein coding RNA 2284 |
| TMEM260 | 14 | transmembrane protein 260 [Source:HGNC Sy |
| AL355103.1 | 14 | novel transcript |
| AL161757.2 | 14 | novel transcript, antisense to C14orf101 |
| AL161757.5 | 14 | novel transcript, antisense to TMEM260 |
| AL161757.1 | 14 | ribosomal protein L36a (RPL36A) pseudogene |
| AL161757.3 | 14 | novel transcript |
| OTX2 | 14 | orthodenticle homeobox 2 [Source:HGNC Sym |
| OTX2-AS1 | 14 | OTX2 antisense RNA 1 (head to head) [Source |
| AL161757.4 | 14 | novel transcript |
| RN7SL461P | 14 | RNA, 7SL, cytoplasmic 461, pseudogene [Sour |
| RNU6-1204P | 14 | RNA, U6 small nuclear 1204, pseudogene [Sou |
| AL137100.3 | 14 | novel transcript |
| RPL3P3 | 14 | ribosomal protein L3 pseudogene 3 [Source:H |
| AL137100.1 | 14 | zinc finger, CCHC domain containing 7 (ZCCHC |
| AL137100.2 | 14 | zinc finger, CCHC domain containing 7 (ZCCHC |
| AL391152.1 | 14 | novel transcript |
| EXOC5 | 14 | exocyst complex component 5 [Source:HGNC |
| AP5M1 | 14 | adaptor related protein complex 5 subunit mu |
| AL355834.2 | 14 | novel transcript |
| AL355834.1 | 14 | novel transcript |
| NAA30 | 14 | N(alpha)-acetyltransferase 30, NatC catalytic [illegible] |

| | | |
|---|---|---|
| CCDC198 | 14 | coiled-coil domain containing 198 [Source:HG |
| SLC35F4 | 14 | solute carrier family 35 member F4 [Source:H |
| AL161804.1 | 14 | novel transcript, antisense to SLC35F4 |
| AL136520.1 | 14 | serine/arginine-rich splicing factor 10 (SRSF10 |
| RN7SKP99 | 14 | RN7SK pseudogene 99 [Source:HGNC Symbol; |
| AL049838.1 | 14 | novel transcript, overlapping C14orf37 |
| ARMH4 | 14 | armadillo like helical domain containing 4 [Sou |
| AL121579.2 | 14 | polycomb group ring finger 3 (PCGF3) pseudo |
| UBA52P3 | 14 | ubiquitin A-52 residue ribosomal protein fusio |
| RN7SL598P | 14 | RNA, 7SL, cytoplasmic 598, pseudogene [Sour |
| AL121579.1 | 14 | 40S ribosomal protein S2 (RPS2) pseudogene |
| ACTR10 | 14 | actin related protein 10 [Source:HGNC Symbo |
| PSMA3 | 14 | proteasome subunit alpha 3 [Source:HGNC Sy |
| AL132989.1 | 14 | novel transcript, sense overlapping PSMA3 |
| PSMA3-AS1 | 14 | PSMA3 antisense RNA 1 [Source:HGNC Symbo |
| HMGB1P14 | 14 | high mobility group box 1 pseudogene 14 [Sou |
| AL132989.2 | 14 | long intergenic non-protein coding RNA 216 |
| RNU6-341P | 14 | RNA, U6 small nuclear 341, pseudogene [Sour |
| ARID4A | 14 | AT-rich interaction domain 4A [Source:HGNC |
| AL139021.2 | 14 | novel transcript, antisense to ARID4A and TON |
| TOMM20L | 14 | translocase of outer mitochondrial membrane |
| AL139021.1 | 14 | novel transcript, antisense to TIMM9 |
| TIMM9 | 14 | translocase of inner mitochondrial membrane |
| KIAA0586 | 14 | KIAA0586 [Source:HGNC Symbol;Acc:HGNC:1 |
| Y_RNA | 14 | Y RNA [Source:RFAM;Acc:RF00019] |
| HSBP1P1 | 14 | heat shock factor binding protein 1 pseudoge |
| HNRNPCP1 | 14 | heterogeneous nuclear ribonucleoprotein C p |
| RPL9P5 | 14 | ribosomal protein L9 pseudogene 5 [Source:H |
| DACT1 | 14 | dishevelled binding antagonist of beta catenin |
| LINC01500 | 14 | long intergenic non-protein coding RNA 1500 |
| AL049873.1 | 14 | ribosomal protein L31 (RPL31) pseudogene |
| AL049873.2 | 14 | transmembrane protein 98 (TMEM98) pseudo |
| AL359219.1 | 14 | aldo-keto reductase family 1, member B1 (ald |
| PPIAP5 | 14 | peptidylprolyl isomerase A pseudogene 5 [Sou |
| DAAM1 | 14 | dishevelled associated activator of morphoge |
| GPR135 | 14 | G protein-coupled receptor 135 [Source:HGN |
| L3HYPDH | 14 | trans-L-3-hydroxyproline dehydratase [Source |
| AL121694.1 | 14 | novel transcript |
| JKAMP | 14 | JNK1/MAPK8 associated membrane protein [ |
| CCDC175 | 14 | coiled-coil domain containing 175 [Source:HG |
| RTN1 | 14 | reticulon 1 [Source:HGNC Symbol;Acc:HGNC:1 |
| MIR5586 | 14 | microRNA 5586 [Source:HGNC Symbol;Acc:HG |
| AL133299.1 | 14 | novel transcript, antisense to LRRC9 |
| LRRC9 | 14 | leucine rich repeat containing 9 [Source:HGN |
| AL157911.1 | 14 | novel transcript, antisense to LRRC9 |
| PCNX4 | 14 | pecanex 4 [Source:HGNC Symbol;Acc:HGNC:2 |
| DHRS7 | 14 | dehydrogenase/reductase 7 [Source:HGNC Sy |
| SCOCP1 | 14 | short coiled-coil protein pseudogene 1 [Sourc |
| PSMA3P | 14 | proteasome subunit alpha 3 pseudogene [Sou |
| RPL17P2 | 14 | ribosomal protein L17 pseudogene 2 [Source: |

| | | |
|---|---|---|
| AL157756.1 | 14 | novel transcript, antisense to PPM1A |
| PPM1A | 14 | protein phosphatase, Mg2+/Mn2+ dependent |
| LINC02322 | 14 | long intergenic non-protein coding RNA 2322 |
| C14orf39 | 14 | chromosome 14 open reading frame 39 [Sour |
| RBM8B | 14 | RNA binding motif protein 8B pseudogene [So |
| GNRHR2P1 | 14 | GNRHR2 pseudogene 1 [Source:HGNC Symbol |
| SALL4P7 | 14 | spalt like transcription factor 4 pseudogene 7 |
| SIX6 | 14 | SIX homeobox 6 [Source:HGNC Symbol;Acc:H( |
| AL049874.3 | 14 | novel transcript, antisense to C14orf39 |
| AL049874.4 | 14 | vomeronasal 1 receptor 4 (VN1R4) pseudoger |
| AL049874.2 | 14 | ribosomal protein L37 (RPL37) pseudogene |
| AL049874.1 | 14 | ribosomal protein S15a (RPS15A) pseudogene |
| SALRNA1 | 14 | senescence associated long non-coding RNA 1 |
| SIX1 | 14 | SIX homeobox 1 [Source:HGNC Symbol;Acc:H( |
| SIX4 | 14 | SIX homeobox 4 [Source:HGNC Symbol;Acc:H( |
| MNAT1 | 14 | MNAT1 component of CDK activating kinase [ |
| AL132777.1 | 14 | MAD2 mitotic arrest deficient-like 1 (yeast) (N |
| AL160236.2 | 14 | novel transcript |
| SRMP2 | 14 | spermidine synthase pseudogene 2 [Source:H |
| TRMT5 | 14 | tRNA methyltransferase 5 [Source:HGNC Syml |
| AL160236.1 | 14 | high mobility group nucleosomal binding dom |
| RNU6-398P | 14 | RNA, U6 small nuclear 398, pseudogene [Sour |
| SLC38A6 | 14 | solute carrier family 38 member 6 [Source:HG |
| PRKCH | 14 | protein kinase C eta [Source:HGNC Symbol;Ac |
| TMEM30B | 14 | transmembrane protein 30B [Source:HGNC Sy |
| AL359220.1 | 14 | novel transcript, antisense to PRKCH |
| AL355916.3 | 14 | novel protein |
| AL355916.2 | 14 | novel transcript, antisense to PRKCH |
| LINC01303 | 14 | long intergenic non-protein coding RNA 1303 |
| AL355916.1 | 14 | novel transcript |
| HIF1A-AS1 | 14 | HIF1A antisense RNA 1 [Source:HGNC Symbol, |
| HIF1A | 14 | hypoxia inducible factor 1 subunit alpha [Sour |
| HIF1A-AS3 | 14 | HIF1A antisense RNA 3 [Source:HGNC Symbol, |
| AL137129.1 | 14 | novel transcript |
| SNAPC1 | 14 | small nuclear RNA activating complex polypep |
| AL137918.1 | 14 | TEC |
| SYT16 | 14 | synaptotagmin 16 [Source:HGNC Symbol;Acc: |
| COX4I1P1 | 14 | cytochrome c oxidase subunit 4I1 pseudogene |
| AL390816.1 | 14 | novel transcript |
| AL390816.2 | 14 | novel transcript |
| LINC00643 | 14 | long intergenic non-protein coding RNA 643 [S |
| LINC00644 | 14 | long intergenic non-protein coding RNA 644 [S |
| AL390816.3 | 14 | novel pseudogene |
| AL389895.1 | 14 | protein tyrosine phosphatase type IVA, memb |
| ATP5F1AP4 | 14 | ATP synthase F1 subunit alpha pseudogene 4 |
| KCNH5 | 14 | potassium voltage-gated channel subfamily H |
| AL137191.1 | 14 | novel transcript |
| PARP1P2 | 14 | poly(ADP-ribose) polymerase 1 pseudogene 2 |
| RHOJ | 14 | ras homolog family member J [Source:HGNC S |
| AL049871.1 | 14 | novel transcript |

| | | |
|---|---|---|
| GPHB5 | 14 | glycoprotein hormone subunit beta 5 [Source: |
| PPP2R5E | 14 | protein phosphatase 2 regulatory subunit B'ep |
| AL132992.1 | 14 | NULL |
| AL136038.3 | 14 | novel transcript, antisense to PPP2R5E |
| AL136038.1 | 14 | ribosomal protein L31 (RPL31) pseudogene |
| AL136038.4 | 14 | glycine C-acetyltransferase (GCAT) pseudogen |
| WDR89 | 14 | WD repeat domain 89 [Source:HGNC Symbol; |
| AL136038.2 | 14 | novel transcript |
| HSPE1P2 | 14 | heat shock protein family E (Hsp10) member : |
| RNU6-597P | 14 | RNA, U6 small nuclear 597, pseudogene [Sour |
| Y_RNA | 14 | Y RNA [Source:RFAM;Acc:RF00019] |
| RNU6-1162P | 14 | RNA, U6 small nuclear 1162, pseudogene [Sou |
| AL136038.5 | 14 | novel transcript |
| U3 | 14 | Small nucleolar RNA U3 [Source:RFAM;Acc:RF |
| SGPP1 | 14 | sphingosine-1-phosphate phosphatase 1 [Sou |
| AL161670.2 | 14 | eukaryotic translation initiation factor 2, subu |
| RNU7-116P | 14 | RNA, U7 small nuclear 116 pseudogene [Sourc |
| AL161670.1 | 14 | RAP1B, member of RAS oncogene family (RAP |
| RPS28P1 | 14 | ribosomal protein S28 pseudogene 1 [Source: |
| Y_RNA | 14 | Y RNA [Source:RFAM;Acc:RF00019] |
| SYNE2 | 14 | spectrin repeat containing nuclear envelope p |
| ESR2 | 14 | estrogen receptor 2 [Source:HGNC Symbol;Ac |
| MIR548H1 | 14 | microRNA 548h-1 [Source:HGNC Symbol;Acc:I |
| AL161756.1 | 14 | novel transcript |
| TEX21P | 14 | testis expressed 21, pseudogene [Source:HGN |
| AL161756.3 | 14 | novel transcript |
| AL161756.2 | 14 | translocase of outer mitochondrial membrane |
| MTHFD1 | 14 | methylenetetrahydrofolate dehydrogenase, c |
| AL122035.1 | 14 | novel transcript |
| AL122035.2 | 14 | novel transcript, antisense to MTHFD1 |
| ZBTB25 | 14 | zinc finger and BTB domain containing 25 [Sou |
| AKAP5 | 14 | A-kinase anchoring protein 5 [Source:HGNC Sy |
| ZBTB1 | 14 | zinc finger and BTB domain containing 1 [Sour |
| AL049869.3 | 14 | novel transcript, antisense to HSPA2 & ZBTB1 |
| AL049869.4 | 14 | ribosomal protein L18 (RPL18) pseudogene |
| HSPA2 | 14 | heat shock protein family A (Hsp70) member : |
| PPP1R36 | 14 | protein phosphatase 1 regulatory subunit 36 [ |
| AL049869.2 | 14 | novel transcript, antisense to C14orf50 |
| AL049869.1 | 14 | nucleoporin 50kDa (NUP50) pseudogene |
| PLEKHG3 | 14 | pleckstrin homology and RhoGEF domain cont |
| SPTB | 14 | spectrin beta, erythrocytic [Source:HGNC Sym |
| MIR7855 | 14 | microRNA 7855 [Source:HGNC Symbol;Acc:HG |
| RPPH1-2P | 14 | ribonuclease P RNA component H1, 2 pseudo |
| CHURC1 | 14 | churchill domain containing 1 [Source:HGNC S |
| CHURC1-FNTB | 14 | CHURC1-FNTB readthrough [Source:HGNC Syr |
| GPX2 | 14 | glutathione peroxidase 2 [Source:HGNC Symb |
| RAB15 | 14 | RAB15, member RAS oncogene family [Source |
| FNTB | 14 | farnesyltransferase, CAAX box, beta [Source:H |
| AL139022.2 | 14 | novel transcript, antisense to CHURC1-FNTB |
| MAX | 14 | MYC associated factor X [Source:HGNC Symbo |

| | | |
|---|---|---|
| MIR4706 | 14 | microRNA 4706 [Source:HGNC Symbol;Acc:HG |
| AL139022.1 | 14 | novel transcript, antisense to MAX |
| RNU2-14P | 14 | RNA, U2 small nuclear 14, pseudogene [Sourc |
| LINC02324 | 14 | long intergenic non-protein coding RNA 2324 |
| AL355076.3 | 14 | transmembrane protein 149 (TMEM149) pseu |
| AL355076.2 | 14 | novel transcript |
| RPL21P7 | 14 | ribosomal protein L21 pseudogene 7 [Source: |
| AL355076.1 | 14 | ribosomal protein L36a (RPL36A) pseudogene |
| PTBP1P | 14 | polypyrimidine tract binding protein 1 pseudo |
| MIR4708 | 14 | microRNA 4708 [Source:HGNC Symbol;Acc:HG |
| FUT8 | 14 | fucosyltransferase 8 [Source:HGNC Symbol;Ac |
| FUT8-AS1 | 14 | FUT8 antisense RNA 1 [Source:HGNC Symbol; |
| RPL21P8 | 14 | ribosomal protein L21 pseudogene 8 [Source: |
| MIR625 | 14 | microRNA 625 [Source:HGNC Symbol;Acc:HGN |
| EIF1AXP2 | 14 | EIF1AX pseudogene 2 [Source:HGNC Symbol;A |
| AL391261.1 | 14 | nuclear receptor coactivator 4 (NCOA4) pseud |
| AL391261.3 | 14 | ferritin, heavy polypeptide 1 (FTH1) pseudoge |
| AL391261.2 | 14 | novel transcript |
| AL391261.4 | 14 | novel transcript, sense intronic to CTD-2014B |
| YBX1P1 | 14 | Y-box binding protein 1 pseudogene 1 [Source |
| LINC02290 | 14 | long intergenic non-protein coding RNA 2290 |
| Y_RNA | 14 | Y RNA [Source:RFAM;Acc:RF00019] |
| AL359232.1 | 14 | novel transcript |
| AL157997.1 | 14 | novel transcript |
| CCDC196 | 14 | coiled-coil domain containing 196 [Source:HG |
| GPHN | 14 | gephyrin [Source:HGNC Symbol;Acc:HGNC:15 |
| AL133241.1 | 14 | ribosomal L24 domain containing 1 (RSL24D1) |
| AL049835.1 | 14 | novel transcript, antisense to GPHN |
| FAM71D | 14 | family with sequence similarity 71 member D |
| AL135978.1 | 14 | splicing factor 3b, subunit 4, 49kDa (SF3B4) ps |
| MPP5 | 14 | membrane palmitoylated protein 5 [Source:H |
| AL135978.2 | 14 | novel transcript, antisense to MPP5 |
| ATP6V1D | 14 | ATPase H+ transporting V1 subunit D [Source: |
| Y_RNA | 14 | Y RNA [Source:RFAM;Acc:RF00019] |
| EIF2S1 | 14 | eukaryotic translation initiation factor 2 subur |
| PLEK2 | 14 | pleckstrin 2 [Source:HGNC Symbol;Acc:HGNC: |
| MIR5694 | 14 | microRNA 5694 [Source:HGNC Symbol;Acc:HG |
| TMEM229B | 14 | transmembrane protein 229B [Source:HGNC S |
| PLEKHH1 | 14 | pleckstrin homology, MyTH4 and FERM doma |
| PIGH | 14 | phosphatidylinositol glycan anchor biosynthes |
| AL132640.1 | 14 | protein kinase C, eta (PRKCH) pseudogene |
| AL132640.2 | 14 | high mobility group box 1 (HMGB1) pseudoge |
| AL132640.3 | 14 | NULL |
| Y_RNA | 14 | Y RNA [Source:RFAM;Acc:RF00019] |
| ARG2 | 14 | arginase 2 [Source:HGNC Symbol;Acc:HGNC:6 |
| AL049779.4 | 14 | novel transcript, antisense to ARG2 |
| VTI1B | 14 | vesicle transport through interaction with t-SN |
| COX7A2P1 | 14 | cytochrome c oxidase subunit 7A2 pseudogen |
| RNA5SP386 | 14 | RNA, 5S ribosomal pseudogene 386 [Source:H |
| AL049779.1 | 14 | novel protein |

| | | |
|---|---|---|
| RDH11 | 14 | retinol dehydrogenase 11 [Source:HGNC Symb |
| RDH12 | 14 | retinol dehydrogenase 12 [Source:HGNC Symb |
| RN7SL369P | 14 | RNA, 7SL, cytoplasmic 369, pseudogene [Sour |
| RPL21P9 | 14 | ribosomal protein L21 pseudogene 9 [Source: |
| ZFYVE26 | 14 | zinc finger FYVE-type containing 26 [Source:H |
| AL049779.3 | 14 | gephyrin (GPHN) pseudogene |
| RN7SL213P | 14 | RNA, 7SL, cytoplasmic 213, pseudogene [Sour |
| AL049779.2 | 14 | DnaJ (Hsp40) homolog, subfamily C, member |
| RAD51B | 14 | RAD51 paralog B [Source:HGNC Symbol;Acc:H |
| AL133370.2 | 14 | zinc finger protein 585B (ZNF585B) pseudoger |
| AL133370.1 | 14 | novel transcript, antisense to RAD51L1 |
| RN7SL706P | 14 | RNA, 7SL, cytoplasmic 706, pseudogene [Sour |
| RN7SL108P | 14 | RNA, 7SL, cytoplasmic 108, pseudogene [Sour |
| AL122013.1 | 14 | ribosomal protein L7a (RPL7A) pseudogene |
| PPIAP6 | 14 | peptidylprolyl isomerase A pseudogene 6 [Sou |
| AL121820.3 | 14 | uncharacterized LOC107984016 [Source:NCBI |
| AL121820.2 | 14 | novel transcript, antisense to RAD51L1 |
| AL121820.1 | 14 | novel transcript, antisense to RAD51L1 |
| RPL12P7 | 14 | ribosomal protein L12 pseudogene 7 [Source: |
| AL133313.1 | 14 | novel transcript, antisense to RAD51B |
| AL132986.1 | 14 | novel transcript |
| RNU6-921P | 14 | RNA, U6 small nuclear 921, pseudogene [Sour |
| ZFP36L1 | 14 | ZFP36 ring finger protein like 1 [Source:HGNC |
| MAGOH3P | 14 | mago homolog 3, pseudogene [Source:HGNC |
| BLZF2P | 14 | basic leucine zipper nuclear factor 2 pseudoge |
| ACTN1 | 14 | actinin alpha 1 [Source:HGNC Symbol;Acc:HGI |
| HMGN1P3 | 14 | high mobility group nucleosome binding doma |
| BANF1P1 | 14 | barrier to autointegration factor 1 pseudogen |
| ACTN1-AS1 | 14 | ACTN1 antisense RNA 1 [Source:HGNC Symbo |
| RPS29P1 | 14 | ribosomal protein S29 pseudogene 1 [Source: |
| DCAF5 | 14 | DDB1 and CUL4 associated factor 5 [Source:H |
| DDX18P1 | 14 | DEAD-box helicase 18 pseudogene 1 [Source:I |
| Y_RNA | 14 | Y RNA [Source:RFAM;Acc:RF00019] |
| AL391262.1 | 14 | novel transcript |
| AL359317.2 | 14 | novel transcript |
| EXD2 | 14 | exonuclease 3'-5' domain containing 2 [Source |
| AL359317.1 | 14 | novel transcript |
| GALNT16 | 14 | polypeptide N-acetylgalactosaminyltransferas |
| AL157996.1 | 14 | TEC |
| ERH | 14 | ERH mRNA splicing and mitosis factor [Source |
| SLC39A9 | 14 | solute carrier family 39 member 9 [Source:HG |
| AL157996.2 | 14 | novel transcript |
| PLEKHD1 | 14 | pleckstrin homology and coiled-coil domain co |
| AL133445.1 | 14 | ribosomal protein L24 (RPL24) pseudogene |
| CCDC177 | 14 | coiled-coil domain containing 177 [Source:HG |
| AL133445.3 | 14 | novel transcript |
| SUSD6 | 14 | sushi domain containing 6 [Source:HGNC Sym |
| AL133445.2 | 14 | novel transcript, sense intronic to KIAA0247 |
| SRSF5 | 14 | serine and arginine rich splicing factor 5 [Sour |
| SLC10A1 | 14 | solute carrier family 10 member 1 [Source:HG |

| | | |
|---|---|---|
| AL157789.1 | 14 | NULL |
| SMOC1 | 14 | SPARC related modular calcium binding 1 [Sou |
| RPL7AP6 | 14 | ribosomal protein L7a pseudogene 6 [Source:I |
| SLC8A3 | 14 | solute carrier family 8 member A3 [Source:HG |
| AL160191.1 | 14 | novel transcript, antisense to SLC8A3 |
| AL160191.3 | 14 | ADAM metallopeptidase domain 20 (ADAM20 |
| AL160191.2 | 14 | Sin3A-associated protein, 25kDa (SAP25) pseu |
| ADAM21P1 | 14 | ADAM metallopeptidase domain 21 pseudoge |
| AL356804.1 | 14 | novel transcript |
| COX16 | 14 | cytochrome c oxidase assembly factor COX16 |
| SYNJ2BP-COX16 | 14 | SYNJ2BP-COX16 readthrough [Source:HGNC S |
| RNU2-51P | 14 | RNA, U2 small nuclear 51, pseudogene [Sourc |
| SYNJ2BP | 14 | synaptojanin 2 binding protein [Source:HGNC |
| AL357153.2 | 14 | novel transcript |
| ADAM21 | 14 | ADAM metallopeptidase domain 21 [Source:H |
| ADAM20P1 | 14 | ADAM metallopeptidase domain 20 pseudoge |
| RNU6-659P | 14 | RNA, U6 small nuclear 659, pseudogene [Sour |
| ADAM20 | 14 | ADAM metallopeptidase domain 20 [Source:H |
| AL357153.4 | 14 | ADAM metallopeptidase domain 20 (ADAM20 |
| MED6 | 14 | mediator complex subunit 6 [Source:HGNC Sy |
| AL357153.3 | 14 | novel transcript |
| KRT18P7 | 14 | keratin 18 pseudogene 7 [Source:HGNC Symb |
| AL357153.1 | 14 | novel transcript |
| RN7SL77P | 14 | RNA, 7SL, cytoplasmic 77, pseudogene [Sourc |
| TTC9 | 14 | tetratricopeptide repeat domain 9 [Source:HG |
| LINC01269 | 14 | long intergenic non-protein coding RNA 1269 |
| MAP3K9 | 14 | mitogen-activated protein kinase kinase kinas |
| AC004816.1 | 14 | novel transcript |
| AC004816.2 | 14 | novel transcript |
| AC004825.1 | 14 | transmembrane protein 183A (TMEM183A) ps |
| AC004825.2 | 14 | novel transcript |
| PCNX1 | 14 | pecanex 1 [Source:HGNC Symbol;Acc:HGNC:1 |
| AC004825.3 | 14 | novel transcript, antisense to PCNX1 |
| AC005230.1 | 14 | tetratricopeptide repeat domain 7B (TTC7B) p |
| GTF3AP2 | 14 | general transcription factor IIIA pseudogene 2 |
| PTTG4P | 14 | pituitary tumor-transforming 4 pseudogene [S |
| AC004817.3 | 14 | novel transcript |
| AC004817.5 | 14 | novel transcript |
| AC004817.2 | 14 | novel transcript |
| AC004817.1 | 14 | novel transcript |
| AC004817.4 | 14 | novel transcript |
| AC005476.2 | 14 | novel transcript, antisense to SIPA1L1 |
| SIPA1L1 | 14 | signal induced proliferation associated 1 like 1 |
| AC005476.1 | 14 | PDZ domain-containing protein 11 (PDZD11) p |
| SNORD56B | 14 | small nucleolar RNA, C/D box 56B [Source:HG |
| RN7SL683P | 14 | RNA, 7SL, cytoplasmic 683, pseudogene [Sour |
| AC004974.1 | 14 | novel transcript, antisense to SIPA1L1 |
| AC004900.1 | 14 | novel transcript, antisense to SIPA1L1 |
| AC005993.1 | 14 | novel transcript |
| RGS6 | 14 | regulator of G protein signaling 6 [Source:HGN |

| | | |
|---|---|---|
| Y_RNA | 14 | Y RNA [Source:RFAM;Acc:RF00019] |
| AC004828.2 | 14 | novel transcript, antisense to RGS6 |
| MIR7843 | 14 | microRNA 7843 [Source:HGNC Symbol;Acc:HG |
| AC004828.1 | 14 | novel transcript |
| DPF3 | 14 | double PHD fingers 3 [Source:HGNC Symbol;A |
| AL392024.1 | 14 | novel transcript, antisense to DPF3 |
| Y_RNA | 14 | Y RNA [Source:RFAM;Acc:RF00019] |
| DCAF4 | 14 | DDB1 and CUL4 associated factor 4 [Source:H |
| AC007160.1 | 14 | ribosomal protein L36 (RPL36) pseudogene |
| AL442663.4 | 14 | chromosome 17 open reading frame 98 (C17c |
| AL442663.1 | 14 | ribosomal protein S12 (RPS12) pseudogene |
| ZFYVE1 | 14 | zinc finger FYVE-type containing 1 [Source:HG |
| RN7SL586P | 14 | RNA, 7SL, cytoplasmic 586, pseudogene [Sour |
| AL442663.3 | 14 | novel transcript, antisense to Rbm25 |
| RBM25 | 14 | RNA binding motif protein 25 [Source:HGNC S |
| AL442663.2 | 14 | required for meiotic nuclear division 1 homolo |
| PSEN1 | 14 | presenilin 1 [Source:HGNC Symbol;Acc:HGNC: |
| AC004858.1 | 14 | WD repeat-containing protein 62 (WDR62) ps |
| PAPLN | 14 | papilin, proteoglycan like sulfated glycoprotei |
| AC004846.1 | 14 | novel transcript, antisense to PAPLN |
| RNU6-419P | 14 | RNA, U6 small nuclear 419, pseudogene [Sour |
| AC004846.2 | 14 | novel transcript, antisense to PAPLN |
| NUMB | 14 | NUMB endocytic adaptor protein [Source:HGI |
| AC005280.3 | 14 | novel transcript |
| AC005280.2 | 14 | novel transcript |
| AC005280.1 | 14 | novel transcript, antisense to NUMB and HEA |
| HEATR4 | 14 | HEAT repeat containing 4 [Source:HGNC Symb |
| RIOX1 | 14 | ribosomal oxygenase 1 [Source:HGNC Symbol |
| AC005225.1 | 14 | novel transcript, sense intronic to HEATR4 |
| AC005225.3 | 14 | novel transcript, antisense to HEATR4 |
| ACOT1 | 14 | acyl-CoA thioesterase 1 [Source:HGNC Symbo |
| NT5CP2 | 14 | 5',3'-nucleotidase, cytosolic pseudogene 2 [Sc |
| AC005225.2 | 14 | novel transcript, antisense to ACOT6 |
| ACOT2 | 14 | acyl-CoA thioesterase 2 [Source:HGNC Symbo |
| NT5CP1 | 14 | 5',3'-nucleotidase, cytosolic pseudogene 1 [Sc |
| AC005225.4 | 14 | TEC |
| ACOT4 | 14 | acyl-CoA thioesterase 4 [Source:HGNC Symbo |
| ACOT6 | 14 | acyl-CoA thioesterase 6 [Source:HGNC Symbo |
| NDUFB8P1 | 14 | NADH:ubiquinone oxidoreductase subunit B8 |
| DNAL1 | 14 | dynein axonemal light chain 1 [Source:HGNC S |
| RNU6-240P | 14 | RNA, U6 small nuclear 240, pseudogene [Sour |
| AC006146.1 | 14 | novel transcript, antisense to DNAL1 |
| PNMA1 | 14 | PNMA family member 1 [Source:HGNC Symbc |
| ELMSAN1 | 14 | ELM2 and Myb/SANT domain containing 1 [Sc |
| MIR4505 | 14 | microRNA 4505 [Source:HGNC Symbol;Acc:HG |
| AC005520.2 | 14 | novel transcript, antisense to C14orf43 |
| LINC02274 | 14 | long intergenic non-protein coding RNA 2274 |
| AC005520.4 | 14 | TAF15 RNA polymerase II, TATA box binding p |
| PTGR2 | 14 | prostaglandin reductase 2 [Source:HGNC Sym |
| AC005520.1 | 14 | novel protein |

| | | |
|---|---|---|
| Y_RNA | 14 | Y RNA [Source:RFAM;Acc:RF00019] |
| AC005520.5 | 14 | novel transcript, antisense to PTGR2 |
| ZNF410 | 14 | zinc finger protein 410 [Source:HGNC Symbol; |
| AC005480.1 | 14 | novel transcript, antisense to ZNF410 |
| AC005520.3 | 14 | novel transcript |
| Y_RNA | 14 | Y RNA [Source:RFAM;Acc:RF00019] |
| FAM161B | 14 | FAM161 centrosomal protein B [Source:HGNC |
| COQ6 | 14 | coenzyme Q6, monooxygenase [Source:HGNC |
| ENTPD5 | 14 | ectonucleoside triphosphate diphosphohydro |
| AC005480.2 | 14 | ribosomal protein L41 (RPL41) pseudogene |
| BBOF1 | 14 | basal body orientation factor 1 [Source:HGNC |
| ALDH6A1 | 14 | aldehyde dehydrogenase 6 family member A1 |
| LIN52 | 14 | lin-52 DREAM MuvB core complex componen |
| RN7SL530P | 14 | RNA, 7SL, cytoplasmic 530, pseudogene [Sour |
| AC006349.1 | 14 | SMT3 suppressor of mif two 3 homolog 2 (S. c |
| VSX2 | 14 | visual system homeobox 2 [Source:HGNC Sym |
| ABCD4 | 14 | ATP binding cassette subfamily D member 4 [S |
| AC005519.1 | 14 | novel transcript, sense overlapping ABCD4 |
| VRTN | 14 | vertebrae development associated [Source:HC |
| SUB1P2 | 14 | SUB1 pseudogene 2 [Source:HGNC Symbol;Ac |
| RPS2P2 | 14 | ribosomal protein S2 pseudogene 2 [Source:H |
| SYNDIG1L | 14 | synapse differentiation inducing 1 like [Source |
| AC005479.2 | 14 | novel transcript |
| AC005479.1 | 14 | novel transcript |
| NPC2 | 14 | NPC intracellular cholesterol transporter 2 [So |
| MIR4709 | 14 | microRNA 4709 [Source:HGNC Symbol;Acc:HG |
| RAP1AP | 14 | RAP1A, member of RAS oncogene family pseu |
| ISCA2 | 14 | iron-sulfur cluster assembly 2 [Source:HGNC S |
| LTBP2 | 14 | latent transforming growth factor beta bindin |
| AC013451.1 | 14 | novel transcript, antisense to LTBP2 |
| AC013451.2 | 14 | novel transcript |
| AREL1 | 14 | apoptosis resistant E3 ubiquitin protein ligase |
| AC007956.2 | 14 | ribosomal protein S2 (RPS2) pseudogene |
| AC007956.1 | 14 | NULL |
| FCF1 | 14 | FCF1 rRNA-processing protein [Source:HGNC S |
| YLPM1 | 14 | YLP motif containing 1 [Source:HGNC Symbol; |
| PROX2 | 14 | prospero homeobox 2 [Source:HGNC Symbol; |
| AC006530.1 | 14 | methyltransferase like 5 (METTL5) pseudogen |
| DLST | 14 | dihydrolipoamide S-succinyltransferase [Sourc |
| RPS6KL1 | 14 | ribosomal protein S6 kinase like 1 [Source:HG |
| PGF | 14 | placental growth factor [Source:HGNC Symbo |
| EIF2B2 | 14 | eukaryotic translation initiation factor 2B subu |
| AL049780.1 | 14 | novel transcript, antisense to EIF2B2 |
| AL049780.3 | 14 | TEC |
| MLH3 | 14 | mutL homolog 3 [Source:HGNC Symbol;Acc:H |
| RNU6-689P | 14 | RNA, U6 small nuclear 689, pseudogene [Sour |
| ACYP1 | 14 | acylphosphatase 1 [Source:HGNC Symbol;Acc: |
| ZC2HC1C | 14 | zinc finger C2HC-type containing 1C [Source:H |
| NEK9 | 14 | NIMA related kinase 9 [Source:HGNC Symbol; |
| HIF1AP1 | 14 | HIF1AP pseudogene 1 [Source:HGNC Symbol;A |

| | | |
|---|---|---|
| AL049780.2 | 14 | novel transcript, antisense to NEK9 and TMED |
| TMED10 | 14 | transmembrane p24 trafficking protein 10 [So |
| AL691403.2 | 14 | novel transcript |
| RNU4ATAC14P | 14 | RNA, U4atac small nuclear 14, pseudogene [S |
| AL691403.1 | 14 | novel transcript |
| AF111167.1 | 14 | novel transcript, antisense to a novel orphan |
| U2 | 14 | U2 spliceosomal RNA [Source:RFAM;Acc:RF00 |
| FOS | 14 | Fos proto-oncogene, AP-1 transcription factor |
| DPPA5P4 | 14 | developmental pluripotency associated 5 pse |
| LINC01220 | 14 | long intergenic non-protein coding RNA 1220 |
| AF111167.2 | 14 | novel transcript, antisense to JDP2 |
| JDP2 | 14 | Jun dimerization protein 2 [Source:HGNC Sym |
| BATF | 14 | basic leucine zipper ATF-like transcription fact |
| AC007182.1 | 14 | novel transcript, antisense to FLVCR2 |
| FLVCR2 | 14 | feline leukemia virus subgroup C cellular rece |
| AC007182.2 | 14 | ribosomal protein S24 (RPS24) pseudogene |
| RNA5SP387 | 14 | RNA, 5S ribosomal pseudogene 387 [Source:H |
| AC007182.3 | 14 | NULL |
| TTLL5 | 14 | tubulin tyrosine ligase like 5 [Source:HGNC Sy |
| ERG28 | 14 | ergosterol biosynthesis 28 homolog [Source:H |
| AF107885.2 | 14 | novel transcript, antisense to TTLL5 |
| AF107885.1 | 14 | ribosomal protein S2 (RPS2) pseudogene |
| IFT43 | 14 | intraflagellar transport 43 [Source:HGNC Sym |
| TGFB3 | 14 | transforming growth factor beta 3 [Source:HG |
| TGFB3-AS1 | 14 | TGFB3 antisense RNA 1 [Source:HGNC Symbol |
| AC008015.1 | 14 | ubiquitin-conjugating enzyme E2L 3 (UBE2L3) |
| AC016526.4 | 14 | novel transcript, antisense to GPATCH2L |
| GPATCH2L | 14 | G-patch domain containing 2 like [Source:HGN |
| AC016526.2 | 14 | novel transcript, sense overlapping GPATCH2L |
| AC016526.3 | 14 | TEC |
| AC016526.1 | 14 | novel transcript |
| ESRRB | 14 | estrogen related receptor beta [Source:HGNC |
| AC008050.1 | 14 | novel transcript |
| CYCSP1 | 14 | CYCS pseudogene 1 [Source:HGNC Symbol;Ac |
| RPSAP3 | 14 | ribosomal protein SA pseudogene 3 [Source:H |
| AC007376.1 | 14 | H1 histone family, member O, oocyte-specific |
| AC007376.2 | 14 | novel transcript |
| VASH1 | 14 | vasohibin 1 [Source:HGNC Symbol;Acc:HGNC: |
| AF111169.3 | 14 | novel transcript |
| AF111169.1 | 14 | novel transcript |
| VASH1-AS1 | 14 | VASH1 antisense RNA 1 [Source:HGNC Symbo |
| ANGEL1 | 14 | angel homolog 1 [Source:HGNC Symbol;Acc:H |
| AF111169.4 | 14 | novel transcript, antisense to ANGEL1 |
| LRRC74A | 14 | leucine rich repeat containing 74A [Source:HG |
| RPL22P2 | 14 | ribosomal protein L22 pseudogene 2 [Source: |
| RN7SKP17 | 14 | RN7SK pseudogene 17 [Source:HGNC Symbol; |
| AF111169.2 | 14 | novel transcript |
| AC007686.4 | 14 | novel transcript |
| LINC01629 | 14 | long intergenic non-protein coding RNA 1629 |
| RN7SL356P | 14 | RNA, 7SL, cytoplasmic 356, pseudogene [Sour |

| | | |
|---|---|---|
| AC007686.1 | 14 | ribosomal protein, large, P1 (RPLP1) pseudoge |
| IRF2BPL | 14 | interferon regulatory factor 2 binding protein |
| AC007686.3 | 14 | novel transcript, antisense to IRF2BPL |
| LINC02288 | 14 | long intergenic non-protein coding RNA 2288 |
| AC007686.2 | 14 | novel transcript |
| LINC02289 | 14 | long intergenic non-protein coding RNA 2289 |
| AC007686.5 | 14 | novel transcript |
| CIPC | 14 | CLOCK interacting pacemaker [Source:HGNC S |
| AC007375.2 | 14 | novel protein |
| TMEM63C | 14 | transmembrane protein 63C [Source:HGNC Sy |
| ZDHHC22 | 14 | zinc finger DHHC-type containing 22 [Source:H |
| AC007375.3 | 14 | novel transcript, antisense to ZDHHC22 |
| FAM204DP | 14 | family with sequence similarity 204 member D |
| AC007375.1 | 14 | NULL |
| NGB | 14 | neuroglobin [Source:HGNC Symbol;Acc:HGNC |
| MIR1260A | 14 | microRNA 1260a [Source:HGNC Symbol;Acc:H |
| POMT2 | 14 | protein O-mannosyltransferase 2 [Source:HGN |
| GSTZ1 | 14 | glutathione S-transferase zeta 1 [Source:HGN |
| TMED8 | 14 | transmembrane p24 trafficking protein family |
| AC007954.1 | 14 | ATP-binding cassette, sub-family G (WHITE), m |
| SAMD15 | 14 | sterile alpha motif domain containing 15 [Sou |
| NOXRED1 | 14 | NADP dependent oxidoreductase domain con |
| VIPAS39 | 14 | VPS33B interacting protein, apical-basolateral |
| AHSA1 | 14 | activator of HSP90 ATPase activity 1 [Source:H |
| AF111168.1 | 14 | NULL |
| ISM2 | 14 | isthmin 2 [Source:HGNC Symbol;Acc:HGNC:23 |
| SPTLC2 | 14 | serine palmitoyltransferase long chain base su |
| RN7SL587P | 14 | RNA, 7SL, cytoplasmic 587, pseudogene [Sour |
| COX6CP11 | 14 | cytochrome c oxidase subunit 6C pseudogene |
| ALKBH1 | 14 | alkB homolog 1, histone H2A dioxygenase [So |
| RPL21P10 | 14 | ribosomal protein L21 pseudogene 10 [Source |
| ZMYND19P1 | 14 | zinc finger MYND-type containing 19 pseudog |
| SLIRP | 14 | SRA stem-loop interacting RNA binding protei |
| SNW1 | 14 | SNW domain containing 1 [Source:HGNC Syml |
| C14orf178 | 14 | chromosome 14 open reading frame 178 [Sou |
| ADCK1 | 14 | aarF domain containing kinase 1 [Source:HGN |
| Y_RNA | 14 | Y RNA [Source:RFAM;Acc:RF00019] |
| FXNP1 | 14 | frataxin pseudogene 1 [Source:HGNC Symbol; |
| NRXN3 | 14 | neurexin 3 [Source:HGNC Symbol;Acc:HGNC:8 |
| RNA5SP388 | 14 | RNA, 5S ribosomal pseudogene 388 [Source:H |
| RPS26P48 | 14 | ribosomal protein S26 pseudogene 48 [Source |
| AF099810.1 | 14 | novel transcript, antisense to NRXN3 |
| AC009396.2 | 14 | novel transcript, antisense to NRXN3 |
| AC009396.3 | 14 | novel transcript, antisense to NRXN3 |
| AC009396.1 | 14 | novel transcript, antisense to NRXN3 |
| AC026888.1 | 14 | novel transcript, antisense to NRXN3 |
| AC022469.1 | 14 | novel transcript, sense intronic to NRXN3 |
| AC022469.2 | 14 | novel transcript, antisense to NRXN3 |
| AC008056.2 | 14 | novel transcript, antisense to NRXN3 |
| AC008056.3 | 14 | novel transcript, antisense to NRXN3 |

| | | |
|---|---|---|
| AC008056.1 | 14 | novel transcript, antisense to NRXN3 |
| AF123462.1 | 14 | novel transcript |
| DIO2 | 14 | iodothyronine deiodinase 2 [Source:HGNC Syr |
| DIO2-AS1 | 14 | DIO2 antisense RNA 1 [Source:HGNC Symbol;/ |
| CEP128 | 14 | centrosomal protein 128 [Source:HGNC Symb |
| AC022404.1 | 14 | high mobility group nucleosomal binding dom |
| TSHR | 14 | thyroid stimulating hormone receptor [Source |
| BHLHB9P1 | 14 | basic helix-loop-helix family member b9 pseuc |
| RPL17P3 | 14 | ribosomal protein L17 pseudogene 3 [Source: |
| AC007262.1 | 14 | ribosomal protein L17 (RPL17) pseudogene |
| AC007262.2 | 14 | novel transcript, antisense to TSHR |
| NMNAT1P1 | 14 | NMNAT1 pseudogene 1 [Source:HGNC Symbo |
| AL136040.2 | 14 | novel transcript |
| GTF2A1 | 14 | general transcription factor IIA subunit 1 [Sou |
| SNORA79 | 14 | small nucleolar RNA, H/ACA box 79 [Source:H |
| AL136040.1 | 14 | novel transcript, antisense to GTF2A1 |
| DYNLL1P1 | 14 | dynein light chain LC8-type 1 pseudogene 1 [S |
| UNGP3 | 14 | uracil-DNA glycosylase pseudogene 3 [Source: |
| STON2 | 14 | stonin 2 [Source:HGNC Symbol;Acc:HGNC:306 |
| AL121769.1 | 14 | ribosomal protein S24 (RPS24) pseudogene |
| DYNLL1P2 | 14 | dynein light chain LC8-type 1 pseudogene 2 [S |
| LINC02308 | 14 | long intergenic non-protein coding RNA 2308 |
| SEL1L | 14 | SEL1L adaptor subunit of ERAD E3 ubiquitin lig |
| LINC01467 | 14 | long intergenic non-protein coding RNA 1467 |
| EEF1A1P2 | 14 | eukaryotic translation elongation factor 1 alpl |
| LINC02311 | 14 | long intergenic non-protein coding RNA 2311 |
| AL355838.1 | 14 | novel transcript |
| RPL9P6 | 14 | ribosomal protein L9 pseudogene 6 [Source:H |
| EIF3LP1 | 14 | eukaryotic translation initiation factor 3 subur |
| Metazoa_SRP | 14 | Metazoan signal recognition particle RNA [Sou |
| LINC02301 | 14 | long intergenic non-protein coding RNA 2301 |
| ENSAP2 | 14 | endosulfine alpha pseudogene 2 [Source:HGN |
| AL355095.1 | 14 | novel transcript |
| AL359238.1 | 14 | novel transcript |
| AL162872.1 | 14 | novel transcript |
| RNU7-51P | 14 | RNA, U7 small nuclear 51 pseudogene [Source |
| RNU6ATAC28P | 14 | RNA, U6atac small nuclear 28, pseudogene [S |
| AL161713.1 | 14 | zinc finger protein 268 (ZNF268) pseudogene |
| LINC02305 | 14 | long intergenic non-protein coding RNA 2305 |
| AL356807.1 | 14 | novel transcript |
| MTND4P33 | 14 | MT-ND4 pseudogene 33 [Source:HGNC Symbo |
| MTND5P35 | 14 | MT-ND5 pseudogene 35 [Source:HGNC Symbo |
| MTCYBP27 | 14 | MT-CYB pseudogene 27 [Source:HGNC Symbo |
| RNU6-976P | 14 | RNA, U6 small nuclear 976, pseudogene [Sour |
| AL163642.1 | 14 | ribosomal protein L13a (RPL13A) pseudogene |
| AL163642.2 | 14 | family with sequence similarity 71, member B |
| AL357172.1 | 14 | novel transcript |
| AL049775.2 | 14 | novel transcript |
| LINC02329 | 14 | long intergenic non-protein coding RNA 2329 |
| LINC00911 | 14 | long intergenic non-protein coding RNA 911 [S |

| | | |
|---|---|---|
| AL049775.1 | 14 | novel transcript |
| AL049775.3 | 14 | novel transcript |
| FLRT2 | 14 | fibronectin leucine rich transmembrane prote |
| LINC02328 | 14 | long intergenic non-protein coding RNA 2328 |
| LINC02316 | 14 | long intergenic non-protein coding RNA 2316 |
| AL161753.1 | 14 | novel transcript |
| LINC02309 | 14 | long intergenic non-protein coding RNA 2309 |
| LINC01148 | 14 | long intergenic non-protein coding RNA 1148 |
| AL358292.1 | 14 | novel transcript |
| AL352955.1 | 14 | novel transcript |
| AL157688.1 | 14 | novel transcript |
| LINC02296 | 14 | long intergenic non-protein coding RNA 2296 |
| AL135746.1 | 14 | novel transcript |
| LINC02330 | 14 | long intergenic non-protein coding RNA 2330 |
| AL359237.1 | 14 | novel transcript |
| AL136501.1 | 14 | novel transcript |
| GALC | 14 | galactosylceramidase [Source:HGNC Symbol;A |
| SHLD2P2 | 14 | shieldin complex subunit 2 pseudogene 2 [Sou |
| RNU6-835P | 14 | RNA, U6 small nuclear 835, pseudogene [Sour |
| GPR65 | 14 | G protein-coupled receptor 65 [Source:HGNC |
| AL157955.1 | 14 | novel transcript, antisense to GPR65 |
| LINC01147 | 14 | long intergenic non-protein coding RNA 1147 |
| LINC01146 | 14 | long intergenic non-protein coding RNA 1146 |
| AL133279.2 | 14 | novel transcript |
| AL133279.3 | 14 | novel transcript |
| AL133279.1 | 14 | novel transcript, antisense to KCNK10 |
| KCNK10 | 14 | potassium two pore domain channel subfamil |
| SPATA7 | 14 | spermatogenesis associated 7 [Source:HGNC S |
| PTPN21 | 14 | protein tyrosine phosphatase non-receptor ty |
| AL162171.2 | 14 | novel transcript, antisense to PTPN21 |
| RNU4-22P | 14 | RNA, U4 small nuclear 22, pseudogene [Sourc |
| AL162171.1 | 14 | novel transcript, antisense to PTPN21 |
| ZC3H14 | 14 | zinc finger CCCH-type containing 14 [Source:H |
| AL162171.3 | 14 | novel transcript, sense intronic to ZC3H14 |
| EML5 | 14 | EMAP like 5 [Source:HGNC Symbol;Acc:HGNC: |
| AL121768.1 | 14 | novel transcript |
| RNU4-92P | 14 | RNA, U4 small nuclear 92, pseudogene [Sourc |
| TTC8 | 14 | tetratricopeptide repeat domain 8 [Source:HG |
| Y_RNA | 14 | Y RNA [Source:RFAM;Acc:RF00019] |
| AL137785.1 | 14 | novel transcript |
| MPPE1P1 | 14 | metallophosphoesterase 1 pseudogene 1 [Sou |
| FOXN3 | 14 | forkhead box N3 [Source:HGNC Symbol;Acc:H |
| AL138478.1 | 14 | novel transcript |
| CAP2P1 | 14 | cyclase associated actin cytoskeleton regulato |
| AL357093.2 | 14 | novel transcript, antisense to FOXN3 |
| AL357093.1 | 14 | novel transcript, antisense to FOXN3 |
| AL356805.1 | 14 | novel transcript, antisense to FOXN3 |
| AL137230.2 | 14 | novel transcript |
| FOXN3-AS1 | 14 | FOXN3 antisense RNA 1 [Source:HGNC Symbo |
| FOXN3-AS2 | 14 | FOXN3 antisense RNA 2 [Source:HGNC Symbo |

| | | |
|---|---|---|
| RN7SKP107 | 14 | RN7SK pseudogene 107 [Source:HGNC Symbo |
| AL137230.1 | 14 | uncharacterized LOC105370616 [Source:NCBI |
| Y_RNA | 14 | Y RNA [Source:RFAM;Acc:RF00019] |
| RN7SKP255 | 14 | RN7SK pseudogene 255 [Source:HGNC Symbo |
| CHORDC2P | 14 | cysteine and histidine rich domain containing |
| EFCAB11 | 14 | EF-hand calcium binding domain 11 [Source:H |
| RAB42P1 | 14 | RAB42, member RAS oncogene family, pseudo |
| TDP1 | 14 | tyrosyl-DNA phosphodiesterase 1 [Source:HG |
| KCNK13 | 14 | potassium two pore domain channel subfamil |
| GLRXP2 | 14 | glutaredoxin pseudogene 2 [Source:HGNC Syr |
| Y_RNA | 14 | Y RNA [Source:RFAM;Acc:RF00019] |
| PSMC1 | 14 | proteasome 26S subunit, ATPase 1 [Source:H( |
| NRDE2 | 14 | NRDE-2, necessary for RNA interference, dom |
| AL161662.1 | 14 | novel transcript, antisense to NRDE2 |
| RPL21P11 | 14 | ribosomal protein L21 pseudogene 11 [Source |
| AL512791.2 | 14 | novel transcript |
| CALM1 | 14 | calmodulin 1 [Source:HGNC Symbol;Acc:HGN( |
| AL512791.1 | 14 | novel transcript, antisense to CALM1 |
| LINC02317 | 14 | long intergenic non-protein coding RNA 2317 |
| LINC00642 | 14 | long intergenic non-protein coding RNA 642 [S |
| AL096869.3 | 14 | novel transcript |
| TTC7B | 14 | tetratricopeptide repeat domain 7B [Source:H |
| AL096869.1 | 14 | novel transcript, antisense & intronic to TTC7I |
| AL096869.2 | 14 | novel transcript, antisense to TTC7B |
| AL139193.1 | 14 | novel transcript, antisense to TTC7B |
| AL139193.2 | 14 | novel transcript, antisense to TTC7B |
| AL122020.2 | 14 | novel transcript, antisense to TTC7B |
| AL122020.1 | 14 | ribosomal protein S18 (RPS18) pseudogene |
| LINC02321 | 14 | long intergenic non-protein coding RNA 2321 |
| RPS6KA5 | 14 | ribosomal protein S6 kinase A5 [Source:HGNC |
| DGLUCY | 14 | D-glutamate cyclase [Source:HGNC Symbol;Ac |
| RNU7-30P | 14 | RNA, U7 small nuclear 30 pseudogene [Source |
| SNORA11B | 14 | small nucleolar RNA, H/ACA box 11B [Source:I |
| GPR68 | 14 | G protein-coupled receptor 68 [Source:HGNC |
| AL135818.1 | 14 | novel transcript, antisense to GPR68 |
| AL135818.2 | 14 | novel transcript |
| AL135818.3 | 14 | novel transcript |
| CCDC88C | 14 | coiled-coil domain containing 88C [Source:HG |
| AL133153.2 | 14 | novel transcript, antisense to CCDC88C |
| AL133153.1 | 14 | COX assembly mitochondrial protein homolog |
| PPP4R3A | 14 | protein phosphatase 4 regulatory subunit 3A [ |
| AL133373.1 | 14 | zinc finger protein 257 (ZNF257) pseudogene |
| AL133373.2 | 14 | TEC |
| CATSPERB | 14 | cation channel sperm associated auxiliary sub |
| POLR3GP1 | 14 | RNA polymerase III subunit G pseudogene 1 [S |
| NANOGP7 | 14 | Nanog homeobox pseudogene 7 [Source:HGN |
| AL121839.1 | 14 | pyruvate dehydrogenase kinase, isozyme 3 (P |
| U3 | 14 | Small nucleolar RNA U3 [Source:RFAM;Acc:RF |
| AL121839.2 | 14 | novel transcript, intronic to CATSPERB |
| TC2N | 14 | tandem C2 domains, nuclear [Source:HGNC Sy |

| | | |
|---|---|---|
| FBLN5 | 14 | fibulin 5 [Source:HGNC Symbol;Acc:HGNC:360 |
| TRIP11 | 14 | thyroid hormone receptor interactor 11 [Sour |
| AL049872.1 | 14 | prothymosin, alpha (PTMA) pseudogene |
| ATXN3 | 14 | ataxin 3 [Source:HGNC Symbol;Acc:HGNC:710 |
| NDUFB1 | 14 | NADH:ubiquinone oxidoreductase subunit B1 |
| CPSF2 | 14 | cleavage and polyadenylation specific factor 2 |
| AL133240.1 | 14 | novel transcript |
| RNU6-366P | 14 | RNA, U6 small nuclear 366, pseudogene [Sour |
| AL133240.2 | 14 | novel transcript |
| SLC24A4 | 14 | solute carrier family 24 member 4 [Source:HG |
| RIN3 | 14 | Ras and Rab interactor 3 [Source:HGNC Symb |
| LGMN | 14 | legumain [Source:HGNC Symbol;Acc:HGNC:94 |
| GOLGA5 | 14 | golgin A5 [Source:HGNC Symbol;Acc:HGNC:44 |
| LINC02833 | 14 | long intergenic non-protein coding RNA 2833 |
| LINC02287 | 14 | long intergenic non-protein coding RNA 2287 |
| CHGA | 14 | chromogranin A [Source:HGNC Symbol;Acc:H |
| ITPK1 | 14 | inositol-tetrakisphosphate 1-kinase [Source:H |
| ITPK1-AS1 | 14 | ITPK1 antisense RNA 1 [Source:HGNC Symbol; |
| AL110118.1 | 14 | cytochrome b5 type B (outer mitochondrial m |
| MOAP1 | 14 | modulator of apoptosis 1 [Source:HGNC Symb |
| TMEM251 | 14 | transmembrane protein 251 [Source:HGNC Sy |
| AL110118.2 | 14 | novel protein |
| GON7 | 14 | GON7 subunit of KEOPS complex [Source:HGN |
| UBR7 | 14 | ubiquitin protein ligase E3 component n-recog |
| AL132838.1 | 14 | ribosomal protein L18a (RPL18A) pseudogene |
| BTBD7 | 14 | BTB domain containing 7 [Source:HGNC Symb |
| AL132838.2 | 14 | ribosomal protein L36a-like (RPL36AL) pseudo |
| UNC79 | 14 | unc-79 homolog, NALCN channel complex sub |
| AL122023.1 | 14 | novel transcript, sense intronic UNC79 |
| COX8C | 14 | cytochrome c oxidase subunit 8C [Source:HGN |
| Y_RNA | 14 | Y RNA [Source:RFAM;Acc:RF00019] |
| RNU6-1258P | 14 | RNA, U6 small nuclear 1258, pseudogene [Sou |
| AL157858.2 | 14 | NULL |
| PRIMA1 | 14 | proline rich membrane anchor 1 [Source:HGN |
| AL132642.2 | 14 | TEC |
| FAM181A-AS1 | 14 | FAM181A antisense RNA 1 [Source:HGNC Sym |
| FAM181A | 14 | family with sequence similarity 181 member A |
| ASB2 | 14 | ankyrin repeat and SOCS box containing 2 [So |
| AL132642.1 | 14 | novel transcript, antisense to ASB2 |
| MIR4506 | 14 | microRNA 4506 [Source:HGNC Symbol;Acc:H |
| CCDC197 | 14 | coiled-coil domain containing 197 [Source:HG |
| OTUB2 | 14 | OTU deubiquitinase, ubiquitin aldehyde bindi |
| DDX24 | 14 | DEAD-box helicase 24 [Source:HGNC Symbol; |
| IFI27L1 | 14 | interferon alpha inducible protein 27 like 1 [S |
| IFI27 | 14 | interferon alpha inducible protein 27 [Source: |
| IFI27L2 | 14 | interferon alpha inducible protein 27 like 2 [S |
| PPP4R4 | 14 | protein phosphatase 4 regulatory subunit 4 [S |
| SERPINA10 | 14 | serpin family A member 10 [Source:HGNC Syn |
| SERPINA6 | 14 | serpin family A member 6 [Source:HGNC Symb |
| SERPINA2 | 14 | serpin family A member 2 (gene/pseudogene) |

| | | |
|---|---|---|
| SERPINA1 | 14 | serpin family A member 1 [Source:HGNC Syml |
| AL132708.1 | 14 | novel transcript, antisense to SERPINA9 |
| SERPINA11 | 14 | serpin family A member 11 [Source:HGNC Syn |
| SERPINA9 | 14 | serpin family A member 9 [Source:HGNC Syml |
| SERPINA12 | 14 | serpin family A member 12 [Source:HGNC Syn |
| AL132990.1 | 14 | novel transcript |
| SERPINA4 | 14 | serpin family A member 4 [Source:HGNC Syml |
| SERPINA5 | 14 | serpin family A member 5 [Source:HGNC Syml |
| AL049839.2 | 14 | novel protein |
| AL049839.3 | 14 | novel transcript |
| SERPINA3 | 14 | serpin family A member 3 [Source:HGNC Syml |
| ADIPOR1P2 | 14 | adiponectin receptor 1 pseudogene 2 [Source |
| SERPINA13P | 14 | serpin family A member 13, pseudogene [Sou |
| AL049839.1 | 14 | coiled-coil domain containing 59 (CCDS59) pse |
| RPSAP4 | 14 | ribosomal protein SA pseudogene 4 [Source:H |
| GSC | 14 | goosecoid homeobox [Source:HGNC Symbol;A |
| AL121612.2 | 14 | novel transcript |
| RPL15P2 | 14 | ribosomal protein L15 pseudogene 2 [Source: |
| AL121612.1 | 14 | novel transcript |
| LINC02279 | 14 | long intergenic non-protein coding RNA 2279 |
| AL390254.1 | 14 | novel transcript |
| DICER1 | 14 | dicer 1, ribonuclease III [Source:HGNC Symbol |
| MIR3173 | 14 | microRNA 3173 [Source:HGNC Symbol;Acc:HG |
| DICER1-AS1 | 14 | DICER1 antisense RNA 1 [Source:HGNC Symbo |
| CLMN | 14 | calmin [Source:HGNC Symbol;Acc:HGNC:1997 |
| AL356017.1 | 14 | novel transcript, sense overlapping CLMN |
| AL117187.1 | 14 | mitochondrially encoded ATP synthase 6 (MT- |
| AL117187.2 | 14 | calmin (calponin-like, transmembrane) (CLMN |
| LINC02292 | 14 | long intergenic non-protein coding RNA 2292 |
| SYNE3 | 14 | spectrin repeat containing nuclear envelope fa |
| AL133467.1 | 14 | novel transcript |
| SNHG10 | 14 | small nucleolar RNA host gene 10 [Source:HGI |
| SCARNA13 | 14 | small Cajal body-specific RNA 13 [Source:HGN |
| GLRX5 | 14 | glutaredoxin 5 [Source:HGNC Symbol;Acc:HGI |
| LINC02318 | 14 | long intergenic non-protein coding RNA 2318 |
| AL133467.2 | 14 | novel transcript |
| AL133467.4 | 14 | novel transcript |
| TCL6 | 14 | T cell leukemia/lymphoma 6 [Source:HGNC Sy |
| AL133467.5 | 14 | NULL |
| AL133467.3 | 14 | novel transcript |
| AL133467.6 | 14 | NULL |
| TCL1B | 14 | T cell leukemia/lymphoma 1B [Source:HGNC S |
| TCL1A | 14 | T cell leukemia/lymphoma 1A [Source:HGNC S |
| AL139020.1 | 14 | novel transcript, antisense to TCL1A |
| AL133167.1 | 14 | novel transcript |
| TUNAR | 14 | TCL1 upstream neural differentiation-associat |
| AL137190.1 | 14 | ubiquitin specific peptidase 5 (isopeptidase T) |
| C14orf132 | 14 | chromosome 14 open reading frame 132 [Sou |
| BDKRB2 | 14 | bradykinin receptor B2 [Source:HGNC Symbol |
| AL355102.2 | 14 | novel protein |

| | | |
|---|---|---|
| AL355102.1 | 14 | novel transcript, sense intronic to BDKRB2 |
| CKS1BP1 | 14 | CDC28 protein kinase regulatory subunit 1B ps |
| AL355102.5 | 14 | novel transcript, antisense to BDKRB2 |
| BDKRB1 | 14 | bradykinin receptor B1 [Source:HGNC Symbol |
| AL355102.4 | 14 | novel transcript, antisense to BDKRB1 |
| AL355102.3 | 14 | novel transcript |
| ATG2B | 14 | autophagy related 2B [Source:HGNC Symbol;A |
| GSKIP | 14 | GSK3B interacting protein [Source:HGNC Syml |
| RNU2-33P | 14 | RNA, U2 small nuclear 33, pseudogene [Sourc |
| AK7 | 14 | adenylate kinase 7 [Source:HGNC Symbol;Acc |
| PEBP1P1 | 14 | phosphatidylethanolamine binding protein 1 p |
| RPL23AP10 | 14 | ribosomal protein L23a pseudogene 10 [Sourc |
| AL163051.1 | 14 | novel transcript, antisense to PAPOLA |
| PAPOLA | 14 | poly(A) polymerase alpha [Source:HGNC Symb |
| AL137786.1 | 14 | novel transcript |
| RN7SKP108 | 14 | RN7SK pseudogene 108 [Source:HGNC Symbo |
| AL137786.2 | 14 | novel transcript |
| LINC02299 | 14 | long intergenic non-protein coding RNA 2299 |
| AL137786.3 | 14 | novel transcript |
| VRK1 | 14 | VRK serine/threonine kinase 1 [Source:HGNC |
| LINC00618 | 14 | long intergenic non-protein coding RNA 618 [S |
| AL049833.3 | 14 | TEC |
| AL049833.4 | 14 | novel transcript |
| AL049833.2 | 14 | novel transcript |
| AL049833.1 | 14 | novel transcript |
| LINC02304 | 14 | long intergenic non-protein coding RNA 2304 |
| AL158800.1 | 14 | novel transcript |
| LINC02325 | 14 | long intergenic non-protein coding RNA 2325 |
| LINC02291 | 14 | long intergenic non-protein coding RNA 2291 |
| AL163872.1 | 14 | novel transcript |
| LINC02312 | 14 | long intergenic non-protein coding RNA 2312 |
| AL355097.1 | 14 | novel transcript |
| LINC01550 | 14 | long intergenic non-protein coding RNA 1550 |
| AL163932.1 | 14 | novel transcript |
| LINC02295 | 14 | long intergenic non-protein coding RNA 2295 |
| RN7SL714P | 14 | RNA, 7SL, cytoplasmic 714, pseudogene [Sour |
| AL132719.1 | 14 | novel transcript |
| C14orf177 | 14 | chromosome 14 open reading frame 177 [Sou |
| AL132796.1 | 14 | novel transcript |
| AL132796.3 | 14 | TEC |
| AL132796.2 | 14 | novel transcript |
| RPL3P4 | 14 | ribosomal protein L3 pseudogene 4 [Source:H |
| AL162151.2 | 14 | pseudogene similar to part of ribosomal prote |
| AL162151.3 | 14 | novel transcript |
| AL162151.1 | 14 | novel transcript |
| BCL11B | 14 | BAF chromatin remodeling complex subunit B |
| AL109767.1 | 14 | novel transcript |
| AL132819.1 | 14 | novel transcript |
| SETD3 | 14 | SET domain containing 3, actin histidine meth |
| RNU6-91P | 14 | RNA, U6 small nuclear 91, pseudogene [Sourc |

| | | |
|---|---|---|
| CCNK | 14 | cyclin K [Source:HGNC Symbol;Acc:HGNC:159 |
| CCDC85C | 14 | coiled-coil domain containing 85C [Source:HG |
| AL110504.1 | 14 | novel transcript, antisense to CCDC85C |
| Y_RNA | 14 | Y RNA [Source:RFAM;Acc:RF00019] |
| AL160313.1 | 14 | novel transcript |
| HHIPL1 | 14 | HHIP like 1 [Source:HGNC Symbol;Acc:HGNC:1 |
| CYP46A1 | 14 | cytochrome P450 family 46 subfamily A meml |
| AL160313.2 | 14 | novel transcript |
| EML1 | 14 | EMAP like 1 [Source:HGNC Symbol;Acc:HGNC: |
| RNU1-47P | 14 | RNA, U1 small nuclear 47, pseudogene [Sourc |
| VDAC3P1 | 14 | voltage dependent anion channel 3 pseudoge |
| EVL | 14 | Enah/Vasp-like [Source:HGNC Symbol;Acc:HG |
| AL133368.1 | 14 | ribosomal protein S2 (RPS2) pseudogene |
| AL133368.2 | 14 | novel transcript |
| AL157912.1 | 14 | novel transcript, antisense to EVL |
| MIR151B | 14 | microRNA 151b [Source:HGNC Symbol;Acc:HG |
| MIR342 | 14 | microRNA 342 [Source:HGNC Symbol;Acc:HGI |
| DEGS2 | 14 | delta 4-desaturase, sphingolipid 2 [Source:HG |
| AL133523.1 | 14 | novel transcript, antisense to YY1 |
| YY1 | 14 | YY1 transcription factor [Source:HGNC Symbo |
| AL157871.5 | 14 | novel transcript, intronic to YY1 |
| MIR6764 | 14 | microRNA 6764 [Source:HGNC Symbol;Acc:HG |
| AL157871.6 | 14 | novel transcript, sense overlapping to SLC25A |
| SLC25A29 | 14 | solute carrier family 25 member 29 [Source:H |
| AL157871.1 | 14 | novel transcript |
| MIR345 | 14 | microRNA 345 [Source:HGNC Symbol;Acc:HGI |
| RN7SL523P | 14 | RNA, 7SL, cytoplasmic 523, pseudogene [Sour |
| SLC25A47 | 14 | solute carrier family 25 member 47 [Source:H |
| WARS | 14 | tryptophanyl-tRNA synthetase [Source:HGNC |
| AL157871.4 | 14 | novel transcript, antisense to WARS |
| AL157871.2 | 14 | novel transcript, antisense to WARS |
| AL157871.3 | 14 | NADH dehydrogenase (ubiquinone) 1 beta sul |
| WDR25 | 14 | WD repeat domain 25 [Source:HGNC Symbol; |
| AL135838.1 | 14 | novel transcript, intronic to WDR25 |
| AL845552.1 | 14 | high mobility group nucleosome binding dom |
| RN7SKP92 | 14 | RN7SK pseudogene 92 [Source:HGNC Symbol; |
| BEGAIN | 14 | brain enriched guanylate kinase associated [S |
| AL845552.2 | 14 | novel transcript |
| AL163974.1 | 14 | novel transcript, overlapping to BEGAIN |
| LINC00523 | 14 | long intergenic non-protein coding RNA 523 [ |
| AL132711.1 | 14 | novel transcript |
| DLK1 | 14 | delta like non-canonical Notch ligand 1 [Sourc |
| MEG3 | 14 | maternally expressed 3 [Source:HGNC Symbol |
| MIR2392 | 14 | microRNA 2392 [Source:HGNC Symbol;Acc:HG |
| AL117190.5 | 14 | NULL |
| AL117190.3 | 14 | NULL |
| AL117190.7 | 14 | NULL |
| AL117190.2 | 14 | novel transcript, antisense to MEG3 |
| AL117190.1 | 14 | novel transcript |
| MIR770 | 14 | microRNA 770 [Source:HGNC Symbol;Acc:HGI |

| | | |
|---|---|---|
| MIR493 | 14 | microRNA 493 [Source:HGNC Symbol;Acc:HGI |
| MIR337 | 14 | microRNA 337 [Source:HGNC Symbol;Acc:HGI |
| MIR665 | 14 | microRNA 665 [Source:HGNC Symbol;Acc:HGI |
| RTL1 | 14 | retrotransposon Gag like 1 [Source:HGNC Syr |
| MIR431 | 14 | microRNA 431 [Source:HGNC Symbol;Acc:HGI |
| MIR433 | 14 | microRNA 433 [Source:HGNC Symbol;Acc:HGI |
| MIR127 | 14 | microRNA 127 [Source:HGNC Symbol;Acc:HGI |
| MIR432 | 14 | microRNA 432 [Source:HGNC Symbol;Acc:HGI |
| MIR136 | 14 | microRNA 136 [Source:HGNC Symbol;Acc:HGI |
| MEG8 | 14 | maternally expressed 8, small nucleolar RNA h |
| AL117190.6 | 14 | NULL |
| SNORD112 | 14 | small nucleolar RNA, C/D box 112 [Source:HG |
| AL117190.4 | 14 | NULL |
| MIR370 | 14 | microRNA 370 [Source:HGNC Symbol;Acc:HGI |
| SNORD113-1 | 14 | small nucleolar RNA, C/D box 113-1 [Source:H |
| SNORD113-2 | 14 | small nucleolar RNA, C/D box 113-2 [Source:H |
| SNORD113-3 | 14 | small nucleolar RNA, C/D box 113-3 [Source:H |
| SNORD113-4 | 14 | small nucleolar RNA, C/D box 113-4 [Source:H |
| SNORD113-5 | 14 | small nucleolar RNA, C/D box 113-5 [Source:H |
| SNORD113-6 | 14 | small nucleolar RNA, C/D box 113-6 [Source:H |
| SNORD113-7 | 14 | small nucleolar RNA, C/D box 113-7 [Source:H |
| SNORD113-8 | 14 | small nucleolar RNA, C/D box 113-8 [Source:H |
| SNORD113-9 | 14 | small nucleolar RNA, C/D box 113-9 [Source:H |
| SNORD114-1 | 14 | small nucleolar RNA, C/D box 114-1 [Source:H |
| SNORD114-2 | 14 | small nucleolar RNA, C/D box 114-2 [Source:H |
| SNORD114-3 | 14 | small nucleolar RNA, C/D box 114-3 [Source:H |
| SNORD114-4 | 14 | small nucleolar RNA, C/D box 114-4 [Source:H |
| SNORD114-5 | 14 | small nucleolar RNA, C/D box 114-5 [Source:H |
| AL132709.1 | 14 | NULL |
| SNORD114-6 | 14 | small nucleolar RNA, C/D box 114-6 [Source:H |
| SNORD114-7 | 14 | small nucleolar RNA, C/D box 114-7 [Source:H |
| SNORD114-9 | 14 | small nucleolar RNA, C/D box 114-9 [Source:H |
| SNORD114-10 | 14 | small nucleolar RNA, C/D box 114-10 [Source: |
| SNORD114-11 | 14 | small nucleolar RNA, C/D box 114-11 [Source: |
| SNORD114-12 | 14 | small nucleolar RNA, C/D box 114-12 [Source: |
| SNORD114-13 | 14 | small nucleolar RNA, C/D box 114-13 [Source: |
| SNORD114-14 | 14 | small nucleolar RNA, C/D box 114-14 [Source: |
| SNORD114-15 | 14 | small nucleolar RNA, C/D box 114-15 [Source: |
| SNORD114-16 | 14 | small nucleolar RNA, C/D box 114-16 [Source: |
| SNORD114-17 | 14 | small nucleolar RNA, C/D box 114-17 [Source: |
| SNORD114-18 | 14 | small nucleolar RNA, C/D box 114-18 [Source: |
| SNORD114-19 | 14 | small nucleolar RNA, C/D box 114-19 [Source: |
| AL132709.3 | 14 | NULL |
| AL132709.2 | 14 | NULL |
| AL132709.4 | 14 | NULL |
| SNORD114-20 | 14 | small nucleolar RNA, C/D box 114-20 [Source: |
| SNORD114-21 | 14 | small nucleolar RNA, C/D box 114-21 [Source: |
| SNORD114-22 | 14 | small nucleolar RNA, C/D box 114-22 [Source: |
| SNORD114-23 | 14 | small nucleolar RNA, C/D box 114-23 [Source: |
| SNORD114-24 | 14 | small nucleolar RNA, C/D box 114-24 [Source: |

| | | |
|---|---|---|
| SNORD114-25 | 14 | small nucleolar RNA, C/D box 114-25 [Source: |
| SNORD114-26 | 14 | small nucleolar RNA, C/D box 114-26 [Source: |
| SNORD114-27 | 14 | small nucleolar RNA, C/D box 114-27 [Source: |
| SNORD114-28 | 14 | small nucleolar RNA, C/D box 114-28 [Source: |
| SNORD114-29 | 14 | small nucleolar RNA, C/D box 114-29 [Source: |
| SNORD114-30 | 14 | small nucleolar RNA, C/D box 114-30 [Source: |
| SNORD114-31 | 14 | small nucleolar RNA, C/D box 114-31 [Source: |
| AL132709.5 | 14 | NULL |
| AL132709.6 | 14 | NULL |
| MIR379 | 14 | microRNA 379 [Source:HGNC Symbol;Acc:HGN |
| MIR411 | 14 | microRNA 411 [Source:HGNC Symbol;Acc:HGN |
| MIR299 | 14 | microRNA 299 [Source:HGNC Symbol;Acc:HGN |
| MIR380 | 14 | microRNA 380 [Source:HGNC Symbol;Acc:HGN |
| MIR1197 | 14 | microRNA 1197 [Source:HGNC Symbol;Acc:HG |
| MIR323A | 14 | microRNA 323a [Source:HGNC Symbol;Acc:HG |
| MIR758 | 14 | microRNA 758 [Source:HGNC Symbol;Acc:HGN |
| MIR329-1 | 14 | microRNA 329-1 [Source:HGNC Symbol;Acc:H |
| MIR329-2 | 14 | microRNA 329-2 [Source:HGNC Symbol;Acc:H |
| MIR494 | 14 | microRNA 494 [Source:HGNC Symbol;Acc:HGN |
| MIR1193 | 14 | microRNA 1193 [Source:HGNC Symbol;Acc:HG |
| MIR543 | 14 | microRNA 543 [Source:HGNC Symbol;Acc:HGN |
| MIR495 | 14 | microRNA 495 [Source:HGNC Symbol;Acc:HGN |
| MIR376C | 14 | microRNA 376c [Source:HGNC Symbol;Acc:HG |
| MIR376A2 | 14 | microRNA 376a-2 [Source:HGNC Symbol;Acc:H |
| MIR654 | 14 | microRNA 654 [Source:HGNC Symbol;Acc:HGN |
| MIR376B | 14 | microRNA 376b [Source:HGNC Symbol;Acc:HG |
| MIR376A1 | 14 | microRNA 376a-1 [Source:HGNC Symbol;Acc:H |
| MIR300 | 14 | microRNA 300 [Source:HGNC Symbol;Acc:HGN |
| MIR1185-1 | 14 | microRNA 1185-1 [Source:HGNC Symbol;Acc:H |
| MIR1185-2 | 14 | microRNA 1185-2 [Source:HGNC Symbol;Acc:H |
| MIR381HG | 14 | MIR381 host gene [Source:HGNC Symbol;Acc: |
| MIR381 | 14 | microRNA 381 [Source:HGNC Symbol;Acc:HGN |
| MIR487B | 14 | microRNA 487b [Source:HGNC Symbol;Acc:HG |
| MIR539 | 14 | microRNA 539 [Source:HGNC Symbol;Acc:HGN |
| MIR889 | 14 | microRNA 889 [Source:HGNC Symbol;Acc:HGN |
| MIR544A | 14 | microRNA 544a [Source:HGNC Symbol;Acc:HG |
| MIR655 | 14 | microRNA 655 [Source:HGNC Symbol;Acc:HGN |
| MIR487A | 14 | microRNA 487a [Source:HGNC Symbol;Acc:HG |
| MIR382 | 14 | microRNA 382 [Source:HGNC Symbol;Acc:HGN |
| MIR134 | 14 | microRNA 134 [Source:HGNC Symbol;Acc:HGN |
| MIR668 | 14 | microRNA 668 [Source:HGNC Symbol;Acc:HGN |
| MIR485 | 14 | microRNA 485 [Source:HGNC Symbol;Acc:HGN |
| MIR323B | 14 | microRNA 323b [Source:HGNC Symbol;Acc:HG |
| MIR154 | 14 | microRNA 154 [Source:HGNC Symbol;Acc:HGN |
| MIR496 | 14 | microRNA 496 [Source:HGNC Symbol;Acc:HGN |
| MIR377 | 14 | microRNA 377 [Source:HGNC Symbol;Acc:HGN |
| MIR541 | 14 | microRNA 541 [Source:HGNC Symbol;Acc:HGN |
| MIR409 | 14 | microRNA 409 [Source:HGNC Symbol;Acc:HGN |
| MIR412 | 14 | microRNA 412 [Source:HGNC Symbol;Acc:HGN |
| MIR369 | 14 | microRNA 369 [Source:HGNC Symbol;Acc:HGN |

| | | |
|---|---|---|
| MIR410 | 14 | microRNA 410 [Source:HGNC Symbol;Acc:HGI |
| MIR656 | 14 | microRNA 656 [Source:HGNC Symbol;Acc:HGI |
| MEG9 | 14 | maternally expressed 9 [Source:HGNC Symbol |
| AL132709.7 | 14 | novel transcript |
| AL132709.8 | 14 | novel transcript |
| LINC02285 | 14 | long intergenic non-protein coding RNA 2285 |
| AL355836.2 | 14 | novel transcript |
| U3 | 14 | Small nucleolar RNA U3 [Source:RFAM;Acc:RF |
| AL355836.4 | 14 | novel transcript |
| AL355836.3 | 14 | novel transcript |
| AL355836.1 | 14 | AKT interacting protein (AKTIP) pseudogene |
| AL359682.1 | 14 | novel transcript |
| AL355096.1 | 14 | novel transcript |
| LINC00524 | 14 | long intergenic non-protein coding RNA 524 [S |
| LINC02314 | 14 | long intergenic non-protein coding RNA 2314 |
| AL049836.2 | 14 | novel transcript |
| DIO3OS | 14 | DIO3 opposite strand upstream RNA [Source:I |
| MIR1247 | 14 | microRNA 1247 [Source:HGNC Symbol;Acc:HG |
| DIO3 | 14 | iodothyronine deiodinase 3 [Source:HGNC Syr |
| AL049836.1 | 14 | novel transcript |
| LINC02320 | 14 | long intergenic non-protein coding RNA 2320 |
| AL355032.1 | 14 | ribosomal protein L26-like 1 (RPL26L1) pseud |
| LINC00239 | 14 | long intergenic non-protein coding RNA 239 [S |
| PPP2R5C | 14 | protein phosphatase 2 regulatory subunit B'ga |
| AL137779.2 | 14 | novel transcript, antisense to PPP2R5C |
| RNU6-790P | 14 | RNA, U6 small nuclear 790, pseudogene [Sour |
| AL137779.1 | 14 | novel transcript, antisense to PPP2R5C |
| AL137779.3 | 14 | nucleophosmin (nucleolar phosphoprotein B2 |
| AL118558.3 | 14 | novel transcript |
| AL118558.4 | 14 | novel transcript |
| DYNC1H1 | 14 | dynein cytoplasmic 1 heavy chain 1 [Source:H |
| AL118558.2 | 14 | NADH dehydrogenase (ubiquinone) 1 alpha su |
| AL118558.1 | 14 | novel pseudogene, ortholog of 1700001K19Ri |
| RN7SL472P | 14 | RNA, 7SL, cytoplasmic 472, pseudogene [Sour |
| HSP90AA1 | 14 | heat shock protein 90 alpha family class A me |
| WDR20 | 14 | WD repeat domain 20 [Source:HGNC Symbol;. |
| MOK | 14 | MOK protein kinase [Source:HGNC Symbol;Ac |
| ZNF839 | 14 | zinc finger protein 839 [Source:HGNC Symbol; |
| CINP | 14 | cyclin dependent kinase 2 interacting protein |
| TECPR2 | 14 | tectonin beta-propeller repeat containing 2 [S |
| RNU6-244P | 14 | RNA, U6 small nuclear 244, pseudogene [Sour |
| ANKRD9 | 14 | ankyrin repeat domain 9 [Source:HGNC Symb |
| MIR4309 | 14 | microRNA 4309 [Source:HGNC Symbol;Acc:HG |
| LINC02323 | 14 | long intergenic non-protein coding RNA 2323 |
| RN7SL546P | 14 | RNA, 7SL, cytoplasmic 546, pseudogene [Sour |
| RCOR1 | 14 | REST corepressor 1 [Source:HGNC Symbol;Acc |
| RPL23AP11 | 14 | ribosomal protein L23a pseudogene 11 [Sourc |
| Y_RNA | 14 | Y RNA [Source:RFAM;Acc:RF00019] |
| AL132801.1 | 14 | novel transcript |
| TRAF3 | 14 | TNF receptor associated factor 3 [Source:HGN |

| | | |
|---|---|---|
| RNU6-1316P | 14 | RNA, U6 small nuclear 1316, pseudogene [Sou |
| AMN | 14 | amnion associated transmembrane protein [S |
| CDC42BPB | 14 | CDC42 binding protein kinase beta [Source:H |
| AL117209.1 | 14 | novel transcript, antisense to CDC42BPB |
| RPL13P6 | 14 | ribosomal protein L13 pseudogene 6 [Source: |
| LBHD2 | 14 | LBH domain containing 2 [Source:HGNC Symb |
| AL161669.1 | 14 | novel transcript, sense overlapping EXOC3L4 |
| EXOC3L4 | 14 | exocyst complex component 3 like 4 [Source: |
| LINC00677 | 14 | long intergenic non-protein coding RNA 677 [ |
| TNFAIP2 | 14 | TNF alpha induced protein 2 [Source:HGNC Sy |
| NDUFB4P11 | 14 | NADH:ubiquinone oxidoreductase subunit B4 |
| RPL21P12 | 14 | ribosomal protein L21 pseudogene 12 [Source |
| GCSHP2 | 14 | glycine cleavage system protein H pseudogen |
| LINC00605 | 14 | long intergenic non-protein coding RNA 605 [ |
| AL161669.2 | 14 | collagen, type IV, alpha 1 (COL4A1) pseudoge |
| AL161669.3 | 14 | novel transcript |
| RPL21P13 | 14 | ribosomal protein L21 pseudogene 13 [Source |
| RAP2CP1 | 14 | RAP2C pseudogene 1 [Source:HGNC Symbol;A |
| AL138976.1 | 14 | ribosomal protein L17 (RPL17) pseudogene |
| AL138976.2 | 14 | novel transcript, antisense to EIF5 |
| EIF5 | 14 | eukaryotic translation initiation factor 5 [Sour |
| SNORA28 | 14 | small nucleolar RNA, H/ACA box 28 [Source:H |
| HMGB3P26 | 14 | high mobility group box 3 pseudogene 26 [Sou |
| RPSAP5 | 14 | ribosomal protein SA pseudogene 5 [Source:H |
| MARK3 | 14 | microtubule affinity regulating kinase 3 [Sourc |
| RPL10AP1 | 14 | ribosomal protein L10a pseudogene 1 [Source |
| CKB | 14 | creatine kinase B [Source:HGNC Symbol;Acc:H |
| AL133367.1 | 14 | novel transcript, antisense to TRMT61A |
| TRMT61A | 14 | tRNA methyltransferase 61A [Source:HGNC Sy |
| RNU7-160P | 14 | RNA, U7 small nuclear 160 pseudogene [Sourc |
| AL139300.2 | 14 | novel transcript, antisense to BAG5 |
| BAG5 | 14 | BCL2 associated athanogene 5 [Source:HGNC |
| KLC1 | 14 | kinesin light chain 1 [Source:HGNC Symbol;Ac |
| AL139300.1 | 14 | novel protein |
| COA8 | 14 | cytochrome c oxidase assembly factor 8 [Sour |
| RNU4-68P | 14 | RNA, U4 small nuclear 68, pseudogene [Sourc |
| AL049840.2 | 14 | novel transcript, antisense to KLC1 |
| AL049840.6 | 14 | novel transcript, sense intronic to KLC1 |
| AL049840.3 | 14 | novel transcript, antisense to KLC1 |
| AL049840.4 | 14 | novel transcript, sense intronic to KLC1 |
| AL049840.5 | 14 | novel transcript, sense intronic to KLC1 |
| XRCC3 | 14 | X-ray repair cross complementing 3 [Source:H |
| ZFYVE21 | 14 | zinc finger FYVE-type containing 21 [Source:H |
| Y_RNA | 14 | Y RNA [Source:RFAM;Acc:RF00019] |
| PPP1R13B | 14 | protein phosphatase 1 regulatory subunit 13B |
| AL049840.7 | 14 | protein phosphatase 2, regulatory subunit B', |
| AL049840.1 | 14 | NULL |
| LINC00637 | 14 | long intergenic non-protein coding RNA 637 [ |
| AL132712.1 | 14 | novel transcript |
| AL132712.2 | 14 | novel transcript |

| | | |
|---|---|---|
| ATP5MPL | 14 | ATP synthase membrane subunit 6.8PL [Sourc |
| TDRD9 | 14 | tudor domain containing 9 [Source:HGNC Sym |
| RD3L | 14 | retinal degeneration 3 like [Source:HGNC Sym |
| RN7SL634P | 14 | RNA, 7SL, cytoplasmic 634, pseudogene [Sour |
| ASPG | 14 | asparaginase [Source:HGNC Symbol;Acc:HGN( |
| MIR203A | 14 | microRNA 203a [Source:HGNC Symbol;Acc:HG |
| MIR203B | 14 | microRNA 203b [Source:HGNC Symbol;Acc:HG |
| AL359399.1 | 14 | novel transcript, antisense to KIF26A |
| KIF26A | 14 | kinesin family member 26A [Source:HGNC Syr |
| LINC02691 | 14 | long intergenic non-protein coding RNA 2691 |
| AL512357.2 | 14 | novel transcript |
| AL512357.1 | 14 | novel transcript |
| RNU6-684P | 14 | RNA, U6 small nuclear 684, pseudogene [Sour |
| CEND1P1 | 14 | cell cycle exit and neuronal differentiation 1 p |
| TMEM179 | 14 | transmembrane protein 179 [Source:HGNC Sy |
| C14orf180 | 14 | chromosome 14 open reading frame 180 [Sou |
| BX927359.1 | 14 | novel transcript, antisense to C14orf180 |
| AL583722.3 | 14 | novel transcript |
| LINC02280 | 14 | long intergenic non-protein coding RNA 2280 |
| MIR4710 | 14 | microRNA 4710 [Source:HGNC Symbol;Acc:HG |
| AL583722.1 | 14 | novel transcript |
| INF2 | 14 | inverted formin, FH2 and WH2 domain contai |
| AL583722.4 | 14 | novel transcript, antisense to INF2 |
| ADSSL1 | 14 | adenylosuccinate synthase like 1 [Source:HGN |
| SIVA1 | 14 | SIVA1 apoptosis inducing factor [Source:HGN( |
| AL583722.2 | 14 | novel transcript, antisense to AKT1 |
| AKT1 | 14 | AKT serine/threonine kinase 1 [Source:HGNC : |
| ZBTB42 | 14 | zinc finger and BTB domain containing 42 [Sou |
| AL590326.1 | 14 | novel transcript |
| LINC00638 | 14 | long intergenic non-protein coding RNA 638 [S |
| RPS26P49 | 14 | ribosomal protein S26 pseudogene 49 [Source |
| RPS2P4 | 14 | ribosomal protein S2 pseudogene 4 [Source:H |
| AL583810.2 | 14 | novel transcript |
| AL583810.1 | 14 | novel transcript, antisense to KIAA0284 |
| CEP170B | 14 | centrosomal protein 170B [Source:HGNC Sym |
| AL583810.3 | 14 | novel transcript |
| PLD4 | 14 | phospholipase D family member 4 [Source:HG |
| AHNAK2 | 14 | AHNAK nucleoprotein 2 [Source:HGNC Symbo |
| CLBA1 | 14 | clathrin binding box of aftiphilin containing 1 [ |
| CDCA4 | 14 | cell division cycle associated 4 [Source:HGNC : |
| GPR132 | 14 | G protein-coupled receptor 132 [Source:HGN( |
| LINC02298 | 14 | long intergenic non-protein coding RNA 2298 |
| AL512356.4 | 14 | novel transcript |
| AL512356.3 | 14 | novel transcript |
| AL512356.2 | 14 | microtubule associated serine/threonine kina: |
| AL512356.5 | 14 | novel transcript |
| JAG2 | 14 | jagged canonical Notch ligand 2 [Source:HGN( |
| MIR6765 | 14 | microRNA 6765 [Source:HGNC Symbol;Acc:HG |
| AL512356.1 | 14 | novel transcript |
| NUDT14 | 14 | nudix hydrolase 14 [Source:HGNC Symbol;Acc |

| | | |
|---|---|---|
| BRF1 | 14 | BRF1 RNA polymerase III transcription initiatic |
| BTBD6 | 14 | BTB domain containing 6 [Source:HGNC Symb |
| PACS2 | 14 | phosphofurin acidic cluster sorting protein 2 [ |
| AL928654.4 | 14 | novel transcript |
| RPS20P33 | 14 | ribosomal protein S20 pseudogene 33 [Source |
| TEX22 | 14 | testis expressed 22 [Source:HGNC Symbol;Acc |
| AL928654.1 | 14 | novel transcript, antisense to MTA1 |
| MTA1 | 14 | metastasis associated 1 [Source:HGNC Symbo |
| AL928654.2 | 14 | novel transcript, antisense to MTA1 |
| CRIP2 | 14 | cysteine rich protein 2 [Source:HGNC Symbol; |
| CRIP1 | 14 | cysteine rich protein 1 [Source:HGNC Symbol; |
| AL928654.3 | 14 | novel protein |
| TEDC1 | 14 | tubulin epsilon and delta complex 1 [Source:H |
| TMEM121 | 14 | transmembrane protein 121 [Source:HGNC Sy |
| ATP5MC1P1 | 14 | ATP synthase membrane subunit c locus 1 pse |
| ELK2BP | 14 | ETS transcription factor ELK2B, pseudogene [S |
| IGHA2 | 14 | immunoglobulin heavy constant alpha 2 (A2m |
| IGHE | 14 | immunoglobulin heavy constant epsilon [Sour |
| IGHG4 | 14 | immunoglobulin heavy constant gamma 4 (G4 |
| MIR8071-1 | 14 | microRNA 8071-1 [Source:HGNC Symbol;Acc:I |
| IGHG2 | 14 | immunoglobulin heavy constant gamma 2 (G2 |
| MIR8071-2 | 14 | microRNA 8071-2 [Source:HGNC Symbol;Acc:I |
| COPDA1 | 14 | chronic obstructive pulmonary disease associa |
| IGHGP | 14 | immunoglobulin heavy constant gamma P (no |
| ELK2AP | 14 | ETS transcription factor ELK2A, pseudogene [S |
| IGHA1 | 14 | immunoglobulin heavy constant alpha 1 [Sour |
| IGHEP1 | 14 | immunoglobulin heavy constant epsilon P1 (p |
| IGHG1 | 14 | immunoglobulin heavy constant gamma 1 (G1 |
| IGHG3 | 14 | immunoglobulin heavy constant gamma 3 (G3 |
| AL122127.1 | 14 | ATPase, H+ transporting, lysosomal 13kDa, V1 |
| IGHD | 14 | immunoglobulin heavy constant delta [Source |
| IGHM | 14 | immunoglobulin heavy constant mu [Source:H |
| MIR4539 | 14 | microRNA 4539 [Source:HGNC Symbol;Acc:HG |
| MIR4507 | 14 | microRNA 4507 [Source:HGNC Symbol;Acc:HG |
| MIR4538 | 14 | microRNA 4538 [Source:HGNC Symbol;Acc:HG |
| MIR4537 | 14 | microRNA 4537 [Source:HGNC Symbol;Acc:HG |
| IGHJ6 | 14 | immunoglobulin heavy joining 6 [Source:HGN |
| IGHJ3P | 14 | immunoglobulin heavy joining 3P (pseudogen |
| IGHJ5 | 14 | immunoglobulin heavy joining 5 [Source:HGN |
| IGHJ4 | 14 | immunoglobulin heavy joining 4 [Source:HGN |
| IGHJ3 | 14 | immunoglobulin heavy joining 3 [Source:HGN |
| IGHJ2P | 14 | immunoglobulin heavy joining 2P (pseudogen |
| IGHJ2 | 14 | immunoglobulin heavy joining 2 [Source:HGN |
| IGHJ1 | 14 | immunoglobulin heavy joining 1 [Source:HGN |
| IGHD7-27 | 14 | immunoglobulin heavy diversity 7-27 [Source: |
| IGHJ1P | 14 | immunoglobulin heavy joining 1P (pseudogen |
| IGHD1-26 | 14 | immunoglobulin heavy diversity 1-26 [Source: |
| IGHD6-25 | 14 | immunoglobulin heavy diversity 6-25 [Source: |
| IGHD5-24 | 14 | immunoglobulin heavy diversity 5-24 (non-fur |
| IGHD4-23 | 14 | immunoglobulin heavy diversity 4-23 (non-fur |

| | | |
|---|---|---|
| IGHD3-22 | 14 | immunoglobulin heavy diversity 3-22 [Source: |
| IGHD2-21 | 14 | immunoglobulin heavy diversity 2-21 [Source: |
| AC246787.2 | 14 | novel transcript |
| IGHD1-20 | 14 | immunoglobulin heavy diversity 1-20 [Source: |
| IGHD6-19 | 14 | immunoglobulin heavy diversity 6-19 [Source: |
| IGHD5-18 | 14 | immunoglobulin heavy diversity 5-18 [Source: |
| IGHD4-17 | 14 | immunoglobulin heavy diversity 4-17 [Source: |
| IGHD3-16 | 14 | immunoglobulin heavy diversity 3-16 [Source: |
| IGHD2-15 | 14 | immunoglobulin heavy diversity 2-15 [Source: |
| IGHD1-14 | 14 | immunoglobulin heavy diversity 1-14 (non-fur |
| IGHD6-13 | 14 | immunoglobulin heavy diversity 6-13 [Source: |
| IGHD5-12 | 14 | immunoglobulin heavy diversity 5-12 [Source: |
| IGHD4-11 | 14 | immunoglobulin heavy diversity 4-11 (non-fur |
| IGHD3-10 | 14 | immunoglobulin heavy diversity 3-10 [Source: |
| IGHD3-9 | 14 | immunoglobulin heavy diversity 3-9 [Source:H |
| IGHD2-8 | 14 | immunoglobulin heavy diversity 2-8 [Source:H |
| IGHD1-7 | 14 | immunoglobulin heavy diversity 1-7 [Source:H |
| IGHD6-6 | 14 | immunoglobulin heavy diversity 6-6 [Source:H |
| IGHD5-5 | 14 | immunoglobulin heavy diversity 5-5 [Source:H |
| IGHD4-4 | 14 | immunoglobulin heavy diversity 4-4 [Source:H |
| IGHD3-3 | 14 | immunoglobulin heavy diversity 3-3 [Source:H |
| IGHD2-2 | 14 | immunoglobulin heavy diversity 2-2 [Source:H |
| FAM30A | 14 | family with sequence similarity 30 member A |
| IGHD1-1 | 14 | immunoglobulin heavy diversity 1-1 [Source:H |
| IGHV6-1 | 14 | immunoglobulin heavy variable 6-1 [Source:H |
| IGHVII-1-1 | 14 | immunoglobulin heavy variable (II)-1-1 (pseuc |
| ADAM6 | 14 | ADAM metallopeptidase domain 6 (pseudoge |
| AC246787.1 | 14 | ribosomal protein S8 (RPS8) pseudogene |
| IGHV1-2 | 14 | immunoglobulin heavy variable 1-2 [Source:H |
| IGHVIII-2-1 | 14 | immunoglobulin heavy variable (III)-2-1 (pseu |
| IGHV1-3 | 14 | immunoglobulin heavy variable 1-3 [Source:H |
| IGHV4-4 | 14 | immunoglobulin heavy variable 4-4 [Source:H |
| IGHV7-4-1 | 14 | immunoglobulin heavy variable 7-4-1 [Source: |
| IGHV2-5 | 14 | immunoglobulin heavy variable 2-5 [Source:H |
| IGHVIII-5-1 | 14 | immunoglobulin heavy variable (III)-5-1 (pseu |
| IGHVIII-5-2 | 14 | immunoglobulin heavy variable (III)-5-2 (pseu |
| IGHV3-6 | 14 | immunoglobulin heavy variable 3-6 (pseudoge |
| IGHV3-7 | 14 | immunoglobulin heavy variable 3-7 [Source:H |
| IGHV3-64D | 14 | immunoglobulin heavy variable 3-64D [Source |
| IGHV5-10-1 | 14 | immunoglobulin heavy variable 5-10-1 [Sourc |
| IGHV3-11 | 14 | immunoglobulin heavy variable 3-11 (gene/ps |
| IGHVIII-11-1 | 14 | immunoglobulin heavy variable (III)-11-1 (pse |
| IGHV1-12 | 14 | immunoglobulin heavy variable 1-12 (pseudog |
| IGHV3-13 | 14 | immunoglobulin heavy variable 3-13 [Source: |
| IGHVIII-13-1 | 14 | immunoglobulin heavy variable (III)-13-1 (pse |
| IGHV1-14 | 14 | immunoglobulin heavy variable 1-14 (pseudog |
| IGHV3-15 | 14 | immunoglobulin heavy variable 3-15 [Source: |
| IGHVII-15-1 | 14 | immunoglobulin heavy variable (II)-15-1 (pseu |
| IGHV3-16 | 14 | immunoglobulin heavy variable 3-16 (non-fun |
| IGHVIII-16-1 | 14 | immunoglobulin heavy variable (III)-16-1 (pse |

| | | |
|---|---|---|
| IGHV1-17 | 14 | immunoglobulin heavy variable 1-17 (pseudog |
| SLC20A1P2 | 14 | solute carrier family 20 member 1 pseudogen |
| IGHV1-18 | 14 | immunoglobulin heavy variable 1-18 [Source:I |
| IGHV3-19 | 14 | immunoglobulin heavy variable 3-19 (pseudog |
| AC247036.1 | 14 | novel transcript |
| IGHV3-20 | 14 | immunoglobulin heavy variable 3-20 [Source:I |
| IGHV3-21 | 14 | immunoglobulin heavy variable 3-21 [Source:I |
| IGHV3-22 | 14 | immunoglobulin heavy variable 3-22 (pseudog |
| IGHVII-22-1 | 14 | immunoglobulin heavy variable (II)-22-1 (pseu |
| IGHVIII-22-2 | 14 | immunoglobulin heavy variable (III)-22-2 (pseu |
| IGHV3-23 | 14 | immunoglobulin heavy variable 3-23 [Source:I |
| IGHV1-24 | 14 | immunoglobulin heavy variable 1-24 [Source:I |
| HOMER2P2 | 14 | HOMER2 pseudogene 2 [Source:HGNC Symbo |
| LINC00226 | 14 | long intergenic non-protein coding RNA 226 [S |
| IGHV3-25 | 14 | immunoglobulin heavy variable 3-25 (pseudog |
| IGHVIII-25-1 | 14 | immunoglobulin heavy variable (III)-25-1 (pseu |
| IGHV2-26 | 14 | immunoglobulin heavy variable 2-26 [Source:I |
| IGHVIII-26-1 | 14 | immunoglobulin heavy variable (III)-26-1 (pseu |
| IGHVII-26-2 | 14 | immunoglobulin heavy variable (II)-26-2 (pseu |
| IGHV7-27 | 14 | immunoglobulin heavy variable 7-27 (pseudog |
| IGHV4-28 | 14 | immunoglobulin heavy variable 4-28 [Source:I |
| IGHVII-28-1 | 14 | immunoglobulin heavy variable (II)-28-1 (pseu |
| IGHV3-32 | 14 | immunoglobulin heavy variable 3-32 (pseudog |
| IGHV3-30 | 14 | immunoglobulin heavy variable 3-30 [Source:I |
| IGHVII-30-1 | 14 | immunoglobulin heavy variable (II)-30-1 (pseu |
| IGHV3-30-2 | 14 | immunoglobulin heavy variable 3-30-2 (pseud |
| IGHV4-31 | 14 | immunoglobulin heavy variable 4-31 [Source:I |
| IGHVII-30-21 | 14 | immunoglobulin heavy variable (II)-30-21 (pse |
| IGHV3-29 | 14 | immunoglobulin heavy variable 3-29 (pseudog |
| IGHV3-33 | 14 | immunoglobulin heavy variable 3-33 [Source:I |
| IGHVII-33-1 | 14 | immunoglobulin heavy variable (II)-33-1 (pseu |
| IGHV3-33-2 | 14 | immunoglobulin heavy variable 3-33-2 (pseud |
| IGHV4-34 | 14 | immunoglobulin heavy variable 4-34 [Source:I |
| IGHV7-34-1 | 14 | immunoglobulin heavy variable 7-34-1 (pseud |
| IGHV3-35 | 14 | immunoglobulin heavy variable 3-35 (non-fun |
| IGHV3-36 | 14 | immunoglobulin heavy variable 3-36 (pseudog |
| IGHV3-37 | 14 | immunoglobulin heavy variable 3-37 (pseudog |
| IGHV3-38 | 14 | immunoglobulin heavy variable 3-38 (non-fun |
| IGHVIII-38-1 | 14 | immunoglobulin heavy variable (III)-38-1 (pseu |
| IGHV4-39 | 14 | immunoglobulin heavy variable 4-39 [Source:I |
| IGHV7-40 | 14 | immunoglobulin heavy variable 7-40 (pseudog |
| AC244452.1 | 14 | pseudogene similar to part of homer homolog |
| IGHVII-40-1 | 14 | immunoglobulin heavy variable (II)-40-1 (pseu |
| IGHV3-41 | 14 | immunoglobulin heavy variable 3-41 (pseudog |
| HOMER2P1 | 14 | HOMER2 pseudogene 1 [Source:HGNC Symbo |
| IGHV3-42 | 14 | immunoglobulin heavy variable 3-42 (pseudog |
| IGHV3-43 | 14 | immunoglobulin heavy variable 3-43 [Source:I |
| IGHVII-43-1 | 14 | immunoglobulin heavy variable (II)-43-1 (pseu |
| IGHVIII-44 | 14 | immunoglobulin heavy variable (III)-44 (pseud |
| AC244452.3 | 14 | novel pseudogene identical to IGHVII-44-2D |

| | | |
|---|---|---|
| LINC00221 | 14 | long intergenic non-protein coding RNA 221 [S |
| IGHVIV-44-1 | 14 | immunoglobulin heavy variable (IV)-44-1 (pse |
| IGHVII-44-2 | 14 | immunoglobulin heavy variable (II)-44-2 (pseu |
| IGHV1-45 | 14 | immunoglobulin heavy variable 1-45 [Source:I |
| IGHV1-46 | 14 | immunoglobulin heavy variable 1-46 [Source:I |
| IGHVII-46-1 | 14 | immunoglobulin heavy variable (II)-46-1 (pseu |
| IGHV3-47 | 14 | immunoglobulin heavy variable 3-47 (pseudog |
| IGHVIII-47-1 | 14 | immunoglobulin heavy variable (III)-47-1 (pseu |
| IGHV3-48 | 14 | immunoglobulin heavy variable 3-48 [Source:I |
| IGHV3-49 | 14 | immunoglobulin heavy variable 3-49 [Source:I |
| IGHVII-49-1 | 14 | immunoglobulin heavy variable (II)-49-1 (pseu |
| IGHV3-50 | 14 | immunoglobulin heavy variable 3-50 (pseudog |
| IGHV5-51 | 14 | immunoglobulin heavy variable 5-51 [Source:I |
| IGHVIII-51-1 | 14 | immunoglobulin heavy variable (III)-51-1 (pseu |
| IGHVII-51-2 | 14 | immunoglobulin heavy variable (II)-51-2 (pseu |
| IGHV3-52 | 14 | immunoglobulin heavy variable 3-52 (pseudog |
| IGHV3-53 | 14 | immunoglobulin heavy variable 3-53 [Source:I |
| IGHVII-53-1 | 14 | immunoglobulin heavy variable (II)-53-1 (pseu |
| IGHV3-54 | 14 | immunoglobulin heavy variable 3-54 (pseudog |
| IGHV4-55 | 14 | immunoglobulin heavy variable 4-55 (pseudog |
| IGHV7-56 | 14 | immunoglobulin heavy variable 7-56 (pseudog |
| IGHV3-57 | 14 | immunoglobulin heavy variable 3-57 (pseudog |
| IGHV1-58 | 14 | immunoglobulin heavy variable 1-58 [Source:I |
| IGHV4-59 | 14 | immunoglobulin heavy variable 4-59 [Source:I |
| IGHV3-60 | 14 | immunoglobulin heavy variable 3-60 (pseudog |
| RNA5SP389 | 14 | RNA, 5S ribosomal pseudogene 389 [Source:H |
| IGHVII-60-1 | 14 | immunoglobulin heavy variable (II)-60-1 (pseu |
| IGHV4-61 | 14 | immunoglobulin heavy variable 4-61 [Source:I |
| IGHV3-62 | 14 | immunoglobulin heavy variable 3-62 (pseudog |
| IGHVII-62-1 | 14 | immunoglobulin heavy variable (II)-62-1 (pseu |
| IGHV3-63 | 14 | immunoglobulin heavy variable 3-63 (pseudog |
| IGHV3-64 | 14 | immunoglobulin heavy variable 3-64 [Source:I |
| IGHV3-65 | 14 | immunoglobulin heavy variable 3-65 (pseudog |
| IGHVII-65-1 | 14 | immunoglobulin heavy variable (II)-65-1 (pseu |
| IGHV3-66 | 14 | immunoglobulin heavy variable 3-66 [Source:I |
| IGHV1-67 | 14 | immunoglobulin heavy variable 1-67 (pseudog |
| SLC20A1P1 | 14 | solute carrier family 20 member 1 pseudogen |
| IGHVII-67-1 | 14 | immunoglobulin heavy variable (II)-67-1 (pseu |
| IGHVIII-67-2 | 14 | immunoglobulin heavy variable (III)-67-2 (pseu |
| IGHVIII-67-3 | 14 | immunoglobulin heavy variable (III)-67-3 (pseu |
| IGHVIII-67-4 | 14 | immunoglobulin heavy variable (III)-67-4 (pseu |
| IGHV1-68 | 14 | immunoglobulin heavy variable 1-68 (pseudog |
| IGHV1-69 | 14 | immunoglobulin heavy variable 1-69 [Source:I |
| IGHV2-70D | 14 | immunoglobulin heavy variable 2-70D [Source |
| IGHV3-69-1 | 14 | immunoglobulin heavy variable 3-69-1 (pseud |
| IGHV1-69-2 | 14 | immunoglobulin heavy variable 1-69-2 [Sourc |
| AC245369.1 | 14 | novel IG pseudogene |
| IGHV1-69D | 14 | immunoglobulin heavy variable 1-69D [Source |
| IGHV2-70 | 14 | immunoglobulin heavy variable 2-70 [Source:I |
| IGHV3-71 | 14 | immunoglobulin heavy variable 3-71 (pseudog |

| | | |
|---|---|---|
| IGHV3-72 | 14 | immunoglobulin heavy variable 3-72 [Source:I |
| IGHV3-73 | 14 | immunoglobulin heavy variable 3-73 [Source:I |
| IGHV3-74 | 14 | immunoglobulin heavy variable 3-74 [Source:I |
| IGHVII-74-1 | 14 | immunoglobulin heavy variable (II)-74-1 (pseu |
| IGHV3-75 | 14 | immunoglobulin heavy variable 3-75 (pseudog |
| IGHV3-76 | 14 | immunoglobulin heavy variable 3-76 (pseudog |
| IGHVIII-76-1 | 14 | immunoglobulin heavy variable (III)-76-1 (pseu |
| MIR5195 | 14 | microRNA 5195 [Source:HGNC Symbol;Acc:HG |
| IGHV5-78 | 14 | immunoglobulin heavy variable 5-78 (pseudog |
| IGHVII-78-1 | 14 | immunoglobulin heavy variable (II)-78-1 (pseu |
| IGHV3-79 | 14 | immunoglobulin heavy variable 3-79 (pseudog |
| IGHV4-80 | 14 | immunoglobulin heavy variable 4-80 (pseudog |
| IGHV7-81 | 14 | immunoglobulin heavy variable 7-81 (non-fun |
| IGHVIII-82 | 14 | immunoglobulin heavy variable (III)-82 (pseud |
| AC138701.1 | 15 | ankyrin repeat domain 30B (ANKRD30B) pseu |
| RNU6-978P | 15 | RNA, U6 small nuclear 978, pseudogene [Sour |
| AC138701.2 | 15 | connector enhancer of kinase suppressor of R |
| AC127381.2 | 15 | Ig heavy chain V-II pseudogene |
| IGHV1OR15-9 | 15 | immunoglobulin heavy variable 1/OR15-9 (no |
| SLC20A1P3 | 15 | solute carrier family 20 member 1 pseudogen |
| IGHV1OR15-2 | 15 | immunoglobulin heavy variable 1/OR15-2 (pse |
| IGHV3OR15-7 | 15 | immunoglobulin heavy variable 3/OR15-7 (pse |
| IGHD5OR15-5A | 15 | immunoglobulin heavy diversity 5/OR15-5A (n |
| IGHD4OR15-4A | 15 | immunoglobulin heavy diversity 4/OR15-4A (n |
| IGHD3OR15-3A | 15 | immunoglobulin heavy diversity 3/OR15-3A (n |
| IGHD2OR15-2A | 15 | immunoglobulin heavy diversity 2/OR15-2A (n |
| IGHD1OR15-1A | 15 | immunoglobulin heavy diversity 1/OR15-1A (n |
| FAM30B | 15 | family with sequence similarity 30 member B |
| AC127381.1 | 15 | ADAM metallopeptidase domain 22 (ADAM22 |
| BCAR1P1 | 15 | BCAR1 pseudogene 1 [Source:HGNC Symbol;A |
| RN7SL584P | 15 | RNA, 7SL, cytoplasmic 584, pseudogene [Sour |
| AC087386.1 | 15 | novel transcript |
| AC087386.2 | 15 | novel transcript |
| BMS1P15 | 15 | BMS1 pseudogene 15 [Source:HGNC Symbol;A |
| AC126603.1 | 15 | rhophilin Rho GTPase binding protein 2 pseud |
| CHEK2P2 | 15 | checkpoint kinase 2 pseudogene 2 [Source:HG |
| AC026495.2 | 15 | novel transcript |
| AC026495.1 | 15 | novel transcript |
| HERC2P3 | 15 | hect domain and RLD 2 pseudogene 3 [Source |
| AC023310.1 | 15 | hect domain and RLD 2 (HERC2) pseudogene |
| GOLGA6L6 | 15 | golgin A6 family like 6 [Source:HGNC Symbol;A |
| AC023310.3 | 15 | M-phase phosphoprotein 10 (U3 small nucleo |
| GOLGA8CP | 15 | golgin A8 family member C, pseudogene [Sour |
| RN7SL759P | 15 | RNA, 7SL, cytoplasmic 759, pseudogene [Sour |
| SPATA31E2P | 15 | SPATA31 subfamily E member 2, pseudogene |
| AC023310.2 | 15 | ring finger protein 145 (RNF145) pseudogene |
| IGHV1OR15-6 | 15 | immunoglobulin heavy variable 1/OR15-6 (pse |
| AC023310.4 | 15 | novel transcript |
| NBEAP1 | 15 | neurobeachin pseudogene 1 [Source:HGNC Sy |
| AC131280.1 | 15 | novel transcript |

| | | |
|---|---|---|
| AC012414.5 | 15 | novel transcript |
| RNU6-498P | 15 | RNA, U6 small nuclear 498, pseudogene [Sour |
| AC012414.3 | 15 | novel transcript |
| AC012414.4 | 15 | novel transcript |
| AC012414.1 | 15 | LON peptidase N-terminal domain and ring fir |
| AC012414.7 | 15 | coxsackie virus and adenovirus receptor (CXAI |
| GRAMD4P5 | 15 | GRAM domain containing 4 pseudogene 5 [So |
| AC012414.2 | 15 | ankyrin repeat domain 18B (ANKRD18B) pseu |
| MIR3118-2 | 15 | microRNA 3118-2 [Source:HGNC Symbol;Acc:I |
| POTEB2 | 15 | POTE ankyrin domain family member B2 [Sou |
| RNU6-749P | 15 | RNA, U6 small nuclear 749, pseudogene [Sour |
| AC012414.6 | 15 | vomeronasal type-1 receptor 4 (VN1R4) pseu |
| AC037471.2 | 15 | vomeronasal 1 receptor (VN1R) pseudogene |
| ZNF519P2 | 15 | zinc finger protein 519 pseudogene 2 [Source: |
| NF1P1 | 15 | neurofibromin 1 pseudogene 1 [Source:HGNC |
| MIR5701-1 | 15 | microRNA 5701-1 [Source:HGNC Symbol;Acc:I |
| LINC01193 | 15 | long intergenic non-protein coding RNA 1193 |
| OR11J2P | 15 | olfactory receptor family 11 subfamily J meml |
| OR11J5P | 15 | olfactory receptor family 11 subfamily J meml |
| IGHD5OR15-5B | 15 | immunoglobulin heavy diversity 5/OR15-5B (r |
| IGHD4OR15-4B | 15 | immunoglobulin heavy diversity 4/OR15-4B (r |
| IGHD3OR15-3B | 15 | immunoglobulin heavy diversity 3/OR15-3B (r |
| IGHD2OR15-2B | 15 | immunoglobulin heavy diversity 2/OR15-2B (r |
| IGHD1OR15-1B | 15 | immunoglobulin heavy diversity 1/OR15-1B (r |
| FAM30C | 15 | family with sequence similarity 30 member C |
| AC037471.1 | 15 | ADAM metallopeptidase domain 11 (ADAM11 |
| BCAR1P2 | 15 | BCAR1 pseudogene 2 [Source:HGNC Symbol;A |
| RN7SL400P | 15 | RNA, 7SL, cytoplasmic 400, pseudogene [Sour |
| AC126335.1 | 15 | novel transcript |
| AC126335.2 | 15 | TEC |
| BMS1P16 | 15 | BMS1 pseudogene 16 [Source:HGNC Symbol;A |
| AC068446.3 | 15 | neurobeachin (NBEA) pseudogene |
| AC068446.1 | 15 | p21 protein (Cdc42/Rac)-activated kinase 2 (P |
| AC068446.2 | 15 | novel transcript |
| RNU6-1235P | 15 | RNA, U6 small nuclear 1235, pseudogene [Sou |
| AC060814.5 | 15 | novel transcript |
| AC060814.4 | 15 | novel transcript |
| AC060814.1 | 15 | LON peptidase N-terminal domain and ring fir |
| AC060814.3 | 15 | novel transcript |
| CXADRP2 | 15 | CXADR pseudogene 2 [Source:HGNC Symbol;A |
| AC060814.2 | 15 | GRAM domain containing 4 (GRAMD4) pseud |
| GRAMD4P6 | 15 | GRAM domain containing 4 pseudogene 6 [So |
| MIR3118-3 | 15 | microRNA 3118-3 [Source:HGNC Symbol;Acc:I |
| POTEB3 | 15 | POTE ankyrin domain family member B3 [Sou |
| U6 | 15 | U6 spliceosomal RNA [Source:RFAM;Acc:RF00 |
| NF1P9 | 15 | neurofibromin 1 pseudogene 9 [Source:HGNC |
| MIR5701-2 | 15 | microRNA 5701-2 [Source:HGNC Symbol;Acc:I |
| AC183088.3 | 15 | novel transcript |
| AC183088.2 | 15 | olfactory receptor pseudogene |
| AC183088.1 | 15 | olfactory receptor family 11 subfamily J meml |

| | | |
|---|---|---|
| LINC02203 | 15 | long intergenic non-protein coding RNA 2203 |
| LINC02203 | 15 | long intergenic non-protein coding RNA 2203 |
| AC135068.5 | 15 | olfactory receceptor pseudogene |
| OR11K1P | 15 | olfactory receptor family 11 subfamily K mem |
| AC135068.11 | 15 | olfactory receptor pseudogene |
| AC135068.10 | 15 | olfactory receptor pseudogene |
| AC135068.1 | 15 | novel olfactory receptor protein similar to OR |
| AC135068.3 | 15 | novel olfactory receptor protein similar to OR |
| AC135068.4 | 15 | olfactory receptor pseudogene |
| AC135068.7 | 15 | novel transcript |
| AC135068.6 | 15 | ribosomal protein S8 (RPS8) pseudogene |
| AC135068.8 | 15 | novel gene identicle to IGHV1OR15-1 |
| AC135068.9 | 15 | novel gene identical to IGHV1OR15-2 |
| AC135068.2 | 15 | novel protein, identical to IGHV4-4 |
| AC134981.1 | 15 | novel gene identical to IGHV1OR15-4 |
| MIR3118-4 | 15 | microRNA 3118-4 [Source:HGNC Symbol;Acc: |
| POTEB | 15 | POTE ankyrin domain family member B [Sourc |
| RNU6-631P | 15 | RNA, U6 small nuclear 631, pseudogene [Sour |
| ZNF519P3 | 15 | zinc finger protein 519 pseudogene 3 [Source: |
| NF1P2 | 15 | neurofibromin 1 pseudogene 2 [Source:HGNC |
| MIR5701-3 | 15 | microRNA 5701-3 [Source:HGNC Symbol;Acc: |
| OR11J7P | 15 | olfactory receptor family 11 subfamily J meml |
| OR11J6P | 15 | olfactory receptor family 11 subfamily J meml |
| AC134980.2 | 15 | novel transcript |
| OR11H3P | 15 | olfactory receptor family 11 subfamily H mem |
| AC134980.3 | 15 | novel protein |
| AC134980.1 | 15 | olfactory receptor family 11 subfamily K mem |
| OR4Q1P | 15 | olfactory receptor family 4 subfamily Q memb |
| OR4H6P | 15 | olfactory receptor family 4 subfamily H memb |
| OR4M2 | 15 | olfactory receptor family 4 subfamily M meml |
| OR4N4 | 15 | olfactory receptor family 4 subfamily N memb |
| OR4N3P | 15 | olfactory receptor family 4 subfamily N memb |
| AC010760.1 | 15 | novel transcript |
| RPS8P10 | 15 | ribosomal protein S8 pseudogene 10 [Source: |
| IGHV1OR15-1 | 15 | immunoglobulin heavy variable 1/OR15-1 (no |
| IGHV1OR15-3 | 15 | immunoglobulin heavy variable 1/OR15-3 (ps |
| IGHV4OR15-8 | 15 | immunoglobulin heavy variable 4/OR15-8 (no |
| IGHV1OR15-4 | 15 | immunoglobulin heavy variable 1/OR15-4 (ps |
| AC025884.3 | 15 | novel immunoglobulin heavy variable 1/OR15 |
| MIR1268A | 15 | microRNA 1268a [Source:HGNC Symbol;Acc:H |
| AC025884.2 | 15 | zinc finger, CCHC domain containing 2 (ZCCHC |
| AC025884.1 | 15 | arginine-glutamic acid dipeptide (RE) repeats |
| AC100757.2 | 15 | Ig heavy chain V-I pseudogene |
| AC100757.1 | 15 | ring finger protein 145 (Rnf145) pseudogene |
| SPATA31E3P | 15 | SPATA31 subfamily E member 3, pseudogene |
| GOLGA8DP | 15 | golgin A8 family member D, pseudogene [Sou |
| RN7SL545P | 15 | RNA, 7SL, cytoplasmic 545, pseudogene [Sour |
| AC100757.3 | 15 | M-phase phosphoprotein 10 (U3 small nucleo |
| GOLGA6L22 | 15 | golgin A6 family like 22 [Source:HGNC Symbol |
| HERC2P7 | 15 | hect domain and RLD 2 pseudogene 7 [Source |

| | | |
|---|---|---|
| HERC2P2 | 15 | hect domain and RLD 2 pseudogene 2 [Source |
| GOLGA8IP | 15 | golgin A8 family member I, pseudogene [Sour |
| RN7SL495P | 15 | RNA, 7SL, cytoplasmic 495, pseudogene [Sour |
| WHAMMP3 | 15 | WAS protein homolog associated with actin, g |
| AC138649.2 | 15 | zinc finger, FYVE domain containing 9 (ZFYVE9 |
| AC138649.5 | 15 | taspase 1 (TASP1) pseudogene |
| AC138649.4 | 15 | dexamethasone-induced transcript (DEXI) pse |
| PDCD6IPP1 | 15 | PDCD6IP pseudogene 1 [Source:HGNC Symbo |
| AC138649.3 | 15 | C-x(9)-C motif containing 2 (CMC2) pseudoger |
| AC138649.1 | 15 | novel transcript, sense overlapping NIPA1 |
| AC011767.1 | 15 | Golgin subfamily A member 6 pseudogene |
| NIPA1 | 15 | NIPA magnesium transporter 1 [Source:HGNC |
| NIPA2 | 15 | NIPA magnesium transporter 2 [Source:HGNC |
| CYFIP1 | 15 | cytoplasmic FMR1 interacting protein 1 [Sourc |
| TUBGCP5 | 15 | tubulin gamma complex associated protein 5 |
| ELMO2P1 | 15 | engulfment and cell motility 2 pseudogene 1 [ |
| GOLGA6L1 | 15 | golgin A6 family like 1 [Source:HGNC Symbol;A |
| AC116165.1 | 15 | M-phase phosphoprotein 10 (U3 small nucleo |
| GOLGA8EP | 15 | golgin A8 family member E, pseudogene [Sour |
| RN7SL106P | 15 | RNA, 7SL, cytoplasmic 106, pseudogene [Sour |
| ABCB10P1 | 15 | ABCB10 pseudogene 1 [Source:HGNC Symbol; |
| AC116165.2 | 15 | family with sequence similarity 208, member |
| MIR4509-1 | 15 | microRNA 4509-1 [Source:HGNC Symbol;Acc:H |
| AC116165.3 | 15 | abhydrolase domain containing 17A (ABHD17. |
| AC242376.1 | 15 | hect domain and RLD 2 pseudogene 7 |
| AC242376.2 | 15 | hect domain and RLD 2 (HERC2) pseudogene |
| AC100756.3 | 15 | golgin subfamily A member 6-like protein 6 [S |
| AC100756.1 | 15 | M-phase phosphoprotein 10 (U3 small nucleo |
| GOLGA8S | 15 | golgin A8 family member S [Source:HGNC Sym |
| RN7SL536P | 15 | RNA, 7SL, cytoplasmic 536, pseudogene [Sour |
| AC100756.4 | 15 | novel transcript |
| AC100756.2 | 15 | hect domain and RLD 2 (HERC2) pseudogene |
| GOLGA6L2 | 15 | golgin A6 family like 2 [Source:HGNC Symbol;A |
| MIR4508 | 15 | microRNA 4508 [Source:HGNC Symbol;Acc:HG |
| MKRN3 | 15 | makorin ring finger protein 3 [Source:HGNC Sy |
| AC126407.1 | 15 | MKRN3 antisense RNA 1 |
| MAGEL2 | 15 | MAGE family member L2 [Source:HGNC Symb |
| NDN | 15 | necdin, MAGE family member [Source:HGNC S |
| AC087490.1 | 15 | novel transcript |
| RNU6-741P | 15 | RNA, U6 small nuclear 741, pseudogene [Sour |
| PWRN4 | 15 | Prader-Willi region non-protein coding RNA 4 |
| PWRN1 | 15 | Prader-Willi region non-protein coding RNA 1 |
| PWRN2 | 15 | Prader-Willi region non-protein coding RNA 2 |
| AC087463.1 | 15 | novel transcript |
| AC087463.2 | 15 | novel transcript |
| AC087463.3 | 15 | novel transcript |
| AC090983.2 | 15 | novel transcript |
| NPAP1 | 15 | nuclear pore associated protein 1 [Source:HGI |
| SNRPN | 15 | small nuclear ribonucleoprotein polypeptide N |
| AC090983.1 | 15 | novel transcript |

| | | |
|---|---|---|
| RPL5P1 | 15 | ribosomal protein L5 pseudogene 1 [Source:H |
| AC090602.1 | 15 | ribosomal protein S27 (RPS27) pseudogene |
| SNURF | 15 | SNRPN upstream reading frame [Source:HGN( |
| AC124312.1 | 15 | novel protein |
| SNHG14 | 15 | small nucleolar RNA host gene 14 [Source:HGI |
| SNORD107 | 15 | small nucleolar RNA, C/D box 107 [Source:HGI |
| PWAR5 | 15 | Prader Willi/Angelman region RNA 5 [Source:I |
| SNORD64 | 15 | small nucleolar RNA, C/D box 64 [Source:HGN |
| SNORD108 | 15 | small nucleolar RNA, C/D box 108 [Source:HGI |
| AC124312.2 | 15 | novel transcript |
| SNORD109A | 15 | small nucleolar RNA, C/D box 109A [Source:H( |
| SNORD116-1 | 15 | small nucleolar RNA, C/D box 116-1 [Source:H |
| SNORD116-2 | 15 | small nucleolar RNA, C/D box 116-2 [Source:H |
| SNORD116-3 | 15 | small nucleolar RNA, C/D box 116-3 [Source:H |
| SNORD116-4 | 15 | small nucleolar RNA, C/D box 116-4 [Source:H |
| SNORD116-5 | 15 | small nucleolar RNA, C/D box 116-5 [Source:H |
| SNORD116-6 | 15 | small nucleolar RNA, C/D box 116-6 [Source:H |
| SNORD116-7 | 15 | small nucleolar RNA, C/D box 116-7 [Source:H |
| SNORD116-8 | 15 | small nucleolar RNA, C/D box 116-8 [Source:H |
| SNORD116-9 | 15 | small nucleolar RNA, C/D box 116-9 [Source:H |
| SNORD116-10 | 15 | small nucleolar RNA, C/D box 116-10 [Source: |
| SNORD116-11 | 15 | small nucleolar RNA, C/D box 116-11 [Source: |
| SNORD116-12 | 15 | small nucleolar RNA, C/D box 116-12 [Source: |
| SNORD116-13 | 15 | small nucleolar RNA, C/D box 116-13 [Source: |
| SNORD116-14 | 15 | small nucleolar RNA, C/D box 116-14 [Source: |
| SNORD116-15 | 15 | small nucleolar RNA, C/D box 116-15 [Source: |
| SNORD116-16 | 15 | small nucleolar RNA, C/D box 116-16 [Source: |
| SNORD116-17 | 15 | small nucleolar RNA, C/D box 116-17 [Source: |
| SNORD116-18 | 15 | small nucleolar RNA, C/D box 116-18 [Source: |
| SNORD116-19 | 15 | small nucleolar RNA, C/D box 116-19 [Source: |
| AC124312.3 | 15 | novel transcript, SNORD host |
| SNORD116-20 | 15 | small nucleolar RNA, C/D box 116-20 [Source: |
| SNORD116-21 | 15 | small nucleolar RNA, C/D box 116-21 [Source: |
| SNORD116-22 | 15 | small nucleolar RNA, C/D box 116-22 [Source: |
| SNORD116-23 | 15 | small nucleolar RNA, C/D box 116-23 [Source: |
| AC124312.5 | 15 | novel transcript |
| AC124312.4 | 15 | TEC |
| SNORD116-24 | 15 | small nucleolar RNA, C/D box 116-24 [Source: |
| SNORD116-25 | 15 | small nucleolar RNA, C/D box 116-25 [Source: |
| SNORD116-26 | 15 | small nucleolar RNA, C/D box 116-26 [Source: |
| SNORD116-27 | 15 | small nucleolar RNA, C/D box 116-27 [Source: |
| SNORD116-28 | 15 | small nucleolar RNA, C/D box 116-28 [Source: |
| SNORD116-29 | 15 | small nucleolar RNA, C/D box 116-29 [Source: |
| SNORD116-30 | 15 | small nucleolar RNA, C/D box 116-30 [Source: |
| PWAR1 | 15 | Prader Willi/Angelman region RNA 1 [Source:I |
| SNORD115 | 15 | Small nucleolar RNA SNORD115 [Source:RFAN |
| TMEM261P1 | 15 | transmembrane protein 261 pseudogene 1 [S( |
| SNORD115-1 | 15 | small nucleolar RNA, C/D box 115-1 [Source:H |
| SNORD115-2 | 15 | small nucleolar RNA, C/D box 115-2 [Source:H |
| SNORD115-3 | 15 | small nucleolar RNA, C/D box 115-3 [Source:H |

| | | |
|---|---|---|
| SNORD115-4 | 15 | small nucleolar RNA, C/D box 115-4 [Source:H |
| SNORD115-5 | 15 | small nucleolar RNA, C/D box 115-5 [Source:H |
| SNORD115-6 | 15 | small nucleolar RNA, C/D box 115-6 [Source:H |
| SNORD115-7 | 15 | small nucleolar RNA, C/D box 115-7 [Source:H |
| SNORD115-8 | 15 | small nucleolar RNA, C/D box 115-8 [Source:H |
| SNORD115-9 | 15 | small nucleolar RNA, C/D box 115-9 [Source:H |
| SNORD115-10 | 15 | small nucleolar RNA, C/D box 115-10 [Source: |
| SNORD115-11 | 15 | small nucleolar RNA, C/D box 115-11 [Source: |
| SNORD115-12 | 15 | small nucleolar RNA, C/D box 115-12 [Source: |
| SNORD115-13 | 15 | small nucleolar RNA, C/D box 115-13 [Source: |
| SNORD115-14 | 15 | small nucleolar RNA, C/D box 115-14 [Source: |
| SNORD115-15 | 15 | small nucleolar RNA, C/D box 115-15 [Source: |
| SNORD115-16 | 15 | small nucleolar RNA, C/D box 115-16 [Source: |
| SNORD115-17 | 15 | small nucleolar RNA, C/D box 115-17 [Source: |
| SNORD115-18 | 15 | small nucleolar RNA, C/D box 115-18 [Source: |
| SNORD115-19 | 15 | small nucleolar RNA, C/D box 115-19 [Source: |
| SNORD115-20 | 15 | small nucleolar RNA, C/D box 115-20 [Source: |
| SNORD115-21 | 15 | small nucleolar RNA, C/D box 115-21 [Source: |
| SNORD115-22 | 15 | small nucleolar RNA, C/D box 115-22 [Source: |
| SNORD115-23 | 15 | small nucleolar RNA, C/D box 115-23 [Source: |
| SNORD115-24 | 15 | small nucleolar RNA, C/D box 115-24 [Source: |
| SNORD115-25 | 15 | small nucleolar RNA, C/D box 115-25 [Source: |
| SNORD115-26 | 15 | small nucleolar RNA, C/D box 115-26 [Source: |
| SNORD115-27 | 15 | small nucleolar RNA, C/D box 115-27 [Source: |
| SNORD115-28 | 15 | small nucleolar RNA, C/D box 115-28 [Source: |
| SNORD115-29 | 15 | small nucleolar RNA, C/D box 115-29 [Source: |
| SNORD115-30 | 15 | small nucleolar RNA, C/D box 115-30 [Source: |
| SNORD115-31 | 15 | small nucleolar RNA, C/D box 115-31 [Source: |
| SNORD115-32 | 15 | small nucleolar RNA, C/D box 115-32 [Source: |
| SNORD115-33 | 15 | small nucleolar RNA, C/D box 115-33 [Source: |
| SNORD115-34 | 15 | small nucleolar RNA, C/D box 115-34 [Source: |
| SNORD115-35 | 15 | small nucleolar RNA, C/D box 115-35 [Source: |
| SNORD115-36 | 15 | small nucleolar RNA, C/D box 115-36 [Source: |
| SNORD115-37 | 15 | small nucleolar RNA, C/D box 115-37 [Source: |
| SNORD115-38 | 15 | small nucleolar RNA, C/D box 115-38 [Source: |
| SNORD115-39 | 15 | small nucleolar RNA, C/D box 115-39 [Source: |
| SNORD115-40 | 15 | small nucleolar RNA, C/D box 115-40 [Source: |
| SNORD115-41 | 15 | small nucleolar RNA, C/D box 115-41 [Source: |
| SNORD115-42 | 15 | small nucleolar RNA, C/D box 115-42 [Source: |
| SNORD115-43 | 15 | small nucleolar RNA, C/D box 115-43 [Source: |
| SNORD115-44 | 15 | small nucleolar RNA, C/D box 115-44 [Source: |
| AC124303.1 | 15 | TEC |
| SNORD115-45 | 15 | small nucleolar RNA, C/D box 115-45 [Source: |
| SNORD115-46 | 15 | small nucleolar RNA, C/D box 115-46 [Source: |
| SNORD115-47 | 15 | small nucleolar RNA, C/D box 115-47 [Source: |
| SNORD115-48 | 15 | small nucleolar RNA, C/D box 115-48 [Source: |
| SNORD109B | 15 | small nucleolar RNA, C/D box 109B [Source:H( |
| UBE3A | 15 | ubiquitin protein ligase E3A [Source:HGNC Syr |
| AC100774.1 | 15 | novel transcript, sense intronic to UBE3A |
| LINC02250 | 15 | long intergenic non-protein coding RNA 2250 |

| | | |
|---|---|---|
| AC023449.1 | 15 | ribosomal protein L39 (RPL39) pseudogene |
| ATP10A | 15 | ATPase phospholipid transporting 10A (putati |
| AC023449.2 | 15 | novel transcript, sense intronic to ATP10A |
| RNA5SP390 | 15 | RNA, 5S ribosomal pseudogene 390 [Source:H |
| MIR4715 | 15 | microRNA 4715 [Source:HGNC Symbol;Acc:H |
| AC016266.1 | 15 | novel transcript |
| LINC02346 | 15 | long intergenic non-protein coding RNA 2346 |
| AC044913.2 | 15 | mitochondrial ribosomal protein S18C (MRPS1 |
| AC044913.1 | 15 | NULL |
| AC100836.1 | 15 | novel transcript |
| LINC00929 | 15 | long intergenic non-protein coding RNA 929 [ |
| AC012060.1 | 15 | novel transcript |
| LINC02248 | 15 | long intergenic non-protein coding RNA 2248 |
| AC009878.2 | 15 | novel transcript |
| GABRB3 | 15 | gamma-aminobutyric acid type A receptor be |
| AC009878.1 | 15 | novel transcript, sense intronic to GABRB3 |
| AC011196.1 | 15 | novel transcript |
| AC135999.1 | 15 | ribosomal protein L9 (RPL9) pseudogene |
| GABRA5 | 15 | gamma-aminobutyric acid type A receptor alp |
| TVP23BP1 | 15 | TVP23B pseudogene 1 [Source:HGNC Symbol; |
| GABRG3 | 15 | gamma-aminobutyric acid type A receptor ga |
| GABRG3-AS1 | 15 | GABRG3 antisense RNA 1 [Source:HGNC Symb |
| AC136896.1 | 15 | dual specificity phosphatase 3 (DUSP3) pseud |
| RNA5SP391 | 15 | RNA, 5S ribosomal pseudogene 391 [Source:H |
| SERPINE4P | 15 | serpin family E member 4, pseudogene [Sourc |
| AC144833.1 | 15 | novel transcript, sense intronic to GABRG3 |
| AC104002.1 | 15 | novel transcript, antisense to GABRG3 |
| AC104002.2 | 15 | novel transcript, intronic to GABRG3 |
| AC104002.3 | 15 | novel transcript, antisense to GABRG3 |
| AC021979.1 | 15 | novel transcript |
| AC021979.3 | 15 | novel transcript |
| AC021979.2 | 15 | novel transcript |
| OCA2 | 15 | OCA2 melanosomal transmembrane protein [ |
| AC090696.1 | 15 | novel transcript, antisense to OCA2 |
| RPL5P32 | 15 | ribosomal protein L5 pseudogene 32 [Source: |
| HERC2 | 15 | HECT and RLD domain containing E3 ubiquitin |
| RPL41P2 | 15 | ribosomal protein L41 pseudogene 2 [Source: |
| AC091304.3 | 15 | hect domain and RLD 2 (HERC2) pseudogene |
| AC091304.5 | 15 | cholinergic receptor, nicotinic, beta 4 (CHRNB |
| AC091304.1 | 15 | golgin A6 family-like [Source:NCBI gene;Acc:6 |
| AC091304.4 | 15 | M-phase phosphoprotein 10 (U3 small nucleo |
| GOLGA8F | 15 | golgin A8 family member F [Source:HGNC Sym |
| RN7SL238P | 15 | RNA, 7SL, cytoplasmic 238, pseudogene [Sour |
| ABCB10P3 | 15 | ABCB10 pseudogene 3 [Source:HGNC Symbol; |
| AC091304.9 | 15 | family with sequence similarity 208, member |
| AC091304.6 | 15 | family with sequence similarity 208, member |
| MIR4509-2 | 15 | microRNA 4509-2 [Source:HGNC Symbol;Acc: |
| AC091304.10 | 15 | glypican 5 (GPC5) pseudogene |
| AC091304.2 | 15 | family with sequence similarity 108, member |
| AC138749.4 | 15 | glypican 5 (GPC5) pseudogene |

| | | |
|---|---|---|
| MIR4509-3 | 15 | microRNA 4509-3 [Source:HGNC Symbol;Acc:I |
| AC138749.5 | 15 | family with sequence similarity 208, member |
| ABCB10P4 | 15 | ABCB10 pseudogene 4 [Source:HGNC Symbol; |
| GOLGA8G | 15 | golgin A8 family member G [Source:HGNC Syr |
| Metazoa_SRP | 15 | Metazoan signal recognition particle RNA [Sou |
| AC138749.3 | 15 | Golgin subfamily A member 2 pseudogene |
| AC138749.1 | 15 | golgin subfamily A member 6-like protein 22 [ |
| AC138749.2 | 15 | cholinergic receptor, nicotinic, beta 4 (CHRNB |
| HERC2P11 | 15 | hect domain and RLD 2 pseudogene 11 [Sourc |
| HERC2P9 | 15 | hect domain and RLD 2 pseudogene 9 [Source |
| AC055876.2 | 15 | hect domain and RLD 2 (HERC2) pseudogene |
| GOLGA8M | 15 | golgin A8 family member M [Source:HGNC Syı |
| RN7SL719P | 15 | RNA, 7SL, cytoplasmic 719, pseudogene [Sour |
| GOLGA8M | 15 | golgin A8 family member M [Source:HGNC Syı |
| WHAMMP2 | 15 | WAS protein homolog associated with actin, g |
| AC055876.4 | 15 | zinc finger, FYVE domain containing 9 (ZFYVE9 |
| AC055876.1 | 15 | taspase, threonine aspartase, 1 (TASP1) pseuc |
| AC055876.3 | 15 | dexamethasone-induced transcript (DEXI) pse |
| PDCD6IPP2 | 15 | PDCD6IP pseudogene 2 [Source:HGNC Symbo |
| AC055876.5 | 15 | COX assembly mitochondrial protein 2 homol |
| GOLGA6L7 | 15 | golgin A6 family like 7 [Source:HGNC Symbol;/ |
| AC174071.1 | 15 | ferritin, light polypeptide (FTL) pseudogene |
| AC174469.1 | 15 | novel transcript, antisense to APBA2 |
| APBA2 | 15 | amyloid beta precursor protein binding family |
| AC127522.1 | 15 | novel transcript |
| FAM189A1 | 15 | family with sequence similarity 189 member / |
| AC061965.2 | 15 | DDB1 and CUL4 associated factor 6 (DCAF6) p |
| AC061965.1 | 15 | NULL |
| NSMCE3 | 15 | NSE3 homolog, SMC5-SMC6 complex compon |
| AC107980.1 | 15 | novel transcript |
| AC102941.1 | 15 | novel transcript |
| TUBBP8 | 15 | tubulin beta class I pseudogene 8 [Source:HGI |
| AC102941.2 | 15 | novel transcript |
| AC022613.1 | 15 | novel transcript, antisense to TJP1 |
| TJP1 | 15 | tight junction protein 1 [Source:HGNC Symbol |
| AC022613.3 | 15 | novel transcript |
| HMGN2P5 | 15 | high mobility group nucleosomal binding dom |
| AC022613.2 | 15 | novel transcript, antisense to TJP1 |
| NCAPGP2 | 15 | non-SMC condensin I complex subunit G pseu |
| AC111152.2 | 15 | novel transcript |
| AC111152.3 | 15 | synaptogyrin 2 (SYNGR2) pseudogene |
| AC111152.1 | 15 | tight junction protein 1 (zona occludens 1) (TJ |
| GOLGA8J | 15 | golgin A8 family member J [Source:HGNC Sym |
| RN7SL673P | 15 | RNA, 7SL, cytoplasmic 673, pseudogene [Sour |
| DNM1P28 | 15 | dynamin 1 pseudogene 28 [Source:HGNC Sym |
| ULK4P3 | 15 | ULK4 pseudogene 3 [Source:HGNC Symbol;Ac |
| U8 | 15 | U8 small nucleolar RNA [Source:RFAM;Acc:RF |
| GOLGA8T | 15 | golgin A8 family member T [Source:HGNC Sym |
| RN7SL469P | 15 | RNA, 7SL, cytoplasmic 469, pseudogene [Sour |
| DNM1P30 | 15 | dynamin 1 pseudogene 30 [Source:HGNC Sym |

| | | |
|---|---|---|
| AC120045.2 | 15 | synaptotagmin II (SYT2) pseudogene |
| AC120045.1 | 15 | novel transcript |
| LINC02249 | 15 | long intergenic non-protein coding RNA 2249 |
| AC135731.1 | 15 | hyperpolarization activated cyclic nucleotide-g |
| RNU6-17P | 15 | RNA, U6 small nuclear 17, pseudogene [Sourc |
| AC135731.2 | 15 | NEDD4 binding protein 2 (N4BP2) pseudogene |
| AC019322.4 | 15 | novel transcript |
| Y_RNA | 15 | Y RNA [Source:RFAM;Acc:RF00019] |
| CHRFAM7A | 15 | CHRNA7 (exons 5-10) and FAM7A (exons A-E) |
| AC019322.1 | 15 | unc-51-like kinase 4 (C. elegans) (ULK4) pseud |
| U8 | 15 | U8 small nucleolar RNA [Source:RFAM;Acc:RF |
| AC019322.3 | 15 | dynamin 1 (DNM1) pseudogene |
| GOLGA8R | 15 | golgin A8 family member R [Source:HGNC Sym |
| RN7SL196P | 15 | RNA, 7SL, cytoplasmic 196, pseudogene [Sour |
| AC019322.2 | 15 | novel pseudogene |
| AC127502.1 | 15 | WAS protein homolog associated with actin, g |
| AC127502.2 | 15 | novel transcript |
| AC127502.3 | 15 | M-phase phosphoprotein 10 (MPHOSPH10) ps |
| AC026150.2 | 15 | golgin A6 family-like pseudogene |
| AC026150.1 | 15 | novel transcript |
| GOLGA8Q | 15 | golgin A8 family member Q [Source:HGNC Syr |
| RN7SL796P | 15 | RNA, 7SL, cytoplasmic 796, pseudogene [Sour |
| DNM1P50 | 15 | dynamin 1 pseudogene 50 [Source:HGNC Sym |
| ULK4P2 | 15 | ULK4 pseudogene 2 [Source:HGNC Symbol;Ac |
| U8 | 15 | U8 small nucleolar RNA [Source:RFAM;Acc:RF |
| GOLGA8H | 15 | golgin A8 family member H [Source:HGNC Syn |
| AC026150.3 | 15 | novel transcript, antisense to AC026150.11 |
| RN7SL628P | 15 | RNA, 7SL, cytoplasmic 628, pseudogene [Sour |
| AC091057.3 | 15 | novel transcript |
| AC091057.2 | 15 | novel transcript, antisense to ARHGAP11B |
| AC091057.6 | 15 | Rho GTPase activating protein 11B |
| ARHGAP11B | 15 | Rho GTPase activating protein 11B [Source:HG |
| AC091057.5 | 15 | Serine/threonine-protein kinase ULK4 (ULK4), |
| AC091057.1 | 15 | novel transcript |
| Y_RNA | 15 | Y RNA [Source:RFAM;Acc:RF00019] |
| AC091057.4 | 15 | OTU deubiquitinase 7A (OTUD7A), pseudogen |
| GOLGA8UP | 15 | golgin A8 family member U, pseudogene [Sou |
| RN7SL82P | 15 | RNA, 7SL, cytoplasmic 82, pseudogene [Sourc |
| HERC2P10 | 15 | hect domain and RLD 2 pseudogene 10 [Sourc |
| AC087481.1 | 15 | transmembrane protein 183A (TMEM183A) ps |
| FAN1 | 15 | FANCD2 and FANCI associated nuclease 1 [Sou |
| AC087481.2 | 15 | M-phase phosphoprotein 10 (U3 small nucleo |
| MTMR10 | 15 | myotubularin related protein 10 [Source:HGN |
| RNU6-466P | 15 | RNA, U6 small nuclear 466, pseudogene [Sour |
| AC009562.1 | 15 | novel transcript, antisense to TRPM1 |
| TRPM1 | 15 | transient receptor potential cation channel su |
| MIR211 | 15 | microRNA 211 [Source:HGNC Symbol;Acc:HGI |
| LINC02352 | 15 | long intergenic non-protein coding RNA 2352 |
| AC012236.1 | 15 | uncharacterized LOC283710 [Source:NCBI ger |
| KLF13 | 15 | Kruppel like factor 13 [Source:HGNC Symbol;A |

| | | |
|---|---|---|
| UBE2CP4 | 15 | ubiquitin conjugating enzyme E2 C pseudogen |
| AC104759.1 | 15 | REX4, RNA exonuclease 4 homolog (S. cerevisi |
| OTUD7A | 15 | OTU deubiquitinase 7A [Source:HGNC Symbol |
| AC026951.1 | 15 | DEP domain containing 1 (DEPDC1) pseudoge |
| AC026951.2 | 15 | zinc finger protein 442 (ZNF442) pseudogene |
| CHRNA7 | 15 | cholinergic receptor nicotinic alpha 7 subunit |
| AC079969.1 | 15 | NULL |
| Y_RNA | 15 | Y RNA [Source:RFAM;Acc:RF00019] |
| AC068448.1 | 15 | TEC |
| AC139426.3 | 15 | NEDD4 binding protein 2 (N4BP2) pseudogene |
| RNU6-18P | 15 | RNA, U6 small nuclear 18, pseudogene [Sourc |
| AC139426.1 | 15 | hyperpolarization activated cyclic nucleotide- |
| AC139426.2 | 15 | novel transcript |
| DNM1P31 | 15 | dynamin 1 pseudogene 31 [Source:HGNC Sym |
| GOLGA8K | 15 | golgin A8 family member K [Source:HGNC Syn |
| RN7SL185P | 15 | RNA, 7SL, cytoplasmic 185, pseudogene [Sour |
| AC135983.1 | 15 | novel transcript |
| ULK4P1 | 15 | ULK4 pseudogene 1 [Source:HGNC Symbol;Ac |
| U8 | 15 | U8 small nucleolar RNA [Source:RFAM;Acc:RF |
| DNM1P32 | 15 | dynamin 1 pseudogene 32 [Source:HGNC Sym |
| GOLGA8O | 15 | golgin A8 family member O [Source:HGNC Syr |
| RN7SL539P | 15 | RNA, 7SL, cytoplasmic 539, pseudogene [Sour |
| AC135983.5 | 15 | novel pseudogene |
| AC135983.4 | 15 | golgin subfamily A member 6-like pseudogene |
| AC135983.3 | 15 | M-phase phosphoprotein 10 (U3 small nucleo |
| AC135983.2 | 15 | WAS protein homolog associated with actin, g |
| LINC02256 | 15 | long intergenic non-protein coding RNA 2256 |
| AC123768.4 | 15 | novel transcript |
| AC123768.3 | 15 | novel transcript, antisense to GOLGA8N |
| GOLGA8N | 15 | golgin A8 family member N [Source:HGNC Syr |
| RN7SL286P | 15 | RNA, 7SL, cytoplasmic 286, pseudogene [Sour |
| AC123768.2 | 15 | novel transcript |
| ARHGAP11A | 15 | Rho GTPase activating protein 11A [Source:H |
| AC123768.5 | 15 | novel transcript, antisense to ARHGAP11A |
| SCG5 | 15 | secretogranin V [Source:HGNC Symbol;Acc:H |
| AC123768.1 | 15 | novel transcript |
| AC090877.2 | 15 | novel transcript |
| GREM1 | 15 | gremlin 1, DAN family BMP antagonist [Source |
| FMN1 | 15 | formin 1 [Source:HGNC Symbol;Acc:HGNC:37 |
| AC090877.1 | 15 | microtubule-associated protein 1 light chain 3 |
| AC018515.1 | 15 | NULL |
| HNRNPA1P71 | 15 | heterogeneous nuclear ribonucleoprotein A1 |
| AC019278.1 | 15 | TEC |
| TMCO5B | 15 | transmembrane and coiled-coil domains 5B (p |
| AC055874.1 | 15 | novel transcript, antisense to RYR3 |
| RYR3 | 15 | ryanodine receptor 3 [Source:HGNC Symbol;A |
| Y_RNA | 15 | Y RNA [Source:RFAM;Acc:RF00019] |
| AC010809.3 | 15 | novel transcript, antisense to RYR3 |
| AC010809.2 | 15 | novel transcript, antisense to RYR3 |
| AC010809.1 | 15 | novel transcript, antisense to RYR3 |

| | | |
|---|---|---|
| AVEN | 15 | apoptosis and caspase activation inhibitor [So |
| CHRM5 | 15 | cholinergic receptor muscarinic 5 [Source:HGI |
| AC009268.2 | 15 | novel transcript, sense intronic to AVEN |
| AC009268.3 | 15 | novel transcript, antisense to AVEN |
| AC009268.1 | 15 | Sjogren syndrome antigen B (autoantigen La) |
| EMC7 | 15 | ER membrane protein complex subunit 7 [Sou |
| AC079203.1 | 15 | ribosomal protein L32 (RPL32) pseudogene |
| PGBD4 | 15 | piggyBac transposable element derived 4 [Sou |
| KATNBL1 | 15 | katanin regulatory subunit B1 like 1 [Source:H |
| EMC4 | 15 | ER membrane protein complex subunit 4 [Sou |
| SLC12A6 | 15 | solute carrier family 12 member 6 [Source:HG |
| AC079203.2 | 15 | novel transcript, antisense to SLC12A6 |
| Y_RNA | 15 | Y RNA [Source:RFAM;Acc:RF00019] |
| NOP10 | 15 | NOP10 ribonucleoprotein [Source:HGNC Symb |
| NUTM1 | 15 | NUT midline carcinoma family member 1 [Sou |
| LPCAT4 | 15 | lysophosphatidylcholine acyltransferase 4 [So |
| ACTG1P15 | 15 | actin gamma 1 pseudogene 15 [Source:HGNC |
| GOLGA8A | 15 | golgin A8 family member A [Source:HGNC Syn |
| MIR1233-1 | 15 | microRNA 1233-1 [Source:HGNC Symbol;Acc:I |
| AC025678.1 | 15 | mitochondrially encoded cytochrome c oxidas |
| AC025678.2 | 15 | amyloid beta (A4) precursor protein-binding, f |
| AC025678.3 | 15 | TEC |
| HNRNPLP2 | 15 | heterogeneous nuclear ribonucleoprotein L ps |
| FSCN1P1 | 15 | fascin actin-bundling protein 1 pseudogene 1 |
| AC027139.2 | 15 | dynamin 1 (DNM1) pseudogene |
| GOLGA8B | 15 | golgin A8 family member B [Source:HGNC Syn |
| MIR1233-2 | 15 | microRNA 1233-2 [Source:HGNC Symbol;Acc:I |
| AC027139.1 | 15 | amyloid beta (A4) precursor protein-binding, f |
| AC100834.1 | 15 | NULL |
| LINC02252 | 15 | long intergenic non-protein coding RNA 2252 |
| AC100834.2 | 15 | novel transcript |
| GJD2 | 15 | gap junction protein delta 2 [Source:HGNC Syı |
| AC087457.1 | 15 | novel transcript, antisense to ACTC1 |
| ACTC1 | 15 | actin alpha cardiac muscle 1 [Source:HGNC Sy |
| AC018868.2 | 15 | tubulin, alpha 1b (TUBA1B) pseudogene |
| AQR | 15 | aquarius intron-binding spliceosomal factor [S |
| AC018868.1 | 15 | ribosomal protein L36a (RPL36A) pseudogene |
| ZNF770 | 15 | zinc finger protein 770 [Source:HGNC Symbol; |
| AC114546.3 | 15 | TEC |
| AC114546.1 | 15 | novel transcript |
| AC114546.4 | 15 | TEC |
| AC114546.2 | 15 | novel transcript |
| AC021231.3 | 15 | transmembrane phosphoinositide 3-phosphat |
| NANOGP8 | 15 | Nanog homeobox retrogene P8 [Source:HGN( |
| AC021231.2 | 15 | U2 small nuclear RNA auxiliary factor 1 (U2AF |
| AC021231.1 | 15 | novel transcript |
| PRELID1P4 | 15 | PRELI domain containing 1 pseudogene 4 [Sou |
| ANP32AP1 | 15 | acidic nuclear phosphoprotein 32 family mem |
| DPH6 | 15 | diphthamine biosynthesis 6 [Source:HGNC Syı |
| NUTF2P6 | 15 | nuclear transport factor 2 pseudogene 6 [Sou |

| | | |
|---|---|---|
| RBM17P4 | 15 | RNA binding motif protein 17 pseudogene 4 [S |
| MIR3942 | 15 | microRNA 3942 [Source:HGNC Symbol;Acc:HG |
| AC019288.1 | 15 | mitochondrially encoded NADH dehydrogenas |
| HNRNPA1P45 | 15 | heterogeneous nuclear ribonucleoprotein A1 |
| DPH6-DT | 15 | DPH6 divergent transcript [Source:HGNC Sym |
| AC068867.1 | 15 | castor zinc finger 1 (CASZ1) pseudogene |
| LINC02853 | 15 | long intergenic non-protein coding RNA 2853 |
| MIR4510 | 15 | microRNA 4510 [Source:HGNC Symbol;Acc:HG |
| AC021351.1 | 15 | novel transcript |
| AC087516.1 | 15 | novel transcript |
| AC087516.2 | 15 | novel transcript |
| COX6CP4 | 15 | cytochrome c oxidase subunit 6C pseudogene |
| C15orf41 | 15 | chromosome 15 open reading frame 41 [Sour |
| AC103988.1 | 15 | tropomyosin 3 (TPM3) pseudogene |
| LARP4P | 15 | La ribonucleoprotein domain family member 4 |
| TPST2P1 | 15 | tyrosylprotein sulfotransferase 2 pseudogene |
| AC013640.1 | 15 | novel transcript, antisense to C15orf41 |
| CSNK1A1P1 | 15 | casein kinase 1 alpha 1 pseudogene 1 [Source |
| U3 | 15 | Small nucleolar RNA U3 [Source:RFAM;Acc:RF |
| AC018563.1 | 15 | novel transcript |
| MEIS2 | 15 | Meis homeobox 2 [Source:HGNC Symbol;Acc: |
| MIR8063 | 15 | microRNA 8063 [Source:HGNC Symbol;Acc:HG |
| Y_RNA | 15 | Y RNA [Source:RFAM;Acc:RF00019] |
| AC078909.1 | 15 | novel transcript, antisense to MEIS2 |
| AC078909.2 | 15 | novel transcript |
| Metazoa_SRP | 15 | Metazoan signal recognition particle RNA [Sou |
| AC087283.1 | 15 | partner of NOB1 homolog (S. cerevisiae) (PNC |
| AC068875.1 | 15 | novel transcript |
| AC016304.1 | 15 | ribosomal protein S15 (RPS15) pseudogene |
| TMCO5A | 15 | transmembrane and coiled-coil domains 5A [S |
| LINC02345 | 15 | long intergenic non-protein coding RNA 2345 |
| LINC01852 | 15 | long intergenic non-protein coding RNA 1852 |
| AC087473.1 | 15 | novel transcript |
| SPRED1 | 15 | sprouty related EVH1 domain containing 1 [So |
| FAM98B | 15 | family with sequence similarity 98 member B |
| AC109631.1 | 15 | CTD (carboxy-terminal domain, RNA polymera |
| RASGRP1 | 15 | RAS guanyl releasing protein 1 [Source:HGNC |
| AC116158.3 | 15 | novel transcript, antisense to RASGRP1 |
| AC116158.1 | 15 | novel transcript, antisense to RASGRP1 |
| AC116158.2 | 15 | novel transcript, antisense to RASGRP1 |
| LINC02694 | 15 | long intergenic non-protein coding RNA 2694 |
| AC022929.1 | 15 | novel transcript |
| AC022929.2 | 15 | novel transcript |
| AC013652.1 | 15 | novel transcript |
| AC087878.1 | 15 | novel transcript |
| AC113146.1 | 15 | novel transcript |
| C15orf54 | 15 | chromosome 15 open reading frame 54 (puta |
| AC013652.2 | 15 | novel transcript |
| AC109630.1 | 15 | novel transcript |
| AC022196.1 | 15 | novel transcript |

| | | |
|---|---|---|
| THBS1 | 15 | thrombospondin 1 [Source:HGNC Symbol;Acc |
| AC037198.1 | 15 | novel transcript, sense intronic to THBS1 |
| AC037198.2 | 15 | novel transcript, antisense to THBS1 |
| FSIP1 | 15 | fibrous sheath interacting protein 1 [Source:H |
| AC023908.2 | 15 | novel transcript |
| AC023908.3 | 15 | novel transcript, antisense to FSIP1 |
| GPR176 | 15 | G protein-coupled receptor 176 [Source:HGN( |
| AC023908.1 | 15 | novel transcript, sense overlapping GPR176 |
| GPR176-DT | 15 | GPR176-DT [Source:HGNC Symbol;Acc:HGNC: |
| Y_RNA | 15 | Y RNA [Source:RFAM;Acc:RF00019] |
| EIF2AK4 | 15 | eukaryotic translation initiation factor 2 alpha |
| H3F3AP1 | 15 | H3 histone family member 3A pseudogene 1 [ |
| SRP14 | 15 | signal recognition particle 14 [Source:HGNC Sy |
| SRP14-AS1 | 15 | SRP14 antisense RNA1 (head to head) [Source |
| AC021755.2 | 15 | novel transcript |
| AC021755.3 | 15 | novel transcript |
| BMF | 15 | Bcl2 modifying factor [Source:HGNC Symbol;A |
| BMF-AS1 | 15 | BMF antisense RNA 1 [Source:HGNC Symbol;A |
| AC021755.1 | 15 | mitochondrially encoded NADH dehydrogenas |
| BUB1B | 15 | BUB1 mitotic checkpoint serine/threonine kin |
| PAK6 | 15 | p21 (RAC1) activated kinase 6 [Source:HGNC S |
| BUB1B-PAK6 | 15 | BUB1B-PAK6 readthrough [Source:HGNC Sym |
| AC020658.4 | 15 | novel transcript, sense intronic to novel prote |
| AC020658.3 | 15 | novel transcript, sense intronic to PAK6 |
| AC020658.7 | 15 | TEC |
| C15orf56 | 15 | chromosome 15 open reading frame 56 [Sour |
| AC020658.6 | 15 | TEC |
| PLCB2 | 15 | phospholipase C beta 2 [Source:HGNC Symbol |
| ANKRD63 | 15 | ankyrin repeat domain 63 [Source:HGNC Syml |
| AC020658.5 | 15 | novel transcript, sense intronic to PLCB2 |
| PLCB2-AS1 | 15 | PLCB2 antisense RNA 1 [Source:HGNC Symbol |
| AC020658.1 | 15 | ribosomal protein, large, P0 (RPLP0) pseudoge |
| AC020658.2 | 15 | novel transcript, antisense to PLCB2 |
| INAFM2 | 15 | InaF motif containing 2 [Source:HGNC Symbol |
| CCDC9B | 15 | coiled-coil domain containing 9B [Source:HGN |
| RNA5SP392 | 15 | RNA, 5S ribosomal pseudogene 392 [Source:H |
| PHGR1 | 15 | proline, histidine and glycine rich 1 [Source:H( |
| DISP2 | 15 | dispatched RND transporter family member 2 |
| AC013356.4 | 15 | novel transcript, antisense to DISP2 |
| KNSTRN | 15 | kinetochore localized astrin (SPAG5) binding p |
| IVD | 15 | isovaleryl-CoA dehydrogenase [Source:HGNC : |
| BAHD1 | 15 | bromo adjacent homology domain containing |
| AC013356.3 | 15 | novel transcript, antisense to BAHD1 |
| AC013356.2 | 15 | novel transcript, antisense to BAHD1 |
| CHST14 | 15 | carbohydrate sulfotransferase 14 [Source:HGI |
| AC013356.1 | 15 | ribosomal protein L9 (RPL9) pseudogene |
| AC091045.1 | 15 | novel transcript, antisense to C15orf57 |
| CCDC32 | 15 | coiled-coil domain containing 32 [Source:HGN |
| RNU6-516P | 15 | RNA, U6 small nuclear 516, pseudogene [Sour |
| RPUSD2 | 15 | RNA pseudouridine synthase domain containi |

| | | |
|---|---|---|
| Metazoa_SRP | 15 | Metazoan signal recognition particle RNA [Sou |
| KNL1 | 15 | kinetochore scaffold 1 [Source:HGNC Symbol; |
| RN7SL376P | 15 | RNA, 7SL, cytoplasmic 376, pseudogene [Sour |
| AC022405.1 | 15 | novel transcript, sense intronic to CASC5 |
| RAD51-AS1 | 15 | RAD51 antisense RNA 1 [Source:HGNC Symbo |
| RAD51 | 15 | RAD51 recombinase [Source:HGNC Symbol;Ac |
| RMDN3 | 15 | regulator of microtubule dynamics 3 [Source:I |
| SUMO2P15 | 15 | SUMO2 pseudogene 15 [Source:HGNC Symbo |
| GCHFR | 15 | GTP cyclohydrolase I feedback regulator [Sour |
| DNAJC17 | 15 | DnaJ heat shock protein family (Hsp40) memb |
| C15orf62 | 15 | chromosome 15 open reading frame 62 [Sour |
| ZFYVE19 | 15 | zinc finger FYVE-type containing 19 [Source:H |
| PPP1R14D | 15 | protein phosphatase 1 regulatory inhibitor sul |
| SPINT1-AS1 | 15 | SPINT1 antisense RNA 1 [Source:HGNC Symbo |
| SPINT1 | 15 | serine peptidase inhibitor, Kunitz type 1 [Sour |
| RHOV | 15 | ras homolog family member V [Source:HGNC |
| AC025166.1 | 15 | novel transcript |
| VPS18 | 15 | VPS18 core subunit of CORVET and HOPS com |
| AC020661.1 | 15 | novel transcript |
| DLL4 | 15 | delta like canonical Notch ligand 4 [Source:HG |
| ISCA1P4 | 15 | iron-sulfur cluster assembly 1 pseudogene 4 [ |
| CHAC1 | 15 | ChaC glutathione specific gamma-glutamylcyc |
| INO80 | 15 | INO80 complex ATPase subunit [Source:HGNC |
| AC020661.3 | 15 | novel transcript, antisense to INO80 |
| AC020661.2 | 15 | novel transcript, antisense to INO80 |
| INO80-AS1 | 15 | INO80 antisense RNA 1 [Source:HGNC Symbol |
| AC020661.4 | 15 | novel transcript, sense intronic to INO80 |
| CYCSP2 | 15 | CYCS pseudogene 2 [Source:HGNC Symbol;Ac |
| AC021753.1 | 15 | mitochondrially encoded NADH dehydrogenas |
| FAM92A1P1 | 15 | family with sequence similarity 92 member A |
| EXD1 | 15 | exonuclease 3'-5' domain containing 1 [Sourc |
| RN7SL497P | 15 | RNA, 7SL, cytoplasmic 497, pseudogene [Sour |
| CHP1 | 15 | calcineurin like EF-hand protein 1 [Source:HG |
| OIP5-AS1 | 15 | OIP5 antisense RNA 1 [Source:HGNC Symbol; |
| AC087721.2 | 15 | novel protein |
| OIP5 | 15 | Opa interacting protein 5 [Source:HGNC Symb |
| NUSAP1 | 15 | nucleolar and spindle associated protein 1 [So |
| AC087721.1 | 15 | novel transcript, sense intronic to NUSAP1 |
| NDUFAF1 | 15 | NADH:ubiquinone oxidoreductase complex as |
| RTF1 | 15 | RTF1 homolog, Paf1/RNA polymerase II comp |
| RNU6-1169P | 15 | RNA, U6 small nuclear 1169, pseudogene [Sou |
| ITPKA | 15 | inositol-trisphosphate 3-kinase A [Source:HGN |
| LTK | 15 | leukocyte receptor tyrosine kinase [Source:HG |
| RPAP1 | 15 | RNA polymerase II associated protein 1 [Sourc |
| ELOCP2 | 15 | elongin C pseudogene 2 [Source:HGNC Symbo |
| TYRO3 | 15 | TYRO3 protein tyrosine kinase [Source:HGNC |
| AC016134.1 | 15 | novel transcript |
| MGA | 15 | MAX dimerization protein MGA [Source:HGN |
| MIR626 | 15 | microRNA 626 [Source:HGNC Symbol;Acc:HGI |
| AC073657.2 | 15 | ribosomal protein L6 (RPL6) pseudogene |

| | | |
|---|---|---|
| AC073657.1 | 15 | novel transcript |
| MAPKBP1 | 15 | mitogen-activated protein kinase binding prot |
| AC020659.2 | 15 | novel transcript |
| JMJD7 | 15 | jumonji domain containing 7 [Source:HGNC Sy |
| JMJD7-PLA2G4B | 15 | JMJD7-PLA2G4B readthrough [Source:HGNC S |
| PLA2G4B | 15 | phospholipase A2 group IVB [Source:HGNC Sy |
| SPTBN5 | 15 | spectrin beta, non-erythrocytic 5 [Source:HGN |
| RNA5SP393 | 15 | RNA, 5S ribosomal pseudogene 393 [Source:H |
| MIR4310 | 15 | microRNA 4310 [Source:HGNC Symbol;Acc:HG |
| AC020659.1 | 15 | novel transcript, antisense to SPTBN5 |
| EHD4 | 15 | EH domain containing 4 [Source:HGNC Symbo |
| AC039056.2 | 15 | novel transcript, sense intronic to EHD4 |
| EHD4-AS1 | 15 | EHD4 antisense RNA 1 [Source:HGNC Symbol; |
| PLA2G4E-AS1 | 15 | PLA2G4E antisense RNA 1 [Source:HGNC Syml |
| PLA2G4E | 15 | phospholipase A2 group IVE [Source:HGNC Sy |
| AC039056.1 | 15 | novel transcript, antisense to PLA2G4E |
| PLA2G4D | 15 | phospholipase A2 group IVD [Source:HGNC Sy |
| PLA2G4F | 15 | phospholipase A2 group IVF [Source:HGNC Syı |
| VPS39 | 15 | VPS39 subunit of HOPS complex [Source:HGN |
| MIR627 | 15 | microRNA 627 [Source:HGNC Symbol;Acc:HGI |
| AC036103.1 | 15 | novel transcript, antisense to TMEM87A |
| TMEM87A | 15 | transmembrane protein 87A [Source:HGNC Sy |
| GANC | 15 | glucosidase alpha, neutral C [Source:HGNC Syı |
| BNIP3P5 | 15 | BCL2 interacting protein 3 pseudogene 5 [Sou |
| AC012651.1 | 15 | novel transcript |
| CAPN3 | 15 | calpain 3 [Source:HGNC Symbol;Acc:HGNC:14 |
| ZNF106 | 15 | zinc finger protein 106 [Source:HGNC Symbol; |
| RNU6-188P | 15 | RNA, U6 small nuclear 188, pseudogene [Sour |
| SNAP23 | 15 | synaptosome associated protein 23 [Source:H |
| AC018362.1 | 15 | novel transcript, antisense to SNAP23 |
| AC018362.3 | 15 | novel protein |
| LRRC57 | 15 | leucine rich repeat containing 57 [Source:HGN |
| HAUS2 | 15 | HAUS augmin like complex subunit 2 [Source:I |
| AC018362.2 | 15 | novel transcript, antisense to HAUS2 |
| MYL12BP1 | 15 | myosin light chain 12B pseudogene 1 [Source: |
| STARD9 | 15 | StAR related lipid transfer domain containing |
| EIF4EBP2P2 | 15 | eukaryotic translation initiation factor 4E bind |
| CDAN1 | 15 | codanin 1 [Source:HGNC Symbol;Acc:HGNC:1 |
| AC090510.3 | 15 | novel transcript, antisense to CDAN1 |
| AC090510.2 | 15 | novel transcript, antisense to CDAN1 |
| TTBK2 | 15 | tau tubulin kinase 2 [Source:HGNC Symbol;Ac |
| AC090510.1 | 15 | novel transcript |
| AC090510.4 | 15 | novel transcript, antisense to TTBK2 |
| KRT8P50 | 15 | keratin 8 pseudogene 50 [Source:HGNC Symb |
| FDPSP4 | 15 | farnesyl diphosphate synthase pseudogene 4 |
| UBR1 | 15 | ubiquitin protein ligase E3 component n-recog |
| AC068724.3 | 15 | novel transcript, antisense to TMEM62 |
| AC068724.2 | 15 | NULL |
| RPS3AP47 | 15 | ribosomal protein S3a pseudogene 47 [Source |
| TMEM62 | 15 | transmembrane protein 62 [Source:HGNC Syn |

| | | |
|---|---|---|
| SPCS2P1 | 15 | signal peptidase complex subunit 2 pseudoge |
| AC068724.4 | 15 | novel transcript |
| AC068724.1 | 15 | novel transcript, antisense to TMEM62 |
| CCNDBP1 | 15 | cyclin D1 binding protein 1 [Source:HGNC Sym |
| EPB42 | 15 | erythrocyte membrane protein band 4.2 [Sou |
| TGM5 | 15 | transglutaminase 5 [Source:HGNC Symbol;Acc |
| Y_RNA | 15 | Y RNA [Source:RFAM;Acc:RF00019] |
| ATP5PDP1 | 15 | ATP synthase peripheral stalk subunit d pseud |
| TGM7 | 15 | transglutaminase 7 [Source:HGNC Symbol;Acc |
| AC009852.1 | 15 | novel transcript, antisense to TGM7 |
| LCMT2 | 15 | leucine carboxyl methyltransferase 2 [Source: |
| ADAL | 15 | adenosine deaminase like [Source:HGNC Symb |
| ZSCAN29 | 15 | zinc finger and SCAN domain containing 29 [S |
| TUBGCP4 | 15 | tubulin gamma complex associated protein 4 |
| RN7SL487P | 15 | RNA, 7SL, cytoplasmic 487, pseudogene [Sour |
| TP53BP1 | 15 | tumor protein p53 binding protein 1 [Source: |
| MAP1A | 15 | microtubule associated protein 1A [Source:H |
| PPIP5K1 | 15 | diphosphoinositol pentakisphosphate kinase 1 |
| CKMT1B | 15 | creatine kinase, mitochondrial 1B [Source:HG |
| STRC | 15 | stereocilin [Source:HGNC Symbol;Acc:HGNC:1 |
| RNU6-554P | 15 | RNA, U6 small nuclear 554, pseudogene [Sour |
| AC011330.3 | 15 | novel transcript |
| CATSPER2 | 15 | cation channel sperm associated 2 [Source:H |
| RNU6-610P | 15 | RNA, U6 small nuclear 610, pseudogene [Sour |
| AC011330.2 | 15 | novel transcript, sense intronic to CATSPER2 |
| PDIA3P2 | 15 | protein disulfide isomerase family A member |
| AC011330.1 | 15 | histidine acid phosphatase domain containing |
| CKMT1A | 15 | creatine kinase, mitochondrial 1A [Source:HG |
| STRCP1 | 15 | stereocilin pseudogene 1 [Source:HGNC Symb |
| RNU6-353P | 15 | RNA, U6 small nuclear 353, pseudogene [Sour |
| CATSPER2P1 | 15 | cation channel sperm associated 2 pseudogen |
| RNU6-354P | 15 | RNA, U6 small nuclear 354, pseudogene [Sour |
| PDIA3 | 15 | protein disulfide isomerase family A member |
| ELL3 | 15 | elongation factor for RNA polymerase II 3 [Sou |
| AC018512.1 | 15 | novel protein |
| SERF2 | 15 | small EDRK-rich factor 2 [Source:HGNC Symbo |
| MIR1282 | 15 | microRNA 1282 [Source:HGNC Symbol;Acc:HG |
| SERINC4 | 15 | serine incorporator 4 [Source:HGNC Symbol;A |
| HYPK | 15 | huntingtin interacting protein K [Source:HGN |
| MFAP1 | 15 | microfibril associated protein 1 [Source:HGNC |
| WDR76 | 15 | WD repeat domain 76 [Source:HGNC Symbol; |
| Y_RNA | 15 | Y RNA [Source:RFAM;Acc:RF00019] |
| FRMD5 | 15 | FERM domain containing 5 [Source:HGNC Sym |
| PIN4P1 | 15 | peptidylprolyl cis/trans isomerase, NIMA-inte |
| ACTBP7 | 15 | ACTB pseudogene 7 [Source:HGNC Symbol;Ac |
| GAPDHP55 | 15 | glyceraldehyde 3 phosphate dehydrogenase p |
| CASC4 | 15 | cancer susceptibility 4 [Source:HGNC Symbol; |
| AC090519.1 | 15 | glyceraldehyde-3-phosphate dehydrogenase ( |
| AC090519.2 | 15 | novel transcript |
| AC025043.1 | 15 | novel transcript, antisense to CASC4 |

| | | |
|---|---|---|
| CTDSPL2 | 15 | CTD small phosphatase like 2 [Source:HGNC S |
| HNRNPMP1 | 15 | heterogeneous nuclear ribonucleoprotein M p |
| AC025430.1 | 15 | novel transcript, antisense to CTDSPL2 |
| EIF3J-DT | 15 | EIF3J divergent transcript [Source:HGNC Symb |
| EIF3J | 15 | eukaryotic translation initiation factor 3 subur |
| AC009996.1 | 15 | novel transcript |
| SPG11 | 15 | SPG11 vesicle trafficking associated, spatacsin |
| PATL2 | 15 | PAT1 homolog 2 [Source:HGNC Symbol;Acc:H |
| Y_RNA | 15 | Y RNA [Source:RFAM;Acc:RF00019] |
| B2M | 15 | beta-2-microglobulin [Source:HGNC Symbol;A |
| TRIM69 | 15 | tripartite motif containing 69 [Source:HGNC S |
| AC122108.2 | 15 | novel transcript |
| SORD2P | 15 | sorbitol dehydrogenase 2, pseudogene [Sourc |
| AC122108.1 | 15 | novel transcript |
| Y_RNA | 15 | Y RNA [Source:RFAM;Acc:RF00019] |
| RNU1-119P | 15 | RNA, U1 small nuclear 119, pseudogene [Sour |
| AC090888.3 | 15 | chromosome 15 open reading frame 43 (C15c |
| AC090888.1 | 15 | 5-azacytidine induced 2 (AZI2) pseudogene |
| RNU6-1108P | 15 | RNA, U6 small nuclear 1108, pseudogene [Sou |
| TERB2 | 15 | telomere repeat binding bouquet formation p |
| RNU6-1332P | 15 | RNA, U6 small nuclear 1332, pseudogene [Sou |
| AC090888.2 | 15 | 5-azacytidine induced 2 (AZI2) pseudogene |
| RNU6-966P | 15 | RNA, U6 small nuclear 966, pseudogene [Sour |
| RNU1-78P | 15 | RNA, U1 small nuclear 78, pseudogene [Sourc |
| SORD | 15 | sorbitol dehydrogenase [Source:HGNC Symbo |
| AC091117.2 | 15 | novel transcript, antisense to SORD |
| Y_RNA | 15 | Y RNA [Source:RFAM;Acc:RF00019] |
| AC091117.1 | 15 | novel transcript, antisense to SORD |
| DUOX2 | 15 | dual oxidase 2 [Source:HGNC Symbol;Acc:HGN |
| DUOXA2 | 15 | dual oxidase maturation factor 2 [Source:HGN |
| DUOXA1 | 15 | dual oxidase maturation factor 1 [Source:HGN |
| DUOX1 | 15 | dual oxidase 1 [Source:HGNC Symbol;Acc:HGN |
| AC051619.6 | 15 | novel transcript |
| SHF | 15 | Src homology 2 domain containing F [Source:H |
| AC051619.4 | 15 | novel transcript, antisense to SHF |
| AC051619.7 | 15 | novel transcript |
| AC051619.8 | 15 | novel transcript |
| AC051619.10 | 15 | novel transcript |
| AC051619.9 | 15 | NULL |
| AC051619.2 | 15 | H3 histone, family 3A (H3F3A) pseudogene |
| AC051619.5 | 15 | novel transcript, antisense to SLC28A2 |
| SLC28A2 | 15 | solute carrier family 28 member 2 [Source:HG |
| AC051619.11 | 15 | novel transcript |
| AC051619.1 | 15 | high mobility group nucleosomal binding dom |
| AC051619.3 | 15 | protein-kinase, interferon-inducible double st |
| RNU6-953P | 15 | RNA, U6 small nuclear 953, pseudogene [Sour |
| GATM | 15 | glycine amidinotransferase [Source:HGNC Syn |
| AC025580.3 | 15 | GATM antisense RNA 1 |
| SPATA5L1 | 15 | spermatogenesis associated 5 like 1 [Source:H |
| C15orf48 | 15 | chromosome 15 open reading frame 48 [Sour |

| | | |
|---|---|---|
| AC025580.1 | 15 | novel transcript, antisense to C15orf48 |
| MIR147B | 15 | microRNA 147b [Source:HGNC Symbol;Acc:HG |
| AC025580.2 | 15 | novel transcript, antisense to SLC30A4 |
| SLC30A4 | 15 | solute carrier family 30 member 4 [Source:HG |
| HMGN2P46 | 15 | high mobility group nucleosomal binding dom |
| SNORA41B | 15 | small nucleolar RNA, H/ACA box 41B [Source:I |
| AC090527.1 | 15 | novel transcript |
| DPPA5P2 | 15 | developmental pluripotency associated 5 pseu |
| AC090527.3 | 15 | novel transcript |
| BLOC1S6 | 15 | biogenesis of lysosomal organelles complex 1 |
| AC090527.2 | 15 | novel protein |
| Y_RNA | 15 | Y RNA [Source:RFAM;Acc:RF00019] |
| SQOR | 15 | sulfide quinone oxidoreductase [Source:HGNC |
| AC068722.2 | 15 | novel transcript |
| AC068722.1 | 15 | novel transcript |
| AC068714.1 | 15 | phosphodiesterase 8B (PDE8B) pseudogene |
| AC091074.3 | 15 | novel transcript |
| MTND5P40 | 15 | MT-ND5 pseudogene 40 [Source:HGNC Symbo |
| AC091074.1 | 15 | NULL |
| AC091074.2 | 15 | novel transcript |
| AC012405.1 | 15 | novel transcript |
| RNU6-1014P | 15 | RNA, U6 small nuclear 1014, pseudogene [Sou |
| AC073941.1 | 15 | novel transcript |
| RN7SKP101 | 15 | RN7SK pseudogene 101 [Source:HGNC Symbo |
| AC073941.2 | 15 | cytochrome b5 type B (outer mitochondrial m |
| AC066615.1 | 15 | TEC |
| SEMA6D | 15 | semaphorin 6D [Source:HGNC Symbol;Acc:HG |
| RN7SKP139 | 15 | RN7SK pseudogene 139 [Source:HGNC Symbo |
| AC084882.1 | 15 | novel transcript, intronic to SEMA6D |
| AC023905.1 | 15 | novel transcript, antisense to SEMA6D |
| AC009558.1 | 15 | novel transcript, antisense to SEMA6D |
| AC009558.2 | 15 | novel transcript, intronic to SEMA6D |
| AC012050.1 | 15 | novel transcript, antisense to SEMA6D |
| AC044787.1 | 15 | ribosomal protein L7a (RPL7A) pseudogene |
| LINC01491 | 15 | long intergenic non-protein coding RNA 1491 |
| AC092078.2 | 15 | novel transcript |
| AC092078.1 | 15 | novel transcript |
| AC092078.3 | 15 | novel transcript |
| SLC24A5 | 15 | solute carrier family 24 member 5 [Source:HG |
| MYEF2 | 15 | myelin expression factor 2 [Source:HGNC Sym |
| CTXN2 | 15 | cortexin 2 [Source:HGNC Symbol;Acc:HGNC:3 |
| SLC12A1 | 15 | solute carrier family 12 member 1 [Source:HG |
| AC066612.1 | 15 | novel transcript |
| AC066612.2 | 15 | novel transcript, antisense to CTXN2 |
| AC023355.1 | 15 | novel transcript, antisense to DUT |
| Y_RNA | 15 | Y RNA [Source:RFAM;Acc:RF00019] |
| DUT | 15 | deoxyuridine triphosphatase [Source:HGNC Sy |
| FBN1 | 15 | fibrillin 1 [Source:HGNC Symbol;Acc:HGNC:36 |
| AC022467.2 | 15 | novel transcript, antisense to FBN1 |
| AC022467.1 | 15 | novel transcript, sense intronic to FBN1 |

| | | |
|---|---|---|
| AC084757.3 | 15 | novel transcript |
| AC084757.4 | 15 | novel transcript |
| CEP152 | 15 | centrosomal protein 152 [Source:HGNC Symb |
| AC084757.1 | 15 | oxidation resistance 1 (OXR1) pseudogene |
| AC084757.2 | 15 | novel transcript |
| AC012379.1 | 15 | novel transcript, antisense to CEP152 |
| AC012379.2 | 15 | novel transcript, antisense to CEP152 |
| SHC4 | 15 | SHC adaptor protein 4 [Source:HGNC Symbol; |
| EID1 | 15 | EP300 interacting inhibitor of differentiation 1 |
| AC091073.1 | 15 | novel transcript, antisense to SHC4 |
| KRT8P24 | 15 | keratin 8 pseudogene 24 [Source:HGNC Symb |
| SECISBP2L | 15 | SECIS binding protein 2 like [Source:HGNC Syn |
| Y_RNA | 15 | Y RNA [Source:RFAM;Acc:RF00019] |
| RN7SL577P | 15 | RNA, 7SL, cytoplasmic 577, pseudogene [Sour |
| AC013452.2 | 15 | novel transcript |
| COPS2 | 15 | COP9 signalosome subunit 2 [Source:HGNC Sy |
| Y_RNA | 15 | Y RNA [Source:RFAM;Acc:RF00019] |
| GALK2 | 15 | galactokinase 2 [Source:HGNC Symbol;Acc:HG |
| NDUFAF4P1 | 15 | NADH:ubiquinone oxidoreductase complex as |
| MIR4716 | 15 | microRNA 4716 [Source:HGNC Symbol;Acc:HG |
| AC013452.1 | 15 | ribosomal protein L15 (RPL15) pseudogene |
| RN7SL307P | 15 | RNA, 7SL, cytoplasmic 307, pseudogene [Sour |
| FAM227B | 15 | family with sequence similarity 227 member E |
| AC022306.2 | 15 | novel transcript, sense intronic to GALK2 |
| AC022306.3 | 15 | novel transcript, sense intronic to GALK2 |
| AC022306.1 | 15 | signal peptide peptidase-like 2A (SPPL2A) pseu |
| FGF7 | 15 | fibroblast growth factor 7 [Source:HGNC Symb |
| DTWD1 | 15 | DTW domain containing 1 [Source:HGNC Sym |
| AC025040.2 | 15 | ring finger protein, LIM domain interacting (RI |
| ATP8B4 | 15 | ATPase phospholipid transporting 8B4 (putativ |
| AC025040.1 | 15 | novel transcript, antisense to ATP8B4 |
| RNA5SP394 | 15 | RNA, 5S ribosomal pseudogene 394 [Source:H |
| SLC27A2 | 15 | solute carrier family 27 member 2 [Source:HG |
| Y_RNA | 15 | Y RNA [Source:RFAM;Acc:RF00019] |
| HDC | 15 | histidine decarboxylase [Source:HGNC Symbo |
| RN7SL494P | 15 | RNA, 7SL, cytoplasmic 494, pseudogene [Sour |
| GABPB1 | 15 | GA binding protein transcription factor subuni |
| RNU6-94P | 15 | RNA, U6 small nuclear 94, pseudogene [Sourc |
| GABPB1-IT1 | 15 | GABPB1 intronic transcript [Source:HGNC Sym |
| GABPB1-AS1 | 15 | GABPB1 antisense RNA 1 [Source:HGNC Symb |
| AC022087.1 | 15 | novel transcript |
| MIR4712 | 15 | microRNA 4712 [Source:HGNC Symbol;Acc:HG |
| AHCYP7 | 15 | adenosylhomocysteinase pseudogene 7 [Sour |
| USP8 | 15 | ubiquitin specific peptidase 8 [Source:HGNC S |
| RNA5SP395 | 15 | RNA, 5S ribosomal pseudogene 395 [Source:H |
| AC012170.1 | 15 | ribosomal protein S20 (RPS20) pseudogene |
| AC012170.3 | 15 | novel transcript |
| AC012170.2 | 15 | novel transcript, antisense to USP8 |
| USP50 | 15 | ubiquitin specific peptidase 50 [Source:HGNC |
| TRPM7 | 15 | transient receptor potential cation channel su |

| | | |
|---|---|---|
| AC084756.1 | 15 | novel transcript, antisense to TRPM7 |
| RN7SL354P | 15 | RNA, 7SL, cytoplasmic 354, pseudogene [Sour |
| SPPL2A | 15 | signal peptide peptidase like 2A [Source:HGN( |
| AC012100.3 | 15 | TEC |
| AC012100.2 | 15 | novel transcript, antisense to SPPL2A |
| AC012100.1 | 15 | ribosomal protein L32 (RPL32) pseudogene |
| AC021752.1 | 15 | novel transcript |
| AP4E1 | 15 | adaptor related protein complex 4 subunit ep: |
| DCAF13P3 | 15 | DDB1 and CUL4 associated factor 13 pseudog |
| MIR4713HG | 15 | MIR4713 host gene [Source:HGNC Symbol;Ac |
| TNFAIP8L3 | 15 | TNF alpha induced protein 8 like 3 [Source:HG |
| AC073964.1 | 15 | novel transcript, intronic to TNFAIP8L3 |
| CYP19A1 | 15 | cytochrome P450 family 19 subfamily A meml |
| MIR4713 | 15 | microRNA 4713 [Source:HGNC Symbol;Acc:HG |
| AC020891.3 | 15 | novel transcript, intronic to CYP19A1 |
| MIR7973-2 | 15 | microRNA 7973-2 [Source:HGNC Symbol;Acc:I |
| MIR7973-1 | 15 | microRNA 7973-1 [Source:HGNC Symbol;Acc:I |
| AC020891.2 | 15 | novel transcript |
| GLDN | 15 | gliomedin [Source:HGNC Symbol;Acc:HGNC:2! |
| AC020891.4 | 15 | ribosomal protein L36a-like (RPL36AL) pseudc |
| AC020891.1 | 15 | novel transcript, intronic to GLDN |
| AC066613.2 | 15 | novel transcript, antisense to DMXL2 |
| DMXL2 | 15 | Dmx like 2 [Source:HGNC Symbol;Acc:HGNC:2 |
| AC066613.1 | 15 | novel transcript, antisense to DMXL2 |
| SCG3 | 15 | secretogranin III [Source:HGNC Symbol;Acc:H( |
| AC020892.1 | 15 | novel transcript |
| AC020892.2 | 15 | novel transcript, antisense to SCG3 |
| LYSMD2 | 15 | LysM domain containing 2 [Source:HGNC Sym |
| TMOD2 | 15 | tropomodulin 2 [Source:HGNC Symbol;Acc:HG |
| AC026770.1 | 15 | novel transcript, intronic to TMOD2 |
| TMOD3 | 15 | tropomodulin 3 [Source:HGNC Symbol;Acc:HG |
| AC090971.5 | 15 | novel transcript |
| Metazoa_SRP | 15 | Metazoan signal recognition particle RNA [Sou |
| AC090971.4 | 15 | UTP6, small subunit (SSU) processome compo |
| Y_RNA | 15 | Y RNA [Source:RFAM;Acc:RF00019] |
| AC090971.2 | 15 | novel transcript, antisense to TMOD3 |
| RNU6-90P | 15 | RNA, U6 small nuclear 90, pseudogene [Sourc |
| AC090971.1 | 15 | novel transcript, antisense to TMOD3 |
| AC090971.3 | 15 | Leo1, Paf1/RNA polymerase II complex compo |
| LEO1 | 15 | LEO1 homolog, Paf1/RNA polymerase II comp |
| MAPK6 | 15 | mitogen-activated protein kinase 6 [Source:H( |
| MAPK6-DT | 15 | MAPK6 divergent transcript [Source:HGNC Syr |
| AC090970.2 | 15 | novel transcript |
| AC090970.1 | 15 | novel transcript, intronic to MAPK6 |
| AC023906.5 | 15 | novel transcript, antisense to MAPK6 |
| AC023906.2 | 15 | novel transcript |
| AC023906.1 | 15 | ribosomal protein S13 (RPS13) pseudogene |
| BCL2L10 | 15 | BCL2 like 10 [Source:HGNC Symbol;Acc:HGNC |
| GNB5 | 15 | G protein subunit beta 5 [Source:HGNC Symb( |
| AC023906.3 | 15 | novel transcript |

| | | |
|---|---|---|
| AC023906.4 | 15 | novel transcript, antisense to GNB5 |
| Y_RNA | 15 | Y RNA [Source:RFAM;Acc:RF00019] |
| CERNA1 | 15 | competing endogenous lncRNA 1 for miR-470 |
| MYO5C | 15 | myosin VC [Source:HGNC Symbol;Acc:HGNC:7 |
| MIR1266 | 15 | microRNA 1266 [Source:HGNC Symbol;Acc:HG |
| MYO5A | 15 | myosin VA [Source:HGNC Symbol;Acc:HGNC:7 |
| EEF1B2P1 | 15 | eukaryotic translation elongation factor 1 beta |
| ARPP19 | 15 | cAMP regulated phosphoprotein 19 [Source:H |
| AC025917.1 | 15 | novel transcript, antisense to KIAA1370 |
| FAM214A | 15 | family with sequence similarity 214 member A |
| AC009754.1 | 15 | novel transcript, intronic to KIAA1370 |
| AC009754.2 | 15 | novel transcript, antisense to KIAA1370 |
| ONECUT1 | 15 | one cut homeobox 1 [Source:HGNC Symbol;A |
| AC016044.1 | 15 | novel transcript |
| RPSAP55 | 15 | ribosomal protein SA pseudogene 55 [Source: |
| EEF1A1P22 | 15 | eukaryotic translation elongation factor 1 alph |
| LINC02490 | 15 | long intergenic non-protein coding RNA 2490 |
| AC066614.1 | 15 | novel transcript |
| WDR72 | 15 | WD repeat domain 72 [Source:HGNC Symbol; |
| RNU2-53P | 15 | RNA, U2 small nuclear 53, pseudogene [Sourc |
| RNU6-449P | 15 | RNA, U6 small nuclear 449, pseudogene [Sour |
| AC084759.1 | 15 | SMT3 suppressor of mif two 3 homolog 2 (S. c |
| AC084759.3 | 15 | TEC |
| AC084759.2 | 15 | novel transcript |
| UNC13C | 15 | unc-13 homolog C [Source:HGNC Symbol;Acc: |
| AC010867.1 | 15 | intraflagellar transport 80 homolog (Chlamydo |
| HNRNPA1P74 | 15 | heterogeneous nuclear ribonucleoprotein A1 |
| AC025272.1 | 15 | novel transcript |
| AC011912.1 | 15 | TEC |
| RSL24D1 | 15 | ribosomal L24 domain containing 1 [Source:H |
| RAB27A | 15 | RAB27A, member RAS oncogene family [Sourc |
| AC018926.2 | 15 | novel transcript, sense intronic to RAB27A |
| PIGBOS1 | 15 | PIGB opposite strand 1 [Source:HGNC Symbol |
| PIGB | 15 | phosphatidylinositol glycan anchor biosynthes |
| CCPG1 | 15 | cell cycle progression 1 [Source:HGNC Symbol |
| AC018926.1 | 15 | novel transcript, sense intronic to CCPG1 |
| AC018926.3 | 15 | novel transcript, sense intronic to CCPG1 |
| DNAAF4-CCPG1 | 15 | DNAAF4-CCPG1 readthrough (NMD candidate |
| MIR628 | 15 | microRNA 628 [Source:HGNC Symbol;Acc:HGN |
| C15orf65 | 15 | chromosome 15 open reading frame 65 [Sour |
| DNAAF4 | 15 | dynein axonemal assembly factor 4 [Source:H |
| AC013355.1 | 15 | thioesterase superfamily member 4 (THEM4) |
| PYGO1 | 15 | pygopus family PHD finger 1 [Source:HGNC Sy |
| AC012378.2 | 15 | novel transcript, antisense to PYGO1 |
| PRTG | 15 | protogenin [Source:HGNC Symbol;Acc:HGNC:2 |
| AC012378.1 | 15 | novel transcript, antisense to PRTG |
| AC009997.1 | 15 | ribosomal protein L19 (RPL19) pseudogene |
| NEDD4 | 15 | NEDD4 E3 ubiquitin protein ligase [Source:HG |
| CNOT6LP1 | 15 | CNOT6L pseudogene 1 [Source:HGNC Symbol; |
| RN7SL568P | 15 | RNA, 7SL, cytoplasmic 568, pseudogene [Sour |

| | | |
|---|---|---|
| CD24P2 | 15 | CD24 molecule pseudogene 2 [Source:HGNC S |
| RFX7 | 15 | regulatory factor X7 [Source:HGNC Symbol;Ac |
| AC084783.1 | 15 | hydroxyacyl-CoA dehydrogenase (HADH) pseu |
| TEX9 | 15 | testis expressed 9 [Source:HGNC Symbol;Acc: |
| AC068726.1 | 15 | novel transcript, sense intronic to TEX9 |
| RNU6-1287P | 15 | RNA, U6 small nuclear 1287, pseudogene [Sou |
| HMGB1P33 | 15 | high mobility group box 1 pseudogene 33 [Sou |
| AC084782.1 | 15 | novel transcript |
| AC084782.2 | 15 | novel transcript |
| MNS1 | 15 | meiosis specific nuclear structural 1 [Source:H |
| AC084782.3 | 15 | novel transcript, antisense to MNS1 |
| AC090518.1 | 15 | novel transcript |
| ZNF280D | 15 | zinc finger protein 280D [Source:HGNC Symbo |
| Y_RNA | 15 | Y RNA [Source:RFAM;Acc:RF00019] |
| AC090517.4 | 15 | zinc finger protein 280D [Source:NCBI gene;Ac |
| AC090517.2 | 15 | novel transcript, sense intronic to |
| AC090517.3 | 15 | novel transcript |
| AC090517.5 | 15 | novel transcript |
| AC090517.1 | 15 | acyl-Coenzyme A binding domain containing 7 |
| AC010999.2 | 15 | novel transcript |
| AC010999.1 | 15 | novel transcript, sense intronic to ZNF280D |
| TCF12 | 15 | transcription factor 12 [Source:HGNC Symbol; |
| HNRNPA3P11 | 15 | heterogeneous nuclear ribonucleoprotein A3 |
| SNORD13D | 15 | small nucleolar RNA, C/D box 13D [Source:HG |
| LINC00926 | 15 | long intergenic non-protein coding RNA 926 [S |
| LINC01413 | 15 | long intergenic non-protein coding RNA 1413 |
| NDUFB10P1 | 15 | NADH:ubiquinone oxidoreductase subunit B1( |
| CGNL1 | 15 | cingulin like 1 [Source:HGNC Symbol;Acc:HGN |
| RNU6-844P | 15 | RNA, U6 small nuclear 844, pseudogene [Sour |
| AC025271.3 | 15 | novel transcript |
| AC025271.2 | 15 | novel transcript |
| AC025271.4 | 15 | novel transcript |
| AC025271.1 | 15 | ribosomal protein S13 (RPS13) pseudogene |
| MYZAP | 15 | myocardial zonula adherens protein [Source:H |
| GCOM1 | 15 | GRINL1A complex locus 1 [Source:HGNC Symb |
| POLR2M | 15 | RNA polymerase II subunit M [Source:HGNC S |
| AC090651.1 | 15 | novel transcript |
| ALDH1A2 | 15 | aldehyde dehydrogenase 1 family member A2 |
| AC012653.1 | 15 | calponin 2 (CNN2) pseudogene |
| AC012653.2 | 15 | novel transcript, sense intronic to ALDH1A2 |
| AC025431.1 | 15 | novel transcript, antisense to ALDH1A2 |
| AQP9 | 15 | aquaporin 9 [Source:HGNC Symbol;Acc:HGNC |
| MTCO3P23 | 15 | MT-CO3 pseudogene 23 [Source:HGNC Symbo |
| MTND3P12 | 15 | MT-ND3 pseudogene 12 [Source:HGNC Symbo |
| AC066616.2 | 15 | mitochondrially encoded NADH 4L (MT-ND4L) |
| MTND5P32 | 15 | MT-ND5 pseudogene 32 [Source:HGNC Symbo |
| MTCYBP23 | 15 | MT-CYB pseudogene 23 [Source:HGNC Symbo |
| AC066616.1 | 15 | ribosomal protein L28 (RPL28) pseudogene |
| LIPC | 15 | lipase C, hepatic type [Source:HGNC Symbol;A |
| LIPC-AS1 | 15 | LIPC antisense RNA 1 [Source:HGNC Symbol;A |

| | | |
|---|---|---|
| AC084781.1 | 15 | novel transcript, sense intronic to LIPC |
| AC084781.2 | 15 | novel transcript, sense intronic to LIPC |
| AC018904.2 | 15 | novel transcript, sense intronic to LIPC |
| AC018904.1 | 15 | novel transcript, antisense to ADAM10 |
| ADAM10 | 15 | ADAM metallopeptidase domain 10 [Source:H |
| HMGB1P51 | 15 | high mobility group box 1 pseudogene 51 [Sou |
| AC091046.1 | 15 | guanine nucleotide binding protein (G protein |
| HSP90AB4P | 15 | heat shock protein 90 alpha family class B mer |
| RN7SKP95 | 15 | RN7SK pseudogene 95 [Source:HGNC Symbol; |
| AC091046.2 | 15 | zinc finger protein 551 (ZNF551) pseudogene |
| AC090515.6 | 15 | novel transcript |
| AC090515.3 | 15 | ATP synthase, H+ transporting, mitochondrial |
| U3 | 15 | Small nucleolar RNA U3 [Source:RFAM;Acc:RF |
| AC090515.2 | 15 | novel transcript |
| MINDY2 | 15 | MINDY lysine 48 deubiquitinase 2 [Source:HG |
| AC090515.5 | 15 | novel transcript, sense intronic to FAM63B |
| AC090515.4 | 15 | novel transcript, antisense to FAM63B |
| ZNF444P1 | 15 | zinc finger protein 444 pseudogene 1 [Source: |
| RNF111 | 15 | ring finger protein 111 [Source:HGNC Symbol; |
| AC090515.1 | 15 | ribosomal protein L21 (RPL21) pseudogene |
| SLTM | 15 | SAFB like transcription modulator [Source:HG |
| AC025918.1 | 15 | novel transcript |
| AC025918.2 | 15 | ribosome binding factor A (putative) (RBFA) ps |
| AC092757.1 | 15 | ribosomal protein L21 (RPL21) pseudogene |
| CCNB2 | 15 | cyclin B2 [Source:HGNC Symbol;Acc:HGNC:15 |
| AC092757.3 | 15 | novel transcript, antisense to CCNB2 |
| AC092757.2 | 15 | novel transcript, sense overlapping to CCNB2 |
| MYO1E | 15 | myosin IE [Source:HGNC Symbol;Acc:HGNC:75 |
| MIR2116 | 15 | microRNA 2116 [Source:HGNC Symbol;Acc:HG |
| LDHAL6B | 15 | lactate dehydrogenase A like 6B [Source:HGN |
| RNU4-80P | 15 | RNA, U4 small nuclear 80, pseudogene [Sourc |
| AC092756.1 | 15 | novel transcript, sense intronic to MYO1E |
| RNU6-212P | 15 | RNA, U6 small nuclear 212, pseudogene [Sour |
| AC092868.2 | 15 | novel transcript, sense intronic to MYO1E |
| FAM81A | 15 | family with sequence similarity 81 member A |
| AC092868.3 | 15 | NSA2 ribosome biogenesis homolog (S. cerevi |
| AC092868.1 | 15 | ribosomal protein L7a (RPL7A) pseudogene |
| Y_RNA | 15 | Y RNA [Source:RFAM;Acc:RF00019] |
| AC092754.2 | 15 | ribosomal protein L21 (RPL21) pseudogene |
| AC092754.1 | 15 | ribosomal protein L21 (RPL21) pseudogene |
| RNA5SP396 | 15 | RNA, 5S ribosomal pseudogene 396 [Source:H |
| GCNT3 | 15 | glucosaminyl (N-acetyl) transferase 3, mucin t |
| GTF2A2 | 15 | general transcription factor IIA subunit 2 [Sou |
| AC092755.1 | 15 | novel transcript, sense overlapping BNIP2 and |
| BNIP2 | 15 | BCL2 interacting protein 2 [Source:HGNC Sym |
| PIGHP1 | 15 | phosphatidylinositol glycan anchor biosynthes |
| AC092755.2 | 15 | novel transcript, antisense to BNIP2 |
| RPS3AP6 | 15 | ribosomal protein S3A pseudogene 6 [Source: |
| AC092079.2 | 15 | novel transcript |
| AC092079.1 | 15 | DEAD (Asp-Glu-Ala-Asp) box polypeptide 18 (D |

| | | |
|---|---|---|
| NMNAT1P5 | 15 | NMNAT1 pseudogene 5 [Source:HGNC Symbo |
| FOXB1 | 15 | forkhead box B1 [Source:HGNC Symbol;Acc:H |
| MESTP2 | 15 | mesoderm specific transcript pseudogene 2 [S |
| AC009654.1 | 15 | novel transcript |
| AC037433.1 | 15 | novel transcript |
| ANXA2 | 15 | annexin A2 [Source:HGNC Symbol;Acc:HGNC: |
| AC087385.1 | 15 | ribosomal protein L36a (RPL36A) pseudogene |
| AC087385.2 | 15 | ribosomal protein S10 (RPS10) pseudogene |
| AC087385.3 | 15 | tumor protein p63 regulated 1 (TPRG1) pseud |
| ICE2 | 15 | interactor of little elongation complex ELL sub |
| RORA-AS1 | 15 | RORA antisense RNA 1 [Source:HGNC Symbol; |
| RORA | 15 | RAR related orphan receptor A [Source:HGNC |
| AC107241.1 | 15 | novel transcript, antisense to RORA |
| CYCSP38 | 15 | CYCS pseudogene 38 [Source:HGNC Symbol;A |
| AC009560.1 | 15 | novel transcript |
| AC009560.3 | 15 | novel transcript, antisense to RORA |
| AC009560.4 | 15 | novel transcript, antisense to RORA |
| AC009560.2 | 15 | NULL |
| RORA-AS2 | 15 | RORA antisense RNA 2 [Source:HGNC Symbol; |
| RNA5SP397 | 15 | RNA, 5S ribosomal pseudogene 397 [Source:H |
| AC022898.1 | 15 | novel transcript, antisense to RORA |
| AC022898.2 | 15 | novel transcript, sense intronic to RORA |
| AC012404.1 | 15 | novel transcript, antisense to RORA |
| AC012404.2 | 15 | novel transcript, antisense to RORA |
| AC107905.1 | 15 | novel transcript |
| AC104574.2 | 15 | novel transcript |
| AC104574.1 | 15 | uncharacterized LOC101928850 [Source:NCBI |
| AC018618.1 | 15 | novel transcript |
| AC018618.2 | 15 | ubiquitin-conjugating enzyme E2D 3 (UBE2D3 |
| LINC02349 | 15 | long intergenic non-protein coding RNA 2349 |
| AC009554.1 | 15 | novel transcript, antisense to VPS13C |
| VPS13C | 15 | vacuolar protein sorting 13 homolog C [Sourc |
| AC009554.2 | 15 | novel transcript, antisense to VPS13C |
| AC104590.1 | 15 | novel transcript, antisense to VPS13C |
| C2CD4A | 15 | C2 calcium dependent domain containing 4A [ |
| NPM1P47 | 15 | nucleophosmin 1 pseudogene 47 [Source:HGN |
| C2CD4B | 15 | C2 calcium dependent domain containing 4B [ |
| AC126323.2 | 15 | novel transcript |
| AC126323.1 | 15 | golgin subfamily A pseudogene |
| AC126323.3 | 15 | novel transcript |
| Metazoa_SRP | 15 | Metazoan signal recognition particle RNA [Sou |
| GOLGA2P11 | 15 | GOLGA2 pseudogene 11 [Source:HGNC Symbo |
| AC126323.4 | 15 | golgin subfamily A pseudogene |
| AC126323.6 | 15 | novel transcript |
| AC126323.5 | 15 | novel transcript |
| MIR8067 | 15 | microRNA 8067 [Source:HGNC Symbol;Acc:HG |
| MIR6085 | 15 | microRNA 6085 [Source:HGNC Symbol;Acc:HG |
| HMGN1P26 | 15 | high mobility group nucleosome binding doma |
| AC032011.1 | 15 | novel transcript |
| TLN2 | 15 | talin 2 [Source:HGNC Symbol;Acc:HGNC:1544 |

| | | |
|---|---|---|
| AC100839.1 | 15 | novel transcript, antisense to TLN2 |
| AC100839.2 | 15 | novel transcript, antisense to TLN2 |
| MIR190A | 15 | microRNA 190a [Source:HGNC Symbol;Acc:HG |
| AC103740.1 | 15 | uncharacterized LOC105370854 [Source:NCBI |
| AC103740.2 | 15 | novel transcript |
| AC079328.1 | 15 | Y box binding protein 1 (YBX1) pseudogene |
| TPM1 | 15 | tropomyosin 1 [Source:HGNC Symbol;Acc:HGI |
| TPM1-AS | 15 | TPM1 antisense RNA [Source:HGNC Symbol;A |
| AC079328.2 | 15 | novel transcript, antisense to TPM1 |
| AC087612.1 | 15 | novel transcript |
| LACTB | 15 | lactamase beta [Source:HGNC Symbol;Acc:HG |
| RPS27L | 15 | ribosomal protein S27 like [Source:HGNC Sym |
| RAB8B | 15 | RAB8B, member RAS oncogene family [Source |
| APH1B | 15 | aph-1 homolog B, gamma-secretase subunit [ |
| CA12 | 15 | carbonic anhydrase 12 [Source:HGNC Symbol; |
| LINC02568 | 15 | long intergenic non-protein coding RNA 2568 |
| AC007950.1 | 15 | novel transcript |
| AC007950.2 | 15 | novel transcript |
| USP3 | 15 | ubiquitin specific peptidase 3 [Source:HGNC S |
| USP3-AS1 | 15 | USP3 antisense RNA 1 [Source:HGNC Symbol; |
| FBXL22 | 15 | F-box and leucine rich repeat protein 22 [Sour |
| HERC1 | 15 | HECT and RLD domain containing E3 ubiquitin |
| AC073167.1 | 15 | novel transcript, antisense to HERC1 |
| MIR422A | 15 | microRNA 422a [Source:HGNC Symbol;Acc:HG |
| AC015914.2 | 15 | coiled-coil domain containing 11 (CCDC11) |
| DAPK2 | 15 | death associated protein kinase 2 [Source:HGI |
| AC015914.1 | 15 | novel transcript, antisense to DAPK2 |
| CIAO2A | 15 | cytosolic iron-sulfur assembly component 2A |
| SNX1 | 15 | sorting nexin 1 [Source:HGNC Symbol;Acc:HGI |
| Y_RNA | 15 | Y RNA [Source:RFAM;Acc:RF00019] |
| SNX22 | 15 | sorting nexin 22 [Source:HGNC Symbol;Acc:HG |
| PPIB | 15 | peptidylprolyl isomerase B [Source:HGNC Sym |
| CSNK1G1 | 15 | casein kinase 1 gamma 1 [Source:HGNC Symb |
| AC087632.2 | 15 | novel protein |
| AC087632.1 | 15 | NULL |
| PCLAF | 15 | PCNA clamp associated factor [Source:HGNC S |
| TRIP4 | 15 | thyroid hormone receptor interactor 4 [Sourc |
| Y_RNA | 15 | Y RNA [Source:RFAM;Acc:RF00019] |
| ZNF609 | 15 | zinc finger protein 609 [Source:HGNC Symbol; |
| AC091231.1 | 15 | novel transcript, sense intronic to ZNF609 |
| GAPDHP61 | 15 | glyceraldehyde 3 phosphate dehydrogenase p |
| AC090543.2 | 15 | ribosomal protein S26 (RPS26) pseudogene |
| AC090543.1 | 15 | ribosomal protein L21 (RPL21) pseudogene |
| Y_RNA | 15 | Y RNA [Source:RFAM;Acc:RF00019] |
| RNU6-549P | 15 | RNA, U6 small nuclear 549, pseudogene [Sour |
| OAZ2 | 15 | ornithine decarboxylase antizyme 2 [Source:H |
| AC100830.3 | 15 | novel transcript, antisense to ornithine decarb |
| AC100830.2 | 15 | novel transcript, antisense to ornithine decarb |
| AC100830.1 | 15 | novel transcript, antisense OAZ2 |
| RBPMS2 | 15 | RNA binding protein, mRNA processing factor |

| | | |
|---|---|---|
| MIR1272 | 15 | microRNA 1272 [Source:HGNC Symbol;Acc:HG |
| PIF1 | 15 | PIF1 5'-to-3' DNA helicase [Source:HGNC Symb |
| AC069368.1 | 15 | novel protein |
| PLEKHO2 | 15 | pleckstrin homology domain containing O2 [So |
| AC069368.2 | 15 | novel transcript |
| ANKDD1A | 15 | ankyrin repeat and death domain containing 1 |
| AC103691.2 | 15 | novel transcript, antisense to ANKDD1A |
| AC103691.1 | 15 | novel transcript, antisense to ANKDD1A |
| SPG21 | 15 | SPG21 abhydrolase domain containing, maspa |
| MTFMT | 15 | mitochondrial methionyl-tRNA formyltransfer |
| AC013553.2 | 15 | transmembrane protein 126A (TMEM126A) ps |
| SLC51B | 15 | solute carrier family 51 beta subunit [Source:H |
| AC013553.3 | 15 | novel transcript, antisense to SLC51B |
| RASL12 | 15 | RAS like family 12 [Source:HGNC Symbol;Acc:H |
| KBTBD13 | 15 | kelch repeat and BTB domain containing 13 [S |
| AC013553.4 | 15 | novel transcript |
| UBAP1L | 15 | ubiquitin associated protein 1 like [Source:HG |
| AC013553.1 | 15 | ribosomal protein L17 (RPL17) pseudogene |
| PDCD7 | 15 | programmed cell death 7 [Source:HGNC Symb |
| CLPX | 15 | caseinolytic mitochondrial matrix peptidase cl |
| CILP | 15 | cartilage intermediate layer protein [Source:H |
| AC068213.1 | 15 | tec |
| RNU6-686P | 15 | RNA, U6 small nuclear 686, pseudogene [Sour |
| PARP16 | 15 | poly(ADP-ribose) polymerase family member |
| SNORA24B | 15 | small nucleolar RNA, H/ACA box 24B [Source:H |
| RNU5A-1 | 15 | RNA, U5A small nuclear 1 [Source:HGNC Symb |
| AC105129.2 | 15 | cyclin-dependent kinase 11A (CDK11A) pseudo |
| RNU5B-1 | 15 | RNA, U5B small nuclear 1 [Source:HGNC Symb |
| IGDCC3 | 15 | immunoglobulin superfamily DCC subclass me |
| IGDCC4 | 15 | immunoglobulin superfamily DCC subclass me |
| DPP8 | 15 | dipeptidyl peptidase 8 [Source:HGNC Symbol; |
| AC105129.3 | 15 | peroxiredoxin 3 (PRDX3) pseudogene |
| AC105129.1 | 15 | target of EGR1, member 1 (nuclear) (TOE1) ps |
| Y_RNA | 15 | Y RNA [Source:RFAM;Acc:RF00019] |
| HACD3 | 15 | 3-hydroxyacyl-CoA dehydratase 3 [Source:HGN |
| RNU6-19P | 15 | RNA, U6 small nuclear 19, pseudogene [Sourc |
| INTS14 | 15 | integrator complex subunit 14 [Source:HGNC |
| SLC24A1 | 15 | solute carrier family 24 member 1 [Source:HG |
| AC011939.3 | 15 | novel transcript, antisense to SLC24A1 |
| DENND4A | 15 | DENN domain containing 4A [Source:HGNC Sy |
| SNORD13E | 15 | small nucleolar RNA, C/D box 13E [Source:HGN |
| MIR4511 | 15 | microRNA 4511 [Source:HGNC Symbol;Acc:HG |
| RAB11A | 15 | RAB11A, member RAS oncogene family [Sourc |
| AC011939.2 | 15 | novel transcript, intronic to DENND4A |
| AC011939.1 | 15 | DnaJ (Hsp40) homolog, subfamily B, member |
| HNRNPA1P44 | 15 | heterogeneous nuclear ribonucleoprotein A1 |
| U6 | 15 | U6 spliceosomal RNA [Source:RFAM;Acc:RF00 |
| MEGF11 | 15 | multiple EGF like domains 11 [Source:HGNC Sy |
| MIR4311 | 15 | microRNA 4311 [Source:HGNC Symbol;Acc:HG |
| AC055855.2 | 15 | novel transcript, antisense to DIS3L |

| | | |
|---|---|---|
| DIS3L | 15 | DIS3 like exosome 3'-5' exoribonuclease [Sour |
| AC055855.1 | 15 | novel transcript, antisense to DIS3L |
| TIPIN | 15 | TIMELESS interacting protein [Source:HGNC S |
| SCARNA14 | 15 | small Cajal body-specific RNA 14 [Source:HGN |
| Y_RNA | 15 | Y RNA [Source:RFAM;Acc:RF00019] |
| RPL9P25 | 15 | ribosomal protein L9 pseudogene 25 [Source: |
| MAP2K1 | 15 | mitogen-activated protein kinase kinase 1 [So |
| ATP5MFP6 | 15 | ATP synthase membrane subunit f pseudogen |
| RPL35AP32 | 15 | ribosomal protein L35a pseudogene 32 [Sourc |
| AC116913.1 | 15 | novel transcript, antisense to MAP2K1 |
| SNAPC5 | 15 | small nuclear RNA activating complex polypep |
| MIR4512 | 15 | microRNA 4512 [Source:HGNC Symbol;Acc:HG |
| RPL4 | 15 | ribosomal protein L4 [Source:HGNC Symbol;A |
| SNORD18C | 15 | small nucleolar RNA, C/D box 18C [Source:HG |
| AC116913.2 | 15 | NULL |
| SNORD18B | 15 | small nucleolar RNA, C/D box 18B [Source:HG |
| SNORD16 | 15 | small nucleolar RNA, C/D box 16 [Source:HGN |
| SNORD18A | 15 | small nucleolar RNA, C/D box 18A [Source:HG |
| ZWILCH | 15 | zwilch kinetochore protein [Source:HGNC Sym |
| LCTL | 15 | lactase like [Source:HGNC Symbol;Acc:HGNC: |
| LINC01169 | 15 | long intergenic non-protein coding RNA 1169 |
| SMAD6 | 15 | SMAD family member 6 [Source:HGNC Symbo |
| AC013564.1 | 15 | novel transcript, sense intronic to SMAD6 |
| AC110048.1 | 15 | leucine zipper-EF-hand containing transmemb |
| AC110048.2 | 15 | novel transcript |
| AC093334.1 | 15 | novel transcript |
| LINC02206 | 15 | long intergenic non-protein coding RNA 2206 |
| AC087482.1 | 15 | novel transcript, antisense to SMAD3 |
| HMGN2P47 | 15 | high mobility group nucleosomal binding dom |
| SMAD3 | 15 | SMAD family member 3 [Source:HGNC Symbo |
| AC012568.1 | 15 | novel transcript, antisense to SMAD3 |
| AAGAB | 15 | alpha and gamma adaptin binding protein [So |
| RPS24P16 | 15 | ribosomal protein S24 pseudogene 16 [Source |
| IQCH | 15 | IQ motif containing H [Source:HGNC Symbol;A |
| IQCH-AS1 | 15 | IQCH antisense RNA 1 [Source:HGNC Symbol; |
| C15orf61 | 15 | chromosome 15 open reading frame 61 [Sour |
| AC016355.1 | 15 | novel transcript |
| MAP2K5 | 15 | mitogen-activated protein kinase kinase 5 [So |
| HNRNPA1P5 | 15 | heterogeneous nuclear ribonucleoprotein A1 |
| SKOR1 | 15 | SKI family transcriptional corepressor 1 [Sourc |
| AC009292.2 | 15 | novel transcript |
| AC009292.1 | 15 | novel transcript |
| RNU6-1 | 15 | RNA, U6 small nuclear 1 [Source:HGNC Symbo |
| AC022254.1 | 15 | ribosomal protein S15 (RPS15A) pseudogene |
| AC135628.1 | 15 | ribosomal protein S27a (RPS27A) pseudogene |
| PIAS1 | 15 | protein inhibitor of activated STAT 1 [Source: |
| AC107871.1 | 15 | novel protein |
| CALML4 | 15 | calmodulin like 4 [Source:HGNC Symbol;Acc:H |
| CLN6 | 15 | CLN6 transmembrane ER protein [Source:HGN |
| HMGN2P40 | 15 | high mobility group nucleosomal binding dom |

| | | |
|---|---|---|
| AC107871.2 | 15 | novel transcript |
| FEM1B | 15 | fem-1 homolog B [Source:HGNC Symbol;Acc:H |
| ITGA11 | 15 | integrin subunit alpha 11 [Source:HGNC Symb |
| AC100825.1 | 15 | novel transcript |
| CORO2B | 15 | coronin 2B [Source:HGNC Symbol;Acc:HGNC:2 |
| CARSP1 | 15 | cysteinyl-tRNA synthetase pseudogene 1 [Sou |
| ANP32A | 15 | acidic nuclear phosphoprotein 32 family mem |
| MIR4312 | 15 | microRNA 4312 [Source:HGNC Symbol;Acc:HG |
| SPESP1 | 15 | sperm equatorial segment protein 1 [Source:H |
| AC087639.1 | 15 | cytokine induced apoptosis inhibitor 1 (CIAPIN |
| AC087639.2 | 15 | novel transcript, antisense to SPESP1 |
| AC027088.1 | 15 | novel transcript |
| NOX5 | 15 | NADPH oxidase 5 [Source:HGNC Symbol;Acc:H |
| AC027088.5 | 15 | novel transcript, sense intronic to NOX5 |
| AC027088.4 | 15 | ATP-binding cassette, sub-family G (WHITE), n |
| AC027088.3 | 15 | novel transcript, antisense to NOX5 |
| EWSAT1 | 15 | Ewing sarcoma associated transcript 1 [Source |
| AC027088.2 | 15 | novel transcript |
| GLCE | 15 | glucuronic acid epimerase [Source:HGNC Sym |
| AC026992.2 | 15 | novel transcript, antisense to PAQR5 |
| AC026992.1 | 15 | novel transcript |
| PAQR5 | 15 | progestin and adipoQ receptor family membe |
| AC027237.4 | 15 | novel transcript, sense intronic to PAQR5 |
| AC027237.3 | 15 | novel transcript, antisense to PAQR5 |
| Y_RNA | 15 | Y RNA [Source:RFAM;Acc:RF00019] |
| KIF23 | 15 | kinesin family member 23 [Source:HGNC Syml |
| AC027237.1 | 15 | fumarylacetoacetate hydrolase domain conta |
| RPLP1 | 15 | ribosomal protein lateral stalk subunit P1 [Sou |
| U3 | 15 | Small nucleolar RNA U3 [Source:RFAM;Acc:RF |
| AC027237.2 | 15 | novel transcript |
| DRAIC | 15 | downregulated RNA in cancer, inhibitor of cell |
| AC100826.1 | 15 | novel transcript |
| AC021818.2 | 15 | gem (nuclear organelle) associated protein 8 ( |
| AC021818.1 | 15 | novel transcript |
| TLE3 | 15 | TLE family member 3, transcriptional corepres |
| MIR629 | 15 | microRNA 629 [Source:HGNC Symbol;Acc:HGN |
| RNU6-745P | 15 | RNA, U6 small nuclear 745, pseudogene [Sour |
| AC026583.1 | 15 | novel transcript |
| AC048383.1 | 15 | novel transcript |
| LINC02205 | 15 | long intergenic non-protein coding RNA 2205 |
| AC009434.1 | 15 | novel transcript |
| LINC02204 | 15 | long intergenic non-protein coding RNA 2204 |
| AC087699.1 | 15 | chromosome 3 open reading frame 56 (C3orf5 |
| SALRNA3 | 15 | senescence associated long non-coding RNA 3 |
| SALRNA2 | 15 | senescence associated long non-coding RNA 2 |
| UACA | 15 | uveal autoantigen with coiled-coil domains an |
| AC009269.2 | 15 | novel transcript |
| AC009269.5 | 15 | novel transcript, sense intronic to UACA |
| AC009269.3 | 15 | novel transcript, antisense to novel protein |
| AC009269.1 | 15 | ribosomal protein L29 (RPL29) pseudogene |

| | | |
|---|---|---|
| RPL29P30 | 15 | ribosomal protein L29 pseudogene 30 [Source |
| LARP6 | 15 | La ribonucleoprotein domain family member ( |
| AC009269.4 | 15 | novel transcript, sense intronic to LARP6 |
| LRRC49 | 15 | leucine rich repeat containing 49 [Source:HGN |
| KRT8P9 | 15 | keratin 8 pseudogene 9 [Source:HGNC Symbo |
| THAP10 | 15 | THAP domain containing 10 [Source:HGNC Syı |
| RPL5P3 | 15 | ribosomal protein L5 pseudogene 3 [Source:H |
| THSD4 | 15 | thrombospondin type 1 domain containing 4 [ |
| CT62 | 15 | cancer/testis antigen 62 [Source:HGNC Symbo |
| THSD4-AS1 | 15 | THSD4 antisense RNA 1 [Source:HGNC Symbo |
| HMGB1P6 | 15 | high mobility group box 1 pseudogene 6 [Sour |
| AC064799.2 | 15 | novel transcript, intronic to THSD4 |
| AC064799.1 | 15 | ribosomal protein L17 (RPL17) pseudogene |
| AC108861.1 | 15 | novel transcript, antisense to THSD4 |
| NR2E3 | 15 | nuclear receptor subfamily 2 group E member |
| AC104938.1 | 15 | novel transcript, antisense to MYO9A |
| MYO9A | 15 | myosin IXA [Source:HGNC Symbol;Acc:HGNC:[illegible] |
| RNA5SP399 | 15 | RNA, 5S ribosomal pseudogene 399 [Source:H |
| AC022872.1 | 15 | novel transcript, antisense to MYO9A |
| EIF5A2P1 | 15 | eukaryotic translation initiation factor 5A2 ps |
| RNU2-65P | 15 | RNA, U2 small nuclear 65, pseudogene [Sourc |
| SENP8 | 15 | SUMO peptidase family member, NEDD8 spec |
| AC020779.1 | 15 | ribosomal protein L21 (RPL21) pseudogene |
| AC020779.2 | 15 | novel transcript, antisense to SENP8 |
| GRAMD2A | 15 | GRAM domain containing 2A [Source:HGNC Sy |
| PKM | 15 | pyruvate kinase M1/2 [Source:HGNC Symbol; |
| PARP6 | 15 | poly(ADP-ribose) polymerase family member |
| AC009690.3 | 15 | novel transcript |
| AC009690.2 | 15 | novel transcript, antisense to HEXA and CELF6 |
| CELF6 | 15 | CUGBP Elav-like family member 6 [Source:HG |
| AC009690.1 | 15 | novel protein |
| HEXA | 15 | hexosaminidase subunit alpha [Source:HGNC |
| HEXA-AS1 | 15 | HEXA antisense RNA 1 [Source:HGNC Symbol; |
| RPL12P35 | 15 | ribosomal protein L12 pseudogene 35 [Source |
| TMEM202 | 15 | transmembrane protein 202 [Source:HGNC Sy |
| TMEM202-AS1 | 15 | TMEM202 antisense RNA 1 [Source:HGNC Syr |
| PHBP20 | 15 | prohibitin pseudogene 20 [Source:HGNC Symb |
| AC079322.1 | 15 | novel transcript |
| ARIH1 | 15 | ariadne RBR E3 ubiquitin protein ligase 1 [Sou |
| MIR630 | 15 | microRNA 630 [Source:HGNC Symbol;Acc:HGN |
| AC100827.3 | 15 | novel transcript |
| AC100827.5 | 15 | SWI5 recombination repair homolog (yeast) (S |
| AC100827.2 | 15 | golgin A6 family, member A (GOLGA6A) pseuc |
| LINC02259 | 15 | long intergenic non-protein coding RNA 2259 |
| AC100827.4 | 15 | novel transcript |
| AC100827.1 | 15 | COMM domain containing 4 (COMMD4) pseu |
| GOLGA6B | 15 | golgin A6 family member B [Source:HGNC Syn |
| RN7SL853P | 15 | RNA, 7SL, cytoplasmic 853, pseudogene [Sour |
| HIGD2B | 15 | HIG1 hypoxia inducible domain family membe |
| AC009712.1 | 15 | small nuclear ribonucleoprotein polypeptide F |

| | | |
|---|---|---|
| BBS4 | 15 | Bardet-Biedl syndrome 4 [Source:HGNC Symb |
| ADPGK | 15 | ADP dependent glucokinase [Source:HGNC Sy |
| ADPGK-AS1 | 15 | ADPGK antisense RNA 1 [Source:HGNC Symbo |
| AC103874.1 | 15 | novel transcript |
| NPM1P42 | 15 | nucleophosmin 1 pseudogene 42 [Source:HGN |
| NEO1 | 15 | neogenin 1 [Source:HGNC Symbol;Acc:HGNC: |
| FKBP1AP2 | 15 | FKBP prolyl isomerase 1A pseudogene 2 [Sour |
| NPM1P43 | 15 | nucleophosmin 1 pseudogene 43 [Source:HGN |
| AC068397.1 | 15 | novel transcript, antisense to NEO1 |
| HCN4 | 15 | hyperpolarization activated cyclic nucleotide g |
| AC068397.2 | 15 | novel transcript, antisense to HCN4 |
| REC114 | 15 | REC114 meiotic recombination protein [Sourc |
| MRPS15P1 | 15 | mitochondrial ribosomal protein S15 pseudog |
| NPTN | 15 | neuroplastin [Source:HGNC Symbol;Acc:HGNC |
| NPTN-IT1 | 15 | NPTN intronic transcript 1 [Source:HGNC Sym |
| CD276 | 15 | CD276 molecule [Source:HGNC Symbol;Acc:H |
| AC022188.1 | 15 | novel transcript |
| INSYN1 | 15 | inhibitory synaptic factor 1 [Source:HGNC Syn |
| INSYN1-AS1 | 15 | INSYN1 antisense RNA 1 [Source:HGNC Symbo |
| AC018943.1 | 15 | novel transcript, antisense to TBC1D21 |
| TBC1D21 | 15 | TBC1 domain family member 21 [Source:HGN |
| LOXL1-AS1 | 15 | LOXL1 antisense RNA 1 [Source:HGNC Symbol |
| LOXL1 | 15 | lysyl oxidase like 1 [Source:HGNC Symbol;Acc: |
| STOML1 | 15 | stomatin like 1 [Source:HGNC Symbol;Acc:HG |
| PML | 15 | promyelocytic leukemia [Source:HGNC Symbo |
| AC013486.1 | 15 | novel transcript, antisense to PML |
| DNM1P33 | 15 | dynamin 1 pseudogene 33 [Source:HGNC Sym |
| GOLGA6A | 15 | golgin A6 family member A [Source:HGNC Syn |
| RN7SL429P | 15 | RNA, 7SL, cytoplasmic 429, pseudogene [Sour |
| AC010931.1 | 15 | COMM domain containing 4 (COMMD4) pseu |
| ISLR2 | 15 | immunoglobulin superfamily containing leucir |
| AC010931.2 | 15 | novel transcript |
| AC010931.3 | 15 | novel transcript, antisense to ISLR |
| Metazoa_SRP | 15 | Metazoan signal recognition particle RNA [Sou |
| ISLR | 15 | immunoglobulin superfamily containing leucir |
| STRA6 | 15 | stimulated by retinoic acid 6 [Source:HGNC Sy |
| CCDC33 | 15 | coiled-coil domain containing 33 [Source:HGN |
| AC023300.3 | 15 | TEC |
| AC023300.1 | 15 | novel transcript |
| AC023300.2 | 15 | novel transcript, antisense to CCDC33 |
| CYP11A1 | 15 | cytochrome P450 family 11 subfamily A meml |
| PPIAP46 | 15 | peptidylprolyl isomerase A pseudogene 46 [Sc |
| AC090826.1 | 15 | novel transcript, antisense to CYP11A1 |
| AC090826.2 | 15 | novel transcript |
| LINC02255 | 15 | long intergenic non-protein coding RNA 2255 |
| SEMA7A | 15 | semaphorin 7A (John Milton Hagen blood grou |
| MIR6881 | 15 | microRNA 6881 [Source:HGNC Symbol;Acc:HG |
| AC090826.3 | 15 | novel transcript, antisense to SEMA7A |
| UBL7 | 15 | ubiquitin like 7 [Source:HGNC Symbol;Acc:HG |
| UBL7-AS1 | 15 | UBL7 antisense RNA 1 (head to head) [Source: |

| | | |
|---|---|---|
| AC012435.2 | 15 | coiled-coil-helix-coiled-coil-helix domain-cont |
| AC012435.3 | 15 | novel transcript |
| AC012435.1 | 15 | NULL |
| ARID3B | 15 | AT-rich interaction domain 3B [Source:HGNC S |
| CLK3 | 15 | CDC like kinase 3 [Source:HGNC Symbol;Acc:H |
| AC100835.2 | 15 | novel transcript |
| AC100835.1 | 15 | novel transcript, antisense to CLK3 |
| EDC3 | 15 | enhancer of mRNA decapping 3 [Source:HGN |
| CYP1A1 | 15 | cytochrome P450 family 1 subfamily A memb |
| CYP1A2 | 15 | cytochrome P450 family 1 subfamily A memb |
| CSK | 15 | C-terminal Src kinase [Source:HGNC Symbol;A |
| MIR4513 | 15 | microRNA 4513 [Source:HGNC Symbol;Acc:HG |
| LMAN1L | 15 | lectin, mannose binding 1 like [Source:HGNC S |
| AC091230.1 | 15 | novel transcript |
| CPLX3 | 15 | complexin 3 [Source:HGNC Symbol;Acc:HGNC |
| ULK3 | 15 | unc-51 like kinase 3 [Source:HGNC Symbol;Ac |
| MIR6882 | 15 | microRNA 6882 [Source:HGNC Symbol;Acc:HG |
| SCAMP2 | 15 | secretory carrier membrane protein 2 [Source |
| MPI | 15 | mannose phosphate isomerase [Source:HGNC |
| FAM219B | 15 | family with sequence similarity 219 member B |
| COX5A | 15 | cytochrome c oxidase subunit 5A [Source:HGN |
| RPP25 | 15 | ribonuclease P and MRP subunit p25 [Source: |
| SCAMP5 | 15 | secretory carrier membrane protein 5 [Source |
| AC015720.2 | 15 | ribosomal protein L15 (RPL15) pseudogene |
| Y_RNA | 15 | Y RNA [Source:RFAM;Acc:RF00019] |
| AC015720.1 | 15 | ribosomal protein S17 (RPS17) pseudogene |
| PPCDC | 15 | phosphopantothenoylcysteine decarboxylase |
| AC113208.2 | 15 | stromal cell-derived factor 2 (SDF2) pseudoge |
| Metazoa_SRP | 15 | Metazoan signal recognition particle RNA [Sou |
| AC113208.1 | 15 | NULL |
| RPL36AP45 | 15 | ribosomal protein L36a pseudogene 45 [Sourc |
| C15orf39 | 15 | chromosome 15 open reading frame 39 [Sour |
| AC113208.4 | 15 | novel protein, antisense to C15orf39 |
| AC113208.5 | 15 | novel transcript |
| AC113208.3 | 15 | taste receptor cell protein 1-like [Source:NCBI |
| AC068338.1 | 15 | SIN3 homolog A, transcription regulator (yeas |
| NIFKP4 | 15 | nucleolar protein interacting with the FHA dor |
| GOLGA6C | 15 | golgin A6 family member C [Source:HGNC Sym |
| RN7SL489P | 15 | RNA, 7SL, cytoplasmic 489, pseudogene [Sour |
| GOLGA6D | 15 | golgin A6 family member D [Source:HGNC Sym |
| RN7SL327P | 15 | RNA, 7SL, cytoplasmic 327, pseudogene [Sour |
| DNM1P34 | 15 | dynamin 1 pseudogene 34 [Source:HGNC Sym |
| ANP32BP1 | 15 | acidic nuclear phosphoprotein 32 family mem |
| COMMD4 | 15 | COMM domain containing 4 [Source:HGNC Sy |
| AC068338.3 | 15 | novel transcript, antisense to NEIL1 |
| NEIL1 | 15 | nei like DNA glycosylase 1 [Source:HGNC Syml |
| MIR631 | 15 | microRNA 631 [Source:HGNC Symbol;Acc:HGN |
| MAN2C1 | 15 | mannosidase alpha class 2C member 1 [Sourc |
| AC068338.2 | 15 | novel transcript |
| SIN3A | 15 | SIN3 transcription regulator family member A |

| | | |
|---|---|---|
| RPL13P4 | 15 | ribosomal protein L13 pseudogene 4 [Source:I |
| AC105137.1 | 15 | high mobility group nucleosomal binding dom |
| AC105137.2 | 15 | novel transcript, sense intronic to SIN3A |
| PTPN9 | 15 | protein tyrosine phosphatase non-receptor ty |
| AC105036.2 | 15 | microtubule associated serine/threonine kina: |
| AC105036.1 | 15 | small nuclear ribonucleoprotein polypeptide F |
| AC105036.3 | 15 | novel transcript |
| SNUPN | 15 | snurportin 1 [Source:HGNC Symbol;Acc:HGNC |
| AC105020.3 | 15 | novel transcript, antisense to SNUPN |
| AC105020.2 | 15 | novel transcript, antisense to IMP3 |
| IMP3 | 15 | IMP U3 small nucleolar ribonucleoprotein 3 [S |
| AC105020.6 | 15 | novel transcript, antisense to IMP3 |
| AC105020.5 | 15 | novel transcript, antisense to IMP3 |
| SNX33 | 15 | sorting nexin 33 [Source:HGNC Symbol;Acc:H( |
| CSPG4 | 15 | chondroitin sulfate proteoglycan 4 [Source:H( |
| AC105020.4 | 15 | novel transcript, antisense to CSPG4 |
| AC105020.1 | 15 | novel transcript, antisense to CSPG4 |
| ODF3L1 | 15 | outer dense fiber of sperm tails 3 like 1 [Sourc |
| DNM1P35 | 15 | dynamin 1 pseudogene 35 [Source:HGNC Sym |
| AC019294.2 | 15 | novel transcript |
| PPIAP47 | 15 | peptidylprolyl isomerase A pseudogene 47 [Sc |
| AC019294.3 | 15 | novel transcript |
| MIR4313 | 15 | microRNA 4313 [Source:HGNC Symbol;Acc:H( |
| AC019294.1 | 15 | Golgin subfamily A member 6 pseudogene |
| RN7SL319P | 15 | RNA, 7SL, cytoplasmic 319, pseudogene [Sour |
| DNM1P49 | 15 | dynamin 1 pseudogene 49 [Source:HGNC Sym |
| UBE2Q2 | 15 | ubiquitin conjugating enzyme E2 Q2 [Source:H |
| RN7SL510P | 15 | RNA, 7SL, cytoplasmic 510, pseudogene [Sour |
| FBXO22 | 15 | F-box protein 22 [Source:HGNC Symbol;Acc:H |
| NRG4 | 15 | neuregulin 4 [Source:HGNC Symbol;Acc:HGNC |
| AC027104.1 | 15 | RWD domain containing 1 (RWDD1) pseudoge |
| TMEM266 | 15 | transmembrane protein 266 [Source:HGNC Sy |
| AC091100.1 | 15 | novel transcript, antisense to C15orf27 |
| ETFA | 15 | electron transfer flavoprotein subunit alpha [S |
| Metazoa_SRP | 15 | Metazoan signal recognition particle RNA [Sou |
| TYRO3P | 15 | TYRO3P protein tyrosine kinase pseudogene [ |
| AC027243.3 | 15 | novel transcript, antisense to ETFA |
| ISL2 | 15 | ISL LIM homeobox 2 [Source:HGNC Symbol;Ac |
| AC027243.1 | 15 | novel transcript, antisense to ISL2 |
| AC027243.2 | 15 | novel transcript |
| SCAPER | 15 | S-phase cyclin A associated protein in the ER [ |
| AC090751.1 | 15 | high mobility group nucleosomal binding dom |
| MIR3713 | 15 | microRNA 3713 [Source:HGNC Symbol;Acc:H( |
| RN7SKP217 | 15 | RN7SK pseudogene 217 [Source:HGNC Symbo |
| RCN2 | 15 | reticulocalbin 2 [Source:HGNC Symbol;Acc:HG |
| RN7SL278P | 15 | RNA, 7SL, cytoplasmic 278, pseudogene [Sour |
| KRT8P23 | 15 | keratin 8 pseudogene 23 [Source:HGNC Symb |
| PSTPIP1 | 15 | proline-serine-threonine phosphatase interac |
| TSPAN3 | 15 | tetraspanin 3 [Source:HGNC Symbol;Acc:HGN |
| AC090181.1 | 15 | novel transcript, antisense to TSPAN3 |

| | | |
|---|---|---|
| AC090181.3 | 15 | TEC |
| AC090181.2 | 15 | novel transcript, sense intronic to TSPAN3 |
| PEAK1 | 15 | pseudopodium enriched atypical kinase 1 [Sou |
| AC087465.1 | 15 | high mobility group nucleosome binding doma |
| AC060773.1 | 15 | high mobility group nucleosomal binding dom |
| HMG20A | 15 | high mobility group 20A [Source:HGNC Symbo |
| AC090984.1 | 15 | tec |
| AC046168.1 | 15 | novel transcript |
| AC046168.2 | 15 | novel transcript |
| LINGO1 | 15 | leucine rich repeat and Ig domain containing 1 |
| LINGO1-AS1 | 15 | LINGO1 antisense RNA 1 [Source:HGNC Symbo |
| LINGO1-AS2 | 15 | LINGO1 antisense RNA 2 [Source:HGNC Symbo |
| AC105133.2 | 15 | novel transcript |
| AC105133.1 | 15 | novel transcript, antisense to LINGO1 |
| CSPG4P13 | 15 | chondroitin sulfate proteoglycan 4 pseudogen |
| AC104758.3 | 15 | golgin A6 family, member A (GOLGA6A) pseuc |
| AC104758.4 | 15 | novel transcript |
| RN7SL214P | 15 | RNA, 7SL, cytoplasmic 214, pseudogene [Sour |
| AC104758.1 | 15 | COMM domain containing 4 (COMMD4) pseu |
| AC104758.5 | 15 | novel transcript |
| ADAMTS7P3 | 15 | ADAMTS7 pseudogene 3 [Source:HGNC Symb |
| TBC1D2B | 15 | TBC1 domain family member 2B [Source:HGN |
| AC104758.2 | 15 | novel transcript, antisense to TBC1D2B |
| SH2D7 | 15 | SH2 domain containing 7 [Source:HGNC Symb |
| SNORA63 | 15 | Small nucleolar RNA SNORA63 [Source:RFAM; |
| CIB2 | 15 | calcium and integrin binding family member 2 |
| IDH3A | 15 | isocitrate dehydrogenase (NAD(+)) 3 alpha [Sc |
| AC090260.1 | 15 | novel transcript |
| ACSBG1 | 15 | acyl-CoA synthetase bubblegum family memb |
| AC090607.3 | 15 | novel transcript, antisense to DNAJA4 |
| DNAJA4 | 15 | DnaJ heat shock protein family (Hsp40) memb |
| WDR61 | 15 | WD repeat domain 61 [Source:HGNC Symbol; |
| AC090607.4 | 15 | novel transcript , sense intronic to WDR61 |
| AC090607.2 | 15 | zinc finger CCHC-type and RNA binding motif 1 |
| AC090607.1 | 15 | novel transcript, antisense to WDR61 |
| AC090607.5 | 15 | novel transcript, antisense to WDR61 |
| AC011270.2 | 15 | TEC |
| CRABP1 | 15 | cellular retinoic acid binding protein 1 [Source |
| AC011270.1 | 15 | RWD domain containing 1 (RWDD1) pseudoge |
| IREB2 | 15 | iron responsive element binding protein 2 [So |
| AC027228.1 | 15 | novel transcript, antisense to IREB2 |
| HYKK | 15 | hydroxylysine kinase [Source:HGNC Symbol;A |
| AC027228.3 | 15 | novel transcript |
| PSMA4 | 15 | proteasome subunit alpha 4 [Source:HGNC Sy |
| CHRNA5 | 15 | cholinergic receptor nicotinic alpha 5 subunit |
| AC027228.2 | 15 | novel transcript, antisense to CHRNA5 |
| CHRNA3 | 15 | cholinergic receptor nicotinic alpha 3 subunit |
| AC067863.2 | 15 | TEC |
| CHRNB4 | 15 | cholinergic receptor nicotinic beta 4 subunit [S |
| AC067863.1 | 15 | novel transcript, antisense to CHRNB4 |

| | | |
|---|---|---|
| AC022748.2 | 15 | Golgin subfamily A pseudogene |
| RPL18P11 | 15 | ribosomal protein L18 pseudogene 11 [Source |
| AC022748.3 | 15 | dynamin 1 (DNM1) pseudogene |
| AC022748.1 | 15 | TBC1 domain family, member 2B (TBC1D2B) p |
| ADAMTS7 | 15 | ADAM metallopeptidase with thrombospondi |
| MORF4L1 | 15 | mortality factor 4 like 1 [Source:HGNC Symbo |
| RPL21P116 | 15 | ribosomal protein L21 pseudogene 116 [Sourc |
| RNU6-415P | 15 | RNA, U6 small nuclear 415, pseudogene [Sour |
| AC011944.2 | 15 | tec |
| CTSH | 15 | cathepsin H [Source:HGNC Symbol;Acc:HGNC: |
| RASGRF1 | 15 | Ras protein specific guanine nucleotide releas |
| AC011944.1 | 15 | novel transcript, antisense to RASGRF1 |
| ANKRD34C-AS1 | 15 | ANKRD34C antisense RNA 1 [Source:HGNC Syr |
| MIR184 | 15 | microRNA 184 [Source:HGNC Symbol;Acc:HGN |
| HNRNPCP3 | 15 | heterogeneous nuclear ribonucleoprotein C p |
| ANKRD34C | 15 | ankyrin repeat domain 34C [Source:HGNC Syn |
| TMED3 | 15 | transmembrane p24 trafficking protein 3 [Sou |
| AC027811.1 | 15 | novel transcript, antisense to TMED3 |
| MINAR1 | 15 | membrane integral NOTCH2 associated recep |
| AC023968.2 | 15 | transcription factor Dp-1 (TFDP1) pseudogene |
| AC023968.1 | 15 | ribosomal protein L7-like 1 (RPL7L1) pseudoge |
| AC026826.3 | 15 | chromosome 17 open reading frame 98 (C17o |
| AC026826.1 | 15 | ribosomal protein S12 (RPS12) pseudogene |
| AC026826.2 | 15 | zinc finger, FYVE domain containing 1 (ZFYVE1 |
| RNU6-667P | 15 | RNA, U6 small nuclear 667, pseudogene [Sour |
| AC021483.1 | 15 | novel transcript |
| MTHFS | 15 | methenyltetrahydrofolate synthetase [Source |
| AC021483.2 | 15 | novel transcript |
| ST20-MTHFS | 15 | ST20-MTHFS readthrough [Source:HGNC Syml |
| AC015871.5 | 15 | novel transcript |
| AC015871.7 | 15 | novel transcript, sense intronic to ST20-MTHF |
| AC015871.1 | 15 | Suppressor of tumorigenicity 20 protein [Sou |
| AC015871.4 | 15 | novel transcript, sense intronic to ST20 |
| ST20-AS1 | 15 | ST20 antisense RNA 1 [Source:HGNC Symbol;/ |
| AC015871.2 | 15 | farnesyl diphosphate synthase (FDRS) pseudo |
| BCL2A1 | 15 | BCL2 related protein A1 [Source:HGNC Symbo |
| AC015871.3 | 15 | ribonucleotide reductase M2 (RRM2) pseudog |
| AC015871.6 | 15 | novel transcript |
| AC092701.1 | 15 | novel transcript |
| Metazoa_SRP | 15 | Metazoan signal recognition particle RNA [Sou |
| ZFAND6 | 15 | zinc finger AN1-type containing 6 [Source:HGI |
| FAH | 15 | fumarylacetoacetate hydrolase [Source:HGNC |
| AC087761.1 | 15 | novel transcript |
| CTXND1 | 15 | cortexin domain containing 1 [Source:HGNC S |
| LINC00927 | 15 | long intergenic non-protein coding RNA 927 [S |
| AC016705.2 | 15 | novel transcript |
| ARNT2 | 15 | aryl hydrocarbon receptor nuclear translocato |
| AC016705.1 | 15 | novel transcript, antisense to ARNT2 |
| AC016705.3 | 15 | novel transcript, antisense to ARNT2 |
| AC108451.1 | 15 | novel transcript, antisense to ARNT2 |

| | | |
|---|---|---|
| AC108451.2 | 15 | novel transcript, antisense to ARNT2 |
| MIR5572 | 15 | microRNA 5572 [Source:HGNC Symbol;Acc:HG |
| RNU6-380P | 15 | RNA, U6 small nuclear 380, pseudogene [Sour |
| ABHD17C | 15 | abhydrolase domain containing 17C [Source:H |
| AC023302.1 | 15 | novel transcript |
| AC023302.2 | 15 | novel transcript |
| CEMIP | 15 | cell migration inducing hyaluronidase 1 [Sourc |
| MIR549A | 15 | microRNA 549a [Source:HGNC Symbol;Acc:HG |
| AC027808.1 | 15 | novel transcript, antisense to CEMIP |
| AC027808.2 | 15 | novel transcript, antisense to CEMIP |
| MESD | 15 | mesoderm development LRP chaperone [Sour |
| AC068870.1 | 15 | novel transcript |
| MIR4514 | 15 | microRNA 4514 [Source:HGNC Symbol;Acc:HG |
| AC068870.2 | 15 | novel transcript |
| TLNRD1 | 15 | talin rod domain containing 1 [Source:HGNC S |
| CFAP161 | 15 | cilia and flagella associated protein 161 [Sourc |
| AC068870.3 | 15 | tec |
| ANP32BP3 | 15 | acidic nuclear phosphoprotein 32 family mem |
| IL16 | 15 | interleukin 16 [Source:HGNC Symbol;Acc:HGN |
| AC103858.1 | 15 | novel transcript |
| STARD5 | 15 | StAR related lipid transfer domain containing |
| AC103858.3 | 15 | novel transcript, antisense to STARD5 |
| TMC3-AS1 | 15 | TMC3 antisense RNA 1 [Source:HGNC Symbol; |
| TMC3 | 15 | transmembrane channel like 3 [Source:HGNC |
| AC103858.2 | 15 | novel transcript, antisense to TMC3 |
| KF456110.1 | 15 | StAR related lipid transfer domain containing |
| AC109809.1 | 15 | novel transcript |
| AC060809.1 | 15 | novel transcript |
| AC023034.1 | 15 | novel transcript |
| AC104041.1 | 15 | novel transcript |
| AC026956.1 | 15 | protein (peptidylprolyl cis/trans isomerase) NI |
| MEX3B | 15 | mex-3 RNA binding family member B [Source: |
| LINC01583 | 15 | long intergenic non-protein coding RNA 1583 |
| AC026956.2 | 15 | novel transcript |
| EFL1 | 15 | elongation factor like GTPase 1 [Source:HGNC |
| RNU1-77P | 15 | RNA, U1 small nuclear 77, pseudogene [Sourc |
| AC026624.1 | 15 | novel transcript, antisense to EFL1 |
| SAXO2 | 15 | stabilizer of axonemal microtubules 2 [Source |
| ADAMTS7P1 | 15 | ADAMTS7 pseudogene 1 [Source:HGNC Symb |
| GOLGA6L10 | 15 | golgin A6 family like 10 [Source:HGNC Symbol |
| UBE2Q2P2 | 15 | ubiquitin conjugating enzyme E2 Q2 pseudoge |
| AC243919.1 | 15 | ribosomal protein L9 pseudogene 9 |
| AC243919.2 | 15 | novel transcript |
| GOLGA6L9 | 15 | golgin A6 family like 9 [Source:HGNC Symbol; |
| UBE2Q2P6 | 15 | ubiquitin conjugating enzyme E2 Q2 pseudoge |
| CSPG4P10 | 15 | chondroitin sulfate proteoglycan 4 pseudoger |
| AC245033.3 | 15 | dynamin 1 (DNM1) pseudogene |
| GOLGA2P10 | 15 | GOLGA2 pseudogene 10 [Source:HGNC Symbo |
| GOLGA6L17P | 15 | golgin A6 family like 17, pseudogene [Source:I |
| DNM1P38 | 15 | dynamin 1 pseudogene 38 [Source:HGNC Sym |

| | | |
|---|---|---|
| AC245033.4 | 15 | novel transcript |
| RPS17 | 15 | ribosomal protein S17 [Source:HGNC Symbol;/ |
| AC245033.1 | 15 | novel protein |
| AC245033.2 | 15 | novel transcript, antisense to CPEB1 |
| CPEB1 | 15 | cytoplasmic polyadenylation element binding |
| CPEB1-AS1 | 15 | CPEB1 antisense RNA 1 [Source:HGNC Symbol |
| AP3B2 | 15 | adaptor related protein complex 3 subunit be |
| AC105339.4 | 15 | serine/arginine-rich splicing factor 9 (SRSF9) p |
| AC105339.1 | 15 | novel pseudogene |
| ACTG1P17 | 15 | actin gamma 1 pseudogene 17 [Source:HGNC |
| AC105339.5 | 15 | tec |
| AC105339.3 | 15 | novel transcript, antisense FSD2 |
| AC105339.6 | 15 | novel transcript |
| SNHG21 | 15 | small nucleolar RNA host gene 21 [Source:HG |
| AC105339.2 | 15 | TEC |
| FSD2 | 15 | fibronectin type III and SPRY domain containir |
| SCARNA15 | 15 | small Cajal body-specific RNA 15 [Source:HGN |
| WHAMM | 15 | WASP homolog associated with actin, golgi m |
| HOMER2 | 15 | homer scaffold protein 2 [Source:HGNC Symb |
| AC022558.2 | 15 | novel transcript, antisense to HOMER2, C15or |
| RAMAC | 15 | RNA guanine-7 methyltransferase activating s |
| C15orf40 | 15 | chromosome 15 open reading frame 40 [Sour |
| AC024270.4 | 15 | novel transcript, antisense to BTBD1 |
| BTBD1 | 15 | BTB domain containing 1 [Source:HGNC Symb |
| AC022558.3 | 15 | novel transcript, antisense to BTBD1 |
| AC022558.1 | 15 | novel transcript, sense intronic to BTBD1 |
| AC024270.2 | 15 | novel transcript, antisense to BTBD1 |
| AC024270.1 | 15 | fatty acid binding protein 5 (psoriasis-associat |
| MIR4515 | 15 | microRNA 4515 [Source:HGNC Symbol;Acc:HG |
| AC024270.5 | 15 | novel transcript, antisense to TM6SF1 |
| TM6SF1 | 15 | transmembrane 6 superfamily member 1 [Sou |
| HDGFL3 | 15 | HDGF like 3 [Source:HGNC Symbol;Acc:HGNC: |
| AC024270.3 | 15 | novel transcript, antisense to TM6SF1 |
| AC103876.1 | 15 | novel transcript, antisense to HDGFRP3 |
| BNC1 | 15 | basonuclin 1 [Source:HGNC Symbol;Acc:HGNC |
| SH3GL3 | 15 | SH3 domain containing GRB2 like 3, endophili |
| ADAMTSL3 | 15 | ADAMTS like 3 [Source:HGNC Symbol;Acc:HG |
| RNU6-401P | 15 | RNA, U6 small nuclear 401, pseudogene [Sour |
| TUBAP4 | 15 | tubulin alpha pseudogene 4 [Source:HGNC Syı |
| RNU6-1339P | 15 | RNA, U6 small nuclear 1339, pseudogene [Sou |
| AC027807.2 | 15 | novel transcript, sense intronic to ADAMTSL3 |
| AC027807.1 | 15 | family with sequence similarity 154, member |
| EFL1P1 | 15 | elongation factor like GTPase 1 pseudogene 1 |
| DNM1P41 | 15 | dynamin 1 pseudogene 41 [Source:HGNC Sym |
| AC136698.1 | 15 | novel transcript |
| UBE2Q2L | 15 | ubiquitin conjugating enzyme E2 Q2 like [Sour |
| CSPG4P11 | 15 | chondroitin sulfate proteoglycan 4 pseudoger |
| GOLGA2P7 | 15 | GOLGA2 pseudogene 7 [Source:HGNC Symbol |
| RN7SL331P | 15 | RNA, 7SL, cytoplasmic 331, pseudogene [Sour |
| GOLGA6L4 | 15 | golgin A6 family like 4 [Source:HGNC Symbol;/ |

| | | |
|---|---|---|
| UBE2Q2P8 | 15 | ubiquitin conjugating enzyme E2 Q2 pseudoge |
| AC243562.2 | 15 | golgin A6 family-like 9 (GOLGA6L9) pseudoger |
| RN7SL417P | 15 | RNA, 7SL, cytoplasmic 417, pseudogene [Sour |
| AC243562.1 | 15 | golgin subfamily A member 2-like (AGSK1) pse |
| DNM1P51 | 15 | dynamin 1 pseudogene 51 [Source:HGNC Sym |
| CSPG4P5 | 15 | chondroitin sulfate proteoglycan 4 pseudoger |
| UBE2Q2P11 | 15 | ubiquitin conjugating enzyme E2 Q2 pseudoge |
| AC243562.3 | 15 | novel transcript |
| Metazoa_SRP | 15 | Metazoan signal recognition particle RNA [Sou |
| UBE2Q2P12 | 15 | ubiquitin conjugating enzyme E2 Q2 pseudoge |
| AC048382.4 | 15 | novel transcript |
| AC048382.3 | 15 | SIN3 homolog A, transcription regulator (yeas |
| GOLGA6L5P | 15 | golgin A6 family like 5, pseudogene [Source:H |
| AC048382.2 | 15 | novel transcript |
| UBE2Q2P1 | 15 | ubiquitin conjugating enzyme E2 Q2 pseudoge |
| LINC00933 | 15 | long intergenic non-protein coding RNA 933 [S |
| AC048382.5 | 15 | novel transcript, antisense to ZSCAN2 |
| ZSCAN2 | 15 | zinc finger and SCAN domain containing 2 [Sou |
| AC048382.1 | 15 | novel transcript, antisense to ZSCAN2 |
| AC048382.6 | 15 | novel transcript, antisense to ZSCAN2 |
| SCAND2P | 15 | SCAN domain containing 2 pseudogene [Sourc |
| EGLN1P1 | 15 | egl-9 family hypoxia inducible factor 1 pseudo |
| WDR73 | 15 | WD repeat domain 73 [Source:HGNC Symbol;. |
| Metazoa_SRP | 15 | Metazoan signal recognition particle RNA [Sou |
| NMB | 15 | neuromedin B [Source:HGNC Symbol;Acc:HGN |
| SEC11A | 15 | SEC11 homolog A, signal peptidase complex su |
| AC115102.1 | 15 | novel transcript, antisense to SEC11A |
| AC012291.3 | 15 | novel transcript |
| ZNF592 | 15 | zinc finger protein 592 [Source:HGNC Symbol; |
| AC012291.2 | 15 | novel transcript |
| ALPK3 | 15 | alpha kinase 3 [Source:HGNC Symbol;Acc:HGN |
| AC012291.1 | 15 | NULL |
| SLC28A1 | 15 | solute carrier family 28 member 1 [Source:HG |
| AC087468.2 | 15 | novel transcript, antisense to SLC28A1 |
| RNU6-339P | 15 | RNA, U6 small nuclear 339, pseudogene [Sour |
| RNU6-796P | 15 | RNA, U6 small nuclear 796, pseudogene [Sour |
| AC087468.1 | 15 | high mobility group box 2 (HMGB2) pseudoge |
| PDE8A | 15 | phosphodiesterase 8A [Source:HGNC Symbol; |
| AC044860.2 | 15 | SIN3 homolog A, transcription regulator (yeas |
| AC044860.1 | 15 | golgin subfamily A pseudogene |
| NIFKP8 | 15 | nucleolar protein interacting with the FHA dor |
| CSPG4P12 | 15 | chondroitin sulfate proteoglycan 4 pseudoger |
| RN7SL428P | 15 | RNA, 7SL, cytoplasmic 428, pseudogene [Sour |
| GOLGA6L3 | 15 | golgin A6 family like 3 [Source:HGNC Symbol;/ |
| ADAMTS7P4 | 15 | ADAMTS7 pseudogene 4 [Source:HGNC Symb |
| AC044860.3 | 15 | SNW domain containing 1 (SNW1) pseudogen |
| AKAP13 | 15 | A-kinase anchoring protein 13 [Source:HGNC S |
| MIR7706 | 15 | microRNA 7706 [Source:HGNC Symbol;Acc:HG |
| RNU7-79P | 15 | RNA, U7 small nuclear 79 pseudogene [Source |
| AC104046.1 | 15 | fatty acid binding protein 5 (psoriasis-associat |

| | | |
|---|---|---|
| AC087286.3 | 15 | novel transcript, antisense to AKAP13 |
| AC087286.2 | 15 | novel transcript, antisense to AKAP13 |
| AC087286.1 | 15 | novel transcript, antisense to AKAP13 |
| RNU6-1280P | 15 | RNA, U6 small nuclear 1280, pseudogene [Sou |
| AC087286.4 | 15 | novel transcript, sense intronic to AKAP13 |
| AC021739.4 | 15 | novel transcript, antisense to AKAP13 |
| AC021739.2 | 15 | novel transcript, antisense to AKAP13 |
| AC021739.5 | 15 | novel transcript, antisense to AKAP13 |
| AC021739.3 | 15 | novel transcript, antisense to AKAP13 |
| KLHL25 | 15 | kelch like family member 25 [Source:HGNC Sy |
| MIR1276 | 15 | microRNA 1276 [Source:HGNC Symbol;Acc:HG |
| AC021739.1 | 15 | ribosomal protein S2 (RPS2) pseudogene |
| RNU6-231P | 15 | RNA, U6 small nuclear 231, pseudogene [Sour |
| MIR548AP | 15 | microRNA 548ap [Source:HGNC Symbol;Acc:H |
| AC104229.1 | 15 | ADP-ribosylation factor-like 2 binding protein |
| LINC01584 | 15 | long intergenic non-protein coding RNA 1584 |
| AGBL1 | 15 | ATP/GTP binding protein like 1 [Source:HGNC |
| RNA5SP400 | 15 | RNA, 5S ribosomal pseudogene 400 [Source:H |
| AC016180.2 | 15 | ribosomal protein L18a (RPL18A) pseudogene |
| AC016180.1 | 15 | NULL |
| AGBL1-AS1 | 15 | AGBL1 antisense RNA 1 [Source:HGNC Symbo |
| AC012229.1 | 15 | novel transcript, antisense to AGBL1 |
| AC016987.2 | 15 | novel transcript |
| AC016987.1 | 15 | novel transcript, antisense to AGBL1 |
| AC078905.1 | 15 | novel transcript |
| AC020687.1 | 15 | novel transcript |
| RNU6-185P | 15 | RNA, U6 small nuclear 185, pseudogene [Sour |
| LINC00052 | 15 | long intergenic non-protein coding RNA 52 [So |
| AC103871.1 | 15 | novel transcript |
| NTRK3 | 15 | neurotrophic receptor tyrosine kinase 3 [Sour |
| MED28P6 | 15 | mediator complex subunit 28 pseudogene 6 [S |
| NTRK3-AS1 | 15 | NTRK3 antisense RNA 1 [Source:HGNC Symbo |
| MRPL46 | 15 | mitochondrial ribosomal protein L46 [Source:I |
| AC013489.1 | 15 | novel transcript |
| MRPS11 | 15 | mitochondrial ribosomal protein S11 [Source: |
| AC013489.3 | 15 | novel transcript |
| DET1 | 15 | DET1 partner of COP1 E3 ubiquitin ligase [Sou |
| AC013489.2 | 15 | dolichyl-phosphate mannosyltransferase poly |
| LINC01586 | 15 | long intergenic non-protein coding RNA 1586 |
| MIR1179 | 15 | microRNA 1179 [Source:HGNC Symbol;Acc:HG |
| MIR7-2 | 15 | microRNA 7-2 [Source:HGNC Symbol;Acc:HGN |
| MIR3529 | 15 | microRNA 3529 [Source:HGNC Symbol;Acc:HG |
| AEN | 15 | apoptosis enhancing nuclease [Source:HGNC S |
| ISG20 | 15 | interferon stimulated exonuclease gene 20 [So |
| AC103982.1 | 15 | novel transcript |
| ACAN | 15 | aggrecan [Source:HGNC Symbol;Acc:HGNC:31 |
| HAPLN3 | 15 | hyaluronan and proteoglycan link protein 3 [S |
| MFGE8 | 15 | milk fat globule-EGF factor 8 protein [Source:I |
| AC067805.1 | 15 | high mobility group nucleosome binding doma |
| KRT18P47 | 15 | keratin 18 pseudogene 47 [Source:HGNC Sym |

| | | |
|---|---|---|
| AC107954.1 | 15 | capping protein (actin filament) muscle Z-line, |
| AC013565.2 | 15 | intraflagellar transport 20 homolog (Chlamydo |
| AC013565.1 | 15 | novel transcript |
| AC013565.3 | 15 | novel transcript, antisense to ABHD2 |
| ABHD2 | 15 | abhydrolase domain containing 2 [Source:HGI |
| RNU7-195P | 15 | RNA, U7 small nuclear 195 pseudogene [Sourc |
| HMGB1P8 | 15 | high mobility group box 1 pseudogene 8 [Sour |
| AC124068.1 | 15 | VAMP (vesicle-associated membrane protein) |
| RLBP1 | 15 | retinaldehyde binding protein 1 [Source:HGN |
| FANCI | 15 | FA complementation group I [Source:HGNC Sy |
| POLG | 15 | DNA polymerase gamma, catalytic subunit [Sc |
| MIR6766 | 15 | microRNA 6766 [Source:HGNC Symbol;Acc:HG |
| AC124068.2 | 15 | novel transcript, antisense to POLG |
| AC133637.1 | 15 | tec |
| AC133637.2 | 15 | novel transcript |
| MIR9-3HG | 15 | MIR9-3 host gene [Source:HGNC Symbol;Acc:I |
| MIR9-3 | 15 | microRNA 9-3 [Source:HGNC Symbol;Acc:HGN |
| RHCG | 15 | Rh family C glycoprotein [Source:HGNC Symbo |
| LINC00928 | 15 | long intergenic non-protein coding RNA 928 [S |
| AC013391.3 | 15 | novel transcript |
| AC013391.2 | 15 | novel transcript |
| TICRR | 15 | TOPBP1 interacting checkpoint and replicatior |
| AC013391.1 | 15 | novel transcript, antisense to TICRR |
| KIF7 | 15 | kinesin family member 7 [Source:HGNC Symb |
| PLIN1 | 15 | perilipin 1 [Source:HGNC Symbol;Acc:HGNC:9 |
| PEX11A | 15 | peroxisomal biogenesis factor 11 alpha [Sourc |
| RPL36AP43 | 15 | ribosomal protein L36a pseudogene 43 [Sourc |
| WDR93 | 15 | WD repeat domain 93 [Source:HGNC Symbol; |
| AC079075.1 | 15 | novel transcript |
| RNU6-132P | 15 | RNA, U6 small nuclear 132, pseudogene [Sour |
| MESP1 | 15 | mesoderm posterior bHLH transcription facto |
| MRPL15P1 | 15 | mitochondrial ribosomal protein L15 pseudog |
| MESP2 | 15 | mesoderm posterior bHLH transcription facto |
| ANPEP | 15 | alanyl aminopeptidase, membrane [Source:HC |
| AP3S2 | 15 | adaptor related protein complex 3 subunit sig |
| ARPIN-AP3S2 | 15 | ARPIN-AP3S2 readthrough [Source:HGNC Sym |
| MIR5094 | 15 | microRNA 5094 [Source:HGNC Symbol;Acc:HG |
| AC027176.2 | 15 | Arginine/serine-rich coiled-coil protein (RSRC2 |
| MIR5009 | 15 | microRNA 5009 [Source:HGNC Symbol;Acc:HG |
| RNU6-1111P | 15 | RNA, U6 small nuclear 1111, pseudogene [Sou |
| ARPIN | 15 | actin related protein 2/3 complex inhibitor [Sc |
| RNU7-111P | 15 | RNA, U7 small nuclear 111 pseudogene [Sourc |
| AC027176.1 | 15 | zinc finger and SCAN domain containing 5A (Z: |
| AC027176.3 | 15 | tec |
| ZNF710 | 15 | zinc finger protein 710 [Source:HGNC Symbol; |
| MIR3174 | 15 | microRNA 3174 [Source:HGNC Symbol;Acc:HG |
| AC087284.1 | 15 | novel transcript, sense intronic to ZNF710 |
| ZNF710-AS1 | 15 | ZNF710 antisense RNA 1 [Source:HGNC Symbo |
| U7 | 15 | U7 small nuclear RNA [Source:RFAM;Acc:RF00 |
| IDH2 | 15 | isocitrate dehydrogenase (NADP(+)) 2, mitoch |

| | | |
|---|---|---|
| IDH2-DT | 15 | IDH2 divergent transcript [Source:HGNC Symb |
| SEMA4B | 15 | semaphorin 4B [Source:HGNC Symbol;Acc:HG |
| RN7SL346P | 15 | RNA, 7SL, cytoplasmic 346, pseudogene [Sour |
| RPS12P26 | 15 | ribosomal protein S12 pseudogene 26 [Source |
| CIB1 | 15 | calcium and integrin binding 1 [Source:HGNC S |
| GDPGP1 | 15 | GDP-D-glucose phosphorylase 1 [Source:HGN( |
| AC091167.7 | 15 | novel protein |
| TTLL13P | 15 | tubulin tyrosine ligase like 13, pseudogene [Sc |
| AC091167.2 | 15 | novel protein |
| NGRN | 15 | neugrin, neurite outgrowth associated [Source |
| AC091167.6 | 15 | novel protein |
| AC091167.1 | 15 | PERP, TP53 apoptosis effector (PERP) pseudog |
| AC091167.3 | 15 | SIN3 transcription regulator homolog A (yeast |
| AC091167.5 | 15 | novel transcript |
| AC091167.4 | 15 | MKI67 (FHA domain) interacting nucleolar pho |
| GOLGA2P8 | 15 | GOLGA2 pseudogene 8 [Source:HGNC Symbol |
| RN7SL736P | 15 | RNA, 7SL, cytoplasmic 736, pseudogene [Sour |
| AC018946.2 | 15 | TEC |
| GABARAPL3 | 15 | GABA type A receptor associated protein like |
| ZNF774 | 15 | zinc finger protein 774 [Source:HGNC Symbol; |
| NDUFA3P4 | 15 | NADH:ubiquinone oxidoreductase subunit A3 |
| IQGAP1 | 15 | IQ motif containing GTPase activating protein |
| AC018946.1 | 15 | novel transcript, antisense to IQGAP1 |
| CRTC3 | 15 | CREB regulated transcription coactivator 3 [Sc |
| AC103739.1 | 15 | novel transcript, antisense to CRTC3 |
| AC103739.2 | 15 | novel transcript, antisense to CRTC3 |
| CRTC3-AS1 | 15 | CRTC3 antisense RNA 1 [Source:HGNC Symbol |
| AC103739.3 | 15 | novel transcript, antisense to CRTC3 |
| HSPE1P3 | 15 | heat shock protein family E (Hsp10) member : |
| AC021422.1 | 15 | novel transcript |
| LINC01585 | 15 | long intergenic non-protein coding RNA 1585 |
| BLM | 15 | BLM RecQ like helicase [Source:HGNC Symbol |
| SNORD18 | 15 | Small nucleolar RNA SNORD18 [Source:RFAM; |
| AC124248.2 | 15 | novel transcript, antisense to BLM |
| AC124248.1 | 15 | novel transcript, sense overlapping 3' UTR of I |
| RN7SL363P | 15 | RNA, 7SL, cytoplasmic 363, pseudogene [Sour |
| FURIN | 15 | furin, paired basic amino acid cleaving enzyme |
| FES | 15 | FES proto-oncogene, tyrosine kinase [Source:I |
| MAN2A2 | 15 | mannosidase alpha class 2A member 2 [Sourc |
| AC068831.4 | 15 | novel transcript, antisense to MAN2A2 |
| HDDC3 | 15 | HD domain containing 3 [Source:HGNC Symbo |
| UNC45A | 15 | unc-45 myosin chaperone A [Source:HGNC Syı |
| AC068831.1 | 15 | novel transcript, antisense to UNC45A and RC |
| RCCD1 | 15 | RCC1 domain containing 1 [Source:HGNC Sym |
| PRC1 | 15 | protein regulator of cytokinesis 1 [Source:HGI |
| AC068831.7 | 15 | novel protein |
| PRC1-AS1 | 15 | PRC1 antisense RNA 1 [Source:HGNC Symbol;/ |
| Y_RNA | 15 | Y RNA [Source:RFAM;Acc:RF00019] |
| VPS33B | 15 | VPS33B late endosome and lysosome associat |
| VPS33B-DT | 15 | VPS33B divergent transcript [Source:HGNC Sy |

| | | |
|---|---|---|
| AC068831.6 | 15 | novel transcript |
| AC068831.2 | 15 | chromosome 14 open reading frame 156 (C14 |
| AC068831.5 | 15 | SNW domain containing 1 (SNW1) pseudogen |
| AC068831.3 | 15 | golgin A5 (GOLGA5) pseudogene |
| AC127520.1 | 15 | novel transcript |
| SV2B | 15 | synaptic vesicle glycoprotein 2B [Source:HGN( |
| AC123784.1 | 15 | novel transcript |
| AC104035.1 | 15 | novel transcript |
| AC107958.3 | 15 | novel transcript |
| AC107958.2 | 15 | novel transcript |
| AC107958.1 | 15 | destrin (actin depolymerizing factor) (DSTN) p |
| AC105109.1 | 15 | thyroid hormone receptor associated protein |
| SLCO3A1 | 15 | solute carrier organic anion transporter family |
| AC116903.2 | 15 | novel transcript, antisense to SLCO3A1 |
| AC116903.1 | 15 | novel transcript, antisense to SLCO3A1 |
| DUXAP6 | 15 | double homeobox A pseudogene 6 [Source:H( |
| NPM1P5 | 15 | nucleophosmin 1 pseudogene 5 [Source:HGN( |
| ST8SIA2 | 15 | ST8 alpha-N-acetyl-neuraminide alpha-2,8-sia |
| AC090985.1 | 15 | NULL |
| ENO1P2 | 15 | enolase 1 pseudogene 2 [Source:HGNC Symbo |
| AC090985.2 | 15 | novel transcript, antisense to ST8SIA2 |
| C15orf32 | 15 | chromosome 15 open reading frame 32 [Sour |
| LINC00930 | 15 | long intergenic non-protein coding RNA 930 [S |
| AC091544.2 | 15 | novel transcript |
| AC091544.4 | 15 | novel transcript |
| AC091544.5 | 15 | novel transcript, antisense to FAM174B |
| FAM174B | 15 | family with sequence similarity 174 member E |
| AC091544.3 | 15 | late endosomal/lysosomal adaptor, MAPK and |
| AC091544.7 | 15 | novel transcript, antisense to FAM174B |
| AC091544.6 | 15 | novel transcript |
| HMGN1P38 | 15 | high mobility group nucleosome binding doma |
| H2AFVP1 | 15 | H2A histone family member V pseudogene 1 [ |
| AC091544.1 | 15 | ribosomal protein L31 (RPL31) pseudogene |
| AC106028.1 | 15 | erritin, light polypeptide (FTL) pseudogene |
| AC106028.3 | 15 | novel transcript |
| ASB9P1 | 15 | ankyrin repeat and SOCS box containing 9 pse |
| AC106028.2 | 15 | novel transcript |
| AC106028.4 | 15 | novel transcript |
| LINC01578 | 15 | long intergenic non-protein coding RNA 1578 |
| AC013394.1 | 15 | novel protein |
| CHD2 | 15 | chromodomain helicase DNA binding protein |
| MIR3175 | 15 | microRNA 3175 [Source:HGNC Symbol;Acc:HG |
| RGMA | 15 | repulsive guidance molecule BMP co-receptor |
| AC087641.1 | 15 | Y box binding protein 2 (YBX2) pseudogene |
| AC108457.1 | 15 | novel transcript |
| AC091078.1 | 15 | novel transcript |
| AC112693.1 | 15 | novel transcript |
| AC112693.2 | 15 | spermatogenesis associated 16 (SPATA16) pse |
| AC091078.2 | 15 | novel transcript |
| SEPHS1P2 | 15 | selenophosphate synthetase 1 pseudogene 2 |

| | | |
|---|---|---|
| AC110023.1 | 15 | novel transcript |
| LINC01579 | 15 | long intergenic non-protein coding RNA 1579 |
| AC103996.1 | 15 | ribosomal protein L39 (RPL39) |
| LINC02207 | 15 | long intergenic non-protein coding RNA 2207 |
| AC103996.3 | 15 | novel transcript |
| AC103996.2 | 15 | novel transcript |
| LINC01580 | 15 | long intergenic non-protein coding RNA 1580 |
| LINC01581 | 15 | long intergenic non-protein coding RNA 1581 |
| AC104390.1 | 15 | poly(A) binding protein, cytoplasmic 1 (PABPC |
| AC110027.1 | 15 | H3 histone, family 3A (H3F3A) pseudogene |
| MCTP2 | 15 | multiple C2 and transmembrane domain cont |
| AC135626.1 | 15 | ribosomal protein L26-like 1 (RPL26L1) pseudo |
| AC135626.2 | 15 | multiple C2 domains, transmembrane 21 (MC |
| AC009432.2 | 15 | novel transcript, antisense to MCTP2 |
| LINC02852 | 15 | long intergenic non-protein coding RNA 2852 |
| AC009432.1 | 15 | novel transcript |
| AC022308.1 | 15 | novel transcript |
| AC087633.2 | 15 | glioma tumor suppressor candidate region ge |
| AC107976.1 | 15 | novel transcript |
| AC087633.1 | 15 | novel transcript |
| AC087636.1 | 15 | novel transcript |
| LINC01197 | 15 | long intergenic non-protein coding RNA 1197 |
| AC104260.1 | 15 | novel transcript |
| AC104260.2 | 15 | TEC |
| AC104260.3 | 15 | novel transcript |
| LINC00924 | 15 | long intergenic non-protein coding RNA 924 [S |
| AC027013.1 | 15 | novel transcript |
| AC015574.1 | 15 | novel transcript |
| RNU2-3P | 15 | RNA, U2 small nuclear 3, pseudogene [Source |
| AC012409.2 | 15 | novel transcript |
| AC024337.1 | 15 | novel transcript |
| AC024337.2 | 15 | novel transcript |
| NR2F2-AS1 | 15 | NR2F2 antisense RNA 1 [Source:HGNC Symbo |
| AC012409.5 | 15 | TEC |
| AC012409.3 | 15 | novel transcript |
| AC012409.1 | 15 | novel transcript |
| AC012409.4 | 15 | novel transcript |
| LINC02157 | 15 | long intergenic non-protein coding RNA 2157 |
| AC016251.1 | 15 | TEC |
| NR2F2 | 15 | nuclear receptor subfamily 2 group F member |
| MIR1469 | 15 | microRNA 1469 [Source:HGNC Symbol;Acc:HG |
| AC103746.1 | 15 | novel transcript |
| AC087477.2 | 15 | novel transcript |
| AC087477.6 | 15 | tubulin, alpha 1a (TUBA1A) pseudogene |
| AC087477.5 | 15 | novel transcript |
| AC087477.3 | 15 | novel transcript |
| PGAM1P12 | 15 | phosphoglycerate mutase 1 pseudogene 12 [S |
| AC087477.1 | 15 | ribosomal protein L31 (RPL31) pseudogene |
| AC087477.4 | 15 | novel transcript |
| RN7SKP254 | 15 | RN7SK pseudogene 254 [Source:HGNC Symbo |

| | | |
|---|---|---|
| Metazoa_SRP | 15 | Metazoan signal recognition particle RNA [Sou |
| AC110588.1 | 15 | CDC28 protein kinase regulatory subunit 2 (CK |
| FAM149B1P1 | 15 | family with sequence similarity 149 member E |
| SPATA8-AS1 | 15 | SPATA8 antisense RNA 1 (head to head) [Sour |
| SPATA8 | 15 | spermatogenesis associated 8 [Source:HGNC S |
| RN7SKP181 | 15 | RN7SK pseudogene 181 [Source:HGNC Symbo |
| LINC02253 | 15 | long intergenic non-protein coding RNA 2253 |
| AC020704.1 | 15 | novel transcript |
| LINC02254 | 15 | long intergenic non-protein coding RNA 2254 |
| RN7SL677P | 15 | RNA, 7SL, cytoplasmic 677, pseudogene [Sour |
| RNA5SP401 | 15 | RNA, 5S ribosomal pseudogene 401 [Source:H |
| AC026523.2 | 15 | novel transcript |
| AC026523.1 | 15 | novel transcript |
| AC026523.3 | 15 | novel transcript |
| AC026523.4 | 15 | novel transcript |
| LINC00923 | 15 | long intergenic non-protein coding RNA 923 [S |
| AC024651.2 | 15 | novel transcript |
| AC024651.3 | 15 | novel transcript |
| ARRDC4 | 15 | arrestin domain containing 4 [Source:HGNC Sy |
| AC024651.1 | 15 | novel transcript |
| LINC02251 | 15 | long intergenic non-protein coding RNA 2251 |
| U6 | 15 | U6 spliceosomal RNA [Source:RFAM;Acc:RF00 |
| AC022523.1 | 15 | novel transcript |
| RNU6-1186P | 15 | RNA, U6 small nuclear 1186, pseudogene [Sou |
| LINC01582 | 15 | long intergenic non-protein coding RNA 1582 |
| AC022523.3 | 15 | novel transcript |
| AC022523.2 | 15 | S phase cyclin A-associated protein in the end |
| AC015722.1 | 15 | novel transcript |
| AC015722.2 | 15 | novel transcript |
| LINC02351 | 15 | long intergenic non-protein coding RNA 2351 |
| FAM169B | 15 | family with sequence similarity 169 member E |
| AC103968.1 | 15 | zinc finger CCHC domain-containing protein 9 |
| IRAIN | 15 | IGF1R antisense imprinted non-protein coding |
| IGF1R | 15 | insulin like growth factor 1 receptor [Source:H |
| AC118658.1 | 15 | novel transcript, sense intronic to IGF1R |
| MIR4714 | 15 | microRNA 4714 [Source:HGNC Symbol;Acc:HG |
| AC055807.1 | 15 | novel transcript, antisense to IGF1R |
| AC069029.1 | 15 | novel transcript, antisense to IGF1R |
| AC036108.1 | 15 | novel transcript, antisense to IGF1R and SYNM |
| PGPEP1L | 15 | pyroglutamyl-peptidase I like [Source:HGNC Sy |
| LUNAR1 | 15 | leukemia-associated non-coding IGF1R activat |
| SYNM | 15 | synemin [Source:HGNC Symbol;Acc:HGNC:244 |
| AC036108.2 | 15 | novel transcript, antisense to SYNM |
| TTC23 | 15 | tetratricopeptide repeat domain 23 [Source:H |
| AC036108.3 | 15 | novel transcript, antisense to TTC23 |
| AC022819.1 | 15 | novel transcript, sense intronic to TTC23 |
| LRRC28 | 15 | leucine rich repeat containing 28 [Source:HGN |
| HSP90B2P | 15 | heat shock protein 90 beta family member 2, |
| HNRNPA1P62 | 15 | heterogeneous nuclear ribonucleoprotein A1 |
| AC037479.1 | 15 | Coiled-coil domain-containing protein 117 (CC |

| | | |
|---|---|---|
| LINC02244 | 15 | long intergenic non-protein coding RNA 2244 |
| AC015660.2 | 15 | novel transcript, antisense to LRRC28 |
| AC015660.3 | 15 | novel transcript |
| AC015660.1 | 15 | novel transcript |
| AC015660.5 | 15 | novel transcript |
| AC015660.4 | 15 | TEC |
| RPL7P5 | 15 | ribosomal protein L7 pseudogene 5 [Source:H |
| MEF2A | 15 | myocyte enhancer factor 2A [Source:HGNC Sy |
| Y_RNA | 15 | Y RNA [Source:RFAM;Acc:RF00019] |
| LYSMD4 | 15 | LysM domain containing 4 [Source:HGNC Sym |
| DNM1P46 | 15 | dynamin 1 pseudogene 46 [Source:HGNC Sym |
| AC090825.2 | 15 | golgin subfamily A (GOLGA) pseudogene |
| RN7SL484P | 15 | RNA, 7SL, cytoplasmic 484, pseudogene [Sour |
| AC090825.1 | 15 | novel transcript |
| AC084855.1 | 15 | novel transcript, antisense to ADAMTS17 |
| ADAMTS17 | 15 | ADAM metallopeptidase with thrombospondi |
| AC084855.2 | 15 | novel transcript, antisense to ADAMTS17 |
| RNA5SP402 | 15 | RNA, 5S ribosomal pseudogene 402 [Source:H |
| AC022710.1 | 15 | novel transcript, antisense to ADAMTS17 |
| SPATA41 | 15 | spermatogenesis associated 41 [Source:HGNC |
| CERS3-AS1 | 15 | CERS3 antisense RNA 1 [Source:HGNC Symbol |
| CERS3 | 15 | ceramide synthase 3 [Source:HGNC Symbol;A |
| AC027020.2 | 15 | novel transcript |
| RNU6-322P | 15 | RNA, U6 small nuclear 322, pseudogene [Sour |
| PRKXP1 | 15 | PRKX pseudogene 1 [Source:HGNC Symbol;Ac |
| AC027020.1 | 15 | novel transcript, antisense to LINS1 |
| LINS1 | 15 | lines homolog 1 [Source:HGNC Symbol;Acc:H |
| RNU6-181P | 15 | RNA, U6 small nuclear 181, pseudogene [Sour |
| ASB7 | 15 | ankyrin repeat and SOCS box containing 7 [So |
| Y_RNA | 15 | Y RNA [Source:RFAM;Acc:RF00019] |
| AC090695.1 | 15 | WD repeat domain 12 (WDR12) pseudogene |
| AC087762.1 | 15 | novel transcript |
| AC087762.2 | 15 | TEC |
| AC015712.3 | 15 | PHD finger protein 5A (PHF5A) pseudogene |
| AC015712.1 | 15 | novel transcript |
| AC015712.6 | 15 | novel transcript |
| AC015712.4 | 15 | novel transcript |
| AC015712.5 | 15 | novel transcript |
| ALDH1A3 | 15 | aldehyde dehydrogenase 1 family member A3 |
| AC015712.7 | 15 | novel transcript, antisense to ALDH1A3 |
| AC015712.2 | 15 | novel transcript, antisense to LRRK1 and ALDH |
| LRRK1 | 15 | leucine rich repeat kinase 1 [Source:HGNC Syr |
| AC090907.1 | 15 | novel transcript, antisense to LRRK1 |
| AC090907.2 | 15 | novel transcript, antisense to LRRK1 |
| AC090907.3 | 15 | novel transcript, antisense to LRRK1 |
| AC019254.2 | 15 | novel transcript |
| AC019254.1 | 15 | novel transcript |
| CHSY1 | 15 | chondroitin sulfate synthase 1 [Source:HGNC |
| SELENOS | 15 | selenoprotein S [Source:HGNC Symbol;Acc:HG |
| SNRPA1 | 15 | small nuclear ribonucleoprotein polypeptide |

| | | |
|---|---|---|
| AC023024.1 | 15 | novel transcript, antisense to SNRPA1 and PCS |
| PCSK6 | 15 | proprotein convertase subtilisin/kexin type 6 |
| AC023024.2 | 15 | TEC |
| PCSK6-AS1 | 15 | PCSK6 antisense RNA 1 [Source:HGNC Symbol |
| AC090164.2 | 15 | novel transcript |
| AC090164.3 | 15 | novel transcript |
| AC090164.4 | 15 | novel transcript |
| SNRPCP18 | 15 | small nuclear ribonucleoprotein polypeptide C |
| AC090164.1 | 15 | ribosomal protein L17 (RPL17) pseudogene |
| AC090164.5 | 15 | novel transcript |
| TM2D3 | 15 | TM2 domain containing 3 [Source:HGNC Symb |
| RNU6-807P | 15 | RNA, U6 small nuclear 807, pseudogene [Sour |
| TARSL2 | 15 | threonyl-tRNA synthetase like 2 [Source:HGNC |
| AC027559.1 | 15 | putative ubiquitin-conjugating enzyme E2Q2-l |
| DNM1P47 | 15 | dynamin 1 pseudogene 47 [Source:HGNC Sym |
| GOLGA8VP | 15 | golgin A8 family member V, pseudogene [Sour |
| RN7SL209P | 15 | RNA, 7SL, cytoplasmic 209, pseudogene [Sour |
| OR4F6 | 15 | olfactory receptor family 4 subfamily F memb |
| OR4F15 | 15 | olfactory receptor family 4 subfamily F memb |
| OR4F14P | 15 | olfactory receptor family 4 subfamily F memb |
| OR4F13P | 15 | olfactory receptor family 4 subfamily F memb |
| OR4F28P | 15 | olfactory receptor family 4 subfamily F memb |
| WBP1LP5 | 15 | WBP1L pseudogene 5 [Source:HGNC Symbol;A |
| AC140725.2 | 15 | olfactory receptor pseudogene |
| OR4F4 | 15 | olfactory receptor family 4 subfamily F memb |
| OR4G2P | 15 | olfactory receptor family 4 subfamily G memb |
| OR4G6P | 15 | olfactory receptor family 4 subfamily G memb |
| FAM138E | 15 | family with sequence similarity 138 member E |
| AC140725.1 | 15 | novel transcript |
| MIR1302-10 | 15 | microRNA 1302-10 [Source:HGNC Symbol;Acc |
| WASH3P | 15 | WASP family homolog 3, pseudogene [Source |
| MIR6859-3 | 15 | microRNA 6859-3 [Source:HGNC Symbol;Acc:H |
| DDX11L9 | 15 | DEAD/H-box helicase 11 like 9 [Source:HGNC S |
| DDX11L10 | 16 | DEAD/H-box helicase 11 like 10 [Source:HGNC |
| WASH4P | 16 | WASP family homolog 4, pseudogene [Source |
| MIR6859-4 | 16 | microRNA 6859-4 [Source:HGNC Symbol;Acc:H |
| Z84723.1 | 16 | novel transcript |
| WASIR2 | 16 | WASH and IL9R antisense RNA 2 [Source:HGN |
| IL9RP3 | 16 | interleukin 9 receptor pseudogene 3 [Source:H |
| POLR3K | 16 | RNA polymerase III subunit K [Source:HGNC Sy |
| SNRNP25 | 16 | small nuclear ribonucleoprotein U11/U12 sub |
| RHBDF1 | 16 | rhomboid 5 homolog 1 [Source:HGNC Symbol |
| Z69720.2 | 16 | TEC |
| MPG | 16 | N-methylpurine DNA glycosylase [Source:HGN |
| NPRL3 | 16 | NPR3 like, GATOR1 complex subunit [Source:H |
| Z69720.1 | 16 | novel transcript, antisense to NPRL3 |
| Z69666.1 | 16 | novel transcript |
| HBZ | 16 | hemoglobin subunit zeta [Source:HGNC Symb |
| HBM | 16 | hemoglobin subunit mu [Source:HGNC Symbo |
| HBZP1 | 16 | hemoglobin subunit zeta pseudogene 1 [Sourc |

| | | |
|---|---|---|
| Z84721.1 | 16 | hemoglobin, alpha (HBA) pseudogene |
| HBA2 | 16 | hemoglobin subunit alpha 2 [Source:HGNC Syı |
| HBA1 | 16 | hemoglobin subunit alpha 1 [Source:HGNC Syı |
| Y_RNA | 16 | Y RNA [Source:RFAM;Acc:RF00019] |
| HBQ1 | 16 | hemoglobin subunit theta 1 [Source:HGNC Syı |
| Z69706.1 | 16 | novel transcript |
| LUC7L | 16 | LUC7 like [Source:HGNC Symbol;Acc:HGNC:67 |
| Z69890.1 | 16 | novel pseudogene |
| FAM234A | 16 | family with sequence similarity 234 member / |
| RGS11 | 16 | regulator of G protein signaling 11 [Source:HG |
| Z69667.1 | 16 | novel transcript |
| ARHGDIG | 16 | Rho GDP dissociation inhibitor gamma [Source |
| PDIA2 | 16 | protein disulfide isomerase family A member |
| AXIN1 | 16 | axin 1 [Source:HGNC Symbol;Acc:HGNC:903] |
| MRPL28 | 16 | mitochondrial ribosomal protein L28 [Source:I |
| TMEM8A | 16 | transmembrane protein 8A [Source:HGNC Syr |
| Z97634.1 | 16 | ribosomal protein L23a (RPL23A) pseudogene |
| NME4 | 16 | NME/NM23 nucleoside diphosphate kinase 4 |
| DECR2 | 16 | 2,4-dienoyl-CoA reductase 2 [Source:HGNC Sy |
| AL023881.1 | 16 | novel transcript |
| RAB11FIP3 | 16 | RAB11 family interacting protein 3 [Source:HG |
| Y_RNA | 16 | Y RNA [Source:RFAM;Acc:RF00019] |
| AL049542.1 | 16 | microtubule-associated protein 1 light chain 3 |
| LINC00235 | 16 | long intergenic non-protein coding RNA 235 [S |
| CAPN15 | 16 | calpain 15 [Source:HGNC Symbol;Acc:HGNC:1 |
| MIR5587 | 16 | microRNA 5587 [Source:HGNC Symbol;Acc:HG |
| MIR3176 | 16 | microRNA 3176 [Source:HGNC Symbol;Acc:HG |
| Z97986.1 | 16 | novel transcript, antisense to SOLH |
| PRR35 | 16 | proline rich 35 [Source:HGNC Symbol;Acc:HGI |
| PIGQ | 16 | phosphatidylinositol glycan anchor biosynthes |
| NHLRC4 | 16 | NHL repeat containing 4 [Source:HGNC Symbo |
| Z98883.1 | 16 | novel transcript |
| RAB40C | 16 | RAB40C, member RAS oncogene family [Sourc |
| WFIKKN1 | 16 | WAP, follistatin/kazal, immunoglobulin, kunitz |
| METTL26 | 16 | methyltransferase like 26 [Source:HGNC Symb |
| MCRIP2 | 16 | MAPK regulated corepressor interacting prote |
| AL022341.1 | 16 | novel transcript |
| WDR90 | 16 | WD repeat domain 90 [Source:HGNC Symbol;/ |
| AL022341.2 | 16 | novel transcript, antisense to WDR90 |
| RHOT2 | 16 | ras homolog family member T2 [Source:HGNC |
| RHBDL1 | 16 | rhomboid like 1 [Source:HGNC Symbol;Acc:HG |
| Z92544.1 | 16 | novel transcript, antisense to RHBDL1 |
| Z92544.3 | 16 | TEC |
| STUB1 | 16 | STIP1 homology and U-box containing protein |
| JMJD8 | 16 | jumonji domain containing 8 [Source:HGNC Sy |
| WDR24 | 16 | WD repeat domain 24 [Source:HGNC Symbol;/ |
| Z92544.2 | 16 | novel transcript, antisense to FBXL16 |
| FBXL16 | 16 | F-box and leucine rich repeat protein 16 [Sour |
| Z97653.2 | 16 | TEC |
| Z97653.1 | 16 | novel transcript |

| | | |
|---|---|---|
| METRN | 16 | meteorin, glial cell differentiation regulator [S |
| ANTKMT | 16 | adenine nucleotide translocase lysine methylt |
| CCDC78 | 16 | coiled-coil domain containing 78 [Source:HGN |
| HAGHL | 16 | hydroxyacylglutathione hydrolase like [Source |
| CIAO3 | 16 | cytosolic iron-sulfur assembly component 3 [S |
| MSLN | 16 | mesothelin [Source:HGNC Symbol;Acc:HGNC:: |
| AL031258.1 | 16 | TEC |
| MSLNL | 16 | mesothelin like [Source:HGNC Symbol;Acc:HG |
| MIR662 | 16 | microRNA 662 [Source:HGNC Symbol;Acc:HGN |
| RPUSD1 | 16 | RNA pseudouridine synthase domain containi |
| CHTF18 | 16 | chromosome transmission fidelity factor 18 [S |
| GNG13 | 16 | G protein subunit gamma 13 [Source:HGNC Sy |
| AL023882.1 | 16 | novel transcript, antisense to LMF1 |
| AL031008.1 | 16 | novel transcript |
| LMF1 | 16 | lipase maturation factor 1 [Source:HGNC Sym |
| AL031716.1 | 16 | novel transcript, antisense to LMF1 |
| AL008727.1 | 16 | novel transcript, intronic to LMF1 |
| LMF1-AS1 | 16 | LMF1 antisense RNA 1 [Source:HGNC Symbol; |
| AC009041.4 | 16 | novel transcript, sense intronic to LMF1 |
| AC009041.2 | 16 | novel transcript |
| SOX8 | 16 | SRY-box transcription factor 8 [Source:HGNC S |
| AC009041.1 | 16 | novel transcript |
| SSTR5-AS1 | 16 | SSTR5 antisense RNA 1 [Source:HGNC Symbol |
| AC009041.3 | 16 | novel transcript |
| SSTR5 | 16 | somatostatin receptor 5 [Source:HGNC Symbo |
| C1QTNF8 | 16 | C1q and TNF related 8 [Source:HGNC Symbol; |
| AL031713.1 | 16 | tektin 4 (TEKT4) pseudogene |
| AL031598.1 | 16 | novel transcript |
| AC120498.8 | 16 | novel transcript |
| CACNA1H | 16 | calcium voltage-gated channel subunit alpha1 |
| AC120498.3 | 16 | novel transcript, antisense to CACNA1H |
| AC120498.1 | 16 | novel transcript, antisense to CACNA1H |
| AC120498.6 | 16 | novel transcript, antisense to CACNA1H |
| TPSG1 | 16 | tryptase gamma 1 [Source:HGNC Symbol;Acc: |
| AC120498.10 | 16 | novel transcript, antisense to TPSG1 |
| TPSB2 | 16 | tryptase beta 2 (gene/pseudogene) [Source:H |
| TPSAB1 | 16 | tryptase alpha/beta 1 [Source:HGNC Symbol;A |
| TPSD1 | 16 | tryptase delta 1 [Source:HGNC Symbol;Acc:HG |
| AC120498.2 | 16 | novel transcript, intronic to TPSD1 |
| PRSS29P | 16 | serine protease 29, pseudogene [Source:HGN |
| AC120498.7 | 16 | tryptase delta 1 (TPSD1) pseudogene |
| AC120498.5 | 16 | protease, serine, 42 9 9 (PRSS42) pseudogene |
| TPSP2 | 16 | tryptase pseudogene 2 [Source:HGNC Symbol |
| AC120498.4 | 16 | novel transcript |
| AC120498.9 | 16 | novel transcript, antisense to UBE2I |
| UBE2I | 16 | ubiquitin conjugating enzyme E2 I [Source:HG |
| AL031714.1 | 16 | novel transcript, antisense to UBE2I |
| RPS20P2 | 16 | ribosomal protein S20 pseudogene 2 [Source: |
| BAIAP3 | 16 | BAI1 associated protein 3 [Source:HGNC Symb |
| TSR3 | 16 | TSR3 ribosome maturation factor [Source:HGI |

| | | |
|---|---|---|
| GNPTG | 16 | N-acetylglucosamine-1-phosphate transferase |
| AL031709.1 | 16 | novel transcript, antisense to GNPTG |
| UNKL | 16 | unk like zinc finger [Source:HGNC Symbol;Acc: |
| AL032819.1 | 16 | novel transcript, antisense to UNKL |
| C16orf91 | 16 | chromosome 16 open reading frame 91 [Sour |
| AL032819.3 | 16 | uncharacterized LOC105371045 [Source:NCBI |
| CCDC154 | 16 | coiled-coil domain containing 154 [Source:HG |
| AL032819.2 | 16 | TEC |
| CLCN7 | 16 | chloride voltage-gated channel 7 [Source:HGN |
| AL031600.3 | 16 | novel transcript, antisense to CLCN7 |
| AL031600.1 | 16 | novel transcript, antisense to CLCN7 |
| AL031600.2 | 16 | novel transcript, intronic to CLCN7 |
| RPS3AP2 | 16 | ribosomal protein S3A pseudogene 2 [Source: |
| PTX4 | 16 | pentraxin 4 [Source:HGNC Symbol;Acc:HGNC: |
| TELO2 | 16 | telomere maintenance 2 [Source:HGNC Symb |
| IFT140 | 16 | intraflagellar transport 140 [Source:HGNC Sym |
| AL031705.1 | 16 | novel transcript |
| TMEM204 | 16 | transmembrane protein 204 [Source:HGNC Sy |
| AL031719.1 | 16 | TEC |
| AL031719.2 | 16 | tec |
| Z97633.1 | 16 | tec |
| AL133297.1 | 16 | novel transcript, intronic to IFT140 |
| AL133297.2 | 16 | novel transcript, antisense to IFT140 |
| CRAMP1 | 16 | cramped chromatin regulator homolog 1 [Sou |
| Z97652.1 | 16 | novel transcript, antisense to CRAMP1L |
| AL031708.1 | 16 | novel protein |
| JPT2 | 16 | Jupiter microtubule associated homolog 2 [So |
| AL031009.1 | 16 | TEC |
| MAPK8IP3 | 16 | mitogen-activated protein kinase 8 interacting |
| AL031710.2 | 16 | novel transcript, sense intronic to MAPK8IP3 |
| AL031710.1 | 16 | novel transcript, antisense to MAPK8IP3 |
| MIR3177 | 16 | microRNA 3177 [Source:HGNC Symbol;Acc:HG |
| AL031717.1 | 16 | novel transcript, antisense to MAPK8IP3 |
| NME3 | 16 | NME/NM23 nucleoside diphosphate kinase 3 |
| MRPS34 | 16 | mitochondrial ribosomal protein S34 [Source: |
| EME2 | 16 | essential meiotic structure-specific endonucle |
| SPSB3 | 16 | splA/ryanodine receptor domain and SOCS bo |
| NUBP2 | 16 | nucleotide binding protein 2 [Source:HGNC Sy |
| IGFALS | 16 | insulin like growth factor binding protein acid |
| HAGH | 16 | hydroxyacylglutathione hydrolase [Source:HG |
| FAHD1 | 16 | fumarylacetoacetate hydrolase domain conta |
| MEIOB | 16 | meiosis specific with OB-fold [Source:HGNC Sy |
| AL031722.1 | 16 | novel transcript, overlapping C16orf73 |
| LINC00254 | 16 | long intergenic non-protein coding RNA 254 [S |
| LINC02124 | 16 | long intergenic non-protein coding RNA 2124 |
| HS3ST6 | 16 | heparan sulfate-glucosamine 3-sulfotransfera: |
| MSRB1 | 16 | methionine sulfoxide reductase B1 [Source:H( |
| RPL3L | 16 | ribosomal protein L3 like [Source:HGNC Symb |
| NDUFB10 | 16 | NADH:ubiquinone oxidoreductase subunit B1( |
| RPS2 | 16 | ribosomal protein S2 [Source:HGNC Symbol;A |

| | | |
|---|---|---|
| SNORA10 | 16 | small nucleolar RNA, H/ACA box 10 [Source:H |
| SNORA64 | 16 | small nucleolar RNA, H/ACA box 64 [Source:H |
| AC005363.1 | 16 | ribosomal protein L26 (RPL26) pseudogene |
| SNHG9 | 16 | small nucleolar RNA host gene 9 [Source:HGN |
| SNORA78 | 16 | small nucleolar RNA, H/ACA box 78 [Source:H |
| RNF151 | 16 | ring finger protein 151 [Source:HGNC Symbol; |
| AC005363.2 | 16 | novel transcript |
| TBL3 | 16 | transducin beta like 3 [Source:HGNC Symbol;A |
| NOXO1 | 16 | NADPH oxidase organizer 1 [Source:HGNC Syr |
| GFER | 16 | growth factor, augmenter of liver regeneratio |
| AC005606.2 | 16 | novel transcript |
| SYNGR3 | 16 | synaptogyrin 3 [Source:HGNC Symbol;Acc:HG |
| AC005606.1 | 16 | novel transcript |
| ZNF598 | 16 | zinc finger protein 598 [Source:HGNC Symbol; |
| NPW | 16 | neuropeptide W [Source:HGNC Symbol;Acc:H |
| RN7SL219P | 16 | RNA, 7SL, cytoplasmic 219, pseudogene [Sour |
| SLC9A3R2 | 16 | SLC9A3 regulator 2 [Source:HGNC Symbol;Acc |
| NTHL1 | 16 | nth like DNA glycosylase 1 [Source:HGNC Sym |
| TSC2 | 16 | TSC complex subunit 2 [Source:HGNC Symbol; |
| PKD1 | 16 | polycystin 1, transient receptor potential char |
| MIR1225 | 16 | microRNA 1225 [Source:HGNC Symbol;Acc:HG |
| AC009065.2 | 16 | novel transcript, overlapping TSC2 |
| AC009065.5 | 16 | novel transcript, intronic to PKD1 |
| MIR6511B1 | 16 | microRNA 6511b-1 [Source:HGNC Symbol;Acc |
| AC009065.6 | 16 | novel transcript, antisense to PKD1 |
| AC009065.3 | 16 | novel transcript, antisense to PKD1 |
| Y_RNA | 16 | Y RNA [Source:RFAM;Acc:RF00019] |
| MIR4516 | 16 | microRNA 4516 [Source:HGNC Symbol;Acc:HG |
| MIR3180-5 | 16 | microRNA 3180-5 [Source:HGNC Symbol;Acc:I |
| RAB26 | 16 | RAB26, member RAS oncogene family [Source |
| SNHG19 | 16 | small nucleolar RNA host gene 19 [Source:HGI |
| SNORD60 | 16 | small nucleolar RNA, C/D box 60 [Source:HGN |
| AC009065.9 | 16 | NULL |
| TRAF7 | 16 | TNF receptor associated factor 7 [Source:HGN |
| CASKIN1 | 16 | CASK interacting protein 1 [Source:HGNC Sym |
| MLST8 | 16 | MTOR associated protein, LST8 homolog [Sou |
| BRICD5 | 16 | BRICHOS domain containing 5 [Source:HGNC S |
| PGP | 16 | phosphoglycolate phosphatase [Source:HGNC |
| AC009065.7 | 16 | novel transcript |
| E4F1 | 16 | E4F transcription factor 1 [Source:HGNC Symb |
| AC009065.1 | 16 | novel transcript |
| DNASE1L2 | 16 | deoxyribonuclease 1 like 2 [Source:HGNC Sym |
| ECI1 | 16 | enoyl-CoA delta isomerase 1 [Source:HGNC Sy |
| AC009065.8 | 16 | novel transcript, antisense to DCI |
| RNPS1 | 16 | RNA binding protein with serine rich domain 1 |
| AC009065.4 | 16 | novel transcript |
| MIR3677 | 16 | microRNA 3677 [Source:HGNC Symbol;Acc:HG |
| MIR940 | 16 | microRNA 940 [Source:HGNC Symbol;Acc:HGI |
| MIR4717 | 16 | microRNA 4717 [Source:HGNC Symbol;Acc:HG |
| ABCA3 | 16 | ATP binding cassette subfamily A member 3 [S |

| | | |
|---|---|---|
| AC106820.1 | 16 | ribosomal protein S16 (RPS16) pseudogene |
| ABCA17P | 16 | ATP binding cassette subfamily A member 17, |
| CCNF | 16 | cyclin F [Source:HGNC Symbol;Acc:HGNC:159 |
| AC106820.6 | 16 | novel transcript, antisense to CCNF |
| MIR6767 | 16 | microRNA 6767 [Source:HGNC Symbol;Acc:HG |
| AC106820.3 | 16 | novel transcript, antisense to CCNF |
| AC106820.5 | 16 | novel transcript |
| TEDC2 | 16 | tubulin epsilon and delta complex 2 [Source:H |
| MIR6768 | 16 | microRNA 6768 [Source:HGNC Symbol;Acc:HG |
| AC106820.2 | 16 | novel transcript, antisense to C16orf59 |
| NTN3 | 16 | netrin 3 [Source:HGNC Symbol;Acc:HGNC:803 |
| TBC1D24 | 16 | TBC1 domain family member 24 [Source:HGN |
| AC106820.4 | 16 | novel transcript, intronic to TBC1D24 |
| AC093525.2 | 16 | novel protein |
| ATP6V0C | 16 | ATPase H+ transporting V0 subunit c [Source: |
| AC093525.1 | 16 | novel protein |
| AC093525.8 | 16 | tec |
| AMDHD2 | 16 | amidohydrolase domain containing 2 [Source: |
| CEMP1 | 16 | cementum protein 1 [Source:HGNC Symbol;A |
| MIR3178 | 16 | microRNA 3178 [Source:HGNC Symbol;Acc:HG |
| PDPK1 | 16 | 3-phosphoinositide dependent protein kinase |
| AC093525.6 | 16 | novel transcript, antisense to PDPK1 |
| AC093525.5 | 16 | novel transcript, intronic to PDPK1 |
| AC093525.7 | 16 | novel transcript, antisense to PDPK1 |
| AC093525.4 | 16 | novel transcript, antisense to PDPK1 |
| AC093525.3 | 16 | novel transcript, antisense to PDPK1 |
| AC093525.9 | 16 | tec |
| AC093525.10 | 16 | tec |
| AC141586.5 | 16 | tec |
| AC141586.3 | 16 | novel transcript, antisense to PDPK1 |
| AC141586.1 | 16 | potassium channel tetramerisation domain co |
| PDPK2P | 16 | 3-phosphoinositide dependent protein kinase |
| AC141586.4 | 16 | tec |
| AC141586.2 | 16 | novel transcript |
| ERVK13-1 | 16 | endogenous retrovirus group K13 member 1 [ |
| KCTD5 | 16 | potassium channel tetramerization domain co |
| AC092117.2 | 16 | tec |
| PRSS27 | 16 | serine protease 27 [Source:HGNC Symbol;Acc |
| SRRM2-AS1 | 16 | SRRM2 antisense RNA 1 [Source:HGNC Symbo |
| SRRM2 | 16 | serine/arginine repetitive matrix 2 [Source:HG |
| ELOB | 16 | elongin B [Source:HGNC Symbol;Acc:HGNC:11 |
| AC092117.1 | 16 | novel transcript |
| PRSS33 | 16 | serine protease 33 [Source:HGNC Symbol;Acc |
| SNORA3C | 16 | small nucleolar RNA, H/ACA box 3C [Source:H |
| PRSS41 | 16 | serine protease 41 [Source:HGNC Symbol;Acc |
| PRSS21 | 16 | serine protease 21 [Source:HGNC Symbol;Acc |
| EIF1P4 | 16 | eukaryotic translation initiation factor 1 pseu |
| ZG16B | 16 | zymogen granule protein 16B [Source:HGNC S |
| PRSS30P | 16 | serine protease 30, pseudogene [Source:HGN |
| PRSS22 | 16 | serine protease 22 [Source:HGNC Symbol;Acc |

| | | |
|---|---|---|
| AC003965.2 | 16 | novel transcript, antisense to PRSS22 |
| AC003965.1 | 16 | novel transcript |
| FLYWCH2 | 16 | FLYWCH family member 2 [Source:HGNC Sym |
| FLYWCH1 | 16 | FLYWCH-type zinc finger 1 [Source:HGNC Sym |
| RPL23AP86 | 16 | ribosomal protein L23a pseudogene 86 [Sourc |
| AC004034.1 | 16 | novel transcript, antisense to FLYWCH1 |
| KREMEN2 | 16 | kringle containing transmembrane protein 2 [ |
| PKMYT1 | 16 | protein kinase, membrane associated tyrosine |
| PAQR4 | 16 | progestin and adipoQ receptor family membe |
| AC004233.3 | 16 | novel transcript |
| LINC00514 | 16 | long intergenic non-protein coding RNA 514 [S |
| AC004233.1 | 16 | novel transcript, sense overlapping CLDN6 |
| AC004233.2 | 16 | novel transcript |
| CLDN9 | 16 | claudin 9 [Source:HGNC Symbol;Acc:HGNC:20 |
| CLDN6 | 16 | claudin 6 [Source:HGNC Symbol;Acc:HGNC:20 |
| TNFRSF12A | 16 | TNF receptor superfamily member 12A [Sourc |
| HCFC1R1 | 16 | host cell factor C1 regulator 1 [Source:HGNC S |
| THOC6 | 16 | THO complex 6 [Source:HGNC Symbol;Acc:HG |
| BICDL2 | 16 | BICD family like cargo adaptor 2 [Source:HGN |
| AC108134.1 | 16 | novel transcript, antisense to CCDC64B |
| MMP25-AS1 | 16 | MMP25 antisense RNA 1 [Source:HGNC Symb |
| MMP25 | 16 | matrix metallopeptidase 25 [Source:HGNC Syr |
| IL32 | 16 | interleukin 32 [Source:HGNC Symbol;Acc:HGN |
| RNU1-125P | 16 | RNA, U1 small nuclear 125, pseudogene [Sour |
| AC108134.3 | 16 | novel transcript |
| RNU1-22P | 16 | RNA, U1 small nuclear 22, pseudogene [Sourc |
| ZSCAN10 | 16 | zinc finger and SCAN domain containing 10 [S |
| AC108134.4 | 16 | novel transcript, sense overlapping ZNF205 |
| ZNF213-AS1 | 16 | ZNF213 antisense RNA 1 (head to head) [Sour |
| ZNF205 | 16 | zinc finger protein 205 [Source:HGNC Symbol; |
| ZNF213 | 16 | zinc finger protein 213 [Source:HGNC Symbol; |
| CASP16P | 16 | caspase 16, pseudogene [Source:HGNC Symbo |
| AC108134.2 | 16 | novel transcript |
| AJ003147.1 | 16 | novel transcript |
| AJ003147.2 | 16 | novel transcript, antisense to ZNF200 |
| OR1F1 | 16 | olfactory receptor family 1 subfamily F memb |
| OR1F2P | 16 | olfactory receptor family 1 subfamily F memb |
| ZNF200 | 16 | zinc finger protein 200 [Source:HGNC Symbol; |
| MEFV | 16 | MEFV innate immuity regulator, pyrin [Source |
| LINC00921 | 16 | long intergenic non-protein coding RNA 921 [S |
| ZNF263 | 16 | zinc finger protein 263 [Source:HGNC Symbol; |
| AJ003147.3 | 16 | TEC |
| AC004232.2 | 16 | TEC |
| TIGD7 | 16 | tigger transposable element derived 7 [Source |
| ZNF75A | 16 | zinc finger protein 75a [Source:HGNC Symbol; |
| AC004232.1 | 16 | novel transcript |
| AC025283.1 | 16 | ribosomal protein L7-like 1 (RPL7L1) pseudoge |
| OR2C1 | 16 | olfactory receptor family 2 subfamily C memb |
| AC025283.2 | 16 | novel protein |
| MTCO1P28 | 16 | MT-CO1 pseudogene 28 [Source:HGNC Symbo |

| | | |
|---|---|---|
| MTND1P8 | 16 | MT-ND1 pseudogene 8 [Source:HGNC Symbol |
| MTRNR2L4 | 16 | MT-RNR2 like 4 [Source:HGNC Symbol;Acc:HG |
| ZSCAN32 | 16 | zinc finger and SCAN domain containing 32 [Sc |
| ZNF174 | 16 | zinc finger protein 174 [Source:HGNC Symbol; |
| ZNF597 | 16 | zinc finger protein 597 [Source:HGNC Symbol; |
| NAA60 | 16 | N(alpha)-acetyltransferase 60, NatF catalytic s |
| AC004224.2 | 16 | novel transcript |
| MIR6126 | 16 | microRNA 6126 [Source:HGNC Symbol;Acc:HG |
| C16orf90 | 16 | chromosome 16 open reading frame 90 [Sour |
| AC004224.1 | 16 | ribosomal protein L18 (RPL18) pseudogene |
| CLUAP1 | 16 | clusterin associated protein 1 [Source:HGNC S |
| AC004494.2 | 16 | TEC |
| NLRC3 | 16 | NLR family CARD domain containing 3 [Source |
| AC004494.1 | 16 | novel transcript, antisense to NLRC3 |
| SLX4 | 16 | SLX4 structure-specific endonuclease subunit |
| AC006111.1 | 16 | novel transcript |
| DNASE1 | 16 | deoxyribonuclease 1 [Source:HGNC Symbol;A |
| AC006111.2 | 16 | novel transcript, sense overlapping TRAP1 |
| TRAP1 | 16 | TNF receptor associated protein 1 [Source:HG |
| AC006111.3 | 16 | novel transcript, antisense to TRAP1 |
| CREBBP | 16 | CREB binding protein [Source:HGNC Symbol;A |
| AC007151.1 | 16 | developmental pluripotency associated 3 (DPI |
| AC005736.1 | 16 | novel transcript |
| ADCY9 | 16 | adenylate cyclase 9 [Source:HGNC Symbol;Acc |
| AC005736.2 | 16 | novel transcript, antisense ADCY9 |
| AC009171.1 | 16 | diazepam binding inhibitor (GABA receptor m |
| AC009171.2 | 16 | novel transcript |
| SRL | 16 | sarcalumenin [Source:HGNC Symbol;Acc:HGN |
| LINC01569 | 16 | long intergenic non-protein coding RNA 1569 |
| TFAP4 | 16 | transcription factor AP-4 [Source:HGNC Symb |
| GLIS2 | 16 | GLIS family zinc finger 2 [Source:HGNC Symbo |
| GLIS2-AS1 | 16 | GLIS2 antisense RNA 1 [Source:HGNC Symbol; |
| PAM16 | 16 | presequence translocase associated motor 16 |
| AC012676.1 | 16 | novel transcript |
| CORO7-PAM16 | 16 | CORO7-PAM16 readthrough [Source:HGNC Sy |
| AC012676.5 | 16 | tec |
| CORO7 | 16 | coronin 7 [Source:HGNC Symbol;Acc:HGNC:26 |
| VASN | 16 | vasorin [Source:HGNC Symbol;Acc:HGNC:185 |
| DNAJA3 | 16 | DnaJ heat shock protein family (Hsp40) memb |
| AC012676.3 | 16 | novel transcript, sense intronic with DNAJA3 |
| AC012676.4 | 16 | novel transcript, sense intronic with DNAJA3 |
| AC012676.2 | 16 | cytochrome c oxidase subunit VIIc (COX7C) ps |
| NMRAL1 | 16 | NmrA like redox sensor 1 [Source:HGNC Symb |
| HMOX2 | 16 | heme oxygenase 2 [Source:HGNC Symbol;Acc |
| CDIP1 | 16 | cell death inducing p53 target 1 [Source:HGN |
| AC023830.3 | 16 | novel transcript, sense intronic to CDIP1 |
| C16orf96 | 16 | chromosome 16 open reading frame 96 [Sour |
| AC023830.2 | 16 | novel transcript, antisense to C16orf96 |
| SUB1P3 | 16 | SUB1 pseudogene 3 [Source:HGNC Symbol;Ac |
| UBALD1 | 16 | UBA like domain containing 1 [Source:HGNC S |

| | | |
|---|---|---|
| MGRN1 | 16 | mahogunin ring finger 1 [Source:HGNC Symbo |
| RN7SL850P | 16 | RNA, 7SL, cytoplasmic 850, pseudogene [Sour |
| AC023830.1 | 16 | novel transcript, antisense to MGRN1 |
| Metazoa_SRP | 16 | Metazoan signal recognition particle RNA [Sou |
| MIR6769A | 16 | microRNA 6769a [Source:HGNC Symbol;Acc:H |
| NUDT16L1 | 16 | nudix hydrolase 16 like 1 [Source:HGNC Symb |
| ANKS3 | 16 | ankyrin repeat and sterile alpha motif domain |
| AC020663.1 | 16 | novel transcript |
| C16orf71 | 16 | chromosome 16 open reading frame 71 [Sour |
| ZNF500 | 16 | zinc finger protein 500 [Source:HGNC Symbol; |
| Metazoa_SRP | 16 | Metazoan signal recognition particle RNA [Sou |
| SEPTIN12 | 16 | septin 12 [Source:HGNC Symbol;Acc:HGNC:26 |
| AC020663.4 | 16 | novel transcript |
| SMIM22 | 16 | small integral membrane protein 22 [Source:H |
| AC020663.2 | 16 | novel transcript, antisense to novel protein |
| ROGDI | 16 | rogdi atypical leucine zipper [Source:HGNC Sy |
| GLYR1 | 16 | glyoxylate reductase 1 homolog [Source:HGN |
| AC020663.3 | 16 | novel transcript, sense intronic to GLYR1 |
| UBN1 | 16 | ubinuclein 1 [Source:HGNC Symbol;Acc:HGNC |
| PPL | 16 | periplakin [Source:HGNC Symbol;Acc:HGNC:9 |
| SEC14L5 | 16 | SEC14 like lipid binding 5 [Source:HGNC Symb |
| NAGPA-AS1 | 16 | NAGPA antisense RNA 1 [Source:HGNC Symbo |
| NAGPA | 16 | N-acetylglucosamine-1-phosphodiester alpha- |
| ALG1 | 16 | ALG1 chitobiosyldiphosphodolichol beta-man |
| C16orf89 | 16 | chromosome 16 open reading frame 89 [Sour |
| AC026458.1 | 16 | novel pseudogene |
| EEF2KMT | 16 | eukaryotic elongation factor 2 lysine methyltr |
| AC026458.2 | 16 | novel transcript |
| AC074051.5 | 16 | novel transcript |
| ENPP7P14 | 16 | ectonucleotide pyrophosphatase/phosphodie |
| AC074051.2 | 16 | C11orf2 chromosome 11 open reading frame |
| LINC02164 | 16 | long intergenic non-protein coding RNA 2164 |
| RBFOX1 | 16 | RNA binding fox-1 homolog 1 [Source:HGNC S |
| AC074051.3 | 16 | ribosomal protein S24 (RPS24) pseudogene |
| AC074051.1 | 16 | ribosomal protein S3A (RPS3A) pseudogene |
| SNRPCP20 | 16 | small nuclear ribonucleoprotein polypeptide C |
| AC074051.4 | 16 | nucleophosmin (nucleolar phosphoprotein B2 |
| LINC01570 | 16 | long intergenic non-protein coding RNA 1570 |
| MIR8065 | 16 | microRNA 8065 [Source:HGNC Symbol;Acc:HG |
| AC012175.1 | 16 | F-box and WD repeat domain containing 12 (F |
| AC009135.2 | 16 | TEC |
| AC009135.1 | 16 | novel transcript |
| AC006112.1 | 16 | novel transcript, sense intronic to RBFOX1 |
| AC007223.1 | 16 | novel transcript |
| AC007012.2 | 16 | TEC |
| AC007012.1 | 16 | ring finger protein 39 (RNF39) pseudogene |
| RNU7-99P | 16 | RNA, U7 small nuclear 99 pseudogene [Source |
| RNU6-457P | 16 | RNA, U6 small nuclear 457, pseudogene [Sour |
| RNU6-328P | 16 | RNA, U6 small nuclear 328, pseudogene [Sour |
| AC022206.1 | 16 | ribosomal protein S26 (RPS26) pseudogene |

| | | |
|---|---|---|
| AC007222.2 | 16 | novel transcript, antisense to RBFOX1 |
| AC007222.1 | 16 | NULL |
| AC005774.2 | 16 | novel transcript, antisense to RBFOX1 |
| AC005774.1 | 16 | NULL |
| AC093515.1 | 16 | novel transcript |
| AC018767.2 | 16 | novel transcript |
| LINC02152 | 16 | long intergenic non-protein coding RNA 2152 |
| AC018767.3 | 16 | novel transcript |
| AC018767.1 | 16 | novel transcript |
| AC074052.1 | 16 | ribosomal protein S14 (RPS14) pseudogene |
| AC074052.2 | 16 | novel transcript |
| TMEM114 | 16 | transmembrane protein 114 [Source:HGNC Sy |
| AC138420.1 | 16 | novel transcript |
| METTL22 | 16 | methyltransferase like 22 [Source:HGNC Symb |
| AC007224.2 | 16 | TEC |
| ABAT | 16 | 4-aminobutyrate aminotransferase [Source:H |
| RN7SL743P | 16 | RNA, 7SL, cytoplasmic 743, pseudogene [Sour |
| RNU7-63P | 16 | RNA, U7 small nuclear 63 pseudogene [Source |
| AC007224.1 | 16 | cytochrome c oxidase subunit VIc (COX6C) pse |
| TMEM186 | 16 | transmembrane protein 186 [Source:HGNC Sy |
| PMM2 | 16 | phosphomannomutase 2 [Source:HGNC Symb |
| AC012173.1 | 16 | novel transcript, antisense to PMM2 |
| AC022167.2 | 16 | novel transcript, antisense to CARHSP1 |
| CARHSP1 | 16 | calcium regulated heat stable protein 1 [Sourc |
| AC022167.1 | 16 | novel transcript, antisense to CARHSP1 |
| AC022167.4 | 16 | novel transcript, antisense to CARHSP1 |
| LITAFD | 16 | LITAF domain containing [Source:HGNC Symb |
| USP7 | 16 | ubiquitin specific peptidase 7 [Source:HGNC S |
| AC022167.3 | 16 | novel transcript |
| AC087190.2 | 16 | novel transcript |
| C16orf72 | 16 | chromosome 16 open reading frame 72 [Sour |
| AC087190.3 | 16 | novel transcript, 3' overlapping chromosome |
| AC087190.1 | 16 | novel transcript, antisense to C16orf72 |
| RPL21P119 | 16 | ribosomal protein L21 pseudogene 119 [Sourc |
| LINC02177 | 16 | long intergenic non-protein coding RNA 2177 |
| AC012178.1 | 16 | novel transcript |
| LINC01177 | 16 | long intergenic non-protein coding RNA 1177 |
| LINC01195 | 16 | long intergenic non-protein coding RNA 1195 |
| AC007221.1 | 16 | novel transcript |
| RNA5SP403 | 16 | RNA, 5S ribosomal pseudogene 403 [Source:H |
| RNA5SP404 | 16 | RNA, 5S ribosomal pseudogene 404 [Source:H |
| AC007218.1 | 16 | novel transcript |
| GRIN2A | 16 | glutamate ionotropic receptor NMDA type sul |
| AC007218.2 | 16 | novel transcript, antisense to GRIN2A |
| AC007218.3 | 16 | novel transcript, antisense to GRIN2A |
| AC133565.1 | 16 | novel transcript, antisense to GRIN2A |
| IMPDH1P11 | 16 | inosine monophosphate dehydrogenase 1 pse |
| AC022168.1 | 16 | novel transcript |
| RN7SL493P | 16 | RNA, 7SL, cytoplasmic 493, pseudogene [Sour |
| ATF7IP2 | 16 | activating transcription factor 7 interacting pr |

| | | |
|---|---|---|
| AC131649.1 | 16 | novel transcript |
| AC131649.2 | 16 | TEC |
| RNU6-633P | 16 | RNA, U6 small nuclear 633, pseudogene [Sour |
| LINC01290 | 16 | long intergenic non-protein coding RNA 1290 |
| EMP2 | 16 | epithelial membrane protein 2 [Source:HGNC |
| AC027277.1 | 16 | novel transcript, antisense to EMP2 |
| AC027277.2 | 16 | novel transcript, sense intronic to EMP2 |
| TEKT5 | 16 | tektin 5 [Source:HGNC Symbol;Acc:HGNC:265 |
| AC007595.1 | 16 | novel transcript, antisense to TEKT5 |
| MTCYBP33 | 16 | MT-CYB pseudogene 33 [Source:HGNC Symbo |
| MTND6P33 | 16 | MT-ND6 pseudogene 33 [Source:HGNC Symbo |
| MTND5P33 | 16 | MT-ND5 pseudogene 33 [Source:HGNC Symbo |
| MTND4P34 | 16 | MT-ND4 pseudogene 34 [Source:HGNC Symbo |
| MTND4LP24 | 16 | MT-ND4L pseudogene 24 [Source:HGNC Symb |
| MTND3P13 | 16 | MT-ND3 pseudogene 13 [Source:HGNC Symbo |
| MTCO3P24 | 16 | MT-CO3 pseudogene 24 [Source:HGNC Symbo |
| MTATP6P24 | 16 | MT-ATP6 pseudogene 24 [Source:HGNC Symb |
| NUBP1 | 16 | nucleotide binding protein 1 [Source:HGNC Sy |
| TVP23A | 16 | trans-golgi network vesicle protein 23 homolo |
| AC133065.2 | 16 | Rho GTPase activating protein 21 (ARHGAP21 |
| AC133065.1 | 16 | novel transcript, antisense to CIITA |
| CIITA | 16 | class II major histocompatibility complex trans |
| DEXI | 16 | Dexi homolog [Source:HGNC Symbol;Acc:HGN |
| AC133065.3 | 16 | TEC |
| CLEC16A | 16 | C-type lectin domain containing 16A [Source:H |
| RPL7P46 | 16 | ribosomal protein L7 pseudogene 46 [Source: |
| AC007014.2 | 16 | novel transcript, sense intronic to CLEC16A |
| AC007014.1 | 16 | novel transcript, antisense to CLEC16A |
| AC007220.2 | 16 | novel transcript, antisense to CLEC16A |
| AC007220.1 | 16 | novel transcript |
| HNRNPCP4 | 16 | heterogeneous nuclear ribonucleoprotein C p |
| RMI2 | 16 | RecQ mediated genome instability 2 [Source:H |
| SOCS1 | 16 | suppressor of cytokine signaling 1 [Source:HG |
| TNP2 | 16 | transition protein 2 [Source:HGNC Symbol;Ac |
| PRM3 | 16 | protamine 3 [Source:HGNC Symbol;Acc:HGNC |
| AC009121.1 | 16 | NULL |
| PRM2 | 16 | protamine 2 [Source:HGNC Symbol;Acc:HGNC |
| PRM1 | 16 | protamine 1 [Source:HGNC Symbol;Acc:HGNC |
| AC009121.4 | 16 | novel transcript, sense intronic to RMI2 |
| MIR548H2 | 16 | microRNA 548h-2 [Source:HGNC Symbol;Acc:I |
| Y_RNA | 16 | Y RNA [Source:RFAM;Acc:RF00019] |
| AC009121.3 | 16 | novel transcript, antisense to RMI2 |
| AC009121.2 | 16 | novel transcript, antisense to RMI2 |
| AC099489.1 | 16 | novel lipoprotein amino terminal region conta |
| AC099489.2 | 16 | mitochondrial ribosomal protein L18 (MRPL18 |
| AC099489.3 | 16 | novel transcript, antisense to a novel protein |
| LITAF | 16 | lipopolysaccharide induced TNF factor [Source |
| SNN | 16 | stannin [Source:HGNC Symbol;Acc:HGNC:111 |
| TXNDC11 | 16 | thioredoxin domain containing 11 [Source:HG |
| AC007613.1 | 16 | novel transcript, antisense to TXNDC11 |

| | | |
|---|---|---|
| ZC3H7A | 16 | zinc finger CCCH-type containing 7A [Source:H |
| Metazoa_SRP | 16 | Metazoan signal recognition particle RNA [Sou |
| AC010654.1 | 16 | novel transcript |
| BCAR4 | 16 | breast cancer anti-estrogen resistance 4 [Sour |
| RSL1D1 | 16 | ribosomal L1 domain containing 1 [Source:HG |
| AC007216.4 | 16 | novel transcript, antisense to GSPT1 |
| GSPT1 | 16 | G1 to S phase transition 1 [Source:HGNC Syml |
| AC007216.3 | 16 | novel transcript, sense intronic to GSPT1 |
| COX6CP1 | 16 | cytochrome c oxidase subunit 6C pseudogene |
| AC007216.2 | 16 | novel transcript, antisense to GSPT1 |
| AC007216.5 | 16 | phospholipase A2, group X (PLA2G10) pseudo |
| NPIPB2 | 16 | nuclear pore complex interacting protein fami |
| TNFRSF17 | 16 | TNF receptor superfamily member 17 [Source |
| UBL5P4 | 16 | ubiquitin like 5 pseudogene 4 [Source:HGNC S |
| SNX29 | 16 | sorting nexin 29 [Source:HGNC Symbol;Acc:H( |
| AC007216.1 | 16 | novel transcript, antisense to RUNDC2A |
| RPS23P6 | 16 | ribosomal protein S23 pseudogene 6 [Source: |
| AC007601.1 | 16 | novel transcript, antisense to SNX29 |
| AC007601.2 | 16 | novel transcript, antisense to a novel protein |
| AC092365.1 | 16 | NULL |
| AC007598.2 | 16 | novel transcript, antisense to a novel protein |
| AC007598.1 | 16 | novel transcript, sense intronic to a novel prot |
| AC007598.3 | 16 | TEC |
| AC131391.1 | 16 | novel transcript, sense intronic to a novel prot |
| AC010333.1 | 16 | novel transcript, antisense to a novel protein |
| AC010333.2 | 16 | novel transcript, antisense to a novel protein |
| AC010333.3 | 16 | novel transcript |
| CPPED1 | 16 | calcineurin like phosphoesterase domain cont |
| MIR4718 | 16 | microRNA 4718 [Source:HGNC Symbol;Acc:HG |
| AC109597.1 | 16 | novel transcript, antisense to CPPED1 |
| AC109597.2 | 16 | novel transcript, antisense to CPPED1 |
| AC092324.1 | 16 | NULL |
| AC092324.2 | 16 | ribosomal protein L35a (RPL35A) pseudogene |
| SHISA9 | 16 | shisa family member 9 [Source:HGNC Symbol; |
| AC009134.1 | 16 | novel transcript, antisense to SHISA9 |
| U91319.1 | 16 | novel transcript |
| AC003009.1 | 16 | novel transcript, antisense to a novel gene |
| TMF1P1 | 16 | TATA element modulatory factor 1 pseudoger |
| U95743.1 | 16 | novel transcript |
| ERCC4 | 16 | ERCC excision repair 4, endonuclease catalytic |
| RPS26P52 | 16 | ribosomal protein S26 pseudogene 52 [Source |
| AC010401.1 | 16 | novel transcript, antisense to ERCC4 |
| AC010401.2 | 16 | novel transcript |
| LINC02185 | 16 | long intergenic non-protein coding RNA 2185 |
| LINC02186 | 16 | long intergenic non-protein coding RNA 2186 |
| MRTFB | 16 | myocardin related transcription factor B [Sour |
| AC130650.2 | 16 | novel transcript |
| AC130650.1 | 16 | novel transcript, antisense to MKL2 |
| TVP23CP2 | 16 | TVP23C pseudogene 2 [Source:HGNC Symbol; |
| Y_RNA | 16 | Y RNA [Source:RFAM;Acc:RF00019] |

| | | |
|---|---|---|
| MIR193BHG | 16 | MIR193B host gene [Source:HGNC Symbol;Ac |
| MIR193B | 16 | microRNA 193b [Source:HGNC Symbol;Acc:HG |
| MIR365A | 16 | microRNA 365a [Source:HGNC Symbol;Acc:HG |
| LINC02130 | 16 | long intergenic non-protein coding RNA 2130 |
| AC040173.1 | 16 | novel transcript |
| PARN | 16 | poly(A)-specific ribonuclease [Source:HGNC Sy |
| RN7SL274P | 16 | RNA, 7SL, cytoplasmic 274, pseudogene [Sour |
| BFAR | 16 | bifunctional apoptosis regulator [Source:HGN |
| RNU7-125P | 16 | RNA, U7 small nuclear 125 pseudogene [Sourc |
| PLA2G10 | 16 | phospholipase A2 group X [Source:HGNC Sym |
| AC009167.1 | 16 | novel transcript, antisense to a novel member |
| NPIPA3 | 16 | nuclear pore complex interacting protein fami |
| AC136443.1 | 16 | phospholipase A2, group X (PLA2G10) pseudo |
| AC136443.4 | 16 | novel transcript, antisense to 2 novel member |
| NPIPA2 | 16 | nuclear pore complex interacting protein fami |
| AC136443.2 | 16 | phospholipase A2, group X (PLA2G10) pseudo |
| ABCC6P2 | 16 | ATP binding cassette subfamily C member 6 p |
| NOMO1 | 16 | NODAL modulator 1 [Source:HGNC Symbol;Ac |
| AC136443.5 | 16 | acid phosphatase 1, soluble (ACP1) pseudoger |
| AC136443.3 | 16 | microRNA 3179-1 |
| MIR3179-1 | 16 | microRNA 3179-1 [Source:HGNC Symbol;Acc:I |
| MIR3670-1 | 16 | microRNA 3670-1 [Source:HGNC Symbol;Acc:I |
| AC138932.2 | 16 | microRNA 3180-1 |
| MIR3180-1 | 16 | microRNA 3180-1 [Source:HGNC Symbol;Acc:I |
| AC138932.1 | 16 | polycystic kidney disease 1 (autosomal domin |
| Y_RNA | 16 | Y RNA [Source:RFAM;Acc:RF00019] |
| NPIPA1 | 16 | nuclear pore complex interacting protein fami |
| MIR6511A1 | 16 | microRNA 6511a-1 [Source:HGNC Symbol;Acc |
| MIR6770-1 | 16 | microRNA 6770-1 [Source:HGNC Symbol;Acc:I |
| AC138932.4 | 16 | phospholipase A2, group X (PLA2G10) pseudo |
| PDXDC1 | 16 | pyridoxal dependent decarboxylase domain co |
| AC138932.3 | 16 | novel pseudogene KIAA2013 |
| MIR1972-1 | 16 | microRNA 1972-1 [Source:HGNC Symbol;Acc:I |
| AC138932.5 | 16 | novel transcript, antisense to PDXDC1 |
| NTAN1 | 16 | N-terminal asparagine amidase [Source:HGNC |
| RRN3 | 16 | RRN3 homolog, RNA polymerase I transcriptic |
| AC139256.1 | 16 | phospholipase A2 group X (PLA2G10) pseudog |
| NPIPP1 | 16 | nuclear pore complex interacting protein pseu |
| AC139256.2 | 16 | novel transcript |
| PKD1P6 | 16 | polycystin 1, transient receptor potential char |
| MIR6511B2 | 16 | microRNA 6511b-2 [Source:HGNC Symbol;Acc |
| Y_RNA | 16 | Y RNA [Source:RFAM;Acc:RF00019] |
| MIR3180-4 | 16 | microRNA 3180-4 [Source:HGNC Symbol;Acc:I |
| AC126763.1 | 16 | microRNA 3180-4 |
| AC137803.1 | 16 | phospholipase A2, group X (PLA2G10) pseudo |
| NPIPA5 | 16 | nuclear pore complex interacting protein fami |
| MPV17L | 16 | MPV17 mitochondrial inner membrane protei |
| AC140504.1 | 16 | MPV17 mitochondrial inner membrane protei |
| BMERB1 | 16 | bMERB domain containing 1 [Source:HGNC Sy |
| MARF1 | 16 | meiosis regulator and mRNA stability factor 1 |

| | | |
|---|---|---|
| AC026401.1 | 16 | novel transcript |
| MIR6506 | 16 | microRNA 6506 [Source:HGNC Symbol;Acc:HG |
| NDE1 | 16 | nudE neurodevelopment protein 1 [Source:HG |
| MIR484 | 16 | microRNA 484 [Source:HGNC Symbol;Acc:HGN |
| AC026401.2 | 16 | novel transcript, sense to NDE1 |
| AC026401.3 | 16 | novel transcript |
| MYH11 | 16 | myosin heavy chain 11 [Source:HGNC Symbol; |
| AF001548.2 | 16 | novel transcript, antisense to MYH11 |
| AF001548.1 | 16 | novel transcript, antisense to MYH11 |
| AF001548.3 | 16 | novel transcript |
| FOPNL | 16 | FGFR1OP N-terminal like [Source:HGNC Symb |
| RNU6-213P | 16 | RNA, U6 small nuclear 213, pseudogene [Sour |
| AC130651.1 | 16 | novel transcript, antisense to C16orf63 |
| RPL15P20 | 16 | ribosomal protein L15 pseudogene 20 [Source |
| ABCC1 | 16 | ATP binding cassette subfamily C member 1 [S |
| RPL17P40 | 16 | ribosomal protein L17 pseudogene 40 [Source |
| ABCC6 | 16 | ATP binding cassette subfamily C member 6 [S |
| AC136624.1 | 16 | novel transcript, antisense to ABCC6 |
| NOMO3 | 16 | NODAL modulator 3 [Source:HGNC Symbol;Ac |
| AC136624.3 | 16 | acid phosphatase 1, soluble (ACP1) pseudoger |
| AC136624.2 | 16 | novel transcript, antisense to NOMO3 |
| MIR3179-2 | 16 | microRNA 3179-2 [Source:HGNC Symbol;Acc:I |
| MIR3670-2 | 16 | microRNA 3670-2 [Source:HGNC Symbol;Acc:I |
| AC138969.3 | 16 | microRNA 3180-2 |
| MIR3180-2 | 16 | microRNA 3180-2 [Source:HGNC Symbol;Acc:I |
| PKD1P1 | 16 | polycystin 1, transient receptor potential char |
| Y_RNA | 16 | Y RNA [Source:RFAM;Acc:RF00019] |
| AC138969.1 | 16 | novel member of the nuclear pore complex in |
| MIR6511A2 | 16 | microRNA 6511a-2 [Source:HGNC Symbol;Acc |
| MIR6770-2 | 16 | microRNA 6770-2 [Source:HGNC Symbol;Acc:I |
| AC138969.4 | 16 | phospholipase A2, group X (PLA2G10) pseudo |
| AC138969.2 | 16 | polycystic kidney disease 1 (autosomal domin |
| MIR6511A3 | 16 | microRNA 6511a-3 [Source:HGNC Symbol;Acc |
| NPIPA7 | 16 | nuclear pore complex interacting protein fami |
| AC136619.1 | 16 | phospholipase A2, group X (PLA2G10) pseudo |
| AC136431.1 | 16 | novel transcript |
| AC136619.2 | 16 | geminin, DNA replication inhibitor (GMNN) ps |
| AC136431.2 | 16 | novel transcript |
| AC092326.1 | 16 | novel transcript |
| AC098965.1 | 16 | chromosome 12 open reading frame 12 (C12c |
| AC109446.4 | 16 | novel transcript |
| AC109446.2 | 16 | transcription elongation regulator 1 (TCERG1) |
| AC109446.1 | 16 | LYR motif containing 2 (LYRM2) pseudogene |
| XYLT1 | 16 | xylosyltransferase 1 [Source:HGNC Symbol;Ac |
| AC109446.3 | 16 | novel transcript, antisense to XYLT1 |
| AC099494.1 | 16 | novel transcript |
| AC099494.2 | 16 | novel transcript |
| AC099494.3 | 16 | novel transcript |
| AC009152.6 | 16 | novel transcript, antisense to XYLT1 |
| AC009152.1 | 16 | novel transcript, sense intronic to XYLT1 |

| | | |
|---|---|---|
| AC009152.3 | 16 | novel transcript |
| AC109495.1 | 16 | novel transcript |
| AC025277.1 | 16 | novel transcript |
| RPL7P47 | 16 | ribosomal protein L7 pseudogene 47 [Source: |
| AC010601.1 | 16 | novel transcript |
| AC010601.2 | 16 | TEC |
| AC091489.1 | 16 | novel transcript |
| AC091489.2 | 16 | geminin, DNA replication inhibitor (GMNN) ps |
| AC008785.1 | 16 | novel transcript |
| AC126755.4 | 16 | phospholipase A2, group X (PLA2G10) pseudo |
| NPIPA8 | 16 | nuclear pore complex interacting protein fami |
| AC126755.6 | 16 | novel transcript |
| AC126755.1 | 16 | polycystic kidney disease 1 (autosomal domin |
| MIR6511A4 | 16 | microRNA 6511a-4 [Source:HGNC Symbol;Acc |
| AC126755.5 | 16 | phospholipase A2, group X (PLA2G10) pseudo |
| NPIPA9 | 16 | nuclear pore complex interacting protein fami |
| PKD1P5 | 16 | polycystin 1, transient receptor potential char |
| MIR6770-3 | 16 | microRNA 6770-3 [Source:HGNC Symbol;Acc: |
| Y_RNA | 16 | Y RNA [Source:RFAM;Acc:RF00019] |
| AC126755.3 | 16 | microRNA 3180-3 |
| MIR3180-3 | 16 | microRNA 3180-3 [Source:HGNC Symbol;Acc: |
| MIR3670-3 | 16 | microRNA 3670-3 [Source:HGNC Symbol;Acc: |
| AC126755.2 | 16 | microRNA 3179-3 |
| MIR3179-3 | 16 | microRNA 3179-3 [Source:HGNC Symbol;Acc: |
| NOMO2 | 16 | NODAL modulator 2 [Source:HGNC Symbol;Ac |
| MIR3670-4 | 16 | microRNA 3670-4 [Source:HGNC Symbol;Acc: |
| MIR3179-4 | 16 | microRNA 3179-4 [Source:HGNC Symbol;Acc: |
| AC136618.2 | 16 | acid phosphatase 1, soluble (ACP1) pseudoger |
| AC136618.1 | 16 | novel transcript |
| ABCC6P1 | 16 | ATP binding cassette subfamily C member 6 p |
| RPS15A | 16 | ribosomal protein S15a [Source:HGNC Symbol |
| AC138811.2 | 16 | novel protein |
| ARL6IP1 | 16 | ADP ribosylation factor like GTPase 6 interacti |
| AC138811.1 | 16 | novel transcript, antisense to SMG1 |
| SMG1 | 16 | SMG1 nonsense mediated mRNA decay assoc |
| AC092287.1 | 16 | novel transcript, antisense to SMG1 |
| TMC7 | 16 | transmembrane channel like 7 [Source:HGNC |
| RNU6-1340P | 16 | RNA, U6 small nuclear 1340, pseudogene [Sou |
| AC099518.4 | 16 | novel transcript, antisense to TMC7 and COQ7 |
| AC099518.2 | 16 | novel transcript |
| COQ7 | 16 | coenzyme Q7, hydroxylase [Source:HGNC Sym |
| AC099518.1 | 16 | novel transcript |
| AC099518.6 | 16 | novel transcript |
| ITPRIPL2 | 16 | ITPRIP like 2 [Source:HGNC Symbol;Acc:HGNC |
| AC099518.3 | 16 | novel transcript |
| AC099518.5 | 16 | novel transcript, antisense to ITPRIPL2 |
| SYT17 | 16 | synaptotagmin 17 [Source:HGNC Symbol;Acc: |
| AC010494.1 | 16 | novel transcript, antisense to SYT17 |
| CLEC19A | 16 | C-type lectin domain containing 19A [Source: |
| AC130456.1 | 16 | novel transcript, sense intronic to CLEC19A |

| | | |
|---|---|---|
| LINC02858 | 16 | long intergenic non-protein coding RNA 2858 |
| AC130456.2 | 16 | novel transcript |
| AC130456.7 | 16 | novel transcript |
| TMC5 | 16 | transmembrane channel like 5 [Source:HGNC |
| AC130456.4 | 16 | novel transcript, antisense to TMC5 |
| AC130456.3 | 16 | novel transcript, antisense to TMC5 |
| RNU4-46P | 16 | RNA, U4 small nuclear 46, pseudogene [Sourc |
| AC130456.5 | 16 | novel transcript, antisense to GDE1 |
| GDE1 | 16 | glycerophosphodiester phosphodiesterase 1 [ |
| AC130456.6 | 16 | TEC |
| CCP110 | 16 | centriolar coiled-coil protein 110 [Source:HGN |
| VPS35L | 16 | VPS35 endosomal protein sorting factor like [S |
| AC002550.2 | 16 | novel transcript, antisense to C16orf62 |
| KNOP1 | 16 | lysine rich nucleolar protein 1 [Source:HGNC S |
| AC002550.1 | 16 | novel transcript, antisense to C16orf88 |
| IQCK | 16 | IQ motif containing K [Source:HGNC Symbol;A |
| AC027130.1 | 16 | novel transcript, antisense to IQCK |
| LARP7P2 | 16 | La ribonucleoprotein domain family member 7 |
| GPRC5B | 16 | G protein-coupled receptor class C group 5 m |
| GPR139 | 16 | G protein-coupled receptor 139 [Source:HGN |
| AC092132.2 | 16 | spalt-like transcription factor 1 (SALL1) pseud |
| AC092132.1 | 16 | novel transcript |
| SNRPEP3 | 16 | small nuclear ribonucleoprotein polypeptide E |
| GP2 | 16 | glycoprotein 2 [Source:HGNC Symbol;Acc:HGN |
| UMOD | 16 | uromodulin [Source:HGNC Symbol;Acc:HGNC: |
| PDILT | 16 | protein disulfide isomerase like, testis express |
| AC106796.1 | 16 | novel transcript, antisense to PDILT |
| ACSM5 | 16 | acyl-CoA synthetase medium chain family mer |
| AC137056.1 | 16 | novel transcript, antisense to ACSM5 |
| ACSM2A | 16 | acyl-CoA synthetase medium chain family mer |
| AC137056.2 | 16 | TEC |
| ACSM2B | 16 | acyl-CoA synthetase medium chain family mer |
| AC141273.1 | 16 | novel transcript |
| ACSM5P1 | 16 | acyl-CoA synthetase medium chain family mer |
| ACSM3 | 16 | acyl-CoA synthetase medium chain family mer |
| AC141273.3 | 16 | nuclear pore complex interacting protein fami |
| AC141273.2 | 16 | nuclear pore complex interacting protein fami |
| ACSM1 | 16 | acyl-CoA synthetase medium chain family mer |
| THUMPD1 | 16 | THUMP domain containing 1 [Source:HGNC Sy |
| AC004381.3 | 16 | novel acyl-CoA synthetase medium-chain fam |
| AC004381.4 | 16 | mitochondrially encoded NADH dehydrogenas |
| AC004381.1 | 16 | novel transcript, antisense to ACSM3 |
| ERI2 | 16 | ERI1 exoribonuclease family member 2 [Sourc |
| RNU6-944P | 16 | RNA, U6 small nuclear 944, pseudogene [Sour |
| REXO5 | 16 | RNA exonuclease 5 [Source:HGNC Symbol;Acc |
| AC004381.2 | 16 | NULL |
| DCUN1D3 | 16 | defective in cullin neddylation 1 domain conta |
| LYRM1 | 16 | LYR motif containing 1 [Source:HGNC Symbol; |
| DNAH3 | 16 | dynein axonemal heavy chain 3 [Source:HGNC |
| AC008551.1 | 16 | novel transcript, antisense to DNAH3 |

| | | |
|---|---|---|
| TMEM159 | 16 | transmembrane protein 159 [Source:HGNC Sy |
| AF001550.1 | 16 | novel transcript, antisense to ZP2 |
| ZP2 | 16 | zona pellucida glycoprotein 2 [Source:HGNC S |
| ANKS4B | 16 | ankyrin repeat and sterile alpha motif domain |
| CRYM | 16 | crystallin mu [Source:HGNC Symbol;Acc:HGN( |
| AF001550.2 | 16 | ATP-binding cassette, sub-family A (ABC1), me |
| CRYM-AS1 | 16 | CRYM antisense RNA 1 [Source:HGNC Symbol; |
| AC008740.1 | 16 | BTG3 associated nuclear protein (BNAP) pseu( |
| SNX29P1 | 16 | sorting nexin 29 pseudogene 1 [Source:HGNC |
| AC008740.2 | 16 | phospholipase A2, group X (PLA2G10) pseudo |
| NPIPB3 | 16 | nuclear pore complex interacting protein fami |
| SMG1P3 | 16 | SMG1 pseudogene 3 [Source:HGNC Symbol;A( |
| MIR3680-1 | 16 | microRNA 3680-1 [Source:HGNC Symbol;Acc:I |
| AC005632.6 | 16 | novel transcript |
| SLC7A5P2 | 16 | solute carrier family 7 member 5 pseudogene |
| AC005632.5 | 16 | novel transcript |
| AC005632.2 | 16 | carbonic anhydrase VA, mitochondrial (CA5A) |
| AC005632.4 | 16 | ribosomal protein S10 (RPS10) pseudogene |
| AC005632.1 | 16 | NULL |
| METTL9 | 16 | methyltransferase like 9 [Source:HGNC Symbo |
| AC005632.3 | 16 | novel transcript, sense intronic to METTL9 |
| IGSF6 | 16 | immunoglobulin superfamily member 6 [Sour( |
| RNU6-1005P | 16 | RNA, U6 small nuclear 1005, pseudogene [Sou |
| RNU6-196P | 16 | RNA, U6 small nuclear 196, pseudogene [Sour |
| OTOA | 16 | otoancorin [Source:HGNC Symbol;Acc:HGNC:1 |
| AC092375.2 | 16 | novel transcript |
| RRN3P1 | 16 | RRN3 homolog, RNA polymerase I transcriptic |
| AC092375.3 | 16 | phospholipase A2, group X (PLA2G10) pseudo |
| NPIPB4 | 16 | nuclear pore complex interacting protein fami |
| SMG1P4 | 16 | SMG1 pseudogene 4 [Source:HGNC Symbol;A( |
| AC092375.1 | 16 | ATP-binding cassette, sub-family A (ABC1), me |
| AC092119.2 | 16 | novel transcript |
| UQCRC2 | 16 | ubiquinol-cytochrome c reductase core protei |
| PDZD9 | 16 | PDZ domain containing 9 [Source:HGNC Symb |
| AC092119.1 | 16 | islet cell autoantigen 1, 69kDa (ICA1) pseudog |
| AC092119.3 | 16 | novel transcript, antisense to C16orf52 |
| MOSMO | 16 | modulator of smoothened [Source:HGNC Sym |
| AC009019.1 | 16 | novel transcript, antisense to C16orf52 and V\ |
| VWA3A | 16 | von Willebrand factor A domain containing 3A |
| SDR42E2 | 16 | short chain dehydrogenase/reductase family 4 |
| AC009019.2 | 16 | novel transcript, antisense to SDR42E2 |
| EEF2K | 16 | eukaryotic elongation factor 2 kinase [Source: |
| AC009034.1 | 16 | novel transcript, antisense to EEF2K |
| POLR3E | 16 | RNA polymerase III subunit E [Source:HGNC Sy |
| AC092338.3 | 16 | novel transcript, antisense to POLR3E |
| CDR2 | 16 | cerebellar degeneration related protein 2 [Sou |
| AC092338.1 | 16 | novel transcript, antisense to CDR2 |
| AC092338.2 | 16 | novel transcript, antisense to CDR2 |
| MFSD13B | 16 | major facilitator superfamily domain containi |
| RRN3P3 | 16 | RRN3 homolog, RNA polymerase I transcriptic |

| | | |
|---|---|---|
| SMG1P1 | 16 | SMG1 pseudogene 1 [Source:HGNC Symbol;A |
| NPIPB5 | 16 | nuclear pore complex interacting protein fami |
| AC106788.1 | 16 | phospholipase A2, group X (PLA2G10) pseudo |
| OTOAP1 | 16 | OTOA pseudogene 1 [Source:HGNC Symbol;A |
| AC009021.2 | 16 | novel transcript |
| AC009021.1 | 16 | novel transcript |
| AC130466.1 | 16 | novel transcript |
| HS3ST2 | 16 | heparan sulfate-glucosamine 3-sulfotransfera |
| AC127459.2 | 16 | novel transcript |
| AC127459.1 | 16 | novel transcript |
| USP31 | 16 | ubiquitin specific peptidase 31 [Source:HGNC |
| AC127459.3 | 16 | novel transcript |
| AC099482.2 | 16 | novel transcript |
| SCNN1G | 16 | sodium channel epithelial 1 gamma subunit [S |
| AC099482.1 | 16 | novel transcript |
| SCNN1B | 16 | sodium channel epithelial 1 beta subunit [Sou |
| COG7 | 16 | component of oligomeric golgi complex 7 [Sou |
| RN7SKP23 | 16 | RN7SK pseudogene 23 [Source:HGNC Symbol; |
| SNORA75 | 16 | Small nucleolar RNA SNORA75 [Source:RFAM; |
| AC008915.1 | 16 | novel transcript |
| AC008915.2 | 16 | novel transcript |
| GGA2 | 16 | golgi associated, gamma adaptin ear containir |
| AC002400.1 | 16 | cyclin Y-like 1 (CCNYL1) pseudogene |
| EARS2 | 16 | glutamyl-tRNA synthetase 2, mitochondrial [S |
| SUB1P4 | 16 | SUB1 pseudogene 4 [Source:HGNC Symbol;Ac |
| UBFD1 | 16 | ubiquitin family domain containing 1 [Source: |
| AC008870.2 | 16 | novel transcript |
| NDUFAB1 | 16 | NADH:ubiquinone oxidoreductase subunit AB |
| PALB2 | 16 | partner and localizer of BRCA2 [Source:HGNC |
| AC008870.4 | 16 | novel transcript, antisense to PALB2 |
| DCTN5 | 16 | dynactin subunit 5 [Source:HGNC Symbol;Acc: |
| AC008870.1 | 16 | novel transcript |
| PLK1 | 16 | polo like kinase 1 [Source:HGNC Symbol;Acc:H |
| AC008870.3 | 16 | novel transcript |
| ERN2 | 16 | endoplasmic reticulum to nucleus signaling 2 [ |
| AC008870.5 | 16 | novel transcript |
| AC012317.1 | 16 | novel transcript |
| AC012317.2 | 16 | novel transcript |
| CHP2 | 16 | calcineurin like EF-hand protein 2 [Source:HGI |
| PRKCB | 16 | protein kinase C beta [Source:HGNC Symbol;A |
| AC130454.1 | 16 | novel transcript |
| AC130448.1 | 16 | novel transcript |
| MIR1273H | 16 | microRNA 1273h [Source:HGNC Symbol;Acc:H |
| AC130448.2 | 16 | novel transcript |
| LINC2194 | 16 | long intergenic non-protein coding RNA 2194 |
| CACNG3 | 16 | calcium voltage-gated channel auxiliary subun |
| AC004125.1 | 16 | NULL |
| RNU7-24P | 16 | RNA, U7 small nuclear 24 pseudogene [Source |
| AC008938.1 | 16 | novel transcript |
| RBBP6 | 16 | RB binding protein 6, ubiquitin ligase [Source: |

| | | |
|---|---|---|
| TNRC6A | 16 | trinucleotide repeat containing adaptor 6A [So |
| LINC01567 | 16 | long intergenic non-protein coding RNA 1567 |
| AC008731.1 | 16 | novel transcript, antisense to TNRC6A |
| SLC5A11 | 16 | solute carrier family 5 member 11 [Source:HG |
| ARHGAP17 | 16 | Rho GTPase activating protein 17 [Source:HGI |
| AC133552.1 | 16 | thyroid hormone receptor interactor 11 (TRIP |
| AC133552.2 | 16 | leucine carboxyl methyltransferase 1 (LCMT1) |
| LINC02175 | 16 | long intergenic non-protein coding RNA 2175 |
| SCML2P2 | 16 | SCML2 pseudogene 2 [Source:HGNC Symbol;A |
| AC133552.4 | 16 | thyroid hormone receptor interactor 11 (TRIP |
| LCMT1-AS1 | 16 | LCMT1 antisense RNA 1 [Source:HGNC Symbo |
| AC133552.3 | 16 | thyroid hormone receptor interactor 11 (TRIP |
| AC133552.5 | 16 | novel transcript |
| LCMT1 | 16 | leucine carboxyl methyltransferase 1 [Source: |
| LCMT1-AS2 | 16 | LCMT1 antisense RNA 2 [Source:HGNC Symbo |
| RN7SL557P | 16 | RNA, 7SL, cytoplasmic 557, pseudogene [Sour |
| AQP8 | 16 | aquaporin 8 [Source:HGNC Symbol;Acc:HGNC |
| ZKSCAN2 | 16 | zinc finger with KRAB and SCAN domains 2 [So |
| AC008741.1 | 16 | novel transcript, antisense to ZKSCAN2 |
| ZKSCAN2-DT | 16 | ZKSCAN2 divergent transcript [Source:HGNC S |
| LINC02191 | 16 | long intergenic non-protein coding RNA 2191 |
| CYCSP39 | 16 | CYCS pseudogene 39 [Source:HGNC Symbol;A |
| HS3ST4 | 16 | heparan sulfate-glucosamine 3-sulfotransfera |
| AC093516.1 | 16 | novel transcript, sense intronic to HS3ST4 |
| AC093524.1 | 16 | TNF receptor-associated factor 7 (TRAF7) pseu |
| MIR548W | 16 | microRNA 548w [Source:HGNC Symbol;Acc:H( |
| RNA5SP405 | 16 | RNA, 5S ribosomal pseudogene 405 [Source:H |
| HMGN2P3 | 16 | high mobility group nucleosomal binding dom |
| HSPE1P16 | 16 | heat shock protein family E (Hsp10) member [illegible] |
| AC009158.1 | 16 | novel transcript |
| AC130464.1 | 16 | ER degradation enhancer, mannosidase alpha |
| AC002331.1 | 16 | novel transcript |
| LINC02195 | 16 | long intergenic non-protein coding RNA 2195 |
| AC009035.1 | 16 | novel transcript |
| C16orf82 | 16 | chromosome 16 open reading frame 82 [Sour |
| AC092725.1 | 16 | novel transcript, antisense to C16orf82 |
| EEF1A1P38 | 16 | eukaryotic translation elongation factor 1 alpl |
| LINC02129 | 16 | long intergenic non-protein coding RNA 2129 |
| KDM8 | 16 | lysine demethylase 8 [Source:HGNC Symbol;A |
| AC109449.1 | 16 | novel transcript, antisense to JMJD5 |
| NSMCE1 | 16 | NSE1 homolog, SMC5-SMC6 complex compon |
| NSMCE1-DT | 16 | NSMCE1 divergent transcript [Source:HGNC S |
| AC106739.1 | 16 | novel transcript, antisense to IL4R |
| IL4R | 16 | interleukin 4 receptor [Source:HGNC Symbol;/ |
| IL21R | 16 | interleukin 21 receptor [Source:HGNC Symbol |
| IL21R-AS1 | 16 | IL21R antisense RNA 1 [Source:HGNC Symbol; |
| GTF3C1 | 16 | general transcription factor IIIC subunit 1 [Sou |
| KIAA0556 | 16 | KIAA0556 [Source:HGNC Symbol;Acc:HGNC:2! |
| AC002551.1 | 16 | novel transcript |
| AC016597.1 | 16 | novel transcript, antisense to a novel protein |

| | | |
|---|---|---|
| AC016597.2 | 16 | novel transcript, sense intronic with KIAA0556 |
| Y_RNA | 16 | Y RNA [Source:RFAM;Acc:RF00019] |
| GSG1L | 16 | GSG1 like [Source:HGNC Symbol;Acc:HGNC:28 |
| RNU6-159P | 16 | RNA, U6 small nuclear 159, pseudogene [Sour |
| RNU6-1241P | 16 | RNA, U6 small nuclear 1241, pseudogene [Sou |
| XPO6 | 16 | exportin 6 [Source:HGNC Symbol;Acc:HGNC:1 |
| TPRKBP2 | 16 | TP53RK binding protein pseudogene 2 [Source |
| AC136611.1 | 16 | NULL |
| RNY1P10 | 16 | RNY1 pseudogene 10 [Source:HGNC Symbol;A |
| GAPDHP35 | 16 | glyceraldehyde 3 phosphate dehydrogenase p |
| AC138904.3 | 16 | novel transcript |
| SBK1 | 16 | SH3 domain binding kinase 1 [Source:HGNC Sy |
| AC138904.1 | 16 | novel transcript LOC100506705 |
| AC138904.2 | 16 | phospholipase A2, group X (PLA2G10) pseudo |
| NPIPB6 | 16 | nuclear pore complex interacting protein fami |
| AC138894.3 | 16 | PRKC, apoptosis, WT1, regulator (PAWR) pseu |
| EIF3CL | 16 | eukaryotic translation initiation factor 3 subur |
| MIR6862-1 | 16 | microRNA 6862-1 [Source:HGNC Symbol;Acc:I |
| CDC37P2 | 16 | cell division cycle 37 pseudogene 2 [Source:H( |
| AC138894.4 | 16 | phospholipase A2, group X (PLA2G10) pseudo |
| AC138894.2 | 16 | novel transcript |
| AC138894.1 | 16 | novel protein |
| NPIPB7 | 16 | nuclear pore complex interacting protein fami |
| CLN3 | 16 | CLN3 lysosomal/endosomal transmembrane p |
| APOBR | 16 | apolipoprotein B receptor [Source:HGNC Sym |
| IL27 | 16 | interleukin 27 [Source:HGNC Symbol;Acc:HGN |
| NUPR1 | 16 | nuclear protein 1, transcriptional regulator [Sc |
| AC020765.1 | 16 | ependymin related protein 1 (zebrafish) (EPDI |
| AC020765.2 | 16 | novel transcript, antisense to CCDC101 |
| SGF29 | 16 | SAGA complex associated factor 29 [Source:H |
| SULT1A2 | 16 | sulfotransferase family 1A member 2 [Source: |
| AC020765.3 | 16 | SLX1 structure-specific endonuclease subunit |
| AC020765.4 | 16 | coiled-coil domain containing 101 (CCDC101) |
| SULT1A1 | 16 | sulfotransferase family 1A member 1 [Source: |
| AC020765.5 | 16 | SLX1 structure-specific endonuclease subunit |
| NPIPB8 | 16 | nuclear pore complex interacting protein fami |
| AC145285.4 | 16 | phospholipase A2, group X (PLA2G10) pseudo |
| EIF3C | 16 | eukaryotic translation initiation factor 3 subur |
| CDC37P1 | 16 | cell division cycle 37 pseudogene 1 [Source:H( |
| MIR6862-2 | 16 | microRNA 6862-2 [Source:HGNC Symbol;Acc:I |
| AC145285.5 | 16 | PRKC, apoptosis, WT1, regulator (PAWR) pseu |
| NPIPB9 | 16 | nuclear pore complex interacting protein fami |
| AC145285.7 | 16 | phospholipase A2, group X (PLA2G10) pseudo |
| AC145285.2 | 16 | novel transcript |
| AC145285.1 | 16 | ribosomal protein S15a (RPS15A) pseudogene |
| AC145285.3 | 16 | novel transcript, antisense to ATXN2L |
| AC145285.6 | 16 | novel transcript, antisense to ATXN2L |
| ATXN2L | 16 | ataxin 2 like [Source:HGNC Symbol;Acc:HGNC |
| AC133550.2 | 16 | novel transcript, antisense to ATXN2L |
| TUFM | 16 | Tu translation elongation factor, mitochondria |

| | | |
|---|---|---|
| MIR4721 | 16 | microRNA 4721 [Source:HGNC Symbol;Acc:HG |
| SH2B1 | 16 | SH2B adaptor protein 1 [Source:HGNC Symbo |
| AC133550.3 | 16 | novel transcript, antisense to SH2B1 |
| ATP2A1 | 16 | ATPase sarcoplasmic/endoplasmic reticulum ( |
| ATP2A1-AS1 | 16 | ATP2A1 antisense RNA 1 [Source:HGNC Symb |
| AC133550.1 | 16 | NULL |
| RABEP2 | 16 | rabaptin, RAB GTPase binding effector protei |
| CD19 | 16 | CD19 molecule [Source:HGNC Symbol;Acc:HG |
| NFATC2IP | 16 | nuclear factor of activated T cells 2 interacting |
| AC109460.2 | 16 | novel transcript, antisense to NFATC2IP |
| MIR4517 | 16 | microRNA 4517 [Source:HGNC Symbol;Acc:HG |
| AC109460.1 | 16 | novel transcript, antisense to SPNS1 |
| SPNS1 | 16 | sphingolipid transporter 1 (putative) [Source:I |
| AC109460.3 | 16 | novel transcript |
| LAT | 16 | linker for activation of T cells [Source:HGNC Sy |
| AC109460.4 | 16 | novel transcript, antisense to LAT & SPNS1 |
| NPIPB10P | 16 | nuclear pore complex interacting protein fami |
| AC009093.3 | 16 | phospholipase A2, group X (PLA2G10) pseudo |
| RRN3P2 | 16 | RRN3 homolog, RNA polymerase I transcriptic |
| AC009093.9 | 16 | RRN3 RNA polymerase I transcription factor h |
| AC009093.8 | 16 | BTG3 associated nuclear protein (BANP) pseu |
| AC009093.10 | 16 | novel transcript |
| AC009093.2 | 16 | novel transcript |
| AC009093.7 | 16 | TEC |
| AC009093.1 | 16 | novel transcript |
| AC009093.5 | 16 | novel transcript |
| AC009093.4 | 16 | novel transcript |
| AC009093.6 | 16 | novel transcript |
| AC025279.1 | 16 | novel transcript |
| AC025279.2 | 16 | BTG3 associated nuclear protein (BANP) pseu |
| SNX29P2 | 16 | sorting nexin 29 pseudogene 2 [Source:HGNC |
| AC025279.3 | 16 | phospholipase A2, group X (PLA2G10) pseudo |
| NPIPB11 | 16 | nuclear pore complex interacting protein fami |
| SMG1P6 | 16 | SMG1 pseudogene 6 [Source:HGNC Symbol;A |
| BOLA2 | 16 | bolA family member 2 [Source:HGNC Symbol; |
| BOLA2-SMG1P6 | 16 | BOLA2-SMG1P6 readthrough [Source:HGNC S |
| AC133555.4 | 16 | coronin, actin binding protein, 1A (CORO1A) p |
| SLX1B | 16 | SLX1 homolog B, structure-specific endonucle |
| SLX1B-SULT1A4 | 16 | SLX1B-SULT1A4 readthrough (NMD candidate |
| SULT1A4 | 16 | sulfotransferase family 1A member 4 [Source: |
| AC133555.3 | 16 | coiled-coil domain containing 101 (CCDC101) |
| AC133555.5 | 16 | SLX1 structure-specific endonuclease subunit |
| AC133555.2 | 16 | MPV17 mitochondrial membrane protein-like |
| AC133555.1 | 16 | phospholipase A2, group X (PLA2G10) pseudo |
| NPIPB12 | 16 | nuclear pore complex interacting protein fami |
| SMG1P2 | 16 | SMG1 pseudogene 2 [Source:HGNC Symbol;A |
| AC009086.2 | 16 | novel transcript |
| MIR3680-2 | 16 | microRNA 3680-2 [Source:HGNC Symbol;Acc:I |
| SLC7A5P1 | 16 | solute carrier family 7 member 5 pseudogene |
| CA5AP1 | 16 | carbonic anhydrase 5A pseudogene 1 [Source |

| | | |
|---|---|---|
| AC009086.1 | 16 | ribosomal protein S10 (RPS10) pseudogene |
| SPN | 16 | sialophorin [Source:HGNC Symbol;Acc:HGNC:[illegible] |
| QPRT | 16 | quinolinate phosphoribosyltransferase [Sourc[illegible] |
| RN7SKP127 | 16 | RN7SK pseudogene 127 [Source:HGNC Symbo[illegible] |
| C16orf54 | 16 | chromosome 16 open reading frame 54 [Sour[illegible] |
| AC009133.3 | 16 | novel transcript, antisense to C16orf54 |
| ZG16 | 16 | zymogen granule protein 16 [Source:HGNC Sy[illegible] |
| KIF22 | 16 | kinesin family member 22 [Source:HGNC Sym[illegible] |
| AC009133.4 | 16 | novel transcript, antisense to KIF22 |
| MAZ | 16 | MYC associated zinc finger protein [Source:H[illegible] |
| AC009133.2 | 16 | novel transcript, antisense to MAZ |
| AC009133.1 | 16 | novel transcript, antisense to C16orf53 and M[illegible] |
| AC009133.5 | 16 | novel transcript |
| PRRT2 | 16 | proline rich transmembrane protein 2 [Source[illegible] |
| AC009133.6 | 16 | novel transcript |
| PAGR1 | 16 | PAXIP1 associated glutamate rich protein 1 [S[illegible] |
| AC120114.4 | 16 | novel protein |
| MVP | 16 | major vault protein [Source:HGNC Symbol;Ac[illegible] |
| CDIPT | 16 | CDP-diacylglycerol--inositol 3-phosphatidyltra[illegible] |
| AC120114.2 | 16 | novel transcript, antisense to CDIPT |
| CDIPTOSP | 16 | CDIP transferase opposite strand, pseudogene[illegible] |
| SEZ6L2 | 16 | seizure related 6 homolog like 2 [Source:HGN[illegible] |
| ASPHD1 | 16 | aspartate beta-hydroxylase domain containin[illegible] |
| KCTD13 | 16 | potassium channel tetramerization domain c[illegible] |
| AC120114.3 | 16 | novel transcript, sense intronic to KCTD13 |
| AC120114.1 | 16 | novel transcript, antisense to KCTD13 |
| TMEM219 | 16 | transmembrane protein 219 [Source:HGNC Sy[illegible] |
| TAOK2 | 16 | TAO kinase 2 [Source:HGNC Symbol;Acc:HGN[illegible] |
| HIRIP3 | 16 | HIRA interacting protein 3 [Source:HGNC Sym[illegible] |
| INO80E | 16 | INO80 complex subunit E [Source:HGNC Symb[illegible] |
| DOC2A | 16 | double C2 domain alpha [Source:HGNC Symbo[illegible] |
| C16orf92 | 16 | chromosome 16 open reading frame 92 [Sour[illegible] |
| TLCD3B | 16 | TLC domain containing 3B [Source:HGNC Sym[illegible] |
| AC093512.2 | 16 | uncharaterized LOC112694756 [Source:NCBI [illegible] |
| ALDOA | 16 | aldolase, fructose-bisphosphate A [Source:HG[illegible] |
| AC093512.1 | 16 | novel transcript, antisense to ALDOA |
| PPP4C | 16 | protein phosphatase 4 catalytic subunit [Sour[illegible] |
| TBX6 | 16 | T-box transcription factor 6 [Source:HGNC Sy[illegible] |
| YPEL3 | 16 | yippee like 3 [Source:HGNC Symbol;Acc:HGN[illegible] |
| AC012645.1 | 16 | novel transcript, antisense to YPEL3 |
| GDPD3 | 16 | glycerophosphodiester phosphodiesterase do[illegible] |
| AC012645.2 | 16 | novel transcript, antisense to GDPD3 |
| AC012645.4 | 16 | novel transcript, antisense to GDPD3 |
| MAPK3 | 16 | mitogen-activated protein kinase 3 [Source:H[illegible] |
| CORO1A | 16 | coronin 1A [Source:HGNC Symbol;Acc:HGNC:[illegible] |
| AC012645.3 | 16 | novel transcript, antisense to CORO1A |
| BOLA2B | 16 | bolA family member 2B [Source:HGNC Symbo[illegible] |
| SLX1A | 16 | SLX1 homolog A, structure-specific endonucle[illegible] |
| SLX1A-SULT1A3 | 16 | SLX1A-SULT1A3 readthrough (NMD candidate[illegible] |
| SULT1A3 | 16 | sulfotransferase family 1A member 3 [Source:[illegible] |

| | | |
|---|---|---|
| AC106782.1 | 16 | coiled-coil domain containing 101 (CCDC101) |
| AC106782.6 | 16 | SLX1 structure-specific endonuclease subunit |
| AC106782.3 | 16 | MPV17 mitochondrial membrane protein-like |
| AC106782.2 | 16 | phospholipase A2, group X (PLA2G10) pseudo |
| NPIPB13 | 16 | nuclear pore complex interacting protein fami |
| SMG1P5 | 16 | SMG1 pseudogene 5 [Source:HGNC Symbol;A |
| AC106782.4 | 16 | carbonic anhydrase VA, mitochondrial (CA5A) |
| CD2BP2 | 16 | CD2 cytoplasmic tail binding protein 2 [Source |
| CD2BP2-DT | 16 | CD2BP2 divergent transcript [Source:HGNC Sy |
| TBC1D10B | 16 | TBC1 domain family member 10B [Source:HG |
| AC106782.5 | 16 | novel transcript, antisense to TBC1D10B |
| MYLPF | 16 | myosin light chain, phosphorylatable, fast ske |
| ZNF48 | 16 | zinc finger protein 48 [Source:HGNC Symbol;A |
| SEPTIN1 | 16 | septin 1 [Source:HGNC Symbol;Acc:HGNC:287 |
| AC116348.4 | 16 | novel transcript |
| ZNF771 | 16 | zinc finger protein 771 [Source:HGNC Symbol; |
| SNORA80C | 16 | small nucleolar RNA, H/ACA box 80C [Source: |
| DCTPP1 | 16 | dCTP pyrophosphatase 1 [Source:HGNC Symb |
| SEPHS2 | 16 | selenophosphate synthetase 2 [Source:HGNC |
| Y_RNA | 16 | Y RNA [Source:RFAM;Acc:RF00019] |
| Y_RNA | 16 | Y RNA [Source:RFAM;Acc:RF00019] |
| ITGAL | 16 | integrin subunit alpha L [Source:HGNC Symbo |
| AC116348.3 | 16 | novel transcript, antisense to ITGAL |
| AC116348.1 | 16 | novel transcript, antisense to ITGAL |
| RNU7-61P | 16 | RNA, U7 small nuclear 61 pseudogene [Source |
| AC116348.2 | 16 | novel transcript, antisense to ITGAL |
| MIR4518 | 16 | microRNA 4518 [Source:HGNC Symbol;Acc:HG |
| ZNF768 | 16 | zinc finger protein 768 [Source:HGNC Symbol; |
| AC002310.4 | 16 | novel transcript |
| AC002310.5 | 16 | novel transcript |
| ZNF747 | 16 | zinc finger protein 747 [Source:HGNC Symbol; |
| AC002310.1 | 16 | novel transcript, antisense to ZNF747 |
| AC002310.3 | 16 | ribosomal protein L7 (RPL7) pseudogene |
| ZNF764 | 16 | zinc finger protein 764 [Source:HGNC Symbol; |
| ZNF688 | 16 | zinc finger protein 688 [Source:HGNC Symbol; |
| AC002310.2 | 16 | novel transcript |
| ZNF785 | 16 | zinc finger protein 785 [Source:HGNC Symbol; |
| AC093249.2 | 16 | novel transcript, antisense to ZNF689 |
| AC093249.4 | 16 | zinc finger protein (ZNF) pseudogene |
| ZNF689 | 16 | zinc finger protein 689 [Source:HGNC Symbol; |
| AC093249.3 | 16 | zinc finger protein (ZNF) pseudogene |
| AC093249.5 | 16 | zinc finger protein (ZNF) pseudogene |
| AC093249.1 | 16 | ribosomal protein L5 (RPL5) pseudogene |
| PRR14 | 16 | proline rich 14 [Source:HGNC Symbol;Acc:HGI |
| FBRS | 16 | fibrosin [Source:HGNC Symbol;Acc:HGNC:204 |
| RNU6-416P | 16 | RNA, U6 small nuclear 416, pseudogene [Sour |
| AC093249.6 | 16 | uncharacterized LOC730183 [Source:NCBI ger |
| SRCAP | 16 | Snf2 related CREBBP activator protein [Source |
| RNU6-1043P | 16 | RNA, U6 small nuclear 1043, pseudogene [Sou |
| AC106886.5 | 16 | novel transcript |

| | | |
|---|---|---|
| SNORA30 | 16 | small nucleolar RNA, H/ACA box 30 [Source:H |
| TMEM265 | 16 | transmembrane protein 265 [Source:HGNC Sy |
| AC106886.2 | 16 | novel transcript |
| PHKG2 | 16 | phosphorylase kinase catalytic subunit gamma |
| CCDC189 | 16 | coiled-coil domain containing 189 [Source:HG |
| RNF40 | 16 | ring finger protein 40 [Source:HGNC Symbol;A |
| AC106886.4 | 16 | novel transcript, antisense to RNF40 |
| ZNF629 | 16 | zinc finger protein 629 [Source:HGNC Symbol; |
| Metazoa_SRP | 16 | Metazoan signal recognition particle RNA [Sou |
| AC106886.1 | 16 | chromosome 20 open reading frame 27 (C20o |
| AC106886.3 | 16 | novel transcript |
| BCL7C | 16 | BAF chromatin remodeling complex subunit B |
| MIR762HG | 16 | MIR762 host gene [Source:HGNC Symbol;Acc: |
| MIR4519 | 16 | microRNA 4519 [Source:HGNC Symbol;Acc:HG |
| AC135048.2 | 16 | novel transcript |
| MIR762 | 16 | microRNA 762 [Source:HGNC Symbol;Acc:HGN |
| CTF1 | 16 | cardiotrophin 1 [Source:HGNC Symbol;Acc:HG |
| CTF2P | 16 | cardiotrophin 2, pseudogene [Source:HGNC Sy |
| FBXL19-AS1 | 16 | FBXL19 antisense RNA 1 [Source:HGNC Symbo |
| FBXL19 | 16 | F-box and leucine rich repeat protein 19 [Sour |
| AC135048.1 | 16 | novel transcript |
| ORAI3 | 16 | ORAI calcium release-activated calcium modu |
| AC135048.3 | 16 | novel transcript |
| SETD1A | 16 | SET domain containing 1A, histone lysine meth |
| AC135048.4 | 16 | novel transcript |
| HSD3B7 | 16 | hydroxy-delta-5-steroid dehydrogenase, 3 bet |
| STX1B | 16 | syntaxin 1B [Source:HGNC Symbol;Acc:HGNC: |
| STX4 | 16 | syntaxin 4 [Source:HGNC Symbol;Acc:HGNC:1 |
| AC135050.3 | 16 | novel transcript |
| AC135050.1 | 16 | novel transcript, antisense to ZNF668 |
| ZNF668 | 16 | zinc finger protein 668 [Source:HGNC Symbol; |
| AC135050.4 | 16 | novel transcript, antisense to ZNF668 |
| ZNF646 | 16 | zinc finger protein 646 [Source:HGNC Symbol; |
| PRSS53 | 16 | serine protease 53 [Source:HGNC Symbol;Acc |
| AC135050.2 | 16 | novel protein, VKORC1 and PRSS53 readthrou |
| VKORC1 | 16 | vitamin K epoxide reductase complex subunit |
| AC135050.7 | 16 | novel transcript |
| BCKDK | 16 | branched chain keto acid dehydrogenase kina |
| KAT8 | 16 | lysine acetyltransferase 8 [Source:HGNC Symb |
| AC135050.5 | 16 | novel transcript |
| AC135050.6 | 16 | novel transcript, sense intronic to KAT8 |
| PRSS8 | 16 | serine protease 8 [Source:HGNC Symbol;Acc:H |
| AC009088.3 | 16 | novel transcript, antisense to PRSS8 |
| PRSS36 | 16 | serine protease 36 [Source:HGNC Symbol;Acc |
| AC009088.4 | 16 | filamin B, beta (FLNB) pseudogene |
| NDUFA3P6 | 16 | NADH:ubiquinone oxidoreductase subunit A3 |
| FUS | 16 | FUS RNA binding protein [Source:HGNC Symbo |
| AC009088.2 | 16 | novel transcript, antisense to FUS |
| AC009088.1 | 16 | novel transcript, antisense to PYCARD |
| PYCARD | 16 | PYD and CARD domain containing [Source:HG |

| | | |
|---|---|---|
| PYCARD-AS1 | 16 | PYCARD antisense RNA 1 [Source:HGNC Symb |
| TRIM72 | 16 | tripartite motif containing 72 [Source:HGNC S |
| PYDC1 | 16 | pyrin domain containing 1 [Source:HGNC Sym |
| ITGAM | 16 | integrin subunit alpha M [Source:HGNC Symb |
| ITGAX | 16 | integrin subunit alpha X [Source:HGNC Symbo |
| AC093520.2 | 16 | integrin, alpha X (complement component 3 r |
| ITGAD | 16 | integrin subunit alpha D [Source:HGNC Symbo |
| AC093520.1 | 16 | novel transcript |
| COX6A2 | 16 | cytochrome c oxidase subunit 6A2 [Source:HG |
| AC026471.5 | 16 | novel transcript, antisense to COX6A2 |
| ZNF843 | 16 | zinc finger protein 843 [Source:HGNC Symbol; |
| AC026471.4 | 16 | novel transcript |
| AC026471.1 | 16 | novel transcript, antisense to ARMC5 |
| ARMC5 | 16 | armadillo repeat containing 5 [Source:HGNC S |
| AC026471.6 | 16 | novel transcript |
| TGFB1I1 | 16 | transforming growth factor beta 1 induced tra |
| SLC5A2 | 16 | solute carrier family 5 member 2 [Source:HGN |
| AC026471.3 | 16 | novel transcript, antisense to SLC5A2 |
| C16orf58 | 16 | chromosome 16 open reading frame 58 [Sour |
| AC026471.2 | 16 | novel transcript, antisense to C16orf58 |
| AHSP | 16 | alpha hemoglobin stabilizing protein [Source:H |
| LINC02190 | 16 | long intergenic non-protein coding RNA 2190 |
| AC106730.1 | 16 | novel transcript |
| VN1R64P | 16 | vomeronasal 1 receptor 64 pseudogene [Sour |
| VN1R65P | 16 | vomeronasal 1 receptor 65 pseudogene [Sour |
| FRG2KP | 16 | FSHD region gene 2 family member K, pseudo |
| AC106730.2 | 16 | TEC |
| YBX3P1 | 16 | Y-box binding protein 3 pseudogene 1 [Source |
| ZNF720P1 | 16 | zinc finger protein 720 pseudogene 1 [Source: |
| VN1R66P | 16 | vomeronasal 1 receptor 66 pseudogene [Sour |
| RBM22P12 | 16 | RNA binding motif protein 22 pseudogene 12 |
| AC074050.1 | 16 | SMT3 suppressor of mif two 3 homolog 2 (S. c |
| AC074050.2 | 16 | novel transcript |
| CLUHP3 | 16 | clustered mitochondria homolog pseudogene |
| AC074050.4 | 16 | novel transcript |
| AC074050.3 | 16 | novel transcript, antisense to ZNF720 |
| ZNF720 | 16 | zinc finger protein 720 [Source:HGNC Symbol; |
| AC002519.1 | 16 | novel transcript |
| VN1R67P | 16 | vomeronasal 1 receptor 67 pseudogene [Sour |
| AC002519.2 | 16 | zinc finger protein 267 (ZNF267) pseudogene |
| VN1R3 | 16 | vomeronasal 1 receptor 3 (gene/pseudogene) |
| RBM22P13 | 16 | RNA binding motif protein 22 pseudogene 13 |
| AC034105.2 | 16 | SMT3 suppressor of mif two 3 homolog 3 (S. c |
| ZNF267 | 16 | zinc finger protein 267 [Source:HGNC Symbol; |
| AC034105.3 | 16 | protein tyrosine phosphatase, receptor type, f |
| AC034105.5 | 16 | ig heavy variable (IGHV) pseudogene |
| AC034105.1 | 16 | ig heavy variable (IGHV) pseudogene |
| AC034105.4 | 16 | V-set and immunoglobulin domain containing |
| AC142381.3 | 16 | Rho GTPase activating protein 23 (ARHGAP23 |
| AC142381.4 | 16 | kinase suppressor of ras 1 (KSR1) pseudogene |

| | | |
|---|---|---|
| IGHV1OR16-1 | 16 | immunoglobulin heavy variable 1/OR16-1 (ps |
| AC142381.1 | 16 | immunoglobulin heavy variable 3-23 (IGHV3-2 |
| IGHV1OR16-3 | 16 | immunoglobulin heavy variable 1/OR16-3 (ps |
| IGHV3OR16-9 | 16 | immunoglobulin heavy variable 3/OR16-9 (no |
| HERC2P4 | 16 | hect domain and RLD 2 pseudogene 4 [Source |
| AC142381.2 | 16 | kinase suppressor of Ras 1 (KSR1) pseudogene |
| AC133485.7 | 16 | novel transcript |
| AC133485.6 | 16 | novel transcript |
| AC133485.5 | 16 | novel transcript |
| AC133485.4 | 16 | novel pseudogene |
| ABHD17AP8 | 16 | abhydrolase domain containing 17A pseudoge |
| AC133485.1 | 16 | glypican 5 (GPC5) pseudogene |
| AC133485.3 | 16 | novel transcript, antisense to TP53TG3 |
| TP53TG3D | 16 | TP53 target 3D [Source:HGNC Symbol;Acc:HG |
| AC133485.2 | 16 | novel transcript |
| AC140878.1 | 16 | kinase suppressor of ras 1 (KSR1) pseudogene |
| AC133548.2 | 16 | protein phosphatase 2, regulatory subunit B'', |
| AC133548.1 | 16 | CHK2 checkpoint homolog (S. pombe) (CHEK2 |
| ACTR3BP3 | 16 | ACTR3B pseudogene 3 [Source:HGNC Symbol; |
| AC138915.2 | 16 | solute carrier family 9, subfamily B (cation pro |
| PABPC1P13 | 16 | poly(A) binding protein cytoplasmic 1 pseudog |
| AC138915.3 | 16 | novel transcript |
| AC138915.1 | 16 | CDGSH iron sulfur domain 2 (CISD2) pseudoge |
| ABCD1P3 | 16 | ATP binding cassette subfamily D member 1 p |
| AC137800.1 | 16 | family with sequence similarity 153, member |
| AC137800.2 | 16 | centrosomal protein 192 (CEP192) pseudogen |
| AC133569.1 | 16 | selenophosphate synthetase 1 (SEPHS1) pseu |
| AC138907.7 | 16 | novel transcript |
| AC138907.5 | 16 | kinase suppressor of ras 1 (KSR1) pseudogene |
| AC138907.1 | 16 | novel transcript |
| TP53TG3 | 16 | TP53 target 3 [Source:HGNC Symbol;Acc:HGN |
| AC138907.3 | 16 | novel transcript, antisense to TP53TG3 |
| AC138907.6 | 16 | glypican 5 (GPC5) pseudogene |
| ABHD17AP7 | 16 | abhydrolase domain containing 17A pseudoge |
| AC138907.2 | 16 | novel transcript |
| AC138907.4 | 16 | chromosome 16 open reading frame 61 (C16o |
| HERC2P5 | 16 | hect domain and RLD 2 pseudogene 5 [Source |
| AC138907.8 | 16 | novel transcript |
| AC138907.9 | 16 | novel transcript |
| AC138907.10 | 16 | novel transcript |
| AC137761.1 | 16 | novel transcript |
| AC137761.2 | 16 | integrator complex subunit 4-like (INTS4L2) ps |
| AC142086.2 | 16 | kinase suppressor of ras 1 (KSR1) pseudogene |
| IGHV2OR16-5 | 16 | immunoglobulin heavy variable 2/OR16-5 (no |
| BCAP31P2 | 16 | B cell receptor associated protein 31 pseudog |
| SLC6A10P | 16 | solute carrier family 6 member 10, pseudoger |
| AC142086.1 | 16 | novel transcript |
| IGHV3OR16-15 | 16 | immunoglobulin heavy variable 3/OR16-15 (ps |
| IGHV3OR16-6 | 16 | immunoglobulin heavy variable 3/OR16-6 (ps |
| AC142086.6 | 16 | Rho GTPase activating protein 23 (ARHGAP23 |

| | | |
|---|---|---|
| AC142086.3 | 16 | proline rich 21 (PRR21) pseudogene |
| AC142086.5 | 16 | kinase suppressor of ras 1 (KSR1) pseudogene |
| IGHV1OR16-2 | 16 | immunoglobulin heavy variable 1/OR16-2 (ps |
| IGHV3OR16-10 | 16 | immunoglobulin heavy variable 3/OR16-10 (n |
| AC142086.4 | 16 | novel transcript |
| IGHV1OR16-4 | 16 | immunoglobulin heavy variable 1/OR16-4 (ps |
| IGHV3OR16-8 | 16 | immunoglobulin heavy variable 3/OR16-8 (no |
| AC145350.2 | 16 | novel transcript |
| AC145350.3 | 16 | integrator complex subunit 4-like 2 (INTS4L2) |
| HERC2P8 | 16 | hect domain and RLD 2 pseudogene 8 [Source |
| AC145350.4 | 16 | novel transcript |
| AC145350.1 | 16 | chromosome 16 open reading frame 61 (C16c |
| ABHD17AP9 | 16 | abhydrolase domain containing 17A pseudoge |
| AC138869.1 | 16 | glypican 5 (GPC5) pseudogene |
| AC138869.2 | 16 | novel transcript, antisense to TP53-target gen |
| TP53TG3C | 16 | TP53 target 3C [Source:HGNC Symbol;Acc:HG |
| AC138869.3 | 16 | novel transcript |
| AC141257.2 | 16 | novel transcript |
| TP53TG3E | 16 | TP53 target 3 family member E [Source:HGNC |
| AC141257.3 | 16 | double homeobox protein (DUX) pseudogene |
| TP53TG3B | 16 | TP53 target 3B [Source:HGNC Symbol;Acc:HG |
| AC141257.1 | 16 | novel transcript |
| AC141257.4 | 16 | kinase suppressor of ras 1 (KSR1) pseudogene |
| TP53TG3F | 16 | TP53 target 3 family member F [Source:HGNC |
| AC136944.6 | 16 | kinase suppressor of ras 1 (KSR1) pseudogene |
| AC136944.5 | 16 | protein phosphatase 2, regulatory subunit B'', |
| AC136944.1 | 16 | novel transcript |
| AC136944.2 | 16 | novel transcript |
| AC136944.4 | 16 | novel transcript |
| AC136944.3 | 16 | CHK2 checkpoint homolog (S. pombe) (CHEK2 |
| AC142384.1 | 16 | rhophilin, Rho GTPase binding protein 2 (RHP |
| BMS1P8 | 16 | BMS1 pseudogene 8 [Source:HGNC Symbol;A |
| AC136428.4 | 16 | ankyrin repeat domain 36 (ANKRD36) pseudo |
| ENPP7P13 | 16 | ectonucleotide pyrophosphatase/phosphodie |
| AC136428.2 | 16 | kinase suppressor of ras 1 (KSR1) pseudogene |
| IGHV3OR16-12 | 16 | immunoglobulin heavy variable 3/OR16-12 (n |
| AC136428.3 | 16 | ADAM metallopeptidase domain 21 (ADAM21 |
| IGHV3OR16-13 | 16 | immunoglobulin heavy variable 3/OR16-13 (n |
| AC136428.1 | 16 | novel gene similar to an immunoglobulin hea |
| AC140658.6 | 16 | novel IGHV pseudogene |
| AC140658.7 | 16 | novel IGHV pseudogene |
| AC140658.2 | 16 | novel transcript |
| IGHV3OR16-11 | 16 | immunoglobulin heavy variable 3/OR16-11 (p |
| AC140658.3 | 16 | immunoglobulin heavy variable region (IGHV) |
| ARHGAP23P1 | 16 | Rho GTPase activating protein 23 pseudogene |
| IGHV3OR16-7 | 16 | immunoglobulin heavy variable 3/OR16-7 (ps |
| AC140658.4 | 16 | novel immunoglobulin heavy variable (IGHV) |
| IGHV3OR16-16 | 16 | immunoglobulin heavy variable 3/OR16-16 (p |
| AC140658.1 | 16 | novel transcript |
| AC140658.5 | 16 | novel immunoglobulin heavy variable region ( |

| | | |
|---|---|---|
| AC133561.1 | 16 | solute carrier family 6 (neurotransmitter trans |
| BCAP31P1 | 16 | B cell receptor associated protein 31 pseudog |
| AC133561.2 | 16 | novel immunoglobulin heavy chain variable (I |
| AC133561.4 | 16 | kinase suppressor of ras 1 (KSR1) pseudogene |
| AC133561.3 | 16 | kinase suppressor of ras 1 (KSR1) pseudogene |
| AC136932.1 | 16 | mitochondrial ribosomal protein 63 (MRP63) p |
| DUX4L45 | 16 | double homeobox 4 like 45 (pseudogene) [Sou |
| PCMTD1P2 | 16 | protein-L-isoaspartate (D-aspartate) O-methyl |
| DUX4L46 | 16 | double homeobox 4 like 46 (pseudogene) [Sou |
| DUX4L47 | 16 | double homeobox 4 like 47 (pseudogene) [Sou |
| LINC00273 | 16 | long intergenic non-protein coding RNA 273 [S |
| RNA5-8SP2 | 16 | RNA, 5.8S ribosomal pseudogene 2 [Source:HG |
| BCLAF1P2 | 16 | BCL2 associated transcription factor 1 pseudo |
| FP325313.1 | 16 | cyclin Y (CCNY) pseudogene |
| FP325313.3 | 16 | novel transcript |
| FP325313.2 | 16 | novel pseudogene |
| AC135776.4 | 16 | novel transcript |
| AC135776.5 | 16 | novel pseudogene |
| LINC02184 | 16 | long intergenic non-protein coding RNA 2184 |
| AC135776.1 | 16 | novel transcript |
| AC135776.2 | 16 | novel transcript |
| AC135776.3 | 16 | sal-like 3 (Drosophila) (SALL3) pseudogene |
| CCNYL3 | 16 | cyclin Y like 3 (pseudogene) [Source:HGNC Syr |
| CLUHP11 | 16 | clustered mitochondria homolog pseudogene |
| NAMPTP3 | 16 | nicotinamide phosphoribosyltransferase pseu |
| VPS35P1 | 16 | VPS35 pseudogene 1 [Source:HGNC Symbol;A |
| VN1R68P | 16 | vomeronasal 1 receptor 68 pseudogene [Sour |
| VN1R69P | 16 | vomeronasal 1 receptor 69 pseudogene [Sour |
| AC023824.5 | 16 | novel transcript |
| AC023824.6 | 16 | novel transcript |
| UBE2MP1 | 16 | ubiquitin conjugating enzyme E2 M pseudoge |
| SLC25A1P4 | 16 | solute carrier family 25 member 1 pseudogen |
| AC023824.1 | 16 | novel transcript |
| AC023824.2 | 16 | double homeobox (DUX) pseudogene |
| AC023824.4 | 16 | novel transcript |
| AC023824.3 | 16 | novel transcript |
| TP53TG3GP | 16 | TP53 target 3 family member G, pseudogene [ |
| RARRES2P5 | 16 | retinoic acid receptor responder 2 pseudogen |
| FGFR3P5 | 16 | fibroblast growth factor receptor 3 pseudoger |
| FRG2JP | 16 | FSHD region gene 2 family member J, pseudog |
| RARRES2P6 | 16 | retinoic acid receptor responder 2 pseudogen |
| AGGF1P8 | 16 | angiogenic factor with G-patch and FHA doma |
| FRG2HP | 16 | FSHD region gene 2 family member H, pseudo |
| RARRES2P9 | 16 | retinoic acid receptor responder 2 pseudogen |
| AGGF1P9 | 16 | angiogenic factor with G-patch and FHA doma |
| C2orf69P4 | 16 | chromosome 2 open reading frame 69 pseudo |
| LINC01566 | 16 | long intergenic non-protein coding RNA 1566 |
| FRG2GP | 16 | FSHD region gene 2 family member G, pseudo |
| AGGF1P4 | 16 | angiogenic factor with G-patch and FHA doma |
| FRG2IP | 16 | FSHD region gene 2 family member I, pseudog |

| | | |
|---|---|---|
| RARRES2P7 | 16 | retinoic acid receptor responder 2 pseudogen |
| AGGF1P5 | 16 | angiogenic factor with G-patch and FHA doma |
| C2orf69P3 | 16 | chromosome 2 open reading frame 69 pseudo |
| AC018558.4 | 16 | coiled-coil domain containing 180 (CCDC180) |
| ZNF971P | 16 | zinc finger protein 971, pseudogene [Source:H |
| AC018558.6 | 16 | novel transcript |
| FRG2DP | 16 | FSHD region gene 2 family member D, pseudo |
| RARRES2P10 | 16 | retinoic acid receptor responder 2 pseudogen |
| AGGF1P6 | 16 | angiogenic factor with G-patch and FHA doma |
| AC018558.2 | 16 | novel transcript |
| C2orf69P2 | 16 | chromosome 2 open reading frame 69 pseudo |
| AC018558.3 | 16 | novel transcript |
| AC018558.7 | 16 | novel transcript |
| AC018558.1 | 16 | novel transcript |
| TP53TG3HP | 16 | TP53 target 3 family member H, pseudogene \| |
| AC018558.5 | 16 | double homeobox 4 (DUX4) pseudogene |
| RARRES2P8 | 16 | retinoic acid receptor responder 2 pseudogen |
| AGGF1P7 | 16 | angiogenic factor with G-patch and FHA doma |
| C2orf69P1 | 16 | chromosome 2 open reading frame 69 pseudo |
| C1QL1P1 | 16 | complement C1q like 1 pseudogene 1 [Source |
| KIF18BP1 | 16 | kinesin family member 18B pseudogene 1 [So |
| AC092325.1 | 16 | novel transcript |
| AC106785.1 | 16 | HECT, UBA and WWE domain containing 1 (HU |
| RNA5SP406 | 16 | RNA, 5S ribosomal pseudogene 406 [Source:H |
| RNA5SP407 | 16 | RNA, 5S ribosomal pseudogene 407 [Source:H |
| RNA5SP408 | 16 | RNA, 5S ribosomal pseudogene 408 [Source:H |
| RNA5SP409 | 16 | RNA, 5S ribosomal pseudogene 409 [Source:H |
| LINC02167 | 16 | long intergenic non-protein coding RNA 2167 |
| RNA5SP410 | 16 | RNA, 5S ribosomal pseudogene 410 [Source:H |
| RNA5SP411 | 16 | RNA, 5S ribosomal pseudogene 411 [Source:H |
| RNA5SP412 | 16 | RNA, 5S ribosomal pseudogene 412 [Source:H |
| RNA5SP413 | 16 | RNA, 5S ribosomal pseudogene 413 [Source:H |
| RNA5SP414 | 16 | RNA, 5S ribosomal pseudogene 414 [Source:H |
| RNA5SP415 | 16 | RNA, 5S ribosomal pseudogene 415 [Source:H |
| RNA5SP416 | 16 | RNA, 5S ribosomal pseudogene 416 [Source:H |
| RNA5SP417 | 16 | RNA, 5S ribosomal pseudogene 417 [Source:H |
| RNA5SP418 | 16 | RNA, 5S ribosomal pseudogene 418 [Source:H |
| RNA5SP419 | 16 | RNA, 5S ribosomal pseudogene 419 [Source:H |
| RNA5SP420 | 16 | RNA, 5S ribosomal pseudogene 420 [Source:H |
| RNA5SP421 | 16 | RNA, 5S ribosomal pseudogene 421 [Source:H |
| RNA5SP422 | 16 | RNA, 5S ribosomal pseudogene 422 [Source:H |
| RNA5SP423 | 16 | RNA, 5S ribosomal pseudogene 423 [Source:H |
| AC106785.2 | 16 | novel transcript |
| HMGN2P41 | 16 | high mobility group nucleosomal binding dom |
| VN1R70P | 16 | vomeronasal 1 receptor 70 pseudogene [Sour |
| AC116553.1 | 16 | WD repeat domain phosphoinositide-interacti |
| AC116553.2 | 16 | novel transcript |
| PPP1R1AP2 | 16 | protein phosphatase 1 regulatory inhibitor sul |
| ANKRD26P1 | 16 | ankyrin repeat domain 26 pseudogene 1 [Sou |
| RNU6-845P | 16 | RNA, U6 small nuclear 845, pseudogene [Sour |

| | | |
|---|---|---|
| AC092368.2 | 16 | RAB31, member RAS oncogene family (RAB31 |
| AC092368.1 | 16 | zinc finger protein 161 homolog (mouse) (ZFP |
| SHCBP1 | 16 | SHC binding and spindle associated 1 [Source: |
| AC092368.3 | 16 | novel transcript |
| RAB43P1 | 16 | RAB43 pseudogene 1 [Source:HGNC Symbol;A |
| AC012186.1 | 16 | histamine receptor H3 (HRH3) pseudogene |
| VPS35 | 16 | VPS35 retromer complex component [Source: |
| AC012186.2 | 16 | novel transcript |
| AC012186.3 | 16 | novel transcript |
| ORC6 | 16 | origin recognition complex subunit 6 [Source:I |
| MYLK3 | 16 | myosin light chain kinase 3 [Source:HGNC Sym |
| AC007225.1 | 16 | nuclear factor (erythroid-derived 2)-like 3 (NF |
| C16orf87 | 16 | chromosome 16 open reading frame 87 [Sour |
| CKBP1 | 16 | creatine kinase B pseudogene 1 [Source:HGN( |
| AC007225.2 | 16 | origin recognition complex, subunit 3 (ORC3) p |
| GPT2 | 16 | glutamic--pyruvic transaminase 2 [Source:HGI |
| DNAJA2 | 16 | DnaJ heat shock protein family (Hsp40) memb |
| AC018845.3 | 16 | novel transcript, antisense to DNAJA2 |
| AC018845.1 | 16 | novel transcript |
| AC018845.2 | 16 | chromosome 8 open reading frame 33 (C8orf3 |
| NETO2 | 16 | neuropilin and tolloid like 2 [Source:HGNC Syr |
| ITFG1-AS1 | 16 | ITFG1 antisense RNA 1 [Source:HGNC Symbol; |
| RPL23AP72 | 16 | ribosomal protein L23a pseudogene 72 [Sourc |
| ITFG1 | 16 | integrin alpha FG-GAP repeat containing 1 [So |
| AC007494.2 | 16 | novel transcript, antisense to ITFG1 |
| AC007494.3 | 16 | novel transcript |
| AC007494.1 | 16 | proteasome (prosome, macropain) subunit, b |
| AC007533.1 | 16 | novel transcript, antisense to ITFG1 |
| Y_RNA | 16 | Y RNA [Source:RFAM;Acc:RF00019] |
| PHKB | 16 | phosphorylase kinase regulatory subunit beta |
| RNA5SP425 | 16 | RNA, 5S ribosomal pseudogene 425 [Source:H |
| AC007599.2 | 16 | novel transcript |
| AC007599.1 | 16 | Rho GTPase activating protein 20 (ARHGAP20 |
| EIF4BP5 | 16 | eukaryotic translation initiation factor 4B pseu |
| NDUFA5P11 | 16 | NADH:ubiquinone oxidoreductase subunit A5 |
| LINC02133 | 16 | long intergenic non-protein coding RNA 2133 |
| AC141846.1 | 16 | ATP-binding cassette, sub-family A (ABC1), me |
| LINC02192 | 16 | long intergenic non-protein coding RNA 2192 |
| LINC02134 | 16 | long intergenic non-protein coding RNA 2134 |
| AC096996.3 | 16 | novel transcript |
| AC096996.1 | 16 | PH domain and leucine rich repeat protein ph |
| ABCC12 | 16 | ATP binding cassette subfamily C member 12 |
| AC096996.2 | 16 | novel transcript, sense overlapping ABCC11 |
| ABCC11 | 16 | ATP binding cassette subfamily C member 11 |
| LONP2 | 16 | lon peptidase 2, peroxisomal [Source:HGNC Sy |
| UBA52P8 | 16 | ubiquitin A-52 residue ribosomal protein fusic |
| AC023818.1 | 16 | novel transcript |
| SIAH1 | 16 | siah E3 ubiquitin protein ligase 1 [Source:HGN |
| Metazoa_SRP | 16 | Metazoan signal recognition particle RNA [Sou |
| AC026470.3 | 16 | novel transcript |

| | | |
|---|---|---|
| AC026470.1 | 16 | chromosome 18 open reading frame 32 (C18c |
| MOCS1P1 | 16 | molybdenum cofactor synthesis 1 pseudogen |
| N4BP1 | 16 | NEDD4 binding protein 1 [Source:HGNC Symb |
| AC026470.2 | 16 | novel transcript, antisense to N4BP1 |
| RPS2P44 | 16 | ribosomal protein S2 pseudogene 44 [Source: |
| AC023813.2 | 16 | ribosomal L24 domain containing 1 (RSL24D1) |
| AC023813.1 | 16 | novel transcript |
| AC007611.1 | 16 | novel transcript |
| AC023813.3 | 16 | novel transcript |
| AC023813.4 | 16 | novel transcript |
| KLF8P1 | 16 | Kruppel like factor 8 pseudogene 1 [Source:H |
| RNU6-257P | 16 | RNA, U6 small nuclear 257, pseudogene [Sour |
| AC023827.1 | 16 | general transcription factor IIA, 1-like (GTF2A |
| MTND4LP25 | 16 | MT-ND4L pseudogene 25 [Source:HGNC Symb |
| AC044798.3 | 16 | novel transcript |
| AC044798.1 | 16 | novel transcript |
| AC044798.2 | 16 | novel transcript |
| CBLN1 | 16 | cerebellin 1 precursor [Source:HGNC Symbol; |
| AC007614.1 | 16 | novel transcript, antisense to CBLN1 |
| AC007614.2 | 16 | novel transcript |
| C16orf78 | 16 | chromosome 16 open reading frame 78 [Sour |
| AC007861.1 | 16 | novel transcript |
| AC007861.2 | 16 | novel transcript |
| LINC02179 | 16 | long intergenic non-protein coding RNA 2179 |
| ZNF423 | 16 | zinc finger protein 423 [Source:HGNC Symbol; |
| ADAM3B | 16 | ADAM metallopeptidase domain 3B (pseudog |
| MRPS21P7 | 16 | mitochondrial ribosomal protein S21 pseudog |
| AC027348.2 | 16 | novel transcript |
| MRPS21P8 | 16 | mitochondrial ribosomal protein S21 pseudog |
| AC027348.1 | 16 | novel transcript, sense intronic to ZNF423 |
| AC007603.2 | 16 | novel transcript |
| AC007603.1 | 16 | novel transcript |
| RPL34P29 | 16 | ribosomal protein L34 pseudogene 29 [Source |
| AC007603.3 | 16 | nuclear receptor coactivator 5-like pseudoger |
| CNEP1R1 | 16 | CTD nuclear envelope phosphatase 1 regulato |
| AC007610.1 | 16 | novel transcript, antisense to HEATR3 |
| AC007610.3 | 16 | regulation of nuclear pre-mRNA domain conta |
| Metazoa_SRP | 16 | Metazoan signal recognition particle RNA [Sou |
| ACTG1P16 | 16 | actin gamma 1 pseudogene 16 [Source:HGNC |
| HEATR3 | 16 | HEAT repeat containing 3 [Source:HGNC Symb |
| AC007610.4 | 16 | novel transcript |
| RNY4P3 | 16 | RNY4 pseudogene 3 [Source:HGNC Symbol;Ac |
| AC007610.2 | 16 | novel transcript, antisense to HEATR3 |
| AC007610.5 | 16 | novel transcript |
| RPL10P14 | 16 | ribosomal protein L10 pseudogene 14 [Source |
| SNORA70 | 16 | Small nucleolar RNA SNORA70 [Source:RFAM; |
| TENT4B | 16 | terminal nucleotidyltransferase 4B [Source:H |
| ADCY7 | 16 | adenylate cyclase 7 [Source:HGNC Symbol;Ac |
| MIR6771 | 16 | microRNA 6771 [Source:HGNC Symbol;Acc:H |
| BRD7 | 16 | bromodomain containing 7 [Source:HGNC Syn |

| | | |
|---|---|---|
| AC007493.2 | 16 | novel transcript, antisense to BRD7 |
| LINC02178 | 16 | long intergenic non-protein coding RNA 2178 |
| AC007493.1 | 16 | novel transcript |
| NKD1 | 16 | NKD inhibitor of WNT signaling pathway 1 [So |
| AC007608.2 | 16 | novel transcript, antisense to NKD1 |
| AC007608.4 | 16 | novel transcript |
| AC007608.1 | 16 | novel transcript, antisense to NKD1 |
| AC007608.3 | 16 | novel transcript, antisense to SNX20 |
| SNX20 | 16 | sorting nexin 20 [Source:HGNC Symbol;Acc:H( |
| NOD2 | 16 | nucleotide binding oligomerization domain co |
| AC007728.3 | 16 | novel transcript |
| AC007728.2 | 16 | novel transcript, antisense to CYLD and NOD2 |
| CYLD | 16 | CYLD lysine 63 deubiquitinase [Source:HGNC S |
| MIR3181 | 16 | microRNA 3181 [Source:HGNC Symbol;Acc:HG |
| AC007728.1 | 16 | novel transcript, antisense to CYLD |
| LINC02168 | 16 | long intergenic non-protein coding RNA 2168 |
| LINC02128 | 16 | long intergenic non-protein coding RNA 2128 |
| AC025810.1 | 16 | novel transcript |
| RNA5SP426 | 16 | RNA, 5S ribosomal pseudogene 426 [Source:H |
| LINC02127 | 16 | long intergenic non-protein coding RNA 2127 |
| AC027688.1 | 16 | novel transcript |
| AC027688.2 | 16 | novel transcript |
| AC009166.3 | 16 | novel transcript |
| SOD1P2 | 16 | superoxide dismutase 1 pseudogene 2 [Source |
| SALL1 | 16 | spalt like transcription factor 1 [Source:HGNC |
| AC009166.1 | 16 | novel transcript, antisense to SALL1 |
| AC087564.1 | 16 | novel transcript |
| AC009166.2 | 16 | TEC |
| AC137494.1 | 16 | novel transcript |
| AC137527.1 | 16 | uracil-DNA glycosylase (UNG) pseudogene |
| UNGP1 | 16 | uracil-DNA glycosylase pseudogene 1 [Source: |
| AC007344.1 | 16 | novel transcript |
| HNRNPA1P48 | 16 | heterogeneous nuclear ribonucleoprotein A1 |
| RN7SKP142 | 16 | RN7SK pseudogene 142 [Source:HGNC Symbo |
| LINC01571 | 16 | long intergenic non-protein coding RNA 1571 |
| C16orf97 | 16 | chromosome 16 open reading frame 97 [Sour |
| AC009039.1 | 16 | novel transcript, antisense to novel protein |
| AC009039.2 | 16 | novel transcript |
| LINC00919 | 16 | long intergenic non-protein coding RNA 919 [S |
| LINC02180 | 16 | long intergenic non-protein coding RNA 2180 |
| AC007333.2 | 16 | novel transcript |
| AC007333.1 | 16 | novel transcript |
| CASC22 | 16 | cancer susceptibility 22 [Source:HGNC Symbo |
| TOX3 | 16 | TOX high mobility group box family member 3 |
| AC007490.1 | 16 | novel transcript |
| CASC16 | 16 | cancer susceptibility 16 [Source:HGNC Symbo |
| AC026462.5 | 16 | novel transcript |
| AC026462.3 | 16 | novel transcript |
| AC026462.2 | 16 | novel transcript |
| AC026462.4 | 16 | novel transcript |

| | | |
|---|---|---|
| AC026462.1 | 16 | tropomyosin 1 (alpha) (TPM1) pseudogene |
| PHBP21 | 16 | prohibitin pseudogene 21 [Source:HGNC Symb |
| AC007346.1 | 16 | novel pseudogene (KIAA1191) |
| AC007346.2 | 16 | mitochondrial carrier 2 (MTCH2) pseudogene |
| AC007906.2 | 16 | novel protein |
| CHD9 | 16 | chromodomain helicase DNA binding protein |
| AC007906.1 | 16 | novel transcript |
| AC079416.4 | 16 | novel transcript |
| AC079416.2 | 16 | thyroid hormone receptor interactor 13 (TRIP |
| AC079416.3 | 16 | novel transcript |
| AC079416.1 | 16 | NULL |
| RNA5SP427 | 16 | RNA, 5S ribosomal pseudogene 427 [Source:H |
| AC007342.2 | 16 | golgin A6 family-like 2 (GOLGA6L2) pseudoger |
| AC007342.3 | 16 | u3 small nucleolar ribonucleoprotein protein |
| AC007342.4 | 16 | novel transcript |
| AC007342.5 | 16 | novel transcript |
| AC007342.8 | 16 | novel transcript |
| AC007342.9 | 16 | novel transcript |
| RBL2 | 16 | RB transcriptional corepressor like 2 [Source:H |
| RNU6-1153P | 16 | RNA, U6 small nuclear 1153, pseudogene [Sou |
| AC007342.1 | 16 | novel transcript |
| AC007342.6 | 16 | novel transcript |
| AC007342.7 | 16 | novel transcript |
| AKTIP | 16 | AKT interacting protein [Source:HGNC Symbol |
| AC007497.2 | 16 | novel transcript |
| RPGRIP1L | 16 | RPGRIP1 like [Source:HGNC Symbol;Acc:HGN( |
| AC007497.1 | 16 | novel transcript, sense intronic to RPGRIP1L |
| FTO | 16 | FTO alpha-ketoglutarate dependent dioxygen |
| AC007496.2 | 16 | novel transcript, antisense to FTO |
| AC007496.3 | 16 | novel transcript |
| AC007496.1 | 16 | novel transcript, antisense to FTO |
| AC007347.1 | 16 | novel transcript, antisense to FTO |
| AC007347.2 | 16 | NULL |
| AC007907.1 | 16 | novel transcript |
| LINC02169 | 16 | long intergenic non-protein coding RNA 2169 |
| IRX3 | 16 | iroquois homeobox 3 [Source:HGNC Symbol;A |
| AC018553.2 | 16 | novel transcript, antisense to IRX3 |
| AC018553.1 | 16 | novel transcript |
| LINC02140 | 16 | long intergenic non-protein coding RNA 2140 |
| AC018553.3 | 16 | novel transcript |
| AC007343.1 | 16 | novel transcript |
| LINC02183 | 16 | long intergenic non-protein coding RNA 2183 |
| AC007491.1 | 16 | novel transcript |
| CRNDE | 16 | colorectal neoplasia differentially expressed [ |
| AC106742.1 | 16 | novel transcript |
| IRX5 | 16 | iroquois homeobox 5 [Source:HGNC Symbol;A |
| AC106738.1 | 16 | novel transcript |
| AC106738.2 | 16 | novel transcript |
| MTND5P34 | 16 | MT-ND5 pseudogene 34 [Source:HGNC Symbo |
| AC109136.1 | 16 | novel transcript |

| | | |
|---|---|---|
| AC109462.1 | 16 | novel transcript, antisense to IRX6 |
| RN7SL841P | 16 | RNA, 7SL, cytoplasmic 841, pseudogene [Sour |
| AC109462.3 | 16 | novel transcript |
| IRX6 | 16 | iroquois homeobox 6 [Source:HGNC Symbol;A |
| AC109462.2 | 16 | novel transcript |
| RPL31P56 | 16 | ribosomal protein L31 pseudogene 56 [Source |
| MMP2 | 16 | matrix metallopeptidase 2 [Source:HGNC Sym |
| AC007336.2 | 16 | novel transcript |
| MMP2-AS1 | 16 | MMP2 antisense RNA 1 [Source:HGNC Symbo |
| AC007336.3 | 16 | novel transcript |
| LPCAT2 | 16 | lysophosphatidylcholine acyltransferase 2 [So |
| AC007336.1 | 16 | novel transcript |
| CAPNS2 | 16 | calpain small subunit 2 [Source:HGNC Symbol |
| SLC6A2 | 16 | solute carrier family 6 member 2 [Source:HGN |
| AC136621.1 | 16 | novel transcript |
| CES1P2 | 16 | carboxylesterase 1 pseudogene 2 [Source:HGI |
| CES1P1 | 16 | carboxylesterase 1 pseudogene 1 [Source:HGI |
| CES1 | 16 | carboxylesterase 1 [Source:HGNC Symbol;Acc |
| CES5A | 16 | carboxylesterase 5A [Source:HGNC Symbol;Ac |
| AC040168.1 | 16 | novel transcript |
| AC040168.2 | 16 | novel transcript |
| AC007495.1 | 16 | novel transcript |
| GNAO1 | 16 | G protein subunit alpha o1 [Source:HGNC Sym |
| AC079411.1 | 16 | novel transcript, antisense to GNAO1 |
| MIR3935 | 16 | microRNA 3935 [Source:HGNC Symbol;Acc:HG |
| AC009102.4 | 16 | novel transcript, antisense to GNAO1 |
| AC009102.1 | 16 | novel transcript, sense intronic to GNAO1 |
| AC009102.3 | 16 | TEC |
| AC009102.2 | 16 | novel transcript, antisense to GNAO1 |
| AMFR | 16 | autocrine motility factor receptor [Source:HG |
| AC092140.1 | 16 | novel transcript, antisense to AMFR |
| NUDT21 | 16 | nudix hydrolase 21 [Source:HGNC Symbol;Acc |
| OGFOD1 | 16 | 2-oxoglutarate and iron dependent oxygenase |
| AC092140.2 | 16 | novel transcript, antisense to OGFOD1 |
| BBS2 | 16 | Bardet-Biedl syndrome 2 [Source:HGNC Symb |
| MT4 | 16 | metallothionein 4 [Source:HGNC Symbol;Acc:I |
| MT3 | 16 | metallothionein 3 [Source:HGNC Symbol;Acc:I |
| MT2A | 16 | metallothionein 2A [Source:HGNC Symbol;Acc |
| AC026461.3 | 16 | novel transcript |
| MT1L | 16 | metallothionein 1L, pseudogene [Source:HGN |
| MT1E | 16 | metallothionein 1E [Source:HGNC Symbol;Acc |
| MT1M | 16 | metallothionein 1M [Source:HGNC Symbol;Ac |
| MT1JP | 16 | metallothionein 1J, pseudogene [Source:HGN |
| MT1A | 16 | metallothionein 1A [Source:HGNC Symbol;Acc |
| MT1DP | 16 | metallothionein 1D, pseudogene [Source:HGN |
| MT1CP | 16 | metallothionein 1C, pseudogene [Source:HGN |
| AC026461.2 | 16 | novel transcript |
| MT1B | 16 | metallothionein 1B [Source:HGNC Symbol;Acc |
| MT1F | 16 | metallothionein 1F [Source:HGNC Symbol;Acc |
| MT1G | 16 | metallothionein 1G [Source:HGNC Symbol;Acc |

| | | |
|---|---|---|
| MT1H | 16 | metallothionein 1H [Source:HGNC Symbol;Acc |
| AC026461.4 | 16 | metallothionein family pseudogene |
| MT1X | 16 | metallothionein 1X [Source:HGNC Symbol;Acc |
| AC026461.1 | 16 | novel transcript |
| DPPA2P4 | 16 | developmental pluripotency associated 2 pseu |
| AC106779.1 | 16 | novel transcript, antisense to NUP93 |
| NUP93 | 16 | nucleoporin 93 [Source:HGNC Symbol;Acc:HG |
| MIR138-2 | 16 | microRNA 138-2 [Source:HGNC Symbol;Acc:H |
| SLC12A3 | 16 | solute carrier family 12 member 3 [Source:HG |
| MIR6863 | 16 | microRNA 6863 [Source:HGNC Symbol;Acc:HG |
| RPS24P17 | 16 | ribosomal protein S24 pseudogene 17 [Source |
| HERPUD1 | 16 | homocysteine inducible ER protein with ubiqu |
| AC012181.2 | 16 | novel transcript, sense intronic to HERPUD1 |
| AC012181.1 | 16 | novel transcript, sense intronic to HERPUD1 |
| CETP | 16 | cholesteryl ester transfer protein [Source:HGN |
| NLRC5 | 16 | NLR family CARD domain containing 5 [Source |
| AC023825.1 | 16 | chromosome 7 open reading frame 63 (C7orf6 |
| AC023825.2 | 16 | novel transcript, antisense to NLRC5 |
| AC009090.2 | 16 | novel transcript, antisense to CPNE2 |
| CPNE2 | 16 | copine 2 [Source:HGNC Symbol;Acc:HGNC:23 |
| AC009090.4 | 16 | TEC |
| AC009090.5 | 16 | TEC |
| FAM192A | 16 | family with sequence similarity 192 member A |
| AC009090.6 | 16 | novel transcript, antisense to FAM192A |
| RSPRY1 | 16 | ring finger and SPRY domain containing 1 [Sou |
| ARL2BP | 16 | ADP ribosylation factor like GTPase 2 binding |
| AC009090.3 | 16 | novel transcript, antisense to ARL2BP |
| AC009090.1 | 16 | novel transcript, sense to intronic to ARL2BP |
| PLLP | 16 | plasmolipin [Source:HGNC Symbol;Acc:HGNC: |
| RPL23AP91 | 16 | ribosomal protein L23a pseudogene 91 [Sourc |
| AC108081.1 | 16 | ribosomal protein S2 (RPS2) pseudogene |
| CCL22 | 16 | C-C motif chemokine ligand 22 [Source:HGNC |
| CX3CL1 | 16 | C-X3-C motif chemokine ligand 1 [Source:HGN |
| CCL17 | 16 | C-C motif chemokine ligand 17 [Source:HGNC |
| CIAPIN1 | 16 | cytokine induced apoptosis inhibitor 1 [Source |
| COQ9 | 16 | coenzyme Q9 [Source:HGNC Symbol;Acc:HGN |
| AC009052.1 | 16 | novel transcript |
| POLR2C | 16 | RNA polymerase II subunit C [Source:HGNC Sy |
| DOK4 | 16 | docking protein 4 [Source:HGNC Symbol;Acc:H |
| CCDC102A | 16 | coiled-coil domain containing 102A [Source:H |
| ADGRG5 | 16 | adhesion G protein-coupled receptor G5 [Sour |
| HMGB3P32 | 16 | high mobility group box 3 pseudogene 32 [Sou |
| ADGRG1 | 16 | adhesion G protein-coupled receptor G1 [Sour |
| AC018552.3 | 16 | novel transcript, antisense to GPR56 |
| ADGRG3 | 16 | adhesion G protein-coupled receptor G3 [Sour |
| AC018552.2 | 16 | novel transcript, antisense to CCDC135 and G |
| DRC7 | 16 | dynein regulatory complex subunit 7 [Source: |
| AC018552.1 | 16 | mitochondrial carrier 2 (MTCH2) pseudogene |
| KATNB1 | 16 | katanin regulatory subunit B1 [Source:HGNC S |
| KIFC3 | 16 | kinesin family member C3 [Source:HGNC Sym |

| | | |
|---|---|---|
| AC092118.2 | 16 | novel transcript, antisense to KIFC3 |
| MIR6772 | 16 | microRNA 6772 [Source:HGNC Symbol;Acc:HG |
| AC092118.1 | 16 | uncharacterized LOC388282 [Source:NCBI ger |
| RNU6-20P | 16 | RNA, U6 small nuclear 20, pseudogene [Sourc |
| CNGB1 | 16 | cyclic nucleotide gated channel subunit beta 1 |
| TEPP | 16 | testis, prostate and placenta expressed [Sourc |
| ZNF319 | 16 | zinc finger protein 319 [Source:HGNC Symbol; |
| USB1 | 16 | U6 snRNA biogenesis phosphodiesterase 1 [Sc |
| MMP15 | 16 | matrix metallopeptidase 15 [Source:HGNC Syr |
| Y_RNA | 16 | Y RNA [Source:RFAM;Acc:RF00019] |
| AC012182.1 | 16 | TIMELESS interacting protein (TIPIN) pseudoge |
| CFAP20 | 16 | cilia and flagella associated protein 20 [Source |
| AC026771.1 | 16 | novel transcript, antisense to C16orf80 |
| AC009107.2 | 16 | uncharacterized LOC101927556 [Source:NCBI |
| CSNK2A2 | 16 | casein kinase 2 alpha 2 [Source:HGNC Symbol |
| AC009107.1 | 16 | novel transcript, antisense to CSNK2A2 |
| RN7SL645P | 16 | RNA, 7SL, cytoplasmic 645, pseudogene [Sour |
| CCDC113 | 16 | coiled-coil domain containing 113 [Source:HG |
| PRSS54 | 16 | serine protease 54 [Source:HGNC Symbol;Acc |
| GINS3 | 16 | GINS complex subunit 3 [Source:HGNC Symbo |
| RNU6-269P | 16 | RNA, U6 small nuclear 269, pseudogene [Sour |
| AC009118.1 | 16 | novel transcript, antisense to GINS3 |
| RNU6-1110P | 16 | RNA, U6 small nuclear 1110, pseudogene [Sou |
| AC009118.2 | 16 | novel transcript, antisense to GINS3 |
| LINC02137 | 16 | long intergenic non-protein coding RNA 2137 |
| NDRG4 | 16 | NDRG family member 4 [Source:HGNC Symbo |
| RNU6-103P | 16 | RNA, U6 small nuclear 103, pseudogene [Sour |
| SETD6 | 16 | SET domain containing 6, protein lysine methy |
| CNOT1 | 16 | CCR4-NOT transcription complex subunit 1 [Sc |
| AC009118.3 | 16 | novel transcript, antisense to CNOT1 |
| SNORA46 | 16 | small nucleolar RNA, H/ACA box 46 [Source:H |
| SNORA50A | 16 | small nucleolar RNA, H/ACA box 50A [Source:I |
| SNORA50A | 16 | small nucleolar RNA, H/ACA box 50A [Source:I |
| AC010287.1 | 16 | chromosome 10 open reading frame 57 (C10c |
| SLC38A7 | 16 | solute carrier family 38 member 7 [Source:HG |
| AC012183.1 | 16 | TEC |
| GOT2 | 16 | glutamic-oxaloacetic transaminase 2 [Source:I |
| AC106793.1 | 16 | novel transcript |
| RNU6-1155P | 16 | RNA, U6 small nuclear 1155, pseudogene [Sou |
| RN7SL143P | 16 | RNA, 7SL, cytoplasmic 143, pseudogene [Sour |
| GEMIN8P2 | 16 | gem nuclear organelle associated protein 8 ps |
| RPL12P36 | 16 | ribosomal protein L12 pseudogene 36 [Source |
| AC092378.1 | 16 | novel transcript |
| Metazoa_SRP | 16 | Metazoan signal recognition particle RNA [Sou |
| RPS27P27 | 16 | ribosomal protein S27 pseudogene 27 [Source |
| AC092121.1 | 16 | karyopherin alpha 2 (RAG cohort 1, importin a |
| RNU4-58P | 16 | RNA, U4 small nuclear 58, pseudogene [Sourc |
| DUXAP11 | 16 | double homeobox A pseudogene 11 [Source:H |
| APOOP5 | 16 | apolipoprotein O pseudogene 5 [Source:HGN( |
| LINC02141 | 16 | long intergenic non-protein coding RNA 2141 |

| | | |
|---|---|---|
| AC009094.1 | 16 | NOP58 ribonucleoprotein homolog (yeast) (N( |
| AC009081.2 | 16 | novel transcript |
| AC018554.1 | 16 | serine palmitoyltransferase, long chain base s |
| NPAP1L | 16 | nuclear pore associated protein 1 like [Source |
| AC009081.1 | 16 | novel transcript |
| GNPATP | 16 | glyceronephosphate O-acyltransferase pseud( |
| AC009169.1 | 16 | family with sequence similarity 71, member A |
| RPS27AP16 | 16 | ribosomal protein S27a pseudogene 16 [Sourc |
| AC092125.2 | 16 | novel transcript |
| CDH8 | 16 | cadherin 8 [Source:HGNC Symbol;Acc:HGNC:1 |
| AC092125.1 | 16 | novel transcript |
| RN7SKP76 | 16 | RN7SK pseudogene 76 [Source:HGNC Symbol; |
| AC012174.1 | 16 | novel transcript, antisense to CDH8 |
| RNU6-21P | 16 | RNA, U6 small nuclear 21, pseudogene [Sourc |
| AC009161.2 | 16 | novel transcript |
| AC009161.1 | 16 | ribosomal protein L32 (RPL32) pseudogene |
| AC009110.1 | 16 | novel transcript |
| AC010546.1 | 16 | developmental pluripotency associated 3 (DPI |
| AC040174.1 | 16 | novel transcript |
| AC040174.2 | 16 | novel transcript |
| UBE2FP2 | 16 | UBE2F pseudogene 2 [Source:HGNC Symbol;A |
| AC138305.1 | 16 | novel transcript |
| RPS15AP34 | 16 | ribosomal protein S15a pseudogene 34 [Sourc |
| AC138305.2 | 16 | novel transcript |
| AC138305.3 | 16 | novel transcript |
| AC018846.1 | 16 | phosphatidylinositol-specific phospholipase C, |
| LINC02165 | 16 | long intergenic non-protein coding RNA 2165 |
| AC092337.1 | 16 | novel transcript |
| AC093536.2 | 16 | novel transcript |
| AC012322.1 | 16 | novel transcript |
| AC093536.1 | 16 | emopamil binding protein-like (EBPL) pseudog |
| AC092131.1 | 16 | novel transcript |
| PPIAP48 | 16 | peptidylprolyl isomerase A pseudogene 48 [Sc |
| AC009101.1 | 16 | myosin, light chain 6B, alkali, smooth muscle a |
| CDH11 | 16 | cadherin 11 [Source:HGNC Symbol;Acc:HGNC: |
| AC010533.1 | 16 | novel transcript |
| LINC02126 | 16 | long intergenic non-protein coding RNA 2126 |
| AC009055.2 | 16 | novel transcript |
| AC009055.1 | 16 | novel transcript |
| LINC00922 | 16 | long intergenic non-protein coding RNA 922 [S |
| AC092138.2 | 16 | novel transcript |
| AC092138.1 | 16 | novel transcript |
| AC022164.1 | 16 | novel transcript |
| AC025281.1 | 16 | ribosomal protein L35a (RPL35A) pseudogene |
| RNA5SP428 | 16 | RNA, 5S ribosomal pseudogene 428 [Source:H |
| AC012325.1 | 16 | novel transcript |
| CDH5 | 16 | cadherin 5 [Source:HGNC Symbol;Acc:HGNC:1 |
| LINC00920 | 16 | long intergenic non-protein coding RNA 920 [S |
| BEAN1 | 16 | brain expressed associated with NEDD4 1 [Sou |
| AC132186.1 | 16 | transmembrane protein 188 (TMEM188) pseu |

| | | |
|---|---|---|
| BEAN1-AS1 | 16 | BEAN1 antisense RNA 1 [Source:HGNC Symbo |
| AC010542.3 | 16 | novel protein |
| TK2 | 16 | thymidine kinase 2 [Source:HGNC Symbol;Acc |
| AC010542.2 | 16 | novel transcript |
| AC010542.4 | 16 | novel transcript |
| Y_RNA | 16 | Y RNA [Source:RFAM;Acc:RF00019] |
| CKLF | 16 | chemokine like factor [Source:HGNC Symbol;A |
| CKLF-CMTM1 | 16 | CKLF-CMTM1 readthrough [Source:HGNC Sym |
| AC010542.5 | 16 | novel transcript, antisense to CKLF |
| CMTM1 | 16 | CKLF like MARVEL transmembrane domain co |
| CMTM2 | 16 | CKLF like MARVEL transmembrane domain co |
| AC010542.1 | 16 | novel transcript |
| CMTM3 | 16 | CKLF like MARVEL transmembrane domain co |
| CMTM4 | 16 | CKLF like MARVEL transmembrane domain co |
| DYNC1LI2 | 16 | dynein cytoplasmic 1 light intermediate chain |
| AC018557.1 | 16 | novel transcript |
| AC018557.3 | 16 | novel transcript, antisense to DYNC1LI2 |
| AC018557.2 | 16 | novel transcript |
| AC044802.1 | 16 | novel transcript |
| TERB1 | 16 | telomere repeat binding bouquet formation p |
| NAE1 | 16 | NEDD8 activating enzyme E1 subunit 1 [Sourc |
| AC044802.2 | 16 | novel transcript, antisense to CA7 |
| CA7 | 16 | carbonic anhydrase 7 [Source:HGNC Symbol;A |
| PDP2 | 16 | pyruvate dehyrogenase phosphatase catalytic |
| CDH16 | 16 | cadherin 16 [Source:HGNC Symbol;Acc:HGNC: |
| RRAD | 16 | RRAD, Ras related glycolysis inhibitor and calc |
| CIAO2B | 16 | cytosolic iron-sulfur assembly component 2B [ |
| CES2 | 16 | carboxylesterase 2 [Source:HGNC Symbol;Acc |
| AC009084.1 | 16 | novel transcript |
| AC009084.3 | 16 | novel transcript |
| CES3 | 16 | carboxylesterase 3 [Source:HGNC Symbol;Acc |
| CES4A | 16 | carboxylesterase 4A [Source:HGNC Symbol;Ac |
| AC009084.2 | 16 | TEC |
| RN7SL543P | 16 | RNA, 7SL, cytoplasmic 543, pseudogene [Sour |
| CBFB | 16 | core-binding factor subunit beta [Source:HGN |
| C16orf70 | 16 | chromosome 16 open reading frame 70 [Sour |
| B3GNT9 | 16 | UDP-GlcNAc:betaGal beta-1,3-N-acetylglucosa |
| TRADD | 16 | TNFRSF1A associated via death domain [Sourc |
| FBXL8 | 16 | F-box and leucine rich repeat protein 8 [Sourc |
| AC074143.1 | 16 | novel protein |
| HSF4 | 16 | heat shock transcription factor 4 [Source:HGN |
| NOL3 | 16 | nucleolar protein 3 [Source:HGNC Symbol;Acc |
| KIAA0895L | 16 | KIAA0895 like [Source:HGNC Symbol;Acc:HGN |
| EXOC3L1 | 16 | exocyst complex component 3 like 1 [Source:H |
| E2F4 | 16 | E2F transcription factor 4 [Source:HGNC Symb |
| ELMO3 | 16 | engulfment and cell motility 3 [Source:HGNC S |
| MIR328 | 16 | microRNA 328 [Source:HGNC Symbol;Acc:HGN |
| AC040160.2 | 16 | novel transcript |
| LRRC29 | 16 | leucine rich repeat containing 29 [Source:HGN |
| TMEM208 | 16 | transmembrane protein 208 [Source:HGNC Sy |

| | | |
|---|---|---|
| FHOD1 | 16 | formin homology 2 domain containing 1 [Sour |
| SLC9A5 | 16 | solute carrier family 9 member A5 [Source:HG |
| AC040160.1 | 16 | novel transcript |
| PLEKHG4 | 16 | pleckstrin homology and RhoGEF domain cont |
| KCTD19 | 16 | potassium channel tetramerization domain cc |
| RN7SKP118 | 16 | RN7SK pseudogene 118 [Source:HGNC Symbo |
| LRRC36 | 16 | leucine rich repeat containing 36 [Source:HGN |
| TPPP3 | 16 | tubulin polymerization promoting protein fam |
| RNU1-123P | 16 | RNA, U1 small nuclear 123, pseudogene [Sour |
| ZDHHC1 | 16 | zinc finger DHHC-type containing 1 [Source:H( |
| HSD11B2 | 16 | hydroxysteroid 11-beta dehydrogenase 2 [Sou |
| AC009061.1 | 16 | novel transcript |
| ATP6V0D1 | 16 | ATPase H+ transporting V0 subunit d1 [Source |
| AC009061.2 | 16 | novel transcript, antisense to AGRP |
| AGRP | 16 | agouti related neuropeptide [Source:HGNC Sy |
| AC027682.6 | 16 | novel transcript, antisense to FAM65A |
| RIPOR1 | 16 | RHO family interacting cell polarization regula |
| AC027682.5 | 16 | novel transcript, antisense to FAM65A |
| AC027682.2 | 16 | novel transcript, antisense to FAM65A |
| AC027682.3 | 16 | novel transcript, antisense to FAM65A |
| AC027682.4 | 16 | novel transcript |
| AC027682.7 | 16 | TEC |
| AC027682.1 | 16 | novel transcript, antisense to CTCF |
| CTCF | 16 | CCCTC-binding factor [Source:HGNC Symbol;A |
| AC009095.1 | 16 | novel transcript |
| CARMIL2 | 16 | capping protein regulator and myosin 1 linker |
| ACD | 16 | ACD shelterin complex subunit and telomeras |
| PARD6A | 16 | par-6 family cell polarity regulator alpha [Sour |
| ENKD1 | 16 | enkurin domain containing 1 [Source:HGNC Sy |
| C16orf86 | 16 | chromosome 16 open reading frame 86 [Sour |
| GFOD2 | 16 | glucose-fructose oxidoreductase domain cont |
| RANBP10 | 16 | RAN binding protein 10 [Source:HGNC Symbol |
| AC010530.1 | 16 | novel transcript |
| TSNAXIP1 | 16 | translin associated factor X interacting proteir |
| CENPT | 16 | centromere protein T [Source:HGNC Symbol;/ |
| THAP11 | 16 | THAP domain containing 11 [Source:HGNC Syr |
| NUTF2 | 16 | nuclear transport factor 2 [Source:HGNC Syml |
| AC040162.2 | 16 | ribosomal protein S11 (RPS11) pseudogene |
| EDC4 | 16 | enhancer of mRNA decapping 4 [Source:HGN( |
| AC040162.3 | 16 | novel transcript |
| NRN1L | 16 | neuritin 1 like [Source:HGNC Symbol;Acc:HGN |
| PSKH1 | 16 | protein serine kinase H1 [Source:HGNC Symbo |
| CTRL | 16 | chymotrypsin like [Source:HGNC Symbol;Acc:I |
| AC040162.1 | 16 | novel transcript |
| PSMB10 | 16 | proteasome subunit beta 10 [Source:HGNC Sy |
| LCAT | 16 | lecithin-cholesterol acyltransferase [Source:H( |
| SLC12A4 | 16 | solute carrier family 12 member 4 [Source:HG |
| DPEP3 | 16 | dipeptidase 3 [Source:HGNC Symbol;Acc:HGN |
| DPEP2 | 16 | dipeptidase 2 [Source:HGNC Symbol;Acc:HGN |
| DUS2 | 16 | dihydrouridine synthase 2 [Source:HGNC Sym |

| | | |
|---|---|---|
| KARSP3 | 16 | lysyl-tRNA synthetase pseudogene 3 [Source:H |
| DPEP2NB | 16 | DPEP2 neighbor [Source:HGNC Symbol;Acc:HG |
| RNU6-359P | 16 | RNA, U6 small nuclear 359, pseudogene [Sour |
| DDX28 | 16 | DEAD-box helicase 28 [Source:HGNC Symbol;A |
| AC130462.2 | 16 | novel transcript |
| NFATC3 | 16 | nuclear factor of activated T cells 3 [Source:HG |
| Y_RNA | 16 | Y RNA [Source:RFAM;Acc:RF00019] |
| AC130462.1 | 16 | NULL |
| AC020978.6 | 16 | novel transcript |
| AC020978.8 | 16 | novel transcript |
| RPS12P27 | 16 | ribosomal protein S12 pseudogene 27 [Source |
| AC020978.3 | 16 | novel transcript, intronic to NFATC3 |
| AC020978.7 | 16 | novel transcript |
| AC020978.5 | 16 | novel transcript, antisense to NFATC3 |
| ESRP2 | 16 | epithelial splicing regulatory protein 2 [Source |
| MIR6773 | 16 | microRNA 6773 [Source:HGNC Symbol;Acc:HG |
| AC020978.4 | 16 | novel transcript, sense intronic to ESRP2 |
| PLA2G15 | 16 | phospholipase A2 group XV [Source:HGNC Syn |
| AC020978.2 | 16 | novel transcript, antisense to PLA2G15 |
| SLC7A6 | 16 | solute carrier family 7 member 6 [Source:HGN |
| RNU6-1262P | 16 | RNA, U6 small nuclear 1262, pseudogene [Sou |
| SLC7A6OS | 16 | solute carrier family 7 member 6 opposite stra |
| AC020978.9 | 16 | novel transcript |
| PRMT7 | 16 | protein arginine methyltransferase 7 [Source: |
| AC020978.1 | 16 | novel transcript, intronic to PRMT7 |
| RNU4-30P | 16 | RNA, U4 small nuclear 30, pseudogene [Sourc |
| SMPD3 | 16 | sphingomyelin phosphodiesterase 3 [Source:H |
| AC099521.2 | 16 | novel transcript |
| AC099521.3 | 16 | novel transcript |
| AC099521.1 | 16 | novel transcript |
| Y_RNA | 16 | Y RNA [Source:RFAM;Acc:RF00019] |
| RPL35AP33 | 16 | ribosomal protein L35a pseudogene 33 [Sourc |
| ZFP90 | 16 | ZFP90 zinc finger protein [Source:HGNC Symb |
| RNU4-36P | 16 | RNA, U4 small nuclear 36, pseudogene [Sourc |
| AC126773.3 | 16 | novel transcript, antisense to ZFP90 |
| AC126773.6 | 16 | novel transcript |
| AC126773.1 | 16 | NULL |
| AC126773.2 | 16 | NULL |
| CDH3 | 16 | cadherin 3 [Source:HGNC Symbol;Acc:HGNC:1 |
| AC126773.4 | 16 | novel transcript, antisense to CDH3 |
| AC126773.5 | 16 | calponin 2 (CNN2) pseudogene |
| HSPE1P5 | 16 | heat shock protein family E (Hsp10) member 1 |
| CDH1 | 16 | cadherin 1 [Source:HGNC Symbol;Acc:HGNC:1 |
| RNA5SP429 | 16 | RNA, 5S ribosomal pseudogene 429 [Source:H |
| AC099314.1 | 16 | novel transcript, antisense to CDH1 |
| FTLP14 | 16 | ferritin light chain pseudogene 14 [Source:HG |
| TANGO6 | 16 | transport and golgi organization 6 homolog [S |
| Y_RNA | 16 | Y RNA [Source:RFAM;Acc:RF00019] |
| U7 | 16 | U7 small nuclear RNA [Source:RFAM;Acc:RF0C |
| AC009137.1 | 16 | novel transcript |

| | | |
|---|---|---|
| AC009137.2 | 16 | novel transcript, sense intronic to transmemb |
| RNU6-898P | 16 | RNA, U6 small nuclear 898, pseudogene [Sour |
| AC009027.1 | 16 | deoxyuridine triphosphatase (DUT) pseudoge |
| RPS2P45 | 16 | ribosomal protein S2 pseudogene 45 [Source: |
| HAS3 | 16 | hyaluronan synthase 3 [Source:HGNC Symbol; |
| CHTF8 | 16 | chromosome transmission fidelity factor 8 [Sc |
| DERPC | 16 | DERPC proline and glycine rich nuclear protei |
| UTP4 | 16 | UTP4 small subunit processome component [ |
| RNU6-22P | 16 | RNA, U6 small nuclear 22, pseudogene [Sourc |
| SNTB2 | 16 | syntrophin beta 2 [Source:HGNC Symbol;Acc: |
| Y_RNA | 16 | Y RNA [Source:RFAM;Acc:RF00019] |
| AC026474.1 | 16 | novel transcript, sense intronic to SNTB2 |
| AC026464.4 | 16 | novel protein |
| VPS4A | 16 | vacuolar protein sorting 4 homolog A [Source: |
| COG8 | 16 | component of oligomeric golgi complex 8 [Sou |
| PDF | 16 | peptide deformylase, mitochondrial [Source:H |
| AC026464.6 | 16 | novel protein, COG8-PDF readthrough |
| AC026464.3 | 16 | novel protein |
| AC026464.1 | 16 | novel protein |
| NIP7 | 16 | nucleolar pre-rRNA processing protein NIP7 [S |
| TMED6 | 16 | transmembrane p24 trafficking protein 6 [Sou |
| TERF2 | 16 | telomeric repeat binding factor 2 [Source:HG |
| CYB5B | 16 | cytochrome b5 type B [Source:HGNC Symbol; |
| AC026464.5 | 16 | transmembrane protein 111 (TMEM111) pseu |
| AC026464.2 | 16 | novel transcript |
| NFAT5 | 16 | nuclear factor of activated T cells 5 [Source:H |
| MIR1538 | 16 | microRNA 1538 [Source:HGNC Symbol;Acc:H |
| AC009032.1 | 16 | novel transcript, sense intronic to NFAT5 |
| AC012321.1 | 16 | novel transcript |
| NQO1 | 16 | NAD(P)H quinone dehydrogenase 1 [Source:H |
| AC092115.2 | 16 | novel transcript |
| AC092115.3 | 16 | novel transcript |
| NOB1 | 16 | NIN1 (RPN12) binding protein 1 homolog [Sou |
| AC092115.1 | 16 | non-POU domain containing, octamer-binding |
| WWP2 | 16 | WW domain containing E3 ubiquitin protein li |
| SNORA62 | 16 | Small nucleolar RNA SNORA62/SNORA6 family |
| MIR140 | 16 | microRNA 140 [Source:HGNC Symbol;Acc:HG |
| CLEC18A | 16 | C-type lectin domain family 18 member A [So |
| AC026468.1 | 16 | phospholipase A2, group X (PLA2G10 |
| AC009022.1 | 16 | novel transcript |
| NPIPB14P | 16 | nuclear pore complex interacting protein fam |
| PDXDC2P | 16 | pyridoxal dependent decarboxylase domain c |
| MIR1972-2 | 16 | microRNA 1972-2 [Source:HGNC Symbol;Acc: |
| AC009022.2 | 16 | ribosomal protein S10 (RPS10) pseudogene |
| PDPR | 16 | pyruvate dehydrogenase phosphatase regulat |
| AC009060.1 | 16 | novel transcript, antisense to PDPR & CLEC18 |
| CLEC18C | 16 | C-type lectin domain family 18 member C [So |
| AC009060.2 | 16 | phospholipase A2, group X (PLA2G10) pseudo |
| SMG1P7 | 16 | SMG1 pseudogene 7 [Source:HGNC Symbol;A |
| EXOSC6 | 16 | exosome component 6 [Source:HGNC Symbol |

| | | |
|---|---|---|
| AARS | 16 | alanyl-tRNA synthetase [Source:HGNC Symbol |
| RN7SL279P | 16 | RNA, 7SL, cytoplasmic 279, pseudogene [Sour |
| DDX19B | 16 | DEAD-box helicase 19B [Source:HGNC Symbol |
| AC012184.2 | 16 | novel protein, DDX19B and DDX19A readthrou |
| AC012184.3 | 16 | novel transcript, antisense to DDX19B and DD |
| DDX19A | 16 | DEAD-box helicase 19A [Source:HGNC Symbol |
| AC012184.4 | 16 | novel transcript, antisense to DDX19A |
| ST3GAL2 | 16 | ST3 beta-galactoside alpha-2,3-sialyltransfera |
| AC012184.1 | 16 | novel transcript, antisense to ST3GAL2 |
| RPS27P26 | 16 | ribosomal protein S27 pseudogene 26 [Source |
| RNU6-23P | 16 | RNA, U6 small nuclear 23, pseudogene [Sourc |
| FCSK | 16 | fucose kinase [Source:HGNC Symbol;Acc:HGN |
| COG4 | 16 | component of oligomeric golgi complex 4 [Sou |
| SF3B3 | 16 | splicing factor 3b subunit 3 [Source:HGNC Sym |
| SNORD111B | 16 | small nucleolar RNA, C/D box 111B [Source:H( |
| SNORD111 | 16 | small nucleolar RNA, C/D box 111 [Source:HGI |
| IL34 | 16 | interleukin 34 [Source:HGNC Symbol;Acc:HGN |
| MTSS2 | 16 | MTSS I-BAR domain containing 2 [Source:HGN |
| AC020763.4 | 16 | novel transcript |
| VAC14 | 16 | VAC14 component of PIKFYVE complex [Sourc |
| AC020763.3 | 16 | novel transcript |
| AC020763.1 | 16 | novel transcript, antisense to VAC14 |
| VAC14-AS1 | 16 | VAC14 antisense RNA 1 [Source:HGNC Symbo |
| AC020763.5 | 16 | TEC |
| AC020763.2 | 16 | novel transcript |
| AC027281.2 | 16 | novel transcript |
| AC027281.1 | 16 | novel transcript |
| HYDIN | 16 | HYDIN axonemal central pair apparatus protei |
| RNU6ATAC25P | 16 | RNA, U6atac small nuclear 25, pseudogene [S( |
| AC138625.2 | 16 | novel transcript, antisense to HYDIN |
| AC138625.1 | 16 | novel transcript HYDIN |
| CMTR2 | 16 | cap methyltransferase 2 [Source:HGNC Symbc |
| AC106736.1 | 16 | novel transcript |
| CALB2 | 16 | calbindin 2 [Source:HGNC Symbol;Acc:HGNC:1 |
| LINC02136 | 16 | long intergenic non-protein coding RNA 2136 |
| AC010547.6 | 16 | 60S ribosomal protein pseudogene (RPL) pseu |
| TLE7 | 16 | TLE family member 7 [Source:HGNC Symbol;A |
| AC010547.5 | 16 | chemokine (C-C motif) ligand 17 (CCL17) pseu |
| ZNF23 | 16 | zinc finger protein 23 [Source:HGNC Symbol;A |
| AC010547.4 | 16 | novel transcript |
| AC010547.1 | 16 | novel transcript, antisense to ZNF23 |
| ZNF19 | 16 | zinc finger protein 19 [Source:HGNC Symbol;A |
| AC010547.3 | 16 | novel transcript, antisense to ZNF19 |
| CHST4 | 16 | carbohydrate sulfotransferase 4 [Source:HGN( |
| AC010547.2 | 16 | novel transcript, sense intronic to ZNF19 |
| RNU6-1061P | 16 | RNA, U6 small nuclear 1061, pseudogene [Sou |
| TAT-AS1 | 16 | TAT antisense RNA 1 [Source:HGNC Symbol;A( |
| TAT | 16 | tyrosine aminotransferase [Source:HGNC Sym |
| AC009097.2 | 16 | novel transcript, antisense to MARVELD3 |
| MARVELD3 | 16 | MARVEL domain containing 3 [Source:HGNC S |

| | | |
|---|---|---|
| PHLPP2 | 16 | PH domain and leucine rich repeat protein ph |
| AC009097.1 | 16 | novel transcript, antisense to PHLPP1 |
| RNU6-208P | 16 | RNA, U6 small nuclear 208, pseudogene [Sour |
| SNORA70D | 16 | small nucleolar RNA, H/ACA box 70D [Source: |
| AC009097.4 | 16 | novel transcript, antisense to PHLPP1 |
| AC009097.3 | 16 | novel transcript |
| AP1G1 | 16 | adaptor related protein complex 1 subunit ga |
| SNORD71 | 16 | small nucleolar RNA, C/D box 71 [Source:HGN |
| AC010653.3 | 16 | parathymosin (PTMS) pseudogene |
| AC010653.1 | 16 | tripartite motif containing 13 (TRIM13) pseud |
| ATXN1L | 16 | ataxin 1 like [Source:HGNC Symbol;Acc:HGNC |
| IST1 | 16 | IST1 factor associated with ESCRT-III [Source: |
| AC010653.2 | 16 | novel transcript |
| ZNF821 | 16 | zinc finger protein 821 [Source:HGNC Symbol; |
| AC009127.1 | 16 | novel transcript |
| PKD1L3 | 16 | polycystin 1 like 3, transient receptor potenti |
| RPL39P31 | 16 | ribosomal protein L39 pseudogene 31 [Source |
| AC009127.2 | 16 | coiled-coil domain-containing 25 (CCDC25) ps |
| ATP5F1AP3 | 16 | ATP synthase F1 subunit alpha pseudogene 3 |
| DHODH | 16 | dihydroorotate dehydrogenase (quinone) [So |
| TXNL4B | 16 | thioredoxin like 4B [Source:HGNC Symbol;Acc |
| HP | 16 | haptoglobin [Source:HGNC Symbol;Acc:HGNC |
| HPR | 16 | haptoglobin-related protein [Source:HGNC Sy |
| AC009087.1 | 16 | novel transcript, antisense to TXNL4B |
| DHX38 | 16 | DEAH-box helicase 38 [Source:HGNC Symbol; |
| PMFBP1 | 16 | polyamine modulated factor 1 binding protei |
| AC009075.1 | 16 | novel transcript |
| LINC01572 | 16 | long intergenic non-protein coding RNA 1572 |
| AC009075.2 | 16 | c-myc binding protein (MYCBP) |
| U6 | 16 | U6 spliceosomal RNA [Source:RFAM;Acc:RF00 |
| AC004158.1 | 16 | novel transcript |
| AC004943.2 | 16 | novel transcript, antisense to ZFHX3 |
| RNU7-90P | 16 | RNA, U7 small nuclear 90 pseudogene [Source |
| KRT18P18 | 16 | keratin 18 pseudogene 18 [Source:HGNC Sym |
| AC004943.3 | 16 | novel transcript |
| ZFHX3 | 16 | zinc finger homeobox 3 [Source:HGNC Symbo |
| AC004943.1 | 16 | novel transcript, antisense to ZFHX3 |
| RNU7-71P | 16 | RNA, U7 small nuclear 71 pseudogene [Source |
| AC132068.1 | 16 | novel transcript |
| AC002044.3 | 16 | novel transcript |
| AC002044.1 | 16 | novel transcript |
| AC002044.2 | 16 | novel transcript |
| AC116667.1 | 16 | novel transcript |
| HCCAT5 | 16 | hepatocellular carcinoma associated transcrip |
| AC116667.2 | 16 | novel transcript |
| AC140912.1 | 16 | novel transcript |
| AC009033.1 | 16 | novel transcript |
| LINC01568 | 16 | long intergenic non-protein coding RNA 1568 |
| AC092114.1 | 16 | novel transcript |
| AC087565.2 | 16 | novel transcript |

| | | |
|---|---|---|
| AC087565.1 | 16 | novel transcript |
| AC092128.1 | 16 | ribosomal protein SA pseudogene 56 |
| AC138627.1 | 16 | novel transcript |
| AC138627.2 | 16 | novel transcript |
| AC009120.6 | 16 | novel transcript |
| PPIAP49 | 16 | peptidylprolyl isomerase A pseudogene 49 [Sc |
| AC009120.4 | 16 | novel transcript |
| AC009120.5 | 16 | novel transcript |
| PSMD7 | 16 | proteasome 26S subunit, non-ATPase 7 [Sourc |
| AC009120.2 | 16 | novel transcript, antisense to PSMD7 |
| AC009120.1 | 16 | ribosomal protein L21 (RPL21) pseudogene |
| AC009120.3 | 16 | novel transcript |
| AC009053.1 | 16 | pyruvate dehydrogenase phosphatase regulat |
| AC009053.2 | 16 | novel transcript |
| NPIPB15 | 16 | nuclear pore complex interacting protein fami |
| AC009053.4 | 16 | phospholipase A2, group X (PLA2G10) pseudo |
| CLEC18B | 16 | C-type lectin domain family 18 member B [Sou |
| AC009053.3 | 16 | novel transcript, antisense to CLEC18B |
| GLG1 | 16 | golgi glycoprotein 1 [Source:HGNC Symbol;Ac |
| RNU6-237P | 16 | RNA, U6 small nuclear 237, pseudogene [Sour |
| Y_RNA | 16 | Y RNA [Source:RFAM;Acc:RF00019] |
| AC009153.1 | 16 | ribosomal protein S10 (RPS10) pseudogene |
| HSPE1P7 | 16 | heat shock protein family E (Hsp10) member 1 |
| RFWD3 | 16 | ring finger and WD repeat domain 3 [Source:H |
| TMPOP2 | 16 | thymopoietin pseudogene 2 [Source:HGNC Sy |
| MLKL | 16 | mixed lineage kinase domain like pseudokinas |
| FA2H | 16 | fatty acid 2-hydroxylase [Source:HGNC Symbo |
| AC009132.1 | 16 | WD repeat domain 59 (WDR59) pseudogene |
| WDR59 | 16 | WD repeat domain 59 [Source:HGNC Symbol; |
| ZNRF1 | 16 | zinc and ring finger 1 [Source:HGNC Symbol;A |
| AC099508.1 | 16 | novel transcript, antisense to ZNRF1 |
| LDHD | 16 | lactate dehydrogenase D [Source:HGNC Symb |
| AC099508.2 | 16 | novel transcript |
| ZFP1 | 16 | ZFP1 zinc finger protein [Source:HGNC Symbo |
| AC009078.1 | 16 | ribosomal protein S4, Y-linked 2 (RPS4Y2) pseu |
| CTRB2 | 16 | chymotrypsinogen B2 [Source:HGNC Symbol;/ |
| CTRB1 | 16 | chymotrypsinogen B1 [Source:HGNC Symbol;/ |
| AC009078.2 | 16 | chymotrypsinogen B (CTRB) pseudogene |
| BCAR1 | 16 | BCAR1 scaffold protein, Cas family member [S |
| AC009078.3 | 16 | TEC |
| RNU6-758P | 16 | RNA, U6 small nuclear 758, pseudogene [Sour |
| CFDP1 | 16 | craniofacial development protein 1 [Source:H |
| AC009054.1 | 16 | novel transcript, sense intronic to CFDP1 |
| AC009054.2 | 16 | novel transcript, sense intronic to CFDP1 |
| AC009163.2 | 16 | NULL |
| AC009163.5 | 16 | novel TMEM170A-CFDP1 readthrough proteir |
| AC009163.7 | 16 | novel transcript |
| TMEM170A | 16 | transmembrane protein 170A [Source:HGNC S |
| AC009163.4 | 16 | novel transcript |
| CHST6 | 16 | carbohydrate sulfotransferase 6 [Source:HGN |

| | | |
|---|---|---|
| AC009163.1 | 16 | novel transcript, antisense to CHST6 |
| AC009163.6 | 16 | novel transcript |
| TMEM231P1 | 16 | TMEM231 pseudogene 1 [Source:HGNC Symb |
| AC009163.3 | 16 | TMEM231-CHST5 readthrough protein |
| CHST5 | 16 | carbohydrate sulfotransferase 5 [Source:HGN |
| TMEM231 | 16 | transmembrane protein 231 [Source:HGNC Sy |
| AC025287.1 | 16 | ADAT1 and TMEM231 readthrough transcript |
| AC025287.2 | 16 | novel transcript |
| GABARAPL2 | 16 | GABA type A receptor associated protein like |
| AC025287.3 | 16 | novel transcript, antisense to GABARAPL2 |
| ADAT1 | 16 | adenosine deaminase tRNA specific 1 [Source |
| KARS | 16 | lysyl-tRNA synthetase [Source:HGNC Symbol; |
| AC025287.5 | 16 | telomeric repeat binding factor 2, interacting |
| TERF2IP | 16 | TERF2 interacting protein [Source:HGNC Symb |
| AC025287.4 | 16 | novel protein similar to cytoplasmic polyaden |
| DUXB | 16 | double homeobox B [Source:HGNC Symbol;Ac |
| RN7SL520P | 16 | RNA, 7SL, cytoplasmic 520, pseudogene [Sour |
| ATP5PBP7 | 16 | ATP synthase peripheral stalk-membrane sub |
| CPHXL | 16 | cytoplasmic polyadenylated homeobox like [S |
| AC105430.1 | 16 | novel transcript |
| RNA5SP430 | 16 | RNA, 5S ribosomal pseudogene 430 [Source:H |
| AC099511.1 | 16 | novel transcript |
| AC010528.1 | 16 | novel transcript |
| RPL18P13 | 16 | ribosomal protein L18 pseudogene 13 [Source |
| AC010528.2 | 16 | novel transcript |
| CNTNAP4 | 16 | contactin associated protein like 4 [Source:HG |
| AC106741.1 | 16 | novel protein |
| RN7SKP233 | 16 | RN7SK pseudogene 233 [Source:HGNC Symbo |
| SNORD33 | 16 | Small nucleolar RNA Z195/SNORD33/SNORD3 |
| AC108125.1 | 16 | novel transcript |
| LINC02125 | 16 | long intergenic non-protein coding RNA 2125 |
| AC106729.1 | 16 | novel transcript |
| MIR4719 | 16 | microRNA 4719 [Source:HGNC Symbol;Acc:HG |
| AC106734.1 | 16 | MAD2 mitotic arrest deficient-like 1 (yeast) (M |
| AC106733.1 | 16 | novel transcript |
| MON1B | 16 | MON1 homolog B, secretory trafficking associ |
| SYCE1L | 16 | synaptonemal complex central element prote |
| AC009139.2 | 16 | novel transcript, antisense to MON1B & SYCE |
| VN2R10P | 16 | vomeronasal 2 receptor 10, pseudogene [Sou |
| AC009139.1 | 16 | novel transcript, antisense to ADAMTS18 |
| ADAMTS18 | 16 | ADAM metallopeptidase with thrombospondi |
| AC025284.1 | 16 | novel transcript, antisense to ADAMTS18 |
| LINC02131 | 16 | long intergenic non-protein coding RNA 2131 |
| AC092724.1 | 16 | novel transcript |
| NUDT7 | 16 | nudix hydrolase 7 [Source:HGNC Symbol;Acc: |
| AC092134.1 | 16 | novel transcript, antisense to NUDT7 |
| AC092134.3 | 16 | novel transcript |
| VAT1L | 16 | vesicle amine transport 1 like [Source:HGNC S |
| AC092134.2 | 16 | novel transcript |
| PPIAP50 | 16 | peptidylprolyl isomerase A pseudogene 50 [So |

| | | |
|---|---|---|
| AC079414.1 | 16 | novel transcript, antisense to CLEC3A |
| CLEC3A | 16 | C-type lectin domain family 3 member A [Sour |
| KRT8P22 | 16 | keratin 8 pseudogene 22 [Source:HGNC Symb |
| AC079414.2 | 16 | leucine rich repeat containing 59 (LRRC59) pse |
| WWOX | 16 | WW domain containing oxidoreductase [Sour |
| AC079414.3 | 16 | novel transcript, antisense WWOX |
| AC009044.1 | 16 | TEC |
| WWOX-AS1 | 16 | WWOX antisense RNA 1 [Source:HGNC Symbo |
| AC106743.1 | 16 | novel transcript, antisense to WWOX |
| LSM3P5 | 16 | LSM3 homolog, U6 small nuclear RNA and mR |
| AC046158.1 | 16 | novel transcript, antisense to WWOX |
| AC046158.4 | 16 | TEC |
| AC046158.2 | 16 | novel transcript, sense intronic to WWOX |
| AC046158.3 | 16 | TEC |
| AC009141.1 | 16 | novel transcript, antisense to WWOX |
| RPS3P7 | 16 | ribosomal protein S3 pseudogene 7 [Source:H |
| AC136603.1 | 16 | TEC |
| AC027279.4 | 16 | TEC |
| AC027279.1 | 16 | novel transcript, sense intronic to WWOX |
| AC027279.3 | 16 | TEC |
| AC027279.2 | 16 | novel transcript, sense intronic to WWOX |
| AC009145.4 | 16 | TEC |
| AC009145.3 | 16 | novel transcript, antisense to WWOX |
| AC009145.1 | 16 | novel transcript, sense intronic to WWOX |
| AC009145.2 | 16 | novel transcript, sense intronic to WWOX |
| AC092376.2 | 16 | novel transcript, antisense to WWOX |
| AC092376.3 | 16 | novel transcript, antisense to WWOX |
| AC092376.1 | 16 | novel transcript |
| RNA5SP431 | 16 | RNA, 5S ribosomal pseudogene 431 [Source:H |
| AC084064.1 | 16 | novel transcript |
| MAF | 16 | MAF bZIP transcription factor [Source:HGNC S |
| AC009159.2 | 16 | novel transcript |
| AC009159.3 | 16 | novel transcript |
| AC009159.1 | 16 | novel transcript |
| AC009159.4 | 16 | TEC |
| LINC01229 | 16 | long intergenic non-protein coding RNA 1229 |
| MAFTRR | 16 | MAF transcriptional regulator RNA [Source:HG |
| LINC01228 | 16 | long intergenic non-protein coding RNA 1228 |
| AC092130.1 | 16 | novel transcript |
| AC022166.1 | 16 | novel transcript |
| AC105411.1 | 16 | novel transcript, antisense DYNLRB2 |
| AC022166.2 | 16 | novel transcript |
| DYNLRB2 | 16 | dynein light chain roadblock-type 2 [Source:H |
| AC108097.1 | 16 | novel transcript, antisense DYNLRB2 |
| LINC01227 | 16 | long intergenic non-protein coding RNA 1227 |
| CDYL2 | 16 | chromodomain Y like 2 [Source:HGNC Symbol |
| AC092332.1 | 16 | novel transcript, sense intronic to CDYL2 |
| AC099313.1 | 16 | novel transcript, antisense to CDYL2 |
| ARLNC1 | 16 | androgen receptor regulated long noncoding |
| AC009070.1 | 16 | novel protein |

| | | |
|---|---|---|
| CMC2 | 16 | C-X9-C motif containing 2 [Source:HGNC Symb |
| AC092718.7 | 16 | novel transcript |
| CENPN | 16 | centromere protein N [Source:HGNC Symbol;/ |
| AC092718.2 | 16 | novel transcript, antisense to CENPN |
| AC092718.8 | 16 | novel protein |
| AC092718.4 | 16 | novel transcript |
| ATMIN | 16 | ATM interactor [Source:HGNC Symbol;Acc:HG |
| C16orf46 | 16 | chromosome 16 open reading frame 46 [Sour |
| AC092718.3 | 16 | novel protein |
| AC092718.5 | 16 | novel transcript, antisense to C16orf46 |
| AC092718.6 | 16 | novel transcript, antisense to C16orf46 |
| AC092718.1 | 16 | novel transcript |
| GCSH | 16 | glycine cleavage system protein H [Source:HG |
| PKD1L2 | 16 | polycystin 1 like 2 (gene/pseudogene) [Source |
| RNU6-1191P | 16 | RNA, U6 small nuclear 1191, pseudogene [Sou |
| AC092718.9 | 16 | novel transcript |
| BCO1 | 16 | beta-carotene oxygenase 1 [Source:HGNC Syn |
| AC131888.1 | 16 | novel transcript |
| AC009148.1 | 16 | novel transcript, antisense to GAN |
| GAN | 16 | gigaxonin [Source:HGNC Symbol;Acc:HGNC:41 |
| MIR4720 | 16 | microRNA 4720 [Source:HGNC Symbol;Acc:HG |
| AC092139.1 | 16 | novel transcript |
| CMIP | 16 | c-Maf inducing protein [Source:HGNC Symbol, |
| AC092139.2 | 16 | novel transcript |
| PPIAP51 | 16 | peptidylprolyl isomerase A pseudogene 51 [Sc |
| AC092139.3 | 16 | novel transcript |
| MIR7854 | 16 | microRNA 7854 [Source:HGNC Symbol;Acc:HG |
| MIR6504 | 16 | microRNA 6504 [Source:HGNC Symbol;Acc:HG |
| AC092135.2 | 16 | novel transcript |
| AC092135.1 | 16 | novel transcript |
| AC092135.3 | 16 | novel transcript |
| AC099524.1 | 16 | novel transcript |
| PLCG2 | 16 | phospholipase C gamma 2 [Source:HGNC Sym |
| RN7SKP176 | 16 | RN7SK pseudogene 176 [Source:HGNC Symbo |
| AC092142.2 | 16 | novel transcript |
| SDR42E1 | 16 | short chain dehydrogenase/reductase family 4 |
| HSD17B2 | 16 | hydroxysteroid 17-beta dehydrogenase 2 [Sou |
| AC092142.1 | 16 | novel transcript, antisense to HSD17B2 |
| MTCYBP28 | 16 | MT-CYB pseudogene 28 [Source:HGNC Symbo |
| MPHOSPH6 | 16 | M-phase phosphoprotein 6 [Source:HGNC Syr |
| AC138304.1 | 16 | novel transcript |
| RN7SKP190 | 16 | RN7SK pseudogene 190 [Source:HGNC Symbo |
| AC009117.1 | 16 | ribosomal protein L21 (RPL21) pseudogene |
| AC009117.2 | 16 | novel transcript |
| CDH13 | 16 | cadherin 13 [Source:HGNC Symbol;Acc:HGNC: |
| MIR8058 | 16 | microRNA 8058 [Source:HGNC Symbol;Acc:HG |
| AC099506.2 | 16 | novel transcript |
| AC099506.1 | 16 | novel transcript |
| AC099506.3 | 16 | novel transcript, antisense to CDH13 |
| RN7SL134P | 16 | RNA, 7SL, cytoplasmic 134, pseudogene [Sour |

| | | |
|---|---|---|
| AC125793.1 | 16 | novel transcript, antisense to CDH13 |
| AC009142.1 | 16 | novel transcript, antisense to CDH13 |
| MIR3182 | 16 | microRNA 3182 [Source:HGNC Symbol;Acc:H( |
| AC009063.3 | 16 | novel transcript, antisense to CDH13 |
| AC009063.2 | 16 | novel transcript, antisense to CDH13 |
| AC009063.1 | 16 | novel transcript, antisense to CDH13 |
| AC009119.1 | 16 | novel transcript, antisense to CDH13 |
| HSBP1 | 16 | heat shock factor binding protein 1 [Source:H( |
| MLYCD | 16 | malonyl-CoA decarboxylase [Source:HGNC Syr |
| AC009119.2 | 16 | novel transcript, antisense to MLYCD - OSGIN: |
| OSGIN1 | 16 | oxidative stress induced growth inhibitor 1 [S( |
| NECAB2 | 16 | N-terminal EF-hand calcium binding protein 2 |
| SLC38A8 | 16 | solute carrier family 38 member 8 [Source:HG |
| RNA5SP432 | 16 | RNA, 5S ribosomal pseudogene 432 [Source:H |
| AC040169.4 | 16 | novel transcript, antisense to SLC38A8 |
| MBTPS1 | 16 | membrane bound transcription factor peptida |
| AC040169.3 | 16 | novel transcript, sense intronic to MBTPS1 |
| AC040169.1 | 16 | novel transcript, antisense to MBTPS1 |
| AC040169.2 | 16 | polo-like kinase 1 (PLK1) pseudogene |
| HSDL1 | 16 | hydroxysteroid dehydrogenase like 1 [Source: |
| DNAAF1 | 16 | dynein axonemal assembly factor 1 [Source:H |
| TAF1C | 16 | TATA-box binding protein associated factor, R |
| ADAD2 | 16 | adenosine deaminase domain containing 2 [S( |
| AC009123.1 | 16 | novel transcript, antisense to ADAD2 |
| KCNG4 | 16 | potassium voltage-gated channel modifier suk |
| AC010551.3 | 16 | novel transcript |
| RNA5SP433 | 16 | RNA, 5S ribosomal pseudogene 433 [Source:H |
| WFDC1 | 16 | WAP four-disulfide core domain 1 [Source:HG |
| AC010551.2 | 16 | novel transcript, antisense to WFDC1 |
| AC010551.1 | 16 | novel transcript |
| ATP2C2 | 16 | ATPase secretory pathway Ca2+ transporting : |
| ATP2C2-AS1 | 16 | ATP2C2 antisense RNA 1 [Source:HGNC Symb( |
| MEAK7 | 16 | MTOR associated protein, eak-7 homolog [Sou |
| AC022165.1 | 16 | novel transcript, antisense to KIAA1609 |
| COTL1 | 16 | coactosin like F-actin binding protein 1 [Sourc( |
| AC092145.1 | 16 | novel transcript, antisense to COTL1 |
| KLHL36 | 16 | kelch like family member 36 [Source:HGNC Sy |
| USP10 | 16 | ubiquitin specific peptidase 10 [Source:HGNC |
| AC025280.3 | 16 | novel transcript, antisense to CRISPLD2 |
| CRISPLD2 | 16 | cysteine rich secretory protein LCCL domain c( |
| AC025280.1 | 16 | novel transcript, sense intronic to CRISPLD2 |
| AC025280.2 | 16 | novel transcript, antisense to CRISPLD2 |
| LINC02176 | 16 | long intergenic non-protein coding RNA 2176 |
| ZDHHC7 | 16 | zinc finger DHHC-type containing 7 [Source:H( |
| KIAA0513 | 16 | KIAA0513 [Source:HGNC Symbol;Acc:HGNC:2( |
| FAM92B | 16 | family with sequence similarity 92 member B |
| LINC02139 | 16 | long intergenic non-protein coding RNA 2139 |
| AC026469.1 | 16 | novel transcript |
| GSE1 | 16 | Gse1 coiled-coil protein [Source:HGNC Symbo |
| AC092275.1 | 16 | novel transcript, sense intronic to GSE1 |

| | | |
|---|---|---|
| COX6CP16 | 16 | cytochrome c oxidase subunit 6C pseudogene |
| LINC00311 | 16 | long intergenic non-protein coding RNA 311 [S |
| MIR5093 | 16 | microRNA 5093 [Source:HGNC Symbol;Acc:HG |
| AC092377.1 | 16 | novel transcript, antisense to GSE1 |
| AC133540.1 | 16 | novel transcript |
| AC092127.2 | 16 | novel transcript |
| AC092127.1 | 16 | novel transcript |
| RN7SL381P | 16 | RNA, 7SL, cytoplasmic 381, pseudogene [Sour |
| GINS2 | 16 | GINS complex subunit 2 [Source:HGNC Symbo |
| C16orf74 | 16 | chromosome 16 open reading frame 74 [Sour |
| AC018695.6 | 16 | novel transcript, sense intronic to C16orf74 |
| AC018695.9 | 16 | novel transcript, antisense to C16orf74 |
| MIR1910 | 16 | microRNA 1910 [Source:HGNC Symbol;Acc:HG |
| AC018695.2 | 16 | novel transcript, antisense to C16orf74 |
| EMC8 | 16 | ER membrane protein complex subunit 8 [Sou |
| RNU1-103P | 16 | RNA, U1 small nuclear 103, pseudogene [Sour |
| AC018695.4 | 16 | novel transcript, antisense to COX4NB |
| AC018695.3 | 16 | novel transcript, antisense to COX4NB |
| COX4I1 | 16 | cytochrome c oxidase subunit 4I1 [Source:HGI |
| AC018695.5 | 16 | iron-sulfur cluster assembly 1 homolog (S. cer |
| AC018695.1 | 16 | ribosomal protein L10a (RPL10A) pseudogene |
| AC018695.8 | 16 | novel transcript |
| AC018695.7 | 16 | TEC |
| IRF8 | 16 | interferon regulatory factor 8 [Source:HGNC S |
| MIR6774 | 16 | microRNA 6774 [Source:HGNC Symbol;Acc:HG |
| AC092723.5 | 16 | novel transcript |
| LINC02132 | 16 | long intergenic non-protein coding RNA 2132 |
| AC092723.4 | 16 | novel transcript |
| AC092723.1 | 16 | novel transcript |
| AC092723.3 | 16 | novel transcript |
| AC092723.2 | 16 | novel transcript |
| AC135012.1 | 16 | novel transcript |
| AC135012.2 | 16 | novel transcript |
| AC135012.3 | 16 | novel transcript |
| LINC01082 | 16 | long intergenic non-protein coding RNA 1082 |
| LINC01081 | 16 | long intergenic non-protein coding RNA 1081 |
| Y_RNA | 16 | Y RNA [Source:RFAM;Acc:RF00019] |
| LINC02135 | 16 | long intergenic non-protein coding RNA 2135 |
| LINC00917 | 16 | long intergenic non-protein coding RNA 917 [S |
| AC092327.2 | 16 | novel transcript |
| AC092327.1 | 16 | chloride intracellular channel 1 (CLIC1) pseudo |
| FENDRR | 16 | FOXF1 adjacent non-coding developmental re |
| FOXF1 | 16 | forkhead box F1 [Source:HGNC Symbol;Acc:H |
| AC009108.3 | 16 | novel transcript |
| AC009108.1 | 16 | ribosomal protein L7a (RPL7A) pseudogene |
| MTHFSD | 16 | methenyltetrahydrofolate synthetase domain |
| AC009108.5 | 16 | novel transcript |
| FOXC2-AS1 | 16 | FOXC2 antisense RNA 1 [Source:HGNC Symbo |
| FOXC2 | 16 | forkhead box C2 [Source:HGNC Symbol;Acc:H |
| FOXL1 | 16 | forkhead box L1 [Source:HGNC Symbol;Acc:H |

| | | |
|---|---|---|
| AC009108.4 | 16 | novel transcript |
| AC009108.2 | 16 | novel transcript |
| AC009154.1 | 16 | novel transcript |
| AC009154.2 | 16 | novel transcript |
| LINC02188 | 16 | long intergenic non-protein coding RNA 2188 |
| LINC02189 | 16 | long intergenic non-protein coding RNA 2189 |
| AC130467.1 | 16 | novel transcript |
| AC093519.2 | 16 | novel transcript |
| AC093519.1 | 16 | coiled-coil domain containing 7 (CCDC7) pseu |
| LINC02181 | 16 | long intergenic non-protein coding RNA 2181 |
| AC106745.1 | 16 | novel transcript |
| C16orf95 | 16 | chromosome 16 open reading frame 95 [Sour |
| AC136285.3 | 16 | novel transcript |
| AC136285.1 | 16 | novel transcript, antisense to C16orf95 |
| AC136285.2 | 16 | novel transcript, antisense to C16orf95 |
| AC010531.4 | 16 | novel transcript, sense intronic to C16orf95 |
| AC010531.2 | 16 | ribosomal protein L39 (RPL39) pseudogene |
| AC010531.8 | 16 | novel transcript |
| AC010531.1 | 16 | novel protein |
| AC010531.6 | 16 | novel transcript, antisense to FBXO31 and C1 |
| FBXO31 | 16 | F-box protein 31 [Source:HGNC Symbol;Acc:H |
| AC010531.7 | 16 | novel transcript, antisense to FBXO31 and C1 |
| AC010531.5 | 16 | novel transcript, antisense to FBXO31 |
| MAP1LC3B | 16 | microtubule associated protein 1 light chain 3 |
| AC010531.3 | 16 | novel transcript, antisense to MAP1LC3B |
| ZCCHC14 | 16 | zinc finger CCHC-type containing 14 [Source:H |
| AC105429.2 | 16 | nuclear receptor subfamily 3, group C, memb |
| AC105429.1 | 16 | novel transcript, sense intronic to ZCCHC14 |
| AC092720.1 | 16 | novel transcript |
| AC092720.2 | 16 | novel transcript, antisense to JPH3 |
| JPH3 | 16 | junctophilin 3 [Source:HGNC Symbol;Acc:HGN |
| AC010536.3 | 16 | novel transcript, antisense to JPH3 |
| AC010536.1 | 16 | novel transcript |
| KLHDC4 | 16 | kelch domain containing 4 [Source:HGNC Sym |
| AC010536.2 | 16 | novel transcript, sense intronic to KLHDC4 |
| AC126696.3 | 16 | novel transcript |
| AC126696.1 | 16 | novel transcript |
| SLC7A5 | 16 | solute carrier family 7 member 5 [Source:HGN |
| MIR6775 | 16 | microRNA 6775 [Source:HGNC Symbol;Acc:H |
| AC126696.2 | 16 | novel transcript, antisense to SLC7A5 |
| CA5A | 16 | carbonic anhydrase 5A [Source:HGNC Symbol |
| AC133539.1 | 16 | ribosomal protein S10 (RPS10) pseudogene |
| AC127455.1 | 16 | small EDRL-rich factor 1 protein pseudogene |
| BANP | 16 | BTG3 associated nuclear protein [Source:HGN |
| AC134312.4 | 16 | novel transcript |
| AC134312.3 | 16 | novel transcript |
| AC134312.1 | 16 | novel transcript |
| AC134312.2 | 16 | novel transcript |
| AC134312.5 | 16 | novel transcript |
| LINC02182 | 16 | long intergenic non-protein coding RNA 2182 |

| | | |
|---|---|---|
| AC134312.6 | 16 | novel transcript |
| AC138512.1 | 16 | novel transcript |
| ZNF469 | 16 | zinc finger protein 469 [Source:HGNC Symbol; |
| ZFPM1 | 16 | zinc finger protein, FOG family member 1 [Sou |
| MIR5189 | 16 | microRNA 5189 [Source:HGNC Symbol;Acc:HG |
| AC116552.1 | 16 | novel transcript, antisense to ZFPM1 |
| ZC3H18 | 16 | zinc finger CCCH-type containing 18 [Source:H |
| IL17C | 16 | interleukin 17C [Source:HGNC Symbol;Acc:HG |
| CYBA | 16 | cytochrome b-245 alpha chain [Source:HGNC |
| MVD | 16 | mevalonate diphosphate decarboxylase [Sour |
| SNAI3-AS1 | 16 | SNAI3 antisense RNA 1 [Source:HGNC Symbol; |
| SNAI3 | 16 | snail family transcriptional repressor 3 [Source |
| RNF166 | 16 | ring finger protein 166 [Source:HGNC Symbol; |
| AC138028.3 | 16 | novel transcript, antisense to RNF166 |
| CTU2 | 16 | cytosolic thiouridylase subunit 2 [Source:HGN |
| AC138028.6 | 16 | novel transcript, antisense to CTU2 |
| PIEZO1 | 16 | piezo type mechanosensitive ion channel com |
| MIR4722 | 16 | microRNA 4722 [Source:HGNC Symbol;Acc:HG |
| AC138028.4 | 16 | novel transcript, antisense to FAM38A |
| AC138028.2 | 16 | novel transcript, antisense to FAM38A |
| AC138028.5 | 16 | novel transcript, antisense to FAM38A |
| AC138028.1 | 16 | novel transcript, antisense to FAM38A |
| CDT1 | 16 | chromatin licensing and DNA replication facto |
| APRT | 16 | adenine phosphoribosyltransferase [Source:H |
| GALNS | 16 | galactosamine (N-acetyl)-6-sulfatase [Source:I |
| TRAPPC2L | 16 | trafficking protein particle complex 2 like [Sou |
| PABPN1L | 16 | poly(A) binding protein nuclear 1 like, cytopla |
| CBFA2T3 | 16 | CBFA2/RUNX1 partner transcriptional co-repr |
| AC092384.2 | 16 | novel transcript, antisense to CBFA2T3 |
| AC092384.3 | 16 | novel transcript |
| AC092384.1 | 16 | uncharacterized LOC100129697 [Source:NCBI |
| AC135782.2 | 16 | novel transcript |
| AC135782.1 | 16 | novel transcript |
| ACSF3 | 16 | acyl-CoA synthetase family member 3 [Source |
| AC135782.3 | 16 | novel transcript, antisense to ACSF3 |
| LINC00304 | 16 | long intergenic non-protein coding RNA 304 [ |
| LINC02138 | 16 | long intergenic non-protein coding RNA 2138 |
| CDH15 | 16 | cadherin 15 [Source:HGNC Symbol;Acc:HGNC: |
| SLC22A31 | 16 | solute carrier family 22 member 31 [Source:H |
| AC009113.1 | 16 | novel transcript, antisense to ZNF778 |
| ZNF778 | 16 | zinc finger protein 778 [Source:HGNC Symbol; |
| AC009113.2 | 16 | novel transcript, antisense to ZNF778 |
| ANKRD11 | 16 | ankyrin repeat domain 11 [Source:HGNC Syml |
| AC137932.3 | 16 | novel transcript, antisense to ANKRD11 |
| AC137932.1 | 16 | novel transcript, antisense to ANKRD11 |
| AC137932.2 | 16 | uncharacterized LOC100287036 [Source:NCBI |
| AC092120.1 | 16 | novel transcript |
| AC092120.3 | 16 | novel transcript, antisense to ANKRD11 |
| RNU6-430P | 16 | RNA, U6 small nuclear 430, pseudogene [Sour |
| AC092120.2 | 16 | cytidine monophosphate (UMP-CMP) kinase 1 |

| | | |
|---|---|---|
| SPG7 | 16 | SPG7 matrix AAA peptidase subunit, paraplegi |
| AC092123.1 | 16 | novel transcript, antisense to SPG7 |
| RPL13 | 16 | ribosomal protein L13 [Source:HGNC Symbol;/ |
| SNORD68 | 16 | small nucleolar RNA, C/D box 68 [Source:HGN |
| CPNE7 | 16 | copine 7 [Source:HGNC Symbol;Acc:HGNC:23 |
| DPEP1 | 16 | dipeptidase 1 [Source:HGNC Symbol;Acc:HGN |
| CHMP1A | 16 | charged multivesicular body protein 1A [Sourc |
| SPATA33 | 16 | spermatogenesis associated 33 [Source:HGNC |
| AC010538.1 | 16 | novel transcript, sense overlapping CDK10 |
| CDK10 | 16 | cyclin dependent kinase 10 [Source:HGNC Syn |
| LINC02166 | 16 | long intergenic non-protein coding RNA 2166 |
| SPATA2L | 16 | spermatogenesis associated 2 like [Source:HG |
| VPS9D1 | 16 | VPS9 domain containing 1 [Source:HGNC Sym |
| VPS9D1-AS1 | 16 | VPS9D1 antisense RNA 1 [Source:HGNC Symb |
| ZNF276 | 16 | zinc finger protein 276 [Source:HGNC Symbol; |
| FANCA | 16 | FA complementation group A [Source:HGNC S |
| SPIRE2 | 16 | spire type actin nucleation factor 2 [Source:H |
| TCF25 | 16 | transcription factor 25 [Source:HGNC Symbol; |
| AC092143.3 | 16 | novel transcript (MC1R-TUBB3 readthrough) |
| MC1R | 16 | melanocortin 1 receptor [Source:HGNC Symb |
| AC092143.1 | 16 | novel protein (MC1R-TUBB3 readthrough) |
| AC092143.2 | 16 | novel transcript, antisense to MC1R |
| TUBB3 | 16 | tubulin beta 3 class III [Source:HGNC Symbol;/ |
| DEF8 | 16 | differentially expressed in FDCP 8 homolog [S |
| CENPBD1 | 16 | CENPB DNA-binding domain containing 1 [Sou |
| AFG3L1P | 16 | AFG3 like matrix AAA peptidase subunit 1, pse |
| DBNDD1 | 16 | dysbindin domain containing 1 [Source:HGNC |
| GAS8 | 16 | growth arrest specific 8 [Source:HGNC Symbo |
| GAS8-AS1 | 16 | GAS8 antisense RNA 1 [Source:HGNC Symbol; |
| URAHP | 16 | urate (hydroxyiso-) hydrolase, pseudogene [S |
| PRDM7 | 16 | PR/SET domain 7 [Source:HGNC Symbol;Acc:H |
| TUBB8P7 | 16 | tubulin beta 8 class VIII pseudogene 7 [Source |
| FAM157C | 16 | family with sequence similarity 157 member ( |
| AC133919.1 | 16 | novel transcript |
| AC133919.2 | 16 | novel transcript |
| RNU6-355P | 16 | RNA, U6 small nuclear 355, pseudogene [Sour |
| LINC02193 | 16 | long intergenic non-protein coding RNA 2193 |
| AC133919.3 | 16 | capicua transcriptional repressor (CIC) pseudc |
| AC240565.2 | 17 | novel transcript |
| AC240565.1 | 17 | novel transcript |
| U6 | 17 | U6 spliceosomal RNA [Source:RFAM;Acc:RF00 |
| SCGB1C2 | 17 | secretoglobin family 1C member 2 [Source:HG |
| DOC2B | 17 | double C2 domain beta [Source:HGNC Symbol |
| LINC02091 | 17 | long intergenic non-protein coding RNA 2091 |
| RPH3AL | 17 | rabphilin 3A like (without C2 domains) [Source |
| AC129507.4 | 17 | novel transcript, antisense to RPH3AL |
| AC129507.1 | 17 | uncharacterized LOC100506388 [Source:NCBI |
| AC129507.2 | 17 | novel transcript, antisense to RPH3AL |
| AC129507.3 | 17 | novel transcript, antisense to RPH3AL |
| AC141424.1 | 17 | novel transcript, antisense to C17orf97 |

| | | |
|---|---|---|
| C17orf97 | 17 | chromosome 17 open reading frame 97 [Sour |
| RFLNB | 17 | refilin B [Source:HGNC Symbol;Acc:HGNC:287 |
| AC015853.2 | 17 | novel transcript, antisense to VPS53 |
| VPS53 | 17 | VPS53 subunit of GARP complex [Source:HGN |
| AC015853.3 | 17 | novel transcript, antisense to VPS53 |
| AC015853.1 | 17 | novel transcript, antisense to VPS53 |
| AC027455.1 | 17 | BCL2-associated athanogene (BAG1) pseudog |
| RPS4XP17 | 17 | ribosomal protein S4X pseudogene 17 [Source |
| TLCD3A | 17 | TLC domain containing 3A [Source:HGNC Sym |
| GEMIN4 | 17 | gem nuclear organelle associated protein 4 [S |
| DBIL5P | 17 | diazepam binding inhibitor-like 5, pseudogene |
| GLOD4 | 17 | glyoxalase domain containing 4 [Source:HGN( |
| AC087392.3 | 17 | novel transcript |
| MRM3 | 17 | mitochondrial rRNA methyltransferase 3 [Sou |
| AC087392.4 | 17 | novel transcript, antisense to RNMTL1 |
| AC087392.5 | 17 | novel transcript |
| NXN | 17 | nucleoredoxin [Source:HGNC Symbol;Acc:HGI |
| AC087392.2 | 17 | novel transcript, antisense to NXN |
| AC087392.1 | 17 | novel transcript, antisense to NXN |
| AC036164.1 | 17 | chromosome 19 open reading frame 70 (C19c |
| TIMM22 | 17 | translocase of inner mitochondrial membrane |
| ABR | 17 | ABR activator of RhoGEF and GTPase [Source: |
| MIR3183 | 17 | microRNA 3183 [Source:HGNC Symbol;Acc:H( |
| Metazoa_SRP | 17 | Metazoan signal recognition particle RNA [Sou |
| AC016292.2 | 17 | mitochondrial ribosomal protein L14 (MRPL1 |
| AC016292.1 | 17 | novel transcript, antisense to ABR |
| AC144836.1 | 17 | novel transcript |
| BHLHA9 | 17 | basic helix-loop-helix family member a9 [Sour |
| TRARG1 | 17 | trafficking regulator of GLUT4 (SLC2A4) 1 [Sou |
| YWHAE | 17 | tyrosine 3-monooxygenase/tryptophan 5-mo |
| AC032044.2 | 17 | TEC |
| CRK | 17 | CRK proto-oncogene, adaptor protein [Source |
| AC032044.1 | 17 | novel transcript, antisense to CRK |
| MYO1C | 17 | myosin IC [Source:HGNC Symbol;Acc:HGNC:7! |
| INPP5K | 17 | inositol polyphosphate-5-phosphatase K [Sou |
| PITPNA-AS1 | 17 | PITPNA antisense RNA 1 [Source:HGNC Symbo |
| PITPNA | 17 | phosphatidylinositol transfer protein alpha [S |
| SLC43A2 | 17 | solute carrier family 43 member 2 [Source:HG |
| RN7SL105P | 17 | RNA, 7SL, cytoplasmic 105, pseudogene [Sour |
| SCARF1 | 17 | scavenger receptor class F member 1 [Source: |
| RILP | 17 | Rab interacting lysosomal protein [Source:HG |
| PRPF8 | 17 | pre-mRNA processing factor 8 [Source:HGNC |
| AC130343.2 | 17 | novel transcript, antisense to PRPF8 |
| TLCD2 | 17 | TLC domain containing 2 [Source:HGNC Symb |
| MIR22HG | 17 | MIR22 host gene [Source:HGNC Symbol;Acc:H |
| MIR22 | 17 | microRNA 22 [Source:HGNC Symbol;Acc:HGN |
| WDR81 | 17 | WD repeat domain 81 [Source:HGNC Symbol; |
| AC130343.1 | 17 | novel transcript, antisense to WDR81 |
| SERPINF2 | 17 | serpin family F member 2 [Source:HGNC Symb |
| SERPINF1 | 17 | serpin family F member 1 [Source:HGNC Symb |

| | | |
|---|---|---|
| SMYD4 | 17 | SET and MYND domain containing 4 [Source:H |
| RPA1 | 17 | replication protein A1 [Source:HGNC Symbol;/ |
| AC130689.1 | 17 | succinate dehydrogenase complex, subunit C, |
| RTN4RL1 | 17 | reticulon 4 receptor like 1 [Source:HGNC Symb |
| AC099684.2 | 17 | novel transcript, antisense to RTN4RL1 |
| AC099684.1 | 17 | novel transcript, antisense to RTN4RL1 |
| AC099684.3 | 17 | novel transcript, sense intronic to RTN4RL1 |
| DPH1 | 17 | diphthamide biosynthesis 1 [Source:HGNC Syr |
| AC090617.10 | 17 | novel transcript, antisense to DPH1 |
| OVCA2 | 17 | OVCA2 serine hydrolase domain containing [S |
| AC090617.3 | 17 | novel transcript, antisense to DPH1 and OVCA |
| AC090617.5 | 17 | novel transcript |
| MIR132 | 17 | microRNA 132 [Source:HGNC Symbol;Acc:HGI |
| MIR212 | 17 | microRNA 212 [Source:HGNC Symbol;Acc:HGI |
| HIC1 | 17 | HIC ZBTB transcriptional repressor 1 [Source:H |
| SMG6 | 17 | SMG6 nonsense mediated mRNA decay factor |
| AC090617.9 | 17 | novel transcript |
| AC090617.4 | 17 | novel transcript, antisense to SMG6 |
| AC090617.6 | 17 | novel transcript |
| AC090617.8 | 17 | novel transcript |
| AC090617.7 | 17 | novel transcript |
| MCUR1P1 | 17 | MCUR1 pseudogene 1 [Source:HGNC Symbol; |
| RN7SL624P | 17 | RNA, 7SL, cytoplasmic 624, pseudogene [Sour |
| AC090617.2 | 17 | novel transcript, antisense to SMG6 |
| AC090617.1 | 17 | novel transcript, sense intronic to SMG6 |
| AL450226.1 | 17 | novel transcript, antisense to SMG6 |
| SMG6-IT1 | 17 | SMG6 intronic transcript 1 [Source:HGNC Sym |
| AL450226.2 | 17 | novel transcript |
| SRR | 17 | serine racemase [Source:HGNC Symbol;Acc:H |
| HNRNPA1P16 | 17 | heterogeneous nuclear ribonucleoprotein A1 |
| TSR1 | 17 | TSR1 ribosome maturation factor [Source:HGI |
| SNORD91B | 17 | small nucleolar RNA, C/D box 91B [Source:HG |
| SNORD91A | 17 | small nucleolar RNA, C/D box 91A [Source:HG |
| SGSM2 | 17 | small G protein signaling modulator 2 [Source |
| AC006435.3 | 17 | novel transcript, antisense to SGSM2 |
| AC006435.2 | 17 | novel transcript, antisense to SGSM2 |
| MNT | 17 | MAX network transcriptional repressor [Sourc |
| AC006435.1 | 17 | novel transcript, antisense to MNT |
| METTL16 | 17 | methyltransferase like 16 [Source:HGNC Symb |
| AC006435.4 | 17 | novel transcript |
| AC006435.5 | 17 | novel transcript |
| RN7SL33P | 17 | RNA, 7SL, cytoplasmic 33, pseudogene [Sourc |
| EIF4A1P9 | 17 | eukaryotic translation initiation factor 4A1 ps |
| PAFAH1B1 | 17 | platelet activating factor acetylhydrolase 1b r |
| SAMD11P1 | 17 | sterile alpha motif domain containing 11 pseu |
| AC015799.1 | 17 | novel transcript |
| RN7SL608P | 17 | RNA, 7SL, cytoplasmic 608, pseudogene [Sour |
| AC005696.2 | 17 | novel transcript, antisense to PAFAH1B1 |
| AC005696.3 | 17 | novel transcript |
| CLUH | 17 | clustered mitochondria homolog [Source:HGN |

| | | |
|---|---|---|
| MIR6776 | 17 | microRNA 6776 [Source:HGNC Symbol;Acc:HG |
| AC005696.1 | 17 | novel transcript, antisense to CLUH |
| AC005696.4 | 17 | novel transcript |
| CCDC92B | 17 | coiled-coil domain containing 92B [Source:HG |
| MIR1253 | 17 | microRNA 1253 [Source:HGNC Symbol;Acc:HG |
| hsa-mir-1253 | 17 | hsa-mir-1253 [Source:miRBase;Acc:MI000638 |
| RAP1GAP2 | 17 | RAP1 GTPase activating protein 2 [Source:HGI |
| AC015921.1 | 17 | novel transcript, antisense to RAP1GAP2 |
| OR1D5 | 17 | olfactory receptor family 1 subfamily D memb |
| OR1D2 | 17 | olfactory receptor family 1 subfamily D memb |
| OR1E3 | 17 | olfactory receptor family 1 subfamily E memb |
| OR1G1 | 17 | olfactory receptor family 1 subfamily G memb |
| AC090282.1 | 17 | novel transcript |
| OR1P1 | 17 | olfactory receptor family 1 subfamily P memb |
| OR1A2 | 17 | olfactory receptor family 1 subfamily A memb |
| OR1A1 | 17 | olfactory receptor family 1 subfamily A memb |
| OR1D4 | 17 | olfactory receptor family 1 subfamily D memb |
| OR1D3P | 17 | olfactory receptor family 1 subfamily D memb |
| OR3A2 | 17 | olfactory receptor family 3 subfamily A memb |
| OR3A1 | 17 | olfactory receptor family 3 subfamily A memb |
| OR3A4P | 17 | olfactory receptor family 3 subfamily A memb |
| AC087498.2 | 17 | novel olfactory receptor pseudogene |
| OR1AC1P | 17 | olfactory receptor family 1 subfamily AC mem |
| AC087498.1 | 17 | olfactory receptor, family 1, subfamily R, mem |
| OR1E1 | 17 | olfactory receptor family 1 subfamily E memb |
| OR3A3 | 17 | olfactory receptor family 3 subfamily A memb |
| OR1E2 | 17 | olfactory receptor family 1 subfamily E memb |
| AC025125.1 | 17 | TEC |
| SPATA22 | 17 | spermatogenesis associated 22 [Source:HGNC |
| ASPA | 17 | aspartoacylase [Source:HGNC Symbol;Acc:HG |
| TRPV3 | 17 | transient receptor potential cation channel su |
| TRPV1 | 17 | transient receptor potential cation channel su |
| AC027796.5 | 17 | novel transcript, sense intronic to TRPV1 |
| AC027796.3 | 17 | novel readthrough SHPK-TRPV1 protein |
| AC027796.1 | 17 | novel transcript, sense intronic to readthrougl |
| SHPK | 17 | sedoheptulokinase [Source:HGNC Symbol;Acc |
| AC027796.2 | 17 | pterin-4 alpha-carbinolamine dehydratase/dir |
| CTNS | 17 | cystinosin, lysosomal cystine transporter [Sou |
| AC027796.4 | 17 | novel transcript, antisense to CTNS |
| TAX1BP3 | 17 | Tax1 binding protein 3 [Source:HGNC Symbol; |
| P2RX5-TAX1BP3 | 17 | P2RX5-TAX1BP3 readthrough (NMD candidate |
| AC132942.1 | 17 | ribosomal protein L21 (RPL21)pseudogene |
| EMC6 | 17 | ER membrane protein complex subunit 6 [Sou |
| P2RX5 | 17 | purinergic receptor P2X 5 [Source:HGNC Symb |
| ITGAE | 17 | integrin subunit alpha E [Source:HGNC Symbo |
| AC116914.2 | 17 | novel transcript, antisense to ITGAE |
| HASPIN | 17 | histone H3 associated protein kinase [Source: |
| AC116914.1 | 17 | Novel protein, antisense to ITGAE |
| NCBP3 | 17 | nuclear cap binding subunit 3 [Source:HGNC S |
| CAMKK1 | 17 | calcium/calmodulin dependent protein kinase |

| | | |
|---|---|---|
| P2RX1 | 17 | purinergic receptor P2X 1 [Source:HGNC Symb |
| ATP2A3 | 17 | ATPase sarcoplasmic/endoplasmic reticulum C |
| LINC01975 | 17 | long intergenic non-protein coding RNA 1975 |
| ZZEF1 | 17 | zinc finger ZZ-type and EF-hand domain conta |
| RNA5SP434 | 17 | RNA, 5S ribosomal pseudogene 434 [Source:H |
| CYB5D2 | 17 | cytochrome b5 domain containing 2 [Source:H |
| ANKFY1 | 17 | ankyrin repeat and FYVE domain containing 1 |
| AC087292.1 | 17 | novel transcript, antisense to ANKFY1 |
| Y_RNA | 17 | Y RNA [Source:RFAM;Acc:RF00019] |
| RYKP1 | 17 | RYK pseudogene 1 [Source:HGNC Symbol;Acc: |
| AC087742.1 | 17 | novel transcript |
| UBE2G1 | 17 | ubiquitin conjugating enzyme E2 G1 [Source:H |
| MFSD1P1 | 17 | major facilitator superfamily domain containir |
| RN7SL774P | 17 | RNA, 7SL, cytoplasmic 774, pseudogene [Sour |
| SPNS3 | 17 | sphingolipid transporter 3 (putative) [Source:H |
| AC127521.1 | 17 | novel transcript, antisense to SPNS3 |
| AC118754.1 | 17 | novel transcript |
| SPNS2 | 17 | sphingolipid transporter 2 [Source:HGNC Sym |
| MYBBP1A | 17 | MYB binding protein 1a [Source:HGNC Symbo |
| GGT6 | 17 | gamma-glutamyltransferase 6 [Source:HGNC S |
| AC118754.2 | 17 | novel transcript |
| TXNP4 | 17 | thioredoxin pseudogene 4 [Source:HGNC Sym |
| SMTNL2 | 17 | smoothelin like 2 [Source:HGNC Symbol;Acc:H |
| LINC01996 | 17 | long intergenic non-protein coding RNA 1996 |
| RNU6-955P | 17 | RNA, U6 small nuclear 955, pseudogene [Sour |
| ALOX15 | 17 | arachidonate 15-lipoxygenase [Source:HGNC S |
| PELP1 | 17 | proline, glutamate and leucine rich protein 1 [ |
| AC091153.2 | 17 | novel transcript, antisense to PELP1 |
| AC091153.3 | 17 | novel transcript |
| AC091153.1 | 17 | ribosomal protein S12 (RPS12) pseudogene |
| ARRB2 | 17 | arrestin beta 2 [Source:HGNC Symbol;Acc:HGI |
| MED11 | 17 | mediator complex subunit 11 [Source:HGNC S |
| AC091153.4 | 17 | novel transcript, antisense to MED11 |
| CXCL16 | 17 | C-X-C motif chemokine ligand 16 [Source:HGN |
| ZMYND15 | 17 | zinc finger MYND-type containing 15 [Source:I |
| TM4SF5 | 17 | transmembrane 4 L six family member 5 [Sour |
| VMO1 | 17 | vitelline membrane outer layer 1 homolog [Sc |
| AC233723.2 | 17 | TEC |
| GLTPD2 | 17 | glycolipid transfer protein domain containing |
| PSMB6 | 17 | proteasome subunit beta 6 [Source:HGNC Syn |
| AC233723.1 | 17 | novel protein |
| PLD2 | 17 | phospholipase D2 [Source:HGNC Symbol;Acc:I |
| MINK1 | 17 | misshapen like kinase 1 [Source:HGNC Symbo |
| ATP6V0CP1 | 17 | ATPase H+ transporting V0 subunit c pseudog |
| RN7SL784P | 17 | RNA, 7SL, cytoplasmic 784, pseudogene [Sour |
| CHRNE | 17 | cholinergic receptor nicotinic epsilon subunit |
| C17orf107 | 17 | chromosome 17 open reading frame 107 [Sou |
| GP1BA | 17 | glycoprotein Ib platelet subunit alpha [Source |
| SLC25A11 | 17 | solute carrier family 25 member 11 [Source:H |
| RNF167 | 17 | ring finger protein 167 [Source:HGNC Symbol; |

| | | |
|---|---|---|
| PFN1 | 17 | profilin 1 [Source:HGNC Symbol;Acc:HGNC:88 |
| ENO3 | 17 | enolase 3 [Source:HGNC Symbol;Acc:HGNC:33 |
| SPAG7 | 17 | sperm associated antigen 7 [Source:HGNC Syr |
| CAMTA2 | 17 | calmodulin binding transcription activator 2 [S |
| AC004771.4 | 17 | novel transcript, antisense to CAMTA2 |
| MIR6864 | 17 | microRNA 6864 [Source:HGNC Symbol;Acc:HG |
| MIR6865 | 17 | microRNA 6865 [Source:HGNC Symbol;Acc:HG |
| AC004771.2 | 17 | novel transcript, antisense to CAMTA2 |
| AC004771.5 | 17 | Novel transcript, antisense to CAMTA2 |
| AC004771.3 | 17 | novel transcript, antisense to INCA1 |
| INCA1 | 17 | inhibitor of CDK, cyclin A1 interacting protein |
| KIF1C | 17 | kinesin family member 1C [Source:HGNC Sym |
| AC004771.1 | 17 | novel transcript, antisense to KIF1C |
| SLC52A1 | 17 | solute carrier family 52 member 1 [Source:HG |
| AC012146.2 | 17 | zinc finger protein fiamily pseudogene |
| AC012146.3 | 17 | novel transcript, antisense to ZFP3 |
| ZFP3 | 17 | ZFP3 zinc finger protein [Source:HGNC Symbo |
| ZNF232 | 17 | zinc finger protein 232 [Source:HGNC Symbol; |
| AC012146.1 | 17 | novel transcript, antisense to ZNF232 |
| USP6 | 17 | ubiquitin specific peptidase 6 [Source:HGNC S |
| AC012146.4 | 17 | tumor necrosis factor receptor superfamily, m |
| ZNF594 | 17 | zinc finger protein 594 [Source:HGNC Symbol; |
| AC087500.1 | 17 | novel transcript, antisense to C17orf87 |
| SCIMP | 17 | SLP adaptor and CSK interacting membrane pr |
| AC087500.2 | 17 | novel transcript |
| RABEP1 | 17 | rabaptin, RAB GTPase binding effector protein |
| NUP88 | 17 | nucleoporin 88 [Source:HGNC Symbol;Acc:HG |
| AC015727.1 | 17 | novel transcript, antisense to RABEP1 |
| RPAIN | 17 | RPA interacting protein [Source:HGNC Symbol |
| AC004148.1 | 17 | novel transcript, antisense to RPAIN |
| C1QBP | 17 | complement C1q binding protein [Source:HGN |
| DHX33 | 17 | DEAH-box helicase 33 [Source:HGNC Symbol;A |
| DHX33-DT | 17 | DHX33 divergent transcript [Source:HGNC Syr |
| DERL2 | 17 | derlin 2 [Source:HGNC Symbol;Acc:HGNC:179 |
| MIS12 | 17 | MIS12 kinetochore complex component [Sour |
| NLRP1 | 17 | NLR family pyrin domain containing 1 [Source |
| AC055839.2 | 17 | uncharacterized LOC728392 [Source:NCBI ger |
| AC055839.1 | 17 | zinc finger, CCHC domain containing 9 (ZCCHC |
| AC135726.1 | 17 | novel transcript |
| AC006236.1 | 17 | glioblastoma amplified sequence (GBAS) pseu |
| AC007846.1 | 17 | novel transcript |
| AC007846.2 | 17 | novel protein |
| WSCD1 | 17 | WSC domain containing 1 [Source:HGNC Symb |
| AC104770.1 | 17 | novel transcript |
| RNU6-1264P | 17 | RNA, U6 small nuclear 1264, pseudogene [Sou |
| BTF3P14 | 17 | basic transcription factor 3 pseudogene 14 [Sc |
| AC055872.2 | 17 | novel transcript |
| AIPL1 | 17 | aryl hydrocarbon receptor interacting protein |
| AC055872.1 | 17 | vacuolar protein sorting 13 homolog B (yeast) |
| PIMREG | 17 | PICALM interacting mitotic regulator [Source: |

| | | |
|---|---|---|
| PITPNM3 | 17 | PITPNM family member 3 [Source:HGNC Symb |
| Metazoa_SRP | 17 | Metazoan signal recognition particle RNA [Sou |
| KIAA0753 | 17 | KIAA0753 [Source:HGNC Symbol;Acc:HGNC:29 |
| RNA5SP435 | 17 | RNA, 5S ribosomal pseudogene 435 [Source:H |
| AC004706.3 | 17 | novel protein |
| TXNDC17 | 17 | thioredoxin domain containing 17 [Source:HG |
| MED31 | 17 | mediator complex subunit 31 [Source:HGNC S |
| C17orf100 | 17 | chromosome 17 open reading frame 100 [Sou |
| AC004706.1 | 17 | novel transcript |
| MIR4520-1 | 17 | microRNA 4520-1 [Source:HGNC Symbol;Acc:H |
| MIR4520-2 | 17 | microRNA 4520-2 [Source:HGNC Symbol;Acc:H |
| ALOX15P1 | 17 | arachidonate 15-lipoxygenase pseudogene 1 [ |
| SLC13A5 | 17 | solute carrier family 13 member 5 [Source:HG |
| AC004706.2 | 17 | major histocompatibility complex, class I, F (H |
| RPL23AP73 | 17 | ribosomal protein L23a pseudogene 73 [Sourc |
| XAF1 | 17 | XIAP associated factor 1 [Source:HGNC Symbo |
| PRAL | 17 | p53 regulation associated lncRNA [Source:HGI |
| FBXO39 | 17 | F-box protein 39 [Source:HGNC Symbol;Acc:H |
| TEKT1 | 17 | tektin 1 [Source:HGNC Symbol;Acc:HGNC:155 |
| ALOX12P2 | 17 | arachidonate 12-lipoxygenase pseudogene 2 [ |
| ALOX12-AS1 | 17 | ALOX12 antisense RNA 1 [Source:HGNC Symb |
| AC027763.1 | 17 | Spi-C transcription factor (Spi-1/PU.1 related) |
| AC027763.2 | 17 | novel transcript, antisense to ALOX12P2 |
| AC040977.2 | 17 | novel transcript |
| AC040977.1 | 17 | novel transcript |
| ALOX12 | 17 | arachidonate 12-lipoxygenase, 12S type [Sour |
| RNASEK | 17 | ribonuclease K [Source:HGNC Symbol;Acc:HGI |
| RNASEK-C17orf49 | 17 | RNASEK-C17orf49 readthrough [Source:HGNC |
| C17orf49 | 17 | chromosome 17 open reading frame 49 [Sour |
| MIR497HG | 17 | mir-497-195 cluster host gene [Source:HGNC |
| MIR195 | 17 | microRNA 195 [Source:HGNC Symbol;Acc:HGI |
| MIR497 | 17 | microRNA 497 [Source:HGNC Symbol;Acc:HGI |
| BCL6B | 17 | BCL6B transcription repressor [Source:HGNC S |
| SLC16A13 | 17 | solute carrier family 16 member 13 [Source:H |
| SLC16A11 | 17 | solute carrier family 16 member 11 [Source:H |
| AC120057.1 | 17 | v-crk sarcoma virus CT10 oncogene homolog ( |
| CLEC10A | 17 | C-type lectin domain containing 10A [Source:H |
| ASGR2 | 17 | asialoglycoprotein receptor 2 [Source:HGNC S |
| RPL7AP64 | 17 | ribosomal protein L7a pseudogene 64 [Source |
| ASGR1 | 17 | asialoglycoprotein receptor 1 [Source:HGNC S |
| DLG4 | 17 | discs large MAGUK scaffold protein 4 [Source: |
| AC120057.4 | 17 | TEC |
| ACADVL | 17 | acyl-CoA dehydrogenase very long chain [Sour |
| MIR324 | 17 | microRNA 324 [Source:HGNC Symbol;Acc:HGI |
| DVL2 | 17 | dishevelled segment polarity protein 2 [Sourc |
| PHF23 | 17 | PHD finger protein 23 [Source:HGNC Symbol;A |
| GABARAP | 17 | GABA type A receptor-associated protein [Sou |
| AC120057.2 | 17 | novel protein |
| AC120057.3 | 17 | TEC |
| CTDNEP1 | 17 | CTD nuclear envelope phosphatase 1 [Source: |

| | | |
|---|---|---|
| AC003688.1 | 17 | novel protein |
| ELP5 | 17 | elongator acetyltransferase complex subunit [illegible] |
| CLDN7 | 17 | claudin 7 [Source:HGNC Symbol;Acc:HGNC:20 |
| Y_RNA | 17 | Y RNA [Source:RFAM;Acc:RF00019] |
| SLC2A4 | 17 | solute carrier family 2 member 4 [Source:HGN |
| AC003688.2 | 17 | novel transcript, antisense to SLC2A4 |
| YBX2 | 17 | Y-box binding protein 2 [Source:HGNC Symbol |
| EIF5A | 17 | eukaryotic translation initiation factor 5A [Sou |
| GPS2 | 17 | G protein pathway suppressor 2 [Source:HGN |
| AC026954.2 | 17 | novel protein |
| NEURL4 | 17 | neuralized E3 ubiquitin protein ligase 4 [Sourc |
| AC026954.1 | 17 | novel protein phosphatase inhibitor 2 (IPP-2) |
| ACAP1 | 17 | ArfGAP with coiled-coil, ankyrin repeat and PH |
| KCTD11 | 17 | potassium channel tetramerization domain cc |
| AC026954.3 | 17 | novel transcript, antisense to KCTD11 |
| TMEM95 | 17 | transmembrane protein 95 [Source:HGNC Syn |
| TNK1 | 17 | tyrosine kinase non receptor 1 [Source:HGNC |
| PLSCR3 | 17 | phospholipid scramblase 3 [Source:HGNC Sym |
| TMEM256-PLSCR3 | 17 | TMEM256-PLSCR3 readthrough (NMD candida |
| TMEM256 | 17 | transmembrane protein 256 [Source:HGNC Sy |
| NLGN2 | 17 | neuroligin 2 [Source:HGNC Symbol;Acc:HGNC: |
| AC113189.2 | 17 | novel transcript |
| SPEM1 | 17 | spermatid maturation 1 [Source:HGNC Symbo |
| SPEM2 | 17 | SPEM family member 2 [Source:HGNC Symbo |
| SPEM3 | 17 | SPEM family member 3 [Source:HGNC Symbo |
| AC113189.9 | 17 | novel protein |
| TMEM102 | 17 | transmembrane protein 102 [Source:HGNC Sy |
| AC113189.1 | 17 | novel transcript, antisense to TMEM102 |
| FGF11 | 17 | fibroblast growth factor 11 [Source:HGNC Sym |
| AC113189.3 | 17 | novel transcript, antisense to FGF11 |
| AC113189.4 | 17 | novel transcript |
| CHRNB1 | 17 | cholinergic receptor nicotinic beta 1 subunit [ |
| ZBTB4 | 17 | zinc finger and BTB domain containing 4 [Sour |
| SLC35G6 | 17 | solute carrier family 35 member G6 [Source:H |
| POLR2A | 17 | RNA polymerase II subunit A [Source:HGNC Sy |
| Y_RNA | 17 | Y RNA [Source:RFAM;Acc:RF00019] |
| TNFSF12 | 17 | TNF superfamily member 12 [Source:HGNC Sy |
| TNFSF12-TNFSF13 | 17 | TNFSF12-TNFSF13 readthrough [Source:HGNC |
| AC016876.3 | 17 | novel transcript, antisense to TNFSF12 |
| TNFSF13 | 17 | TNF superfamily member 13 [Source:HGNC Sy |
| SENP3 | 17 | SUMO specific peptidase 3 [Source:HGNC Sym |
| SENP3-EIF4A1 | 17 | SENP3-EIF4A1 readthrough (NMD candidate) |
| EIF4A1 | 17 | eukaryotic translation initiation factor 4A1 [Sc |
| AC016876.2 | 17 | novel transcript |
| SNORA48 | 17 | small nucleolar RNA, H/ACA box 48 [Source:H |
| SNORD10 | 17 | small nucleolar RNA, C/D box 10 [Source:HGN |
| SNORA67 | 17 | small nucleolar RNA, H/ACA box 67 [Source:H |
| CD68 | 17 | CD68 molecule [Source:HGNC Symbol;Acc:HG |
| AC016876.1 | 17 | novel transcript, antisense to MPDU1 and CD6 |
| MPDU1 | 17 | mannose-P-dolichol utilization defect 1 [Sourc |

| | | |
|---|---|---|
| SOX15 | 17 | SRY-box transcription factor 15 [Source:HGNC |
| FXR2 | 17 | FMR1 autosomal homolog 2 [Source:HGNC Sy |
| SHBG | 17 | sex hormone binding globulin [Source:HGNC S |
| SAT2 | 17 | spermidine/spermine N1-acetyltransferase fa |
| ATP1B2 | 17 | ATPase Na+/K+ transporting subunit beta 2 [S |
| TP53 | 17 | tumor protein p53 [Source:HGNC Symbol;Acc: |
| WRAP53 | 17 | WD repeat containing antisense to TP53 [Sour |
| EFNB3 | 17 | ephrin B3 [Source:HGNC Symbol;Acc:HGNC:32 |
| DNAH2 | 17 | dynein axonemal heavy chain 2 [Source:HGNC |
| RPL29P2 | 17 | ribosomal protein L29 pseudogene 2 [Source: |
| KDM6B | 17 | lysine demethylase 6B [Source:HGNC Symbol; |
| TMEM88 | 17 | transmembrane protein 88 [Source:HGNC Syn |
| NAA38 | 17 | N(alpha)-acetyltransferase 38, NatC auxiliary s |
| CYB5D1 | 17 | cytochrome b5 domain containing 1 [Source:H |
| AC104581.4 | 17 | TEC |
| CHD3 | 17 | chromodomain helicase DNA binding protein |
| SCARNA21 | 17 | small Cajal body-specific RNA 21 [Source:HGN |
| RNF227 | 17 | ring finger protein 227 [Source:HGNC Symbol; |
| KCNAB3 | 17 | potassium voltage-gated channel subfamily A |
| AC104581.2 | 17 | novel transcript |
| TRAPPC1 | 17 | trafficking protein particle complex 1 [Source: |
| CNTROB | 17 | centrobin, centriole duplication and spindle as |
| AC104581.1 | 17 | ribosomal protein L21 (RPL21) pseudogene |
| GUCY2D | 17 | guanylate cyclase 2D, retinal [Source:HGNC Sy |
| AC104581.3 | 17 | TEC |
| ALOX15B | 17 | arachidonate 15-lipoxygenase type B [Source: |
| AC129492.2 | 17 | novel transcript |
| AC129492.6 | 17 | arachidonate 15-lipoxygenase, type B (ALOX15 |
| ALOX12B | 17 | arachidonate 12-lipoxygenase, 12R type [Sour |
| AC129492.1 | 17 | novel transcript, antisense to ALOX12B |
| MIR4314 | 17 | microRNA 4314 [Source:HGNC Symbol;Acc:HG |
| ALOXE3 | 17 | arachidonate lipoxygenase 3 [Source:HGNC Sy |
| HES7 | 17 | hes family bHLH transcription factor 7 [Source |
| PER1 | 17 | period circadian regulator 1 [Source:HGNC Syn |
| MIR6883 | 17 | microRNA 6883 [Source:HGNC Symbol;Acc:HG |
| AC129492.3 | 17 | novel protein |
| VAMP2 | 17 | vesicle associated membrane protein 2 [Sourc |
| TMEM107 | 17 | transmembrane protein 107 [Source:HGNC Sy |
| SNORD118 | 17 | small nucleolar RNA, C/D box 118 [Source:HGI |
| AC129492.4 | 17 | novel transcript |
| MIR4521 | 17 | microRNA 4521 [Source:HGNC Symbol;Acc:HG |
| BORCS6 | 17 | BLOC-1 related complex subunit 6 [Source:HG |
| AC129492.7 | 17 | TEC |
| AC129492.5 | 17 | ribosomal protein L11 (RPL11) pseudogene |
| AURKB | 17 | aurora kinase B [Source:HGNC Symbol;Acc:HG |
| LINC00324 | 17 | long intergenic non-protein coding RNA 324 [S |
| CTC1 | 17 | CST telomere replication complex component |
| PFAS | 17 | phosphoribosylformylglycinamidine synthase |
| AC135178.6 | 17 | novel transcript |
| SLC25A35 | 17 | solute carrier family 25 member 35 [Source:H |

| | | |
|---|---|---|
| RANGRF | 17 | RAN guanine nucleotide release factor [Sourc |
| AC135178.7 | 17 | ribosomal protein S24 (RPS24) pseudogene |
| ARHGEF15 | 17 | Rho guanine nucleotide exchange factor 15 [S |
| AC135178.2 | 17 | novel transcript |
| AC135178.1 | 17 | NULL |
| AC135178.5 | 17 | novel transcript |
| ODF4 | 17 | outer dense fiber of sperm tails 4 [Source:HG |
| KRBA2 | 17 | KRAB-A domain containing 2 [Source:HGNC Sy |
| AC135178.4 | 17 | novel transcript, antisense to KRBA2 and RPL2 |
| AC135178.3 | 17 | novel protein |
| RPL26 | 17 | ribosomal protein L26 [Source:HGNC Symbol; |
| RNF222 | 17 | ring finger protein 222 [Source:HGNC Symbol; |
| NDEL1 | 17 | nudE neurodevelopment protein 1 like 1 [Sou |
| MYH10 | 17 | myosin heavy chain 10 [Source:HGNC Symbol |
| RNU7-43P | 17 | RNA, U7 small nuclear 43 pseudogene [Source |
| PPIAP52 | 17 | peptidylprolyl isomerase A pseudogene 52 [Sc |
| AC025518.1 | 17 | ribosomal protein S26 (RPS26) pseudogene |
| CCDC42 | 17 | coiled-coil domain containing 42 [Source:HGN |
| SPDYE4 | 17 | speedy/RINGO cell cycle regulator family men |
| MFSD6L | 17 | major facilitator superfamily domain containi |
| PIK3R6 | 17 | phosphoinositide-3-kinase regulatory subunit |
| PIK3R5 | 17 | phosphoinositide-3-kinase regulatory subunit |
| AC002091.2 | 17 | novel transcript |
| AC002091.1 | 17 | novel transcript |
| NTN1 | 17 | netrin 1 [Source:HGNC Symbol;Acc:HGNC:802 |
| AC005695.1 | 17 | novel transcript, antisense to NTN1 |
| AC005695.3 | 17 | novel transcript |
| STX8 | 17 | syntaxin 8 [Source:HGNC Symbol;Acc:HGNC:1 |
| AC005695.2 | 17 | novel transcript, antisense to STX8 |
| AC087501.1 | 17 | novel transcript, antisense to STX8 |
| AC087501.2 | 17 | novel transcript, antisense to STX8 |
| AC087501.3 | 17 | novel transcript, antisense to STX8 |
| AC087501.4 | 17 | novel transcript, sense intronic to STX8 |
| CFAP52 | 17 | cilia and flagella associated protein 52 [Source |
| RPL19P18 | 17 | ribosomal protein L19 pseudogene 18 [Source |
| AC118755.2 | 17 | novel transcript, antisense to WDR16 and USI |
| USP43 | 17 | ubiquitin specific peptidase 43 [Source:HGNC |
| AC118755.1 | 17 | novel transcript, antisense to USP43 |
| AC027045.2 | 17 | novel transcript |
| DHRS7C | 17 | dehydrogenase/reductase 7C [Source:HGNC S |
| AC027045.1 | 17 | novel transcript |
| GSG1L2 | 17 | GSG1 like 2 [Source:HGNC Symbol;Acc:HGNC: |
| AC027045.3 | 17 | novel transcript, antisense to novel protein |
| GLP2R | 17 | glucagon like peptide 2 receptor [Source:HGN |
| NPM1P45 | 17 | nucleophosmin 1 pseudogene 45 [Source:HGI |
| RCVRN | 17 | recoverin [Source:HGNC Symbol;Acc:HGNC:9 |
| GAS7 | 17 | growth arrest specific 7 [Source:HGNC Symbo |
| RPS27AP1 | 17 | ribosomal protein S27a pseudogene 1 [Source |
| AC005291.2 | 17 | novel transcript, antisense to MYH13 |
| MYH13 | 17 | myosin heavy chain 13 [Source:HGNC Symbol |

| | | |
|---|---|---|
| AC005291.1 | 17 | novel transcript, antisense to MYH13 |
| AC005323.2 | 17 | novel transcript |
| MYHAS | 17 | myosin heavy chain gene cluster antisense RN |
| MYH8 | 17 | myosin heavy chain 8 [Source:HGNC Symbol;A |
| MYH4 | 17 | myosin heavy chain 4 [Source:HGNC Symbol;A |
| MYH1 | 17 | myosin heavy chain 1 [Source:HGNC Symbol;A |
| MYH2 | 17 | myosin heavy chain 2 [Source:HGNC Symbol;A |
| AC005323.1 | 17 | novel transcript, antisense to MYH2, MYH8, M |
| MYH3 | 17 | myosin heavy chain 3 [Source:HGNC Symbol;A |
| AC002347.1 | 17 | novel transcript |
| SCO1 | 17 | SCO cytochrome c oxidase assembly protein 1 |
| AC002347.2 | 17 | novel transcript, antisense to SCO1 |
| ADPRM | 17 | ADP-ribose/CDP-alcohol diphosphatase, mang |
| TMEM220 | 17 | transmembrane protein 220 [Source:HGNC Sy |
| MAGOH2P | 17 | mago homolog 2, pseudogene [Source:HGNC |
| TMEM220-AS1 | 17 | TMEM220 antisense RNA 1 [Source:HGNC Syn |
| AC015908.7 | 17 | novel transcript |
| AC015908.3 | 17 | novel transcript |
| AC015908.1 | 17 | ribosomal protein S27a (RPS27A) pseudogene |
| AC015908.4 | 17 | novel transcript |
| AC015908.6 | 17 | novel transcript |
| TMEM238L | 17 | transmembrane protein 238 like [Source:HGN |
| AC015908.5 | 17 | novel transcript |
| PIRT | 17 | phosphoinositide interacting regulator of tran |
| AC015908.2 | 17 | novel transcript |
| RNU6-1065P | 17 | RNA, U6 small nuclear 1065, pseudogene [Sou |
| RPL15P21 | 17 | ribosomal protein L15 pseudogene 21 [Source |
| RN7SL601P | 17 | RNA, 7SL, cytoplasmic 601, pseudogene [Sour |
| AC005548.1 | 17 | novel transcript, antisense to SHISA6 |
| SHISA6 | 17 | shisa family member 6 [Source:HGNC Symbol; |
| AC005725.1 | 17 | novel transcript |
| DNAH9 | 17 | dynein axonemal heavy chain 9 [Source:HGNC |
| AC005209.1 | 17 | novel transcript, antisense to DNAH9 |
| AC005410.2 | 17 | novel transcript, antisense to DNAH9 |
| ZNF18 | 17 | zinc finger protein 18 [Source:HGNC Symbol;A |
| AC005410.1 | 17 | ribosomal protein L21 pseudogene 122 |
| MAP2K4 | 17 | mitogen-activated protein kinase kinase 4 [So |
| RNU11-2P | 17 | RNA, U11 small nuclear 2, pseudogene [Sourc |
| MIR744 | 17 | microRNA 744 [Source:HGNC Symbol;Acc:HGN |
| AC005244.1 | 17 | dynein, light chain, Tctex-type 1 (DYNLT1) pse |
| AC005244.2 | 17 | novel transcript |
| AC084167.1 | 17 | novel transcript |
| LINC00670 | 17 | long intergenic non-protein coding RNA 670 [S |
| AC068442.1 | 17 | novel transcript |
| MYOCD | 17 | myocardin [Source:HGNC Symbol;Acc:HGNC:1 |
| AC005358.1 | 17 | novel transcript, antisense to MYOCD |
| AC005358.2 | 17 | novel transcript, antisense to MYOCD |
| ARHGAP44 | 17 | Rho GTPase activating protein 44 [Source:HGN |
| MIR1269B | 17 | microRNA 1269b [Source:HGNC Symbol;Acc:H |
| AC005274.1 | 17 | TEC |

| | | |
|---|---|---|
| AC005277.2 | 17 | novel transcript, antisense to ARHGAP44 |
| AC005277.1 | 17 | novel transcript, antisense to ARHGAP44 |
| ELAC2 | 17 | elaC ribonuclease Z 2 [Source:HGNC Symbol;A |
| AC005303.1 | 17 | novel transcript |
| LINC02093 | 17 | long intergenic non-protein coding RNA 2093 |
| HS3ST3A1 | 17 | heparan sulfate-glucosamine 3-sulfotransfera: |
| MIR548H3 | 17 | microRNA 548h-3 [Source:HGNC Symbol;Acc:I |
| COX10-AS1 | 17 | COX10 antisense RNA 1 [Source:HGNC Symbo |
| AC005304.2 | 17 | novel transcript |
| AC005304.1 | 17 | novel transcript |
| AC005304.3 | 17 | novel transcript |
| CDRT15P1 | 17 | CMT1A duplicated region transcript 15 pseud |
| COX10 | 17 | cytochrome c oxidase assembly factor heme / |
| SNORA74 | 17 | Small nucleolar RNA SNORA74 [Source:RFAM; |
| CDRT15 | 17 | CMT1A duplicated region transcript 15 [Sourc |
| AC005224.1 | 17 | novel transcript |
| HS3ST3B1 | 17 | heparan sulfate-glucosamine 3-sulfotransfera: |
| AC005224.3 | 17 | novel transcript |
| AC005224.2 | 17 | novel transcript |
| AC022816.1 | 17 | novel transcript |
| RPS18P12 | 17 | ribosomal protein S18 pseudogene 12 [Source |
| AC013248.1 | 17 | novel transcript |
| AC005863.1 | 17 | novel transcript |
| LINC02096 | 17 | long intergenic non-protein coding RNA 2096 |
| RPL23AP76 | 17 | ribosomal protein L23a pseudogene 76 [Sourc |
| AC005772.1 | 17 | uncharacterized LOC101928475 [Source:NCBI |
| CDRT7 | 17 | CMT1A duplicated region transcript 7 [Source |
| AC005772.2 | 17 | novel transcript |
| CDRT8 | 17 | CMT1A duplicated region transcript 8 [Source |
| PMP22 | 17 | peripheral myelin protein 22 [Source:HGNC Sy |
| MIR4731 | 17 | microRNA 4731 [Source:HGNC Symbol;Acc:HG |
| AC005703.6 | 17 | novel transcript |
| AC005703.3 | 17 | novel transcript |
| AC005703.7 | 17 | novel transcript, antisense to TEKT3 |
| AC005703.1 | 17 | novel transcript |
| AC005703.2 | 17 | novel transcript |
| TEKT3 | 17 | tektin 3 [Source:HGNC Symbol;Acc:HGNC:142 |
| RNU6-799P | 17 | RNA, U6 small nuclear 799, pseudogene [Sour |
| AC005703.5 | 17 | TEC |
| AC005703.4 | 17 | novel transcript |
| RN7SL792P | 17 | RNA, 7SL, cytoplasmic 792, pseudogene [Sour |
| CDRT4 | 17 | CMT1A duplicated region transcript 4 [Source |
| TVP23C-CDRT4 | 17 | TVP23C-CDRT4 readthrough [Source:HGNC Sy |
| RPL9P2 | 17 | ribosomal protein L9 pseudogene 2 [Source:H |
| TVP23C | 17 | trans-golgi network vesicle protein 23 homolc |
| PPIAP53 | 17 | peptidylprolyl isomerase A pseudogene 53 [Sc |
| AC005838.2 | 17 | novel transcript |
| CDRT1 | 17 | CMT1A duplicated region transcript 1 [Source |
| AC005324.3 | 17 | novel tripartite motif-containing 16 (TRIM16) |
| SNORA74 | 17 | Small nucleolar RNA SNORA74 [Source:RFAM; |

| | | |
|---|---|---|
| AC005838.1 | 17 | pseudogene fragment similar to COX10 homo |
| TRIM16 | 17 | tripartite motif containing 16 [Source:HGNC S |
| AC005324.5 | 17 | novel transcript |
| ZNF29P | 17 | zinc finger protein 29, pseudogene [Source:HG |
| ZNF286A | 17 | zinc finger protein 286A [Source:HGNC Symbo |
| AC005324.4 | 17 | novel transcript |
| UBE2SP1 | 17 | ubiquitin conjugating enzyme E2 S pseudogen |
| TBC1D26 | 17 | TBC1 domain family member 26 [Source:HGN |
| AC005324.1 | 17 | novel transcript |
| AC005324.6 | 17 | NIMA (never in mitosis gene a)-related kinase |
| AC005324.2 | 17 | novel transcript |
| CDRT15P2 | 17 | CMT1A duplicated region transcript 15 pseud |
| ZSWIM5P1 | 17 | zinc finger SWIM-type containing 5 pseudoger |
| RNA5SP436 | 17 | RNA, 5S ribosomal pseudogene 436 [Source:H |
| IL6STP1 | 17 | interleukin 6 signal transducer pseudogene 1 [ |
| MEIS3P1 | 17 | Meis homeobox 3 pseudogene 1 [Source:HGN |
| AC015922.2 | 17 | novel transcript |
| AC015922.3 | 17 | novel transcript |
| AC015922.1 | 17 | membrane targeting (tandem) C2 domain con |
| LINC02087 | 17 | long intergenic non-protein coding RNA 2087 |
| SPECC1P1 | 17 | SPECC1 pseudogene 1 [Source:HGNC Symbol; |
| ADORA2B | 17 | adenosine A2b receptor [Source:HGNC Symbo |
| ZSWIM7 | 17 | zinc finger SWIM-type containing 7 [Source:HG |
| TTC19 | 17 | tetratricopeptide repeat domain 19 [Source:H |
| AC002553.2 | 17 | novel transcript, antisense to TTC19 |
| NCOR1 | 17 | nuclear receptor corepressor 1 [Source:HGNC |
| AC002553.1 | 17 | novel transcript, antisense to NCOR1 |
| RPLP1P11 | 17 | ribosomal protein lateral stalk subunit P1 pseu |
| RPL22P21 | 17 | ribosomal protein L22 pseudogene 21 [Source |
| RNU6-314P | 17 | RNA, U6 small nuclear 314, pseudogene [Sour |
| RNU6-862P | 17 | RNA, U6 small nuclear 862, pseudogene [Sour |
| RN7SL442P | 17 | RNA, 7SL, cytoplasmic 442, pseudogene [Sour |
| PIGL | 17 | phosphatidylinositol glycan anchor biosynthes |
| MIR1288 | 17 | microRNA 1288 [Source:HGNC Symbol;Acc:HG |
| CENPV | 17 | centromere protein V [Source:HGNC Symbol;A |
| UBB | 17 | ubiquitin B [Source:HGNC Symbol;Acc:HGNC:1 |
| AC093484.3 | 17 | novel transcript |
| FTLP12 | 17 | ferritin light chain pseudogene 12 [Source:HG |
| AC093484.2 | 17 | novel transcript |
| TRPV2 | 17 | transient receptor potential cation channel su |
| AC093484.1 | 17 | novel transcript |
| SNHG29 | 17 | small nucleolar RNA host gene 29 [Source:HGI |
| SNORD49B | 17 | small nucleolar RNA, C/D box 49B [Source:HG |
| SNORD49A | 17 | small nucleolar RNA, C/D box 49A [Source:HG |
| AC093484.4 | 17 | novel transcript |
| SNORD65 | 17 | small nucleolar RNA, C/D box 65 [Source:HGN |
| LRRC75A | 17 | leucine rich repeat containing 75A [Source:HG |
| AC127540.2 | 17 | novel transcript |
| ZNF287 | 17 | zinc finger protein 287 [Source:HGNC Symbol; |
| AC127540.1 | 17 | novel transcript |

| | | |
|---|---|---|
| AC005822.1 | 17 | splicing factor, arginine/serine-rich 6 (SFRS6) p |
| ZNF624 | 17 | zinc finger protein 624 [Source:HGNC Symbol; |
| AC098850.2 | 17 | novel transcript |
| NEK4P2 | 17 | NIMA-related kinase 4 pseudogene 2 [Source: |
| RNASEH1P2 | 17 | ribonuclease H1 pseudogene 2 [Source:HGNC |
| CCDC144A | 17 | coiled-coil domain containing 144A [Source:H |
| AC098850.3 | 17 | novel transcript |
| RNU6-405P | 17 | RNA, U6 small nuclear 405, pseudogene [Sour |
| RN7SL620P | 17 | RNA, 7SL, cytoplasmic 620, pseudogene [Sour |
| UPF3AP1 | 17 | UPF3A pseudogene 1 [Source:HGNC Symbol;A |
| USP32P1 | 17 | ubiquitin specific peptidase 32 pseudogene 1 |
| SRP68P1 | 17 | signal recognition particle 68 pseudogene 1 [S |
| AC098850.4 | 17 | novel transcript |
| AC098850.1 | 17 | novel transcript |
| FAM106C | 17 | family with sequence similarity 106 member C |
| AC022596.1 | 17 | novel transcript |
| NOS2P4 | 17 | nitric oxide synthase 2 pseudogene 4 [Source: |
| KRT16P6 | 17 | keratin 16 pseudogene 6 [Source:HGNC Symb |
| KRT16P2 | 17 | keratin 16 pseudogene 2 [Source:HGNC Symb |
| KRT17P1 | 17 | keratin 17 pseudogene 1 [Source:HGNC Symb |
| KRT17P4 | 17 | keratin 17 pseudogene 4 [Source:HGNC Symb |
| COTL1P1 | 17 | COTL1 pseudogene 1 [Source:HGNC Symbol;A |
| AC022596.2 | 17 | novel transcript |
| AL353996.1 | 17 | novel pseudogene |
| TBC1D27P | 17 | TBC1 domain family member 27, pseudogene |
| TNFRSF13B | 17 | TNF receptor superfamily member 13B [Sourc |
| AC104024.1 | 17 | novel transcript |
| LINC02090 | 17 | long intergenic non-protein coding RNA 2090 |
| AC104024.2 | 17 | novel transcript |
| AC104024.3 | 17 | novel transcript |
| AC104024.4 | 17 | novel transcript |
| MPRIP | 17 | myosin phosphatase Rho interacting protein [ |
| MPRIP-AS1 | 17 | MPRIP antisense RNA 1 [Source:HGNC Symbo |
| AC055811.1 | 17 | novel transcript |
| RNU6-767P | 17 | RNA, U6 small nuclear 767, pseudogene [Sour |
| RN7SL775P | 17 | RNA, 7SL, cytoplasmic 775, pseudogene [Sour |
| AC055811.4 | 17 | novel transcript |
| PLD6 | 17 | phospholipase D family member 6 [Source:HG |
| AC055811.2 | 17 | novel protein |
| FLCN | 17 | folliculin [Source:HGNC Symbol;Acc:HGNC:27 |
| AC055811.3 | 17 | novel transcript |
| ACTG1P24 | 17 | actin gamma 1 pseudogene 24 [Source:HGNC |
| COPS3 | 17 | COP9 signalosome subunit 3 [Source:HGNC Sy |
| NT5M | 17 | 5',3'-nucleotidase, mitochondrial [Source:HGN |
| RPL13P12 | 17 | ribosomal protein L13 pseudogene 12 [Source |
| AC073621.1 | 17 | ligand of numb-protein X 1, E3 ubiquitin prote |
| TSEN15P1 | 17 | tRNA splicing endonuclease subunit 15 pseud |
| Y_RNA | 17 | Y RNA [Source:RFAM;Acc:RF00019] |
| MED9 | 17 | mediator complex subunit 9 [Source:HGNC Sy |
| RASD1 | 17 | ras related dexamethasone induced 1 [Source |

| | | |
|---|---|---|
| AC020558.6 | 17 | novel transcript, antisense to PEMT |
| PEMT | 17 | phosphatidylethanolamine N-methyltransfera |
| AC020558.2 | 17 | novel transcript |
| RNU6-468P | 17 | RNA, U6 small nuclear 468, pseudogene [Sour |
| AC020558.1 | 17 | novel transcript |
| AC020558.5 | 17 | novel transcript |
| SMCR2 | 17 | Smith-Magenis syndrome chromosome regior |
| AC020558.3 | 17 | NULL |
| AC020558.4 | 17 | NULL |
| RAI1 | 17 | retinoic acid induced 1 [Source:HGNC Symbol; |
| RAI1-AS1 | 17 | RAI1 antisense RNA 1 [Source:HGNC Symbol;A |
| SMCR5 | 17 | Smith-Magenis syndrome chromosome regior |
| AC122129.2 | 17 | novel transcript |
| SREBF1 | 17 | sterol regulatory element binding transcriptio |
| MIR6777 | 17 | microRNA 6777 [Source:HGNC Symbol;Acc:HG |
| MIR33B | 17 | microRNA 33b [Source:HGNC Symbol;Acc:HGI |
| TOM1L2 | 17 | target of myb1 like 2 membrane trafficking pr |
| AC122129.1 | 17 | novel transcript, antisense to TOM1L2 |
| DRC3 | 17 | dynein regulatory complex subunit 3 [Source:I |
| ATPAF2 | 17 | ATP synthase mitochondrial F1 complex assen |
| AC087163.1 | 17 | brain protein I3 (BRI3) pseudogene |
| AC087163.2 | 17 | ribosomal protein S2 (RPS2) pseudogene |
| AC087163.3 | 17 | novel transcript |
| GID4 | 17 | GID complex subunit 4 homolog [Source:HGN |
| DRG2 | 17 | developmentally regulated GTP binding prote |
| AC087164.2 | 17 | TEC |
| MYO15A | 17 | myosin XVA [Source:HGNC Symbol;Acc:HGNC: |
| AC087164.1 | 17 | novel transcript |
| ALKBH5 | 17 | alkB homolog 5, RNA demethylase [Source:HG |
| LLGL1 | 17 | LLGL scribble cell polarity complex componen |
| FLII | 17 | FLII actin remodeling protein [Source:HGNC Sy |
| MIEF2 | 17 | mitochondrial elongation factor 2 [Source:HG |
| AC127537.1 | 17 | novel transcript, antisense to TOP3A |
| TOP3A | 17 | DNA topoisomerase III alpha [Source:HGNC Sy |
| RPL7AP65 | 17 | ribosomal protein L7a pseudogene 65 [Source |
| RPL21P121 | 17 | ribosomal protein L21 pseudogene 121 [Sourc |
| SMCR8 | 17 | SMCR8-C9orf72 complex subunit [Source:HGN |
| SHMT1 | 17 | serine hydroxymethyltransferase 1 [Source:H( |
| MIR6778 | 17 | microRNA 6778 [Source:HGNC Symbol;Acc:HG |
| EVPLL | 17 | envoplakin like [Source:HGNC Symbol;Acc:HG |
| AL353997.2 | 17 | novel transcript |
| AL353997.3 | 17 | CMT1A duplicated region transcript 15-like 2 ( |
| AL353997.5 | 17 | tumor necrosis factor receptor superfamily, m |
| TBC1D3P4 | 17 | TBC1 domain family member 3 pseudogene 4 |
| LINC02076 | 17 | long intergenic non-protein coding RNA 2076 |
| KRT17P5 | 17 | keratin 17 pseudogene 5 [Source:HGNC Symb |
| YWHAEP2 | 17 | tyrosine 3-monooxygenase/tryptophan 5-mor |
| KRT17P2 | 17 | keratin 17 pseudogene 2 [Source:HGNC Symb |
| KRT16P1 | 17 | keratin 16 pseudogene 1 [Source:HGNC Symb |
| KRT16P4 | 17 | keratin 16 pseudogene 4 [Source:HGNC Symb |

| | | |
|---|---|---|
| AL353997.1 | 17 | tyrosine 3-monooxygenase/tryptophan 5-mor |
| AL353997.4 | 17 | tyrosine 3-monooxygenase/tryptophan 5-mor |
| TNPO1P2 | 17 | transportin 1 pseudogene 2 [Source:HGNC Syr |
| LGALS9C | 17 | galectin 9C [Source:HGNC Symbol;Acc:HGNC:3 |
| NOS2P2 | 17 | nitric oxide synthase 2 pseudogene 2 [Source: |
| FAM106A | 17 | family with sequence similarity 106 member A |
| USP32P2 | 17 | ubiquitin specific peptidase 32 pseudogene 2 |
| SRP68P2 | 17 | signal recognition particle 68 pseudogene 2 [S |
| CCDC144B | 17 | coiled-coil domain containing 144B (pseudoge |
| AC107983.1 | 17 | ribosomal protein S28 (RPS28) pseudogene |
| Metazoa_SRP | 17 | Metazoan signal recognition particle RNA [Sou |
| TBC1D28 | 17 | TBC1 domain family member 28 [Source:HGN |
| AC026271.2 | 17 | novel pseudogene |
| AC026271.4 | 17 | novel transcript |
| AC026271.1 | 17 | poly(A) binding protein interacting protein 1 ( |
| ZNF286B | 17 | zinc finger protein 286B (pseudogene) [Source |
| FOXO3B | 17 | forkhead box O3B [Source:HGNC Symbol;Acc: |
| UBE2SP2 | 17 | ubiquitin conjugating enzyme E2 S pseudogen |
| TRIM16L | 17 | tripartite motif containing 16 like [Source:HGI |
| AC026271.3 | 17 | zinc finger protein 695 (ZNF695) pseudogene |
| FBXW10 | 17 | F-box and WD repeat domain containing 10 [S |
| TVP23B | 17 | trans-golgi network vesicle protein 23 homolo |
| AC107982.3 | 17 | novel transcript |
| AC107982.1 | 17 | coiled-coil domain-containing protein 144B-lik |
| PRPSAP2 | 17 | phosphoribosyl pyrophosphate synthetase ass |
| RN7SL627P | 17 | RNA, 7SL, cytoplasmic 627, pseudogene [Sour |
| AC107982.2 | 17 | novel transcript, antisense to PRPSAP2 |
| AC090286.3 | 17 | RAN binding protein 1 (RANBP1) pseudogene |
| AC090286.2 | 17 | NULL |
| SLC5A10 | 17 | solute carrier family 5 member 10 [Source:HG |
| AC090286.1 | 17 | novel transcript, antisense to SLC5A10 |
| FAM83G | 17 | family with sequence similarity 83 member G |
| AC090286.4 | 17 | ATP synthase, H+ transporting, mitochondrial |
| GRAP | 17 | GRB2 related adaptor protein [Source:HGNC S |
| AC003957.1 | 17 | novel transcript |
| SNORD3B-1 | 17 | small nucleolar RNA, C/D box 3B-1 [Source:HG |
| SNORD3B-2 | 17 | small nucleolar RNA, C/D box 3B-2 [Source:HG |
| AC007952.3 | 17 | novel transcript |
| KYNUP2 | 17 | kynureninase pseudogene 2 [Source:HGNC Sy |
| AC007952.7 | 17 | novel transcript |
| AC007952.2 | 17 | novel transcript |
| AC007952.4 | 17 | novel transcript |
| SNORD3D | 17 | small nucleolar RNA, C/D box 3D [Source:HGN |
| SNORD3D | 17 | small nucleolar RNA, C/D box 3D [Source:HGN |
| GRAPL | 17 | GRB2 related adaptor protein like [Source:HG |
| AC007952.6 | 17 | novel transcript, antisense to GRAPL |
| AC007952.9 | 17 | novel transcript |
| AC007952.1 | 17 | novel transcript, antisense to GRAPL |
| AC007952.5 | 17 | novel transcript |
| AC007952.8 | 17 | novel transcript |

| | | |
|---|---|---|
| KYNUP1 | 17 | kynureninase pseudogene 1 [Source:HGNC Sy |
| SNORD3A | 17 | small nucleolar RNA, C/D box 3A [Source:HGN |
| SNORD3C | 17 | small nucleolar RNA, C/D box 3C [Source:HGN |
| AC106017.1 | 17 | novel transcript |
| KYNUP3 | 17 | kynureninase pseudogene 3 [Source:HGNC Sy |
| EPN2 | 17 | epsin 2 [Source:HGNC Symbol;Acc:HGNC:186 |
| AC106017.2 | 17 | novel transcript |
| EPN2-AS1 | 17 | EPN2 antisense RNA 1 [Source:HGNC Symbol; |
| B9D1 | 17 | B9 domain containing 1 [Source:HGNC Symbo |
| MIR1180 | 17 | microRNA 1180 [Source:HGNC Symbol;Acc:H |
| AC124066.1 | 17 | novel transcript, antisense to B9D1 |
| MAPK7 | 17 | mitogen-activated protein kinase 7 [Source:H |
| MFAP4 | 17 | microfibril associated protein 4 [Source:HGNC |
| AC004448.5 | 17 | novel transcript |
| RNF112 | 17 | ring finger protein 112 [Source:HGNC Symbol; |
| AC004448.3 | 17 | novel transcript, antisense to RNF112 |
| AC004448.2 | 17 | novel transcript |
| LINC02094 | 17 | long intergenic non-protein coding RNA 2094 |
| AC004448.4 | 17 | NULL |
| RPS2P46 | 17 | ribosomal protein S2 pseudogene 46 [Source: |
| AC004448.1 | 17 | novel transcript |
| SLC47A1 | 17 | solute carrier family 47 member 1 [Source:HG |
| RPL17P43 | 17 | ribosomal protein L17 pseudogene 43 [Source |
| SNORA59B | 17 | small nucleolar RNA, H/ACA box 59B [Source: |
| AC025627.1 | 17 | novel transcript, antisense to SLC47A1 |
| SLC47A1P1 | 17 | SLC47A1 pseudogene 1 [Source:HGNC Symbol |
| AC025627.2 | 17 | solute carrier family 47, member 1 (SLC47A1) |
| AC025627.3 | 17 | LSU rRNA pseudogene |
| AC025627.4 | 17 | transfer RNA leucine (anticodon TTA) pseudog |
| MTND1P14 | 17 | MT-ND1 pseudogene 14 [Source:HGNC Symb |
| NMTRQ-TTG12-1 | 17 | nuclear-encoded mitochondrial tRNA-Gln (TT |
| MTND2P12 | 17 | MT-ND2 pseudogene 12 [Source:HGNC Symb |
| MTCO1P39 | 17 | MT-CO1 pseudogene 39 [Source:HGNC Symb |
| SLC47A1P2 | 17 | SLC47A1 pseudogene 2 [Source:HGNC Symbol |
| ALDH3A2 | 17 | aldehyde dehydrogenase 3 family member A2 |
| AC115989.1 | 17 | novel transcript, antisense to AL3A2 |
| Y_RNA | 17 | Y RNA [Source:RFAM;Acc:RF00019] |
| AC005722.2 | 17 | NULL |
| SLC47A2 | 17 | solute carrier family 47 member 2 [Source:HG |
| AC005722.3 | 17 | novel transcript |
| AC005722.4 | 17 | novel transcript, antisense to ALDH3A1 |
| ALDH3A1 | 17 | aldehyde dehydrogenase 3 family member A1 |
| AC005722.1 | 17 | TBC1 domain family, member 25 (TBC1D25) p |
| ULK2 | 17 | unc-51 like autophagy activating kinase 2 [Sou |
| NDUFB4P3 | 17 | NADH:ubiquinone oxidoreductase subunit B4 |
| AC015726.3 | 17 | TEC |
| AC015726.2 | 17 | novel transcript |
| AC015726.1 | 17 | novel transcript |
| AKAP10 | 17 | A-kinase anchoring protein 10 [Source:HGNC |
| AC005730.3 | 17 | novel transcript |

| | | |
|---|---|---|
| AC005730.2 | 17 | novel transcript, antisense to SPECC1 |
| SPECC1 | 17 | sperm antigen with calponin homology and cc |
| AC005730.1 | 17 | coiled-coil-helix-coiled-coil-helix domain conta |
| KCTD9P1 | 17 | potassium channel tetramerization domain cc |
| RNU6-258P | 17 | RNA, U6 small nuclear 258, pseudogene [Sour |
| AC004702.1 | 17 | novel transcript |
| RNU6-1057P | 17 | RNA, U6 small nuclear 1057, pseudogene [Sou |
| CCDC144CP | 17 | coiled-coil domain containing 144C, pseudoge |
| RNU6-467P | 17 | RNA, U6 small nuclear 467, pseudogene [Sour |
| AC008088.1 | 17 | novel transcript |
| Metazoa_SRP | 17 | Metazoan signal recognition particle RNA [Sou |
| UPF3AP2 | 17 | UPF3A pseudogene 2 [Source:HGNC Symbol;A |
| USP32P3 | 17 | ubiquitin specific peptidase 32 pseudogene 3 |
| SRP68P3 | 17 | signal recognition particle 68 pseudogene 3 [S |
| AC015818.9 | 17 | novel transcript |
| AC015818.10 | 17 | novel family with sequence similarity 106 pseu |
| NOS2P3 | 17 | nitric oxide synthase 2 pseudogene 3 [Source: |
| LGALS9B | 17 | galectin 9B [Source:HGNC Symbol;Acc:HGNC:2 |
| TNPO1P3 | 17 | transportin 1 pseudogene 3 [Source:HGNC Syr |
| YWHAEP3 | 17 | tyrosine 3-monooxygenase/tryptophan 5-mor |
| KRT16P5 | 17 | keratin 16 pseudogene 5 [Source:HGNC Symb |
| KRT16P3 | 17 | keratin 16 pseudogene 3 [Source:HGNC Symb |
| AC015818.7 | 17 | novel transcript |
| KRT17P6 | 17 | keratin 17 pseudogene 6 [Source:HGNC Symb |
| KRT17P7 | 17 | keratin 17 pseudogene 7 [Source:HGNC Symb |
| AC015818.6 | 17 | tyrosine 3-monooxygenase/tryptophan 5-mor |
| AC015818.2 | 17 | novel transcript |
| AC015818.5 | 17 | novel pseudogene |
| TBC1D3P3 | 17 | TBC1 domain family member 3 pseudogene 3 |
| AC015818.8 | 17 | novel transcript, piRNA host |
| AC015818.1 | 17 | tumor necrosis factor receptor superfamily, m |
| AC015818.3 | 17 | chromosome 2 open reading frame 27B (C2or |
| COTL1P2 | 17 | COTL1 pseudogene 2 [Source:HGNC Symbol;A |
| AC015818.4 | 17 | novel transcript |
| CDRT15L2 | 17 | CMT1A duplicated region transcript 15 like 2 [ |
| ZSWIM5P2 | 17 | zinc finger SWIM-type containing 5 pseudoger |
| MEIS3P2 | 17 | Meis homeobox 3 pseudogene 2 [Source:HGN |
| AC087499.3 | 17 | tandem C2 domains, nuclear (TC2N) pseudoge |
| AC087499.1 | 17 | contactin associated protein-like 3B (CNTNAP: |
| AC087499.4 | 17 | differentially expressed in FDCP 8 homolog (D |
| LINC02088 | 17 | long intergenic non-protein coding RNA 2088 |
| AC087499.7 | 17 | novel pseudogene |
| AC087499.5 | 17 | sperm antigen with calponin homology and cc |
| AC087499.6 | 17 | tyrosine 3-monooxygenase/tryptophan 5-mor |
| AC087499.2 | 17 | ribosomal protein S6 kinase, 70kDa, polypepti |
| RNFT1P3 | 17 | ring finger protein, transmembrane 1 pseudog |
| HNRNPA1P19 | 17 | heterogeneous nuclear ribonucleoprotein A1 |
| OLA1P2 | 17 | OLA1 pseudogene 2 [Source:HGNC Symbol;Ac |
| SCDP1 | 17 | stearoyl-CoA desaturase pseudogene 1 [Sourc |
| AC126365.1 | 17 | novel transcript |

| | | |
|---|---|---|
| AC126365.2 | 17 | novel pseudogene |
| ABHD17AP6 | 17 | abhydrolase domain containing 17A pseudoge |
| CCDC144NL | 17 | coiled-coil domain containing 144 family, N-te |
| AC090774.2 | 17 | novel transcript, antisense to CCDC144NL |
| CCDC144NL-AS1 | 17 | CCDC144NL antisense RNA 1 [Source:HGNC Sy |
| RNU6-1178P | 17 | RNA, U6 small nuclear 1178, pseudogene [Sou |
| RNASEH1P1 | 17 | ribonuclease H1 pseudogene 1 [Source:HGNC |
| AC090774.1 | 17 | novel pseudogene |
| SPECC1P2 | 17 | SPECC1 pseudogene 2 [Source:HGNC Symbol; |
| AC107926.1 | 17 | novel transcript |
| AC087393.3 | 17 | tumor necrosis factor receptor superfamily, m |
| AC087393.1 | 17 | microtubule associated serine/threonine kina |
| USP22 | 17 | ubiquitin specific peptidase 22 [Source:HGNC |
| AC087393.2 | 17 | novel transcript, antisense to USP22 |
| LINC01563 | 17 | long intergenic non-protein coding RNA 1563 |
| DHRS7B | 17 | dehydrogenase/reductase 7B [Source:HGNC S |
| TMEM11 | 17 | transmembrane protein 11 [Source:HGNC Syn |
| AC087294.1 | 17 | novel transcript, antisense to TMEM11 |
| RN7SL426P | 17 | RNA, 7SL, cytoplasmic 426, pseudogene [Sour |
| NATD1 | 17 | N-acetyltransferase domain containing 1 [Sou |
| EIF1P5 | 17 | eukaryotic translation initiation factor 1 pseu |
| MAP2K3 | 17 | mitogen-activated protein kinase kinase 3 [So |
| KCNJ12 | 17 | potassium voltage-gated channel subfamily J r |
| LINC02693 | 17 | long intergenic non-protein coding RNA 2693 |
| AC068418.3 | 17 | novel transcript |
| PDLIM1P2 | 17 | PDZ and LIM domain 1 pseudogene 2 [Source: |
| AC068418.2 | 17 | novel transcript |
| AC068418.1 | 17 | novel transcript |
| RPL21P120 | 17 | ribosomal protein L21 pseudogene 120 [Sourc |
| AC233702.5 | 17 | novel transcript |
| AC233702.7 | 17 | novel transcript, antisense to C17orf51 |
| AC233702.8 | 17 | ribonuclease P/MRP 40kDa subunit (RPP40) p |
| AC233702.6 | 17 | family with sequence similarity 27-like (FAM2 |
| AC233702.3 | 17 | family with sequence similarity 27, member E |
| AC233702.2 | 17 | fermitin family member 3 (FERMT3) pseudoge |
| AC233702.1 | 17 | nuclear receptor corepressor 1 (NCOR1) pseu |
| AC233702.4 | 17 | family with sequence similarity 182 (FAM182 |
| AC233702.9 | 17 | family with sequence similarity 27, member E |
| KCNJ18 | 17 | potassium voltage-gated channel subfamily J r |
| AC243725.1 | 17 | dual specificity mitogen-activated protein kina |
| ABBA01006766.1 | 17 | Bartter syndrome, infantile, with sensorineura |
| NCOR1P2 | 17 | nuclear receptor corepressor 1 pseudogene 2 |
| UBBP4 | 17 | ubiquitin B pseudogene 4 [Source:HGNC Symb |
| FTLP13 | 17 | ferritin light chain pseudogene 13 [Source:HG |
| LINC02002 | 17 | long intergenic non-protein coding RNA 2002 |
| AC138761.1 | 17 | novel transcript |
| AC138761.2 | 17 | fermitin family member 3 (FERMT3) pseudoge |
| FAM27E5 | 17 | family with sequence similarity E5 [Source:HG |
| AC087575.1 | 17 | zinc finger protein 850 (ZNF850) pseudogene |
| AC209154.1 | 17 | novel transcript, piRNA host |

| | | |
|---|---|---|
| AC132825.3 | 17 | novel transcript |
| AC132825.4 | 17 | chromosome 17 open reading frame 51 (C17c |
| AC132825.5 | 17 | mitochondrially encoded NADH dehydrogenas |
| MTCYBP13 | 17 | MT-CYB pseudogene 13 [Source:HGNC Symbo |
| MTRNR2L1 | 17 | MT-RNR2 like 1 [Source:HGNC Symbol;Acc:HG |
| AC132825.2 | 17 | novel transcript, antisense to MTRNR2L1 |
| MTND1P15 | 17 | MT-ND1 pseudogene 15 [Source:HGNC Symbo |
| MTND2P13 | 17 | MT-ND2 pseudogene 13 [Source:HGNC Symbo |
| AC132825.1 | 17 | mitochondrially encoded cytochrome c oxidas |
| NMTRS-TGA3-1 | 17 | nuclear-encoded mitochondrial tRNA-Ser (TG/ |
| MTATP6P3 | 17 | MT-ATP6 pseudogene 3 [Source:HGNC Symbo |
| MTCO3P13 | 17 | MT-CO3 pseudogene 13 [Source:HGNC Symbo |
| AC132825.7 | 17 | mitochondrially encoded NADH dehydrogenas |
| AC132825.8 | 17 | mitochondrially encoded NADH 4L dehydroge |
| AC132825.6 | 17 | mitochondrially encoded NADH dehydrogenas |
| AC131055.2 | 17 | coiled-coil domain containing 144 family, N-te |
| AC131055.1 | 17 | zinc finger, MYM-type 4 (ZMYM4) pseudogene |
| AC131274.3 | 17 | WEE1 homolog (S. pombe) (WEE1) pseudoge |
| AC131274.1 | 17 | novel transcript |
| AC131274.2 | 17 | WEE1 homolog (S. pombe) (WEE1) pseudoger |
| AC069061.1 | 17 | potassium inwardly-rectifying channel, subfan |
| AC069061.2 | 17 | novel transcript |
| PDLIM1P3 | 17 | PDZ and LIM domain 1 pseudogene 3 [Source: |
| GTF2IP6 | 17 | general transcription factor IIi pseudogene 6 [ |
| VN1R71P | 17 | vomeronasal 1 receptor 71 pseudogene [Sour |
| TUFMP1 | 17 | Tu translation elongation factor, mitochondria |
| AC129926.2 | 17 | novel transcript |
| AC129926.1 | 17 | novel transcript |
| RPL34P31 | 17 | ribosomal protein L34 pseudogene 31 [Source |
| RPS16P8 | 17 | ribosomal protein S16 pseudogene 8 [Source: |
| AC026254.1 | 17 | fission 1 (mitochondrial outer membrane) hor |
| MIR4522 | 17 | microRNA 4522 [Source:HGNC Symbol;Acc:HG |
| WSB1 | 17 | WD repeat and SOCS box containing 1 [Source |
| AC026254.2 | 17 | novel transcript |
| SYPL1P2 | 17 | synaptophysin like 1 pseudogene 2 [Source:H( |
| AC090615.1 | 17 | tyrosine 3-monooxygenase/tryptophan 5-mor |
| TBC1D3P5 | 17 | TBC1 domain family member 3 pseudogene 5 |
| KSR1 | 17 | kinase suppressor of ras 1 [Source:HGNC Syml |
| AC069366.2 | 17 | novel transcript |
| AC069366.1 | 17 | chromosome 9 open reading frame 30 (C9orf3 |
| ITM2BP1 | 17 | integral membrane protein 2B pseudogene 1 |
| AC015688.4 | 17 | novel protein |
| AC015688.6 | 17 | novel transcript, antisense to KSR1 |
| LGALS9 | 17 | galectin 9 [Source:HGNC Symbol;Acc:HGNC:65 |
| NOS2P1 | 17 | nitric oxide synthase 2 pseudogene 1 [Source: |
| AC015688.3 | 17 | TBC1 domain family, member 29 (TBC1D29) p |
| AC015688.5 | 17 | nitric oxide synthase 3 (endothelial cell) (NOS: |
| CPDP1 | 17 | carboxypeptidase D pseudogene 1 [Source:HG |
| AC015688.2 | 17 | golgi SNAP receptor complex member 1 (GOS |
| AC015688.1 | 17 | nitric oxide synthase 2, inducible (NOS2) pseu |

| | | |
|---|---|---|
| AC130289.1 | 17 | golgi SNAP receptor complex member 1 (GOS |
| AC130289.2 | 17 | lectin, galactoside-binding, soluble, 9 (LGALS9 |
| LGALS9DP | 17 | galectin 9D, pseudogene [Source:HGNC Symb |
| NOS2 | 17 | nitric oxide synthase 2 [Source:HGNC Symbol; |
| AC005697.1 | 17 | novel protein |
| AC005697.2 | 17 | novel transcript, antisense to C17orf108 |
| LYRM9 | 17 | LYR motif containing 9 [Source:HGNC Symbol; |
| LINC01992 | 17 | long intergenic non-protein coding RNA 1992 |
| AC005697.3 | 17 | novel transcript |
| AC090287.1 | 17 | NULL |
| AC090287.2 | 17 | ribosomal protein L10 (RPL10) pseudogene |
| SNORA70 | 17 | Small nucleolar RNA SNORA70 [Source:RFAM; |
| NLK | 17 | nemo like kinase [Source:HGNC Symbol;Acc:H |
| Vault | 17 | Vault RNA [Source:RFAM;Acc:RF00006] |
| RPS29P22 | 17 | ribosomal protein S29 pseudogene 22 [Source |
| AC061975.8 | 17 | novel transcript |
| AC061975.7 | 17 | novel transcript |
| PYY2 | 17 | peptide YY 2 (pseudogene) [Source:HGNC Sym |
| ERVE-1 | 17 | endogenous retrovirus group E member 1 [So |
| AC061975.6 | 17 | novel transcript |
| PPY2P | 17 | pancreatic polypeptide 2, pseudogene [Source |
| AC061975.2 | 17 | wingless-type MMTV integration site family, n |
| AC061975.4 | 17 | novel transcript |
| AC061975.1 | 17 | novel transcript |
| AC061975.5 | 17 | leucine rich repeat containing 37, member A3 |
| AC061975.3 | 17 | golgi SNAP receptor complex member 2 (GOS |
| KRT18P55 | 17 | keratin 18 pseudogene 55 [Source:HGNC Sym |
| TMEM97 | 17 | transmembrane protein 97 [Source:HGNC Syn |
| IFT20 | 17 | intraflagellar transport 20 [Source:HGNC Syml |
| TNFAIP1 | 17 | TNF alpha induced protein 1 [Source:HGNC Sy |
| POLDIP2 | 17 | DNA polymerase delta interacting protein 2 [S |
| TMEM199 | 17 | transmembrane protein 199 [Source:HGNC Sy |
| AC002094.1 | 17 | novel transcript readthrough between TMEM |
| MIR4723 | 17 | microRNA 4723 [Source:HGNC Symbol;Acc:HG |
| AC002094.2 | 17 | novel transcript, antisense to TMEM199 |
| SEBOX | 17 | SEBOX homeobox [Source:HGNC Symbol;Acc: |
| AC002094.3 | 17 | novel protein, readthrough between VTN and |
| SARM1 | 17 | sterile alpha and TIR motif containing 1 [Sourc |
| VTN | 17 | vitronectin [Source:HGNC Symbol;Acc:HGNC:1 |
| AC002094.4 | 17 | novel transcript, sense intronic to transcript re |
| AC002094.5 | 17 | novel transcript |
| SLC46A1 | 17 | solute carrier family 46 member 1 [Source:HG |
| AC015917.2 | 17 | novel transcript, antisense to SLC46A1 |
| H3F3BP2 | 17 | H3 histone, family 3B pseudogene 2 [Source:H |
| AC015917.1 | 17 | RAP1A, member of RAS oncogene family (RAP |
| AC005726.1 | 17 | novel protein |
| RPS7P1 | 17 | ribosomal protein S7 pseudogene 1 [Source:H |
| SLC13A2 | 17 | solute carrier family 13 member 2 [Source:HG |
| FOXN1 | 17 | forkhead box N1 [Source:HGNC Symbol;Acc:H |
| UNC119 | 17 | unc-119 lipid binding chaperone [Source:HGN |

| | | |
|---|---|---|
| PIGS | 17 | phosphatidylinositol glycan anchor biosynthes |
| ALDOC | 17 | aldolase, fructose-bisphosphate C [Source:HG |
| AC005726.4 | 17 | novel transcript, antisense to ALDOC |
| SPAG5 | 17 | sperm associated antigen 5 [Source:HGNC Syr |
| SPAG5-AS1 | 17 | SPAG5 antisense RNA 1 [Source:HGNC Symbo |
| AC005726.3 | 17 | novel transcript, sense intronic to novel prote |
| AC005726.5 | 17 | novel transcript, antisense to novel protein |
| RSKR | 17 | ribosomal protein S6 kinase related [Source:H |
| KIAA0100 | 17 | KIAA0100 [Source:HGNC Symbol;Acc:HGNC:2ɛ |
| AC005726.2 | 17 | novel transcript, antisense to KIAA0100 |
| SDF2 | 17 | stromal cell derived factor 2 [Source:HGNC Sy |
| RPS12P28 | 17 | ribosomal protein S12 pseudogene 28 [Source |
| SUPT6H | 17 | SPT6 homolog, histone chaperone and transcr |
| AC010761.3 | 17 | novel transcript, antisense to SUPT6H |
| PROCA1 | 17 | protein interacting with cyclin A1 [Source:HGI |
| RAB34 | 17 | RAB34, member RAS oncogene family [Source |
| RPL23A | 17 | ribosomal protein L23a [Source:HGNC Symbol |
| SNORD42B | 17 | small nucleolar RNA, C/D box 42B [Source:HG |
| AC010761.1 | 17 | novel transcript, antisense to RPL23A |
| SNORD4A | 17 | small nucleolar RNA, C/D box 4A [Source:HGN |
| SNORD42A | 17 | small nucleolar RNA, C/D box 42A [Source:HG |
| SNORD4B | 17 | small nucleolar RNA, C/D box 4B [Source:HGN |
| TLCD1 | 17 | TLC domain containing 1 [Source:HGNC Symb |
| NEK8 | 17 | NIMA related kinase 8 [Source:HGNC Symbol; |
| AC010761.6 | 17 | novel transcript, antisense to TLCD1 |
| AC010761.2 | 17 | novel transcript, antisense to NEK8 |
| TRAF4 | 17 | TNF receptor associated factor 4 [Source:HGN |
| AC010761.4 | 17 | novel transcript, antisense to TRAF4 |
| AC010761.5 | 17 | novel transcript, antisense to TRAF4 |
| FAM222B | 17 | family with sequence similarity 222 member E |
| RPL31P58 | 17 | ribosomal protein L31 pseudogene 58 [Source |
| Y_RNA | 17 | Y RNA [Source:RFAM;Acc:RF00019] |
| AC024267.2 | 17 | novel pseudogene |
| ERAL1 | 17 | Era like 12S mitochondrial rRNA chaperone 1 |
| AC024267.1 | 17 | novel transcript |
| MIR451A | 17 | microRNA 451a [Source:HGNC Symbol;Acc:HG |
| MIR451B | 17 | microRNA 451b [Source:HGNC Symbol;Acc:HG |
| MIR144 | 17 | microRNA 144 [Source:HGNC Symbol;Acc:HGI |
| MIR4732 | 17 | microRNA 4732 [Source:HGNC Symbol;Acc:HG |
| FLOT2 | 17 | flotillin 2 [Source:HGNC Symbol;Acc:HGNC:37 |
| AC024267.3 | 17 | novel transcript, antisense to FLOT2 |
| AC024267.6 | 17 | novel transcript, antisense to DHRS13 |
| DHRS13 | 17 | dehydrogenase/reductase 13 [Source:HGNC S |
| PHF12 | 17 | PHD finger protein 12 [Source:HGNC Symbol; |
| AC024267.4 | 17 | novel transcript, antisense to PHF12 |
| AC024267.5 | 17 | novel transcript, antisense to PHF12 |
| PIPOX | 17 | pipecolic acid and sarcosine oxidase [Source:H |
| SEZ6 | 17 | seizure related 6 homolog [Source:HGNC Sym |
| AC024619.2 | 17 | novel transcript, antisense to PIPOX |
| AC024619.1 | 17 | novel transcript |

| | | |
|---|---|---|
| AC024619.4 | 17 | NADH dehydrogenase (ubiquinone) Fe-S prote |
| MYO18A | 17 | myosin XVIIIA [Source:HGNC Symbol;Acc:HGN |
| TIAF1 | 17 | TGFB1-induced anti-apoptotic factor 1 [Source |
| AC024619.3 | 17 | novel transcript, antisense to MYO18A |
| AC005412.1 | 17 | gap junction protein, alpha 4, 37kDa (GJA4) ps |
| TWF1P1 | 17 | twinfilin 1 pseudogene 1 [Source:HGNC Symb |
| CRYBA1 | 17 | crystallin beta A1 [Source:HGNC Symbol;Acc:H |
| NUFIP2 | 17 | nuclear FMR1 interacting protein 2 [Source:H |
| RPL35AP35 | 17 | ribosomal protein L35a pseudogene 35 [Sourc |
| AC068025.2 | 17 | novel transcript |
| AC068025.1 | 17 | novel transcript, antisense to TAOK1 |
| RNU4-34P | 17 | RNA, U4 small nuclear 34, pseudogene [Sourc |
| TAOK1 | 17 | TAO kinase 1 [Source:HGNC Symbol;Acc:HGN |
| MIR4523 | 17 | microRNA 4523 [Source:HGNC Symbol;Acc:HG |
| RNU6-711P | 17 | RNA, U6 small nuclear 711, pseudogene [Sour |
| RNU6-1034P | 17 | RNA, U6 small nuclear 1034, pseudogene [Sou |
| ABHD15-AS1 | 17 | ABHD15 antisense RNA 1 [Source:HGNC Symb |
| ABHD15 | 17 | abhydrolase domain containing 15 [Source:HG |
| TP53I13 | 17 | tumor protein p53 inducible protein 13 [Sourc |
| AC104564.3 | 17 | novel transcript |
| GIT1 | 17 | GIT ArfGAP 1 [Source:HGNC Symbol;Acc:HGN |
| ANKRD13B | 17 | ankyrin repeat domain 13B [Source:HGNC Syn |
| AC104564.4 | 17 | novel transcript, antisense to ANKRD13B |
| CORO6 | 17 | coronin 6 [Source:HGNC Symbol;Acc:HGNC:21 |
| AC104564.2 | 17 | novel transcript |
| SSH2 | 17 | slingshot protein phosphatase 2 [Source:HGN |
| AC104564.1 | 17 | novel transcript, antisense to SSH2 |
| RNU6-920P | 17 | RNA, U6 small nuclear 920, pseudogene [Sour |
| AC104564.5 | 17 | novel transcript, sense intronic to SSH2 |
| RPL21P123 | 17 | ribosomal protein L21 pseudogene 123 [Sourc |
| AC023389.1 | 17 | novel transcript, antisense to SSH2 |
| AC023389.2 | 17 | RNA terminal phosphate cyclase-like 1 (RCL1) |
| SNORA70 | 17 | Small nucleolar RNA SNORA70 [Source:RFAM; |
| RPL9P30 | 17 | ribosomal protein L9 pseudogene 30 [Source: |
| AC104982.2 | 17 | novel transcript, antisense to SSH2 |
| AC104982.1 | 17 | NULL |
| EFCAB5 | 17 | EF-hand calcium binding domain 5 [Source:HG |
| AC104982.3 | 17 | non-metastatic cells 4, protein expressed in (N |
| RNY4P13 | 17 | RNY4 pseudogene 13 [Source:HGNC Symbol;A |
| AC104996.3 | 17 | RNA, Ro-associated Y4 pseudogene 13 |
| AC104996.1 | 17 | novel transcript, antisense to EFCAB5 |
| AC104996.2 | 17 | syntaxin 18 (STX18) pseudogene |
| AC104984.2 | 17 | novel transcript, antisense to EFCAB5 |
| NSRP1 | 17 | nuclear speckle splicing regulatory protein 1 [S |
| MIR423 | 17 | microRNA 423 [Source:HGNC Symbol;Acc:HGN |
| hsa-mir-423 | 17 | hsa-mir-423 [Source:miRBase;Acc:MI0001445 |
| MIR3184 | 17 | microRNA 3184 [Source:HGNC Symbol;Acc:HG |
| AC104984.3 | 17 | novel transcript, antisense to CCDC55 |
| AC104984.5 | 17 | novel transcript, antisense to NSRP1 |
| SLC6A4 | 17 | solute carrier family 6 member 4 [Source:HGN |

| | | |
|---|---|---|
| AC104984.1 | 17 | novel transcript, antisense to SLC6A4 |
| AC104984.4 | 17 | novel transcript, antisense to BLMH |
| SNORD63 | 17 | Small nucleolar RNA SNORD63 [Source:RFAM; |
| BLMH | 17 | bleomycin hydrolase [Source:HGNC Symbol;A |
| RNU6-1267P | 17 | RNA, U6 small nuclear 1267, pseudogene [Sou |
| TMIGD1 | 17 | transmembrane and immunoglobulin domain |
| Y_RNA | 17 | Y RNA [Source:RFAM;Acc:RF00019] |
| RNU6-990P | 17 | RNA, U6 small nuclear 990, pseudogene [Sour |
| CPD | 17 | carboxypeptidase D [Source:HGNC Symbol;Ac |
| GOSR1 | 17 | golgi SNAP receptor complex member 1 [Sour |
| ALOX12P1 | 17 | arachidonate 12-lipoxygenase pseudogene 1 [ |
| AC011840.3 | 17 | novel transcript |
| AC011840.5 | 17 | TEC |
| TBC1D29P | 17 | TBC1 domain family member 29, pseudogene |
| AC011840.4 | 17 | novel transcript, antisense to TBC1D29 |
| AC011840.2 | 17 | novel transcript |
| KRT17P3 | 17 | keratin 17 pseudogene 3 [Source:HGNC Symb |
| AC011840.1 | 17 | novel transcript |
| AC005562.1 | 17 | novel transcript |
| SMURF2P1 | 17 | SMAD specific E3 ubiquitin protein ligase 2 ps |
| AC005562.2 | 17 | leucine rich repeat containing 37B (LRRC37B) |
| SH3GL1P2 | 17 | SH3 domain containing GRB2 like 1, endophili |
| LRRC37BP1 | 17 | leucine rich repeat containing 37B pseudogen |
| RN7SL316P | 17 | RNA, 7SL, cytoplasmic 316, pseudogene [Sour |
| SUZ12P1 | 17 | SUZ12 pseudogene 1 [Source:HGNC Symbol;A |
| AC127024.7 | 17 | TEC |
| AC127024.3 | 17 | novel transcript |
| AC127024.2 | 17 | novel transcript |
| AC127024.8 | 17 | TEC |
| CRLF3 | 17 | cytokine receptor like factor 3 [Source:HGNC |
| AC127024.4 | 17 | novel transcript, sense intronic CRLF3 |
| AC127024.5 | 17 | novel transcript |
| AC127024.6 | 17 | novel transcript, antisense to CRLF3 |
| AC127024.1 | 17 | ribosomal protein S17 (RPS17) pseudogene |
| ATAD5 | 17 | ATPase family AAA domain containing 5 [Sour |
| AC130324.2 | 17 | novel transcript, antisense ATAD5 |
| RNU6-298P | 17 | RNA, U6 small nuclear 298, pseudogene [Sour |
| AC130324.1 | 17 | novel transcript, antisense ATAD5 |
| TEFM | 17 | transcription elongation factor, mitochondrial |
| AC130324.3 | 17 | novel transcript, antisense to TEFM |
| ADAP2 | 17 | ArfGAP with dual PH domains 2 [Source:HGN( |
| AC138207.1 | 17 | novel transcript |
| RN7SL138P | 17 | RNA, 7SL, cytoplasmic 138, pseudogene [Sour |
| AC138207.5 | 17 | novel transcript |
| AC138207.9 | 17 | Novel transcript |
| RNF135 | 17 | ring finger protein 135 [Source:HGNC Symbol; |
| AC138207.4 | 17 | novel transcript, sense intronic RNF135 |
| DPRXP4 | 17 | divergent-paired related homeobox pseudoge |
| AC138207.7 | 17 | novel transcript, antisense RNF135 |
| AC138207.3 | 17 | tyrosine 3-monooxygenase/tryptophan 5-mor |

| | | |
|---|---|---|
| AC138207.8 | 17 | SMAD specific E3 ubiquitin protein ligase 2 (SI |
| AC138207.6 | 17 | leucine rich repeat containing 37B (LRRC37B) |
| AC138207.2 | 17 | novel transcript, antisense to NF1 |
| MIR4733 | 17 | microRNA 4733 [Source:HGNC Symbol;Acc:HG |
| NF1 | 17 | neurofibromin 1 [Source:HGNC Symbol;Acc:H |
| AC079915.1 | 17 | novel transcript, sense intronic NF1 |
| OMG | 17 | oligodendrocyte myelin glycoprotein [Source: |
| EVI2B | 17 | ecotropic viral integration site 2B [Source:HGI |
| AC134669.1 | 17 | novel protein |
| EVI2A | 17 | ecotropic viral integration site 2A [Source:HGI |
| AK4P1 | 17 | adenylate kinase 4 pseudogene 1 [Source:HGI |
| RAB11FIP4 | 17 | RAB11 family interacting protein 4 [Source:HG |
| RN7SL79P | 17 | RNA, 7SL, cytoplasmic 79, pseudogene [Sourc |
| RN7SL45P | 17 | RNA, 7SL, cytoplasmic 45, pseudogene [Sourc |
| AC003101.3 | 17 | TEC |
| MIR4724 | 17 | microRNA 4724 [Source:HGNC Symbol;Acc:HG |
| MIR193A | 17 | microRNA 193a [Source:HGNC Symbol;Acc:HG |
| AC003101.2 | 17 | novel transcript |
| RNU6ATAC7P | 17 | RNA, U6atac small nuclear 7, pseudogene [So |
| AC003101.1 | 17 | novel transcript |
| MIR4725 | 17 | microRNA 4725 [Source:HGNC Symbol;Acc:HG |
| MIR365B | 17 | microRNA 365b [Source:HGNC Symbol;Acc:HG |
| AC007923.4 | 17 | novel transcript |
| AC007923.3 | 17 | myelin transcription factor 1-like (MYT1L) pse |
| RNU6-1134P | 17 | RNA, U6 small nuclear 1134, pseudogene [Sou |
| AC007923.2 | 17 | G protein-coupled receptor 160 (GPR160) pse |
| AC007923.1 | 17 | novel transcript |
| AC004253.2 | 17 | novel transcript |
| COPRS | 17 | coordinator of PRMT5 and differentiation stin |
| UTP6 | 17 | UTP6 small subunit processome component [ |
| AC004253.1 | 17 | novel transcript |
| SUZ12 | 17 | SUZ12 polycomb repressive complex 2 subuni |
| RNA5SP437 | 17 | RNA, 5S ribosomal pseudogene 437 [Source:H |
| AC090616.3 | 17 | bromodomain PHD finger transcription factor |
| LRRC37B | 17 | leucine rich repeat containing 37B [Source:HG |
| AC090616.4 | 17 | ADP-ribosylation factor-like 17B (ARL17B) pse |
| SH3GL1P1 | 17 | SH3 domain containing GRB2 like 1, endophili |
| AC090616.6 | 17 | TEC |
| AC090616.2 | 17 | bromodomain PHD finger transcription factor |
| AC090616.5 | 17 | leucine rich repeat containing 37B (LRRC37B) |
| AC090616.1 | 17 | bromodomain PHD finger transcription factor |
| WDR45BP1 | 17 | WD repeat domain 45B pseudogene 1 [Source |
| AC116407.2 | 17 | novel transcript |
| AC116407.1 | 17 | novel transcript, antisense to RHOT1 |
| RHOT1 | 17 | ras homolog family member T1 [Source:HGNC |
| ARGFXP2 | 17 | arginine-fifty homeobox pseudogene 2 [Sourc |
| AC116407.4 | 17 | TEC |
| AC116407.3 | 17 | TEC |
| UBL5P2 | 17 | ubiquitin like 5 pseudogene 2 [Source:HGNC S |
| AC026620.2 | 17 | novel transcript |

| | | |
|---|---|---|
| RHBDL3 | 17 | rhomboid like 3 [Source:HGNC Symbol;Acc:H( |
| AC026620.1 | 17 | novel transcript, antisense RHBDL3 |
| AC005899.4 | 17 | novel transcript |
| C17orf75 | 17 | chromosome 17 open reading frame 75 [Sour |
| AC005899.3 | 17 | novel transcript |
| OOSP1P2 | 17 | oocyte secreted protein 1 pseudogene 2 [Sou |
| MIR632 | 17 | microRNA 632 [Source:HGNC Symbol;Acc:HGI |
| ZNF207 | 17 | zinc finger protein 207 [Source:HGNC Symbol; |
| AC005899.6 | 17 | novel transcript |
| AC005899.7 | 17 | novel transcript |
| AC005899.5 | 17 | novel transcript |
| AC005899.8 | 17 | novel transcript, antisense to PSMD11 |
| AC005899.1 | 17 | novel transcript |
| AC005899.2 | 17 | nudix (nucleoside diphosphate linked moiety ) |
| PSMD11 | 17 | proteasome 26S subunit, non-ATPase 11 [Sou |
| Y_RNA | 17 | Y RNA [Source:RFAM;Acc:RF00019] |
| CDK5R1 | 17 | cyclin dependent kinase 5 regulatory subunit : |
| MYO1D | 17 | myosin ID [Source:HGNC Symbol;Acc:HGNC:7! |
| AC079336.5 | 17 | novel transcript, antisense MYO1D |
| AC079336.7 | 17 | TEC |
| AC079336.2 | 17 | novel transcript, sense intronic MYO1D |
| AC079336.1 | 17 | novel transcript, sense intronic MYO1D |
| AC079336.6 | 17 | TEC |
| AC079336.4 | 17 | novel transcript, antisense MYO1D |
| AC079336.3 | 17 | novel transcript, sense intronic MYO1D |
| AC025211.1 | 17 | novel transcript, antisense MYO1D |
| Y_RNA | 17 | Y RNA [Source:RFAM;Acc:RF00019] |
| AC084809.1 | 17 | novel transcript |
| AC084809.2 | 17 | novel transcript |
| TMEM98 | 17 | transmembrane protein 98 [Source:HGNC Syn |
| SPACA3 | 17 | sperm acrosome associated 3 [Source:HGNC S |
| ASIC2 | 17 | acid sensing ion channel subunit 2 [Source:HG |
| AC003687.1 | 17 | novel transcript, sense intronic ASIC2 |
| AC008133.2 | 17 | TEC |
| AC008133.1 | 17 | novel transcript, antisense ASIC2 |
| AC011824.3 | 17 | novel transcript, sense intronic ASIC2 |
| AC011824.4 | 17 | novel transcript, antisense ASIC2 |
| AC011824.2 | 17 | novel transcript, antisense ASIC2 |
| AC011824.1 | 17 | novel transcript, antisense ASIC2 |
| AC024614.1 | 17 | novel transcript |
| AC024614.2 | 17 | novel transcript |
| AC024614.3 | 17 | mitochondrially encoded NADH dehydrogenas |
| AC024614.4 | 17 | novel transcript |
| AC024610.2 | 17 | novel transcript |
| AC004147.4 | 17 | novel transcript |
| AC024610.1 | 17 | novel transcript, antisense ASIC2 |
| AC123769.1 | 17 | TEC |
| TLK2P1 | 17 | tousled like kinase 2 pseudogene 1 [Source:H( |
| AC004147.1 | 17 | novel transcript, sense intronic ASIC2 |
| AC004147.5 | 17 | novel transcript, sense intronic ASIC2 |

| | | |
|---|---|---|
| RNA5SP438 | 17 | RNA, 5S ribosomal pseudogene 438 [Source:H |
| AC004147.3 | 17 | novel transcript, antisense ASIC2 |
| AC004147.2 | 17 | novel transcript |
| LINC01989 | 17 | long intergenic non-protein coding RNA 1989 |
| AC005549.1 | 17 | novel transcript |
| CCL2 | 17 | C-C motif chemokine ligand 2 [Source:HGNC S |
| CCL7 | 17 | C-C motif chemokine ligand 7 [Source:HGNC S |
| CCL11 | 17 | C-C motif chemokine ligand 11 [Source:HGNC |
| CCL8 | 17 | C-C motif chemokine ligand 8 [Source:HGNC S |
| AC011193.2 | 17 | progesterone receptor membrane componen |
| CCL13 | 17 | C-C motif chemokine ligand 13 [Source:HGNC |
| CCL1 | 17 | C-C motif chemokine ligand 1 [Source:HGNC S |
| AC011193.1 | 17 | novel transcript |
| C17orf102 | 17 | chromosome 17 open reading frame 102 [Sou |
| TMEM132E | 17 | transmembrane protein 132E [Source:HGNC S |
| AC005691.1 | 17 | TEC |
| AC005552.1 | 17 | novel transcript |
| AC022903.1 | 17 | novel transcript |
| CCT6B | 17 | chaperonin containing TCP1 subunit 6B [Sourc |
| AC022903.2 | 17 | novel transcript, antisense to CCT6B |
| ZNF830 | 17 | zinc finger protein 830 [Source:HGNC Symbol; |
| LIG3 | 17 | DNA ligase 3 [Source:HGNC Symbol;Acc:HGNC |
| RFFL | 17 | ring finger and FYVE like domain containing E3 |
| AC004223.3 | 17 | RAD51L3-RFFL readthrough |
| AC004223.4 | 17 | novel transcript, antisense to RFFL |
| AC004223.2 | 17 | novel transcript |
| AC004223.1 | 17 | ribosomal protein L37 (RPL37) pseudogene |
| RAD51D | 17 | RAD51 paralog D [Source:HGNC Symbol;Acc:H |
| RNU6-840P | 17 | RNA, U6 small nuclear 840, pseudogene [Sour |
| Vault | 17 | Vault RNA [Source:RFAM;Acc:RF00006] |
| FNDC8 | 17 | fibronectin type III domain containing 8 [Sourc |
| NLE1 | 17 | notchless homolog 1 [Source:HGNC Symbol;A |
| UNC45B | 17 | unc-45 myosin chaperone B [Source:HGNC Syı |
| AC022916.4 | 17 | novel transcript, antisense UNC45B |
| AC022916.2 | 17 | novel transcript |
| AC022916.3 | 17 | Spi-C transcription factor (Spi-1/PU.1 related) |
| SLC35G3 | 17 | solute carrier family 35 member G3 [Source:H |
| AC022916.1 | 17 | novel transcript, antisense SLFN5 |
| SLFN5 | 17 | schlafen family member 5 [Source:HGNC Sym |
| AC022706.1 | 17 | novel transcript |
| AC060766.2 | 17 | ring finger protein 7 (RNF7) pseudogene |
| SLFN11 | 17 | schlafen family member 11 [Source:HGNC Syn |
| AC060766.3 | 17 | ubiquitin-fold modifier 1 (UFM1) pseudogene |
| AC060766.6 | 17 | novel transcript |
| AC060766.4 | 17 | novel transcript |
| AC060766.7 | 17 | novel transcript |
| AC060766.5 | 17 | DTW domain containing 2 (DTWD2) pseudoge |
| AC060766.1 | 17 | RAB32, member RAS oncogene family (RAB32 |
| SLFN12 | 17 | schlafen family member 12 [Source:HGNC Syn |
| SLFN13 | 17 | schlafen family member 13 [Source:HGNC Syn |

| | | |
|---|---|---|
| AC015911.4 | 17 | adenylosuccinate synthase (ADSS) pseudogen |
| SLFN12L | 17 | schlafen family member 12 like [Source:HGNC |
| AC015911.8 | 17 | RAB32, member RAS oncogene family (RAB32 |
| AC015911.11 | 17 | novel transcript |
| AC015911.5 | 17 | novel transcript |
| E2F3P1 | 17 | E2F transcription factor 3 pseudogene 1 [Sour |
| AC015911.7 | 17 | TAF5-like RNA polymerase II, p300/CBP-assoc |
| AC015911.3 | 17 | novel transcript, sense intronic SLFN12L |
| TOMM20P2 | 17 | TOMM20 pseudogene 2 [Source:HGNC Symbo |
| AC015911.10 | 17 | schlafen family member pseudogene |
| AC015911.2 | 17 | novel transcript |
| SLFN14 | 17 | schlafen family member 14 [Source:HGNC Syr |
| AC015911.6 | 17 | novel transcript, antisense SLFN14 |
| AC015911.9 | 17 | novel transcript |
| AC015911.1 | 17 | ribosomal protein L39 (RPL39) pseudogene |
| SNHG30 | 17 | small nucleolar RNA host gene 30 [Source:HGI |
| SNORD7 | 17 | small nucleolar RNA, C/D box 7 [Source:HGNC |
| PEX12 | 17 | peroxisomal biogenesis factor 12 [Source:HGN |
| AP2B1 | 17 | adaptor related protein complex 2 subunit be |
| AC004134.1 | 17 | transmembrane protein 160 (TMEM160) pseu |
| TAF15 | 17 | TATA-box binding protein associated factor 15 |
| RASL10B | 17 | RAS like family 10 member B [Source:HGNC Sy |
| GAS2L2 | 17 | growth arrest specific 2 like 2 [Source:HGNC S |
| MMP28 | 17 | matrix metallopeptidase 28 [Source:HGNC Syr |
| AC006237.1 | 17 | novel transcript, sense intronic to MMP28 |
| C17orf50 | 17 | chromosome 17 open reading frame 50 [Sour |
| AC015849.2 | 17 | novel transcript |
| AC015849.3 | 17 | novel transcript, antisense TAF15 |
| AC015849.4 | 17 | novel transcript, sense overlapping TAF15 |
| HEATR9 | 17 | HEAT repeat containing 9 [Source:HGNC Symb |
| AC015849.6 | 17 | ribosomal L24 domain containing 1 (RSL24D1) |
| AC015849.1 | 17 | novel transcript, antisense to CCL5 |
| CCL5 | 17 | C-C motif chemokine ligand 5 [Source:HGNC S |
| 7SK | 17 | 7SK RNA [Source:RFAM;Acc:RF00100] |
| AC015849.5 | 17 | leucine rich repeat containing 37 member A3 |
| LRRC37A9P | 17 | leucine rich repeat containing 37 member A9, |
| RDM1 | 17 | RAD52 motif containing 1 [Source:HGNC Symb |
| LYZL6 | 17 | lysozyme like 6 [Source:HGNC Symbol;Acc:HG |
| AC244100.1 | 17 | tetratricopeptide repeat, ankyrin repeat and c |
| CCL16 | 17 | C-C motif chemokine ligand 16 [Source:HGNC |
| CCL14 | 17 | C-C motif chemokine ligand 14 [Source:HGNC |
| AC244100.2 | 17 | novel transcript, antisense to CCL14 |
| CCL15-CCL14 | 17 | CCL15-CCL14 readthrough (NMD candidate) [S |
| CCL15 | 17 | C-C motif chemokine ligand 15 [Source:HGNC |
| AC244100.3 | 17 | novel transcript, antisense CCL15 |
| AC244100.4 | 17 | novel transcript |
| CCL23 | 17 | C-C motif chemokine ligand 23 [Source:HGNC |
| CCL18 | 17 | C-C motif chemokine ligand 18 [Source:HGNC |
| AC243829.4 | 17 | novel transcript, antisense CCL3 |
| CCL3 | 17 | C-C motif chemokine ligand 3 [Source:HGNC S |

| | | |
|---|---|---|
| CCL4 | 17 | C-C motif chemokine ligand 4 [Source:HGNC S |
| RN7SL301P | 17 | RNA, 7SL, cytoplasmic 301, pseudogene [Sour |
| AC243829.2 | 17 | novel transcript, antisense TBC1D3B |
| AC243829.7 | 17 | nitric oxide synthase 2, inducible (NOS2) pseu |
| TBC1D3B | 17 | TBC1 domain family member 3B [Source:HGN |
| AC243829.1 | 17 | novel transcript, antisense CCL3L3 |
| CCL3L1 | 17 | C-C motif chemokine ligand 3 like 1 [Source:H |
| CCL4L2 | 17 | C-C motif chemokine ligand 4 like 2 [Source:H |
| AC243829.5 | 17 | novel transcript |
| AC243829.3 | 17 | ribosomal protein S6 kinase, 70kDa, polypepti |
| AC243829.6 | 17 | tyrosine 3-monooxygenase/tryptophan 5-mor |
| TBC1D3I | 17 | TBC1 domain family member 3I [Source:HGN( |
| AC233699.1 | 17 | aminopeptidase puromycin sensitive (NPEPPS |
| Y_RNA | 17 | Y RNA [Source:RFAM;Acc:RF00019] |
| TBC1D3G | 17 | TBC1 domain family member 3G [Source:HGN |
| TBC1D3H | 17 | TBC1 domain family member 3H [Source:HGN |
| RNU6-1192P | 17 | RNA, U6 small nuclear 1192, pseudogene [Sou |
| TBC1D3F | 17 | TBC1 domain family member 3F [Source:HGN |
| TBC1D3J | 17 | TBC1 domain family member 3J [Source:HGN( |
| ZNHIT3 | 17 | zinc finger HIT-type containing 3 [Source:HGN |
| RNA5SP439 | 17 | RNA, 5S ribosomal pseudogene 439 [Source:H |
| MYO19 | 17 | myosin XIX [Source:HGNC Symbol;Acc:HGNC:2 |
| PIGW | 17 | phosphatidylinositol glycan anchor biosynthes |
| GGNBP2 | 17 | gametogenetin binding protein 2 [Source:HGN |
| AC243732.1 | 17 | ribosomal protein S2 (RPS2) pseudogene |
| DHRS11 | 17 | dehydrogenase/reductase 11 [Source:HGNC S |
| MRM1 | 17 | mitochondrial rRNA methyltransferase 1 [Sou |
| AC243830.2 | 17 | novel transcript |
| AC243830.3 | 17 | novel transcript |
| AC243830.1 | 17 | novel transcript |
| LHX1-DT | 17 | LHX1 divergent transcript [Source:HGNC Symb |
| LHX1 | 17 | LIM homeobox 1 [Source:HGNC Symbol;Acc:H |
| AC243773.1 | 17 | novel transcript, antisense to LHX1 |
| AATF | 17 | apoptosis antagonizing transcription factor [S |
| AC243773.2 | 17 | novel transcript, sense intronic AATF |
| AC244093.4 | 17 | novel transcript, sense intronic AATF |
| MIR2909 | 17 | microRNA 2909 [Source:HGNC Symbol;Acc:HG |
| AC244093.3 | 17 | novel transcript, sense intronic AATF |
| AC244093.5 | 17 | novel transcript, antisense AATF |
| ACACA | 17 | acetyl-CoA carboxylase alpha [Source:HGNC S |
| AC244093.1 | 17 | ribosomal protein S2 (RPS2) pseudogene |
| HMGB1P24 | 17 | high mobility group box 1 pseudogene 24 [Sou |
| AC244093.2 | 17 | ribosomal protein L18a (RPL18A) pseudogene |
| C17orf78 | 17 | chromosome 17 open reading frame 78 [Sour |
| AC243654.3 | 17 | novel transcript, sense intronic to C17orf78 |
| TADA2A | 17 | transcriptional adaptor 2A [Source:HGNC Sym |
| AC243654.1 | 17 | novel transcript, antisense TADA2A |
| AC243654.2 | 17 | ribosomal protein L24 (RPL24) pseudogene |
| DUSP14 | 17 | dual specificity phosphatase 14 [Source:HGNC |
| SYNRG | 17 | synergin gamma [Source:HGNC Symbol;Acc:H |

| | | |
|---|---|---|
| DDX52 | 17 | DExD-box helicase 52 [Source:HGNC Symbol;A |
| AC243585.2 | 17 | novel transcript, antisense DDX52 |
| MIR378J | 17 | microRNA 378j [Source:HGNC Symbol;Acc:HG |
| AC243571.2 | 17 | novel transcript, antisense DDX52 |
| AC243585.1 | 17 | novel transcript |
| HNF1B | 17 | HNF1 homeobox B [Source:HGNC Symbol;Acc |
| AC243571.1 | 17 | novel transcript |
| YWHAEP7 | 17 | tyrosine 3-monooxygenase/tryptophan 5-mor |
| RNU6-489P | 17 | RNA, U6 small nuclear 489, pseudogene [Sour |
| TBC1D3K | 17 | TBC1 domain family member 3K [Source:HGN |
| 5S_rRNA | 17 | 5S ribosomal RNA [Source:RFAM;Acc:RF00001 |
| TBC1D3L | 17 | TBC1 domain family member 3L [Source:HGN |
| TBC1D3D | 17 | TBC1 domain family member 3D [Source:HGN |
| TBC1D3C | 17 | TBC1 domain family member 3C [Source:HGN |
| AC233968.1 | 17 | aminopeptidase puromycin sensitive (NPEPPS |
| Y_RNA | 17 | Y RNA [Source:RFAM;Acc:RF00019] |
| TBC1D3E | 17 | TBC1 domain family member 3E [Source:HGN |
| RNA5SP526 | 17 | RNA, 5S ribosomal pseudogene 526 [Source:H |
| TBC1D3 | 17 | TBC1 domain family member 3 [Source:HGNC |
| AC244154.1 | 17 | aminopeptidase puromycin sensitive (NPEPPS |
| Y_RNA | 17 | Y RNA [Source:RFAM;Acc:RF00019] |
| MRPL45 | 17 | mitochondrial ribosomal protein L45 [Source: |
| GPR179 | 17 | G protein-coupled receptor 179 [Source:HGN |
| SOCS7 | 17 | suppressor of cytokine signaling 7 [Source:HG |
| ARHGAP23 | 17 | Rho GTPase activating protein 23 [Source:HGI |
| AC244153.1 | 17 | novel transcript, antisense to ARHGAP23 |
| SRCIN1 | 17 | SRC kinase signaling inhibitor 1 [Source:HGNC |
| AC006449.1 | 17 | novel transcript, antisense to SRCIN1 |
| EPOP | 17 | elongin BC and polycomb repressive complex |
| MIR4734 | 17 | microRNA 4734 [Source:HGNC Symbol;Acc:HG |
| AC006449.6 | 17 | novel transcript |
| AC006449.2 | 17 | novel transcript, antisense to MLLT6 |
| MLLT6 | 17 | MLLT6, PHD finger containing [Source:HGNC S |
| AC006449.3 | 17 | novel transcript, antisense to MLLT6 |
| MIR4726 | 17 | microRNA 4726 [Source:HGNC Symbol;Acc:HG |
| AC006449.7 | 17 | TEC |
| CISD3 | 17 | CDGSH iron sulfur domain 3 [Source:HGNC Syı |
| RNA5SP440 | 17 | RNA, 5S ribosomal pseudogene 440 [Source:H |
| PCGF2 | 17 | polycomb group ring finger 2 [Source:HGNC S |
| AC006449.5 | 17 | novel transcript, antisense to PCGF2 |
| PSMB3 | 17 | proteasome subunit beta 3 [Source:HGNC Syn |
| RNU6-866P | 17 | RNA, U6 small nuclear 866, pseudogene [Sour |
| PIP4K2B | 17 | phosphatidylinositol-5-phosphate 4-kinase typ |
| CWC25 | 17 | CWC25 spliceosome associated protein homo |
| MIR4727 | 17 | microRNA 4727 [Source:HGNC Symbol;Acc:HG |
| C17orf98 | 17 | chromosome 17 open reading frame 98 [Sour |
| RPL23 | 17 | ribosomal protein L23 [Source:HGNC Symbol;/ |
| SNORA21B | 17 | small nucleolar RNA, H/ACA box 21B [Source: |
| SNORA21 | 17 | small nucleolar RNA, H/ACA box 21 [Source:H |
| AC110749.1 | 17 | ribosomal protein S20 pseudogene 35 |

| | | |
|---|---|---|
| LASP1 | 17 | LIM and SH3 protein 1 [Source:HGNC Symbol; |
| AC006441.1 | 17 | novel transcript, antisense to LASP1 |
| MIR6779 | 17 | microRNA 6779 [Source:HGNC Symbol;Acc:HG |
| AC006441.3 | 17 | novel transcript, antisense to LASP1 |
| LINC00672 | 17 | long intergenic non-protein coding RNA 672 [S |
| FBXO47 | 17 | F-box protein 47 [Source:HGNC Symbol;Acc:H |
| AC006441.2 | 17 | ribosomal protein L36 (RPL36) pseudogene |
| AC006441.4 | 17 | novel transcript |
| AC006441.5 | 17 | RAD52 motif 1 (RDM1) pseudogene |
| LINC02079 | 17 | long intergenic non-protein coding RNA 2079 |
| LRRC37A11P | 17 | leucine rich repeat containing 37 member A1 |
| AC091178.1 | 17 | ADP-ribosylation factor-like 17B (ARL17B) pse |
| RDM1P5 | 17 | RDM1 pseudogene 5 [Source:HGNC Symbol;A |
| PLXDC1 | 17 | plexin domain containing 1 [Source:HGNC Syn |
| AC091178.2 | 17 | novel transcript, antisense to PLXDC1 |
| ARL5C | 17 | ADP ribosylation factor like GTPase 5C [Source |
| AC004408.2 | 17 | TEC |
| AC004408.1 | 17 | novel transcript, antisense to CACNB1 |
| CACNB1 | 17 | calcium voltage-gated channel auxiliary subun |
| RPL19 | 17 | ribosomal protein L19 [Source:HGNC Symbol; |
| STAC2 | 17 | SH3 and cysteine rich domain 2 [Source:HGNC |
| AC015910.1 | 17 | novel pseudogene |
| FBXL20 | 17 | F-box and leucine rich repeat protein 20 [Sour |
| AC005288.1 | 17 | novel transcript, antisense to MED1 & FBXL20 |
| MED1 | 17 | mediator complex subunit 1 [Source:HGNC Sy |
| CDK12 | 17 | cyclin dependent kinase 12 [Source:HGNC Syn |
| RNU6-233P | 17 | RNA, U6 small nuclear 233, pseudogene [Sour |
| AC009283.1 | 17 | novel transcript |
| NEUROD2 | 17 | neuronal differentiation 2 [Source:HGNC Syml |
| AC087491.1 | 17 | novel transcript |
| PPP1R1B | 17 | protein phosphatase 1 regulatory inhibitor sul |
| STARD3 | 17 | StAR related lipid transfer domain containing |
| TCAP | 17 | titin-cap [Source:HGNC Symbol;Acc:HGNC:116 |
| PNMT | 17 | phenylethanolamine N-methyltransferase [So |
| PGAP3 | 17 | post-GPI attachment to proteins 3 [Source:HG |
| ERBB2 | 17 | erb-b2 receptor tyrosine kinase 2 [Source:HGI |
| MIR4728 | 17 | microRNA 4728 [Source:HGNC Symbol;Acc:HG |
| MIEN1 | 17 | migration and invasion enhancer 1 [Source:HC |
| GRB7 | 17 | growth factor receptor bound protein 7 [Sour |
| IKZF3 | 17 | IKAROS family zinc finger 3 [Source:HGNC Sym |
| KRT8P34 | 17 | keratin 8 pseudogene 34 [Source:HGNC Symb |
| AC090844.1 | 17 | ribosomal protein L39 (RPL39) pseudogene |
| ZPBP2 | 17 | zona pellucida binding protein 2 [Source:HGN |
| GSDMB | 17 | gasdermin B [Source:HGNC Symbol;Acc:HGNC |
| ORMDL3 | 17 | ORMDL sphingolipid biosynthesis regulator 3 |
| AC090844.2 | 17 | novel transcript |
| LRRC3C | 17 | leucine rich repeat containing 3C [Source:HGN |
| GSDMA | 17 | gasdermin A [Source:HGNC Symbol;Acc:HGNC |
| PSMD3 | 17 | proteasome 26S subunit, non-ATPase 3 [Sourc |
| AC090844.3 | 17 | novel transcript |

| | | |
|---|---|---|
| CSF3 | 17 | colony stimulating factor 3 [Source:HGNC Syn |
| MED24 | 17 | mediator complex subunit 24 [Source:HGNC S |
| MIR6884 | 17 | microRNA 6884 [Source:HGNC Symbol;Acc:HG |
| SNORD124 | 17 | small nucleolar RNA, C/D box 124 [Source:HGI |
| THRA | 17 | thyroid hormone receptor alpha [Source:HGN |
| NR1D1 | 17 | nuclear receptor subfamily 1 group D membe |
| AC068669.1 | 17 | novel transcript |
| MSL1 | 17 | MSL complex subunit 1 [Source:HGNC Symbol |
| CASC3 | 17 | CASC3 exon junction complex subunit [Source |
| MIR6866 | 17 | microRNA 6866 [Source:HGNC Symbol;Acc:HG |
| RAPGEFL1 | 17 | Rap guanine nucleotide exchange factor like 1 |
| MIR6867 | 17 | microRNA 6867 [Source:HGNC Symbol;Acc:HG |
| WIPF2 | 17 | WAS/WASL interacting protein family membe |
| CDC6 | 17 | cell division cycle 6 [Source:HGNC Symbol;Acc |
| RARA | 17 | retinoic acid receptor alpha [Source:HGNC Syr |
| AC080112.4 | 17 | novel transcript, antisense to RARA |
| RARA-AS1 | 17 | RARA antisense RNA 1 [Source:HGNC Symbol; |
| AC080112.3 | 17 | novel transcript |
| GJD3 | 17 | gap junction protein delta 3 [Source:HGNC Syr |
| AC080112.1 | 17 | novel transcript, antisense to GJD3 |
| PPIAP54 | 17 | peptidylprolyl isomerase A pseudogene 54 [Sc |
| RNA5SP441 | 17 | RNA, 5S ribosomal pseudogene 441 [Source:H |
| AC080112.2 | 17 | kelch repeat and BTB (POZ) domain containin |
| TOP2A | 17 | DNA topoisomerase II alpha [Source:HGNC Sy |
| Y_RNA | 17 | Y RNA [Source:RFAM;Acc:RF00019] |
| RPL23AP75 | 17 | ribosomal protein L23a pseudogene 75 [Sourc |
| IGFBP4 | 17 | insulin like growth factor binding protein 4 [Sc |
| AC018629.1 | 17 | TEC |
| TNS4 | 17 | tensin 4 [Source:HGNC Symbol;Acc:HGNC:243 |
| AC004585.1 | 17 | novel transcript |
| CCR7 | 17 | C-C motif chemokine receptor 7 [Source:HGN |
| AC004585.2 | 17 | TEC |
| SMARCE1 | 17 | SWI/SNF related, matrix associated, actin dep |
| AC073508.2 | 17 | novel transcript |
| AC073508.3 | 17 | novel transcript, antisense to SMARCE1 |
| KRT222 | 17 | keratin 222 [Source:HGNC Symbol;Acc:HGNC: |
| AC073508.1 | 17 | novel keratin (KRT) family pseudogene |
| KRT24 | 17 | keratin 24 [Source:HGNC Symbol;Acc:HGNC:1 |
| KRT223P | 17 | keratin 223 pseudogene [Source:HGNC Symbo |
| KRT25 | 17 | keratin 25 [Source:HGNC Symbol;Acc:HGNC:3 |
| KRT26 | 17 | keratin 26 [Source:HGNC Symbol;Acc:HGNC:3 |
| KRT27 | 17 | keratin 27 [Source:HGNC Symbol;Acc:HGNC:3 |
| KRT28 | 17 | keratin 28 [Source:HGNC Symbol;Acc:HGNC:3 |
| AC090283.1 | 17 | novel transcript |
| KRT10 | 17 | keratin 10 [Source:HGNC Symbol;Acc:HGNC:6 |
| TMEM99 | 17 | transmembrane protein 99 (putative) [Source |
| AC004231.3 | 17 | novel transcript, antisense to KRT12 |
| KRT12 | 17 | keratin 12 [Source:HGNC Symbol;Acc:HGNC:6 |
| KRT20 | 17 | keratin 20 [Source:HGNC Symbol;Acc:HGNC:2 |
| AC004231.1 | 17 | novel transcript |

| | | |
|---|---|---|
| KRT23 | 17 | keratin 23 [Source:HGNC Symbol;Acc:HGNC:6 |
| KRT39 | 17 | keratin 39 [Source:HGNC Symbol;Acc:HGNC:3 |
| KRT40 | 17 | keratin 40 [Source:HGNC Symbol;Acc:HGNC:2 |
| KRTAP3-3 | 17 | keratin associated protein 3-3 [Source:HGNC S |
| KRTAP3-2 | 17 | keratin associated protein 3-2 [Source:HGNC S |
| KRTAP3-4P | 17 | keratin associated protein 3-4, pseudogene [S |
| KRTAP3-1 | 17 | keratin associated protein 3-1 [Source:HGNC S |
| KRTAP1-5 | 17 | keratin associated protein 1-5 [Source:HGNC S |
| KRTAP1-4 | 17 | keratin associated protein 1-4 [Source:HGNC S |
| KRTAP1-3 | 17 | keratin associated protein 1-3 [Source:HGNC S |
| KRTAP1-1 | 17 | keratin associated protein 1-1 [Source:HGNC S |
| KRTAP2-1 | 17 | keratin associated protein 2-1 [Source:HGNC S |
| KRTAP2-2 | 17 | keratin associated protein 2-2 [Source:HGNC S |
| KRTAP2-3 | 17 | keratin associated protein 2-3 [Source:HGNC S |
| KRTAP2-4 | 17 | keratin associated protein 2-4 [Source:HGNC S |
| KRTAP2-5P | 17 | keratin associated protein 2-5, pseudogene [S |
| KRTAP4-7 | 17 | keratin associated protein 4-7 [Source:HGNC S |
| AC100808.1 | 17 | novel transcript |
| KRTAP4-8 | 17 | keratin associated protein 4-8 [Source:HGNC S |
| KRTAP4-16 | 17 | keratin associated protein 4-16 [Source:HGNC |
| KRTAP4-9 | 17 | keratin associated protein 4-9 [Source:HGNC S |
| KRTAP4-11 | 17 | keratin associated protein 4-11 [Source:HGNC |
| KRTAP4-12 | 17 | keratin associated protein 4-12 [Source:HGNC |
| KRTAP4-6 | 17 | keratin associated protein 4-6 [Source:HGNC S |
| KRTAP4-5 | 17 | keratin associated protein 4-5 [Source:HGNC S |
| KRTAP4-4 | 17 | keratin associated protein 4-4 [Source:HGNC S |
| KRTAP4-3 | 17 | keratin associated protein 4-3 [Source:HGNC S |
| KRTAP4-2 | 17 | keratin associated protein 4-2 [Source:HGNC S |
| KRTAP4-1 | 17 | keratin associated protein 4-1 [Source:HGNC S |
| KRTAP4-17P | 17 | keratin associated protein 4-17, pseudogene [ |
| KRTAP9-1 | 17 | keratin associated protein 9-1 [Source:HGNC S |
| KRTAP9-12P | 17 | keratin associated protein 9-12, pseudogene [ |
| KRTAP9-2 | 17 | keratin associated protein 9-2 [Source:HGNC S |
| KRTAP9-3 | 17 | keratin associated protein 9-3 [Source:HGNC S |
| KRTAP9-8 | 17 | keratin associated protein 9-8 [Source:HGNC S |
| KRTAP9-4 | 17 | keratin associated protein 9-4 [Source:HGNC S |
| KRTAP9-9 | 17 | keratin associated protein 9-9 [Source:HGNC S |
| KRTAP9-6 | 17 | keratin associated protein 9-6 [Source:HGNC S |
| KRTAP9-11P | 17 | keratin associated protein 9-11, pseudogene [ |
| KRTAP9-7 | 17 | keratin associated protein 9-7 [Source:HGNC S |
| KRTAP9-10P | 17 | keratin associated protein 9-10, pseudogene [ |
| KRTAP29-1 | 17 | keratin associated like protein 29-1 [Source:H |
| KRTAP16-1 | 17 | keratin associated protein 16-1 [Source:HGNC |
| KRTAP17-1 | 17 | keratin associated protein 17-1 [Source:HGNC |
| TBC1D3P7 | 17 | TBC1 domain family member 3 pseudogene 7 |
| AC003958.1 | 17 | tumor necrosis factor receptor superfamily, m |
| KRT33A | 17 | keratin 33A [Source:HGNC Symbol;Acc:HGNC: |
| KRT33B | 17 | keratin 33B [Source:HGNC Symbol;Acc:HGNC: |
| KRT34 | 17 | keratin 34 [Source:HGNC Symbol;Acc:HGNC:6 |
| KRT31 | 17 | keratin 31 [Source:HGNC Symbol;Acc:HGNC:6 |

| | | |
|---|---|---|
| AC003958.2 | 17 | novel transcript |
| KRT41P | 17 | keratin 41 pseudogene [Source:HGNC Symbol |
| KRT37 | 17 | keratin 37 [Source:HGNC Symbol;Acc:HGNC:6 |
| KRT38 | 17 | keratin 38 [Source:HGNC Symbol;Acc:HGNC:6 |
| KRT43P | 17 | keratin 43 pseudogene [Source:HGNC Symbol |
| KRT32 | 17 | keratin 32 [Source:HGNC Symbol;Acc:HGNC:6 |
| RNU2-32P | 17 | RNA, U2 small nuclear 32, pseudogene [Sourc |
| KRT35 | 17 | keratin 35 [Source:HGNC Symbol;Acc:HGNC:6 |
| KRT36 | 17 | keratin 36 [Source:HGNC Symbol;Acc:HGNC:6 |
| KRT13 | 17 | keratin 13 [Source:HGNC Symbol;Acc:HGNC:6 |
| AC019349.1 | 17 | novel transcript |
| KRT15 | 17 | keratin 15 [Source:HGNC Symbol;Acc:HGNC:6 |
| MIR6510 | 17 | microRNA 6510 [Source:HGNC Symbol;Acc:HG |
| KRT19 | 17 | keratin 19 [Source:HGNC Symbol;Acc:HGNC:6 |
| LINC00974 | 17 | long intergenic non-protein coding RNA 974 [S |
| KRT9 | 17 | keratin 9 [Source:HGNC Symbol;Acc:HGNC:64 |
| KRT14 | 17 | keratin 14 [Source:HGNC Symbol;Acc:HGNC:6 |
| KRT16 | 17 | keratin 16 [Source:HGNC Symbol;Acc:HGNC:6 |
| KRT17 | 17 | keratin 17 [Source:HGNC Symbol;Acc:HGNC:6 |
| KRT42P | 17 | keratin 42 pseudogene [Source:HGNC Symbol |
| AC130686.1 | 17 | novel transcript |
| EIF1 | 17 | eukaryotic translation initiation factor 1 [Sour |
| GAST | 17 | gastrin [Source:HGNC Symbol;Acc:HGNC:4164 |
| HAP1 | 17 | huntingtin associated protein 1 [Source:HGNC |
| RNA5SP442 | 17 | RNA, 5S ribosomal pseudogene 442 [Source:H |
| RN7SL399P | 17 | RNA, 7SL, cytoplasmic 399, pseudogene [Sour |
| JUP | 17 | junction plakoglobin [Source:HGNC Symbol;Ac |
| P3H4 | 17 | prolyl 3-hydroxylase family member 4 (non-er |
| FKBP10 | 17 | FKBP prolyl isomerase 10 [Source:HGNC Symb |
| NT5C3B | 17 | 5'-nucleotidase, cytosolic IIIB [Source:HGNC Sy |
| KLHL10 | 17 | kelch like family member 10 [Source:HGNC Sy |
| AC125257.1 | 17 | novel transcript, sense overlapping to KLHL11 |
| KLHL11 | 17 | kelch like family member 11 [Source:HGNC Sy |
| ACLY | 17 | ATP citrate lyase [Source:HGNC Symbol;Acc:H |
| AC125257.2 | 17 | novel transcript, antisense to ACLY |
| TTC25 | 17 | tetratricopeptide repeat domain 25 [Source:H |
| CNP | 17 | 2',3'-cyclic nucleotide 3' phosphodiesterase [S |
| DNAJC7 | 17 | DnaJ heat shock protein family (Hsp40) memb |
| NKIRAS2 | 17 | NFKB inhibitor interacting Ras like 2 [Source:H |
| ZNF385C | 17 | zinc finger protein 385C [Source:HGNC Symbo |
| C17orf113 | 17 | chromosome 17 open reading frame 113 [Sou |
| DHX58 | 17 | DExH-box helicase 58 [Source:HGNC Symbol;A |
| KAT2A | 17 | lysine acetyltransferase 2A [Source:HGNC Sym |
| AC099811.2 | 17 | novel protein |
| HSPB9 | 17 | heat shock protein family B (small) member 9 |
| RAB5C | 17 | RAB5C, member RAS oncogene family [Source |
| AC099811.3 | 17 | novel transcript |
| KCNH4 | 17 | potassium voltage-gated channel subfamily H |
| HCRT | 17 | hypocretin neuropeptide precursor [Source:H |
| GHDC | 17 | GH3 domain containing [Source:HGNC Symbo |

| | | |
|---|---|---|
| STAT5B | 17 | signal transducer and activator of transcriptio |
| AC099811.6 | 17 | novel transcript |
| AC099811.4 | 17 | novel transcript |
| AC099811.1 | 17 | novel transcript, sense intronic to STAT5B |
| AC099811.5 | 17 | novel transcript, sense intronic to STAT5B |
| STAT5A | 17 | signal transducer and activator of transcriptio |
| STAT3 | 17 | signal transducer and activator of transcriptio |
| CAVIN1 | 17 | caveolae associated protein 1 [Source:HGNC S |
| RNU7-97P | 17 | RNA, U7 small nuclear 97 pseudogene [Source |
| ATP6V0A1 | 17 | ATPase H+ transporting V0 subunit a1 [Source |
| MIR548AT | 17 | microRNA 548at [Source:HGNC Symbol;Acc:H |
| AC107993.1 | 17 | novel transcript |
| AC067852.3 | 17 | novel transcript |
| MIR5010 | 17 | microRNA 5010 [Source:HGNC Symbol;Acc:HG |
| PTP4A2P1 | 17 | PTP4A2 pseudogene 1 [Source:HGNC Symbol; |
| NAGLU | 17 | N-acetyl-alpha-glucosaminidase [Source:HGN( |
| AC067852.1 | 17 | novel transcript |
| HSD17B1P1 | 17 | hydroxysteroid 17-beta dehydrogenase 1 pseu |
| HSD17B1 | 17 | hydroxysteroid 17-beta dehydrogenase 1 [Sou |
| AC067852.2 | 17 | novel transcript, antisense to HSD17B1 |
| COASY | 17 | Coenzyme A synthase [Source:HGNC Symbol;A |
| MLX | 17 | MAX dimerization protein MLX [Source:HGNC |
| PSMC3IP | 17 | PSMC3 interacting protein [Source:HGNC Sym |
| AC067852.5 | 17 | novel transcript, antisense to PSMC3IP |
| RETREG3 | 17 | reticulophagy regulator family member 3 [Sou |
| AC067852.4 | 17 | novel transcript |
| TUBG1 | 17 | tubulin gamma 1 [Source:HGNC Symbol;Acc:H |
| ATP5MGP7 | 17 | ATP synthase membrane subunit g pseudoger |
| HMGB3P27 | 17 | high mobility group box 3 pseudogene 27 [Sou |
| TUBG2 | 17 | tubulin gamma 2 [Source:HGNC Symbol;Acc:H |
| PLEKHH3 | 17 | pleckstrin homology, MyTH4 and FERM doma |
| CCR10 | 17 | C-C motif chemokine receptor 10 [Source:HGI |
| AC100793.2 | 17 | novel transcript, antisense to CCR10 |
| CNTNAP1 | 17 | contactin associated protein 1 [Source:HGNC |
| AC100793.3 | 17 | novel transcript, antisense to CNTNAP1 |
| EZH1 | 17 | enhancer of zeste 1 polycomb repressive com |
| MIR6780A | 17 | microRNA 6780a [Source:HGNC Symbol;Acc:H |
| AC100793.1 | 17 | ribosomal protein L34 (RPL34) pseudogene |
| AC100793.4 | 17 | novel transcript |
| HMGN2P15 | 17 | high mobility group nucleosomal binding dom |
| RAMP2-AS1 | 17 | RAMP2 antisense RNA 1 [Source:HGNC Symbo |
| RAMP2 | 17 | receptor activity modifying protein 2 [Source: |
| VPS25 | 17 | vacuolar protein sorting 25 homolog [Source:I |
| WNK4 | 17 | WNK lysine deficient protein kinase 4 [Source |
| COA3 | 17 | cytochrome c oxidase assembly factor 3 [Sour |
| CNTD1 | 17 | cyclin N-terminal domain containing 1 [Source |
| BECN1 | 17 | beclin 1 [Source:HGNC Symbol;Acc:HGNC:103 |
| MIR6781 | 17 | microRNA 6781 [Source:HGNC Symbol;Acc:HG |
| PSME3 | 17 | proteasome activator subunit 3 [Source:HGN( |
| AOC2 | 17 | amine oxidase copper containing 2 [Source:H( |

| | | |
|---|---|---|
| AOC3 | 17 | amine oxidase copper containing 3 [Source:H( |
| AOC4P | 17 | amine oxidase copper containing 4, pseudoge |
| LINC00671 | 17 | long intergenic non-protein coding RNA 671 [S |
| RNU6-287P | 17 | RNA, U6 small nuclear 287, pseudogene [Sour |
| G6PC | 17 | glucose-6-phosphatase catalytic subunit [Sour |
| AC055866.1 | 17 | NULL |
| PTGES3L-AARSD1 | 17 | PTGES3L-AARSD1 readthrough [Source:HGNC |
| AARSD1 | 17 | alanyl-tRNA synthetase domain containing 1 [ |
| PTGES3L | 17 | prostaglandin E synthase 3 like [Source:HGNC |
| RUNDC1 | 17 | RUN domain containing 1 [Source:HGNC Symb |
| RPL27 | 17 | ribosomal protein L27 [Source:HGNC Symbol;/ |
| IFI35 | 17 | interferon induced protein 35 [Source:HGNC S |
| VAT1 | 17 | vesicle amine transport 1 [Source:HGNC Symb |
| RND2 | 17 | Rho family GTPase 2 [Source:HGNC Symbol;A( |
| BRCA1 | 17 | BRCA1 DNA repair associated [Source:HGNC S |
| RPL21P4 | 17 | ribosomal protein L21 pseudogene 4 [Source: |
| NBR2 | 17 | neighbor of BRCA1 lncRNA 2 [Source:HGNC Sy |
| AC135721.2 | 17 | neighbor of BRCA1 gene 1 (NBR1) pseudogen |
| AC135721.1 | 17 | high mobility group nucleosome binding dom |
| AC060780.1 | 17 | novel transcript |
| AC060780.3 | 17 | breast cancer 1, early onset (BRCA1) pseudog |
| AC060780.2 | 17 | ribosomal protein, large, P1 (RPLP1) pseudoge |
| NBR1 | 17 | NBR1 autophagy cargo receptor [Source:HGN |
| TMEM106A | 17 | transmembrane protein 106A [Source:HGNC S |
| CCDC200 | 17 | coiled-coil domain containing 200 [Source:HG |
| RNU2-1 | 17 | RNA, U2 small nuclear 1 [Source:HGNC Symbo |
| U2 | 17 | U2 spliceosomal RNA [Source:RFAM;Acc:RF00 |
| U2 | 17 | U2 spliceosomal RNA [Source:RFAM;Acc:RF00 |
| U2 | 17 | U2 spliceosomal RNA [Source:RFAM;Acc:RF00 |
| U2 | 17 | U2 spliceosomal RNA [Source:RFAM;Acc:RF00 |
| U2 | 17 | U2 spliceosomal RNA [Source:RFAM;Acc:RF00 |
| U2 | 17 | U2 spliceosomal RNA [Source:RFAM;Acc:RF00 |
| U2 | 17 | U2 spliceosomal RNA [Source:RFAM;Acc:RF00 |
| U2 | 17 | U2 spliceosomal RNA [Source:RFAM;Acc:RF00 |
| U2 | 17 | U2 spliceosomal RNA [Source:RFAM;Acc:RF00 |
| U2 | 17 | U2 spliceosomal RNA [Source:RFAM;Acc:RF00 |
| U2 | 17 | U2 spliceosomal RNA [Source:RFAM;Acc:RF00 |
| U2 | 17 | U2 spliceosomal RNA [Source:RFAM;Acc:RF00 |
| LINC00910 | 17 | long intergenic non-protein coding RNA 910 [S |
| AC109326.1 | 17 | TEC |
| RNU2-4P | 17 | RNA, U2 small nuclear 4, pseudogene [Source |
| RNU6-1137P | 17 | RNA, U6 small nuclear 1137, pseudogene [Sou |
| ARL4D | 17 | ADP ribosylation factor like GTPase 4D [Sourc |
| RNU6-470P | 17 | RNA, U6 small nuclear 470, pseudogene [Sour |
| MIR2117HG | 17 | MIR2117 host gene [Source:HGNC Symbol;Ac |
| MIR2117 | 17 | microRNA 2117 [Source:HGNC Symbol;Acc:HG |
| RNU6-406P | 17 | RNA, U6 small nuclear 406, pseudogene [Sour |
| RNU6-971P | 17 | RNA, U6 small nuclear 971, pseudogene [Sour |
| AC087650.1 | 17 | ribosomal protein L29(RPL29) pseudogene |
| DHX8 | 17 | DEAH-box helicase 8 [Source:HGNC Symbol;A( |

| | | |
|---|---|---|
| ETV4 | 17 | ETS variant 4 [Source:HGNC Symbol;Acc:HGN( |
| Metazoa_SRP | 17 | Metazoan signal recognition particle RNA [Sou |
| MEOX1 | 17 | mesenchyme homeobox 1 [Source:HGNC Sym |
| LINC02594 | 17 | long intergenic non-protein coding RNA 2594 |
| WHSC1L2P | 17 | Wolf-Hirschhorn syndrome candidate 1-like 2, |
| SOST | 17 | sclerostin [Source:HGNC Symbol;Acc:HGNC:1: |
| DUSP3 | 17 | dual specificity phosphatase 3 [Source:HGNC : |
| CFAP97D1 | 17 | CFAP97 domain containing 1 [Source:HGNC Sy |
| AC003098.1 | 17 | novel transcript, antisense to chromosome 17 |
| MPP3 | 17 | membrane palmitoylated protein 3 [Source:H |
| CD300LG | 17 | CD300 molecule like family member g [Source |
| MPP2 | 17 | membrane palmitoylated protein 2 [Source:H |
| AC007993.3 | 17 | novel transcript, antisense to FAM215A |
| FAM215A | 17 | family with sequence similarity 215 member / |
| AC007993.1 | 17 | leucine rich repeat containing 37A (LRRC37A) |
| AC007993.2 | 17 | novel transcript |
| LINC01976 | 17 | long intergenic non-protein coding RNA 1976 |
| PPY | 17 | pancreatic polypeptide [Source:HGNC Symbol |
| PYY | 17 | peptide YY [Source:HGNC Symbol;Acc:HGNC:9 |
| NAGS | 17 | N-acetylglutamate synthase [Source:HGNC Sy |
| TMEM101 | 17 | transmembrane protein 101 [Source:HGNC Sy |
| U3 | 17 | Small nucleolar RNA U3 [Source:RFAM;Acc:RF |
| LSM12 | 17 | LSM12 homolog [Source:HGNC Symbol;Acc:H( |
| G6PC3 | 17 | glucose-6-phosphatase catalytic subunit 3 [So |
| HDAC5 | 17 | histone deacetylase 5 [Source:HGNC Symbol;/ |
| AC023855.1 | 17 | novel transcript, antisense to HDAC5 |
| RNU6-131P | 17 | RNA, U6 small nuclear 131, pseudogene [Sour |
| C17orf53 | 17 | chromosome 17 open reading frame 53 [Sour |
| ASB16 | 17 | ankyrin repeat and SOCS box containing 16 [S |
| ASB16-AS1 | 17 | ASB16 antisense RNA 1 [Source:HGNC Symbol |
| TMUB2 | 17 | transmembrane and ubiquitin like domain cor |
| ATXN7L3 | 17 | ataxin 7 like 3 [Source:HGNC Symbol;Acc:HGN |
| AC004596.1 | 17 | novel transcript, antisense to ATXN7L3 |
| UBTF | 17 | upstream binding transcription factor [Source |
| MIR6782 | 17 | microRNA 6782 [Source:HGNC Symbol;Acc:HG |
| AC003102.1 | 17 | novel transcript, antisense to UBTF |
| SHC1P2 | 17 | SHC adaptor protein 1 pseudogene 2 [Source: |
| SLC4A1 | 17 | solute carrier family 4 member 1 (Diego blooc |
| AC003043.2 | 17 | novel transcript |
| RN7SL507P | 17 | RNA, 7SL, cytoplasmic 507, pseudogene [Sour |
| RUNDC3A-AS1 | 17 | RUNDC3A antisense RNA 1 [Source:HGNC Sym |
| U3 | 17 | Small nucleolar RNA U3 [Source:RFAM;Acc:RF |
| RNU6-453P | 17 | RNA, U6 small nuclear 453, pseudogene [Sour |
| RUNDC3A | 17 | RUN domain containing 3A [Source:HGNC Sym |
| SLC25A39 | 17 | solute carrier family 25 member 39 [Source:H |
| AC003043.1 | 17 | novel transcript |
| GRN | 17 | granulin precursor [Source:HGNC Symbol;Acc: |
| FAM171A2 | 17 | family with sequence similarity 171 member / |
| RPL7L1P5 | 17 | ribosomal protein L7 like 1 pseudogene 5 [Sou |
| ITGA2B | 17 | integrin subunit alpha 2b [Source:HGNC Symb |

| | | |
|---|---|---|
| GPATCH8 | 17 | G-patch domain containing 8 [Source:HGNC S |
| RN7SL258P | 17 | RNA, 7SL, cytoplasmic 258, pseudogene [Sour |
| AC103703.1 | 17 | novel transcript, sense intronic to GPATCH8 |
| FZD2 | 17 | frizzled class receptor 2 [Source:HGNC Symbo |
| AC091152.3 | 17 | ribosomal protein L7 (RPL7) pseudogene |
| SMCO4P1 | 17 | single-pass membrane protein with coiled-coi |
| LINC01180 | 17 | long intergenic non-protein coding RNA 1180 |
| MEIOC | 17 | meiosis specific with coiled-coil domain [Sour |
| CCDC43 | 17 | coiled-coil domain containing 43 [Source:HGN |
| AC091152.2 | 17 | novel transcript, antisense to C17orf104 |
| AC091152.4 | 17 | TEC |
| DBF4B | 17 | DBF4 zinc finger B [Source:HGNC Symbol;Acc: |
| RN7SL819P | 17 | RNA, 7SL, cytoplasmic 819, pseudogene [Sour |
| AC091152.1 | 17 | ribosomal protein L6 (RPL6) pseudogene |
| ADAM11 | 17 | ADAM metallopeptidase domain 11 [Source:H |
| AC005180.2 | 17 | novel transcript |
| AC005180.1 | 17 | novel transcript |
| GJC1 | 17 | gap junction protein gamma 1 [Source:HGNC |
| HIGD1B | 17 | HIG1 hypoxia inducible domain family membe |
| EFTUD2 | 17 | elongation factor Tu GTP binding domain cont |
| RN7SL405P | 17 | RNA, 7SL, cytoplasmic 405, pseudogene [Sour |
| CCDC103 | 17 | coiled-coil domain containing 103 [Source:HG |
| FAM187A | 17 | family with sequence similarity 187 member |
| GFAP | 17 | glial fibrillary acidic protein [Source:HGNC Syn |
| KIF18B | 17 | kinesin family member 18B [Source:HGNC Syr |
| MIR6783 | 17 | microRNA 6783 [Source:HGNC Symbol;Acc:HG |
| AC015936.1 | 17 | novel transcript |
| C1QL1 | 17 | complement C1q like 1 [Source:HGNC Symbol |
| AC015936.2 | 17 | novel transcript |
| DCAKD | 17 | dephospho-CoA kinase domain containing [So |
| NMT1 | 17 | N-myristoyltransferase 1 [Source:HGNC Symb |
| Y_RNA | 17 | Y RNA [Source:RFAM;Acc:RF00019] |
| PLCD3 | 17 | phospholipase C delta 3 [Source:HGNC Symbo |
| MIR6784 | 17 | microRNA 6784 [Source:HGNC Symbol;Acc:HG |
| ACBD4 | 17 | acyl-CoA binding domain containing 4 [Source |
| AC142472.1 | 17 | novel transcript |
| HEXIM1 | 17 | HEXIM P-TEFb complex subunit 1 [Source:HGN |
| AC138150.1 | 17 | novel transcript, antisense to HEXIM1 and HE |
| HEXIM2 | 17 | HEXIM P-TEFb complex subunit 2 [Source:HGN |
| AC138150.2 | 17 | novel transcript, antisense to HEXIM2 |
| AC008105.3 | 17 | novel transcript, antisense to FMNL1 |
| FMNL1 | 17 | formin like 1 [Source:HGNC Symbol;Acc:HGNC |
| AC008105.1 | 17 | novel transcript, antisense to FMNL1 |
| AC008105.2 | 17 | novel pseudogene, ortholog of 1700023F06Ri |
| MAP3K14-AS1 | 17 | MAP3K14 antisense RNA 1 [Source:HGNC Sym |
| SPATA32 | 17 | spermatogenesis associated 32 [Source:HGNC |
| MAP3K14 | 17 | mitogen-activated protein kinase kinase kinas |
| AC003070.7 | 17 | NULL |
| RNA5SP443 | 17 | RNA, 5S ribosomal pseudogene 443 [Source:H |
| AC003070.2 | 17 | novel transcript |

| | | |
|---|---|---|
| ARHGAP27 | 17 | Rho GTPase activating protein 27 [Source:HGN |
| AC003070.1 | 17 | novel transcript, antisense to ARHGAP27 |
| PLEKHM1 | 17 | pleckstrin homology and RUN domain contain |
| AC091132.2 | 17 | novel transcript, antisense to PLEKHM1 |
| MIR4315-1 | 17 | microRNA 4315-1 [Source:HGNC Symbol;Acc:I |
| LRRC37A4P | 17 | leucine rich repeat containing 37 member A4, |
| Metazoa_SRP | 17 | Metazoan signal recognition particle RNA [Sou |
| AC091132.3 | 17 | TEC |
| AC091132.1 | 17 | novel transcript |
| AC091132.4 | 17 | novel transcript |
| AC091132.5 | 17 | novel transcript |
| RDM1P1 | 17 | RDM1 pseudogene 1 [Source:HGNC Symbol;A |
| DND1P1 | 17 | DND microRNA-mediated repression inhibitor |
| AC126544.2 | 17 | novel transcript |
| AC126544.1 | 17 | TEC |
| MAPK8IP1P2 | 17 | mitogen-activated protein kinase 8 interacting |
| RPS26P8 | 17 | ribosomal protein S26 pseudogene 8 [Source: |
| LINC02210 | 17 | long intergenic non-protein coding RNA 2210 |
| LINC02210-CRHR1 | 17 | LINC02210-CRHR1 readthrough [Source:HGNC |
| AC217774.1 | 17 | novel transcript |
| AC217774.2 | 17 | novel transcript |
| CRHR1 | 17 | corticotropin releasing hormone receptor 1 [S |
| MAPT-AS1 | 17 | MAPT antisense RNA 1 [Source:HGNC Symbol |
| SPPL2C | 17 | signal peptide peptidase like 2C [Source:HGNC |
| MAPT | 17 | microtubule associated protein tau [Source:H |
| MAPT-IT1 | 17 | MAPT intronic transcript 1 [Source:HGNC Sym |
| CR936218.2 | 17 | novel transcript, antisense to MAPT |
| Metazoa_SRP | 17 | Metazoan signal recognition particle RNA [Sou |
| STH | 17 | saitohin [Source:HGNC Symbol;Acc:HGNC:188 |
| KANSL1 | 17 | KAT8 regulatory NSL complex subunit 1 [Sourc |
| CR936218.1 | 17 | novel transcript, antisense to a novel protein |
| KANSL1-AS1 | 17 | KANSL1 antisense RNA 1 [Source:HGNC Symb |
| Y_RNA | 17 | Y RNA [Source:RFAM;Acc:RF00019] |
| MAPK8IP1P1 | 17 | mitogen-activated protein kinase 8 interacting |
| AC005829.2 | 17 | dead end homolog 1 (zebrafish) (DND1) pseu |
| AC005829.1 | 17 | C17orf58 pseudogene |
| ARL17B | 17 | ADP ribosylation factor like GTPase 17B [Sour |
| LRRC37A | 17 | leucine rich repeat containing 37A [Source:HG |
| RN7SL656P | 17 | RNA, 7SL, cytoplasmic 656, pseudogene [Sour |
| NSFP1 | 17 | N-ethylmaleimide-sensitive factor pseudogen |
| RDM1P2 | 17 | RDM1 pseudogene 2 [Source:HGNC Symbol;A |
| LRRC37A2 | 17 | leucine rich repeat containing 37 member A2 |
| ARL17A | 17 | ADP ribosylation factor like GTPase 17A [Sour |
| RN7SL199P | 17 | RNA, 7SL, cytoplasmic 199, pseudogene [Sour |
| FAM215B | 17 | family with sequence similarity 215 member E |
| NSF | 17 | N-ethylmaleimide sensitive factor, vesicle fusi |
| RPS7P11 | 17 | ribosomal protein S7 pseudogene 11 [Source: |
| WNT3 | 17 | Wnt family member 3 [Source:HGNC Symbol; |
| WNT9B | 17 | Wnt family member 9B [Source:HGNC Symbol |
| LINC01974 | 17 | long intergenic non-protein coding RNA 1974 |

| | | |
|---|---|---|
| AC005670.1 | 17 | novel transcript |
| RNU6ATAC3P | 17 | RNA, U6atac small nuclear 3, pseudogene [Sou |
| GOSR2 | 17 | golgi SNAP receptor complex member 2 [Sour |
| AC005670.2 | 17 | novel protein |
| MIR5089 | 17 | microRNA 5089 [Source:HGNC Symbol;Acc:HG |
| RPRML | 17 | reprimo like [Source:HGNC Symbol;Acc:HGNC |
| LRRC37A17P | 17 | leucine rich repeat containing 37 member A17 |
| AC005670.3 | 17 | novel transcript |
| RN7SL270P | 17 | RNA, 7SL, cytoplasmic 270, pseudogene [Sour |
| CDC27 | 17 | cell division cycle 27 [Source:HGNC Symbol;Ac |
| AC002558.1 | 17 | ribosomal protein S2 (RPS2) pseudogene |
| AC002558.3 | 17 | novel transcript, sense intronic to CDC27 |
| AC002558.2 | 17 | novel transcript, antisense to CDC27 |
| MYL4 | 17 | myosin light chain 4 [Source:HGNC Symbol;Ac |
| ITGB3 | 17 | integrin subunit beta 3 [Source:HGNC Symbol; |
| AC068234.1 | 17 | novel protein |
| RNU7-186P | 17 | RNA, U7 small nuclear 186 pseudogene [Sourc |
| AC068234.3 | 17 | novel transcript, antisense to ITGB3 |
| AC068234.2 | 17 | novel transcript, antisense to ITGB3 |
| EFCAB13 | 17 | EF-hand calcium binding domain 13 [Source:H |
| AC040934.1 | 17 | novel transcript |
| MRPL45P2 | 17 | mitochondrial ribosomal protein L45 pseudog |
| NPEPPS | 17 | aminopeptidase puromycin sensitive [Source: |
| Y_RNA | 17 | Y RNA [Source:RFAM;Acc:RF00019] |
| AC025682.1 | 17 | novel transcript, antisense to NPEPPS |
| KPNB1 | 17 | karyopherin subunit beta 1 [Source:HGNC Syn |
| AC015674.1 | 17 | novel transcript, antisense to KPNB1 |
| TBKBP1 | 17 | TBK1 binding protein 1 [Source:HGNC Symbol; |
| TBX21 | 17 | T-box transcription factor 21 [Source:HGNC Sy |
| OSBPL7 | 17 | oxysterol binding protein like 7 [Source:HGNC |
| MRPL10 | 17 | mitochondrial ribosomal protein L10 [Source:I |
| LRRC46 | 17 | leucine rich repeat containing 46 [Source:HGN |
| SCRN2 | 17 | secernin 2 [Source:HGNC Symbol;Acc:HGNC:3 |
| SP6 | 17 | Sp6 transcription factor [Source:HGNC Symbo |
| AC018521.3 | 17 | novel transcript |
| AC018521.7 | 17 | novel transcript |
| AC018521.5 | 17 | novel transcript |
| SP2 | 17 | Sp2 transcription factor [Source:HGNC Symbo |
| SP2-AS1 | 17 | SP2 antisense RNA 1 [Source:HGNC Symbol;Ac |
| AC018521.6 | 17 | novel transcript, antisense to SP2 |
| PNPO | 17 | pyridoxamine 5'-phosphate oxidase [Source:H |
| AC018521.1 | 17 | novel transcript |
| AC018521.2 | 17 | novel transcript, antisense to PNPO |
| PRR15L | 17 | proline rich 15 like [Source:HGNC Symbol;Acc: |
| CDK5RAP3 | 17 | CDK5 regulatory subunit associated protein 3 |
| AC018521.4 | 17 | novel transcript, antisense to CDK5RAP3 |
| COPZ2 | 17 | coatomer protein complex subunit zeta 2 [Sou |
| MIR152 | 17 | microRNA 152 [Source:HGNC Symbol;Acc:HGI |
| AC004477.1 | 17 | novel transcript, antisense to NFE2L1 |
| NFE2L1 | 17 | nuclear factor, erythroid 2 like 1 [Source:HGN |

| | | |
|---|---|---|
| AC004477.2 | 17 | novel transcript, antisense to NFE2L1 |
| AC004477.3 | 17 | novel transcript |
| CBX1 | 17 | chromobox 1 [Source:HGNC Symbol;Acc:HGN |
| RNU6-1201P | 17 | RNA, U6 small nuclear 1201, pseudogene [Sou |
| SNX11 | 17 | sorting nexin 11 [Source:HGNC Symbol;Acc:HG |
| SKAP1 | 17 | src kinase associated phosphoprotein 1 [Sourc |
| MIR1203 | 17 | microRNA 1203 [Source:HGNC Symbol;Acc:HG |
| SKAP1-AS1 | 17 | SKAP1 antisense RNA 1 [Source:HGNC Symbol |
| RNU6-1152P | 17 | RNA, U6 small nuclear 1152, pseudogene [Sou |
| THRA1/BTR | 17 | uncharacterized LOC105371807 [Source:NCBI |
| AC036222.2 | 17 | novel transcript, sense intronic to SKAP1 |
| U3 | 17 | Small nucleolar RNA U3 [Source:RFAM;Acc:RF |
| AC036222.1 | 17 | novel transcript |
| Y_RNA | 17 | Y RNA [Source:RFAM;Acc:RF00019] |
| HOXB1 | 17 | homeobox B1 [Source:HGNC Symbol;Acc:HGN |
| HOXB2 | 17 | homeobox B2 [Source:HGNC Symbol;Acc:HGN |
| HOXB-AS1 | 17 | HOXB cluster antisense RNA 1 [Source:HGNC S |
| HOXB3 | 17 | homeobox B3 [Source:HGNC Symbol;Acc:HGN |
| HOXB-AS3 | 17 | HOXB cluster antisense RNA 3 [Source:HGNC S |
| HOXB-AS2 | 17 | HOXB cluster antisense RNA 2 [Source:HGNC S |
| HOXB4 | 17 | homeobox B4 [Source:HGNC Symbol;Acc:HGN |
| AC103702.1 | 17 | novel transcript, sense intronic to HOXB3 |
| MIR10A | 17 | microRNA 10a [Source:HGNC Symbol;Acc:HGN |
| HOXB5 | 17 | homeobox B5 [Source:HGNC Symbol;Acc:HGN |
| HOXB6 | 17 | homeobox B6 [Source:HGNC Symbol;Acc:HGN |
| HOXB7 | 17 | homeobox B7 [Source:HGNC Symbol;Acc:HGN |
| HOXB8 | 17 | homeobox B8 [Source:HGNC Symbol;Acc:HGN |
| HOXB9 | 17 | homeobox B9 [Source:HGNC Symbol;Acc:HGN |
| HOXB-AS4 | 17 | HOXB cluster antisense RNA 4 [Source:HGNC S |
| MIR196A1 | 17 | microRNA 196a-1 [Source:HGNC Symbol;Acc:H |
| AC103702.2 | 17 | novel transcript |
| LINC02086 | 17 | long intergenic non-protein coding RNA 2086 |
| AC091179.2 | 17 | ribosomal protein L9 (RPL9) pseudogene |
| RPL9P28 | 17 | ribosomal protein L9 pseudogene 28 [Source: |
| COX6B1P2 | 17 | cytochrome c oxidase subunit 6B1 pseudogen |
| PRAC1 | 17 | PRAC1 small nuclear protein [Source:HGNC Sy |
| PRAC2 | 17 | PRAC2 small nuclear protein [Source:HGNC Sy |
| AC091179.3 | 17 | NULL |
| AC091179.4 | 17 | NULL |
| MIR3185 | 17 | microRNA 3185 [Source:HGNC Symbol;Acc:HG |
| HOXB13 | 17 | homeobox B13 [Source:HGNC Symbol;Acc:HG |
| AC091179.1 | 17 | novel transcript |
| TTLL6 | 17 | tubulin tyrosine ligase like 6 [Source:HGNC Sy |
| RN7SL125P | 17 | RNA, 7SL, cytoplasmic 125, pseudogene [Sour |
| CALCOCO2 | 17 | calcium binding and coiled-coil domain 2 [Sou |
| SUMO2P17 | 17 | SUMO2 pseudogene 17 [Source:HGNC Symbo |
| AC068531.1 | 17 | NULL |
| AC068531.2 | 17 | pseudogene similar to part of ribosomal prote |
| ATP5MC1 | 17 | ATP synthase membrane subunit c locus 1 [So |
| AC091133.6 | 17 | ribosomal protein L37 (RPL37) pseudogene |

| | | |
|---|---|---|
| UBE2Z | 17 | ubiquitin conjugating enzyme E2 Z [Source:HG |
| SNF8 | 17 | SNF8 subunit of ESCRT-II [Source:HGNC Symbo |
| AC091133.1 | 17 | novel transcript |
| RNU1-42P | 17 | RNA, U1 small nuclear 42, pseudogene [Sourc |
| AC091133.2 | 17 | NULL |
| GIP | 17 | gastric inhibitory polypeptide [Source:HGNC S |
| AC091133.3 | 17 | novel transcript |
| IGF2BP1 | 17 | insulin like growth factor 2 mRNA binding pro |
| AC091133.4 | 17 | novel transcript, antisense to IGF2BP1 |
| Metazoa_SRP | 17 | Metazoan signal recognition particle RNA [Sou |
| RNU6-826P | 17 | RNA, U6 small nuclear 826, pseudogene [Sour |
| AC091133.5 | 17 | COX assembly mitochondrial protein homolog |
| B4GALNT2P1 | 17 | B4GALNT2 pseudogene 1 [Source:HGNC Symb |
| B4GALNT2 | 17 | beta-1,4-N-acetyl-galactosaminyltransferase 2 |
| AC069454.1 | 17 | ribosomal protein S10 (RPS10) pseudogene |
| GNGT2 | 17 | G protein subunit gamma transducin 2 [Sourc |
| ABI3 | 17 | ABI family member 3 [Source:HGNC Symbol;A |
| PHOSPHO1 | 17 | phosphoethanolamine/phosphocholine phosp |
| AC004797.1 | 17 | novel transcript FLJ40194 |
| MIR6129 | 17 | microRNA 6129 [Source:HGNC Symbol;Acc:HG |
| ZNF652 | 17 | zinc finger protein 652 [Source:HGNC Symbol; |
| AC091180.2 | 17 | novel transcript |
| AC091180.5 | 17 | novel transcript, antisense to PHB |
| AC091180.4 | 17 | novel transcript |
| AC091180.1 | 17 | ribosomal protein L21 (RPL21) pseudogene |
| PHB | 17 | prohibitin [Source:HGNC Symbol;Acc:HGNC:8 |
| AC091180.3 | 17 | novel transcript, antisense to PHB |
| Y_RNA | 17 | Y RNA [Source:RFAM;Acc:RF00019] |
| EIF4EP2 | 17 | eukaryotic translation initiation factor 4E pseu |
| LINC02075 | 17 | long intergenic non-protein coding RNA 2075 |
| AC015656.1 | 17 | TEC |
| NGFR | 17 | nerve growth factor receptor [Source:HGNC S |
| AC006487.1 | 17 | novel transcript |
| MIR6165 | 17 | microRNA 6165 [Source:HGNC Symbol;Acc:HG |
| AC006487.2 | 17 | novel transcript |
| NXPH3 | 17 | neurexophilin 3 [Source:HGNC Symbol;Acc:HG |
| SPOP | 17 | speckle type BTB/POZ protein [Source:HGNC S |
| SLC35B1 | 17 | solute carrier family 35 member B1 [Source:H |
| AC015795.1 | 17 | novel transcript |
| FAM117A | 17 | family with sequence similarity 117 member A |
| KAT7 | 17 | lysine acetyltransferase 7 [Source:HGNC Symb |
| SRP14P3 | 17 | signal recognition particle 14 pseudogene 3 [S |
| AC027801.4 | 17 | novel transcript |
| TAC4 | 17 | tachykinin 4 [Source:HGNC Symbol;Acc:HGNC |
| FLJ45513 | 17 | uncharacterized LOC729220 [Source:NCBI ger |
| AC027801.1 | 17 | novel transcript |
| AC027801.3 | 17 | novel transcript |
| AC027801.2 | 17 | mRNA turnover 4 homolog (S. cerevisiae) (MR |
| RNU6-1313P | 17 | RNA, U6 small nuclear 1313, pseudogene [Sou |
| DLX4 | 17 | distal-less homeobox 4 [Source:HGNC Symbol |

| | | |
|---|---|---|
| DLX3 | 17 | distal-less homeobox 3 [Source:HGNC Symbol |
| AC009720.1 | 17 | novel transcript |
| PICART1 | 17 | p53 inducible cancer associated RNA transcrip |
| ITGA3 | 17 | integrin subunit alpha 3 [Source:HGNC Symbo |
| AC002401.4 | 17 | novel transcript |
| PDK2 | 17 | pyruvate dehydrogenase kinase 2 [Source:HG |
| AC002401.3 | 17 | novel transcript, antisense to PDK2 |
| AC002401.2 | 17 | novel transcript, antisense to PDK2 |
| SAMD14 | 17 | sterile alpha motif domain containing 14 [Sou |
| PPP1R9B | 17 | protein phosphatase 1 regulatory subunit 9B [ |
| AC002401.1 | 17 | novel transcript, antisense to PPP1R9B |
| AC015909.2 | 17 | novel transcript |
| AC015909.3 | 17 | novel transcript, antisense to SGCA |
| SGCA | 17 | sarcoglycan alpha [Source:HGNC Symbol;Acc:I |
| HILS1 | 17 | histone linker H1 domain, spermatid-specific : |
| COL1A1 | 17 | collagen type I alpha 1 chain [Source:HGNC Sy |
| AC015909.1 | 17 | novel transcript |
| AC015909.4 | 17 | novel transcript |
| SUMO2P7 | 17 | SUMO2 pseudogene 7 [Source:HGNC Symbol; |
| TMEM92 | 17 | transmembrane protein 92 [Source:HGNC Syn |
| TMEM92-AS1 | 17 | TMEM92 antisense RNA 1 [Source:HGNC Sym |
| XYLT2 | 17 | xylosyltransferase 2 [Source:HGNC Symbol;Ac |
| AC015909.5 | 17 | TEC |
| MRPL27 | 17 | mitochondrial ribosomal protein L27 [Source:I |
| EME1 | 17 | essential meiotic structure-specific endonucle |
| LRRC59 | 17 | leucine rich repeat containing 59 [Source:HGN |
| Y_RNA | 17 | Y RNA [Source:RFAM;Acc:RF00019] |
| AC004707.1 | 17 | novel transcript, antisense to LRRC59 |
| ACSF2 | 17 | acyl-CoA synthetase family member 2 [Source |
| MRPS21P9 | 17 | mitochondrial ribosomal protein S21 pseudog |
| CHAD | 17 | chondroadherin [Source:HGNC Symbol;Acc:H( |
| AC021491.4 | 17 | novel transcript |
| RSAD1 | 17 | radical S-adenosyl methionine domain contair |
| AC021491.1 | 17 | novel transcript |
| MYCBPAP | 17 | MYCBP associated protein [Source:HGNC Sym |
| AC021491.2 | 17 | novel transcript, antisense to EPN3 |
| EPN3 | 17 | epsin 3 [Source:HGNC Symbol;Acc:HGNC:182: |
| SPATA20 | 17 | spermatogenesis associated 20 [Source:HGNC |
| CACNA1G-AS1 | 17 | CACNA1G antisense RNA 1 [Source:HGNC Syn |
| CACNA1G | 17 | calcium voltage-gated channel subunit alpha1 |
| AC021491.3 | 17 | novel transcript |
| AC004590.1 | 17 | novel transcript, antisense to CACNA1G |
| ABCC3 | 17 | ATP binding cassette subfamily C member 3 [S |
| ANKRD40 | 17 | ankyrin repeat domain 40 [Source:HGNC Syml |
| AC005921.2 | 17 | novel transcript, antisense to ANKRD40 |
| Y_RNA | 17 | Y RNA [Source:RFAM;Acc:RF00019] |
| LUC7L3 | 17 | LUC7 like 3 pre-mRNA splicing factor [Source:I |
| AC005921.3 | 17 | TEC |
| ANKRD40CL | 17 | ANKRD40 C-terminal like [Source:HGNC Symb |
| MIR8059 | 17 | microRNA 8059 [Source:HGNC Symbol;Acc:H( |

| | | |
|---|---|---|
| AC005921.4 | 17 | novel transcript |
| AC005921.1 | 17 | S100 calcium binding protein A10 (S100A10) p |
| WFIKKN2 | 17 | WAP, follistatin/kazal, immunoglobulin, kunitz |
| AC091062.1 | 17 | novel transcript, antisense to WFIKKN2 |
| RPL5P33 | 17 | ribosomal protein L5 pseudogene 33 [Source: |
| TOB1 | 17 | transducer of ERBB2, 1 [Source:HGNC Symbol |
| TOB1-AS1 | 17 | TOB1 antisense RNA 1 [Source:HGNC Symbol; |
| AC005920.4 | 17 | novel transcript |
| AC005920.1 | 17 | novel transcript |
| AC005920.3 | 17 | novel transcript |
| SPAG9 | 17 | sperm associated antigen 9 [Source:HGNC Syr |
| AC005920.2 | 17 | novel transcript |
| NME1 | 17 | NME/NM23 nucleoside diphosphate kinase 1 |
| NME1-NME2 | 17 | NME1-NME2 readthrough [Source:HGNC Sym |
| NME2 | 17 | NME/NM23 nucleoside diphosphate kinase 2 |
| MBTD1 | 17 | mbt domain containing 1 [Source:HGNC Symb |
| AC005839.1 | 17 | TEC |
| AC006141.1 | 17 | novel transcript, sense intronic to MBTD1 |
| UTP18 | 17 | UTP18 small subunit processome component |
| LINC02071 | 17 | long intergenic non-protein coding RNA 2071 |
| AC005823.2 | 17 | novel transcript |
| AC005823.1 | 17 | novel transcript |
| LINC02073 | 17 | long intergenic non-protein coding RNA 2073 |
| RPL7P48 | 17 | ribosomal protein L7 pseudogene 48 [Source: |
| CA10 | 17 | carbonic anhydrase 10 [Source:HGNC Symbol; |
| AC002090.1 | 17 | TEC |
| LINC01982 | 17 | long intergenic non-protein coding RNA 1982 |
| LINC02089 | 17 | long intergenic non-protein coding RNA 2089 |
| C17orf112 | 17 | chromosome 17 open reading frame 112 [Sou |
| MTCO1P40 | 17 | MT-CO1 pseudogene 40 [Source:HGNC Symbo |
| AC034268.2 | 17 | novel transcript, antisense to KIF2B |
| AC005341.1 | 17 | mitogen-activated protein kinase 1 interacting |
| AC090079.1 | 17 | phosphatidylinositol 4-kinase type 2 alpha (PI4 |
| AC090079.2 | 17 | tubulin, beta class I (TUBB) pseudogene |
| AC034268.1 | 17 | proteasome (prosome, macropain) 26S subun |
| RPS2P48 | 17 | ribosomal protein S2 pseudogene 48 [Source: |
| KIF2B | 17 | kinesin family member 2B [Source:HGNC Sym |
| AC023934.1 | 17 | Zinc finger protein pseudogene |
| ISCA1P3 | 17 | iron-sulfur cluster assembly 1 pseudogene 3 [S |
| AC015943.1 | 17 | novel transcript |
| AC005951.1 | 17 | myosin, light chain 12-like, regulatory pseudog |
| ARL2BPP8 | 17 | ADP ribosylation factor like GTPase 2 binding p |
| RN7SKP14 | 17 | RN7SK pseudogene 14 [Source:HGNC Symbol; |
| TOM1L1 | 17 | target of myb1 like 1 membrane trafficking pr |
| COX11 | 17 | cytochrome c oxidase copper chaperone COX1 |
| AC007485.2 | 17 | TEC |
| AC007485.1 | 17 | novel transcript, antisense to COX11 |
| STXBP4 | 17 | syntaxin binding protein 4 [Source:HGNC Sym |
| HLF | 17 | HLF transcription factor, PAR bZIP family mem |
| AC007638.2 | 17 | novel transcript, antisense to HLF |

| | | |
|---|---|---|
| AC007638.1 | 17 | novel transcript |
| MMD | 17 | monocyte to macrophage differentiation asso |
| SMIM36 | 17 | small integral membrane protein 36 [Source:H |
| AC105021.1 | 17 | peptidase D (PEPD) pseudogene |
| GARSP1 | 17 | glycyl-tRNA synthetase pseudogene 1 [Source |
| RNU6-1249P | 17 | RNA, U6 small nuclear 1249, pseudogene [Sou |
| TMEM100 | 17 | transmembrane protein 100 [Source:HGNC Sy |
| PCTP | 17 | phosphatidylcholine transfer protein [Source: |
| AC090618.1 | 17 | novel transcript |
| ANKFN1 | 17 | ankyrin repeat and fibronectin type III domain |
| AC006600.1 | 17 | phosphatase and tensin homologphosphatase |
| AC006600.2 | 17 | TEC |
| RPL39P33 | 17 | ribosomal protein L39 pseudogene 33 [Source |
| NOG | 17 | noggin [Source:HGNC Symbol;Acc:HGNC:7866 |
| C17orf67 | 17 | chromosome 17 open reading frame 67 [Sour |
| AC106858.1 | 17 | novel transcript, sense intronic to C17orf67 |
| DGKE | 17 | diacylglycerol kinase epsilon [Source:HGNC Sy |
| TRIM25 | 17 | tripartite motif containing 25 [Source:HGNC S |
| RNU6-1158P | 17 | RNA, U6 small nuclear 1158, pseudogene [Sou |
| AC015912.2 | 17 | family with sequence similarity 89, member B |
| AC015912.1 | 17 | novel transcript, antisense to TRIM25 |
| MIR3614 | 17 | microRNA 3614 [Source:HGNC Symbol;Acc:HG |
| AC015912.3 | 17 | novel transcript |
| COIL | 17 | coilin [Source:HGNC Symbol;Acc:HGNC:2184] |
| AC004584.1 | 17 | novel transcript, antisense to COIL |
| AC004584.2 | 17 | ribosomal protein S15a (RPS15A) pseudogene |
| SCPEP1 | 17 | serine carboxypeptidase 1 [Source:HGNC Sym |
| AC004584.3 | 17 | novel transcript, sense intronic to SCPEP1 |
| RNF126P1 | 17 | ring finger protein 126 pseudogene 1 [Source: |
| AC007114.1 | 17 | novel transcript |
| AKAP1 | 17 | A-kinase anchoring protein 1 [Source:HGNC Sy |
| AC007114.2 | 17 | novel transcript, antisense to AKAP1 |
| AC003950.1 | 17 | novel transcript |
| AC091181.2 | 17 | novel transcript, antisense to MSI2 |
| MSI2 | 17 | musashi RNA binding protein 2 [Source:HGNC |
| AC091181.1 | 17 | TEC |
| AC015883.1 | 17 | TEC |
| AC007431.2 | 17 | novel transcript, antisense to MSI2 |
| AC007431.3 | 17 | TEC |
| AC007431.1 | 17 | novel transcript, antisense to MSI2 |
| RN7SL449P | 17 | RNA, 7SL, cytoplasmic 449, pseudogene [Sour |
| RNU7-134P | 17 | RNA, U7 small nuclear 134 pseudogene [Sourc |
| CCDC182 | 17 | coiled-coil domain containing 182 [Source:HG |
| AC015845.2 | 17 | novel transcript |
| RN7SKP94 | 17 | RN7SK pseudogene 94 [Source:HGNC Symbol; |
| MRPS23 | 17 | mitochondrial ribosomal protein S23 [Source: |
| CUEDC1 | 17 | CUE domain containing 1 [Source:HGNC Symb |
| AC015845.1 | 17 | novel transcript, antisense to CUEDC1 |
| AC015813.4 | 17 | novel transcript |
| VEZF1 | 17 | vascular endothelial zinc finger 1 [Source:HGN |

| | | |
|---|---|---|
| AC015813.7 | 17 | novel transcript |
| AC015813.2 | 17 | novel transcript |
| AC015813.1 | 17 | novel transcript |
| SRSF1 | 17 | serine and arginine rich splicing factor 1 [Sour |
| AC015813.5 | 17 | novel transcript |
| AC015813.8 | 17 | novel transcript |
| AC015813.3 | 17 | novel transcript |
| DYNLL2 | 17 | dynein light chain LC8-type 2 [Source:HGNC Sy |
| AC015813.6 | 17 | novel transcript |
| OR4D1 | 17 | olfactory receptor family 4 subfamily D memb |
| MSX2P1 | 17 | msh homeobox 2 pseudogene 1 [Source:HGN |
| OR4D2 | 17 | olfactory receptor family 4 subfamily D memb |
| EPX | 17 | eosinophil peroxidase [Source:HGNC Symbol; |
| AC005962.1 | 17 | novel transcript, antisense to EPX |
| MKS1 | 17 | MKS transition zone complex subunit 1 [Sourc |
| LPO | 17 | lactoperoxidase [Source:HGNC Symbol;Acc:HC |
| AC005962.2 | 17 | novel transcript, antisense to MKS1 |
| AC004687.3 | 17 | novel transcript, antisense to LPO |
| MPO | 17 | myeloperoxidase [Source:HGNC Symbol;Acc:H |
| TSPOAP1 | 17 | TSPO associated protein 1 [Source:HGNC Sym |
| TSPOAP1-AS1 | 17 | TSPOAP1, SUPT4H1 and RNF43 antisense RNA |
| AC004687.1 | 17 | novel transcript, MIR142 host |
| MIR142 | 17 | microRNA 142 [Source:HGNC Symbol;Acc:HGI |
| MIR4736 | 17 | microRNA 4736 [Source:HGNC Symbol;Acc:HG |
| SUPT4H1 | 17 | SPT4 homolog, DSIF elongation factor subunit |
| AC004687.2 | 17 | novel transcript |
| RNF43 | 17 | ring finger protein 43 [Source:HGNC Symbol;A |
| HSF5 | 17 | heat shock transcription factor 5 [Source:HGN |
| SETP3 | 17 | SET pseudogene 3 [Source:HGNC Symbol;Acc: |
| MTMR4 | 17 | myotubularin related protein 4 [Source:HGNC |
| SEPTIN4-AS1 | 17 | SEPT4 antisense RNA 1 [Source:HGNC Symbol |
| SEPTIN4 | 17 | septin 4 [Source:HGNC Symbol;Acc:HGNC:916 |
| TEX14 | 17 | testis expressed 14, intercellular bridge formi |
| U3 | 17 | Small nucleolar RNA U3 [Source:RFAM;Acc:RF |
| AC011195.2 | 17 | novel protein similar to immunoglobulin (CD7 |
| RNU1-108P | 17 | RNA, U1 small nuclear 108, pseudogene [Sour |
| AC011195.1 | 17 | ribosomal protein L36a-like (RPL36AL) pseudo |
| RNU1-52P | 17 | RNA, U1 small nuclear 52, pseudogene [Sourc |
| RNU1-85P | 17 | RNA, U1 small nuclear 85, pseudogene [Sourc |
| RAD51C | 17 | RAD51 paralog C [Source:HGNC Symbol;Acc:H |
| AC025521.1 | 17 | novel transcript |
| PPM1E | 17 | protein phosphatase, Mg2+/Mn2+ dependent |
| Metazoa_SRP | 17 | Metazoan signal recognition particle RNA [Sou |
| RNU6-518P | 17 | RNA, U6 small nuclear 518, pseudogene [Sour |
| AC100832.1 | 17 | ribosomal protein S12 (RPS12) pseudogene |
| Metazoa_SRP | 17 | Metazoan signal recognition particle RNA [Sou |
| AC100832.2 | 17 | novel transcript, sense intronic to PPM1E |
| TRIM37 | 17 | tripartite motif containing 37 [Source:HGNC S |
| RN7SL716P | 17 | RNA, 7SL, cytoplasmic 716, pseudogene [Sour |
| AC099850.1 | 17 | novel transcript |

| | | |
|---|---|---|
| SKA2 | 17 | spindle and kinetochore associated complex s |
| RNU2-58P | 17 | RNA, U2 small nuclear 58, pseudogene [Sourc |
| MIR454 | 17 | microRNA 454 [Source:HGNC Symbol;Acc:HGI |
| MIR301A | 17 | microRNA 301a [Source:HGNC Symbol;Acc:HG |
| PRR11 | 17 | proline rich 11 [Source:HGNC Symbol;Acc:HGI |
| SPDYE22P | 17 | speedy/RINGO cell cycle regulator family men |
| AC099850.2 | 17 | novel protein |
| AC099850.3 | 17 | novel transcript, antisense to PRR11 |
| SMG8 | 17 | SMG8 nonsense mediated mRNA decay factor |
| GDPD1 | 17 | glycerophosphodiester phosphodiesterase do |
| Y_RNA | 17 | Y RNA [Source:RFAM;Acc:RF00019] |
| SNRPGP17 | 17 | small nuclear ribonucleoprotein polypeptide ( |
| YPEL2 | 17 | yippee like 2 [Source:HGNC Symbol;Acc:HGNC |
| AC091059.2 | 17 | TEC |
| MIR4729 | 17 | microRNA 4729 [Source:HGNC Symbol;Acc:HG |
| AC091059.1 | 17 | novel transcript, antisense to YPEL2 |
| LINC01476 | 17 | long intergenic non-protein coding RNA 1476 |
| RNU4-13P | 17 | RNA, U4 small nuclear 13, pseudogene [Sourc |
| DHX40 | 17 | DEAH-box helicase 40 [Source:HGNC Symbol;/ |
| AC091271.1 | 17 | novel transcript |
| CLTC | 17 | clathrin heavy chain [Source:HGNC Symbol;Ac |
| PTRH2 | 17 | peptidyl-tRNA hydrolase 2 [Source:HGNC Sym |
| VMP1 | 17 | vacuole membrane protein 1 [Source:HGNC S |
| AC040904.1 | 17 | novel transcript, sense intronic to VMP1 |
| MIR21 | 17 | microRNA 21 [Source:HGNC Symbol;Acc:HGN |
| RNU6-450P | 17 | RNA, U6 small nuclear 450, pseudogene [Sour |
| TUBD1 | 17 | tubulin delta 1 [Source:HGNC Symbol;Acc:HGI |
| NDUFB8P2 | 17 | NADH:ubiquinone oxidoreductase subunit B8 |
| RPS6KB1 | 17 | ribosomal protein S6 kinase B1 [Source:HGNC |
| RPS29P21 | 17 | ribosomal protein S29 pseudogene 21 [Source |
| AC005702.1 | 17 | novel protein |
| RNFT1 | 17 | ring finger protein, transmembrane 1 [Source: |
| TBC1D3P1-DHX40P1 | 17 | TBC1D3P1-DHX40P1 readthrough, transcribec |
| RNFT1-DT | 17 | RNFT1 divergent transcript [Source:HGNC Sym |
| DHX40P1 | 17 | DEAH-box helicase 40 pseudogene 1 [Source:I |
| TBC1D3P1 | 17 | TBC1 domain family member 3 pseudogene 1 |
| AC005702.5 | 17 | uncharacterized LOC102723851 [Source:NCBI |
| AC005702.4 | 17 | tumor necrosis factor receptor superfamily, m |
| AC005702.3 | 17 | double C2-like domains, beta (DOC2B) pseudo |
| MIR4737 | 17 | microRNA 4737 [Source:HGNC Symbol;Acc:HG |
| HEATR6 | 17 | HEAT repeat containing 6 [Source:HGNC Symb |
| AC005702.2 | 17 | UDP-N-acetyl-alpha-D-galactosamine:polypep |
| AC025048.4 | 17 | novel transcript |
| WFDC21P | 17 | WAP four-disulfide core domain 21, pseudoge |
| AC025048.6 | 17 | adaptor-related protein complex 1, sigma 2 su |
| AC025048.3 | 17 | double C2-like domains, beta (DOC2B) pseudo |
| AC025048.2 | 17 | novel transcript |
| AC025048.5 | 17 | tumor necrosis factor receptor superfamily, m |
| AC025048.7 | 17 | novel transcript |
| AC025048.1 | 17 | novel transcript |

| | | |
|---|---|---|
| CA4 | 17 | carbonic anhydrase 4 [Source:HGNC Symbol;A |
| USP32 | 17 | ubiquitin specific peptidase 32 [Source:HGNC |
| SCARNA20 | 17 | small Cajal body-specific RNA 20 [Source:HGN |
| AC104763.1 | 17 | ribosomal protein L32 (RPL32) pseudogene |
| AC104763.3 | 17 | septin 7 (SEPT7) pseudogene |
| AC104763.2 | 17 | H3 histone, family 3B (H3.3B) (H3F3B) pseudo |
| C17orf64 | 17 | chromosome 17 open reading frame 64 [Sour |
| RPL12P38 | 17 | ribosomal protein L12 pseudogene 38 [Source |
| APPBP2 | 17 | amyloid beta precursor protein binding protei |
| AC011921.3 | 17 | splicing factor 3B, 14 kDa subunit (SF3B14) ps |
| AC011921.1 | 17 | novel transcript, antisense to APPBP2 |
| AC011921.2 | 17 | sema domain, immunoglobulin domain (Ig), sh |
| LINC01999 | 17 | long intergenic non-protein coding RNA 1999 |
| RPSAP66 | 17 | ribosomal protein SA pseudogene 66 [Source: |
| PPM1D | 17 | protein phosphatase, Mg2+/Mn2+ dependent |
| RNU6-623P | 17 | RNA, U6 small nuclear 623, pseudogene [Sour |
| BCAS3 | 17 | BCAS3 microtubule associated cell migration f |
| RN7SL606P | 17 | RNA, 7SL, cytoplasmic 606, pseudogene [Sour |
| AC110602.1 | 17 | ribosomal protein L36a (RPL36A) pseudogene |
| Y_RNA | 17 | Y RNA [Source:RFAM;Acc:RF00019] |
| KRT18P61 | 17 | keratin 18 pseudogene 61 [Source:HGNC Sym |
| HMGN1P28 | 17 | high mobility group nucleosome binding doma |
| RN7SL448P | 17 | RNA, 7SL, cytoplasmic 448, pseudogene [Sour |
| AC005884.1 | 17 | novel transcript, antisense to BCAS3 |
| AC005884.2 | 17 | novel transcript, antisense to BCAS3 |
| RPL23AP74 | 17 | ribosomal protein L23a pseudogene 74 [Sourc |
| AC005856.1 | 17 | novel transcript, antisense to BCAS3 |
| AC005746.3 | 17 | novel transcript, antisense to BCAS3 |
| AC005746.2 | 17 | novel transcript, antisense to BCAS3 |
| AC005746.1 | 17 | novel transcript, sense intronic to BCAS3 |
| TBX2-AS1 | 17 | TBX2 antisense RNA 1 [Source:HGNC Symbol;/ |
| TBX2 | 17 | T-box transcription factor 2 [Source:HGNC Syr |
| C17orf82 | 17 | chromosome 17 open reading frame 82 [Sour |
| TBX4 | 17 | T-box transcription factor 4 [Source:HGNC Syr |
| AC005901.1 | 17 | novel transcript, antisense to TBX4 |
| NACA2 | 17 | nascent polypeptide associated complex subu |
| BRIP1 | 17 | BRCA1 interacting protein C-terminal helicase |
| INTS2 | 17 | integrator complex subunit 2 [Source:HGNC S |
| Y_RNA | 17 | Y RNA [Source:RFAM;Acc:RF00019] |
| MED13 | 17 | mediator complex subunit 13 [Source:HGNC S |
| AC018628.1 | 17 | TEC |
| RN7SL800P | 17 | RNA, 7SL, cytoplasmic 800, pseudogene [Sour |
| AC018628.2 | 17 | novel transcript, antisense to MED13 |
| Y_RNA | 17 | Y RNA [Source:RFAM;Acc:RF00019] |
| Y_RNA | 17 | Y RNA [Source:RFAM;Acc:RF00019] |
| POLRMTP1 | 17 | RNA polymerase mitochondrial pseudogene 1 |
| AC053481.3 | 17 | ribosomal protein L36a pseudogene 47 |
| AC053481.4 | 17 | nitric oxide synthase 2, inducible (NOS2) pseu |
| TBC1D3P2 | 17 | TBC1 domain family member 3 pseudogene 2 |
| AC053481.2 | 17 | ubiquitin specific peptidase 32 (USP32) pseud |

| | | |
|---|---|---|
| AC053481.1 | 17 | ribosomal protein L36a (RPL36A) pseudogene |
| EFCAB3 | 17 | EF-hand calcium binding domain 3 [Source:HG |
| AC008026.2 | 17 | ribosomal protein, large, P2 (RPLP2) pseudoge |
| RNU7-52P | 17 | RNA, U7 small nuclear 52 pseudogene [Source |
| RPS23P7 | 17 | ribosomal protein S23 pseudogene 7 [Source: |
| METTL2A | 17 | methyltransferase like 2A [Source:HGNC Symb |
| TLK2 | 17 | tousled like kinase 2 [Source:HGNC Symbol;Ac |
| AC008026.1 | 17 | ribosomal protein S10 (RPS10) pseudogene |
| AC008026.3 | 17 | novel transcript |
| AC080038.2 | 17 | novel transcript |
| AC080038.1 | 17 | novel transcript |
| MRC2 | 17 | mannose receptor C type 2 [Source:HGNC Syn |
| Y_RNA | 17 | Y RNA [Source:RFAM;Acc:RF00019] |
| AC080038.3 | 17 | TEC |
| RNU6-446P | 17 | RNA, U6 small nuclear 446, pseudogene [Sour |
| AC005821.1 | 17 | novel transcript, antisense to MARCH10 |
| MARCH10 | 17 | membrane associated ring-CH-type finger 10 |
| AC005821.2 | 17 | novel transcript, antisense to MARCH10 |
| MARCH10-DT | 17 | MARCH10 divergent transcript [Source:HGNC |
| PRELID3BP3 | 17 | PRELI domain containing 3B pseudogene 3 [Sc |
| AC005972.3 | 17 | novel transcript |
| MIR633 | 17 | microRNA 633 [Source:HGNC Symbol;Acc:HGI |
| TRMT112P3 | 17 | tRNA methyltransferase subunit 11-2 pseudog |
| AC005972.1 | 17 | ribosomal protein L29 (RPL29) pseudogene |
| AC005972.2 | 17 | high mobility group nucleosomal binding dom |
| TANC2 | 17 | tetratricopeptide repeat, ankyrin repeat and c |
| AC006270.3 | 17 | zinc finger protein 329 (ZNF329) pseudogene |
| AC006270.1 | 17 | novel transcript, antisense to TANC2 |
| AC006270.2 | 17 | DnaJ (Hsp40) homolog, subfamily B, member |
| AC015923.1 | 17 | novel transcript, antisense to TANC2 |
| AC005828.2 | 17 | novel transcript, antisense to TANC2 |
| AC005828.7 | 17 | density-regulated protein (DENR) pseudogene |
| AC005828.1 | 17 | novel transcript, antisense to TANC2 |
| AC005828.3 | 17 | novel transcript, antisense to TANC2 |
| AC005828.6 | 17 | mitochondrially encoded cytochrome c oxidas |
| AC005828.4 | 17 | novel transcript |
| CYB561 | 17 | cytochrome b561 [Source:HGNC Symbol;Acc:I |
| AC005828.5 | 17 | novel transcript |
| PPIAP55 | 17 | peptidylprolyl isomerase A pseudogene 55 [Sc |
| ACE | 17 | angiotensin I converting enzyme [Source:HGN |
| AC113554.1 | 17 | novel protein |
| ACE3P | 17 | angiotensin I converting enzyme 3, pseudoger |
| KCNH6 | 17 | potassium voltage-gated channel subfamily H |
| DCAF7 | 17 | DDB1 and CUL4 associated factor 7 [Source:H |
| RNU6-288P | 17 | RNA, U6 small nuclear 288, pseudogene [Sour |
| TACO1 | 17 | translational activator of cytochrome c oxidas |
| AC046185.2 | 17 | family with sequence similarity 136, member |
| MAP3K3 | 17 | mitogen-activated protein kinase kinase kinas |
| EEF1DP7 | 17 | eukaryotic translation elongation factor 1 delt |
| STRADA | 17 | STE20 related adaptor alpha [Source:HGNC Sy |

| | | |
|---|---|---|
| LIMD2 | 17 | LIM domain containing 2 [Source:HGNC Symb |
| AC046185.3 | 17 | TEC |
| AC046185.1 | 17 | novel transcript |
| CCDC47 | 17 | coiled-coil domain containing 47 [Source:HGN |
| DDX42 | 17 | DEAD-box helicase 42 [Source:HGNC Symbol;/ |
| FTSJ3 | 17 | FtsJ RNA 2'-O-methyltransferase 3 [Source:HG |
| PSMC5 | 17 | proteasome 26S subunit, ATPase 5 [Source:H( |
| SMARCD2 | 17 | SWI/SNF related, matrix associated, actin dep |
| TCAM1P | 17 | testicular cell adhesion molecule 1, pseudoge |
| CSH2 | 17 | chorionic somatomammotropin hormone 2 [S |
| GH2 | 17 | growth hormone 2 [Source:HGNC Symbol;Acc |
| AC040958.1 | 17 | intercellular adhesion molecule (ICAM) pseud |
| CSH1 | 17 | chorionic somatomammotropin hormone 1 [S |
| AC127029.1 | 17 | intercellular adhesion molecule (ICAM) pseud |
| CSHL1 | 17 | chorionic somatomammotropin hormone like |
| GH1 | 17 | growth hormone 1 [Source:HGNC Symbol;Acc |
| AC127029.3 | 17 | novel protein |
| CD79B | 17 | CD79b molecule [Source:HGNC Symbol;Acc:H |
| SCN4A | 17 | sodium voltage-gated channel alpha subunit 4 |
| AC127029.2 | 17 | novel transcript |
| PRR29-AS1 | 17 | PRR29 antisense RNA 1 [Source:HGNC Symbol |
| PRR29 | 17 | proline rich 29 [Source:HGNC Symbol;Acc:HGI |
| ICAM2 | 17 | intercellular adhesion molecule 2 [Source:HGI |
| ERN1 | 17 | endoplasmic reticulum to nucleus signaling 1 |
| AC005803.1 | 17 | novel transcript, antisense to ERN1 |
| SNHG25 | 17 | small nucleolar RNA host gene 25 [Source:HGI |
| SNORD104 | 17 | small nucleolar RNA, C/D box 104 [Source:HGI |
| AC025362.1 | 17 | NULL |
| SNORA50C | 17 | small nucleolar RNA, H/ACA box 50C [Source:I |
| TEX2 | 17 | testis expressed 2 [Source:HGNC Symbol;Acc:I |
| RPL31P57 | 17 | ribosomal protein L31 pseudogene 57 [Source |
| PECAM1 | 17 | platelet and endothelial cell adhesion molecul |
| Y_RNA | 17 | Y RNA [Source:RFAM;Acc:RF00019] |
| AC234063.1 | 17 | NULL |
| MILR1 | 17 | mast cell immunoglobulin like receptor 1 [Sou |
| POLG2 | 17 | DNA polymerase gamma 2, accessory subunit |
| ATP5MFP4 | 17 | ATP synthase membrane subunit f pseudogen |
| DDX5 | 17 | DEAD-box helicase 5 [Source:HGNC Symbol;A |
| MIR3064 | 17 | microRNA 3064 [Source:HGNC Symbol;Acc:HG |
| MIR5047 | 17 | microRNA 5047 [Source:HGNC Symbol;Acc:HG |
| CEP95 | 17 | centrosomal protein 95 [Source:HGNC Symbo |
| SMURF2 | 17 | SMAD specific E3 ubiquitin protein ligase 2 [S |
| AC009994.1 | 17 | G protein-coupled receptor 128 (GPR128) pse |
| MICOS10P2 | 17 | MICOS10 pseudogene 2 [Source:HGNC Symbo |
| ARHGAP27P1-BPTFP1-KPI | 17 | ARHGAP27P1-BPTFP1-KPNA2P3 readthrough, |
| KPNA2P3 | 17 | karyopherin subunit alpha 2 pseudogene 3 [S |
| AC132812.1 | 17 | small nuclear ribonucleoprotein 200kDa (U5) |
| BPTFP1 | 17 | bromodomain PHD finger transcription factor |
| AC103810.3 | 17 | Rho GTPase-activating protein 27-like protein |
| PLEKHM1P1 | 17 | pleckstrin homology and RUN domain contain |

| | | |
|---|---|---|
| MIR6080 | 17 | microRNA 6080 [Source:HGNC Symbol;Acc:HG |
| MIR4315-2 | 17 | microRNA 4315-2 [Source:HGNC Symbol;Acc:H |
| RNU7-115P | 17 | RNA, U7 small nuclear 115 pseudogene [Sourc |
| AC103810.9 | 17 | family with sequence similarity 215, member |
| LRRC37A3 | 17 | leucine rich repeat containing 37 member A3 |
| RN7SL404P | 17 | RNA, 7SL, cytoplasmic 404, pseudogene [Sour |
| AC103810.6 | 17 | neighbor of BRCA1 gene 2 (non-protein coding |
| AC103810.7 | 17 | ADP-ribosylation factor-like 17B (ARL17B) pse |
| AC103810.2 | 17 | novel transcript, antisense to LRRC37A3 |
| AC103810.5 | 17 | novel transcript, antisense to LRRC37A3 |
| RDM1P4 | 17 | RDM1 pseudogene 4 [Source:HGNC Symbol;A |
| AC103810.1 | 17 | bromodomain PHD finger transcription factor |
| AC103810.4 | 17 | pleckstrin homology domain containing, famil |
| AC037487.4 | 17 | novel transcript |
| AC103810.8 | 17 | pleckstrin homology domain containing, famil |
| AC037487.1 | 17 | novel transcript |
| AC037487.3 | 17 | eyes shut homolog (Drosophila) 9 (EYS) pseud |
| SLC16A6P1 | 17 | SLC16A6 pseudogene 1 [Source:HGNC Symbol |
| AMZ2P1 | 17 | archaelysin family metallopeptidase 2 pseudo |
| GNA13 | 17 | G protein subunit alpha 13 [Source:HGNC Sym |
| AC037487.2 | 17 | novel transcript, antisense to GNA13 |
| RGS9 | 17 | regulator of G protein signaling 9 [Source:HGN |
| AC060771.1 | 17 | Up-regulated during skeletal muscle growth p |
| LINC02563 | 17 | long intergenic non-protein coding RNA 2563 |
| AXIN2 | 17 | axin 2 [Source:HGNC Symbol;Acc:HGNC:904] |
| AC004805.1 | 17 | novel transcript |
| CEP112 | 17 | centrosomal protein 112 [Source:HGNC Symb |
| AC004805.3 | 17 | ribosomal protein S7 (RPS7) pseudogene |
| AC004805.2 | 17 | dynactin 6 (DCTN6) pseudogene |
| PSMD7P1 | 17 | proteasome 26S subunit, non-ATPase 7 pseud |
| APOH | 17 | apolipoprotein H [Source:HGNC Symbol;Acc:H |
| RNA5SP444 | 17 | RNA, 5S ribosomal pseudogene 444 [Source:H |
| AC009452.1 | 17 | mitochondrially encoded ATP synthase 6 (MT- |
| PRKCA | 17 | protein kinase C alpha [Source:HGNC Symbol; |
| RN7SL735P | 17 | RNA, 7SL, cytoplasmic 735, pseudogene [Sour |
| PRKCA-AS1 | 17 | PRKCA antisense RNA 1 [Source:HGNC Symbo |
| RNU6-928P | 17 | RNA, U6 small nuclear 928, pseudogene [Sour |
| RNA5SP445 | 17 | RNA, 5S ribosomal pseudogene 445 [Source:H |
| AC006947.1 | 17 | novel transcript |
| MIR634 | 17 | microRNA 634 [Source:HGNC Symbol;Acc:HGN |
| CACNG5 | 17 | calcium voltage-gated channel auxiliary subur |
| RNA5SP446 | 17 | RNA, 5S ribosomal pseudogene 446 [Source:H |
| CACNG4 | 17 | calcium voltage-gated channel auxiliary subur |
| AC005544.1 | 17 | novel transcript |
| AC005544.2 | 17 | novel transcript |
| CACNG1 | 17 | calcium voltage-gated channel auxiliary subur |
| HELZ | 17 | helicase with zinc finger [Source:HGNC Symbo |
| AC007448.3 | 17 | novel transcript |
| RPL36AP48 | 17 | ribosomal protein L36a pseudogene 48 [Sourc |
| AC007448.5 | 17 | novel transcript, antisense to HELZ |

| | | |
|---|---|---|
| AC007448.4 | 17 | novel transcript |
| RNA5SP447 | 17 | RNA, 5S ribosomal pseudogene 447 [Source:H |
| AC007448.1 | 17 | NULL |
| AC007448.2 | 17 | keratin 18 (KRT18) pseudogene |
| PSMD12 | 17 | proteasome 26S subunit, non-ATPase 12 [Sou |
| PITPNC1 | 17 | phosphatidylinositol transfer protein cytoplas |
| Y_RNA | 17 | Y RNA [Source:RFAM;Acc:RF00019] |
| RN7SL756P | 17 | RNA, 7SL, cytoplasmic 756, pseudogene [Sour |
| MIR548D2 | 17 | microRNA 548d-2 [Source:HGNC Symbol;Acc:I |
| MIR548AA2 | 17 | microRNA 548aa-2 [Source:HGNC Symbol;Acc |
| AC110921.1 | 17 | novel transcript |
| AC079331.2 | 17 | TEC |
| AC079331.1 | 17 | TEC |
| AC079331.3 | 17 | novel transcript, antisense to PITPNC1 |
| NOL11 | 17 | nucleolar protein 11 [Source:HGNC Symbol;Ac |
| SNORA38B | 17 | small nucleolar RNA, H/ACA box 38B [Source:I |
| RPSAP67 | 17 | ribosomal protein SA pseudogene 67 [Source: |
| RPL17P41 | 17 | ribosomal protein L17 pseudogene 41 [Source |
| BPTF | 17 | bromodomain PHD finger transcription factor |
| AC134407.1 | 17 | novel transcript, antisense to BPTF |
| AC134407.2 | 17 | TEC |
| AC134407.3 | 17 | TEC |
| RN7SL622P | 17 | RNA, 7SL, cytoplasmic 622, pseudogene [Sour |
| C17orf58 | 17 | chromosome 17 open reading frame 58 [Sour |
| KPNA2 | 17 | karyopherin subunit alpha 2 [Source:HGNC Sy |
| FBXO36P1 | 17 | F-box protein 36 pseudogene 1 [Source:HGNC |
| AC145343.1 | 17 | novel transcript |
| LINC00674 | 17 | long intergenic non-protein coding RNA 674 [S |
| LRRC37A16P | 17 | leucine rich repeat containing 37 member A1( |
| AC005332.6 | 17 | novel transcript |
| AC005332.3 | 17 | novel transcript |
| AC005332.5 | 17 | novel transcript |
| SH3GL1P3 | 17 | SH3 domain containing GRB2 like 1, endophili |
| RDM1P3 | 17 | RDM1 pseudogene 3 [Source:HGNC Symbol;A |
| AC005332.7 | 17 | novel transcript |
| AC005332.2 | 17 | novel transcript |
| ARHGAP27P2 | 17 | Rho GTPase activating protein 27 pseudogene |
| AC005332.4 | 17 | novel transcript |
| AC005332.1 | 17 | novel transcript, antisense to AMZ2 |
| AMZ2 | 17 | archaelysin family metallopeptidase 2 [Source |
| ARSG | 17 | arylsulfatase G [Source:HGNC Symbol;Acc:HGI |
| SLC16A6 | 17 | solute carrier family 16 member 6 [Source:HG |
| AC007780.1 | 17 | novel transcript |
| WIPI1 | 17 | WD repeat domain, phosphoinositide interact |
| MIR635 | 17 | microRNA 635 [Source:HGNC Symbol;Acc:HGI |
| PRKAR1A | 17 | protein kinase cAMP-dependent type I regulat |
| FAM20A | 17 | FAM20A golgi associated secretory pathway p |
| AC079210.1 | 17 | novel transcript |
| LINC01482 | 17 | long intergenic non-protein coding RNA 1482 |
| AC011591.2 | 17 | novel pseudogene |

| | | |
|---|---|---|
| AC011591.1 | 17 | novel transcript |
| ABCA8 | 17 | ATP binding cassette subfamily A member 8 [S |
| ABCA9-AS1 | 17 | ABCA9 antisense RNA 1 [Source:HGNC Symbo |
| ABCA9 | 17 | ATP binding cassette subfamily A member 9 [S |
| ABCA6 | 17 | ATP binding cassette subfamily A member 6 [S |
| SEC24AP1 | 17 | SEC24 homolog A pseudogene 1 [Source:HGN |
| MIR4524B | 17 | microRNA 4524b [Source:HGNC Symbol;Acc:H |
| MIR4524A | 17 | microRNA 4524a [Source:HGNC Symbol;Acc:H |
| ABCA10 | 17 | ATP binding cassette subfamily A member 10 |
| AC005495.1 | 17 | TEC |
| AC005495.2 | 17 | novel transcript, antisense to ABCA10 |
| ABCA5 | 17 | ATP binding cassette subfamily A member 5 [S |
| SNRPGP4 | 17 | small nuclear ribonucleoprotein polypeptide G |
| AC115085.1 | 17 | NULL |
| MAP2K6 | 17 | mitogen-activated protein kinase kinase 6 [So |
| AC015920.1 | 17 | TEC |
| AC002546.1 | 17 | novel transcript |
| LINC01483 | 17 | long intergenic non-protein coding RNA 1483 |
| AC002545.1 | 17 | IMP1 inner mitochondrial membrane peptidas |
| LINC01497 | 17 | long intergenic non-protein coding RNA 1497 |
| Y_RNA | 17 | Y RNA [Source:RFAM;Acc:RF00019] |
| AC005208.1 | 17 | novel transcript |
| LINC01028 | 17 | long intergenic non-protein coding RNA 1028 |
| KCNJ16 | 17 | potassium voltage-gated channel subfamily J r |
| KCNJ2-AS1 | 17 | KCNJ2 antisense RNA 1 [Source:HGNC Symbol |
| KCNJ2 | 17 | potassium voltage-gated channel subfamily J r |
| CALM2P1 | 17 | calmodulin 2 pseudogene 1 [Source:HGNC Syr |
| AC011990.1 | 17 | novel transcript |
| AC005771.1 | 17 | member RAS oncogene family (RAB9) pseudog |
| AC007423.1 | 17 | novel transcript |
| AC005181.1 | 17 | SERPINE1 mRNA binding protein 1 (SERBP1) p |
| CASC17 | 17 | cancer susceptibility 17 [Source:HGNC Symbo |
| RNU7-155P | 17 | RNA, U7 small nuclear 155 pseudogene [Sourc |
| AC118653.1 | 17 | novel transcript |
| MYL6P5 | 17 | myosin light chain 6 pseudogene 5 [Source:HG |
| AC007432.1 | 17 | novel transcript |
| AC005144.1 | 17 | novel transcript |
| ROCR | 17 | regulator of chondrogenesis RNA [Source:HGN |
| LINC01152 | 17 | long intergenic non-protein coding RNA 1152 |
| SOX9-AS1 | 17 | SOX9 antisense RNA 1 [Source:HGNC Symbol;A |
| LINC02097 | 17 | long intergenic non-protein coding RNA 2097 |
| SOX9 | 17 | SRY-box transcription factor 9 [Source:HGNC S |
| LINC00511 | 17 | long intergenic non-protein coding RNA 511 [S |
| LINC02003 | 17 | long intergenic non-protein coding RNA 2003 |
| AC007639.1 | 17 | novel transcript |
| RPL32P33 | 17 | ribosomal protein L32 pseudogene 33 [Source |
| AC080037.2 | 17 | novel transcript |
| SLC39A11 | 17 | solute carrier family 39 member 11 [Source:H |
| RN7SKP180 | 17 | RN7SK pseudogene 180 [Source:HGNC Symbo |
| AC080037.1 | 17 | NULL |

| | | |
|---|---|---|
| AC011120.1 | 17 | novel transcript, antisense to SLC39A11 |
| AC025198.1 | 17 | ATG12 autophagy related 12 homolog (S. cere |
| POLR3KP2 | 17 | RNA polymerase III subunit K pseudogene 2 [S |
| AC097641.1 | 17 | novel transcript |
| SSTR2 | 17 | somatostatin receptor 2 [Source:HGNC Symbo |
| COG1 | 17 | component of oligomeric golgi complex 1 [Sou |
| AC097641.2 | 17 | novel transcript, antisense to COG1 |
| FAM104A | 17 | family with sequence similarity 104 member A |
| C17orf80 | 17 | chromosome 17 open reading frame 80 [Sour |
| AC087301.1 | 17 | novel transcript, sense intronic to C17orf80 |
| CPSF4L | 17 | cleavage and polyadenylation specific factor 4 |
| CDC42EP4 | 17 | CDC42 effector protein 4 [Source:HGNC Symb |
| SDK2 | 17 | sidekick cell adhesion molecule 2 [Source:HGN |
| AC124804.1 | 17 | novel transcript, antisense to SDK2 |
| AC032019.1 | 17 | novel transcript, antisense to SDK2 |
| AC032019.2 | 17 | novel transcript |
| AC125421.1 | 17 | uncharacterized LOC100134391 [Source:NCBI |
| LINC00469 | 17 | long intergenic non-protein coding RNA 469 [S |
| LINC02092 | 17 | long intergenic non-protein coding RNA 2092 |
| AC125421.2 | 17 | novel transcript |
| AC137735.1 | 17 | novel transcript |
| AC137735.3 | 17 | novel transcript |
| AC137735.2 | 17 | lincRNA |
| LINC02074 | 17 | long intergenic non-protein coding RNA 2074 |
| RPL38 | 17 | ribosomal protein L38 [Source:HGNC Symbol;A |
| AC100786.1 | 17 | novel transcript, antisense to RPL38 |
| TTYH2 | 17 | tweety family member 2 [Source:HGNC Symbo |
| AC100786.2 | 17 | novel transcript, antisense to TTYH2 |
| DNAI2 | 17 | dynein axonemal intermediate chain 2 [Sourc |
| AC103809.1 | 17 | novel transcript, antisense to DNAI2 |
| KIF19 | 17 | kinesin family member 19 [Source:HGNC Sym |
| BTBD17 | 17 | BTB domain containing 17 [Source:HGNC Sym |
| GPR142 | 17 | G protein-coupled receptor 142 [Source:HGN |
| RNA5SP448 | 17 | RNA, 5S ribosomal pseudogene 448 [Source:H |
| GPRC5C | 17 | G protein-coupled receptor class C group 5 m |
| CD300A | 17 | CD300a molecule [Source:HGNC Symbol;Acc:H |
| CD300LB | 17 | CD300 molecule like family member b [Source |
| CD300C | 17 | CD300c molecule [Source:HGNC Symbol;Acc:H |
| AC079325.1 | 17 | CD300 antigen pseudogene |
| CD300H | 17 | CD300H molecule (gene/pseudogene) [Source |
| CD300LD | 17 | CD300 molecule like family member d [Source |
| C17orf77 | 17 | chromosome 17 open reading frame 77 [Sour |
| AC064805.1 | 17 | uncharacterized LOC101928343 [Source:NCBI |
| CD300E | 17 | CD300e molecule [Source:HGNC Symbol;Acc:H |
| AC064805.2 | 17 | novel transcript, antisense to RAB37 |
| RAB37 | 17 | RAB37, member RAS oncogene family [Source |
| CD300LF | 17 | CD300 molecule like family member f [Source |
| AC016888.1 | 17 | novel transcript, antisense to SLC9A3R1 |
| MIR3615 | 17 | microRNA 3615 [Source:HGNC Symbol;Acc:HG |
| SLC9A3R1 | 17 | SLC9A3 regulator 1 [Source:HGNC Symbol;Acc |

| | | |
|---|---|---|
| NAT9 | 17 | N-acetyltransferase 9 (putative) [Source:HGN( |
| TMEM104 | 17 | transmembrane protein 104 [Source:HGNC Sy |
| GRIN2C | 17 | glutamate ionotropic receptor NMDA type sul |
| FDXR | 17 | ferredoxin reductase [Source:HGNC Symbol;A |
| FADS6 | 17 | fatty acid desaturase 6 [Source:HGNC Symbol; |
| USH1G | 17 | USH1 protein network component sans [Sour( |
| OTOP2 | 17 | otopetrin 2 [Source:HGNC Symbol;Acc:HGNC: |
| OTOP3 | 17 | otopetrin 3 [Source:HGNC Symbol;Acc:HGNC: |
| HID1 | 17 | HID1 domain containing [Source:HGNC Symbo |
| HID1-AS1 | 17 | HID1 antisense RNA 1 [Source:HGNC Symbol;/ |
| CDR2L | 17 | cerebellar degeneration related protein 2 like |
| MRPL58 | 17 | mitochondrial ribosomal protein L58 [Source:I |
| RNU6-362P | 17 | RNA, U6 small nuclear 362, pseudogene [Sour |
| KCTD2 | 17 | potassium channel tetramerization domain cc |
| ATP5PD | 17 | ATP synthase peripheral stalk subunit d [Sour( |
| RN7SL573P | 17 | RNA, 7SL, cytoplasmic 573, pseudogene [Sour |
| AC111186.1 | 17 | WD repeat domain, phosphoinositide interact |
| TRIM80P | 17 | tripartite motif containing 80, pseudogene [Sc |
| SLC16A5 | 17 | solute carrier family 16 member 5 [Source:HG |
| Y_RNA | 17 | Y RNA [Source:RFAM;Acc:RF00019] |
| ARMC7 | 17 | armadillo repeat containing 7 [Source:HGNC S |
| NT5C | 17 | 5', 3'-nucleotidase, cytosolic [Source:HGNC Sy |
| AC022211.4 | 17 | TEC |
| JPT1 | 17 | Jupiter microtubule associated homolog 1 [So |
| AC022211.3 | 17 | novel transcript, sense intronic to HN1 |
| AC022211.1 | 17 | novel transcript, sense intronic to HN1 |
| Y_RNA | 17 | Y RNA [Source:RFAM;Acc:RF00019] |
| SUMO2 | 17 | small ubiquitin like modifier 2 [Source:HGNC S |
| NUP85 | 17 | nucleoporin 85 [Source:HGNC Symbol;Acc:HG |
| GGA3 | 17 | golgi associated, gamma adaptin ear containir |
| MRPS7 | 17 | mitochondrial ribosomal protein S7 [Source:H |
| MIF4GD | 17 | MIF4G domain containing [Source:HGNC Syml |
| AC022211.2 | 17 | novel transcript, antisense to SLC25A19 |
| SLC25A19 | 17 | solute carrier family 25 member 19 [Source:H |
| GRB2 | 17 | growth factor receptor bound protein 2 [Sour( |
| AC011933.1 | 17 | ribosomal protein L36a (RPL36A) pseudogene |
| AC011933.4 | 17 | novel transcript, antisense to GRB2 |
| AC011933.2 | 17 | novel transcript, sense intronic to GRB2 |
| AC011933.3 | 17 | novel transcript, antisense to GRB2 |
| MIR3678 | 17 | microRNA 3678 [Source:HGNC Symbol;Acc:HG |
| RNU6-938P | 17 | RNA, U6 small nuclear 938, pseudogene [Sour |
| Y_RNA | 17 | Y RNA [Source:RFAM;Acc:RF00019] |
| TMEM94 | 17 | transmembrane protein 94 [Source:HGNC Syn |
| MIR6785 | 17 | microRNA 6785 [Source:HGNC Symbol;Acc:HG |
| CASKIN2 | 17 | CASK interacting protein 2 [Source:HGNC Sym |
| TSEN54 | 17 | tRNA splicing endonuclease subunit 54 [Sourc |
| LLGL2 | 17 | LLGL scribble cell polarity complex componen |
| MYO15B | 17 | myosin XVB [Source:HGNC Symbol;Acc:HGNC: |
| RECQL5 | 17 | RecQ like helicase 5 [Source:HGNC Symbol;Ac |
| SMIM5 | 17 | small integral membrane protein 5 [Source:H( |

| | | |
|---|---|---|
| SMIM6 | 17 | small integral membrane protein 6 [Source:H( |
| SAP30BP | 17 | SAP30 binding protein [Source:HGNC Symbol; |
| AC087749.2 | 17 | novel transcript, antisense to SAP30BP |
| AC087749.1 | 17 | novel transcript, antisense to SAP30BP |
| ITGB4 | 17 | integrin subunit beta 4 [Source:HGNC Symbol; |
| GALK1 | 17 | galactokinase 1 [Source:HGNC Symbol;Acc:HG |
| H3F3B | 17 | H3 histone family member 3B [Source:HGNC S |
| MIR4738 | 17 | microRNA 4738 [Source:HGNC Symbol;Acc:HG |
| UNK | 17 | unk zinc finger [Source:HGNC Symbol;Acc:HGI |
| AC087289.1 | 17 | novel transcript, antisense to UNK |
| UNC13D | 17 | unc-13 homolog D [Source:HGNC Symbol;Acc: |
| WBP2 | 17 | WW domain binding protein 2 [Source:HGNC |
| TRIM47 | 17 | tripartite motif containing 47 [Source:HGNC S |
| AC087289.5 | 17 | novel transcript, antisense to TRIM47 |
| TRIM65 | 17 | tripartite motif containing 65 [Source:HGNC S |
| AC087289.2 | 17 | novel transcript, antisense to MRPL38 |
| MRPL38 | 17 | mitochondrial ribosomal protein L38 [Source: |
| AC087289.3 | 17 | novel protein |
| FBF1 | 17 | Fas binding factor 1 [Source:HGNC Symbol;Ac |
| ACOX1 | 17 | acyl-CoA oxidase 1 [Source:HGNC Symbol;Acc |
| AC087289.4 | 17 | novel transcript, antisense to ACOX1 |
| TEN1-CDK3 | 17 | TEN1-CDK3 readthrough (NMD candidate) [So |
| TEN1 | 17 | TEN1 subunit of CST complex [Source:HGNC S |
| CDK3 | 17 | cyclin dependent kinase 3 [Source:HGNC Syml |
| EVPL | 17 | envoplakin [Source:HGNC Symbol;Acc:HGNC:3 |
| SRP68 | 17 | signal recognition particle 68 [Source:HGNC Sy |
| ZACN | 17 | zinc activated ion channel [Source:HGNC Syml |
| GALR2 | 17 | galanin receptor 2 [Source:HGNC Symbol;Acc: |
| EXOC7 | 17 | exocyst complex component 7 [Source:HGNC |
| AC018665.1 | 17 | TEC |
| MIR6868 | 17 | microRNA 6868 [Source:HGNC Symbol;Acc:HG |
| FOXJ1 | 17 | forkhead box J1 [Source:HGNC Symbol;Acc:HG |
| RNF157-AS1 | 17 | RNF157 antisense RNA 1 [Source:HGNC Symb |
| RNF157 | 17 | ring finger protein 157 [Source:HGNC Symbol; |
| ATP5MGP6 | 17 | ATP synthase membrane subunit g pseudoger |
| ATF4P3 | 17 | activating transcription factor 4 pseudogene 3 |
| UBALD2 | 17 | UBA like domain containing 2 [Source:HGNC S |
| QRICH2 | 17 | glutamine rich 2 [Source:HGNC Symbol;Acc:H( |
| PRPSAP1 | 17 | phosphoribosyl pyrophosphate synthetase ass |
| Y_RNA | 17 | Y RNA [Source:RFAM;Acc:RF00019] |
| RNU6-24P | 17 | RNA, U6 small nuclear 24, pseudogene [Sourc |
| AC090699.1 | 17 | high mobility group nucleosomal binding dom |
| SPHK1 | 17 | sphingosine kinase 1 [Source:HGNC Symbol;A( |
| UBE2O | 17 | ubiquitin conjugating enzyme E2 O [Source:H( |
| RPL7P49 | 17 | ribosomal protein L7 pseudogene 49 [Source: |
| AANAT | 17 | aralkylamine N-acetyltransferase [Source:HGN |
| RHBDF2 | 17 | rhomboid 5 homolog 2 [Source:HGNC Symbol |
| CYGB | 17 | cytoglobin [Source:HGNC Symbol;Acc:HGNC:1 |
| PRCD | 17 | photoreceptor disc component [Source:HGNC |
| AC015802.4 | 17 | novel transcript, antisense to PRCD |

| | | |
|---|---|---|
| AC015802.3 | 17 | novel transcript, sense intronic to PRCD |
| AC015802.5 | 17 | novel transcript, sense intronic to PRCD |
| SNHG16 | 17 | small nucleolar RNA host gene 16 [Source:HGI |
| SNORD1C | 17 | small nucleolar RNA, C/D box 1C [Source:HGN |
| SNORD1B | 17 | small nucleolar RNA, C/D box 1B [Source:HGN |
| SNORD1A | 17 | small nucleolar RNA, C/D box 1A [Source:HGN |
| AC015802.6 | 17 | novel protein |
| ST6GALNAC2 | 17 | ST6 N-acetylgalactosaminide alpha-2,6-sialyltr |
| AC015802.1 | 17 | novel transcript, antisense to ST6GALNAC2 |
| AC015802.2 | 17 | COX20 Cox2 chaperone homolog (S. cerevisia |
| AC015802.7 | 17 | novel transcript |
| ST6GALNAC1 | 17 | ST6 N-acetylgalactosaminide alpha-2,6-sialyltr |
| RNU6-227P | 17 | RNA, U6 small nuclear 227, pseudogene [Sour |
| AC005837.1 | 17 | novel transcript, antisense to MXRA7 |
| MXRA7 | 17 | matrix remodeling associated 7 [Source:HGN( |
| RNY4P36 | 17 | RNY4 pseudogene 36 [Source:HGNC Symbol;A |
| AC005837.4 | 17 | novel transcript, antisense to MXRA7 |
| JMJD6 | 17 | jumonji domain containing 6, arginine demeth |
| AC005837.3 | 17 | NULL |
| METTL23 | 17 | methyltransferase like 23 [Source:HGNC Symb |
| AC005837.2 | 17 | novel protein |
| SRSF2 | 17 | serine and arginine rich splicing factor 2 [Sour |
| MFSD11 | 17 | major facilitator superfamily domain containi |
| MIR636 | 17 | microRNA 636 [Source:HGNC Symbol;Acc:HGI |
| RNU6-97P | 17 | RNA, U6 small nuclear 97, pseudogene [Sourc |
| LINC02080 | 17 | long intergenic non-protein coding RNA 2080 |
| AC016168.4 | 17 | novel transcript |
| LINC00868 | 17 | long intergenic non-protein coding RNA 868 [S |
| MGAT5B | 17 | alpha-1,6-mannosylglycoprotein 6-beta-N-ace |
| AC016168.2 | 17 | novel transcript |
| AC016168.1 | 17 | novel transcript |
| AC016168.3 | 17 | TEC |
| Metazoa_SRP | 17 | Metazoan signal recognition particle RNA [Sou |
| SNHG20 | 17 | small nucleolar RNA host gene 20 [Source:HGI |
| SEC14L1 | 17 | SEC14 like lipid binding 1 [Source:HGNC Symb |
| SCARNA16 | 17 | small Cajal body-specific RNA 16 [Source:HGN |
| MIR6516 | 17 | microRNA 6516 [Source:HGNC Symbol;Acc:HG |
| RNU4-47P | 17 | RNA, U4 small nuclear 47, pseudogene [Sourc |
| CYCSP40 | 17 | CYCS pseudogene 40 [Source:HGNC Symbol;A |
| SEPTIN9-DT | 17 | SEPTIN9 divergent transcript [Source:HGNC Sy |
| AC068594.1 | 17 | novel transcript |
| SEPTIN9 | 17 | septin 9 [Source:HGNC Symbol;Acc:HGNC:732 |
| AC111182.1 | 17 | novel transcript, antisense to SEPT9 |
| AC111182.2 | 17 | novel transcript, antisense to SEPT9 |
| MIR4316 | 17 | microRNA 4316 [Source:HGNC Symbol;Acc:HG |
| Y_RNA | 17 | Y RNA [Source:RFAM;Acc:RF00019] |
| AC111170.3 | 17 | TEC |
| AC111170.2 | 17 | novel transcript, sense intronic to SEPT9 |
| AC111170.1 | 17 | novel transcript, antisense to SEPT9 |
| AC021683.3 | 17 | novel transcript |

| | | |
|---|---|---|
| AC021683.2 | 17 | novel transcript |
| AC021683.6 | 17 | novel transcript |
| AC021683.1 | 17 | novel transcript |
| AC021683.4 | 17 | TEC |
| AC021683.5 | 17 | novel transcript |
| LINC01987 | 17 | long intergenic non-protein coding RNA 1987 |
| LINC01973 | 17 | long intergenic non-protein coding RNA 1973 |
| RNU1-80P | 17 | RNA, U1 small nuclear 80, pseudogene [Sourc |
| TNRC6C | 17 | trinucleotide repeat containing adaptor 6C [Sc |
| AC015804.1 | 17 | TEC |
| UBE2V2P2 | 17 | ubiquitin conjugating enzyme E2 V2 pseudoge |
| RNU6-625P | 17 | RNA, U6 small nuclear 625, pseudogene [Sour |
| TNRC6C-AS1 | 17 | TNRC6C antisense RNA 1 [Source:HGNC Symb |
| TMC6 | 17 | transmembrane channel like 6 [Source:HGNC |
| TMC8 | 17 | transmembrane channel like 8 [Source:HGNC |
| C17orf99 | 17 | chromosome 17 open reading frame 99 [Sour |
| EIF5AP2 | 17 | eukaryotic translation initiation factor 5A pseu |
| SYNGR2 | 17 | synaptogyrin 2 [Source:HGNC Symbol;Acc:HGI |
| TK1 | 17 | thymidine kinase 1 [Source:HGNC Symbol;Acc |
| AFMID | 17 | arylformamidase [Source:HGNC Symbol;Acc:H |
| AC087645.1 | 17 | MED9_HUMAN Mediator of RNA polymerase |
| BIRC5 | 17 | baculoviral IAP repeat containing 5 [Source:H( |
| TMEM235 | 17 | transmembrane protein 235 [Source:HGNC Sy |
| AC087645.4 | 17 | novel transcript |
| THA1P | 17 | threonine aldolase 1, pseudogene [Source:HG |
| AC087645.3 | 17 | TEC |
| LINC01993 | 17 | long intergenic non-protein coding RNA 1993 |
| AC087645.2 | 17 | novel transcript |
| SOCS3 | 17 | suppressor of cytokine signaling 3 [Source:HG |
| AC061992.2 | 17 | novel transcript |
| RN7SL236P | 17 | RNA, 7SL, cytoplasmic 236, pseudogene [Sour |
| PGS1 | 17 | phosphatidylglycerophosphate synthase 1 [So |
| AC061992.1 | 17 | NULL |
| DNAH17 | 17 | dynein axonemal heavy chain 17 [Source:HGN |
| DNAH17-AS1 | 17 | DNAH17 antisense RNA 1 [Source:HGNC Symb |
| AC016182.1 | 17 | Ran-specific GTPase-activating protein pseudo |
| RN7SL454P | 17 | RNA, 7SL, cytoplasmic 454, pseudogene [Sour |
| SCAT1 | 17 | S-phase cancer associated transcript 1 [Source |
| CYTH1 | 17 | cytohesin 1 [Source:HGNC Symbol;Acc:HGNC: |
| AC099804.1 | 17 | TEC |
| RNU6-638P | 17 | RNA, U6 small nuclear 638, pseudogene [Sour |
| USP36 | 17 | ubiquitin specific peptidase 36 [Source:HGNC |
| AC022966.2 | 17 | novel transcript |
| RPL9P29 | 17 | ribosomal protein L9 pseudogene 29 [Source: |
| TIMP2 | 17 | TIMP metallopeptidase inhibitor 2 [Source:HG |
| AC022966.1 | 17 | novel transcript, antisense to TIMP2 |
| CEP295NL | 17 | CEP295 N-terminal like [Source:HGNC Symbol |
| AC100788.2 | 17 | TEC |
| AC100788.1 | 17 | novel transcript, antisense to TIMP2 |
| LGALS3BP | 17 | galectin 3 binding protein [Source:HGNC Symb |

| | | |
|---|---|---|
| CANT1 | 17 | calcium activated nucleotidase 1 [Source:HGN |
| C1QTNF1-AS1 | 17 | C1QTNF1 antisense RNA 1 [Source:HGNC Sym |
| C1QTNF1 | 17 | C1q and TNF related 1 [Source:HGNC Symbol; |
| ENGASE | 17 | endo-beta-N-acetylglucosaminidase [Source:H |
| RBFOX3 | 17 | RNA binding fox-1 homolog 3 [Source:HGNC S |
| AC021534.1 | 17 | novel transcript, antisense to RBFOX3 |
| AC233701.1 | 17 | novel transcript, sense intronic to RBFOX3 |
| MIR4739 | 17 | microRNA 4739 [Source:HGNC Symbol;Acc:HG |
| LINC02078 | 17 | long intergenic non-protein coding RNA 2078 |
| ENPP7 | 17 | ectonucleotide pyrophosphatase/phosphodie |
| CBX2 | 17 | chromobox 2 [Source:HGNC Symbol;Acc:HGN |
| CBX8 | 17 | chromobox 8 [Source:HGNC Symbol;Acc:HGN |
| AC100791.2 | 17 | novel transcript, antisense to CBX8 |
| LINC01977 | 17 | long intergenic non-protein coding RNA 1977 |
| CBX4 | 17 | chromobox 4 [Source:HGNC Symbol;Acc:HGN |
| AC100791.1 | 17 | novel transcript |
| LINC01979 | 17 | long intergenic non-protein coding RNA 1979 |
| LINC01978 | 17 | long intergenic non-protein coding RNA 1978 |
| TBC1D16 | 17 | TBC1 domain family member 16 [Source:HGN |
| AC100791.3 | 17 | novel transcript, antisense to TBC1D16 |
| AC116025.2 | 17 | novel transcript |
| AC116025.1 | 17 | novel transcript, antisense to TBC1D16 |
| CCDC40 | 17 | coiled-coil domain containing 40 [Source:HGN |
| MIR1268B | 17 | microRNA 1268b [Source:HGNC Symbol;Acc:H |
| GAA | 17 | glucosidase alpha, acid [Source:HGNC Symbol |
| EIF4A3 | 17 | eukaryotic translation initiation factor 4A3 [Sc |
| AC087741.3 | 17 | TEC |
| AC087741.2 | 17 | novel transcript |
| CARD14 | 17 | caspase recruitment domain family member 1 |
| AC087741.1 | 17 | novel transcript, antisense to CARD14 |
| SGSH | 17 | N-sulfoglucosamine sulfohydrolase [Source:H |
| SLC26A11 | 17 | solute carrier family 26 member 11 [Source:H |
| RNF213 | 17 | ring finger protein 213 [Source:HGNC Symbol; |
| AC124319.1 | 17 | novel transcript |
| AC124319.2 | 17 | novel transcript, antisense to RNF213 |
| AC124319.3 | 17 | TEC |
| RNF213-AS1 | 17 | RNF213 antisense RNA 1 [Source:HGNC Symb |
| ENDOV | 17 | endonuclease V [Source:HGNC Symbol;Acc:H |
| MIR4730 | 17 | microRNA 4730 [Source:HGNC Symbol;Acc:HG |
| AC120024.1 | 17 | novel transcript |
| AC120024.4 | 17 | TEC |
| NPTX1 | 17 | neuronal pentraxin 1 [Source:HGNC Symbol;A |
| AC120024.2 | 17 | REV1 homolog (S. cerevisiae) (REV1) pseudog |
| AC120024.3 | 17 | COX assembly mitochondrial protein 2 homol |
| RPL32P31 | 17 | ribosomal protein L32 pseudogene 31 [Source |
| RPTOR | 17 | regulatory associated protein of MTOR compl |
| RPL31P7 | 17 | ribosomal protein L31 pseudogene 7 [Source: |
| AC016245.1 | 17 | novel transcript, antisense to RPTOR |
| AC016245.2 | 17 | novel transcript, antisense to RPTOR |
| AC127496.3 | 17 | novel transcript, antisense to RPTOR |

| | | |
|---|---|---|
| AC127496.4 | 17 | novel transcript, antisense to RPTOR |
| AC127496.1 | 17 | novel transcript, antisense to RPTOR |
| RPL12P37 | 17 | ribosomal protein L12 pseudogene 37 [Source |
| CHMP6 | 17 | charged multivesicular body protein 6 [Source |
| AC127496.6 | 17 | novel transcript, antisense to CHMP6 |
| AC127496.7 | 17 | TEC |
| AC127496.5 | 17 | novel transcript |
| AC127496.2 | 17 | novel transcript |
| BAIAP2-DT | 17 | BAIAP2 divergent transcript [Source:HGNC Sy |
| BAIAP2 | 17 | BAI1 associated protein 2 [Source:HGNC Symb |
| AATK | 17 | apoptosis associated tyrosine kinase [Source: |
| MIR657 | 17 | microRNA 657 [Source:HGNC Symbol;Acc:HGI |
| MIR3065 | 17 | microRNA 3065 [Source:HGNC Symbol;Acc:HC |
| MIR338 | 17 | microRNA 338 [Source:HGNC Symbol;Acc:HGI |
| MIR1250 | 17 | microRNA 1250 [Source:HGNC Symbol;Acc:HC |
| AC115099.1 | 17 | novel transcript, antisense to AATK |
| PVALEF | 17 | parvalbumin like EF-hand containing [Source: |
| CEP131 | 17 | centrosomal protein 131 [Source:HGNC Symb |
| AC027601.3 | 17 | novel transcript, antisense to AZI1 |
| TEPSIN | 17 | TEPSIN adaptor related protein complex 4 acc |
| AC027601.2 | 17 | novel transcript, antisense to C17orf56 |
| NDUFAF8 | 17 | NADH:ubiquinone oxidoreductase complex as |
| SLC38A10 | 17 | solute carrier family 38 member 10 [Source:H |
| AC027601.4 | 17 | novel transcript, antisense to SLC38A10 |
| AC027601.5 | 17 | TEC |
| LINC00482 | 17 | long intergenic non-protein coding RNA 482 [ |
| AC027601.1 | 17 | novel transcript |
| AC027601.6 | 17 | novel transcript |
| AC110285.3 | 17 | novel transcript |
| AC110285.1 | 17 | uncharacterized LOC100130370 [Source:NCBI |
| AC110285.5 | 17 | novel transcript |
| AC110285.2 | 17 | novel transcript |
| BAHCC1 | 17 | BAH domain and coiled-coil containing 1 [Sou |
| AC110285.6 | 17 | TEC |
| MIR4740 | 17 | microRNA 4740 [Source:HGNC Symbol;Acc:HC |
| MIR3186 | 17 | microRNA 3186 [Source:HGNC Symbol;Acc:HC |
| AC110285.4 | 17 | novel transcript, antisense to BAHCC1 |
| LINC01971 | 17 | long intergenic non-protein coding RNA 1971 |
| ACTG1 | 17 | actin gamma 1 [Source:HGNC Symbol;Acc:HGI |
| AC139149.1 | 17 | novel transcript, antisense to ACTG1 |
| FSCN2 | 17 | fascin actin-bundling protein 2, retinal [Source |
| FAAP100 | 17 | FA core complex associated protein 100 [Sour |
| NPLOC4 | 17 | NPL4 homolog, ubiquitin recognition factor [S |
| Y_RNA | 17 | Y RNA [Source:RFAM;Acc:RF00019] |
| TSPAN10 | 17 | tetraspanin 10 [Source:HGNC Symbol;Acc:HGI |
| PDE6G | 17 | phosphodiesterase 6G [Source:HGNC Symbol; |
| OXLD1 | 17 | oxidoreductase like domain containing 1 [Sou |
| CCDC137 | 17 | coiled-coil domain containing 137 [Source:HG |
| ARL16 | 17 | ADP ribosylation factor like GTPase 16 [Source |
| HGS | 17 | hepatocyte growth factor-regulated tyrosine |

| | | |
|---|---|---|
| MIR6786 | 17 | microRNA 6786 [Source:HGNC Symbol;Acc:HG |
| AC139530.3 | 17 | novel transcript, antisense to HGS |
| AC139530.1 | 17 | novel transcript, antisense to HGS |
| MRPL12 | 17 | mitochondrial ribosomal protein L12 [Source: |
| AC139530.2 | 17 | novel protein |
| SLC25A10 | 17 | solute carrier family 25 member 10 [Source:H |
| GCGR | 17 | glucagon receptor [Source:HGNC Symbol;Acc: |
| MCRIP1 | 17 | MAPK regulated corepressor interacting prote |
| PPP1R27 | 17 | protein phosphatase 1 regulatory subunit 27 [ |
| P4HB | 17 | prolyl 4-hydroxylase subunit beta [Source:HGI |
| AC145207.3 | 17 | novel transcript, antisense P4HB |
| AC145207.9 | 17 | TEC |
| ARHGDIA | 17 | Rho GDP dissociation inhibitor alpha [Source:I |
| AC145207.2 | 17 | novel transcript, antisense ARHGDIA |
| AC145207.5 | 17 | novel transcript |
| AC145207.6 | 17 | novel transcript |
| ALYREF | 17 | Aly/REF export factor [Source:HGNC Symbol;A |
| ANAPC11 | 17 | anaphase promoting complex subunit 11 [Sou |
| NPB | 17 | neuropeptide B [Source:HGNC Symbol;Acc:HG |
| PCYT2 | 17 | phosphate cytidylyltransferase 2, ethanolamir |
| SIRT7 | 17 | sirtuin 7 [Source:HGNC Symbol;Acc:HGNC:149 |
| MAFG | 17 | MAF bZIP transcription factor G [Source:HGN( |
| AC145207.8 | 17 | novel transcript, antisense MAFG |
| MAFG-DT | 17 | MAFG divergent transcript [Source:HGNC Sym |
| PYCR1 | 17 | pyrroline-5-carboxylate reductase 1 [Source:H |
| AC145207.4 | 17 | novel transcript, antisense PYCR1 |
| MYADML2 | 17 | myeloid associated differentiation marker like |
| AC145207.1 | 17 | novel transcript |
| NOTUM | 17 | notum, palmitoleoyl-protein carboxylesterase |
| AC145207.7 | 17 | novel transcript |
| ASPSCR1 | 17 | ASPSCR1 tether for SLC2A4, UBX domain cont |
| CENPX | 17 | centromere protein X [Source:HGNC Symbol;A |
| LRRC45 | 17 | leucine rich repeat containing 45 [Source:HGN |
| RAC3 | 17 | Rac family small GTPase 3 [Source:HGNC Syml |
| DCXR | 17 | dicarbonyl and L-xylulose reductase [Source:H |
| AC137723.1 | 17 | TEC |
| DCXR-DT | 17 | DCXR divergent transcript [Source:HGNC Syml |
| RFNG | 17 | RFNG O-fucosylpeptide 3-beta-N-acetylglucos |
| GPS1 | 17 | G protein pathway suppressor 1 [Source:HGN |
| DUS1L | 17 | dihydrouridine synthase 1 like [Source:HGNC S |
| FASN | 17 | fatty acid synthase [Source:HGNC Symbol;Acc |
| CCDC57 | 17 | coiled-coil domain containing 57 [Source:HGN |
| AC129510.1 | 17 | novel transcript, antisense CCDC57 |
| AC129510.2 | 17 | novel transcript, antisense CCDC57 |
| AC132872.2 | 17 | novel transcript, antisense CCDC57 |
| SLC16A3 | 17 | solute carrier family 16 member 3 [Source:HG |
| MIR6787 | 17 | microRNA 6787 [Source:HGNC Symbol;Acc:HG |
| CSNK1D | 17 | casein kinase 1 delta [Source:HGNC Symbol;A |
| AC132872.3 | 17 | novel transcript, antisense to CSNK1D |
| AC132872.4 | 17 | TEC |

| | | |
|---|---|---|
| AC132872.5 | 17 | novel transcript |
| LINC01970 | 17 | long intergenic non-protein coding RNA 1970 |
| AC132872.1 | 17 | novel transcript |
| CD7 | 17 | CD7 molecule [Source:HGNC Symbol;Acc:HGN |
| SECTM1 | 17 | secreted and transmembrane 1 [Source:HGNC |
| TEX19 | 17 | testis expressed 19 [Source:HGNC Symbol;Acc |
| AC132938.4 | 17 | TEC |
| UTS2R | 17 | urotensin 2 receptor [Source:HGNC Symbol;A |
| AC132938.1 | 17 | novel transcript |
| OGFOD3 | 17 | 2-oxoglutarate and iron dependent oxygenase |
| AC132938.2 | 17 | novel transcript, sense intronic C17orf101 |
| Y_RNA | 17 | Y RNA [Source:RFAM;Acc:RF00019] |
| HEXD | 17 | hexosaminidase D [Source:HGNC Symbol;Acc: |
| HEXD-IT1 | 17 | HEXD intronic transcript 1 [Source:HGNC Sym |
| CYBC1 | 17 | cytochrome b-245 chaperone 1 [Source:HGNC |
| AC132938.6 | 17 | novel transcript |
| AC132938.3 | 17 | novel transcript, antisense NARF |
| NARF | 17 | nuclear prelamin A recognition factor [Source |
| AC132938.5 | 17 | TEC |
| NARF-AS1 | 17 | NARF antisense RNA 1 [Source:HGNC Symbol; |
| NARF-IT1 | 17 | NARF intronic transcript 1 [Source:HGNC Sym |
| FOXK2 | 17 | forkhead box K2 [Source:HGNC Symbol;Acc:H |
| AC124283.4 | 17 | ADP-ribosylation factor-like 2 binding protein |
| AC124283.3 | 17 | novel transcript, sense intronic FOXK2 |
| AC124283.2 | 17 | novel transcript, antisense FOXK2 |
| AC124283.1 | 17 | novel transcript, antisense FOXK2 |
| WDR45B | 17 | WD repeat domain 45B [Source:HGNC Symbo |
| AC124283.5 | 17 | TEC |
| RAB40B | 17 | RAB40B, member RAS oncogene family [Sourc |
| MIR4525 | 17 | microRNA 4525 [Source:HGNC Symbol;Acc:HG |
| AC024361.2 | 17 | novel transcript |
| FN3KRP | 17 | fructosamine 3 kinase related protein [Source |
| AC024361.3 | 17 | novel transcript |
| FN3K | 17 | fructosamine 3 kinase [Source:HGNC Symbol; |
| AC024361.1 | 17 | novel transcript, sense intronic FN3K |
| TBCD | 17 | tubulin folding cofactor D [Source:HGNC Symb |
| AC068014.1 | 17 | novel transcript, antisense TBCD |
| ZNF750 | 17 | zinc finger protein 750 [Source:HGNC Symbol; |
| AC087222.1 | 17 | novel transcript, sense intronic TBCD |
| B3GNTL1 | 17 | UDP-GlcNAc:betaGal beta-1,3-N-acetylglucosa |
| AC130371.1 | 17 | novel transcript, antisense to B3GNTL1 |
| METRNL | 17 | meteorin like, glial cell differentiation regulato |
| AC130371.2 | 17 | novel transcript |
| AC144831.1 | 17 | novel transcript |
| AC139099.3 | 17 | TEC |
| AC139099.2 | 17 | novel transcript |
| AC139099.1 | 17 | novel transcript |
| RPL23AP87 | 17 | ribosomal protein L23a pseudogene 87 [Sourc |
| LINC02564 | 18 | long intergenic non-protein coding RNA 2564 |
| AP005530.1 | 18 | novel transcript |

| | | |
|---|---|---|
| AP001005.1 | 18 | FAM166A, pseudogene 6 |
| TUBB8P12 | 18 | tubulin beta 8 pseudogene 12 [Source:HGNC S |
| AP001005.3 | 18 | novel transcript |
| IL9RP4 | 18 | interleukin 9 receptor pseudogene 4 [Source:I |
| AP001005.2 | 18 | polymerase (RNA) III (DNA directed) polypepti |
| ROCK1P1 | 18 | Rho associated coiled-coil containing protein k |
| MIR8078 | 18 | microRNA 8078 [Source:HGNC Symbol;Acc:HG |
| USP14 | 18 | ubiquitin specific peptidase 14 [Source:HGNC |
| THOC1 | 18 | THO complex 1 [Source:HGNC Symbol;Acc:HG |
| AP000845.1 | 18 | novel transcript |
| COLEC12 | 18 | collectin subfamily member 12 [Source:HGNC |
| AP000915.1 | 18 | novel transcript |
| AP000915.2 | 18 | novel transcript |
| LINC01925 | 18 | long intergenic non-protein coding RNA 1925 |
| CETN1 | 18 | centrin 1 [Source:HGNC Symbol;Acc:HGNC:18 |
| RN7SKP146 | 18 | RN7SK pseudogene 146 [Source:HGNC Symbo |
| CLUL1 | 18 | clusterin like 1 [Source:HGNC Symbol;Acc:HGN |
| TYMSOS | 18 | TYMS opposite strand [Source:HGNC Symbol;A |
| AP001178.3 | 18 | novel transcript, antisense to C18orf56 |
| AP001178.1 | 18 | novel transcript |
| TYMS | 18 | thymidylate synthetase [Source:HGNC Symbo |
| ENOSF1 | 18 | enolase superfamily member 1 [Source:HGNC |
| AP001178.2 | 18 | novel transcript, antisense to ENOSF1 |
| YES1 | 18 | YES proto-oncogene 1, Src family tyrosine kina |
| AP001020.3 | 18 | novel transcript, antisense to YES1 |
| AP001020.2 | 18 | novel transcript, sense intronic toYES1 |
| AP001020.1 | 18 | novel transcript, antisense to YES1 |
| RNU1-109P | 18 | RNA, U1 small nuclear 109, pseudogene [Sour |
| AP000894.4 | 18 | novel transcript |
| BOLA2P1 | 18 | bolA family member 2 pseudogene 1 [Source: |
| AP000894.1 | 18 | proteasome (prosome, macropain) 26S subun |
| AP000894.3 | 18 | novel transcript, antisense to ADCYAP1 |
| AP000894.2 | 18 | novel transcript, antisense to ADCYAP1 |
| ADCYAP1 | 18 | adenylate cyclase activating polypeptide 1 [So |
| LINC01904 | 18 | long intergenic non-protein coding RNA 1904 |
| AP005328.1 | 18 | novel transcript |
| AP000829.1 | 18 | novel transcript |
| AP005265.1 | 18 | solute carrier family 25 (mitochondrial carrier |
| COX6CP3 | 18 | cytochrome c oxidase subunit 6C pseudogene |
| LINC00470 | 18 | long intergenic non-protein coding RNA 470 [S |
| RN7SKP72 | 18 | RN7SK pseudogene 72 [Source:HGNC Symbol; |
| AP005119.1 | 18 | LIM domain kinase 1 (LIMK1) pseudogene |
| AP005119.2 | 18 | ATPase family, AAA domain containing (ATD1) |
| AP005262.2 | 18 | novel transcript |
| AC019183.1 | 18 | novel transcript |
| AP005057.1 | 18 | novel transcript |
| AP005230.1 | 18 | novel transcript |
| AP005262.1 | 18 | axin interactor, dorsalization associated (AIDA |
| AP005136.4 | 18 | novel transcript |
| METTL4 | 18 | methyltransferase like 4 [Source:HGNC Symbo |

| | | |
|---|---|---|
| AP005136.1 | 18 | NULL |
| AP005136.3 | 18 | novel transcript, sense intronic METTL4 |
| AP005136.2 | 18 | ubiquitin-conjugating enzyme E2 variant 1 (UE |
| NDC80 | 18 | NDC80 kinetochore complex component [Sou |
| KATNBL1P3 | 18 | katanin regulatory subunit B1 like 1 pseudoge |
| Y_RNA | 18 | Y RNA [Source:RFAM;Acc:RF00019] |
| RNU6-340P | 18 | RNA, U6 small nuclear 340, pseudogene [Sour |
| AP005061.1 | 18 | Chromobox protein homolog 3 (CBX3) pseudo |
| SMCHD1 | 18 | structural maintenance of chromosomes flexi |
| AP001011.1 | 18 | novel transcript, antisense to SMCHD1 |
| Y_RNA | 18 | Y RNA [Source:RFAM;Acc:RF00019] |
| EMILIN2 | 18 | elastin microfibril interfacer 2 [Source:HGNC S |
| LPIN2 | 18 | lipin 2 [Source:HGNC Symbol;Acc:HGNC:1445 |
| AP000919.4 | 18 | novel transcript, antisense to LPIN2 |
| AP000919.1 | 18 | cysteine and histidine-rich domain (CHORD) c |
| AP000919.2 | 18 | novel transcript, antisense to LPIN2 |
| AP000919.3 | 18 | novel transcript, antisense to LPIN2 |
| SNRPCP4 | 18 | small nuclear ribonucleoprotein polypeptide C |
| AP005431.1 | 18 | ribosomal protein L10 (RPL10) pseudogene |
| SNORA70 | 18 | Small nucleolar RNA SNORA70 [Source:RFAM; |
| MYOM1 | 18 | myomesin 1 [Source:HGNC Symbol;Acc:HGNC |
| RNU7-25P | 18 | RNA, U7 small nuclear 25 pseudogene [Source |
| AP005329.2 | 18 | novel transcript, antisense to MYOM1 |
| AP005329.3 | 18 | novel transcript |
| MYL12A | 18 | myosin light chain 12A [Source:HGNC Symbol; |
| AP005329.1 | 18 | novel transcript, antisense to MYL12A |
| MYL12B | 18 | myosin light chain 12B [Source:HGNC Symbol; |
| AP001025.1 | 18 | novel transcript |
| LINC01895 | 18 | long intergenic non-protein coding RNA 1895 |
| RPL31P59 | 18 | ribosomal protein L31 pseudogene 59 [Source |
| IGLJCOR18 | 18 | immunoglobulin lambda joining-constant/OR1 |
| RPL21P127 | 18 | ribosomal protein L21 pseudogene 127 [Sourc |
| TGIF1 | 18 | TGFB induced factor homeobox 1 [Source:HG |
| BOD1P1 | 18 | biorientation of chromosomes in cell division |
| GAPLINC | 18 | gastric adenocarcinoma associated, positive C |
| DLGAP1 | 18 | DLG associated protein 1 [Source:HGNC Symb |
| RN7SL39P | 18 | RNA, 7SL, cytoplasmic 39, pseudogene [Sourc |
| AP002472.1 | 18 | zinc finger protein 592 (ZNF592) pseudogene |
| DLGAP1-AS1 | 18 | DLGAP1 antisense RNA 1 [Source:HGNC Symb |
| DLGAP1-AS2 | 18 | DLGAP1 antisense RNA 2 [Source:HGNC Symb |
| AP002478.1 | 18 | novel transcript, antisense to DLGAP1 |
| AP002478.2 | 18 | novel transcript, overlapping DLGAP1 |
| DLGAP1-AS3 | 18 | DLGAP1 antisense RNA 3 [Source:HGNC Symb |
| MIR6718 | 18 | microRNA 6718 [Source:HGNC Symbol;Acc:HG |
| RNU6-831P | 18 | RNA, U6 small nuclear 831, pseudogene [Sour |
| DLGAP1-AS4 | 18 | DLGAP1 antisense RNA 4 [Source:HGNC Symb |
| GAPDHP66 | 18 | glyceraldehyde-3-phosphate dehydrogenase p |
| DLGAP1-AS5 | 18 | DLGAP1 antisense RNA 5 [Source:HGNC Symb |
| AP005208.1 | 18 | chromosome 5 open reading frame 43 (C5orf4 |
| AP005403.1 | 18 | transcription elongation factor B (SIII), polype |

| | | |
|---|---|---|
| AP005203.1 | 18 | novel transcript |
| PPIAP14 | 18 | peptidylprolyl isomerase A pseudogene 14 [Sc |
| LINC01892 | 18 | long intergenic non-protein coding RNA 1892 |
| BOD1P2 | 18 | biorientation of chromosomes in cell division |
| AKAIN1 | 18 | A-kinase anchor inhibitor 1 [Source:HGNC Syn |
| AP005380.1 | 18 | novel transcript |
| AP001496.4 | 18 | novel transcript, antisense to C18orf42 |
| AP001496.2 | 18 | novel transcript |
| LINC00526 | 18 | long intergenic non-protein coding RNA 526 [S |
| LINC00667 | 18 | long intergenic non-protein coding RNA 667 [S |
| AP001496.3 | 18 | transforming, acidic coiled-coil containing pro |
| ZBTB14 | 18 | zinc finger and BTB domain containing 14 [Sou |
| AP001496.1 | 18 | novel transcript |
| AP005671.1 | 18 | novel transcript |
| EPB41L3 | 18 | erythrocyte membrane protein band 4.1 like 3 |
| AP005059.2 | 18 | novel transcript, antisense to EPB41L3 |
| AP005059.1 | 18 | novel transcript, antisense to EPB41L3 |
| MIR3976HG | 18 | MIR3976 host gene [Source:HGNC Symbol;Acc |
| MIR3976 | 18 | microRNA 3976 [Source:HGNC Symbol;Acc:HG |
| TMEM200C | 18 | transmembrane protein 200C [Source:HGNC S |
| AP001021.1 | 18 | novel transcript, antisense to TMEM200C |
| AP001021.3 | 18 | novel transcript, antisense to L3MBTL4 |
| L3MBTL4 | 18 | L3MBTL histone methyl-lysine binding protein |
| AP001021.2 | 18 | novel transcript, antisense to L3MBTL4 |
| RNU5F-3P | 18 | RNA, U5F small nuclear 3, pseudogene [Sourc |
| L3MBTL4-AS1 | 18 | L3MBTL4 antisense RNA 1 [Source:HGNC Sym |
| MIR4317 | 18 | microRNA 4317 [Source:HGNC Symbol;Acc:HG |
| AP005060.1 | 18 | novel transcript |
| RNU6-349P | 18 | RNA, U6 small nuclear 349, pseudogene [Sour |
| RPL6P27 | 18 | ribosomal protein L6 pseudogene 27 [Source: |
| AP005202.2 | 18 | novel transcript |
| LINC01387 | 18 | long intergenic non-protein coding RNA 1387 |
| AP005202.1 | 18 | novel transcript |
| RN7SL282P | 18 | RNA, 7SL, cytoplasmic 282, pseudogene [Sour |
| AP005205.1 | 18 | novel transcript |
| AP005205.3 | 18 | novel transcript |
| AP005205.2 | 18 | novel transcript, antisense to ARHGAP28. |
| ARHGAP28 | 18 | Rho GTPase activating protein 28 [Source:HGN |
| AP005210.2 | 18 | novel transcript, antisense to ARHGAP28 |
| AP005210.1 | 18 | novel transcript, antisense to ARHGAP28 |
| LINC00668 | 18 | long intergenic non-protein coding RNA 668 [S |
| SCML2P1 | 18 | SCML2 pseudogene 1 [Source:HGNC Symbol;A |
| RNU6-916P | 18 | RNA, U6 small nuclear 916, pseudogene [Sour |
| LAMA1 | 18 | laminin subunit alpha 1 [Source:HGNC Symbol |
| AP002409.1 | 18 | novel transcript, antisense to LAMA1 |
| AP005062.1 | 18 | novel transcript, antisense to LAMA1 |
| SLC25A51P2 | 18 | SLC25A51 pseudogene 2 [Source:HGNC Symb |
| LRRC30 | 18 | leucine rich repeat containing 30 [Source:HGN |
| AP005270.1 | 18 | NULL |
| AP001095.1 | 18 | novel transcript |

| | | |
|---|---|---|
| PTPRM | 18 | protein tyrosine phosphatase receptor type M |
| AP000897.2 | 18 | novel transcript, antisense to PTPRM |
| AP000897.1 | 18 | novel transcript, sense intronic to PTPRM |
| U3 | 18 | Small nucleolar RNA U3 [Source:RFAM;Acc:RF |
| AP005900.1 | 18 | novel transcript, antisense to PTPRM |
| AC006566.2 | 18 | novel transcript, antisense to PTPRM |
| AC006566.1 | 18 | novel transcript, sense intronic to PTPRM |
| AP001094.3 | 18 | novel transcript, antisense to PTPRM |
| AP001094.2 | 18 | novel transcript, sense intronic to PTPRM |
| AP001094.1 | 18 | novel transcript, antisense to PTPRM |
| COP1P1 | 18 | COP1 pseudogene 1 [Source:HGNC Symbol;Ac |
| RN7SL50P | 18 | RNA, 7SL, cytoplasmic 50, pseudogene [Source |
| THEMIS3P | 18 | thymocyte selection associated family membe |
| AKR1B1P6 | 18 | aldo-keto reductase family 1 member B1 pseu |
| RAB12 | 18 | RAB12, member RAS oncogene family [Source |
| AP001793.1 | 18 | novel transcript, antisense to RAB12 |
| TOMM20P3 | 18 | TOMM20 pseudogene 3 [Source:HGNC Symbo |
| GACAT2 | 18 | gastric cancer associated transcript 2 [Source: |
| MTCL1 | 18 | microtubule crosslinking factor 1 [Source:HGN |
| Y_RNA | 18 | Y RNA [Source:RFAM;Acc:RF00019] |
| AP000864.1 | 18 | novel transcript, sense intronic to SOGA2 |
| AP001531.1 | 18 | novel transcript |
| AP005899.2 | 18 | novel transcript |
| RPS4XP19 | 18 | ribosomal protein S4X pseudogene 19 [Source |
| NDUFV2 | 18 | NADH:ubiquinone oxidoreductase core subun |
| AP005263.1 | 18 | novel transcript |
| AP005899.1 | 18 | novel transcript, antisense to NDUFV2 |
| NDUFV2-AS1 | 18 | NDUFV2 antisense RNA 1 [Source:HGNC Symb |
| ANKRD12 | 18 | ankyrin repeat domain 12 [Source:HGNC Syml |
| Y_RNA | 18 | Y RNA [Source:RFAM;Acc:RF00019] |
| Y_RNA | 18 | Y RNA [Source:RFAM;Acc:RF00019] |
| AP001033.2 | 18 | novel transcript, sense intronic to ANKRD12 |
| AP001033.1 | 18 | novel transcript |
| TWSG1 | 18 | twisted gastrulation BMP signaling modulator |
| AP001033.3 | 18 | novel transcript, sense intronic to TWSG1 |
| AP005432.2 | 18 | novel transcript |
| RALBP1 | 18 | ralA binding protein 1 [Source:HGNC Symbol;A |
| AP005432.1 | 18 | novel transcript, sense intronic to RALBP1 |
| RNU2-27P | 18 | RNA, U2 small nuclear 27, pseudogene [Sourc |
| PPP4R1 | 18 | protein phosphatase 4 regulatory subunit 1 [S |
| AP001381.1 | 18 | novel transcript, antisense to PPP4R1 |
| PPP4R1-AS1 | 18 | PPP4R1 antisense RNA 1 [Source:HGNC Symbo |
| RNU6-903P | 18 | RNA, U6 small nuclear 903, pseudogene [Sour |
| LINC02856 | 18 | long intergenic non-protein coding RNA 2856 |
| KRT18P8 | 18 | keratin 18 pseudogene 8 [Source:HGNC Symb |
| RAB31 | 18 | RAB31, member RAS oncogene family [Source |
| AP000902.1 | 18 | ribosomal protein L27a (RPL27A) pseudogene |
| ZNF415P1 | 18 | zinc finger protein 415 pseudogene 1 [Source: |
| RN7SL862P | 18 | RNA, 7SL, cytoplasmic 862, pseudogene [Sour |
| AC006238.2 | 18 | novel transcript, antisense to RAB31 |

| | | |
|---|---|---|
| RNA5SP449 | 18 | RNA, 5S ribosomal pseudogene 449 [Source:H |
| PIGPP4 | 18 | phosphatidylinositol glycan anchor biosynthes |
| TXNDC2 | 18 | thioredoxin domain containing 2 [Source:HGN |
| AC006238.1 | 18 | novel transcript |
| VAPA | 18 | VAMP associated protein A [Source:HGNC Syn |
| RNA5SP450 | 18 | RNA, 5S ribosomal pseudogene 450 [Source:H |
| AP005271.1 | 18 | novel transcript |
| AP005209.2 | 18 | novel transcript |
| AP005209.1 | 18 | novel transcript |
| AP006219.1 | 18 | novel transcript |
| AP006219.2 | 18 | novel transcript |
| LINC01254 | 18 | long intergenic non-protein coding RNA 1254 |
| AP006219.3 | 18 | novel transcript |
| APCDD1 | 18 | APC down-regulated 1 [Source:HGNC Symbol; |
| NAPG | 18 | NSF attachment protein gamma [Source:HGN |
| AP001099.1 | 18 | novel transcript |
| LINC01887 | 18 | long intergenic non-protein coding RNA 1887 |
| AP001180.3 | 18 | Low affinity sodium-glucose cotransporter (SL |
| CCDC58P1 | 18 | coiled-coil domain containing 58 pseudogene |
| AP001180.2 | 18 | novel transcript |
| AP001180.5 | 18 | ubiquitin specific peptidase 41 (USP41) pseud |
| PMM2P1 | 18 | phosphomannomutase 2 pseudogene 1 [Sour |
| AP001180.1 | 18 | novel transcript, antisense to PIEZO2 |
| PIEZO2 | 18 | piezo type mechanosensitive ion channel com |
| AP001180.4 | 18 | novel transcript, antisense to PIEZO2 |
| KIAA0895LP1 | 18 | KIAA00895L pseudogene 1 [Source:HGNC Sym |
| MIR6788 | 18 | microRNA 6788 [Source:HGNC Symbol;Acc:HG |
| AP005120.1 | 18 | novel transcript, antisense to PIEZO2 |
| AP005139.1 | 18 | solute carrier family 41, member 2 (SLC41A2) |
| AP005229.1 | 18 | novel transcript |
| AP005229.2 | 18 | novel transcript |
| AP001109.1 | 18 | novel transcript |
| LINC01928 | 18 | long intergenic non-protein coding RNA 1928 |
| LINC01255 | 18 | long intergenic non-protein coding RNA 1255 |
| AP001017.1 | 18 | nuclear factor of kappa light polypeptide gene |
| SLC35G4 | 18 | solute carrier family 35 member G4 [Source:H |
| AP001120.4 | 18 | phospholipase A2, group X (PLA2G10) pseudo |
| NPIPB1P | 18 | nuclear pore complex interacting protein fami |
| AP001120.2 | 18 | novel transcript |
| AP001120.3 | 18 | solute carrier family 35, member G4 (SLC35G4 |
| MIR7153 | 18 | microRNA 7153 [Source:HGNC Symbol;Acc:HG |
| AP001120.1 | 18 | novel transcript |
| GNAL | 18 | G protein subunit alpha L [Source:HGNC Symb |
| ASNSP6 | 18 | asparagine synthetase pseudogene 6 [Source: |
| AP001120.5 | 18 | novel transcript, antisense to GNAL |
| AP005137.1 | 18 | novel transcript, antisense GNAL |
| CHMP1B | 18 | charged multivesicular body protein 1B [Sourc |
| CHMP1B-AS1 | 18 | CHMP1B antisense RNA 1 [Source:HGNC Symb |
| AP005137.2 | 18 | novel transcript |
| MPPE1 | 18 | metallophosphoesterase 1 [Source:HGNC Sym |

| | | |
|---|---|---|
| AP001269.4 | 18 | novel transcript, antisense to MPPE1 |
| AP001269.2 | 18 | novel transcript |
| AP001269.3 | 18 | ribosomal protein S20 (RPS20) pseudogene |
| IMPA2 | 18 | inositol monophosphatase 2 [Source:HGNC Sy |
| AP001269.1 | 18 | novel transcript, antisense IMPA2 |
| AP001542.3 | 18 | novel transcript, antisense IMPA2 |
| AP001542.1 | 18 | telomeric repeat binding factor (NIMA-interac |
| AP001542.2 | 18 | WD repeat domain, phosphoinositide interact |
| AP002414.1 | 18 | kinesin family member 1C (KIF1C) pseudogene |
| AP002414.4 | 18 | single stranded DNA binding protein 4 (SSBP4) |
| AP002414.2 | 18 | G protein-coupled receptor 125 (GPR125) pse |
| AP002414.5 | 18 | novel transcript |
| AP002414.3 | 18 | nuclear factor of kappa light polypeptide gene |
| ANKRD62 | 18 | ankyrin repeat domain 62 [Source:HGNC Symb |
| RNU6-324P | 18 | RNA, U6 small nuclear 324, pseudogene [Sour |
| SDHDP1 | 18 | succinate dehydrogenase complex subunit D p |
| AP005264.2 | 18 | phosphomannomutase 2 (PMM2) pseudogene |
| AP005264.6 | 18 | TEC |
| AP005264.4 | 18 | novel transcript |
| AP005264.5 | 18 | solute carrier family 5 (low affinity glucose co |
| CCDC58P3 | 18 | coiled-coil domain containing 58 pseudogene |
| AP005264.3 | 18 | novel transcript |
| PPIAP56 | 18 | peptidylprolyl isomerase A pseudogene 56 [Sc |
| CIDEA | 18 | cell death inducing DFFA like effector a [Sourc |
| AP005264.7 | 18 | TEC |
| AP005264.1 | 18 | novel transcript |
| RNU6-170P | 18 | RNA, U6 small nuclear 170, pseudogene [Sour |
| TUBB6 | 18 | tubulin beta 6 class V [Source:HGNC Symbol;A |
| AFG3L2 | 18 | AFG3 like matrix AAA peptidase subunit 2 [Sou |
| RNU7-129P | 18 | RNA, U7 small nuclear 129 pseudogene [Sourc |
| PRELID3A | 18 | PRELI domain containing 3A [Source:HGNC Sy |
| AP001029.1 | 18 | novel transcript |
| AP001029.2 | 18 | novel transcript, antisense SPIRE1 |
| SPIRE1 | 18 | spire type actin nucleation factor 1 [Source:H |
| AP001029.3 | 18 | high mobility group box 3 (HMGB3) pseudoge |
| PSMG2 | 18 | proteasome assembly chaperone 2 [Source:H |
| CEP76 | 18 | centrosomal protein 76 [Source:HGNC Symbo |
| AP005482.3 | 18 | novel transcript, sense intronic CEP76 |
| AP005482.2 | 18 | novel transcript |
| AP005482.1 | 18 | novel transcript |
| Y_RNA | 18 | Y RNA [Source:RFAM;Acc:RF00019] |
| PTPN2 | 18 | protein tyrosine phosphatase non-receptor ty |
| STK25P1 | 18 | serine/threonine kinase 25 pseudogene 1 [Sou |
| EIF4A2P1 | 18 | eukaryotic translation initiation factor 4A2 ps |
| SEH1L | 18 | SEH1 like nucleoporin [Source:HGNC Symbol;A |
| AP002449.1 | 18 | novel transcript, antisense SEH1L |
| CEP192 | 18 | centrosomal protein 192 [Source:HGNC Symb |
| AP001357.2 | 18 | mRNA turnover 4 homolog (S. cerevisiae) (MR |
| AP001357.1 | 18 | WD repeat domain 89 (WDR89) pseudogene |
| AP001198.2 | 18 | novel transcript |

| | | |
|---|---|---|
| AP001198.1 | 18 | novel transcript, antisense C18orf1 |
| LDLRAD4 | 18 | low density lipoprotein receptor class A doma |
| AP002505.1 | 18 | novel transcript, antisense C18orf1 |
| C18orf15 | 18 | chromosome 18 open reading frame 15 (puta |
| AP002505.3 | 18 | novel transcript, antisense to LDLRAD4 |
| AP002505.2 | 18 | TEC |
| AP002439.1 | 18 | novel transcript, antisense C18orf1 |
| LDLRAD4-AS1 | 18 | LDLRAD4 antisense RNA 1 [Source:HGNC Sym |
| MIR5190 | 18 | microRNA 5190 [Source:HGNC Symbol;Acc:HG |
| AP005131.5 | 18 | novel transcript, antisense C18orf1 |
| AP005131.4 | 18 | novel transcript, antisense C18orf1 |
| AP005131.6 | 18 | novel transcript, sense intronic C18orf1 |
| AP005131.2 | 18 | novel transcript, antisense C18orf1 |
| AP005131.1 | 18 | novel transcript, antisense C18orf1 |
| AP005131.7 | 18 | novel transcript, sense intronic to LDLRAD4 |
| AP005131.3 | 18 | novel transcript, antisense C18orf1 |
| MIR4526 | 18 | microRNA 4526 [Source:HGNC Symbol;Acc:HG |
| AP001010.1 | 18 | novel transcript, antisense C18orf1 |
| FAM210A | 18 | family with sequence similarity 210 member A |
| AP001525.1 | 18 | TEC |
| RN7SL362P | 18 | RNA, 7SL, cytoplasmic 362, pseudogene [Sour |
| RNMT | 18 | RNA guanine-7 methyltransferase [Source:HG |
| MC5R | 18 | melanocortin 5 receptor [Source:HGNC Symbo |
| MC2R | 18 | melanocortin 2 receptor [Source:HGNC Symbo |
| AP001086.1 | 18 | ribosomal protein L36a-like (RPL36AL) pseudo |
| AC006557.6 | 18 | TEC |
| AC006557.5 | 18 | non-SMC element 4 homolog A (S. cerevisiae) |
| ZNF519 | 18 | zinc finger protein 519 [Source:HGNC Symbol; |
| AC006557.4 | 18 | vomeronasal 1 receptor 4 (VN1R4) pseudoger |
| AC006557.1 | 18 | novel transcript, antisense ZNF519 |
| AC006557.3 | 18 | novel transcript, antisense ZNF519 |
| FRG2LP | 18 | FSHD region gene 2 family member L, pseudo |
| AC006557.2 | 18 | protein phosphatase 6, regulatory subunit 2 (I |
| NF1P5 | 18 | neurofibromin 1 pseudogene 5 [Source:HGNC |
| ANKRD20A5P | 18 | ankyrin repeat domain 20 family member A5, |
| RNU6-316P | 18 | RNA, U6 small nuclear 316, pseudogene [Sour |
| RHOT1P1 | 18 | ras homolog family member T1 pseudogene 1 |
| AP005212.4 | 18 | novel transcript |
| AP005212.2 | 18 | sorting nexin 18 (SNX18) pseudogene |
| CYP4F35P | 18 | cytochrome P450 family 4 subfamily F membe |
| AP005212.3 | 18 | zinc finger protein 114 (ZNF114) pseudogene |
| AP005212.1 | 18 | family with sequence similarity 207, member |
| TERF1P2 | 18 | telomeric repeat binding factor 1 pseudogene |
| FEM1AP2 | 18 | fem-1 homolog A pseudogene 2 [Source:HGN |
| AP006261.1 | 18 | novel transcript |
| LONRF2P1 | 18 | LON peptidase N-terminal domain and ring fir |
| CXADRP3 | 18 | CXADR pseudogene 3 [Source:HGNC Symbol;A |
| GRAMD4P7 | 18 | GRAM domain containing 4 pseudogene 7 [So |
| POTEC | 18 | POTE ankyrin domain family member C [Sourc |
| RNU6-1021P | 18 | RNA, U6 small nuclear 1021, pseudogene [Sou |

| | | |
|---|---|---|
| OR4K7P | 18 | olfactory receptor family 4 subfamily K memb |
| OR4K8P | 18 | olfactory receptor family 4 subfamily K memb |
| SNX19P3 | 18 | sorting nexin 19 pseudogene 3 [Source:HGNC |
| VN1R74P | 18 | vomeronasal 1 receptor 74 pseudogene [Sour |
| GTF2IP8 | 18 | general transcription factor IIi pseudogene 8 [ |
| ANKRD30B | 18 | ankyrin repeat domain 30B [Source:HGNC Syn |
| RNU6-1210P | 18 | RNA, U6 small nuclear 1210, pseudogene [Sou |
| AP006565.1 | 18 | novel transcript, antisense to ANKRD30B |
| MIR3156-2 | 18 | microRNA 3156-2 [Source:HGNC Symbol;Acc:I |
| LINC01906 | 18 | long intergenic non-protein coding RNA 1906 |
| FGF7P1 | 18 | fibroblast growth factor 7 pseudogene 1 [Sour |
| AP005121.1 | 18 | novel transcript |
| LINC01443 | 18 | long intergenic non-protein coding RNA 1443 |
| LINC01444 | 18 | long intergenic non-protein coding RNA 1444 |
| AP005121.2 | 18 | novel transcript |
| AP005242.4 | 18 | TEC |
| AP005242.2 | 18 | chromosome 9 open reading frame 86 pseudo |
| AP005242.3 | 18 | novel transcript |
| AP005242.1 | 18 | novel transcript |
| AP005242.5 | 18 | novel transcript |
| AP005901.2 | 18 | novel transcript |
| AP005901.6 | 18 | novel transcript |
| AP005901.5 | 18 | novel transcript |
| AP005901.3 | 18 | kinase suppressor of ras 1 pseudogene |
| BNIP3P3 | 18 | BCL2 interacting protein 3 pseudogene 3 [Sou |
| AP005901.4 | 18 | zinc finger pseudogene |
| AP005901.1 | 18 | ankyrin repeat domain 30A (ANKRD30A) pseu |
| RNU6-721P | 18 | RNA, U6 small nuclear 721, pseudogene [Sour |
| ROCK1 | 18 | Rho associated coiled-coil containing protein I |
| AC022795.1 | 18 | ribosomal protein L21 (RPL21) pseudogene |
| RNU6-120P | 18 | RNA, U6 small nuclear 120, pseudogene [Sour |
| EXOGP1 | 18 | EXOG pseudogene 1 [Source:HGNC Symbol;Ac |
| AC022809.1 | 18 | novel transcript |
| GREB1L | 18 | GREB1 like retinoic acid receptor coactivator [ |
| AC015878.2 | 18 | novel transcript |
| AC015878.1 | 18 | novel transcript, antisense to GREB1L |
| AC011774.1 | 18 | NULL |
| ESCO1 | 18 | establishment of sister chromatid cohesion N- |
| SNRPD1 | 18 | small nuclear ribonucleoprotein D1 polypeptid |
| ABHD3 | 18 | abhydrolase domain containing 3 [Source:HGI |
| AC106037.2 | 18 | novel transcript |
| AC106037.3 | 18 | novel transcript, antisense to ABHD3 |
| MIR320C1 | 18 | microRNA 320c-1 [Source:HGNC Symbol;Acc:I |
| MIB1 | 18 | mindbomb E3 ubiquitin protein ligase 1 [Sourc |
| AC106037.1 | 18 | NULL |
| Y_RNA | 18 | Y RNA [Source:RFAM;Acc:RF00019] |
| RN7SL233P | 18 | RNA, 7SL, cytoplasmic 233, pseudogene [Sour |
| SNORA73 | 18 | Small nucleolar RNA SNORA73 family [Source: |
| MIR133A1HG | 18 | MIR133A1 host gene [Source:HGNC Symbol;A |
| MIR133A1 | 18 | microRNA 133a-1 [Source:HGNC Symbol;Acc:I |

| | | |
|---|---|---|
| MIR1-2 | 18 | microRNA 1-2 [Source:HGNC Symbol;Acc:HGN |
| AC103987.1 | 18 | ribosomal protein L34 (RPL34) pseudogene |
| AC103987.2 | 18 | novel transcript |
| RNU6-1038P | 18 | RNA, U6 small nuclear 1038, pseudogene [Sou |
| AC103987.3 | 18 | prefoldin subunit 4 (PFD4) pseudogene |
| RNA5SP451 | 18 | RNA, 5S ribosomal pseudogene 451 [Source:H |
| LINC01900 | 18 | long intergenic non-protein coding RNA 1900 |
| AC091043.1 | 18 | novel transcript |
| RNU6ATAC20P | 18 | RNA, U6atac small nuclear 20, pseudogene [So |
| GATA6-AS1 | 18 | GATA6 antisense RNA 1 (head to head) [Sourc |
| GATA6 | 18 | GATA binding protein 6 [Source:HGNC Symbol |
| AC091588.1 | 18 | novel transcript, antisense to GATA6 |
| RNU6-702P | 18 | RNA, U6 small nuclear 702, pseudogene [Sour |
| AC091588.3 | 18 | novel transcript, antisense to GATA6 |
| AC091588.2 | 18 | novel transcript |
| AC027449.1 | 18 | novel transcript |
| CTAGE1 | 18 | cutaneous T cell lymphoma-associated antige |
| AC099849.1 | 18 | ATPase, Cu++ transporting, beta polypeptide, |
| RPS4XP18 | 18 | ribosomal protein S4X pseudogene 18 [Source |
| RNU6-1032P | 18 | RNA, U6 small nuclear 1032, pseudogene [Sou |
| AC090912.1 | 18 | novel transcript, antisense to RBBP8 |
| AC090912.2 | 18 | novel transcript, antisense to RBBP8 |
| RBBP8 | 18 | RB binding protein 8, endonuclease [Source:H |
| AC090912.3 | 18 | novel transcript, sense intronic to RBBP8 |
| UBE2CP2 | 18 | ubiquitin conjugating enzyme E2 C pseudogen |
| MIR4741 | 18 | microRNA 4741 [Source:HGNC Symbol;Acc:HG |
| RN7SL745P | 18 | RNA, 7SL, cytoplasmic 745, pseudogene [Sour |
| Y_RNA | 18 | Y RNA [Source:RFAM;Acc:RF00019] |
| CABLES1 | 18 | Cdk5 and Abl enzyme substrate 1 [Source:HGI |
| TMEM241 | 18 | transmembrane protein 241 [Source:HGNC Sy |
| AC011731.1 | 18 | novel transcript, antisense to CABLES1 |
| RIOK3 | 18 | RIO kinase 3 [Source:HGNC Symbol;Acc:HGNC |
| Y_RNA | 18 | Y RNA [Source:RFAM;Acc:RF00019] |
| AC026634.1 | 18 | high mobility group nucleosome binding doma |
| RMC1 | 18 | regulator of MON1-CCZ1 [Source:HGNC Symb |
| NPC1 | 18 | NPC intracellular cholesterol transporter 1 [Sc |
| Y_RNA | 18 | Y RNA [Source:RFAM;Acc:RF00019] |
| ANKRD29 | 18 | ankyrin repeat domain 29 [Source:HGNC Syml |
| RPS10P27 | 18 | ribosomal protein S10 pseudogene 27 [Source |
| LAMA3 | 18 | laminin subunit alpha 3 [Source:HGNC Symbol |
| RPL23AP77 | 18 | ribosomal protein L23a pseudogene 77 [Sourc |
| AC010754.1 | 18 | novel transcript |
| TTC39C | 18 | tetratricopeptide repeat domain 39C [Source: |
| TTC39C-AS1 | 18 | TTC39C antisense RNA 1 [Source:HGNC Symbo |
| AC090772.2 | 18 | novel transcript, antisense to TTC39C |
| AC090772.1 | 18 | novel transcript, antisense to TTC39C |
| RNU5A-6P | 18 | RNA, U5A small nuclear 6, pseudogene [Sourc |
| AC090772.4 | 18 | TEC |
| CABYR | 18 | calcium binding tyrosine phosphorylation regu |
| AC090772.3 | 18 | novel transcript |

| | | |
|---|---|---|
| OSBPL1A | 18 | oxysterol binding protein like 1A [Source:HGN |
| RNA5SP452 | 18 | RNA, 5S ribosomal pseudogene 452 [Source:H |
| Metazoa_SRP | 18 | Metazoan signal recognition particle RNA [Sou |
| RNU6-435P | 18 | RNA, U6 small nuclear 435, pseudogene [Sour |
| MIR320C2 | 18 | microRNA 320c-2 [Source:HGNC Symbol;Acc:H |
| AC023983.1 | 18 | novel transcript, antisense to OSBPL1A |
| AC023983.2 | 18 | novel transcript |
| IMPACT | 18 | impact RWD domain protein [Source:HGNC Sy |
| Y_RNA | 18 | Y RNA [Source:RFAM;Acc:RF00019] |
| AC007922.2 | 18 | novel transcript |
| HRH4 | 18 | histamine receptor H4 [Source:HGNC Symbol; |
| AC007922.1 | 18 | novel transcript |
| AC007922.4 | 18 | novel transcript |
| AC007922.3 | 18 | novel transcript |
| EIF4A3P1 | 18 | eukaryotic translation initiation factor 4A3 ps |
| LINC01915 | 18 | long intergenic non-protein coding RNA 1915 |
| RAC1P1 | 18 | Rac family small GTPase 1 pseudogene 1 [Sou |
| PPIAP57 | 18 | peptidylprolyl isomerase A pseudogene 57 [Sc |
| AC018697.1 | 18 | novel transcript |
| LINC01894 | 18 | long intergenic non-protein coding RNA 1894 |
| WBP2P1 | 18 | WW domain binding protein 2 pseudogene 1 [ |
| AC104961.1 | 18 | novel transcript |
| ZNF521 | 18 | zinc finger protein 521 [Source:HGNC Symbol; |
| AC105114.2 | 18 | novel transcript, antisense to ZNF521 |
| AC105114.1 | 18 | TEC |
| AC110603.1 | 18 | novel transcript, antisense to ZNF521 |
| RN7SL97P | 18 | RNA, 7SL, cytoplasmic 97, pseudogene [Sourc |
| AC027229.1 | 18 | vacuolar protein sorting 53 (VPS53) homolog ( |
| SS18 | 18 | SS18 subunit of BAF chromatin remodeling co |
| RPS24P18 | 18 | ribosomal protein S24 pseudogene 18 [Source |
| PSMA8 | 18 | proteasome subunit alpha 8 [Source:HGNC Sy |
| DHFRP1 | 18 | dihydrofolate reductase pseudogene 1 [Sourc |
| NPM1P2 | 18 | nucleophosmin 1 pseudogene 2 [Source:HGN( |
| TAF4B | 18 | TATA-box binding protein associated factor 4b |
| SINHCAFP1 | 18 | SINHCAF pseudogene 1 [Source:HGNC Symbol |
| AC121320.1 | 18 | novel transcript, sense intronic to TAF4B |
| AC022069.1 | 18 | zinc finger protein pseudogene |
| RNU6-1289P | 18 | RNA, U6 small nuclear 1289, pseudogene [Sou |
| LINC01543 | 18 | long intergenic non-protein coding RNA 1543 |
| KCTD1 | 18 | potassium channel tetramerization domain cc |
| AC007996.1 | 18 | novel transcript, sense intronic to KCTD1 |
| AC090206.1 | 18 | novel transcript, sense intronic to KCTD1 |
| MIR8057 | 18 | microRNA 8057 [Source:HGNC Symbol;Acc:HG |
| CIAPIN1P | 18 | cytokine induced apoptosis inhibitor 1 pseudc |
| AQP4-AS1 | 18 | AQP4 antisense RNA 1 [Source:HGNC Symbol; |
| AC012588.1 | 18 | novel transcript |
| PCAT18 | 18 | prostate cancer associated transcript 18 [Sour |
| U3 | 18 | Small nucleolar RNA U3 [Source:RFAM;Acc:RF |
| AC018371.1 | 18 | novel transcript |
| AC018371.2 | 18 | ubiquitin fusion degradation 1 like (yeast) (UF |

| | | |
|---|---|---|
| AQP4 | 18 | aquaporin 4 [Source:HGNC Symbol;Acc:HGNC |
| CHST9 | 18 | carbohydrate sulfotransferase 9 [Source:HGN |
| AC023575.1 | 18 | ATPase, H+ transporting, lysosomal 31kDa, V1 |
| LINC01908 | 18 | long intergenic non-protein coding RNA 1908 |
| AC090403.1 | 18 | novel transcript |
| AC021382.1 | 18 | novel transcript |
| UBA52P9 | 18 | ubiquitin A-52 residue ribosomal protein fusic |
| AC068408.1 | 18 | novel transcript |
| RBM22P1 | 18 | RNA binding motif protein 22 pseudogene 1 [S |
| PA2G4P3 | 18 | proliferation-associated 2G4 pseudogene 3 [S |
| AC068408.2 | 18 | novel transcript |
| AC090936.1 | 18 | novel transcript |
| CDH2 | 18 | cadherin 2 [Source:HGNC Symbol;Acc:HGNC:1 |
| AC110015.1 | 18 | novel transcript, antisense to CDH2 |
| AC006249.1 | 18 | novel transcript, sense intronic to CDH2 |
| AC090365.1 | 18 | novel transcript |
| ARIH2P1 | 18 | ariadne RBR E3 ubiquitin protein ligase 2 pseu |
| AC023932.1 | 18 | ALS2 amyotrophic lateral sclerosis 2 (juvenile) |
| RNU6-408P | 18 | RNA, U6 small nuclear 408, pseudogene [Sour |
| AC105245.1 | 18 | heat shock 70kDa protein 9 (mortalin) (HSPA9 |
| AC074237.1 | 18 | novel transcript |
| AC117569.1 | 18 | novel transcript |
| AC117569.2 | 18 | novel transcript |
| AC115100.1 | 18 | novel transcript |
| MIR302F | 18 | microRNA 302f [Source:HGNC Symbol;Acc:HG |
| AC090506.1 | 18 | novel transcript |
| AC090506.2 | 18 | TEC |
| AC016382.1 | 18 | novel transcript |
| RNU6-857P | 18 | RNA, U6 small nuclear 857, pseudogene [Sour |
| DSC3 | 18 | desmocollin 3 [Source:HGNC Symbol;Acc:HGN |
| DSC2 | 18 | desmocollin 2 [Source:HGNC Symbol;Acc:HGN |
| DSCAS | 18 | DSC1/DSC2 antisense RNA [Source:HGNC Sym |
| DSC1 | 18 | desmocollin 1 [Source:HGNC Symbol;Acc:HGN |
| AC012417.1 | 18 | novel transcript, antisense to DSC1 |
| DSG1 | 18 | desmoglein 1 [Source:HGNC Symbol;Acc:HGN |
| DSG1-AS1 | 18 | DSG1 antisense RNA 1 [Source:HGNC Symbol; |
| RNU6-167P | 18 | RNA, U6 small nuclear 167, pseudogene [Sour |
| DSG4 | 18 | desmoglein 4 [Source:HGNC Symbol;Acc:HGN |
| Y_RNA | 18 | Y RNA [Source:RFAM;Acc:RF00019] |
| DSG3 | 18 | desmoglein 3 [Source:HGNC Symbol;Acc:HGN |
| AC021549.1 | 18 | novel transcript |
| DSG2 | 18 | desmoglein 2 [Source:HGNC Symbol;Acc:HGN |
| DSG2-AS1 | 18 | DSG2 antisense RNA 1 [Source:HGNC Symbol; |
| TTR | 18 | transthyretin [Source:HGNC Symbol;Acc:HGN |
| B4GALT6 | 18 | beta-1,4-galactosyltransferase 6 [Source:HGN |
| AC017100.1 | 18 | novel transcript, antisense to B4GALT6 |
| RN7SKP44 | 18 | RN7SK pseudogene 44 [Source:HGNC Symbol; |
| LRRC37A7P | 18 | leucine rich repeat containing 37 member A7, |
| AC017100.2 | 18 | golgi SNAP receptor complex member 2 (GOS |
| SLC25A52 | 18 | solute carrier family 25 member 52 [Source:H |

| | | |
|---|---|---|
| TRAPPC8 | 18 | trafficking protein particle complex 8 [Source: |
| AC022960.1 | 18 | novel transcript, antisense to KIAA1012 |
| AC022960.3 | 18 | family with sequence similarity 76, member A |
| AC022960.2 | 18 | novel transcript, sense intronic to TRAPPC8 |
| RNU6-1050P | 18 | RNA, U6 small nuclear 1050, pseudogene [Sou |
| AC009831.1 | 18 | novel transcript, antisense to KIAA1012 |
| PGDP1 | 18 | phosphogluconate dehydrogenase pseudoger |
| AC009831.2 | 18 | epsin- like pseudogene |
| AC009831.4 | 18 | TEC |
| RNF125 | 18 | ring finger protein 125 [Source:HGNC Symbol; |
| AC011825.3 | 18 | novel transcript |
| AC009831.3 | 18 | novel transcript, antisense to RNF125 |
| AC011825.1 | 18 | ribosomal protein S15a (RPS15A) pseudogene |
| AC011825.4 | 18 | novel transcript, antisense to RNF138 |
| RNF138 | 18 | ring finger protein 138 [Source:HGNC Symbol; |
| AC011825.2 | 18 | novel transcript, antisense to RNF138 |
| GAREM1 | 18 | GRB2 associated regulator of MAPK1 subtype |
| RNA5SP453 | 18 | RNA, 5S ribosomal pseudogene 453 [Source:H |
| MEP1B | 18 | meprin A subunit beta [Source:HGNC Symbol; |
| CLUHP6 | 18 | clustered mitochondria homolog pseudogene |
| AC015563.1 | 18 | novel transcript, sense intronic to FAM59A |
| Y_RNA | 18 | Y RNA [Source:RFAM;Acc:RF00019] |
| AC015563.2 | 18 | novel transcript, antisense to FAM59A |
| HNRNPA1P7 | 18 | heterogeneous nuclear ribonucleoprotein A1 |
| AC025887.2 | 18 | novel transcript |
| WBP11P1 | 18 | WW domain binding protein 11 pseudogene 1 |
| AC009835.1 | 18 | novel transcript |
| AC025887.1 | 18 | novel transcript |
| KLHL14 | 18 | kelch like family member 14 [Source:HGNC Sy |
| AC012123.1 | 18 | novel transcript, antisense to KLHL14 |
| AC012123.2 | 18 | ribosomal protein S27 (RPS27) pseudogene |
| AC090371.2 | 18 | novel transcript, antisense to C18orf34 |
| CCDC178 | 18 | coiled-coil domain containing 178 [Source:HG |
| AC090371.1 | 18 | novel transcript, antisense to C18orf34 |
| AC091198.1 | 18 | novel transcript |
| ASXL3 | 18 | ASXL transcriptional regulator 3 [Source:HGN( |
| NOL4 | 18 | nucleolar protein 4 [Source:HGNC Symbol;Acc |
| Y_RNA | 18 | Y RNA [Source:RFAM;Acc:RF00019] |
| AC104985.3 | 18 | TEC |
| AC104985.1 | 18 | novel transcript, antisense to NOL4 |
| AC104985.2 | 18 | novel transcript |
| DTNA | 18 | dystrobrevin alpha [Source:HGNC Symbol;Acc |
| AC013290.1 | 18 | ribosomal protein L10a (RPL10A) pseudogene |
| AC022601.1 | 18 | novel transcript, antisense to DTNA |
| AC068506.1 | 18 | novel transcript |
| MAPRE2 | 18 | microtubule associated protein RP/EB family r |
| AC015967.2 | 18 | TEC |
| AC015967.1 | 18 | novel transcript |
| ZNF397 | 18 | zinc finger protein 397 [Source:HGNC Symbol; |
| ZSCAN30 | 18 | zinc finger and SCAN domain containing 30 [S( |

| | | |
|---|---|---|
| AC011815.3 | 18 | TEC |
| AC011815.2 | 18 | novel transcript, antisense to ZSCAN30 |
| AC011815.1 | 18 | novel transcript |
| ZNF271P | 18 | zinc finger protein 271, pseudogene [Source:H |
| AC116447.1 | 18 | novel transcript |
| ZNF24 | 18 | zinc finger protein 24 [Source:HGNC Symbol;A |
| ZNF396 | 18 | zinc finger protein 396 [Source:HGNC Symbol; |
| AC007998.1 | 18 | ribosomal protein L7a (RPL7A) pseudogene |
| AC007998.4 | 18 | novel transcript |
| AC007998.3 | 18 | novel transcript |
| AC007998.2 | 18 | novel protein |
| INO80C | 18 | INO80 complex subunit C [Source:HGNC Symb |
| AC007998.5 | 18 | TEC |
| GALNT1 | 18 | polypeptide N-acetylgalactosaminyltransferas |
| MIR3975 | 18 | microRNA 3975 [Source:HGNC Symbol;Acc:HG |
| AC090220.1 | 18 | novel transcript |
| AC090229.1 | 18 | novel transcript |
| NRBF2P1 | 18 | nuclear receptor binding factor 2 pseudogene |
| MIR187 | 18 | microRNA 187 [Source:HGNC Symbol;Acc:HGN |
| MIR3929 | 18 | microRNA 3929 [Source:HGNC Symbol;Acc:HG |
| AC118757.1 | 18 | novel transcript |
| AC091060.1 | 18 | TEC |
| C18orf21 | 18 | chromosome 18 open reading frame 21 [Sour |
| RPRD1A | 18 | regulation of nuclear pre-mRNA domain conta |
| SLC39A6 | 18 | solute carrier family 39 member 6 [Source:HG |
| ELP2 | 18 | elongator acetyltransferase complex subunit 2 |
| AC023043.1 | 18 | novel transcript, antisense to MOCOS |
| MOCOS | 18 | molybdenum cofactor sulfurase [Source:HGN( |
| AC023043.4 | 18 | novel transcript, sense intronic to MOCOS |
| AC023043.2 | 18 | WD repeat domain phosphoinositide-interacti |
| AC023043.3 | 18 | related RAS viral (r-ras) oncogene homolog 2, |
| FHOD3 | 18 | formin homology 2 domain containing 3 [Sour |
| AC055840.1 | 18 | novel transcript, antisense to FHOD3 |
| TPGS2 | 18 | tubulin polyglutamylase complex subunit 2 [S |
| KIAA1328 | 18 | KIAA1328 [Source:HGNC Symbol;Acc:HGNC:2 |
| AC016493.1 | 18 | novel transcript, antisense to KIAA1328 |
| AC015961.1 | 18 | novel transcript |
| CELF4 | 18 | CUGBP Elav-like family member 4 [Source:HG |
| AC015961.2 | 18 | novel transcript, antisense to CELF4 |
| AC090386.1 | 18 | novel transcript, antisense to CELF4 |
| AC090386.2 | 18 | TEC |
| AC129908.1 | 18 | TEC |
| AC009899.1 | 18 | novel transcript, antisense to CELF4 |
| MIR4318 | 18 | microRNA 4318 [Source:HGNC Symbol;Acc:HG |
| RPL12P40 | 18 | ribosomal protein L12 pseudogene 40 [Source |
| AC108452.1 | 18 | novel transcript |
| AC100781.1 | 18 | novel transcript |
| RN7SKP182 | 18 | RN7SK pseudogene 182 [Source:HGNC Symbo |
| RNU6-706P | 18 | RNA, U6 small nuclear 706, pseudogene [Sour |
| MIR924HG | 18 | MIR924 host gene [Source:HGNC Symbol;Acc: |

| | | |
|---|---|---|
| AC011139.1 | 18 | novel transcript |
| RPL7AP66 | 18 | ribosomal protein L7a pseudogene 66 [Source |
| MIR924 | 18 | microRNA 924 [Source:HGNC Symbol;Acc:HGN |
| AC099845.1 | 18 | novel transcript |
| MIR5583-2 | 18 | microRNA 5583-2 [Source:HGNC Symbol;Acc:H |
| MIR5583-1 | 18 | microRNA 5583-1 [Source:HGNC Symbol;Acc:H |
| RNU6-1242P | 18 | RNA, U6 small nuclear 1242, pseudogene [Sou |
| LINC01901 | 18 | long intergenic non-protein coding RNA 1901 |
| LINC01902 | 18 | long intergenic non-protein coding RNA 1902 |
| AC011266.1 | 18 | TEC |
| LINC01477 | 18 | long intergenic non-protein coding RNA 1477 |
| RPL17P45 | 18 | ribosomal protein L17 pseudogene 45 [Source |
| AC068688.1 | 18 | novel transcript |
| AC079052.1 | 18 | novel transcript |
| AC021504.1 | 18 | KC6, keratoconus gene 6 |
| NPM1P1 | 18 | nucleophosmin 1 pseudogene 1 [Source:HGNC |
| AC011225.1 | 18 | C15orf40 pseudogene |
| PIK3C3 | 18 | phosphatidylinositol 3-kinase catalytic subunit |
| AC087683.2 | 18 | novel transcript, sense intronic to PIK3C3 |
| AC087683.1 | 18 | Nucleosome assembly protein 1-like 2, pseudo |
| AC087683.3 | 18 | novel transcript |
| LINC00907 | 18 | long intergenic non-protein coding RNA 907 [S |
| RNA5SP454 | 18 | RNA, 5S ribosomal pseudogene 454 [Source:H |
| AC084335.1 | 18 | novel transcript |
| RIT2 | 18 | Ras like without CAAX 2 [Source:HGNC Symbo |
| AC100779.1 | 18 | novel transcript |
| SYT4 | 18 | synaptotagmin 4 [Source:HGNC Symbol;Acc:H |
| AC022743.1 | 18 | TEC |
| AC083760.1 | 18 | inhibitor of Bruton agammaglobulinemia tyros |
| RNA5SP455 | 18 | RNA, 5S ribosomal pseudogene 455 [Source:H |
| AC110014.1 | 18 | novel transcript |
| LINC01478 | 18 | long intergenic non-protein coding RNA 1478 |
| KRT8P5 | 18 | keratin 8 pseudogene 5 [Source:HGNC Symbo |
| MLECP1 | 18 | malectin pseudogene 1 [Source:HGNC Symbol |
| AC120049.1 | 18 | uncharacterized LOC101927943 [Source:NCBI |
| SETBP1 | 18 | SET binding protein 1 [Source:HGNC Symbol;A |
| AC021766.1 | 18 | TEC |
| MIR4319 | 18 | microRNA 4319 [Source:HGNC Symbol;Acc:HG |
| AC105074.1 | 18 | TEC |
| AC090376.1 | 18 | novel transcript |
| SLC14A2 | 18 | solute carrier family 14 member 2 [Source:HG |
| SLC14A2-AS1 | 18 | SLC14A2 antisense RNA 1 [Source:HGNC Symb |
| AC021517.1 | 18 | uncharacterized LOC112268408 [Source:NCBI |
| AC021517.2 | 18 | novel transcript |
| AC091151.1 | 18 | novel transcript |
| AC091151.5 | 18 | novel transcript, antisense to SLC14A2 |
| AC023421.2 | 18 | novel transcript, antisense to SLC14A1 and SL |
| AC023421.1 | 18 | novel transcript |
| SLC14A1 | 18 | solute carrier family 14 member 1 (Kidd blood |
| AC087685.1 | 18 | ribosomal protein S24 (RPS24) pseudogene |

| | | |
|---|---|---|
| SIGLEC15 | 18 | sialic acid binding Ig like lectin 15 [Source:HGN |
| EPG5 | 18 | ectopic P-granules autophagy protein 5 homo |
| PSTPIP2 | 18 | proline-serine-threonine phosphatase interac |
| RN7SKP26 | 18 | RN7SK pseudogene 26 [Source:HGNC Symbol; |
| AC012569.1 | 18 | U3 small nucleolar RNA-associated protein 18 |
| RNY4P37 | 18 | RNY4 pseudogene 37 [Source:HGNC Symbol;A |
| ATP5F1A | 18 | ATP synthase F1 subunit alpha [Source:HGNC |
| HAUS1 | 18 | HAUS augmin like complex subunit 1 [Source:I |
| RNU6-1278P | 18 | RNA, U6 small nuclear 1278, pseudogene [Sou |
| C18orf25 | 18 | chromosome 18 open reading frame 25 [Sour |
| RNF165 | 18 | ring finger protein 165 [Source:HGNC Symbol; |
| AC015959.1 | 18 | TEC |
| AC018931.1 | 18 | novel transcript, antisense to RNF165 |
| LOXHD1 | 18 | lipoxygenase homology domains 1 [Source:HG |
| ST8SIA5 | 18 | ST8 alpha-N-acetyl-neuraminide alpha-2,8-sia |
| AC090311.1 | 18 | ribosomal protein S21 (RPS21) pseudogene |
| AC090241.2 | 18 | novel transcript, antisense to ST8SIA5 |
| AC090241.3 | 18 | novel transcript |
| PIAS2 | 18 | protein inhibitor of activated STAT 2 [Source:H |
| AC090241.1 | 18 | Ubiquitin carboxyl-terminal hydrolase 4, pseu |
| AC090373.1 | 18 | TEC |
| KATNAL2 | 18 | katanin catalytic subunit A1 like 2 [Source:HG |
| ELOA3 | 18 | elongin A3 [Source:HGNC Symbol;Acc:HGNC:2 |
| ELOA3C | 18 | elongin A3 family member C [Source:HGNC Sy |
| ELOA3B | 18 | elongin A3 family member B [Source:HGNC Sy |
| ELOA3D | 18 | elongin A3 family member D [Source:HGNC Sy |
| ELOA2 | 18 | elongin A2 [Source:HGNC Symbol;Acc:HGNC:3 |
| AC012254.5 | 18 | TEC |
| AC012254.1 | 18 | novel transcript, antisense to KATNAL2 |
| AC012254.3 | 18 | novel transcript, antisense to HDHD2 |
| HDHD2 | 18 | haloacid dehalogenase like hydrolase domain |
| AC012254.2 | 18 | novel transcript |
| AC012254.4 | 18 | SUMO2 SMT3 suppressor of mif two 3 homol |
| IER3IP1 | 18 | immediate early response 3 interacting protei |
| SKOR2 | 18 | SKI family transcriptional corepressor 2 [Sourc |
| RNU6-1131P | 18 | RNA, U6 small nuclear 1131, pseudogene [Sou |
| AC051635.1 | 18 | novel transcript |
| MIR4527HG | 18 | MIR4527 host gene [Source:HGNC Symbol;Acc |
| MIR4527 | 18 | microRNA 4527 [Source:HGNC Symbol;Acc:HG |
| AC102797.1 | 18 | novel transcript |
| AC102797.2 | 18 | TEC |
| TPMTP1 | 18 | thiopurine S-methyltransferase pseudogene 1 |
| AC026898.1 | 18 | sal-like 2 (Drosophila), pseudogene |
| SMAD2 | 18 | SMAD family member 2 [Source:HGNC Symbo |
| MTCO2P2 | 18 | MT-CO2 pseudogene 2 [Source:HGNC Symbol |
| AC120349.3 | 18 | TEC |
| AC120349.1 | 18 | novel transcript |
| AC120349.2 | 18 | TEC |
| ZBTB7C | 18 | zinc finger and BTB domain containing 7C [Sou |
| AC105101.1 | 18 | TEC |

| | | |
|---|---|---|
| AC105101.2 | 18 | novel transcript, antisense to ZBTB7C |
| RNU6-708P | 18 | RNA, U6 small nuclear 708, pseudogene [Sour |
| C18orf12 | 18 | chromosome 18 open reading frame 12 [Sour |
| RNA5SP456 | 18 | RNA, 5S ribosomal pseudogene 456 [Source:H |
| AC024288.1 | 18 | metallo-beta-lactamase domain containing 2 ( |
| POLR3GP2 | 18 | RNA polymerase III subunit G pseudogene 2 [S |
| CTIF | 18 | cap binding complex dependent translation in |
| AC048380.2 | 18 | TEC |
| AC048380.3 | 18 | TEC |
| MIR4743 | 18 | microRNA 4743 [Source:HGNC Symbol;Acc:HG |
| AC048380.1 | 18 | novel transcript, antisense to CTIF |
| AC022919.1 | 18 | TEC |
| AC093567.1 | 18 | novel transcript, antisense to CTIF |
| SMAD7 | 18 | SMAD family member 7 [Source:HGNC Symbo |
| AC114684.1 | 18 | novel transcript |
| AC016866.3 | 18 | TEC |
| AC016866.1 | 18 | novel transcript, antisense to DYM |
| DYM | 18 | dymeclin [Source:HGNC Symbol;Acc:HGNC:21 |
| MIR4744 | 18 | microRNA 4744 [Source:HGNC Symbol;Acc:HG |
| AC016866.2 | 18 | novel transcript, antisense to DYM |
| AC044840.1 | 18 | novel transcript, antisense to DYM |
| AC100778.1 | 18 | novel transcript, sense intronic to DYM |
| AC100778.4 | 18 | novel transcript, antisense to DYM |
| PRR13P4 | 18 | proline rich 13 pseudogene 4 [Source:HGNC S |
| C18orf32 | 18 | chromosome 18 open reading frame 32 [Sour |
| RPL17-C18orf32 | 18 | RPL17-C18orf32 readthrough [Source:HGNC S |
| AC100778.2 | 18 | novel transcript, sense intronic to C18orf32 |
| AC100778.3 | 18 | microRNA 1539 |
| MIR1539 | 18 | microRNA 1539 [Source:HGNC Symbol;Acc:HG |
| RPL17 | 18 | ribosomal protein L17 [Source:HGNC Symbol;A |
| SNORD58C | 18 | small nucleolar RNA, C/D box 58C [Source:HG |
| SNORD58A | 18 | small nucleolar RNA, C/D box 58A [Source:HG |
| SNORD58B | 18 | small nucleolar RNA, C/D box 58B [Source:HG |
| SRP72P1 | 18 | signal recognition particle 72 pseudogene 1 [S |
| LINC02837 | 18 | long intergenic non-protein coding RNA 2837 |
| LIPG | 18 | lipase G, endothelial type [Source:HGNC Symb |
| SMUG1P1 | 18 | single-strand-selective monofunctional uracil- |
| AC090227.3 | 18 | novel transcript |
| ACAA2 | 18 | acetyl-CoA acyltransferase 2 [Source:HGNC Sy |
| AC090227.2 | 18 | novel protein |
| SNHG22 | 18 | small nucleolar RNA host gene 22 [Source:HGI |
| SCARNA17 | 18 | small Cajal body-specific RNA 17 [Source:HGN |
| AC090227.1 | 18 | NULL |
| MYO5B | 18 | myosin VB [Source:HGNC Symbol;Acc:HGNC:7 |
| Y_RNA | 18 | Y RNA [Source:RFAM;Acc:RF00019] |
| RNA5SP457 | 18 | RNA, 5S ribosomal pseudogene 457 [Source:H |
| AC091044.1 | 18 | novel transcript, sense intronic to MYO5B |
| ADAD1P2 | 18 | adenosine deaminase domain containing 1 ps |
| AC105224.1 | 18 | myosin VB (MYO5B) pseudogene |
| MIR4320 | 18 | microRNA 4320 [Source:HGNC Symbol;Acc:HG |

| | | |
|---|---|---|
| RN7SL310P | 18 | RNA, 7SL, cytoplasmic 310, pseudogene [Sour |
| CFAP53 | 18 | cilia and flagella associated protein 53 [Source |
| AC090246.1 | 18 | novel transcript, sense intronic to CCDC11 |
| MBD1 | 18 | methyl-CpG binding domain protein 1 [Source |
| CXXC1 | 18 | CXXC finger protein 1 [Source:HGNC Symbol;A |
| AC105227.1 | 18 | novel transcript |
| AC105227.2 | 18 | ankyrin repeat domain 26 (ANKRD26) pseudo |
| RNA5SP458 | 18 | RNA, 5S ribosomal pseudogene 458 [Source:H |
| SKA1 | 18 | spindle and kinetochore associated complex s |
| RPLP0P11 | 18 | ribosomal protein lateral stalk subunit P0 pse |
| MAPK4 | 18 | mitogen-activated protein kinase 4 [Source:H |
| AC012433.2 | 18 | novel transcript, antisense to MAPK4 |
| AC012433.1 | 18 | NADH dehydrogenase pseudogene |
| MRO | 18 | maestro [Source:HGNC Symbol;Acc:HGNC:241 |
| HNRNPA3P16 | 18 | heterogeneous nuclear ribonucleoprotein A3 |
| RPL17P46 | 18 | ribosomal protein L17 pseudogene 46 [Source |
| ME2 | 18 | malic enzyme 2 [Source:HGNC Symbol;Acc:HG |
| Y_RNA | 18 | Y RNA [Source:RFAM;Acc:RF00019] |
| ELAC1 | 18 | elaC ribonuclease Z 1 [Source:HGNC Symbol;A |
| AC091551.1 | 18 | novel protein |
| SMAD4 | 18 | SMAD family member 4 [Source:HGNC Symbo |
| SRSF10P1 | 18 | serine and arginine rich splicing factor 10 pse |
| MEX3C | 18 | mex-3 RNA binding family member C [Source: |
| RNU1-46P | 18 | RNA, U1 small nuclear 46, pseudogene [Sourc |
| LINC01630 | 18 | long intergenic non-protein coding RNA 1630 |
| SS18L2P2 | 18 | SS18 like 2 pseudogene 2 [Source:HGNC Symb |
| AC109315.1 | 18 | Rho guanine nucleotide exchange factor (GEF |
| RSL24D1P9 | 18 | ribosomal L24 domain containing 1 pseudoge |
| AC027216.1 | 18 | intraflagellar transport 57 homolog (Chlamyd |
| RPS8P3 | 18 | ribosomal protein S8 pseudogene 3 [Source:H |
| AC016383.3 | 18 | novel transcript |
| AC016383.2 | 18 | novel transcript |
| AC016383.1 | 18 | novel transcript |
| DCC | 18 | DCC netrin 1 receptor [Source:HGNC Symbol; |
| AC011155.1 | 18 | TEC |
| VN1R76P | 18 | vomeronasal 1 receptor 76 pseudogene [Sour |
| MIR4528 | 18 | microRNA 4528 [Source:HGNC Symbol;Acc:HG |
| LINC01917 | 18 | long intergenic non-protein coding RNA 1917 |
| LINC01919 | 18 | long intergenic non-protein coding RNA 1919 |
| RPL29P32 | 18 | ribosomal protein L29 pseudogene 32 [Source |
| AC090666.1 | 18 | zinc finger protein (ZNF) pseudogene |
| MBD2 | 18 | methyl-CpG binding domain protein 2 [Source |
| SNORA37 | 18 | small nucleolar RNA, H/ACA box 37 [Source:H |
| AC093462.1 | 18 | novel transcript, antisense to POLI |
| POLI | 18 | DNA polymerase iota [Source:HGNC Symbol;A |
| STARD6 | 18 | StAR related lipid transfer domain containing |
| C18orf54 | 18 | chromosome 18 open reading frame 54 [Sour |
| SNRPGP2 | 18 | small nuclear ribonucleoprotein polypeptide |
| AC090897.2 | 18 | novel pseudogene |
| AC090897.1 | 18 | chromosome 18 open reading frame 26 (C18o |

| | | |
|---|---|---|
| AC091135.1 | 18 | dynactin associated protein (DYNAP) pseudog |
| AC091135.2 | 18 | dynactin associated protein (DYNAP) pseudog |
| DYNAP | 18 | dynactin associated protein [Source:HGNC Syr |
| RAB27B | 18 | RAB27B, member RAS oncogene family [Sourc |
| RPSAP57 | 18 | ribosomal protein SA pseudogene 57 [Source: |
| AC007673.1 | 18 | novel transcript, sense intronic to RAB27B |
| AC098848.1 | 18 | novel transcript, antisense to RAB27B |
| CCDC68 | 18 | coiled-coil domain containing 68 [Source:HGN |
| MAP1LC3P | 18 | microtubule associated protein 1 light chain 3 |
| LINC01929 | 18 | long intergenic non-protein coding RNA 1929 |
| RNA5SP459 | 18 | RNA, 5S ribosomal pseudogene 459 [Source:H |
| TCF4 | 18 | transcription factor 4 [Source:HGNC Symbol;A |
| TCF4-AS1 | 18 | TCF4 antisense RNA 1 [Source:HGNC Symbol;A |
| MIR4529 | 18 | microRNA 4529 [Source:HGNC Symbol;Acc:HG |
| TCF4-AS2 | 18 | TCF4 antisense RNA 2 [Source:HGNC Symbol;A |
| AC022031.2 | 18 | novel transcript |
| RPL21P126 | 18 | ribosomal protein L21 pseudogene 126 [Sourc |
| LINC01415 | 18 | long intergenic non-protein coding RNA 1415 |
| AC022031.1 | 18 | novel transcript |
| AC027584.1 | 18 | novel transcript |
| LINC01416 | 18 | long intergenic non-protein coding RNA 1416 |
| AC009271.1 | 18 | novel transcript |
| AC006305.1 | 18 | novel transcript |
| AC009271.2 | 18 | novel transcript |
| LINC01905 | 18 | long intergenic non-protein coding RNA 1905 |
| AC006305.2 | 18 | novel transcript |
| SNORA73 | 18 | Small nucleolar RNA SNORA73 family [Source: |
| LINC01539 | 18 | long intergenic non-protein coding RNA 1539 |
| AC006305.3 | 18 | novel transcript |
| TXNL1 | 18 | thioredoxin like 1 [Source:HGNC Symbol;Acc:H |
| WDR7 | 18 | WD repeat domain 7 [Source:HGNC Symbol;A |
| Y_RNA | 18 | Y RNA [Source:RFAM;Acc:RF00019] |
| AC012301.1 | 18 | novel transcript, antisense to WDR7 |
| U3 | 18 | Small nucleolar RNA U3 [Source:RFAM;Acc:RF |
| WDR7-OT1 | 18 | WDR7 overlapping transcript 1 [Source:HGNC |
| AC012301.2 | 18 | novel transcript |
| LINC-ROR | 18 | long intergenic non-protein coding RNA, regul |
| AC100775.1 | 18 | novel transcript, antisense to BOD1L2 |
| BOD1L2 | 18 | biorientation of chromosomes in cell division |
| LINC02565 | 18 | long intergenic non-protein coding RNA 2565 |
| RNU6-737P | 18 | RNA, U6 small nuclear 737, pseudogene [Sour |
| ST8SIA3 | 18 | ST8 alpha-N-acetyl-neuraminide alpha-2,8-sia |
| AC090340.1 | 18 | novel transcript |
| ONECUT2 | 18 | one cut homeobox 2 [Source:HGNC Symbol;A |
| FECH | 18 | ferrochelatase [Source:HGNC Symbol;Acc:HGI |
| AC100847.1 | 18 | novel transcript |
| NARS | 18 | asparaginyl-tRNA synthetase [Source:HGNC Sy |
| AC027097.1 | 18 | uncharacterized LOC100505549 [Source:NCBI |
| AC027097.2 | 18 | novel transcript, antisense to ATP8B1 |
| ATP8B1 | 18 | ATPase phospholipid transporting 8B1 [Source |

| | | |
|---|---|---|
| RNU6-742P | 18 | RNA, U6 small nuclear 742, pseudogene [Sour |
| RSL24D1P11 | 18 | ribosomal L24 domain containing 1 pseudoge |
| LINC01897 | 18 | long intergenic non-protein coding RNA 1897 |
| HMGN1P30 | 18 | high mobility group nucleosome binding dom |
| NEDD4L | 18 | NEDD4 like E3 ubiquitin protein ligase [Source |
| AC107896.1 | 18 | uncharacterized LOC105372143 [Source:NCBI |
| AC090236.2 | 18 | novel transcript, sense intronic to NEDD4L |
| AC090236.1 | 18 | novel transcript, antisense to NEDD4L |
| AC105105.2 | 18 | novel transcript |
| MIR122HG | 18 | MIR122 host gene [Source:HGNC Symbol;Acc: |
| MIR122 | 18 | microRNA 122 [Source:HGNC Symbol;Acc:HG |
| MIR3591 | 18 | microRNA 3591 [Source:HGNC Symbol;Acc:H |
| AC105105.4 | 18 | novel transcript |
| ALPK2 | 18 | alpha kinase 2 [Source:HGNC Symbol;Acc:HG |
| AC105105.3 | 18 | novel transcript, sense intronic to ALPK2 |
| AC105105.1 | 18 | novel transcript, antisense to ALPK2 |
| AC104971.2 | 18 | novel transcript, antisense to ALPK2 |
| SNORA108 | 18 | small nucleolar RNA, H/ACA box 108 [Source: |
| RPL9P31 | 18 | ribosomal protein L9 pseudogene 31 [Source: |
| AC104971.3 | 18 | novel transcript |
| AC104971.1 | 18 | uncharacterized LOC101927322 [Source:NCBI |
| MALT1 | 18 | MALT1 paracaspase [Source:HGNC Symbol;Ac |
| AC104365.1 | 18 | novel transcript, antisense to MALT1 |
| MRPL37P1 | 18 | mitochondrial ribosomal protein L37 pseudog |
| AC104365.3 | 18 | novel transcript |
| AC104365.2 | 18 | novel transcript |
| RNU6-219P | 18 | RNA, U6 small nuclear 219, pseudogene [Sour |
| LINC01926 | 18 | long intergenic non-protein coding RNA 1926 |
| U8 | 18 | U8 small nucleolar RNA [Source:RFAM;Acc:RF |
| ZNF532 | 18 | zinc finger protein 532 [Source:HGNC Symbol; |
| RNU2-69P | 18 | RNA, U2 small nuclear 69, pseudogene [Sourc |
| OACYLP | 18 | O-acyltransferase like, pseudogene [Source:H |
| AC040963.1 | 18 | novel transcript |
| SEC11C | 18 | SEC11 homolog C, signal peptidase complex s |
| GRP | 18 | gastrin releasing peptide [Source:HGNC Symb |
| RAX | 18 | retina and anterior neural fold homeobox [So |
| CPLX4 | 18 | complexin 4 [Source:HGNC Symbol;Acc:HGNC |
| LMAN1 | 18 | lectin, mannose binding 1 [Source:HGNC Sym |
| AC016229.2 | 18 | novel transcript |
| CCBE1 | 18 | collagen and calcium binding EGF domains 1 [ |
| AC016229.1 | 18 | novel transcript, antisense to CCBE1 |
| AC090213.1 | 18 | novel transcript, antisense to CCBE1 |
| AC010776.1 | 18 | novel transcript, antisense to CCBE1 |
| AC010776.2 | 18 | novel transcript, antisense to CCBE1 |
| AC010776.3 | 18 | novel transcript |
| RPS26P54 | 18 | ribosomal protein S26 pseudogene 54 [Source |
| GLUD1P4 | 18 | glutamate dehydrogenase 1 pseudogene 4 [So |
| AC098847.1 | 18 | ribosomal protein, large, P0 pseudogene |
| PMAIP1 | 18 | phorbol-12-myristate-13-acetate-induced pro |
| AC107990.1 | 18 | pleckstrin homology domain containing, famil |

| | | |
|---|---|---|
| NFE2L3P1 | 18 | nuclear factor, erythroid 2 like 3 pseudogene |
| RN7SL342P | 18 | RNA, 7SL, cytoplasmic 342, pseudogene [Sour |
| ENTR1P1 | 18 | ENTR1 pseudogene 1 [Source:HGNC Symbol;A |
| AC090377.1 | 18 | novel transcript |
| SINHCAFP2 | 18 | SINHCAF pseudogene 2 [Source:HGNC Symbol |
| RNU6-567P | 18 | RNA, U6 small nuclear 567, pseudogene [Sour |
| AC090771.2 | 18 | novel transcript |
| RPS3AP49 | 18 | ribosomal protein S3a pseudogene 49 [Source |
| RNU4-17P | 18 | RNA, U4 small nuclear 17, pseudogene [Sourc |
| AC090771.1 | 18 | general transcription factor IIH, polypeptide 1 |
| AC090621.1 | 18 | novel transcript |
| AC091576.1 | 18 | novel transcript, antisense to MC4R |
| MC4R | 18 | melanocortin 4 receptor [Source:HGNC Symbo |
| AC010928.2 | 18 | dynein, light chain, roadblock-type 2 (DYNLRB |
| MRPS5P4 | 18 | mitochondrial ribosomal protein S5 pseudoge |
| AC010928.3 | 18 | testis derived transcript (3 LIM domains) pseu |
| AC010928.1 | 18 | glutamate decarboxylase 1-like pseudogene |
| AC113137.1 | 18 | novel transcript |
| CTBP2P3 | 18 | C-terminal binding protein 2 pseudogene 3 [S |
| HMGN1P31 | 18 | high mobility group nucleosome binding doma |
| CDH20 | 18 | cadherin 20 [Source:HGNC Symbol;Acc:HGNC: |
| RNU6-116P | 18 | RNA, U6 small nuclear 116, pseudogene [Sour |
| RPL30P14 | 18 | ribosomal protein L30 pseudogene 14 [Source |
| AC090409.2 | 18 | novel transcript |
| AC090409.1 | 18 | novel transcript |
| AC105094.2 | 18 | novel transcript |
| AC105094.1 | 18 | novel transcript |
| LINC01544 | 18 | long intergenic non-protein coding RNA 1544 |
| RNF152 | 18 | ring finger protein 152 [Source:HGNC Symbol; |
| PIGN | 18 | phosphatidylinositol glycan anchor biosynthes |
| RPIAP1 | 18 | ribose 5-phosphate isomerase A pseudogene |
| RELCH | 18 | RAB11 binding and LisH domain, coiled-coil ar |
| AC027514.1 | 18 | novel transcript, sense intronic to KIAA1468 |
| AC027514.2 | 18 | novel transcript, antisense to KIAA1468 |
| TNFRSF11A | 18 | TNF receptor superfamily member 11a [Sourc |
| Y_RNA | 18 | Y RNA [Source:RFAM;Acc:RF00019] |
| AC100843.1 | 18 | novel transcript, antisense to TNFRSF11A |
| RPL17P44 | 18 | ribosomal protein L17 pseudogene 44 [Source |
| AC100843.2 | 18 | ATP synthase, H+ transporting, mitochondrial |
| ACTBP9 | 18 | ACTB pseudogene 9 [Source:HGNC Symbol;Ac |
| ZCCHC2 | 18 | zinc finger CCHC-type containing 2 [Source:HG |
| AC064801.1 | 18 | novel transcript, antisense to ZCCHC2 |
| AC064801.2 | 18 | novel transcript |
| RN7SL705P | 18 | RNA, 7SL, cytoplasmic 705, pseudogene [Sour |
| PHLPP1 | 18 | PH domain and leucine rich repeat protein ph |
| RNU6-142P | 18 | RNA, U6 small nuclear 142, pseudogene [Sour |
| BCL2 | 18 | BCL2 apoptosis regulator [Source:HGNC Symb |
| AC022726.1 | 18 | novel transcript, sense intronic to BCL2 |
| AC022726.2 | 18 | novel transcript, antisense to BCL2 |
| KDSR | 18 | 3-ketodihydrosphingosine reductase [Source:I |

| | | |
|---|---|---|
| AC036176.1 | 18 | novel transcript, antisense to KDSR |
| VPS4B | 18 | vacuolar protein sorting 4 homolog B [Source: |
| AC036176.2 | 18 | proteasome (prosome, macropain) 26S subun |
| SERPINB5 | 18 | serpin family B member 5 [Source:HGNC Syml |
| AC036176.3 | 18 | novel transcript |
| ATP5MC1P6 | 18 | ATP synthase membrane subunit c locus 1 pse |
| SERPINB12 | 18 | serpin family B member 12 [Source:HGNC Syn |
| SERPINB13 | 18 | serpin family B member 13 [Source:HGNC Syn |
| SERPINB4 | 18 | serpin family B member 4 [Source:HGNC Syml |
| SERPINB11 | 18 | serpin family B member 11 (gene/pseudogene |
| SERPINB3 | 18 | serpin family B member 3 [Source:HGNC Syml |
| SERPINB7 | 18 | serpin family B member 7 [Source:HGNC Syml |
| SERPINB2 | 18 | serpin family B member 2 [Source:HGNC Syml |
| SERPINB10 | 18 | serpin family B member 10 [Source:HGNC Syn |
| HMSD | 18 | histocompatibility minor serpin domain conta |
| AC009802.1 | 18 | novel transcript, antisense to SERPINB8 |
| SERPINB8 | 18 | serpin family B member 8 [Source:HGNC Syml |
| LINC01924 | 18 | long intergenic non-protein coding RNA 1924 |
| RPL12P39 | 18 | ribosomal protein L12 pseudogene 39 [Source |
| LINC00305 | 18 | long intergenic non-protein coding RNA 305 [S |
| LINC01538 | 18 | long intergenic non-protein coding RNA 1538 |
| AC027506.1 | 18 | H3 histone, family 3B (H3.3B) (H3F3B) pseudo |
| AC090348.1 | 18 | novel transcript |
| AC007948.1 | 18 | novel transcript |
| AC007631.1 | 18 | novel transcript |
| LINC01916 | 18 | long intergenic non-protein coding RNA 1916 |
| AC090358.1 | 18 | novel transcript |
| CDH7 | 18 | cadherin 7 [Source:HGNC Symbol;Acc:HGNC:1 |
| AC023394.2 | 18 | novel transcript, sense intronic to CDH7 |
| AC023394.1 | 18 | novel transcript |
| PRPF19P1 | 18 | PRPF19 pseudogene 1 [Source:HGNC Symbol; |
| CDH19 | 18 | cadherin 19 [Source:HGNC Symbol;Acc:HGNC: |
| RNU6-1037P | 18 | RNA, U6 small nuclear 1037, pseudogene [Sou |
| MIR5011 | 18 | microRNA 5011 [Source:HGNC Symbol;Acc:HG |
| RPL31P9 | 18 | ribosomal protein L31 pseudogene 9 [Source: |
| AC091042.1 | 18 | ribosomal protein L10a (RPL10A) pseudogene |
| AC110597.1 | 18 | novel transcript |
| DSEL | 18 | dermatan sulfate epimerase like [Source:HGN |
| AC110597.3 | 18 | novel transcript, antisense to DSEL |
| AC114689.3 | 18 | novel transcript |
| AC022662.1 | 18 | fibrous sheath CABYR binding protein (FSCB) p |
| AC022655.1 | 18 | novel transcript |
| AC022655.2 | 18 | family with sequence similarity 32, member A |
| AC114689.1 | 18 | SECIS binding protein 2 (SECISBP2) pseudogen |
| LINC01903 | 18 | long intergenic non-protein coding RNA 1903 |
| AC114689.2 | 18 | myotubularin related protein 14 (MTMR14) ps |
| AC100844.1 | 18 | novel transcript |
| AC068112.1 | 18 | novel transcript |
| AC005909.2 | 18 | novel transcript |
| LINC01912 | 18 | long intergenic non-protein coding RNA 1912 |

| | | |
|---|---|---|
| AC022968.1 | 18 | ATP synthase, H+ transporting, mitochondrial |
| AC005909.1 | 18 | novel transcript |
| AKR1B10P2 | 18 | aldo-keto reductase family 1 member B10 pse |
| AC067819.1 | 18 | metallothionein 1E (MT1E) pseudogene |
| TMX3 | 18 | thioredoxin related transmembrane protein 3 |
| CCDC102B | 18 | coiled-coil domain containing 102B [Source:H |
| AC040896.1 | 18 | novel transcript, sense intronic to CCDC102B |
| AC022035.1 | 18 | novel transcript, antisense to CCDC102B |
| RNU6-39P | 18 | RNA, U6 small nuclear 39, pseudogene [Sourc |
| SDHCP1 | 18 | succinate dehydrogenase complex subunit C p |
| AC096708.2 | 18 | novel transcript, antisense to CCDC102B |
| AC096708.3 | 18 | novel transcript, sense intronic to CCDC102B |
| AC096708.1 | 18 | aminoacylase 1 (ACY1) pseudogene |
| AC090337.1 | 18 | novel transcript |
| AC090337.2 | 18 | novel transcript |
| AC026585.1 | 18 | novel transcript |
| DOK6 | 18 | docking protein 6 [Source:HGNC Symbol;Acc:H |
| AC119868.1 | 18 | cholinergic receptor, nicotinic, epsilon (CHRNE |
| AC119868.2 | 18 | novel transcript, antisense to DOK6 |
| AC068254.2 | 18 | TEC |
| AC068254.1 | 18 | novel transcript, antisense to DOK6 |
| CD226 | 18 | CD226 molecule [Source:HGNC Symbol;Acc:H |
| RTTN | 18 | rotatin [Source:HGNC Symbol;Acc:HGNC:1865 |
| AC021701.1 | 18 | novel transcript, antisense to RTTN |
| SOCS6 | 18 | suppressor of cytokine signaling 6 [Source:HG |
| LINC01909 | 18 | long intergenic non-protein coding RNA 1909 |
| LIVAR | 18 | liver cell viability associated lncRNA [Source:H |
| LINC01910 | 18 | long intergenic non-protein coding RNA 1910 |
| LARP7P3 | 18 | La ribonucleoprotein domain family member |
| RPS2P6 | 18 | ribosomal protein S2 pseudogene 6 [Source:H |
| AC091305.1 | 18 | TEC |
| AC091646.1 | 18 | novel transcript |
| RN7SL795P | 18 | RNA, 7SL, cytoplasmic 795, pseudogene [Sour |
| GTSCR1 | 18 | Gilles de la Tourette syndrome chromosome r |
| AC090415.2 | 18 | novel transcript |
| AC090415.1 | 18 | lysosomal-associated membrane protein 1 (LA |
| AC091691.3 | 18 | novel transcript |
| AC091691.2 | 18 | novel transcript |
| AC091691.1 | 18 | signal peptide peptidase-like 3 (SPPL3) pseudo |
| LINC01541 | 18 | long intergenic non-protein coding RNA 1541 |
| AC090312.1 | 18 | microtubule-associated protein 7 (MAP7) pseu |
| AC090312.2 | 18 | SRSF protein kinase 1 (SRPK1) pseudogene |
| LINC01899 | 18 | long intergenic non-protein coding RNA 1899 |
| AC027458.1 | 18 | novel transcript |
| AC069114.1 | 18 | novel transcript |
| AC069114.2 | 18 | BCL2-associated transcription factor 1, pseudo |
| CBLN2 | 18 | cerebellin 2 precursor [Source:HGNC Symbol;A |
| HNRNPA1P11 | 18 | heterogeneous nuclear ribonucleoprotein A1 |
| NETO1 | 18 | neuropilin and tolloid like 1 [Source:HGNC Syn |
| AC023301.1 | 18 | novel transcript , sense intronic to NETO1 |

| | | |
|---|---|---|
| RNA5SP460 | 18 | RNA, 5S ribosomal pseudogene 460 [Source:H |
| MIR548AV | 18 | microRNA 548av [Source:HGNC Symbol;Acc:H |
| AC091138.1 | 18 | novel transcript |
| AC079062.1 | 18 | novel transcript |
| LINC02582 | 18 | long intergenic non-protein coding RNA 2582 |
| AC079070.1 | 18 | novel transcript |
| RN7SL401P | 18 | RNA, 7SL, cytoplasmic 401, pseudogene [Sour |
| AC090125.1 | 18 | novel transcript |
| AC010947.1 | 18 | TEC |
| AC090125.2 | 18 | novel transcript |
| FBXO15 | 18 | F-box protein 15 [Source:HGNC Symbol;Acc:H |
| TIMM21 | 18 | translocase of inner mitochondrial membrane |
| AC090398.1 | 18 | novel transcript |
| RN7SL551P | 18 | RNA, 7SL, cytoplasmic 551, pseudogene [Sour |
| CYB5A | 18 | cytochrome b5 type A [Source:HGNC Symbol;. |
| AC090398.2 | 18 | novel transcript |
| C18orf63 | 18 | chromosome 18 open reading frame 63 [Sour |
| FAUP1 | 18 | FAU pseudogene 1 [Source:HGNC Symbol;Acc |
| LINC01922 | 18 | long intergenic non-protein coding RNA 1922 |
| DIPK1C | 18 | divergent protein kinase domain 1C [Source:H |
| AC009704.3 | 18 | novel transcript |
| CNDP2 | 18 | carnosine dipeptidase 2 [Source:HGNC Symbo |
| AC009704.2 | 18 | novel transcript, sense intronic to CNDP2 |
| CNDP1 | 18 | carnosine dipeptidase 1 [Source:HGNC Symbo |
| AC009704.1 | 18 | novel transcript, antisense to CNDP1 |
| LINC00909 | 18 | long intergenic non-protein coding RNA 909 [S |
| ZNF407 | 18 | zinc finger protein 407 [Source:HGNC Symbol; |
| AC015819.2 | 18 | novel transcript |
| AC015819.1 | 18 | novel transcript |
| AC015819.4 | 18 | NULL |
| AC015819.3 | 18 | novel transcript |
| AC015819.5 | 18 | novel transcript |
| ZADH2 | 18 | zinc binding alcohol dehydrogenase domain c |
| TSHZ1 | 18 | teashirt zinc finger homeobox 1 [Source:HGN( |
| AC116003.3 | 18 | novel transcript |
| AC116003.1 | 18 | novel transcript, antisense to SMIM21 |
| SMIM21 | 18 | small integral membrane protein 21 [Source:H |
| AC116003.2 | 18 | novel transcript |
| AC012101.2 | 18 | novel transcript |
| AC012101.1 | 18 | novel transcript |
| AC090812.1 | 18 | novel transcript |
| AC090338.1 | 18 | novel transcript |
| LINC01898 | 18 | long intergenic non-protein coding RNA 1898 |
| AC021506.1 | 18 | novel transcript |
| AC021506.2 | 18 | novel pseudogene |
| AC090457.1 | 18 | novel transcript |
| AC011095.1 | 18 | novel transcript |
| AC103808.6 | 18 | novel transcript |
| LINC01893 | 18 | long intergenic non-protein coding RNA 1893 |
| AC103808.2 | 18 | novel transcript |

| | | |
|---|---|---|
| AC103808.4 | 18 | novel transcript |
| AC103808.3 | 18 | novel transcript |
| AC103808.1 | 18 | novel transcript |
| AC103808.5 | 18 | novel transcript |
| ZNF516 | 18 | zinc finger protein 516 [Source:HGNC Symbol; |
| AC009716.1 | 18 | novel transcript, antisense to ZFN156 |
| AC009716.2 | 18 | TEC |
| AC018413.1 | 18 | novel transcript, antisense to ZFN156 |
| C18orf65 | 18 | chromosome 18 open reading frame 65 [Sour |
| LINC00683 | 18 | long intergenic non-protein coding RNA 683 [S |
| AC034110.1 | 18 | novel transcript |
| LINC01927 | 18 | long intergenic non-protein coding RNA 1927 |
| LINC01879 | 18 | long intergenic non-protein coding RNA 1879 |
| ARL2BPP1 | 18 | ADP ribosylation factor like GTPase 2 binding |
| CCND3P2 | 18 | cyclin D3 pseudogene 2 [Source:HGNC Symbo |
| AC139085.1 | 18 | ribosomal protein S3 (RPS3) pseudogene |
| ZNF236-DT | 18 | ZNF236 divergent transcript [Source:HGNC Sy |
| RNU6-346P | 18 | RNA, U6 small nuclear 346, pseudogene [Sour |
| AC027575.2 | 18 | novel transcript |
| ZNF236 | 18 | zinc finger protein 236 [Source:HGNC Symbol; |
| RPL26P35 | 18 | ribosomal protein L26 pseudogene 35 [Source |
| Metazoa_SRP | 18 | Metazoan signal recognition particle RNA [Sou |
| AC027575.1 | 18 | exportin, tRNA (nuclear export receptor for tR |
| AC093330.1 | 18 | novel transcript, antisense to MBP |
| MBP | 18 | myelin basic protein [Source:HGNC Symbol;Ac |
| AC093330.2 | 18 | TEC |
| AC093330.3 | 18 | novel transcript, antisense to MBP |
| AC018529.1 | 18 | novel transcript, antisense to MBP |
| AC018529.3 | 18 | TEC |
| AC018529.2 | 18 | novel transcript, antisense to MBP |
| AC100863.1 | 18 | novel transcript |
| GALR1 | 18 | galanin receptor 1 [Source:HGNC Symbol;Acc: |
| AC124254.2 | 18 | novel transcript |
| AC124254.1 | 18 | NULL |
| BDP1P | 18 | B double prime 1 pseudogene [Source:HGNC S |
| AC123786.1 | 18 | novel transcript |
| AC123786.2 | 18 | DnaJ (Hsp40) homolog, subfamily C, member |
| AC123786.3 | 18 | novel transcript |
| RNA5SP461 | 18 | RNA, 5S ribosomal pseudogene 461 [Source:H |
| LINC01029 | 18 | long intergenic non-protein coding RNA 1029 |
| AC134978.1 | 18 | novel transcript |
| RNU6-655P | 18 | RNA, U6 small nuclear 655, pseudogene [Sour |
| AC107892.1 | 18 | novel transcript |
| AC087399.1 | 18 | novel transcript |
| AC012572.1 | 18 | novel transcript |
| Y_RNA | 18 | Y RNA [Source:RFAM;Acc:RF00019] |
| AC044873.1 | 18 | novel transcript |
| AC091027.2 | 18 | novel transcript |
| AC091027.1 | 18 | novel transcript |
| AC091027.3 | 18 | novel transcript |

| | | |
|---|---|---|
| LINC01896 | 18 | long intergenic non-protein coding RNA 1896 |
| SALL3 | 18 | spalt like transcription factor 3 [Source:HGNC |
| ATP9B | 18 | ATPase phospholipid transporting 9B (putative |
| AC099689.1 | 18 | TEC |
| AC125437.1 | 18 | novel transcript, sense intronic to ATP9B |
| AC125437.2 | 18 | TEC |
| AC125437.3 | 18 | novel transcript, antisense to ATP9B |
| AC104423.1 | 18 | novel transcript, antisense to ATP9B |
| AC023090.1 | 18 | novel transcript, antisense to ATP9B |
| AC023090.2 | 18 | TEC |
| NFATC1 | 18 | nuclear factor of activated T cells 1 [Source:H |
| AC018445.3 | 18 | TEC |
| AC018445.5 | 18 | novel transcript, antisense to NFATC1 |
| AC018445.1 | 18 | novel transcript, antisense to NFATC1 |
| AC018445.4 | 18 | TEC |
| AC018445.2 | 18 | TEC |
| AC018445.6 | 18 | novel transcript |
| AC068473.3 | 18 | novel transcript |
| AC068473.4 | 18 | novel transcript |
| AC068473.1 | 18 | novel transcript |
| AC068473.5 | 18 | novel transcript |
| CTDP1 | 18 | CTD phosphatase subunit 1 [Source:HGNC Syr |
| AC068473.2 | 18 | novel transcript, antisense to CTDP1 |
| AC021594.1 | 18 | novel transcript |
| AC021594.2 | 18 | novel transcript |
| KCNG2 | 18 | potassium voltage-gated channel modifier sub |
| AC114341.1 | 18 | novel transcript |
| SLC66A2 | 18 | solute carrier family 66 member 2 [Source:HG |
| AC114341.2 | 18 | novel transcript |
| HSBP1L1 | 18 | heat shock factor binding protein 1 like 1 [Sou |
| TXNL4A | 18 | thioredoxin like 4A [Source:HGNC Symbol;Acc |
| AC090360.1 | 18 | novel protein |
| RBFA | 18 | ribosome binding factor A [Source:HGNC Sym |
| RBFADN | 18 | RBFA downstream neighbor [Source:HGNC Sy |
| SLC25A6P4 | 18 | solute carrier family 25 member 6 pseudogen |
| ADNP2 | 18 | ADNP homeobox 2 [Source:HGNC Symbol;Acc |
| PARD6G-AS1 | 18 | PARD6G antisense RNA 1 [Source:HGNC Symb |
| PARD6G | 18 | par-6 family cell polarity regulator gamma [So |
| AC139100.2 | 18 | novel transcript |
| AC139100.1 | 18 | novel transcript, antisense to PARD6G |
| WASH5P | 19 | WASP family homolog 5, pseudogene [Source |
| AC008993.1 | 19 | TEC |
| AC008993.2 | 19 | TEC |
| AC008993.3 | 19 | novel transcript |
| MIR1302-11 | 19 | microRNA 1302-11 [Source:HGNC Symbol;Acc |
| FAM138F | 19 | family with sequence similarity 138 member F |
| OR4G3P | 19 | olfactory receptor family 4 subfamily G memb |
| OR4G1P | 19 | olfactory receptor family 4 subfamily G memb |
| OR4F17 | 19 | olfactory receptor family 4 subfamily F memb |
| WBP1LP11 | 19 | WBP1L pseudogene 11 [Source:HGNC Symbol |

| | | |
|---|---|---|
| OR4F8P | 19 | olfactory receptor family 4 subfamily F memb |
| AC092192.1 | 19 | novel transcript |
| SEPTIN14P19 | 19 | septin 14 pseudogene 19 [Source:HGNC Symb |
| CICP19 | 19 | capicua transcriptional repressor pseudogene |
| AC092299.1 | 19 | novel pseudogene |
| AC010507.1 | 19 | novel pseudogene |
| LINC01002 | 19 | long intergenic non-protein coding RNA 1002 |
| AC010507.2 | 19 | novel pseudogene |
| RNU6-1076P | 19 | RNA, U6 small nuclear 1076, pseudogene [Sou |
| AC016588.2 | 19 | novel transcript |
| PLPP2 | 19 | phospholipid phosphatase 2 [Source:HGNC Sy |
| MIER2 | 19 | MIER family member 2 [Source:HGNC Symbol |
| AC016588.1 | 19 | novel transcript, antisense to MIER2 |
| THEG | 19 | theg spermatid protein [Source:HGNC Symbol |
| AC010641.2 | 19 | novel transcript |
| AC010641.1 | 19 | novel transcript |
| C2CD4C | 19 | C2 calcium dependent domain containing 4C [ |
| SHC2 | 19 | SHC adaptor protein 2 [Source:HGNC Symbol; |
| RNA5SP462 | 19 | RNA, 5S ribosomal pseudogene 462 [Source:H |
| ODF3L2 | 19 | outer dense fiber of sperm tails 3 like 2 [Sourc |
| MADCAM1 | 19 | mucosal vascular addressin cell adhesion mol |
| AC005775.1 | 19 | novel transcript, antisense to MADCAM1 and |
| TPGS1 | 19 | tubulin polyglutamylase complex subunit 1 [S |
| CDC34 | 19 | cell division cycle 34 [Source:HGNC Symbol;Ac |
| GZMM | 19 | granzyme M [Source:HGNC Symbol;Acc:HGNC |
| AC009005.1 | 19 | novel transcript, antisense to BSG |
| BSG | 19 | basigin (Ok blood group) [Source:HGNC Symb |
| HCN2 | 19 | hyperpolarization activated cyclic nucleotide g |
| AC005559.1 | 19 | Rho GTPase activating protein 33 (ARHGAP33 |
| POLRMT | 19 | RNA polymerase mitochondrial [Source:HGNC |
| AC004449.1 | 19 | novel transcript |
| FGF22 | 19 | fibroblast growth factor 22 [Source:HGNC Sym |
| RNF126 | 19 | ring finger protein 126 [Source:HGNC Symbol; |
| AC004156.1 | 19 | novel transcript |
| FSTL3 | 19 | follistatin like 3 [Source:HGNC Symbol;Acc:HG |
| PRSS57 | 19 | serine protease 57 [Source:HGNC Symbol;Acc |
| RPS2P52 | 19 | ribosomal protein S2 pseudogene 52 [Source: |
| PALM | 19 | paralemmin [Source:HGNC Symbol;Acc:HGNC |
| MISP | 19 | mitotic spindle positioning [Source:HGNC Sym |
| AC006273.1 | 19 | novel transcript |
| LINC01836 | 19 | long intergenic non-protein coding RNA 1836 |
| PTBP1 | 19 | polypyrimidine tract binding protein 1 [Source |
| MIR4745 | 19 | microRNA 4745 [Source:HGNC Symbol;Acc:HG |
| PLPPR3 | 19 | phospholipid phosphatase related 3 [Source:H |
| MIR3187 | 19 | microRNA 3187 [Source:HGNC Symbol;Acc:HG |
| AZU1 | 19 | azurocidin 1 [Source:HGNC Symbol;Acc:HGNC |
| PRTN3 | 19 | proteinase 3 [Source:HGNC Symbol;Acc:HGNC |
| ELANE | 19 | elastase, neutrophil expressed [Source:HGNC |
| CFD | 19 | complement factor D [Source:HGNC Symbol;A |
| MED16 | 19 | mediator complex subunit 16 [Source:HGNC S |

| | | |
|---|---|---|
| RNU6-9 | 19 | RNA, U6 small nuclear 9 [Source:HGNC Symbo |
| R3HDM4 | 19 | R3H domain containing 4 [Source:HGNC Symb |
| KISS1R | 19 | KISS1 receptor [Source:HGNC Symbol;Acc:HG |
| ARID3A | 19 | AT-rich interaction domain 3A [Source:HGNC |
| AC005379.1 | 19 | novel transcript, antisense to ARID3A |
| AC005391.1 | 19 | novel transcript, sense intronic to ARID3A |
| WDR18 | 19 | WD repeat domain 18 [Source:HGNC Symbol; |
| AC004528.1 | 19 | novel transcript, antisense to GRIN3B |
| GRIN3B | 19 | glutamate ionotropic receptor NMDA type su |
| TMEM259 | 19 | transmembrane protein 259 [Source:HGNC Sy |
| AC004528.2 | 19 | novel transcript, antisense to TMEM259 |
| RNU6-2 | 19 | RNA, U6 small nuclear 2 [Source:HGNC Symbo |
| CNN2 | 19 | calponin 2 [Source:HGNC Symbol;Acc:HGNC:2 |
| AC011558.1 | 19 | TEC |
| ABCA7 | 19 | ATP binding cassette subfamily A member 7 [ |
| ARHGAP45 | 19 | Rho GTPase activating protein 45 [Source:HG |
| POLR2E | 19 | RNA polymerase II subunit E [Source:HGNC Sy |
| GPX4 | 19 | glutathione peroxidase 4 [Source:HGNC Symb |
| SBNO2 | 19 | strawberry notch homolog 2 [Source:HGNC Sy |
| STK11 | 19 | serine/threonine kinase 11 [Source:HGNC Sym |
| HMGB2P1 | 19 | high mobility group box 2 pseudogene 1 [Sour |
| CBARP | 19 | CACN subunit beta associated regulatory prot |
| AC004221.1 | 19 | novel transcript |
| ATP5F1D | 19 | ATP synthase F1 subunit delta [Source:HGNC |
| MIDN | 19 | midnolin [Source:HGNC Symbol;Acc:HGNC:16 |
| CIRBP | 19 | cold inducible RNA binding protein [Source:H |
| CIRBP-AS1 | 19 | CIRBP antisense RNA 1 [Source:HGNC Symbol; |
| FAM174C | 19 | family with sequence similarity 174 member |
| EFNA2 | 19 | ephrin A2 [Source:HGNC Symbol;Acc:HGNC:3 |
| RPS15P9 | 19 | ribosomal protein S15 pseudogene 9 [Source: |
| AC005330.1 | 19 | novel transcript |
| PWWP3A | 19 | PWWP domain containing 3A, DNA repair fact |
| AC004623.1 | 19 | TEC |
| AC005329.2 | 19 | novel transcript, antisense to MUM1 |
| NDUFS7 | 19 | NADH:ubiquinone oxidoreductase core subun |
| AC005329.3 | 19 | novel transcript, antisense to NDUFS7 |
| AC005329.1 | 19 | novel transcript, antisense to NDUFS7 |
| GAMT | 19 | guanidinoacetate N-methyltransferase [Sourc |
| DAZAP1 | 19 | DAZ associated protein 1 [Source:HGNC Symb |
| RPS15 | 19 | ribosomal protein S15 [Source:HGNC Symbol; |
| AC027307.3 | 19 | novel transcript |
| APC2 | 19 | APC regulator of WNT signaling pathway 2 [So |
| AC027307.2 | 19 | novel transcript, antisense to APC2 |
| C19orf25 | 19 | chromosome 19 open reading frame 25 [Sour |
| PCSK4 | 19 | proprotein convertase subtilisin/kexin type 4 |
| REEP6 | 19 | receptor accessory protein 6 [Source:HGNC Sy |
| ADAMTSL5 | 19 | ADAMTS like 5 [Source:HGNC Symbol;Acc:HG |
| AC027307.1 | 19 | novel transcript, antisense to ADAMTSL5 |
| PLK5 | 19 | polo like kinase 5 [Source:HGNC Symbol;Acc:H |
| MEX3D | 19 | mex-3 RNA binding family member D [Source: |

| | | |
|---|---|---|
| RN7SL477P | 19 | RNA, 7SL, cytoplasmic 477, pseudogene [Sour |
| MBD3 | 19 | methyl-CpG binding domain protein 3 [Source |
| AC005943.2 | 19 | TEC |
| AC005943.1 | 19 | novel transcript, readthrough between UQCR: |
| UQCR11 | 19 | ubiquinol-cytochrome c reductase, complex II |
| TCF3 | 19 | transcription factor 3 [Source:HGNC Symbol;A |
| RNU6-1223P | 19 | RNA, U6 small nuclear 1223, pseudogene [Sou |
| Metazoa_SRP | 19 | Metazoan signal recognition particle RNA [Sou |
| AC005256.1 | 19 | novel transcript |
| ONECUT3 | 19 | one cut homeobox 3 [Source:HGNC Symbol;A |
| ATP8B3 | 19 | ATPase phospholipid transporting 8B3 [Source |
| REXO1 | 19 | RNA exonuclease 1 homolog [Source:HGNC Sy |
| AC012615.2 | 19 | novel transcript, antisense to REXO1 |
| MIR1909 | 19 | microRNA 1909 [Source:HGNC Symbol;Acc:HG |
| AC012615.6 | 19 | novel transcript, antisense to REXO1 |
| AC012615.3 | 19 | novel transcript, antisense to KLF16 |
| KLF16 | 19 | Kruppel like factor 16 [Source:HGNC Symbol;A |
| AC012615.4 | 19 | novel transcript, antisense to KLF16 |
| AC012615.1 | 19 | novel transcript |
| AC012615.5 | 19 | novel transcript |
| ABHD17A | 19 | abhydrolase domain containing 17A [Source:H |
| SCAMP4 | 19 | secretory carrier membrane protein 4 [Source |
| ADAT3 | 19 | adenosine deaminase tRNA specific 3 [Source |
| CSNK1G2 | 19 | casein kinase 1 gamma 2 [Source:HGNC Symb |
| CSNK1G2-AS1 | 19 | CSNK1G2 antisense RNA 1 [Source:HGNC Sym |
| BTBD2 | 19 | BTB domain containing 2 [Source:HGNC Symb |
| AC005306.1 | 19 | novel transcript, antisense to BTBD2 |
| AC004678.2 | 19 | TEC |
| AC004678.1 | 19 | novel pseudogene |
| AC007136.1 | 19 | ribosomal protein L36a (RPL36A) pseudogene |
| MKNK2 | 19 | MAPK interacting serine/threonine kinase 2 [S |
| MOB3A | 19 | MOB kinase activator 3A [Source:HGNC Symb |
| IZUMO4 | 19 | IZUMO family member 4 [Source:HGNC Symb |
| AP3D1 | 19 | adaptor related protein complex 3 subunit de |
| DOT1L | 19 | DOT1 like histone lysine methyltransferase [Sc |
| AC004490.1 | 19 | novel transcript, antisense to DOT1L |
| PLEKHJ1 | 19 | pleckstrin homology domain containing J1 [So |
| MIR1227 | 19 | microRNA 1227 [Source:HGNC Symbol;Acc:HG |
| MIR6789 | 19 | microRNA 6789 [Source:HGNC Symbol;Acc:HG |
| SF3A2 | 19 | splicing factor 3a subunit 2 [Source:HGNC Sym |
| AMH | 19 | anti-Mullerian hormone [Source:HGNC Symbo |
| MIR4321 | 19 | microRNA 4321 [Source:HGNC Symbol;Acc:HG |
| JSRP1 | 19 | junctional sarcoplasmic reticulum protein 1 [S |
| OAZ1 | 19 | ornithine decarboxylase antizyme 1 [Source:H |
| AC005258.1 | 19 | novel protein, readthrough between OAZ1 an |
| PEAK3 | 19 | PEAK family member 3 [Source:HGNC Symbol |
| LINGO3 | 19 | leucine rich repeat and Ig domain containing 3 |
| AC104530.1 | 19 | novel transcript |
| AC005258.2 | 19 | novel transcript |
| LSM7 | 19 | LSM7 homolog, U6 small nuclear RNA and mR |

| | | |
|---|---|---|
| SPPL2B | 19 | signal peptide peptidase like 2B [Source:HGN( |
| TMPRSS9 | 19 | transmembrane serine protease 9 [Source:HG |
| TIMM13 | 19 | translocase of inner mitochondrial membrane |
| LMNB2 | 19 | lamin B2 [Source:HGNC Symbol;Acc:HGNC:66: |
| MIR7108 | 19 | microRNA 7108 [Source:HGNC Symbol;Acc:HG |
| LINC01775 | 19 | long intergenic non-protein coding RNA 1775 |
| GADD45B | 19 | growth arrest and DNA damage inducible beta |
| RNU6-993P | 19 | RNA, U6 small nuclear 993, pseudogene [Sour |
| GNG7 | 19 | G protein subunit gamma 7 [Source:HGNC Syr |
| ELOCP28 | 19 | elongin C pseudogene 28 [Source:HGNC Symb |
| RN7SL121P | 19 | RNA, 7SL, cytoplasmic 121, pseudogene [Sour |
| AC092068.2 | 19 | novel transcript, sense intronic to GNG7 |
| MIR7850 | 19 | microRNA 7850 [Source:HGNC Symbol;Acc:HG |
| AC092068.1 | 19 | novel transcript, antisense to GNG7 |
| AC092068.3 | 19 | novel transcript, sense intronic to GNG7 |
| AC006538.3 | 19 | novel transcript, sense intronic to GNG7 |
| DIRAS1 | 19 | DIRAS family GTPase 1 [Source:HGNC Symbol; |
| AC006538.2 | 19 | novel protein |
| AC006538.4 | 19 | novel transcript |
| AC006538.1 | 19 | novel transcript |
| SLC39A3 | 19 | solute carrier family 39 member 3 [Source:HG |
| SGTA | 19 | small glutamine rich tetratricopeptide repeat |
| THOP1 | 19 | thimet oligopeptidase 1 [Source:HGNC Symbo |
| ZNF554 | 19 | zinc finger protein 554 [Source:HGNC Symbol; |
| ZNF555 | 19 | zinc finger protein 555 [Source:HGNC Symbol; |
| AC006130.1 | 19 | novel transcript, antisense to ZNF555 |
| AC006130.3 | 19 | novel transcript |
| ZNF556 | 19 | zinc finger protein 556 [Source:HGNC Symbol; |
| AC006130.2 | 19 | ribosomal protein L35a (RPL35A) pseudogene |
| AC119403.2 | 19 | zinc finger protein pseudogene |
| ZNF57 | 19 | zinc finger protein 57 [Source:HGNC Symbol;A |
| AC119403.1 | 19 | novel transcript, antisense to ZNF57 |
| ZNF77 | 19 | zinc finger protein 77 [Source:HGNC Symbol;A |
| AC006277.1 | 19 | TEC |
| TLE6 | 19 | TLE family member 6, subcortical maternal co |
| TLE2 | 19 | TLE family member 2, transcriptional corepres |
| AC005944.1 | 19 | novel transcript, antisense to AES |
| TLE5 | 19 | TLE family member 5, transcriptional modulat |
| GNA11 | 19 | G protein subunit alpha 11 [Source:HGNC Sym |
| AC005262.2 | 19 | novel transcript, antisense to GNA11 |
| KF456478.1 | 19 | novel transcript, antisense to GNA11 |
| AC005262.1 | 19 | ribosomal protein L35 (RPL35) pseudogene |
| GNA15 | 19 | G protein subunit alpha 15 [Source:HGNC Sym |
| AC005264.1 | 19 | novel transcript, antisense to GNA15 |
| S1PR4 | 19 | sphingosine-1-phosphate receptor 4 [Source:I |
| NCLN | 19 | nicalin [Source:HGNC Symbol;Acc:HGNC:2692 |
| CELF5 | 19 | CUGBP Elav-like family member 5 [Source:HGI |
| AC010649.1 | 19 | novel transcript, antisense to CELF5 |
| NFIC | 19 | nuclear factor I C [Source:HGNC Symbol;Acc:H |
| AC007792.1 | 19 | TEC |

| | | |
|---|---|---|
| AC005514.1 | 19 | TEC |
| SMIM24 | 19 | small integral membrane protein 24 [Source:H |
| AC005551.1 | 19 | novel protein |
| DOHH | 19 | deoxyhypusine hydroxylase [Source:HGNC Syr |
| FZR1 | 19 | fizzy and cell division cycle 20 related 1 [Sourc |
| AC005787.1 | 19 | NULL |
| MFSD12 | 19 | major facilitator superfamily domain containir |
| C19orf71 | 19 | chromosome 19 open reading frame 71 [Sour |
| AC005786.3 | 19 | novel transcript, antisense to MFSD12 |
| AC005786.2 | 19 | novel transcript |
| AC005786.4 | 19 | TEC |
| AC005786.1 | 19 | ribosomal protein S7 (RPS7) pseudogene |
| HMG20B | 19 | high mobility group 20B [Source:HGNC Symbc |
| GIPC3 | 19 | GIPC PDZ domain containing family member 3 |
| TBXA2R | 19 | thromboxane A2 receptor [Source:HGNC Sym |
| CACTIN-AS1 | 19 | CACTIN antisense RNA 1 [Source:HGNC Symbc |
| CACTIN | 19 | cactin, spliceosome C complex subunit [Sourc |
| PIP5K1C | 19 | phosphatidylinositol-4-phosphate 5-kinase ty |
| AC004637.1 | 19 | novel transcript, antisense to PIP5K1C |
| TJP3 | 19 | tight junction protein 3 [Source:HGNC Symbol |
| APBA3 | 19 | amyloid beta precursor protein binding family |
| AC005954.2 | 19 | novel transcript, antisense to APBA3 |
| AC005954.1 | 19 | novel transcript, sense intronic to APBA3 |
| MRPL54 | 19 | mitochondrial ribosomal protein L54 [Source: |
| RAX2 | 19 | retina and anterior neural fold homeobox 2 [S |
| AC005777.1 | 19 | megakaryocyte-associated tyrosine kinase (M |
| MATK | 19 | megakaryocyte-associated tyrosine kinase [So |
| ZFR2 | 19 | zinc finger RNA binding protein 2 [Source:HGN |
| FTLP5 | 19 | ferritin light chain pseudogene 5 [Source:HGN |
| ATCAY | 19 | ATCAY kinesin light chain interacting caytaxin |
| RN7SL202P | 19 | RNA, 7SL, cytoplasmic 202, pseudogene [Sour |
| NMRK2 | 19 | nicotinamide riboside kinase 2 [Source:HGNC |
| DAPK3 | 19 | death associated protein kinase 3 [Source:HGI |
| MIR637 | 19 | microRNA 637 [Source:HGNC Symbol;Acc:HGI |
| EEF2 | 19 | eukaryotic translation elongation factor 2 [Sou |
| SNORD37 | 19 | small nucleolar RNA, C/D box 37 [Source:HGN |
| PIAS4 | 19 | protein inhibitor of activated STAT 4 [Source:H |
| ZBTB7A | 19 | zinc finger and BTB domain containing 7A [Sou |
| AC016586.1 | 19 | novel transcript, sense intronic to ZBTB7A |
| MAP2K2 | 19 | mitogen-activated protein kinase kinase 2 [So |
| Metazoa_SRP | 19 | Metazoan signal recognition particle RNA [Sou |
| CREB3L3 | 19 | cAMP responsive element binding protein 3 li |
| SIRT6 | 19 | sirtuin 6 [Source:HGNC Symbol;Acc:HGNC:14 |
| ANKRD24 | 19 | ankyrin repeat domain 24 [Source:HGNC Syml |
| EBI3 | 19 | Epstein-Barr virus induced 3 [Source:HGNC Sy |
| AC005578.1 | 19 | TEC |
| YJU2 | 19 | YJU2 splicing factor homolog [Source:HGNC S |
| SHD | 19 | Src homology 2 domain containing transformi |
| TMIGD2 | 19 | transmembrane and immunoglobulin domain |
| FSD1 | 19 | fibronectin type III and SPRY domain containir |

| | | |
|---|---|---|
| STAP2 | 19 | signal transducing adaptor family member 2 [S |
| AC104521.1 | 19 | novel transcript, antisense to STAP2 |
| MPND | 19 | MPN domain containing [Source:HGNC Symbo |
| AC007292.3 | 19 | novel transcript, antisense to MPND |
| EIF1P6 | 19 | eukaryotic translation initiation factor 1 pseu |
| AC007292.2 | 19 | novel transcript, antisense to MPND |
| SH3GL1 | 19 | SH3 domain containing GRB2 like 1, endophili |
| AC007292.1 | 19 | novel transcript, antisense to SH3GL1 |
| CHAF1A | 19 | chromatin assembly factor 1 subunit A [Sourc |
| AC011498.3 | 19 | novel transcript, antisense to CHAF1A |
| UBXN6 | 19 | UBX domain protein 6 [Source:HGNC Symbol; |
| MIR4746 | 19 | microRNA 4746 [Source:HGNC Symbol;Acc:HG |
| AC011498.2 | 19 | novel transcript, antisense to UBXN6 |
| AC011498.7 | 19 | TEC |
| AC011498.6 | 19 | novel transcript, antisense to UBX6 |
| AC011498.1 | 19 | novel transcript |
| HDGFL2 | 19 | HDGF like 2 [Source:HGNC Symbol;Acc:HGNC: |
| PLIN4 | 19 | perilipin 4 [Source:HGNC Symbol;Acc:HGNC:2 |
| PLIN5 | 19 | perilipin 5 [Source:HGNC Symbol;Acc:HGNC:3 |
| AC011498.4 | 19 | novel protein |
| LRG1 | 19 | leucine rich alpha-2-glycoprotein 1 [Source:H |
| SEMA6B | 19 | semaphorin 6B [Source:HGNC Symbol;Acc:HG |
| AC011498.5 | 19 | ribosomal protein S10 (RPS10) pseudogene |
| TNFAIP8L1 | 19 | TNF alpha induced protein 8 like 1 [Source:HG |
| MYDGF | 19 | myeloid derived growth factor [Source:HGNC |
| AC005339.1 | 19 | novel transcript, antisense to TNFAIP8L1 |
| DPP9 | 19 | dipeptidyl peptidase 9 [Source:HGNC Symbol; |
| DPP9-AS1 | 19 | DPP9 antisense RNA 1 [Source:HGNC Symbol; |
| AC005594.1 | 19 | novel transcript, antisense to DPP9 |
| MIR7-3HG | 19 | MIR7-3 host gene [Source:HGNC Symbol;Acc: |
| MIR7-3 | 19 | microRNA 7-3 [Source:HGNC Symbol;Acc:HGN |
| AC005523.1 | 19 | novel transcript |
| FEM1A | 19 | fem-1 homolog A [Source:HGNC Symbol;Acc:H |
| AC005523.2 | 19 | novel transcript, antisense to FEM1A |
| TICAM1 | 19 | toll like receptor adaptor molecule 1 [Source: |
| AC027319.1 | 19 | novel transcript, antisense to PLIN3 |
| PLIN3 | 19 | perilipin 3 [Source:HGNC Symbol;Acc:HGNC:1 |
| ARRDC5 | 19 | arrestin domain containing 5 [Source:HGNC Sy |
| UHRF1 | 19 | ubiquitin like with PHD and ring finger domair |
| MIR4747 | 19 | microRNA 4747 [Source:HGNC Symbol;Acc:HG |
| KDM4B | 19 | lysine demethylase 4B [Source:HGNC Symbol; |
| PTPRS | 19 | protein tyrosine phosphatase receptor type S |
| AC022517.1 | 19 | novel transcript, antisense to PTPRS |
| RPL32P34 | 19 | ribosomal protein L32 pseudogene 34 [Source |
| AC005790.1 | 19 | novel transcript, antisense to PTPRS |
| ZNRF4 | 19 | zinc and ring finger 4 [Source:HGNC Symbol;A |
| TINCR | 19 | TINCR ubiquitin domain containing [Source:H |
| SNRPEP4 | 19 | small nuclear ribonucleoprotein polypeptide E |
| SAFB2 | 19 | scaffold attachment factor B2 [Source:HGNC S |
| SAFB | 19 | scaffold attachment factor B [Source:HGNC Sy |

| | | |
|---|---|---|
| RPL36 | 19 | ribosomal protein L36 [Source:HGNC Symbol;/ |
| MICOS13 | 19 | mitochondrial contact site and cristae organiz |
| HSD11B1L | 19 | hydroxysteroid 11-beta dehydrogenase 1 like |
| LONP1 | 19 | lon peptidase 1, mitochondrial [Source:HGNC |
| CATSPERD | 19 | cation channel sperm associated auxiliary sub |
| PRR22 | 19 | proline rich 22 [Source:HGNC Symbol;Acc:HGI |
| AC011499.1 | 19 | novel protein |
| DUS3L | 19 | dihydrouridine synthase 3 like [Source:HGNC S |
| NRTN | 19 | neurturin [Source:HGNC Symbol;Acc:HGNC:80 |
| FUT6 | 19 | fucosyltransferase 6 [Source:HGNC Symbol;Ac |
| FUT3 | 19 | fucosyltransferase 3 (Lewis blood group) [Sou |
| AC024592.2 | 19 | novel transcript, antisense to FUT3 |
| FUT5 | 19 | fucosyltransferase 5 [Source:HGNC Symbol;Ac |
| AC024592.3 | 19 | novel protein |
| NDUFA11 | 19 | NADH:ubiquinone oxidoreductase subunit A1: |
| AC024592.1 | 19 | novel transcript, antisense to NDUFA11 |
| AC104532.1 | 19 | novel protein, readthrough VMAC-CAPS |
| VMAC | 19 | vimentin type intermediate filament associate |
| AC104532.2 | 19 | novel transcript, antisense to CAPS |
| CAPS | 19 | calcyphosine [Source:HGNC Symbol;Acc:HGN( |
| RANBP3 | 19 | RAN binding protein 3 [Source:HGNC Symbol;/ |
| AC011444.1 | 19 | novel transcript, antisense to RFX2 |
| RFX2 | 19 | regulatory factor X2 [Source:HGNC Symbol;Ac |
| AC011444.2 | 19 | novel transcript, antisense to RFX2 |
| AC011444.4 | 19 | ribosomal protein L32 (RPL32) pseudogene |
| AC011444.3 | 19 | novel transcript, antisense to RFX2 |
| ACSBG2 | 19 | acyl-CoA synthetase bubblegum family memb |
| AC011471.1 | 19 | acyl-CoA synthetase bubblegum family memb |
| AC011471.4 | 19 | acyl-CoA synthetase bubblegum family memb |
| MLLT1 | 19 | MLLT1 super elongation complex subunit [Sou |
| AC011471.2 | 19 | novel transcript, antisense to MLLT1 |
| AC011471.3 | 19 | novel transcript, sense intronic to MLLT1 |
| ACER1 | 19 | alkaline ceramidase 1 [Source:HGNC Symbol;/ |
| AC011491.3 | 19 | novel transcript, antisense to CLPP |
| CLPP | 19 | caseinolytic mitochondrial matrix peptidase p |
| ALKBH7 | 19 | alkB homolog 7 [Source:HGNC Symbol;Acc:HG |
| PSPN | 19 | persephin [Source:HGNC Symbol;Acc:HGNC:9! |
| GTF2F1 | 19 | general transcription factor IIF subunit 1 [Sour |
| AC011491.2 | 19 | novel transcript, sense overlapping GTF2F1 |
| MIR6885 | 19 | microRNA 6885 [Source:HGNC Symbol;Acc:HG |
| MIR6790 | 19 | microRNA 6790 [Source:HGNC Symbol;Acc:HG |
| AC011491.1 | 19 | adenylate kinase isoenzyme 1-like [Source:NC |
| KHSRP | 19 | KH-type splicing regulatory protein [Source:H( |
| MIR3940 | 19 | microRNA 3940 [Source:HGNC Symbol;Acc:HG |
| SLC25A41 | 19 | solute carrier family 25 member 41 [Source:H |
| SLC25A23 | 19 | solute carrier family 25 member 23 [Source:H |
| CRB3 | 19 | crumbs cell polarity complex component 3 [Sc |
| DENND1C | 19 | DENN domain containing 1C [Source:HGNC Sy |
| AC010503.4 | 19 | novel transcript, antisense to DENND1C |
| TUBB4A | 19 | tubulin beta 4A class IVa [Source:HGNC Symbo |

| | | |
|---|---|---|
| AC010503.1 | 19 | novel transcript, antisense to TUBB4A |
| AC010503.2 | 19 | novel transcript, antisense to TUBB4A |
| TNFSF9 | 19 | TNF superfamily member 9 [Source:HGNC Syn |
| AC010503.3 | 19 | transforming growth factor, beta receptor 1 (T |
| CLIC4P2 | 19 | chloride intracellular channel 4 pseudogene 2 |
| Y_RNA | 19 | Y RNA [Source:RFAM;Acc:RF00019] |
| CD70 | 19 | CD70 molecule [Source:HGNC Symbol;Acc:HG |
| RPL7P50 | 19 | ribosomal protein L7 pseudogene 50 [Source: |
| TNFSF14 | 19 | TNF superfamily member 14 [Source:HGNC Sy |
| C3 | 19 | complement C3 [Source:HGNC Symbol;Acc:HC |
| AC008760.2 | 19 | novel transcript, sense intronic C3 |
| GPR108 | 19 | G protein-coupled receptor 108 [Source:HGN( |
| MIR6791 | 19 | microRNA 6791 [Source:HGNC Symbol;Acc:HC |
| TRIP10 | 19 | thyroid hormone receptor interactor 10 [Sour |
| AC008760.1 | 19 | novel transcript, antisense to TRIP10 |
| SH2D3A | 19 | SH2 domain containing 3A [Source:HGNC Sym |
| VAV1 | 19 | vav guanine nucleotide exchange factor 1 [Sou |
| ADGRE1 | 19 | adhesion G protein-coupled receptor E1 [Sour |
| AC020895.2 | 19 | novel transcript |
| AC020895.1 | 19 | egf-like module containing, mucin-like, hormo |
| Metazoa_SRP | 19 | Metazoan signal recognition particle RNA [Sou |
| ADGRE4P | 19 | adhesion G protein-coupled receptor E4, pseu |
| AC025278.1 | 19 | novel transcript, antisense to EMR4P |
| AC025278.2 | 19 | MAP/microtubule affinity-regulating kinase (M |
| AC025278.3 | 19 | MAP/microtubule affinity-regulating kinase 1 |
| MBD3L2B | 19 | methyl-CpG binding domain protein 3 like 2B |
| MBD3L5 | 19 | methyl-CpG binding domain protein 3 like 5 [S |
| MBD3L4 | 19 | methyl-CpG binding domain protein 3 like 4 [S |
| MBD3L2 | 19 | methyl-CpG binding domain protein 3 like 2 [S |
| MBD3L3 | 19 | methyl-CpG binding domain protein 3 like 3 [S |
| ZNF557 | 19 | zinc finger protein 557 [Source:HGNC Symbol; |
| INSR | 19 | insulin receptor [Source:HGNC Symbol;Acc:HC |
| AC119396.1 | 19 | novel protein |
| AC008878.3 | 19 | Rho/Rac guanine nucleotide exchange factor : |
| AC119396.2 | 19 | novel transcript, antisense to ARHGEF18 |
| ARHGEF18 | 19 | Rho/Rac guanine nucleotide exchange factor : |
| PEX11G | 19 | peroxisomal biogenesis factor 11 gamma [Sou |
| TEX45 | 19 | testis expressed 45 [Source:HGNC Symbol;Acc |
| AC008878.1 | 19 | novel transcript |
| ZNF358 | 19 | zinc finger protein 358 [Source:HGNC Symbol; |
| AC008878.4 | 19 | novel transcript, antisense to ZNF358 |
| MCOLN1 | 19 | mucolipin 1 [Source:HGNC Symbol;Acc:HGNC: |
| AC008878.2 | 19 | novel transcript |
| PNPLA6 | 19 | patatin like phospholipase domain containing |
| CAMSAP3 | 19 | calmodulin regulated spectrin associated prot |
| MIR6792 | 19 | microRNA 6792 [Source:HGNC Symbol;Acc:HC |
| XAB2 | 19 | XPA binding protein 2 [Source:HGNC Symbol;A |
| PET100 | 19 | PET100 cytochrome c oxidase chaperone [Sou |
| AC008763.2 | 19 | novel transcript |
| PCP2 | 19 | Purkinje cell protein 2 [Source:HGNC Symbol;A |

| | | |
|---|---|---|
| AC008763.1 | 19 | novel transcript, antisense to STXBP2 |
| STXBP2 | 19 | syntaxin binding protein 2 [Source:HGNC Sym |
| RPS27AP19 | 19 | ribosomal protein S27a pseudogene 19 [Sourc |
| RETN | 19 | resistin [Source:HGNC Symbol;Acc:HGNC:2038 |
| MCEMP1 | 19 | mast cell expressed membrane protein 1 [Sou |
| AC008763.3 | 19 | novel protein |
| TRAPPC5 | 19 | trafficking protein particle complex 5 [Source: |
| FCER2 | 19 | Fc fragment of IgE receptor II [Source:HGNC S |
| CLEC4G | 19 | C-type lectin domain family 4 member G [Sou |
| CD209 | 19 | CD209 molecule [Source:HGNC Symbol;Acc:H |
| RPL21P129 | 19 | ribosomal protein L21 pseudogene 129 [Sourc |
| CLEC4M | 19 | C-type lectin domain family 4 member M [Sou |
| CLEC4GP1 | 19 | C-type lectin domain family 4 member G pseu |
| EXOSC3P2 | 19 | exosome component 3 pseudogene 2 [Source |
| EVI5L | 19 | ecotropic viral integration site 5 like [Source:H |
| PRR36 | 19 | proline rich 36 [Source:HGNC Symbol;Acc:HGN |
| AC010336.3 | 19 | novel transcript |
| LYPLA2P2 | 19 | LYPLA2 pseudogene 2 [Source:HGNC Symbol;A |
| AC010336.7 | 19 | NULL |
| RNA5SP463 | 19 | RNA, 5S ribosomal pseudogene 463 [Source:H |
| LRRC8E | 19 | leucine rich repeat containing 8 VRAC subunit |
| AC010336.1 | 19 | novel transcript, antisense to LRRC8E |
| MAP2K7 | 19 | mitogen-activated protein kinase kinase 7 [So |
| AC010336.4 | 19 | novel transcript, antisense to MAP2K7 |
| TGFBR3L | 19 | transforming growth factor beta receptor 3 lik |
| AC010336.2 | 19 | novel transcript, antisense to CTD-3193O13.2 |
| SNAPC2 | 19 | small nuclear RNA activating complex polypep |
| CTXN1 | 19 | cortexin 1 [Source:HGNC Symbol;Acc:HGNC:3 |
| AC010336.5 | 19 | novel transcript, antisense to CTXN1 and TIMI |
| TIMM44 | 19 | translocase of inner mitochondrial membrane |
| AC010336.8 | 19 | novel transcript |
| ELAVL1 | 19 | ELAV like RNA binding protein 1 [Source:HGN( |
| AC010336.6 | 19 | novel transcript, antisense to ELAVL1 |
| AC008946.1 | 19 | novel transcript |
| CCL25 | 19 | C-C motif chemokine ligand 25 [Source:HGNC |
| FBN3 | 19 | fibrillin 3 [Source:HGNC Symbol;Acc:HGNC:18 |
| AC022146.1 | 19 | asparagine-linked glycosylation 11, alpha-1,2- |
| CERS4 | 19 | ceramide synthase 4 [Source:HGNC Symbol;A( |
| AC022146.2 | 19 | novel transcript, antisense to CERS4 |
| CD320 | 19 | CD320 molecule [Source:HGNC Symbol;Acc:H |
| AC010323.1 | 19 | novel transcript |
| NDUFA7 | 19 | NADH:ubiquinone oxidoreductase subunit A7 |
| RPS28 | 19 | ribosomal protein S28 [Source:HGNC Symbol;A |
| KANK3 | 19 | KN motif and ankyrin repeat domains 3 [Sourc |
| AC010323.2 | 19 | ribosomal protein L17 (RPL17) pseudogene |
| ANGPTL4 | 19 | angiopoietin like 4 [Source:HGNC Symbol;Acc: |
| RAB11B-AS1 | 19 | RAB11B antisense RNA 1 [Source:HGNC Symb |
| MIR4999 | 19 | microRNA 4999 [Source:HGNC Symbol;Acc:HG |
| RAB11B | 19 | RAB11B, member RAS oncogene family [Sourc |
| MARCH2 | 19 | membrane associated ring-CH-type finger 2 [S |

| | | |
|---|---|---|
| AC136469.1 | 19 | novel transcript, antisense to MARCH2 |
| AC136469.2 | 19 | novel transcript |
| HNRNPM | 19 | heterogeneous nuclear ribonucleoprotein M [ |
| PRAM1 | 19 | PML-RARA regulated adaptor molecule 1 [Sou |
| ZNF414 | 19 | zinc finger protein 414 [Source:HGNC Symbol; |
| MYO1F | 19 | myosin IF [Source:HGNC Symbol;Acc:HGNC:76 |
| AC092316.1 | 19 | novel transcript, antisense to MYO1F |
| ADAMTS10 | 19 | ADAM metallopeptidase with thrombospondi |
| AC130469.1 | 19 | novel transcript, antisense to ADAMTS10 |
| LINC01862 | 19 | long intergenic non-protein coding RNA 1862 |
| AC243312.1 | 19 | TEC |
| ACTL9 | 19 | actin like 9 [Source:HGNC Symbol;Acc:HGNC:2 |
| OR2Z1 | 19 | olfactory receptor family 2 subfamily Z memb |
| RPL23AP78 | 19 | ribosomal protein L23a pseudogene 78 [Sourc |
| AC012616.1 | 19 | proline-rich protein pseudogene |
| ZNF558 | 19 | zinc finger protein 558 [Source:HGNC Symbol; |
| AC008734.1 | 19 | novel transcript, antisense to ZNF558 |
| MBD3L1 | 19 | methyl-CpG binding domain protein 3 like 1 [S |
| AC008734.2 | 19 | novel transcript, sense intronic to MBD3L1 |
| MUC16 | 19 | mucin 16, cell surface associated [Source:HGN |
| BOLA3P2 | 19 | bolA family member 3 pseudogene 2 [Source: |
| OR1M4P | 19 | olfactory receptor family 1 subfamily M meml |
| OR1M1 | 19 | olfactory receptor family 1 subfamily M meml |
| OR7G2 | 19 | olfactory receptor family 7 subfamily G memb |
| OR7G1 | 19 | olfactory receptor family 7 subfamily G memb |
| OR7G15P | 19 | olfactory receptor family 7 subfamily G memb |
| OR7G3 | 19 | olfactory receptor family 7 subfamily G memb |
| ZNF317 | 19 | zinc finger protein 317 [Source:HGNC Symbol; |
| OR7D2 | 19 | olfactory receptor family 7 subfamily D memb |
| ELOCP29 | 19 | elongin C pseudogene 29 [Source:HGNC Symb |
| OR7E16P | 19 | olfactory receptor family 7 subfamily E memb |
| OR7E25P | 19 | olfactory receptor family 7 subfamily E memb |
| OR7D4 | 19 | olfactory receptor family 7 subfamily D memb |
| OR7D1P | 19 | olfactory receptor family 7 subfamily D memb |
| OR7E24 | 19 | olfactory receptor family 7 subfamily E memb |
| OR7E18P | 19 | olfactory receptor family 7 subfamily E memb |
| OR7E19P | 19 | olfactory receptor family 7 subfamily E memb |
| OR7H1P | 19 | olfactory receptor family 7 subfamily H memb |
| ZNF699 | 19 | zinc finger protein 699 [Source:HGNC Symbol; |
| AC011451.1 | 19 | novel transcript, antisense to ZNF699 |
| ZNF559 | 19 | zinc finger protein 559 [Source:HGNC Symbol; |
| ZNF559-ZNF177 | 19 | ZNF559-ZNF177 readthrough [Source:HGNC S |
| AC011451.2 | 19 | UBX domain protein 2A (UBXN2A) pseudogen |
| ZNF177 | 19 | zinc finger protein 177 [Source:HGNC Symbol; |
| AC011451.3 | 19 | novel transcript |
| ZNF266 | 19 | zinc finger protein 266 [Source:HGNC Symbol; |
| AC011451.4 | 19 | novel transcript |
| ZNF560 | 19 | zinc finger protein 560 [Source:HGNC Symbol; |
| AC008567.2 | 19 | novel transcript |
| AC008567.3 | 19 | novel transcript |

| | | |
|---|---|---|
| AC008567.1 | 19 | actin, beta (ACTB) pseudogene |
| ZNF426 | 19 | zinc finger protein 426 [Source:HGNC Symbol; |
| ZNF426-DT | 19 | ZNF426 divergent transcript [Source:HGNC Sy |
| ZNF121 | 19 | zinc finger protein 121 [Source:HGNC Symbol; |
| ZNF561 | 19 | zinc finger protein 561 [Source:HGNC Symbol; |
| ZNF561-AS1 | 19 | ZNF561 antisense RNA 1 (head to head) [Sour |
| ZNF562 | 19 | zinc finger protein 562 [Source:HGNC Symbol; |
| RPS4XP22 | 19 | ribosomal protein S4X pseudogene 22 [Source |
| ZNF812P | 19 | zinc finger protein 812, pseudogene [Source:H |
| AC008759.3 | 19 | novel transcript |
| AC008759.1 | 19 | zinc finger protein 562 (ZNF562) pseudogene |
| AC008759.2 | 19 | novel transcript |
| ZNF846 | 19 | zinc finger protein 846 [Source:HGNC Symbol; |
| AC008752.3 | 19 | zinc finger protein 420 (ZNF420) pseudogene |
| UBE2L4 | 19 | ubiquitin conjugating enzyme E2 L4 (pseudoge |
| AC008752.1 | 19 | novel transcript |
| FBXL12 | 19 | F-box and leucine rich repeat protein 12 [Sour |
| SNORA70 | 19 | Small nucleolar RNA SNORA70 [Source:RFAM; |
| RPL10P15 | 19 | ribosomal protein L10 pseudogene 15 [Source |
| UBL5 | 19 | ubiquitin like 5 [Source:HGNC Symbol;Acc:HGI |
| AC008752.2 | 19 | novel transcript |
| PIN1 | 19 | peptidylprolyl cis/trans isomerase, NIMA-inte |
| OLFM2 | 19 | olfactomedin 2 [Source:HGNC Symbol;Acc:HG |
| COL5A3 | 19 | collagen type V alpha 3 chain [Source:HGNC S |
| AC008742.1 | 19 | novel transcript, antisense to COL5A3 |
| RDH8 | 19 | retinol dehydrogenase 8 [Source:HGNC Symbo |
| C3P1 | 19 | complement component 3 precursor pseudog |
| MIR5589 | 19 | microRNA 5589 [Source:HGNC Symbol;Acc:HG |
| SHFL | 19 | shiftless antiviral inhibitor of ribosomal frame |
| AC020931.1 | 19 | novel transcript, antisense to C19orf66 |
| ANGPTL6 | 19 | angiopoietin like 6 [Source:HGNC Symbol;Acc: |
| PPAN-P2RY11 | 19 | PPAN-P2RY11 readthrough [Source:HGNC Syn |
| PPAN | 19 | peter pan homolog [Source:HGNC Symbol;Acc |
| SNORD105 | 19 | small nucleolar RNA, C/D box 105 [Source:HGI |
| SNORD105B | 19 | small nucleolar RNA, C/D box 105B [Source:H( |
| P2RY11 | 19 | purinergic receptor P2Y11 [Source:HGNC Sym |
| EIF3G | 19 | eukaryotic translation initiation factor 3 subur |
| DNMT1 | 19 | DNA methyltransferase 1 [Source:HGNC Symb |
| S1PR2 | 19 | sphingosine-1-phosphate receptor 2 [Source:I |
| MIR4322 | 19 | microRNA 4322 [Source:HGNC Symbol;Acc:HG |
| MRPL4 | 19 | mitochondrial ribosomal protein L4 [Source:H |
| AC011511.2 | 19 | novel transcript, antisense to MRPL4 and ICAN |
| AC011511.3 | 19 | novel transcript, antisense to MRPL4 |
| ICAM1 | 19 | intercellular adhesion molecule 1 [Source:HGI |
| AC011511.5 | 19 | novel transcript, antisense to ICAM4 and ICAN |
| ICAM4 | 19 | intercellular adhesion molecule 4 (Landsteine |
| ICAM5 | 19 | intercellular adhesion molecule 5 [Source:HGI |
| ZGLP1 | 19 | zinc finger GATA like protein 1 [Source:HGNC : |
| AC011511.1 | 19 | novel transcript |
| FDX2 | 19 | ferredoxin 2 [Source:HGNC Symbol;Acc:HGNC |

| | | |
|---|---|---|
| AC011511.4 | 19 | novel transcript |
| RAVER1 | 19 | ribonucleoprotein, PTB binding 1 [Source:HGN |
| AC114271.1 | 19 | novel transcript, antisense to ICAM3 |
| ICAM3 | 19 | intercellular adhesion molecule 3 [Source:HGI |
| TYK2 | 19 | tyrosine kinase 2 [Source:HGNC Symbol;Acc:H |
| CDC37 | 19 | cell division cycle 37 [Source:HGNC Symbol;Ac |
| MIR1181 | 19 | microRNA 1181 [Source:HGNC Symbol;Acc:HG |
| PDE4A | 19 | phosphodiesterase 4A [Source:HGNC Symbol; |
| KEAP1 | 19 | kelch like ECH associated protein 1 [Source:HG |
| AC011461.1 | 19 | novel transcript, sense intronic with KEAP1 |
| S1PR5 | 19 | sphingosine-1-phosphate receptor 5 [Source:H |
| ATG4D | 19 | autophagy related 4D cysteine peptidase [Sou |
| RNU7-140P | 19 | RNA, U7 small nuclear 140 pseudogene [Sourc |
| MIR1238 | 19 | microRNA 1238 [Source:HGNC Symbol;Acc:HG |
| KRI1 | 19 | KRI1 homolog [Source:HGNC Symbol;Acc:HGN |
| CDKN2D | 19 | cyclin dependent kinase inhibitor 2D [Source:I |
| AP1M2 | 19 | adaptor related protein complex 1 subunit mu |
| AC011475.1 | 19 | LSM2 homolog, U6 small nuclear RNA associa |
| SLC44A2 | 19 | solute carrier family 44 member 2 [Source:HG |
| ILF3-DT | 19 | ILF3 divergent transcript [Source:HGNC Symbo |
| ILF3 | 19 | interleukin enhancer binding factor 3 [Source: |
| QTRT1 | 19 | queuine tRNA-ribosyltransferase catalytic sub |
| DNM2 | 19 | dynamin 2 [Source:HGNC Symbol;Acc:HGNC:2 |
| MIR638 | 19 | microRNA 638 [Source:HGNC Symbol;Acc:HGN |
| MIR4748 | 19 | microRNA 4748 [Source:HGNC Symbol;Acc:HG |
| MIR199A1 | 19 | microRNA 199a-1 [Source:HGNC Symbol;Acc:H |
| MIR6793 | 19 | microRNA 6793 [Source:HGNC Symbol;Acc:HG |
| TMED1 | 19 | transmembrane p24 trafficking protein 1 [Sou |
| C19orf38 | 19 | chromosome 19 open reading frame 38 [Sour |
| AC007229.1 | 19 | chromosome 11 open reading frame 73 (C11c |
| CARM1 | 19 | coactivator associated arginine methyltransfe |
| YIPF2 | 19 | Yip1 domain family member 2 [Source:HGNC S |
| TIMM29 | 19 | translocase of inner mitochondrial membrane |
| SMARCA4 | 19 | SWI/SNF related, matrix associated, actin dep |
| AC011442.1 | 19 | novel transcript, antisense to SMARCA4 |
| RN7SL192P | 19 | RNA, 7SL, cytoplasmic 192, pseudogene [Sour |
| AC006127.1 | 19 | NULL |
| LDLR | 19 | low density lipoprotein receptor [Source:HGN |
| MIR6886 | 19 | microRNA 6886 [Source:HGNC Symbol;Acc:HG |
| SPC24 | 19 | SPC24 component of NDC80 kinetochore com |
| KANK2 | 19 | KN motif and ankyrin repeat domains 2 [Sourc |
| DOCK6 | 19 | dedicator of cytokinesis 6 [Source:HGNC Symb |
| AC011472.1 | 19 | novel transcript, antisense to DOCK6 |
| AC011472.5 | 19 | TEC |
| AC011472.4 | 19 | novel transcript, antisense to DOCK6 |
| ANGPTL8 | 19 | angiopoietin like 8 [Source:HGNC Symbol;Acc: |
| TSPAN16 | 19 | tetraspanin 16 [Source:HGNC Symbol;Acc:HGI |
| AC011472.2 | 19 | novel transcript |
| RAB3D | 19 | RAB3D, member RAS oncogene family [Source |
| AC011472.3 | 19 | novel transcript, sense intronic to TSPAN16 |

| | | |
|---|---|---|
| TMEM205 | 19 | transmembrane protein 205 [Source:HGNC Sy |
| CCDC159 | 19 | coiled-coil domain containing 159 [Source:HG |
| PLPPR2 | 19 | phospholipid phosphatase related 2 [Source:H |
| AC024575.1 | 19 | novel transcript, antisense to SWSAP1 |
| SWSAP1 | 19 | SWIM-type zinc finger 7 associated protein 1 [ |
| EPOR | 19 | erythropoietin receptor [Source:HGNC Symbo |
| RGL3 | 19 | ral guanine nucleotide dissociation stimulator |
| Y_RNA | 19 | Y RNA [Source:RFAM;Acc:RF00019] |
| CCDC151 | 19 | coiled-coil domain containing 151 [Source:HG |
| PRKCSH | 19 | protein kinase C substrate 80K-H [Source:HGN |
| ELAVL3 | 19 | ELAV like RNA binding protein 3 [Source:HGN( |
| AC008481.3 | 19 | novel protein |
| ZNF653 | 19 | zinc finger protein 653 [Source:HGNC Symbol; |
| MIR7974 | 19 | microRNA 7974 [Source:HGNC Symbol;Acc:HG |
| ECSIT | 19 | ECSIT signalling integrator [Source:HGNC Syml |
| RN7SL833P | 19 | RNA, 7SL, cytoplasmic 833, pseudogene [Sour |
| AC008481.2 | 19 | ribosomal protein L18a (RPL18A) pseudogene |
| AC008481.1 | 19 | ribosomal protein S12 (RPS12) pseudogene |
| CNN1 | 19 | calponin 1 [Source:HGNC Symbol;Acc:HGNC:2 |
| ELOF1 | 19 | elongation factor 1 homolog [Source:HGNC Sy |
| ZNF627 | 19 | zinc finger protein 627 [Source:HGNC Symbol; |
| ACP5 | 19 | acid phosphatase 5, tartrate resistant [Source |
| GAPDHP76 | 19 | glyceraldehyde-3-phosphate dehydrogenase p |
| AC008543.1 | 19 | novel transcript |
| ZNF887P | 19 | zinc finger protein 887, pseudogene [Source:H |
| HNRNPA1P10 | 19 | heterogeneous nuclear ribonucleoprotein A1 |
| ZNF833P | 19 | zinc finger protein 833, pseudogene [Source:H |
| ZNF823 | 19 | zinc finger protein 823 [Source:HGNC Symbol; |
| AC008543.3 | 19 | novel transcript, sense intronic to ZNF823 |
| AC008543.2 | 19 | novel zinc finger protein pseudogene |
| ZNF441 | 19 | zinc finger protein 441 [Source:HGNC Symbol; |
| AC008543.5 | 19 | novel transcript, antisense to ZNF491 |
| ZNF491 | 19 | zinc finger protein 491 [Source:HGNC Symbol; |
| ZNF440 | 19 | zinc finger protein 440 [Source:HGNC Symbol; |
| AC008543.4 | 19 | novel transcript |
| ZNF439 | 19 | zinc finger protein 439 [Source:HGNC Symbol; |
| AC020951.1 | 19 | TEC |
| ZNF69 | 19 | zinc finger protein 69 [Source:HGNC Symbol;A |
| ZNF700 | 19 | zinc finger protein 700 [Source:HGNC Symbol; |
| AC008770.2 | 19 | novel protein |
| AC008770.3 | 19 | novel transcript, sense overlapping with ZNF7 |
| ZNF763 | 19 | zinc finger protein 763 [Source:HGNC Symbol; |
| ZNF433-AS1 | 19 | ZNF433 and ZNF878 antisense RNA 1 [Source: |
| AC008770.4 | 19 | zinc finger protein 844 [Source:NCBI gene;Acc |
| RNA5SP464 | 19 | RNA, 5S ribosomal pseudogene 464 [Source:H |
| ZNF433 | 19 | zinc finger protein 433 [Source:HGNC Symbol; |
| AC008770.1 | 19 | novel protein |
| RNA5SP465 | 19 | RNA, 5S ribosomal pseudogene 465 [Source:H |
| ZNF878 | 19 | zinc finger protein 878 [Source:HGNC Symbol; |
| ZNF844 | 19 | zinc finger protein 844 [Source:HGNC Symbol; |

| | | |
|---|---|---|
| RNA5SP466 | 19 | RNA, 5S ribosomal pseudogene 466 [Source:H |
| AC022415.2 | 19 | novel protein |
| Y_RNA | 19 | Y RNA [Source:RFAM;Acc:RF00019] |
| ZNF788P | 19 | zinc finger family member 788, pseudogene [S |
| ZNF20 | 19 | zinc finger protein 20 [Source:HGNC Symbol;A |
| RSL24D1P8 | 19 | ribosomal L24 domain containing 1 pseudoger |
| RNA5SP467 | 19 | RNA, 5S ribosomal pseudogene 467 [Source:H |
| ZNF625-ZNF20 | 19 | ZNF625-ZNF20 readthrough (NMD candidate) |
| AC022415.1 | 19 | ribosomal protein L17 (RPL17) pseudogene |
| ZNF625 | 19 | zinc finger protein 625 [Source:HGNC Symbol; |
| ZNF136 | 19 | zinc finger protein 136 [Source:HGNC Symbol; |
| AC012618.3 | 19 | novel zinc finger protein pseudogene |
| ZNF44 | 19 | zinc finger protein 44 [Source:HGNC Symbol;A |
| AC012618.1 | 19 | ribosomal protein L28 (RPL28) pseudogene |
| AC012618.2 | 19 | ribosomal protein S2 (RPS2) pseudogene |
| ZNF563 | 19 | zinc finger protein 563 [Source:HGNC Symbol; |
| ZNF442 | 19 | zinc finger protein 442 [Source:HGNC Symbol; |
| RPS29P23 | 19 | ribosomal protein S29 pseudogene 23 [Source |
| AC008758.4 | 19 | novel transcript |
| AC008758.3 | 19 | novel pseudogene similar to zinc finger protei |
| ZNF799 | 19 | zinc finger protein 799 [Source:HGNC Symbol; |
| AC008758.5 | 19 | novel zinc finger protein |
| ZNF443 | 19 | zinc finger protein 443 [Source:HGNC Symbol; |
| AC008758.6 | 19 | novel transcript |
| AC008758.2 | 19 | novel transcript, antisense to ZNF709 |
| ZNF709 | 19 | zinc finger protein 709 [Source:HGNC Symbol; |
| AC008758.1 | 19 | novel zinc finger protein |
| MTND2P40 | 19 | MT-ND2 pseudogene 40 [Source:HGNC Symbo |
| MTCO1P27 | 19 | MT-CO1 pseudogene 27 [Source:HGNC Symbo |
| MTCO2P27 | 19 | MT-CO2 pseudogene 27 [Source:HGNC Symbo |
| MTATP6P27 | 19 | MT-ATP6 pseudogene 27 [Source:HGNC Symb |
| PPIAP20 | 19 | peptidylprolyl isomerase A pseudogene 20 [Sc |
| ZNF564 | 19 | zinc finger protein 564 [Source:HGNC Symbol; |
| AC010422.6 | 19 | novel transcript |
| AC010422.4 | 19 | novel transcript, sense intronic to ZNF564 |
| AC010422.1 | 19 | novel transcript |
| PGK1P2 | 19 | phosphoglycerate kinase 1, pseudogene 2 [So |
| ZNF490 | 19 | zinc finger protein 490 [Source:HGNC Symbol; |
| ZNF791 | 19 | zinc finger protein 791 [Source:HGNC Symbol; |
| RPL10P16 | 19 | ribosomal protein L10 pseudogene 16 [Source |
| AC010422.3 | 19 | novel transcript |
| MAN2B1 | 19 | mannosidase alpha class 2B member 1 [Sourc |
| AC010422.5 | 19 | novel protein |
| WDR83 | 19 | WD repeat domain 83 [Source:HGNC Symbol; |
| AC010422.8 | 19 | novel transcript |
| WDR83OS | 19 | WD repeat domain 83 opposite strand [Source |
| AC010422.7 | 19 | novel transcript, antisense to WDR83 |
| DHPS | 19 | deoxyhypusine synthase [Source:HGNC Symbo |
| AC010422.2 | 19 | novel transcript |
| GNG14 | 19 | G protein subunit gamma 14 [Source:HGNC Sy |

| | | |
|---|---|---|
| FBXW9 | 19 | F-box and WD repeat domain containing 9 [Sc |
| AC018761.4 | 19 | novel transcript, antisense to FBXW9 |
| TNPO2 | 19 | transportin 2 [Source:HGNC Symbol;Acc:HGN( |
| SNORD41 | 19 | small nucleolar RNA, C/D box 41 [Source:HGN |
| AC018761.1 | 19 | novel transcript |
| TRIR | 19 | telomerase RNA component interacting RNas |
| GET3 | 19 | guided entry of tail-anchored proteins factor 3 |
| BEST2 | 19 | bestrophin 2 [Source:HGNC Symbol;Acc:HGN( |
| HOOK2 | 19 | hook microtubule tethering protein 2 [Source |
| AC018761.3 | 19 | novel transcript |
| MIR5684 | 19 | microRNA 5684 [Source:HGNC Symbol;Acc:HG |
| JUNB | 19 | JunB proto-oncogene, AP-1 transcription facto |
| PRDX2 | 19 | peroxiredoxin 2 [Source:HGNC Symbol;Acc:HG |
| AC018761.2 | 19 | novel transcript, antisense to PRDX2 |
| THSD8 | 19 | thrombospondin type 1 domain containing 8 [ |
| RNASEH2A | 19 | ribonuclease H2 subunit A [Source:HGNC Sym |
| RTBDN | 19 | retbindin [Source:HGNC Symbol;Acc:HGNC:30 |
| AC020934.1 | 19 | novel transcript, antisense to RTBDN |
| MAST1 | 19 | microtubule associated serine/threonine kina |
| MIR6794 | 19 | microRNA 6794 [Source:HGNC Symbol;Acc:HG |
| DNASE2 | 19 | deoxyribonuclease 2, lysosomal [Source:HGN( |
| AC020934.2 | 19 | novel transcript, antisense to DNASE2 |
| KLF1 | 19 | Kruppel like factor 1 [Source:HGNC Symbol;Ac |
| GCDH | 19 | glutaryl-CoA dehydrogenase [Source:HGNC Sy |
| RPS6P25 | 19 | ribosomal protein S6 pseudogene 25 [Source: |
| SYCE2 | 19 | synaptonemal complex central element prote |
| MIR5695 | 19 | microRNA 5695 [Source:HGNC Symbol;Acc:HG |
| FARSA | 19 | phenylalanyl-tRNA synthetase subunit alpha [ |
| FARSA-AS1 | 19 | FARSA antisense RNA 1 [Source:HGNC Symbol |
| CALR | 19 | calreticulin [Source:HGNC Symbol;Acc:HGNC:1 |
| MIR6515 | 19 | microRNA 6515 [Source:HGNC Symbol;Acc:HG |
| AC092069.1 | 19 | novel transcript, antisense to CALR |
| RAD23A | 19 | RAD23 homolog A, nucleotide excision repair |
| GADD45GIP1 | 19 | GADD45G interacting protein 1 [Source:HGNC |
| DAND5 | 19 | DAN domain BMP antagonist family member |
| NFIX | 19 | nuclear factor I X [Source:HGNC Symbol;Acc:H |
| AC138474.1 | 19 | novel transcript, antisense to NFIX |
| AC007787.1 | 19 | novel transcript, antisense to NFIX |
| AC007787.2 | 19 | TEC |
| LYL1 | 19 | LYL1 basic helix-loop-helix family member [So |
| TRMT1 | 19 | tRNA methyltransferase 1 [Source:HGNC Syml |
| NACC1 | 19 | nucleus accumbens associated 1 [Source:HGN |
| AC005546.1 | 19 | novel transcript |
| AC011446.1 | 19 | novel transcript, antisense to NACC1 |
| STX10 | 19 | syntaxin 10 [Source:HGNC Symbol;Acc:HGNC: |
| IER2 | 19 | immediate early response 2 [Source:HGNC Sy |
| AC011446.3 | 19 | novel transcript, antisense to IER2 |
| AC011446.2 | 19 | novel transcript, antisense to IER2 |
| RPL12P42 | 19 | ribosomal protein L12 pseudogene 42 [Source |
| CACNA1A | 19 | calcium voltage-gated channel subunit alpha1 |

| | | |
|---|---|---|
| CCDC130 | 19 | coiled-coil domain containing 130 [Source:HG |
| MRI1 | 19 | methylthioribose-1-phosphate isomerase 1 [S |
| AC008686.1 | 19 | novel transcript, antisense to MRI1 |
| C19orf53 | 19 | chromosome 19 open reading frame 53 [Sour |
| ZSWIM4 | 19 | zinc finger SWIM-type containing 4 [Source:H( |
| AC020916.2 | 19 | novel transcript, antisense to ZSWIM4 |
| RN7SL619P | 19 | RNA, 7SL, cytoplasmic 619, pseudogene [Sour |
| AC020916.1 | 19 | novel transcript |
| MIR24-2 | 19 | microRNA 24-2 [Source:HGNC Symbol;Acc:HG |
| MIR27A | 19 | microRNA 27a [Source:HGNC Symbol;Acc:HGN |
| MIR23A | 19 | microRNA 23a [Source:HGNC Symbol;Acc:HGN |
| NANOS3 | 19 | nanos C2HC-type zinc finger 3 [Source:HGNC S |
| MIR181C | 19 | microRNA 181c [Source:HGNC Symbol;Acc:HG |
| MIR181D | 19 | microRNA 181d [Source:HGNC Symbol;Acc:HG |
| C19orf57 | 19 | chromosome 19 open reading frame 57 [Sour |
| CC2D1A | 19 | coiled-coil and C2 domain containing 1A [Sour |
| PODNL1 | 19 | podocan like 1 [Source:HGNC Symbol;Acc:HGI |
| DCAF15 | 19 | DDB1 and CUL4 associated factor 15 [Source:I |
| RFX1 | 19 | regulatory factor X1 [Source:HGNC Symbol;Ac |
| AC022098.4 | 19 | novel transcript, antisense to RFX1 |
| RLN3 | 19 | relaxin 3 [Source:HGNC Symbol;Acc:HGNC:17: |
| AC022098.2 | 19 | novel transcript, antisense to RLN3 |
| IL27RA | 19 | interleukin 27 receptor subunit alpha [Source: |
| PALM3 | 19 | paralemmin 3 [Source:HGNC Symbol;Acc:HGN |
| EEF1DP1 | 19 | eukaryotic translation elongation factor 1 delt |
| MISP3 | 19 | MISP family member 3 [Source:HGNC Symbol; |
| MIR1199 | 19 | microRNA 1199 [Source:HGNC Symbol;Acc:HG |
| C19orf67 | 19 | chromosome 19 open reading frame 67 [Sour |
| SAMD1 | 19 | sterile alpha motif domain containing 1 [Sourc |
| PRKACA | 19 | protein kinase cAMP-activated catalytic subur |
| AC022098.3 | 19 | novel transcript, antisense to ASF1B |
| ASF1B | 19 | anti-silencing function 1B histone chaperone [ |
| AC022098.1 | 19 | novel transcript, antisense to LPHN1 |
| ADGRL1 | 19 | adhesion G protein-coupled receptor L1 [Sour |
| RN7SL231P | 19 | RNA, 7SL, cytoplasmic 231, pseudogene [Sour |
| AC011509.3 | 19 | novel transcript |
| AC011509.2 | 19 | novel transcript |
| AC011509.1 | 19 | novel transcript |
| LINC01841 | 19 | long intergenic non-protein coding RNA 1841 |
| LINC01842 | 19 | long intergenic non-protein coding RNA 1842 |
| ADGRE5 | 19 | adhesion G protein-coupled receptor E5 [Sour |
| AC008569.1 | 19 | novel transcript, antisense to CD97 |
| DDX39A | 19 | DExD-box helicase 39A [Source:HGNC Symbol; |
| PKN1 | 19 | protein kinase N1 [Source:HGNC Symbol;Acc:I |
| AC008569.2 | 19 | novel transcript, sense intronic to PKN1 |
| PTGER1 | 19 | prostaglandin E receptor 1 [Source:HGNC Sym |
| GIPC1 | 19 | GIPC PDZ domain containing family member 1 |
| SNRPGP15 | 19 | small nuclear ribonucleoprotein polypeptide ( |
| DNAJB1 | 19 | DnaJ heat shock protein family (Hsp40) memb |
| TECR | 19 | trans-2,3-enoyl-CoA reductase [Source:HGNC |

| | | |
|---|---|---|
| MIR639 | 19 | microRNA 639 [Source:HGNC Symbol;Acc:HGN |
| NDUFB7 | 19 | NADH:ubiquinone oxidoreductase subunit B7 |
| CLEC17A | 19 | C-type lectin domain containing 17A [Source:H |
| RN7SL337P | 19 | RNA, 7SL, cytoplasmic 337, pseudogene [Sour |
| RN7SL842P | 19 | RNA, 7SL, cytoplasmic 842, pseudogene [Sour |
| ADGRE3 | 19 | adhesion G protein-coupled receptor E3 [Sour |
| AC022149.1 | 19 | integrin beta 1 pseudogene 1 |
| AC022149.2 | 19 | yippee-like 5 (Drosophila) (YPEL5) pseudogene |
| ZNF333 | 19 | zinc finger protein 333 [Source:HGNC Symbol; |
| ADGRE2 | 19 | adhesion G protein-coupled receptor E2 [Sour |
| OR7C1 | 19 | olfactory receptor family 7 subfamily C memb |
| AC005255.1 | 19 | transmembrane protein 167A (TMEM167A) ps |
| OR7A5 | 19 | olfactory receptor family 7 subfamily A memb |
| OR7A10 | 19 | olfactory receptor family 7 subfamily A memb |
| OR7A8P | 19 | olfactory receptor family 7 subfamily A memb |
| OR7A2P | 19 | olfactory receptor family 7 subfamily A memb |
| OR7A17 | 19 | olfactory receptor family 7 subfamily A memb |
| OR7A18P | 19 | olfactory receptor family 7 subfamily A memb |
| OR7A1P | 19 | olfactory receptor family 7 subfamily A memb |
| OR7A3P | 19 | olfactory receptor family 7 subfamily A memb |
| OR7A11P | 19 | olfactory receptor family 7 subfamily A memb |
| OR7A15P | 19 | olfactory receptor family 7 subfamily A memb |
| OR7C2 | 19 | olfactory receptor family 7 subfamily C memb |
| SLC1A6 | 19 | solute carrier family 1 member 6 [Source:HGN |
| CCDC105 | 19 | coiled-coil domain containing 105 [Source:HG |
| CASP14 | 19 | caspase 14 [Source:HGNC Symbol;Acc:HGNC:1 |
| OR1I1 | 19 | olfactory receptor family 1 subfamily I membe |
| SYDE1 | 19 | synapse defective Rho GTPase homolog 1 [Sou |
| ILVBL | 19 | ilvB acetolactate synthase like [Source:HGNC S |
| AC003956.1 | 19 | novel transcript, antisense to ILVBL |
| OR10B1P | 19 | olfactory receptor family 10 subfamily B mem |
| RNU6-782P | 19 | RNA, U6 small nuclear 782, pseudogene [Sour |
| NOTCH3 | 19 | notch receptor 3 [Source:HGNC Symbol;Acc:H |
| MIR6795 | 19 | microRNA 6795 [Source:HGNC Symbol;Acc:HG |
| AC004257.1 | 19 | novel transcript |
| EPHX3 | 19 | epoxide hydrolase 3 [Source:HGNC Symbol;Ac |
| BRD4 | 19 | bromodomain containing 4 [Source:HGNC Syn |
| AC005776.1 | 19 | ribosomal protein L21 (RPL21) pseudogene |
| AC005776.2 | 19 | novel transcript |
| AKAP8 | 19 | A-kinase anchoring protein 8 [Source:HGNC Sy |
| AC005785.1 | 19 | novel transcript, antisense to AKAP8 and AKAI |
| AKAP8L | 19 | A-kinase anchoring protein 8 like [Source:HGN |
| AC005785.2 | 19 | TEC |
| AC006128.1 | 19 | TEC |
| WIZ | 19 | WIZ zinc finger [Source:HGNC Symbol;Acc:HGI |
| MIR1470 | 19 | microRNA 1470 [Source:HGNC Symbol;Acc:HG |
| RASAL3 | 19 | RAS protein activator like 3 [Source:HGNC Syn |
| PGLYRP2 | 19 | peptidoglycan recognition protein 2 [Source:H |
| CYP4F22 | 19 | cytochrome P450 family 4 subfamily F membe |
| AC011492.1 | 19 | staufen, RNA binding protein, homolog 1 (Dro |

| | | |
|---|---|---|
| CYP4F23P | 19 | cytochrome P450 family 4 subfamily F membe |
| AD000091.1 | 19 | major histocompatibility complex, class I, F (H |
| RPL23AP2 | 19 | ribosomal protein L23a pseudogene 2 [Source |
| AC093072.1 | 19 | novel transcript |
| CYP4F8 | 19 | cytochrome P450 family 4 subfamily F membe |
| CYP4F3 | 19 | cytochrome P450 family 4 subfamily F membe |
| CYP4F10P | 19 | cytochrome P450 family 4 subfamily F membe |
| CYP4F12 | 19 | cytochrome P450 family 4 subfamily F membe |
| OR10H2 | 19 | olfactory receptor family 10 subfamily H mem |
| OR10H3 | 19 | olfactory receptor family 10 subfamily H mem |
| CYP4F24P | 19 | cytochrome P450 family 4 subfamily F membe |
| AC004597.1 | 19 | novel transcript |
| OR10H5 | 19 | olfactory receptor family 10 subfamily H mem |
| OR10H1 | 19 | olfactory receptor family 10 subfamily H mem |
| ZNF861P | 19 | zinc finger protein 861, pseudogene [Source:H |
| LINC01764 | 19 | long intergenic non-protein coding RNA 1764 |
| UCA1 | 19 | urothelial cancer associated 1 [Source:HGNC S |
| AC004510.1 | 19 | NULL |
| CLEC4O | 19 | C-type lectin domain family 4 member O [Sou |
| CYP4F36P | 19 | cytochrome P450 family 4 subfamily F membe |
| CYP4F2 | 19 | cytochrome P450 family 4 subfamily F membe |
| AC005336.2 | 19 | polymerase (RNA) mitochondrial (DNA directe |
| AC005336.3 | 19 | TEC |
| AC005336.1 | 19 | inositol polyphosphate multikinase (IPMK) pse |
| CYP4F11 | 19 | cytochrome P450 family 4 subfamily F membe |
| OR10H4 | 19 | olfactory receptor family 10 subfamily H mem |
| CYP4F9P | 19 | cytochrome P450 family 4 subfamily F membe |
| LINC00661 | 19 | long intergenic non-protein coding RNA 661 [S |
| AC004790.2 | 19 | TEC |
| LINC00905 | 19 | long intergenic non-protein coding RNA 905 [S |
| SNX33P1 | 19 | sorting nexin 33 pseudogene 1 [Source:HGNC |
| AC004790.1 | 19 | ubiquitin specific peptidase 6 (Tre-2 oncogene |
| OR1AB1P | 19 | olfactory receptor family 1 subfamily AB mem |
| LINC01855 | 19 | long intergenic non-protein coding RNA 1855 |
| TPM4 | 19 | tropomyosin 4 [Source:HGNC Symbol;Acc:HGI |
| AC008894.3 | 19 | TEC |
| AC008894.1 | 19 | novel transcript |
| RAB8A | 19 | RAB8A, member RAS oncogene family [Source |
| AC008894.2 | 19 | novel transcript, antisense RAB8A |
| HSH2D | 19 | hematopoietic SH2 domain containing [Source |
| CIB3 | 19 | calcium and integrin binding family member 3 |
| FAM32A | 19 | family with sequence similarity 32 member A |
| AC020911.1 | 19 | novel transcript, antisense FAM32A |
| Metazoa_SRP | 19 | Metazoan signal recognition particle RNA [Sou |
| AC020911.3 | 19 | novel transcript, antisense to AP1M1 |
| AP1M1 | 19 | adaptor related protein complex 1 subunit mu |
| AC020911.2 | 19 | novel transcript, antisense KLF2 |
| KLF2 | 19 | Kruppel like factor 2 [Source:HGNC Symbol;Ac |
| AC020917.3 | 19 | novel transcript |
| EPS15L1 | 19 | epidermal growth factor receptor pathway su |

| | | |
|---|---|---|
| AC020917.4 | 19 | TEC |
| AC020917.1 | 19 | ribosomal protein S2 (RPS2) pseudogene |
| AC020917.2 | 19 | novel transcript, antisense EPS15L1 |
| CALR3 | 19 | calreticulin 3 [Source:HGNC Symbol;Acc:HGN( |
| AC008764.1 | 19 | novel protein |
| C19orf44 | 19 | chromosome 19 open reading frame 44 [Sour |
| CHERP | 19 | calcium homeostasis endoplasmic reticulum p |
| RN7SL146P | 19 | RNA, 7SL, cytoplasmic 146, pseudogene [Sour |
| AC008764.6 | 19 | novel transcript |
| SLC35E1 | 19 | solute carrier family 35 member E1 [Source:H |
| AC008764.3 | 19 | novel transcript, antisense SLC35E1 |
| AC008764.5 | 19 | novel transcript, antisense SLC35E1 |
| AC008764.8 | 19 | novel transcript, antisense to MED26 |
| MED26 | 19 | mediator complex subunit 26 [Source:HGNC S |
| AC008764.4 | 19 | novel protein |
| AC008764.7 | 19 | novel transcript, sense intronic MED26 |
| AC008764.9 | 19 | TEC |
| AC008764.10 | 19 | TEC |
| AC008764.2 | 19 | novel transcript |
| AC024075.3 | 19 | novel transcript, antisense C19orf42 |
| SMIM7 | 19 | small integral membrane protein 7 [Source:H( |
| AC024075.2 | 19 | novel transcript, sense intronic C19orf42 |
| AC024075.1 | 19 | novel transcript, sense intronic C19orf42 |
| TMEM38A | 19 | transmembrane protein 38A [Source:HGNC Sy |
| AC008737.2 | 19 | transmembrane protein 38A (TMEM38A) pseu |
| NWD1 | 19 | NACHT and WD repeat domain containing 1 [S |
| AC008737.3 | 19 | novel transcript, sense intronic NWD1 |
| SIN3B | 19 | SIN3 transcription regulator family member B |
| AC008737.1 | 19 | novel transcript, sense intronic SIN3B |
| F2RL3 | 19 | F2R like thrombin or trypsin receptor 3 [Sourc |
| CPAMD8 | 19 | C3 and PZP like alpha-2-macroglobulin domair |
| RN7SL835P | 19 | RNA, 7SL, cytoplasmic 835, pseudogene [Sour |
| RN7SL823P | 19 | RNA, 7SL, cytoplasmic 823, pseudogene [Sour |
| AC020908.2 | 19 | novel transcript, antisense CPAMD8 |
| HAUS8 | 19 | HAUS augmin like complex subunit 8 [Source:I |
| AC020908.1 | 19 | novel transcript, antisense HAUS8 |
| MYO9B | 19 | myosin IXB [Source:HGNC Symbol;Acc:HGNC:7 |
| AC020908.3 | 19 | novel transcript, antisense MYO9B |
| AC020913.1 | 19 | novel transcript, antisense MYO9B |
| AC020913.3 | 19 | novel transcript, antisense MYO9B |
| AC020913.2 | 19 | novel transcript, antisense MYO9B |
| USE1 | 19 | unconventional SNARE in the ER 1 [Source:HG |
| OCEL1 | 19 | occludin/ELL domain containing 1 [Source:HG |
| NR2F6 | 19 | nuclear receptor subfamily 2 group F member |
| AC010646.1 | 19 | novel protein |
| USHBP1 | 19 | USH1 protein network component harmonin I |
| BABAM1 | 19 | BRISC and BRCA1 A complex member 1 [Sourc |
| AC010463.1 | 19 | novel transcript |
| ANKLE1 | 19 | ankyrin repeat and LEM domain containing 1 [ |
| ABHD8 | 19 | abhydrolase domain containing 8 [Source:HGI |

| | | |
|---|---|---|
| MRPL34 | 19 | mitochondrial ribosomal protein L34 [Source:I |
| AC010463.3 | 19 | novel transcript, sense intronic ABHD8 |
| DDA1 | 19 | DET1 and DDB1 associated 1 [Source:HGNC Sy |
| ANO8 | 19 | anoctamin 8 [Source:HGNC Symbol;Acc:HGNC |
| GTPBP3 | 19 | GTP binding protein 3, mitochondrial [Source: |
| PLVAP | 19 | plasmalemma vesicle associated protein [Sour |
| AC010463.2 | 19 | novel transcript, antisense PLVAP |
| CCDC194 | 19 | coiled-coil domain containing 194 [Source:HG |
| BST2 | 19 | bone marrow stromal cell antigen 2 [Source:H |
| BISPR | 19 | BST2 interferon stimulated positive regulator |
| MVB12A | 19 | multivesicular body subunit 12A [Source:HGN |
| AC010319.4 | 19 | novel transcript, antisense FAM125A |
| AC010319.1 | 19 | high-mobility group box 3 (HMGB3) pseudoge |
| AC010319.3 | 19 | novel transcript, antisense FAM125A |
| TMEM221 | 19 | transmembrane protein 221 [Source:HGNC Sy |
| AC010319.2 | 19 | novel protein |
| AC010319.5 | 19 | novel transcript |
| NXNL1 | 19 | nucleoredoxin like 1 [Source:HGNC Symbol;Ac |
| SLC27A1 | 19 | solute carrier family 27 member 1 [Source:HG |
| AC010618.3 | 19 | novel transcript |
| PGLS | 19 | 6-phosphogluconolactonase [Source:HGNC Sy |
| AC010618.2 | 19 | novel transcript, antisense PGLS |
| NIBAN3 | 19 | niban apoptosis regulator 3 [Source:HGNC Syr |
| COLGALT1 | 19 | collagen beta(1-O)galactosyltransferase 1 [Sou |
| AC010618.1 | 19 | ribosomal protein L21 (RPL21) pseudogene |
| UNC13A | 19 | unc-13 homolog A [Source:HGNC Symbol;Acc: |
| AC008761.2 | 19 | novel transcript, antisense UNC13A |
| AC008761.3 | 19 | TEC |
| MAP1S | 19 | microtubule associated protein 1S [Source:HG |
| AC008761.1 | 19 | novel transcript, antisense MAP1S |
| FCHO1 | 19 | FCH domain only 1 [Source:HGNC Symbol;Acc |
| B3GNT3 | 19 | UDP-GlcNAc:betaGal beta-1,3-N-acetylglucosa |
| INSL3 | 19 | insulin like 3 [Source:HGNC Symbol;Acc:HGNC |
| JAK3 | 19 | Janus kinase 3 [Source:HGNC Symbol;Acc:HGN |
| RPL18A | 19 | ribosomal protein L18a [Source:HGNC Symbol |
| SNORA68 | 19 | small nucleolar RNA, H/ACA box 68 [Source:H |
| AC005796.1 | 19 | TEC |
| SLC5A5 | 19 | solute carrier family 5 member 5 [Source:HGN |
| CCDC124 | 19 | coiled-coil domain containing 124 [Source:HG |
| KCNN1 | 19 | potassium calcium-activated channel subfami |
| RNA5SP468 | 19 | RNA, 5S ribosomal pseudogene 468 [Source:H |
| ARRDC2 | 19 | arrestin domain containing 2 [Source:HGNC Sy |
| AC020904.3 | 19 | TEC |
| AC020904.2 | 19 | citrate synthase (CS) pseudogene |
| AC020904.1 | 19 | apolipoprotein A-I binding protein (APOA1BP) |
| RPS18P13 | 19 | ribosomal protein S18 pseudogene 13 [Source |
| IL12RB1 | 19 | interleukin 12 receptor subunit beta 1 [Source |
| MAST3 | 19 | microtubule associated serine/threonine kina: |
| AC007192.2 | 19 | novel transcript, antisense to MAST3 |
| AC007192.1 | 19 | novel protein, readthrough between PIK3R2 a |

| | | |
|---|---|---|
| PIK3R2 | 19 | phosphoinositide-3-kinase regulatory subunit |
| IFI30 | 19 | IFI30 lysosomal thiol reductase [Source:HGNC |
| MPV17L2 | 19 | MPV17 mitochondrial inner membrane protei |
| RAB3A | 19 | RAB3A, member RAS oncogene family [Source |
| AC005759.1 | 19 | novel transcript, antisense to PDE4C |
| AC008397.2 | 19 | novel protein |
| PDE4C | 19 | phosphodiesterase 4C [Source:HGNC Symbol; |
| AC008397.1 | 19 | novel protein |
| IQCN | 19 | IQ motif containing N [Source:HGNC Symbol;A |
| JUND | 19 | JunD proto-oncogene, AP-1 transcription fact |
| MIR3188 | 19 | microRNA 3188 [Source:HGNC Symbol;Acc:HG |
| RPL39P38 | 19 | ribosomal protein L39 pseudogene 38 [Source |
| LSM4 | 19 | LSM4 homolog, U6 small nuclear RNA and mR |
| RN7SL513P | 19 | RNA, 7SL, cytoplasmic 513, pseudogene [Sour |
| PGPEP1 | 19 | pyroglutamyl-peptidase I [Source:HGNC Symb |
| GDF15 | 19 | growth differentiation factor 15 [Source:HGN |
| MIR3189 | 19 | microRNA 3189 [Source:HGNC Symbol;Acc:HG |
| LRRC25 | 19 | leucine rich repeat containing 25 [Source:HGN |
| SSBP4 | 19 | single stranded DNA binding protein 4 [Source |
| ISYNA1 | 19 | inositol-3-phosphate synthase 1 [Source:HGN |
| AC010335.1 | 19 | novel transcript, antisense to ELL |
| ELL | 19 | elongation factor for RNA polymerase II [Sour |
| AC010335.2 | 19 | TEC |
| AC010335.3 | 19 | TEC |
| AC005387.1 | 19 | novel transcript, antisense to FKBP8 |
| FKBP8 | 19 | FKBP prolyl isomerase 8 [Source:HGNC Symbo |
| AC005387.2 | 19 | novel transcript, antisense to FKBP8 |
| KXD1 | 19 | KxDL motif containing 1 [Source:HGNC Symbo |
| AC005253.1 | 19 | novel transcript, antisense to KXD1 |
| AC005253.2 | 19 | novel transcript, antisense to KXD1 |
| UBA52 | 19 | ubiquitin A-52 residue ribosomal protein fusic |
| CRLF1 | 19 | cytokine receptor like factor 1 [Source:HGNC S |
| REX1BD | 19 | required for excision 1-B domain containing [S |
| TMEM59L | 19 | transmembrane protein 59 like [Source:HGNC |
| RN7SL155P | 19 | RNA, 7SL, cytoplasmic 155, pseudogene [Sour |
| KLHL26 | 19 | kelch like family member 26 [Source:HGNC Sy |
| CRTC1 | 19 | CREB regulated transcription coactivator 1 [Sc |
| COMP | 19 | cartilage oligomeric matrix protein [Source:HG |
| UPF1 | 19 | UPF1 RNA helicase and ATPase [Source:HGNC |
| CERS1 | 19 | ceramide synthase 1 [Source:HGNC Symbol;Ac |
| GDF1 | 19 | growth differentiation factor 1 [Source:HGNC |
| AC005197.1 | 19 | novel transcript, antisense to CERS1 |
| COPE | 19 | coatomer protein complex subunit epsilon [Sc |
| AC002985.1 | 19 | novel transcript |
| DDX49 | 19 | DEAD-box helicase 49 [Source:HGNC Symbol;A |
| HOMER3 | 19 | homer scaffold protein 3 [Source:HGNC Symb |
| HOMER3-AS1 | 19 | HOMER3 antisense RNA 1 [Source:HGNC Syml |
| RN7SL70P | 19 | RNA, 7SL, cytoplasmic 70, pseudogene [Sourc |
| SUGP2 | 19 | SURP and G-patch domain containing 2 [Sourc |
| ARMC6 | 19 | armadillo repeat containing 6 [Source:HGNC S |

| | | |
|---|---|---|
| SLC25A42 | 19 | solute carrier family 25 member 42 [Source:H |
| TMEM161A | 19 | transmembrane protein 161A [Source:HGNC S |
| MEF2B | 19 | myocyte enhancer factor 2B [Source:HGNC Sy |
| BORCS8-MEF2B | 19 | BORCS8-MEF2B readthrough [Source:HGNC Sy |
| BORCS8 | 19 | BLOC-1 related complex subunit 8 [Source:HG |
| RFXANK | 19 | regulatory factor X associated ankyrin contain |
| NR2C2AP | 19 | nuclear receptor 2C2 associated protein [Sour |
| NCAN | 19 | neurocan [Source:HGNC Symbol;Acc:HGNC:24 |
| RNU6-1028P | 19 | RNA, U6 small nuclear 1028, pseudogene [Sou |
| AC138430.1 | 19 | novel transcript, readthrough between TM6SF |
| HAPLN4 | 19 | hyaluronan and proteoglycan link protein 4 [S |
| AC138430.2 | 19 | TEC |
| TM6SF2 | 19 | transmembrane 6 superfamily member 2 [Sou |
| SUGP1 | 19 | SURP and G-patch domain containing 1 [Sourc |
| MAU2 | 19 | MAU2 sister chromatid cohesion factor [Sourc |
| GATAD2A | 19 | GATA zinc finger domain containing 2A [Sourc |
| AC092067.1 | 19 | novel transcript, antisense to GATAD2A |
| MIR640 | 19 | microRNA 640 [Source:HGNC Symbol;Acc:HGN |
| TSSK6 | 19 | testis specific serine kinase 6 [Source:HGNC Sy |
| NDUFA13 | 19 | NADH:ubiquinone oxidoreductase subunit A1 |
| AC011448.1 | 19 | novel protein |
| YJEFN3 | 19 | YjeF N-terminal domain containing 3 [Source:I |
| CILP2 | 19 | cartilage intermediate layer protein 2 [Source |
| PBX4 | 19 | PBX homeobox 4 [Source:HGNC Symbol;Acc:H |
| PHF5CP | 19 | PHD finger protein 5C pseudogene [Source:HC |
| AC002306.1 | 19 | novel transcript, antisense to PBX4 |
| LPAR2 | 19 | lysophosphatidic acid receptor 2 [Source:HGN |
| GMIP | 19 | GEM interacting protein [Source:HGNC Symbo |
| ATP13A1 | 19 | ATPase 13A1 [Source:HGNC Symbol;Acc:HGN |
| ZNF101 | 19 | zinc finger protein 101 [Source:HGNC Symbol; |
| AC011458.2 | 19 | iron-sulfur cluster assembly 1 homolog (S. cer |
| AC011458.1 | 19 | ankyrin repeat domain 26 (ANKRD26) pseudo |
| ZNF14 | 19 | zinc finger protein 14 [Source:HGNC Symbol;A |
| AC011477.7 | 19 | ubiquitin specific peptidase 6 (Tre-2 oncogene |
| AC011477.8 | 19 | novel transcript |
| AC011477.6 | 19 | sorting nexin 18 (SNX18) pseudogene |
| LINC00663 | 19 | long intergenic non-protein coding RNA 663 [S |
| AC011477.9 | 19 | novel transcript |
| AC011477.4 | 19 | novel transcript |
| AC011477.5 | 19 | ribosomal protein L10a (RPL10A) pseudogene |
| AC011477.1 | 19 | zinc finger protein 506 (ZNF506) pseudogene |
| ZNF506 | 19 | zinc finger protein 506 [Source:HGNC Symbol; |
| AC011477.2 | 19 | novel transcript, sense intronic to ZNF506 |
| BNIP3P9 | 19 | BCL2 interacting protein 3 pseudogene 9 [Sou |
| ZNF253 | 19 | zinc finger protein 253 [Source:HGNC Symbol; |
| BNIP3P10 | 19 | BCL2 interacting protein 3 pseudogene 10 [So |
| AC011477.3 | 19 | novel transcript, antisense to ZNF253 |
| ZNF93 | 19 | zinc finger protein 93 [Source:HGNC Symbol;A |
| AC007204.1 | 19 | BCL2/adenovirus E1B 19kDa interacting prote |
| AC007204.2 | 19 | vomeronasal 1 receptor 3 (VN1R3) pseudoger |

| | | |
|---|---|---|
| ZNF682 | 19 | zinc finger protein 682 [Source:HGNC Symbol; |
| AC006539.2 | 19 | BCL2/adenovirus E1B 19kDa interacting prote |
| AC006539.3 | 19 | novel zinc finger protein pseudogene |
| BNIP3P13 | 19 | BCL2 interacting protein 3 pseudogene 13 [So |
| ZNF90 | 19 | zinc finger protein 90 [Source:HGNC Symbol;A |
| AC006539.1 | 19 | ribosomal protein S8 (RPS8) pseudogene |
| AC011447.5 | 19 | BCL2/adenovirus E1B 19kDa interacting prote |
| AC011447.3 | 19 | novel transcript, antisense to ZNF486 and ZNF |
| AC011447.1 | 19 | ribosomal protein S16 (RPS16) pseudogene |
| AC011447.2 | 19 | zinc finger (ZNF) pseudogene |
| BNIP3P15 | 19 | BCL2 interacting protein 3 pseudogene 15 [So |
| ZNF486 | 19 | zinc finger protein 486 [Source:HGNC Symbol; |
| BNIP3P16 | 19 | BCL2 interacting protein 3 pseudogene 16 [So |
| AC011447.4 | 19 | zinc finger protein (ZNF) pseudogene |
| AC011447.6 | 19 | TEC |
| AC011447.7 | 19 | TEC |
| BNIP3P17 | 19 | BCL2 interacting protein 3 pseudogene 17 [So |
| AC078899.3 | 19 | novel zinc finger protein pseudogene |
| AC078899.1 | 19 | ARP2 actin-related protein 2 homolog (yeast) |
| BNIP3P18 | 19 | BCL2 interacting protein 3 pseudogene 18 [So |
| AC078899.4 | 19 | novel zinc finger protein pseudogene |
| BNIP3P19 | 19 | BCL2 interacting protein 3 pseudogene 19 [So |
| AC078899.5 | 19 | novel zinc finger protein pseudogene |
| ZNF826P | 19 | zinc finger protein 826, pseudogene [Source:H |
| BNIP3P20 | 19 | BCL2 interacting protein 3 pseudogene 20 [So |
| AC078899.2 | 19 | novel zinc finger protein pseudogene |
| MIR1270 | 19 | microRNA 1270 [Source:HGNC Symbol;Acc:HG |
| BNIP3P22 | 19 | BCL2 interacting protein 3 pseudogene 22 [So |
| AC008554.1 | 19 | novel transcript |
| BNIP3P23 | 19 | BCL2 interacting protein 3 pseudogene 23 [So |
| ZNF737 | 19 | zinc finger protein 737 [Source:HGNC Symbol; |
| AC010636.2 | 19 | novel transcript |
| VN1R78P | 19 | vomeronasal 1 receptor 78 pseudogene [Sour |
| AC010636.1 | 19 | novel transcript |
| ZNF626 | 19 | zinc finger protein 626 [Source:HGNC Symbol; |
| AC010329.5 | 19 | novel zinc finger pseudogene |
| AC010329.2 | 19 | zinc finger protein 85 pseudogene |
| AC010329.1 | 19 | novel transcript |
| VN1R79P | 19 | vomeronasal 1 receptor 79 pseudogene [Sour |
| ZNF66 | 19 | zinc finger protein 66 [Source:HGNC Symbol;A |
| BNIP3P24 | 19 | BCL2 interacting protein 3 pseudogene 24 [So |
| AC008739.2 | 19 | novel zinc finger protein pseudogene |
| ZNF85 | 19 | zinc finger protein 85 [Source:HGNC Symbol;A |
| AC008739.5 | 19 | TEC |
| KRT18P40 | 19 | keratin 18 pseudogene 40 [Source:HGNC Sym |
| AC008739.1 | 19 | novel transcript |
| ZNF430 | 19 | zinc finger protein 430 [Source:HGNC Symbol; |
| AC008739.3 | 19 | novel pseudogene |
| VN1R80P | 19 | vomeronasal 1 receptor 80 pseudogene [Sour |
| ZNF714 | 19 | zinc finger protein 714 [Source:HGNC Symbol; |

| | | |
|---|---|---|
| RNA5SP469 | 19 | RNA, 5S ribosomal pseudogene 469 [Source:H |
| VN1R81P | 19 | vomeronasal 1 receptor 81 pseudogene [Sour |
| ZNF431 | 19 | zinc finger protein 431 [Source:HGNC Symbol; |
| RPL7AP10 | 19 | ribosomal protein L7a pseudogene 10 [Source |
| VN1R82P | 19 | vomeronasal 1 receptor 82 pseudogene [Sour |
| AC010620.1 | 19 | novel zinc finger protein pseudogene |
| AC010620.2 | 19 | zinc finger protein 738 (ZNF738) pseudogene |
| RPL36AP51 | 19 | ribosomal protein L36a pseudogene 51 [Sourc |
| AC022432.1 | 19 | COX16 cytochrome c oxidase assembly homol |
| VN1R83P | 19 | vomeronasal 1 receptor 83 pseudogene [Sour |
| ZNF708 | 19 | zinc finger protein 708 [Source:HGNC Symbol; |
| BNIP3P25 | 19 | BCL2 interacting protein 3 pseudogene 25 [So |
| ZNF738 | 19 | zinc finger protein 738 [Source:HGNC Symbol; |
| AC010615.3 | 19 | iron-sulfur cluster assembly 1 homolog (S. cer |
| AC010615.1 | 19 | zinc finger protein pseudogene |
| ZNF493 | 19 | zinc finger protein 493 [Source:HGNC Symbol; |
| AC010615.4 | 19 | novel transcript |
| AC010615.2 | 19 | novel transcript |
| LINC00664 | 19 | long intergenic non-protein coding RNA 664 [S |
| ZNF429 | 19 | zinc finger protein 429 [Source:HGNC Symbol; |
| BNIP3P26 | 19 | BCL2 interacting protein 3 pseudogene 26 [So |
| AC123912.1 | 19 | novel transcript, antisense to ZNF429 |
| AC123912.4 | 19 | novel transcript |
| AC123912.2 | 19 | novel transcript |
| AC123912.6 | 19 | transmembrane protein 61 (TMEM61) pseudo |
| AC123912.5 | 19 | cleavage and polyadenylation specific factor 6 |
| AC123912.3 | 19 | zinc finger protein 100 (ZNF100) pseudogene |
| MTDHP3 | 19 | metadherin pseudogene 3 [Source:HGNC Sym |
| MTDHP4 | 19 | metadherin pseudogene 4 [Source:HGNC Sym |
| VN1R84P | 19 | vomeronasal 1 receptor 84 pseudogene [Sour |
| ZNF100 | 19 | zinc finger protein 100 [Source:HGNC Symbol; |
| AC092364.2 | 19 | cyclin Y-like 1 (CCNYL1) pseudogene |
| AC092364.1 | 19 | novel transcript |
| BRI3BPP1 | 19 | BRI3 binding protein pseudogene 1 [Source:H |
| ZNF43 | 19 | zinc finger protein 43 [Source:HGNC Symbol;A |
| BNIP3P27 | 19 | BCL2 interacting protein 3 pseudogene 27 [So |
| ZNF208 | 19 | zinc finger protein 208 [Source:HGNC Symbol; |
| MTDHP2 | 19 | metadherin pseudogene 2 [Source:HGNC Sym |
| AC003973.2 | 19 | TEC |
| BNIP3P28 | 19 | BCL2 interacting protein 3 pseudogene 28 [So |
| AC003973.1 | 19 | novel transcript |
| ZNF257 | 19 | zinc finger protein 257 [Source:HGNC Symbol; |
| BNIP3P29 | 19 | BCL2 interacting protein 3 pseudogene 29 [So |
| ZNF92P2 | 19 | zinc finger protein 92 pseudogene 2 [Source:H |
| AC073539.2 | 19 | ribosomal protein L17 (RPL17) pseudogene |
| PCGF7P | 19 | polycomb group ring finger 7 pseudogene [So |
| MTDHP5 | 19 | metadherin pseudogene 5 [Source:HGNC Sym |
| VN1R85P | 19 | vomeronasal 1 receptor 85 pseudogene [Sour |
| ZNF676 | 19 | zinc finger protein 676 [Source:HGNC Symbol; |
| BNIP3P30 | 19 | BCL2 interacting protein 3 pseudogene 30 [So |

| | | |
|---|---|---|
| AC073539.7 | 19 | novel pseudogene similar to part of zinc finge |
| AC073539.1 | 19 | novel transcript |
| AC073539.3 | 19 | novel transcript |
| ZNF729 | 19 | zinc finger protein 729 [Source:HGNC Symbol; |
| BNIP3P31 | 19 | BCL2 interacting protein 3 pseudogene 31 [So |
| AC011494.1 | 19 | metadherin (MTDH) pseudogene |
| RPL34P34 | 19 | ribosomal protein L34 pseudogene 34 [Source |
| ZNF98 | 19 | zinc finger protein 98 [Source:HGNC Symbol;A |
| AC011516.1 | 19 | novel transcript |
| BNIP3P32 | 19 | BCL2 interacting protein 3 pseudogene 32 [So |
| AC025811.1 | 19 | novel transcript |
| BNIP3P33 | 19 | BCL2 interacting protein 3 pseudogene 33 [So |
| ZNF209P | 19 | zinc finger protein 209, pseudogene [Source:H |
| AC004004.1 | 19 | folliculogenesis specific basic helix-loop-helix ( |
| AC011467.7 | 19 | novel transcript, antisense to ZNF98 |
| AC011467.1 | 19 | novel transcript |
| LINC01233 | 19 | long intergenic non-protein coding RNA 1233 |
| RNU6-1179P | 19 | RNA, U6 small nuclear 1179, pseudogene [Sou |
| AC011467.6 | 19 | SWI5 recombination repair homolog (yeast) (S |
| GOLGA2P9 | 19 | GOLGA2 pseudogene 9 [Source:HGNC Symbol |
| RN7SL860P | 19 | RNA, 7SL, cytoplasmic 860, pseudogene [Sour |
| AC011467.3 | 19 | novel transcript |
| AC011467.4 | 19 | testis expressed 264 (TEX264) pseudogene |
| AC011467.5 | 19 | RAD54-like 2 (S. cerevisiae) (RAD54L2) pseudo |
| LINC01785 | 19 | long intergenic non-protein coding RNA 1785 |
| AC011467.2 | 19 | testis expressed 264 (TEX264) pseudogene |
| ZNF492 | 19 | zinc finger protein 492 [Source:HGNC Symbol; |
| AC024563.1 | 19 | novel transcript, antisense to ZNF492 |
| ZNF849P | 19 | zinc finger protein 849, pseudogene [Source:H |
| RPL34P33 | 19 | ribosomal protein L34 pseudogene 33 [Source |
| AC024563.2 | 19 | metadherin (MTDH) pseudogene |
| VN1R87P | 19 | vomeronasal 1 receptor 87 pseudogene [Sour |
| ZNF99 | 19 | zinc finger protein 99 [Source:HGNC Symbol;A |
| AC008626.2 | 19 | zinc finger protein 99 (ZNF99) pseudogene |
| AC008626.1 | 19 | zinc finger protein 99 (ZNF99) pseudogene |
| BNIP3P34 | 19 | BCL2 interacting protein 3 pseudogene 34 [So |
| ZNF723 | 19 | zinc finger protein 723 [Source:HGNC Symbol; |
| BNIP3P35 | 19 | BCL2 interacting protein 3 pseudogene 35 [So |
| VN1R88P | 19 | vomeronasal 1 receptor 88 pseudogene [Sour |
| AC022145.1 | 19 | novel transcript |
| AC022145.2 | 19 | microtubule associated serine/threonine kina |
| ZNF728 | 19 | zinc finger protein 728 [Source:HGNC Symbol; |
| BNIP3P36 | 19 | BCL2 interacting protein 3 pseudogene 36 [So |
| LINC01859 | 19 | long intergenic non-protein coding RNA 1859 |
| LINC01858 | 19 | long intergenic non-protein coding RNA 1858 |
| AC074135.2 | 19 | novel transcript |
| AC074135.1 | 19 | novel transcript |
| ZNF730 | 19 | zinc finger protein 730 [Source:HGNC Symbol; |
| BNIP3P37 | 19 | BCL2 interacting protein 3 pseudogene 37 [So |
| AC124856.1 | 19 | ribosomal protein L5 (RPL5) pseudogene |

| | | |
|---|---|---|
| SNX6P1 | 19 | sorting nexin 6 pseudogene 1 [Source:HGNC S |
| AC092329.2 | 19 | metadherin (MTDH) pseudogene |
| ZNF724 | 19 | zinc finger protein 724 [Source:HGNC Symbol; |
| AC092329.3 | 19 | zinc finger protein 724 [Source:NCBI gene;Acc |
| BNIP3P38 | 19 | BCL2 interacting protein 3 pseudogene 38 [So |
| IPO5P1 | 19 | importin 5 pseudogene 1 [Source:HGNC Symb |
| AC092329.1 | 19 | novel transcript |
| AC092329.4 | 19 | novel transcript |
| VN1R90P | 19 | vomeronasal 1 receptor 90 pseudogene [Sour |
| VN1R91P | 19 | vomeronasal 1 receptor 91 pseudogene [Sour |
| ZNF91 | 19 | zinc finger protein 91 [Source:HGNC Symbol;A |
| AC010300.1 | 19 | novel transcript, antisense to ZNF91 |
| BNIP3P8 | 19 | BCL2 interacting protein 3 pseudogene 8 [Sou |
| CDC42EP3P1 | 19 | CDC42 effector protein 3 pseudogene 1 [Sour |
| LINC01224 | 19 | long intergenic non-protein coding RNA 1224 |
| VN1R92P | 19 | vomeronasal 1 receptor 92 pseudogene [Sour |
| AC074140.1 | 19 | novel zinc finger protein pseudogene |
| ZNF725P | 19 | zinc finger protein 725, pseudogene [Source:H |
| ZNF675 | 19 | zinc finger protein 675 [Source:HGNC Symbol; |
| AC073544.1 | 19 | zinc finger protein 430 (ZNF430) pseudogene |
| RPS27P29 | 19 | ribosomal protein S27 pseudogene 29 [Source |
| VN1R93P | 19 | vomeronasal 1 receptor 93 pseudogene [Sour |
| ZNF681 | 19 | zinc finger protein 681 [Source:HGNC Symbol; |
| AC139769.1 | 19 | zinc finger protein 726 pseudogene 1 |
| BNIP3P39 | 19 | BCL2 interacting protein 3 pseudogene 39 [So |
| AC139769.2 | 19 | novel transcript |
| RPSAP58 | 19 | ribosomal protein SA pseudogene 58 [Source: |
| AC139769.3 | 19 | novel zinc finger protein pseudogene |
| AC011503.4 | 19 | novel transcript |
| ZNF726 | 19 | zinc finger protein 726 [Source:HGNC Symbol; |
| AC011503.1 | 19 | novel transcript, antisense to ZNF726 |
| AC011503.2 | 19 | novel transcript, sense intronic to ZNF726 |
| AC011503.3 | 19 | zinc finger protein 92 pseudogene 3 |
| RNA5-8SP4 | 19 | RNA, 5.8S ribosomal pseudogene 4 [Source:H( |
| AC092279.1 | 19 | novel transcript |
| ZNF254 | 19 | zinc finger protein 254 [Source:HGNC Symbol; |
| AC092279.2 | 19 | TEC |
| BNIP3P40 | 19 | BCL2 interacting protein 3 pseudogene 40 [So |
| AC073534.2 | 19 | novel transcript |
| AC073534.1 | 19 | hepatitis A virus cellular receptor 1 (HAVCR1) |
| ERVK-28 | 19 | endogenous retrovirus group K member 28 [S |
| AC112702.1 | 19 | TEC |
| LINC00662 | 19 | long intergenic non-protein coding RNA 662 [S |
| AC006504.1 | 19 | novel transcript |
| AC006504.5 | 19 | novel transcript |
| AC006504.7 | 19 | novel transcript, sense intronic to novel transc |
| AC006504.3 | 19 | vomeronasal receptor pseudogene |
| SLC25A1P5 | 19 | solute carrier family 25 member 1 pseudogen |
| AC006504.2 | 19 | novel transcript |
| AC006504.4 | 19 | ladinin 1 (LAD1) pseudogene |

| | | |
|---|---|---|
| AC005357.2 | 19 | cyclin Y-like pseudogene |
| AC006504.6 | 19 | ubiquitin specific peptidase 24 (USP24) pseud |
| AC005758.1 | 19 | ATPase, aminophospholipid transporter, class |
| AC005357.1 | 19 | zinc finger protein pseudogene |
| AC010511.1 | 19 | novel transcript |
| AC020905.1 | 19 | novel transcript |
| AC005580.1 | 19 | novel transcript |
| AC093074.1 | 19 | novel transcript |
| AC005307.1 | 19 | novel transcript |
| AC005394.2 | 19 | novel transcript |
| AC005381.1 | 19 | novel transcript |
| AC005394.1 | 19 | novel transcript |
| AC005616.1 | 19 | novel transcript |
| AC079466.2 | 19 | novel transcript |
| AC079466.1 | 19 | novel transcript |
| AC005524.1 | 19 | SHC SH2-domain binding protein 1 (SHCBP1) p |
| MAN1A2P1 | 19 | mannosidase alpha class 1A member 2 pseud |
| AC007795.1 | 19 | novel transcript |
| LINC00906 | 19 | long intergenic non-protein coding RNA 906 [S |
| AC008991.1 | 19 | novel transcript |
| AC011524.1 | 19 | novel transcript |
| AC011524.2 | 19 | novel transcript |
| RNA5SP470 | 19 | RNA, 5S ribosomal pseudogene 470 [Source:H |
| LINC01532 | 19 | long intergenic non-protein coding RNA 1532 |
| AC011505.1 | 19 | novel transcript |
| UQCRFS1 | 19 | ubiquinol-cytochrome c reductase, Rieske iror |
| AC007786.1 | 19 | uncharacterized LOC105372352 [Source:NCBI |
| AC007786.2 | 19 | novel transcript |
| AC007786.3 | 19 | novel transcript |
| RN7SL340P | 19 | RNA, 7SL, cytoplasmic 340, pseudogene [Sour |
| AC007786.4 | 19 | novel transcript |
| AC011474.1 | 19 | novel transcript |
| AC011474.3 | 19 | novel transcript |
| AC011474.2 | 19 | novel transcript, antisense to VSTM2B |
| AC011474.4 | 19 | TEC |
| VSTM2B | 19 | V-set and transmembrane domain containing |
| POP4 | 19 | POP4 homolog, ribonuclease P/MRP subunit [ |
| PLEKHF1 | 19 | pleckstrin homology and FYVE domain contair |
| C19orf12 | 19 | chromosome 19 open reading frame 12 [Sour |
| AC008798.3 | 19 | novel transcript |
| CCNE1 | 19 | cyclin E1 [Source:HGNC Symbol;Acc:HGNC:15 |
| AC008798.2 | 19 | novel transcript |
| AC008798.1 | 19 | microtubule associated serine/threonine kina |
| AC008507.3 | 19 | novel transcript |
| PPIAP58 | 19 | peptidylprolyl isomerase A pseudogene 58 [Sc |
| URI1 | 19 | URI1 prefoldin like chaperone [Source:HGNC S |
| RPL9P32 | 19 | ribosomal protein L9 pseudogene 32 [Source: |
| AC008507.2 | 19 | novel transcript |
| TAF9P3 | 19 | TATA-box binding protein associated factor 9 |
| AC008507.1 | 19 | TAF9 RNA polymerase II, TATA box binding pro |

| | | |
|---|---|---|
| AC008507.4 | 19 | novel transcript |
| AC008507.5 | 19 | novel transcript |
| AC005597.1 | 19 | novel transcript, antisense to ZNF536 |
| ZNF536 | 19 | zinc finger protein 536 [Source:HGNC Symbol; |
| AC011504.1 | 19 | TEC |
| AC011478.1 | 19 | novel transcript |
| AC011507.1 | 19 | novel transcript |
| LINC01834 | 19 | long intergenic non-protein coding RNA 1834 |
| AC020897.1 | 19 | novel transcript |
| AC020912.1 | 19 | novel transcript |
| TSHZ3 | 19 | teashirt zinc finger homeobox 3 [Source:HGN( |
| LINC01791 | 19 | long intergenic non-protein coding RNA 1791 |
| AC025809.1 | 19 | novel transcript, antisense to TSHZ3 |
| AC025809.2 | 19 | TEC |
| AC008992.1 | 19 | novel transcript |
| AC008794.1 | 19 | novel transcript |
| LINC02841 | 19 | long intergenic non-protein coding RNA 2841 |
| AC011525.1 | 19 | novel transcript |
| AC011525.2 | 19 | testis highly expressed protein 5 [Source:NCBI |
| RNA5SP471 | 19 | RNA, 5S ribosomal pseudogene 471 [Source:H |
| RNU6-967P | 19 | RNA, U6 small nuclear 967, pseudogene [Sour |
| LINC01837 | 19 | long intergenic non-protein coding RNA 1837 |
| LINC01533 | 19 | long intergenic non-protein coding RNA 1533 |
| LINC01782 | 19 | long intergenic non-protein coding RNA 1782 |
| AC011518.1 | 19 | novel transcript |
| AC074139.1 | 19 | novel transcript |
| RNA5SP472 | 19 | RNA, 5S ribosomal pseudogene 472 [Source:H |
| ZNF507 | 19 | zinc finger protein 507 [Source:HGNC Symbol; |
| AC007773.1 | 19 | novel transcript, antisense to DPY19L3 |
| DPY19L3 | 19 | dpy-19 like C-mannosyltransferase 3 [Source:I |
| PDCD5 | 19 | programmed cell death 5 [Source:HGNC Symb |
| ANKRD27 | 19 | ankyrin repeat domain 27 [Source:HGNC Syml |
| SNORA68B | 19 | small nucleolar RNA, H/ACA box 68B [Source:I |
| RPS12P31 | 19 | ribosomal protein S12 pseudogene 31 [Source |
| RN7SL789P | 19 | RNA, 7SL, cytoplasmic 789, pseudogene [Sour |
| AC008474.1 | 19 | novel transcript, antisense to ANKRD27 |
| RGS9BP | 19 | regulator of G protein signaling 9 binding prot |
| AC008736.1 | 19 | novel transcript |
| NUDT19 | 19 | nudix hydrolase 19 [Source:HGNC Symbol;Acc |
| AC008736.2 | 19 | novel transcript |
| TDRD12 | 19 | tudor domain containing 12 [Source:HGNC Sy |
| AC008805.1 | 19 | glial cells missing homolog 1 (Drosophila) pseu |
| SLC7A9 | 19 | solute carrier family 7 member 9 [Source:HGN |
| AC008805.2 | 19 | novel transcript, antisense to SLC7A9 |
| RN7SKP22 | 19 | RN7SK pseudogene 22 [Source:HGNC Symbol; |
| CEP89 | 19 | centrosomal protein 89 [Source:HGNC Symbo |
| RPL31P60 | 19 | ribosomal protein L31 pseudogene 60 [Source |
| FAAP24 | 19 | FA core complex associated protein 24 [Sourc |
| RHPN2 | 19 | rhophilin Rho GTPase binding protein 2 [Sourc |
| AC008521.1 | 19 | MAD2 mitotic arrest deficient-like 1 (yeast) (N |

| | | |
|---|---|---|
| AC008521.2 | 19 | ribosomal protein S26 (RPS26) pseudogene |
| GPATCH1 | 19 | G-patch domain containing 1 [Source:HGNC S |
| WDR88 | 19 | WD repeat domain 88 [Source:HGNC Symbol; |
| LRP3 | 19 | LDL receptor related protein 3 [Source:HGNC |
| AC008738.1 | 19 | novel transcript, sense intronic to LRP3 |
| AC008738.7 | 19 | novel transcript |
| SLC7A10 | 19 | solute carrier family 7 member 10 [Source:HG |
| AC008738.4 | 19 | novel transcript, antisense to SLC7A10 |
| AC008738.6 | 19 | novel transcript |
| AC008738.3 | 19 | novel transcript |
| CEBPA | 19 | CCAAT enhancer binding protein alpha [Sourc |
| AC008738.5 | 19 | novel transcript |
| CEBPA-DT | 19 | CEBPA divergent transcript [Source:HGNC Syn |
| AC008738.2 | 19 | novel transcript |
| RPS3AP50 | 19 | ribosomal protein S3a pseudogene 50 [Source |
| AKR1B1P7 | 19 | aldo-keto reductase family 1 member B1 pseu |
| CEBPG | 19 | CCAAT enhancer binding protein gamma [Sou |
| PEPD | 19 | peptidase D [Source:HGNC Symbol;Acc:HGNC |
| AC010485.1 | 19 | novel transcript |
| RPL21P131 | 19 | ribosomal protein L21 pseudogene 131 [Sourc |
| CHST8 | 19 | carbohydrate sulfotransferase 8 [Source:HGN |
| KCTD15 | 19 | potassium channel tetramerization domain cc |
| AC008556.1 | 19 | novel transcript |
| RN7SL150P | 19 | RNA, 7SL, cytoplasmic 150, pseudogene [Sour |
| AC016587.1 | 19 | novel transcript |
| RPS4XP23 | 19 | ribosomal protein S4X pseudogene 23 [Source |
| RPS4XP20 | 19 | ribosomal protein S4X pseudogene 20 [Source |
| RPS4XP21 | 19 | ribosomal protein S4X pseudogene 21 [Source |
| CHCHD2P3 | 19 | coiled-coil-helix-coiled-coil-helix domain cont |
| LSM14A | 19 | LSM14A mRNA processing body assembly fact |
| KIAA0355 | 19 | KIAA0355 [Source:HGNC Symbol;Acc:HGNC:2 |
| RPL29P33 | 19 | ribosomal protein L29 pseudogene 33 [Source |
| AC010504.1 | 19 | novel transcript, antisense to KIAA0355 |
| GPI | 19 | glucose-6-phosphate isomerase [Source:HGN |
| AC092073.1 | 19 | novel protein, readthrough between GPI and |
| PDCD2L | 19 | programmed cell death 2 like [Source:HGNC S |
| RN7SL154P | 19 | RNA, 7SL, cytoplasmic 154, pseudogene [Sour |
| AC008747.1 | 19 | novel transcript, antisense to PDCD2 |
| UBA2 | 19 | ubiquitin like modifier activating enzyme 2 [Sc |
| WTIP | 19 | WT1 interacting protein [Source:HGNC Symbc |
| RPS26P55 | 19 | ribosomal protein S26 pseudogene 55 [Source |
| ZNF807 | 19 | zinc finger protein 807 [Source:HGNC Symbol; |
| SCGB1B2P | 19 | secretoglobin family 1B member 2, pseudoge |
| SCGB2B2 | 19 | secretoglobin family 2B member 2 [Source:HG |
| AC020910.4 | 19 | small nuclear ribonucleoprotein polypeptide E |
| AC020910.1 | 19 | NULL |
| AC020910.2 | 19 | novel transcript, sense intronic to SCGB2B2 |
| AC020910.7 | 19 | novel transcript |
| SCGB2B3P | 19 | secretoglobin family 2B member 3, pseudoge |
| AC020910.6 | 19 | novel transcript |

| | | |
|---|---|---|
| ZNF302 | 19 | zinc finger protein 302 [Source:HGNC Symbol; |
| AC020910.5 | 19 | novel transcript |
| ZNF181 | 19 | zinc finger protein 181 [Source:HGNC Symbol; |
| ZNF599 | 19 | zinc finger protein 599 [Source:HGNC Symbol; |
| AC020910.3 | 19 | zinc finger protein 134 pseudogene |
| LINC01801 | 19 | long intergenic non-protein coding RNA 1801 |
| AC008555.1 | 19 | novel transcript |
| AC008555.6 | 19 | zinc finger protein 396 (ZNF396) pseudogene |
| AC008555.2 | 19 | novel transcript |
| AC008555.4 | 19 | novel transcript |
| AC008555.3 | 19 | ABO blood group (transferase A, alpha 1-3-N-a |
| AC008555.5 | 19 | novel transcript |
| LINC01838 | 19 | long intergenic non-protein coding RNA 1838 |
| ZNF30-AS1 | 19 | ZNF30 antisense RNA 1 [Source:HGNC Symbol |
| ZNF30 | 19 | zinc finger protein 30 [Source:HGNC Symbol;A |
| AC008555.7 | 19 | TEC |
| ZNF792 | 19 | zinc finger protein 792 [Source:HGNC Symbol; |
| GRAMD1A | 19 | GRAM domain containing 1A [Source:HGNC Sy |
| AC020907.3 | 19 | novel transcript |
| AC020907.4 | 19 | novel transcript |
| SCN1B | 19 | sodium voltage-gated channel beta subunit 1 |
| HPN | 19 | hepsin [Source:HGNC Symbol;Acc:HGNC:5155 |
| HPN-AS1 | 19 | HPN antisense RNA 1 [Source:HGNC Symbol;A |
| AC020907.1 | 19 | novel transcript |
| AC020907.6 | 19 | novel protein |
| FXYD3 | 19 | FXYD domain containing ion transport regulat |
| MIR6887 | 19 | microRNA 6887 [Source:HGNC Symbol;Acc:HG |
| LGI4 | 19 | leucine rich repeat LGI family member 4 [Sour |
| AC020907.5 | 19 | novel transcript |
| FXYD1 | 19 | FXYD domain containing ion transport regulat |
| AC020907.2 | 19 | novel transcript |
| FXYD7 | 19 | FXYD domain containing ion transport regulat |
| FXYD5 | 19 | FXYD domain containing ion transport regulat |
| FAM187B | 19 | family with sequence similarity 187 member B |
| FAM187B2P | 19 | family with sequence similarity 187 member B |
| LSR | 19 | lipolysis stimulated lipoprotein receptor [Sour |
| AC002128.2 | 19 | novel transcript |
| AC002128.1 | 19 | novel transcript |
| USF2 | 19 | upstream transcription factor 2, c-fos interact |
| HAMP | 19 | hepcidin antimicrobial peptide [Source:HGNC |
| MAG | 19 | myelin associated glycoprotein [Source:HGNC |
| CD22 | 19 | CD22 molecule [Source:HGNC Symbol;Acc:HG |
| U62631.1 | 19 | novel transcript |
| MIR5196 | 19 | microRNA 5196 [Source:HGNC Symbol;Acc:HG |
| FFAR1 | 19 | free fatty acid receptor 1 [Source:HGNC Symb |
| FFAR3 | 19 | free fatty acid receptor 3 [Source:HGNC Symb |
| GPR42 | 19 | G protein-coupled receptor 42 (gene/pseudog |
| EEF1A1P7 | 19 | eukaryotic translation elongation factor 1 alph |
| LINC01531 | 19 | long intergenic non-protein coding RNA 1531 |
| AC002511.3 | 19 | free fatty acid receptor 2 (FFAR2) pseudogene |

| | | |
|---|---|---|
| RN7SL491P | 19 | RNA, 7SL, cytoplasmic 491, pseudogene [Sour |
| AC002511.2 | 19 | novel transcript |
| AC002511.1 | 19 | novel transcript |
| FFAR2 | 19 | free fatty acid receptor 2 [Source:HGNC Symb |
| KRTDAP | 19 | keratinocyte differentiation associated protei |
| DMKN | 19 | dermokine [Source:HGNC Symbol;Acc:HGNC:2 |
| SBSN | 19 | suprabasin [Source:HGNC Symbol;Acc:HGNC:2 |
| GAPDHS | 19 | glyceraldehyde-3-phosphate dehydrogenase, |
| TMEM147-AS1 | 19 | TMEM147 antisense RNA 1 [Source:HGNC Syn |
| TMEM147 | 19 | transmembrane protein 147 [Source:HGNC Sy |
| ATP4A | 19 | ATPase H+/K+ transporting subunit alpha [Sou |
| AD000090.1 | 19 | novel transcript, antisense to ATP4A |
| PMIS2 | 19 | PMIS2 transmembrane protein [Source:HGNC |
| LINC01766 | 19 | long intergenic non-protein coding RNA 1766 |
| AC002115.1 | 19 | novel transcript |
| HAUS5 | 19 | HAUS augmin like complex subunit 5 [Source:I |
| RBM42 | 19 | RNA binding motif protein 42 [Source:HGNC S |
| ETV2 | 19 | ETS variant 2 [Source:HGNC Symbol;Acc:HGN( |
| COX6B1 | 19 | cytochrome c oxidase subunit 6B1 [Source:HG |
| UPK1A | 19 | uroplakin 1A [Source:HGNC Symbol;Acc:HGN( |
| UPK1A-AS1 | 19 | UPK1A antisense RNA 1 [Source:HGNC Symbo |
| TYMSP2 | 19 | thymidylate synthetase pseudogene 2 [Source |
| ZBTB32 | 19 | zinc finger and BTB domain containing 32 [Sou |
| KMT2B | 19 | lysine methyltransferase 2B [Source:HGNC Syı |
| IGFLR1 | 19 | IGF like family receptor 1 [Source:HGNC Symb |
| AD000671.2 | 19 | novel transcript |
| U2AF1L4 | 19 | U2 small nuclear RNA auxiliary factor 1 like 4 [ |
| PSENEN | 19 | presenilin enhancer, gamma-secretase subuni |
| AD000671.1 | 19 | novel protein |
| AD000671.3 | 19 | novel transcript |
| LIN37 | 19 | lin-37 DREAM MuvB core complex componen |
| AC002398.2 | 19 | novel transcript |
| HSPB6 | 19 | heat shock protein family B (small) member 6 |
| PROSER3 | 19 | proline and serine rich 3 [Source:HGNC Symbo |
| AC002398.1 | 19 | novel transcript |
| ARHGAP33 | 19 | Rho GTPase activating protein 33 [Source:HGI |
| LINC01529 | 19 | long intergenic non-protein coding RNA 1529 |
| PRODH2 | 19 | proline dehydrogenase 2 [Source:HGNC Symb |
| NPHS1 | 19 | NPHS1 adhesion molecule, nephrin [Source:H |
| KIRREL2 | 19 | kirre like nephrin family adhesion molecule 2 |
| APLP1 | 19 | amyloid beta precursor like protein 1 [Source: |
| RN7SL402P | 19 | RNA, 7SL, cytoplasmic 402, pseudogene [Sour |
| NFKBID | 19 | NFKB inhibitor delta [Source:HGNC Symbol;Ac |
| AD000864.1 | 19 | novel transcript |
| HCST | 19 | hematopoietic cell signal transducer [Source:I |
| TYROBP | 19 | TYRO protein tyrosine kinase binding protein |
| LRFN3 | 19 | leucine rich repeat and fibronectin type III dor |
| AF038458.3 | 19 | novel transcript |
| AF038458.1 | 19 | LYR motif containing 2 (LYRM2) pseudogene |
| AF038458.2 | 19 | novel transcript |

| | | |
|---|---|---|
| SDHAF1 | 19 | succinate dehydrogenase complex assembly f |
| SYNE4 | 19 | spectrin repeat containing nuclear envelope fa |
| AC002116.1 | 19 | novel transcript |
| ALKBH6 | 19 | alkB homolog 6 [Source:HGNC Symbol;Acc:HG |
| AC002116.2 | 19 | uncharacterized LOC101927572 [Source:NCBI |
| CLIP3 | 19 | CAP-Gly domain containing linker protein 3 [S |
| THAP8 | 19 | THAP domain containing 8 [Source:HGNC Sym |
| WDR62 | 19 | WD repeat domain 62 [Source:HGNC Symbol; |
| AD000813.1 | 19 | TEC |
| OVOL3 | 19 | ovo like zinc finger 3 [Source:HGNC Symbol;Ac |
| POLR2I | 19 | RNA polymerase II subunit I [Source:HGNC Syr |
| TBCB | 19 | tubulin folding cofactor B [Source:HGNC Symb |
| AD001527.2 | 19 | TEC |
| CAPNS1 | 19 | calpain small subunit 1 [Source:HGNC Symbol |
| AD001527.1 | 19 | novel transcript, antisense to CAPNS1 |
| COX7A1 | 19 | cytochrome c oxidase subunit 7A1 [Source:HG |
| AC002984.1 | 19 | ectonucleoside triphosphate diphosphohydro |
| ZNF565 | 19 | zinc finger protein 565 [Source:HGNC Symbol; |
| Y_RNA | 19 | Y RNA [Source:RFAM;Acc:RF00019] |
| ZNF146 | 19 | zinc finger protein 146 [Source:HGNC Symbol; |
| AC092296.2 | 19 | novel transcript |
| LINC00665 | 19 | long intergenic non-protein coding RNA 665 [S |
| ZFP14 | 19 | ZFP14 zinc finger protein [Source:HGNC Symb |
| AC092296.1 | 19 | novel transcript, antisense to ZFP14 |
| AC092296.4 | 19 | novel transcript |
| ZFP82 | 19 | ZFP82 zinc finger protein [Source:HGNC Symb |
| AC092296.3 | 19 | acyl-CoA thioesterase 2 (ACOT2) pseudogene |
| ZNF566 | 19 | zinc finger protein 566 [Source:HGNC Symbol; |
| AC092295.2 | 19 | novel transcript, antisense to ZNF566 |
| CTBP2P7 | 19 | C-terminal binding protein 2 pseudogene 7 [S |
| ZNF260 | 19 | zinc finger protein 260 [Source:HGNC Symbol; |
| AC092295.1 | 19 | novel transcript, antisense to ZNF260 |
| ZNF529 | 19 | zinc finger protein 529 [Source:HGNC Symbol; |
| ZNF529-AS1 | 19 | ZNF529 antisense RNA 1 [Source:HGNC Symbo |
| ZNF382 | 19 | zinc finger protein 382 [Source:HGNC Symbol; |
| ZNF461 | 19 | zinc finger protein 461 [Source:HGNC Symbol; |
| AC074138.1 | 19 | novel transcript |
| LINC01534 | 19 | long intergenic non-protein coding RNA 1534 |
| ZNF567 | 19 | zinc finger protein 567 [Source:HGNC Symbol; |
| ZNF850 | 19 | zinc finger protein 850 [Source:HGNC Symbol; |
| AC020928.1 | 19 | uncharacterized LOC728485 [Source:NCBI ger |
| AC020928.2 | 19 | novel transcript |
| ZNF790-AS1 | 19 | ZNF790 antisense RNA 1 [Source:HGNC Symbo |
| ZNF790 | 19 | zinc finger protein 790 [Source:HGNC Symbol; |
| ZNF345 | 19 | zinc finger protein 345 [Source:HGNC Symbol; |
| Y_RNA | 19 | Y RNA [Source:RFAM;Acc:RF00019] |
| ZNF829 | 19 | zinc finger protein 829 [Source:HGNC Symbol; |
| RPL31P61 | 19 | ribosomal protein L31 pseudogene 61 [Source |
| ZNF568 | 19 | zinc finger protein 568 [Source:HGNC Symbol; |
| ZNF420 | 19 | zinc finger protein 420 [Source:HGNC Symbol; |

| | | |
|---|---|---|
| AC010632.3 | 19 | novel zinc finger pseudogene |
| AC010632.1 | 19 | novel transcript, antisense to ZNF420 |
| AC010632.2 | 19 | novel transcript |
| AC012309.1 | 19 | novel protein |
| ZNF585A | 19 | zinc finger protein 585A [Source:HGNC Symbo |
| AC012309.2 | 19 | novel transcript, antisense to ZNF420 |
| ZNF585B | 19 | zinc finger protein 585B [Source:HGNC Symbo |
| ZNF383 | 19 | zinc finger protein 383 [Source:HGNC Symbol; |
| AC016590.1 | 19 | novel transcript, antisense to ZNF383 |
| LINC01535 | 19 | long intergenic non-protein coding RNA 1535 |
| AC016590.4 | 19 | novel transcript |
| ZNF875 | 19 | zinc finger protein 875 [Source:HGNC Symbol; |
| AC016590.3 | 19 | novel transcript, antisense to HKR1 |
| AC016590.2 | 19 | novel transcript, sense intronic to HKR1 |
| ZNF527 | 19 | zinc finger protein 527 [Source:HGNC Symbol; |
| AC008806.1 | 19 | TEC |
| ZNF569 | 19 | zinc finger protein 569 [Source:HGNC Symbol; |
| ZNF570 | 19 | zinc finger protein 570 [Source:HGNC Symbol; |
| ZNF793-AS1 | 19 | ZNF793 antisense RNA 1 (head to head) [Sour |
| AC022148.2 | 19 | novel transcript |
| ZNF793 | 19 | zinc finger protein 793 [Source:HGNC Symbol; |
| AC022148.1 | 19 | novel transcript, antisense to ZNF793 |
| ZNF571-AS1 | 19 | ZNF571 antisense RNA 1 [Source:HGNC Symbo |
| ZNF540 | 19 | zinc finger protein 540 [Source:HGNC Symbol; |
| ZNF571 | 19 | zinc finger protein 571 [Source:HGNC Symbol; |
| ZFP30 | 19 | ZFP30 zinc finger protein [Source:HGNC Symb |
| ZNF781 | 19 | zinc finger protein 781 [Source:HGNC Symbol; |
| AC093227.2 | 19 | novel transcript |
| SELENOKP1 | 19 | selenoprotein K pseudogene 1 [Source:HGNC |
| ZNF607 | 19 | zinc finger protein 607 [Source:HGNC Symbol; |
| AC093227.1 | 19 | novel transcript |
| AC093227.3 | 19 | novel transcript |
| ZNF573 | 19 | zinc finger protein 573 [Source:HGNC Symbol; |
| AC016582.2 | 19 | increased sodium tolerance 1 homolog (yeast |
| AC016582.3 | 19 | novel zinc finger protein pseudogene |
| AC016582.1 | 19 | WD repeat domain 87 (WDR87) pseudogene |
| WDR87 | 19 | WD repeat domain 87 [Source:HGNC Symbol; |
| SIPA1L3 | 19 | signal induced proliferation associated 1 like 3 |
| AC008395.1 | 19 | novel transcript, sense intronic to SIPA1L3 |
| AC011465.1 | 19 | novel transcript, antisense to SIPA1L3 |
| AC011479.3 | 19 | novel transcript, sense intronic to SIPA1L3 |
| AC011479.2 | 19 | novel transcript, sense intronic to SIPA1L3 |
| RN7SL663P | 19 | RNA, 7SL, cytoplasmic 663, pseudogene [Sour |
| DPF1 | 19 | double PHD fingers 1 [Source:HGNC Symbol;A |
| SPINT2 | 19 | serine peptidase inhibitor, Kunitz type 2 [Sour |
| PPP1R14A | 19 | protein phosphatase 1 regulatory inhibitor sul |
| AC011479.1 | 19 | novel protein |
| AC011479.4 | 19 | novel transcript, antisense to SPINT2 |
| Y_RNA | 19 | Y RNA [Source:RFAM;Acc:RF00019] |
| YIF1B | 19 | Yip1 interacting factor homolog B, membrane |

| | | |
|---|---|---|
| C19orf33 | 19 | chromosome 19 open reading frame 33 [Sour |
| KCNK6 | 19 | potassium two pore domain channel subfamil |
| CATSPERG | 19 | cation channel sperm associated auxiliary sub |
| AC005625.1 | 19 | novel transcript, antisense to CATSPERG |
| PSMD8 | 19 | proteasome 26S subunit, non-ATPase 8 [Sourc |
| GGN | 19 | gametogenetin [Source:HGNC Symbol;Acc:HG |
| AC005789.1 | 19 | novel transcript, antisense to GGN |
| SPRED3 | 19 | sprouty related EVH1 domain containing 3 [Sc |
| FAM98C | 19 | family with sequence similarity 98 member C |
| RASGRP4 | 19 | RAS guanyl releasing protein 4 [Source:HGNC |
| RYR1 | 19 | ryanodine receptor 1 [Source:HGNC Symbol;A |
| AC067969.2 | 19 | novel transcript, antisense to RYR1 |
| AC067969.1 | 19 | novel transcript, antisense to RYR1 |
| MAP4K1 | 19 | mitogen-activated protein kinase kinase kinas |
| AC008649.1 | 19 | novel transcript |
| EIF3K | 19 | eukaryotic translation initiation factor 3 subur |
| ACTN4 | 19 | actinin alpha 4 [Source:HGNC Symbol;Acc:HGI |
| AC008649.2 | 19 | novel transcript, antisense to ACTN4 |
| CAPN12 | 19 | calpain 12 [Source:HGNC Symbol;Acc:HGNC:1 |
| AC022144.1 | 19 | novel transcript, antisense to CAPN12 |
| LGALS7 | 19 | galectin 7 [Source:HGNC Symbol;Acc:HGNC:6 |
| LGALS7B | 19 | galectin 7B [Source:HGNC Symbol;Acc:HGNC: |
| RNU6-140P | 19 | RNA, U6 small nuclear 140, pseudogene [Sour |
| LGALS4 | 19 | galectin 4 [Source:HGNC Symbol;Acc:HGNC:6 |
| ECH1 | 19 | enoyl-CoA hydratase 1 [Source:HGNC Symbol; |
| AC008982.1 | 19 | novel protein |
| AC104534.1 | 19 | novel transcript, antisense to ECH1 |
| HNRNPL | 19 | heterogeneous nuclear ribonucleoprotein L [S |
| AC008982.2 | 19 | novel transcript, sense intronic to HNRNPL |
| RINL | 19 | Ras and Rab interactor like [Source:HGNC Sym |
| AC011455.1 | 19 | novel transcript, antisense to RINL |
| SIRT2 | 19 | sirtuin 2 [Source:HGNC Symbol;Acc:HGNC:108 |
| NFKBIB | 19 | NFKB inhibitor beta [Source:HGNC Symbol;Acc |
| CCER2 | 19 | coiled-coil glutamate rich protein 2 [Source:H |
| SARS2 | 19 | seryl-tRNA synthetase 2, mitochondrial [Sourc |
| AC011455.2 | 19 | novel protein |
| MRPS12 | 19 | mitochondrial ribosomal protein S12 [Source: |
| ERVK9-11 | 19 | endogenous retrovirus group K9, member 11 |
| FBXO17 | 19 | F-box protein 17 [Source:HGNC Symbol;Acc:H |
| AC011455.6 | 19 | novel transcript, antisense to FBXO17and FBX |
| FBXO27 | 19 | F-box protein 27 [Source:HGNC Symbol;Acc:H |
| AC010605.1 | 19 | novel transcript, antisense to FBXO27 |
| ACP7 | 19 | acid phosphatase 7, tartrate resistant (putativ |
| PAK4 | 19 | p21 (RAC1) activated kinase 4 [Source:HGNC S |
| AC011443.1 | 19 | novel transcript, antisense to PAK4 |
| NCCRP1 | 19 | non-specific cytotoxic cell receptor protein 1 h |
| SYCN | 19 | syncollin [Source:HGNC Symbol;Acc:HGNC:184 |
| IFNL3P1 | 19 | interferon lambda 3 pseudogene 1 [Source:HC |
| IFNL3 | 19 | interferon lambda 3 [Source:HGNC Symbol;Ac |
| IFNL4 | 19 | interferon lambda 4 (gene/pseudogene) [Sour |

| | | |
|---|---|---|
| MSRB1P1 | 19 | methionine sulfoxide reductase B1 pseudoger |
| IFNL4P1 | 19 | interferon lambda 4 pseudogene 1 [Source:HG |
| IFNL2 | 19 | interferon lambda 2 [Source:HGNC Symbol;Ac |
| IFNL1 | 19 | interferon lambda 1 [Source:HGNC Symbol;Ac |
| LRFN1 | 19 | leucine rich repeat and fibronectin type III dor |
| AC011445.1 | 19 | novel transcript, sense overlapping LRFN1 |
| GMFG | 19 | glia maturation factor gamma [Source:HGNC S |
| AC011445.2 | 19 | novel transcript |
| SAMD4B | 19 | sterile alpha motif domain containing 4B [Sou |
| RN7SL566P | 19 | RNA, 7SL, cytoplasmic 566, pseudogene [Sour |
| PAF1 | 19 | PAF1 homolog, Paf1/RNA polymerase II comp |
| MED29 | 19 | mediator complex subunit 29 [Source:HGNC S |
| ZFP36 | 19 | ZFP36 ring finger protein [Source:HGNC Symb |
| MIR4530 | 19 | microRNA 4530 [Source:HGNC Symbol;Acc:HG |
| PLEKHG2 | 19 | pleckstrin homology and RhoGEF domain cont |
| RPS16 | 19 | ribosomal protein S16 [Source:HGNC Symbol; |
| SUPT5H | 19 | SPT5 homolog, DSIF elongation factor subunit |
| TIMM50 | 19 | translocase of inner mitochondrial membrane |
| DLL3 | 19 | delta like canonical Notch ligand 3 [Source:HG |
| SELENOV | 19 | selenoprotein V [Source:HGNC Symbol;Acc:HG |
| EID2B | 19 | EP300 interacting inhibitor of differentiation 2 |
| AC011500.3 | 19 | novel transcript, antisense to EID2B |
| TDGF1P7 | 19 | teratocarcinoma-derived growth factor 1 pseu |
| EID2 | 19 | EP300 interacting inhibitor of differentiation 2 |
| AC011500.2 | 19 | lectin, galactoside-binding, soluble, 13 (LGALS |
| AC011500.1 | 19 | lectin, galactoside-binding, soluble, 13 (LGALS |
| RPS29P24 | 19 | ribosomal protein S29 pseudogene 24 [Source |
| RPS29P25 | 19 | ribosomal protein S29 pseudogene 25 [Source |
| RPS29P26 | 19 | ribosomal protein S29 pseudogene 26 [Source |
| LGALS13 | 19 | galectin 13 [Source:HGNC Symbol;Acc:HGNC:1 |
| AC005176.2 | 19 | transmembrane protein 128 (TMEM128) pseu |
| AC005176.1 | 19 | lectin, galactoside-binding, soluble, 14 (LGALS |
| RPS29P30 | 19 | ribosomal protein S29 pseudogene 30 [Source |
| LGALS16 | 19 | galectin 16 [Source:HGNC Symbol;Acc:HGNC:4 |
| AC005515.1 | 19 | Charcot-Leyden crystal protein pseudogene |
| AC005515.2 | 19 | novel transcript |
| AC093063.1 | 19 | lectin, galactoside-binding, soluble, 14 (LGALS |
| RPS29P27 | 19 | ribosomal protein S29 pseudogene 27 [Source |
| LGALS14 | 19 | galectin 14 [Source:HGNC Symbol;Acc:HGNC:3 |
| AC006133.1 | 19 | Charcot-Leyden crystal galectin (CLC) pseudog |
| CLC | 19 | Charcot-Leyden crystal galectin [Source:HGNC |
| LEUTX | 19 | leucine twenty homeobox [Source:HGNC Sym |
| AC005393.1 | 19 | novel transcript |
| DYRK1B | 19 | dual specificity tyrosine phosphorylation regu |
| MIR6719 | 19 | microRNA 6719 [Source:HGNC Symbol;Acc:HG |
| FBL | 19 | fibrillarin [Source:HGNC Symbol;Acc:HGNC:35 |
| FCGBP | 19 | Fc fragment of IgG binding protein [Source:HG |
| PRR13P5 | 19 | proline rich 13 pseudogene 5 [Source:HGNC S |
| PSMC4 | 19 | proteasome 26S subunit, ATPase 4 [Source:HG |
| ZNF546 | 19 | zinc finger protein 546 [Source:HGNC Symbol; |

| | | |
|---|---|---|
| AC007842.1 | 19 | protein associated with topoisomerase II hom |
| ZNF780B | 19 | zinc finger protein 780B [Source:HGNC Symbo |
| AC005614.2 | 19 | novel transcript |
| ZNF780A | 19 | zinc finger protein 780A [Source:HGNC Symbo |
| AC005614.1 | 19 | novel transcript, antisense to ZNF780A |
| VN1R96P | 19 | vomeronasal 1 receptor 96 pseudogene [Sour |
| MAP3K10 | 19 | mitogen-activated protein kinase kinase kinas |
| TTC9B | 19 | tetratricopeptide repeat domain 9B [Source:H |
| CNTD2 | 19 | cyclin N-terminal domain containing 2 [Source |
| AKT2 | 19 | AKT serine/threonine kinase 2 [Source:HGNC : |
| AC118344.1 | 19 | novel transcript, sense intronic to AKT2 |
| MIR641 | 19 | microRNA 641 [Source:HGNC Symbol;Acc:HGI |
| RNU6-945P | 19 | RNA, U6 small nuclear 945, pseudogene [Sour |
| C19orf47 | 19 | chromosome 19 open reading frame 47 [Sour |
| Y_RNA | 19 | Y RNA [Source:RFAM;Acc:RF00019] |
| Y_RNA | 19 | Y RNA [Source:RFAM;Acc:RF00019] |
| PLD3 | 19 | phospholipase D family member 3 [Source:HG |
| AC118344.2 | 19 | TEC |
| MIR6796 | 19 | microRNA 6796 [Source:HGNC Symbol;Acc:HG |
| HIPK4 | 19 | homeodomain interacting protein kinase 4 [So |
| PRX | 19 | periaxin [Source:HGNC Symbol;Acc:HGNC:137 |
| SERTAD1 | 19 | SERTA domain containing 1 [Source:HGNC Syr |
| AC010271.2 | 19 | novel transcript |
| SERTAD3 | 19 | SERTA domain containing 3 [Source:HGNC Syr |
| AC010271.1 | 19 | novel transcript, antisense to SERTAD3 |
| BLVRB | 19 | biliverdin reductase B [Source:HGNC Symbol;/ |
| Metazoa_SRP | 19 | Metazoan signal recognition particle RNA [Sou |
| SPTBN4 | 19 | spectrin beta, non-erythrocytic 4 [Source:HGI |
| SHKBP1 | 19 | SH3KBP1 binding protein 1 [Source:HGNC Sym |
| LTBP4 | 19 | latent transforming growth factor beta bindin |
| NUMBL | 19 | NUMB like endocytic adaptor protein [Source |
| COQ8B | 19 | coenzyme Q8B [Source:HGNC Symbol;Acc:HG |
| RNU6-195P | 19 | RNA, U6 small nuclear 195, pseudogene [Sour |
| ITPKC | 19 | inositol-trisphosphate 3-kinase C [Source:HGI |
| C19orf54 | 19 | chromosome 19 open reading frame 54 [Sour |
| SNRPA | 19 | small nuclear ribonucleoprotein polypeptide / |
| MIA-RAB4B | 19 | MIA-RAB4B readthrough (NMD candidate) [Sc |
| MIA | 19 | MIA SH3 domain containing [Source:HGNC Syr |
| RAB4B | 19 | RAB4B, member RAS oncogene family [Source |
| RAB4B-EGLN2 | 19 | RAB4B-EGLN2 readthrough (NMD candidate) [ |
| AC008537.4 | 19 | novel transcript, antisense to RAB4B |
| EGLN2 | 19 | egl-9 family hypoxia inducible factor 2 [Source |
| AC008537.1 | 19 | novel protein |
| CYP2T1P | 19 | cytochrome P450 family 2 subfamily T membe |
| CYP2F2P | 19 | cytochrome P450 family 2 subfamily F membe |
| AC008537.2 | 19 | novel transcript |
| AC008537.3 | 19 | TEC |
| CYP2A6 | 19 | cytochrome P450 family 2 subfamily A membe |
| CYP2A7 | 19 | cytochrome P450 family 2 subfamily A membe |
| CYP2G1P | 19 | cytochrome P450 family 2 subfamily G membe |

| | | |
|---|---|---|
| CYP2A7P2 | 19 | cytochrome P450 family 2 subfamily A memb |
| CYP2B7P | 19 | cytochrome P450 family 2 subfamily B membe |
| AC092071.1 | 19 | novel transcript |
| CYP2B6 | 19 | cytochrome P450 family 2 subfamily B membe |
| CYP2A7P1 | 19 | cytochrome P450 family 2 subfamily A memb |
| CYP2G2P | 19 | cytochrome P450 family 2 subfamily G memb |
| AC008962.1 | 19 | adenosylmethionine decarboxylase 1 (AMD1) |
| CYP2A13 | 19 | cytochrome P450 family 2 subfamily A memb |
| CYP2F1 | 19 | cytochrome P450 family 2 subfamily F membe |
| CYP2T3P | 19 | cytochrome P450 family 2 subfamily T membe |
| RPL36P16 | 19 | ribosomal protein L36 pseudogene 16 [Source |
| RN7SL718P | 19 | RNA, 7SL, cytoplasmic 718, pseudogene [Sour |
| CYP2S1 | 19 | cytochrome P450 family 2 subfamily S membe |
| AXL | 19 | AXL receptor tyrosine kinase [Source:HGNC Sy |
| AC011510.1 | 19 | novel transcript, antisense to AXL |
| HNRNPUL1 | 19 | heterogeneous nuclear ribonucleoprotein U li |
| RN7SL34P | 19 | RNA, 7SL, cytoplasmic 34, pseudogene [Sourc |
| TGFB1 | 19 | transforming growth factor beta 1 [Source:HG |
| AC011462.5 | 19 | novel transcript, antisense to CCDC97 and HN |
| CCDC97 | 19 | coiled-coil domain containing 97 [Source:HGN |
| AC011462.1 | 19 | novel protein |
| TMEM91 | 19 | transmembrane protein 91 [Source:HGNC Syn |
| B9D2 | 19 | B9 domain containing 2 [Source:HGNC Symbo |
| AC011462.4 | 19 | novel transcript, sense intronic TMEM91 |
| EXOSC5 | 19 | exosome component 5 [Source:HGNC Symbol |
| BCKDHA | 19 | branched chain keto acid dehydrogenase E1 s |
| AC011462.3 | 19 | novel transcript, sense intronic to BCKDHA |
| AC011462.2 | 19 | novel transcript, antisense to B3GNT8 |
| B3GNT8 | 19 | UDP-GlcNAc:betaGal beta-1,3-N-acetylglucosa |
| DMAC2 | 19 | distal membrane arm assembly complex 2 [So |
| Metazoa_SRP | 19 | Metazoan signal recognition particle RNA [Sou |
| ERICH4 | 19 | glutamate rich 4 [Source:HGNC Symbol;Acc:H |
| PCAT19 | 19 | prostate cancer associated transcript 19 [Sour |
| AC243960.2 | 19 | tropomyosin pseudogene |
| PLEKHA3P1 | 19 | pleckstrin homology domain containing A3 ps |
| AC243960.3 | 19 | novel transcript |
| LINC01480 | 19 | long intergenic non-protein coding RNA 1480 |
| AC243960.1 | 19 | novel transcript, sense intronic to CEACAM21 |
| CEACAM21 | 19 | carcinoembryonic antigen related cell adhesic |
| AC243960.7 | 19 | TEC |
| CEACAMP3 | 19 | carcinoembryonic antigen related cell adhesic |
| DNAJC19P2 | 19 | DnaJ heat shock protein family (Hsp40) memb |
| CEACAM4 | 19 | carcinoembryonic antigen related cell adhesic |
| DNAJC19P3 | 19 | DnaJ heat shock protein family (Hsp40) memb |
| AC243960.5 | 19 | pregnancy-specific glycoprotein pseudogene |
| AC243960.4 | 19 | phosphodiesterase 7 pseudogene |
| CEACAM7 | 19 | carcinoembryonic antigen related cell adhesic |
| CEACAM5 | 19 | carcinoembryonic antigen related cell adhesic |
| AC243967.1 | 19 | novel protein, readthrough between CEACAM |
| CEACAM6 | 19 | carcinoembryonic antigen related cell adhesic |

| | | |
|---|---|---|
| AC243967.2 | 19 | novel transcript, antisense to CEACAM6 |
| CEACAM3 | 19 | carcinoembryonic antigen related cell adhesic |
| HNRNPA1P52 | 19 | heterogeneous nuclear ribonucleoprotein A1 |
| AC243967.3 | 19 | novel transcript |
| LYPD4 | 19 | LY6/PLAUR domain containing 4 [Source:HGN |
| DMRTC2 | 19 | DMRT like family C2 [Source:HGNC Symbol;Ac |
| RPS19 | 19 | ribosomal protein S19 [Source:HGNC Symbol;. |
| MIR6797 | 19 | microRNA 6797 [Source:HGNC Symbol;Acc:HG |
| CD79A | 19 | CD79a molecule [Source:HGNC Symbol;Acc:H( |
| ARHGEF1 | 19 | Rho guanine nucleotide exchange factor 1 [So |
| ERFL | 19 | ETS repressor factor like [Source:HGNC Symbc |
| RABAC1 | 19 | Rab acceptor 1 [Source:HGNC Symbol;Acc:HG |
| AC010616.1 | 19 | novel protein, ATP1A3-RABAC1 readthrough |
| ATP1A3 | 19 | ATPase Na+/K+ transporting subunit alpha 3 [! |
| GRIK5 | 19 | glutamate ionotropic receptor kainate type su |
| ZNF574 | 19 | zinc finger protein 574 [Source:HGNC Symbol; |
| POU2F2 | 19 | POU class 2 homeobox 2 [Source:HGNC Symb |
| AC010247.1 | 19 | novel transcript, antisense to POU2F2 |
| MIR4323 | 19 | microRNA 4323 [Source:HGNC Symbol;Acc:HG |
| AC010247.2 | 19 | novel transcript, antisense to POU2F2 |
| DEDD2 | 19 | death effector domain containing 2 [Source:H |
| ZNF526 | 19 | zinc finger protein 526 [Source:HGNC Symbol; |
| GSK3A | 19 | glycogen synthase kinase 3 alpha [Source:HGN |
| AC006486.1 | 19 | novel protein |
| AC006486.2 | 19 | TEC |
| ERF | 19 | ETS2 repressor factor [Source:HGNC Symbol;A |
| CIC | 19 | capicua transcriptional repressor [Source:HGN |
| PAFAH1B3 | 19 | platelet activating factor acetylhydrolase 1b c. |
| PRR19 | 19 | proline rich 19 [Source:HGNC Symbol;Acc:HGI |
| TMEM145 | 19 | transmembrane protein 145 [Source:HGNC Sy |
| MEGF8 | 19 | multiple EGF like domains 8 [Source:HGNC Syı |
| MIR8077 | 19 | microRNA 8077 [Source:HGNC Symbol;Acc:HG |
| CNFN | 19 | cornifelin [Source:HGNC Symbol;Acc:HGNC:3C |
| LIPE-AS1 | 19 | LIPE antisense RNA 1 [Source:HGNC Symbol;A |
| LIPE | 19 | lipase E, hormone sensitive type [Source:HGN |
| AC011497.1 | 19 | novel transcript |
| CXCL17 | 19 | C-X-C motif chemokine ligand 17 [Source:HGN |
| RNU4-60P | 19 | RNA, U4 small nuclear 60, pseudogene [Sourc |
| AC011497.2 | 19 | zinc finger, AN1-type domain 5 (ZFAND5) pseu |
| CEACAM1 | 19 | carcinoembryonic antigen related cell adhesic |
| CEACAMP2 | 19 | carcinoembryonic antigen related cell adhesic |
| CEACAM8 | 19 | carcinoembryonic antigen related cell adhesic |
| CEACAMP1 | 19 | carcinoembryonic antigen related cell adhesic |
| CEACAMP5 | 19 | carcinoembryonic antigen related cell adhesic |
| RPS10P28 | 19 | ribosomal protein S10 pseudogene 28 [Source |
| PSG3 | 19 | pregnancy specific beta-1-glycoprotein 3 [Sou |
| PSG8 | 19 | pregnancy specific beta-1-glycoprotein 8 [Sou |
| CEACAMP6 | 19 | carcinoembryonic antigen related cell adhesic |
| PSG8-AS1 | 19 | PSG8 antisense RNA 1 [Source:HGNC Symbol;. |
| PSG10P | 19 | pregnancy specific beta-1-glycoprotein 10, psε |

| | | |
|---|---|---|
| PSG1 | 19 | pregnancy specific beta-1-glycoprotein 1 [Sou |
| PSG6 | 19 | pregnancy specific beta-1-glycoprotein 6 [Sou |
| PSG7 | 19 | pregnancy specific beta-1-glycoprotein 7 (gen |
| CEACAMP7 | 19 | carcinoembryonic antigen related cell adhesic |
| AC004784.1 | 19 | novel transcript, antisense to PSG11, PSG2 an |
| PSG11 | 19 | pregnancy specific beta-1-glycoprotein 11 [So |
| CEACAMP8 | 19 | carcinoembryonic antigen related cell adhesic |
| PSG2 | 19 | pregnancy specific beta-1-glycoprotein 2 [Sou |
| CEACAMP9 | 19 | carcinoembryonic antigen related cell adhesic |
| AC005392.1 | 19 | novel transcript |
| PSG5 | 19 | pregnancy specific beta-1-glycoprotein 5 [Sou |
| PSG4 | 19 | pregnancy specific beta-1-glycoprotein 4 [Sou |
| CEACAMP10 | 19 | carcinoembryonic antigen related cell adhesic |
| PSG9 | 19 | pregnancy specific beta-1-glycoprotein 9 [Sou |
| CEACAMP11 | 19 | carcinoembryonic antigen related cell adhesic |
| CEACAMP4 | 19 | carcinoembryonic antigen related cell adhesic |
| AC005392.2 | 19 | novel transcript |
| CD177 | 19 | CD177 molecule [Source:HGNC Symbol;Acc:H |
| AC005392.3 | 19 | novel transcript, antisense to CD177 |
| CD177P1 | 19 | CD177 molecule pseudogene 1 [Source:HGNC |
| TEX101 | 19 | testis expressed 101 [Source:HGNC Symbol;Ac |
| LYPD3 | 19 | LY6/PLAUR domain containing 3 [Source:HGN |
| PHLDB3 | 19 | pleckstrin homology like domain family B mer |
| ETHE1 | 19 | ETHE1 persulfide dioxygenase [Source:HGNC S |
| ZNF575 | 19 | zinc finger protein 575 [Source:HGNC Symbol; |
| XRCC1 | 19 | X-ray repair cross complementing 1 [Source:H |
| L34079.2 | 19 | novel transcript, antisense to XRCC1 |
| L34079.1 | 19 | novel protein |
| L34079.3 | 19 | novel transcript, antisense to XRCC1 |
| PINLYP | 19 | phospholipase A2 inhibitor and LY6/PLAUR do |
| IRGQ | 19 | immunity related GTPase Q [Source:HGNC Syr |
| ZNF576 | 19 | zinc finger protein 576 [Source:HGNC Symbol; |
| SRRM5 | 19 | serine/arginine repetitive matrix 5 [Source:HG |
| ZNF428 | 19 | zinc finger protein 428 [Source:HGNC Symbol; |
| CADM4 | 19 | cell adhesion molecule 4 [Source:HGNC Symb |
| PLAUR | 19 | plasminogen activator, urokinase receptor [Sc |
| RN7SL368P | 19 | RNA, 7SL, cytoplasmic 368, pseudogene [Sour |
| AC005622.1 | 19 | phospholipase A2 inhibitor and Ly6/PLAUR do |
| IRGC | 19 | immunity related GTPase cinema [Source:HGN |
| SMG9 | 19 | SMG9 nonsense mediated mRNA decay factor |
| KCNN4 | 19 | potassium calcium-activated channel subfami |
| LYPD5 | 19 | LY6/PLAUR domain containing 5 [Source:HGN |
| AC115522.1 | 19 | novel transcript |
| ZNF283 | 19 | zinc finger protein 283 [Source:HGNC Symbol; |
| ZNF404 | 19 | zinc finger protein 404 [Source:HGNC Symbol; |
| AC006213.6 | 19 | TEC |
| AC006213.2 | 19 | novel transcript |
| AC006213.7 | 19 | novel transcript, sense intronic to ZNF404 |
| AC006213.3 | 19 | novel transcript, antisense to ZNF45 |
| ZNF45 | 19 | zinc finger protein 45 [Source:HGNC Symbol;A |

| | | |
|---|---|---|
| ZNF221 | 19 | zinc finger protein 221 [Source:HGNC Symbol; |
| ZNF155 | 19 | zinc finger protein 155 [Source:HGNC Symbol; |
| AC006213.4 | 19 | novel transcript, sense intronic to ZNF155 |
| AC006213.5 | 19 | novel transcript, sense intronic to ZNF155 |
| AC006213.1 | 19 | novel transcript, antisense to ZNF155 |
| ZNF230 | 19 | zinc finger protein 230 [Source:HGNC Symbol; |
| AC067968.2 | 19 | novel transcript |
| ZNF222 | 19 | zinc finger protein 222 [Source:HGNC Symbol; |
| AC067968.1 | 19 | novel protein |
| ZNF223 | 19 | zinc finger protein 223 [Source:HGNC Symbol; |
| ZNF284 | 19 | zinc finger protein 284 [Source:HGNC Symbol; |
| RNU6-902P | 19 | RNA, U6 small nuclear 902, pseudogene [Sour |
| ZNF224 | 19 | zinc finger protein 224 [Source:HGNC Symbol; |
| AC021092.1 | 19 | novel transcript, antisense to ZNF225 and ZNF |
| ZNF225 | 19 | zinc finger protein 225 [Source:HGNC Symbol; |
| ZNF234 | 19 | zinc finger protein 234 [Source:HGNC Symbol; |
| AC138470.2 | 19 | novel transcript |
| ZNF226 | 19 | zinc finger protein 226 [Source:HGNC Symbol; |
| ZNF227 | 19 | zinc finger protein 227 [Source:HGNC Symbol; |
| ZNF235 | 19 | zinc finger protein 235 [Source:HGNC Symbol; |
| AC138470.1 | 19 | TEC |
| ZNF233 | 19 | zinc finger protein 233 [Source:HGNC Symbol; |
| AC138473.1 | 19 | mitochondrial ribosomal protein S15 (MRPS15 |
| NDUFA3P1 | 19 | NADH:ubiquinone oxidoreductase subunit A3 |
| ZNF112 | 19 | zinc finger protein 112 [Source:HGNC Symbol; |
| AC245748.1 | 19 | novel protein |
| ZNF285 | 19 | zinc finger protein 285 [Source:HGNC Symbol; |
| AC245748.2 | 19 | novel transcript |
| ZNF229 | 19 | zinc finger protein 229 [Source:HGNC Symbol; |
| ZNF285B | 19 | zinc finger protein 285B (pseudogene) [Source |
| ZNF180 | 19 | zinc finger protein 180 [Source:HGNC Symbol; |
| CEACAM20 | 19 | carcinoembryonic antigen related cell adhesio |
| AC245748.3 | 19 | novel transcript |
| CEACAM22P | 19 | carcinoembryonic antigen related cell adhesio |
| AC243964.1 | 19 | carcinoembryonic antigen-related cell adhesio |
| IGSF23 | 19 | immunoglobulin superfamily member 23 [Sou |
| AC243964.3 | 19 | novel transcript, antisense to CEACAM16, CEA |
| PVR | 19 | PVR cell adhesion molecule [Source:HGNC Syn |
| MIR4531 | 19 | microRNA 4531 [Source:HGNC Symbol;Acc:HG |
| CEACAM19 | 19 | carcinoembryonic antigen related cell adhesio |
| AC243964.4 | 19 | TEC |
| CEACAM16 | 19 | carcinoembryonic antigen related cell adhesio |
| AC243964.2 | 19 | NULL |
| BCL3 | 19 | BCL3 transcription coactivator [Source:HGNC S |
| MIR8085 | 19 | microRNA 8085 [Source:HGNC Symbol;Acc:HG |
| SNORA70 | 19 | Small nucleolar RNA SNORA70 [Source:RFAM; |
| AC092066.1 | 19 | ribosomal protein L10 (RPL10) pseudogene |
| CBLC | 19 | Cbl proto-oncogene C [Source:HGNC Symbol;A |
| AC092306.1 | 19 | novel transcript |
| BCAM | 19 | basal cell adhesion molecule (Lutheran blood |

| | | |
|---|---|---|
| NECTIN2 | 19 | nectin cell adhesion molecule 2 [Source:HGN( |
| AC011481.2 | 19 | novel transcript, antisense to PVRL2 |
| TOMM40 | 19 | translocase of outer mitochondrial membrane |
| APOE | 19 | apolipoprotein E [Source:HGNC Symbol;Acc:H |
| AC011481.3 | 19 | TEC |
| APOC1 | 19 | apolipoprotein C1 [Source:HGNC Symbol;Acc: |
| APOC1P1 | 19 | apolipoprotein C1 pseudogene 1 [Source:HGN |
| APOC4 | 19 | apolipoprotein C4 [Source:HGNC Symbol;Acc: |
| APOC4-APOC2 | 19 | APOC4-APOC2 readthrough (NMD candidate) |
| APOC2 | 19 | apolipoprotein C2 [Source:HGNC Symbol;Acc: |
| AC011481.1 | 19 | novel transcript |
| CLPTM1 | 19 | CLPTM1 regulator of GABA type A receptor fo |
| RELB | 19 | RELB proto-oncogene, NF-kB subunit [Source: |
| CLASRP | 19 | CLK4 associating serine/arginine rich protein [ |
| RNU6-611P | 19 | RNA, U6 small nuclear 611, pseudogene [Sour |
| ZNF296 | 19 | zinc finger protein 296 [Source:HGNC Symbol; |
| GEMIN7-AS1 | 19 | GEMIN7 antisense RNA 1 [Source:HGNC Symb |
| GEMIN7 | 19 | gem nuclear organelle associated protein 7 [S |
| MARK4 | 19 | microtubule affinity regulating kinase 4 [Sourc |
| PPP1R37 | 19 | protein phosphatase 1 regulatory subunit 37 [ |
| EIF5AP3 | 19 | eukaryotic translation initiation factor 5A pseu |
| AC005757.1 | 19 | novel transcript, antisense to PPP1R37 |
| NKPD1 | 19 | NTPase KAP family P-loop domain containing |
| TRAPPC6A | 19 | trafficking protein particle complex 6A [Sourc |
| BLOC1S3 | 19 | biogenesis of lysosomal organelles complex 1 |
| EXOC3L2 | 19 | exocyst complex component 3 like 2 [Source:I |
| CKM | 19 | creatine kinase, M-type [Source:HGNC Symbo |
| RPS16P9 | 19 | ribosomal protein S16 pseudogene 9 [Source: |
| KLC3 | 19 | kinesin light chain 3 [Source:HGNC Symbol;Ac |
| ERCC2 | 19 | ERCC excision repair 2, TFIIH core complex hel |
| PPP1R13L | 19 | protein phosphatase 1 regulatory subunit 13 l |
| CD3EAP | 19 | CD3e molecule associated protein [Source:HG |
| ERCC1 | 19 | ERCC excision repair 1, endonuclease non-cat |
| MIR6088 | 19 | microRNA 6088 [Source:HGNC Symbol;Acc:HG |
| FOSB | 19 | FosB proto-oncogene, AP-1 transcription facto |
| RTN2 | 19 | reticulon 2 [Source:HGNC Symbol;Acc:HGNC:1 |
| PPM1N | 19 | protein phosphatase, Mg2+/Mn2+ dependent |
| VASP | 19 | vasodilator stimulated phosphoprotein [Sourc |
| OPA3 | 19 | outer mitochondrial membrane lipid metaboli |
| GPR4 | 19 | G protein-coupled receptor 4 [Source:HGNC S |
| EML2 | 19 | EMAP like 2 [Source:HGNC Symbol;Acc:HGNC: |
| MIR330 | 19 | microRNA 330 [Source:HGNC Symbol;Acc:HGN |
| EML2-AS1 | 19 | EML2 antisense RNA 1 [Source:HGNC Symbol; |
| RN7SL836P | 19 | RNA, 7SL, cytoplasmic 836, pseudogene [Sour |
| GIPR | 19 | gastric inhibitory polypeptide receptor [Sourc |
| MIR642A | 19 | microRNA 642a [Source:HGNC Symbol;Acc:HG |
| MIR642B | 19 | microRNA 642b [Source:HGNC Symbol;Acc:HG |
| SNRPD2 | 19 | small nuclear ribonucleoprotein D2 polypeptic |
| QPCTL | 19 | glutaminyl-peptide cyclotransferase like [Sour |
| FBXO46 | 19 | F-box protein 46 [Source:HGNC Symbol;Acc:H |

| | | |
|---|---|---|
| AC007191.1 | 19 | TEC |
| BHMG1 | 19 | basic helix-loop-helix and HMG-box containin |
| AC074212.1 | 19 | novel transcript, antisense to SIX5 |
| SIX5 | 19 | SIX homeobox 5 [Source:HGNC Symbol;Acc:H |
| DM1-AS | 19 | DM1 locus antisense RNA [Source:HGNC Symb |
| DMPK | 19 | DM1 protein kinase [Source:HGNC Symbol;Ac |
| AC011530.1 | 19 | novel protein |
| DMWD | 19 | DM1 locus, WD repeat containing [Source:HG |
| RSPH6A | 19 | radial spoke head 6 homolog A [Source:HGNC |
| SYMPK | 19 | symplekin [Source:HGNC Symbol;Acc:HGNC:2 |
| AC092301.1 | 19 | novel transcript, antisense to SYMPK |
| FOXA3 | 19 | forkhead box A3 [Source:HGNC Symbol;Acc:H |
| IRF2BP1 | 19 | interferon regulatory factor 2 binding protein |
| MYPOP | 19 | Myb related transcription factor, partner of p |
| NANOS2 | 19 | nanos C2HC-type zinc finger 2 [Source:HGNC |
| NOVA2 | 19 | NOVA alternative splicing regulator 2 [Source: |
| CCDC61 | 19 | coiled-coil domain containing 61 [Source:HGN |
| MIR769 | 19 | microRNA 769 [Source:HGNC Symbol;Acc:HG |
| PGLYRP1 | 19 | peptidoglycan recognition protein 1 [Source:H |
| AC007785.3 | 19 | novel transcript |
| IGFL4 | 19 | IGF like family member 4 [Source:HGNC Symb |
| AC007785.1 | 19 | novel transcript |
| AC007785.2 | 19 | solute carrier family 30 (zinc transporter), me |
| IGFL3 | 19 | IGF like family member 3 [Source:HGNC Symb |
| TGIF1P1 | 19 | TGFB induced factor homeobox 1 pseudogene |
| IGFL2 | 19 | IGF like family member 2 [Source:HGNC Symb |
| AC006262.2 | 19 | novel transcript |
| IGFL2-AS1 | 19 | IGFL2 antisense RNA 1 [Source:HGNC Symbol; |
| IGFL1P1 | 19 | IGF like family member 1 pseudogene 1 [Sour |
| AC006262.1 | 19 | novel transcript |
| RPL12P41 | 19 | ribosomal protein L12 pseudogene 41 [Source |
| AC006262.3 | 19 | novel transcript, antisense to IGFL1 |
| IGFL1 | 19 | IGF like family member 1 [Source:HGNC Symb |
| AC006262.4 | 19 | IGF-like family member 4 (IGFL4) pseudogene |
| IGFL1P2 | 19 | IGF like family member 1 pseudogene 2 [Sour |
| HIF3A | 19 | hypoxia inducible factor 3 subunit alpha [Sour |
| RNU6-924P | 19 | RNA, U6 small nuclear 924, pseudogene [Sour |
| AC007193.2 | 19 | novel transcript, antisense to HIF3A |
| PPP5C | 19 | protein phosphatase 5 catalytic subunit [Sour |
| AC007193.1 | 19 | novel transcript, antisense to PPP5C |
| AC007193.3 | 19 | novel transcript, antisense to PPP5C |
| CCDC8 | 19 | coiled-coil domain containing 8 [Source:HGNC |
| PNMA8C | 19 | PNMA family member 8C [Source:HGNC Symb |
| PNMA8A | 19 | PNMA family member 8A [Source:HGNC Symb |
| PPP5D1 | 19 | PPP5 tetratricopeptide repeat domain contai |
| PNMA8B | 19 | PNMA family member 8B [Source:HGNC Symb |
| AC011484.1 | 19 | novel transcript, antisense to PPP5D1 |
| AC011551.1 | 19 | paraneoplastic Ma antigen family-like 2 (PNM |
| AC093503.1 | 19 | novel transcript, sense intronic to PPP5D1 |
| GAPDHP38 | 19 | glyceraldehyde 3 phosphate dehydrogenase p |

| | | |
|---|---|---|
| CALM3 | 19 | calmodulin 3 [Source:HGNC Symbol;Acc:HGN( |
| AC093503.2 | 19 | novel transcript, antisense to CALM3 |
| PTGIR | 19 | prostaglandin I2 receptor [Source:HGNC Symb |
| GNG8 | 19 | G protein subunit gamma 8 [Source:HGNC Syr |
| AC093503.3 | 19 | TEC |
| DACT3 | 19 | dishevelled binding antagonist of beta catenir |
| DACT3-AS1 | 19 | DACT3 antisense RNA 1 [Source:HGNC Symbo |
| PRKD2 | 19 | protein kinase D2 [Source:HGNC Symbol;Acc:I |
| RN7SL364P | 19 | RNA, 7SL, cytoplasmic 364, pseudogene [Sour |
| MIR320E | 19 | microRNA 320e [Source:HGNC Symbol;Acc:HG |
| STRN4 | 19 | striatin 4 [Source:HGNC Symbol;Acc:HGNC:15 |
| AC008635.1 | 19 | novel transcript, antisense to STRN4 |
| Y_RNA | 19 | Y RNA [Source:RFAM;Acc:RF00019] |
| FKRP | 19 | fukutin related protein [Source:HGNC Symbol |
| SLC1A5 | 19 | solute carrier family 1 member 5 [Source:HGN |
| AC008622.2 | 19 | novel transcript, antisense to SLC1A5 |
| AC008622.3 | 19 | heterogeneous nuclear ribonucleoprotein M ( |
| HNRNPMP2 | 19 | heterogeneous nuclear ribonucleoprotein M p |
| AC008622.1 | 19 | heterogeneous nuclear ribonucleoprotein A1- |
| AP2S1 | 19 | adaptor related protein complex 2 subunit sig |
| ARHGAP35 | 19 | Rho GTPase activating protein 35 [Source:HGI |
| AC008895.1 | 19 | TEC |
| NPAS1 | 19 | neuronal PAS domain protein 1 [Source:HGNC |
| TMEM160 | 19 | transmembrane protein 160 [Source:HGNC Sy |
| ZC3H4 | 19 | zinc finger CCCH-type containing 4 [Source:HG |
| SAE1 | 19 | SUMO1 activating enzyme subunit 1 [Source:I |
| BBC3 | 19 | BCL2 binding component 3 [Source:HGNC Syn |
| MIR3190 | 19 | microRNA 3190 [Source:HGNC Symbol;Acc:HG |
| MIR3191 | 19 | microRNA 3191 [Source:HGNC Symbol;Acc:HG |
| AC008532.1 | 19 | novel transcript |
| CCDC9 | 19 | coiled-coil domain containing 9 [Source:HGNC |
| INAFM1 | 19 | InaF motif containing 1 [Source:HGNC Symbol |
| C5AR1 | 19 | complement C5a receptor 1 [Source:HGNC Sy |
| C5AR2 | 19 | complement component 5a receptor 2 [Sourc |
| DHX34 | 19 | DExH-box helicase 34 [Source:HGNC Symbol;A |
| MEIS3 | 19 | Meis homeobox 3 [Source:HGNC Symbol;Acc: |
| SLC8A2 | 19 | solute carrier family 8 member A2 [Source:HG |
| AC073548.2 | 19 | novel transcript, antisense to KPTNand SLC8A |
| KPTN | 19 | kaptin, actin binding protein [Source:HGNC Sy |
| NAPA-AS1 | 19 | NAPA antisense RNA 1 [Source:HGNC Symbol; |
| NAPA | 19 | NSF attachment protein alpha [Source:HGNC ! |
| AC073548.1 | 19 | TEC |
| ZNF541 | 19 | zinc finger protein 541 [Source:HGNC Symbol; |
| RN7SL322P | 19 | RNA, 7SL, cytoplasmic 322, pseudogene [Sour |
| AC010519.1 | 19 | novel transcript |
| AC010331.1 | 19 | novel transcript, antisense to GLTSCR1 |
| BICRA | 19 | BRD4 interacting chromatin remodeling comp |
| BICRA-AS1 | 19 | BICRA antisense RNA 1 [Source:HGNC Symbol |
| AC008985.1 | 19 | TEC |
| EHD2 | 19 | EH domain containing 2 [Source:HGNC Symbo |

| | | |
|---|---|---|
| NOP53 | 19 | NOP53 ribosome biogenesis factor [Source:H( |
| SNORD23 | 19 | small nucleolar RNA, C/D box 23 [Source:HGN |
| NOP53-AS1 | 19 | NOP53 antisense RNA 1 [Source:HGNC Symbo |
| SELENOW | 19 | selenoprotein W [Source:HGNC Symbol;Acc:H |
| RPL23AP80 | 19 | ribosomal protein L23a pseudogene 80 [Sourc |
| AC008745.1 | 19 | major histocompatibility complex, class I, F (H |
| TPRX1 | 19 | tetrapeptide repeat homeobox 1 [Source:HGN |
| CRX | 19 | cone-rod homeobox [Source:HGNC Symbol;Ac |
| TPRX2P | 19 | tetrapeptide repeat homeobox 2, pseudogene |
| LINC01595 | 19 | long intergenic non-protein coding RNA 1595 |
| SULT2A1 | 19 | sulfotransferase family 2A member 1 [Source: |
| BSPH1 | 19 | binder of sperm protein homolog 1 [Source:H |
| ELSPBP1 | 19 | epididymal sperm binding protein 1 [Source:H |
| CABP5 | 19 | calcium binding protein 5 [Source:HGNC Symb |
| PLA2G4C | 19 | phospholipase A2 group IVC [Source:HGNC Sy |
| PLA2G4C-AS1 | 19 | PLA2G4C antisense RNA 1 [Source:HGNC Sym |
| LIG1 | 19 | DNA ligase 1 [Source:HGNC Symbol;Acc:HGNC |
| AC011466.3 | 19 | novel transcript, antisense to LIG1 |
| AC011466.2 | 19 | novel transcript, antisense to LIG1 |
| ZSWIM9 | 19 | zinc finger SWIM-type containing 9 [Source:H( |
| CARD8 | 19 | caspase recruitment domain family member 8 |
| AC011466.4 | 19 | TEC |
| AC011466.1 | 19 | novel transcript, antisense to CARD8 |
| CARD8-AS1 | 19 | CARD8 antisense RNA 1 [Source:HGNC Symbo |
| ZNF114-AS1 | 19 | ZNF114 antisense RNA 1 [Source:HGNC Symb |
| ZNF114 | 19 | zinc finger protein 114 [Source:HGNC Symbol; |
| CCDC114 | 19 | coiled-coil domain containing 114 [Source:HG |
| EMP3 | 19 | epithelial membrane protein 3 [Source:HGNC |
| TMEM143 | 19 | transmembrane protein 143 [Source:HGNC Sy |
| SYNGR4 | 19 | synaptogyrin 4 [Source:HGNC Symbol;Acc:HG |
| KDELR1 | 19 | KDEL endoplasmic reticulum protein retentior |
| GRIN2D | 19 | glutamate ionotropic receptor NMDA type sul |
| GRWD1 | 19 | glutamate rich WD repeat containing 1 [Sourc |
| KCNJ14 | 19 | potassium voltage-gated channel subfamily J r |
| AC008403.2 | 19 | novel transcript, antisense to KCNJ14 |
| AC008403.1 | 19 | novel protein |
| CYTH2 | 19 | cytohesin 2 [Source:HGNC Symbol;Acc:HGNC: |
| LMTK3 | 19 | lemur tyrosine kinase 3 [Source:HGNC Symbo |
| AC008403.3 | 19 | novel transcript, antisense to SULT2B1 |
| SULT2B1 | 19 | sulfotransferase family 2B member 1 [Source: |
| FAM83E | 19 | family with sequence similarity 83 member E |
| SPACA4 | 19 | sperm acrosome associated 4 [Source:HGNC S |
| RPL18 | 19 | ribosomal protein L18 [Source:HGNC Symbol;/ |
| AC022154.1 | 19 | novel transcript, antisense to SPHK2 |
| SPHK2 | 19 | sphingosine kinase 2 [Source:HGNC Symbol;A( |
| DBP | 19 | D-box binding PAR bZIP transcription factor [S |
| CA11 | 19 | carbonic anhydrase 11 [Source:HGNC Symbol; |
| SEC1P | 19 | secretory blood group 1, pseudogene [Source |
| NTN5 | 19 | netrin 5 [Source:HGNC Symbol;Acc:HGNC:252 |
| FUT2 | 19 | fucosyltransferase 2 [Source:HGNC Symbol;Ac |

| | | |
|---|---|---|
| MAMSTR | 19 | MEF2 activating motif and SAP domain contai |
| RASIP1 | 19 | Ras interacting protein 1 [Source:HGNC Symb |
| IZUMO1 | 19 | izumo sperm-egg fusion 1 [Source:HGNC Syml |
| FUT1 | 19 | fucosyltransferase 1 (H blood group) [Source: |
| FGF21 | 19 | fibroblast growth factor 21 [Source:HGNC Syn |
| RNU6-317P | 19 | RNA, U6 small nuclear 317, pseudogene [Sour |
| BCAT2 | 19 | branched chain amino acid transaminase 2 [So |
| AC026803.3 | 19 | novel transcript, antisense to BCAT2 |
| HSD17B14 | 19 | hydroxysteroid 17-beta dehydrogenase 14 [So |
| PLEKHA4 | 19 | pleckstrin homology domain containing A4 [So |
| PPP1R15A | 19 | protein phosphatase 1 regulatory subunit 15A |
| TULP2 | 19 | TUB like protein 2 [Source:HGNC Symbol;Acc: |
| NUCB1 | 19 | nucleobindin 1 [Source:HGNC Symbol;Acc:HG |
| NUCB1-AS1 | 19 | NUCB1 antisense RNA 1 [Source:HGNC Symbo |
| AC026803.1 | 19 | novel transcript, overlapping NUCB1 |
| DHDH | 19 | dihydrodiol dehydrogenase [Source:HGNC Syr |
| BAX | 19 | BCL2 associated X, apoptosis regulator [Sourc |
| AC026803.2 | 19 | novel transcript |
| FTL | 19 | ferritin light chain [Source:HGNC Symbol;Acc: |
| GYS1 | 19 | glycogen synthase 1 [Source:HGNC Symbol;Ac |
| RUVBL2 | 19 | RuvB like AAA ATPase 2 [Source:HGNC Symbo |
| MIR6798 | 19 | microRNA 6798 [Source:HGNC Symbol;Acc:HG |
| LHB | 19 | luteinizing hormone beta polypeptide [Source |
| AC008687.4 | 19 | novel protein |
| CGB3 | 19 | chorionic gonadotropin subunit beta 3 [Sourc |
| AC008687.1 | 19 | novel protein |
| NTF6A | 19 | neurotrophin 6 alpha (pseudogene) [Source:H |
| NTF6G | 19 | neurotrophin 6 gamma (pseudogene) [Source |
| CGB2 | 19 | chorionic gonadotropin subunit beta 2 [Sourc |
| CGB1 | 19 | chorionic gonadotropin subunit beta 1 [Sourc |
| NTF6B | 19 | neurotrophin 6 beta (pseudogene) [Source:HG |
| AC008687.5 | 19 | neurotrophin 4 (NTF4) pseudogene |
| CGB5 | 19 | chorionic gonadotropin subunit beta 5 [Sourc |
| CGB8 | 19 | chorionic gonadotropin subunit beta 8 [Sourc |
| AC008687.6 | 19 | neurotrophin 4 (NTF4) pseudogene |
| CGB7 | 19 | chorionic gonadotropin subunit beta 7 [Sourc |
| AC008687.8 | 19 | novel transcript |
| AC008687.7 | 19 | novel pseudogene |
| AC008687.2 | 19 | novel transcript, antisense to NTF4 |
| NTF4 | 19 | neurotrophin 4 [Source:HGNC Symbol;Acc:HG |
| AC008687.3 | 19 | novel transcript |
| KCNA7 | 19 | potassium voltage-gated channel subfamily A |
| RN7SL708P | 19 | RNA, 7SL, cytoplasmic 708, pseudogene [Sour |
| SNRNP70 | 19 | small nuclear ribonucleoprotein U1 subunit 70 |
| LIN7B | 19 | lin-7 homolog B, crumbs cell polarity complex |
| C19orf73 | 19 | chromosome 19 open reading frame 73 [Sour |
| PPFIA3 | 19 | PTPRF interacting protein alpha 3 [Source:HG |
| HRC | 19 | histidine rich calcium binding protein [Source: |
| TRPM4 | 19 | transient receptor potential cation channel su |
| SLC6A21P | 19 | solute carrier family 6 member 21, pseudoger |

| | | |
|---|---|---|
| SUMO1P4 | 19 | SUMO1 pseudogene 4 [Source:HGNC Symbol; |
| SLC6A16 | 19 | solute carrier family 6 member 16 [Source:HG |
| MIR4324 | 19 | microRNA 4324 [Source:HGNC Symbol;Acc:HG |
| AC011450.1 | 19 | novel transcript |
| CD37 | 19 | CD37 molecule [Source:HGNC Symbol;Acc:HG |
| TEAD2 | 19 | TEA domain transcription factor 2 [Source:HG |
| AC010524.1 | 19 | novel transcript, antisense to TEAD2 |
| DKKL1 | 19 | dickkopf like acrosomal protein 1 [Source:HGI |
| AC010643.1 | 19 | novel transcript, antisense to DKKL1 |
| CCDC155 | 19 | coiled-coil domain containing 155 [Source:HG |
| PTH2 | 19 | parathyroid hormone 2 [Source:HGNC Symbo |
| GFY | 19 | golgi associated olfactory signaling regulator [ |
| SLC17A7 | 19 | solute carrier family 17 member 7 [Source:HG |
| PIH1D1 | 19 | PIH1 domain containing 1 [Source:HGNC Syml |
| ALDH16A1 | 19 | aldehyde dehydrogenase 16 family member A |
| AC010619.1 | 19 | novel protein |
| FLT3LG | 19 | fms related tyrosine kinase 3 ligand [Source:H |
| AC010619.2 | 19 | novel transcript |
| RPL13A | 19 | ribosomal protein L13a [Source:HGNC Symbol |
| SNORD32A | 19 | small nucleolar RNA, C/D box 32A [Source:HG |
| SNORD33 | 19 | small nucleolar RNA, C/D box 33 [Source:HGN |
| SNORD34 | 19 | small nucleolar RNA, C/D box 34 [Source:HGN |
| SNORD35A | 19 | small nucleolar RNA, C/D box 35A [Source:HG |
| RPS11 | 19 | ribosomal protein S11 [Source:HGNC Symbol;. |
| SNORD35B | 19 | small nucleolar RNA, C/D box 35B [Source:HG |
| MIR150 | 19 | microRNA 150 [Source:HGNC Symbol;Acc:HGI |
| COX6CP7 | 19 | cytochrome c oxidase subunit 6C pseudogene |
| FCGRT | 19 | Fc fragment of IgG receptor and transporter [! |
| RCN3 | 19 | reticulocalbin 3 [Source:HGNC Symbol;Acc:HG |
| NOSIP | 19 | nitric oxide synthase interacting protein [Sour |
| Y_RNA | 19 | Y RNA [Source:RFAM;Acc:RF00019] |
| PRRG2 | 19 | proline rich and Gla domain 2 [Source:HGNC S |
| PRR12 | 19 | proline rich 12 [Source:HGNC Symbol;Acc:HGI |
| AC011495.2 | 19 | novel transcript, antisense to PRR12 |
| RRAS | 19 | RAS related [Source:HGNC Symbol;Acc:HGNC: |
| SCAF1 | 19 | SR-related CTD associated factor 1 [Source:HG |
| IRF3 | 19 | interferon regulatory factor 3 [Source:HGNC S |
| BCL2L12 | 19 | BCL2 like 12 [Source:HGNC Symbol;Acc:HGNC |
| PRMT1 | 19 | protein arginine methyltransferase 1 [Source: |
| MIR5088 | 19 | microRNA 5088 [Source:HGNC Symbol;Acc:HG |
| ADM5 | 19 | adrenomedullin 5 (putative) [Source:HGNC Sy |
| AC011495.3 | 19 | novel transcript, antisense to C19orf76 |
| CPT1C | 19 | carnitine palmitoyltransferase 1C [Source:HGI |
| AC011495.1 | 19 | ribosomal protein S9 (RPS9) pseudogene |
| TSKS | 19 | testis specific serine kinase substrate [Source: |
| RNU6-841P | 19 | RNA, U6 small nuclear 841, pseudogene [Sour |
| AP2A1 | 19 | adaptor related protein complex 2 subunit alp |
| MIR6799 | 19 | microRNA 6799 [Source:HGNC Symbol;Acc:HG |
| FUZ | 19 | fuzzy planar cell polarity protein [Source:HGN |
| AC006942.1 | 19 | novel transcript, antisense to FUZ |

| | | |
|---|---|---|
| MED25 | 19 | mediator complex subunit 25 [Source:HGNC S |
| MIR6800 | 19 | microRNA 6800 [Source:HGNC Symbol;Acc:HG |
| PTOV1-AS1 | 19 | PTOV1 antisense RNA 1 [Source:HGNC Symbo |
| PTOV1 | 19 | PTOV1 extended AT-hook containing adaptor |
| AC018766.1 | 19 | novel transcript, antisense to PTOV1 |
| MIR4749 | 19 | microRNA 4749 [Source:HGNC Symbol;Acc:HG |
| PTOV1-AS2 | 19 | PTOV1 antisense RNA 2 [Source:HGNC Symbo |
| PNKP | 19 | polynucleotide kinase 3'-phosphatase [Source |
| AKT1S1 | 19 | AKT1 substrate 1 [Source:HGNC Symbol;Acc:H |
| TBC1D17 | 19 | TBC1 domain family member 17 [Source:HGN |
| MIR4750 | 19 | microRNA 4750 [Source:HGNC Symbol;Acc:HG |
| IL4I1 | 19 | interleukin 4 induced 1 [Source:HGNC Symbol |
| NUP62 | 19 | nucleoporin 62 [Source:HGNC Symbol;Acc:HG |
| AC011452.1 | 19 | novel protein |
| ATF5 | 19 | activating transcription factor 5 [Source:HGNC |
| MIR4751 | 19 | microRNA 4751 [Source:HGNC Symbol;Acc:HG |
| SIGLEC11 | 19 | sialic acid binding Ig like lectin 11 [Source:HGN |
| U3 | 19 | Small nucleolar RNA U3 [Source:RFAM;Acc:RF |
| SIGLEC16 | 19 | sialic acid binding Ig like lectin 16 (gene/pseuc |
| VRK3 | 19 | VRK serine/threonine kinase 3 [Source:HGNC |
| Metazoa_SRP | 19 | Metazoan signal recognition particle RNA [Sou |
| ZNF473 | 19 | zinc finger protein 473 [Source:HGNC Symbol; |
| AC010624.3 | 19 | novel transcript, antisense to ZNF473 |
| AC010624.1 | 19 | novel transcript, sense overlapping ZNF473 |
| AC010624.4 | 19 | novel zinc finger protein pseudogene |
| AC010624.2 | 19 | novel transcript, similar to SYT3 |
| AC010624.5 | 19 | novel transcript |
| IZUMO2 | 19 | IZUMO family member 2 [Source:HGNC Symb |
| MYH14 | 19 | myosin heavy chain 14 [Source:HGNC Symbol; |
| Metazoa_SRP | 19 | Metazoan signal recognition particle RNA [Sou |
| AC008655.1 | 19 | novel transcript, antisense to MYH14 |
| KCNC3 | 19 | potassium voltage-gated channel subfamily C |
| NR1H2 | 19 | nuclear receptor subfamily 1 group H membe |
| NAPSB | 19 | napsin B aspartic peptidase, pseudogene [Sou |
| NAPSA | 19 | napsin A aspartic peptidase [Source:HGNC Syr |
| AC008655.2 | 19 | novel transcript, antisense to NR1H2 |
| POLD1 | 19 | DNA polymerase delta 1, catalytic subunit [So |
| AC020909.1 | 19 | novel protein |
| SPIB | 19 | Spi-B transcription factor [Source:HGNC Symb |
| MYBPC2 | 19 | myosin binding protein C, fast type [Source:H( |
| FAM71E1 | 19 | family with sequence similarity 71 member E1 |
| EMC10 | 19 | ER membrane protein complex subunit 10 [So |
| AC020909.3 | 19 | novel transcript, antisense to EMC10 |
| AC020909.2 | 19 | novel transcript |
| JOSD2 | 19 | Josephin domain containing 2 [Source:HGNC S |
| ASPDH | 19 | aspartate dehydrogenase domain containing [ |
| LRRC4B | 19 | leucine rich repeat containing 4B [Source:HGN |
| AC008743.1 | 19 | novel transcript, sense intronic to LRRC4B |
| SYT3 | 19 | synaptotagmin 3 [Source:HGNC Symbol;Acc:H |
| C19orf81 | 19 | chromosome 19 open reading frame 81 [Sour |

| | | |
|---|---|---|
| SHANK1 | 19 | SH3 and multiple ankyrin repeat domains 1 [S |
| CLEC11A | 19 | C-type lectin domain containing 11A [Source:I |
| GPR32P1 | 19 | GPR32 pseudogene 1 [Source:HGNC Symbol;A |
| GPR32 | 19 | G protein-coupled receptor 32 [Source:HGNC |
| AC010325.1 | 19 | novel protein |
| ACP4 | 19 | acid phosphatase 4 [Source:HGNC Symbol;Acc |
| AC010325.2 | 19 | novel transcript, antisense to ACPT |
| C19orf48 | 19 | chromosome 19 open reading frame 48 [Sour |
| SNORD88B | 19 | small nucleolar RNA, C/D box 88B [Source:HG |
| SNORD88A | 19 | small nucleolar RNA, C/D box 88A [Source:HG |
| SNORD88C | 19 | small nucleolar RNA, C/D box 88C [Source:HG |
| AC010325.3 | 19 | novel pseudogene |
| LINC01869 | 19 | long intergenic non-protein coding RNA 1869 |
| KLK1 | 19 | kallikrein 1 [Source:HGNC Symbol;Acc:HGNC:6 |
| KLK15 | 19 | kallikrein related peptidase 15 [Source:HGNC |
| AC011523.1 | 19 | novel transcript, antisense to KLK15 |
| KLK3 | 19 | kallikrein related peptidase 3 [Source:HGNC Sy |
| KLK2 | 19 | kallikrein related peptidase 2 [Source:HGNC Sy |
| KLKP1 | 19 | kallikrein pseudogene 1 [Source:HGNC Symbo |
| KLK4 | 19 | kallikrein related peptidase 4 [Source:HGNC Sy |
| PPIAP59 | 19 | peptidylprolyl isomerase A pseudogene 59 [Sc |
| KLK5 | 19 | kallikrein related peptidase 5 [Source:HGNC Sy |
| AC011483.2 | 19 | novel transcript, antisense to KLK5 and KLK6 |
| KLK6 | 19 | kallikrein related peptidase 6 [Source:HGNC Sy |
| AC011483.3 | 19 | TEC |
| AC011483.1 | 19 | novel transcript, antisense to KLK6, KLK7, KLK |
| KLK7 | 19 | kallikrein related peptidase 7 [Source:HGNC Sy |
| KLK8 | 19 | kallikrein related peptidase 8 [Source:HGNC Sy |
| AC011473.4 | 19 | novel transcript |
| KLK9 | 19 | kallikrein related peptidase 9 [Source:HGNC Sy |
| KLK10 | 19 | kallikrein related peptidase 10 [Source:HGNC |
| AC011473.2 | 19 | novel transcript, antisense to KLK10 |
| KLK11 | 19 | kallikrein related peptidase 11 [Source:HGNC |
| KLK12 | 19 | kallikrein related peptidase 12 [Source:HGNC |
| AC011473.3 | 19 | novel transcript, antisense to KLK12 |
| AC011473.1 | 19 | novel transcript |
| KLK13 | 19 | kallikrein related peptidase 13 [Source:HGNC |
| KLK14 | 19 | kallikrein related peptidase 14 [Source:HGNC |
| CTU1 | 19 | cytosolic thiouridylase subunit 1 [Source:HGN |
| SIGLEC18P | 19 | sialic acid binding Ig like lectin 18, pseudogene |
| SIGLEC9 | 19 | sialic acid binding Ig like lectin 9 [Source:HGN( |
| AC063977.2 | 19 | novel sialic acid binding Ig-like lectin (SIGLEC) |
| SIGLEC7 | 19 | sialic acid binding Ig like lectin 7 [Source:HGN( |
| AC063977.1 | 19 | sialic acid binding Ig-like lectin 19, pseudogen |
| AC063977.6 | 19 | novel transcript, antisense to SIGLEC7 and SIG |
| AC063977.7 | 19 | carcinoembryonic antigen-related cell adhesic |
| SIGLEC17P | 19 | sialic acid binding Ig like lectin 17, pseudogene |
| SIGLEC20P | 19 | sialic acid binding Ig like lectin 20, pseudogene |
| AC063977.4 | 19 | zinc finger, DHHC-type containing 7 (ZDHHC7) |
| SIGLEC21P | 19 | sialic acid binding Ig like lectin 21, pseudogene |

| | | |
|---|---|---|
| MIR8074 | 19 | microRNA 8074 [Source:HGNC Symbol;Acc:HG |
| SIGLEC22P | 19 | sialic acid binding Ig like lectin 22, pseudogene |
| CD33 | 19 | CD33 molecule [Source:HGNC Symbol;Acc:HG |
| SIGLECL1 | 19 | SIGLEC family like 1 [Source:HGNC Symbol;Acc |
| AC063977.3 | 19 | novel transcript, antisense to SIGLECL1 |
| LINC01872 | 19 | long intergenic non-protein coding RNA 1872 |
| AC063977.5 | 19 | SIGLEC family like 1 (SIGLECL1) pseudogene |
| IGLON5 | 19 | IgLON family member 5 [Source:HGNC Symbo |
| VSIG10L | 19 | V-set and immunoglobulin domain containing |
| AC008750.3 | 19 | novel transcript, antisense to VSIG10L |
| AC008750.4 | 19 | novel transcript, antisense to ETFB |
| ETFB | 19 | electron transfer flavoprotein subunit beta [So |
| AC008750.8 | 19 | novel protein |
| AC008750.5 | 19 | novel transcript, sense intronic to ETFB |
| CLDND2 | 19 | claudin domain containing 2 [Source:HGNC Sy |
| NKG7 | 19 | natural killer cell granule protein 7 [Source:HG |
| LIM2 | 19 | lens intrinsic membrane protein 2 [Source:HG |
| C19orf84 | 19 | chromosome 19 open reading frame 84 [Sour |
| AC008750.7 | 19 | novel transcript |
| NIFKP6 | 19 | nucleolar protein interacting with the FHA dor |
| SIGLEC10 | 19 | sialic acid binding Ig like lectin 10 [Source:HGN |
| AC008750.1 | 19 | novel transcript, antisense to SIGLEC10 |
| AC008750.2 | 19 | novel transcript, antisense to SIGLEC10 |
| AC008750.6 | 19 | novel sialic acid binding Ig-like lectin (SIGLEC) |
| AC020914.5 | 19 | ZDHHC7 zinc finger, DHHC-type containing 7 ( |
| SIGLEC8 | 19 | sialic acid binding Ig like lectin 8 [Source:HGN( |
| AC020914.3 | 19 | sialic acid binding Ig-like lectin 26, pseudogene |
| CEACAM18 | 19 | carcinoembryonic antigen related cell adhesio |
| SIGLEC12 | 19 | sialic acid binding Ig like lectin 12 (gene/pseuc |
| AC020914.2 | 19 | novel pseudogene |
| SIGLEC27P | 19 | sialic acid binding Ig like lectin 27, pseudogene |
| SIGLEC6 | 19 | sialic acid binding Ig like lectin 6 [Source:HGN( |
| AC020914.1 | 19 | novel transcript, antisense to SIGLEC6 |
| AC020914.4 | 19 | sialic acid binding Ig-like lectin 28, pseudogene |
| SIGLEC29P | 19 | sialic acid binding Ig like lectin 29, pseudogene |
| ZNF175 | 19 | zinc finger protein 175 [Source:HGNC Symbol; |
| AC018755.3 | 19 | putative ATP-binding domain-containing prote |
| AC018755.1 | 19 | novel transcript |
| SIGLEC5 | 19 | sialic acid binding Ig like lectin 5 [Source:HGN( |
| AC018755.2 | 19 | novel transcript, sense intronic to AC018755.1 |
| RPL9P33 | 19 | ribosomal protein L9 pseudogene 33 [Source: |
| AC018755.4 | 19 | novel transcript, sense intronic to AC018755.1 |
| SIGLEC14 | 19 | sialic acid binding Ig like lectin 14 [Source:HGN |
| RPL7P51 | 19 | ribosomal protein L7 pseudogene 51 [Source: |
| SPACA6P-AS | 19 | SPACA6P antisense RNA [Source:HGNC Symbo |
| SPACA6 | 19 | sperm acrosome associated 6 [Source:HGNC S |
| MIR99B | 19 | microRNA 99b [Source:HGNC Symbol;Acc:HGN |
| MIRLET7E | 19 | microRNA let-7e [Source:HGNC Symbol;Acc:H |
| MIR125A | 19 | microRNA 125a [Source:HGNC Symbol;Acc:HG |
| HAS1 | 19 | hyaluronan synthase 1 [Source:HGNC Symbol; |

| | | |
|---|---|---|
| FPR1 | 19 | formyl peptide receptor 1 [Source:HGNC Syml |
| FPR2 | 19 | formyl peptide receptor 2 [Source:HGNC Syml |
| AC018755.5 | 19 | novel transcript, sense intronic to FPR1 |
| FPR3 | 19 | formyl peptide receptor 3 [Source:HGNC Syml |
| ZNF577 | 19 | zinc finger protein 577 [Source:HGNC Symbol; |
| AC006272.1 | 19 | novel transcript |
| AC006272.2 | 19 | novel zinc finger protein pseudogene |
| AC074141.1 | 19 | novel zinc finger protein pseudogene |
| ZNF649-AS1 | 19 | ZNF649 antisense RNA 1 [Source:HGNC Symbo |
| ZNF649 | 19 | zinc finger protein 649 [Source:HGNC Symbol; |
| AC011460.1 | 19 | novel zinc finger protein pseudogene |
| ZNF613 | 19 | zinc finger protein 613 [Source:HGNC Symbol; |
| ZNF350-AS1 | 19 | ZNF350 antisense RNA 1 [Source:HGNC Symbo |
| ZNF350 | 19 | zinc finger protein 350 [Source:HGNC Symbol; |
| ZNF615 | 19 | zinc finger protein 615 [Source:HGNC Symbol; |
| ZNF614 | 19 | zinc finger protein 614 [Source:HGNC Symbol; |
| ZNF432 | 19 | zinc finger protein 432 [Source:HGNC Symbol; |
| AC011468.5 | 19 | novel transcript, antisense to ZNF432 |
| AC011468.1 | 19 | novel transcript |
| ZNF841 | 19 | zinc finger protein 841 [Source:HGNC Symbol; |
| AC011468.4 | 19 | transmembrane protein 50B (TMEM50B) pseu |
| ZNF616 | 19 | zinc finger protein 616 [Source:HGNC Symbol; |
| AC011468.2 | 19 | novel transcript |
| AC011468.3 | 19 | novel transcript |
| RPL37P23 | 19 | ribosomal protein L37 pseudogene 23 [Source |
| ZNF836 | 19 | zinc finger protein 836 [Source:HGNC Symbol; |
| AC010320.1 | 19 | novel transcript, antisense to ZNF836 |
| PPP2R1A | 19 | protein phosphatase 2 scaffold subunit Aalpha |
| MIR6801 | 19 | microRNA 6801 [Source:HGNC Symbol;Acc:HG |
| AC010320.2 | 19 | novel transcript, antisense to PPP2R1A |
| AC010320.5 | 19 | vomeronasal 1 receptor 2 (VN1R2) pseudoger |
| Y_RNA | 19 | Y RNA [Source:RFAM;Acc:RF00019] |
| ZNF766 | 19 | zinc finger protein 766 [Source:HGNC Symbol; |
| MIR643 | 19 | microRNA 643 [Source:HGNC Symbol;Acc:HGN |
| AC010320.3 | 19 | novel transcript, antisense to ZNF766 and ZNF |
| ZNF480 | 19 | zinc finger protein 480 [Source:HGNC Symbol; |
| AC010320.4 | 19 | novel transcript, antisense to ZNF480 and ZNF |
| ZNF610 | 19 | zinc finger protein 610 [Source:HGNC Symbol; |
| ZNF880 | 19 | zinc finger protein 880 [Source:HGNC Symbol; |
| ZNF528-AS1 | 19 | ZNF528 antisense RNA 1 [Source:HGNC Symbo |
| AC010332.1 | 19 | novel zinc finger pseudogene |
| ZNF528 | 19 | zinc finger protein 528 [Source:HGNC Symbol; |
| DPPA5P1 | 19 | developmental pluripotency associated 5 pseu |
| ZNF534 | 19 | zinc finger protein 534 [Source:HGNC Symbol; |
| ZNF578 | 19 | zinc finger protein 578 [Source:HGNC Symbol; |
| AC010332.3 | 19 | novel transcript |
| AC010332.2 | 19 | novel zinc finger pseudogene |
| AC022150.3 | 19 | novel transcript, antisense to ZNF578 |
| ZNF808 | 19 | zinc finger protein 808 [Source:HGNC Symbol; |
| ZNF701 | 19 | zinc finger protein 701 [Source:HGNC Symbol; |

| | | |
|---|---|---|
| RPL39P34 | 19 | ribosomal protein L39 pseudogene 34 [Source |
| AC022150.1 | 19 | novel transcript, antisense to ZNF701 |
| ZNF137P | 19 | zinc finger protein 137, pseudogene [Source:H |
| ZNF83 | 19 | zinc finger protein 83 [Source:HGNC Symbol;A |
| AC022150.2 | 19 | novel transcript, sense intronic to ZNF83 |
| AC022150.4 | 19 | novel transcript, sense intronic to ZNF83 |
| ZNF611 | 19 | zinc finger protein 611 [Source:HGNC Symbol; |
| ZNF600 | 19 | zinc finger protein 600 [Source:HGNC Symbol; |
| ZNF28 | 19 | zinc finger protein 28 [Source:HGNC Symbol;A |
| AC008813.1 | 19 | poly(A) binding protein, nuclear 1 (PABPN1) p |
| ZNF468 | 19 | zinc finger protein 468 [Source:HGNC Symbol; |
| AC010487.1 | 19 | zinc finger protein pseudogene |
| ZNF320 | 19 | zinc finger protein 320 [Source:HGNC Symbol; |
| AC010487.3 | 19 | novel zinc finger protein pseudogene |
| ZNF888 | 19 | zinc finger protein 888 [Source:HGNC Symbol; |
| AC010487.2 | 19 | novel transcript |
| ZNF321P | 19 | zinc finger protein 321, pseudogene [Source:H |
| ZNF816-ZNF321P | 19 | ZNF816-ZNF321P readthrough [Source:HGNC |
| ZNF816 | 19 | zinc finger protein 816 [Source:HGNC Symbol; |
| AC010328.3 | 19 | novel transcript |
| ZNF702P | 19 | zinc finger protein 702, pseudogene [Source:H |
| AC010328.2 | 19 | actin related protein 2/3 complex, subunit 1A |
| AC010328.1 | 19 | novel transcript |
| ERVV-1 | 19 | endogenous retrovirus group V member 1, en |
| ERVV-2 | 19 | endogenous retrovirus group V member 2, en |
| ZNF160 | 19 | zinc finger protein 160 [Source:HGNC Symbol; |
| ZNF415 | 19 | zinc finger protein 415 [Source:HGNC Symbol; |
| ZNF347 | 19 | zinc finger protein 347 [Source:HGNC Symbol; |
| ZNF665 | 19 | zinc finger protein 665 [Source:HGNC Symbol; |
| AC092070.1 | 19 | novel transcript, antisense to ZNF665 |
| AC092070.2 | 19 | zinc finger protein 818, pseudogene |
| NDUFV2P1 | 19 | NADH:ubiquinone oxidoreductase core subun |
| ZNF677 | 19 | zinc finger protein 677 [Source:HGNC Symbol; |
| AC092070.3 | 19 | novel transcript, antisense to ZNF677 |
| VN1R2 | 19 | vomeronasal 1 receptor 2 [Source:HGNC Syml |
| VN1R4 | 19 | vomeronasal 1 receptor 4 [Source:HGNC Syml |
| AC092070.4 | 19 | novel transcript |
| FAM90A27P | 19 | family with sequence similarity 90 member A |
| BIRC8 | 19 | baculoviral IAP repeat containing 8 [Source:H |
| FAM90A28P | 19 | family with sequence similarity 90 member A |
| VN1R6P | 19 | vomeronasal 1 receptor 6 pseudogene [Sourc |
| ZNF845 | 19 | zinc finger protein 845 [Source:HGNC Symbol; |
| ZNF525 | 19 | zinc finger protein 525 [Source:HGNC Symbol; |
| AC125388.1 | 19 | ribosomal protein L39 (RPL39) pseudogene |
| ZNF765 | 19 | zinc finger protein 765 [Source:HGNC Symbol; |
| RPL39P36 | 19 | ribosomal protein L39 pseudogene 36 [Source |
| AC022137.2 | 19 | FSHD region gene 2 (FRG2) pseudogene |
| AC022137.3 | 19 | novel protein |
| TPM3P9 | 19 | tropomyosin 3 pseudogene 9 [Source:HGNC S |
| AC022137.4 | 19 | novel transcript, antisense to ZNF761and ZNF |

| | | |
|---|---|---|
| ZNF761 | 19 | zinc finger protein 761 [Source:HGNC Symbol; |
| ZNF813 | 19 | zinc finger protein 813 [Source:HGNC Symbol; |
| TPM3P6 | 19 | tropomyosin 3 pseudogene 6 [Source:HGNC S |
| AC022137.1 | 19 | ribosomal protein L39 (RPL39) pseudogene |
| AC011487.1 | 19 | vomeronasal 1 receptor (VN1R) pseudogene |
| AC011487.4 | 19 | FSHD region gene 2 family, member C (FRG2C |
| ZNF331 | 19 | zinc finger protein 331 [Source:HGNC Symbol; |
| AC011487.2 | 19 | solute carrier family 7 (cationic amino acid tra |
| AC011487.3 | 19 | oligosaccharyltransferase complex subunit (O |
| RN7SL317P | 19 | RNA, 7SL, cytoplasmic 317, pseudogene [Sour |
| DPRX | 19 | divergent-paired related homeobox [Source:H |
| RNU6-698P | 19 | RNA, U6 small nuclear 698, pseudogene [Sour |
| AC011453.2 | 19 | nuclear RNA export factor 1 (NXF1) pseudoge |
| MIR512-1 | 19 | microRNA 512-1 [Source:HGNC Symbol;Acc:H |
| MIR512-2 | 19 | microRNA 512-2 [Source:HGNC Symbol;Acc:H |
| MIR1323 | 19 | microRNA 1323 [Source:HGNC Symbol;Acc:HG |
| MIR498 | 19 | microRNA 498 [Source:HGNC Symbol;Acc:HGN |
| MIR520E | 19 | microRNA 520e [Source:HGNC Symbol;Acc:HG |
| MIR515-1 | 19 | microRNA 515-1 [Source:HGNC Symbol;Acc:H |
| MIR519E | 19 | microRNA 519e [Source:HGNC Symbol;Acc:HG |
| MIR520F | 19 | microRNA 520f [Source:HGNC Symbol;Acc:HG |
| MIR515-2 | 19 | microRNA 515-2 [Source:HGNC Symbol;Acc:H |
| MIR519C | 19 | microRNA 519c [Source:HGNC Symbol;Acc:HG |
| MIR1283-1 | 19 | microRNA 1283-1 [Source:HGNC Symbol;Acc: |
| MIR520A | 19 | microRNA 520a [Source:HGNC Symbol;Acc:HG |
| MIR526B | 19 | microRNA 526b [Source:HGNC Symbol;Acc:HG |
| MIR519B | 19 | microRNA 519b [Source:HGNC Symbol;Acc:HG |
| MIR525 | 19 | microRNA 525 [Source:HGNC Symbol;Acc:HGN |
| MIR523 | 19 | microRNA 523 [Source:HGNC Symbol;Acc:HGN |
| MIR518F | 19 | microRNA 518f [Source:HGNC Symbol;Acc:HG |
| MIR520B | 19 | microRNA 520b [Source:HGNC Symbol;Acc:HG |
| MIR518B | 19 | microRNA 518b [Source:HGNC Symbol;Acc:HG |
| MIR526A1 | 19 | microRNA 526a-1 [Source:HGNC Symbol;Acc: |
| MIR520C | 19 | microRNA 520c [Source:HGNC Symbol;Acc:HG |
| MIR518C | 19 | microRNA 518c [Source:HGNC Symbol;Acc:HG |
| MIR524 | 19 | microRNA 524 [Source:HGNC Symbol;Acc:HGN |
| MIR517A | 19 | microRNA 517a [Source:HGNC Symbol;Acc:HG |
| MIR519D | 19 | microRNA 519d [Source:HGNC Symbol;Acc:HG |
| MIR521-2 | 19 | microRNA 521-2 [Source:HGNC Symbol;Acc:H |
| RNU6-803P | 19 | RNA, U6 small nuclear 803, pseudogene [Sour |
| MIR520D | 19 | microRNA 520d [Source:HGNC Symbol;Acc:HG |
| MIR517B | 19 | microRNA 517b [Source:HGNC Symbol;Acc:HG |
| MIR520G | 19 | microRNA 520g [Source:HGNC Symbol;Acc:HG |
| MIR516B2 | 19 | microRNA 516b-2 [Source:HGNC Symbol;Acc: |
| MIR526A2 | 19 | microRNA 526a-2 [Source:HGNC Symbol;Acc: |
| MIR518E | 19 | microRNA 518e [Source:HGNC Symbol;Acc:HG |
| MIR518A1 | 19 | microRNA 518a-1 [Source:HGNC Symbol;Acc: |
| RNU6-980P | 19 | RNA, U6 small nuclear 980, pseudogene [Sour |
| MIR518D | 19 | microRNA 518d [Source:HGNC Symbol;Acc:HG |
| MIR516B1 | 19 | microRNA 516b-1 [Source:HGNC Symbol;Acc: |

| | | |
|---|---|---|
| MIR518A2 | 19 | microRNA 518a-2 [Source:HGNC Symbol;Acc:H |
| MIR517C | 19 | microRNA 517c [Source:HGNC Symbol;Acc:HG |
| MIR520H | 19 | microRNA 520h [Source:HGNC Symbol;Acc:HG |
| RNU6-982P | 19 | RNA, U6 small nuclear 982, pseudogene [Sour |
| MIR521-1 | 19 | microRNA 521-1 [Source:HGNC Symbol;Acc:H |
| RNU6-751P | 19 | RNA, U6 small nuclear 751, pseudogene [Sour |
| MIR522 | 19 | microRNA 522 [Source:HGNC Symbol;Acc:HGN |
| MIR519A1 | 19 | microRNA 519a-1 [Source:HGNC Symbol;Acc:H |
| MIR527 | 19 | microRNA 527 [Source:HGNC Symbol;Acc:HGN |
| MIR516A1 | 19 | microRNA 516a-1 [Source:HGNC Symbol;Acc:H |
| MIR1283-2 | 19 | microRNA 1283-2 [Source:HGNC Symbol;Acc:H |
| RNU6-1041P | 19 | RNA, U6 small nuclear 1041, pseudogene [Sou |
| MIR516A2 | 19 | microRNA 516a-2 [Source:HGNC Symbol;Acc:H |
| AC011453.1 | 19 | novel transcript, host to microRNA 519a-2 |
| MIR519A2 | 19 | microRNA 519a-2 [Source:HGNC Symbol;Acc:H |
| RNU6-165P | 19 | RNA, U6 small nuclear 165, pseudogene [Sour |
| HMGN1P32 | 19 | high mobility group nucleosome binding doma |
| SEPTIN7P8 | 19 | septin 7 pseudogene 8 [Source:HGNC Symbol; |
| AC008753.1 | 19 | ribosomal protein, large, P1 (RPLP1) pseudoge |
| AC008753.3 | 19 | novel transcript |
| MIR371A | 19 | microRNA 371a [Source:HGNC Symbol;Acc:HG |
| MIR371B | 19 | microRNA 371b [Source:HGNC Symbol;Acc:HG |
| MIR372 | 19 | microRNA 372 [Source:HGNC Symbol;Acc:HGN |
| MIR373 | 19 | microRNA 373 [Source:HGNC Symbol;Acc:HGN |
| AC008753.2 | 19 | novel transcript |
| NLRP12 | 19 | NLR family pyrin domain containing 12 [Sourc |
| AC008440.1 | 19 | novel transcript, antisense to MYADM |
| MYADM | 19 | myeloid associated differentiation marker [So |
| AC008440.2 | 19 | novel transcript |
| PRKCG | 19 | protein kinase C gamma [Source:HGNC Symbo |
| CACNG7 | 19 | calcium voltage-gated channel auxiliary subun |
| CACNG8 | 19 | calcium voltage-gated channel auxiliary subun |
| MIR935 | 19 | microRNA 935 [Source:HGNC Symbol;Acc:HGN |
| CACNG6 | 19 | calcium voltage-gated channel auxiliary subun |
| AC012314.13 | 19 | heterogeneous nuclear ribonucleoprotein A3 |
| VSTM1 | 19 | V-set and transmembrane domain containing |
| TARM1 | 19 | T cell-interacting, activating receptor on myel |
| OSCAR | 19 | osteoclast associated Ig-like receptor [Source: |
| NDUFA3 | 19 | NADH:ubiquinone oxidoreductase subunit A3 |
| TFPT | 19 | TCF3 fusion partner [Source:HGNC Symbol;Ac |
| PRPF31 | 19 | pre-mRNA processing factor 31 [Source:HGNC |
| AC245052.4 | 19 | novel transcript |
| CNOT3 | 19 | CCR4-NOT transcription complex subunit 3 [So |
| LENG1 | 19 | leukocyte receptor cluster member 1 [Source: |
| TMC4 | 19 | transmembrane channel like 4 [Source:HGNC |
| MBOAT7 | 19 | membrane bound O-acyltransferase domain c |
| TSEN34 | 19 | tRNA splicing endonuclease subunit 34 [Sourc |
| AC245052.1 | 19 | novel transcript |
| RPS9 | 19 | ribosomal protein S9 [Source:HGNC Symbol;A |
| AC245052.2 | 19 | leukocyte immunoglobulin-like receptor, subf |

| | | |
|---|---|---|
| LILRB3 | 19 | leukocyte immunoglobulin like receptor B3 [S |
| AC245052.7 | 19 | novel transcript |
| AC245052.3 | 19 | novel pseudogene |
| LILRA6 | 19 | leukocyte immunoglobulin like receptor A6 [S |
| AC245052.6 | 19 | pseudogene similar to part of leukocyte immu |
| LILRB5 | 19 | leukocyte immunoglobulin like receptor B5 [S |
| RNU6-1307P | 19 | RNA, U6 small nuclear 1307, pseudogene [Sou |
| AC245052.5 | 19 | YME1-like 1 (S. cerevisiae) (YME1L1) pseudog |
| LILRB2 | 19 | leukocyte immunoglobulin like receptor B2 [S |
| MIR4752 | 19 | microRNA 4752 [Source:HGNC Symbol;Acc:H |
| AC245884.5 | 19 | leukocyte immunoglobulin-like receptor, subf |
| AC245884.12 | 19 | leukocyte immunoglobulin-like receptor, subf |
| AC245884.7 | 19 | pseudogene similar to part of leukocyte immu |
| AC245884.2 | 19 | leukocyte immunoglobulin-like receptor, subf |
| LILRA5 | 19 | leukocyte immunoglobulin like receptor A5 [S |
| AC245884.11 | 19 | novel transcript |
| VN1R104P | 19 | vomeronasal 1 receptor 104 pseudogene [Sou |
| AC245884.6 | 19 | pseudogene similar to part of leukocyte immu |
| LILRA4 | 19 | leukocyte immunoglobulin like receptor A4 [S |
| LAIR1 | 19 | leukocyte associated immunoglobulin like rec |
| AC245884.4 | 19 | mitochondrial ribosomal protein L40 (MRPL40 |
| TTYH1 | 19 | tweety family member 1 [Source:HGNC Symb |
| AC245884.1 | 19 | novel transcript |
| AC245884.9 | 19 | novel transcript, sense intronic to TTYH1 |
| AC245884.10 | 19 | novel transcript |
| AC245884.8 | 19 | novel transcript |
| LENG8-AS1 | 19 | LENG8 antisense RNA 1 [Source:HGNC Symbo |
| AC245884.3 | 19 | LSM6 homolog, U6 small nuclear RNA associa |
| LENG8 | 19 | leukocyte receptor cluster member 8 [Source: |
| LENG9 | 19 | leukocyte receptor cluster member 9 [Source: |
| CDC42EP5 | 19 | CDC42 effector protein 5 [Source:HGNC Symb |
| LAIR2 | 19 | leukocyte associated immunoglobulin like rec |
| AC245036.1 | 19 | pseudogene similar to part of killer cell immu |
| KIR3DX1 | 19 | killer cell immunoglobulin like receptor, three |
| AC245036.2 | 19 | pseudogene similar to part of leukocyte immu |
| LILRA2 | 19 | leukocyte immunoglobulin like receptor A2 [S |
| AC245036.5 | 19 | novel transcript |
| LILRA1 | 19 | leukocyte immunoglobulin like receptor A1 [S |
| AC245036.4 | 19 | pseudogene similar to part of leukocyte immu |
| LILRB1 | 19 | leukocyte immunoglobulin like receptor B1 [S |
| LILRB1-AS1 | 19 | LILRB1 antisense RNA 1 [Source:HGNC Symbo |
| LILRB4 | 19 | leukocyte immunoglobulin like receptor B4 [S |
| MIR8061 | 19 | microRNA 8061 [Source:HGNC Symbol;Acc:H |
| VN1R105P | 19 | vomeronasal 1 receptor 105 pseudogene [Sou |
| AC245036.3 | 19 | pseudogene similar to part of leukocyte-assoc |
| AC243962.1 | 19 | pseudogene similar to leukocyte immunoglob |
| LILRP1 | 19 | leukocyte immunoglobulin-like receptor pseu |
| AC245128.2 | 19 | killer cell immunoglobulin-like receptor family |
| LILRP2 | 19 | leukocyte immunoglobulin-like receptor pseu |
| KIR3DL3 | 19 | killer cell immunoglobulin like receptor, three |

| | | |
|---|---|---|
| AC245128.1 | 19 | novel transcript |
| KIR2DL3 | 19 | killer cell immunoglobulin like receptor, two Ig |
| KIR2DP1 | 19 | killer cell immunoglobulin like receptor, two Ig |
| KIR2DL1 | 19 | killer cell immunoglobulin like receptor, two Ig |
| KIR3DP1 | 19 | killer cell immunoglobulin like receptor, three |
| KIR2DL4 | 19 | killer cell immunoglobulin like receptor, two Ig |
| KIR3DL1 | 19 | killer cell immunoglobulin like receptor, three |
| KIR2DS4 | 19 | killer cell immunoglobulin like receptor, two Ig |
| KIR3DL2 | 19 | killer cell immunoglobulin like receptor, three |
| RNU6-222P | 19 | RNA, U6 small nuclear 222, pseudogene [Sour |
| FCAR | 19 | Fc fragment of IgA receptor [Source:HGNC Syr |
| AC245128.3 | 19 | novel transcript |
| NCR1 | 19 | natural cytotoxicity triggering receptor 1 [Sou |
| NLRP7 | 19 | NLR family pyrin domain containing 7 [Source |
| AC011476.8 | 19 | 3-hydroxy-3-methylglutaryl-CoA reductase (H |
| NLRP2 | 19 | NLR family pyrin domain containing 2 [Source |
| AC011476.1 | 19 | ribosomal protein L36a (RPL36A) pseudogene |
| AC011476.3 | 19 | novel transcript, antisense to RDH13 and GP6 |
| GP6 | 19 | glycoprotein VI platelet [Source:HGNC Symbo |
| AC011476.2 | 19 | novel transcript |
| RDH13 | 19 | retinol dehydrogenase 13 [Source:HGNC Symb |
| EPS8L1 | 19 | EPS8 like 1 [Source:HGNC Symbol;Acc:HGNC:2 |
| PPP1R12C | 19 | protein phosphatase 1 regulatory subunit 12C |
| AC010327.8 | 19 | novel transcript, antisense to PPP1R12C |
| AC010327.3 | 19 | nuclear transport factor 2, pseudogene |
| MIR7975 | 19 | microRNA 7975 [Source:HGNC Symbol;Acc:HG |
| TNNT1 | 19 | troponin T1, slow skeletal type [Source:HGNC |
| TNNI3 | 19 | troponin I3, cardiac type [Source:HGNC Symbo |
| AC010327.2 | 19 | novel transcript |
| DNAAF3 | 19 | dynein axonemal assembly factor 3 [Source:H |
| AC010327.4 | 19 | novel transcript, antisense to C19orf51 and SY |
| AC010327.1 | 19 | NULL |
| SYT5 | 19 | synaptotagmin 5 [Source:HGNC Symbol;Acc:H |
| PTPRH | 19 | protein tyrosine phosphatase receptor type H |
| AC010327.7 | 19 | novel transcript |
| AC010327.5 | 19 | novel transcript |
| TMEM86B | 19 | transmembrane protein 86B [Source:HGNC Sy |
| AC010327.6 | 19 | novel transcript |
| PPP6R1 | 19 | protein phosphatase 6 regulatory subunit 1 [S |
| MIR6804 | 19 | microRNA 6804 [Source:HGNC Symbol;Acc:HG |
| MIR6802 | 19 | microRNA 6802 [Source:HGNC Symbol;Acc:HG |
| MIR6803 | 19 | microRNA 6803 [Source:HGNC Symbol;Acc:HG |
| HSPBP1 | 19 | HSPA (Hsp70) binding protein 1 [Source:HGNC |
| BRSK1 | 19 | BR serine/threonine kinase 1 [Source:HGNC Sy |
| AC020922.2 | 19 | novel transcript, antisense to BRSK1 |
| TMEM150B | 19 | transmembrane protein 150B [Source:HGNC S |
| AC020922.4 | 19 | novel transcript |
| KMT5C | 19 | lysine methyltransferase 5C [Source:HGNC Syr |
| COX6B2 | 19 | cytochrome c oxidase subunit 6B2 [Source:HG |
| AC020922.1 | 19 | novel transcript |

| | | |
|---|---|---|
| FAM71E2 | 19 | family with sequence similarity 71 member E2 |
| IL11 | 19 | interleukin 11 [Source:HGNC Symbol;Acc:HGN |
| TMEM190 | 19 | transmembrane protein 190 [Source:HGNC Sy |
| AC020922.3 | 19 | novel transcript, antisense to TMEM190 |
| TMEM238 | 19 | transmembrane protein 238 [Source:HGNC Sy |
| RPL28 | 19 | ribosomal protein L28 [Source:HGNC Symbol;A |
| MIR6805 | 19 | microRNA 6805 [Source:HGNC Symbol;Acc:HG |
| UBE2S | 19 | ubiquitin conjugating enzyme E2 S [Source:HG |
| SHISA7 | 19 | shisa family member 7 [Source:HGNC Symbol; |
| ISOC2 | 19 | isochorismatase domain containing 2 [Source: |
| C19orf85 | 19 | chromosome 19 open reading frame 85 [Sour |
| AC008735.5 | 19 | TEC |
| AC008735.3 | 19 | novel transcript, antisense to ZNF628 |
| ZNF628 | 19 | zinc finger protein 628 [Source:HGNC Symbol; |
| NAT14 | 19 | N-acetyltransferase 14 (putative) [Source:HGN |
| SSC5D | 19 | scavenger receptor cysteine rich family memb |
| SBK2 | 19 | SH3 domain binding kinase family member 2 [ |
| SBK3 | 19 | SH3 domain binding kinase family member 3 [ |
| ZNF579 | 19 | zinc finger protein 579 [Source:HGNC Symbol; |
| FIZ1 | 19 | FLT3 interacting zinc finger 1 [Source:HGNC Sy |
| ZNF524 | 19 | zinc finger protein 524 [Source:HGNC Symbol; |
| ZNF865 | 19 | zinc finger protein 865 [Source:HGNC Symbol; |
| AC008735.4 | 19 | novel transcript, antisense to ZNF865 |
| ZNF784 | 19 | zinc finger protein 784 [Source:HGNC Symbol; |
| ZNF580 | 19 | zinc finger protein 580 [Source:HGNC Symbol; |
| ZNF581 | 19 | zinc finger protein 581 [Source:HGNC Symbol; |
| CCDC106 | 19 | coiled-coil domain containing 106 [Source:HG |
| U2AF2 | 19 | U2 small nuclear RNA auxiliary factor 2 [Sourc |
| AC008735.2 | 19 | novel transcript, antisense to U2AF2A |
| AC008735.1 | 19 | novel transcript |
| EPN1 | 19 | epsin 1 [Source:HGNC Symbol;Acc:HGNC:2160 |
| AC010525.1 | 19 | novel transcript, antisense to EPN1 |
| AC010525.2 | 19 | Diphosphoinositol polyphosphate phosphohyd |
| AC008749.1 | 19 | Vomeronasal type-1 receptor 1, pseudogene |
| NLRP9 | 19 | NLR family pyrin domain containing 9 [Source |
| RN7SKP109 | 19 | RN7SK pseudogene 109 [Source:HGNC Symbo |
| RFPL4A | 19 | ret finger protein like 4A [Source:HGNC Symb |
| RFPL4AL1 | 19 | ret finger protein like 4A like 1 [Source:HGNC |
| RFPL4AP1 | 19 | ret finger protein like 4A pseudogene 1 [Sourc |
| NLRP11 | 19 | NLR family pyrin domain containing 11 [Sourc |
| NLRP4 | 19 | NLR family pyrin domain containing 4 [Source |
| NLRP13 | 19 | NLR family pyrin domain containing 13 [Sourc |
| NLRP8 | 19 | NLR family pyrin domain containing 8 [Source |
| NLRP5 | 19 | NLR family pyrin domain containing 5 [Source |
| LINC01864 | 19 | long intergenic non-protein coding RNA 1864 |
| ZNF787 | 19 | zinc finger protein 787 [Source:HGNC Symbol; |
| Y_RNA | 19 | Y RNA [Source:RFAM;Acc:RF00019] |
| AC024580.2 | 19 | TEC |
| ZNF444 | 19 | zinc finger protein 444 [Source:HGNC Symbol; |
| AC024580.1 | 19 | TEC |

| | | |
|---|---|---|
| GALP | 19 | galanin like peptide [Source:HGNC Symbol;Acc |
| ZSCAN5B | 19 | zinc finger and SCAN domain containing 5B [So |
| ZSCAN5C | 19 | zinc finger and SCAN domain containing 5C [So |
| AC011506.1 | 19 | ribosomal protein L13 (RPL13) pseudogene |
| ZSCAN5A | 19 | zinc finger and SCAN domain containing 5A [So |
| ZSCAN5DP | 19 | zinc finger and SCAN domain containing 5D ps |
| EDDM13 | 19 | epididymal protein 13 [Source:HGNC Symbol;A |
| AC006116.5 | 19 | novel transcript, sense intronic to ZSCAN5A |
| AC006116.8 | 19 | novel transcript, antisense to ZSCAN5A |
| Y_RNA | 19 | Y RNA [Source:RFAM;Acc:RF00019] |
| VN2R17P | 19 | vomeronasal 2 receptor 17, pseudogene [Sou |
| AC006116.1 | 19 | glycerol-3-phosphate dehydrogenase 2 (mitoc |
| AC006116.3 | 19 | Vomeronasal type-2 receptor, pseudogene |
| AC006116.2 | 19 | NADH dehydrogenase (ubiquinone) complex I |
| AC006116.6 | 19 | novel transcript, sense intronic to ZSCAN5A |
| AC006116.9 | 19 | novel transcript |
| ZNF542P | 19 | zinc finger protein 542, pseudogene [Source:H |
| ZNF582 | 19 | zinc finger protein 582 [Source:HGNC Symbol; |
| AC006116.10 | 19 | novel transcript |
| AC006116.7 | 19 | unamed protein product, pseudogene |
| AC006116.11 | 19 | novel transcript, antisense to ZNF582 |
| SLC25A36P1 | 19 | SLC25A36 pseudogene 1 [Source:HGNC Symbo |
| AC006116.4 | 19 | novel transcript, antisense to ZNF582 |
| ZNF582-AS1 | 19 | ZNF582 antisense RNA 1 (head to head) [Sour |
| ZNF583 | 19 | zinc finger protein 583 [Source:HGNC Symbol; |
| ZNF667 | 19 | zinc finger protein 667 [Source:HGNC Symbol; |
| ZNF667-AS1 | 19 | ZNF667 antisense RNA 1 (head to head) [Sour |
| AC004696.2 | 19 | novel transcript |
| AC004696.1 | 19 | sulfotransferase family, cytosolic, 2A, dehydro |
| ZNF471 | 19 | zinc finger protein 471 [Source:HGNC Symbol; |
| AC005498.1 | 19 | novel transcript |
| AC005498.2 | 19 | novel transcript |
| ZFP28 | 19 | ZFP28 zinc finger protein [Source:HGNC Symb |
| AC005498.3 | 19 | novel transcript, antisense to ZFP28 |
| ZNF470 | 19 | zinc finger protein 470 [Source:HGNC Symbol; |
| SIGLEC31P | 19 | sialic acid binding Ig like lectin 31, pseudogene |
| ZNF71 | 19 | zinc finger protein 71 [Source:HGNC Symbol;A |
| ZIM2-AS1 | 19 | ZIM2 antisense RNA 1 [Source:HGNC Symbol;A |
| SMIM17 | 19 | small integral membrane protein 17 [Source:H |
| ZNF835 | 19 | zinc finger protein 835 [Source:HGNC Symbol; |
| AC006115.2 | 19 | novel transcript (contains ZIM2-AS1 and PEG3 |
| OR5AH1P | 19 | olfactory receptor family 5 subfamily AH mem |
| ZIM2 | 19 | zinc finger imprinted 2 [Source:HGNC Symbol; |
| AC006115.1 | 19 | zinc finger protein pseudogene |
| PEG3 | 19 | paternally expressed 3 [Source:HGNC Symbol; |
| MIMT1 | 19 | MER1 repeat containing imprinted transcript 1 |
| AC004792.1 | 19 | NULL |
| AC044792.1 | 19 | NADH dehydrogenase pseudogene |
| RPL7AP69 | 19 | ribosomal protein L7a pseudogene 69 [Source |
| AC025588.4 | 19 | novel zinc finger protein pseudogene |

| | | |
|---|---|---|
| USP29 | 19 | ubiquitin specific peptidase 29 [Source:HGNC |
| ZIM3 | 19 | zinc finger imprinted 3 [Source:HGNC Symbol; |
| AC025588.2 | 19 | developmental pluripotency associated 3 (DPI |
| AC025588.3 | 19 | developmental pluripotency associated 3 (DPI |
| DUXA | 19 | double homeobox A [Source:HGNC Symbol;Ac |
| AC025588.1 | 19 | pentatricopeptide repeat domain 3 pseudoge |
| ZNF264 | 19 | zinc finger protein 264 [Source:HGNC Symbol; |
| AURKC | 19 | aurora kinase C [Source:HGNC Symbol;Acc:HG |
| ZNF805 | 19 | zinc finger protein 805 [Source:HGNC Symbol; |
| AC005261.3 | 19 | novel transcript |
| ZNF460-AS1 | 19 | ZNF460 antisense RNA 1 [Source:HGNC Symb |
| ZNF460 | 19 | zinc finger protein 460 [Source:HGNC Symbol; |
| AC005261.5 | 19 | TEC |
| AC005261.1 | 19 | novel transcript |
| AC005261.4 | 19 | PPPDE peptidase domain containing 1 pseudo |
| ZNF543 | 19 | zinc finger protein 543 [Source:HGNC Symbol; |
| AC005261.2 | 19 | novel transcript, antisense to ZNF304 |
| ZNF304 | 19 | zinc finger protein 304 [Source:HGNC Symbol; |
| ZNF547 | 19 | zinc finger protein 547 [Source:HGNC Symbol; |
| TRAPPC2B | 19 | trafficking protein particle complex 2B [Sourc |
| AC003002.1 | 19 | novel protein |
| ZNF548 | 19 | zinc finger protein 548 [Source:HGNC Symbol; |
| AC003002.3 | 19 | novel transcript |
| AC003002.2 | 19 | novel protein |
| ZNF17 | 19 | zinc finger protein 17 [Source:HGNC Symbol;A |
| ZNF749 | 19 | zinc finger protein 749 [Source:HGNC Symbol; |
| AC004076.1 | 19 | novel transcript |
| AC004076.2 | 19 | novel transcript |
| VN1R1 | 19 | vomeronasal 1 receptor 1 [Source:HGNC Syml |
| VN1R107P | 19 | vomeronasal 1 receptor 107 pseudogene [Sou |
| ZNF772 | 19 | zinc finger protein 772 [Source:HGNC Symbol; |
| AC003005.2 | 19 | novel transcript |
| ZNF419 | 19 | zinc finger protein 419 [Source:HGNC Symbol; |
| AC003005.1 | 19 | novel transcript |
| ZNF773 | 19 | zinc finger protein 773 [Source:HGNC Symbol; |
| ZNF549 | 19 | zinc finger protein 549 [Source:HGNC Symbol; |
| ZNF550 | 19 | zinc finger protein 550 [Source:HGNC Symbol; |
| ZNF416 | 19 | zinc finger protein 416 [Source:HGNC Symbol; |
| ZIK1 | 19 | zinc finger protein interacting with K protein 1 |
| ZNF530 | 19 | zinc finger protein 530 [Source:HGNC Symbol; |
| ZNF134 | 19 | zinc finger protein 134 [Source:HGNC Symbol; |
| AC003682.1 | 19 | novel transcript |
| ZNF211 | 19 | zinc finger protein 211 [Source:HGNC Symbol; |
| TPRG1LP1 | 19 | tumor protein p63 regulated 1 like pseudoger |
| ZSCAN4 | 19 | zinc finger and SCAN domain containing 4 [Sou |
| ZNF551 | 19 | zinc finger protein 551 [Source:HGNC Symbol; |
| AC003006.1 | 19 | novel protein |
| ZNF154 | 19 | zinc finger protein 154 [Source:HGNC Symbol; |
| ZNF671 | 19 | zinc finger protein 671 [Source:HGNC Symbol; |
| ZNF776 | 19 | zinc finger protein 776 [Source:HGNC Symbol; |

| | | |
|---|---|---|
| ZNF586 | 19 | zinc finger protein 586 [Source:HGNC Symbol; |
| ZNF552 | 19 | zinc finger protein 552 [Source:HGNC Symbol; |
| AC010522.1 | 19 | novel protein |
| ZNF587B | 19 | zinc finger protein 587B [Source:HGNC Symbo |
| FKBP1AP1 | 19 | FKBP prolyl isomerase 1A pseudogene 1 [Sour |
| ZNF814 | 19 | zinc finger protein 814 [Source:HGNC Symbol; |
| ZNF587 | 19 | zinc finger protein 587 [Source:HGNC Symbol; |
| AC010326.3 | 19 | novel transcript, sense intronic to ZNF814 |
| AC010326.4 | 19 | ubiquitin-conjugating enzyme E2C (UBE2C) ps |
| AC010326.2 | 19 | novel transcript |
| ZNF417 | 19 | zinc finger protein 417 [Source:HGNC Symbol; |
| ZNF418 | 19 | zinc finger protein 418 [Source:HGNC Symbol; |
| ZNF256 | 19 | zinc finger protein 256 [Source:HGNC Symbol; |
| AC010326.1 | 19 | ribosomal protein L19 (RPL19) pseudogene |
| C19orf18 | 19 | chromosome 19 open reading frame 18 [Sour |
| ZNF606 | 19 | zinc finger protein 606 [Source:HGNC Symbol; |
| AC008969.1 | 19 | novel transcript, antisense to ZNF606 |
| VN2R19P | 19 | vomeronasal 2 receptor 19, pseudogene [Sou |
| AC008969.2 | 19 | novel transcript |
| ZSCAN1 | 19 | zinc finger and SCAN domain containing 1 [So |
| LETM1P2 | 19 | leucine zipper and EF-hand containing transm |
| HNRNPDLP4 | 19 | HNRNPDL pseudogene 4 [Source:HGNC Symb |
| ZNF135 | 19 | zinc finger protein 135 [Source:HGNC Symbol; |
| RN7SL526P | 19 | RNA, 7SL, cytoplasmic 526, pseudogene [Sour |
| ZSCAN18 | 19 | zinc finger and SCAN domain containing 18 [S |
| ZNF329 | 19 | zinc finger protein 329 [Source:HGNC Symbol; |
| AC008751.1 | 19 | NADH dehydrogenase (ubiquinone) 1 alpha su |
| AC008751.3 | 19 | novel transcript |
| AC008751.2 | 19 | zinc finger protein family pseudogene |
| ZNF274 | 19 | zinc finger protein 274 [Source:HGNC Symbol; |
| RPS15AP36 | 19 | ribosomal protein S15a pseudogene 36 [Sourc |
| AC020915.3 | 19 | zinc finger protein family pseudogene |
| ZNF544 | 19 | zinc finger protein 544 [Source:HGNC Symbol; |
| AC020915.4 | 19 | novel protein |
| AC020915.2 | 19 | novel transcript, antisense to ZNF544 |
| AC020915.1 | 19 | novel transcript, sense intronic to ZNF544 |
| ZNF8 | 19 | zinc finger protein 8 [Source:HGNC Symbol;Ac |
| ZNF8-ERVK3-1 | 19 | ZNF8-ERVK3-1 readthrough [Source:NCBI gen |
| ERVK3-1 | 19 | endogenous retrovirus group K3 member 1 [S |
| AC010642.2 | 19 | novel transcript, antisense to ERVK3-1 |
| AC010642.1 | 19 | zinc finger protein family pseudogene |
| ZSCAN22 | 19 | zinc finger and SCAN domain containing 22 [S |
| MIR6806 | 19 | microRNA 6806 [Source:HGNC Symbol;Acc:HG |
| A1BG | 19 | alpha-1-B glycoprotein [Source:HGNC Symbol; |
| AC012313.3 | 19 | novel transcript |
| A1BG-AS1 | 19 | A1BG antisense RNA 1 [Source:HGNC Symbol; |
| AC012313.10 | 19 | novel transcript |
| ZNF497 | 19 | zinc finger protein 497 [Source:HGNC Symbol; |
| AC012313.2 | 19 | novel transcript, antisense to ZNF497 |
| AC012313.6 | 19 | novel transcript, antisense to ZNF497 |

| | | |
|---|---|---|
| RNA5SP473 | 19 | RNA, 5S ribosomal pseudogene 473 [Source:H |
| ZNF837 | 19 | zinc finger protein 837 [Source:HGNC Symbol; |
| RPS5 | 19 | ribosomal protein S5 [Source:HGNC Symbol;A |
| MIR4754 | 19 | microRNA 4754 [Source:HGNC Symbol;Acc:HG |
| RNF225 | 19 | ring finger protein 225 [Source:HGNC Symbol; |
| LINC02560 | 19 | long intergenic non-protein coding RNA 2560 |
| ZNF584 | 19 | zinc finger protein 584 [Source:HGNC Symbol; |
| AC012313.1 | 19 | novel transcript |
| AC012313.4 | 19 | novel transcript |
| AC012313.5 | 19 | novel transcript |
| ZNF132 | 19 | zinc finger protein 132 [Source:HGNC Symbol; |
| AC012313.8 | 19 | novel transcript |
| ZNF324B | 19 | zinc finger protein 324B [Source:HGNC Symbo |
| ZNF324 | 19 | zinc finger protein 324 [Source:HGNC Symbol; |
| ZNF446 | 19 | zinc finger protein 446 [Source:HGNC Symbol; |
| AC012313.7 | 19 | novel transcript, antisense to ZNF446 |
| SLC27A5 | 19 | solute carrier family 27 member 5 [Source:HG |
| RNU6-1337P | 19 | RNA, U6 small nuclear 1337, pseudogene [Sou |
| RN7SL693P | 19 | RNA, 7SL, cytoplasmic 693, pseudogene [Sour |
| AC012313.9 | 19 | novel transcript, antisense to SLC27A5 |
| ZBTB45 | 19 | zinc finger and BTB domain containing 45 [Sou |
| RN7SL525P | 19 | RNA, 7SL, cytoplasmic 525, pseudogene [Sour |
| TRIM28 | 19 | tripartite motif containing 28 [Source:HGNC S |
| MIR6807 | 19 | microRNA 6807 [Source:HGNC Symbol;Acc:HG |
| CHMP2A | 19 | charged multivesicular body protein 2A [Sourc |
| UBE2M | 19 | ubiquitin conjugating enzyme E2 M [Source:H |
| MZF1-AS1 | 19 | MZF1 antisense RNA 1 [Source:HGNC Symbol; |
| MZF1 | 19 | myeloid zinc finger 1 [Source:HGNC Symbol;A |
| CENPBD1P1 | 19 | CENPB DNA-binding domains containing 1 pse |
| AC016629.3 | 19 | Novel transcript |
| AC016629.1 | 19 | interleukin 9 receptor (IL9R) pseudogene |
| AC016629.2 | 19 | novel transcript, antisense to RPL23AP79 |
| DEFB125 | 20 | defensin beta 125 [Source:HGNC Symbol;Acc: |
| DEFB126 | 20 | defensin beta 126 [Source:HGNC Symbol;Acc: |
| DEFB127 | 20 | defensin beta 127 [Source:HGNC Symbol;Acc: |
| DEFB128 | 20 | defensin beta 128 [Source:HGNC Symbol;Acc: |
| DEFB129 | 20 | defensin beta 129 [Source:HGNC Symbol;Acc: |
| DEFB132 | 20 | defensin beta 132 [Source:HGNC Symbol;Acc: |
| AL034548.1 | 20 | novel transcript |
| C20orf96 | 20 | chromosome 20 open reading frame 96 [Sour |
| ZCCHC3 | 20 | zinc finger CCHC-type containing 3 [Source:HG |
| AL034548.2 | 20 | novel transcript |
| NRSN2-AS1 | 20 | NRSN2 antisense RNA 1 [Source:HGNC Symbo |
| SOX12 | 20 | SRY-box transcription factor 12 [Source:HGNC |
| NRSN2 | 20 | neurensin 2 [Source:HGNC Symbol;Acc:HGNC: |
| TRIB3 | 20 | tribbles pseudokinase 3 [Source:HGNC Symbo |
| RBCK1 | 20 | RANBP2-type and C3HC4-type zinc finger cont |
| TBC1D20 | 20 | TBC1 domain family member 20 [Source:HGN |
| Y_RNA | 20 | Y RNA [Source:RFAM;Acc:RF00019] |
| CSNK2A1 | 20 | casein kinase 2 alpha 1 [Source:HGNC Symbol |

| | | |
|---|---|---|
| TCF15 | 20 | transcription factor 15 [Source:HGNC Symbol; |
| SRXN1 | 20 | sulfiredoxin 1 [Source:HGNC Symbol;Acc:HGN |
| AL121758.1 | 20 | novel protein |
| SCRT2 | 20 | scratch family transcriptional repressor 2 [Sou |
| SLC52A3 | 20 | solute carrier family 52 member 3 [Source:HG |
| FAM110A | 20 | family with sequence similarity 110 member A |
| RPS10P5 | 20 | ribosomal protein S10 pseudogene 5 [Source: |
| ANGPT4 | 20 | angiopoietin 4 [Source:HGNC Symbol;Acc:HGN |
| RSPO4 | 20 | R-spondin 4 [Source:HGNC Symbol;Acc:HGNC: |
| AL110114.1 | 20 | novel transcript, antisense to PSMF1 |
| PSMF1 | 20 | proteasome inhibitor subunit 1 [Source:HGNC |
| ACTG1P3 | 20 | actin gamma 1 pseudogene 3 [Source:HGNC S |
| TMEM74B | 20 | transmembrane protein 74B [Source:HGNC Sy |
| AL031665.1 | 20 | novel transcript |
| C20orf202 | 20 | chromosome 20 open reading frame 202 [Sou |
| RAD21L1 | 20 | RAD21 cohesin complex component like 1 [So |
| SNPH | 20 | syntaphilin [Source:HGNC Symbol;Acc:HGNC:1 |
| SDCBP2 | 20 | syndecan binding protein 2 [Source:HGNC Sym |
| AL136531.2 | 20 | novel protein |
| AL136531.3 | 20 | TEC |
| SDCBP2-AS1 | 20 | SDCBP2 antisense RNA 1 [Source:HGNC Symbo |
| AL136531.1 | 20 | novel transcript |
| FKBP1A | 20 | FKBP prolyl isomerase 1A [Source:HGNC Symb |
| MIR6869 | 20 | microRNA 6869 [Source:HGNC Symbol;Acc:HG |
| NSFL1C | 20 | NSFL1 cofactor [Source:HGNC Symbol;Acc:HG |
| SIRPB2 | 20 | signal regulatory protein beta 2 [Source:HGNC |
| AL049634.1 | 20 | PTPNS (protein tyrosine phosphatase, non-rec |
| RNU6-917P | 20 | RNA, U6 small nuclear 917, pseudogene [Sour |
| SIRPD | 20 | signal regulatory protein delta [Source:HGNC |
| AL049634.2 | 20 | novel protein, SIRPB1-SIRPD readthrough |
| SIRPB1 | 20 | signal regulatory protein beta 1 [Source:HGNC |
| AL049634.3 | 20 | trans-golgi network vesicle protein 23 homolo |
| SIRPG | 20 | signal regulatory protein gamma [Source:HGN |
| SIRPG-AS1 | 20 | SIRPG antisense RNA 1 [Source:HGNC Symbol; |
| SIRPB3P | 20 | signal regulatory protein beta 3, pseudogene |
| RN7SL561P | 20 | RNA, 7SL, cytoplasmic 561, pseudogene [Sour |
| AL109809.3 | 20 | pseudogene similar to part of PTPNS (protein |
| AL109809.2 | 20 | pseudogene similar to part of PTPNS (protein |
| AL109809.5 | 20 | novel transcript |
| AL109809.4 | 20 | cytoskeleton associated protein 2-like (CKAP2 |
| AL109809.1 | 20 | pseudogene similar to F830045P16Rik |
| AL117335.1 | 20 | novel transcript, antisense to SIRPA |
| SIRPA | 20 | signal regulatory protein alpha [Source:HGNC |
| PDYN-AS1 | 20 | PDYN antisense RNA 1 [Source:HGNC Symbol; |
| PDYN | 20 | prodynorphin [Source:HGNC Symbol;Acc:HGN |
| RPL7P2 | 20 | ribosomal protein L7 pseudogene 2 [Source:H |
| STK35 | 20 | serine/threonine kinase 35 [Source:HGNC Sym |
| AL359916.1 | 20 | novel transcript, antisense to STK35 |
| AL121899.3 | 20 | novel transcript |
| AL121899.4 | 20 | novel transcript |

| | | |
|---|---|---|
| AL121899.1 | 20 | novel transcript |
| AL121899.2 | 20 | novel protein |
| TGM3 | 20 | transglutaminase 3 [Source:HGNC Symbol;Acc |
| TGM6 | 20 | transglutaminase 6 [Source:HGNC Symbol;Acc |
| Y_RNA | 20 | Y RNA [Source:RFAM;Acc:RF00019] |
| SNRPB | 20 | small nuclear ribonucleoprotein polypeptides |
| AL049650.1 | 20 | novel protein |
| ZNF343 | 20 | zinc finger protein 343 [Source:HGNC Symbol; |
| TMC2 | 20 | transmembrane channel like 2 [Source:HGNC |
| NOP56 | 20 | NOP56 ribonucleoprotein [Source:HGNC Symb |
| MIR1292 | 20 | microRNA 1292 [Source:HGNC Symbol;Acc:HG |
| SNORD110 | 20 | small nucleolar RNA, C/D box 110 [Source:HG |
| SNORA51 | 20 | small nucleolar RNA, H/ACA box 51 [Source:H |
| SNORD86 | 20 | small nucleolar RNA, C/D box 86 [Source:HGN |
| SNORD56 | 20 | small nucleolar RNA, C/D box 56 [Source:HGN |
| SNORD57 | 20 | small nucleolar RNA, C/D box 57 [Source:HGN |
| IDH3B | 20 | isocitrate dehydrogenase (NAD(+)) 3 beta [Sou |
| AL049712.1 | 20 | novel transcript |
| EBF4 | 20 | EBF family member 4 [Source:HGNC Symbol;A |
| RPL19P1 | 20 | ribosomal protein L19 pseudogene 1 [Source: |
| CPXM1 | 20 | carboxypeptidase X, M14 family member 1 [S |
| AL035460.1 | 20 | novel protein |
| C20orf141 | 20 | chromosome 20 open reading frame 141 [Sou |
| TMEM239 | 20 | transmembrane protein 239 [Source:HGNC Sy |
| PCED1A | 20 | PC-esterase domain containing 1A [Source:HG |
| VPS16 | 20 | VPS16 core subunit of CORVET and HOPS com |
| PTPRA | 20 | protein tyrosine phosphatase receptor type A |
| GNRH2 | 20 | gonadotropin releasing hormone 2 [Source:H |
| MRPS26 | 20 | mitochondrial ribosomal protein S26 [Source: |
| OXT | 20 | oxytocin/neurophysin I prepropeptide [Source |
| AVP | 20 | arginine vasopressin [Source:HGNC Symbol;Ac |
| RN7SL555P | 20 | RNA, 7SL, cytoplasmic 555, pseudogene [Sour |
| UBOX5-AS1 | 20 | UBOX5 antisense RNA 1 [Source:HGNC Symbo |
| UBOX5 | 20 | U-box domain containing 5 [Source:HGNC Sym |
| FASTKD5 | 20 | FAST kinase domains 5 [Source:HGNC Symbol; |
| LZTS3 | 20 | leucine zipper tumor suppressor family memb |
| DDRGK1 | 20 | DDRGK domain containing 1 [Source:HGNC Sy |
| ITPA | 20 | inosine triphosphatase [Source:HGNC Symbol; |
| SLC4A11 | 20 | solute carrier family 4 member 11 [Source:HG |
| AL109976.1 | 20 | novel transcript |
| C20orf194 | 20 | chromosome 20 open reading frame 194 [Sou |
| RNU6-1019P | 20 | RNA, U6 small nuclear 1019, pseudogene [Sou |
| UBE2V1P1 | 20 | ubiquitin conjugating enzyme E2 V1 pseudoge |
| U3 | 20 | Small nucleolar RNA U3 [Source:RFAM;Acc:RF |
| AL353193.1 | 20 | pseudogene similar to part of transcription el |
| ATRN | 20 | attractin [Source:HGNC Symbol;Acc:HGNC:88 |
| SF3A3P1 | 20 | splicing factor 3a, subunit 3 pseudogene 1 [So |
| GFRA4 | 20 | GDNF family receptor alpha 4 [Source:HGNC S |
| ADAM33 | 20 | ADAM metallopeptidase domain 33 [Source:H |
| SIGLEC1 | 20 | sialic acid binding Ig like lectin 1 [Source:HGN |

| | | |
|---|---|---|
| HSPA12B | 20 | heat shock protein family A (Hsp70) member [illegible] |
| C20orf27 | 20 | chromosome 20 open reading frame 27 [Sour |
| SPEF1 | 20 | sperm flagellar 1 [Source:HGNC Symbol;Acc:H |
| CENPB | 20 | centromere protein B [Source:HGNC Symbol;[illegible] |
| CDC25B | 20 | cell division cycle 25B [Source:HGNC Symbol;[illegible] |
| LINC01730 | 20 | long intergenic non-protein coding RNA 1730 |
| AL109804.1 | 20 | novel transcript |
| AP5S1 | 20 | adaptor related protein complex 5 subunit sig |
| MAVS | 20 | mitochondrial antiviral signaling protein [Sour |
| AL353194.1 | 20 | novel transcript |
| PANK2 | 20 | pantothenate kinase 2 [Source:HGNC Symbol; |
| MIR103A2 | 20 | microRNA 103a-2 [Source:HGNC Symbol;Acc:[illegible] |
| MIR103B2 | 20 | microRNA 103b-2 [Source:HGNC Symbol;Acc:[illegible] |
| AL031670.1 | 20 | novel transcript, antisense to PANK2 |
| RNF24 | 20 | ring finger protein 24 [Source:HGNC Symbol;[illegible] |
| FTLP3 | 20 | ferritin light chain pseudogene 3 [Source:HGN |
| RPL21P2 | 20 | ribosomal protein L21 pseudogene 2 [Source:[illegible] |
| AL356414.1 | 20 | novel transcript |
| SMOX | 20 | spermine oxidase [Source:HGNC Symbol;Acc:[illegible] |
| LINC01433 | 20 | long intergenic non-protein coding RNA 1433 |
| ADRA1D | 20 | adrenoceptor alpha 1D [Source:HGNC Symbol |
| AL139350.1 | 20 | novel transcript |
| AL121781.1 | 20 | novel transcript |
| AL121916.1 | 20 | ribosomal protein L7a-like 2 |
| RPS4XP2 | 20 | ribosomal protein S4X pseudogene 2 [Source: |
| PRNP | 20 | prion protein [Source:HGNC Symbol;Acc:HGN[illegible] |
| PRND | 20 | prion like protein doppel [Source:HGNC Symb |
| PRNT | 20 | prion locus lncRNA, testis expressed [Source:[illegible] |
| AL133396.1 | 20 | isopentenyl-diphosphate delta isomerase 1 (I[illegible] |
| AL133396.2 | 20 | novel transcript |
| RASSF2 | 20 | Ras association domain family member 2 [Sou |
| SLC23A2 | 20 | solute carrier family 23 member 2 [Source:HG |
| AL389886.1 | 20 | NULL |
| AL121890.1 | 20 | RPS21 (ribosomal protein S21) pseudogene |
| AL121890.3 | 20 | novel transcript |
| AL121890.2 | 20 | novel transcript |
| AL121890.5 | 20 | novel transcript |
| AL121890.4 | 20 | novel transcript |
| TMEM230 | 20 | transmembrane protein 230 [Source:HGNC Sy |
| RNA5SP474 | 20 | RNA, 5S ribosomal pseudogene 474 [Source:H |
| Y_RNA | 20 | Y RNA [Source:RFAM;Acc:RF00019] |
| PCNA | 20 | proliferating cell nuclear antigen [Source:HGN |
| AL121924.1 | 20 | NULL |
| CDS2 | 20 | CDP-diacylglycerol synthase 2 [Source:HGNC [illegible] |
| UBE2D3P1 | 20 | ubiquitin conjugating enzyme E2 D3 pseudoge |
| PROKR2 | 20 | prokineticin receptor 2 [Source:HGNC Symbol |
| AL121757.2 | 20 | novel transcript |
| RNA5-8SP7 | 20 | RNA, 5.8S ribosomal pseudogene 7 [Source:H[illegible] |
| AL121757.3 | 20 | novel transcript |
| LINC00658 | 20 | long intergenic non-protein coding RNA 658 [[illegible] |

| | | |
|---|---|---|
| AL121757.1 | 20 | novel transcript |
| LINC00654 | 20 | long intergenic non-protein coding RNA 654 [S |
| LINC01729 | 20 | long intergenic non-protein coding RNA 1729 |
| RPS18P1 | 20 | ribosomal protein S18 pseudogene 1 [Source: |
| AL109935.1 | 20 | novel transcript |
| GPCPD1 | 20 | glycerophosphocholine phosphodiesterase 1 [ |
| EIF4EP1 | 20 | eukaryotic translation initiation factor 4E pseu |
| SHLD1 | 20 | shieldin complex subunit 1 [Source:HGNC Sym |
| RNU1-55P | 20 | RNA, U1 small nuclear 55, pseudogene [Sourc |
| CHGB | 20 | chromogranin B [Source:HGNC Symbol;Acc:H |
| KANK1P1 | 20 | KN motif and ankyrin repeat domains 1 pseud |
| TRMT6 | 20 | tRNA methyltransferase 6 [Source:HGNC Sym |
| MCM8 | 20 | minichromosome maintenance 8 homologous |
| AL035461.3 | 20 | novel protein |
| Y_RNA | 20 | Y RNA [Source:RFAM;Acc:RF00019] |
| AL035461.1 | 20 | pseudogene similar to part of cytochrome c o |
| MCM8-AS1 | 20 | MCM8 antisense RNA 1 [Source:HGNC Symbo |
| AL035461.2 | 20 | novel transcript |
| RN7SL498P | 20 | RNA, 7SL, cytoplasmic 498, pseudogene [Sour |
| CRLS1 | 20 | cardiolipin synthase 1 [Source:HGNC Symbol; |
| LRRN4 | 20 | leucine rich repeat neuronal 4 [Source:HGNC |
| AL118505.1 | 20 | novel transcript |
| FERMT1 | 20 | fermitin family member 1 [Source:HGNC Sym |
| TARDBPP1 | 20 | TAR DNA binding protein pseudogene 1 [Sour |
| AL109618.1 | 20 | NS1-associated protein 1 (NSAP1) pseudogen |
| AL109618.2 | 20 | ribonuclease P/MRP 40kDa subunit (RPP40) p |
| AL109618.3 | 20 | fibroblast growth factor receptor 3 (FGFR3) ps |
| CASC20 | 20 | cancer susceptibility 20 [Source:HGNC Symbo |
| LINC01713 | 20 | long intergenic non-protein coding RNA 1713 |
| BMP2 | 20 | bone morphogenetic protein 2 [Source:HGNC |
| AL096799.1 | 20 | novel transcript |
| LINC01428 | 20 | long intergenic non-protein coding RNA 1428 |
| AL080248.1 | 20 | novel transcript |
| LINC01751 | 20 | long intergenic non-protein coding RNA 1751 |
| LINC01706 | 20 | long intergenic non-protein coding RNA 1706 |
| MIR8062 | 20 | microRNA 8062 [Source:HGNC Symbol;Acc:H |
| RN7SL547P | 20 | RNA, 7SL, cytoplasmic 547, pseudogene [Sour |
| AL031679.1 | 20 | FUS interacting protein (serine/arginine-rich) |
| AL021879.1 | 20 | HSPB (heat shock 27kDa) associated protein 1 |
| HAO1 | 20 | hydroxyacid oxidase 1 [Source:HGNC Symbol; |
| TMX4 | 20 | thioredoxin related transmembrane protein 4 |
| AL021396.1 | 20 | novel transcript, antisense to TMX4 |
| PLCB1 | 20 | phospholipase C beta 1 [Source:HGNC Symbol |
| PHKBP1 | 20 | phosphorylase kinase regulatory subunit beta |
| PLCB1-IT1 | 20 | PLCB1 intronic transcript 1 [Source:HGNC Sym |
| RNU105B | 20 | RNA, U105B small nucleolar [Source:HGNC Sy |
| AL445567.1 | 20 | ribosomal protein L31 (RPL31) pseudogene |
| AL445567.2 | 20 | CTAGE family, member 5 (CTAGE5) pseudoge |
| Metazoa_SRP | 20 | Metazoan signal recognition particle RNA [Sou |
| PLCB4 | 20 | phospholipase C beta 4 [Source:HGNC Symbol |

| | | |
|---|---|---|
| LAMP5-AS1 | 20 | LAMP5 antisense RNA 1 [Source:HGNC Symbo |
| LAMP5 | 20 | lysosomal associated membrane protein fami |
| PAK5 | 20 | p21 (RAC1) activated kinase 5 [Source:HGNC S |
| AL353612.2 | 20 | novel transcript, antisense to PAK7 |
| AL353612.1 | 20 | novel transcript |
| AL034427.1 | 20 | novel transcript, antisense to PAK7 |
| PARAL1 | 20 | PPARG activating RBM14 associated lncRNA 1 |
| ANKEF1 | 20 | ankyrin repeat and EF-hand domain containin |
| SNAP25-AS1 | 20 | SNAP25 antisense RNA 1 [Source:HGNC Symb |
| Metazoa_SRP | 20 | Metazoan signal recognition particle RNA [Sou |
| AL354824.1 | 20 | novel transcript |
| AL354824.2 | 20 | novel transcript |
| SNAP25 | 20 | synaptosome associated protein 25 [Source:H |
| AL023913.1 | 20 | novel transcript |
| HIGD1AP15 | 20 | HIG1 hypoxia inducible domain family membe |
| RPL23AP6 | 20 | ribosomal protein L23a pseudogene 6 [Source |
| SDAD1P2 | 20 | SDA1 domain containing 1 pseudogene 2 [Sou |
| MKKS | 20 | McKusick-Kaufman syndrome [Source:HGNC S |
| AL034430.1 | 20 | novel protein |
| AL034430.2 | 20 | novel protein |
| SLX4IP | 20 | SLX4 interacting protein [Source:HGNC Symbo |
| AL035456.1 | 20 | pseudogene similar to DNAJ protein |
| JAG1 | 20 | jagged canonical Notch ligand 1 [Source:HGN( |
| MIR6870 | 20 | microRNA 6870 [Source:HGNC Symbol;Acc:HG |
| AL050403.2 | 20 | novel transcript, antisense to JAG1 |
| LINC01752 | 20 | long intergenic non-protein coding RNA 1752 |
| AL135937.1 | 20 | novel transcript |
| AL050403.1 | 20 | novel transcript |
| FAT1P1 | 20 | FAT atypical cadherin 1 pseudogene 1 [Source |
| C20orf187 | 20 | chromosome 20 open reading frame 187 (put |
| AL158042.1 | 20 | novel transcript |
| AL049649.1 | 20 | putative novel transcript. |
| RPS11P1 | 20 | ribosomal protein S11 pseudogene 1 [Source: |
| AL109837.2 | 20 | novel transcript |
| AL109837.3 | 20 | novel transcript |
| AL109837.1 | 20 | lin-28 homolog A (C. elegans) (LIN28A) pseudo |
| PGAM3P | 20 | phosphoglycerate mutase 3, pseudogene [Sou |
| AL161938.1 | 20 | novel transcript |
| LINC00687 | 20 | long intergenic non-protein coding RNA 687 [S |
| RN7SKP111 | 20 | RN7SK pseudogene 111 [Source:HGNC Symbo |
| AL080274.1 | 20 | anaphase promoting complex subunit 13 (ANA |
| BTBD3 | 20 | BTB domain containing 3 [Source:HGNC Symb |
| AL035448.1 | 20 | novel transcript |
| AL109838.1 | 20 | novel transcript |
| AL136460.1 | 20 | novel transcript |
| PA2G4P2 | 20 | proliferation-associated 2G4 pseudogene 2 [S |
| LINC01722 | 20 | long intergenic non-protein coding RNA 1722 |
| AL078623.1 | 20 | novel transcript |
| AL133331.1 | 20 | novel transcript |
| LINC01723 | 20 | long intergenic non-protein coding RNA 1723 |

| | | |
|---|---|---|
| SPTLC3 | 20 | serine palmitoyltransferase long chain base su |
| AL109983.1 | 20 | coiled-coil domain containing 84 (CCDC84) pse |
| ISM1 | 20 | isthmin 1 [Source:HGNC Symbol;Acc:HGNC:16 |
| ISM1-AS1 | 20 | ISM1 antisense RNA 1 [Source:HGNC Symbol;A |
| AL050320.1 | 20 | novel transcript |
| AL121782.1 | 20 | novel transcript |
| TASP1 | 20 | taspase 1 [Source:HGNC Symbol;Acc:HGNC:15 |
| GAPDHP2 | 20 | glyceraldehyde-3-phosphate dehydrogenase p |
| ESF1 | 20 | ESF1 nucleolar pre-rRNA processing protein ho |
| NDUFAF5 | 20 | NADH:ubiquinone oxidoreductase complex as |
| SEL1L2 | 20 | SEL1L2 adaptor subunit of ERAD E3 ligase [Sou |
| RNU6-278P | 20 | RNA, U6 small nuclear 278, pseudogene [Sour |
| MACROD2 | 20 | mono-ADP ribosylhydrolase 2 [Source:HGNC S |
| RPS3P1 | 20 | ribosomal protein S3 pseudogene 1 [Source:H |
| AIMP1P1 | 20 | aminoacyl tRNA synthetase complex interacti |
| RN7SL864P | 20 | RNA, 7SL, cytoplasmic 864, pseudogene [Sour |
| FLRT3 | 20 | fibronectin leucine rich transmembrane prote |
| RNU6-228P | 20 | RNA, U6 small nuclear 228, pseudogene [Sour |
| AL121582.1 | 20 | novel transcript, antisense to MACROD2 |
| RNF11P2 | 20 | ring finger protein 11 pseudogene 2 [Source:H |
| MACROD2-IT1 | 20 | MACROD2 intronic transcript 1 [Source:HGNC |
| RPS10P2 | 20 | ribosomal protein S10 pseudogene 2 [Source: |
| MACROD2-AS1 | 20 | MACROD2 antisense RNA 1 [Source:HGNC Syr |
| AL138808.1 | 20 | novel transcript |
| RNU6-1159P | 20 | RNA, U6 small nuclear 1159, pseudogene [Sou |
| RNU6-115P | 20 | RNA, U6 small nuclear 115, pseudogene [Sour |
| RNA5SP475 | 20 | RNA, 5S ribosomal pseudogene 475 [Source:H |
| AL121584.1 | 20 | novel transcript |
| ENSAP1 | 20 | endosulfine alpha pseudogene 1 [Source:HGN |
| AL079338.1 | 20 | novel transcript, antisense to MACROD2 |
| AL161941.1 | 20 | novel transcript |
| PPIAP17 | 20 | peptidylprolyl isomerase A pseudogene 17 [So |
| KIF16B | 20 | kinesin family member 16B [Source:HGNC Syr |
| AL049794.1 | 20 | novel transcript |
| AL049794.2 | 20 | novel transcript |
| RPLP0P1 | 20 | ribosomal protein lateral stalk subunit P0 pseu |
| Y_RNA | 20 | Y RNA [Source:RFAM;Acc:RF00019] |
| AL135938.1 | 20 | ribosomal protein L7a-like 3 |
| AL034428.1 | 20 | novel transcript, antisense to SNRPB2 |
| SNRPB2 | 20 | small nuclear ribonucleoprotein polypeptide E |
| OTOR | 20 | otoraplin [Source:HGNC Symbol;Acc:HGNC:85 |
| AL121892.1 | 20 | novel transcript |
| U3 | 20 | Small nucleolar RNA U3 [Source:RFAM;Acc:RF |
| RNU6-27P | 20 | RNA, U6 small nuclear 27, pseudogene [Sourc |
| AL359511.1 | 20 | zinc finger protein (KRAB box-containing) pseu |
| RNU1-131P | 20 | RNA, U1 small nuclear 131, pseudogene [Sour |
| AL359511.2 | 20 | novel transcript |
| AL359511.3 | 20 | novel transcript, antisense to PCSK2 |
| PCSK2 | 20 | proprotein convertase subtilisin/kexin type 2 |
| DYNLT3P1 | 20 | dynein light chain Tctex-type 3 pseudogene 1 |

| | | |
|---|---|---|
| BFSP1 | 20 | beaded filament structural protein 1 [Source:I |
| RPS27AP2 | 20 | ribosomal protein S27a pseudogene 2 [Source |
| AL132765.2 | 20 | novel transcript, sense intronic to BFSP1 |
| DSTN | 20 | destrin, actin depolymerizing factor [Source:H |
| RN7SKP69 | 20 | RN7SK pseudogene 69 [Source:HGNC Symbol; |
| AL132765.1 | 20 | ras-related pseudogene |
| RRBP1 | 20 | ribosome binding protein 1 [Source:HGNC Syr |
| BANF2 | 20 | barrier to autointegration factor 2 [Source:HG |
| AL035045.1 | 20 | novel transcript |
| RNU6-192P | 20 | RNA, U6 small nuclear 192, pseudogene [Sour |
| SNX5 | 20 | sorting nexin 5 [Source:HGNC Symbol;Acc:HGI |
| OVOL2 | 20 | ovo like zinc finger 2 [Source:HGNC Symbol;Ac |
| SNORD17 | 20 | small nucleolar RNA, C/D box 17 [Source:HGN |
| MGME1 | 20 | mitochondrial genome maintenance exonucle |
| PTMAP3 | 20 | prothymosin alpha pseudogene 3 [Source:HGI |
| AL160411.1 | 20 | novel transcript |
| RPL15P1 | 20 | ribosomal protein L15 pseudogene 1 [Source:I |
| RNU7-137P | 20 | RNA, U7 small nuclear 137 pseudogene [Sourc |
| Y_RNA | 20 | Y RNA [Source:RFAM;Acc:RF00019] |
| PET117 | 20 | PET117 cytochrome c oxidase chaperone [Sou |
| KAT14 | 20 | lysine acetyltransferase 14 [Source:HGNC Sym |
| AL050321.1 | 20 | chromosome 7 open reading frame 24 (C7orf2 |
| RN7SL14P | 20 | RNA, 7SL, cytoplasmic 14, pseudogene [Sourc |
| RNU2-56P | 20 | RNA, U2 small nuclear 56, pseudogene [Sourc |
| AL049646.3 | 20 | novel transcript |
| ZNF133 | 20 | zinc finger protein 133 [Source:HGNC Symbol; |
| AL049646.1 | 20 | novel transcript |
| RN7SKP74 | 20 | RN7SK pseudogene 74 [Source:HGNC Symbol; |
| AL049646.2 | 20 | novel transcript |
| LINC00851 | 20 | long intergenic non-protein coding RNA 851 [S |
| DZANK1 | 20 | double zinc ribbon and ankyrin repeat domair |
| GCNT1P1 | 20 | glucosaminyl (N-acetyl) transferase 1 pseudog |
| RNA5SP476 | 20 | RNA, 5S ribosomal pseudogene 476 [Source:H |
| POLR3F | 20 | RNA polymerase III subunit F [Source:HGNC Sy |
| MIR3192 | 20 | microRNA 3192 [Source:HGNC Symbol;Acc:HG |
| RPL21P3 | 20 | ribosomal protein L21 pseudogene 3 [Source:I |
| RBBP9 | 20 | RB binding protein 9, serine hydrolase [Source |
| RPS19P1 | 20 | ribosomal protein S19 pseudogene 1 [Source: |
| SEC23B | 20 | SEC23 homolog B, coat complex II component |
| SMIM26 | 20 | small integral membrane protein 26 [Source:H |
| AL121900.2 | 20 | novel protein |
| DTD1 | 20 | D-aminoacyl-tRNA deacylase 1 [Source:HGNC |
| RN7SL638P | 20 | RNA, 7SL, cytoplasmic 638, pseudogene [Sour |
| RNU6ATAC34P | 20 | RNA, U6atac small nuclear 34, pseudogene [So |
| DUXAP7 | 20 | double homeobox A pseudogene 7 [Source:H( |
| AL121900.1 | 20 | novel transcript |
| EEF1A1P34 | 20 | eukaryotic translation elongation factor 1 alpl |
| LINC00652 | 20 | long intergenic non-protein coding RNA 652 [S |
| LINC00653 | 20 | long intergenic non-protein coding RNA 653 [S |
| C20orf78 | 20 | chromosome 20 open reading frame 78 [Sour |

| | | |
|---|---|---|
| SCP2D1 | 20 | SCP2 sterol binding domain containing 1 [Sour |
| AL135936.1 | 20 | novel transcript |
| AL136090.2 | 20 | novel transcript |
| SLC24A3 | 20 | solute carrier family 24 member 3 [Source:HG |
| AL136090.1 | 20 | novel transcript |
| AL049647.1 | 20 | novel transcript |
| AL121830.1 | 20 | novel transcript, antisense to SLC24A3 |
| AL121761.1 | 20 | novel transcript |
| RIN2 | 20 | Ras and Rab interactor 2 [Source:HGNC Symb |
| RPL12P12 | 20 | ribosomal protein L12 pseudogene 12 [Source |
| NAA20 | 20 | N(alpha)-acetyltransferase 20, NatB catalytic s |
| CRNKL1 | 20 | crooked neck pre-mRNA splicing factor 1 [Sou |
| CFAP61 | 20 | cilia and flagella associated protein 61 [Source |
| AL035454.1 | 20 | novel transcript |
| AL049648.1 | 20 | novel transcript |
| RPL17P1 | 20 | ribosomal protein L17 pseudogene 1 [Source: |
| AL121721.1 | 20 | novel transcript |
| RN7SL690P | 20 | RNA, 7SL, cytoplasmic 690, pseudogene [Sour |
| AL161658.1 | 20 | TEC |
| INSM1 | 20 | INSM transcriptional repressor 1 [Source:HGN |
| RALGAPA2 | 20 | Ral GTPase activating protein catalytic alpha s |
| EIF4E2P1 | 20 | eukaryotic translation initiation factor 4E fami |
| LLPHP1 | 20 | LLP homolog, long-term synaptic facilitation p |
| RN7SL607P | 20 | RNA, 7SL, cytoplasmic 607, pseudogene [Sour |
| MRPS11P1 | 20 | mitochondrial ribosomal protein S11 pseudog |
| AL133465.1 | 20 | novel transcript |
| LINC00237 | 20 | long intergenic non-protein coding RNA 237 [S |
| AL121759.2 | 20 | novel transcript |
| RPL24P2 | 20 | ribosomal protein L24 pseudogene 2 [Source: |
| KIZ | 20 | kizuna centrosomal protein [Source:HGNC Syr |
| RNA5SP477 | 20 | RNA, 5S ribosomal pseudogene 477 [Source:H |
| AL121759.1 | 20 | novel transcript |
| KIZ-AS1 | 20 | KIZ antisense RNA 1 [Source:HGNC Symbol;Ac |
| RPS15AP1 | 20 | ribosomal protein S15a pseudogene 1 [Source |
| ZNF877P | 20 | zinc finger protein 877, pseudogene [Source:H |
| AL117332.1 | 20 | novel transcript, antisense to XRN2 |
| XRN2 | 20 | 5'-3' exoribonuclease 2 [Source:HGNC Symbol |
| NKX2-4 | 20 | NK2 homeobox 4 [Source:HGNC Symbol;Acc:H |
| AL158013.1 | 20 | novel transcript, antisense to NKX2-4 |
| RN7SKP140 | 20 | RN7SK pseudogene 140 [Source:HGNC Symbo |
| AL133325.1 | 20 | hydroxysteroid (17-beta) dehydrogenase 12 (I |
| AL133325.3 | 20 | novel transcript |
| GSTM3P1 | 20 | glutathione S-transferase mu 3 pseudogene 1 |
| NKX2-2 | 20 | NK2 homeobox 2 [Source:HGNC Symbol;Acc:H |
| NKX2-2-AS1 | 20 | NKX2-2 antisense RNA 1 [Source:HGNC Symbo |
| AL133325.2 | 20 | novel transcript |
| LINC01727 | 20 | long intergenic non-protein coding RNA 1727 |
| LINC01726 | 20 | long intergenic non-protein coding RNA 1726 |
| AL121590.1 | 20 | NULL |
| PAX1 | 20 | paired box 1 [Source:HGNC Symbol;Acc:HGNC |

| | | |
|---|---|---|
| SLC25A6P1 | 20 | solute carrier family 25 member 6 pseudogen |
| RPL41P1 | 20 | ribosomal protein L41 pseudogene 1 [Source: |
| AL109807.1 | 20 | novel transcript |
| LINC01432 | 20 | long intergenic non-protein coding RNA 1432 |
| AL035258.1 | 20 | novel transcript |
| LINC01427 | 20 | long intergenic non-protein coding RNA 1427 |
| AL049737.1 | 20 | suppression of tumorigenicity 13 (colon carcir |
| AL121912.1 | 20 | chromosome 9 open reading frame 16 (C9orf1 |
| AL133464.1 | 20 | novel transcript |
| LINC00261 | 20 | long intergenic non-protein coding RNA 261 [S |
| FOXA2 | 20 | forkhead box A2 [Source:HGNC Symbol;Acc:H |
| LNCNEF | 20 | lncRNA neighboring enhancer of FOXA2 [Sour |
| LINC01747 | 20 | long intergenic non-protein coding RNA 1747 |
| AL158175.1 | 20 | novel transcript |
| KRT18P3 | 20 | keratin 18 pseudogene 3 [Source:HGNC Symb |
| CYB5AP4 | 20 | cytochrome b5 type A pseudogene 4 [Source: |
| AL353132.1 | 20 | kinesin-like 7 (KNSL7) pseudogene |
| AL049651.1 | 20 | novel transcript |
| AL049651.2 | 20 | novel transcript, sense overlapping SSTR4 |
| SSTR4 | 20 | somatostatin receptor 4 [Source:HGNC Symbo |
| THBD | 20 | thrombomodulin [Source:HGNC Symbol;Acc:H |
| AL118508.3 | 20 | TEC |
| CD93 | 20 | CD93 molecule [Source:HGNC Symbol;Acc:HG |
| LINC00656 | 20 | long intergenic non-protein coding RNA 656 [S |
| AL118508.2 | 20 | TEC |
| RNA5SP478 | 20 | RNA, 5S ribosomal pseudogene 478 [Source:H |
| AL118508.1 | 20 | novel transcript |
| AL390037.1 | 20 | novel transcript |
| AL096677.1 | 20 | novel transcript |
| NXT1 | 20 | nuclear transport factor 2 like export factor 1 |
| LINC01431 | 20 | long intergenic non-protein coding RNA 1431 |
| GZF1 | 20 | GDNF inducible zinc finger protein 1 [Source:H |
| NAPB | 20 | NSF attachment protein beta [Source:HGNC Sy |
| RNA5SP479 | 20 | RNA, 5S ribosomal pseudogene 479 [Source:H |
| CSTL1 | 20 | cystatin like 1 [Source:HGNC Symbol;Acc:HGN |
| CST11 | 20 | cystatin 11 [Source:HGNC Symbol;Acc:HGNC:1 |
| AL096677.2 | 20 | novel transcript |
| Y_RNA | 20 | Y RNA [Source:RFAM;Acc:RF00019] |
| CST12P | 20 | cystatin 12, pseudogene [Source:HGNC Symbo |
| AL109954.1 | 20 | novel transcript |
| AL109954.2 | 20 | novel transcript |
| CST8 | 20 | cystatin 8 [Source:HGNC Symbol;Acc:HGNC:24 |
| CST13P | 20 | cystatin 13, pseudogene [Source:HGNC Symbo |
| CST9LP1 | 20 | cystatin 9-like pseudogene 1 [Source:HGNC Sy |
| CST9L | 20 | cystatin 9 like [Source:HGNC Symbol;Acc:HGN |
| CST9LP2 | 20 | cystatin 9-like pseudogene 2 [Source:HGNC Sy |
| CST9 | 20 | cystatin 9 [Source:HGNC Symbol;Acc:HGNC:13 |
| CST3 | 20 | cystatin C [Source:HGNC Symbol;Acc:HGNC:24 |
| AL121894.2 | 20 | novel transcript |
| AL121894.1 | 20 | novel transcript |

| | | |
|---|---|---|
| AL121894.3 | 20 | novel transcript |
| CST4 | 20 | cystatin S [Source:HGNC Symbol;Acc:HGNC:24 |
| CST2P1 | 20 | cystatin SA pseudogene 1 [Source:HGNC Symb |
| AL591074.2 | 20 | novel transcript |
| CST1 | 20 | cystatin SN [Source:HGNC Symbol;Acc:HGNC:2 |
| CSTP2 | 20 | cystatin pseudogene 2 [Source:HGNC Symbol; |
| AL591074.1 | 20 | novel transcript |
| CST2 | 20 | cystatin SA [Source:HGNC Symbol;Acc:HGNC:2 |
| CST5 | 20 | cystatin D [Source:HGNC Symbol;Acc:HGNC:24 |
| CSTP1 | 20 | cystatin pseudogene 1 [Source:HGNC Symbol; |
| AL133466.1 | 20 | gamma-glutamyltransferase family pseudoger |
| GGTLC1 | 20 | gamma-glutamyltransferase light chain 1 [Sou |
| POM121L3P | 20 | POM121 transmembrane nucleoporin like 3, p |
| LINC01721 | 20 | long intergenic non-protein coding RNA 1721 |
| AL110503.1 | 20 | novel transcript |
| RNU1-23P | 20 | RNA, U1 small nuclear 23, pseudogene [Sourc |
| AL158090.1 | 20 | novel transcript |
| GAPDHP53 | 20 | glyceraldehyde 3 phosphate dehydrogenase p |
| SYNDIG1 | 20 | synapse differentiation inducing 1 [Source:HG |
| AL157413.1 | 20 | novel transcript, antisense to SYNDIG1 |
| AL049594.1 | 20 | novel transcript |
| AL035661.1 | 20 | novel transcript |
| CST7 | 20 | cystatin F [Source:HGNC Symbol;Acc:HGNC:24 |
| APMAP | 20 | adipocyte plasma membrane associated prote |
| RNU6-1257P | 20 | RNA, U6 small nuclear 1257, pseudogene [Sou |
| AL035661.2 | 20 | novel transcript, antisense to APMAP |
| ACSS1 | 20 | acyl-CoA synthetase short chain family memb |
| AL080312.2 | 20 | novel transcript |
| VSX1 | 20 | visual system homeobox 1 [Source:HGNC Sym |
| AL080312.1 | 20 | V-set domain containing T cell activation inhib |
| AL035252.2 | 20 | novel transcript |
| AL035252.1 | 20 | novel pseudogene |
| AL035252.5 | 20 | novel transcript |
| ENTPD6 | 20 | ectonucleoside triphosphate diphosphohydro |
| Y_RNA | 20 | Y RNA [Source:RFAM;Acc:RF00019] |
| AL035252.4 | 20 | TEC |
| AL035252.3 | 20 | novel transcript |
| AL121772.1 | 20 | novel transcript |
| PYGB | 20 | glycogen phosphorylase B [Source:HGNC Sym |
| AL121772.2 | 20 | novel transcript, antisense to PYGB |
| AL121772.3 | 20 | novel transcript, antisense to PYGB |
| ABHD12 | 20 | abhydrolase domain containing 12 [Source:HG |
| PPIAP2 | 20 | peptidylprolyl isomerase A pseudogene 2 [Sou |
| GINS1 | 20 | GINS complex subunit 1 [Source:HGNC Symbo |
| NINL | 20 | ninein like [Source:HGNC Symbol;Acc:HGNC:2 |
| RNU6ATAC17P | 20 | RNA, U6atac small nuclear 17, pseudogene [So |
| NANP | 20 | N-acetylneuraminic acid phosphatase [Source |
| ZNF337-AS1 | 20 | ZNF337 antisense RNA 1 [Source:HGNC Symbo |
| RN7SL594P | 20 | RNA, 7SL, cytoplasmic 594, pseudogene [Sour |
| MED28P7 | 20 | mediator complex subunit 28 pseudogene 7 [S |

| | | |
|---|---|---|
| ZNF337 | 20 | zinc finger protein 337 [Source:HGNC Symbol; |
| AL031673.1 | 20 | novel transcript, antisense to ZNF337 |
| AL390198.1 | 20 | novel transcript |
| VN1R108P | 20 | vomeronasal 1 receptor 108 pseudogene [Sou |
| FAM182B | 20 | family with sequence similarity 182 member B |
| BSNDP1 | 20 | barttin CLCNK type accessory beta subunit pse |
| AL390198.2 | 20 | novel zinc finger pseudogene |
| BSNDP2 | 20 | barttin CLCNK type accessory beta subunit pse |
| CFTRP1 | 20 | CFTR pseudogene 1 [Source:HGNC Symbol;Acc |
| LINC01733 | 20 | long intergenic non-protein coding RNA 1733 |
| FAM182A | 20 | family with sequence similarity 182 member A |
| AL078587.1 | 20 | novel transcript |
| AL078587.2 | 20 | novel transcript |
| AL450124.1 | 20 | zinc-finger pseudogene |
| BSNDP3 | 20 | barttin CLCNK type accessory beta subunit pse |
| NCOR1P1 | 20 | nuclear receptor corepressor 1 pseudogene 1 |
| AL391119.1 | 20 | novel transcript |
| AL121904.2 | 20 | novel transcript |
| MIR663AHG | 20 | MIR663A host gene [Source:HGNC Symbol;Acc |
| AL121904.1 | 20 | novel transcript |
| MIR663A | 20 | microRNA 663a [Source:HGNC Symbol;Acc:HG |
| FRG1CP | 20 | FSHD region gene 1 family member C, pseudo |
| FRG1DP | 20 | FSHD region gene 1 family member D, pseudo |
| 5_8S_rRNA | 20 | 5.8S ribosomal RNA [Source:RFAM;Acc:RF000 |
| DUX4L32 | 20 | double homeobox 4 like 32 (pseudogene) [Sou |
| PCMTD1P7 | 20 | protein-L-isoaspartate (D-aspartate) O-methyl |
| DUX4L33 | 20 | double homeobox 4 like 33 (pseudogene) [Sou |
| DUX4L34 | 20 | double homeobox 4 like 34 (pseudogene) [Sou |
| FRG2EP | 20 | FSHD region gene 2 family member E, pseudo |
| RARRES2P11 | 20 | retinoic acid receptor responder 2 pseudogen |
| AGGF1P10 | 20 | angiogenic factor with G-patch and FHA doma |
| DUX4L35 | 20 | double homeobox 4 like 35 (pseudogene) [Sou |
| FRG1EP | 20 | FSHD region gene 1 family member E, pseudo |
| FAM242B | 20 | family with sequence similarity 242 member B |
| Y_RNA | 20 | Y RNA [Source:RFAM;Acc:RF00019] |
| CFTRP2 | 20 | CFTR pseudogene 2 [Source:HGNC Symbol;Acc |
| 5_8S_rRNA | 20 | 5.8S ribosomal RNA [Source:RFAM;Acc:RF000 |
| CDC27P3 | 20 | cell division cycle 27 pseudogene 3 [Source:HG |
| RNA5SP532 | 20 | RNA, 5S ribosomal pseudogene 532 [Source:H |
| DUX4L37 | 20 | double homeobox 4 like 37 (pseudogene) [Sou |
| CFTRP3 | 20 | CFTR pseudogene 3 [Source:HGNC Symbol;Acc |
| LINC01597 | 20 | long intergenic non-protein coding RNA 1597 |
| AL121762.1 | 20 | novel transcript |
| AL121762.2 | 20 | novel transcript |
| Y_RNA | 20 | Y RNA [Source:RFAM;Acc:RF00019] |
| FAM242A | 20 | family with sequence similarity 242 member A |
| FRG1BP | 20 | FSHD region gene 1 family member B, pseudo |
| MLLT10P1 | 20 | MLLT10 pseudogene 1 [Source:HGNC Symbol; |
| 5_8S_rRNA | 20 | 5.8S ribosomal RNA [Source:RFAM;Acc:RF000 |
| AC018688.1 | 20 | zinc finger protein 285 (ZNF285) pseudogene |

| | | |
|---|---|---|
| ANKRD20A21P | 20 | ankyrin repeat domain 20 family member A21 |
| U6 | 20 | U6 spliceosomal RNA [Source:RFAM;Acc:RF00 |
| RNA5SP528 | 20 | RNA, 5S ribosomal pseudogene 528 [Source:H |
| DEFB115 | 20 | defensin beta 115 [Source:HGNC Symbol;Acc: |
| DKKL1P1 | 20 | DKKL1 pseudogene 1 [Source:HGNC Symbol;A |
| AL121723.1 | 20 | CDK5 regulatory subunit associated protein 3 |
| DEFB116 | 20 | defensin beta 116 [Source:HGNC Symbol;Acc: |
| RPL31P3 | 20 | ribosomal protein L31 pseudogene 3 [Source: |
| HAUS6P2 | 20 | HAUS augmin like complex subunit 6 pseudog |
| RNA5SP480 | 20 | RNA, 5S ribosomal pseudogene 480 [Source:H |
| DEFB117 | 20 | defensin beta 117 (pseudogene) [Source:HGN |
| DEFB118 | 20 | defensin beta 118 [Source:HGNC Symbol;Acc: |
| AL031650.1 | 20 | NULL |
| DEFB119 | 20 | defensin beta 119 [Source:HGNC Symbol;Acc: |
| DEFB121 | 20 | defensin beta 121 [Source:HGNC Symbol;Acc: |
| DEFB122 | 20 | defensin beta 122 (pseudogene) [Source:HGN |
| DEFB123 | 20 | defensin beta 123 [Source:HGNC Symbol;Acc: |
| DEFB124 | 20 | defensin beta 124 [Source:HGNC Symbol;Acc: |
| REM1 | 20 | RRAD and GEM like GTPase 1 [Source:HGNC S |
| LINC00028 | 20 | long intergenic non-protein coding RNA 28 [So |
| HM13 | 20 | histocompatibility minor 13 [Source:HGNC Syr |
| AL110115.1 | 20 | novel transcript, sense intronic to HM13 |
| MCTS2P | 20 | MCTS family member 2, pseudogene [Source: |
| HM13-IT1 | 20 | HM13 intronic transcript 1 [Source:HGNC Sym |
| HM13-AS1 | 20 | HM13 antisense RNA 1 [Source:HGNC Symbol |
| AL110115.2 | 20 | novel transcript |
| RNU6-384P | 20 | RNA, U6 small nuclear 384, pseudogene [Sour |
| ID1 | 20 | inhibitor of DNA binding 1, HLH protein [Sourc |
| MIR3193 | 20 | microRNA 3193 [Source:HGNC Symbol;Acc:HG |
| COX4I2 | 20 | cytochrome c oxidase subunit 4I2 [Source:HG |
| BCL2L1 | 20 | BCL2 like 1 [Source:HGNC Symbol;Acc:HGNC:9 |
| AL117381.1 | 20 | novel transcript |
| ABALON | 20 | apoptotic BCL2L1-antisense long non-coding F |
| TPX2 | 20 | TPX2 microtubule nucleation factor [Source:H |
| AL160175.1 | 20 | ATP synthase, H+ transporting, mitochondrial |
| MYLK2 | 20 | myosin light chain kinase 2 [Source:HGNC Sym |
| FOXS1 | 20 | forkhead box S1 [Source:HGNC Symbol;Acc:HG |
| DUSP15 | 20 | dual specificity phosphatase 15 [Source:HGNC |
| TTLL9 | 20 | tubulin tyrosine ligase like 9 [Source:HGNC Syr |
| RNU1-94P | 20 | RNA, U1 small nuclear 94, pseudogene [Sourc |
| PDRG1 | 20 | p53 and DNA damage regulated 1 [Source:HG |
| XKR7 | 20 | XK related 7 [Source:HGNC Symbol;Acc:HGNC |
| AL031658.2 | 20 | novel transcript, sense intronic to XKR7 |
| AL031658.1 | 20 | novel transcript |
| RNA5SP481 | 20 | RNA, 5S ribosomal pseudogene 481 [Source:H |
| CCM2L | 20 | CCM2 like scaffold protein [Source:HGNC Sym |
| RNA5SP482 | 20 | RNA, 5S ribosomal pseudogene 482 [Source:H |
| HCK | 20 | HCK proto-oncogene, Src family tyrosine kinas |
| TM9SF4 | 20 | transmembrane 9 superfamily member 4 [Sou |
| AL049539.1 | 20 | novel transcript, sense intronic to TM9SF4 |

| | | |
|---|---|---|
| RSL24D1P6 | 20 | ribosomal L24 domain containing 1 pseudoger |
| TSPY26P | 20 | testis specific protein Y-linked 26, pseudogene |
| PLAGL2 | 20 | PLAG1 like zinc finger 2 [Source:HGNC Symbol |
| POFUT1 | 20 | protein O-fucosyltransferase 1 [Source:HGNC |
| MIR1825 | 20 | microRNA 1825 [Source:HGNC Symbol;Acc:HG |
| AL121897.1 | 20 | novel transcript |
| KIF3B | 20 | kinesin family member 3B [Source:HGNC Sym |
| AL121583.1 | 20 | novel transcript |
| ASXL1 | 20 | ASXL transcriptional regulator 1 [Source:HGN( |
| NOL4L | 20 | nucleolar protein 4 like [Source:HGNC Symbol |
| AL034550.1 | 20 | novel transcript, antisense to NOL4L |
| AL034550.3 | 20 | novel transcript, antisense to NOL4L |
| AL034550.2 | 20 | novel transcript, antisense to NOL4L |
| AL133343.2 | 20 | novel transcript, antisense to NOL4L |
| NOL4L-DT | 20 | NOL4L divergent transcript [Source:HGNC Syn |
| AL133343.1 | 20 | novel transcript, antisense to NOL4L |
| C20orf203 | 20 | chromosome 20 open reading frame 203 [Sou |
| FO393400.1 | 20 | COMM domain containing 7 [Source:NCBI ger |
| BAK1P1 | 20 | BCL2 antagonist/killer 1 pseudogene 1 [Sourc |
| COMMD7 | 20 | COMM domain containing 7 [Source:HGNC Sy |
| DNMT3B | 20 | DNA methyltransferase 3 beta [Source:HGNC |
| MAPRE1 | 20 | microtubule associated protein RP/EB family r |
| AL035071.2 | 20 | novel transcript, antisense to MAPRE1 |
| AL035071.1 | 20 | novel transcript |
| EFCAB8 | 20 | EF-hand calcium binding domain 8 [Source:HG |
| SUN5 | 20 | Sad1 and UNC84 domain containing 5 [Source |
| BPIFB2 | 20 | BPI fold containing family B member 2 [Sourc |
| BPIFB6 | 20 | BPI fold containing family B member 6 [Sourc |
| PUDPP3 | 20 | pseudouridine 5'-phosphatase pseudogene 3 |
| BPIFB3 | 20 | BPI fold containing family B member 3 [Sourc |
| BPIFB4 | 20 | BPI fold containing family B member 4 [Sourc |
| BPIFA2 | 20 | BPI fold containing family A member 2 [Sourc |
| SOCS2P1 | 20 | suppressor of cytokine signaling 2 pseudogen |
| BPIFA4P | 20 | BPI fold containing family A member 4, pseud |
| AL121901.1 | 20 | novel transcript |
| BPIFA3 | 20 | BPI fold containing family A member 3 [Sourc |
| BPIFA1 | 20 | BPI fold containing family A member 1 [Sourc |
| RPL12P3 | 20 | ribosomal protein L12 pseudogene 3 [Source: |
| BPIFB1 | 20 | BPI fold containing family B member 1 [Sourc |
| BPIFB5P | 20 | BPI fold containing family B member 5, pseud |
| BPIFB9P | 20 | BPI fold containing family B member 9, pseud |
| CDK5RAP1 | 20 | CDK5 regulatory subunit associated protein 1 |
| SNTA1 | 20 | syntrophin alpha 1 [Source:HGNC Symbol;Acc |
| CBFA2T2 | 20 | CBFA2/RUNX1 partner transcriptional co-repr |
| AL121906.2 | 20 | novel transcript |
| NECAB3 | 20 | N-terminal EF-hand calcium binding protein 3 |
| C20orf144 | 20 | chromosome 20 open reading frame 144 [Sou |
| ACTL10 | 20 | actin like 10 [Source:HGNC Symbol;Acc:HGNC |
| AL121906.1 | 20 | novel transcript, antisense to NECAB3 |
| E2F1 | 20 | E2F transcription factor 1 [Source:HGNC Symb |

| | | |
|---|---|---|
| PXMP4 | 20 | peroxisomal membrane protein 4 [Source:HG |
| AL050349.1 | 20 | novel transcript |
| ZNF341 | 20 | zinc finger protein 341 [Source:HGNC Symbol; |
| RPL31P2 | 20 | ribosomal protein L31 pseudogene 2 [Source: |
| ZNF341-AS1 | 20 | ZNF341 antisense RNA 1 [Source:HGNC Symbo |
| CHMP4B | 20 | charged multivesicular body protein 4B [Sourc |
| TPM3P2 | 20 | tropomyosin 3 pseudogene 2 [Source:HGNC S |
| PIGPP3 | 20 | phosphatidylinositol glycan anchor biosynthes |
| RALY-AS1 | 20 | RALY antisense RNA 1 [Source:HGNC Symbol;A |
| RALY | 20 | RALY heterogeneous nuclear ribonucleoprote |
| AL031668.1 | 20 | novel transcript |
| MIR4755 | 20 | microRNA 4755 [Source:HGNC Symbol;Acc:HG |
| EIF2S2 | 20 | eukaryotic translation initiation factor 2 subur |
| RPS2P1 | 20 | ribosomal protein S2 pseudogene 1 [Source:H |
| AL031668.2 | 20 | novel transcript |
| ASIP | 20 | agouti signaling protein [Source:HGNC Symbo |
| XPOTP1 | 20 | exportin for tRNA pseudogene 1 [Source:HGN |
| AL035458.2 | 20 | novel transcript, overlapping AHCY |
| AL035458.1 | 20 | density-regulated protein (DENR) pseudogene |
| AHCY | 20 | adenosylhomocysteinase [Source:HGNC Symb |
| AL356299.2 | 20 | novel transcript |
| AL356299.1 | 20 | pseudogene similar to part of laminin recepto |
| ITCH | 20 | itchy E3 ubiquitin protein ligase [Source:HGNC |
| Y_RNA | 20 | Y RNA [Source:RFAM;Acc:RF00019] |
| CDC42P1 | 20 | cell division cycle 42 pseudogene 1 [Source:HG |
| ITCH-AS1 | 20 | ITCH antisense RNA 1 [Source:HGNC Symbol;A |
| ITCH-IT1 | 20 | ITCH intronic transcript 1 [Source:HGNC Symb |
| MIR644A | 20 | microRNA 644a [Source:HGNC Symbol;Acc:HG |
| FDX1P1 | 20 | ferredoxin 1 pseudogene 1 [Source:HGNC Sym |
| AL109923.1 | 20 | novel transcript, sense intronic to ITCH |
| DYNLRB1 | 20 | dynein light chain roadblock-type 1 [Source:H |
| Y_RNA | 20 | Y RNA [Source:RFAM;Acc:RF00019] |
| MAP1LC3A | 20 | microtubule associated protein 1 light chain 3 |
| PIGU | 20 | phosphatidylinositol glycan anchor biosynthes |
| AL118520.1 | 20 | cytochrome c oxidase III (MTCO3) pseudogene |
| NCOA6 | 20 | nuclear receptor coactivator 6 [Source:HGNC |
| TP53INP2 | 20 | tumor protein p53 inducible nuclear protein 2 |
| AL109824.1 | 20 | novel transcript, antisense to NCOA6 |
| HMGB3P1 | 20 | high mobility group box 3 pseudogene 1 [Sour |
| GGT7 | 20 | gamma-glutamyltransferase 7 [Source:HGNC S |
| ACSS2 | 20 | acyl-CoA synthetase short chain family memb |
| GSS | 20 | glutathione synthetase [Source:HGNC Symbol |
| MYH7B | 20 | myosin heavy chain 7B [Source:HGNC Symbol; |
| MIR499A | 20 | microRNA 499a [Source:HGNC Symbol;Acc:HG |
| MIR499B | 20 | microRNA 499b [Source:HGNC Symbol;Acc:HG |
| TRPC4AP | 20 | transient receptor potential cation channel su |
| RNU6-407P | 20 | RNA, U6 small nuclear 407, pseudogene [Sour |
| EDEM2 | 20 | ER degradation enhancing alpha-mannosidase |
| AL135844.1 | 20 | NULL |
| PROCR | 20 | protein C receptor [Source:HGNC Symbol;Acc: |

| | | |
|---|---|---|
| AL356652.1 | 20 | novel transcript, antisense to PROCR |
| RNA5SP483 | 20 | RNA, 5S ribosomal pseudogene 483 [Source:H |
| MMP24OS | 20 | MMP24 opposite strand [Source:HGNC Symbo |
| MT1P3 | 20 | metallothionein 1 pseudogene 3 [Source:HGN |
| MMP24 | 20 | matrix metallopeptidase 24 [Source:HGNC Syr |
| AL121753.2 | 20 | novel transcript |
| AL121753.1 | 20 | novel transcript |
| EIF6 | 20 | eukaryotic translation initiation factor 6 [Sour |
| FAM83C-AS1 | 20 | FAM83C antisense RNA 1 [Source:HGNC Symb |
| FAM83C | 20 | family with sequence similarity 83 member C |
| UQCC1 | 20 | ubiquinol-cytochrome c reductase complex as |
| GDF5OS | 20 | growth differentiation factor 5 opposite stran |
| GDF5 | 20 | growth differentiation factor 5 [Source:HGNC |
| MIR1289-1 | 20 | microRNA 1289-1 [Source:HGNC Symbol;Acc:I |
| CEP250 | 20 | centrosomal protein 250 [Source:HGNC Symb |
| FO393401.1 | 20 | novel transcript |
| FO393401.2 | 20 | pseudogene similar to sialyltransferase protei |
| C20orf173 | 20 | chromosome 20 open reading frame 173 [Sou |
| ERGIC3 | 20 | ERGIC and golgi 3 [Source:HGNC Symbol;Acc:H |
| RPL36P4 | 20 | ribosomal protein L36 pseudogene 4 [Source: |
| FER1L4 | 20 | fer-1 like family member 4 (pseudogene) [Sou |
| RPL37P1 | 20 | ribosomal protein L37 pseudogene 1 [Source: |
| SPAG4 | 20 | sperm associated antigen 4 [Source:HGNC Syr |
| CPNE1 | 20 | copine 1 [Source:HGNC Symbol;Acc:HGNC:23 |
| AL109827.1 | 20 | novel protein |
| RN7SKP271 | 20 | RN7SK pseudogene 271 [Source:HGNC Symbo |
| RNU6-759P | 20 | RNA, U6 small nuclear 759, pseudogene [Sour |
| RBM12 | 20 | RNA binding motif protein 12 [Source:HGNC S |
| NFS1 | 20 | NFS1 cysteine desulfurase [Source:HGNC Sym |
| ROMO1 | 20 | reactive oxygen species modulator 1 [Source: |
| RBM39 | 20 | RNA binding motif protein 39 [Source:HGNC S |
| RPF2P1 | 20 | ribosome production factor 2 homolog pseud |
| U6 | 20 | U6 spliceosomal RNA [Source:RFAM;Acc:RF00 |
| PHF20 | 20 | PHD finger protein 20 [Source:HGNC Symbol;A |
| COX7BP2 | 20 | cytochrome c oxidase subunit 7B pseudogene |
| Y_RNA | 20 | Y RNA [Source:RFAM;Acc:RF00019] |
| HIGD1AP16 | 20 | HIG1 hypoxia inducible domain family membe |
| RNU4-40P | 20 | RNA, U4 small nuclear 40, pseudogene [Sourc |
| RNU6-937P | 20 | RNA, U6 small nuclear 937, pseudogene [Sour |
| SCAND1 | 20 | SCAN domain containing 1 [Source:HGNC Sym |
| CNBD2 | 20 | cyclic nucleotide binding domain containing 2 |
| NORAD | 20 | non-coding RNA activated by DNA damage [So |
| AL035420.1 | 20 | novel transcript |
| AL035420.3 | 20 | novel transcript |
| HMGB3P2 | 20 | high mobility group box 3 pseudogene 2 [Sour |
| AL035420.2 | 20 | novel transcript |
| EPB41L1 | 20 | erythrocyte membrane protein band 4.1 like 1 |
| AL121895.1 | 20 | novel transcript, antisense to EPB41L1 |
| AL121895.2 | 20 | novel transcript |
| AAR2 | 20 | AAR2 splicing factor [Source:HGNC Symbol;Ac |

| | | |
|---|---|---|
| DLGAP4 | 20 | DLG associated protein 4 [Source:HGNC Symb |
| DLGAP4-AS1 | 20 | DLGAP4 antisense RNA 1 [Source:HGNC Symb |
| MYL9 | 20 | myosin light chain 9 [Source:HGNC Symbol;Ac |
| Metazoa_SRP | 20 | Metazoan signal recognition particle RNA [Sou |
| TGIF2 | 20 | TGFB induced factor homeobox 2 [Source:HGI |
| TGIF2-RAB5IF | 20 | TGIF2-RAB5IF readthrough [Source:HGNC Sym |
| RAB5IF | 20 | RAB5 interacting factor [Source:HGNC Symbol |
| SLA2 | 20 | Src like adaptor 2 [Source:HGNC Symbol;Acc:I |
| HNRNPA3P2 | 20 | heterogeneous nuclear ribonucleoprotein A3 |
| NDRG3 | 20 | NDRG family member 3 [Source:HGNC Symbo |
| Y_RNA | 20 | Y RNA [Source:RFAM;Acc:RF00019] |
| DSN1 | 20 | DSN1 component of MIS12 kinetochore comp |
| SOGA1 | 20 | suppressor of glucose, autophagy associated : |
| RN7SL156P | 20 | RNA, 7SL, cytoplasmic 156, pseudogene [Sour |
| TLDC2 | 20 | TBC/LysM-associated domain containing 2 [Sc |
| SAMHD1 | 20 | SAM and HD domain containing deoxynucleos |
| RBL1 | 20 | RB transcriptional corepressor like 1 [Source:I |
| RPS3AP3 | 20 | ribosomal protein S3A pseudogene 3 [Source: |
| RPS27AP3 | 20 | ribosomal protein S27a pseudogene 3 [Source |
| AL136172.1 | 20 | novel transcript, antisense to RBL1 |
| MROH8 | 20 | maestro heat like repeat family member 8 [Sc |
| AL031659.1 | 20 | ribosomal protein L13 (RPL13) pseudogene |
| RPN2 | 20 | ribophorin II [Source:HGNC Symbol;Acc:HGNC |
| GHRH | 20 | growth hormone releasing hormone [Source:I |
| MANBAL | 20 | mannosidase beta like [Source:HGNC Symbol; |
| AL034422.1 | 20 | novel transcript |
| SRC | 20 | SRC proto-oncogene, non-receptor tyrosine ki |
| RPL7AP14 | 20 | ribosomal protein L7a pseudogene 14 [Source |
| BLCAP | 20 | BLCAP apoptosis inducing factor [Source:HGN |
| NNAT | 20 | neuronatin [Source:HGNC Symbol;Acc:HGNC: |
| AL109614.1 | 20 | novel transcript, sense intronic to BLCAP |
| PPIAP3 | 20 | peptidylprolyl isomerase A pseudogene 3 [Sou |
| LINC01746 | 20 | long intergenic non-protein coding RNA 1746 |
| GLRXP1 | 20 | glutaredoxin pseudogene 1 [Source:HGNC Syr |
| LINC00489 | 20 | long intergenic non-protein coding RNA 489 [S |
| AL162293.1 | 20 | novel transcript |
| CTNNBL1 | 20 | catenin beta like 1 [Source:HGNC Symbol;Acc: |
| VSTM2L | 20 | V-set and transmembrane domain containing |
| RN7SKP185 | 20 | RN7SK pseudogene 185 [Source:HGNC Symbo |
| TTI1 | 20 | TELO2 interacting protein 1 [Source:HGNC Syr |
| RPRD1B | 20 | regulation of nuclear pre-mRNA domain conta |
| RN7SL237P | 20 | RNA, 7SL, cytoplasmic 237, pseudogene [Sour |
| AL031651.1 | 20 | novel transcript |
| TGM2 | 20 | transglutaminase 2 [Source:HGNC Symbol;Acc |
| KIAA1755 | 20 | KIAA1755 [Source:HGNC Symbol;Acc:HGNC:2! |
| AL031651.2 | 20 | novel transcript, sense intronic to KIAA1755 |
| AL359555.3 | 20 | novel transcript |
| AL359555.4 | 20 | novel transcript |
| AL359555.2 | 20 | novel transcript |
| AL359555.1 | 20 | novel transcript |

| | | |
|---|---|---|
| BPI | 20 | bactericidal permeability increasing protein [S |
| LBP | 20 | lipopolysaccharide binding protein [Source:HG |
| AL391095.2 | 20 | novel transcript |
| AL391095.1 | 20 | novel transcript |
| AL391095.3 | 20 | novel transcript |
| SNHG17 | 20 | small nucleolar RNA host gene 17 [Source:HGI |
| SNORA71 | 20 | Small nucleolar RNA SNORA71 [Source:RFAM; |
| SNORA71B | 20 | small nucleolar RNA, H/ACA box 71B [Source:I |
| SNORA71A | 20 | small nucleolar RNA, H/ACA box 71A [Source:I |
| SNORA71C | 20 | small nucleolar RNA, H/ACA box 71C [Source:I |
| SNORA71D | 20 | small nucleolar RNA, H/ACA box 71D [Source:I |
| SNORA71 | 20 | Small nucleolar RNA SNORA71 [Source:RFAM; |
| SNHG11 | 20 | small nucleolar RNA host gene 11 [Source:HGI |
| SNORA71E | 20 | small nucleolar RNA, H/ACA box 71E [Source:I |
| SNORA60 | 20 | small nucleolar RNA, H/ACA box 60 [Source:H |
| RALGAPB | 20 | Ral GTPase activating protein non-catalytic be |
| MIR548O2 | 20 | microRNA 548o-2 [Source:HGNC Symbol;Acc:I |
| RPS3P2 | 20 | ribosomal protein S3 pseudogene 2 [Source:H |
| ADIG | 20 | adipogenin [Source:HGNC Symbol;Acc:HGNC:. |
| ARHGAP40 | 20 | Rho GTPase activating protein 40 [Source:HGI |
| AL035419.1 | 20 | GINS complex subunit 2 (Psf2 homolog) (GINS |
| Metazoa_SRP | 20 | Metazoan signal recognition particle RNA [Sou |
| SLC32A1 | 20 | solute carrier family 32 member 1 [Source:HG |
| ACTR5 | 20 | actin related protein 5 [Source:HGNC Symbol; |
| RN7SKP173 | 20 | RN7SK pseudogene 173 [Source:HGNC Symbo |
| PPP1R16B | 20 | protein phosphatase 1 regulatory subunit 16B |
| RN7SL116P | 20 | RNA, 7SL, cytoplasmic 116, pseudogene [Sour |
| FAM83D | 20 | family with sequence similarity 83 member D |
| AL023803.2 | 20 | novel transcript |
| AL023803.1 | 20 | novel transcript, antisense to DHX35 |
| DHX35 | 20 | DEAH-box helicase 35 [Source:HGNC Symbol;/ |
| AL023803.3 | 20 | novel transcript, sense intronic to DHX35 |
| NPM1P19 | 20 | nucleophosmin 1 pseudogene 19 [Source:HGI |
| LINC01734 | 20 | long intergenic non-protein coding RNA 1734 |
| AL035251.1 | 20 | novel transcript |
| ATG3P1 | 20 | autophagy related 3 pseudogene 1 [Source:H |
| RN7SL680P | 20 | RNA, 7SL, cytoplasmic 680, pseudogene [Sour |
| AL132981.1 | 20 | novel transcript |
| AL118523.1 | 20 | novel transcript |
| HSPE1P1 | 20 | heat shock protein family E (Hsp10) member : |
| LINC01370 | 20 | long intergenic non-protein coding RNA 1370 |
| AL133419.1 | 20 | novel transcript |
| AL009050.1 | 20 | novel transcript |
| MAFB | 20 | MAF bZIP transcription factor B [Source:HGN( |
| AL035665.1 | 20 | novel transcript |
| AL035665.2 | 20 | novel transcript |
| RNA5SP484 | 20 | RNA, 5S ribosomal pseudogene 484 [Source:H |
| RNU2-52P | 20 | RNA, U2 small nuclear 52, pseudogene [Sourc |
| TOP1 | 20 | DNA topoisomerase I [Source:HGNC Symbol;A |
| PLCG1-AS1 | 20 | PLCG1 antisense RNA 1 [Source:HGNC Symbol |

| | | |
|---|---|---|
| PLCG1 | 20 | phospholipase C gamma 1 [Source:HGNC Sym |
| MIR6871 | 20 | microRNA 6871 [Source:HGNC Symbol;Acc:HG |
| ZHX3 | 20 | zinc fingers and homeoboxes 3 [Source:HGNC |
| AL022394.1 | 20 | major histocompatibility complex, class I (HLA |
| RPL23AP81 | 20 | ribosomal protein L23a pseudogene 81 [Sourc |
| RN7SL615P | 20 | RNA, 7SL, cytoplasmic 615, pseudogene [Sour |
| ADI1P1 | 20 | acireductone dioxygenase 1 pseudogene 1 [Sc |
| LPIN3 | 20 | lipin 3 [Source:HGNC Symbol;Acc:HGNC:1445 |
| EMILIN3 | 20 | elastin microfibril interfacer 3 [Source:HGNC S |
| AL031667.1 | 20 | novel pseudogene |
| CHD6 | 20 | chromodomain helicase DNA binding protein |
| AL031667.2 | 20 | lymphocyte antigen 6 complex, locus H (LY6H) |
| AL031667.3 | 20 | novel transcript, sense intronic to CHD6 |
| RPL12P11 | 20 | ribosomal protein L12 pseudogene 11 [Source |
| RNU6-1018P | 20 | RNA, U6 small nuclear 1018, pseudogene [Sou |
| AL133229.1 | 20 | NULL |
| AL049812.2 | 20 | novel transcript |
| AL049812.1 | 20 | neurofilament heavy polypeptide 200kDa (NE |
| AL049812.3 | 20 | novel transcript |
| PTPRT | 20 | protein tyrosine phosphatase receptor type T |
| AL035666.1 | 20 | pseudogene similar to part of ATP synthase, H |
| AL031656.1 | 20 | novel transcript, antisense to PTPRT |
| AL031676.1 | 20 | novel transcript |
| RN7SKP100 | 20 | RN7SK pseudogene 100 [Source:HGNC Symbo |
| AL021395.1 | 20 | novel transcript |
| RN7SL666P | 20 | RNA, 7SL, cytoplasmic 666, pseudogene [Sour |
| PPIAP21 | 20 | peptidylprolyl isomerase A pseudogene 21 [Sc |
| AL021395.2 | 20 | NULL |
| AL031681.1 | 20 | pseudogene similar to part of nucleotide bind |
| RNU6-743P | 20 | RNA, U6 small nuclear 743, pseudogene [Sour |
| EIF4EBP2P1 | 20 | eukaryotic translation initiation factor 4E bind |
| Y_RNA | 20 | Y RNA [Source:RFAM;Acc:RF00019] |
| SRSF6 | 20 | serine and arginine rich splicing factor 6 [Sour |
| AL031681.2 | 20 | novel protein |
| RNU6-1251P | 20 | RNA, U6 small nuclear 1251, pseudogene [Sou |
| L3MBTL1 | 20 | L3MBTL histone methyl-lysine binding protein |
| Z98752.2 | 20 | transmembrane protein 14C (TMEM14C) pseu |
| Z98752.3 | 20 | novel protein |
| Z98752.1 | 20 | novel transcript |
| SGK2 | 20 | serum/glucocorticoid regulated kinase 2 [Sour |
| Z98752.4 | 20 | novel transcript |
| IFT52 | 20 | intraflagellar transport 52 [Source:HGNC Syml |
| RPL27AP | 20 | ribosomal protein L27a pseudogene [Source:H |
| MYBL2 | 20 | MYB proto-oncogene like 2 [Source:HGNC Syr |
| GTSF1L | 20 | gametocyte specific factor 1 like [Source:HGN |
| RNU6-639P | 20 | RNA, U6 small nuclear 639, pseudogene [Sour |
| LINC01728 | 20 | long intergenic non-protein coding RNA 1728 |
| TOX2 | 20 | TOX high mobility group box family member 2 |
| RN7SL443P | 20 | RNA, 7SL, cytoplasmic 443, pseudogene [Sour |
| JPH2 | 20 | junctophilin 2 [Source:HGNC Symbol;Acc:HGN |

| | | |
|---|---|---|
| AL035447.1 | 20 | novel transcript, antisense to JPH2 |
| OSER1 | 20 | oxidative stress responsive serine rich 1 [Sour |
| OSER1-DT | 20 | OSER1 divergent transcript [Source:HGNC Syn |
| GDAP1L1 | 20 | ganglioside induced differentiation associated |
| FITM2 | 20 | fat storage inducing transmembrane protein 2 |
| R3HDML | 20 | R3H domain containing like [Source:HGNC Syr |
| Y_RNA | 20 | Y RNA [Source:RFAM;Acc:RF00019] |
| AL117382.1 | 20 | novel transcript, antisense to R3HDML |
| HNF4A | 20 | hepatocyte nuclear factor 4 alpha [Source:HG |
| HNF4A-AS1 | 20 | HNF4A antisense RNA 1 [Source:HGNC Symbo |
| AL117382.2 | 20 | novel transcript, antisense to HNF4A |
| MIR3646 | 20 | microRNA 3646 [Source:HGNC Symbol;Acc:HG |
| LINC01620 | 20 | long intergenic non-protein coding RNA 1620 |
| RPL37AP1 | 20 | ribosomal protein L37a pseudogene 1 [Source |
| TTPAL | 20 | alpha tocopherol transfer protein like [Source |
| SERINC3 | 20 | serine incorporator 3 [Source:HGNC Symbol;A |
| PKIG | 20 | cAMP-dependent protein kinase inhibitor gam |
| ADA | 20 | adenosine deaminase [Source:HGNC Symbol;A |
| LINC01260 | 20 | long intergenic non-protein coding RNA 1260 |
| AL139352.1 | 20 | novel transcript |
| KCNK15-AS1 | 20 | KCNK15 and WISP2 antisense RNA 1 [Source:H |
| CCN5 | 20 | cellular communication network factor 5 [Sou |
| KCNK15 | 20 | potassium two pore domain channel subfamil |
| AL118522.1 | 20 | novel transcript, sense intronic to KCNK15 |
| RIMS4 | 20 | regulating synaptic membrane exocytosis 4 [S |
| PGBD4P2 | 20 | piggyBac transposable element derived 4 pseu |
| AL008725.1 | 20 | novel transcript |
| YWHAB | 20 | tyrosine 3-monooxygenase/tryptophan 5-mon |
| PABPC1L | 20 | poly(A) binding protein cytoplasmic 1 like [Sou |
| TOMM34 | 20 | translocase of outer mitochondrial membrane |
| STK4-AS1 | 20 | STK4 antisense RNA 1 (head to head) [Source: |
| STK4 | 20 | serine/threonine kinase 4 [Source:HGNC Symb |
| Y_RNA | 20 | Y RNA [Source:RFAM;Acc:RF00019] |
| KCNS1 | 20 | potassium voltage-gated channel modifier sub |
| WFDC5 | 20 | WAP four-disulfide core domain 5 [Source:HG |
| WFDC12 | 20 | WAP four-disulfide core domain 12 [Source:H |
| AL049767.1 | 20 | WAP four-disulfide core domain 15 family pse |
| PI3 | 20 | peptidase inhibitor 3 [Source:HGNC Symbol;A |
| SEMG1 | 20 | semenogelin 1 [Source:HGNC Symbol;Acc:HGI |
| SEMG2 | 20 | semenogelin 2 [Source:HGNC Symbol;Acc:HGI |
| AL035660.1 | 20 | WAP four-disulfide core domain 15B (Wfdc15 |
| AL035660.2 | 20 | WAP four-disulfide core domain 15A (Wfdc15 |
| SLPI | 20 | secretory leukocyte peptidase inhibitor [Sourc |
| MATN4 | 20 | matrilin 4 [Source:HGNC Symbol;Acc:HGNC:69 |
| RBPJL | 20 | recombination signal binding protein for immu |
| SDC4 | 20 | syndecan 4 [Source:HGNC Symbol;Acc:HGNC:1 |
| AL021578.1 | 20 | novel transcript, sense intronic to SDC4 |
| SYS1 | 20 | SYS1 golgi trafficking protein [Source:HGNC Sy |
| SYS1-DBNDD2 | 20 | SYS1-DBNDD2 readthrough (NMD candidate) |
| TP53TG5 | 20 | TP53 target 5 [Source:HGNC Symbol;Acc:HGN |

| | | |
|---|---|---|
| NDUFB4P10 | 20 | NADH:ubiquinone oxidoreductase subunit B4 |
| DBNDD2 | 20 | dysbindin domain containing 2 [Source:HGNC |
| PIGT | 20 | phosphatidylinositol glycan anchor biosynthes |
| MIR6812 | 20 | microRNA 6812 [Source:HGNC Symbol;Acc:HG |
| AL031663.3 | 20 | novel transcript |
| WFDC2 | 20 | WAP four-disulfide core domain 2 [Source:HG |
| AL031663.2 | 20 | novel transcript |
| RPL5P2 | 20 | ribosomal protein L5 pseudogene 2 [Source:H |
| SPINT3 | 20 | serine peptidase inhibitor, Kunitz type 3 [Sour |
| WFDC6 | 20 | WAP four-disulfide core domain 6 [Source:HG |
| EPPIN-WFDC6 | 20 | EPPIN-WFDC6 readthrough [Source:HGNC Syr |
| AL031663.1 | 20 | cytochrome C oxidase subunit VIc (COX6C) ps |
| HSPD1P21 | 20 | heat shock protein family D (Hsp60) member |
| EPPIN | 20 | epididymal peptidase inhibitor [Source:HGNC |
| WFDC8 | 20 | WAP four-disulfide core domain 8 [Source:HG |
| RNA5SP485 | 20 | RNA, 5S ribosomal pseudogene 485 [Source:H |
| CCNB1IP1P2 | 20 | cyclin B1 interacting protein 1 pseudogene 2 [ |
| WFDC9 | 20 | WAP four-disulfide core domain 9 [Source:HG |
| WFDC10A | 20 | WAP four-disulfide core domain 10A [Source:I |
| RPS2P7 | 20 | ribosomal protein S2 pseudogene 7 [Source:H |
| WFDC11 | 20 | WAP four-disulfide core domain 11 [Source:H |
| WFDC10B | 20 | WAP four-disulfide core domain 10B [Source:I |
| WFDC13 | 20 | WAP four-disulfide core domain 13 [Source:H |
| MIR3617 | 20 | microRNA 3617 [Source:HGNC Symbol;Acc:HG |
| SPINT4 | 20 | serine peptidase inhibitor, Kunitz type 4 [Sour |
| HNRNPA1P3 | 20 | heterogeneous nuclear ribonucleoprotein A1 |
| WFDC3 | 20 | WAP four-disulfide core domain 3 [Source:HG |
| SPINT5P | 20 | serine peptidase inhibitor, Kunitz type 5, pseu |
| RNU6ATAC38P | 20 | RNA, U6atac small nuclear 38, pseudogene [S |
| DNTTIP1 | 20 | deoxynucleotidyltransferase terminal interact |
| UBE2C | 20 | ubiquitin conjugating enzyme E2 C [Source:HG |
| TNNC2 | 20 | troponin C2, fast skeletal type [Source:HGNC S |
| SNX21 | 20 | sorting nexin family member 21 [Source:HGN |
| ACOT8 | 20 | acyl-CoA thioesterase 8 [Source:HGNC Symbo |
| Metazoa_SRP | 20 | Metazoan signal recognition particle RNA [Sou |
| ZSWIM3 | 20 | zinc finger SWIM-type containing 3 [Source:H |
| ZSWIM1 | 20 | zinc finger SWIM-type containing 1 [Source:H |
| SPATA25 | 20 | spermatogenesis associated 25 [Source:HGNC |
| NEURL2 | 20 | neuralized E3 ubiquitin protein ligase 2 [Sourc |
| CTSA | 20 | cathepsin A [Source:HGNC Symbol;Acc:HGNC: |
| AL008726.1 | 20 | novel transcript, antisense to CTSA |
| PLTP | 20 | phospholipid transfer protein [Source:HGNC S |
| AL162458.1 | 20 | novel transcript |
| PCIF1 | 20 | PDX1 C-terminal inhibiting factor 1 [Source:H |
| ZNF335 | 20 | zinc finger protein 335 [Source:HGNC Symbol; |
| FTLP1 | 20 | ferritin light chain pseudogene 1 [Source:HGN |
| MMP9 | 20 | matrix metallopeptidase 9 [Source:HGNC Sym |
| SLC12A5-AS1 | 20 | SLC12A5 and MMP9 antisense RNA 1 [Source: |
| SLC12A5 | 20 | solute carrier family 12 member 5 [Source:HG |
| NCOA5 | 20 | nuclear receptor coactivator 5 [Source:HGNC |

| | | |
|---|---|---|
| RPL13P2 | 20 | ribosomal protein L13 pseudogene 2 [Source: |
| CD40 | 20 | CD40 molecule [Source:HGNC Symbol;Acc:HG |
| AL031687.1 | 20 | novel transcript |
| CDH22 | 20 | cadherin 22 [Source:HGNC Symbol;Acc:HGNC: |
| SLC35C2 | 20 | solute carrier family 35 member C2 [Source:H |
| AL133227.1 | 20 | novel transcript, antisense to ELMO2 |
| ELMO2 | 20 | engulfment and cell motility 2 [Source:HGNC S |
| ZNF663P | 20 | zinc finger protein 663, pseudogene [Source:H |
| AL031686.1 | 20 | novel transcript |
| MKRN7P | 20 | makorin ring finger protein 7, pseudogene [So |
| ZNF840P | 20 | zinc finger protein 840, pseudogene [Source:H |
| ZNF334 | 20 | zinc finger protein 334 [Source:HGNC Symbol; |
| OCSTAMP | 20 | osteoclast stimulatory transmembrane protei |
| SLC13A3 | 20 | solute carrier family 13 member 3 [Source:HG |
| AL133520.1 | 20 | novel transcript, sense intronic to SLC13A3 |
| TP53RK | 20 | TP53 regulating kinase [Source:HGNC Symbol; |
| AL031055.1 | 20 | novel transcript, antisense to TP53RK |
| SLC2A10 | 20 | solute carrier family 2 member 10 [Source:HG |
| RN7SKP33 | 20 | RN7SK pseudogene 33 [Source:HGNC Symbol; |
| AL354766.1 | 20 | ribosomal protein L13 (RPL13) pseudogene |
| EYA2 | 20 | EYA transcriptional coactivator and phosphata |
| AL354766.2 | 20 | novel transcript |
| GAPDHP54 | 20 | glyceraldehyde 3 phosphate dehydrogenase p |
| AL121776.1 | 20 | ribosomal protein L27A (RPL27A) pseudogene |
| AL022342.1 | 20 | ribosomal protein S2 (RPS2) pseudogene |
| MIR3616 | 20 | microRNA 3616 [Source:HGNC Symbol;Acc:HG |
| ZMYND8 | 20 | zinc finger MYND-type containing 8 [Source:H |
| AL031666.2 | 20 | novel transcript, antisense to ZMYND8 |
| AL031666.3 | 20 | novel transcript, sense intronic to ZMYND8 |
| AL031666.1 | 20 | novel transcript, antisense to ZMYND8 |
| LINC01754 | 20 | long intergenic non-protein coding RNA 1754 |
| RPL35AP | 20 | ribosomal protein L35a pseudogene [Source:H |
| RNU6-497P | 20 | RNA, U6 small nuclear 497, pseudogene [Sour |
| RNU6-563P | 20 | RNA, U6 small nuclear 563, pseudogene [Sour |
| NCOA3 | 20 | nuclear receptor coactivator 3 [Source:HGNC |
| AL034418.1 | 20 | ribosomal protein S3A (RPS3A) pseudogene |
| SULF2 | 20 | sulfatase 2 [Source:HGNC Symbol;Acc:HGNC:2 |
| AL354813.1 | 20 | novel transcript, antisense to SULF2 |
| RNU7-173P | 20 | RNA, U7 small nuclear 173 pseudogene [Sourc |
| SRMP1 | 20 | spermidine synthase pseudogene 1 [Source:H |
| RNA5SP486 | 20 | RNA, 5S ribosomal pseudogene 486 [Source:H |
| RNU7-92P | 20 | RNA, U7 small nuclear 92 pseudogene [Source |
| AL357558.2 | 20 | novel transcript |
| AL357558.1 | 20 | novel transcript |
| AL357558.3 | 20 | novel transcript |
| LINC01522 | 20 | long intergenic non-protein coding RNA 1522 |
| LINC01523 | 20 | long intergenic non-protein coding RNA 1523 |
| AL139351.2 | 20 | novel transcript |
| AL139351.3 | 20 | novel transcript |
| AL139351.1 | 20 | novel transcript |

| | | |
|---|---|---|
| AL136102.1 | 20 | novel transcript |
| AL121888.2 | 20 | novel transcript |
| AL121888.1 | 20 | novel transcript |
| LINC00494 | 20 | long intergenic non-protein coding RNA 494 [S |
| AL137078.1 | 20 | novel transcript |
| AL137078.2 | 20 | novel transcript |
| AL049541.1 | 20 | novel transcript |
| AL049541.2 | 20 | novel transcript |
| RNU7-144P | 20 | RNA, U7 small nuclear 144 pseudogene [Sourc |
| PREX1 | 20 | phosphatidylinositol-3,4,5-trisphosphate depe |
| AL035106.1 | 20 | novel transcript, antisense to PREX1 |
| AL133342.1 | 20 | novel transcript |
| ARFGEF2 | 20 | ADP ribosylation factor guanine nucleotide ex |
| SNAP23P1 | 20 | synaptosome associated protein 23 pseudoge |
| CSE1L-AS1 | 20 | CSE1L antisense RNA 1 [Source:HGNC Symbol, |
| CSE1L | 20 | chromosome segregation 1 like [Source:HGNC |
| STAU1 | 20 | staufen double-stranded RNA binding protein |
| ARPC3P1 | 20 | actin related protein 2/3 complex subunit 3 ps |
| DDX27 | 20 | DEAD-box helicase 27 [Source:HGNC Symbol;A |
| ZNFX1 | 20 | zinc finger NFX1-type containing 1 [Source:HG |
| ZFAS1 | 20 | ZNFX1 antisense RNA 1 [Source:HGNC Symbol |
| AL049766.2 | 20 | NULL |
| SNORD12C | 20 | small nucleolar RNA, C/D box 12C [Source:HG |
| AL049766.1 | 20 | NULL |
| SNORD12B | 20 | small nucleolar RNA, C/D box 12B [Source:HG |
| AL049766.3 | 20 | NULL |
| SNORD12 | 20 | small nucleolar RNA, C/D box 12 [Source:HGN |
| KCNB1 | 20 | potassium voltage-gated channel subfamily B |
| AL035685.1 | 20 | somatomedin B and thrombospondin type 1 c |
| PTGIS | 20 | prostaglandin I2 synthase [Source:HGNC Symb |
| B4GALT5 | 20 | beta-1,4-galactosyltransferase 5 [Source:HGN |
| RN7SL197P | 20 | RNA, 7SL, cytoplasmic 197, pseudogene [Sour |
| SNRPFP1 | 20 | small nuclear ribonucleoprotein polypeptide F |
| RNU6-919P | 20 | RNA, U6 small nuclear 919, pseudogene [Sour |
| SLC9A8 | 20 | solute carrier family 9 member A8 [Source:HG |
| AL162615.1 | 20 | cytokine-like nuclear factor n-pac (N-PAC) pse |
| SPATA2 | 20 | spermatogenesis associated 2 [Source:HGNC S |
| Y_RNA | 20 | Y RNA [Source:RFAM;Acc:RF00019] |
| Metazoa_SRP | 20 | Metazoan signal recognition particle RNA [Sou |
| RNF114 | 20 | ring finger protein 114 [Source:HGNC Symbol; |
| KRT18P4 | 20 | keratin 18 pseudogene 4 [Source:HGNC Symb |
| RNU6-147P | 20 | RNA, U6 small nuclear 147, pseudogene [Sour |
| SNAI1 | 20 | snail family transcriptional repressor 1 [Sourc |
| TRERNA1 | 20 | translation regulatory long non-coding RNA 1 |
| UBE2V1 | 20 | ubiquitin conjugating enzyme E2 V1 [Source:H |
| TMEM189-UBE2V1 | 20 | TMEM189-UBE2V1 readthrough [Source:HGN |
| TMEM189 | 20 | transmembrane protein 189 [Source:HGNC Sy |
| LINC01275 | 20 | long intergenic non-protein coding RNA 1275 |
| AL161937.1 | 20 | novel transcript |
| LINC01273 | 20 | long intergenic non-protein coding RNA 1273 |

| | | |
|---|---|---|
| CEBPB-AS1 | 20 | CEBPB antisense RNA 1 [Source:HGNC Symbol |
| CEBPB | 20 | CCAAT enhancer binding protein beta [Source |
| SMIM25 | 20 | small integral membrane protein 25 [Source:H |
| LINC01270 | 20 | long intergenic non-protein coding RNA 1270 |
| LINC01271 | 20 | long intergenic non-protein coding RNA 1271 |
| RN7SL636P | 20 | RNA, 7SL, cytoplasmic 636, pseudogene [Sour |
| COX6CP2 | 20 | cytochrome c oxidase subunit 6C pseudogene |
| PTPN1 | 20 | protein tyrosine phosphatase non-receptor ty |
| RN7SL672P | 20 | RNA, 7SL, cytoplasmic 672, pseudogene [Sour |
| Y_RNA | 20 | Y RNA [Source:RFAM;Acc:RF00019] |
| AL133230.1 | 20 | novel transcript, antisense to PTPN1 |
| MIR645 | 20 | microRNA 645 [Source:HGNC Symbol;Acc:HGI |
| RIPOR3 | 20 | RIPOR family member 3 [Source:HGNC Symbo |
| MIR1302-5 | 20 | microRNA 1302-5 [Source:HGNC Symbol;Acc:I |
| RPL36P2 | 20 | ribosomal protein L36 pseudogene 2 [Source: |
| AL353653.1 | 20 | novel transcript |
| PARD6B | 20 | par-6 family cell polarity regulator beta [Sourc |
| BCAS4 | 20 | breast carcinoma amplified sequence 4 [Sourc |
| TMSB4XP6 | 20 | TMSB4X pseudogene 6 [Source:HGNC Symbol |
| ADNP | 20 | activity dependent neuroprotector homeobox |
| PSMD10P1 | 20 | proteasome 26S subunit, non-ATPase, 10 pseu |
| ADNP-AS1 | 20 | ADNP antisense RNA 1 [Source:HGNC Symbol; |
| DPM1 | 20 | dolichyl-phosphate mannosyltransferase subu |
| MOCS3 | 20 | molybdenum cofactor synthesis 3 [Source:HG |
| AL050404.1 | 20 | novel transcript |
| KCNG1 | 20 | potassium voltage-gated channel modifier sub |
| AL121785.1 | 20 | TEC |
| RPSAP1 | 20 | ribosomal protein SA pseudogene 1 [Source:H |
| NFATC2 | 20 | nuclear factor of activated T cells 2 [Source:H( |
| MIR3194 | 20 | microRNA 3194 [Source:HGNC Symbol;Acc:HC |
| ATP9A | 20 | ATPase phospholipid transporting 9A (putativ |
| AL138807.1 | 20 | RPL29 (ribosomal protein L29) pseudogene |
| SALL4 | 20 | spalt like transcription factor 4 [Source:HGNC |
| LINC01429 | 20 | long intergenic non-protein coding RNA 1429 |
| RNU6-347P | 20 | RNA, U6 small nuclear 347, pseudogene [Sour |
| RNU7-6P | 20 | RNA, U7 small nuclear 6 pseudogene [Source: |
| ZFP64 | 20 | ZFP64 zinc finger protein [Source:HGNC Symb |
| AL109984.1 | 20 | novel transcript, antisense to ZFP64 |
| ERP29P1 | 20 | endoplasmic reticulum lumenal protein 29 pse |
| LINC01524 | 20 | long intergenic non-protein coding RNA 1524 |
| AL109610.1 | 20 | novel transcript |
| AL049765.2 | 20 | NULL |
| AL049765.1 | 20 | novel transcript |
| MRPS33P4 | 20 | mitochondrial ribosomal protein S33 pseudog |
| AL031674.1 | 20 | novel transcript |
| AL161945.1 | 20 | novel transcript |
| AL121902.1 | 20 | novel transcript |
| RPL36P1 | 20 | ribosomal protein L36 pseudogene 1 [Source: |
| TSHZ2 | 20 | teashirt zinc finger homeobox 2 [Source:HGN( |
| RN7SKP184 | 20 | RN7SK pseudogene 184 [Source:HGNC Symbo |

| | | |
|---|---|---|
| AL391097.3 | 20 | novel transcript, antisense to TSHZ2 |
| AL391097.1 | 20 | novel transcript, antisense to TSHZ2 |
| AL391097.2 | 20 | novel transcript, antisense to TSHZ2 |
| AL109930.1 | 20 | novel transcript, antisense to TSHZ2 |
| AL354993.1 | 20 | novel transcript, antisense to TSHZ2 |
| PPIAP10 | 20 | peptidylprolyl isomerase A pseudogene 10 [Sc |
| AL354993.2 | 20 | novel transcript, antisense to TSHZ2 |
| Y_RNA | 20 | Y RNA [Source:RFAM;Acc:RF00019] |
| AL157838.1 | 20 | novel transcript, antisense to ZNF217 |
| ZNF217 | 20 | zinc finger protein 217 [Source:HGNC Symbol; |
| AC005808.1 | 20 | novel transcript |
| RNU7-14P | 20 | RNA, U7 small nuclear 14 pseudogene [Source |
| AC005914.1 | 20 | novel transcript |
| AC006076.1 | 20 | novel transcript |
| SUMO1P1 | 20 | SUMO1 pseudogene 1 [Source:HGNC Symbol; |
| BCAS1 | 20 | breast carcinoma amplified sequence 1 [Sourc |
| AC005220.1 | 20 | novel transcript |
| MIR4756 | 20 | microRNA 4756 [Source:HGNC Symbol;Acc:HG |
| CYP24A1 | 20 | cytochrome P450 family 24 subfamily A meml |
| AL133335.2 | 20 | novel transcript |
| PFDN4 | 20 | prefoldin subunit 4 [Source:HGNC Symbol;Acc |
| AL133335.1 | 20 | novel transcript |
| DOK5 | 20 | docking protein 5 [Source:HGNC Symbol;Acc:H |
| RNU4ATAC7P | 20 | RNA, U4atac small nuclear 7, pseudogene [Sou |
| RPL12P4 | 20 | ribosomal protein L12 pseudogene 4 [Source: |
| LINC01440 | 20 | long intergenic non-protein coding RNA 1440 |
| LINC01441 | 20 | long intergenic non-protein coding RNA 1441 |
| AL160413.1 | 20 | novel transcript |
| AL109829.1 | 20 | novel transcript |
| AL109828.1 | 20 | novel transcript |
| CBLN4 | 20 | cerebellin 4 precursor [Source:HGNC Symbol;A |
| RNA5SP487 | 20 | RNA, 5S ribosomal pseudogene 487 [Source:H |
| MC3R | 20 | melanocortin 3 receptor [Source:HGNC Symbo |
| AL139824.2 | 20 | zinc finger protein pseudogene |
| AL139824.1 | 20 | novel transcript |
| FAM210B | 20 | family with sequence similarity 210 member E |
| AURKA | 20 | aurora kinase A [Source:HGNC Symbol;Acc:HG |
| CSTF1 | 20 | cleavage stimulation factor subunit 1 [Source: |
| CASS4 | 20 | Cas scaffold protein family member 4 [Source |
| RPL39P | 20 | ribosomal protein L39 pseudogene [Source:H( |
| RTF2 | 20 | replication termination factor 2 [Source:HGN( |
| GCNT7 | 20 | glucosaminyl (N-acetyl) transferase family me |
| FAM209A | 20 | family with sequence similarity 209 member A |
| AL109806.1 | 20 | TEC |
| FAM209B | 20 | family with sequence similarity 209 member E |
| AL109806.2 | 20 | novel transcript |
| LINC01716 | 20 | long intergenic non-protein coding RNA 1716 |
| RPS4XP3 | 20 | ribosomal protein S4X pseudogene 3 [Source: |
| TFAP2C | 20 | transcription factor AP-2 gamma [Source:HGN |
| RNU6-1146P | 20 | RNA, U6 small nuclear 1146, pseudogene [Sou |

| | | |
|---|---|---|
| RN7SL170P | 20 | RNA, 7SL, cytoplasmic 170, pseudogene [Sour |
| PTMAP6 | 20 | prothymosin alpha pseudogene 6 [Source:HGI |
| AL133232.1 | 20 | novel transcript |
| RNU6-929P | 20 | RNA, U6 small nuclear 929, pseudogene [Sour |
| AL157414.2 | 20 | novel transcript |
| AL157414.1 | 20 | novel transcript |
| AL157414.4 | 20 | novel transcript |
| AL157414.3 | 20 | TEC |
| BMP7 | 20 | bone morphogenetic protein 7 [Source:HGNC |
| BMP7-AS1 | 20 | BMP7 antisense RNA 1 [Source:HGNC Symbol; |
| AL122058.1 | 20 | novel transcript |
| Y_RNA | 20 | Y RNA [Source:RFAM;Acc:RF00019] |
| MIR4325 | 20 | microRNA 4325 [Source:HGNC Symbol;Acc:HG |
| RPL39P39 | 20 | ribosomal protein L39 pseudogene 39 [Source |
| SPO11 | 20 | SPO11 initiator of meiotic double stranded br |
| RAE1 | 20 | ribonucleic acid export 1 [Source:HGNC Symb |
| MTND1P9 | 20 | MT-ND1 pseudogene 9 [Source:HGNC Symbol |
| MTRNR2L3 | 20 | MT-RNR2 like 3 [Source:HGNC Symbol;Acc:HG |
| AL109955.1 | 20 | novel transcript |
| RBM38 | 20 | RNA binding motif protein 38 [Source:HGNC S |
| AL109955.2 | 20 | novel transcript |
| HMGB1P1 | 20 | high mobility group box 1 pseudogene 1 [Sour |
| CTCFL | 20 | CCCTC-binding factor like [Source:HGNC Symb |
| PCK1 | 20 | phosphoenolpyruvate carboxykinase 1 [Sourc |
| AL035541.1 | 20 | novel transcript |
| ZBP1 | 20 | Z-DNA binding protein 1 [Source:HGNC Symbo |
| PMEPA1 | 20 | prostate transmembrane protein, androgen ir |
| NKILA | 20 | NF-kappaB interacting lncRNA [Source:HGNC S |
| AL162291.1 | 20 | NULL |
| AL354984.1 | 20 | novel transcript |
| LINC01742 | 20 | long intergenic non-protein coding RNA 1742 |
| AL354984.2 | 20 | novel transcript |
| AL354984.3 | 20 | heat shock 60kDa protein 1 (chaperonin) (HSP |
| C20orf85 | 20 | chromosome 20 open reading frame 85 [Sour |
| ANKRD60 | 20 | ankyrin repeat domain 60 [Source:HGNC Syml |
| PPP4R1L | 20 | protein phosphatase 4 regulatory subunit 1 lik |
| RAB22A | 20 | RAB22A, member RAS oncogene family [Sourc |
| VAPB | 20 | VAMP associated protein B and C [Source:HGI |
| APCDD1L | 20 | APC down-regulated 1 like [Source:HGNC Sym |
| APCDD1L-DT | 20 | APCDD1L divergent transcript [Source:HGNC S |
| AL050327.1 | 20 | novel transcript |
| LINC01711 | 20 | long intergenic non-protein coding RNA 1711 |
| STX16 | 20 | syntaxin 16 [Source:HGNC Symbol;Acc:HGNC: |
| STX16-NPEPL1 | 20 | STX16-NPEPL1 readthrough (NMD candidate) |
| NPEPL1 | 20 | aminopeptidase like 1 [Source:HGNC Symbol;/ |
| AL139349.1 | 20 | novel transcript |
| PIEZO1P2 | 20 | piezo type mechanosensitive ion channel com |
| AL132655.3 | 20 | novel transcript |
| AL132655.2 | 20 | novel transcript |
| MIR296 | 20 | microRNA 296 [Source:HGNC Symbol;Acc:HGI |

| | | |
|---|---|---|
| MIR298 | 20 | microRNA 298 [Source:HGNC Symbol;Acc:HGN |
| GNAS-AS1 | 20 | GNAS antisense RNA 1 [Source:HGNC Symbol; |
| AL132655.1 | 20 | novel transcript |
| GNAS | 20 | GNAS complex locus [Source:HGNC Symbol;Ac |
| AL132655.7 | 20 | NULL |
| AL132655.6 | 20 | NULL |
| AL132655.4 | 20 | NULL |
| AL132655.5 | 20 | NULL |
| AL121917.1 | 20 | novel transcript, antisense to GNAS |
| AL121917.2 | 20 | novel transcript, antisense to GNAS |
| AL109840.2 | 20 | novel transcript |
| AL109840.1 | 20 | novel pseudogene |
| NELFCD | 20 | negative elongation factor complex member C |
| CTSZ | 20 | cathepsin Z [Source:HGNC Symbol;Acc:HGNC: |
| TUBB1 | 20 | tubulin beta 1 class VI [Source:HGNC Symbol;A |
| ATP5F1E | 20 | ATP synthase F1 subunit epsilon [Source:HGN |
| PRELID3B | 20 | PRELI domain containing 3B [Source:HGNC Sy |
| MRPS16P2 | 20 | mitochondrial ribosomal protein S16 pseudog |
| ZNF831 | 20 | zinc finger protein 831 [Source:HGNC Symbol; |
| EDN3 | 20 | endothelin 3 [Source:HGNC Symbol;Acc:HGNC |
| AL035250.1 | 20 | novel transcript |
| AL035250.2 | 20 | novel transcript |
| AL160410.1 | 20 | novel transcript |
| PIEZO1P1 | 20 | piezo type mechanosensitive ion channel com |
| AL389889.2 | 20 | novel transcript |
| AL389889.1 | 20 | novel transcript |
| PHACTR3 | 20 | phosphatase and actin regulator 3 [Source:HG |
| AL121908.1 | 20 | novel transcript |
| RNU7-141P | 20 | RNA, U7 small nuclear 141 pseudogene [Sourc |
| SYCP2 | 20 | synaptonemal complex protein 2 [Source:HGN |
| FAM217B | 20 | family with sequence similarity 217 member B |
| PPP1R3D | 20 | protein phosphatase 1 regulatory subunit 3D [ |
| CDH26 | 20 | cadherin 26 [Source:HGNC Symbol;Acc:HGNC: |
| C20orf197 | 20 | chromosome 20 open reading frame 197 [Sou |
| AL132822.1 | 20 | novel transcript |
| MIR646HG | 20 | MIR646 host gene [Source:HGNC Symbol;Acc: |
| MIR646 | 20 | microRNA 646 [Source:HGNC Symbol;Acc:HGN |
| MTCO2P1 | 20 | MT-CO2 pseudogene 1 [Source:HGNC Symbol |
| MIR4533 | 20 | microRNA 4533 [Source:HGNC Symbol;Acc:HG |
| MIR548AG2 | 20 | microRNA 548ag-2 [Source:HGNC Symbol;Acc |
| AL117372.1 | 20 | novel transcript |
| AL139348.1 | 20 | novel transcript |
| AL121910.1 | 20 | novel transcript |
| LINC01718 | 20 | long intergenic non-protein coding RNA 1718 |
| CDH4 | 20 | cadherin 4 [Source:HGNC Symbol;Acc:HGNC:1 |
| AL365229.1 | 20 | novel transcript |
| BX640515.1 | 20 | novel transcript |
| AL160412.1 | 20 | novel transcript |
| AL162457.1 | 20 | novel transcript, antisense to CDH4 |
| AL162457.2 | 20 | novel transcript, antisense to CDH4 |

| | | |
|---|---|---|
| TAF4 | 20 | TATA-box binding protein associated factor 4 |
| MIR1257 | 20 | microRNA 1257 [Source:HGNC Symbol;Acc:HG |
| AL137077.1 | 20 | ribosomal protein L17 (RPL17) pseudogene |
| MIR3195 | 20 | microRNA 3195 [Source:HGNC Symbol;Acc:HG |
| AL137077.2 | 20 | novel transcript |
| LSM14B | 20 | LSM family member 14B [Source:HGNC Symb |
| PSMA7 | 20 | proteasome subunit alpha 7 [Source:HGNC Sy |
| SS18L1 | 20 | SS18L1 subunit of BAF chromatin remodeling |
| MTG2 | 20 | mitochondrial ribosome associated GTPase 2 |
| HRH3 | 20 | histamine receptor H3 [Source:HGNC Symbol; |
| OSBPL2 | 20 | oxysterol binding protein like 2 [Source:HGNC |
| ADRM1 | 20 | adhesion regulating molecule 1 [Source:HGNC |
| AL354836.1 | 20 | novel transcript |
| LAMA5 | 20 | laminin subunit alpha 5 [Source:HGNC Symbol |
| MIR4758 | 20 | microRNA 4758 [Source:HGNC Symbol;Acc:HG |
| LAMA5-AS1 | 20 | LAMA5 antisense RNA 1 [Source:HGNC Symbo |
| AL121832.2 | 20 | novel transcript, antisense to RPS21 |
| RPS21 | 20 | ribosomal protein S21 [Source:HGNC Symbol; |
| CABLES2 | 20 | Cdk5 and Abl enzyme substrate 2 [Source:HGI |
| AL121832.3 | 20 | novel transcript, sense intronic to CABLES2 |
| RBBP8NL | 20 | RBBP8 N-terminal like [Source:HGNC Symbol; |
| AL121832.1 | 20 | novel transcript, antisense to RBBP8NL |
| GATA5 | 20 | GATA binding protein 5 [Source:HGNC Symbol |
| AL499627.2 | 20 | novel transcript |
| AL499627.1 | 20 | novel transcript |
| MIR1-1HG-AS1 | 20 | MIR1-1HG antisense RNA 1 [Source:HGNC Syn |
| MIR1-1HG | 20 | MIR1-1 host gene [Source:HGNC Symbol;Acc:I |
| MIR1-1 | 20 | microRNA 1-1 [Source:HGNC Symbol;Acc:HGN |
| MIR133A2 | 20 | microRNA 133a-2 [Source:HGNC Symbol;Acc:I |
| RPL7P3 | 20 | ribosomal protein L7 pseudogene 3 [Source:H |
| BX640514.1 | 20 | novel transcript |
| BX640514.2 | 20 | novel transcript |
| AL450469.2 | 20 | novel transcript |
| AL450469.1 | 20 | novel transcript |
| AL357033.1 | 20 | novel transcript, antisense to SLCO4A1 |
| SLCO4A1 | 20 | solute carrier organic anion transporter family |
| AL357033.4 | 20 | novel transcript, antisense to SLCO4A1 |
| AL357033.3 | 20 | novel transcript, antisense to SLCO4A1 |
| SLCO4A1-AS1 | 20 | SLCO4A1 antisense RNA 1 [Source:HGNC Sym |
| LINC00686 | 20 | long intergenic non-protein coding RNA 686 [S |
| NTSR1 | 20 | neurotensin receptor 1 [Source:HGNC Symbol |
| AL357033.2 | 20 | novel transcript |
| LINC00659 | 20 | long intergenic non-protein coding RNA 659 [S |
| MRGBP | 20 | MRG domain binding protein [Source:HGNC S |
| OGFR-AS1 | 20 | OGFR antisense RNA 1 [Source:HGNC Symbol; |
| OGFR | 20 | opioid growth factor receptor [Source:HGNC S |
| COL9A3 | 20 | collagen type IX alpha 3 chain [Source:HGNC S |
| TCFL5 | 20 | transcription factor like 5 [Source:HGNC Symb |
| DPH3P1 | 20 | diphthamide biosynthesis 3 pseudogene 1 [So |
| ARF4P2 | 20 | ADP ribosylation factor 4 pseudogene 2 [Sour |

| | | |
|---|---|---|
| DIDO1 | 20 | death inducer-obliterator 1 [Source:HGNC Syr |
| AL117379.1 | 20 | novel transcript |
| GID8 | 20 | GID complex subunit 8 homolog [Source:HGN |
| SLC17A9 | 20 | solute carrier family 17 member 9 [Source:HG |
| BHLHE23 | 20 | basic helix-loop-helix family member e23 [Sou |
| LINC01749 | 20 | long intergenic non-protein coding RNA 1749 |
| LINC00029 | 20 | long intergenic non-protein coding RNA 29 [Sc |
| LINC01056 | 20 | long intergenic non-protein coding RNA 1056 |
| HAR1B | 20 | highly accelerated region 1B [Source:HGNC Sy |
| AL096828.4 | 20 | NULL |
| HAR1A | 20 | highly accelerated region 1A [Source:HGNC Sy |
| AL096828.6 | 20 | novel transcript |
| AL096828.2 | 20 | novel transcript |
| AL096828.5 | 20 | novel transcript |
| AL096828.1 | 20 | novel transcript |
| MIR124-3 | 20 | microRNA 124-3 [Source:HGNC Symbol;Acc:H |
| YTHDF1 | 20 | YTH N6-methyladenosine RNA binding proteir |
| AL096828.3 | 20 | novel transcript |
| BIRC7 | 20 | baculoviral IAP repeat containing 7 [Source:H |
| MIR3196 | 20 | microRNA 3196 [Source:HGNC Symbol;Acc:HG |
| NKAIN4 | 20 | sodium/potassium transporting ATPase intera |
| AL121827.2 | 20 | novel transcript, antisense to NKAIN4 |
| ARFGAP1 | 20 | ADP ribosylation factor GTPase activating prot |
| MIR4326 | 20 | microRNA 4326 [Source:HGNC Symbol;Acc:HG |
| COL20A1 | 20 | collagen type XX alpha 1 chain [Source:HGNC |
| RNU6-994P | 20 | RNA, U6 small nuclear 994, pseudogene [Sour |
| CHRNA4 | 20 | cholinergic receptor nicotinic alpha 4 subunit |
| AL121827.1 | 20 | novel transcript, antisense to CHRNA4 |
| KCNQ2 | 20 | potassium voltage-gated channel subfamily Q |
| AL353658.2 | 20 | novel transcript, antisense to KCNQ2 |
| AL353658.1 | 20 | novel transcript |
| EEF1A2 | 20 | eukaryotic translation elongation factor 1 alpl |
| AL121829.1 | 20 | novel transcript |
| PPDPF | 20 | pancreatic progenitor cell differentiation and |
| PTK6 | 20 | protein tyrosine kinase 6 [Source:HGNC Symb |
| SRMS | 20 | src-related kinase lacking C-terminal regulator |
| AL121829.2 | 20 | novel transcript, antisense to SRMS |
| FNDC11 | 20 | fibronectin type III domain containing 11 [Sou |
| HELZ2 | 20 | helicase with zinc finger 2 [Source:HGNC Syml |
| GMEB2 | 20 | glucocorticoid modulatory element binding pr |
| MHENCR | 20 | melanoma highly expressed competing endog |
| STMN3 | 20 | stathmin 3 [Source:HGNC Symbol;Acc:HGNC:1 |
| RTEL1 | 20 | regulator of telomere elongation helicase 1 [S |
| RTEL1-TNFRSF6B | 20 | RTEL1-TNFRSF6B readthrough (NMD candidat |
| TNFRSF6B | 20 | TNF receptor superfamily member 6b [Source |
| ARFRP1 | 20 | ADP ribosylation factor related protein 1 [Sou |
| ZGPAT | 20 | zinc finger CCCH-type and G-patch domain co |
| AL121845.3 | 20 | novel protein, ZGPAT-LIME1 readthrough |
| AL121845.4 | 20 | novel transcript, antisense to ZGPAT |
| LIME1 | 20 | Lck interacting transmembrane adaptor 1 [Sou |

| | | |
|---|---|---|
| AL121845.2 | 20 | novel transcript, LIME1-SLC2A4RG readthroug |
| SLC2A4RG | 20 | SLC2A4 regulator [Source:HGNC Symbol;Acc:H |
| ZBTB46 | 20 | zinc finger and BTB domain containing 46 [Sou |
| AL121845.1 | 20 | novel transcript |
| ZBTB46-AS1 | 20 | ZBTB46 antisense RNA 1 [Source:HGNC Symbo |
| C20orf181 | 20 | chromosome 20 open reading frame 181 [Sou |
| AL118506.1 | 20 | novel transcript, antisense to ABHD16B |
| ABHD16B | 20 | abhydrolase domain containing 16B [Source:H |
| TPD52L2 | 20 | TPD52 like 2 [Source:HGNC Symbol;Acc:HGNC |
| DNAJC5 | 20 | DnaJ heat shock protein family (Hsp40) memb |
| RNU1-134P | 20 | RNA, U1 small nuclear 134, pseudogene [Sour |
| MIR941-1 | 20 | microRNA 941-1 [Source:HGNC Symbol;Acc:H |
| MIR941-2 | 20 | microRNA 941-2 [Source:HGNC Symbol;Acc:H |
| MIR941-3 | 20 | microRNA 941-3 [Source:HGNC Symbol;Acc:H |
| MIR941-4 | 20 | microRNA 941-4 [Source:HGNC Symbol;Acc:H |
| MIR941-5 | 20 | microRNA 941-5 [Source:HGNC Symbol;Acc:H |
| UCKL1 | 20 | uridine-cytidine kinase 1 like 1 [Source:HGNC |
| MIR1914 | 20 | microRNA 1914 [Source:HGNC Symbol;Acc:HG |
| MIR647 | 20 | microRNA 647 [Source:HGNC Symbol;Acc:HGN |
| UCKL1-AS1 | 20 | UCKL1 antisense RNA 1 [Source:HGNC Symbol |
| ZNF512B | 20 | zinc finger protein 512B [Source:HGNC Symbo |
| SAMD10 | 20 | sterile alpha motif domain containing 10 [Sou |
| PRPF6 | 20 | pre-mRNA processing factor 6 [Source:HGNC S |
| C20orf204 | 20 | chromosome 20 open reading frame 204 [Sou |
| SOX18 | 20 | SRY-box transcription factor 18 [Source:HGNC |
| TCEA2 | 20 | transcription elongation factor A2 [Source:HG |
| AL355803.1 | 20 | novel transcript |
| RGS19 | 20 | regulator of G protein signaling 19 [Source:HG |
| MIR6813 | 20 | microRNA 6813 [Source:HGNC Symbol;Acc:HG |
| OPRL1 | 20 | opioid related nociceptin receptor 1 [Source:H |
| LKAAEAR1 | 20 | LKAAEAR motif containing 1 [Source:HGNC Sy |
| AL121581.2 | 20 | novel transcript, antisense to OPRL1 |
| MYT1 | 20 | myelin transcription factor 1 [Source:HGNC Sy |
| NPBWR2 | 20 | neuropeptides B and W receptor 2 [Source:HG |
| PCMTD2 | 20 | protein-L-isoaspartate (D-aspartate) O-methyl |
| AL137028.2 | 20 | interleukin 9 receptor (IL9R) pseudogene |
| CICP4 | 20 | capicua transcriptional repressor pseudogene |
| LINC00266-1 | 20 | long intergenic non-protein coding RNA 266-1 |
| AL137028.1 | 20 | IQ motif and Sec7 domain 3 (IQSEC3) pseudog |
| FP565260.4 | 21 | novel protein, similar to DNA (cytosine-5-)-me |
| FP565260.3 | 21 | ICOS ligand [Source:NCBI gene;Acc:10272399 |
| FP565260.5 | 21 | novel transcript, antisense to CH507-9B2.3 |
| GATD3B | 21 | glutamine amidotransferase like class 1 doma |
| FP565260.2 | 21 | novel protein |
| FP565260.1 | 21 | periodic tryptophan protein 2 homolog [Sourc |
| FP565260.6 | 21 | novel protein, similar to trafficking protein pa |
| AC079801.1 | 21 | novel transcript |
| LINC01670 | 21 | long intergenic non-protein coding RNA 1670 |
| CU633967.1 | 21 | uncharacterized LOC102724219 [Source:NCBI |
| Y_RNA | 21 | Y RNA [Source:RFAM;Acc:RF00019] |

| | | |
|---|---|---|
| FP236315.2 | 21 | TEC |
| FP236315.3 | 21 | novel transcript |
| FP236315.1 | 21 | novel transcript |
| CU639417.1 | 21 | histone H2B type F-S-like [Source:NCBI gene;A |
| CU639417.3 | 21 | heat shock transcription factor 2 binding prot |
| CU639417.2 | 21 | TEC |
| LINC01669 | 21 | long intergenic non-protein coding RNA 1669 |
| CU639417.4 | 21 | novel transcript |
| CU639417.5 | 21 | uncharacterized CH507-42P11.6 [Source:NCBI |
| SIK1B | 21 | salt inducible kinase 1B (putative) [Source:HG |
| CU633906.5 | 21 | novel transcript |
| Y_RNA | 21 | Y RNA [Source:RFAM;Acc:RF00019] |
| CU633906.1 | 21 | uncharacterized LOC102724701 [Source:NCBI |
| CU633906.4 | 21 | clustered mitochondria (cluA/CLU1) homolog |
| CU633906.3 | 21 | clustered mitochondria (cluA/CLU1) homolog |
| CU633906.2 | 21 | novel transcript |
| Y_RNA | 21 | Y RNA [Source:RFAM;Acc:RF00019] |
| CBSL | 21 | cystathionine-beta-synthase like [Source:HGN |
| U2AF1L5 | 21 | U2 small nuclear RNA auxiliary factor 1 like 5 [ |
| FP236240.3 | 21 | mitochondrial ribosomal protein L51 (MRPL51 |
| FP236240.2 | 21 | TEC |
| FP236240.1 | 21 | novel transcript |
| CRYAA2 | 21 | crystallin alpha A2 [Source:HGNC Symbol;Acc: |
| CU638689.4 | 21 | novel transcript |
| CU638689.1 | 21 | novel transcript |
| CU638689.3 | 21 | clustered mitochondria (cluA/CLU1) homolog |
| CU638689.2 | 21 | clustered mitochondria (cluA/CLU1) homolog |
| CU638689.5 | 21 | novel transcript |
| FP475955.3 | 21 | novel transcript |
| FP475955.1 | 21 | novel transcript |
| MIR8069-1 | 21 | microRNA 8069-1 [Source:HGNC Symbol;Acc:I |
| FP475955.4 | 21 | fem-1 homolog a (C. elegans) (FEM1A) pseudo |
| FP475955.2 | 21 | novel transcript |
| CU634019.5 | 21 | novel transcript |
| CU634019.4 | 21 | clustered mitochondria (cluA/CLU1) homolog |
| CU634019.2 | 21 | novel transcript |
| CU634019.1 | 21 | uncharacterized LOC102724843 [Source:NCBI |
| Y_RNA | 21 | Y RNA [Source:RFAM;Acc:RF00019] |
| CU634019.6 | 21 | C-terminal binding protein 2 (Ctbp2) pseudog |
| CU634019.3 | 21 | TEC |
| CU633904.2 | 21 | clustered mitochondria (cluA/CLU1) homolog |
| CU633904.3 | 21 | clustered mitochondria (cluA/CLU1) homolog |
| CU633904.1 | 21 | uncharacterized LOC102724951 [Source:NCBI |
| Y_RNA | 21 | Y RNA [Source:RFAM;Acc:RF00019] |
| FP236241.2 | 21 | C-terminal binding protein 2 (CTBP2) pseudog |
| Y_RNA | 21 | Y RNA [Source:RFAM;Acc:RF00019] |
| FP236241.1 | 21 | uncharacterized LOC102723360 [Source:NCBI |
| SMIM11B | 21 | small integral membrane protein 11B [Source |
| FAM243B | 21 | family with sequence similarity 243 member B |
| SMIM34B | 21 | small integral membrane protein 34B [Source |

| | | |
|---|---|---|
| KCNE1B | 21 | potassium voltage-gated channel subfamily E |
| CU633980.1 | 21 | NULL |
| FP671120.5 | 21 | TEC |
| FP671120.4 | 21 | novel transcript |
| MIR6724-1 | 21 | microRNA 6724-1 [Source:HGNC Symbol;Acc:I |
| FP671120.1 | 21 | NULL |
| MIR3648-1 | 21 | microRNA 3648-1 [Source:HGNC Symbol;Acc:I |
| FP671120.2 | 21 | NULL |
| FP671120.6 | 21 | novel transcript, similar to YY1 associated myo |
| RNA5-8SN2 | 21 | RNA, 5.8S ribosomal N2 [Source:HGNC Symbo |
| FP671120.10 | 21 | TEC |
| FP671120.11 | 21 | TEC |
| FP671120.9 | 21 | TEC |
| FP671120.8 | 21 | TEC |
| MIR6724-2 | 21 | microRNA 6724-2 [Source:HGNC Symbol;Acc:I |
| FP671120.3 | 21 | NULL |
| FP671120.7 | 21 | novel transcript, similar to YY1 associated myo |
| 5_8S_rRNA | 21 | 5.8S ribosomal RNA [Source:RFAM;Acc:RF000 |
| FP236383.3 | 21 | novel transcript |
| MIR6724-3 | 21 | microRNA 6724-3 [Source:HGNC Symbol;Acc:I |
| FP236383.1 | 21 | NULL |
| FP236383.4 | 21 | novel transcript, similar to YY1 associated myo |
| RNA5-8SN3 | 21 | RNA, 5.8S ribosomal N3 [Source:HGNC Symbo |
| FP236383.13 | 21 | TEC |
| FP236383.9 | 21 | TEC |
| FP236383.7 | 21 | TEC |
| FP236383.12 | 21 | TEC |
| MIR6724-4 | 21 | microRNA 6724-4 [Source:HGNC Symbol;Acc:I |
| FP236383.2 | 21 | NULL |
| FP236383.5 | 21 | novel transcript, similar to YY1 associated myo |
| RNA5-8SN1 | 21 | RNA, 5.8S ribosomal N1 [Source:HGNC Symbo |
| FP236383.11 | 21 | TEC |
| FP236383.6 | 21 | TEC |
| FP236383.8 | 21 | TEC |
| FP236383.10 | 21 | TEC |
| RPSAP68 | 21 | ribosomal protein SA pseudogene 68 [Source: |
| CR381572.2 | 21 | tripartite motif containing 43 (TRIM43) pseud |
| CR381572.1 | 21 | N-acetylated alpha-linked acidic dipeptidase 2 |
| LINC01666 | 21 | long intergenic non-protein coding RNA 1666 |
| CR381670.2 | 21 | novel transcript |
| SNX18P10 | 21 | sorting nexin 18 pseudogene 10 [Source:HGN |
| MTCO1P1 | 21 | MT-CO1 pseudogene 1 [Source:HGNC Symbol |
| CR381670.1 | 21 | ankyrin repeat domain 20 family, member A3 |
| MIR3648-2 | 21 | microRNA 3648-2 [Source:HGNC Symbol;Acc:I |
| CR392039.1 | 21 | NULL |
| CR392039.4 | 21 | novel transcript |
| CR392039.3 | 21 | chromosome 2 open reading frame 27A (C2or |
| CR392039.6 | 21 | asparagine synthetase (glutamine-hydrolyzing |
| TEKT4P2 | 21 | tektin 4 pseudogene 2 [Source:HGNC Symbol; |
| CR392039.5 | 21 | sosondowah ankyrin repeat domain family me |

| | | |
|---|---|---|
| CR392039.2 | 21 | ankyrin repeat domain 20 family, member A2 |
| CR381653.1 | 21 | novel transcript |
| SNX18P12 | 21 | sorting nexin 18 pseudogene 12 [Source:HGN |
| CR381653.2 | 21 | ankyrin repeat domain 18A (ANKRD18A) pseu |
| AC124864.2 | 21 | nuclear receptor corepressor 1 (NCOR1) pseu |
| AC124864.1 | 21 | family with sequence similarity 182, member |
| AC124864.3 | 21 | novel transcript |
| U1 | 21 | U1 spliceosomal RNA [Source:RFAM;Acc:RF00 |
| LINC01667 | 21 | long intergenic non-protein coding RNA 1667 |
| AC018692.2 | 21 | family with sequence similarity 27, member D |
| RN7SL52P | 21 | RNA, 7SL, cytoplasmic 52, pseudogene [Sourc |
| SNORA70 | 21 | Small nucleolar RNA SNORA70 [Source:RFAM; |
| AC018692.1 | 21 | novel transcript |
| CR382285.1 | 21 | ribosomal protein L7a (RPL7A) pseudogene |
| CR382285.2 | 21 | TEC |
| CR382287.2 | 21 | novel transcript |
| CR382287.1 | 21 | chromosome 2 open reading frame 27A (C2or |
| AP003900.1 | 21 | novel transcript |
| EIF3FP1 | 21 | eukaryotic translation initiation factor 3 subur |
| VN1R7P | 21 | vomeronasal 1 receptor 7 pseudogene [Sourc |
| AF254983.2 | 21 | tubulin tyrosine ligase-like family, member 1 ( |
| BAGE2 | 21 | BAGE family member 2 [Source:HGNC Symbol |
| AF254983.1 | 21 | novel transcript |
| TPTE | 21 | transmembrane phosphatase with tensin hom |
| CYCSP41 | 21 | CYCS pseudogene 41 [Source:HGNC Symbol;A |
| SLC25A15P4 | 21 | solute carrier family 25 member 15 pseudoge |
| AF254982.1 | 21 | pseudogene similar to part of immunoglobulir |
| IGHV1OR21-1 | 21 | immunoglobulin heavy variable 1/OR21-1 (no |
| AP001464.1 | 21 | novel transcript |
| ANKRD30BP2 | 21 | ankyrin repeat domain 30B pseudogene 2 [So |
| RNU6-614P | 21 | RNA, U6 small nuclear 614, pseudogene [Sour |
| ZNF355P | 21 | zinc finger protein 355, pseudogene [Source:H |
| AJ239321.1 | 21 | BCL2/adenovirus E1B 19kDa interacting prote |
| AJ239318.1 | 21 | novel transcript |
| FGF7P2 | 21 | fibroblast growth factor 7 pseudogene 2 [Sour |
| ANKRD30BP1 | 21 | ankyrin repeat domain 30B pseudogene 1 [So |
| MIR3156-3 | 21 | microRNA 3156-3 [Source:HGNC Symbol;Acc:I |
| GTF2IP2 | 21 | general transcription factor IIi pseudogene 2 [ |
| VN1R8P | 21 | vomeronasal 1 receptor 8 pseudogene [Sourc |
| LINC01674 | 21 | long intergenic non-protein coding RNA 1674 |
| SNX19P1 | 21 | sorting nexin 19 pseudogene 1 [Source:HGNC |
| OR4K11P | 21 | olfactory receptor family 4 subfamily K memb |
| OR4K12P | 21 | olfactory receptor family 4 subfamily K memb |
| POTED | 21 | POTE ankyrin domain family member D [Sourc |
| RNU6-286P | 21 | RNA, U6 small nuclear 286, pseudogene [Sour |
| MIR3118-1 | 21 | microRNA 3118-1 [Source:HGNC Symbol;Acc:I |
| AL050303.3 | 21 | novel pseudogene |
| GRAMD4P1 | 21 | GRAM domain containing 4 pseudogene 1 [So |
| CXADRP1 | 21 | CXADR pseudogene 1 [Source:HGNC Symbol;A |
| AL050303.4 | 21 | TEC |

| | | |
|---|---|---|
| AL050303.2 | 21 | LON peptidase N-terminal domain and ring fir |
| MIR8069-2 | 21 | microRNA 8069-2 [Source:HGNC Symbol;Acc:I |
| FEM1AP1 | 21 | fem-1 homolog A pseudogene 1 [Source:HGN( |
| AL050303.1 | 21 | novel transcript |
| TERF1P1 | 21 | telomeric repeat binding factor 1 pseudogene |
| FAM207CP | 21 | family with sequence similarity 207 member ( |
| GXYLT1P2 | 21 | glucoside xylosyltransferase 1 pseudogene 2 [ |
| CNN2P7 | 21 | calponin 2 pseudogene 7 [Source:HGNC Symb |
| ZNF114P1 | 21 | zinc finger protein 114 pseudogene 1 [Source: |
| CYP4F29P | 21 | cytochrome P450 family 4 subfamily F membe |
| SNX18P13 | 21 | sorting nexin 18 pseudogene 13 [Source:HGN( |
| ANKRD20A11P | 21 | ankyrin repeat domain 20 family member A11 |
| RHOT1P2 | 21 | ras homolog family member T1 pseudogene 2 |
| RNU6-954P | 21 | RNA, U6 small nuclear 954, pseudogene [Sour |
| NF1P3 | 21 | neurofibromin 1 pseudogene 3 [Source:HGNC |
| PPP6R2P1 | 21 | protein phosphatase 6 regulatory subunit 2 ps |
| FRG2MP | 21 | FSHD region gene 2 family member M, pseud( |
| AP001347.1 | 21 | novel transcript, antisense to LIPI |
| ANKRD20A18P | 21 | ankyrin repeat domain 20 family member A18 |
| RNA5SP488 | 21 | RNA, 5S ribosomal pseudogene 488 [Source:H |
| AP001347.2 | 21 | NULL |
| LIPI | 21 | lipase I [Source:HGNC Symbol;Acc:HGNC:1882 |
| ERLEC1P1 | 21 | endoplasmic reticulum lectin 1 pseudogene 1 |
| RBM11 | 21 | RNA binding motif protein 11 [Source:HGNC S |
| ABCC13 | 21 | ATP binding cassette subfamily C member 13 |
| HSPA13 | 21 | heat shock protein family A (Hsp70) member : |
| SAMSN1 | 21 | SAM domain, SH3 domain and nuclear localiza |
| SAMSN1-AS1 | 21 | SAMSN1 antisense RNA 1 [Source:HGNC Symb |
| AF127936.2 | 21 | novel transcript |
| POLR2CP1 | 21 | RNA polymerase II subunit C pseudogene 1 [S( |
| AF127936.1 | 21 | novel transcript |
| GAPDHP16 | 21 | glyceraldehyde-3-phosphate dehydrogenase p |
| AF127577.4 | 21 | novel transcript |
| LINC02246 | 21 | long intergenic non-protein coding RNA 2246 |
| RBMX2P1 | 21 | RBMX2 pseudogene 1 [Source:HGNC Symbol;/ |
| AF127577.3 | 21 | novel transcript |
| NRIP1 | 21 | nuclear receptor interacting protein 1 [Source |
| AF127577.1 | 21 | novel transcript |
| AF127577.2 | 21 | novel transcript |
| AF127577.6 | 21 | TEC |
| AF127577.5 | 21 | novel transcript |
| AF246928.1 | 21 | TEC |
| AJ009632.2 | 21 | novel transcript |
| CYCSP42 | 21 | CYCS pseudogene 42 [Source:HGNC Symbol;A |
| AJ009632.1 | 21 | novel transcript |
| RNU6-1326P | 21 | RNA, U6 small nuclear 1326, pseudogene [Sou |
| RAD23BP3 | 21 | RAD23B pseudogene 3 [Source:HGNC Symbol, |
| USP25 | 21 | ubiquitin specific peptidase 25 [Source:HGNC |
| RBPMSLP | 21 | RNA binding protein with multiple splicing-like |
| MIR99AHG | 21 | mir-99a-let-7c cluster host gene [Source:HGN( |

| | | |
|---|---|---|
| RNU6-426P | 21 | RNA, U6 small nuclear 426, pseudogene [Sour |
| VDAC2P1 | 21 | voltage dependent anion channel 2 pseudoge |
| RPS26P5 | 21 | ribosomal protein S26 pseudogene 5 [Source: |
| SNORD74B | 21 | small nucleolar RNA, C/D box 74B [Source:HG |
| AP001172.1 | 21 | novel transcript |
| AP001172.2 | 21 | novel transcript |
| AP000962.3 | 21 | novel transcript |
| AP000962.2 | 21 | TEC |
| MIR99A | 21 | microRNA 99a [Source:HGNC Symbol;Acc:HGI |
| MIRLET7C | 21 | microRNA let-7c [Source:HGNC Symbol;Acc:H |
| AP000962.1 | 21 | novel transcript |
| MIR125B2 | 21 | microRNA 125b-2 [Source:HGNC Symbol;Acc:I |
| RNU1-98P | 21 | RNA, U1 small nuclear 98, pseudogene [Sourc |
| AF130359.1 | 21 | novel transcript |
| AF212831.1 | 21 | novel transcript |
| NEK4P1 | 21 | NIMA-related kinase 4 pseudogene 1 [Source: |
| AP000457.1 | 21 | G protein-coupled receptor 157 (GPR157) pse |
| RNU6-113P | 21 | RNA, U6 small nuclear 113, pseudogene [Sour |
| LINC01549 | 21 | long intergenic non-protein coding RNA 1549 |
| RPL39P40 | 21 | ribosomal protein L39 pseudogene 40 [Source |
| RN7SL163P | 21 | RNA, 7SL, cytoplasmic 163, pseudogene [Sour |
| CXADR | 21 | CXADR Ig-like cell adhesion molecule [Source: |
| BTF3L4P1 | 21 | basic transcription factor 3 like 4 pseudogene |
| Y_RNA | 21 | Y RNA [Source:RFAM;Acc:RF00019] |
| Y_RNA | 21 | Y RNA [Source:RFAM;Acc:RF00019] |
| BTG3 | 21 | BTG anti-proliferation factor 3 [Source:HGNC |
| BTG3-AS1 | 21 | BTG3 antisense RNA 1 [Source:HGNC Symbol; |
| AP000432.1 | 21 | novel transcript |
| AP000432.2 | 21 | TEC |
| C21orf91-OT1 | 21 | C21orf91 overlapping transcript 1 [Source:HG |
| C21orf91 | 21 | chromosome 21 open reading frame 91 [Sour |
| AL109761.1 | 21 | novel transcript, antisense to C21orf91 |
| CHODL-AS1 | 21 | CHODL antisense RNA 1 [Source:HGNC Symbo |
| RPL37P3 | 21 | ribosomal protein L37 pseudogene 3 [Source: |
| CHODL | 21 | chondrolectin [Source:HGNC Symbol;Acc:HGN |
| AF130417.1 | 21 | novel transcript |
| TMPRSS15 | 21 | transmembrane serine protease 15 [Source:H |
| AL109763.1 | 21 | novel transcript |
| MIR548XHG | 21 | MIR548X host gene [Source:HGNC Symbol;Ac |
| AF240627.1 | 21 | novel transcript |
| MIR548X | 21 | microRNA 548x [Source:HGNC Symbol;Acc:HG |
| PPIAP22 | 21 | peptidylprolyl isomerase A pseudogene 22 [Sc |
| AL157359.3 | 21 | TEC |
| AL157359.2 | 21 | novel transcript |
| AL157359.1 | 21 | novel transcript |
| AP000431.2 | 21 | novel transcript |
| AP000431.1 | 21 | novel transcript |
| SLC6A6P1 | 21 | solute carrier family 6 member 6 pseudogene |
| AP000855.1 | 21 | novel transcript |
| RNU1-139P | 21 | RNA, U1 small nuclear 139, pseudogene [Sour |

| | | |
|---|---|---|
| RPL37P4 | 21 | ribosomal protein L37 pseudogene 4 [Source: |
| AP000403.1 | 21 | non imprinted in Prader-Willi/Angelman syndr |
| AP000568.1 | 21 | novel pseudogene |
| C1QBPP1 | 21 | complement C1q binding protein pseudogene |
| LINC01683 | 21 | long intergenic non-protein coding RNA 1683 |
| LINC02573 | 21 | long intergenic non-protein coding RNA 2573 |
| RNU6-772P | 21 | RNA, U6 small nuclear 772, pseudogene [Sour |
| RN7SKP147 | 21 | RN7SK pseudogene 147 [Source:HGNC Symbo |
| FDPSP6 | 21 | farnesyl diphosphate synthase pseudogene 6 |
| KRT18P2 | 21 | keratin 18 pseudogene 2 [Source:HGNC Symb |
| RPS3AP1 | 21 | ribosomal protein S3A pseudogene 1 [Source: |
| AP001506.1 | 21 | zinc finger protein 799 (ZNF799) pseudogene |
| LINC00320 | 21 | long intergenic non-protein coding RNA 320 [S |
| PPIAP1 | 21 | peptidylprolyl isomerase A pseudogene 1 [Sou |
| NCAM2 | 21 | neural cell adhesion molecule 2 [Source:HGN( |
| AP001136.1 | 21 | novel transcript |
| AP001117.1 | 21 | novel transcript |
| AF241725.1 | 21 | novel transcript |
| LINC00317 | 21 | long intergenic non-protein coding RNA 317 [S |
| LINC01425 | 21 | long intergenic non-protein coding RNA 1425 |
| AP000472.1 | 21 | novel transcript |
| LINC01687 | 21 | long intergenic non-protein coding RNA 1687 |
| LINC00308 | 21 | long intergenic non-protein coding RNA 308 [S |
| AP000705.2 | 21 | phosphoglycerate kinase 1 (PGK1) pseudogen |
| AP000705.1 | 21 | IBR domain containing 3 (IBRDC3) pseudogen |
| RNU4-45P | 21 | RNA, U4 small nuclear 45, pseudogene [Sourc |
| AP000561.1 | 21 | novel transcript |
| MAPK6P2 | 21 | mitogen-activated protein kinase 6 pseudoger |
| AP000959.1 | 21 | novel transcript |
| AP000949.1 | 21 | glomulin, FKBP associated protein (GLMN) pse |
| AP001116.1 | 21 | novel transcript |
| MIR6130 | 21 | microRNA 6130 [Source:HGNC Symbol;Acc:HG |
| ZNF299P | 21 | zinc finger protein 299, pseudogene [Source:H |
| MSANTD2P1 | 21 | Myb/SANT DNA binding domain containing 2 p |
| AP001255.1 | 21 | novel transcript |
| RNU2-55P | 21 | RNA, U2 small nuclear 55, pseudogene [Sourc |
| AP000459.2 | 21 | novel transcript |
| EEF1A1P1 | 21 | eukaryotic translation elongation factor 1 alpl |
| TUBAP | 21 | tubulin alpha pseudogene [Source:HGNC Sym |
| Y_RNA | 21 | Y RNA [Source:RFAM;Acc:RF00019] |
| AP000459.1 | 21 | novel transcript |
| AP000961.1 | 21 | novel transcript |
| AP000474.2 | 21 | novel transcript |
| AP000474.1 | 21 | novel transcript |
| AP000477.2 | 21 | novel transcript |
| AP000477.3 | 21 | novel transcript |
| AP000477.1 | 21 | novel transcript |
| AP000470.1 | 21 | novel transcript |
| LINC01689 | 21 | long intergenic non-protein coding RNA 1689 |
| LINC01684 | 21 | long intergenic non-protein coding RNA 1684 |

| | | |
|---|---|---|
| LINC01692 | 21 | long intergenic non-protein coding RNA 1692 |
| AP000402.1 | 21 | novel transcript |
| AP000146.1 | 21 | novel transcript |
| AP001342.1 | 21 | CTAGE family, member 4 (CTAGE4) pseudoger |
| AP000235.1 | 21 | novel transcript |
| AP000233.2 | 21 | novel transcript |
| RN7SKP236 | 21 | RN7SK pseudogene 236 [Source:HGNC Symbo |
| AP000233.1 | 21 | novel transcript |
| AP001341.1 | 21 | novel transcript |
| RNA5SP489 | 21 | RNA, 5S ribosomal pseudogene 489 [Source:H |
| RPL13AP7 | 21 | ribosomal protein L13a pseudogene 7 [Source |
| AP001340.1 | 21 | TEC |
| LINC00158 | 21 | long intergenic non-protein coding RNA 158 [S |
| AP000221.1 | 21 | novel transcript |
| MIR155HG | 21 | MIR155 host gene [Source:HGNC Symbol;Acc: |
| MIR155 | 21 | microRNA 155 [Source:HGNC Symbol;Acc:HGI |
| AP000223.1 | 21 | novel transcript, antisense to MRPL39 |
| MRPL39 | 21 | mitochondrial ribosomal protein L39 [Source:I |
| JAM2 | 21 | junctional adhesion molecule 2 [Source:HGNC |
| RNGTTP1 | 21 | RNA guanylyltransferase and 5'-phosphatase p |
| AP000224.1 | 21 | novel transcript, antisense to JAM2 |
| FDX1P2 | 21 | ferredoxin 1 pseudogene 2 [Source:HGNC Syn |
| ATP5PF | 21 | ATP synthase peripheral stalk subunit F6 [Sour |
| GABPA | 21 | GA binding protein transcription factor subuni |
| LLPHP2 | 21 | LLP homolog, long-term synaptic facilitation p |
| APP | 21 | amyloid beta precursor protein [Source:HGNC |
| AP001442.1 | 21 | novel transcript |
| RNU6-123P | 21 | RNA, U6 small nuclear 123, pseudogene [Sour |
| AP001439.1 | 21 | novel transcript |
| AP000229.1 | 21 | novel transcript |
| RNU6-926P | 21 | RNA, U6 small nuclear 926, pseudogene [Sour |
| AP001595.2 | 21 | novel transcript |
| AP001595.1 | 21 | novel transcript |
| AP001596.1 | 21 | novel transcript |
| CYYR1-AS1 | 21 | cysteine and tyrosine rich 1 antisense RNA 1 [S |
| AP001596.2 | 21 | novel transcript |
| CYYR1 | 21 | cysteine and tyrosine rich 1 [Source:HGNC Syr |
| ADAMTS1 | 21 | ADAM metallopeptidase with thrombospondi |
| AP001599.1 | 21 | novel transcript |
| ADAMTS5 | 21 | ADAM metallopeptidase with thrombospondi |
| MIR4759 | 21 | microRNA 4759 [Source:HGNC Symbol;Acc:HG |
| GPX1P2 | 21 | glutathione peroxidase pseudogene 2 [Source |
| AP001605.1 | 21 | novel transcript |
| AP001604.1 | 21 | novel transcript |
| EIF4A1P1 | 21 | eukaryotic translation initiation factor 4A1 pse |
| RPL10P1 | 21 | ribosomal protein L10 pseudogene 1 [Source:I |
| NCSTNP1 | 21 | nicastrin pseudogene 1 [Source:HGNC Symbol |
| LINC01673 | 21 | long intergenic non-protein coding RNA 1673 |
| LINC00113 | 21 | long intergenic non-protein coding RNA 113 [S |
| AJ006995.1 | 21 | novel transcript |

| | | |
|---|---|---|
| LINC00314 | 21 | long intergenic non-protein coding RNA 314 [S |
| LINC01697 | 21 | long intergenic non-protein coding RNA 1697 |
| LINC01695 | 21 | long intergenic non-protein coding RNA 1695 |
| AF165147.1 | 21 | novel transcript |
| LINC00161 | 21 | long intergenic non-protein coding RNA 161 [S |
| U6 | 21 | U6 spliceosomal RNA [Source:RFAM;Acc:RF00 |
| N6AMT1 | 21 | N-6 adenine-specific DNA methyltransferase 1 |
| HSPD1P7 | 21 | heat shock protein family D (Hsp60) member |
| THUMPD1P1 | 21 | THUMP domain containing 1 pseudogene 1 [S |
| LTN1 | 21 | listerin E3 ubiquitin protein ligase 1 [Source:H |
| RPL23P2 | 21 | ribosomal protein L23 pseudogene 2 [Source: |
| RWDD2B | 21 | RWD domain containing 2B [Source:HGNC Syr |
| AF129075.2 | 21 | novel transcript, antisense to USP16 |
| USP16 | 21 | ubiquitin specific peptidase 16 [Source:HGNC |
| CCT8 | 21 | chaperonin containing TCP1 subunit 8 [Source |
| AF129075.1 | 21 | novel transcript, sense intronic to CCT8 |
| AF129075.3 | 21 | novel transcript, antisense to CCT8 |
| MAP3K7CL | 21 | MAP3K7 C-terminal like [Source:HGNC Symbo |
| RPL12P9 | 21 | ribosomal protein L12 pseudogene 9 [Source: |
| Y_RNA | 21 | Y RNA [Source:RFAM;Acc:RF00019] |
| U3 | 21 | Small nucleolar RNA U3 [Source:RFAM;Acc:RF |
| AF124730.1 | 21 | novel transcript, antisense to BACH1 |
| LINC00189 | 21 | long intergenic non-protein coding RNA 189 [S |
| BACH1 | 21 | BTB domain and CNC homolog 1 [Source:HGN |
| GAPDHP14 | 21 | glyceraldehyde-3-phosphate dehydrogenase p |
| BACH1-IT1 | 21 | BACH1 intronic transcript 1 [Source:HGNC Syr |
| AP000240.1 | 21 | novel transcript, sense intronic BACH1 |
| BACH1-AS1 | 21 | BACH1 antisense RNA 1 [Source:HGNC Symbo |
| BACH1-IT2 | 21 | BACH1 intronic transcript 2 [Source:HGNC Syr |
| BACH1-IT3 | 21 | BACH1 intronic transcript 3 [Source:HGNC Syr |
| GRIK1 | 21 | glutamate ionotropic receptor kainate type su |
| AP000238.1 | 21 | novel transcript, antisense to GRIK1 |
| GRIK1-AS1 | 21 | GRIK1 antisense RNA 1 [Source:HGNC Symbol |
| AF096876.1 | 21 | novel transcript |
| CLDN17 | 21 | claudin 17 [Source:HGNC Symbol;Acc:HGNC:2 |
| LINC00307 | 21 | long intergenic non-protein coding RNA 307 [S |
| CLDN8 | 21 | claudin 8 [Source:HGNC Symbol;Acc:HGNC:20 |
| RPL8P2 | 21 | ribosomal protein L8 pseudogene 2 [Source:H |
| KRTAP24-1 | 21 | keratin associated protein 24-1 [Source:HGNC |
| KRTAP25-1 | 21 | keratin associated protein 25-1 [Source:HGNC |
| KRTAP26-1 | 21 | keratin associated protein 26-1 [Source:HGNC |
| KRTAP27-1 | 21 | keratin associated protein 27-1 [Source:HGNC |
| KRTAP23-1 | 21 | keratin associated protein 23-1 [Source:HGNC |
| KRTAP13-6P | 21 | keratin associated protein 13-6, pseudogene [ |
| KRTAP13-2 | 21 | keratin associated protein 13-2 [Source:HGNC |
| MIR4327 | 21 | microRNA 4327 [Source:HGNC Symbol;Acc:HG |
| KRTAP13-1 | 21 | keratin associated protein 13-1 [Source:HGNC |
| KRTAP13-3 | 21 | keratin associated protein 13-3 [Source:HGNC |
| KRTAP13-4 | 21 | keratin associated protein 13-4 [Source:HGNC |
| KRTAP13-5P | 21 | keratin associated protein 13-5, pseudogene [ |

| | | |
|---|---|---|
| KRTAP15-1 | 21 | keratin associated protein 15-1 [Source:HGNC |
| KRTAP19-1 | 21 | keratin associated protein 19-1 [Source:HGNC |
| KRTAP19-2 | 21 | keratin associated protein 19-2 [Source:HGNC |
| KRTAP19-3 | 21 | keratin associated protein 19-3 [Source:HGNC |
| KRTAP19-4 | 21 | keratin associated protein 19-4 [Source:HGNC |
| KRTAP19-5 | 21 | keratin associated protein 19-5 [Source:HGNC |
| KRTAP19-9P | 21 | keratin associated protein 19-9, pseudogene [ |
| KRTAP19-10P | 21 | keratin associated protein 19-10, pseudogene |
| AP000567.1 | 21 | novel keratin associated protein (KRTAP) fami |
| KRTAP19-11P | 21 | keratin associated protein 19-11, pseudogene |
| KRTAP19-6 | 21 | keratin associated protein 19-6 [Source:HGNC |
| KRTAP19-7 | 21 | keratin associated protein 19-7 [Source:HGNC |
| KRTAP22-2 | 21 | keratin associated protein 22-2 [Source:HGNC |
| KRTAP6-3 | 21 | keratin associated protein 6-3 [Source:HGNC S |
| KRTAP6-2 | 21 | keratin associated protein 6-2 [Source:HGNC S |
| KRTAP22-1 | 21 | keratin associated protein 22-1 [Source:HGNC |
| KRTAP6-1 | 21 | keratin associated protein 6-1 [Source:HGNC S |
| KRTAP20-1 | 21 | keratin associated protein 20-1 [Source:HGNC |
| KRTAP20-4 | 21 | keratin associated protein 20-4 [Source:HGNC |
| KRTAP20-2 | 21 | keratin associated protein 20-2 [Source:HGNC |
| KRTAP20-3 | 21 | keratin associated protein 20-3 [Source:HGNC |
| KRTAP21-3 | 21 | keratin associated protein 21-3 [Source:HGNC |
| KRTAP21-4P | 21 | keratin associated protein 21-4, pseudogene [ |
| KRTAP21-2 | 21 | keratin associated protein 21-2 [Source:HGNC |
| KRTAP21-1 | 21 | keratin associated protein 21-1 [Source:HGNC |
| AP000244.1 | 21 | novel keratin associated protein 21 (KRTAP21 |
| KRTAP8-2P | 21 | keratin associated protein 8-2, pseudogene [S |
| KRTAP8-3P | 21 | keratin associated protein 8-3, pseudogene [S |
| KRTAP8-1 | 21 | keratin associated protein 8-1 [Source:HGNC S |
| KRTAP7-1 | 21 | keratin associated protein 7-1 (gene/pseudog |
| KRTAP11-1 | 21 | keratin associated protein 11-1 [Source:HGNC |
| KRTAP19-8 | 21 | keratin associated protein 19-8 [Source:HGNC |
| UBE3AP2 | 21 | ubiquitin protein ligase E3A pseudogene 2 [So |
| TIAM1 | 21 | T cell lymphoma invasion and metastasis 1 [So |
| AP000248.1 | 21 | novel transcript, antisense to TIAM1 |
| AP000563.1 | 21 | chromosome 6 open reading frame 203 (C6or |
| AP000251.1 | 21 | uncharacterized LOC150051 [Source:NCBI ger |
| FBXW11P1 | 21 | F-box and WD repeat domain containing 11 ps |
| AP000253.1 | 21 | novel transcript |
| SOD1 | 21 | superoxide dismutase 1 [Source:HGNC Symbo |
| AP000254.1 | 21 | NULL |
| AP000254.2 | 21 | novel transcript, antisense to SOD1 |
| SCAF4 | 21 | SR-related CTD associated factor 4 [Source:HG |
| HMGN1P2 | 21 | high mobility group nucleosome binding doma |
| AP000255.1 | 21 | novel transcript |
| TPT1P1 | 21 | tumor protein, translationally-controlled 1 pse |
| HUNK | 21 | hormonally up-regulated Neu-associated kina |
| HUNK-AS1 | 21 | HUNK antisense RNA 1 [Source:HGNC Symbol, |
| LINC00159 | 21 | long intergenic non-protein coding RNA 159 [S |
| AP000265.1 | 21 | novel transcript |

| | | |
|---|---|---|
| MIS18A | 21 | MIS18 kinetochore protein A [Source:HGNC Sy |
| MIS18A-AS1 | 21 | MIS18A antisense RNA 1 [Source:HGNC Symb |
| MRAP | 21 | melanocortin 2 receptor accessory protein [Sc |
| AP000266.1 | 21 | novel transcript |
| URB1 | 21 | URB1 ribosome biogenesis homolog [Source:H |
| SNORA80A | 21 | small nucleolar RNA, H/ACA box 80A [Source:I |
| URB1-AS1 | 21 | URB1 antisense RNA 1 (head to head) [Source |
| EVA1C | 21 | eva-1 homolog C [Source:HGNC Symbol;Acc:H |
| EXOSC3P1 | 21 | exosome component 3 pseudogene 1 [Source |
| AP000272.1 | 21 | NULL |
| RNA5SP490 | 21 | RNA, 5S ribosomal pseudogene 490 [Source:H |
| TCP10L | 21 | t-complex 10 like [Source:HGNC Symbol;Acc:H |
| C21orf59-TCP10L | 21 | C21orf59-TCP10L readthrough [Source:NCBI g |
| CFAP298 | 21 | cilia and flagella associated protein 298 [Sourc |
| OR7E23P | 21 | olfactory receptor family 7 subfamily E memb |
| AP000275.1 | 21 | ribosomal protein S24 (RPS24) pseudogene |
| SYNJ1 | 21 | synaptojanin 1 [Source:HGNC Symbol;Acc:HGI |
| PAXBP1-AS1 | 21 | PAXBP1 antisense RNA 1 [Source:HGNC Symb |
| PAXBP1 | 21 | PAX3 and PAX7 binding protein 1 [Source:HGN |
| C21orf62-AS1 | 21 | C21orf62 antisense RNA 1 [Source:HGNC Sym |
| AP000280.1 | 21 | TEC |
| C21orf62 | 21 | chromosome 21 open reading frame 62 [Sour |
| AP000281.1 | 21 | pseudogene similar to part of ribosomal prote |
| SNORA70 | 21 | Small nucleolar RNA SNORA70 [Source:RFAM; |
| AP000281.2 | 21 | novel transcript |
| AP000282.1 | 21 | novel transcript |
| LINC01690 | 21 | long intergenic non-protein coding RNA 1690 |
| OLIG2 | 21 | oligodendrocyte transcription factor 2 [Source |
| LINC00945 | 21 | long intergenic non-protein coding RNA 945 [S |
| OLIG1 | 21 | oligodendrocyte transcription factor 1 [Source |
| AP000289.1 | 21 | novel transcript |
| AP000290.1 | 21 | novel transcript |
| LINC01548 | 21 | long intergenic non-protein coding RNA 1548 |
| AP000295.1 | 21 | novel protein |
| IFNAR2 | 21 | interferon alpha and beta receptor subunit 2 [ |
| IL10RB-DT | 21 | IL10RB divergent transcript [Source:HGNC Syn |
| IL10RB | 21 | interleukin 10 receptor subunit beta [Source:H |
| IFNAR1 | 21 | interferon alpha and beta receptor subunit 1 [ |
| USF1P1 | 21 | upstream transcription factor 1 pseudogene 1 |
| IFNGR2 | 21 | interferon gamma receptor 2 [Source:HGNC S |
| TMEM50B | 21 | transmembrane protein 50B [Source:HGNC Sy |
| RPS5P3 | 21 | ribosomal protein S5 pseudogene 3 [Source:H |
| AP000302.1 | 21 | novel transcript |
| DNAJC28 | 21 | DnaJ heat shock protein family (Hsp40) memb |
| GART | 21 | phosphoribosylglycinamide formyltransferase |
| BTF3P6 | 21 | basic transcription factor 3 pseudogene 6 [Sou |
| SON | 21 | SON DNA binding protein [Source:HGNC Symb |
| MIR6501 | 21 | microRNA 6501 [Source:HGNC Symbol;Acc:HG |
| DONSON | 21 | downstream neighbor of SON [Source:HGNC S |
| AP000311.1 | 21 | novel protein similar to ATP synthase delta (O |

| | | |
|---|---|---|
| CRYZL1 | 21 | crystallin zeta like 1 [Source:HGNC Symbol;Acc |
| ITSN1 | 21 | intersectin 1 [Source:HGNC Symbol;Acc:HGNC |
| ATP5PO | 21 | ATP synthase peripheral stalk subunit OSCP [S |
| LINC00649 | 21 | long intergenic non-protein coding RNA 649 [S |
| RN7SL740P | 21 | RNA, 7SL, cytoplasmic 740, pseudogene [Sour |
| AP000569.1 | 21 | novel transcript |
| MRPS6 | 21 | mitochondrial ribosomal protein S6 [Source:H |
| SLC5A3 | 21 | solute carrier family 5 member 3 [Source:HGN |
| AP000317.2 | 21 | novel transcript |
| RPS5P2 | 21 | ribosomal protein S5 pseudogene 2 [Source:H |
| LINC00310 | 21 | long intergenic non-protein coding RNA 310 [S |
| AP000317.1 | 21 | novel transcript |
| AP000320.1 | 21 | novel transcript |
| KCNE2 | 21 | potassium voltage-gated channel subfamily E |
| SMIM11A | 21 | small integral membrane protein 11A [Source |
| FAM243A | 21 | family with sequence similarity 243 member A |
| AP000322.2 | 21 | novel transcript |
| SMIM34A | 21 | small integral membrane protein 34A [Source |
| AP000322.1 | 21 | novel transcript |
| KCNE1 | 21 | potassium voltage-gated channel subfamily E |
| AP000324.1 | 21 | NULL |
| RCAN1 | 21 | regulator of calcineurin 1 [Source:HGNC Symb |
| CLIC6 | 21 | chloride intracellular channel 6 [Source:HGNC |
| LINC00160 | 21 | long intergenic non-protein coding RNA 160 [S |
| LINC01426 | 21 | long intergenic non-protein coding RNA 1426 |
| RUNX1 | 21 | RUNX family transcription factor 1 [Source:HG |
| AP000331.1 | 21 | novel transcript, antisense to RUNX1 |
| AF015262.1 | 21 | novel transcript |
| RPL34P3 | 21 | ribosomal protein L34 pseudogene 3 [Source: |
| EZH2P1 | 21 | enhancer of zeste 2 polycomb repressive com |
| AF015720.1 | 21 | novel transcript |
| MIR802 | 21 | microRNA 802 [Source:HGNC Symbol;Acc:HGN |
| RPS20P1 | 21 | ribosomal protein S20 pseudogene 1 [Source: |
| PPP1R2P2 | 21 | protein phosphatase 1 regulatory inhibitor sul |
| LINC01436 | 21 | long intergenic non-protein coding RNA 1436 |
| RPL23AP3 | 21 | ribosomal protein L23a pseudogene 3 [Source |
| SETD4 | 21 | SET domain containing 4 [Source:HGNC Symbo |
| RIMKLBP1 | 21 | ribosomal modification protein rimK like famil |
| AP000688.4 | 21 | novel transcript |
| RNU6-992P | 21 | RNA, U6 small nuclear 992, pseudogene [Sour |
| AP000688.1 | 21 | novel transcript |
| CBR1 | 21 | carbonyl reductase 1 [Source:HGNC Symbol;A |
| AP000688.3 | 21 | novel transcript |
| AP000688.2 | 21 | novel transcript |
| MEMO1P1 | 21 | mediator of cell motility 1 pseudogene 1 [Sou |
| CBR3-AS1 | 21 | CBR3 antisense RNA 1 [Source:HGNC Symbol;A |
| RPS9P1 | 21 | ribosomal protein S9 pseudogene 1 [Source:H |
| CBR3 | 21 | carbonyl reductase 3 [Source:HGNC Symbol;A |
| DOP1B | 21 | DOP1 leucine zipper like protein B [Source:HG |
| RPL3P1 | 21 | ribosomal protein L3 pseudogene 1 [Source:H |

| | | |
|---|---|---|
| SRSF9P1 | 21 | serine and arginine rich splicing factor 9 pseuc |
| AP000692.2 | 21 | novel transcript, antisense to MORC3 |
| MORC3 | 21 | MORC family CW-type zinc finger 3 [Source:H |
| AP000692.1 | 21 | novel transcript, sense overlapping MORC3 |
| CHAF1B | 21 | chromatin assembly factor 1 subunit B [Sourc |
| ATP5MFP1 | 21 | ATP synthase membrane subunit f pseudogen |
| AP000695.1 | 21 | novel transcript |
| AP000695.2 | 21 | novel transcript |
| CLDN14 | 21 | claudin 14 [Source:HGNC Symbol;Acc:HGNC:2 |
| AP000695.3 | 21 | novel transcript |
| PSMD4P1 | 21 | proteasome 26S subunit, non-ATPase 4 pseud |
| AP000696.2 | 21 | novel transcript |
| AP000696.3 | 21 | novel transcript |
| AP000696.1 | 21 | novel transcript |
| AP000697.1 | 21 | novel transcript |
| SIM2 | 21 | SIM bHLH transcription factor 2 [Source:HGN |
| HLCS | 21 | holocarboxylase synthetase [Source:HGNC Sy |
| HLCS-IT1 | 21 | HLCS intronic transcript 1 [Source:HGNC Symb |
| RNA5SP491 | 21 | RNA, 5S ribosomal pseudogene 491 [Source:H |
| DPRXP5 | 21 | divergent-paired related homeobox pseudoge |
| AP000704.1 | 21 | novel transcript |
| Y_RNA | 21 | Y RNA [Source:RFAM;Acc:RF00019] |
| MRPL20P1 | 21 | mitochondrial ribosomal protein L20 pseudog |
| RIPPLY3 | 21 | ripply transcriptional repressor 3 [Source:HGN |
| RNU6-696P | 21 | RNA, U6 small nuclear 696, pseudogene [Sour |
| PIGP | 21 | phosphatidylinositol glycan anchor biosynthes |
| TTC3 | 21 | tetratricopeptide repeat domain 3 [Source:HG |
| AP001429.1 | 21 | novel transcript, sense intronic to TTC3 |
| TTC3-AS1 | 21 | TTC3 antisense RNA 1 [Source:HGNC Symbol;/ |
| DSCR9 | 21 | Down syndrome critical region 9 [Source:HGN |
| RN7SL678P | 21 | RNA, 7SL, cytoplasmic 678, pseudogene [Sour |
| AP001432.1 | 21 | novel transcript, antisense to DSCR3 |
| VPS26C | 21 | VPS26 endosomal protein sorting factor C [So |
| AP001412.1 | 21 | novel transcript, antisense to DSCR3 |
| AP001437.2 | 21 | novel transcript, antisense to DYRK1A |
| AP001437.1 | 21 | novel transcript, antisense to DYRK1A |
| DYRK1A | 21 | dual specificity tyrosine phosphorylation regu |
| AP001407.1 | 21 | novel transcript, antisense to KCNJ6 |
| KCNJ6 | 21 | potassium voltage-gated channel subfamily J r |
| KCNJ6-AS1 | 21 | KCNJ6 antisense RNA 1 [Source:HGNC Symbol |
| DSCR4 | 21 | Down syndrome critical region 4 [Source:HGN |
| DSCR4-IT1 | 21 | DSCR4 intronic transcript 1 [Source:HGNC Syn |
| DSCR8 | 21 | Down syndrome critical region 8 [Source:HGN |
| KCNJ15 | 21 | potassium voltage-gated channel subfamily J r |
| AP001425.1 | 21 | CTAGE family, member 5 (CTAGE5) pseudoger |
| DSCR10 | 21 | Down syndrome critical region 10 [Source:HG |
| AP001434.1 | 21 | novel transcript |
| SPATA20P1 | 21 | spermatogenesis associated 20 pseudogene 1 |
| LINC01423 | 21 | long intergenic non-protein coding RNA 1423 |
| ERG | 21 | ETS transcription factor ERG [Source:HGNC Sy |

| | | |
|---|---|---|
| SNRPGP13 | 21 | small nuclear ribonucleoprotein polypeptide ( |
| AP001037.1 | 21 | novel transcript, antisense to ERG |
| LINC00114 | 21 | long intergenic non-protein coding RNA 114 [S |
| ETS2 | 21 | ETS proto-oncogene 2, transcription factor [So |
| AP001042.1 | 21 | novel transcript |
| AP001042.2 | 21 | novel transcript |
| AP001042.3 | 21 | novel transcript |
| AP001043.1 | 21 | novel transcript |
| SNORA62 | 21 | Small nucleolar RNA SNORA62/SNORA6 family |
| RPSAP64 | 21 | ribosomal protein SA pseudogene 64 [Source: |
| LINC01700 | 21 | long intergenic non-protein coding RNA 1700 |
| AF064858.1 | 21 | novel transcript |
| AF064858.3 | 21 | TEC |
| AF064858.2 | 21 | novel transcript |
| RPL23AP12 | 21 | ribosomal protein L23a pseudogene 12 [Sourc |
| PCBP2P1 | 21 | poly(rC) binding protein 2 pseudogene 1 [Sou |
| AF129408.2 | 21 | NULL |
| PSMG1 | 21 | proteasome assembly chaperone 1 [Source:H |
| BRWD1 | 21 | bromodomain and WD repeat domain contair |
| AF129408.1 | 21 | novel transcript, antisense to BRWD1 |
| TIMM9P2 | 21 | TIMM9 pseudogene 2 [Source:HGNC Symbol; |
| BRWD1-IT1 | 21 | BRWD1 intronic transcript 1 [Source:HGNC Sy |
| METTL21AP1 | 21 | methyltransferase like 21A pseudogene 1 [Sou |
| BRWD1-AS2 | 21 | BRWD1 antisense RNA 2 [Source:HGNC Symbo |
| BRWD1-AS1 | 21 | BRWD1 antisense RNA 1 [Source:HGNC Symbo |
| HMGN1 | 21 | high mobility group nucleosome binding doma |
| Y_RNA | 21 | Y RNA [Source:RFAM;Acc:RF00019] |
| RNF6P1 | 21 | ring finger protein 6 pseudogene 1 [Source:HG |
| GET1 | 21 | guided entry of tail-anchored proteins factor 1 |
| WRB-SH3BGR | 21 | WRB-SH3BGR readthrough [Source:NCBI gene |
| LCA5L | 21 | lebercilin LCA5 like [Source:HGNC Symbol;Acc |
| SH3BGR | 21 | SH3 domain binding glutamate rich protein [So |
| MIR6508 | 21 | microRNA 6508 [Source:HGNC Symbol;Acc:HG |
| MYL6P2 | 21 | myosin light chain 6 pseudogene 2 [Source:HG |
| RPS26P4 | 21 | ribosomal protein S26 pseudogene 4 [Source: |
| AF121897.1 | 21 | KIAA1462 pseudogene |
| B3GALT5 | 21 | beta-1,3-galactosyltransferase 5 [Source:HGN |
| B3GALT5-AS1 | 21 | B3GALT5 antisense RNA 1 [Source:HGNC Syml |
| AF064860.1 | 21 | novel transcript, sense overlapping B3GALT5 |
| AF064860.2 | 21 | novel transcript |
| IGSF5 | 21 | immunoglobulin superfamily member 5 [Sour |
| PCP4 | 21 | Purkinje cell protein 4 [Source:HGNC Symbol; |
| DSCAM | 21 | DS cell adhesion molecule [Source:HGNC Sym |
| MIR4760 | 21 | microRNA 4760 [Source:HGNC Symbol;Acc:HG |
| DSCAM-AS1 | 21 | DSCAM antisense RNA 1 [Source:HGNC Symbo |
| AF064863.1 | 21 | NULL |
| DSCAM-IT1 | 21 | DSCAM intronic transcript 1 [Source:HGNC Sy |
| AF064866.1 | 21 | novel transcript, antisense to DSCAM |
| YRDCP3 | 21 | yrdC N(6)-threonylcarbamoyltransferase dom |
| AL773573.1 | 21 | novel transcript |

| | | |
|---|---|---|
| LINC00323 | 21 | long intergenic non-protein coding RNA 323 [S |
| MIR3197 | 21 | microRNA 3197 [Source:HGNC Symbol;Acc:HG |
| BACE2 | 21 | beta-secretase 2 [Source:HGNC Symbol;Acc:H |
| PLAC4 | 21 | placenta enriched 4 [Source:HGNC Symbol;Ac |
| BACE2-IT1 | 21 | BACE2 intronic transcript 1 [Source:HGNC Sym |
| FAM3B | 21 | family with sequence similarity 3 member B [S |
| MX2 | 21 | MX dynamin like GTPase 2 [Source:HGNC Sym |
| MX1 | 21 | MX dynamin like GTPase 1 [Source:HGNC Sym |
| AP001610.2 | 21 | novel transcript |
| TMPRSS2 | 21 | transmembrane serine protease 2 [Source:HG |
| AP001610.1 | 21 | NULL |
| AP001610.3 | 21 | novel transcript |
| AP006748.1 | 21 | novel transcript |
| AP006748.2 | 21 | novel transcript |
| LINC00111 | 21 | long intergenic non-protein coding RNA 111 [S |
| LINC00479 | 21 | long intergenic non-protein coding RNA 479 [S |
| LINC00112 | 21 | long intergenic non-protein coding RNA 112 [S |
| RIPK4 | 21 | receptor interacting serine/threonine kinase 4 |
| AP001615.1 | 21 | novel transcript |
| MIR6814 | 21 | microRNA 6814 [Source:HGNC Symbol;Acc:HG |
| PRDM15 | 21 | PR/SET domain 15 [Source:HGNC Symbol;Acc: |
| AP001619.2 | 21 | novel transcript |
| AP001619.1 | 21 | novel transcript |
| AP001619.3 | 21 | NULL |
| C2CD2 | 21 | C2 calcium dependent domain containing 2 [S |
| ZBTB21 | 21 | zinc finger and BTB domain containing 21 [Sou |
| ZNF295-AS1 | 21 | ZNF295 antisense RNA 1 [Source:HGNC Symbo |
| UMODL1 | 21 | uromodulin like 1 [Source:HGNC Symbol;Acc:H |
| UMODL1-AS1 | 21 | UMODL1 antisense RNA 1 [Source:HGNC Sym |
| ABCG1 | 21 | ATP binding cassette subfamily G member 1 [S |
| RNA5SP492 | 21 | RNA, 5S ribosomal pseudogene 492 [Source:H |
| TFF3 | 21 | trefoil factor 3 [Source:HGNC Symbol;Acc:HGN |
| TFF2 | 21 | trefoil factor 2 [Source:HGNC Symbol;Acc:HGN |
| TFF1 | 21 | trefoil factor 1 [Source:HGNC Symbol;Acc:HGN |
| TMPRSS3 | 21 | transmembrane serine protease 3 [Source:HG |
| UBASH3A | 21 | ubiquitin associated and SH3 domain containi |
| RNU6-1149P | 21 | RNA, U6 small nuclear 1149, pseudogene [Sou |
| RSPH1 | 21 | radial spoke head component 1 [Source:HGNC |
| SLC37A1 | 21 | solute carrier family 37 member 1 [Source:HG |
| AP001625.3 | 21 | novel transcript |
| AP001625.1 | 21 | novel transcript |
| AP001625.2 | 21 | novel transcript |
| LINC01671 | 21 | long intergenic non-protein coding RNA 1671 |
| AP001626.1 | 21 | novel transcript |
| PDE9A | 21 | phosphodiesterase 9A [Source:HGNC Symbol; |
| AP001627.1 | 21 | novel transcript |
| LINC01668 | 21 | long intergenic non-protein coding RNA 1668 |
| AP001628.2 | 21 | novel transcript |
| AP001628.1 | 21 | novel transcript |
| WDR4 | 21 | WD repeat domain 4 [Source:HGNC Symbol;A |

| | | |
|---|---|---|
| NDUFV3 | 21 | NADH:ubiquinone oxidoreductase subunit V3 |
| ERVH48-1 | 21 | endogenous retrovirus group 48 member 1 [S |
| MIR5692B | 21 | microRNA 5692b [Source:HGNC Symbol;Acc:H |
| AP001630.1 | 21 | novel transcript, antisense to PKNOX1 |
| PKNOX1 | 21 | PBX/knotted 1 homeobox 1 [Source:HGNC Syı |
| CBS | 21 | cystathionine-beta-synthase [Source:HGNC Sy |
| U2AF1 | 21 | U2 small nuclear RNA auxiliary factor 1 [Sourc |
| AP001631.2 | 21 | novel transcript, sense intronic to MX2 |
| MRPL51P2 | 21 | mitochondrial ribosomal protein L51 pseudog |
| FRGCA | 21 | FOXM1-regulated, gastric cancer associated [S |
| AP001631.1 | 21 | novel transcript |
| CRYAA | 21 | crystallin alpha A [Source:HGNC Symbol;Acc:H |
| LINC00322 | 21 | long intergenic non-protein coding RNA 322 [S |
| LINC01679 | 21 | long intergenic non-protein coding RNA 1679 |
| AP001046.1 | 21 | novel transcript |
| SIK1 | 21 | salt inducible kinase 1 [Source:HGNC Symbol;/ |
| LINC00319 | 21 | long intergenic non-protein coding RNA 319 [S |
| LINC00313 | 21 | long intergenic non-protein coding RNA 313 [S |
| AP001048.1 | 21 | novel transcript |
| HSF2BP | 21 | heat shock transcription factor 2 binding prot |
| RPL31P1 | 21 | ribosomal protein L31 pseudogene 1 [Source: |
| H2BFS | 21 | H2B histone family member S [Source:HGNC S |
| MIR6070 | 21 | microRNA 6070 [Source:HGNC Symbol;Acc:HG |
| RRP1B | 21 | ribosomal RNA processing 1B [Source:HGNC S |
| PDXK | 21 | pyridoxal kinase [Source:HGNC Symbol;Acc:H( |
| AP001052.1 | 21 | NULL |
| AP001053.1 | 21 | novel transcript, sense intronic to C21orf33an |
| CSTB | 21 | cystatin B [Source:HGNC Symbol;Acc:HGNC:24 |
| TMEM97P1 | 21 | transmembrane protein 97 pseudogene 1 [Sou |
| RRP1 | 21 | ribosomal RNA processing 1 [Source:HGNC Sy |
| AATBC | 21 | apoptosis associated transcript in bladder can |
| MYL6P1 | 21 | myosin light chain 6 pseudogene 1 [Source:HG |
| AGPAT3 | 21 | 1-acylglycerol-3-phosphate O-acyltransferase |
| RNU6-859P | 21 | RNA, U6 small nuclear 859, pseudogene [Sour |
| RNU6-1150P | 21 | RNA, U6 small nuclear 1150, pseudogene [Sou |
| TRAPPC10 | 21 | trafficking protein particle complex 10 [Source |
| H2AFZP1 | 21 | H2A histone family member Z pseudogene 1 [ |
| PWP2 | 21 | PWP2 small subunit processome component [ |
| GATD3A | 21 | glutamine amidotransferase like class 1 doma |
| LINC01678 | 21 | long intergenic non-protein coding RNA 1678 |
| AP001056.2 | 21 | novel transcript |
| AP001056.1 | 21 | novel transcript |
| AP001057.1 | 21 | novel transcript |
| ICOSLG | 21 | inducible T cell costimulator ligand [Source:H( |
| AP001059.3 | 21 | novel transcript |
| AP001059.2 | 21 | novel transcript |
| DNMT3L | 21 | DNA methyltransferase 3 like [Source:HGNC S |
| AP001059.1 | 21 | novel transcript |
| AIRE | 21 | autoimmune regulator [Source:HGNC Symbol, |
| PFKL | 21 | phosphofructokinase, liver type [Source:HGN( |

| | | |
|---|---|---|
| CFAP410 | 21 | cilia and flagella associated protein 410 [Sourc |
| AP001062.3 | 21 | novel transcript, sense overlapping C21orf2 |
| AP001062.1 | 21 | novel transcript, antisense to C21orf2 |
| AP001062.2 | 21 | novel transcript |
| AP001063.1 | 21 | novel transcript |
| TRPM2 | 21 | transient receptor potential cation channel su |
| TRPM2-AS | 21 | TRPM2 antisense RNA [Source:HGNC Symbol; |
| AP001065.2 | 21 | NULL |
| AP001065.1 | 21 | NULL |
| LRRC3-DT | 21 | LRRC3 divergent transcript [Source:HGNC Sym |
| LRRC3 | 21 | leucine rich repeat containing 3 [Source:HGNC |
| MTCYBP21 | 21 | MT-CYB pseudogene 21 [Source:HGNC Symbo |
| MTND6P21 | 21 | MT-ND6 pseudogene 21 [Source:HGNC Symbo |
| MTND5P1 | 21 | MT-ND5 pseudogene 1 [Source:HGNC Symbol |
| AP001065.3 | 21 | novel transcript |
| LINC02575 | 21 | long intergenic non-protein coding RNA 2575 |
| AP001065.4 | 21 | novel transcript |
| TSPEAR | 21 | thrombospondin type laminin G domain and E |
| TSPEAR-AS1 | 21 | TSPEAR antisense RNA 1 [Source:HGNC Symbo |
| TSPEAR-AS2 | 21 | TSPEAR antisense RNA 2 [Source:HGNC Symbo |
| AP001066.1 | 21 | novel transcript, antisense to TSPEAR |
| KRTAP10-1 | 21 | keratin associated protein 10-1 [Source:HGNC |
| KRTAP10-2 | 21 | keratin associated protein 10-2 [Source:HGNC |
| KRTAP10-3 | 21 | keratin associated protein 10-3 [Source:HGNC |
| KRTAP10-4 | 21 | keratin associated protein 10-4 [Source:HGNC |
| KRTAP10-5 | 21 | keratin associated protein 10-5 [Source:HGNC |
| KRTAP10-6 | 21 | keratin associated protein 10-6 [Source:HGNC |
| KRTAP10-7 | 21 | keratin associated protein 10-7 [Source:HGNC |
| KRTAP10-8 | 21 | keratin associated protein 10-8 [Source:HGNC |
| KRTAP10-9 | 21 | keratin associated protein 10-9 [Source:HGNC |
| KRTAP10-10 | 21 | keratin associated protein 10-10 [Source:HGN |
| KRTAP10-11 | 21 | keratin associated protein 10-11 [Source:HGN |
| KRTAP12-4 | 21 | keratin associated protein 12-4 [Source:HGNC |
| KRTAP12-3 | 21 | keratin associated protein 12-3 [Source:HGNC |
| KRTAP12-2 | 21 | keratin associated protein 12-2 [Source:HGNC |
| IMMTP1 | 21 | inner membrane mitochondrial protein pseud |
| KRTAP12-1 | 21 | keratin associated protein 12-1 [Source:HGNC |
| KRTAP12-6P | 21 | keratin associated protein 12-6, pseudogene [ |
| KRTAP10-12 | 21 | keratin associated protein 10-12 [Source:HGN |
| KRTAP10-13P | 21 | keratin associated protein 10-13, pseudogene |
| UBE2G2 | 21 | ubiquitin conjugating enzyme E2 G2 [Source:H |
| LINC01424 | 21 | long intergenic non-protein coding RNA 1424 |
| SUMO3 | 21 | small ubiquitin like modifier 3 [Source:HGNC S |
| PTTG1IP | 21 | PTTG1 interacting protein [Source:HGNC Syml |
| ITGB2 | 21 | integrin subunit beta 2 [Source:HGNC Symbol; |
| ITGB2-AS1 | 21 | ITGB2 antisense RNA 1 [Source:HGNC Symbol, |
| AL844908.2 | 21 | novel transcript, sense intronic to ITGB2 |
| LINC01547 | 21 | long intergenic non-protein coding RNA 1547 |
| AL844908.1 | 21 | novel transcript |
| FAM207A | 21 | family with sequence similarity 207 member A |

| | | |
|---|---|---|
| AP001505.1 | 21 | novel transcript |
| LINC00163 | 21 | long intergenic non-protein coding RNA 163 [S |
| LINC00165 | 21 | long intergenic non-protein coding RNA 165 [S |
| PICSAR | 21 | P38 inhibited cutaneous squamous cell carcin |
| SSR4P1 | 21 | signal sequence receptor subunit 4 pseudoger |
| ADARB1 | 21 | adenosine deaminase RNA specific B1 [Source |
| AL133499.1 | 21 | novel transcript |
| LINC00334 | 21 | long intergenic non-protein coding RNA 334 [S |
| POFUT2 | 21 | protein O-fucosyltransferase 2 [Source:HGNC |
| LINC00205 | 21 | long intergenic non-protein coding RNA 205 [S |
| BX322557.2 | 21 | novel transcript |
| LINC00315 | 21 | long intergenic non-protein coding RNA 315 [S |
| BX322557.1 | 21 | novel transcript |
| LINC00316 | 21 | long intergenic non-protein coding RNA 316 [S |
| MTCO1P3 | 21 | MT-CO1 pseudogene 3 [Source:HGNC Symbol |
| BX322559.1 | 21 | novel transcript |
| BX322562.1 | 21 | novel transcript |
| COL18A1 | 21 | collagen type XVIII alpha 1 chain [Source:HGN |
| COL18A1-AS2 | 21 | COL18A1 antisense RNA 2 [Source:HGNC Sym |
| COL18A1-AS1 | 21 | COL18A1 antisense RNA 1 [Source:HGNC Sym |
| MIR6815 | 21 | microRNA 6815 [Source:HGNC Symbol;Acc:HG |
| SLC19A1 | 21 | solute carrier family 19 member 1 [Source:HG |
| LINC01694 | 21 | long intergenic non-protein coding RNA 1694 |
| AL133493.1 | 21 | novel transcript |
| AL133493.2 | 21 | novel transcript |
| PCBP3 | 21 | poly(rC) binding protein 3 [Source:HGNC Symb |
| AL133492.1 | 21 | novel transcript, antisense to PCBP3 |
| AL592528.1 | 21 | novel transcript |
| AJ011931.2 | 21 | novel transcript, antisense to PCBP3 |
| AJ011931.1 | 21 | novel transcript |
| AJ239328.1 | 21 | PCBP3 overlapping transcript 1 |
| AJ011932.1 | 21 | novel transcript |
| COL6A1 | 21 | collagen type VI alpha 1 chain [Source:HGNC S |
| AP001476.1 | 21 | novel transcript |
| AP001476.3 | 21 | novel transcript |
| AP001476.2 | 21 | novel transcript |
| PSMA6P3 | 21 | proteasome subunit alpha 6 pseudogene 3 [So |
| AP001471.1 | 21 | novel transcript |
| COL6A2 | 21 | collagen type VI alpha 2 chain [Source:HGNC S |
| FTCD | 21 | formimidoyltransferase cyclodeaminase [Sour |
| FTCD-AS1 | 21 | FTCD antisense RNA 1 [Source:HGNC Symbol;A |
| SPATC1L | 21 | spermatogenesis and centriole associated 1 lil |
| AP001468.1 | 21 | novel transcript |
| LSS | 21 | lanosterol synthase [Source:HGNC Symbol;Acc |
| AP001469.1 | 21 | novel transcript |
| MCM3AP-AS1 | 21 | MCM3AP antisense RNA 1 [Source:HGNC Sym |
| MCM3AP | 21 | minichromosome maintenance complex comp |
| AP001469.2 | 21 | novel transcript |
| AP001469.3 | 21 | novel transcript, antisense to MCM3APAS |
| YBEY | 21 | ybeY metalloendoribonuclease [Source:HGNC |

| | | |
|---|---|---|
| C21orf58 | 21 | chromosome 21 open reading frame 58 [Sour |
| PCNT | 21 | pericentrin [Source:HGNC Symbol;Acc:HGNC:: |
| AP000471.1 | 21 | novel protein |
| RPL18AP2 | 21 | ribosomal protein L18a pseudogene 2 [Source |
| DIP2A | 21 | disco interacting protein 2 homolog A [Source |
| DIP2A-IT1 | 21 | DIP2A intronic transcript 1 [Source:HGNC Sym |
| RNU6-396P | 21 | RNA, U6 small nuclear 396, pseudogene [Sour |
| S100B | 21 | S100 calcium binding protein B [Source:HGNC |
| PRMT2 | 21 | protein arginine methyltransferase 2 [Source: |
| DSTNP1 | 21 | DSTN pseudogene 1 [Source:HGNC Symbol;Ac |
| RPL23AP4 | 21 | ribosomal protein L23a pseudogene 4 [Source |
| U2 | 22 | U2 spliceosomal RNA [Source:RFAM;Acc:RF00 |
| FRG1FP | 22 | FSHD region gene 1 family member F, pseudo |
| CU104787.1 | 22 | novel transcript |
| ACTR3BP6 | 22 | ACTR3B pseudogene 6 [Source:HGNC Symbol; |
| 5_8S_rRNA | 22 | 5.8S ribosomal RNA [Source:RFAM;Acc:RF000 |
| CT867976.1 | 22 | novel zinc finger protein pseudogene |
| AC138776.1 | 22 | novel ankyrin repeat domain family pseudoge |
| FRG1GP | 22 | FSHD region gene 1 family member G, pseudo |
| U6 | 22 | U6 spliceosomal RNA [Source:RFAM;Acc:RF00 |
| AP000542.3 | 22 | novel transcript |
| AP000542.1 | 22 | Ankyrin repeat domain 30A (ANKRD30A) pseu |
| AP000542.2 | 22 | novel transcript |
| ZNF72P | 22 | zinc finger protein 72, pseudogene [Source:H( |
| BNIP3P2 | 22 | BCL2 interacting protein 3 pseudogene 2 [Sou |
| YME1L1P1 | 22 | YME1L1 pseudogene 1 [Source:HGNC Symbol; |
| OR11H1 | 22 | olfactory receptor family 11 subfamily H mem |
| AP000534.2 | 22 | male sterility domain containing 1 (MLSTD1) p |
| ARHGAP42P3 | 22 | Rho GTPase activating protein 42 pseudogene |
| AP000534.1 | 22 | novel transcript |
| AP000533.2 | 22 | ubiquitin specific peptidase 10 (USP10) pseud |
| AP000533.1 | 22 | single stranded DNA binding protein 3 (SSBP3) |
| AP000533.3 | 22 | chromosome 10 open reading frame 57 (C10c |
| AP000532.2 | 22 | novel transcript |
| NEK2P2 | 22 | NEK2 pseudogene 2 [Source:HGNC Symbol;Ac |
| AP000532.1 | 22 | nitric oxide synthase 2A (inducible, hepatocyt |
| NF1P6 | 22 | neurofibromin 1 pseudogene 6 [Source:HGNC |
| MED15P7 | 22 | mediator complex subunit 15 pseudogene 7 [S |
| POTEH | 22 | POTE ankyrin domain family member H [Sourc |
| POTEH-AS1 | 22 | POTEH antisense RNA 1 [Source:HGNC Symbo |
| RNU6-816P | 22 | RNA, U6 small nuclear 816, pseudogene [Sour |
| AP000529.1 | 22 | actin, beta (ACTB) pseudogene |
| AP000527.1 | 22 | novel transcript |
| GRAMD4P2 | 22 | GRAM domain containing 4 pseudogene 2 [So |
| DUXAP8 | 22 | double homeobox A pseudogene 8 [Source:H( |
| AP000526.1 | 22 | novel pseudogene |
| BMS1P22 | 22 | BMS1 pseudogene 22 [Source:HGNC Symbol;/ |
| AP000525.1 | 22 | novel transcript, sense intronic to novel transc |
| DUXAP8 | 22 | double homeobox A pseudogene 8 [Source:H( |
| NBEAP3 | 22 | neurobeachin pseudogene 3 [Source:HGNC Sy |

| | | |
|---|---|---|
| AP000523.1 | 22 | translocase of outer mitochondrial membrane |
| AP000522.1 | 22 | pseudogene similar to TBC1 domain family, m |
| AC145543.1 | 22 | novel transcript |
| U6 | 22 | U6 spliceosomal RNA [Source:RFAM;Acc:RF00 |
| ABCD1P4 | 22 | ATP binding cassette subfamily D member 1 p |
| AC137499.1 | 22 | zinc finger, CDGSH-type domain 2 (ZCD2) pseu |
| PABPC1P9 | 22 | poly(A) binding protein cytoplasmic 1 pseudog |
| SLC9B1P4 | 22 | solute carrier family 9 member B1 pseudogen |
| AC137499.2 | 22 | NULL |
| CHEK2P4 | 22 | checkpoint kinase 2 pseudogene 4 [Source:HG |
| AP000547.2 | 22 | novel transcript |
| AP000547.1 | 22 | phosphoinositide-3-kinase, catalytic, alpha po |
| KCNMB3P1 | 22 | potassium calcium-activated channel subfami |
| CCT8L2 | 22 | chaperonin containing TCP1 subunit 8 like 2 [S |
| AP000547.4 | 22 | novel transcript |
| FABP5P11 | 22 | fatty acid binding protein 5 pseudogene 11 [S |
| TPTEP1 | 22 | TPTE pseudogene 1 [Source:HGNC Symbol;Acc |
| AP000547.3 | 22 | novel transcript |
| AP000365.1 | 22 | novel transcript |
| SLC25A15P5 | 22 | solute carrier family 25 member 15 pseudoge |
| PARP4P3 | 22 | poly(ADP-ribose) polymerase family member |
| ANKRD62P1-PARP4P3 | 22 | ANKRD62P1-PARP4P3 readthrough, transcribe |
| ANKRD62P1 | 22 | ankyrin repeat domain 62 pseudogene 1 [Sour |
| AC005301.1 | 22 | VWFP1-ANKRD62P1 readthrough transcript |
| VWFP1 | 22 | von Willebrand factor pseudogene 1 [Source:I |
| AC005301.2 | 22 | novel transcript |
| LINC01665 | 22 | long intergenic non-protein coding RNA 1665 |
| XKR3 | 22 | XK related 3 [Source:HGNC Symbol;Acc:HGNC |
| HSFY1P1 | 22 | HSFY1 pseudogene 1 [Source:HGNC Symbol;A |
| GPM6BP3 | 22 | glycoprotein M6B pseudogene 3 [Source:HGN |
| AC007064.4 | 22 | novel pseudogene |
| AC007064.3 | 22 | Zinc finger protein pseudogene |
| ZNF402P | 22 | zinc finger protein 402, pseudogene [Source:H |
| AC007064.2 | 22 | novel transcript |
| MTND1P17 | 22 | MT-ND1 pseudogene 17 [Source:HGNC Symbo |
| AC007064.1 | 22 | tripartite motif-containing 43 (TRIM43) pseud |
| AC006548.1 | 22 | N-acetylated alpha-linked acidic dipeptidase 2 |
| IGKV1OR22-5 | 22 | immunoglobulin kappa variable 1/OR22-5 (ps |
| IGKV2OR22-4 | 22 | immunoglobulin kappa variable 2/OR22-4 (ps |
| IGKV2OR22-3 | 22 | immunoglobulin kappa variable 2/OR22-3 (ps |
| IGKV3OR22-2 | 22 | immunoglobulin kappa variable 3/OR22-2 (ps |
| IGKV1OR22-1 | 22 | immunoglobulin kappa variable 1/OR22-1 (ps |
| AC006548.3 | 22 | ubiquitin specific peptidase 35 (USP35) pseud |
| GAB4 | 22 | GRB2 associated binding protein family memb |
| AC006548.2 | 22 | interleukin 9 receptor (IL9R) pseudogene |
| VN1R9P | 22 | vomeronasal 1 receptor 9 pseudogene [Sourc |
| CECR7 | 22 | cat eye syndrome chromosome region, candid |
| AC006946.2 | 22 | novel transcript |
| AC006946.3 | 22 | novel transcript |
| IL17RA | 22 | interleukin 17 receptor A [Source:HGNC Symb |

| | | |
|---|---|---|
| AC006946.1 | 22 | ribosomal protein L31 (RPL31) pseudogene |
| TMEM121B | 22 | transmembrane protein 121B [Source:HGNC S |
| LINC01664 | 22 | long intergenic non-protein coding RNA 1664 |
| HDHD5 | 22 | haloacid dehalogenase like hydrolase domain |
| HDHD5-AS1 | 22 | HDHD5 antisense RNA 1 [Source:HGNC Symbo |
| ADA2 | 22 | adenosine deaminase 2 [Source:HGNC Symbo |
| AC005300.1 | 22 | 60S ribosomal protein L32 (RPL32) pseudogen |
| FAM32BP | 22 | family with sequence similarity 32 member B, |
| CECR3 | 22 | cat eye syndrome chromosome region, candic |
| CECR9 | 22 | cat eye syndrome chromosome region, candic |
| RN7SL843P | 22 | RNA, 7SL, cytoplasmic 843, pseudogene [Sour |
| CECR2 | 22 | CECR2 histone acetyl-lysine reader [Source:HC |
| FO681548.1 | 22 | Telomeric repeat-binding factor 2-interacting |
| CLCP1 | 22 | Charcot-Leyden crystal protein pseudogene 1 |
| DNAJA1P6 | 22 | DnaJ heat shock protein family (Hsp40) memb |
| AC007666.2 | 22 | novel transcript, antisense to CECR2 and SLC2 |
| SLC25A18 | 22 | solute carrier family 25 member 18 [Source:H |
| AC007666.1 | 22 | novel transcript |
| ATP6V1E1 | 22 | ATPase H+ transporting V1 subunit E1 [Source |
| BCL2L13 | 22 | BCL2 like 13 [Source:HGNC Symbol;Acc:HGNC |
| BID | 22 | BH3 interacting domain death agonist [Source |
| MIR3198-1 | 22 | microRNA 3198-1 [Source:HGNC Symbol;Acc:I |
| LINC00528 | 22 | long intergenic non-protein coding RNA 528 [S |
| MICAL3 | 22 | microtubule associated monooxygenase, calp |
| AC016026.1 | 22 | novel transcript |
| AC016026.2 | 22 | novel transcript, antisense to MICAL3 |
| MIR648 | 22 | microRNA 648 [Source:HGNC Symbol;Acc:HGI |
| AC016027.3 | 22 | novel transcript |
| AC016027.2 | 22 | novel transcript |
| RHEBP3 | 22 | RHEB pseudogene 3 [Source:HGNC Symbol;Ac |
| LINC01634 | 22 | long intergenic non-protein coding RNA 1634 |
| AC016027.1 | 22 | novel transcript |
| PEX26 | 22 | peroxisomal biogenesis factor 26 [Source:HGI |
| AC016027.5 | 22 | ribosomal protein L18a (RPL18A) pseudogene |
| AC016027.4 | 22 | neurexin (NRXN) pseudogene |
| ARL2BPP10 | 22 | ADP ribosylation factor like GTPase 2 binding |
| TUBA8 | 22 | tubulin alpha 8 [Source:HGNC Symbol;Acc:HG |
| AC008079.1 | 22 | novel transcript, antisense to TUBA8 |
| USP18 | 22 | ubiquitin specific peptidase 18 [Source:HGNC |
| FAM230D | 22 | family with sequence similarity 230 member I |
| GGTLC5P | 22 | gamma-glutamyltransferase light chain 5 pseu |
| FAM230J | 22 | family with sequence similarity 230 member J |
| AC011718.1 | 22 | novel transcript |
| PPP1R26P3 | 22 | protein phosphatase 1 regulatory subunit 26 |
| FAM230A | 22 | family with sequence similarity 230 member / |
| AC023490.2 | 22 | novel transcript |
| AC023490.4 | 22 | novel transcript |
| GGTLC3 | 22 | gamma-glutamyltransferase light chain family |
| Metazoa_SRP | 22 | Metazoan signal recognition particle RNA [Sou |
| AC023490.3 | 22 | transmembrane protein 191A (pseudogene) T |

| | | |
|---|---|---|
| TMEM191B | 22 | transmembrane protein 191B [Source:HGNC S |
| PI4KAP1 | 22 | phosphatidylinositol 4-kinase alpha pseudoge |
| AC023490.1 | 22 | NULL |
| RN7SKP131 | 22 | RN7SK pseudogene 131 [Source:HGNC Symbo |
| RIMBP3 | 22 | RIMS binding protein 3 [Source:HGNC Symbol |
| Metazoa_SRP | 22 | Metazoan signal recognition particle RNA [Sou |
| SUSD2P2 | 22 | sushi domain containing 2 pseudogene 2 [Sou |
| AC023490.5 | 22 | novel protein, similar to B830017H08Rik (M. r |
| CA15P2 | 22 | CA15 pseudogene 2 [Source:HGNC Symbol;Ac |
| PPP1R26P2 | 22 | protein phosphatase 1 regulatory subunit 26 p |
| PPP1R26P4 | 22 | protein phosphatase 1 regulatory subunit 26 p |
| FAM230E | 22 | family with sequence similarity 230 member E |
| GGT3P | 22 | gamma-glutamyltransferase 3 pseudogene [So |
| AC008132.2 | 22 | immunoglobulin superfamily, member 3 (IGSF |
| E2F6P1 | 22 | E2F transcription factor 6 pseudogene 1 [Sour |
| AC008132.1 | 22 | POM121 transmembrane nucleoporin (POM1 |
| BCRP7 | 22 | BCR pseudogene 7 [Source:HGNC Symbol;Acc |
| FAM230F | 22 | family with sequence similarity 230 member F |
| AC008103.1 | 22 | novel transcript |
| DGCR6 | 22 | DiGeorge syndrome critical region gene 6 [Sou |
| AC007326.4 | 22 | novel protein |
| PRODH | 22 | proline dehydrogenase 1 [Source:HGNC Symb |
| AC007326.5 | 22 | novel transcript |
| AC007326.2 | 22 | novel transcript |
| DGCR5 | 22 | DiGeorge syndrome critical region gene 5 [Sou |
| DGCR9 | 22 | DiGeorge syndrome critical region gene 9 [Sou |
| AC007326.3 | 22 | fructosamine 3 kinase (FN3K) pseudogene |
| AC007326.1 | 22 | POM121 transmembrane nucleoporin (POM1 |
| AC000095.2 | 22 | sperm antigen with calponin homology and co |
| AC000095.1 | 22 | POM121 transmembrane nucleoporin (POM1 |
| DGCR10 | 22 | DiGeorge syndrome critical region gene 10 [So |
| AC000095.3 | 22 | novel protein, similar to B830017H08Rik (M. r |
| CA15P1 | 22 | CA15 pseudogene 1 [Source:HGNC Symbol;Ac |
| DGCR2 | 22 | DiGeorge syndrome critical region gene 2 [Sou |
| Y_RNA | 22 | Y RNA [Source:RFAM;Acc:RF00019] |
| DGCR11 | 22 | DiGeorge syndrome critical region gene 11 [So |
| AC004461.1 | 22 | 60S ribosomal protein L28 (RPL28) pseudogen |
| AC004461.2 | 22 | DiGeorge syndrome critical region gene 12 (no |
| AC004471.1 | 22 | novel transcript |
| AC004471.2 | 22 | novel transcript |
| TSSK1A | 22 | testis specific serine kinase 1A, pseudogene [S |
| ESS2 | 22 | ess-2 splicing factor homolog [Source:HGNC S |
| TSSK2 | 22 | testis specific serine kinase 2 [Source:HGNC Sy |
| GSC2 | 22 | goosecoid homeobox 2 [Source:HGNC Symbol |
| LINC01311 | 22 | long intergenic non-protein coding RNA 1311 |
| SLC25A1 | 22 | solute carrier family 25 member 1 [Source:HG |
| CLTCL1 | 22 | clathrin heavy chain like 1 [Source:HGNC Syml |
| AC000072.1 | 22 | novel transcript, antisense to CLTCL1 |
| AC000081.1 | 22 | ribosomal protein L34 (RPL34) pseudogene |
| AC000081.2 | 22 | NULL |

| | | |
|---|---|---|
| KRT18P62 | 22 | keratin 18 pseudogene 62 [Source:HGNC Sym |
| AC000085.1 | 22 | novel transcript |
| HIRA | 22 | histone cell cycle regulator [Source:HGNC Syn |
| C22orf39 | 22 | chromosome 22 open reading frame 39 [Sour |
| RN7SL168P | 22 | RNA, 7SL, cytoplasmic 168, pseudogene [Sour |
| MRPL40 | 22 | mitochondrial ribosomal protein L40 [Source:I |
| AC000068.1 | 22 | novel transcript, antisense to C22orf39 |
| UFD1 | 22 | ubiquitin recognition factor in ER associated d |
| AC000068.3 | 22 | novel transcript, antisense to UFD1L |
| AC000068.2 | 22 | novel transcript, antisense to UFD1L |
| CDC45 | 22 | cell division cycle 45 [Source:HGNC Symbol;Ac |
| AC000082.1 | 22 | novel transcript, antisense to DGCR8and CDC4 |
| CLDN5 | 22 | claudin 5 [Source:HGNC Symbol;Acc:HGNC:20 |
| LINC00895 | 22 | long intergenic non-protein coding RNA 895 [S |
| AC000077.1 | 22 | chromosome 3 open reading frame 38 (C3orf3 |
| AC000067.1 | 22 | novel transcript |
| SEPTIN5 | 22 | septin 5 [Source:HGNC Symbol;Acc:HGNC:916 |
| AC000093.1 | 22 | novel transcript |
| GP1BB | 22 | glycoprotein Ib platelet subunit beta [Source:I |
| TBX1 | 22 | T-box transcription factor 1 [Source:HGNC Syr |
| GNB1L | 22 | G protein subunit beta 1 like [Source:HGNC Sy |
| AC000089.1 | 22 | ribosomal protein L7a (RPL7A) pseudogene |
| RTL10 | 22 | retrotransposon Gag like 10 [Source:HGNC Syi |
| TXNRD2 | 22 | thioredoxin reductase 2 [Source:HGNC Symbo |
| AC000078.1 | 22 | ribosomal protein L8 (RPL8) pseudogene |
| COMT | 22 | catechol-O-methyltransferase [Source:HGNC S |
| MIR4761 | 22 | microRNA 4761 [Source:HGNC Symbol;Acc:HG |
| ARVCF | 22 | ARVCF delta catenin family member [Source:H |
| TANGO2 | 22 | transport and golgi organization 2 homolog [S |
| MIR185 | 22 | microRNA 185 [Source:HGNC Symbol;Acc:HGI |
| AC006547.1 | 22 | novel transcript, antisense to C22orf25 |
| AC006547.3 | 22 | novel transcript, antisense to C22orf25 |
| Metazoa_SRP | 22 | Metazoan signal recognition particle RNA [Sou |
| DGCR8 | 22 | DGCR8 microprocessor complex subunit [Sour |
| MIR3618 | 22 | microRNA 3618 [Source:HGNC Symbol;Acc:HG |
| MIR1306 | 22 | microRNA 1306 [Source:HGNC Symbol;Acc:HG |
| AC006547.2 | 22 | novel transcript, antisense to DGCR8 |
| TRMT2A | 22 | tRNA methyltransferase 2 homolog A [Source: |
| MIR6816 | 22 | microRNA 6816 [Source:HGNC Symbol;Acc:HG |
| RANBP1 | 22 | RAN binding protein 1 [Source:HGNC Symbol;. |
| SNORA77B | 22 | small nucleolar RNA, H/ACA box 77B [Source:I |
| ZDHHC8 | 22 | zinc finger DHHC-type containing 8 [Source:HG |
| CCDC188 | 22 | coiled-coil domain containing 188 [Source:HG |
| AC007663.2 | 22 | novel transcript |
| LINC00896 | 22 | long intergenic non-protein coding RNA 896 [S |
| RTN4R | 22 | reticulon 4 receptor [Source:HGNC Symbol;Ac |
| MIR1286 | 22 | microRNA 1286 [Source:HGNC Symbol;Acc:HG |
| AC007663.1 | 22 | proline dehydrogenase (oxidase) 1 (PRODH) p |
| DGCR6L | 22 | DiGeorge syndrome critical region gene 6 like |
| AC007663.4 | 22 | novel transcript, sense intronic to DGCR6L |

| | | |
|---|---|---|
| AC007663.3 | 22 | novel transcript |
| FAM230G | 22 | family with sequence similarity 230 member ( |
| AC007731.2 | 22 | novel transcript |
| AC007731.5 | 22 | novel transcript |
| USP41 | 22 | ubiquitin specific peptidase 41 [Source:HGNC |
| ZNF74 | 22 | zinc finger protein 74 [Source:HGNC Symbol;A |
| RNU6-225P | 22 | RNA, U6 small nuclear 225, pseudogene [Sour |
| SCARF2 | 22 | scavenger receptor class F member 2 [Source: |
| AC007731.4 | 22 | novel protein |
| KLHL22 | 22 | kelch like family member 22 [Source:HGNC Sy |
| AC007731.1 | 22 | kelch-like 12 (Drosophila) (KLH12) pseudogen |
| RNY1P9 | 22 | RNY1 pseudogene 9 [Source:HGNC Symbol;Ac |
| Y_RNA | 22 | Y RNA [Source:RFAM;Acc:RF00019] |
| RN7SL812P | 22 | RNA, 7SL, cytoplasmic 812, pseudogene [Sour |
| KRT18P5 | 22 | keratin 18 pseudogene 5 [Source:HGNC Symb |
| MED15 | 22 | mediator complex subunit 15 [Source:HGNC S |
| AC007731.3 | 22 | novel transcript |
| CCDC74BP1 | 22 | coiled-coil domain containing 74B pseudogen |
| SMPD4P1 | 22 | sphingomyelin phosphodiesterase 4 pseudoge |
| IGLL4P | 22 | immunoglobulin lambda like polypeptide 4, ps |
| SLC9A3P2 | 22 | solute carrier family 9 member 3 pseudogene |
| ABHD17AP4 | 22 | abhydrolase domain containing 17A pseudoge |
| POM121L4P | 22 | POM121 transmembrane nucleoporin like 4, p |
| BCRP5 | 22 | BCR pseudogene 5 [Source:HGNC Symbol;Acc |
| TMEM191A | 22 | transmembrane protein 191A (pseudogene) [S |
| PI4KA | 22 | phosphatidylinositol 4-kinase alpha [Source:H |
| SERPIND1 | 22 | serpin family D member 1 [Source:HGNC Syml |
| SNAP29 | 22 | synaptosome associated protein 29 [Source:H |
| AC007308.1 | 22 | novel transcript, antisense to SNAP29 |
| CRKL | 22 | CRK like proto-oncogene, adaptor protein [Sou |
| LINC01637 | 22 | long intergenic non-protein coding RNA 1637 |
| AIFM3 | 22 | apoptosis inducing factor mitochondria associ |
| AC002470.2 | 22 | novel transcript |
| AC002470.1 | 22 | novel transcript |
| LZTR1 | 22 | leucine zipper like transcription regulator 1 [S |
| THAP7 | 22 | THAP domain containing 7 [Source:HGNC Sym |
| THAP7-AS1 | 22 | THAP7 antisense RNA 1 [Source:HGNC Symbo |
| AC002472.2 | 22 | pseudogene similar to part of tubulin, alpha 3 |
| TUBA3FP | 22 | tubulin alpha 3f pseudogene [Source:HGNC Sy |
| P2RX6 | 22 | purinergic receptor P2X 6 [Source:HGNC Symb |
| AC002472.3 | 22 | leucine rich repeat containing 74B (LRRC74B) |
| SLC7A4 | 22 | solute carrier family 7 member 4 [Source:HGN |
| AC002472.1 | 22 | novel transcript |
| MIR649 | 22 | microRNA 649 [Source:HGNC Symbol;Acc:HGN |
| P2RX6P | 22 | purinergic receptor P2X 6 pseudogene [Sourc |
| LRRC74B | 22 | leucine rich repeat containing 74B [Source:HG |
| TUBA3GP | 22 | tubulin alpha 3g pseudogene [Source:HGNC S |
| AC002472.4 | 22 | mitotic spindle organizing protein 2A (MZT2A) |
| BCRP2 | 22 | BCR pseudogene 2 [Source:HGNC Symbol;Acc |
| AP000550.1 | 22 | novel transcript |

| | | |
|---|---|---|
| POM121L7P | 22 | POM121 transmembrane nucleoporin like 7 p |
| E2F6P2 | 22 | E2F transcription factor 6 pseudogene 2 [Sour |
| FAM230B | 22 | family with sequence similarity 230 member E |
| AP000550.2 | 22 | novel transcript |
| AP000550.4 | 22 | gamma-glutamyltransferase 5 (GGT5) pseudo |
| GGT2 | 22 | gamma-glutamyltransferase 2 [Source:HGNC S |
| AP000550.3 | 22 | immunoglobulin superfamily member 3 (IGSF3 |
| E2F6P3 | 22 | E2F transcription factor 6 pseudogene 3 [Sour |
| POM121L8P | 22 | POM121 transmembrane nucleoporin like 8, p |
| BCRP6 | 22 | BCR pseudogene 6 [Source:HGNC Symbol;Acc |
| FAM230H | 22 | family with sequence similarity 230 member H |
| AP000552.2 | 22 | novel transcript |
| PPP1R26P5 | 22 | protein phosphatase 1 regulatory subunit 26 p |
| LINC01651 | 22 | long intergenic non-protein coding RNA 1651 |
| AP000552.4 | 22 | novel protein, similar to B830017H08Rik (M. m |
| AP000552.1 | 22 | pseudogene similar to part of sushi domain co |
| Metazoa_SRP | 22 | Metazoan signal recognition particle RNA [Sou |
| RIMBP3B | 22 | RIMS binding protein 3B [Source:HGNC Symbo |
| RN7SKP63 | 22 | RN7SK pseudogene 63 [Source:HGNC Symbol; |
| AP000552.3 | 22 | NULL |
| Metazoa_SRP | 22 | Metazoan signal recognition particle RNA [Sou |
| HIC2 | 22 | HIC ZBTB transcriptional repressor 2 [Source:H |
| Metazoa_SRP | 22 | Metazoan signal recognition particle RNA [Sou |
| TMEM191C | 22 | transmembrane protein 191C [Source:HGNC S |
| PI4KAP2 | 22 | phosphatidylinositol 4-kinase alpha pseudoge |
| AP000557.1 | 22 | NULL |
| RN7SKP221 | 22 | RN7SK pseudogene 221 [Source:HGNC Symbo |
| RIMBP3C | 22 | RIMS binding protein 3C [Source:HGNC Symbo |
| UBE2L3 | 22 | ubiquitin conjugating enzyme E2 L3 [Source:H |
| Metazoa_SRP | 22 | Metazoan signal recognition particle RNA [Sou |
| YDJC | 22 | YdjC chitooligosaccharide deacetylase homolo |
| CCDC116 | 22 | coiled-coil domain containing 116 [Source:HG |
| AP000553.3 | 22 | novel transcript |
| SDF2L1 | 22 | stromal cell derived factor 2 like 1 [Source:HG |
| AP000553.1 | 22 | novel transcript |
| MIR301B | 22 | microRNA 301b [Source:HGNC Symbol;Acc:HG |
| MIR130B | 22 | microRNA 130b [Source:HGNC Symbol;Acc:HG |
| AP000553.5 | 22 | novel pseudogene |
| AP000553.6 | 22 | novel pseudogene |
| AP000553.7 | 22 | novel pseudogene |
| AP000553.4 | 22 | novel pseudogene |
| AP000553.2 | 22 | novel transcript, sense intronic to PPIL2 |
| PPIL2 | 22 | peptidylprolyl isomerase like 2 [Source:HGNC |
| YPEL1 | 22 | yippee like 1 [Source:HGNC Symbol;Acc:HGNC |
| AP000553.9 | 22 | novel transcript, antisense to YPEL1 |
| RN7SL280P | 22 | RNA, 7SL, cytoplasmic 280, pseudogene [Sour |
| AP000553.8 | 22 | novel transcript |
| MAPK1 | 22 | mitogen-activated protein kinase 1 [Source:HG |
| RNA5SP493 | 22 | RNA, 5S ribosomal pseudogene 493 [Source:H |
| AC245452.2 | 22 | pseudogene similar to part of metallophospho |

| | | |
|---|---|---|
| PPM1F | 22 | protein phosphatase, Mg2+/Mn2+ dependent |
| AC245452.1 | 22 | novel transcript, antisense to PPM1F and TOP |
| TOP3B | 22 | DNA topoisomerase III beta [Source:HGNC Syr |
| PRAMENP | 22 | PRAME N-terminal like, pseudogene [Source:H |
| IGLVI-70 | 22 | immunoglobulin lambda variable (I)-70 (pseuc |
| IGLV4-69 | 22 | immunoglobulin lambda variable 4-69 [Source |
| IGLVI-68 | 22 | immunoglobulin lambda variable (I)-68 (pseuc |
| AC245452.5 | 22 | novel transcript |
| AC245452.4 | 22 | Ig lambda chain V-I region pseudogene |
| IGLV10-54 | 22 | immunoglobulin lambda variable 10-54 [Sourc |
| IGLV10-67 | 22 | immunoglobulin lambda variable 10-67 (pseu |
| AC245452.3 | 22 | Ig lambda chain V-VI region pseudogene |
| IGLVIV-66-1 | 22 | immunoglobulin lambda variable (IV)-66-1 (ps |
| IGLVV-66 | 22 | immunoglobulin lambda variable (V)-66 (pseu |
| IGLVIV-65 | 22 | immunoglobulin lambda variable (IV)-65 (pseu |
| IGLVIV-64 | 22 | immunoglobulin lambda variable (IV)-64 (pseu |
| IGLVI-63 | 22 | immunoglobulin lambda variable (I)-63 (pseuc |
| IGLV1-62 | 22 | immunoglobulin lambda variable 1-62 (pseudo |
| AC245517.5 | 22 | Ig Lambda chain V-IV region pseudogene |
| IGLV8-61 | 22 | immunoglobulin lambda variable 8-61 [Source |
| AC245517.1 | 22 | novel pseudogene |
| ABHD17AP5 | 22 | abhydrolase domain containing 17A pseudoge |
| AC245517.4 | 22 | Ig lambda-1 chain V region pseudogene |
| AC245517.3 | 22 | Ig lambda chain V region pseudogene |
| IGLV4-60 | 22 | immunoglobulin lambda variable 4-60 [Source |
| AC245517.2 | 22 | Ig lambda chain V region pseudogene |
| SOCS2P2 | 22 | suppressor of cytokine signaling 2 pseudogene |
| IGLVIV-59 | 22 | immunoglobulin lambda variable (IV)-59 (pseu |
| IGLVV-58 | 22 | immunoglobulin lambda variable (V)-58 (pseu |
| BMP6P1 | 22 | bone morphogenetic protein 6 pseudogene 1 |
| IGLV6-57 | 22 | immunoglobulin lambda variable 6-57 [Source |
| IGLVI-56 | 22 | immunoglobulin lambda variable (I)-56 (pseuc |
| IGLV11-55 | 22 | immunoglobulin lambda variable 11-55 (non-f |
| IGLVIV-53 | 22 | immunoglobulin lambda variable (IV)-53 (pseu |
| TOP3BP1 | 22 | DNA topoisomerase III beta pseudogene 1 [So |
| AC245060.3 | 22 | chromosome 19 open reading frame 2 (C19or |
| VPREB1 | 22 | V-set pre-B cell surrogate light chain 1 [Source |
| AC245060.6 | 22 | novel transcript |
| AC245060.5 | 22 | novel transcript |
| AC245060.2 | 22 | novel transcript |
| AC245060.7 | 22 | novel transcript |
| AC245060.4 | 22 | BMS1, ribosome biogenesis factor (BSM1) pse |
| IGLV5-52 | 22 | immunoglobulin lambda variable 5-52 [Source |
| IGLV1-51 | 22 | immunoglobulin lambda variable 1-51 [Source |
| IGLV1-50 | 22 | immunoglobulin lambda variable 1-50 (non-fu |
| ASH2LP3 | 22 | ASH2L pseudogene 3 [Source:HGNC Symbol;A |
| IGLV9-49 | 22 | immunoglobulin lambda variable 9-49 [Source |
| AC245060.1 | 22 | Ig lambda chain V-I region pseudogene |
| IGLV5-48 | 22 | immunoglobulin lambda variable 5-48 (non-fu |
| IGLV1-47 | 22 | immunoglobulin lambda variable 1-47 [Source |

| | | |
|---|---|---|
| ASH2LP2 | 22 | ASH2L pseudogene 2 [Source:HGNC Symbol;A |
| IGLV7-46 | 22 | immunoglobulin lambda variable 7-46 (gene/p |
| AC245291.3 | 22 | Ig lambda chain V-11 pseudogene |
| IGLV5-45 | 22 | immunoglobulin lambda variable 5-45 [Source |
| IGLV1-44 | 22 | immunoglobulin lambda variable 1-44 [Source |
| IGLV7-43 | 22 | immunoglobulin lambda variable 7-43 [Source |
| IGLVI-42 | 22 | immunoglobulin lambda variable (I)-42 (pseuc |
| IGLVVII-41-1 | 22 | immunoglobulin lambda variable (VII)-41-1 (ps |
| IGLV1-41 | 22 | immunoglobulin lambda variable 1-41 (pseudo |
| IGLV1-40 | 22 | immunoglobulin lambda variable 1-40 [Source |
| ASH2LP1 | 22 | ASH2L pseudogene 1 [Source:HGNC Symbol;A |
| IGLVI-38 | 22 | immunoglobulin lambda variable (I)-38 (pseuc |
| IGLV5-37 | 22 | immunoglobulin lambda variable 5-37 [Source |
| IGLV1-36 | 22 | immunoglobulin lambda variable 1-36 [Source |
| AC245291.2 | 22 | ash2 (absent, small, or homeotic)-like (Drosop |
| IGLV7-35 | 22 | immunoglobulin lambda variable 7-35 (pseudo |
| AC245291.1 | 22 | Ig lambda chain V-I region pseudogene |
| ZNF280B | 22 | zinc finger protein 280B [Source:HGNC Symbo |
| ZNF280A | 22 | zinc finger protein 280A [Source:HGNC Symbo |
| PRAME | 22 | preferentially expressed antigen in melanoma |
| AC246793.1 | 22 | novel transcript, antisense to PRAME |
| IGLV2-34 | 22 | immunoglobulin lambda variable 2-34 (pseudo |
| IGLV2-33 | 22 | immunoglobulin lambda variable 2-33 (non-fu |
| IGLV3-32 | 22 | immunoglobulin lambda variable 3-32 (non-fu |
| IGLV3-31 | 22 | immunoglobulin lambda variable 3-31 (pseudo |
| IGLV3-30 | 22 | immunoglobulin lambda variable 3-30 (pseudo |
| BCRP4 | 22 | BCR pseudogene 4 [Source:HGNC Symbol;Acc |
| AC244250.4 | 22 | novel transcript |
| POM121L1P | 22 | POM121 transmembrane nucleoporin like 1, p |
| GGTLC2 | 22 | gamma-glutamyltransferase light chain 2 [Sou |
| IGLV3-29 | 22 | immunoglobulin lambda variable 3-29 (pseudo |
| IGLV2-28 | 22 | immunoglobulin lambda variable 2-28 (pseudo |
| IGLV3-27 | 22 | immunoglobulin lambda variable 3-27 [Source |
| IGLV3-26 | 22 | immunoglobulin lambda variable 3-26 (pseudo |
| IGLVVI-25-1 | 22 | immunoglobulin lambda variable (VI)-25-1 (ps |
| IGLV3-25 | 22 | immunoglobulin lambda variable 3-25 [Source |
| AC244250.2 | 22 | Ig lambda chain V-VI region pseudogene |
| AC244250.1 | 22 | novel transcript |
| IGLV3-24 | 22 | immunoglobulin lambda variable 3-24 (pseudo |
| IGLV2-23 | 22 | immunoglobulin lambda variable 2-23 [Source |
| IGLVVI-22-1 | 22 | immunoglobulin lambda variable (VI)-22-1 (ps |
| IGLV3-22 | 22 | immunoglobulin lambda variable 3-22 (gene/p |
| IGLV3-21 | 22 | immunoglobulin lambda variable 3-21 [Source |
| IGLVI-20 | 22 | immunoglobulin lambda variable (I)-20 (pseuc |
| IGLV3-19 | 22 | immunoglobulin lambda variable 3-19 [Source |
| AC244250.3 | 22 | chromosome 2 open reading frame 27A (C2or |
| IGLV2-18 | 22 | immunoglobulin lambda variable 2-18 [Source |
| IGLV3-17 | 22 | immunoglobulin lambda variable 3-17 (pseudo |
| IGLV3-16 | 22 | immunoglobulin lambda variable 3-16 [Source |
| IGLV3-15 | 22 | immunoglobulin lambda variable 3-15 (pseudo |

| | | |
|---|---|---|
| IGLV2-14 | 22 | immunoglobulin lambda variable 2-14 [Source |
| IGLV3-13 | 22 | immunoglobulin lambda variable 3-13 (pseud |
| IGLV3-12 | 22 | immunoglobulin lambda variable 3-12 [Source |
| AC244157.1 | 22 | Ig lambda chain V-II region pseudogene |
| AC244157.2 | 22 | CMT1A duplicated region transcript 15-like 2 ( |
| IGLV2-11 | 22 | immunoglobulin lambda variable 2-11 [Source |
| AC245028.3 | 22 | Ig Lambda chain V-III region pseudogene |
| IGLV3-10 | 22 | immunoglobulin lambda variable 3-10 [Source |
| AC245028.1 | 22 | Ig Lambda chain V-IV region pseudogene |
| IGLV3-9 | 22 | immunoglobulin lambda variable 3-9 (gene/ps |
| IGLV2-8 | 22 | immunoglobulin lambda variable 2-8 [Source: |
| MIR650 | 22 | microRNA 650 [Source:HGNC Symbol;Acc:HGI |
| AC245028.2 | 22 | Ig lambda chain V-IV region pseudogene |
| IGLV3-7 | 22 | immunoglobulin lambda variable 3-7 (pseudo |
| IGLV3-6 | 22 | immunoglobulin lambda variable 3-6 (pseudo |
| IGLV2-5 | 22 | immunoglobulin lambda variable 2-5 (pseudo |
| IGLV3-4 | 22 | immunoglobulin lambda variable 3-4 (pseudo |
| IGLV4-3 | 22 | immunoglobulin lambda variable 4-3 [Source: |
| IGLV3-2 | 22 | immunoglobulin lambda variable 3-2 (pseudo |
| IGLV3-1 | 22 | immunoglobulin lambda variable 3-1 [Source: |
| MIR5571 | 22 | microRNA 5571 [Source:HGNC Symbol;Acc:HG |
| IGLL5 | 22 | immunoglobulin lambda like polypeptide 5 [So |
| IGLJ1 | 22 | immunoglobulin lambda joining 1 [Source:HGI |
| IGLC1 | 22 | immunoglobulin lambda constant 1 [Source:H |
| IGLJ2 | 22 | immunoglobulin lambda joining 2 [Source:HGI |
| IGLC2 | 22 | immunoglobulin lambda constant 2 [Source:H |
| IGLJ3 | 22 | immunoglobulin lambda joining 3 [Source:HGI |
| IGLC3 | 22 | immunoglobulin lambda constant 3 (Kern-Oz+ |
| IGLJ4 | 22 | immunoglobulin lambda joining 4 (non-functio |
| IGLC4 | 22 | immunoglobulin lambda constant 4 (pseudoge |
| IGLJ5 | 22 | immunoglobulin lambda joining 5 (non-functio |
| IGLC5 | 22 | immunoglobulin lambda constant 5 (pseudoge |
| IGLJ6 | 22 | immunoglobulin lambda joining 6 [Source:HGI |
| IGLC6 | 22 | immunoglobulin lambda constant 6 (gene/pse |
| IGLJ7 | 22 | immunoglobulin lambda joining 7 [Source:HGI |
| IGLC7 | 22 | immunoglobulin lambda constant 7 [Source:H |
| AC245054.2 | 22 | ATP synthase, H+ transporting, mitochondrial |
| AC245054.1 | 22 | solute carrier family 5 (sodium/glucose cotrar |
| RSPH14 | 22 | radial spoke head 14 homolog [Source:HGNC |
| GNAZ | 22 | G protein subunit alpha z [Source:HGNC Symb |
| Metazoa_SRP | 22 | Metazoan signal recognition particle RNA [Sou |
| U7 | 22 | U7 small nuclear RNA [Source:RFAM;Acc:RF0C |
| RAB36 | 22 | RAB36, member RAS oncogene family [Source |
| BCR | 22 | BCR activator of RhoGEF and GTPase [Source: |
| BCRP8 | 22 | BCR pseudogene 8 [Source:HGNC Symbol;Acc |
| RN7SL263P | 22 | RNA, 7SL, cytoplasmic 263, pseudogene [Sour |
| FBXW4P1 | 22 | F-box and WD repeat domain containing 4 pse |
| AP000343.1 | 22 | ribosomal protein S10 (RPS10) pseudogene |
| LINC02556 | 22 | long intergenic non-protein coding RNA 2556 |
| CES5AP1 | 22 | carboxylesterase 5A pseudogene 1 [Source:HC |

| | | |
|---|---|---|
| ZDHHC8P1 | 22 | zinc finger DHHC-type containing 8 pseudoger |
| AP000344.2 | 22 | coiled-coil domain containing 188 (CCDC188) |
| LINC01659 | 22 | long intergenic non-protein coding RNA 1659 |
| AP000344.1 | 22 | MAD1 mitotic arrest deficient-like 1 (yeast)(M |
| FAM230I | 22 | family with sequence similarity 230 member I |
| AP000345.3 | 22 | novel transcript |
| LINC02557 | 22 | long intergenic non-protein coding RNA 2557 |
| PCAT14 | 22 | prostate cancer associated transcript 14 [Sour |
| AP000345.1 | 22 | novel transcript |
| IGLL1 | 22 | immunoglobulin lambda like polypeptide 1 [So |
| AP000345.2 | 22 | novel transcript |
| AP000346.4 | 22 | rhabdoid tumor deletion region gene 1 (RTDR |
| DRICH1 | 22 | aspartate rich 1 [Source:HGNC Symbol;Acc:HG |
| AP000346.3 | 22 | solute carrier family 5 (sodium/glucose cotrar |
| AP000346.1 | 22 | novel transcript |
| GUSBP11 | 22 | GUSB pseudogene 11 [Source:HGNC Symbol;A |
| AP000346.2 | 22 | immunoglobulin lambda-like polypeptide 1 (IG |
| ASLP1 | 22 | argininosuccinate lyase pseudogene 1 [Source |
| AP000347.1 | 22 | glucuronidase, beta (GUSB) pseudogene |
| RGL4 | 22 | ral guanine nucleotide dissociation stimulator |
| ZNF70 | 22 | zinc finger protein 70 [Source:HGNC Symbol;A |
| VPREB3 | 22 | V-set pre-B cell surrogate light chain 3 [Source |
| C22orf15 | 22 | chromosome 22 open reading frame 15 [Sour |
| CHCHD10 | 22 | coiled-coil-helix-coiled-coil-helix domain conta |
| MMP11 | 22 | matrix metallopeptidase 11 [Source:HGNC Syr |
| SMARCB1 | 22 | SWI/SNF related, matrix associated, actin dep |
| DERL3 | 22 | derlin 3 [Source:HGNC Symbol;Acc:HGNC:142 |
| AP000350.5 | 22 | novel transcript, antisense to SLC2A11 |
| SLC2A11 | 22 | solute carrier family 2 member 11 [Source:HG |
| AP000350.4 | 22 | novel protein |
| AP000350.7 | 22 | novel transcript, antisense to SLC2A11 |
| RN7SL268P | 22 | RNA, 7SL, cytoplasmic 268, pseudogene [Sour |
| MIF | 22 | macrophage migration inhibitory factor [Sourc |
| MIF-AS1 | 22 | MIF antisense RNA 1 [Source:HGNC Symbol;Ac |
| AP000350.6 | 22 | novel transcript |
| AP000350.2 | 22 | kelch-like 5 (Drosophila) (KLHL5) pseudogene |
| AP000350.1 | 22 | glutathione S-transferase (GST) theta pseudog |
| AP000350.3 | 22 | glutathione S-transferase (GST) theta pseudog |
| GSTT2B | 22 | glutathione S-transferase theta 2B (gene/pseu |
| AC253536.2 | 22 | novel pseudogene |
| DDTL | 22 | D-dopachrome tautomerase like [Source:HGN |
| AC253536.6 | 22 | novel transcript |
| DDT | 22 | D-dopachrome tautomerase [Source:HGNC Sy |
| AC253536.7 | 22 | novel protein, AP000351.4-DDT readtrhough |
| AC253536.1 | 22 | glutathione S-transferase (GST) theta pseudog |
| AC253536.5 | 22 | glutathione S-transferase (GST) theta pseudog |
| GSTT2 | 22 | glutathione S-transferase theta 2 (gene/pseuc |
| GSTT4 | 22 | glutathione S-transferase theta 4 [Source:HGN |
| AC253536.4 | 22 | glutathione S-transferase (GST) theta pseudog |
| CABIN1 | 22 | calcineurin binding protein 1 [Source:HGNC Sy |

| | | |
|---|---|---|
| AC253536.3 | 22 | novel transcript |
| SUSD2 | 22 | sushi domain containing 2 [Source:HGNC Sym |
| GGT5 | 22 | gamma-glutamyltransferase 5 [Source:HGNC S |
| GGTLC4P | 22 | gamma-glutamyltransferase light chain 4 pseu |
| POM121L9P | 22 | POM121 transmembrane nucleoporin like 9, p |
| BCRP1 | 22 | BCR pseudogene 1 [Source:HGNC Symbol;Acc |
| AP000354.1 | 22 | 40S ribosomal protein S10 (RPS10) pseudoger |
| SPECC1L | 22 | sperm antigen with calponin homology and cc |
| SPECC1L-ADORA2A | 22 | SPECC1L-ADORA2A readthrough (NMD candic |
| ADORA2A | 22 | adenosine A2a receptor [Source:HGNC Symbc |
| ADORA2A-AS1 | 22 | ADORA2A antisense RNA 1 [Source:HGNC Syn |
| UPB1 | 22 | beta-ureidopropionase 1 [Source:HGNC Symb |
| AP000355.1 | 22 | novel transcript |
| GUCD1 | 22 | guanylyl cyclase domain containing 1 [Source: |
| SNRPD3 | 22 | small nuclear ribonucleoprotein D3 polypeptic |
| GGT1 | 22 | gamma-glutamyltransferase 1 [Source:HGNC S |
| LRRC75B | 22 | leucine rich repeat containing 75B [Source:HG |
| GGT1 | 22 | gamma-glutamyltransferase 1 [Source:HGNC S |
| AP000356.1 | 22 | novel transcript |
| BCRP3 | 22 | BCR pseudogene 3 [Source:HGNC Symbol;Acc |
| AP000356.4 | 22 | gamma-glutamyltransferase 5 (GGT5) pseudo |
| AP000356.3 | 22 | breakpoint cluster region pseudogene 3 |
| AP000356.2 | 22 | novel transcript |
| POM121L10P | 22 | POM121 transmembrane nucleoporin like 10, |
| ARL5AP4 | 22 | ADP ribosylation factor like GTPase 5A pseudc |
| AP000357.1 | 22 | ARP2 actin-related protein 2 homolog (yeast) |
| AP000357.2 | 22 | novel transcript |
| AP000358.1 | 22 | pseudogene similar to part of fructosamine 3 |
| CRIP1P4 | 22 | cysteine rich protein 1 pseudogene 4 [Source: |
| PIWIL3 | 22 | piwi like RNA-mediated gene silencing 3 [Sour |
| SGSM1 | 22 | small G protein signaling modulator 1 [Source |
| AL049759.1 | 22 | NULL |
| TMEM211 | 22 | transmembrane protein 211 [Source:HGNC Sy |
| KIAA1671 | 22 | KIAA1671 [Source:HGNC Symbol;Acc:HGNC:2 |
| AL022323.3 | 22 | novel transcript |
| AL022323.5 | 22 | novel transcript |
| AL022323.2 | 22 | novel transcript |
| AL022323.4 | 22 | novel transcript |
| AL022323.1 | 22 | novel transcript |
| Z99916.2 | 22 | stathmin-like 3 (STMN3) pseudogene |
| CRYBB3 | 22 | crystallin beta B3 [Source:HGNC Symbol;Acc:H |
| Z99916.1 | 22 | novel transcript |
| CRYBB2 | 22 | crystallin beta B2 [Source:HGNC Symbol;Acc:H |
| Z99916.3 | 22 | novel transcript |
| AL022332.1 | 22 | novel transcript |
| AL022324.4 | 22 | novel transcript |
| AL022332.2 | 22 | G protein-coupled receptor kinase 3 (GRK3) p |
| IGLL3P | 22 | immunoglobulin lambda like polypeptide 3, ps |
| AL022324.1 | 22 | novel pseudogene |
| AL022324.2 | 22 | PHD finger protein 10 (PHF10) pseudogene |

| | | |
|---|---|---|
| LRP5L | 22 | LDL receptor related protein 5 like [Source:HG |
| AL022324.3 | 22 | novel transcript |
| AL008721.1 | 22 | novel transcript |
| IGLVIVOR22-1 | 22 | immunoglobulin lambda variable (IV)/OR22-1 |
| CRYBB2P1 | 22 | crystallin beta B2 pseudogene 1 [Source:HGN |
| MIR6817 | 22 | microRNA 6817 [Source:HGNC Symbol;Acc:HG |
| AL008721.2 | 22 | novel transcript |
| AL022329.1 | 22 | novel transcript |
| GRK3 | 22 | G protein-coupled receptor kinase 3 [Source:H |
| AL022329.2 | 22 | novel transcript |
| YES1P1 | 22 | YES1 pseudogene 1 [Source:HGNC Symbol;Acc |
| RNA5SP494 | 22 | RNA, 5S ribosomal pseudogene 494 [Source:H |
| MYO18B | 22 | myosin XVIIIB [Source:HGNC Symbol;Acc:HGN |
| AL022329.3 | 22 | novel transcript, antisense to MYO18B |
| Z98949.1 | 22 | novel transcript, antisense to MYO18B |
| Z98949.3 | 22 | novel transcript |
| RN7SKP169 | 22 | RN7SK pseudogene 169 [Source:HGNC Symbo |
| Z98949.2 | 22 | pseudogene similar to part of ubiquitin-conju |
| LINC02559 | 22 | long intergenic non-protein coding RNA 2559 |
| AL022337.1 | 22 | novel transcript |
| SEZ6L | 22 | seizure related 6 homolog like [Source:HGNC S |
| AL080273.1 | 22 | novel transcript |
| RNA5SP495 | 22 | RNA, 5S ribosomal pseudogene 495 [Source:H |
| Z99714.1 | 22 | ribosomal protein S3A (RPS3A) pseudogene |
| ASPHD2 | 22 | aspartate beta-hydroxylase domain containin |
| HPS4 | 22 | HPS4 biogenesis of lysosomal organelles com |
| SRRD | 22 | SRR1 domain containing [Source:HGNC Symbo |
| TFIP11 | 22 | tuftelin interacting protein 11 [Source:HGNC S |
| Z95115.1 | 22 | novel transcript, antisense to TFIP11 |
| TPST2 | 22 | tyrosylprotein sulfotransferase 2 [Source:HGN |
| MIR548J | 22 | microRNA 548j [Source:HGNC Symbol;Acc:HG |
| HMGB1P10 | 22 | high mobility group box 1 pseudogene 10 [Sou |
| Z95115.2 | 22 | novel transcript, antisense to CRYBB1 |
| CRYBB1 | 22 | crystallin beta B1 [Source:HGNC Symbol;Acc:H |
| CRYBA4 | 22 | crystallin beta A4 [Source:HGNC Symbol;Acc:H |
| ISCA2P1 | 22 | iron-sulfur cluster assembly 2 pseudogene 1 [S |
| MIAT | 22 | myocardial infarction associated transcript [So |
| Z99774.5 | 22 | NULL |
| Z99774.1 | 22 | novel transcript |
| Z99774.4 | 22 | NULL |
| Z99774.2 | 22 | NULL |
| MIATNB | 22 | MIAT neighbor [Source:HGNC Symbol;Acc:HG |
| Z99774.3 | 22 | NULL |
| LINC01422 | 22 | long intergenic non-protein coding RNA 1422 |
| Z97353.1 | 22 | ribosomal protein L15 (RPL15) pseudogene |
| Z97353.2 | 22 | novel transcript |
| AL008638.3 | 22 | novel transcript |
| RNU6-1066P | 22 | RNA, U6 small nuclear 1066, pseudogene [Sou |
| AL008638.5 | 22 | novel transcript |
| AL008638.4 | 22 | novel transcript |

| | | |
|---|---|---|
| AL008638.2 | 22 | uncharacterized LOC284898 [Source:NCBI ger |
| AL008638.6 | 22 | novel transcript |
| AL008638.1 | 22 | novel transcript |
| AL021153.1 | 22 | novel transcript |
| LINC01638 | 22 | long intergenic non-protein coding RNA 1638 |
| AL020994.3 | 22 | novel transcript |
| AL020994.1 | 22 | novel transcript |
| LINC02554 | 22 | long intergenic non-protein coding RNA 2554 |
| AL020994.2 | 22 | novel transcript |
| AL049536.1 | 22 | novel transcript |
| AL050402.2 | 22 | uncharacterized LOC105372980 [Source:NCBI |
| AL050402.1 | 22 | novel transcript |
| AL133456.1 | 22 | novel transcript |
| AL121885.2 | 22 | novel transcript |
| AL121885.3 | 22 | novel transcript |
| AL121885.1 | 22 | novel transcript |
| MN1 | 22 | MN1 proto-oncogene, transcriptional regulato |
| PITPNB | 22 | phosphatidylinositol transfer protein beta [So |
| TTC28-AS1 | 22 | TTC28 antisense RNA 1 [Source:HGNC Symbol |
| AL031591.2 | 22 | NULL |
| MIR3199-1 | 22 | microRNA 3199-1 [Source:HGNC Symbol;Acc:I |
| MIR3199-2 | 22 | microRNA 3199-2 [Source:HGNC Symbol;Acc:I |
| AL031591.1 | 22 | NULL |
| TTC28 | 22 | tetratricopeptide repeat domain 28 [Source:H |
| AL033538.1 | 22 | NULL |
| AL033538.2 | 22 | NULL |
| RN7SL757P | 22 | RNA, 7SL, cytoplasmic 757, pseudogene [Sour |
| SNORD42 | 22 | Small nucleolar RNA SNORD42 [Source:RFAM; |
| Y_RNA | 22 | Y RNA [Source:RFAM;Acc:RF00019] |
| MIR5739 | 22 | microRNA 5739 [Source:HGNC Symbol;Acc:HG |
| RN7SL162P | 22 | RNA, 7SL, cytoplasmic 162, pseudogene [Sour |
| CHEK2 | 22 | checkpoint kinase 2 [Source:HGNC Symbol;Ac |
| HSCB | 22 | HscB mitochondrial iron-sulfur cluster cochap |
| CCDC117 | 22 | coiled-coil domain containing 117 [Source:HG |
| XBP1 | 22 | X-box binding protein 1 [Source:HGNC Symbo |
| Z93930.2 | 22 | novel transcript |
| Z93930.3 | 22 | novel transcript |
| Z93930.1 | 22 | chromosome 9 open reading frame 16 (C9orf1 |
| ZNRF3 | 22 | zinc and ring finger 3 [Source:HGNC Symbol;A |
| ZNRF3-IT1 | 22 | ZNRF3 intronic transcript 1 [Source:HGNC Syn |
| ZNRF3-AS1 | 22 | ZNRF3 antisense RNA 1 [Source:HGNC Symbol |
| C22orf31 | 22 | chromosome 22 open reading frame 31 [Sour |
| KREMEN1 | 22 | kringle containing transmembrane protein 1 [ |
| AL021393.1 | 22 | novel transcript |
| RNU6-810P | 22 | RNA, U6 small nuclear 810, pseudogene [Sour |
| RNU6-1219P | 22 | RNA, U6 small nuclear 1219, pseudogene [Sou |
| EMID1 | 22 | EMI domain containing 1 [Source:HGNC Symb |
| RHBDD3 | 22 | rhomboid domain containing 3 [Source:HGNC |
| AL031186.1 | 22 | novel transcript |
| EWSR1 | 22 | EWS RNA binding protein 1 [Source:HGNC Syn |

| | | |
|---|---|---|
| GAS2L1 | 22 | growth arrest specific 2 like 1 [Source:HGNC S |
| RASL10A | 22 | RAS like family 10 member A [Source:HGNC Sy |
| AC002059.1 | 22 | novel transcript, novel transcript, sense intror |
| AP1B1 | 22 | adaptor related protein complex 1 subunit be |
| AC002059.2 | 22 | NULL |
| SNORD125 | 22 | small nucleolar RNA, C/D box 125 [Source:HG |
| AC002059.3 | 22 | chromosome 22 open reading frame 42 (C22c |
| AC000041.1 | 22 | adaptor-related protein complex 1, beta 1 sub |
| RFPL4AP6 | 22 | ret finger protein like 4A pseudogene 6 [Sourc |
| RFPL1S | 22 | RFPL1 antisense RNA 1 [Source:HGNC Symbol |
| RFPL1 | 22 | ret finger protein like 1 [Source:HGNC Symbol |
| NEFH | 22 | neurofilament heavy [Source:HGNC Symbol;A |
| AC000035.1 | 22 | novel transcript, antisense to NEFH |
| THOC5 | 22 | THO complex 5 [Source:HGNC Symbol;Acc:HG |
| AC005529.1 | 22 | novel transcript, sense intronic to THOC5 |
| NIPSNAP1 | 22 | nipsnap homolog 1 [Source:HGNC Symbol;Acc |
| NF2 | 22 | neurofibromin 2 [Source:HGNC Symbol;Acc:H |
| RPEP4 | 22 | ribulose-5-phosphate-3-epimerase pseudoger |
| AC004882.1 | 22 | novel transcript |
| AC004882.2 | 22 | novel transcript |
| CABP7 | 22 | calcium binding protein 7 [Source:HGNC Symb |
| ZMAT5 | 22 | zinc finger matrin-type 5 [Source:HGNC Symb |
| Y_RNA | 22 | Y RNA [Source:RFAM;Acc:RF00019] |
| AC004882.3 | 22 | novel transcript, antisense to UQCR10 |
| UQCR10 | 22 | ubiquinol-cytochrome c reductase, complex II |
| ASCC2 | 22 | activating signal cointegrator 1 complex subu |
| MTMR3 | 22 | myotubularin related protein 3 [Source:HGNC |
| RNU6-331P | 22 | RNA, U6 small nuclear 331, pseudogene [Sour |
| AC003681.1 | 22 | novel transcript, antisense to MTMR3 |
| MIR6818 | 22 | microRNA 6818 [Source:HGNC Symbol;Acc:HG |
| HORMAD2-AS1 | 22 | HORMAD2 and MTMR3 antisense RNA 1 [Sou |
| CNN2P1 | 22 | calponin 2 pseudogene 1 [Source:HGNC Symb |
| HORMAD2 | 22 | HORMA domain containing 2 [Source:HGNC Sy |
| AC003071.2 | 22 | ribosomal protein L36 (RPL36) pseudogene |
| AC003071.1 | 22 | ribosomal protein S3A (RPS3A) pseudogene |
| AC002378.1 | 22 | novel transcript |
| LIF-AS1 | 22 | LIF antisense RNA 1 [Source:HGNC Symbol;Ac |
| LIF | 22 | LIF interleukin 6 family cytokine [Source:HGN |
| AC004264.1 | 22 | novel transcript, antisense to LIF |
| OSM | 22 | oncostatin M [Source:HGNC Symbol;Acc:HGN |
| AC004264.2 | 22 | TEC |
| CASTOR1 | 22 | cytosolic arginine sensor for mTORC1 subunit |
| AC004997.1 | 22 | novel protein |
| TBC1D10A | 22 | TBC1 domain family member 10A [Source:HG |
| SF3A1 | 22 | splicing factor 3a subunit 1 [Source:HGNC Sym |
| CCDC157 | 22 | coiled-coil domain containing 157 [Source:HG |
| RNF215 | 22 | ring finger protein 215 [Source:HGNC Symbol; |
| SEC14L2 | 22 | SEC14 like lipid binding 2 [Source:HGNC Symb |
| AC004832.3 | 22 | novel SEC14-like 2 (S. cerevisiae) (SEC14L2) ar |
| AC004832.5 | 22 | novel transcript, sense intronic to SEC14L2 |

| | | |
|---|---|---|
| RNU6-564P | 22 | RNA, U6 small nuclear 564, pseudogene [Sour |
| AC004832.4 | 22 | novel transcript, sense intonic to |
| MTFP1 | 22 | mitochondrial fission process 1 [Source:HGNC |
| AC004832.6 | 22 | novel transcript |
| AC004832.2 | 22 | solute carrier family 39 (zinc transporter), mer |
| SEC14L3 | 22 | SEC14 like lipid binding 3 [Source:HGNC Symb |
| AC004832.1 | 22 | novel transcript, antisense to SEC14L4 |
| SDC4P | 22 | syndecan 4 pseudogene [Source:HGNC Symbo |
| SEC14L4 | 22 | SEC14 like lipid binding 4 [Source:HGNC Symb |
| SEC14L6 | 22 | SEC14 like lipid binding 6 [Source:HGNC Symb |
| SIRPAP1 | 22 | signal regulatory protein alpha pseudogene 1 |
| GAL3ST1 | 22 | galactose-3-O-sulfotransferase 1 [Source:HGN |
| PES1 | 22 | pescadillo ribosomal biogenesis factor 1 [Sour |
| AC005006.1 | 22 | novel transcript |
| TCN2 | 22 | transcobalamin 2 [Source:HGNC Symbol;Acc:H |
| SLC35E4 | 22 | solute carrier family 35 member E4 [Source:H |
| DUSP18 | 22 | dual specificity phosphatase 18 [Source:HGNC |
| AC003072.1 | 22 | ribosomal protein L13a (RPL13A) pseudogene |
| AC003072.2 | 22 | chromosome 5 open reading frame 52 (C5orf5 |
| OSBP2 | 22 | oxysterol binding protein 2 [Source:HGNC Sym |
| MIR3200 | 22 | microRNA 3200 [Source:HGNC Symbol;Acc:HG |
| EIF4HP2 | 22 | eukaryotic translation initiation factor 4H pse |
| MORC2-AS1 | 22 | MORC2 antisense RNA 1 [Source:HGNC Symbo |
| MORC2 | 22 | MORC family CW-type zinc finger 2 [Source:HG |
| TUG1 | 22 | taurine up-regulated 1 [Source:HGNC Symbol; |
| AC004542.5 | 22 | NULL |
| AC004542.4 | 22 | NULL |
| AC004542.6 | 22 | NULL |
| AC004542.3 | 22 | NULL |
| AC004542.2 | 22 | novel transcript |
| AC004542.1 | 22 | novel transcript |
| AC005005.4 | 22 | novel transcript, sense overlapping SMTN |
| RN7SL633P | 22 | RNA, 7SL, cytoplasmic 633, pseudogene [Sour |
| SMTN | 22 | smoothelin [Source:HGNC Symbol;Acc:HGNC: |
| AC005005.3 | 22 | novel transcript, antisense to SMTN |
| SELENOM | 22 | selenoprotein M [Source:HGNC Symbol;Acc:H |
| AC005005.2 | 22 | ribosomal protein S15a (RPS15A) pseudogene |
| INPP5J | 22 | inositol polyphosphate-5-phosphatase J [Sour |
| PLA2G3 | 22 | phospholipase A2 group III [Source:HGNC Sym |
| AC005005.1 | 22 | small nuclear ribonucleoprotein polypeptide N |
| MIR3928 | 22 | microRNA 3928 [Source:HGNC Symbol;Acc:HG |
| RNF185 | 22 | ring finger protein 185 [Source:HGNC Symbol; |
| RNF185-AS1 | 22 | RNF185 antisense RNA 1 [Source:HGNC Symbo |
| LIMK2 | 22 | LIM domain kinase 2 [Source:HGNC Symbol;Ac |
| RNU6-1128P | 22 | RNA, U6 small nuclear 1128, pseudogene [Sou |
| Y_RNA | 22 | Y RNA [Source:RFAM;Acc:RF00019] |
| PIK3IP1 | 22 | phosphoinositide-3-kinase interacting protein |
| PIK3IP1-AS1 | 22 | PIK3IP1 antisense RNA 1 (head to head) [Sour |
| RNA5SP496 | 22 | RNA, 5S ribosomal pseudogene 496 [Source:H |
| PATZ1 | 22 | POZ/BTB and AT hook containing zinc finger 1 |

| | | |
|---|---|---|
| LINC01521 | 22 | long intergenic non-protein coding RNA 1521 |
| RNU6-338P | 22 | RNA, U6 small nuclear 338, pseudogene [Sour |
| DRG1 | 22 | developmentally regulated GTP binding prote |
| EIF4ENIF1 | 22 | eukaryotic translation initiation factor 4E nucl |
| AL096701.3 | 22 | novel transcript |
| RNU6-28P | 22 | RNA, U6 small nuclear 28, pseudogene [Sourc |
| SFI1 | 22 | SFI1 centrin binding protein [Source:HGNC Syr |
| AL096701.4 | 22 | novel transcript, antisense to EIF4ENIF1 |
| H2AFZP6 | 22 | H2A histone family member Z pseudogene 6 [ |
| AL096701.2 | 22 | ribosomal protein S26 (RPS260 pseudogene |
| AL096701.1 | 22 | ribosomal protein S18 (RPS18) pseudogene |
| PISD | 22 | phosphatidylserine decarboxylase [Source:HG |
| MIR7109 | 22 | microRNA 7109 [Source:HGNC Symbol;Acc:HG |
| PRR14L | 22 | proline rich 14 like [Source:HGNC Symbol;Acc: |
| DEPDC5 | 22 | DEP domain containing 5 [Source:HGNC Symb |
| Z82190.2 | 22 | novel protein, DEPDC5-YWHAH readthrough |
| RN7SL20P | 22 | RNA, 7SL, cytoplasmic 20, pseudogene [Sourc |
| AL022331.1 | 22 | ribosomal protein L17 (RPL17) pseudogene |
| RNU6-201P | 22 | RNA, U6 small nuclear 201, pseudogene [Sour |
| Z82190.1 | 22 | novel pseudogene similar to e(y)2 protein |
| C22orf24 | 22 | chromosome 22 open reading frame 24 [Sour |
| YWHAH | 22 | tyrosine 3-monooxygenase/tryptophan 5-mor |
| AL008719.1 | 22 | novel transcript |
| LINC02558 | 22 | long intergenic non-protein coding RNA 2558 |
| RN7SL305P | 22 | RNA, 7SL, cytoplasmic 305, pseudogene [Sour |
| Z74021.1 | 22 | ribosomal protein S17 (RPS17) pseudogene |
| SLC5A1 | 22 | solute carrier family 5 member 1 [Source:HGN |
| AP1B1P1 | 22 | AP1B1 pseudogene 1 [Source:HGNC Symbol;A |
| Z83839.1 | 22 | novel pseudogene |
| AP1B1P2 | 22 | AP1B1 pseudogene 2 [Source:HGNC Symbol;A |
| Z83839.2 | 22 | novel transcript |
| C22orf42 | 22 | chromosome 22 open reading frame 42 [Sour |
| AL008723.1 | 22 | novel transcript |
| AL008723.3 | 22 | NULL |
| RFPL2 | 22 | ret finger protein like 2 [Source:HGNC Symbol |
| IGLCOR22-1 | 22 | immunoglobulin lambda constant/OR22-1 (ps |
| SLC5A4-AS1 | 22 | SLC5A4 antisense RNA 1 [Source:HGNC Symbo |
| SLC5A4 | 22 | solute carrier family 5 member 4 [Source:HGN |
| CPSF1P1 | 22 | cleavage and polyadenylation specific factor 1 |
| AL008723.2 | 22 | novel transcript |
| AL021937.3 | 22 | novel transcript |
| RFPL3 | 22 | ret finger protein like 3 [Source:HGNC Symbol |
| IGLCOR22-2 | 22 | immunoglobulin lambda constant/OR22-2 (ps |
| RFPL3S | 22 | RFPL3 antisense [Source:HGNC Symbol;Acc:H |
| AL021937.6 | 22 | NULL |
| AL021937.5 | 22 | NULL |
| AL021937.1 | 22 | novel transcript |
| IGLVIVOR22-2 | 22 | immunoglobulin lambda variable (IV)/OR22-2 |
| AL021937.4 | 22 | novel transcript |
| AL021937.2 | 22 | Ral-GDS related protein Rgr (Rgr) pseudogene |

| | | |
|---|---|---|
| RTCB | 22 | RNA 2',3'-cyclic phosphate and 5'-OH ligase [S |
| BPIFC | 22 | BPI fold containing family C [Source:HGNC Syr |
| FBXO7 | 22 | F-box protein 7 [Source:HGNC Symbol;Acc:HG |
| SYN3 | 22 | synapsin III [Source:HGNC Symbol;Acc:HGNC:: |
| Z82246.1 | 22 | novel transcript |
| RNA5SP497 | 22 | RNA, 5S ribosomal pseudogene 497 [Source:H |
| Z73495.1 | 22 | novel transcript |
| TIMP3 | 22 | TIMP metallopeptidase inhibitor 3 [Source:HG |
| Z83846.1 | 22 | novel pseudogene |
| LINC01640 | 22 | long intergenic non-protein coding RNA 1640 |
| Z82198.1 | 22 | novel transcript |
| Z82198.3 | 22 | ibernator-specific blood complex, 20 kDa subu |
| LARGE1 | 22 | LARGE xylosyl- and glucuronyltransferase 1 [S |
| Z82198.2 | 22 | novel transcript, antisense to LARGE |
| Z82173.1 | 22 | novel transcript |
| MIR4764 | 22 | microRNA 4764 [Source:HGNC Symbol;Acc:HG |
| SNORA50B | 22 | small nucleolar RNA, H/ACA box 50B [Source:I |
| Z69713.1 | 22 | NULL |
| LARGE-IT1 | 22 | LARGE intronic transcript 1 [Source:HGNC Sym |
| LARGE-AS1 | 22 | LARGE antisense RNA 1 [Source:HGNC Symbo |
| Z73429.1 | 22 | novel transcript, antisense to LARGE |
| LINC01643 | 22 | long intergenic non-protein coding RNA 1643 |
| Z68323.1 | 22 | novel transcript |
| AL021877.1 | 22 | aconitase 2, mitochondrial (ACO2) pseudogen |
| AL021877.2 | 22 | novel transcript |
| Z82196.1 | 22 | novel transcript |
| Z82196.2 | 22 | novel transcript |
| AL024495.1 | 22 | novel transcript |
| ISX | 22 | intestine specific homeobox [Source:HGNC Sy |
| LINC01399 | 22 | long intergenic non-protein coding RNA 1399 |
| Z99755.2 | 22 | novel transcript |
| Z99755.1 | 22 | nerve growth factor receptor (TNFRSF16) assc |
| RNU7-167P | 22 | RNA, U7 small nuclear 167 pseudogene [Sourc |
| HMGXB4 | 22 | HMG-box containing 4 [Source:HGNC Symbol; |
| AL008635.1 | 22 | novel transcript, antisense to TOM1 |
| TOM1 | 22 | target of myb1 membrane trafficking protein |
| MIR3909 | 22 | microRNA 3909 [Source:HGNC Symbol;Acc:HG |
| MIR6069 | 22 | microRNA 6069 [Source:HGNC Symbol;Acc:HG |
| Z82244.2 | 22 | novel transcript |
| Z82244.1 | 22 | novel transcript |
| HMOX1 | 22 | heme oxygenase 1 [Source:HGNC Symbol;Acc |
| MCM5 | 22 | minichromosome maintenance complex comp |
| AL022334.1 | 22 | tRNA methyltransferase 11 homolog (S. cerev |
| AL022334.2 | 22 | novel transcript |
| RASD2 | 22 | RASD family member 2 [Source:HGNC Symbol |
| MB | 22 | myoglobin [Source:HGNC Symbol;Acc:HGNC:6 |
| AL049747.1 | 22 | novel transcript |
| APOL6 | 22 | apolipoprotein L6 [Source:HGNC Symbol;Acc:I |
| AL049748.1 | 22 | novel transcript |
| MRPS16P3 | 22 | mitochondrial ribosomal protein S16 pseudog |

| | | |
|---|---|---|
| APOL5 | 22 | apolipoprotein L5 [Source:HGNC Symbol;Acc:I |
| RBFOX2 | 22 | RNA binding fox-1 homolog 2 [Source:HGNC S |
| NDUFA9P1 | 22 | NADH:ubiquinone oxidoreductase subunit A9 |
| Z82217.1 | 22 | novel transcript |
| Z95114.4 | 22 | novel transcript |
| Z95114.1 | 22 | novel transcript |
| Z95114.2 | 22 | novel transcript |
| Z95114.3 | 22 | TEC |
| APOL3 | 22 | apolipoprotein L3 [Source:HGNC Symbol;Acc:I |
| MTCO1P20 | 22 | MT-CO1 pseudogene 20 [Source:HGNC Symbo |
| MTCO2P20 | 22 | MT-CO2 pseudogene 20 [Source:HGNC Symbo |
| MTATP6P20 | 22 | MT-ATP6 pseudogene 20 [Source:HGNC Symb |
| MTCO3P20 | 22 | MT-CO3 pseudogene 20 [Source:HGNC Symbo |
| MTCYBP34 | 22 | MT-CYB pseudogene 34 [Source:HGNC Symbo |
| MTND1P10 | 22 | MT-ND1 pseudogene 10 [Source:HGNC Symbo |
| APOL4 | 22 | apolipoprotein L4 [Source:HGNC Symbol;Acc:I |
| APOL2 | 22 | apolipoprotein L2 [Source:HGNC Symbol;Acc:I |
| APOL1 | 22 | apolipoprotein L1 [Source:HGNC Symbol;Acc:I |
| MYH9 | 22 | myosin heavy chain 9 [Source:HGNC Symbol;A |
| MIR6819 | 22 | microRNA 6819 [Source:HGNC Symbol;Acc:HG |
| Z82215.1 | 22 | novel transcript, antisense to ANKRD54and M |
| FO393418.1 | 22 | novel transcript |
| RPS15AP38 | 22 | ribosomal protein S15a pseudogene 38 [Sourc |
| Y_RNA | 22 | Y RNA [Source:RFAM;Acc:RF00019] |
| AL022313.2 | 22 | novel transcript |
| TXN2 | 22 | thioredoxin 2 [Source:HGNC Symbol;Acc:HGN |
| FOXRED2 | 22 | FAD dependent oxidoreductase domain conta |
| EIF3D | 22 | eukaryotic translation initiation factor 3 subur |
| AL022313.1 | 22 | Hermansky-Pudlak syndrome 1 (HPS1) pseudo |
| AL022313.3 | 22 | novel transcript |
| AL022313.4 | 22 | novel transcript |
| CACNG2 | 22 | calcium voltage-gated channel auxiliary subun |
| AL049749.1 | 22 | novel transcript |
| Z80897.2 | 22 | novel transcript, antisense to TRIOBP |
| IFT27 | 22 | intraflagellar transport 27 [Source:HGNC Syml |
| Z80897.1 | 22 | novel transcript, antisense to ANKRD54and IF |
| PVALB | 22 | parvalbumin [Source:HGNC Symbol;Acc:HGNC |
| Z82185.1 | 22 | novel transcript |
| NCF4-AS1 | 22 | NCF4 antisense RNA 1 [Source:HGNC Symbol;. |
| NCF4 | 22 | neutrophil cytosolic factor 4 [Source:HGNC Sy |
| CSF2RB | 22 | colony stimulating factor 2 receptor beta com |
| CSF2RBP1 | 22 | colony stimulating factor 2 receptor beta com |
| Z82180.1 | 22 | novel transcript |
| TEX33 | 22 | testis expressed 33 [Source:HGNC Symbol;Acc |
| TST | 22 | thiosulfate sulfurtransferase [Source:HGNC Sy |
| Y_RNA | 22 | Y RNA [Source:RFAM;Acc:RF00019] |
| MPST | 22 | mercaptopyruvate sulfurtransferase [Source:H |
| Z73420.1 | 22 | novel transcript, antisense to ANKRD54 |
| KCTD17 | 22 | potassium channel tetramerization domain cc |
| RN7SKP214 | 22 | RN7SK pseudogene 214 [Source:HGNC Symbo |

| | | |
|---|---|---|
| TMPRSS6 | 22 | transmembrane serine protease 6 [Source:HG |
| AL022314.1 | 22 | novel transcript |
| IL2RB | 22 | interleukin 2 receptor subunit beta [Source:H |
| Z82188.2 | 22 | novel transcript |
| C1QTNF6 | 22 | C1q and TNF related 6 [Source:HGNC Symbol; |
| SSTR3 | 22 | somatostatin receptor 3 [Source:HGNC Symbo |
| Z82188.1 | 22 | ribosomal protein L39 (RPL39) pseudogene |
| RAC2 | 22 | Rac family small GTPase 2 [Source:HGNC Sym |
| CYTH4 | 22 | cytohesin 4 [Source:HGNC Symbol;Acc:HGNC: |
| FP325335.1 | 22 | novel transcript |
| Z94160.1 | 22 | novel transcript |
| ELFN2 | 22 | extracellular leucine rich repeat and fibronect |
| Z94160.2 | 22 | novel transcript, antisense to ELFN2 |
| MFNG | 22 | MFNG O-fucosylpeptide 3-beta-N-acetylgluco |
| CARD10 | 22 | caspase recruitment domain family member 1 |
| AL022315.1 | 22 | novel transcript |
| CDC42EP1 | 22 | CDC42 effector protein 1 [Source:HGNC Symb |
| LGALS2 | 22 | galectin 2 [Source:HGNC Symbol;Acc:HGNC:6 |
| Metazoa_SRP | 22 | Metazoan signal recognition particle RNA [Sou |
| GGA1 | 22 | golgi associated, gamma adaptin ear containi |
| SH3BP1 | 22 | SH3 domain binding protein 1 [Source:HGNC |
| Z83844.3 | 22 | novel protein |
| Z83844.2 | 22 | novel transcript |
| PDXP | 22 | pyridoxal phosphatase [Source:HGNC Symbol; |
| RN7SL385P | 22 | RNA, 7SL, cytoplasmic 385, pseudogene [Sour |
| LGALS1 | 22 | galectin 1 [Source:HGNC Symbol;Acc:HGNC:6 |
| NOL12 | 22 | nucleolar protein 12 [Source:HGNC Symbol;A |
| Z83844.1 | 22 | novel transcript |
| TRIOBP | 22 | TRIO and F-actin binding protein [Source:HGN |
| H1F0 | 22 | H1 histone family member 0 [Source:HGNC S |
| GCAT | 22 | glycine C-acetyltransferase [Source:HGNC Sym |
| GALR3 | 22 | galanin receptor 3 [Source:HGNC Symbol;Acc: |
| ANKRD54 | 22 | ankyrin repeat domain 54 [Source:HGNC Sym |
| MIR658 | 22 | microRNA 658 [Source:HGNC Symbol;Acc:HG |
| MIR659 | 22 | microRNA 659 [Source:HGNC Symbol;Acc:HG |
| EIF3L | 22 | eukaryotic translation initiation factor 3 subu |
| AL022311.1 | 22 | novel transcript, sense overlapping to EIF3L |
| RNU6-900P | 22 | RNA, U6 small nuclear 900, pseudogene [Sour |
| MICALL1 | 22 | MICAL like 1 [Source:HGNC Symbol;Acc:HGNC |
| C22orf23 | 22 | chromosome 22 open reading frame 23 [Sour |
| AL031587.5 | 22 | TEC |
| AL031587.3 | 22 | novel transcript, antisense to C22orf23 |
| POLR2F | 22 | RNA polymerase II subunit F [Source:HGNC Sy |
| MIR6820 | 22 | microRNA 6820 [Source:HGNC Symbol;Acc:H |
| SOX10 | 22 | SRY-box transcription factor 10 [Source:HGNC |
| MIR4534 | 22 | microRNA 4534 [Source:HGNC Symbol;Acc:H |
| AL031587.1 | 22 | novel transcript |
| AL031587.4 | 22 | novel transcript |
| PICK1 | 22 | protein interacting with PRKCA 1 [Source:HG |
| AL031587.2 | 22 | novel transcript |

| | | |
|---|---|---|
| SLC16A8 | 22 | solute carrier family 16 member 8 [Source:HG |
| BAIAP2L2 | 22 | BAI1 associated protein 2 like 2 [Source:HGNC |
| AL022322.1 | 22 | novel transcript, antisense to BAIAP2L2 |
| PLA2G6 | 22 | phospholipase A2 group VI [Source:HGNC Sym |
| AL022322.2 | 22 | novel transcript, antisense to PLA2G6 |
| MAFF | 22 | MAF bZIP transcription factor F [Source:HGNC |
| TMEM184B | 22 | transmembrane protein 184B [Source:HGNC S |
| AL020993.1 | 22 | novel transcript, antisense to TMEM184B |
| RN7SL704P | 22 | RNA, 7SL, cytoplasmic 704, pseudogene [Sour |
| Metazoa_SRP | 22 | Metazoan signal recognition particle RNA [Sou |
| CSNK1E | 22 | casein kinase 1 epsilon [Source:HGNC Symbol; |
| TPTEP2-CSNK1E | 22 | TPTEP2-CSNK1E readthrough [Source:HGNC Sy |
| TPTEP2 | 22 | TPTE pseudogene 2 [Source:HGNC Symbol;Acc |
| Z98749.1 | 22 | novel transcript, sense intronic to CSNK1E |
| Z97056.2 | 22 | novel transcript |
| Z97056.1 | 22 | putaitve novel transcript |
| KCNJ4 | 22 | potassium voltage-gated channel subfamily J r |
| Z97056.3 | 22 | novel transcript, antisense to KCNJ4 |
| KDELR3 | 22 | KDEL endoplasmic reticulum protein retentior |
| DDX17 | 22 | DEAD-box helicase 17 [Source:HGNC Symbol;A |
| DMC1 | 22 | DNA meiotic recombinase 1 [Source:HGNC Syr |
| RPS29P31 | 22 | ribosomal protein S29 pseudogene 31 [Source |
| FAM227A | 22 | family with sequence similarity 227 member A |
| CBY1 | 22 | chibby family member 1, beta catenin antagor |
| AL021707.4 | 22 | novel transcript |
| AL021707.2 | 22 | novel transcript, antisense to CBY1 |
| AL021707.9 | 22 | novel transcript |
| TOMM22 | 22 | translocase of outer mitochondrial membrane |
| JOSD1 | 22 | Josephin domain containing 1 [Source:HGNC S |
| GTPBP1 | 22 | GTP binding protein 1 [Source:HGNC Symbol;A |
| PRDX3P1 | 22 | peroxiredoxin 3 pseudogene 1 [Source:HGNC |
| AL021707.5 | 22 | novel transcript, antisense to SUN2 |
| SUN2 | 22 | Sad1 and UNC84 domain containing 2 [Source |
| AL021707.3 | 22 | novel transcript, antisense to SUN2 |
| AL021707.8 | 22 | novel transcript, sense intronic to SUN2 |
| AL021707.1 | 22 | novel transcript |
| AL021707.6 | 22 | novel transcript, antisense to SUN2 |
| AL021707.7 | 22 | novel transcript, antisense to SUN2 |
| DNAL4 | 22 | dynein axonemal light chain 4 [Source:HGNC S |
| NPTXR | 22 | neuronal pentraxin receptor [Source:HGNC Sy |
| CBX6 | 22 | chromobox 6 [Source:HGNC Symbol;Acc:HGN |
| AL022318.5 | 22 | novel transcript |
| AL022318.1 | 22 | novel transcript |
| APOBEC3A | 22 | apolipoprotein B mRNA editing enzyme cataly |
| APOBEC3B | 22 | apolipoprotein B mRNA editing enzyme cataly |
| APOBEC3B-AS1 | 22 | APOBEC3B antisense RNA 1 [Source:HGNC Syr |
| APOBEC3C | 22 | apolipoprotein B mRNA editing enzyme cataly |
| AL022318.4 | 22 | novel protein |
| APOBEC3D | 22 | apolipoprotein B mRNA editing enzyme cataly |
| APOBEC3F | 22 | apolipoprotein B mRNA editing enzyme cataly |

| | | |
|---|---|---|
| AL022318.2 | 22 | apolipoprotein B mRNA editing enzyme, catal |
| APOBEC3G | 22 | apolipoprotein B mRNA editing enzyme catal |
| AL031846.1 | 22 | apolipoprotein B mRNA editing enzyme, catal |
| APOBEC3H | 22 | apolipoprotein B mRNA editing enzyme catal |
| CBX7 | 22 | chromobox 7 [Source:HGNC Symbol;Acc:HGN |
| COX5BP7 | 22 | cytochrome c oxidase subunit 5B pseudogene |
| AL031846.2 | 22 | novel transcript, antisense to CBX7 |
| FUNDC2P4 | 22 | FUN14 domain containing 2 pseudogene 4 [Sc |
| PDGFB | 22 | platelet derived growth factor subunit B [Sour |
| AL031590.1 | 22 | novel transcript, antisense to PDGFB |
| AL022326.1 | 22 | novel transcript |
| RPL3 | 22 | ribosomal protein L3 [Source:HGNC Symbol;A |
| SNORD83B | 22 | small nucleolar RNA, C/D box 83B [Source:HG |
| SNORD83A | 22 | small nucleolar RNA, C/D box 83A [Source:HG |
| SNORD43 | 22 | small nucleolar RNA, C/D box 43 [Source:HGN |
| SYNGR1 | 22 | synaptogyrin 1 [Source:HGNC Symbol;Acc:HG |
| AL022326.2 | 22 | TEC |
| TAB1 | 22 | TGF-beta activated kinase 1 (MAP3K7) binding |
| MGAT3 | 22 | mannosyl (beta-1,4-)-glycoprotein beta-1,4-N |
| MGAT3-AS1 | 22 | MGAT3 antisense RNA 1 [Source:HGNC Symbo |
| MIEF1 | 22 | mitochondrial elongation factor 1 [Source:HG |
| AL022312.1 | 22 | MIEF1 upstream open reading frame protein |
| ATF4 | 22 | activating transcription factor 4 [Source:HGN |
| RPS19BP1 | 22 | ribosomal protein S19 binding protein 1 [Sour |
| CACNA1I | 22 | calcium voltage-gated channel subunit alpha1 |
| ENTHD1 | 22 | ENTH domain containing 1 [Source:HGNC Sym |
| AL022319.1 | 22 | family with sequence similarity 54, member A |
| RN7SKP210 | 22 | RN7SK pseudogene 210 [Source:HGNC Symbo |
| UQCRFS1P1 | 22 | ubiquinol-cytochrome c reductase, Rieske iron |
| GRAP2 | 22 | GRB2 related adaptor protein 2 [Source:HGN |
| Z82206.1 | 22 | novel protein similar to translation machinery |
| FAM83F | 22 | family with sequence similarity 83 member F |
| Z83847.1 | 22 | novel transcript, antisense to FAM83F |
| TNRC6B | 22 | trinucleotide repeat containing adaptor 6B [Sc |
| RPL7P52 | 22 | ribosomal protein L7 pseudogene 52 [Source: |
| ADSL | 22 | adenylosuccinate lyase [Source:HGNC Symbol |
| AL022238.4 | 22 | novel protein |
| SGSM3 | 22 | small G protein signaling modulator 3 [Source |
| AL022238.2 | 22 | novel transcript, antisense to SGSM3 and ADS |
| MRTFA | 22 | myocardin related transcription factor A [Sour |
| AL022238.3 | 22 | novel transcript, antisense to MKL1 |
| AL022238.1 | 22 | novel pseudogene |
| MRTFA-AS1 | 22 | MRTFA antisense RNA 1 [Source:HGNC Symbo |
| COX6B1P3 | 22 | cytochrome c oxidase subunit 6B1 pseudogen |
| RPL4P6 | 22 | ribosomal protein L4 pseudogene 6 [Source:H |
| GAPDHP37 | 22 | glyceraldehyde 3 phosphate dehydrogenase p |
| MCHR1 | 22 | melanin concentrating hormone receptor 1 [S |
| SLC25A17 | 22 | solute carrier family 25 member 17 [Source:H |
| Z98048.1 | 22 | riboflavin kinase (RFK) pseudogene |
| JTBP1 | 22 | jumping translocation breakpoint pseudogene |

| | | |
|---|---|---|
| MIR4766 | 22 | microRNA 4766 [Source:HGNC Symbol;Acc:HG |
| ST13 | 22 | ST13 Hsp70 interacting protein [Source:HGNC |
| XPNPEP3 | 22 | X-prolyl aminopeptidase 3 [Source:HGNC Sym |
| DNAJB7 | 22 | DnaJ heat shock protein family (Hsp40) memb |
| Metazoa_SRP | 22 | Metazoan signal recognition particle RNA [Sou |
| RNU6-379P | 22 | RNA, U6 small nuclear 379, pseudogene [Sour |
| RBX1 | 22 | ring-box 1 [Source:HGNC Symbol;Acc:HGNC:9 |
| Y_RNA | 22 | Y RNA [Source:RFAM;Acc:RF00019] |
| AL080243.2 | 22 | ribosomal protein S9 (RPS9) pseudogene |
| Y_RNA | 22 | Y RNA [Source:RFAM;Acc:RF00019] |
| AL080243.1 | 22 | actin, beta (ACTB) pseudogene |
| MIR1281 | 22 | microRNA 1281 [Source:HGNC Symbol;Acc:HG |
| EP300 | 22 | E1A binding protein p300 [Source:HGNC Symb |
| RNU6-375P | 22 | RNA, U6 small nuclear 375, pseudogene [Sour |
| AL035658.1 | 22 | novel transcript, antisense to EP300 |
| EP300-AS1 | 22 | EP300 antisense RNA 1 [Source:HGNC Symbol |
| LRRC37A14P | 22 | leucine rich repeat containing 37 member A14 |
| L3MBTL2 | 22 | L3MBTL histone methyl-lysine binding protein |
| AL035681.1 | 22 | novel transcript, antisense to L3MBTL2 |
| CHADL | 22 | chondroadherin like [Source:HGNC Symbol;Ac |
| RANGAP1 | 22 | Ran GTPase activating protein 1 [Source:HGN |
| MIR6889 | 22 | microRNA 6889 [Source:HGNC Symbol;Acc:HG |
| ZC3H7B | 22 | zinc finger CCCH-type containing 7B [Source:H |
| TEF | 22 | TEF transcription factor, PAR bZIP family mem |
| RNU6-495P | 22 | RNA, U6 small nuclear 495, pseudogene [Sour |
| AL008582.1 | 22 | novel transcript |
| TOB2 | 22 | transducer of ERBB2, 2 [Source:HGNC Symbol |
| PHF5A | 22 | PHD finger protein 5A [Source:HGNC Symbol;A |
| ACO2 | 22 | aconitase 2 [Source:HGNC Symbol;Acc:HGNC: |
| POLR3H | 22 | RNA polymerase III subunit H [Source:HGNC S |
| AL023553.1 | 22 | novel transcript, antisense to TSPO |
| CSDC2 | 22 | cold shock domain containing C2 [Source:HGN |
| PMM1 | 22 | phosphomannomutase 1 [Source:HGNC Symb |
| DESI1 | 22 | desumoylating isopeptidase 1 [Source:HGNC S |
| XRCC6 | 22 | X-ray repair cross complementing 6 [Source:H |
| Y_RNA | 22 | Y RNA [Source:RFAM;Acc:RF00019] |
| SNU13 | 22 | small nuclear ribonucleoprotein 13 [Source:H |
| Z83840.1 | 22 | LLP homolog, long-term synaptic facilitation ( |
| RNU6-476P | 22 | RNA, U6 small nuclear 476, pseudogene [Sour |
| C22orf46 | 22 | chromosome 22 open reading frame 46 [Sour |
| MEI1 | 22 | meiotic double-stranded break formation pro |
| HMGN2P10 | 22 | high mobility group nucleosomal binding dom |
| RNU6ATAC22P | 22 | RNA, U6atac small nuclear 22, pseudogene [S |
| Y_RNA | 22 | Y RNA [Source:RFAM;Acc:RF00019] |
| CCDC134 | 22 | coiled-coil domain containing 134 [Source:HG |
| SREBF2-AS1 | 22 | SREBF2 antisense RNA 1 [Source:HGNC Symbo |
| SREBF2 | 22 | sterol regulatory element binding transcriptio |
| MIR33A | 22 | microRNA 33a [Source:HGNC Symbol;Acc:HGN |
| SHISA8 | 22 | shisa family member 8 [Source:HGNC Symbol; |
| TNFRSF13C | 22 | TNF receptor superfamily member 13C [Sourc |

| | | |
|---|---|---|
| MIR378I | 22 | microRNA 378i [Source:HGNC Symbol;Acc:HG |
| CENPM | 22 | centromere protein M [Source:HGNC Symbol; |
| LINC00634 | 22 | long intergenic non-protein coding RNA 634 [S |
| SEPTIN3 | 22 | septin 3 [Source:HGNC Symbol;Acc:HGNC:107 |
| Z99716.1 | 22 | novel transcript, antisense to SEPT3 |
| WBP2NL | 22 | WBP2 N-terminal like [Source:HGNC Symbol;A |
| SLC25A5P1 | 22 | solute carrier family 25 member 5 pseudogen |
| NAGA | 22 | alpha-N-acetylgalactosaminidase [Source:HGN |
| Z99716.2 | 22 | novel transcript, antisense to POLDIP3 |
| PHETA2 | 22 | PH domain containing endocytic trafficking ad |
| SNORD13P1 | 22 | small nucleolar RNA, C/D box 13 pseudogene |
| SMDT1 | 22 | single-pass membrane protein with aspartate |
| NDUFA6 | 22 | NADH:ubiquinone oxidoreductase subunit A6 |
| AL021878.2 | 22 | novel transcript, sense intronic to NDUFA6 |
| NDUFA6-DT | 22 | NDUFA6 divergent transcript [Source:HGNC Sy |
| OLA1P1 | 22 | OLA1 pseudogene 1 [Source:HGNC Symbol;Ac |
| CYP2D6 | 22 | cytochrome P450 family 2 subfamily D memb |
| AC254562.1 | 22 | chromosome 6 open reading frame 108 (C6or |
| AC254562.2 | 22 | novel transcript |
| AC254562.3 | 22 | novel transcript |
| CYP2D7 | 22 | cytochrome P450 family 2 subfamily D memb |
| CYP2D8P | 22 | ccytochrome P450 family 2 subfamily D meml |
| TCF20 | 22 | transcription factor 20 [Source:HGNC Symbol; |
| OGFRP1 | 22 | opioid growth factor receptor pseudogene 1 [ |
| LINC01315 | 22 | long intergenic non-protein coding RNA 1315 |
| NFAM1 | 22 | NFAT activating protein with ITAM motif 1 [Sc |
| AL022316.1 | 22 | novel transcript |
| SERHL | 22 | serine hydrolase like (pseudogene) [Source:H( |
| RRP7A | 22 | ribosomal RNA processing 7 homolog A [Sourc |
| SERHL2 | 22 | serine hydrolase like 2 [Source:HGNC Symbol; |
| RRP7BP | 22 | ribosomal RNA processing 7 homolog B, pseuc |
| RN7SKP80 | 22 | RN7SK pseudogene 80 [Source:HGNC Symbol; |
| RNU6-513P | 22 | RNA, U6 small nuclear 513, pseudogene [Sour |
| POLDIP3 | 22 | DNA polymerase delta interacting protein 3 [S |
| RNU12 | 22 | RNA, U12 small nuclear [Source:HGNC Symbo |
| Z93241.1 | 22 | RNA, U12 small nuclear |
| CYB5R3 | 22 | cytochrome b5 reductase 3 [Source:HGNC Syr |
| ATP5MGL | 22 | ATP synthase membrane subunit g like [Sourc |
| A4GALT | 22 | alpha 1,4-galactosyltransferase (P blood grou |
| Y_RNA | 22 | Y RNA [Source:RFAM;Acc:RF00019] |
| RPL5P34 | 22 | ribosomal protein L5 pseudogene 34 [Source: |
| GOLGA2P4 | 22 | GOLGA2 pseudogene 4 [Source:HGNC Symbol |
| AL049757.1 | 22 | ADAM metallopeptidase with thrombospondi |
| ARFGAP3 | 22 | ADP ribosylation factor GTPase activating pro |
| PACSIN2 | 22 | protein kinase C and casein kinase substrate i |
| AL049758.1 | 22 | chromosome 7 open reading frame 44 (C7orf |
| AL022476.1 | 22 | novel transcript |
| TTLL1 | 22 | tubulin tyrosine ligase like 1 [Source:HGNC Syr |
| AL022237.1 | 22 | ribosomal protein S25 (RPS25) pseudogene |
| BIK | 22 | BCL2 interacting killer [Source:HGNC Symbol;/ |

| | | |
|---|---|---|
| MCAT | 22 | malonyl-CoA-acyl carrier protein transacylase |
| TSPO | 22 | translocator protein [Source:HGNC Symbol;Ac |
| TTLL12 | 22 | tubulin tyrosine ligase like 12 [Source:HGNC S |
| SCUBE1 | 22 | signal peptide, CUB domain and EGF like doma |
| Z82214.2 | 22 | novel transcript |
| Z82214.1 | 22 | novel transcript |
| Z99756.1 | 22 | novel transcript |
| LINC01639 | 22 | long intergenic non-protein coding RNA 1639 |
| MPPED1 | 22 | metallophosphoesterase domain containing 1 |
| BX546033.1 | 22 | novel transcript, antisense to TSPOand MPPEI |
| EFCAB6-AS1 | 22 | EFCAB6 antisense RNA 1 [Source:HGNC Symbo |
| EFCAB6 | 22 | EF-hand calcium binding domain 6 [Source:HG |
| Z97055.1 | 22 | ribosomal protein L4 (RPL4) pseudogene |
| HMGN2P9 | 22 | high mobility group nucleosomal binding dom |
| Z97055.2 | 22 | novel transcript, antisense to EFCAB6 |
| SULT4A1 | 22 | sulfotransferase family 4A member 1 [Source: |
| PNPLA5 | 22 | patatin like phospholipase domain containing |
| PNPLA3 | 22 | patatin like phospholipase domain containing |
| SAMM50 | 22 | SAMM50 sorting and assembly machinery cor |
| AL035398.1 | 22 | ribosomal protein L35a (RPL35A) pseudogene |
| PARVB | 22 | parvin beta [Source:HGNC Symbol;Acc:HGNC: |
| AL033543.1 | 22 | novel transcript, antisense to PARVB |
| AL031595.3 | 22 | novel transcript, sense overlapping PARVB |
| PARVG | 22 | parvin gamma [Source:HGNC Symbol;Acc:HGN |
| AL031595.1 | 22 | TEC |
| AL031595.2 | 22 | novel transcript, antisense to PARVG |
| SHISAL1 | 22 | shisa like 1 [Source:HGNC Symbol;Acc:HGNC:2 |
| Z85994.1 | 22 | novel transcript |
| Z85994.2 | 22 | S-phase kinase-associated protein 1A (p19A) ( |
| LINC01656 | 22 | long intergenic non-protein coding RNA 1656 |
| MRPS18CP6 | 22 | mitochondrial ribosomal protein S18C pseudo |
| AL023973.1 | 22 | TEC |
| RTL6 | 22 | retrotransposon Gag like 6 [Source:HGNC Sym |
| KRT18P23 | 22 | keratin 18 pseudogene 23 [Source:HGNC Sym |
| LINC00207 | 22 | long intergenic non-protein coding RNA 207 [S |
| LINC00229 | 22 | long intergenic non-protein coding RNA 229 [S |
| ANP32BP2 | 22 | acidic nuclear phosphoprotein 32 family mem |
| PRR5 | 22 | proline rich 5 [Source:HGNC Symbol;Acc:HGN |
| PRR5-ARHGAP8 | 22 | PRR5-ARHGAP8 readthrough [Source:HGNC Sy |
| ARHGAP8 | 22 | Rho GTPase activating protein 8 [Source:HGN |
| PHF21B | 22 | PHD finger protein 21B [Source:HGNC Symbol |
| Z83838.1 | 22 | ribosomal protein L6 (RPL6) pseudogene |
| AL079301.1 | 22 | novel transcript |
| NUP50-DT | 22 | NUP50 divergent transcript [Source:HGNC Syr |
| Z82243.1 | 22 | novel transcript |
| NUP50 | 22 | nucleoporin 50 [Source:HGNC Symbol;Acc:HG |
| AL008718.3 | 22 | novel transcript, sense intronic to NUP50 |
| KIAA0930 | 22 | KIAA0930 [Source:HGNC Symbol;Acc:HGNC:1: |
| MIR1249 | 22 | microRNA 1249 [Source:HGNC Symbol;Acc:HG |
| AL008718.2 | 22 | novel transcript |

| | | |
|---|---|---|
| AL008718.1 | 22 | ribosomal protein S10 (RPS10) pseudogene |
| UPK3A | 22 | uroplakin 3A [Source:HGNC Symbol;Acc:HGN( |
| FAM118A | 22 | family with sequence similarity 118 member / |
| SMC1B | 22 | structural maintenance of chromosomes 1B [S |
| RIBC2 | 22 | RIB43A domain with coiled-coils 2 [Source:HG |
| AL021391.1 | 22 | novel transcript |
| FBLN1 | 22 | fibulin 1 [Source:HGNC Symbol;Acc:HGNC:360 |
| LINC01589 | 22 | long intergenic non-protein coding RNA 1589 |
| RNU6-1161P | 22 | RNA, U6 small nuclear 1161, pseudogene [Sou |
| Z95331.1 | 22 | novel transcript, sense overlapping ATXN10 |
| ATXN10 | 22 | ataxin 10 [Source:HGNC Symbol;Acc:HGNC:10 |
| MIR4762 | 22 | microRNA 4762 [Source:HGNC Symbol;Acc:HG |
| Z84478.1 | 22 | tRNA nucleotidyl transferase, CCA-adding, 1 (T |
| BX324167.2 | 22 | novel transcript |
| BX324167.1 | 22 | novel transcript |
| WNT7B | 22 | Wnt family member 7B [Source:HGNC Symbol |
| BX537318.2 | 22 | TEC |
| BX537318.1 | 22 | novel transcript |
| LINC00899 | 22 | long intergenic non-protein coding RNA 899 [S |
| PRR34 | 22 | PRR34 long non-coding RNA [Source:HGNC Sy |
| AL121672.3 | 22 | novel transcript, antisense to C22orf26 |
| PRR34-AS1 | 22 | PRR34 antisense RNA 1 [Source:HGNC Symbol |
| MIRLET7BHG | 22 | MIRLET7B host gene [Source:HGNC Symbol;A( |
| AL121672.1 | 22 | novel transcript |
| AL121672.2 | 22 | novel transcript |
| MIR3619 | 22 | microRNA 3619 [Source:HGNC Symbol;Acc:HG |
| MIRLET7A3 | 22 | microRNA let-7a-3 [Source:HGNC Symbol;Acc: |
| MIR4763 | 22 | microRNA 4763 [Source:HGNC Symbol;Acc:HG |
| MIRLET7B | 22 | microRNA let-7b [Source:HGNC Symbol;Acc:H |
| PPARA | 22 | peroxisome proliferator activated receptor al |
| FP325332.1 | 22 | novel transcript, sense intronic to PPARA |
| CDPF1 | 22 | cysteine rich DPF motif domain containing 1 [S |
| PKDREJ | 22 | polycystin family receptor for egg jelly [Source |
| TTC38 | 22 | tetratricopeptide repeat domain 38 [Source:H |
| GTSE1-DT | 22 | GTSE1 divergent transcript [Source:HGNC Sym |
| GTSE1 | 22 | G2 and S-phase expressed 1 [Source:HGNC Sy |
| TRMU | 22 | tRNA 5-methylaminomethyl-2-thiouridylate m |
| CELSR1 | 22 | cadherin EGF LAG seven-pass G-type receptor |
| AL031597.1 | 22 | novel pseudogene |
| AL021392.1 | 22 | novel transcript |
| GRAMD4 | 22 | GRAM domain containing 4 [Source:HGNC Syr |
| CERK | 22 | ceramide kinase [Source:HGNC Symbol;Acc:H( |
| AL118516.1 | 22 | novel transcript, antisense to TBC1D22A |
| TBC1D22A | 22 | TBC1 domain family member 22A [Source:HG |
| TBC1D22A-AS1 | 22 | TBC1D22A antisense RNA 1 [Source:HGNC Syr |
| FP325331.1 | 22 | TEC |
| Z83836.1 | 22 | TEC |
| Z82186.1 | 22 | novel transcript |
| LINC01644 | 22 | long intergenic non-protein coding RNA 1644 |
| LINC00898 | 22 | long intergenic non-protein coding RNA 898 [S |

| | | |
|---|---|---|
| AL117329.1 | 22 | novel transcript |
| AL117329.2 | 22 | ribosomal protein L7 (RPL7) pseudogene |
| FP325330.3 | 22 | novel transcript |
| FP325330.1 | 22 | novel transcript |
| FP325330.2 | 22 | novel transcript |
| BX284656.2 | 22 | novel transcript |
| BX284656.1 | 22 | novel transcript |
| AL008720.1 | 22 | novel transcript |
| MIR3201 | 22 | microRNA 3201 [Source:HGNC Symbol;Acc:HG |
| Z72006.1 | 22 | novel transcript |
| Z82249.1 | 22 | novel transcript |
| Z84468.2 | 22 | novel transcript |
| TAFA5 | 22 | TAFA chemokine like family member 5 [Sourc |
| Z84468.1 | 22 | novel transcript |
| MIR4535 | 22 | microRNA 4535 [Source:HGNC Symbol;Acc:HG |
| LINC01310 | 22 | long intergenic non-protein coding RNA 1310 |
| AL078622.1 | 22 | novel transcript |
| Z82202.2 | 22 | novel transcript |
| RPL35P8 | 22 | ribosomal protein L35 pseudogene 8 [Source: |
| Z82202.1 | 22 | novel transcript |
| AC207130.1 | 22 | novel transcript |
| C22orf34 | 22 | chromosome 22 open reading frame 34 [Sour |
| AL008636.1 | 22 | novel transcript |
| AL031593.1 | 22 | novel transcript |
| MIR3667 | 22 | microRNA 3667 [Source:HGNC Symbol;Acc:HG |
| Z97192.1 | 22 | novel transcript |
| Z97192.2 | 22 | novel transcript |
| Z97192.3 | 22 | novel transcript |
| Z97192.4 | 22 | novel transcript |
| AL023802.1 | 22 | novel transcript |
| RN7SKP252 | 22 | RN7SK pseudogene 252 [Source:HGNC Symbo |
| RPL5P35 | 22 | ribosomal protein L5 pseudogene 35 [Source: |
| BRD1 | 22 | bromodomain containing 1 [Source:HGNC Syn |
| Z98885.3 | 22 | novel transcript, antisense to BRD1 |
| Z98885.1 | 22 | suppressor of cytokine signaling 2 (SOCS2) pse |
| Z98885.2 | 22 | novel transcript |
| AL117328.1 | 22 | chromosome 6 open reading frame 49 (C6orf4 |
| Metazoa_SRP | 22 | Metazoan signal recognition particle RNA [Sou |
| AL117328.2 | 22 | novel transcript |
| ZBED4 | 22 | zinc finger BED-type containing 4 [Source:HGN |
| ALG12 | 22 | ALG12 alpha-1,6-mannosyltransferase [Source |
| AL671710.1 | 22 | novel transcript, antisense to ALG12 |
| CRELD2 | 22 | cysteine rich with EGF like domains 2 [Source: |
| BX539320.1 | 22 | novel transcript |
| PIM3 | 22 | Pim-3 proto-oncogene, serine/threonine kinas |
| MIR6821 | 22 | microRNA 6821 [Source:HGNC Symbol;Acc:HG |
| IL17REL | 22 | interleukin 17 receptor E like [Source:HGNC S |
| TTLL8 | 22 | tubulin tyrosine ligase like 8 [Source:HGNC Syn |
| MLC1 | 22 | modulator of VRAC current 1 [Source:HGNC Sy |
| MOV10L1 | 22 | Mov10 like RISC complex RNA helicase 1 [Sour |

| | | |
|---|---|---|
| AL034546.1 | 22 | novel transcript, sense intronic to MOV10L1 |
| PANX2 | 22 | pannexin 2 [Source:HGNC Symbol;Acc:HGNC:8 |
| TRABD | 22 | TraB domain containing [Source:HGNC Symbo |
| AL022328.3 | 22 | novel transcript, antisense to TRABD |
| AL022328.4 | 22 | novel transcript, antisense TRABD |
| SELENOO | 22 | selenoprotein O [Source:HGNC Symbol;Acc:H( |
| AL022328.1 | 22 | novel transcript, antisense to SELO |
| AL022328.2 | 22 | novel transcript, antisense to SELO |
| TUBGCP6 | 22 | tubulin gamma complex associated protein 6 |
| HDAC10 | 22 | histone deacetylase 10 [Source:HGNC Symbol |
| MAPK12 | 22 | mitogen-activated protein kinase 12 [Source:I |
| MAPK11 | 22 | mitogen-activated protein kinase 11 [Source:I |
| PLXNB2 | 22 | plexin B2 [Source:HGNC Symbol;Acc:HGNC:91 |
| DENND6B | 22 | DENN domain containing 6B [Source:HGNC Sy |
| CR559946.1 | 22 | novel transcript |
| CR559946.2 | 22 | novel transcript, antisense to DENND6B |
| PPP6R2 | 22 | protein phosphatase 6 regulatory subunit 2 [S |
| SBF1 | 22 | SET binding factor 1 [Source:HGNC Symbol;Ac |
| ADM2 | 22 | adrenomedullin 2 [Source:HGNC Symbol;Acc:I |
| MIOX | 22 | myo-inositol oxygenase [Source:HGNC Symbo |
| LMF2 | 22 | lipase maturation factor 2 [Source:HGNC Sym |
| NCAPH2 | 22 | non-SMC condensin II complex subunit H2 [So |
| SCO2 | 22 | SCO cytochrome c oxidase assembly protein 2 |
| U62317.2 | 22 | novel transcript, antisense to SCO2 |
| TYMP | 22 | thymidine phosphorylase [Source:HGNC Symb |
| ODF3B | 22 | outer dense fiber of sperm tails 3B [Source:H( |
| U62317.4 | 22 | novel transcript |
| U62317.1 | 22 | uncharacterized LOC105373098 [Source:NCBI |
| KLHDC7B | 22 | kelch domain containing 7B [Source:HGNC Syr |
| SYCE3 | 22 | synaptonemal complex central element prote |
| CPT1B | 22 | carnitine palmitoyltransferase 1B [Source:HGI |
| CHKB-CPT1B | 22 | CHKB-CPT1B readthrough (NMD candidate) [S |
| CHKB | 22 | choline kinase beta [Source:HGNC Symbol;Acc |
| CHKB-DT | 22 | CHKB divergent transcript [Source:HGNC Syml |
| U62317.3 | 22 | novel transcript |
| MAPK8IP2 | 22 | mitogen-activated protein kinase 8 interacting |
| ARSA | 22 | arylsulfatase A [Source:HGNC Symbol;Acc:HGI |
| Y_RNA | 22 | Y RNA [Source:RFAM;Acc:RF00019] |
| AC000050.1 | 22 | novel transcript |
| SHANK3 | 22 | SH3 and multiple ankyrin repeat domains 3 [S |
| RNU6-409P | 22 | RNA, U6 small nuclear 409, pseudogene [Sour |
| AC000036.1 | 22 | novel transcript |
| ACR | 22 | acrosin [Source:HGNC Symbol;Acc:HGNC:126] |
| AC002056.2 | 22 | novel transcript |
| AC002056.1 | 22 | small nuclear ribonucleoprotein polypeptide A |
| RPL23AP82 | 22 | ribosomal protein L23a pseudogene 82 [Sourc |
| RABL2B | 22 | RAB, member of RAS oncogene family like 2B |

| seq_region_start | seq_region_end | seq_region_strand | biotype | stable_id |
|---|---|---|---|---|
| 1012823 | 1036718 | -1 | protein_coding | ENSG0000C |
| 1049876 | 1055745 | 1 | lncRNA | ENSG0000C |
| 1075018 | 1112365 | 1 | protein_coding | ENSG0000C |
| 1159911 | 1203106 | 1 | protein_coding | ENSG0000C |
| 13795460 | 13832230 | 1 | protein_coding | ENSG0000C |
| 13796574 | 13796933 | -1 | lncRNA | ENSG0000C |
| 13815062 | 13815319 | -1 | misc_RNA | ENSG0000C |
| 13823880 | 13842928 | -1 | lncRNA | ENSG0000C |
| 13836287 | 13836359 | -1 | miRNA | ENSG0000C |
| 13836440 | 13836517 | -1 | miRNA | ENSG0000C |
| 13836587 | 13836659 | -1 | miRNA | ENSG0000C |
| 13862063 | 13880757 | 1 | protein_coding | ENSG0000C |
| 13874699 | 13874808 | 1 | miRNA | ENSG0000C |
| 13874875 | 13875011 | 1 | miRNA | ENSG0000C |
| 13882348 | 13906452 | -1 | protein_coding | ENSG0000C |
| 13906201 | 13930879 | 1 | protein_coding | ENSG0000C |
| 13931187 | 13953392 | -1 | protein_coding | ENSG0000C |
| 13952492 | 13961449 | 1 | protein_coding | ENSG0000C |
| 13961530 | 14007039 | -1 | protein_coding | ENSG0000C |
| 14006422 | 14006773 | 1 | lncRNA | ENSG0000C |
| 14028148 | 14031551 | 1 | protein_coding | ENSG0000C |
| 14030855 | 14031557 | -1 | lncRNA | ENSG0000C |
| 14031762 | 14053218 | 1 | protein_coding | ENSG0000C |
| 14053363 | 14062076 | -1 | protein_coding | ENSG0000C |
| 14070342 | 14071237 | 1 | processed_pseudoge | ENSG0000C |
| 14072536 | 14075062 | 1 | protein_coding | ENSG0000C |
| 14073361 | 14073479 | 1 | miRNA | ENSG0000C |
| 14081624 | 14085875 | -1 | protein_coding | ENSG0000C |
| 14088420 | 14090738 | -1 | protein_coding | ENSG0000C |
| 14092006 | 14118402 | -1 | protein_coding | ENSG0000C |
| 14119424 | 14119855 | 1 | lncRNA | ENSG0000C |
| 14119830 | 14136931 | -1 | protein_coding | ENSG0000C |
| 14137464 | 14171585 | 1 | lncRNA | ENSG0000C |
| 14148061 | 14206505 | -1 | protein_coding | ENSG0000C |
| 14213370 | 14213641 | 1 | misc_RNA | ENSG0000C |
| 14217811 | 14219923 | 1 | lncRNA | ENSG0000C |
| 14254507 | 14293205 | -1 | lncRNA | ENSG0000C |
| 14268125 | 14269695 | -1 | lncRNA | ENSG0000C |
| 72378406 | 72447301 | -1 | protein_coding | ENSG0000C |
| 72505075 | 72550889 | -1 | lncRNA | ENSG0000C |
| 72577040 | 72577203 | 1 | snRNA | ENSG0000C |
| 72583719 | 72583884 | 1 | processed_pseudoge | ENSG0000C |
| 72692054 | 72692160 | -1 | rRNA_pseudogene | ENSG0000C |
| 47750864 | 47763592 | -1 | protein_coding | ENSG0000C |
| 47791021 | 47792394 | 1 | pseudogene | ENSG0000C |
| 47865495 | 47949308 | -1 | lncRNA | ENSG0000C |
| 47876170 | 47880955 | 1 | protein_coding | ENSG0000C |
| 47927474 | 47930927 | -1 | lncRNA | ENSG0000C |
| 47952753 | 47957222 | -1 | protein_coding | ENSG0000C |

| | | | | |
|---|---|---|---|---|
| 47967304 | 47989071 | -1 | transcribed_unproce | ENSG00000 |
| 47997893 | 47998008 | 1 | rRNA_pseudogene | ENSG00000 |
| 48007713 | 48017447 | -1 | transcribed_unproce | ENSG00000 |
| 48021739 | 48119455 | -1 | unprocessed_pseudc | ENSG00000 |
| 8745391 | 8747727 | -1 | lncRNA | ENSG00000 |
| 8827806 | 8853569 | -1 | protein_coding | ENSG00000 |
| 8840445 | 8842557 | -1 | lncRNA | ENSG00000 |
| 90561 | 115272 | -1 | lncRNA | ENSG00000 |
| 96627 | 100767 | 1 | lncRNA | ENSG00000 |
| 134775 | 151623 | -1 | lncRNA | ENSG00000 |
| 155442 | 169116 | -1 | lncRNA | ENSG00000 |
| 169138 | 175054 | 1 | lncRNA | ENSG00000 |
| 181354 | 184561 | -1 | lncRNA | ENSG00000 |
| 194509 | 195602 | 1 | lncRNA | ENSG00000 |
| 202524 | 213769 | -1 | lncRNA | ENSG00000 |
| 212160 | 279890 | 1 | protein_coding | ENSG00000 |
| 247143 | 249146 | -1 | TEC | ENSG00000 |
| 281927 | 282363 | 1 | protein_coding | ENSG00000 |
| 50669804 | 50669901 | 1 | misc_RNA | ENSG00000 |
| 50672714 | 50674174 | 1 | lncRNA | ENSG00000 |
| 50691077 | 50691180 | -1 | snRNA | ENSG00000 |
| 50721436 | 50733232 | 1 | protein_coding | ENSG00000 |
| 50738219 | 50745356 | 1 | protein_coding | ENSG00000 |
| 50740616 | 50743537 | -1 | lncRNA | ENSG00000 |
| 50754693 | 50755452 | -1 | processed_pseudoge | ENSG00000 |
| 50756966 | 50801309 | 1 | transcribed_unproce | ENSG00000 |
| 81754364 | 81764240 | 1 | protein_coding | ENSG00000 |
| 81772593 | 81783534 | -1 | protein_coding | ENSG00000 |
| 81783724 | 81785282 | -1 | protein_coding | ENSG00000 |
| 12822686 | 12823159 | -1 | lncRNA | ENSG00000 |
| 12824605 | 12831410 | -1 | protein_coding | ENSG00000 |
| 12838125 | 12841357 | -1 | unprocessed_pseudc | ENSG00000 |
| 12847408 | 12848725 | -1 | protein_coding | ENSG00000 |
| 12857086 | 12861909 | 1 | protein_coding | ENSG00000 |
| 12879212 | 12886201 | -1 | protein_coding | ENSG00000 |
| 12892896 | 12898270 | -1 | protein_coding | ENSG00000 |
| 12916610 | 12920482 | 1 | protein_coding | ENSG00000 |
| 12922554 | 12922660 | 1 | snRNA | ENSG00000 |
| 12926162 | 12928253 | 1 | unprocessed_pseudc | ENSG00000 |
| 12938472 | 12947580 | -1 | protein_coding | ENSG00000 |
| 13002166 | 13004957 | 1 | unprocessed_pseudc | ENSG00000 |
| 13022303 | 13029318 | -1 | protein_coding | ENSG00000 |
| 13033696 | 13035056 | 1 | protein_coding | ENSG00000 |
| 13041504 | 13050711 | 1 | protein_coding | ENSG00000 |
| 13068006 | 13070798 | -1 | unprocessed_pseudc | ENSG00000 |
| 13077230 | 13078516 | -1 | unprocessed_pseudc | ENSG00000 |
| 13088315 | 13089681 | -1 | protein_coding | ENSG00000 |
| 13101645 | 13104442 | 1 | transcribed_unproce | ENSG00000 |
| 13119808 | 13120287 | -1 | lncRNA | ENSG00000 |
| 13121732 | 13128788 | -1 | protein_coding | ENSG00000 |

| | | | | |
|---|---|---|---|---|
| 13137413 | 13138294 | -1 | protein_coding | ENSG00000( |
| 13145282 | 13152291 | 1 | protein_coding | ENSG00000( |
| 13169015 | 13174236 | -1 | protein_coding | ENSG00000( |
| 13195532 | 13198933 | -1 | protein_coding | ENSG00000( |
| 13218690 | 13221479 | 1 | unprocessed_pseudc | ENSG00000( |
| 13227039 | 13236141 | 1 | protein_coding | ENSG00000( |
| 13246366 | 13248454 | -1 | unprocessed_pseudc | ENSG00000( |
| 13251952 | 13252058 | -1 | snRNA | ENSG00000( |
| 13253862 | 13258001 | -1 | protein_coding | ENSG00000( |
| 13276366 | 13281734 | 1 | protein_coding | ENSG00000( |
| 12049844 | 12211084 | 1 | protein_coding | ENSG00000( |
| 12111163 | 12111489 | 1 | processed_pseudoge | ENSG00000( |
| 12111952 | 12112036 | 1 | miRNA | ENSG00000( |
| 12116025 | 12267044 | -1 | protein_coding | ENSG00000( |
| 12155262 | 12155772 | 1 | processed_pseudoge | ENSG00000( |
| 12226674 | 12226777 | 1 | snRNA | ENSG00000( |
| 12248991 | 12249427 | 1 | processed_pseudoge | ENSG00000( |
| 12289227 | 12289329 | -1 | snRNA | ENSG00000( |
| 12326056 | 12350242 | -1 | protein_coding | ENSG00000( |
| 12338776 | 12339254 | 1 | processed_pseudoge | ENSG00000( |
| 12355406 | 12357067 | -1 | lncRNA | ENSG00000( |
| 12357078 | 12471238 | 1 | protein_coding | ENSG00000( |
| 12452108 | 12452486 | -1 | processed_pseudoge | ENSG00000( |
| 12473287 | 12562868 | -1 | protein_coding | ENSG00000( |
| 12485358 | 12491586 | 1 | lncRNA | ENSG00000( |
| 12568206 | 12568781 | -1 | processed_pseudoge | ENSG00000( |
| 16085222 | 16105841 | -1 | lncRNA | ENSG00000( |
| 16151181 | 16151324 | 1 | processed_pseudoge | ENSG00000( |
| 139408588 | 139444491 | -1 | protein_coding | ENSG00000( |
| 139442316 | 139461973 | -1 | protein_coding | ENSG00000( |
| 139447907 | 139448017 | 1 | snRNA | ENSG00000( |
| 139448560 | 139462009 | -1 | protein_coding | ENSG00000( |
| 139470044 | 139474038 | 1 | protein_coding | ENSG00000( |
| 139474799 | 139482201 | -1 | protein_coding | ENSG00000( |
| 139491804 | 139492371 | 1 | processed_pseudoge | ENSG00000( |
| 139493522 | 139495382 | 1 | processed_pseudoge | ENSG00000( |
| 7080214 | 7091771 | -1 | protein_coding | ENSG00000( |
| 7088818 | 7108469 | -1 | protein_coding | ENSG00000( |
| 7107283 | 7121732 | 1 | lncRNA | ENSG00000( |
| 7114967 | 7128120 | -1 | protein_coding | ENSG00000( |
| 7118016 | 7118122 | 1 | snRNA | ENSG00000( |
| 7128310 | 7130429 | -1 | lncRNA | ENSG00000( |
| 7128929 | 7158176 | 1 | protein_coding | ENSG00000( |
| 7145713 | 7147048 | -1 | lncRNA | ENSG00000( |
| 7165905 | 7188300 | -1 | lncRNA | ENSG00000( |
| 7187916 | 7217805 | 1 | protein_coding | ENSG00000( |
| 7236663 | 7240217 | 1 | processed_pseudoge | ENSG00000( |
| 56205125 | 56205625 | -1 | processed_pseudoge | ENSG00000( |
| 56210873 | 56211820 | -1 | polymorphic_pseudc | ENSG00000( |
| 56232106 | 56233185 | -1 | protein_coding | ENSG00000( |

| | | | | |
|---|---|---|---|---|
| 56252200 | 56253222 | 1 | protein_coding | ENSG00000( |
| 56275639 | 56276619 | 1 | protein_coding | ENSG00000( |
| 56290053 | 56291090 | -1 | protein_coding | ENSG00000( |
| 56296541 | 56297471 | -1 | unprocessed_pseudc | ENSG00000( |
| 56318307 | 56319245 | 1 | polymorphic_pseudc | ENSG00000( |
| 56331347 | 56332488 | -1 | unprocessed_pseudc | ENSG00000( |
| 56335142 | 56336068 | 1 | unprocessed_pseudc | ENSG00000( |
| 56346039 | 56346998 | 1 | protein_coding | ENSG00000( |
| 56357328 | 56358221 | 1 | processed_pseudoge | ENSG00000( |
| 56360215 | 56361288 | 1 | protein_coding | ENSG00000( |
| 56375625 | 56376584 | 1 | protein_coding | ENSG00000( |
| 56383230 | 56384159 | 1 | protein_coding | ENSG00000( |
| 56389241 | 56390195 | 1 | unprocessed_pseudc | ENSG00000( |
| 87847259 | 88022145 | -1 | lncRNA | ENSG00000( |
| 88026643 | 88038647 | 1 | lncRNA | ENSG00000( |
| 88038311 | 88074684 | -1 | lncRNA | ENSG00000( |
| 88075128 | 88092372 | 1 | lncRNA | ENSG00000( |
| 88076532 | 88076642 | 1 | misc_RNA | ENSG00000( |
| 16132469 | 16145958 | 1 | protein_coding | ENSG00000( |
| 16141767 | 16141872 | 1 | snRNA | ENSG00000( |
| 16484487 | 16487680 | 1 | unprocessed_pseudc | ENSG00000( |
| 16502123 | 16502210 | 1 | processed_pseudoge | ENSG00000( |
| 16520736 | 16522824 | 1 | unprocessed_pseudc | ENSG00000( |
| 16525740 | 16573663 | -1 | unprocessed_pseudc | ENSG00000( |
| 16583778 | 16585115 | 1 | processed_pseudoge | ENSG00000( |
| 56700386 | 56701303 | 1 | protein_coding | ENSG00000( |
| 56711466 | 56712383 | 1 | protein_coding | ENSG00000( |
| 56747390 | 56747536 | -1 | processed_pseudoge | ENSG00000( |
| 56751253 | 56752170 | 1 | unprocessed_pseudc | ENSG00000( |
| 56753901 | 56754885 | -1 | protein_coding | ENSG00000( |
| 56762097 | 56762976 | -1 | unprocessed_pseudc | ENSG00000( |
| 56786335 | 56787280 | -1 | unprocessed_pseudc | ENSG00000( |
| 56801280 | 56802039 | -1 | unprocessed_pseudc | ENSG00000( |
| 56812930 | 56831496 | -1 | transcribed_unproce | ENSG00000( |
| 56830655 | 56831590 | -1 | polymorphic_pseudc | ENSG00000( |
| 56859642 | 56882325 | 1 | lncRNA | ENSG00000( |
| 113648804 | 113658198 | -1 | protein_coding | ENSG00000( |
| 113667219 | 113688512 | 1 | protein_coding | ENSG00000( |
| 113699287 | 113701865 | -1 | TEC | ENSG00000( |
| 9030594 | 9039988 | 1 | unprocessed_pseudc | ENSG00000( |
| 21565092 | 21605081 | -1 | lncRNA | ENSG00000( |
| 21631683 | 21632015 | -1 | processed_pseudoge | ENSG00000( |
| 21637475 | 21639041 | -1 | unprocessed_pseudc | ENSG00000( |
| 21661926 | 21674008 | 1 | unprocessed_pseudc | ENSG00000( |
| 21677246 | 21681118 | -1 | unprocessed_pseudc | ENSG00000( |
| 21696570 | 21698220 | 1 | unprocessed_pseudc | ENSG00000( |
| 21777583 | 21778222 | -1 | unprocessed_pseudc | ENSG00000( |
| 21785445 | 21846721 | 1 | lncRNA | ENSG00000( |
| 61240898 | 61241311 | 1 | processed_pseudoge | ENSG00000( |
| 49981473 | 49982535 | -1 | protein_coding | ENSG00000( |

| | | | | |
|---|---|---|---|---|
| 50018798 | 50019695 | 1 | unprocessed_pseudc | ENSG0000C |
| 50053408 | 50054307 | -1 | unprocessed_pseudc | ENSG0000C |
| 1660315 | 1660380 | -1 | miRNA | ENSG0000C |
| 1688574 | 1788674 | 1 | protein_coding | ENSG0000C |
| 1695830 | 1698044 | -1 | processed_pseudoge | ENSG0000C |
| 1741197 | 1747099 | -1 | lncRNA | ENSG0000C |
| 1757231 | 1758011 | -1 | processed_pseudoge | ENSG0000C |
| 1791963 | 1918666 | -1 | protein_coding | ENSG0000C |
| 1800862 | 1807486 | 1 | lncRNA | ENSG0000C |
| 1820267 | 1836753 | 1 | protein_coding | ENSG0000C |
| 1917951 | 1922867 | 1 | lncRNA | ENSG0000C |
| 1929202 | 1936574 | -1 | lncRNA | ENSG0000C |
| 1946053 | 2004535 | -1 | protein_coding | ENSG0000C |
| 1970786 | 2703922 | 1 | protein_coding | ENSG0000C |
| 2004666 | 2011392 | 1 | lncRNA | ENSG0000C |
| 2018213 | 2020532 | 1 | lncRNA | ENSG0000C |
| 2048352 | 2049463 | 1 | lncRNA | ENSG0000C |
| 2217462 | 2217920 | -1 | lncRNA | ENSG0000C |
| 2219485 | 2223479 | -1 | lncRNA | ENSG0000C |
| 2275748 | 2294909 | 1 | lncRNA | ENSG0000C |
| 2338833 | 2341674 | -1 | lncRNA | ENSG0000C |
| 2446161 | 2463364 | -1 | lncRNA | ENSG0000C |
| 2609322 | 2613412 | -1 | lncRNA | ENSG0000C |
| 2665909 | 2666941 | 1 | processed_pseudoge | ENSG0000C |
| 2674472 | 2678192 | -1 | lncRNA | ENSG0000C |
| 2681973 | 2697172 | -1 | lncRNA | ENSG0000C |
| 2701737 | 2812902 | -1 | lncRNA | ENSG0000C |
| 210231456 | 210234047 | -1 | lncRNA | ENSG0000C |
| 210232796 | 210246631 | 1 | protein_coding | ENSG0000C |
| 210265636 | 210267241 | -1 | processed_pseudoge | ENSG0000C |
| 210303684 | 210304540 | 1 | processed_pseudoge | ENSG0000C |
| 210328252 | 210679653 | 1 | protein_coding | ENSG0000C |
| 210362861 | 210366948 | -1 | lncRNA | ENSG0000C |
| 210374154 | 210374267 | -1 | snRNA | ENSG0000C |
| 210386657 | 210394112 | -1 | lncRNA | ENSG0000C |
| 210462345 | 210463012 | -1 | processed_pseudoge | ENSG0000C |
| 23699797 | 23729757 | -1 | lncRNA | ENSG0000C |
| 23749236 | 23751150 | 1 | processed_pseudoge | ENSG0000C |
| 23767662 | 23768998 | 1 | unprocessed_pseudc | ENSG0000C |
| 23791930 | 23792236 | 1 | misc_RNA | ENSG0000C |
| 23817281 | 23827396 | -1 | unprocessed_pseudc | ENSG0000C |
| 23875938 | 23882428 | -1 | lncRNA | ENSG0000C |
| 23922247 | 23934568 | -1 | lncRNA | ENSG0000C |
| 23938219 | 24052554 | -1 | lncRNA | ENSG0000C |
| 23953734 | 24011760 | 1 | protein_coding | ENSG0000C |
| 23970058 | 24027781 | 1 | lncRNA | ENSG0000C |
| 24012412 | 24051376 | 1 | lncRNA | ENSG0000C |
| 24036452 | 24051027 | 1 | transcribed_unproce | ENSG0000C |
| 24052008 | 24069728 | 1 | protein_coding | ENSG0000C |
| 24070836 | 24078099 | 1 | protein_coding | ENSG0000C |

| | | | | |
|---|---|---|---|---|
| 24080106 | 24115013 | -1 | protein_coding | ENSG00000 |
| 24094052 | 24110597 | 1 | protein_coding | ENSG00000 |
| 24114194 | 24125241 | 1 | protein_coding | ENSG00000 |
| 24120955 | 24131828 | 1 | protein_coding | ENSG00000 |
| 24131274 | 24132848 | 1 | protein_coding | ENSG00000 |
| 24136162 | 24138966 | 1 | protein_coding | ENSG00000 |
| 24138958 | 24141590 | -1 | protein_coding | ENSG00000 |
| 24139444 | 24140443 | 1 | lncRNA | ENSG00000 |
| 24143361 | 24147569 | -1 | protein_coding | ENSG00000 |
| 24143488 | 24143564 | -1 | miRNA | ENSG00000 |
| 24146682 | 24160659 | 1 | protein_coding | ENSG00000 |
| 24147547 | 24166451 | 1 | protein_coding | ENSG00000 |
| 24158602 | 24158721 | 1 | rRNA_pseudogene | ENSG00000 |
| 24161264 | 24166564 | 1 | protein_coding | ENSG00000 |
| 24171852 | 24180256 | 1 | protein_coding | ENSG00000 |
| 24180218 | 24188868 | -1 | protein_coding | ENSG00000 |
| 24180394 | 24190415 | -1 | protein_coding | ENSG00000 |
| 24188913 | 24193105 | 1 | lncRNA | ENSG00000 |
| 24189148 | 24195686 | -1 | protein_coding | ENSG00000 |
| 24189156 | 24213472 | -1 | protein_coding | ENSG00000 |
| 24198432 | 24199089 | 1 | lncRNA | ENSG00000 |
| 24201611 | 24202810 | -1 | lncRNA | ENSG00000 |
| 24205695 | 24208361 | 1 | protein_coding | ENSG00000 |
| 24209582 | 24213868 | -1 | protein_coding | ENSG00000 |
| 24209645 | 24215986 | -1 | lncRNA | ENSG00000 |
| 24213942 | 24216069 | -1 | protein_coding | ENSG00000 |
| 24213954 | 24232351 | -1 | protein_coding | ENSG00000 |
| 24216856 | 24232366 | -1 | protein_coding | ENSG00000 |
| 24232421 | 24239241 | 1 | protein_coding | ENSG00000 |
| 24239642 | 24242673 | -1 | protein_coding | ENSG00000 |
| 24249113 | 24264431 | -1 | protein_coding | ENSG00000 |
| 24265537 | 24271738 | -1 | protein_coding | ENSG00000 |
| 24271212 | 24298845 | 1 | lncRNA | ENSG00000 |
| 24271725 | 24279088 | 1 | lncRNA | ENSG00000 |
| 24290597 | 24299779 | -1 | protein_coding | ENSG00000 |
| 24299849 | 24309123 | 1 | protein_coding | ENSG00000 |
| 24305095 | 24311429 | -1 | protein_coding | ENSG00000 |
| 24305733 | 24312052 | 1 | protein_coding | ENSG00000 |
| 24311449 | 24318035 | 1 | protein_coding | ENSG00000 |
| 24318348 | 24335092 | -1 | protein_coding | ENSG00000 |
| 24332309 | 24337381 | -1 | lncRNA | ENSG00000 |
| 24336024 | 24340021 | -1 | protein_coding | ENSG00000 |
| 24352056 | 24352156 | -1 | snRNA | ENSG00000 |
| 24365672 | 24379603 | 1 | protein_coding | ENSG00000 |
| 228407365 | 228410870 | 1 | lncRNA | ENSG00000 |
| 228407935 | 228416861 | -1 | protein_coding | ENSG00000 |
| 228424845 | 228425325 | -1 | protein_coding | ENSG00000 |
| 228449163 | 228449621 | -1 | processed_pseudoge | ENSG00000 |
| 228456979 | 228457873 | -1 | protein_coding | ENSG00000 |
| 228458107 | 228460470 | 1 | protein_coding | ENSG00000 |

| | | | | |
|---|---|---|---|---|
| 228462074 | 228462152 | 1 | miRNA | ENSG0000C |
| 228464103 | 228464626 | -1 | unitary_pseudogene | ENSG0000C |
| 228486188 | 228487120 | -1 | TEC | ENSG0000C |
| 228487382 | 228496188 | 1 | protein_coding | ENSG0000C |
| 228510425 | 228512305 | -1 | transcribed_unitary_ | ENSG0000C |
| 228558286 | 228558404 | -1 | rRNA | ENSG0000C |
| 228560525 | 228560643 | -1 | rRNA | ENSG0000C |
| 228620966 | 228621084 | -1 | rRNA | ENSG0000C |
| 228629928 | 228630046 | -1 | rRNA | ENSG0000C |
| 228632170 | 228632288 | -1 | rRNA | ENSG0000C |
| 228634040 | 228634160 | -1 | rRNA_pseudogene | ENSG0000C |
| 228636369 | 228637507 | 1 | processed_pseudoge | ENSG0000C |
| 228638564 | 228638688 | 1 | snoRNA | ENSG0000C |
| 228673218 | 228673629 | 1 | processed_pseudoge | ENSG0000C |
| 228721284 | 228732469 | 1 | protein_coding | ENSG0000C |
| 39776595 | 39786167 | 1 | protein_coding | ENSG0000C |
| 39812972 | 39813555 | -1 | lncRNA | ENSG0000C |
| 39825350 | 39834201 | -1 | protein_coding | ENSG0000C |
| 39829716 | 39829802 | -1 | miRNA | ENSG0000C |
| 39834458 | 39846379 | -1 | protein_coding | ENSG0000C |
| 39863324 | 39949893 | -1 | protein_coding | ENSG0000C |
| 39958506 | 39958947 | -1 | processed_pseudoge | ENSG0000C |
| 39986432 | 39997031 | 1 | protein_coding | ENSG0000C |
| 39999401 | 40036305 | 1 | protein_coding | ENSG0000C |
| 40038651 | 40040769 | -1 | processed_pseudoge | ENSG0000C |
| 40043527 | 40071498 | -1 | protein_coding | ENSG0000C |
| 40064024 | 40106192 | -1 | lncRNA | ENSG0000C |
| 40084419 | 40106205 | -1 | protein_coding | ENSG0000C |
| 40106021 | 40109673 | 1 | lncRNA | ENSG0000C |
| 91356877 | 91357036 | 1 | sRNA | ENSG0000C |
| 91374331 | 91514157 | -1 | unprocessed_pseudc | ENSG0000C |
| 189764391 | 189940733 | -1 | lncRNA | ENSG0000C |
| 189940870 | 189963202 | 1 | protein_coding | ENSG0000C |
| 189973599 | 189973847 | -1 | processed_pseudoge | ENSG0000C |
| 189979902 | 189982379 | 1 | processed_pseudoge | ENSG0000C |
| 189982523 | 189984871 | -1 | unprocessed_pseudc | ENSG0000C |
| 190014800 | 190014917 | 1 | rRNA_pseudogene | ENSG0000C |
| 190017360 | 190017478 | 1 | rRNA_pseudogene | ENSG0000C |
| 190021078 | 190022336 | -1 | unprocessed_pseudc | ENSG0000C |
| 190024025 | 190026913 | -1 | protein_coding | ENSG0000C |
| 190037743 | 190038226 | -1 | processed_pseudoge | ENSG0000C |
| 190067620 | 190068894 | 1 | unprocessed_pseudc | ENSG0000C |
| 190070926 | 190072197 | 1 | protein_coding | ENSG0000C |
| 190074232 | 190075503 | 1 | protein_coding | ENSG0000C |
| 190077538 | 190078809 | 1 | protein_coding | ENSG0000C |
| 190080844 | 190082115 | 1 | protein_coding | ENSG0000C |
| 190084150 | 190085421 | 1 | protein_coding | ENSG0000C |
| 190087456 | 190088727 | 1 | protein_coding | ENSG0000C |
| 190090762 | 190092033 | 1 | protein_coding | ENSG0000C |
| 190094068 | 190095339 | 1 | protein_coding | ENSG0000C |

| | | | | |
|---|---|---|---|---|
| 190097374 | 190098645 | 1 | protein_coding | ENSG00000( |
| 190100680 | 190101951 | 1 | protein_coding | ENSG00000( |
| 190103986 | 190105257 | 1 | protein_coding | ENSG00000( |
| 190107292 | 190108563 | 1 | protein_coding | ENSG00000( |
| 190128676 | 190129139 | 1 | processed_pseudoge | ENSG00000( |
| 190128687 | 190128927 | -1 | processed_pseudoge | ENSG00000( |
| 133205277 | 133209250 | -1 | protein_coding | ENSG00000( |
| 133224905 | 133228591 | -1 | unprocessed_pseudc | ENSG00000( |
| 133255585 | 133275349 | -1 | protein_coding | ENSG00000( |
| 133302618 | 133302711 | 1 | misc_RNA | ENSG00000( |
| 133317942 | 133318617 | 1 | unprocessed_pseudc | ENSG00000( |
| 133328924 | 133336336 | -1 | protein_coding | ENSG00000( |
| 133337876 | 133337972 | -1 | misc_RNA | ENSG00000( |
| 133338460 | 133348279 | -1 | protein_coding | ENSG00000( |
| 133339138 | 133339613 | 1 | processed_pseudoge | ENSG00000( |
| 133348366 | 133351574 | 1 | protein_coding | ENSG00000( |
| 133349544 | 133349618 | 1 | snoRNA | ENSG00000( |
| 133350243 | 133350316 | 1 | snoRNA | ENSG00000( |
| 133350604 | 133350676 | 1 | snoRNA | ENSG00000( |
| 133350829 | 133351541 | -1 | protein_coding | ENSG00000( |
| 133350995 | 133351060 | 1 | snoRNA | ENSG00000( |
| 133351906 | 133356824 | -1 | protein_coding | ENSG00000( |
| 133356698 | 133361306 | 1 | protein_coding | ENSG00000( |
| 133361598 | 133376314 | -1 | protein_coding | ENSG00000( |
| 133376514 | 133406244 | 1 | protein_coding | ENSG00000( |
| 133406207 | 133418244 | -1 | protein_coding | ENSG00000( |
| 133414506 | 133459550 | 1 | protein_coding | ENSG00000( |
| 133460113 | 133470996 | 1 | protein_coding | ENSG00000( |
| 133471242 | 133479275 | -1 | protein_coding | ENSG00000( |
| 7380534 | 7389774 | 1 | protein_coding | ENSG00000( |
| 7389727 | 7394842 | -1 | protein_coding | ENSG00000( |
| 7389734 | 7404097 | -1 | protein_coding | ENSG00000( |
| 7402975 | 7404097 | -1 | protein_coding | ENSG00000( |
| 7404874 | 7419860 | 1 | protein_coding | ENSG00000( |
| 7420103 | 7444081 | 1 | lncRNA | ENSG00000( |
| 7420324 | 7421632 | 1 | protein_coding | ENSG00000( |
| 7425616 | 7427568 | 1 | protein_coding | ENSG00000( |
| 7428857 | 7432820 | 1 | protein_coding | ENSG00000( |
| 7428861 | 7436278 | 1 | protein_coding | ENSG00000( |
| 7435435 | 7437679 | 1 | protein_coding | ENSG00000( |
| 7436557 | 7437523 | -1 | lncRNA | ENSG00000( |
| 7438273 | 7444937 | 1 | protein_coding | ENSG00000( |
| 7439159 | 7443327 | -1 | lncRNA | ENSG00000( |
| 7439506 | 7445966 | 1 | lncRNA | ENSG00000( |
| 7445061 | 7457710 | 1 | protein_coding | ENSG00000( |
| 7459366 | 7484263 | -1 | protein_coding | ENSG00000( |
| 7481332 | 7483496 | 1 | protein_coding | ENSG00000( |
| 7484366 | 7514614 | 1 | protein_coding | ENSG00000( |
| 121604902 | 121605012 | 1 | snRNA | ENSG00000( |
| 121652732 | 121669478 | -1 | protein_coding | ENSG00000( |

| | | | | |
|---|---|---|---|---|
| 10492223 | 10629368 | 1 | protein_coding | ENSG00000( |
| 10511036 | 10514546 | -1 | lncRNA | ENSG00000( |
| 10514282 | 10514973 | 1 | processed_pseudoge | ENSG00000( |
| 10574749 | 10575227 | -1 | processed_pseudoge | ENSG00000( |
| 10633762 | 10634695 | 1 | processed_pseudoge | ENSG00000( |
| 10659498 | 10692860 | 1 | lncRNA | ENSG00000( |
| 10671418 | 10694797 | -1 | protein_coding | ENSG00000( |
| 10674658 | 10674768 | -1 | misc_RNA | ENSG00000( |
| 10694972 | 10709782 | 1 | protein_coding | ENSG00000( |
| 10722915 | 10731129 | 1 | protein_coding | ENSG00000( |
| 26519354 | 26520250 | -1 | processed_pseudoge | ENSG00000( |
| 26529761 | 26575030 | 1 | protein_coding | ENSG00000( |
| 26554542 | 26554593 | 1 | miRNA | ENSG00000( |
| 26593243 | 26593546 | -1 | misc_RNA | ENSG00000( |
| 26593940 | 26594041 | -1 | misc_RNA | ENSG00000( |
| 26602369 | 26602432 | -1 | snRNA | ENSG00000( |
| 26640377 | 26640614 | -1 | processed_pseudoge | ENSG00000( |
| 26645671 | 26645774 | 1 | snoRNA | ENSG00000( |
| 26647447 | 26647681 | 1 | processed_pseudoge | ENSG00000( |
| 49558546 | 49810390 | 1 | transcribed_unproce | ENSG00000( |
| 49817749 | 49825524 | 1 | unprocessed_pseudc | ENSG00000( |
| 49824810 | 49826695 | -1 | unprocessed_pseudc | ENSG00000( |
| 49833111 | 49839291 | -1 | unprocessed_pseudc | ENSG00000( |
| 56890001 | 56890245 | 1 | processed_pseudoge | ENSG00000( |
| 85955666 | 86838884 | -1 | protein_coding | ENSG00000( |
| 86599578 | 86838978 | -1 | lncRNA | ENSG00000( |
| 87041747 | 87042254 | 1 | TEC | ENSG00000( |
| 44431705 | 44477670 | -1 | protein_coding | ENSG00000( |
| 44494847 | 44510636 | -1 | protein_coding | ENSG00000( |
| 44524744 | 44525805 | -1 | processed_pseudoge | ENSG00000( |
| 44528955 | 44529391 | -1 | processed_pseudoge | ENSG00000( |
| 44555193 | 44594173 | 1 | protein_coding | ENSG00000( |
| 44557357 | 44685653 | -1 | lncRNA | ENSG00000( |
| 44578223 | 44599694 | 1 | protein_coding | ENSG00000( |
| 44579938 | 44581026 | -1 | processed_pseudoge | ENSG00000( |
| 44584888 | 44648471 | 1 | protein_coding | ENSG00000( |
| 44617128 | 44624797 | -1 | lncRNA | ENSG00000( |
| 44648732 | 44660791 | 1 | protein_coding | ENSG00000( |
| 44667412 | 44669364 | 1 | lncRNA | ENSG00000( |
| 44709307 | 44710882 | -1 | processed_pseudoge | ENSG00000( |
| 44712639 | 44723827 | 1 | protein_coding | ENSG00000( |
| 44729592 | 44737079 | 1 | protein_coding | ENSG00000( |
| 44737657 | 44761615 | -1 | protein_coding | ENSG00000( |
| 44761717 | 44873372 | 1 | protein_coding | ENSG00000( |
| 44861884 | 44861977 | 1 | miRNA | ENSG00000( |
| 44861900 | 44865666 | 1 | protein_coding | ENSG00000( |
| 44874604 | 44914986 | 1 | protein_coding | ENSG00000( |
| 44899174 | 44900921 | 1 | lncRNA | ENSG00000( |
| 57132453 | 57132560 | -1 | rRNA_pseudogene | ENSG00000( |
| 57148457 | 57244793 | 1 | lncRNA | ENSG00000( |

| | | | | |
|---|---|---|---|---|
| 81844602 | 82169391 | -1 | lncRNA | ENSG00000 |
| 82169986 | 82247754 | -1 | protein_coding | ENSG00000 |
| 82210338 | 82210440 | -1 | snRNA | ENSG00000 |
| 82274199 | 82275177 | -1 | processed_pseudoge | ENSG00000 |
| 82364064 | 82373860 | -1 | lncRNA | ENSG00000 |
| 82373409 | 82381031 | 1 | protein_coding | ENSG00000 |
| 7243591 | 7252491 | -1 | protein_coding | ENSG00000 |
| 7246829 | 7262089 | -1 | protein_coding | ENSG00000 |
| 7251416 | 7259940 | 1 | protein_coding | ENSG00000 |
| 7259903 | 7263983 | -1 | protein_coding | ENSG00000 |
| 7266055 | 7266166 | -1 | misc_RNA | ENSG00000 |
| 7281718 | 7288257 | 1 | protein_coding | ENSG00000 |
| 7282947 | 7284071 | -1 | lncRNA | ENSG00000 |
| 7288263 | 7294639 | -1 | protein_coding | ENSG00000 |
| 7310036 | 7315500 | 1 | protein_coding | ENSG00000 |
| 7314361 | 7318601 | -1 | protein_coding | ENSG00000 |
| 7315698 | 7322211 | -1 | protein_coding | ENSG00000 |
| 7318665 | 7332430 | -1 | protein_coding | ENSG00000 |
| 7333489 | 7338450 | 1 | unitary_pseudogene | ENSG00000 |
| 7339566 | 7354514 | 1 | protein_coding | ENSG00000 |
| 7354926 | 7357981 | 1 | protein_coding | ENSG00000 |
| 7355724 | 7357981 | -1 | lncRNA | ENSG00000 |
| 7358160 | 7360256 | 1 | protein_coding | ENSG00000 |
| 99027438 | 99144100 | -1 | protein_coding | ENSG00000 |
| 99173574 | 99207506 | -1 | protein_coding | ENSG00000 |
| 17189783 | 17197617 | 1 | lncRNA | ENSG00000 |
| 17193232 | 17201733 | -1 | lncRNA | ENSG00000 |
| 17205128 | 17246007 | 1 | protein_coding | ENSG00000 |
| 17249098 | 17284233 | 1 | protein_coding | ENSG00000 |
| 17277889 | 17277975 | 1 | miRNA | ENSG00000 |
| 17310905 | 17366713 | 1 | protein_coding | ENSG00000 |
| 17374905 | 17404408 | 1 | protein_coding | ENSG00000 |
| 17409469 | 17410091 | 1 | lncRNA | ENSG00000 |
| 17409469 | 17442386 | -1 | protein_coding | ENSG00000 |
| 112911847 | 112957013 | -1 | protein_coding | ENSG00000 |
| 112923423 | 112924337 | 1 | processed_pseudoge | ENSG00000 |
| 112956601 | 112956752 | 1 | pseudogene | ENSG00000 |
| 113013524 | 113073113 | -1 | lncRNA | ENSG00000 |
| 18261982 | 18263091 | 1 | processed_pseudoge | ENSG00000 |
| 18266260 | 18269977 | 1 | protein_coding | ENSG00000 |
| 18266393 | 18266510 | 1 | rRNA_pseudogene | ENSG00000 |
| 18278668 | 18322198 | -1 | protein_coding | ENSG00000 |
| 18322295 | 18367045 | 1 | protein_coding | ENSG00000 |
| 18390413 | 18390486 | 1 | miRNA | ENSG00000 |
| 18397186 | 18411051 | 1 | protein_coding | ENSG00000 |
| 18408235 | 18409357 | 1 | lncRNA | ENSG00000 |
| 47565336 | 47584562 | 1 | protein_coding | ENSG00000 |
| 47565430 | 47573461 | 1 | protein_coding | ENSG00000 |
| 47572197 | 47578976 | -1 | protein_coding | ENSG00000 |
| 47576471 | 47576588 | -1 | snRNA | ENSG00000 |

| | | | | |
|---|---|---|---|---|
| 47577725 | 47578277 | 1 | lncRNA | ENSG0000C |
| 47586693 | 47589194 | 1 | protein_coding | ENSG0000C |
| 47589667 | 47594411 | -1 | protein_coding | ENSG0000C |
| 47617315 | 47647147 | -1 | protein_coding | ENSG0000C |
| 47664132 | 47719930 | -1 | protein_coding | ENSG0000C |
| 47721035 | 47771984 | -1 | protein_coding | ENSG0000C |
| 47731435 | 47731533 | -1 | misc_RNA | ENSG0000C |
| 47800710 | 47800920 | 1 | processed_pseudoge | ENSG0000C |
| 47808837 | 47808943 | -1 | rRNA_pseudogene | ENSG0000C |
| 47824741 | 47824830 | -1 | misc_RNA | ENSG0000C |
| 47845865 | 47846223 | 1 | processed_pseudoge | ENSG0000C |
| 70272173 | 70387610 | 1 | protein_coding | ENSG0000C |
| 70278921 | 70279297 | -1 | processed_pseudoge | ENSG0000C |
| 70282367 | 70363368 | -1 | lncRNA | ENSG0000C |
| 70283955 | 70284070 | 1 | miRNA | ENSG0000C |
| 70319928 | 70321415 | 1 | lncRNA | ENSG0000C |
| 70324871 | 70327209 | -1 | lncRNA | ENSG0000C |
| 70358198 | 70358677 | -1 | lncRNA | ENSG0000C |
| 70377713 | 70403955 | -1 | lncRNA | ENSG0000C |
| 70403871 | 70442051 | 1 | protein_coding | ENSG0000C |
| 92340108 | 92340572 | -1 | processed_pseudoge | ENSG0000C |
| 92370572 | 92371326 | -1 | processed_pseudoge | ENSG0000C |
| 89642498 | 89819794 | -1 | protein_coding | ENSG0000C |
| 89814709 | 89815463 | 1 | processed_pseudoge | ENSG0000C |
| 89829894 | 89876779 | 1 | protein_coding | ENSG0000C |
| 89841329 | 89841424 | 1 | misc_RNA | ENSG0000C |
| 89878887 | 89879221 | 1 | processed_pseudoge | ENSG0000C |
| 89888499 | 89890191 | 1 | processed_pseudoge | ENSG0000C |
| 89896812 | 89924103 | 1 | protein_coding | ENSG0000C |
| 89900626 | 89900735 | -1 | misc_RNA | ENSG0000C |
| 67546651 | 67546754 | 1 | snRNA | ENSG0000C |
| 67631019 | 67631127 | 1 | misc_RNA | ENSG0000C |
| 94939812 | 94940172 | -1 | processed_pseudoge | ENSG0000C |
| 94941138 | 94942076 | -1 | processed_pseudoge | ENSG0000C |
| 94942032 | 95168351 | 1 | lncRNA | ENSG0000C |
| 74823667 | 74824187 | -1 | lncRNA | ENSG0000C |
| 74826984 | 75034681 | 1 | protein_coding | ENSG0000C |
| 75005112 | 75027231 | -1 | lncRNA | ENSG0000C |
| 75025532 | 75026308 | -1 | lncRNA | ENSG0000C |
| 75039893 | 75060571 | -1 | protein_coding | ENSG0000C |
| 75043933 | 75057811 | 1 | lncRNA | ENSG0000C |
| 74152800 | 74179226 | -1 | lncRNA | ENSG0000C |
| 74173710 | 74176872 | 1 | protein_coding | ENSG0000C |
| 74179466 | 74212267 | -1 | protein_coding | ENSG0000C |
| 74202705 | 74338316 | 1 | protein_coding | ENSG0000C |
| 74301347 | 74306371 | 1 | TEC | ENSG0000C |
| 74304849 | 74306187 | 1 | lncRNA | ENSG0000C |
| 74313360 | 74321532 | -1 | lncRNA | ENSG0000C |
| 74339603 | 74369490 | -1 | protein_coding | ENSG0000C |
| 74352612 | 74366464 | 1 | transcribed_processe | ENSG0000C |

| | | | | |
|---|---|---|---|---|
| 74367279 | 74373055 | 1 | lncRNA | ENSG0000C |
| 74376522 | 74377355 | 1 | lncRNA | ENSG0000C |
| 74380927 | 74392379 | 1 | lncRNA | ENSG0000C |
| 74411133 | 74435802 | -1 | protein_coding | ENSG0000C |
| 74413201 | 74413276 | -1 | miRNA | ENSG0000C |
| 74431289 | 74433293 | 1 | lncRNA | ENSG0000C |
| 74447821 | 74463026 | -1 | protein_coding | ENSG0000C |
| 74463109 | 74515480 | 1 | lncRNA | ENSG0000C |
| 74479914 | 74492130 | -1 | transcribed_processe | ENSG0000C |
| 74491446 | 74518803 | 1 | lncRNA | ENSG0000C |
| 74492803 | 74492872 | -1 | snoRNA | ENSG0000C |
| 48539031 | 48867468 | 1 | protein_coding | ENSG0000C |
| 48564795 | 48568342 | 1 | TEC | ENSG0000C |
| 48670632 | 48670700 | 1 | miRNA | ENSG0000C |
| 48677826 | 48692670 | -1 | lncRNA | ENSG0000C |
| 48759944 | 48760569 | 1 | TEC | ENSG0000C |
| 48830302 | 48839021 | -1 | lncRNA | ENSG0000C |
| 118971712 | 118981287 | 1 | protein_coding | ENSG0000C |
| 118994334 | 118994630 | 1 | misc_RNA | ENSG0000C |
| 118994560 | 118994655 | 1 | misc_RNA | ENSG0000C |
| 118994824 | 118998004 | -1 | lncRNA | ENSG0000C |
| 118998138 | 119015793 | 1 | protein_coding | ENSG0000C |
| 119003012 | 119003446 | -1 | processed_pseudoge | ENSG0000C |
| 119003742 | 119004893 | -1 | lncRNA | ENSG0000C |
| 119005727 | 119005934 | -1 | processed_pseudoge | ENSG0000C |
| 119015712 | 119018691 | -1 | protein_coding | ENSG0000C |
| 119018763 | 119025454 | 1 | protein_coding | ENSG0000C |
| 119018944 | 119019012 | 1 | miRNA | ENSG0000C |
| 119024112 | 119030905 | -1 | protein_coding | ENSG0000C |
| 119044187 | 119045492 | 1 | lncRNA | ENSG0000C |
| 119044187 | 119057226 | -1 | protein_coding | ENSG0000C |
| 119065262 | 119065676 | -1 | lncRNA | ENSG0000C |
| 119067373 | 119067697 | -1 | lncRNA | ENSG0000C |
| 119067691 | 119081977 | 1 | protein_coding | ENSG0000C |
| 233059967 | 233207903 | 1 | protein_coding | ENSG0000C |
| 233205199 | 233205479 | 1 | misc_RNA | ENSG0000C |
| 233210051 | 233295674 | 1 | protein_coding | ENSG0000C |
| 233275727 | 233276002 | 1 | scaRNA | ENSG0000C |
| 233288676 | 233288940 | 1 | scaRNA | ENSG0000C |
| 233307816 | 233347055 | 1 | protein_coding | ENSG0000C |
| 233351132 | 233353416 | -1 | lncRNA | ENSG0000C |
| 233354507 | 233472104 | 1 | protein_coding | ENSG0000C |
| 233379843 | 233379955 | -1 | misc_RNA | ENSG0000C |
| 239734956 | 239735538 | 1 | lncRNA | ENSG0000C |
| 239762860 | 239824764 | 1 | lncRNA | ENSG0000C |
| 65353525 | 66038834 | -1 | protein_coding | ENSG0000C |
| 66133610 | 66388212 | 1 | protein_coding | ENSG0000C |
| 66293210 | 66293316 | -1 | snRNA | ENSG0000C |
| 66323951 | 66324248 | 1 | misc_RNA | ENSG0000C |
| 16687986 | 16697479 | -1 | lncRNA | ENSG0000C |

| | | | | |
|---|---|---|---|---|
| 16692745 | 16694202 | -1 | unprocessed_pseudc | ENSG00000 |
| 16802651 | 17090582 | 1 | protein_coding | ENSG00000 |
| 16933174 | 16933238 | 1 | miRNA | ENSG00000 |
| 17042720 | 17044170 | -1 | lncRNA | ENSG00000 |
| 154544169 | 154547285 | -1 | lncRNA | ENSG00000 |
| 132084283 | 132131639 | 1 | transcribed_unproce | ENSG00000 |
| 132100013 | 132100820 | -1 | TEC | ENSG00000 |
| 132126461 | 132126764 | -1 | processed_pseudoge | ENSG00000 |
| 132136594 | 132144319 | -1 | protein_coding | ENSG00000 |
| 132144457 | 132152473 | 1 | protein_coding | ENSG00000 |
| 132169823 | 132170043 | 1 | lncRNA | ENSG00000 |
| 132186735 | 132189695 | -1 | lncRNA | ENSG00000 |
| 132190213 | 132190997 | 1 | lncRNA | ENSG00000 |
| 132196372 | 132328920 | -1 | protein_coding | ENSG00000 |
| 132200436 | 132201287 | 1 | TEC | ENSG00000 |
| 132274722 | 132277660 | 1 | lncRNA | ENSG00000 |
| 132276680 | 132280231 | 1 | lncRNA | ENSG00000 |
| 132278587 | 132280231 | -1 | TEC | ENSG00000 |
| 132282145 | 132283937 | 1 | lncRNA | ENSG00000 |
| 132329181 | 132330906 | 1 | lncRNA | ENSG00000 |
| 132331336 | 132331763 | 1 | TEC | ENSG00000 |
| 132343808 | 132344683 | 1 | lncRNA | ENSG00000 |
| 132351186 | 132351653 | 1 | TEC | ENSG00000 |
| 132360065 | 132360529 | 1 | TEC | ENSG00000 |
| 132362141 | 132362679 | 1 | TEC | ENSG00000 |
| 122019422 | 122062044 | 1 | protein_coding | ENSG00000 |
| 122063306 | 122068616 | 1 | lncRNA | ENSG00000 |
| 122078746 | 122143107 | 1 | protein_coding | ENSG00000 |
| 100708325 | 101059286 | 1 | lncRNA | ENSG00000 |
| 100934023 | 100944553 | -1 | lncRNA | ENSG00000 |
| 17082295 | 17085049 | -1 | TEC | ENSG00000 |
| 17094351 | 17094927 | -1 | unprocessed_pseudc | ENSG00000 |
| 17096202 | 17096421 | -1 | processed_pseudoge | ENSG00000 |
| 17108698 | 17259501 | -1 | protein_coding | ENSG00000 |
| 17134504 | 17138736 | 1 | lncRNA | ENSG00000 |
| 17192863 | 17194232 | -1 | TEC | ENSG00000 |
| 17227023 | 17228000 | -1 | TEC | ENSG00000 |
| 17249674 | 17251395 | -1 | TEC | ENSG00000 |
| 17405678 | 17408073 | 1 | lncRNA | ENSG00000 |
| 17445825 | 17446380 | -1 | lncRNA | ENSG00000 |
| 17453470 | 17456236 | 1 | lncRNA | ENSG00000 |
| 107044705 | 107161811 | 1 | protein_coding | ENSG00000 |
| 107168961 | 107202522 | 1 | protein_coding | ENSG00000 |
| 107192559 | 107193300 | -1 | lncRNA | ENSG00000 |
| 107201555 | 107564647 | -1 | protein_coding | ENSG00000 |
| 107450200 | 107450479 | 1 | processed_pseudoge | ENSG00000 |
| 107470143 | 107475810 | 1 | protein_coding | ENSG00000 |
| 107563617 | 107578656 | 1 | protein_coding | ENSG00000 |
| 107579690 | 107580190 | -1 | lncRNA | ENSG00000 |
| 107580906 | 107618493 | 1 | protein_coding | ENSG00000 |

| | | | | |
|---|---|---|---|---|
| 107603496 | 107603640 | 1 | snoRNA | ENSG0000C |
| 404468 | 414023 | -1 | lncRNA | ENSG0000C |
| 410325 | 431062 | 1 | protein_coding | ENSG0000C |
| 439978 | 445939 | -1 | protein_coding | ENSG0000C |
| 465285 | 473461 | 1 | lncRNA | ENSG0000C |
| 470782 | 683831 | -1 | protein_coding | ENSG0000C |
| 485254 | 502690 | 1 | lncRNA | ENSG0000C |
| 514680 | 515531 | 1 | lncRNA | ENSG0000C |
| 578116 | 578709 | 1 | processed_pseudoge | ENSG0000C |
| 618908 | 619661 | -1 | processed_pseudoge | ENSG0000C |
| 113953705 | 113974077 | -1 | protein_coding | ENSG0000C |
| 113977784 | 114128689 | -1 | protein_coding | ENSG0000C |
| 114103635 | 114104886 | -1 | lncRNA | ENSG0000C |
| 88897232 | 88897784 | 1 | IG_V_gene | ENSG0000C |
| 88915081 | 88915378 | -1 | IG_V_pseudogene | ENSG0000C |
| 88922328 | 88922753 | -1 | processed_pseudoge | ENSG0000C |
| 88931666 | 88932380 | -1 | IG_V_pseudogene | ENSG0000C |
| 88947301 | 88947957 | -1 | IG_V_gene | ENSG0000C |
| 88966262 | 88966767 | -1 | IG_V_gene | ENSG0000C |
| 88978468 | 88979081 | -1 | IG_V_gene | ENSG0000C |
| 88992409 | 88992931 | -1 | IG_V_gene | ENSG0000C |
| 89009982 | 89010515 | -1 | IG_V_gene | ENSG0000C |
| 89019992 | 89020686 | -1 | IG_V_pseudogene | ENSG0000C |
| 89027171 | 89027731 | -1 | IG_V_gene | ENSG0000C |
| 89040224 | 89040745 | -1 | IG_V_gene | ENSG0000C |
| 89045995 | 89046466 | -1 | IG_V_pseudogene | ENSG0000C |
| 89078010 | 89078784 | -1 | IG_V_pseudogene | ENSG0000C |
| 89085177 | 89085787 | -1 | IG_V_gene | ENSG0000C |
| 89099859 | 89100361 | -1 | IG_V_gene | ENSG0000C |
| 89117342 | 89117844 | -1 | IG_V_gene | ENSG0000C |
| 89128724 | 89129483 | -1 | IG_V_pseudogene | ENSG0000C |
| 89134975 | 89135257 | -1 | IG_V_pseudogene | ENSG0000C |
| 89142574 | 89143160 | -1 | IG_V_gene | ENSG0000C |
| 89159751 | 89160366 | -1 | IG_V_gene | ENSG0000C |
| 89170775 | 89171212 | -1 | IG_V_pseudogene | ENSG0000C |
| 89172022 | 89172553 | -1 | IG_V_pseudogene | ENSG0000C |
| 89176328 | 89177160 | -1 | IG_V_gene | ENSG0000C |
| 89192500 | 89192752 | -1 | IG_V_pseudogene | ENSG0000C |
| 89196096 | 89196829 | -1 | IG_V_pseudogene | ENSG0000C |
| 89203509 | 89203906 | -1 | unprocessed_pseudc | ENSG0000C |
| 89213423 | 89213928 | -1 | IG_V_gene | ENSG0000C |
| 89221698 | 89222461 | -1 | IG_V_gene | ENSG0000C |
| 89234174 | 89234912 | -1 | IG_V_pseudogene | ENSG0000C |
| 89244781 | 89245596 | -1 | IG_V_gene | ENSG0000C |
| 89252211 | 89252736 | -1 | IG_V_pseudogene | ENSG0000C |
| 89253571 | 89254015 | -1 | IG_V_pseudogene | ENSG0000C |
| 89266494 | 89268506 | -1 | IG_V_gene | ENSG0000C |
| 89275298 | 89275787 | -1 | IG_V_pseudogene | ENSG0000C |
| 89286689 | 89286973 | -1 | IG_V_pseudogene | ENSG0000C |
| 89295233 | 89295455 | -1 | IG_V_pseudogene | ENSG0000C |

| | | | | |
|---|---|---|---|---|
| 89297264 | 89297785 | -1 | IG_V_gene | ENSG00000 |
| 89309898 | 89310144 | -1 | IG_V_pseudogene | ENSG00000 |
| 89319625 | 89320146 | -1 | IG_V_gene | ENSG00000 |
| 89330110 | 89330429 | -1 | IG_V_gene | ENSG00000 |
| 18343918 | 18348373 | -1 | unprocessed_pseudo | ENSG00000 |
| 18365205 | 18401932 | 1 | lncRNA | ENSG00000 |
| 18376478 | 18377374 | -1 | processed_pseudoge | ENSG00000 |
| 18405408 | 18407072 | 1 | protein_coding | ENSG00000 |
| 18418821 | 18418943 | -1 | processed_pseudoge | ENSG00000 |
| 18423434 | 18440730 | -1 | unprocessed_pseudo | ENSG00000 |
| 18449228 | 18479842 | -1 | unprocessed_pseudo | ENSG00000 |
| 18467462 | 18468641 | 1 | processed_pseudoge | ENSG00000 |
| 18501841 | 18518504 | 1 | unprocessed_pseudo | ENSG00000 |
| 87751512 | 87841343 | -1 | protein_coding | ENSG00000 |
| 87817928 | 87845612 | 1 | transcribed_unproce | ENSG00000 |
| 87832818 | 87833098 | -1 | misc_RNA | ENSG00000 |
| 87861956 | 87862489 | -1 | protein_coding | ENSG00000 |
| 87863440 | 87966341 | 1 | protein_coding | ENSG00000 |
| 87878694 | 87880428 | 1 | lncRNA | ENSG00000 |
| 87945469 | 87945991 | 1 | processed_pseudoge | ENSG00000 |
| 87994585 | 87994662 | -1 | snoRNA | ENSG00000 |
| 88048102 | 88049823 | -1 | processed_pseudoge | ENSG00000 |
| 19878555 | 19887814 | 1 | unprocessed_pseudo | ENSG00000 |
| 19883614 | 19883716 | 1 | snRNA | ENSG00000 |
| 19899334 | 19899559 | 1 | processed_pseudoge | ENSG00000 |
| 19955300 | 19955602 | -1 | unprocessed_pseudo | ENSG00000 |
| 19964668 | 19965101 | -1 | protein_coding | ENSG00000 |
| 19987663 | 19988122 | -1 | protein_coding | ENSG00000 |
| 20012746 | 20014191 | 1 | protein_coding | ENSG00000 |
| 20122821 | 20123707 | 1 | protein_coding | ENSG00000 |
| 20321344 | 20321616 | 1 | protein_coding | ENSG00000 |
| 20331892 | 20332782 | -1 | protein_coding | ENSG00000 |
| 20374618 | 20374849 | -1 | pseudogene | ENSG00000 |
| 20469564 | 20470996 | -1 | protein_coding | ENSG00000 |
| 20589394 | 20589682 | 1 | protein_coding | ENSG00000 |
| 20643749 | 20644108 | -1 | protein_coding | ENSG00000 |
| 20741608 | 20742444 | -1 | protein_coding | ENSG00000 |
| 20816371 | 20817048 | 1 | protein_coding | ENSG00000 |
| 20820670 | 20821762 | -1 | pseudogene | ENSG00000 |
| 20855523 | 20886468 | -1 | protein_coding | ENSG00000 |
| 21084472 | 21085410 | 1 | protein_coding | ENSG00000 |
| 21098400 | 21099347 | 1 | protein_coding | ENSG00000 |
| 21156147 | 21156770 | -1 | protein_coding | ENSG00000 |
| 21164308 | 21164743 | -1 | protein_coding | ENSG00000 |
| 21181946 | 21189242 | -1 | protein_coding | ENSG00000 |
| 21376529 | 21376760 | 1 | pseudogene | ENSG00000 |
| 21406249 | 21407139 | 1 | protein_coding | ENSG00000 |
| 21417407 | 21417679 | -1 | protein_coding | ENSG00000 |
| 21641276 | 21642848 | -1 | pseudogene | ENSG00000 |
| 21733035 | 21733712 | 1 | protein_coding | ENSG00000 |

| | | | | |
|---|---|---|---|---|
| 21737334 | 21738426 | -1 | pseudogene | ENSG0000( |
| 21773471 | 21804450 | -1 | protein_coding | ENSG0000( |
| 22002424 | 22003362 | 1 | protein_coding | ENSG0000( |
| 22016348 | 22017295 | 1 | protein_coding | ENSG0000( |
| 22082221 | 22082656 | -1 | protein_coding | ENSG0000( |
| 22106780 | 22107213 | -1 | protein_coding | ENSG0000( |
| 22287263 | 22287494 | 1 | pseudogene | ENSG0000( |
| 22324750 | 22325640 | 1 | protein_coding | ENSG0000( |
| 22335908 | 22336180 | -1 | protein_coding | ENSG0000( |
| 22539500 | 22540386 | -1 | protein_coding | ENSG0000( |
| 22799188 | 22830039 | 1 | protein_coding | ENSG0000( |
| 22865088 | 22866180 | 1 | pseudogene | ENSG0000( |
| 22869800 | 22870477 | -1 | protein_coding | ENSG0000( |
| 22943679 | 22945251 | 1 | pseudogene | ENSG0000( |
| 23042969 | 23043328 | 1 | protein_coding | ENSG0000( |
| 23097381 | 23107645 | -1 | protein_coding | ENSG0000( |
| 23215931 | 23217363 | 1 | protein_coding | ENSG0000( |
| 23277139 | 23287363 | -1 | protein_coding | ENSG0000( |
| 23455943 | 23493534 | 1 | protein_coding | ENSG0000( |
| 23539361 | 23616629 | -1 | protein_coding | ENSG0000( |
| 23758443 | 23760479 | 1 | protein_coding | ENSG0000( |
| 23798790 | 23806603 | -1 | protein_coding | ENSG0000( |
| 23827481 | 23837765 | 1 | protein_coding | ENSG0000( |
| 23961862 | 23972149 | -1 | protein_coding | ENSG0000( |
| 23992969 | 24000781 | 1 | protein_coding | ENSG0000( |
| 24050655 | 24058495 | -1 | protein_coding | ENSG0000( |
| 24089314 | 24089592 | 1 | protein_coding | ENSG0000( |
| 24160220 | 24168592 | -1 | protein_coding | ENSG0000( |
| 24288509 | 24298144 | 1 | protein_coding | ENSG0000( |
| 24366714 | 24370460 | -1 | protein_coding | ENSG0000( |
| 24409558 | 24410281 | -1 | protein_coding | ENSG0000( |
| 25064722 | 25066457 | -1 | lncRNA | ENSG0000( |
| 45423764 | 45483218 | -1 | lncRNA | ENSG0000( |
| 45483343 | 45529854 | 1 | lncRNA | ENSG0000( |
| 45491581 | 45503824 | -1 | lncRNA | ENSG0000( |
| 45507220 | 45550182 | 1 | lncRNA | ENSG0000( |
| 45571405 | 45611540 | -1 | lncRNA | ENSG0000( |
| 45573083 | 45747215 | -1 | lncRNA | ENSG0000( |
| 144262045 | 144291438 | -1 | protein_coding | ENSG0000( |
| 144291591 | 144314720 | 1 | protein_coding | ENSG0000( |
| 144314584 | 144326910 | -1 | protein_coding | ENSG0000( |
| 144314590 | 144315138 | -1 | lncRNA | ENSG0000( |
| 144317246 | 144317315 | -1 | miRNA | ENSG0000( |
| 144330565 | 144336482 | -1 | protein_coding | ENSG0000( |
| 144352219 | 144354914 | -1 | protein_coding | ENSG0000( |
| 144353228 | 144355609 | -1 | protein_coding | ENSG0000( |
| 144354135 | 144361272 | 1 | protein_coding | ENSG0000( |
| 144355431 | 144359376 | -1 | protein_coding | ENSG0000( |
| 144373101 | 144393242 | 1 | protein_coding | ENSG0000( |
| 144393229 | 144409335 | -1 | protein_coding | ENSG0000( |

| | | | | |
|---|---|---|---|---|
| 144394149 | 144394230 | -1 | miRNA | ENSG0000C |
| 144400086 | 144400165 | -1 | miRNA | ENSG0000C |
| 144400277 | 144400345 | -1 | miRNA | ENSG0000C |
| 144409492 | 144409976 | 1 | lncRNA | ENSG0000C |
| 144409742 | 144416895 | -1 | protein_coding | ENSG0000C |
| 144423600 | 144428562 | -1 | protein_coding | ENSG0000C |
| 70430231 | 70431125 | 1 | processed_pseudoge | ENSG0000C |
| 70495527 | 70497102 | 1 | TEC | ENSG0000C |
| 70503820 | 70554100 | -1 | lncRNA | ENSG0000C |
| 24610209 | 24827632 | 1 | protein_coding | ENSG0000C |
| 24803451 | 24819739 | -1 | lncRNA | ENSG0000C |
| 24845841 | 24914411 | 1 | protein_coding | ENSG0000C |
| 24922172 | 25018449 | -1 | protein_coding | ENSG0000C |
| 101903154 | 101903204 | 1 | unprocessed_pseudc | ENSG0000C |
| 101922042 | 101923113 | -1 | protein_coding | ENSG0000C |
| 101926164 | 101933350 | -1 | transcribed_unproce | ENSG0000C |
| 101937591 | 101938470 | -1 | unprocessed_pseudc | ENSG0000C |
| 101954885 | 101956412 | 1 | lncRNA | ENSG0000C |
| 101959539 | 101961408 | -1 | lncRNA | ENSG0000C |
| 101960459 | 101960596 | -1 | miRNA | ENSG0000C |
| 101961603 | 101976543 | 1 | transcribed_unproce | ENSG0000C |
| 101973524 | 101973591 | 1 | miRNA | ENSG0000C |
| 101976558 | 101979093 | -1 | transcribed_unproce | ENSG0000C |
| 101984125 | 101986741 | 1 | protein_coding | ENSG0000C |
| 101987809 | 101988823 | -1 | protein_coding | ENSG0000C |
| 15675621 | 15675927 | -1 | pseudogene | ENSG0000C |
| 15719866 | 15724275 | -1 | protein_coding | ENSG0000C |
| 15846439 | 15850834 | 1 | protein_coding | ENSG0000C |
| 15671165 | 15671474 | -1 | pseudogene | ENSG0000C |
| 15715696 | 15720088 | -1 | protein_coding | ENSG0000C |
| 15844938 | 15849328 | 1 | protein_coding | ENSG0000C |
| 16709202 | 16709476 | -1 | pseudogene | ENSG0000C |
| 16752660 | 16757051 | -1 | protein_coding | ENSG0000C |
| 16929570 | 16929879 | 1 | pseudogene | ENSG0000C |
| 10186517 | 10186826 | -1 | pseudogene | ENSG0000C |
| 10354850 | 10359245 | 1 | protein_coding | ENSG0000C |
| 8415714 | 8416023 | -1 | pseudogene | ENSG0000C |
| 8459946 | 8464342 | -1 | protein_coding | ENSG0000C |
| 8589051 | 8593445 | 1 | protein_coding | ENSG0000C |
| 8746458 | 8746549 | 1 | miRNA | ENSG0000C |
| 8746994 | 8747083 | 1 | misc_RNA | ENSG0000C |
| 8751515 | 8752437 | -1 | lncRNA | ENSG0000C |
| 8753703 | 8753855 | 1 | rRNA | ENSG0000C |
| 8775853 | 8776331 | 1 | TEC | ENSG0000C |
| 8778771 | 8779099 | 1 | TEC | ENSG0000C |
| 8779979 | 8780212 | 1 | TEC | ENSG0000C |
| 8781856 | 8782167 | 1 | TEC | ENSG0000C |
| 8790626 | 8790717 | 1 | miRNA | ENSG0000C |
| 8791181 | 8791270 | 1 | misc_RNA | ENSG0000C |
| 8795725 | 8796647 | -1 | lncRNA | ENSG0000C |

| | | | | |
|---|---|---|---|---|
| 8797919 | 8798071 | 1 | rRNA | ENSG00000 |
| 8820416 | 8820894 | 1 | TEC | ENSG00000 |
| 8823350 | 8823682 | 1 | TEC | ENSG00000 |
| 8824562 | 8824795 | 1 | TEC | ENSG00000 |
| 8826416 | 8826727 | 1 | TEC | ENSG00000 |
| 49203171 | 49203726 | -1 | processed_pseudoge | ENSG00000 |
| 49212251 | 49212624 | -1 | processed_pseudoge | ENSG00000 |
| 49229573 | 49231284 | 1 | TEC | ENSG00000 |
| 49233289 | 49233965 | -1 | processed_pseudoge | ENSG00000 |
| 49238032 | 49245419 | -1 | processed_pseudoge | ENSG00000 |
| 49246021 | 49246538 | 1 | processed_pseudoge | ENSG00000 |
| 49246601 | 49246915 | 1 | processed_pseudoge | ENSG00000 |
| 49270489 | 49271208 | -1 | protein_coding | ENSG00000 |
| 49339472 | 49340191 | 1 | protein_coding | ENSG00000 |
| 38587641 | 38618882 | -1 | protein_coding | ENSG00000 |
| 38596346 | 38601694 | 1 | lncRNA | ENSG00000 |
| 38619082 | 38636955 | 1 | protein_coding | ENSG00000 |
| 38647649 | 38731589 | 1 | protein_coding | ENSG00000 |
| 38683873 | 38693606 | -1 | lncRNA | ENSG00000 |
| 38738251 | 38739830 | 1 | lncRNA | ENSG00000 |
| 38770969 | 38773523 | -1 | protein_coding | ENSG00000 |
| 38789280 | 38791794 | 1 | protein_coding | ENSG00000 |
| 38797048 | 38797155 | -1 | snRNA | ENSG00000 |
| 38801718 | 38812981 | -1 | protein_coding | ENSG00000 |
| 38815458 | 38831877 | -1 | protein_coding | ENSG00000 |
| 38817507 | 38840214 | -1 | protein_coding | ENSG00000 |
| 38820202 | 38823259 | 1 | lncRNA | ENSG00000 |
| 38836424 | 38852383 | -1 | protein_coding | ENSG00000 |
| 38844765 | 38845535 | -1 | lncRNA | ENSG00000 |
| 38867866 | 38878311 | -1 | protein_coding | ENSG00000 |
| 38870195 | 38873799 | 1 | lncRNA | ENSG00000 |
| 38878591 | 38899898 | -1 | protein_coding | ENSG00000 |
| 38899736 | 38908929 | 1 | protein_coding | ENSG00000 |
| 38909016 | 38912194 | -1 | protein_coding | ENSG00000 |
| 38915302 | 38930932 | -1 | protein_coding | ENSG00000 |
| 38915440 | 38949891 | -1 | protein_coding | ENSG00000 |
| 38930584 | 38933198 | 1 | protein_coding | ENSG00000 |
| 38935333 | 38938668 | -1 | lncRNA | ENSG00000 |
| 38941437 | 38975778 | -1 | protein_coding | ENSG00000 |
| 38975197 | 39014527 | 1 | lncRNA | ENSG00000 |
| 1828307 | 1837521 | 1 | protein_coding | ENSG00000 |
| 1838981 | 1841680 | 1 | protein_coding | ENSG00000 |
| 1852970 | 1892267 | 1 | protein_coding | ENSG00000 |
| 1859464 | 1859536 | -1 | miRNA | ENSG00000 |
| 1864177 | 1866667 | 1 | lncRNA | ENSG00000 |
| 1880045 | 1880147 | 1 | miRNA | ENSG00000 |
| 1888577 | 1891895 | -1 | protein_coding | ENSG00000 |
| 1898024 | 1917031 | 1 | protein_coding | ENSG00000 |
| 1925868 | 1937511 | -1 | lncRNA | ENSG00000 |
| 1949297 | 1986541 | 1 | protein_coding | ENSG00000 |

| | | | | |
|---|---|---|---|---|
| 1985256 | 1991939 | -1 | lncRNA | ENSG00000 |
| 1993115 | 1995689 | 1 | lncRNA | ENSG00000 |
| 1997195 | 2003489 | -1 | lncRNA | ENSG00000 |
| 1998559 | 1998629 | 1 | misc_RNA | ENSG00000 |
| 1998778 | 1998850 | -1 | miRNA | ENSG00000 |
| 1999503 | 1999571 | 1 | misc_RNA | ENSG00000 |
| 179110853 | 179345461 | -1 | protein_coding | ENSG00000 |
| 179377505 | 179378865 | 1 | lncRNA | ENSG00000 |
| 179377531 | 179378761 | -1 | lncRNA | ENSG00000 |
| 179385819 | 179385992 | -1 | processed_pseudoge | ENSG00000 |
| 179455294 | 179455974 | 1 | lncRNA | ENSG00000 |
| 179492015 | 179492096 | -1 | snRNA | ENSG00000 |
| 179503254 | 179515530 | -1 | lncRNA | ENSG00000 |
| 179522252 | 179524868 | 1 | unprocessed_pseudo | ENSG00000 |
| 179527020 | 179527309 | -1 | processed_pseudoge | ENSG00000 |
| 179550489 | 179611566 | 1 | protein_coding | ENSG00000 |
| 179572092 | 179572677 | 1 | processed_pseudoge | ENSG00000 |
| 179597471 | 179605295 | -1 | lncRNA | ENSG00000 |
| 179615760 | 179636304 | -1 | protein_coding | ENSG00000 |
| 179630732 | 179630991 | 1 | misc_RNA | ENSG00000 |
| 179641610 | 179642783 | -1 | unprocessed_pseudo | ENSG00000 |
| 179649036 | 179651563 | 1 | protein_coding | ENSG00000 |
| 179679544 | 179682019 | -1 | protein_coding | ENSG00000 |
| 179679612 | 179732514 | 1 | protein_coding | ENSG00000 |
| 179680016 | 179695621 | 1 | transcribed_process | ENSG00000 |
| 179733695 | 179778170 | 1 | protein_coding | ENSG00000 |
| 179794850 | 179797517 | 1 | protein_coding | ENSG00000 |
| 179798463 | 179807814 | -1 | protein_coding | ENSG00000 |
| 179799144 | 179799212 | -1 | miRNA | ENSG00000 |
| 179821758 | 179838931 | 1 | protein_coding | ENSG00000 |
| 179835986 | 179863026 | -1 | protein_coding | ENSG00000 |
| 179846685 | 179847005 | 1 | misc_RNA | ENSG00000 |
| 179847888 | 179847983 | -1 | misc_RNA | ENSG00000 |
| 179859866 | 179862136 | 1 | lncRNA | ENSG00000 |
| 179862919 | 179907859 | -1 | protein_coding | ENSG00000 |
| 120594010 | 120621159 | 1 | protein_coding | ENSG00000 |
| 120625674 | 120630054 | -1 | protein_coding | ENSG00000 |
| 120709061 | 120709392 | -1 | processed_pseudoge | ENSG00000 |
| 120733094 | 120733396 | 1 | processed_pseudoge | ENSG00000 |
| 120872597 | 120875937 | -1 | protein_coding | ENSG00000 |
| 120876560 | 120876813 | 1 | processed_pseudoge | ENSG00000 |
| 120877498 | 120880820 | -1 | protein_coding | ENSG00000 |
| 120887218 | 120890540 | -1 | protein_coding | ENSG00000 |
| 120891140 | 120891391 | 1 | processed_pseudoge | ENSG00000 |
| 120901808 | 120905120 | -1 | protein_coding | ENSG00000 |
| 120905720 | 120905971 | 1 | processed_pseudoge | ENSG00000 |
| 120944299 | 120945898 | -1 | processed_pseudoge | ENSG00000 |
| 120966945 | 120969429 | 1 | protein_coding | ENSG00000 |
| 120985217 | 120997873 | 1 | lncRNA | ENSG00000 |
| 38356380 | 38378505 | 1 | transcribed_unproce | ENSG00000 |

| | | | | |
|---|---|---|---|---|
| 38383069 | 38402916 | -1 | transcribed_unproce | ENSG00000( |
| 38403189 | 38452153 | 1 | transcribed_unproce | ENSG00000( |
| 38423625 | 38423728 | 1 | snRNA | ENSG00000( |
| 38452987 | 38455365 | -1 | processed_pseudoge | ENSG00000( |
| 38453181 | 38466176 | 1 | lncRNA | ENSG00000( |
| 38637657 | 38641848 | 1 | unprocessed_pseudc | ENSG00000( |
| 38653494 | 38653583 | 1 | processed_pseudoge | ENSG00000( |
| 38672147 | 38674321 | 1 | processed_pseudoge | ENSG00000( |
| 38677210 | 38725307 | -1 | unprocessed_pseudc | ENSG00000( |
| 38732663 | 38734141 | 1 | processed_pseudoge | ENSG00000( |
| 38749434 | 38755468 | 1 | unprocessed_pseudc | ENSG00000( |
| 38785406 | 38785678 | 1 | protein_coding | ENSG00000( |
| 38798334 | 38798730 | -1 | processed_pseudoge | ENSG00000( |
| 1546949 | 1547237 | 1 | lncRNA | ENSG00000( |
| 1570727 | 1576918 | -1 | lncRNA | ENSG00000( |
| 1638229 | 1665406 | -1 | protein_coding | ENSG00000( |
| 58780626 | 58942190 | -1 | lncRNA | ENSG00000( |
| 58897284 | 58988182 | -1 | lncRNA | ENSG00000( |
| 58942211 | 59046959 | 1 | lncRNA | ENSG00000( |
| 143381295 | 143391111 | 1 | protein_coding | ENSG00000( |
| 143382686 | 143382800 | 1 | miRNA | ENSG00000( |
| 143390289 | 143408856 | -1 | protein_coding | ENSG00000( |
| 143407813 | 143523449 | 1 | lncRNA | ENSG00000( |
| 143437034 | 143437973 | 1 | unitary_pseudogene | ENSG00000( |
| 143443453 | 143444409 | 1 | protein_coding | ENSG00000( |
| 143477873 | 143478796 | 1 | protein_coding | ENSG00000( |
| 143488462 | 143489402 | -1 | unitary_pseudogene | ENSG00000( |
| 143509256 | 143519469 | -1 | polymorphic_pseudc | ENSG00000( |
| 143532749 | 143533164 | -1 | processed_pseudoge | ENSG00000( |
| 143545186 | 143546735 | -1 | processed_pseudoge | ENSG00000( |
| 143571801 | 143574387 | 1 | protein_coding | ENSG00000( |
| 143679585 | 143679687 | -1 | snRNA | ENSG00000( |
| 143680046 | 143682653 | -1 | protein_coding | ENSG00000( |
| 143686747 | 143761873 | -1 | transcribed_unproce | ENSG00000( |
| 143721228 | 143721653 | 1 | processed_pseudoge | ENSG00000( |
| 143725664 | 143742911 | 1 | transcribed_unproce | ENSG00000( |
| 143860031 | 143861144 | 1 | protein_coding | ENSG00000( |
| 143879710 | 143922178 | 1 | protein_coding | ENSG00000( |
| 143903987 | 143905091 | -1 | processed_pseudoge | ENSG00000( |
| 143905771 | 143906686 | 1 | unprocessed_pseudc | ENSG00000( |
| 143910008 | 143911517 | 1 | processed_pseudoge | ENSG00000( |
| 143925186 | 143933659 | 1 | protein_coding | ENSG00000( |
| 238771768 | 238784764 | 1 | lncRNA | ENSG00000( |
| 238788648 | 238789716 | -1 | lncRNA | ENSG00000( |
| 238848032 | 238911221 | 1 | protein_coding | ENSG00000( |
| 6406596 | 6648508 | 1 | protein_coding | ENSG00000( |
| 6615604 | 6617198 | -1 | lncRNA | ENSG00000( |
| 6617310 | 6708224 | -1 | lncRNA | ENSG00000( |
| 6622124 | 6622220 | -1 | miRNA | ENSG00000( |
| 6708642 | 6761503 | 1 | protein_coding | ENSG00000( |

| | | | | |
|---|---|---|---|---|
| 6745164 | 6745244 | 1 | miRNA | ENSG0000C |
| 6745168 | 6745240 | -1 | miRNA | ENSG0000C |
| 6780873 | 6789020 | 1 | lncRNA | ENSG0000C |
| 6811203 | 6835487 | -1 | protein_coding | ENSG0000C |
| 6835529 | 6885235 | 1 | lncRNA | ENSG0000C |
| 6870569 | 6877898 | -1 | protein_coding | ENSG0000C |
| 6924551 | 6925929 | -1 | protein_coding | ENSG0000C |
| 6928461 | 6930326 | 1 | lncRNA | ENSG0000C |
| 6935673 | 6938159 | -1 | protein_coding | ENSG0000C |
| 6950568 | 6951441 | -1 | unprocessed_pseudc | ENSG0000C |
| 6959129 | 6960001 | -1 | unprocessed_pseudc | ENSG0000C |
| 6967980 | 6968952 | -1 | unprocessed_pseudc | ENSG0000C |
| 6977491 | 6979922 | -1 | protein_coding | ENSG0000C |
| 6987258 | 6988100 | -1 | protein_coding | ENSG0000C |
| 6996733 | 6997594 | -1 | protein_coding | ENSG0000C |
| 7006376 | 7007219 | -1 | protein_coding | ENSG0000C |
| 7015719 | 7018151 | -1 | protein_coding | ENSG0000C |
| 7028495 | 7029343 | -1 | unprocessed_pseudc | ENSG0000C |
| 7038426 | 7039294 | -1 | unprocessed_pseudc | ENSG0000C |
| 7055160 | 7056595 | -1 | protein_coding | ENSG0000C |
| 7082825 | 7083182 | 1 | processed_pseudoge | ENSG0000C |
| 7112189 | 7113603 | -1 | processed_pseudoge | ENSG0000C |
| 7122734 | 7123581 | 1 | processed_pseudoge | ENSG0000C |
| 7143520 | 7143729 | -1 | processed_pseudoge | ENSG0000C |
| 7170420 | 7170624 | 1 | processed_pseudoge | ENSG0000C |
| 7190772 | 7191434 | -1 | processed_pseudoge | ENSG0000C |
| 7200615 | 7200716 | 1 | processed_pseudoge | ENSG0000C |
| 7221552 | 7221769 | -1 | unprocessed_pseudc | ENSG0000C |
| 7227682 | 7227902 | -1 | unprocessed_pseudc | ENSG0000C |
| 7238096 | 7238359 | -1 | processed_pseudoge | ENSG0000C |
| 7246441 | 7247372 | -1 | unprocessed_pseudc | ENSG0000C |
| 7264368 | 7267379 | 1 | protein_coding | ENSG0000C |
| 7278782 | 7279406 | -1 | unprocessed_pseudc | ENSG0000C |
| 7281632 | 7284454 | -1 | protein_coding | ENSG0000C |
| 7296789 | 7298943 | -1 | unprocessed_pseudc | ENSG0000C |
| 7314775 | 7328172 | -1 | protein_coding | ENSG0000C |
| 7334489 | 7336403 | -1 | protein_coding | ENSG0000C |
| 7337700 | 7341475 | 1 | protein_coding | ENSG0000C |
| 7348998 | 7353771 | 1 | protein_coding | ENSG0000C |
| 7360575 | 7376076 | 1 | protein_coding | ENSG0000C |
| 7393965 | 7395109 | 1 | protein_coding | ENSG0000C |
| 7403550 | 7405105 | -1 | processed_pseudoge | ENSG0000C |
| 7407233 | 7409249 | 1 | protein_coding | ENSG0000C |
| 7446613 | 7450965 | -1 | protein_coding | ENSG0000C |
| 7462876 | 7466132 | 1 | protein_coding | ENSG0000C |
| 7481168 | 7482284 | -1 | unprocessed_pseudc | ENSG0000C |
| 7485404 | 7486996 | -1 | protein_coding | ENSG0000C |
| 7490130 | 7491722 | -1 | protein_coding | ENSG0000C |
| 7504143 | 7511247 | -1 | unprocessed_pseudc | ENSG0000C |
| 7526067 | 7529077 | -1 | unprocessed_pseudc | ENSG0000C |

| | | | | |
|---|---|---|---|---|
| 7533689 | 7536699 | -1 | unprocessed_pseudo | ENSG00000 |
| 7545459 | 7548469 | -1 | protein_coding | ENSG00000 |
| 7715430 | 7715600 | -1 | protein_coding | ENSG00000 |
| 7743638 | 7744268 | 1 | processed_pseudoge | ENSG00000 |
| 7755882 | 7766294 | 1 | protein_coding | ENSG00000 |
| 7792863 | 7799990 | 1 | unprocessed_pseudo | ENSG00000 |
| 7812941 | 7814527 | 1 | processed_pseudoge | ENSG00000 |
| 7817656 | 7818836 | 1 | processed_pseudoge | ENSG00000 |
| 7833826 | 7837080 | -1 | protein_coding | ENSG00000 |
| 7847445 | 7851803 | 1 | unprocessed_pseudo | ENSG00000 |
| 7859537 | 7859586 | 1 | rRNA_pseudogene | ENSG00000 |
| 7874916 | 7882269 | 1 | protein_coding | ENSG00000 |
| 7903002 | 7903166 | 1 | processed_pseudoge | ENSG00000 |
| 7912058 | 7913024 | 1 | unprocessed_pseudo | ENSG00000 |
| 7937724 | 7937941 | 1 | transcribed_unproce | ENSG00000 |
| 7948684 | 7949366 | 1 | transcribed_processe | ENSG00000 |
| 7953280 | 7953661 | -1 | processed_pseudoge | ENSG00000 |
| 7956435 | 7959574 | 1 | protein_coding | ENSG00000 |
| 7968158 | 7970427 | -1 | protein_coding | ENSG00000 |
| 7978152 | 7978814 | 1 | protein_coding | ENSG00000 |
| 8002008 | 8006300 | 1 | lncRNA | ENSG00000 |
| 8011487 | 8016729 | 1 | lncRNA | ENSG00000 |
| 8019559 | 8024165 | 1 | transcribed_unproce | ENSG00000 |
| 8055913 | 8056933 | 1 | unprocessed_pseudo | ENSG00000 |
| 8084702 | 8110299 | 1 | protein_coding | ENSG00000 |
| 8106433 | 8106670 | 1 | lncRNA | ENSG00000 |
| 8113504 | 8150080 | -1 | protein_coding | ENSG00000 |
| 8150253 | 8154895 | 1 | lncRNA | ENSG00000 |
| 8162440 | 8162821 | -1 | processed_pseudoge | ENSG00000 |
| 8166558 | 8166788 | -1 | protein_coding | ENSG00000 |
| 8190710 | 8191399 | -1 | protein_coding | ENSG00000 |
| 8192811 | 8275840 | -1 | protein_coding | ENSG00000 |
| 8304852 | 8309394 | -1 | lncRNA | ENSG00000 |
| 8388152 | 8400340 | 1 | lncRNA | ENSG00000 |
| 8388247 | 8409603 | -1 | protein_coding | ENSG00000 |
| 8471756 | 8473621 | 1 | lncRNA | ENSG00000 |
| 8508887 | 8509267 | 1 | protein_coding | ENSG00000 |
| 8514640 | 8583179 | -1 | lncRNA | ENSG00000 |
| 8528676 | 8528930 | 1 | protein_coding | ENSG00000 |
| 8560716 | 8560822 | 1 | snRNA | ENSG00000 |
| 8571907 | 8572187 | -1 | misc_RNA | ENSG00000 |
| 8583129 | 8611938 | -1 | transcribed_unitary_ | ENSG00000 |
| 8619367 | 8620687 | -1 | protein_coding | ENSG00000 |
| 8626149 | 8635719 | 1 | lncRNA | ENSG00000 |
| 8628793 | 8666765 | -1 | protein_coding | ENSG00000 |
| 8666513 | 8668100 | 1 | lncRNA | ENSG00000 |
| 8669080 | 8670217 | 1 | unprocessed_pseudo | ENSG00000 |
| 8683726 | 8703973 | 1 | lncRNA | ENSG00000 |
| 8694985 | 8695567 | -1 | processed_pseudoge | ENSG00000 |
| 8747990 | 8748696 | -1 | lncRNA | ENSG00000 |

| | | | | |
|---|---|---|---|---|
| 8750316 | 8756526 | 1 | protein_coding | ENSG00000 |
| 8776061 | 8776580 | -1 | processed_pseudoge | ENSG00000 |
| 8813027 | 8814736 | 1 | lncRNA | ENSG00000 |
| 8825581 | 8828264 | 1 | lncRNA | ENSG00000 |
| 8841928 | 8843866 | 1 | lncRNA | ENSG00000 |
| 8844965 | 8846978 | 1 | lncRNA | ENSG00000 |
| 8884661 | 8889109 | 1 | lncRNA | ENSG00000 |
| 8916436 | 8916532 | 1 | miRNA | ENSG00000 |
| 9092550 | 9095901 | 1 | TEC | ENSG00000 |
| 9112146 | 9226509 | 1 | protein_coding | ENSG00000 |
| 9126419 | 9126489 | 1 | miRNA | ENSG00000 |
| 9178905 | 9222594 | -1 | lncRNA | ENSG00000 |
| 9221397 | 9228140 | 1 | protein_coding | ENSG00000 |
| 9251333 | 9279211 | -1 | protein_coding | ENSG00000 |
| 9265241 | 9265359 | -1 | rRNA_pseudogene | ENSG00000 |
| 9284761 | 9284853 | -1 | miRNA | ENSG00000 |
| 9397615 | 9426245 | -1 | protein_coding | ENSG00000 |
| 9462088 | 9464947 | -1 | lncRNA | ENSG00000 |
| 9469366 | 9477174 | -1 | lncRNA | ENSG00000 |
| 9472308 | 9475148 | 1 | lncRNA | ENSG00000 |
| 9475410 | 9475516 | -1 | snRNA | ENSG00000 |
| 9476694 | 9476950 | 1 | unprocessed_pseudo | ENSG00000 |
| 9522867 | 9898846 | -1 | protein_coding | ENSG00000 |
| 9614095 | 9614990 | 1 | processed_pseudoge | ENSG00000 |
| 9672054 | 9676292 | 1 | lncRNA | ENSG00000 |
| 9805716 | 9835135 | 1 | lncRNA | ENSG00000 |
| 10046775 | 10053583 | 1 | lncRNA | ENSG00000 |
| 10049982 | 10050066 | 1 | miRNA | ENSG00000 |
| 10177060 | 10400259 | -1 | protein_coding | ENSG00000 |
| 10214536 | 10214632 | -1 | miRNA | ENSG00000 |
| 10297849 | 10297992 | 1 | unprocessed_pseudo | ENSG00000 |
| 10498801 | 10502522 | -1 | processed_pseudoge | ENSG00000 |
| 10519986 | 10588362 | 1 | lncRNA | ENSG00000 |
| 10530661 | 10804664 | -1 | lncRNA | ENSG00000 |
| 10585066 | 10585171 | 1 | snRNA | ENSG00000 |
| 10597817 | 10605016 | 1 | lncRNA | ENSG00000 |
| 10668715 | 10668821 | 1 | snRNA | ENSG00000 |
| 10696074 | 10700966 | 1 | lncRNA | ENSG00000 |
| 10722855 | 10723291 | -1 | processed_pseudoge | ENSG00000 |
| 10773783 | 10776052 | 1 | lncRNA | ENSG00000 |
| 10797689 | 10798247 | -1 | lncRNA | ENSG00000 |
| 10804785 | 10820105 | 1 | protein_coding | ENSG00000 |
| 10810856 | 10814936 | -1 | lncRNA | ENSG00000 |
| 10864871 | 10865139 | -1 | misc_RNA | ENSG00000 |
| 10883871 | 10883932 | 1 | snRNA | ENSG00000 |
| 10908190 | 10908263 | -1 | miRNA | ENSG00000 |
| 10977657 | 10995482 | -1 | transcribed_process | ENSG00000 |
| 10994974 | 10995107 | -1 | snoRNA | ENSG00000 |
| 11022678 | 11022963 | 1 | processed_pseudoge | ENSG00000 |
| 11060783 | 11060888 | -1 | snRNA | ENSG00000 |

| | | | | |
|---|---|---|---|---|
| 11063643 | 11170702 | 1 | protein_coding | ENSG00000 |
| 11174247 | 11233162 | 1 | lncRNA | ENSG00000 |
| 11237319 | 11242800 | 1 | unitary_pseudogene | ENSG00000 |
| 11252761 | 11254920 | -1 | protein_coding | ENSG00000 |
| 11287437 | 11291344 | -1 | lncRNA | ENSG00000 |
| 11374196 | 11374499 | 1 | misc_RNA | ENSG00000 |
| 11394731 | 11396963 | -1 | lncRNA | ENSG00000 |
| 11395890 | 11401112 | 1 | lncRNA | ENSG00000 |
| 11495510 | 11499939 | 1 | lncRNA | ENSG00000 |
| 11707730 | 11724000 | -1 | transcribed_unproce | ENSG00000 |
| 11707928 | 11716592 | 1 | transcribed_unproce | ENSG00000 |
| 11763403 | 11764063 | 1 | lncRNA | ENSG00000 |
| 11898795 | 11899143 | -1 | processed_pseudoge | ENSG00000 |
| 11925041 | 11928051 | 1 | protein_coding | ENSG00000 |
| 12013695 | 12017701 | -1 | protein_coding | ENSG00000 |
| 12023992 | 12025826 | -1 | protein_coding | ENSG00000 |
| 12027161 | 12030974 | 1 | protein_coding | ENSG00000 |
| 12038494 | 12043266 | 1 | protein_coding | ENSG00000 |
| 12050031 | 12065549 | 1 | protein_coding | ENSG00000 |
| 12083221 | 12084682 | 1 | protein_coding | ENSG00000 |
| 12093021 | 12094573 | -1 | processed_pseudoge | ENSG00000 |
| 12096646 | 12098724 | 1 | protein_coding | ENSG00000 |
| 12136093 | 12140452 | -1 | protein_coding | ENSG00000 |
| 12152311 | 12155567 | 1 | protein_coding | ENSG00000 |
| 12170522 | 12171713 | -1 | processed_pseudoge | ENSG00000 |
| 12174834 | 12176855 | -1 | protein_coding | ENSG00000 |
| 12179565 | 12181474 | -1 | protein_coding | ENSG00000 |
| 12193295 | 12193534 | -1 | processed_pseudoge | ENSG00000 |
| 12225889 | 12236336 | -1 | protein_coding | ENSG00000 |
| 12276938 | 12277108 | 1 | protein_coding | ENSG00000 |
| 12370482 | 12371262 | -1 | processed_pseudoge | ENSG00000 |
| 12380410 | 12381186 | 1 | processed_pseudoge | ENSG00000 |
| 12401301 | 12401519 | -1 | transcribed_unproce | ENSG00000 |
| 12407435 | 12407653 | -1 | transcribed_unproce | ENSG00000 |
| 12428616 | 12429396 | -1 | processed_pseudoge | ENSG00000 |
| 12438569 | 12438743 | 1 | processed_pseudoge | ENSG00000 |
| 12459568 | 12459786 | -1 | transcribed_unproce | ENSG00000 |
| 12484243 | 12485242 | -1 | unprocessed_pseudo | ENSG00000 |
| 12496396 | 12497362 | -1 | unprocessed_pseudo | ENSG00000 |
| 12503144 | 12504134 | -1 | unprocessed_pseudo | ENSG00000 |
| 12519216 | 12519274 | 1 | miRNA | ENSG00000 |
| 12521983 | 12556166 | -1 | protein_coding | ENSG00000 |
| 12527310 | 12527382 | -1 | miRNA | ENSG00000 |
| 12527315 | 12527377 | 1 | miRNA | ENSG00000 |
| 12565917 | 12611581 | -1 | lncRNA | ENSG00000 |
| 12594340 | 12618397 | 1 | lncRNA | ENSG00000 |
| 12758554 | 12762363 | 1 | unprocessed_pseudo | ENSG00000 |
| 12806341 | 12822192 | 1 | protein_coding | ENSG00000 |
| 12844527 | 12844633 | 1 | snRNA | ENSG00000 |
| 12886135 | 12915978 | -1 | protein_coding | ENSG00000 |

| | | | | |
|---|---|---|---|---|
| 12947742 | 13207125 | 1 | lncRNA | ENSG0000C |
| 12960531 | 12960632 | 1 | misc_RNA | ENSG0000C |
| 21158377 | 21180616 | 1 | protein_coding | ENSG0000C |
| 21194847 | 21205977 | 1 | lncRNA | ENSG0000C |
| 21197450 | 21214510 | -1 | protein_coding | ENSG0000C |
| 21233699 | 21253850 | 1 | protein_coding | ENSG0000C |
| 21402110 | 21424449 | -1 | protein_coding | ENSG0000C |
| 21472115 | 21478634 | -1 | protein_coding | ENSG0000C |
| 21744964 | 21767544 | -1 | protein_coding | ENSG0000C |
| 21815465 | 21821981 | -1 | protein_coding | ENSG0000C |
| 21946243 | 21983930 | -1 | protein_coding | ENSG0000C |
| 21992621 | 22075070 | -1 | protein_coding | ENSG0000C |
| 22200470 | 22273045 | -1 | protein_coding | ENSG0000C |
| 22280232 | 22297482 | 1 | protein_coding | ENSG0000C |
| 22321945 | 22327924 | -1 | protein_coding | ENSG0000C |
| 22369984 | 22376491 | 1 | protein_coding | ENSG0000C |
| 22423969 | 22446363 | 1 | protein_coding | ENSG0000C |
| 22520552 | 22559704 | -1 | protein_coding | ENSG0000C |
| 22625708 | 22681332 | -1 | protein_coding | ENSG0000C |
| 22628529 | 22641587 | 1 | protein_coding | ENSG0000C |
| 22801555 | 22808047 | 1 | protein_coding | ENSG0000C |
| 22855983 | 22878913 | 1 | protein_coding | ENSG0000C |
| 41241772 | 41338644 | 1 | lncRNA | ENSG0000C |
| 41242373 | 41284861 | 1 | lncRNA | ENSG0000C |
| 41264550 | 41264862 | -1 | processed_pseudoge | ENSG0000C |
| 41302932 | 41306148 | -1 | transcribed_unproce | ENSG0000C |
| 41361434 | 41383813 | 1 | protein_coding | ENSG0000C |
| 41375060 | 41375701 | -1 | lncRNA | ENSG0000C |
| 27288669 | 27288857 | 1 | unprocessed_pseudc | ENSG0000C |
| 27315120 | 27315226 | 1 | snRNA | ENSG0000C |
| 27378657 | 27388552 | -1 | unprocessed_pseudc | ENSG0000C |
| 27382801 | 27382907 | -1 | snRNA | ENSG0000C |
| 27397017 | 27397590 | -1 | processed_pseudoge | ENSG0000C |
| 27425438 | 27425662 | -1 | processed_pseudoge | ENSG0000C |
| 27445487 | 27461668 | -1 | protein_coding | ENSG0000C |
| 45176724 | 45176925 | 1 | unprocessed_pseudc | ENSG0000C |
| 45181082 | 45181427 | -1 | lncRNA | ENSG0000C |
| 45187292 | 45198323 | -1 | unprocessed_pseudc | ENSG0000C |
| 45188790 | 45188896 | -1 | snRNA | ENSG0000C |
| 45232026 | 45247361 | -1 | unprocessed_pseudc | ENSG0000C |
| 45258185 | 45259123 | 1 | unitary_pseudogene | ENSG0000C |
| 45302654 | 45315608 | -1 | protein_coding | ENSG0000C |
| 45390239 | 45462182 | 1 | protein_coding | ENSG0000C |
| 45460633 | 45469184 | -1 | lncRNA | ENSG0000C |
| 45470650 | 45480390 | -1 | protein_coding | ENSG0000C |
| 42149310 | 42149549 | -1 | processed_pseudoge | ENSG0000C |
| 42186432 | 42186884 | 1 | IG_V_pseudogene | ENSG0000C |
| 42228517 | 42262638 | 1 | transcribed_unproce | ENSG0000C |
| 42265530 | 42267615 | -1 | processed_pseudoge | ENSG0000C |
| 77776951 | 77777055 | -1 | snRNA | ENSG0000C |

| | | | | |
|---|---|---|---|---|
| 77780337 | 77781876 | -1 | processed_pseudoge | ENSG00000 |
| 77782866 | 77793176 | 1 | lncRNA | ENSG00000 |
| 77790791 | 77926526 | -1 | protein_coding | ENSG00000 |
| 77866875 | 77869610 | 1 | lncRNA | ENSG00000 |
| 77912271 | 77912545 | -1 | misc_RNA | ENSG00000 |
| 77927372 | 77929824 | 1 | lncRNA | ENSG00000 |
| 65574155 | 65880289 | 1 | lncRNA | ENSG00000 |
| 7704628 | 7893073 | -1 | lncRNA | ENSG00000 |
| 7754393 | 7916081 | -1 | lncRNA | ENSG00000 |
| 7783697 | 7784407 | 1 | processed_pseudoge | ENSG00000 |
| 7784186 | 7784434 | -1 | protein_coding | ENSG00000 |
| 7795905 | 7796973 | -1 | protein_coding | ENSG00000 |
| 7834940 | 7840960 | -1 | protein_coding | ENSG00000 |
| 7858826 | 7859752 | 1 | unprocessed_pseudo | ENSG00000 |
| 7899308 | 7899405 | -1 | snRNA | ENSG00000 |
| 7934718 | 7941394 | -1 | protein_coding | ENSG00000 |
| 7947297 | 7951834 | -1 | protein_coding | ENSG00000 |
| 7967663 | 7975552 | -1 | protein_coding | ENSG00000 |
| 59938432 | 59957480 | 1 | protein_coding | ENSG00000 |
| 59942879 | 59970146 | 1 | lncRNA | ENSG00000 |
| 59964033 | 59970146 | 1 | protein_coding | ENSG00000 |
| 59978020 | 60031077 | 1 | protein_coding | ENSG00000 |
| 60040409 | 60048047 | 1 | protein_coding | ENSG00000 |
| 60060882 | 60075349 | 1 | protein_coding | ENSG00000 |
| 60081584 | 60082623 | 1 | processed_pseudoge | ENSG00000 |
| 7783697 | 7784407 | 1 | processed_pseudoge | ENSG00000 |
| 7795905 | 7796973 | -1 | protein_coding | ENSG00000 |
| 7825037 | 7825972 | -1 | protein_coding | ENSG00000 |
| 7859351 | 7860277 | 1 | unprocessed_pseudo | ENSG00000 |
| 7861399 | 7916588 | -1 | lncRNA | ENSG00000 |
| 7899815 | 7899912 | -1 | snRNA | ENSG00000 |
| 7938266 | 7939295 | -1 | polymorphic_pseudo | ENSG00000 |
| 7949167 | 7950227 | -1 | protein_coding | ENSG00000 |
| 7970057 | 7976059 | -1 | protein_coding | ENSG00000 |
| 1559754 | 1577630 | -1 | protein_coding | ENSG00000 |
| 1577063 | 1604894 | 1 | lncRNA | ENSG00000 |
| 1586202 | 1591699 | 1 | protein_coding | ENSG00000 |
| 1590052 | 1590993 | -1 | protein_coding | ENSG00000 |
| 1602887 | 1604004 | -1 | protein_coding | ENSG00000 |
| 1613275 | 1614173 | -1 | protein_coding | ENSG00000 |
| 1626668 | 1627848 | -1 | protein_coding | ENSG00000 |
| 1635485 | 1636417 | 1 | protein_coding | ENSG00000 |
| 1668294 | 1669053 | -1 | lncRNA | ENSG00000 |
| 1671307 | 1673566 | 1 | lncRNA | ENSG00000 |
| 1688979 | 1691339 | -1 | lncRNA | ENSG00000 |
| 1694007 | 1694766 | 1 | lncRNA | ENSG00000 |
| 2768844 | 2849088 | 1 | protein_coding | ENSG00000 |
| 2771356 | 2771577 | 1 | processed_pseudoge | ENSG00000 |
| 2840112 | 2861539 | -1 | lncRNA | ENSG00000 |
| 2870047 | 2872091 | 1 | lncRNA | ENSG00000 |

| | | | | |
|---|---|---|---|---|
| 2871423 | 2871484 | 1 | misc_RNA | ENSG0000C |
| 2883205 | 2885904 | -1 | protein_coding | ENSG0000C |
| 2887804 | 2903769 | -1 | lncRNA | ENSG0000C |
| 2899745 | 2925175 | 1 | protein_coding | ENSG0000C |
| 2928201 | 2929383 | -1 | protein_coding | ENSG0000C |
| 2943292 | 2991260 | -1 | protein_coding | ENSG0000C |
| 2962636 | 2962758 | -1 | snoRNA | ENSG0000C |
| 2988746 | 2990227 | 1 | lncRNA | ENSG0000C |
| 2999797 | 3056458 | -1 | protein_coding | ENSG0000C |
| 3027879 | 3040098 | 1 | lncRNA | ENSG0000C |
| 3047202 | 3047364 | -1 | snRNA | ENSG0000C |
| 473895 | 491399 | 1 | protein_coding | ENSG0000C |
| 494512 | 507300 | -1 | protein_coding | ENSG0000C |
| 528907 | 532261 | 1 | lncRNA | ENSG0000C |
| 532242 | 537287 | -1 | protein_coding | ENSG0000C |
| 537527 | 554915 | 1 | protein_coding | ENSG0000C |
| 554854 | 560793 | -1 | protein_coding | ENSG0000C |
| 557613 | 560121 | 1 | lncRNA | ENSG0000C |
| 560418 | 564036 | 1 | protein_coding | ENSG0000C |
| 565674 | 568471 | -1 | lncRNA | ENSG0000C |
| 576501 | 612474 | 1 | protein_coding | ENSG0000C |
| 612805 | 616257 | -1 | protein_coding | ENSG0000C |
| 616823 | 626336 | -1 | protein_coding | ENSG0000C |
| 626689 | 627401 | -1 | protein_coding | ENSG0000C |
| 637525 | 640936 | 1 | protein_coding | ENSG0000C |
| 646472 | 677067 | -1 | protein_coding | ENSG0000C |
| 668204 | 680202 | 1 | lncRNA | ENSG0000C |
| 940735 | 1012239 | 1 | protein_coding | ENSG0000C |
| 948421 | 948524 | 1 | snRNA | ENSG0000C |
| 1012821 | 1043528 | -1 | polymorphic_pseudc | ENSG0000C |
| 1554044 | 1571920 | -1 | protein_coding | ENSG0000C |
| 1571353 | 1599200 | 1 | lncRNA | ENSG0000C |
| 1584352 | 1585293 | -1 | protein_coding | ENSG0000C |
| 1597193 | 1598310 | -1 | protein_coding | ENSG0000C |
| 1607591 | 1608489 | -1 | protein_coding | ENSG0000C |
| 1620988 | 1622346 | -1 | protein_coding | ENSG0000C |
| 1629984 | 1630868 | 1 | protein_coding | ENSG0000C |
| 1659817 | 1660576 | -1 | lncRNA | ENSG0000C |
| 1663142 | 1665432 | 1 | lncRNA | ENSG0000C |
| 1680530 | 1682598 | -1 | lncRNA | ENSG0000C |
| 1685548 | 1686307 | 1 | lncRNA | ENSG0000C |
| 1694414 | 1694974 | 1 | protein_coding | ENSG0000C |
| 1729629 | 1732648 | -1 | protein_coding | ENSG0000C |
| 90098421 | 90098817 | -1 | processed_pseudoge | ENSG0000C |
| 90110638 | 90112022 | -1 | lncRNA | ENSG0000C |
| 90118391 | 90118461 | 1 | snoRNA | ENSG0000C |
| 90125658 | 90126061 | 1 | TEC | ENSG0000C |
| 90131515 | 90192894 | 1 | protein_coding | ENSG0000C |
| 90193614 | 90198120 | 1 | TEC | ENSG0000C |
| 90200429 | 90223077 | -1 | protein_coding | ENSG0000C |

| | | | | |
|---|---|---|---|---|
| 90223153 | 90226538 | 1 | TEC | ENSG00000 |
| 90313371 | 90314983 | 1 | processed_pseudoge | ENSG00000 |
| 940735 | 1019087 | 1 | protein_coding | ENSG00000 |
| 948421 | 948524 | 1 | snRNA | ENSG00000 |
| 1019669 | 1048356 | -1 | polymorphic_pseudo | ENSG00000 |
| 1061530 | 1067399 | 1 | lncRNA | ENSG00000 |
| 1086525 | 1116069 | 1 | protein_coding | ENSG00000 |
| 12310 | 13501 | 1 | unprocessed_pseudo | ENSG00000 |
| 14522 | 32015 | -1 | unprocessed_pseudo | ENSG00000 |
| 36667 | 38073 | -1 | lncRNA | ENSG00000 |
| 70780 | 73513 | 1 | protein_coding | ENSG00000 |
| 70780 | 175481 | 1 | protein_coding | ENSG00000 |
| 110539 | 115865 | -1 | lncRNA | ENSG00000 |
| 141426 | 153184 | -1 | lncRNA | ENSG00000 |
| 168679 | 170336 | -1 | lncRNA | ENSG00000 |
| 170871 | 171722 | -1 | lncRNA | ENSG00000 |
| 57935074 | 57936175 | -1 | lncRNA | ENSG00000 |
| 57941541 | 57957269 | 1 | protein_coding | ENSG00000 |
| 57967058 | 57968399 | 1 | processed_pseudoge | ENSG00000 |
| 57977965 | 57978277 | -1 | misc_RNA | ENSG00000 |
| 57983795 | 57984761 | -1 | processed_pseudoge | ENSG00000 |
| 58097051 | 58097709 | 1 | lncRNA | ENSG00000 |
| 10801532 | 10802627 | -1 | protein_coding | ENSG00000 |
| 10806051 | 10807293 | -1 | protein_coding | ENSG00000 |
| 10809137 | 10810168 | -1 | protein_coding | ENSG00000 |
| 10824960 | 11203561 | -1 | protein_coding | ENSG00000 |
| 10825317 | 10826358 | -1 | protein_coding | ENSG00000 |
| 10831074 | 10832016 | -1 | processed_pseudoge | ENSG00000 |
| 10894943 | 10895882 | -1 | processed_pseudoge | ENSG00000 |
| 10907926 | 10909562 | -1 | protein_coding | ENSG00000 |
| 10929236 | 10932341 | 1 | protein_coding | ENSG00000 |
| 10937406 | 11203521 | -1 | protein_coding | ENSG00000 |
| 10964425 | 10965352 | -1 | processed_pseudoge | ENSG00000 |
| 10985913 | 10986912 | -1 | protein_coding | ENSG00000 |
| 10996495 | 10997875 | -1 | protein_coding | ENSG00000 |
| 11021619 | 11022620 | -1 | protein_coding | ENSG00000 |
| 11030387 | 11031407 | -1 | protein_coding | ENSG00000 |
| 11048332 | 11049256 | -1 | processed_pseudoge | ENSG00000 |
| 11061365 | 11062294 | -1 | protein_coding | ENSG00000 |
| 11076769 | 11079171 | -1 | transcribed_processe | ENSG00000 |
| 11091308 | 11092334 | -1 | protein_coding | ENSG00000 |
| 11097196 | 11097291 | -1 | processed_pseudoge | ENSG00000 |
| 11115354 | 11116274 | -1 | unprocessed_pseudo | ENSG00000 |
| 11128789 | 11129718 | -1 | unprocessed_pseudo | ENSG00000 |
| 11164906 | 11166592 | -1 | protein_coding | ENSG00000 |
| 11190733 | 11191642 | -1 | processed_pseudoge | ENSG00000 |
| 11198038 | 11203301 | -1 | lncRNA | ENSG00000 |
| 11203312 | 11203513 | 1 | processed_pseudoge | ENSG00000 |
| 59027218 | 59027943 | 1 | processed_pseudoge | ENSG00000 |
| 59060029 | 59061233 | -1 | processed_pseudoge | ENSG00000 |

| | | | | |
|---|---|---|---|---|
| 59063055 | 59063157 | 1 | lncRNA | ENSG00000 |
| 10801532 | 10802627 | -1 | protein_coding | ENSG00000 |
| 10806051 | 10807293 | -1 | protein_coding | ENSG00000 |
| 10809137 | 10810168 | -1 | protein_coding | ENSG00000 |
| 10824960 | 11167778 | -1 | protein_coding | ENSG00000 |
| 10825317 | 10826358 | -1 | protein_coding | ENSG00000 |
| 10831074 | 10832016 | -1 | processed_pseudoge | ENSG00000 |
| 10845849 | 11167847 | -1 | protein_coding | ENSG00000 |
| 10845849 | 11167832 | -1 | protein_coding | ENSG00000 |
| 10881011 | 10884255 | -1 | protein_coding | ENSG00000 |
| 10894943 | 10896952 | -1 | transcribed_processe | ENSG00000 |
| 10907926 | 10909562 | -1 | protein_coding | ENSG00000 |
| 10929236 | 10932341 | 1 | protein_coding | ENSG00000 |
| 10937405 | 11167807 | -1 | protein_coding | ENSG00000 |
| 10964425 | 10965352 | -1 | processed_pseudoge | ENSG00000 |
| 10985900 | 10986899 | -1 | protein_coding | ENSG00000 |
| 10996484 | 10997864 | -1 | protein_coding | ENSG00000 |
| 11021607 | 11022608 | -1 | protein_coding | ENSG00000 |
| 11030686 | 11031706 | -1 | protein_coding | ENSG00000 |
| 11048621 | 11049550 | -1 | processed_pseudoge | ENSG00000 |
| 11063523 | 11064452 | -1 | protein_coding | ENSG00000 |
| 11079729 | 11080651 | -1 | unprocessed_pseudo | ENSG00000 |
| 11093151 | 11094080 | -1 | protein_coding | ENSG00000 |
| 11129613 | 11130572 | -1 | protein_coding | ENSG00000 |
| 11155045 | 11155955 | -1 | processed_pseudoge | ENSG00000 |
| 11162354 | 11167587 | -1 | lncRNA | ENSG00000 |
| 11167456 | 11172249 | 1 | lncRNA | ENSG00000 |
| 11182250 | 11183194 | -1 | protein_coding | ENSG00000 |
| 11192332 | 11193253 | -1 | processed_pseudoge | ENSG00000 |
| 11208476 | 11247646 | 1 | lncRNA | ENSG00000 |
| 11262181 | 11266062 | -1 | protein_coding | ENSG00000 |
| 11303340 | 11306692 | -1 | protein_coding | ENSG00000 |
| 11348080 | 11351848 | -1 | protein_coding | ENSG00000 |
| 101111599 | 101128641 | 1 | protein_coding | ENSG00000 |
| 101161669 | 101216388 | -1 | protein_coding | ENSG00000 |
| 101205448 | 101205554 | -1 | snRNA | ENSG00000 |
| 101234318 | 101235053 | -1 | processed_pseudoge | ENSG00000 |
| 31111652 | 31208427 | -1 | protein_coding | ENSG00000 |
| 31246713 | 31246958 | 1 | processed_pseudoge | ENSG00000 |
| 31254265 | 31254870 | -1 | processed_pseudoge | ENSG00000 |
| 31256634 | 31256718 | -1 | processed_pseudoge | ENSG00000 |
| 31282857 | 31329331 | -1 | protein_coding | ENSG00000 |
| 31313890 | 31314302 | 1 | processed_pseudoge | ENSG00000 |
| 31326589 | 31328102 | 1 | lncRNA | ENSG00000 |
| 126364076 | 126364938 | 1 | lncRNA | ENSG00000 |
| 126365344 | 126367107 | 1 | lncRNA | ENSG00000 |
| 126406834 | 126411516 | -1 | lncRNA | ENSG00000 |
| 11351822 | 11356402 | -1 | protein_coding | ENSG00000 |
| 130337887 | 130372618 | 1 | protein_coding | ENSG00000 |
| 130350996 | 130351707 | -1 | TEC | ENSG00000 |

| | | | | |
|---|---|---|---|---|
| 130383653 | 130384179 | 1 | TEC | ENSG00000 |
| 123826687 | 123935766 | 1 | protein_coding | ENSG00000 |
| 123918723 | 123946006 | -1 | protein_coding | ENSG00000 |
| 123927107 | 123935036 | -1 | lncRNA | ENSG00000 |
| 85955666 | 86838904 | -1 | protein_coding | ENSG00000 |
| 126915199 | 127060565 | -1 | lncRNA | ENSG00000 |
| 127130005 | 127207706 | 1 | lncRNA | ENSG00000 |
| 127146555 | 127151342 | 1 | lncRNA | ENSG00000 |
| 127149967 | 127175607 | -1 | lncRNA | ENSG00000 |
| 112197350 | 112201000 | 1 | lncRNA | ENSG00000 |
| 112297337 | 112305608 | 1 | lncRNA | ENSG00000 |
| 112306417 | 112315069 | -1 | lncRNA | ENSG00000 |
| 112360163 | 112418533 | 1 | protein_coding | ENSG00000 |
| 112468853 | 112488558 | -1 | protein_coding | ENSG00000 |
| 18724415 | 18724904 | -1 | processed_pseudoge | ENSG00000 |
| 18726706 | 18727414 | 1 | processed_pseudoge | ENSG00000 |
| 18734080 | 18734896 | 1 | processed_pseudoge | ENSG00000 |
| 18741730 | 18742718 | -1 | unprocessed_pseudo | ENSG00000 |
| 18807279 | 18808018 | -1 | processed_pseudoge | ENSG00000 |
| 18809830 | 18810048 | 1 | processed_pseudoge | ENSG00000 |
| 18816718 | 18880683 | -1 | unprocessed_pseudo | ENSG00000 |
| 18869185 | 18869291 | -1 | snRNA | ENSG00000 |
| 18885950 | 18886053 | 1 | snRNA | ENSG00000 |
| 113731864 | 113738248 | 1 | protein_coding | ENSG00000 |
| 113749533 | 113752109 | -1 | TEC | ENSG00000 |
| 113760262 | 113760337 | 1 | protein_coding | ENSG00000 |
| 91554410 | 91554503 | 1 | unprocessed_pseudo | ENSG00000 |
| 91572444 | 91575271 | -1 | TEC | ENSG00000 |
| 91580696 | 91784366 | -1 | protein_coding | ENSG00000 |
| 91603173 | 91603433 | -1 | transcribed_process | ENSG00000 |
| 91604402 | 91605319 | -1 | transcribed_process | ENSG00000 |
| 91712770 | 91714276 | 1 | processed_pseudoge | ENSG00000 |
| 91718574 | 91718789 | -1 | snoRNA | ENSG00000 |
| 91756515 | 91763457 | -1 | lncRNA | ENSG00000 |
| 91783410 | 91840185 | -1 | protein_coding | ENSG00000 |
| 104474682 | 104612791 | -1 | protein_coding | ENSG00000 |
| 104586828 | 104597659 | 1 | protein_coding | ENSG00000 |
| 104596165 | 104596991 | -1 | lncRNA | ENSG00000 |
| 105536861 | 105537239 | -1 | processed_pseudoge | ENSG00000 |
| 105546610 | 105549694 | -1 | unprocessed_pseudo | ENSG00000 |
| 105588247 | 105589753 | -1 | IG_C_gene | ENSG00000 |
| 105600717 | 105602504 | -1 | IG_C_gene | ENSG00000 |
| 105625112 | 105626837 | -1 | IG_C_gene | ENSG00000 |
| 105643841 | 105645561 | -1 | IG_C_gene | ENSG00000 |
| 105648695 | 105649828 | -1 | lncRNA | ENSG00000 |
| 105665404 | 105670614 | -1 | IG_C_pseudogene | ENSG00000 |
| 105673079 | 105674085 | -1 | processed_pseudoge | ENSG00000 |
| 105704732 | 105705373 | -1 | lncRNA | ENSG00000 |
| 105707889 | 105709436 | -1 | IG_C_gene | ENSG00000 |
| 105722565 | 105723298 | -1 | IG_C_pseudogene | ENSG00000 |

| | | | | |
|---|---|---|---|---|
| 105737114 | 105743842 | -1 | IG_C_gene | ENSG0000C |
| 105769875 | 105772176 | -1 | IG_C_gene | ENSG0000C |
| 105786276 | 105786602 | -1 | processed_pseudoge | ENSG0000C |
| 105837536 | 105846448 | -1 | IG_C_gene | ENSG0000C |
| 105854991 | 105856989 | -1 | IG_C_gene | ENSG0000C |
| 105863969 | 105864031 | -1 | IG_J_gene | ENSG0000C |
| 105864187 | 105864236 | -1 | IG_J_pseudogene | ENSG0000C |
| 105864585 | 105864635 | -1 | IG_J_gene | ENSG0000C |
| 105864986 | 105865033 | -1 | IG_J_gene | ENSG0000C |
| 105865358 | 105865407 | -1 | IG_J_gene | ENSG0000C |
| 105865564 | 105865623 | -1 | IG_J_pseudogene | ENSG0000C |
| 105865970 | 105866022 | -1 | IG_J_gene | ENSG0000C |
| 105866178 | 105866229 | -1 | IG_J_gene | ENSG0000C |
| 105866322 | 105866332 | -1 | IG_D_gene | ENSG0000C |
| 105866395 | 105866449 | -1 | IG_J_pseudogene | ENSG0000C |
| 105881805 | 105881824 | -1 | IG_D_gene | ENSG0000C |
| 105882310 | 105882327 | -1 | IG_D_gene | ENSG0000C |
| 105884674 | 105884693 | -1 | IG_D_gene | ENSG0000C |
| 105885641 | 105885659 | -1 | IG_D_gene | ENSG0000C |
| 105886802 | 105886832 | -1 | IG_D_gene | ENSG0000C |
| 105889322 | 105889349 | -1 | IG_D_gene | ENSG0000C |
| 105890855 | 105897348 | 1 | lncRNA | ENSG0000C |
| 105891962 | 105891978 | -1 | IG_D_gene | ENSG0000C |
| 105892470 | 105892490 | -1 | IG_D_gene | ENSG0000C |
| 105894313 | 105894332 | -1 | IG_D_gene | ENSG0000C |
| 105895279 | 105895294 | -1 | IG_D_gene | ENSG0000C |
| 105896405 | 105896441 | -1 | IG_D_gene | ENSG0000C |
| 105898728 | 105898758 | -1 | IG_D_gene | ENSG0000C |
| 105901409 | 105901425 | -1 | IG_D_gene | ENSG0000C |
| 105901913 | 105901933 | -1 | IG_D_gene | ENSG0000C |
| 105903420 | 105903442 | -1 | IG_D_gene | ENSG0000C |
| 105904387 | 105904402 | -1 | IG_D_gene | ENSG0000C |
| 105905268 | 105905298 | -1 | IG_D_gene | ENSG0000C |
| 105905452 | 105905482 | -1 | IG_D_gene | ENSG0000C |
| 105907982 | 105908012 | -1 | IG_D_gene | ENSG0000C |
| 105910678 | 105910694 | -1 | IG_D_gene | ENSG0000C |
| 105911181 | 105911198 | -1 | IG_D_gene | ENSG0000C |
| 105913028 | 105913047 | -1 | IG_D_gene | ENSG0000C |
| 105913993 | 105914008 | -1 | IG_D_gene | ENSG0000C |
| 105915130 | 105915160 | -1 | IG_D_gene | ENSG0000C |
| 105917597 | 105917627 | -1 | IG_D_gene | ENSG0000C |
| 105918750 | 105933413 | 1 | lncRNA | ENSG0000C |
| 105920273 | 105920289 | -1 | IG_D_gene | ENSG0000C |
| 105940525 | 105941024 | -1 | IG_V_gene | ENSG0000C |
| 105945981 | 105946160 | -1 | IG_V_pseudogene | ENSG0000C |
| 105970500 | 105973040 | -1 | transcribed_unproce | ENSG0000C |
| 105979291 | 105979917 | -1 | processed_pseudoge | ENSG0000C |
| 105987353 | 105987854 | -1 | IG_V_gene | ENSG0000C |
| 106002305 | 106002595 | -1 | IG_V_pseudogene | ENSG0000C |
| 106005866 | 106006345 | -1 | IG_V_gene | ENSG0000C |

| | | | | |
|---|---|---|---|---|
| 106012693 | 106013191 | -1 | IG_V_gene | ENSG0000C |
| 106025916 | 106026401 | -1 | IG_V_gene | ENSG0000C |
| 106038673 | 106039136 | -1 | IG_V_gene | ENSG0000C |
| 106040437 | 106040535 | -1 | IG_V_pseudogene | ENSG0000C |
| 106046371 | 106046631 | -1 | IG_V_pseudogene | ENSG0000C |
| 106056320 | 106056769 | -1 | IG_V_pseudogene | ENSG0000C |
| 106062920 | 106063454 | -1 | IG_V_gene | ENSG0000C |
| 106088893 | 106089344 | -1 | IG_V_gene | ENSG0000C |
| 106108743 | 106109178 | -1 | IG_V_gene | ENSG0000C |
| 106117406 | 106117975 | -1 | IG_V_gene | ENSG0000C |
| 106120978 | 106121244 | -1 | IG_V_pseudogene | ENSG0000C |
| 106123191 | 106123480 | -1 | IG_V_pseudogene | ENSG0000C |
| 106130311 | 106130843 | -1 | IG_V_gene | ENSG0000C |
| 106143058 | 106143348 | -1 | IG_V_pseudogene | ENSG0000C |
| 106146662 | 106146902 | -1 | IG_V_pseudogene | ENSG0000C |
| 106154393 | 106154934 | -1 | IG_V_gene | ENSG0000C |
| 106164354 | 106164573 | -1 | IG_V_pseudogene | ENSG0000C |
| 106165976 | 106166501 | -1 | IG_V_gene | ENSG0000C |
| 106175113 | 106175531 | -1 | IG_V_pseudogene | ENSG0000C |
| 106180325 | 106181290 | -1 | processed_pseudoge | ENSG0000C |
| 106185670 | 106186165 | -1 | IG_V_gene | ENSG0000C |
| 106197471 | 106197761 | -1 | IG_V_pseudogene | ENSG0000C |
| 106207374 | 106209342 | 1 | lncRNA | ENSG0000C |
| 106211707 | 106212224 | -1 | IG_V_gene | ENSG0000C |
| 106235833 | 106236365 | -1 | IG_V_gene | ENSG0000C |
| 106258533 | 106258994 | -1 | IG_V_pseudogene | ENSG0000C |
| 106264133 | 106264397 | -1 | IG_V_pseudogene | ENSG0000C |
| 106265459 | 106265486 | -1 | IG_V_pseudogene | ENSG0000C |
| 106269377 | 106269911 | -1 | IG_V_gene | ENSG0000C |
| 106277317 | 106277814 | -1 | IG_V_gene | ENSG0000C |
| 106287582 | 106288428 | -1 | processed_pseudoge | ENSG0000C |
| 106289800 | 106290250 | -1 | IG_V_pseudogene | ENSG0000C |
| 106294339 | 106294579 | -1 | IG_V_pseudogene | ENSG0000C |
| 106302166 | 106302633 | -1 | IG_V_gene | ENSG0000C |
| 106310188 | 106310437 | -1 | IG_V_pseudogene | ENSG0000C |
| 106315422 | 106315746 | -1 | IG_V_pseudogene | ENSG0000C |
| 106318594 | 106319007 | -1 | IG_V_pseudogene | ENSG0000C |
| 106325025 | 106325531 | -1 | IG_V_gene | ENSG0000C |
| 106331887 | 106332323 | -1 | IG_V_pseudogene | ENSG0000C |
| 106335853 | 106336384 | -1 | IG_V_gene | ENSG0000C |
| 106343495 | 106343726 | -1 | IG_V_pseudogene | ENSG0000C |
| 106345166 | 106345604 | -1 | IG_V_pseudogene | ENSG0000C |
| 106350054 | 106350526 | -1 | IG_V_gene | ENSG0000C |
| 106355443 | 106355479 | -1 | IG_V_pseudogene | ENSG0000C |
| 106356982 | 106357409 | -1 | IG_V_pseudogene | ENSG0000C |
| 106359784 | 106360304 | -1 | IG_V_gene | ENSG0000C |
| 106369091 | 106369528 | -1 | IG_V_pseudogene | ENSG0000C |
| 106373990 | 106374420 | -1 | IG_V_gene | ENSG0000C |
| 106380821 | 106381287 | -1 | IG_V_pseudogene | ENSG0000C |
| 106384472 | 106384925 | -1 | IG_V_gene | ENSG0000C |

| | | | | |
|---|---|---|---|---|
| 106392066 | 106392345 | -1 | IG_V_pseudogene | ENSG00000 |
| 106393788 | 106394227 | -1 | IG_V_pseudogene | ENSG00000 |
| 106398342 | 106398826 | -1 | IG_V_gene | ENSG00000 |
| 106401981 | 106402414 | -1 | IG_V_pseudogene | ENSG00000 |
| 106414071 | 106414539 | -1 | IG_V_gene | ENSG00000 |
| 106417455 | 106417912 | -1 | IG_V_pseudogene | ENSG00000 |
| 106421357 | 106421795 | -1 | IG_V_pseudogene | ENSG00000 |
| 106435173 | 106435702 | -1 | IG_V_gene | ENSG00000 |
| 106442706 | 106442759 | -1 | IG_V_pseudogene | ENSG00000 |
| 106446390 | 106446899 | -1 | IG_V_gene | ENSG00000 |
| 106450046 | 106450244 | -1 | IG_V_pseudogene | ENSG00000 |
| 106461096 | 106461236 | -1 | unprocessed_pseudo | ENSG00000 |
| 106465631 | 106465688 | -1 | IG_V_pseudogene | ENSG00000 |
| 106467814 | 106468265 | -1 | IG_V_pseudogene | ENSG00000 |
| 106484773 | 106485736 | -1 | processed_pseudoge | ENSG00000 |
| 106487939 | 106488239 | -1 | IG_V_pseudogene | ENSG00000 |
| 106494944 | 106495481 | -1 | IG_V_gene | ENSG00000 |
| 106502901 | 106503074 | -1 | IG_V_pseudogene | ENSG00000 |
| 106514026 | 106514437 | -1 | IG_V_pseudogene | ENSG00000 |
| 106518815 | 106519064 | -1 | IG_V_pseudogene | ENSG00000 |
| 106531677 | 106532172 | -1 | IG_V_gene | ENSG00000 |
| 106535796 | 106536537 | -1 | IG_V_gene | ENSG00000 |
| 106540499 | 106540535 | -1 | IG_V_pseudogene | ENSG00000 |
| 106543263 | 106543708 | -1 | IG_V_pseudogene | ENSG00000 |
| 106555822 | 106556152 | -1 | IG_V_pseudogene | ENSG00000 |
| 106562491 | 106563025 | -1 | IG_V_gene | ENSG00000 |
| 106581617 | 106582158 | -1 | IG_V_gene | ENSG00000 |
| 106589056 | 106589279 | -1 | IG_V_pseudogene | ENSG00000 |
| 106590798 | 106591236 | -1 | IG_V_pseudogene | ENSG00000 |
| 106603423 | 106603917 | -1 | IG_V_gene | ENSG00000 |
| 106608187 | 106608488 | -1 | IG_V_pseudogene | ENSG00000 |
| 106609399 | 106609657 | -1 | IG_V_pseudogene | ENSG00000 |
| 106611057 | 106611507 | -1 | IG_V_pseudogene | ENSG00000 |
| 106617357 | 106618028 | -1 | IG_V_gene | ENSG00000 |
| 106624351 | 106624605 | -1 | IG_V_pseudogene | ENSG00000 |
| 106626027 | 106626473 | -1 | IG_V_pseudogene | ENSG00000 |
| 106630782 | 106631232 | -1 | IG_V_pseudogene | ENSG00000 |
| 106634443 | 106634877 | -1 | IG_V_pseudogene | ENSG00000 |
| 106643575 | 106643854 | -1 | IG_V_pseudogene | ENSG00000 |
| 106647038 | 106647536 | -1 | IG_V_gene | ENSG00000 |
| 106651930 | 106652506 | -1 | IG_V_gene | ENSG00000 |
| 106655878 | 106656334 | -1 | IG_V_pseudogene | ENSG00000 |
| 106661353 | 106661467 | 1 | rRNA | ENSG00000 |
| 106662399 | 106662654 | -1 | IG_V_pseudogene | ENSG00000 |
| 106663800 | 106664338 | -1 | IG_V_gene | ENSG00000 |
| 106667823 | 106668266 | -1 | IG_V_pseudogene | ENSG00000 |
| 106675232 | 106675466 | -1 | IG_V_pseudogene | ENSG00000 |
| 106676918 | 106677362 | -1 | IG_V_pseudogene | ENSG00000 |
| 106682404 | 106682939 | -1 | IG_V_gene | ENSG00000 |
| 106690773 | 106691213 | -1 | IG_V_pseudogene | ENSG00000 |

| | | | | |
|---|---|---|---|---|
| 106696478 | 106696725 | -1 | IG_V_pseudogene | ENSG00000( |
| 106699696 | 106700225 | -1 | IG_V_gene | ENSG00000( |
| 106705284 | 106705723 | -1 | IG_V_pseudogene | ENSG00000( |
| 106708076 | 106709031 | -1 | processed_pseudoge | ENSG00000( |
| 106711198 | 106711243 | -1 | IG_V_pseudogene | ENSG00000( |
| 106711794 | 106711881 | -1 | IG_V_pseudogene | ENSG00000( |
| 106717338 | 106717569 | -1 | IG_V_pseudogene | ENSG00000( |
| 106719783 | 106720078 | -1 | IG_V_pseudogene | ENSG00000( |
| 106728533 | 106728827 | -1 | IG_V_pseudogene | ENSG00000( |
| 106739363 | 106739862 | -1 | IG_V_gene | ENSG00000( |
| 106748255 | 106748774 | -1 | IG_V_gene | ENSG00000( |
| 106752844 | 106753288 | -1 | IG_V_pseudogene | ENSG00000( |
| 106761791 | 106762228 | -1 | IG_V_gene | ENSG00000( |
| 106774973 | 106775426 | -1 | IG_V_gene | ENSG00000( |
| 106786363 | 106786859 | -1 | IG_V_gene | ENSG00000( |
| 106794747 | 106795266 | -1 | IG_V_gene | ENSG00000( |
| 106799336 | 106799794 | -1 | IG_V_pseudogene | ENSG00000( |
| 106814867 | 106815409 | -1 | IG_V_gene | ENSG00000( |
| 106826868 | 106827409 | -1 | IG_V_gene | ENSG00000( |
| 106834616 | 106835307 | -1 | IG_V_gene | ENSG00000( |
| 106845350 | 106845590 | -1 | IG_V_pseudogene | ENSG00000( |
| 106847863 | 106848323 | -1 | IG_V_pseudogene | ENSG00000( |
| 106852045 | 106852339 | -1 | IG_V_pseudogene | ENSG00000( |
| 106855943 | 106856228 | -1 | IG_V_pseudogene | ENSG00000( |
| 106875299 | 106875593 | -1 | IG_V_pseudogene | ENSG00000( |
| 92886352 | 92893506 | 1 | lncRNA | ENSG00000( |
| 92905697 | 92907482 | -1 | lncRNA | ENSG00000( |
| 92923080 | 92935293 | 1 | protein_coding | ENSG00000( |
| 92936914 | 93116320 | -1 | protein_coding | ENSG00000( |
| 93067452 | 93072152 | 1 | lncRNA | ENSG00000( |
| 93138066 | 93138465 | 1 | processed_pseudoge | ENSG00000( |
| 93182196 | 93184928 | -1 | protein_coding | ENSG00000( |
| 93184951 | 93187089 | 1 | protein_coding | ENSG00000( |
| 93184973 | 93218586 | 1 | protein_coding | ENSG00000( |
| 93202894 | 93207094 | -1 | protein_coding | ENSG00000( |
| 93207056 | 93229215 | 1 | protein_coding | ENSG00000( |
| 93223598 | 93224139 | -1 | processed_pseudoge | ENSG00000( |
| 93237550 | 93333092 | -1 | protein_coding | ENSG00000( |
| 93244010 | 93244323 | 1 | processed_pseudoge | ENSG00000( |
| 93333219 | 93707876 | 1 | protein_coding | ENSG00000( |
| 93334528 | 93335057 | 1 | lncRNA | ENSG00000( |
| 93347191 | 93348356 | 1 | protein_coding | ENSG00000( |
| 93397139 | 93397234 | -1 | misc_RNA | ENSG00000( |
| 93437400 | 93437510 | -1 | snRNA | ENSG00000( |
| 93637179 | 93637338 | 1 | misc_RNA | ENSG00000( |
| 93718298 | 93788481 | -1 | protein_coding | ENSG00000( |
| 93895762 | 93897759 | 1 | TEC | ENSG00000( |
| 93904730 | 93927066 | -1 | lncRNA | ENSG00000( |
| 93918894 | 93929608 | 1 | protein_coding | ENSG00000( |
| 93934153 | 93976791 | -1 | protein_coding | ENSG00000( |

| | | | | |
|---|---|---|---|---|
| 93939555 | 93944129 | 1 | lncRNA | ENSG0000C |
| 93997270 | 94008863 | 1 | lncRNA | ENSG0000C |
| 94026329 | 94049624 | 1 | protein_coding | ENSG0000C |
| 94050920 | 94081245 | -1 | protein_coding | ENSG0000C |
| 94081282 | 94103846 | 1 | protein_coding | ENSG0000C |
| 94104836 | 94116698 | 1 | protein_coding | ENSG0000C |
| 94127779 | 94130253 | -1 | protein_coding | ENSG0000C |
| 94146128 | 94279735 | 1 | protein_coding | ENSG0000C |
| 94280460 | 94293271 | -1 | protein_coding | ENSG0000C |
| 94304248 | 94323394 | -1 | protein_coding | ENSG0000C |
| 94363640 | 94374417 | -1 | protein_coding | ENSG0000C |
| 94378757 | 94392703 | -1 | protein_coding | ENSG0000C |
| 63642035 | 63665593 | 1 | lncRNA | ENSG0000C |
| 63684216 | 63728065 | -1 | protein_coding | ENSG0000C |
| 63734697 | 63735923 | 1 | processed_pseudoge | ENSG0000C |
| 63736444 | 63736505 | 1 | snRNA | ENSG0000C |
| 63762013 | 63762149 | 1 | processed_pseudoge | ENSG0000C |
| 63815247 | 63815433 | 1 | processed_pseudoge | ENSG0000C |
| 45891087 | 46508547 | 1 | lncRNA | ENSG0000C |
| 21250180 | 21260147 | -1 | unprocessed_pseudc | ENSG0000C |
| 21293635 | 21295183 | -1 | processed_pseudoge | ENSG0000C |
| 21298216 | 21325353 | -1 | lncRNA | ENSG0000C |
| 21307731 | 21307837 | 1 | snRNA | ENSG0000C |
| 21328492 | 21343937 | -1 | lncRNA | ENSG0000C |
| 21342227 | 21344014 | 1 | lncRNA | ENSG0000C |
| 21344605 | 21345142 | -1 | processed_pseudoge | ENSG0000C |
| 21368518 | 21370717 | 1 | lncRNA | ENSG0000C |
| 21372682 | 21387280 | -1 | transcribed_processe | ENSG0000C |
| 21382467 | 21382926 | 1 | processed_pseudoge | ENSG0000C |
| 21383905 | 21384148 | 1 | processed_pseudoge | ENSG0000C |
| 21408280 | 21440481 | -1 | protein_coding | ENSG0000C |
| 21428930 | 21429036 | -1 | snRNA | ENSG0000C |
| 21490371 | 21509391 | -1 | transcribed_unproce | ENSG0000C |
| 21514120 | 21566989 | 1 | lncRNA | ENSG0000C |
| 21534392 | 21535271 | 1 | unprocessed_pseudc | ENSG0000C |
| 21552724 | 21660461 | 1 | lncRNA | ENSG0000C |
| 21645163 | 21646246 | 1 | protein_coding | ENSG0000C |
| 21659028 | 21660224 | 1 | protein_coding | ENSG0000C |
| 21690124 | 21691048 | 1 | unprocessed_pseudc | ENSG0000C |
| 21716735 | 21717356 | -1 | processed_pseudoge | ENSG0000C |
| 22903706 | 22981018 | -1 | protein_coding | ENSG0000C |
| 22983146 | 23039654 | -1 | protein_coding | ENSG0000C |
| 23086215 | 23124800 | 1 | transcribed_unproce | ENSG0000C |
| 23131736 | 23140290 | -1 | protein_coding | ENSG0000C |
| 23144031 | 23147890 | -1 | unprocessed_pseudc | ENSG0000C |
| 23160896 | 23171217 | 1 | unprocessed_pseudc | ENSG0000C |
| 23169365 | 23169654 | 1 | misc_RNA | ENSG0000C |
| 23183337 | 23185517 | 1 | processed_pseudoge | ENSG0000C |
| 23196959 | 23197164 | 1 | processed_pseudoge | ENSG0000C |
| 23229344 | 23229859 | -1 | unprocessed_pseudc | ENSG0000C |

| | | | | |
|---|---|---|---|---|
| 28315693 | 28315770 | 1 | protein_coding | ENSG00000 |
| 28320122 | 28322152 | -1 | protein_coding | ENSG00000 |
| 28332230 | 28336599 | 1 | unprocessed_pseudc | ENSG00000 |
| 28346477 | 28346643 | 1 | processed_pseudoge | ENSG00000 |
| 28349050 | 28358014 | -1 | unprocessed_pseudc | ENSG00000 |
| 28361822 | 28365684 | -1 | unprocessed_pseudc | ENSG00000 |
| 28378587 | 28391990 | 1 | protein_coding | ENSG00000 |
| 28387160 | 28387417 | 1 | misc_RNA | ENSG00000 |
| 28402766 | 28404923 | 1 | processed_pseudoge | ENSG00000 |
| 28416342 | 28416567 | 1 | processed_pseudoge | ENSG00000 |
| 28422060 | 28422264 | 1 | processed_pseudoge | ENSG00000 |
| 28427760 | 28427964 | 1 | processed_pseudoge | ENSG00000 |
| 28455755 | 28455878 | 1 | processed_pseudoge | ENSG00000 |
| 28466463 | 28469402 | -1 | unprocessed_pseudc | ENSG00000 |
| 28484946 | 28485066 | -1 | processed_pseudoge | ENSG00000 |
| 28506596 | 28506797 | -1 | unprocessed_pseudc | ENSG00000 |
| 28518244 | 28520426 | -1 | processed_pseudoge | ENSG00000 |
| 28531231 | 28544591 | -1 | protein_coding | ENSG00000 |
| 28535771 | 28536060 | -1 | misc_RNA | ENSG00000 |
| 28557532 | 28561397 | 1 | unprocessed_pseudc | ENSG00000 |
| 28565258 | 28572999 | 1 | unprocessed_pseudc | ENSG00000 |
| 28576608 | 28576845 | -1 | processed_pseudoge | ENSG00000 |
| 28586639 | 28591020 | -1 | unprocessed_pseudc | ENSG00000 |
| 28601112 | 28602743 | 1 | transcribed_unproce | ENSG00000 |
| 23036897 | 23039673 | -1 | protein_coding | ENSG00000 |
| 23087871 | 23126101 | 1 | transcribed_unproce | ENSG00000 |
| 23133045 | 23141599 | -1 | protein_coding | ENSG00000 |
| 23145340 | 23149199 | -1 | unprocessed_pseudc | ENSG00000 |
| 23162205 | 23173124 | 1 | transcribed_unproce | ENSG00000 |
| 23170674 | 23170963 | 1 | misc_RNA | ENSG00000 |
| 23184646 | 23186826 | 1 | processed_pseudoge | ENSG00000 |
| 23198268 | 23198473 | 1 | processed_pseudoge | ENSG00000 |
| 23219948 | 23220085 | 1 | processed_pseudoge | ENSG00000 |
| 23230653 | 23231168 | -1 | unprocessed_pseudc | ENSG00000 |
| 65909030 | 66265884 | -1 | protein_coding | ENSG00000 |
| 66270636 | 66273940 | 1 | lncRNA | ENSG00000 |
| 66290635 | 66291180 | -1 | lncRNA | ENSG00000 |
| 22363371 | 22363650 | 1 | unprocessed_pseudc | ENSG00000 |
| 22374469 | 22376303 | 1 | processed_pseudoge | ENSG00000 |
| 22403919 | 22409655 | 1 | unprocessed_pseudc | ENSG00000 |
| 22429756 | 22440620 | -1 | transcribed_unproce | ENSG00000 |
| 22431917 | 22432206 | -1 | misc_RNA | ENSG00000 |
| 22453604 | 22457549 | 1 | unprocessed_pseudc | ENSG00000 |
| 22461390 | 22471066 | 1 | protein_coding | ENSG00000 |
| 22482318 | 22486719 | -1 | unprocessed_pseudc | ENSG00000 |
| 22497423 | 22593430 | 1 | transcribed_unproce | ENSG00000 |
| 22610118 | 22620428 | -1 | transcribed_unproce | ENSG00000 |
| 22611719 | 22611955 | -1 | misc_RNA | ENSG00000 |
| 44679746 | 44684155 | -1 | TEC | ENSG00000 |
| 44697333 | 44697430 | -1 | misc_RNA | ENSG00000 |

| | | | | |
|---|---|---|---|---|
| 44711032 | 44711328 | 1 | lncRNA | ENSG0000C |
| 44711489 | 44718888 | 1 | protein_coding | ENSG0000C |
| 44729020 | 44767863 | 1 | protein_coding | ENSG0000C |
| 44827352 | 44893221 | -1 | transcribed_unproce | ENSG0000C |
| 44844600 | 44844712 | 1 | misc_RNA | ENSG0000C |
| 44893143 | 44893307 | 1 | snRNA | ENSG0000C |
| 27871154 | 28099370 | -1 | protein_coding | ENSG0000C |
| 27895704 | 27896582 | 1 | processed_pseudoge | ENSG0000C |
| 28110217 | 28435080 | -1 | protein_coding | ENSG0000C |
| 28428588 | 28428996 | 1 | processed_pseudoge | ENSG0000C |
| 28445158 | 28447993 | 1 | unprocessed_pseudc | ENSG0000C |
| 28459418 | 28459575 | 1 | processed_pseudoge | ENSG0000C |
| 28461979 | 28470930 | -1 | unprocessed_pseudc | ENSG0000C |
| 28475678 | 28478612 | -1 | unprocessed_pseudc | ENSG0000C |
| 28491516 | 28504915 | 1 | transcribed_unproce | ENSG0000C |
| 28500089 | 28500346 | 1 | misc_RNA | ENSG0000C |
| 28515677 | 28517839 | 1 | processed_pseudoge | ENSG0000C |
| 28529259 | 28529484 | 1 | processed_pseudoge | ENSG0000C |
| 28535031 | 28535183 | 1 | processed_pseudoge | ENSG0000C |
| 28540731 | 28540883 | 1 | processed_pseudoge | ENSG0000C |
| 28546431 | 28546583 | 1 | processed_pseudoge | ENSG0000C |
| 28552131 | 28552283 | 1 | processed_pseudoge | ENSG0000C |
| 28557827 | 28557979 | 1 | processed_pseudoge | ENSG0000C |
| 28563475 | 28563679 | 1 | processed_pseudoge | ENSG0000C |
| 28585644 | 28585769 | 1 | unprocessed_pseudc | ENSG0000C |
| 28596172 | 28599296 | -1 | unprocessed_pseudc | ENSG0000C |
| 28614840 | 28614963 | -1 | processed_pseudoge | ENSG0000C |
| 28636485 | 28636692 | -1 | processed_pseudoge | ENSG0000C |
| 28648137 | 28650318 | -1 | processed_pseudoge | ENSG0000C |
| 28661119 | 28674479 | -1 | protein_coding | ENSG0000C |
| 28665659 | 28665948 | -1 | misc_RNA | ENSG0000C |
| 28687420 | 28691285 | 1 | unprocessed_pseudc | ENSG0000C |
| 28695142 | 28702879 | 1 | unprocessed_pseudc | ENSG0000C |
| 28706490 | 28706648 | -1 | processed_pseudoge | ENSG0000C |
| 28716523 | 28720895 | -1 | unprocessed_pseudc | ENSG0000C |
| 28731693 | 28826770 | 1 | transcribed_unproce | ENSG0000C |
| 28843455 | 28853824 | -1 | unprocessed_pseudc | ENSG0000C |
| 28845055 | 28845291 | -1 | misc_RNA | ENSG0000C |
| 28879087 | 28900706 | 1 | transcribed_unproce | ENSG0000C |
| 28904945 | 28905656 | -1 | processed_pseudoge | ENSG0000C |
| 28929748 | 28933507 | 1 | transcribed_unproce | ENSG0000C |
| 28930704 | 28930991 | 1 | transcribed_processe | ENSG0000C |
| 28931169 | 28943001 | 1 | transcribed_unproce | ENSG0000C |
| 29060035 | 29259075 | 1 | protein_coding | ENSG0000C |
| 29260596 | 29720617 | -1 | protein_coding | ENSG0000C |
| 29374919 | 29376383 | -1 | processed_pseudoge | ENSG0000C |
| 29375114 | 29375225 | 1 | snoRNA | ENSG0000C |
| 29409413 | 29411093 | -1 | protein_coding | ENSG0000C |
| 29514804 | 29518158 | -1 | lncRNA | ENSG0000C |
| 29850002 | 29972346 | -1 | protein_coding | ENSG0000C |

| | | | | |
|---|---|---|---|---|
| 29880557 | 29880829 | 1 | processed_pseudoge | ENSG00000 |
| 30233872 | 30247618 | 1 | protein_coding | ENSG00000 |
| 30242579 | 30242815 | 1 | misc_RNA | ENSG00000 |
| 30263132 | 30263264 | 1 | snoRNA | ENSG00000 |
| 30294670 | 30294906 | 1 | misc_RNA | ENSG00000 |
| 30361040 | 30361438 | 1 | protein_coding | ENSG00000 |
| 30405073 | 30405179 | -1 | snRNA | ENSG00000 |
| 30506091 | 30506195 | 1 | misc_RNA | ENSG00000 |
| 30511927 | 30544252 | -1 | protein_coding | ENSG00000 |
| 30535548 | 30535680 | -1 | snoRNA | ENSG00000 |
| 30555912 | 30556202 | -1 | misc_RNA | ENSG00000 |
| 30711324 | 30711614 | 1 | misc_RNA | ENSG00000 |
| 30731838 | 30731970 | 1 | snoRNA | ENSG00000 |
| 30763377 | 30763613 | 1 | misc_RNA | ENSG00000 |
| 30776860 | 30837039 | 1 | protein_coding | ENSG00000 |
| 30825003 | 30825096 | 1 | misc_RNA | ENSG00000 |
| 30951585 | 30951821 | 1 | misc_RNA | ENSG00000 |
| 31055318 | 31094547 | 1 | protein_coding | ENSG00000 |
| 31092912 | 31142971 | -1 | protein_coding | ENSG00000 |
| 31107692 | 31107794 | -1 | snRNA | ENSG00000 |
| 31152534 | 31312750 | -1 | protein_coding | ENSG00000 |
| 31478417 | 31529455 | 1 | protein_coding | ENSG00000 |
| 32079158 | 32079289 | 1 | snoRNA | ENSG00000 |
| 32181324 | 32319966 | 1 | protein_coding | ENSG00000 |
| 32326292 | 32326396 | -1 | misc_RNA | ENSG00000 |
| 32435572 | 32435678 | 1 | snRNA | ENSG00000 |
| 32682415 | 32682705 | 1 | misc_RNA | ENSG00000 |
| 32695437 | 32720246 | 1 | protein_coding | ENSG00000 |
| 32721966 | 32777405 | 1 | protein_coding | ENSG00000 |
| 32798311 | 32814964 | 1 | protein_coding | ENSG00000 |
| 82298559 | 82334628 | 1 | transcribed_unproce | ENSG00000 |
| 82342783 | 82349497 | -1 | protein_coding | ENSG00000 |
| 82380817 | 82395723 | -1 | transcribed_unproce | ENSG00000 |
| 82403250 | 82412947 | -1 | protein_coding | ENSG00000 |
| 82418974 | 82428498 | -1 | protein_coding | ENSG00000 |
| 82434179 | 82499143 | 1 | transcribed_unproce | ENSG00000 |
| 82451282 | 82451992 | -1 | transcribed_processe | ENSG00000 |
| 82497719 | 82513298 | 1 | lncRNA | ENSG00000 |
| 82509086 | 82518216 | 1 | protein_coding | ENSG00000 |
| 82524734 | 82533748 | 1 | unprocessed_pseudo | ENSG00000 |
| 82538538 | 82556321 | 1 | transcribed_unproce | ENSG00000 |
| 82551263 | 82552010 | 1 | unprocessed_pseudo | ENSG00000 |
| 82552056 | 82593266 | -1 | transcribed_unproce | ENSG00000 |
| 82598917 | 82605538 | 1 | transcribed_unproce | ENSG00000 |
| 82610742 | 82613228 | -1 | unprocessed_pseudo | ENSG00000 |
| 82615842 | 82619653 | -1 | protein_coding | ENSG00000 |
| 82619121 | 82635020 | -1 | protein_coding | ENSG00000 |
| 82622289 | 82720771 | -1 | protein_coding | ENSG00000 |
| 30005443 | 30045854 | -1 | lncRNA | ENSG00000 |
| 30044461 | 30045302 | 1 | processed_pseudoge | ENSG00000 |

| | | | | |
|---|---|---|---|---|
| 30052948 | 30073074 | -1 | transcribed_unproce | ENSG00000 |
| 30085788 | 30090449 | 1 | protein_coding | ENSG00000 |
| 30091657 | 30091893 | 1 | misc_RNA | ENSG00000 |
| 30098819 | 30102086 | -1 | unprocessed_pseudo | ENSG00000 |
| 30103719 | 30131750 | 1 | transcribed_unproce | ENSG00000 |
| 30112190 | 30112322 | 1 | snoRNA | ENSG00000 |
| 30135046 | 30148832 | 1 | protein_coding | ENSG00000 |
| 30143737 | 30143973 | 1 | misc_RNA | ENSG00000 |
| 30150880 | 30153410 | -1 | unprocessed_pseudo | ENSG00000 |
| 30153934 | 30154111 | -1 | unprocessed_pseudo | ENSG00000 |
| 30224299 | 30226846 | 1 | processed_pseudoge | ENSG00000 |
| 30255704 | 30255810 | -1 | snRNA | ENSG00000 |
| 30353939 | 30357863 | -1 | TEC | ENSG00000 |
| 30365063 | 30365167 | 1 | misc_RNA | ENSG00000 |
| 30371954 | 30510022 | -1 | protein_coding | ENSG00000 |
| 31891116 | 31914762 | -1 | protein_coding | ENSG00000 |
| 31915115 | 31923892 | 1 | lncRNA | ENSG00000 |
| 31923141 | 31923932 | -1 | lncRNA | ENSG00000 |
| 31927931 | 31928167 | -1 | misc_RNA | ENSG00000 |
| 31930192 | 31930690 | 1 | lncRNA | ENSG00000 |
| 31937745 | 31965690 | -1 | transcribed_unproce | ENSG00000 |
| 31957111 | 31957243 | -1 | snoRNA | ENSG00000 |
| 31967338 | 31970605 | 1 | unprocessed_pseudo | ENSG00000 |
| 31975928 | 31986353 | -1 | protein_coding | ENSG00000 |
| 31977496 | 31977786 | -1 | misc_RNA | ENSG00000 |
| 31992464 | 31998340 | -1 | lncRNA | ENSG00000 |
| 32024724 | 32031065 | -1 | unprocessed_pseudo | ENSG00000 |
| 32117941 | 32118087 | 1 | unprocessed_pseudo | ENSG00000 |
| 32124112 | 32137984 | 1 | protein_coding | ENSG00000 |
| 32132796 | 32133086 | 1 | misc_RNA | ENSG00000 |
| 32146220 | 32159477 | 1 | protein_coding | ENSG00000 |
| 32154795 | 32172305 | -1 | lncRNA | ENSG00000 |
| 32172366 | 32227841 | 1 | protein_coding | ENSG00000 |
| 32186776 | 32203791 | -1 | lncRNA | ENSG00000 |
| 32248010 | 32249747 | -1 | lncRNA | ENSG00000 |
| 32248744 | 32275846 | 1 | protein_coding | ENSG00000 |
| 32367723 | 32368066 | -1 | transcribed_processe | ENSG00000 |
| 14435701 | 14632727 | -1 | protein_coding | ENSG00000 |
| 14601823 | 14602094 | -1 | misc_RNA | ENSG00000 |
| 14632814 | 14669237 | 1 | protein_coding | ENSG00000 |
| 14643787 | 14643848 | 1 | snRNA | ENSG00000 |
| 14672547 | 14694674 | -1 | protein_coding | ENSG00000 |
| 14695572 | 14707060 | -1 | lncRNA | ENSG00000 |
| 14708949 | 14726344 | 1 | protein_coding | ENSG00000 |
| 14938852 | 14938951 | -1 | misc_RNA | ENSG00000 |
| 14952495 | 14992495 | 1 | transcribed_unproce | ENSG00000 |
| 14987680 | 15002408 | -1 | lncRNA | ENSG00000 |
| 15002502 | 15036801 | 1 | protein_coding | ENSG00000 |
| 15040743 | 15058977 | 1 | protein_coding | ENSG00000 |
| 15058498 | 15122225 | -1 | protein_coding | ENSG00000 |

| | | | | |
|---|---|---|---|---|
| 15080438 | 15081000 | 1 | lncRNA | ENSG00000 |
| 15106659 | 15108560 | 1 | processed_pseudoge | ENSG00000 |
| 15144756 | 15171447 | -1 | protein_coding | ENSG00000 |
| 15144760 | 15175928 | -1 | lncRNA | ENSG00000 |
| 15172578 | 15187178 | -1 | transcribed_unproce | ENSG00000 |
| 15183174 | 15183273 | 1 | misc_RNA | ENSG00000 |
| 15241230 | 15297417 | 1 | protein_coding | ENSG00000 |
| 15298518 | 15298617 | 1 | misc_RNA | ENSG00000 |
| 15317897 | 15380303 | -1 | protein_coding | ENSG00000 |
| 15351236 | 15351605 | 1 | processed_pseudoge | ENSG00000 |
| 15389326 | 15393236 | 1 | transcribed_unproce | ENSG00000 |
| 15448713 | 15463457 | -1 | protein_coding | ENSG00000 |
| 15480860 | 15498390 | 1 | protein_coding | ENSG00000 |
| 15480860 | 15600484 | 1 | protein_coding | ENSG00000 |
| 15519415 | 15710184 | 1 | protein_coding | ENSG00000 |
| 15679520 | 15728320 | -1 | protein_coding | ENSG00000 |
| 15693630 | 15695719 | 1 | lncRNA | ENSG00000 |
| 15728421 | 15811507 | 1 | protein_coding | ENSG00000 |
| 15768444 | 15769724 | 1 | lncRNA | ENSG00000 |
| 15786391 | 15787272 | 1 | lncRNA | ENSG00000 |
| 15788326 | 15942169 | -1 | protein_coding | ENSG00000 |
| 15811828 | 15818147 | 1 | lncRNA | ENSG00000 |
| 15826305 | 15826945 | 1 | lncRNA | ENSG00000 |
| 15950862 | 15973769 | -1 | protein_coding | ENSG00000 |
| 15962302 | 15962405 | -1 | snRNA | ENSG00000 |
| 15970173 | 15971302 | 1 | lncRNA | ENSG00000 |
| 16016803 | 16017400 | -1 | processed_pseudoge | ENSG00000 |
| 16034690 | 16228109 | 1 | protein_coding | ENSG00000 |
| 16179193 | 16180211 | -1 | processed_pseudoge | ENSG00000 |
| 16233991 | 16308660 | -1 | protein_coding | ENSG00000 |
| 16308165 | 16309399 | 1 | lncRNA | ENSG00000 |
| 16317633 | 16379962 | 1 | protein_coding | ENSG00000 |
| 16346007 | 16346376 | -1 | processed_pseudoge | ENSG00000 |
| 16375283 | 16377390 | -1 | lncRNA | ENSG00000 |
| 16395246 | 16465697 | 1 | transcribed_unproce | ENSG00000 |
| 16399228 | 16399327 | -1 | misc_RNA | ENSG00000 |
| 16418201 | 16436927 | 1 | protein_coding | ENSG00000 |
| 16467435 | 16482129 | 1 | protein_coding | ENSG00000 |
| 16629420 | 16630050 | -1 | processed_pseudoge | ENSG00000 |
| 16746309 | 16752220 | 1 | lncRNA | ENSG00000 |
| 16848984 | 16850544 | 1 | TEC | ENSG00000 |
| 17040606 | 17041572 | -1 | processed_pseudoge | ENSG00000 |
| 55773402 | 55790834 | 1 | unprocessed_pseudo | ENSG00000 |
| 55799704 | 55830035 | -1 | protein_coding | ENSG00000 |
| 55843031 | 55952925 | -1 | protein_coding | ENSG00000 |
| 69142116 | 69169025 | 1 | protein_coding | ENSG00000 |
| 69175662 | 69175768 | -1 | snRNA | ENSG00000 |
| 69187122 | 69187739 | 1 | protein_coding | ENSG00000 |
| 69191115 | 69191215 | 1 | misc_RNA | ENSG00000 |
| 4314761 | 4339597 | 1 | protein_coding | ENSG00000 |

| | | | | |
|---|---|---|---|---|
| 4324667 | 4328340 | -1 | lncRNA | ENSG00000 |
| 4331549 | 4356579 | -1 | protein_coding | ENSG00000 |
| 4340251 | 4422453 | -1 | protein_coding | ENSG00000 |
| 4348654 | 4350608 | -1 | TEC | ENSG00000 |
| 4356503 | 4427664 | -1 | protein_coding | ENSG00000 |
| 4373807 | 4385487 | 1 | protein_coding | ENSG00000 |
| 4427764 | 4458734 | 1 | protein_coding | ENSG00000 |
| 4428861 | 4429339 | 1 | lncRNA | ENSG00000 |
| 4432481 | 4433062 | 1 | lncRNA | ENSG00000 |
| 4435161 | 4435346 | -1 | processed_pseudoge | ENSG00000 |
| 4463639 | 4497722 | -1 | protein_coding | ENSG00000 |
| 4476649 | 4512306 | 1 | protein_coding | ENSG00000 |
| 4512634 | 4526261 | -1 | protein_coding | ENSG00000 |
| 1874025 | 1886319 | -1 | lncRNA | ENSG00000 |
| 1880310 | 1886269 | -1 | lncRNA | ENSG00000 |
| 1891136 | 1892455 | 1 | lncRNA | ENSG00000 |
| 1913487 | 1920464 | -1 | protein_coding | ENSG00000 |
| 70802084 | 71230718 | -1 | protein_coding | ENSG00000 |
| 71288746 | 71314716 | 1 | lncRNA | ENSG00000 |
| 71293954 | 71294528 | 1 | pseudogene | ENSG00000 |
| 71363481 | 71395224 | 1 | protein_coding | ENSG00000 |
| 71371721 | 71426464 | -1 | lncRNA | ENSG00000 |
| 2817180 | 2826304 | 1 | protein_coding | ENSG00000 |
| 2822659 | 2822993 | -1 | processed_pseudoge | ENSG00000 |
| 2830169 | 2839582 | 1 | protein_coding | ENSG00000 |
| 2839565 | 2842741 | -1 | transcribed_unitary_ | ENSG00000 |
| 2852743 | 2858163 | -1 | protein_coding | ENSG00000 |
| 2857891 | 2859719 | 1 | lncRNA | ENSG00000 |
| 2872255 | 2874181 | -1 | lncRNA | ENSG00000 |
| 75148494 | 75172236 | 1 | protein_coding | ENSG00000 |
| 75192487 | 75193241 | -1 | pseudogene | ENSG00000 |
| 75204116 | 75209969 | -1 | protein_coding | ENSG00000 |
| 75221828 | 75224885 | 1 | protein_coding | ENSG00000 |
| 75226100 | 75228223 | 1 | transcribed_unproce | ENSG00000 |
| 75228207 | 75268085 | -1 | protein_coding | ENSG00000 |
| 75246026 | 75250109 | -1 | TEC | ENSG00000 |
| 401826 | 412487 | 1 | protein_coding | ENSG00000 |
| 424243 | 424543 | -1 | lncRNA | ENSG00000 |
| 425619 | 526533 | 1 | protein_coding | ENSG00000 |
| 439297 | 439397 | -1 | misc_RNA | ENSG00000 |
| 451401 | 453138 | 1 | transcribed_unproce | ENSG00000 |
| 528677 | 530929 | -1 | lncRNA | ENSG00000 |
| 531239 | 531551 | 1 | lncRNA | ENSG00000 |
| 36064280 | 36072032 | 1 | protein_coding | ENSG00000 |
| 36072866 | 36090134 | 1 | lncRNA | ENSG00000 |
| 36088256 | 36090169 | -1 | protein_coding | ENSG00000 |
| 36103827 | 36105621 | 1 | protein_coding | ENSG00000 |
| 36110150 | 36110447 | 1 | misc_RNA | ENSG00000 |
| 36116177 | 36177516 | 1 | lncRNA | ENSG00000 |
| 36161606 | 36162932 | 1 | unprocessed_pseudc | ENSG00000 |

| | | | | |
|---|---|---|---|---|
| 36165687 | 36176601 | -1 | unprocessed_pseudc | ENSG00000( |
| 36183253 | 36196508 | 1 | lncRNA | ENSG00000( |
| 36194906 | 36196795 | -1 | protein_coding | ENSG00000( |
| 36210948 | 36212918 | 1 | protein_coding | ENSG00000( |
| 36249651 | 36250970 | 1 | unprocessed_pseudc | ENSG00000( |
| 36253723 | 36264678 | -1 | protein_coding | ENSG00000( |
| 36282949 | 36284006 | -1 | unprocessed_pseudc | ENSG00000( |
| 36296480 | 36298369 | -1 | protein_coding | ENSG00000( |
| 36312669 | 36314484 | 1 | protein_coding | ENSG00000( |
| 36326893 | 36328453 | -1 | lncRNA | ENSG00000( |
| 36336091 | 36336195 | -1 | unprocessed_pseudc | ENSG00000( |
| 36343345 | 36343444 | 1 | unprocessed_pseudc | ENSG00000( |
| 36355285 | 36366216 | -1 | protein_coding | ENSG00000( |
| 52390515 | 52535701 | 1 | lncRNA | ENSG00000( |
| 72839039 | 72839718 | 1 | lncRNA | ENSG00000( |
| 64099 | 76866 | -1 | lncRNA | ENSG00000( |
| 90796 | 90899 | -1 | snRNA | ENSG00000( |
| 97494 | 118578 | -1 | lncRNA | ENSG00000( |
| 137526 | 139067 | 1 | protein_coding | ENSG00000( |
| 142789 | 180568 | -1 | protein_coding | ENSG00000( |
| 182756 | 190502 | -1 | lncRNA | ENSG00000( |
| 217541 | 419308 | -1 | protein_coding | ENSG00000( |
| 352806 | 353888 | 1 | lncRNA | ENSG00000( |
| 362640 | 364923 | 1 | lncRNA | ENSG00000( |
| 384081 | 390528 | 1 | lncRNA | ENSG00000( |
| 386179 | 389829 | 1 | lncRNA | ENSG00000( |
| 1569973 | 1628886 | -1 | protein_coding | ENSG00000( |
| 1604374 | 1604672 | -1 | misc_RNA | ENSG00000( |
| 1633858 | 1645747 | -1 | protein_coding | ENSG00000( |
| 1646145 | 1650077 | -1 | protein_coding | ENSG00000( |
| 1650629 | 1684882 | -1 | protein_coding | ENSG00000( |
| 1702790 | 1710438 | -1 | protein_coding | ENSG00000( |
| 1711493 | 1717174 | -1 | lncRNA | ENSG00000( |
| 1716523 | 1738610 | 1 | protein_coding | ENSG00000( |
| 1741760 | 1754189 | 1 | protein_coding | ENSG00000( |
| 1760880 | 1765915 | 1 | protein_coding | ENSG00000( |
| 41307700 | 41309253 | -1 | protein_coding | ENSG00000( |
| 41314917 | 41315695 | -1 | protein_coding | ENSG00000( |
| 41325648 | 41329544 | 1 | unprocessed_pseudc | ENSG00000( |
| 41334632 | 41335733 | 1 | unprocessed_pseudc | ENSG00000( |
| 41346092 | 41350812 | -1 | protein_coding | ENSG00000( |
| 41363494 | 41369800 | -1 | protein_coding | ENSG00000( |
| 41377665 | 41382400 | -1 | protein_coding | ENSG00000( |
| 41384972 | 41389707 | -1 | protein_coding | ENSG00000( |
| 41401024 | 41404892 | -1 | protein_coding | ENSG00000( |
| 41409741 | 41432374 | 1 | lncRNA | ENSG00000( |
| 41413837 | 41419100 | -1 | unprocessed_pseudc | ENSG00000( |
| 41428078 | 41431848 | -1 | protein_coding | ENSG00000( |
| 45327366 | 45327497 | -1 | rRNA_pseudogene | ENSG00000( |
| 45371402 | 45372057 | -1 | lncRNA | ENSG00000( |

| | | | | |
|---|---|---|---|---|
| 45393920 | 45434353 | -1 | protein_coding | ENSG00000 |
| 45396950 | 45397495 | 1 | lncRNA | ENSG00000 |
| 45435832 | 45490687 | -1 | protein_coding | ENSG00000 |
| 45505209 | 45550562 | -1 | protein_coding | ENSG00000 |
| 45523061 | 45523341 | -1 | misc_RNA | ENSG00000 |
| 45551530 | 45556925 | -1 | unprocessed_pseudo | ENSG00000 |
| 45573716 | 45688015 | -1 | unprocessed_pseudo | ENSG00000 |
| 45699087 | 45820912 | 1 | protein_coding | ENSG00000 |
| 45723122 | 45768499 | -1 | protein_coding | ENSG00000 |
| 45740978 | 45741258 | -1 | misc_RNA | ENSG00000 |
| 45835461 | 45836529 | 1 | processed_pseudoge | ENSG00000 |
| 45851975 | 45852968 | -1 | processed_pseudoge | ENSG00000 |
| 45871009 | 46068204 | 1 | protein_coding | ENSG00000 |
| 45881211 | 45881312 | 1 | misc_RNA | ENSG00000 |
| 45899740 | 45902887 | -1 | lncRNA | ENSG00000 |
| 46062345 | 46062802 | -1 | lncRNA | ENSG00000 |
| 46069784 | 46203150 | -1 | protein_coding | ENSG00000 |
| 46098193 | 46098637 | -1 | protein_coding | ENSG00000 |
| 46165745 | 46166022 | -1 | misc_RNA | ENSG00000 |
| 46187101 | 46190209 | 1 | lncRNA | ENSG00000 |
| 46198938 | 46201928 | -1 | lncRNA | ENSG00000 |
| 46202111 | 46254322 | 1 | lncRNA | ENSG00000 |
| 46250606 | 46252843 | -1 | protein_coding | ENSG00000 |
| 46261820 | 46313387 | -1 | protein_coding | ENSG00000 |
| 46495985 | 46497453 | 1 | processed_pseudoge | ENSG00000 |
| 46518953 | 46520589 | 1 | transcribed_unproce | ENSG00000 |
| 46660217 | 46660318 | 1 | misc_RNA | ENSG00000 |
| 46706769 | 46708058 | 1 | processed_pseudoge | ENSG00000 |
| 46769907 | 46770900 | 1 | processed_pseudoge | ENSG00000 |
| 46784842 | 46785913 | -1 | processed_pseudoge | ENSG00000 |
| 46799540 | 46804933 | 1 | unprocessed_pseudo | ENSG00000 |
| 46805899 | 46851260 | 1 | protein_coding | ENSG00000 |
| 46812294 | 46875243 | -1 | protein_coding | ENSG00000 |
| 46833127 | 46833407 | 1 | misc_RNA | ENSG00000 |
| 46854433 | 46858400 | -1 | lncRNA | ENSG00000 |
| 46886236 | 47052838 | 1 | protein_coding | ENSG00000 |
| 47057885 | 47113984 | -1 | protein_coding | ENSG00000 |
| 82923652 | 82946138 | 1 | protein_coding | ENSG00000 |
| 82942371 | 83050530 | -1 | protein_coding | ENSG00000 |
| 82978708 | 82981921 | 1 | lncRNA | ENSG00000 |
| 83077507 | 83097050 | 1 | protein_coding | ENSG00000 |
| 281768 | 388309 | -1 | protein_coding | ENSG00000 |
| 323447 | 324508 | 1 | lncRNA | ENSG00000 |
| 333260 | 335543 | 1 | lncRNA | ENSG00000 |
| 354693 | 359636 | 1 | lncRNA | ENSG00000 |
| 356792 | 358937 | 1 | lncRNA | ENSG00000 |
| 978299 | 979770 | -1 | protein_coding | ENSG00000 |
| 997117 | 1003671 | 1 | protein_coding | ENSG00000 |
| 1003519 | 1192547 | -1 | protein_coding | ENSG00000 |
| 1105307 | 1105573 | -1 | misc_RNA | ENSG00000 |

| | | | | |
|---|---|---|---|---|
| 1160565 | 1160774 | 1 | processed_pseudoge | ENSG00000( |
| 1187118 | 1190311 | 1 | lncRNA | ENSG00000( |
| 1201029 | 1203765 | 1 | polymorphic_pseudc | ENSG00000( |
| 1247054 | 1303157 | -1 | protein_coding | ENSG00000( |
| 70312831 | 70374969 | -1 | lncRNA | ENSG00000( |
| 45316960 | 45317040 | -1 | lncRNA | ENSG00000( |
| 45327366 | 45327497 | -1 | rRNA_pseudogene | ENSG00000( |
| 45371402 | 45372057 | -1 | lncRNA | ENSG00000( |
| 45393902 | 45434423 | -1 | protein_coding | ENSG00000( |
| 45396932 | 45397477 | 1 | lncRNA | ENSG00000( |
| 45435902 | 45490401 | -1 | protein_coding | ENSG00000( |
| 45452499 | 45463720 | 1 | lncRNA | ENSG00000( |
| 45506393 | 45561970 | -1 | transcribed_unproce | ENSG00000( |
| 45538099 | 45538379 | -1 | misc_RNA | ENSG00000( |
| 45543212 | 45545473 | 1 | lncRNA | ENSG00000( |
| 45557439 | 45574865 | 1 | lncRNA | ENSG00000( |
| 45561416 | 45570373 | -1 | lncRNA | ENSG00000( |
| 45597507 | 45598565 | 1 | processed_pseudoge | ENSG00000( |
| 45612507 | 45613978 | -1 | processed_pseudoge | ENSG00000( |
| 45620204 | 45620551 | 1 | processed_pseudoge | ENSG00000( |
| 45631949 | 45659841 | 1 | transcribed_unproce | ENSG00000( |
| 45633529 | 45828162 | 1 | lncRNA | ENSG00000( |
| 45743300 | 45744574 | -1 | lncRNA | ENSG00000( |
| 45744950 | 45746266 | -1 | lncRNA | ENSG00000( |
| 45795888 | 45847447 | 1 | protein_coding | ENSG00000( |
| 45854974 | 45907225 | -1 | lncRNA | ENSG00000( |
| 45856453 | 45858690 | 1 | protein_coding | ENSG00000( |
| 45906010 | 46039943 | 1 | protein_coding | ENSG00000( |
| 45907408 | 45910423 | 1 | lncRNA | ENSG00000( |
| 45919274 | 45922376 | -1 | lncRNA | ENSG00000( |
| 45943436 | 45943713 | 1 | misc_RNA | ENSG00000( |
| 46041525 | 46237041 | -1 | protein_coding | ENSG00000( |
| 46046922 | 46047379 | 1 | lncRNA | ENSG00000( |
| 46205177 | 46208324 | 1 | lncRNA | ENSG00000( |
| 46226740 | 46226841 | -1 | misc_RNA | ENSG00000( |
| 46255525 | 46256963 | 1 | processed_pseudoge | ENSG00000( |
| 46271462 | 46272509 | -1 | processed_pseudoge | ENSG00000( |
| 46278924 | 46280579 | 1 | transcribed_unproce | ENSG00000( |
| 46286670 | 46408885 | -1 | protein_coding | ENSG00000( |
| 46339497 | 46384852 | 1 | protein_coding | ENSG00000( |
| 46366710 | 46366990 | 1 | misc_RNA | ENSG00000( |
| 46419843 | 46586373 | 1 | protein_coding | ENSG00000( |
| 46419950 | 46533777 | 1 | unprocessed_pseudc | ENSG00000( |
| 46550508 | 46551093 | -1 | processed_pseudoge | ENSG00000( |
| 46591415 | 46662074 | -1 | protein_coding | ENSG00000( |
| 46662121 | 46715644 | 1 | protein_coding | ENSG00000( |
| 18225635 | 18244875 | 1 | protein_coding | ENSG00000( |
| 18244815 | 18258738 | -1 | protein_coding | ENSG00000( |
| 18260597 | 18266552 | 1 | protein_coding | ENSG00000( |
| 18268080 | 18268828 | 1 | lncRNA | ENSG00000( |

| | | | | |
|---|---|---|---|---|
| 18271428 | 18315007 | -1 | protein_coding | ENSG00000( |
| 18311904 | 18312651 | 1 | processed_pseudoge | ENSG00000( |
| 18312760 | 18313533 | 1 | processed_pseudoge | ENSG00000( |
| 18315293 | 18328056 | 1 | protein_coding | ENSG00000( |
| 18327881 | 18363567 | -1 | protein_coding | ENSG00000( |
| 18340814 | 18340886 | -1 | miRNA | ENSG00000( |
| 18387383 | 18399252 | 1 | protein_coding | ENSG00000( |
| 18389460 | 18398589 | -1 | lncRNA | ENSG00000( |
| 18398476 | 18399064 | 1 | processed_pseudoge | ENSG00000( |
| 18399758 | 18399928 | -1 | unprocessed_pseudc | ENSG00000( |
| 18403085 | 18410566 | 1 | unprocessed_pseudc | ENSG00000( |
| 18420764 | 18423985 | 1 | lncRNA | ENSG00000( |
| 18425333 | 18434702 | 1 | transcribed_unproce | ENSG00000( |
| 18427800 | 18427911 | -1 | processed_pseudoge | ENSG00000( |
| 18436466 | 18441453 | 1 | transcribed_unproce | ENSG00000( |
| 18441656 | 18452500 | 1 | transcribed_unproce | ENSG00000( |
| 18459849 | 18485525 | 1 | transcribed_process | ENSG00000( |
| 18462560 | 18462667 | 1 | unprocessed_pseudc | ENSG00000( |
| 18463179 | 18463344 | -1 | processed_pseudoge | ENSG00000( |
| 18469996 | 18472660 | 1 | processed_pseudoge | ENSG00000( |
| 18486342 | 18504550 | 1 | protein_coding | ENSG00000( |
| 18506700 | 18515918 | -1 | transcribed_unproce | ENSG00000( |
| 978299 | 979770 | -1 | protein_coding | ENSG00000( |
| 997117 | 1003671 | 1 | protein_coding | ENSG00000( |
| 1003519 | 1116206 | -1 | protein_coding | ENSG00000( |
| 1101777 | 1102043 | -1 | misc_RNA | ENSG00000( |
| 77269359 | 77293519 | -1 | lncRNA | ENSG00000( |
| 77275802 | 77276398 | -1 | lncRNA | ENSG00000( |
| 77292191 | 77306525 | 1 | protein_coding | ENSG00000( |
| 40850800 | 40863282 | 1 | lncRNA | ENSG00000( |
| 40861303 | 40867210 | -1 | protein_coding | ENSG00000( |
| 40875941 | 40885227 | -1 | protein_coding | ENSG00000( |
| 40921430 | 40976053 | 1 | lncRNA | ENSG00000( |
| 40922696 | 40937634 | -1 | protein_coding | ENSG00000( |
| 40958544 | 40967019 | -1 | protein_coding | ENSG00000( |
| 40977843 | 40987262 | -1 | protein_coding | ENSG00000( |
| 40993557 | 40994260 | -1 | protein_coding | ENSG00000( |
| 40999320 | 41000021 | -1 | protein_coding | ENSG00000( |
| 41004608 | 41004887 | -1 | unprocessed_pseudc | ENSG00000( |
| 41008648 | 41019451 | -1 | protein_coding | ENSG00000( |
| 41026153 | 41027329 | -1 | protein_coding | ENSG00000( |
| 41029824 | 41030231 | -1 | protein_coding | ENSG00000( |
| 41034011 | 41034982 | -1 | protein_coding | ENSG00000( |
| 41040684 | 41041586 | -1 | protein_coding | ENSG00000( |
| 41046666 | 41047441 | -1 | protein_coding | ENSG00000( |
| 41054623 | 41055489 | -1 | protein_coding | ENSG00000( |
| 41059470 | 41060344 | -1 | protein_coding | ENSG00000( |
| 41065346 | 41066109 | -1 | protein_coding | ENSG00000( |
| 41072178 | 41072573 | 1 | unprocessed_pseudc | ENSG00000( |
| 41084386 | 41085542 | 1 | protein_coding | ENSG00000( |

| | | | | |
|---|---|---|---|---|
| 41097371 | 41098544 | -1 | protein_coding | ENSG00000( |
| 41101921 | 41102628 | -1 | protein_coding | ENSG00000( |
| 41105751 | 41106907 | 1 | protein_coding | ENSG00000( |
| 41117663 | 41118842 | -1 | protein_coding | ENSG00000( |
| 41123559 | 41124635 | -1 | protein_coding | ENSG00000( |
| 41226648 | 41227652 | 1 | protein_coding | ENSG00000( |
| 41232462 | 41233453 | 1 | protein_coding | ENSG00000( |
| 41238001 | 41238465 | 1 | unprocessed_pseudc | ENSG00000( |
| 41249619 | 41250585 | 1 | protein_coding | ENSG00000( |
| 41255319 | 41256311 | 1 | protein_coding | ENSG00000( |
| 41265358 | 41265882 | 1 | protein_coding | ENSG00000( |
| 41270282 | 41270488 | 1 | unprocessed_pseudc | ENSG00000( |
| 41291504 | 41292529 | -1 | protein_coding | ENSG00000( |
| 41212710 | 41213241 | 1 | unprocessed_pseudc | ENSG00000( |
| 41226627 | 41227631 | 1 | protein_coding | ENSG00000( |
| 41238317 | 41239283 | 1 | protein_coding | ENSG00000( |
| 41244014 | 41244994 | 1 | protein_coding | ENSG00000( |
| 41253969 | 41255271 | 1 | protein_coding | ENSG00000( |
| 41259941 | 41260465 | 1 | unprocessed_pseudc | ENSG00000( |
| 41264289 | 41265327 | 1 | protein_coding | ENSG00000( |
| 41269234 | 41269440 | 1 | unprocessed_pseudc | ENSG00000( |
| 41290456 | 41291481 | -1 | protein_coding | ENSG00000( |
| 35765873 | 35795707 | -1 | protein_coding | ENSG00000( |
| 35808489 | 35808897 | -1 | lncRNA | ENSG00000( |
| 35809455 | 35864615 | 1 | protein_coding | ENSG00000( |
| 35816717 | 35830293 | -1 | lncRNA | ENSG00000( |
| 35818399 | 35823713 | 1 | lncRNA | ENSG00000( |
| 35854946 | 35868891 | -1 | protein_coding | ENSG00000( |
| 35861093 | 35861566 | -1 | processed_pseudoge | ENSG00000( |
| 35868967 | 35885863 | 1 | lncRNA | ENSG00000( |
| 35871491 | 35880793 | -1 | protein_coding | ENSG00000( |
| 35889558 | 35889804 | 1 | misc_RNA | ENSG00000( |
| 35893707 | 35911023 | -1 | lncRNA | ENSG00000( |
| 35912635 | 35918010 | -1 | processed_pseudoge | ENSG00000( |
| 35918066 | 35930773 | -1 | protein_coding | ENSG00000( |
| 35934518 | 35943699 | -1 | protein_coding | ENSG00000( |
| 35963502 | 35965476 | -1 | transcribed_unproce | ENSG00000( |
| 35976525 | 35981528 | -1 | protein_coding | ENSG00000( |
| 35983323 | 35987036 | -1 | protein_coding | ENSG00000( |
| 35983688 | 36001656 | -1 | protein_coding | ENSG00000( |
| 35983688 | 35990302 | 1 | lncRNA | ENSG00000( |
| 35996472 | 36002080 | -1 | protein_coding | ENSG00000( |
| 36009598 | 36011654 | 1 | lncRNA | ENSG00000( |
| 36012540 | 36012927 | 1 | lncRNA | ENSG00000( |
| 36013092 | 36018001 | -1 | protein_coding | ENSG00000( |
| 36064636 | 36072388 | 1 | protein_coding | ENSG00000( |
| 36073222 | 36090476 | 1 | lncRNA | ENSG00000( |
| 36088598 | 36090511 | -1 | protein_coding | ENSG00000( |
| 36103979 | 36106010 | 1 | protein_coding | ENSG00000( |
| 36110539 | 36110836 | 1 | misc_RNA | ENSG00000( |

| | | | | |
|---|---|---|---|---|
| 36116566 | 36177866 | 1 | lncRNA | ENSG0000C |
| 36163208 | 36163292 | 1 | processed_pseudoge | ENSG0000C |
| 36166057 | 36176992 | -1 | protein_coding | ENSG0000C |
| 36183607 | 36196843 | 1 | lncRNA | ENSG0000C |
| 36195241 | 36197130 | -1 | protein_coding | ENSG0000C |
| 36211296 | 36213250 | 1 | protein_coding | ENSG0000C |
| 36225618 | 36227179 | -1 | lncRNA | ENSG0000C |
| 36234819 | 36234926 | -1 | unprocessed_pseudc | ENSG0000C |
| 36242073 | 36242178 | 1 | processed_pseudoge | ENSG0000C |
| 36254011 | 36264925 | -1 | protein_coding | ENSG0000C |
| 36274563 | 36320919 | 1 | unprocessed_pseudc | ENSG0000C |
| 36285684 | 36285785 | 1 | misc_RNA | ENSG0000C |
| 36324256 | 36335131 | 1 | protein_coding | ENSG0000C |
| 36377903 | 36388795 | -1 | protein_coding | ENSG0000C |
| 36405198 | 36405304 | -1 | snRNA | ENSG0000C |
| 36428990 | 36439938 | -1 | protein_coding | ENSG0000C |
| 36450330 | 36457907 | -1 | unprocessed_pseudc | ENSG0000C |
| 36487001 | 36499682 | 1 | protein_coding | ENSG0000C |
| 36491571 | 36491694 | -1 | rRNA | ENSG0000C |
| 36496005 | 36543807 | -1 | protein_coding | ENSG0000C |
| 36535392 | 36539682 | 1 | protein_coding | ENSG0000C |
| 36545260 | 36590811 | 1 | protein_coding | ENSG0000C |
| 36574834 | 36575697 | -1 | processed_pseudoge | ENSG0000C |
| 36592761 | 36601768 | 1 | protein_coding | ENSG0000C |
| 36602534 | 36609940 | 1 | protein_coding | ENSG0000C |
| 36635060 | 36635689 | 1 | lncRNA | ENSG0000C |
| 36685736 | 36686111 | 1 | lncRNA | ENSG0000C |
| 36723427 | 36727184 | 1 | lncRNA | ENSG0000C |
| 36863468 | 36938493 | -1 | lncRNA | ENSG0000C |
| 36938617 | 36946445 | 1 | protein_coding | ENSG0000C |
| 36941879 | 36945288 | -1 | lncRNA | ENSG0000C |
| 36950708 | 37058704 | 1 | protein_coding | ENSG0000C |
| 36981999 | 36982254 | 1 | lncRNA | ENSG0000C |
| 37000430 | 37001866 | 1 | lncRNA | ENSG0000C |
| 37036498 | 37045083 | 1 | lncRNA | ENSG0000C |
| 37047293 | 37054608 | -1 | lncRNA | ENSG0000C |
| 37086456 | 37411442 | -1 | protein_coding | ENSG0000C |
| 37142069 | 37142882 | -1 | processed_pseudoge | ENSG0000C |
| 37145064 | 37145920 | -1 | transcribed_processe | ENSG0000C |
| 37164063 | 37164586 | 1 | processed_pseudoge | ENSG0000C |
| 37377461 | 37394195 | 1 | protein_coding | ENSG0000C |
| 37388354 | 37389394 | 1 | lncRNA | ENSG0000C |
| 37411498 | 37484368 | 1 | protein_coding | ENSG0000C |
| 37412556 | 37413214 | -1 | lncRNA | ENSG0000C |
| 37458806 | 37459566 | 1 | processed_pseudoge | ENSG0000C |
| 37494470 | 37518136 | 1 | protein_coding | ENSG0000C |
| 37519433 | 37614077 | -1 | protein_coding | ENSG0000C |
| 37614320 | 37648026 | -1 | protein_coding | ENSG0000C |
| 37690967 | 37749771 | -1 | protein_coding | ENSG0000C |
| 37803402 | 37807292 | -1 | lncRNA | ENSG0000C |

| | | | | |
|---|---|---|---|---|
| 37832458 | 37888896 | -1 | transcribed_unproce | ENSG00000 |
| 37912034 | 37912140 | 1 | snRNA | ENSG00000 |
| 37928493 | 37939442 | 1 | protein_coding | ENSG00000 |
| 37982250 | 37993199 | -1 | protein_coding | ENSG00000 |
| 37996476 | 38057692 | -1 | unprocessed_pseudo | ENSG00000 |
| 38031608 | 38031709 | -1 | misc_RNA | ENSG00000 |
| 38097522 | 38123634 | 1 | protein_coding | ENSG00000 |
| 38126026 | 38144226 | -1 | protein_coding | ENSG00000 |
| 38152644 | 38206380 | 1 | protein_coding | ENSG00000 |
| 38229196 | 38313162 | 1 | protein_coding | ENSG00000 |
| 38251172 | 38253222 | -1 | lncRNA | ENSG00000 |
| 38330785 | 38406717 | -1 | protein_coding | ENSG00000 |
| 38401836 | 38403488 | 1 | lncRNA | ENSG00000 |
| 38472490 | 38476208 | -1 | protein_coding | ENSG00000 |
| 38503239 | 38505534 | 1 | lncRNA | ENSG00000 |
| 38506329 | 38530590 | 1 | protein_coding | ENSG00000 |
| 38528620 | 38528985 | -1 | protein_coding | ENSG00000 |
| 38531022 | 38535831 | 1 | protein_coding | ENSG00000 |
| 38532589 | 38532707 | -1 | rRNA | ENSG00000 |
| 38534684 | 38550604 | -1 | protein_coding | ENSG00000 |
| 38550147 | 38552244 | 1 | lncRNA | ENSG00000 |
| 38553523 | 38565018 | 1 | protein_coding | ENSG00000 |
| 38556066 | 38556187 | -1 | snRNA | ENSG00000 |
| 38566476 | 38600913 | -1 | protein_coding | ENSG00000 |
| 38601221 | 38626268 | -1 | protein_coding | ENSG00000 |
| 38635875 | 38642242 | -1 | protein_coding | ENSG00000 |
| 71596338 | 71597279 | 1 | lncRNA | ENSG00000 |
| 71620978 | 71621436 | 1 | processed_pseudoge | ENSG00000 |
| 72837800 | 72849806 | -1 | protein_coding | ENSG00000 |
| 72839039 | 72839718 | 1 | lncRNA | ENSG00000 |
| 1655177 | 1779955 | 1 | lncRNA | ENSG00000 |
| 1780329 | 1782064 | 1 | lncRNA | ENSG00000 |
| 50302190 | 50313595 | 1 | lncRNA | ENSG00000 |
| 50318420 | 50319024 | 1 | processed_pseudoge | ENSG00000 |
| 50319002 | 50319120 | 1 | rRNA_pseudogene | ENSG00000 |
| 50374995 | 50394173 | 1 | protein_coding | ENSG00000 |
| 50465102 | 50466010 | 1 | processed_pseudoge | ENSG00000 |
| 50567310 | 50567991 | 1 | lncRNA | ENSG00000 |
| 78505165 | 78506451 | 1 | lncRNA | ENSG00000 |
| 78544095 | 78544188 | -1 | misc_RNA | ENSG00000 |
| 51687527 | 51688210 | -1 | processed_pseudoge | ENSG00000 |
| 67457499 | 67458554 | 1 | TEC | ENSG00000 |
| 67551243 | 67554415 | -1 | lncRNA | ENSG00000 |
| 67576031 | 67586429 | -1 | protein_coding | ENSG00000 |
| 67576038 | 67583479 | 1 | lncRNA | ENSG00000 |
| 67585995 | 67586198 | 1 | lncRNA | ENSG00000 |
| 78976555 | 78979074 | -1 | lncRNA | ENSG00000 |
| 78980275 | 79002678 | 1 | protein_coding | ENSG00000 |
| 79074837 | 79074964 | 1 | protein_coding | ENSG00000 |
| 79576460 | 79589010 | 1 | lncRNA | ENSG00000 |

| | | | | |
|---|---|---|---|---|
| 79610642 | 79612198 | 1 | lncRNA | ENSG0000C |
| 79644682 | 79685499 | -1 | lncRNA | ENSG0000C |
| 79683041 | 79685112 | -1 | lncRNA | ENSG0000C |
| 79685555 | 79723770 | 1 | protein_coding | ENSG0000C |
| 79708475 | 79709405 | -1 | lncRNA | ENSG0000C |
| 22413599 | 22417915 | -1 | protein_coding | ENSG0000C |
| 22418829 | 22419294 | -1 | processed_pseudoge | ENSG0000C |
| 78505165 | 78506410 | 1 | lncRNA | ENSG0000C |
| 78544095 | 78544188 | -1 | misc_RNA | ENSG0000C |
| 78976555 | 78979074 | -1 | lncRNA | ENSG0000C |
| 78980275 | 79002677 | 1 | protein_coding | ENSG0000C |
| 79072883 | 79091276 | 1 | protein_coding | ENSG0000C |
| 79088732 | 79090033 | 1 | TEC | ENSG0000C |
| 54696604 | 54700905 | 1 | unprocessed_pseudc | ENSG0000C |
| 54703952 | 54704235 | 1 | unprocessed_pseudc | ENSG0000C |
| 54707916 | 54713453 | 1 | transcribed_processe | ENSG0000C |
| 54724485 | 54736499 | 1 | protein_coding | ENSG0000C |
| 54738476 | 54753016 | 1 | protein_coding | ENSG0000C |
| 54754987 | 54767335 | 1 | unprocessed_pseudc | ENSG0000C |
| 54786172 | 54787996 | 1 | protein_coding | ENSG0000C |
| 54789705 | 54789811 | -1 | snRNA | ENSG0000C |
| 54794996 | 54812181 | 1 | protein_coding | ENSG0000C |
| 54696604 | 54700905 | 1 | unprocessed_pseudc | ENSG0000C |
| 54703952 | 54704235 | 1 | unprocessed_pseudc | ENSG0000C |
| 54707916 | 54713453 | 1 | transcribed_processe | ENSG0000C |
| 54724479 | 54736542 | 1 | protein_coding | ENSG0000C |
| 54724496 | 54798294 | 1 | lncRNA | ENSG0000C |
| 54738518 | 54753046 | 1 | protein_coding | ENSG0000C |
| 54769802 | 54784331 | 1 | protein_coding | ENSG0000C |
| 54786364 | 54790327 | 1 | transcribed_unproce | ENSG0000C |
| 54803543 | 54814528 | 1 | protein_coding | ENSG0000C |
| 54816479 | 54830810 | 1 | protein_coding | ENSG0000C |
| 54832708 | 54848388 | 1 | polymorphic_pseudc | ENSG0000C |
| 54850262 | 54867032 | 1 | protein_coding | ENSG0000C |
| 54696604 | 54700905 | 1 | unprocessed_pseudc | ENSG0000C |
| 54703952 | 54704235 | 1 | unprocessed_pseudc | ENSG0000C |
| 54707916 | 54713453 | 1 | transcribed_processe | ENSG0000C |
| 54724500 | 54738318 | 1 | protein_coding | ENSG0000C |
| 54744096 | 54746206 | 1 | protein_coding | ENSG0000C |
| 54757172 | 54766181 | 1 | protein_coding | ENSG0000C |
| 54768588 | 54782745 | 1 | protein_coding | ENSG0000C |
| 54785376 | 54797726 | 1 | unprocessed_pseudc | ENSG0000C |
| 54800209 | 54814178 | 1 | protein_coding | ENSG0000C |
| 54816720 | 54820679 | 1 | transcribed_unproce | ENSG0000C |
| 54833970 | 54844886 | 1 | protein_coding | ENSG0000C |
| 54846871 | 54860739 | 1 | protein_coding | ENSG0000C |
| 54863509 | 54889060 | 1 | protein_coding | ENSG0000C |
| 54891695 | 54905523 | 1 | protein_coding | ENSG0000C |
| 54908052 | 54913516 | 1 | protein_coding | ENSG0000C |
| 54696604 | 54700905 | 1 | unprocessed_pseudc | ENSG0000C |

| | | | | |
|---|---|---|---|---|
| 54703952 | 54704235 | 1 | unprocessed_pseudc | ENSG00000( |
| 54707916 | 54713453 | 1 | transcribed_processe | ENSG00000( |
| 54724485 | 54736500 | 1 | protein_coding | ENSG00000( |
| 54724502 | 54798229 | 1 | lncRNA | ENSG00000( |
| 54738475 | 54752995 | 1 | protein_coding | ENSG00000( |
| 54754966 | 54767314 | 1 | unprocessed_pseudc | ENSG00000( |
| 54769736 | 54784266 | 1 | protein_coding | ENSG00000( |
| 54786299 | 54790262 | 1 | transcribed_unproce | ENSG00000( |
| 54803469 | 54814492 | 1 | protein_coding | ENSG00000( |
| 54816443 | 54830754 | 1 | protein_coding | ENSG00000( |
| 54832652 | 54848544 | 1 | polymorphic_pseudc | ENSG00000( |
| 54850414 | 54867200 | 1 | protein_coding | ENSG00000( |
| 54696604 | 54700905 | 1 | unprocessed_pseudc | ENSG00000( |
| 54703952 | 54704235 | 1 | unprocessed_pseudc | ENSG00000( |
| 54707916 | 54713453 | 1 | transcribed_processe | ENSG00000( |
| 54724496 | 54798077 | 1 | lncRNA | ENSG00000( |
| 54754806 | 54767156 | 1 | unprocessed_pseudc | ENSG00000( |
| 54769582 | 54784118 | 1 | protein_coding | ENSG00000( |
| 54786151 | 54790110 | 1 | transcribed_unproce | ENSG00000( |
| 54803392 | 54814342 | 1 | protein_coding | ENSG00000( |
| 54816294 | 54830607 | 1 | protein_coding | ENSG00000( |
| 54832536 | 54847547 | 1 | protein_coding | ENSG00000( |
| 54849415 | 54866185 | 1 | protein_coding | ENSG00000( |
| 54867917 | 54868023 | -1 | snRNA | ENSG00000( |
| 54873209 | 54890394 | 1 | protein_coding | ENSG00000( |
| 54696604 | 54700905 | 1 | unprocessed_pseudc | ENSG00000( |
| 54703952 | 54704235 | 1 | unprocessed_pseudc | ENSG00000( |
| 54707916 | 54713453 | 1 | transcribed_processe | ENSG00000( |
| 54724479 | 54736535 | 1 | protein_coding | ENSG00000( |
| 54724496 | 54798268 | 1 | lncRNA | ENSG00000( |
| 54738511 | 54753035 | 1 | protein_coding | ENSG00000( |
| 54755006 | 54767354 | 1 | unprocessed_pseudc | ENSG00000( |
| 54769776 | 54784305 | 1 | protein_coding | ENSG00000( |
| 54786338 | 54790301 | 1 | transcribed_unproce | ENSG00000( |
| 54803507 | 54814530 | 1 | protein_coding | ENSG00000( |
| 54816481 | 54830792 | 1 | protein_coding | ENSG00000( |
| 54832690 | 54848582 | 1 | polymorphic_pseudc | ENSG00000( |
| 54850452 | 54867240 | 1 | protein_coding | ENSG00000( |
| 54696604 | 54700905 | 1 | unprocessed_pseudc | ENSG00000( |
| 54703952 | 54704235 | 1 | unprocessed_pseudc | ENSG00000( |
| 54707916 | 54713453 | 1 | transcribed_processe | ENSG00000( |
| 54724479 | 54736532 | 1 | protein_coding | ENSG00000( |
| 54724496 | 54798231 | 1 | lncRNA | ENSG00000( |
| 54738509 | 54753057 | 1 | protein_coding | ENSG00000( |
| 54755028 | 54767374 | 1 | unprocessed_pseudc | ENSG00000( |
| 54773339 | 54775455 | 1 | protein_coding | ENSG00000( |
| 54786301 | 54790264 | 1 | transcribed_unproce | ENSG00000( |
| 54803544 | 54814494 | 1 | protein_coding | ENSG00000( |
| 54816446 | 54830759 | 1 | protein_coding | ENSG00000( |
| 54832688 | 54848578 | 1 | protein_coding | ENSG00000( |

| | | | | |
|---|---|---|---|---|
| 54850445 | 54867195 | 1 | protein_coding | ENSG00000 |
| 54868927 | 54869033 | -1 | snRNA | ENSG00000 |
| 54724507 | 54736522 | 1 | protein_coding | ENSG00000 |
| 54738499 | 54753027 | 1 | protein_coding | ENSG00000 |
| 54754998 | 54767346 | 1 | unprocessed_pseudo | ENSG00000 |
| 54769794 | 54784109 | 1 | protein_coding | ENSG00000 |
| 54786437 | 54798259 | 1 | transcribed_processe | ENSG00000 |
| 54803411 | 54814521 | 1 | protein_coding | ENSG00000 |
| 54816473 | 54830785 | 1 | protein_coding | ENSG00000 |
| 54832684 | 54848599 | 1 | protein_coding | ENSG00000 |
| 54850485 | 54867216 | 1 | protein_coding | ENSG00000 |
| 54868948 | 54869054 | -1 | snRNA | ENSG00000 |
| 54874257 | 54890481 | 1 | protein_coding | ENSG00000 |
| 54906159 | 54910074 | 1 | protein_coding | ENSG00000 |
| 54700593 | 54700905 | 1 | unprocessed_pseudo | ENSG00000 |
| 54703952 | 54704235 | 1 | unprocessed_pseudo | ENSG00000 |
| 54707916 | 54713453 | 1 | transcribed_processe | ENSG00000 |
| 54724453 | 54767462 | 1 | protein_coding | ENSG00000 |
| 54738471 | 54753005 | 1 | protein_coding | ENSG00000 |
| 54754976 | 54767325 | 1 | unprocessed_pseudo | ENSG00000 |
| 54769765 | 54784000 | 1 | protein_coding | ENSG00000 |
| 54786274 | 54790272 | 1 | unprocessed_pseudo | ENSG00000 |
| 54803541 | 54814525 | 1 | protein_coding | ENSG00000 |
| 54816496 | 54820397 | 1 | protein_coding | ENSG00000 |
| 54700593 | 54700905 | 1 | unprocessed_pseudo | ENSG00000 |
| 54703952 | 54704235 | 1 | unprocessed_pseudo | ENSG00000 |
| 54707916 | 54713453 | 1 | transcribed_processe | ENSG00000 |
| 54724479 | 54736536 | 1 | protein_coding | ENSG00000 |
| 54738506 | 54752764 | 1 | protein_coding | ENSG00000 |
| 54755023 | 54767371 | 1 | unprocessed_pseudo | ENSG00000 |
| 54769814 | 54784135 | 1 | protein_coding | ENSG00000 |
| 54786355 | 54790318 | 1 | unprocessed_pseudo | ENSG00000 |
| 54803535 | 54814517 | 1 | protein_coding | ENSG00000 |
| 54816438 | 54830737 | 1 | protein_coding | ENSG00000 |
| 54832765 | 54848078 | 1 | unprocessed_pseudo | ENSG00000 |
| 54850439 | 54867235 | 1 | protein_coding | ENSG00000 |
| 54868959 | 54869065 | -1 | snRNA | ENSG00000 |
| 54696604 | 54700905 | 1 | unprocessed_pseudo | ENSG00000 |
| 54703952 | 54704235 | 1 | unprocessed_pseudo | ENSG00000 |
| 54707916 | 54713453 | 1 | transcribed_processe | ENSG00000 |
| 54724479 | 54736533 | 1 | protein_coding | ENSG00000 |
| 54724496 | 54798302 | 1 | lncRNA | ENSG00000 |
| 54738510 | 54753069 | 1 | protein_coding | ENSG00000 |
| 54755039 | 54767385 | 1 | unprocessed_pseudo | ENSG00000 |
| 54769810 | 54784338 | 1 | protein_coding | ENSG00000 |
| 54786371 | 54790334 | 1 | transcribed_unproce | ENSG00000 |
| 54803554 | 54814536 | 1 | protein_coding | ENSG00000 |
| 54816487 | 54830797 | 1 | protein_coding | ENSG00000 |
| 54832695 | 54848588 | 1 | polymorphic_pseudo | ENSG00000 |
| 54850462 | 54867223 | 1 | protein_coding | ENSG00000 |

| | | | | |
|---|---|---|---|---|
| 54696604 | 54700905 | 1 | unprocessed_pseudc | ENSG0000C |
| 54703952 | 54704235 | 1 | unprocessed_pseudc | ENSG0000C |
| 54707916 | 54713453 | 1 | transcribed_processe | ENSG0000C |
| 54724488 | 54736506 | 1 | protein_coding | ENSG0000C |
| 54724505 | 54798268 | 1 | lncRNA | ENSG0000C |
| 54738483 | 54753037 | 1 | protein_coding | ENSG0000C |
| 54755008 | 54767354 | 1 | unprocessed_pseudc | ENSG0000C |
| 54769779 | 54784306 | 1 | protein_coding | ENSG0000C |
| 54786338 | 54790301 | 1 | transcribed_unproce | ENSG0000C |
| 54803507 | 54814530 | 1 | protein_coding | ENSG0000C |
| 54816481 | 54830792 | 1 | protein_coding | ENSG0000C |
| 54832690 | 54848582 | 1 | polymorphic_pseudc | ENSG0000C |
| 54850452 | 54867238 | 1 | protein_coding | ENSG0000C |
| 54696604 | 54700905 | 1 | unprocessed_pseudc | ENSG0000C |
| 54703952 | 54704235 | 1 | unprocessed_pseudc | ENSG0000C |
| 54707916 | 54713453 | 1 | transcribed_processe | ENSG0000C |
| 54724496 | 54798270 | 1 | lncRNA | ENSG0000C |
| 54738510 | 54753036 | 1 | protein_coding | ENSG0000C |
| 54758593 | 54767076 | 1 | protein_coding | ENSG0000C |
| 54769779 | 54784308 | 1 | protein_coding | ENSG0000C |
| 54786341 | 54790304 | 1 | transcribed_unproce | ENSG0000C |
| 54803603 | 54814519 | 1 | protein_coding | ENSG0000C |
| 54816504 | 54830372 | 1 | protein_coding | ENSG0000C |
| 54833142 | 54858873 | 1 | protein_coding | ENSG0000C |
| 54861549 | 54875377 | 1 | protein_coding | ENSG0000C |
| 54877873 | 54894659 | 1 | protein_coding | ENSG0000C |
| 54896391 | 54896497 | -1 | snRNA | ENSG0000C |
| 54901684 | 54918869 | 1 | protein_coding | ENSG0000C |
| 54933637 | 54940317 | 1 | protein_coding | ENSG0000C |
| 54696604 | 54700905 | 1 | unprocessed_pseudc | ENSG0000C |
| 54703952 | 54704235 | 1 | unprocessed_pseudc | ENSG0000C |
| 54707916 | 54713453 | 1 | transcribed_processe | ENSG0000C |
| 54724479 | 54736502 | 1 | protein_coding | ENSG0000C |
| 54738477 | 54753006 | 1 | protein_coding | ENSG0000C |
| 54754977 | 54767325 | 1 | unprocessed_pseudc | ENSG0000C |
| 54788172 | 54790295 | 1 | protein_coding | ENSG0000C |
| 54803584 | 54814500 | 1 | protein_coding | ENSG0000C |
| 54816484 | 54830352 | 1 | protein_coding | ENSG0000C |
| 54833122 | 54858674 | 1 | protein_coding | ENSG0000C |
| 54876141 | 54890110 | 1 | protein_coding | ENSG0000C |
| 54892652 | 54896615 | 1 | transcribed_unproce | ENSG0000C |
| 54908995 | 54919911 | 1 | protein_coding | ENSG0000C |
| 54921895 | 54935763 | 1 | protein_coding | ENSG0000C |
| 54938533 | 54964264 | 1 | protein_coding | ENSG0000C |
| 54966899 | 54980727 | 1 | protein_coding | ENSG0000C |
| 54983256 | 54988720 | 1 | protein_coding | ENSG0000C |
| 54696604 | 54700905 | 1 | unprocessed_pseudc | ENSG0000C |
| 54703952 | 54704235 | 1 | unprocessed_pseudc | ENSG0000C |
| 54707916 | 54713453 | 1 | transcribed_processe | ENSG0000C |
| 54738494 | 54753018 | 1 | protein_coding | ENSG0000C |

| | | | | |
|---|---|---|---|---|
| 54754989 | 54767345 | 1 | unprocessed_pseudc | ENSG00000( |
| 54769770 | 54786707 | 1 | protein_coding | ENSG00000( |
| 54788439 | 54788545 | -1 | snRNA | ENSG00000( |
| 54793733 | 54810918 | 1 | protein_coding | ENSG00000( |
| 54724486 | 54736500 | 1 | protein_coding | ENSG00000( |
| 54738479 | 54753050 | 1 | protein_coding | ENSG00000( |
| 54754958 | 54767824 | 1 | unprocessed_pseudc | ENSG00000( |
| 54769810 | 54784323 | 1 | protein_coding | ENSG00000( |
| 54786460 | 54798282 | 1 | transcribed_unproce | ENSG00000( |
| 54803627 | 54814543 | 1 | protein_coding | ENSG00000( |
| 54816499 | 54830812 | 1 | protein_coding | ENSG00000( |
| 54832710 | 54847746 | 1 | protein_coding | ENSG00000( |
| 54849620 | 54866390 | 1 | protein_coding | ENSG00000( |
| 54868122 | 54868228 | -1 | snRNA | ENSG00000( |
| 54873427 | 54889651 | 1 | protein_coding | ENSG00000( |
| 54905329 | 54909244 | 1 | protein_coding | ENSG00000( |
| 54724203 | 54736221 | 1 | protein_coding | ENSG00000( |
| 54738243 | 54752150 | 1 | protein_coding | ENSG00000( |
| 54754397 | 54768727 | 1 | protein_coding | ENSG00000( |
| 54770973 | 54780252 | 1 | protein_coding | ENSG00000( |
| 54782354 | 54796789 | 1 | protein_coding | ENSG00000( |
| 54799171 | 54811528 | 1 | unprocessed_pseudc | ENSG00000( |
| 54813971 | 54828213 | 1 | protein_coding | ENSG00000( |
| 54830630 | 54842447 | 1 | transcribed_unproce | ENSG00000( |
| 54847612 | 54858688 | 1 | protein_coding | ENSG00000( |
| 54860636 | 54875223 | 1 | protein_coding | ENSG00000( |
| 54877311 | 54886807 | 1 | protein_coding | ENSG00000( |
| 54888671 | 54903107 | 1 | protein_coding | ENSG00000( |
| 54905478 | 54919948 | 1 | protein_coding | ENSG00000( |
| 54921821 | 54938615 | 1 | protein_coding | ENSG00000( |
| 54940339 | 54940445 | -1 | snRNA | ENSG00000( |
| 54945648 | 54961872 | 1 | protein_coding | ENSG00000( |
| 54977550 | 54981465 | 1 | protein_coding | ENSG00000( |
| 54724479 | 54736494 | 1 | protein_coding | ENSG00000( |
| 54738481 | 54752986 | 1 | protein_coding | ENSG00000( |
| 54754958 | 54767306 | 1 | unprocessed_pseudc | ENSG00000( |
| 54769746 | 54783981 | 1 | protein_coding | ENSG00000( |
| 54786398 | 54798220 | 1 | transcribed_unproce | ENSG00000( |
| 54803572 | 54814476 | 1 | protein_coding | ENSG00000( |
| 54816432 | 54830744 | 1 | protein_coding | ENSG00000( |
| 54832686 | 54848263 | 1 | polymorphic_pseudc | ENSG00000( |
| 54850408 | 54867186 | 1 | protein_coding | ENSG00000( |
| 54868910 | 54869016 | -1 | snRNA | ENSG00000( |
| 54874219 | 54890443 | 1 | protein_coding | ENSG00000( |
| 54906121 | 54910036 | 1 | protein_coding | ENSG00000( |
| 54709037 | 54713341 | 1 | transcribed_unproce | ENSG00000( |
| 54724486 | 54736514 | 1 | protein_coding | ENSG00000( |
| 54738501 | 54769226 | 1 | protein_coding | ENSG00000( |
| 54754632 | 54769000 | 1 | protein_coding | ENSG00000( |
| 54771258 | 54784662 | 1 | protein_coding | ENSG00000( |

| | | | | |
|---|---|---|---|---|
| 54789912 | 54800893 | 1 | protein_coding | ENSG00000( |
| 54802859 | 54817155 | 1 | protein_coding | ENSG00000( |
| 54819131 | 54834528 | 1 | polymorphic_pseudo | ENSG00000( |
| 54836820 | 54853592 | 1 | protein_coding | ENSG00000( |
| 54855316 | 54855422 | -1 | snRNA | ENSG00000( |
| 54860626 | 54876850 | 1 | protein_coding | ENSG00000( |
| 54892528 | 54896443 | 1 | protein_coding | ENSG00000( |
| 54726054 | 54736537 | 1 | protein_coding | ENSG00000( |
| 54738538 | 54753053 | 1 | protein_coding | ENSG00000( |
| 54755024 | 54767369 | 1 | unprocessed_pseudo | ENSG00000( |
| 54769838 | 54784324 | 1 | protein_coding | ENSG00000( |
| 54786463 | 54790318 | 1 | unprocessed_pseudo | ENSG00000( |
| 54803599 | 54814548 | 1 | protein_coding | ENSG00000( |
| 54816470 | 54830813 | 1 | protein_coding | ENSG00000( |
| 54832771 | 54848624 | 1 | protein_coding | ENSG00000( |
| 54850500 | 54867258 | 1 | protein_coding | ENSG00000( |
| 54868982 | 54869088 | -1 | snRNA | ENSG00000( |
| 54874290 | 54891462 | 1 | protein_coding | ENSG00000( |
| 54906192 | 54910107 | 1 | protein_coding | ENSG00000( |
| 54700593 | 54700905 | 1 | unprocessed_pseudo | ENSG00000( |
| 54703952 | 54704235 | 1 | unprocessed_pseudo | ENSG00000( |
| 54707916 | 54713453 | 1 | transcribed_processe | ENSG00000( |
| 54727609 | 54736462 | 1 | protein_coding | ENSG00000( |
| 54738530 | 54753070 | 1 | protein_coding | ENSG00000( |
| 54755098 | 54764695 | 1 | protein_coding | ENSG00000( |
| 54766539 | 54781397 | 1 | protein_coding | ENSG00000( |
| 54783578 | 54795926 | 1 | unprocessed_pseudo | ENSG00000( |
| 54798381 | 54812886 | 1 | protein_coding | ENSG00000( |
| 54814901 | 54878560 | 1 | protein_coding | ENSG00000( |
| 54814919 | 54820584 | 1 | unprocessed_pseudo | ENSG00000( |
| 54833799 | 54844782 | 1 | protein_coding | ENSG00000( |
| 54863003 | 54878657 | 1 | polymorphic_pseudo | ENSG00000( |
| 54880532 | 54897308 | 1 | protein_coding | ENSG00000( |
| 54899032 | 54899138 | -1 | snRNA | ENSG00000( |
| 54700593 | 54700905 | 1 | unprocessed_pseudo | ENSG00000( |
| 54703952 | 54704235 | 1 | unprocessed_pseudo | ENSG00000( |
| 54707916 | 54713453 | 1 | transcribed_processe | ENSG00000( |
| 54724478 | 54736594 | 1 | protein_coding | ENSG00000( |
| 54738484 | 54752990 | 1 | protein_coding | ENSG00000( |
| 54754924 | 54767305 | 1 | unprocessed_pseudo | ENSG00000( |
| 54769748 | 54784069 | 1 | protein_coding | ENSG00000( |
| 54786397 | 54790252 | 1 | unprocessed_pseudo | ENSG00000( |
| 54803533 | 54814482 | 1 | protein_coding | ENSG00000( |
| 54816442 | 54830745 | 1 | protein_coding | ENSG00000( |
| 54832694 | 54847510 | 1 | protein_coding | ENSG00000( |
| 54849551 | 54866326 | 1 | protein_coding | ENSG00000( |
| 54868050 | 54868156 | -1 | snRNA | ENSG00000( |
| 54698377 | 54701106 | 1 | transcribed_unproce | ENSG00000( |
| 54707916 | 54713341 | 1 | transcribed_unproce | ENSG00000( |
| 54724448 | 54736134 | 1 | protein_coding | ENSG00000( |

| | | | | |
|---|---|---|---|---|
| 54738568 | 54752475 | 1 | protein_coding | ENSG00000( |
| 54754722 | 54769049 | 1 | protein_coding | ENSG00000( |
| 54771241 | 54780725 | 1 | protein_coding | ENSG00000( |
| 54782658 | 54797195 | 1 | protein_coding | ENSG00000( |
| 54799680 | 54812030 | 1 | unprocessed_pseudo | ENSG00000( |
| 54814475 | 54828769 | 1 | protein_coding | ENSG00000( |
| 54831023 | 54842954 | 1 | transcribed_unproce | ENSG00000( |
| 54848193 | 54859216 | 1 | protein_coding | ENSG00000( |
| 54861173 | 54875478 | 1 | protein_coding | ENSG00000( |
| 54877376 | 54892982 | 1 | polymorphic_pseudo | ENSG00000( |
| 54895138 | 54911932 | 1 | protein_coding | ENSG00000( |
| 54913656 | 54913762 | -1 | snRNA | ENSG00000( |
| 54700593 | 54700905 | 1 | unprocessed_pseudo | ENSG00000( |
| 54703952 | 54704235 | 1 | unprocessed_pseudo | ENSG00000( |
| 54707916 | 54713453 | 1 | transcribed_processe | ENSG00000( |
| 54724447 | 54736132 | 1 | protein_coding | ENSG00000( |
| 54738514 | 54753065 | 1 | protein_coding | ENSG00000( |
| 54754987 | 54767361 | 1 | unprocessed_pseudo | ENSG00000( |
| 54769825 | 54784337 | 1 | protein_coding | ENSG00000( |
| 54786474 | 54798295 | 1 | transcribed_unproce | ENSG00000( |
| 54803379 | 54814527 | 1 | protein_coding | ENSG00000( |
| 54816448 | 54830767 | 1 | protein_coding | ENSG00000( |
| 54832788 | 54848156 | 1 | polymorphic_pseudo | ENSG00000( |
| 54850300 | 54867064 | 1 | protein_coding | ENSG00000( |
| 54868796 | 54868902 | -1 | snRNA | ENSG00000( |
| 54700593 | 54700905 | 1 | unprocessed_pseudo | ENSG00000( |
| 54703952 | 54704235 | 1 | unprocessed_pseudo | ENSG00000( |
| 54707916 | 54713453 | 1 | transcribed_processe | ENSG00000( |
| 54724479 | 54736497 | 1 | protein_coding | ENSG00000( |
| 54738502 | 54752797 | 1 | protein_coding | ENSG00000( |
| 54754688 | 54769216 | 1 | protein_coding | ENSG00000( |
| 54771247 | 54780763 | 1 | protein_coding | ENSG00000( |
| 54782627 | 54797062 | 1 | protein_coding | ENSG00000( |
| 54799416 | 54812309 | 1 | unprocessed_pseudo | ENSG00000( |
| 54814247 | 54828575 | 1 | protein_coding | ENSG00000( |
| 54830903 | 54842719 | 1 | transcribed_unproce | ENSG00000( |
| 54837484 | 54858960 | 1 | protein_coding | ENSG00000( |
| 54860882 | 54875495 | 1 | protein_coding | ENSG00000( |
| 54877583 | 54887079 | 1 | protein_coding | ENSG00000( |
| 54888943 | 54903379 | 1 | protein_coding | ENSG00000( |
| 54905771 | 54920219 | 1 | protein_coding | ENSG00000( |
| 54922093 | 54938887 | 1 | protein_coding | ENSG00000( |
| 54940611 | 54940717 | -1 | snRNA | ENSG00000( |
| 54700593 | 54700905 | 1 | unprocessed_pseudo | ENSG00000( |
| 54703952 | 54704235 | 1 | unprocessed_pseudo | ENSG00000( |
| 54707916 | 54713453 | 1 | transcribed_processe | ENSG00000( |
| 54724447 | 54736085 | 1 | protein_coding | ENSG00000( |
| 54738465 | 54752989 | 1 | protein_coding | ENSG00000( |
| 54754923 | 54767817 | 1 | unprocessed_pseudo | ENSG00000( |
| 54769756 | 54784259 | 1 | protein_coding | ENSG00000( |

| | | | | |
|---|---|---|---|---|
| 54786292 | 54798222 | 1 | transcribed_unproce | ENSG0000C |
| 54803296 | 54814484 | 1 | protein_coding | ENSG0000C |
| 54816441 | 54830746 | 1 | protein_coding | ENSG0000C |
| 54832726 | 54848250 | 1 | polymorphic_pseudc | ENSG0000C |
| 54850406 | 54867200 | 1 | protein_coding | ENSG0000C |
| 54700593 | 54700905 | 1 | unprocessed_pseudc | ENSG0000C |
| 54703952 | 54704235 | 1 | unprocessed_pseudc | ENSG0000C |
| 54707923 | 54713345 | 1 | transcribed_processe | ENSG0000C |
| 54724454 | 54736547 | 1 | protein_coding | ENSG0000C |
| 54738560 | 54752721 | 1 | protein_coding | ENSG0000C |
| 54754663 | 54769262 | 1 | protein_coding | ENSG0000C |
| 54771283 | 54784696 | 1 | transcribed_unproce | ENSG0000C |
| 54789863 | 54800933 | 1 | protein_coding | ENSG0000C |
| 54802891 | 54817199 | 1 | protein_coding | ENSG0000C |
| 54819179 | 54834168 | 1 | polymorphic_pseudc | ENSG0000C |
| 54836668 | 54853397 | 1 | protein_coding | ENSG0000C |
| 54855129 | 54855235 | -1 | snRNA | ENSG0000C |
| 54700593 | 54700905 | 1 | unprocessed_pseudc | ENSG0000C |
| 54703952 | 54704235 | 1 | unprocessed_pseudc | ENSG0000C |
| 54707916 | 54713453 | 1 | transcribed_processe | ENSG0000C |
| 54724452 | 54736131 | 1 | protein_coding | ENSG0000C |
| 54738461 | 54752744 | 1 | protein_coding | ENSG0000C |
| 54755019 | 54767464 | 1 | unprocessed_pseudc | ENSG0000C |
| 54769815 | 54784318 | 1 | protein_coding | ENSG0000C |
| 54786351 | 54798281 | 1 | transcribed_unproce | ENSG0000C |
| 54803354 | 54814544 | 1 | protein_coding | ENSG0000C |
| 54816466 | 54830808 | 1 | protein_coding | ENSG0000C |
| 54832707 | 54848622 | 1 | protein_coding | ENSG0000C |
| 54850495 | 54867253 | 1 | protein_coding | ENSG0000C |
| 54868977 | 54869083 | -1 | snRNA | ENSG0000C |
| 54724485 | 54736502 | 1 | protein_coding | ENSG0000C |
| 54738507 | 54752802 | 1 | protein_coding | ENSG0000C |
| 54754693 | 54769187 | 1 | protein_coding | ENSG0000C |
| 54771163 | 54780643 | 1 | protein_coding | ENSG0000C |
| 54782575 | 54796808 | 1 | protein_coding | ENSG0000C |
| 54799454 | 54811769 | 1 | unprocessed_pseudc | ENSG0000C |
| 54814235 | 54828734 | 1 | protein_coding | ENSG0000C |
| 54830833 | 54834726 | 1 | unprocessed_pseudc | ENSG0000C |
| 54848033 | 54858928 | 1 | protein_coding | ENSG0000C |
| 54860886 | 54875194 | 1 | protein_coding | ENSG0000C |
| 54877174 | 54891588 | 1 | unprocessed_pseudc | ENSG0000C |
| 54707923 | 54712937 | 1 | transcribed_unproce | ENSG0000C |
| 54723963 | 54736023 | 1 | protein_coding | ENSG0000C |
| 54738018 | 54752516 | 1 | protein_coding | ENSG0000C |
| 54754487 | 54766832 | 1 | unprocessed_pseudc | ENSG0000C |
| 54769275 | 54783790 | 1 | protein_coding | ENSG0000C |
| 54785929 | 54797435 | 1 | unprocessed_pseudc | ENSG0000C |
| 54802913 | 54813989 | 1 | protein_coding | ENSG0000C |
| 54815911 | 54830525 | 1 | protein_coding | ENSG0000C |
| 54832558 | 54842109 | 1 | protein_coding | ENSG0000C |

| | | | | |
|---|---|---|---|---|
| 54843950 | 54858409 | 1 | protein_coding | ENSG00000 |
| 54860710 | 54875253 | 1 | protein_coding | ENSG00000 |
| 54877123 | 54893917 | 1 | protein_coding | ENSG00000 |
| 54895641 | 54895747 | -1 | snRNA | ENSG00000 |
| 54707913 | 54713335 | 1 | transcribed_unproce | ENSG00000 |
| 54724478 | 54736626 | 1 | protein_coding | ENSG00000 |
| 54738498 | 54752702 | 1 | protein_coding | ENSG00000 |
| 54754680 | 54769221 | 1 | protein_coding | ENSG00000 |
| 54771253 | 54784657 | 1 | protein_coding | ENSG00000 |
| 54789822 | 54800920 | 1 | protein_coding | ENSG00000 |
| 54802842 | 54817184 | 1 | protein_coding | ENSG00000 |
| 54819083 | 54833565 | 1 | protein_coding | ENSG00000 |
| 54835435 | 54852194 | 1 | protein_coding | ENSG00000 |
| 54724503 | 54752350 | 1 | protein_coding | ENSG00000 |
| 54758126 | 54760236 | 1 | protein_coding | ENSG00000 |
| 54771254 | 54780267 | 1 | protein_coding | ENSG00000 |
| 54782674 | 54797009 | 1 | protein_coding | ENSG00000 |
| 54799641 | 54813469 | 1 | protein_coding | ENSG00000 |
| 54815973 | 54832763 | 1 | protein_coding | ENSG00000 |
| 54834495 | 54834601 | -1 | snRNA | ENSG00000 |
| 54839791 | 54856976 | 1 | protein_coding | ENSG00000 |
| 54856229 | 54856976 | 1 | lncRNA | ENSG00000 |
| 54871744 | 54878424 | 1 | protein_coding | ENSG00000 |
| 54696604 | 54700905 | 1 | unprocessed_pseudo | ENSG00000 |
| 54703952 | 54704235 | 1 | unprocessed_pseudo | ENSG00000 |
| 54707916 | 54713453 | 1 | transcribed_processe | ENSG00000 |
| 54724485 | 54736499 | 1 | protein_coding | ENSG00000 |
| 54724502 | 54798256 | 1 | lncRNA | ENSG00000 |
| 54738477 | 54753024 | 1 | protein_coding | ENSG00000 |
| 54754995 | 54767341 | 1 | unprocessed_pseudo | ENSG00000 |
| 54769766 | 54784293 | 1 | protein_coding | ENSG00000 |
| 54786326 | 54790289 | 1 | transcribed_unproce | ENSG00000 |
| 54803569 | 54814519 | 1 | protein_coding | ENSG00000 |
| 54816470 | 54830783 | 1 | protein_coding | ENSG00000 |
| 54832712 | 54848602 | 1 | protein_coding | ENSG00000 |
| 54850469 | 54867221 | 1 | protein_coding | ENSG00000 |
| 54696604 | 54700905 | 1 | unprocessed_pseudo | ENSG00000 |
| 54703952 | 54704235 | 1 | unprocessed_pseudo | ENSG00000 |
| 54707916 | 54713453 | 1 | transcribed_processe | ENSG00000 |
| 54724502 | 54784651 | 1 | lncRNA | ENSG00000 |
| 54758122 | 54760232 | 1 | protein_coding | ENSG00000 |
| 54789901 | 54800883 | 1 | protein_coding | ENSG00000 |
| 54802834 | 54817144 | 1 | protein_coding | ENSG00000 |
| 54819042 | 54834935 | 1 | polymorphic_pseudo | ENSG00000 |
| 54836809 | 54853573 | 1 | protein_coding | ENSG00000 |
| 54724479 | 54736534 | 1 | protein_coding | ENSG00000 |
| 54738512 | 54753028 | 1 | protein_coding | ENSG00000 |
| 54754999 | 54767347 | 1 | unprocessed_pseudo | ENSG00000 |
| 54769801 | 54784302 | 1 | protein_coding | ENSG00000 |
| 54786437 | 54797952 | 1 | unprocessed_pseudo | ENSG00000 |

| | | | | |
|---|---|---|---|---|
| 54803616 | 54814523 | 1 | protein_coding | ENSG00000( |
| 54816482 | 54830789 | 1 | protein_coding | ENSG00000( |
| 54832687 | 54848602 | 1 | protein_coding | ENSG00000( |
| 54850475 | 54867233 | 1 | protein_coding | ENSG00000( |
| 54868957 | 54869063 | -1 | snRNA | ENSG00000( |
| 54700593 | 54700905 | 1 | unprocessed_pseudo | ENSG00000( |
| 54703952 | 54704235 | 1 | unprocessed_pseudo | ENSG00000( |
| 54707916 | 54713453 | 1 | transcribed_processe | ENSG00000( |
| 54724481 | 54736671 | 1 | protein_coding | ENSG00000( |
| 54738547 | 54752855 | 1 | protein_coding | ENSG00000( |
| 54754729 | 54769262 | 1 | protein_coding | ENSG00000( |
| 54771294 | 54784697 | 1 | protein_coding | ENSG00000( |
| 54789862 | 54800938 | 1 | protein_coding | ENSG00000( |
| 54802886 | 54817473 | 1 | protein_coding | ENSG00000( |
| 54819506 | 54828970 | 1 | protein_coding | ENSG00000( |
| 54830927 | 54845947 | 1 | protein_coding | ENSG00000( |
| 54847887 | 54862123 | 1 | protein_coding | ENSG00000( |
| 54864251 | 54881045 | 1 | protein_coding | ENSG00000( |
| 54882769 | 54882875 | -1 | snRNA | ENSG00000( |
| 54700593 | 54700905 | 1 | unprocessed_pseudo | ENSG00000( |
| 54707916 | 54713453 | 1 | transcribed_processe | ENSG00000( |
| 54724498 | 54736060 | 1 | protein_coding | ENSG00000( |
| 54738439 | 54769215 | 1 | protein_coding | ENSG00000( |
| 54754679 | 54769001 | 1 | protein_coding | ENSG00000( |
| 54771247 | 54784650 | 1 | protein_coding | ENSG00000( |
| 54789815 | 54800891 | 1 | protein_coding | ENSG00000( |
| 54802839 | 54817200 | 1 | protein_coding | ENSG00000( |
| 54819459 | 54828923 | 1 | protein_coding | ENSG00000( |
| 54830903 | 54845777 | 1 | protein_coding | ENSG00000( |
| 54847867 | 54861881 | 1 | protein_coding | ENSG00000( |
| 54864204 | 54872914 | 1 | protein_coding | ENSG00000( |
| 54700593 | 54700905 | 1 | unprocessed_pseudo | ENSG00000( |
| 54703952 | 54704235 | 1 | unprocessed_pseudo | ENSG00000( |
| 54707916 | 54713453 | 1 | transcribed_processe | ENSG00000( |
| 54724453 | 54736091 | 1 | protein_coding | ENSG00000( |
| 54738470 | 54752993 | 1 | protein_coding | ENSG00000( |
| 54754928 | 54766523 | 1 | unprocessed_pseudo | ENSG00000( |
| 54769704 | 54783960 | 1 | protein_coding | ENSG00000( |
| 54786260 | 54790260 | 1 | unprocessed_pseudo | ENSG00000( |
| 54803565 | 54814279 | 1 | protein_coding | ENSG00000( |
| 54816451 | 54830756 | 1 | protein_coding | ENSG00000( |
| 54832654 | 54848284 | 1 | protein_coding | ENSG00000( |
| 54850443 | 54867215 | 1 | protein_coding | ENSG00000( |
| 54868939 | 54869045 | -1 | snRNA | ENSG00000( |
| 54700593 | 54700905 | 1 | unprocessed_pseudo | ENSG00000( |
| 54703952 | 54704235 | 1 | unprocessed_pseudo | ENSG00000( |
| 54707916 | 54713453 | 1 | transcribed_processe | ENSG00000( |
| 54724450 | 54736092 | 1 | protein_coding | ENSG00000( |
| 54738519 | 54752638 | 1 | protein_coding | ENSG00000( |
| 54754673 | 54769220 | 1 | protein_coding | ENSG00000( |

| | | | | |
|---|---|---|---|---|
| 54771198 | 54780682 | 1 | protein_coding | ENSG0000C |
| 54782617 | 54797414 | 1 | protein_coding | ENSG0000C |
| 54799595 | 54812147 | 1 | unprocessed_pseudc | ENSG0000C |
| 54814388 | 54828892 | 1 | protein_coding | ENSG0000C |
| 54848094 | 54859121 | 1 | protein_coding | ENSG0000C |
| 54861074 | 54875379 | 1 | protein_coding | ENSG0000C |
| 54877320 | 54892899 | 1 | polymorphic_pseudc | ENSG0000C |
| 54700593 | 54700905 | 1 | unprocessed_pseudc | ENSG0000C |
| 54703952 | 54704235 | 1 | unprocessed_pseudc | ENSG0000C |
| 54707916 | 54713453 | 1 | transcribed_processe | ENSG0000C |
| 54724451 | 54736098 | 1 | protein_coding | ENSG0000C |
| 54738496 | 54752881 | 1 | protein_coding | ENSG0000C |
| 54754966 | 54767314 | 1 | unprocessed_pseudc | ENSG0000C |
| 54769705 | 54783961 | 1 | protein_coding | ENSG0000C |
| 54786406 | 54790261 | 1 | unprocessed_pseudc | ENSG0000C |
| 54803571 | 54814487 | 1 | protein_coding | ENSG0000C |
| 54816451 | 54830756 | 1 | protein_coding | ENSG0000C |
| 54832654 | 54848283 | 1 | protein_coding | ENSG0000C |
| 54850442 | 54867200 | 1 | protein_coding | ENSG0000C |
| 54868924 | 54869030 | -1 | snRNA | ENSG0000C |
| 54698377 | 54700905 | -1 | unprocessed_pseudc | ENSG0000C |
| 54707916 | 54713453 | 1 | transcribed_processe | ENSG0000C |
| 54724485 | 54736504 | 1 | protein_coding | ENSG0000C |
| 54738476 | 54752501 | 1 | protein_coding | ENSG0000C |
| 54754600 | 54768961 | 1 | protein_coding | ENSG0000C |
| 54771219 | 54784641 | 1 | transcribed_unproce | ENSG0000C |
| 54789806 | 54800882 | 1 | protein_coding | ENSG0000C |
| 54802804 | 54817417 | 1 | protein_coding | ENSG0000C |
| 54819505 | 54829001 | 1 | protein_coding | ENSG0000C |
| 54830865 | 54845301 | 1 | protein_coding | ENSG0000C |
| 54847678 | 54861692 | 1 | protein_coding | ENSG0000C |
| 54864015 | 54880809 | 1 | protein_coding | ENSG0000C |
| 54882533 | 54882639 | -1 | snRNA | ENSG0000C |
| 54040833 | 54063953 | -1 | protein_coding | ENSG0000C |
| 54069947 | 54081380 | -1 | protein_coding | ENSG0000C |
| 54094681 | 54102740 | -1 | protein_coding | ENSG0000C |
| 54102906 | 54107026 | 1 | protein_coding | ENSG0000C |
| 54107066 | 54115801 | -1 | protein_coding | ENSG0000C |
| 54115536 | 54131896 | 1 | protein_coding | ENSG0000C |
| 54138182 | 54156190 | 1 | protein_coding | ENSG0000C |
| 54156104 | 54160226 | -1 | protein_coding | ENSG0000C |
| 54160591 | 54173655 | -1 | protein_coding | ENSG0000C |
| 54173818 | 54190312 | -1 | protein_coding | ENSG0000C |
| 54190368 | 54194965 | 1 | protein_coding | ENSG0000C |
| 54201241 | 54249371 | 1 | protein_coding | ENSG0000C |
| 54217107 | 54223520 | -1 | protein_coding | ENSG0000C |
| 54229470 | 54230647 | 1 | processed_pseudoge | ENSG0000C |
| 54238892 | 54243172 | -1 | protein_coding | ENSG0000C |
| 54250781 | 54257977 | -1 | protein_coding | ENSG0000C |
| 54264323 | 54264425 | -1 | snRNA | ENSG0000C |

| | | | | |
|---|---|---|---|---|
| 54268383 | 54269416 | 1 | processed_pseudoge | ENSG0000C |
| 54274195 | 54281689 | -1 | protein_coding | ENSG0000C |
| 54294163 | 54306461 | -1 | protein_coding | ENSG0000C |
| 54314131 | 54327636 | -1 | protein_coding | ENSG0000C |
| 54347370 | 54369057 | 1 | protein_coding | ENSG0000C |
| 54362810 | 54366855 | -1 | lncRNA | ENSG0000C |
| 54380864 | 54394163 | 1 | protein_coding | ENSG0000C |
| 54393938 | 54395858 | -1 | protein_coding | ENSG0000C |
| 54397172 | 54405337 | -1 | protein_coding | ENSG0000C |
| 54429706 | 54442942 | 1 | protein_coding | ENSG0000C |
| 54464632 | 54477600 | 1 | protein_coding | ENSG0000C |
| 54504943 | 54519583 | 1 | protein_coding | ENSG0000C |
| 54525581 | 54534404 | 1 | protein_coding | ENSG0000C |
| 54548949 | 54569575 | 1 | protein_coding | ENSG0000C |
| 54575908 | 54581617 | 1 | transcribed_unproce | ENSG0000C |
| 54594151 | 54602395 | 1 | protein_coding | ENSG0000C |
| 54628233 | 54632455 | 1 | unprocessed_pseudc | ENSG0000C |
| 54639543 | 54644657 | 1 | transcribed_unproce | ENSG0000C |
| 54655800 | 54667814 | 1 | protein_coding | ENSG0000C |
| 54669793 | 54684345 | 1 | protein_coding | ENSG0000C |
| 54686316 | 54698662 | 1 | unprocessed_pseudc | ENSG0000C |
| 54701102 | 54715618 | 1 | protein_coding | ENSG0000C |
| 54717647 | 54721610 | 1 | unprocessed_pseudc | ENSG0000C |
| 54835171 | 54842110 | 1 | protein_coding | ENSG0000C |
| 54852540 | 54876536 | -1 | protein_coding | ENSG0000C |
| 54894315 | 54930171 | 1 | protein_coding | ENSG0000C |
| 54942736 | 54967295 | -1 | protein_coding | ENSG0000C |
| 54973355 | 54992248 | -1 | protein_coding | ENSG0000C |
| 54040833 | 54063953 | -1 | protein_coding | ENSG0000C |
| 54069947 | 54081380 | -1 | protein_coding | ENSG0000C |
| 54094681 | 54102740 | -1 | protein_coding | ENSG0000C |
| 54102906 | 54107026 | 1 | protein_coding | ENSG0000C |
| 54107066 | 54115801 | -1 | protein_coding | ENSG0000C |
| 54115536 | 54131896 | 1 | protein_coding | ENSG0000C |
| 54138182 | 54156191 | 1 | protein_coding | ENSG0000C |
| 54156127 | 54160192 | -1 | protein_coding | ENSG0000C |
| 54160649 | 54173691 | -1 | protein_coding | ENSG0000C |
| 54173854 | 54190480 | -1 | protein_coding | ENSG0000C |
| 54190866 | 54195141 | 1 | protein_coding | ENSG0000C |
| 54201473 | 54208260 | 1 | protein_coding | ENSG0000C |
| 54294043 | 54306352 | -1 | protein_coding | ENSG0000C |
| 54314055 | 54320118 | -1 | protein_coding | ENSG0000C |
| 54315894 | 54346149 | -1 | protein_coding | ENSG0000C |
| 54360692 | 54377679 | -1 | protein_coding | ENSG0000C |
| 54426396 | 54427817 | 1 | protein_coding | ENSG0000C |
| 54429845 | 54441468 | 1 | unprocessed_pseudc | ENSG0000C |
| 54446939 | 54458015 | 1 | protein_coding | ENSG0000C |
| 54459987 | 54474463 | 1 | protein_coding | ENSG0000C |
| 54476583 | 54485924 | 1 | protein_coding | ENSG0000C |
| 54488004 | 54503024 | 1 | protein_coding | ENSG0000C |

| | | | | |
|---|---|---|---|---|
| 54504993 | 54518921 | 1 | protein_coding | ENSG00000 |
| 54521328 | 54538122 | 1 | protein_coding | ENSG00000 |
| 54539846 | 54539952 | -1 | snRNA | ENSG00000 |
| 54545136 | 54562282 | 1 | protein_coding | ENSG00000 |
| 54576966 | 54586961 | 1 | protein_coding | ENSG00000 |
| 54594345 | 54618340 | -1 | protein_coding | ENSG00000 |
| 54636119 | 54671975 | 1 | protein_coding | ENSG00000 |
| 54684540 | 54709099 | -1 | protein_coding | ENSG00000 |
| 54715159 | 54734052 | -1 | protein_coding | ENSG00000 |
| 54040833 | 54063953 | -1 | protein_coding | ENSG00000 |
| 54069947 | 54081380 | -1 | protein_coding | ENSG00000 |
| 54094681 | 54102740 | -1 | protein_coding | ENSG00000 |
| 54102906 | 54107026 | 1 | protein_coding | ENSG00000 |
| 54107066 | 54115801 | -1 | protein_coding | ENSG00000 |
| 54115536 | 54131896 | 1 | protein_coding | ENSG00000 |
| 54138182 | 54156191 | 1 | protein_coding | ENSG00000 |
| 54156127 | 54160192 | -1 | protein_coding | ENSG00000 |
| 54160649 | 54173691 | -1 | protein_coding | ENSG00000 |
| 54173854 | 54190480 | -1 | protein_coding | ENSG00000 |
| 54190866 | 54195141 | 1 | protein_coding | ENSG00000 |
| 54201473 | 54208260 | 1 | protein_coding | ENSG00000 |
| 54746804 | 54761173 | 1 | protein_coding | ENSG00000 |
| 54763061 | 54777585 | 1 | protein_coding | ENSG00000 |
| 54763075 | 54854081 | 1 | protein_coding | ENSG00000 |
| 54779616 | 54785052 | 1 | unprocessed_pseudc | ENSG00000 |
| 54855924 | 54864640 | 1 | protein_coding | ENSG00000 |
| 54911759 | 54918698 | 1 | protein_coding | ENSG00000 |
| 54929128 | 54953124 | -1 | protein_coding | ENSG00000 |
| 54970903 | 55006759 | 1 | protein_coding | ENSG00000 |
| 55019324 | 55043883 | -1 | protein_coding | ENSG00000 |
| 55049943 | 55068836 | -1 | protein_coding | ENSG00000 |
| 54040833 | 54063953 | -1 | protein_coding | ENSG00000 |
| 54069947 | 54081380 | -1 | protein_coding | ENSG00000 |
| 54094681 | 54102740 | -1 | protein_coding | ENSG00000 |
| 54102906 | 54107026 | 1 | protein_coding | ENSG00000 |
| 54107066 | 54115801 | -1 | protein_coding | ENSG00000 |
| 54115536 | 54131896 | 1 | protein_coding | ENSG00000 |
| 54138182 | 54156191 | 1 | protein_coding | ENSG00000 |
| 54156127 | 54160192 | -1 | protein_coding | ENSG00000 |
| 54160649 | 54173691 | -1 | protein_coding | ENSG00000 |
| 54173854 | 54190480 | -1 | protein_coding | ENSG00000 |
| 54190866 | 54195141 | 1 | protein_coding | ENSG00000 |
| 54201473 | 54208260 | 1 | protein_coding | ENSG00000 |
| 54838897 | 54850914 | 1 | protein_coding | ENSG00000 |
| 54852887 | 54865626 | 1 | protein_coding | ENSG00000 |
| 54867554 | 54882047 | 1 | protein_coding | ENSG00000 |
| 54883911 | 54885779 | 1 | protein_coding | ENSG00000 |
| 54939296 | 54946235 | 1 | protein_coding | ENSG00000 |
| 54956665 | 54980661 | -1 | protein_coding | ENSG00000 |
| 54998440 | 55034296 | 1 | protein_coding | ENSG00000 |

| | | | | |
|---|---|---|---|---|
| 55046861 | 55071420 | -1 | protein_coding | ENSG00000 |
| 55077480 | 55096373 | -1 | protein_coding | ENSG00000 |
| 54040833 | 54063953 | -1 | protein_coding | ENSG00000 |
| 54069947 | 54081380 | -1 | protein_coding | ENSG00000 |
| 54094681 | 54102740 | -1 | protein_coding | ENSG00000 |
| 54102906 | 54107026 | 1 | protein_coding | ENSG00000 |
| 54107066 | 54115801 | -1 | protein_coding | ENSG00000 |
| 54115536 | 54131896 | 1 | protein_coding | ENSG00000 |
| 54138182 | 54156191 | 1 | protein_coding | ENSG00000 |
| 54156127 | 54160192 | -1 | protein_coding | ENSG00000 |
| 54160649 | 54173691 | -1 | protein_coding | ENSG00000 |
| 54173854 | 54190480 | -1 | protein_coding | ENSG00000 |
| 54190866 | 54195141 | 1 | protein_coding | ENSG00000 |
| 54201473 | 54208260 | 1 | protein_coding | ENSG00000 |
| 54845266 | 54856124 | 1 | protein_coding | ENSG00000 |
| 54857994 | 54874788 | 1 | protein_coding | ENSG00000 |
| 54876512 | 54876618 | -1 | snRNA | ENSG00000 |
| 54913845 | 54920784 | 1 | protein_coding | ENSG00000 |
| 54931214 | 54955210 | -1 | protein_coding | ENSG00000 |
| 54972989 | 55008845 | 1 | protein_coding | ENSG00000 |
| 55021410 | 55045969 | -1 | protein_coding | ENSG00000 |
| 55052029 | 55070922 | -1 | protein_coding | ENSG00000 |
| 54040833 | 54063953 | -1 | protein_coding | ENSG00000 |
| 54069947 | 54081380 | -1 | protein_coding | ENSG00000 |
| 54094681 | 54102740 | -1 | protein_coding | ENSG00000 |
| 54102906 | 54107026 | 1 | protein_coding | ENSG00000 |
| 54107066 | 54115801 | -1 | protein_coding | ENSG00000 |
| 54115536 | 54131896 | 1 | protein_coding | ENSG00000 |
| 54138182 | 54156191 | 1 | protein_coding | ENSG00000 |
| 54156127 | 54160192 | -1 | protein_coding | ENSG00000 |
| 54160649 | 54173691 | -1 | protein_coding | ENSG00000 |
| 54173854 | 54190480 | -1 | protein_coding | ENSG00000 |
| 54190866 | 54195141 | 1 | protein_coding | ENSG00000 |
| 54201473 | 54208260 | 1 | protein_coding | ENSG00000 |
| 54733008 | 54744567 | 1 | protein_coding | ENSG00000 |
| 54746977 | 54761521 | 1 | protein_coding | ENSG00000 |
| 54763492 | 54775840 | 1 | unprocessed_pseudo | ENSG00000 |
| 54778277 | 54792578 | 1 | protein_coding | ENSG00000 |
| 54794614 | 54811438 | 1 | protein_coding | ENSG00000 |
| 54813162 | 54813268 | -1 | snRNA | ENSG00000 |
| 54850138 | 54857077 | 1 | protein_coding | ENSG00000 |
| 54867507 | 54891503 | -1 | protein_coding | ENSG00000 |
| 54909282 | 54945138 | 1 | protein_coding | ENSG00000 |
| 54957703 | 54982262 | -1 | protein_coding | ENSG00000 |
| 54988322 | 55007215 | -1 | protein_coding | ENSG00000 |
| 54040833 | 54063953 | -1 | protein_coding | ENSG00000 |
| 54069947 | 54081380 | -1 | protein_coding | ENSG00000 |
| 54094681 | 54102740 | -1 | protein_coding | ENSG00000 |
| 54102906 | 54107026 | 1 | protein_coding | ENSG00000 |
| 54107066 | 54115801 | -1 | protein_coding | ENSG00000 |

| | | | | |
|---|---|---|---|---|
| 54115536 | 54131896 | 1 | protein_coding | ENSG0000C |
| 54138182 | 54156191 | 1 | protein_coding | ENSG0000C |
| 54156107 | 54160227 | -1 | protein_coding | ENSG0000C |
| 54160592 | 54173661 | -1 | protein_coding | ENSG0000C |
| 54173824 | 54190479 | -1 | protein_coding | ENSG0000C |
| 54190535 | 54195132 | 1 | protein_coding | ENSG0000C |
| 54201424 | 54249604 | 1 | protein_coding | ENSG0000C |
| 54217293 | 54223789 | -1 | protein_coding | ENSG0000C |
| 54229605 | 54230979 | 1 | processed_pseudoge | ENSG0000C |
| 54239108 | 54243457 | -1 | protein_coding | ENSG0000C |
| 54251026 | 54258210 | -1 | protein_coding | ENSG0000C |
| 54264566 | 54264668 | -1 | snRNA | ENSG0000C |
| 54268619 | 54269662 | 1 | processed_pseudoge | ENSG0000C |
| 54274438 | 54278995 | -1 | protein_coding | ENSG0000C |
| 54283996 | 54300924 | -1 | protein_coding | ENSG0000C |
| 54345546 | 54367221 | 1 | protein_coding | ENSG0000C |
| 54360973 | 54365019 | -1 | lncRNA | ENSG0000C |
| 54379038 | 54392337 | 1 | protein_coding | ENSG0000C |
| 54392112 | 54394035 | -1 | protein_coding | ENSG0000C |
| 54395351 | 54403578 | -1 | protein_coding | ENSG0000C |
| 54428232 | 54441459 | 1 | protein_coding | ENSG0000C |
| 54463149 | 54476118 | 1 | protein_coding | ENSG0000C |
| 54503437 | 54518077 | 1 | protein_coding | ENSG0000C |
| 54524093 | 54532898 | 1 | protein_coding | ENSG0000C |
| 54547681 | 54568080 | 1 | protein_coding | ENSG0000C |
| 54574401 | 54580112 | 1 | transcribed_unproce | ENSG0000C |
| 54592642 | 54600251 | 1 | protein_coding | ENSG0000C |
| 54626866 | 54631219 | 1 | unprocessed_pseudc | ENSG0000C |
| 54638174 | 54643363 | 1 | transcribed_unproce | ENSG0000C |
| 54654506 | 54666521 | 1 | protein_coding | ENSG0000C |
| 54668491 | 54683019 | 1 | protein_coding | ENSG0000C |
| 54684990 | 54697367 | 1 | unprocessed_pseudc | ENSG0000C |
| 54699775 | 54714233 | 1 | protein_coding | ENSG0000C |
| 54716322 | 54720285 | 1 | unprocessed_pseudc | ENSG0000C |
| 54733566 | 54744515 | 1 | protein_coding | ENSG0000C |
| 54746462 | 54760779 | 1 | protein_coding | ENSG0000C |
| 54762712 | 54778593 | 1 | protein_coding | ENSG0000C |
| 54780466 | 54797223 | 1 | protein_coding | ENSG0000C |
| 54798947 | 54799053 | -1 | snRNA | ENSG0000C |
| 54804243 | 54821411 | 1 | protein_coding | ENSG0000C |
| 54836140 | 54846123 | 1 | protein_coding | ENSG0000C |
| 54853492 | 54877496 | -1 | protein_coding | ENSG0000C |
| 54894717 | 54929555 | 1 | protein_coding | ENSG0000C |
| 54942120 | 54966679 | -1 | protein_coding | ENSG0000C |
| 54972739 | 54991632 | -1 | protein_coding | ENSG0000C |
| 54040833 | 54063953 | -1 | protein_coding | ENSG0000C |
| 54069947 | 54081380 | -1 | protein_coding | ENSG0000C |
| 54094681 | 54102740 | -1 | protein_coding | ENSG0000C |
| 54102906 | 54107026 | 1 | protein_coding | ENSG0000C |
| 54107066 | 54115801 | -1 | protein_coding | ENSG0000C |

| | | | | |
|---|---|---|---|---|
| 54115536 | 54131896 | 1 | protein_coding | ENSG00000( |
| 54138182 | 54156191 | 1 | protein_coding | ENSG00000( |
| 54156127 | 54160192 | -1 | protein_coding | ENSG00000( |
| 54160649 | 54173691 | -1 | protein_coding | ENSG00000( |
| 54173854 | 54190480 | -1 | protein_coding | ENSG00000( |
| 54190866 | 54195141 | 1 | protein_coding | ENSG00000( |
| 54201473 | 54208260 | 1 | protein_coding | ENSG00000( |
| 54216894 | 54243483 | -1 | protein_coding | ENSG00000( |
| 54251046 | 54258245 | -1 | protein_coding | ENSG00000( |
| 54264578 | 54264680 | -1 | snRNA | ENSG00000( |
| 54268639 | 54269669 | 1 | processed_pseudoge | ENSG00000( |
| 54274450 | 54281944 | -1 | protein_coding | ENSG00000( |
| 54294418 | 54306726 | -1 | protein_coding | ENSG00000( |
| 54314236 | 54320273 | -1 | protein_coding | ENSG00000( |
| 54316049 | 54346292 | -1 | protein_coding | ENSG00000( |
| 54361220 | 54378234 | -1 | protein_coding | ENSG00000( |
| 54388712 | 54389685 | 1 | protein_coding | ENSG00000( |
| 54396006 | 54401717 | 1 | transcribed_unproce | ENSG00000( |
| 54414251 | 54422496 | 1 | protein_coding | ENSG00000( |
| 54448346 | 54452936 | 1 | unprocessed_pseudc | ENSG00000( |
| 54459891 | 54465035 | 1 | transcribed_unproce | ENSG00000( |
| 54476158 | 54520914 | 1 | protein_coding | ENSG00000( |
| 54476178 | 54488196 | 1 | protein_coding | ENSG00000( |
| 54476196 | 54520909 | 1 | protein_coding | ENSG00000( |
| 54522942 | 54528379 | 1 | unprocessed_pseudc | ENSG00000( |
| 54643934 | 54650873 | 1 | protein_coding | ENSG00000( |
| 54661303 | 54685299 | -1 | protein_coding | ENSG00000( |
| 54703078 | 54738934 | 1 | protein_coding | ENSG00000( |
| 54751499 | 54776058 | -1 | protein_coding | ENSG00000( |
| 54782118 | 54801011 | -1 | protein_coding | ENSG00000( |
| 20674923 | 20696533 | 1 | unprocessed_pseudc | ENSG00000( |
| 20746923 | 20755250 | -1 | lncRNA | ENSG00000( |
| 20776304 | 20807420 | 1 | protein_coding | ENSG00000( |
| 20784596 | 20785152 | -1 | processed_pseudoge | ENSG00000( |
| 20815244 | 20822244 | 1 | lncRNA | ENSG00000( |
| 20844829 | 20853768 | -1 | unprocessed_pseudc | ENSG00000( |
| 20929638 | 20957113 | 1 | protein_coding | ENSG00000( |
| 20933201 | 20935130 | -1 | TEC | ENSG00000( |
| 20967473 | 20971556 | 1 | unprocessed_pseudc | ENSG00000( |
| 20978678 | 20989626 | 1 | lncRNA | ENSG00000( |
| 21027036 | 21043218 | 1 | protein_coding | ENSG00000( |
| 34172447 | 34229322 | 1 | protein_coding | ENSG00000( |
| 34261710 | 34308123 | 1 | protein_coding | ENSG00000( |
| 34303449 | 34303913 | -1 | processed_pseudoge | ENSG00000( |
| 21655154 | 21656539 | 1 | unprocessed_pseudc | ENSG00000( |
| 21687036 | 21687706 | -1 | processed_pseudoge | ENSG00000( |
| 21719001 | 21719670 | 1 | processed_pseudoge | ENSG00000( |
| 21729280 | 21729514 | -1 | unprocessed_pseudc | ENSG00000( |
| 21732245 | 21777028 | -1 | protein_coding | ENSG00000( |
| 21760226 | 21762917 | 1 | transcribed_processe | ENSG00000( |

| | | | | |
|---|---|---|---|---|
| 21778251 | 21803339 | -1 | lncRNA | ENSG00000 |
| 21809967 | 21810673 | 1 | processed_pseudoge | ENSG00000 |
| 21814429 | 21818511 | -1 | protein_coding | ENSG00000 |
| 34351514 | 34355586 | 1 | protein_coding | ENSG00000 |
| 34358485 | 34359412 | -1 | lncRNA | ENSG00000 |
| 34359480 | 34381539 | 1 | protein_coding | ENSG00000 |
| 22128748 | 22130323 | -1 | processed_pseudoge | ENSG00000 |
| 22130325 | 22130489 | 1 | processed_pseudoge | ENSG00000 |
| 22137931 | 22138981 | 1 | processed_pseudoge | ENSG00000 |
| 22140532 | 22141117 | 1 | processed_pseudoge | ENSG00000 |
| 22174766 | 22175191 | -1 | unprocessed_pseudo | ENSG00000 |
| 22179091 | 22215686 | -1 | protein_coding | ENSG00000 |
| 22203120 | 22203995 | 1 | processed_pseudoge | ENSG00000 |
| 22251461 | 22253554 | -1 | lncRNA | ENSG00000 |
| 22267900 | 22269542 | 1 | lncRNA | ENSG00000 |
| 22294615 | 22294762 | 1 | protein_coding | ENSG00000 |
| 22307184 | 22308075 | -1 | processed_pseudoge | ENSG00000 |
| 41479081 | 41500644 | -1 | lncRNA | ENSG00000 |
| 41506152 | 41506898 | 1 | processed_pseudoge | ENSG00000 |
| 41521043 | 41521989 | -1 | processed_pseudoge | ENSG00000 |
| 41531206 | 41532026 | 1 | lncRNA | ENSG00000 |
| 41543524 | 41545246 | 1 | lncRNA | ENSG00000 |
| 41557860 | 41595186 | 1 | protein_coding | ENSG00000 |
| 41557862 | 41563804 | 1 | lncRNA | ENSG00000 |
| 41559910 | 41560616 | -1 | TEC | ENSG00000 |
| 41608077 | 41614326 | 1 | transcribed_unproce | ENSG00000 |
| 41613683 | 41614013 | -1 | processed_pseudoge | ENSG00000 |
| 41627313 | 41635416 | -1 | protein_coding | ENSG00000 |
| 41646911 | 41647241 | -1 | processed_pseudoge | ENSG00000 |
| 41648001 | 41648944 | -1 | processed_pseudoge | ENSG00000 |
| 41661018 | 41661305 | 1 | processed_pseudoge | ENSG00000 |
| 41681649 | 41685845 | -1 | protein_coding | ENSG00000 |
| 1107635 | 1156979 | -1 | protein_coding | ENSG00000 |
| 54040829 | 54063953 | -1 | protein_coding | ENSG00000 |
| 54069895 | 54081380 | -1 | protein_coding | ENSG00000 |
| 54094679 | 54102746 | -1 | protein_coding | ENSG00000 |
| 54102782 | 54108779 | 1 | protein_coding | ENSG00000 |
| 54107066 | 54115801 | -1 | protein_coding | ENSG00000 |
| 54115536 | 54131896 | 1 | protein_coding | ENSG00000 |
| 54119637 | 54125468 | -1 | lncRNA | ENSG00000 |
| 54138190 | 54156165 | 1 | protein_coding | ENSG00000 |
| 54156123 | 54160192 | -1 | protein_coding | ENSG00000 |
| 54160592 | 54173691 | -1 | protein_coding | ENSG00000 |
| 54173853 | 54190413 | -1 | protein_coding | ENSG00000 |
| 54190536 | 54194332 | 1 | protein_coding | ENSG00000 |
| 54200068 | 54200585 | -1 | lncRNA | ENSG00000 |
| 54201357 | 54249609 | 1 | protein_coding | ENSG00000 |
| 54209707 | 54210033 | -1 | unprocessed_pseudo | ENSG00000 |
| 54217271 | 54223706 | -1 | protein_coding | ENSG00000 |
| 54225142 | 54225500 | 1 | lncRNA | ENSG00000 |

| | | | | |
|---|---|---|---|---|
| 54229658 | 54230835 | 1 | processed_pseudoge | ENSG00000 |
| 54237215 | 54243458 | -1 | protein_coding | ENSG00000 |
| 54246700 | 54247928 | -1 | unprocessed_pseudo | ENSG00000 |
| 54251016 | 54257914 | -1 | protein_coding | ENSG00000 |
| 54264552 | 54264654 | -1 | snRNA | ENSG00000 |
| 54268606 | 54269646 | 1 | processed_pseudoge | ENSG00000 |
| 54274422 | 54281786 | -1 | protein_coding | ENSG00000 |
| 54294391 | 54301485 | -1 | protein_coding | ENSG00000 |
| 54294649 | 54294966 | -1 | unprocessed_pseudo | ENSG00000 |
| 54305461 | 54306170 | -1 | unprocessed_pseudo | ENSG00000 |
| 54309794 | 54314056 | -1 | unprocessed_pseudo | ENSG00000 |
| 54315100 | 54321156 | -1 | protein_coding | ENSG00000 |
| 54316929 | 54345186 | -1 | lncRNA | ENSG00000 |
| 54328223 | 54329236 | -1 | transcribed_unproce | ENSG00000 |
| 54336291 | 54339453 | -1 | transcribed_unproce | ENSG00000 |
| 54341203 | 54347168 | -1 | protein_coding | ENSG00000 |
| 54359738 | 54378912 | -1 | protein_coding | ENSG00000 |
| 54387840 | 54388311 | 1 | processed_pseudoge | ENSG00000 |
| 54423352 | 54444827 | 1 | protein_coding | ENSG00000 |
| 54438579 | 54442625 | -1 | lncRNA | ENSG00000 |
| 54445939 | 54446275 | -1 | lncRNA | ENSG00000 |
| 54446594 | 54447473 | -1 | lncRNA | ENSG00000 |
| 54452738 | 54456970 | -1 | lncRNA | ENSG00000 |
| 54456812 | 54469943 | 1 | protein_coding | ENSG00000 |
| 54469723 | 54471641 | -1 | protein_coding | ENSG00000 |
| 54472957 | 54481158 | -1 | protein_coding | ENSG00000 |
| 54505847 | 54518644 | 1 | protein_coding | ENSG00000 |
| 54529653 | 54529961 | 1 | unprocessed_pseudo | ENSG00000 |
| 54540656 | 54553413 | 1 | protein_coding | ENSG00000 |
| 54560000 | 54560286 | 1 | unprocessed_pseudo | ENSG00000 |
| 54581134 | 54595774 | 1 | protein_coding | ENSG00000 |
| 54597653 | 54598499 | 1 | lncRNA | ENSG00000 |
| 54601794 | 54610302 | 1 | protein_coding | ENSG00000 |
| 54616083 | 54616589 | 1 | unprocessed_pseudo | ENSG00000 |
| 54638682 | 54645767 | 1 | protein_coding | ENSG00000 |
| 54643920 | 54647090 | -1 | lncRNA | ENSG00000 |
| 54671018 | 54678557 | 1 | protein_coding | ENSG00000 |
| 54679429 | 54680789 | 1 | unprocessed_pseudo | ENSG00000 |
| 54705130 | 54709430 | 1 | unprocessed_pseudo | ENSG00000 |
| 54712477 | 54712760 | 1 | unprocessed_pseudo | ENSG00000 |
| 54716447 | 54721701 | 1 | transcribed_processe | ENSG00000 |
| 54732699 | 54744341 | 1 | protein_coding | ENSG00000 |
| 54746721 | 54761251 | 1 | protein_coding | ENSG00000 |
| 54763222 | 54775570 | 1 | transcribed_processe | ENSG00000 |
| 54778013 | 54792334 | 1 | protein_coding | ENSG00000 |
| 54794554 | 54798517 | 1 | unprocessed_pseudo | ENSG00000 |
| 54811737 | 54822719 | 1 | protein_coding | ENSG00000 |
| 54824676 | 54838980 | 1 | protein_coding | ENSG00000 |
| 54840878 | 54856485 | 1 | polymorphic_pseudo | ENSG00000 |
| 54858645 | 54875417 | 1 | protein_coding | ENSG00000 |

| | | | | |
|---|---|---|---|---|
| 54877141 | 54877247 | -1 | snRNA | ENSG0000C |
| 54882451 | 54899533 | 1 | protein_coding | ENSG0000C |
| 54914255 | 54924255 | 1 | protein_coding | ENSG0000C |
| 54931624 | 54974427 | -1 | protein_coding | ENSG0000C |
| 54961245 | 55009257 | 1 | protein_coding | ENSG0000C |
| 55021820 | 55046379 | -1 | protein_coding | ENSG0000C |
| 55035708 | 55062043 | -1 | lncRNA | ENSG0000C |
| 55047223 | 55079406 | -1 | protein_coding | ENSG0000C |
| 821485 | 821977 | -1 | protein_coding | ENSG0000C |
| 825097 | 832018 | 1 | protein_coding | ENSG0000C |
| 840960 | 848175 | 1 | protein_coding | ENSG0000C |
| 851006 | 856290 | 1 | protein_coding | ENSG0000C |
| 859680 | 863666 | 1 | protein_coding | ENSG0000C |
| 868013 | 881626 | -1 | protein_coding | ENSG0000C |
| 30406695 | 30413531 | -1 | lncRNA | ENSG0000C |
| 30414392 | 30416755 | -1 | lncRNA | ENSG0000C |
| 30420942 | 30426225 | 1 | lncRNA | ENSG0000C |
| 2467459 | 2505530 | 1 | protein_coding | ENSG0000C |
| 2492300 | 2493258 | -1 | lncRNA | ENSG0000C |
| 2493437 | 2494479 | -1 | lncRNA | ENSG0000C |
| 2508533 | 2526600 | -1 | protein_coding | ENSG0000C |
| 2528745 | 2530245 | -1 | protein_coding | ENSG0000C |
| 2546465 | 2547460 | 1 | lncRNA | ENSG0000C |
| 2549920 | 2557031 | -1 | lncRNA | ENSG0000C |
| 2555639 | 2565622 | 1 | protein_coding | ENSG0000C |
| 2566410 | 2569888 | 1 | TEC | ENSG0000C |
| 2581560 | 2584534 | 1 | lncRNA | ENSG0000C |
| 2586491 | 2591469 | 1 | protein_coding | ENSG0000C |
| 2590639 | 2632987 | -1 | protein_coding | ENSG0000C |
| 2632565 | 2636215 | 1 | lncRNA | ENSG0000C |
| 2635571 | 2801699 | -1 | protein_coding | ENSG0000C |
| 153700659 | 153700956 | 1 | misc_RNA | ENSG0000C |
| 153713833 | 153714877 | -1 | processed_pseudoge | ENSG0000C |
| 153721328 | 153721625 | 1 | misc_RNA | ENSG0000C |
| 153734523 | 153735546 | -1 | processed_pseudoge | ENSG0000C |
| 153743483 | 153743584 | -1 | misc_RNA | ENSG0000C |
| 153745298 | 153792039 | 1 | protein_coding | ENSG0000C |
| 153764082 | 153768458 | -1 | lncRNA | ENSG0000C |
| 153768214 | 153769407 | -1 | lncRNA | ENSG0000C |
| 153789602 | 153791310 | -1 | protein_coding | ENSG0000C |
| 153791585 | 153797388 | 1 | protein_coding | ENSG0000C |
| 153802951 | 153803052 | -1 | misc_RNA | ENSG0000C |
| 153812404 | 153819322 | 1 | unprocessed_pseudc | ENSG0000C |
| 153821956 | 153845579 | -1 | protein_coding | ENSG0000C |
| 153869337 | 153870645 | -1 | processed_pseudoge | ENSG0000C |
| 245353678 | 245713153 | 1 | protein_coding | ENSG0000C |
| 13076471 | 13079295 | -1 | unprocessed_pseudc | ENSG0000C |
| 13093633 | 13098515 | 1 | unprocessed_pseudc | ENSG0000C |
| 13115828 | 13118626 | -1 | unprocessed_pseudc | ENSG0000C |
| 13125206 | 13126314 | -1 | unprocessed_pseudc | ENSG0000C |

| | | | | |
|---|---|---|---|---|
| 13136151 | 13137509 | -1 | protein_coding | ENSG0000C |
| 13149474 | 13152272 | 1 | unprocessed_pseudc | ENSG0000C |
| 13169610 | 13178799 | -1 | protein_coding | ENSG0000C |
| 13185268 | 13186628 | -1 | protein_coding | ENSG0000C |
| 13191011 | 13198026 | 1 | protein_coding | ENSG0000C |
| 13215394 | 13218160 | -1 | unprocessed_pseudc | ENSG0000C |
| 13241912 | 13244686 | -1 | protein_coding | ENSG0000C |
| 13264443 | 13267241 | 1 | unprocessed_pseudc | ENSG0000C |
| 13272778 | 13282015 | 1 | protein_coding | ENSG0000C |
| 13292116 | 13294207 | -1 | unprocessed_pseudc | ENSG0000C |
| 13297705 | 13297811 | -1 | snRNA | ENSG0000C |
| 13299884 | 13303717 | -1 | protein_coding | ENSG0000C |
| 13322112 | 13327566 | 1 | protein_coding | ENSG0000C |
| 155211179 | 155213671 | -1 | unprocessed_pseudc | ENSG0000C |
| 155225758 | 155229107 | 1 | unprocessed_pseudc | ENSG0000C |
| 155229777 | 155234285 | -1 | unprocessed_pseudc | ENSG0000C |
| 155246379 | 155249729 | 1 | unprocessed_pseudc | ENSG0000C |
| 155249852 | 155260266 | -1 | protein_coding | ENSG0000C |
| 155262609 | 155270887 | -1 | protein_coding | ENSG0000C |
| 155271383 | 155277834 | -1 | protein_coding | ENSG0000C |
| 155278272 | 155293895 | -1 | protein_coding | ENSG0000C |
| 155292987 | 155305252 | 1 | protein_coding | ENSG0000C |
| 155305243 | 155316838 | -1 | protein_coding | ENSG0000C |
| 248361581 | 248362627 | 1 | protein_coding | ENSG0000C |
| 248387950 | 248388876 | 1 | protein_coding | ENSG0000C |
| 248412061 | 248413170 | 1 | protein_coding | ENSG0000C |
| 248447273 | 248448199 | 1 | protein_coding | ENSG0000C |
| 248458842 | 248459895 | 1 | protein_coding | ENSG0000C |
| 248479392 | 248480399 | 1 | protein_coding | ENSG0000C |
| 248494630 | 248495663 | 1 | protein_coding | ENSG0000C |
| 103926563 | 103927802 | 1 | processed_pseudoge | ENSG0000C |
| 198534710 | 198552432 | -1 | protein_coding | ENSG0000C |
| 198609211 | 198610355 | -1 | lncRNA | ENSG0000C |
| 198650158 | 198733963 | 1 | protein_coding | ENSG0000C |
| 198669040 | 198678548 | -1 | lncRNA | ENSG0000C |
| 198690621 | 198691520 | -1 | processed_pseudoge | ENSG0000C |
| 243005845 | 243007468 | -1 | lncRNA | ENSG0000C |
| 243029512 | 243047869 | -1 | lncRNA | ENSG0000C |
| 243056307 | 243101744 | -1 | lncRNA | ENSG0000C |
| 243081156 | 243081259 | -1 | snRNA | ENSG0000C |
| 243124428 | 243255321 | -1 | protein_coding | ENSG0000C |
| 243256056 | 243500091 | 1 | protein_coding | ENSG0000C |
| 243488233 | 243855434 | -1 | protein_coding | ENSG0000C |
| 243855738 | 243856491 | 1 | lncRNA | ENSG0000C |
| 23244765 | 23245224 | 1 | processed_pseudoge | ENSG0000C |
| 23288771 | 23294952 | -1 | lncRNA | ENSG0000C |
| 23305261 | 23310330 | 1 | lncRNA | ENSG0000C |
| 23311235 | 23351800 | -1 | protein_coding | ENSG0000C |
| 23366912 | 23377300 | -1 | protein_coding | ENSG0000C |
| 23376461 | 23379303 | 1 | lncRNA | ENSG0000C |

| | | | | |
|---|---|---|---|---|
| 23377718 | 23377810 | 1 | misc_RNA | ENSG00000( |
| 23385844 | 23386493 | -1 | lncRNA | ENSG00000( |
| 23388373 | 23432212 | -1 | protein_coding | ENSG00000( |
| 23418296 | 23419610 | 1 | processed_pseudoge | ENSG00000( |
| 23436027 | 23492032 | -1 | protein_coding | ENSG00000( |
| 23513902 | 23538697 | -1 | protein_coding | ENSG00000( |
| 23556603 | 23558379 | -1 | lncRNA | ENSG00000( |
| 23565390 | 23566965 | -1 | protein_coding | ENSG00000( |
| 23583900 | 23593621 | -1 | lncRNA | ENSG00000( |
| 144523860 | 144524021 | -1 | snRNA | ENSG00000( |
| 144551779 | 144551943 | -1 | snRNA | ENSG00000( |
| 144560668 | 144560831 | -1 | snRNA | ENSG00000( |
| 144561389 | 144568027 | -1 | lncRNA | ENSG00000( |
| 144567601 | 144568743 | 1 | lncRNA | ENSG00000( |
| 144592865 | 144593524 | 1 | protein_coding | ENSG00000( |
| 144627218 | 144641637 | -1 | lncRNA | ENSG00000( |
| 144660404 | 144669313 | 1 | TEC | ENSG00000( |
| 13029339 | 13032130 | 1 | unprocessed_pseudc | ENSG00000( |
| 13049476 | 13056491 | -1 | protein_coding | ENSG00000( |
| 13060869 | 13062229 | 1 | protein_coding | ENSG00000( |
| 13068677 | 13077884 | 1 | protein_coding | ENSG00000( |
| 13095179 | 13097971 | -1 | unprocessed_pseudc | ENSG00000( |
| 13104403 | 13105689 | -1 | unprocessed_pseudc | ENSG00000( |
| 13115488 | 13116854 | -1 | protein_coding | ENSG00000( |
| 13128818 | 13131615 | 1 | transcribed_unproce | ENSG00000( |
| 13146981 | 13147460 | -1 | lncRNA | ENSG00000( |
| 13148905 | 13155961 | -1 | protein_coding | ENSG00000( |
| 13164586 | 13165467 | -1 | protein_coding | ENSG00000( |
| 13172455 | 13179464 | 1 | protein_coding | ENSG00000( |
| 13216103 | 13218892 | 1 | unprocessed_pseudc | ENSG00000( |
| 13224452 | 13233554 | 1 | protein_coding | ENSG00000( |
| 13243779 | 13245867 | -1 | unprocessed_pseudc | ENSG00000( |
| 13249365 | 13249471 | -1 | snRNA | ENSG00000( |
| 13251275 | 13255414 | -1 | protein_coding | ENSG00000( |
| 13273779 | 13279147 | 1 | protein_coding | ENSG00000( |
| 235328570 | 235452743 | 1 | protein_coding | ENSG00000( |
| 235361153 | 235362540 | 1 | processed_pseudoge | ENSG00000( |
| 235366353 | 235367366 | -1 | lncRNA | ENSG00000( |
| 235436776 | 235437022 | 1 | transcribed_processe | ENSG00000( |
| 235450981 | 235508243 | -1 | protein_coding | ENSG00000( |
| 235523762 | 235524476 | 1 | processed_pseudoge | ENSG00000( |
| 235541311 | 235541832 | 1 | processed_pseudoge | ENSG00000( |
| 235541843 | 235543636 | -1 | processed_pseudoge | ENSG00000( |
| 235543844 | 235545206 | -1 | processed_pseudoge | ENSG00000( |
| 235545203 | 235545497 | -1 | processed_pseudoge | ENSG00000( |
| 235545550 | 235545888 | -1 | processed_pseudoge | ENSG00000( |
| 235545950 | 235546181 | -1 | processed_pseudoge | ENSG00000( |
| 235551476 | 235650754 | -1 | protein_coding | ENSG00000( |
| 235569552 | 235570728 | 1 | processed_pseudoge | ENSG00000( |
| 107964443 | 107994607 | 1 | lncRNA | ENSG00000( |

| | | | | |
|---|---|---|---|---|
| 108040263 | 108076020 | 1 | lncRNA | ENSG00000 |
| 108134010 | 108200843 | -1 | protein_coding | ENSG00000 |
| 108199920 | 108201485 | 1 | lncRNA | ENSG00000 |
| 108200407 | 108202737 | 1 | TEC | ENSG00000 |
| 108494877 | 108500545 | -1 | unprocessed_pseudo | ENSG00000 |
| 108501780 | 108502877 | 1 | processed_pseudoge | ENSG00000 |
| 108507994 | 108508663 | 1 | processed_pseudoge | ENSG00000 |
| 28199456 | 28233029 | -1 | protein_coding | ENSG00000 |
| 28236109 | 28246906 | 1 | protein_coding | ENSG00000 |
| 28239509 | 28241453 | -1 | lncRNA | ENSG00000 |
| 28247144 | 28247568 | 1 | lncRNA | ENSG00000 |
| 28259518 | 28282491 | 1 | protein_coding | ENSG00000 |
| 28329002 | 28335965 | 1 | protein_coding | ENSG00000 |
| 72283170 | 72753772 | 1 | lncRNA | ENSG00000 |
| 72301472 | 72301829 | 1 | processed_pseudoge | ENSG00000 |
| 248361581 | 248362627 | 1 | protein_coding | ENSG00000 |
| 248387949 | 248388875 | 1 | protein_coding | ENSG00000 |
| 248408227 | 248409336 | 1 | protein_coding | ENSG00000 |
| 248443651 | 248444577 | 1 | protein_coding | ENSG00000 |
| 248455242 | 248456216 | 1 | protein_coding | ENSG00000 |
| 248475795 | 248476751 | 1 | protein_coding | ENSG00000 |
| 248486689 | 248487928 | -1 | lncRNA | ENSG00000 |
| 248491033 | 248491980 | 1 | protein_coding | ENSG00000 |
| 248500752 | 248501072 | 1 | unprocessed_pseudo | ENSG00000 |
| 17814608 | 17815038 | -1 | lncRNA | ENSG00000 |
| 17872581 | 17878345 | 1 | lncRNA | ENSG00000 |
| 17893379 | 17894176 | -1 | lncRNA | ENSG00000 |
| 61894910 | 61940574 | 1 | protein_coding | ENSG00000 |
| 61950345 | 62005079 | -1 | protein_coding | ENSG00000 |
| 64097802 | 64100643 | 1 | protein_coding | ENSG00000 |
| 64105820 | 64107171 | -1 | protein_coding | ENSG00000 |
| 64143780 | 64147290 | -1 | lncRNA | ENSG00000 |
| 64155437 | 64245898 | 1 | protein_coding | ENSG00000 |
| 64211718 | 64278487 | 1 | protein_coding | ENSG00000 |
| 64259386 | 64278487 | 1 | protein_coding | ENSG00000 |
| 63414903 | 63466150 | -1 | protein_coding | ENSG00000 |
| 63422269 | 63424555 | 1 | lncRNA | ENSG00000 |
| 63449173 | 63450451 | 1 | lncRNA | ENSG00000 |
| 22109451 | 22116528 | 1 | lncRNA | ENSG00000 |
| 22134698 | 22136083 | -1 | processed_pseudoge | ENSG00000 |
| 22153950 | 22154299 | -1 | processed_pseudoge | ENSG00000 |
| 22205192 | 22205332 | 1 | snRNA | ENSG00000 |
| 22209939 | 22212813 | 1 | lncRNA | ENSG00000 |
| 32769814 | 32771858 | -1 | protein_coding | ENSG00000 |
| 32772100 | 32798969 | 1 | lncRNA | ENSG00000 |
| 32790673 | 32791504 | -1 | TEC | ENSG00000 |
| 32793564 | 32813728 | -1 | protein_coding | ENSG00000 |
| 33410836 | 33488905 | 1 | protein_coding | ENSG00000 |
| 33432485 | 33489568 | -1 | protein_coding | ENSG00000 |
| 33491137 | 33491752 | -1 | processed_pseudoge | ENSG00000 |

| | | | | |
|---|---|---|---|---|
| 33492056 | 33493815 | -1 | lncRNA | ENSG00000 |
| 33495087 | 33501278 | -1 | protein_coding | ENSG00000 |
| 33508081 | 33521016 | -1 | protein_coding | ENSG00000 |
| 46142109 | 46155567 | -1 | protein_coding | ENSG00000 |
| 46151614 | 46152647 | 1 | lncRNA | ENSG00000 |
| 46161526 | 46186787 | -1 | protein_coding | ENSG00000 |
| 46187390 | 46191252 | 1 | lncRNA | ENSG00000 |
| 46190452 | 46221883 | -1 | protein_coding | ENSG00000 |
| 17604145 | 17612947 | -1 | protein_coding | ENSG00000 |
| 17611744 | 17633199 | 1 | lncRNA | ENSG00000 |
| 17659276 | 17660384 | -1 | lncRNA | ENSG00000 |
| 17705408 | 17707455 | 1 | TEC | ENSG00000 |
| 17732440 | 17742196 | -1 | lncRNA | ENSG00000 |
| 17210469 | 17211347 | -1 | processed_pseudoge | ENSG00000 |
| 17299421 | 17299798 | 1 | processed_pseudoge | ENSG00000 |
| 42079691 | 42124100 | 1 | protein_coding | ENSG00000 |
| 42085525 | 42090955 | -1 | protein_coding | ENSG00000 |
| 42090956 | 42125351 | 1 | lncRNA | ENSG00000 |
| 42107765 | 42108953 | -1 | processed_pseudoge | ENSG00000 |
| 42124281 | 42127775 | -1 | lncRNA | ENSG00000 |
| 42128051 | 42132425 | -1 | polymorphic_pseudo | ENSG00000 |
| 42137734 | 42142850 | -1 | unprocessed_pseudo | ENSG00000 |
| 42147859 | 42203293 | -1 | protein_coding | ENSG00000 |
| 38921227 | 38924708 | 1 | lncRNA | ENSG00000 |
| 38949113 | 38949127 | 1 | unprocessed_pseudo | ENSG00000 |
| 38952742 | 38962839 | 1 | protein_coding | ENSG00000 |
| 38961611 | 38968270 | -1 | lncRNA | ENSG00000 |
| 22375985 | 22376504 | 1 | IG_V_gene | ENSG00000 |
| 22380766 | 22381346 | 1 | IG_V_gene | ENSG00000 |
| 22395019 | 22395490 | 1 | IG_V_gene | ENSG00000 |
| 22397030 | 22397522 | 1 | IG_V_pseudogene | ENSG00000 |
| 22402588 | 22403079 | 1 | IG_V_gene | ENSG00000 |
| 22407947 | 22408097 | 1 | IG_V_pseudogene | ENSG00000 |
| 22413303 | 22413817 | 1 | IG_V_pseudogene | ENSG00000 |
| 22418863 | 22419379 | 1 | IG_V_gene | ENSG00000 |
| 22429412 | 22430396 | 1 | processed_pseudoge | ENSG00000 |
| 22434907 | 22435400 | 1 | IG_V_pseudogene | ENSG00000 |
| 22436625 | 22437120 | 1 | IG_V_gene | ENSG00000 |
| 22441045 | 22441552 | 1 | IG_V_gene | ENSG00000 |
| 22455773 | 22457423 | 1 | processed_pseudoge | ENSG00000 |
| 22459289 | 22459739 | 1 | IG_V_pseudogene | ENSG00000 |
| 22474747 | 22475074 | 1 | unprocessed_pseudo | ENSG00000 |
| 22493526 | 22518259 | -1 | protein_coding | ENSG00000 |
| 22522841 | 22529375 | -1 | protein_coding | ENSG00000 |
| 22556806 | 22568466 | -1 | protein_coding | ENSG00000 |
| 22568448 | 22575707 | 1 | lncRNA | ENSG00000 |
| 22588947 | 22589401 | 1 | IG_V_pseudogene | ENSG00000 |
| 22597260 | 22597793 | 1 | IG_V_gene | ENSG00000 |
| 22603633 | 22604161 | 1 | IG_V_gene | ENSG00000 |
| 22613550 | 22614270 | 1 | IG_V_pseudogene | ENSG00000 |

| | | | | |
|---|---|---|---|---|
| 22627677 | 22628268 | 1 | IG_V_pseudogene | ENSG00000 |
| 35685940 | 35689373 | 1 | lncRNA | ENSG00000 |
| 35703803 | 35704176 | 1 | processed_pseudoge | ENSG00000 |
| 35717872 | 35729483 | 1 | protein_coding | ENSG00000 |
| 35744967 | 35945128 | -1 | protein_coding | ENSG00000 |
| 36800684 | 36819479 | -1 | protein_coding | ENSG00000 |
| 36816326 | 36819736 | 1 | lncRNA | ENSG00000 |
| 36847372 | 36873605 | -1 | lncRNA | ENSG00000 |
| 36860995 | 36880544 | 1 | protein_coding | ENSG00000 |
| 23359603 | 23387731 | -1 | transcribed_unproce | ENSG00000 |
| 23390606 | 23402726 | -1 | transcribed_unproce | ENSG00000 |
| 23392771 | 23393839 | 1 | unprocessed_pseudo | ENSG00000 |
| 23433564 | 23435071 | 1 | lncRNA | ENSG00000 |
| 23446598 | 23460833 | 1 | transcribed_process | ENSG00000 |
| 23462086 | 23486981 | -1 | lncRNA | ENSG00000 |
| 23536736 | 23537844 | -1 | lncRNA | ENSG00000 |
| 23764087 | 23765861 | 1 | protein_coding | ENSG00000 |
| 23765834 | 23768443 | -1 | protein_coding | ENSG00000 |
| 23768226 | 23784316 | 1 | protein_coding | ENSG00000 |
| 23782283 | 23783958 | -1 | protein_coding | ENSG00000 |
| 23786963 | 23834516 | 1 | protein_coding | ENSG00000 |
| 23834503 | 23839128 | -1 | protein_coding | ENSG00000 |
| 23856427 | 23857039 | -1 | lncRNA | ENSG00000 |
| 23856703 | 23886309 | 1 | protein_coding | ENSG00000 |
| 23862188 | 23895223 | 1 | protein_coding | ENSG00000 |
| 23865248 | 23873277 | -1 | lncRNA | ENSG00000 |
| 23879493 | 23879790 | -1 | misc_RNA | ENSG00000 |
| 23894004 | 23895227 | 1 | protein_coding | ENSG00000 |
| 23894426 | 23898930 | -1 | lncRNA | ENSG00000 |
| 23901432 | 23907068 | -1 | lncRNA | ENSG00000 |
| 23926900 | 23929574 | 1 | processed_pseudoge | ENSG00000 |
| 23939998 | 23942798 | 1 | unprocessed_pseudo | ENSG00000 |
| 23949918 | 23954042 | 1 | unprocessed_pseudo | ENSG00000 |
| 23957418 | 23961206 | -1 | protein_coding | ENSG00000 |
| 23963780 | 23964374 | 1 | processed_pseudoge | ENSG00000 |
| 23966902 | 23972534 | 1 | protein_coding | ENSG00000 |
| 23969213 | 23969875 | 1 | lncRNA | ENSG00000 |
| 23971367 | 23980473 | -1 | protein_coding | ENSG00000 |
| 23976905 | 23977589 | -1 | unprocessed_pseudo | ENSG00000 |
| 23980152 | 23983919 | 1 | protein_coding | ENSG00000 |
| 23987324 | 23991428 | -1 | unprocessed_pseudo | ENSG00000 |
| 23998411 | 24005163 | -1 | protein_coding | ENSG00000 |
| 24008537 | 24009893 | -1 | unprocessed_pseudo | ENSG00000 |
| 24022930 | 24023387 | -1 | protein_coding | ENSG00000 |
| 24024565 | 24028697 | 1 | unprocessed_pseudo | ENSG00000 |
| 24030925 | 24031599 | 1 | unprocessed_pseudo | ENSG00000 |
| 24033952 | 24042493 | -1 | protein_coding | ENSG00000 |
| 24052465 | 24059615 | -1 | protein_coding | ENSG00000 |
| 24060497 | 24062557 | -1 | unprocessed_pseudo | ENSG00000 |
| 24065455 | 24065991 | 1 | lncRNA | ENSG00000 |

| | | | | |
|---|---|---|---|---|
| 42079691 | 42084284 | 1 | protein_coding | ENSG00000 |
| 42085525 | 42090955 | -1 | protein_coding | ENSG00000 |
| 42089630 | 42090028 | -1 | lncRNA | ENSG00000 |
| 42090931 | 42148836 | 1 | lncRNA | ENSG00000 |
| 42107765 | 42108953 | -1 | processed_pseudoge | ENSG00000 |
| 42126481 | 42130888 | -1 | polymorphic_pseudc | ENSG00000 |
| 42132528 | 42132983 | 1 | processed_pseudoge | ENSG00000 |
| 42138640 | 42143020 | -1 | protein_coding | ENSG00000 |
| 42148576 | 42150238 | -1 | lncRNA | ENSG00000 |
| 42079691 | 42084284 | 1 | protein_coding | ENSG00000 |
| 42085525 | 42090955 | -1 | protein_coding | ENSG00000 |
| 42089630 | 42090028 | -1 | lncRNA | ENSG00000 |
| 42090933 | 42137053 | 1 | lncRNA | ENSG00000 |
| 42107765 | 42108953 | -1 | processed_pseudoge | ENSG00000 |
| 42126495 | 42130902 | -1 | protein_coding | ENSG00000 |
| 42136436 | 42139932 | -1 | lncRNA | ENSG00000 |
| 42138063 | 42139731 | 1 | lncRNA | ENSG00000 |
| 42140208 | 42144572 | -1 | polymorphic_pseudc | ENSG00000 |
| 42149871 | 42155001 | -1 | unprocessed_pseudc | ENSG00000 |
| 42160013 | 42215442 | -1 | protein_coding | ENSG00000 |
| 42085526 | 42090894 | -1 | protein_coding | ENSG00000 |
| 42089640 | 42090038 | -1 | lncRNA | ENSG00000 |
| 42107776 | 42108964 | -1 | processed_pseudoge | ENSG00000 |
| 42124292 | 42127786 | -1 | lncRNA | ENSG00000 |
| 42125919 | 42127585 | 1 | lncRNA | ENSG00000 |
| 42129815 | 42131435 | -1 | protein_coding | ENSG00000 |
| 42137724 | 42142854 | -1 | unprocessed_pseudc | ENSG00000 |
| 42147866 | 42343616 | -1 | protein_coding | ENSG00000 |
| 42126481 | 42130863 | -1 | protein_coding | ENSG00000 |
| 42132527 | 42132982 | 1 | processed_pseudoge | ENSG00000 |
| 42150129 | 42153625 | -1 | lncRNA | ENSG00000 |
| 42151756 | 42153424 | 1 | lncRNA | ENSG00000 |
| 42163591 | 42168706 | -1 | unprocessed_pseudc | ENSG00000 |
| 42173715 | 42343616 | -1 | protein_coding | ENSG00000 |
| 42107763 | 42108951 | -1 | processed_pseudoge | ENSG00000 |
| 42124279 | 42127773 | -1 | lncRNA | ENSG00000 |
| 42125906 | 42127572 | 1 | lncRNA | ENSG00000 |
| 42128127 | 42141929 | -1 | protein_coding | ENSG00000 |
| 42147351 | 42152466 | -1 | unprocessed_pseudc | ENSG00000 |
| 42157475 | 42343616 | -1 | protein_coding | ENSG00000 |
| 42107765 | 42108953 | -1 | processed_pseudoge | ENSG00000 |
| 42126481 | 42130863 | -1 | protein_coding | ENSG00000 |
| 42132527 | 42132982 | 1 | processed_pseudoge | ENSG00000 |
| 42163834 | 42167328 | -1 | lncRNA | ENSG00000 |
| 42165461 | 42167127 | 1 | lncRNA | ENSG00000 |
| 42167604 | 42176857 | -1 | polymorphic_pseudc | ENSG00000 |
| 42177288 | 42182405 | -1 | unprocessed_pseudc | ENSG00000 |
| 42187414 | 42343616 | -1 | protein_coding | ENSG00000 |
| 42126498 | 42130880 | -1 | protein_coding | ENSG00000 |
| 42132544 | 42132999 | 1 | processed_pseudoge | ENSG00000 |

| | | | | |
|---|---|---|---|---|
| 42136436 | 42139932 | -1 | lncRNA | ENSG00000( |
| 42138063 | 42139731 | 1 | lncRNA | ENSG00000( |
| 42153945 | 42158119 | -1 | protein_coding | ENSG00000( |
| 42163563 | 42168680 | -1 | unprocessed_pseudc | ENSG00000( |
| 42173689 | 42343616 | -1 | protein_coding | ENSG00000( |
| 3401777 | 3486461 | 1 | protein_coding | ENSG00000( |
| 3427276 | 3427885 | 1 | lncRNA | ENSG00000( |
| 3498320 | 3498795 | 1 | lncRNA | ENSG00000( |
| 3498728 | 3502360 | -1 | protein_coding | ENSG00000( |
| 241970683 | 241977276 | 1 | lncRNA | ENSG00000( |
| 242003691 | 242006013 | 1 | lncRNA | ENSG00000( |
| 242030558 | 242031529 | -1 | lncRNA | ENSG00000( |
| 242053059 | 242089513 | -1 | lncRNA | ENSG00000( |
| 242100269 | 242121607 | 1 | unprocessed_pseudc | ENSG00000( |
| 36299388 | 36299830 | -1 | processed_pseudoge | ENSG00000( |
| 36360778 | 36361147 | -1 | lncRNA | ENSG00000( |
| 36361959 | 36362656 | 1 | protein_coding | ENSG00000( |
| 90309230 | 90309333 | 1 | unprocessed_pseudc | ENSG00000( |
| 90315366 | 90315836 | -1 | IG_V_pseudogene | ENSG00000( |
| 90359653 | 90359876 | -1 | processed_pseudoge | ENSG00000( |
| 90365762 | 90367722 | 1 | lncRNA | ENSG00000( |
| 168831472 | 168865516 | 1 | protein_coding | ENSG00000( |
| 168834132 | 168930360 | -1 | protein_coding | ENSG00000( |
| 168918229 | 168926984 | 1 | protein_coding | ENSG00000( |
| 168939927 | 168953422 | -1 | protein_coding | ENSG00000( |
| 745961 | 754657 | 1 | lncRNA | ENSG00000( |
| 753629 | 754224 | 1 | lncRNA | ENSG00000( |
| 784657 | 859032 | -1 | lncRNA | ENSG00000( |
| 241970683 | 241977276 | 1 | lncRNA | ENSG00000( |
| 242003691 | 242006013 | 1 | lncRNA | ENSG00000( |
| 242030199 | 242031170 | -1 | lncRNA | ENSG00000( |
| 242052699 | 242089153 | -1 | lncRNA | ENSG00000( |
| 242099909 | 242121247 | 1 | unprocessed_pseudc | ENSG00000( |
| 90220727 | 90221384 | 1 | IG_V_gene | ENSG00000( |
| 90245744 | 90246220 | 1 | IG_V_gene | ENSG00000( |
| 90259509 | 90260067 | 1 | IG_V_gene | ENSG00000( |
| 90340444 | 90340547 | 1 | unprocessed_pseudc | ENSG00000( |
| 90346572 | 90347042 | -1 | IG_V_pseudogene | ENSG00000( |
| 90389792 | 90390015 | -1 | processed_pseudoge | ENSG00000( |
| 90395720 | 90397682 | 1 | lncRNA | ENSG00000( |
| 148937284 | 149030199 | 1 | protein_coding | ENSG00000( |
| 1209102 | 1269668 | 1 | protein_coding | ENSG00000( |
| 241799063 | 241804208 | 1 | protein_coding | ENSG00000( |
| 241800916 | 241801907 | -1 | lncRNA | ENSG00000( |
| 241808312 | 241812016 | -1 | lncRNA | ENSG00000( |
| 241808825 | 241817413 | 1 | protein_coding | ENSG00000( |
| 241844380 | 241845036 | 1 | lncRNA | ENSG00000( |
| 241849881 | 241858908 | -1 | protein_coding | ENSG00000( |
| 241869600 | 241873823 | 1 | protein_coding | ENSG00000( |
| 241881363 | 241959696 | 1 | lncRNA | ENSG00000( |

| | | | | |
|---|---|---|---|---|
| 241893989 | 241902551 | -1 | lncRNA | ENSG00000 |
| 131722418 | 131768326 | 1 | transcribed_unproce | ENSG00000 |
| 131723253 | 131724279 | -1 | processed_pseudoge | ENSG00000 |
| 131795990 | 131802689 | -1 | lncRNA | ENSG00000 |
| 1316265 | 1365970 | 1 | protein_coding | ENSG00000 |
| 1340838 | 1344763 | -1 | lncRNA | ENSG00000 |
| 1372594 | 1540021 | 1 | protein_coding | ENSG00000 |
| 205900630 | 205900958 | 1 | misc_RNA | ENSG00000 |
| 205949091 | 205950145 | 1 | processed_pseudoge | ENSG00000 |
| 205989585 | 206000858 | 1 | lncRNA | ENSG00000 |
| 205993721 | 206086303 | -1 | protein_coding | ENSG00000 |
| 206007880 | 206007974 | -1 | misc_RNA | ENSG00000 |
| 206025593 | 206025697 | 1 | misc_RNA | ENSG00000 |
| 206040432 | 206040845 | 1 | processed_pseudoge | ENSG00000 |
| 206084605 | 206086564 | 1 | lncRNA | ENSG00000 |
| 206114817 | 206159509 | -1 | protein_coding | ENSG00000 |
| 206115547 | 206122323 | 1 | lncRNA | ENSG00000 |
| 206116110 | 206116500 | -1 | processed_pseudoge | ENSG00000 |
| 206159585 | 206162928 | 1 | protein_coding | ENSG00000 |
| 206160843 | 206161024 | 1 | misc_RNA | ENSG00000 |
| 206161841 | 206161929 | 1 | snoRNA | ENSG00000 |
| 206161878 | 206161957 | 1 | snoRNA | ENSG00000 |
| 206162228 | 206162359 | 1 | snoRNA | ENSG00000 |
| 206175316 | 206218047 | -1 | protein_coding | ENSG00000 |
| 206203543 | 206269946 | 1 | lncRNA | ENSG00000 |
| 206221316 | 206221635 | 1 | misc_RNA | ENSG00000 |
| 206264248 | 206264538 | -1 | processed_pseudoge | ENSG00000 |
| 206264686 | 206264834 | -1 | misc_RNA | ENSG00000 |
| 206279283 | 206319047 | 1 | protein_coding | ENSG00000 |
| 206414707 | 206415529 | -1 | processed_pseudoge | ENSG00000 |
| 206423912 | 206424858 | -1 | processed_pseudoge | ENSG00000 |
| 126154583 | 126203372 | 1 | lncRNA | ENSG00000 |
| 126311469 | 126311930 | -1 | processed_pseudoge | ENSG00000 |
| 207821288 | 208025527 | -1 | protein_coding | ENSG00000 |
| 208019638 | 208019741 | -1 | snRNA | ENSG00000 |
| 208031383 | 208031953 | -1 | processed_pseudoge | ENSG00000 |
| 208054161 | 208054258 | -1 | rRNA_pseudogene | ENSG00000 |
| 208057248 | 208057745 | -1 | processed_pseudoge | ENSG00000 |
| 208114374 | 208118652 | -1 | transcribed_unproce | ENSG00000 |
| 208127748 | 208130665 | -1 | protein_coding | ENSG00000 |
| 208134278 | 208135969 | -1 | protein_coding | ENSG00000 |
| 11817080 | 11846919 | -1 | protein_coding | ENSG00000 |
| 11844908 | 11846845 | -1 | protein_coding | ENSG00000 |
| 11859673 | 11893500 | 1 | unprocessed_pseudc | ENSG00000 |
| 11880784 | 11881057 | -1 | processed_pseudoge | ENSG00000 |
| 11901932 | 11903166 | 1 | unprocessed_pseudc | ENSG00000 |
| 11908448 | 11914234 | 1 | unprocessed_pseudc | ENSG00000 |
| 11911555 | 11912686 | -1 | unprocessed_pseudc | ENSG00000 |
| 11954709 | 11955006 | -1 | misc_RNA | ENSG00000 |
| 151637174 | 151657956 | 1 | lncRNA | ENSG00000 |

| | | | | |
|---|---|---|---|---|
| 151744454 | 151770036 | 1 | protein_coding | ENSG00000( |
| 151761981 | 151765669 | -1 | lncRNA | ENSG00000( |
| 195639990 | 195640460 | 1 | lncRNA | ENSG00000( |
| 195650146 | 195651472 | -1 | processed_pseudoge | ENSG00000( |
| 195655565 | 195657927 | -1 | lncRNA | ENSG00000( |
| 195658062 | 195739964 | 1 | lncRNA | ENSG00000( |
| 195658096 | 195685904 | 1 | transcribed_unproce | ENSG00000( |
| 195720882 | 195741123 | 1 | protein_coding | ENSG00000( |
| 195746765 | 195819600 | -1 | protein_coding | ENSG00000( |
| 154119686 | 154121332 | -1 | lncRNA | ENSG00000( |
| 154121003 | 154261004 | 1 | protein_coding | ENSG00000( |
| 154275723 | 154284039 | -1 | protein_coding | ENSG00000( |
| 197445061 | 197458322 | -1 | lncRNA | ENSG00000( |
| 197450327 | 197456654 | -1 | lncRNA | ENSG00000( |
| 197458404 | 197468515 | 1 | lncRNA | ENSG00000( |
| 197506664 | 197557450 | -1 | protein_coding | ENSG00000( |
| 20347019 | 20355479 | 1 | lncRNA | ENSG00000( |
| 195650146 | 195651472 | -1 | processed_pseudoge | ENSG00000( |
| 195655565 | 195657927 | -1 | lncRNA | ENSG00000( |
| 195658062 | 195776180 | 1 | lncRNA | ENSG00000( |
| 195658096 | 195685904 | 1 | transcribed_unproce | ENSG00000( |
| 195719081 | 195738792 | 1 | protein_coding | ENSG00000( |
| 195744429 | 195789161 | -1 | protein_coding | ENSG00000( |
| 84638405 | 84881924 | -1 | lncRNA | ENSG00000( |
| 84881984 | 84893375 | 1 | lncRNA | ENSG00000( |
| 84955809 | 84962251 | 1 | lncRNA | ENSG00000( |
| 125703375 | 125704277 | 1 | unprocessed_pseudc | ENSG00000( |
| 125712140 | 125713051 | 1 | unprocessed_pseudc | ENSG00000( |
| 125724567 | 125725463 | 1 | unprocessed_pseudc | ENSG00000( |
| 125734326 | 125735208 | 1 | unprocessed_pseudc | ENSG00000( |
| 125747135 | 125748058 | 1 | unprocessed_pseudc | ENSG00000( |
| 125756086 | 125756441 | 1 | unprocessed_pseudc | ENSG00000( |
| 125800930 | 125801709 | 1 | unprocessed_pseudc | ENSG00000( |
| 195650146 | 195651472 | -1 | processed_pseudoge | ENSG00000( |
| 195655565 | 195657927 | -1 | lncRNA | ENSG00000( |
| 195658062 | 195739964 | 1 | lncRNA | ENSG00000( |
| 195658096 | 195685904 | 1 | transcribed_unproce | ENSG00000( |
| 195720882 | 195741123 | 1 | protein_coding | ENSG00000( |
| 195746774 | 195792809 | -1 | protein_coding | ENSG00000( |
| 75370448 | 75371342 | -1 | unprocessed_pseudc | ENSG00000( |
| 75384059 | 75391810 | -1 | unprocessed_pseudc | ENSG00000( |
| 75395188 | 75395320 | 1 | processed_pseudoge | ENSG00000( |
| 75395404 | 75395536 | -1 | processed_pseudoge | ENSG00000( |
| 75425349 | 75432428 | 1 | unprocessed_pseudc | ENSG00000( |
| 75425608 | 75426048 | -1 | protein_coding | ENSG00000( |
| 75431837 | 75445395 | -1 | transcribed_unproce | ENSG00000( |
| 75445517 | 75548314 | 1 | lncRNA | ENSG00000( |
| 75457054 | 75501487 | -1 | unprocessed_pseudc | ENSG00000( |
| 75532088 | 75532297 | 1 | processed_pseudoge | ENSG00000( |
| 75550334 | 75580199 | -1 | transcribed_unproce | ENSG00000( |

| | | | | |
|---|---|---|---|---|
| 75552434 | 75553214 | -1 | processed_pseudoge | ENSG00000( |
| 184426592 | 184444693 | 1 | lncRNA | ENSG00000( |
| 184510148 | 184513434 | 1 | lncRNA | ENSG00000( |
| 195639990 | 195640460 | 1 | lncRNA | ENSG00000( |
| 195650146 | 195651472 | -1 | processed_pseudoge | ENSG00000( |
| 195655565 | 195657927 | -1 | lncRNA | ENSG00000( |
| 195658062 | 195776742 | 1 | lncRNA | ENSG00000( |
| 195658096 | 195685904 | 1 | transcribed_unproce | ENSG00000( |
| 195719644 | 195739354 | 1 | protein_coding | ENSG00000( |
| 195744993 | 195789727 | -1 | protein_coding | ENSG00000( |
| 112086335 | 112131004 | 1 | protein_coding | ENSG00000( |
| 112120839 | 112133270 | -1 | protein_coding | ENSG00000( |
| 112133423 | 112135359 | 1 | lncRNA | ENSG00000( |
| 112140898 | 112303675 | -1 | protein_coding | ENSG00000( |
| 112194928 | 112195446 | -1 | processed_pseudoge | ENSG00000( |
| 195650146 | 195651472 | -1 | processed_pseudoge | ENSG00000( |
| 195655565 | 195657927 | -1 | lncRNA | ENSG00000( |
| 195658062 | 195739964 | 1 | lncRNA | ENSG00000( |
| 195658096 | 195685904 | 1 | transcribed_unproce | ENSG00000( |
| 195720882 | 195741123 | 1 | protein_coding | ENSG00000( |
| 195747176 | 195811179 | -1 | unprocessed_pseudc | ENSG00000( |
| 195650146 | 195651472 | -1 | processed_pseudoge | ENSG00000( |
| 195655565 | 195657927 | -1 | lncRNA | ENSG00000( |
| 195658062 | 195738903 | 1 | lncRNA | ENSG00000( |
| 195658096 | 195685904 | 1 | transcribed_unproce | ENSG00000( |
| 195720361 | 195740062 | 1 | protein_coding | ENSG00000( |
| 195745705 | 195790439 | -1 | protein_coding | ENSG00000( |
| 175369545 | 175369626 | 1 | miRNA | ENSG00000( |
| 175403968 | 175404069 | -1 | snRNA | ENSG00000( |
| 175480640 | 175484323 | -1 | lncRNA | ENSG00000( |
| 195650146 | 195651472 | -1 | processed_pseudoge | ENSG00000( |
| 195655565 | 195657927 | -1 | lncRNA | ENSG00000( |
| 195658062 | 195740223 | 1 | lncRNA | ENSG00000( |
| 195658096 | 195685904 | 1 | transcribed_unproce | ENSG00000( |
| 195721575 | 195741387 | 1 | protein_coding | ENSG00000( |
| 195747140 | 195791872 | -1 | protein_coding | ENSG00000( |
| 128064778 | 128160821 | -1 | protein_coding | ENSG00000( |
| 128148917 | 128149019 | -1 | snRNA | ENSG00000( |
| 128160388 | 128415576 | 1 | protein_coding | ENSG00000( |
| 128188319 | 128201362 | -1 | lncRNA | ENSG00000( |
| 198053522 | 198080720 | -1 | transcribed_unproce | ENSG00000( |
| 198071311 | 198071417 | -1 | snRNA | ENSG00000( |
| 198106449 | 198108214 | 1 | transcribed_unproce | ENSG00000( |
| 198116056 | 198117113 | 1 | unprocessed_pseudc | ENSG00000( |
| 198117723 | 198119668 | -1 | processed_pseudoge | ENSG00000( |
| 198148759 | 198166452 | 1 | lncRNA | ENSG00000( |
| 198177148 | 198186438 | 1 | unprocessed_pseudc | ENSG00000( |
| 198193139 | 198193242 | 1 | snRNA | ENSG00000( |
| 198219890 | 198222703 | -1 | processed_pseudoge | ENSG00000( |
| 198224587 | 198224769 | 1 | unprocessed_pseudc | ENSG00000( |

| | | | | |
|---|---|---|---|---|
| 189764391 | 189940733 | -1 | lncRNA | ENSG0000C |
| 189940870 | 189963202 | 1 | protein_coding | ENSG0000C |
| 189973599 | 189973847 | -1 | processed_pseudoge | ENSG0000C |
| 189979902 | 189982379 | 1 | processed_pseudoge | ENSG0000C |
| 189982523 | 189984871 | -1 | unprocessed_pseudc | ENSG0000C |
| 190014800 | 190014917 | 1 | rRNA_pseudogene | ENSG0000C |
| 190017360 | 190017478 | 1 | rRNA_pseudogene | ENSG0000C |
| 190021078 | 190022336 | -1 | unprocessed_pseudc | ENSG0000C |
| 190024025 | 190026913 | -1 | protein_coding | ENSG0000C |
| 190037743 | 190038226 | -1 | processed_pseudoge | ENSG0000C |
| 190067620 | 190068894 | 1 | unprocessed_pseudc | ENSG0000C |
| 190070926 | 190072197 | 1 | protein_coding | ENSG0000C |
| 190074232 | 190075503 | 1 | protein_coding | ENSG0000C |
| 190077538 | 190078809 | 1 | protein_coding | ENSG0000C |
| 190080844 | 190082115 | 1 | protein_coding | ENSG0000C |
| 190084150 | 190085421 | 1 | protein_coding | ENSG0000C |
| 190087456 | 190088727 | 1 | protein_coding | ENSG0000C |
| 190090762 | 190092033 | 1 | protein_coding | ENSG0000C |
| 190094068 | 190095339 | 1 | protein_coding | ENSG0000C |
| 190097374 | 190098645 | 1 | protein_coding | ENSG0000C |
| 190100680 | 190101951 | 1 | protein_coding | ENSG0000C |
| 190103986 | 190105260 | 1 | unprocessed_pseudc | ENSG0000C |
| 190107292 | 190108563 | 1 | protein_coding | ENSG0000C |
| 153948718 | 154307593 | 1 | lncRNA | ENSG0000C |
| 154238835 | 154498892 | -1 | protein_coding | ENSG0000C |
| 154243073 | 154244691 | 1 | lncRNA | ENSG0000C |
| 154268828 | 154269286 | 1 | lncRNA | ENSG0000C |
| 154380979 | 154381440 | 1 | processed_pseudoge | ENSG0000C |
| 155836689 | 155866273 | -1 | protein_coding | ENSG0000C |
| 155854738 | 155920406 | 1 | protein_coding | ENSG0000C |
| 155924118 | 155953911 | -1 | protein_coding | ENSG0000C |
| 21544390 | 21737314 | -1 | protein_coding | ENSG0000C |
| 21582138 | 21620924 | 1 | lncRNA | ENSG0000C |
| 21663714 | 21664071 | -1 | processed_pseudoge | ENSG0000C |
| 21704652 | 21726228 | 1 | lncRNA | ENSG0000C |
| 34657606 | 34669432 | 1 | lncRNA | ENSG0000C |
| 68310387 | 68350089 | -1 | protein_coding | ENSG0000C |
| 68376327 | 68376509 | 1 | processed_pseudoge | ENSG0000C |
| 68447498 | 68497649 | 1 | protein_coding | ENSG0000C |
| 68509485 | 68511653 | -1 | unprocessed_pseudc | ENSG0000C |
| 68529322 | 68553341 | -1 | protein_coding | ENSG0000C |
| 68587305 | 68595199 | -1 | unprocessed_pseudc | ENSG0000C |
| 68667002 | 68669858 | 1 | unprocessed_pseudc | ENSG0000C |
| 68696372 | 68697104 | 1 | unprocessed_pseudc | ENSG0000C |
| 68698370 | 68713573 | 1 | protein_coding | ENSG0000C |
| 68716590 | 68716745 | -1 | unprocessed_pseudc | ENSG0000C |
| 68743709 | 68743987 | -1 | unprocessed_pseudc | ENSG0000C |
| 68760003 | 68762508 | -1 | unprocessed_pseudc | ENSG0000C |
| 68766262 | 68766817 | -1 | processed_pseudoge | ENSG0000C |
| 68783028 | 68783287 | -1 | unprocessed_pseudc | ENSG0000C |

| | | | | |
|---|---|---|---|---|
| 68783385 | 68789360 | -1 | lncRNA | ENSG00000 |
| 68790295 | 68790411 | -1 | unprocessed_pseudc | ENSG00000 |
| 68797305 | 68807118 | -1 | unprocessed_pseudc | ENSG00000 |
| 68810840 | 68834168 | -1 | protein_coding | ENSG00000 |
| 68855376 | 68858370 | -1 | unprocessed_pseudc | ENSG00000 |
| 68867386 | 68870910 | -1 | unprocessed_pseudc | ENSG00000 |
| 68879312 | 68880086 | -1 | processed_pseudoge | ENSG00000 |
| 68887233 | 68892380 | -1 | unprocessed_pseudc | ENSG00000 |
| 186971648 | 187014215 | -1 | lncRNA | ENSG00000 |
| 14909961 | 15005388 | -1 | lncRNA | ENSG00000 |
| 189703881 | 189704490 | -1 | lncRNA | ENSG00000 |
| 189709539 | 189709714 | -1 | snRNA | ENSG00000 |
| 189780336 | 189899116 | -1 | lncRNA | ENSG00000 |
| 189884988 | 189888341 | 1 | lncRNA | ENSG00000 |
| 120165490 | 120165892 | 1 | lncRNA | ENSG00000 |
| 188349376 | 188350024 | -1 | processed_pseudoge | ENSG00000 |
| 188387471 | 188389666 | -1 | processed_pseudoge | ENSG00000 |
| 188400736 | 188543559 | 1 | lncRNA | ENSG00000 |
| 189703881 | 189704490 | -1 | lncRNA | ENSG00000 |
| 189709539 | 189709714 | -1 | snRNA | ENSG00000 |
| 189780336 | 189944248 | -1 | lncRNA | ENSG00000 |
| 189885470 | 189888818 | 1 | lncRNA | ENSG00000 |
| 189944765 | 189967182 | 1 | protein_coding | ENSG00000 |
| 189977577 | 189977825 | -1 | processed_pseudoge | ENSG00000 |
| 189983880 | 189984585 | 1 | processed_pseudoge | ENSG00000 |
| 189986501 | 189988849 | -1 | unprocessed_pseudc | ENSG00000 |
| 190019116 | 190019233 | 1 | rRNA | ENSG00000 |
| 190021674 | 190021792 | 1 | rRNA | ENSG00000 |
| 190025385 | 190026643 | -1 | unprocessed_pseudc | ENSG00000 |
| 190028329 | 190031235 | -1 | protein_coding | ENSG00000 |
| 190041894 | 190042377 | -1 | processed_pseudoge | ENSG00000 |
| 189472965 | 189475192 | 1 | transcribed_processe | ENSG00000 |
| 189669074 | 189670954 | 1 | lncRNA | ENSG00000 |
| 189714467 | 189715076 | -1 | lncRNA | ENSG00000 |
| 189720125 | 189720300 | -1 | snRNA | ENSG00000 |
| 189790922 | 189906229 | -1 | lncRNA | ENSG00000 |
| 189892101 | 189895454 | 1 | lncRNA | ENSG00000 |
| 66002771 | 66150012 | 1 | lncRNA | ENSG00000 |
| 66007206 | 66020717 | 1 | lncRNA | ENSG00000 |
| 69955256 | 69961007 | -1 | protein_coding | ENSG00000 |
| 69995931 | 70002570 | 1 | protein_coding | ENSG00000 |
| 70028413 | 70036538 | 1 | protein_coding | ENSG00000 |
| 70050438 | 70058848 | 1 | protein_coding | ENSG00000 |
| 70067389 | 70085270 | 1 | transcribed_unitary_ | ENSG00000 |
| 70138066 | 70181249 | 1 | protein_coding | ENSG00000 |
| 12686551 | 12731442 | 1 | lncRNA | ENSG00000 |
| 69910279 | 69916203 | 1 | unprocessed_pseudc | ENSG00000 |
| 69930714 | 69941207 | 1 | unprocessed_pseudc | ENSG00000 |
| 70005540 | 70005841 | -1 | unprocessed_pseudc | ENSG00000 |
| 70045915 | 70063796 | 1 | protein_coding | ENSG00000 |

| | | | | |
|---|---|---|---|---|
| 70070287 | 70099184 | 1 | protein_coding | ENSG00000( |
| 70076476 | 70078088 | -1 | lncRNA | ENSG00000( |
| 70115323 | 70149120 | -1 | unprocessed_pseudc | ENSG00000( |
| 70153366 | 70153660 | 1 | unprocessed_pseudc | ENSG00000( |
| 70244409 | 70262299 | 1 | protein_coding | ENSG00000( |
| 70268827 | 70296783 | 1 | protein_coding | ENSG00000( |
| 70274990 | 70276602 | -1 | lncRNA | ENSG00000( |
| 70312214 | 70368830 | -1 | protein_coding | ENSG00000( |
| 70376388 | 70411404 | -1 | protein_coding | ENSG00000( |
| 70425738 | 70449761 | -1 | unprocessed_pseudc | ENSG00000( |
| 70448301 | 70556133 | -1 | lncRNA | ENSG00000( |
| 70518623 | 70529103 | -1 | transcribed_unproce | ENSG00000( |
| 70543654 | 70549533 | -1 | transcribed_unproce | ENSG00000( |
| 70677171 | 70771675 | -1 | lncRNA | ENSG00000( |
| 70697990 | 70700212 | -1 | lncRNA | ENSG00000( |
| 70701100 | 70711308 | 1 | transcribed_unproce | ENSG00000( |
| 70770949 | 70771902 | 1 | lncRNA | ENSG00000( |
| 70780935 | 70893131 | 1 | protein_coding | ENSG00000( |
| 70912599 | 70929779 | 1 | protein_coding | ENSG00000( |
| 162424378 | 162427345 | -1 | lncRNA | ENSG00000( |
| 162477891 | 162477993 | 1 | snRNA | ENSG00000( |
| 29079384 | 29079498 | 1 | snoRNA | ENSG00000( |
| 29113759 | 29114317 | 1 | processed_pseudoge | ENSG00000( |
| 29152383 | 29181238 | 1 | lncRNA | ENSG00000( |
| 29200579 | 29225491 | 1 | lncRNA | ENSG00000( |
| 69217760 | 69230127 | 1 | protein_coding | ENSG00000( |
| 69234793 | 69277413 | 1 | protein_coding | ENSG00000( |
| 69281282 | 69332741 | -1 | protein_coding | ENSG00000( |
| 69313353 | 69313854 | -1 | processed_pseudoge | ENSG00000( |
| 69333861 | 69334181 | 1 | processed_pseudoge | ENSG00000( |
| 69349947 | 69350456 | 1 | processed_pseudoge | ENSG00000( |
| 69350995 | 69370027 | -1 | protein_coding | ENSG00000( |
| 69364756 | 69370027 | -1 | protein_coding | ENSG00000( |
| 69369307 | 69415042 | 1 | protein_coding | ENSG00000( |
| 69415353 | 69444141 | 1 | protein_coding | ENSG00000( |
| 69469588 | 69469841 | 1 | processed_pseudoge | ENSG00000( |
| 69477193 | 69502181 | -1 | lncRNA | ENSG00000( |
| 69478716 | 69478984 | -1 | misc_RNA | ENSG00000( |
| 69492007 | 69556424 | 1 | protein_coding | ENSG00000( |
| 69530371 | 69530475 | -1 | snRNA | ENSG00000( |
| 69557495 | 69559073 | -1 | protein_coding | ENSG00000( |
| 69559974 | 69592641 | 1 | protein_coding | ENSG00000( |
| 69593256 | 69594998 | 1 | lncRNA | ENSG00000( |
| 69602550 | 69659159 | 1 | protein_coding | ENSG00000( |
| 69605896 | 69607737 | -1 | TEC | ENSG00000( |
| 69659867 | 69661133 | -1 | TEC | ENSG00000( |
| 69673667 | 69702549 | -1 | protein_coding | ENSG00000( |
| 69694773 | 69696385 | 1 | lncRNA | ENSG00000( |
| 69709074 | 69726961 | -1 | protein_coding | ENSG00000( |
| 69755077 | 69848798 | -1 | transcribed_unproce | ENSG00000( |

| | | | | |
|---|---|---|---|---|
| 69758658 | 69759641 | -1 | lncRNA | ENSG0000C |
| 69761566 | 69762272 | -1 | lncRNA | ENSG0000C |
| 69766913 | 69768768 | -1 | lncRNA | ENSG0000C |
| 69767062 | 69767356 | 1 | unprocessed_pseudc | ENSG0000C |
| 69814552 | 69814989 | -1 | lncRNA | ENSG0000C |
| 69826364 | 69826946 | -1 | lncRNA | ENSG0000C |
| 69856054 | 69898246 | -1 | transcribed_unproce | ENSG0000C |
| 70064043 | 70134190 | 1 | transcribed_unproce | ENSG0000C |
| 70150924 | 70167868 | -1 | transcribed_unproce | ENSG0000C |
| 70179769 | 70181513 | -1 | lncRNA | ENSG0000C |
| 70182128 | 70214752 | -1 | protein_coding | ENSG0000C |
| 70221290 | 70240276 | -1 | unprocessed_pseudc | ENSG0000C |
| 70249980 | 70275822 | -1 | unprocessed_pseudc | ENSG0000C |
| 70280062 | 70280356 | 1 | unprocessed_pseudc | ENSG0000C |
| 70344694 | 70355160 | -1 | unprocessed_pseudc | ENSG0000C |
| 70567180 | 70661645 | -1 | lncRNA | ENSG0000C |
| 70587997 | 70590219 | -1 | lncRNA | ENSG0000C |
| 70591107 | 70601315 | 1 | transcribed_unproce | ENSG0000C |
| 70660919 | 70661872 | 1 | lncRNA | ENSG0000C |
| 70670918 | 70783123 | 1 | protein_coding | ENSG0000C |
| 70691064 | 70691619 | -1 | processed_pseudoge | ENSG0000C |
| 70752955 | 70753149 | -1 | processed_pseudoge | ENSG0000C |
| 70802600 | 70819771 | 1 | protein_coding | ENSG0000C |
| 177870853 | 177871194 | -1 | lncRNA | ENSG0000C |
| 177875522 | 177955274 | 1 | transcribed_unproce | ENSG0000C |
| 177944158 | 177987099 | 1 | lncRNA | ENSG0000C |
| 177954877 | 177967034 | 1 | lncRNA | ENSG0000C |
| 177970078 | 177970531 | 1 | lncRNA | ENSG0000C |
| 177974393 | 177975358 | 1 | processed_pseudoge | ENSG0000C |
| 177996182 | 177996994 | -1 | protein_coding | ENSG0000C |
| 1318299 | 1345099 | -1 | protein_coding | ENSG0000C |
| 1363582 | 1380067 | -1 | lncRNA | ENSG0000C |
| 1370543 | 1370945 | -1 | processed_pseudoge | ENSG0000C |
| 1383159 | 1386409 | -1 | TEC | ENSG0000C |
| 1392790 | 1449666 | -1 | protein_coding | ENSG0000C |
| 1460716 | 1505839 | -1 | protein_coding | ENSG0000C |
| 177870853 | 177871194 | -1 | lncRNA | ENSG0000C |
| 177875522 | 177950732 | 1 | transcribed_unproce | ENSG0000C |
| 177939622 | 177983334 | 1 | lncRNA | ENSG0000C |
| 177950335 | 177963960 | 1 | lncRNA | ENSG0000C |
| 177967004 | 177967457 | 1 | lncRNA | ENSG0000C |
| 177971313 | 177972279 | 1 | processed_pseudoge | ENSG0000C |
| 177992545 | 177996421 | -1 | protein_coding | ENSG0000C |
| 987180 | 997308 | -1 | lncRNA | ENSG0000C |
| 1005039 | 1006623 | 1 | lncRNA | ENSG0000C |
| 1008829 | 1038943 | 1 | protein_coding | ENSG0000C |
| 1050378 | 1111210 | -1 | protein_coding | ENSG0000C |
| 125325350 | 125367734 | 1 | lncRNA | ENSG0000C |
| 125350868 | 125351131 | -1 | misc_RNA | ENSG0000C |
| 125376828 | 125377439 | -1 | lncRNA | ENSG0000C |

| | | | | |
|---|---|---|---|---|
| 524112 | 524332 | -1 | protein_coding | ENSG00000( |
| 524705 | 526594 | 1 | lncRNA | ENSG00000( |
| 604294 | 613884 | -1 | lncRNA | ENSG00000( |
| 614020 | 626253 | 1 | protein_coding | ENSG00000( |
| 33842143 | 33894178 | -1 | protein_coding | ENSG00000( |
| 33890040 | 33890146 | -1 | snRNA | ENSG00000( |
| 33938372 | 33940223 | 1 | protein_coding | ENSG00000( |
| 33946602 | 33956490 | -1 | protein_coding | ENSG00000( |
| 21196720 | 21196860 | 1 | processed_pseudoge | ENSG00000( |
| 21522076 | 21527875 | 1 | protein_coding | ENSG00000( |
| 21542357 | 21548155 | 1 | protein_coding | ENSG00000( |
| 21544470 | 21545630 | 1 | TEC | ENSG00000( |
| 21561834 | 21567636 | 1 | protein_coding | ENSG00000( |
| 21582121 | 21587921 | 1 | protein_coding | ENSG00000( |
| 21592308 | 21594398 | -1 | protein_coding | ENSG00000( |
| 21601596 | 21607404 | 1 | protein_coding | ENSG00000( |
| 21621899 | 21627692 | 1 | protein_coding | ENSG00000( |
| 21700313 | 21700603 | 1 | lncRNA | ENSG00000( |
| 21746820 | 21779522 | 1 | lncRNA | ENSG00000( |
| 79354443 | 79354713 | -1 | processed_pseudoge | ENSG00000( |
| 79384784 | 79385673 | 1 | processed_pseudoge | ENSG00000( |
| 79434072 | 79434606 | -1 | processed_pseudoge | ENSG00000( |
| 79448129 | 79449137 | 1 | lncRNA | ENSG00000( |
| 79464740 | 79464999 | -1 | processed_pseudoge | ENSG00000( |
| 168820423 | 168820656 | 1 | TEC | ENSG00000( |
| 169725091 | 169725854 | -1 | lncRNA | ENSG00000( |
| 169725504 | 169738992 | 1 | lncRNA | ENSG00000( |
| 169740114 | 169751587 | -1 | protein_coding | ENSG00000( |
| 169751622 | 169781584 | 1 | protein_coding | ENSG00000( |
| 169770413 | 169772042 | 1 | lncRNA | ENSG00000( |
| 169788790 | 169798825 | -1 | lncRNA | ENSG00000( |
| 169790321 | 169802873 | 1 | lncRNA | ENSG00000( |
| 169809318 | 169810102 | -1 | lncRNA | ENSG00000( |
| 170272763 | 170279887 | 1 | lncRNA | ENSG00000( |
| 170275331 | 170276762 | -1 | lncRNA | ENSG00000( |
| 170282206 | 170306565 | -1 | protein_coding | ENSG00000( |
| 170290703 | 170416388 | 1 | protein_coding | ENSG00000( |
| 170297876 | 170298375 | -1 | lncRNA | ENSG00000( |
| 170423462 | 170428648 | 1 | lncRNA | ENSG00000( |
| 170544440 | 170544544 | -1 | lncRNA | ENSG00000( |
| 68266500 | 68342717 | -1 | lncRNA | ENSG00000( |
| 65788417 | 65789287 | 1 | processed_pseudoge | ENSG00000( |
| 65836930 | 65838039 | -1 | processed_pseudoge | ENSG00000( |
| 127654860 | 127655278 | -1 | processed_pseudoge | ENSG00000( |
| 127659425 | 127660338 | 1 | unprocessed_pseudc | ENSG00000( |
| 127659690 | 127660579 | -1 | unprocessed_pseudc | ENSG00000( |
| 127664554 | 127681555 | 1 | lncRNA | ENSG00000( |
| 127708072 | 127918597 | -1 | protein_coding | ENSG00000( |
| 127909833 | 127910052 | 1 | processed_pseudoge | ENSG00000( |
| 127968779 | 128524810 | -1 | protein_coding | ENSG00000( |

| | | | | |
|---|---|---|---|---|
| 128032022 | 128089384 | 1 | lncRNA | ENSG00000 |
| 128504663 | 128505312 | 1 | lncRNA | ENSG00000 |
| 166229334 | 166229554 | 1 | unprocessed_pseudc | ENSG00000 |
| 166236969 | 166256947 | 1 | lncRNA | ENSG00000 |
| 166240290 | 166240493 | -1 | processed_pseudoge | ENSG00000 |
| 166275457 | 166277068 | -1 | lncRNA | ENSG00000 |
| 32402558 | 32414973 | -1 | protein_coding | ENSG00000 |
| 32446915 | 32452117 | 1 | protein_coding | ENSG00000 |
| 28783625 | 28783996 | -1 | processed_pseudoge | ENSG00000 |
| 28837861 | 28838998 | -1 | lncRNA | ENSG00000 |
| 28859616 | 28863341 | -1 | lncRNA | ENSG00000 |
| 28861407 | 28862038 | 1 | processed_pseudoge | ENSG00000 |
| 28888823 | 28889897 | -1 | processed_pseudoge | ENSG00000 |
| 28896521 | 28897313 | 1 | lncRNA | ENSG00000 |
| 28902993 | 28923981 | -1 | protein_coding | ENSG00000 |
| 28915637 | 28915736 | 1 | snRNA | ENSG00000 |
| 28986195 | 28987476 | 1 | lncRNA | ENSG00000 |
| 28986792 | 28988567 | 1 | lncRNA | ENSG00000 |
| 28994816 | 29005348 | -1 | protein_coding | ENSG00000 |
| 29026712 | 29027639 | -1 | unprocessed_pseudc | ENSG00000 |
| 29044245 | 29045272 | -1 | protein_coding | ENSG00000 |
| 29086337 | 29087343 | -1 | protein_coding | ENSG00000 |
| 29100980 | 29101918 | 1 | unprocessed_pseudc | ENSG00000 |
| 29111918 | 29112853 | 1 | protein_coding | ENSG00000 |
| 29124236 | 29128934 | 1 | lncRNA | ENSG00000 |
| 29137903 | 29138839 | -1 | unprocessed_pseudc | ENSG00000 |
| 29173336 | 29174607 | 1 | protein_coding | ENSG00000 |
| 29266972 | 29267126 | 1 | lncRNA | ENSG00000 |
| 29268503 | 29269462 | -1 | unprocessed_pseudc | ENSG00000 |
| 29290770 | 29291712 | 1 | unprocessed_pseudc | ENSG00000 |
| 29306763 | 29307728 | 1 | protein_coding | ENSG00000 |
| 29422681 | 29432038 | -1 | lncRNA | ENSG00000 |
| 29425576 | 29457138 | -1 | protein_coding | ENSG00000 |
| 29439377 | 29441048 | 1 | protein_coding | ENSG00000 |
| 29444942 | 29450170 | -1 | lncRNA | ENSG00000 |
| 29457248 | 29464396 | 1 | protein_coding | ENSG00000 |
| 29464664 | 29469874 | -1 | unprocessed_pseudc | ENSG00000 |
| 29475093 | 29476139 | -1 | unprocessed_pseudc | ENSG00000 |
| 29486834 | 29487970 | -1 | protein_coding | ENSG00000 |
| 29489340 | 29489745 | -1 | processed_pseudoge | ENSG00000 |
| 29497578 | 29510624 | 1 | lncRNA | ENSG00000 |
| 29529494 | 29533642 | 1 | lncRNA | ENSG00000 |
| 29537778 | 29538861 | -1 | unprocessed_pseudc | ENSG00000 |
| 29553320 | 29554267 | 1 | unprocessed_pseudc | ENSG00000 |
| 29555714 | 29633284 | -1 | protein_coding | ENSG00000 |
| 29555714 | 29560028 | -1 | protein_coding | ENSG00000 |
| 29574010 | 29574937 | 1 | unprocessed_pseudc | ENSG00000 |
| 29577561 | 29577850 | 1 | processed_pseudoge | ENSG00000 |
| 29582350 | 29582433 | 1 | snoRNA | ENSG00000 |
| 29582609 | 29583126 | 1 | processed_pseudoge | ENSG00000 |

| | | | | |
|---|---|---|---|---|
| 29587555 | 29589138 | 1 | protein_coding | ENSG00000 |
| 29636180 | 29636463 | -1 | processed_pseudoge | ENSG00000 |
| 29657097 | 29672487 | 1 | protein_coding | ENSG00000 |
| 29672598 | 29673248 | -1 | protein_coding | ENSG00000 |
| 29791100 | 29797819 | 1 | transcribed_unproce | ENSG00000 |
| 29791993 | 29792967 | -1 | unprocessed_pseudo | ENSG00000 |
| 29800445 | 29802452 | 1 | unprocessed_pseudo | ENSG00000 |
| 29803224 | 29804020 | -1 | processed_pseudoge | ENSG00000 |
| 29812592 | 29812723 | -1 | unprocessed_pseudo | ENSG00000 |
| 29826986 | 29831158 | 1 | protein_coding | ENSG00000 |
| 29827404 | 29828384 | -1 | unprocessed_pseudo | ENSG00000 |
| 29852167 | 29852298 | -1 | unprocessed_pseudo | ENSG00000 |
| 29881442 | 29882408 | -1 | unprocessed_pseudo | ENSG00000 |
| 29881755 | 30002076 | 1 | protein_coding | ENSG00000 |
| 29881898 | 29884620 | 1 | unprocessed_pseudo | ENSG00000 |
| 29890694 | 29891825 | 1 | unprocessed_pseudo | ENSG00000 |
| 29900659 | 29901025 | 1 | unprocessed_pseudo | ENSG00000 |
| 29901898 | 29902760 | -1 | unprocessed_pseudo | ENSG00000 |
| 29920067 | 29921057 | -1 | unprocessed_pseudo | ENSG00000 |
| 29920543 | 29923320 | 1 | unprocessed_pseudo | ENSG00000 |
| 29926431 | 29926616 | 1 | unprocessed_pseudo | ENSG00000 |
| 29934229 | 29935221 | -1 | unprocessed_pseudo | ENSG00000 |
| 29948716 | 29950690 | 1 | unprocessed_pseudo | ENSG00000 |
| 29985328 | 29985724 | 1 | unprocessed_pseudo | ENSG00000 |
| 29986556 | 29987433 | -1 | unprocessed_pseudo | ENSG00000 |
| 29993130 | 30054145 | -1 | lncRNA | ENSG00000 |
| 29998237 | 29999232 | -1 | unprocessed_pseudo | ENSG00000 |
| 29998699 | 30002072 | 1 | transcribed_unproce | ENSG00000 |
| 30024180 | 30025470 | 1 | processed_pseudoge | ENSG00000 |
| 30028383 | 30028450 | 1 | misc_RNA | ENSG00000 |
| 30050620 | 30050695 | 1 | misc_RNA | ENSG00000 |
| 30051404 | 30057408 | 1 | protein_coding | ENSG00000 |
| 30053585 | 30053688 | 1 | misc_RNA | ENSG00000 |
| 30059208 | 30062832 | 1 | protein_coding | ENSG00000 |
| 30062765 | 30068386 | -1 | protein_coding | ENSG00000 |
| 30102946 | 30105603 | -1 | protein_coding | ENSG00000 |
| 30104215 | 30107221 | 1 | lncRNA | ENSG00000 |
| 30125305 | 30125466 | -1 | snoRNA | ENSG00000 |
| 30128631 | 30141238 | 1 | protein_coding | ENSG00000 |
| 30144448 | 30153436 | -1 | protein_coding | ENSG00000 |
| 30155693 | 30165183 | 1 | protein_coding | ENSG00000 |
| 30176913 | 30205889 | -1 | protein_coding | ENSG00000 |
| 30179256 | 30181079 | -1 | processed_pseudoge | ENSG00000 |
| 30226495 | 30390031 | -1 | lncRNA | ENSG00000 |
| 30230757 | 30234733 | 1 | unprocessed_pseudo | ENSG00000 |
| 30354705 | 30391047 | -1 | lncRNA | ENSG00000 |
| 30390376 | 30407632 | 1 | protein_coding | ENSG00000 |
| 30393209 | 30410787 | 1 | protein_coding | ENSG00000 |
| 30415319 | 30415453 | 1 | unprocessed_pseudo | ENSG00000 |
| 30423182 | 30423815 | -1 | processed_pseudoge | ENSG00000 |

| | | | | |
|---|---|---|---|---|
| 30427014 | 30428182 | -1 | processed_pseudoge | ENSG00000( |
| 30478618 | 30483222 | 1 | unprocessed_pseudc | ENSG00000( |
| 30580177 | 30583131 | 1 | lncRNA | ENSG00000( |
| 30605288 | 30617469 | -1 | protein_coding | ENSG00000( |
| 30723385 | 30736950 | -1 | protein_coding | ENSG00000( |
| 30740303 | 30751809 | -1 | protein_coding | ENSG00000( |
| 30751961 | 30755334 | -1 | protein_coding | ENSG00000( |
| 30760718 | 30761447 | -1 | processed_pseudoge | ENSG00000( |
| 30763719 | 30781800 | -1 | protein_coding | ENSG00000( |
| 30766979 | 30777095 | 1 | lncRNA | ENSG00000( |
| 30784111 | 30789336 | 1 | protein_coding | ENSG00000( |
| 30791619 | 30794413 | -1 | protein_coding | ENSG00000( |
| 30949136 | 30953340 | 1 | protein_coding | ENSG00000( |
| 31047625 | 31053805 | 1 | protein_coding | ENSG00000( |
| 31094157 | 31098956 | 1 | lncRNA | ENSG00000( |
| 31120427 | 31126868 | 1 | lncRNA | ENSG00000( |
| 31561856 | 31578081 | 1 | protein_coding | ENSG00000( |
| 31563664 | 31563765 | -1 | misc_RNA | ENSG00000( |
| 31586437 | 31587248 | -1 | processed_pseudoge | ENSG00000( |
| 31595070 | 31595649 | 1 | processed_pseudoge | ENSG00000( |
| 31595673 | 31597188 | 1 | protein_coding | ENSG00000( |
| 31597175 | 31606140 | -1 | protein_coding | ENSG00000( |
| 31603330 | 31603405 | -1 | snoRNA | ENSG00000( |
| 31643520 | 31645322 | 1 | protein_coding | ENSG00000( |
| 31647541 | 31649508 | -1 | protein_coding | ENSG00000( |
| 31653110 | 31655895 | 1 | protein_coding | ENSG00000( |
| 31655881 | 31659971 | -1 | protein_coding | ENSG00000( |
| 31678099 | 31678372 | -1 | processed_pseudoge | ENSG00000( |
| 31823611 | 31830163 | 1 | protein_coding | ENSG00000( |
| 31823970 | 31824072 | -1 | snRNA | ENSG00000( |
| 31825184 | 31831870 | 1 | protein_coding | ENSG00000( |
| 31829818 | 31831870 | 1 | protein_coding | ENSG00000( |
| 31831329 | 31832607 | -1 | lncRNA | ENSG00000( |
| 31832609 | 31844312 | -1 | protein_coding | ENSG00000( |
| 31844536 | 31862971 | -1 | protein_coding | ENSG00000( |
| 31845835 | 31845923 | -1 | misc_RNA | ENSG00000( |
| 31864414 | 31874003 | -1 | protein_coding | ENSG00000( |
| 31876648 | 31882094 | -1 | protein_coding | ENSG00000( |
| 31882493 | 31884975 | 1 | protein_coding | ENSG00000( |
| 31894767 | 31897281 | 1 | protein_coding | ENSG00000( |
| 31901630 | 31906788 | 1 | protein_coding | ENSG00000( |
| 31902285 | 31902347 | 1 | snoRNA | ENSG00000( |
| 31904099 | 31904165 | 1 | snoRNA | ENSG00000( |
| 31924682 | 31929929 | -1 | protein_coding | ENSG00000( |
| 31930215 | 31946068 | -1 | protein_coding | ENSG00000( |
| 31946781 | 31964716 | -1 | protein_coding | ENSG00000( |
| 31950783 | 31951076 | 1 | lncRNA | ENSG00000( |
| 32140675 | 32153125 | -1 | protein_coding | ENSG00000( |
| 32145516 | 32145636 | -1 | rRNA | ENSG00000( |
| 32215364 | 32221379 | -1 | protein_coding | ENSG00000( |

| | | | | |
|---|---|---|---|---|
| 32220447 | 32238987 | 1 | protein_coding | ENSG00000 |
| 32231591 | 32235290 | 1 | protein_coding | ENSG00000 |
| 32235221 | 32245105 | -1 | protein_coding | ENSG00000 |
| 32245363 | 32247802 | 1 | protein_coding | ENSG00000 |
| 32247977 | 32251333 | -1 | protein_coding | ENSG00000 |
| 32251742 | 32257192 | -1 | protein_coding | ENSG00000 |
| 32257772 | 32262529 | -1 | protein_coding | ENSG00000 |
| 32261849 | 32271406 | -1 | protein_coding | ENSG00000 |
| 32383385 | 32438560 | -1 | protein_coding | ENSG00000 |
| 32392216 | 32393175 | 1 | processed_pseudoge | ENSG00000 |
| 32419522 | 32419628 | 1 | snRNA | ENSG00000 |
| 32457148 | 32460324 | 1 | lncRNA | ENSG00000 |
| 32460601 | 32473766 | -1 | protein_coding | ENSG00000 |
| 32526470 | 32540151 | -1 | unprocessed_pseudo | ENSG00000 |
| 32556348 | 32556496 | -1 | snRNA | ENSG00000 |
| 32558296 | 32572041 | -1 | protein_coding | ENSG00000 |
| 32758260 | 32759090 | 1 | lncRNA | ENSG00000 |
| 32770986 | 32771923 | -1 | unprocessed_pseudo | ENSG00000 |
| 32781927 | 32787758 | 1 | protein_coding | ENSG00000 |
| 32796644 | 32804081 | -1 | protein_coding | ENSG00000 |
| 32852194 | 32856479 | -1 | protein_coding | ENSG00000 |
| 32861254 | 32878050 | -1 | protein_coding | ENSG00000 |
| 32879989 | 32883975 | -1 | protein_coding | ENSG00000 |
| 32883358 | 32885767 | 1 | lncRNA | ENSG00000 |
| 32883408 | 32918836 | 1 | protein_coding | ENSG00000 |
| 32884481 | 32893250 | -1 | protein_coding | ENSG00000 |
| 32918409 | 32919085 | -1 | transcribed_processe | ENSG00000 |
| 32935645 | 32935719 | 1 | unprocessed_pseudo | ENSG00000 |
| 32973881 | 32992379 | -1 | protein_coding | ENSG00000 |
| 32987867 | 32992379 | -1 | protein_coding | ENSG00000 |
| 33103656 | 33119850 | -1 | protein_coding | ENSG00000 |
| 33114995 | 33126260 | 1 | protein_coding | ENSG00000 |
| 33118526 | 33118935 | 1 | processed_pseudoge | ENSG00000 |
| 33130812 | 33136354 | -1 | unprocessed_pseudo | ENSG00000 |
| 33142826 | 33146362 | -1 | unprocessed_pseudo | ENSG00000 |
| 33151483 | 33168112 | 1 | transcribed_unproce | ENSG00000 |
| 33170244 | 33170370 | -1 | unprocessed_pseudo | ENSG00000 |
| 33404609 | 33405304 | -1 | processed_pseudoge | ENSG00000 |
| 28510773 | 28515743 | -1 | protein_coding | ENSG00000 |
| 28526012 | 28534951 | 1 | protein_coding | ENSG00000 |
| 28573507 | 28587330 | -1 | protein_coding | ENSG00000 |
| 28587373 | 28591739 | 1 | lncRNA | ENSG00000 |
| 28648243 | 28654475 | -1 | lncRNA | ENSG00000 |
| 28732809 | 28733696 | 1 | processed_pseudoge | ENSG00000 |
| 28784450 | 28784821 | -1 | processed_pseudoge | ENSG00000 |
| 28838507 | 28839644 | -1 | lncRNA | ENSG00000 |
| 28860261 | 28864009 | -1 | lncRNA | ENSG00000 |
| 28862047 | 28862678 | 1 | processed_pseudoge | ENSG00000 |
| 28889657 | 28890731 | -1 | processed_pseudoge | ENSG00000 |
| 28897348 | 28898138 | 1 | lncRNA | ENSG00000 |

| | | | | |
|---|---|---|---|---|
| 28903816 | 28924807 | -1 | protein_coding | ENSG00000 |
| 28916461 | 28916560 | 1 | snRNA | ENSG00000 |
| 28940401 | 28941061 | 1 | protein_coding | ENSG00000 |
| 28965658 | 28966988 | -1 | processed_pseudoge | ENSG00000 |
| 28974068 | 28974366 | 1 | misc_RNA | ENSG00000 |
| 28982795 | 28984076 | 1 | lncRNA | ENSG00000 |
| 28983392 | 28985128 | 1 | lncRNA | ENSG00000 |
| 28991375 | 29001915 | -1 | protein_coding | ENSG00000 |
| 29023286 | 29024213 | -1 | unprocessed_pseudo | ENSG00000 |
| 29040820 | 29041847 | -1 | protein_coding | ENSG00000 |
| 29060593 | 29073129 | 1 | protein_coding | ENSG00000 |
| 29068429 | 29069201 | -1 | unprocessed_pseudo | ENSG00000 |
| 29073178 | 29073773 | -1 | processed_pseudoge | ENSG00000 |
| 29082970 | 29083976 | -1 | protein_coding | ENSG00000 |
| 29097608 | 29098546 | 1 | unprocessed_pseudo | ENSG00000 |
| 29108558 | 29109493 | 1 | protein_coding | ENSG00000 |
| 29120878 | 29125574 | 1 | lncRNA | ENSG00000 |
| 29134543 | 29135479 | -1 | unprocessed_pseudo | ENSG00000 |
| 29169965 | 29171236 | 1 | protein_coding | ENSG00000 |
| 29178191 | 29179126 | 1 | unprocessed_pseudo | ENSG00000 |
| 29211927 | 29212864 | 1 | unprocessed_pseudo | ENSG00000 |
| 29220658 | 29263772 | 1 | lncRNA | ENSG00000 |
| 29225912 | 29226842 | -1 | unprocessed_pseudo | ENSG00000 |
| 29259390 | 29260766 | -1 | unprocessed_pseudo | ENSG00000 |
| 29265149 | 29266108 | -1 | unprocessed_pseudo | ENSG00000 |
| 29287419 | 29288361 | 1 | unprocessed_pseudo | ENSG00000 |
| 29303399 | 29304364 | 1 | protein_coding | ENSG00000 |
| 29326337 | 29327771 | -1 | processed_pseudoge | ENSG00000 |
| 29351939 | 29428535 | -1 | protein_coding | ENSG00000 |
| 29370147 | 29446668 | -1 | protein_coding | ENSG00000 |
| 29393135 | 29394203 | 1 | protein_coding | ENSG00000 |
| 29413850 | 29414796 | 1 | unprocessed_pseudo | ENSG00000 |
| 29422072 | 29453643 | -1 | protein_coding | ENSG00000 |
| 29435874 | 29437545 | 1 | protein_coding | ENSG00000 |
| 29453753 | 29460899 | 1 | protein_coding | ENSG00000 |
| 29461167 | 29466379 | -1 | unprocessed_pseudo | ENSG00000 |
| 29471596 | 29472642 | -1 | unprocessed_pseudo | ENSG00000 |
| 29483337 | 29484473 | -1 | protein_coding | ENSG00000 |
| 29485842 | 29486247 | -1 | processed_pseudoge | ENSG00000 |
| 29494080 | 29507127 | 1 | lncRNA | ENSG00000 |
| 29526002 | 29530150 | 1 | lncRNA | ENSG00000 |
| 29534286 | 29535369 | -1 | processed_pseudoge | ENSG00000 |
| 29549819 | 29550766 | 1 | unprocessed_pseudo | ENSG00000 |
| 29552212 | 29556525 | -1 | protein_coding | ENSG00000 |
| 29552212 | 29629760 | -1 | protein_coding | ENSG00000 |
| 29570508 | 29571435 | 1 | unprocessed_pseudo | ENSG00000 |
| 29574058 | 29574347 | 1 | processed_pseudoge | ENSG00000 |
| 29578848 | 29578931 | 1 | snoRNA | ENSG00000 |
| 29579107 | 29579624 | 1 | processed_pseudoge | ENSG00000 |
| 29584054 | 29585637 | 1 | protein_coding | ENSG00000 |

| | | | | |
|---|---|---|---|---|
| 29632655 | 29632938 | -1 | processed_pseudoge | ENSG00000( |
| 29653572 | 29668964 | 1 | protein_coding | ENSG00000( |
| 29668984 | 29677737 | -1 | protein_coding | ENSG00000( |
| 29704757 | 29705179 | -1 | processed_pseudoge | ENSG00000( |
| 29716754 | 29732497 | 1 | protein_coding | ENSG00000( |
| 29716960 | 29717950 | -1 | unprocessed_pseudc | ENSG00000( |
| 29720576 | 29743014 | -1 | lncRNA | ENSG00000( |
| 29720644 | 29721114 | -1 | transcribed_process | ENSG00000( |
| 29735695 | 29742934 | -1 | transcribed_unproce | ENSG00000( |
| 29742254 | 29742478 | 1 | unprocessed_pseudc | ENSG00000( |
| 29744335 | 29744436 | 1 | misc_RNA | ENSG00000( |
| 29744691 | 29745111 | -1 | processed_pseudoge | ENSG00000( |
| 29785254 | 29791477 | 1 | transcribed_unproce | ENSG00000( |
| 29785655 | 29786629 | -1 | unprocessed_pseudc | ENSG00000( |
| 29794107 | 29796116 | 1 | unprocessed_pseudc | ENSG00000( |
| 29796886 | 29797682 | -1 | processed_pseudoge | ENSG00000( |
| 29806254 | 29806385 | -1 | processed_pseudoge | ENSG00000( |
| 29820648 | 29824817 | 1 | protein_coding | ENSG00000( |
| 29821066 | 29822046 | -1 | unprocessed_pseudc | ENSG00000( |
| 29849539 | 29849688 | -1 | unprocessed_pseudc | ENSG00000( |
| 29878828 | 29879794 | -1 | unprocessed_pseudc | ENSG00000( |
| 29879141 | 29999464 | 1 | protein_coding | ENSG00000( |
| 29879284 | 29882006 | 1 | unprocessed_pseudc | ENSG00000( |
| 29888080 | 29889211 | 1 | unprocessed_pseudc | ENSG00000( |
| 29898045 | 29898411 | 1 | unprocessed_pseudc | ENSG00000( |
| 29899284 | 29900148 | -1 | unprocessed_pseudc | ENSG00000( |
| 29917455 | 29918445 | -1 | unprocessed_pseudc | ENSG00000( |
| 29917929 | 29920706 | 1 | unprocessed_pseudc | ENSG00000( |
| 29923817 | 29924002 | 1 | unprocessed_pseudc | ENSG00000( |
| 29931616 | 29932608 | -1 | unprocessed_pseudc | ENSG00000( |
| 29946103 | 29948077 | 1 | unprocessed_pseudc | ENSG00000( |
| 29960309 | 29961971 | -1 | unprocessed_pseudc | ENSG00000( |
| 29964608 | 29967904 | 1 | lncRNA | ENSG00000( |
| 29982716 | 29983112 | 1 | unprocessed_pseudc | ENSG00000( |
| 29983944 | 29984821 | -1 | unprocessed_pseudc | ENSG00000( |
| 29990518 | 30051536 | -1 | lncRNA | ENSG00000( |
| 29995625 | 29996628 | -1 | unprocessed_pseudc | ENSG00000( |
| 29996087 | 29999460 | 1 | transcribed_unproce | ENSG00000( |
| 30021567 | 30022857 | 1 | processed_pseudoge | ENSG00000( |
| 30025770 | 30025837 | 1 | misc_RNA | ENSG00000( |
| 30048011 | 30048086 | 1 | misc_RNA | ENSG00000( |
| 30048795 | 30054799 | 1 | protein_coding | ENSG00000( |
| 30050976 | 30051079 | 1 | misc_RNA | ENSG00000( |
| 30056599 | 30060223 | 1 | protein_coding | ENSG00000( |
| 30060156 | 30065777 | -1 | protein_coding | ENSG00000( |
| 30092809 | 30103024 | -1 | protein_coding | ENSG00000( |
| 30095152 | 30104642 | 1 | lncRNA | ENSG00000( |
| 30122730 | 30122889 | -1 | snoRNA | ENSG00000( |
| 30126032 | 30138662 | 1 | protein_coding | ENSG00000( |
| 30141872 | 30150859 | -1 | protein_coding | ENSG00000( |

| | | | | |
|---|---|---|---|---|
| 30153123 | 30162613 | 1 | protein_coding | ENSG0000C |
| 30174354 | 30203317 | -1 | protein_coding | ENSG0000C |
| 30176697 | 30178517 | -1 | processed_pseudoge | ENSG0000C |
| 30223924 | 30315930 | -1 | lncRNA | ENSG0000C |
| 30228173 | 30232162 | 1 | unprocessed_pseudc | ENSG0000C |
| 30249483 | 30256765 | 1 | transcribed_unproce | ENSG0000C |
| 30280607 | 30316946 | -1 | lncRNA | ENSG0000C |
| 30316275 | 30333529 | 1 | protein_coding | ENSG0000C |
| 30319108 | 30336684 | 1 | protein_coding | ENSG0000C |
| 30341216 | 30341350 | 1 | unprocessed_pseudc | ENSG0000C |
| 30349077 | 30349710 | -1 | processed_pseudoge | ENSG0000C |
| 30352921 | 30354089 | -1 | processed_pseudoge | ENSG0000C |
| 30404517 | 30409122 | 1 | unprocessed_pseudc | ENSG0000C |
| 30456371 | 30457898 | 1 | processed_pseudoge | ENSG0000C |
| 30458739 | 30460108 | 1 | processed_pseudoge | ENSG0000C |
| 30475795 | 30476445 | 1 | processed_pseudoge | ENSG0000C |
| 30479321 | 30484059 | 1 | protein_coding | ENSG0000C |
| 30506108 | 30509062 | 1 | lncRNA | ENSG0000C |
| 30531215 | 30547013 | -1 | protein_coding | ENSG0000C |
| 30546725 | 30553562 | 1 | protein_coding | ENSG0000C |
| 30561217 | 30587022 | 1 | protein_coding | ENSG0000C |
| 30590243 | 30608455 | -1 | protein_coding | ENSG0000C |
| 30607552 | 30616238 | 1 | protein_coding | ENSG0000C |
| 30616685 | 30636664 | 1 | protein_coding | ENSG0000C |
| 30623474 | 30623740 | 1 | processed_pseudoge | ENSG0000C |
| 30636880 | 30642942 | 1 | protein_coding | ENSG0000C |
| 30642952 | 30662900 | -1 | protein_coding | ENSG0000C |
| 30666252 | 30677758 | -1 | protein_coding | ENSG0000C |
| 30677910 | 30681283 | -1 | protein_coding | ENSG0000C |
| 30686667 | 30687396 | -1 | processed_pseudoge | ENSG0000C |
| 30689666 | 30707748 | -1 | protein_coding | ENSG0000C |
| 30692927 | 30703043 | 1 | lncRNA | ENSG0000C |
| 30710059 | 30715284 | 1 | protein_coding | ENSG0000C |
| 30717565 | 30732592 | -1 | protein_coding | ENSG0000C |
| 30726158 | 30726268 | -1 | misc_RNA | ENSG0000C |
| 30733058 | 30734413 | -1 | protein_coding | ENSG0000C |
| 30740863 | 30741114 | 1 | misc_RNA | ENSG0000C |
| 30756658 | 30782086 | 1 | lncRNA | ENSG0000C |
| 30788497 | 30820509 | -1 | lncRNA | ENSG0000C |
| 30829379 | 30838017 | -1 | lncRNA | ENSG0000C |
| 30854117 | 30854419 | 1 | misc_RNA | ENSG0000C |
| 30856848 | 30865810 | -1 | lncRNA | ENSG0000C |
| 30870863 | 30890046 | 1 | protein_coding | ENSG0000C |
| 30898049 | 30903971 | 1 | protein_coding | ENSG0000C |
| 30898107 | 30916176 | 1 | lncRNA | ENSG0000C |
| 30904070 | 30916176 | 1 | protein_coding | ENSG0000C |
| 30921066 | 30921888 | -1 | protein_coding | ENSG0000C |
| 30922332 | 30922420 | -1 | misc_RNA | ENSG0000C |
| 30930703 | 30943914 | 1 | protein_coding | ENSG0000C |
| 30935673 | 30944555 | -1 | lncRNA | ENSG0000C |

| | | | | |
|---|---|---|---|---|
| 30973408 | 30979603 | 1 | protein_coding | ENSG00000( |
| 31000184 | 31025215 | 1 | lncRNA | ENSG00000( |
| 31045946 | 31052387 | 1 | transcribed_unproce | ENSG00000( |
| 31075504 | 31075603 | 1 | snRNA | ENSG00000( |
| 31103717 | 31105053 | -1 | protein_coding | ENSG00000( |
| 31107244 | 31132550 | 1 | protein_coding | ENSG00000( |
| 31107584 | 31112941 | -1 | protein_coding | ENSG00000( |
| 31130046 | 31131862 | -1 | protein_coding | ENSG00000( |
| 31133236 | 31133422 | 1 | processed_pseudoge | ENSG00000( |
| 31134989 | 31150760 | -1 | protein_coding | ENSG00000( |
| 31151089 | 31156758 | 1 | protein_coding | ENSG00000( |
| 31156880 | 31163254 | -1 | protein_coding | ENSG00000( |
| 31166295 | 31170460 | -1 | lncRNA | ENSG00000( |
| 31190323 | 31196527 | 1 | lncRNA | ENSG00000( |
| 31192728 | 31194480 | -1 | lncRNA | ENSG00000( |
| 31259794 | 31263175 | -1 | protein_coding | ENSG00000( |
| 31260536 | 31347723 | -1 | protein_coding | ENSG00000( |
| 31266612 | 31269783 | 1 | processed_pseudoge | ENSG00000( |
| 31271346 | 31272548 | 1 | processed_pseudoge | ENSG00000( |
| 31278540 | 31279994 | -1 | processed_pseudoge | ENSG00000( |
| 31298844 | 31299578 | -1 | unprocessed_pseudc | ENSG00000( |
| 31356825 | 31357371 | -1 | processed_pseudoge | ENSG00000( |
| 31360589 | 31360695 | 1 | snRNA | ENSG00000( |
| 31367866 | 31368468 | 1 | processed_pseudoge | ENSG00000( |
| 31370871 | 31371297 | 1 | processed_pseudoge | ENSG00000( |
| 31372541 | 31372755 | -1 | unprocessed_pseudc | ENSG00000( |
| 31390270 | 31406729 | 1 | protein_coding | ENSG00000( |
| 31392641 | 31392739 | -1 | misc_RNA | ENSG00000( |
| 31432091 | 31437391 | 1 | lncRNA | ENSG00000( |
| 31450619 | 31453253 | 1 | lncRNA | ENSG00000( |
| 31484150 | 31484251 | -1 | misc_RNA | ENSG00000( |
| 31485545 | 31498583 | 1 | protein_coding | ENSG00000( |
| 31506943 | 31507754 | -1 | processed_pseudoge | ENSG00000( |
| 31515566 | 31516145 | 1 | processed_pseudoge | ENSG00000( |
| 31516169 | 31517684 | 1 | protein_coding | ENSG00000( |
| 31517671 | 31534027 | -1 | protein_coding | ENSG00000( |
| 31523826 | 31523901 | -1 | snoRNA | ENSG00000( |
| 31528550 | 31528627 | -1 | snoRNA | ENSG00000( |
| 31529753 | 31530587 | 1 | lncRNA | ENSG00000( |
| 31531911 | 31534293 | -1 | protein_coding | ENSG00000( |
| 31534315 | 31546261 | 1 | protein_coding | ENSG00000( |
| 31559460 | 31561730 | 1 | protein_coding | ENSG00000( |
| 31562973 | 31565742 | 1 | protein_coding | ENSG00000( |
| 31567961 | 31569928 | -1 | protein_coding | ENSG00000( |
| 31573530 | 31576311 | 1 | protein_coding | ENSG00000( |
| 31576297 | 31580387 | -1 | protein_coding | ENSG00000( |
| 31598491 | 31598764 | -1 | processed_pseudoge | ENSG00000( |
| 31602592 | 31604429 | 1 | protein_coding | ENSG00000( |
| 31608127 | 31625177 | 1 | protein_coding | ENSG00000( |
| 31610483 | 31610614 | 1 | snoRNA | ENSG00000( |

| | | | | |
|---|---|---|---|---|
| 31626429 | 31640114 | -1 | protein_coding | ENSG0000C |
| 31639825 | 31645618 | 1 | protein_coding | ENSG0000C |
| 31645706 | 31648178 | -1 | protein_coding | ENSG0000C |
| 31645737 | 31648128 | 1 | lncRNA | ENSG0000C |
| 31648638 | 31653693 | -1 | protein_coding | ENSG0000C |
| 31650698 | 31650811 | 1 | misc_RNA | ENSG0000C |
| 31652646 | 31660956 | 1 | protein_coding | ENSG0000C |
| 31656528 | 31660956 | 1 | protein_coding | ENSG0000C |
| 31664094 | 31671449 | -1 | protein_coding | ENSG0000C |
| 31674358 | 31690824 | -1 | protein_coding | ENSG0000C |
| 31694294 | 31698004 | 1 | protein_coding | ENSG0000C |
| 31694313 | 31705325 | 1 | protein_coding | ENSG0000C |
| 31699739 | 31701215 | -1 | protein_coding | ENSG0000C |
| 31706001 | 31714117 | 1 | protein_coding | ENSG0000C |
| 31706055 | 31709252 | -1 | protein_coding | ENSG0000C |
| 31714445 | 31718024 | -1 | protein_coding | ENSG0000C |
| 31717988 | 31727162 | -1 | protein_coding | ENSG0000C |
| 31727390 | 31750052 | 1 | protein_coding | ENSG0000C |
| 31727419 | 31752228 | 1 | protein_coding | ENSG0000C |
| 31744327 | 31744429 | -1 | snRNA | ENSG0000C |
| 31750175 | 31752228 | 1 | protein_coding | ENSG0000C |
| 31751686 | 31752965 | -1 | lncRNA | ENSG0000C |
| 31752967 | 31764655 | -1 | protein_coding | ENSG0000C |
| 31764879 | 31783312 | -1 | protein_coding | ENSG0000C |
| 31766178 | 31766266 | -1 | misc_RNA | ENSG0000C |
| 31784754 | 31794345 | -1 | protein_coding | ENSG0000C |
| 31796990 | 31802435 | -1 | protein_coding | ENSG0000C |
| 31802834 | 31805316 | 1 | protein_coding | ENSG0000C |
| 31815107 | 31817630 | 1 | protein_coding | ENSG0000C |
| 31821974 | 31827129 | 1 | protein_coding | ENSG0000C |
| 31822629 | 31822691 | 1 | snoRNA | ENSG0000C |
| 31824440 | 31824506 | 1 | snoRNA | ENSG0000C |
| 31845195 | 31850443 | -1 | protein_coding | ENSG0000C |
| 31850729 | 31866590 | -1 | protein_coding | ENSG0000C |
| 31867303 | 31885231 | -1 | protein_coding | ENSG0000C |
| 31871305 | 31871598 | 1 | lncRNA | ENSG0000C |
| 31885329 | 31933220 | 1 | protein_coding | ENSG0000C |
| 31887151 | 31889536 | -1 | protein_coding | ENSG0000C |
| 31921995 | 31929270 | -1 | lncRNA | ENSG0000C |
| 31933196 | 31939630 | 1 | protein_coding | ENSG0000C |
| 31939633 | 31946638 | -1 | protein_coding | ENSG0000C |
| 31946626 | 31957297 | 1 | protein_coding | ENSG0000C |
| 31957356 | 31959838 | -1 | protein_coding | ENSG0000C |
| 31958721 | 31968992 | 1 | protein_coding | ENSG0000C |
| 31969570 | 31983859 | 1 | protein_coding | ENSG0000C |
| 31981170 | 31984699 | -1 | lncRNA | ENSG0000C |
| 31986739 | 31990111 | 1 | protein_coding | ENSG0000C |
| 31989596 | 32066062 | -1 | protein_coding | ENSG0000C |
| 32026965 | 32027085 | -1 | rRNA | ENSG0000C |
| 32063716 | 32076703 | -1 | protein_coding | ENSG0000C |

| | | | | |
|---|---|---|---|---|
| 32077157 | 32078741 | -1 | protein_coding | ENSG00000( |
| 32096815 | 32102831 | -1 | protein_coding | ENSG00000( |
| 32101899 | 32112151 | 1 | protein_coding | ENSG00000( |
| 32102303 | 32120449 | 1 | protein_coding | ENSG00000( |
| 32113054 | 32116752 | 1 | protein_coding | ENSG00000( |
| 32116683 | 32126567 | -1 | protein_coding | ENSG00000( |
| 32126831 | 32129264 | 1 | protein_coding | ENSG00000( |
| 32129439 | 32132795 | -1 | protein_coding | ENSG00000( |
| 32133204 | 32138654 | -1 | protein_coding | ENSG00000( |
| 32139234 | 32143991 | -1 | protein_coding | ENSG00000( |
| 32143311 | 32172533 | -1 | protein_coding | ENSG00000( |
| 32241266 | 32320157 | -1 | protein_coding | ENSG00000( |
| 32273814 | 32274773 | 1 | processed_pseudoge | ENSG00000( |
| 32301118 | 32301224 | 1 | snRNA | ENSG00000( |
| 32338729 | 32341897 | 1 | lncRNA | ENSG00000( |
| 32342174 | 32355327 | -1 | protein_coding | ENSG00000( |
| 32388087 | 32393254 | 1 | protein_coding | ENSG00000( |
| 32408020 | 32421718 | -1 | unprocessed_pseudc | ENSG00000( |
| 32441551 | 32441699 | -1 | snRNA | ENSG00000( |
| 32444128 | 32457245 | -1 | protein_coding | ENSG00000( |
| 32479741 | 32479887 | -1 | snRNA | ENSG00000( |
| 32483164 | 32497705 | -1 | unprocessed_pseudc | ENSG00000( |
| 32505701 | 32505847 | -1 | snRNA | ENSG00000( |
| 32508164 | 32538771 | -1 | protein_coding | ENSG00000( |
| 32565603 | 32671108 | 1 | protein_coding | ENSG00000( |
| 32582490 | 32590101 | -1 | protein_coding | ENSG00000( |
| 32582940 | 32583789 | 1 | lncRNA | ENSG00000( |
| 32582946 | 32585541 | -1 | lncRNA | ENSG00000( |
| 32629554 | 32630385 | -1 | unprocessed_pseudc | ENSG00000( |
| 32641404 | 32642234 | 1 | lncRNA | ENSG00000( |
| 32654132 | 32655069 | -1 | unprocessed_pseudc | ENSG00000( |
| 32679790 | 32687227 | -1 | protein_coding | ENSG00000( |
| 32735083 | 32739368 | -1 | protein_coding | ENSG00000( |
| 32744153 | 32761098 | -1 | protein_coding | ENSG00000( |
| 32763034 | 32767021 | -1 | protein_coding | ENSG00000( |
| 32766404 | 32768813 | 1 | lncRNA | ENSG00000( |
| 32766454 | 32801905 | 1 | protein_coding | ENSG00000( |
| 32767527 | 32776296 | -1 | protein_coding | ENSG00000( |
| 32801489 | 32802166 | -1 | transcribed_processe | ENSG00000( |
| 32818730 | 32818804 | 1 | unprocessed_pseudc | ENSG00000( |
| 32856943 | 32875441 | -1 | protein_coding | ENSG00000( |
| 32870929 | 32893044 | -1 | protein_coding | ENSG00000( |
| 32890988 | 32903839 | 1 | protein_coding | ENSG00000( |
| 32892631 | 32893145 | 1 | lncRNA | ENSG00000( |
| 32926465 | 32931899 | -1 | protein_coding | ENSG00000( |
| 32986539 | 33002744 | -1 | protein_coding | ENSG00000( |
| 32997896 | 33009106 | 1 | protein_coding | ENSG00000( |
| 33001420 | 33001829 | 1 | processed_pseudoge | ENSG00000( |
| 33013645 | 33019231 | -1 | unprocessed_pseudc | ENSG00000( |
| 33025668 | 33029223 | -1 | unprocessed_pseudc | ENSG00000( |

| | | | | |
|---|---|---|---|---|
| 33034341 | 33050662 | 1 | transcribed_unproce | ENSG0000C |
| 33052792 | 33064929 | -1 | unprocessed_pseudc | ENSG0000C |
| 33066394 | 33069387 | 1 | lncRNA | ENSG0000C |
| 33084308 | 33114125 | -1 | protein_coding | ENSG0000C |
| 33115214 | 33122477 | -1 | protein_coding | ENSG0000C |
| 33121225 | 33121320 | 1 | misc_RNA | ENSG0000C |
| 33122069 | 33126063 | 1 | protein_coding | ENSG0000C |
| 33126266 | 33128451 | 1 | protein_coding | ENSG0000C |
| 33130117 | 33134342 | 1 | protein_coding | ENSG0000C |
| 33137325 | 33137948 | 1 | processed_pseudoge | ENSG0000C |
| 33159435 | 33161326 | 1 | processed_pseudoge | ENSG0000C |
| 33171150 | 33176605 | 1 | lncRNA | ENSG0000C |
| 33171888 | 33193647 | -1 | protein_coding | ENSG0000C |
| 33193610 | 33198110 | 1 | protein_coding | ENSG0000C |
| 33198732 | 33200425 | 1 | protein_coding | ENSG0000C |
| 33200708 | 33211131 | -1 | protein_coding | ENSG0000C |
| 33210906 | 33220005 | 1 | protein_coding | ENSG0000C |
| 33213258 | 33220928 | -1 | protein_coding | ENSG0000C |
| 33221298 | 33235989 | -1 | protein_coding | ENSG0000C |
| 33236020 | 33239546 | -1 | protein_coding | ENSG0000C |
| 33240162 | 33250873 | -1 | protein_coding | ENSG0000C |
| 33260584 | 33261101 | 1 | processed_pseudoge | ENSG0000C |
| 33287152 | 33287847 | -1 | processed_pseudoge | ENSG0000C |
| 28783633 | 28784004 | -1 | processed_pseudoge | ENSG0000C |
| 28837878 | 28839015 | -1 | lncRNA | ENSG0000C |
| 28859634 | 28863359 | -1 | lncRNA | ENSG0000C |
| 28861425 | 28862056 | 1 | processed_pseudoge | ENSG0000C |
| 28977485 | 28977783 | 1 | misc_RNA | ENSG0000C |
| 28986213 | 28987494 | 1 | lncRNA | ENSG0000C |
| 28986810 | 28988546 | 1 | lncRNA | ENSG0000C |
| 28994795 | 29005326 | -1 | protein_coding | ENSG0000C |
| 29026690 | 29027617 | -1 | unprocessed_pseudc | ENSG0000C |
| 29044223 | 29045250 | -1 | protein_coding | ENSG0000C |
| 29064006 | 29076532 | 1 | protein_coding | ENSG0000C |
| 29071832 | 29072607 | -1 | unprocessed_pseudc | ENSG0000C |
| 29076581 | 29077176 | -1 | processed_pseudoge | ENSG0000C |
| 29086315 | 29087321 | -1 | protein_coding | ENSG0000C |
| 29100958 | 29101896 | 1 | unprocessed_pseudc | ENSG0000C |
| 29111900 | 29112835 | 1 | protein_coding | ENSG0000C |
| 29124218 | 29128916 | 1 | lncRNA | ENSG0000C |
| 29137885 | 29138821 | -1 | unprocessed_pseudc | ENSG0000C |
| 29173317 | 29174588 | 1 | protein_coding | ENSG0000C |
| 29181532 | 29182467 | 1 | unprocessed_pseudc | ENSG0000C |
| 29215281 | 29216224 | 1 | unprocessed_pseudc | ENSG0000C |
| 29224018 | 29267104 | 1 | lncRNA | ENSG0000C |
| 29229272 | 29230202 | -1 | unprocessed_pseudc | ENSG0000C |
| 29262722 | 29264098 | -1 | unprocessed_pseudc | ENSG0000C |
| 29268481 | 29269440 | -1 | unprocessed_pseudc | ENSG0000C |
| 29290747 | 29291689 | 1 | unprocessed_pseudc | ENSG0000C |
| 29306740 | 29307705 | 1 | protein_coding | ENSG0000C |

| | | | | |
|---|---|---|---|---|
| 29329677 | 29331112 | -1 | processed_pseudoge | ENSG00000( |
| 29373478 | 29450027 | -1 | protein_coding | ENSG00000( |
| 29396471 | 29397539 | 1 | protein_coding | ENSG00000( |
| 29417201 | 29418163 | 1 | unprocessed_pseudc | ENSG00000( |
| 29422544 | 29431896 | -1 | lncRNA | ENSG00000( |
| 29425439 | 29457001 | -1 | protein_coding | ENSG00000( |
| 29439235 | 29440906 | 1 | protein_coding | ENSG00000( |
| 29457111 | 29464258 | 1 | protein_coding | ENSG00000( |
| 29464526 | 29469736 | -1 | unprocessed_pseudc | ENSG00000( |
| 29474955 | 29476001 | -1 | unprocessed_pseudc | ENSG00000( |
| 29486696 | 29487832 | -1 | protein_coding | ENSG00000( |
| 29489201 | 29489606 | -1 | processed_pseudoge | ENSG00000( |
| 29497439 | 29510486 | 1 | lncRNA | ENSG00000( |
| 29529354 | 29533502 | 1 | lncRNA | ENSG00000( |
| 29537638 | 29538721 | -1 | unprocessed_pseudc | ENSG00000( |
| 29553169 | 29554116 | 1 | unprocessed_pseudc | ENSG00000( |
| 29555562 | 29559875 | -1 | protein_coding | ENSG00000( |
| 29555562 | 29633117 | -1 | protein_coding | ENSG00000( |
| 29573865 | 29574792 | 1 | unprocessed_pseudc | ENSG00000( |
| 29577415 | 29577704 | 1 | processed_pseudoge | ENSG00000( |
| 29582205 | 29582288 | 1 | snoRNA | ENSG00000( |
| 29582464 | 29582981 | 1 | processed_pseudoge | ENSG00000( |
| 29587411 | 29588994 | 1 | protein_coding | ENSG00000( |
| 29636014 | 29636297 | -1 | processed_pseudoge | ENSG00000( |
| 29656933 | 29672325 | 1 | protein_coding | ENSG00000( |
| 29672345 | 29681055 | -1 | protein_coding | ENSG00000( |
| 29708062 | 29708484 | -1 | processed_pseudoge | ENSG00000( |
| 29720256 | 29736008 | 1 | protein_coding | ENSG00000( |
| 29720462 | 29721452 | -1 | unprocessed_pseudc | ENSG00000( |
| 29724084 | 29746526 | -1 | lncRNA | ENSG00000( |
| 29724152 | 29724622 | -1 | transcribed_processe | ENSG00000( |
| 29739210 | 29746446 | -1 | transcribed_unproce | ENSG00000( |
| 29745766 | 29745990 | 1 | unprocessed_pseudc | ENSG00000( |
| 29747847 | 29747948 | 1 | misc_RNA | ENSG00000( |
| 29748203 | 29748623 | -1 | processed_pseudoge | ENSG00000( |
| 29788252 | 29795033 | 1 | transcribed_unproce | ENSG00000( |
| 29789145 | 29790142 | -1 | unprocessed_pseudc | ENSG00000( |
| 29797637 | 29799645 | 1 | unprocessed_pseudc | ENSG00000( |
| 29800417 | 29801213 | -1 | processed_pseudoge | ENSG00000( |
| 29809789 | 29809920 | -1 | unprocessed_pseudc | ENSG00000( |
| 29824187 | 29828345 | 1 | protein_coding | ENSG00000( |
| 29824605 | 29825584 | -1 | unprocessed_pseudc | ENSG00000( |
| 29849566 | 29849694 | -1 | unprocessed_pseudc | ENSG00000( |
| 29878909 | 29879877 | -1 | unprocessed_pseudc | ENSG00000( |
| 29879362 | 29999797 | 1 | protein_coding | ENSG00000( |
| 29879367 | 29882089 | 1 | unprocessed_pseudc | ENSG00000( |
| 29888173 | 29889304 | 1 | unprocessed_pseudc | ENSG00000( |
| 29898144 | 29898510 | 1 | unprocessed_pseudc | ENSG00000( |
| 29899383 | 29900245 | -1 | unprocessed_pseudc | ENSG00000( |
| 29917544 | 29918534 | -1 | unprocessed_pseudc | ENSG00000( |

| | | | | |
|---|---|---|---|---|
| 29918020 | 29920795 | 1 | unprocessed_pseudc | ENSG00000( |
| 29923905 | 29924090 | 1 | unprocessed_pseudc | ENSG00000( |
| 29932035 | 29933027 | -1 | unprocessed_pseudc | ENSG00000( |
| 29946543 | 29948517 | 1 | unprocessed_pseudc | ENSG00000( |
| 29960749 | 29962411 | -1 | unprocessed_pseudc | ENSG00000( |
| 29965045 | 29968344 | 1 | lncRNA | ENSG00000( |
| 29983154 | 29983550 | 1 | unprocessed_pseudc | ENSG00000( |
| 29984382 | 29985259 | -1 | unprocessed_pseudc | ENSG00000( |
| 29990956 | 30051969 | -1 | lncRNA | ENSG00000( |
| 29996063 | 29997058 | -1 | unprocessed_pseudc | ENSG00000( |
| 29996525 | 29999898 | 1 | transcribed_unproce | ENSG00000( |
| 30022006 | 30023296 | 1 | processed_pseudoge | ENSG00000( |
| 30026209 | 30026276 | 1 | misc_RNA | ENSG00000( |
| 30048444 | 30048519 | 1 | misc_RNA | ENSG00000( |
| 30049228 | 30055232 | 1 | protein_coding | ENSG00000( |
| 30051409 | 30051512 | 1 | misc_RNA | ENSG00000( |
| 30057032 | 30060656 | 1 | protein_coding | ENSG00000( |
| 30060589 | 30066210 | -1 | protein_coding | ENSG00000( |
| 30093221 | 30103423 | -1 | protein_coding | ENSG00000( |
| 30095564 | 30105041 | 1 | lncRNA | ENSG00000( |
| 30123125 | 30123286 | -1 | snoRNA | ENSG00000( |
| 30126433 | 30139059 | 1 | protein_coding | ENSG00000( |
| 30142269 | 30151257 | -1 | protein_coding | ENSG00000( |
| 30153514 | 30163004 | 1 | protein_coding | ENSG00000( |
| 30174734 | 30203710 | -1 | protein_coding | ENSG00000( |
| 30177077 | 30178900 | -1 | processed_pseudoge | ENSG00000( |
| 30224316 | 30316381 | -1 | lncRNA | ENSG00000( |
| 30228578 | 30232554 | 1 | unprocessed_pseudc | ENSG00000( |
| 30249873 | 30257227 | 1 | transcribed_unproce | ENSG00000( |
| 30281062 | 30317397 | -1 | lncRNA | ENSG00000( |
| 30316726 | 30333982 | 1 | protein_coding | ENSG00000( |
| 30319559 | 30337137 | 1 | protein_coding | ENSG00000( |
| 30341669 | 30341803 | 1 | unprocessed_pseudc | ENSG00000( |
| 30349532 | 30350165 | -1 | processed_pseudoge | ENSG00000( |
| 30353364 | 30354532 | -1 | processed_pseudoge | ENSG00000( |
| 30404972 | 30409576 | 1 | unprocessed_pseudc | ENSG00000( |
| 30456729 | 30458271 | 1 | processed_pseudoge | ENSG00000( |
| 30459163 | 30460532 | 1 | processed_pseudoge | ENSG00000( |
| 30476183 | 30476833 | 1 | processed_pseudoge | ENSG00000( |
| 30479706 | 30484444 | 1 | protein_coding | ENSG00000( |
| 30506497 | 30509451 | 1 | lncRNA | ENSG00000( |
| 30531607 | 30547405 | -1 | protein_coding | ENSG00000( |
| 30547117 | 30553954 | 1 | protein_coding | ENSG00000( |
| 30561610 | 30587416 | 1 | protein_coding | ENSG00000( |
| 30590637 | 30608849 | -1 | protein_coding | ENSG00000( |
| 30607946 | 30616632 | 1 | protein_coding | ENSG00000( |
| 30617079 | 30637059 | 1 | protein_coding | ENSG00000( |
| 30623868 | 30624134 | 1 | processed_pseudoge | ENSG00000( |
| 30637275 | 30643339 | 1 | protein_coding | ENSG00000( |
| 30643349 | 30663298 | -1 | protein_coding | ENSG00000( |

| | | | | |
|---|---|---|---|---|
| 30666650 | 30678156 | -1 | protein_coding | ENSG00000C |
| 30678308 | 30681681 | -1 | protein_coding | ENSG00000C |
| 30687065 | 30687794 | -1 | processed_pseudoge | ENSG00000C |
| 30690066 | 30708147 | -1 | protein_coding | ENSG00000C |
| 30693326 | 30703442 | 1 | lncRNA | ENSG00000C |
| 30710458 | 30715683 | 1 | protein_coding | ENSG00000C |
| 30717966 | 30732991 | -1 | protein_coding | ENSG00000C |
| 30726562 | 30726672 | -1 | misc_RNA | ENSG00000C |
| 30733457 | 30734812 | -1 | protein_coding | ENSG00000C |
| 30741416 | 30741667 | 1 | misc_RNA | ENSG00000C |
| 30757206 | 30782937 | 1 | lncRNA | ENSG00000C |
| 30789330 | 30821330 | -1 | lncRNA | ENSG00000C |
| 30830194 | 30838828 | -1 | lncRNA | ENSG00000C |
| 30854930 | 30855232 | 1 | misc_RNA | ENSG00000C |
| 30857662 | 30866606 | -1 | lncRNA | ENSG00000C |
| 30871677 | 30890873 | 1 | protein_coding | ENSG00000C |
| 30898895 | 30904817 | 1 | protein_coding | ENSG00000C |
| 30898953 | 30917183 | 1 | lncRNA | ENSG00000C |
| 30904916 | 30917183 | 1 | protein_coding | ENSG00000C |
| 30922062 | 30922884 | -1 | protein_coding | ENSG00000C |
| 30923328 | 30923419 | -1 | misc_RNA | ENSG00000C |
| 30931715 | 30944934 | 1 | protein_coding | ENSG00000C |
| 30936693 | 30945575 | -1 | lncRNA | ENSG00000C |
| 30974428 | 30980608 | 1 | protein_coding | ENSG00000C |
| 31020962 | 31025761 | 1 | lncRNA | ENSG00000C |
| 31047228 | 31053669 | 1 | transcribed_processe | ENSG00000C |
| 31076788 | 31076887 | 1 | snRNA | ENSG00000C |
| 31104994 | 31106330 | -1 | protein_coding | ENSG00000C |
| 31108521 | 31133775 | 1 | protein_coding | ENSG00000C |
| 31108861 | 31114200 | -1 | protein_coding | ENSG00000C |
| 31131271 | 31133087 | -1 | protein_coding | ENSG00000C |
| 31134461 | 31134647 | 1 | processed_pseudoge | ENSG00000C |
| 31136168 | 31151940 | -1 | protein_coding | ENSG00000C |
| 31152269 | 31157938 | 1 | protein_coding | ENSG00000C |
| 31158060 | 31164434 | -1 | protein_coding | ENSG00000C |
| 31167475 | 31171640 | -1 | lncRNA | ENSG00000C |
| 31191499 | 31197704 | 1 | lncRNA | ENSG00000C |
| 31193904 | 31195656 | -1 | lncRNA | ENSG00000C |
| 31260956 | 31264327 | -1 | protein_coding | ENSG00000C |
| 31267736 | 31270908 | 1 | processed_pseudoge | ENSG00000C |
| 31272464 | 31273666 | 1 | processed_pseudoge | ENSG00000C |
| 31279657 | 31281111 | -1 | processed_pseudoge | ENSG00000C |
| 31299956 | 31300690 | -1 | unprocessed_pseudc | ENSG00000C |
| 31478704 | 31478805 | -1 | misc_RNA | ENSG00000C |
| 31480101 | 31493317 | 1 | protein_coding | ENSG00000C |
| 31501676 | 31502487 | -1 | processed_pseudoge | ENSG00000C |
| 31510294 | 31510873 | 1 | processed_pseudoge | ENSG00000C |
| 31510897 | 31512412 | 1 | protein_coding | ENSG00000C |
| 31512399 | 31528755 | -1 | protein_coding | ENSG00000C |
| 31518553 | 31518628 | -1 | snoRNA | ENSG00000C |

| | | | | |
|---|---|---|---|---|
| 31523278 | 31523355 | -1 | snoRNA | ENSG00000 |
| 31524481 | 31525315 | 1 | lncRNA | ENSG00000 |
| 31526639 | 31529021 | -1 | protein_coding | ENSG00000 |
| 31529043 | 31540993 | 1 | protein_coding | ENSG00000 |
| 31554192 | 31556464 | 1 | protein_coding | ENSG00000 |
| 31557707 | 31560476 | 1 | protein_coding | ENSG00000 |
| 31562695 | 31564662 | -1 | protein_coding | ENSG00000 |
| 31568266 | 31571053 | 1 | protein_coding | ENSG00000 |
| 31571039 | 31575129 | -1 | protein_coding | ENSG00000 |
| 31593261 | 31593534 | -1 | processed_pseudoge | ENSG00000 |
| 31597358 | 31599195 | 1 | protein_coding | ENSG00000 |
| 31602891 | 31619942 | 1 | protein_coding | ENSG00000 |
| 31605253 | 31605384 | 1 | snoRNA | ENSG00000 |
| 31621198 | 31634883 | -1 | protein_coding | ENSG00000 |
| 31634594 | 31640388 | 1 | protein_coding | ENSG00000 |
| 31640476 | 31642950 | -1 | protein_coding | ENSG00000 |
| 31640507 | 31642899 | 1 | lncRNA | ENSG00000 |
| 31643407 | 31648462 | -1 | protein_coding | ENSG00000 |
| 31645466 | 31645579 | 1 | misc_RNA | ENSG00000 |
| 31647414 | 31654525 | 1 | protein_coding | ENSG00000 |
| 31648281 | 31654629 | 1 | protein_coding | ENSG00000 |
| 31652558 | 31655955 | 1 | protein_coding | ENSG00000 |
| 31658863 | 31666220 | -1 | protein_coding | ENSG00000 |
| 31669120 | 31685587 | -1 | protein_coding | ENSG00000 |
| 31689056 | 31692766 | 1 | protein_coding | ENSG00000 |
| 31689075 | 31700087 | 1 | protein_coding | ENSG00000 |
| 31694505 | 31695981 | -1 | protein_coding | ENSG00000 |
| 31697525 | 31699973 | 1 | protein_coding | ENSG00000 |
| 31700763 | 31708879 | 1 | protein_coding | ENSG00000 |
| 31700817 | 31704014 | -1 | protein_coding | ENSG00000 |
| 31709206 | 31712785 | -1 | protein_coding | ENSG00000 |
| 31712749 | 31721936 | -1 | protein_coding | ENSG00000 |
| 31722164 | 31744857 | 1 | protein_coding | ENSG00000 |
| 31722193 | 31747032 | 1 | protein_coding | ENSG00000 |
| 31739132 | 31739234 | -1 | snRNA | ENSG00000 |
| 31744980 | 31747032 | 1 | protein_coding | ENSG00000 |
| 31746491 | 31747769 | -1 | lncRNA | ENSG00000 |
| 31747771 | 31759475 | -1 | protein_coding | ENSG00000 |
| 31759699 | 31778134 | -1 | protein_coding | ENSG00000 |
| 31760998 | 31761086 | -1 | misc_RNA | ENSG00000 |
| 31779577 | 31789166 | -1 | protein_coding | ENSG00000 |
| 31791805 | 31797251 | -1 | protein_coding | ENSG00000 |
| 31797650 | 31800132 | 1 | protein_coding | ENSG00000 |
| 31809921 | 31812441 | 1 | protein_coding | ENSG00000 |
| 31816794 | 31821951 | 1 | protein_coding | ENSG00000 |
| 31817449 | 31817511 | 1 | snoRNA | ENSG00000 |
| 31819261 | 31819327 | 1 | snoRNA | ENSG00000 |
| 31839844 | 31845091 | -1 | protein_coding | ENSG00000 |
| 31845377 | 31861227 | -1 | protein_coding | ENSG00000 |
| 31861940 | 31879868 | -1 | protein_coding | ENSG00000 |

| | | | | |
|---|---|---|---|---|
| 31865942 | 31866235 | 1 | lncRNA | ENSG0000( |
| 31879966 | 31927861 | 1 | protein_coding | ENSG0000( |
| 31881788 | 31884173 | -1 | protein_coding | ENSG0000( |
| 31916635 | 31923909 | -1 | lncRNA | ENSG0000( |
| 31927837 | 31934271 | 1 | protein_coding | ENSG0000( |
| 31934274 | 31941274 | -1 | protein_coding | ENSG0000( |
| 31941262 | 31951933 | 1 | protein_coding | ENSG0000( |
| 31951992 | 31954474 | -1 | protein_coding | ENSG0000( |
| 31953357 | 31965001 | 1 | protein_coding | ENSG0000( |
| 31964204 | 31984862 | 1 | protein_coding | ENSG0000( |
| 31982173 | 31985702 | -1 | lncRNA | ENSG0000( |
| 31987818 | 31990633 | 1 | transcribed_unproce | ENSG0000( |
| 31988350 | 31989286 | -1 | protein_coding | ENSG0000( |
| 31990796 | 31994654 | -1 | unprocessed_pseudo | ENSG0000( |
| 31995452 | 31995969 | 1 | unprocessed_pseudo | ENSG0000( |
| 31996944 | 32011233 | 1 | protein_coding | ENSG0000( |
| 32008544 | 32012073 | -1 | lncRNA | ENSG0000( |
| 32014113 | 32017485 | 1 | protein_coding | ENSG0000( |
| 32016970 | 32093383 | -1 | protein_coding | ENSG0000( |
| 32054322 | 32054442 | -1 | rRNA | ENSG0000( |
| 32091037 | 32104024 | -1 | protein_coding | ENSG0000( |
| 32104478 | 32106062 | -1 | protein_coding | ENSG0000( |
| 32124129 | 32130145 | -1 | protein_coding | ENSG0000( |
| 32129213 | 32134104 | 1 | protein_coding | ENSG0000( |
| 32268544 | 32347428 | -1 | protein_coding | ENSG0000( |
| 32301091 | 32302050 | 1 | processed_pseudoge | ENSG0000( |
| 32328397 | 32328503 | 1 | snRNA | ENSG0000( |
| 32366008 | 32369184 | 1 | lncRNA | ENSG0000( |
| 32369461 | 32382615 | -1 | protein_coding | ENSG0000( |
| 32414554 | 32419764 | 1 | protein_coding | ENSG0000( |
| 32434517 | 32448172 | -1 | unprocessed_pseudo | ENSG0000( |
| 32460410 | 32460505 | -1 | unprocessed_pseudo | ENSG0000( |
| 32501548 | 32501955 | -1 | processed_pseudoge | ENSG0000( |
| 32504263 | 32523409 | -1 | unprocessed_pseudo | ENSG0000( |
| 32507110 | 32508241 | 1 | processed_pseudoge | ENSG0000( |
| 32546409 | 32546557 | -1 | snRNA | ENSG0000( |
| 32548950 | 32583052 | -1 | protein_coding | ENSG0000( |
| 32606510 | 32628863 | 1 | protein_coding | ENSG0000( |
| 32634703 | 32644644 | -1 | protein_coding | ENSG0000( |
| 32635116 | 32635966 | 1 | lncRNA | ENSG0000( |
| 32635122 | 32637710 | -1 | lncRNA | ENSG0000( |
| 32685134 | 32685966 | -1 | unprocessed_pseudo | ENSG0000( |
| 32696518 | 32697352 | 1 | lncRNA | ENSG0000( |
| 32709925 | 32710861 | -1 | unprocessed_pseudo | ENSG0000( |
| 32720520 | 32726372 | 1 | protein_coding | ENSG0000( |
| 32735261 | 32742700 | -1 | protein_coding | ENSG0000( |
| 32790888 | 32795173 | -1 | protein_coding | ENSG0000( |
| 32799958 | 32816755 | -1 | protein_coding | ENSG0000( |
| 32818694 | 32822680 | -1 | protein_coding | ENSG0000( |
| 32822063 | 32824472 | 1 | lncRNA | ENSG0000( |

| | | | | |
|---|---|---|---|---|
| 32822113 | 32857517 | 1 | protein_coding | ENSG00000( |
| 32823186 | 32831955 | -1 | protein_coding | ENSG00000( |
| 32857101 | 32857778 | -1 | transcribed_processe | ENSG00000( |
| 32874341 | 32874415 | 1 | unprocessed_pseudc | ENSG00000( |
| 32912565 | 32931062 | -1 | protein_coding | ENSG00000( |
| 32926550 | 32931062 | -1 | protein_coding | ENSG00000( |
| 32955787 | 32959446 | 1 | protein_coding | ENSG00000( |
| 32982126 | 32987559 | -1 | protein_coding | ENSG00000( |
| 33042507 | 33058713 | -1 | protein_coding | ENSG00000( |
| 33053864 | 33065131 | 1 | protein_coding | ENSG00000( |
| 33057389 | 33057798 | 1 | processed_pseudoge | ENSG00000( |
| 33069683 | 33075225 | -1 | unprocessed_pseudc | ENSG00000( |
| 33081698 | 33081901 | -1 | unprocessed_pseudc | ENSG00000( |
| 33105723 | 33106989 | 1 | unprocessed_pseudc | ENSG00000( |
| 33109121 | 33121230 | -1 | unprocessed_pseudc | ENSG00000( |
| 33122692 | 33125676 | 1 | lncRNA | ENSG00000( |
| 33140587 | 33170405 | -1 | protein_coding | ENSG00000( |
| 33171494 | 33178594 | -1 | protein_coding | ENSG00000( |
| 33177507 | 33177602 | 1 | misc_RNA | ENSG00000( |
| 33178351 | 33182345 | 1 | protein_coding | ENSG00000( |
| 33182548 | 33184737 | 1 | protein_coding | ENSG00000( |
| 33186401 | 33190628 | 1 | protein_coding | ENSG00000( |
| 33193611 | 33194234 | 1 | processed_pseudoge | ENSG00000( |
| 33215710 | 33217601 | 1 | processed_pseudoge | ENSG00000( |
| 33227453 | 33232908 | 1 | lncRNA | ENSG00000( |
| 33228191 | 33249968 | -1 | protein_coding | ENSG00000( |
| 33249931 | 33254431 | 1 | protein_coding | ENSG00000( |
| 33255053 | 33256746 | 1 | protein_coding | ENSG00000( |
| 33257029 | 33267452 | -1 | protein_coding | ENSG00000( |
| 33267227 | 33276326 | 1 | protein_coding | ENSG00000( |
| 33269579 | 33277249 | -1 | protein_coding | ENSG00000( |
| 33277619 | 33292309 | -1 | protein_coding | ENSG00000( |
| 33292340 | 33295864 | -1 | protein_coding | ENSG00000( |
| 33296480 | 33307190 | -1 | protein_coding | ENSG00000( |
| 33316901 | 33317418 | 1 | processed_pseudoge | ENSG00000( |
| 28783623 | 28783994 | -1 | processed_pseudoge | ENSG00000( |
| 28837838 | 28838975 | -1 | lncRNA | ENSG00000( |
| 28859590 | 28863315 | -1 | lncRNA | ENSG00000( |
| 28861381 | 28862012 | 1 | processed_pseudoge | ENSG00000( |
| 28914742 | 28923955 | -1 | protein_coding | ENSG00000( |
| 28915611 | 28915710 | 1 | snRNA | ENSG00000( |
| 28943846 | 28944506 | 1 | protein_coding | ENSG00000( |
| 28969099 | 28970429 | -1 | processed_pseudoge | ENSG00000( |
| 28977442 | 28977740 | 1 | misc_RNA | ENSG00000( |
| 28986169 | 28987450 | 1 | lncRNA | ENSG00000( |
| 28986766 | 28988502 | 1 | lncRNA | ENSG00000( |
| 28994750 | 29005281 | -1 | protein_coding | ENSG00000( |
| 29026645 | 29027572 | -1 | unprocessed_pseudc | ENSG00000( |
| 29044178 | 29045205 | -1 | protein_coding | ENSG00000( |
| 29063961 | 29076488 | 1 | protein_coding | ENSG00000( |

| | | | | |
|---|---|---|---|---|
| 29071788 | 29072563 | -1 | unprocessed_pseudc | ENSG00000C |
| 29076537 | 29077132 | -1 | processed_pseudoge | ENSG00000C |
| 29086272 | 29087278 | -1 | protein_coding | ENSG00000C |
| 29100915 | 29101853 | 1 | unprocessed_pseudc | ENSG00000C |
| 29111857 | 29112792 | 1 | protein_coding | ENSG00000C |
| 29124176 | 29128873 | 1 | lncRNA | ENSG00000C |
| 29137841 | 29138777 | -1 | unprocessed_pseudc | ENSG00000C |
| 29173273 | 29174544 | 1 | protein_coding | ENSG00000C |
| 29181492 | 29182427 | 1 | unprocessed_pseudc | ENSG00000C |
| 29215243 | 29216186 | 1 | unprocessed_pseudc | ENSG00000C |
| 29223980 | 29267066 | 1 | lncRNA | ENSG00000C |
| 29229234 | 29230164 | -1 | unprocessed_pseudc | ENSG00000C |
| 29262684 | 29264060 | -1 | unprocessed_pseudc | ENSG00000C |
| 29268443 | 29269402 | -1 | unprocessed_pseudc | ENSG00000C |
| 29290710 | 29291652 | 1 | unprocessed_pseudc | ENSG00000C |
| 29306699 | 29307664 | 1 | protein_coding | ENSG00000C |
| 29329636 | 29331071 | -1 | processed_pseudoge | ENSG00000C |
| 29355241 | 29431509 | -1 | protein_coding | ENSG00000C |
| 29373120 | 29449634 | -1 | protein_coding | ENSG00000C |
| 29396107 | 29397175 | 1 | protein_coding | ENSG00000C |
| 29416825 | 29417771 | 1 | unprocessed_pseudc | ENSG00000C |
| 29425047 | 29456602 | -1 | protein_coding | ENSG00000C |
| 29438848 | 29440519 | 1 | protein_coding | ENSG00000C |
| 29456712 | 29463860 | 1 | protein_coding | ENSG00000C |
| 29464128 | 29469338 | -1 | unprocessed_pseudc | ENSG00000C |
| 29474557 | 29475603 | -1 | unprocessed_pseudc | ENSG00000C |
| 29486298 | 29487434 | -1 | protein_coding | ENSG00000C |
| 29488804 | 29489209 | -1 | processed_pseudoge | ENSG00000C |
| 29497042 | 29510088 | 1 | lncRNA | ENSG00000C |
| 29528958 | 29533106 | 1 | lncRNA | ENSG00000C |
| 29537242 | 29538325 | -1 | unprocessed_pseudc | ENSG00000C |
| 29552784 | 29553731 | 1 | unprocessed_pseudc | ENSG00000C |
| 29555178 | 29559492 | -1 | protein_coding | ENSG00000C |
| 29555178 | 29632716 | -1 | protein_coding | ENSG00000C |
| 29573499 | 29574426 | 1 | unprocessed_pseudc | ENSG00000C |
| 29577050 | 29577339 | 1 | processed_pseudoge | ENSG00000C |
| 29581840 | 29581923 | 1 | snoRNA | ENSG00000C |
| 29582099 | 29582616 | 1 | processed_pseudoge | ENSG00000C |
| 29587046 | 29588629 | 1 | protein_coding | ENSG00000C |
| 29635589 | 29635872 | -1 | processed_pseudoge | ENSG00000C |
| 29656505 | 29671893 | 1 | protein_coding | ENSG00000C |
| 29671913 | 29680622 | -1 | protein_coding | ENSG00000C |
| 29707631 | 29708053 | -1 | processed_pseudoge | ENSG00000C |
| 29719844 | 29735599 | 1 | protein_coding | ENSG00000C |
| 29720050 | 29721040 | -1 | unprocessed_pseudc | ENSG00000C |
| 29723672 | 29746121 | -1 | lncRNA | ENSG00000C |
| 29723740 | 29724210 | -1 | transcribed_processe | ENSG00000C |
| 29738801 | 29746041 | -1 | transcribed_unproce | ENSG00000C |
| 29745361 | 29745585 | 1 | unprocessed_pseudc | ENSG00000C |
| 29747442 | 29747543 | 1 | misc_RNA | ENSG00000C |

| | | | | |
|---|---|---|---|---|
| 29747798 | 29748218 | -1 | processed_pseudoge | ENSG00000 |
| 29787843 | 29794717 | 1 | transcribed_unproce | ENSG00000 |
| 29788736 | 29789733 | -1 | unprocessed_pseudc | ENSG00000 |
| 29797321 | 29799329 | 1 | unprocessed_pseudc | ENSG00000 |
| 29800101 | 29800897 | -1 | processed_pseudoge | ENSG00000 |
| 29809470 | 29809601 | -1 | unprocessed_pseudc | ENSG00000 |
| 29823861 | 29828018 | 1 | protein_coding | ENSG00000 |
| 29824278 | 29825257 | -1 | unprocessed_pseudc | ENSG00000 |
| 29848896 | 29849024 | -1 | unprocessed_pseudc | ENSG00000 |
| 29878153 | 29879115 | -1 | unprocessed_pseudc | ENSG00000 |
| 29878611 | 29881327 | 1 | unprocessed_pseudc | ENSG00000 |
| 29887395 | 29888526 | 1 | unprocessed_pseudc | ENSG00000 |
| 29897284 | 29897650 | 1 | unprocessed_pseudc | ENSG00000 |
| 29898524 | 29899386 | -1 | unprocessed_pseudc | ENSG00000 |
| 29916754 | 29917744 | -1 | unprocessed_pseudc | ENSG00000 |
| 29917230 | 29920003 | 1 | unprocessed_pseudc | ENSG00000 |
| 29923115 | 29923300 | 1 | unprocessed_pseudc | ENSG00000 |
| 29930208 | 29934849 | 1 | protein_coding | ENSG00000 |
| 29931024 | 29932015 | -1 | unprocessed_pseudc | ENSG00000 |
| 29951762 | 29953727 | 1 | unprocessed_pseudc | ENSG00000 |
| 29965942 | 29967605 | -1 | unprocessed_pseudc | ENSG00000 |
| 29970259 | 29973553 | 1 | lncRNA | ENSG00000 |
| 29988342 | 29988738 | 1 | unprocessed_pseudc | ENSG00000 |
| 29989570 | 29990447 | -1 | unprocessed_pseudc | ENSG00000 |
| 29996148 | 30056801 | -1 | lncRNA | ENSG00000 |
| 30001259 | 30002254 | -1 | unprocessed_pseudc | ENSG00000 |
| 30001721 | 30005094 | 1 | transcribed_unproce | ENSG00000 |
| 30026870 | 30028160 | 1 | processed_pseudoge | ENSG00000 |
| 30031073 | 30031140 | 1 | misc_RNA | ENSG00000 |
| 30053279 | 30053354 | 1 | misc_RNA | ENSG00000 |
| 30054063 | 30060065 | 1 | protein_coding | ENSG00000 |
| 30056241 | 30056344 | 1 | misc_RNA | ENSG00000 |
| 30061865 | 30065490 | 1 | protein_coding | ENSG00000 |
| 30065423 | 30071044 | -1 | protein_coding | ENSG00000 |
| 30098021 | 30108223 | -1 | protein_coding | ENSG00000 |
| 30100364 | 30109841 | 1 | lncRNA | ENSG00000 |
| 30177929 | 30178090 | -1 | snoRNA | ENSG00000 |
| 30181237 | 30193864 | 1 | protein_coding | ENSG00000 |
| 30197074 | 30206062 | -1 | protein_coding | ENSG00000 |
| 30208315 | 30217805 | 1 | protein_coding | ENSG00000 |
| 30229518 | 30258487 | -1 | protein_coding | ENSG00000 |
| 30231863 | 30233684 | -1 | processed_pseudoge | ENSG00000 |
| 30279091 | 30370766 | -1 | lncRNA | ENSG00000 |
| 30283353 | 30286914 | 1 | unprocessed_pseudc | ENSG00000 |
| 30304241 | 30311591 | 1 | transcribed_unproce | ENSG00000 |
| 30335445 | 30371782 | -1 | lncRNA | ENSG00000 |
| 30371111 | 30388367 | 1 | protein_coding | ENSG00000 |
| 30373944 | 30391525 | 1 | protein_coding | ENSG00000 |
| 30396057 | 30396191 | 1 | unprocessed_pseudc | ENSG00000 |
| 30403920 | 30404553 | -1 | processed_pseudoge | ENSG00000 |

| | | | | |
|---|---|---|---|---|
| 30407752 | 30408920 | -1 | processed_pseudoge | ENSG00000 |
| 30459359 | 30463963 | 1 | unprocessed_pseudo | ENSG00000 |
| 30511094 | 30512636 | 1 | processed_pseudoge | ENSG00000 |
| 30513524 | 30514893 | 1 | processed_pseudoge | ENSG00000 |
| 30530582 | 30531232 | 1 | processed_pseudoge | ENSG00000 |
| 30534109 | 30538847 | 1 | protein_coding | ENSG00000 |
| 30560894 | 30563845 | 1 | lncRNA | ENSG00000 |
| 30586013 | 30601810 | -1 | protein_coding | ENSG00000 |
| 30601522 | 30608359 | 1 | protein_coding | ENSG00000 |
| 30616012 | 30641816 | 1 | protein_coding | ENSG00000 |
| 30645036 | 30663248 | -1 | protein_coding | ENSG00000 |
| 30662345 | 30671031 | 1 | protein_coding | ENSG00000 |
| 30671478 | 30691459 | 1 | protein_coding | ENSG00000 |
| 30678268 | 30678534 | 1 | processed_pseudoge | ENSG00000 |
| 30691675 | 30697740 | 1 | protein_coding | ENSG00000 |
| 30697750 | 30717697 | -1 | protein_coding | ENSG00000 |
| 30721049 | 30732555 | -1 | protein_coding | ENSG00000 |
| 30732707 | 30736080 | -1 | protein_coding | ENSG00000 |
| 30741465 | 30742194 | -1 | processed_pseudoge | ENSG00000 |
| 30744466 | 30762547 | -1 | protein_coding | ENSG00000 |
| 30747726 | 30757842 | 1 | lncRNA | ENSG00000 |
| 30764859 | 30770084 | 1 | protein_coding | ENSG00000 |
| 30772367 | 30787391 | -1 | protein_coding | ENSG00000 |
| 30787857 | 30789212 | -1 | protein_coding | ENSG00000 |
| 30795686 | 30795937 | 1 | misc_RNA | ENSG00000 |
| 30811474 | 30836899 | 1 | lncRNA | ENSG00000 |
| 30843303 | 30875311 | -1 | lncRNA | ENSG00000 |
| 30884177 | 30892811 | -1 | lncRNA | ENSG00000 |
| 30908902 | 30909204 | 1 | misc_RNA | ENSG00000 |
| 30911634 | 30920567 | -1 | lncRNA | ENSG00000 |
| 30925624 | 30944801 | 1 | protein_coding | ENSG00000 |
| 30952827 | 30958749 | 1 | protein_coding | ENSG00000 |
| 30952885 | 30971104 | 1 | lncRNA | ENSG00000 |
| 30958848 | 30971104 | 1 | protein_coding | ENSG00000 |
| 30975998 | 30976820 | -1 | protein_coding | ENSG00000 |
| 30977263 | 30977354 | -1 | misc_RNA | ENSG00000 |
| 31028368 | 31034648 | 1 | protein_coding | ENSG00000 |
| 31055372 | 31080324 | 1 | lncRNA | ENSG00000 |
| 31098369 | 31104804 | 1 | transcribed_unproce | ENSG00000 |
| 31127920 | 31128019 | 1 | snRNA | ENSG00000 |
| 31156133 | 31157469 | -1 | protein_coding | ENSG00000 |
| 31159671 | 31184970 | 1 | protein_coding | ENSG00000 |
| 31160057 | 31165363 | -1 | protein_coding | ENSG00000 |
| 31182466 | 31184282 | -1 | protein_coding | ENSG00000 |
| 31185651 | 31185837 | 1 | processed_pseudoge | ENSG00000 |
| 31187361 | 31203155 | -1 | protein_coding | ENSG00000 |
| 31203494 | 31209123 | 1 | protein_coding | ENSG00000 |
| 31209245 | 31215620 | -1 | protein_coding | ENSG00000 |
| 31215489 | 31217512 | 1 | protein_coding | ENSG00000 |
| 31218662 | 31222831 | -1 | lncRNA | ENSG00000 |

| | | | | |
|---|---|---|---|---|
| 31242712 | 31243133 | 1 | lncRNA | ENSG00000 |
| 31245121 | 31245594 | -1 | lncRNA | ENSG00000 |
| 31312208 | 31315579 | -1 | protein_coding | ENSG00000 |
| 31312948 | 31400264 | -1 | protein_coding | ENSG00000 |
| 31318990 | 31322162 | 1 | processed_pseudoge | ENSG00000 |
| 31323725 | 31324927 | 1 | processed_pseudoge | ENSG00000 |
| 31330914 | 31332365 | -1 | processed_pseudoge | ENSG00000 |
| 31351206 | 31351939 | -1 | unprocessed_pseudo | ENSG00000 |
| 31409367 | 31409913 | -1 | processed_pseudoge | ENSG00000 |
| 31413150 | 31413256 | 1 | snRNA | ENSG00000 |
| 31420428 | 31421029 | 1 | processed_pseudoge | ENSG00000 |
| 31423425 | 31423853 | 1 | processed_pseudoge | ENSG00000 |
| 31425088 | 31425302 | -1 | unprocessed_pseudo | ENSG00000 |
| 31442784 | 31459233 | 1 | protein_coding | ENSG00000 |
| 31445150 | 31445248 | -1 | misc_RNA | ENSG00000 |
| 31484113 | 31489410 | 1 | lncRNA | ENSG00000 |
| 31545102 | 31550607 | 1 | protein_coding | ENSG00000 |
| 31558969 | 31559780 | -1 | processed_pseudoge | ENSG00000 |
| 31567594 | 31568173 | 1 | processed_pseudoge | ENSG00000 |
| 31568197 | 31569712 | 1 | protein_coding | ENSG00000 |
| 31569699 | 31586038 | -1 | protein_coding | ENSG00000 |
| 31575852 | 31575927 | -1 | snoRNA | ENSG00000 |
| 31580561 | 31580638 | -1 | snoRNA | ENSG00000 |
| 31581764 | 31582598 | 1 | lncRNA | ENSG00000 |
| 31583922 | 31586304 | -1 | protein_coding | ENSG00000 |
| 31586326 | 31598283 | 1 | protein_coding | ENSG00000 |
| 31611502 | 31613772 | 1 | protein_coding | ENSG00000 |
| 31615015 | 31617784 | 1 | protein_coding | ENSG00000 |
| 31620003 | 31621970 | -1 | protein_coding | ENSG00000 |
| 31625572 | 31628357 | 1 | protein_coding | ENSG00000 |
| 31628343 | 31632433 | -1 | protein_coding | ENSG00000 |
| 31660168 | 31677219 | 1 | protein_coding | ENSG00000 |
| 31662524 | 31662655 | 1 | snoRNA | ENSG00000 |
| 31678475 | 31692158 | -1 | protein_coding | ENSG00000 |
| 31691869 | 31697663 | 1 | protein_coding | ENSG00000 |
| 31697751 | 31700226 | -1 | protein_coding | ENSG00000 |
| 31697782 | 31700175 | 1 | lncRNA | ENSG00000 |
| 31700683 | 31705737 | -1 | protein_coding | ENSG00000 |
| 31702742 | 31702855 | 1 | misc_RNA | ENSG00000 |
| 31704690 | 31711800 | 1 | protein_coding | ENSG00000 |
| 31705556 | 31711904 | 1 | protein_coding | ENSG00000 |
| 31709833 | 31713000 | 1 | protein_coding | ENSG00000 |
| 31716138 | 31723494 | -1 | protein_coding | ENSG00000 |
| 31726403 | 31742871 | -1 | protein_coding | ENSG00000 |
| 31746341 | 31750054 | 1 | protein_coding | ENSG00000 |
| 31746360 | 31757371 | 1 | protein_coding | ENSG00000 |
| 31751793 | 31753269 | -1 | protein_coding | ENSG00000 |
| 31754813 | 31757257 | 1 | protein_coding | ENSG00000 |
| 31758047 | 31766164 | 1 | protein_coding | ENSG00000 |
| 31758101 | 31761298 | -1 | protein_coding | ENSG00000 |

| | | | | |
|---|---|---|---|---|
| 31766492 | 31770071 | -1 | protein_coding | ENSG00000( |
| 31770035 | 31779217 | -1 | protein_coding | ENSG00000( |
| 31779445 | 31786880 | 1 | protein_coding | ENSG00000( |
| 31897282 | 31902529 | -1 | protein_coding | ENSG00000( |
| 31902815 | 31903058 | -1 | lncRNA | ENSG00000( |
| 31959563 | 31985058 | 1 | protein_coding | ENSG00000( |
| 31973822 | 31981098 | -1 | lncRNA | ENSG00000( |
| 31985034 | 31990148 | 1 | protein_coding | ENSG00000( |
| 32104206 | 32137613 | -1 | protein_coding | ENSG00000( |
| 32117921 | 32118041 | -1 | rRNA | ENSG00000( |
| 32187767 | 32193781 | -1 | protein_coding | ENSG00000( |
| 32192849 | 32211366 | 1 | protein_coding | ENSG00000( |
| 32203996 | 32207669 | 1 | protein_coding | ENSG00000( |
| 32207600 | 32217484 | -1 | protein_coding | ENSG00000( |
| 32217742 | 32220181 | 1 | protein_coding | ENSG00000( |
| 32220356 | 32223712 | -1 | protein_coding | ENSG00000( |
| 32224129 | 32229571 | -1 | protein_coding | ENSG00000( |
| 32230151 | 32234908 | -1 | protein_coding | ENSG00000( |
| 32234228 | 32263476 | -1 | protein_coding | ENSG00000( |
| 32332098 | 32411315 | -1 | protein_coding | ENSG00000( |
| 32364622 | 32365581 | 1 | processed_pseudoge | ENSG00000( |
| 32392288 | 32392394 | 1 | snRNA | ENSG00000( |
| 32429909 | 32433086 | 1 | lncRNA | ENSG00000( |
| 32433363 | 32446554 | -1 | protein_coding | ENSG00000( |
| 32478466 | 32483677 | 1 | protein_coding | ENSG00000( |
| 32498480 | 32512136 | -1 | unprocessed_pseudc | ENSG00000( |
| 32524368 | 32524463 | -1 | unprocessed_pseudc | ENSG00000( |
| 32549943 | 32550090 | -1 | snRNA | ENSG00000( |
| 32555366 | 32555461 | -1 | unprocessed_pseudc | ENSG00000( |
| 32572064 | 32572207 | -1 | snRNA | ENSG00000( |
| 32574846 | 32589794 | -1 | protein_coding | ENSG00000( |
| 32611491 | 32611638 | -1 | snRNA | ENSG00000( |
| 32621417 | 32623878 | -1 | unprocessed_pseudc | ENSG00000( |
| 32672452 | 32672615 | -1 | snRNA | ENSG00000( |
| 32674660 | 32675599 | 1 | processed_pseudoge | ENSG00000( |
| 32677106 | 32677513 | -1 | processed_pseudoge | ENSG00000( |
| 32679833 | 32698994 | -1 | unprocessed_pseudc | ENSG00000( |
| 32682668 | 32683798 | 1 | processed_pseudoge | ENSG00000( |
| 32720817 | 32720965 | -1 | snRNA | ENSG00000( |
| 32723350 | 32757450 | -1 | protein_coding | ENSG00000( |
| 32780908 | 32803273 | 1 | protein_coding | ENSG00000( |
| 32811324 | 32820618 | -1 | protein_coding | ENSG00000( |
| 32811737 | 32812588 | 1 | lncRNA | ENSG00000( |
| 32811743 | 32814338 | -1 | lncRNA | ENSG00000( |
| 32860339 | 32861169 | -1 | unprocessed_pseudc | ENSG00000( |
| 32871715 | 32872545 | 1 | lncRNA | ENSG00000( |
| 32884471 | 32885407 | -1 | unprocessed_pseudc | ENSG00000( |
| 32895064 | 32900625 | 1 | protein_coding | ENSG00000( |
| 32909857 | 32917296 | -1 | protein_coding | ENSG00000( |
| 32975593 | 32992541 | -1 | protein_coding | ENSG00000( |

| | | | | |
|---|---|---|---|---|
| 32994477 | 32998463 | -1 | protein_coding | ENSG00000 |
| 32997846 | 33000255 | 1 | lncRNA | ENSG00000 |
| 32997896 | 33033347 | 1 | protein_coding | ENSG00000 |
| 32998969 | 33007738 | -1 | protein_coding | ENSG00000 |
| 33032931 | 33033608 | -1 | transcribed_processe | ENSG00000 |
| 33050172 | 33050246 | 1 | unprocessed_pseudo | ENSG00000 |
| 33088404 | 33106898 | -1 | protein_coding | ENSG00000 |
| 33102388 | 33124507 | -1 | protein_coding | ENSG00000 |
| 33122451 | 33135290 | 1 | protein_coding | ENSG00000 |
| 33124094 | 33124608 | 1 | lncRNA | ENSG00000 |
| 33157934 | 33163367 | -1 | protein_coding | ENSG00000 |
| 33218673 | 33234885 | -1 | protein_coding | ENSG00000 |
| 33230030 | 33241311 | 1 | protein_coding | ENSG00000 |
| 33233561 | 33233970 | 1 | processed_pseudoge | ENSG00000 |
| 33245863 | 33251405 | -1 | unprocessed_pseudo | ENSG00000 |
| 33257906 | 33261442 | -1 | unprocessed_pseudo | ENSG00000 |
| 33266563 | 33282911 | 1 | transcribed_unproce | ENSG00000 |
| 33285042 | 33297177 | -1 | unprocessed_pseudo | ENSG00000 |
| 33298642 | 33301626 | 1 | lncRNA | ENSG00000 |
| 33316561 | 33346378 | -1 | protein_coding | ENSG00000 |
| 33347482 | 33354568 | -1 | protein_coding | ENSG00000 |
| 33353481 | 33353576 | 1 | misc_RNA | ENSG00000 |
| 33354325 | 33358319 | 1 | protein_coding | ENSG00000 |
| 33358522 | 33360715 | 1 | protein_coding | ENSG00000 |
| 33362379 | 33366606 | 1 | protein_coding | ENSG00000 |
| 33369589 | 33370212 | 1 | processed_pseudoge | ENSG00000 |
| 33391691 | 33393582 | 1 | processed_pseudoge | ENSG00000 |
| 33403402 | 33408857 | 1 | lncRNA | ENSG00000 |
| 33404140 | 33405839 | -1 | protein_coding | ENSG00000 |
| 28837874 | 28839011 | -1 | lncRNA | ENSG00000 |
| 28859629 | 28863351 | -1 | lncRNA | ENSG00000 |
| 28861420 | 28862051 | 1 | processed_pseudoge | ENSG00000 |
| 28888999 | 28890073 | -1 | processed_pseudoge | ENSG00000 |
| 28896693 | 28897483 | 1 | lncRNA | ENSG00000 |
| 28903164 | 28924151 | -1 | protein_coding | ENSG00000 |
| 28915807 | 28915906 | 1 | snRNA | ENSG00000 |
| 28944039 | 28944699 | 1 | protein_coding | ENSG00000 |
| 29026834 | 29027761 | -1 | unprocessed_pseudo | ENSG00000 |
| 29044362 | 29045389 | -1 | protein_coding | ENSG00000 |
| 29064147 | 29076673 | 1 | protein_coding | ENSG00000 |
| 29071973 | 29072748 | -1 | unprocessed_pseudo | ENSG00000 |
| 29076722 | 29077317 | -1 | processed_pseudoge | ENSG00000 |
| 29086444 | 29087450 | -1 | protein_coding | ENSG00000 |
| 29100753 | 29102025 | 1 | protein_coding | ENSG00000 |
| 29112037 | 29112810 | 1 | protein_coding | ENSG00000 |
| 29138023 | 29138959 | -1 | unprocessed_pseudo | ENSG00000 |
| 29173447 | 29174718 | 1 | protein_coding | ENSG00000 |
| 29181658 | 29182593 | 1 | unprocessed_pseudo | ENSG00000 |
| 29215392 | 29216335 | 1 | unprocessed_pseudo | ENSG00000 |
| 29290867 | 29291809 | 1 | unprocessed_pseudo | ENSG00000 |

| | | | | |
|---|---|---|---|---|
| 29306920 | 29307885 | 1 | protein_coding | ENSG0000C |
| 29329857 | 29331292 | -1 | processed_pseudoge | ENSG0000C |
| 29355463 | 29431743 | -1 | protein_coding | ENSG0000C |
| 29373343 | 29449876 | -1 | protein_coding | ENSG0000C |
| 29396336 | 29397404 | 1 | protein_coding | ENSG0000C |
| 29417055 | 29418001 | 1 | unprocessed_pseudc | ENSG0000C |
| 29425279 | 29456852 | -1 | protein_coding | ENSG0000C |
| 29439082 | 29440753 | 1 | protein_coding | ENSG0000C |
| 29456962 | 29464109 | 1 | protein_coding | ENSG0000C |
| 29464377 | 29469587 | -1 | unprocessed_pseudc | ENSG0000C |
| 29474806 | 29475852 | -1 | unprocessed_pseudc | ENSG0000C |
| 29486547 | 29487683 | -1 | protein_coding | ENSG0000C |
| 29489052 | 29489457 | -1 | processed_pseudoge | ENSG0000C |
| 29497291 | 29510338 | 1 | lncRNA | ENSG0000C |
| 29529209 | 29533357 | 1 | lncRNA | ENSG0000C |
| 29537493 | 29538576 | -1 | unprocessed_pseudc | ENSG0000C |
| 29553030 | 29553977 | 1 | unprocessed_pseudc | ENSG0000C |
| 29555423 | 29559736 | -1 | protein_coding | ENSG0000C |
| 29555423 | 29632960 | -1 | protein_coding | ENSG0000C |
| 29573709 | 29574636 | 1 | unprocessed_pseudc | ENSG0000C |
| 29577260 | 29577549 | 1 | processed_pseudoge | ENSG0000C |
| 29582051 | 29582134 | 1 | snoRNA | ENSG0000C |
| 29582310 | 29582827 | 1 | processed_pseudoge | ENSG0000C |
| 29587257 | 29588840 | 1 | protein_coding | ENSG0000C |
| 29635855 | 29636138 | -1 | processed_pseudoge | ENSG0000C |
| 29656787 | 29672153 | 1 | protein_coding | ENSG0000C |
| 29672173 | 29680888 | -1 | protein_coding | ENSG0000C |
| 29707877 | 29708299 | -1 | processed_pseudoge | ENSG0000C |
| 29719887 | 29722948 | 1 | protein_coding | ENSG0000C |
| 29720114 | 29721104 | -1 | unprocessed_pseudc | ENSG0000C |
| 29738863 | 29746102 | -1 | unprocessed_pseudc | ENSG0000C |
| 29742604 | 29746182 | -1 | lncRNA | ENSG0000C |
| 29745422 | 29745646 | 1 | unprocessed_pseudc | ENSG0000C |
| 29747504 | 29747605 | 1 | misc_RNA | ENSG0000C |
| 29747862 | 29748281 | -1 | processed_pseudoge | ENSG0000C |
| 29787918 | 29794690 | 1 | transcribed_unproce | ENSG0000C |
| 29788811 | 29789808 | -1 | unprocessed_pseudc | ENSG0000C |
| 29797294 | 29799302 | 1 | unprocessed_pseudc | ENSG0000C |
| 29800074 | 29800870 | -1 | processed_pseudoge | ENSG0000C |
| 29809446 | 29809577 | -1 | unprocessed_pseudc | ENSG0000C |
| 29823845 | 29828003 | 1 | protein_coding | ENSG0000C |
| 29824263 | 29825242 | -1 | unprocessed_pseudc | ENSG0000C |
| 29849214 | 29849342 | -1 | unprocessed_pseudc | ENSG0000C |
| 29878546 | 29879514 | -1 | unprocessed_pseudc | ENSG0000C |
| 29879004 | 29881726 | 1 | unprocessed_pseudc | ENSG0000C |
| 29887810 | 29888941 | 1 | unprocessed_pseudc | ENSG0000C |
| 29897781 | 29898147 | 1 | unprocessed_pseudc | ENSG0000C |
| 29899020 | 29899882 | -1 | unprocessed_pseudc | ENSG0000C |
| 30021043 | 30024386 | 1 | protein_coding | ENSG0000C |
| 30021043 | 30021554 | -1 | unprocessed_pseudc | ENSG0000C |

| | | | | |
|---|---|---|---|---|
| 30035070 | 30037031 | 1 | unprocessed_pseudc | ENSG0000C |
| 30049251 | 30050914 | -1 | unprocessed_pseudc | ENSG0000C |
| 30053572 | 30056871 | 1 | lncRNA | ENSG0000C |
| 30071672 | 30072068 | 1 | unprocessed_pseudc | ENSG0000C |
| 30072900 | 30073777 | -1 | unprocessed_pseudc | ENSG0000C |
| 30079474 | 30140140 | -1 | lncRNA | ENSG0000C |
| 30084584 | 30085579 | -1 | unprocessed_pseudc | ENSG0000C |
| 30085046 | 30088419 | 1 | transcribed_unproce | ENSG0000C |
| 30110194 | 30111484 | 1 | processed_pseudoge | ENSG0000C |
| 30114397 | 30114464 | 1 | misc_RNA | ENSG0000C |
| 30136615 | 30136690 | 1 | misc_RNA | ENSG0000C |
| 30137399 | 30143406 | 1 | protein_coding | ENSG0000C |
| 30139580 | 30139683 | 1 | misc_RNA | ENSG0000C |
| 30145206 | 30148830 | 1 | protein_coding | ENSG0000C |
| 30148763 | 30154384 | -1 | protein_coding | ENSG0000C |
| 30181379 | 30191586 | -1 | protein_coding | ENSG0000C |
| 30183722 | 30193204 | 1 | lncRNA | ENSG0000C |
| 30211301 | 30211463 | -1 | snoRNA | ENSG0000C |
| 30214604 | 30227224 | 1 | protein_coding | ENSG0000C |
| 30230427 | 30239418 | -1 | protein_coding | ENSG0000C |
| 30241686 | 30251176 | 1 | protein_coding | ENSG0000C |
| 30262891 | 30291774 | -1 | protein_coding | ENSG0000C |
| 30265236 | 30267045 | -1 | processed_pseudoge | ENSG0000C |
| 30493160 | 30497764 | 1 | unprocessed_pseudc | ENSG0000C |
| 30567930 | 30572668 | 1 | protein_coding | ENSG0000C |
| 30594716 | 30597667 | 1 | lncRNA | ENSG0000C |
| 30619830 | 30635627 | -1 | protein_coding | ENSG0000C |
| 30635339 | 30642176 | 1 | protein_coding | ENSG0000C |
| 30649829 | 30675633 | 1 | protein_coding | ENSG0000C |
| 30678854 | 30697067 | -1 | protein_coding | ENSG0000C |
| 30696164 | 30704850 | 1 | protein_coding | ENSG0000C |
| 30705297 | 30725280 | 1 | protein_coding | ENSG0000C |
| 30712087 | 30712353 | 1 | processed_pseudoge | ENSG0000C |
| 30725496 | 30731566 | 1 | protein_coding | ENSG0000C |
| 30731576 | 30751521 | -1 | protein_coding | ENSG0000C |
| 30754873 | 30766379 | -1 | protein_coding | ENSG0000C |
| 30766531 | 30769904 | -1 | protein_coding | ENSG0000C |
| 30775289 | 30776018 | -1 | processed_pseudoge | ENSG0000C |
| 30778290 | 30796371 | -1 | protein_coding | ENSG0000C |
| 30781550 | 30791666 | 1 | lncRNA | ENSG0000C |
| 30798683 | 30803908 | 1 | protein_coding | ENSG0000C |
| 30806191 | 30820021 | -1 | protein_coding | ENSG0000C |
| 30814786 | 30814896 | -1 | misc_RNA | ENSG0000C |
| 30918008 | 30926645 | -1 | lncRNA | ENSG0000C |
| 30942815 | 30943117 | 1 | misc_RNA | ENSG0000C |
| 30945549 | 30954477 | -1 | lncRNA | ENSG0000C |
| 30959531 | 30970646 | 1 | protein_coding | ENSG0000C |
| 30986738 | 30992660 | 1 | protein_coding | ENSG0000C |
| 30986796 | 31005022 | 1 | lncRNA | ENSG0000C |
| 30992759 | 31005022 | 1 | protein_coding | ENSG0000C |

| | | | | |
|---|---|---|---|---|
| 31009891 | 31010713 | -1 | protein_coding | ENSG0000C |
| 31011157 | 31011248 | -1 | misc_RNA | ENSG0000C |
| 31019512 | 31029672 | 1 | protein_coding | ENSG0000C |
| 31024520 | 31024900 | -1 | lncRNA | ENSG0000C |
| 31062263 | 31068453 | 1 | protein_coding | ENSG0000C |
| 31089020 | 31113948 | 1 | lncRNA | ENSG0000C |
| 31131982 | 31138413 | 1 | transcribed_unproce | ENSG0000C |
| 31161534 | 31161633 | 1 | snRNA | ENSG0000C |
| 31189747 | 31191083 | -1 | protein_coding | ENSG0000C |
| 31193274 | 31218582 | 1 | protein_coding | ENSG0000C |
| 31193614 | 31198966 | -1 | protein_coding | ENSG0000C |
| 31216078 | 31217894 | -1 | protein_coding | ENSG0000C |
| 31219263 | 31219449 | 1 | processed_pseudoge | ENSG0000C |
| 31220974 | 31236769 | -1 | protein_coding | ENSG0000C |
| 31237108 | 31242737 | 1 | protein_coding | ENSG0000C |
| 31242859 | 31249234 | -1 | protein_coding | ENSG0000C |
| 31249103 | 31251125 | 1 | protein_coding | ENSG0000C |
| 31252275 | 31256445 | -1 | lncRNA | ENSG0000C |
| 31276325 | 31282581 | 1 | lncRNA | ENSG0000C |
| 31278735 | 31280532 | -1 | lncRNA | ENSG0000C |
| 31345885 | 31349256 | -1 | protein_coding | ENSG0000C |
| 31346626 | 31433637 | -1 | protein_coding | ENSG0000C |
| 31352684 | 31355856 | 1 | processed_pseudoge | ENSG0000C |
| 31357419 | 31358618 | 1 | processed_pseudoge | ENSG0000C |
| 31364601 | 31366055 | -1 | processed_pseudoge | ENSG0000C |
| 31384925 | 31385659 | -1 | unprocessed_pseudc | ENSG0000C |
| 31540726 | 31542189 | 1 | lncRNA | ENSG0000C |
| 31573052 | 31573153 | -1 | misc_RNA | ENSG0000C |
| 31574447 | 31587478 | 1 | protein_coding | ENSG0000C |
| 31595839 | 31596650 | -1 | processed_pseudoge | ENSG0000C |
| 31604464 | 31605043 | 1 | processed_pseudoge | ENSG0000C |
| 31605067 | 31606582 | 1 | protein_coding | ENSG0000C |
| 31606569 | 31622924 | -1 | protein_coding | ENSG0000C |
| 31612723 | 31612798 | -1 | snoRNA | ENSG0000C |
| 31617447 | 31617524 | -1 | snoRNA | ENSG0000C |
| 31618650 | 31619484 | 1 | lncRNA | ENSG0000C |
| 31620808 | 31623190 | -1 | protein_coding | ENSG0000C |
| 31623212 | 31635161 | 1 | protein_coding | ENSG0000C |
| 31648357 | 31650629 | 1 | protein_coding | ENSG0000C |
| 31651872 | 31654641 | 1 | protein_coding | ENSG0000C |
| 31656860 | 31658827 | -1 | protein_coding | ENSG0000C |
| 31662426 | 31665209 | 1 | protein_coding | ENSG0000C |
| 31665195 | 31669285 | -1 | protein_coding | ENSG0000C |
| 31691477 | 31693314 | 1 | protein_coding | ENSG0000C |
| 31697009 | 31714058 | 1 | protein_coding | ENSG0000C |
| 31699371 | 31699502 | 1 | snoRNA | ENSG0000C |
| 31715314 | 31728984 | -1 | protein_coding | ENSG0000C |
| 31728695 | 31734489 | 1 | protein_coding | ENSG0000C |
| 31734577 | 31737051 | -1 | protein_coding | ENSG0000C |
| 31734608 | 31737000 | 1 | lncRNA | ENSG0000C |

| | | | | |
|---|---|---|---|---|
| 31737508 | 31742562 | -1 | protein_coding | ENSG00000 |
| 31739567 | 31739680 | 1 | misc_RNA | ENSG00000 |
| 31741515 | 31749825 | 1 | protein_coding | ENSG00000 |
| 31745397 | 31749825 | 1 | protein_coding | ENSG00000 |
| 31752963 | 31760319 | -1 | protein_coding | ENSG00000 |
| 31763228 | 31779696 | -1 | protein_coding | ENSG00000 |
| 31783165 | 31786875 | 1 | protein_coding | ENSG00000 |
| 31783184 | 31794200 | 1 | protein_coding | ENSG00000 |
| 31788620 | 31790096 | -1 | protein_coding | ENSG00000 |
| 31794876 | 31802992 | 1 | protein_coding | ENSG00000 |
| 31794930 | 31798127 | -1 | protein_coding | ENSG00000 |
| 31803320 | 31806899 | -1 | protein_coding | ENSG00000 |
| 31806863 | 31816046 | -1 | protein_coding | ENSG00000 |
| 31816274 | 31839440 | 1 | protein_coding | ENSG00000 |
| 31816303 | 31841147 | 1 | protein_coding | ENSG00000 |
| 31833247 | 31833349 | -1 | snRNA | ENSG00000 |
| 31839095 | 31841147 | 1 | protein_coding | ENSG00000 |
| 31840606 | 31841884 | -1 | lncRNA | ENSG00000 |
| 31841886 | 31853590 | -1 | protein_coding | ENSG00000 |
| 31853814 | 31872249 | -1 | protein_coding | ENSG00000 |
| 31855113 | 31855201 | -1 | misc_RNA | ENSG00000 |
| 31873692 | 31883281 | -1 | protein_coding | ENSG00000 |
| 31934136 | 31939383 | -1 | protein_coding | ENSG00000 |
| 31939669 | 31955518 | -1 | protein_coding | ENSG00000 |
| 31956231 | 31974166 | -1 | protein_coding | ENSG00000 |
| 31960233 | 31960526 | 1 | lncRNA | ENSG00000 |
| 31974264 | 32022117 | 1 | protein_coding | ENSG00000 |
| 31976089 | 31978474 | -1 | protein_coding | ENSG00000 |
| 32010892 | 32018166 | -1 | lncRNA | ENSG00000 |
| 32022093 | 32028527 | 1 | protein_coding | ENSG00000 |
| 32028530 | 32035535 | -1 | protein_coding | ENSG00000 |
| 32035523 | 32046194 | 1 | protein_coding | ENSG00000 |
| 32046253 | 32048735 | -1 | protein_coding | ENSG00000 |
| 32047618 | 32057890 | 1 | protein_coding | ENSG00000 |
| 32058468 | 32058494 | 1 | lncRNA | ENSG00000 |
| 32084376 | 32084894 | 1 | transcribed_unproce | ENSG00000 |
| 32085057 | 32088915 | -1 | unprocessed_pseudo | ENSG00000 |
| 32089713 | 32090230 | 1 | unprocessed_pseudo | ENSG00000 |
| 32091205 | 32111862 | 1 | protein_coding | ENSG00000 |
| 32109173 | 32112702 | -1 | lncRNA | ENSG00000 |
| 32114742 | 32118110 | 1 | protein_coding | ENSG00000 |
| 32117595 | 32194037 | -1 | protein_coding | ENSG00000 |
| 32154947 | 32155067 | -1 | rRNA | ENSG00000 |
| 32191691 | 32204680 | -1 | protein_coding | ENSG00000 |
| 32205134 | 32206718 | -1 | protein_coding | ENSG00000 |
| 32224781 | 32230795 | -1 | protein_coding | ENSG00000 |
| 32229863 | 32248406 | 1 | protein_coding | ENSG00000 |
| 32241011 | 32244709 | 1 | protein_coding | ENSG00000 |
| 32244640 | 32254524 | -1 | protein_coding | ENSG00000 |
| 32254782 | 32257219 | 1 | protein_coding | ENSG00000 |

| | | | | |
|---|---|---|---|---|
| 32257394 | 32260750 | -1 | protein_coding | ENSG0000C |
| 32261167 | 32266609 | -1 | protein_coding | ENSG0000C |
| 32267189 | 32271946 | -1 | protein_coding | ENSG0000C |
| 32271266 | 32300513 | -1 | protein_coding | ENSG0000C |
| 32392894 | 32448091 | -1 | protein_coding | ENSG0000C |
| 32401725 | 32402684 | 1 | processed_pseudoge | ENSG0000C |
| 32429038 | 32429144 | 1 | snRNA | ENSG0000C |
| 32466666 | 32469843 | 1 | lncRNA | ENSG0000C |
| 32470120 | 32483292 | -1 | protein_coding | ENSG0000C |
| 32515215 | 32520425 | 1 | protein_coding | ENSG0000C |
| 32535112 | 32548767 | -1 | unprocessed_pseudc | ENSG0000C |
| 32561005 | 32561100 | -1 | unprocessed_pseudc | ENSG0000C |
| 32586571 | 32586718 | -1 | snRNA | ENSG0000C |
| 32591998 | 32592093 | -1 | unprocessed_pseudc | ENSG0000C |
| 32608365 | 32608516 | -1 | snRNA | ENSG0000C |
| 32611143 | 32626106 | -1 | protein_coding | ENSG0000C |
| 32663658 | 32684403 | 1 | protein_coding | ENSG0000C |
| 32690241 | 32699289 | -1 | protein_coding | ENSG0000C |
| 32690654 | 32691504 | 1 | lncRNA | ENSG0000C |
| 32690660 | 32693255 | -1 | lncRNA | ENSG0000C |
| 32739769 | 32740601 | -1 | unprocessed_pseudc | ENSG0000C |
| 32751146 | 32751980 | 1 | lncRNA | ENSG0000C |
| 32764544 | 32765480 | -1 | unprocessed_pseudc | ENSG0000C |
| 32775123 | 32780991 | 1 | protein_coding | ENSG0000C |
| 32789880 | 32797319 | -1 | protein_coding | ENSG0000C |
| 32846263 | 32850548 | -1 | protein_coding | ENSG0000C |
| 32855328 | 32872254 | -1 | protein_coding | ENSG0000C |
| 32874193 | 32878179 | -1 | protein_coding | ENSG0000C |
| 32877562 | 32879971 | 1 | lncRNA | ENSG0000C |
| 32877612 | 32913027 | 1 | protein_coding | ENSG0000C |
| 32878685 | 32887454 | -1 | protein_coding | ENSG0000C |
| 32912612 | 32913288 | -1 | transcribed_processe | ENSG0000C |
| 32929849 | 32929923 | 1 | unprocessed_pseudc | ENSG0000C |
| 32968079 | 32986577 | -1 | protein_coding | ENSG0000C |
| 32982065 | 33004180 | -1 | protein_coding | ENSG0000C |
| 33002124 | 33012662 | 1 | protein_coding | ENSG0000C |
| 33003767 | 33004281 | 1 | lncRNA | ENSG0000C |
| 33037640 | 33043074 | -1 | protein_coding | ENSG0000C |
| 33098041 | 33114235 | -1 | protein_coding | ENSG0000C |
| 33109380 | 33120645 | 1 | protein_coding | ENSG0000C |
| 33112911 | 33113320 | 1 | processed_pseudoge | ENSG0000C |
| 33125197 | 33130739 | -1 | unprocessed_pseudc | ENSG0000C |
| 33137211 | 33140747 | -1 | unprocessed_pseudc | ENSG0000C |
| 33145868 | 33150636 | 1 | transcribed_unproce | ENSG0000C |
| 33333046 | 33362862 | -1 | protein_coding | ENSG0000C |
| 33363951 | 33371048 | -1 | protein_coding | ENSG0000C |
| 33369961 | 33370056 | 1 | misc_RNA | ENSG0000C |
| 33370805 | 33374799 | 1 | protein_coding | ENSG0000C |
| 33375002 | 33377191 | 1 | protein_coding | ENSG0000C |
| 33378855 | 33383082 | 1 | protein_coding | ENSG0000C |

| | | | | |
|---|---|---|---|---|
| 33386065 | 33386688 | 1 | processed_pseudoge | ENSG00000 |
| 33408140 | 33410038 | 1 | processed_pseudoge | ENSG00000 |
| 33419923 | 33425374 | 1 | lncRNA | ENSG00000 |
| 33420661 | 33442416 | -1 | protein_coding | ENSG00000 |
| 33442379 | 33446883 | 1 | protein_coding | ENSG00000 |
| 33447505 | 33449198 | 1 | protein_coding | ENSG00000 |
| 33449481 | 33459870 | -1 | protein_coding | ENSG00000 |
| 33459645 | 33468744 | 1 | protein_coding | ENSG00000 |
| 33461997 | 33469667 | -1 | protein_coding | ENSG00000 |
| 33470037 | 33484728 | -1 | protein_coding | ENSG00000 |
| 33484759 | 33488283 | -1 | protein_coding | ENSG00000 |
| 33488900 | 33499609 | -1 | protein_coding | ENSG00000 |
| 33509323 | 33509840 | 1 | processed_pseudoge | ENSG00000 |
| 33535893 | 33536588 | -1 | processed_pseudoge | ENSG00000 |
| 33559730 | 33559831 | -1 | processed_pseudoge | ENSG00000 |
| 33561891 | 33562220 | 1 | lncRNA | ENSG00000 |
| 28783631 | 28784002 | -1 | processed_pseudoge | ENSG00000 |
| 28837857 | 28838994 | -1 | lncRNA | ENSG00000 |
| 28859613 | 28863339 | -1 | lncRNA | ENSG00000 |
| 28861404 | 28862035 | 1 | processed_pseudoge | ENSG00000 |
| 28888821 | 28889895 | -1 | processed_pseudoge | ENSG00000 |
| 28896519 | 28897311 | 1 | lncRNA | ENSG00000 |
| 28902991 | 28923978 | -1 | protein_coding | ENSG00000 |
| 28915634 | 28915733 | 1 | snRNA | ENSG00000 |
| 28943865 | 28944525 | 1 | protein_coding | ENSG00000 |
| 28969118 | 28970448 | -1 | processed_pseudoge | ENSG00000 |
| 28977461 | 28977759 | 1 | misc_RNA | ENSG00000 |
| 28986188 | 28987469 | 1 | lncRNA | ENSG00000 |
| 28986785 | 28988521 | 1 | lncRNA | ENSG00000 |
| 28994770 | 29005301 | -1 | protein_coding | ENSG00000 |
| 29026665 | 29027592 | -1 | unprocessed_pseudc | ENSG00000 |
| 29044198 | 29045225 | -1 | protein_coding | ENSG00000 |
| 29063984 | 29076511 | 1 | protein_coding | ENSG00000 |
| 29071811 | 29072586 | -1 | unprocessed_pseudc | ENSG00000 |
| 29076560 | 29077155 | -1 | processed_pseudoge | ENSG00000 |
| 29086195 | 29087300 | -1 | protein_coding | ENSG00000 |
| 29100937 | 29101875 | 1 | unprocessed_pseudc | ENSG00000 |
| 29111883 | 29112818 | 1 | protein_coding | ENSG00000 |
| 29124201 | 29128899 | 1 | lncRNA | ENSG00000 |
| 29137868 | 29138804 | -1 | unprocessed_pseudc | ENSG00000 |
| 29173300 | 29174571 | 1 | protein_coding | ENSG00000 |
| 29181515 | 29182450 | 1 | unprocessed_pseudc | ENSG00000 |
| 29215267 | 29216210 | 1 | unprocessed_pseudc | ENSG00000 |
| 29229258 | 29230188 | -1 | unprocessed_pseudc | ENSG00000 |
| 29417159 | 29418105 | 1 | unprocessed_pseudc | ENSG00000 |
| 29422486 | 29431843 | -1 | lncRNA | ENSG00000 |
| 29425381 | 29456943 | -1 | protein_coding | ENSG00000 |
| 29439182 | 29440853 | 1 | protein_coding | ENSG00000 |
| 29457053 | 29464201 | 1 | protein_coding | ENSG00000 |
| 29464469 | 29469679 | -1 | unprocessed_pseudc | ENSG00000 |

| | | | | |
|---|---|---|---|---|
| 29474898 | 29475944 | -1 | unprocessed_pseudc | ENSG00000( |
| 29486639 | 29487775 | -1 | protein_coding | ENSG00000( |
| 29489145 | 29489550 | -1 | processed_pseudoge | ENSG00000( |
| 29497383 | 29510429 | 1 | lncRNA | ENSG00000( |
| 29529298 | 29533446 | 1 | lncRNA | ENSG00000( |
| 29537652 | 29538442 | -1 | unprocessed_pseudc | ENSG00000( |
| 29553124 | 29554071 | 1 | unprocessed_pseudc | ENSG00000( |
| 29555518 | 29633080 | -1 | protein_coding | ENSG00000( |
| 29555518 | 29559832 | -1 | protein_coding | ENSG00000( |
| 29573828 | 29574755 | 1 | unprocessed_pseudc | ENSG00000( |
| 29577378 | 29577667 | 1 | processed_pseudoge | ENSG00000( |
| 29582168 | 29582251 | 1 | snoRNA | ENSG00000( |
| 29582427 | 29582944 | 1 | processed_pseudoge | ENSG00000( |
| 29587374 | 29588957 | 1 | protein_coding | ENSG00000( |
| 29635969 | 29636252 | -1 | processed_pseudoge | ENSG00000( |
| 29656888 | 29672280 | 1 | protein_coding | ENSG00000( |
| 29672300 | 29681010 | -1 | protein_coding | ENSG00000( |
| 29708017 | 29708439 | -1 | processed_pseudoge | ENSG00000( |
| 29720222 | 29735978 | 1 | protein_coding | ENSG00000( |
| 29720428 | 29721418 | -1 | unprocessed_pseudc | ENSG00000( |
| 29724050 | 29746500 | -1 | lncRNA | ENSG00000( |
| 29724118 | 29724588 | -1 | transcribed_processe | ENSG00000( |
| 29739180 | 29746420 | -1 | transcribed_unproce | ENSG00000( |
| 29745740 | 29745964 | 1 | unprocessed_pseudc | ENSG00000( |
| 29747821 | 29747922 | 1 | misc_RNA | ENSG00000( |
| 29748177 | 29748597 | -1 | processed_pseudoge | ENSG00000( |
| 29788252 | 29794972 | 1 | transcribed_unproce | ENSG00000( |
| 29789145 | 29790119 | -1 | unprocessed_pseudc | ENSG00000( |
| 29797597 | 29799601 | 1 | unprocessed_pseudc | ENSG00000( |
| 29800376 | 29801172 | -1 | processed_pseudoge | ENSG00000( |
| 29809741 | 29809872 | -1 | unprocessed_pseudc | ENSG00000( |
| 29824140 | 29828314 | 1 | protein_coding | ENSG00000( |
| 29824558 | 29825536 | -1 | unprocessed_pseudc | ENSG00000( |
| 29849309 | 29849437 | -1 | unprocessed_pseudc | ENSG00000( |
| 29878575 | 29879543 | -1 | unprocessed_pseudc | ENSG00000( |
| 29879033 | 29881754 | 1 | unprocessed_pseudc | ENSG00000( |
| 29887829 | 29888960 | 1 | unprocessed_pseudc | ENSG00000( |
| 29897789 | 29898155 | 1 | unprocessed_pseudc | ENSG00000( |
| 29899031 | 29899890 | -1 | unprocessed_pseudc | ENSG00000( |
| 29917179 | 29918169 | -1 | unprocessed_pseudc | ENSG00000( |
| 29917655 | 29920429 | 1 | unprocessed_pseudc | ENSG00000( |
| 29923540 | 29923724 | 1 | unprocessed_pseudc | ENSG00000( |
| 29930625 | 29999286 | 1 | protein_coding | ENSG00000( |
| 29931441 | 29932433 | -1 | unprocessed_pseudc | ENSG00000( |
| 29945986 | 29947960 | 1 | unprocessed_pseudc | ENSG00000( |
| 29960191 | 29961854 | -1 | unprocessed_pseudc | ENSG00000( |
| 29964492 | 29967793 | 1 | lncRNA | ENSG00000( |
| 29982616 | 29983011 | 1 | unprocessed_pseudc | ENSG00000( |
| 29983843 | 29984720 | -1 | unprocessed_pseudc | ENSG00000( |
| 29990513 | 30051230 | -1 | lncRNA | ENSG00000( |

| | | | | |
|---|---|---|---|---|
| 29995625 | 29996621 | -1 | unprocessed_pseudc | ENSG00000C |
| 29996088 | 29999447 | 1 | transcribed_unproce | ENSG00000C |
| 30021246 | 30022535 | 1 | processed_pseudoge | ENSG00000C |
| 30025452 | 30025519 | 1 | misc_RNA | ENSG00000C |
| 30047700 | 30047775 | 1 | misc_RNA | ENSG00000C |
| 30048487 | 30054477 | 1 | protein_coding | ENSG00000C |
| 30050669 | 30050772 | 1 | misc_RNA | ENSG00000C |
| 30056276 | 30059900 | 1 | protein_coding | ENSG00000C |
| 30059837 | 30065413 | -1 | protein_coding | ENSG00000C |
| 30092451 | 30102654 | -1 | protein_coding | ENSG00000C |
| 30094794 | 30104272 | 1 | lncRNA | ENSG00000C |
| 30122351 | 30122511 | -1 | snoRNA | ENSG00000C |
| 30125643 | 30138271 | 1 | protein_coding | ENSG00000C |
| 30141481 | 30150469 | -1 | protein_coding | ENSG00000C |
| 30152741 | 30162231 | 1 | protein_coding | ENSG00000C |
| 30173958 | 30202936 | -1 | protein_coding | ENSG00000C |
| 30176301 | 30178125 | -1 | processed_pseudoge | ENSG00000C |
| 30223542 | 30315616 | -1 | lncRNA | ENSG00000C |
| 30227804 | 30231780 | 1 | unprocessed_pseudc | ENSG00000C |
| 30249099 | 30256416 | 1 | transcribed_unproce | ENSG00000C |
| 30280294 | 30316632 | -1 | lncRNA | ENSG00000C |
| 30315961 | 30333218 | 1 | protein_coding | ENSG00000C |
| 30318794 | 30336373 | 1 | protein_coding | ENSG00000C |
| 30340905 | 30341039 | 1 | unprocessed_pseudc | ENSG00000C |
| 30348767 | 30349400 | -1 | processed_pseudoge | ENSG00000C |
| 30352597 | 30353765 | -1 | processed_pseudoge | ENSG00000C |
| 30404207 | 30408812 | 1 | unprocessed_pseudc | ENSG00000C |
| 30456041 | 30457570 | 1 | processed_pseudoge | ENSG00000C |
| 30458389 | 30459758 | 1 | processed_pseudoge | ENSG00000C |
| 30475446 | 30476096 | 1 | processed_pseudoge | ENSG00000C |
| 30478970 | 30483708 | 1 | protein_coding | ENSG00000C |
| 30505760 | 30508714 | 1 | lncRNA | ENSG00000C |
| 30530868 | 30546666 | -1 | protein_coding | ENSG00000C |
| 30546378 | 30553215 | 1 | protein_coding | ENSG00000C |
| 30560870 | 30586674 | 1 | protein_coding | ENSG00000C |
| 30589896 | 30608107 | -1 | protein_coding | ENSG00000C |
| 30607204 | 30615890 | 1 | protein_coding | ENSG00000C |
| 30616337 | 30636318 | 1 | protein_coding | ENSG00000C |
| 30623124 | 30623393 | 1 | processed_pseudoge | ENSG00000C |
| 30636534 | 30642598 | 1 | protein_coding | ENSG00000C |
| 30642608 | 30662557 | -1 | protein_coding | ENSG00000C |
| 30665910 | 30677416 | -1 | protein_coding | ENSG00000C |
| 30677568 | 30680941 | -1 | protein_coding | ENSG00000C |
| 30686326 | 30687055 | -1 | processed_pseudoge | ENSG00000C |
| 30689327 | 30707408 | -1 | protein_coding | ENSG00000C |
| 30692588 | 30702703 | 1 | lncRNA | ENSG00000C |
| 30709720 | 30714946 | 1 | protein_coding | ENSG00000C |
| 30717229 | 30732250 | -1 | protein_coding | ENSG00000C |
| 30725828 | 30725938 | -1 | misc_RNA | ENSG00000C |
| 30732716 | 30734071 | -1 | protein_coding | ENSG00000C |

| | | | | |
|---|---|---|---|---|
| 30740537 | 30740788 | 1 | misc_RNA | ENSG00000 |
| 30756325 | 30781741 | 1 | lncRNA | ENSG00000 |
| 30788147 | 30820172 | -1 | lncRNA | ENSG00000 |
| 30829032 | 30837670 | -1 | lncRNA | ENSG00000 |
| 30853833 | 30854135 | 1 | misc_RNA | ENSG00000 |
| 30856563 | 30865496 | -1 | lncRNA | ENSG00000 |
| 30870550 | 30889733 | 1 | protein_coding | ENSG00000 |
| 30897757 | 30903679 | 1 | protein_coding | ENSG00000 |
| 30897815 | 30916043 | 1 | lncRNA | ENSG00000 |
| 30903778 | 30916043 | 1 | protein_coding | ENSG00000 |
| 30920912 | 30921734 | -1 | protein_coding | ENSG00000 |
| 30922178 | 30922269 | -1 | misc_RNA | ENSG00000 |
| 30930561 | 30943780 | 1 | protein_coding | ENSG00000 |
| 30935539 | 30944421 | -1 | lncRNA | ENSG00000 |
| 30973281 | 30979676 | 1 | protein_coding | ENSG00000 |
| 31000494 | 31025446 | 1 | lncRNA | ENSG00000 |
| 31046189 | 31052260 | 1 | transcribed_unproce | ENSG00000 |
| 31075424 | 31075523 | 1 | snRNA | ENSG00000 |
| 31103629 | 31104965 | -1 | protein_coding | ENSG00000 |
| 31107156 | 31132470 | 1 | protein_coding | ENSG00000 |
| 31107496 | 31112855 | -1 | protein_coding | ENSG00000 |
| 31129969 | 31131783 | -1 | protein_coding | ENSG00000 |
| 31133166 | 31133352 | 1 | processed_pseudoge | ENSG00000 |
| 31134861 | 31150652 | -1 | protein_coding | ENSG00000 |
| 31150981 | 31156655 | 1 | protein_coding | ENSG00000 |
| 31156777 | 31163150 | -1 | protein_coding | ENSG00000 |
| 31166192 | 31170357 | -1 | lncRNA | ENSG00000 |
| 31190191 | 31196403 | 1 | lncRNA | ENSG00000 |
| 31192602 | 31194361 | -1 | lncRNA | ENSG00000 |
| 31258588 | 31261959 | -1 | protein_coding | ENSG00000 |
| 31259328 | 31347305 | -1 | protein_coding | ENSG00000 |
| 31265372 | 31268544 | 1 | processed_pseudoge | ENSG00000 |
| 31270107 | 31271306 | 1 | processed_pseudoge | ENSG00000 |
| 31277583 | 31279034 | -1 | processed_pseudoge | ENSG00000 |
| 31297920 | 31298654 | -1 | unprocessed_pseudo | ENSG00000 |
| 31356448 | 31356994 | -1 | processed_pseudoge | ENSG00000 |
| 31360234 | 31360340 | 1 | snRNA | ENSG00000 |
| 31367516 | 31368113 | 1 | processed_pseudoge | ENSG00000 |
| 31370510 | 31370936 | 1 | processed_pseudoge | ENSG00000 |
| 31372160 | 31372374 | -1 | unprocessed_pseudo | ENSG00000 |
| 31389818 | 31406373 | 1 | protein_coding | ENSG00000 |
| 31392186 | 31392284 | -1 | misc_RNA | ENSG00000 |
| 31486996 | 31487097 | -1 | misc_RNA | ENSG00000 |
| 31488393 | 31501409 | 1 | protein_coding | ENSG00000 |
| 31509763 | 31510574 | -1 | processed_pseudoge | ENSG00000 |
| 31518383 | 31518962 | 1 | processed_pseudoge | ENSG00000 |
| 31518986 | 31520501 | 1 | protein_coding | ENSG00000 |
| 31520488 | 31536841 | -1 | protein_coding | ENSG00000 |
| 31526642 | 31526717 | -1 | snoRNA | ENSG00000 |
| 31531364 | 31531441 | -1 | snoRNA | ENSG00000 |

| | | | | |
|---|---|---|---|---|
| 31532567 | 31533401 | 1 | lncRNA | ENSG00000 |
| 31534725 | 31537107 | -1 | protein_coding | ENSG00000 |
| 31537129 | 31549080 | 1 | protein_coding | ENSG00000 |
| 31562278 | 31564550 | 1 | protein_coding | ENSG00000 |
| 31565793 | 31568564 | 1 | protein_coding | ENSG00000 |
| 31570783 | 31572750 | -1 | protein_coding | ENSG00000 |
| 31576352 | 31579139 | 1 | protein_coding | ENSG00000 |
| 31579125 | 31583213 | -1 | protein_coding | ENSG00000 |
| 31601313 | 31601586 | -1 | processed_pseudoge | ENSG00000 |
| 31605413 | 31607250 | 1 | protein_coding | ENSG00000 |
| 31610949 | 31627996 | 1 | protein_coding | ENSG00000 |
| 31613305 | 31613436 | 1 | snoRNA | ENSG00000 |
| 31629248 | 31642931 | -1 | protein_coding | ENSG00000 |
| 31642642 | 31648435 | 1 | protein_coding | ENSG00000 |
| 31648523 | 31650997 | -1 | protein_coding | ENSG00000 |
| 31648554 | 31650946 | 1 | lncRNA | ENSG00000 |
| 31651454 | 31656514 | -1 | protein_coding | ENSG00000 |
| 31653519 | 31653632 | 1 | misc_RNA | ENSG00000 |
| 31655467 | 31663777 | 1 | protein_coding | ENSG00000 |
| 31659349 | 31663777 | 1 | protein_coding | ENSG00000 |
| 31666915 | 31674267 | -1 | protein_coding | ENSG00000 |
| 31677180 | 31693655 | -1 | protein_coding | ENSG00000 |
| 31697109 | 31700820 | 1 | protein_coding | ENSG00000 |
| 31697128 | 31708139 | 1 | protein_coding | ENSG00000 |
| 31702563 | 31704039 | -1 | protein_coding | ENSG00000 |
| 31708815 | 31716928 | 1 | protein_coding | ENSG00000 |
| 31708869 | 31712066 | -1 | protein_coding | ENSG00000 |
| 31717256 | 31720835 | -1 | protein_coding | ENSG00000 |
| 31720799 | 31729989 | -1 | protein_coding | ENSG00000 |
| 31730217 | 31751987 | 1 | protein_coding | ENSG00000 |
| 31747180 | 31747282 | -1 | snRNA | ENSG00000 |
| 31781844 | 31786187 | -1 | protein_coding | ENSG00000 |
| 31787629 | 31797209 | -1 | protein_coding | ENSG00000 |
| 31799855 | 31805300 | -1 | protein_coding | ENSG00000 |
| 31805699 | 31808181 | 1 | protein_coding | ENSG00000 |
| 31817974 | 31820498 | 1 | protein_coding | ENSG00000 |
| 31824855 | 31830009 | 1 | protein_coding | ENSG00000 |
| 31825510 | 31825572 | 1 | snoRNA | ENSG00000 |
| 31827320 | 31827386 | 1 | snoRNA | ENSG00000 |
| 31848038 | 31853285 | -1 | protein_coding | ENSG00000 |
| 31850614 | 31999312 | 1 | protein_coding | ENSG00000 |
| 31853571 | 31869426 | -1 | protein_coding | ENSG00000 |
| 31870139 | 31888078 | -1 | protein_coding | ENSG00000 |
| 31874140 | 31874433 | 1 | lncRNA | ENSG00000 |
| 31888164 | 31936056 | 1 | protein_coding | ENSG00000 |
| 31889987 | 31892374 | -1 | protein_coding | ENSG00000 |
| 31924830 | 31932095 | -1 | lncRNA | ENSG00000 |
| 31936032 | 31942466 | 1 | protein_coding | ENSG00000 |
| 31942469 | 31949469 | -1 | protein_coding | ENSG00000 |
| 31949457 | 31960128 | 1 | protein_coding | ENSG00000 |

| | | | | |
|---|---|---|---|---|
| 31960187 | 31962669 | -1 | protein_coding | ENSG00000( |
| 31961552 | 31971824 | 1 | protein_coding | ENSG00000( |
| 31962758 | 31973199 | 1 | protein_coding | ENSG00000( |
| 31972402 | 31993060 | 1 | protein_coding | ENSG00000( |
| 31990371 | 31993900 | -1 | lncRNA | ENSG00000( |
| 31998796 | 32043729 | -1 | protein_coding | ENSG00000( |
| 32036120 | 32036240 | -1 | rRNA | ENSG00000( |
| 32105963 | 32111974 | -1 | protein_coding | ENSG00000( |
| 32111042 | 32121288 | 1 | protein_coding | ENSG00000( |
| 32111446 | 32129586 | 1 | protein_coding | ENSG00000( |
| 32122191 | 32125889 | 1 | protein_coding | ENSG00000( |
| 32125820 | 32135704 | -1 | protein_coding | ENSG00000( |
| 32135962 | 32138401 | 1 | protein_coding | ENSG00000( |
| 32138576 | 32141932 | -1 | protein_coding | ENSG00000( |
| 32142341 | 32147792 | -1 | protein_coding | ENSG00000( |
| 32148372 | 32153129 | -1 | protein_coding | ENSG00000( |
| 32152449 | 32181676 | -1 | protein_coding | ENSG00000( |
| 32250505 | 32329363 | -1 | protein_coding | ENSG00000( |
| 32283008 | 32283967 | 1 | processed_pseudoge | ENSG00000( |
| 32310327 | 32310433 | 1 | snRNA | ENSG00000( |
| 32347957 | 32351132 | 1 | lncRNA | ENSG00000( |
| 32351409 | 32364583 | -1 | protein_coding | ENSG00000( |
| 32397298 | 32402508 | 1 | protein_coding | ENSG00000( |
| 32417316 | 32430998 | -1 | unprocessed_pseudo | ENSG00000( |
| 32447192 | 32447340 | -1 | snRNA | ENSG00000( |
| 32449765 | 32462852 | -1 | protein_coding | ENSG00000( |
| 32485042 | 32485188 | -1 | snRNA | ENSG00000( |
| 32488465 | 32502952 | -1 | unprocessed_pseudo | ENSG00000( |
| 32510948 | 32511094 | -1 | snRNA | ENSG00000( |
| 32513411 | 32544028 | -1 | protein_coding | ENSG00000( |
| 32570792 | 32577071 | 1 | protein_coding | ENSG00000( |
| 32587742 | 32595353 | -1 | protein_coding | ENSG00000( |
| 32588193 | 32589041 | 1 | lncRNA | ENSG00000( |
| 32588198 | 32590793 | -1 | lncRNA | ENSG00000( |
| 32634806 | 32635637 | -1 | unprocessed_pseudo | ENSG00000( |
| 32646664 | 32647494 | 1 | lncRNA | ENSG00000( |
| 32659420 | 32660356 | -1 | unprocessed_pseudo | ENSG00000( |
| 32670013 | 32675872 | 1 | protein_coding | ENSG00000( |
| 32684806 | 32692245 | -1 | protein_coding | ENSG00000( |
| 32750528 | 32767476 | -1 | protein_coding | ENSG00000( |
| 32769412 | 32773398 | -1 | protein_coding | ENSG00000( |
| 32772781 | 32775190 | 1 | lncRNA | ENSG00000( |
| 32772831 | 32808281 | 1 | protein_coding | ENSG00000( |
| 32773904 | 32782673 | -1 | protein_coding | ENSG00000( |
| 32807865 | 32808542 | -1 | transcribed_process | ENSG00000( |
| 32825106 | 32825180 | 1 | unprocessed_pseudo | ENSG00000( |
| 32863344 | 32881839 | -1 | protein_coding | ENSG00000( |
| 32877327 | 32899448 | -1 | protein_coding | ENSG00000( |
| 32897392 | 32910230 | 1 | protein_coding | ENSG00000( |
| 32899035 | 32899549 | 1 | lncRNA | ENSG00000( |

| | | | | |
|---|---|---|---|---|
| 32932859 | 32938293 | -1 | protein_coding | ENSG00000( |
| 32993608 | 33009820 | -1 | protein_coding | ENSG00000( |
| 33004965 | 33016242 | 1 | protein_coding | ENSG00000( |
| 33008496 | 33008905 | 1 | processed_pseudoge | ENSG00000( |
| 33020794 | 33026336 | -1 | unprocessed_pseudc | ENSG00000( |
| 33032837 | 33036373 | -1 | unprocessed_pseudc | ENSG00000( |
| 33041494 | 33057842 | 1 | transcribed_unproce | ENSG00000( |
| 33059973 | 33072108 | -1 | unprocessed_pseudc | ENSG00000( |
| 33073573 | 33076557 | 1 | lncRNA | ENSG00000( |
| 33091492 | 33121309 | -1 | protein_coding | ENSG00000( |
| 33122398 | 33129499 | -1 | protein_coding | ENSG00000( |
| 33128412 | 33128507 | 1 | misc_RNA | ENSG00000( |
| 33129256 | 33133250 | 1 | protein_coding | ENSG00000( |
| 33133453 | 33135646 | 1 | protein_coding | ENSG00000( |
| 33137310 | 33141537 | 1 | protein_coding | ENSG00000( |
| 33144520 | 33145143 | 1 | processed_pseudoge | ENSG00000( |
| 33166626 | 33168528 | 1 | processed_pseudoge | ENSG00000( |
| 33178341 | 33183793 | 1 | lncRNA | ENSG00000( |
| 33179079 | 33200871 | -1 | protein_coding | ENSG00000( |
| 33200834 | 33205337 | 1 | protein_coding | ENSG00000( |
| 33205959 | 33207652 | 1 | protein_coding | ENSG00000( |
| 33207935 | 33218358 | -1 | protein_coding | ENSG00000( |
| 33218133 | 33227229 | 1 | protein_coding | ENSG00000( |
| 33220485 | 33228152 | -1 | protein_coding | ENSG00000( |
| 33228522 | 33243221 | -1 | protein_coding | ENSG00000( |
| 33243252 | 33246776 | -1 | protein_coding | ENSG00000( |
| 33247394 | 33258105 | -1 | protein_coding | ENSG00000( |
| 33267813 | 33268330 | 1 | processed_pseudoge | ENSG00000( |
| 33294377 | 33295072 | -1 | processed_pseudoge | ENSG00000( |
| 33318189 | 33318290 | -1 | processed_pseudoge | ENSG00000( |
| 33320351 | 33338717 | 1 | protein_coding | ENSG00000( |
| 33328865 | 33329362 | -1 | processed_pseudoge | ENSG00000( |
| 33339192 | 33340037 | 1 | lncRNA | ENSG00000( |
| 28732818 | 28733705 | 1 | processed_pseudoge | ENSG00000( |
| 28784437 | 28784808 | -1 | processed_pseudoge | ENSG00000( |
| 28838582 | 28839719 | -1 | lncRNA | ENSG00000( |
| 28860338 | 28864060 | -1 | lncRNA | ENSG00000( |
| 28862129 | 28862760 | 1 | processed_pseudoge | ENSG00000( |
| 28889708 | 28890782 | -1 | processed_pseudoge | ENSG00000( |
| 28897398 | 28898188 | 1 | lncRNA | ENSG00000( |
| 28903869 | 28924846 | -1 | protein_coding | ENSG00000( |
| 28916503 | 28916602 | 1 | snRNA | ENSG00000( |
| 28944738 | 28945398 | 1 | protein_coding | ENSG00000( |
| 28969980 | 28971310 | -1 | processed_pseudoge | ENSG00000( |
| 28978391 | 28978689 | 1 | misc_RNA | ENSG00000( |
| 28987124 | 28988405 | 1 | lncRNA | ENSG00000( |
| 28987721 | 28989460 | 1 | lncRNA | ENSG00000( |
| 28995715 | 29006139 | -1 | protein_coding | ENSG00000( |
| 29027504 | 29028431 | -1 | unprocessed_pseudc | ENSG00000( |
| 29045035 | 29046062 | -1 | protein_coding | ENSG00000( |

| | | | | |
|---|---|---|---|---|
| 29064823 | 29077352 | 1 | protein_coding | ENSG00000( |
| 29072652 | 29073427 | -1 | unprocessed_pseudc | ENSG00000( |
| 29077401 | 29077996 | -1 | processed_pseudoge | ENSG00000( |
| 29087135 | 29088141 | -1 | protein_coding | ENSG00000( |
| 29101778 | 29102716 | 1 | unprocessed_pseudc | ENSG00000( |
| 29112720 | 29113655 | 1 | protein_coding | ENSG00000( |
| 29125038 | 29129735 | 1 | lncRNA | ENSG00000( |
| 29138704 | 29139640 | -1 | unprocessed_pseudc | ENSG00000( |
| 29174136 | 29175407 | 1 | protein_coding | ENSG00000( |
| 29182359 | 29183294 | 1 | unprocessed_pseudc | ENSG00000( |
| 29216117 | 29217054 | 1 | unprocessed_pseudc | ENSG00000( |
| 29224848 | 29267951 | 1 | lncRNA | ENSG00000( |
| 29230102 | 29231032 | -1 | unprocessed_pseudc | ENSG00000( |
| 29263569 | 29264945 | -1 | unprocessed_pseudc | ENSG00000( |
| 29269328 | 29270287 | -1 | unprocessed_pseudc | ENSG00000( |
| 29291595 | 29292537 | 1 | unprocessed_pseudc | ENSG00000( |
| 29307588 | 29308553 | 1 | protein_coding | ENSG00000( |
| 29330530 | 29331965 | -1 | processed_pseudoge | ENSG00000( |
| 29356146 | 29432443 | -1 | protein_coding | ENSG00000( |
| 29374013 | 29450575 | -1 | protein_coding | ENSG00000( |
| 29397025 | 29398093 | 1 | protein_coding | ENSG00000( |
| 29417758 | 29418704 | 1 | unprocessed_pseudc | ENSG00000( |
| 29425980 | 29457549 | -1 | protein_coding | ENSG00000( |
| 29439782 | 29441453 | 1 | protein_coding | ENSG00000( |
| 29457659 | 29464806 | 1 | protein_coding | ENSG00000( |
| 29465074 | 29470282 | -1 | unprocessed_pseudc | ENSG00000( |
| 29475501 | 29476547 | -1 | unprocessed_pseudc | ENSG00000( |
| 29487242 | 29488378 | -1 | protein_coding | ENSG00000( |
| 29489747 | 29490152 | -1 | processed_pseudoge | ENSG00000( |
| 29497985 | 29511032 | 1 | lncRNA | ENSG00000( |
| 29529901 | 29534049 | 1 | lncRNA | ENSG00000( |
| 29538185 | 29539268 | -1 | unprocessed_pseudc | ENSG00000( |
| 29636562 | 29636845 | -1 | processed_pseudoge | ENSG00000( |
| 29657481 | 29672869 | 1 | protein_coding | ENSG00000( |
| 29672889 | 29681605 | -1 | protein_coding | ENSG00000( |
| 29708627 | 29709049 | -1 | processed_pseudoge | ENSG00000( |
| 29720539 | 29736235 | 1 | protein_coding | ENSG00000( |
| 29720705 | 29721695 | -1 | unprocessed_pseudc | ENSG00000( |
| 29724295 | 29746743 | -1 | lncRNA | ENSG00000( |
| 29724363 | 29724833 | -1 | transcribed_processe | ENSG00000( |
| 29739435 | 29746663 | -1 | transcribed_unproce | ENSG00000( |
| 29745983 | 29746207 | 1 | unprocessed_pseudc | ENSG00000( |
| 29748420 | 29748840 | -1 | processed_pseudoge | ENSG00000( |
| 29788968 | 29795307 | 1 | transcribed_unproce | ENSG00000( |
| 29789369 | 29790366 | -1 | unprocessed_pseudc | ENSG00000( |
| 29797915 | 29799945 | 1 | unprocessed_pseudc | ENSG00000( |
| 29800695 | 29801491 | -1 | processed_pseudoge | ENSG00000( |
| 29810065 | 29810196 | -1 | unprocessed_pseudc | ENSG00000( |
| 29824463 | 29828635 | 1 | protein_coding | ENSG00000( |
| 29824881 | 29825861 | -1 | unprocessed_pseudc | ENSG00000( |

| | | | | |
|---|---|---|---|---|
| 29849350 | 29849494 | -1 | unprocessed_pseudc | ENSG00000C |
| 29878595 | 29879557 | -1 | unprocessed_pseudc | ENSG00000C |
| 29879053 | 29881769 | 1 | unprocessed_pseudc | ENSG00000C |
| 29887842 | 29888973 | 1 | unprocessed_pseudc | ENSG00000C |
| 29897735 | 29898101 | 1 | unprocessed_pseudc | ENSG00000C |
| 29898975 | 29899837 | -1 | unprocessed_pseudc | ENSG00000C |
| 29917138 | 29918128 | -1 | unprocessed_pseudc | ENSG00000C |
| 29917614 | 29920391 | 1 | unprocessed_pseudc | ENSG00000C |
| 29923501 | 29923685 | 1 | unprocessed_pseudc | ENSG00000C |
| 29930538 | 29999232 | 1 | protein_coding | ENSG00000C |
| 29931354 | 29932345 | -1 | unprocessed_pseudc | ENSG00000C |
| 29945879 | 29947844 | 1 | unprocessed_pseudc | ENSG00000C |
| 29960061 | 29961724 | -1 | unprocessed_pseudc | ENSG00000C |
| 29964383 | 29967676 | 1 | lncRNA | ENSG00000C |
| 29982480 | 29982876 | 1 | unprocessed_pseudc | ENSG00000C |
| 29983708 | 29984585 | -1 | unprocessed_pseudc | ENSG00000C |
| 29990282 | 30052027 | -1 | lncRNA | ENSG00000C |
| 29995393 | 29996388 | -1 | unprocessed_pseudc | ENSG00000C |
| 29995855 | 29999228 | 1 | transcribed_unproce | ENSG00000C |
| 30022074 | 30023364 | 1 | processed_pseudoge | ENSG00000C |
| 30026277 | 30026344 | 1 | misc_RNA | ENSG00000C |
| 30048502 | 30048577 | 1 | misc_RNA | ENSG00000C |
| 30049286 | 30055293 | 1 | protein_coding | ENSG00000C |
| 30051467 | 30051570 | 1 | misc_RNA | ENSG00000C |
| 30057093 | 30060717 | 1 | protein_coding | ENSG00000C |
| 30060650 | 30066271 | -1 | protein_coding | ENSG00000C |
| 30093294 | 30103494 | -1 | protein_coding | ENSG00000C |
| 30095634 | 30105111 | 1 | lncRNA | ENSG00000C |
| 30123209 | 30123371 | -1 | snoRNA | ENSG00000C |
| 30126512 | 30139132 | 1 | protein_coding | ENSG00000C |
| 30142335 | 30151326 | -1 | protein_coding | ENSG00000C |
| 30153602 | 30163092 | 1 | protein_coding | ENSG00000C |
| 30174807 | 30203783 | -1 | protein_coding | ENSG00000C |
| 30177152 | 30178961 | -1 | processed_pseudoge | ENSG00000C |
| 30224388 | 30316053 | -1 | lncRNA | ENSG00000C |
| 30228650 | 30232212 | 1 | unprocessed_pseudc | ENSG00000C |
| 30249542 | 30256896 | 1 | transcribed_unproce | ENSG00000C |
| 30280729 | 30317069 | -1 | lncRNA | ENSG00000C |
| 30316398 | 30333651 | 1 | protein_coding | ENSG00000C |
| 30319231 | 30336806 | 1 | protein_coding | ENSG00000C |
| 30341338 | 30341472 | 1 | unprocessed_pseudc | ENSG00000C |
| 30349200 | 30349833 | -1 | processed_pseudoge | ENSG00000C |
| 30353022 | 30354190 | -1 | processed_pseudoge | ENSG00000C |
| 30404626 | 30409231 | 1 | unprocessed_pseudc | ENSG00000C |
| 30456357 | 30457894 | 1 | processed_pseudoge | ENSG00000C |
| 30458704 | 30460073 | 1 | processed_pseudoge | ENSG00000C |
| 30478177 | 30478827 | 1 | processed_pseudoge | ENSG00000C |
| 30481701 | 30486439 | 1 | protein_coding | ENSG00000C |
| 30508488 | 30511442 | 1 | lncRNA | ENSG00000C |
| 30533598 | 30549394 | -1 | protein_coding | ENSG00000C |

| | | | | |
|---|---|---|---|---|
| 30549106 | 30555943 | 1 | protein_coding | ENSG00000 |
| 30563598 | 30589402 | 1 | protein_coding | ENSG00000 |
| 30592623 | 30610835 | -1 | protein_coding | ENSG00000 |
| 30609932 | 30618618 | 1 | protein_coding | ENSG00000 |
| 30619065 | 30639047 | 1 | protein_coding | ENSG00000 |
| 30625856 | 30626122 | 1 | processed_pseudoge | ENSG00000 |
| 30639263 | 30645325 | 1 | protein_coding | ENSG00000 |
| 30645335 | 30665285 | -1 | protein_coding | ENSG00000 |
| 30668637 | 30671637 | -1 | protein_coding | ENSG00000 |
| 30692058 | 30710140 | -1 | protein_coding | ENSG00000 |
| 30695318 | 30705434 | 1 | lncRNA | ENSG00000 |
| 30712452 | 30717677 | 1 | protein_coding | ENSG00000 |
| 30719960 | 30734994 | -1 | protein_coding | ENSG00000 |
| 30728556 | 30728666 | -1 | misc_RNA | ENSG00000 |
| 30735460 | 30736815 | -1 | protein_coding | ENSG00000 |
| 30743271 | 30743522 | 1 | misc_RNA | ENSG00000 |
| 30780908 | 30784476 | 1 | lncRNA | ENSG00000 |
| 30790867 | 30822832 | -1 | lncRNA | ENSG00000 |
| 31002649 | 31027602 | 1 | lncRNA | ENSG00000 |
| 31048375 | 31054804 | 1 | transcribed_unproce | ENSG00000 |
| 31077965 | 31078064 | 1 | snRNA | ENSG00000 |
| 31153422 | 31159096 | 1 | protein_coding | ENSG00000 |
| 31159218 | 31165590 | -1 | protein_coding | ENSG00000 |
| 31168631 | 31172797 | -1 | lncRNA | ENSG00000 |
| 31192649 | 31198853 | 1 | lncRNA | ENSG00000 |
| 31195054 | 31196805 | -1 | lncRNA | ENSG00000 |
| 31262172 | 31265564 | -1 | protein_coding | ENSG00000 |
| 31262912 | 31350932 | -1 | protein_coding | ENSG00000 |
| 31268956 | 31272128 | 1 | processed_pseudoge | ENSG00000 |
| 31273691 | 31274890 | 1 | processed_pseudoge | ENSG00000 |
| 31281167 | 31282618 | -1 | processed_pseudoge | ENSG00000 |
| 31301504 | 31302238 | -1 | unprocessed_pseudo | ENSG00000 |
| 31360052 | 31360598 | -1 | processed_pseudoge | ENSG00000 |
| 31363816 | 31363922 | 1 | snRNA | ENSG00000 |
| 31371102 | 31371704 | 1 | processed_pseudoge | ENSG00000 |
| 31374124 | 31374550 | 1 | processed_pseudoge | ENSG00000 |
| 31375802 | 31376019 | -1 | unprocessed_pseudo | ENSG00000 |
| 31393526 | 31409981 | 1 | protein_coding | ENSG00000 |
| 31395893 | 31395991 | -1 | misc_RNA | ENSG00000 |
| 31435351 | 31440654 | 1 | lncRNA | ENSG00000 |
| 31453889 | 31456523 | 1 | lncRNA | ENSG00000 |
| 31487449 | 31487550 | -1 | misc_RNA | ENSG00000 |
| 31488846 | 31501875 | 1 | protein_coding | ENSG00000 |
| 31510228 | 31511039 | -1 | processed_pseudoge | ENSG00000 |
| 31518863 | 31519442 | 1 | processed_pseudoge | ENSG00000 |
| 31519466 | 31520981 | 1 | protein_coding | ENSG00000 |
| 31520968 | 31533197 | -1 | protein_coding | ENSG00000 |
| 31520968 | 31537331 | -1 | protein_coding | ENSG00000 |
| 31527123 | 31527198 | -1 | snoRNA | ENSG00000 |
| 31531850 | 31531927 | -1 | snoRNA | ENSG00000 |

| | | | | |
|---|---|---|---|---|
| 31533053 | 31533887 | 1 | lncRNA | ENSG0000C |
| 31535211 | 31537597 | -1 | protein_coding | ENSG0000C |
| 31537619 | 31549575 | 1 | protein_coding | ENSG0000C |
| 31562799 | 31565069 | 1 | protein_coding | ENSG0000C |
| 31566312 | 31569081 | 1 | protein_coding | ENSG0000C |
| 31571300 | 31573267 | -1 | protein_coding | ENSG0000C |
| 31576870 | 31579655 | 1 | protein_coding | ENSG0000C |
| 31579641 | 31583730 | -1 | protein_coding | ENSG0000C |
| 31601835 | 31602108 | -1 | processed_pseudoge | ENSG0000C |
| 31605935 | 31607772 | 1 | protein_coding | ENSG0000C |
| 31611467 | 31628517 | 1 | protein_coding | ENSG0000C |
| 31613823 | 31613954 | 1 | snoRNA | ENSG0000C |
| 31629769 | 31643448 | -1 | protein_coding | ENSG0000C |
| 31643159 | 31648952 | 1 | protein_coding | ENSG0000C |
| 31649040 | 31651514 | -1 | protein_coding | ENSG0000C |
| 31649071 | 31651461 | 1 | lncRNA | ENSG0000C |
| 31651971 | 31657032 | -1 | protein_coding | ENSG0000C |
| 31654037 | 31654150 | 1 | misc_RNA | ENSG0000C |
| 31655985 | 31664295 | 1 | protein_coding | ENSG0000C |
| 31659867 | 31664295 | 1 | protein_coding | ENSG0000C |
| 31667433 | 31674786 | -1 | protein_coding | ENSG0000C |
| 31677699 | 31694178 | -1 | protein_coding | ENSG0000C |
| 31697642 | 31701354 | 1 | protein_coding | ENSG0000C |
| 31697661 | 31708679 | 1 | protein_coding | ENSG0000C |
| 31703097 | 31704573 | -1 | protein_coding | ENSG0000C |
| 31706117 | 31708565 | 1 | protein_coding | ENSG0000C |
| 31709355 | 31717471 | 1 | protein_coding | ENSG0000C |
| 31709409 | 31712606 | -1 | protein_coding | ENSG0000C |
| 31717799 | 31721378 | -1 | protein_coding | ENSG0000C |
| 31721342 | 31730525 | -1 | protein_coding | ENSG0000C |
| 31730753 | 31755602 | 1 | protein_coding | ENSG0000C |
| 31747708 | 31747810 | -1 | snRNA | ENSG0000C |
| 31748922 | 31755608 | 1 | protein_coding | ENSG0000C |
| 31753556 | 31755608 | 1 | protein_coding | ENSG0000C |
| 31756347 | 31768050 | -1 | protein_coding | ENSG0000C |
| 31768274 | 31785735 | -1 | protein_coding | ENSG0000C |
| 31769573 | 31769661 | -1 | misc_RNA | ENSG0000C |
| 31850351 | 31855624 | -1 | protein_coding | ENSG0000C |
| 31855884 | 31871733 | -1 | protein_coding | ENSG0000C |
| 31872446 | 31882289 | -1 | protein_coding | ENSG0000C |
| 31876448 | 31876741 | 1 | lncRNA | ENSG0000C |
| 31912644 | 31938328 | 1 | protein_coding | ENSG0000C |
| 31927103 | 31934377 | -1 | lncRNA | ENSG0000C |
| 31938304 | 31944738 | 1 | protein_coding | ENSG0000C |
| 31944741 | 31951746 | -1 | protein_coding | ENSG0000C |
| 31951734 | 31962405 | 1 | protein_coding | ENSG0000C |
| 31962464 | 31964946 | -1 | protein_coding | ENSG0000C |
| 31963829 | 31974100 | 1 | protein_coding | ENSG0000C |
| 31974678 | 31995335 | 1 | protein_coding | ENSG0000C |
| 31992630 | 31996175 | -1 | lncRNA | ENSG0000C |

| | | | | |
|---|---|---|---|---|
| 31998290 | 32001104 | 1 | transcribed_unproce | ENSG0000C |
| 32001268 | 32005126 | -1 | unprocessed_pseudc | ENSG0000C |
| 32005925 | 32006441 | 1 | unprocessed_pseudc | ENSG0000C |
| 32007416 | 32028073 | 1 | protein_coding | ENSG0000C |
| 32025384 | 32028913 | -1 | lncRNA | ENSG0000C |
| 32030953 | 32034322 | 1 | protein_coding | ENSG0000C |
| 32033807 | 32060653 | -1 | protein_coding | ENSG0000C |
| 32156028 | 32162042 | -1 | protein_coding | ENSG0000C |
| 32160569 | 32179650 | 1 | protein_coding | ENSG0000C |
| 32172254 | 32175953 | 1 | protein_coding | ENSG0000C |
| 32175884 | 32185768 | -1 | protein_coding | ENSG0000C |
| 32186026 | 32188465 | 1 | protein_coding | ENSG0000C |
| 32188640 | 32191996 | -1 | protein_coding | ENSG0000C |
| 32192405 | 32197856 | -1 | protein_coding | ENSG0000C |
| 32198436 | 32203193 | -1 | protein_coding | ENSG0000C |
| 32202513 | 32231740 | -1 | protein_coding | ENSG0000C |
| 32300437 | 32379326 | -1 | protein_coding | ENSG0000C |
| 32332954 | 32333913 | 1 | processed_pseudoge | ENSG0000C |
| 32360267 | 32360373 | 1 | snRNA | ENSG0000C |
| 32397896 | 32401073 | 1 | lncRNA | ENSG0000C |
| 32401346 | 32414522 | -1 | protein_coding | ENSG0000C |
| 32446450 | 32451660 | 1 | protein_coding | ENSG0000C |
| 32466569 | 32480224 | -1 | unprocessed_pseudc | ENSG0000C |
| 32492462 | 32492557 | -1 | unprocessed_pseudc | ENSG0000C |
| 32518027 | 32518174 | -1 | snRNA | ENSG0000C |
| 32523454 | 32523549 | -1 | unprocessed_pseudc | ENSG0000C |
| 32539820 | 32539971 | -1 | snRNA | ENSG0000C |
| 32542598 | 32557561 | -1 | protein_coding | ENSG0000C |
| 32579248 | 32579395 | -1 | snRNA | ENSG0000C |
| 32589215 | 32591752 | -1 | unprocessed_pseudc | ENSG0000C |
| 32619811 | 32619974 | -1 | snRNA | ENSG0000C |
| 32621929 | 32622868 | 1 | processed_pseudoge | ENSG0000C |
| 32624375 | 32624782 | -1 | processed_pseudoge | ENSG0000C |
| 32627102 | 32646281 | -1 | unprocessed_pseudc | ENSG0000C |
| 32629937 | 32631068 | 1 | processed_pseudoge | ENSG0000C |
| 32668064 | 32668212 | -1 | snRNA | ENSG0000C |
| 32670593 | 32703817 | -1 | protein_coding | ENSG0000C |
| 32730753 | 32736617 | 1 | protein_coding | ENSG0000C |
| 32748163 | 32755410 | -1 | protein_coding | ENSG0000C |
| 32748576 | 32749425 | 1 | lncRNA | ENSG0000C |
| 32748582 | 32751171 | -1 | lncRNA | ENSG0000C |
| 32798219 | 32799050 | -1 | unprocessed_pseudc | ENSG0000C |
| 32809613 | 32810443 | 1 | lncRNA | ENSG0000C |
| 32822370 | 32823307 | -1 | unprocessed_pseudc | ENSG0000C |
| 32832954 | 32838824 | 1 | protein_coding | ENSG0000C |
| 32847708 | 32855144 | -1 | protein_coding | ENSG0000C |
| 32903640 | 32907925 | -1 | protein_coding | ENSG0000C |
| 32912700 | 32929495 | -1 | protein_coding | ENSG0000C |
| 32931434 | 32935368 | -1 | protein_coding | ENSG0000C |
| 32934803 | 32937212 | 1 | lncRNA | ENSG0000C |

| | | | | |
|---|---|---|---|---|
| 32934853 | 32970271 | 1 | protein_coding | ENSG0000C |
| 32935926 | 32944695 | -1 | protein_coding | ENSG0000C |
| 32969856 | 32970532 | -1 | transcribed_processe | ENSG0000C |
| 32987088 | 32987162 | 1 | unprocessed_pseudc | ENSG0000C |
| 33025327 | 33043825 | -1 | protein_coding | ENSG0000C |
| 33039313 | 33061428 | -1 | protein_coding | ENSG0000C |
| 33059372 | 33122227 | 1 | protein_coding | ENSG0000C |
| 33061015 | 33061529 | 1 | lncRNA | ENSG0000C |
| 33144838 | 33150271 | -1 | protein_coding | ENSG0000C |
| 33205017 | 33221223 | -1 | protein_coding | ENSG0000C |
| 33216374 | 33227649 | 1 | protein_coding | ENSG0000C |
| 33219899 | 33220308 | 1 | processed_pseudoge | ENSG0000C |
| 33232201 | 33237743 | -1 | unprocessed_pseudc | ENSG0000C |
| 33244216 | 33247752 | -1 | unprocessed_pseudc | ENSG0000C |
| 33252873 | 33269193 | 1 | transcribed_unproce | ENSG0000C |
| 33271325 | 33271451 | -1 | unprocessed_pseudc | ENSG0000C |
| 33302792 | 33329578 | -1 | protein_coding | ENSG0000C |
| 33355816 | 33356439 | 1 | processed_pseudoge | ENSG0000C |
| 33377903 | 33379802 | 1 | processed_pseudoge | ENSG0000C |
| 33389703 | 33395155 | 1 | lncRNA | ENSG0000C |
| 33390441 | 33412193 | -1 | protein_coding | ENSG0000C |
| 33412156 | 33415676 | 1 | protein_coding | ENSG0000C |
| 33469139 | 33469421 | -1 | lncRNA | ENSG0000C |
| 33529535 | 33529636 | -1 | processed_pseudoge | ENSG0000C |
| 33531696 | 33550087 | 1 | protein_coding | ENSG0000C |
| 33540223 | 33540720 | -1 | processed_pseudoge | ENSG0000C |
| 33550562 | 33556643 | 1 | protein_coding | ENSG0000C |
| 33556632 | 33558507 | -1 | protein_coding | ENSG0000C |
| 33560260 | 33593877 | 1 | protein_coding | ENSG0000C |
| 33594767 | 33597736 | 1 | protein_coding | ENSG0000C |
| 70972 | 95683 | -1 | lncRNA | ENSG0000C |
| 77038 | 80418 | 1 | lncRNA | ENSG0000C |
| 135676 | 149167 | -1 | lncRNA | ENSG0000C |
| 149303 | 155171 | 1 | lncRNA | ENSG0000C |
| 165082 | 168277 | -1 | lncRNA | ENSG0000C |
| 178225 | 179318 | 1 | lncRNA | ENSG0000C |
| 186240 | 197485 | -1 | lncRNA | ENSG0000C |
| 196274 | 199037 | 1 | protein_coding | ENSG0000C |
| 103086630 | 103099764 | 1 | protein_coding | ENSG0000C |
| 103088664 | 103089610 | -1 | processed_pseudoge | ENSG0000C |
| 103099776 | 103149564 | -1 | protein_coding | ENSG0000C |
| 103141340 | 103141930 | -1 | processed_pseudoge | ENSG0000C |
| 103152009 | 103152327 | -1 | processed_pseudoge | ENSG0000C |
| 103164581 | 103165294 | -1 | TEC | ENSG0000C |
| 103177764 | 103243284 | -1 | transcribed_unproce | ENSG0000C |
| 141633901 | 141655244 | 1 | protein_coding | ENSG0000C |
| 141652381 | 141656810 | -1 | lncRNA | ENSG0000C |
| 141656728 | 141702156 | -1 | protein_coding | ENSG0000C |
| 141710346 | 141744354 | -1 | lncRNA | ENSG0000C |
| 141714361 | 141737279 | 1 | protein_coding | ENSG0000C |

| | | | | |
|---|---|---|---|---|
| 141733992 | 141734156 | 1 | snRNA | ENSG00000 |
| 141744329 | 141752534 | 1 | protein_coding | ENSG00000 |
| 158291152 | 158424386 | -1 | protein_coding | ENSG00000 |
| 19790 | 37822 | -1 | lncRNA | ENSG00000 |
| 39354 | 39799 | 1 | processed_pseudoge | ENSG00000 |
| 73293 | 98004 | -1 | lncRNA | ENSG00000 |
| 79359 | 82739 | 1 | lncRNA | ENSG00000 |
| 138174 | 151787 | -1 | lncRNA | ENSG00000 |
| 151918 | 157786 | 1 | lncRNA | ENSG00000 |
| 164086 | 167293 | -1 | lncRNA | ENSG00000 |
| 177241 | 178334 | 1 | lncRNA | ENSG00000 |
| 185256 | 196501 | -1 | lncRNA | ENSG00000 |
| 195290 | 199896 | 1 | protein_coding | ENSG00000 |
| 76501267 | 76505995 | 1 | protein_coding | ENSG00000 |
| 76510496 | 76515744 | 1 | protein_coding | ENSG00000 |
| 76515627 | 76520574 | 1 | unprocessed_pseudc | ENSG00000 |
| 76519194 | 76529340 | -1 | protein_coding | ENSG00000 |
| 76537175 | 76615798 | 1 | transcribed_unproce | ENSG00000 |
| 76597802 | 76615077 | -1 | protein_coding | ENSG00000 |
| 76638217 | 76638713 | -1 | processed_pseudoge | ENSG00000 |
| 76646507 | 76684889 | -1 | lncRNA | ENSG00000 |
| 142038524 | 142106747 | 1 | protein_coding | ENSG00000 |
| 142111749 | 142222324 | 1 | protein_coding | ENSG00000 |
| 142240759 | 142247086 | -1 | transcribed_unproce | ENSG00000 |
| 142252163 | 142258078 | -1 | protein_coding | ENSG00000 |
| 142265850 | 142272254 | -1 | transcribed_unproce | ENSG00000 |
| 142285779 | 142288898 | 1 | lncRNA | ENSG00000 |
| 142287280 | 142289809 | -1 | transcribed_unproce | ENSG00000 |
| 142299217 | 142299500 | 1 | TR_V_pseudogene | ENSG00000 |
| 142300947 | 142301455 | 1 | TR_V_gene | ENSG00000 |
| 142308565 | 142309071 | 1 | TR_V_gene | ENSG00000 |
| 142313207 | 142313689 | 1 | TR_V_gene | ENSG00000 |
| 142320696 | 142321563 | 1 | TR_V_gene | ENSG00000 |
| 142328321 | 142328810 | 1 | TR_V_gene | ENSG00000 |
| 142332206 | 142332727 | 1 | TR_V_gene | ENSG00000 |
| 142345452 | 142346016 | 1 | TR_V_gene | ENSG00000 |
| 142349215 | 142349695 | 1 | TR_V_gene | ENSG00000 |
| 142359222 | 142359681 | 1 | TR_V_pseudogene | ENSG00000 |
| 142362570 | 142363134 | 1 | TR_V_gene | ENSG00000 |
| 142370836 | 142371348 | 1 | TR_V_gene | ENSG00000 |
| 142374511 | 142375050 | 1 | TR_V_gene | ENSG00000 |
| 142380331 | 142380609 | 1 | TR_V_pseudogene | ENSG00000 |
| 142394342 | 142394614 | 1 | TR_V_pseudogene | ENSG00000 |
| 142402503 | 142402958 | 1 | TR_V_gene | ENSG00000 |
| 142406045 | 142406551 | 1 | TR_V_gene | ENSG00000 |
| 142408088 | 142408357 | 1 | TR_V_pseudogene | ENSG00000 |
| 142410913 | 142411379 | 1 | TR_V_gene | ENSG00000 |
| 142413602 | 142414123 | 1 | TR_V_gene | ENSG00000 |
| 142421573 | 142422090 | 1 | TR_V_gene | ENSG00000 |
| 142429379 | 142429843 | 1 | TR_V_gene | ENSG00000 |

| | | | | |
|---|---|---|---|---|
| 142436931 | 142437373 | 1 | TR_V_pseudogene | ENSG00000 |
| 142446652 | 142447152 | 1 | TR_V_gene | ENSG00000 |
| 142455594 | 142456093 | 1 | lncRNA | ENSG00000 |
| 142462583 | 142463025 | 1 | TR_V_pseudogene | ENSG00000 |
| 142472647 | 142473148 | 1 | TR_V_gene | ENSG00000 |
| 142476873 | 142477334 | 1 | TR_V_gene | ENSG00000 |
| 142484618 | 142485283 | 1 | TR_V_gene | ENSG00000 |
| 142491254 | 142491732 | 1 | TR_V_gene | ENSG00000 |
| 142495814 | 142496285 | 1 | TR_V_pseudogene | ENSG00000 |
| 142504264 | 142504735 | 1 | TR_V_gene | ENSG00000 |
| 142509537 | 142510011 | 1 | TR_V_gene | ENSG00000 |
| 142513848 | 142514385 | 1 | TR_V_gene | ENSG00000 |
| 142521745 | 142522251 | 1 | TR_V_gene | ENSG00000 |
| 142529032 | 142529526 | 1 | TR_V_gene | ENSG00000 |
| 142533342 | 142533843 | 1 | TR_V_gene | ENSG00000 |
| 142541804 | 142542270 | 1 | TR_V_gene | ENSG00000 |
| 142549110 | 142549542 | 1 | TR_V_gene | ENSG00000 |
| 142553725 | 142554208 | 1 | TR_V_gene | ENSG00000 |
| 142561449 | 142562408 | 1 | TR_V_gene | ENSG00000 |
| 142571143 | 142571615 | 1 | TR_V_gene | ENSG00000 |
| 142577660 | 142578143 | 1 | TR_V_gene | ENSG00000 |
| 142586067 | 142586516 | 1 | TR_V_gene | ENSG00000 |
| 142596665 | 142597147 | 1 | TR_V_gene | ENSG00000 |
| 142602235 | 142602743 | 1 | TR_V_gene | ENSG00000 |
| 142605549 | 142606054 | 1 | TR_V_gene | ENSG00000 |
| 142622755 | 142623265 | 1 | TR_V_gene | ENSG00000 |
| 142629704 | 142630195 | 1 | TR_V_gene | ENSG00000 |
| 142634764 | 142635309 | 1 | TR_V_gene | ENSG00000 |
| 142639852 | 142640305 | 1 | TR_V_gene | ENSG00000 |
| 142643462 | 142644194 | 1 | TR_V_gene | ENSG00000 |
| 142657499 | 142658198 | 1 | TR_V_gene | ENSG00000 |
| 142660632 | 142661315 | 1 | TR_V_gene | ENSG00000 |
| 142668431 | 142669181 | 1 | TR_V_gene | ENSG00000 |
| 142678692 | 142679152 | 1 | TR_V_pseudogene | ENSG00000 |
| 142683514 | 142683964 | 1 | TR_V_pseudogene | ENSG00000 |
| 142687733 | 142688229 | 1 | TR_V_gene | ENSG00000 |
| 142697038 | 142697550 | 1 | TR_V_gene | ENSG00000 |
| 142706945 | 142708391 | 1 | protein_coding | ENSG00000 |
| 142711413 | 142711917 | 1 | TR_V_gene | ENSG00000 |
| 142722076 | 142722530 | 1 | TR_V_pseudogene | ENSG00000 |
| 142736369 | 142736853 | 1 | TR_V_pseudogene | ENSG00000 |
| 142752085 | 142752625 | 1 | TR_V_pseudogene | ENSG00000 |
| 142756048 | 142756563 | 1 | TR_V_gene | ENSG00000 |
| 142761362 | 142761862 | 1 | TR_V_gene | ENSG00000 |
| 142763060 | 142763466 | 1 | processed_pseudoge | ENSG00000 |
| 142780919 | 142781607 | 1 | TR_V_gene | ENSG00000 |
| 142790185 | 142793793 | 1 | protein_coding | ENSG00000 |
| 142801130 | 142804658 | 1 | unprocessed_pseudc | ENSG00000 |
| 142811706 | 142815230 | 1 | unprocessed_pseudc | ENSG00000 |
| 142821798 | 142825331 | 1 | unprocessed_pseudc | ENSG00000 |

| | | | | |
|---|---|---|---|---|
| 142832063 | 142835652 | 1 | protein_coding | ENSG00000( |
| 142839736 | 142840629 | -1 | processed_pseudoge | ENSG00000( |
| 142847306 | 142847317 | 1 | TR_D_gene | ENSG00000( |
| 142847973 | 142848020 | 1 | TR_J_gene | ENSG00000( |
| 142848110 | 142848157 | 1 | TR_J_gene | ENSG00000( |
| 142848723 | 142848772 | 1 | TR_J_gene | ENSG00000( |
| 142849318 | 142849368 | 1 | TR_J_gene | ENSG00000( |
| 142849591 | 142849640 | 1 | TR_J_gene | ENSG00000( |
| 142850081 | 142850133 | 1 | TR_J_gene | ENSG00000( |
| 142852787 | 142854461 | 1 | TR_C_gene | ENSG00000( |
| 142857458 | 142857507 | 1 | TR_J_gene | ENSG00000( |
| 142857653 | 142857703 | 1 | TR_J_gene | ENSG00000( |
| 142857790 | 142857835 | 1 | TR_J_gene | ENSG00000( |
| 142857940 | 142857988 | 1 | TR_J_gene | ENSG00000( |
| 142858091 | 142858140 | 1 | TR_J_gene | ENSG00000( |
| 142858212 | 142858259 | 1 | TR_J_gene | ENSG00000( |
| 142858332 | 142858384 | 1 | TR_J_gene | ENSG00000( |
| 142858549 | 142858595 | 1 | TR_J_gene | ENSG00000( |
| 142862134 | 142863838 | 1 | TR_C_gene | ENSG00000( |
| 142873679 | 142874492 | -1 | TR_V_gene | ENSG00000( |
| 142916237 | 142932255 | 1 | protein_coding | ENSG00000( |
| 142932365 | 142946922 | -1 | protein_coding | ENSG00000( |
| 142936996 | 142953930 | 1 | lncRNA | ENSG00000( |
| 142969288 | 142994995 | -1 | protein_coding | ENSG00000( |
| 143000530 | 143002052 | 1 | protein_coding | ENSG00000( |
| 143002301 | 143023868 | -1 | protein_coding | ENSG00000( |
| 143087345 | 143088382 | -1 | protein_coding | ENSG00000( |
| 143108400 | 143108767 | -1 | unprocessed_pseudo | ENSG00000( |
| 143113507 | 143114534 | 1 | protein_coding | ENSG00000( |
| 143123517 | 143124772 | -1 | transcribed_unproce | ENSG00000( |
| 157741015 | 157742461 | -1 | lncRNA | ENSG00000( |
| 9089110 | 9196410 | -1 | lncRNA | ENSG00000( |
| 9090575 | 9090728 | 1 | processed_pseudoge | ENSG00000( |
| 9177501 | 9177826 | -1 | processed_pseudoge | ENSG00000( |
| 144048948 | 144058845 | 1 | protein_coding | ENSG00000( |
| 144074223 | 144075152 | 1 | protein_coding | ENSG00000( |
| 144095100 | 144096029 | 1 | protein_coding | ENSG00000( |
| 144109569 | 144110522 | 1 | protein_coding | ENSG00000( |
| 144129097 | 144130023 | 1 | protein_coding | ENSG00000( |
| 144183495 | 144185825 | 1 | protein_coding | ENSG00000( |
| 144186923 | 144188374 | -1 | protein_coding | ENSG00000( |
| 144231943 | 144232863 | -1 | protein_coding | ENSG00000( |
| 144251330 | 144251623 | -1 | protein_coding | ENSG00000( |
| 144258719 | 144259648 | -1 | protein_coding | ENSG00000( |
| 144266940 | 144269270 | -1 | protein_coding | ENSG00000( |
| 144285553 | 144286986 | -1 | protein_coding | ENSG00000( |
| 144299403 | 144299696 | 1 | protein_coding | ENSG00000( |
| 144318162 | 144319082 | 1 | protein_coding | ENSG00000( |
| 144360650 | 144373656 | 1 | protein_coding | ENSG00000( |
| 144378365 | 144380695 | 1 | protein_coding | ENSG00000( |

| | | | | |
|---|---|---|---|---|
| 144387992 | 144388921 | 1 | protein_coding | ENSG00000 |
| 144396023 | 144396316 | 1 | protein_coding | ENSG00000 |
| 144414782 | 144415702 | 1 | protein_coding | ENSG00000 |
| 144450317 | 144475530 | 1 | protein_coding | ENSG00000 |
| 144492136 | 144505111 | -1 | protein_coding | ENSG00000 |
| 144718185 | 144718358 | -1 | protein_coding | ENSG00000 |
| 148217288 | 148419466 | 1 | protein_coding | ENSG00000 |
| 148386174 | 148386384 | 1 | snoRNA | ENSG00000 |
| 148436777 | 148437045 | 1 | misc_RNA | ENSG00000 |
| 148439467 | 148475983 | -1 | lncRNA | ENSG00000 |
| 148444254 | 148444767 | 1 | unprocessed_pseudo | ENSG00000 |
| 1296034 | 1302607 | -1 | lncRNA | ENSG00000 |
| 1368648 | 1369833 | -1 | lncRNA | ENSG00000 |
| 1373685 | 1381062 | 1 | lncRNA | ENSG00000 |
| 1377533 | 1379973 | 1 | lncRNA | ENSG00000 |
| 1487682 | 1552210 | 1 | protein_coding | ENSG00000 |
| 1551825 | 1552054 | -1 | lncRNA | ENSG00000 |
| 57193590 | 57252058 | 1 | lncRNA | ENSG00000 |
| 57251821 | 57252305 | 1 | processed_pseudoge | ENSG00000 |
| 57272990 | 57278377 | 1 | lncRNA | ENSG00000 |
| 57291307 | 57296495 | 1 | lncRNA | ENSG00000 |
| 57301316 | 57301413 | 1 | snRNA | ENSG00000 |
| 128559028 | 128564679 | -1 | lncRNA | ENSG00000 |
| 128908170 | 128924280 | -1 | lncRNA | ENSG00000 |
| 883198 | 1140708 | 1 | lncRNA | ENSG00000 |
| 1021688 | 1022635 | 1 | lncRNA | ENSG00000 |
| 1107813 | 1107997 | 1 | lncRNA | ENSG00000 |
| 144701846 | 144702417 | -1 | lncRNA | ENSG00000 |
| 144703585 | 144711749 | 1 | transcribed_unproce | ENSG00000 |
| 144724139 | 144759649 | -1 | protein_coding | ENSG00000 |
| 144739024 | 144760993 | 1 | lncRNA | ENSG00000 |
| 7318568 | 7348494 | -1 | lncRNA | ENSG00000 |
| 7332922 | 7334514 | 1 | unprocessed_pseudo | ENSG00000 |
| 7334978 | 7336158 | 1 | unprocessed_pseudo | ENSG00000 |
| 7348669 | 7377169 | -1 | protein_coding | ENSG00000 |
| 7364723 | 7369092 | 1 | unprocessed_pseudo | ENSG00000 |
| 7406786 | 7408809 | -1 | protein_coding | ENSG00000 |
| 7410936 | 7412491 | 1 | processed_pseudoge | ENSG00000 |
| 7420832 | 7422292 | -1 | protein_coding | ENSG00000 |
| 7434631 | 7455657 | -1 | protein_coding | ENSG00000 |
| 7462290 | 7467065 | -1 | protein_coding | ENSG00000 |
| 7474484 | 7478378 | -1 | protein_coding | ENSG00000 |
| 7479647 | 7481569 | 1 | protein_coding | ENSG00000 |
| 7487822 | 7501287 | 1 | protein_coding | ENSG00000 |
| 7531604 | 7534426 | 1 | protein_coding | ENSG00000 |
| 7541029 | 7544038 | -1 | unprocessed_pseudo | ENSG00000 |
| 7548676 | 7551688 | -1 | unprocessed_pseudo | ENSG00000 |
| 7556326 | 7559336 | -1 | unprocessed_pseudo | ENSG00000 |
| 7563971 | 7566982 | -1 | unprocessed_pseudo | ENSG00000 |
| 7571620 | 7574629 | -1 | unprocessed_pseudo | ENSG00000 |

| | | | | |
|---|---|---|---|---|
| 7583982 | 7584892 | 1 | unprocessed_pseudc | ENSG0000C |
| 7592980 | 7593243 | 1 | processed_pseudoge | ENSG0000C |
| 143536257 | 143541453 | -1 | protein_coding | ENSG0000C |
| 143542110 | 143542398 | 1 | misc_RNA | ENSG0000C |
| 143553207 | 143563062 | 1 | protein_coding | ENSG0000C |
| 143566187 | 143572971 | -1 | protein_coding | ENSG0000C |
| 143573490 | 143577397 | 1 | lncRNA | ENSG0000C |
| 143574785 | 143578649 | -1 | protein_coding | ENSG0000C |
| 143578358 | 143583305 | 1 | lncRNA | ENSG0000C |
| 143579697 | 143597744 | -1 | protein_coding | ENSG0000C |
| 143597835 | 143603224 | 1 | protein_coding | ENSG0000C |
| 143603913 | 143609773 | -1 | protein_coding | ENSG0000C |
| 143612618 | 143618048 | -1 | protein_coding | ENSG0000C |
| 143621803 | 143641845 | 1 | protein_coding | ENSG0000C |
| 143671459 | 143700905 | 1 | protein_coding | ENSG0000C |
| 143683161 | 143685420 | 1 | lncRNA | ENSG0000C |
| 143693701 | 143695116 | -1 | protein_coding | ENSG0000C |
| 143696014 | 143700591 | -1 | lncRNA | ENSG0000C |
| 143703266 | 143709465 | 1 | protein_coding | ENSG0000C |
| 143705253 | 143705898 | -1 | lncRNA | ENSG0000C |
| 143710940 | 143720808 | -1 | protein_coding | ENSG0000C |
| 143721147 | 143733344 | 1 | lncRNA | ENSG0000C |
| 143775927 | 143777466 | 1 | lncRNA | ENSG0000C |
| 143777927 | 143802386 | -1 | protein_coding | ENSG0000C |
| 143803351 | 143816310 | -1 | protein_coding | ENSG0000C |
| 725188 | 725877 | -1 | lncRNA | ENSG0000C |
| 731140 | 738106 | -1 | protein_coding | ENSG0000C |
| 737651 | 824587 | 1 | lncRNA | ENSG0000C |
| 738548 | 740374 | 1 | lncRNA | ENSG0000C |
| 143012485 | 143018437 | -1 | lncRNA | ENSG0000C |
| 143017982 | 143023832 | 1 | protein_coding | ENSG0000C |
| 143039209 | 143059942 | 1 | lncRNA | ENSG0000C |
| 143057059 | 143057816 | -1 | processed_pseudoge | ENSG0000C |
| 143065916 | 143066603 | -1 | processed_pseudoge | ENSG0000C |
| 143085735 | 143088278 | 1 | protein_coding | ENSG0000C |
| 143163211 | 143166008 | -1 | protein_coding | ENSG0000C |
| 1501879 | 1707457 | 1 | protein_coding | ENSG0000C |
| 1565035 | 1621861 | -1 | lncRNA | ENSG0000C |
| 141340549 | 141344621 | 1 | lncRNA | ENSG0000C |
| 141353403 | 141355365 | -1 | lncRNA | ENSG0000C |
| 141356500 | 141367267 | -1 | protein_coding | ENSG0000C |
| 141391991 | 141438356 | 1 | protein_coding | ENSG0000C |
| 141439731 | 141455651 | -1 | polymorphic_pseudc | ENSG0000C |
| 141443230 | 141443856 | 1 | lncRNA | ENSG0000C |
| 141453377 | 141453451 | 1 | snoRNA | ENSG0000C |
| 141464977 | 141465933 | 1 | processed_pseudoge | ENSG0000C |
| 1160740 | 1186922 | 1 | lncRNA | ENSG0000C |
| 1247928 | 1260201 | 1 | lncRNA | ENSG0000C |
| 1604655 | 1622707 | -1 | lncRNA | ENSG0000C |
| 1627028 | 1708042 | 1 | protein_coding | ENSG0000C |

| | | | | |
|---|---|---|---|---|
| 1729770 | 1731018 | 1 | processed_pseudoge | ENSG00000 |
| 1755340 | 1786132 | 1 | protein_coding | ENSG00000 |
| 1757770 | 1760009 | -1 | lncRNA | ENSG00000 |
| 1761552 | 1764146 | -1 | lncRNA | ENSG00000 |
| 129728744 | 129730445 | 1 | processed_pseudoge | ENSG00000 |
| 129757225 | 129795674 | -1 | protein_coding | ENSG00000 |
| 1823976 | 1958641 | 1 | protein_coding | ENSG00000 |
| 1847451 | 1847916 | 1 | lncRNA | ENSG00000 |
| 1934747 | 1938216 | -1 | lncRNA | ENSG00000 |
| 1971397 | 1976478 | 1 | lncRNA | ENSG00000 |
| 1972570 | 1974637 | -1 | lncRNA | ENSG00000 |
| 1973878 | 2006936 | 1 | protein_coding | ENSG00000 |
| 1974818 | 1975314 | 1 | lncRNA | ENSG00000 |
| 2033484 | 2039347 | 1 | lncRNA | ENSG00000 |
| 2044989 | 2165303 | 1 | protein_coding | ENSG00000 |
| 2199057 | 2201817 | 1 | lncRNA | ENSG00000 |
| 2478186 | 2496887 | -1 | lncRNA | ENSG00000 |
| 2503318 | 2506502 | 1 | lncRNA | ENSG00000 |
| 2544294 | 2607662 | -1 | lncRNA | ENSG00000 |
| 2649201 | 2651108 | 1 | lncRNA | ENSG00000 |
| 2650769 | 2713141 | -1 | lncRNA | ENSG00000 |
| 2712008 | 2793124 | 1 | lncRNA | ENSG00000 |
| 39101863 | 39105144 | 1 | protein_coding | ENSG00000 |
| 39106990 | 39284943 | 1 | protein_coding | ENSG00000 |
| 39157538 | 39158701 | -1 | processed_pseudoge | ENSG00000 |
| 39309931 | 39310465 | -1 | processed_pseudoge | ENSG00000 |
| 39314619 | 39346102 | 1 | transcribed_unitary_ | ENSG00000 |
| 39429440 | 39575001 | 1 | protein_coding | ENSG00000 |
| 39588691 | 39683242 | -1 | protein_coding | ENSG00000 |
| 39705172 | 39705484 | 1 | processed_pseudoge | ENSG00000 |
| 39712389 | 39712765 | -1 | processed_pseudoge | ENSG00000 |
| 7491857 | 7493132 | -1 | processed_pseudoge | ENSG00000 |
| 69418516 | 69419862 | -1 | unprocessed_pseudo | ENSG00000 |
| 69435727 | 69484339 | -1 | protein_coding | ENSG00000 |
| 69480662 | 69482549 | 1 | lncRNA | ENSG00000 |
| 70043581 | 70176862 | 1 | protein_coding | ENSG00000 |
| 70087297 | 70113866 | -1 | lncRNA | ENSG00000 |
| 70153120 | 70181968 | -1 | lncRNA | ENSG00000 |
| 70194651 | 70194830 | -1 | snRNA | ENSG00000 |
| 88180669 | 88181409 | 1 | processed_pseudoge | ENSG00000 |
| 88187735 | 88190202 | -1 | processed_pseudoge | ENSG00000 |
| 100975668 | 100976682 | 1 | processed_pseudoge | ENSG00000 |
| 101028709 | 101325135 | 1 | protein_coding | ENSG00000 |
| 101125436 | 101125921 | -1 | processed_pseudoge | ENSG00000 |
| 101322477 | 101322533 | 1 | snoRNA | ENSG00000 |
| 101334084 | 101339373 | -1 | unitary_pseudogene | ENSG00000 |
| 101356497 | 101356875 | -1 | processed_pseudoge | ENSG00000 |
| 101368525 | 101370831 | 1 | processed_pseudoge | ENSG00000 |
| 101371093 | 101381063 | -1 | protein_coding | ENSG00000 |
| 101384932 | 101386191 | -1 | processed_pseudoge | ENSG00000 |

| | | | | |
|---|---|---|---|---|
| 8314246 | 10612723 | -1 | protein_coding | ENSG00000( |
| 9095713 | 9096060 | 1 | processed_pseudoge | ENSG00000( |
| 320990 | 321711 | 1 | lncRNA | ENSG00000( |
| 336496 | 389645 | -1 | protein_coding | ENSG00000( |
| 389670 | 408269 | 1 | lncRNA | ENSG00000( |
| 390790 | 391079 | -1 | misc_RNA | ENSG00000( |
| 421847 | 424670 | 1 | lncRNA | ENSG00000( |
| 526465 | 526792 | -1 | processed_pseudoge | ENSG00000( |
| 547926 | 548042 | -1 | processed_pseudoge | ENSG00000( |
| 80027033 | 80044046 | 1 | protein_coding | ENSG00000( |
| 333240 | 385935 | -1 | protein_coding | ENSG00000( |
| 385960 | 404559 | 1 | lncRNA | ENSG00000( |
| 418137 | 420960 | 1 | lncRNA | ENSG00000( |
| 522755 | 523082 | -1 | processed_pseudoge | ENSG00000( |
| 544216 | 544332 | -1 | processed_pseudoge | ENSG00000( |
| 56140 | 58376 | -1 | protein_coding | ENSG00000( |
| 53590 | 115018 | -1 | protein_coding | ENSG00000( |
| 53594 | 115055 | -1 | protein_coding | ENSG00000( |
| 37434 | 37534 | -1 | misc_RNA | ENSG00000( |
| 42939 | 49164 | -1 | protein_coding | ENSG00000( |
| 173872 | 179372 | 1 | protein_coding | ENSG00000( |
| 93050 | 93150 | -1 | misc_RNA | ENSG00000( |
| 99351 | 104855 | -1 | protein_coding | ENSG00000( |
| 108007 | 139659 | -1 | protein_coding | ENSG00000( |
| 127840 | 127946 | -1 | snRNA | ENSG00000( |
| 93197 | 93309 | -1 | snRNA | ENSG00000( |
| 51867 | 54893 | -1 | protein_coding | ENSG00000( |
| 54224 | 83311 | -1 | protein_coding | ENSG00000( |
| 105287 | 105376 | 1 | misc_RNA | ENSG00000( |
| 112025 | 112177 | 1 | rRNA | ENSG00000( |
| 149259 | 149348 | 1 | misc_RNA | ENSG00000( |
| 155997 | 156149 | 1 | rRNA | ENSG00000( |
| 183252 | 183364 | 1 | snRNA | ENSG00000( |
| 217250 | 217401 | -1 | rRNA | ENSG00000( |
| 380608 | 380726 | 1 | rRNA | ENSG00000( |
| 4612 | 29626 | -1 | protein_coding | ENSG00000( |
| 21861 | 22024 | -1 | snRNA | ENSG00000( |
| 30437 | 30580 | -1 | snRNA | ENSG00000( |
| 31698 | 32528 | -1 | protein_coding | ENSG00000( |
| 35407 | 35916 | 1 | protein_coding | ENSG00000( |
| 2585 | 11802 | 1 | protein_coding | ENSG00000( |
| 51722 | 51792 | 1 | misc_RNA | ENSG00000( |
| 52666 | 52734 | 1 | misc_RNA | ENSG00000( |
| 52895 | 53010 | 1 | misc_RNA | ENSG00000( |
| 26241 | 26534 | 1 | protein_coding | ENSG00000( |
| 41444 | 41876 | 1 | protein_coding | ENSG00000( |
| 20741 | 22392 | -1 | pseudogene | ENSG00000( |
| 34890 | 34996 | 1 | snRNA | ENSG00000( |
| 100123 | 101141 | -1 | protein_coding | ENSG00000( |
| 136160 | 167571 | -1 | protein_coding | ENSG00000( |

| | | | | |
|---|---|---|---|---|
| 156089 | 156195 | -1 | snRNA | ENSG0000( |
| 372322 | 373405 | 1 | protein_coding | ENSG0000( |
| 386278 | 387620 | 1 | protein_coding | ENSG0000( |
| 417283 | 418206 | 1 | pseudogene | ENSG0000( |
| 17234 | 19833 | 1 | protein_coding | ENSG0000( |
| 137798 | 139135 | 1 | processed_pseudoge | ENSG0000( |
| 933862 | 936467 | -1 | protein_coding | ENSG0000( |
| 1109945 | 1110235 | 1 | processed_pseudoge | ENSG0000( |
| 1139577 | 1147868 | -1 | protein_coding | ENSG0000( |
| 1157687 | 1240349 | 1 | protein_coding | ENSG0000( |
| 1167457 | 1257196 | 1 | protein_coding | ENSG0000( |
| 1270984 | 1271271 | 1 | protein_coding | ENSG0000( |
| 10598 | 13001 | -1 | protein_coding | ENSG0000( |
| 26533 | 27138 | -1 | pseudogene | ENSG0000( |
| 122136 | 122225 | 1 | misc_RNA | ENSG0000( |
| 128877 | 129029 | 1 | rRNA | ENSG0000( |
| 167218 | 167307 | 1 | misc_RNA | ENSG0000( |
| 173956 | 174108 | 1 | rRNA | ENSG0000( |
| 59711 | 60316 | 1 | pseudogene | ENSG0000( |
| 72411 | 74814 | 1 | protein_coding | ENSG0000( |
| 131494 | 137392 | 1 | protein_coding | ENSG0000( |
| 138082 | 161852 | -1 | protein_coding | ENSG0000( |
| 51009 | 51114 | -1 | snRNA | ENSG0000( |
| 148668 | 148843 | 1 | snRNA | ENSG0000( |
| 5001 | 22544 | 1 | LRG_gene | LRG_1 |
| 5001 | 12341 | 1 | LRG_gene | LRG_10 |
| 5001 | 23546 | 1 | LRG_gene | LRG_100 |
| 4959 | 70493 | 1 | LRG_gene | LRG_1008 |
| 5001 | 11654 | 1 | LRG_gene | LRG_101 |
| 5001 | 33236 | 1 | LRG_gene | LRG_1010 |
| 5001 | 482060 | 1 | LRG_gene | LRG_1012 |
| 6444 | 65748 | 1 | LRG_gene | LRG_1018 |
| 5001 | 14671 | 1 | LRG_gene | LRG_102 |
| 5001 | 33709 | 1 | LRG_gene | LRG_1020 |
| 5001 | 20561 | 1 | LRG_gene | LRG_1021 |
| 5001 | 113603 | 1 | LRG_gene | LRG_1022 |
| 5001 | 70083 | 1 | LRG_gene | LRG_1026 |
| 5001 | 28029 | 1 | LRG_gene | LRG_1028 |
| 5001 | 15402 | 1 | LRG_gene | LRG_103 |
| 5001 | 12899 | 1 | LRG_gene | LRG_104 |
| 5001 | 22300 | 1 | LRG_gene | LRG_105 |
| 4983 | 31861 | 1 | LRG_gene | LRG_106 |
| 6019 | 13945 | 1 | LRG_gene | LRG_107 |
| 5005 | 13673 | 1 | LRG_gene | LRG_1076 |
| 5001 | 19166 | 1 | LRG_gene | LRG_1077 |
| 5097 | 8418 | 1 | LRG_gene | LRG_1079 |
| 5001 | 75640 | 1 | LRG_gene | LRG_108 |
| 5001 | 56194 | 1 | LRG_gene | LRG_109 |
| 5001 | 31740 | 1 | LRG_gene | LRG_11 |
| 5001 | 78391 | 1 | LRG_gene | LRG_110 |

| | | | | |
|---|---|---|---|---|
| 5001 | 50215 | 1 | LRG_gene | LRG_111 |
| 5001 | 80171 | 1 | LRG_gene | LRG_112 |
| 5001 | 9450 | 1 | LRG_gene | LRG_1120 |
| 4739 | 56354 | 1 | LRG_gene | LRG_1122 |
| 5040 | 28281 | 1 | LRG_gene | LRG_1124 |
| 5055 | 10732 | 1 | LRG_gene | LRG_1125 |
| 5008 | 463642 | 1 | LRG_gene | LRG_1127 |
| 5010 | 11430 | 1 | LRG_gene | LRG_1129 |
| 5001 | 46423 | 1 | LRG_gene | LRG_113 |
| 4986 | 138610 | 1 | LRG_gene | LRG_1130 |
| 5002 | 81011 | 1 | LRG_gene | LRG_1131 |
| 5309 | 38802 | 1 | LRG_gene | LRG_1132 |
| 5047 | 19028 | 1 | LRG_gene | LRG_1133 |
| 5459 | 58934 | 1 | LRG_gene | LRG_1134 |
| 4738 | 1312200 | 1 | LRG_gene | LRG_1135 |
| 4997 | 17142 | 1 | LRG_gene | LRG_1136 |
| 5001 | 74883 | 1 | LRG_gene | LRG_1137 |
| 5031 | 13646 | 1 | LRG_gene | LRG_1138 |
| 4826 | 19519 | 1 | LRG_gene | LRG_114 |
| 5028 | 10057 | 1 | LRG_gene | LRG_1140 |
| 5782 | 221330 | 1 | LRG_gene | LRG_1141 |
| 5088 | 8795 | 1 | LRG_gene | LRG_1143 |
| 5017 | 7796 | 1 | LRG_gene | LRG_1146 |
| 4998 | 10658 | 1 | LRG_gene | LRG_1148 |
| 5001 | 16884 | 1 | LRG_gene | LRG_115 |
| 5716 | 393338 | 1 | LRG_gene | LRG_1150 |
| 5022 | 41086 | 1 | LRG_gene | LRG_1151 |
| 5588 | 9445 | 1 | LRG_gene | LRG_1152 |
| 5054 | 286392 | 1 | LRG_gene | LRG_1153 |
| 5195 | 21645 | 1 | LRG_gene | LRG_1154 |
| 5055 | 14939 | 1 | LRG_gene | LRG_1155 |
| 5147 | 9687 | 1 | LRG_gene | LRG_1156 |
| 5008 | 6401 | 1 | LRG_gene | LRG_1158 |
| 5001 | 24887 | 1 | LRG_gene | LRG_116 |
| 5036 | 27798 | 1 | LRG_gene | LRG_1161 |
| 5001 | 110236 | 1 | LRG_gene | LRG_1162 |
| 5088 | 27010 | 1 | LRG_gene | LRG_1163 |
| 5092 | 18676 | 1 | LRG_gene | LRG_1164 |
| 5407 | 23093 | 1 | LRG_gene | LRG_1165 |
| 5029 | 52789 | 1 | LRG_gene | LRG_1166 |
| 5222 | 32596 | 1 | LRG_gene | LRG_1168 |
| 5031 | 15310 | 1 | LRG_gene | LRG_1169 |
| 5001 | 20944 | 1 | LRG_gene | LRG_117 |
| 5484 | 83037 | 1 | LRG_gene | LRG_1170 |
| 9839 | 33630 | 1 | LRG_gene | LRG_1171 |
| 5104 | 39182 | 1 | LRG_gene | LRG_1173 |
| 10605 | 47822 | 1 | LRG_gene | LRG_1178 |
| 5541 | 219092 | 1 | LRG_gene | LRG_1179 |
| 5001 | 24490 | 1 | LRG_gene | LRG_118 |
| 5001 | 32649 | 1 | LRG_gene | LRG_1186 |

| | | | | |
|---|---|---|---|---|
| 5056 | 25142 | 1 | LRG_gene | LRG_1188 |
| 5001 | 17191 | 1 | LRG_gene | LRG_119 |
| 4743 | 10770 | 1 | LRG_gene | LRG_1190 |
| 5048 | 41363 | 1 | LRG_gene | LRG_1191 |
| 5501 | 56934 | 1 | LRG_gene | LRG_1195 |
| 5068 | 44548 | 1 | LRG_gene | LRG_1196 |
| 5073 | 115616 | 1 | LRG_gene | LRG_1199 |
| 5001 | 15508 | 1 | LRG_gene | LRG_12 |
| 5001 | 38005 | 1 | LRG_gene | LRG_120 |
| 78371 | 159302 | 1 | LRG_gene | LRG_1201 |
| 4959 | 72303 | 1 | LRG_gene | LRG_1203 |
| 5508 | 162397 | 1 | LRG_gene | LRG_1205 |
| 977 | 33665 | 1 | LRG_gene | LRG_1206 |
| 17903 | 68481 | 1 | LRG_gene | LRG_1207 |
| 5043 | 74997 | 1 | LRG_gene | LRG_1208 |
| 4990 | 271474 | 1 | LRG_gene | LRG_1209 |
| 5001 | 35045 | 1 | LRG_gene | LRG_121 |
| 5273 | 16129 | 1 | LRG_gene | LRG_1210 |
| 4998 | 6730 | 1 | LRG_gene | LRG_1211 |
| 28375 | 54504 | 1 | LRG_gene | LRG_1212 |
| 4992 | 287009 | 1 | LRG_gene | LRG_1213 |
| 5020 | 34425 | 1 | LRG_gene | LRG_1214 |
| 5041 | 12401 | 1 | LRG_gene | LRG_1215 |
| 4971 | 20855 | 1 | LRG_gene | LRG_1216 |
| 7420 | 48065 | 1 | LRG_gene | LRG_1217 |
| 6342 | 9327 | 1 | LRG_gene | LRG_1218 |
| 5708 | 14468 | 1 | LRG_gene | LRG_1219 |
| 5001 | 22493 | 1 | LRG_gene | LRG_122 |
| 4986 | 11180 | 1 | LRG_gene | LRG_1220 |
| 5080 | 88092 | 1 | LRG_gene | LRG_1221 |
| 5029 | 58930 | 1 | LRG_gene | LRG_1222 |
| 4989 | 9252 | 1 | LRG_gene | LRG_1223 |
| 5039 | 35620 | 1 | LRG_gene | LRG_1224 |
| 5029 | 9354 | 1 | LRG_gene | LRG_1226 |
| 5029 | 451136 | 1 | LRG_gene | LRG_1227 |
| 5024 | 8763 | 1 | LRG_gene | LRG_1228 |
| 5398 | 38267 | 1 | LRG_gene | LRG_1229 |
| 5001 | 18020 | 1 | LRG_gene | LRG_123 |
| 5001 | 9893 | 1 | LRG_gene | LRG_1230 |
| 17360 | 18403 | 1 | LRG_gene | LRG_1231 |
| 6499 | 143899 | 1 | LRG_gene | LRG_1233 |
| 5011 | 124731 | 1 | LRG_gene | LRG_1234 |
| 4846 | 56456 | 1 | LRG_gene | LRG_1235 |
| 5556 | 8304 | 1 | LRG_gene | LRG_1237 |
| 5788 | 13768 | 1 | LRG_gene | LRG_1238 |
| 5739 | 22958 | 1 | LRG_gene | LRG_1239 |
| 5001 | 18384 | 1 | LRG_gene | LRG_124 |
| 5001 | 66159 | 1 | LRG_gene | LRG_1241 |
| 5001 | 13345 | 1 | LRG_gene | LRG_1242 |
| 5313 | 76985 | 1 | LRG_gene | LRG_1243 |

| | | | | |
|---|---|---|---|---|
| 5007 | 20221 | 1 | LRG_gene | LRG_1244 |
| 5071 | 15530 | 1 | LRG_gene | LRG_1246 |
| 4993 | 23767 | 1 | LRG_gene | LRG_1247 |
| 5001 | 10363 | 1 | LRG_gene | LRG_1248 |
| 5001 | 59595 | 1 | LRG_gene | LRG_1249 |
| 5001 | 12633 | 1 | LRG_gene | LRG_125 |
| 5005 | 12913 | 1 | LRG_gene | LRG_1250 |
| 5133 | 108967 | 1 | LRG_gene | LRG_1251 |
| 4989 | 29098 | 1 | LRG_gene | LRG_1253 |
| 4978 | 10305 | 1 | LRG_gene | LRG_1254 |
| 5028 | 8969 | 1 | LRG_gene | LRG_1255 |
| 5028 | 54727 | 1 | LRG_gene | LRG_1256 |
| 5001 | 31293 | 1 | LRG_gene | LRG_126 |
| 5038 | 75964 | 1 | LRG_gene | LRG_1260 |
| 5001 | 10987 | 1 | LRG_gene | LRG_1262 |
| 5001 | 15959 | 1 | LRG_gene | LRG_1263 |
| 5001 | 20081 | 1 | LRG_gene | LRG_1264 |
| 5029 | 15831 | 1 | LRG_gene | LRG_1265 |
| 4984 | 32100 | 1 | LRG_gene | LRG_1267 |
| 5001 | 44495 | 1 | LRG_gene | LRG_127 |
| 5001 | 12763 | 1 | LRG_gene | LRG_1274 |
| 5001 | 41778 | 1 | LRG_gene | LRG_128 |
| 5217 | 44176 | 1 | LRG_gene | LRG_1285 |
| 5012 | 199571 | 1 | LRG_gene | LRG_1286 |
| 5025 | 290926 | 1 | LRG_gene | LRG_1287 |
| 5001 | 11093 | 1 | LRG_gene | LRG_129 |
| 5001 | 10618 | 1 | LRG_gene | LRG_13 |
| 20001 | 158719 | 1 | LRG_gene | LRG_130 |
| 5061 | 55055 | 1 | LRG_gene | LRG_1306 |
| 5033 | 12258 | 1 | LRG_gene | LRG_1308 |
| 5001 | 21524 | 1 | LRG_gene | LRG_1311 |
| 5636 | 34028 | 1 | LRG_gene | LRG_1312 |
| 5050 | 8445 | 1 | LRG_gene | LRG_1313 |
| 5025 | 27108 | 1 | LRG_gene | LRG_1314 |
| 5624 | 18781 | 1 | LRG_gene | LRG_1315 |
| 5031 | 120974 | 1 | LRG_gene | LRG_1316 |
| 5858 | 46291 | 1 | LRG_gene | LRG_1317 |
| 5001 | 49127 | 1 | LRG_gene | LRG_1318 |
| 5486 | 8299 | 1 | LRG_gene | LRG_1319 |
| 5001 | 8454 | 1 | LRG_gene | LRG_132 |
| 5001 | 13824 | 1 | LRG_gene | LRG_1320 |
| 5071 | 30802 | 1 | LRG_gene | LRG_1322 |
| 30096 | 164995 | 1 | LRG_gene | LRG_1323 |
| 5214 | 756063 | 1 | LRG_gene | LRG_1324 |
| 4932 | 48255 | 1 | LRG_gene | LRG_1325 |
| 10201 | 41709 | 1 | LRG_gene | LRG_1327 |
| 5257 | 9219 | 1 | LRG_gene | LRG_1328 |
| 5147 | 23657 | 1 | LRG_gene | LRG_1329 |
| 5001 | 33907 | 1 | LRG_gene | LRG_133 |
| 5039 | 44340 | 1 | LRG_gene | LRG_1330 |

| | | | | |
|---|---|---|---|---|
| 5873 | 144561 | 1 | LRG_gene | LRG_1332 |
| 5009 | 77911 | 1 | LRG_gene | LRG_1333 |
| 5063 | 53074 | 1 | LRG_gene | LRG_1334 |
| 5315 | 59971 | 1 | LRG_gene | LRG_1336 |
| 5001 | 55498 | 1 | LRG_gene | LRG_1337 |
| 5154 | 154190 | 1 | LRG_gene | LRG_1338 |
| 5001 | 30255 | 1 | LRG_gene | LRG_134 |
| 5001 | 87973 | 1 | LRG_gene | LRG_1341 |
| 6596 | 13415 | 1 | LRG_gene | LRG_1347 |
| 5001 | 151268 | 1 | LRG_gene | LRG_135 |
| 5001 | 11141 | 1 | LRG_gene | LRG_136 |
| 5003 | 76912 | 1 | LRG_gene | LRG_1364 |
| 5001 | 32103 | 1 | LRG_gene | LRG_1365 |
| 5033 | 13047 | 1 | LRG_gene | LRG_1366 |
| 5032 | 28682 | 1 | LRG_gene | LRG_1367 |
| 27959 | 71812 | 1 | LRG_gene | LRG_1368 |
| 25778 | 41732 | 1 | LRG_gene | LRG_1369 |
| 5001 | 25624 | 1 | LRG_gene | LRG_137 |
| 5054 | 18595 | 1 | LRG_gene | LRG_1372 |
| 5039 | 45709 | 1 | LRG_gene | LRG_1375 |
| 5002 | 25625 | 1 | LRG_gene | LRG_138 |
| 4993 | 114070 | 1 | LRG_gene | LRG_1383 |
| 5212 | 206211 | 1 | LRG_gene | LRG_1384 |
| 5001 | 10385 | 1 | LRG_gene | LRG_1386 |
| 5057 | 193932 | 1 | LRG_gene | LRG_1387 |
| 6276 | 227907 | 1 | LRG_gene | LRG_1389 |
| 5001 | 68452 | 1 | LRG_gene | LRG_139 |
| 6252 | 161193 | 1 | LRG_gene | LRG_1390 |
| 5014 | 60042 | 1 | LRG_gene | LRG_1391 |
| 4986 | 22355 | 1 | LRG_gene | LRG_1392 |
| 4762 | 29674 | 1 | LRG_gene | LRG_1393 |
| 5837 | 356356 | 1 | LRG_gene | LRG_1396 |
| 106198 | 239518 | 1 | LRG_gene | LRG_1398 |
| 5030 | 70524 | 1 | LRG_gene | LRG_1399 |
| 5001 | 19944 | 1 | LRG_gene | LRG_140 |
| 9863 | 35899 | 1 | LRG_gene | LRG_1400 |
| 5001 | 17214 | 1 | LRG_gene | LRG_141 |
| 5001 | 25103 | 1 | LRG_gene | LRG_142 |
| 4933 | 227610 | 1 | LRG_gene | LRG_143 |
| 4595 | 21866 | 1 | LRG_gene | LRG_144 |
| 5001 | 12439 | 1 | LRG_gene | LRG_145 |
| 5001 | 11125 | 1 | LRG_gene | LRG_146 |
| 5001 | 17572 | 1 | LRG_gene | LRG_147 |
| 5555 | 21182 | 1 | LRG_gene | LRG_148 |
| 5001 | 17459 | 1 | LRG_gene | LRG_149 |
| 38476 | 106733 | 1 | LRG_gene | LRG_15 |
| 4923 | 9150 | 1 | LRG_gene | LRG_150 |
| 5001 | 20093 | 1 | LRG_gene | LRG_151 |
| 5001 | 35849 | 1 | LRG_gene | LRG_152 |
| 5001 | 11321 | 1 | LRG_gene | LRG_154 |

| | | | | |
|---|---|---|---|---|
| 4982 | 48460 | 1 | LRG_gene | LRG_155 |
| 5001 | 28572 | 1 | LRG_gene | LRG_156 |
| 5001 | 9545 | 1 | LRG_gene | LRG_157 |
| 5001 | 56336 | 1 | LRG_gene | LRG_158 |
| 5001 | 22030 | 1 | LRG_gene | LRG_159 |
| 5001 | 37214 | 1 | LRG_gene | LRG_16 |
| 4985 | 21088 | 1 | LRG_gene | LRG_160 |
| 5001 | 40868 | 1 | LRG_gene | LRG_161 |
| 5001 | 192076 | 1 | LRG_gene | LRG_162 |
| 5001 | 5268 | 1 | LRG_gene | LRG_163 |
| 5001 | 242508 | 1 | LRG_gene | LRG_164 |
| 5001 | 15769 | 1 | LRG_gene | LRG_165 |
| 5001 | 9032 | 1 | LRG_gene | LRG_168 |
| 5001 | 19461 | 1 | LRG_gene | LRG_169 |
| 5001 | 15681 | 1 | LRG_gene | LRG_17 |
| 4950 | 297379 | 1 | LRG_gene | LRG_170 |
| 5001 | 10715 | 1 | LRG_gene | LRG_171 |
| 5001 | 47207 | 1 | LRG_gene | LRG_172 |
| 5001 | 96111 | 1 | LRG_gene | LRG_173 |
| 5001 | 99954 | 1 | LRG_gene | LRG_174 |
| 5001 | 24274 | 1 | LRG_gene | LRG_175 |
| 5001 | 193127 | 1 | LRG_gene | LRG_176 |
| 5001 | 40938 | 1 | LRG_gene | LRG_177 |
| 5001 | 14726 | 1 | LRG_gene | LRG_178 |
| 5001 | 18493 | 1 | LRG_gene | LRG_179 |
| 5001 | 17340 | 1 | LRG_gene | LRG_18 |
| 5001 | 22212 | 1 | LRG_gene | LRG_180 |
| 5001 | 10615 | 1 | LRG_gene | LRG_182 |
| 5001 | 16567 | 1 | LRG_gene | LRG_183 |
| 5001 | 6786 | 1 | LRG_gene | LRG_184 |
| 5001 | 39986 | 1 | LRG_gene | LRG_185 |
| 5001 | 46136 | 1 | LRG_gene | LRG_186 |
| 5001 | 11555 | 1 | LRG_gene | LRG_187 |
| 5001 | 21124 | 1 | LRG_gene | LRG_188 |
| 5001 | 79203 | 1 | LRG_gene | LRG_189 |
| 4970 | 58782 | 1 | LRG_gene | LRG_19 |
| 5001 | 19600 | 1 | LRG_gene | LRG_190 |
| 5001 | 82383 | 1 | LRG_gene | LRG_191 |
| 5001 | 82230 | 1 | LRG_gene | LRG_192 |
| 4979 | 18339 | 1 | LRG_gene | LRG_193 |
| 5001 | 65957 | 1 | LRG_gene | LRG_194 |
| 5001 | 10472 | 1 | LRG_gene | LRG_195 |
| 5001 | 255395 | 1 | LRG_gene | LRG_196 |
| 6877 | 37936 | 1 | LRG_gene | LRG_197 |
| 5001 | 40990 | 1 | LRG_gene | LRG_198 |
| 133054 | 2225382 | 1 | LRG_gene | LRG_199 |
| 5001 | 41672 | 1 | LRG_gene | LRG_2 |
| 4978 | 103085 | 1 | LRG_gene | LRG_20 |
| 5001 | 12030 | 1 | LRG_gene | LRG_200 |
| 5001 | 24996 | 1 | LRG_gene | LRG_201 |

| | | | | |
|---|---|---|---|---|
| 5001 | 254149 | 1 | LRG_gene | LRG_202 |
| 5001 | 14928 | 1 | LRG_gene | LRG_203 |
| 5001 | 22625 | 1 | LRG_gene | LRG_204 |
| 461414 | 902446 | 1 | LRG_gene | LRG_205 |
| 5001 | 149245 | 1 | LRG_gene | LRG_207 |
| 5001 | 1153420 | 1 | LRG_gene | LRG_208 |
| 78346 | 117787 | 1 | LRG_gene | LRG_209 |
| 9994 | 89872 | 1 | LRG_gene | LRG_21 |
| 5001 | 6209 | 1 | LRG_gene | LRG_210 |
| 5001 | 18999 | 1 | LRG_gene | LRG_211 |
| 5001 | 41124 | 1 | LRG_gene | LRG_212 |
| 5001 | 9442 | 1 | LRG_gene | LRG_213 |
| 4951 | 287701 | 1 | LRG_gene | LRG_214 |
| 28986 | 46866 | 1 | LRG_gene | LRG_215 |
| 5001 | 62497 | 1 | LRG_gene | LRG_216 |
| 5001 | 42769 | 1 | LRG_gene | LRG_217 |
| 4944 | 85105 | 1 | LRG_gene | LRG_218 |
| 4936 | 28807 | 1 | LRG_gene | LRG_219 |
| 5001 | 8058 | 1 | LRG_gene | LRG_22 |
| 5001 | 16229 | 1 | LRG_gene | LRG_220 |
| 5001 | 98545 | 1 | LRG_gene | LRG_221 |
| 5001 | 15630 | 1 | LRG_gene | LRG_226 |
| 5001 | 37138 | 1 | LRG_gene | LRG_227 |
| 5001 | 9240 | 1 | LRG_gene | LRG_228 |
| 5001 | 139022 | 1 | LRG_gene | LRG_229 |
| 5001 | 13348 | 1 | LRG_gene | LRG_23 |
| 5001 | 155228 | 1 | LRG_gene | LRG_230 |
| 5001 | 166849 | 1 | LRG_gene | LRG_231 |
| 5001 | 139659 | 1 | LRG_gene | LRG_234 |
| 6868 | 24313 | 1 | LRG_gene | LRG_235 |
| 5001 | 79710 | 1 | LRG_gene | LRG_236 |
| 5001 | 35934 | 1 | LRG_gene | LRG_237 |
| 4975 | 118832 | 1 | LRG_gene | LRG_238 |
| 108535 | 112904 | 1 | LRG_gene | LRG_239 |
| 5001 | 9486 | 1 | LRG_gene | LRG_24 |
| 91444 | 251522 | 1 | LRG_gene | LRG_240 |
| 5001 | 139211 | 1 | LRG_gene | LRG_241 |
| 5001 | 70233 | 1 | LRG_gene | LRG_242 |
| 5001 | 44469 | 1 | LRG_gene | LRG_243 |
| 34254 | 50981 | 1 | LRG_gene | LRG_244 |
| 5001 | 15305 | 1 | LRG_gene | LRG_245 |
| 10001 | 13179 | 1 | LRG_gene | LRG_246 |
| 5001 | 6740 | 1 | LRG_gene | LRG_248 |
| 5001 | 20957 | 1 | LRG_gene | LRG_249 |
| 5001 | 15357 | 1 | LRG_gene | LRG_25 |
| 5001 | 41751 | 1 | LRG_gene | LRG_250 |
| 5001 | 71809 | 1 | LRG_gene | LRG_251 |
| 5001 | 175898 | 1 | LRG_gene | LRG_252 |
| 5001 | 44229 | 1 | LRG_gene | LRG_253 |
| 37098 | 62515 | 1 | LRG_gene | LRG_254 |

| | | | |
|---|---|---|---|
| 5001 | 38335 | 1 LRG_gene | LRG_255 |
| 5001 | 10238 | 1 LRG_gene | LRG_256 |
| 5001 | 96328 | 1 LRG_gene | LRG_257 |
| 4952 | 65085 | 1 LRG_gene | LRG_258 |
| 4749 | 10663 | 1 LRG_gene | LRG_259 |
| 5001 | 23184 | 1 LRG_gene | LRG_26 |
| 5001 | 57321 | 1 LRG_gene | LRG_260 |
| 5001 | 71101 | 1 LRG_gene | LRG_261 |
| 33310 | 58918 | 1 LRG_gene | LRG_262 |
| 5001 | 40548 | 1 LRG_gene | LRG_263 |
| 5001 | 27603 | 1 LRG_gene | LRG_264 |
| 5001 | 24601 | 1 LRG_gene | LRG_265 |
| 5001 | 93663 | 1 LRG_gene | LRG_266 |
| 5001 | 520541 | 1 LRG_gene | LRG_267 |
| 5001 | 98546 | 1 LRG_gene | LRG_268 |
| 5001 | 86025 | 1 LRG_gene | LRG_269 |
| 5001 | 47817 | 1 LRG_gene | LRG_27 |
| 6221 | 113069 | 1 LRG_gene | LRG_270 |
| 5001 | 17221 | 1 LRG_gene | LRG_271 |
| 5001 | 89264 | 1 LRG_gene | LRG_272 |
| 5001 | 47794 | 1 LRG_gene | LRG_273 |
| 4982 | 49450 | 1 LRG_gene | LRG_274 |
| 4930 | 30307 | 1 LRG_gene | LRG_275 |
| 5001 | 30302 | 1 LRG_gene | LRG_276 |
| 5001 | 12035 | 1 LRG_gene | LRG_278 |
| 4923 | 65705 | 1 LRG_gene | LRG_279 |
| 5001 | 102942 | 1 LRG_gene | LRG_28 |
| 5001 | 8266 | 1 LRG_gene | LRG_280 |
| 5001 | 65020 | 1 LRG_gene | LRG_281 |
| 5088 | 221929 | 1 LRG_gene | LRG_283 |
| 5001 | 16855 | 1 LRG_gene | LRG_284 |
| 5001 | 28668 | 1 LRG_gene | LRG_285 |
| 5093 | 36180 | 1 LRG_gene | LRG_286 |
| 5001 | 409120 | 1 LRG_gene | LRG_287 |
| 4613 | 37971 | 1 LRG_gene | LRG_288 |
| 5001 | 106611 | 1 LRG_gene | LRG_289 |
| 52874 | 124205 | 1 LRG_gene | LRG_29 |
| 5001 | 69626 | 1 LRG_gene | LRG_290 |
| 5001 | 12118 | 1 LRG_gene | LRG_291 |
| 92501 | 173689 | 1 LRG_gene | LRG_292 |
| 5001 | 89193 | 1 LRG_gene | LRG_293 |
| 5001 | 28439 | 1 LRG_gene | LRG_294 |
| 5001 | 18766 | 1 LRG_gene | LRG_295 |
| 5001 | 38060 | 1 LRG_gene | LRG_296 |
| 5029 | 89066 | 1 LRG_gene | LRG_297 |
| 5001 | 173550 | 1 LRG_gene | LRG_298 |
| 5001 | 195753 | 1 LRG_gene | LRG_299 |
| 5001 | 43374 | 1 LRG_gene | LRG_3 |
| 5001 | 78443 | 1 LRG_gene | LRG_30 |
| 5001 | 189374 | 1 LRG_gene | LRG_300 |

| | | | | |
|---|---|---|---|---|
| 5001 | 103250 | 1 | LRG_gene | LRG_301 |
| 5001 | 9312 | 1 | LRG_gene | LRG_303 |
| 5001 | 193307 | 1 | LRG_gene | LRG_304 |
| 5001 | 80502 | 1 | LRG_gene | LRG_306 |
| 4935 | 87721 | 1 | LRG_gene | LRG_307 |
| 5001 | 43196 | 1 | LRG_gene | LRG_308 |
| 5001 | 74149 | 1 | LRG_gene | LRG_309 |
| 5001 | 41806 | 1 | LRG_gene | LRG_31 |
| 5001 | 91190 | 1 | LRG_gene | LRG_310 |
| 5001 | 110337 | 1 | LRG_gene | LRG_311 |
| 5001 | 46730 | 1 | LRG_gene | LRG_314 |
| 5001 | 40449 | 1 | LRG_gene | LRG_316 |
| 5001 | 55376 | 1 | LRG_gene | LRG_317 |
| 67174 | 122003 | 1 | LRG_gene | LRG_318 |
| 21393 | 44029 | 1 | LRG_gene | LRG_319 |
| 5001 | 85279 | 1 | LRG_gene | LRG_32 |
| 5001 | 24149 | 1 | LRG_gene | LRG_321 |
| 5001 | 17036 | 1 | LRG_gene | LRG_322 |
| 5001 | 29977 | 1 | LRG_gene | LRG_325 |
| 5001 | 25722 | 1 | LRG_gene | LRG_326 |
| 5001 | 570658 | 1 | LRG_gene | LRG_327 |
| 5001 | 15510 | 1 | LRG_gene | LRG_328 |
| 4991 | 17956 | 1 | LRG_gene | LRG_329 |
| 4767 | 43529 | 1 | LRG_gene | LRG_33 |
| 5001 | 24539 | 1 | LRG_gene | LRG_330 |
| 5001 | 174799 | 1 | LRG_gene | LRG_331 |
| 5001 | 40936 | 1 | LRG_gene | LRG_332 |
| 5001 | 31652 | 1 | LRG_gene | LRG_333 |
| 87465 | 732164 | 1 | LRG_gene | LRG_334 |
| 5214 | 50794 | 1 | LRG_gene | LRG_335 |
| 83915 | 206423 | 1 | LRG_gene | LRG_336 |
| 5001 | 109668 | 1 | LRG_gene | LRG_337 |
| 22990 | 72325 | 1 | LRG_gene | LRG_338 |
| 4986 | 59254 | 1 | LRG_gene | LRG_34 |
| 5001 | 19021 | 1 | LRG_gene | LRG_340 |
| 5001 | 10922 | 1 | LRG_gene | LRG_341 |
| 5001 | 46881 | 1 | LRG_gene | LRG_343 |
| 5075 | 50758 | 1 | LRG_gene | LRG_344 |
| 5001 | 6446 | 1 | LRG_gene | LRG_345 |
| 5001 | 9498 | 1 | LRG_gene | LRG_346 |
| 5001 | 5451 | 1 | LRG_gene | LRG_347 |
| 5001 | 40596 | 1 | LRG_gene | LRG_348 |
| 5001 | 80938 | 1 | LRG_gene | LRG_349 |
| 5001 | 12409 | 1 | LRG_gene | LRG_35 |
| 5001 | 54813 | 1 | LRG_gene | LRG_350 |
| 5001 | 869321 | 1 | LRG_gene | LRG_351 |
| 5001 | 25251 | 1 | LRG_gene | LRG_352 |
| 5001 | 74205 | 1 | LRG_gene | LRG_353 |
| 5001 | 153136 | 1 | LRG_gene | LRG_354 |
| 4999 | 35734 | 1 | LRG_gene | LRG_355 |

| | | | | |
|---|---|---|---|---|
| 5001 | 12815 | 1 | LRG_gene | LRG_356 |
| 5001 | 11834 | 1 | LRG_gene | LRG_357 |
| 5001 | 20819 | 1 | LRG_gene | LRG_358 |
| 5001 | 42116 | 1 | LRG_gene | LRG_359 |
| 5000 | 92970 | 1 | LRG_gene | LRG_36 |
| 5001 | 104987 | 1 | LRG_gene | LRG_360 |
| 5001 | 21220 | 1 | LRG_gene | LRG_361 |
| 41208 | 102941 | 1 | LRG_gene | LRG_362 |
| 4952 | 148972 | 1 | LRG_gene | LRG_363 |
| 5001 | 83292 | 1 | LRG_gene | LRG_364 |
| 5001 | 136883 | 1 | LRG_gene | LRG_365 |
| 5001 | 24964 | 1 | LRG_gene | LRG_366 |
| 4900 | 111444 | 1 | LRG_gene | LRG_367 |
| 5001 | 23702 | 1 | LRG_gene | LRG_368 |
| 5001 | 185803 | 1 | LRG_gene | LRG_369 |
| 5001 | 8671 | 1 | LRG_gene | LRG_37 |
| 5001 | 115771 | 1 | LRG_gene | LRG_371 |
| 5001 | 55321 | 1 | LRG_gene | LRG_372 |
| 5001 | 57016 | 1 | LRG_gene | LRG_373 |
| 5001 | 128326 | 1 | LRG_gene | LRG_374 |
| 7279 | 22432 | 1 | LRG_gene | LRG_375 |
| 45198 | 224198 | 1 | LRG_gene | LRG_376 |
| 5001 | 144306 | 1 | LRG_gene | LRG_377 |
| 5001 | 7951 | 1 | LRG_gene | LRG_378 |
| 5001 | 14176 | 1 | LRG_gene | LRG_379 |
| 5001 | 16596 | 1 | LRG_gene | LRG_38 |
| 5001 | 13363 | 1 | LRG_gene | LRG_380 |
| 5001 | 200073 | 1 | LRG_gene | LRG_382 |
| 5001 | 127047 | 1 | LRG_gene | LRG_383 |
| 5001 | 27924 | 1 | LRG_gene | LRG_384 |
| 5001 | 72505 | 1 | LRG_gene | LRG_385 |
| 5001 | 26297 | 1 | LRG_gene | LRG_386 |
| 5001 | 12631 | 1 | LRG_gene | LRG_388 |
| 5001 | 31288 | 1 | LRG_gene | LRG_389 |
| 5001 | 14439 | 1 | LRG_gene | LRG_39 |
| 5001 | 17146 | 1 | LRG_gene | LRG_390 |
| 28380 | 309814 | 1 | LRG_gene | LRG_391 |
| 5001 | 20323 | 1 | LRG_gene | LRG_392 |
| 4978 | 14759 | 1 | LRG_gene | LRG_393 |
| 5001 | 80882 | 1 | LRG_gene | LRG_394 |
| 23687 | 29303 | 1 | LRG_gene | LRG_395 |
| 5001 | 57006 | 1 | LRG_gene | LRG_396 |
| 4823 | 55610 | 1 | LRG_gene | LRG_397 |
| 5001 | 111101 | 1 | LRG_gene | LRG_398 |
| 5001 | 28651 | 1 | LRG_gene | LRG_399 |
| 5001 | 38816 | 1 | LRG_gene | LRG_4 |
| 5001 | 16479 | 1 | LRG_gene | LRG_40 |
| 5001 | 41450 | 1 | LRG_gene | LRG_400 |
| 5001 | 37106 | 1 | LRG_gene | LRG_401 |
| 5001 | 796805 | 1 | LRG_gene | LRG_402 |

| | | | | |
|---|---|---|---|---|
| 5001 | 11037 | 1 | LRG_gene | LRG_403 |
| 5001 | 73803 | 1 | LRG_gene | LRG_404 |
| 5001 | 153373 | 1 | LRG_gene | LRG_405 |
| 5001 | 28919 | 1 | LRG_gene | LRG_406 |
| 5001 | 11954 | 1 | LRG_gene | LRG_408 |
| 5001 | 638426 | 1 | LRG_gene | LRG_409 |
| 5000 | 38469 | 1 | LRG_gene | LRG_41 |
| 8317 | 110901 | 1 | LRG_gene | LRG_410 |
| 5001 | 399215 | 1 | LRG_gene | LRG_411 |
| 4979 | 85579 | 1 | LRG_gene | LRG_413 |
| 11910 | 26877 | 1 | LRG_gene | LRG_414 |
| 11867 | 42382 | 1 | LRG_gene | LRG_415 |
| 5001 | 12258 | 1 | LRG_gene | LRG_416 |
| 5001 | 71320 | 1 | LRG_gene | LRG_417 |
| 5001 | 67205 | 1 | LRG_gene | LRG_419 |
| 5001 | 9250 | 1 | LRG_gene | LRG_42 |
| 5001 | 30421 | 1 | LRG_gene | LRG_421 |
| 137013 | 154148 | 1 | LRG_gene | LRG_422 |
| 5001 | 50077 | 1 | LRG_gene | LRG_423 |
| 64855 | 69547 | 1 | LRG_gene | LRG_424 |
| 5001 | 158302 | 1 | LRG_gene | LRG_426 |
| 5001 | 7852 | 1 | LRG_gene | LRG_429 |
| 5001 | 8607 | 1 | LRG_gene | LRG_43 |
| 5001 | 10966 | 1 | LRG_gene | LRG_432 |
| 5001 | 151695 | 1 | LRG_gene | LRG_433 |
| 5001 | 87989 | 1 | LRG_gene | LRG_434 |
| 5001 | 23741 | 1 | LRG_gene | LRG_435 |
| 5001 | 498614 | 1 | LRG_gene | LRG_437 |
| 5001 | 17715 | 1 | LRG_gene | LRG_439 |
| 21683 | 28792 | 1 | LRG_gene | LRG_44 |
| 13532 | 33543 | 1 | LRG_gene | LRG_440 |
| 5001 | 60381 | 1 | LRG_gene | LRG_442 |
| 5001 | 39807 | 1 | LRG_gene | LRG_443 |
| 5001 | 12138 | 1 | LRG_gene | LRG_444 |
| 5001 | 218324 | 1 | LRG_gene | LRG_445 |
| 5002 | 8137 | 1 | LRG_gene | LRG_446 |
| 5001 | 25937 | 1 | LRG_gene | LRG_447 |
| 5001 | 17316 | 1 | LRG_gene | LRG_448 |
| 5001 | 153110 | 1 | LRG_gene | LRG_449 |
| 5001 | 6960 | 1 | LRG_gene | LRG_45 |
| 5001 | 8118 | 1 | LRG_gene | LRG_450 |
| 5001 | 54378 | 1 | LRG_gene | LRG_451 |
| 5001 | 7187 | 1 | LRG_gene | LRG_452 |
| 5001 | 21219 | 1 | LRG_gene | LRG_454 |
| 5001 | 10989 | 1 | LRG_gene | LRG_455 |
| 5001 | 7591 | 1 | LRG_gene | LRG_456 |
| 4977 | 102295 | 1 | LRG_gene | LRG_457 |
| 5001 | 28181 | 1 | LRG_gene | LRG_458 |
| 5001 | 114630 | 1 | LRG_gene | LRG_459 |
| 5001 | 8960 | 1 | LRG_gene | LRG_46 |

| | | | | |
|---|---|---|---|---|
| 5001 | 13075 | 1 | LRG_gene | LRG_460 |
| 5001 | 24197 | 1 | LRG_gene | LRG_461 |
| 5001 | 41887 | 1 | LRG_gene | LRG_462 |
| 5001 | 37192 | 1 | LRG_gene | LRG_463 |
| 5018 | 35178 | 1 | LRG_gene | LRG_464 |
| 5001 | 87657 | 1 | LRG_gene | LRG_465 |
| 5001 | 76247 | 1 | LRG_gene | LRG_466 |
| 5001 | 29277 | 1 | LRG_gene | LRG_467 |
| 5001 | 35998 | 1 | LRG_gene | LRG_469 |
| 4868 | 100494 | 1 | LRG_gene | LRG_47 |
| 5001 | 49383 | 1 | LRG_gene | LRG_470 |
| 5001 | 27501 | 1 | LRG_gene | LRG_471 |
| 5001 | 38526 | 1 | LRG_gene | LRG_472 |
| 5001 | 95196 | 1 | LRG_gene | LRG_473 |
| 14385 | 48150 | 1 | LRG_gene | LRG_474 |
| 5001 | 28301 | 1 | LRG_gene | LRG_475 |
| 5001 | 39731 | 1 | LRG_gene | LRG_476 |
| 4966 | 6610 | 1 | LRG_gene | LRG_477 |
| 5001 | 6232 | 1 | LRG_gene | LRG_478 |
| 5001 | 22324 | 1 | LRG_gene | LRG_479 |
| 4807 | 66294 | 1 | LRG_gene | LRG_48 |
| 5001 | 63870 | 1 | LRG_gene | LRG_481 |
| 940414 | 1201911 | 1 | LRG_gene | LRG_482 |
| 5001 | 82045 | 1 | LRG_gene | LRG_483 |
| 5001 | 42610 | 1 | LRG_gene | LRG_484 |
| 5004 | 7714 | 1 | LRG_gene | LRG_485 |
| 5001 | 58286 | 1 | LRG_gene | LRG_486 |
| 3684 | 44407 | 1 | LRG_gene | LRG_487 |
| 5001 | 733793 | 1 | LRG_gene | LRG_488 |
| 5001 | 65128 | 1 | LRG_gene | LRG_489 |
| 5001 | 52786 | 1 | LRG_gene | LRG_49 |
| 5000 | 9162 | 1 | LRG_gene | LRG_490 |
| 5001 | 64886 | 1 | LRG_gene | LRG_491 |
| 5001 | 76195 | 1 | LRG_gene | LRG_492 |
| 5001 | 317457 | 1 | LRG_gene | LRG_493 |
| 5001 | 154882 | 1 | LRG_gene | LRG_494 |
| 5001 | 84107 | 1 | LRG_gene | LRG_495 |
| 5001 | 34656 | 1 | LRG_gene | LRG_496 |
| 5001 | 223656 | 1 | LRG_gene | LRG_497 |
| 5001 | 19744 | 1 | LRG_gene | LRG_498 |
| 5001 | 11179 | 1 | LRG_gene | LRG_499 |
| 5001 | 25750 | 1 | LRG_gene | LRG_5 |
| 5001 | 49182 | 1 | LRG_gene | LRG_50 |
| 5001 | 78169 | 1 | LRG_gene | LRG_500 |
| 5000 | 87137 | 1 | LRG_gene | LRG_501 |
| 5001 | 69958 | 1 | LRG_gene | LRG_502 |
| 5001 | 35404 | 1 | LRG_gene | LRG_503 |
| 4970 | 27198 | 1 | LRG_gene | LRG_504 |
| 4994 | 454894 | 1 | LRG_gene | LRG_505 |
| 5001 | 137858 | 1 | LRG_gene | LRG_507 |

| | | | | |
|---|---|---|---|---|
| 5579 | 12781 | 1 | LRG_gene | LRG_509 |
| 5001 | 8807 | 1 | LRG_gene | LRG_51 |
| 5001 | 21661 | 1 | LRG_gene | LRG_510 |
| 5001 | 100045 | 1 | LRG_gene | LRG_511 |
| 5754 | 172135 | 1 | LRG_gene | LRG_512 |
| 5001 | 9889 | 1 | LRG_gene | LRG_513 |
| 103757 | 124809 | 1 | LRG_gene | LRG_514 |
| 5001 | 78984 | 1 | LRG_gene | LRG_515 |
| 5001 | 25078 | 1 | LRG_gene | LRG_516 |
| 5001 | 183144 | 1 | LRG_gene | LRG_517 |
| 5001 | 58283 | 1 | LRG_gene | LRG_518 |
| 5001 | 21643 | 1 | LRG_gene | LRG_519 |
| 5001 | 12761 | 1 | LRG_gene | LRG_52 |
| 5001 | 52556 | 1 | LRG_gene | LRG_520 |
| 5001 | 134496 | 1 | LRG_gene | LRG_521 |
| 5001 | 28767 | 1 | LRG_gene | LRG_522 |
| 5001 | 195778 | 1 | LRG_gene | LRG_523 |
| 5001 | 145500 | 1 | LRG_gene | LRG_524 |
| 5001 | 52767 | 1 | LRG_gene | LRG_525 |
| 5018 | 47230 | 1 | LRG_gene | LRG_526 |
| 5001 | 8309 | 1 | LRG_gene | LRG_527 |
| 5001 | 20806 | 1 | LRG_gene | LRG_528 |
| 5001 | 13986 | 1 | LRG_gene | LRG_529 |
| 5001 | 38445 | 1 | LRG_gene | LRG_53 |
| 5001 | 32383 | 1 | LRG_gene | LRG_530 |
| 5001 | 81978 | 1 | LRG_gene | LRG_531 |
| 5001 | 35535 | 1 | LRG_gene | LRG_532 |
| 5001 | 7548 | 1 | LRG_gene | LRG_533 |
| 5001 | 380616 | 1 | LRG_gene | LRG_534 |
| 5001 | 118475 | 1 | LRG_gene | LRG_535 |
| 4965 | 197335 | 1 | LRG_gene | LRG_536 |
| 5001 | 10901 | 1 | LRG_gene | LRG_537 |
| 5001 | 12864 | 1 | LRG_gene | LRG_538 |
| 5001 | 8788 | 1 | LRG_gene | LRG_539 |
| 5001 | 52225 | 1 | LRG_gene | LRG_54 |
| 5001 | 310852 | 1 | LRG_gene | LRG_540 |
| 5001 | 19429 | 1 | LRG_gene | LRG_541 |
| 5001 | 20944 | 1 | LRG_gene | LRG_543 |
| 12662 | 50065 | 1 | LRG_gene | LRG_544 |
| 5001 | 8057 | 1 | LRG_gene | LRG_546 |
| 5001 | 31731 | 1 | LRG_gene | LRG_548 |
| 5001 | 181614 | 1 | LRG_gene | LRG_549 |
| 5001 | 19934 | 1 | LRG_gene | LRG_55 |
| 5001 | 33077 | 1 | LRG_gene | LRG_550 |
| 4974 | 25316 | 1 | LRG_gene | LRG_551 |
| 5001 | 79578 | 1 | LRG_gene | LRG_553 |
| 4998 | 19891 | 1 | LRG_gene | LRG_554 |
| 5001 | 37723 | 1 | LRG_gene | LRG_556 |
| 5001 | 12618 | 1 | LRG_gene | LRG_557 |
| 5001 | 14784 | 1 | LRG_gene | LRG_558 |

| | | | | |
|---|---|---|---|---|
| 5001 | 12738 | 1 | LRG_gene | LRG_559 |
| 5001 | 51972 | 1 | LRG_gene | LRG_56 |
| 5001 | 29560 | 1 | LRG_gene | LRG_560 |
| 5001 | 9060 | 1 | LRG_gene | LRG_561 |
| 4978 | 7649 | 1 | LRG_gene | LRG_564 |
| 5001 | 44986 | 1 | LRG_gene | LRG_566 |
| 4952 | 111742 | 1 | LRG_gene | LRG_567 |
| 5001 | 35022 | 1 | LRG_gene | LRG_568 |
| 4945 | 52914 | 1 | LRG_gene | LRG_569 |
| 5001 | 8956 | 1 | LRG_gene | LRG_57 |
| 5001 | 37959 | 1 | LRG_gene | LRG_570 |
| 5001 | 56862 | 1 | LRG_gene | LRG_571 |
| 5001 | 106054 | 1 | LRG_gene | LRG_572 |
| 5001 | 10981 | 1 | LRG_gene | LRG_574 |
| 5001 | 18575 | 1 | LRG_gene | LRG_577 |
| 5001 | 17328 | 1 | LRG_gene | LRG_578 |
| 4867 | 246944 | 1 | LRG_gene | LRG_579 |
| 5001 | 12829 | 1 | LRG_gene | LRG_58 |
| 4938 | 11210 | 1 | LRG_gene | LRG_580 |
| 5001 | 12401 | 1 | LRG_gene | LRG_581 |
| 5001 | 28718 | 1 | LRG_gene | LRG_583 |
| 5001 | 48505 | 1 | LRG_gene | LRG_586 |
| 4983 | 145240 | 1 | LRG_gene | LRG_588 |
| 4996 | 44757 | 1 | LRG_gene | LRG_589 |
| 4963 | 14787 | 1 | LRG_gene | LRG_59 |
| 5001 | 37982 | 1 | LRG_gene | LRG_590 |
| 5001 | 11398 | 1 | LRG_gene | LRG_593 |
| 4992 | 17169 | 1 | LRG_gene | LRG_597 |
| 5051 | 32102 | 1 | LRG_gene | LRG_598 |
| 5001 | 15827 | 1 | LRG_gene | LRG_599 |
| 5001 | 32834 | 1 | LRG_gene | LRG_6 |
| 5001 | 13865 | 1 | LRG_gene | LRG_60 |
| 5001 | 7198 | 1 | LRG_gene | LRG_600 |
| 10737 | 23025 | 1 | LRG_gene | LRG_601 |
| 5001 | 101383 | 1 | LRG_gene | LRG_605 |
| 4976 | 8884 | 1 | LRG_gene | LRG_607 |
| 4997 | 106870 | 1 | LRG_gene | LRG_608 |
| 5001 | 250549 | 1 | LRG_gene | LRG_609 |
| 5001 | 19217 | 1 | LRG_gene | LRG_61 |
| 15932 | 34848 | 1 | LRG_gene | LRG_610 |
| 4923 | 23499 | 1 | LRG_gene | LRG_611 |
| 5001 | 147939 | 1 | LRG_gene | LRG_612 |
| 5001 | 95335 | 1 | LRG_gene | LRG_613 |
| 5001 | 96182 | 1 | LRG_gene | LRG_614 |
| 5001 | 19623 | 1 | LRG_gene | LRG_615 |
| 4999 | 244435 | 1 | LRG_gene | LRG_616 |
| 5001 | 19241 | 1 | LRG_gene | LRG_617 |
| 5001 | 37810 | 1 | LRG_gene | LRG_618 |
| 4999 | 8301 | 1 | LRG_gene | LRG_619 |
| 5001 | 19392 | 1 | LRG_gene | LRG_62 |

| | | | | |
|---|---|---|---|---|
| 5001 | 52532 | 1 | LRG_gene | LRG_620 |
| 5001 | 50676 | 1 | LRG_gene | LRG_621 |
| 5001 | 29937 | 1 | LRG_gene | LRG_622 |
| 5001 | 25667 | 1 | LRG_gene | LRG_623 |
| 4955 | 48074 | 1 | LRG_gene | LRG_624 |
| 60197 | 81452 | 1 | LRG_gene | LRG_625 |
| 5001 | 138929 | 1 | LRG_gene | LRG_626 |
| 5001 | 131084 | 1 | LRG_gene | LRG_627 |
| 27496 | 80390 | 1 | LRG_gene | LRG_628 |
| 5001 | 60481 | 1 | LRG_gene | LRG_629 |
| 5001 | 17116 | 1 | LRG_gene | LRG_63 |
| 5001 | 85976 | 1 | LRG_gene | LRG_630 |
| 4999 | 9679 | 1 | LRG_gene | LRG_631 |
| 5001 | 8313 | 1 | LRG_gene | LRG_64 |
| 5001 | 8297 | 1 | LRG_gene | LRG_640 |
| 5001 | 13187 | 1 | LRG_gene | LRG_642 |
| 5001 | 118229 | 1 | LRG_gene | LRG_643 |
| 4969 | 29798 | 1 | LRG_gene | LRG_65 |
| 5001 | 52132 | 1 | LRG_gene | LRG_650 |
| 5001 | 14310 | 1 | LRG_gene | LRG_652 |
| 5001 | 15010 | 1 | LRG_gene | LRG_653 |
| 4979 | 19740 | 1 | LRG_gene | LRG_655 |
| 4931 | 66913 | 1 | LRG_gene | LRG_656 |
| 5001 | 21675 | 1 | LRG_gene | LRG_657 |
| 5001 | 17871 | 1 | LRG_gene | LRG_659 |
| 5001 | 26947 | 1 | LRG_gene | LRG_66 |
| 5001 | 12980 | 1 | LRG_gene | LRG_661 |
| 5001 | 130982 | 1 | LRG_gene | LRG_662 |
| 19180 | 207882 | 1 | LRG_gene | LRG_663 |
| 5001 | 81283 | 1 | LRG_gene | LRG_664 |
| 5001 | 8418 | 1 | LRG_gene | LRG_665 |
| 5001 | 53333 | 1 | LRG_gene | LRG_666 |
| 5001 | 30435 | 1 | LRG_gene | LRG_669 |
| 22904 | 57530 | 1 | LRG_gene | LRG_67 |
| 5001 | 59513 | 1 | LRG_gene | LRG_670 |
| 4951 | 15173 | 1 | LRG_gene | LRG_672 |
| 5001 | 23325 | 1 | LRG_gene | LRG_673 |
| 5001 | 20298 | 1 | LRG_gene | LRG_674 |
| 5001 | 23468 | 1 | LRG_gene | LRG_675 |
| 5001 | 33072 | 1 | LRG_gene | LRG_676 |
| 5002 | 13065 | 1 | LRG_gene | LRG_680 |
| 5001 | 8123 | 1 | LRG_gene | LRG_682 |
| 5001 | 42767 | 1 | LRG_gene | LRG_683 |
| 1855 | 28518 | 1 | LRG_gene | LRG_684 |
| 4975 | 26648 | 1 | LRG_gene | LRG_685 |
| 5001 | 19170 | 1 | LRG_gene | LRG_687 |
| 5001 | 68276 | 1 | LRG_gene | LRG_688 |
| 5001 | 20024 | 1 | LRG_gene | LRG_689 |
| 5001 | 12184 | 1 | LRG_gene | LRG_69 |
| 5001 | 29761 | 1 | LRG_gene | LRG_690 |

| | | | | |
|---|---|---|---|---|
| 12927 | 35793 | 1 | LRG_gene | LRG_691 |
| 5001 | 15594 | 1 | LRG_gene | LRG_692 |
| 22295 | 48916 | 1 | LRG_gene | LRG_693 |
| 15010 | 61599 | 1 | LRG_gene | LRG_695 |
| 5001 | 136692 | 1 | LRG_gene | LRG_697 |
| 5001 | 191383 | 1 | LRG_gene | LRG_699 |
| 5001 | 305019 | 1 | LRG_gene | LRG_7 |
| 10604 | 27803 | 1 | LRG_gene | LRG_70 |
| 5001 | 51837 | 1 | LRG_gene | LRG_700 |
| 10346 | 42761 | 1 | LRG_gene | LRG_702 |
| 5001 | 80442 | 1 | LRG_gene | LRG_703 |
| 5001 | 20691 | 1 | LRG_gene | LRG_71 |
| 5001 | 196425 | 1 | LRG_gene | LRG_712 |
| 5001 | 129091 | 1 | LRG_gene | LRG_713 |
| 5001 | 99027 | 1 | LRG_gene | LRG_714 |
| 5001 | 7169 | 1 | LRG_gene | LRG_715 |
| 5001 | 13141 | 1 | LRG_gene | LRG_717 |
| 5001 | 7503 | 1 | LRG_gene | LRG_719 |
| 17063 | 44389 | 1 | LRG_gene | LRG_72 |
| 11631 | 38170 | 1 | LRG_gene | LRG_720 |
| 5001 | 31396 | 1 | LRG_gene | LRG_721 |
| 5001 | 848317 | 1 | LRG_gene | LRG_722 |
| 5001 | 8059 | 1 | LRG_gene | LRG_723 |
| 4945 | 45523 | 1 | LRG_gene | LRG_724 |
| 5001 | 109672 | 1 | LRG_gene | LRG_725 |
| 5001 | 25374 | 1 | LRG_gene | LRG_726 |
| 5001 | 9334 | 1 | LRG_gene | LRG_727 |
| 4940 | 56616 | 1 | LRG_gene | LRG_73 |
| 5001 | 11107 | 1 | LRG_gene | LRG_730 |
| 5001 | 18027 | 1 | LRG_gene | LRG_733 |
| 5001 | 161021 | 1 | LRG_gene | LRG_734 |
| 5000 | 208038 | 1 | LRG_gene | LRG_737 |
| 5001 | 24934 | 1 | LRG_gene | LRG_74 |
| 5001 | 229167 | 1 | LRG_gene | LRG_741 |
| 5001 | 31450 | 1 | LRG_gene | LRG_742 |
| 5001 | 7204 | 1 | LRG_gene | LRG_744 |
| 9802 | 12088 | 1 | LRG_gene | LRG_745 |
| 5001 | 12961 | 1 | LRG_gene | LRG_748 |
| 5001 | 48202 | 1 | LRG_gene | LRG_749 |
| 5001 | 35600 | 1 | LRG_gene | LRG_75 |
| 5001 | 38808 | 1 | LRG_gene | LRG_750 |
| 5001 | 7075 | 1 | LRG_gene | LRG_751 |
| 5001 | 99125 | 1 | LRG_gene | LRG_753 |
| 5001 | 143915 | 1 | LRG_gene | LRG_754 |
| 5001 | 56670 | 1 | LRG_gene | LRG_755 |
| 5001 | 21331 | 1 | LRG_gene | LRG_757 |
| 41373 | 171001 | 1 | LRG_gene | LRG_759 |
| 12789 | 47886 | 1 | LRG_gene | LRG_76 |
| 5001 | 15311 | 1 | LRG_gene | LRG_760 |
| 4961 | 44137 | 1 | LRG_gene | LRG_762 |

| | | | | |
|---|---|---|---|---|
| 5001 | 174286 | 1 | LRG_gene | LRG_763 |
| 5001 | 23491 | 1 | LRG_gene | LRG_765 |
| 5001 | 158865 | 1 | LRG_gene | LRG_766 |
| 5001 | 6870 | 1 | LRG_gene | LRG_767 |
| 5001 | 10880 | 1 | LRG_gene | LRG_768 |
| 4960 | 28210 | 1 | LRG_gene | LRG_77 |
| 4991 | 71088 | 1 | LRG_gene | LRG_770 |
| 5001 | 8987 | 1 | LRG_gene | LRG_771 |
| 5001 | 33933 | 1 | LRG_gene | LRG_772 |
| 4943 | 53522 | 1 | LRG_gene | LRG_773 |
| 5001 | 41944 | 1 | LRG_gene | LRG_774 |
| 5001 | 152570 | 1 | LRG_gene | LRG_775 |
| 202119 | 233856 | 1 | LRG_gene | LRG_776 |
| 5001 | 27740 | 1 | LRG_gene | LRG_777 |
| 5001 | 242483 | 1 | LRG_gene | LRG_778 |
| 5001 | 92641 | 1 | LRG_gene | LRG_779 |
| 5001 | 59858 | 1 | LRG_gene | LRG_78 |
| 5001 | 215663 | 1 | LRG_gene | LRG_780 |
| 5001 | 61317 | 1 | LRG_gene | LRG_781 |
| 5001 | 147097 | 1 | LRG_gene | LRG_782 |
| 5001 | 20896 | 1 | LRG_gene | LRG_783 |
| 5001 | 37267 | 1 | LRG_gene | LRG_784 |
| 5001 | 38696 | 1 | LRG_gene | LRG_785 |
| 5001 | 113024 | 1 | LRG_gene | LRG_786 |
| 4997 | 72482 | 1 | LRG_gene | LRG_787 |
| 5001 | 39618 | 1 | LRG_gene | LRG_788 |
| 5001 | 68763 | 1 | LRG_gene | LRG_789 |
| 5753 | 13093 | 1 | LRG_gene | LRG_79 |
| 4979 | 17341 | 1 | LRG_gene | LRG_798 |
| 5001 | 89480 | 1 | LRG_gene | LRG_8 |
| 5001 | 40928 | 1 | LRG_gene | LRG_80 |
| 63301 | 77124 | 1 | LRG_gene | LRG_808 |
| 5001 | 8785 | 1 | LRG_gene | LRG_81 |
| 5001 | 29670 | 1 | LRG_gene | LRG_818 |
| 5001 | 36700 | 1 | LRG_gene | LRG_822 |
| 5001 | 7782 | 1 | LRG_gene | LRG_825 |
| 5001 | 10891 | 1 | LRG_gene | LRG_828 |
| 4949 | 73032 | 1 | LRG_gene | LRG_83 |
| 5001 | 25222 | 1 | LRG_gene | LRG_837 |
| 4990 | 44322 | 1 | LRG_gene | LRG_838 |
| 5001 | 109570 | 1 | LRG_gene | LRG_839 |
| 5001 | 16080 | 1 | LRG_gene | LRG_84 |
| 5001 | 50927 | 1 | LRG_gene | LRG_844 |
| 5001 | 19483 | 1 | LRG_gene | LRG_847 |
| 5001 | 81575 | 1 | LRG_gene | LRG_85 |
| 5001 | 9042 | 1 | LRG_gene | LRG_855 |
| 5001 | 226768 | 1 | LRG_gene | LRG_86 |
| 5001 | 17180 | 1 | LRG_gene | LRG_860 |
| 5001 | 10259 | 1 | LRG_gene | LRG_862 |
| 5001 | 467379 | 1 | LRG_gene | LRG_863 |

| | | | | |
|---|---|---|---|---|
| 5001 | 56030 | 1 | LRG_gene | LRG_867 |
| 5001 | 95192 | 1 | LRG_gene | LRG_868 |
| 5001 | 33846 | 1 | LRG_gene | LRG_870 |
| 5001 | 91080 | 1 | LRG_gene | LRG_875 |
| 5001 | 11711 | 1 | LRG_gene | LRG_876 |
| 5001 | 51143 | 1 | LRG_gene | LRG_879 |
| 4978 | 40020 | 1 | LRG_gene | LRG_88 |
| 5001 | 26146 | 1 | LRG_gene | LRG_880 |
| 4905 | 27498 | 1 | LRG_gene | LRG_883 |
| 4983 | 29188 | 1 | LRG_gene | LRG_884 |
| 5001 | 17430 | 1 | LRG_gene | LRG_885 |
| 5001 | 110244 | 1 | LRG_gene | LRG_886 |
| 4998 | 30411 | 1 | LRG_gene | LRG_887 |
| 4985 | 8229 | 1 | LRG_gene | LRG_89 |
| 5001 | 90542 | 1 | LRG_gene | LRG_90 |
| 4997 | 13624 | 1 | LRG_gene | LRG_91 |
| 5001 | 17438 | 1 | LRG_gene | LRG_92 |
| 4981 | 20466 | 1 | LRG_gene | LRG_93 |
| 5001 | 10428 | 1 | LRG_gene | LRG_94 |
| 5001 | 91210 | 1 | LRG_gene | LRG_96 |
| 5001 | 23996 | 1 | LRG_gene | LRG_97 |
| 5001 | 16750 | 1 | LRG_gene | LRG_98 |
| 4975 | 11294 | 1 | LRG_gene | LRG_99 |
| 5001 | 18370 | 1 | LRG_gene | LRG_990 |
| 5001 | 236707 | 1 | LRG_gene | LRG_991 |
| 155989 | 451584 | 1 | LRG_gene | LRG_992 |
| 5001 | 62697 | 1 | LRG_gene | LRG_993 |
| 5001 | 125129 | 1 | LRG_gene | LRG_994 |
| 4889 | 35692 | 1 | LRG_gene | LRG_995 |
| 4918 | 28400 | 1 | LRG_gene | LRG_996 |
| 5001 | 142489 | 1 | LRG_gene | LRG_997 |
| 5001 | 118962 | 1 | LRG_gene | LRG_998 |
| 5001 | 32260 | 1 | LRG_gene | LRG_999 |
| 577 | 647 | 1 | Mt_tRNA | ENSG00000 |
| 648 | 1601 | 1 | Mt_rRNA | ENSG00000 |
| 1602 | 1670 | 1 | Mt_tRNA | ENSG00000 |
| 1671 | 3229 | 1 | Mt_rRNA | ENSG00000 |
| 3230 | 3304 | 1 | Mt_tRNA | ENSG00000 |
| 3307 | 4262 | 1 | protein_coding | ENSG00000 |
| 4263 | 4331 | 1 | Mt_tRNA | ENSG00000 |
| 4329 | 4400 | -1 | Mt_tRNA | ENSG00000 |
| 4402 | 4469 | 1 | Mt_tRNA | ENSG00000 |
| 4470 | 5511 | 1 | protein_coding | ENSG00000 |
| 5512 | 5579 | 1 | Mt_tRNA | ENSG00000 |
| 5587 | 5655 | -1 | Mt_tRNA | ENSG00000 |
| 5657 | 5729 | -1 | Mt_tRNA | ENSG00000 |
| 5761 | 5826 | -1 | Mt_tRNA | ENSG00000 |
| 5826 | 5891 | -1 | Mt_tRNA | ENSG00000 |
| 5904 | 7445 | 1 | protein_coding | ENSG00000 |
| 7446 | 7514 | -1 | Mt_tRNA | ENSG00000 |

| | | | | |
|---|---|---|---|---|
| 7518 | 7585 | 1 | Mt_tRNA | ENSG00000 |
| 7586 | 8269 | 1 | protein_coding | ENSG00000 |
| 8295 | 8364 | 1 | Mt_tRNA | ENSG00000 |
| 8366 | 8572 | 1 | protein_coding | ENSG00000 |
| 8527 | 9207 | 1 | protein_coding | ENSG00000 |
| 9207 | 9990 | 1 | protein_coding | ENSG00000 |
| 9991 | 10058 | 1 | Mt_tRNA | ENSG00000 |
| 10059 | 10404 | 1 | protein_coding | ENSG00000 |
| 10405 | 10469 | 1 | Mt_tRNA | ENSG00000 |
| 10470 | 10766 | 1 | protein_coding | ENSG00000 |
| 10760 | 12137 | 1 | protein_coding | ENSG00000 |
| 12138 | 12206 | 1 | Mt_tRNA | ENSG00000 |
| 12207 | 12265 | 1 | Mt_tRNA | ENSG00000 |
| 12266 | 12336 | 1 | Mt_tRNA | ENSG00000 |
| 12337 | 14148 | 1 | protein_coding | ENSG00000 |
| 14149 | 14673 | -1 | protein_coding | ENSG00000 |
| 14674 | 14742 | -1 | Mt_tRNA | ENSG00000 |
| 14747 | 15887 | 1 | protein_coding | ENSG00000 |
| 15888 | 15953 | 1 | Mt_tRNA | ENSG00000 |
| 15956 | 16023 | -1 | Mt_tRNA | ENSG00000 |
| 253743 | 255091 | 1 | unprocessed_pseudo | ENSG00000 |
| 276322 | 303356 | 1 | protein_coding | ENSG00000 |
| 304529 | 318819 | -1 | protein_coding | ENSG00000 |
| 320990 | 321851 | 1 | lncRNA | ENSG00000 |
| 333933 | 386955 | -1 | protein_coding | ENSG00000 |
| 386980 | 405579 | 1 | lncRNA | ENSG00000 |
| 388100 | 388389 | -1 | misc_RNA | ENSG00000 |
| 419157 | 421980 | 1 | lncRNA | ENSG00000 |
| 523775 | 524102 | -1 | processed_pseudoge | ENSG00000 |
| 545236 | 545352 | -1 | processed_pseudoge | ENSG00000 |
| 624344 | 659411 | 1 | protein_coding | ENSG00000 |
| 990221 | 994365 | 1 | lncRNA | ENSG00000 |
| 1008503 | 1010101 | -1 | processed_pseudoge | ENSG00000 |
| 1187549 | 1212723 | -1 | protein_coding | ENSG00000 |
| 1268800 | 1310381 | 1 | protein_coding | ENSG00000 |
| 1293918 | 1293992 | 1 | miRNA | ENSG00000 |
| 1300256 | 1300375 | -1 | rRNA_pseudogene | ENSG00000 |
| 1336616 | 1382689 | 1 | protein_coding | ENSG00000 |
| 1386152 | 1392113 | -1 | protein_coding | ENSG00000 |
| 1396427 | 1399402 | 1 | lncRNA | ENSG00000 |
| 1401769 | 1414028 | 1 | lncRNA | ENSG00000 |
| 1403139 | 1453762 | -1 | protein_coding | ENSG00000 |
| 1462581 | 1537185 | -1 | protein_coding | ENSG00000 |
| 1591604 | 1602520 | 1 | protein_coding | ENSG00000 |
| 1615059 | 1643081 | 1 | protein_coding | ENSG00000 |
| 1732584 | 1763212 | 1 | lncRNA | ENSG00000 |
| 1767347 | 1768776 | 1 | lncRNA | ENSG00000 |
| 2219506 | 2502805 | -1 | protein_coding | ENSG00000 |
| 2334295 | 2336410 | -1 | lncRNA | ENSG00000 |
| 2486414 | 2500976 | -1 | protein_coding | ENSG00000 |

| | | | | |
|---|---|---|---|---|
| 2609191 | 2609254 | 1 | miRNA | ENSG0000( |
| 2609348 | 2657229 | 1 | transcribed_unproce | ENSG0000( |
| 2612988 | 2615347 | -1 | lncRNA | ENSG0000( |
| 2691187 | 2741309 | 1 | protein_coding | ENSG0000( |
| 2752048 | 2816500 | 1 | protein_coding | ENSG0000( |
| 2828822 | 2882820 | 1 | protein_coding | ENSG0000( |
| 2852740 | 2853760 | -1 | lncRNA | ENSG0000( |
| 2903972 | 2929349 | -1 | protein_coding | ENSG0000( |
| 2904904 | 2906081 | 1 | lncRNA | ENSG0000( |
| 2934521 | 2968475 | -1 | protein_coding | ENSG0000( |
| 3006613 | 3033571 | 1 | protein_coding | ENSG0000( |
| 3041471 | 3112727 | 1 | protein_coding | ENSG0000( |
| 3058220 | 3058502 | -1 | misc_RNA | ENSG0000( |
| 3112900 | 3113596 | -1 | processed_pseudoge | ENSG0000( |
| 3271544 | 3286138 | 1 | lncRNA | ENSG0000( |
| 3308565 | 3346652 | -1 | protein_coding | ENSG0000( |
| 3367149 | 3368382 | 1 | processed_pseudoge | ENSG0000( |
| 3532117 | 3532251 | -1 | snoRNA | ENSG0000( |
| 3572857 | 3572961 | 1 | snRNA | ENSG0000( |
| 3604340 | 3713649 | -1 | protein_coding | ENSG0000( |
| 3607258 | 3607365 | -1 | snRNA | ENSG0000( |
| 3641443 | 3641555 | -1 | misc_RNA | ENSG0000( |
| 3659487 | 3668192 | 1 | lncRNA | ENSG0000( |
| 3717149 | 3717600 | 1 | processed_pseudoge | ENSG0000( |
| 3817528 | 3937855 | -1 | lncRNA | ENSG0000( |
| 3819924 | 3823898 | 1 | lncRNA | ENSG0000( |
| 3953056 | 3953561 | 1 | processed_pseudoge | ENSG0000( |
| 4627200 | 4633572 | -1 | lncRNA | ENSG0000( |
| 5168918 | 5169439 | -1 | processed_pseudoge | ENSG0000( |
| 5169512 | 5170645 | 1 | processed_pseudoge | ENSG0000( |
| 5305714 | 5307128 | -1 | processed_pseudoge | ENSG0000( |
| 5410122 | 5410251 | 1 | snoRNA | ENSG0000( |
| 5653418 | 5726326 | -1 | lncRNA | ENSG0000( |
| 5840637 | 6228867 | -1 | protein_coding | ENSG0000( |
| 6061644 | 6061749 | -1 | snoRNA | ENSG0000( |
| 6383906 | 6383963 | -1 | miRNA | ENSG0000( |
| 6422092 | 6422701 | -1 | processed_pseudoge | ENSG0000( |
| 6533618 | 6535118 | -1 | protein_coding | ENSG0000( |
| 6667865 | 7148190 | -1 | protein_coding | ENSG0000( |
| 6989329 | 6989783 | 1 | processed_pseudoge | ENSG0000( |
| 7113005 | 7113404 | -1 | processed_pseudoge | ENSG0000( |
| 7147237 | 7804358 | 1 | protein_coding | ENSG0000( |
| 7147860 | 7147937 | 1 | miRNA | ENSG0000( |
| 7489659 | 7491199 | 1 | TEC | ENSG0000( |
| 7842262 | 7844143 | 1 | protein_coding | ENSG0000( |
| 7896514 | 7896722 | -1 | unprocessed_pseudc | ENSG0000( |
| 7898247 | 7927739 | -1 | protein_coding | ENSG0000( |
| 7928443 | 8374386 | 1 | lncRNA | ENSG0000( |
| 8126965 | 8127061 | 1 | miRNA | ENSG0000( |
| 8169944 | 8171267 | -1 | protein_coding | ENSG0000( |

| | | | | |
|---|---|---|---|---|
| 8464830 | 8466510 | 1 | protein_coding | ENSG00000( |
| 8487165 | 8504564 | -1 | lncRNA | ENSG00000( |
| 8528874 | 8732137 | -1 | protein_coding | ENSG00000( |
| 8783514 | 8784461 | -1 | processed_pseudoge | ENSG00000( |
| 8790795 | 8801383 | -1 | protein_coding | ENSG00000( |
| 8856665 | 8857059 | -1 | processed_pseudoge | ENSG00000( |
| 8863861 | 8867601 | -1 | lncRNA | ENSG00000( |
| 8897642 | 8927153 | -1 | lncRNA | ENSG00000( |
| 8980425 | 8982505 | -1 | processed_pseudoge | ENSG00000( |
| 9024232 | 9295682 | -1 | protein_coding | ENSG00000( |
| 9133828 | 9133946 | 1 | rRNA_pseudogene | ENSG00000( |
| 9179385 | 9179606 | 1 | processed_pseudoge | ENSG00000( |
| 9249920 | 9275206 | 1 | lncRNA | ENSG00000( |
| 9402751 | 9404802 | 1 | processed_pseudoge | ENSG00000( |
| 9404954 | 9405731 | 1 | processed_pseudoge | ENSG00000( |
| 9406025 | 9406778 | 1 | processed_pseudoge | ENSG00000( |
| 9406952 | 9407708 | 1 | processed_pseudoge | ENSG00000( |
| 9407861 | 9408638 | 1 | processed_pseudoge | ENSG00000( |
| 9408818 | 9409571 | 1 | processed_pseudoge | ENSG00000( |
| 9409748 | 9410501 | 1 | processed_pseudoge | ENSG00000( |
| 9410789 | 9411545 | 1 | processed_pseudoge | ENSG00000( |
| 9411701 | 9412478 | 1 | processed_pseudoge | ENSG00000( |
| 9412772 | 9413522 | 1 | processed_pseudoge | ENSG00000( |
| 9413816 | 9414569 | 1 | processed_pseudoge | ENSG00000( |
| 9463295 | 9741037 | 1 | protein_coding | ENSG00000( |
| 9725346 | 9786297 | -1 | protein_coding | ENSG00000( |
| 9786429 | 9949443 | 1 | protein_coding | ENSG00000( |
| 9818239 | 9819469 | -1 | processed_pseudoge | ENSG00000( |
| 9865093 | 9865543 | -1 | processed_pseudoge | ENSG00000( |
| 9874601 | 9874701 | -1 | misc_RNA | ENSG00000( |
| 9967358 | 9968352 | 1 | protein_coding | ENSG00000( |
| 10015562 | 10144478 | 1 | protein_coding | ENSG00000( |
| 10024842 | 10038654 | -1 | lncRNA | ENSG00000( |
| 10061222 | 10061323 | 1 | misc_RNA | ENSG00000( |
| 10156945 | 10237660 | 1 | protein_coding | ENSG00000( |
| 10242339 | 10365323 | 1 | lncRNA | ENSG00000( |
| 10445310 | 10833654 | -1 | protein_coding | ENSG00000( |
| 10582817 | 10582919 | 1 | snRNA | ENSG00000( |
| 10847578 | 11111220 | -1 | lncRNA | ENSG00000( |
| 11111301 | 11123086 | 1 | protein_coding | ENSG00000( |
| 11116092 | 11116193 | -1 | misc_RNA | ENSG00000( |
| 11117651 | 11665920 | -1 | protein_coding | ENSG00000( |
| 11148584 | 11148769 | -1 | misc_RNA | ENSG00000( |
| 11221348 | 11225731 | 1 | unprocessed_pseudc | ENSG00000( |
| 11293413 | 11300761 | 1 | protein_coding | ENSG00000( |
| 11318614 | 11318686 | -1 | miRNA | ENSG00000( |
| 11334233 | 11335901 | -1 | processed_pseudoge | ENSG00000( |
| 11668401 | 11757718 | -1 | lncRNA | ENSG00000( |
| 11704123 | 11705590 | -1 | processed_pseudoge | ENSG00000( |
| 11758159 | 11775772 | 1 | protein_coding | ENSG00000( |

| | | | | |
|---|---|---|---|---|
| 11822423 | 12724523 | 1 | protein_coding | ENSG00000 |
| 12054824 | 12054926 | -1 | misc_RNA | ENSG00000 |
| 12149126 | 12149238 | 1 | misc_RNA | ENSG00000 |
| 12373167 | 12375133 | -1 | lncRNA | ENSG00000 |
| 12395757 | 12396167 | -1 | processed_pseudoge | ENSG00000 |
| 12583415 | 12583526 | -1 | unprocessed_pseudo | ENSG00000 |
| 12614003 | 12614197 | -1 | misc_RNA | ENSG00000 |
| 12791355 | 12824222 | 1 | protein_coding | ENSG00000 |
| 12825840 | 12826833 | 1 | processed_pseudoge | ENSG00000 |
| 12844865 | 12845430 | -1 | processed_pseudoge | ENSG00000 |
| 12867072 | 12890361 | 1 | protein_coding | ENSG00000 |
| 12902817 | 12908333 | -1 | lncRNA | ENSG00000 |
| 12906620 | 12923169 | 1 | protein_coding | ENSG00000 |
| 12975110 | 12977227 | 1 | protein_coding | ENSG00000 |
| 13035617 | 13044682 | -1 | protein_coding | ENSG00000 |
| 13093660 | 13094573 | 1 | lncRNA | ENSG00000 |
| 13266048 | 13303452 | -1 | lncRNA | ENSG00000 |
| 13310652 | 13319933 | 1 | lncRNA | ENSG00000 |
| 13318236 | 13320399 | -1 | protein_coding | ENSG00000 |
| 13327937 | 13329867 | 1 | unprocessed_pseudo | ENSG00000 |
| 13333952 | 13403035 | 1 | lncRNA | ENSG00000 |
| 13378735 | 13379340 | -1 | processed_pseudoge | ENSG00000 |
| 13569601 | 13633575 | 1 | protein_coding | ENSG00000 |
| 13574746 | 13575066 | 1 | processed_pseudoge | ENSG00000 |
| 13590292 | 13590346 | 1 | miRNA | ENSG00000 |
| 13596236 | 13596548 | 1 | misc_RNA | ENSG00000 |
| 13653189 | 13681964 | 1 | protein_coding | ENSG00000 |
| 13689128 | 13710504 | 1 | protein_coding | ENSG00000 |
| 13712244 | 13734635 | -1 | protein_coding | ENSG00000 |
| 13734745 | 13769357 | 1 | protein_coding | ENSG00000 |
| 13770939 | 13938638 | -1 | protein_coding | ENSG00000 |
| 13955393 | 13963904 | 1 | lncRNA | ENSG00000 |
| 13980066 | 13988820 | -1 | lncRNA | ENSG00000 |
| 14008279 | 14029893 | -1 | protein_coding | ENSG00000 |
| 14244382 | 14244928 | 1 | processed_pseudoge | ENSG00000 |
| 14529298 | 14731812 | 1 | protein_coding | ENSG00000 |
| 14834476 | 14835319 | -1 | processed_pseudoge | ENSG00000 |
| 14835961 | 14873069 | -1 | protein_coding | ENSG00000 |
| 14873421 | 14922327 | 1 | protein_coding | ENSG00000 |
| 14977959 | 14978463 | -1 | processed_pseudoge | ENSG00000 |
| 15117437 | 15117768 | -1 | processed_pseudoge | ENSG00000 |
| 15235288 | 15270467 | -1 | protein_coding | ENSG00000 |
| 15281697 | 15315640 | -1 | protein_coding | ENSG00000 |
| 15319451 | 15335580 | -1 | protein_coding | ENSG00000 |
| 15345596 | 15384413 | -1 | protein_coding | ENSG00000 |
| 15384799 | 15493564 | -1 | protein_coding | ENSG00000 |
| 15464246 | 15556529 | 1 | protein_coding | ENSG00000 |
| 15561033 | 15602148 | -1 | protein_coding | ENSG00000 |
| 15561047 | 15675608 | -1 | protein_coding | ENSG00000 |
| 15602881 | 15621484 | 1 | lncRNA | ENSG00000 |

| | | | | |
|---|---|---|---|---|
| 15627318 | 15675012 | -1 | protein_coding | ENSG00000 |
| 15646167 | 15646875 | -1 | processed_pseudoge | ENSG00000 |
| 15674932 | 15703724 | 1 | transcribed_unproce | ENSG00000 |
| 15688830 | 15788411 | 1 | protein_coding | ENSG00000 |
| 15702602 | 15703243 | -1 | pseudogene | ENSG00000 |
| 15716208 | 15716346 | 1 | snoRNA | ENSG00000 |
| 15785716 | 15787589 | -1 | lncRNA | ENSG00000 |
| 15790472 | 15823260 | 1 | protein_coding | ENSG00000 |
| 15825806 | 15854931 | -1 | protein_coding | ENSG00000 |
| 15916189 | 15916303 | -1 | snRNA | ENSG00000 |
| 15917548 | 15918631 | 1 | processed_pseudoge | ENSG00000 |
| 16123565 | 16153518 | 1 | protein_coding | ENSG00000 |
| 16152941 | 16170869 | -1 | lncRNA | ENSG00000 |
| 16167481 | 16171464 | 1 | protein_coding | ENSG00000 |
| 16198428 | 16199289 | -1 | processed_pseudoge | ENSG00000 |
| 16474342 | 16475340 | -1 | processed_pseudoge | ENSG00000 |
| 16539137 | 16539439 | -1 | misc_RNA | ENSG00000 |
| 16587999 | 16712936 | -1 | protein_coding | ENSG00000 |
| 16627012 | 16627085 | -1 | miRNA | ENSG00000 |
| 16650158 | 16654670 | 1 | protein_coding | ENSG00000 |
| 16719612 | 16765340 | 1 | protein_coding | ENSG00000 |
| 16767844 | 16767905 | 1 | snRNA | ENSG00000 |
| 16786432 | 16844519 | 1 | protein_coding | ENSG00000 |
| 16803233 | 16803353 | 1 | processed_pseudoge | ENSG00000 |
| 16839283 | 16870414 | -1 | protein_coding | ENSG00000 |
| 16875146 | 16875267 | -1 | snRNA | ENSG00000 |
| 16897651 | 16898521 | -1 | processed_pseudoge | ENSG00000 |
| 16912215 | 16912317 | -1 | misc_RNA | ENSG00000 |
| 16946658 | 17153272 | 1 | protein_coding | ENSG00000 |
| 17044380 | 17044511 | -1 | snoRNA | ENSG00000 |
| 17099782 | 17100239 | -1 | processed_pseudoge | ENSG00000 |
| 17282637 | 17283093 | 1 | processed_pseudoge | ENSG00000 |
| 17357930 | 17358033 | 1 | snoRNA | ENSG00000 |
| 17375420 | 17735994 | 1 | protein_coding | ENSG00000 |
| 17425881 | 17425954 | 1 | miRNA | ENSG00000 |
| 17528435 | 17587160 | 1 | lncRNA | ENSG00000 |
| 17552349 | 17557387 | -1 | lncRNA | ENSG00000 |
| 17705339 | 17705691 | -1 | processed_pseudoge | ENSG00000 |
| 17737449 | 17754988 | 1 | protein_coding | ENSG00000 |
| 17769005 | 17769398 | -1 | processed_pseudoge | ENSG00000 |
| 17800049 | 17861337 | -1 | protein_coding | ENSG00000 |
| 17897711 | 17900333 | -1 | unprocessed_pseudc | ENSG00000 |
| 17933349 | 17934823 | 1 | processed_pseudoge | ENSG00000 |
| 17970197 | 18104644 | -1 | lncRNA | ENSG00000 |
| 18162931 | 18220886 | -1 | protein_coding | ENSG00000 |
| 18239313 | 18354688 | -1 | protein_coding | ENSG00000 |
| 18341216 | 18341346 | 1 | processed_pseudoge | ENSG00000 |
| 18395994 | 18396095 | -1 | misc_RNA | ENSG00000 |
| 18425583 | 18653629 | 1 | protein_coding | ENSG00000 |
| 18535079 | 18536120 | 1 | processed_pseudoge | ENSG00000 |

| | | | | |
|---|---|---|---|---|
| 18639910 | 18672109 | -1 | protein_coding | ENSG00000( |
| 18675909 | 18827921 | 1 | protein_coding | ENSG00000( |
| 18688643 | 18691604 | -1 | lncRNA | ENSG00000( |
| 18838709 | 18839452 | -1 | processed_pseudoge | ENSG00000( |
| 18865771 | 18866554 | 1 | processed_pseudoge | ENSG00000( |
| 18890296 | 18894974 | 1 | lncRNA | ENSG00000( |
| 18892298 | 18984114 | -1 | protein_coding | ENSG00000( |
| 18915243 | 18915535 | -1 | misc_RNA | ENSG00000( |
| 18984291 | 19060329 | 1 | lncRNA | ENSG00000( |
| 18989307 | 19122637 | -1 | protein_coding | ENSG00000( |
| 19343893 | 19361718 | 1 | protein_coding | ENSG00000( |
| 19360056 | 19515261 | -1 | protein_coding | ENSG00000( |
| 19376774 | 19376875 | 1 | misc_RNA | ENSG00000( |
| 19496378 | 19497721 | -1 | unprocessed_pseudc | ENSG00000( |
| 19533977 | 19887600 | -1 | protein_coding | ENSG00000( |
| 19912860 | 19970298 | -1 | protein_coding | ENSG00000( |
| 19985058 | 19985159 | -1 | misc_RNA | ENSG00000( |
| 20006713 | 20116907 | -1 | protein_coding | ENSG00000( |
| 20017088 | 20017187 | -1 | miRNA | ENSG00000( |
| 20124525 | 20141838 | -1 | protein_coding | ENSG00000( |
| 20136066 | 20136135 | -1 | snoRNA | ENSG00000( |
| 20136306 | 20136385 | -1 | snoRNA | ENSG00000( |
| 20139968 | 20140444 | 1 | lncRNA | ENSG00000( |
| 20149911 | 20267519 | -1 | protein_coding | ENSG00000( |
| 20311838 | 20317783 | -1 | lncRNA | ENSG00000( |
| 20443326 | 20492349 | 1 | lncRNA | ENSG00000( |
| 20452108 | 20452431 | 1 | misc_RNA | ENSG00000( |
| 20769718 | 21374265 | -1 | lncRNA | ENSG00000( |
| 21131496 | 21163726 | 1 | lncRNA | ENSG00000( |
| 21214637 | 21214743 | 1 | snRNA | ENSG00000( |
| 21372801 | 21654695 | 1 | protein_coding | ENSG00000( |
| 21421822 | 21422294 | -1 | processed_pseudoge | ENSG00000( |
| 21654690 | 21658330 | -1 | protein_coding | ENSG00000( |
| 21705978 | 21758116 | -1 | protein_coding | ENSG00000( |
| 21839617 | 21885423 | 1 | protein_coding | ENSG00000( |
| 21841794 | 21841961 | -1 | unprocessed_pseudc | ENSG00000( |
| 21855987 | 21858740 | 1 | protein_coding | ENSG00000( |
| 21872707 | 21872808 | 1 | misc_RNA | ENSG00000( |
| 21939461 | 21940160 | 1 | processed_pseudoge | ENSG00000( |
| 21940709 | 21994837 | 1 | protein_coding | ENSG00000( |
| 22032325 | 22251310 | 1 | protein_coding | ENSG00000( |
| 22162733 | 22172983 | -1 | lncRNA | ENSG00000( |
| 22191895 | 23293146 | -1 | lncRNA | ENSG00000( |
| 22272943 | 22274461 | 1 | protein_coding | ENSG00000( |
| 22428132 | 22494713 | 1 | lncRNA | ENSG00000( |
| 22435598 | 22436818 | 1 | processed_pseudoge | ENSG00000( |
| 22527835 | 22527941 | 1 | snRNA | ENSG00000( |
| 22616744 | 22617143 | -1 | processed_pseudoge | ENSG00000( |
| 22698457 | 22768966 | 1 | lncRNA | ENSG00000( |
| 22999960 | 23003589 | 1 | protein_coding | ENSG00000( |

| | | | | |
|---|---|---|---|---|
| 23075758 | 23076767 | 1 | processed_pseudoge | ENSG00000 |
| 23198339 | 23198690 | 1 | processed_pseudoge | ENSG00000 |
| 23334849 | 23404374 | 1 | protein_coding | ENSG00000 |
| 23351013 | 23351450 | 1 | processed_pseudoge | ENSG00000 |
| 23664262 | 23686397 | 1 | protein_coding | ENSG00000 |
| 23701055 | 23766475 | -1 | protein_coding | ENSG00000 |
| 23772992 | 23782956 | -1 | lncRNA | ENSG00000 |
| 23782210 | 23782310 | 1 | misc_RNA | ENSG00000 |
| 23783173 | 23786226 | 1 | protein_coding | ENSG00000 |
| 23833353 | 23907938 | -1 | protein_coding | ENSG00000 |
| 23836742 | 23837321 | -1 | processed_pseudoge | ENSG00000 |
| 23907801 | 23939509 | 1 | protein_coding | ENSG00000 |
| 23983720 | 24027186 | -1 | protein_coding | ENSG00000 |
| 24054946 | 24078810 | 1 | protein_coding | ENSG00000 |
| 24133186 | 24133317 | 1 | snoRNA | ENSG00000 |
| 24146225 | 24149654 | -1 | lncRNA | ENSG00000 |
| 24149173 | 24216255 | 1 | protein_coding | ENSG00000 |
| 24310907 | 24313315 | -1 | processed_pseudoge | ENSG00000 |
| 24362761 | 24365378 | 1 | processed_pseudoge | ENSG00000 |
| 24429573 | 24429920 | -1 | processed_pseudoge | ENSG00000 |
| 24465221 | 24550466 | 1 | protein_coding | ENSG00000 |
| 24513450 | 24513687 | 1 | processed_pseudoge | ENSG00000 |
| 24558087 | 24672677 | -1 | protein_coding | ENSG00000 |
| 24632257 | 24632601 | 1 | processed_pseudoge | ENSG00000 |
| 24650073 | 24658237 | 1 | lncRNA | ENSG00000 |
| 24681909 | 24682634 | 1 | processed_pseudoge | ENSG00000 |
| 24693909 | 24996986 | 1 | protein_coding | ENSG00000 |
| 24744441 | 24744570 | 1 | scaRNA | ENSG00000 |
| 24788392 | 24789069 | -1 | processed_pseudoge | ENSG00000 |
| 25003694 | 25016420 | -1 | protein_coding | ENSG00000 |
| 25029545 | 25030235 | 1 | processed_pseudoge | ENSG00000 |
| 25060649 | 25060939 | -1 | misc_RNA | ENSG00000 |
| 25194381 | 25195207 | -1 | processed_pseudoge | ENSG00000 |
| 25591353 | 25591808 | -1 | processed_pseudoge | ENSG00000 |
| 25645722 | 25646327 | -1 | processed_pseudoge | ENSG00000 |
| 25878339 | 25893544 | 1 | lncRNA | ENSG00000 |
| 26138343 | 26140736 | 1 | protein_coding | ENSG00000 |
| 26147774 | 26148281 | 1 | unprocessed_pseudo | ENSG00000 |
| 26160601 | 26161824 | 1 | protein_coding | ENSG00000 |
| 26170052 | 26170913 | 1 | unprocessed_pseudo | ENSG00000 |
| 26192440 | 26195646 | 1 | protein_coding | ENSG00000 |
| 26200042 | 26200647 | 1 | unprocessed_pseudo | ENSG00000 |
| 26216169 | 26218270 | 1 | protein_coding | ENSG00000 |
| 26223903 | 26224658 | 1 | unprocessed_pseudo | ENSG00000 |
| 26345236 | 26346242 | 1 | processed_pseudoge | ENSG00000 |
| 26496936 | 26505603 | 1 | processed_pseudoge | ENSG00000 |
| 26557675 | 26558330 | 1 | processed_pseudoge | ENSG00000 |
| 26558337 | 26561052 | 1 | lncRNA | ENSG00000 |
| 26657291 | 26657507 | -1 | processed_pseudoge | ENSG00000 |
| 26686026 | 26687955 | 1 | processed_pseudoge | ENSG00000 |

| | | | | |
|---|---|---|---|---|
| 26748029 | 26749090 | 1 | processed_pseudoge | ENSG00000 |
| 26790142 | 26790286 | 1 | snRNA | ENSG00000 |
| 27042907 | 27176298 | -1 | lncRNA | ENSG00000 |
| 27174920 | 27399006 | -1 | lncRNA | ENSG00000 |
| 27218131 | 27218678 | -1 | processed_pseudoge | ENSG00000 |
| 27252119 | 27252235 | -1 | misc_RNA | ENSG00000 |
| 27427777 | 27428231 | -1 | processed_pseudoge | ENSG00000 |
| 27460207 | 27463341 | -1 | protein_coding | ENSG00000 |
| 27517771 | 27517881 | -1 | processed_pseudoge | ENSG00000 |
| 27517884 | 27519759 | 1 | processed_pseudoge | ENSG00000 |
| 27590382 | 27748821 | 1 | protein_coding | ENSG00000 |
| 27807990 | 27823014 | 1 | protein_coding | ENSG00000 |
| 27811885 | 27812902 | 1 | processed_pseudoge | ENSG00000 |
| 27832175 | 27833204 | 1 | processed_pseudoge | ENSG00000 |
| 27847588 | 27847939 | -1 | processed_pseudoge | ENSG00000 |
| 27854323 | 27855353 | 1 | processed_pseudoge | ENSG00000 |
| 27863893 | 27864349 | 1 | processed_pseudoge | ENSG00000 |
| 27893348 | 27894059 | -1 | processed_pseudoge | ENSG00000 |
| 27977992 | 27981449 | -1 | protein_coding | ENSG00000 |
| 28495555 | 28495663 | -1 | miRNA | ENSG00000 |
| 28571532 | 28586395 | -1 | lncRNA | ENSG00000 |
| 28587446 | 29956718 | 1 | protein_coding | ENSG00000 |
| 28659065 | 28659469 | -1 | processed_pseudoge | ENSG00000 |
| 28924069 | 28924138 | 1 | snoRNA | ENSG00000 |
| 28942050 | 28942568 | 1 | lncRNA | ENSG00000 |
| 28982852 | 28982970 | -1 | rRNA_pseudogene | ENSG00000 |
| 29369001 | 29369845 | -1 | processed_pseudoge | ENSG00000 |
| 29574278 | 29574358 | 1 | miRNA | ENSG00000 |
| 30017160 | 30017881 | -1 | processed_pseudoge | ENSG00000 |
| 30209423 | 30210209 | 1 | processed_pseudoge | ENSG00000 |
| 30215563 | 30220089 | 1 | protein_coding | ENSG00000 |
| 30230436 | 30237492 | 1 | protein_coding | ENSG00000 |
| 30242000 | 30244187 | 1 | protein_coding | ENSG00000 |
| 30243730 | 30252040 | 1 | protein_coding | ENSG00000 |
| 30290047 | 30290351 | -1 | unprocessed_pseudc | ENSG00000 |
| 30304206 | 30309390 | -1 | protein_coding | ENSG00000 |
| 30558809 | 30577766 | -1 | protein_coding | ENSG00000 |
| 30617454 | 30617693 | 1 | processed_pseudoge | ENSG00000 |
| 30630298 | 30630825 | 1 | processed_pseudoge | ENSG00000 |
| 30653359 | 30731456 | 1 | protein_coding | ENSG00000 |
| 30671635 | 30672166 | 1 | lncRNA | ENSG00000 |
| 30698207 | 30721932 | 1 | lncRNA | ENSG00000 |
| 30699998 | 30724174 | -1 | lncRNA | ENSG00000 |
| 30821815 | 30823407 | -1 | unprocessed_pseudc | ENSG00000 |
| 30827442 | 30975084 | -1 | protein_coding | ENSG00000 |
| 30834623 | 30835300 | 1 | lncRNA | ENSG00000 |
| 30854321 | 30854707 | 1 | lncRNA | ENSG00000 |
| 31071233 | 31072041 | -1 | protein_coding | ENSG00000 |
| 31097677 | 33339441 | -1 | protein_coding | ENSG00000 |
| 31348192 | 31348296 | 1 | snRNA | ENSG00000 |

| | | | | |
|---|---|---|---|---|
| 31411033 | 31413701 | 1 | lncRNA | ENSG0000C |
| 32102065 | 32102176 | 1 | rRNA_pseudogene | ENSG0000C |
| 32205902 | 32206738 | 1 | processed_pseudoge | ENSG0000C |
| 32583656 | 32583752 | -1 | miRNA | ENSG0000C |
| 32641474 | 32641559 | -1 | miRNA | ENSG0000C |
| 32754940 | 32756400 | 1 | lncRNA | ENSG0000C |
| 33041395 | 33041721 | 1 | processed_pseudoge | ENSG0000C |
| 33410944 | 33412173 | 1 | processed_pseudoge | ENSG0000C |
| 33718958 | 33726013 | -1 | lncRNA | ENSG0000C |
| 33726337 | 34346527 | 1 | lncRNA | ENSG0000C |
| 34129756 | 34132314 | -1 | protein_coding | ENSG0000C |
| 34147036 | 34147586 | 1 | processed_pseudoge | ENSG0000C |
| 34206725 | 34415537 | -1 | lncRNA | ENSG0000C |
| 34387722 | 34388464 | 1 | processed_pseudoge | ENSG0000C |
| 34516920 | 34517358 | 1 | processed_pseudoge | ENSG0000C |
| 34627075 | 34657285 | -1 | protein_coding | ENSG0000C |
| 34717440 | 34717992 | -1 | processed_pseudoge | ENSG0000C |
| 34768327 | 34768688 | -1 | unprocessed_pseudc | ENSG0000C |
| 34942796 | 34944915 | 1 | protein_coding | ENSG0000C |
| 35058927 | 35059974 | 1 | processed_pseudoge | ENSG0000C |
| 35594173 | 35594273 | 1 | snRNA | ENSG0000C |
| 35626142 | 35627064 | 1 | processed_pseudoge | ENSG0000C |
| 35710911 | 35711942 | 1 | processed_pseudoge | ENSG0000C |
| 35741600 | 35744582 | 1 | processed_pseudoge | ENSG0000C |
| 35761499 | 35762479 | 1 | processed_pseudoge | ENSG0000C |
| 35798342 | 35803735 | 1 | protein_coding | ENSG0000C |
| 35864858 | 35869631 | 1 | processed_pseudoge | ENSG0000C |
| 35919734 | 36385317 | 1 | protein_coding | ENSG0000C |
| 36021939 | 36022045 | -1 | snRNA | ENSG0000C |
| 36211096 | 36211635 | 1 | processed_pseudoge | ENSG0000C |
| 36365626 | 36440292 | -1 | lncRNA | ENSG0000C |
| 36374793 | 36375183 | -1 | processed_pseudoge | ENSG0000C |
| 36678678 | 36866689 | -1 | lncRNA | ENSG0000C |
| 36719918 | 36720221 | 1 | processed_pseudoge | ENSG0000C |
| 36794500 | 36795460 | -1 | unprocessed_pseudc | ENSG0000C |
| 36957913 | 36958139 | 1 | unprocessed_pseudc | ENSG0000C |
| 36962196 | 36963425 | -1 | processed_pseudoge | ENSG0000C |
| 36985523 | 36986093 | -1 | unprocessed_pseudc | ENSG0000C |
| 37008397 | 37011666 | 1 | protein_coding | ENSG0000C |
| 37043023 | 37043571 | -1 | unprocessed_pseudc | ENSG0000C |
| 37077851 | 37078401 | -1 | unprocessed_pseudc | ENSG0000C |
| 37291168 | 37292227 | 1 | processed_pseudoge | ENSG0000C |
| 37349275 | 37457295 | 1 | protein_coding | ENSG0000C |
| 37349330 | 38687674 | 1 | protein_coding | ENSG0000C |
| 37441523 | 37442068 | 1 | unprocessed_pseudc | ENSG0000C |
| 37492021 | 37492548 | 1 | unprocessed_pseudc | ENSG0000C |
| 37505310 | 37505860 | 1 | unprocessed_pseudc | ENSG0000C |
| 37541077 | 37545262 | -1 | processed_pseudoge | ENSG0000C |
| 37571569 | 37684463 | 1 | protein_coding | ENSG0000C |
| 37685791 | 37732130 | 1 | protein_coding | ENSG0000C |

| | | | | |
|---|---|---|---|---|
| 37700005 | 37700277 | 1 | unprocessed_pseudc | ENSG00000( |
| 37780011 | 37813461 | 1 | protein_coding | ENSG00000( |
| 37836757 | 37847571 | -1 | protein_coding | ENSG00000( |
| 37870109 | 37870215 | 1 | snRNA | ENSG00000( |
| 37906147 | 37949405 | 1 | lncRNA | ENSG00000( |
| 37945884 | 37946775 | 1 | processed_pseudoge | ENSG00000( |
| 37990779 | 37991314 | 1 | protein_coding | ENSG00000( |
| 37994252 | 37994918 | 1 | protein_coding | ENSG00000( |
| 38006553 | 38128819 | 1 | protein_coding | ENSG00000( |
| 38023895 | 38023986 | -1 | miRNA | ENSG00000( |
| 38149336 | 38220924 | -1 | protein_coding | ENSG00000( |
| 38221391 | 38223667 | 1 | lncRNA | ENSG00000( |
| 38269170 | 38327544 | -1 | protein_coding | ENSG00000( |
| 38308112 | 38308241 | -1 | snoRNA | ENSG00000( |
| 38352586 | 38421446 | 1 | protein_coding | ENSG00000( |
| 38395369 | 38395895 | -1 | processed_pseudoge | ENSG00000( |
| 38479215 | 38479913 | 1 | processed_pseudoge | ENSG00000( |
| 38484752 | 38485264 | -1 | processed_pseudoge | ENSG00000( |
| 38531923 | 38534174 | -1 | processed_pseudoge | ENSG00000( |
| 38561370 | 38688920 | 1 | protein_coding | ENSG00000( |
| 38608215 | 38608276 | 1 | snRNA | ENSG00000( |
| 38770331 | 38799658 | 1 | lncRNA | ENSG00000( |
| 38779584 | 38800678 | -1 | lncRNA | ENSG00000( |
| 38801432 | 38806537 | 1 | protein_coding | ENSG00000( |
| 38801568 | 38803883 | -1 | lncRNA | ENSG00000( |
| 39081180 | 39081285 | -1 | snRNA | ENSG00000( |
| 39226103 | 39299786 | -1 | lncRNA | ENSG00000( |
| 39304956 | 39327362 | -1 | lncRNA | ENSG00000( |
| 39367285 | 39391807 | -1 | lncRNA | ENSG00000( |
| 39401252 | 39434824 | -1 | lncRNA | ENSG00000( |
| 39661216 | 39661321 | 1 | miRNA | ENSG00000( |
| 39786524 | 39786823 | -1 | misc_RNA | ENSG00000( |
| 39787132 | 39788136 | -1 | processed_pseudoge | ENSG00000( |
| 39791580 | 39791904 | -1 | processed_pseudoge | ENSG00000( |
| 39837536 | 39848358 | -1 | lncRNA | ENSG00000( |
| 39837561 | 39837613 | 1 | miRNA | ENSG00000( |
| 39865424 | 39865888 | 1 | processed_pseudoge | ENSG00000( |
| 39908309 | 39914419 | 1 | lncRNA | ENSG00000( |
| 40009276 | 40012182 | 1 | lncRNA | ENSG00000( |
| 40049815 | 40177329 | -1 | protein_coding | ENSG00000( |
| 40262917 | 40299061 | 1 | lncRNA | ENSG00000( |
| 40359183 | 40360726 | -1 | processed_pseudoge | ENSG00000( |
| 40492844 | 40492941 | 1 | misc_RNA | ENSG00000( |
| 40514895 | 40515508 | 1 | processed_pseudoge | ENSG00000( |
| 40535502 | 40535563 | -1 | snRNA | ENSG00000( |
| 40579372 | 40606848 | 1 | protein_coding | ENSG00000( |
| 40620701 | 40622109 | -1 | lncRNA | ENSG00000( |
| 40623566 | 40624136 | 1 | protein_coding | ENSG00000( |
| 40626921 | 40647561 | -1 | protein_coding | ENSG00000( |
| 40648305 | 40735858 | -1 | protein_coding | ENSG00000( |

| | | | | |
|---|---|---|---|---|
| 40735400 | 40738701 | 1 | lncRNA | ENSG00000( |
| 40758483 | 40759050 | 1 | processed_pseudoge | ENSG00000( |
| 40760104 | 40761270 | -1 | processed_pseudoge | ENSG00000( |
| 40831844 | 40832562 | -1 | processed_pseudoge | ENSG00000( |
| 40834485 | 40836005 | 1 | processed_pseudoge | ENSG00000( |
| 40890647 | 40891537 | -1 | processed_pseudoge | ENSG00000( |
| 40907987 | 40908391 | 1 | processed_pseudoge | ENSG00000( |
| 40934982 | 40935864 | -1 | processed_pseudoge | ENSG00000( |
| 41008427 | 41029306 | -1 | lncRNA | ENSG00000( |
| 41076999 | 41077750 | 1 | processed_pseudoge | ENSG00000( |
| 41085445 | 41236579 | 1 | protein_coding | ENSG00000( |
| 41275739 | 41276778 | 1 | lncRNA | ENSG00000( |
| 41292798 | 41292916 | 1 | rRNA_pseudogene | ENSG00000( |
| 41316488 | 41316589 | 1 | misc_RNA | ENSG00000( |
| 41322601 | 41323111 | -1 | processed_pseudoge | ENSG00000( |
| 41333284 | 41364472 | 1 | protein_coding | ENSG00000( |
| 41345576 | 41345876 | -1 | misc_RNA | ENSG00000( |
| 41447434 | 41475710 | 1 | protein_coding | ENSG00000( |
| 41472264 | 41472653 | -1 | processed_pseudoge | ENSG00000( |
| 41514934 | 41923645 | -1 | protein_coding | ENSG00000( |
| 41520036 | 41522336 | 1 | lncRNA | ENSG00000( |
| 41570923 | 41571029 | -1 | snRNA | ENSG00000( |
| 41675760 | 41676494 | -1 | processed_pseudoge | ENSG00000( |
| 41683168 | 41683462 | -1 | misc_RNA | ENSG00000( |
| 41688973 | 41697275 | 1 | protein_coding | ENSG00000( |
| 41724181 | 41730130 | 1 | protein_coding | ENSG00000( |
| 41748222 | 41748608 | 1 | processed_pseudoge | ENSG00000( |
| 41790192 | 41790250 | -1 | processed_pseudoge | ENSG00000( |
| 41919254 | 41919349 | -1 | snRNA | ENSG00000( |
| 42047883 | 42054836 | -1 | lncRNA | ENSG00000( |
| 42098754 | 42099173 | -1 | processed_pseudoge | ENSG00000( |
| 42139425 | 42139530 | -1 | snRNA | ENSG00000( |
| 42150511 | 42150744 | -1 | processed_pseudoge | ENSG00000( |
| 42174279 | 42174385 | -1 | snRNA | ENSG00000( |
| 42252459 | 42699385 | 1 | lncRNA | ENSG00000( |
| 42285130 | 42285925 | 1 | TEC | ENSG00000( |
| 42723209 | 42755892 | -1 | lncRNA | ENSG00000( |
| 42777366 | 42778163 | -1 | protein_coding | ENSG00000( |
| 43000174 | 43000396 | 1 | processed_pseudoge | ENSG00000( |
| 43176994 | 43226598 | 1 | lncRNA | ENSG00000( |
| 43278125 | 43279658 | -1 | processed_pseudoge | ENSG00000( |
| 43279560 | 43438657 | 1 | lncRNA | ENSG00000( |
| 43407665 | 43408559 | -1 | processed_pseudoge | ENSG00000( |
| 43654907 | 43746824 | 1 | protein_coding | ENSG00000( |
| 43766610 | 43882450 | -1 | protein_coding | ENSG00000( |
| 43948776 | 43973395 | -1 | protein_coding | ENSG00000( |
| 43949732 | 43971582 | 1 | lncRNA | ENSG00000( |
| 44029332 | 44030921 | 1 | processed_pseudoge | ENSG00000( |
| 44147872 | 44343672 | -1 | protein_coding | ENSG00000( |
| 44283916 | 44286246 | 1 | processed_pseudoge | ENSG00000( |

| | | | | |
|---|---|---|---|---|
| 44309314 | 44310476 | -1 | processed_pseudoge | ENSG0000C |
| 44401603 | 44401704 | 1 | misc_RNA | ENSG0000C |
| 44466691 | 44468955 | 1 | processed_pseudoge | ENSG0000C |
| 44476027 | 44477422 | -1 | processed_pseudoge | ENSG0000C |
| 44523639 | 44542859 | -1 | protein_coding | ENSG0000C |
| 44633692 | 44635232 | 1 | processed_pseudoge | ENSG0000C |
| 44649129 | 44649719 | 1 | processed_pseudoge | ENSG0000C |
| 44741087 | 44741964 | 1 | processed_pseudoge | ENSG0000C |
| 44772773 | 44773061 | -1 | misc_RNA | ENSG0000C |
| 44844021 | 44844888 | 1 | protein_coding | ENSG0000C |
| 44873177 | 45112602 | 1 | protein_coding | ENSG0000C |
| 45148373 | 45200901 | -1 | protein_coding | ENSG0000C |
| 45183251 | 45333917 | 1 | lncRNA | ENSG0000C |
| 45381527 | 45381995 | -1 | processed_pseudoge | ENSG0000C |
| 45505388 | 45632121 | 1 | lncRNA | ENSG0000C |
| 45632292 | 45633733 | -1 | processed_pseudoge | ENSG0000C |
| 45730752 | 45731688 | 1 | processed_pseudoge | ENSG0000C |
| 45745211 | 45770274 | -1 | lncRNA | ENSG0000C |
| 45746157 | 45746266 | -1 | miRNA | ENSG0000C |
| 45747015 | 45747124 | -1 | miRNA | ENSG0000C |
| 45764772 | 45765299 | -1 | lncRNA | ENSG0000C |
| 45847042 | 45851541 | -1 | lncRNA | ENSG0000C |
| 45892912 | 45892979 | -1 | snoRNA | ENSG0000C |
| 45913118 | 45913871 | 1 | processed_pseudoge | ENSG0000C |
| 45913267 | 45917040 | -1 | lncRNA | ENSG0000C |
| 45951762 | 45952333 | -1 | processed_pseudoge | ENSG0000C |
| 45983365 | 45984364 | 1 | processed_pseudoge | ENSG0000C |
| 46058751 | 46059528 | -1 | processed_pseudoge | ENSG0000C |
| 46073651 | 46073935 | -1 | misc_RNA | ENSG0000C |
| 46256759 | 46327680 | -1 | lncRNA | ENSG0000C |
| 46288016 | 46289128 | -1 | processed_pseudoge | ENSG0000C |
| 46400686 | 46402957 | -1 | lncRNA | ENSG0000C |
| 46418450 | 46420079 | -1 | processed_pseudoge | ENSG0000C |
| 46439709 | 46440714 | -1 | processed_pseudoge | ENSG0000C |
| 46447292 | 46497422 | 1 | protein_coding | ENSG0000C |
| 46454097 | 46457199 | -1 | processed_pseudoge | ENSG0000C |
| 46497727 | 46545457 | -1 | protein_coding | ENSG0000C |
| 46517766 | 46517868 | 1 | snRNA | ENSG0000C |
| 46545438 | 46548841 | 1 | lncRNA | ENSG0000C |
| 46553522 | 46553775 | -1 | misc_RNA | ENSG0000C |
| 46560928 | 46562749 | -1 | processed_pseudoge | ENSG0000C |
| 46573765 | 46598496 | 1 | protein_coding | ENSG0000C |
| 46599251 | 46759118 | -1 | protein_coding | ENSG0000C |
| 46646594 | 46647330 | -1 | processed_pseudoge | ENSG0000C |
| 46684765 | 46685707 | -1 | processed_pseudoge | ENSG0000C |
| 46725164 | 46725818 | -1 | processed_pseudoge | ENSG0000C |
| 46837043 | 46882358 | 1 | protein_coding | ENSG0000C |
| 46887417 | 46899703 | -1 | lncRNA | ENSG0000C |
| 46912276 | 47061242 | 1 | protein_coding | ENSG0000C |
| 47078355 | 47093314 | 1 | protein_coding | ENSG0000C |

| | | | | |
|---|---|---|---|---|
| 47087506 | 47087612 | 1 | snRNA | ENSG0000C |
| 47132671 | 47132826 | 1 | snRNA | ENSG0000C |
| 47142216 | 47145504 | -1 | protein_coding | ENSG0000C |
| 47145221 | 47186813 | 1 | protein_coding | ENSG0000C |
| 47156638 | 47158199 | -1 | processed_pseudoge | ENSG0000C |
| 47190861 | 47215128 | 1 | protein_coding | ENSG0000C |
| 47204921 | 47205865 | 1 | lncRNA | ENSG0000C |
| 47217860 | 47229997 | 1 | protein_coding | ENSG0000C |
| 47232690 | 47248328 | 1 | protein_coding | ENSG0000C |
| 47273366 | 47274390 | -1 | processed_pseudoge | ENSG0000C |
| 47280811 | 47281063 | -1 | misc_RNA | ENSG0000C |
| 47297852 | 47298721 | 1 | lncRNA | ENSG0000C |
| 47370583 | 47414305 | 1 | protein_coding | ENSG0000C |
| 47388649 | 47388777 | 1 | snoRNA | ENSG0000C |
| 47389961 | 47392300 | -1 | processed_pseudoge | ENSG0000C |
| 47438736 | 47439774 | -1 | processed_pseudoge | ENSG0000C |
| 47445879 | 47482946 | -1 | protein_coding | ENSG0000C |
| 47476822 | 47477152 | -1 | misc_RNA | ENSG0000C |
| 47483571 | 47484823 | 1 | lncRNA | ENSG0000C |
| 47512602 | 47513483 | -1 | processed_pseudoge | ENSG0000C |
| 47561100 | 47571920 | 1 | protein_coding | ENSG0000C |
| 47571901 | 47619857 | -1 | protein_coding | ENSG0000C |
| 47575128 | 47628742 | -1 | lncRNA | ENSG0000C |
| 47582408 | 47586789 | 1 | protein_coding | ENSG0000C |
| 47587429 | 47587505 | 1 | miRNA | ENSG0000C |
| 47623172 | 47630305 | -1 | protein_coding | ENSG0000C |
| 47635521 | 47650604 | -1 | protein_coding | ENSG0000C |
| 47651796 | 47659180 | -1 | protein_coding | ENSG0000C |
| 47658833 | 47692326 | 1 | lncRNA | ENSG0000C |
| 47707191 | 47753456 | 1 | transcribed_unproce | ENSG0000C |
| 47718002 | 47719099 | -1 | processed_pseudoge | ENSG0000C |
| 47776827 | 47778697 | -1 | processed_pseudoge | ENSG0000C |
| 47803296 | 47804789 | 1 | processed_pseudoge | ENSG0000C |
| 47836902 | 48002561 | 1 | protein_coding | ENSG0000C |
| 47840963 | 47841702 | -1 | processed_pseudoge | ENSG0000C |
| 47974851 | 48003978 | -1 | protein_coding | ENSG0000C |
| 47983356 | 48071658 | -1 | protein_coding | ENSG0000C |
| 48004336 | 48009729 | 1 | protein_coding | ENSG0000C |
| 48056310 | 48066583 | 1 | lncRNA | ENSG0000C |
| 48071226 | 48078721 | 1 | lncRNA | ENSG0000C |
| 48107992 | 48120210 | 1 | transcribed_unproce | ENSG0000C |
| 48130657 | 48132613 | 1 | protein_coding | ENSG0000C |
| 48135658 | 48136159 | -1 | processed_pseudoge | ENSG0000C |
| 48153868 | 48155941 | -1 | processed_pseudoge | ENSG0000C |
| 48153980 | 48154074 | -1 | snRNA | ENSG0000C |
| 48156367 | 48166350 | 1 | unprocessed_pseudc | ENSG0000C |
| 48177731 | 48178023 | -1 | processed_pseudoge | ENSG0000C |
| 48186220 | 48196763 | -1 | protein_coding | ENSG0000C |
| 48197221 | 48199431 | 1 | processed_pseudoge | ENSG0000C |
| 48206258 | 48206374 | 1 | rRNA_pseudogene | ENSG0000C |

| | | | | |
|---|---|---|---|---|
| 48212791 | 48218010 | 1 | processed_pseudoge | ENSG00000 |
| 48228833 | 48228959 | -1 | processed_pseudoge | ENSG00000 |
| 48244894 | 48245745 | 1 | processed_pseudoge | ENSG00000 |
| 48255317 | 48267444 | 1 | protein_coding | ENSG00000 |
| 48274904 | 48275047 | -1 | processed_pseudoge | ENSG00000 |
| 48276704 | 48277578 | -1 | unprocessed_pseudo | ENSG00000 |
| 48279423 | 48281792 | 1 | processed_pseudoge | ENSG00000 |
| 48296816 | 48306179 | -1 | transcribed_unproce | ENSG00000 |
| 48306769 | 48309055 | 1 | processed_pseudoge | ENSG00000 |
| 48322349 | 48326683 | 1 | unprocessed_pseudo | ENSG00000 |
| 48333675 | 48334439 | -1 | lncRNA | ENSG00000 |
| 48336899 | 48337169 | -1 | processed_pseudoge | ENSG00000 |
| 48346427 | 48356707 | -1 | protein_coding | ENSG00000 |
| 48357125 | 48359495 | 1 | processed_pseudoge | ENSG00000 |
| 48371905 | 48372095 | 1 | processed_pseudoge | ENSG00000 |
| 48383516 | 48393347 | 1 | protein_coding | ENSG00000 |
| 48402078 | 48411910 | -1 | protein_coding | ENSG00000 |
| 48412303 | 48415285 | 1 | processed_pseudoge | ENSG00000 |
| 48423342 | 48423468 | -1 | processed_pseudoge | ENSG00000 |
| 48433201 | 48441619 | 1 | unprocessed_pseudo | ENSG00000 |
| 48446893 | 48448879 | -1 | processed_pseudoge | ENSG00000 |
| 48458537 | 48470260 | -1 | protein_coding | ENSG00000 |
| 48476021 | 48486364 | 1 | protein_coding | ENSG00000 |
| 48506523 | 48508838 | -1 | lncRNA | ENSG00000 |
| 48508962 | 48520814 | 1 | protein_coding | ENSG00000 |
| 48521799 | 48528716 | 1 | protein_coding | ENSG00000 |
| 48521806 | 48545469 | 1 | protein_coding | ENSG00000 |
| 48524480 | 48524806 | 1 | misc_RNA | ENSG00000 |
| 48539714 | 48562609 | 1 | protein_coding | ENSG00000 |
| 48568014 | 48574860 | -1 | lncRNA | ENSG00000 |
| 48574449 | 48581162 | 1 | protein_coding | ENSG00000 |
| 48579774 | 48581157 | -1 | lncRNA | ENSG00000 |
| 48580741 | 48581165 | 1 | TEC | ENSG00000 |
| 48583093 | 48583339 | -1 | processed_pseudoge | ENSG00000 |
| 48590042 | 48608869 | 1 | protein_coding | ENSG00000 |
| 48636165 | 48637135 | 1 | processed_pseudoge | ENSG00000 |
| 48676596 | 48691427 | 1 | protein_coding | ENSG00000 |
| 48695554 | 48709012 | 1 | protein_coding | ENSG00000 |
| 48698963 | 48737163 | -1 | lncRNA | ENSG00000 |
| 48724455 | 48724561 | -1 | snRNA | ENSG00000 |
| 48761747 | 48773648 | 1 | protein_coding | ENSG00000 |
| 48775644 | 48776970 | -1 | transcribed_processe | ENSG00000 |
| 48776965 | 48777071 | -1 | snRNA | ENSG00000 |
| 48786554 | 48794311 | 1 | protein_coding | ENSG00000 |
| 48801377 | 48824982 | 1 | protein_coding | ENSG00000 |
| 48826513 | 48830138 | 1 | protein_coding | ENSG00000 |
| 48831096 | 48835610 | -1 | protein_coding | ENSG00000 |
| 48890197 | 48903143 | 1 | protein_coding | ENSG00000 |
| 48893447 | 48898143 | -1 | protein_coding | ENSG00000 |
| 48903180 | 48911958 | -1 | protein_coding | ENSG00000 |

| | | | | |
|---|---|---|---|---|
| 48913182 | 48919024 | -1 | protein_coding | ENSG0000C |
| 48922028 | 48958386 | -1 | protein_coding | ENSG0000C |
| 48939992 | 48940808 | 1 | TEC | ENSG0000C |
| 48959179 | 48959282 | -1 | snRNA | ENSG0000C |
| 48961378 | 48971844 | -1 | protein_coding | ENSG0000C |
| 48973720 | 49002264 | -1 | protein_coding | ENSG0000C |
| 49028726 | 49043410 | -1 | protein_coding | ENSG0000C |
| 49053572 | 49069857 | 1 | protein_coding | ENSG0000C |
| 49071161 | 49074002 | -1 | protein_coding | ENSG0000C |
| 49071470 | 49079887 | -1 | protein_coding | ENSG0000C |
| 49074433 | 49101170 | -1 | protein_coding | ENSG0000C |
| 49082028 | 49082165 | 1 | snRNA | ENSG0000C |
| 49113407 | 49123735 | -1 | protein_coding | ENSG0000C |
| 49152651 | 49152944 | 1 | misc_RNA | ENSG0000C |
| 49155242 | 49157351 | -1 | unprocessed_pseudc | ENSG0000C |
| 49162564 | 49168483 | 1 | protein_coding | ENSG0000C |
| 49165576 | 49165972 | -1 | TEC | ENSG0000C |
| 49171898 | 49175235 | 1 | protein_coding | ENSG0000C |
| 49174802 | 49186528 | -1 | protein_coding | ENSG0000C |
| 49187804 | 49200259 | -1 | protein_coding | ENSG0000C |
| 49198966 | 49202454 | 1 | lncRNA | ENSG0000C |
| 49205063 | 49233371 | -1 | protein_coding | ENSG0000C |
| 49233956 | 49234501 | -1 | processed_pseudoge | ENSG0000C |
| 49235470 | 49250520 | 1 | protein_coding | ENSG0000C |
| 49250436 | 49270477 | -1 | protein_coding | ENSG0000C |
| 49262866 | 49270521 | -1 | lncRNA | ENSG0000C |
| 49269793 | 49301461 | 1 | protein_coding | ENSG0000C |
| 49273054 | 49275768 | 1 | lncRNA | ENSG0000C |
| 49279677 | 49281440 | -1 | lncRNA | ENSG0000C |
| 49303669 | 49319844 | 1 | protein_coding | ENSG0000C |
| 49322057 | 49329384 | 1 | protein_coding | ENSG0000C |
| 49331616 | 49338952 | 1 | protein_coding | ENSG0000C |
| 49341192 | 49345922 | 1 | protein_coding | ENSG0000C |
| 49529869 | 49529985 | 1 | protein_coding | ENSG0000C |
| 49532211 | 49539538 | 1 | protein_coding | ENSG0000C |
| 49541767 | 49549094 | 1 | protein_coding | ENSG0000C |
| 49551278 | 49568218 | 1 | protein_coding | ENSG0000C |
| 49551333 | 49558649 | 1 | protein_coding | ENSG0000C |
| 49570434 | 49577754 | 1 | protein_coding | ENSG0000C |
| 49579983 | 49587301 | 1 | protein_coding | ENSG0000C |
| 49589496 | 49608536 | 1 | protein_coding | ENSG0000C |
| 49589529 | 49596827 | 1 | protein_coding | ENSG0000C |
| 49632500 | 49633349 | -1 | processed_pseudoge | ENSG0000C |
| 49664844 | 49668680 | -1 | processed_pseudoge | ENSG0000C |
| 49687447 | 49695984 | -1 | protein_coding | ENSG0000C |
| 49829260 | 49834264 | 1 | protein_coding | ENSG0000C |
| 49876724 | 49879241 | -1 | lncRNA | ENSG0000C |
| 49879484 | 49882558 | 1 | protein_coding | ENSG0000C |
| 49911620 | 49912362 | -1 | processed_pseudoge | ENSG0000C |
| 49922596 | 50099235 | 1 | protein_coding | ENSG0000C |

| | | | | |
|---|---|---|---|---|
| 49945336 | 49945442 | -1 | snRNA | ENSG0000C |
| 50003148 | 50003238 | 1 | miRNA | ENSG0000C |
| 50003503 | 50003588 | 1 | miRNA | ENSG0000C |
| 50008431 | 50008514 | 1 | miRNA | ENSG0000C |
| 50008964 | 50009028 | 1 | miRNA | ENSG0000C |
| 50009722 | 50009805 | 1 | miRNA | ENSG0000C |
| 50010672 | 50010750 | 1 | miRNA | ENSG0000C |
| 50013241 | 50013337 | 1 | miRNA | ENSG0000C |
| 50014598 | 50014683 | 1 | miRNA | ENSG0000C |
| 50156159 | 50156371 | 1 | processed_pseudoge | ENSG0000C |
| 50171197 | 50171297 | -1 | misc_RNA | ENSG0000C |
| 50190777 | 50201007 | -1 | protein_coding | ENSG0000C |
| 50202713 | 50351910 | 1 | protein_coding | ENSG0000C |
| 50365409 | 50470825 | -1 | protein_coding | ENSG0000C |
| 50591647 | 50814302 | -1 | protein_coding | ENSG0000C |
| 50649641 | 50649740 | 1 | snRNA | ENSG0000C |
| 50905438 | 50905866 | 1 | processed_pseudoge | ENSG0000C |
| 50910784 | 50916607 | 1 | protein_coding | ENSG0000C |
| 50931114 | 50931548 | -1 | processed_pseudoge | ENSG0000C |
| 51030422 | 51031357 | -1 | unprocessed_pseudc | ENSG0000C |
| 51082818 | 51091228 | 1 | unprocessed_pseudc | ENSG0000C |
| 51095836 | 51224432 | -1 | lncRNA | ENSG0000C |
| 51162864 | 51163677 | 1 | unprocessed_pseudc | ENSG0000C |
| 51190598 | 51396462 | -1 | lncRNA | ENSG0000C |
| 51325790 | 51330662 | 1 | lncRNA | ENSG0000C |
| 51332231 | 51337525 | 1 | protein_coding | ENSG0000C |
| 51395959 | 51465661 | 1 | lncRNA | ENSG0000C |
| 51406948 | 51408843 | 1 | protein_coding | ENSG0000C |
| 51490011 | 51496592 | -1 | protein_coding | ENSG0000C |
| 51498490 | 51511005 | -1 | lncRNA | ENSG0000C |
| 51617024 | 51618968 | -1 | protein_coding | ENSG0000C |
| 51681212 | 51682831 | -1 | protein_coding | ENSG0000C |
| 51710512 | 51712131 | 1 | protein_coding | ENSG0000C |
| 51743442 | 51746232 | 1 | protein_coding | ENSG0000C |
| 51803007 | 51902357 | 1 | protein_coding | ENSG0000C |
| 51856968 | 51858066 | 1 | processed_pseudoge | ENSG0000C |
| 51870706 | 51870809 | -1 | snRNA | ENSG0000C |
| 51903338 | 51903850 | -1 | unprocessed_pseudc | ENSG0000C |
| 51921864 | 51922499 | -1 | processed_pseudoge | ENSG0000C |
| 51923552 | 51923740 | 1 | processed_pseudoge | ENSG0000C |
| 51979223 | 51980012 | -1 | processed_pseudoge | ENSG0000C |
| 52050860 | 52054255 | -1 | lncRNA | ENSG0000C |
| 52053176 | 52055684 | 1 | lncRNA | ENSG0000C |
| 52061827 | 52069248 | -1 | protein_coding | ENSG0000C |
| 52063347 | 52063474 | -1 | snoRNA | ENSG0000C |
| 52184823 | 52192268 | 1 | protein_coding | ENSG0000C |
| 52190621 | 52190748 | 1 | snoRNA | ENSG0000C |
| 52195836 | 52265919 | -1 | lncRNA | ENSG0000C |
| 52199840 | 52203235 | 1 | lncRNA | ENSG0000C |
| 52326166 | 52326762 | 1 | processed_pseudoge | ENSG0000C |

| | | | | |
|---|---|---|---|---|
| 52336557 | 52336642 | -1 | miRNA | ENSG0000C |
| 52369021 | 52375680 | 1 | protein_coding | ENSG0000C |
| 52382053 | 52385614 | 1 | unprocessed_pseudc | ENSG0000C |
| 52409772 | 52410432 | -1 | unprocessed_pseudc | ENSG0000C |
| 52422069 | 52459066 | 1 | unprocessed_pseudc | ENSG0000C |
| 52448587 | 52451138 | -1 | unprocessed_pseudc | ENSG0000C |
| 52480836 | 52481131 | 1 | processed_pseudoge | ENSG0000C |
| 52481515 | 52483950 | 1 | unprocessed_pseudc | ENSG0000C |
| 52485046 | 52485912 | -1 | processed_pseudoge | ENSG0000C |
| 52495808 | 52500812 | 1 | protein_coding | ENSG0000C |
| 52509903 | 52510182 | 1 | processed_pseudoge | ENSG0000C |
| 52512077 | 52520803 | -1 | protein_coding | ENSG0000C |
| 52531709 | 52532051 | -1 | processed_pseudoge | ENSG0000C |
| 52545151 | 52547695 | 1 | unprocessed_pseudc | ENSG0000C |
| 52561769 | 52562301 | -1 | unprocessed_pseudc | ENSG0000C |
| 52583989 | 52586312 | 1 | unprocessed_pseudc | ENSG0000C |
| 52598494 | 52603289 | 1 | unprocessed_pseudc | ENSG0000C |
| 52606535 | 52611758 | -1 | unprocessed_pseudc | ENSG0000C |
| 52612656 | 52613523 | 1 | processed_pseudoge | ENSG0000C |
| 52624998 | 52632895 | 1 | unprocessed_pseudc | ENSG0000C |
| 52644061 | 52654900 | -1 | protein_coding | ENSG0000C |
| 52655207 | 52657254 | 1 | processed_pseudoge | ENSG0000C |
| 52665231 | 52665347 | 1 | rRNA_pseudogene | ENSG0000C |
| 52672718 | 52677051 | 1 | unprocessed_pseudc | ENSG0000C |
| 52687363 | 52687633 | -1 | processed_pseudoge | ENSG0000C |
| 52696896 | 52707189 | -1 | protein_coding | ENSG0000C |
| 52707602 | 52709933 | 1 | processed_pseudoge | ENSG0000C |
| 52722338 | 52722524 | 1 | processed_pseudoge | ENSG0000C |
| 52735901 | 52736107 | -1 | processed_pseudoge | ENSG0000C |
| 52751132 | 52790305 | 1 | protein_coding | ENSG0000C |
| 52770809 | 52771080 | 1 | processed_pseudoge | ENSG0000C |
| 52781452 | 52785786 | -1 | unprocessed_pseudc | ENSG0000C |
| 52796144 | 52797427 | -1 | protein_coding | ENSG0000C |
| 52801400 | 52801529 | -1 | processed_pseudoge | ENSG0000C |
| 52812204 | 52818301 | 1 | protein_coding | ENSG0000C |
| 52824269 | 52829267 | -1 | unprocessed_pseudc | ENSG0000C |
| 52832704 | 52833899 | -1 | processed_pseudoge | ENSG0000C |
| 52862525 | 52868083 | -1 | protein_coding | ENSG0000C |
| 52891306 | 52908560 | 1 | protein_coding | ENSG0000C |
| 52925956 | 52929574 | 1 | lncRNA | ENSG0000C |
| 52926402 | 52995472 | -1 | protein_coding | ENSG0000C |
| 53049091 | 53080615 | 1 | protein_coding | ENSG0000C |
| 53065053 | 53065548 | -1 | processed_pseudoge | ENSG0000C |
| 53082367 | 53088540 | 1 | protein_coding | ENSG0000C |
| 53093710 | 53094858 | -1 | lncRNA | ENSG0000C |
| 53094145 | 53170914 | 1 | protein_coding | ENSG0000C |
| 53113018 | 53113274 | -1 | processed_pseudoge | ENSG0000C |
| 53142832 | 53143913 | -1 | processed_pseudoge | ENSG0000C |
| 53164391 | 53164557 | -1 | processed_pseudoge | ENSG0000C |
| 53191321 | 53225422 | -1 | protein_coding | ENSG0000C |

| | | | | |
|---|---|---|---|---|
| 53195411 | 53195488 | -1 | miRNA | ENSG0000C |
| 53198889 | 53198945 | -1 | miRNA | ENSG0000C |
| 53212408 | 53214679 | -1 | lncRNA | ENSG0000C |
| 53225828 | 53321350 | -1 | protein_coding | ENSG0000C |
| 53322990 | 53323822 | 1 | processed_pseudoge | ENSG0000C |
| 53324562 | 53324674 | 1 | misc_RNA | ENSG0000C |
| 53337783 | 53337973 | -1 | processed_pseudoge | ENSG0000C |
| 53340986 | 53341505 | 1 | processed_pseudoge | ENSG0000C |
| 53374149 | 53422728 | -1 | protein_coding | ENSG0000C |
| 53405673 | 53405765 | -1 | miRNA | ENSG0000C |
| 53422690 | 53431120 | 1 | protein_coding | ENSG0000C |
| 53431258 | 53434373 | -1 | protein_coding | ENSG0000C |
| 53432722 | 53433032 | -1 | lncRNA | ENSG0000C |
| 53456273 | 53457029 | -1 | processed_pseudoge | ENSG0000C |
| 53462209 | 53462310 | 1 | misc_RNA | ENSG0000C |
| 53532096 | 53686728 | -1 | protein_coding | ENSG0000C |
| 53556223 | 53556341 | -1 | miRNA | ENSG0000C |
| 53557192 | 53557274 | -1 | miRNA | ENSG0000C |
| 53759026 | 53759204 | 1 | processed_pseudoge | ENSG0000C |
| 53768986 | 53769219 | -1 | processed_pseudoge | ENSG0000C |
| 53807776 | 53808623 | 1 | processed_pseudoge | ENSG0000C |
| 53825887 | 53826309 | 1 | processed_pseudoge | ENSG0000C |
| 53909054 | 53909161 | 1 | rRNA_pseudogene | ENSG0000C |
| 53936676 | 54048958 | -1 | protein_coding | ENSG0000C |
| 54068324 | 54183281 | -1 | protein_coding | ENSG0000C |
| 54147372 | 54147659 | 1 | processed_pseudoge | ENSG0000C |
| 54192823 | 54358642 | -1 | protein_coding | ENSG0000C |
| 54223324 | 54224062 | 1 | processed_pseudoge | ENSG0000C |
| 54343546 | 54343652 | 1 | snRNA | ENSG0000C |
| 54440404 | 54448032 | 1 | protein_coding | ENSG0000C |
| 54445454 | 54496234 | -1 | protein_coding | ENSG0000C |
| 54530211 | 54567289 | 1 | protein_coding | ENSG0000C |
| 54671985 | 54672638 | 1 | processed_pseudoge | ENSG0000C |
| 54748918 | 54798255 | -1 | protein_coding | ENSG0000C |
| 54807599 | 54816015 | 1 | protein_coding | ENSG0000C |
| 54814370 | 54814497 | 1 | snoRNA | ENSG0000C |
| 54842014 | 54842134 | 1 | unprocessed_pseudc | ENSG0000C |
| 54920462 | 54931431 | 1 | protein_coding | ENSG0000C |
| 54927305 | 54927433 | 1 | snoRNA | ENSG0000C |
| 54932961 | 54998534 | -1 | protein_coding | ENSG0000C |
| 55000363 | 55009057 | 1 | protein_coding | ENSG0000C |
| 55009055 | 55030977 | -1 | protein_coding | ENSG0000C |
| 55044749 | 55045121 | -1 | unprocessed_pseudc | ENSG0000C |
| 55054945 | 55055838 | -1 | processed_pseudoge | ENSG0000C |
| 55075030 | 55078909 | 1 | protein_coding | ENSG0000C |
| 55089018 | 55092842 | 1 | protein_coding | ENSG0000C |
| 55101637 | 55102026 | -1 | unprocessed_pseudc | ENSG0000C |
| 55137930 | 55142433 | 1 | unprocessed_pseudc | ENSG0000C |
| 55143102 | 55161310 | -1 | protein_coding | ENSG0000C |
| 55172717 | 55176026 | 1 | unprocessed_pseudc | ENSG0000C |

| | | | | |
|---|---|---|---|---|
| 55178207 | 55178737 | -1 | processed_pseudoge | ENSG00000 |
| 55179194 | 55179999 | -1 | processed_pseudoge | ENSG00000 |
| 55180377 | 55181277 | -1 | processed_pseudoge | ENSG00000 |
| 55181392 | 55182442 | -1 | protein_coding | ENSG00000 |
| 55220355 | 55224108 | 1 | protein_coding | ENSG00000 |
| 55258412 | 55264846 | -1 | protein_coding | ENSG00000 |
| 55279839 | 55288785 | 1 | transcribed_unproce | ENSG00000 |
| 55451495 | 55451582 | 1 | miRNA | ENSG00000 |
| 55451495 | 55451582 | -1 | miRNA | ENSG00000 |
| 55452127 | 55453566 | 1 | protein_coding | ENSG00000 |
| 55484616 | 55489202 | -1 | protein_coding | ENSG00000 |
| 55520382 | 55520714 | -1 | unprocessed_pseudo | ENSG00000 |
| 55582733 | 55582834 | 1 | misc_RNA | ENSG00000 |
| 55599663 | 55600106 | -1 | processed_pseudoge | ENSG00000 |
| 55623400 | 55626192 | 1 | protein_coding | ENSG00000 |
| 55633031 | 55635656 | -1 | unprocessed_pseudo | ENSG00000 |
| 55654709 | 55657014 | -1 | transcribed_unproce | ENSG00000 |
| 55717739 | 55758774 | 1 | protein_coding | ENSG00000 |
| 55908123 | 56209271 | 1 | lncRNA | ENSG00000 |
| 55961208 | 55962482 | 1 | processed_pseudoge | ENSG00000 |
| 56232356 | 56291531 | 1 | protein_coding | ENSG00000 |
| 56242937 | 56243409 | -1 | processed_pseudoge | ENSG00000 |
| 56563639 | 56567868 | 1 | protein_coding | ENSG00000 |
| 56618391 | 56624458 | 1 | lncRNA | ENSG00000 |
| 56729241 | 56819179 | 1 | protein_coding | ENSG00000 |
| 56737242 | 56737577 | -1 | processed_pseudoge | ENSG00000 |
| 56818298 | 56995827 | -1 | protein_coding | ENSG00000 |
| 56973510 | 56974796 | 1 | lncRNA | ENSG00000 |
| 57068914 | 57069685 | -1 | processed_pseudoge | ENSG00000 |
| 57118551 | 57121546 | -1 | protein_coding | ENSG00000 |
| 57121662 | 57127243 | 1 | lncRNA | ENSG00000 |
| 57134530 | 57137625 | -1 | protein_coding | ENSG00000 |
| 57222706 | 57224791 | 1 | lncRNA | ENSG00000 |
| 57229034 | 57229415 | 1 | processed_pseudoge | ENSG00000 |
| 57286706 | 57489196 | 1 | protein_coding | ENSG00000 |
| 57392646 | 57395409 | -1 | processed_pseudoge | ENSG00000 |
| 57591652 | 57597545 | 1 | protein_coding | ENSG00000 |
| 57677067 | 57680260 | -1 | protein_coding | ENSG00000 |
| 57766320 | 57767292 | 1 | processed_pseudoge | ENSG00000 |
| 57906708 | 57910820 | -1 | protein_coding | ENSG00000 |
| 57933837 | 57934803 | -1 | processed_pseudoge | ENSG00000 |
| 57984686 | 57986133 | 1 | processed_pseudoge | ENSG00000 |
| 62779250 | 62780326 | 1 | processed_pseudoge | ENSG00000 |
| 62842542 | 62843502 | 1 | processed_pseudoge | ENSG00000 |
| 62843697 | 62844306 | 1 | processed_pseudoge | ENSG00000 |
| 62878811 | 62884842 | -1 | processed_pseudoge | ENSG00000 |
| 63087071 | 63088664 | 1 | processed_pseudoge | ENSG00000 |
| 63222993 | 63561095 | -1 | lncRNA | ENSG00000 |
| 63299247 | 63299673 | -1 | processed_pseudoge | ENSG00000 |
| 63347228 | 63351332 | -1 | protein_coding | ENSG00000 |

| | | | | |
|---|---|---|---|---|
| 63349646 | 63352178 | 1 | lncRNA | ENSG0000C |
| 63634967 | 63809274 | -1 | protein_coding | ENSG0000C |
| 63670196 | 63671502 | -1 | lncRNA | ENSG0000C |
| 63766875 | 63767400 | 1 | processed_pseudoge | ENSG0000C |
| 63786002 | 63786087 | -1 | miRNA | ENSG0000C |
| 64044305 | 64045058 | 1 | processed_pseudoge | ENSG0000C |
| 64144872 | 64147052 | -1 | processed_pseudoge | ENSG0000C |
| 64185117 | 64205708 | -1 | protein_coding | ENSG0000C |
| 64205974 | 64233855 | 1 | lncRNA | ENSG0000C |
| 64210691 | 64210988 | 1 | misc_RNA | ENSG0000C |
| 64224194 | 64230607 | -1 | protein_coding | ENSG0000C |
| 64268081 | 64395452 | -1 | protein_coding | ENSG0000C |
| 64305047 | 64306733 | -1 | processed_pseudoge | ENSG0000C |
| 64405473 | 64406136 | 1 | transcribed_unitary_ | ENSG0000C |
| 64432401 | 64434145 | 1 | processed_pseudoge | ENSG0000C |
| 64612632 | 64613262 | -1 | processed_pseudoge | ENSG0000C |
| 64623117 | 64624573 | 1 | processed_pseudoge | ENSG0000C |
| 64697878 | 64698547 | -1 | processed_pseudoge | ENSG0000C |
| 64726748 | 64727048 | -1 | processed_pseudoge | ENSG0000C |
| 64915802 | 65034713 | -1 | protein_coding | ENSG0000C |
| 65185018 | 65187265 | -1 | processed_pseudoge | ENSG0000C |
| 65270913 | 65273019 | 1 | processed_pseudoge | ENSG0000C |
| 65327988 | 65331233 | -1 | processed_pseudoge | ENSG0000C |
| 65356890 | 65357547 | -1 | processed_pseudoge | ENSG0000C |
| 65366638 | 65507887 | 1 | protein_coding | ENSG0000C |
| 65408084 | 65408672 | 1 | processed_pseudoge | ENSG0000C |
| 65469087 | 65470380 | 1 | processed_pseudoge | ENSG0000C |
| 65512582 | 65534775 | -1 | protein_coding | ENSG0000C |
| 65550898 | 65552421 | -1 | processed_pseudoge | ENSG0000C |
| 65588377 | 65741931 | 1 | protein_coding | ENSG0000C |
| 65772741 | 65773632 | -1 | processed_pseudoge | ENSG0000C |
| 65786763 | 65787169 | 1 | processed_pseudoge | ENSG0000C |
| 65794345 | 65795835 | 1 | processed_pseudoge | ENSG0000C |
| 65821255 | 65822249 | -1 | processed_pseudoge | ENSG0000C |
| 65925836 | 66001436 | 1 | lncRNA | ENSG0000C |
| 65956289 | 65957196 | -1 | processed_pseudoge | ENSG0000C |
| 66015461 | 66020422 | 1 | lncRNA | ENSG0000C |
| 66018870 | 66018979 | 1 | miRNA | ENSG0000C |
| 66021738 | 66040125 | -1 | protein_coding | ENSG0000C |
| 66055587 | 66056587 | -1 | processed_pseudoge | ENSG0000C |
| 66074835 | 66076347 | -1 | processed_pseudoge | ENSG0000C |
| 66162549 | 66268867 | 1 | protein_coding | ENSG0000C |
| 66360766 | 66361776 | -1 | processed_pseudoge | ENSG0000C |
| 66437345 | 66440668 | 1 | unprocessed_pseudc | ENSG0000C |
| 66497748 | 66499424 | 1 | processed_pseudoge | ENSG0000C |
| 66595637 | 66639298 | -1 | protein_coding | ENSG0000C |
| 66676258 | 66676364 | -1 | snRNA | ENSG0000C |
| 67373573 | 67375520 | 1 | processed_pseudoge | ENSG0000C |
| 67533163 | 67533987 | 1 | processed_pseudoge | ENSG0000C |
| 67544021 | 67730619 | 1 | protein_coding | ENSG0000C |

| | | | | |
|---|---|---|---|---|
| 67791955 | 67792895 | -1 | processed_pseudoge | ENSG00000 |
| 68013470 | 68014901 | -1 | lncRNA | ENSG00000 |
| 68042344 | 68433913 | -1 | protein_coding | ENSG00000 |
| 68070520 | 68071767 | -1 | processed_pseudoge | ENSG00000 |
| 68102068 | 68102132 | -1 | snRNA | ENSG00000 |
| 68132868 | 68133437 | 1 | processed_pseudoge | ENSG00000 |
| 68361075 | 68361187 | 1 | misc_RNA | ENSG00000 |
| 68474710 | 68476874 | 1 | processed_pseudoge | ENSG00000 |
| 68498562 | 68537282 | 1 | protein_coding | ENSG00000 |
| 68539443 | 68539549 | -1 | snRNA | ENSG00000 |
| 68600099 | 68600473 | 1 | processed_pseudoge | ENSG00000 |
| 68645326 | 68645527 | -1 | processed_pseudoge | ENSG00000 |
| 68647666 | 68725842 | 1 | protein_coding | ENSG00000 |
| 68771322 | 68772578 | 1 | processed_pseudoge | ENSG00000 |
| 68783472 | 68785066 | 1 | processed_pseudoge | ENSG00000 |
| 68829021 | 68842160 | 1 | protein_coding | ENSG00000 |
| 69160851 | 69165793 | -1 | protein_coding | ENSG00000 |
| 69174124 | 69174717 | 1 | processed_pseudoge | ENSG00000 |
| 69179557 | 69209924 | 1 | lncRNA | ENSG00000 |
| 69485343 | 69485652 | -1 | processed_pseudoge | ENSG00000 |
| 69505241 | 69532508 | 1 | protein_coding | ENSG00000 |
| 69569635 | 69576337 | -1 | lncRNA | ENSG00000 |
| 69616067 | 70039472 | 1 | protein_coding | ENSG00000 |
| 69672479 | 69672597 | 1 | rRNA_pseudogene | ENSG00000 |
| 69672479 | 69672597 | 1 | rRNA | ENSG00000 |
| 69692956 | 69693155 | 1 | snoRNA | ENSG00000 |
| 69937166 | 69938017 | -1 | processed_pseudoge | ENSG00000 |
| 70022857 | 70022923 | 1 | miRNA | ENSG00000 |
| 70037840 | 70039325 | -1 | lncRNA | ENSG00000 |
| 70040542 | 70049938 | -1 | protein_coding | ENSG00000 |
| 70062457 | 70064179 | 1 | protein_coding | ENSG00000 |
| 70120353 | 70121474 | -1 | processed_pseudoge | ENSG00000 |
| 70124972 | 70129071 | -1 | processed_pseudoge | ENSG00000 |
| 70133447 | 70166324 | 1 | protein_coding | ENSG00000 |
| 70148582 | 70149654 | -1 | lncRNA | ENSG00000 |
| 70163842 | 70165206 | -1 | lncRNA | ENSG00000 |
| 70177483 | 70205704 | 1 | protein_coding | ENSG00000 |
| 70222006 | 70222314 | -1 | misc_RNA | ENSG00000 |
| 70234655 | 70240659 | 1 | protein_coding | ENSG00000 |
| 70253042 | 70253162 | 1 | rRNA_pseudogene | ENSG00000 |
| 70258170 | 70259764 | -1 | protein_coding | ENSG00000 |
| 70268305 | 70281840 | 1 | protein_coding | ENSG00000 |
| 70282093 | 70285002 | 1 | protein_coding | ENSG00000 |
| 70286595 | 70290514 | -1 | protein_coding | ENSG00000 |
| 70290104 | 70420886 | 1 | protein_coding | ENSG00000 |
| 70361486 | 70362374 | -1 | processed_pseudoge | ENSG00000 |
| 70396279 | 70396375 | 1 | misc_RNA | ENSG00000 |
| 70423031 | 70433390 | 1 | protein_coding | ENSG00000 |
| 70427450 | 70435350 | -1 | lncRNA | ENSG00000 |
| 70444835 | 70505490 | 1 | protein_coding | ENSG00000 |

| | | | | |
|---|---|---|---|---|
| 70450879 | 70451022 | 1 | snRNA | ENSG0000C |
| 70452958 | 70455994 | -1 | lncRNA | ENSG0000C |
| 70528940 | 70908731 | -1 | protein_coding | ENSG0000C |
| 70672895 | 70673515 | -1 | processed_pseudoge | ENSG0000C |
| 70846079 | 70846295 | -1 | snoRNA | ENSG0000C |
| 70846080 | 70846295 | -1 | snoRNA | ENSG0000C |
| 70857163 | 70857502 | 1 | processed_pseudoge | ENSG0000C |
| 70925579 | 70931125 | -1 | protein_coding | ENSG0000C |
| 70962964 | 70963293 | 1 | processed_pseudoge | ENSG0000C |
| 71016531 | 71016905 | -1 | processed_pseudoge | ENSG0000C |
| 71020275 | 71020438 | -1 | snRNA | ENSG0000C |
| 71043214 | 71045470 | -1 | processed_pseudoge | ENSG0000C |
| 71056332 | 71073426 | -1 | protein_coding | ENSG0000C |
| 71095851 | 71103532 | 1 | protein_coding | ENSG0000C |
| 71103889 | 71106788 | -1 | protein_coding | ENSG0000C |
| 71103987 | 71111575 | -1 | protein_coding | ENSG0000C |
| 71107404 | 71112108 | -1 | protein_coding | ENSG0000C |
| 71118556 | 71142454 | 1 | protein_coding | ENSG0000C |
| 71144831 | 71171201 | 1 | protein_coding | ENSG0000C |
| 71183384 | 71198208 | -1 | lncRNA | ENSG0000C |
| 71215194 | 71225516 | 1 | protein_coding | ENSG0000C |
| 71239624 | 71255146 | -1 | protein_coding | ENSG0000C |
| 71283192 | 71301168 | 1 | protein_coding | ENSG0000C |
| 71301750 | 71305371 | 1 | protein_coding | ENSG0000C |
| 71314912 | 71315879 | -1 | processed_pseudoge | ENSG0000C |
| 71352418 | 71352988 | 1 | processed_pseudoge | ENSG0000C |
| 71366239 | 71532374 | 1 | protein_coding | ENSG0000C |
| 71368666 | 71368968 | 1 | misc_RNA | ENSG0000C |
| 71413834 | 71414086 | -1 | processed_pseudoge | ENSG0000C |
| 71420083 | 71420514 | -1 | processed_pseudoge | ENSG0000C |
| 71491066 | 71491167 | -1 | misc_RNA | ENSG0000C |
| 71491682 | 71492928 | -1 | transcribed_processe | ENSG0000C |
| 71527814 | 71530225 | -1 | processed_pseudoge | ENSG0000C |
| 71533104 | 71575892 | 1 | protein_coding | ENSG0000C |
| 71578411 | 71613583 | 1 | protein_coding | ENSG0000C |
| 71615916 | 71618511 | -1 | protein_coding | ENSG0000C |
| 71662998 | 71666767 | -1 | transcribed_unproce | ENSG0000C |
| 71667542 | 71671524 | 1 | protein_coding | ENSG0000C |
| 71691932 | 71692179 | 1 | processed_pseudoge | ENSG0000C |
| 71697196 | 71706455 | 1 | lncRNA | ENSG0000C |
| 71714371 | 71718151 | -1 | protein_coding | ENSG0000C |
| 71719142 | 71721278 | 1 | transcribed_unproce | ENSG0000C |
| 71736454 | 71736703 | -1 | processed_pseudoge | ENSG0000C |
| 71744828 | 71745077 | 1 | processed_pseudoge | ENSG0000C |
| 71760764 | 71763075 | -1 | transcribed_unproce | ENSG0000C |
| 71763424 | 71767204 | 1 | protein_coding | ENSG0000C |
| 71789386 | 71789636 | -1 | processed_pseudoge | ENSG0000C |
| 71848775 | 71849004 | 1 | processed_pseudoge | ENSG0000C |
| 71869763 | 71873295 | -1 | unprocessed_pseudc | ENSG0000C |
| 71910818 | 72161750 | 1 | protein_coding | ENSG0000C |

| | | | | |
|---|---|---|---|---|
| 71965972 | 71966074 | -1 | snRNA | ENSG0000C |
| 72044545 | 72044892 | 1 | processed_pseudoge | ENSG0000C |
| 72127110 | 72131901 | -1 | protein_coding | ENSG0000C |
| 72159845 | 72160346 | 1 | transcribed_processe | ENSG0000C |
| 72181353 | 72302926 | 1 | protein_coding | ENSG0000C |
| 72198712 | 72199050 | 1 | misc_RNA | ENSG0000C |
| 72204657 | 72239027 | -1 | protein_coding | ENSG0000C |
| 72255679 | 72277248 | -1 | protein_coding | ENSG0000C |
| 72284945 | 72285045 | -1 | misc_RNA | ENSG0000C |
| 72301638 | 72307187 | -1 | protein_coding | ENSG0000C |
| 72301691 | 72572843 | -1 | protein_coding | ENSG0000C |
| 72329516 | 72573101 | -1 | protein_coding | ENSG0000C |
| 72376979 | 72377169 | -1 | snRNA | ENSG0000C |
| 72578814 | 72714319 | -1 | protein_coding | ENSG0000C |
| 72660470 | 72660719 | 1 | processed_pseudoge | ENSG0000C |
| 72688950 | 72712348 | 1 | lncRNA | ENSG0000C |
| 72726122 | 72726337 | -1 | snoRNA | ENSG0000C |
| 72727977 | 72728821 | -1 | processed_pseudoge | ENSG0000C |
| 72740706 | 72740869 | -1 | snRNA | ENSG0000C |
| 72777009 | 72848802 | 1 | protein_coding | ENSG0000C |
| 72777608 | 72779097 | 1 | lncRNA | ENSG0000C |
| 72781865 | 72782660 | -1 | protein_coding | ENSG0000C |
| 72807425 | 72808210 | -1 | protein_coding | ENSG0000C |
| 72872025 | 72943814 | -1 | protein_coding | ENSG0000C |
| 72912615 | 72913401 | 1 | protein_coding | ENSG0000C |
| 72938163 | 72938958 | 1 | protein_coding | ENSG0000C |
| 72998388 | 73002856 | -1 | lncRNA | ENSG0000C |
| 73003517 | 73005713 | 1 | protein_coding | ENSG0000C |
| 73077276 | 73079512 | -1 | protein_coding | ENSG0000C |
| 73080167 | 73084635 | 1 | lncRNA | ENSG0000C |
| 73126037 | 73128080 | -1 | transcribed_processe | ENSG0000C |
| 73212299 | 73214851 | -1 | protein_coding | ENSG0000C |
| 73223124 | 73223558 | 1 | processed_pseudoge | ENSG0000C |
| 73237647 | 73238709 | -1 | processed_pseudoge | ENSG0000C |
| 73274785 | 73275515 | 1 | processed_pseudoge | ENSG0000C |
| 73397531 | 73397637 | -1 | snRNA | ENSG0000C |
| 73447254 | 73455245 | 1 | protein_coding | ENSG0000C |
| 73474563 | 73474830 | 1 | processed_pseudoge | ENSG0000C |
| 73524275 | 73563085 | -1 | transcribed_processe | ENSG0000C |
| 73535503 | 73536241 | 1 | processed_pseudoge | ENSG0000C |
| 73563200 | 73687102 | 1 | protein_coding | ENSG0000C |
| 73769248 | 73770423 | 1 | processed_pseudoge | ENSG0000C |
| 73792205 | 73829231 | 1 | lncRNA | ENSG0000C |
| 73820649 | 73852723 | -1 | lncRNA | ENSG0000C |
| 73821657 | 73821724 | -1 | misc_RNA | ENSG0000C |
| 73831145 | 73831270 | -1 | misc_RNA | ENSG0000C |
| 73850487 | 73850571 | 1 | misc_RNA | ENSG0000C |
| 73875068 | 73875635 | 1 | processed_pseudoge | ENSG0000C |
| 73940435 | 74293574 | -1 | lncRNA | ENSG0000C |
| 73944182 | 74070408 | 1 | lncRNA | ENSG0000C |

| | | | | |
|---|---|---|---|---|
| 73944332 | 73944466 | 1 | misc_RNA | ENSG00000 |
| 73944595 | 73944663 | 1 | misc_RNA | ENSG00000 |
| 73948973 | 73949558 | 1 | lncRNA | ENSG00000 |
| 73958059 | 73969485 | -1 | lncRNA | ENSG00000 |
| 74031462 | 74031820 | 1 | processed_pseudoge | ENSG00000 |
| 74066083 | 74066592 | 1 | processed_pseudoge | ENSG00000 |
| 74069276 | 74070832 | 1 | processed_pseudoge | ENSG00000 |
| 74105572 | 74107958 | 1 | processed_pseudoge | ENSG00000 |
| 74121012 | 74131923 | 1 | processed_pseudoge | ENSG00000 |
| 74122134 | 74122725 | -1 | TEC | ENSG00000 |
| 74133004 | 74133114 | -1 | processed_pseudoge | ENSG00000 |
| 74161062 | 74162528 | -1 | processed_pseudoge | ENSG00000 |
| 74173890 | 74174066 | 1 | processed_pseudoge | ENSG00000 |
| 74185929 | 74187153 | 1 | processed_pseudoge | ENSG00000 |
| 74202834 | 74204595 | 1 | processed_pseudoge | ENSG00000 |
| 74209976 | 74213660 | -1 | lncRNA | ENSG00000 |
| 74218377 | 74218461 | -1 | miRNA | ENSG00000 |
| 74218547 | 74218618 | -1 | miRNA | ENSG00000 |
| 74218549 | 74218618 | 1 | miRNA | ENSG00000 |
| 74242608 | 74242908 | -1 | misc_RNA | ENSG00000 |
| 74274341 | 74274411 | 1 | misc_RNA | ENSG00000 |
| 74280402 | 74280495 | 1 | misc_RNA | ENSG00000 |
| 74280936 | 74281082 | 1 | misc_RNA | ENSG00000 |
| 74281703 | 74281849 | 1 | misc_RNA | ENSG00000 |
| 74287104 | 74287209 | -1 | miRNA | ENSG00000 |
| 74287286 | 74287357 | -1 | miRNA | ENSG00000 |
| 74293357 | 74293574 | 1 | misc_RNA | ENSG00000 |
| 74304180 | 74305034 | 1 | protein_coding | ENSG00000 |
| 74342713 | 74342860 | -1 | processed_pseudoge | ENSG00000 |
| 74376125 | 74376885 | -1 | processed_pseudoge | ENSG00000 |
| 74390692 | 74390989 | 1 | misc_RNA | ENSG00000 |
| 74396871 | 74399550 | -1 | processed_pseudoge | ENSG00000 |
| 74409518 | 74410100 | 1 | processed_pseudoge | ENSG00000 |
| 74421493 | 74533917 | 1 | protein_coding | ENSG00000 |
| 74473017 | 74473953 | -1 | processed_pseudoge | ENSG00000 |
| 74582976 | 74614624 | -1 | protein_coding | ENSG00000 |
| 74583088 | 74583546 | -1 | processed_pseudoge | ENSG00000 |
| 74680053 | 74680159 | 1 | snRNA | ENSG00000 |
| 74732856 | 74925485 | -1 | protein_coding | ENSG00000 |
| 74769639 | 74769798 | 1 | processed_pseudoge | ENSG00000 |
| 75051048 | 75156732 | -1 | protein_coding | ENSG00000 |
| 75156388 | 75304885 | 1 | protein_coding | ENSG00000 |
| 75201491 | 75201790 | 1 | processed_pseudoge | ENSG00000 |
| 75202703 | 75202809 | 1 | snRNA | ENSG00000 |
| 75326772 | 75327999 | -1 | processed_pseudoge | ENSG00000 |
| 75368427 | 75523502 | -1 | protein_coding | ENSG00000 |
| 75384346 | 75384828 | 1 | processed_pseudoge | ENSG00000 |
| 75537547 | 75538702 | -1 | processed_pseudoge | ENSG00000 |
| 75582676 | 75585506 | 1 | unprocessed_pseudo | ENSG00000 |
| 75740831 | 75746911 | -1 | processed_pseudoge | ENSG00000 |

| | | | | |
|---|---|---|---|---|
| 75782987 | 75785254 | -1 | protein_coding | ENSG00000( |
| 75884546 | 75885136 | 1 | processed_pseudoge | ENSG00000( |
| 75888700 | 75888888 | 1 | processed_pseudoge | ENSG00000( |
| 75903105 | 75904858 | 1 | lncRNA | ENSG00000( |
| 76029895 | 76030330 | 1 | processed_pseudoge | ENSG00000( |
| 76173040 | 76178314 | 1 | protein_coding | ENSG00000( |
| 76222637 | 76223097 | 1 | processed_pseudoge | ENSG00000( |
| 76244968 | 76245074 | 1 | snRNA | ENSG00000( |
| 76250183 | 76251796 | -1 | processed_pseudoge | ENSG00000( |
| 76334841 | 76335845 | -1 | processed_pseudoge | ENSG00000( |
| 76427710 | 76431342 | 1 | protein_coding | ENSG00000( |
| 76655009 | 76655127 | -1 | rRNA_pseudogene | ENSG00000( |
| 76655498 | 76656559 | -1 | processed_pseudoge | ENSG00000( |
| 76656866 | 77014532 | -1 | lncRNA | ENSG00000( |
| 76919273 | 76919360 | -1 | miRNA | ENSG00000( |
| 77005404 | 77005501 | -1 | miRNA | ENSG00000( |
| 77066709 | 77066827 | 1 | rRNA_pseudogene | ENSG00000( |
| 77374212 | 77381319 | 1 | processed_pseudoge | ENSG00000( |
| 77447389 | 77457278 | 1 | protein_coding | ENSG00000( |
| 77504878 | 77786233 | -1 | protein_coding | ENSG00000( |
| 77727867 | 77728080 | 1 | processed_pseudoge | ENSG00000( |
| 77826364 | 77895593 | -1 | protein_coding | ENSG00000( |
| 77837289 | 77837395 | 1 | snRNA | ENSG00000( |
| 77885377 | 77885634 | 1 | misc_RNA | ENSG00000( |
| 77899440 | 77907376 | 1 | protein_coding | ENSG00000( |
| 77910656 | 78050395 | 1 | protein_coding | ENSG00000( |
| 77910739 | 78129295 | 1 | protein_coding | ENSG00000( |
| 77910741 | 77964890 | 1 | transcribed_processe | ENSG00000( |
| 77967949 | 77969638 | -1 | protein_coding | ENSG00000( |
| 78129748 | 78139650 | -1 | protein_coding | ENSG00000( |
| 78165696 | 78172422 | -1 | unitary_pseudogene | ENSG00000( |
| 78271468 | 78327691 | -1 | protein_coding | ENSG00000( |
| 78519593 | 78520186 | -1 | processed_pseudoge | ENSG00000( |
| 78554412 | 78554751 | -1 | processed_pseudoge | ENSG00000( |
| 78656068 | 78659328 | -1 | protein_coding | ENSG00000( |
| 78699384 | 78699891 | -1 | processed_pseudoge | ENSG00000( |
| 78747709 | 78757094 | 1 | protein_coding | ENSG00000( |
| 78763226 | 78763899 | 1 | processed_pseudoge | ENSG00000( |
| 78901194 | 78901249 | -1 | miRNA | ENSG00000( |
| 78945332 | 78963727 | 1 | protein_coding | ENSG00000( |
| 79084936 | 79085848 | 1 | unitary_pseudogene | ENSG00000( |
| 79144663 | 79175315 | 1 | protein_coding | ENSG00000( |
| 79177200 | 79177668 | -1 | processed_pseudoge | ENSG00000( |
| 79323446 | 79325217 | 1 | processed_pseudoge | ENSG00000( |
| 79360384 | 79367667 | -1 | protein_coding | ENSG00000( |
| 79369364 | 79370165 | -1 | processed_pseudoge | ENSG00000( |
| 80014753 | 80031774 | 1 | protein_coding | ENSG00000( |
| 80228489 | 80335364 | -1 | transcribed_unitary_ | ENSG00000( |
| 80335504 | 80445311 | 1 | protein_coding | ENSG00000( |
| 80529037 | 80530027 | 1 | processed_pseudoge | ENSG00000( |

| | | | | |
|---|---|---|---|---|
| 80560146 | 80562054 | 1 | processed_pseudoge | ENSG00000( |
| 80571871 | 80574607 | -1 | processed_pseudoge | ENSG00000( |
| 80669503 | 80809877 | -1 | protein_coding | ENSG00000( |
| 80709928 | 80710705 | 1 | processed_pseudoge | ENSG00000( |
| 80857076 | 80857202 | -1 | snoRNA | ENSG00000( |
| 80900757 | 80900864 | 1 | snRNA | ENSG00000( |
| 80929500 | 80930347 | 1 | processed_pseudoge | ENSG00000( |
| 80936434 | 80936540 | 1 | snRNA | ENSG00000( |
| 81000150 | 81004217 | -1 | lncRNA | ENSG00000( |
| 81113699 | 81201913 | -1 | protein_coding | ENSG00000( |
| 81202102 | 81298547 | 1 | protein_coding | ENSG00000( |
| 81363268 | 81364170 | 1 | processed_pseudoge | ENSG00000( |
| 81408938 | 81427276 | 1 | unprocessed_pseudc | ENSG00000( |
| 81475532 | 81498083 | 1 | lncRNA | ENSG00000( |
| 81600072 | 81601306 | 1 | processed_pseudoge | ENSG00000( |
| 82001220 | 82001327 | -1 | snRNA | ENSG00000( |
| 82497669 | 82497924 | -1 | processed_pseudoge | ENSG00000( |
| 82506434 | 82506812 | -1 | processed_pseudoge | ENSG00000( |
| 82561201 | 82561347 | 1 | snoRNA | ENSG00000( |
| 82875649 | 82875885 | 1 | processed_pseudoge | ENSG00000( |
| 82998699 | 82999236 | 1 | processed_pseudoge | ENSG00000( |
| 83506023 | 83509214 | -1 | TEC | ENSG00000( |
| 83508290 | 83512127 | 1 | protein_coding | ENSG00000( |
| 83559896 | 83560312 | -1 | processed_pseudoge | ENSG00000( |
| 83748825 | 83749939 | -1 | processed_pseudoge | ENSG00000( |
| 83861126 | 83886699 | 1 | protein_coding | ENSG00000( |
| 84058346 | 84188199 | -1 | protein_coding | ENSG00000( |
| 84225752 | 84225828 | -1 | miRNA | ENSG00000( |
| 84317874 | 84502479 | -1 | protein_coding | ENSG00000( |
| 84403898 | 84404024 | -1 | transcribed_process( | ENSG00000( |
| 84755136 | 84756003 | 1 | processed_pseudoge | ENSG00000( |
| 84934151 | 84934890 | 1 | transcribed_process( | ENSG00000( |
| 84973217 | 84977829 | 1 | unitary_pseudogene | ENSG00000( |
| 85003877 | 85093316 | 1 | protein_coding | ENSG00000( |
| 85092287 | 85243961 | -1 | protein_coding | ENSG00000( |
| 85143427 | 85156444 | 1 | unprocessed_pseudc | ENSG00000( |
| 85207187 | 85210649 | 1 | processed_pseudoge | ENSG00000( |
| 85210684 | 85220021 | 1 | lncRNA | ENSG00000( |
| 85244032 | 85273362 | 1 | protein_coding | ENSG00000( |
| 85277396 | 85379717 | -1 | protein_coding | ENSG00000( |
| 85835780 | 85835858 | 1 | miRNA | ENSG00000( |
| 85849410 | 85850139 | -1 | processed_pseudoge | ENSG00000( |
| 85851575 | 85851682 | -1 | processed_pseudoge | ENSG00000( |
| 85861180 | 86047561 | -1 | protein_coding | ENSG00000( |
| 85903636 | 85903707 | -1 | miRNA | ENSG00000( |
| 86019608 | 86019715 | 1 | misc_RNA | ENSG00000( |
| 86066227 | 86066468 | -1 | processed_pseudoge | ENSG00000( |
| 86084716 | 86086339 | -1 | processed_pseudoge | ENSG00000( |
| 86128893 | 86129570 | -1 | processed_pseudoge | ENSG00000( |
| 86148451 | 86832604 | 1 | protein_coding | ENSG00000( |

| | | | | |
|---|---|---|---|---|
| 86160264 | 86161359 | -1 | processed_pseudoge | ENSG00000( |
| 86304367 | 86305086 | 1 | processed_pseudoge | ENSG00000( |
| 86481028 | 86481130 | 1 | snRNA | ENSG00000( |
| 86481532 | 86481813 | 1 | transcribed_processe | ENSG00000( |
| 86798598 | 86799226 | -1 | processed_pseudoge | ENSG00000( |
| 86888586 | 86888816 | 1 | processed_pseudoge | ENSG00000( |
| 87143854 | 87144748 | -1 | processed_pseudoge | ENSG00000( |
| 87146733 | 87146804 | 1 | snoRNA | ENSG00000( |
| 87425583 | 87425690 | -1 | processed_pseudoge | ENSG00000( |
| 87517409 | 87670050 | 1 | protein_coding | ENSG00000( |
| 87703343 | 87704203 | -1 | processed_pseudoge | ENSG00000( |
| 87707579 | 87750883 | -1 | lncRNA | ENSG00000( |
| 87723089 | 87723238 | -1 | processed_pseudoge | ENSG00000( |
| 87806433 | 87806769 | -1 | processed_pseudoge | ENSG00000( |
| 87807549 | 87809883 | -1 | processed_pseudoge | ENSG00000( |
| 88061590 | 88062844 | -1 | processed_pseudoge | ENSG00000( |
| 88096393 | 88097225 | -1 | processed_pseudoge | ENSG00000( |
| 88148792 | 88148918 | 1 | snoRNA | ENSG00000( |
| 88369343 | 88369459 | 1 | processed_pseudoge | ENSG00000( |
| 88747225 | 88754785 | 1 | protein_coding | ENSG00000( |
| 88827554 | 88829858 | -1 | processed_pseudoge | ENSG00000( |
| 88908403 | 88908763 | 1 | processed_pseudoge | ENSG00000( |
| 88982215 | 88982353 | -1 | processed_pseudoge | ENSG00000( |
| 89367435 | 89367600 | -1 | processed_pseudoge | ENSG00000( |
| 89423738 | 89446983 | 1 | lncRNA | ENSG00000( |
| 89921908 | 89922883 | 1 | protein_coding | ENSG00000( |
| 90039213 | 90039834 | -1 | processed_pseudoge | ENSG00000( |
| 90112650 | 90113706 | 1 | processed_pseudoge | ENSG00000( |
| 90208895 | 90209418 | -1 | processed_pseudoge | ENSG00000( |
| 90209796 | 90210420 | -1 | processed_pseudoge | ENSG00000( |
| 90289536 | 90289903 | -1 | processed_pseudoge | ENSG00000( |
| 90597235 | 90597340 | -1 | snRNA | ENSG00000( |
| 91307781 | 91308878 | 1 | lncRNA | ENSG00000( |
| 91414878 | 91434999 | -1 | lncRNA | ENSG00000( |
| 91422309 | 91422538 | 1 | processed_pseudoge | ENSG00000( |
| 91434595 | 91438584 | 1 | protein_coding | ENSG00000( |
| 91560436 | 91561597 | -1 | processed_pseudoge | ENSG00000( |
| 91779261 | 92623230 | 1 | protein_coding | ENSG00000( |
| 91798093 | 91798283 | 1 | snRNA | ENSG00000( |
| 91982655 | 91983505 | -1 | processed_pseudoge | ENSG00000( |
| 92113246 | 92114461 | -1 | processed_pseudoge | ENSG00000( |
| 92459670 | 92460952 | 1 | processed_pseudoge | ENSG00000( |
| 92515323 | 92515497 | -1 | processed_pseudoge | ENSG00000( |
| 92676519 | 92676954 | -1 | processed_pseudoge | ENSG00000( |
| 92770855 | 92772091 | 1 | processed_pseudoge | ENSG00000( |
| 93042490 | 93042942 | 1 | processed_pseudoge | ENSG00000( |
| 93222220 | 93225015 | -1 | processed_pseudoge | ENSG00000( |
| 93287733 | 93288759 | -1 | processed_pseudoge | ENSG00000( |
| 93388075 | 93389497 | -1 | processed_pseudoge | ENSG00000( |
| 93573834 | 93574579 | 1 | processed_pseudoge | ENSG00000( |

| | | | | |
|---|---|---|---|---|
| 93670930 | 93673578 | -1 | protein_coding | ENSG0000C |
| 93674013 | 93712274 | 1 | protein_coding | ENSG0000C |
| 93776911 | 93777384 | -1 | processed_pseudoge | ENSG0000C |
| 94003815 | 94003922 | -1 | snRNA | ENSG0000C |
| 94178005 | 94178106 | 1 | misc_RNA | ENSG0000C |
| 94319038 | 94320025 | 1 | processed_pseudoge | ENSG0000C |
| 94636663 | 94639570 | -1 | processed_pseudoge | ENSG0000C |
| 94776739 | 94777567 | -1 | processed_pseudoge | ENSG0000C |
| 95063141 | 95063226 | -1 | miRNA | ENSG0000C |
| 95453369 | 95453581 | 1 | processed_pseudoge | ENSG0000C |
| 95499604 | 95500036 | 1 | processed_pseudoge | ENSG0000C |
| 95582064 | 95582162 | 1 | rRNA_pseudogene | ENSG0000C |
| 95584303 | 95584942 | 1 | processed_pseudoge | ENSG0000C |
| 95610578 | 95610957 | 1 | processed_pseudoge | ENSG0000C |
| 95638375 | 95638924 | 1 | processed_pseudoge | ENSG0000C |
| 95679130 | 95679627 | -1 | processed_pseudoge | ENSG0000C |
| 95720243 | 95721144 | 1 | processed_pseudoge | ENSG0000C |
| 95973328 | 95974315 | -1 | processed_pseudoge | ENSG0000C |
| 96023360 | 96023652 | 1 | misc_RNA | ENSG0000C |
| 96070669 | 96071778 | -1 | processed_pseudoge | ENSG0000C |
| 96127747 | 96372889 | -1 | lncRNA | ENSG0000C |
| 96195477 | 96195617 | 1 | processed_pseudoge | ENSG0000C |
| 96337236 | 96337912 | -1 | processed_pseudoge | ENSG0000C |
| 96410432 | 96410782 | -1 | misc_RNA | ENSG0000C |
| 96620654 | 96621903 | 1 | processed_pseudoge | ENSG0000C |
| 96684663 | 97604997 | 1 | protein_coding | ENSG0000C |
| 96701507 | 96701619 | -1 | misc_RNA | ENSG0000C |
| 96883908 | 96885467 | 1 | protein_coding | ENSG0000C |
| 96953630 | 96954195 | -1 | processed_pseudoge | ENSG0000C |
| 97081674 | 97081956 | 1 | misc_RNA | ENSG0000C |
| 97341046 | 97341344 | 1 | misc_RNA | ENSG0000C |
| 97431286 | 97642589 | -1 | lncRNA | ENSG0000C |
| 97847263 | 97849017 | 1 | processed_pseudoge | ENSG0000C |
| 97906644 | 97908090 | 1 | processed_pseudoge | ENSG0000C |
| 97975878 | 97978641 | -1 | processed_pseudoge | ENSG0000C |
| 98251679 | 98252543 | 1 | processed_pseudoge | ENSG0000C |
| 98348451 | 98348748 | -1 | processed_pseudoge | ENSG0000C |
| 98389512 | 98390919 | 1 | processed_pseudoge | ENSG0000C |
| 98422138 | 98422809 | 1 | processed_pseudoge | ENSG0000C |
| 98450688 | 98892061 | 1 | lncRNA | ENSG0000C |
| 99656997 | 99657381 | 1 | processed_pseudoge | ENSG0000C |
| 99719364 | 99721158 | -1 | processed_pseudoge | ENSG0000C |
| 99886740 | 99887691 | -1 | processed_pseudoge | ENSG0000C |
| 100100084 | 100100172 | 1 | snoRNA | ENSG0000C |
| 100149783 | 100151340 | -1 | processed_pseudoge | ENSG0000C |
| 100155885 | 100156773 | -1 | processed_pseudoge | ENSG0000C |
| 100291644 | 100410273 | -1 | protein_coding | ENSG0000C |
| 100584936 | 100599885 | 1 | protein_coding | ENSG0000C |
| 100627108 | 100639991 | -1 | protein_coding | ENSG0000C |
| 100644195 | 100675788 | 1 | protein_coding | ENSG0000C |

| | | | | |
|---|---|---|---|---|
| 100674491 | 100732123 | -1 | protein_coding | ENSG00000 |
| 100800303 | 100800784 | -1 | processed_pseudoge | ENSG00000 |
| 100801188 | 100803060 | 1 | processed_pseudoge | ENSG00000 |
| 100815865 | 100817174 | -1 | processed_pseudoge | ENSG00000 |
| 100820359 | 100840932 | 1 | protein_coding | ENSG00000 |
| 100822620 | 100822752 | -1 | snoRNA | ENSG00000 |
| 100843324 | 100874359 | -1 | protein_coding | ENSG00000 |
| 100855288 | 100856379 | -1 | processed_pseudoge | ENSG00000 |
| 100887636 | 100888590 | -1 | processed_pseudoge | ENSG00000 |
| 100910712 | 100911489 | 1 | processed_pseudoge | ENSG00000 |
| 100913445 | 100929433 | -1 | protein_coding | ENSG00000 |
| 100956593 | 100957538 | 1 | processed_pseudoge | ENSG00000 |
| 100969708 | 100990831 | 1 | protein_coding | ENSG00000 |
| 101009346 | 101052116 | -1 | protein_coding | ENSG00000 |
| 101043564 | 101094476 | -1 | lncRNA | ENSG00000 |
| 101078879 | 101096367 | 1 | protein_coding | ENSG00000 |
| 101098218 | 101163681 | 1 | protein_coding | ENSG00000 |
| 101191115 | 101191409 | -1 | processed_pseudoge | ENSG00000 |
| 101219769 | 101264497 | 1 | protein_coding | ENSG00000 |
| 101268253 | 101293057 | -1 | protein_coding | ENSG00000 |
| 101325404 | 101325510 | 1 | snRNA | ENSG00000 |
| 101337725 | 101338289 | -1 | processed_pseudoge | ENSG00000 |
| 101339917 | 101340934 | 1 | processed_pseudoge | ENSG00000 |
| 101345661 | 101348742 | -1 | protein_coding | ENSG00000 |
| 101349447 | 101390796 | -1 | protein_coding | ENSG00000 |
| 101390824 | 101396154 | 1 | protein_coding | ENSG00000 |
| 101391011 | 101412297 | 1 | protein_coding | ENSG00000 |
| 101397803 | 101408012 | -1 | protein_coding | ENSG00000 |
| 101408222 | 101414133 | 1 | protein_coding | ENSG00000 |
| 101418287 | 101533459 | 1 | protein_coding | ENSG00000 |
| 101443707 | 101444103 | -1 | processed_pseudoge | ENSG00000 |
| 101537485 | 101539276 | -1 | processed_pseudoge | ENSG00000 |
| 101550547 | 101554700 | 1 | protein_coding | ENSG00000 |
| 101597510 | 101598671 | 1 | transcribed_unproce | ENSG00000 |
| 101615118 | 101618001 | -1 | protein_coding | ENSG00000 |
| 101622797 | 101627843 | 1 | protein_coding | ENSG00000 |
| 101622983 | 101624164 | -1 | lncRNA | ENSG00000 |
| 101627868 | 101628523 | 1 | lncRNA | ENSG00000 |
| 101634006 | 101634112 | -1 | snRNA | ENSG00000 |
| 101640421 | 101644352 | 1 | lncRNA | ENSG00000 |
| 101655281 | 101659850 | -1 | protein_coding | ENSG00000 |
| 101683780 | 101684449 | 1 | processed_pseudoge | ENSG00000 |
| 101691580 | 101691685 | 1 | snRNA | ENSG00000 |
| 101720340 | 101721709 | -1 | processed_pseudoge | ENSG00000 |
| 101778969 | 101780538 | -1 | processed_pseudoge | ENSG00000 |
| 101832112 | 101857577 | -1 | protein_coding | ENSG00000 |
| 101882288 | 101932079 | -1 | protein_coding | ENSG00000 |
| 101911529 | 101912072 | 1 | processed_pseudoge | ENSG00000 |
| 101915589 | 101915695 | 1 | snRNA | ENSG00000 |
| 101989481 | 101989872 | -1 | processed_pseudoge | ENSG00000 |

| | | | | |
|---|---|---|---|---|
| 102006883 | 102007294 | 1 | processed_pseudoge | ENSG00000 |
| 102008524 | 102009046 | -1 | processed_pseudoge | ENSG00000 |
| 102125679 | 102140426 | 1 | protein_coding | ENSG00000 |
| 102140476 | 102142970 | -1 | protein_coding | ENSG00000 |
| 102142588 | 102143971 | 1 | lncRNA | ENSG00000 |
| 102153708 | 102155977 | -1 | protein_coding | ENSG00000 |
| 102173100 | 102185021 | 1 | transcribed_unproce | ENSG00000 |
| 102215301 | 102229532 | 1 | protein_coding | ENSG00000 |
| 102247161 | 102326719 | 1 | protein_coding | ENSG00000 |
| 102259043 | 102261692 | 1 | unprocessed_pseudo | ENSG00000 |
| 102360395 | 102471812 | -1 | protein_coding | ENSG00000 |
| 102360396 | 102439932 | -1 | protein_coding | ENSG00000 |
| 102425343 | 102428080 | -1 | unitary_pseudogene | ENSG00000 |
| 102456862 | 102471842 | -1 | protein_coding | ENSG00000 |
| 102513682 | 102516739 | -1 | protein_coding | ENSG00000 |
| 102546734 | 102548213 | -1 | processed_pseudoge | ENSG00000 |
| 102549965 | 102571695 | 1 | transcribed_unproce | ENSG00000 |
| 102599168 | 102604159 | 1 | protein_coding | ENSG00000 |
| 102599512 | 102714671 | 1 | lncRNA | ENSG00000 |
| 102599657 | 102659712 | -1 | lncRNA | ENSG00000 |
| 102615473 | 102615820 | -1 | processed_pseudoge | ENSG00000 |
| 102651092 | 102659083 | 1 | protein_coding | ENSG00000 |
| 102659983 | 102663455 | 1 | lncRNA | ENSG00000 |
| 102678707 | 102678813 | -1 | snRNA | ENSG00000 |
| 102705784 | 102706240 | 1 | processed_pseudoge | ENSG00000 |
| 102712176 | 102717733 | 1 | protein_coding | ENSG00000 |
| 102712495 | 102753530 | 1 | protein_coding | ENSG00000 |
| 102720688 | 102753540 | 1 | protein_coding | ENSG00000 |
| 102769158 | 102885406 | 1 | lncRNA | ENSG00000 |
| 102785667 | 102786604 | 1 | processed_pseudoge | ENSG00000 |
| 102786813 | 102788387 | 1 | processed_pseudoge | ENSG00000 |
| 102796251 | 102797093 | 1 | processed_pseudoge | ENSG00000 |
| 102797531 | 102798128 | 1 | processed_pseudoge | ENSG00000 |
| 102798396 | 102799560 | 1 | processed_pseudoge | ENSG00000 |
| 102799563 | 102799883 | 1 | processed_pseudoge | ENSG00000 |
| 102799942 | 102801399 | 1 | processed_pseudoge | ENSG00000 |
| 102803041 | 102806138 | 1 | processed_pseudoge | ENSG00000 |
| 102806164 | 102806646 | -1 | processed_pseudoge | ENSG00000 |
| 102806741 | 102807824 | 1 | processed_pseudoge | ENSG00000 |
| 102884414 | 102906158 | 1 | lncRNA | ENSG00000 |
| 102937272 | 102938300 | 1 | protein_coding | ENSG00000 |
| 102937770 | 102940561 | -1 | lncRNA | ENSG00000 |
| 102945971 | 102952039 | 1 | lncRNA | ENSG00000 |
| 103007461 | 103008072 | 1 | processed_pseudoge | ENSG00000 |
| 103062651 | 103064171 | -1 | protein_coding | ENSG00000 |
| 103075810 | 103093143 | -1 | protein_coding | ENSG00000 |
| 103087198 | 103089277 | 1 | processed_pseudoge | ENSG00000 |
| 103206010 | 103206899 | -1 | processed_pseudoge | ENSG00000 |
| 103215108 | 103217246 | 1 | protein_coding | ENSG00000 |
| 103252995 | 103255192 | -1 | protein_coding | ENSG00000 |

| | | | | |
|---|---|---|---|---|
| 103258572 | 103259581 | -1 | processed_pseudoge | ENSG00000 |
| 103273691 | 103276750 | -1 | protein_coding | ENSG00000 |
| 103309346 | 103311046 | -1 | protein_coding | ENSG00000 |
| 103330229 | 103332326 | 1 | protein_coding | ENSG00000 |
| 103347462 | 103347729 | -1 | processed_pseudoge | ENSG00000 |
| 103356489 | 103358462 | 1 | protein_coding | ENSG00000 |
| 103376340 | 103378077 | 1 | protein_coding | ENSG00000 |
| 103404777 | 103452754 | 1 | lncRNA | ENSG00000 |
| 103486461 | 103517919 | 1 | lncRNA | ENSG00000 |
| 103499130 | 103519489 | -1 | protein_coding | ENSG00000 |
| 103563917 | 103573278 | -1 | lncRNA | ENSG00000 |
| 103576231 | 103587736 | 1 | protein_coding | ENSG00000 |
| 103596608 | 103598904 | 1 | TEC | ENSG00000 |
| 103607451 | 103629690 | 1 | protein_coding | ENSG00000 |
| 103626076 | 103626492 | -1 | lncRNA | ENSG00000 |
| 103628704 | 103630953 | 1 | protein_coding | ENSG00000 |
| 103651735 | 103652106 | -1 | processed_pseudoge | ENSG00000 |
| 103675496 | 103688158 | -1 | protein_coding | ENSG00000 |
| 103687284 | 103691772 | 1 | lncRNA | ENSG00000 |
| 103707224 | 103728655 | -1 | protein_coding | ENSG00000 |
| 103710909 | 103714032 | 1 | protein_coding | ENSG00000 |
| 103773718 | 103792619 | 1 | protein_coding | ENSG00000 |
| 103822327 | 103832257 | -1 | protein_coding | ENSG00000 |
| 103850797 | 103850911 | -1 | rRNA_pseudogene | ENSG00000 |
| 103880668 | 103919548 | -1 | lncRNA | ENSG00000 |
| 103904322 | 103904754 | -1 | processed_pseudoge | ENSG00000 |
| 103909126 | 103909449 | 1 | unprocessed_pseudc | ENSG00000 |
| 103910305 | 103911352 | -1 | processed_pseudoge | ENSG00000 |
| 103918896 | 103966712 | 1 | protein_coding | ENSG00000 |
| 103928099 | 103929017 | 1 | processed_pseudoge | ENSG00000 |
| 103961811 | 103962279 | -1 | processed_pseudoge | ENSG00000 |
| 104011147 | 104013687 | -1 | protein_coding | ENSG00000 |
| 104039949 | 104042454 | 1 | protein_coding | ENSG00000 |
| 104062542 | 104063037 | -1 | processed_pseudoge | ENSG00000 |
| 104063871 | 104076212 | -1 | protein_coding | ENSG00000 |
| 104099214 | 104157009 | -1 | protein_coding | ENSG00000 |
| 104112131 | 104115846 | 1 | protein_coding | ENSG00000 |
| 104166453 | 104195902 | 1 | protein_coding | ENSG00000 |
| 104250038 | 104254933 | -1 | protein_coding | ENSG00000 |
| 104566199 | 105767829 | 1 | protein_coding | ENSG00000 |
| 104647216 | 104647999 | -1 | processed_pseudoge | ENSG00000 |
| 104803915 | 104804440 | 1 | processed_pseudoge | ENSG00000 |
| 105218929 | 105220674 | -1 | protein_coding | ENSG00000 |
| 105405647 | 105407039 | 1 | processed_pseudoge | ENSG00000 |
| 105676071 | 105676177 | -1 | snRNA | ENSG00000 |
| 105822539 | 105958610 | 1 | protein_coding | ENSG00000 |
| 106032435 | 106038727 | -1 | protein_coding | ENSG00000 |
| 106070481 | 106073202 | 1 | processed_pseudoge | ENSG00000 |
| 106168305 | 106208956 | 1 | protein_coding | ENSG00000 |
| 106327534 | 106327824 | 1 | processed_pseudoge | ENSG00000 |

| | | | | |
|---|---|---|---|---|
| 106426568 | 106426669 | 1 | misc_RNA | ENSG0000C |
| 106466542 | 106467440 | -1 | processed_pseudoge | ENSG0000C |
| 106562774 | 106563404 | -1 | unprocessed_pseudc | ENSG0000C |
| 106566021 | 106566161 | 1 | unprocessed_pseudc | ENSG0000C |
| 106611930 | 106679439 | 1 | protein_coding | ENSG0000C |
| 106639814 | 106639896 | 1 | miRNA | ENSG0000C |
| 106693794 | 106797016 | 1 | protein_coding | ENSG0000C |
| 106802673 | 106876150 | 1 | protein_coding | ENSG0000C |
| 106813871 | 107000212 | -1 | protein_coding | ENSG0000C |
| 106900063 | 106903341 | -1 | protein_coding | ENSG0000C |
| 106900164 | 106930861 | 1 | protein_coding | ENSG0000C |
| 107032248 | 107032820 | 1 | processed_pseudoge | ENSG0000C |
| 107064420 | 107118823 | -1 | protein_coding | ENSG0000C |
| 107123427 | 107206433 | -1 | protein_coding | ENSG0000C |
| 107131659 | 107132269 | 1 | processed_pseudoge | ENSG0000C |
| 107202577 | 107202683 | 1 | misc_RNA | ENSG0000C |
| 107206632 | 107244243 | 1 | protein_coding | ENSG0000C |
| 107272582 | 107273656 | 1 | processed_pseudoge | ENSG0000C |
| 107351650 | 107352843 | -1 | processed_pseudoge | ENSG0000C |
| 107428630 | 107429461 | 1 | processed_pseudoge | ENSG0000C |
| 107512983 | 107545821 | -1 | lncRNA | ENSG0000C |
| 107522450 | 107605251 | 1 | protein_coding | ENSG0000C |
| 107628424 | 107651026 | 1 | protein_coding | ENSG0000C |
| 107713221 | 107777342 | -1 | protein_coding | ENSG0000C |
| 107774899 | 107795829 | 1 | protein_coding | ENSG0000C |
| 107825755 | 107927193 | 1 | protein_coding | ENSG0000C |
| 107857364 | 107936230 | -1 | lncRNA | ENSG0000C |
| 107901145 | 107901246 | -1 | misc_RNA | ENSG0000C |
| 107939263 | 107939642 | -1 | processed_pseudoge | ENSG0000C |
| 107974078 | 107985008 | 1 | unprocessed_pseudc | ENSG0000C |
| 107980864 | 107982370 | -1 | protein_coding | ENSG0000C |
| 108044970 | 108079184 | 1 | protein_coding | ENSG0000C |
| 108084207 | 108091549 | -1 | protein_coding | ENSG0000C |
| 108091668 | 108154671 | 1 | protein_coding | ENSG0000C |
| 108155607 | 108439497 | -1 | protein_coding | ENSG0000C |
| 108383398 | 108383993 | 1 | unprocessed_pseudc | ENSG0000C |
| 108439844 | 108697545 | 1 | protein_coding | ENSG0000C |
| 108676951 | 108677057 | -1 | misc_RNA | ENSG0000C |
| 108719949 | 108724944 | -1 | lncRNA | ENSG0000C |
| 108722070 | 108732687 | -1 | protein_coding | ENSG0000C |
| 108725886 | 108726948 | -1 | TEC | ENSG0000C |
| 108732482 | 108736409 | -1 | protein_coding | ENSG0000C |
| 108736011 | 108739223 | 1 | lncRNA | ENSG0000C |
| 108826764 | 108826869 | 1 | snRNA | ENSG0000C |
| 109054542 | 109054590 | 1 | miRNA | ENSG0000C |
| 109341813 | 109342697 | 1 | processed_pseudoge | ENSG0000C |
| 109372906 | 109482086 | -1 | protein_coding | ENSG0000C |
| 109535781 | 109544690 | 1 | protein_coding | ENSG0000C |
| 109568478 | 109570289 | 1 | processed_pseudoge | ENSG0000C |
| 109623700 | 109625172 | -1 | protein_coding | ENSG0000C |

| | | | | |
|---|---|---|---|---|
| 109624244 | 109733403 | -1 | protein_coding | ENSG00000( |
| 109636834 | 109637063 | 1 | processed_pseudoge | ENSG00000( |
| 109744643 | 109744749 | -1 | misc_RNA | ENSG00000( |
| 109853312 | 109853919 | 1 | processed_pseudoge | ENSG00000( |
| 110002631 | 110182734 | 1 | protein_coding | ENSG00000( |
| 110055329 | 110055426 | 1 | miRNA | ENSG00000( |
| 110082118 | 110082218 | 1 | miRNA | ENSG00000( |
| 110194186 | 110440233 | -1 | protein_coding | ENSG00000( |
| 110224985 | 110225063 | -1 | snoRNA | ENSG00000( |
| 110305420 | 110307221 | -1 | lncRNA | ENSG00000( |
| 110346703 | 110346909 | -1 | processed_pseudoge | ENSG00000( |
| 110358816 | 110456334 | 1 | protein_coding | ENSG00000( |
| 110472692 | 110524320 | 1 | lncRNA | ENSG00000( |
| 110520312 | 110523021 | 1 | transcribed_processe | ENSG00000( |
| 110636841 | 110637672 | 1 | processed_pseudoge | ENSG00000( |
| 110673856 | 110795819 | -1 | protein_coding | ENSG00000( |
| 110944285 | 111227361 | 1 | protein_coding | ENSG00000( |
| 111169221 | 111169403 | 1 | processed_pseudoge | ENSG00000( |
| 111229889 | 111231061 | -1 | processed_pseudoge | ENSG00000( |
| 111245099 | 111270483 | -1 | protein_coding | ENSG00000( |
| 111293779 | 111412429 | -1 | protein_coding | ENSG00000( |
| 111421510 | 111421608 | -1 | snRNA | ENSG00000( |
| 111434427 | 111435024 | 1 | processed_pseudoge | ENSG00000( |
| 111511662 | 111522399 | 1 | protein_coding | ENSG00000( |
| 111618808 | 111619094 | 1 | processed_pseudoge | ENSG00000( |
| 111619677 | 111621489 | -1 | processed_pseudoge | ENSG00000( |
| 111624210 | 111624726 | -1 | processed_pseudoge | ENSG00000( |
| 111665811 | 111760649 | 1 | protein_coding | ENSG00000( |
| 111669829 | 111669948 | -1 | rRNA_pseudogene | ENSG00000( |
| 111706649 | 111711101 | -1 | lncRNA | ENSG00000( |
| 111774315 | 112082776 | -1 | protein_coding | ENSG00000( |
| 111876051 | 111903990 | 1 | protein_coding | ENSG00000( |
| 111930735 | 111931092 | 1 | processed_pseudoge | ENSG00000( |
| 112319952 | 112320417 | -1 | processed_pseudoge | ENSG00000( |
| 112381392 | 112382220 | 1 | processed_pseudoge | ENSG00000( |
| 112454500 | 112457112 | 1 | protein_coding | ENSG00000( |
| 112630648 | 112680054 | -1 | protein_coding | ENSG00000( |
| 112661657 | 112661768 | 1 | misc_RNA | ENSG00000( |
| 112689915 | 112690520 | 1 | processed_pseudoge | ENSG00000( |
| 112774503 | 112840815 | -1 | protein_coding | ENSG00000( |
| 112780718 | 112780788 | -1 | miRNA | ENSG00000( |
| 112855874 | 112856162 | 1 | misc_RNA | ENSG00000( |
| 112880563 | 113212186 | 1 | lncRNA | ENSG00000( |
| 112919132 | 112919230 | 1 | snRNA | ENSG00000( |
| 113157873 | 113188018 | 1 | lncRNA | ENSG00000( |
| 113347413 | 113347619 | 1 | processed_pseudoge | ENSG00000( |
| 113522482 | 113522989 | 1 | processed_pseudoge | ENSG00000( |
| 113616300 | 114059289 | -1 | lncRNA | ENSG00000( |
| 113882590 | 113883667 | 1 | processed_pseudoge | ENSG00000( |
| 113944186 | 113944399 | -1 | snoRNA | ENSG00000( |

| | | | | |
|---|---|---|---|---|
| 113949917 | 113950082 | 1 | snRNA | ENSG0000C |
| 113976341 | 113976633 | -1 | misc_RNA | ENSG0000C |
| 114017033 | 114017102 | 1 | snoRNA | ENSG0000C |
| 114388988 | 114389155 | -1 | processed_pseudoge | ENSG0000C |
| 114389135 | 114389954 | 1 | processed_pseudoge | ENSG0000C |
| 114390674 | 114391256 | 1 | processed_pseudoge | ENSG0000C |
| 114584078 | 114910061 | 1 | protein_coding | ENSG0000C |
| 114589620 | 114589935 | 1 | processed_pseudoge | ENSG0000C |
| 114630797 | 114630924 | 1 | snoRNA | ENSG0000C |
| 114639435 | 114639519 | 1 | miRNA | ENSG0000C |
| 114651544 | 114651623 | 1 | miRNA | ENSG0000C |
| 114652655 | 114652723 | 1 | miRNA | ENSG0000C |
| 114715233 | 114715344 | 1 | miRNA | ENSG0000C |
| 114763184 | 114763263 | 1 | miRNA | ENSG0000C |
| 114774828 | 114776775 | -1 | processed_pseudoge | ENSG0000C |
| 114823454 | 114823564 | 1 | miRNA | ENSG0000C |
| 114900359 | 114900574 | -1 | processed_pseudoge | ENSG0000C |
| 115003975 | 115019977 | -1 | protein_coding | ENSG0000C |
| 115032427 | 115032533 | 1 | snRNA | ENSG0000C |
| 115063936 | 115064794 | -1 | processed_pseudoge | ENSG0000C |
| 115069237 | 115069846 | 1 | processed_pseudoge | ENSG0000C |
| 115110616 | 115234096 | -1 | protein_coding | ENSG0000C |
| 115126316 | 115126444 | -1 | snoRNA | ENSG0000C |
| 115139030 | 115139123 | -1 | misc_RNA | ENSG0000C |
| 115189427 | 115192868 | 1 | protein_coding | ENSG0000C |
| 115198432 | 115198749 | 1 | processed_pseudoge | ENSG0000C |
| 115289715 | 115307563 | 1 | protein_coding | ENSG0000C |
| 115424064 | 115425600 | -1 | processed_pseudoge | ENSG0000C |
| 115454733 | 115455194 | -1 | processed_pseudoge | ENSG0000C |
| 115518182 | 115562731 | -1 | lncRNA | ENSG0000C |
| 115545637 | 115545717 | 1 | snoRNA | ENSG0000C |
| 115561174 | 115650861 | 1 | protein_coding | ENSG0000C |
| 115659713 | 115660022 | 1 | processed_pseudoge | ENSG0000C |
| 115664895 | 115666133 | 1 | processed_pseudoge | ENSG0000C |
| 115694880 | 115969130 | -1 | lncRNA | ENSG0000C |
| 115702811 | 115704120 | 1 | processed_pseudoge | ENSG0000C |
| 115718942 | 115719229 | -1 | processed_pseudoge | ENSG0000C |
| 115840964 | 115843050 | 1 | lncRNA | ENSG0000C |
| 115992466 | 115992773 | -1 | misc_RNA | ENSG0000C |
| 116043483 | 116044418 | 1 | processed_pseudoge | ENSG0000C |
| 116106921 | 116108430 | -1 | processed_pseudoge | ENSG0000C |
| 116170744 | 116174974 | 1 | protein_coding | ENSG0000C |
| 116400848 | 116400953 | -1 | snRNA | ENSG0000C |
| 116436606 | 116461458 | 1 | protein_coding | ENSG0000C |
| 116461686 | 116462976 | -1 | protein_coding | ENSG0000C |
| 116669810 | 116680303 | -1 | unprocessed_pseudc | ENSG0000C |
| 116688870 | 116690185 | 1 | unprocessed_pseudc | ENSG0000C |
| 116692910 | 116694188 | -1 | lncRNA | ENSG0000C |
| 116748530 | 116748877 | -1 | processed_pseudoge | ENSG0000C |
| 116850839 | 116850945 | -1 | snRNA | ENSG0000C |

| | | | | |
|---|---|---|---|---|
| 116963049 | 116963676 | 1 | processed_pseudoge | ENSG00000( |
| 117397938 | 117398206 | 1 | processed_pseudoge | ENSG00000( |
| 117399452 | 117400081 | 1 | processed_pseudoge | ENSG00000( |
| 117894632 | 117895683 | -1 | TEC | ENSG00000( |
| 117897813 | 118117340 | -1 | protein_coding | ENSG00000( |
| 117914244 | 117914345 | -1 | misc_RNA | ENSG00000( |
| 118128748 | 118129018 | 1 | processed_pseudoge | ENSG00000( |
| 118146063 | 118146561 | 1 | processed_pseudoge | ENSG00000( |
| 118278159 | 118283367 | 1 | processed_pseudoge | ENSG00000( |
| 118346073 | 118449961 | 1 | protein_coding | ENSG00000( |
| 118386394 | 118386471 | 1 | miRNA | ENSG00000( |
| 118495898 | 118686163 | 1 | protein_coding | ENSG00000( |
| 118569405 | 118569515 | -1 | misc_RNA | ENSG00000( |
| 118632375 | 118634218 | -1 | processed_pseudoge | ENSG00000( |
| 118727133 | 118794535 | 1 | protein_coding | ENSG00000( |
| 118738055 | 118738182 | 1 | snoRNA | ENSG00000( |
| 118769796 | 118770857 | 1 | processed_pseudoge | ENSG00000( |
| 118823824 | 118826968 | 1 | protein_coding | ENSG00000( |
| 118839539 | 118919669 | 1 | lncRNA | ENSG00000( |
| 118877886 | 118879185 | -1 | processed_pseudoge | ENSG00000( |
| 118938153 | 118939134 | 1 | processed_pseudoge | ENSG00000( |
| 118974614 | 119018355 | 1 | protein_coding | ENSG00000( |
| 119073226 | 119076373 | 1 | protein_coding | ENSG00000( |
| 119078635 | 119150579 | -1 | protein_coding | ENSG00000( |
| 119186442 | 119186829 | -1 | processed_pseudoge | ENSG00000( |
| 119220782 | 119221742 | 1 | processed_pseudoge | ENSG00000( |
| 119222564 | 119223245 | -1 | processed_pseudoge | ENSG00000( |
| 119236245 | 119244466 | 1 | protein_coding | ENSG00000( |
| 119249346 | 119257667 | -1 | unprocessed_pseudc | ENSG00000( |
| 119291529 | 119335610 | 1 | lncRNA | ENSG00000( |
| 119356301 | 119393595 | -1 | lncRNA | ENSG00000( |
| 119379712 | 119380104 | 1 | processed_pseudoge | ENSG00000( |
| 119399336 | 119454478 | 1 | protein_coding | ENSG00000( |
| 119422917 | 119423547 | 1 | lncRNA | ENSG00000( |
| 119423742 | 119423905 | 1 | snRNA | ENSG00000( |
| 119434519 | 119434820 | 1 | misc_RNA | ENSG00000( |
| 119456558 | 119456658 | 1 | misc_RNA | ENSG00000( |
| 119465986 | 119469128 | -1 | lncRNA | ENSG00000( |
| 119468422 | 119471396 | 1 | protein_coding | ENSG00000( |
| 119538149 | 119565408 | -1 | protein_coding | ENSG00000( |
| 119574536 | 119591083 | 1 | protein_coding | ENSG00000( |
| 119588337 | 119606443 | -1 | protein_coding | ENSG00000( |
| 119615724 | 119693370 | -1 | protein_coding | ENSG00000( |
| 119646738 | 119646848 | -1 | miRNA | ENSG00000( |
| 119693568 | 119696059 | 1 | TEC | ENSG00000( |
| 119758588 | 119760202 | 1 | protein_coding | ENSG00000( |
| 119786504 | 119791630 | -1 | protein_coding | ENSG00000( |
| 119787353 | 119787484 | -1 | snoRNA | ENSG00000( |
| 119791971 | 119810949 | 1 | lncRNA | ENSG00000( |
| 119805311 | 119852998 | -1 | protein_coding | ENSG00000( |

| | | | | |
|---|---|---|---|---|
| 119870475 | 119871733 | -1 | protein_coding | ENSG00000 |
| 119871832 | 119876662 | 1 | protein_coding | ENSG00000 |
| 119895837 | 119920716 | 1 | protein_coding | ENSG00000 |
| 119920672 | 119943751 | -1 | protein_coding | ENSG00000 |
| 119942834 | 119992810 | 1 | transcribed_unproce | ENSG00000 |
| 119978713 | 120025138 | -1 | transcribed_unproce | ENSG00000 |
| 119998087 | 120000088 | -1 | unprocessed_pseudo | ENSG00000 |
| 120036236 | 120146854 | 1 | lncRNA | ENSG00000 |
| 120070672 | 120077705 | -1 | protein_coding | ENSG00000 |
| 120109051 | 120115913 | -1 | protein_coding | ENSG00000 |
| 120117642 | 120119700 | -1 | lncRNA | ENSG00000 |
| 120120953 | 120245267 | -1 | transcribed_process | ENSG00000 |
| 120158613 | 120165630 | 1 | protein_coding | ENSG00000 |
| 120202061 | 120202748 | -1 | processed_pseudoge | ENSG00000 |
| 120210461 | 120212680 | -1 | processed_pseudoge | ENSG00000 |
| 120250752 | 120258398 | 1 | protein_coding | ENSG00000 |
| 120258650 | 120311556 | -1 | protein_coding | ENSG00000 |
| 120273127 | 120273526 | 1 | processed_pseudoge | ENSG00000 |
| 120362085 | 120383249 | 1 | protein_coding | ENSG00000 |
| 120427827 | 120469365 | -1 | protein_coding | ENSG00000 |
| 120503037 | 120503096 | -1 | snRNA | ENSG00000 |
| 120524609 | 120575794 | -1 | protein_coding | ENSG00000 |
| 120594010 | 120621159 | 1 | protein_coding | ENSG00000 |
| 120625674 | 120630054 | -1 | protein_coding | ENSG00000 |
| 120709061 | 120709392 | -1 | processed_pseudoge | ENSG00000 |
| 120733094 | 120733396 | 1 | processed_pseudoge | ENSG00000 |
| 120868611 | 120869807 | -1 | processed_pseudoge | ENSG00000 |
| 120872603 | 120875929 | -1 | protein_coding | ENSG00000 |
| 120876552 | 120876805 | 1 | processed_pseudoge | ENSG00000 |
| 120877496 | 120878924 | -1 | protein_coding | ENSG00000 |
| 120930250 | 120932301 | -1 | protein_coding | ENSG00000 |
| 120932901 | 120933152 | 1 | processed_pseudoge | ENSG00000 |
| 120933840 | 120937158 | -1 | protein_coding | ENSG00000 |
| 120937762 | 120938015 | 1 | processed_pseudoge | ENSG00000 |
| 120938701 | 120942023 | -1 | protein_coding | ENSG00000 |
| 120942623 | 120942875 | 1 | processed_pseudoge | ENSG00000 |
| 120943561 | 120946883 | -1 | protein_coding | ENSG00000 |
| 120947483 | 120947735 | 1 | processed_pseudoge | ENSG00000 |
| 120948422 | 120951744 | -1 | protein_coding | ENSG00000 |
| 120952344 | 120952595 | 1 | processed_pseudoge | ENSG00000 |
| 120953282 | 120956600 | -1 | protein_coding | ENSG00000 |
| 120957227 | 120957479 | 1 | processed_pseudoge | ENSG00000 |
| 120958165 | 120961487 | -1 | protein_coding | ENSG00000 |
| 120962087 | 120962339 | 1 | processed_pseudoge | ENSG00000 |
| 120963026 | 120966348 | -1 | protein_coding | ENSG00000 |
| 120966948 | 120967199 | 1 | processed_pseudoge | ENSG00000 |
| 120967886 | 120971208 | -1 | protein_coding | ENSG00000 |
| 120971808 | 120972059 | 1 | processed_pseudoge | ENSG00000 |
| 120972746 | 120976068 | -1 | protein_coding | ENSG00000 |
| 120976668 | 120976919 | 1 | processed_pseudoge | ENSG00000 |

| | | | | |
|---|---|---|---|---|
| 120977606 | 120980928 | -1 | protein_coding | ENSG00000 |
| 120981529 | 120981779 | 1 | processed_pseudoge | ENSG00000 |
| 120982476 | 120985788 | -1 | protein_coding | ENSG00000 |
| 120986388 | 120986639 | 1 | processed_pseudoge | ENSG00000 |
| 121024964 | 121026563 | -1 | processed_pseudoge | ENSG00000 |
| 121047610 | 121050094 | 1 | protein_coding | ENSG00000 |
| 121065882 | 121078538 | 1 | lncRNA | ENSG00000 |
| 121093392 | 121298035 | 1 | lncRNA | ENSG00000 |
| 121203182 | 121205014 | -1 | processed_pseudoge | ENSG00000 |
| 121370972 | 121371053 | 1 | miRNA | ENSG00000 |
| 121863698 | 121864196 | -1 | processed_pseudoge | ENSG00000 |
| 121871869 | 121876865 | 1 | lncRNA | ENSG00000 |
| 122010976 | 122011612 | -1 | processed_pseudoge | ENSG00000 |
| 122242008 | 122242616 | 1 | TEC | ENSG00000 |
| 122422006 | 122478043 | 1 | lncRNA | ENSG00000 |
| 122538253 | 122539460 | 1 | processed_pseudoge | ENSG00000 |
| 122840426 | 122840645 | 1 | snoRNA | ENSG00000 |
| 123116107 | 123117781 | -1 | processed_pseudoge | ENSG00000 |
| 123184153 | 123490915 | 1 | protein_coding | ENSG00000 |
| 123514473 | 123514872 | 1 | processed_pseudoge | ENSG00000 |
| 123558350 | 123558584 | 1 | processed_pseudoge | ENSG00000 |
| 123561642 | 123562966 | 1 | processed_pseudoge | ENSG00000 |
| 123600561 | 123733056 | -1 | protein_coding | ENSG00000 |
| 123603139 | 123603931 | 1 | processed_pseudoge | ENSG00000 |
| 123765301 | 123765686 | 1 | processed_pseudoge | ENSG00000 |
| 123772078 | 123772139 | -1 | snRNA | ENSG00000 |
| 123785009 | 123785451 | 1 | processed_pseudoge | ENSG00000 |
| 123811331 | 123811587 | -1 | misc_RNA | ENSG00000 |
| 123857062 | 123857163 | -1 | misc_RNA | ENSG00000 |
| 123859724 | 123913979 | 1 | protein_coding | ENSG00000 |
| 123872626 | 123873932 | -1 | lncRNA | ENSG00000 |
| 123874459 | 123874565 | 1 | snRNA | ENSG00000 |
| 123878590 | 123879296 | -1 | processed_pseudoge | ENSG00000 |
| 123959233 | 123961341 | -1 | lncRNA | ENSG00000 |
| 123960212 | 124422664 | 1 | protein_coding | ENSG00000 |
| 124173001 | 124174459 | -1 | processed_pseudoge | ENSG00000 |
| 124197743 | 124197849 | 1 | snoRNA | ENSG00000 |
| 124204499 | 124205806 | -1 | processed_pseudoge | ENSG00000 |
| 124227868 | 124373197 | 1 | protein_coding | ENSG00000 |
| 124246249 | 124336863 | 1 | protein_coding | ENSG00000 |
| 124280530 | 124281422 | 1 | processed_pseudoge | ENSG00000 |
| 124326175 | 124326258 | -1 | processed_pseudoge | ENSG00000 |
| 124375903 | 124963817 | -1 | protein_coding | ENSG00000 |
| 124688504 | 124688838 | -1 | processed_pseudoge | ENSG00000 |
| 125203805 | 125204338 | -1 | processed_pseudoge | ENSG00000 |
| 125320120 | 125325214 | 1 | protein_coding | ENSG00000 |
| 125326897 | 125328021 | -1 | processed_pseudoge | ENSG00000 |
| 126004446 | 126006118 | -1 | TEC | ENSG00000 |
| 126109762 | 126115562 | 1 | lncRNA | ENSG00000 |
| 126163499 | 126166289 | -1 | protein_coding | ENSG00000 |

| | | | | |
|---|---|---|---|---|
| 126471705 | 126472403 | 1 | processed_pseudoge | ENSG00000 |
| 126472728 | 126472884 | 1 | processed_pseudoge | ENSG00000 |
| 126472885 | 126473281 | 1 | processed_pseudoge | ENSG00000 |
| 126502867 | 126503155 | -1 | unprocessed_pseudc | ENSG00000 |
| 126549383 | 126552814 | -1 | protein_coding | ENSG00000 |
| 126729104 | 126730226 | 1 | processed_pseudoge | ENSG00000 |
| 126819729 | 126821786 | 1 | protein_coding | ENSG00000 |
| 127390342 | 127392526 | 1 | processed_pseudoge | ENSG00000 |
| 127392760 | 127394771 | 1 | TEC | ENSG00000 |
| 127623410 | 127623530 | -1 | processed_pseudoge | ENSG00000 |
| 127660631 | 127662530 | -1 | lncRNA | ENSG00000 |
| 127728734 | 127730858 | -1 | processed_pseudoge | ENSG00000 |
| 127796124 | 127796733 | 1 | processed_pseudoge | ENSG00000 |
| 128027322 | 129326707 | -1 | lncRNA | ENSG00000 |
| 128050962 | 128052398 | -1 | protein_coding | ENSG00000 |
| 128098032 | 128098529 | -1 | processed_pseudoge | ENSG00000 |
| 128122734 | 128123496 | 1 | processed_pseudoge | ENSG00000 |
| 128127090 | 128128212 | -1 | processed_pseudoge | ENSG00000 |
| 128313281 | 128314048 | 1 | processed_pseudoge | ENSG00000 |
| 128440929 | 128441785 | 1 | processed_pseudoge | ENSG00000 |
| 128714506 | 128715790 | 1 | processed_pseudoge | ENSG00000 |
| 128838441 | 128838834 | -1 | processed_pseudoge | ENSG00000 |
| 129041536 | 129043183 | -1 | processed_pseudoge | ENSG00000 |
| 129408382 | 129408621 | 1 | processed_pseudoge | ENSG00000 |
| 129436530 | 129436647 | -1 | rRNA_pseudogene | ENSG00000 |
| 129446501 | 129523500 | -1 | protein_coding | ENSG00000 |
| 129518576 | 129518647 | 1 | snoRNA | ENSG00000 |
| 129539849 | 129592561 | 1 | protein_coding | ENSG00000 |
| 129645259 | 129654956 | -1 | protein_coding | ENSG00000 |
| 129675888 | 129678348 | -1 | lncRNA | ENSG00000 |
| 129738974 | 129769536 | 1 | protein_coding | ENSG00000 |
| 129779949 | 129795201 | 1 | protein_coding | ENSG00000 |
| 129794994 | 129796963 | 1 | lncRNA | ENSG00000 |
| 129796491 | 129796783 | -1 | misc_RNA | ENSG00000 |
| 129803288 | 129843909 | -1 | protein_coding | ENSG00000 |
| 129869064 | 129957528 | -1 | lncRNA | ENSG00000 |
| 129871309 | 129877567 | 1 | processed_pseudoge | ENSG00000 |
| 129906121 | 129929761 | 1 | protein_coding | ENSG00000 |
| 129981107 | 130058083 | 1 | protein_coding | ENSG00000 |
| 130064874 | 130110716 | -1 | protein_coding | ENSG00000 |
| 130129362 | 130165887 | -1 | protein_coding | ENSG00000 |
| 130165954 | 130166905 | 1 | lncRNA | ENSG00000 |
| 130171962 | 130184870 | 1 | protein_coding | ENSG00000 |
| 130202707 | 130268899 | -1 | protein_coding | ENSG00000 |
| 130310072 | 130310178 | -1 | snRNA | ENSG00000 |
| 130330857 | 130331069 | 1 | processed_pseudoge | ENSG00000 |
| 130339888 | 130373361 | 1 | protein_coding | ENSG00000 |
| 130384345 | 130385660 | -1 | protein_coding | ENSG00000 |
| 130401987 | 130413656 | 1 | protein_coding | ENSG00000 |
| 130445085 | 130524257 | -1 | transcribed_unproce | ENSG00000 |

| | | | | |
|---|---|---|---|---|
| 130494965 | 130496588 | 1 | transcribed_processe | ENSG00000 |
| 130497785 | 130520309 | 1 | lncRNA | ENSG00000 |
| 130545586 | 130546474 | 1 | unprocessed_pseudc | ENSG00000 |
| 130623369 | 130903317 | -1 | protein_coding | ENSG00000 |
| 130645723 | 130646325 | 1 | processed_pseudoge | ENSG00000 |
| 130981590 | 131058146 | -1 | lncRNA | ENSG00000 |
| 131058346 | 131089885 | 1 | protein_coding | ENSG00000 |
| 131134086 | 131134390 | -1 | misc_RNA | ENSG00000 |
| 131204068 | 131205010 | 1 | unprocessed_pseudc | ENSG00000 |
| 131221196 | 131221436 | -1 | unprocessed_pseudc | ENSG00000 |
| 131273506 | 131578899 | -1 | protein_coding | ENSG00000 |
| 131324391 | 131325313 | -1 | unprocessed_pseudc | ENSG00000 |
| 131341570 | 131342425 | -1 | unprocessed_pseudc | ENSG00000 |
| 131385453 | 131407083 | 1 | lncRNA | ENSG00000 |
| 131398072 | 131398957 | -1 | unprocessed_pseudc | ENSG00000 |
| 131424301 | 131425239 | 1 | unprocessed_pseudc | ENSG00000 |
| 131437055 | 131437670 | -1 | processed_pseudoge | ENSG00000 |
| 131492279 | 131492886 | -1 | processed_pseudoge | ENSG00000 |
| 131543976 | 131545056 | 1 | protein_coding | ENSG00000 |
| 131562729 | 131563846 | 1 | unprocessed_pseudc | ENSG00000 |
| 131578681 | 131611684 | 1 | lncRNA | ENSG00000 |
| 131596121 | 131596987 | 1 | unprocessed_pseudc | ENSG00000 |
| 131619013 | 131619951 | 1 | unprocessed_pseudc | ENSG00000 |
| 131646639 | 131646890 | 1 | processed_pseudoge | ENSG00000 |
| 131688779 | 131830862 | -1 | lncRNA | ENSG00000 |
| 131795657 | 131796167 | -1 | processed_pseudoge | ENSG00000 |
| 131803312 | 131803741 | -1 | processed_pseudoge | ENSG00000 |
| 131937670 | 131937853 | 1 | processed_pseudoge | ENSG00000 |
| 131938846 | 131938956 | -1 | rRNA_pseudogene | ENSG00000 |
| 132023302 | 132075943 | 1 | protein_coding | ENSG00000 |
| 132076993 | 132128020 | -1 | protein_coding | ENSG00000 |
| 132130091 | 132131352 | -1 | processed_pseudoge | ENSG00000 |
| 132203024 | 132219480 | -1 | protein_coding | ENSG00000 |
| 132217053 | 132432862 | 1 | lncRNA | ENSG00000 |
| 132369317 | 132489968 | -1 | protein_coding | ENSG00000 |
| 132580117 | 132580223 | -1 | snRNA | ENSG00000 |
| 132626016 | 132961395 | -1 | protein_coding | ENSG00000 |
| 132667642 | 132669888 | 1 | lncRNA | ENSG00000 |
| 132907279 | 132907575 | -1 | processed_pseudoge | ENSG00000 |
| 133024631 | 133097109 | -1 | protein_coding | ENSG00000 |
| 133141300 | 133143940 | -1 | processed_pseudoge | ENSG00000 |
| 133152754 | 133152892 | 1 | snoRNA | ENSG00000 |
| 133216662 | 133218354 | -1 | protein_coding | ENSG00000 |
| 133238657 | 133238816 | -1 | snRNA | ENSG00000 |
| 133300103 | 133415489 | -1 | protein_coding | ENSG00000 |
| 133531723 | 133531826 | -1 | snRNA | ENSG00000 |
| 133535745 | 133985895 | -1 | protein_coding | ENSG00000 |
| 133670971 | 133671386 | 1 | processed_pseudoge | ENSG00000 |
| 133762796 | 133763683 | 1 | processed_pseudoge | ENSG00000 |
| 133897443 | 133897934 | 1 | TEC | ENSG00000 |

| | | | | |
|---|---|---|---|---|
| 133989443 | 133990086 | 1 | unprocessed_pseudc | ENSG0000C |
| 134142228 | 134142340 | -1 | misc_RNA | ENSG0000C |
| 134166682 | 134166932 | 1 | processed_pseudoge | ENSG0000C |
| 134168911 | 134174089 | -1 | lncRNA | ENSG0000C |
| 134169378 | 134169452 | -1 | miRNA | ENSG0000C |
| 134169538 | 134169612 | -1 | miRNA | ENSG0000C |
| 134169671 | 134169766 | -1 | miRNA | ENSG0000C |
| 134169809 | 134169877 | -1 | miRNA | ENSG0000C |
| 134170041 | 134170111 | -1 | miRNA | ENSG0000C |
| 134170198 | 134170278 | -1 | miRNA | ENSG0000C |
| 134204353 | 134205916 | 1 | processed_pseudoge | ENSG0000C |
| 134237047 | 134246207 | 1 | protein_coding | ENSG0000C |
| 134272287 | 134273658 | 1 | processed_pseudoge | ENSG0000C |
| 134373253 | 134428791 | 1 | protein_coding | ENSG0000C |
| 134460165 | 134520513 | 1 | protein_coding | ENSG0000C |
| 134522702 | 134522990 | 1 | processed_pseudoge | ENSG0000C |
| 134540185 | 134540262 | -1 | miRNA | ENSG0000C |
| 134540341 | 134540431 | -1 | miRNA | ENSG0000C |
| 134540508 | 134540607 | -1 | miRNA | ENSG0000C |
| 134541341 | 134541437 | -1 | miRNA | ENSG0000C |
| 134543119 | 134546642 | -1 | lncRNA | ENSG0000C |
| 134546328 | 134546398 | -1 | miRNA | ENSG0000C |
| 134546614 | 134546711 | -1 | miRNA | ENSG0000C |
| 134549770 | 134560399 | 1 | lncRNA | ENSG0000C |
| 134565838 | 134764322 | -1 | protein_coding | ENSG0000C |
| 134599457 | 134606254 | 1 | lncRNA | ENSG0000C |
| 134607157 | 134607632 | 1 | processed_pseudoge | ENSG0000C |
| 134650830 | 134651411 | 1 | processed_pseudoge | ENSG0000C |
| 134760208 | 134760348 | -1 | snRNA | ENSG0000C |
| 134769566 | 134797232 | -1 | protein_coding | ENSG0000C |
| 134796395 | 134854835 | 1 | protein_coding | ENSG0000C |
| 134799782 | 134799945 | -1 | pseudogene | ENSG0000C |
| 134887632 | 134915257 | -1 | protein_coding | ENSG0000C |
| 134918310 | 134918416 | -1 | snRNA | ENSG0000C |
| 134949549 | 134953382 | 1 | lncRNA | ENSG0000C |
| 134961711 | 134962305 | -1 | processed_pseudoge | ENSG0000C |
| 134990991 | 134992473 | 1 | protein_coding | ENSG0000C |
| 135020513 | 135022542 | -1 | protein_coding | ENSG0000C |
| 135032355 | 135033546 | 1 | protein_coding | ENSG0000C |
| 135037281 | 135040355 | 1 | lncRNA | ENSG0000C |
| 135050932 | 135052196 | -1 | protein_coding | ENSG0000C |
| 135059685 | 135123952 | 1 | lncRNA | ENSG0000C |
| 135075136 | 135076597 | -1 | processed_pseudoge | ENSG0000C |
| 135094985 | 135098712 | -1 | protein_coding | ENSG0000C |
| 135118955 | 135120526 | -1 | protein_coding | ENSG0000C |
| 135123239 | 135123604 | -1 | lncRNA | ENSG0000C |
| 135135149 | 135135770 | -1 | processed_pseudoge | ENSG0000C |
| 135156536 | 135171398 | -1 | protein_coding | ENSG0000C |
| 135180255 | 135194679 | -1 | unprocessed_pseudc | ENSG0000C |
| 135201731 | 135215232 | 1 | unprocessed_pseudc | ENSG0000C |

| | | | | |
|---|---|---|---|---|
| 135236895 | 135237408 | 1 | processed_pseudoge | ENSG00000 |
| 135248920 | 135344109 | -1 | protein_coding | ENSG00000 |
| 135252061 | 135253583 | 1 | protein_coding | ENSG00000 |
| 135297046 | 135298585 | 1 | processed_pseudoge | ENSG00000 |
| 135309480 | 135309659 | 1 | protein_coding | ENSG00000 |
| 135327758 | 135330743 | 1 | lncRNA | ENSG00000 |
| 135344796 | 135363413 | 1 | protein_coding | ENSG00000 |
| 135396429 | 135397764 | 1 | transcribed_unproce | ENSG00000 |
| 135421944 | 135428075 | 1 | protein_coding | ENSG00000 |
| 135428168 | 135428285 | -1 | rRNA_pseudogene | ENSG00000 |
| 135432960 | 135438634 | 1 | processed_pseudoge | ENSG00000 |
| 135445897 | 135447368 | -1 | unprocessed_pseudo | ENSG00000 |
| 135464329 | 135465484 | -1 | processed_pseudoge | ENSG00000 |
| 135520083 | 135520674 | -1 | lncRNA | ENSG00000 |
| 135520643 | 135582510 | 1 | protein_coding | ENSG00000 |
| 135583651 | 135585160 | -1 | processed_pseudoge | ENSG00000 |
| 135598927 | 135615431 | 1 | unprocessed_pseudo | ENSG00000 |
| 135637623 | 135642756 | -1 | unprocessed_pseudo | ENSG00000 |
| 135673591 | 135686151 | -1 | unprocessed_pseudo | ENSG00000 |
| 135710095 | 135710377 | -1 | unprocessed_pseudo | ENSG00000 |
| 135713453 | 135723539 | 1 | protein_coding | ENSG00000 |
| 135730733 | 135732197 | -1 | processed_pseudoge | ENSG00000 |
| 135751119 | 135752583 | 1 | processed_pseudoge | ENSG00000 |
| 135759846 | 135768191 | -1 | protein_coding | ENSG00000 |
| 135777130 | 135785298 | -1 | protein_coding | ENSG00000 |
| 135794706 | 135802656 | -1 | protein_coding | ENSG00000 |
| 135811668 | 135820012 | -1 | protein_coding | ENSG00000 |
| 135829247 | 135837268 | -1 | protein_coding | ENSG00000 |
| 135846497 | 135854538 | -1 | protein_coding | ENSG00000 |
| 135863418 | 135871812 | -1 | protein_coding | ENSG00000 |
| 135881063 | 135889086 | -1 | protein_coding | ENSG00000 |
| 135889205 | 135913061 | 1 | protein_coding | ENSG00000 |
| 135942478 | 135947139 | -1 | unprocessed_pseudo | ENSG00000 |
| 135962070 | 135974063 | -1 | protein_coding | ENSG00000 |
| 135973841 | 136047269 | 1 | protein_coding | ENSG00000 |
| 136077844 | 136078631 | 1 | processed_pseudoge | ENSG00000 |
| 136097835 | 136098348 | -1 | processed_pseudoge | ENSG00000 |
| 136134317 | 136134420 | -1 | snoRNA | ENSG00000 |
| 136146702 | 136211359 | 1 | protein_coding | ENSG00000 |
| 136213220 | 136256482 | -1 | protein_coding | ENSG00000 |
| 136295690 | 136300298 | -1 | processed_pseudoge | ENSG00000 |
| 136300963 | 136416890 | 1 | protein_coding | ENSG00000 |
| 136487947 | 136493780 | 1 | protein_coding | ENSG00000 |
| 136497079 | 136512346 | 1 | protein_coding | ENSG00000 |
| 136532215 | 136556807 | 1 | protein_coding | ENSG00000 |
| 136550878 | 136550960 | 1 | miRNA | ENSG00000 |
| 136571705 | 136571815 | -1 | misc_RNA | ENSG00000 |
| 136583240 | 136583345 | 1 | snRNA | ENSG00000 |
| 136639539 | 136642429 | 1 | lncRNA | ENSG00000 |
| 136648193 | 136660390 | 1 | protein_coding | ENSG00000 |

| | | | | |
|---|---|---|---|---|
| 136665547 | 136780932 | -1 | protein_coding | ENSG00000 |
| 136773999 | 136774105 | 1 | snRNA | ENSG00000 |
| 136788286 | 136788787 | 1 | processed_pseudoge | ENSG00000 |
| 136791064 | 136791780 | 1 | processed_pseudoge | ENSG00000 |
| 136807223 | 136807877 | -1 | processed_pseudoge | ENSG00000 |
| 136816477 | 136816574 | -1 | misc_RNA | ENSG00000 |
| 136840931 | 136847797 | -1 | lncRNA | ENSG00000 |
| 136847384 | 136848034 | 1 | processed_pseudoge | ENSG00000 |
| 136848004 | 136880764 | -1 | protein_coding | ENSG00000 |
| 136879199 | 136879271 | -1 | snoRNA | ENSG00000 |
| 136909369 | 137022143 | 1 | transcribed_unproce | ENSG00000 |
| 136947748 | 136948133 | -1 | processed_pseudoge | ENSG00000 |
| 136976417 | 136985705 | -1 | processed_pseudoge | ENSG00000 |
| 137027447 | 137033847 | -1 | protein_coding | ENSG00000 |
| 137324972 | 137325049 | -1 | processed_pseudoge | ENSG00000 |
| 137441258 | 137441836 | -1 | processed_pseudoge | ENSG00000 |
| 137556191 | 137564387 | -1 | lncRNA | ENSG00000 |
| 137566127 | 137577691 | 1 | protein_coding | ENSG00000 |
| 137573752 | 137981780 | 1 | lncRNA | ENSG00000 |
| 137595265 | 137595547 | 1 | misc_RNA | ENSG00000 |
| 137660519 | 137662242 | 1 | processed_pseudoge | ENSG00000 |
| 137687891 | 137687994 | -1 | snRNA | ENSG00000 |
| 138397351 | 138397628 | 1 | misc_RNA | ENSG00000 |
| 138482745 | 138483695 | 1 | processed_pseudoge | ENSG00000 |
| 138487665 | 138488871 | -1 | processed_pseudoge | ENSG00000 |
| 138558996 | 138559298 | 1 | unitary_pseudogene | ENSG00000 |
| 138614727 | 139222777 | -1 | protein_coding | ENSG00000 |
| 138627077 | 138627343 | 1 | lncRNA | ENSG00000 |
| 138667711 | 138667793 | -1 | miRNA | ENSG00000 |
| 138711452 | 138716617 | 1 | lncRNA | ENSG00000 |
| 138806374 | 138807091 | 1 | processed_pseudoge | ENSG00000 |
| 139362040 | 139362382 | 1 | processed_pseudoge | ENSG00000 |
| 139446891 | 139447676 | 1 | processed_pseudoge | ENSG00000 |
| 139530758 | 139563458 | 1 | protein_coding | ENSG00000 |
| 139581770 | 139708227 | -1 | protein_coding | ENSG00000 |
| 139702287 | 139702390 | -1 | snoRNA | ENSG00000 |
| 139726346 | 139945276 | -1 | protein_coding | ENSG00000 |
| 139885481 | 139885603 | 1 | snRNA | ENSG00000 |
| 139909904 | 139910034 | 1 | snoRNA | ENSG00000 |
| 139924148 | 139924231 | -1 | miRNA | ENSG00000 |
| 139955728 | 139965521 | -1 | protein_coding | ENSG00000 |
| 140017268 | 140018037 | 1 | lncRNA | ENSG00000 |
| 140032344 | 140033480 | 1 | processed_pseudoge | ENSG00000 |
| 140085855 | 140086150 | 1 | misc_RNA | ENSG00000 |
| 140091743 | 140092561 | 1 | processed_pseudoge | ENSG00000 |
| 140216035 | 140216804 | -1 | lncRNA | ENSG00000 |
| 140397062 | 140398311 | 1 | processed_pseudoge | ENSG00000 |
| 140451148 | 140451208 | 1 | snRNA | ENSG00000 |
| 140470866 | 140471274 | -1 | processed_pseudoge | ENSG00000 |
| 140502985 | 140505116 | -1 | protein_coding | ENSG00000 |

| | | | | |
|---|---|---|---|---|
| 140681106 | 140689819 | -1 | lncRNA | ENSG0000C |
| 140709562 | 140793215 | 1 | lncRNA | ENSG0000C |
| 140783578 | 140784366 | -1 | protein_coding | ENSG0000C |
| 140926172 | 140926219 | -1 | miRNA | ENSG0000C |
| 140931738 | 140997448 | 1 | lncRNA | ENSG0000C |
| 141002594 | 141003706 | 1 | protein_coding | ENSG0000C |
| 141111605 | 141177129 | -1 | protein_coding | ENSG0000C |
| 141118030 | 141118136 | -1 | snRNA | ENSG0000C |
| 141138478 | 141139174 | 1 | processed_pseudoge | ENSG0000C |
| 141177284 | 141649927 | 1 | lncRNA | ENSG0000C |
| 141241463 | 141242517 | -1 | protein_coding | ENSG0000C |
| 141261119 | 141264213 | -1 | processed_pseudoge | ENSG0000C |
| 141380823 | 141381107 | 1 | processed_pseudoge | ENSG0000C |
| 141583674 | 141585011 | -1 | protein_coding | ENSG0000C |
| 141589708 | 141590762 | 1 | protein_coding | ENSG0000C |
| 141625864 | 141626733 | 1 | lncRNA | ENSG0000C |
| 141697411 | 141698739 | -1 | protein_coding | ENSG0000C |
| 141838316 | 141897832 | 1 | protein_coding | ENSG0000C |
| 141903894 | 141909374 | 1 | protein_coding | ENSG0000C |
| 141934496 | 142740581 | 1 | lncRNA | ENSG0000C |
| 142147862 | 142148866 | -1 | unprocessed_pseudc | ENSG0000C |
| 142172866 | 142173989 | -1 | unprocessed_pseudc | ENSG0000C |
| 142194352 | 142195490 | -1 | unprocessed_pseudc | ENSG0000C |
| 142202342 | 142205290 | -1 | protein_coding | ENSG0000C |
| 142372276 | 142373990 | 1 | processed_pseudoge | ENSG0000C |
| 142411053 | 142411171 | 1 | rRNA_pseudogene | ENSG0000C |
| 143025918 | 143034702 | 1 | protein_coding | ENSG0000C |
| 143090347 | 143090677 | 1 | misc_RNA | ENSG0000C |
| 143197248 | 143197591 | -1 | misc_RNA | ENSG0000C |
| 143284977 | 143516803 | 1 | lncRNA | ENSG0000C |
| 143343976 | 143344585 | 1 | processed_pseudoge | ENSG0000C |
| 143433042 | 143433958 | 1 | processed_pseudoge | ENSG0000C |
| 143434517 | 143435280 | 1 | processed_pseudoge | ENSG0000C |
| 143508735 | 143517475 | -1 | protein_coding | ENSG0000C |
| 143622790 | 143636101 | -1 | protein_coding | ENSG0000C |
| 143677153 | 143678362 | -1 | processed_pseudoge | ENSG0000C |
| 143711955 | 143721423 | -1 | protein_coding | ENSG0000C |
| 143757737 | 143828133 | 1 | lncRNA | ENSG0000C |
| 143806391 | 143806733 | 1 | processed_pseudoge | ENSG0000C |
| 143884071 | 143885255 | 1 | polymorphic_pseudc | ENSG0000C |
| 144308834 | 144309080 | -1 | processed_pseudoge | ENSG0000C |
| 144309132 | 144309693 | 1 | processed_pseudoge | ENSG0000C |
| 144840777 | 144841660 | 1 | processed_pseudoge | ENSG0000C |
| 145057108 | 145057437 | 1 | misc_RNA | ENSG0000C |
| 145176582 | 145176894 | 1 | processed_pseudoge | ENSG0000C |
| 145232929 | 145233619 | -1 | lncRNA | ENSG0000C |
| 145247503 | 145256208 | 1 | protein_coding | ENSG0000C |
| 145630218 | 145632375 | 1 | processed_pseudoge | ENSG0000C |
| 145699056 | 145820632 | -1 | lncRNA | ENSG0000C |
| 145768095 | 145768342 | 1 | processed_pseudoge | ENSG0000C |

| | | | | |
|---|---|---|---|---|
| 145797375 | 145798439 | 1 | TEC | ENSG0000( |
| 145817832 | 145829856 | 1 | protein_coding | ENSG0000( |
| 145992750 | 145992826 | -1 | miRNA | ENSG0000( |
| 145994275 | 145994351 | -1 | miRNA | ENSG0000( |
| 145994784 | 145994860 | -1 | miRNA | ENSG0000( |
| 145996669 | 145996743 | -1 | miRNA | ENSG0000( |
| 145997198 | 145997274 | -1 | miRNA | ENSG0000( |
| 146001053 | 146001131 | -1 | miRNA | ENSG0000( |
| 146027794 | 146027872 | -1 | miRNA | ENSG0000( |
| 146427456 | 146427572 | -1 | rRNA_pseudogene | ENSG0000( |
| 146537535 | 146538048 | 1 | processed_pseudoge | ENSG0000( |
| 146553728 | 146555765 | 1 | processed_pseudoge | ENSG0000( |
| 146619597 | 146620584 | -1 | processed_pseudoge | ENSG0000( |
| 146809771 | 146810411 | 1 | protein_coding | ENSG0000( |
| 146814106 | 146814726 | -1 | protein_coding | ENSG0000( |
| 146955075 | 146955209 | 1 | processed_pseudoge | ENSG0000( |
| 146964176 | 146964485 | -1 | processed_pseudoge | ENSG0000( |
| 147181064 | 147271894 | 1 | lncRNA | ENSG0000( |
| 147189704 | 147189787 | -1 | miRNA | ENSG0000( |
| 147199044 | 147199127 | -1 | miRNA | ENSG0000( |
| 147213463 | 147213591 | -1 | miRNA | ENSG0000( |
| 147225826 | 147225952 | -1 | miRNA | ENSG0000( |
| 147230720 | 147230843 | -1 | miRNA | ENSG0000( |
| 147230984 | 147231077 | -1 | miRNA | ENSG0000( |
| 147236913 | 147237027 | -1 | miRNA | ENSG0000( |
| 147250151 | 147250230 | -1 | miRNA | ENSG0000( |
| 147258760 | 147258850 | -1 | miRNA | ENSG0000( |
| 147259652 | 147259726 | -1 | miRNA | ENSG0000( |
| 147260532 | 147260625 | -1 | miRNA | ENSG0000( |
| 147272335 | 147272408 | -1 | miRNA | ENSG0000( |
| 147279247 | 147279344 | -1 | miRNA | ENSG0000( |
| 147279481 | 147286481 | -1 | lncRNA | ENSG0000( |
| 147281943 | 147282030 | -1 | miRNA | ENSG0000( |
| 147284641 | 147284728 | -1 | miRNA | ENSG0000( |
| 147476324 | 147476446 | 1 | processed_pseudoge | ENSG0000( |
| 147591612 | 147591909 | -1 | processed_pseudoge | ENSG0000( |
| 147801005 | 147801302 | 1 | processed_pseudoge | ENSG0000( |
| 147812733 | 147812974 | -1 | processed_pseudoge | ENSG0000( |
| 147827279 | 147827384 | 1 | snRNA | ENSG0000( |
| 147879361 | 147885249 | 1 | lncRNA | ENSG0000( |
| 147900178 | 147901459 | -1 | lncRNA | ENSG0000( |
| 147909431 | 147911817 | -1 | lncRNA | ENSG0000( |
| 147911822 | 147911948 | 1 | misc_RNA | ENSG0000( |
| 147911951 | 147951125 | 1 | protein_coding | ENSG0000( |
| 147946941 | 147947583 | 1 | lncRNA | ENSG0000( |
| 147981337 | 148026665 | 1 | protein_coding | ENSG0000( |
| 148008100 | 148008215 | -1 | rRNA | ENSG0000( |
| 148052233 | 148052746 | 1 | processed_pseudoge | ENSG0000( |
| 148198014 | 148198878 | -1 | processed_pseudoge | ENSG0000( |
| 148464708 | 148465428 | 1 | processed_pseudoge | ENSG0000( |

| | | | | |
|---|---|---|---|---|
| 148498113 | 148500615 | -1 | lncRNA | ENSG00000 |
| 148500617 | 149000663 | 1 | protein_coding | ENSG00000 |
| 148546878 | 148547397 | 1 | lncRNA | ENSG00000 |
| 148822164 | 148822448 | -1 | misc_RNA | ENSG00000 |
| 149056716 | 149057166 | 1 | unprocessed_pseudo | ENSG00000 |
| 149414886 | 149415495 | -1 | protein_coding | ENSG00000 |
| 149476988 | 149521096 | -1 | protein_coding | ENSG00000 |
| 149477104 | 149540926 | -1 | protein_coding | ENSG00000 |
| 149511509 | 149526264 | -1 | lncRNA | ENSG00000 |
| 149525001 | 149526043 | 1 | unprocessed_pseudo | ENSG00000 |
| 149527591 | 149540959 | -1 | lncRNA | ENSG00000 |
| 149540355 | 149550510 | 1 | protein_coding | ENSG00000 |
| 149548210 | 149549932 | -1 | protein_coding | ENSG00000 |
| 149581653 | 149587453 | -1 | protein_coding | ENSG00000 |
| 149592512 | 149595314 | -1 | protein_coding | ENSG00000 |
| 149596556 | 149631912 | -1 | protein_coding | ENSG00000 |
| 149649174 | 149650124 | -1 | processed_pseudoge | ENSG00000 |
| 149688228 | 149717268 | 1 | protein_coding | ENSG00000 |
| 149757797 | 149757895 | 1 | unprocessed_pseudo | ENSG00000 |
| 149758021 | 149758295 | -1 | unprocessed_pseudo | ENSG00000 |
| 149760779 | 149760874 | 1 | unprocessed_pseudo | ENSG00000 |
| 149768897 | 149771368 | 1 | unprocessed_pseudo | ENSG00000 |
| 149774068 | 149776867 | 1 | protein_coding | ENSG00000 |
| 149781930 | 149787737 | 1 | protein_coding | ENSG00000 |
| 149808214 | 149809155 | 1 | unprocessed_pseudo | ENSG00000 |
| 149817170 | 149817644 | 1 | processed_pseudoge | ENSG00000 |
| 149825708 | 149879799 | 1 | lncRNA | ENSG00000 |
| 149878836 | 149881070 | -1 | lncRNA | ENSG00000 |
| 149881141 | 149885835 | 1 | protein_coding | ENSG00000 |
| 149929527 | 149938700 | -1 | protein_coding | ENSG00000 |
| 149929645 | 149931287 | 1 | protein_coding | ENSG00000 |
| 149938613 | 150224580 | 1 | lncRNA | ENSG00000 |
| 150114378 | 150116520 | 1 | processed_pseudoge | ENSG00000 |
| 150228004 | 150228083 | 1 | miRNA | ENSG00000 |
| 150231075 | 150232896 | -1 | processed_pseudoge | ENSG00000 |
| 150361422 | 150514178 | 1 | protein_coding | ENSG00000 |
| 150568619 | 150673322 | 1 | protein_coding | ENSG00000 |
| 150692971 | 150765108 | 1 | protein_coding | ENSG00000 |
| 150766336 | 150898816 | -1 | protein_coding | ENSG00000 |
| 150818753 | 150819246 | 1 | processed_pseudoge | ENSG00000 |
| 150915813 | 150920938 | 1 | lncRNA | ENSG00000 |
| 150955494 | 150955587 | -1 | snRNA | ENSG00000 |
| 150980509 | 150990771 | 1 | protein_coding | ENSG00000 |
| 150993501 | 150994091 | 1 | processed_pseudoge | ENSG00000 |
| 151023607 | 151023722 | -1 | processed_pseudoge | ENSG00000 |
| 151026295 | 151026406 | -1 | rRNA | ENSG00000 |
| 151168222 | 151168326 | 1 | miRNA | ENSG00000 |
| 151175192 | 151177836 | -1 | lncRNA | ENSG00000 |
| 151176584 | 151181465 | 1 | protein_coding | ENSG00000 |
| 151182386 | 151182855 | 1 | lncRNA | ENSG00000 |

| | | | | |
|---|---|---|---|---|
| 151226229 | 151227715 | -1 | processed_pseudoge | ENSG00000 |
| 151275509 | 151275789 | 1 | misc_RNA | ENSG00000 |
| 151303384 | 151397142 | -1 | lncRNA | ENSG00000 |
| 151396515 | 151409364 | 1 | protein_coding | ENSG00000 |
| 151497726 | 151498354 | 1 | lncRNA | ENSG00000 |
| 151515551 | 151516245 | 1 | lncRNA | ENSG00000 |
| 151563675 | 151676739 | 1 | protein_coding | ENSG00000 |
| 151694658 | 151705924 | 1 | protein_coding | ENSG00000 |
| 151716035 | 151723194 | 1 | protein_coding | ENSG00000 |
| 151738451 | 151745304 | 1 | protein_coding | ENSG00000 |
| 151904431 | 151913968 | -1 | lncRNA | ENSG00000 |
| 151912509 | 151925170 | 1 | protein_coding | ENSG00000 |
| 151953124 | 151974680 | -1 | protein_coding | ENSG00000 |
| 151958578 | 151958658 | -1 | miRNA | ENSG00000 |
| 151959628 | 151959712 | -1 | miRNA | ENSG00000 |
| 152045314 | 152046193 | -1 | processed_pseudoge | ENSG00000 |
| 152114599 | 152115540 | -1 | transcribed_unproce | ENSG00000 |
| 152114994 | 152138556 | -1 | lncRNA | ENSG00000 |
| 152133310 | 152138578 | -1 | protein_coding | ENSG00000 |
| 152138883 | 152186857 | 1 | lncRNA | ENSG00000 |
| 152166234 | 152451315 | -1 | protein_coding | ENSG00000 |
| 152250572 | 152250676 | -1 | snRNA | ENSG00000 |
| 152392219 | 152392299 | -1 | miRNA | ENSG00000 |
| 152393421 | 152393529 | -1 | miRNA | ENSG00000 |
| 152394412 | 152394492 | -1 | miRNA | ENSG00000 |
| 152479577 | 152481024 | 1 | processed_pseudoge | ENSG00000 |
| 152574677 | 152698700 | -1 | lncRNA | ENSG00000 |
| 152637895 | 152657542 | 1 | protein_coding | ENSG00000 |
| 152682810 | 152683842 | 1 | unprocessed_pseudo | ENSG00000 |
| 152698767 | 152702347 | 1 | protein_coding | ENSG00000 |
| 152708261 | 152714549 | -1 | protein_coding | ENSG00000 |
| 152714586 | 152718607 | 1 | protein_coding | ENSG00000 |
| 152727484 | 152733735 | -1 | protein_coding | ENSG00000 |
| 152733757 | 152737669 | 1 | protein_coding | ENSG00000 |
| 152733827 | 152740985 | 1 | transcribed_unproce | ENSG00000 |
| 152746817 | 152747762 | 1 | lncRNA | ENSG00000 |
| 152749863 | 152753884 | -1 | protein_coding | ENSG00000 |
| 152753921 | 152760222 | 1 | protein_coding | ENSG00000 |
| 152766136 | 152769747 | -1 | protein_coding | ENSG00000 |
| 152769764 | 152792268 | 1 | lncRNA | ENSG00000 |
| 152826979 | 152830757 | -1 | protein_coding | ENSG00000 |
| 152830967 | 152869729 | 1 | protein_coding | ENSG00000 |
| 152914442 | 152973480 | 1 | protein_coding | ENSG00000 |
| 152925356 | 152927043 | -1 | lncRNA | ENSG00000 |
| 152988824 | 152994116 | -1 | protein_coding | ENSG00000 |
| 153028785 | 153030148 | 1 | unprocessed_pseudo | ENSG00000 |
| 153052150 | 153052413 | -1 | processed_pseudoge | ENSG00000 |
| 153056409 | 153060467 | 1 | protein_coding | ENSG00000 |
| 153072482 | 153075018 | 1 | protein_coding | ENSG00000 |
| 153075769 | 153076968 | -1 | unprocessed_pseudo | ENSG00000 |

| | | | | |
|---|---|---|---|---|
| 153122145 | 153128323 | -1 | processed_pseudoge | ENSG00000( |
| 153179284 | 153183880 | 1 | protein_coding | ENSG00000( |
| 153203044 | 153205534 | 1 | unprocessed_pseudc | ENSG00000( |
| 153225649 | 153230357 | 1 | lncRNA | ENSG00000( |
| 153249115 | 153249691 | -1 | unprocessed_pseudc | ENSG00000( |
| 153271618 | 153277803 | 1 | protein_coding | ENSG00000( |
| 153317680 | 153321767 | -1 | protein_coding | ENSG00000( |
| 153334147 | 153360110 | 1 | protein_coding | ENSG00000( |
| 153355733 | 153355956 | 1 | unprocessed_pseudc | ENSG00000( |
| 153395639 | 153401392 | -1 | protein_coding | ENSG00000( |
| 153403844 | 153404141 | -1 | misc_RNA | ENSG00000( |
| 153418322 | 153426481 | 1 | protein_coding | ENSG00000( |
| 153444720 | 153470587 | -1 | protein_coding | ENSG00000( |
| 153447666 | 153495516 | -1 | protein_coding | ENSG00000( |
| 153479266 | 153487088 | 1 | transcribed_unitary_ | ENSG00000( |
| 153494980 | 153509546 | 1 | protein_coding | ENSG00000( |
| 153517676 | 153582939 | 1 | protein_coding | ENSG00000( |
| 153587925 | 153600045 | -1 | protein_coding | ENSG00000( |
| 153599315 | 153613937 | 1 | transcribed_processe | ENSG00000( |
| 153604497 | 153605348 | -1 | processed_pseudoge | ENSG00000( |
| 153642492 | 153651326 | 1 | protein_coding | ENSG00000( |
| 153662162 | 153662692 | 1 | processed_pseudoge | ENSG00000( |
| 153666226 | 153666510 | 1 | misc_RNA | ENSG00000( |
| 153669730 | 153689010 | -1 | protein_coding | ENSG00000( |
| 153687926 | 153696588 | 1 | protein_coding | ENSG00000( |
| 153700492 | 153724565 | -1 | protein_coding | ENSG00000( |
| 153724856 | 153744755 | 1 | protein_coding | ENSG00000( |
| 153735626 | 153766478 | -1 | lncRNA | ENSG00000( |
| 153764196 | 153779346 | 1 | protein_coding | ENSG00000( |
| 153776412 | 153785732 | 1 | protein_coding | ENSG00000( |
| 153785766 | 153794523 | -1 | protein_coding | ENSG00000( |
| 153793516 | 153798499 | 1 | protein_coding | ENSG00000( |
| 153802166 | 153830565 | -1 | protein_coding | ENSG00000( |
| 153841251 | 153841565 | 1 | processed_pseudoge | ENSG00000( |
| 153861514 | 153886173 | -1 | protein_coding | ENSG00000( |
| 153872221 | 153909223 | -1 | protein_coding | ENSG00000( |
| 153880672 | 153888990 | 1 | lncRNA | ENSG00000( |
| 153902531 | 153907166 | 1 | protein_coding | ENSG00000( |
| 153907367 | 153934999 | -1 | protein_coding | ENSG00000( |
| 153929225 | 153935080 | -1 | protein_coding | ENSG00000( |
| 153935269 | 153944687 | -1 | protein_coding | ENSG00000( |
| 153947557 | 153971818 | -1 | protein_coding | ENSG00000( |
| 153969325 | 153970087 | 1 | lncRNA | ENSG00000( |
| 153972754 | 153983194 | 1 | protein_coding | ENSG00000( |
| 153981097 | 153981177 | 1 | miRNA | ENSG00000( |
| 153981098 | 153981176 | -1 | miRNA | ENSG00000( |
| 154010506 | 154019902 | -1 | protein_coding | ENSG00000( |
| 154019920 | 154019989 | -1 | miRNA | ENSG00000( |
| 154021573 | 154137103 | -1 | protein_coding | ENSG00000( |
| 154144243 | 154159032 | 1 | protein_coding | ENSG00000( |

| | | | | |
|---|---|---|---|---|
| 154159357 | 154176349 | -1 | unprocessed_pseudc | ENSG00000( |
| 154182596 | 154196861 | 1 | protein_coding | ENSG00000( |
| 154196471 | 154213487 | -1 | unprocessed_pseudc | ENSG00000( |
| 154219756 | 154233286 | 1 | protein_coding | ENSG00000( |
| 154233615 | 154251288 | -1 | unprocessed_pseudc | ENSG00000( |
| 154257538 | 154271805 | 1 | protein_coding | ENSG00000( |
| 154271265 | 154295853 | -1 | protein_coding | ENSG00000( |
| 154295795 | 154330350 | 1 | protein_coding | ENSG00000( |
| 154333960 | 154335037 | -1 | lncRNA | ENSG00000( |
| 154348524 | 154374638 | -1 | protein_coding | ENSG00000( |
| 154374595 | 154376934 | 1 | lncRNA | ENSG00000( |
| 154379197 | 154381523 | 1 | protein_coding | ENSG00000( |
| 154389955 | 154409168 | 1 | protein_coding | ENSG00000( |
| 154396878 | 154398816 | -1 | lncRNA | ENSG00000( |
| 154400281 | 154400415 | 1 | snoRNA | ENSG00000( |
| 154401236 | 154412112 | -1 | protein_coding | ENSG00000( |
| 154411524 | 154421726 | 1 | protein_coding | ENSG00000( |
| 154424380 | 154428479 | -1 | lncRNA | ENSG00000( |
| 154428645 | 154436516 | 1 | protein_coding | ENSG00000( |
| 154436913 | 154443467 | 1 | protein_coding | ENSG00000( |
| 154444141 | 154450654 | 1 | protein_coding | ENSG00000( |
| 154450320 | 154450386 | 1 | miRNA | ENSG00000( |
| 154458281 | 154477779 | 1 | protein_coding | ENSG00000( |
| 154477769 | 154479257 | -1 | protein_coding | ENSG00000( |
| 154483717 | 154486615 | -1 | protein_coding | ENSG00000( |
| 154487306 | 154490690 | -1 | protein_coding | ENSG00000( |
| 154495815 | 154496113 | -1 | misc_RNA | ENSG00000( |
| 154506159 | 154516242 | -1 | protein_coding | ENSG00000( |
| 154517840 | 154518631 | -1 | lncRNA | ENSG00000( |
| 154531391 | 154547572 | -1 | protein_coding | ENSG00000( |
| 154541199 | 154565046 | 1 | protein_coding | ENSG00000( |
| 154571248 | 154571960 | 1 | lncRNA | ENSG00000( |
| 154580926 | 154581998 | 1 | processed_pseudoge | ENSG00000( |
| 154585133 | 154586821 | 1 | protein_coding | ENSG00000( |
| 154605858 | 154605950 | 1 | processed_pseudoge | ENSG00000( |
| 154617609 | 154619282 | -1 | protein_coding | ENSG00000( |
| 154620096 | 154620171 | 1 | unprocessed_pseudc | ENSG00000( |
| 154622428 | 154623500 | -1 | processed_pseudoge | ENSG00000( |
| 154632470 | 154633182 | -1 | lncRNA | ENSG00000( |
| 154639978 | 154648275 | -1 | unprocessed_pseudc | ENSG00000( |
| 154651972 | 154653579 | -1 | protein_coding | ENSG00000( |
| 154654546 | 154655325 | 1 | unprocessed_pseudc | ENSG00000( |
| 154658741 | 154659643 | -1 | unprocessed_pseudc | ENSG00000( |
| 154675249 | 154751583 | -1 | protein_coding | ENSG00000( |
| 154762742 | 154777689 | 1 | protein_coding | ENSG00000( |
| 154768528 | 154768659 | 1 | snoRNA | ENSG00000( |
| 154768596 | 154768656 | 1 | miRNA | ENSG00000( |
| 154774998 | 154775126 | 1 | snoRNA | ENSG00000( |
| 154778684 | 154821007 | -1 | protein_coding | ENSG00000( |
| 154817099 | 154817401 | -1 | processed_pseudoge | ENSG00000( |

| | | | | |
|---|---|---|---|---|
| 154823348 | 154834662 | -1 | protein_coding | ENSG00000 |
| 154835788 | 155026940 | -1 | protein_coding | ENSG00000 |
| 154884972 | 154885558 | 1 | protein_coding | ENSG00000 |
| 154886349 | 154888061 | 1 | protein_coding | ENSG00000 |
| 154887360 | 154887458 | -1 | miRNA | ENSG00000 |
| 154897561 | 154898092 | -1 | processed_pseudoge | ENSG00000 |
| 154908885 | 154919103 | -1 | unprocessed_pseudc | ENSG00000 |
| 154919578 | 154920182 | -1 | processed_pseudoge | ENSG00000 |
| 155025980 | 155060304 | 1 | protein_coding | ENSG00000 |
| 155061622 | 155065519 | -1 | protein_coding | ENSG00000 |
| 155061622 | 155071136 | -1 | protein_coding | ENSG00000 |
| 155064034 | 155147937 | -1 | protein_coding | ENSG00000 |
| 155071420 | 155123074 | 1 | protein_coding | ENSG00000 |
| 155137111 | 155156503 | 1 | unprocessed_pseudc | ENSG00000 |
| 155197007 | 155239841 | 1 | protein_coding | ENSG00000 |
| 155212921 | 155213638 | -1 | processed_pseudoge | ENSG00000 |
| 155258235 | 155264491 | -1 | protein_coding | ENSG00000 |
| 155276211 | 155334657 | -1 | protein_coding | ENSG00000 |
| 155290360 | 155291353 | -1 | processed_pseudoge | ENSG00000 |
| 155292169 | 155293432 | 1 | processed_pseudoge | ENSG00000 |
| 155310247 | 155310490 | 1 | processed_pseudoge | ENSG00000 |
| 155334858 | 155351957 | -1 | lncRNA | ENSG00000 |
| 155348752 | 155349799 | 1 | transcribed_unproce | ENSG00000 |
| 155380787 | 155381134 | 1 | protein_coding | ENSG00000 |
| 155382115 | 155383230 | 1 | protein_coding | ENSG00000 |
| 155383100 | 155383198 | -1 | miRNA | ENSG00000 |
| 155456914 | 155458672 | -1 | protein_coding | ENSG00000 |
| 155457517 | 155457615 | 1 | miRNA | ENSG00000 |
| 155459415 | 155460005 | -1 | protein_coding | ENSG00000 |
| 155466540 | 155611616 | 1 | lncRNA | ENSG00000 |
| 155468286 | 155487046 | 1 | lncRNA | ENSG00000 |
| 155489011 | 155669944 | -1 | protein_coding | ENSG00000 |
| 155767812 | 155782459 | 1 | protein_coding | ENSG00000 |
| 155828585 | 155829576 | -1 | processed_pseudoge | ENSG00000 |
| 155875636 | 155875885 | 1 | processed_pseudoge | ENSG00000 |
| 155881345 | 155943769 | 1 | protein_coding | ENSG00000 |
| 155978992 | 155979325 | -1 | processed_pseudoge | ENSG00000 |
| 155985370 | 155986249 | -1 | processed_pseudoge | ENSG00000 |
| 155997581 | 156010817 | 1 | protein_coding | ENSG00000 |
| 156004218 | 156022236 | 1 | lncRNA | ENSG00000 |
| 156014623 | 156016837 | -1 | lncRNA | ENSG00000 |
| 156020826 | 156025710 | 1 | transcribed_unproce | ENSG00000 |
| 156025664 | 156027877 | -1 | unprocessed_pseudc | ENSG00000 |
| 2784749 | 2784853 | 1 | snRNA | ENSG00000 |
| 2786855 | 2787682 | -1 | protein_coding | ENSG00000 |
| 2789827 | 2790328 | 1 | processed_pseudoge | ENSG00000 |
| 2827982 | 2828218 | 1 | processed_pseudoge | ENSG00000 |
| 2828192 | 2840851 | -1 | lncRNA | ENSG00000 |
| 2841602 | 2932000 | 1 | protein_coding | ENSG00000 |
| 2881683 | 2890551 | -1 | transcribed_unproce | ENSG00000 |

| | | | | |
|---|---|---|---|---|
| 2929001 | 2931120 | -1 | processed_pseudoge | ENSG00000 |
| 2934406 | 2934771 | -1 | lncRNA | ENSG00000 |
| 2935281 | 2982506 | 1 | protein_coding | ENSG00000 |
| 2966844 | 3002626 | -1 | lncRNA | ENSG00000 |
| 2995067 | 2995273 | -1 | processed_pseudoge | ENSG00000 |
| 3002887 | 3200509 | 1 | lncRNA | ENSG00000 |
| 3293808 | 3294826 | 1 | processed_pseudoge | ENSG00000 |
| 3549652 | 3549810 | -1 | processed_pseudoge | ENSG00000 |
| 3579041 | 3580041 | 1 | protein_coding | ENSG00000 |
| 3682805 | 3683868 | 1 | processed_pseudoge | ENSG00000 |
| 3777997 | 3779546 | -1 | processed_pseudoge | ENSG00000 |
| 3786282 | 3864893 | 1 | lncRNA | ENSG00000 |
| 3851224 | 3852869 | -1 | processed_pseudoge | ENSG00000 |
| 3866306 | 3866722 | -1 | processed_pseudoge | ENSG00000 |
| 4036497 | 4100320 | 1 | lncRNA | ENSG00000 |
| 4174985 | 4175090 | -1 | snRNA | ENSG00000 |
| 4801685 | 4802848 | -1 | processed_pseudoge | ENSG00000 |
| 4993858 | 4999650 | -1 | lncRNA | ENSG00000 |
| 5000226 | 5742224 | 1 | protein_coding | ENSG00000 |
| 5019076 | 5019266 | 1 | snRNA | ENSG00000 |
| 5207215 | 5208069 | -1 | processed_pseudoge | ENSG00000 |
| 5337745 | 5338964 | -1 | processed_pseudoge | ENSG00000 |
| 5573145 | 5574431 | 1 | processed_pseudoge | ENSG00000 |
| 5634480 | 5634655 | -1 | processed_pseudoge | ENSG00000 |
| 5793300 | 5793737 | -1 | processed_pseudoge | ENSG00000 |
| 5887504 | 5888740 | 1 | processed_pseudoge | ENSG00000 |
| 6158802 | 6159265 | 1 | processed_pseudoge | ENSG00000 |
| 6242446 | 6245283 | -1 | transcribed_unproce | ENSG00000 |
| 6246223 | 6249019 | 1 | protein_coding | ENSG00000 |
| 6256267 | 6265595 | -1 | transcribed_unproce | ENSG00000 |
| 6266593 | 6269275 | 1 | unprocessed_pseudo | ENSG00000 |
| 6273579 | 6274065 | -1 | unprocessed_pseudo | ENSG00000 |
| 6303957 | 6305074 | -1 | unprocessed_pseudo | ENSG00000 |
| 6306042 | 6308202 | 1 | unprocessed_pseudo | ENSG00000 |
| 6328052 | 6343323 | 1 | unprocessed_pseudo | ENSG00000 |
| 6357219 | 6369921 | -1 | lncRNA | ENSG00000 |
| 6390431 | 6411564 | 1 | lncRNA | ENSG00000 |
| 6406244 | 6462091 | -1 | lncRNA | ENSG00000 |
| 6443434 | 6450685 | 1 | lncRNA | ENSG00000 |
| 6449468 | 6457906 | 1 | lncRNA | ENSG00000 |
| 6470773 | 6473630 | -1 | lncRNA | ENSG00000 |
| 6492976 | 6496757 | -1 | unprocessed_pseudo | ENSG00000 |
| 6520463 | 6520929 | 1 | unprocessed_pseudo | ENSG00000 |
| 6718962 | 6719180 | -1 | processed_pseudoge | ENSG00000 |
| 6789153 | 6789344 | 1 | processed_pseudoge | ENSG00000 |
| 6837707 | 6838252 | 1 | processed_pseudoge | ENSG00000 |
| 6865918 | 6911774 | -1 | protein_coding | ENSG00000 |
| 6900753 | 6901372 | 1 | processed_pseudoge | ENSG00000 |
| 6910686 | 7101970 | 1 | protein_coding | ENSG00000 |
| 6918682 | 6920791 | 1 | lncRNA | ENSG00000 |

| | | | | |
|---|---|---|---|---|
| 7100733 | 7106502 | -1 | unprocessed_pseudc | ENSG00000C |
| 7273972 | 7381548 | 1 | protein_coding | ENSG00000C |
| 7324297 | 7324595 | -1 | misc_RNA | ENSG00000C |
| 7341531 | 7341642 | 1 | misc_RNA | ENSG00000C |
| 7378672 | 7378779 | 1 | snRNA | ENSG00000C |
| 7423054 | 7423158 | -1 | snRNA | ENSG00000C |
| 7656968 | 7657352 | 1 | processed_pseudoge | ENSG00000C |
| 7671743 | 7676024 | -1 | unprocessed_pseudc | ENSG00000C |
| 7678899 | 7681028 | 1 | unprocessed_pseudc | ENSG00000C |
| 7687739 | 7689548 | -1 | unprocessed_pseudc | ENSG00000C |
| 7690511 | 7692678 | 1 | unprocessed_pseudc | ENSG00000C |
| 7694707 | 7810305 | -1 | transcribed_unproce | ENSG00000C |
| 7803996 | 7810683 | 1 | lncRNA | ENSG00000C |
| 7845485 | 7846454 | -1 | processed_pseudoge | ENSG00000C |
| 7853686 | 7854876 | -1 | processed_pseudoge | ENSG00000C |
| 7905536 | 7907303 | -1 | processed_pseudoge | ENSG00000C |
| 7913422 | 7914090 | -1 | processed_pseudoge | ENSG00000C |
| 7932997 | 7937640 | -1 | unprocessed_pseudc | ENSG00000C |
| 7990987 | 7991806 | -1 | processed_pseudoge | ENSG00000C |
| 8068445 | 8126293 | -1 | transcribed_processe | ENSG00000C |
| 8127130 | 8144203 | 1 | unprocessed_pseudc | ENSG00000C |
| 8264449 | 8277188 | -1 | unprocessed_pseudc | ENSG00000C |
| 8280198 | 8282209 | 1 | unprocessed_pseudc | ENSG00000C |
| 8350931 | 8352143 | 1 | processed_pseudoge | ENSG00000C |
| 8363536 | 8363959 | -1 | processed_pseudoge | ENSG00000C |
| 8364032 | 8365150 | 1 | processed_pseudoge | ENSG00000C |
| 8371663 | 8372030 | 1 | processed_pseudoge | ENSG00000C |
| 8372241 | 8372710 | 1 | processed_pseudoge | ENSG00000C |
| 8413037 | 8414054 | -1 | processed_pseudoge | ENSG00000C |
| 8418791 | 8419900 | -1 | processed_pseudoge | ENSG00000C |
| 8517364 | 8517748 | -1 | processed_pseudoge | ENSG00000C |
| 8550518 | 8713825 | 1 | lncRNA | ENSG00000C |
| 8682594 | 8700977 | -1 | lncRNA | ENSG00000C |
| 8704472 | 8705283 | 1 | lncRNA | ENSG00000C |
| 8783310 | 8817382 | -1 | transcribed_unproce | ENSG00000C |
| 8901203 | 8903276 | -1 | unprocessed_pseudc | ENSG00000C |
| 8906224 | 8916073 | 1 | transcribed_unproce | ENSG00000C |
| 8931069 | 8931454 | -1 | processed_pseudoge | ENSG00000C |
| 8971751 | 8974095 | -1 | unprocessed_pseudc | ENSG00000C |
| 9030594 | 9039988 | 1 | unprocessed_pseudc | ENSG00000C |
| 9166085 | 9167702 | -1 | processed_pseudoge | ENSG00000C |
| 9183694 | 9185803 | 1 | unprocessed_pseudc | ENSG00000C |
| 9189629 | 9204611 | -1 | unprocessed_pseudc | ENSG00000C |
| 9230915 | 9231247 | 1 | processed_pseudoge | ENSG00000C |
| 9311130 | 9322869 | -1 | unprocessed_pseudc | ENSG00000C |
| 9325799 | 9326047 | 1 | processed_pseudoge | ENSG00000C |
| 9329880 | 9334832 | -1 | lncRNA | ENSG00000C |
| 9337464 | 9340284 | 1 | protein_coding | ENSG00000C |
| 9347511 | 9356814 | -1 | transcribed_processe | ENSG00000C |
| 9357797 | 9360599 | 1 | protein_coding | ENSG00000C |

| | | | | |
|---|---|---|---|---|
| 9367803 | 9375681 | -1 | lncRNA | ENSG00000( |
| 9375129 | 9377121 | -1 | transcribed_unproce | ENSG00000( |
| 9378122 | 9380871 | 1 | unprocessed_pseudc | ENSG00000( |
| 9388122 | 9396027 | -1 | lncRNA | ENSG00000( |
| 9395475 | 9397467 | -1 | transcribed_unproce | ENSG00000( |
| 9398421 | 9401223 | 1 | protein_coding | ENSG00000( |
| 9458613 | 9466003 | -1 | transcribed_unproce | ENSG00000( |
| 9466955 | 9469749 | 1 | protein_coding | ENSG00000( |
| 9479053 | 9486313 | -1 | transcribed_unproce | ENSG00000( |
| 9487313 | 9489893 | 1 | protein_coding | ENSG00000( |
| 9497287 | 9506596 | -1 | transcribed_unproce | ENSG00000( |
| 9507596 | 9510175 | 1 | unprocessed_pseudc | ENSG00000( |
| 9517599 | 9526926 | -1 | transcribed_unproce | ENSG00000( |
| 9527880 | 9530682 | 1 | protein_coding | ENSG00000( |
| 9544917 | 9547147 | -1 | unprocessed_pseudc | ENSG00000( |
| 9548108 | 9550681 | 1 | unprocessed_pseudc | ENSG00000( |
| 9610571 | 9621276 | -1 | unprocessed_pseudc | ENSG00000( |
| 9624183 | 9626352 | 1 | unprocessed_pseudc | ENSG00000( |
| 9640249 | 9640716 | -1 | unprocessed_pseudc | ENSG00000( |
| 9665229 | 9669010 | 1 | unprocessed_pseudc | ENSG00000( |
| 9691100 | 9693957 | 1 | lncRNA | ENSG00000( |
| 9706824 | 9715262 | -1 | lncRNA | ENSG00000( |
| 9717653 | 9721296 | -1 | lncRNA | ENSG00000( |
| 9740584 | 9758476 | 1 | lncRNA | ENSG00000( |
| 9753156 | 9774289 | -1 | lncRNA | ENSG00000( |
| 9784372 | 9833895 | 1 | lncRNA | ENSG00000( |
| 9813315 | 9817513 | 1 | lncRNA | ENSG00000( |
| 9831418 | 9846696 | -1 | unprocessed_pseudc | ENSG00000( |
| 9868462 | 9868702 | -1 | processed_pseudoge | ENSG00000( |
| 9869664 | 9870781 | 1 | unprocessed_pseudc | ENSG00000( |
| 9905595 | 9908139 | -1 | unprocessed_pseudc | ENSG00000( |
| 9910798 | 9911962 | 1 | lncRNA | ENSG00000( |
| 9951553 | 9959423 | 1 | unprocessed_pseudc | ENSG00000( |
| 9954532 | 10035299 | -1 | transcribed_unproce | ENSG00000( |
| 10036195 | 10038375 | 1 | unprocessed_pseudc | ENSG00000( |
| 10066554 | 10069151 | -1 | transcribed_unproce | ENSG00000( |
| 10070131 | 10070409 | 1 | processed_pseudoge | ENSG00000( |
| 10090410 | 10090528 | -1 | rRNA_pseudogene | ENSG00000( |
| 10092875 | 10092993 | -1 | rRNA_pseudogene | ENSG00000( |
| 10171590 | 10172725 | -1 | unprocessed_pseudc | ENSG00000( |
| 10173853 | 10174207 | 1 | unprocessed_pseudc | ENSG00000( |
| 10190377 | 10192298 | -1 | processed_pseudoge | ENSG00000( |
| 10197256 | 10199103 | -1 | processed_pseudoge | ENSG00000( |
| 10200155 | 10200306 | 1 | rRNA_pseudogene | ENSG00000( |
| 11042722 | 11042943 | -1 | processed_pseudoge | ENSG00000( |
| 11085476 | 11086061 | -1 | processed_pseudoge | ENSG00000( |
| 11107065 | 11107263 | -1 | processed_pseudoge | ENSG00000( |
| 11107389 | 11107596 | -1 | processed_pseudoge | ENSG00000( |
| 11107719 | 11107887 | -1 | processed_pseudoge | ENSG00000( |
| 11144250 | 11144492 | 1 | processed_pseudoge | ENSG00000( |

| | | | | |
|---|---|---|---|---|
| 11153858 | 11215026 | -1 | transcribed_unproce | ENSG00000( |
| 11173618 | 11173821 | 1 | processed_pseudoge | ENSG00000( |
| 11178306 | 11178816 | 1 | processed_pseudoge | ENSG00000( |
| 11306918 | 11308181 | 1 | unprocessed_pseudc | ENSG00000( |
| 11314921 | 11316187 | 1 | unprocessed_pseudc | ENSG00000( |
| 11321557 | 11322823 | 1 | unprocessed_pseudc | ENSG00000( |
| 11332329 | 11333595 | 1 | unprocessed_pseudc | ENSG00000( |
| 11335627 | 11337693 | 1 | processed_pseudoge | ENSG00000( |
| 11340579 | 11385687 | -1 | unprocessed_pseudc | ENSG00000( |
| 11395699 | 11397076 | 1 | processed_pseudoge | ENSG00000( |
| 11412548 | 11418716 | 1 | unprocessed_pseudc | ENSG00000( |
| 11445418 | 11445564 | -1 | unprocessed_pseudc | ENSG00000( |
| 11445708 | 11446101 | -1 | processed_pseudoge | ENSG00000( |
| 11473727 | 11474237 | -1 | processed_pseudoge | ENSG00000( |
| 11781052 | 11782527 | -1 | processed_pseudoge | ENSG00000( |
| 11922536 | 11923769 | -1 | processed_pseudoge | ENSG00000( |
| 11957208 | 11987386 | 1 | unprocessed_pseudc | ENSG00000( |
| 12088010 | 12421587 | -1 | transcribed_unproce | ENSG00000( |
| 12244752 | 12245457 | 1 | processed_pseudoge | ENSG00000( |
| 12252321 | 12258033 | -1 | transcribed_unproce | ENSG00000( |
| 12273473 | 12273764 | 1 | misc_RNA | ENSG00000( |
| 12321623 | 12322835 | 1 | processed_pseudoge | ENSG00000( |
| 12339813 | 12347495 | 1 | transcribed_unproce | ENSG00000( |
| 12354096 | 12387328 | 1 | unprocessed_pseudc | ENSG00000( |
| 12440046 | 12507369 | -1 | unprocessed_pseudc | ENSG00000( |
| 12537650 | 12860839 | 1 | protein_coding | ENSG00000( |
| 12537773 | 12544883 | 1 | transcribed_unproce | ENSG00000( |
| 12618983 | 12634402 | -1 | unprocessed_pseudc | ENSG00000( |
| 12661309 | 12663478 | -1 | lncRNA | ENSG00000( |
| 12801973 | 12803830 | -1 | processed_pseudoge | ENSG00000( |
| 12904108 | 12920478 | 1 | protein_coding | ENSG00000( |
| 12930165 | 12948182 | -1 | unprocessed_pseudc | ENSG00000( |
| 13083790 | 13095805 | 1 | processed_pseudoge | ENSG00000( |
| 13153931 | 13162214 | -1 | transcribed_unproce | ENSG00000( |
| 13230884 | 13231439 | 1 | processed_pseudoge | ENSG00000( |
| 13248379 | 13480673 | -1 | protein_coding | ENSG00000( |
| 13286638 | 13287378 | 1 | processed_pseudoge | ENSG00000( |
| 13703899 | 13706024 | 1 | protein_coding | ENSG00000( |
| 13703902 | 13915824 | 1 | transcribed_unproce | ENSG00000( |
| 13985772 | 13986513 | -1 | protein_coding | ENSG00000( |
| 14056227 | 14056958 | 1 | protein_coding | ENSG00000( |
| 14081075 | 14086600 | 1 | unprocessed_pseudc | ENSG00000( |
| 14276212 | 14277489 | -1 | lncRNA | ENSG00000( |
| 14522573 | 14845650 | 1 | protein_coding | ENSG00000( |
| 14639096 | 14640358 | 1 | processed_pseudoge | ENSG00000( |
| 14793642 | 14804033 | -1 | lncRNA | ENSG00000( |
| 14907898 | 14908068 | 1 | processed_pseudoge | ENSG00000( |
| 14941547 | 14941750 | -1 | processed_pseudoge | ENSG00000( |
| 14941746 | 14942715 | 1 | processed_pseudoge | ENSG00000( |
| 15348662 | 15456074 | -1 | unprocessed_pseudc | ENSG00000( |

| | | | | |
|---|---|---|---|---|
| 15547216 | 15593331 | 1 | unprocessed_pseudo | ENSG00000 |
| 16138248 | 16138379 | -1 | snoRNA | ENSG00000 |
| 16248935 | 16249041 | 1 | snRNA | ENSG00000 |
| 16336283 | 16336389 | -1 | snRNA | ENSG00000 |
| 16599730 | 16644539 | -1 | lncRNA | ENSG00000 |
| 17192826 | 17193652 | 1 | processed_pseudoge | ENSG00000 |
| 17464879 | 17465214 | -1 | processed_pseudoge | ENSG00000 |
| 17500958 | 17515018 | 1 | lncRNA | ENSG00000 |
| 17514502 | 17516715 | 1 | transcribed_unproce | ENSG00000 |
| 17516836 | 17519038 | -1 | unprocessed_pseudo | ENSG00000 |
| 17552800 | 17579754 | -1 | lncRNA | ENSG00000 |
| 17557864 | 17557976 | 1 | rRNA_pseudogene | ENSG00000 |
| 17559774 | 17559889 | 1 | rRNA_pseudogene | ENSG00000 |
| 17568962 | 17569051 | -1 | misc_RNA | ENSG00000 |
| 17621259 | 17625832 | -1 | unprocessed_pseudo | ENSG00000 |
| 17628513 | 17629402 | 1 | processed_pseudoge | ENSG00000 |
| 17705968 | 17708185 | -1 | unprocessed_pseudo | ENSG00000 |
| 17721598 | 17723177 | 1 | unprocessed_pseudo | ENSG00000 |
| 17726752 | 17727080 | 1 | processed_pseudoge | ENSG00000 |
| 17729528 | 17744596 | -1 | unprocessed_pseudo | ENSG00000 |
| 17757001 | 17758125 | -1 | processed_pseudoge | ENSG00000 |
| 17768980 | 17777400 | -1 | transcribed_unproce | ENSG00000 |
| 17782210 | 17784815 | 1 | unprocessed_pseudo | ENSG00000 |
| 17788315 | 17789484 | -1 | unprocessed_pseudo | ENSG00000 |
| 17789003 | 17789162 | -1 | snRNA | ENSG00000 |
| 17797983 | 17800696 | -1 | unprocessed_pseudo | ENSG00000 |
| 17809807 | 17825329 | 1 | unprocessed_pseudo | ENSG00000 |
| 17837201 | 17837528 | -1 | processed_pseudoge | ENSG00000 |
| 17877410 | 17880220 | -1 | protein_coding | ENSG00000 |
| 17882099 | 17883708 | -1 | unprocessed_pseudo | ENSG00000 |
| 17908755 | 17910806 | 1 | unprocessed_pseudo | ENSG00000 |
| 17912699 | 17921630 | -1 | unprocessed_pseudo | ENSG00000 |
| 17951589 | 17953500 | 1 | unprocessed_pseudo | ENSG00000 |
| 17984349 | 17993260 | 1 | unprocessed_pseudo | ENSG00000 |
| 17995201 | 17997252 | -1 | unprocessed_pseudo | ENSG00000 |
| 18022299 | 18023908 | 1 | unprocessed_pseudo | ENSG00000 |
| 18025787 | 18028597 | 1 | protein_coding | ENSG00000 |
| 18068483 | 18068810 | 1 | processed_pseudoge | ENSG00000 |
| 18080681 | 18096202 | -1 | unprocessed_pseudo | ENSG00000 |
| 18105314 | 18108102 | 1 | unprocessed_pseudo | ENSG00000 |
| 18116527 | 18117695 | 1 | unprocessed_pseudo | ENSG00000 |
| 18116848 | 18117007 | 1 | snRNA | ENSG00000 |
| 18121195 | 18123799 | -1 | unprocessed_pseudo | ENSG00000 |
| 18128610 | 18137029 | 1 | transcribed_unproce | ENSG00000 |
| 18147884 | 18149008 | 1 | processed_pseudoge | ENSG00000 |
| 18161366 | 18176484 | 1 | unprocessed_pseudo | ENSG00000 |
| 18178932 | 18179260 | -1 | processed_pseudoge | ENSG00000 |
| 18182835 | 18184414 | -1 | unprocessed_pseudo | ENSG00000 |
| 18276607 | 18277495 | -1 | processed_pseudoge | ENSG00000 |
| 18280178 | 18284761 | 1 | unprocessed_pseudo | ENSG00000 |

| | | | | |
|---|---|---|---|---|
| 18326253 | 18353210 | 1 | lncRNA | ENSG00000 |
| 18336960 | 18337049 | 1 | misc_RNA | ENSG00000 |
| 18346123 | 18346238 | -1 | rRNA_pseudogene | ENSG00000 |
| 18348035 | 18348147 | -1 | rRNA_pseudogene | ENSG00000 |
| 18386974 | 18389151 | 1 | unprocessed_pseudo | ENSG00000 |
| 18389297 | 18405046 | -1 | transcribed_unproce | ENSG00000 |
| 18390994 | 18405046 | -1 | lncRNA | ENSG00000 |
| 18440797 | 18441125 | 1 | processed_pseudoge | ENSG00000 |
| 18453150 | 18472137 | -1 | unprocessed_pseudo | ENSG00000 |
| 18481087 | 18487400 | 1 | unprocessed_pseudo | ENSG00000 |
| 18486511 | 18486672 | 1 | snRNA | ENSG00000 |
| 18490915 | 18494295 | -1 | unprocessed_pseudo | ENSG00000 |
| 18491740 | 18547698 | 1 | lncRNA | ENSG00000 |
| 18508577 | 18509068 | 1 | unprocessed_pseudo | ENSG00000 |
| 18529358 | 18529623 | 1 | unprocessed_pseudo | ENSG00000 |
| 18532326 | 18532656 | -1 | processed_pseudoge | ENSG00000 |
| 18540980 | 18544128 | 1 | unprocessed_pseudo | ENSG00000 |
| 18546671 | 18588963 | 1 | protein_coding | ENSG00000 |
| 18578417 | 18578560 | 1 | unprocessed_pseudo | ENSG00000 |
| 18581206 | 18590521 | -1 | lncRNA | ENSG00000 |
| 18582063 | 18629077 | -1 | unprocessed_pseudo | ENSG00000 |
| 18649671 | 18650279 | 1 | unprocessed_pseudo | ENSG00000 |
| 18655520 | 18656190 | -1 | unprocessed_pseudo | ENSG00000 |
| 18674033 | 18739197 | 1 | transcribed_unproce | ENSG00000 |
| 18731440 | 18773735 | -1 | protein_coding | ENSG00000 |
| 18741843 | 18741986 | -1 | unprocessed_pseudo | ENSG00000 |
| 18772706 | 19077563 | -1 | lncRNA | ENSG00000 |
| 18776279 | 18779427 | -1 | unprocessed_pseudo | ENSG00000 |
| 18787750 | 18788080 | 1 | processed_pseudoge | ENSG00000 |
| 18790709 | 18791051 | -1 | transcribed_unproce | ENSG00000 |
| 18811338 | 18811829 | -1 | unprocessed_pseudo | ENSG00000 |
| 18826107 | 18829487 | 1 | unprocessed_pseudo | ENSG00000 |
| 18833002 | 18839313 | -1 | unprocessed_pseudo | ENSG00000 |
| 18833729 | 18833890 | -1 | snRNA | ENSG00000 |
| 18848264 | 18867280 | 1 | unprocessed_pseudo | ENSG00000 |
| 18872102 | 18872272 | 1 | processed_pseudoge | ENSG00000 |
| 18985175 | 18986398 | 1 | processed_pseudoge | ENSG00000 |
| 18992467 | 18992709 | -1 | processed_pseudoge | ENSG00000 |
| 19018983 | 19019087 | -1 | snRNA | ENSG00000 |
| 19076946 | 19077416 | -1 | protein_coding | ENSG00000 |
| 19253960 | 19260875 | -1 | processed_pseudoge | ENSG00000 |
| 19327569 | 19328573 | 1 | processed_pseudoge | ENSG00000 |
| 19455431 | 19567092 | -1 | transcribed_unproce | ENSG00000 |
| 19567313 | 19606274 | 1 | transcribed_unproce | ENSG00000 |
| 19598188 | 19598757 | 1 | transcribed_processe | ENSG00000 |
| 19691941 | 19694606 | -1 | lncRNA | ENSG00000 |
| 19703865 | 19744939 | -1 | protein_coding | ENSG00000 |
| 19744756 | 19759978 | 1 | lncRNA | ENSG00000 |
| 19932130 | 19932500 | -1 | processed_pseudoge | ENSG00000 |
| 19986575 | 19987852 | -1 | processed_pseudoge | ENSG00000 |

| | | | | |
|---|---|---|---|---|
| 20001185 | 20012204 | -1 | unprocessed_pseudc | ENSG00000( |
| 20390051 | 20396796 | 1 | processed_pseudoge | ENSG00000( |
| 20464916 | 20575519 | -1 | lncRNA | ENSG00000( |
| 20570222 | 20570519 | -1 | processed_pseudoge | ENSG00000( |
| 20575776 | 20593154 | 1 | protein_coding | ENSG00000( |
| 20725331 | 20734526 | -1 | lncRNA | ENSG00000( |
| 20727254 | 20742900 | 1 | unprocessed_pseudc | ENSG00000( |
| 20756108 | 20781032 | 1 | protein_coding | ENSG00000( |
| 20808776 | 20811809 | -1 | unprocessed_pseudc | ENSG00000( |
| 20843256 | 20843710 | -1 | processed_pseudoge | ENSG00000( |
| 20861337 | 20862337 | 1 | processed_pseudoge | ENSG00000( |
| 20907344 | 20907562 | -1 | processed_pseudoge | ENSG00000( |
| 21038289 | 21044724 | -1 | lncRNA | ENSG00000( |
| 21130870 | 21131181 | 1 | processed_pseudoge | ENSG00000( |
| 21221106 | 21222089 | -1 | processed_pseudoge | ENSG00000( |
| 21311268 | 21318895 | 1 | unprocessed_pseudc | ENSG00000( |
| 21382954 | 21386360 | -1 | lncRNA | ENSG00000( |
| 21394991 | 21401585 | -1 | transcribed_unproce | ENSG00000( |
| 21405770 | 21405995 | 1 | unprocessed_pseudc | ENSG00000( |
| 21424028 | 21456216 | -1 | lncRNA | ENSG00000( |
| 21430697 | 21437245 | -1 | unprocessed_pseudc | ENSG00000( |
| 21468158 | 21470683 | 1 | unprocessed_pseudc | ENSG00000( |
| 21493519 | 21501968 | 1 | unprocessed_pseudc | ENSG00000( |
| 21508299 | 21510470 | -1 | unprocessed_pseudc | ENSG00000( |
| 21511338 | 21549094 | 1 | protein_coding | ENSG00000( |
| 21511372 | 21549326 | 1 | protein_coding | ENSG00000( |
| 21531839 | 21534011 | -1 | unprocessed_pseudc | ENSG00000( |
| 21565092 | 21605081 | -1 | lncRNA | ENSG00000( |
| 21631683 | 21632015 | -1 | processed_pseudoge | ENSG00000( |
| 21637475 | 21639041 | -1 | unprocessed_pseudc | ENSG00000( |
| 21661926 | 21674008 | 1 | unprocessed_pseudc | ENSG00000( |
| 21677246 | 21681118 | -1 | unprocessed_pseudc | ENSG00000( |
| 21696570 | 21698220 | 1 | unprocessed_pseudc | ENSG00000( |
| 21817579 | 21818218 | -1 | unprocessed_pseudc | ENSG00000( |
| 21825441 | 21846721 | 1 | lncRNA | ENSG00000( |
| 21871331 | 21873550 | 1 | unprocessed_pseudc | ENSG00000( |
| 21880076 | 21894526 | -1 | protein_coding | ENSG00000( |
| 21895394 | 21897559 | 1 | unprocessed_pseudc | ENSG00000( |
| 21903618 | 21918067 | -1 | protein_coding | ENSG00000( |
| 21918935 | 21921106 | 1 | unprocessed_pseudc | ENSG00000( |
| 21927521 | 21937150 | -1 | unprocessed_pseudc | ENSG00000( |
| 21940012 | 21941593 | 1 | unprocessed_pseudc | ENSG00000( |
| 21972313 | 22005349 | 1 | unprocessed_pseudc | ENSG00000( |
| 22048554 | 22049347 | -1 | unprocessed_pseudc | ENSG00000( |
| 22049755 | 22058113 | -1 | unprocessed_pseudc | ENSG00000( |
| 22064698 | 22065712 | 1 | unprocessed_pseudc | ENSG00000( |
| 22068820 | 22069151 | 1 | processed_pseudoge | ENSG00000( |
| 22071756 | 22096007 | -1 | protein_coding | ENSG00000( |
| 22100814 | 22147484 | -1 | lncRNA | ENSG00000( |
| 22144966 | 22146831 | 1 | lncRNA | ENSG00000( |

| | | | | |
|---|---|---|---|---|
| 22168542 | 22182982 | -1 | protein_coding | ENSG00000 |
| 22183850 | 22186021 | 1 | unprocessed_pseudo | ENSG00000 |
| 22198429 | 22203712 | 1 | unprocessed_pseudo | ENSG00000 |
| 22209331 | 22216580 | 1 | unprocessed_pseudo | ENSG00000 |
| 22296798 | 22298876 | -1 | lncRNA | ENSG00000 |
| 22305925 | 22307951 | -1 | unprocessed_pseudo | ENSG00000 |
| 22308859 | 22321825 | 1 | transcribed_unproce | ENSG00000 |
| 22330452 | 22332500 | 1 | unprocessed_pseudo | ENSG00000 |
| 22400403 | 22402568 | -1 | unprocessed_pseudo | ENSG00000 |
| 22403461 | 22417881 | 1 | protein_coding | ENSG00000 |
| 22438940 | 22485592 | 1 | lncRNA | ENSG00000 |
| 22439593 | 22441458 | -1 | lncRNA | ENSG00000 |
| 22490397 | 22514637 | 1 | protein_coding | ENSG00000 |
| 22517242 | 22517573 | -1 | processed_pseudoge | ENSG00000 |
| 22520681 | 22521695 | -1 | unprocessed_pseudo | ENSG00000 |
| 22528281 | 22536639 | 1 | unprocessed_pseudo | ENSG00000 |
| 22537047 | 22537849 | 1 | unprocessed_pseudo | ENSG00000 |
| 22580989 | 22614081 | -1 | unprocessed_pseudo | ENSG00000 |
| 22646031 | 22646327 | -1 | unprocessed_pseudo | ENSG00000 |
| 22649245 | 22658878 | 1 | transcribed_unproce | ENSG00000 |
| 22670616 | 22672859 | -1 | unprocessed_pseudo | ENSG00000 |
| 22762851 | 22769787 | -1 | unprocessed_pseudo | ENSG00000 |
| 22851584 | 22852715 | 1 | lncRNA | ENSG00000 |
| 22861112 | 22861834 | -1 | unprocessed_pseudo | ENSG00000 |
| 22866544 | 22867756 | -1 | processed_pseudoge | ENSG00000 |
| 22936455 | 22973284 | 1 | lncRNA | ENSG00000 |
| 22973819 | 23005465 | 1 | protein_coding | ENSG00000 |
| 23017063 | 23017821 | 1 | processed_pseudoge | ENSG00000 |
| 23025594 | 23026663 | 1 | processed_pseudoge | ENSG00000 |
| 23049247 | 23051195 | 1 | processed_pseudoge | ENSG00000 |
| 23058030 | 23059186 | 1 | processed_pseudoge | ENSG00000 |
| 23129355 | 23199094 | -1 | protein_coding | ENSG00000 |
| 23219434 | 23291356 | 1 | protein_coding | ENSG00000 |
| 23379484 | 23392697 | 1 | unprocessed_pseudo | ENSG00000 |
| 23423099 | 23423236 | -1 | snoRNA | ENSG00000 |
| 23435905 | 23436006 | 1 | unprocessed_pseudo | ENSG00000 |
| 23462465 | 23472280 | 1 | unprocessed_pseudo | ENSG00000 |
| 23523321 | 23525369 | 1 | unprocessed_pseudo | ENSG00000 |
| 23537308 | 23546942 | -1 | transcribed_processe | ENSG00000 |
| 23549809 | 23551385 | 1 | unprocessed_pseudo | ENSG00000 |
| 23581595 | 23614640 | 1 | unprocessed_pseudo | ENSG00000 |
| 23657839 | 23659058 | -1 | unprocessed_pseudo | ENSG00000 |
| 23659466 | 23667822 | -1 | unprocessed_pseudo | ENSG00000 |
| 23674379 | 23675392 | 1 | unprocessed_pseudo | ENSG00000 |
| 23678504 | 23678835 | 1 | unprocessed_pseudo | ENSG00000 |
| 23681440 | 23694579 | -1 | protein_coding | ENSG00000 |
| 23715922 | 23726808 | -1 | unprocessed_pseudo | ENSG00000 |
| 23734193 | 23736249 | 1 | unprocessed_pseudo | ENSG00000 |
| 23740258 | 23746693 | -1 | unprocessed_pseudo | ENSG00000 |
| 23740942 | 23741105 | -1 | snRNA | ENSG00000 |

| | | | | |
|---|---|---|---|---|
| 23751188 | 23751760 | 1 | unprocessed_pseudc | ENSG00000( |
| 23762838 | 23765583 | -1 | unprocessed_pseudc | ENSG00000( |
| 23771232 | 23798150 | 1 | unprocessed_pseudc | ENSG00000( |
| 23808821 | 23809059 | -1 | processed_pseudoge | ENSG00000( |
| 23855522 | 23857091 | -1 | unprocessed_pseudc | ENSG00000( |
| 23917241 | 23917723 | -1 | unprocessed_pseudc | ENSG00000( |
| 23919905 | 23921510 | 1 | unprocessed_pseudc | ENSG00000( |
| 23934953 | 23940101 | 1 | unprocessed_pseudc | ENSG00000( |
| 23936727 | 23941622 | -1 | lncRNA | ENSG00000( |
| 23945927 | 23952293 | -1 | unprocessed_pseudc | ENSG00000( |
| 23946618 | 23946771 | -1 | snRNA | ENSG00000( |
| 23956569 | 23960218 | 1 | unprocessed_pseudc | ENSG00000( |
| 23967564 | 23970694 | -1 | unprocessed_pseudc | ENSG00000( |
| 23972780 | 23995081 | 1 | unprocessed_pseudc | ENSG00000( |
| 24006911 | 24007237 | -1 | processed_pseudoge | ENSG00000( |
| 24045229 | 24048019 | -1 | protein_coding | ENSG00000( |
| 24049878 | 24051487 | -1 | unprocessed_pseudc | ENSG00000( |
| 24077799 | 24079812 | 1 | unprocessed_pseudc | ENSG00000( |
| 24081704 | 24090346 | -1 | unprocessed_pseudc | ENSG00000( |
| 24104107 | 24106018 | 1 | unprocessed_pseudc | ENSG00000( |
| 24142358 | 24157843 | 1 | unprocessed_pseudc | ENSG00000( |
| 24168187 | 24174494 | 1 | unprocessed_pseudc | ENSG00000( |
| 24179316 | 24183505 | -1 | unprocessed_pseudc | ENSG00000( |
| 24182477 | 24183071 | 1 | unprocessed_pseudc | ENSG00000( |
| 24183434 | 24187231 | -1 | lncRNA | ENSG00000( |
| 24186509 | 24191870 | -1 | unprocessed_pseudc | ENSG00000( |
| 24196518 | 24206132 | -1 | unprocessed_pseudc | ENSG00000( |
| 24209567 | 24215581 | -1 | transcribed_unproce | ENSG00000( |
| 24210958 | 24211216 | -1 | misc_RNA | ENSG00000( |
| 24232826 | 24238984 | -1 | unprocessed_pseudc | ENSG00000( |
| 24276245 | 24277026 | 1 | processed_pseudoge | ENSG00000( |
| 24278337 | 24281140 | -1 | processed_pseudoge | ENSG00000( |
| 24283062 | 24283230 | 1 | unprocessed_pseudc | ENSG00000( |
| 24362335 | 24362409 | -1 | processed_pseudoge | ENSG00000( |
| 24396604 | 24403526 | -1 | unprocessed_pseudc | ENSG00000( |
| 24485332 | 24486463 | 1 | lncRNA | ENSG00000( |
| 24494709 | 24495583 | -1 | processed_pseudoge | ENSG00000( |
| 24500263 | 24501505 | -1 | processed_pseudoge | ENSG00000( |
| 24570202 | 24607025 | 1 | lncRNA | ENSG00000( |
| 24607560 | 24639207 | 1 | protein_coding | ENSG00000( |
| 24650808 | 24651534 | 1 | processed_pseudoge | ENSG00000( |
| 24659337 | 24660406 | 1 | processed_pseudoge | ENSG00000( |
| 24683657 | 24684935 | 1 | processed_pseudoge | ENSG00000( |
| 24691791 | 24692932 | 1 | processed_pseudoge | ENSG00000( |
| 24763069 | 24813492 | -1 | protein_coding | ENSG00000( |
| 24833843 | 24907040 | 1 | protein_coding | ENSG00000( |
| 24977176 | 24978317 | -1 | unprocessed_pseudc | ENSG00000( |
| 24985201 | 24986476 | -1 | processed_pseudoge | ENSG00000( |
| 25009700 | 25010769 | -1 | unprocessed_pseudc | ENSG00000( |
| 25018499 | 25019303 | -1 | processed_pseudoge | ENSG00000( |

| | | | | |
|---|---|---|---|---|
| 25030901 | 25062548 | -1 | protein_coding | ENSG00000( |
| 25063083 | 25099892 | -1 | lncRNA | ENSG00000( |
| 25168599 | 25169841 | 1 | processed_pseudoge | ENSG00000( |
| 25174520 | 25175237 | 1 | processed_pseudoge | ENSG00000( |
| 25182277 | 25213389 | -1 | lncRNA | ENSG00000( |
| 25266584 | 25273531 | 1 | unprocessed_pseudo | ENSG00000( |
| 25307729 | 25307803 | 1 | processed_pseudoge | ENSG00000( |
| 25378300 | 25394719 | -1 | lncRNA | ENSG00000( |
| 25388992 | 25391811 | 1 | processed_pseudoge | ENSG00000( |
| 25393154 | 25394056 | -1 | processed_pseudoge | ENSG00000( |
| 25431156 | 25437315 | 1 | unprocessed_pseudo | ENSG00000( |
| 25454561 | 25460572 | 1 | unprocessed_pseudo | ENSG00000( |
| 25458926 | 25459184 | 1 | misc_RNA | ENSG00000( |
| 25464006 | 25473621 | 1 | unprocessed_pseudo | ENSG00000( |
| 25478269 | 25483630 | 1 | unprocessed_pseudo | ENSG00000( |
| 25482908 | 25486705 | 1 | lncRNA | ENSG00000( |
| 25486640 | 25487322 | 1 | processed_pseudoge | ENSG00000( |
| 25487284 | 25487662 | -1 | processed_pseudoge | ENSG00000( |
| 25495651 | 25501958 | -1 | unprocessed_pseudo | ENSG00000( |
| 25512301 | 25527786 | -1 | unprocessed_pseudo | ENSG00000( |
| 25564122 | 25566033 | -1 | unprocessed_pseudo | ENSG00000( |
| 25579790 | 25588444 | 1 | unprocessed_pseudo | ENSG00000( |
| 25590323 | 25592345 | -1 | unprocessed_pseudo | ENSG00000( |
| 25618644 | 25620253 | 1 | unprocessed_pseudo | ENSG00000( |
| 25622162 | 25624902 | 1 | protein_coding | ENSG00000( |
| 25662900 | 25663226 | 1 | processed_pseudoge | ENSG00000( |
| 25675060 | 25697346 | -1 | unprocessed_pseudo | ENSG00000( |
| 25699451 | 25702562 | 1 | unprocessed_pseudo | ENSG00000( |
| 25709908 | 25713557 | -1 | unprocessed_pseudo | ENSG00000( |
| 25723016 | 25724186 | 1 | unprocessed_pseudo | ENSG00000( |
| 25723342 | 25723495 | 1 | snRNA | ENSG00000( |
| 25728490 | 25733388 | 1 | lncRNA | ENSG00000( |
| 25730011 | 25735160 | -1 | unprocessed_pseudo | ENSG00000( |
| 25748596 | 25750206 | -1 | unprocessed_pseudo | ENSG00000( |
| 25752388 | 25752870 | 1 | unprocessed_pseudo | ENSG00000( |
| 25813013 | 25814566 | 1 | unprocessed_pseudo | ENSG00000( |
| 25860943 | 25861268 | 1 | processed_pseudoge | ENSG00000( |
| 25871931 | 25898653 | -1 | unprocessed_pseudo | ENSG00000( |
| 25904518 | 25907250 | 1 | unprocessed_pseudo | ENSG00000( |
| 25918323 | 25918895 | -1 | processed_pseudoge | ENSG00000( |
| 25923388 | 25929826 | 1 | unprocessed_pseudo | ENSG00000( |
| 25928979 | 25929142 | 1 | snRNA | ENSG00000( |
| 25933833 | 25935881 | -1 | unprocessed_pseudo | ENSG00000( |
| 25943268 | 25954152 | 1 | unprocessed_pseudo | ENSG00000( |
| 25975549 | 25988069 | 1 | unprocessed_pseudo | ENSG00000( |
| 25991241 | 25991572 | -1 | processed_pseudoge | ENSG00000( |
| 25994684 | 25995697 | -1 | unprocessed_pseudo | ENSG00000( |
| 26002254 | 26008856 | 1 | unprocessed_pseudo | ENSG00000( |
| 26011435 | 26012237 | 1 | unprocessed_pseudo | ENSG00000( |
| 26055779 | 26088493 | -1 | unprocessed_pseudo | ENSG00000( |

| | | | | |
|---|---|---|---|---|
| 26123674 | 26133308 | 1 | transcribed_processe | ENSG00000 |
| 26198331 | 26208148 | -1 | unprocessed_pseudo | ENSG00000 |
| 26244351 | 26244573 | -1 | unprocessed_pseudo | ENSG00000 |
| 26247384 | 26247521 | 1 | snoRNA | ENSG00000 |
| 26277923 | 26354418 | -1 | unprocessed_pseudo | ENSG00000 |
| 26360989 | 26361092 | 1 | snRNA | ENSG00000 |
| 26400611 | 26401230 | 1 | processed_pseudoge | ENSG00000 |
| 26409815 | 26420535 | 1 | unprocessed_pseudo | ENSG00000 |
| 26453982 | 26454092 | 1 | processed_pseudoge | ENSG00000 |
| 26508213 | 26579690 | 1 | unprocessed_pseudo | ENSG00000 |
| 26549425 | 26549743 | 1 | processed_pseudoge | ENSG00000 |
| 26586642 | 26591601 | -1 | unprocessed_pseudo | ENSG00000 |
| 26594851 | 26634652 | -1 | unprocessed_pseudo | ENSG00000 |
| 26626520 | 26627159 | -1 | processed_pseudoge | ENSG00000 |
| 56855244 | 56855488 | 1 | processed_pseudoge | ENSG00000 |
| 11869 | 14409 | 1 | transcribed_unproce | ENSG00000 |
| 14404 | 29570 | -1 | unprocessed_pseudo | ENSG00000 |
| 17369 | 17436 | -1 | miRNA | ENSG00000 |
| 29554 | 31109 | 1 | lncRNA | ENSG00000 |
| 30366 | 30503 | 1 | miRNA | ENSG00000 |
| 34554 | 36081 | -1 | lncRNA | ENSG00000 |
| 52473 | 53312 | 1 | unprocessed_pseudo | ENSG00000 |
| 57598 | 64116 | 1 | transcribed_unproce | ENSG00000 |
| 65419 | 71585 | 1 | protein_coding | ENSG00000 |
| 89295 | 133723 | -1 | lncRNA | ENSG00000 |
| 89551 | 91105 | -1 | lncRNA | ENSG00000 |
| 131025 | 134836 | 1 | processed_pseudoge | ENSG00000 |
| 135141 | 135895 | -1 | processed_pseudoge | ENSG00000 |
| 137682 | 137965 | -1 | processed_pseudoge | ENSG00000 |
| 139790 | 140339 | -1 | lncRNA | ENSG00000 |
| 141474 | 173862 | -1 | lncRNA | ENSG00000 |
| 157784 | 157887 | -1 | snRNA | ENSG00000 |
| 160446 | 161525 | 1 | lncRNA | ENSG00000 |
| 182696 | 184174 | 1 | unprocessed_pseudo | ENSG00000 |
| 185217 | 195411 | -1 | unprocessed_pseudo | ENSG00000 |
| 187891 | 187958 | -1 | miRNA | ENSG00000 |
| 257864 | 359681 | -1 | transcribed_processe | ENSG00000 |
| 266855 | 268655 | 1 | lncRNA | ENSG00000 |
| 347982 | 348366 | -1 | processed_pseudoge | ENSG00000 |
| 358857 | 366052 | 1 | lncRNA | ENSG00000 |
| 365389 | 522928 | -1 | transcribed_unproce | ENSG00000 |
| 439870 | 440232 | 1 | processed_pseudoge | ENSG00000 |
| 450703 | 451697 | -1 | protein_coding | ENSG00000 |
| 487101 | 489906 | 1 | processed_pseudoge | ENSG00000 |
| 491225 | 493241 | -1 | unprocessed_pseudo | ENSG00000 |
| 516376 | 516479 | -1 | snRNA | ENSG00000 |
| 586071 | 827796 | -1 | transcribed_processe | ENSG00000 |
| 587629 | 594768 | 1 | lncRNA | ENSG00000 |
| 629062 | 629433 | 1 | unprocessed_pseudo | ENSG00000 |
| 629640 | 630683 | 1 | unprocessed_pseudo | ENSG00000 |

| | | | | |
|---|---|---|---|---|
| 631074 | 632616 | 1 | unprocessed_pseudc | ENSG00000 |
| 632325 | 632413 | -1 | miRNA | ENSG00000 |
| 632757 | 633438 | 1 | unprocessed_pseudc | ENSG00000 |
| 633535 | 633741 | 1 | unprocessed_pseudc | ENSG00000 |
| 633696 | 634376 | 1 | unprocessed_pseudc | ENSG00000 |
| 634376 | 634922 | 1 | unprocessed_pseudc | ENSG00000 |
| 674842 | 675265 | 1 | processed_pseudoge | ENSG00000 |
| 685679 | 686673 | -1 | protein_coding | ENSG00000 |
| 722092 | 724903 | 1 | processed_pseudoge | ENSG00000 |
| 725885 | 778626 | -1 | transcribed_unproce | ENSG00000 |
| 758233 | 758336 | -1 | snRNA | ENSG00000 |
| 760911 | 761989 | 1 | lncRNA | ENSG00000 |
| 778747 | 810065 | 1 | lncRNA | ENSG00000 |
| 781937 | 782050 | 1 | transcribed_processe | ENSG00000 |
| 800879 | 817712 | -1 | transcribed_unproce | ENSG00000 |
| 817371 | 819837 | 1 | lncRNA | ENSG00000 |
| 825138 | 868202 | 1 | lncRNA | ENSG00000 |
| 826206 | 827522 | -1 | lncRNA | ENSG00000 |
| 868071 | 876903 | -1 | lncRNA | ENSG00000 |
| 873292 | 874349 | 1 | unprocessed_pseudc | ENSG00000 |
| 874529 | 877234 | -1 | transcribed_unproce | ENSG00000 |
| 904834 | 915976 | 1 | lncRNA | ENSG00000 |
| 911435 | 914948 | 1 | lncRNA | ENSG00000 |
| 914171 | 914971 | 1 | lncRNA | ENSG00000 |
| 916865 | 921016 | -1 | lncRNA | ENSG00000 |
| 923928 | 944581 | 1 | protein_coding | ENSG00000 |
| 944203 | 959309 | -1 | protein_coding | ENSG00000 |
| 960584 | 965719 | 1 | protein_coding | ENSG00000 |
| 966482 | 975865 | 1 | protein_coding | ENSG00000 |
| 975204 | 982093 | -1 | protein_coding | ENSG00000 |
| 995966 | 998051 | -1 | lncRNA | ENSG00000 |
| 998962 | 1000172 | -1 | protein_coding | ENSG00000 |
| 1001138 | 1014540 | 1 | protein_coding | ENSG00000 |
| 1008076 | 1008229 | -1 | processed_pseudoge | ENSG00000 |
| 1011997 | 1013193 | -1 | lncRNA | ENSG00000 |
| 1020120 | 1056118 | 1 | protein_coding | ENSG00000 |
| 1055033 | 1056116 | 1 | lncRNA | ENSG00000 |
| 1059708 | 1069355 | 1 | transcribed_unproce | ENSG00000 |
| 1062208 | 1063288 | -1 | lncRNA | ENSG00000 |
| 1070967 | 1074306 | -1 | protein_coding | ENSG00000 |
| 1081818 | 1116361 | -1 | protein_coding | ENSG00000 |
| 1097585 | 1104598 | 1 | lncRNA | ENSG00000 |
| 1137017 | 1144056 | 1 | lncRNA | ENSG00000 |
| 1167104 | 1167198 | 1 | miRNA | ENSG00000 |
| 1167863 | 1167952 | 1 | miRNA | ENSG00000 |
| 1169005 | 1169087 | 1 | miRNA | ENSG00000 |
| 1169357 | 1170343 | 1 | lncRNA | ENSG00000 |
| 1173056 | 1179555 | -1 | lncRNA | ENSG00000 |
| 1173884 | 1197935 | 1 | protein_coding | ENSG00000 |
| 1203508 | 1206592 | -1 | protein_coding | ENSG00000 |

| | | | | |
|---|---|---|---|---|
| 1211340 | 1214153 | -1 | protein_coding | ENSG00000( |
| 1216908 | 1232067 | -1 | protein_coding | ENSG00000( |
| 1232237 | 1235041 | 1 | protein_coding | ENSG00000( |
| 1242446 | 1246722 | -1 | protein_coding | ENSG00000( |
| 1249777 | 1251334 | -1 | lncRNA | ENSG00000( |
| 1253909 | 1273885 | -1 | protein_coding | ENSG00000( |
| 1275223 | 1280420 | 1 | lncRNA | ENSG00000( |
| 1280436 | 1292029 | 1 | protein_coding | ENSG00000( |
| 1292390 | 1309609 | -1 | protein_coding | ENSG00000( |
| 1296110 | 1296170 | -1 | miRNA | ENSG00000( |
| 1308597 | 1311677 | 1 | protein_coding | ENSG00000( |
| 1311585 | 1324691 | -1 | protein_coding | ENSG00000( |
| 1312502 | 1312566 | -1 | miRNA | ENSG00000( |
| 1317581 | 1318689 | -1 | lncRNA | ENSG00000( |
| 1324756 | 1328896 | 1 | protein_coding | ENSG00000( |
| 1331280 | 1335314 | 1 | protein_coding | ENSG00000( |
| 1335276 | 1349418 | -1 | protein_coding | ENSG00000( |
| 1339650 | 1339708 | -1 | miRNA | ENSG00000( |
| 1352689 | 1361777 | -1 | protein_coding | ENSG00000( |
| 1373730 | 1375495 | -1 | protein_coding | ENSG00000( |
| 1378666 | 1379032 | 1 | processed_pseudoge | ENSG00000( |
| 1385711 | 1399335 | -1 | protein_coding | ENSG00000( |
| 1399520 | 1402046 | 1 | lncRNA | ENSG00000( |
| 1401909 | 1407293 | -1 | protein_coding | ENSG00000( |
| 1405460 | 1405752 | -1 | misc_RNA | ENSG00000( |
| 1409096 | 1410618 | 1 | lncRNA | ENSG00000( |
| 1418420 | 1421769 | -1 | protein_coding | ENSG00000( |
| 1420245 | 1422691 | 1 | lncRNA | ENSG00000( |
| 1426128 | 1427787 | 1 | protein_coding | ENSG00000( |
| 1430539 | 1434573 | -1 | lncRNA | ENSG00000( |
| 1434861 | 1442882 | 1 | protein_coding | ENSG00000( |
| 1449689 | 1470163 | 1 | protein_coding | ENSG00000( |
| 1471765 | 1497848 | 1 | protein_coding | ENSG00000( |
| 1503250 | 1509452 | 1 | transcribed_unproce | ENSG00000( |
| 1512162 | 1534685 | 1 | protein_coding | ENSG00000( |
| 1534778 | 1540624 | -1 | protein_coding | ENSG00000( |
| 1541673 | 1574863 | -1 | protein_coding | ENSG00000( |
| 1574102 | 1577075 | 1 | lncRNA | ENSG00000( |
| 1583510 | 1583909 | -1 | processed_pseudoge | ENSG00000( |
| 1598012 | 1600135 | -1 | protein_coding | ENSG00000( |
| 1600314 | 1602703 | 1 | lncRNA | ENSG00000( |
| 1613758 | 1615795 | -1 | lncRNA | ENSG00000( |
| 1615415 | 1630610 | 1 | protein_coding | ENSG00000( |
| 1632163 | 1635263 | 1 | protein_coding | ENSG00000( |
| 1635227 | 1659012 | -1 | protein_coding | ENSG00000( |
| 1659325 | 1662602 | 1 | lncRNA | ENSG00000( |
| 1659529 | 1692795 | -1 | protein_coding | ENSG00000( |
| 1671990 | 1674397 | 1 | transcribed_processe | ENSG00000( |
| 1699942 | 1701782 | 1 | unprocessed_pseudc | ENSG00000( |
| 1702379 | 1724357 | -1 | protein_coding | ENSG00000( |

| | | | | |
|---|---|---|---|---|
| 1702736 | 1737688 | -1 | lncRNA | ENSG0000C |
| 1724512 | 1737251 | 1 | transcribed_processe | ENSG0000C |
| 1724838 | 1745999 | -1 | protein_coding | ENSG0000C |
| 1751232 | 1780457 | -1 | protein_coding | ENSG0000C |
| 1785285 | 1891117 | -1 | protein_coding | ENSG0000C |
| 1891471 | 1892658 | 1 | lncRNA | ENSG0000C |
| 1915108 | 1917296 | 1 | protein_coding | ENSG0000C |
| 1917590 | 1919279 | -1 | protein_coding | ENSG0000C |
| 1921951 | 2003837 | -1 | protein_coding | ENSG0000C |
| 2013213 | 2015639 | -1 | lncRNA | ENSG0000C |
| 2019329 | 2030758 | 1 | protein_coding | ENSG0000C |
| 2049203 | 2050070 | -1 | lncRNA | ENSG0000C |
| 2050411 | 2185395 | 1 | protein_coding | ENSG0000C |
| 2141084 | 2145279 | -1 | lncRNA | ENSG0000C |
| 2181794 | 2184389 | -1 | lncRNA | ENSG0000C |
| 2184461 | 2212720 | -1 | protein_coding | ENSG0000C |
| 2212523 | 2220738 | 1 | lncRNA | ENSG0000C |
| 2228319 | 2310213 | 1 | protein_coding | ENSG0000C |
| 2315040 | 2323085 | 1 | lncRNA | ENSG0000C |
| 2321253 | 2391707 | -1 | protein_coding | ENSG0000C |
| 2326201 | 2326693 | -1 | lncRNA | ENSG0000C |
| 2350414 | 2352820 | -1 | transcribed_processe | ENSG0000C |
| 2363061 | 2363628 | -1 | lncRNA | ENSG0000C |
| 2391775 | 2405442 | 1 | protein_coding | ENSG0000C |
| 2403964 | 2413797 | -1 | protein_coding | ENSG0000C |
| 2425980 | 2505532 | 1 | protein_coding | ENSG0000C |
| 2492300 | 2493258 | -1 | lncRNA | ENSG0000C |
| 2493437 | 2494479 | -1 | lncRNA | ENSG0000C |
| 2508537 | 2526611 | -1 | protein_coding | ENSG0000C |
| 2528745 | 2530263 | -1 | protein_coding | ENSG0000C |
| 2530064 | 2547460 | 1 | lncRNA | ENSG0000C |
| 2549920 | 2557031 | -1 | lncRNA | ENSG0000C |
| 2555639 | 2565382 | 1 | protein_coding | ENSG0000C |
| 2566410 | 2569888 | 1 | TEC | ENSG0000C |
| 2581560 | 2584533 | 1 | lncRNA | ENSG0000C |
| 2586491 | 2591469 | 1 | protein_coding | ENSG0000C |
| 2590639 | 2633016 | -1 | protein_coding | ENSG0000C |
| 2632568 | 2636620 | 1 | lncRNA | ENSG0000C |
| 2635976 | 2801717 | -1 | protein_coding | ENSG0000C |
| 2768091 | 2784733 | 1 | lncRNA | ENSG0000C |
| 2773603 | 2776473 | 1 | lncRNA | ENSG0000C |
| 2811850 | 2812692 | -1 | lncRNA | ENSG0000C |
| 2814056 | 2817767 | 1 | lncRNA | ENSG0000C |
| 2814432 | 2814998 | -1 | lncRNA | ENSG0000C |
| 2960658 | 2968707 | -1 | lncRNA | ENSG0000C |
| 3021467 | 3022903 | 1 | protein_coding | ENSG0000C |
| 3059615 | 3068437 | -1 | lncRNA | ENSG0000C |
| 3069168 | 3438621 | 1 | protein_coding | ENSG0000C |
| 3127975 | 3128035 | 1 | miRNA | ENSG0000C |
| 3132927 | 3133709 | -1 | lncRNA | ENSG0000C |

| | | | | |
|---|---|---|---|---|
| 3205988 | 3208664 | 1 | TEC | ENSG0000C |
| 3306636 | 3310096 | -1 | lncRNA | ENSG0000C |
| 3313052 | 3323149 | -1 | lncRNA | ENSG0000C |
| 3367815 | 3373928 | -1 | lncRNA | ENSG0000C |
| 3454665 | 3481113 | 1 | protein_coding | ENSG0000C |
| 3487246 | 3487627 | 1 | lncRNA | ENSG0000C |
| 3487951 | 3611508 | -1 | protein_coding | ENSG0000C |
| 3560695 | 3560790 | -1 | miRNA | ENSG0000C |
| 3623190 | 3624743 | -1 | lncRNA | ENSG0000C |
| 3625015 | 3630127 | 1 | protein_coding | ENSG0000C |
| 3630767 | 3652761 | -1 | protein_coding | ENSG0000C |
| 3652516 | 3736201 | 1 | protein_coding | ENSG0000C |
| 3658938 | 3668772 | -1 | lncRNA | ENSG0000C |
| 3712200 | 3714298 | -1 | lncRNA | ENSG0000C |
| 3735511 | 3747373 | -1 | transcribed_unitary_ | ENSG0000C |
| 3736943 | 3737103 | 1 | misc_RNA | ENSG0000C |
| 3746460 | 3771645 | 1 | protein_coding | ENSG0000C |
| 3772761 | 3775982 | 1 | protein_coding | ENSG0000C |
| 3778559 | 3796498 | -1 | protein_coding | ENSG0000C |
| 3782815 | 3783111 | 1 | misc_RNA | ENSG0000C |
| 3785008 | 3785538 | 1 | lncRNA | ENSG0000C |
| 3812086 | 3857214 | -1 | protein_coding | ENSG0000C |
| 3857267 | 3885429 | 1 | protein_coding | ENSG0000C |
| 3889125 | 3900293 | -1 | protein_coding | ENSG0000C |
| 3900352 | 3917225 | 1 | lncRNA | ENSG0000C |
| 3940486 | 3955262 | 1 | lncRNA | ENSG0000C |
| 3944547 | 3949024 | -1 | lncRNA | ENSG0000C |
| 3976133 | 4012704 | 1 | lncRNA | ENSG0000C |
| 4012921 | 4019508 | -1 | lncRNA | ENSG0000C |
| 4175528 | 4175899 | -1 | processed_pseudoge | ENSG0000C |
| 4412027 | 4424689 | 1 | lncRNA | ENSG0000C |
| 4479131 | 4484375 | -1 | lncRNA | ENSG0000C |
| 4551735 | 4552145 | -1 | lncRNA | ENSG0000C |
| 4571481 | 4594016 | 1 | lncRNA | ENSG0000C |
| 4654609 | 4792534 | 1 | protein_coding | ENSG0000C |
| 4730211 | 4734992 | -1 | lncRNA | ENSG0000C |
| 4963954 | 4973298 | -1 | lncRNA | ENSG0000C |
| 4973381 | 4981568 | 1 | lncRNA | ENSG0000C |
| 5086459 | 5090899 | -1 | lncRNA | ENSG0000C |
| 5301928 | 5307394 | -1 | lncRNA | ENSG0000C |
| 5478736 | 5493057 | -1 | lncRNA | ENSG0000C |
| 5480787 | 5482028 | -1 | lncRNA | ENSG0000C |
| 5492978 | 5494674 | 1 | lncRNA | ENSG0000C |
| 5554747 | 5554881 | 1 | unprocessed_pseudc | ENSG0000C |
| 5561709 | 5668295 | -1 | lncRNA | ENSG0000C |
| 5564071 | 5564143 | 1 | miRNA | ENSG0000C |
| 5862672 | 5862741 | -1 | miRNA | ENSG0000C |
| 5862811 | 5992473 | -1 | protein_coding | ENSG0000C |
| 5990927 | 6101193 | 1 | protein_coding | ENSG0000C |
| 6101787 | 6180321 | -1 | protein_coding | ENSG0000C |

| | | | | |
|---|---|---|---|---|
| 6159430 | 6197757 | -1 | protein_coding | ENSG00000 |
| 6185020 | 6209389 | -1 | protein_coding | ENSG00000 |
| 6204840 | 6205780 | -1 | lncRNA | ENSG00000 |
| 6205475 | 6221299 | 1 | protein_coding | ENSG00000 |
| 6221193 | 6235972 | -1 | protein_coding | ENSG00000 |
| 6234692 | 6239444 | 1 | lncRNA | ENSG00000 |
| 6244179 | 6245578 | 1 | protein_coding | ENSG00000 |
| 6247353 | 6261098 | -1 | protein_coding | ENSG00000 |
| 6264269 | 6394391 | -1 | protein_coding | ENSG00000 |
| 6393555 | 6394391 | 1 | lncRNA | ENSG00000 |
| 6412418 | 6424670 | -1 | protein_coding | ENSG00000 |
| 6424776 | 6461367 | 1 | protein_coding | ENSG00000 |
| 6429834 | 6429896 | -1 | miRNA | ENSG00000 |
| 6443034 | 6447006 | -1 | lncRNA | ENSG00000 |
| 6460786 | 6466195 | -1 | protein_coding | ENSG00000 |
| 6466092 | 6520061 | -1 | protein_coding | ENSG00000 |
| 6521347 | 6554513 | -1 | protein_coding | ENSG00000 |
| 6540854 | 6540964 | 1 | snRNA | ENSG00000 |
| 6547905 | 6548619 | 1 | processed_pseudoge | ENSG00000 |
| 6555307 | 6579755 | 1 | protein_coding | ENSG00000 |
| 6579994 | 6589280 | 1 | protein_coding | ENSG00000 |
| 6590724 | 6614607 | -1 | protein_coding | ENSG00000 |
| 6613731 | 6624030 | 1 | protein_coding | ENSG00000 |
| 6624866 | 6635586 | 1 | protein_coding | ENSG00000 |
| 6634168 | 6701924 | -1 | protein_coding | ENSG00000 |
| 6724637 | 6730012 | 1 | lncRNA | ENSG00000 |
| 6767954 | 6770038 | 1 | lncRNA | ENSG00000 |
| 6783892 | 6784843 | -1 | lncRNA | ENSG00000 |
| 6785454 | 7769706 | 1 | protein_coding | ENSG00000 |
| 6834333 | 6834618 | -1 | processed_pseudoge | ENSG00000 |
| 7008376 | 7014279 | -1 | lncRNA | ENSG00000 |
| 7219360 | 7219524 | 1 | snRNA | ENSG00000 |
| 7368942 | 7370270 | 1 | lncRNA | ENSG00000 |
| 7382487 | 7389754 | -1 | lncRNA | ENSG00000 |
| 7441096 | 7441554 | -1 | lncRNA | ENSG00000 |
| 7693124 | 7694844 | -1 | lncRNA | ENSG00000 |
| 7698303 | 7698872 | -1 | lncRNA | ENSG00000 |
| 7700704 | 7700970 | -1 | lncRNA | ENSG00000 |
| 7771296 | 7781432 | 1 | protein_coding | ENSG00000 |
| 7776383 | 7776775 | 1 | lncRNA | ENSG00000 |
| 7784320 | 7845177 | 1 | protein_coding | ENSG00000 |
| 7810242 | 7827342 | -1 | lncRNA | ENSG00000 |
| 7843083 | 7853512 | -1 | protein_coding | ENSG00000 |
| 7915894 | 7943165 | -1 | protein_coding | ENSG00000 |
| 7942372 | 7942726 | 1 | processed_pseudoge | ENSG00000 |
| 7954291 | 7985505 | 1 | protein_coding | ENSG00000 |
| 7982881 | 7982983 | -1 | misc_RNA | ENSG00000 |
| 7991134 | 8005312 | 1 | lncRNA | ENSG00000 |
| 7998187 | 7999934 | -1 | lncRNA | ENSG00000 |
| 8004404 | 8026309 | -1 | protein_coding | ENSG00000 |

| | | | | |
|---|---|---|---|---|
| 8026738 | 8122702 | 1 | lncRNA | ENSG00000 |
| 8057245 | 8058463 | -1 | processed_pseudoge | ENSG00000 |
| 8189824 | 8192770 | 1 | processed_pseudoge | ENSG00000 |
| 8201518 | 8215210 | 1 | lncRNA | ENSG00000 |
| 8206434 | 8206597 | -1 | snRNA | ENSG00000 |
| 8218190 | 8220529 | -1 | lncRNA | ENSG00000 |
| 8292157 | 8292263 | -1 | snRNA | ENSG00000 |
| 8317826 | 8344167 | 1 | protein_coding | ENSG00000 |
| 8328067 | 8328179 | 1 | misc_RNA | ENSG00000 |
| 8352397 | 8848921 | -1 | protein_coding | ENSG00000 |
| 8424645 | 8435013 | 1 | lncRNA | ENSG00000 |
| 8511795 | 8511919 | -1 | snoRNA | ENSG00000 |
| 8512653 | 8513021 | 1 | processed_pseudoge | ENSG00000 |
| 8750430 | 8751087 | 1 | processed_pseudoge | ENSG00000 |
| 8786211 | 8786913 | -1 | processed_pseudoge | ENSG00000 |
| 8796571 | 8796673 | -1 | misc_RNA | ENSG00000 |
| 8798475 | 8799506 | 1 | unprocessed_pseudo | ENSG00000 |
| 8805860 | 8807051 | -1 | lncRNA | ENSG00000 |
| 8831007 | 8831375 | 1 | processed_pseudoge | ENSG00000 |
| 8861000 | 8879190 | -1 | protein_coding | ENSG00000 |
| 8866502 | 8866590 | -1 | miRNA | ENSG00000 |
| 8878835 | 8879894 | 1 | lncRNA | ENSG00000 |
| 8883427 | 8883533 | -1 | snRNA | ENSG00000 |
| 8893409 | 8894151 | 1 | processed_pseudoge | ENSG00000 |
| 8907393 | 8907744 | -1 | processed_pseudoge | ENSG00000 |
| 8909742 | 8909904 | 1 | processed_pseudoge | ENSG00000 |
| 8945867 | 8975092 | 1 | protein_coding | ENSG00000 |
| 8979576 | 8979874 | -1 | misc_RNA | ENSG00000 |
| 9003300 | 9026345 | -1 | protein_coding | ENSG00000 |
| 9035106 | 9088478 | -1 | protein_coding | ENSG00000 |
| 9082696 | 9082879 | 1 | scaRNA | ENSG00000 |
| 9100305 | 9129102 | -1 | protein_coding | ENSG00000 |
| 9148011 | 9198906 | -1 | lncRNA | ENSG00000 |
| 9151668 | 9151777 | -1 | miRNA | ENSG00000 |
| 9182004 | 9196284 | 1 | lncRNA | ENSG00000 |
| 9234775 | 9271337 | 1 | protein_coding | ENSG00000 |
| 9292894 | 9369532 | 1 | protein_coding | ENSG00000 |
| 9421098 | 9422862 | -1 | lncRNA | ENSG00000 |
| 9425094 | 9440564 | -1 | lncRNA | ENSG00000 |
| 9437669 | 9437778 | -1 | rRNA_pseudogene | ENSG00000 |
| 9501092 | 9503471 | -1 | lncRNA | ENSG00000 |
| 9539465 | 9585173 | 1 | protein_coding | ENSG00000 |
| 9576427 | 9576985 | 1 | processed_pseudoge | ENSG00000 |
| 9588911 | 9614877 | 1 | protein_coding | ENSG00000 |
| 9651731 | 9729114 | 1 | protein_coding | ENSG00000 |
| 9652610 | 9654586 | -1 | lncRNA | ENSG00000 |
| 9660828 | 9662626 | -1 | processed_pseudoge | ENSG00000 |
| 9672426 | 9687555 | -1 | lncRNA | ENSG00000 |
| 9729026 | 9824526 | -1 | protein_coding | ENSG00000 |
| 9780822 | 9781240 | 1 | processed_pseudoge | ENSG00000 |

| | | | |
|---|---|---|---|
| 9826289 | 9828271 | -1 TEC | ENSG0000C |
| 9848276 | 9910336 | -1 protein_coding | ENSG0000C |
| 9848318 | 9850154 | 1 lncRNA | ENSG0000C |
| 9900614 | 9908034 | 1 lncRNA | ENSG0000C |
| 9922113 | 9943407 | -1 protein_coding | ENSG0000C |
| 9942923 | 9949974 | 1 lncRNA | ENSG0000C |
| 9943428 | 9985501 | 1 protein_coding | ENSG0000C |
| 9947318 | 9947636 | 1 misc_RNA | ENSG0000C |
| 9950572 | 9960787 | -1 unitary_pseudogene | ENSG0000C |
| 9967381 | 9967458 | 1 miRNA | ENSG0000C |
| 9983141 | 9984568 | 1 lncRNA | ENSG0000C |
| 9997206 | 10016021 | 1 protein_coding | ENSG0000C |
| 10032832 | 10181239 | 1 protein_coding | ENSG0000C |
| 10054445 | 10054781 | -1 processed_pseudoge | ENSG0000C |
| 10058671 | 10059648 | 1 processed_pseudoge | ENSG0000C |
| 10163268 | 10163374 | 1 snRNA | ENSG0000C |
| 10210805 | 10381603 | 1 protein_coding | ENSG0000C |
| 10227718 | 10227803 | 1 miRNA | ENSG0000C |
| 10298966 | 10299072 | 1 snRNA | ENSG0000C |
| 10306465 | 10306757 | 1 misc_RNA | ENSG0000C |
| 10381906 | 10387150 | -1 lncRNA | ENSG0000C |
| 10395416 | 10397432 | 1 lncRNA | ENSG0000C |
| 10398592 | 10420511 | 1 protein_coding | ENSG0000C |
| 10429881 | 10430677 | -1 lncRNA | ENSG0000C |
| 10430102 | 10452153 | 1 protein_coding | ENSG0000C |
| 10430433 | 10442808 | 1 protein_coding | ENSG0000C |
| 10449719 | 10451902 | 1 protein_coding | ENSG0000C |
| 10456522 | 10472529 | -1 protein_coding | ENSG0000C |
| 10458555 | 10459338 | 1 lncRNA | ENSG0000C |
| 10472288 | 10630758 | 1 protein_coding | ENSG0000C |
| 10612222 | 10616452 | -1 lncRNA | ENSG0000C |
| 10616836 | 10617115 | 1 misc_RNA | ENSG0000C |
| 10636604 | 10796650 | -1 protein_coding | ENSG0000C |
| 10639241 | 10654333 | 1 lncRNA | ENSG0000C |
| 10895761 | 10895998 | 1 processed_pseudoge | ENSG0000C |
| 10946471 | 10982037 | -1 protein_coding | ENSG0000C |
| 10990978 | 10991797 | -1 processed_pseudoge | ENSG0000C |
| 11012344 | 11026420 | 1 protein_coding | ENSG0000C |
| 11012662 | 11030528 | 1 protein_coding | ENSG0000C |
| 11026523 | 11047239 | -1 protein_coding | ENSG0000C |
| 11029659 | 11030528 | 1 lncRNA | ENSG0000C |
| 11054584 | 11060020 | -1 protein_coding | ENSG0000C |
| 11066618 | 11099869 | -1 protein_coding | ENSG0000C |
| 11068471 | 11073097 | 1 lncRNA | ENSG0000C |
| 11099675 | 11102100 | 1 lncRNA | ENSG0000C |
| 11106535 | 11262551 | -1 protein_coding | ENSG0000C |
| 11143898 | 11149537 | 1 lncRNA | ENSG0000C |
| 11152350 | 11152452 | 1 snRNA | ENSG0000C |
| 11189355 | 11195981 | 1 protein_coding | ENSG0000C |
| 11226254 | 11226360 | 1 snRNA | ENSG0000C |

| | | | | |
|---|---|---|---|---|
| 11232963 | 11233112 | 1 | processed_pseudoge | ENSG00000 |
| 11273206 | 11296049 | 1 | protein_coding | ENSG00000 |
| 11278616 | 11279351 | -1 | processed_pseudoge | ENSG00000 |
| 11311734 | 11319093 | -1 | lncRNA | ENSG00000 |
| 11425257 | 11425821 | -1 | processed_pseudoge | ENSG00000 |
| 11479155 | 11537551 | 1 | protein_coding | ENSG00000 |
| 11500803 | 11502016 | -1 | lncRNA | ENSG00000 |
| 11609468 | 11613358 | 1 | lncRNA | ENSG00000 |
| 11623558 | 11643416 | 1 | lncRNA | ENSG00000 |
| 11637018 | 11655785 | -1 | protein_coding | ENSG00000 |
| 11654375 | 11663327 | 1 | protein_coding | ENSG00000 |
| 11664200 | 11674354 | 1 | protein_coding | ENSG00000 |
| 11674480 | 11691650 | -1 | protein_coding | ENSG00000 |
| 11691710 | 11725857 | 1 | protein_coding | ENSG00000 |
| 11736084 | 11754802 | 1 | protein_coding | ENSG00000 |
| 11761787 | 11789585 | 1 | protein_coding | ENSG00000 |
| 11777077 | 11779619 | -1 | lncRNA | ENSG00000 |
| 11785723 | 11806920 | -1 | protein_coding | ENSG00000 |
| 11806096 | 11848079 | 1 | protein_coding | ENSG00000 |
| 11843812 | 11843984 | 1 | misc_RNA | ENSG00000 |
| 11845549 | 11845697 | 1 | misc_RNA | ENSG00000 |
| 11845709 | 11848345 | -1 | protein_coding | ENSG00000 |
| 11847442 | 11847549 | 1 | misc_RNA | ENSG00000 |
| 11857464 | 11858945 | -1 | protein_coding | ENSG00000 |
| 11877770 | 11880406 | 1 | processed_pseudoge | ENSG00000 |
| 11907940 | 11914298 | 1 | lncRNA | ENSG00000 |
| 11908152 | 11908271 | 1 | snRNA | ENSG00000 |
| 11909808 | 11909927 | -1 | snRNA | ENSG00000 |
| 11919591 | 11926428 | -1 | protein_coding | ENSG00000 |
| 11934205 | 11975538 | 1 | protein_coding | ENSG00000 |
| 11979533 | 11980127 | -1 | lncRNA | ENSG00000 |
| 11980181 | 12013514 | 1 | protein_coding | ENSG00000 |
| 12017216 | 12017308 | -1 | processed_pseudoge | ENSG00000 |
| 12019466 | 12032045 | 1 | protein_coding | ENSG00000 |
| 12024012 | 12024124 | -1 | misc_RNA | ENSG00000 |
| 12029158 | 12029222 | 1 | miRNA | ENSG00000 |
| 12036740 | 12037014 | -1 | misc_RNA | ENSG00000 |
| 12063303 | 12144207 | 1 | protein_coding | ENSG00000 |
| 12077881 | 12077984 | -1 | snRNA | ENSG00000 |
| 12080293 | 12080547 | -1 | processed_pseudoge | ENSG00000 |
| 12088441 | 12092264 | -1 | lncRNA | ENSG00000 |
| 12166943 | 12167038 | 1 | miRNA | ENSG00000 |
| 12166991 | 12209228 | 1 | protein_coding | ENSG00000 |
| 12191713 | 12191773 | 1 | miRNA | ENSG00000 |
| 12220794 | 12221109 | -1 | processed_pseudoge | ENSG00000 |
| 12221148 | 12221271 | -1 | snoRNA | ENSG00000 |
| 12230030 | 12512047 | 1 | protein_coding | ENSG00000 |
| 12507246 | 12507397 | 1 | snoRNA | ENSG00000 |
| 12527724 | 12528420 | 1 | lncRNA | ENSG00000 |
| 12567910 | 12618210 | -1 | protein_coding | ENSG00000 |

| | | | | |
|---|---|---|---|---|
| 12569972 | 12570072 | -1 | snRNA | ENSG00000 |
| 12578957 | 12579023 | -1 | miRNA | ENSG00000 |
| 12618900 | 12619244 | -1 | lncRNA | ENSG00000 |
| 12644547 | 12667086 | 1 | protein_coding | ENSG00000 |
| 12692909 | 12693020 | 1 | processed_pseudoge | ENSG00000 |
| 12716110 | 12728760 | 1 | protein_coding | ENSG00000 |
| 12746200 | 12763699 | 1 | protein_coding | ENSG00000 |
| 12774841 | 12777906 | 1 | protein_coding | ENSG00000 |
| 12791397 | 12796628 | 1 | protein_coding | ENSG00000 |
| 12822686 | 12823159 | -1 | lncRNA | ENSG00000 |
| 12824605 | 12831410 | -1 | protein_coding | ENSG00000 |
| 12838125 | 12841357 | -1 | unprocessed_pseudo | ENSG00000 |
| 12847408 | 12848725 | -1 | protein_coding | ENSG00000 |
| 12857086 | 12861909 | 1 | protein_coding | ENSG00000 |
| 12879212 | 12886201 | -1 | protein_coding | ENSG00000 |
| 12892896 | 12898270 | -1 | protein_coding | ENSG00000 |
| 12916610 | 12920482 | 1 | protein_coding | ENSG00000 |
| 12922554 | 12922660 | 1 | snRNA | ENSG00000 |
| 12926162 | 12928253 | 1 | unprocessed_pseudo | ENSG00000 |
| 12938472 | 12947580 | -1 | protein_coding | ENSG00000 |
| 13029339 | 13032130 | 1 | unprocessed_pseudo | ENSG00000 |
| 13049476 | 13056491 | -1 | protein_coding | ENSG00000 |
| 13060869 | 13062229 | 1 | protein_coding | ENSG00000 |
| 13068677 | 13077884 | 1 | protein_coding | ENSG00000 |
| 13095179 | 13097971 | -1 | unprocessed_pseudo | ENSG00000 |
| 13104403 | 13105689 | -1 | unprocessed_pseudo | ENSG00000 |
| 13115488 | 13116854 | -1 | protein_coding | ENSG00000 |
| 13128818 | 13131615 | 1 | transcribed_unproce | ENSG00000 |
| 13146981 | 13147460 | -1 | lncRNA | ENSG00000 |
| 13148905 | 13155961 | -1 | protein_coding | ENSG00000 |
| 13164586 | 13165467 | -1 | protein_coding | ENSG00000 |
| 13172455 | 13179464 | 1 | protein_coding | ENSG00000 |
| 13196188 | 13201409 | -1 | protein_coding | ENSG00000 |
| 13222705 | 13226106 | -1 | protein_coding | ENSG00000 |
| 13245863 | 13248652 | 1 | unprocessed_pseudo | ENSG00000 |
| 13254212 | 13263314 | 1 | protein_coding | ENSG00000 |
| 13273539 | 13275627 | -1 | unprocessed_pseudo | ENSG00000 |
| 13279125 | 13279231 | -1 | snRNA | ENSG00000 |
| 13281035 | 13285174 | -1 | protein_coding | ENSG00000 |
| 13303539 | 13308907 | 1 | protein_coding | ENSG00000 |
| 13315581 | 13322598 | 1 | protein_coding | ENSG00000 |
| 13324039 | 13324518 | 1 | lncRNA | ENSG00000 |
| 13341892 | 13347134 | -1 | protein_coding | ENSG00000 |
| 13369067 | 13371900 | -1 | protein_coding | ENSG00000 |
| 13389632 | 13392629 | 1 | protein_coding | ENSG00000 |
| 13410450 | 13421328 | 1 | protein_coding | ENSG00000 |
| 13474973 | 13514003 | -1 | protein_coding | ENSG00000 |
| 13513220 | 13516270 | 1 | lncRNA | ENSG00000 |
| 13555001 | 13555694 | 1 | processed_pseudoge | ENSG00000 |
| 13583465 | 13617957 | 1 | protein_coding | ENSG00000 |

| | | | | |
|---|---|---|---|---|
| 13623184 | 13623284 | -1 | rRNA_pseudogene | ENSG00000 |
| 13657311 | 13659910 | -1 | lncRNA | ENSG00000 |
| 13696070 | 13696199 | -1 | scaRNA | ENSG00000 |
| 13700198 | 13825079 | 1 | protein_coding | ENSG00000 |
| 13892792 | 15118043 | 1 | protein_coding | ENSG00000 |
| 14124233 | 14124335 | -1 | snRNA | ENSG00000 |
| 14221887 | 14304445 | -1 | lncRNA | ENSG00000 |
| 14338825 | 14419973 | -1 | lncRNA | ENSG00000 |
| 14692129 | 14692613 | -1 | processed_pseudoge | ENSG00000 |
| 14774469 | 14777727 | -1 | lncRNA | ENSG00000 |
| 15111815 | 15153618 | -1 | lncRNA | ENSG00000 |
| 15152532 | 15220478 | 1 | protein_coding | ENSG00000 |
| 15164344 | 15171317 | -1 | lncRNA | ENSG00000 |
| 15191828 | 15192845 | -1 | processed_pseudoge | ENSG00000 |
| 15226373 | 15227549 | 1 | processed_pseudoge | ENSG00000 |
| 15247272 | 15400283 | 1 | protein_coding | ENSG00000 |
| 15326680 | 15343876 | -1 | lncRNA | ENSG00000 |
| 15334166 | 15335464 | -1 | lncRNA | ENSG00000 |
| 15402979 | 15409433 | -1 | lncRNA | ENSG00000 |
| 15409888 | 15430339 | 1 | protein_coding | ENSG00000 |
| 15438442 | 15449242 | 1 | protein_coding | ENSG00000 |
| 15456728 | 15472091 | 1 | protein_coding | ENSG00000 |
| 15465909 | 15491395 | 1 | protein_coding | ENSG00000 |
| 15490832 | 15526534 | -1 | protein_coding | ENSG00000 |
| 15526813 | 15592379 | 1 | protein_coding | ENSG00000 |
| 15542165 | 15542304 | 1 | scaRNA | ENSG00000 |
| 15565611 | 15565956 | -1 | lncRNA | ENSG00000 |
| 15571699 | 15585051 | -1 | protein_coding | ENSG00000 |
| 15586136 | 15603626 | -1 | lncRNA | ENSG00000 |
| 15604597 | 15605043 | 1 | processed_pseudoge | ENSG00000 |
| 15608078 | 15608145 | -1 | snRNA | ENSG00000 |
| 15614643 | 15614867 | -1 | processed_pseudoge | ENSG00000 |
| 15617458 | 15669044 | 1 | protein_coding | ENSG00000 |
| 15659869 | 15661722 | 1 | protein_coding | ENSG00000 |
| 15682873 | 15683128 | -1 | lncRNA | ENSG00000 |
| 15684320 | 15734769 | 1 | protein_coding | ENSG00000 |
| 15720312 | 15736896 | -1 | lncRNA | ENSG00000 |
| 15736258 | 15741392 | 1 | protein_coding | ENSG00000 |
| 15740051 | 15749896 | -1 | lncRNA | ENSG00000 |
| 15742499 | 15747982 | 1 | protein_coding | ENSG00000 |
| 15756607 | 15786594 | 1 | protein_coding | ENSG00000 |
| 15792796 | 15793285 | 1 | processed_pseudoge | ENSG00000 |
| 15807169 | 15809348 | -1 | protein_coding | ENSG00000 |
| 15828232 | 15828669 | 1 | processed_pseudoge | ENSG00000 |
| 15834474 | 15848147 | -1 | lncRNA | ENSG00000 |
| 15847707 | 15940456 | 1 | protein_coding | ENSG00000 |
| 15917698 | 15917908 | -1 | unprocessed_pseudc | ENSG00000 |
| 15941869 | 15976132 | -1 | protein_coding | ENSG00000 |
| 15969632 | 15970194 | 1 | processed_pseudoge | ENSG00000 |
| 15988182 | 15988441 | 1 | unprocessed_pseudc | ENSG00000 |

| | | | | |
|---|---|---|---|---|
| 15989140 | 15989375 | -1 | processed_pseudoge | ENSG00000 |
| 15989871 | 15994751 | 1 | unprocessed_pseudo | ENSG00000 |
| 16004236 | 16008807 | 1 | protein_coding | ENSG00000 |
| 16006160 | 16006671 | -1 | lncRNA | ENSG00000 |
| 16014028 | 16019594 | -1 | protein_coding | ENSG00000 |
| 16018875 | 16034050 | 1 | protein_coding | ENSG00000 |
| 16035178 | 16040029 | -1 | unprocessed_pseudo | ENSG00000 |
| 16043736 | 16057308 | 1 | protein_coding | ENSG00000 |
| 16057769 | 16073651 | -1 | protein_coding | ENSG00000 |
| 16124337 | 16156069 | -1 | protein_coding | ENSG00000 |
| 16155211 | 16157329 | 1 | lncRNA | ENSG00000 |
| 16159266 | 16161883 | 1 | lncRNA | ENSG00000 |
| 16197854 | 16198357 | 1 | lncRNA | ENSG00000 |
| 16197854 | 16212652 | -1 | protein_coding | ENSG00000 |
| 16215909 | 16228027 | -1 | transcribed_unproce | ENSG00000 |
| 16228674 | 16231335 | -1 | lncRNA | ENSG00000 |
| 16231692 | 16237183 | -1 | protein_coding | ENSG00000 |
| 16241213 | 16241398 | -1 | processed_pseudoge | ENSG00000 |
| 16246840 | 16352480 | -1 | protein_coding | ENSG00000 |
| 16352575 | 16398145 | 1 | protein_coding | ENSG00000 |
| 16369150 | 16369530 | -1 | processed_pseudoge | ENSG00000 |
| 16387117 | 16437424 | -1 | protein_coding | ENSG00000 |
| 16440721 | 16460078 | 1 | protein_coding | ENSG00000 |
| 16460948 | 16468481 | 1 | lncRNA | ENSG00000 |
| 16467436 | 16499257 | -1 | transcribed_unproce | ENSG00000 |
| 16514122 | 16514285 | -1 | snRNA | ENSG00000 |
| 16514645 | 16515754 | 1 | lncRNA | ENSG00000 |
| 16515034 | 16521788 | 1 | lncRNA | ENSG00000 |
| 16520694 | 16521796 | 1 | lncRNA | ENSG00000 |
| 16533886 | 16536172 | -1 | lncRNA | ENSG00000 |
| 16534081 | 16534224 | -1 | snRNA | ENSG00000 |
| 16548651 | 16552531 | -1 | unprocessed_pseudo | ENSG00000 |
| 16562319 | 16613562 | -1 | protein_coding | ENSG00000 |
| 16617391 | 16617729 | 1 | lncRNA | ENSG00000 |
| 16618253 | 16657232 | -1 | transcribed_unproce | ENSG00000 |
| 16642767 | 16643109 | -1 | transcribed_unproce | ENSG00000 |
| 16645622 | 16650289 | 1 | unprocessed_pseudo | ENSG00000 |
| 16656879 | 16667524 | 1 | lncRNA | ENSG00000 |
| 16666785 | 16666948 | -1 | snRNA | ENSG00000 |
| 16681097 | 16688749 | -1 | unprocessed_pseudo | ENSG00000 |
| 16685621 | 16686055 | -1 | processed_pseudoge | ENSG00000 |
| 16687339 | 16720157 | -1 | transcribed_unproce | ENSG00000 |
| 16701546 | 16705594 | 1 | lncRNA | ENSG00000 |
| 16739938 | 16750589 | -1 | lncRNA | ENSG00000 |
| 16740273 | 16972964 | 1 | protein_coding | ENSG00000 |
| 16740280 | 16750968 | 1 | transcribed_unproce | ENSG00000 |
| 16740516 | 16740679 | 1 | snRNA | ENSG00000 |
| 16754910 | 16770237 | -1 | transcribed_unproce | ENSG00000 |
| 16851257 | 16853129 | -1 | lncRNA | ENSG00000 |
| 16855407 | 16859284 | 1 | unprocessed_pseudo | ENSG00000 |

| | | | | |
|---|---|---|---|---|
| 16858949 | 16859021 | -1 | miRNA | ENSG00000 |
| 16870945 | 16883659 | 1 | lncRNA | ENSG00000 |
| 16873708 | 16873851 | 1 | snRNA | ENSG00000 |
| 16887577 | 16889706 | -1 | lncRNA | ENSG00000 |
| 16889095 | 16889602 | 1 | processed_pseudoge | ENSG00000 |
| 16895980 | 16896143 | 1 | snRNA | ENSG00000 |
| 16904339 | 16904776 | -1 | lncRNA | ENSG00000 |
| 16974502 | 16980632 | -1 | protein_coding | ENSG00000 |
| 16976302 | 16977792 | 1 | lncRNA | ENSG00000 |
| 16978926 | 17005091 | 1 | lncRNA | ENSG00000 |
| 16985958 | 17011928 | -1 | protein_coding | ENSG00000 |
| 17018722 | 17054170 | -1 | protein_coding | ENSG00000 |
| 17066761 | 17119451 | -1 | protein_coding | ENSG00000 |
| 17189783 | 17197617 | 1 | lncRNA | ENSG00000 |
| 17193232 | 17201733 | -1 | lncRNA | ENSG00000 |
| 17205128 | 17246007 | 1 | protein_coding | ENSG00000 |
| 17249098 | 17284233 | 1 | protein_coding | ENSG00000 |
| 17277889 | 17277975 | 1 | miRNA | ENSG00000 |
| 17308195 | 17364004 | 1 | protein_coding | ENSG00000 |
| 17372196 | 17401699 | 1 | protein_coding | ENSG00000 |
| 17406760 | 17439677 | -1 | protein_coding | ENSG00000 |
| 17406760 | 17407382 | 1 | lncRNA | ENSG00000 |
| 17539698 | 17697874 | 1 | protein_coding | ENSG00000 |
| 17717625 | 17749978 | 1 | lncRNA | ENSG00000 |
| 17755333 | 17827063 | 1 | protein_coding | ENSG00000 |
| 18015712 | 18045612 | 1 | lncRNA | ENSG00000 |
| 18065657 | 18074412 | 1 | lncRNA | ENSG00000 |
| 18107798 | 18378483 | 1 | protein_coding | ENSG00000 |
| 18109389 | 18115861 | 1 | TEC | ENSG00000 |
| 18166929 | 18179346 | -1 | lncRNA | ENSG00000 |
| 18385829 | 18388514 | 1 | lncRNA | ENSG00000 |
| 18480982 | 18486126 | 1 | protein_coding | ENSG00000 |
| 18513118 | 18513383 | 1 | processed_pseudoge | ENSG00000 |
| 18595414 | 18595636 | 1 | processed_pseudoge | ENSG00000 |
| 18630846 | 18748866 | 1 | protein_coding | ENSG00000 |
| 18839599 | 18859682 | -1 | protein_coding | ENSG00000 |
| 18871430 | 18902724 | -1 | protein_coding | ENSG00000 |
| 18883202 | 18883275 | -1 | miRNA | ENSG00000 |
| 18897071 | 18897148 | -1 | miRNA | ENSG00000 |
| 18904280 | 18956676 | -1 | protein_coding | ENSG00000 |
| 19072110 | 19075511 | -1 | lncRNA | ENSG00000 |
| 19074510 | 19210266 | -1 | protein_coding | ENSG00000 |
| 19210501 | 19240704 | 1 | lncRNA | ENSG00000 |
| 19215660 | 19251527 | -1 | protein_coding | ENSG00000 |
| 19251805 | 19260128 | 1 | protein_coding | ENSG00000 |
| 19260521 | 19262307 | -1 | unprocessed_pseudo | ENSG00000 |
| 19265982 | 19274194 | -1 | polymorphic_pseudo | ENSG00000 |
| 19282573 | 19288770 | -1 | protein_coding | ENSG00000 |
| 19297080 | 19297903 | 1 | processed_pseudoge | ENSG00000 |
| 19303965 | 19312146 | -1 | protein_coding | ENSG00000 |

| | | | | |
|---|---|---|---|---|
| 19305076 | 19305182 | -1 | snRNA | ENSG00000 |
| 19312326 | 19329300 | 1 | protein_coding | ENSG00000 |
| 19338775 | 19485539 | -1 | protein_coding | ENSG00000 |
| 19424384 | 19424671 | 1 | misc_RNA | ENSG00000 |
| 19484403 | 19629821 | 1 | protein_coding | ENSG00000 |
| 19510593 | 19510733 | 1 | snRNA | ENSG00000 |
| 19591802 | 19596832 | -1 | lncRNA | ENSG00000 |
| 19596979 | 19658456 | 1 | protein_coding | ENSG00000 |
| 19597067 | 19656927 | 1 | protein_coding | ENSG00000 |
| 19608114 | 19608568 | 1 | processed_pseudoge | ENSG00000 |
| 19664875 | 19680966 | 1 | protein_coding | ENSG00000 |
| 19682213 | 19799945 | -1 | protein_coding | ENSG00000 |
| 19814029 | 19815283 | -1 | protein_coding | ENSG00000 |
| 19814367 | 19819502 | 1 | lncRNA | ENSG00000 |
| 19882395 | 19912945 | 1 | protein_coding | ENSG00000 |
| 19920009 | 19923617 | -1 | protein_coding | ENSG00000 |
| 19970969 | 19971269 | 1 | misc_RNA | ENSG00000 |
| 19975431 | 19980416 | -1 | protein_coding | ENSG00000 |
| 20028179 | 20091911 | 1 | protein_coding | ENSG00000 |
| 20111939 | 20119566 | -1 | protein_coding | ENSG00000 |
| 20139326 | 20150386 | 1 | protein_coding | ENSG00000 |
| 20154171 | 20160568 | 1 | lncRNA | ENSG00000 |
| 20161253 | 20177424 | -1 | protein_coding | ENSG00000 |
| 20184242 | 20186486 | -1 | lncRNA | ENSG00000 |
| 20186096 | 20196050 | 1 | protein_coding | ENSG00000 |
| 20243095 | 20244664 | 1 | lncRNA | ENSG00000 |
| 20272018 | 20275005 | 1 | lncRNA | ENSG00000 |
| 20290919 | 20354894 | 1 | protein_coding | ENSG00000 |
| 20294211 | 20323187 | -1 | lncRNA | ENSG00000 |
| 20360579 | 20431234 | -1 | lncRNA | ENSG00000 |
| 20412304 | 20413253 | -1 | lncRNA | ENSG00000 |
| 20476222 | 20478937 | 1 | lncRNA | ENSG00000 |
| 20478779 | 20480397 | -1 | lncRNA | ENSG00000 |
| 20482391 | 20486210 | -1 | protein_coding | ENSG00000 |
| 20499448 | 20508151 | -1 | protein_coding | ENSG00000 |
| 20525989 | 20526751 | 1 | processed_pseudoge | ENSG00000 |
| 20552573 | 20555020 | 1 | protein_coding | ENSG00000 |
| 20589086 | 20618903 | 1 | protein_coding | ENSG00000 |
| 20633458 | 20651511 | 1 | protein_coding | ENSG00000 |
| 20633679 | 20633788 | 1 | miRNA | ENSG00000 |
| 20642657 | 20652193 | -1 | lncRNA | ENSG00000 |
| 20651767 | 20661544 | -1 | protein_coding | ENSG00000 |
| 20664014 | 20718017 | -1 | protein_coding | ENSG00000 |
| 20692734 | 20694013 | -1 | unprocessed_pseudo | ENSG00000 |
| 20719731 | 20732837 | -1 | protein_coding | ENSG00000 |
| 20732880 | 20733952 | 1 | lncRNA | ENSG00000 |
| 20742679 | 20787323 | -1 | protein_coding | ENSG00000 |
| 20806292 | 21176888 | -1 | protein_coding | ENSG00000 |
| 20841241 | 20841302 | 1 | snRNA | ENSG00000 |
| 20988314 | 20988432 | -1 | miRNA | ENSG00000 |

| | | | | |
|---|---|---|---|---|
| 21003550 | 21003934 | -1 | processed_pseudoge | ENSG00000 |
| 21073639 | 21073753 | -1 | processed_pseudoge | ENSG00000 |
| 21161475 | 21162081 | 1 | processed_pseudoge | ENSG00000 |
| 21177054 | 21178164 | 1 | lncRNA | ENSG00000 |
| 21217247 | 21345572 | -1 | protein_coding | ENSG00000 |
| 21266082 | 21267251 | 1 | lncRNA | ENSG00000 |
| 21293290 | 21299874 | 1 | lncRNA | ENSG00000 |
| 21397987 | 21398362 | 1 | processed_pseudoge | ENSG00000 |
| 21411460 | 21413293 | -1 | unprocessed_pseudo | ENSG00000 |
| 21415898 | 21417492 | 1 | lncRNA | ENSG00000 |
| 21424625 | 21427967 | -1 | unprocessed_pseudo | ENSG00000 |
| 21428303 | 21429536 | 1 | processed_pseudoge | ENSG00000 |
| 21434318 | 21437558 | 1 | unprocessed_pseudo | ENSG00000 |
| 21440128 | 21485005 | 1 | protein_coding | ENSG00000 |
| 21459756 | 21460082 | -1 | processed_pseudoge | ENSG00000 |
| 21509397 | 21578410 | 1 | protein_coding | ENSG00000 |
| 21586472 | 21591187 | 1 | lncRNA | ENSG00000 |
| 21596215 | 21669363 | -1 | protein_coding | ENSG00000 |
| 21678298 | 21783606 | -1 | protein_coding | ENSG00000 |
| 21768269 | 21768575 | 1 | processed_pseudoge | ENSG00000 |
| 21812265 | 21825225 | 1 | protein_coding | ENSG00000 |
| 21822244 | 21937310 | -1 | protein_coding | ENSG00000 |
| 21908098 | 21908580 | 1 | processed_pseudoge | ENSG00000 |
| 21950679 | 21956871 | -1 | transcribed_unitary_ | ENSG00000 |
| 21977022 | 21998642 | 1 | protein_coding | ENSG00000 |
| 21983606 | 22030617 | 1 | lncRNA | ENSG00000 |
| 21987481 | 21987777 | 1 | misc_RNA | ENSG00000 |
| 21987816 | 21987919 | 1 | snRNA | ENSG00000 |
| 22001657 | 22012542 | 1 | protein_coding | ENSG00000 |
| 22010650 | 22010946 | 1 | misc_RNA | ENSG00000 |
| 22010985 | 22011088 | 1 | snRNA | ENSG00000 |
| 22023990 | 22026048 | -1 | lncRNA | ENSG00000 |
| 22024558 | 22031223 | 1 | lncRNA | ENSG00000 |
| 22025511 | 22101360 | 1 | protein_coding | ENSG00000 |
| 22028317 | 22052927 | -1 | lncRNA | ENSG00000 |
| 22030527 | 22030748 | 1 | transcribed_processe | ENSG00000 |
| 22059197 | 22064199 | 1 | lncRNA | ENSG00000 |
| 22068340 | 22068827 | -1 | processed_pseudoge | ENSG00000 |
| 22117313 | 22143969 | -1 | protein_coding | ENSG00000 |
| 22142850 | 22157401 | 1 | lncRNA | ENSG00000 |
| 22266239 | 22266300 | 1 | miRNA | ENSG00000 |
| 22322840 | 22323331 | -1 | processed_pseudoge | ENSG00000 |
| 22364630 | 22366482 | -1 | TEC | ENSG00000 |
| 22428838 | 22531157 | 1 | protein_coding | ENSG00000 |
| 22517474 | 22519708 | 1 | lncRNA | ENSG00000 |
| 22563489 | 22603595 | 1 | protein_coding | ENSG00000 |
| 22633258 | 22633366 | -1 | miRNA | ENSG00000 |
| 22636628 | 22639678 | 1 | protein_coding | ENSG00000 |
| 22643633 | 22648110 | 1 | protein_coding | ENSG00000 |
| 22652762 | 22661637 | 1 | protein_coding | ENSG00000 |

| | | | | |
|---|---|---|---|---|
| 22710839 | 22921500 | 1 | protein_coding | ENSG0000C |
| 22719517 | 22719598 | 1 | miRNA | ENSG0000C |
| 22835713 | 22836849 | -1 | lncRNA | ENSG0000C |
| 22863159 | 22863226 | -1 | miRNA | ENSG0000C |
| 22953043 | 22965338 | -1 | protein_coding | ENSG0000C |
| 23010834 | 23015852 | -1 | protein_coding | ENSG0000C |
| 23019443 | 23083689 | 1 | protein_coding | ENSG0000C |
| 23020147 | 23088058 | -1 | lncRNA | ENSG0000C |
| 23044305 | 23044372 | 1 | miRNA | ENSG0000C |
| 23057858 | 23057934 | -1 | miRNA | ENSG0000C |
| 23084023 | 23177808 | -1 | protein_coding | ENSG0000C |
| 23140325 | 23141142 | 1 | processed_pseudoge | ENSG0000C |
| 23162704 | 23162810 | 1 | snRNA | ENSG0000C |
| 23163953 | 23164053 | 1 | snRNA | ENSG0000C |
| 23191895 | 23194729 | -1 | protein_coding | ENSG0000C |
| 23244765 | 23245224 | 1 | processed_pseudoge | ENSG0000C |
| 23281307 | 23287488 | -1 | lncRNA | ENSG0000C |
| 23297797 | 23302866 | 1 | lncRNA | ENSG0000C |
| 23303771 | 23344336 | -1 | protein_coding | ENSG0000C |
| 23359448 | 23369836 | -1 | protein_coding | ENSG0000C |
| 23368997 | 23371839 | 1 | lncRNA | ENSG0000C |
| 23370254 | 23370346 | 1 | misc_RNA | ENSG0000C |
| 23378380 | 23379029 | -1 | lncRNA | ENSG0000C |
| 23380909 | 23424748 | -1 | protein_coding | ENSG0000C |
| 23410832 | 23412146 | 1 | processed_pseudoge | ENSG0000C |
| 23428563 | 23484568 | -1 | protein_coding | ENSG0000C |
| 23506438 | 23531233 | -1 | protein_coding | ENSG0000C |
| 23549139 | 23550915 | -1 | lncRNA | ENSG0000C |
| 23557926 | 23559501 | -1 | protein_coding | ENSG0000C |
| 23576436 | 23593621 | -1 | lncRNA | ENSG0000C |
| 23581495 | 23640568 | 1 | lncRNA | ENSG0000C |
| 23670294 | 23680259 | 1 | unprocessed_pseudc | ENSG0000C |
| 23691742 | 23696835 | 1 | protein_coding | ENSG0000C |
| 23706901 | 23778296 | -1 | lncRNA | ENSG0000C |
| 23743155 | 23762059 | 1 | protein_coding | ENSG0000C |
| 23778418 | 23788232 | 1 | protein_coding | ENSG0000C |
| 23791145 | 23795539 | 1 | protein_coding | ENSG0000C |
| 23795599 | 23800781 | -1 | protein_coding | ENSG0000C |
| 23801885 | 23838620 | -1 | protein_coding | ENSG0000C |
| 23845077 | 23868294 | -1 | protein_coding | ENSG0000C |
| 23870515 | 23913362 | -1 | protein_coding | ENSG0000C |
| 23901471 | 23902737 | -1 | processed_pseudoge | ENSG0000C |
| 23907111 | 23907885 | 1 | lncRNA | ENSG0000C |
| 23929070 | 23929147 | 1 | miRNA | ENSG0000C |
| 23949016 | 23949383 | -1 | processed_pseudoge | ENSG0000C |
| 23956839 | 23963462 | 1 | protein_coding | ENSG0000C |
| 23964347 | 23980927 | -1 | protein_coding | ENSG0000C |
| 24007881 | 24008519 | -1 | processed_pseudoge | ENSG0000C |
| 24040835 | 24086799 | 1 | lncRNA | ENSG0000C |
| 24056035 | 24112135 | -1 | protein_coding | ENSG0000C |

| | | | | |
|---|---|---|---|---|
| 24066774 | 24083565 | 1 | lncRNA | ENSG0000C |
| 24119771 | 24143140 | -1 | protein_coding | ENSG0000C |
| 24154168 | 24187959 | -1 | protein_coding | ENSG0000C |
| 24200240 | 24211693 | 1 | lncRNA | ENSG0000C |
| 24307556 | 24321901 | -1 | lncRNA | ENSG0000C |
| 24319322 | 24364482 | 1 | protein_coding | ENSG0000C |
| 24356999 | 24416934 | -1 | protein_coding | ENSG0000C |
| 24415802 | 24472976 | 1 | protein_coding | ENSG0000C |
| 24496254 | 24535400 | -1 | lncRNA | ENSG0000C |
| 24502351 | 24541040 | 1 | protein_coding | ENSG0000C |
| 24538802 | 24556024 | -1 | lncRNA | ENSG0000C |
| 24556087 | 24609328 | 1 | protein_coding | ENSG0000C |
| 24563627 | 24563974 | 1 | processed_pseudoge | ENSG0000C |
| 24625411 | 24625513 | -1 | misc_RNA | ENSG0000C |
| 24631716 | 24673281 | 1 | protein_coding | ENSG0000C |
| 24704894 | 24717596 | 1 | lncRNA | ENSG0000C |
| 24745382 | 24844321 | 1 | protein_coding | ENSG0000C |
| 24777873 | 24777979 | -1 | snRNA | ENSG0000C |
| 24899511 | 24965121 | -1 | protein_coding | ENSG0000C |
| 24919345 | 24919416 | -1 | miRNA | ENSG0000C |
| 24961345 | 24963097 | 1 | lncRNA | ENSG0000C |
| 24968423 | 24970865 | -1 | lncRNA | ENSG0000C |
| 25023503 | 25023586 | 1 | miRNA | ENSG0000C |
| 25041136 | 25047404 | 1 | lncRNA | ENSG0000C |
| 25043707 | 25113120 | -1 | lncRNA | ENSG0000C |
| 25125053 | 25125454 | -1 | processed_pseudoge | ENSG0000C |
| 25208139 | 25209437 | 1 | lncRNA | ENSG0000C |
| 25222276 | 25232502 | -1 | protein_coding | ENSG0000C |
| 25232586 | 25234775 | 1 | lncRNA | ENSG0000C |
| 25239494 | 25240253 | 1 | lncRNA | ENSG0000C |
| 25242249 | 25338213 | -1 | protein_coding | ENSG0000C |
| 25247837 | 25248321 | 1 | lncRNA | ENSG0000C |
| 25266102 | 25267136 | -1 | unprocessed_pseudc | ENSG0000C |
| 25272393 | 25330445 | 1 | protein_coding | ENSG0000C |
| 25294164 | 25294643 | -1 | processed_pseudoge | ENSG0000C |
| 25336429 | 25337465 | -1 | unprocessed_pseudc | ENSG0000C |
| 25338317 | 25362361 | 1 | protein_coding | ENSG0000C |
| 25362249 | 25430192 | -1 | protein_coding | ENSG0000C |
| 25398721 | 25399198 | 1 | processed_pseudoge | ENSG0000C |
| 25430858 | 25500209 | 1 | protein_coding | ENSG0000C |
| 25543580 | 25568886 | 1 | protein_coding | ENSG0000C |
| 25581478 | 25590356 | 1 | lncRNA | ENSG0000C |
| 25617468 | 25786207 | 1 | protein_coding | ENSG0000C |
| 25644544 | 25659111 | -1 | lncRNA | ENSG0000C |
| 25800176 | 25818221 | 1 | protein_coding | ENSG0000C |
| 25811470 | 25823744 | 1 | protein_coding | ENSG0000C |
| 25816749 | 25820797 | -1 | lncRNA | ENSG0000C |
| 25818640 | 25832942 | 1 | protein_coding | ENSG0000C |
| 25831913 | 25832134 | -1 | lncRNA | ENSG0000C |
| 25831913 | 25859458 | -1 | protein_coding | ENSG0000C |

| | | | | |
|---|---|---|---|---|
| 25859613 | 25863420 | 1 | lncRNA | ENSG0000C |
| 25861210 | 25871253 | -1 | protein_coding | ENSG0000C |
| 25877496 | 25877605 | 1 | misc_RNA | ENSG0000C |
| 25884181 | 25906991 | -1 | protein_coding | ENSG0000C |
| 25887360 | 25887610 | 1 | processed_pseudoge | ENSG0000C |
| 25888970 | 25889109 | 1 | processed_pseudoge | ENSG0000C |
| 25906362 | 25906454 | -1 | miRNA | ENSG0000C |
| 25959767 | 25998117 | -1 | protein_coding | ENSG0000C |
| 25964197 | 25964300 | 1 | snRNA | ENSG0000C |
| 26006006 | 26006130 | 1 | scaRNA | ENSG0000C |
| 26006216 | 26006299 | 1 | scaRNA | ENSG0000C |
| 26019884 | 26036464 | 1 | protein_coding | ENSG0000C |
| 26037252 | 26046118 | -1 | protein_coding | ENSG0000C |
| 26046665 | 26049099 | 1 | lncRNA | ENSG0000C |
| 26051304 | 26068436 | -1 | protein_coding | ENSG0000C |
| 26111165 | 26125555 | 1 | protein_coding | ENSG0000C |
| 26159079 | 26163962 | 1 | protein_coding | ENSG0000C |
| 26164161 | 26168581 | -1 | protein_coding | ENSG0000C |
| 26169516 | 26171831 | -1 | protein_coding | ENSG0000C |
| 26169908 | 26170873 | 1 | protein_coding | ENSG0000C |
| 26177484 | 26189884 | 1 | protein_coding | ENSG0000C |
| 26190561 | 26202968 | 1 | protein_coding | ENSG0000C |
| 26209741 | 26229840 | 1 | transcribed_unproce | ENSG0000C |
| 26218581 | 26220476 | -1 | unprocessed_pseudc | ENSG0000C |
| 26234200 | 26278808 | 1 | protein_coding | ENSG0000C |
| 26263041 | 26263398 | -1 | processed_pseudoge | ENSG0000C |
| 26280086 | 26281522 | 1 | protein_coding | ENSG0000C |
| 26281328 | 26318363 | -1 | protein_coding | ENSG0000C |
| 26317958 | 26320523 | 1 | protein_coding | ENSG0000C |
| 26321698 | 26360080 | -1 | protein_coding | ENSG0000C |
| 26326688 | 26327033 | 1 | processed_pseudoge | ENSG0000C |
| 26361634 | 26374522 | -1 | protein_coding | ENSG0000C |
| 26410817 | 26429728 | 1 | protein_coding | ENSG0000C |
| 26432282 | 26471306 | 1 | protein_coding | ENSG0000C |
| 26454653 | 26455187 | -1 | processed_pseudoge | ENSG0000C |
| 26462756 | 26468014 | -1 | lncRNA | ENSG0000C |
| 26472440 | 26476642 | 1 | protein_coding | ENSG0000C |
| 26519354 | 26520250 | -1 | processed_pseudoge | ENSG0000C |
| 26529761 | 26575030 | 1 | protein_coding | ENSG0000C |
| 26554542 | 26554593 | 1 | miRNA | ENSG0000C |
| 26593243 | 26593546 | -1 | misc_RNA | ENSG0000C |
| 26593940 | 26594041 | -1 | misc_RNA | ENSG0000C |
| 26602369 | 26602432 | -1 | snRNA | ENSG0000C |
| 26640377 | 26640614 | -1 | processed_pseudoge | ENSG0000C |
| 26642286 | 26642389 | -1 | snoRNA | ENSG0000C |
| 26647447 | 26647681 | 1 | processed_pseudoge | ENSG0000C |
| 26692132 | 26694131 | -1 | lncRNA | ENSG0000C |
| 26693236 | 26782104 | 1 | protein_coding | ENSG0000C |
| 26787472 | 26798398 | 1 | protein_coding | ENSG0000C |
| 26814822 | 26815112 | 1 | misc_RNA | ENSG0000C |

| | | | | |
|---|---|---|---|---|
| 26817300 | 26817476 | 1 | unprocessed_pseudo | ENSG00000 |
| 26826688 | 26857604 | 1 | protein_coding | ENSG00000 |
| 26863149 | 26864456 | 1 | protein_coding | ENSG00000 |
| 26876132 | 26890283 | -1 | protein_coding | ENSG00000 |
| 26876133 | 26878245 | 1 | lncRNA | ENSG00000 |
| 26890488 | 26900467 | -1 | protein_coding | ENSG00000 |
| 26900238 | 26946871 | 1 | protein_coding | ENSG00000 |
| 26911489 | 26913975 | -1 | protein_coding | ENSG00000 |
| 26949562 | 26960468 | -1 | protein_coding | ENSG00000 |
| 26980165 | 26980975 | 1 | unprocessed_pseudo | ENSG00000 |
| 26983628 | 26984024 | 1 | processed_pseudoge | ENSG00000 |
| 26985897 | 26986340 | -1 | processed_pseudoge | ENSG00000 |
| 26993692 | 27000886 | 1 | protein_coding | ENSG00000 |
| 27005020 | 27012850 | -1 | protein_coding | ENSG00000 |
| 27098809 | 27166981 | -1 | protein_coding | ENSG00000 |
| 27176751 | 27177277 | 1 | processed_pseudoge | ENSG00000 |
| 27200834 | 27201473 | -1 | processed_pseudoge | ENSG00000 |
| 27206930 | 27207796 | 1 | processed_pseudoge | ENSG00000 |
| 27211265 | 27211484 | 1 | processed_pseudoge | ENSG00000 |
| 27229106 | 27234352 | -1 | lncRNA | ENSG00000 |
| 27234632 | 27308636 | 1 | protein_coding | ENSG00000 |
| 27255464 | 27255567 | -1 | misc_RNA | ENSG00000 |
| 27311240 | 27311838 | 1 | processed_pseudoge | ENSG00000 |
| 27322145 | 27336400 | 1 | protein_coding | ENSG00000 |
| 27325219 | 27325325 | 1 | snRNA | ENSG00000 |
| 27325329 | 27325796 | 1 | processed_pseudoge | ENSG00000 |
| 27342020 | 27353937 | 1 | protein_coding | ENSG00000 |
| 27354067 | 27366961 | -1 | protein_coding | ENSG00000 |
| 27369112 | 27374824 | -1 | protein_coding | ENSG00000 |
| 27379176 | 27383333 | -1 | protein_coding | ENSG00000 |
| 27389468 | 27390145 | 1 | processed_pseudoge | ENSG00000 |
| 27392622 | 27395814 | 1 | protein_coding | ENSG00000 |
| 27404230 | 27490167 | -1 | protein_coding | ENSG00000 |
| 27457198 | 27459582 | 1 | lncRNA | ENSG00000 |
| 27525805 | 27530561 | 1 | lncRNA | ENSG00000 |
| 27534035 | 27604431 | -1 | protein_coding | ENSG00000 |
| 27612064 | 27635185 | -1 | protein_coding | ENSG00000 |
| 27649419 | 27649610 | 1 | processed_pseudoge | ENSG00000 |
| 27660328 | 27662722 | -1 | lncRNA | ENSG00000 |
| 27666064 | 27672198 | -1 | protein_coding | ENSG00000 |
| 27669468 | 27703063 | 1 | lncRNA | ENSG00000 |
| 27675603 | 27675709 | -1 | snRNA | ENSG00000 |
| 27686810 | 27687471 | -1 | processed_pseudoge | ENSG00000 |
| 27693731 | 27693835 | 1 | snRNA | ENSG00000 |
| 27724822 | 27725756 | -1 | lncRNA | ENSG00000 |
| 27725979 | 27763122 | 1 | protein_coding | ENSG00000 |
| 27739091 | 27739439 | 1 | processed_pseudoge | ENSG00000 |
| 27739954 | 27740629 | 1 | processed_pseudoge | ENSG00000 |
| 27773219 | 27824443 | 1 | protein_coding | ENSG00000 |
| 27773858 | 27774041 | 1 | lncRNA | ENSG00000 |

| | | | | |
|---|---|---|---|---|
| 27819983 | 27820341 | 1 | lncRNA | ENSG0000C |
| 27824538 | 27824643 | 1 | snRNA | ENSG0000C |
| 27827812 | 27831209 | -1 | lncRNA | ENSG0000C |
| 27830782 | 27851676 | 1 | protein_coding | ENSG0000C |
| 27834401 | 27834566 | 1 | scaRNA | ENSG0000C |
| 27872543 | 27886685 | 1 | protein_coding | ENSG0000C |
| 27891524 | 27914746 | -1 | protein_coding | ENSG0000C |
| 27935000 | 27959157 | 1 | protein_coding | ENSG0000C |
| 27938875 | 27960193 | -1 | lncRNA | ENSG0000C |
| 27959462 | 27968096 | 1 | protein_coding | ENSG0000C |
| 27970344 | 28088637 | -1 | protein_coding | ENSG0000C |
| 27990158 | 27990524 | -1 | processed_pseudoge | ENSG0000C |
| 28031884 | 28032180 | -1 | misc_RNA | ENSG0000C |
| 28095742 | 28096422 | -1 | processed_pseudoge | ENSG0000C |
| 28109739 | 28110298 | 1 | processed_pseudoge | ENSG0000C |
| 28120449 | 28121321 | -1 | processed_pseudoge | ENSG0000C |
| 28142737 | 28142841 | -1 | snRNA | ENSG0000C |
| 28147166 | 28193936 | -1 | protein_coding | ENSG0000C |
| 28199456 | 28233029 | -1 | protein_coding | ENSG0000C |
| 28236109 | 28246906 | 1 | protein_coding | ENSG0000C |
| 28239509 | 28241453 | -1 | lncRNA | ENSG0000C |
| 28247144 | 28247568 | 1 | lncRNA | ENSG0000C |
| 28259518 | 28282491 | 1 | protein_coding | ENSG0000C |
| 28329002 | 28335965 | 1 | protein_coding | ENSG0000C |
| 28369582 | 28500364 | 1 | protein_coding | ENSG0000C |
| 28422555 | 28422655 | -1 | misc_RNA | ENSG0000C |
| 28453541 | 28453934 | 1 | processed_pseudoge | ENSG0000C |
| 28481362 | 28481480 | 1 | snRNA | ENSG0000C |
| 28505943 | 28539300 | 1 | protein_coding | ENSG0000C |
| 28505980 | 28510892 | 1 | lncRNA | ENSG0000C |
| 28507366 | 28507571 | 1 | snoRNA | ENSG0000C |
| 28508559 | 28508762 | 1 | snoRNA | ENSG0000C |
| 28526318 | 28527227 | -1 | processed_pseudoge | ENSG0000C |
| 28544460 | 28546542 | 1 | TEC | ENSG0000C |
| 28553085 | 28578545 | 1 | protein_coding | ENSG0000C |
| 28578538 | 28583132 | -1 | lncRNA | ENSG0000C |
| 28578749 | 28578822 | -1 | snoRNA | ENSG0000C |
| 28579764 | 28579893 | -1 | snoRNA | ENSG0000C |
| 28580381 | 28580512 | -1 | snoRNA | ENSG0000C |
| 28580920 | 28581054 | -1 | snoRNA | ENSG0000C |
| 28580920 | 28581056 | -1 | snoRNA | ENSG0000C |
| 28589323 | 28643085 | -1 | protein_coding | ENSG0000C |
| 28592200 | 28595443 | 1 | protein_coding | ENSG0000C |
| 28643228 | 28648581 | 1 | lncRNA | ENSG0000C |
| 28648600 | 28648733 | 1 | snRNA | ENSG0000C |
| 28648600 | 28648730 | 1 | lncRNA | ENSG0000C |
| 28668732 | 28719353 | 1 | protein_coding | ENSG0000C |
| 28689665 | 28689794 | 1 | scaRNA | ENSG0000C |
| 28736621 | 28769775 | 1 | protein_coding | ENSG0000C |
| 28812142 | 28871267 | 1 | protein_coding | ENSG0000C |

| | | | | |
|---|---|---|---|---|
| 28870483 | 28877336 | -1 | lncRNA | ENSG00000 |
| 28881726 | 28881835 | -1 | misc_RNA | ENSG00000 |
| 28887091 | 29120046 | 1 | protein_coding | ENSG00000 |
| 28982278 | 28982608 | -1 | processed_pseudoge | ENSG00000 |
| 28985710 | 28985810 | 1 | misc_RNA | ENSG00000 |
| 29041199 | 29041480 | 1 | misc_RNA | ENSG00000 |
| 29119429 | 29123903 | -1 | protein_coding | ENSG00000 |
| 29144494 | 29146994 | -1 | lncRNA | ENSG00000 |
| 29147743 | 29181900 | -1 | protein_coding | ENSG00000 |
| 29171285 | 29171453 | -1 | misc_RNA | ENSG00000 |
| 29192657 | 29230942 | -1 | protein_coding | ENSG00000 |
| 29223933 | 29224816 | 1 | lncRNA | ENSG00000 |
| 29236516 | 29326813 | 1 | protein_coding | ENSG00000 |
| 29329620 | 29350114 | -1 | lncRNA | ENSG00000 |
| 29488193 | 29496893 | -1 | unprocessed_pseudo | ENSG00000 |
| 29708851 | 29709547 | -1 | lncRNA | ENSG00000 |
| 29755175 | 29790597 | 1 | lncRNA | ENSG00000 |
| 29904865 | 29905357 | -1 | processed_pseudoge | ENSG00000 |
| 30013952 | 30037612 | -1 | lncRNA | ENSG00000 |
| 30140263 | 30141607 | 1 | lncRNA | ENSG00000 |
| 30226523 | 30226615 | 1 | processed_pseudoge | ENSG00000 |
| 30409560 | 30411638 | -1 | lncRNA | ENSG00000 |
| 30415825 | 30421108 | 1 | lncRNA | ENSG00000 |
| 30711277 | 30723585 | -1 | protein_coding | ENSG00000 |
| 30718504 | 30726827 | 1 | lncRNA | ENSG00000 |
| 30731693 | 30733427 | 1 | lncRNA | ENSG00000 |
| 30732469 | 30757774 | -1 | protein_coding | ENSG00000 |
| 30739156 | 30739232 | -1 | miRNA | ENSG00000 |
| 30810378 | 30815553 | -1 | lncRNA | ENSG00000 |
| 30824217 | 30834864 | 1 | lncRNA | ENSG00000 |
| 30843823 | 30844110 | 1 | misc_RNA | ENSG00000 |
| 30858158 | 30860254 | 1 | lncRNA | ENSG00000 |
| 30869466 | 30908758 | -1 | protein_coding | ENSG00000 |
| 30918469 | 30918735 | -1 | misc_RNA | ENSG00000 |
| 30931506 | 31065991 | -1 | protein_coding | ENSG00000 |
| 31036734 | 31037240 | 1 | processed_pseudoge | ENSG00000 |
| 31050872 | 31051034 | 1 | processed_pseudoge | ENSG00000 |
| 31094987 | 31095558 | -1 | processed_pseudoge | ENSG00000 |
| 31108188 | 31108623 | -1 | processed_pseudoge | ENSG00000 |
| 31179745 | 31239887 | -1 | protein_coding | ENSG00000 |
| 31259568 | 31296788 | -1 | protein_coding | ENSG00000 |
| 31263245 | 31263681 | -1 | processed_pseudoge | ENSG00000 |
| 31296982 | 31364953 | 1 | protein_coding | ENSG00000 |
| 31333067 | 31346799 | -1 | lncRNA | ENSG00000 |
| 31365253 | 31376850 | -1 | protein_coding | ENSG00000 |
| 31409565 | 31434680 | 1 | protein_coding | ENSG00000 |
| 31487589 | 31488930 | -1 | processed_pseudoge | ENSG00000 |
| 31497577 | 31497683 | -1 | snRNA | ENSG00000 |
| 31500085 | 31509648 | 1 | transcribed_unitary_ | ENSG00000 |
| 31503631 | 31503744 | -1 | misc_RNA | ENSG00000 |

| | | | | |
|---|---|---|---|---|
| 31506226 | 31583306 | 1 | lncRNA | ENSG00000( |
| 31571585 | 31577898 | -1 | lncRNA | ENSG00000( |
| 31576485 | 31587686 | 1 | protein_coding | ENSG00000( |
| 31617686 | 31632518 | 1 | protein_coding | ENSG00000( |
| 31629866 | 31644896 | -1 | protein_coding | ENSG00000( |
| 31644049 | 31660162 | 1 | lncRNA | ENSG00000( |
| 31644694 | 31649371 | 1 | lncRNA | ENSG00000( |
| 31652263 | 31704319 | -1 | protein_coding | ENSG00000( |
| 31727117 | 31764893 | -1 | protein_coding | ENSG00000( |
| 31758660 | 31758735 | -1 | miRNA | ENSG00000( |
| 31789130 | 31791322 | 1 | lncRNA | ENSG00000( |
| 31790422 | 31816051 | -1 | protein_coding | ENSG00000( |
| 31842019 | 31855568 | -1 | lncRNA | ENSG00000( |
| 31851913 | 31921841 | -1 | lncRNA | ENSG00000( |
| 31906421 | 31944856 | -1 | protein_coding | ENSG00000( |
| 31933020 | 31933975 | 1 | lncRNA | ENSG00000( |
| 31972189 | 31987034 | -1 | lncRNA | ENSG00000( |
| 32013868 | 32060850 | 1 | protein_coding | ENSG00000( |
| 32052291 | 32073474 | -1 | lncRNA | ENSG00000( |
| 32072031 | 32102866 | 1 | protein_coding | ENSG00000( |
| 32086949 | 32087007 | 1 | miRNA | ENSG00000( |
| 32087523 | 32087813 | -1 | misc_RNA | ENSG00000( |
| 32108056 | 32176563 | 1 | protein_coding | ENSG00000( |
| 32170733 | 32176568 | 1 | lncRNA | ENSG00000( |
| 32179675 | 32198285 | 1 | protein_coding | ENSG00000( |
| 32200386 | 32205387 | 1 | protein_coding | ENSG00000( |
| 32204769 | 32206814 | -1 | lncRNA | ENSG00000( |
| 32205661 | 32208687 | 1 | protein_coding | ENSG00000( |
| 32209094 | 32216196 | 1 | protein_coding | ENSG00000( |
| 32214472 | 32222359 | -1 | protein_coding | ENSG00000( |
| 32221928 | 32231604 | 1 | protein_coding | ENSG00000( |
| 32231658 | 32241620 | -1 | transcribed_unproce | ENSG00000( |
| 32247222 | 32248856 | 1 | protein_coding | ENSG00000( |
| 32251239 | 32286165 | 1 | protein_coding | ENSG00000( |
| 32269338 | 32269597 | -1 | misc_RNA | ENSG00000( |
| 32286452 | 32286561 | -1 | misc_RNA | ENSG00000( |
| 32292083 | 32333635 | 1 | protein_coding | ENSG00000( |
| 32333839 | 32336233 | -1 | protein_coding | ENSG00000( |
| 32349194 | 32350663 | -1 | lncRNA | ENSG00000( |
| 32351521 | 32364312 | 1 | protein_coding | ENSG00000( |
| 32361270 | 32364278 | -1 | protein_coding | ENSG00000( |
| 32365103 | 32394731 | -1 | protein_coding | ENSG00000( |
| 32402109 | 32403115 | -1 | processed_pseudoge | ENSG00000( |
| 32421979 | 32422319 | 1 | processed_pseudoge | ENSG00000( |
| 32423214 | 32426789 | -1 | unprocessed_pseudc | ENSG00000( |
| 32457835 | 32458135 | 1 | misc_RNA | ENSG00000( |
| 32465057 | 32600792 | 1 | protein_coding | ENSG00000( |
| 32465072 | 32496686 | 1 | protein_coding | ENSG00000( |
| 32539427 | 32605941 | 1 | protein_coding | ENSG00000( |
| 32600172 | 32650903 | -1 | protein_coding | ENSG00000( |

| | | | | |
|---|---|---|---|---|
| 32639951 | 32640052 | 1 | misc_RNA | ENSG00000 |
| 32651142 | 32686211 | 1 | protein_coding | ENSG00000 |
| 32679906 | 32703596 | -1 | protein_coding | ENSG00000 |
| 32717734 | 32725237 | -1 | lncRNA | ENSG00000 |
| 32741830 | 32774970 | 1 | protein_coding | ENSG00000 |
| 32775237 | 32818153 | -1 | protein_coding | ENSG00000 |
| 32816767 | 32858875 | 1 | protein_coding | ENSG00000 |
| 32862268 | 32872482 | -1 | protein_coding | ENSG00000 |
| 32885994 | 32898441 | 1 | protein_coding | ENSG00000 |
| 32894594 | 32901438 | -1 | protein_coding | ENSG00000 |
| 32925454 | 32951638 | 1 | lncRNA | ENSG00000 |
| 32936445 | 32964685 | -1 | protein_coding | ENSG00000 |
| 32973553 | 32974463 | 1 | TEC | ENSG00000 |
| 32979947 | 32980472 | 1 | processed_pseudoge | ENSG00000 |
| 32986952 | 32988233 | 1 | unprocessed_pseudo | ENSG00000 |
| 32987075 | 33032469 | 1 | lncRNA | ENSG00000 |
| 33007940 | 33080996 | -1 | protein_coding | ENSG00000 |
| 33029232 | 33029576 | -1 | misc_RNA | ENSG00000 |
| 33081104 | 33123492 | 1 | protein_coding | ENSG00000 |
| 33141871 | 33143230 | -1 | TEC | ENSG00000 |
| 33145399 | 33182059 | -1 | protein_coding | ENSG00000 |
| 33162851 | 33166298 | 1 | TEC | ENSG00000 |
| 33194788 | 33200353 | 1 | lncRNA | ENSG00000 |
| 33256492 | 33300719 | 1 | protein_coding | ENSG00000 |
| 33261212 | 33261680 | 1 | lncRNA | ENSG00000 |
| 33306766 | 33321098 | -1 | protein_coding | ENSG00000 |
| 33307348 | 33349245 | 1 | lncRNA | ENSG00000 |
| 33323623 | 33431052 | -1 | protein_coding | ENSG00000 |
| 33332393 | 33332492 | -1 | miRNA | ENSG00000 |
| 33336566 | 33336864 | -1 | misc_RNA | ENSG00000 |
| 33350352 | 33363245 | 1 | lncRNA | ENSG00000 |
| 33466249 | 33468954 | 1 | unitary_pseudogene | ENSG00000 |
| 33472645 | 33496507 | 1 | protein_coding | ENSG00000 |
| 33512008 | 33512743 | -1 | processed_pseudoge | ENSG00000 |
| 33513999 | 34165842 | -1 | protein_coding | ENSG00000 |
| 33838523 | 33840140 | 1 | processed_pseudoge | ENSG00000 |
| 33860475 | 33864791 | 1 | protein_coding | ENSG00000 |
| 33868953 | 33893726 | 1 | lncRNA | ENSG00000 |
| 34112949 | 34113063 | 1 | rRNA_pseudogene | ENSG00000 |
| 34166883 | 34219131 | 1 | protein_coding | ENSG00000 |
| 34276859 | 34277292 | -1 | processed_pseudoge | ENSG00000 |
| 34640157 | 34685732 | -1 | lncRNA | ENSG00000 |
| 34669599 | 34669694 | -1 | miRNA | ENSG00000 |
| 34712737 | 34859755 | -1 | protein_coding | ENSG00000 |
| 34755047 | 34758512 | 1 | protein_coding | ENSG00000 |
| 34759740 | 34762327 | 1 | protein_coding | ENSG00000 |
| 34761426 | 34788097 | -1 | lncRNA | ENSG00000 |
| 34781214 | 34786369 | 1 | protein_coding | ENSG00000 |
| 34792999 | 34795747 | 1 | protein_coding | ENSG00000 |
| 34850694 | 34851555 | 1 | lncRNA | ENSG00000 |

| | | | | |
|---|---|---|---|---|
| 34865436 | 34929650 | -1 | protein_coding | ENSG00000( |
| 34974356 | 34985313 | -1 | protein_coding | ENSG00000( |
| 34975699 | 34978706 | -1 | transcribed_unitary_ | ENSG00000( |
| 34981533 | 35031741 | -1 | protein_coding | ENSG00000( |
| 34981535 | 34985353 | -1 | protein_coding | ENSG00000( |
| 34986165 | 35031945 | -1 | protein_coding | ENSG00000( |
| 35032172 | 35115859 | 1 | protein_coding | ENSG00000( |
| 35053468 | 35059271 | 1 | processed_pseudoge | ENSG00000( |
| 35122022 | 35122484 | 1 | processed_pseudoge | ENSG00000( |
| 35141515 | 35146435 | -1 | lncRNA | ENSG00000( |
| 35176378 | 35193145 | -1 | protein_coding | ENSG00000( |
| 35195969 | 35196062 | 1 | misc_RNA | ENSG00000( |
| 35268709 | 35422058 | 1 | protein_coding | ENSG00000( |
| 35292196 | 35292498 | -1 | misc_RNA | ENSG00000( |
| 35310274 | 35310425 | 1 | snoRNA | ENSG00000( |
| 35350722 | 35351607 | 1 | processed_pseudoge | ENSG00000( |
| 35358822 | 35366077 | -1 | lncRNA | ENSG00000( |
| 35399997 | 35400319 | 1 | misc_RNA | ENSG00000( |
| 35433492 | 35557950 | -1 | protein_coding | ENSG00000( |
| 35509742 | 35511438 | 1 | processed_pseudoge | ENSG00000( |
| 35557473 | 35567274 | 1 | protein_coding | ENSG00000( |
| 35569813 | 35577729 | -1 | lncRNA | ENSG00000( |
| 35573314 | 35595328 | 1 | protein_coding | ENSG00000( |
| 35599541 | 35641526 | -1 | protein_coding | ENSG00000( |
| 35713877 | 35718894 | -1 | protein_coding | ENSG00000( |
| 35720218 | 35769978 | -1 | protein_coding | ENSG00000( |
| 35739389 | 35743576 | 1 | lncRNA | ENSG00000( |
| 35808016 | 35857890 | 1 | protein_coding | ENSG00000( |
| 35869808 | 35930532 | 1 | protein_coding | ENSG00000( |
| 35873270 | 35873752 | -1 | processed_pseudoge | ENSG00000( |
| 35908980 | 35910872 | -1 | lncRNA | ENSG00000( |
| 35929720 | 35930115 | -1 | lncRNA | ENSG00000( |
| 35930718 | 36072500 | 1 | protein_coding | ENSG00000( |
| 35992109 | 36013630 | -1 | lncRNA | ENSG00000( |
| 36080066 | 36084026 | -1 | transcribed_process( | ENSG00000( |
| 36084094 | 36088275 | 1 | protein_coding | ENSG00000( |
| 36088892 | 36093932 | 1 | protein_coding | ENSG00000( |
| 36095239 | 36125222 | -1 | protein_coding | ENSG00000( |
| 36136570 | 36156053 | -1 | protein_coding | ENSG00000( |
| 36155579 | 36180849 | 1 | protein_coding | ENSG00000( |
| 36191915 | 36192170 | 1 | misc_RNA | ENSG00000( |
| 36224432 | 36305357 | 1 | protein_coding | ENSG00000( |
| 36241898 | 36242156 | -1 | processed_pseudoge | ENSG00000( |
| 36306368 | 36329340 | 1 | protein_coding | ENSG00000( |
| 36322031 | 36324154 | -1 | protein_coding | ENSG00000( |
| 36329630 | 36335406 | 1 | lncRNA | ENSG00000( |
| 36339624 | 36385896 | -1 | protein_coding | ENSG00000( |
| 36391238 | 36397908 | -1 | protein_coding | ENSG00000( |
| 36402721 | 36402859 | -1 | snRNA | ENSG00000( |
| 36415827 | 36450451 | -1 | protein_coding | ENSG00000( |

| | | | | |
|---|---|---|---|---|
| 36418450 | 36418578 | -1 | snoRNA | ENSG00000 |
| 36455718 | 36464384 | -1 | protein_coding | ENSG00000 |
| 36466043 | 36483278 | -1 | protein_coding | ENSG00000 |
| 36630335 | 36630857 | 1 | processed_pseudoge | ENSG00000 |
| 36703953 | 36710582 | 1 | lncRNA | ENSG00000 |
| 36768122 | 36769725 | -1 | lncRNA | ENSG00000 |
| 36769812 | 36775768 | 1 | lncRNA | ENSG00000 |
| 36795527 | 37034515 | -1 | protein_coding | ENSG00000 |
| 37133489 | 37136146 | 1 | lncRNA | ENSG00000 |
| 37154761 | 37325100 | -1 | lncRNA | ENSG00000 |
| 37161563 | 37161634 | 1 | miRNA | ENSG00000 |
| 37203610 | 37203716 | 1 | snRNA | ENSG00000 |
| 37264677 | 37264786 | -1 | rRNA_pseudogene | ENSG00000 |
| 37330852 | 37330996 | 1 | processed_pseudoge | ENSG00000 |
| 37350934 | 37474411 | -1 | lncRNA | ENSG00000 |
| 37474580 | 37484377 | 1 | protein_coding | ENSG00000 |
| 37480230 | 37480289 | 1 | miRNA | ENSG00000 |
| 37492575 | 37514774 | -1 | protein_coding | ENSG00000 |
| 37500935 | 37500994 | -1 | miRNA | ENSG00000 |
| 37534449 | 37554293 | -1 | protein_coding | ENSG00000 |
| 37544763 | 37545294 | 1 | processed_pseudoge | ENSG00000 |
| 37556247 | 37556499 | 1 | processed_pseudoge | ENSG00000 |
| 37556919 | 37566857 | 1 | protein_coding | ENSG00000 |
| 37566816 | 37595937 | -1 | protein_coding | ENSG00000 |
| 37596126 | 37607336 | 1 | lncRNA | ENSG00000 |
| 37611350 | 37634892 | -1 | protein_coding | ENSG00000 |
| 37681570 | 37692249 | -1 | protein_coding | ENSG00000 |
| 37692481 | 37709719 | 1 | protein_coding | ENSG00000 |
| 37713880 | 37765133 | -1 | protein_coding | ENSG00000 |
| 37737955 | 37738054 | -1 | misc_RNA | ENSG00000 |
| 37776670 | 37779418 | 1 | processed_pseudoge | ENSG00000 |
| 37782457 | 37782885 | -1 | processed_pseudoge | ENSG00000 |
| 37793802 | 37801137 | 1 | protein_coding | ENSG00000 |
| 37799720 | 37800879 | -1 | lncRNA | ENSG00000 |
| 37802945 | 37808208 | -1 | protein_coding | ENSG00000 |
| 37806979 | 37809454 | 1 | protein_coding | ENSG00000 |
| 37809574 | 37859592 | -1 | protein_coding | ENSG00000 |
| 37840986 | 37841676 | 1 | processed_pseudoge | ENSG00000 |
| 37860697 | 37947057 | -1 | protein_coding | ENSG00000 |
| 37860697 | 37861580 | 1 | lncRNA | ENSG00000 |
| 37884237 | 37884317 | 1 | snoRNA | ENSG00000 |
| 37885023 | 37885117 | 1 | snRNA | ENSG00000 |
| 37953508 | 37953780 | 1 | misc_RNA | ENSG00000 |
| 37956975 | 37990075 | -1 | protein_coding | ENSG00000 |
| 37991462 | 37991569 | 1 | snRNA | ENSG00000 |
| 37996770 | 38005606 | -1 | protein_coding | ENSG00000 |
| 38009258 | 38024820 | 1 | protein_coding | ENSG00000 |
| 38043829 | 38046793 | -1 | protein_coding | ENSG00000 |
| 38047314 | 38119025 | 1 | lncRNA | ENSG00000 |
| 38080572 | 38080911 | -1 | processed_pseudoge | ENSG00000 |

| | | | | |
|---|---|---|---|---|
| 38089231 | 38089329 | 1 | miRNA | ENSG0000C |
| 38129464 | 38140593 | -1 | lncRNA | ENSG0000C |
| 38149544 | 38162545 | 1 | lncRNA | ENSG0000C |
| 38193619 | 38210340 | 1 | lncRNA | ENSG0000C |
| 38209034 | 38214806 | -1 | lncRNA | ENSG0000C |
| 38383838 | 38384422 | -1 | processed_pseudoge | ENSG0000C |
| 38396659 | 38396765 | 1 | snRNA | ENSG0000C |
| 38474825 | 38516520 | 1 | lncRNA | ENSG0000C |
| 38708931 | 38710890 | 1 | processed_pseudoge | ENSG0000C |
| 38754216 | 38815979 | -1 | lncRNA | ENSG0000C |
| 38838198 | 38859772 | -1 | protein_coding | ENSG0000C |
| 38839333 | 38874494 | -1 | lncRNA | ENSG0000C |
| 38859912 | 38965038 | 1 | lncRNA | ENSG0000C |
| 38862493 | 38873368 | -1 | protein_coding | ENSG0000C |
| 38864501 | 38881617 | -1 | protein_coding | ENSG0000C |
| 38874069 | 38881587 | -1 | protein_coding | ENSG0000C |
| 38885807 | 38941830 | -1 | protein_coding | ENSG0000C |
| 38926870 | 38926977 | 1 | snRNA | ENSG0000C |
| 38950825 | 38950925 | -1 | misc_RNA | ENSG0000C |
| 38958275 | 38958925 | -1 | processed_pseudoge | ENSG0000C |
| 38991276 | 39006059 | 1 | protein_coding | ENSG0000C |
| 39026318 | 39034636 | 1 | protein_coding | ENSG0000C |
| 39081316 | 39487177 | 1 | protein_coding | ENSG0000C |
| 39120940 | 39121043 | -1 | snRNA | ENSG0000C |
| 39154164 | 39154296 | -1 | rRNA_pseudogene | ENSG0000C |
| 39206512 | 39206957 | 1 | lncRNA | ENSG0000C |
| 39226670 | 39238620 | -1 | lncRNA | ENSG0000C |
| 39249838 | 39257649 | -1 | lncRNA | ENSG0000C |
| 39304294 | 39304898 | -1 | processed_pseudoge | ENSG0000C |
| 39491636 | 39529869 | 1 | protein_coding | ENSG0000C |
| 39514956 | 39516490 | 1 | unprocessed_pseudc | ENSG0000C |
| 39522280 | 39559671 | -1 | transcribed_unproce | ENSG0000C |
| 39560816 | 39576790 | -1 | protein_coding | ENSG0000C |
| 39565052 | 39573860 | 1 | lncRNA | ENSG0000C |
| 39567374 | 39567508 | -1 | snoRNA | ENSG0000C |
| 39623435 | 39639643 | -1 | protein_coding | ENSG0000C |
| 39633416 | 39633903 | 1 | lncRNA | ENSG0000C |
| 39659121 | 39672038 | -1 | protein_coding | ENSG0000C |
| 39678648 | 39691485 | -1 | protein_coding | ENSG0000C |
| 39692182 | 39763914 | 1 | protein_coding | ENSG0000C |
| 39718028 | 39718595 | 1 | processed_pseudoge | ENSG0000C |
| 39723566 | 39723627 | -1 | snRNA | ENSG0000C |
| 39757182 | 39788865 | -1 | protein_coding | ENSG0000C |
| 39769523 | 39771348 | -1 | protein_coding | ENSG0000C |
| 39779969 | 39781418 | 1 | lncRNA | ENSG0000C |
| 39788976 | 39790171 | 1 | lncRNA | ENSG0000C |
| 39795843 | 39795996 | -1 | unprocessed_pseudc | ENSG0000C |
| 39799422 | 39809450 | 1 | lncRNA | ENSG0000C |
| 39801414 | 39817460 | -1 | lncRNA | ENSG0000C |
| 39838110 | 39883511 | -1 | protein_coding | ENSG0000C |

| | | | | |
|---|---|---|---|---|
| 39881566 | 39881678 | -1 | misc_RNA | ENSG00000 |
| 39895426 | 39902256 | -1 | protein_coding | ENSG00000 |
| 39897745 | 39899098 | 1 | lncRNA | ENSG00000 |
| 39944890 | 39944998 | -1 | misc_RNA | ENSG00000 |
| 39955112 | 39969968 | 1 | protein_coding | ENSG00000 |
| 39962680 | 39963404 | 1 | processed_pseudoge | ENSG00000 |
| 40040233 | 40072649 | 1 | protein_coding | ENSG00000 |
| 40071461 | 40097727 | -1 | protein_coding | ENSG00000 |
| 40132764 | 40133448 | 1 | processed_pseudoge | ENSG00000 |
| 40161387 | 40240921 | 1 | protein_coding | ENSG00000 |
| 40177843 | 40177949 | 1 | snRNA | ENSG00000 |
| 40245947 | 40251684 | 1 | protein_coding | ENSG00000 |
| 40256427 | 40257967 | -1 | lncRNA | ENSG00000 |
| 40258236 | 40294180 | 1 | protein_coding | ENSG00000 |
| 40262672 | 40262984 | -1 | processed_pseudoge | ENSG00000 |
| 40300489 | 40317813 | -1 | protein_coding | ENSG00000 |
| 40333078 | 40333557 | -1 | processed_pseudoge | ENSG00000 |
| 40344850 | 40423326 | 1 | protein_coding | ENSG00000 |
| 40364766 | 40365183 | 1 | processed_pseudoge | ENSG00000 |
| 40436199 | 40450039 | -1 | lncRNA | ENSG00000 |
| 40450102 | 40463718 | 1 | protein_coding | ENSG00000 |
| 40464319 | 40466767 | 1 | lncRNA | ENSG00000 |
| 40473055 | 40474059 | 1 | TEC | ENSG00000 |
| 40477290 | 40496343 | 1 | protein_coding | ENSG00000 |
| 40493157 | 40508661 | -1 | lncRNA | ENSG00000 |
| 40508741 | 40516556 | 1 | protein_coding | ENSG00000 |
| 40514461 | 40515057 | -1 | lncRNA | ENSG00000 |
| 40515754 | 40517174 | -1 | lncRNA | ENSG00000 |
| 40531573 | 40548167 | 1 | protein_coding | ENSG00000 |
| 40559666 | 40584335 | -1 | lncRNA | ENSG00000 |
| 40563534 | 40563659 | 1 | unprocessed_pseudc | ENSG00000 |
| 40593633 | 40594353 | -1 | processed_pseudoge | ENSG00000 |
| 40620680 | 40665682 | -1 | protein_coding | ENSG00000 |
| 40659848 | 40667480 | 1 | lncRNA | ENSG00000 |
| 40669089 | 40687588 | -1 | lncRNA | ENSG00000 |
| 40690380 | 40692066 | -1 | lncRNA | ENSG00000 |
| 40691648 | 40771603 | 1 | protein_coding | ENSG00000 |
| 40754355 | 40754446 | 1 | miRNA | ENSG00000 |
| 40757284 | 40757372 | 1 | miRNA | ENSG00000 |
| 40783787 | 40840452 | 1 | protein_coding | ENSG00000 |
| 40804846 | 40805141 | 1 | misc_RNA | ENSG00000 |
| 40861054 | 40862363 | -1 | protein_coding | ENSG00000 |
| 40863914 | 40876670 | 1 | lncRNA | ENSG00000 |
| 40938104 | 40939763 | -1 | unprocessed_pseudc | ENSG00000 |
| 40939294 | 40945524 | 1 | lncRNA | ENSG00000 |
| 40942251 | 40942515 | -1 | processed_pseudoge | ENSG00000 |
| 40979688 | 41012565 | 1 | protein_coding | ENSG00000 |
| 41014590 | 41043890 | 1 | lncRNA | ENSG00000 |
| 41015597 | 41023237 | -1 | protein_coding | ENSG00000 |
| 41027200 | 41242154 | -1 | protein_coding | ENSG00000 |

| | | | | |
|---|---|---|---|---|
| 41098638 | 41099136 | 1 | processed_pseudoge | ENSG00000 |
| 41241772 | 41338644 | 1 | lncRNA | ENSG00000 |
| 41242373 | 41284861 | 1 | lncRNA | ENSG00000 |
| 41264550 | 41264862 | -1 | processed_pseudoge | ENSG00000 |
| 41302932 | 41306148 | -1 | transcribed_unproce | ENSG00000 |
| 41361922 | 41383590 | 1 | protein_coding | ENSG00000 |
| 41375004 | 41375669 | -1 | lncRNA | ENSG00000 |
| 41466937 | 41467028 | -1 | rRNA_pseudogene | ENSG00000 |
| 41478775 | 41484683 | -1 | protein_coding | ENSG00000 |
| 41506365 | 42035925 | -1 | protein_coding | ENSG00000 |
| 41535443 | 41535868 | 1 | lncRNA | ENSG00000 |
| 41542069 | 41544310 | 1 | processed_pseudoge | ENSG00000 |
| 41585306 | 41628816 | -1 | protein_coding | ENSG00000 |
| 42036143 | 42051784 | 1 | lncRNA | ENSG00000 |
| 42040597 | 42041729 | 1 | processed_pseudoge | ENSG00000 |
| 42140635 | 42141391 | 1 | processed_pseudoge | ENSG00000 |
| 42153410 | 42155820 | 1 | protein_coding | ENSG00000 |
| 42162690 | 42164745 | -1 | protein_coding | ENSG00000 |
| 42176539 | 42335877 | -1 | protein_coding | ENSG00000 |
| 42335386 | 42338376 | -1 | lncRNA | ENSG00000 |
| 42380792 | 42424232 | 1 | protein_coding | ENSG00000 |
| 42412398 | 42412794 | -1 | processed_pseudoge | ENSG00000 |
| 42430329 | 42456253 | -1 | protein_coding | ENSG00000 |
| 42456117 | 42473385 | 1 | protein_coding | ENSG00000 |
| 42463221 | 42657190 | 1 | protein_coding | ENSG00000 |
| 42491739 | 42492493 | 1 | processed_pseudoge | ENSG00000 |
| 42500205 | 42500339 | -1 | processed_pseudoge | ENSG00000 |
| 42569033 | 42569139 | 1 | snRNA | ENSG00000 |
| 42570970 | 42571390 | 1 | processed_pseudoge | ENSG00000 |
| 42658423 | 42676758 | 1 | protein_coding | ENSG00000 |
| 42658687 | 42682160 | -1 | lncRNA | ENSG00000 |
| 42678735 | 42681659 | 1 | lncRNA | ENSG00000 |
| 42682418 | 42703805 | 1 | protein_coding | ENSG00000 |
| 42733093 | 42740254 | -1 | protein_coding | ENSG00000 |
| 42746335 | 42767084 | -1 | protein_coding | ENSG00000 |
| 42767249 | 42779491 | 1 | protein_coding | ENSG00000 |
| 42767292 | 42794493 | 1 | protein_coding | ENSG00000 |
| 42775813 | 42776790 | -1 | lncRNA | ENSG00000 |
| 42785007 | 42816619 | 1 | protein_coding | ENSG00000 |
| 42807052 | 42817397 | -1 | protein_coding | ENSG00000 |
| 42817122 | 42844991 | 1 | protein_coding | ENSG00000 |
| 42832522 | 42846422 | -1 | lncRNA | ENSG00000 |
| 42846573 | 42852477 | 1 | protein_coding | ENSG00000 |
| 42886597 | 42888767 | -1 | processed_pseudoge | ENSG00000 |
| 42891094 | 42892430 | -1 | processed_pseudoge | ENSG00000 |
| 42903232 | 42903912 | 1 | processed_pseudoge | ENSG00000 |
| 42925375 | 42958868 | -1 | protein_coding | ENSG00000 |
| 42952202 | 42952641 | -1 | processed_pseudoge | ENSG00000 |
| 42959049 | 42996814 | 1 | lncRNA | ENSG00000 |
| 42959065 | 42961864 | -1 | lncRNA | ENSG00000 |

| | | | | |
|---|---|---|---|---|
| 42991438 | 42991544 | 1 | snRNA | ENSG0000C |
| 43023549 | 43023637 | -1 | snRNA | ENSG0000C |
| 43104086 | 43104497 | 1 | processed_pseudoge | ENSG0000C |
| 43145153 | 43156396 | 1 | protein_coding | ENSG0000C |
| 43164175 | 43270936 | -1 | protein_coding | ENSG0000C |
| 43171652 | 43171712 | -1 | miRNA | ENSG0000C |
| 43172149 | 43254358 | 1 | protein_coding | ENSG0000C |
| 43196417 | 43196536 | 1 | rRNA_pseudogene | ENSG0000C |
| 43269983 | 43274002 | 1 | protein_coding | ENSG0000C |
| 43281877 | 43285617 | -1 | protein_coding | ENSG0000C |
| 43300982 | 43323108 | 1 | protein_coding | ENSG0000C |
| 43337849 | 43352772 | 1 | protein_coding | ENSG0000C |
| 43348288 | 43348844 | -1 | lncRNA | ENSG0000C |
| 43354684 | 43358658 | -1 | lncRNA | ENSG0000C |
| 43358981 | 43363203 | 1 | protein_coding | ENSG0000C |
| 43363398 | 43368074 | -1 | protein_coding | ENSG0000C |
| 43364648 | 43364715 | -1 | miRNA | ENSG0000C |
| 43383917 | 43389808 | -1 | protein_coding | ENSG0000C |
| 43385113 | 43389155 | 1 | lncRNA | ENSG0000C |
| 43389882 | 43454247 | 1 | protein_coding | ENSG0000C |
| 43447776 | 43448644 | -1 | lncRNA | ENSG0000C |
| 43448539 | 43448611 | 1 | miRNA | ENSG0000C |
| 43451003 | 43453989 | -1 | protein_coding | ENSG0000C |
| 43453927 | 43456995 | 1 | lncRNA | ENSG0000C |
| 43525187 | 43623666 | 1 | protein_coding | ENSG0000C |
| 43650149 | 43931014 | 1 | protein_coding | ENSG0000C |
| 43650149 | 43705518 | 1 | protein_coding | ENSG0000C |
| 43685123 | 43708138 | -1 | lncRNA | ENSG0000C |
| 43705824 | 43931165 | 1 | protein_coding | ENSG0000C |
| 43709392 | 43727343 | -1 | lncRNA | ENSG0000C |
| 43716467 | 43716556 | 1 | snRNA | ENSG0000C |
| 43743471 | 43743741 | 1 | processed_pseudoge | ENSG0000C |
| 43838622 | 43838683 | 1 | miRNA | ENSG0000C |
| 43850300 | 43851611 | -1 | processed_pseudoge | ENSG0000C |
| 43933320 | 43937240 | 1 | protein_coding | ENSG0000C |
| 43937918 | 43946551 | -1 | lncRNA | ENSG0000C |
| 43946950 | 43968022 | 1 | protein_coding | ENSG0000C |
| 43968351 | 43974821 | -1 | lncRNA | ENSG0000C |
| 43970000 | 43973369 | 1 | protein_coding | ENSG0000C |
| 43974487 | 43978295 | 1 | protein_coding | ENSG0000C |
| 43978943 | 43991170 | 1 | protein_coding | ENSG0000C |
| 43991359 | 43996528 | 1 | protein_coding | ENSG0000C |
| 43991500 | 44031467 | -1 | protein_coding | ENSG0000C |
| 44030437 | 44115913 | 1 | lncRNA | ENSG0000C |
| 44087958 | 44088477 | -1 | processed_pseudoge | ENSG0000C |
| 44103306 | 44104703 | -1 | processed_pseudoge | ENSG0000C |
| 44117098 | 44117397 | -1 | misc_RNA | ENSG0000C |
| 44118821 | 44135140 | 1 | protein_coding | ENSG0000C |
| 44122153 | 44122512 | -1 | processed_pseudoge | ENSG0000C |
| 44137821 | 44141631 | -1 | protein_coding | ENSG0000C |

| | | | | |
|---|---|---|---|---|
| 44150594 | 44151753 | -1 | processed_pseudoge | ENSG0000C |
| 44153385 | 44153480 | -1 | misc_RNA | ENSG0000C |
| 44155028 | 44155301 | -1 | processed_pseudoge | ENSG0000C |
| 44172506 | 44172719 | -1 | processed_pseudoge | ENSG0000C |
| 44187943 | 44189049 | -1 | processed_pseudoge | ENSG0000C |
| 44213455 | 44220681 | 1 | protein_coding | ENSG0000C |
| 44221070 | 44355260 | -1 | protein_coding | ENSG0000C |
| 44243408 | 44244273 | -1 | lncRNA | ENSG0000C |
| 44390722 | 44390823 | 1 | snRNA | ENSG0000C |
| 44405194 | 44651724 | 1 | protein_coding | ENSG0000C |
| 44545493 | 44545552 | 1 | miRNA | ENSG0000C |
| 44635238 | 44674481 | -1 | protein_coding | ENSG0000C |
| 44674692 | 44725591 | 1 | protein_coding | ENSG0000C |
| 44721786 | 44721902 | -1 | snRNA | ENSG0000C |
| 44731055 | 44731170 | -1 | snRNA | ENSG0000C |
| 44739818 | 44767767 | 1 | protein_coding | ENSG0000C |
| 44759037 | 44775810 | -1 | lncRNA | ENSG0000C |
| 44775251 | 44778779 | 1 | protein_coding | ENSG0000C |
| 44775864 | 44775943 | 1 | snoRNA | ENSG0000C |
| 44776490 | 44776593 | 1 | snoRNA | ENSG0000C |
| 44777843 | 44777912 | 1 | snoRNA | ENSG0000C |
| 44778390 | 44778458 | 1 | snoRNA | ENSG0000C |
| 44778390 | 44778456 | 1 | snoRNA | ENSG0000C |
| 44780331 | 44780715 | -1 | processed_pseudoge | ENSG0000C |
| 44783585 | 44788170 | -1 | protein_coding | ENSG0000C |
| 44800377 | 44805990 | 1 | protein_coding | ENSG0000C |
| 44805913 | 44806675 | -1 | protein_coding | ENSG0000C |
| 44808482 | 44815585 | 1 | protein_coding | ENSG0000C |
| 44819844 | 44843253 | -1 | protein_coding | ENSG0000C |
| 44819883 | 44819997 | -1 | snRNA | ENSG0000C |
| 44843921 | 44844082 | -1 | processed_pseudoge | ENSG0000C |
| 44850522 | 44986722 | -1 | protein_coding | ENSG0000C |
| 44932323 | 44932431 | 1 | rRNA_pseudogene | ENSG0000C |
| 44958557 | 44959395 | 1 | processed_pseudoge | ENSG0000C |
| 44988234 | 44988725 | -1 | processed_pseudoge | ENSG0000C |
| 44988705 | 44991834 | -1 | processed_pseudoge | ENSG0000C |
| 45002540 | 45011324 | -1 | protein_coding | ENSG0000C |
| 45010950 | 45015575 | 1 | protein_coding | ENSG0000C |
| 45016399 | 45306209 | -1 | protein_coding | ENSG0000C |
| 45069977 | 45070429 | -1 | processed_pseudoge | ENSG0000C |
| 45303910 | 45305619 | 1 | lncRNA | ENSG0000C |
| 45326895 | 45328679 | 1 | protein_coding | ENSG0000C |
| 45329163 | 45340893 | -1 | protein_coding | ENSG0000C |
| 45329262 | 45491155 | -1 | protein_coding | ENSG0000C |
| 45340052 | 45343973 | 1 | protein_coding | ENSG0000C |
| 45343883 | 45491163 | -1 | protein_coding | ENSG0000C |
| 45415020 | 45415344 | 1 | processed_pseudoge | ENSG0000C |
| 45493866 | 45500079 | -1 | protein_coding | ENSG0000C |
| 45500300 | 45513382 | 1 | protein_coding | ENSG0000C |
| 45511036 | 45523047 | -1 | protein_coding | ENSG0000C |

| | | | | |
|---|---|---|---|---|
| 45530927 | 45532139 | 1 | unprocessed_pseudo | ENSG00000 |
| 45550543 | 45570049 | 1 | protein_coding | ENSG00000 |
| 45580892 | 45580996 | -1 | pseudogene | ENSG00000 |
| 45581219 | 45581321 | -1 | pseudogene | ENSG00000 |
| 45583846 | 45618904 | 1 | protein_coding | ENSG00000 |
| 45592722 | 45592814 | -1 | pseudogene | ENSG00000 |
| 45605657 | 45605730 | -1 | pseudogene | ENSG00000 |
| 45620044 | 45624057 | -1 | protein_coding | ENSG00000 |
| 45627304 | 45688113 | -1 | protein_coding | ENSG00000 |
| 45645816 | 45646197 | -1 | processed_pseudoge | ENSG00000 |
| 45651039 | 45651826 | -1 | processed_pseudoge | ENSG00000 |
| 45688181 | 45694436 | 1 | protein_coding | ENSG00000 |
| 45694324 | 45750653 | -1 | protein_coding | ENSG00000 |
| 45694684 | 45697075 | -1 | lncRNA | ENSG00000 |
| 45781277 | 45781957 | 1 | processed_pseudoge | ENSG00000 |
| 45786987 | 46036122 | 1 | protein_coding | ENSG00000 |
| 45846994 | 45847594 | -1 | processed_pseudoge | ENSG00000 |
| 46040140 | 46133036 | -1 | protein_coding | ENSG00000 |
| 46043661 | 46176488 | -1 | protein_coding | ENSG00000 |
| 46046818 | 46048368 | 1 | lncRNA | ENSG00000 |
| 46104950 | 46105175 | 1 | processed_pseudoge | ENSG00000 |
| 46134531 | 46139081 | 1 | lncRNA | ENSG00000 |
| 46175073 | 46185962 | 1 | protein_coding | ENSG00000 |
| 46175486 | 46176478 | -1 | protein_coding | ENSG00000 |
| 46188682 | 46220305 | -1 | protein_coding | ENSG00000 |
| 46203334 | 46221256 | 1 | protein_coding | ENSG00000 |
| 46246461 | 46278480 | 1 | protein_coding | ENSG00000 |
| 46261196 | 46303608 | -1 | protein_coding | ENSG00000 |
| 46303698 | 46316776 | 1 | protein_coding | ENSG00000 |
| 46340177 | 46365152 | 1 | protein_coding | ENSG00000 |
| 46370586 | 46370955 | 1 | processed_pseudoge | ENSG00000 |
| 46394317 | 46413848 | 1 | protein_coding | ENSG00000 |
| 46432129 | 46482493 | 1 | transcribed_unproce | ENSG00000 |
| 46446600 | 46451317 | -1 | lncRNA | ENSG00000 |
| 46506996 | 46514226 | 1 | protein_coding | ENSG00000 |
| 46532166 | 46543969 | -1 | protein_coding | ENSG00000 |
| 46538611 | 46570255 | 1 | lncRNA | ENSG00000 |
| 46545644 | 46551527 | -1 | protein_coding | ENSG00000 |
| 46557408 | 46616843 | -1 | protein_coding | ENSG00000 |
| 46607715 | 46616891 | -1 | protein_coding | ENSG00000 |
| 46632737 | 46673867 | -1 | protein_coding | ENSG00000 |
| 46665910 | 46666375 | -1 | processed_pseudoge | ENSG00000 |
| 46668855 | 46673594 | 1 | protein_coding | ENSG00000 |
| 46674036 | 46692098 | 1 | lncRNA | ENSG00000 |
| 46675159 | 46719114 | -1 | protein_coding | ENSG00000 |
| 46742329 | 46743133 | 1 | processed_pseudoge | ENSG00000 |
| 46757838 | 46819413 | 1 | protein_coding | ENSG00000 |
| 46843095 | 46900437 | -1 | transcribed_unproce | ENSG00000 |
| 46891639 | 46892830 | 1 | processed_pseudoge | ENSG00000 |
| 46929177 | 46941484 | -1 | protein_coding | ENSG00000 |

| | | | | |
|---|---|---|---|---|
| 46967679 | 46967874 | -1 | unprocessed_pseudo | ENSG00000 |
| 46994382 | 46994502 | -1 | unprocessed_pseudo | ENSG00000 |
| 47000898 | 47001087 | -1 | unprocessed_pseudo | ENSG00000 |
| 47002995 | 47003148 | -1 | unprocessed_pseudo | ENSG00000 |
| 47023669 | 47050751 | 1 | protein_coding | ENSG00000 |
| 47067231 | 47118318 | 1 | protein_coding | ENSG00000 |
| 47074778 | 47075979 | -1 | processed_pseudoge | ENSG00000 |
| 47096653 | 47179271 | -1 | lncRNA | ENSG00000 |
| 47137435 | 47149735 | 1 | protein_coding | ENSG00000 |
| 47164510 | 47165920 | -1 | transcribed_processe | ENSG00000 |
| 47179250 | 47180339 | 1 | lncRNA | ENSG00000 |
| 47183582 | 47191044 | -1 | protein_coding | ENSG00000 |
| 47216290 | 47232220 | -1 | protein_coding | ENSG00000 |
| 47225797 | 47230750 | 1 | lncRNA | ENSG00000 |
| 47250139 | 47314147 | -1 | protein_coding | ENSG00000 |
| 47333797 | 47378839 | 1 | protein_coding | ENSG00000 |
| 47380928 | 47408477 | -1 | lncRNA | ENSG00000 |
| 47416285 | 47418052 | 1 | protein_coding | ENSG00000 |
| 47432133 | 47434641 | -1 | lncRNA | ENSG00000 |
| 47436017 | 47440691 | 1 | protein_coding | ENSG00000 |
| 47483698 | 47484162 | -1 | processed_pseudoge | ENSG00000 |
| 47497894 | 47498341 | -1 | processed_pseudoge | ENSG00000 |
| 47550196 | 47550753 | 1 | processed_pseudoge | ENSG00000 |
| 47688463 | 47704029 | 1 | lncRNA | ENSG00000 |
| 47760528 | 47997385 | -1 | protein_coding | ENSG00000 |
| 47761132 | 47765547 | -1 | lncRNA | ENSG00000 |
| 47818066 | 47820237 | 1 | lncRNA | ENSG00000 |
| 48050659 | 48091736 | 1 | lncRNA | ENSG00000 |
| 48078787 | 48082196 | -1 | lncRNA | ENSG00000 |
| 48089368 | 48089634 | 1 | processed_pseudoge | ENSG00000 |
| 48096092 | 48182428 | -1 | transcribed_unproce | ENSG00000 |
| 48102068 | 48104736 | 1 | TEC | ENSG00000 |
| 48172972 | 48206857 | 1 | lncRNA | ENSG00000 |
| 48222685 | 48248644 | 1 | protein_coding | ENSG00000 |
| 48227888 | 48229561 | -1 | lncRNA | ENSG00000 |
| 48262230 | 48263179 | -1 | TEC | ENSG00000 |
| 48295373 | 48472208 | -1 | protein_coding | ENSG00000 |
| 48325080 | 48326123 | -1 | processed_pseudoge | ENSG00000 |
| 48344209 | 48344315 | 1 | snRNA | ENSG00000 |
| 48435967 | 48437223 | 1 | processed_pseudoge | ENSG00000 |
| 48447936 | 48448042 | 1 | snRNA | ENSG00000 |
| 48497263 | 48497736 | -1 | processed_pseudoge | ENSG00000 |
| 48532854 | 50023954 | -1 | protein_coding | ENSG00000 |
| 48552991 | 48558106 | 1 | lncRNA | ENSG00000 |
| 48727519 | 48776969 | -1 | protein_coding | ENSG00000 |
| 48926021 | 48936734 | 1 | lncRNA | ENSG00000 |
| 49025595 | 49187585 | 1 | lncRNA | ENSG00000 |
| 49257411 | 49269285 | 1 | lncRNA | ENSG00000 |
| 49374201 | 49472085 | -1 | lncRNA | ENSG00000 |
| 49691262 | 49691523 | -1 | processed_pseudoge | ENSG00000 |

| | | | | |
|---|---|---|---|---|
| 49841821 | 49846400 | 1 | unprocessed_pseudc | ENSG0000C |
| 49994318 | 49996218 | 1 | TEC | ENSG0000C |
| 50017092 | 50017503 | 1 | processed_pseudoge | ENSG0000C |
| 50024029 | 50203786 | 1 | protein_coding | ENSG0000C |
| 50114937 | 50115282 | 1 | processed_pseudoge | ENSG0000C |
| 50174306 | 50175868 | -1 | lncRNA | ENSG0000C |
| 50206084 | 50223127 | -1 | lncRNA | ENSG0000C |
| 50229662 | 50321192 | -1 | lncRNA | ENSG0000C |
| 50252569 | 50258994 | 1 | lncRNA | ENSG0000C |
| 50326131 | 50327007 | 1 | processed_pseudoge | ENSG0000C |
| 50398825 | 50399773 | -1 | processed_pseudoge | ENSG0000C |
| 50405430 | 50406024 | -1 | processed_pseudoge | ENSG0000C |
| 50417550 | 50423443 | -1 | protein_coding | ENSG0000C |
| 50423609 | 50425316 | 1 | lncRNA | ENSG0000C |
| 50437028 | 50960267 | -1 | protein_coding | ENSG0000C |
| 50461469 | 50471150 | 1 | lncRNA | ENSG0000C |
| 50582404 | 50582511 | 1 | snRNA | ENSG0000C |
| 50780340 | 50781150 | 1 | processed_pseudoge | ENSG0000C |
| 50846468 | 50846837 | 1 | processed_pseudoge | ENSG0000C |
| 50960745 | 50974634 | 1 | protein_coding | ENSG0000C |
| 51059837 | 51059905 | 1 | miRNA | ENSG0000C |
| 51060018 | 51060103 | 1 | miRNA | ENSG0000C |
| 51102221 | 51148086 | 1 | protein_coding | ENSG0000C |
| 51107222 | 51107331 | 1 | misc_RNA | ENSG0000C |
| 51157788 | 51158471 | -1 | processed_pseudoge | ENSG0000C |
| 51195095 | 51235096 | -1 | lncRNA | ENSG0000C |
| 51236273 | 51273447 | 1 | protein_coding | ENSG0000C |
| 51250603 | 51251472 | -1 | processed_pseudoge | ENSG0000C |
| 51264916 | 51266074 | -1 | lncRNA | ENSG0000C |
| 51287258 | 51345116 | -1 | protein_coding | ENSG0000C |
| 51329654 | 51335324 | 1 | lncRNA | ENSG0000C |
| 51354263 | 51519266 | -1 | protein_coding | ENSG0000C |
| 51372270 | 51373224 | 1 | processed_pseudoge | ENSG0000C |
| 51382308 | 51382414 | 1 | snRNA | ENSG0000C |
| 51461721 | 51463416 | 1 | lncRNA | ENSG0000C |
| 51518309 | 51561629 | 1 | lncRNA | ENSG0000C |
| 51538625 | 51538729 | -1 | snRNA | ENSG0000C |
| 51561866 | 51594897 | -1 | transcribed_unitary_ | ENSG0000C |
| 51577179 | 51798427 | 1 | protein_coding | ENSG0000C |
| 51707138 | 51708137 | -1 | processed_pseudoge | ENSG0000C |
| 51709062 | 51709940 | 1 | processed_pseudoge | ENSG0000C |
| 51724775 | 51724896 | -1 | snoRNA | ENSG0000C |
| 51789191 | 51878937 | -1 | protein_coding | ENSG0000C |
| 51793934 | 51799154 | 1 | lncRNA | ENSG0000C |
| 51801028 | 51801307 | 1 | lncRNA | ENSG0000C |
| 51836344 | 51836402 | -1 | miRNA | ENSG0000C |
| 51859778 | 51860135 | 1 | processed_pseudoge | ENSG0000C |
| 51907956 | 51990700 | -1 | protein_coding | ENSG0000C |
| 51973410 | 51973535 | 1 | rRNA_pseudogene | ENSG0000C |
| 51980473 | 51980814 | 1 | processed_pseudoge | ENSG0000C |

| | | | | |
|---|---|---|---|---|
| 51995740 | 51996039 | 1 | misc_RNA | ENSG00000( |
| 52020131 | 52056171 | -1 | protein_coding | ENSG00000( |
| 52020153 | 52033350 | -1 | protein_coding | ENSG00000( |
| 52032103 | 52033810 | -1 | protein_coding | ENSG00000( |
| 52033391 | 52044279 | 1 | lncRNA | ENSG00000( |
| 52050918 | 52052683 | 1 | lncRNA | ENSG00000( |
| 52056199 | 52090716 | 1 | protein_coding | ENSG00000( |
| 52142094 | 52346686 | 1 | protein_coding | ENSG00000( |
| 52150104 | 52150419 | 1 | misc_RNA | ENSG00000( |
| 52160261 | 52160600 | -1 | processed_pseudoge | ENSG00000( |
| 52162186 | 52163026 | -1 | processed_pseudoge | ENSG00000( |
| 52179848 | 52180553 | 1 | processed_pseudoge | ENSG00000( |
| 52189916 | 52190309 | -1 | processed_pseudoge | ENSG00000( |
| 52252062 | 52252773 | -1 | processed_pseudoge | ENSG00000( |
| 52253621 | 52254176 | 1 | processed_pseudoge | ENSG00000( |
| 52345723 | 52366193 | -1 | protein_coding | ENSG00000( |
| 52353487 | 52353877 | 1 | lncRNA | ENSG00000( |
| 52365443 | 52367865 | 1 | lncRNA | ENSG00000( |
| 52368677 | 52369244 | 1 | processed_pseudoge | ENSG00000( |
| 52372829 | 52404423 | -1 | protein_coding | ENSG00000( |
| 52404602 | 52420836 | 1 | protein_coding | ENSG00000( |
| 52408282 | 52553487 | -1 | protein_coding | ENSG00000( |
| 52554818 | 52555273 | 1 | lncRNA | ENSG00000( |
| 52573114 | 52573816 | 1 | processed_pseudoge | ENSG00000( |
| 52602371 | 52609051 | 1 | protein_coding | ENSG00000( |
| 52633168 | 52669683 | 1 | protein_coding | ENSG00000( |
| 52684449 | 52698347 | -1 | protein_coding | ENSG00000( |
| 52709122 | 52709442 | 1 | processed_pseudoge | ENSG00000( |
| 52714399 | 52714695 | 1 | misc_RNA | ENSG00000( |
| 52726453 | 52827336 | 1 | protein_coding | ENSG00000( |
| 52754322 | 52754462 | 1 | snRNA | ENSG00000( |
| 52772194 | 52772648 | -1 | processed_pseudoge | ENSG00000( |
| 52805108 | 52805212 | -1 | snRNA | ENSG00000( |
| 52842511 | 52894998 | 1 | protein_coding | ENSG00000( |
| 52881216 | 52881730 | -1 | lncRNA | ENSG00000( |
| 52895910 | 52927212 | -1 | protein_coding | ENSG00000( |
| 52920422 | 52920596 | 1 | processed_pseudoge | ENSG00000( |
| 52925249 | 52925423 | -1 | processed_pseudoge | ENSG00000( |
| 52927276 | 53051698 | 1 | protein_coding | ENSG00000( |
| 52928674 | 52928772 | 1 | miRNA | ENSG00000( |
| 52934930 | 52935017 | 1 | miRNA | ENSG00000( |
| 52940314 | 52940413 | 1 | miRNA | ENSG00000( |
| 52943536 | 52943930 | -1 | processed_pseudoge | ENSG00000( |
| 52993201 | 52993702 | -1 | processed_pseudoge | ENSG00000( |
| 52994726 | 52996026 | 1 | processed_pseudoge | ENSG00000( |
| 53062052 | 53085502 | 1 | protein_coding | ENSG00000( |
| 53069938 | 53085502 | -1 | lncRNA | ENSG00000( |
| 53073110 | 53073389 | -1 | processed_pseudoge | ENSG00000( |
| 53087179 | 53142632 | -1 | protein_coding | ENSG00000( |
| 53114576 | 53118609 | 1 | lncRNA | ENSG00000( |

| | | | | |
|---|---|---|---|---|
| 53180921 | 53186900 | 1 | unprocessed_pseudo | ENSG00000 |
| 53196792 | 53214197 | 1 | protein_coding | ENSG00000 |
| 53209783 | 53213775 | -1 | lncRNA | ENSG00000 |
| 53214099 | 53220634 | -1 | protein_coding | ENSG00000 |
| 53220663 | 53224253 | 1 | lncRNA | ENSG00000 |
| 53226900 | 53238518 | -1 | protein_coding | ENSG00000 |
| 53238550 | 53246482 | 1 | lncRNA | ENSG00000 |
| 53242364 | 53328469 | -1 | protein_coding | ENSG00000 |
| 53267935 | 53268601 | 1 | lncRNA | ENSG00000 |
| 53288024 | 53289706 | 1 | lncRNA | ENSG00000 |
| 53304536 | 53307462 | 1 | lncRNA | ENSG00000 |
| 53328233 | 53336509 | 1 | lncRNA | ENSG00000 |
| 53344031 | 53365473 | 1 | lncRNA | ENSG00000 |
| 53348488 | 53349233 | -1 | lncRNA | ENSG00000 |
| 53366656 | 53368245 | -1 | lncRNA | ENSG00000 |
| 53413149 | 53440020 | -1 | transcribed_processe | ENSG00000 |
| 53459399 | 53467488 | 1 | protein_coding | ENSG00000 |
| 53506237 | 53738106 | -1 | protein_coding | ENSG00000 |
| 53688749 | 53688812 | -1 | snRNA | ENSG00000 |
| 53765478 | 53838463 | -1 | protein_coding | ENSG00000 |
| 53771018 | 53771153 | -1 | snoRNA | ENSG00000 |
| 53841547 | 53841832 | 1 | processed_pseudoge | ENSG00000 |
| 53851719 | 53889798 | -1 | protein_coding | ENSG00000 |
| 53891239 | 53911086 | 1 | protein_coding | ENSG00000 |
| 53916574 | 53945929 | -1 | protein_coding | ENSG00000 |
| 53946085 | 53968168 | 1 | protein_coding | ENSG00000 |
| 53974969 | 53976031 | -1 | processed_pseudoge | ENSG00000 |
| 54007298 | 54018186 | -1 | protein_coding | ENSG00000 |
| 54026681 | 54053504 | -1 | protein_coding | ENSG00000 |
| 54033126 | 54034880 | -1 | TEC | ENSG00000 |
| 54053584 | 54112519 | 1 | protein_coding | ENSG00000 |
| 54054079 | 54054154 | 1 | miRNA | ENSG00000 |
| 54089856 | 54090093 | 1 | processed_pseudoge | ENSG00000 |
| 54099968 | 54100224 | -1 | pseudogene | ENSG00000 |
| 54132686 | 54200073 | -1 | protein_coding | ENSG00000 |
| 54132968 | 54153770 | -1 | protein_coding | ENSG00000 |
| 54137746 | 54142980 | 1 | lncRNA | ENSG00000 |
| 54169651 | 54200036 | -1 | protein_coding | ENSG00000 |
| 54184041 | 54225464 | 1 | protein_coding | ENSG00000 |
| 54225432 | 54413479 | -1 | protein_coding | ENSG00000 |
| 54236440 | 54239063 | 1 | lncRNA | ENSG00000 |
| 54285405 | 54287371 | 1 | lncRNA | ENSG00000 |
| 54416256 | 54420860 | 1 | lncRNA | ENSG00000 |
| 54514417 | 54517383 | 1 | lncRNA | ENSG00000 |
| 54516412 | 54517259 | -1 | lncRNA | ENSG00000 |
| 54524824 | 54524931 | 1 | processed_pseudoge | ENSG00000 |
| 54536796 | 54537758 | 1 | processed_pseudoge | ENSG00000 |
| 54542257 | 54639192 | 1 | protein_coding | ENSG00000 |
| 54609181 | 54623556 | -1 | protein_coding | ENSG00000 |
| 54621477 | 54628438 | -1 | lncRNA | ENSG00000 |

| | | | | |
|---|---|---|---|---|
| 54641754 | 54710266 | 1 | protein_coding | ENSG00000 |
| 54641786 | 54742308 | 1 | protein_coding | ENSG00000 |
| 54715861 | 54742657 | 1 | protein_coding | ENSG00000 |
| 54756898 | 54764523 | -1 | protein_coding | ENSG00000 |
| 54779712 | 54801323 | -1 | protein_coding | ENSG00000 |
| 54792885 | 54794905 | 1 | lncRNA | ENSG00000 |
| 54806063 | 54842252 | 1 | protein_coding | ENSG00000 |
| 54849627 | 54887195 | -1 | protein_coding | ENSG00000 |
| 54874672 | 54883664 | 1 | lncRNA | ENSG00000 |
| 54886875 | 54888001 | 1 | lncRNA | ENSG00000 |
| 54887563 | 54888850 | 1 | lncRNA | ENSG00000 |
| 54974900 | 54980464 | -1 | lncRNA | ENSG00000 |
| 54980628 | 54992288 | 1 | protein_coding | ENSG00000 |
| 54980950 | 54992274 | -1 | lncRNA | ENSG00000 |
| 54998933 | 55017172 | 1 | protein_coding | ENSG00000 |
| 55039548 | 55064852 | 1 | protein_coding | ENSG00000 |
| 55066359 | 55215364 | -1 | protein_coding | ENSG00000 |
| 55217645 | 55324691 | 1 | lncRNA | ENSG00000 |
| 55222379 | 55223372 | 1 | processed_pseudoge | ENSG00000 |
| 55225641 | 55225723 | 1 | miRNA | ENSG00000 |
| 55329288 | 56070513 | 1 | lncRNA | ENSG00000 |
| 55367466 | 55368774 | 1 | processed_pseudoge | ENSG00000 |
| 55372710 | 55373316 | -1 | processed_pseudoge | ENSG00000 |
| 55376526 | 55376852 | -1 | misc_RNA | ENSG00000 |
| 55398514 | 55398620 | 1 | snRNA | ENSG00000 |
| 55484871 | 55484972 | -1 | misc_RNA | ENSG00000 |
| 55868254 | 55950119 | 1 | lncRNA | ENSG00000 |
| 55915603 | 55944980 | -1 | lncRNA | ENSG00000 |
| 55938714 | 55939280 | -1 | transcribed_unproce | ENSG00000 |
| 56145721 | 56155224 | 1 | lncRNA | ENSG00000 |
| 56154545 | 56477687 | -1 | lncRNA | ENSG00000 |
| 56173433 | 56524495 | -1 | protein_coding | ENSG00000 |
| 56207567 | 56208456 | -1 | processed_pseudoge | ENSG00000 |
| 56414935 | 56417137 | 1 | lncRNA | ENSG00000 |
| 56494761 | 56645301 | -1 | protein_coding | ENSG00000 |
| 56538452 | 56538911 | 1 | processed_pseudoge | ENSG00000 |
| 56585612 | 56586242 | -1 | processed_pseudoge | ENSG00000 |
| 56619409 | 56619549 | -1 | processed_pseudoge | ENSG00000 |
| 56645314 | 56715335 | 1 | protein_coding | ENSG00000 |
| 56718789 | 56819696 | -1 | protein_coding | ENSG00000 |
| 56823679 | 56826920 | -1 | lncRNA | ENSG00000 |
| 56854806 | 56918221 | 1 | protein_coding | ENSG00000 |
| 56929207 | 56966140 | -1 | protein_coding | ENSG00000 |
| 56963886 | 56996757 | 1 | lncRNA | ENSG00000 |
| 56994778 | 58546734 | -1 | protein_coding | ENSG00000 |
| 57386576 | 57387450 | 1 | lncRNA | ENSG00000 |
| 57605847 | 57606206 | -1 | processed_pseudoge | ENSG00000 |
| 57860532 | 57880905 | 1 | lncRNA | ENSG00000 |
| 58047889 | 58049335 | -1 | transcribed_unproce | ENSG00000 |
| 58056133 | 58056480 | -1 | processed_pseudoge | ENSG00000 |

| | | | | |
|---|---|---|---|---|
| 58060139 | 58080274 | 1 | lncRNA | ENSG0000C |
| 58084419 | 58084559 | 1 | unprocessed_pseudc | ENSG0000C |
| 58228682 | 58229003 | -1 | processed_pseudoge | ENSG0000C |
| 58415384 | 58546802 | -1 | protein_coding | ENSG0000C |
| 58546168 | 58563897 | 1 | lncRNA | ENSG0000C |
| 58552913 | 58553333 | -1 | processed_pseudoge | ENSG0000C |
| 58565631 | 58565932 | -1 | misc_RNA | ENSG0000C |
| 58575433 | 58577252 | -1 | protein_coding | ENSG0000C |
| 58630841 | 58631530 | -1 | processed_pseudoge | ENSG0000C |
| 58643440 | 58700077 | -1 | protein_coding | ENSG0000C |
| 58715609 | 58771295 | 1 | lncRNA | ENSG0000C |
| 58780791 | 58784047 | -1 | protein_coding | ENSG0000C |
| 58785128 | 58901109 | 1 | lncRNA | ENSG0000C |
| 58812808 | 58813342 | -1 | lncRNA | ENSG0000C |
| 58838448 | 58851254 | -1 | lncRNA | ENSG0000C |
| 58882868 | 58931897 | -1 | lncRNA | ENSG0000C |
| 58933643 | 59240555 | 1 | lncRNA | ENSG0000C |
| 58999676 | 59000494 | 1 | processed_pseudoge | ENSG0000C |
| 59054397 | 59056049 | -1 | lncRNA | ENSG0000C |
| 59055999 | 59078116 | -1 | lncRNA | ENSG0000C |
| 59131932 | 59296491 | -1 | lncRNA | ENSG0000C |
| 59289303 | 59289640 | -1 | lncRNA | ENSG0000C |
| 59296638 | 59810647 | 1 | protein_coding | ENSG0000C |
| 59733227 | 59733296 | -1 | miRNA | ENSG0000C |
| 59754747 | 59789182 | -1 | lncRNA | ENSG0000C |
| 59814786 | 59876370 | 1 | protein_coding | ENSG0000C |
| 59893308 | 59926773 | -1 | protein_coding | ENSG0000C |
| 59974757 | 59975054 | -1 | misc_RNA | ENSG0000C |
| 59987269 | 60073770 | -1 | protein_coding | ENSG0000C |
| 60097415 | 60097976 | 1 | processed_pseudoge | ENSG0000C |
| 60114875 | 60149784 | 1 | lncRNA | ENSG0000C |
| 60515716 | 60640507 | -1 | lncRNA | ENSG0000C |
| 60659631 | 60868009 | -1 | lncRNA | ENSG0000C |
| 60865259 | 61462788 | 1 | protein_coding | ENSG0000C |
| 60912675 | 61056885 | -1 | lncRNA | ENSG0000C |
| 61248945 | 61253510 | -1 | lncRNA | ENSG0000C |
| 61481087 | 61533271 | 1 | lncRNA | ENSG0000C |
| 61588049 | 61589904 | -1 | lncRNA | ENSG0000C |
| 61654194 | 61654908 | 1 | processed_pseudoge | ENSG0000C |
| 61681046 | 61725423 | -1 | protein_coding | ENSG0000C |
| 61741998 | 61742424 | -1 | lncRNA | ENSG0000C |
| 61742406 | 62178675 | 1 | protein_coding | ENSG0000C |
| 61816419 | 61816522 | 1 | snRNA | ENSG0000C |
| 61852499 | 61852602 | 1 | snRNA | ENSG0000C |
| 62038842 | 62039111 | 1 | processed_pseudoge | ENSG0000C |
| 62072448 | 62072741 | 1 | misc_RNA | ENSG0000C |
| 62078786 | 62078859 | 1 | miRNA | ENSG0000C |
| 62078789 | 62078856 | -1 | miRNA | ENSG0000C |
| 62189131 | 62189953 | 1 | processed_pseudoge | ENSG0000C |
| 62190522 | 62190926 | 1 | processed_pseudoge | ENSG0000C |

| | | | | |
|---|---|---|---|---|
| 62194849 | 62212328 | 1 | protein_coding | ENSG00000 |
| 62208136 | 62208442 | -1 | processed_pseudoge | ENSG00000 |
| 62211557 | 62211666 | 1 | misc_RNA | ENSG00000 |
| 62236165 | 62319434 | -1 | protein_coding | ENSG00000 |
| 62298149 | 62298256 | -1 | snRNA | ENSG00000 |
| 62436297 | 62451804 | 1 | protein_coding | ENSG00000 |
| 62454298 | 62688386 | -1 | protein_coding | ENSG00000 |
| 62530636 | 62530873 | 1 | processed_pseudoge | ENSG00000 |
| 62597520 | 62606313 | 1 | protein_coding | ENSG00000 |
| 62607766 | 62609584 | 1 | TEC | ENSG00000 |
| 62641122 | 62641667 | -1 | processed_pseudoge | ENSG00000 |
| 62688482 | 62710694 | 1 | lncRNA | ENSG00000 |
| 62784132 | 62865516 | 1 | protein_coding | ENSG00000 |
| 62896009 | 62901639 | 1 | lncRNA | ENSG00000 |
| 62905180 | 62905482 | 1 | processed_pseudoge | ENSG00000 |
| 62975751 | 63022504 | -1 | lncRNA | ENSG00000 |
| 63011197 | 63018614 | 1 | lncRNA | ENSG00000 |
| 63024207 | 63025658 | 1 | lncRNA | ENSG00000 |
| 63078081 | 63079031 | -1 | lncRNA | ENSG00000 |
| 63139250 | 63163153 | 1 | lncRNA | ENSG00000 |
| 63159083 | 63317274 | -1 | lncRNA | ENSG00000 |
| 63186336 | 63186450 | -1 | rRNA_pseudogene | ENSG00000 |
| 63249920 | 63251771 | 1 | lncRNA | ENSG00000 |
| 63262113 | 63262707 | 1 | processed_pseudoge | ENSG00000 |
| 63320884 | 63324441 | -1 | lncRNA | ENSG00000 |
| 63322567 | 63325128 | 1 | protein_coding | ENSG00000 |
| 63326925 | 63326984 | -1 | miRNA | ENSG00000 |
| 63338263 | 63338372 | 1 | misc_RNA | ENSG00000 |
| 63359823 | 63361179 | -1 | processed_pseudoge | ENSG00000 |
| 63367575 | 63438553 | 1 | protein_coding | ENSG00000 |
| 63440770 | 63593721 | -1 | protein_coding | ENSG00000 |
| 63487957 | 63508204 | 1 | lncRNA | ENSG00000 |
| 63523372 | 63572693 | 1 | protein_coding | ENSG00000 |
| 63529613 | 63529898 | -1 | misc_RNA | ENSG00000 |
| 63536711 | 63536770 | -1 | snRNA | ENSG00000 |
| 63547082 | 63550636 | 1 | lncRNA | ENSG00000 |
| 63548977 | 63549112 | -1 | misc_RNA | ENSG00000 |
| 63549133 | 63549216 | -1 | misc_RNA | ENSG00000 |
| 63549227 | 63549284 | -1 | misc_RNA | ENSG00000 |
| 63549294 | 63549366 | -1 | misc_RNA | ENSG00000 |
| 63549374 | 63549463 | -1 | misc_RNA | ENSG00000 |
| 63549489 | 63549608 | -1 | misc_RNA | ENSG00000 |
| 63593411 | 63660245 | 1 | protein_coding | ENSG00000 |
| 63655743 | 63656047 | 1 | misc_RNA | ENSG00000 |
| 63774017 | 64181498 | 1 | protein_coding | ENSG00000 |
| 63788721 | 63789304 | -1 | processed_pseudoge | ENSG00000 |
| 63843196 | 63843685 | -1 | processed_pseudoge | ENSG00000 |
| 64028894 | 64029000 | -1 | snRNA | ENSG00000 |
| 64066640 | 64066741 | -1 | misc_RNA | ENSG00000 |
| 64094442 | 64171297 | -1 | lncRNA | ENSG00000 |

| | | | | |
|---|---|---|---|---|
| 64186791 | 64192685 | 1 | lncRNA | ENSG0000C |
| 64203627 | 64267368 | 1 | protein_coding | ENSG0000C |
| 64384398 | 64384459 | -1 | snRNA | ENSG0000C |
| 64470129 | 64693058 | 1 | protein_coding | ENSG0000C |
| 64579847 | 64579923 | 1 | miRNA | ENSG0000C |
| 64745095 | 64833232 | 1 | protein_coding | ENSG0000C |
| 64833229 | 65067754 | -1 | protein_coding | ENSG0000C |
| 64918443 | 64918683 | 1 | processed_pseudoge | ENSG0000C |
| 64941979 | 64942365 | 1 | processed_pseudoge | ENSG0000C |
| 64972225 | 65002489 | -1 | lncRNA | ENSG0000C |
| 64984608 | 64986087 | -1 | processed_pseudoge | ENSG0000C |
| 65003470 | 65004087 | -1 | lncRNA | ENSG0000C |
| 65022968 | 65023074 | -1 | snRNA | ENSG0000C |
| 65057755 | 65057842 | -1 | miRNA | ENSG0000C |
| 65058434 | 65058508 | -1 | miRNA | ENSG0000C |
| 65077413 | 65078131 | -1 | processed_pseudoge | ENSG0000C |
| 65092392 | 65092614 | -1 | processed_pseudoge | ENSG0000C |
| 65147549 | 65232145 | 1 | protein_coding | ENSG0000C |
| 65154480 | 65154650 | 1 | processed_pseudoge | ENSG0000C |
| 65248219 | 65415871 | 1 | protein_coding | ENSG0000C |
| 65279456 | 65306521 | -1 | lncRNA | ENSG0000C |
| 65298755 | 65298980 | -1 | processed_pseudoge | ENSG0000C |
| 65415816 | 65416006 | 1 | snRNA | ENSG0000C |
| 65420587 | 65436007 | 1 | protein_coding | ENSG0000C |
| 65420652 | 65641559 | 1 | protein_coding | ENSG0000C |
| 65486406 | 65494188 | 1 | lncRNA | ENSG0000C |
| 65576129 | 65578380 | -1 | processed_pseudoge | ENSG0000C |
| 65703962 | 65714847 | 1 | lncRNA | ENSG0000C |
| 65761058 | 65761352 | -1 | misc_RNA | ENSG0000C |
| 65792514 | 66374579 | 1 | protein_coding | ENSG0000C |
| 66042500 | 66050718 | -1 | lncRNA | ENSG0000C |
| 66094461 | 66094546 | -1 | snRNA | ENSG0000C |
| 66533383 | 66751139 | 1 | protein_coding | ENSG0000C |
| 66628440 | 66628517 | 1 | miRNA | ENSG0000C |
| 66665864 | 66677027 | -1 | lncRNA | ENSG0000C |
| 66752459 | 66779047 | 1 | protein_coding | ENSG0000C |
| 66797741 | 66801256 | -1 | protein_coding | ENSG0000C |
| 66812885 | 66924856 | -1 | protein_coding | ENSG0000C |
| 66826942 | 66828558 | 1 | lncRNA | ENSG0000C |
| 66924895 | 66988619 | 1 | protein_coding | ENSG0000C |
| 66999350 | 67054148 | -1 | protein_coding | ENSG0000C |
| 67092165 | 67231853 | -1 | protein_coding | ENSG0000C |
| 67102645 | 67102779 | -1 | snoRNA | ENSG0000C |
| 67121605 | 67123956 | -1 | lncRNA | ENSG0000C |
| 67138907 | 67259979 | 1 | protein_coding | ENSG0000C |
| 67196140 | 67196243 | 1 | snRNA | ENSG0000C |
| 67267601 | 67267726 | -1 | snRNA | ENSG0000C |
| 67278052 | 67278779 | 1 | processed_pseudoge | ENSG0000C |
| 67307364 | 67397090 | 1 | protein_coding | ENSG0000C |
| 67407810 | 67430415 | -1 | protein_coding | ENSG0000C |

| | | | | |
|---|---|---|---|---|
| 67417214 | 67417318 | 1 | snRNA | ENSG0000C |
| 67522299 | 67532612 | 1 | lncRNA | ENSG0000C |
| 67541127 | 67541233 | 1 | snRNA | ENSG0000C |
| 67561073 | 67561446 | 1 | processed_pseudoge | ENSG0000C |
| 67656831 | 67657129 | -1 | misc_RNA | ENSG0000C |
| 67660155 | 67661013 | -1 | processed_pseudoge | ENSG0000C |
| 67685201 | 67688334 | 1 | protein_coding | ENSG0000C |
| 67701475 | 67833467 | -1 | protein_coding | ENSG0000C |
| 67772593 | 67772653 | -1 | snRNA | ENSG0000C |
| 67832303 | 68202987 | 1 | lncRNA | ENSG0000C |
| 68043330 | 68044233 | 1 | processed_pseudoge | ENSG0000C |
| 68045886 | 68051631 | -1 | protein_coding | ENSG0000C |
| 68049360 | 68049745 | -1 | processed_pseudoge | ENSG0000C |
| 68098473 | 68233120 | -1 | protein_coding | ENSG0000C |
| 68161761 | 68163090 | -1 | processed_pseudoge | ENSG0000C |
| 68183518 | 68183610 | -1 | miRNA | ENSG0000C |
| 68242474 | 68243055 | -1 | processed_pseudoge | ENSG0000C |
| 68282388 | 68282622 | 1 | processed_pseudoge | ENSG0000C |
| 68375327 | 68375663 | -1 | processed_pseudoge | ENSG0000C |
| 68381441 | 68381657 | -1 | processed_pseudoge | ENSG0000C |
| 68428822 | 68449954 | -1 | protein_coding | ENSG0000C |
| 68474152 | 68497221 | -1 | protein_coding | ENSG0000C |
| 68479129 | 68483539 | 1 | lncRNA | ENSG0000C |
| 68496676 | 68538627 | 1 | lncRNA | ENSG0000C |
| 68514375 | 68514669 | -1 | processed_pseudoge | ENSG0000C |
| 68633701 | 68640425 | 1 | lncRNA | ENSG0000C |
| 68679202 | 68943452 | 1 | lncRNA | ENSG0000C |
| 68974010 | 69023558 | 1 | lncRNA | ENSG0000C |
| 69055838 | 69229607 | 1 | lncRNA | ENSG0000C |
| 69215835 | 69249477 | -1 | lncRNA | ENSG0000C |
| 69433255 | 69435409 | 1 | lncRNA | ENSG0000C |
| 69551848 | 69563715 | 1 | lncRNA | ENSG0000C |
| 69568261 | 70151945 | 1 | protein_coding | ENSG0000C |
| 69606855 | 69608301 | -1 | processed_pseudoge | ENSG0000C |
| 69706950 | 69710274 | -1 | lncRNA | ENSG0000C |
| 69879592 | 69879895 | 1 | misc_RNA | ENSG0000C |
| 69919322 | 69920317 | 1 | transcribed_processe | ENSG0000C |
| 70013982 | 70031222 | -1 | lncRNA | ENSG0000C |
| 70144805 | 70205579 | -1 | protein_coding | ENSG0000C |
| 70180144 | 70180440 | -1 | misc_RNA | ENSG0000C |
| 70205682 | 70253052 | 1 | protein_coding | ENSG0000C |
| 70218589 | 70221023 | -1 | lncRNA | ENSG0000C |
| 70258999 | 70354734 | -1 | protein_coding | ENSG0000C |
| 70354805 | 70385339 | 1 | protein_coding | ENSG0000C |
| 70359562 | 70360437 | -1 | lncRNA | ENSG0000C |
| 70411218 | 70439851 | 1 | protein_coding | ENSG0000C |
| 70430896 | 70431178 | 1 | misc_RNA | ENSG0000C |
| 70445071 | 70445536 | 1 | lncRNA | ENSG0000C |
| 70530526 | 70531492 | 1 | processed_pseudoge | ENSG0000C |
| 70660657 | 70661265 | -1 | processed_pseudoge | ENSG0000C |

| | | | | |
|---|---|---|---|---|
| 70706441 | 70786468 | 1 | lncRNA | ENSG0000C |
| 70715933 | 70720289 | -1 | lncRNA | ENSG0000C |
| 70852353 | 71047808 | -1 | protein_coding | ENSG0000C |
| 70947379 | 70951493 | 1 | lncRNA | ENSG0000C |
| 71005854 | 71006502 | -1 | lncRNA | ENSG0000C |
| 71048855 | 71067184 | 1 | lncRNA | ENSG0000C |
| 71063291 | 71081289 | -1 | protein_coding | ENSG0000C |
| 71067631 | 71067716 | -1 | miRNA | ENSG0000C |
| 71081324 | 71489976 | 1 | lncRNA | ENSG0000C |
| 71367054 | 71367303 | -1 | processed_pseudoge | ENSG0000C |
| 71395943 | 72282539 | -1 | protein_coding | ENSG0000C |
| 71570956 | 71573558 | 1 | lncRNA | ENSG0000C |
| 71738173 | 71738354 | 1 | processed_pseudoge | ENSG0000C |
| 71794232 | 71837012 | -1 | lncRNA | ENSG0000C |
| 72274552 | 72275159 | 1 | processed_pseudoge | ENSG0000C |
| 72283170 | 72753772 | 1 | lncRNA | ENSG0000C |
| 72301472 | 72301829 | 1 | processed_pseudoge | ENSG0000C |
| 72636547 | 72899240 | -1 | lncRNA | ENSG0000C |
| 72717663 | 72717769 | -1 | snRNA | ENSG0000C |
| 72765031 | 72791282 | 1 | lncRNA | ENSG0000C |
| 72793104 | 72854475 | 1 | lncRNA | ENSG0000C |
| 72979014 | 72979314 | -1 | TEC | ENSG0000C |
| 73104792 | 73106282 | 1 | processed_pseudoge | ENSG0000C |
| 73191604 | 73191855 | -1 | misc_RNA | ENSG0000C |
| 73305609 | 73355253 | 1 | lncRNA | ENSG0000C |
| 73635216 | 73715214 | 1 | lncRNA | ENSG0000C |
| 73749517 | 73749623 | -1 | rRNA_pseudogene | ENSG0000C |
| 73787370 | 73915340 | -1 | lncRNA | ENSG0000C |
| 73883714 | 73883822 | 1 | snRNA | ENSG0000C |
| 74026015 | 74198187 | -1 | protein_coding | ENSG0000C |
| 74198212 | 74234086 | 1 | protein_coding | ENSG0000C |
| 74198235 | 74544393 | 1 | protein_coding | ENSG0000C |
| 74235387 | 74544428 | 1 | protein_coding | ENSG0000C |
| 74341579 | 74378802 | -1 | lncRNA | ENSG0000C |
| 74468195 | 74469543 | 1 | lncRNA | ENSG0000C |
| 74469878 | 74512614 | -1 | protein_coding | ENSG0000C |
| 74568117 | 74673792 | -1 | protein_coding | ENSG0000C |
| 74577430 | 74626098 | 1 | lncRNA | ENSG0000C |
| 74698769 | 74699333 | -1 | lncRNA | ENSG0000C |
| 74705482 | 74733408 | -1 | protein_coding | ENSG0000C |
| 74733152 | 74766678 | 1 | protein_coding | ENSG0000C |
| 74963314 | 74965118 | 1 | lncRNA | ENSG0000C |
| 75122518 | 75123927 | -1 | lncRNA | ENSG0000C |
| 75127830 | 75133058 | -1 | lncRNA | ENSG0000C |
| 75128434 | 75161533 | 1 | protein_coding | ENSG0000C |
| 75129974 | 75132576 | -1 | lncRNA | ENSG0000C |
| 75183045 | 75183147 | 1 | snRNA | ENSG0000C |
| 75202131 | 75611116 | -1 | protein_coding | ENSG0000C |
| 75521562 | 75522231 | 1 | processed_pseudoge | ENSG0000C |
| 75538015 | 75538119 | -1 | snRNA | ENSG0000C |

| | | | | |
|---|---|---|---|---|
| 75582099 | 75582555 | 1 | processed_pseudoge | ENSG00000( |
| 75641178 | 75724049 | -1 | transcribed_processe | ENSG00000( |
| 75724347 | 75787575 | 1 | protein_coding | ENSG00000( |
| 75743423 | 75744776 | -1 | processed_pseudoge | ENSG00000( |
| 75786197 | 75795086 | 1 | protein_coding | ENSG00000( |
| 75787072 | 75787150 | 1 | snoRNA | ENSG00000( |
| 75787889 | 75787972 | 1 | snoRNA | ENSG00000( |
| 75789477 | 75789548 | 1 | snoRNA | ENSG00000( |
| 75796882 | 75913242 | 1 | protein_coding | ENSG00000( |
| 75918873 | 75932404 | -1 | protein_coding | ENSG00000( |
| 75926454 | 76019356 | 1 | lncRNA | ENSG00000( |
| 76041691 | 76066349 | 1 | lncRNA | ENSG00000( |
| 76074746 | 76634603 | 1 | protein_coding | ENSG00000( |
| 76353583 | 76354246 | -1 | processed_pseudoge | ENSG00000( |
| 76636877 | 76637339 | 1 | lncRNA | ENSG00000( |
| 76699789 | 76700538 | 1 | processed_pseudoge | ENSG00000( |
| 76753135 | 76753240 | 1 | snRNA | ENSG00000( |
| 76758124 | 76779267 | -1 | lncRNA | ENSG00000( |
| 76867480 | 77067546 | 1 | protein_coding | ENSG00000( |
| 77060143 | 77060202 | 1 | miRNA | ENSG00000( |
| 77067920 | 77086402 | 1 | lncRNA | ENSG00000( |
| 77088989 | 77219430 | -1 | protein_coding | ENSG00000( |
| 77129114 | 77129668 | 1 | processed_pseudoge | ENSG00000( |
| 77194825 | 77195395 | 1 | processed_pseudoge | ENSG00000( |
| 77248633 | 77250826 | 1 | lncRNA | ENSG00000( |
| 77282019 | 77559966 | 1 | protein_coding | ENSG00000( |
| 77346046 | 77349585 | -1 | lncRNA | ENSG00000( |
| 77420325 | 77420386 | 1 | snRNA | ENSG00000( |
| 77431314 | 77437604 | -1 | lncRNA | ENSG00000( |
| 77562416 | 77683419 | -1 | protein_coding | ENSG00000( |
| 77614869 | 77614952 | 1 | rRNA_pseudogene | ENSG00000( |
| 77645324 | 77645618 | 1 | misc_RNA | ENSG00000( |
| 77695987 | 77759852 | -1 | protein_coding | ENSG00000( |
| 77773865 | 77774864 | -1 | processed_pseudoge | ENSG00000( |
| 77779624 | 77879539 | 1 | protein_coding | ENSG00000( |
| 77779904 | 77779999 | 1 | rRNA_pseudogene | ENSG00000( |
| 77810861 | 77811781 | -1 | processed_pseudoge | ENSG00000( |
| 77847110 | 77848802 | 1 | processed_pseudoge | ENSG00000( |
| 77853355 | 77853893 | 1 | processed_pseudoge | ENSG00000( |
| 77881348 | 77889539 | -1 | lncRNA | ENSG00000( |
| 77888513 | 77943895 | 1 | protein_coding | ENSG00000( |
| 77944055 | 77979110 | -1 | protein_coding | ENSG00000( |
| 77979175 | 78017964 | 1 | protein_coding | ENSG00000( |
| 77979542 | 78138444 | 1 | protein_coding | ENSG00000( |
| 78004346 | 78004554 | -1 | lncRNA | ENSG00000( |
| 78022565 | 78024861 | -1 | lncRNA | ENSG00000( |
| 78043383 | 78044148 | -1 | processed_pseudoge | ENSG00000( |
| 78088988 | 78089083 | -1 | snRNA | ENSG00000( |
| 78091499 | 78092054 | 1 | processed_pseudoge | ENSG00000( |
| 78094807 | 78094915 | -1 | rRNA_pseudogene | ENSG00000( |

| | | | | |
|---|---|---|---|---|
| 78170481 | 78171237 | -1 | processed_pseudoge | ENSG00000( |
| 78229599 | 78369464 | 1 | lncRNA | ENSG00000( |
| 78303884 | 78540701 | 1 | protein_coding | ENSG00000( |
| 78317157 | 78317263 | -1 | unprocessed_pseudc | ENSG00000( |
| 78375164 | 78375268 | -1 | rRNA_pseudogene | ENSG00000( |
| 78619922 | 78646145 | 1 | protein_coding | ENSG00000( |
| 78649796 | 78664078 | 1 | protein_coding | ENSG00000( |
| 78666272 | 78666695 | 1 | processed_pseudoge | ENSG00000( |
| 78749073 | 78750659 | 1 | processed_pseudoge | ENSG00000( |
| 78889764 | 79282124 | -1 | protein_coding | ENSG00000( |
| 79054945 | 79056055 | -1 | processed_pseudoge | ENSG00000( |
| 79323769 | 79324852 | -1 | lncRNA | ENSG00000( |
| 79323769 | 79324863 | 1 | lncRNA | ENSG00000( |
| 79325008 | 79346445 | 1 | lncRNA | ENSG00000( |
| 79521080 | 79522203 | 1 | processed_pseudoge | ENSG00000( |
| 79967733 | 80051404 | -1 | lncRNA | ENSG00000( |
| 80092103 | 80093237 | -1 | processed_pseudoge | ENSG00000( |
| 80114943 | 80116918 | 1 | lncRNA | ENSG00000( |
| 80124004 | 80124361 | 1 | unprocessed_pseudc | ENSG00000( |
| 80283352 | 80283970 | 1 | processed_pseudoge | ENSG00000( |
| 80373364 | 80374621 | -1 | lncRNA | ENSG00000( |
| 80451083 | 80452004 | -1 | processed_pseudoge | ENSG00000( |
| 80464301 | 80464718 | -1 | processed_pseudoge | ENSG00000( |
| 80495903 | 80496233 | 1 | processed_pseudoge | ENSG00000( |
| 80534978 | 80584363 | -1 | lncRNA | ENSG00000( |
| 80535755 | 80646791 | 1 | lncRNA | ENSG00000( |
| 81080790 | 81081171 | 1 | processed_pseudoge | ENSG00000( |
| 81098267 | 81099005 | -1 | processed_pseudoge | ENSG00000( |
| 81208568 | 81209210 | 1 | processed_pseudoge | ENSG00000( |
| 81209834 | 81222076 | 1 | lncRNA | ENSG00000( |
| 81251789 | 81252096 | 1 | misc_RNA | ENSG00000( |
| 81306160 | 81992436 | 1 | protein_coding | ENSG00000( |
| 81426456 | 81427605 | 1 | processed_pseudoge | ENSG00000( |
| 81501794 | 81503468 | 1 | processed_pseudoge | ENSG00000( |
| 81505099 | 81506296 | 1 | lncRNA | ENSG00000( |
| 81513880 | 81557702 | -1 | lncRNA | ENSG00000( |
| 81557121 | 81557832 | -1 | processed_pseudoge | ENSG00000( |
| 81585941 | 81625713 | -1 | lncRNA | ENSG00000( |
| 81596157 | 81596506 | -1 | processed_pseudoge | ENSG00000( |
| 81721693 | 81723068 | 1 | transcribed_processe | ENSG00000( |
| 82212413 | 82848205 | -1 | lncRNA | ENSG00000( |
| 82903183 | 83166815 | 1 | lncRNA | ENSG00000( |
| 82970820 | 82986208 | -1 | lncRNA | ENSG00000( |
| 83397555 | 83425353 | -1 | lncRNA | ENSG00000( |
| 83445967 | 83480024 | 1 | lncRNA | ENSG00000( |
| 83575776 | 83861023 | -1 | lncRNA | ENSG00000( |
| 83790280 | 83801465 | 1 | lncRNA | ENSG00000( |
| 83865024 | 83999150 | -1 | protein_coding | ENSG00000( |
| 83979118 | 83984300 | -1 | lncRNA | ENSG00000( |
| 84015865 | 84018371 | -1 | processed_pseudoge | ENSG00000( |

| | | | | |
|---|---|---|---|---|
| 84038529 | 84065031 | 1 | lncRNA | ENSG00000 |
| 84076331 | 84077931 | -1 | lncRNA | ENSG00000 |
| 84078062 | 84238498 | 1 | protein_coding | ENSG00000 |
| 84085741 | 84086233 | 1 | processed_pseudoge | ENSG00000 |
| 84244334 | 84244577 | 1 | processed_pseudoge | ENSG00000 |
| 84277321 | 84277457 | 1 | snoRNA | ENSG00000 |
| 84298366 | 84389957 | 1 | protein_coding | ENSG00000 |
| 84344678 | 84349939 | 1 | transcribed_unitary_ | ENSG00000 |
| 84363706 | 84397831 | -1 | transcribed_unitary_ | ENSG00000 |
| 84398532 | 84415018 | 1 | protein_coding | ENSG00000 |
| 84477039 | 84479157 | 1 | lncRNA | ENSG00000 |
| 84479259 | 84498352 | 1 | protein_coding | ENSG00000 |
| 84498325 | 84506581 | -1 | protein_coding | ENSG00000 |
| 84498350 | 84556688 | -1 | lncRNA | ENSG00000 |
| 84506300 | 84566194 | 1 | protein_coding | ENSG00000 |
| 84549611 | 84574480 | -1 | protein_coding | ENSG00000 |
| 84574129 | 84583620 | 1 | lncRNA | ENSG00000 |
| 84607099 | 84610697 | 1 | lncRNA | ENSG00000 |
| 84628230 | 84635020 | -1 | lncRNA | ENSG00000 |
| 84636158 | 84637106 | -1 | processed_pseudoge | ENSG00000 |
| 84643707 | 84690803 | -1 | protein_coding | ENSG00000 |
| 84785427 | 84786714 | 1 | lncRNA | ENSG00000 |
| 84811602 | 84893206 | -1 | protein_coding | ENSG00000 |
| 84925583 | 84997113 | -1 | protein_coding | ENSG00000 |
| 84999147 | 85133138 | 1 | protein_coding | ENSG00000 |
| 85018082 | 85048500 | -1 | protein_coding | ENSG00000 |
| 85133794 | 85133873 | 1 | miRNA | ENSG00000 |
| 85152487 | 85154203 | -1 | TEC | ENSG00000 |
| 85156889 | 85201016 | -1 | protein_coding | ENSG00000 |
| 85249953 | 85259672 | -1 | protein_coding | ENSG00000 |
| 85264296 | 85264402 | 1 | misc_RNA | ENSG00000 |
| 85265776 | 85276632 | -1 | protein_coding | ENSG00000 |
| 85276388 | 85448124 | 1 | lncRNA | ENSG00000 |
| 85318481 | 85578363 | -1 | protein_coding | ENSG00000 |
| 85435175 | 85435284 | -1 | misc_RNA | ENSG00000 |
| 85467295 | 85467660 | 1 | lncRNA | ENSG00000 |
| 85482281 | 85578250 | -1 | lncRNA | ENSG00000 |
| 85578500 | 85578742 | -1 | lncRNA | ENSG00000 |
| 85580761 | 85583950 | 1 | protein_coding | ENSG00000 |
| 85592280 | 85592356 | 1 | snoRNA | ENSG00000 |
| 85599131 | 85600734 | 1 | lncRNA | ENSG00000 |
| 85649417 | 85708433 | -1 | protein_coding | ENSG00000 |
| 85729233 | 86156943 | -1 | protein_coding | ENSG00000 |
| 85883680 | 85883798 | -1 | rRNA_pseudogene | ENSG00000 |
| 86029854 | 86030589 | 1 | processed_pseudoge | ENSG00000 |
| 86288704 | 86320711 | -1 | lncRNA | ENSG00000 |
| 86346824 | 86396342 | -1 | protein_coding | ENSG00000 |
| 86357632 | 86357687 | -1 | miRNA | ENSG00000 |
| 86404176 | 86404866 | -1 | processed_pseudoge | ENSG00000 |
| 86424171 | 86456553 | 1 | protein_coding | ENSG00000 |

| | | | | |
|---|---|---|---|---|
| 86468368 | 86500289 | 1 | protein_coding | ENSG00000( |
| 86547078 | 86580754 | 1 | protein_coding | ENSG00000( |
| 86552625 | 86553342 | -1 | processed_pseudoge | ENSG00000( |
| 86571181 | 86696311 | -1 | lncRNA | ENSG00000( |
| 86634273 | 86655376 | 1 | transcribed_unproce | ENSG00000( |
| 86703502 | 86704462 | -1 | lncRNA | ENSG00000( |
| 86704570 | 86748184 | 1 | protein_coding | ENSG00000( |
| 86784913 | 86785467 | 1 | processed_pseudoge | ENSG00000( |
| 86821558 | 86834169 | -1 | lncRNA | ENSG00000( |
| 86862445 | 86914424 | -1 | protein_coding | ENSG00000( |
| 86914635 | 87109982 | 1 | protein_coding | ENSG00000( |
| 86932199 | 86934891 | -1 | lncRNA | ENSG00000( |
| 86993009 | 87169204 | 1 | protein_coding | ENSG00000( |
| 87044935 | 87045871 | -1 | processed_pseudoge | ENSG00000( |
| 87045875 | 87046700 | 1 | processed_pseudoge | ENSG00000( |
| 87129765 | 87169198 | 1 | lncRNA | ENSG00000( |
| 87212669 | 87264741 | 1 | lncRNA | ENSG00000( |
| 87328880 | 87348923 | 1 | protein_coding | ENSG00000( |
| 87353521 | 87371683 | -1 | lncRNA | ENSG00000( |
| 87453240 | 87453373 | -1 | rRNA_pseudogene | ENSG00000( |
| 87620803 | 88685204 | -1 | lncRNA | ENSG00000( |
| 87805286 | 87808372 | -1 | TEC | ENSG00000( |
| 88313153 | 88313302 | 1 | processed_pseudoge | ENSG00000( |
| 88462936 | 88464079 | 1 | lncRNA | ENSG00000( |
| 88477828 | 88478114 | -1 | misc_RNA | ENSG00000( |
| 88498309 | 88498676 | -1 | processed_pseudoge | ENSG00000( |
| 88520000 | 88520170 | 1 | misc_RNA | ENSG00000( |
| 88577880 | 88578201 | 1 | processed_pseudoge | ENSG00000( |
| 88684222 | 88836255 | 1 | protein_coding | ENSG00000( |
| 88816779 | 88816885 | 1 | snRNA | ENSG00000( |
| 88829102 | 88829419 | 1 | processed_pseudoge | ENSG00000( |
| 88852633 | 88891944 | -1 | protein_coding | ENSG00000( |
| 88923370 | 88923844 | 1 | processed_pseudoge | ENSG00000( |
| 88935773 | 88992953 | -1 | protein_coding | ENSG00000( |
| 88979456 | 88992960 | -1 | protein_coding | ENSG00000( |
| 89006679 | 89022894 | -1 | protein_coding | ENSG00000( |
| 89020246 | 89020365 | -1 | misc_RNA | ENSG00000( |
| 89052319 | 89065230 | -1 | protein_coding | ENSG00000( |
| 89104285 | 89104767 | -1 | processed_pseudoge | ENSG00000( |
| 89106132 | 89150456 | -1 | protein_coding | ENSG00000( |
| 89128432 | 89144695 | 1 | lncRNA | ENSG00000( |
| 89131742 | 89176009 | -1 | protein_coding | ENSG00000( |
| 89181144 | 89198942 | -1 | protein_coding | ENSG00000( |
| 89198714 | 89207040 | 1 | lncRNA | ENSG00000( |
| 89203280 | 89203464 | -1 | unprocessed_pseudc | ENSG00000( |
| 89258950 | 89272804 | -1 | protein_coding | ENSG00000( |
| 89260582 | 89269754 | 1 | lncRNA | ENSG00000( |
| 89289676 | 89290337 | -1 | processed_pseudoge | ENSG00000( |
| 89324522 | 89334880 | 1 | unprocessed_pseudc | ENSG00000( |
| 89364058 | 89386461 | 1 | protein_coding | ENSG00000( |

| | | | | |
|---|---|---|---|---|
| 89394033 | 89394665 | 1 | processed_pseudoge | ENSG00000( |
| 89407679 | 89426243 | 1 | transcribed_unproce | ENSG00000( |
| 89427533 | 89525472 | -1 | lncRNA | ENSG00000( |
| 89524836 | 89597864 | 1 | protein_coding | ENSG00000( |
| 89581291 | 89632916 | -1 | lncRNA | ENSG00000( |
| 89629725 | 89676386 | 1 | lncRNA | ENSG00000( |
| 89633072 | 89769903 | 1 | protein_coding | ENSG00000( |
| 89633140 | 89933250 | 1 | protein_coding | ENSG00000( |
| 89661212 | 89661353 | -1 | processed_pseudoge | ENSG00000( |
| 89788914 | 89790492 | 1 | lncRNA | ENSG00000( |
| 89820174 | 89820868 | -1 | lncRNA | ENSG00000( |
| 89821014 | 89936611 | 1 | protein_coding | ENSG00000( |
| 89939601 | 89940147 | 1 | lncRNA | ENSG00000( |
| 89987713 | 89987944 | -1 | misc_RNA | ENSG00000( |
| 89993593 | 89994321 | -1 | processed_pseudoge | ENSG00000( |
| 89995110 | 90035533 | 1 | protein_coding | ENSG00000( |
| 90045998 | 90047109 | 1 | lncRNA | ENSG00000( |
| 90242088 | 90284188 | 1 | lncRNA | ENSG00000( |
| 90253456 | 90253559 | -1 | snRNA | ENSG00000( |
| 90388193 | 90420396 | -1 | lncRNA | ENSG00000( |
| 90510910 | 90533472 | -1 | lncRNA | ENSG00000( |
| 90657750 | 90657964 | 1 | snoRNA | ENSG00000( |
| 90711539 | 90717302 | -1 | protein_coding | ENSG00000( |
| 90719576 | 90725620 | 1 | lncRNA | ENSG00000( |
| 90769086 | 90851658 | -1 | lncRNA | ENSG00000( |
| 90835660 | 90849472 | 1 | lncRNA | ENSG00000( |
| 90851122 | 90855253 | 1 | lncRNA | ENSG00000( |
| 90860550 | 90862920 | -1 | lncRNA | ENSG00000( |
| 90892992 | 90893612 | -1 | processed_pseudoge | ENSG00000( |
| 90915298 | 91022272 | -1 | protein_coding | ENSG00000( |
| 91023919 | 91024804 | 1 | processed_pseudoge | ENSG00000( |
| 91260766 | 91404856 | -1 | protein_coding | ENSG00000( |
| 91261625 | 91261724 | 1 | misc_RNA | ENSG00000( |
| 91328369 | 91329513 | 1 | processed_pseudoge | ENSG00000( |
| 91500851 | 91525764 | 1 | protein_coding | ENSG00000( |
| 91534666 | 91535593 | -1 | processed_pseudoge | ENSG00000( |
| 91600171 | 91600321 | 1 | processed_pseudoge | ENSG00000( |
| 91642516 | 91644082 | 1 | processed_pseudoge | ENSG00000( |
| 91680343 | 91906335 | -1 | protein_coding | ENSG00000( |
| 91829774 | 91830066 | -1 | misc_RNA | ENSG00000( |
| 91949371 | 92014426 | 1 | protein_coding | ENSG00000( |
| 92029985 | 92063538 | 1 | protein_coding | ENSG00000( |
| 92066306 | 92067839 | 1 | unitary_pseudogene | ENSG00000( |
| 92074533 | 92075441 | -1 | protein_coding | ENSG00000( |
| 92080305 | 92184725 | 1 | protein_coding | ENSG00000( |
| 92114803 | 92115796 | -1 | processed_pseudoge | ENSG00000( |
| 92125301 | 92126462 | 1 | processed_pseudoge | ENSG00000( |
| 92189237 | 92190707 | 1 | lncRNA | ENSG00000( |
| 92203148 | 92203991 | 1 | processed_pseudoge | ENSG00000( |
| 92217915 | 92245813 | 1 | protein_coding | ENSG00000( |

| | | | | |
|---|---|---|---|---|
| 92229018 | 92229435 | -1 | processed_pseudoge | ENSG00000 |
| 92246402 | 92298987 | -1 | protein_coding | ENSG00000 |
| 92299059 | 92402056 | 1 | protein_coding | ENSG00000 |
| 92402389 | 92402685 | -1 | misc_RNA | ENSG00000 |
| 92473043 | 92486925 | -1 | protein_coding | ENSG00000 |
| 92508696 | 92792404 | -1 | protein_coding | ENSG00000 |
| 92580476 | 92580821 | -1 | processed_pseudoge | ENSG00000 |
| 92647048 | 92647630 | -1 | processed_pseudoge | ENSG00000 |
| 92700819 | 92700934 | -1 | snRNA | ENSG00000 |
| 92732000 | 92733257 | 1 | processed_pseudoge | ENSG00000 |
| 92749175 | 92749330 | -1 | processed_pseudoge | ENSG00000 |
| 92755794 | 92756956 | 1 | processed_pseudoge | ENSG00000 |
| 92832013 | 92841924 | 1 | protein_coding | ENSG00000 |
| 92832737 | 92961522 | -1 | protein_coding | ENSG00000 |
| 92837289 | 92837383 | 1 | snoRNA | ENSG00000 |
| 92840719 | 92840851 | 1 | snoRNA | ENSG00000 |
| 92846059 | 92846182 | 1 | snoRNA | ENSG00000 |
| 92930696 | 92934098 | 1 | transcribed_processe | ENSG00000 |
| 92969604 | 92969710 | -1 | snRNA | ENSG00000 |
| 92974827 | 92975114 | -1 | misc_RNA | ENSG00000 |
| 92978265 | 92987367 | 1 | lncRNA | ENSG00000 |
| 93010257 | 93010351 | 1 | snRNA | ENSG00000 |
| 93026252 | 93026542 | 1 | misc_RNA | ENSG00000 |
| 93027410 | 93027506 | -1 | misc_RNA | ENSG00000 |
| 93079235 | 93139079 | 1 | protein_coding | ENSG00000 |
| 93149742 | 93180516 | -1 | protein_coding | ENSG00000 |
| 93179919 | 93278730 | 1 | protein_coding | ENSG00000 |
| 93190740 | 93191060 | -1 | processed_pseudoge | ENSG00000 |
| 93199755 | 93199939 | -1 | processed_pseudoge | ENSG00000 |
| 93262186 | 93346025 | -1 | lncRNA | ENSG00000 |
| 93278961 | 93279129 | -1 | processed_pseudoge | ENSG00000 |
| 93345907 | 93369493 | 1 | protein_coding | ENSG00000 |
| 93384487 | 93384998 | -1 | processed_pseudoge | ENSG00000 |
| 93385711 | 93385817 | 1 | misc_RNA | ENSG00000 |
| 93448131 | 93554661 | 1 | protein_coding | ENSG00000 |
| 93488333 | 93488452 | -1 | rRNA_pseudogene | ENSG00000 |
| 93561741 | 93847150 | -1 | protein_coding | ENSG00000 |
| 93592199 | 93605573 | 1 | lncRNA | ENSG00000 |
| 93846832 | 93846911 | 1 | miRNA | ENSG00000 |
| 93847174 | 93848939 | 1 | lncRNA | ENSG00000 |
| 93866284 | 93879918 | -1 | protein_coding | ENSG00000 |
| 93885199 | 93909456 | -1 | protein_coding | ENSG00000 |
| 93921268 | 93921667 | -1 | processed_pseudoge | ENSG00000 |
| 93922574 | 93923945 | -1 | unprocessed_pseudc | ENSG00000 |
| 93924386 | 93924716 | -1 | unprocessed_pseudc | ENSG00000 |
| 93924743 | 93925385 | -1 | unprocessed_pseudc | ENSG00000 |
| 93925406 | 93926072 | -1 | unprocessed_pseudc | ENSG00000 |
| 93926032 | 93926188 | -1 | unprocessed_pseudc | ENSG00000 |
| 93926615 | 93927569 | -1 | unprocessed_pseudc | ENSG00000 |
| 93927714 | 93928302 | -1 | unprocessed_pseudc | ENSG00000 |

| | | | | |
|---|---|---|---|---|
| 93934479 | 93935244 | -1 | processed_pseudoge | ENSG00000 |
| 93992834 | 94121148 | -1 | protein_coding | ENSG00000 |
| 94145111 | 94145619 | 1 | lncRNA | ENSG00000 |
| 94148988 | 94275068 | -1 | protein_coding | ENSG00000 |
| 94150738 | 94151080 | 1 | misc_RNA | ENSG00000 |
| 94247819 | 94411036 | 1 | lncRNA | ENSG00000 |
| 94302038 | 94303022 | 1 | processed_pseudoge | ENSG00000 |
| 94318479 | 94324569 | -1 | lncRNA | ENSG00000 |
| 94406395 | 94406882 | -1 | processed_pseudoge | ENSG00000 |
| 94417743 | 94418228 | -1 | lncRNA | ENSG00000 |
| 94418389 | 94518666 | 1 | protein_coding | ENSG00000 |
| 94529173 | 94541759 | -1 | protein_coding | ENSG00000 |
| 94585556 | 94592297 | -1 | lncRNA | ENSG00000 |
| 94613814 | 94855426 | -1 | transcribed_processe | ENSG00000 |
| 94650544 | 94651378 | -1 | processed_pseudoge | ENSG00000 |
| 94745860 | 94745900 | -1 | miRNA | ENSG00000 |
| 94820342 | 94895246 | 1 | protein_coding | ENSG00000 |
| 94896949 | 94927223 | -1 | protein_coding | ENSG00000 |
| 94927361 | 94963616 | 1 | lncRNA | ENSG00000 |
| 94974405 | 95072951 | -1 | protein_coding | ENSG00000 |
| 95061596 | 95067545 | 1 | lncRNA | ENSG00000 |
| 95117355 | 95197607 | 1 | protein_coding | ENSG00000 |
| 95117923 | 95247225 | 1 | protein_coding | ENSG00000 |
| 95120147 | 95138554 | 1 | lncRNA | ENSG00000 |
| 95125511 | 95125609 | -1 | misc_RNA | ENSG00000 |
| 95161676 | 95233982 | -1 | lncRNA | ENSG00000 |
| 95234210 | 95247225 | 1 | protein_coding | ENSG00000 |
| 95243167 | 95278940 | -1 | lncRNA | ENSG00000 |
| 95282233 | 95287702 | 1 | lncRNA | ENSG00000 |
| 95310928 | 95318263 | -1 | lncRNA | ENSG00000 |
| 95351251 | 95352676 | -1 | lncRNA | ENSG00000 |
| 95356229 | 95381000 | -1 | lncRNA | ENSG00000 |
| 95474737 | 95479356 | 1 | lncRNA | ENSG00000 |
| 95510059 | 95782342 | 1 | lncRNA | ENSG00000 |
| 95743096 | 95759470 | 1 | lncRNA | ENSG00000 |
| 95937901 | 96022890 | 1 | lncRNA | ENSG00000 |
| 96225901 | 96226060 | 1 | snRNA | ENSG00000 |
| 96254069 | 96374125 | -1 | lncRNA | ENSG00000 |
| 96390652 | 96391393 | 1 | processed_pseudoge | ENSG00000 |
| 96418594 | 96418933 | 1 | processed_pseudoge | ENSG00000 |
| 96446930 | 96448318 | 1 | processed_pseudoge | ENSG00000 |
| 96583209 | 96583527 | 1 | misc_RNA | ENSG00000 |
| 96584422 | 96584727 | -1 | processed_pseudoge | ENSG00000 |
| 96678874 | 96679620 | -1 | processed_pseudoge | ENSG00000 |
| 96695856 | 96696167 | -1 | misc_RNA | ENSG00000 |
| 96721665 | 96823738 | 1 | protein_coding | ENSG00000 |
| 97077743 | 97995000 | -1 | protein_coding | ENSG00000 |
| 97095923 | 97322955 | 1 | lncRNA | ENSG00000 |
| 97394154 | 97420141 | -1 | lncRNA | ENSG00000 |
| 97545701 | 97546958 | 1 | processed_pseudoge | ENSG00000 |

| | | | | |
|---|---|---|---|---|
| 97585862 | 97586294 | 1 | processed_pseudoge | ENSG0000( |
| 97774669 | 97775026 | 1 | processed_pseudoge | ENSG0000( |
| 97796921 | 97798066 | 1 | lncRNA | ENSG0000( |
| 97855575 | 97856825 | 1 | processed_pseudoge | ENSG0000( |
| 97933474 | 98049863 | -1 | lncRNA | ENSG0000( |
| 97967005 | 97968814 | -1 | lncRNA | ENSG0000( |
| 98045242 | 98045351 | -1 | miRNA | ENSG0000( |
| 98046070 | 98046171 | -1 | miRNA | ENSG0000( |
| 98052077 | 98054592 | -1 | lncRNA | ENSG0000( |
| 98077000 | 98077468 | 1 | processed_pseudoge | ENSG0000( |
| 98210747 | 98272658 | 1 | lncRNA | ENSG0000( |
| 98660388 | 98661059 | -1 | TEC | ENSG0000( |
| 98661701 | 98760500 | 1 | protein_coding | ENSG0000( |
| 98890245 | 99005032 | -1 | protein_coding | ENSG0000( |
| 99004276 | 99244794 | 1 | lncRNA | ENSG0000( |
| 99008218 | 99008474 | 1 | processed_pseudoge | ENSG0000( |
| 99263953 | 99309590 | 1 | protein_coding | ENSG0000( |
| 99464378 | 99464776 | 1 | processed_pseudoge | ENSG0000( |
| 99472332 | 99600995 | -1 | lncRNA | ENSG0000( |
| 99646113 | 99694541 | 1 | protein_coding | ENSG0000( |
| 99698242 | 99699178 | 1 | processed_pseudoge | ENSG0000( |
| 99708632 | 99766631 | -1 | protein_coding | ENSG0000( |
| 99784740 | 99784885 | -1 | snRNA | ENSG0000( |
| 99791662 | 99791753 | 1 | misc_RNA | ENSG0000( |
| 99842610 | 99842761 | 1 | processed_pseudoge | ENSG0000( |
| 99850361 | 99924023 | 1 | protein_coding | ENSG0000( |
| 99968383 | 99969864 | -1 | lncRNA | ENSG0000( |
| 99969351 | 100035634 | 1 | protein_coding | ENSG0000( |
| 99970011 | 100083321 | 1 | protein_coding | ENSG0000( |
| 99978939 | 99979045 | 1 | snRNA | ENSG0000( |
| 100000637 | 100000739 | -1 | snRNA | ENSG0000( |
| 100038095 | 100083377 | 1 | protein_coding | ENSG0000( |
| 100057990 | 100084471 | -1 | lncRNA | ENSG0000( |
| 100083563 | 100132955 | -1 | protein_coding | ENSG0000( |
| 100099239 | 100099490 | 1 | processed_pseudoge | ENSG0000( |
| 100133150 | 100150498 | 1 | protein_coding | ENSG0000( |
| 100148449 | 100178273 | -1 | protein_coding | ENSG0000( |
| 100186919 | 100249834 | -1 | protein_coding | ENSG0000( |
| 100196816 | 100197051 | 1 | processed_pseudoge | ENSG0000( |
| 100213293 | 100213670 | 1 | processed_pseudoge | ENSG0000( |
| 100220488 | 100253414 | 1 | lncRNA | ENSG0000( |
| 100249090 | 100249311 | 1 | processed_pseudoge | ENSG0000( |
| 100251528 | 100266179 | -1 | lncRNA | ENSG0000( |
| 100266207 | 100292769 | 1 | protein_coding | ENSG0000( |
| 100281241 | 100281308 | 1 | miRNA | ENSG0000( |
| 100331804 | 100332617 | -1 | processed_pseudoge | ENSG0000( |
| 100344477 | 100345242 | 1 | lncRNA | ENSG0000( |
| 100345001 | 100520277 | 1 | protein_coding | ENSG0000( |
| 100393033 | 100394026 | -1 | processed_pseudoge | ENSG0000( |
| 100462399 | 100485997 | -1 | lncRNA | ENSG0000( |

| | | | | |
|---|---|---|---|---|
| 100538139 | 100542021 | 1 | protein_coding | ENSG00000 |
| 100586649 | 100587431 | -1 | processed_pseudoge | ENSG00000 |
| 100627049 | 100640613 | -1 | lncRNA | ENSG00000 |
| 100628230 | 100641723 | 1 | lncRNA | ENSG00000 |
| 100651947 | 100652267 | 1 | processed_pseudoge | ENSG00000 |
| 100719742 | 100739045 | 1 | protein_coding | ENSG00000 |
| 100872372 | 100895179 | -1 | protein_coding | ENSG00000 |
| 100894928 | 100895356 | 1 | lncRNA | ENSG00000 |
| 100896076 | 100981757 | 1 | protein_coding | ENSG00000 |
| 100941017 | 100941995 | -1 | processed_pseudoge | ENSG00000 |
| 100989623 | 101026088 | -1 | protein_coding | ENSG00000 |
| 100995473 | 100996260 | 1 | lncRNA | ENSG00000 |
| 101025844 | 101090513 | 1 | lncRNA | ENSG00000 |
| 101133153 | 101133339 | 1 | scaRNA | ENSG00000 |
| 101190520 | 101190994 | 1 | processed_pseudoge | ENSG00000 |
| 101235683 | 101236528 | -1 | lncRNA | ENSG00000 |
| 101236865 | 101243713 | 1 | protein_coding | ENSG00000 |
| 101256274 | 101256616 | 1 | processed_pseudoge | ENSG00000 |
| 101270875 | 101271363 | -1 | processed_pseudoge | ENSG00000 |
| 101323337 | 101390303 | 1 | lncRNA | ENSG00000 |
| 101639509 | 101787572 | 1 | lncRNA | ENSG00000 |
| 101728642 | 101728737 | -1 | snRNA | ENSG00000 |
| 101786340 | 101787219 | -1 | processed_pseudoge | ENSG00000 |
| 101802574 | 101997030 | -1 | protein_coding | ENSG00000 |
| 101859851 | 101859957 | 1 | snRNA | ENSG00000 |
| 101882516 | 101882894 | 1 | processed_pseudoge | ENSG00000 |
| 101893105 | 101894047 | 1 | processed_pseudoge | ENSG00000 |
| 102199739 | 102389630 | -1 | lncRNA | ENSG00000 |
| 102763322 | 102853842 | -1 | lncRNA | ENSG00000 |
| 102876467 | 103108872 | -1 | protein_coding | ENSG00000 |
| 103100143 | 103100704 | -1 | processed_pseudoge | ENSG00000 |
| 103414879 | 103425465 | 1 | lncRNA | ENSG00000 |
| 103415980 | 103525511 | -1 | lncRNA | ENSG00000 |
| 103523562 | 103523879 | 1 | misc_RNA | ENSG00000 |
| 103525691 | 103555239 | 1 | protein_coding | ENSG00000 |
| 103553815 | 103579534 | 1 | protein_coding | ENSG00000 |
| 103569553 | 103570674 | 1 | processed_pseudoge | ENSG00000 |
| 103616811 | 103625780 | 1 | protein_coding | ENSG00000 |
| 103655290 | 103664554 | 1 | protein_coding | ENSG00000 |
| 103668071 | 103668268 | -1 | unprocessed_pseudo | ENSG00000 |
| 103687415 | 103696680 | -1 | protein_coding | ENSG00000 |
| 103713723 | 103719871 | 1 | unprocessed_pseudo | ENSG00000 |
| 103750406 | 103758690 | 1 | protein_coding | ENSG00000 |
| 103926567 | 103927806 | 1 | processed_pseudoge | ENSG00000 |
| 104072983 | 104226678 | 1 | transcribed_processe | ENSG00000 |
| 104153306 | 104153830 | -1 | processed_pseudoge | ENSG00000 |
| 104998406 | 105032365 | -1 | lncRNA | ENSG00000 |
| 105433994 | 105434821 | 1 | processed_pseudoge | ENSG00000 |
| 105587575 | 105618991 | -1 | lncRNA | ENSG00000 |
| 105698039 | 105698218 | 1 | processed_pseudoge | ENSG00000 |

| | | | | |
|---|---|---|---|---|
| 105890693 | 105891430 | -1 | processed_pseudoge | ENSG00000( |
| 105891739 | 105893517 | 1 | processed_pseudoge | ENSG00000( |
| 105927620 | 106028277 | -1 | lncRNA | ENSG00000( |
| 105956694 | 106124564 | 1 | lncRNA | ENSG00000( |
| 106080801 | 106081379 | -1 | processed_pseudoge | ENSG00000( |
| 106544342 | 106544744 | -1 | processed_pseudoge | ENSG00000( |
| 106780223 | 106780435 | 1 | processed_pseudoge | ENSG00000( |
| 106802755 | 106803427 | -1 | processed_pseudoge | ENSG00000( |
| 106804474 | 106806032 | -1 | processed_pseudoge | ENSG00000( |
| 106818224 | 106865316 | 1 | lncRNA | ENSG00000( |
| 107056674 | 107067636 | 1 | protein_coding | ENSG00000( |
| 107140007 | 107483458 | 1 | protein_coding | ENSG00000( |
| 107505203 | 107505448 | 1 | processed_pseudoge | ENSG00000( |
| 107571161 | 107965180 | -1 | protein_coding | ENSG00000( |
| 107897223 | 107897304 | 1 | miRNA | ENSG00000( |
| 107964443 | 107994607 | 1 | lncRNA | ENSG00000( |
| 108040263 | 108076020 | 1 | lncRNA | ENSG00000( |
| 108134043 | 108200849 | -1 | protein_coding | ENSG00000( |
| 108199926 | 108201491 | 1 | lncRNA | ENSG00000( |
| 108200413 | 108202743 | 1 | TEC | ENSG00000( |
| 108223341 | 108244081 | -1 | protein_coding | ENSG00000( |
| 108261196 | 108273689 | -1 | lncRNA | ENSG00000( |
| 108272943 | 108337853 | 1 | transcribed_unproce | ENSG00000( |
| 108375838 | 108410814 | 1 | transcribed_unproce | ENSG00000( |
| 108383736 | 108421295 | -1 | unprocessed_pseudc | ENSG00000( |
| 108420689 | 108433184 | 1 | lncRNA | ENSG00000( |
| 108450282 | 108471002 | 1 | protein_coding | ENSG00000( |
| 108495475 | 108501144 | -1 | unprocessed_pseudc | ENSG00000( |
| 108502358 | 108503455 | 1 | processed_pseudoge | ENSG00000( |
| 108508574 | 108509243 | 1 | processed_pseudoge | ENSG00000( |
| 108560089 | 108644900 | 1 | protein_coding | ENSG00000( |
| 108648290 | 108661526 | -1 | protein_coding | ENSG00000( |
| 108661533 | 108666246 | 1 | lncRNA | ENSG00000( |
| 108692310 | 108702928 | 1 | protein_coding | ENSG00000( |
| 108712908 | 108742749 | 1 | protein_coding | ENSG00000( |
| 108734256 | 108734725 | -1 | lncRNA | ENSG00000( |
| 108746674 | 108809523 | 1 | protein_coding | ENSG00000( |
| 108766841 | 108767584 | -1 | processed_pseudoge | ENSG00000( |
| 108815898 | 108963484 | -1 | protein_coding | ENSG00000( |
| 108857217 | 108858524 | 1 | lncRNA | ENSG00000( |
| 108875350 | 108934545 | 1 | protein_coding | ENSG00000( |
| 108929508 | 108963457 | -1 | protein_coding | ENSG00000( |
| 108970214 | 109042113 | -1 | protein_coding | ENSG00000( |
| 108986963 | 108987245 | -1 | processed_pseudoge | ENSG00000( |
| 108992282 | 108992836 | 1 | processed_pseudoge | ENSG00000( |
| 109030067 | 109030319 | 1 | processed_pseudoge | ENSG00000( |
| 109046828 | 109047482 | 1 | processed_pseudoge | ENSG00000( |
| 109062486 | 109076003 | -1 | protein_coding | ENSG00000( |
| 109087971 | 109090858 | -1 | lncRNA | ENSG00000( |
| 109089803 | 109096934 | 1 | protein_coding | ENSG00000( |

| | | | | |
|---|---|---|---|---|
| 109100193 | 109100612 | 1 | scaRNA | ENSG00000 |
| 109100193 | 109100619 | 1 | lncRNA | ENSG00000 |
| 109103535 | 109104766 | -1 | processed_pseudoge | ENSG00000 |
| 109105951 | 109113857 | -1 | protein_coding | ENSG00000 |
| 109113679 | 109206781 | 1 | protein_coding | ENSG00000 |
| 109213918 | 109238182 | 1 | protein_coding | ENSG00000 |
| 109249539 | 109275751 | 1 | protein_coding | ENSG00000 |
| 109279556 | 109283186 | -1 | protein_coding | ENSG00000 |
| 109292365 | 109307011 | -1 | protein_coding | ENSG00000 |
| 109309568 | 109397918 | -1 | protein_coding | ENSG00000 |
| 109399042 | 109426448 | -1 | protein_coding | ENSG00000 |
| 109466546 | 109482134 | 1 | protein_coding | ENSG00000 |
| 109483479 | 109492804 | 1 | protein_coding | ENSG00000 |
| 109494052 | 109502932 | 1 | protein_coding | ENSG00000 |
| 109504178 | 109509738 | -1 | protein_coding | ENSG00000 |
| 109539872 | 109548406 | 1 | protein_coding | ENSG00000 |
| 109539906 | 109543837 | -1 | lncRNA | ENSG00000 |
| 109548615 | 109618324 | 1 | protein_coding | ENSG00000 |
| 109591534 | 109591640 | -1 | snRNA | ENSG00000 |
| 109596225 | 109597781 | 1 | lncRNA | ENSG00000 |
| 109598893 | 109598967 | 1 | miRNA | ENSG00000 |
| 109603254 | 109619929 | -1 | protein_coding | ENSG00000 |
| 109616104 | 109632053 | 1 | protein_coding | ENSG00000 |
| 109628417 | 109630305 | -1 | lncRNA | ENSG00000 |
| 109651370 | 109652099 | 1 | processed_pseudoge | ENSG00000 |
| 109656099 | 109674836 | 1 | protein_coding | ENSG00000 |
| 109668022 | 109709551 | 1 | protein_coding | ENSG00000 |
| 109687814 | 109709039 | 1 | protein_coding | ENSG00000 |
| 109693117 | 109693742 | -1 | lncRNA | ENSG00000 |
| 109711780 | 109775428 | 1 | protein_coding | ENSG00000 |
| 109725820 | 109775252 | 1 | lncRNA | ENSG00000 |
| 109733932 | 109741038 | -1 | protein_coding | ENSG00000 |
| 109750080 | 109764027 | -1 | protein_coding | ENSG00000 |
| 109810642 | 109810998 | -1 | processed_pseudoge | ENSG00000 |
| 109828355 | 109871436 | 1 | lncRNA | ENSG00000 |
| 109884176 | 109886264 | -1 | lncRNA | ENSG00000 |
| 109895973 | 109897861 | 1 | lncRNA | ENSG00000 |
| 109910242 | 109930992 | 1 | protein_coding | ENSG00000 |
| 109984765 | 110023742 | 1 | protein_coding | ENSG00000 |
| 110031577 | 110074641 | 1 | protein_coding | ENSG00000 |
| 110058340 | 110062555 | 1 | lncRNA | ENSG00000 |
| 110059870 | 110070672 | -1 | protein_coding | ENSG00000 |
| 110082688 | 110109719 | 1 | lncRNA | ENSG00000 |
| 110112443 | 110113947 | 1 | protein_coding | ENSG00000 |
| 110150494 | 110202202 | 1 | protein_coding | ENSG00000 |
| 110165948 | 110172489 | -1 | lncRNA | ENSG00000 |
| 110177643 | 110178719 | -1 | lncRNA | ENSG00000 |
| 110208834 | 110209987 | -1 | lncRNA | ENSG00000 |
| 110211343 | 110283100 | 1 | protein_coding | ENSG00000 |
| 110256754 | 110256930 | 1 | processed_pseudoge | ENSG00000 |

| | | | | |
|---|---|---|---|---|
| 110272484 | 110272607 | 1 | snoRNA | ENSG00000 |
| 110286375 | 110339171 | -1 | lncRNA | ENSG00000 |
| 110338506 | 110346681 | 1 | protein_coding | ENSG00000 |
| 110347116 | 110443817 | 1 | lncRNA | ENSG00000 |
| 110362851 | 110391082 | -1 | protein_coding | ENSG00000 |
| 110370154 | 110373003 | 1 | lncRNA | ENSG00000 |
| 110401249 | 110407942 | -1 | protein_coding | ENSG00000 |
| 110451149 | 110457358 | 1 | protein_coding | ENSG00000 |
| 110456505 | 110457354 | -1 | lncRNA | ENSG00000 |
| 110472543 | 110474980 | 1 | lncRNA | ENSG00000 |
| 110473756 | 110488663 | 1 | lncRNA | ENSG00000 |
| 110480752 | 110491277 | 1 | transcribed_unitary_ | ENSG00000 |
| 110487680 | 110490258 | -1 | lncRNA | ENSG00000 |
| 110517217 | 110519175 | -1 | protein_coding | ENSG00000 |
| 110519837 | 110631474 | -1 | protein_coding | ENSG00000 |
| 110653560 | 110657040 | -1 | lncRNA | ENSG00000 |
| 110672465 | 110674940 | -1 | protein_coding | ENSG00000 |
| 110764408 | 110764506 | -1 | misc_RNA | ENSG00000 |
| 110848077 | 110848941 | -1 | processed_pseudoge | ENSG00000 |
| 110853939 | 110854892 | 1 | unitary_pseudogene | ENSG00000 |
| 110871188 | 110899922 | 1 | protein_coding | ENSG00000 |
| 110936369 | 110942353 | -1 | lncRNA | ENSG00000 |
| 110943467 | 110952848 | 1 | lncRNA | ENSG00000 |
| 110947190 | 110963965 | -1 | protein_coding | ENSG00000 |
| 110963302 | 110964649 | 1 | lncRNA | ENSG00000 |
| 111007700 | 111015530 | -1 | processed_pseudoge | ENSG00000 |
| 111041834 | 111041925 | 1 | rRNA_pseudogene | ENSG00000 |
| 111117333 | 111140216 | -1 | protein_coding | ENSG00000 |
| 111139627 | 111185104 | 1 | protein_coding | ENSG00000 |
| 111181374 | 111181491 | -1 | lncRNA | ENSG00000 |
| 111184415 | 111185061 | -1 | lncRNA | ENSG00000 |
| 111185969 | 111204535 | -1 | protein_coding | ENSG00000 |
| 111200771 | 111243440 | 1 | protein_coding | ENSG00000 |
| 111250254 | 111256715 | 1 | unprocessed_pseudc | ENSG00000 |
| 111280059 | 111286116 | 1 | transcribed_unproce | ENSG00000 |
| 111290851 | 111320566 | 1 | protein_coding | ENSG00000 |
| 111317600 | 111323981 | -1 | lncRNA | ENSG00000 |
| 111323833 | 111324075 | 1 | processed_pseudoge | ENSG00000 |
| 111346600 | 111353013 | 1 | protein_coding | ENSG00000 |
| 111353275 | 111367409 | -1 | unprocessed_pseudc | ENSG00000 |
| 111380291 | 111380571 | 1 | processed_pseudoge | ENSG00000 |
| 111382860 | 111391819 | -1 | transcribed_unproce | ENSG00000 |
| 111414319 | 111427735 | -1 | protein_coding | ENSG00000 |
| 111431046 | 111433068 | -1 | lncRNA | ENSG00000 |
| 111437514 | 111438037 | 1 | processed_pseudoge | ENSG00000 |
| 111438638 | 111441364 | -1 | lncRNA | ENSG00000 |
| 111439890 | 111449256 | -1 | protein_coding | ENSG00000 |
| 111446798 | 111446908 | 1 | misc_RNA | ENSG00000 |
| 111448864 | 111462773 | 1 | protein_coding | ENSG00000 |
| 111473792 | 111478512 | 1 | protein_coding | ENSG00000 |

| | | | | |
|---|---|---|---|---|
| 111483348 | 111563962 | -1 | protein_coding | ENSG00000( |
| 111490317 | 111490423 | -1 | snRNA | ENSG00000( |
| 111499429 | 111503760 | -1 | protein_coding | ENSG00000( |
| 111542218 | 111716691 | 1 | protein_coding | ENSG00000( |
| 111599655 | 111608723 | -1 | lncRNA | ENSG00000( |
| 111648291 | 111649544 | -1 | processed_pseudoge | ENSG00000( |
| 111650431 | 111650537 | -1 | snRNA | ENSG00000( |
| 111680630 | 111755824 | -1 | protein_coding | ENSG00000( |
| 111739579 | 111747798 | 1 | lncRNA | ENSG00000( |
| 111745299 | 111755537 | -1 | lncRNA | ENSG00000( |
| 111755245 | 111768000 | 1 | protein_coding | ENSG00000( |
| 111770662 | 111989155 | -1 | protein_coding | ENSG00000( |
| 111853762 | 111856905 | -1 | lncRNA | ENSG00000( |
| 111909336 | 111910931 | 1 | lncRNA | ENSG00000( |
| 111989770 | 111998842 | 1 | lncRNA | ENSG00000( |
| 112177234 | 112360625 | -1 | lncRNA | ENSG00000( |
| 112363281 | 112363581 | 1 | processed_pseudoge | ENSG00000( |
| 112371004 | 112371107 | 1 | snoRNA | ENSG00000( |
| 112396214 | 112463456 | 1 | protein_coding | ENSG00000( |
| 112461770 | 112461833 | -1 | miRNA | ENSG00000( |
| 112466541 | 112530165 | 1 | protein_coding | ENSG00000( |
| 112517799 | 112518441 | -1 | lncRNA | ENSG00000( |
| 112523514 | 112620825 | -1 | protein_coding | ENSG00000( |
| 112619805 | 112671616 | 1 | protein_coding | ENSG00000( |
| 112625906 | 112626224 | -1 | processed_pseudoge | ENSG00000( |
| 112634719 | 112634784 | -1 | snRNA | ENSG00000( |
| 112673141 | 112700746 | 1 | protein_coding | ENSG00000( |
| 112693688 | 112696621 | -1 | lncRNA | ENSG00000( |
| 112701106 | 112707434 | -1 | protein_coding | ENSG00000( |
| 112702614 | 112711433 | -1 | protein_coding | ENSG00000( |
| 112709994 | 112715477 | -1 | protein_coding | ENSG00000( |
| 112715672 | 112717796 | 1 | lncRNA | ENSG00000( |
| 112720419 | 112727235 | 1 | protein_coding | ENSG00000( |
| 112748095 | 112748475 | 1 | processed_pseudoge | ENSG00000( |
| 112820170 | 112850643 | -1 | lncRNA | ENSG00000( |
| 112849821 | 112877871 | 1 | lncRNA | ENSG00000( |
| 112890767 | 112890922 | -1 | processed_pseudoge | ENSG00000( |
| 112911847 | 112957013 | -1 | protein_coding | ENSG00000( |
| 112923423 | 112924337 | 1 | processed_pseudoge | ENSG00000( |
| 112956415 | 113047055 | 1 | lncRNA | ENSG00000( |
| 112978610 | 112999636 | -1 | lncRNA | ENSG00000( |
| 113011687 | 113073113 | -1 | lncRNA | ENSG00000( |
| 113073198 | 113132260 | 1 | protein_coding | ENSG00000( |
| 113125321 | 113126883 | -1 | processed_pseudoge | ENSG00000( |
| 113168994 | 113169403 | 1 | processed_pseudoge | ENSG00000( |
| 113198825 | 113199258 | 1 | processed_pseudoge | ENSG00000( |
| 113390515 | 113685923 | 1 | protein_coding | ENSG00000( |
| 113449700 | 113450728 | 1 | processed_pseudoge | ENSG00000( |
| 113576757 | 113576899 | -1 | processed_pseudoge | ENSG00000( |
| 113696831 | 113759489 | -1 | protein_coding | ENSG00000( |

| | | | | |
|---|---|---|---|---|
| 113698884 | 113699631 | -1 | transcribed_processe | ENSG00000 |
| 113761832 | 113812476 | -1 | protein_coding | ENSG00000 |
| 113812379 | 113870159 | 1 | lncRNA | ENSG00000 |
| 113813811 | 113871753 | -1 | protein_coding | ENSG00000 |
| 113856635 | 113901237 | 1 | lncRNA | ENSG00000 |
| 113876816 | 113887581 | -1 | protein_coding | ENSG00000 |
| 113894194 | 113905201 | -1 | protein_coding | ENSG00000 |
| 113905213 | 113914086 | 1 | protein_coding | ENSG00000 |
| 113924000 | 113929495 | -1 | lncRNA | ENSG00000 |
| 113929324 | 113977869 | 1 | protein_coding | ENSG00000 |
| 113979391 | 114035572 | 1 | protein_coding | ENSG00000 |
| 114001433 | 114002235 | -1 | processed_pseudoge | ENSG00000 |
| 114032393 | 114034511 | 1 | lncRNA | ENSG00000 |
| 114089291 | 114153919 | -1 | protein_coding | ENSG00000 |
| 114206427 | 114262278 | -1 | lncRNA | ENSG00000 |
| 114279011 | 114279275 | -1 | processed_pseudoge | ENSG00000 |
| 114353986 | 114355178 | 1 | processed_pseudoge | ENSG00000 |
| 114392790 | 114511203 | -1 | protein_coding | ENSG00000 |
| 114459934 | 114460360 | 1 | processed_pseudoge | ENSG00000 |
| 114468315 | 114469282 | 1 | processed_pseudoge | ENSG00000 |
| 114490724 | 114490832 | 1 | misc_RNA | ENSG00000 |
| 114535995 | 114537840 | 1 | processed_pseudoge | ENSG00000 |
| 114567557 | 114581629 | -1 | protein_coding | ENSG00000 |
| 114582848 | 114670422 | -1 | protein_coding | ENSG00000 |
| 114673090 | 114695618 | -1 | protein_coding | ENSG00000 |
| 114697627 | 114697924 | -1 | misc_RNA | ENSG00000 |
| 114704469 | 114716771 | -1 | protein_coding | ENSG00000 |
| 114716913 | 114758676 | -1 | protein_coding | ENSG00000 |
| 114727720 | 114727824 | 1 | misc_RNA | ENSG00000 |
| 114769479 | 114780685 | -1 | protein_coding | ENSG00000 |
| 114837227 | 114851453 | 1 | unitary_pseudogene | ENSG00000 |
| 114854863 | 114995370 | 1 | protein_coding | ENSG00000 |
| 115029824 | 115034309 | 1 | protein_coding | ENSG00000 |
| 115048011 | 115089503 | -1 | protein_coding | ENSG00000 |
| 115099672 | 115102658 | 1 | lncRNA | ENSG00000 |
| 115270767 | 115283763 | -1 | lncRNA | ENSG00000 |
| 115283034 | 115368072 | 1 | lncRNA | ENSG00000 |
| 115285918 | 115338236 | -1 | protein_coding | ENSG00000 |
| 115356664 | 115364897 | -1 | lncRNA | ENSG00000 |
| 115471941 | 115476027 | 1 | unprocessed_pseudo | ENSG00000 |
| 115479428 | 115481572 | 1 | unprocessed_pseudo | ENSG00000 |
| 115556826 | 115557475 | 1 | unprocessed_pseudo | ENSG00000 |
| 115564601 | 115565432 | 1 | processed_pseudoge | ENSG00000 |
| 115577229 | 115577786 | -1 | processed_pseudoge | ENSG00000 |
| 115606471 | 115606768 | 1 | misc_RNA | ENSG00000 |
| 115621872 | 115622005 | -1 | snoRNA | ENSG00000 |
| 115641970 | 115698224 | 1 | protein_coding | ENSG00000 |
| 115700007 | 115768781 | -1 | protein_coding | ENSG00000 |
| 115836377 | 115843917 | -1 | protein_coding | ENSG00000 |
| 115856910 | 115857819 | -1 | processed_pseudoge | ENSG00000 |

| | | | | |
|---|---|---|---|---|
| 115904855 | 115931616 | 1 | lncRNA | ENSG00000 |
| 115916497 | 115916699 | 1 | processed_pseudoge | ENSG00000 |
| 115976513 | 116070054 | 1 | protein_coding | ENSG00000 |
| 116013813 | 116017705 | -1 | lncRNA | ENSG00000 |
| 116111755 | 116135240 | 1 | protein_coding | ENSG00000 |
| 116164284 | 116166241 | 1 | lncRNA | ENSG00000 |
| 116278606 | 116278821 | 1 | snoRNA | ENSG00000 |
| 116289237 | 116297621 | -1 | lncRNA | ENSG00000 |
| 116372668 | 116410261 | 1 | protein_coding | ENSG00000 |
| 116390575 | 116418635 | -1 | lncRNA | ENSG00000 |
| 116413766 | 116413869 | -1 | snRNA | ENSG00000 |
| 116423724 | 116478842 | -1 | lncRNA | ENSG00000 |
| 116429049 | 116433368 | -1 | lncRNA | ENSG00000 |
| 116452536 | 116452627 | 1 | misc_RNA | ENSG00000 |
| 116493016 | 116499212 | -1 | lncRNA | ENSG00000 |
| 116493350 | 116502634 | 1 | lncRNA | ENSG00000 |
| 116514534 | 116571039 | -1 | protein_coding | ENSG00000 |
| 116532936 | 116534092 | -1 | processed_pseudoge | ENSG00000 |
| 116560024 | 116560111 | -1 | miRNA | ENSG00000 |
| 116574399 | 116667755 | -1 | protein_coding | ENSG00000 |
| 116671749 | 116671827 | 1 | miRNA | ENSG00000 |
| 116694112 | 116706603 | 1 | lncRNA | ENSG00000 |
| 116713833 | 116714835 | -1 | processed_pseudoge | ENSG00000 |
| 116739981 | 116742609 | -1 | processed_pseudoge | ENSG00000 |
| 116754430 | 116769229 | 1 | protein_coding | ENSG00000 |
| 116775104 | 116775623 | -1 | processed_pseudoge | ENSG00000 |
| 116909149 | 116909531 | -1 | lncRNA | ENSG00000 |
| 116909916 | 116990353 | 1 | protein_coding | ENSG00000 |
| 116962347 | 116962471 | -1 | rRNA_pseudogene | ENSG00000 |
| 117001750 | 117036476 | 1 | protein_coding | ENSG00000 |
| 117025482 | 117059490 | -1 | lncRNA | ENSG00000 |
| 117060326 | 117107453 | 1 | protein_coding | ENSG00000 |
| 117094643 | 117094728 | 1 | miRNA | ENSG00000 |
| 117111060 | 117122587 | -1 | protein_coding | ENSG00000 |
| 117138229 | 117138609 | 1 | processed_pseudoge | ENSG00000 |
| 117143587 | 117210960 | -1 | protein_coding | ENSG00000 |
| 117243212 | 117243508 | -1 | misc_RNA | ENSG00000 |
| 117272182 | 117321336 | 1 | lncRNA | ENSG00000 |
| 117364899 | 117365473 | -1 | lncRNA | ENSG00000 |
| 117367449 | 117528872 | 1 | protein_coding | ENSG00000 |
| 117493515 | 117495006 | 1 | TEC | ENSG00000 |
| 117549415 | 117549922 | 1 | processed_pseudoge | ENSG00000 |
| 117596832 | 117605770 | -1 | lncRNA | ENSG00000 |
| 117606048 | 117628389 | 1 | protein_coding | ENSG00000 |
| 117640812 | 117641639 | -1 | processed_pseudoge | ENSG00000 |
| 117688621 | 117688742 | -1 | snoRNA | ENSG00000 |
| 117700029 | 117700242 | -1 | processed_pseudoge | ENSG00000 |
| 117778087 | 117778506 | -1 | processed_pseudoge | ENSG00000 |
| 117863485 | 117929621 | -1 | protein_coding | ENSG00000 |
| 117929720 | 117966543 | 1 | protein_coding | ENSG00000 |

| | | | | |
|---|---|---|---|---|
| 117953590 | 118185228 | -1 | protein_coding | ENSG00000( |
| 118185349 | 118207051 | 1 | lncRNA | ENSG00000( |
| 118264372 | 118264489 | -1 | rRNA_pseudogene | ENSG00000( |
| 118614333 | 118615646 | -1 | processed_pseudoge | ENSG00000( |
| 118712999 | 118713678 | 1 | processed_pseudoge | ENSG00000( |
| 118883046 | 118989556 | -1 | protein_coding | ENSG00000( |
| 119000344 | 119001392 | -1 | lncRNA | ENSG00000( |
| 119000618 | 119001405 | -1 | lncRNA | ENSG00000( |
| 119031216 | 119140671 | -1 | protein_coding | ENSG00000( |
| 119047405 | 119064785 | 1 | lncRNA | ENSG00000( |
| 119084998 | 119085954 | -1 | processed_pseudoge | ENSG00000( |
| 119126539 | 119127291 | -1 | processed_pseudoge | ENSG00000( |
| 119140391 | 119275973 | 1 | lncRNA | ENSG00000( |
| 119219314 | 119220096 | -1 | processed_pseudoge | ENSG00000( |
| 119230313 | 119327915 | -1 | lncRNA | ENSG00000( |
| 119327399 | 119356182 | 1 | lncRNA | ENSG00000( |
| 119331397 | 119331525 | -1 | snRNA | ENSG00000( |
| 119368779 | 119394130 | 1 | protein_coding | ENSG00000( |
| 119368946 | 119370331 | 1 | lncRNA | ENSG00000( |
| 119414931 | 119423035 | 1 | protein_coding | ENSG00000( |
| 119434166 | 119435080 | -1 | transcribed_process | ENSG00000( |
| 119439001 | 119446086 | 1 | unprocessed_pseudc | ENSG00000( |
| 119462029 | 119463021 | -1 | processed_pseudoge | ENSG00000( |
| 119467221 | 119473803 | 1 | unprocessed_pseudc | ENSG00000( |
| 119495836 | 119496841 | -1 | processed_pseudoge | ENSG00000( |
| 119507198 | 119515054 | 1 | protein_coding | ENSG00000( |
| 119533749 | 119534746 | -1 | processed_pseudoge | ENSG00000( |
| 119538509 | 119545913 | 1 | unprocessed_pseudc | ENSG00000( |
| 119558755 | 119559774 | -1 | processed_pseudoge | ENSG00000( |
| 119564066 | 119572067 | 1 | unprocessed_pseudc | ENSG00000( |
| 119596167 | 119597179 | -1 | processed_pseudoge | ENSG00000( |
| 119597702 | 119599271 | -1 | lncRNA | ENSG00000( |
| 119601340 | 119609250 | 1 | transcribed_unproce | ENSG00000( |
| 119619377 | 119648266 | -1 | protein_coding | ENSG00000( |
| 119648411 | 119744226 | 1 | protein_coding | ENSG00000( |
| 119747996 | 119768905 | -1 | protein_coding | ENSG00000( |
| 119794017 | 119811580 | -1 | protein_coding | ENSG00000( |
| 119834870 | 119844514 | -1 | unprocessed_pseudc | ENSG00000( |
| 119853316 | 119853748 | 1 | processed_pseudoge | ENSG00000( |
| 119886304 | 119886927 | 1 | processed_pseudoge | ENSG00000( |
| 119893533 | 119896515 | -1 | protein_coding | ENSG00000( |
| 119911553 | 120100779 | -1 | protein_coding | ENSG00000( |
| 120076616 | 120077091 | 1 | lncRNA | ENSG00000( |
| 120126974 | 120127074 | 1 | snRNA | ENSG00000( |
| 120150898 | 120176520 | -1 | protein_coding | ENSG00000( |
| 120197085 | 120319680 | 1 | unprocessed_pseudc | ENSG00000( |
| 120228046 | 120228295 | -1 | misc_RNA | ENSG00000( |
| 120267334 | 120341871 | -1 | transcribed_unproce | ENSG00000( |
| 120354987 | 120355148 | 1 | snRNA | ENSG00000( |
| 120419850 | 120467739 | 1 | transcribed_unproce | ENSG00000( |

| | | | | |
|---|---|---|---|---|
| 120432204 | 120434052 | -1 | transcribed_processe | ENSG00000( |
| 120489860 | 120701289 | -1 | transcribed_unproce | ENSG00000( |
| 120571120 | 120571264 | 1 | snRNA | ENSG00000( |
| 120670127 | 120670376 | 1 | misc_RNA | ENSG00000( |
| 120723946 | 120793874 | 1 | protein_coding | ENSG00000( |
| 120723949 | 120842110 | 1 | protein_coding | ENSG00000( |
| 120849674 | 120850985 | -1 | lncRNA | ENSG00000( |
| 120850819 | 120850985 | -1 | snRNA | ENSG00000( |
| 120889746 | 120890405 | 1 | protein_coding | ENSG00000( |
| 120913151 | 120914238 | -1 | TEC | ENSG00000( |
| 120913184 | 121009291 | 1 | lncRNA | ENSG00000( |
| 120942600 | 120942763 | 1 | snRNA | ENSG00000( |
| 120963645 | 120977585 | 1 | transcribed_unproce | ENSG00000( |
| 120985692 | 121052167 | -1 | transcribed_unproce | ENSG00000( |
| 121008362 | 121008507 | -1 | snRNA | ENSG00000( |
| 121058948 | 121059353 | -1 | processed_pseudoge | ENSG00000( |
| 121087345 | 121097161 | 1 | protein_coding | ENSG00000( |
| 121087528 | 121116676 | -1 | lncRNA | ENSG00000( |
| 121090289 | 121097655 | -1 | lncRNA | ENSG00000( |
| 121108210 | 121117257 | -1 | transcribed_unproce | ENSG00000( |
| 121118126 | 121146826 | 1 | lncRNA | ENSG00000( |
| 121118195 | 121118610 | -1 | unprocessed_pseudc | ENSG00000( |
| 121142051 | 121142438 | -1 | processed_pseudoge | ENSG00000( |
| 121167646 | 121185539 | -1 | protein_coding | ENSG00000( |
| 121184811 | 121392874 | 1 | protein_coding | ENSG00000( |
| 121314947 | 121315055 | -1 | processed_pseudoge | ENSG00000( |
| 121360156 | 121398806 | -1 | lncRNA | ENSG00000( |
| 121396754 | 121463508 | 1 | lncRNA | ENSG00000( |
| 121494329 | 121511588 | -1 | lncRNA | ENSG00000( |
| 121502344 | 121503891 | 1 | processed_pseudoge | ENSG00000( |
| 121518366 | 121518829 | -1 | lncRNA | ENSG00000( |
| 121519112 | 121575702 | 1 | transcribed_unproce | ENSG00000( |
| 121573946 | 121580524 | -1 | lncRNA | ENSG00000( |
| 143343180 | 143343275 | 1 | unprocessed_pseudc | ENSG00000( |
| 143354191 | 143386122 | 1 | unprocessed_pseudc | ENSG00000( |
| 143419625 | 143420207 | -1 | unprocessed_pseudc | ENSG00000( |
| 143439605 | 143439714 | 1 | rRNA | ENSG00000( |
| 143449571 | 143450245 | -1 | processed_pseudoge | ENSG00000( |
| 143461221 | 143498767 | -1 | unprocessed_pseudc | ENSG00000( |
| 143487772 | 143488353 | 1 | processed_pseudoge | ENSG00000( |
| 143498785 | 143499102 | 1 | processed_pseudoge | ENSG00000( |
| 143499187 | 143500512 | -1 | lncRNA | ENSG00000( |
| 143541768 | 143572834 | 1 | unprocessed_pseudc | ENSG00000( |
| 143595216 | 143635641 | -1 | unprocessed_pseudc | ENSG00000( |
| 143619994 | 143620527 | 1 | processed_pseudoge | ENSG00000( |
| 143652050 | 143652215 | -1 | snRNA | ENSG00000( |
| 143699456 | 143699619 | -1 | snRNA | ENSG00000( |
| 143720510 | 143720644 | -1 | snRNA | ENSG00000( |
| 143729407 | 143729570 | -1 | snRNA | ENSG00000( |
| 143736066 | 143739506 | 1 | lncRNA | ENSG00000( |

| | | | | |
|---|---|---|---|---|
| 143745249 | 143760669 | 1 | unprocessed_pseudc | ENSG0000C |
| 143766540 | 143769083 | 1 | transcribed_processe | ENSG0000C |
| 143784376 | 143784671 | 1 | processed_pseudoge | ENSG0000C |
| 143790010 | 143797108 | 1 | lncRNA | ENSG0000C |
| 143791542 | 143791687 | -1 | snRNA | ENSG0000C |
| 143811359 | 143825837 | -1 | unprocessed_pseudc | ENSG0000C |
| 143846097 | 143846504 | -1 | processed_pseudoge | ENSG0000C |
| 143874793 | 143883575 | 1 | unprocessed_pseudc | ENSG0000C |
| 143875171 | 143904650 | -1 | transcribed_unproce | ENSG0000C |
| 143894544 | 143905966 | -1 | protein_coding | ENSG0000C |
| 143905487 | 143934776 | 1 | lncRNA | ENSG0000C |
| 143929994 | 143930382 | -1 | processed_pseudoge | ENSG0000C |
| 143955364 | 143971965 | -1 | protein_coding | ENSG0000C |
| 143975087 | 144068350 | 1 | unprocessed_pseudc | ENSG0000C |
| 144085628 | 144086098 | 1 | IG_V_pseudogene | ENSG0000C |
| 144168752 | 144168850 | 1 | processed_pseudoge | ENSG0000C |
| 144179777 | 144211729 | 1 | unprocessed_pseudc | ENSG0000C |
| 144227029 | 144250326 | -1 | lncRNA | ENSG0000C |
| 144265217 | 144265326 | 1 | rRNA | ENSG0000C |
| 144274892 | 144277993 | -1 | processed_pseudoge | ENSG0000C |
| 144276484 | 144305180 | -1 | transcribed_unproce | ENSG0000C |
| 144372875 | 144373659 | -1 | protein_coding | ENSG0000C |
| 144401069 | 144401355 | -1 | processed_pseudoge | ENSG0000C |
| 144412576 | 144412740 | 1 | snRNA | ENSG0000C |
| 144418113 | 144419192 | -1 | lncRNA | ENSG0000C |
| 144418678 | 144419190 | 1 | processed_pseudoge | ENSG0000C |
| 144421386 | 144461674 | -1 | protein_coding | ENSG0000C |
| 144442606 | 144443004 | 1 | processed_pseudoge | ENSG0000C |
| 144472535 | 144474790 | -1 | unprocessed_pseudc | ENSG0000C |
| 144523860 | 144524021 | -1 | snRNA | ENSG0000C |
| 144551779 | 144551943 | -1 | snRNA | ENSG0000C |
| 144560666 | 144560829 | -1 | snRNA | ENSG0000C |
| 144592868 | 144593652 | 1 | protein_coding | ENSG0000C |
| 144627192 | 144642372 | -1 | lncRNA | ENSG0000C |
| 144641371 | 144919902 | 1 | lncRNA | ENSG0000C |
| 144661200 | 144679513 | 1 | unprocessed_pseudc | ENSG0000C |
| 144701050 | 144701159 | -1 | rRNA_pseudogene | ENSG0000C |
| 144754572 | 144786606 | -1 | unprocessed_pseudc | ENSG0000C |
| 144797534 | 144797632 | -1 | processed_pseudoge | ENSG0000C |
| 144887265 | 145095528 | -1 | protein_coding | ENSG0000C |
| 144965025 | 144965135 | 1 | processed_pseudoge | ENSG0000C |
| 145095974 | 145112696 | 1 | protein_coding | ENSG0000C |
| 145139986 | 145148375 | 1 | unprocessed_pseudc | ENSG0000C |
| 145164099 | 145216058 | -1 | transcribed_unproce | ENSG0000C |
| 145169519 | 145169664 | 1 | snRNA | ENSG0000C |
| 145214851 | 145215938 | 1 | TEC | ENSG0000C |
| 145233001 | 145233519 | 1 | processed_pseudoge | ENSG0000C |
| 145241630 | 145242124 | -1 | protein_coding | ENSG0000C |
| 145269604 | 145269890 | -1 | processed_pseudoge | ENSG0000C |
| 145281116 | 145281279 | 1 | snRNA | ENSG0000C |

| | | | | |
|---|---|---|---|---|
| 145281116 | 145281462 | 1 | lncRNA | ENSG0000C |
| 145286610 | 145287755 | -1 | lncRNA | ENSG0000C |
| 145289900 | 145405778 | -1 | protein_coding | ENSG0000C |
| 145407772 | 145408173 | 1 | processed_pseudoge | ENSG0000C |
| 145431527 | 145431693 | -1 | snRNA | ENSG0000C |
| 145459798 | 145462007 | 1 | TEC | ENSG0000C |
| 145465617 | 145465780 | -1 | snRNA | ENSG0000C |
| 145475606 | 145480999 | 1 | lncRNA | ENSG0000C |
| 145507444 | 145508607 | 1 | unprocessed_pseudc | ENSG0000C |
| 145509047 | 145520594 | 1 | unprocessed_pseudc | ENSG0000C |
| 145572345 | 145607858 | -1 | transcribed_unproce | ENSG0000C |
| 145607988 | 145670650 | 1 | protein_coding | ENSG0000C |
| 145670852 | 145708148 | -1 | protein_coding | ENSG0000C |
| 145719471 | 145739288 | 1 | protein_coding | ENSG0000C |
| 145738868 | 145824095 | -1 | protein_coding | ENSG0000C |
| 145824088 | 145844402 | 1 | protein_coding | ENSG0000C |
| 145845630 | 145848954 | 1 | protein_coding | ENSG0000C |
| 145848522 | 145859836 | -1 | protein_coding | ENSG0000C |
| 145850587 | 145850645 | -1 | miRNA | ENSG0000C |
| 145866560 | 145885866 | -1 | protein_coding | ENSG0000C |
| 145891208 | 145910111 | -1 | protein_coding | ENSG0000C |
| 145892847 | 145893483 | 1 | lncRNA | ENSG0000C |
| 145911350 | 145918717 | -1 | protein_coding | ENSG0000C |
| 145917714 | 145927678 | -1 | protein_coding | ENSG0000C |
| 145919013 | 145925341 | 1 | transcribed_unitary_ | ENSG0000C |
| 145926590 | 145959179 | 1 | lncRNA | ENSG0000C |
| 145927258 | 145977811 | 1 | protein_coding | ENSG0000C |
| 145933423 | 145958017 | -1 | protein_coding | ENSG0000C |
| 145959441 | 145964575 | -1 | protein_coding | ENSG0000C |
| 145961388 | 145964422 | 1 | lncRNA | ENSG0000C |
| 145964690 | 145978848 | 1 | protein_coding | ENSG0000C |
| 145992435 | 145996579 | -1 | protein_coding | ENSG0000C |
| 146017468 | 146036746 | -1 | protein_coding | ENSG0000C |
| 146050441 | 146052244 | -1 | lncRNA | ENSG0000C |
| 146052081 | 146052244 | -1 | snRNA | ENSG0000C |
| 146052566 | 146061948 | 1 | lncRNA | ENSG0000C |
| 146064711 | 146229000 | -1 | protein_coding | ENSG0000C |
| 146146203 | 146229026 | -1 | protein_coding | ENSG0000C |
| 146235806 | 146237807 | 1 | lncRNA | ENSG0000C |
| 146288596 | 146288696 | 1 | snRNA | ENSG0000C |
| 146298170 | 146298715 | 1 | processed_pseudoge | ENSG0000C |
| 146321214 | 146401647 | -1 | transcribed_unproce | ENSG0000C |
| 146344131 | 146344790 | -1 | protein_coding | ENSG0000C |
| 146376807 | 146376970 | 1 | snRNA | ENSG0000C |
| 146387202 | 146388092 | 1 | lncRNA | ENSG0000C |
| 146409903 | 146410062 | 1 | snRNA | ENSG0000C |
| 146459449 | 146461859 | 1 | unprocessed_pseudc | ENSG0000C |
| 146472566 | 146914294 | 1 | transcribed_unproce | ENSG0000C |
| 146539541 | 146539663 | -1 | rRNA | ENSG0000C |
| 146862729 | 146862845 | -1 | snRNA | ENSG0000C |

| | | | | |
|---|---|---|---|---|
| 146938744 | 146996202 | 1 | protein_coding | ENSG00000( |
| 146957117 | 146957659 | -1 | processed_pseudoge | ENSG00000( |
| 147001931 | 147003618 | -1 | lncRNA | ENSG00000( |
| 147019656 | 147056593 | -1 | unprocessed_pseudc | ENSG00000( |
| 147050817 | 147052481 | 1 | processed_pseudoge | ENSG00000( |
| 147079746 | 147079870 | 1 | snRNA | ENSG00000( |
| 147082338 | 147083578 | -1 | processed_pseudoge | ENSG00000( |
| 147084616 | 147084756 | -1 | snRNA | ENSG00000( |
| 147099482 | 147124285 | -1 | unprocessed_pseudc | ENSG00000( |
| 147155106 | 147172550 | -1 | protein_coding | ENSG00000( |
| 147172744 | 147179622 | 1 | transcribed_processe | ENSG00000( |
| 147172755 | 147295734 | 1 | lncRNA | ENSG00000( |
| 147175351 | 147243050 | -1 | protein_coding | ENSG00000( |
| 147203276 | 147204932 | -1 | processed_pseudoge | ENSG00000( |
| 147223554 | 147224348 | 1 | processed_pseudoge | ENSG00000( |
| 147242654 | 147295765 | 1 | protein_coding | ENSG00000( |
| 147258885 | 147517875 | -1 | lncRNA | ENSG00000( |
| 147319110 | 147320224 | 1 | processed_pseudoge | ENSG00000( |
| 147419053 | 147419900 | 1 | unitary_pseudogene | ENSG00000( |
| 147420199 | 147420300 | -1 | misc_RNA | ENSG00000( |
| 147445579 | 147446133 | 1 | unprocessed_pseudc | ENSG00000( |
| 147482238 | 147482628 | 1 | unprocessed_pseudc | ENSG00000( |
| 147541501 | 147626216 | 1 | protein_coding | ENSG00000( |
| 147629652 | 147670524 | -1 | protein_coding | ENSG00000( |
| 147689256 | 147689537 | 1 | misc_RNA | ENSG00000( |
| 147697794 | 147699335 | 1 | lncRNA | ENSG00000( |
| 147756199 | 147773362 | -1 | protein_coding | ENSG00000( |
| 147757185 | 147758434 | 1 | lncRNA | ENSG00000( |
| 147777590 | 147788953 | 1 | lncRNA | ENSG00000( |
| 147840962 | 147842359 | 1 | processed_pseudoge | ENSG00000( |
| 147907956 | 147909257 | 1 | protein_coding | ENSG00000( |
| 147928393 | 147993592 | 1 | protein_coding | ENSG00000( |
| 147993862 | 148014956 | -1 | transcribed_unproce | ENSG00000( |
| 148011799 | 148012228 | 1 | lncRNA | ENSG00000( |
| 148013203 | 148025934 | 1 | lncRNA | ENSG00000( |
| 148014417 | 148014583 | -1 | snRNA | ENSG00000( |
| 148038753 | 148038916 | -1 | snRNA | ENSG00000( |
| 148080598 | 148093883 | 1 | unprocessed_pseudc | ENSG00000( |
| 148102046 | 148152322 | -1 | protein_coding | ENSG00000( |
| 148129497 | 148129898 | 1 | processed_pseudoge | ENSG00000( |
| 148146395 | 148149566 | 1 | unprocessed_pseudc | ENSG00000( |
| 148156139 | 148186980 | 1 | lncRNA | ENSG00000( |
| 148159213 | 148255012 | -1 | lncRNA | ENSG00000( |
| 148193716 | 148193838 | 1 | rRNA_pseudogene | ENSG00000( |
| 148234273 | 148234489 | 1 | unprocessed_pseudc | ENSG00000( |
| 148246169 | 148246685 | 1 | processed_pseudoge | ENSG00000( |
| 148263476 | 148263632 | 1 | snRNA | ENSG00000( |
| 148271884 | 148280749 | -1 | lncRNA | ENSG00000( |
| 148288001 | 148288951 | -1 | unprocessed_pseudc | ENSG00000( |
| 148290889 | 148519604 | -1 | lncRNA | ENSG00000( |

| | | | | |
|---|---|---|---|---|
| 148295180 | 148297556 | 1 | lncRNA | ENSG0000C |
| 148295895 | 148344638 | 1 | lncRNA | ENSG0000C |
| 148317683 | 148318013 | -1 | processed_pseudoge | ENSG0000C |
| 148330271 | 148330394 | 1 | misc_RNA | ENSG0000C |
| 148334479 | 148334554 | -1 | miRNA | ENSG0000C |
| 148334612 | 148334767 | -1 | snRNA | ENSG0000C |
| 148358245 | 148358686 | -1 | lncRNA | ENSG0000C |
| 148362370 | 148362533 | -1 | snRNA | ENSG0000C |
| 148385829 | 148385963 | -1 | snRNA | ENSG0000C |
| 148388282 | 148388363 | 1 | miRNA | ENSG0000C |
| 148388490 | 148388651 | -1 | snRNA | ENSG0000C |
| 148402516 | 148432545 | 1 | transcribed_unproce | ENSG0000C |
| 148402715 | 148402875 | 1 | snRNA | ENSG0000C |
| 148435105 | 148436926 | 1 | processed_pseudoge | ENSG0000C |
| 148458643 | 148459689 | -1 | transcribed_unproce | ENSG0000C |
| 148458814 | 148459871 | 1 | TEC | ENSG0000C |
| 148479824 | 148483679 | -1 | protein_coding | ENSG0000C |
| 148522601 | 148522765 | 1 | snRNA | ENSG0000C |
| 148531385 | 148679742 | -1 | protein_coding | ENSG0000C |
| 148607040 | 148679779 | -1 | protein_coding | ENSG0000C |
| 148739379 | 148739479 | 1 | snRNA | ENSG0000C |
| 148748894 | 148750099 | 1 | protein_coding | ENSG0000C |
| 148772640 | 148784594 | -1 | unprocessed_pseudc | ENSG0000C |
| 148795797 | 148796325 | 1 | processed_pseudoge | ENSG0000C |
| 148808181 | 149048286 | 1 | protein_coding | ENSG0000C |
| 148839483 | 148839732 | -1 | misc_RNA | ENSG0000C |
| 148939739 | 148939817 | -1 | snRNA | ENSG0000C |
| 149006309 | 149009527 | -1 | lncRNA | ENSG0000C |
| 149018671 | 149026269 | -1 | lncRNA | ENSG0000C |
| 149048576 | 149051273 | 1 | lncRNA | ENSG0000C |
| 149054027 | 149103561 | -1 | protein_coding | ENSG0000C |
| 149111304 | 149121866 | -1 | unprocessed_pseudc | ENSG0000C |
| 149162783 | 149162944 | -1 | snRNA | ENSG0000C |
| 149175748 | 149259987 | 1 | lncRNA | ENSG0000C |
| 149197992 | 149321732 | -1 | transcribed_unproce | ENSG0000C |
| 149264252 | 149264816 | 1 | lncRNA | ENSG0000C |
| 149290513 | 149290762 | 1 | misc_RNA | ENSG0000C |
| 149345661 | 149357612 | 1 | unprocessed_pseudc | ENSG0000C |
| 149390614 | 149468030 | 1 | protein_coding | ENSG0000C |
| 149390623 | 149556361 | 1 | protein_coding | ENSG0000C |
| 149475338 | 149556361 | 1 | protein_coding | ENSG0000C |
| 149583865 | 149584464 | 1 | protein_coding | ENSG0000C |
| 149606334 | 149705716 | 1 | transcribed_unproce | ENSG0000C |
| 149607467 | 149608516 | 1 | transcribed_unproce | ENSG0000C |
| 149607765 | 149612402 | 1 | lncRNA | ENSG0000C |
| 149636766 | 149636929 | 1 | snRNA | ENSG0000C |
| 149675205 | 149677740 | 1 | transcribed_processe | ENSG0000C |
| 149676851 | 149747818 | -1 | transcribed_unproce | ENSG0000C |
| 149700151 | 149700296 | -1 | snRNA | ENSG0000C |
| 149754301 | 149754710 | -1 | processed_pseudoge | ENSG0000C |

| | | | | |
|---|---|---|---|---|
| 149782671 | 149792518 | 1 | protein_coding | ENSG00000( |
| 149782689 | 149812373 | -1 | protein_coding | ENSG00000( |
| 149785659 | 149793020 | -1 | lncRNA | ENSG00000( |
| 149813271 | 149813681 | -1 | protein_coding | ENSG00000( |
| 149832659 | 149839767 | 1 | protein_coding | ENSG00000( |
| 149839538 | 149841193 | -1 | protein_coding | ENSG00000( |
| 149842188 | 149842736 | -1 | protein_coding | ENSG00000( |
| 149842875 | 149846486 | 1 | transcribed_unproce | ENSG00000( |
| 149844498 | 149850937 | -1 | transcribed_unproce | ENSG00000( |
| 149851061 | 149851624 | 1 | protein_coding | ENSG00000( |
| 149852619 | 149854274 | 1 | protein_coding | ENSG00000( |
| 149854045 | 149861210 | -1 | protein_coding | ENSG00000( |
| 149861271 | 149862504 | 1 | lncRNA | ENSG00000( |
| 149884459 | 149886652 | -1 | protein_coding | ENSG00000( |
| 149886975 | 149887364 | 1 | protein_coding | ENSG00000( |
| 149887484 | 149888013 | -1 | protein_coding | ENSG00000( |
| 149887890 | 149900798 | 1 | protein_coding | ENSG00000( |
| 149903318 | 149917844 | -1 | protein_coding | ENSG00000( |
| 149923317 | 149927803 | -1 | protein_coding | ENSG00000( |
| 149928651 | 149936879 | -1 | protein_coding | ENSG00000( |
| 149937812 | 150010726 | -1 | protein_coding | ENSG00000( |
| 150045660 | 150067701 | -1 | lncRNA | ENSG00000( |
| 150053864 | 150055034 | 1 | processed_pseudoge | ENSG00000( |
| 150067279 | 150145329 | 1 | protein_coding | ENSG00000( |
| 150149183 | 150164720 | 1 | protein_coding | ENSG00000( |
| 150173049 | 150181429 | 1 | lncRNA | ENSG00000( |
| 150211632 | 150211925 | 1 | misc_RNA | ENSG00000( |
| 150218417 | 150236156 | -1 | protein_coding | ENSG00000( |
| 150236967 | 150237156 | 1 | snRNA | ENSG00000( |
| 150255095 | 150257286 | -1 | lncRNA | ENSG00000( |
| 150257251 | 150265078 | 1 | protein_coding | ENSG00000( |
| 150265399 | 150269580 | -1 | protein_coding | ENSG00000( |
| 150268200 | 150280916 | 1 | protein_coding | ENSG00000( |
| 150282543 | 150287093 | 1 | protein_coding | ENSG00000( |
| 150293720 | 150308979 | 1 | protein_coding | ENSG00000( |
| 150321479 | 150353233 | 1 | protein_coding | ENSG00000( |
| 150363091 | 150476566 | 1 | protein_coding | ENSG00000( |
| 150487364 | 150507609 | 1 | protein_coding | ENSG00000( |
| 150492345 | 150492410 | 1 | miRNA | ENSG00000( |
| 150508062 | 150513789 | 1 | protein_coding | ENSG00000( |
| 150515757 | 150518032 | 1 | lncRNA | ENSG00000( |
| 150548562 | 150557724 | -1 | lncRNA | ENSG00000( |
| 150549369 | 150560937 | 1 | protein_coding | ENSG00000( |
| 150551929 | 150552014 | 1 | miRNA | ENSG00000( |
| 150560202 | 150574552 | -1 | lncRNA | ENSG00000( |
| 150566562 | 150566860 | -1 | misc_RNA | ENSG00000( |
| 150568971 | 150569269 | -1 | misc_RNA | ENSG00000( |
| 150574551 | 150579738 | -1 | protein_coding | ENSG00000( |
| 150600539 | 150600659 | -1 | snoRNA | ENSG00000( |
| 150600851 | 150629612 | -1 | protein_coding | ENSG00000( |

| | | | | |
|---|---|---|---|---|
| 150608507 | 150608623 | -1 | snRNA | ENSG00000 |
| 150646230 | 150697154 | -1 | protein_coding | ENSG00000 |
| 150698060 | 150720895 | -1 | protein_coding | ENSG00000 |
| 150701866 | 150701972 | 1 | snRNA | ENSG00000 |
| 150730188 | 150765792 | -1 | protein_coding | ENSG00000 |
| 150780272 | 150780644 | 1 | processed_pseudoge | ENSG00000 |
| 150796208 | 150808260 | -1 | protein_coding | ENSG00000 |
| 150800473 | 150801031 | 1 | processed_pseudoge | ENSG00000 |
| 150809713 | 150876708 | -1 | protein_coding | ENSG00000 |
| 150812591 | 150812698 | -1 | snRNA | ENSG00000 |
| 150881236 | 150881683 | -1 | processed_pseudoge | ENSG00000 |
| 150887136 | 150913292 | 1 | protein_coding | ENSG00000 |
| 150903896 | 150904213 | 1 | processed_pseudoge | ENSG00000 |
| 150926263 | 150964744 | 1 | protein_coding | ENSG00000 |
| 150960583 | 150975004 | -1 | protein_coding | ENSG00000 |
| 150965245 | 150966256 | 1 | lncRNA | ENSG00000 |
| 150973123 | 150975534 | 1 | lncRNA | ENSG00000 |
| 150982249 | 150995634 | 1 | protein_coding | ENSG00000 |
| 150996086 | 151008375 | -1 | protein_coding | ENSG00000 |
| 151008420 | 151035713 | 1 | protein_coding | ENSG00000 |
| 151022746 | 151022852 | -1 | snRNA | ENSG00000 |
| 151036321 | 151047720 | 1 | protein_coding | ENSG00000 |
| 151047751 | 151051986 | 1 | protein_coding | ENSG00000 |
| 151050971 | 151070325 | -1 | protein_coding | ENSG00000 |
| 151057758 | 151068497 | 1 | protein_coding | ENSG00000 |
| 151070578 | 151125542 | 1 | protein_coding | ENSG00000 |
| 151111912 | 151112728 | -1 | processed_pseudoge | ENSG00000 |
| 151130075 | 151131610 | -1 | lncRNA | ENSG00000 |
| 151131685 | 151146664 | -1 | protein_coding | ENSG00000 |
| 151156649 | 151159749 | 1 | protein_coding | ENSG00000 |
| 151156664 | 151170297 | 1 | protein_coding | ENSG00000 |
| 151159748 | 151165948 | -1 | protein_coding | ENSG00000 |
| 151169986 | 151176284 | -1 | protein_coding | ENSG00000 |
| 151169987 | 151195321 | -1 | protein_coding | ENSG00000 |
| 151197949 | 151249536 | 1 | protein_coding | ENSG00000 |
| 151254703 | 151267479 | 1 | protein_coding | ENSG00000 |
| 151279678 | 151281950 | -1 | lncRNA | ENSG00000 |
| 151281618 | 151292176 | 1 | protein_coding | ENSG00000 |
| 151291797 | 151327715 | -1 | protein_coding | ENSG00000 |
| 151300665 | 151300964 | -1 | misc_RNA | ENSG00000 |
| 151327949 | 151328429 | 1 | lncRNA | ENSG00000 |
| 151340640 | 151347357 | -1 | protein_coding | ENSG00000 |
| 151340648 | 151341966 | 1 | lncRNA | ENSG00000 |
| 151346967 | 151348027 | 1 | lncRNA | ENSG00000 |
| 151364304 | 151372707 | -1 | protein_coding | ENSG00000 |
| 151399560 | 151401937 | 1 | protein_coding | ENSG00000 |
| 151402724 | 151459494 | -1 | protein_coding | ENSG00000 |
| 151439000 | 151439095 | 1 | misc_RNA | ENSG00000 |
| 151510510 | 151538692 | 1 | protein_coding | ENSG00000 |
| 151527831 | 151527938 | -1 | snoRNA | ENSG00000 |

| | | | | |
|---|---|---|---|---|
| 151540305 | 151583583 | 1 | protein_coding | ENSG00000 |
| 151540516 | 151561855 | 1 | lncRNA | ENSG00000 |
| 151545796 | 151545891 | 1 | miRNA | ENSG00000 |
| 151557446 | 151557940 | 1 | processed_pseudoge | ENSG00000 |
| 151612006 | 151699091 | 1 | protein_coding | ENSG00000 |
| 151612038 | 151613363 | -1 | lncRNA | ENSG00000 |
| 151629324 | 151629430 | 1 | snRNA | ENSG00000 |
| 151700058 | 151716803 | -1 | protein_coding | ENSG00000 |
| 151701026 | 151708386 | 1 | lncRNA | ENSG00000 |
| 151710433 | 151729805 | 1 | protein_coding | ENSG00000 |
| 151747597 | 151747700 | -1 | snRNA | ENSG00000 |
| 151755541 | 151759911 | 1 | lncRNA | ENSG00000 |
| 151757659 | 151758442 | 1 | processed_pseudoge | ENSG00000 |
| 151759647 | 151763496 | -1 | protein_coding | ENSG00000 |
| 151762899 | 151771334 | 1 | protein_coding | ENSG00000 |
| 151763384 | 151769501 | -1 | lncRNA | ENSG00000 |
| 151765709 | 151766389 | 1 | lncRNA | ENSG00000 |
| 151766486 | 151767000 | -1 | lncRNA | ENSG00000 |
| 151770107 | 151791416 | -1 | protein_coding | ENSG00000 |
| 151790804 | 151794402 | 1 | lncRNA | ENSG00000 |
| 151798054 | 151798602 | 1 | lncRNA | ENSG00000 |
| 151800264 | 151806154 | -1 | protein_coding | ENSG00000 |
| 151806071 | 151831845 | -1 | protein_coding | ENSG00000 |
| 151837819 | 151840557 | -1 | protein_coding | ENSG00000 |
| 151841736 | 151841837 | -1 | misc_RNA | ENSG00000 |
| 151841877 | 151850385 | 1 | lncRNA | ENSG00000 |
| 151847101 | 151853712 | -1 | protein_coding | ENSG00000 |
| 151870866 | 151909637 | -1 | protein_coding | ENSG00000 |
| 151885251 | 151899333 | 1 | lncRNA | ENSG00000 |
| 151944890 | 151953985 | 1 | lncRNA | ENSG00000 |
| 151949523 | 151950912 | -1 | processed_pseudoge | ENSG00000 |
| 151982915 | 151993859 | -1 | protein_coding | ENSG00000 |
| 151994531 | 152042774 | 1 | lncRNA | ENSG00000 |
| 152018662 | 152022508 | 1 | unprocessed_pseudc | ENSG00000 |
| 152032506 | 152047907 | -1 | protein_coding | ENSG00000 |
| 152077952 | 152078211 | 1 | processed_pseudoge | ENSG00000 |
| 152079557 | 152080903 | -1 | unprocessed_pseudc | ENSG00000 |
| 152084141 | 152089064 | -1 | protein_coding | ENSG00000 |
| 152106317 | 152115454 | -1 | protein_coding | ENSG00000 |
| 152122534 | 152125065 | 1 | lncRNA | ENSG00000 |
| 152124016 | 152196699 | -1 | transcribed_processe | ENSG00000 |
| 152153595 | 152159228 | -1 | protein_coding | ENSG00000 |
| 152168125 | 152445456 | 1 | lncRNA | ENSG00000 |
| 152205858 | 152207057 | 1 | lncRNA | ENSG00000 |
| 152212076 | 152224193 | -1 | protein_coding | ENSG00000 |
| 152302175 | 152325203 | -1 | protein_coding | ENSG00000 |
| 152348735 | 152360006 | -1 | protein_coding | ENSG00000 |
| 152399577 | 152399769 | 1 | processed_pseudoge | ENSG00000 |
| 152409243 | 152414263 | -1 | protein_coding | ENSG00000 |
| 152510803 | 152512177 | 1 | protein_coding | ENSG00000 |

| | | | | |
|---|---|---|---|---|
| 152514482 | 152516008 | 1 | protein_coding | ENSG00000 |
| 152565654 | 152566780 | -1 | protein_coding | ENSG00000 |
| 152579381 | 152580516 | -1 | protein_coding | ENSG00000 |
| 152600662 | 152601086 | 1 | protein_coding | ENSG00000 |
| 152613811 | 152614098 | 1 | protein_coding | ENSG00000 |
| 152622834 | 152623103 | -1 | protein_coding | ENSG00000 |
| 152644393 | 152644717 | -1 | unprocessed_pseudc | ENSG00000 |
| 152656332 | 152656501 | 1 | unprocessed_pseudc | ENSG00000 |
| 152663380 | 152664659 | 1 | protein_coding | ENSG00000 |
| 152675295 | 152676574 | 1 | protein_coding | ENSG00000 |
| 152686123 | 152687397 | 1 | protein_coding | ENSG00000 |
| 152698345 | 152699442 | 1 | protein_coding | ENSG00000 |
| 152708160 | 152709491 | 1 | protein_coding | ENSG00000 |
| 152719522 | 152720470 | 1 | protein_coding | ENSG00000 |
| 152737518 | 152737868 | 1 | unprocessed_pseudc | ENSG00000 |
| 152744299 | 152744606 | 1 | unprocessed_pseudc | ENSG00000 |
| 152759561 | 152762052 | 1 | protein_coding | ENSG00000 |
| 152776372 | 152776728 | 1 | protein_coding | ENSG00000 |
| 152786214 | 152788426 | 1 | protein_coding | ENSG00000 |
| 152796721 | 152798181 | 1 | protein_coding | ENSG00000 |
| 152804835 | 152806628 | -1 | protein_coding | ENSG00000 |
| 152811971 | 152813108 | 1 | protein_coding | ENSG00000 |
| 152827473 | 152828097 | 1 | protein_coding | ENSG00000 |
| 152842856 | 152843983 | 1 | protein_coding | ENSG00000 |
| 152859996 | 152860985 | 1 | protein_coding | ENSG00000 |
| 152878322 | 152885047 | 1 | protein_coding | ENSG00000 |
| 152908546 | 152911886 | 1 | protein_coding | ENSG00000 |
| 152930040 | 152949210 | -1 | lncRNA | ENSG00000 |
| 152947154 | 152949258 | 1 | protein_coding | ENSG00000 |
| 152970648 | 152972574 | 1 | protein_coding | ENSG00000 |
| 152985231 | 152985500 | 1 | protein_coding | ENSG00000 |
| 153001747 | 153003856 | 1 | protein_coding | ENSG00000 |
| 153012482 | 153012614 | -1 | snoRNA | ENSG00000 |
| 153023962 | 153024260 | 1 | unprocessed_pseudc | ENSG00000 |
| 153031203 | 153032900 | 1 | protein_coding | ENSG00000 |
| 153039732 | 153041931 | -1 | protein_coding | ENSG00000 |
| 153056120 | 153057512 | -1 | protein_coding | ENSG00000 |
| 153070224 | 153070840 | -1 | protein_coding | ENSG00000 |
| 153093135 | 153106184 | -1 | protein_coding | ENSG00000 |
| 153112121 | 153113516 | -1 | protein_coding | ENSG00000 |
| 153140491 | 153140709 | -1 | unprocessed_pseudc | ENSG00000 |
| 153149582 | 153150890 | -1 | protein_coding | ENSG00000 |
| 153174518 | 153191676 | 1 | lncRNA | ENSG00000 |
| 153203430 | 153205120 | 1 | protein_coding | ENSG00000 |
| 153217584 | 153219310 | 1 | protein_coding | ENSG00000 |
| 153225080 | 153226006 | 1 | processed_pseudoge | ENSG00000 |
| 153259687 | 153262124 | 1 | protein_coding | ENSG00000 |
| 153297862 | 153310718 | -1 | protein_coding | ENSG00000 |
| 153330120 | 153348841 | -1 | protein_coding | ENSG00000 |
| 153331622 | 153331728 | 1 | snRNA | ENSG00000 |

| | | | | |
|---|---|---|---|---|
| 153357854 | 153361023 | 1 | protein_coding | ENSG0000C |
| 153373711 | 153375621 | -1 | protein_coding | ENSG0000C |
| 153379821 | 153380799 | -1 | processed_pseudoge | ENSG0000C |
| 153390032 | 153391073 | -1 | protein_coding | ENSG0000C |
| 153396591 | 153396771 | -1 | processed_pseudoge | ENSG0000C |
| 153416520 | 153423222 | 1 | protein_coding | ENSG0000C |
| 153427020 | 153428401 | -1 | unprocessed_pseudc | ENSG0000C |
| 153437058 | 153439949 | -1 | protein_coding | ENSG0000C |
| 153457744 | 153460651 | -1 | protein_coding | ENSG0000C |
| 153500463 | 153500764 | -1 | misc_RNA | ENSG0000C |
| 153533603 | 153535115 | 1 | lncRNA | ENSG0000C |
| 153534599 | 153536244 | -1 | protein_coding | ENSG0000C |
| 153537147 | 153541765 | -1 | protein_coding | ENSG0000C |
| 153543613 | 153550136 | -1 | protein_coding | ENSG0000C |
| 153547329 | 153549258 | -1 | protein_coding | ENSG0000C |
| 153561108 | 153567890 | -1 | protein_coding | ENSG0000C |
| 153586813 | 153606283 | -1 | lncRNA | ENSG0000C |
| 153606886 | 153613145 | -1 | protein_coding | ENSG0000C |
| 153614255 | 153616986 | -1 | protein_coding | ENSG0000C |
| 153618787 | 153631360 | -1 | protein_coding | ENSG0000C |
| 153626332 | 153634340 | -1 | lncRNA | ENSG0000C |
| 153627926 | 153632039 | 1 | protein_coding | ENSG0000C |
| 153631438 | 153634397 | -1 | lncRNA | ENSG0000C |
| 153633982 | 153646306 | 1 | protein_coding | ENSG0000C |
| 153658703 | 153661852 | 1 | protein_coding | ENSG0000C |
| 153661788 | 153671028 | -1 | protein_coding | ENSG0000C |
| 153678688 | 153693992 | 1 | protein_coding | ENSG0000C |
| 153689705 | 153689793 | -1 | miRNA | ENSG0000C |
| 153704088 | 153704385 | 1 | misc_RNA | ENSG0000C |
| 153717303 | 153718347 | -1 | processed_pseudoge | ENSG0000C |
| 153726252 | 153726353 | -1 | misc_RNA | ENSG0000C |
| 153728050 | 153774808 | 1 | protein_coding | ENSG0000C |
| 153746851 | 153751227 | -1 | lncRNA | ENSG0000C |
| 153750983 | 153752176 | -1 | lncRNA | ENSG0000C |
| 153774354 | 153780157 | 1 | protein_coding | ENSG0000C |
| 153785720 | 153785821 | -1 | misc_RNA | ENSG0000C |
| 153789030 | 153923360 | -1 | protein_coding | ENSG0000C |
| 153793947 | 153802091 | 1 | transcribed_unproce | ENSG0000C |
| 153852106 | 153853414 | -1 | processed_pseudoge | ENSG0000C |
| 153890595 | 153890865 | 1 | processed_pseudoge | ENSG0000C |
| 153923337 | 153935240 | 1 | lncRNA | ENSG0000C |
| 153929501 | 153946718 | -1 | protein_coding | ENSG0000C |
| 153947669 | 153958615 | -1 | protein_coding | ENSG0000C |
| 153959099 | 153968184 | -1 | protein_coding | ENSG0000C |
| 153959151 | 153975887 | -1 | protein_coding | ENSG0000C |
| 153962351 | 153962420 | -1 | miRNA | ENSG0000C |
| 153964361 | 153965070 | 1 | lncRNA | ENSG0000C |
| 153966516 | 153966930 | 1 | lncRNA | ENSG0000C |
| 153967534 | 153974361 | 1 | protein_coding | ENSG0000C |
| 153974269 | 153977674 | -1 | protein_coding | ENSG0000C |

| | | | | |
|---|---|---|---|---|
| 153975850 | 153986246 | -1 | protein_coding | ENSG0000C |
| 153977743 | 153979160 | 1 | lncRNA | ENSG0000C |
| 153981605 | 153986358 | -1 | protein_coding | ENSG0000C |
| 153990762 | 153992155 | 1 | protein_coding | ENSG0000C |
| 153992685 | 154155116 | -1 | protein_coding | ENSG0000C |
| 153995632 | 153995960 | 1 | processed_pseudoge | ENSG0000C |
| 154039916 | 154040022 | -1 | snRNA | ENSG0000C |
| 154078866 | 154079963 | 1 | processed_pseudoge | ENSG0000C |
| 154104521 | 154104592 | -1 | miRNA | ENSG0000C |
| 154155304 | 154194648 | -1 | protein_coding | ENSG0000C |
| 154166245 | 154166549 | -1 | misc_RNA | ENSG0000C |
| 154193665 | 154193743 | -1 | miRNA | ENSG0000C |
| 154199085 | 154206333 | -1 | protein_coding | ENSG0000C |
| 154206696 | 154220637 | -1 | protein_coding | ENSG0000C |
| 154220179 | 154271510 | 1 | protein_coding | ENSG0000C |
| 154259727 | 154259862 | 1 | snoRNA | ENSG0000C |
| 154272511 | 154275875 | 1 | protein_coding | ENSG0000C |
| 154288460 | 154288527 | 1 | snoRNA | ENSG0000C |
| 154295503 | 154295601 | 1 | snRNA | ENSG0000C |
| 154297650 | 154297748 | 1 | snRNA | ENSG0000C |
| 154312462 | 154313084 | 1 | unprocessed_pseudc | ENSG0000C |
| 154321090 | 154325325 | 1 | protein_coding | ENSG0000C |
| 154325553 | 154351304 | 1 | protein_coding | ENSG0000C |
| 154338743 | 154338802 | 1 | snRNA | ENSG0000C |
| 154376966 | 154379273 | -1 | unprocessed_pseudc | ENSG0000C |
| 154378207 | 154379089 | -1 | processed_pseudoge | ENSG0000C |
| 154402328 | 154406564 | -1 | lncRNA | ENSG0000C |
| 154405193 | 154469450 | 1 | protein_coding | ENSG0000C |
| 154414369 | 154415137 | -1 | processed_pseudoge | ENSG0000C |
| 154469772 | 154502412 | -1 | protein_coding | ENSG0000C |
| 154480012 | 154481501 | 1 | lncRNA | ENSG0000C |
| 154502219 | 154548147 | 1 | protein_coding | ENSG0000C |
| 154548577 | 154559028 | -1 | protein_coding | ENSG0000C |
| 154553609 | 154555017 | 1 | lncRNA | ENSG0000C |
| 154564855 | 154567748 | -1 | lncRNA | ENSG0000C |
| 154567778 | 154580013 | 1 | protein_coding | ENSG0000C |
| 154579065 | 154579663 | -1 | lncRNA | ENSG0000C |
| 154582057 | 154628013 | -1 | protein_coding | ENSG0000C |
| 154671593 | 154678345 | 1 | lncRNA | ENSG0000C |
| 154697455 | 154870281 | -1 | protein_coding | ENSG0000C |
| 154924740 | 154936719 | -1 | protein_coding | ENSG0000C |
| 154937370 | 154938059 | 1 | lncRNA | ENSG0000C |
| 154944076 | 154956123 | -1 | protein_coding | ENSG0000C |
| 154950210 | 154950492 | -1 | misc_RNA | ENSG0000C |
| 154957026 | 154963853 | -1 | protein_coding | ENSG0000C |
| 154961825 | 154962623 | 1 | lncRNA | ENSG0000C |
| 154962298 | 154974395 | -1 | protein_coding | ENSG0000C |
| 154974653 | 154979251 | 1 | protein_coding | ENSG0000C |
| 154975693 | 154975783 | 1 | miRNA | ENSG0000C |
| 154983338 | 154993111 | 1 | protein_coding | ENSG0000C |

| | | | | |
|---|---|---|---|---|
| 154993586 | 154994315 | 1 | protein_coding | ENSG00000 |
| 155002630 | 155018522 | 1 | protein_coding | ENSG00000 |
| 155018520 | 155033781 | -1 | protein_coding | ENSG00000 |
| 155033824 | 155050930 | 1 | protein_coding | ENSG00000 |
| 155045191 | 155046118 | -1 | lncRNA | ENSG00000 |
| 155050566 | 155062775 | 1 | protein_coding | ENSG00000 |
| 155063737 | 155069553 | 1 | protein_coding | ENSG00000 |
| 155063748 | 155086807 | 1 | protein_coding | ENSG00000 |
| 155078837 | 155087538 | 1 | protein_coding | ENSG00000 |
| 155120490 | 155120598 | 1 | misc_RNA | ENSG00000 |
| 155127876 | 155134899 | 1 | protein_coding | ENSG00000 |
| 155135344 | 155138857 | 1 | protein_coding | ENSG00000 |
| 155139891 | 155140595 | -1 | protein_coding | ENSG00000 |
| 155148544 | 155148813 | 1 | processed_pseudoge | ENSG00000 |
| 155169408 | 155173475 | -1 | protein_coding | ENSG00000 |
| 155169409 | 155187272 | -1 | protein_coding | ENSG00000 |
| 155173787 | 155184971 | 1 | protein_coding | ENSG00000 |
| 155185824 | 155192916 | -1 | protein_coding | ENSG00000 |
| 155195004 | 155205495 | 1 | lncRNA | ENSG00000 |
| 155195177 | 155195272 | 1 | miRNA | ENSG00000 |
| 155195588 | 155209051 | -1 | protein_coding | ENSG00000 |
| 155208699 | 155213824 | 1 | protein_coding | ENSG00000 |
| 155211151 | 155213819 | -1 | lncRNA | ENSG00000 |
| 155213821 | 155227422 | -1 | transcribed_unproce | ENSG00000 |
| 155230975 | 155234325 | 1 | unprocessed_pseudo | ENSG00000 |
| 155234452 | 155244699 | -1 | protein_coding | ENSG00000 |
| 155247205 | 155255483 | -1 | protein_coding | ENSG00000 |
| 155255979 | 155262430 | -1 | protein_coding | ENSG00000 |
| 155262868 | 155278491 | -1 | protein_coding | ENSG00000 |
| 155277463 | 155289848 | 1 | protein_coding | ENSG00000 |
| 155289293 | 155301438 | -1 | protein_coding | ENSG00000 |
| 155308748 | 155320666 | 1 | protein_coding | ENSG00000 |
| 155316863 | 155324176 | -1 | lncRNA | ENSG00000 |
| 155320894 | 155331114 | 1 | protein_coding | ENSG00000 |
| 155335268 | 155562807 | -1 | protein_coding | ENSG00000 |
| 155346350 | 155346445 | -1 | miRNA | ENSG00000 |
| 155358712 | 155358818 | -1 | snRNA | ENSG00000 |
| 155396010 | 155396978 | -1 | lncRNA | ENSG00000 |
| 155419397 | 155419500 | 1 | snRNA | ENSG00000 |
| 155433178 | 155434262 | 1 | processed_pseudoge | ENSG00000 |
| 155562042 | 155563944 | 1 | lncRNA | ENSG00000 |
| 155566050 | 155566306 | -1 | processed_pseudoge | ENSG00000 |
| 155586644 | 155602197 | 1 | unprocessed_pseudo | ENSG00000 |
| 155590601 | 155591262 | -1 | processed_pseudoge | ENSG00000 |
| 155609776 | 155610380 | -1 | lncRNA | ENSG00000 |
| 155610205 | 155614967 | 1 | protein_coding | ENSG00000 |
| 155614726 | 155660245 | -1 | unprocessed_pseudo | ENSG00000 |
| 155626757 | 155637604 | -1 | lncRNA | ENSG00000 |
| 155659443 | 155689000 | -1 | protein_coding | ENSG00000 |
| 155687960 | 155739010 | 1 | protein_coding | ENSG00000 |

| | | | | |
|---|---|---|---|---|
| 155710098 | 155710563 | -1 | lncRNA | ENSG00000 |
| 155745829 | 155750137 | 1 | unprocessed_pseudo | ENSG00000 |
| 155749662 | 155859400 | -1 | protein_coding | ENSG00000 |
| 155845367 | 155846624 | 1 | processed_pseudoge | ENSG00000 |
| 155859567 | 155885199 | 1 | protein_coding | ENSG00000 |
| 155894281 | 155894386 | 1 | snRNA | ENSG00000 |
| 155897808 | 155911404 | -1 | protein_coding | ENSG00000 |
| 155913045 | 155934413 | -1 | protein_coding | ENSG00000 |
| 155919909 | 155920045 | -1 | snoRNA | ENSG00000 |
| 155925958 | 155926086 | -1 | scaRNA | ENSG00000 |
| 155925958 | 155926085 | -1 | scaRNA | ENSG00000 |
| 155941710 | 155942949 | 1 | protein_coding | ENSG00000 |
| 155946851 | 156007070 | -1 | protein_coding | ENSG00000 |
| 155951273 | 155951336 | -1 | miRNA | ENSG00000 |
| 155978799 | 155982986 | 1 | lncRNA | ENSG00000 |
| 155991390 | 156001787 | 1 | lncRNA | ENSG00000 |
| 156001953 | 156033342 | 1 | lncRNA | ENSG00000 |
| 156009048 | 156020951 | -1 | protein_coding | ENSG00000 |
| 156035299 | 156053798 | -1 | protein_coding | ENSG00000 |
| 156054752 | 156058510 | 1 | protein_coding | ENSG00000 |
| 156061160 | 156070504 | 1 | protein_coding | ENSG00000 |
| 156072013 | 156082465 | -1 | protein_coding | ENSG00000 |
| 156082573 | 156140089 | 1 | protein_coding | ENSG00000 |
| 156147366 | 156177752 | 1 | protein_coding | ENSG00000 |
| 156192063 | 156192203 | -1 | snoRNA | ENSG00000 |
| 156193932 | 156212796 | 1 | protein_coding | ENSG00000 |
| 156212982 | 156243332 | 1 | protein_coding | ENSG00000 |
| 156212993 | 156240042 | 1 | protein_coding | ENSG00000 |
| 156242184 | 156243317 | 1 | protein_coding | ENSG00000 |
| 156243320 | 156248117 | -1 | protein_coding | ENSG00000 |
| 156249224 | 156282825 | -1 | protein_coding | ENSG00000 |
| 156282935 | 156293185 | 1 | protein_coding | ENSG00000 |
| 156290089 | 156295689 | -1 | protein_coding | ENSG00000 |
| 156298624 | 156299299 | -1 | protein_coding | ENSG00000 |
| 156308968 | 156367873 | -1 | protein_coding | ENSG00000 |
| 156337314 | 156346995 | 1 | protein_coding | ENSG00000 |
| 156369211 | 156385219 | 1 | protein_coding | ENSG00000 |
| 156388226 | 156395609 | 1 | lncRNA | ENSG00000 |
| 156404250 | 156456763 | -1 | protein_coding | ENSG00000 |
| 156420341 | 156420429 | -1 | miRNA | ENSG00000 |
| 156463727 | 156500779 | -1 | protein_coding | ENSG00000 |
| 156484098 | 156484210 | -1 | misc_RNA | ENSG00000 |
| 156509854 | 156511681 | 1 | lncRNA | ENSG00000 |
| 156525405 | 156572604 | -1 | protein_coding | ENSG00000 |
| 156579727 | 156586770 | 1 | protein_coding | ENSG00000 |
| 156591762 | 156594299 | 1 | protein_coding | ENSG00000 |
| 156594487 | 156601496 | -1 | protein_coding | ENSG00000 |
| 156614742 | 156615288 | -1 | lncRNA | ENSG00000 |
| 156619331 | 156625725 | 1 | protein_coding | ENSG00000 |
| 156637783 | 156641004 | -1 | lncRNA | ENSG00000 |

| | | | | |
|---|---|---|---|---|
| 156641390 | 156659532 | 1 | protein_coding | ENSG0000C |
| 156641666 | 156644887 | -1 | lncRNA | ENSG0000C |
| 156646507 | 156661424 | -1 | lncRNA | ENSG0000C |
| 156668763 | 156677407 | -1 | protein_coding | ENSG0000C |
| 156687695 | 156691997 | -1 | lncRNA | ENSG0000C |
| 156689676 | 156704757 | 1 | lncRNA | ENSG0000C |
| 156699606 | 156705816 | -1 | protein_coding | ENSG0000C |
| 156712212 | 156713174 | -1 | lncRNA | ENSG0000C |
| 156721891 | 156728799 | -1 | protein_coding | ENSG0000C |
| 156728442 | 156736960 | 1 | protein_coding | ENSG0000C |
| 156737303 | 156741590 | -1 | protein_coding | ENSG0000C |
| 156742109 | 156766925 | -1 | protein_coding | ENSG0000C |
| 156750610 | 156800815 | 1 | protein_coding | ENSG0000C |
| 156768105 | 156769233 | -1 | lncRNA | ENSG0000C |
| 156806243 | 156816862 | -1 | protein_coding | ENSG0000C |
| 156815640 | 156881850 | 1 | protein_coding | ENSG0000C |
| 156840063 | 156859117 | -1 | protein_coding | ENSG0000C |
| 156893698 | 156916434 | 1 | protein_coding | ENSG0000C |
| 156920632 | 156933094 | 1 | protein_coding | ENSG0000C |
| 156934840 | 157045370 | -1 | protein_coding | ENSG0000C |
| 156936131 | 156936244 | -1 | miRNA | ENSG0000C |
| 156985755 | 156986049 | -1 | misc_RNA | ENSG0000C |
| 157059232 | 157060762 | -1 | processed_pseudoge | ENSG0000C |
| 157073257 | 157074703 | 1 | processed_pseudoge | ENSG0000C |
| 157092043 | 157112412 | -1 | protein_coding | ENSG0000C |
| 157121191 | 157138474 | -1 | protein_coding | ENSG0000C |
| 157128362 | 157128671 | 1 | processed_pseudoge | ENSG0000C |
| 157171116 | 157191963 | 1 | transcribed_unproce | ENSG0000C |
| 157204779 | 157205040 | -1 | unprocessed_pseudc | ENSG0000C |
| 157232231 | 157237136 | -1 | lncRNA | ENSG0000C |
| 157273760 | 157283968 | 1 | lncRNA | ENSG0000C |
| 157287267 | 157310655 | -1 | unprocessed_pseudc | ENSG0000C |
| 157400927 | 157417259 | 1 | unprocessed_pseudc | ENSG0000C |
| 157437484 | 157437754 | 1 | unprocessed_pseudc | ENSG0000C |
| 157513377 | 157552515 | -1 | protein_coding | ENSG0000C |
| 157573747 | 157598085 | -1 | protein_coding | ENSG0000C |
| 157629939 | 157630728 | -1 | unprocessed_pseudc | ENSG0000C |
| 157636300 | 157640301 | -1 | unprocessed_pseudc | ENSG0000C |
| 157674321 | 157700769 | -1 | protein_coding | ENSG0000C |
| 157691762 | 157696459 | 1 | lncRNA | ENSG0000C |
| 157709086 | 157709643 | 1 | processed_pseudoge | ENSG0000C |
| 157724180 | 157725020 | -1 | processed_pseudoge | ENSG0000C |
| 157745733 | 157777132 | -1 | protein_coding | ENSG0000C |
| 157794403 | 157820120 | -1 | protein_coding | ENSG0000C |
| 157830911 | 157898256 | -1 | protein_coding | ENSG0000C |
| 157861197 | 157861460 | 1 | processed_pseudoge | ENSG0000C |
| 157925065 | 157949071 | 1 | transcribed_unproce | ENSG0000C |
| 157993273 | 158100262 | 1 | protein_coding | ENSG0000C |
| 158025550 | 158031166 | 1 | lncRNA | ENSG0000C |
| 158127287 | 158128301 | -1 | protein_coding | ENSG0000C |

| | | | | |
|---|---|---|---|---|
| 158131983 | 158149810 | -1 | lncRNA | ENSG0000C |
| 158175850 | 158177755 | -1 | processed_pseudoge | ENSG0000C |
| 158179947 | 158184896 | 1 | protein_coding | ENSG0000C |
| 158195633 | 158196131 | -1 | processed_pseudoge | ENSG0000C |
| 158197922 | 158203877 | -1 | lncRNA | ENSG0000C |
| 158254424 | 158258269 | 1 | protein_coding | ENSG0000C |
| 158266753 | 158267320 | -1 | processed_pseudoge | ENSG0000C |
| 158289923 | 158294774 | 1 | protein_coding | ENSG0000C |
| 158327951 | 158331531 | -1 | protein_coding | ENSG0000C |
| 158353696 | 158357553 | 1 | protein_coding | ENSG0000C |
| 158398522 | 158399466 | -1 | protein_coding | ENSG0000C |
| 158418210 | 158426237 | -1 | protein_coding | ENSG0000C |
| 158445068 | 158446014 | -1 | unprocessed_pseudc | ENSG0000C |
| 158454198 | 158455273 | 1 | processed_pseudoge | ENSG0000C |
| 158461574 | 158470857 | 1 | protein_coding | ENSG0000C |
| 158472220 | 158480936 | 1 | protein_coding | ENSG0000C |
| 158474454 | 158494886 | -1 | lncRNA | ENSG0000C |
| 158491219 | 158492160 | 1 | unprocessed_pseudc | ENSG0000C |
| 158514785 | 158515925 | 1 | unprocessed_pseudc | ENSG0000C |
| 158523672 | 158525838 | 1 | processed_pseudoge | ENSG0000C |
| 158544550 | 158554405 | -1 | protein_coding | ENSG0000C |
| 158560606 | 158570580 | -1 | protein_coding | ENSG0000C |
| 158578919 | 158579899 | -1 | protein_coding | ENSG0000C |
| 158605268 | 158612514 | 1 | protein_coding | ENSG0000C |
| 158610704 | 158686715 | -1 | protein_coding | ENSG0000C |
| 158694539 | 158695509 | -1 | unprocessed_pseudc | ENSG0000C |
| 158699678 | 158700652 | -1 | protein_coding | ENSG0000C |
| 158716327 | 158720720 | -1 | protein_coding | ENSG0000C |
| 158724113 | 158725067 | -1 | unprocessed_pseudc | ENSG0000C |
| 158742146 | 158743415 | -1 | unprocessed_pseudc | ENSG0000C |
| 158747814 | 158772195 | -1 | protein_coding | ENSG0000C |
| 158754720 | 158755891 | 1 | protein_coding | ENSG0000C |
| 158774222 | 158781204 | -1 | protein_coding | ENSG0000C |
| 158796014 | 158796213 | 1 | unprocessed_pseudc | ENSG0000C |
| 158808399 | 158809335 | -1 | unprocessed_pseudc | ENSG0000C |
| 158831351 | 158849506 | 1 | protein_coding | ENSG0000C |
| 158878746 | 158883500 | 1 | unprocessed_pseudc | ENSG0000C |
| 158930796 | 158977059 | 1 | protein_coding | ENSG0000C |
| 158999968 | 159055155 | 1 | protein_coding | ENSG0000C |
| 159062484 | 159147096 | -1 | protein_coding | ENSG0000C |
| 159081133 | 159081795 | -1 | processed_pseudoge | ENSG0000C |
| 159171609 | 159203313 | 1 | protein_coding | ENSG0000C |
| 159178473 | 159178559 | 1 | rRNA_pseudogene | ENSG0000C |
| 159194325 | 159207973 | -1 | lncRNA | ENSG0000C |
| 159203307 | 159206500 | 1 | protein_coding | ENSG0000C |
| 159276503 | 159277111 | 1 | processed_pseudoge | ENSG0000C |
| 159279041 | 159280001 | -1 | unprocessed_pseudc | ENSG0000C |
| 159289714 | 159308224 | 1 | protein_coding | ENSG0000C |
| 159313670 | 159314659 | -1 | polymorphic_pseudc | ENSG0000C |
| 159346166 | 159469068 | -1 | lncRNA | ENSG0000C |

| | | | | |
|---|---|---|---|---|
| 159351093 | 159352026 | 1 | unitary_pseudogene | ENSG00000 |
| 159366161 | 159367089 | 1 | unprocessed_pseudo | ENSG00000 |
| 159405423 | 159406409 | -1 | unprocessed_pseudo | ENSG00000 |
| 159432204 | 159433138 | 1 | polymorphic_pseudo | ENSG00000 |
| 159437845 | 159443078 | 1 | protein_coding | ENSG00000 |
| 159466321 | 159483376 | 1 | lncRNA | ENSG00000 |
| 159535078 | 159536007 | -1 | protein_coding | ENSG00000 |
| 159557368 | 159558200 | 1 | unprocessed_pseudo | ENSG00000 |
| 159581620 | 159582648 | -1 | unprocessed_pseudo | ENSG00000 |
| 159587826 | 159588865 | 1 | protein_coding | ENSG00000 |
| 159598298 | 159599227 | 1 | unprocessed_pseudo | ENSG00000 |
| 159649151 | 159649620 | 1 | processed_pseudoge | ENSG00000 |
| 159704983 | 159705607 | -1 | unprocessed_pseudo | ENSG00000 |
| 159712289 | 159714589 | -1 | protein_coding | ENSG00000 |
| 159759170 | 159759589 | 1 | processed_pseudoge | ENSG00000 |
| 159780932 | 159782543 | 1 | protein_coding | ENSG00000 |
| 159800511 | 159816257 | 1 | protein_coding | ENSG00000 |
| 159826811 | 159837492 | 1 | protein_coding | ENSG00000 |
| 159834474 | 159873053 | -1 | protein_coding | ENSG00000 |
| 159834495 | 159855237 | -1 | lncRNA | ENSG00000 |
| 159851906 | 159851972 | -1 | snoRNA | ENSG00000 |
| 159854316 | 159862657 | -1 | protein_coding | ENSG00000 |
| 159854870 | 159867685 | 1 | lncRNA | ENSG00000 |
| 159872364 | 159900165 | -1 | protein_coding | ENSG00000 |
| 159899979 | 159900079 | -1 | miRNA | ENSG00000 |
| 159900475 | 159903431 | 1 | lncRNA | ENSG00000 |
| 159910094 | 159910554 | -1 | TEC | ENSG00000 |
| 159918107 | 159925732 | -1 | protein_coding | ENSG00000 |
| 159927039 | 159945613 | -1 | protein_coding | ENSG00000 |
| 159951492 | 159954237 | -1 | protein_coding | ENSG00000 |
| 159958035 | 159984750 | 1 | lncRNA | ENSG00000 |
| 159972548 | 159974064 | -1 | unprocessed_pseudo | ENSG00000 |
| 159998651 | 160070160 | -1 | protein_coding | ENSG00000 |
| 160020300 | 160020727 | 1 | processed_pseudoge | ENSG00000 |
| 160024953 | 160031990 | -1 | protein_coding | ENSG00000 |
| 160062461 | 160080769 | 1 | lncRNA | ENSG00000 |
| 160081538 | 160090563 | 1 | protein_coding | ENSG00000 |
| 160091340 | 160098943 | -1 | protein_coding | ENSG00000 |
| 160115759 | 160143591 | 1 | protein_coding | ENSG00000 |
| 160151570 | 160186977 | 1 | protein_coding | ENSG00000 |
| 160190575 | 160201886 | 1 | protein_coding | ENSG00000 |
| 160202199 | 160208869 | -1 | lncRNA | ENSG00000 |
| 160205337 | 160215376 | 1 | protein_coding | ENSG00000 |
| 160215715 | 160262531 | -1 | protein_coding | ENSG00000 |
| 160216800 | 160285130 | -1 | protein_coding | ENSG00000 |
| 160261741 | 160263328 | 1 | lncRNA | ENSG00000 |
| 160266340 | 160267225 | -1 | processed_pseudoge | ENSG00000 |
| 160276812 | 160286348 | -1 | protein_coding | ENSG00000 |
| 160288594 | 160343273 | -1 | protein_coding | ENSG00000 |
| 160317403 | 160317706 | 1 | processed_pseudoge | ENSG00000 |

| | | | | |
|---|---|---|---|---|
| 160326104 | 160326216 | -1 | misc_RNA | ENSG0000C |
| 160343316 | 160358952 | 1 | protein_coding | ENSG0000C |
| 160367071 | 160372846 | 1 | protein_coding | ENSG0000C |
| 160392768 | 160392909 | -1 | snRNA | ENSG0000C |
| 160400564 | 160428670 | 1 | protein_coding | ENSG0000C |
| 160485030 | 160523262 | -1 | protein_coding | ENSG0000C |
| 160537073 | 160571458 | 1 | lncRNA | ENSG0000C |
| 160541095 | 160579516 | -1 | protein_coding | ENSG0000C |
| 160608106 | 160647295 | -1 | protein_coding | ENSG0000C |
| 160670148 | 160670979 | 1 | transcribed_processe | ENSG0000C |
| 160670778 | 160699761 | 1 | lncRNA | ENSG0000C |
| 160678746 | 160711831 | -1 | protein_coding | ENSG0000C |
| 160739057 | 160754821 | 1 | protein_coding | ENSG0000C |
| 160775954 | 160776608 | -1 | unprocessed_pseudc | ENSG0000C |
| 160776975 | 160780140 | -1 | unprocessed_pseudc | ENSG0000C |
| 160796074 | 160828261 | 1 | protein_coding | ENSG0000C |
| 160830160 | 160862887 | -1 | protein_coding | ENSG0000C |
| 160848010 | 160848502 | 1 | processed_pseudoge | ENSG0000C |
| 160876540 | 160885180 | -1 | protein_coding | ENSG0000C |
| 160894980 | 160896076 | 1 | processed_pseudoge | ENSG0000C |
| 160931739 | 160934380 | -1 | lncRNA | ENSG0000C |
| 160932465 | 160949922 | 1 | lncRNA | ENSG0000C |
| 160935537 | 160936126 | 1 | processed_pseudoge | ENSG0000C |
| 160945025 | 160954809 | -1 | protein_coding | ENSG0000C |
| 160995211 | 161021343 | -1 | protein_coding | ENSG0000C |
| 160997957 | 161038962 | -1 | protein_coding | ENSG0000C |
| 161034834 | 161035006 | 1 | processed_pseudoge | ENSG0000C |
| 161037631 | 161038977 | -1 | protein_coding | ENSG0000C |
| 161039251 | 161045977 | -1 | protein_coding | ENSG0000C |
| 161046946 | 161069970 | -1 | protein_coding | ENSG0000C |
| 161070998 | 161089558 | -1 | protein_coding | ENSG0000C |
| 161084465 | 161087571 | 1 | lncRNA | ENSG0000C |
| 161098361 | 161100346 | 1 | protein_coding | ENSG0000C |
| 161100556 | 161118055 | -1 | protein_coding | ENSG0000C |
| 161118086 | 161125445 | 1 | protein_coding | ENSG0000C |
| 161120974 | 161132688 | -1 | protein_coding | ENSG0000C |
| 161141208 | 161141336 | -1 | snoRNA | ENSG0000C |
| 161152776 | 161158856 | 1 | protein_coding | ENSG0000C |
| 161153760 | 161159349 | -1 | lncRNA | ENSG0000C |
| 161159450 | 161165723 | 1 | protein_coding | ENSG0000C |
| 161166056 | 161178013 | 1 | protein_coding | ENSG0000C |
| 161171310 | 161177968 | -1 | protein_coding | ENSG0000C |
| 161184302 | 161199054 | -1 | protein_coding | ENSG0000C |
| 161197104 | 161214395 | 1 | protein_coding | ENSG0000C |
| 161215234 | 161220699 | 1 | protein_coding | ENSG0000C |
| 161222292 | 161223631 | -1 | protein_coding | ENSG0000C |
| 161225939 | 161230746 | 1 | protein_coding | ENSG0000C |
| 161227186 | 161227261 | 1 | miRNA | ENSG0000C |
| 161229666 | 161238302 | -1 | protein_coding | ENSG0000C |
| 161258745 | 161285450 | 1 | protein_coding | ENSG0000C |

| | | | | |
|---|---|---|---|---|
| 161304735 | 161309972 | -1 | protein_coding | ENSG0000C |
| 161314257 | 161375340 | 1 | protein_coding | ENSG0000C |
| 161364731 | 161367874 | -1 | protein_coding | ENSG0000C |
| 161378707 | 161379358 | -1 | processed_pseudoge | ENSG0000C |
| 161399409 | 161422424 | 1 | lncRNA | ENSG0000C |
| 161399998 | 161401868 | 1 | lncRNA | ENSG0000C |
| 161401289 | 161401395 | 1 | snRNA | ENSG0000C |
| 161403409 | 161470523 | -1 | lncRNA | ENSG0000C |
| 161406068 | 161407082 | 1 | processed_pseudoge | ENSG0000C |
| 161411597 | 161411802 | -1 | processed_pseudoge | ENSG0000C |
| 161433444 | 161440996 | -1 | lncRNA | ENSG0000C |
| 161505430 | 161524013 | 1 | protein_coding | ENSG0000C |
| 161513176 | 161605099 | 1 | lncRNA | ENSG0000C |
| 161524540 | 161526894 | 1 | protein_coding | ENSG0000C |
| 161536571 | 161538158 | -1 | unprocessed_pseudc | ENSG0000C |
| 161541759 | 161550737 | -1 | protein_coding | ENSG0000C |
| 161556290 | 161557078 | -1 | lncRNA | ENSG0000C |
| 161581339 | 161605662 | 1 | polymorphic_pseudc | ENSG0000C |
| 161606291 | 161608217 | 1 | unprocessed_pseudc | ENSG0000C |
| 161617992 | 161619585 | -1 | unprocessed_pseudc | ENSG0000C |
| 161623196 | 161631963 | -1 | protein_coding | ENSG0000C |
| 161663147 | 161678654 | 1 | protein_coding | ENSG0000C |
| 161671978 | 161674824 | -1 | lncRNA | ENSG0000C |
| 161683695 | 161685252 | -1 | transcribed_processe | ENSG0000C |
| 161699506 | 161699607 | 1 | misc_RNA | ENSG0000C |
| 161706972 | 161714352 | 1 | protein_coding | ENSG0000C |
| 161721563 | 161728143 | 1 | protein_coding | ENSG0000C |
| 161735808 | 161736102 | 1 | misc_RNA | ENSG0000C |
| 161749758 | 161757238 | 1 | protein_coding | ENSG0000C |
| 161765325 | 161766227 | -1 | lncRNA | ENSG0000C |
| 161766320 | 161964070 | 1 | protein_coding | ENSG0000C |
| 161890833 | 161892196 | 1 | transcribed_processe | ENSG0000C |
| 161983192 | 162023854 | -1 | protein_coding | ENSG0000C |
| 162039016 | 162039567 | 1 | lncRNA | ENSG0000C |
| 162069691 | 162370475 | 1 | protein_coding | ENSG0000C |
| 162146709 | 162175243 | -1 | lncRNA | ENSG0000C |
| 162157107 | 162157182 | 1 | miRNA | ENSG0000C |
| 162316852 | 162317794 | -1 | lncRNA | ENSG0000C |
| 162338643 | 162338742 | 1 | rRNA_pseudogene | ENSG0000C |
| 162342546 | 162342640 | 1 | miRNA | ENSG0000C |
| 162365407 | 162383531 | 1 | protein_coding | ENSG0000C |
| 162373203 | 162376854 | -1 | protein_coding | ENSG0000C |
| 162378841 | 162386812 | 1 | protein_coding | ENSG0000C |
| 162395268 | 162412138 | -1 | protein_coding | ENSG0000C |
| 162445549 | 162446566 | 1 | unprocessed_pseudc | ENSG0000C |
| 162497251 | 162529631 | 1 | protein_coding | ENSG0000C |
| 162541332 | 162541628 | -1 | processed_pseudoge | ENSG0000C |
| 162560227 | 162561308 | -1 | lncRNA | ENSG0000C |
| 162561506 | 162599842 | 1 | protein_coding | ENSG0000C |
| 162593103 | 162593754 | 1 | lncRNA | ENSG0000C |

| | | | | |
|---|---|---|---|---|
| 162631373 | 162787405 | 1 | protein_coding | ENSG00000( |
| 162777730 | 162778014 | 1 | misc_RNA | ENSG00000( |
| 162790702 | 162812817 | 1 | protein_coding | ENSG00000( |
| 162824458 | 162868815 | -1 | protein_coding | ENSG00000( |
| 162824795 | 162849467 | 1 | unprocessed_pseudc | ENSG00000( |
| 162979551 | 162979793 | -1 | processed_pseudoge | ENSG00000( |
| 163068775 | 163076802 | 1 | protein_coding | ENSG00000( |
| 163111121 | 163321791 | -1 | protein_coding | ENSG00000( |
| 163161675 | 163213023 | 1 | lncRNA | ENSG00000( |
| 163237214 | 163239197 | 1 | processed_pseudoge | ENSG00000( |
| 163244505 | 163321894 | -1 | lncRNA | ENSG00000( |
| 163259850 | 163260659 | 1 | lncRNA | ENSG00000( |
| 163266576 | 163355764 | 1 | protein_coding | ENSG00000( |
| 163422405 | 163423164 | -1 | unprocessed_pseudc | ENSG00000( |
| 163468496 | 163468605 | -1 | rRNA_pseudogene | ENSG00000( |
| 163509484 | 163509595 | 1 | rRNA_pseudogene | ENSG00000( |
| 163769339 | 163769691 | -1 | processed_pseudoge | ENSG00000( |
| 163923670 | 163923873 | 1 | snoRNA | ENSG00000( |
| 164343005 | 164343864 | 1 | processed_pseudoge | ENSG00000( |
| 164351273 | 164351388 | 1 | snRNA | ENSG00000( |
| 164356767 | 164357364 | 1 | processed_pseudoge | ENSG00000( |
| 164555584 | 164899296 | 1 | protein_coding | ENSG00000( |
| 164639565 | 164639671 | 1 | snRNA | ENSG00000( |
| 164680085 | 164680799 | 1 | lncRNA | ENSG00000( |
| 164703026 | 164703309 | -1 | misc_RNA | ENSG00000( |
| 164769116 | 164774641 | -1 | lncRNA | ENSG00000( |
| 164828436 | 164829952 | 1 | lncRNA | ENSG00000( |
| 164854231 | 164854332 | 1 | misc_RNA | ENSG00000( |
| 164921318 | 164921629 | -1 | processed_pseudoge | ENSG00000( |
| 164980035 | 164980137 | -1 | snRNA | ENSG00000( |
| 165201867 | 165356715 | -1 | protein_coding | ENSG00000( |
| 165210627 | 165213090 | 1 | lncRNA | ENSG00000( |
| 165215951 | 165219104 | 1 | lncRNA | ENSG00000( |
| 165400922 | 165445355 | -1 | protein_coding | ENSG00000( |
| 165476838 | 165582155 | -1 | lncRNA | ENSG00000( |
| 165497724 | 165498348 | 1 | processed_pseudoge | ENSG00000( |
| 165544007 | 165563961 | 1 | protein_coding | ENSG00000( |
| 165581613 | 165582476 | 1 | processed_pseudoge | ENSG00000( |
| 165598356 | 165624084 | 1 | lncRNA | ENSG00000( |
| 165631213 | 165661796 | 1 | protein_coding | ENSG00000( |
| 165662216 | 165698863 | -1 | protein_coding | ENSG00000( |
| 165662585 | 165662687 | -1 | misc_RNA | ENSG00000( |
| 165671256 | 165671725 | 1 | processed_pseudoge | ENSG00000( |
| 165676310 | 165676781 | 1 | processed_pseudoge | ENSG00000( |
| 165698750 | 165709968 | 1 | transcribed_processe | ENSG00000( |
| 165706556 | 165707284 | -1 | lncRNA | ENSG00000( |
| 165724293 | 165827755 | -1 | protein_coding | ENSG00000( |
| 165768929 | 165775176 | 1 | lncRNA | ENSG00000( |
| 165819353 | 165819787 | 1 | processed_pseudoge | ENSG00000( |
| 165827614 | 165911618 | 1 | protein_coding | ENSG00000( |

| | | | | |
|---|---|---|---|---|
| 165889725 | 165900995 | -1 | lncRNA | ENSG00000( |
| 165907921 | 165907976 | 1 | miRNA | ENSG00000( |
| 165941235 | 165941769 | -1 | processed_pseudoge | ENSG00000( |
| 166022215 | 166023027 | 1 | processed_pseudoge | ENSG00000( |
| 166042244 | 166042350 | 1 | rRNA_pseudogene | ENSG00000( |
| 166057426 | 166166969 | -1 | protein_coding | ENSG00000( |
| 166081183 | 166087483 | 1 | lncRNA | ENSG00000( |
| 166147782 | 166165095 | -1 | lncRNA | ENSG00000( |
| 166154743 | 166154798 | -1 | miRNA | ENSG00000( |
| 166165911 | 166339090 | 1 | lncRNA | ENSG00000( |
| 166275629 | 166277597 | -1 | processed_pseudoge | ENSG00000( |
| 166387727 | 166452632 | -1 | lncRNA | ENSG00000( |
| 166474745 | 166481565 | 1 | unprocessed_pseudc | ENSG00000( |
| 166474879 | 166490307 | -1 | lncRNA | ENSG00000( |
| 166566178 | 166580705 | 1 | unprocessed_pseudc | ENSG00000( |
| 166603916 | 166631400 | 1 | transcribed_unproce | ENSG00000( |
| 166665885 | 166682196 | 1 | unprocessed_pseudc | ENSG00000( |
| 166747379 | 166748660 | 1 | processed_pseudoge | ENSG00000( |
| 166763334 | 166792767 | 1 | unprocessed_pseudc | ENSG00000( |
| 166796266 | 166797183 | 1 | processed_pseudoge | ENSG00000( |
| 166839447 | 166856344 | 1 | protein_coding | ENSG00000( |
| 166856510 | 166876264 | -1 | protein_coding | ENSG00000( |
| 166868748 | 166869209 | 1 | processed_pseudoge | ENSG00000( |
| 166895711 | 166975540 | -1 | protein_coding | ENSG00000( |
| 166975582 | 167022214 | 1 | protein_coding | ENSG00000( |
| 167005959 | 167006077 | 1 | rRNA_pseudogene | ENSG00000( |
| 167052551 | 167058542 | 1 | lncRNA | ENSG00000( |
| 167052836 | 167166479 | -1 | protein_coding | ENSG00000( |
| 167094075 | 167129165 | 1 | protein_coding | ENSG00000( |
| 167162423 | 167162837 | -1 | processed_pseudoge | ENSG00000( |
| 167175363 | 167195792 | -1 | lncRNA | ENSG00000( |
| 167219831 | 167220512 | -1 | lncRNA | ENSG00000( |
| 167220876 | 167427345 | 1 | protein_coding | ENSG00000( |
| 167379108 | 167381000 | 1 | lncRNA | ENSG00000( |
| 167430640 | 167518610 | -1 | protein_coding | ENSG00000( |
| 167455195 | 167458646 | 1 | lncRNA | ENSG00000( |
| 167457383 | 167458661 | -1 | lncRNA | ENSG00000( |
| 167457742 | 167459891 | -1 | lncRNA | ENSG00000( |
| 167519536 | 167520586 | 1 | processed_pseudoge | ENSG00000( |
| 167529117 | 167553805 | -1 | protein_coding | ENSG00000( |
| 167591392 | 167591846 | 1 | processed_pseudoge | ENSG00000( |
| 167627385 | 167630674 | -1 | lncRNA | ENSG00000( |
| 167630093 | 167708696 | 1 | protein_coding | ENSG00000( |
| 167721192 | 167791919 | 1 | protein_coding | ENSG00000( |
| 167809386 | 167914215 | -1 | protein_coding | ENSG00000( |
| 167819898 | 167820248 | -1 | unprocessed_pseudc | ENSG00000( |
| 167820406 | 167821224 | 1 | lncRNA | ENSG00000( |
| 167916675 | 167937072 | -1 | protein_coding | ENSG00000( |
| 167935783 | 168075843 | 1 | protein_coding | ENSG00000( |
| 167998660 | 167998726 | 1 | miRNA | ENSG00000( |

| | | | | |
|---|---|---|---|---|
| 168055901 | 168056422 | -1 | processed_pseudoge | ENSG00000( |
| 168079543 | 168137667 | -1 | protein_coding | ENSG00000( |
| 168178962 | 168202109 | 1 | protein_coding | ENSG00000( |
| 168210616 | 168210969 | 1 | processed_pseudoge | ENSG00000( |
| 168215405 | 168215652 | 1 | processed_pseudoge | ENSG00000( |
| 168225938 | 168253021 | 1 | protein_coding | ENSG00000( |
| 168245565 | 168247343 | -1 | transcribed_processe | ENSG00000( |
| 168263375 | 168263481 | -1 | snRNA | ENSG00000( |
| 168280877 | 168314426 | 1 | protein_coding | ENSG00000( |
| 168317497 | 168376876 | 1 | transcribed_processe | ENSG00000( |
| 168375524 | 168375621 | 1 | miRNA | ENSG00000( |
| 168400829 | 168495685 | -1 | lncRNA | ENSG00000( |
| 168401483 | 168407274 | -1 | lncRNA | ENSG00000( |
| 168449672 | 168451211 | -1 | processed_pseudoge | ENSG00000( |
| 168540768 | 168543997 | -1 | protein_coding | ENSG00000( |
| 168542737 | 168543354 | 1 | processed_pseudoge | ENSG00000( |
| 168576605 | 168582069 | 1 | protein_coding | ENSG00000( |
| 168578653 | 168579439 | -1 | processed_pseudoge | ENSG00000( |
| 168695468 | 168729206 | -1 | protein_coding | ENSG00000( |
| 168695874 | 169103276 | -1 | lncRNA | ENSG00000( |
| 168763365 | 168774490 | -1 | lncRNA | ENSG00000( |
| 168784012 | 168792886 | 1 | lncRNA | ENSG00000( |
| 168898136 | 168898439 | 1 | processed_pseudoge | ENSG00000( |
| 168898633 | 168902836 | 1 | lncRNA | ENSG00000( |
| 168903905 | 169087005 | -1 | lncRNA | ENSG00000( |
| 168938467 | 168938923 | 1 | processed_pseudoge | ENSG00000( |
| 169059576 | 169062441 | 1 | lncRNA | ENSG00000( |
| 169067264 | 169067364 | 1 | rRNA_pseudogene | ENSG00000( |
| 169104124 | 169104907 | 1 | lncRNA | ENSG00000( |
| 169105697 | 169132722 | 1 | protein_coding | ENSG00000( |
| 169132531 | 169367948 | -1 | protein_coding | ENSG00000( |
| 169310665 | 169322479 | 1 | lncRNA | ENSG00000( |
| 169367970 | 169396540 | 1 | protein_coding | ENSG00000( |
| 169394870 | 169460669 | -1 | protein_coding | ENSG00000( |
| 169463909 | 169485944 | -1 | protein_coding | ENSG00000( |
| 169474060 | 169474159 | -1 | processed_pseudoge | ENSG00000( |
| 169486076 | 169500182 | 1 | lncRNA | ENSG00000( |
| 169511953 | 169586588 | -1 | protein_coding | ENSG00000( |
| 169588849 | 169630193 | -1 | protein_coding | ENSG00000( |
| 169662007 | 169854080 | 1 | protein_coding | ENSG00000( |
| 169690665 | 169711702 | -1 | protein_coding | ENSG00000( |
| 169722640 | 169764705 | -1 | protein_coding | ENSG00000( |
| 169762929 | 169764648 | -1 | lncRNA | ENSG00000( |
| 169792529 | 169794963 | -1 | protein_coding | ENSG00000( |
| 169849631 | 169894267 | -1 | protein_coding | ENSG00000( |
| 169859756 | 169860052 | 1 | misc_RNA | ENSG00000( |
| 169915004 | 169915198 | -1 | processed_pseudoge | ENSG00000( |
| 169921326 | 170085208 | -1 | protein_coding | ENSG00000( |
| 169957942 | 169958246 | -1 | misc_RNA | ENSG00000( |
| 169990067 | 169990196 | 1 | processed_pseudoge | ENSG00000( |

| | | | | |
|---|---|---|---|---|
| 170024077 | 170024683 | 1 | lncRNA | ENSG0000C |
| 170115636 | 170116576 | 1 | processed_pseudoge | ENSG0000C |
| 170146001 | 170167790 | 1 | protein_coding | ENSG0000C |
| 170151378 | 170151462 | -1 | miRNA | ENSG0000C |
| 170151378 | 170151462 | 1 | miRNA | ENSG0000C |
| 170173865 | 170304358 | 1 | lncRNA | ENSG0000C |
| 170211010 | 170211457 | -1 | processed_pseudoge | ENSG0000C |
| 170271395 | 170284277 | -1 | lncRNA | ENSG0000C |
| 170369223 | 170370367 | -1 | processed_pseudoge | ENSG0000C |
| 170460453 | 170532647 | -1 | lncRNA | ENSG0000C |
| 170532129 | 170553446 | 1 | protein_coding | ENSG0000C |
| 170587249 | 170588236 | -1 | lncRNA | ENSG0000C |
| 170598854 | 170647339 | 1 | lncRNA | ENSG0000C |
| 170662728 | 170739421 | 1 | protein_coding | ENSG0000C |
| 170667381 | 170669425 | 1 | lncRNA | ENSG0000C |
| 170748573 | 171251857 | -1 | lncRNA | ENSG0000C |
| 170935471 | 171064765 | 1 | protein_coding | ENSG0000C |
| 171083565 | 171085205 | 1 | processed_pseudoge | ENSG0000C |
| 171090901 | 171117819 | 1 | protein_coding | ENSG0000C |
| 171101728 | 171101806 | -1 | miRNA | ENSG0000C |
| 171101739 | 171101798 | 1 | miRNA | ENSG0000C |
| 171137740 | 171161568 | 1 | transcribed_unitary_ | ENSG0000C |
| 171185249 | 171212686 | 1 | protein_coding | ENSG0000C |
| 171199244 | 171227788 | -1 | lncRNA | ENSG0000C |
| 171248471 | 171285978 | 1 | protein_coding | ENSG0000C |
| 171253906 | 171254022 | -1 | misc_RNA | ENSG0000C |
| 171270954 | 171271590 | -1 | processed_pseudoge | ENSG0000C |
| 171314183 | 171342084 | 1 | protein_coding | ENSG0000C |
| 171339299 | 171339959 | 1 | processed_pseudoge | ENSG0000C |
| 171345105 | 171345307 | -1 | processed_pseudoge | ENSG0000C |
| 171392229 | 171392790 | 1 | processed_pseudoge | ENSG0000C |
| 171444699 | 171444963 | -1 | processed_pseudoge | ENSG0000C |
| 171485551 | 171593511 | 1 | protein_coding | ENSG0000C |
| 171600621 | 171638799 | 1 | protein_coding | ENSG0000C |
| 171635417 | 171652688 | -1 | protein_coding | ENSG0000C |
| 171670517 | 171670939 | -1 | processed_pseudoge | ENSG0000C |
| 171683128 | 171684438 | -1 | processed_pseudoge | ENSG0000C |
| 171700160 | 171742074 | -1 | protein_coding | ENSG0000C |
| 171751543 | 171752025 | -1 | unprocessed_pseudc | ENSG0000C |
| 171755803 | 171756323 | -1 | processed_pseudoge | ENSG0000C |
| 171762074 | 171762524 | 1 | processed_pseudoge | ENSG0000C |
| 171768070 | 171768175 | 1 | scaRNA | ENSG0000C |
| 171781660 | 171814023 | 1 | protein_coding | ENSG0000C |
| 171803517 | 171803939 | 1 | processed_pseudoge | ENSG0000C |
| 171824610 | 171824959 | -1 | processed_pseudoge | ENSG0000C |
| 171841498 | 172418466 | 1 | protein_coding | ENSG0000C |
| 171864187 | 171864687 | 1 | lncRNA | ENSG0000C |
| 172138397 | 172144840 | -1 | lncRNA | ENSG0000C |
| 172138798 | 172138907 | -1 | miRNA | ENSG0000C |
| 172138808 | 172138888 | 1 | miRNA | ENSG0000C |

| | | | | |
|---|---|---|---|---|
| 172144535 | 172144644 | -1 | miRNA | ENSG0000C |
| 172210711 | 172213499 | -1 | lncRNA | ENSG0000C |
| 172366540 | 172366646 | 1 | snRNA | ENSG0000C |
| 172370189 | 172444086 | -1 | protein_coding | ENSG0000C |
| 172420685 | 172468831 | 1 | protein_coding | ENSG0000C |
| 172532349 | 172611833 | 1 | protein_coding | ENSG0000C |
| 172613428 | 172613534 | -1 | snRNA | ENSG0000C |
| 172659103 | 172666876 | 1 | protein_coding | ENSG0000C |
| 172748560 | 172749502 | 1 | processed_pseudoge | ENSG0000C |
| 172752586 | 172752924 | 1 | TEC | ENSG0000C |
| 172775905 | 173064015 | 1 | lncRNA | ENSG0000C |
| 172885947 | 172886258 | -1 | processed_pseudoge | ENSG0000C |
| 172906900 | 172907684 | 1 | lncRNA | ENSG0000C |
| 173039960 | 173050963 | -1 | protein_coding | ENSG0000C |
| 173141100 | 173142350 | -1 | processed_pseudoge | ENSG0000C |
| 173174300 | 173175503 | -1 | lncRNA | ENSG0000C |
| 173183731 | 173207331 | -1 | protein_coding | ENSG0000C |
| 173351689 | 173352004 | 1 | processed_pseudoge | ENSG0000C |
| 173362397 | 173363733 | -1 | processed_pseudoge | ENSG0000C |
| 173417793 | 173489407 | -1 | lncRNA | ENSG0000C |
| 173477330 | 173488815 | 1 | protein_coding | ENSG0000C |
| 173500460 | 173603072 | -1 | protein_coding | ENSG0000C |
| 173555251 | 173612772 | 1 | lncRNA | ENSG0000C |
| 173596060 | 173669837 | -1 | protein_coding | ENSG0000C |
| 173608336 | 173669851 | -1 | protein_coding | ENSG0000C |
| 173635338 | 173637130 | 1 | protein_coding | ENSG0000C |
| 173714941 | 173786692 | 1 | protein_coding | ENSG0000C |
| 173741674 | 173741928 | 1 | processed_pseudoge | ENSG0000C |
| 173791548 | 173791887 | 1 | misc_RNA | ENSG0000C |
| 173799550 | 173824720 | -1 | protein_coding | ENSG0000C |
| 173808489 | 173808622 | 1 | misc_RNA | ENSG0000C |
| 173824653 | 173858808 | 1 | protein_coding | ENSG0000C |
| 173858559 | 173868882 | -1 | lncRNA | ENSG0000C |
| 173862473 | 173863941 | 1 | lncRNA | ENSG0000C |
| 173868082 | 173903549 | 1 | protein_coding | ENSG0000C |
| 173903804 | 173917378 | -1 | protein_coding | ENSG0000C |
| 173921070 | 173921200 | 1 | rRNA_pseudogene | ENSG0000C |
| 173931214 | 174022297 | -1 | protein_coding | ENSG0000C |
| 173969318 | 173969436 | -1 | rRNA_pseudogene | ENSG0000C |
| 174009267 | 174016206 | -1 | lncRNA | ENSG0000C |
| 174022509 | 174022985 | 1 | lncRNA | ENSG0000C |
| 174090136 | 174090474 | -1 | processed_pseudoge | ENSG0000C |
| 174110268 | 174160165 | -1 | lncRNA | ENSG0000C |
| 174159410 | 174995308 | 1 | protein_coding | ENSG0000C |
| 174367105 | 174368039 | -1 | processed_pseudoge | ENSG0000C |
| 174447964 | 174449545 | 1 | protein_coding | ENSG0000C |
| 174756850 | 174757119 | -1 | processed_pseudoge | ENSG0000C |
| 174849667 | 174850168 | -1 | processed_pseudoge | ENSG0000C |
| 174892417 | 174893155 | 1 | processed_pseudoge | ENSG0000C |
| 174896958 | 174897996 | 1 | lncRNA | ENSG0000C |

| | | | | |
|---|---|---|---|---|
| 174922107 | 174922440 | -1 | processed_pseudoge | ENSG0000C |
| 174934947 | 174954261 | -1 | lncRNA | ENSG0000C |
| 174996524 | 174996629 | 1 | snRNA | ENSG0000C |
| 174998353 | 174999623 | -1 | lncRNA | ENSG0000C |
| 174999163 | 175012027 | 1 | protein_coding | ENSG0000C |
| 175010789 | 175023425 | -1 | protein_coding | ENSG0000C |
| 175022479 | 175022580 | -1 | misc_RNA | ENSG0000C |
| 175044626 | 175045648 | -1 | processed_pseudoge | ENSG0000C |
| 175067833 | 175148075 | 1 | protein_coding | ENSG0000C |
| 175156986 | 175192999 | -1 | protein_coding | ENSG0000C |
| 175203228 | 175204849 | -1 | lncRNA | ENSG0000C |
| 175297080 | 175297240 | 1 | processed_pseudoge | ENSG0000C |
| 175307218 | 175335459 | 1 | lncRNA | ENSG0000C |
| 175315194 | 175743616 | -1 | protein_coding | ENSG0000C |
| 175538775 | 175556818 | -1 | lncRNA | ENSG0000C |
| 175877343 | 175880468 | 1 | lncRNA | ENSG0000C |
| 175904762 | 175921057 | -1 | lncRNA | ENSG0000C |
| 175921975 | 175922145 | 1 | processed_pseudoge | ENSG0000C |
| 175944831 | 176207286 | -1 | protein_coding | ENSG0000C |
| 175968398 | 175968540 | -1 | scaRNA | ENSG0000C |
| 176017277 | 176018760 | 1 | lncRNA | ENSG0000C |
| 176207648 | 176447919 | 1 | lncRNA | ENSG0000C |
| 176217541 | 176217602 | 1 | snRNA | ENSG0000C |
| 176231200 | 176231830 | -1 | processed_pseudoge | ENSG0000C |
| 176243862 | 176244029 | -1 | snRNA | ENSG0000C |
| 176272483 | 176273402 | 1 | unprocessed_pseudc | ENSG0000C |
| 176305672 | 176306095 | -1 | processed_pseudoge | ENSG0000C |
| 176367699 | 176368401 | -1 | processed_pseudoge | ENSG0000C |
| 176463171 | 176845601 | 1 | protein_coding | ENSG0000C |
| 176616273 | 176616786 | -1 | processed_pseudoge | ENSG0000C |
| 176829128 | 176831841 | -1 | lncRNA | ENSG0000C |
| 176857302 | 177164973 | -1 | protein_coding | ENSG0000C |
| 177029363 | 177029445 | -1 | miRNA | ENSG0000C |
| 177170958 | 177282422 | 1 | protein_coding | ENSG0000C |
| 177351562 | 177437880 | 1 | lncRNA | ENSG0000C |
| 177392667 | 177920152 | -1 | lncRNA | ENSG0000C |
| 177700524 | 177712275 | 1 | lncRNA | ENSG0000C |
| 177923956 | 177984303 | -1 | protein_coding | ENSG0000C |
| 177928788 | 178038007 | -1 | lncRNA | ENSG0000C |
| 178006136 | 178037950 | -1 | transcribed_unitary_ | ENSG0000C |
| 178017127 | 178017343 | 1 | processed_pseudoge | ENSG0000C |
| 178090677 | 178093984 | -1 | lncRNA | ENSG0000C |
| 178094141 | 178484147 | 1 | protein_coding | ENSG0000C |
| 178194342 | 178194709 | -1 | processed_pseudoge | ENSG0000C |
| 178411616 | 178411972 | 1 | processed_pseudoge | ENSG0000C |
| 178479247 | 178499634 | -1 | protein_coding | ENSG0000C |
| 178484408 | 178484682 | 1 | misc_RNA | ENSG0000C |
| 178511563 | 178513051 | -1 | lncRNA | ENSG0000C |
| 178513109 | 178548602 | 1 | protein_coding | ENSG0000C |
| 178542752 | 178548889 | 1 | lncRNA | ENSG0000C |

| | | | | |
|---|---|---|---|---|
| 178560913 | 178561029 | 1 | rRNA_pseudogene | ENSG00000 |
| 178651706 | 178652282 | 1 | lncRNA | ENSG00000 |
| 178677749 | 178677834 | 1 | miRNA | ENSG00000 |
| 178724306 | 178726285 | -1 | lncRNA | ENSG00000 |
| 178725147 | 178921841 | 1 | protein_coding | ENSG00000 |
| 178746683 | 178747754 | -1 | processed_pseudoge | ENSG00000 |
| 178753654 | 178753772 | -1 | snoRNA | ENSG00000 |
| 178849535 | 178871077 | -1 | protein_coding | ENSG00000 |
| 179025804 | 179076567 | 1 | protein_coding | ENSG00000 |
| 179035309 | 179035652 | -1 | processed_pseudoge | ENSG00000 |
| 179082070 | 179098023 | 1 | protein_coding | ENSG00000 |
| 179099330 | 179229684 | -1 | protein_coding | ENSG00000 |
| 179137764 | 179137934 | 1 | processed_pseudoge | ENSG00000 |
| 179183734 | 179184547 | 1 | processed_pseudoge | ENSG00000 |
| 179196473 | 179196579 | 1 | snoRNA | ENSG00000 |
| 179201487 | 179201627 | 1 | snoRNA | ENSG00000 |
| 179201705 | 179201926 | 1 | processed_pseudoge | ENSG00000 |
| 179220938 | 179222125 | 1 | processed_pseudoge | ENSG00000 |
| 179255733 | 179256081 | 1 | processed_pseudoge | ENSG00000 |
| 179271116 | 179271266 | -1 | processed_pseudoge | ENSG00000 |
| 179293714 | 179358680 | 1 | protein_coding | ENSG00000 |
| 179365720 | 179554735 | 1 | protein_coding | ENSG00000 |
| 179447578 | 179448440 | -1 | processed_pseudoge | ENSG00000 |
| 179447602 | 179457315 | -1 | processed_pseudoge | ENSG00000 |
| 179543201 | 179548922 | -1 | lncRNA | ENSG00000 |
| 179550539 | 179575952 | -1 | protein_coding | ENSG00000 |
| 179576268 | 179576349 | 1 | snRNA | ENSG00000 |
| 179586705 | 179589175 | 1 | processed_pseudoge | ENSG00000 |
| 179590372 | 179591305 | 1 | lncRNA | ENSG00000 |
| 179591613 | 179691272 | 1 | protein_coding | ENSG00000 |
| 179730191 | 179742697 | 1 | lncRNA | ENSG00000 |
| 179743163 | 179816198 | 1 | protein_coding | ENSG00000 |
| 179816184 | 179818191 | -1 | lncRNA | ENSG00000 |
| 179829609 | 179836124 | -1 | lncRNA | ENSG00000 |
| 179839967 | 179877803 | -1 | protein_coding | ENSG00000 |
| 179881607 | 179882595 | -1 | lncRNA | ENSG00000 |
| 179882042 | 179925000 | 1 | protein_coding | ENSG00000 |
| 179900262 | 179900564 | 1 | misc_RNA | ENSG00000 |
| 179926641 | 179941796 | -1 | lncRNA | ENSG00000 |
| 179953184 | 179954440 | -1 | lncRNA | ENSG00000 |
| 179954773 | 180114875 | 1 | protein_coding | ENSG00000 |
| 179968686 | 179969506 | 1 | processed_pseudoge | ENSG00000 |
| 180000438 | 180000808 | -1 | processed_pseudoge | ENSG00000 |
| 180117140 | 180123890 | -1 | lncRNA | ENSG00000 |
| 180154869 | 180204030 | 1 | protein_coding | ENSG00000 |
| 180230264 | 180278984 | 1 | protein_coding | ENSG00000 |
| 180269653 | 180502954 | -1 | protein_coding | ENSG00000 |
| 180434800 | 180435936 | 1 | processed_pseudoge | ENSG00000 |
| 180438314 | 180438390 | -1 | miRNA | ENSG00000 |
| 180519016 | 180519119 | -1 | misc_RNA | ENSG00000 |

| | | | | |
|---|---|---|---|---|
| 180547436 | 180566520 | 1 | lncRNA | ENSG0000C |
| 180632022 | 180890279 | 1 | protein_coding | ENSG0000C |
| 180758722 | 180758817 | 1 | snRNA | ENSG0000C |
| 180906651 | 180909166 | 1 | lncRNA | ENSG0000C |
| 180912897 | 180951614 | 1 | protein_coding | ENSG0000C |
| 180944042 | 180976482 | -1 | lncRNA | ENSG0000C |
| 180949699 | 180954887 | -1 | lncRNA | ENSG0000C |
| 180964511 | 180965909 | 1 | unprocessed_pseudc | ENSG0000C |
| 180970837 | 180971247 | 1 | processed_pseudoge | ENSG0000C |
| 180972712 | 181023121 | -1 | protein_coding | ENSG0000C |
| 181033425 | 181061938 | 1 | protein_coding | ENSG0000C |
| 181088700 | 181092900 | 1 | protein_coding | ENSG0000C |
| 181174484 | 181182208 | 1 | lncRNA | ENSG0000C |
| 181190471 | 181191149 | -1 | lncRNA | ENSG0000C |
| 181236388 | 181238604 | 1 | lncRNA | ENSG0000C |
| 181317690 | 181808084 | 1 | protein_coding | ENSG0000C |
| 181668749 | 181669028 | 1 | misc_RNA | ENSG0000C |
| 181771566 | 181771644 | 1 | rRNA_pseudogene | ENSG0000C |
| 181808927 | 181998985 | -1 | lncRNA | ENSG0000C |
| 181812705 | 181813261 | 1 | lncRNA | ENSG0000C |
| 181839473 | 181839774 | -1 | misc_RNA | ENSG0000C |
| 182054570 | 182061712 | -1 | protein_coding | ENSG0000C |
| 182062677 | 182069253 | -1 | lncRNA | ENSG0000C |
| 182086551 | 182090112 | 1 | lncRNA | ENSG0000C |
| 182096338 | 182314061 | -1 | lncRNA | ENSG0000C |
| 182127297 | 182128682 | -1 | lncRNA | ENSG0000C |
| 182182730 | 182183096 | -1 | processed_pseudoge | ENSG0000C |
| 182327068 | 182327174 | 1 | snRNA | ENSG0000C |
| 182328497 | 182329007 | 1 | processed_pseudoge | ENSG0000C |
| 182336001 | 182336334 | 1 | processed_pseudoge | ENSG0000C |
| 182381704 | 182392206 | -1 | protein_coding | ENSG0000C |
| 182398117 | 182400667 | -1 | protein_coding | ENSG0000C |
| 182407621 | 182414815 | 1 | lncRNA | ENSG0000C |
| 182409192 | 182560599 | 1 | protein_coding | ENSG0000C |
| 182433893 | 182434461 | -1 | processed_pseudoge | ENSG0000C |
| 182441577 | 182442362 | 1 | unitary_pseudogene | ENSG0000C |
| 182573634 | 182589256 | -1 | protein_coding | ENSG0000C |
| 182576772 | 182577040 | -1 | misc_RNA | ENSG0000C |
| 182598623 | 182604389 | -1 | protein_coding | ENSG0000C |
| 182615254 | 182616629 | 1 | lncRNA | ENSG0000C |
| 182641816 | 182684576 | -1 | protein_coding | ENSG0000C |
| 182712862 | 182714544 | 1 | lncRNA | ENSG0000C |
| 182733792 | 182734779 | 1 | processed_pseudoge | ENSG0000C |
| 182789293 | 182830384 | 1 | protein_coding | ENSG0000C |
| 182837185 | 182839561 | -1 | lncRNA | ENSG0000C |
| 182839347 | 182887982 | 1 | protein_coding | ENSG0000C |
| 182899865 | 182953525 | -1 | protein_coding | ENSG0000C |
| 182942115 | 182942404 | -1 | processed_pseudoge | ENSG0000C |
| 182944365 | 182944490 | -1 | rRNA_pseudogene | ENSG0000C |
| 182944365 | 182944490 | -1 | rRNA | ENSG0000C |

| | | | | |
|---|---|---|---|---|
| 182955390 | 182956148 | -1 | processed_pseudoge | ENSG00000 |
| 182959074 | 182960405 | -1 | processed_pseudoge | ENSG00000 |
| 182982212 | 182982318 | -1 | snRNA | ENSG00000 |
| 183023420 | 183145592 | 1 | protein_coding | ENSG00000 |
| 183138402 | 183141282 | -1 | lncRNA | ENSG00000 |
| 183186238 | 183245127 | 1 | protein_coding | ENSG00000 |
| 183248237 | 183418380 | -1 | protein_coding | ENSG00000 |
| 183252263 | 183258968 | 1 | lncRNA | ENSG00000 |
| 183266602 | 183267285 | 1 | processed_pseudoge | ENSG00000 |
| 183372870 | 183374556 | 1 | lncRNA | ENSG00000 |
| 183460874 | 183472265 | -1 | lncRNA | ENSG00000 |
| 183472216 | 183598246 | 1 | protein_coding | ENSG00000 |
| 183555563 | 183590876 | -1 | protein_coding | ENSG00000 |
| 183613537 | 183619335 | 1 | lncRNA | ENSG00000 |
| 183620846 | 183635783 | -1 | protein_coding | ENSG00000 |
| 183636085 | 183928531 | 1 | protein_coding | ENSG00000 |
| 183646275 | 183653316 | -1 | protein_coding | ENSG00000 |
| 183709305 | 183710240 | -1 | processed_pseudoge | ENSG00000 |
| 183754418 | 183754937 | -1 | lncRNA | ENSG00000 |
| 183929854 | 184037729 | -1 | protein_coding | ENSG00000 |
| 184051651 | 184123978 | 1 | protein_coding | ENSG00000 |
| 184080657 | 184149652 | -1 | lncRNA | ENSG00000 |
| 184171714 | 184171823 | -1 | misc_RNA | ENSG00000 |
| 184315658 | 184315765 | 1 | misc_RNA | ENSG00000 |
| 184329071 | 184332826 | 1 | lncRNA | ENSG00000 |
| 184335658 | 184335949 | 1 | misc_RNA | ENSG00000 |
| 184385753 | 184386704 | -1 | lncRNA | ENSG00000 |
| 184387029 | 184629019 | 1 | protein_coding | ENSG00000 |
| 184408337 | 184412360 | 1 | lncRNA | ENSG00000 |
| 184566511 | 184567835 | 1 | processed_pseudoge | ENSG00000 |
| 184607599 | 184672166 | -1 | lncRNA | ENSG00000 |
| 184664282 | 184668535 | 1 | lncRNA | ENSG00000 |
| 184690237 | 184754907 | -1 | protein_coding | ENSG00000 |
| 184790724 | 184974508 | -1 | protein_coding | ENSG00000 |
| 184821428 | 184821489 | -1 | snRNA | ENSG00000 |
| 184999710 | 185008789 | 1 | lncRNA | ENSG00000 |
| 185014951 | 185015067 | -1 | rRNA_pseudogene | ENSG00000 |
| 185045526 | 185102603 | 1 | protein_coding | ENSG00000 |
| 185071567 | 185072035 | -1 | processed_pseudoge | ENSG00000 |
| 185118101 | 185157072 | -1 | protein_coding | ENSG00000 |
| 185157080 | 185291781 | 1 | protein_coding | ENSG00000 |
| 185171335 | 185171710 | -1 | processed_pseudoge | ENSG00000 |
| 185226808 | 185227700 | -1 | processed_pseudoge | ENSG00000 |
| 185251313 | 185251411 | 1 | misc_RNA | ENSG00000 |
| 185257911 | 185258015 | -1 | misc_RNA | ENSG00000 |
| 185262286 | 185264356 | -1 | processed_pseudoge | ENSG00000 |
| 185266535 | 185266627 | 1 | misc_RNA | ENSG00000 |
| 185280844 | 185281816 | 1 | processed_pseudoge | ENSG00000 |
| 185292384 | 185294372 | -1 | TEC | ENSG00000 |
| 185296388 | 185317273 | -1 | protein_coding | ENSG00000 |

| | | | | |
|---|---|---|---|---|
| 185317779 | 185318530 | 1 | lncRNA | ENSG0000C |
| 185321157 | 185335039 | -1 | transcribed_processe | ENSG0000C |
| 185434244 | 185434305 | 1 | snRNA | ENSG0000C |
| 185435839 | 185436047 | 1 | processed_pseudoge | ENSG0000C |
| 185518651 | 185520986 | 1 | TEC | ENSG0000C |
| 185558371 | 185628944 | -1 | lncRNA | ENSG0000C |
| 185630428 | 185630528 | -1 | misc_RNA | ENSG0000C |
| 185634073 | 185634182 | -1 | misc_RNA | ENSG0000C |
| 185646463 | 185657168 | 1 | lncRNA | ENSG0000C |
| 185734391 | 186190949 | 1 | protein_coding | ENSG0000C |
| 186176814 | 186177302 | -1 | lncRNA | ENSG0000C |
| 186224472 | 186228191 | 1 | lncRNA | ENSG0000C |
| 186296279 | 186314562 | 1 | protein_coding | ENSG0000C |
| 186311652 | 186375693 | -1 | protein_coding | ENSG0000C |
| 186311825 | 186311928 | -1 | snRNA | ENSG0000C |
| 186375838 | 186421378 | 1 | protein_coding | ENSG0000C |
| 186401046 | 186401180 | 1 | protein_coding | ENSG0000C |
| 186423481 | 186626619 | 1 | lncRNA | ENSG0000C |
| 186443566 | 186461114 | -1 | protein_coding | ENSG0000C |
| 186521773 | 186522304 | 1 | lncRNA | ENSG0000C |
| 186578279 | 186579299 | 1 | processed_pseudoge | ENSG0000C |
| 186580515 | 186581191 | -1 | processed_pseudoge | ENSG0000C |
| 186671791 | 186680423 | -1 | protein_coding | ENSG0000C |
| 186680622 | 186681446 | 1 | lncRNA | ENSG0000C |
| 186828949 | 186988981 | 1 | protein_coding | ENSG0000C |
| 187070700 | 187686628 | 1 | lncRNA | ENSG0000C |
| 187506838 | 187507119 | -1 | processed_pseudoge | ENSG0000C |
| 187563061 | 187564131 | 1 | processed_pseudoge | ENSG0000C |
| 187632166 | 187632439 | 1 | processed_pseudoge | ENSG0000C |
| 187706561 | 187708898 | 1 | unprocessed_pseudc | ENSG0000C |
| 187714243 | 187714735 | -1 | processed_pseudoge | ENSG0000C |
| 188013642 | 188144768 | 1 | lncRNA | ENSG0000C |
| 188067298 | 188067525 | -1 | processed_pseudoge | ENSG0000C |
| 188155839 | 188156180 | -1 | misc_RNA | ENSG0000C |
| 188218400 | 188220676 | 1 | lncRNA | ENSG0000C |
| 188242139 | 188242678 | -1 | processed_pseudoge | ENSG0000C |
| 188508538 | 188536465 | 1 | lncRNA | ENSG0000C |
| 188671353 | 188671759 | 1 | processed_pseudoge | ENSG0000C |
| 188694320 | 188695111 | 1 | processed_pseudoge | ENSG0000C |
| 188705623 | 188732208 | -1 | lncRNA | ENSG0000C |
| 188869474 | 188888540 | -1 | lncRNA | ENSG0000C |
| 188905688 | 189037564 | 1 | lncRNA | ENSG0000C |
| 189035961 | 189036444 | -1 | processed_pseudoge | ENSG0000C |
| 189132350 | 189133292 | -1 | processed_pseudoge | ENSG0000C |
| 189666149 | 189666256 | -1 | rRNA_pseudogene | ENSG0000C |
| 189775465 | 189814918 | 1 | lncRNA | ENSG0000C |
| 189868001 | 189868528 | -1 | lncRNA | ENSG0000C |
| 189868381 | 189876160 | 1 | lncRNA | ENSG0000C |
| 189989570 | 189993097 | -1 | processed_pseudoge | ENSG0000C |
| 190097658 | 190478404 | -1 | protein_coding | ENSG0000C |

| | | | | |
|---|---|---|---|---|
| 190264898 | 190362270 | 1 | lncRNA | ENSG0000C |
| 190478551 | 190480735 | 1 | lncRNA | ENSG0000C |
| 190480379 | 190494297 | 1 | lncRNA | ENSG0000C |
| 190624890 | 190801658 | 1 | lncRNA | ENSG0000C |
| 190781787 | 190781995 | 1 | processed_pseudoge | ENSG0000C |
| 190797524 | 190797956 | 1 | processed_pseudoge | ENSG0000C |
| 190878145 | 191151410 | -1 | lncRNA | ENSG0000C |
| 191146025 | 191146961 | -1 | processed_pseudoge | ENSG0000C |
| 191151510 | 191154634 | 1 | lncRNA | ENSG0000C |
| 191179521 | 191179739 | -1 | processed_pseudoge | ENSG0000C |
| 191221159 | 191228467 | -1 | lncRNA | ENSG0000C |
| 191823432 | 192011260 | 1 | lncRNA | ENSG0000C |
| 191858707 | 191890229 | -1 | lncRNA | ENSG0000C |
| 192158462 | 192185815 | 1 | protein_coding | ENSG0000C |
| 192167786 | 192957713 | -1 | lncRNA | ENSG0000C |
| 192246708 | 192247487 | -1 | processed_pseudoge | ENSG0000C |
| 192247505 | 192247947 | 1 | processed_pseudoge | ENSG0000C |
| 192316992 | 192367285 | 1 | protein_coding | ENSG0000C |
| 192517190 | 192538102 | 1 | lncRNA | ENSG0000C |
| 192575763 | 192580024 | 1 | protein_coding | ENSG0000C |
| 192636138 | 192660306 | 1 | protein_coding | ENSG0000C |
| 192716132 | 192716653 | 1 | protein_coding | ENSG0000C |
| 192716328 | 192716390 | 1 | miRNA | ENSG0000C |
| 192796533 | 192797205 | 1 | unprocessed_pseudc | ENSG0000C |
| 192800571 | 192801141 | 1 | processed_pseudoge | ENSG0000C |
| 192809039 | 192812275 | 1 | protein_coding | ENSG0000C |
| 192875686 | 192875985 | -1 | misc_RNA | ENSG0000C |
| 192993449 | 192994677 | -1 | processed_pseudoge | ENSG0000C |
| 193012250 | 193060080 | -1 | protein_coding | ENSG0000C |
| 193057281 | 193057415 | -1 | snoRNA | ENSG0000C |
| 193059422 | 193091777 | 1 | protein_coding | ENSG0000C |
| 193090866 | 193106114 | -1 | protein_coding | ENSG0000C |
| 193121958 | 193254815 | 1 | protein_coding | ENSG0000C |
| 193136503 | 193136583 | 1 | miRNA | ENSG0000C |
| 193178730 | 193186613 | -1 | protein_coding | ENSG0000C |
| 193236794 | 193237107 | -1 | misc_RNA | ENSG0000C |
| 193304745 | 193365953 | 1 | lncRNA | ENSG0000C |
| 193457422 | 193465198 | 1 | lncRNA | ENSG0000C |
| 193473224 | 194198708 | 1 | lncRNA | ENSG0000C |
| 193678894 | 193727035 | 1 | lncRNA | ENSG0000C |
| 193684246 | 193688429 | -1 | lncRNA | ENSG0000C |
| 193756815 | 193757267 | -1 | processed_pseudoge | ENSG0000C |
| 194188967 | 194190342 | 1 | processed_pseudoge | ENSG0000C |
| 194350943 | 194352426 | 1 | lncRNA | ENSG0000C |
| 194488103 | 194488205 | -1 | snRNA | ENSG0000C |
| 194718795 | 194719236 | 1 | processed_pseudoge | ENSG0000C |
| 194785517 | 194931310 | -1 | lncRNA | ENSG0000C |
| 195747024 | 195763501 | -1 | lncRNA | ENSG0000C |
| 196044883 | 196065235 | -1 | lncRNA | ENSG0000C |
| 196225779 | 196609225 | -1 | protein_coding | ENSG0000C |

| | | | | |
|---|---|---|---|---|
| 196582413 | 196582481 | -1 | miRNA | ENSG00000 |
| 196651878 | 196747504 | 1 | protein_coding | ENSG00000 |
| 196774795 | 196795406 | 1 | protein_coding | ENSG00000 |
| 196819757 | 196832189 | 1 | protein_coding | ENSG00000 |
| 196850283 | 196884793 | 1 | unprocessed_pseudo | ENSG00000 |
| 196888014 | 196918713 | 1 | protein_coding | ENSG00000 |
| 196943738 | 196959622 | 1 | protein_coding | ENSG00000 |
| 196970495 | 196970671 | 1 | unprocessed_pseudo | ENSG00000 |
| 196977556 | 197009674 | 1 | protein_coding | ENSG00000 |
| 197038741 | 197067267 | -1 | protein_coding | ENSG00000 |
| 197084127 | 197146694 | -1 | protein_coding | ENSG00000 |
| 197138748 | 197139307 | -1 | processed_pseudoge | ENSG00000 |
| 197153680 | 197200542 | -1 | protein_coding | ENSG00000 |
| 197222222 | 197223255 | 1 | processed_pseudoge | ENSG00000 |
| 197268204 | 197478455 | 1 | protein_coding | ENSG00000 |
| 197363817 | 197364078 | 1 | processed_pseudoge | ENSG00000 |
| 197437976 | 197447469 | -1 | lncRNA | ENSG00000 |
| 197504748 | 197775696 | -1 | protein_coding | ENSG00000 |
| 197685640 | 197685740 | -1 | misc_RNA | ENSG00000 |
| 197688760 | 197690627 | 1 | unprocessed_pseudo | ENSG00000 |
| 197735636 | 197736107 | 1 | processed_pseudoge | ENSG00000 |
| 197746751 | 197747451 | -1 | processed_pseudoge | ENSG00000 |
| 197757319 | 197761965 | 1 | lncRNA | ENSG00000 |
| 197902630 | 197907367 | 1 | protein_coding | ENSG00000 |
| 197911902 | 197935478 | 1 | protein_coding | ENSG00000 |
| 198156994 | 198322420 | 1 | protein_coding | ENSG00000 |
| 198197779 | 198198186 | -1 | processed_pseudoge | ENSG00000 |
| 198450666 | 198519877 | -1 | lncRNA | ENSG00000 |
| 198523222 | 198540945 | -1 | protein_coding | ENSG00000 |
| 198597724 | 198598868 | -1 | lncRNA | ENSG00000 |
| 198638457 | 198757476 | 1 | protein_coding | ENSG00000 |
| 198657553 | 198667061 | -1 | lncRNA | ENSG00000 |
| 198679139 | 198680033 | -1 | processed_pseudoge | ENSG00000 |
| 198777861 | 198937471 | -1 | lncRNA | ENSG00000 |
| 198858873 | 198858982 | -1 | miRNA | ENSG00000 |
| 198859044 | 198859153 | -1 | miRNA | ENSG00000 |
| 198949016 | 198949805 | 1 | processed_pseudoge | ENSG00000 |
| 198973379 | 198985506 | 1 | lncRNA | ENSG00000 |
| 199006040 | 199021037 | -1 | lncRNA | ENSG00000 |
| 199016133 | 199076735 | 1 | lncRNA | ENSG00000 |
| 199040501 | 199080975 | -1 | lncRNA | ENSG00000 |
| 199148598 | 199393307 | 1 | lncRNA | ENSG00000 |
| 199371877 | 199372297 | 1 | processed_pseudoge | ENSG00000 |
| 199387141 | 199388475 | 1 | processed_pseudoge | ENSG00000 |
| 199752491 | 199752890 | 1 | processed_pseudoge | ENSG00000 |
| 199876978 | 199877558 | 1 | processed_pseudoge | ENSG00000 |
| 199888159 | 199888261 | 1 | snRNA | ENSG00000 |
| 199908921 | 199909648 | 1 | processed_pseudoge | ENSG00000 |
| 200008505 | 200008607 | -1 | snRNA | ENSG00000 |
| 200014689 | 200014795 | -1 | snRNA | ENSG00000 |

| | | | | |
|---|---|---|---|---|
| 200027614 | 200177420 | 1 | protein_coding | ENSG00000 |
| 200054061 | 200054165 | 1 | snRNA | ENSG00000 |
| 200147531 | 200148279 | -1 | processed_pseudoge | ENSG00000 |
| 200213678 | 200214322 | 1 | processed_pseudoge | ENSG00000 |
| 200253419 | 200400705 | -1 | lncRNA | ENSG00000 |
| 200315435 | 200315967 | -1 | lncRNA | ENSG00000 |
| 200329161 | 200329677 | 1 | processed_pseudoge | ENSG00000 |
| 200333193 | 200478669 | 1 | lncRNA | ENSG00000 |
| 200404940 | 200410056 | -1 | protein_coding | ENSG00000 |
| 200478020 | 200483604 | -1 | lncRNA | ENSG00000 |
| 200551497 | 200620751 | -1 | protein_coding | ENSG00000 |
| 200623896 | 200669907 | -1 | protein_coding | ENSG00000 |
| 200669507 | 200694250 | 1 | lncRNA | ENSG00000 |
| 200739558 | 200860704 | 1 | protein_coding | ENSG00000 |
| 200863808 | 200864152 | 1 | processed_pseudoge | ENSG00000 |
| 200872981 | 200874178 | 1 | protein_coding | ENSG00000 |
| 200891048 | 200915742 | 1 | protein_coding | ENSG00000 |
| 200917460 | 200966668 | 1 | transcribed_unitary_ | ENSG00000 |
| 200933345 | 200933447 | -1 | snRNA | ENSG00000 |
| 200969390 | 201023700 | -1 | protein_coding | ENSG00000 |
| 201023949 | 201028792 | 1 | lncRNA | ENSG00000 |
| 201031136 | 201043073 | 1 | lncRNA | ENSG00000 |
| 201039512 | 201112451 | -1 | protein_coding | ENSG00000 |
| 201113953 | 201127184 | -1 | protein_coding | ENSG00000 |
| 201134772 | 201171574 | -1 | protein_coding | ENSG00000 |
| 201190825 | 201228952 | 1 | protein_coding | ENSG00000 |
| 201222113 | 201223123 | -1 | lncRNA | ENSG00000 |
| 201283452 | 201332993 | 1 | protein_coding | ENSG00000 |
| 201359008 | 201377764 | -1 | protein_coding | ENSG00000 |
| 201359018 | 201385784 | -1 | lncRNA | ENSG00000 |
| 201380833 | 201399915 | -1 | protein_coding | ENSG00000 |
| 201399633 | 201401190 | 1 | lncRNA | ENSG00000 |
| 201403768 | 201429866 | -1 | protein_coding | ENSG00000 |
| 201428975 | 201429361 | 1 | processed_pseudoge | ENSG00000 |
| 201464278 | 201469237 | -1 | protein_coding | ENSG00000 |
| 201464383 | 201465146 | -1 | lncRNA | ENSG00000 |
| 201483530 | 201509456 | -1 | protein_coding | ENSG00000 |
| 201507241 | 201534784 | 1 | lncRNA | ENSG00000 |
| 201518703 | 201584367 | 1 | transcribed_processe | ENSG00000 |
| 201622885 | 201826969 | 1 | protein_coding | ENSG00000 |
| 201673105 | 201674144 | -1 | lncRNA | ENSG00000 |
| 201688259 | 201829559 | -1 | lncRNA | ENSG00000 |
| 201719508 | 201719627 | 1 | miRNA | ENSG00000 |
| 201723294 | 201737506 | -1 | lncRNA | ENSG00000 |
| 201733406 | 201733505 | 1 | snRNA | ENSG00000 |
| 201808611 | 201808702 | 1 | miRNA | ENSG00000 |
| 201829149 | 201884291 | 1 | protein_coding | ENSG00000 |
| 201863373 | 201863447 | 1 | miRNA | ENSG00000 |
| 201888680 | 201892306 | 1 | protein_coding | ENSG00000 |
| 201893842 | 201899978 | 1 | lncRNA | ENSG00000 |

| | | | | |
|---|---|---|---|---|
| 201896456 | 201946588 | -1 | protein_coding | ENSG00000 |
| 201955503 | 201970664 | 1 | protein_coding | ENSG00000 |
| 201978461 | 201978595 | 1 | snoRNA | ENSG00000 |
| 201978642 | 201978949 | 1 | processed_pseudoge | ENSG00000 |
| 201982372 | 202006147 | 1 | protein_coding | ENSG00000 |
| 201995696 | 202010463 | -1 | lncRNA | ENSG00000 |
| 202003124 | 202003236 | 1 | miRNA | ENSG00000 |
| 202007945 | 202017183 | 1 | protein_coding | ENSG00000 |
| 202011370 | 202015657 | 1 | lncRNA | ENSG00000 |
| 202028606 | 202028692 | 1 | processed_pseudoge | ENSG00000 |
| 202039851 | 202039934 | 1 | processed_pseudoge | ENSG00000 |
| 202070841 | 202070943 | 1 | misc_RNA | ENSG00000 |
| 202096759 | 202096978 | 1 | processed_pseudoge | ENSG00000 |
| 202122917 | 202133592 | 1 | protein_coding | ENSG00000 |
| 202133404 | 202144743 | -1 | protein_coding | ENSG00000 |
| 202147013 | 202161588 | -1 | protein_coding | ENSG00000 |
| 202168051 | 202189455 | 1 | transcribed_unitary_ | ENSG00000 |
| 202193901 | 202319781 | 1 | protein_coding | ENSG00000 |
| 202331657 | 202341980 | -1 | protein_coding | ENSG00000 |
| 202348699 | 202592706 | 1 | protein_coding | ENSG00000 |
| 202369526 | 202370145 | -1 | processed_pseudoge | ENSG00000 |
| 202410108 | 202410207 | -1 | snRNA | ENSG00000 |
| 202438396 | 202439957 | 1 | transcribed_processe | ENSG00000 |
| 202471864 | 202472117 | -1 | processed_pseudoge | ENSG00000 |
| 202527310 | 202527427 | -1 | snoRNA | ENSG00000 |
| 202590596 | 202710454 | -1 | protein_coding | ENSG00000 |
| 202604268 | 202605293 | 1 | lncRNA | ENSG00000 |
| 202632428 | 202632911 | 1 | lncRNA | ENSG00000 |
| 202724495 | 202809470 | -1 | protein_coding | ENSG00000 |
| 202767229 | 202767418 | -1 | processed_pseudoge | ENSG00000 |
| 202796030 | 202797094 | 1 | processed_pseudoge | ENSG00000 |
| 202810238 | 202810829 | -1 | lncRNA | ENSG00000 |
| 202810954 | 202812156 | 1 | lncRNA | ENSG00000 |
| 202820266 | 202827225 | 1 | transcribed_unitary_ | ENSG00000 |
| 202851828 | 202861620 | -1 | transcribed_unproce | ENSG00000 |
| 202861754 | 202875241 | 1 | transcribed_processe | ENSG00000 |
| 202878282 | 202889149 | -1 | protein_coding | ENSG00000 |
| 202891116 | 202928636 | -1 | protein_coding | ENSG00000 |
| 202911812 | 202912729 | 1 | processed_pseudoge | ENSG00000 |
| 202914880 | 202914980 | -1 | misc_RNA | ENSG00000 |
| 202940826 | 202958572 | -1 | protein_coding | ENSG00000 |
| 202961873 | 202967275 | -1 | protein_coding | ENSG00000 |
| 202986557 | 202988582 | -1 | transcribed_unproce | ENSG00000 |
| 202999738 | 203000352 | 1 | unprocessed_pseudc | ENSG00000 |
| 203007374 | 203024848 | 1 | protein_coding | ENSG00000 |
| 203026498 | 203078740 | 1 | protein_coding | ENSG00000 |
| 203083129 | 203086012 | -1 | protein_coding | ENSG00000 |
| 203090654 | 203167405 | 1 | protein_coding | ENSG00000 |
| 203115468 | 203128367 | -1 | lncRNA | ENSG00000 |
| 203144694 | 203152579 | -1 | lncRNA | ENSG00000 |

| | | | | |
|---|---|---|---|---|
| 203167811 | 203175826 | -1 | protein_coding | ENSG00000 |
| 203178931 | 203186704 | -1 | protein_coding | ENSG00000 |
| 203212827 | 203273641 | -1 | protein_coding | ENSG00000 |
| 203255743 | 203256475 | 1 | processed_pseudoge | ENSG00000 |
| 203273221 | 203288804 | 1 | lncRNA | ENSG00000 |
| 203298758 | 203305309 | -1 | lncRNA | ENSG00000 |
| 203305491 | 203309602 | 1 | protein_coding | ENSG00000 |
| 203318996 | 203319099 | -1 | snRNA | ENSG00000 |
| 203340628 | 203351758 | -1 | protein_coding | ENSG00000 |
| 203353365 | 203353651 | -1 | processed_pseudoge | ENSG00000 |
| 203400266 | 203400581 | -1 | processed_pseudoge | ENSG00000 |
| 203475806 | 203491352 | 1 | protein_coding | ENSG00000 |
| 203494153 | 203508949 | 1 | protein_coding | ENSG00000 |
| 203626561 | 203744081 | 1 | protein_coding | ENSG00000 |
| 203656969 | 203657760 | -1 | processed_pseudoge | ENSG00000 |
| 203729581 | 203729705 | 1 | snoRNA | ENSG00000 |
| 203765177 | 203776372 | 1 | protein_coding | ENSG00000 |
| 203795654 | 203854999 | 1 | protein_coding | ENSG00000 |
| 203795714 | 203854999 | 1 | protein_coding | ENSG00000 |
| 203805621 | 203806263 | -1 | processed_pseudoge | ENSG00000 |
| 203835585 | 203835912 | -1 | processed_pseudoge | ENSG00000 |
| 203861599 | 203871152 | 1 | protein_coding | ENSG00000 |
| 203872574 | 203874187 | -1 | processed_pseudoge | ENSG00000 |
| 203903723 | 203904013 | -1 | processed_pseudoge | ENSG00000 |
| 203954640 | 203955780 | 1 | processed_pseudoge | ENSG00000 |
| 203996532 | 204008357 | 1 | lncRNA | ENSG00000 |
| 204032447 | 204041265 | -1 | lncRNA | ENSG00000 |
| 204064533 | 204073965 | -1 | lncRNA | ENSG00000 |
| 204073115 | 204127743 | 1 | protein_coding | ENSG00000 |
| 204131062 | 204152044 | -1 | protein_coding | ENSG00000 |
| 204131062 | 204131966 | 1 | lncRNA | ENSG00000 |
| 204141404 | 204143327 | 1 | lncRNA | ENSG00000 |
| 204154819 | 204190324 | -1 | protein_coding | ENSG00000 |
| 204183006 | 204183537 | 1 | processed_pseudoge | ENSG00000 |
| 204190341 | 204196491 | -1 | protein_coding | ENSG00000 |
| 204198163 | 204213988 | -1 | protein_coding | ENSG00000 |
| 204218853 | 204377665 | -1 | protein_coding | ENSG00000 |
| 204276901 | 204280363 | 1 | lncRNA | ENSG00000 |
| 204346776 | 204347258 | -1 | processed_pseudoge | ENSG00000 |
| 204368431 | 204369719 | -1 | lncRNA | ENSG00000 |
| 204377850 | 204435846 | 1 | lncRNA | ENSG00000 |
| 204394541 | 204394774 | 1 | processed_pseudoge | ENSG00000 |
| 204403381 | 204411817 | -1 | protein_coding | ENSG00000 |
| 204422628 | 204494724 | -1 | protein_coding | ENSG00000 |
| 204516379 | 204558120 | 1 | protein_coding | ENSG00000 |
| 204528845 | 204529692 | -1 | processed_pseudoge | ENSG00000 |
| 204562413 | 204562521 | -1 | rRNA_pseudogene | ENSG00000 |
| 204603035 | 204616565 | 1 | lncRNA | ENSG00000 |
| 204617170 | 204685738 | -1 | protein_coding | ENSG00000 |
| 204626775 | 204629712 | 1 | lncRNA | ENSG00000 |

| | | | | |
|---|---|---|---|---|
| 204663872 | 204664613 | 1 | lncRNA | ENSG0000C |
| 204707320 | 204707430 | -1 | rRNA_pseudogene | ENSG0000C |
| 204727991 | 204728106 | 1 | scaRNA | ENSG0000C |
| 204822664 | 204829274 | -1 | lncRNA | ENSG0000C |
| 204828651 | 205022822 | 1 | protein_coding | ENSG0000C |
| 204946608 | 204947169 | 1 | processed_pseudoge | ENSG0000C |
| 205042937 | 205078289 | 1 | protein_coding | ENSG0000C |
| 205083129 | 205084460 | -1 | protein_coding | ENSG0000C |
| 205086142 | 205122015 | -1 | protein_coding | ENSG0000C |
| 205091163 | 205091946 | 1 | processed_pseudoge | ENSG0000C |
| 205134646 | 205134950 | 1 | processed_pseudoge | ENSG0000C |
| 205142505 | 205211702 | -1 | protein_coding | ENSG0000C |
| 205202191 | 205202742 | 1 | processed_pseudoge | ENSG0000C |
| 205227946 | 205285632 | 1 | protein_coding | ENSG0000C |
| 205233821 | 205236867 | -1 | lncRNA | ENSG0000C |
| 205302063 | 205321760 | -1 | protein_coding | ENSG0000C |
| 205336065 | 205357090 | -1 | protein_coding | ENSG0000C |
| 205351247 | 205351471 | 1 | processed_pseudoge | ENSG0000C |
| 205373252 | 205387440 | 1 | lncRNA | ENSG0000C |
| 205381378 | 205455954 | -1 | protein_coding | ENSG0000C |
| 205434885 | 205457091 | -1 | protein_coding | ENSG0000C |
| 205448302 | 205448398 | -1 | miRNA | ENSG0000C |
| 205455929 | 205469024 | 1 | lncRNA | ENSG0000C |
| 205504595 | 205532793 | 1 | protein_coding | ENSG0000C |
| 205566716 | 205566799 | 1 | snRNA | ENSG0000C |
| 205568885 | 205602918 | 1 | protein_coding | ENSG0000C |
| 205595041 | 205595147 | 1 | snRNA | ENSG0000C |
| 205597556 | 205631962 | -1 | protein_coding | ENSG0000C |
| 205625483 | 205626153 | 1 | processed_pseudoge | ENSG0000C |
| 205657851 | 205680509 | -1 | protein_coding | ENSG0000C |
| 205684018 | 205684307 | 1 | misc_RNA | ENSG0000C |
| 205712822 | 205750182 | -1 | protein_coding | ENSG0000C |
| 205731221 | 205731352 | -1 | snoRNA | ENSG0000C |
| 205767986 | 205775482 | -1 | protein_coding | ENSG0000C |
| 205775559 | 205783623 | 1 | lncRNA | ENSG0000C |
| 205789094 | 205813748 | -1 | protein_coding | ENSG0000C |
| 205813322 | 205896082 | 1 | lncRNA | ENSG0000C |
| 205828025 | 205850132 | -1 | protein_coding | ENSG0000C |
| 205913048 | 205943460 | -1 | protein_coding | ENSG0000C |
| 205935128 | 205935768 | 1 | lncRNA | ENSG0000C |
| 205976740 | 206003461 | -1 | protein_coding | ENSG0000C |
| 206009264 | 206023895 | -1 | protein_coding | ENSG0000C |
| 206035253 | 206062257 | 1 | transcribed_unitary_ | ENSG0000C |
| 206053173 | 206102449 | 1 | protein_coding | ENSG0000C |
| 206080239 | 206080324 | 1 | snoRNA | ENSG0000C |
| 206106936 | 206117699 | -1 | protein_coding | ENSG0000C |
| 206117783 | 206126805 | 1 | lncRNA | ENSG0000C |
| 206147164 | 206166149 | 1 | lncRNA | ENSG0000C |
| 206160887 | 206161276 | -1 | processed_pseudoge | ENSG0000C |
| 206175060 | 206177246 | -1 | lncRNA | ENSG0000C |

| | | | | |
|---|---|---|---|---|
| 206186179 | 206204414 | -1 | protein_coding | ENSG00000 |
| 206203345 | 206464443 | 1 | protein_coding | ENSG00000 |
| 206333327 | 206333438 | -1 | unprocessed_pseudo | ENSG00000 |
| 206470476 | 206496889 | 1 | protein_coding | ENSG00000 |
| 206474803 | 206474864 | 1 | miRNA | ENSG00000 |
| 206491116 | 206497728 | -1 | lncRNA | ENSG00000 |
| 206503948 | 206504456 | 1 | lncRNA | ENSG00000 |
| 206507531 | 206589448 | 1 | protein_coding | ENSG00000 |
| 206528915 | 206529706 | 1 | processed_pseudoge | ENSG00000 |
| 206541758 | 206544892 | 1 | TEC | ENSG00000 |
| 206571292 | 206612465 | -1 | protein_coding | ENSG00000 |
| 206634196 | 206635714 | -1 | lncRNA | ENSG00000 |
| 206635536 | 206684419 | 1 | protein_coding | ENSG00000 |
| 206684944 | 206734283 | 1 | protein_coding | ENSG00000 |
| 206695837 | 206696269 | -1 | processed_pseudoge | ENSG00000 |
| 206747980 | 206748092 | -1 | misc_RNA | ENSG00000 |
| 206767602 | 206774541 | -1 | protein_coding | ENSG00000 |
| 206770764 | 206842981 | 1 | protein_coding | ENSG00000 |
| 206865354 | 206869223 | 1 | protein_coding | ENSG00000 |
| 206897443 | 206904139 | 1 | protein_coding | ENSG00000 |
| 206903317 | 206923247 | -1 | protein_coding | ENSG00000 |
| 206905928 | 206906393 | 1 | processed_pseudoge | ENSG00000 |
| 206907619 | 206908229 | 1 | processed_pseudoge | ENSG00000 |
| 206928522 | 206946466 | -1 | protein_coding | ENSG00000 |
| 206957965 | 206970625 | -1 | protein_coding | ENSG00000 |
| 207018522 | 207032756 | -1 | protein_coding | ENSG00000 |
| 207034366 | 207081024 | 1 | protein_coding | ENSG00000 |
| 207043849 | 207052980 | -1 | protein_coding | ENSG00000 |
| 207088860 | 207099993 | 1 | protein_coding | ENSG00000 |
| 207104233 | 207144972 | 1 | protein_coding | ENSG00000 |
| 207127010 | 207127486 | -1 | lncRNA | ENSG00000 |
| 207150205 | 207150281 | -1 | processed_pseudoge | ENSG00000 |
| 207165496 | 207183910 | 1 | unprocessed_pseudo | ENSG00000 |
| 207179296 | 207184062 | 1 | lncRNA | ENSG00000 |
| 207225798 | 207229202 | 1 | unprocessed_pseudo | ENSG00000 |
| 207240122 | 207309292 | 1 | lncRNA | ENSG00000 |
| 207321376 | 207386804 | 1 | protein_coding | ENSG00000 |
| 207401691 | 207416236 | -1 | lncRNA | ENSG00000 |
| 207454230 | 207489895 | 1 | protein_coding | ENSG00000 |
| 207496147 | 207641765 | 1 | protein_coding | ENSG00000 |
| 207551925 | 207606555 | -1 | lncRNA | ENSG00000 |
| 207645113 | 207738416 | 1 | protein_coding | ENSG00000 |
| 207645234 | 207657410 | 1 | unprocessed_pseudo | ENSG00000 |
| 207658454 | 207659142 | 1 | unprocessed_pseudo | ENSG00000 |
| 207708898 | 207708981 | -1 | rRNA | ENSG00000 |
| 207709024 | 207711508 | -1 | lncRNA | ENSG00000 |
| 207752057 | 207795513 | 1 | protein_coding | ENSG00000 |
| 207762584 | 207763263 | 1 | unprocessed_pseudo | ENSG00000 |
| 207801518 | 207879115 | -1 | lncRNA | ENSG00000 |
| 207801852 | 207801939 | -1 | miRNA | ENSG00000 |

| | | | | |
|---|---|---|---|---|
| 207802443 | 207802523 | -1 | miRNA | ENSG0000( |
| 207880972 | 207911402 | -1 | protein_coding | ENSG0000( |
| 207909992 | 207917088 | 1 | lncRNA | ENSG0000( |
| 207959292 | 207969329 | 1 | lncRNA | ENSG0000( |
| 208022242 | 208244384 | -1 | protein_coding | ENSG0000( |
| 208106102 | 208107655 | 1 | lncRNA | ENSG0000( |
| 208255290 | 208255376 | -1 | processed_pseudoge | ENSG0000( |
| 208266059 | 208270667 | -1 | lncRNA | ENSG0000( |
| 208606564 | 208614295 | -1 | lncRNA | ENSG0000( |
| 208626741 | 208628085 | -1 | lncRNA | ENSG0000( |
| 208697369 | 208697698 | -1 | processed_pseudoge | ENSG0000( |
| 208728665 | 208736286 | 1 | lncRNA | ENSG0000( |
| 208972454 | 208975307 | 1 | lncRNA | ENSG0000( |
| 209001338 | 209001476 | 1 | unprocessed_pseudc | ENSG0000( |
| 209107503 | 209114419 | 1 | lncRNA | ENSG0000( |
| 209147220 | 209266758 | -1 | lncRNA | ENSG0000( |
| 209173014 | 209173375 | 1 | processed_pseudoge | ENSG0000( |
| 209232196 | 209233419 | 1 | processed_pseudoge | ENSG0000( |
| 209267798 | 209268223 | -1 | processed_pseudoge | ENSG0000( |
| 209325392 | 209328535 | 1 | lncRNA | ENSG0000( |
| 209367662 | 209380518 | 1 | lncRNA | ENSG0000( |
| 209428817 | 209432838 | 1 | lncRNA | ENSG0000( |
| 209432133 | 209432242 | 1 | miRNA | ENSG0000( |
| 209528455 | 209567673 | -1 | lncRNA | ENSG0000( |
| 209531236 | 209536351 | 1 | lncRNA | ENSG0000( |
| 209583714 | 209613939 | 1 | protein_coding | ENSG0000( |
| 209614870 | 209652425 | -1 | protein_coding | ENSG0000( |
| 209623444 | 209623510 | -1 | miRNA | ENSG0000( |
| 209661364 | 209724125 | -1 | lncRNA | ENSG0000( |
| 209675412 | 209676390 | 1 | protein_coding | ENSG0000( |
| 209686178 | 209734949 | 1 | protein_coding | ENSG0000( |
| 209744373 | 209745214 | 1 | processed_pseudoge | ENSG0000( |
| 209756032 | 209782320 | 1 | protein_coding | ENSG0000( |
| 209779208 | 209784559 | -1 | protein_coding | ENSG0000( |
| 209785617 | 209806175 | -1 | protein_coding | ENSG0000( |
| 209827972 | 209857565 | 1 | protein_coding | ENSG0000( |
| 209900923 | 210171389 | 1 | protein_coding | ENSG0000( |
| 209987333 | 209997092 | -1 | lncRNA | ENSG0000( |
| 210231456 | 210234047 | -1 | lncRNA | ENSG0000( |
| 210232796 | 210246631 | 1 | protein_coding | ENSG0000( |
| 210265636 | 210267241 | -1 | processed_pseudoge | ENSG0000( |
| 210303684 | 210304540 | 1 | processed_pseudoge | ENSG0000( |
| 210328252 | 210676296 | 1 | protein_coding | ENSG0000( |
| 210362861 | 210366948 | -1 | lncRNA | ENSG0000( |
| 210374154 | 210374267 | -1 | snRNA | ENSG0000( |
| 210386657 | 210394112 | -1 | lncRNA | ENSG0000( |
| 210462345 | 210463012 | -1 | processed_pseudoge | ENSG0000( |
| 210676823 | 211134165 | -1 | protein_coding | ENSG0000( |
| 210678315 | 210681932 | 1 | TEC | ENSG0000( |
| 210858125 | 211134146 | -1 | protein_coding | ENSG0000( |

| | | | | |
|---|---|---|---|---|
| 210859177 | 210862285 | -1 | processed_pseudoge | ENSG00000 |
| 211082872 | 211188533 | -1 | protein_coding | ENSG00000 |
| 211108445 | 211207138 | -1 | lncRNA | ENSG00000 |
| 211132588 | 211133374 | -1 | lncRNA | ENSG00000 |
| 211173488 | 211173849 | 1 | processed_pseudoge | ENSG00000 |
| 211207239 | 211207897 | 1 | processed_pseudoge | ENSG00000 |
| 211258377 | 211316385 | 1 | protein_coding | ENSG00000 |
| 211326615 | 211374946 | 1 | protein_coding | ENSG00000 |
| 211376834 | 211382717 | -1 | lncRNA | ENSG00000 |
| 211382736 | 211435570 | 1 | lncRNA | ENSG00000 |
| 211417025 | 211417934 | 1 | unprocessed_pseudo | ENSG00000 |
| 211442274 | 211442806 | 1 | processed_pseudoge | ENSG00000 |
| 211476522 | 211492917 | -1 | protein_coding | ENSG00000 |
| 211492255 | 211492917 | 1 | lncRNA | ENSG00000 |
| 211571568 | 211579161 | -1 | protein_coding | ENSG00000 |
| 211583015 | 211583725 | -1 | lncRNA | ENSG00000 |
| 211635865 | 211644114 | -1 | lncRNA | ENSG00000 |
| 211639440 | 211654627 | 1 | lncRNA | ENSG00000 |
| 211658657 | 211675630 | -1 | protein_coding | ENSG00000 |
| 211660189 | 211661277 | -1 | processed_pseudoge | ENSG00000 |
| 211675749 | 211690103 | 1 | lncRNA | ENSG00000 |
| 211715928 | 211725402 | -1 | lncRNA | ENSG00000 |
| 211743457 | 211830763 | -1 | protein_coding | ENSG00000 |
| 211792111 | 211792406 | -1 | misc_RNA | ENSG00000 |
| 211803017 | 211803104 | 1 | misc_RNA | ENSG00000 |
| 211829636 | 211853703 | 1 | lncRNA | ENSG00000 |
| 211936249 | 211936634 | 1 | processed_pseudoge | ENSG00000 |
| 211940399 | 212035557 | -1 | protein_coding | ENSG00000 |
| 212025561 | 212025682 | 1 | snoRNA | ENSG00000 |
| 212035553 | 212107400 | 1 | protein_coding | ENSG00000 |
| 212051524 | 212052006 | -1 | processed_pseudoge | ENSG00000 |
| 212077613 | 212077685 | 1 | miRNA | ENSG00000 |
| 212099521 | 212099676 | -1 | misc_RNA | ENSG00000 |
| 212168207 | 212190259 | 1 | lncRNA | ENSG00000 |
| 212225278 | 212238977 | -1 | lncRNA | ENSG00000 |
| 212285410 | 212361853 | 1 | protein_coding | ENSG00000 |
| 212297448 | 212299579 | 1 | lncRNA | ENSG00000 |
| 212299495 | 212331713 | -1 | lncRNA | ENSG00000 |
| 212309051 | 212309518 | -1 | processed_pseudoge | ENSG00000 |
| 212352816 | 212352950 | 1 | snoRNA | ENSG00000 |
| 212357418 | 212358353 | 1 | lncRNA | ENSG00000 |
| 212363931 | 212414901 | -1 | protein_coding | ENSG00000 |
| 212430269 | 212432775 | -1 | lncRNA | ENSG00000 |
| 212432920 | 212446379 | 1 | protein_coding | ENSG00000 |
| 212466699 | 212472905 | -1 | lncRNA | ENSG00000 |
| 212504178 | 212534060 | 1 | lncRNA | ENSG00000 |
| 212545694 | 212556065 | 1 | lncRNA | ENSG00000 |
| 212557833 | 212559731 | -1 | lncRNA | ENSG00000 |
| 212559363 | 212561399 | 1 | lncRNA | ENSG00000 |
| 212565334 | 212620777 | 1 | protein_coding | ENSG00000 |

| | | | | |
|---|---|---|---|---|
| 212624284 | 212626771 | -1 | lncRNA | ENSG0000C |
| 212624474 | 212626775 | 1 | protein_coding | ENSG0000C |
| 212647296 | 212647755 | 1 | processed_pseudoge | ENSG0000C |
| 212653305 | 212665638 | -1 | lncRNA | ENSG0000C |
| 212686417 | 212699840 | -1 | protein_coding | ENSG0000C |
| 212726153 | 212791782 | -1 | protein_coding | ENSG0000C |
| 212791828 | 212816830 | 1 | protein_coding | ENSG0000C |
| 212824027 | 212824505 | 1 | processed_pseudoge | ENSG0000C |
| 212830141 | 212847649 | -1 | protein_coding | ENSG0000C |
| 212852105 | 212858126 | -1 | lncRNA | ENSG0000C |
| 212853280 | 212853426 | 1 | processed_pseudoge | ENSG0000C |
| 212855175 | 212855366 | -1 | processed_pseudoge | ENSG0000C |
| 212858275 | 212899363 | 1 | protein_coding | ENSG0000C |
| 212916787 | 212917577 | 1 | lncRNA | ENSG0000C |
| 212950520 | 212992037 | 1 | protein_coding | ENSG0000C |
| 212992182 | 213015867 | -1 | protein_coding | ENSG0000C |
| 213051233 | 213274774 | 1 | protein_coding | ENSG0000C |
| 213428708 | 213429077 | -1 | processed_pseudoge | ENSG0000C |
| 213492288 | 213579267 | 1 | lncRNA | ENSG0000C |
| 213731416 | 213794587 | 1 | lncRNA | ENSG0000C |
| 213817751 | 213988508 | -1 | lncRNA | ENSG0000C |
| 213924749 | 213926654 | -1 | lncRNA | ENSG0000C |
| 213983181 | 214041510 | 1 | protein_coding | ENSG0000C |
| 213983793 | 213986419 | -1 | lncRNA | ENSG0000C |
| 214028891 | 214030901 | -1 | lncRNA | ENSG0000C |
| 214051194 | 214187773 | -1 | lncRNA | ENSG0000C |
| 214281102 | 214337131 | 1 | protein_coding | ENSG0000C |
| 214344172 | 214357615 | -1 | lncRNA | ENSG0000C |
| 214348700 | 214552449 | -1 | protein_coding | ENSG0000C |
| 214482813 | 214483387 | -1 | processed_pseudoge | ENSG0000C |
| 214532195 | 214533467 | 1 | processed_pseudoge | ENSG0000C |
| 214603195 | 214664571 | 1 | protein_coding | ENSG0000C |
| 214605470 | 214608840 | -1 | unprocessed_pseudc | ENSG0000C |
| 214612960 | 214613398 | -1 | processed_pseudoge | ENSG0000C |
| 214870734 | 214871716 | 1 | processed_pseudoge | ENSG0000C |
| 214943123 | 214943540 | 1 | processed_pseudoge | ENSG0000C |
| 214946909 | 214988040 | 1 | lncRNA | ENSG0000C |
| 215005775 | 215237093 | 1 | protein_coding | ENSG0000C |
| 215376484 | 215377374 | 1 | processed_pseudoge | ENSG0000C |
| 215393646 | 215394418 | -1 | lncRNA | ENSG0000C |
| 215567304 | 215621807 | 1 | protein_coding | ENSG0000C |
| 215622891 | 216423448 | -1 | protein_coding | ENSG0000C |
| 215630026 | 215630117 | -1 | snoRNA | ENSG0000C |
| 215886582 | 215901464 | 1 | lncRNA | ENSG0000C |
| 216072465 | 216086917 | 1 | lncRNA | ENSG0000C |
| 216194051 | 216204366 | 1 | lncRNA | ENSG0000C |
| 216201635 | 216201826 | -1 | processed_pseudoge | ENSG0000C |
| 216503246 | 217137755 | -1 | protein_coding | ENSG0000C |
| 217426992 | 217631090 | -1 | protein_coding | ENSG0000C |
| 217631324 | 217871696 | 1 | protein_coding | ENSG0000C |

| | | | | |
|---|---|---|---|---|
| 217781198 | 217785120 | -1 | lncRNA | ENSG0000C |
| 217850403 | 217850633 | -1 | processed_pseudoge | ENSG0000C |
| 217892900 | 217920804 | 1 | lncRNA | ENSG0000C |
| 218031835 | 218033660 | 1 | lncRNA | ENSG0000C |
| 218043505 | 218059140 | 1 | lncRNA | ENSG0000C |
| 218046943 | 218049221 | -1 | lncRNA | ENSG0000C |
| 218129795 | 218129952 | 1 | snRNA | ENSG0000C |
| 218285293 | 218337983 | 1 | protein_coding | ENSG0000C |
| 218301262 | 218301972 | -1 | processed_pseudoge | ENSG0000C |
| 218338541 | 218338756 | 1 | processed_pseudoge | ENSG0000C |
| 218344196 | 218345678 | -1 | lncRNA | ENSG0000C |
| 218345336 | 218444619 | 1 | protein_coding | ENSG0000C |
| 218442626 | 218443996 | 1 | lncRNA | ENSG0000C |
| 218459265 | 218525978 | 1 | lncRNA | ENSG0000C |
| 218541690 | 218541899 | -1 | snoRNA | ENSG0000C |
| 218881600 | 218882650 | -1 | processed_pseudoge | ENSG0000C |
| 218912757 | 218918714 | -1 | lncRNA | ENSG0000C |
| 218983023 | 219173961 | -1 | lncRNA | ENSG0000C |
| 219173869 | 219212865 | 1 | protein_coding | ENSG0000C |
| 219199914 | 219200567 | -1 | processed_pseudoge | ENSG0000C |
| 219222248 | 219225497 | -1 | lncRNA | ENSG0000C |
| 219270774 | 219273387 | 1 | lncRNA | ENSG0000C |
| 219294982 | 219297668 | 1 | lncRNA | ENSG0000C |
| 219409039 | 219459369 | -1 | lncRNA | ENSG0000C |
| 219557192 | 219557701 | 1 | lncRNA | ENSG0000C |
| 219608010 | 219613145 | -1 | protein_coding | ENSG0000C |
| 219685427 | 219958647 | -1 | protein_coding | ENSG0000C |
| 219761789 | 219761875 | 1 | rRNA_pseudogene | ENSG0000C |
| 219962652 | 219962865 | 1 | snoRNA | ENSG0000C |
| 219968600 | 220046530 | -1 | protein_coding | ENSG0000C |
| 220057482 | 220090462 | -1 | protein_coding | ENSG0000C |
| 220094132 | 220148041 | 1 | protein_coding | ENSG0000C |
| 220117853 | 220117962 | -1 | miRNA | ENSG0000C |
| 220118157 | 220118241 | -1 | miRNA | ENSG0000C |
| 220143964 | 220144351 | -1 | processed_pseudoge | ENSG0000C |
| 220148293 | 220272453 | -1 | protein_coding | ENSG0000C |
| 220200538 | 220200619 | -1 | miRNA | ENSG0000C |
| 220200546 | 220200676 | -1 | snoRNA | ENSG0000C |
| 220207618 | 220208282 | 1 | processed_pseudoge | ENSG0000C |
| 220220291 | 220220593 | 1 | processed_pseudoge | ENSG0000C |
| 220253570 | 220254536 | 1 | processed_pseudoge | ENSG0000C |
| 220266706 | 220267917 | -1 | transcribed_processe | ENSG0000C |
| 220313945 | 220315565 | -1 | processed_pseudoge | ENSG0000C |
| 220316667 | 220317599 | -1 | processed_pseudoge | ENSG0000C |
| 220359731 | 220360183 | -1 | lncRNA | ENSG0000C |
| 220401122 | 220404033 | 1 | lncRNA | ENSG0000C |
| 220448516 | 220448853 | 1 | processed_pseudoge | ENSG0000C |
| 220455154 | 220455312 | -1 | processed_pseudoge | ENSG0000C |
| 220467954 | 220468808 | 1 | processed_pseudoge | ENSG0000C |
| 220485104 | 220487558 | -1 | lncRNA | ENSG0000C |

| | | | | |
|---|---|---|---|---|
| 220528183 | 220664461 | 1 | protein_coding | ENSG00000( |
| 220571739 | 220572035 | -1 | misc_RNA | ENSG00000( |
| 220625740 | 220628056 | 1 | transcribed_processe | ENSG00000( |
| 220690363 | 220699153 | 1 | protein_coding | ENSG00000( |
| 220748225 | 220784815 | 1 | protein_coding | ENSG00000( |
| 220786913 | 220819659 | 1 | protein_coding | ENSG00000( |
| 220786990 | 220884503 | 1 | protein_coding | ENSG00000( |
| 220825620 | 220826063 | 1 | lncRNA | ENSG00000( |
| 220825776 | 220825893 | 1 | snRNA | ENSG00000( |
| 220828676 | 220829211 | -1 | lncRNA | ENSG00000( |
| 220829255 | 220832429 | 1 | lncRNA | ENSG00000( |
| 220832763 | 220880140 | -1 | lncRNA | ENSG00000( |
| 220879431 | 220885059 | 1 | protein_coding | ENSG00000( |
| 221133865 | 221134221 | -1 | processed_pseudoge | ENSG00000( |
| 221330080 | 221336489 | -1 | lncRNA | ENSG00000( |
| 221508559 | 221510979 | 1 | lncRNA | ENSG00000( |
| 221549362 | 221554314 | 1 | lncRNA | ENSG00000( |
| 221549786 | 221550643 | -1 | processed_pseudoge | ENSG00000( |
| 221555550 | 221563653 | -1 | lncRNA | ENSG00000( |
| 221701424 | 221742089 | -1 | protein_coding | ENSG00000( |
| 221819842 | 221840717 | -1 | lncRNA | ENSG00000( |
| 221837334 | 221837437 | -1 | snRNA | ENSG00000( |
| 221880981 | 221978523 | -1 | lncRNA | ENSG00000( |
| 221966341 | 221984964 | 1 | lncRNA | ENSG00000( |
| 222010825 | 222064773 | -1 | lncRNA | ENSG00000( |
| 222088806 | 222389103 | -1 | lncRNA | ENSG00000( |
| 222261833 | 222262796 | -1 | processed_pseudoge | ENSG00000( |
| 222452738 | 222454705 | -1 | lncRNA | ENSG00000( |
| 222468094 | 222470882 | 1 | processed_pseudoge | ENSG00000( |
| 222477252 | 222504622 | -1 | lncRNA | ENSG00000( |
| 222503632 | 222503735 | -1 | snRNA | ENSG00000( |
| 222510169 | 222519732 | -1 | unprocessed_pseudc | ENSG00000( |
| 222522258 | 222548104 | -1 | protein_coding | ENSG00000( |
| 222557902 | 222589933 | -1 | protein_coding | ENSG00000( |
| 222589825 | 222593843 | 1 | lncRNA | ENSG00000( |
| 222618097 | 222668007 | 1 | protein_coding | ENSG00000( |
| 222641414 | 222641858 | -1 | processed_pseudoge | ENSG00000( |
| 222658867 | 222661512 | -1 | lncRNA | ENSG00000( |
| 222668013 | 222713210 | -1 | protein_coding | ENSG00000( |
| 222712553 | 222735196 | 1 | protein_coding | ENSG00000( |
| 222737207 | 222750805 | 1 | protein_coding | ENSG00000( |
| 222743356 | 222773149 | -1 | lncRNA | ENSG00000( |
| 222815022 | 222837384 | 1 | lncRNA | ENSG00000( |
| 222815064 | 222994906 | 1 | lncRNA | ENSG00000( |
| 222872271 | 223005995 | 1 | protein_coding | ENSG00000( |
| 222945725 | 222945900 | 1 | processed_pseudoge | ENSG00000( |
| 223091872 | 223103335 | -1 | lncRNA | ENSG00000( |
| 223109404 | 223143248 | -1 | protein_coding | ENSG00000( |
| 223144049 | 223144954 | 1 | lncRNA | ENSG00000( |
| 223181144 | 223188154 | 1 | lncRNA | ENSG00000( |

| | | | | |
|---|---|---|---|---|
| 223220819 | 223364178 | -1 | protein_coding | ENSG0000( |
| 223373822 | 223373956 | 1 | snRNA | ENSG0000( |
| 223393415 | 223395465 | 1 | protein_coding | ENSG0000( |
| 223538007 | 223665734 | -1 | protein_coding | ENSG0000( |
| 223644110 | 223644383 | -1 | processed_pseudoge | ENSG0000( |
| 223690051 | 223690114 | 1 | snRNA | ENSG0000( |
| 223701593 | 223776018 | 1 | protein_coding | ENSG0000( |
| 223779893 | 223845954 | -1 | protein_coding | ENSG0000( |
| 223856579 | 223857387 | -1 | processed_pseudoge | ENSG0000( |
| 223863726 | 223864851 | -1 | processed_pseudoge | ENSG0000( |
| 223947605 | 223950416 | 1 | unprocessed_pseudc | ENSG0000( |
| 223951394 | 223992594 | -1 | transcribed_unproce | ENSG0000( |
| 223976146 | 223976249 | -1 | snRNA | ENSG0000( |
| 223992743 | 224010612 | 1 | transcribed_unproce | ENSG0000( |
| 223994262 | 223995196 | 1 | lncRNA | ENSG0000( |
| 224030704 | 224035056 | -1 | unprocessed_pseudc | ENSG0000( |
| 224107282 | 224107541 | -1 | misc_RNA | ENSG0000( |
| 224114111 | 224162047 | 1 | protein_coding | ENSG0000( |
| 224175476 | 224175706 | -1 | processed_pseudoge | ENSG0000( |
| 224175756 | 224193441 | 1 | protein_coding | ENSG0000( |
| 224179641 | 224179767 | 1 | snoRNA | ENSG0000( |
| 224208741 | 224213625 | -1 | lncRNA | ENSG0000( |
| 224219613 | 224228043 | 1 | lncRNA | ENSG0000( |
| 224227334 | 224330189 | -1 | protein_coding | ENSG0000( |
| 224257004 | 224257141 | -1 | miRNA | ENSG0000( |
| 224297646 | 224298600 | 1 | processed_pseudoge | ENSG0000( |
| 224305380 | 224305483 | 1 | snRNA | ENSG0000( |
| 224356858 | 224379459 | 1 | protein_coding | ENSG0000( |
| 224385143 | 224437033 | -1 | protein_coding | ENSG0000( |
| 224398227 | 224398311 | -1 | miRNA | ENSG0000( |
| 224434660 | 224740554 | 1 | protein_coding | ENSG0000( |
| 224574434 | 224575389 | 1 | processed_pseudoge | ENSG0000( |
| 224608130 | 224616220 | -1 | lncRNA | ENSG0000( |
| 224661173 | 224661886 | -1 | processed_pseudoge | ENSG0000( |
| 224703272 | 224712849 | -1 | lncRNA | ENSG0000( |
| 224717504 | 224730662 | -1 | lncRNA | ENSG0000( |
| 224766324 | 224779335 | 1 | lncRNA | ENSG0000( |
| 224802959 | 224881355 | -1 | lncRNA | ENSG0000( |
| 224896262 | 225399292 | 1 | protein_coding | ENSG0000( |
| 225401502 | 225428925 | -1 | protein_coding | ENSG0000( |
| 225447233 | 225465400 | -1 | lncRNA | ENSG0000( |
| 225465021 | 225473837 | 1 | lncRNA | ENSG0000( |
| 225486835 | 225653142 | -1 | protein_coding | ENSG0000( |
| 225700264 | 225752258 | 1 | lncRNA | ENSG0000( |
| 225710968 | 225736274 | -1 | lncRNA | ENSG0000( |
| 225741275 | 225741380 | 1 | snRNA | ENSG0000( |
| 225777813 | 225790468 | 1 | protein_coding | ENSG0000( |
| 225803148 | 225803448 | 1 | processed_pseudoge | ENSG0000( |
| 225810092 | 225845563 | 1 | protein_coding | ENSG0000( |
| 225840883 | 225846522 | -1 | lncRNA | ENSG0000( |

| | | | | |
|---|---|---|---|---|
| 225845536 | 225882380 | -1 | protein_coding | ENSG0000C |
| 225886282 | 225911382 | -1 | protein_coding | ENSG0000C |
| 225886696 | 225924278 | -1 | protein_coding | ENSG0000C |
| 225919877 | 225924340 | -1 | protein_coding | ENSG0000C |
| 225922080 | 225922142 | -1 | miRNA | ENSG0000C |
| 225936411 | 225937557 | -1 | lncRNA | ENSG0000C |
| 225936598 | 225941383 | -1 | protein_coding | ENSG0000C |
| 225964179 | 225964428 | 1 | processed_pseudoge | ENSG0000C |
| 225982702 | 225999343 | -1 | protein_coding | ENSG0000C |
| 226045561 | 226061898 | -1 | lncRNA | ENSG0000C |
| 226061851 | 226072007 | 1 | protein_coding | ENSG0000C |
| 226083590 | 226090465 | 1 | lncRNA | ENSG0000C |
| 226127178 | 226127346 | 1 | processed_pseudoge | ENSG0000C |
| 226144679 | 226186741 | -1 | protein_coding | ENSG0000C |
| 226148003 | 226155071 | 1 | lncRNA | ENSG0000C |
| 226188870 | 226189221 | -1 | processed_pseudoge | ENSG0000C |
| 226223618 | 226227054 | 1 | protein_coding | ENSG0000C |
| 226231149 | 226309869 | -1 | protein_coding | ENSG0000C |
| 226318015 | 226318693 | -1 | processed_pseudoge | ENSG0000C |
| 226331999 | 226333655 | -1 | unprocessed_pseudc | ENSG0000C |
| 226360691 | 226408093 | -1 | protein_coding | ENSG0000C |
| 226411615 | 226412135 | -1 | processed_pseudoge | ENSG0000C |
| 226438564 | 226439344 | 1 | processed_pseudoge | ENSG0000C |
| 226445937 | 226446292 | 1 | misc_RNA | ENSG0000C |
| 226493188 | 226493489 | -1 | processed_pseudoge | ENSG0000C |
| 226538305 | 226542729 | -1 | lncRNA | ENSG0000C |
| 226548764 | 226609230 | 1 | protein_coding | ENSG0000C |
| 226631690 | 226739323 | -1 | protein_coding | ENSG0000C |
| 226656640 | 226675067 | -1 | lncRNA | ENSG0000C |
| 226668897 | 226676345 | 1 | lncRNA | ENSG0000C |
| 226781501 | 226781744 | 1 | processed_pseudoge | ENSG0000C |
| 226827711 | 226833906 | 1 | lncRNA | ENSG0000C |
| 226870184 | 226896105 | 1 | protein_coding | ENSG0000C |
| 226897536 | 226987545 | 1 | protein_coding | ENSG0000C |
| 226958069 | 226958267 | -1 | processed_pseudoge | ENSG0000C |
| 226989865 | 227318474 | -1 | protein_coding | ENSG0000C |
| 226992140 | 226993206 | 1 | lncRNA | ENSG0000C |
| 227123895 | 227132394 | 1 | lncRNA | ENSG0000C |
| 227178333 | 227183444 | 1 | lncRNA | ENSG0000C |
| 227234269 | 227235355 | 1 | processed_pseudoge | ENSG0000C |
| 227264776 | 227265207 | -1 | processed_pseudoge | ENSG0000C |
| 227280449 | 227281967 | 1 | lncRNA | ENSG0000C |
| 227393554 | 227431035 | 1 | lncRNA | ENSG0000C |
| 227410617 | 227411319 | -1 | processed_pseudoge | ENSG0000C |
| 227430526 | 227430976 | 1 | processed_pseudoge | ENSG0000C |
| 227434064 | 227434346 | 1 | processed_pseudoge | ENSG0000C |
| 227482253 | 227482454 | 1 | processed_pseudoge | ENSG0000C |
| 227490691 | 227492296 | 1 | unprocessed_pseudc | ENSG0000C |
| 227493029 | 227495142 | -1 | unprocessed_pseudc | ENSG0000C |
| 227506182 | 227508292 | 1 | unprocessed_pseudc | ENSG0000C |

| | | | | |
|---|---|---|---|---|
| 227509028 | 227510618 | -1 | unprocessed_pseudc | ENSG00000( |
| 227518738 | 227520619 | -1 | lncRNA | ENSG00000( |
| 227561181 | 227561300 | 1 | rRNA_pseudogene | ENSG00000( |
| 227563543 | 227677443 | 1 | protein_coding | ENSG00000( |
| 227598424 | 227599094 | -1 | processed_pseudoge | ENSG00000( |
| 227696892 | 227706699 | -1 | unprocessed_pseudc | ENSG00000( |
| 227728539 | 227781231 | 1 | protein_coding | ENSG00000( |
| 227730425 | 227735411 | -1 | protein_coding | ENSG00000( |
| 227743831 | 227747191 | -1 | lncRNA | ENSG00000( |
| 227786753 | 227792179 | 1 | lncRNA | ENSG00000( |
| 227815693 | 227846470 | 1 | protein_coding | ENSG00000( |
| 227918656 | 227947932 | -1 | protein_coding | ENSG00000( |
| 227941590 | 227941683 | -1 | miRNA | ENSG00000( |
| 227975390 | 227978193 | -1 | unprocessed_pseudc | ENSG00000( |
| 227980051 | 227980227 | 1 | unprocessed_pseudc | ENSG00000( |
| 228006998 | 228061271 | 1 | protein_coding | ENSG00000( |
| 228073909 | 228076550 | -1 | lncRNA | ENSG00000( |
| 228082660 | 228099212 | 1 | protein_coding | ENSG00000( |
| 228097263 | 228097341 | 1 | miRNA | ENSG00000( |
| 228100726 | 228105411 | -1 | protein_coding | ENSG00000( |
| 228106679 | 228109312 | -1 | protein_coding | ENSG00000( |
| 228114997 | 228115473 | -1 | processed_pseudoge | ENSG00000( |
| 228119149 | 228121312 | -1 | lncRNA | ENSG00000( |
| 228121523 | 228123771 | 1 | TEC | ENSG00000( |
| 228134785 | 228134992 | -1 | processed_pseudoge | ENSG00000( |
| 228139962 | 228148984 | 1 | protein_coding | ENSG00000( |
| 228149930 | 228159826 | 1 | protein_coding | ENSG00000( |
| 228164086 | 228165512 | -1 | lncRNA | ENSG00000( |
| 228165804 | 228182257 | 1 | protein_coding | ENSG00000( |
| 228203503 | 228213664 | -1 | lncRNA | ENSG00000( |
| 228208063 | 228378876 | 1 | protein_coding | ENSG00000( |
| 228238241 | 228276066 | -1 | lncRNA | ENSG00000( |
| 228295549 | 228303002 | -1 | lncRNA | ENSG00000( |
| 228329467 | 228331191 | 1 | lncRNA | ENSG00000( |
| 228357012 | 228368554 | -1 | lncRNA | ENSG00000( |
| 228384114 | 228385016 | -1 | lncRNA | ENSG00000( |
| 228393673 | 228406835 | -1 | protein_coding | ENSG00000( |
| 228394290 | 228396967 | 1 | lncRNA | ENSG00000( |
| 228397048 | 228397109 | -1 | miRNA | ENSG00000( |
| 228407365 | 228410870 | 1 | lncRNA | ENSG00000( |
| 228407935 | 228416861 | -1 | protein_coding | ENSG00000( |
| 228424845 | 228425325 | -1 | protein_coding | ENSG00000( |
| 228449163 | 228449621 | -1 | processed_pseudoge | ENSG00000( |
| 228456979 | 228457873 | -1 | protein_coding | ENSG00000( |
| 228458107 | 228460470 | 1 | protein_coding | ENSG00000( |
| 228462074 | 228462152 | 1 | miRNA | ENSG00000( |
| 228464103 | 228464626 | -1 | unitary_pseudogene | ENSG00000( |
| 228486188 | 228487120 | -1 | TEC | ENSG00000( |
| 228487382 | 228496188 | 1 | protein_coding | ENSG00000( |
| 228510425 | 228512305 | -1 | transcribed_unitary_ | ENSG00000( |

| | | | | |
|---|---|---|---|---|
| 228555793 | 228555901 | 1 | rRNA_pseudogene | ENSG0000C |
| 228558296 | 228558339 | -1 | rRNA_pseudogene | ENSG0000C |
| 228610268 | 228610386 | -1 | rRNA | ENSG0000C |
| 228612509 | 228612627 | -1 | rRNA | ENSG0000C |
| 228614750 | 228614868 | -1 | rRNA | ENSG0000C |
| 228616991 | 228617109 | -1 | rRNA | ENSG0000C |
| 228619232 | 228619350 | -1 | rRNA | ENSG0000C |
| 228621447 | 228621565 | -1 | rRNA | ENSG0000C |
| 228623667 | 228623785 | -1 | rRNA | ENSG0000C |
| 228625909 | 228626027 | -1 | rRNA | ENSG0000C |
| 228628148 | 228628266 | -1 | rRNA | ENSG0000C |
| 228630390 | 228630508 | -1 | rRNA | ENSG0000C |
| 228632631 | 228632749 | -1 | rRNA | ENSG0000C |
| 228634871 | 228634989 | -1 | rRNA | ENSG0000C |
| 228637096 | 228637214 | -1 | rRNA | ENSG0000C |
| 228639337 | 228639455 | -1 | rRNA | ENSG0000C |
| 228641568 | 228641686 | -1 | rRNA | ENSG0000C |
| 228643809 | 228643927 | -1 | rRNA | ENSG0000C |
| 228646040 | 228646158 | -1 | rRNA | ENSG0000C |
| 228647912 | 228648032 | -1 | rRNA_pseudogene | ENSG0000C |
| 228650241 | 228651379 | 1 | processed_pseudoge | ENSG0000C |
| 228652436 | 228652560 | 1 | snoRNA | ENSG0000C |
| 228687415 | 228687826 | 1 | processed_pseudoge | ENSG0000C |
| 228735479 | 228746664 | 1 | protein_coding | ENSG0000C |
| 228776312 | 228776865 | 1 | processed_pseudoge | ENSG0000C |
| 228858010 | 228858556 | 1 | processed_pseudoge | ENSG0000C |
| 229022773 | 229038274 | -1 | lncRNA | ENSG0000C |
| 229042171 | 229042591 | 1 | processed_pseudoge | ENSG0000C |
| 229087114 | 229205428 | -1 | lncRNA | ENSG0000C |
| 229223457 | 229242902 | -1 | lncRNA | ENSG0000C |
| 229249636 | 229251810 | 1 | lncRNA | ENSG0000C |
| 229256892 | 229271242 | -1 | lncRNA | ENSG0000C |
| 229271062 | 229305894 | 1 | protein_coding | ENSG0000C |
| 229305135 | 229305326 | 1 | protein_coding | ENSG0000C |
| 229319403 | 229323087 | 1 | lncRNA | ENSG0000C |
| 229321011 | 229343294 | -1 | protein_coding | ENSG0000C |
| 229383128 | 229383188 | 1 | snRNA | ENSG0000C |
| 229410500 | 229410767 | -1 | misc_RNA | ENSG0000C |
| 229425020 | 229425650 | 1 | processed_pseudoge | ENSG0000C |
| 229431245 | 229434098 | -1 | protein_coding | ENSG0000C |
| 229440259 | 229508341 | -1 | protein_coding | ENSG0000C |
| 229440284 | 229441020 | -1 | lncRNA | ENSG0000C |
| 229508369 | 229514272 | 1 | lncRNA | ENSG0000C |
| 229516582 | 229558707 | -1 | protein_coding | ENSG0000C |
| 229535064 | 229535189 | -1 | snRNA | ENSG0000C |
| 229549905 | 229550022 | 1 | rRNA_pseudogene | ENSG0000C |
| 229570532 | 229570796 | 1 | processed_pseudoge | ENSG0000C |
| 229593111 | 229626047 | -1 | protein_coding | ENSG0000C |
| 229626247 | 229660200 | 1 | protein_coding | ENSG0000C |
| 229688999 | 229689904 | -1 | processed_pseudoge | ENSG0000C |

| | | | | |
|---|---|---|---|---|
| 229705234 | 229705861 | -1 | processed_pseudoge | ENSG00000 |
| 229812917 | 229892046 | -1 | lncRNA | ENSG00000 |
| 230002372 | 230007195 | 1 | lncRNA | ENSG00000 |
| 230057990 | 230282122 | 1 | protein_coding | ENSG00000 |
| 230258694 | 230272100 | -1 | lncRNA | ENSG00000 |
| 230280312 | 230281893 | -1 | lncRNA | ENSG00000 |
| 230314490 | 230426332 | -1 | protein_coding | ENSG00000 |
| 230426491 | 230436822 | 1 | lncRNA | ENSG00000 |
| 230592660 | 230595583 | -1 | lncRNA | ENSG00000 |
| 230612009 | 230612412 | -1 | processed_pseudoge | ENSG00000 |
| 230642481 | 230693982 | 1 | protein_coding | ENSG00000 |
| 230692533 | 230696888 | -1 | lncRNA | ENSG00000 |
| 230702523 | 230714122 | -1 | protein_coding | ENSG00000 |
| 230710698 | 230795492 | -1 | lncRNA | ENSG00000 |
| 230747384 | 230802003 | 1 | protein_coding | ENSG00000 |
| 230820250 | 230820358 | 1 | rRNA_pseudogene | ENSG00000 |
| 230823641 | 230824707 | -1 | lncRNA | ENSG00000 |
| 230837119 | 230869589 | -1 | protein_coding | ENSG00000 |
| 230868259 | 230879141 | 1 | lncRNA | ENSG00000 |
| 230878662 | 230890169 | -1 | lncRNA | ENSG00000 |
| 230889949 | 230895788 | 1 | lncRNA | ENSG00000 |
| 230894141 | 230894427 | 1 | misc_RNA | ENSG00000 |
| 230906243 | 230978875 | -1 | protein_coding | ENSG00000 |
| 230978981 | 231000733 | 1 | protein_coding | ENSG00000 |
| 231018958 | 231040254 | -1 | protein_coding | ENSG00000 |
| 231019828 | 231019924 | -1 | miRNA | ENSG00000 |
| 231021611 | 231022183 | 1 | processed_pseudoge | ENSG00000 |
| 231117831 | 231118325 | -1 | processed_pseudoge | ENSG00000 |
| 231162112 | 231221556 | 1 | protein_coding | ENSG00000 |
| 231184098 | 231187627 | -1 | protein_coding | ENSG00000 |
| 231223763 | 231241187 | -1 | protein_coding | ENSG00000 |
| 231241207 | 231277973 | 1 | protein_coding | ENSG00000 |
| 231281414 | 231281524 | -1 | rRNA_pseudogene | ENSG00000 |
| 231332753 | 231337852 | -1 | protein_coding | ENSG00000 |
| 231337104 | 231355023 | 1 | protein_coding | ENSG00000 |
| 231339421 | 231339558 | -1 | unprocessed_pseudo | ENSG00000 |
| 231363751 | 231422287 | -1 | protein_coding | ENSG00000 |
| 231363797 | 231528603 | -1 | protein_coding | ENSG00000 |
| 231475956 | 231476288 | -1 | processed_pseudoge | ENSG00000 |
| 231520729 | 231528618 | -1 | lncRNA | ENSG00000 |
| 231528541 | 231566524 | 1 | protein_coding | ENSG00000 |
| 231528653 | 231819244 | 1 | protein_coding | ENSG00000 |
| 231591292 | 231612090 | -1 | lncRNA | ENSG00000 |
| 231626815 | 232041272 | 1 | protein_coding | ENSG00000 |
| 231670635 | 231670750 | 1 | snRNA | ENSG00000 |
| 231767464 | 231847706 | -1 | lncRNA | ENSG00000 |
| 231814982 | 231815186 | 1 | misc_RNA | ENSG00000 |
| 231925834 | 231945233 | 1 | lncRNA | ENSG00000 |
| 232160091 | 232185643 | 1 | lncRNA | ENSG00000 |
| 232174932 | 232180113 | -1 | lncRNA | ENSG00000 |

| | | | | |
|---|---|---|---|---|
| 232221938 | 232222174 | -1 | processed_pseudoge | ENSG0000C |
| 232222866 | 232223145 | 1 | misc_RNA | ENSG0000C |
| 232397965 | 232561558 | -1 | protein_coding | ENSG0000C |
| 232700204 | 232700308 | 1 | snRNA | ENSG0000C |
| 232718071 | 232722250 | 1 | lncRNA | ENSG0000C |
| 232727251 | 232751299 | -1 | lncRNA | ENSG0000C |
| 232804892 | 232808407 | 1 | protein_coding | ENSG0000C |
| 232832017 | 232832187 | -1 | snRNA | ENSG0000C |
| 232843386 | 232897660 | 1 | lncRNA | ENSG0000C |
| 232917235 | 232918018 | 1 | processed_pseudoge | ENSG0000C |
| 232950605 | 232983882 | 1 | protein_coding | ENSG0000C |
| 232983435 | 233295725 | -1 | protein_coding | ENSG0000C |
| 233288868 | 233289447 | -1 | processed_pseudoge | ENSG0000C |
| 233295325 | 233295727 | -1 | unprocessed_pseudc | ENSG0000C |
| 233327724 | 233385148 | 1 | protein_coding | ENSG0000C |
| 233448626 | 233448781 | -1 | snRNA | ENSG0000C |
| 233527544 | 233543633 | -1 | lncRNA | ENSG0000C |
| 233614106 | 233672514 | 1 | protein_coding | ENSG0000C |
| 233624152 | 233624219 | 1 | miRNA | ENSG0000C |
| 233724454 | 233730746 | 1 | lncRNA | ENSG0000C |
| 233836080 | 233836432 | -1 | processed_pseudoge | ENSG0000C |
| 233844621 | 233865370 | -1 | lncRNA | ENSG0000C |
| 233904104 | 233905931 | -1 | lncRNA | ENSG0000C |
| 233904676 | 234324511 | 1 | protein_coding | ENSG0000C |
| 234026851 | 234027227 | -1 | processed_pseudoge | ENSG0000C |
| 234212606 | 234215088 | -1 | lncRNA | ENSG0000C |
| 234261706 | 234274465 | 1 | lncRNA | ENSG0000C |
| 234268583 | 234272500 | -1 | lncRNA | ENSG0000C |
| 234284972 | 234285541 | 1 | processed_pseudoge | ENSG0000C |
| 234306467 | 234306539 | 1 | miRNA | ENSG0000C |
| 234356704 | 234357141 | -1 | processed_pseudoge | ENSG0000C |
| 234357006 | 234365828 | 1 | lncRNA | ENSG0000C |
| 234372186 | 234372811 | 1 | lncRNA | ENSG0000C |
| 234372807 | 234373593 | -1 | lncRNA | ENSG0000C |
| 234373456 | 234385068 | 1 | protein_coding | ENSG0000C |
| 234391313 | 234479179 | -1 | protein_coding | ENSG0000C |
| 234527887 | 234531803 | -1 | lncRNA | ENSG0000C |
| 234535963 | 234539054 | -1 | lncRNA | ENSG0000C |
| 234550542 | 234554838 | 1 | lncRNA | ENSG0000C |
| 234565298 | 234570088 | 1 | lncRNA | ENSG0000C |
| 234593275 | 234593402 | 1 | snoRNA | ENSG0000C |
| 234604269 | 234609525 | -1 | protein_coding | ENSG0000C |
| 234607008 | 234609483 | 1 | lncRNA | ENSG0000C |
| 234629311 | 234634780 | 1 | lncRNA | ENSG0000C |
| 234644666 | 234647571 | 1 | lncRNA | ENSG0000C |
| 234646289 | 234683176 | 1 | lncRNA | ENSG0000C |
| 234660271 | 234667104 | 1 | lncRNA | ENSG0000C |
| 234669523 | 234696153 | -1 | lncRNA | ENSG0000C |
| 234709383 | 234720220 | 1 | lncRNA | ENSG0000C |
| 234724042 | 234736720 | 1 | lncRNA | ENSG0000C |

| | | | | |
|---|---|---|---|---|
| 234757619 | 234760056 | -1 | lncRNA | ENSG0000C |
| 234811052 | 234932653 | -1 | lncRNA | ENSG0000C |
| 234837976 | 234838069 | 1 | misc_RNA | ENSG0000C |
| 234904186 | 234904488 | 1 | misc_RNA | ENSG0000C |
| 234957231 | 234959989 | 1 | lncRNA | ENSG0000C |
| 234957342 | 234970062 | -1 | lncRNA | ENSG0000C |
| 234979647 | 234980804 | -1 | lncRNA | ENSG0000C |
| 235065479 | 235074220 | -1 | lncRNA | ENSG0000C |
| 235097187 | 235097412 | -1 | processed_pseudoge | ENSG0000C |
| 235104180 | 235104609 | -1 | lncRNA | ENSG0000C |
| 235109341 | 235128837 | -1 | protein_coding | ENSG0000C |
| 235127803 | 235127937 | -1 | snoRNA | ENSG0000C |
| 235131183 | 235161283 | -1 | protein_coding | ENSG0000C |
| 235131634 | 235328219 | -1 | protein_coding | ENSG0000C |
| 235190034 | 235190116 | -1 | miRNA | ENSG0000C |
| 235295865 | 235296335 | 1 | processed_pseudoge | ENSG0000C |
| 235327350 | 235344532 | 1 | protein_coding | ENSG0000C |
| 235328570 | 235448952 | 1 | protein_coding | ENSG0000C |
| 235336806 | 235337087 | 1 | processed_pseudoge | ENSG0000C |
| 235361153 | 235362540 | 1 | processed_pseudoge | ENSG0000C |
| 235366353 | 235367366 | -1 | lncRNA | ENSG0000C |
| 235367360 | 235452443 | 1 | protein_coding | ENSG0000C |
| 235432985 | 235433231 | 1 | transcribed_processe | ENSG0000C |
| 235447190 | 235504452 | -1 | protein_coding | ENSG0000C |
| 235519971 | 235520685 | 1 | processed_pseudoge | ENSG0000C |
| 235537520 | 235538041 | 1 | processed_pseudoge | ENSG0000C |
| 235538052 | 235539845 | -1 | processed_pseudoge | ENSG0000C |
| 235540053 | 235541415 | -1 | processed_pseudoge | ENSG0000C |
| 235541412 | 235541706 | -1 | processed_pseudoge | ENSG0000C |
| 235541759 | 235542097 | -1 | processed_pseudoge | ENSG0000C |
| 235542159 | 235542390 | -1 | processed_pseudoge | ENSG0000C |
| 235547685 | 235650754 | -1 | protein_coding | ENSG0000C |
| 235565761 | 235566937 | 1 | processed_pseudoge | ENSG0000C |
| 235614674 | 235616397 | 1 | processed_pseudoge | ENSG0000C |
| 235661041 | 235883640 | -1 | protein_coding | ENSG0000C |
| 235738005 | 235738989 | 1 | processed_pseudoge | ENSG0000C |
| 235839483 | 235840182 | 1 | lncRNA | ENSG0000C |
| 235853000 | 235853060 | -1 | miRNA | ENSG0000C |
| 235915415 | 235915521 | 1 | snRNA | ENSG0000C |
| 235942553 | 235943805 | -1 | lncRNA | ENSG0000C |
| 235957879 | 235971825 | -1 | lncRNA | ENSG0000C |
| 235975830 | 236065109 | -1 | protein_coding | ENSG0000C |
| 236060677 | 236060772 | 1 | misc_RNA | ENSG0000C |
| 236110061 | 236112370 | 1 | processed_pseudoge | ENSG0000C |
| 236123667 | 236130415 | 1 | lncRNA | ENSG0000C |
| 236142505 | 236221865 | 1 | protein_coding | ENSG0000C |
| 236215101 | 236281958 | -1 | protein_coding | ENSG0000C |
| 236267780 | 236267958 | 1 | snRNA | ENSG0000C |
| 236285976 | 236286960 | -1 | processed_pseudoge | ENSG0000C |
| 236348257 | 236502915 | 1 | protein_coding | ENSG0000C |

| | | | | |
|---|---|---|---|---|
| 236483165 | 236484468 | 1 | transcribed_processe | ENSG0000C |
| 236518000 | 236552981 | 1 | protein_coding | ENSG0000C |
| 236523052 | 236524508 | -1 | lncRNA | ENSG0000C |
| 236536162 | 236536704 | -1 | lncRNA | ENSG0000C |
| 236540094 | 236550280 | -1 | lncRNA | ENSG0000C |
| 236549005 | 236604516 | -1 | protein_coding | ENSG0000C |
| 236664141 | 236764631 | 1 | protein_coding | ENSG0000C |
| 236795292 | 236921278 | 1 | protein_coding | ENSG0000C |
| 236819634 | 236820518 | 1 | processed_pseudoge | ENSG0000C |
| 236981339 | 236981708 | 1 | processed_pseudoge | ENSG0000C |
| 237004103 | 237004441 | -1 | protein_coding | ENSG0000C |
| 237013373 | 237014200 | 1 | lncRNA | ENSG0000C |
| 237042184 | 237833988 | 1 | protein_coding | ENSG0000C |
| 237120807 | 237121109 | 1 | misc_RNA | ENSG0000C |
| 237471119 | 237471191 | 1 | miRNA | ENSG0000C |
| 237555040 | 237555171 | -1 | snoRNA | ENSG0000C |
| 237862175 | 237928321 | 1 | lncRNA | ENSG0000C |
| 237877864 | 237890922 | -1 | protein_coding | ENSG0000C |
| 237926831 | 237927605 | 1 | transcribed_processe | ENSG0000C |
| 237940762 | 237941061 | -1 | processed_pseudoge | ENSG0000C |
| 237941452 | 237942492 | -1 | processed_pseudoge | ENSG0000C |
| 237942698 | 237943653 | -1 | processed_pseudoge | ENSG0000C |
| 237943724 | 237945275 | -1 | protein_coding | ENSG0000C |
| 237948017 | 237949143 | -1 | processed_pseudoge | ENSG0000C |
| 237949214 | 237949734 | 1 | processed_pseudoge | ENSG0000C |
| 237949736 | 237951536 | -1 | processed_pseudoge | ENSG0000C |
| 238107736 | 238108567 | -1 | processed_pseudoge | ENSG0000C |
| 238238943 | 238404906 | 1 | lncRNA | ENSG0000C |
| 238268494 | 238268595 | -1 | processed_pseudoge | ENSG0000C |
| 238325687 | 238325788 | -1 | snRNA | ENSG0000C |
| 238476542 | 238486060 | -1 | lncRNA | ENSG0000C |
| 238485445 | 238538305 | 1 | lncRNA | ENSG0000C |
| 238491358 | 238492644 | 1 | processed_pseudoge | ENSG0000C |
| 238777049 | 238779200 | -1 | processed_pseudoge | ENSG0000C |
| 238842767 | 238848115 | 1 | lncRNA | ENSG0000C |
| 239052748 | 239052903 | -1 | processed_pseudoge | ENSG0000C |
| 239205138 | 239212284 | 1 | lncRNA | ENSG0000C |
| 239247808 | 239250818 | -1 | lncRNA | ENSG0000C |
| 239386565 | 239915452 | 1 | protein_coding | ENSG0000C |
| 239703381 | 239770130 | -1 | lncRNA | ENSG0000C |
| 239898016 | 239899872 | -1 | lncRNA | ENSG0000C |
| 239915439 | 239916955 | -1 | lncRNA | ENSG0000C |
| 239972787 | 239974245 | 1 | processed_pseudoge | ENSG0000C |
| 240007524 | 240008381 | 1 | lncRNA | ENSG0000C |
| 240012646 | 240013226 | 1 | processed_pseudoge | ENSG0000C |
| 240014348 | 240475187 | 1 | protein_coding | ENSG0000C |
| 240142670 | 240145389 | -1 | processed_pseudoge | ENSG0000C |
| 240154651 | 240154772 | -1 | misc_RNA | ENSG0000C |
| 240170155 | 240171291 | 1 | processed_pseudoge | ENSG0000C |
| 240177839 | 240179644 | -1 | lncRNA | ENSG0000C |

| | | | | |
|---|---|---|---|---|
| 240341385 | 240341477 | 1 | misc_RNA | ENSG00000 |
| 240400671 | 240401123 | -1 | lncRNA | ENSG00000 |
| 240489573 | 240612155 | -1 | protein_coding | ENSG00000 |
| 240530452 | 240530862 | 1 | lncRNA | ENSG00000 |
| 240549867 | 240550378 | -1 | processed_pseudoge | ENSG00000 |
| 240588522 | 240590748 | -1 | lncRNA | ENSG00000 |
| 240636599 | 240637052 | -1 | processed_pseudoge | ENSG00000 |
| 240653367 | 240653477 | 1 | snRNA | ENSG00000 |
| 240654241 | 240654625 | 1 | processed_pseudoge | ENSG00000 |
| 240698320 | 240698429 | 1 | misc_RNA | ENSG00000 |
| 240739419 | 240739853 | -1 | lncRNA | ENSG00000 |
| 240763334 | 240776824 | 1 | lncRNA | ENSG00000 |
| 240769420 | 240771534 | -1 | processed_pseudoge | ENSG00000 |
| 240775515 | 241357230 | -1 | protein_coding | ENSG00000 |
| 240823006 | 240823433 | -1 | transcribed_processe | ENSG00000 |
| 240919653 | 240920609 | 1 | processed_pseudoge | ENSG00000 |
| 240998451 | 240998794 | -1 | processed_pseudoge | ENSG00000 |
| 241132272 | 241132346 | 1 | miRNA | ENSG00000 |
| 241305580 | 241305882 | -1 | processed_pseudoge | ENSG00000 |
| 241357343 | 241364054 | 1 | lncRNA | ENSG00000 |
| 241413716 | 241468661 | 1 | lncRNA | ENSG00000 |
| 241453751 | 241484995 | -1 | lncRNA | ENSG00000 |
| 241497603 | 241519761 | -1 | protein_coding | ENSG00000 |
| 241532134 | 241595642 | 1 | protein_coding | ENSG00000 |
| 241590102 | 241677376 | -1 | protein_coding | ENSG00000 |
| 241628853 | 241640254 | -1 | protein_coding | ENSG00000 |
| 241640555 | 241641835 | 1 | lncRNA | ENSG00000 |
| 241652278 | 241802133 | 1 | protein_coding | ENSG00000 |
| 241831935 | 241832787 | 1 | processed_pseudoge | ENSG00000 |
| 241847967 | 241895148 | 1 | protein_coding | ENSG00000 |
| 241916123 | 241916543 | 1 | processed_pseudoge | ENSG00000 |
| 241957767 | 241959062 | 1 | protein_coding | ENSG00000 |
| 241993185 | 241993682 | 1 | processed_pseudoge | ENSG00000 |
| 241995490 | 241999098 | -1 | protein_coding | ENSG00000 |
| 242057085 | 242060242 | 1 | unprocessed_pseudc | ENSG00000 |
| 242060490 | 242062595 | -1 | unprocessed_pseudc | ENSG00000 |
| 242082986 | 242524696 | -1 | protein_coding | ENSG00000 |
| 242147230 | 242209129 | 1 | lncRNA | ENSG00000 |
| 242188647 | 242188789 | -1 | misc_RNA | ENSG00000 |
| 242203555 | 242210827 | 1 | lncRNA | ENSG00000 |
| 242345558 | 242345718 | 1 | unprocessed_pseudc | ENSG00000 |
| 242365189 | 242365822 | 1 | processed_pseudoge | ENSG00000 |
| 242376923 | 242377432 | -1 | processed_pseudoge | ENSG00000 |
| 242671140 | 242671576 | 1 | processed_pseudoge | ENSG00000 |
| 242772620 | 242773103 | 1 | processed_pseudoge | ENSG00000 |
| 242882066 | 242883475 | -1 | processed_pseudoge | ENSG00000 |
| 242890247 | 242890803 | 1 | processed_pseudoge | ENSG00000 |
| 242975005 | 242975084 | 1 | processed_pseudoge | ENSG00000 |
| 243005845 | 243007468 | -1 | lncRNA | ENSG00000 |
| 243029512 | 243052252 | -1 | transcribed_unproce | ENSG00000 |

| | | | | |
|---|---|---|---|---|
| 243049782 | 243051619 | 1 | processed_pseudoge | ENSG00000 |
| 243054861 | 243056394 | -1 | processed_pseudoge | ENSG00000 |
| 243056307 | 243101744 | -1 | transcribed_unproce | ENSG00000 |
| 243081156 | 243081259 | -1 | snRNA | ENSG00000 |
| 243124428 | 243255348 | -1 | protein_coding | ENSG00000 |
| 243135898 | 243140588 | 1 | lncRNA | ENSG00000 |
| 243164638 | 243169445 | 1 | lncRNA | ENSG00000 |
| 243256034 | 243500091 | 1 | protein_coding | ENSG00000 |
| 243267257 | 243268119 | -1 | processed_pseudoge | ENSG00000 |
| 243346176 | 243346255 | 1 | miRNA | ENSG00000 |
| 243488233 | 243851079 | -1 | protein_coding | ENSG00000 |
| 243545532 | 243548329 | 1 | lncRNA | ENSG00000 |
| 243624666 | 243625056 | -1 | processed_pseudoge | ENSG00000 |
| 243702857 | 243740821 | -1 | lncRNA | ENSG00000 |
| 243793205 | 243794400 | -1 | lncRNA | ENSG00000 |
| 243917402 | 244047317 | 1 | lncRNA | ENSG00000 |
| 244048939 | 244057476 | 1 | protein_coding | ENSG00000 |
| 244064330 | 244067909 | 1 | TEC | ENSG00000 |
| 244068820 | 244093026 | -1 | lncRNA | ENSG00000 |
| 244087306 | 244090295 | 1 | lncRNA | ENSG00000 |
| 244103932 | 244104210 | 1 | misc_RNA | ENSG00000 |
| 244107365 | 244109011 | 1 | lncRNA | ENSG00000 |
| 244184953 | 244187559 | -1 | lncRNA | ENSG00000 |
| 244352635 | 244389663 | 1 | protein_coding | ENSG00000 |
| 244375100 | 244409592 | -1 | lncRNA | ENSG00000 |
| 244394976 | 244395660 | -1 | processed_pseudoge | ENSG00000 |
| 244408494 | 244451909 | -1 | protein_coding | ENSG00000 |
| 244454377 | 244641177 | 1 | protein_coding | ENSG00000 |
| 244598391 | 244598687 | -1 | processed_pseudoge | ENSG00000 |
| 244653103 | 244709033 | 1 | protein_coding | ENSG00000 |
| 244694432 | 244694720 | -1 | processed_pseudoge | ENSG00000 |
| 244729898 | 244730962 | -1 | protein_coding | ENSG00000 |
| 244731024 | 244731586 | 1 | lncRNA | ENSG00000 |
| 244834374 | 244835011 | -1 | lncRNA | ENSG00000 |
| 244835616 | 244845057 | 1 | protein_coding | ENSG00000 |
| 244840638 | 244864560 | -1 | protein_coding | ENSG00000 |
| 244864738 | 244865272 | 1 | lncRNA | ENSG00000 |
| 244943910 | 244944216 | 1 | misc_RNA | ENSG00000 |
| 244954858 | 244954963 | 1 | snRNA | ENSG00000 |
| 244969350 | 244971088 | -1 | lncRNA | ENSG00000 |
| 244969705 | 245127164 | 1 | protein_coding | ENSG00000 |
| 245023015 | 245023118 | 1 | snRNA | ENSG00000 |
| 245043928 | 245044032 | 1 | snRNA | ENSG00000 |
| 245123471 | 245124450 | -1 | processed_pseudoge | ENSG00000 |
| 245133927 | 245134090 | -1 | snRNA | ENSG00000 |
| 245154985 | 245709432 | 1 | protein_coding | ENSG00000 |
| 245206444 | 245234501 | -1 | lncRNA | ENSG00000 |
| 245282831 | 245283089 | -1 | processed_pseudoge | ENSG00000 |
| 245614773 | 245615145 | -1 | lncRNA | ENSG00000 |
| 245673732 | 245676478 | -1 | lncRNA | ENSG00000 |

| | | | | |
|---|---|---|---|---|
| 245749342 | 246507312 | -1 | protein_coding | ENSG0000C |
| 246025897 | 246035603 | 1 | lncRNA | ENSG0000C |
| 246108626 | 246113866 | 1 | lncRNA | ENSG0000C |
| 246178778 | 246179204 | 1 | processed_pseudoge | ENSG0000C |
| 246189014 | 246189118 | -1 | snRNA | ENSG0000C |
| 246321663 | 246322438 | -1 | lncRNA | ENSG0000C |
| 246426006 | 246426119 | -1 | misc_RNA | ENSG0000C |
| 246509038 | 246522592 | -1 | unprocessed_pseudc | ENSG0000C |
| 246516039 | 246524287 | -1 | lncRNA | ENSG0000C |
| 246527030 | 246528155 | 1 | unprocessed_pseudc | ENSG0000C |
| 246528562 | 246528841 | -1 | unprocessed_pseudc | ENSG0000C |
| 246540561 | 246566261 | -1 | protein_coding | ENSG0000C |
| 246566444 | 246668595 | 1 | protein_coding | ENSG0000C |
| 246605873 | 246608102 | 1 | lncRNA | ENSG0000C |
| 246632251 | 246632478 | -1 | processed_pseudoge | ENSG0000C |
| 246682108 | 246685075 | 1 | lncRNA | ENSG0000C |
| 246690047 | 246691821 | 1 | lncRNA | ENSG0000C |
| 246724409 | 246768137 | 1 | protein_coding | ENSG0000C |
| 246755511 | 246755832 | -1 | processed_pseudoge | ENSG0000C |
| 246771837 | 246839611 | -1 | transcribed_unitary_ | ENSG0000C |
| 246772301 | 246775772 | 1 | lncRNA | ENSG0000C |
| 246775880 | 246792385 | 1 | lncRNA | ENSG0000C |
| 246818076 | 246818180 | 1 | unprocessed_pseudc | ENSG0000C |
| 246833448 | 246833529 | -1 | snoRNA | ENSG0000C |
| 246839098 | 246931967 | -1 | protein_coding | ENSG0000C |
| 246945547 | 247078811 | -1 | protein_coding | ENSG0000C |
| 246945547 | 247008093 | -1 | protein_coding | ENSG0000C |
| 247015068 | 247017291 | 1 | processed_pseudoge | ENSG0000C |
| 247034637 | 247078811 | -1 | protein_coding | ENSG0000C |
| 247042791 | 247044021 | 1 | processed_pseudoge | ENSG0000C |
| 247085389 | 247086067 | -1 | unprocessed_pseudc | ENSG0000C |
| 247099962 | 247104372 | -1 | protein_coding | ENSG0000C |
| 247105405 | 247108929 | 1 | lncRNA | ENSG0000C |
| 247110160 | 247112417 | -1 | lncRNA | ENSG0000C |
| 247119669 | 247120278 | 1 | processed_pseudoge | ENSG0000C |
| 247121975 | 247172016 | -1 | protein_coding | ENSG0000C |
| 247175256 | 247175525 | 1 | processed_pseudoge | ENSG0000C |
| 247183813 | 247185482 | -1 | processed_pseudoge | ENSG0000C |
| 247187281 | 247188526 | -1 | lncRNA | ENSG0000C |
| 247189851 | 247210856 | -1 | transcribed_unproce | ENSG0000C |
| 247201967 | 247202060 | -1 | miRNA | ENSG0000C |
| 247204570 | 247204661 | -1 | rRNA_pseudogene | ENSG0000C |
| 247210691 | 247241823 | -1 | lncRNA | ENSG0000C |
| 247229940 | 247231880 | 1 | processed_pseudoge | ENSG0000C |
| 247236868 | 247237957 | -1 | processed_pseudoge | ENSG0000C |
| 247255972 | 247257210 | 1 | polymorphic_pseudc | ENSG0000C |
| 247294835 | 247294941 | 1 | misc_RNA | ENSG0000C |
| 247297412 | 247331846 | -1 | protein_coding | ENSG0000C |
| 247337886 | 247348076 | -1 | lncRNA | ENSG0000C |
| 247390243 | 247393839 | 1 | lncRNA | ENSG0000C |

| | | | | |
|---|---|---|---|---|
| 247416156 | 247449108 | 1 | protein_coding | ENSG00000 |
| 247449118 | 247458105 | -1 | protein_coding | ENSG00000 |
| 247488093 | 247492415 | 1 | transcribed_unproce | ENSG00000 |
| 247507058 | 247577690 | 1 | protein_coding | ENSG00000 |
| 247524677 | 247536440 | -1 | protein_coding | ENSG00000 |
| 247524679 | 247532613 | -1 | protein_coding | ENSG00000 |
| 247565639 | 247640854 | -1 | lncRNA | ENSG00000 |
| 247588360 | 247589313 | 1 | protein_coding | ENSG00000 |
| 247605586 | 247606515 | 1 | protein_coding | ENSG00000 |
| 247607126 | 247607201 | -1 | snRNA | ENSG00000 |
| 247617143 | 247622372 | 1 | transcribed_unproce | ENSG00000 |
| 247639630 | 247761255 | 1 | lncRNA | ENSG00000 |
| 247666887 | 247667841 | 1 | unprocessed_pseudo | ENSG00000 |
| 247670812 | 247679739 | -1 | protein_coding | ENSG00000 |
| 247711436 | 247716646 | -1 | protein_coding | ENSG00000 |
| 247711813 | 247723764 | -1 | protein_coding | ENSG00000 |
| 247722957 | 247724043 | -1 | protein_coding | ENSG00000 |
| 247738615 | 247739559 | 1 | protein_coding | ENSG00000 |
| 247754846 | 247760556 | -1 | protein_coding | ENSG00000 |
| 247770169 | 247779524 | 1 | polymorphic_pseudo | ENSG00000 |
| 247814660 | 247824119 | -1 | protein_coding | ENSG00000 |
| 247827486 | 247828502 | 1 | processed_pseudoge | ENSG00000 |
| 247833349 | 247834288 | -1 | unprocessed_pseudo | ENSG00000 |
| 247840928 | 247841896 | -1 | protein_coding | ENSG00000 |
| 247857187 | 247880138 | 1 | protein_coding | ENSG00000 |
| 247895587 | 247896531 | 1 | protein_coding | ENSG00000 |
| 247920252 | 247921956 | 1 | protein_coding | ENSG00000 |
| 247924889 | 247935339 | 1 | protein_coding | ENSG00000 |
| 247936061 | 247937513 | -1 | processed_pseudoge | ENSG00000 |
| 247938973 | 247939659 | 1 | unprocessed_pseudo | ENSG00000 |
| 247948858 | 247949796 | 1 | protein_coding | ENSG00000 |
| 247965233 | 247966386 | 1 | protein_coding | ENSG00000 |
| 247974741 | 247975653 | 1 | unprocessed_pseudo | ENSG00000 |
| 247980872 | 247991204 | 1 | transcribed_unproce | ENSG00000 |
| 247997000 | 247997096 | 1 | misc_RNA | ENSG00000 |
| 248003129 | 248004068 | 1 | unprocessed_pseudo | ENSG00000 |
| 248013660 | 248024276 | 1 | protein_coding | ENSG00000 |
| 248029120 | 248029221 | 1 | misc_RNA | ENSG00000 |
| 248030070 | 248042305 | 1 | protein_coding | ENSG00000 |
| 248045788 | 248045889 | 1 | misc_RNA | ENSG00000 |
| 248046836 | 248063407 | 1 | protein_coding | ENSG00000 |
| 248083522 | 248087522 | -1 | transcribed_unproce | ENSG00000 |
| 248095184 | 248101103 | 1 | protein_coding | ENSG00000 |
| 248121932 | 248122882 | 1 | unprocessed_pseudo | ENSG00000 |
| 248145148 | 248146086 | 1 | protein_coding | ENSG00000 |
| 248174821 | 248181067 | 1 | protein_coding | ENSG00000 |
| 248197265 | 248212925 | 1 | protein_coding | ENSG00000 |
| 248231417 | 248244679 | 1 | protein_coding | ENSG00000 |
| 248269917 | 248277976 | -1 | protein_coding | ENSG00000 |
| 248290139 | 248303424 | -1 | protein_coding | ENSG00000 |

| | | | | |
|---|---|---|---|---|
| 248323630 | 248324568 | -1 | protein_coding | ENSG00000 |
| 248348775 | 248349713 | 1 | protein_coding | ENSG00000 |
| 248361581 | 248362627 | 1 | protein_coding | ENSG00000 |
| 248375746 | 248391811 | 1 | protein_coding | ENSG00000 |
| 248403048 | 248408020 | 1 | protein_coding | ENSG00000 |
| 248436359 | 248444316 | 1 | polymorphic_pseudo | ENSG00000 |
| 248445512 | 248455725 | 1 | protein_coding | ENSG00000 |
| 248473351 | 248474307 | 1 | protein_coding | ENSG00000 |
| 248484245 | 248485484 | -1 | lncRNA | ENSG00000 |
| 248488589 | 248491113 | 1 | protein_coding | ENSG00000 |
| 248498308 | 248498649 | 1 | unprocessed_pseudo | ENSG00000 |
| 248508073 | 248527337 | 1 | protein_coding | ENSG00000 |
| 248548756 | 248565299 | 1 | lncRNA | ENSG00000 |
| 248549444 | 248549762 | -1 | unprocessed_pseudo | ENSG00000 |
| 248556912 | 248562772 | -1 | protein_coding | ENSG00000 |
| 248573801 | 248574757 | -1 | protein_coding | ENSG00000 |
| 248590487 | 248597700 | -1 | protein_coding | ENSG00000 |
| 248594925 | 248595033 | -1 | misc_RNA | ENSG00000 |
| 248601416 | 248601869 | 1 | processed_pseudoge | ENSG00000 |
| 248623557 | 248635091 | -1 | protein_coding | ENSG00000 |
| 248636356 | 248645278 | -1 | protein_coding | ENSG00000 |
| 248649838 | 248655528 | -1 | protein_coding | ENSG00000 |
| 248681249 | 248748349 | -1 | protein_coding | ENSG00000 |
| 248681322 | 248682328 | -1 | protein_coding | ENSG00000 |
| 248691760 | 248716755 | 1 | transcribed_unitary_ | ENSG00000 |
| 248708202 | 248708641 | 1 | processed_pseudoge | ENSG00000 |
| 248721993 | 248722201 | -1 | transcribed_processe | ENSG00000 |
| 248739415 | 248755759 | -1 | protein_coding | ENSG00000 |
| 248771635 | 248773792 | -1 | processed_pseudoge | ENSG00000 |
| 248810446 | 248825915 | -1 | protein_coding | ENSG00000 |
| 248826377 | 248826443 | 1 | miRNA | ENSG00000 |
| 248838210 | 248849517 | 1 | protein_coding | ENSG00000 |
| 248850006 | 248859144 | -1 | protein_coding | ENSG00000 |
| 248859164 | 248864796 | 1 | lncRNA | ENSG00000 |
| 248906196 | 248919946 | 1 | protein_coding | ENSG00000 |
| 248912690 | 248912795 | -1 | snRNA | ENSG00000 |
| 248936581 | 248937043 | 1 | processed_pseudoge | ENSG00000 |
| 38814 | 46870 | -1 | protein_coding | ENSG00000 |
| 197569 | 209892 | 1 | lncRNA | ENSG00000 |
| 217730 | 266398 | -1 | protein_coding | ENSG00000 |
| 264140 | 278283 | 1 | protein_coding | ENSG00000 |
| 279558 | 288851 | -1 | protein_coding | ENSG00000 |
| 286419 | 301515 | 1 | lncRNA | ENSG00000 |
| 290941 | 314373 | -1 | lncRNA | ENSG00000 |
| 307683 | 309424 | 1 | lncRNA | ENSG00000 |
| 317912 | 342118 | 1 | lncRNA | ENSG00000 |
| 388412 | 416885 | 1 | lncRNA | ENSG00000 |
| 421057 | 424059 | 1 | lncRNA | ENSG00000 |
| 490944 | 503905 | -1 | lncRNA | ENSG00000 |
| 495731 | 496633 | 1 | lncRNA | ENSG00000 |

| | | | | |
|---|---|---|---|---|
| 545805 | 546667 | 1 | lncRNA | ENSG0000C |
| 558153 | 578145 | 1 | lncRNA | ENSG0000C |
| 663877 | 677406 | -1 | protein_coding | ENSG0000C |
| 677186 | 697382 | 1 | lncRNA | ENSG0000C |
| 687799 | 688348 | 1 | TEC | ENSG0000C |
| 692083 | 693235 | -1 | lncRNA | ENSG0000C |
| 724194 | 731224 | -1 | lncRNA | ENSG0000C |
| 739588 | 740164 | -1 | lncRNA | ENSG0000C |
| 741977 | 749856 | 1 | lncRNA | ENSG0000C |
| 742488 | 747767 | 1 | lncRNA | ENSG0000C |
| 779840 | 868608 | -1 | lncRNA | ENSG0000C |
| 899462 | 905539 | -1 | lncRNA | ENSG0000C |
| 910926 | 921124 | -1 | lncRNA | ENSG0000C |
| 949627 | 950274 | -1 | lncRNA | ENSG0000C |
| 950849 | 1367613 | 1 | protein_coding | ENSG0000C |
| 1059133 | 1068341 | -1 | lncRNA | ENSG0000C |
| 1158310 | 1160424 | -1 | lncRNA | ENSG0000C |
| 1341044 | 1346568 | -1 | lncRNA | ENSG0000C |
| 1366234 | 1580747 | -1 | lncRNA | ENSG0000C |
| 1374066 | 1543711 | 1 | protein_coding | ENSG0000C |
| 1546665 | 1620113 | -1 | lncRNA | ENSG0000C |
| 1552445 | 1554701 | 1 | lncRNA | ENSG0000C |
| 1620510 | 1625419 | -1 | lncRNA | ENSG0000C |
| 1631887 | 1744852 | -1 | protein_coding | ENSG0000C |
| 1789113 | 2331664 | -1 | protein_coding | ENSG0000C |
| 1795525 | 1811526 | 1 | lncRNA | ENSG0000C |
| 1824412 | 1828667 | 1 | lncRNA | ENSG0000C |
| 2319232 | 2327110 | 1 | lncRNA | ENSG0000C |
| 2638178 | 2696285 | -1 | lncRNA | ENSG0000C |
| 2697490 | 2701812 | -1 | lncRNA | ENSG0000C |
| 2707368 | 2840512 | -1 | lncRNA | ENSG0000C |
| 2729908 | 2730957 | -1 | lncRNA | ENSG0000C |
| 2870558 | 2871231 | -1 | lncRNA | ENSG0000C |
| 2895048 | 3126026 | -1 | lncRNA | ENSG0000C |
| 3131581 | 3145780 | -1 | lncRNA | ENSG0000C |
| 3156756 | 3157797 | 1 | lncRNA | ENSG0000C |
| 3188925 | 3377882 | -1 | protein_coding | ENSG0000C |
| 3298341 | 3301465 | -1 | lncRNA | ENSG0000C |
| 3379675 | 3485094 | 1 | protein_coding | ENSG0000C |
| 3481242 | 3482409 | -1 | lncRNA | ENSG0000C |
| 3496956 | 3497428 | 1 | lncRNA | ENSG0000C |
| 3497366 | 3519531 | -1 | protein_coding | ENSG0000C |
| 3519275 | 3523197 | 1 | lncRNA | ENSG0000C |
| 3531813 | 3536873 | -1 | lncRNA | ENSG0000C |
| 3532008 | 3558333 | -1 | protein_coding | ENSG0000C |
| 3541430 | 3558333 | -1 | protein_coding | ENSG0000C |
| 3558474 | 3564842 | 1 | lncRNA | ENSG0000C |
| 3568505 | 3575100 | -1 | lncRNA | ENSG0000C |
| 3575260 | 3580920 | 1 | protein_coding | ENSG0000C |
| 3594832 | 3644644 | 1 | protein_coding | ENSG0000C |

| | | | | |
|---|---|---|---|---|
| 3603397 | 3604242 | -1 | lncRNA | ENSG0000C |
| 3617548 | 3617757 | -1 | unprocessed_pseudc | ENSG0000C |
| 3658200 | 3702671 | 1 | protein_coding | ENSG0000C |
| 3688021 | 3688981 | -1 | processed_pseudoge | ENSG0000C |
| 3702585 | 3704153 | -1 | lncRNA | ENSG0000C |
| 3703575 | 3848008 | 1 | protein_coding | ENSG0000C |
| 3957652 | 3974124 | -1 | lncRNA | ENSG0000C |
| 4136644 | 4140731 | -1 | lncRNA | ENSG0000C |
| 4514204 | 4515056 | -1 | processed_pseudoge | ENSG0000C |
| 4628216 | 4656498 | -1 | lncRNA | ENSG0000C |
| 4827001 | 4827084 | -1 | snoRNA | ENSG0000C |
| 4945277 | 4945383 | 1 | snRNA | ENSG0000C |
| 5313740 | 5314134 | -1 | processed_pseudoge | ENSG0000C |
| 5549780 | 5556031 | -1 | lncRNA | ENSG0000C |
| 5602505 | 5691488 | -1 | lncRNA | ENSG0000C |
| 5618327 | 5691118 | -1 | lncRNA | ENSG0000C |
| 5692384 | 5701385 | 1 | protein_coding | ENSG0000C |
| 5696220 | 5719670 | 1 | lncRNA | ENSG0000C |
| 5726253 | 5730355 | 1 | lncRNA | ENSG0000C |
| 5810641 | 5812670 | -1 | lncRNA | ENSG0000C |
| 5932543 | 6003510 | 1 | lncRNA | ENSG0000C |
| 5974662 | 5974732 | 1 | miRNA | ENSG0000C |
| 6258348 | 6341788 | -1 | lncRNA | ENSG0000C |
| 6366010 | 6375422 | -1 | lncRNA | ENSG0000C |
| 6495651 | 6511839 | 1 | lncRNA | ENSG0000C |
| 6495877 | 6496798 | -1 | processed_pseudoge | ENSG0000C |
| 6615389 | 6651104 | -1 | lncRNA | ENSG0000C |
| 6649706 | 6650897 | 1 | lncRNA | ENSG0000C |
| 6650373 | 6650439 | 1 | miRNA | ENSG0000C |
| 6728177 | 6770311 | -1 | lncRNA | ENSG0000C |
| 6819463 | 6840464 | -1 | lncRNA | ENSG0000C |
| 6840570 | 6866635 | -1 | protein_coding | ENSG0000C |
| 6865806 | 6898239 | 1 | protein_coding | ENSG0000C |
| 6905724 | 6906301 | -1 | lncRNA | ENSG0000C |
| 6911754 | 6918734 | -1 | lncRNA | ENSG0000C |
| 6917412 | 7068286 | 1 | protein_coding | ENSG0000C |
| 7045329 | 7077916 | -1 | lncRNA | ENSG0000C |
| 7141227 | 7141544 | -1 | misc_RNA | ENSG0000C |
| 7260871 | 7261504 | 1 | lncRNA | ENSG0000C |
| 7324748 | 7325062 | -1 | processed_pseudoge | ENSG0000C |
| 7383227 | 7450544 | 1 | lncRNA | ENSG0000C |
| 7576841 | 7576971 | 1 | snRNA | ENSG0000C |
| 7671330 | 7675736 | 1 | lncRNA | ENSG0000C |
| 7671604 | 7672246 | -1 | lncRNA | ENSG0000C |
| 7725801 | 7730705 | 1 | lncRNA | ENSG0000C |
| 7735645 | 7735783 | -1 | processed_pseudoge | ENSG0000C |
| 7736438 | 7737095 | -1 | processed_pseudoge | ENSG0000C |
| 7886767 | 7899725 | -1 | lncRNA | ENSG0000C |
| 7922425 | 8278186 | -1 | lncRNA | ENSG0000C |
| 7937282 | 7953402 | 1 | lncRNA | ENSG0000C |

| | | | | |
|---|---|---|---|---|
| 7988683 | 8488090 | -1 | lncRNA | ENSG00000( |
| 8139335 | 8144950 | 1 | lncRNA | ENSG00000( |
| 8461703 | 8592903 | -1 | lncRNA | ENSG00000( |
| 8488420 | 8505027 | 1 | lncRNA | ENSG00000( |
| 8600892 | 8622942 | -1 | lncRNA | ENSG00000( |
| 8603082 | 8603359 | 1 | processed_pseudoge | ENSG00000( |
| 8666636 | 8681864 | -1 | lncRNA | ENSG00000( |
| 8678845 | 8684461 | 1 | protein_coding | ENSG00000( |
| 8721081 | 8837630 | -1 | protein_coding | ENSG00000( |
| 8852690 | 9003709 | -1 | protein_coding | ENSG00000( |
| 9018293 | 9018899 | -1 | processed_pseudoge | ENSG00000( |
| 9081395 | 9081727 | -1 | processed_pseudoge | ENSG00000( |
| 9103440 | 9105114 | 1 | lncRNA | ENSG00000( |
| 9106593 | 9109865 | 1 | lncRNA | ENSG00000( |
| 9110457 | 9116947 | 1 | lncRNA | ENSG00000( |
| 9143166 | 9146106 | 1 | lncRNA | ENSG00000( |
| 9206765 | 9405683 | 1 | protein_coding | ENSG00000( |
| 9271292 | 9271628 | -1 | processed_pseudoge | ENSG00000( |
| 9403475 | 9423547 | -1 | protein_coding | ENSG00000( |
| 9423651 | 9473101 | 1 | protein_coding | ENSG00000( |
| 9473658 | 9496543 | 1 | protein_coding | ENSG00000( |
| 9488486 | 9556732 | -1 | protein_coding | ENSG00000( |
| 9501466 | 9512413 | 1 | lncRNA | ENSG00000( |
| 9555899 | 9556775 | 1 | lncRNA | ENSG00000( |
| 9583967 | 9630997 | -1 | protein_coding | ENSG00000( |
| 9615267 | 9615373 | 1 | misc_RNA | ENSG00000( |
| 9638745 | 9712920 | 1 | lncRNA | ENSG00000( |
| 9671875 | 9708413 | 1 | lncRNA | ENSG00000( |
| 9740643 | 9740783 | -1 | snRNA | ENSG00000( |
| 9746704 | 9746869 | 1 | processed_pseudoge | ENSG00000( |
| 9757496 | 9770341 | -1 | lncRNA | ENSG00000( |
| 9843443 | 9934416 | 1 | protein_coding | ENSG00000( |
| 9936360 | 9939590 | 1 | lncRNA | ENSG00000( |
| 9939088 | 9950817 | -1 | lncRNA | ENSG00000( |
| 9951693 | 10002277 | 1 | protein_coding | ENSG00000( |
| 10001757 | 10006030 | -1 | lncRNA | ENSG00000( |
| 10021578 | 10022825 | 1 | lncRNA | ENSG00000( |
| 10039092 | 10040663 | -1 | lncRNA | ENSG00000( |
| 10042849 | 10054836 | 1 | protein_coding | ENSG00000( |
| 10054421 | 10054866 | -1 | lncRNA | ENSG00000( |
| 10056780 | 10080944 | -1 | protein_coding | ENSG00000( |
| 10083781 | 10086101 | 1 | lncRNA | ENSG00000( |
| 10090205 | 10090319 | -1 | snoRNA | ENSG00000( |
| 10120698 | 10211725 | 1 | protein_coding | ENSG00000( |
| 10155072 | 10155206 | -1 | snoRNA | ENSG00000( |
| 10192614 | 10192671 | -1 | miRNA | ENSG00000( |
| 10280628 | 10280891 | 1 | misc_RNA | ENSG00000( |
| 10287800 | 10302485 | -1 | lncRNA | ENSG00000( |
| 10296047 | 10296180 | -1 | snoRNA | ENSG00000( |
| 10302889 | 10427617 | 1 | protein_coding | ENSG00000( |

| | | | | |
|---|---|---|---|---|
| 10439968 | 10448327 | -1 | protein_coding | ENSG0000C |
| 10446714 | 10446849 | -1 | snoRNA | ENSG0000C |
| 10448689 | 10457693 | 1 | lncRNA | ENSG0000C |
| 10570754 | 10689987 | -1 | protein_coding | ENSG0000C |
| 10589166 | 10604830 | 1 | lncRNA | ENSG0000C |
| 10690344 | 10692099 | 1 | lncRNA | ENSG0000C |
| 10691538 | 10691831 | 1 | misc_RNA | ENSG0000C |
| 10717773 | 10720952 | 1 | lncRNA | ENSG0000C |
| 10721100 | 10785110 | 1 | protein_coding | ENSG0000C |
| 10744186 | 10744249 | 1 | snRNA | ENSG0000C |
| 10767875 | 10770058 | -1 | lncRNA | ENSG0000C |
| 10783391 | 10837977 | -1 | protein_coding | ENSG0000C |
| 10815485 | 10815546 | -1 | snRNA | ENSG0000C |
| 10844477 | 10885118 | 1 | lncRNA | ENSG0000C |
| 10847577 | 10854955 | 1 | lncRNA | ENSG0000C |
| 10874291 | 10874567 | -1 | processed_pseudoge | ENSG0000C |
| 10911934 | 10914225 | 1 | protein_coding | ENSG0000C |
| 11102142 | 11102997 | 1 | processed_pseudoge | ENSG0000C |
| 11105317 | 11132821 | -1 | lncRNA | ENSG0000C |
| 11133053 | 11146790 | 1 | protein_coding | ENSG0000C |
| 11155198 | 11178870 | 1 | protein_coding | ENSG0000C |
| 11179759 | 11348330 | -1 | protein_coding | ENSG0000C |
| 11233988 | 11234094 | 1 | snRNA | ENSG0000C |
| 11308025 | 11308941 | 1 | processed_pseudoge | ENSG0000C |
| 11351627 | 11352118 | 1 | processed_pseudoge | ENSG0000C |
| 11357515 | 11365636 | -1 | lncRNA | ENSG0000C |
| 11388023 | 11390367 | -1 | lncRNA | ENSG0000C |
| 11391810 | 11403080 | 1 | lncRNA | ENSG0000C |
| 11405508 | 11427186 | 1 | lncRNA | ENSG0000C |
| 11444375 | 11466177 | -1 | protein_coding | ENSG0000C |
| 11482341 | 11642788 | 1 | protein_coding | ENSG0000C |
| 11517397 | 11517503 | 1 | rRNA_pseudogene | ENSG0000C |
| 11540605 | 11540677 | -1 | miRNA | ENSG0000C |
| 11561194 | 11561306 | 1 | snRNA | ENSG0000C |
| 11561661 | 11561779 | -1 | rRNA_pseudogene | ENSG0000C |
| 11561661 | 11561779 | -1 | rRNA | ENSG0000C |
| 11584773 | 11585047 | 1 | misc_RNA | ENSG0000C |
| 11658178 | 11670195 | -1 | protein_coding | ENSG0000C |
| 11665369 | 11665969 | -1 | processed_pseudoge | ENSG0000C |
| 11673964 | 11676925 | -1 | lncRNA | ENSG0000C |
| 11677595 | 11827409 | 1 | protein_coding | ENSG0000C |
| 11681460 | 11683328 | 1 | lncRNA | ENSG0000C |
| 11721619 | 11724222 | -1 | lncRNA | ENSG0000C |
| 11740997 | 11745301 | -1 | lncRNA | ENSG0000C |
| 11767444 | 11767525 | 1 | miRNA | ENSG0000C |
| 11833926 | 12702913 | 1 | lncRNA | ENSG0000C |
| 11836933 | 11836986 | -1 | miRNA | ENSG0000C |
| 12030303 | 12030372 | -1 | snoRNA | ENSG0000C |
| 12199130 | 12199201 | 1 | miRNA | ENSG0000C |
| 12411398 | 12411501 | -1 | snRNA | ENSG0000C |

| | | | | |
|---|---|---|---|---|
| 12598987 | 12605145 | 1 | lncRNA | ENSG0000C |
| 12702764 | 12716557 | -1 | lncRNA | ENSG0000C |
| 12715415 | 12716227 | 1 | lncRNA | ENSG0000C |
| 12716889 | 12742734 | 1 | protein_coding | ENSG0000C |
| 12737367 | 12737444 | 1 | miRNA | ENSG0000C |
| 12780593 | 13007029 | -1 | lncRNA | ENSG0000C |
| 13000953 | 13331683 | 1 | lncRNA | ENSG0000C |
| 13537673 | 13609168 | 1 | lncRNA | ENSG0000C |
| 13710531 | 14400963 | -1 | lncRNA | ENSG0000C |
| 13723048 | 13758152 | 1 | lncRNA | ENSG0000C |
| 14274622 | 14274725 | -1 | snRNA | ENSG0000C |
| 14530941 | 14531223 | 1 | misc_RNA | ENSG0000C |
| 14616428 | 14630192 | 1 | lncRNA | ENSG0000C |
| 14632700 | 14650814 | 1 | protein_coding | ENSG0000C |
| 14705668 | 14992066 | -1 | lncRNA | ENSG0000C |
| 14886647 | 14907664 | -1 | lncRNA | ENSG0000C |
| 15166914 | 15561334 | -1 | protein_coding | ENSG0000C |
| 15397435 | 15397782 | -1 | processed_pseudoge | ENSG0000C |
| 15564170 | 15573868 | 1 | lncRNA | ENSG0000C |
| 15591178 | 15631111 | 1 | protein_coding | ENSG0000C |
| 15668684 | 15680484 | -1 | lncRNA | ENSG0000C |
| 15690782 | 15744339 | 1 | lncRNA | ENSG0000C |
| 15801747 | 15810877 | -1 | lncRNA | ENSG0000C |
| 15864935 | 15865051 | 1 | snRNA | ENSG0000C |
| 15869939 | 15870243 | 1 | processed_pseudoge | ENSG0000C |
| 15903669 | 15903772 | 1 | snoRNA | ENSG0000C |
| 15918350 | 15942249 | -1 | lncRNA | ENSG0000C |
| 15920399 | 15936017 | 1 | lncRNA | ENSG0000C |
| 15940550 | 15947007 | 1 | protein_coding | ENSG0000C |
| 15950689 | 15950981 | -1 | misc_RNA | ENSG0000C |
| 15997049 | 16062885 | -1 | lncRNA | ENSG0000C |
| 16013928 | 16087201 | 1 | lncRNA | ENSG0000C |
| 16085222 | 16105841 | -1 | lncRNA | ENSG0000C |
| 16149251 | 16149394 | 1 | processed_pseudoge | ENSG0000C |
| 16199203 | 16199295 | 1 | snoRNA | ENSG0000C |
| 16202430 | 16204226 | 1 | lncRNA | ENSG0000C |
| 16224047 | 16333978 | 1 | lncRNA | ENSG0000C |
| 16227027 | 16228194 | -1 | lncRNA | ENSG0000C |
| 16294445 | 16302290 | -1 | lncRNA | ENSG0000C |
| 16316324 | 16319566 | -1 | lncRNA | ENSG0000C |
| 16354256 | 16432702 | 1 | lncRNA | ENSG0000C |
| 16523176 | 16555564 | 1 | lncRNA | ENSG0000C |
| 16541690 | 16545695 | -1 | TEC | ENSG0000C |
| 16549459 | 16666331 | -1 | protein_coding | ENSG0000C |
| 16728124 | 16769567 | 1 | lncRNA | ENSG0000C |
| 16853820 | 16855284 | -1 | processed_pseudoge | ENSG0000C |
| 16970034 | 16974497 | 1 | lncRNA | ENSG0000C |
| 17122841 | 17123140 | -1 | misc_RNA | ENSG0000C |
| 17284292 | 17284895 | 1 | processed_pseudoge | ENSG0000C |
| 17385091 | 17386417 | 1 | processed_pseudoge | ENSG0000C |

| | | | | |
|---|---|---|---|---|
| 17510584 | 17518439 | -1 | protein_coding | ENSG00000( |
| 17539126 | 17657018 | 1 | protein_coding | ENSG00000( |
| 17663812 | 17800242 | -1 | protein_coding | ENSG00000( |
| 17753858 | 17788946 | 1 | protein_coding | ENSG00000( |
| 17816460 | 17817798 | 1 | protein_coding | ENSG00000( |
| 17877847 | 18361616 | 1 | protein_coding | ENSG00000( |
| 18040606 | 18040733 | -1 | snoRNA | ENSG00000( |
| 18403967 | 18504484 | 1 | lncRNA | ENSG00000( |
| 18505729 | 18514332 | -1 | lncRNA | ENSG00000( |
| 18547386 | 18548204 | -1 | lncRNA | ENSG00000( |
| 18554723 | 18560679 | -1 | protein_coding | ENSG00000( |
| 18555545 | 18589564 | -1 | protein_coding | ENSG00000( |
| 18562872 | 18589572 | -1 | protein_coding | ENSG00000( |
| 18583367 | 18583468 | -1 | snRNA | ENSG00000( |
| 18784807 | 18787752 | 1 | lncRNA | ENSG00000( |
| 18939697 | 18946650 | 1 | lncRNA | ENSG00000( |
| 18986451 | 19348067 | -1 | lncRNA | ENSG00000( |
| 19348429 | 19348505 | 1 | miRNA | ENSG00000( |
| 19351485 | 19358623 | -1 | protein_coding | ENSG00000( |
| 19458220 | 19468961 | -1 | lncRNA | ENSG00000( |
| 19468997 | 19520095 | 1 | lncRNA | ENSG00000( |
| 19711715 | 19717576 | 1 | lncRNA | ENSG00000( |
| 19826249 | 19826575 | 1 | processed_pseudoge | ENSG00000( |
| 19868860 | 19885047 | 1 | lncRNA | ENSG00000( |
| 19896631 | 19901983 | -1 | protein_coding | ENSG00000( |
| 19902025 | 19902569 | 1 | lncRNA | ENSG00000( |
| 19910260 | 19990131 | -1 | protein_coding | ENSG00000( |
| 19990209 | 20004795 | 1 | lncRNA | ENSG00000( |
| 19992052 | 20012668 | -1 | protein_coding | ENSG00000( |
| 20032650 | 20051628 | -1 | protein_coding | ENSG00000( |
| 20052134 | 20054496 | 1 | lncRNA | ENSG00000( |
| 20063856 | 20106829 | -1 | lncRNA | ENSG00000( |
| 20155618 | 20156057 | -1 | processed_pseudoge | ENSG00000( |
| 20175346 | 20175639 | 1 | misc_RNA | ENSG00000( |
| 20175805 | 20175909 | 1 | snRNA | ENSG00000( |
| 20200797 | 20225433 | -1 | protein_coding | ENSG00000( |
| 20239892 | 20241046 | -1 | processed_pseudoge | ENSG00000( |
| 20246326 | 20246387 | -1 | snRNA | ENSG00000( |
| 20248691 | 20352234 | -1 | protein_coding | ENSG00000( |
| 20386386 | 20396588 | 1 | unitary_pseudogene | ENSG00000( |
| 20401627 | 20401724 | -1 | rRNA_pseudogene | ENSG00000( |
| 20447074 | 20449440 | 1 | protein_coding | ENSG00000( |
| 20451042 | 20452947 | 1 | lncRNA | ENSG00000( |
| 20480844 | 20480955 | 1 | misc_RNA | ENSG00000( |
| 20499571 | 20501311 | -1 | lncRNA | ENSG00000( |
| 20529467 | 20529976 | -1 | processed_pseudoge | ENSG00000( |
| 20560448 | 20651130 | -1 | protein_coding | ENSG00000( |
| 20586248 | 20586686 | -1 | lncRNA | ENSG00000( |
| 20590775 | 20592548 | -1 | lncRNA | ENSG00000( |
| 20606280 | 20606654 | -1 | processed_pseudoge | ENSG00000( |

| | | | | |
|---|---|---|---|---|
| 20667144 | 20679243 | 1 | protein_coding | ENSG00000( |
| 20678254 | 20678932 | -1 | lncRNA | ENSG00000( |
| 20684014 | 20823130 | -1 | protein_coding | ENSG00000( |
| 20859771 | 20861661 | 1 | lncRNA | ENSG00000( |
| 20999313 | 21000917 | -1 | lncRNA | ENSG00000( |
| 21001429 | 21044073 | -1 | protein_coding | ENSG00000( |
| 21023496 | 21024170 | 1 | TEC | ENSG00000( |
| 21094101 | 21096543 | -1 | lncRNA | ENSG00000( |
| 21123917 | 21143272 | 1 | protein_coding | ENSG00000( |
| 21198518 | 21198896 | -1 | processed_pseudoge | ENSG00000( |
| 21221169 | 21970959 | 1 | lncRNA | ENSG00000( |
| 21317660 | 21561313 | 1 | lncRNA | ENSG00000( |
| 21362292 | 21364025 | -1 | processed_pseudoge | ENSG00000( |
| 21607465 | 21637197 | -1 | lncRNA | ENSG00000( |
| 21638068 | 21649572 | -1 | lncRNA | ENSG00000( |
| 21687430 | 21710656 | -1 | lncRNA | ENSG00000( |
| 21922993 | 21923288 | 1 | misc_RNA | ENSG00000( |
| 21932662 | 22536322 | -1 | lncRNA | ENSG00000( |
| 22188336 | 22190313 | 1 | processed_pseudoge | ENSG00000( |
| 22317053 | 22317244 | 1 | processed_pseudoge | ENSG00000( |
| 22338886 | 22339003 | -1 | rRNA_pseudogene | ENSG00000( |
| 22377594 | 22482004 | 1 | lncRNA | ENSG00000( |
| 22508129 | 22548800 | 1 | lncRNA | ENSG00000( |
| 22861552 | 22861884 | 1 | misc_RNA | ENSG00000( |
| 23018125 | 23199056 | -1 | lncRNA | ENSG00000( |
| 23330664 | 23332044 | -1 | lncRNA | ENSG00000( |
| 23347654 | 23351864 | 1 | lncRNA | ENSG00000( |
| 23357516 | 23360376 | -1 | lncRNA | ENSG00000( |
| 23375229 | 23381299 | -1 | lncRNA | ENSG00000( |
| 23385179 | 23708611 | 1 | protein_coding | ENSG00000( |
| 23507043 | 23524344 | -1 | lncRNA | ENSG00000( |
| 23616499 | 23617987 | -1 | TEC | ENSG00000( |
| 23667208 | 23685453 | -1 | lncRNA | ENSG00000( |
| 23748664 | 23927123 | -1 | protein_coding | ENSG00000( |
| 23872268 | 23873014 | -1 | processed_pseudoge | ENSG00000( |
| 23900693 | 23901149 | 1 | processed_pseudoge | ENSG00000( |
| 23927285 | 24004909 | 1 | protein_coding | ENSG00000( |
| 23943846 | 23944351 | 1 | processed_pseudoge | ENSG00000( |
| 23996795 | 23997085 | -1 | misc_RNA | ENSG00000( |
| 24010081 | 24063321 | 1 | protein_coding | ENSG00000( |
| 24029347 | 24049575 | -1 | protein_coding | ENSG00000( |
| 24045835 | 24045938 | 1 | snRNA | ENSG00000( |
| 24049701 | 24063681 | 1 | protein_coding | ENSG00000( |
| 24067586 | 24076373 | -1 | protein_coding | ENSG00000( |
| 24076526 | 24169640 | 1 | protein_coding | ENSG00000( |
| 24077433 | 24085861 | -1 | protein_coding | ENSG00000( |
| 24114809 | 24123464 | -1 | protein_coding | ENSG00000( |
| 24124366 | 24190436 | 1 | protein_coding | ENSG00000( |
| 24165884 | 24175005 | -1 | lncRNA | ENSG00000( |
| 24175053 | 24200849 | 1 | protein_coding | ENSG00000( |

| | | | | |
|---|---|---|---|---|
| 24199839 | 24201698 | -1 | lncRNA | ENSG0000C |
| 24202864 | 24360714 | -1 | protein_coding | ENSG0000C |
| 24210650 | 24222201 | 1 | lncRNA | ENSG0000C |
| 24273614 | 24273741 | -1 | scaRNA | ENSG0000C |
| 24330402 | 24330674 | 1 | processed_pseudoge | ENSG0000C |
| 24334512 | 24335152 | 1 | processed_pseudoge | ENSG0000C |
| 24402995 | 24413292 | 1 | lncRNA | ENSG0000C |
| 24491914 | 24770702 | 1 | protein_coding | ENSG0000C |
| 24564630 | 24564738 | -1 | rRNA_pseudogene | ENSG0000C |
| 24676309 | 24676415 | 1 | snRNA | ENSG0000C |
| 24789734 | 24793382 | -1 | protein_coding | ENSG0000C |
| 24793136 | 24822376 | 1 | protein_coding | ENSG0000C |
| 24819169 | 24919839 | -1 | protein_coding | ENSG0000C |
| 24825610 | 24826717 | 1 | lncRNA | ENSG0000C |
| 24943636 | 24972094 | -1 | protein_coding | ENSG0000C |
| 24968958 | 24969224 | -1 | processed_pseudoge | ENSG0000C |
| 24969388 | 24969477 | -1 | snoRNA | ENSG0000C |
| 24971390 | 25039716 | 1 | lncRNA | ENSG0000C |
| 25000147 | 25010274 | -1 | lncRNA | ENSG0000C |
| 25008845 | 25008946 | -1 | misc_RNA | ENSG0000C |
| 25042076 | 25159135 | 1 | protein_coding | ENSG0000C |
| 25058032 | 25058329 | 1 | misc_RNA | ENSG0000C |
| 25079901 | 25081689 | -1 | processed_pseudoge | ENSG0000C |
| 25160853 | 25168903 | -1 | protein_coding | ENSG0000C |
| 25204313 | 25209202 | 1 | lncRNA | ENSG0000C |
| 25227855 | 25342590 | -1 | protein_coding | ENSG0000C |
| 25328640 | 25328721 | -1 | miRNA | ENSG0000C |
| 25369136 | 25375845 | -1 | lncRNA | ENSG0000C |
| 25377198 | 25673647 | -1 | protein_coding | ENSG0000C |
| 25421117 | 25427643 | 1 | lncRNA | ENSG0000C |
| 25697076 | 25697188 | -1 | misc_RNA | ENSG0000C |
| 25733753 | 25878487 | -1 | protein_coding | ENSG0000C |
| 25809925 | 25810656 | -1 | processed_pseudoge | ENSG0000C |
| 25822469 | 25822950 | 1 | processed_pseudoge | ENSG0000C |
| 25856461 | 25856966 | -1 | processed_pseudoge | ENSG0000C |
| 25886964 | 25887206 | -1 | processed_pseudoge | ENSG0000C |
| 25926598 | 25982749 | -1 | protein_coding | ENSG0000C |
| 25994794 | 25995081 | 1 | processed_pseudoge | ENSG0000C |
| 26028208 | 26028612 | -1 | processed_pseudoge | ENSG0000C |
| 26034084 | 26137454 | 1 | protein_coding | ENSG0000C |
| 26042675 | 26042781 | 1 | snRNA | ENSG0000C |
| 26101317 | 26102136 | -1 | processed_pseudoge | ENSG0000C |
| 26140263 | 26141264 | -1 | processed_pseudoge | ENSG0000C |
| 26149204 | 26150731 | 1 | processed_pseudoge | ENSG0000C |
| 26160023 | 26161136 | 1 | processed_pseudoge | ENSG0000C |
| 26162056 | 26162554 | 1 | processed_pseudoge | ENSG0000C |
| 26173088 | 26189663 | 1 | protein_coding | ENSG0000C |
| 26190635 | 26244672 | -1 | protein_coding | ENSG0000C |
| 26243170 | 26290465 | 1 | protein_coding | ENSG0000C |
| 26298570 | 26306340 | 1 | lncRNA | ENSG0000C |

| | | | | |
|---|---|---|---|---|
| 26308173 | 26346817 | -1 | protein_coding | ENSG00000 |
| 26308547 | 26395891 | 1 | protein_coding | ENSG00000 |
| 26401920 | 26456711 | 1 | protein_coding | ENSG00000 |
| 26457203 | 26558698 | -1 | protein_coding | ENSG00000 |
| 26562585 | 26579532 | 1 | protein_coding | ENSG00000 |
| 26581205 | 26641366 | -1 | protein_coding | ENSG00000 |
| 26671254 | 26674464 | -1 | lncRNA | ENSG00000 |
| 26678219 | 26678510 | -1 | processed_pseudoge | ENSG00000 |
| 26692690 | 26733420 | 1 | protein_coding | ENSG00000 |
| 26764284 | 26781231 | 1 | protein_coding | ENSG00000 |
| 26764289 | 26801067 | 1 | protein_coding | ENSG00000 |
| 26827169 | 26827520 | -1 | processed_pseudoge | ENSG00000 |
| 26847747 | 26950351 | 1 | protein_coding | ENSG00000 |
| 26950308 | 27009807 | -1 | lncRNA | ENSG00000 |
| 26970637 | 27027219 | 1 | protein_coding | ENSG00000 |
| 26984776 | 27014612 | -1 | lncRNA | ENSG00000 |
| 27032910 | 27041694 | 1 | protein_coding | ENSG00000 |
| 27042364 | 27070622 | 1 | protein_coding | ENSG00000 |
| 27049683 | 27050264 | -1 | lncRNA | ENSG00000 |
| 27053618 | 27054276 | -1 | lncRNA | ENSG00000 |
| 27061038 | 27061815 | 1 | lncRNA | ENSG00000 |
| 27062428 | 27062907 | -1 | lncRNA | ENSG00000 |
| 27070472 | 27071654 | -1 | protein_coding | ENSG00000 |
| 27078615 | 27086403 | 1 | protein_coding | ENSG00000 |
| 27086747 | 27100762 | 1 | protein_coding | ENSG00000 |
| 27098889 | 27119128 | -1 | protein_coding | ENSG00000 |
| 27123789 | 27130812 | 1 | protein_coding | ENSG00000 |
| 27130756 | 27134666 | -1 | protein_coding | ENSG00000 |
| 27136848 | 27139410 | -1 | protein_coding | ENSG00000 |
| 27149004 | 27156974 | 1 | protein_coding | ENSG00000 |
| 27158563 | 27159229 | -1 | unprocessed_pseudc | ENSG00000 |
| 27199587 | 27212958 | -1 | protein_coding | ENSG00000 |
| 27212027 | 27217178 | 1 | protein_coding | ENSG00000 |
| 27217369 | 27243943 | 1 | protein_coding | ENSG00000 |
| 27253684 | 27275817 | -1 | protein_coding | ENSG00000 |
| 27275433 | 27281499 | 1 | protein_coding | ENSG00000 |
| 27282392 | 27307439 | 1 | protein_coding | ENSG00000 |
| 27307400 | 27308445 | -1 | protein_coding | ENSG00000 |
| 27309492 | 27325680 | -1 | protein_coding | ENSG00000 |
| 27325849 | 27357034 | -1 | protein_coding | ENSG00000 |
| 27335520 | 27342599 | 1 | lncRNA | ENSG00000 |
| 27356246 | 27367622 | 1 | lncRNA | ENSG00000 |
| 27364352 | 27370486 | -1 | protein_coding | ENSG00000 |
| 27370496 | 27377535 | 1 | protein_coding | ENSG00000 |
| 27377235 | 27380790 | -1 | protein_coding | ENSG00000 |
| 27381195 | 27409591 | -1 | protein_coding | ENSG00000 |
| 27392784 | 27393367 | -1 | processed_pseudoge | ENSG00000 |
| 27427790 | 27442259 | 1 | protein_coding | ENSG00000 |
| 27442366 | 27446481 | 1 | protein_coding | ENSG00000 |
| 27444377 | 27489789 | -1 | protein_coding | ENSG00000 |

| | | | | |
|---|---|---|---|---|
| 27455156 | 27455862 | 1 | processed_pseudoge | ENSG00000 |
| 27475494 | 27475595 | -1 | snRNA | ENSG00000 |
| 27491883 | 27495200 | -1 | protein_coding | ENSG00000 |
| 27496839 | 27523684 | 1 | protein_coding | ENSG00000 |
| 27537386 | 27582721 | 1 | protein_coding | ENSG00000 |
| 27582969 | 27623217 | 1 | protein_coding | ENSG00000 |
| 27583046 | 27635174 | 1 | lncRNA | ENSG00000 |
| 27625639 | 27629012 | -1 | protein_coding | ENSG00000 |
| 27628247 | 27651511 | 1 | protein_coding | ENSG00000 |
| 27642043 | 27642169 | -1 | snoRNA | ENSG00000 |
| 27650809 | 27663840 | -1 | protein_coding | ENSG00000 |
| 27663471 | 27694976 | 1 | protein_coding | ENSG00000 |
| 27705786 | 27715732 | -1 | lncRNA | ENSG00000 |
| 27771717 | 27988087 | 1 | protein_coding | ENSG00000 |
| 27781379 | 27890681 | -1 | protein_coding | ENSG00000 |
| 27889941 | 28338901 | 1 | protein_coding | ENSG00000 |
| 27896116 | 27897195 | -1 | processed_pseudoge | ENSG00000 |
| 27996367 | 27996449 | 1 | miRNA | ENSG00000 |
| 28015777 | 28016482 | -1 | processed_pseudoge | ENSG00000 |
| 28134408 | 28134585 | -1 | misc_RNA | ENSG00000 |
| 28307063 | 28310560 | -1 | lncRNA | ENSG00000 |
| 28308161 | 28308570 | 1 | processed_pseudoge | ENSG00000 |
| 28384409 | 28394672 | -1 | lncRNA | ENSG00000 |
| 28392448 | 28417317 | 1 | protein_coding | ENSG00000 |
| 28396815 | 28397110 | 1 | lncRNA | ENSG00000 |
| 28425945 | 28426719 | 1 | lncRNA | ENSG00000 |
| 28448167 | 28450184 | -1 | lncRNA | ENSG00000 |
| 28457145 | 28644142 | 1 | protein_coding | ENSG00000 |
| 28460256 | 28460361 | -1 | processed_pseudoge | ENSG00000 |
| 28633282 | 28664540 | -1 | lncRNA | ENSG00000 |
| 28683976 | 28684083 | 1 | rRNA_pseudogene | ENSG00000 |
| 28707511 | 28751722 | -1 | lncRNA | ENSG00000 |
| 28751640 | 28802940 | 1 | protein_coding | ENSG00000 |
| 28782517 | 28850611 | 1 | protein_coding | ENSG00000 |
| 28810281 | 28810706 | -1 | lncRNA | ENSG00000 |
| 28849821 | 28870309 | -1 | protein_coding | ENSG00000 |
| 28894667 | 28948219 | 1 | protein_coding | ENSG00000 |
| 28913664 | 28913748 | 1 | snoRNA | ENSG00000 |
| 28927067 | 28927144 | 1 | snoRNA | ENSG00000 |
| 28927243 | 28927340 | -1 | misc_RNA | ENSG00000 |
| 28927983 | 28928060 | 1 | snoRNA | ENSG00000 |
| 28956611 | 29061373 | 1 | protein_coding | ENSG00000 |
| 28972414 | 28972521 | -1 | misc_RNA | ENSG00000 |
| 29060976 | 29074523 | -1 | protein_coding | ENSG00000 |
| 29088649 | 29097586 | -1 | lncRNA | ENSG00000 |
| 29097705 | 29189643 | 1 | protein_coding | ENSG00000 |
| 29192774 | 29921586 | -1 | protein_coding | ENSG00000 |
| 29319554 | 29352214 | 1 | lncRNA | ENSG00000 |
| 29681027 | 29681323 | -1 | misc_RNA | ENSG00000 |
| 29841187 | 30143804 | -1 | lncRNA | ENSG00000 |

| | | | | |
|---|---|---|---|---|
| 29890371 | 29892354 | 1 | lncRNA | ENSG00000( |
| 29899597 | 29907199 | 1 | lncRNA | ENSG00000( |
| 30077093 | 30078272 | 1 | processed_pseudoge | ENSG00000( |
| 30146941 | 30160533 | 1 | protein_coding | ENSG00000( |
| 30187434 | 30187566 | 1 | snoRNA | ENSG00000( |
| 30209995 | 30210307 | 1 | processed_pseudoge | ENSG00000( |
| 30231534 | 30323730 | 1 | protein_coding | ENSG00000( |
| 30343222 | 30348298 | -1 | lncRNA | ENSG00000( |
| 30346623 | 30361010 | 1 | lncRNA | ENSG00000( |
| 30408170 | 30408346 | 1 | TEC | ENSG00000( |
| 30447226 | 30644225 | 1 | protein_coding | ENSG00000( |
| 30677762 | 30679895 | 1 | TEC | ENSG00000( |
| 30722771 | 30820542 | -1 | protein_coding | ENSG00000( |
| 30746444 | 30746701 | -1 | processed_pseudoge | ENSG00000( |
| 30887626 | 30897387 | -1 | lncRNA | ENSG00000( |
| 30910467 | 31155202 | -1 | protein_coding | ENSG00000( |
| 30986939 | 30991426 | 1 | lncRNA | ENSG00000( |
| 31173056 | 31233858 | -1 | protein_coding | ENSG00000( |
| 31228312 | 31228410 | 1 | rRNA_pseudogene | ENSG00000( |
| 31234152 | 31269451 | 1 | protein_coding | ENSG00000( |
| 31290762 | 31291604 | 1 | processed_pseudoge | ENSG00000( |
| 31334321 | 31414742 | -1 | protein_coding | ENSG00000( |
| 31522480 | 31580938 | -1 | protein_coding | ENSG00000( |
| 31526942 | 31563464 | 1 | lncRNA | ENSG00000( |
| 31651381 | 31651862 | 1 | processed_pseudoge | ENSG00000( |
| 31793823 | 31803980 | -1 | lncRNA | ENSG00000( |
| 31822591 | 31823436 | -1 | processed_pseudoge | ENSG00000( |
| 31823413 | 31823817 | -1 | processed_pseudoge | ENSG00000( |
| 31852976 | 31853423 | -1 | lncRNA | ENSG00000( |
| 31865060 | 32011230 | -1 | protein_coding | ENSG00000( |
| 31867809 | 32039805 | -1 | protein_coding | ENSG00000( |
| 32013061 | 32013368 | 1 | lncRNA | ENSG00000( |
| 32063547 | 32157637 | 1 | protein_coding | ENSG00000( |
| 32165046 | 32165757 | -1 | lncRNA | ENSG00000( |
| 32165841 | 32224379 | 1 | protein_coding | ENSG00000( |
| 32201600 | 32203999 | 1 | processed_pseudoge | ENSG00000( |
| 32214456 | 32214562 | -1 | snRNA | ENSG00000( |
| 32224453 | 32265732 | -1 | protein_coding | ENSG00000( |
| 32277904 | 32316594 | 1 | protein_coding | ENSG00000( |
| 32321638 | 32323002 | 1 | lncRNA | ENSG00000( |
| 32357028 | 32618899 | 1 | protein_coding | ENSG00000( |
| 32377631 | 32379599 | -1 | lncRNA | ENSG00000( |
| 32521927 | 32523547 | 1 | lncRNA | ENSG00000( |
| 32526504 | 32529507 | 1 | lncRNA | ENSG00000( |
| 32532153 | 32532246 | 1 | miRNA | ENSG00000( |
| 32548675 | 32549040 | 1 | lncRNA | ENSG00000( |
| 32557273 | 32574818 | -1 | transcribed_unproce | ENSG00000( |
| 32563328 | 32566436 | 1 | TEC | ENSG00000( |
| 32628032 | 32821051 | 1 | protein_coding | ENSG00000( |
| 32635255 | 32635331 | 1 | miRNA | ENSG00000( |

| | | | | |
|---|---|---|---|---|
| 32825359 | 32926693 | 1 | lncRNA | ENSG0000C |
| 32927085 | 32946149 | 1 | lncRNA | ENSG0000C |
| 32945339 | 32945451 | -1 | misc_RNA | ENSG0000C |
| 32946953 | 33399509 | 1 | protein_coding | ENSG0000C |
| 33056333 | 33063113 | -1 | processed_pseudoge | ENSG0000C |
| 33274465 | 33281261 | -1 | lncRNA | ENSG0000C |
| 33285769 | 33285887 | -1 | rRNA_pseudogene | ENSG0000C |
| 33332898 | 33333007 | 1 | rRNA_pseudogene | ENSG0000C |
| 33418516 | 33418564 | 1 | miRNA | ENSG0000C |
| 33436324 | 33564750 | 1 | protein_coding | ENSG0000C |
| 33555047 | 33563547 | -1 | lncRNA | ENSG0000C |
| 33583660 | 33599382 | -1 | protein_coding | ENSG0000C |
| 33599442 | 33657460 | 1 | lncRNA | ENSG0000C |
| 33602041 | 33602286 | 1 | processed_pseudoge | ENSG0000C |
| 33636502 | 33637425 | 1 | processed_pseudoge | ENSG0000C |
| 33706886 | 34738231 | 1 | lncRNA | ENSG0000C |
| 33722721 | 33728207 | -1 | transcribed_processe | ENSG0000C |
| 33839782 | 33840678 | -1 | processed_pseudoge | ENSG0000C |
| 34067226 | 34069550 | 1 | lncRNA | ENSG0000C |
| 34134371 | 34220575 | -1 | lncRNA | ENSG0000C |
| 34692290 | 34703606 | -1 | lncRNA | ENSG0000C |
| 34732287 | 34822726 | -1 | lncRNA | ENSG0000C |
| 34799850 | 35284997 | 1 | lncRNA | ENSG0000C |
| 34809233 | 34809556 | -1 | misc_RNA | ENSG0000C |
| 34998273 | 35000160 | -1 | lncRNA | ENSG0000C |
| 35219377 | 35219604 | -1 | processed_pseudoge | ENSG0000C |
| 35471405 | 35471486 | 1 | miRNA | ENSG0000C |
| 35471605 | 35471708 | 1 | snRNA | ENSG0000C |
| 35471716 | 35471841 | 1 | processed_pseudoge | ENSG0000C |
| 35724759 | 35725142 | -1 | processed_pseudoge | ENSG0000C |
| 36299388 | 36299830 | -1 | processed_pseudoge | ENSG0000C |
| 36354749 | 36355114 | -1 | lncRNA | ENSG0000C |
| 36355778 | 36551135 | 1 | protein_coding | ENSG0000C |
| 36513255 | 36513732 | -1 | lncRNA | ENSG0000C |
| 36531805 | 36646087 | -1 | protein_coding | ENSG0000C |
| 36656322 | 36656941 | -1 | processed_pseudoge | ENSG0000C |
| 36696690 | 36814792 | 1 | protein_coding | ENSG0000C |
| 36837698 | 36966536 | -1 | protein_coding | ENSG0000C |
| 36839922 | 36842539 | -1 | TEC | ENSG0000C |
| 36867398 | 36867495 | -1 | snRNA | ENSG0000C |
| 36968383 | 37084372 | -1 | protein_coding | ENSG0000C |
| 37084451 | 37099244 | 1 | protein_coding | ENSG0000C |
| 37099210 | 37157065 | -1 | protein_coding | ENSG0000C |
| 37148530 | 37149304 | 1 | processed_pseudoge | ENSG0000C |
| 37167820 | 37196598 | -1 | protein_coding | ENSG0000C |
| 37194382 | 37194737 | -1 | processed_pseudoge | ENSG0000C |
| 37196488 | 37216193 | 1 | protein_coding | ENSG0000C |
| 37201612 | 37231596 | -1 | protein_coding | ENSG0000C |
| 37208875 | 37212677 | 1 | lncRNA | ENSG0000C |
| 37231631 | 37253403 | 1 | protein_coding | ENSG0000C |

| | | | | |
|---|---|---|---|---|
| 37250502 | 37324808 | -1 | protein_coding | ENSG0000C |
| 37325340 | 37326797 | 1 | lncRNA | ENSG0000C |
| 37331770 | 37331876 | -1 | snRNA | ENSG0000C |
| 37339957 | 37343796 | -1 | lncRNA | ENSG0000C |
| 37342827 | 37373322 | 1 | protein_coding | ENSG0000C |
| 37435510 | 37435608 | -1 | snRNA | ENSG0000C |
| 37466781 | 37524807 | 1 | lncRNA | ENSG0000C |
| 37562486 | 37646698 | 1 | lncRNA | ENSG0000C |
| 37641882 | 37738468 | -1 | protein_coding | ENSG0000C |
| 37744333 | 37747046 | 1 | lncRNA | ENSG0000C |
| 37816915 | 37817961 | 1 | processed_pseudoge | ENSG0000C |
| 37820498 | 37876274 | -1 | lncRNA | ENSG0000C |
| 37923187 | 38067142 | 1 | protein_coding | ENSG0000C |
| 37949911 | 38067041 | -1 | lncRNA | ENSG0000C |
| 38066973 | 38109902 | -1 | protein_coding | ENSG0000C |
| 38073447 | 38231651 | 1 | lncRNA | ENSG0000C |
| 38121935 | 38123881 | -1 | TEC | ENSG0000C |
| 38132637 | 38138946 | -1 | lncRNA | ENSG0000C |
| 38147415 | 38147521 | 1 | snRNA | ENSG0000C |
| 38193348 | 38193629 | 1 | lncRNA | ENSG0000C |
| 38203363 | 38239590 | -1 | lncRNA | ENSG0000C |
| 38231568 | 38232317 | 1 | processed_pseudoge | ENSG0000C |
| 38285410 | 38286410 | 1 | processed_pseudoge | ENSG0000C |
| 38293954 | 38377285 | -1 | protein_coding | ENSG0000C |
| 38406527 | 38515934 | -1 | lncRNA | ENSG0000C |
| 38408719 | 38412627 | 1 | lncRNA | ENSG0000C |
| 38431294 | 38433573 | 1 | lncRNA | ENSG0000C |
| 38481851 | 38482804 | 1 | processed_pseudoge | ENSG0000C |
| 38535258 | 38536249 | -1 | transcribed_processe | ENSG0000C |
| 38561969 | 38603586 | -1 | protein_coding | ENSG0000C |
| 38601598 | 38602178 | 1 | lncRNA | ENSG0000C |
| 38666081 | 38741237 | 1 | protein_coding | ENSG0000C |
| 38668202 | 38671421 | -1 | lncRNA | ENSG0000C |
| 38743599 | 38751494 | -1 | protein_coding | ENSG0000C |
| 38751534 | 38785002 | 1 | protein_coding | ENSG0000C |
| 38763534 | 38765221 | 1 | unitary_pseudogene | ENSG0000C |
| 38769265 | 38770124 | 1 | processed_pseudoge | ENSG0000C |
| 38797729 | 38875934 | -1 | protein_coding | ENSG0000C |
| 38810432 | 38811654 | -1 | processed_pseudoge | ENSG0000C |
| 38861720 | 38863133 | 1 | lncRNA | ENSG0000C |
| 38875962 | 38929072 | 1 | protein_coding | ENSG0000C |
| 38884560 | 38884666 | -1 | snRNA | ENSG0000C |
| 38889841 | 38975449 | 1 | protein_coding | ENSG0000C |
| 38936876 | 38937158 | -1 | misc_RNA | ENSG0000C |
| 38959287 | 38960342 | -1 | lncRNA | ENSG0000C |
| 38981396 | 39124345 | -1 | protein_coding | ENSG0000C |
| 38992279 | 38993857 | -1 | lncRNA | ENSG0000C |
| 39082589 | 39082692 | 1 | snRNA | ENSG0000C |
| 39098149 | 39098445 | -1 | processed_pseudoge | ENSG0000C |
| 39128826 | 39128927 | -1 | misc_RNA | ENSG0000C |

| | | | | |
|---|---|---|---|---|
| 39175646 | 39229588 | -1 | protein_coding | ENSG00000 |
| 39249266 | 39437301 | -1 | protein_coding | ENSG00000 |
| 39283657 | 39283791 | -1 | snoRNA | ENSG00000 |
| 39323328 | 39323804 | -1 | lncRNA | ENSG00000 |
| 39436530 | 39665343 | 1 | lncRNA | ENSG00000 |
| 39664982 | 39717963 | 1 | protein_coding | ENSG00000 |
| 39736060 | 39779267 | -1 | protein_coding | ENSG00000 |
| 39786453 | 40255209 | 1 | lncRNA | ENSG00000 |
| 39929110 | 39929998 | 1 | processed_pseudoge | ENSG00000 |
| 40097270 | 40611053 | -1 | protein_coding | ENSG00000 |
| 40534806 | 40545781 | 1 | lncRNA | ENSG00000 |
| 40591285 | 40679604 | 1 | lncRNA | ENSG00000 |
| 40620667 | 40639632 | -1 | lncRNA | ENSG00000 |
| 40746481 | 40767452 | 1 | lncRNA | ENSG00000 |
| 41143780 | 41157555 | 1 | transcribed_processe | ENSG00000 |
| 41716133 | 41731611 | 1 | lncRNA | ENSG00000 |
| 41819747 | 41820754 | 1 | processed_pseudoge | ENSG00000 |
| 41850203 | 41850601 | -1 | processed_pseudoge | ENSG00000 |
| 41857271 | 41857372 | -1 | misc_RNA | ENSG00000 |
| 41860155 | 41894050 | 1 | lncRNA | ENSG00000 |
| 41871271 | 41871388 | 1 | snRNA | ENSG00000 |
| 41931599 | 41933909 | -1 | lncRNA | ENSG00000 |
| 41935368 | 41956806 | -1 | protein_coding | ENSG00000 |
| 42015625 | 42025302 | -1 | lncRNA | ENSG00000 |
| 42048021 | 42058517 | 1 | protein_coding | ENSG00000 |
| 42143238 | 42170301 | -1 | lncRNA | ENSG00000 |
| 42169353 | 42332548 | 1 | protein_coding | ENSG00000 |
| 42333546 | 42425088 | -1 | protein_coding | ENSG00000 |
| 42440377 | 42440436 | 1 | snoRNA | ENSG00000 |
| 42442017 | 42493982 | -1 | protein_coding | ENSG00000 |
| 42463139 | 42463997 | -1 | processed_pseudoge | ENSG00000 |
| 42469817 | 42470266 | 1 | processed_pseudoge | ENSG00000 |
| 42494569 | 42756947 | 1 | protein_coding | ENSG00000 |
| 42511570 | 42511851 | 1 | misc_RNA | ENSG00000 |
| 42532766 | 42533442 | -1 | processed_pseudoge | ENSG00000 |
| 42680088 | 42680279 | -1 | processed_pseudoge | ENSG00000 |
| 42712740 | 42712847 | 1 | snRNA | ENSG00000 |
| 42762481 | 42764261 | -1 | protein_coding | ENSG00000 |
| 42767089 | 42792593 | -1 | protein_coding | ENSG00000 |
| 42797225 | 42798712 | -1 | processed_pseudoge | ENSG00000 |
| 42826322 | 42827617 | -1 | processed_pseudoge | ENSG00000 |
| 43001355 | 43006060 | -1 | lncRNA | ENSG00000 |
| 43027823 | 43040662 | -1 | lncRNA | ENSG00000 |
| 43041193 | 43043782 | -1 | lncRNA | ENSG00000 |
| 43091388 | 43091494 | -1 | snRNA | ENSG00000 |
| 43092530 | 43132482 | -1 | lncRNA | ENSG00000 |
| 43128819 | 43131999 | 1 | lncRNA | ENSG00000 |
| 43219849 | 43223227 | 1 | lncRNA | ENSG00000 |
| 43222402 | 43226606 | -1 | protein_coding | ENSG00000 |
| 43227210 | 43228855 | 1 | lncRNA | ENSG00000 |

| | | | | |
|---|---|---|---|---|
| 43229573 | 43233394 | 1 | lncRNA | ENSG0000C |
| 43230836 | 43596046 | -1 | protein_coding | ENSG0000C |
| 43408307 | 43408412 | 1 | snRNA | ENSG0000C |
| 43492032 | 43492313 | 1 | misc_RNA | ENSG0000C |
| 43620878 | 43620984 | -1 | misc_RNA | ENSG0000C |
| 43637260 | 43767987 | 1 | protein_coding | ENSG0000C |
| 43675151 | 43676429 | -1 | protein_coding | ENSG0000C |
| 43680465 | 43681622 | -1 | processed_pseudoge | ENSG0000C |
| 43772120 | 43772350 | -1 | misc_RNA | ENSG0000C |
| 43774039 | 43810010 | 1 | protein_coding | ENSG0000C |
| 43812472 | 43838865 | -1 | protein_coding | ENSG0000C |
| 43831942 | 43882988 | 1 | protein_coding | ENSG0000C |
| 43886224 | 43995989 | -1 | protein_coding | ENSG0000C |
| 43892690 | 43892796 | 1 | snRNA | ENSG0000C |
| 43904412 | 43904693 | 1 | misc_RNA | ENSG0000C |
| 44065894 | 44066154 | -1 | processed_pseudoge | ENSG0000C |
| 44154789 | 44154895 | -1 | snRNA | ENSG0000C |
| 44166266 | 44166943 | 1 | unprocessed_pseudc | ENSG0000C |
| 44167625 | 44168859 | -1 | lncRNA | ENSG0000C |
| 44167969 | 44244384 | 1 | protein_coding | ENSG0000C |
| 44168851 | 44320310 | 1 | protein_coding | ENSG0000C |
| 44270621 | 44271106 | 1 | processed_pseudoge | ENSG0000C |
| 44275458 | 44321494 | 1 | protein_coding | ENSG0000C |
| 44316281 | 44361862 | -1 | protein_coding | ENSG0000C |
| 44361947 | 44772592 | 1 | protein_coding | ENSG0000C |
| 44921077 | 44939199 | -1 | lncRNA | ENSG0000C |
| 44927914 | 44928396 | -1 | TEC | ENSG0000C |
| 44932320 | 44932406 | 1 | misc_RNA | ENSG0000C |
| 44934055 | 44934271 | 1 | misc_RNA | ENSG0000C |
| 44935294 | 44935368 | 1 | misc_RNA | ENSG0000C |
| 44935585 | 44935908 | 1 | misc_RNA | ENSG0000C |
| 44937426 | 44937696 | 1 | misc_RNA | ENSG0000C |
| 44938833 | 44939028 | 1 | misc_RNA | ENSG0000C |
| 44940045 | 44940245 | 1 | misc_RNA | ENSG0000C |
| 44940154 | 44941873 | -1 | lncRNA | ENSG0000C |
| 44941702 | 44946071 | 1 | protein_coding | ENSG0000C |
| 44954664 | 44968762 | 1 | lncRNA | ENSG0000C |
| 44996413 | 44996873 | 1 | processed_pseudoge | ENSG0000C |
| 45005182 | 45009452 | -1 | protein_coding | ENSG0000C |
| 45013214 | 45013668 | -1 | lncRNA | ENSG0000C |
| 45164816 | 45323397 | -1 | lncRNA | ENSG0000C |
| 45168583 | 45169414 | 1 | lncRNA | ENSG0000C |
| 45169616 | 45211673 | 1 | lncRNA | ENSG0000C |
| 45173722 | 45175935 | 1 | lncRNA | ENSG0000C |
| 45388680 | 45612165 | -1 | protein_coding | ENSG0000C |
| 45569199 | 45569492 | -1 | misc_RNA | ENSG0000C |
| 45651345 | 46187990 | 1 | protein_coding | ENSG0000C |
| 45674701 | 45675476 | -1 | lncRNA | ENSG0000C |
| 46003942 | 46004309 | -1 | processed_pseudoge | ENSG0000C |
| 46078015 | 46078828 | -1 | lncRNA | ENSG0000C |

| | | | | |
|---|---|---|---|---|
| 46166789 | 46167978 | -1 | lncRNA | ENSG00000 |
| 46256860 | 46257177 | -1 | processed_pseudoge | ENSG00000 |
| 46266690 | 46266971 | 1 | misc_RNA | ENSG00000 |
| 46293667 | 46386697 | 1 | protein_coding | ENSG00000 |
| 46392291 | 46394228 | -1 | lncRNA | ENSG00000 |
| 46429190 | 46441833 | 1 | lncRNA | ENSG00000 |
| 46429195 | 46484424 | 1 | lncRNA | ENSG00000 |
| 46448224 | 46448511 | -1 | misc_RNA | ENSG00000 |
| 46479565 | 46484425 | 1 | protein_coding | ENSG00000 |
| 46490750 | 46542577 | -1 | protein_coding | ENSG00000 |
| 46499731 | 46501278 | 1 | lncRNA | ENSG00000 |
| 46541806 | 46584688 | 1 | protein_coding | ENSG00000 |
| 46568256 | 46580238 | -1 | lncRNA | ENSG00000 |
| 46580937 | 46617055 | -1 | protein_coding | ENSG00000 |
| 46616416 | 46630176 | 1 | protein_coding | ENSG00000 |
| 46668870 | 46670778 | 1 | TEC | ENSG00000 |
| 46698940 | 46822804 | 1 | lncRNA | ENSG00000 |
| 46698952 | 46780245 | 1 | protein_coding | ENSG00000 |
| 46816697 | 46859007 | 1 | lncRNA | ENSG00000 |
| 46852020 | 46853496 | -1 | lncRNA | ENSG00000 |
| 46899275 | 46908678 | 1 | lncRNA | ENSG00000 |
| 46901870 | 46941855 | -1 | protein_coding | ENSG00000 |
| 46916157 | 47076137 | 1 | protein_coding | ENSG00000 |
| 46956615 | 46956888 | -1 | lncRNA | ENSG00000 |
| 47035279 | 47040524 | -1 | lncRNA | ENSG00000 |
| 47045538 | 47155308 | -1 | protein_coding | ENSG00000 |
| 47065941 | 47176511 | -1 | protein_coding | ENSG00000 |
| 47067822 | 47071204 | 1 | lncRNA | ENSG00000 |
| 47160083 | 47176921 | -1 | protein_coding | ENSG00000 |
| 47192405 | 47345074 | -1 | lncRNA | ENSG00000 |
| 47225781 | 47240886 | 1 | lncRNA | ENSG00000 |
| 47335315 | 47335514 | 1 | scRNA | ENSG00000 |
| 47345158 | 47387601 | 1 | protein_coding | ENSG00000 |
| 47359505 | 47359777 | -1 | misc_RNA | ENSG00000 |
| 47377675 | 47377770 | 1 | miRNA | ENSG00000 |
| 47383160 | 47383421 | 1 | misc_RNA | ENSG00000 |
| 47402969 | 47663146 | 1 | protein_coding | ENSG00000 |
| 47516581 | 47570939 | -1 | protein_coding | ENSG00000 |
| 47527008 | 47535199 | 1 | lncRNA | ENSG00000 |
| 47690716 | 47691246 | -1 | processed_pseudoge | ENSG00000 |
| 47695530 | 47810101 | 1 | protein_coding | ENSG00000 |
| 47705468 | 47705979 | 1 | processed_pseudoge | ENSG00000 |
| 47731402 | 47732035 | -1 | processed_pseudoge | ENSG00000 |
| 47781379 | 47781465 | 1 | snRNA | ENSG00000 |
| 47789316 | 47905793 | -1 | protein_coding | ENSG00000 |
| 47797826 | 47798091 | -1 | processed_pseudoge | ENSG00000 |
| 47883455 | 47883909 | 1 | processed_pseudoge | ENSG00000 |
| 47905678 | 47907810 | 1 | lncRNA | ENSG00000 |
| 47924181 | 48314224 | -1 | lncRNA | ENSG00000 |
| 47939738 | 47940218 | -1 | processed_pseudoge | ENSG00000 |

| | | | | |
|---|---|---|---|---|
| 47989625 | 47990526 | 1 | processed_pseudoge | ENSG00000 |
| 48217575 | 48217899 | 1 | misc_RNA | ENSG00000 |
| 48314637 | 48379295 | 1 | protein_coding | ENSG00000 |
| 48340687 | 48340826 | -1 | snRNA | ENSG00000 |
| 48440043 | 48440597 | -1 | lncRNA | ENSG00000 |
| 48440598 | 48515391 | 1 | protein_coding | ENSG00000 |
| 48501922 | 48502024 | -1 | snRNA | ENSG00000 |
| 48529383 | 48598513 | 1 | protein_coding | ENSG00000 |
| 48529925 | 48776517 | 1 | protein_coding | ENSG00000 |
| 48617798 | 48733148 | 1 | protein_coding | ENSG00000 |
| 48632291 | 48755724 | -1 | protein_coding | ENSG00000 |
| 48632856 | 48633353 | -1 | transcribed_processe | ENSG00000 |
| 48686774 | 48755730 | -1 | protein_coding | ENSG00000 |
| 48780602 | 48780895 | 1 | processed_pseudoge | ENSG00000 |
| 48809340 | 49419013 | 1 | lncRNA | ENSG00000 |
| 48915267 | 48916530 | -1 | processed_pseudoge | ENSG00000 |
| 48962157 | 49154537 | -1 | protein_coding | ENSG00000 |
| 49059603 | 49059658 | 1 | miRNA | ENSG00000 |
| 49202126 | 49451896 | -1 | lncRNA | ENSG00000 |
| 49233794 | 49233899 | -1 | snRNA | ENSG00000 |
| 49563388 | 49595126 | 1 | lncRNA | ENSG00000 |
| 49878623 | 49879363 | 1 | processed_pseudoge | ENSG00000 |
| 49888868 | 49888986 | -1 | snoRNA | ENSG00000 |
| 49918503 | 51225575 | -1 | protein_coding | ENSG00000 |
| 50324643 | 50345598 | 1 | lncRNA | ENSG00000 |
| 50588690 | 50589415 | -1 | processed_pseudoge | ENSG00000 |
| 50620963 | 50632993 | 1 | lncRNA | ENSG00000 |
| 50696172 | 50696262 | -1 | miRNA | ENSG00000 |
| 50829442 | 50829678 | -1 | processed_pseudoge | ENSG00000 |
| 51011777 | 51016950 | 1 | lncRNA | ENSG00000 |
| 51032601 | 52407917 | 1 | lncRNA | ENSG00000 |
| 51441959 | 51455913 | 1 | lncRNA | ENSG00000 |
| 51511058 | 51511483 | 1 | processed_pseudoge | ENSG00000 |
| 51775258 | 51775402 | -1 | unprocessed_pseudc | ENSG00000 |
| 51925692 | 51925833 | -1 | processed_pseudoge | ENSG00000 |
| 51926882 | 51927362 | -1 | processed_pseudoge | ENSG00000 |
| 51977787 | 52022435 | -1 | lncRNA | ENSG00000 |
| 52071355 | 52077197 | 1 | processed_pseudoge | ENSG00000 |
| 52297995 | 52298096 | 1 | misc_RNA | ENSG00000 |
| 52370602 | 52390056 | 1 | lncRNA | ENSG00000 |
| 52474073 | 52474411 | -1 | processed_pseudoge | ENSG00000 |
| 52494688 | 52500752 | -1 | lncRNA | ENSG00000 |
| 52570648 | 52572522 | 1 | processed_pseudoge | ENSG00000 |
| 52629743 | 52630078 | 1 | processed_pseudoge | ENSG00000 |
| 52702522 | 52702615 | -1 | miRNA | ENSG00000 |
| 52722671 | 52961452 | -1 | lncRNA | ENSG00000 |
| 52864235 | 52866631 | 1 | lncRNA | ENSG00000 |
| 52883243 | 52883455 | 1 | processed_pseudoge | ENSG00000 |
| 53470447 | 53470629 | -1 | scaRNA | ENSG00000 |
| 53486144 | 53486280 | -1 | processed_pseudoge | ENSG00000 |

| | | | | |
|---|---|---|---|---|
| 53532672 | 53860160 | -1 | protein_coding | ENSG00000 |
| 53570374 | 53570480 | -1 | snRNA | ENSG00000 |
| 53767804 | 53775196 | 1 | protein_coding | ENSG00000 |
| 53787044 | 53818819 | 1 | protein_coding | ENSG00000 |
| 53849122 | 53849205 | -1 | miRNA | ENSG00000 |
| 53852912 | 53859967 | -1 | protein_coding | ENSG00000 |
| 53864069 | 53970993 | -1 | protein_coding | ENSG00000 |
| 53970838 | 54305300 | 1 | protein_coding | ENSG00000 |
| 54029552 | 54030682 | -1 | processed_pseudoge | ENSG00000 |
| 54051334 | 54051760 | 1 | processed_pseudoge | ENSG00000 |
| 54079974 | 54080280 | -1 | processed_pseudoge | ENSG00000 |
| 54082554 | 54085066 | 1 | lncRNA | ENSG00000 |
| 54166944 | 54167007 | -1 | snRNA | ENSG00000 |
| 54253178 | 54256229 | -1 | protein_coding | ENSG00000 |
| 54330034 | 54383742 | 1 | protein_coding | ENSG00000 |
| 54456317 | 54671446 | 1 | protein_coding | ENSG00000 |
| 54516048 | 54540697 | -1 | lncRNA | ENSG00000 |
| 54529343 | 54529801 | 1 | processed_pseudoge | ENSG00000 |
| 54545368 | 54546677 | -1 | lncRNA | ENSG00000 |
| 54661011 | 54680045 | -1 | lncRNA | ENSG00000 |
| 54723499 | 54972025 | 1 | protein_coding | ENSG00000 |
| 54747103 | 54750473 | -1 | lncRNA | ENSG00000 |
| 54768492 | 54801540 | -1 | lncRNA | ENSG00000 |
| 54850289 | 54850352 | 1 | snRNA | ENSG00000 |
| 54972187 | 55112621 | -1 | protein_coding | ENSG00000 |
| 55014418 | 55014521 | -1 | snRNA | ENSG00000 |
| 55172547 | 55232563 | -1 | protein_coding | ENSG00000 |
| 55214387 | 55216126 | -1 | lncRNA | ENSG00000 |
| 55224280 | 55225908 | 1 | processed_pseudoge | ENSG00000 |
| 55231903 | 55235853 | 1 | protein_coding | ENSG00000 |
| 55236595 | 55269347 | -1 | protein_coding | ENSG00000 |
| 55282319 | 55284522 | 1 | transcribed_unitary_ | ENSG00000 |
| 55282350 | 55346049 | 1 | lncRNA | ENSG00000 |
| 55286018 | 55286128 | 1 | misc_RNA | ENSG00000 |
| 55287842 | 55419895 | -1 | protein_coding | ENSG00000 |
| 55435156 | 55435622 | -1 | processed_pseudoge | ENSG00000 |
| 55451004 | 55451099 | -1 | snRNA | ENSG00000 |
| 55456106 | 55456204 | -1 | snRNA | ENSG00000 |
| 55499950 | 55500055 | 1 | snRNA | ENSG00000 |
| 55519604 | 55545879 | 1 | protein_coding | ENSG00000 |
| 55547292 | 55618880 | -1 | protein_coding | ENSG00000 |
| 55565703 | 55565850 | 1 | snoRNA | ENSG00000 |
| 55617909 | 55618373 | 1 | lncRNA | ENSG00000 |
| 55634061 | 55693863 | -1 | protein_coding | ENSG00000 |
| 55865967 | 55924139 | -1 | protein_coding | ENSG00000 |
| 55951654 | 55952014 | -1 | misc_RNA | ENSG00000 |
| 55952158 | 56181652 | 1 | lncRNA | ENSG00000 |
| 55963191 | 56047326 | -1 | lncRNA | ENSG00000 |
| 55982967 | 55983076 | -1 | miRNA | ENSG00000 |
| 55988950 | 55989059 | -1 | miRNA | ENSG00000 |

| | | | | |
|---|---|---|---|---|
| 56000714 | 56000795 | -1 | miRNA | ENSG00000 |
| 56077417 | 56090382 | -1 | lncRNA | ENSG00000 |
| 56147630 | 56386171 | 1 | lncRNA | ENSG00000 |
| 56173534 | 56185870 | -1 | lncRNA | ENSG00000 |
| 56183990 | 56386172 | 1 | protein_coding | ENSG00000 |
| 56235217 | 56235330 | -1 | rRNA_pseudogene | ENSG00000 |
| 56750300 | 56750563 | 1 | processed_pseudoge | ENSG00000 |
| 57048350 | 57049336 | -1 | processed_pseudoge | ENSG00000 |
| 57289648 | 57380132 | -1 | lncRNA | ENSG00000 |
| 57429548 | 57430693 | -1 | processed_pseudoge | ENSG00000 |
| 57544535 | 57544599 | 1 | snoRNA | ENSG00000 |
| 57755428 | 57766052 | 1 | transcribed_processe | ENSG00000 |
| 57907629 | 58159920 | 1 | protein_coding | ENSG00000 |
| 58040211 | 58046700 | -1 | lncRNA | ENSG00000 |
| 58159243 | 58241372 | -1 | protein_coding | ENSG00000 |
| 58241349 | 58241686 | 1 | lncRNA | ENSG00000 |
| 58251440 | 58252525 | 1 | processed_pseudoge | ENSG00000 |
| 58275532 | 58296817 | 1 | lncRNA | ENSG00000 |
| 58427799 | 59063766 | 1 | lncRNA | ENSG00000 |
| 58460292 | 58462032 | -1 | processed_pseudoge | ENSG00000 |
| 59014354 | 59277688 | 1 | lncRNA | ENSG00000 |
| 59217708 | 59279400 | 1 | lncRNA | ENSG00000 |
| 59218680 | 60100200 | -1 | lncRNA | ENSG00000 |
| 59238703 | 59733396 | -1 | lncRNA | ENSG00000 |
| 59434552 | 59442261 | 1 | lncRNA | ENSG00000 |
| 59514890 | 59515124 | 1 | processed_pseudoge | ENSG00000 |
| 59647621 | 59647723 | 1 | snRNA | ENSG00000 |
| 59694762 | 59694869 | 1 | rRNA_pseudogene | ENSG00000 |
| 59778685 | 59791656 | 1 | lncRNA | ENSG00000 |
| 60057601 | 60071127 | 1 | lncRNA | ENSG00000 |
| 60336446 | 60439828 | -1 | lncRNA | ENSG00000 |
| 60383141 | 60385794 | 1 | lncRNA | ENSG00000 |
| 60384605 | 60384763 | 1 | snRNA | ENSG00000 |
| 60387362 | 60387445 | -1 | miRNA | ENSG00000 |
| 60450520 | 60554467 | -1 | protein_coding | ENSG00000 |
| 60495686 | 60499964 | 1 | lncRNA | ENSG00000 |
| 60640705 | 60641002 | 1 | misc_RNA | ENSG00000 |
| 60682873 | 60683244 | -1 | processed_pseudoge | ENSG00000 |
| 60711484 | 60711920 | 1 | processed_pseudoge | ENSG00000 |
| 60719640 | 60719741 | 1 | snRNA | ENSG00000 |
| 60734895 | 60735715 | -1 | processed_pseudoge | ENSG00000 |
| 60756253 | 60802086 | 1 | protein_coding | ENSG00000 |
| 60823069 | 60881317 | -1 | lncRNA | ENSG00000 |
| 60831664 | 60831929 | -1 | misc_RNA | ENSG00000 |
| 60852260 | 60852743 | 1 | processed_pseudoge | ENSG00000 |
| 60881521 | 60931610 | 1 | protein_coding | ENSG00000 |
| 60911303 | 60911442 | 1 | snRNA | ENSG00000 |
| 60925909 | 60931610 | 1 | lncRNA | ENSG00000 |
| 60936819 | 60938049 | -1 | processed_pseudoge | ENSG00000 |
| 60938204 | 60938604 | -1 | processed_pseudoge | ENSG00000 |

| | | | | |
|---|---|---|---|---|
| 60940222 | 61018259 | -1 | protein_coding | ENSG00000( |
| 60998752 | 60998888 | 1 | rRNA_pseudogene | ENSG00000( |
| 61017225 | 61051990 | 1 | protein_coding | ENSG00000( |
| 61065871 | 61138034 | 1 | protein_coding | ENSG00000( |
| 61115787 | 61164825 | 1 | lncRNA | ENSG00000( |
| 61141592 | 61144969 | -1 | lncRNA | ENSG00000( |
| 61145068 | 61164829 | 1 | protein_coding | ENSG00000( |
| 61151433 | 61162105 | -1 | lncRNA | ENSG00000( |
| 61177418 | 61191203 | 1 | transcribed_unitary_ | ENSG00000( |
| 61187463 | 61471087 | -1 | protein_coding | ENSG00000( |
| 61199979 | 61200769 | 1 | lncRNA | ENSG00000( |
| 61249780 | 61250621 | 1 | processed_pseudoge | ENSG00000( |
| 61416887 | 61417197 | -1 | processed_pseudoge | ENSG00000( |
| 61417244 | 61417378 | -1 | snoRNA | ENSG00000( |
| 61471188 | 61484130 | 1 | lncRNA | ENSG00000( |
| 61477849 | 61538626 | -1 | protein_coding | ENSG00000( |
| 61527340 | 61529160 | 1 | lncRNA | ENSG00000( |
| 61575774 | 61576394 | 1 | processed_pseudoge | ENSG00000( |
| 61589498 | 61589663 | -1 | processed_pseudoge | ENSG00000( |
| 61605616 | 61605722 | 1 | snRNA | ENSG00000( |
| 61710076 | 61710978 | -1 | processed_pseudoge | ENSG00000( |
| 61803143 | 61803388 | 1 | processed_pseudoge | ENSG00000( |
| 61820208 | 61820566 | 1 | processed_pseudoge | ENSG00000( |
| 61824848 | 61854143 | -1 | protein_coding | ENSG00000( |
| 61856695 | 61857055 | 1 | processed_pseudoge | ENSG00000( |
| 61858137 | 61860298 | 1 | unprocessed_pseudc | ENSG00000( |
| 61868085 | 61888671 | -1 | protein_coding | ENSG00000( |
| 61868432 | 61886082 | 1 | lncRNA | ENSG00000( |
| 61878940 | 61881353 | 1 | lncRNA | ENSG00000( |
| 61888724 | 62147247 | 1 | protein_coding | ENSG00000( |
| 61928634 | 61928704 | -1 | misc_RNA | ENSG00000( |
| 62069447 | 62146881 | -1 | lncRNA | ENSG00000( |
| 62146413 | 62147153 | 1 | processed_pseudoge | ENSG00000( |
| 62168862 | 62171163 | 1 | processed_pseudoge | ENSG00000( |
| 62196115 | 62224731 | 1 | protein_coding | ENSG00000( |
| 62205826 | 62205917 | 1 | miRNA | ENSG00000( |
| 62262387 | 62262671 | -1 | misc_RNA | ENSG00000( |
| 62348948 | 62350739 | -1 | processed_pseudoge | ENSG00000( |
| 62463127 | 62464070 | 1 | lncRNA | ENSG00000( |
| 62491178 | 62491468 | 1 | misc_RNA | ENSG00000( |
| 62500218 | 62511894 | -1 | protein_coding | ENSG00000( |
| 62507545 | 62507823 | -1 | processed_pseudoge | ENSG00000( |
| 62532583 | 62533059 | -1 | processed_pseudoge | ENSG00000( |
| 62533681 | 62662829 | -1 | lncRNA | ENSG00000( |
| 62552463 | 62553434 | -1 | processed_pseudoge | ENSG00000( |
| 62561058 | 62561423 | -1 | processed_pseudoge | ENSG00000( |
| 62673851 | 63046487 | 1 | protein_coding | ENSG00000( |
| 62726636 | 62726740 | -1 | misc_RNA | ENSG00000( |
| 62817764 | 62819706 | -1 | lncRNA | ENSG00000( |
| 62826064 | 62858438 | -1 | lncRNA | ENSG00000( |

| | | | | |
|---|---|---|---|---|
| 62939916 | 62940282 | 1 | processed_pseudoge | ENSG00000 |
| 62957326 | 63048640 | -1 | lncRNA | ENSG00000 |
| 63050057 | 63057836 | 1 | protein_coding | ENSG00000 |
| 63083008 | 63087322 | 1 | processed_pseudoge | ENSG00000 |
| 63106879 | 63197037 | 1 | lncRNA | ENSG00000 |
| 63108118 | 63108429 | -1 | processed_pseudoge | ENSG00000 |
| 63117851 | 63119542 | -1 | transcribed_processe | ENSG00000 |
| 63119559 | 63827843 | -1 | protein_coding | ENSG00000 |
| 63168600 | 63168847 | 1 | misc_RNA | ENSG00000 |
| 63232453 | 63233577 | -1 | processed_pseudoge | ENSG00000 |
| 63517892 | 63519268 | -1 | TEC | ENSG00000 |
| 63588609 | 63607197 | 1 | protein_coding | ENSG00000 |
| 63622178 | 63622831 | -1 | processed_pseudoge | ENSG00000 |
| 63642455 | 63644038 | -1 | transcribed_processe | ENSG00000 |
| 63684305 | 63684710 | 1 | processed_pseudoge | ENSG00000 |
| 63717122 | 63717899 | 1 | processed_pseudoge | ENSG00000 |
| 63751697 | 63752630 | -1 | processed_pseudoge | ENSG00000 |
| 63840940 | 63891562 | 1 | protein_coding | ENSG00000 |
| 63883249 | 63883391 | -1 | snoRNA | ENSG00000 |
| 63892146 | 64019428 | -1 | protein_coding | ENSG00000 |
| 64086353 | 64088246 | -1 | lncRNA | ENSG00000 |
| 64092652 | 64144420 | -1 | protein_coding | ENSG00000 |
| 64143239 | 64252859 | 1 | lncRNA | ENSG00000 |
| 64185078 | 64205485 | -1 | lncRNA | ENSG00000 |
| 64208498 | 64208716 | -1 | processed_pseudoge | ENSG00000 |
| 64330481 | 64332801 | -1 | lncRNA | ENSG00000 |
| 64338067 | 64341647 | -1 | lncRNA | ENSG00000 |
| 64340747 | 64340848 | -1 | miRNA | ENSG00000 |
| 64340759 | 64340839 | 1 | miRNA | ENSG00000 |
| 64347193 | 64347741 | -1 | processed_pseudoge | ENSG00000 |
| 64395220 | 64453967 | -1 | lncRNA | ENSG00000 |
| 64450096 | 64450524 | 1 | processed_pseudoge | ENSG00000 |
| 64453969 | 64461381 | 1 | protein_coding | ENSG00000 |
| 64486353 | 64500904 | 1 | lncRNA | ENSG00000 |
| 64522187 | 64524093 | -1 | lncRNA | ENSG00000 |
| 64524305 | 64593005 | 1 | protein_coding | ENSG00000 |
| 64525513 | 64525565 | 1 | miRNA | ENSG00000 |
| 64578892 | 64578997 | 1 | snRNA | ENSG00000 |
| 64606975 | 64616556 | 1 | lncRNA | ENSG00000 |
| 64631621 | 64751005 | -1 | protein_coding | ENSG00000 |
| 64644612 | 64646698 | 1 | lncRNA | ENSG00000 |
| 64665607 | 64666064 | 1 | processed_pseudoge | ENSG00000 |
| 64817378 | 64817675 | 1 | misc_RNA | ENSG00000 |
| 64834056 | 64834157 | -1 | misc_RNA | ENSG00000 |
| 64836985 | 64848264 | 1 | lncRNA | ENSG00000 |
| 64846130 | 64863626 | -1 | lncRNA | ENSG00000 |
| 64901840 | 65056168 | -1 | lncRNA | ENSG00000 |
| 64906863 | 64907138 | -1 | misc_RNA | ENSG00000 |
| 64960053 | 64960634 | -1 | processed_pseudoge | ENSG00000 |
| 64988477 | 65023865 | 1 | protein_coding | ENSG00000 |

| | | | | |
|---|---|---|---|---|
| 64994746 | 64994860 | 1 | snRNA | ENSG0000C |
| 65030727 | 65053017 | 1 | lncRNA | ENSG0000C |
| 65056366 | 65087004 | 1 | protein_coding | ENSG0000C |
| 65070696 | 65130331 | -1 | protein_coding | ENSG0000C |
| 65158662 | 65158860 | 1 | snoRNA | ENSG0000C |
| 65203502 | 65227605 | -1 | transcribed_processe | ENSG0000C |
| 65205108 | 65205988 | 1 | processed_pseudoge | ENSG0000C |
| 65227753 | 65271253 | 1 | protein_coding | ENSG0000C |
| 65310851 | 65432637 | -1 | protein_coding | ENSG0000C |
| 65373700 | 65380685 | -1 | lncRNA | ENSG0000C |
| 65436711 | 66200373 | 1 | lncRNA | ENSG0000C |
| 65439838 | 65456571 | -1 | lncRNA | ENSG0000C |
| 65500993 | 65502138 | -1 | processed_pseudoge | ENSG0000C |
| 65511771 | 65512076 | -1 | processed_pseudoge | ENSG0000C |
| 65545401 | 65545696 | -1 | misc_RNA | ENSG0000C |
| 65555432 | 65555534 | -1 | misc_RNA | ENSG0000C |
| 65589566 | 65640177 | -1 | lncRNA | ENSG0000C |
| 65623272 | 65628424 | 1 | lncRNA | ENSG0000C |
| 65666695 | 65667737 | 1 | processed_pseudoge | ENSG0000C |
| 65790039 | 65793533 | 1 | TEC | ENSG0000C |
| 66235377 | 66236213 | -1 | lncRNA | ENSG0000C |
| 66327349 | 66328984 | 1 | lncRNA | ENSG0000C |
| 66358249 | 66358328 | -1 | miRNA | ENSG0000C |
| 66383306 | 66392450 | 1 | lncRNA | ENSG0000C |
| 66426735 | 66433470 | -1 | lncRNA | ENSG0000C |
| 66433452 | 66573869 | 1 | protein_coding | ENSG0000C |
| 66439088 | 66444221 | -1 | lncRNA | ENSG0000C |
| 66574030 | 66731223 | 1 | lncRNA | ENSG0000C |
| 66696190 | 66703252 | -1 | lncRNA | ENSG0000C |
| 66820684 | 66822570 | -1 | processed_pseudoge | ENSG0000C |
| 66881087 | 66881222 | -1 | processed_pseudoge | ENSG0000C |
| 66904436 | 66971462 | 1 | lncRNA | ENSG0000C |
| 66921510 | 66922482 | -1 | lncRNA | ENSG0000C |
| 67040546 | 67041966 | 1 | lncRNA | ENSG0000C |
| 67086446 | 67311439 | 1 | lncRNA | ENSG0000C |
| 67123357 | 67397980 | -1 | lncRNA | ENSG0000C |
| 67324627 | 67325304 | 1 | lncRNA | ENSG0000C |
| 67397322 | 67412089 | 1 | protein_coding | ENSG0000C |
| 67562067 | 67620545 | 1 | lncRNA | ENSG0000C |
| 67565604 | 67684077 | -1 | lncRNA | ENSG0000C |
| 67677499 | 67680798 | 1 | lncRNA | ENSG0000C |
| 67796054 | 67825562 | -1 | lncRNA | ENSG0000C |
| 68023694 | 68024766 | -1 | processed_pseudoge | ENSG0000C |
| 68041130 | 68110948 | -1 | protein_coding | ENSG0000C |
| 68117026 | 68117200 | 1 | unprocessed_pseudc | ENSG0000C |
| 68122936 | 68157549 | -1 | protein_coding | ENSG0000C |
| 68125265 | 68125747 | 1 | processed_pseudoge | ENSG0000C |
| 68131238 | 68261230 | -1 | protein_coding | ENSG0000C |
| 68157888 | 68176238 | 1 | protein_coding | ENSG0000C |
| 68178857 | 68256237 | -1 | protein_coding | ENSG0000C |

| | | | | |
|---|---|---|---|---|
| 68179833 | 68180532 | 1 | lncRNA | ENSG00000 |
| 68252870 | 68253848 | 1 | lncRNA | ENSG00000 |
| 68284171 | 68320051 | -1 | protein_coding | ENSG00000 |
| 68361214 | 68365584 | -1 | lncRNA | ENSG00000 |
| 68365282 | 68397453 | 1 | protein_coding | ENSG00000 |
| 68445710 | 68449174 | -1 | unprocessed_pseudo | ENSG00000 |
| 68459422 | 68467294 | -1 | protein_coding | ENSG00000 |
| 68467572 | 68655862 | 1 | protein_coding | ENSG00000 |
| 68528241 | 68529452 | -1 | processed_pseudoge | ENSG00000 |
| 68643589 | 68658247 | 1 | protein_coding | ENSG00000 |
| 68679601 | 68826833 | 1 | protein_coding | ENSG00000 |
| 68822855 | 68837235 | -1 | lncRNA | ENSG00000 |
| 68832014 | 68837726 | 1 | lncRNA | ENSG00000 |
| 68860909 | 68871397 | -1 | protein_coding | ENSG00000 |
| 68921248 | 68926834 | 1 | polymorphic_pseudo | ENSG00000 |
| 68945232 | 68952893 | -1 | protein_coding | ENSG00000 |
| 68974573 | 68980974 | 1 | protein_coding | ENSG00000 |
| 69013178 | 69249327 | 1 | protein_coding | ENSG00000 |
| 69030042 | 69033637 | -1 | lncRNA | ENSG00000 |
| 69103682 | 69103755 | 1 | miRNA | ENSG00000 |
| 69181897 | 69182003 | -1 | rRNA_pseudogene | ENSG00000 |
| 69182877 | 69182983 | 1 | snRNA | ENSG00000 |
| 69251818 | 69252364 | -1 | processed_pseudoge | ENSG00000 |
| 69319769 | 69387254 | -1 | protein_coding | ENSG00000 |
| 69334600 | 69334692 | 1 | misc_RNA | ENSG00000 |
| 69395750 | 69437628 | -1 | protein_coding | ENSG00000 |
| 69457997 | 69674349 | -1 | protein_coding | ENSG00000 |
| 69516751 | 69517046 | 1 | misc_RNA | ENSG00000 |
| 69520043 | 69520174 | -1 | snoRNA | ENSG00000 |
| 69594741 | 69595058 | -1 | processed_pseudoge | ENSG00000 |
| 69597353 | 69598951 | 1 | processed_pseudoge | ENSG00000 |
| 69644425 | 69827112 | 1 | protein_coding | ENSG00000 |
| 69700192 | 69713847 | -1 | lncRNA | ENSG00000 |
| 69829660 | 69881384 | 1 | protein_coding | ENSG00000 |
| 69844509 | 69844933 | -1 | processed_pseudoge | ENSG00000 |
| 69873565 | 69873819 | -1 | processed_pseudoge | ENSG00000 |
| 69893956 | 69905575 | 1 | protein_coding | ENSG00000 |
| 69897688 | 69942945 | 1 | protein_coding | ENSG00000 |
| 69960089 | 69962265 | -1 | protein_coding | ENSG00000 |
| 69962263 | 70103220 | -1 | lncRNA | ENSG00000 |
| 70075014 | 70075272 | -1 | misc_RNA | ENSG00000 |
| 70087477 | 70089203 | 1 | protein_coding | ENSG00000 |
| 70089721 | 70096100 | 1 | lncRNA | ENSG00000 |
| 70102551 | 70102862 | 1 | processed_pseudoge | ENSG00000 |
| 70124036 | 70125317 | -1 | lncRNA | ENSG00000 |
| 70149880 | 70248615 | -1 | protein_coding | ENSG00000 |
| 70209444 | 70248660 | -1 | protein_coding | ENSG00000 |
| 70252918 | 70253005 | -1 | miRNA | ENSG00000 |
| 70253855 | 70254010 | -1 | processed_pseudoge | ENSG00000 |
| 70257386 | 70281185 | 1 | protein_coding | ENSG00000 |

| | | | | |
|---|---|---|---|---|
| 70281362 | 70293740 | -1 | protein_coding | ENSG00000 |
| 70295975 | 70302090 | -1 | protein_coding | ENSG00000 |
| 70301451 | 70302072 | 1 | lncRNA | ENSG00000 |
| 70353010 | 70355915 | 1 | processed_pseudoge | ENSG00000 |
| 70402934 | 70422678 | 1 | lncRNA | ENSG00000 |
| 70447284 | 70554193 | -1 | protein_coding | ENSG00000 |
| 70467385 | 70468495 | -1 | lncRNA | ENSG00000 |
| 70607618 | 70768225 | -1 | protein_coding | ENSG00000 |
| 70629657 | 70648761 | 1 | lncRNA | ENSG00000 |
| 70687142 | 70691824 | 1 | lncRNA | ENSG00000 |
| 70777310 | 70790643 | -1 | protein_coding | ENSG00000 |
| 70803157 | 70803426 | 1 | processed_pseudoge | ENSG00000 |
| 70808643 | 70820599 | -1 | protein_coding | ENSG00000 |
| 70810798 | 70811474 | -1 | processed_pseudoge | ENSG00000 |
| 70830211 | 70835816 | -1 | protein_coding | ENSG00000 |
| 70887871 | 70889959 | 1 | lncRNA | ENSG00000 |
| 70900155 | 70900351 | 1 | misc_RNA | ENSG00000 |
| 70900576 | 70965373 | 1 | protein_coding | ENSG00000 |
| 70935900 | 70965431 | 1 | protein_coding | ENSG00000 |
| 70939318 | 70995336 | -1 | protein_coding | ENSG00000 |
| 70941817 | 70948610 | -1 | lncRNA | ENSG00000 |
| 70955899 | 70956234 | -1 | processed_pseudoge | ENSG00000 |
| 70965584 | 70965867 | 1 | misc_RNA | ENSG00000 |
| 70978380 | 70985499 | 1 | protein_coding | ENSG00000 |
| 70985942 | 70994873 | -1 | protein_coding | ENSG00000 |
| 70994510 | 71002754 | 1 | lncRNA | ENSG00000 |
| 71002531 | 71064743 | -1 | lncRNA | ENSG00000 |
| 71024127 | 71029893 | 1 | transcribed_unproce | ENSG00000 |
| 71037685 | 71038703 | 1 | unprocessed_pseudc | ENSG00000 |
| 71055527 | 71056003 | 1 | unprocessed_pseudc | ENSG00000 |
| 71064344 | 71079808 | 1 | protein_coding | ENSG00000 |
| 71067519 | 71068125 | -1 | lncRNA | ENSG00000 |
| 71109684 | 71130239 | -1 | protein_coding | ENSG00000 |
| 71112877 | 71117406 | 1 | lncRNA | ENSG00000 |
| 71130632 | 71150101 | 1 | protein_coding | ENSG00000 |
| 71178587 | 71178773 | 1 | unprocessed_pseudc | ENSG00000 |
| 71182738 | 71227103 | -1 | protein_coding | ENSG00000 |
| 71276561 | 71435069 | 1 | protein_coding | ENSG00000 |
| 71373938 | 71376320 | 1 | lncRNA | ENSG00000 |
| 71379755 | 71379858 | -1 | snRNA | ENSG00000 |
| 71453722 | 71686768 | 1 | protein_coding | ENSG00000 |
| 71984182 | 71984434 | -1 | processed_pseudoge | ENSG00000 |
| 72129238 | 72148038 | -1 | protein_coding | ENSG00000 |
| 72175984 | 72826041 | -1 | protein_coding | ENSG00000 |
| 72288785 | 72289110 | 1 | processed_pseudoge | ENSG00000 |
| 72721806 | 72722003 | 1 | snRNA | ENSG00000 |
| 72760534 | 72760602 | 1 | snoRNA | ENSG00000 |
| 72887382 | 72892158 | 1 | protein_coding | ENSG00000 |
| 72916260 | 72936071 | 1 | protein_coding | ENSG00000 |
| 72932974 | 72934355 | -1 | lncRNA | ENSG00000 |

| | | | | |
|---|---|---|---|---|
| 72942036 | 73075619 | -1 | protein_coding | ENSG00000( |
| 73073382 | 73156721 | -1 | protein_coding | ENSG00000( |
| 73113018 | 73115907 | 1 | lncRNA | ENSG00000( |
| 73202574 | 73212513 | 1 | protein_coding | ENSG00000( |
| 73214222 | 73227221 | 1 | protein_coding | ENSG00000( |
| 73228010 | 73233239 | -1 | protein_coding | ENSG00000( |
| 73233420 | 73253021 | 1 | protein_coding | ENSG00000( |
| 73254682 | 73284431 | -1 | protein_coding | ENSG00000( |
| 73290929 | 73293701 | -1 | protein_coding | ENSG00000( |
| 73298683 | 73298787 | 1 | snRNA | ENSG00000( |
| 73305941 | 73307890 | -1 | lncRNA | ENSG00000( |
| 73352610 | 73385677 | -1 | lncRNA | ENSG00000( |
| 73370019 | 73370892 | 1 | processed_pseudoge | ENSG00000( |
| 73385758 | 73625166 | 1 | protein_coding | ENSG00000( |
| 73456764 | 73459482 | 1 | lncRNA | ENSG00000( |
| 73469525 | 73471030 | 1 | lncRNA | ENSG00000( |
| 73640723 | 73642422 | -1 | protein_coding | ENSG00000( |
| 73641083 | 73697509 | 1 | transcribed_unproce | ENSG00000( |
| 73700576 | 73701340 | -1 | polymorphic_pseudo | ENSG00000( |
| 73729104 | 73737400 | -1 | protein_coding | ENSG00000( |
| 73750256 | 73750786 | -1 | lncRNA | ENSG00000( |
| 73761269 | 73761781 | -1 | TEC | ENSG00000( |
| 73762184 | 73780157 | -1 | protein_coding | ENSG00000( |
| 73784189 | 73817147 | 1 | protein_coding | ENSG00000( |
| 73823154 | 73824593 | -1 | lncRNA | ENSG00000( |
| 73828916 | 73873659 | 1 | protein_coding | ENSG00000( |
| 73834422 | 73856965 | -1 | lncRNA | ENSG00000( |
| 73877566 | 73878862 | 1 | processed_pseudoge | ENSG00000( |
| 73892314 | 73919865 | 1 | protein_coding | ENSG00000( |
| 73926826 | 73958961 | 1 | protein_coding | ENSG00000( |
| 73947322 | 73981441 | -1 | lncRNA | ENSG00000( |
| 73968123 | 73968206 | -1 | rRNA_pseudogene | ENSG00000( |
| 73985132 | 73986343 | 1 | lncRNA | ENSG00000( |
| 73986404 | 74108176 | 1 | protein_coding | ENSG00000( |
| 73998337 | 73998548 | -1 | TEC | ENSG00000( |
| 74120680 | 74123218 | 1 | processed_pseudoge | ENSG00000( |
| 74123965 | 74135640 | 1 | lncRNA | ENSG00000( |
| 74135400 | 74147912 | -1 | protein_coding | ENSG00000( |
| 74148007 | 74151952 | 1 | lncRNA | ENSG00000( |
| 74152528 | 74178898 | -1 | protein_coding | ENSG00000( |
| 74191388 | 74192245 | 1 | TEC | ENSG00000( |
| 74196698 | 74198560 | -1 | lncRNA | ENSG00000( |
| 74198610 | 74217565 | 1 | protein_coding | ENSG00000( |
| 74211604 | 74363377 | -1 | protein_coding | ENSG00000( |
| 74216242 | 74343414 | -1 | protein_coding | ENSG00000( |
| 74306728 | 74308008 | -1 | processed_pseudoge | ENSG00000( |
| 74319399 | 74319501 | 1 | snRNA | ENSG00000( |
| 74350069 | 74351009 | 1 | processed_pseudoge | ENSG00000( |
| 74351134 | 74351419 | 1 | processed_pseudoge | ENSG00000( |
| 74361154 | 74392087 | -1 | protein_coding | ENSG00000( |

| | | | | |
|---|---|---|---|---|
| 74385474 | 74393882 | 1 | lncRNA | ENSG00000 |
| 74393836 | 74421662 | -1 | protein_coding | ENSG00000 |
| 74407797 | 74413603 | -1 | transcribed_unitary_ | ENSG00000 |
| 74414176 | 74421591 | -1 | protein_coding | ENSG00000 |
| 74415422 | 74416781 | 1 | lncRNA | ENSG00000 |
| 74418122 | 74418442 | -1 | processed_pseudoge | ENSG00000 |
| 74421678 | 74425755 | 1 | protein_coding | ENSG00000 |
| 74425835 | 74442422 | -1 | protein_coding | ENSG00000 |
| 74455087 | 74457944 | 1 | protein_coding | ENSG00000 |
| 74455088 | 74460884 | 1 | protein_coding | ENSG00000 |
| 74458400 | 74460891 | 1 | protein_coding | ENSG00000 |
| 74461057 | 74465410 | -1 | protein_coding | ENSG00000 |
| 74465339 | 74472961 | 1 | lncRNA | ENSG00000 |
| 74471982 | 74472687 | -1 | protein_coding | ENSG00000 |
| 74471986 | 74483408 | -1 | protein_coding | ENSG00000 |
| 74483073 | 74494886 | 1 | protein_coding | ENSG00000 |
| 74497517 | 74503316 | -1 | protein_coding | ENSG00000 |
| 74501717 | 74502365 | 1 | lncRNA | ENSG00000 |
| 74502595 | 74504678 | 1 | lncRNA | ENSG00000 |
| 74505043 | 74507695 | -1 | protein_coding | ENSG00000 |
| 74513463 | 74517148 | 1 | protein_coding | ENSG00000 |
| 74518131 | 74526281 | -1 | protein_coding | ENSG00000 |
| 74526645 | 74529760 | -1 | protein_coding | ENSG00000 |
| 74529377 | 74533348 | 1 | protein_coding | ENSG00000 |
| 74532258 | 74555690 | -1 | protein_coding | ENSG00000 |
| 74549026 | 74557554 | 1 | protein_coding | ENSG00000 |
| 74557883 | 74648338 | -1 | protein_coding | ENSG00000 |
| 74618856 | 74620979 | -1 | processed_pseudoge | ENSG00000 |
| 74628328 | 74628942 | -1 | processed_pseudoge | ENSG00000 |
| 74654228 | 74683853 | 1 | protein_coding | ENSG00000 |
| 74750173 | 74778742 | -1 | lncRNA | ENSG00000 |
| 74754670 | 74754819 | -1 | processed_pseudoge | ENSG00000 |
| 74754773 | 74760227 | 1 | lncRNA | ENSG00000 |
| 74832655 | 74833987 | -1 | lncRNA | ENSG00000 |
| 74833981 | 74893359 | 1 | protein_coding | ENSG00000 |
| 74918148 | 74938418 | 1 | lncRNA | ENSG00000 |
| 74919555 | 74924846 | -1 | lncRNA | ENSG00000 |
| 74940258 | 74942670 | 1 | lncRNA | ENSG00000 |
| 74958643 | 74970128 | 1 | protein_coding | ENSG00000 |
| 75046463 | 75199520 | -1 | protein_coding | ENSG00000 |
| 75090812 | 75090914 | 1 | miRNA | ENSG00000 |
| 75154366 | 75189949 | 1 | lncRNA | ENSG00000 |
| 75455994 | 75456578 | -1 | processed_pseudoge | ENSG00000 |
| 75469302 | 75569722 | -1 | protein_coding | ENSG00000 |
| 75474453 | 75482579 | 1 | lncRNA | ENSG00000 |
| 75489576 | 75489710 | 1 | snoRNA | ENSG00000 |
| 75524068 | 75542706 | 1 | lncRNA | ENSG00000 |
| 75598071 | 75598559 | 1 | processed_pseudoge | ENSG00000 |
| 75627950 | 75628168 | 1 | snoRNA | ENSG00000 |
| 75646783 | 75690851 | 1 | protein_coding | ENSG00000 |

| | | | | |
|---|---|---|---|---|
| 75651288 | 75651627 | -1 | processed_pseudoge | ENSG00000 |
| 75652000 | 75710985 | -1 | protein_coding | ENSG00000 |
| 75660462 | 75662208 | 1 | lncRNA | ENSG00000 |
| 75669989 | 75670454 | 1 | lncRNA | ENSG00000 |
| 75697583 | 75697996 | 1 | lncRNA | ENSG00000 |
| 75710782 | 75728123 | 1 | lncRNA | ENSG00000 |
| 75719120 | 75720018 | 1 | lncRNA | ENSG00000 |
| 75799974 | 76208703 | 1 | lncRNA | ENSG00000 |
| 76106016 | 76107324 | -1 | processed_pseudoge | ENSG00000 |
| 76185020 | 76399490 | 1 | lncRNA | ENSG00000 |
| 76197855 | 76252927 | -1 | lncRNA | ENSG00000 |
| 76258034 | 76258897 | -1 | processed_pseudoge | ENSG00000 |
| 76260755 | 76261661 | -1 | processed_pseudoge | ENSG00000 |
| 76445079 | 76445410 | -1 | misc_RNA | ENSG00000 |
| 76595413 | 76595678 | -1 | misc_RNA | ENSG00000 |
| 76633365 | 76673346 | -1 | lncRNA | ENSG00000 |
| 76684975 | 76698996 | 1 | lncRNA | ENSG00000 |
| 76691007 | 76692489 | -1 | lncRNA | ENSG00000 |
| 76747719 | 77593319 | -1 | protein_coding | ENSG00000 |
| 76772909 | 76773045 | 1 | rRNA_pseudogene | ENSG00000 |
| 76893894 | 76897247 | -1 | lncRNA | ENSG00000 |
| 76985965 | 77009791 | 1 | lncRNA | ENSG00000 |
| 77210587 | 77327103 | -1 | lncRNA | ENSG00000 |
| 77652025 | 78290469 | -1 | lncRNA | ENSG00000 |
| 77672215 | 77673792 | 1 | lncRNA | ENSG00000 |
| 77788382 | 77793087 | 1 | transcribed_processe | ENSG00000 |
| 77915870 | 77918011 | 1 | lncRNA | ENSG00000 |
| 78088729 | 78127806 | 1 | lncRNA | ENSG00000 |
| 78412793 | 78413094 | -1 | processed_pseudoge | ENSG00000 |
| 78437850 | 78438921 | 1 | processed_pseudoge | ENSG00000 |
| 78597911 | 78599406 | 1 | lncRNA | ENSG00000 |
| 78880713 | 78931146 | -1 | lncRNA | ENSG00000 |
| 78882447 | 78882552 | 1 | snRNA | ENSG00000 |
| 78882628 | 78882734 | 1 | snRNA | ENSG00000 |
| 79025686 | 79028505 | 1 | protein_coding | ENSG00000 |
| 79085023 | 79088019 | -1 | protein_coding | ENSG00000 |
| 79120362 | 79123409 | 1 | protein_coding | ENSG00000 |
| 79135503 | 79138427 | -1 | transcribed_unproce | ENSG00000 |
| 79157003 | 79159753 | -1 | protein_coding | ENSG00000 |
| 79158374 | 79185523 | 1 | lncRNA | ENSG00000 |
| 79185231 | 80648861 | 1 | protein_coding | ENSG00000 |
| 79269905 | 79292878 | -1 | lncRNA | ENSG00000 |
| 79492704 | 79513900 | -1 | lncRNA | ENSG00000 |
| 79547151 | 79548128 | -1 | unprocessed_pseudo | ENSG00000 |
| 79573764 | 79575593 | -1 | processed_pseudoge | ENSG00000 |
| 79636862 | 79636963 | -1 | snRNA | ENSG00000 |
| 79649294 | 79649359 | -1 | miRNA | ENSG00000 |
| 79866495 | 79866583 | -1 | miRNA | ENSG00000 |
| 80028012 | 80030551 | -1 | lncRNA | ENSG00000 |
| 80162428 | 80163135 | -1 | processed_pseudoge | ENSG00000 |

| | | | | |
|---|---|---|---|---|
| 80288351 | 80304752 | -1 | protein_coding | ENSG00000( |
| 80572681 | 80618777 | -1 | lncRNA | ENSG00000( |
| 80699388 | 80870190 | -1 | lncRNA | ENSG00000( |
| 81194337 | 81201184 | -1 | processed_pseudoge | ENSG00000( |
| 81418723 | 81419302 | -1 | processed_pseudoge | ENSG00000( |
| 81461358 | 81467468 | -1 | lncRNA | ENSG00000( |
| 81496214 | 81496327 | -1 | rRNA_pseudogene | ENSG00000( |
| 81666477 | 81666728 | -1 | processed_pseudoge | ENSG00000( |
| 81967079 | 81967374 | 1 | misc_RNA | ENSG00000( |
| 81983272 | 82005700 | -1 | lncRNA | ENSG00000( |
| 82268427 | 82269927 | -1 | processed_pseudoge | ENSG00000( |
| 82268612 | 82268706 | 1 | snRNA | ENSG00000( |
| 82307067 | 82307168 | -1 | misc_RNA | ENSG00000( |
| 82476825 | 82538711 | -1 | lncRNA | ENSG00000( |
| 82609652 | 82609977 | 1 | processed_pseudoge | ENSG00000( |
| 82814984 | 82815516 | 1 | processed_pseudoge | ENSG00000( |
| 82815809 | 82817528 | 1 | processed_pseudoge | ENSG00000( |
| 82817538 | 82818061 | -1 | processed_pseudoge | ENSG00000( |
| 82818131 | 82819264 | 1 | processed_pseudoge | ENSG00000( |
| 82831234 | 82866508 | 1 | lncRNA | ENSG00000( |
| 82856826 | 82857384 | 1 | processed_pseudoge | ENSG00000( |
| 83218890 | 83219105 | -1 | processed_pseudoge | ENSG00000( |
| 83522814 | 83523502 | 1 | lncRNA | ENSG00000( |
| 83594956 | 83595243 | 1 | processed_pseudoge | ENSG00000( |
| 83657735 | 83657842 | 1 | snRNA | ENSG00000( |
| 84031140 | 84032358 | 1 | processed_pseudoge | ENSG00000( |
| 84039885 | 84040720 | -1 | processed_pseudoge | ENSG00000( |
| 84290683 | 84292202 | 1 | transcribed_processe | ENSG00000( |
| 84315108 | 84350774 | 1 | lncRNA | ENSG00000( |
| 84423528 | 84460045 | -1 | protein_coding | ENSG00000( |
| 84516455 | 84819589 | 1 | protein_coding | ENSG00000( |
| 84750769 | 84751231 | 1 | transcribed_unproce | ENSG00000( |
| 84777259 | 84778223 | -1 | processed_pseudoge | ENSG00000( |
| 84821650 | 84907008 | -1 | protein_coding | ENSG00000( |
| 84850711 | 84851148 | 1 | unprocessed_pseudc | ENSG00000( |
| 84874696 | 84875020 | 1 | processed_pseudoge | ENSG00000( |
| 84905656 | 84906671 | 1 | protein_coding | ENSG00000( |
| 84915868 | 84916660 | 1 | processed_pseudoge | ENSG00000( |
| 84926019 | 84966685 | 1 | lncRNA | ENSG00000( |
| 84971093 | 85059472 | 1 | protein_coding | ENSG00000( |
| 85061213 | 85067347 | -1 | lncRNA | ENSG00000( |
| 85068809 | 85069031 | -1 | processed_pseudoge | ENSG00000( |
| 85102389 | 85102694 | -1 | processed_pseudoge | ENSG00000( |
| 85133392 | 85310387 | 1 | protein_coding | ENSG00000( |
| 85186409 | 85187253 | 1 | lncRNA | ENSG00000( |
| 85204926 | 85205032 | 1 | snRNA | ENSG00000( |
| 85262144 | 85262267 | -1 | processed_pseudoge | ENSG00000( |
| 85315041 | 85316529 | 1 | unprocessed_pseudc | ENSG00000( |
| 85318020 | 85328296 | -1 | protein_coding | ENSG00000( |
| 85332895 | 85332996 | -1 | misc_RNA | ENSG00000( |

| | | | | |
|---|---|---|---|---|
| 85341281 | 85341838 | -1 | processed_pseudoge | ENSG00000( |
| 85341955 | 85354531 | -1 | protein_coding | ENSG00000( |
| 85354394 | 85391752 | 1 | protein_coding | ENSG00000( |
| 85360798 | 85360925 | -1 | processed_pseudoge | ENSG00000( |
| 85367585 | 85367686 | 1 | misc_RNA | ENSG00000( |
| 85368282 | 85368580 | 1 | misc_RNA | ENSG00000( |
| 85387074 | 85387146 | -1 | lncRNA | ENSG00000( |
| 85394753 | 85418432 | -1 | protein_coding | ENSG00000( |
| 85418504 | 85426724 | 1 | lncRNA | ENSG00000( |
| 85418931 | 85437029 | 1 | protein_coding | ENSG00000( |
| 85434507 | 85434619 | 1 | misc_RNA | ENSG00000( |
| 85442495 | 85442791 | 1 | misc_RNA | ENSG00000( |
| 85460144 | 85460252 | -1 | misc_RNA | ENSG00000( |
| 85490930 | 85491780 | -1 | processed_pseudoge | ENSG00000( |
| 85537462 | 85538666 | -1 | lncRNA | ENSG00000( |
| 85539168 | 85545281 | 1 | protein_coding | ENSG00000( |
| 85544720 | 85561547 | -1 | protein_coding | ENSG00000( |
| 85561562 | 85582031 | 1 | protein_coding | ENSG00000( |
| 85567662 | 85567960 | -1 | misc_RNA | ENSG00000( |
| 85584431 | 85593406 | 1 | protein_coding | ENSG00000( |
| 85595725 | 85597708 | 1 | protein_coding | ENSG00000( |
| 85598547 | 85603196 | -1 | protein_coding | ENSG00000( |
| 85602856 | 85649283 | 1 | protein_coding | ENSG00000( |
| 85605254 | 85612066 | -1 | protein_coding | ENSG00000( |
| 85657314 | 85668741 | -1 | protein_coding | ENSG00000( |
| 85685175 | 85698854 | 1 | protein_coding | ENSG00000( |
| 85686053 | 85687047 | -1 | processed_pseudoge | ENSG00000( |
| 85728194 | 85728337 | -1 | snRNA | ENSG00000( |
| 85751344 | 85791383 | 1 | protein_coding | ENSG00000( |
| 85783600 | 85783677 | -1 | miRNA | ENSG00000( |
| 85815130 | 85825391 | 1 | lncRNA | ENSG00000( |
| 85820435 | 85820612 | -1 | misc_RNA | ENSG00000( |
| 85837120 | 85905199 | -1 | protein_coding | ENSG00000( |
| 85889151 | 85891136 | 1 | lncRNA | ENSG00000( |
| 85904279 | 85904727 | 1 | lncRNA | ENSG00000( |
| 85934535 | 85937548 | -1 | lncRNA | ENSG00000( |
| 86020216 | 86106155 | -1 | protein_coding | ENSG00000( |
| 86106223 | 86142157 | 1 | protein_coding | ENSG00000( |
| 86135870 | 86136006 | 1 | snoRNA | ENSG00000( |
| 86143932 | 86195770 | -1 | protein_coding | ENSG00000( |
| 86159956 | 86160066 | -1 | misc_RNA | ENSG00000( |
| 86193026 | 86193108 | -1 | miRNA | ENSG00000( |
| 86195590 | 86196049 | 1 | lncRNA | ENSG00000( |
| 86199355 | 86213790 | 1 | protein_coding | ENSG00000( |
| 86213993 | 86338083 | -1 | protein_coding | ENSG00000( |
| 86347062 | 86347195 | -1 | snoRNA | ENSG00000( |
| 86364136 | 86364265 | 1 | snoRNA | ENSG00000( |
| 86440647 | 86492716 | 1 | protein_coding | ENSG00000( |
| 86503430 | 86563479 | -1 | protein_coding | ENSG00000( |
| 86505668 | 86721122 | -1 | protein_coding | ENSG00000( |

| | | | | |
|---|---|---|---|---|
| 86515204 | 86515308 | -1 | snRNA | ENSG0000C |
| 86562070 | 86618766 | 1 | lncRNA | ENSG0000C |
| 86603398 | 86623866 | -1 | protein_coding | ENSG0000C |
| 86720291 | 86778041 | 1 | protein_coding | ENSG0000C |
| 86784610 | 86808396 | -1 | protein_coding | ENSG0000C |
| 86815339 | 86861924 | -1 | protein_coding | ENSG0000C |
| 86861825 | 86912978 | 1 | transcribed_unproce | ENSG0000C |
| 86885075 | 86885731 | -1 | processed_pseudoge | ENSG0000C |
| 86907953 | 87013976 | 1 | protein_coding | ENSG0000C |
| 86930250 | 86930754 | -1 | processed_pseudoge | ENSG0000C |
| 86934462 | 86934822 | -1 | processed_pseudoge | ENSG0000C |
| 87002559 | 87021852 | -1 | protein_coding | ENSG0000C |
| 87030675 | 87076429 | -1 | lncRNA | ENSG0000C |
| 87031815 | 87052992 | -1 | unprocessed_pseudc | ENSG0000C |
| 87055658 | 87076060 | -1 | unprocessed_pseudc | ENSG0000C |
| 87075653 | 87118267 | 1 | lncRNA | ENSG0000C |
| 87118534 | 87120980 | -1 | unprocessed_pseudc | ENSG0000C |
| 87125198 | 87196629 | 1 | unprocessed_pseudc | ENSG0000C |
| 87194786 | 87194859 | 1 | miRNA | ENSG0000C |
| 87221113 | 87222488 | -1 | processed_pseudoge | ENSG0000C |
| 87249095 | 87253750 | -1 | lncRNA | ENSG0000C |
| 87301658 | 87302251 | 1 | processed_pseudoge | ENSG0000C |
| 87311460 | 87348728 | -1 | lncRNA | ENSG0000C |
| 87338511 | 87339035 | 1 | IG_V_gene | ENSG0000C |
| 87379880 | 87380767 | -1 | processed_pseudoge | ENSG0000C |
| 87439523 | 87459044 | -1 | lncRNA | ENSG0000C |
| 87454781 | 87636740 | 1 | lncRNA | ENSG0000C |
| 87455476 | 87767359 | 1 | lncRNA | ENSG0000C |
| 87477495 | 87478034 | -1 | lncRNA | ENSG0000C |
| 87565828 | 87566764 | -1 | processed_pseudoge | ENSG0000C |
| 87629755 | 87629834 | 1 | miRNA | ENSG0000C |
| 87654890 | 87655243 | 1 | processed_pseudoge | ENSG0000C |
| 87700984 | 87738105 | 1 | transcribed_unproce | ENSG0000C |
| 87748087 | 87759476 | 1 | protein_coding | ENSG0000C |
| 87755960 | 87825952 | -1 | protein_coding | ENSG0000C |
| 87824942 | 87825143 | 1 | processed_pseudoge | ENSG0000C |
| 87968609 | 87968971 | 1 | processed_pseudoge | ENSG0000C |
| 87972656 | 87973082 | 1 | processed_pseudoge | ENSG0000C |
| 87980714 | 87992858 | -1 | unprocessed_pseudc | ENSG0000C |
| 88003008 | 88003743 | -1 | unprocessed_pseudc | ENSG0000C |
| 88016354 | 88016547 | -1 | snRNA | ENSG0000C |
| 88016780 | 88021451 | 1 | lncRNA | ENSG0000C |
| 88027205 | 88064252 | -1 | protein_coding | ENSG0000C |
| 88067825 | 88113384 | 1 | protein_coding | ENSG0000C |
| 88082519 | 88082599 | -1 | miRNA | ENSG0000C |
| 88122982 | 88128062 | -1 | protein_coding | ENSG0000C |
| 88170295 | 88186636 | 1 | protein_coding | ENSG0000C |
| 88364695 | 88365608 | -1 | processed_pseudoge | ENSG0000C |
| 88367793 | 88367862 | -1 | snRNA | ENSG0000C |
| 88383494 | 88383597 | -1 | snRNA | ENSG0000C |

| | | | | |
|---|---|---|---|---|
| 88414898 | 88415004 | -1 | snRNA | ENSG00000 |
| 88428078 | 88428615 | 1 | processed_pseudoge | ENSG00000 |
| 88446787 | 88452693 | -1 | protein_coding | ENSG00000 |
| 88524649 | 88529585 | 1 | protein_coding | ENSG00000 |
| 88538720 | 88575610 | 1 | lncRNA | ENSG00000 |
| 88556741 | 88627464 | -1 | protein_coding | ENSG00000 |
| 88627539 | 88631821 | 1 | lncRNA | ENSG00000 |
| 88690177 | 88691464 | -1 | lncRNA | ENSG00000 |
| 88691673 | 88750929 | 1 | protein_coding | ENSG00000 |
| 88765807 | 88806612 | 1 | transcribed_unproce | ENSG00000 |
| 88811186 | 88861563 | -1 | lncRNA | ENSG00000 |
| 88811186 | 88825207 | -1 | transcribed_unproce | ENSG00000 |
| 88812370 | 88812454 | 1 | miRNA | ENSG00000 |
| 88857161 | 88857683 | -1 | IG_C_gene | ENSG00000 |
| 88860568 | 88860605 | -1 | IG_J_gene | ENSG00000 |
| 88860886 | 88860922 | -1 | IG_J_gene | ENSG00000 |
| 88861221 | 88861258 | -1 | IG_J_gene | ENSG00000 |
| 88861525 | 88861563 | -1 | IG_J_gene | ENSG00000 |
| 88861886 | 88861923 | -1 | IG_J_gene | ENSG00000 |
| 88885397 | 88886153 | 1 | IG_V_gene | ENSG00000 |
| 88897232 | 88897784 | 1 | IG_V_gene | ENSG00000 |
| 88915081 | 88915378 | -1 | IG_V_pseudogene | ENSG00000 |
| 88922328 | 88922753 | -1 | processed_pseudoge | ENSG00000 |
| 88931666 | 88932380 | -1 | IG_V_pseudogene | ENSG00000 |
| 88947301 | 88947957 | -1 | IG_V_gene | ENSG00000 |
| 88966262 | 88966767 | -1 | IG_V_gene | ENSG00000 |
| 88978468 | 88979081 | -1 | IG_V_gene | ENSG00000 |
| 88992409 | 88992931 | -1 | IG_V_gene | ENSG00000 |
| 89009982 | 89010515 | -1 | IG_V_gene | ENSG00000 |
| 89019992 | 89020686 | -1 | IG_V_pseudogene | ENSG00000 |
| 89027171 | 89027731 | -1 | IG_V_gene | ENSG00000 |
| 89040224 | 89040745 | -1 | IG_V_gene | ENSG00000 |
| 89045995 | 89046466 | -1 | IG_V_pseudogene | ENSG00000 |
| 89078010 | 89078784 | -1 | IG_V_pseudogene | ENSG00000 |
| 89085177 | 89085787 | -1 | IG_V_gene | ENSG00000 |
| 89099859 | 89100361 | -1 | IG_V_gene | ENSG00000 |
| 89117342 | 89117844 | -1 | IG_V_gene | ENSG00000 |
| 89128724 | 89129483 | -1 | IG_V_pseudogene | ENSG00000 |
| 89134975 | 89135257 | -1 | IG_V_pseudogene | ENSG00000 |
| 89142574 | 89143160 | -1 | IG_V_gene | ENSG00000 |
| 89159751 | 89160366 | -1 | IG_V_gene | ENSG00000 |
| 89170775 | 89171212 | -1 | IG_V_pseudogene | ENSG00000 |
| 89172022 | 89172553 | -1 | IG_V_pseudogene | ENSG00000 |
| 89176328 | 89177160 | -1 | IG_V_gene | ENSG00000 |
| 89192500 | 89192752 | -1 | IG_V_pseudogene | ENSG00000 |
| 89196096 | 89196829 | -1 | IG_V_pseudogene | ENSG00000 |
| 89203509 | 89203906 | -1 | unprocessed_pseudc | ENSG00000 |
| 89213423 | 89213928 | -1 | IG_V_gene | ENSG00000 |
| 89221698 | 89222461 | -1 | IG_V_gene | ENSG00000 |
| 89234174 | 89234912 | -1 | IG_V_pseudogene | ENSG00000 |

| | | | | |
|---|---|---|---|---|
| 89244781 | 89245596 | -1 | IG_V_gene | ENSG0000C |
| 89252211 | 89252736 | -1 | IG_V_pseudogene | ENSG0000C |
| 89253571 | 89254015 | -1 | IG_V_pseudogene | ENSG0000C |
| 89266494 | 89268506 | -1 | IG_V_gene | ENSG0000C |
| 89275298 | 89275787 | -1 | IG_V_pseudogene | ENSG0000C |
| 89286689 | 89286973 | -1 | IG_V_pseudogene | ENSG0000C |
| 89295233 | 89295455 | -1 | IG_V_pseudogene | ENSG0000C |
| 89297264 | 89297785 | -1 | IG_V_gene | ENSG0000C |
| 89309898 | 89310144 | -1 | IG_V_pseudogene | ENSG0000C |
| 89319625 | 89320146 | -1 | IG_V_gene | ENSG0000C |
| 89330110 | 89330429 | -1 | IG_V_gene | ENSG0000C |
| 89533185 | 89547106 | 1 | unprocessed_pseudc | ENSG0000C |
| 89542842 | 89585652 | -1 | transcribed_unproce | ENSG0000C |
| 89600571 | 89600680 | 1 | rRNA | ENSG0000C |
| 89622213 | 89640635 | -1 | unprocessed_pseudc | ENSG0000C |
| 89662601 | 89662977 | 1 | unprocessed_pseudc | ENSG0000C |
| 89666033 | 89683204 | 1 | unprocessed_pseudc | ENSG0000C |
| 89851791 | 89852497 | 1 | IG_V_gene | ENSG0000C |
| 89862482 | 89862981 | 1 | IG_V_gene | ENSG0000C |
| 89872463 | 89872702 | 1 | IG_V_pseudogene | ENSG0000C |
| 89884740 | 89885216 | 1 | IG_V_gene | ENSG0000C |
| 89887022 | 89887247 | 1 | IG_V_pseudogene | ENSG0000C |
| 89895502 | 89895772 | 1 | IG_V_pseudogene | ENSG0000C |
| 89906757 | 89907246 | 1 | IG_V_pseudogene | ENSG0000C |
| 89913982 | 89916052 | 1 | IG_V_gene | ENSG0000C |
| 89928422 | 89928867 | 1 | IG_V_pseudogene | ENSG0000C |
| 89929701 | 89930202 | 1 | IG_V_pseudogene | ENSG0000C |
| 89936859 | 89937679 | 1 | IG_V_gene | ENSG0000C |
| 89947512 | 89948279 | 1 | IG_V_gene | ENSG0000C |
| 89959979 | 89960754 | 1 | IG_V_gene | ENSG0000C |
| 89968867 | 89969335 | 1 | IG_V_pseudogene | ENSG0000C |
| 89985922 | 89986702 | 1 | IG_V_gene | ENSG0000C |
| 89989987 | 89990221 | 1 | IG_V_pseudogene | ENSG0000C |
| 90004797 | 90005629 | 1 | IG_V_gene | ENSG0000C |
| 90009402 | 90009927 | 1 | IG_V_pseudogene | ENSG0000C |
| 90010741 | 90011184 | 1 | IG_V_pseudogene | ENSG0000C |
| 90021567 | 90022185 | 1 | IG_V_gene | ENSG0000C |
| 90038848 | 90039479 | 1 | IG_V_gene | ENSG0000C |
| 90046796 | 90047057 | 1 | IG_V_pseudogene | ENSG0000C |
| 90052581 | 90053372 | 1 | IG_V_pseudogene | ENSG0000C |
| 90069662 | 90070238 | 1 | IG_V_gene | ENSG0000C |
| 90082635 | 90083291 | 1 | IG_V_gene | ENSG0000C |
| 90100236 | 90100738 | 1 | IG_V_gene | ENSG0000C |
| 90114838 | 90115402 | 1 | IG_V_gene | ENSG0000C |
| 90121786 | 90122564 | 1 | IG_V_pseudogene | ENSG0000C |
| 90154073 | 90154574 | 1 | IG_V_gene | ENSG0000C |
| 90159840 | 90160335 | 1 | IG_V_gene | ENSG0000C |
| 90172802 | 90173414 | 1 | IG_V_gene | ENSG0000C |
| 90179889 | 90180587 | 1 | IG_V_pseudogene | ENSG0000C |
| 90190193 | 90190681 | 1 | IG_V_gene | ENSG0000C |

| | | | | |
|---|---|---|---|---|
| 90209873 | 90210529 | 1 | IG_V_gene | ENSG00000( |
| 90220727 | 90221384 | 1 | IG_V_gene | ENSG00000( |
| 90234812 | 90235370 | 1 | IG_V_gene | ENSG00000( |
| 90309230 | 90309333 | 1 | unprocessed_pseudc | ENSG00000( |
| 90315366 | 90315836 | -1 | IG_V_pseudogene | ENSG00000( |
| 90359809 | 90360032 | -1 | processed_pseudoge | ENSG00000( |
| 90365737 | 90367699 | 1 | lncRNA | ENSG00000( |
| 91578478 | 91578573 | 1 | processed_pseudoge | ENSG00000( |
| 91580336 | 91580863 | 1 | lncRNA | ENSG00000( |
| 91589494 | 91621421 | 1 | unprocessed_pseudc | ENSG00000( |
| 91617160 | 91659972 | -1 | transcribed_unproce | ENSG00000( |
| 91674857 | 91674966 | 1 | rRNA_pseudogene | ENSG00000( |
| 91684447 | 91685884 | -1 | processed_pseudoge | ENSG00000( |
| 91686102 | 91714745 | -1 | transcribed_unproce | ENSG00000( |
| 91723023 | 91723605 | 1 | processed_pseudoge | ENSG00000( |
| 91736726 | 91767439 | 1 | unprocessed_pseudc | ENSG00000( |
| 91747940 | 91748986 | -1 | processed_pseudoge | ENSG00000( |
| 91759462 | 91767701 | -1 | lncRNA | ENSG00000( |
| 91775944 | 91781169 | 1 | unprocessed_pseudc | ENSG00000( |
| 91817771 | 91818248 | 1 | IG_V_pseudogene | ENSG00000( |
| 91840601 | 91843802 | 1 | unprocessed_pseudc | ENSG00000( |
| 91859384 | 91859481 | 1 | processed_pseudoge | ENSG00000( |
| 91877969 | 91880195 | 1 | processed_pseudoge | ENSG00000( |
| 91883076 | 91931714 | -1 | transcribed_unproce | ENSG00000( |
| 91940668 | 91942040 | 1 | processed_pseudoge | ENSG00000( |
| 91957436 | 91963610 | 1 | unprocessed_pseudc | ENSG00000( |
| 92006680 | 92007121 | -1 | processed_pseudoge | ENSG00000( |
| 92034522 | 92035000 | -1 | unprocessed_pseudc | ENSG00000( |
| 94195806 | 94208379 | -1 | transcribed_unproce | ENSG00000( |
| 94575976 | 94587183 | -1 | transcribed_unproce | ENSG00000( |
| 94588522 | 94604573 | 1 | protein_coding | ENSG00000( |
| 94725674 | 94725935 | -1 | processed_pseudoge | ENSG00000( |
| 94734655 | 94735844 | -1 | processed_pseudoge | ENSG00000( |
| 94737337 | 94738249 | 1 | processed_pseudoge | ENSG00000( |
| 94750582 | 94751293 | 1 | processed_pseudoge | ENSG00000( |
| 94759259 | 94760279 | -1 | unprocessed_pseudc | ENSG00000( |
| 94760774 | 94761116 | -1 | processed_pseudoge | ENSG00000( |
| 94786688 | 94787261 | -1 | processed_pseudoge | ENSG00000( |
| 94791103 | 94857282 | -1 | transcribed_unproce | ENSG00000( |
| 94846533 | 94846639 | -1 | snRNA | ENSG00000( |
| 94861362 | 94862667 | 1 | processed_pseudoge | ENSG00000( |
| 94867486 | 94947341 | -1 | transcribed_unproce | ENSG00000( |
| 94871430 | 94876823 | 1 | protein_coding | ENSG00000( |
| 94899566 | 94900661 | -1 | processed_pseudoge | ENSG00000( |
| 94900990 | 94901602 | 1 | processed_pseudoge | ENSG00000( |
| 94912432 | 94912831 | 1 | processed_pseudoge | ENSG00000( |
| 94953161 | 94953593 | -1 | unprocessed_pseudc | ENSG00000( |
| 94961047 | 94961503 | -1 | unprocessed_pseudc | ENSG00000( |
| 94964473 | 94964836 | -1 | unprocessed_pseudc | ENSG00000( |
| 94971954 | 94972167 | -1 | unprocessed_pseudc | ENSG00000( |

| | | | | |
|---|---|---|---|---|
| 94974544 | 94974838 | -1 | unprocessed_pseudc | ENSG00000( |
| 94991880 | 94994362 | 1 | unprocessed_pseudc | ENSG00000( |
| 95003545 | 95003843 | -1 | misc_RNA | ENSG00000( |
| 95025193 | 95026709 | -1 | lncRNA | ENSG00000( |
| 95025677 | 95053992 | 1 | protein_coding | ENSG00000( |
| 95051395 | 95053176 | -1 | lncRNA | ENSG00000( |
| 95067074 | 95076900 | -1 | lncRNA | ENSG00000( |
| 95085369 | 95149434 | -1 | protein_coding | ENSG00000( |
| 95145000 | 95165413 | -1 | protein_coding | ENSG00000( |
| 95165432 | 95184317 | 1 | protein_coding | ENSG00000( |
| 95196449 | 95197357 | -1 | unprocessed_pseudc | ENSG00000( |
| 95207535 | 95259774 | 1 | protein_coding | ENSG00000( |
| 95274449 | 95291308 | 1 | protein_coding | ENSG00000( |
| 95297327 | 95386077 | 1 | protein_coding | ENSG00000( |
| 95368507 | 95368885 | -1 | processed_pseudoge | ENSG00000( |
| 95402708 | 95416616 | 1 | protein_coding | ENSG00000( |
| 95413456 | 95414296 | -1 | unprocessed_pseudc | ENSG00000( |
| 95434759 | 95435113 | 1 | processed_pseudoge | ENSG00000( |
| 95436133 | 95436556 | 1 | processed_pseudoge | ENSG00000( |
| 95450310 | 95451500 | -1 | unprocessed_pseudc | ENSG00000( |
| 95460144 | 95460336 | -1 | unprocessed_pseudc | ENSG00000( |
| 95468863 | 95469778 | -1 | unprocessed_pseudc | ENSG00000( |
| 95476967 | 95484731 | -1 | protein_coding | ENSG00000( |
| 95485541 | 95485805 | 1 | processed_pseudoge | ENSG00000( |
| 95486480 | 95488627 | -1 | unprocessed_pseudc | ENSG00000( |
| 95496022 | 95496531 | -1 | processed_pseudoge | ENSG00000( |
| 95502791 | 95505059 | 1 | unprocessed_pseudc | ENSG00000( |
| 95514827 | 95520107 | 1 | unprocessed_pseudc | ENSG00000( |
| 95525109 | 95526702 | -1 | lncRNA | ENSG00000( |
| 95525345 | 95532405 | 1 | transcribed_unproce | ENSG00000( |
| 95536117 | 95536495 | -1 | processed_pseudoge | ENSG00000( |
| 95537969 | 95538469 | 1 | lncRNA | ENSG00000( |
| 95542730 | 95543230 | -1 | processed_pseudoge | ENSG00000( |
| 95546531 | 95547545 | 1 | unprocessed_pseudc | ENSG00000( |
| 95574901 | 95580769 | 1 | unprocessed_pseudc | ENSG00000( |
| 95588149 | 95589253 | 1 | unprocessed_pseudc | ENSG00000( |
| 95590969 | 95591206 | -1 | processed_pseudoge | ENSG00000( |
| 95592001 | 95599778 | 1 | protein_coding | ENSG00000( |
| 95606924 | 95607152 | 1 | processed_pseudoge | ENSG00000( |
| 95607073 | 95616592 | 1 | unprocessed_pseudc | ENSG00000( |
| 95616492 | 95617233 | 1 | processed_pseudoge | ENSG00000( |
| 95625213 | 95626236 | 1 | unprocessed_pseudc | ENSG00000( |
| 95633850 | 95636106 | 1 | unprocessed_pseudc | ENSG00000( |
| 95640181 | 95640604 | -1 | processed_pseudoge | ENSG00000( |
| 95641634 | 95641980 | -1 | processed_pseudoge | ENSG00000( |
| 95660588 | 95668727 | 1 | lncRNA | ENSG00000( |
| 95789654 | 95800166 | 1 | transcribed_unproce | ENSG00000( |
| 95807052 | 95835003 | -1 | lncRNA | ENSG00000( |
| 95836919 | 95991831 | -1 | protein_coding | ENSG00000( |
| 96004563 | 96004850 | -1 | misc_RNA | ENSG00000( |

| | | | | |
|---|---|---|---|---|
| 96010551 | 96023380 | 1 | transcribed_unproce | ENSG00000 |
| 96021946 | 96039451 | -1 | protein_coding | ENSG00000 |
| 96112876 | 96116571 | -1 | protein_coding | ENSG00000 |
| 96123850 | 96138436 | -1 | protein_coding | ENSG00000 |
| 96143169 | 96145440 | -1 | protein_coding | ENSG00000 |
| 96145602 | 96147012 | -1 | lncRNA | ENSG00000 |
| 96184859 | 96208825 | -1 | protein_coding | ENSG00000 |
| 96208403 | 96243353 | 1 | lncRNA | ENSG00000 |
| 96248516 | 96265994 | -1 | protein_coding | ENSG00000 |
| 96266159 | 96274173 | 1 | protein_coding | ENSG00000 |
| 96274338 | 96321271 | -1 | protein_coding | ENSG00000 |
| 96307263 | 96321731 | -1 | lncRNA | ENSG00000 |
| 96325331 | 96330517 | 1 | protein_coding | ENSG00000 |
| 96335766 | 96377091 | 1 | protein_coding | ENSG00000 |
| 96497646 | 96508109 | -1 | protein_coding | ENSG00000 |
| 96527940 | 96532306 | -1 | lncRNA | ENSG00000 |
| 96536743 | 96552634 | 1 | protein_coding | ENSG00000 |
| 96593170 | 96642787 | -1 | protein_coding | ENSG00000 |
| 96642737 | 96704887 | 1 | protein_coding | ENSG00000 |
| 96705929 | 96740064 | -1 | protein_coding | ENSG00000 |
| 96760902 | 96811874 | 1 | protein_coding | ENSG00000 |
| 96798278 | 96798353 | 1 | miRNA | ENSG00000 |
| 96812239 | 96813657 | -1 | lncRNA | ENSG00000 |
| 96816245 | 96835382 | 1 | protein_coding | ENSG00000 |
| 96824526 | 96857934 | -1 | protein_coding | ENSG00000 |
| 96836611 | 96858016 | -1 | protein_coding | ENSG00000 |
| 96859718 | 96870757 | -1 | protein_coding | ENSG00000 |
| 96875882 | 96986580 | -1 | protein_coding | ENSG00000 |
| 96912549 | 96917272 | 1 | lncRNA | ENSG00000 |
| 96938923 | 96939203 | 1 | misc_RNA | ENSG00000 |
| 96956708 | 96956815 | -1 | rRNA_pseudogene | ENSG00000 |
| 97000436 | 97001593 | -1 | unprocessed_pseudo | ENSG00000 |
| 97008368 | 97009104 | -1 | processed_pseudoge | ENSG00000 |
| 97018343 | 97018604 | -1 | unprocessed_pseudo | ENSG00000 |
| 97025981 | 97031636 | -1 | unprocessed_pseudo | ENSG00000 |
| 97034442 | 97034706 | 1 | unprocessed_pseudo | ENSG00000 |
| 97035461 | 97037549 | -1 | unprocessed_pseudo | ENSG00000 |
| 97046588 | 97046891 | 1 | IG_V_pseudogene | ENSG00000 |
| 97050729 | 97051028 | 1 | IG_V_pseudogene | ENSG00000 |
| 97060128 | 97060415 | 1 | IG_V_pseudogene | ENSG00000 |
| 97081098 | 97083249 | 1 | unprocessed_pseudo | ENSG00000 |
| 97083583 | 97094882 | -1 | protein_coding | ENSG00000 |
| 97094935 | 97097433 | 1 | TEC | ENSG00000 |
| 97100587 | 97100874 | 1 | misc_RNA | ENSG00000 |
| 97113218 | 97264434 | 1 | protein_coding | ENSG00000 |
| 97281356 | 97291849 | 1 | lncRNA | ENSG00000 |
| 97301729 | 97304151 | 1 | lncRNA | ENSG00000 |
| 97322137 | 97322423 | -1 | IG_V_pseudogene | ENSG00000 |
| 97331533 | 97331843 | -1 | IG_V_pseudogene | ENSG00000 |
| 97335671 | 97335974 | -1 | IG_V_pseudogene | ENSG00000 |

| | | | | |
|---|---|---|---|---|
| 97348899 | 97349179 | 1 | IG_V_pseudogene | ENSG0000C |
| 97355059 | 97355335 | 1 | IG_V_pseudogene | ENSG0000C |
| 97372532 | 97372835 | 1 | IG_V_pseudogene | ENSG0000C |
| 97376674 | 97376973 | 1 | IG_V_pseudogene | ENSG0000C |
| 97386082 | 97386368 | 1 | IG_V_pseudogene | ENSG0000C |
| 97416165 | 97433527 | -1 | lncRNA | ENSG0000C |
| 97421075 | 97434847 | 1 | lncRNA | ENSG0000C |
| 97492663 | 97589965 | -1 | protein_coding | ENSG0000C |
| 97523949 | 97524976 | -1 | processed_pseudoge | ENSG0000C |
| 97618638 | 97638417 | 1 | lncRNA | ENSG0000C |
| 97636780 | 97637803 | -1 | unprocessed_pseudc | ENSG0000C |
| 97646062 | 97648383 | 1 | protein_coding | ENSG0000C |
| 97655939 | 97664044 | -1 | protein_coding | ENSG0000C |
| 97664217 | 97703064 | 1 | protein_coding | ENSG0000C |
| 97664591 | 97664731 | -1 | snRNA | ENSG0000C |
| 97713560 | 97739862 | 1 | protein_coding | ENSG0000C |
| 97756333 | 97995948 | -1 | protein_coding | ENSG0000C |
| 97827248 | 97827545 | 1 | processed_pseudoge | ENSG0000C |
| 97913054 | 97913117 | -1 | snRNA | ENSG0000C |
| 98087116 | 98313299 | 1 | protein_coding | ENSG0000C |
| 98206151 | 98207724 | 1 | processed_pseudoge | ENSG0000C |
| 98331389 | 98356005 | -1 | lncRNA | ENSG0000C |
| 98346155 | 98398601 | 1 | protein_coding | ENSG0000C |
| 98346995 | 98351140 | 1 | lncRNA | ENSG0000C |
| 98444854 | 98594390 | 1 | protein_coding | ENSG0000C |
| 98599314 | 98608515 | -1 | protein_coding | ENSG0000C |
| 98608579 | 98618515 | 1 | protein_coding | ENSG0000C |
| 98619106 | 98731132 | -1 | protein_coding | ENSG0000C |
| 98694109 | 98695490 | 1 | processed_pseudoge | ENSG0000C |
| 98761098 | 98775361 | -1 | lncRNA | ENSG0000C |
| 98782410 | 98782548 | 1 | snRNA | ENSG0000C |
| 98793846 | 98936259 | -1 | protein_coding | ENSG0000C |
| 98840675 | 98840736 | -1 | snRNA | ENSG0000C |
| 98997261 | 99154964 | -1 | protein_coding | ENSG0000C |
| 99102018 | 99102752 | 1 | processed_pseudoge | ENSG0000C |
| 99141485 | 99151487 | 1 | protein_coding | ENSG0000C |
| 99141485 | 99322741 | 1 | protein_coding | ENSG0000C |
| 99154955 | 99163157 | 1 | protein_coding | ENSG0000C |
| 99154998 | 99195298 | 1 | protein_coding | ENSG0000C |
| 99161427 | 99181058 | -1 | protein_coding | ENSG0000C |
| 99181152 | 99199561 | 1 | protein_coding | ENSG0000C |
| 99242246 | 99255282 | -1 | protein_coding | ENSG0000C |
| 99284238 | 99304742 | -1 | protein_coding | ENSG0000C |
| 99318982 | 99340702 | -1 | protein_coding | ENSG0000C |
| 99337371 | 99401326 | 1 | protein_coding | ENSG0000C |
| 99400475 | 99490035 | -1 | protein_coding | ENSG0000C |
| 99405218 | 99405843 | 1 | lncRNA | ENSG0000C |
| 99545419 | 100192428 | -1 | protein_coding | ENSG0000C |
| 100104919 | 100107504 | 1 | lncRNA | ENSG0000C |
| 100208254 | 100251484 | 1 | lncRNA | ENSG0000C |

| | | | | |
|---|---|---|---|---|
| 100273291 | 100322733 | -1 | protein_coding | ENSG0000C |
| 100370255 | 100370545 | -1 | processed_pseudoge | ENSG0000C |
| 100377421 | 100377532 | 1 | processed_pseudoge | ENSG0000C |
| 100391860 | 100417668 | -1 | protein_coding | ENSG0000C |
| 100425348 | 100426107 | 1 | processed_pseudoge | ENSG0000C |
| 100470482 | 100483280 | 1 | protein_coding | ENSG0000C |
| 100509277 | 100509612 | 1 | processed_pseudoge | ENSG0000C |
| 100562993 | 100576739 | 1 | protein_coding | ENSG0000C |
| 100591150 | 100591324 | -1 | processed_pseudoge | ENSG0000C |
| 100603752 | 100611542 | 1 | lncRNA | ENSG0000C |
| 100642444 | 100652567 | -1 | lncRNA | ENSG0000C |
| 100660199 | 100660678 | 1 | processed_pseudoge | ENSG0000C |
| 100669892 | 100676038 | -1 | lncRNA | ENSG0000C |
| 100722221 | 100722882 | -1 | lncRNA | ENSG0000C |
| 100739958 | 100747391 | 1 | lncRNA | ENSG0000C |
| 100820139 | 100996829 | 1 | protein_coding | ENSG0000C |
| 100822661 | 100847220 | 1 | lncRNA | ENSG0000C |
| 100970767 | 100977766 | -1 | lncRNA | ENSG0000C |
| 100993676 | 101002244 | -1 | lncRNA | ENSG0000C |
| 101002229 | 101024032 | 1 | protein_coding | ENSG0000C |
| 101007617 | 101252866 | -1 | protein_coding | ENSG0000C |
| 101068378 | 101068656 | 1 | processed_pseudoge | ENSG0000C |
| 101151660 | 101155412 | 1 | lncRNA | ENSG0000C |
| 101252886 | 101270316 | 1 | protein_coding | ENSG0000C |
| 101271219 | 101308701 | -1 | protein_coding | ENSG0000C |
| 101272936 | 101273049 | -1 | snoRNA | ENSG0000C |
| 101309450 | 101309534 | 1 | miRNA | ENSG0000C |
| 101345550 | 101387503 | -1 | protein_coding | ENSG0000C |
| 101375587 | 101376055 | 1 | processed_pseudoge | ENSG0000C |
| 101397359 | 101475112 | -1 | protein_coding | ENSG0000C |
| 101479390 | 101486822 | 1 | lncRNA | ENSG0000C |
| 101509530 | 101510222 | -1 | processed_pseudoge | ENSG0000C |
| 101520834 | 101521976 | -1 | processed_pseudoge | ENSG0000C |
| 101535601 | 101539077 | 1 | lncRNA | ENSG0000C |
| 101551592 | 101567901 | -1 | lncRNA | ENSG0000C |
| 101696850 | 101894689 | 1 | protein_coding | ENSG0000C |
| 101962056 | 101987167 | 1 | lncRNA | ENSG0000C |
| 101991960 | 102028544 | 1 | protein_coding | ENSG0000C |
| 102037963 | 102058190 | 1 | lncRNA | ENSG0000C |
| 102064544 | 102179874 | 1 | protein_coding | ENSG0000C |
| 102172621 | 102182108 | -1 | lncRNA | ENSG0000C |
| 102187006 | 102240002 | 1 | protein_coding | ENSG0000C |
| 102249857 | 102250255 | -1 | processed_pseudoge | ENSG0000C |
| 102311502 | 102352037 | 1 | protein_coding | ENSG0000C |
| 102311529 | 102398775 | 1 | protein_coding | ENSG0000C |
| 102411243 | 102412108 | 1 | processed_pseudoge | ENSG0000C |
| 102418689 | 102452565 | 1 | protein_coding | ENSG0000C |
| 102432289 | 102432366 | 1 | miRNA | ENSG0000C |
| 102433957 | 102435340 | 1 | lncRNA | ENSG0000C |
| 102438713 | 102440475 | 1 | lncRNA | ENSG0000C |

| | | | | |
|---|---|---|---|---|
| 102473226 | 102533972 | 1 | protein_coding | ENSG00000 |
| 102619553 | 102711355 | 1 | protein_coding | ENSG00000 |
| 102714630 | 102736888 | -1 | protein_coding | ENSG00000 |
| 102736905 | 103019900 | 1 | protein_coding | ENSG00000 |
| 102873183 | 102895780 | -1 | lncRNA | ENSG00000 |
| 102967408 | 102984429 | -1 | lncRNA | ENSG00000 |
| 102987323 | 102988565 | 1 | lncRNA | ENSG00000 |
| 103109759 | 103176260 | -1 | lncRNA | ENSG00000 |
| 103275904 | 103276138 | 1 | processed_pseudoge | ENSG00000 |
| 103362082 | 103403712 | 1 | lncRNA | ENSG00000 |
| 103494241 | 103495842 | 1 | processed_pseudoge | ENSG00000 |
| 103527620 | 103529261 | 1 | processed_pseudoge | ENSG00000 |
| 103623630 | 103623830 | 1 | processed_pseudoge | ENSG00000 |
| 103855829 | 103869651 | -1 | lncRNA | ENSG00000 |
| 103856970 | 103880209 | -1 | lncRNA | ENSG00000 |
| 103874310 | 104077778 | 1 | lncRNA | ENSG00000 |
| 103953098 | 103953284 | -1 | misc_RNA | ENSG00000 |
| 104125268 | 104147229 | -1 | lncRNA | ENSG00000 |
| 104395002 | 104396280 | -1 | processed_pseudoge | ENSG00000 |
| 104406471 | 104416720 | 1 | lncRNA | ENSG00000 |
| 104412146 | 104414195 | -1 | lncRNA | ENSG00000 |
| 104423039 | 104429454 | 1 | lncRNA | ENSG00000 |
| 104430130 | 104532747 | 1 | lncRNA | ENSG00000 |
| 104488458 | 104510509 | -1 | lncRNA | ENSG00000 |
| 104580821 | 104581890 | -1 | lncRNA | ENSG00000 |
| 104583138 | 104621767 | 1 | lncRNA | ENSG00000 |
| 104656135 | 104853183 | -1 | lncRNA | ENSG00000 |
| 104659337 | 104666858 | 1 | lncRNA | ENSG00000 |
| 104678223 | 104678673 | 1 | processed_pseudoge | ENSG00000 |
| 104702650 | 104705813 | 1 | lncRNA | ENSG00000 |
| 104706612 | 104707506 | 1 | TEC | ENSG00000 |
| 104733234 | 104733362 | 1 | snoRNA | ENSG00000 |
| 104746638 | 104757719 | -1 | lncRNA | ENSG00000 |
| 104771282 | 104771885 | 1 | processed_pseudoge | ENSG00000 |
| 104853285 | 104927722 | 1 | lncRNA | ENSG00000 |
| 104854115 | 104858574 | 1 | protein_coding | ENSG00000 |
| 104865407 | 104872595 | -1 | lncRNA | ENSG00000 |
| 104928803 | 104936467 | 1 | lncRNA | ENSG00000 |
| 104936241 | 105038496 | -1 | lncRNA | ENSG00000 |
| 105038069 | 105099960 | 1 | protein_coding | ENSG00000 |
| 105092368 | 105103036 | -1 | lncRNA | ENSG00000 |
| 105144113 | 105145424 | 1 | lncRNA | ENSG00000 |
| 105241743 | 105243467 | 1 | protein_coding | ENSG00000 |
| 105249404 | 105249794 | -1 | lncRNA | ENSG00000 |
| 105264391 | 105329735 | -1 | protein_coding | ENSG00000 |
| 105324210 | 105330529 | 1 | lncRNA | ENSG00000 |
| 105334027 | 105337475 | -1 | lncRNA | ENSG00000 |
| 105337532 | 105349211 | 1 | protein_coding | ENSG00000 |
| 105357712 | 105438513 | -1 | protein_coding | ENSG00000 |
| 105363038 | 105378839 | 1 | lncRNA | ENSG00000 |

| | | | | |
|---|---|---|---|---|
| 105600703 | 105610487 | -1 | lncRNA | ENSG00000( |
| 105601146 | 105636637 | 1 | lncRNA | ENSG00000( |
| 105744912 | 105894274 | 1 | protein_coding | ENSG00000( |
| 105846534 | 105857177 | -1 | lncRNA | ENSG00000( |
| 105874509 | 105962386 | -1 | lncRNA | ENSG00000( |
| 106063246 | 106078155 | 1 | protein_coding | ENSG00000( |
| 106093303 | 106194339 | -1 | protein_coding | ENSG00000( |
| 106179542 | 106183797 | 1 | lncRNA | ENSG00000( |
| 106207394 | 106208115 | 1 | processed_pseudoge | ENSG00000( |
| 106247369 | 106247741 | 1 | processed_pseudoge | ENSG00000( |
| 106253231 | 106253717 | -1 | processed_pseudoge | ENSG00000( |
| 106288803 | 106289053 | -1 | processed_pseudoge | ENSG00000( |
| 106304755 | 106305197 | 1 | processed_pseudoge | ENSG00000( |
| 106344566 | 106345433 | 1 | processed_pseudoge | ENSG00000( |
| 106368209 | 106368876 | -1 | processed_pseudoge | ENSG00000( |
| 106369724 | 106378247 | 1 | unprocessed_pseudc | ENSG00000( |
| 106382171 | 106393031 | 1 | transcribed_unproce | ENSG00000( |
| 106391290 | 106468376 | -1 | protein_coding | ENSG00000( |
| 106470799 | 106477669 | -1 | unprocessed_pseudc | ENSG00000( |
| 106487364 | 106544297 | 1 | protein_coding | ENSG00000( |
| 106521563 | 106544297 | 1 | lncRNA | ENSG00000( |
| 106521903 | 106538079 | -1 | lncRNA | ENSG00000( |
| 106697331 | 106698676 | -1 | processed_pseudoge | ENSG00000( |
| 106801600 | 106887108 | -1 | protein_coding | ENSG00000( |
| 106822923 | 106825031 | -1 | lncRNA | ENSG00000( |
| 106834464 | 106841207 | 1 | lncRNA | ENSG00000( |
| 106933739 | 106996534 | -1 | lncRNA | ENSG00000( |
| 106940880 | 106941482 | -1 | processed_pseudoge | ENSG00000( |
| 106963193 | 106963674 | 1 | processed_pseudoge | ENSG00000( |
| 107254691 | 107365873 | -1 | lncRNA | ENSG00000( |
| 107362282 | 107407329 | 1 | lncRNA | ENSG00000( |
| 107382715 | 107748270 | -1 | lncRNA | ENSG00000( |
| 107529292 | 107556499 | 1 | lncRNA | ENSG00000( |
| 107698737 | 107746941 | -1 | lncRNA | ENSG00000( |
| 107754112 | 107822129 | 1 | lncRNA | ENSG00000( |
| 107823063 | 107826891 | -1 | lncRNA | ENSG00000( |
| 107826937 | 107890841 | 1 | protein_coding | ENSG00000( |
| 107902997 | 107903134 | -1 | processed_pseudoge | ENSG00000( |
| 107903964 | 107904241 | 1 | unprocessed_pseudc | ENSG00000( |
| 107914948 | 107915286 | 1 | processed_pseudoge | ENSG00000( |
| 107920763 | 107921248 | -1 | processed_pseudoge | ENSG00000( |
| 107986523 | 108013994 | 1 | protein_coding | ENSG00000( |
| 108049200 | 108052755 | -1 | lncRNA | ENSG00000( |
| 108099110 | 108099492 | 1 | processed_pseudoge | ENSG00000( |
| 108167125 | 108217886 | -1 | lncRNA | ENSG00000( |
| 108226015 | 108226331 | -1 | processed_pseudoge | ENSG00000( |
| 108247195 | 108265351 | 1 | protein_coding | ENSG00000( |
| 108276812 | 108277942 | 1 | processed_pseudoge | ENSG00000( |
| 108288639 | 108309915 | 1 | protein_coding | ENSG00000( |
| 108317725 | 108318611 | 1 | unprocessed_pseudc | ENSG00000( |

| | | | | |
|---|---|---|---|---|
| 108322238 | 108353799 | 1 | transcribed_unproce | ENSG00000 |
| 108367116 | 108367938 | 1 | processed_pseudoge | ENSG00000 |
| 108377911 | 108388989 | 1 | protein_coding | ENSG00000 |
| 108423612 | 108423885 | 1 | processed_pseudoge | ENSG00000 |
| 108448561 | 108509415 | 1 | protein_coding | ENSG00000 |
| 108507515 | 108534196 | -1 | lncRNA | ENSG00000 |
| 108534355 | 108687246 | 1 | protein_coding | ENSG00000 |
| 108676795 | 108678601 | -1 | lncRNA | ENSG00000 |
| 108694750 | 108694893 | -1 | processed_pseudoge | ENSG00000 |
| 108712262 | 108712900 | -1 | processed_pseudoge | ENSG00000 |
| 108719482 | 108785809 | 1 | protein_coding | ENSG00000 |
| 108786757 | 108885477 | 1 | protein_coding | ENSG00000 |
| 108878308 | 108878460 | 1 | processed_pseudoge | ENSG00000 |
| 108894471 | 108989372 | -1 | protein_coding | ENSG00000 |
| 109124357 | 109124609 | -1 | misc_RNA | ENSG00000 |
| 109127327 | 109128930 | -1 | lncRNA | ENSG00000 |
| 109129205 | 109504634 | 1 | protein_coding | ENSG00000 |
| 109141490 | 109141588 | -1 | miRNA | ENSG00000 |
| 109251422 | 109251646 | 1 | processed_pseudoge | ENSG00000 |
| 109313571 | 109313625 | -1 | miRNA | ENSG00000 |
| 109542799 | 109614143 | -1 | protein_coding | ENSG00000 |
| 109552536 | 109552846 | 1 | processed_pseudoge | ENSG00000 |
| 109594693 | 109605698 | -1 | lncRNA | ENSG00000 |
| 109614364 | 109618990 | 1 | protein_coding | ENSG00000 |
| 109665209 | 109665742 | 1 | processed_pseudoge | ENSG00000 |
| 109667128 | 109668002 | 1 | processed_pseudoge | ENSG00000 |
| 109732731 | 109733136 | 1 | processed_pseudoge | ENSG00000 |
| 109737139 | 109737523 | -1 | processed_pseudoge | ENSG00000 |
| 109758799 | 109760364 | -1 | lncRNA | ENSG00000 |
| 109792758 | 109857705 | 1 | protein_coding | ENSG00000 |
| 109898428 | 109924868 | 1 | protein_coding | ENSG00000 |
| 109898432 | 109968557 | 1 | protein_coding | ENSG00000 |
| 109933161 | 109974085 | 1 | transcribed_unproce | ENSG00000 |
| 109937438 | 109985124 | 1 | lncRNA | ENSG00000 |
| 109984058 | 109984988 | 1 | transcribed_processe | ENSG00000 |
| 109986939 | 109988394 | -1 | processed_pseudoge | ENSG00000 |
| 109987063 | 109996140 | 1 | lncRNA | ENSG00000 |
| 110007675 | 110010783 | 1 | processed_pseudoge | ENSG00000 |
| 110012265 | 110012601 | -1 | processed_pseudoge | ENSG00000 |
| 110022447 | 110050064 | -1 | unprocessed_pseudo | ENSG00000 |
| 110069961 | 110070042 | -1 | miRNA | ENSG00000 |
| 110083870 | 110116566 | -1 | protein_coding | ENSG00000 |
| 110086433 | 110086523 | -1 | miRNA | ENSG00000 |
| 110122311 | 110205066 | -1 | protein_coding | ENSG00000 |
| 110184054 | 110185179 | 1 | processed_pseudoge | ENSG00000 |
| 110211529 | 110245420 | -1 | protein_coding | ENSG00000 |
| 110231106 | 110272050 | 1 | transcribed_unproce | ENSG00000 |
| 110245591 | 110285856 | 1 | lncRNA | ENSG00000 |
| 110279606 | 110285681 | 1 | transcribed_unproce | ENSG00000 |
| 110284853 | 110284943 | 1 | miRNA | ENSG00000 |

| | | | | |
|---|---|---|---|---|
| 110321402 | 110348941 | 1 | unprocessed_pseudc | ENSG00000C |
| 110358790 | 110359126 | 1 | processed_pseudoge | ENSG00000C |
| 110360608 | 110363737 | -1 | processed_pseudoge | ENSG00000C |
| 110375138 | 110384442 | -1 | lncRNA | ENSG00000C |
| 110383112 | 110384642 | 1 | processed_pseudoge | ENSG00000C |
| 110386377 | 110388163 | -1 | transcribed_processe | ENSG00000C |
| 110386411 | 110433673 | -1 | lncRNA | ENSG00000C |
| 110397406 | 110435166 | -1 | transcribed_unproce | ENSG00000C |
| 110402934 | 110473075 | -1 | protein_coding | ENSG00000C |
| 110446640 | 110473075 | -1 | protein_coding | ENSG00000C |
| 110449242 | 110451064 | 1 | lncRNA | ENSG00000C |
| 110500100 | 110500235 | 1 | unprocessed_pseudc | ENSG00000C |
| 110513812 | 110577185 | -1 | protein_coding | ENSG00000C |
| 110610916 | 110612404 | 1 | lncRNA | ENSG00000C |
| 110633947 | 110634333 | 1 | processed_pseudoge | ENSG00000C |
| 110637528 | 110678063 | -1 | protein_coding | ENSG00000C |
| 110709575 | 110716595 | -1 | lncRNA | ENSG00000C |
| 110732573 | 111118222 | 1 | protein_coding | ENSG00000C |
| 110967834 | 110968719 | 1 | processed_pseudoge | ENSG00000C |
| 111036776 | 111523376 | -1 | lncRNA | ENSG00000C |
| 111098345 | 111116407 | -1 | lncRNA | ENSG00000C |
| 111119378 | 111168445 | 1 | protein_coding | ENSG00000C |
| 111195963 | 111206494 | 1 | lncRNA | ENSG00000C |
| 111203964 | 111206215 | -1 | processed_pseudoge | ENSG00000C |
| 111210995 | 111212476 | -1 | lncRNA | ENSG00000C |
| 111265283 | 111279880 | 1 | lncRNA | ENSG00000C |
| 111266868 | 111267473 | -1 | lncRNA | ENSG00000C |
| 111295629 | 111296046 | -1 | processed_pseudoge | ENSG00000C |
| 111321013 | 111321086 | -1 | miRNA | ENSG00000C |
| 111321433 | 111324812 | -1 | TEC | ENSG00000C |
| 111383752 | 111384992 | 1 | transcribed_processe | ENSG00000C |
| 111429324 | 111699033 | -1 | lncRNA | ENSG00000C |
| 111468810 | 111637967 | 1 | lncRNA | ENSG00000C |
| 111491273 | 111570974 | 1 | lncRNA | ENSG00000C |
| 111491943 | 111494811 | -1 | lncRNA | ENSG00000C |
| 111558413 | 111559937 | -1 | processed_pseudoge | ENSG00000C |
| 111607841 | 111612518 | 1 | lncRNA | ENSG00000C |
| 111611639 | 111884690 | -1 | protein_coding | ENSG00000C |
| 111675026 | 111675343 | 1 | processed_pseudoge | ENSG00000C |
| 111717508 | 111717837 | 1 | processed_pseudoge | ENSG00000C |
| 111742586 | 111744662 | 1 | processed_pseudoge | ENSG00000C |
| 111771061 | 111771134 | -1 | miRNA | ENSG00000C |
| 111865798 | 111866405 | 1 | processed_pseudoge | ENSG00000C |
| 111887914 | 111888741 | 1 | processed_pseudoge | ENSG00000C |
| 111898607 | 112029561 | 1 | protein_coding | ENSG00000C |
| 111930175 | 111930467 | 1 | misc_RNA | ENSG00000C |
| 111940302 | 111941036 | -1 | processed_pseudoge | ENSG00000C |
| 111952144 | 111952671 | -1 | processed_pseudoge | ENSG00000C |
| 112039378 | 112055337 | -1 | lncRNA | ENSG00000C |
| 112055269 | 112119318 | 1 | protein_coding | ENSG00000C |

| | | | | |
|---|---|---|---|---|
| 112061084 | 112063709 | -1 | lncRNA | ENSG0000( |
| 112138385 | 112188218 | 1 | protein_coding | ENSG0000( |
| 112188364 | 112190635 | 1 | TEC | ENSG0000( |
| 112211529 | 112255136 | -1 | protein_coding | ENSG0000( |
| 112239621 | 112244375 | 1 | lncRNA | ENSG0000( |
| 112275597 | 112340063 | 1 | protein_coding | ENSG0000( |
| 112286057 | 112286422 | -1 | processed_pseudoge | ENSG0000( |
| 112346011 | 112346258 | 1 | processed_pseudoge | ENSG0000( |
| 112368369 | 112434488 | -1 | protein_coding | ENSG0000( |
| 112439312 | 112469687 | -1 | transcribed_unitary_ | ENSG0000( |
| 112482156 | 112541739 | 1 | protein_coding | ENSG0000( |
| 112541915 | 112577150 | 1 | protein_coding | ENSG0000( |
| 112579484 | 112579584 | -1 | misc_RNA | ENSG0000( |
| 112584240 | 112589275 | 1 | protein_coding | ENSG0000( |
| 112589040 | 112614431 | 1 | lncRNA | ENSG0000( |
| 112590796 | 112591939 | 1 | lncRNA | ENSG0000( |
| 112621809 | 112622167 | -1 | processed_pseudoge | ENSG0000( |
| 112641832 | 112645720 | -1 | lncRNA | ENSG0000( |
| 112645939 | 112663825 | 1 | protein_coding | ENSG0000( |
| 112721486 | 112742879 | 1 | protein_coding | ENSG0000( |
| 112736349 | 112764664 | -1 | protein_coding | ENSG0000( |
| 112773915 | 112784590 | -1 | protein_coding | ENSG0000( |
| 112817076 | 112824457 | -1 | lncRNA | ENSG0000( |
| 112829751 | 112836816 | -1 | protein_coding | ENSG0000( |
| 112840328 | 112844195 | 1 | TEC | ENSG0000( |
| 112853328 | 112854122 | 1 | processed_pseudoge | ENSG0000( |
| 112912971 | 112918882 | 1 | protein_coding | ENSG0000( |
| 112932625 | 112933171 | 1 | processed_pseudoge | ENSG0000( |
| 112973203 | 112985658 | 1 | protein_coding | ENSG0000( |
| 112981166 | 112981434 | -1 | processed_pseudoge | ENSG0000( |
| 112995517 | 112995937 | 1 | processed_pseudoge | ENSG0000( |
| 113005459 | 113008044 | 1 | protein_coding | ENSG0000( |
| 113022089 | 113052867 | -1 | protein_coding | ENSG0000( |
| 113058638 | 113065382 | 1 | protein_coding | ENSG0000( |
| 113067970 | 113075843 | 1 | protein_coding | ENSG0000( |
| 113086805 | 113086911 | 1 | snRNA | ENSG0000( |
| 113107214 | 113134016 | 1 | protein_coding | ENSG0000( |
| 113157325 | 113209396 | 1 | protein_coding | ENSG0000( |
| 113171535 | 113175360 | -1 | lncRNA | ENSG0000( |
| 113211421 | 113276581 | 1 | lncRNA | ENSG0000( |
| 113215997 | 113278921 | -1 | protein_coding | ENSG0000( |
| 113325372 | 113425548 | 1 | lncRNA | ENSG0000( |
| 113406396 | 113406872 | 1 | IG_V_gene | ENSG0000( |
| 113432600 | 113436042 | 1 | lncRNA | ENSG0000( |
| 113437691 | 113496204 | 1 | protein_coding | ENSG0000( |
| 113498665 | 113501155 | 1 | protein_coding | ENSG0000( |
| 113512222 | 113515488 | 1 | lncRNA | ENSG0000( |
| 113526836 | 113542694 | -1 | lncRNA | ENSG0000( |
| 113541937 | 113563298 | 1 | unprocessed_pseudc | ENSG0000( |
| 113577382 | 113578852 | 1 | lncRNA | ENSG0000( |

| | | | | |
|---|---|---|---|---|
| 113582354 | 113583152 | -1 | lncRNA | ENSG0000( |
| 113582959 | 113583096 | -1 | miRNA | ENSG0000( |
| 113588550 | 113599043 | 1 | transcribed_unproce | ENSG0000( |
| 113599036 | 113601261 | -1 | unprocessed_pseudc | ENSG0000( |
| 113605867 | 113607908 | -1 | TEC | ENSG0000( |
| 113610502 | 113627090 | -1 | transcribed_processe | ENSG0000( |
| 113627229 | 113643396 | 1 | protein_coding | ENSG0000( |
| 113657908 | 113658675 | 1 | processed_pseudoge | ENSG0000( |
| 113667478 | 113668621 | -1 | processed_pseudoge | ENSG0000( |
| 113669166 | 113673191 | 1 | lncRNA | ENSG0000( |
| 113677702 | 113704078 | -1 | lncRNA | ENSG0000( |
| 113681111 | 113681217 | 1 | snRNA | ENSG0000( |
| 113705011 | 113756693 | -1 | protein_coding | ENSG0000( |
| 113721290 | 113721368 | -1 | miRNA | ENSG0000( |
| 113829390 | 113831119 | -1 | lncRNA | ENSG0000( |
| 113831049 | 113843356 | -1 | lncRNA | ENSG0000( |
| 113888203 | 113889750 | -1 | lncRNA | ENSG0000( |
| 113890063 | 113962596 | 1 | protein_coding | ENSG0000( |
| 113904086 | 113905258 | 1 | TEC | ENSG0000( |
| 113970719 | 114007304 | 1 | lncRNA | ENSG0000( |
| 113979909 | 113983258 | 1 | lncRNA | ENSG0000( |
| 114005440 | 114005655 | -1 | snoRNA | ENSG0000( |
| 114114679 | 114117390 | -1 | lncRNA | ENSG0000( |
| 114122274 | 114123293 | -1 | lncRNA | ENSG0000( |
| 114260050 | 114261226 | -1 | processed_pseudoge | ENSG0000( |
| 114420125 | 114420302 | 1 | snRNA | ENSG0000( |
| 114442299 | 115845752 | 1 | protein_coding | ENSG0000( |
| 114828432 | 114833548 | -1 | lncRNA | ENSG0000( |
| 114833830 | 114835588 | -1 | lncRNA | ENSG0000( |
| 115079354 | 115099092 | -1 | lncRNA | ENSG0000( |
| 115126622 | 115161384 | -1 | lncRNA | ENSG0000( |
| 115468056 | 115468938 | 1 | processed_pseudoge | ENSG0000( |
| 115835129 | 115940291 | -1 | lncRNA | ENSG0000( |
| 116687218 | 116687387 | -1 | processed_pseudoge | ENSG0000( |
| 116726453 | 116726771 | -1 | processed_pseudoge | ENSG0000( |
| 116751158 | 116751648 | 1 | processed_pseudoge | ENSG0000( |
| 116818086 | 116818761 | 1 | processed_pseudoge | ENSG0000( |
| 117023918 | 117024867 | 1 | unprocessed_pseudc | ENSG0000( |
| 117025077 | 117026118 | 1 | unprocessed_pseudc | ENSG0000( |
| 117026537 | 117027024 | 1 | unprocessed_pseudc | ENSG0000( |
| 117139715 | 117139775 | -1 | snRNA | ENSG0000( |
| 117180892 | 117181562 | -1 | processed_pseudoge | ENSG0000( |
| 117536387 | 117539464 | 1 | lncRNA | ENSG0000( |
| 117754139 | 117804174 | -1 | lncRNA | ENSG0000( |
| 117814691 | 117832377 | 1 | protein_coding | ENSG0000( |
| 117833937 | 117841658 | 1 | lncRNA | ENSG0000( |
| 117859427 | 117934942 | 1 | transcribed_unproce | ENSG0000( |
| 117915478 | 118014133 | -1 | protein_coding | ENSG0000( |
| 117934025 | 117934789 | -1 | processed_pseudoge | ENSG0000( |
| 117995397 | 118055033 | 1 | lncRNA | ENSG0000( |

| | | | | |
|---|---|---|---|---|
| 117998745 | 117999480 | 1 | lncRNA | ENSG0000( |
| 118014174 | 118015673 | 1 | TEC | ENSG0000( |
| 118016385 | 118016669 | 1 | misc_RNA | ENSG0000( |
| 118088452 | 118110997 | 1 | protein_coding | ENSG0000( |
| 118132128 | 118222250 | -1 | lncRNA | ENSG0000( |
| 118766965 | 118835110 | -1 | lncRNA | ENSG0000( |
| 118842171 | 118847678 | -1 | protein_coding | ENSG0000( |
| 118942166 | 118994660 | 1 | protein_coding | ENSG0000( |
| 118949306 | 118952983 | -1 | lncRNA | ENSG0000( |
| 119156243 | 119158751 | -1 | protein_coding | ENSG0000( |
| 119169822 | 119170115 | 1 | misc_RNA | ENSG0000( |
| 119223831 | 119265652 | 1 | protein_coding | ENSG0000( |
| 119244422 | 119270714 | -1 | lncRNA | ENSG0000( |
| 119302225 | 119366834 | -1 | protein_coding | ENSG0000( |
| 119366921 | 119372560 | 1 | protein_coding | ENSG0000( |
| 119429901 | 119438504 | 1 | protein_coding | ENSG0000( |
| 119439843 | 119525301 | -1 | protein_coding | ENSG0000( |
| 119476428 | 119487346 | 1 | lncRNA | ENSG0000( |
| 119544432 | 119662251 | 1 | protein_coding | ENSG0000( |
| 119679167 | 119686507 | 1 | protein_coding | ENSG0000( |
| 119698623 | 119700151 | 1 | processed_pseudoge | ENSG0000( |
| 119759922 | 119984899 | 1 | protein_coding | ENSG0000( |
| 119993199 | 119993611 | -1 | processed_pseudoge | ENSG0000( |
| 120013077 | 120179119 | 1 | protein_coding | ENSG0000( |
| 120174885 | 120216544 | -1 | lncRNA | ENSG0000( |
| 120192245 | 120192357 | 1 | misc_RNA | ENSG0000( |
| 120211054 | 120211715 | 1 | processed_pseudoge | ENSG0000( |
| 120211727 | 120212862 | 1 | processed_pseudoge | ENSG0000( |
| 120212935 | 120213275 | 1 | processed_pseudoge | ENSG0000( |
| 120213344 | 120213631 | 1 | processed_pseudoge | ENSG0000( |
| 120213631 | 120214989 | 1 | processed_pseudoge | ENSG0000( |
| 120215181 | 120217279 | 1 | processed_pseudoge | ENSG0000( |
| 120217479 | 120223400 | -1 | protein_coding | ENSG0000( |
| 120240064 | 120294710 | 1 | protein_coding | ENSG0000( |
| 120319007 | 120326298 | 1 | lncRNA | ENSG0000( |
| 120346136 | 120351803 | 1 | protein_coding | ENSG0000( |
| 120464335 | 120466218 | -1 | TEC | ENSG0000( |
| 120542905 | 120545249 | -1 | lncRNA | ENSG0000( |
| 120552481 | 120558119 | -1 | lncRNA | ENSG0000( |
| 120651273 | 120651374 | 1 | misc_RNA | ENSG0000( |
| 120686635 | 120710517 | -1 | lncRNA | ENSG0000( |
| 120735623 | 120992653 | 1 | protein_coding | ENSG0000( |
| 120866378 | 120867403 | -1 | lncRNA | ENSG0000( |
| 121040610 | 121040710 | -1 | misc_RNA | ENSG0000( |
| 121081437 | 121088726 | -1 | lncRNA | ENSG0000( |
| 121178327 | 121182580 | 1 | lncRNA | ENSG0000( |
| 121216587 | 121285202 | -1 | protein_coding | ENSG0000( |
| 121309995 | 121310412 | 1 | processed_pseudoge | ENSG0000( |
| 121337776 | 121649587 | -1 | protein_coding | ENSG0000( |
| 121530422 | 121532705 | 1 | lncRNA | ENSG0000( |

| | | | | |
|---|---|---|---|---|
| 121530881 | 121531007 | 1 | snRNA | ENSG0000C |
| 121603073 | 121603164 | -1 | misc_RNA | ENSG0000C |
| 121649320 | 121728563 | 1 | lncRNA | ENSG0000C |
| 121658475 | 121658973 | -1 | processed_pseudoge | ENSG0000C |
| 121688203 | 121689714 | 1 | TEC | ENSG0000C |
| 121708512 | 121709115 | -1 | processed_pseudoge | ENSG0000C |
| 121726945 | 121736911 | -1 | protein_coding | ENSG0000C |
| 121737103 | 121767853 | 1 | protein_coding | ENSG0000C |
| 121743479 | 121745921 | -1 | processed_pseudoge | ENSG0000C |
| 121779133 | 121809962 | 1 | lncRNA | ENSG0000C |
| 121902501 | 122887691 | 1 | lncRNA | ENSG0000C |
| 122503366 | 122505118 | -1 | lncRNA | ENSG0000C |
| 123065321 | 123067722 | 1 | lncRNA | ENSG0000C |
| 123066151 | 123073481 | -1 | lncRNA | ENSG0000C |
| 123421486 | 123438693 | 1 | lncRNA | ENSG0000C |
| 123443266 | 123444445 | -1 | lncRNA | ENSG0000C |
| 123695302 | 123697396 | 1 | processed_pseudoge | ENSG0000C |
| 123762171 | 123763071 | 1 | processed_pseudoge | ENSG0000C |
| 123850522 | 123850695 | -1 | TEC | ENSG0000C |
| 123869256 | 123869570 | -1 | misc_RNA | ENSG0000C |
| 124011984 | 124025173 | -1 | lncRNA | ENSG0000C |
| 124025287 | 124915287 | 1 | protein_coding | ENSG0000C |
| 124680722 | 124680916 | -1 | processed_pseudoge | ENSG0000C |
| 124695242 | 124696530 | -1 | lncRNA | ENSG0000C |
| 124778454 | 124778640 | -1 | processed_pseudoge | ENSG0000C |
| 124932915 | 124933020 | 1 | rRNA_pseudogene | ENSG0000C |
| 124989219 | 124989325 | 1 | snRNA | ENSG0000C |
| 125665331 | 125717150 | 1 | lncRNA | ENSG0000C |
| 125710969 | 125765842 | 1 | lncRNA | ENSG0000C |
| 125823108 | 125823315 | 1 | processed_pseudoge | ENSG0000C |
| 126109761 | 126118023 | -1 | lncRNA | ENSG0000C |
| 126154583 | 126203372 | 1 | lncRNA | ENSG0000C |
| 126308494 | 126343992 | 1 | lncRNA | ENSG0000C |
| 126311466 | 126311927 | -1 | processed_pseudoge | ENSG0000C |
| 126557435 | 126558162 | -1 | processed_pseudoge | ENSG0000C |
| 126656133 | 126696667 | 1 | protein_coding | ENSG0000C |
| 126679622 | 126680420 | 1 | processed_pseudoge | ENSG0000C |
| 126702839 | 126702942 | -1 | snRNA | ENSG0000C |
| 126898864 | 126902097 | 1 | protein_coding | ENSG0000C |
| 127017421 | 127017482 | 1 | snRNA | ENSG0000C |
| 127024252 | 127025723 | -1 | lncRNA | ENSG0000C |
| 127024866 | 127033948 | 1 | lncRNA | ENSG0000C |
| 127025211 | 127029686 | 1 | lncRNA | ENSG0000C |
| 127048027 | 127107288 | -1 | protein_coding | ENSG0000C |
| 127141976 | 127143793 | -1 | lncRNA | ENSG0000C |
| 127170011 | 127170231 | 1 | processed_pseudoge | ENSG0000C |
| 127183832 | 127220313 | -1 | protein_coding | ENSG0000C |
| 127247258 | 127250082 | 1 | processed_pseudoge | ENSG0000C |
| 127257290 | 127294176 | -1 | protein_coding | ENSG0000C |
| 127294212 | 127375075 | 1 | lncRNA | ENSG0000C |

| | | | | |
|---|---|---|---|---|
| 127298730 | 127388465 | -1 | protein_coding | ENSG00000( |
| 127316873 | 127316979 | 1 | snRNA | ENSG00000( |
| 127388184 | 127400580 | 1 | lncRNA | ENSG00000( |
| 127406731 | 127408221 | -1 | lncRNA | ENSG00000( |
| 127418427 | 127429246 | 1 | protein_coding | ENSG00000( |
| 127423537 | 127423618 | -1 | miRNA | ENSG00000( |
| 127436207 | 127526886 | -1 | protein_coding | ENSG00000( |
| 127455394 | 127514623 | 1 | lncRNA | ENSG00000( |
| 127472234 | 127472415 | 1 | snRNA | ENSG00000( |
| 127535802 | 127637729 | 1 | protein_coding | ENSG00000( |
| 127625997 | 127626848 | -1 | lncRNA | ENSG00000( |
| 127638381 | 127681786 | -1 | protein_coding | ENSG00000( |
| 127645864 | 127652639 | 1 | protein_coding | ENSG00000( |
| 127701027 | 127811187 | -1 | protein_coding | ENSG00000( |
| 127701497 | 127705242 | 1 | protein_coding | ENSG00000( |
| 127798903 | 127798998 | -1 | misc_RNA | ENSG00000( |
| 127829747 | 127829852 | -1 | misc_RNA | ENSG00000( |
| 127840606 | 127841763 | -1 | processed_pseudoge | ENSG00000( |
| 127843553 | 127858155 | -1 | protein_coding | ENSG00000( |
| 127845236 | 127845341 | 1 | snRNA | ENSG00000( |
| 127846736 | 127847084 | -1 | processed_pseudoge | ENSG00000( |
| 127856890 | 127857147 | 1 | misc_RNA | ENSG00000( |
| 127861630 | 127885922 | -1 | protein_coding | ENSG00000( |
| 127886556 | 127887185 | 1 | lncRNA | ENSG00000( |
| 127888829 | 127892732 | 1 | lncRNA | ENSG00000( |
| 127931327 | 127931440 | 1 | processed_pseudoge | ENSG00000( |
| 127941217 | 128028120 | -1 | protein_coding | ENSG00000( |
| 128067194 | 128067407 | 1 | processed_pseudoge | ENSG00000( |
| 128067800 | 128067901 | 1 | misc_RNA | ENSG00000( |
| 128091200 | 128195677 | 1 | protein_coding | ENSG00000( |
| 128139792 | 128140516 | -1 | processed_pseudoge | ENSG00000( |
| 128151154 | 128151304 | -1 | processed_pseudoge | ENSG00000( |
| 128199901 | 128200195 | 1 | processed_pseudoge | ENSG00000( |
| 128203361 | 128203775 | 1 | processed_pseudoge | ENSG00000( |
| 128217588 | 128218072 | 1 | processed_pseudoge | ENSG00000( |
| 128236716 | 128318868 | -1 | protein_coding | ENSG00000( |
| 128350829 | 128350927 | -1 | misc_RNA | ENSG00000( |
| 128445081 | 128445198 | -1 | rRNA_pseudogene | ENSG00000( |
| 128518788 | 128519177 | -1 | processed_pseudoge | ENSG00000( |
| 128580491 | 128581576 | -1 | lncRNA | ENSG00000( |
| 129063846 | 129071725 | -1 | lncRNA | ENSG00000( |
| 129242173 | 129273848 | -1 | lncRNA | ENSG00000( |
| 129496297 | 129496788 | 1 | processed_pseudoge | ENSG00000( |
| 129825126 | 129877664 | -1 | lncRNA | ENSG00000( |
| 129877896 | 129884441 | 1 | lncRNA | ENSG00000( |
| 129922863 | 129934317 | -1 | transcribed_unproce | ENSG00000( |
| 129923177 | 129946703 | 1 | lncRNA | ENSG00000( |
| 129939848 | 129940217 | -1 | processed_pseudoge | ENSG00000( |
| 129966592 | 129980466 | -1 | transcribed_unproce | ENSG00000( |
| 129979666 | 129982738 | 1 | protein_coding | ENSG00000( |

| | | | | |
|---|---|---|---|---|
| 129988158 | 129990530 | -1 | transcribed_unproce | ENSG00000 |
| 129990456 | 130028412 | -1 | lncRNA | ENSG00000 |
| 129992602 | 129993105 | -1 | unprocessed_pseudo | ENSG00000 |
| 129992889 | 129994040 | 1 | processed_pseudoge | ENSG00000 |
| 130006200 | 130009123 | 1 | unprocessed_pseudo | ENSG00000 |
| 130012065 | 130040560 | -1 | transcribed_unproce | ENSG00000 |
| 130026845 | 130051131 | -1 | lncRNA | ENSG00000 |
| 130036958 | 130038422 | 1 | processed_pseudoge | ENSG00000 |
| 130048426 | 130049440 | -1 | unprocessed_pseudo | ENSG00000 |
| 130073535 | 130129222 | -1 | protein_coding | ENSG00000 |
| 130082092 | 130082220 | -1 | processed_pseudoge | ENSG00000 |
| 130107684 | 130109741 | 1 | lncRNA | ENSG00000 |
| 130109204 | 130109310 | -1 | snRNA | ENSG00000 |
| 130129621 | 130139417 | 1 | transcribed_unproce | ENSG00000 |
| 130139287 | 130145134 | -1 | protein_coding | ENSG00000 |
| 130151392 | 130182750 | -1 | protein_coding | ENSG00000 |
| 130181737 | 130190729 | 1 | protein_coding | ENSG00000 |
| 130191745 | 130198439 | -1 | protein_coding | ENSG00000 |
| 130202312 | 130212628 | 1 | unprocessed_pseudo | ENSG00000 |
| 130212870 | 130213490 | -1 | processed_pseudoge | ENSG00000 |
| 130229379 | 130232106 | 1 | transcribed_processe | ENSG00000 |
| 130245971 | 130248770 | 1 | unprocessed_pseudo | ENSG00000 |
| 130251568 | 130252559 | 1 | processed_pseudoge | ENSG00000 |
| 130270440 | 130272272 | 1 | unprocessed_pseudo | ENSG00000 |
| 130272479 | 130273511 | 1 | unprocessed_pseudo | ENSG00000 |
| 130273908 | 130275449 | 1 | unprocessed_pseudo | ENSG00000 |
| 130275592 | 130276257 | 1 | unprocessed_pseudo | ENSG00000 |
| 130276507 | 130277187 | 1 | unprocessed_pseudo | ENSG00000 |
| 130277184 | 130277957 | 1 | unprocessed_pseudo | ENSG00000 |
| 130278036 | 130278376 | 1 | unprocessed_pseudo | ENSG00000 |
| 130278444 | 130278739 | 1 | unprocessed_pseudo | ENSG00000 |
| 130278757 | 130280110 | 1 | unprocessed_pseudo | ENSG00000 |
| 130280299 | 130282105 | 1 | unprocessed_pseudo | ENSG00000 |
| 130282085 | 130282606 | -1 | unprocessed_pseudo | ENSG00000 |
| 130282679 | 130283329 | 1 | unprocessed_pseudo | ENSG00000 |
| 130293822 | 130294405 | 1 | processed_pseudoge | ENSG00000 |
| 130313934 | 130314411 | -1 | processed_pseudoge | ENSG00000 |
| 130330142 | 130330285 | -1 | unprocessed_pseudo | ENSG00000 |
| 130337933 | 130342699 | -1 | protein_coding | ENSG00000 |
| 130342225 | 130347810 | 1 | protein_coding | ENSG00000 |
| 130356045 | 130375405 | 1 | protein_coding | ENSG00000 |
| 130416755 | 130431363 | -1 | transcribed_unproce | ENSG00000 |
| 130427731 | 130429225 | 1 | processed_pseudoge | ENSG00000 |
| 130429827 | 130441413 | -1 | transcribed_unproce | ENSG00000 |
| 130459455 | 130509707 | -1 | protein_coding | ENSG00000 |
| 130497743 | 130497849 | -1 | snRNA | ENSG00000 |
| 130515997 | 130526691 | -1 | lncRNA | ENSG00000 |
| 130517049 | 130517312 | -1 | misc_RNA | ENSG00000 |
| 130521197 | 130528604 | 1 | protein_coding | ENSG00000 |
| 130530275 | 130537182 | 1 | lncRNA | ENSG00000 |

| | | | | |
|---|---|---|---|---|
| 130539095 | 130549883 | -1 | transcribed_unproce | ENSG00000 |
| 130570829 | 130584161 | 1 | transcribed_unproce | ENSG00000 |
| 130583549 | 130590463 | -1 | lncRNA | ENSG00000 |
| 130592168 | 130599575 | -1 | protein_coding | ENSG00000 |
| 130603461 | 130603724 | 1 | misc_RNA | ENSG00000 |
| 130611481 | 130658091 | 1 | protein_coding | ENSG00000 |
| 130622838 | 130622944 | 1 | snRNA | ENSG00000 |
| 130680050 | 130685863 | 1 | transcribed_unproce | ENSG00000 |
| 130681228 | 130691691 | 1 | lncRNA | ENSG00000 |
| 130690133 | 130718831 | 1 | transcribed_unproce | ENSG00000 |
| 130692243 | 130693770 | -1 | processed_pseudoge | ENSG00000 |
| 130729070 | 130730336 | 1 | protein_coding | ENSG00000 |
| 130742959 | 130755879 | -1 | lncRNA | ENSG00000 |
| 130755435 | 130768134 | 1 | protein_coding | ENSG00000 |
| 130776646 | 130776957 | 1 | processed_pseudoge | ENSG00000 |
| 130830978 | 130836994 | -1 | lncRNA | ENSG00000 |
| 130836916 | 131047263 | 1 | protein_coding | ENSG00000 |
| 130929762 | 130929883 | -1 | scaRNA | ENSG00000 |
| 130929762 | 130929884 | -1 | scaRNA | ENSG00000 |
| 131035092 | 131035262 | 1 | protein_coding | ENSG00000 |
| 131047876 | 131093460 | -1 | protein_coding | ENSG00000 |
| 131104847 | 131353709 | 1 | protein_coding | ENSG00000 |
| 131177618 | 131178946 | -1 | processed_pseudoge | ENSG00000 |
| 131181153 | 131181369 | -1 | processed_pseudoge | ENSG00000 |
| 131189834 | 131198007 | -1 | unprocessed_pseudc | ENSG00000 |
| 131191641 | 131209211 | -1 | lncRNA | ENSG00000 |
| 131201142 | 131202941 | -1 | unprocessed_pseudc | ENSG00000 |
| 131218067 | 131265278 | 1 | protein_coding | ENSG00000 |
| 131229607 | 131229713 | 1 | snRNA | ENSG00000 |
| 131279290 | 131309068 | 1 | transcribed_unproce | ENSG00000 |
| 131299220 | 131300444 | -1 | processed_pseudoge | ENSG00000 |
| 131303037 | 131355683 | 1 | lncRNA | ENSG00000 |
| 131328176 | 131330857 | -1 | unprocessed_pseudc | ENSG00000 |
| 131335502 | 131355769 | 1 | unprocessed_pseudc | ENSG00000 |
| 131352214 | 131353239 | -1 | processed_pseudoge | ENSG00000 |
| 131361111 | 131364142 | -1 | protein_coding | ENSG00000 |
| 131363364 | 131364881 | 1 | lncRNA | ENSG00000 |
| 131369062 | 131369711 | -1 | unprocessed_pseudc | ENSG00000 |
| 131369785 | 131370301 | 1 | unprocessed_pseudc | ENSG00000 |
| 131370306 | 131372075 | -1 | unprocessed_pseudc | ENSG00000 |
| 131372294 | 131373650 | -1 | unprocessed_pseudc | ENSG00000 |
| 131373647 | 131379675 | -1 | unprocessed_pseudc | ENSG00000 |
| 131379743 | 131380083 | -1 | unprocessed_pseudc | ENSG00000 |
| 131380153 | 131380913 | -1 | unprocessed_pseudc | ENSG00000 |
| 131380916 | 131381590 | -1 | unprocessed_pseudc | ENSG00000 |
| 131381840 | 131382497 | -1 | unprocessed_pseudc | ENSG00000 |
| 131382643 | 131384061 | -1 | unprocessed_pseudc | ENSG00000 |
| 131384461 | 131385490 | -1 | unprocessed_pseudc | ENSG00000 |
| 131385700 | 131386314 | -1 | unprocessed_pseudc | ENSG00000 |
| 131395217 | 131395542 | -1 | processed_pseudoge | ENSG00000 |

| | | | | |
|---|---|---|---|---|
| 131398439 | 131401250 | -1 | unprocessed_pseudc | ENSG00000C |
| 131402778 | 131410050 | 1 | lncRNA | ENSG00000C |
| 131419665 | 131419784 | -1 | processed_pseudoge | ENSG00000C |
| 131423801 | 131424867 | -1 | processed_pseudoge | ENSG00000C |
| 131442644 | 131444587 | -1 | processed_pseudoge | ENSG00000C |
| 131460999 | 131461577 | 1 | processed_pseudoge | ENSG00000C |
| 131461821 | 131463615 | -1 | lncRNA | ENSG00000C |
| 131464900 | 131492743 | -1 | protein_coding | ENSG00000C |
| 131476119 | 131482934 | 1 | protein_coding | ENSG00000C |
| 131491160 | 131491236 | -1 | miRNA | ENSG00000C |
| 131492813 | 131521573 | 1 | transcribed_processe | ENSG00000C |
| 131527675 | 131533666 | 1 | protein_coding | ENSG00000C |
| 131535187 | 131536988 | -1 | unprocessed_pseudc | ENSG00000C |
| 131555007 | 131568599 | 1 | lncRNA | ENSG00000C |
| 131591752 | 131628147 | 1 | unprocessed_pseudc | ENSG00000C |
| 131602790 | 131602896 | 1 | snRNA | ENSG00000C |
| 131637025 | 131649615 | 1 | lncRNA | ENSG00000C |
| 131646704 | 131648372 | -1 | processed_pseudoge | ENSG00000C |
| 131647990 | 131767404 | 1 | lncRNA | ENSG00000C |
| 131692392 | 131697954 | 1 | transcribed_unproce | ENSG00000C |
| 131722417 | 131751669 | 1 | transcribed_unproce | ENSG00000C |
| 131723253 | 131724279 | -1 | processed_pseudoge | ENSG00000C |
| 131758031 | 131767221 | 1 | transcribed_unproce | ENSG00000C |
| 131794961 | 131801661 | -1 | transcribed_unproce | ENSG00000C |
| 131821987 | 131822920 | -1 | unprocessed_pseudc | ENSG00000C |
| 131829801 | 131832031 | -1 | lncRNA | ENSG00000C |
| 131833783 | 131836532 | -1 | unprocessed_pseudc | ENSG00000C |
| 131958277 | 131965535 | 1 | lncRNA | ENSG00000C |
| 131964731 | 131979873 | -1 | lncRNA | ENSG00000C |
| 131983937 | 131994161 | -1 | lncRNA | ENSG00000C |
| 132010195 | 132017000 | 1 | lncRNA | ENSG00000C |
| 132037787 | 132037993 | -1 | processed_pseudoge | ENSG00000C |
| 132080290 | 132080755 | -1 | processed_pseudoge | ENSG00000C |
| 132087289 | 132102564 | 1 | unprocessed_pseudc | ENSG00000C |
| 132132856 | 132139429 | -1 | unprocessed_pseudc | ENSG00000C |
| 132147591 | 132257969 | -1 | protein_coding | ENSG00000C |
| 132152243 | 132152349 | -1 | snRNA | ENSG00000C |
| 132253154 | 132253305 | -1 | rRNA_pseudogene | ENSG00000C |
| 132256966 | 132257080 | -1 | miRNA | ENSG00000C |
| 132262328 | 132263042 | 1 | processed_pseudoge | ENSG00000C |
| 132285795 | 132294377 | -1 | lncRNA | ENSG00000C |
| 132309309 | 132318736 | 1 | unprocessed_pseudc | ENSG00000C |
| 132345616 | 132347297 | -1 | lncRNA | ENSG00000C |
| 132347473 | 132365130 | 1 | lncRNA | ENSG00000C |
| 132352722 | 132353318 | 1 | processed_pseudoge | ENSG00000C |
| 132406752 | 132406858 | -1 | snRNA | ENSG00000C |
| 132416805 | 132646582 | 1 | protein_coding | ENSG00000C |
| 132488554 | 132489088 | -1 | processed_pseudoge | ENSG00000C |
| 132523617 | 132523936 | 1 | misc_RNA | ENSG00000C |
| 132643286 | 132671579 | -1 | protein_coding | ENSG00000C |

| | | | | |
|---|---|---|---|---|
| 132660526 | 132666990 | 1 | lncRNA | ENSG00000 |
| 132671788 | 133568463 | -1 | protein_coding | ENSG00000 |
| 132685224 | 132816896 | 1 | lncRNA | ENSG00000 |
| 132911096 | 132911261 | -1 | processed_pseudoge | ENSG00000 |
| 132915557 | 132931494 | 1 | lncRNA | ENSG00000 |
| 133117004 | 133119052 | 1 | TEC | ENSG00000 |
| 133264968 | 133285445 | 1 | lncRNA | ENSG00000 |
| 133333829 | 133333930 | 1 | snRNA | ENSG00000 |
| 133391109 | 133391398 | -1 | misc_RNA | ENSG00000 |
| 133431666 | 133433897 | -1 | lncRNA | ENSG00000 |
| 133554315 | 133558227 | 1 | lncRNA | ENSG00000 |
| 133596091 | 133596399 | 1 | misc_RNA | ENSG00000 |
| 134028608 | 134032597 | 1 | lncRNA | ENSG00000 |
| 134119983 | 134454621 | 1 | protein_coding | ENSG00000 |
| 134127125 | 134127192 | 1 | miRNA | ENSG00000 |
| 134231195 | 134231312 | 1 | rRNA_pseudogene | ENSG00000 |
| 134319654 | 134320078 | -1 | processed_pseudoge | ENSG00000 |
| 134455759 | 134719000 | -1 | protein_coding | ENSG00000 |
| 134718188 | 134725474 | 1 | lncRNA | ENSG00000 |
| 134735464 | 134918710 | -1 | lncRNA | ENSG00000 |
| 134797169 | 134797879 | 1 | processed_pseudoge | ENSG00000 |
| 134816362 | 134816589 | 1 | processed_pseudoge | ENSG00000 |
| 134838616 | 134902034 | 1 | protein_coding | ENSG00000 |
| 134857820 | 134857873 | 1 | miRNA | ENSG00000 |
| 134918235 | 134959342 | 1 | protein_coding | ENSG00000 |
| 134964485 | 135047468 | -1 | protein_coding | ENSG00000 |
| 135052265 | 135176394 | 1 | protein_coding | ENSG00000 |
| 135136628 | 135136755 | 1 | snoRNA | ENSG00000 |
| 135136916 | 135531218 | -1 | protein_coding | ENSG00000 |
| 135510546 | 135511943 | -1 | processed_pseudoge | ENSG00000 |
| 135531455 | 135725270 | 1 | protein_coding | ENSG00000 |
| 135656477 | 135656579 | -1 | snRNA | ENSG00000 |
| 135665397 | 135665478 | 1 | miRNA | ENSG00000 |
| 135741734 | 135785056 | 1 | protein_coding | ENSG00000 |
| 135787850 | 135837184 | -1 | protein_coding | ENSG00000 |
| 135810169 | 135810279 | -1 | misc_RNA | ENSG00000 |
| 135820191 | 135823087 | 1 | lncRNA | ENSG00000 |
| 135839626 | 135876443 | -1 | protein_coding | ENSG00000 |
| 135897955 | 135898978 | -1 | processed_pseudoge | ENSG00000 |
| 135905881 | 135986100 | -1 | protein_coding | ENSG00000 |
| 135985176 | 136022593 | 1 | lncRNA | ENSG00000 |
| 136077892 | 136078513 | 1 | lncRNA | ENSG00000 |
| 136114349 | 136118165 | -1 | protein_coding | ENSG00000 |
| 136199114 | 136200503 | -1 | processed_pseudoge | ENSG00000 |
| 136230753 | 136233245 | 1 | TEC | ENSG00000 |
| 136329441 | 136329913 | -1 | processed_pseudoge | ENSG00000 |
| 136390332 | 136390669 | 1 | misc_RNA | ENSG00000 |
| 136544712 | 136545542 | 1 | unprocessed_pseudc | ENSG00000 |
| 136765545 | 137677718 | 1 | protein_coding | ENSG00000 |
| 137512098 | 137512210 | -1 | rRNA_pseudogene | ENSG00000 |

| | | | | |
|---|---|---|---|---|
| 137715332 | 137715440 | 1 | misc_RNA | ENSG0000C |
| 137964020 | 138016364 | 1 | protein_coding | ENSG0000C |
| 138101878 | 138107178 | 1 | lncRNA | ENSG0000C |
| 138278574 | 138279210 | 1 | processed_pseudoge | ENSG0000C |
| 138288029 | 138288793 | -1 | processed_pseudoge | ENSG0000C |
| 138307319 | 138307970 | 1 | processed_pseudoge | ENSG0000C |
| 138412066 | 138413318 | -1 | processed_pseudoge | ENSG0000C |
| 138418284 | 138501698 | -1 | lncRNA | ENSG0000C |
| 138470656 | 138473932 | 1 | lncRNA | ENSG0000C |
| 138501770 | 138573547 | 1 | protein_coding | ENSG0000C |
| 138569090 | 138574458 | 1 | lncRNA | ENSG0000C |
| 138599663 | 138602426 | -1 | lncRNA | ENSG0000C |
| 138601599 | 138617469 | 1 | lncRNA | ENSG0000C |
| 138669157 | 138780390 | -1 | protein_coding | ENSG0000C |
| 138863597 | 138863895 | -1 | misc_RNA | ENSG0000C |
| 138897151 | 138898098 | 1 | processed_pseudoge | ENSG0000C |
| 138901433 | 138902747 | -1 | processed_pseudoge | ENSG0000C |
| 138917204 | 138975006 | 1 | lncRNA | ENSG0000C |
| 139323921 | 139364583 | 1 | lncRNA | ENSG0000C |
| 139366165 | 139379147 | 1 | lncRNA | ENSG0000C |
| 139469775 | 139477747 | 1 | lncRNA | ENSG0000C |
| 139511424 | 139511555 | 1 | snoRNA | ENSG0000C |
| 139668547 | 139669305 | 1 | processed_pseudoge | ENSG0000C |
| 139732708 | 139733282 | -1 | processed_pseudoge | ENSG0000C |
| 139824896 | 139825877 | -1 | lncRNA | ENSG0000C |
| 139982684 | 139982986 | 1 | misc_RNA | ENSG0000C |
| 140103583 | 140131780 | -1 | lncRNA | ENSG0000C |
| 140216997 | 140217239 | -1 | processed_pseudoge | ENSG0000C |
| 140217651 | 140220368 | -1 | unprocessed_pseudc | ENSG0000C |
| 140220584 | 140221485 | -1 | processed_pseudoge | ENSG0000C |
| 140231423 | 142131701 | -1 | protein_coding | ENSG0000C |
| 140234737 | 140234923 | -1 | misc_RNA | ENSG0000C |
| 140586626 | 140586685 | -1 | miRNA | ENSG0000C |
| 140683360 | 140683888 | -1 | processed_pseudoge | ENSG0000C |
| 140898423 | 140899106 | 1 | processed_pseudoge | ENSG0000C |
| 140992543 | 140992659 | 1 | misc_RNA | ENSG0000C |
| 141167257 | 141167357 | -1 | snRNA | ENSG0000C |
| 141208082 | 141208376 | -1 | processed_pseudoge | ENSG0000C |
| 142131178 | 142137830 | -1 | lncRNA | ENSG0000C |
| 142543418 | 142543975 | 1 | processed_pseudoge | ENSG0000C |
| 142854095 | 142855998 | 1 | processed_pseudoge | ENSG0000C |
| 142877657 | 143055833 | 1 | protein_coding | ENSG0000C |
| 142947300 | 142948086 | -1 | processed_pseudoge | ENSG0000C |
| 143091362 | 143768352 | 1 | protein_coding | ENSG0000C |
| 143093056 | 143094164 | -1 | processed_pseudoge | ENSG0000C |
| 143094233 | 143094715 | 1 | processed_pseudoge | ENSG0000C |
| 143094789 | 143096555 | -1 | processed_pseudoge | ENSG0000C |
| 143096839 | 143098007 | -1 | processed_pseudoge | ENSG0000C |
| 143098123 | 143098417 | -1 | processed_pseudoge | ENSG0000C |
| 143098484 | 143098829 | -1 | processed_pseudoge | ENSG0000C |

| | | | | |
|---|---|---|---|---|
| 143098898 | 143099933 | -1 | processed_pseudoge | ENSG00000 |
| 143099936 | 143100612 | -1 | processed_pseudoge | ENSG00000 |
| 143100930 | 143101355 | -1 | processed_pseudoge | ENSG00000 |
| 143162078 | 143172172 | -1 | lncRNA | ENSG00000 |
| 143295421 | 143598002 | -1 | lncRNA | ENSG00000 |
| 143601517 | 143608759 | 1 | lncRNA | ENSG00000 |
| 143629133 | 143656179 | -1 | lncRNA | ENSG00000 |
| 143674315 | 143741294 | -1 | lncRNA | ENSG00000 |
| 143765940 | 143776119 | -1 | lncRNA | ENSG00000 |
| 143799467 | 143819599 | 1 | lncRNA | ENSG00000 |
| 143897263 | 143897361 | 1 | misc_RNA | ENSG00000 |
| 143937073 | 143964156 | 1 | lncRNA | ENSG00000 |
| 143938068 | 144332568 | -1 | protein_coding | ENSG00000 |
| 144364364 | 144521057 | -1 | protein_coding | ENSG00000 |
| 144444848 | 144450450 | 1 | lncRNA | ENSG00000 |
| 144494265 | 144496878 | -1 | TEC | ENSG00000 |
| 144517978 | 144521477 | 1 | lncRNA | ENSG00000 |
| 144518447 | 144518574 | 1 | misc_RNA | ENSG00000 |
| 144520456 | 144520555 | 1 | misc_RNA | ENSG00000 |
| 144521039 | 144521116 | 1 | misc_RNA | ENSG00000 |
| 144523552 | 144524651 | -1 | lncRNA | ENSG00000 |
| 144523912 | 144579434 | 1 | lncRNA | ENSG00000 |
| 144666312 | 145262988 | 1 | lncRNA | ENSG00000 |
| 144688413 | 144690185 | -1 | lncRNA | ENSG00000 |
| 144705217 | 144705419 | 1 | misc_RNA | ENSG00000 |
| 144754776 | 144755998 | -1 | processed_pseudoge | ENSG00000 |
| 144877734 | 144882033 | -1 | lncRNA | ENSG00000 |
| 145294277 | 145331847 | -1 | lncRNA | ENSG00000 |
| 145337230 | 145338057 | 1 | processed_pseudoge | ENSG00000 |
| 145569294 | 145588024 | -1 | lncRNA | ENSG00000 |
| 145572662 | 145765726 | 1 | lncRNA | ENSG00000 |
| 145600907 | 145603423 | -1 | lncRNA | ENSG00000 |
| 145826285 | 145827830 | -1 | processed_pseudoge | ENSG00000 |
| 145872209 | 145931146 | 1 | lncRNA | ENSG00000 |
| 145874520 | 145874845 | -1 | TEC | ENSG00000 |
| 146145156 | 146145217 | 1 | snRNA | ENSG00000 |
| 146194296 | 146194821 | 1 | processed_pseudoge | ENSG00000 |
| 146477427 | 146478113 | 1 | processed_pseudoge | ENSG00000 |
| 146587506 | 146589310 | 1 | processed_pseudoge | ENSG00000 |
| 146833501 | 146834038 | -1 | processed_pseudoge | ENSG00000 |
| 146837727 | 146878220 | 1 | lncRNA | ENSG00000 |
| 146879909 | 146898218 | 1 | lncRNA | ENSG00000 |
| 147150707 | 147152047 | 1 | lncRNA | ENSG00000 |
| 147408342 | 147408448 | 1 | snRNA | ENSG00000 |
| 147472133 | 147472243 | -1 | misc_RNA | ENSG00000 |
| 147495438 | 147495540 | -1 | snRNA | ENSG00000 |
| 147713455 | 147713554 | -1 | rRNA_pseudogene | ENSG00000 |
| 147810346 | 147810777 | 1 | processed_pseudoge | ENSG00000 |
| 147844517 | 147930826 | 1 | protein_coding | ENSG00000 |
| 147877422 | 147877528 | 1 | snRNA | ENSG00000 |

| | | | | |
|---|---|---|---|---|
| 147899401 | 147902956 | -1 | lncRNA | ENSG00000 |
| 147930397 | 148021604 | -1 | protein_coding | ENSG00000 |
| 148021011 | 148516971 | 1 | protein_coding | ENSG00000 |
| 148044380 | 148044894 | 1 | TEC | ENSG00000 |
| 148062154 | 148062577 | 1 | TEC | ENSG00000 |
| 148260574 | 148260708 | -1 | snoRNA | ENSG00000 |
| 148295656 | 148296679 | -1 | processed_pseudoge | ENSG00000 |
| 148595158 | 148595328 | 1 | processed_pseudoge | ENSG00000 |
| 148644440 | 148787569 | 1 | protein_coding | ENSG00000 |
| 148825691 | 148825835 | -1 | snRNA | ENSG00000 |
| 148863564 | 148863804 | -1 | processed_pseudoge | ENSG00000 |
| 148866470 | 148888823 | -1 | lncRNA | ENSG00000 |
| 148870024 | 148870381 | -1 | processed_pseudoge | ENSG00000 |
| 148872253 | 148872797 | -1 | processed_pseudoge | ENSG00000 |
| 148875227 | 149026759 | 1 | protein_coding | ENSG00000 |
| 148909406 | 148927254 | -1 | lncRNA | ENSG00000 |
| 149038107 | 149215262 | 1 | protein_coding | ENSG00000 |
| 149068596 | 149068910 | -1 | processed_pseudoge | ENSG00000 |
| 149170561 | 149184570 | -1 | lncRNA | ENSG00000 |
| 149320593 | 149321436 | 1 | processed_pseudoge | ENSG00000 |
| 149329985 | 149474138 | 1 | protein_coding | ENSG00000 |
| 149439897 | 149440427 | 1 | processed_pseudoge | ENSG00000 |
| 149569637 | 149587778 | -1 | protein_coding | ENSG00000 |
| 149587196 | 150047447 | 1 | lncRNA | ENSG00000 |
| 149610668 | 149610774 | -1 | snRNA | ENSG00000 |
| 149745648 | 149859191 | -1 | lncRNA | ENSG00000 |
| 150150633 | 150166266 | -1 | lncRNA | ENSG00000 |
| 150169474 | 150356460 | 1 | lncRNA | ENSG00000 |
| 150234892 | 150257007 | -1 | lncRNA | ENSG00000 |
| 150468195 | 150539011 | -1 | protein_coding | ENSG00000 |
| 150552532 | 150572221 | -1 | lncRNA | ENSG00000 |
| 150566134 | 150568080 | 1 | lncRNA | ENSG00000 |
| 150595102 | 150620067 | -1 | lncRNA | ENSG00000 |
| 150612381 | 150635818 | 1 | lncRNA | ENSG00000 |
| 150982506 | 151003474 | 1 | lncRNA | ENSG00000 |
| 151001220 | 151048774 | -1 | lncRNA | ENSG00000 |
| 151114811 | 151186209 | -1 | lncRNA | ENSG00000 |
| 151186188 | 151186595 | 1 | processed_pseudoge | ENSG00000 |
| 151247940 | 151261863 | -1 | protein_coding | ENSG00000 |
| 151270470 | 151289894 | -1 | protein_coding | ENSG00000 |
| 151357592 | 151380046 | 1 | protein_coding | ENSG00000 |
| 151368334 | 151368411 | -1 | miRNA | ENSG00000 |
| 151368334 | 151368411 | 1 | miRNA | ENSG00000 |
| 151372003 | 151372287 | 1 | misc_RNA | ENSG00000 |
| 151409883 | 151508013 | 1 | protein_coding | ENSG00000 |
| 151485336 | 151734487 | -1 | protein_coding | ENSG00000 |
| 151788984 | 151828492 | -1 | protein_coding | ENSG00000 |
| 151796688 | 151798688 | 1 | lncRNA | ENSG00000 |
| 151802651 | 151973949 | -1 | protein_coding | ENSG00000 |
| 151832768 | 152099475 | -1 | protein_coding | ENSG00000 |

| | | | | |
|---|---|---|---|---|
| 152098328 | 152109202 | 1 | lncRNA | ENSG00000 |
| 152116801 | 152175763 | -1 | protein_coding | ENSG00000 |
| 152131555 | 152131845 | 1 | processed_pseudoge | ENSG00000 |
| 152176020 | 152225211 | 1 | lncRNA | ENSG00000 |
| 152284350 | 152284460 | -1 | misc_RNA | ENSG00000 |
| 152335174 | 152649826 | 1 | protein_coding | ENSG00000 |
| 152389937 | 152390630 | 1 | processed_pseudoge | ENSG00000 |
| 152450162 | 152452891 | -1 | lncRNA | ENSG00000 |
| 152539197 | 152539427 | -1 | processed_pseudoge | ENSG00000 |
| 152635268 | 152641275 | -1 | lncRNA | ENSG00000 |
| 152651593 | 152717997 | -1 | protein_coding | ENSG00000 |
| 152717893 | 152761253 | 1 | protein_coding | ENSG00000 |
| 152850653 | 152850824 | -1 | processed_pseudoge | ENSG00000 |
| 152876820 | 152877767 | 1 | processed_pseudoge | ENSG00000 |
| 153158938 | 153159683 | 1 | processed_pseudoge | ENSG00000 |
| 153168399 | 153201762 | -1 | lncRNA | ENSG00000 |
| 153208086 | 153208163 | -1 | processed_pseudoge | ENSG00000 |
| 153337705 | 153357422 | 1 | lncRNA | ENSG00000 |
| 153370612 | 153371064 | 1 | processed_pseudoge | ENSG00000 |
| 153420677 | 153421957 | -1 | transcribed_process | ENSG00000 |
| 153421616 | 153593288 | -1 | lncRNA | ENSG00000 |
| 153446813 | 153446883 | -1 | snoRNA | ENSG00000 |
| 153477338 | 153478762 | -1 | protein_coding | ENSG00000 |
| 153761450 | 153762646 | -1 | processed_pseudoge | ENSG00000 |
| 153871922 | 154453979 | 1 | protein_coding | ENSG00000 |
| 154286589 | 154287425 | 1 | processed_pseudoge | ENSG00000 |
| 154435853 | 154457438 | -1 | lncRNA | ENSG00000 |
| 154459857 | 154460820 | 1 | lncRNA | ENSG00000 |
| 154590407 | 154590528 | 1 | rRNA_pseudogene | ENSG00000 |
| 154688296 | 154697817 | -1 | lncRNA | ENSG00000 |
| 154697855 | 154858354 | 1 | protein_coding | ENSG00000 |
| 154940541 | 154941067 | -1 | processed_pseudoge | ENSG00000 |
| 154964308 | 154969101 | -1 | lncRNA | ENSG00000 |
| 155089439 | 155089538 | -1 | snRNA | ENSG00000 |
| 155263458 | 155263724 | -1 | processed_pseudoge | ENSG00000 |
| 155264163 | 155264823 | -1 | processed_pseudoge | ENSG00000 |
| 155269633 | 155270134 | 1 | transcribed_process | ENSG00000 |
| 155311023 | 155311409 | 1 | processed_pseudoge | ENSG00000 |
| 155311697 | 155313427 | 1 | processed_pseudoge | ENSG00000 |
| 155313440 | 155313958 | -1 | processed_pseudoge | ENSG00000 |
| 155314066 | 155314368 | 1 | processed_pseudoge | ENSG00000 |
| 155330421 | 155330527 | 1 | snRNA | ENSG00000 |
| 155525424 | 155664668 | 1 | lncRNA | ENSG00000 |
| 156011530 | 156254950 | -1 | lncRNA | ENSG00000 |
| 156033247 | 156034080 | -1 | processed_pseudoge | ENSG00000 |
| 156305108 | 156320255 | -1 | lncRNA | ENSG00000 |
| 156324437 | 156342348 | -1 | protein_coding | ENSG00000 |
| 156435290 | 156613735 | 1 | protein_coding | ENSG00000 |
| 156655224 | 156695659 | 1 | lncRNA | ENSG00000 |
| 156777706 | 156778372 | -1 | processed_pseudoge | ENSG00000 |

| | | | | |
|---|---|---|---|---|
| 156788731 | 156865701 | 1 | lncRNA | ENSG0000C |
| 156803339 | 156891160 | 1 | lncRNA | ENSG0000C |
| 157033837 | 157049716 | 1 | lncRNA | ENSG0000C |
| 157210847 | 157211539 | -1 | processed_pseudoge | ENSG0000C |
| 157257705 | 157318491 | 1 | protein_coding | ENSG0000C |
| 157298807 | 157299158 | -1 | misc_RNA | ENSG0000C |
| 157318625 | 157327713 | -1 | protein_coding | ENSG0000C |
| 157379724 | 157380414 | -1 | processed_pseudoge | ENSG0000C |
| 157414619 | 157488961 | -1 | protein_coding | ENSG0000C |
| 157526767 | 157628864 | -1 | protein_coding | ENSG0000C |
| 157725708 | 157736005 | -1 | lncRNA | ENSG0000C |
| 157736444 | 157875862 | -1 | protein_coding | ENSG0000C |
| 157876702 | 158136154 | 1 | protein_coding | ENSG0000C |
| 157877683 | 157878565 | 1 | lncRNA | ENSG0000C |
| 157912772 | 157912885 | 1 | snRNA | ENSG0000C |
| 157917712 | 157918371 | 1 | lncRNA | ENSG0000C |
| 157961410 | 157962888 | -1 | processed_pseudoge | ENSG0000C |
| 157985789 | 157985894 | -1 | snRNA | ENSG0000C |
| 158065545 | 158066059 | 1 | processed_pseudoge | ENSG0000C |
| 158127957 | 158133988 | 1 | lncRNA | ENSG0000C |
| 158166650 | 158236169 | 1 | lncRNA | ENSG0000C |
| 158171073 | 158456753 | -1 | protein_coding | ENSG0000C |
| 158419955 | 158425303 | 1 | unprocessed_pseudc | ENSG0000C |
| 158447166 | 158447461 | -1 | misc_RNA | ENSG0000C |
| 158456964 | 158682879 | 1 | protein_coding | ENSG0000C |
| 158548425 | 158549224 | 1 | processed_pseudoge | ENSG0000C |
| 158658337 | 158735002 | -1 | lncRNA | ENSG0000C |
| 158685903 | 158753775 | 1 | lncRNA | ENSG0000C |
| 158703287 | 158703460 | 1 | processed_pseudoge | ENSG0000C |
| 158795317 | 158862781 | 1 | protein_coding | ENSG0000C |
| 158847619 | 158870393 | -1 | lncRNA | ENSG0000C |
| 158853755 | 158854576 | 1 | transcribed_unproce | ENSG0000C |
| 158862311 | 158863285 | 1 | transcribed_unproce | ENSG0000C |
| 158874642 | 158875670 | 1 | unprocessed_pseudc | ENSG0000C |
| 158935798 | 158935985 | -1 | snRNA | ENSG0000C |
| 158968640 | 159232659 | 1 | protein_coding | ENSG0000C |
| 159003975 | 159004320 | 1 | processed_pseudoge | ENSG0000C |
| 159027036 | 159027142 | -1 | snRNA | ENSG0000C |
| 159056005 | 159056859 | -1 | processed_pseudoge | ENSG0000C |
| 159186835 | 159186901 | 1 | miRNA | ENSG0000C |
| 159235793 | 159286799 | -1 | protein_coding | ENSG0000C |
| 159318979 | 159616569 | -1 | protein_coding | ENSG0000C |
| 159386367 | 159404636 | 1 | lncRNA | ENSG0000C |
| 159444117 | 159444913 | 1 | processed_pseudoge | ENSG0000C |
| 159462417 | 159463256 | -1 | processed_pseudoge | ENSG0000C |
| 159536244 | 159536346 | -1 | misc_RNA | ENSG0000C |
| 159615296 | 159617082 | 1 | lncRNA | ENSG0000C |
| 159670708 | 159712435 | -1 | lncRNA | ENSG0000C |
| 159689217 | 159690291 | -1 | processed_pseudoge | ENSG0000C |
| 159712457 | 159771027 | 1 | protein_coding | ENSG0000C |

| | | | | |
|---|---|---|---|---|
| 159768628 | 159798255 | -1 | protein_coding | ENSG00000( |
| 159771851 | 159904710 | -1 | protein_coding | ENSG00000( |
| 159797288 | 159798043 | -1 | lncRNA | ENSG00000( |
| 159803355 | 159904756 | -1 | protein_coding | ENSG00000( |
| 159812234 | 159814009 | -1 | processed_pseudoge | ENSG00000( |
| 159932006 | 160062615 | -1 | protein_coding | ENSG00000( |
| 160099667 | 160200313 | -1 | protein_coding | ENSG00000( |
| 160141981 | 160271888 | -1 | lncRNA | ENSG00000( |
| 160257718 | 160271880 | -1 | lncRNA | ENSG00000( |
| 160272151 | 160493807 | -1 | protein_coding | ENSG00000( |
| 160407810 | 160407882 | -1 | miRNA | ENSG00000( |
| 160849313 | 160850676 | -1 | TEC | ENSG00000( |
| 160877547 | 160877850 | 1 | misc_RNA | ENSG00000( |
| 161048606 | 161050546 | -1 | processed_pseudoge | ENSG00000( |
| 161096231 | 161160224 | -1 | lncRNA | ENSG00000( |
| 161136908 | 161236230 | 1 | protein_coding | ENSG00000( |
| 161222785 | 161308389 | -1 | lncRNA | ENSG00000( |
| 161244730 | 161249050 | 1 | lncRNA | ENSG00000( |
| 161281902 | 161282618 | 1 | processed_pseudoge | ENSG00000( |
| 161308425 | 161411717 | 1 | protein_coding | ENSG00000( |
| 161340816 | 161341326 | -1 | processed_pseudoge | ENSG00000( |
| 161409554 | 161409670 | 1 | rRNA_pseudogene | ENSG00000( |
| 161416297 | 161425870 | 1 | protein_coding | ENSG00000( |
| 161422659 | 161423577 | -1 | lncRNA | ENSG00000( |
| 161424015 | 161428774 | 1 | lncRNA | ENSG00000( |
| 161424332 | 161985282 | 1 | protein_coding | ENSG00000( |
| 161433874 | 161437458 | 1 | TEC | ENSG00000( |
| 161500885 | 161507705 | -1 | processed_pseudoge | ENSG00000( |
| 161558544 | 161558646 | 1 | snoRNA | ENSG00000( |
| 161568502 | 161570185 | -1 | processed_pseudoge | ENSG00000( |
| 161577192 | 161625752 | -1 | lncRNA | ENSG00000( |
| 161580072 | 161581350 | 1 | processed_pseudoge | ENSG00000( |
| 161931199 | 161931466 | -1 | processed_pseudoge | ENSG00000( |
| 161992245 | 162074215 | -1 | protein_coding | ENSG00000( |
| 162073256 | 162077911 | 1 | lncRNA | ENSG00000( |
| 162077357 | 162077651 | -1 | processed_pseudoge | ENSG00000( |
| 162088259 | 162090072 | 1 | processed_pseudoge | ENSG00000( |
| 162092232 | 162095032 | 1 | lncRNA | ENSG00000( |
| 162142882 | 162152404 | -1 | protein_coding | ENSG00000( |
| 162159762 | 162173223 | 1 | lncRNA | ENSG00000( |
| 162170684 | 162245151 | -1 | protein_coding | ENSG00000( |
| 162267074 | 162318684 | -1 | protein_coding | ENSG00000( |
| 162318840 | 162371595 | 1 | protein_coding | ENSG00000( |
| 162371407 | 162838767 | -1 | protein_coding | ENSG00000( |
| 162496936 | 162497051 | 1 | rRNA_pseudogene | ENSG00000( |
| 162768936 | 162797972 | 1 | lncRNA | ENSG00000( |
| 163152251 | 163153307 | -1 | processed_pseudoge | ENSG00000( |
| 163288995 | 163289096 | 1 | snRNA | ENSG00000( |
| 163593396 | 163736012 | -1 | protein_coding | ENSG00000( |
| 163743913 | 164352223 | -1 | lncRNA | ENSG00000( |

| | | | | |
|---|---|---|---|---|
| 164213539 | 164214414 | 1 | processed_pseudoge | ENSG00000( |
| 164321933 | 164322223 | 1 | processed_pseudoge | ENSG00000( |
| 164492417 | 164621482 | -1 | protein_coding | ENSG00000( |
| 164573741 | 164574054 | 1 | unprocessed_pseudc | ENSG00000( |
| 164653624 | 164843679 | -1 | protein_coding | ENSG00000( |
| 164687287 | 164687596 | -1 | processed_pseudoge | ENSG00000( |
| 164687643 | 164687777 | -1 | snoRNA | ENSG00000( |
| 164840661 | 165208261 | 1 | lncRNA | ENSG00000( |
| 164858432 | 164858546 | 1 | rRNA_pseudogene | ENSG00000( |
| 164895677 | 164895777 | -1 | rRNA_pseudogene | ENSG00000( |
| 164896186 | 164955525 | -1 | protein_coding | ENSG00000( |
| 165087522 | 165204067 | -1 | protein_coding | ENSG00000( |
| 165194993 | 165392310 | 1 | protein_coding | ENSG00000( |
| 165285253 | 165286048 | 1 | processed_pseudoge | ENSG00000( |
| 165433438 | 165436614 | -1 | lncRNA | ENSG00000( |
| 165469647 | 165689407 | 1 | protein_coding | ENSG00000( |
| 165747588 | 165794659 | -1 | protein_coding | ENSG00000( |
| 165794851 | 165810010 | 1 | lncRNA | ENSG00000( |
| 165794857 | 165846091 | -1 | lncRNA | ENSG00000( |
| 165833048 | 165839098 | -1 | lncRNA | ENSG00000( |
| 165857475 | 165953816 | -1 | protein_coding | ENSG00000( |
| 165933857 | 165949891 | 1 | lncRNA | ENSG00000( |
| 165957188 | 166390771 | 1 | lncRNA | ENSG00000( |
| 165984641 | 166149214 | -1 | protein_coding | ENSG00000( |
| 166123494 | 166123597 | -1 | misc_RNA | ENSG00000( |
| 166180652 | 166180966 | -1 | misc_RNA | ENSG00000( |
| 166195185 | 166376001 | -1 | protein_coding | ENSG00000( |
| 166403573 | 166611482 | -1 | protein_coding | ENSG00000( |
| 166414489 | 166414729 | 1 | unprocessed_pseudc | ENSG00000( |
| 166888487 | 167259753 | 1 | protein_coding | ENSG00000( |
| 166889502 | 166889884 | -1 | processed_pseudoge | ENSG00000( |
| 167123904 | 167140955 | -1 | lncRNA | ENSG00000( |
| 167293171 | 167558333 | 1 | lncRNA | ENSG00000( |
| 167631480 | 167631541 | -1 | snRNA | ENSG00000( |
| 167713663 | 167716065 | -1 | processed_pseudoge | ENSG00000( |
| 167814757 | 167941180 | -1 | lncRNA | ENSG00000( |
| 167868948 | 167874041 | 1 | protein_coding | ENSG00000( |
| 167954020 | 168247595 | -1 | protein_coding | ENSG00000( |
| 168231921 | 168232220 | -1 | processed_pseudoge | ENSG00000( |
| 168451324 | 168451606 | 1 | misc_RNA | ENSG00000( |
| 168455862 | 168775137 | 1 | protein_coding | ENSG00000( |
| 168582943 | 168583018 | 1 | miRNA | ENSG00000( |
| 168606390 | 168606496 | -1 | snRNA | ENSG00000( |
| 168771951 | 168786961 | -1 | lncRNA | ENSG00000( |
| 168786539 | 168865514 | 1 | protein_coding | ENSG00000( |
| 168834132 | 168913371 | -1 | protein_coding | ENSG00000( |
| 168901291 | 168910000 | 1 | protein_coding | ENSG00000( |
| 168915498 | 169031324 | -1 | protein_coding | ENSG00000( |
| 169064789 | 169096167 | 1 | protein_coding | ENSG00000( |
| 169100743 | 169101210 | 1 | lncRNA | ENSG00000( |

| | | | | |
|---|---|---|---|---|
| 169115195 | 169115680 | 1 | processed_pseudoge | ENSG00000 |
| 169127109 | 169362534 | -1 | protein_coding | ENSG00000 |
| 169479480 | 169525922 | 1 | protein_coding | ENSG00000 |
| 169479480 | 169506655 | 1 | protein_coding | ENSG00000 |
| 169509702 | 169526258 | 1 | protein_coding | ENSG00000 |
| 169528508 | 169573875 | -1 | protein_coding | ENSG00000 |
| 169577371 | 169577746 | -1 | processed_pseudoge | ENSG00000 |
| 169584344 | 169641406 | 1 | protein_coding | ENSG00000 |
| 169645425 | 169694405 | -1 | protein_coding | ENSG00000 |
| 169658132 | 169661357 | -1 | processed_pseudoge | ENSG00000 |
| 169694454 | 169701708 | 1 | protein_coding | ENSG00000 |
| 169694488 | 169776989 | 1 | protein_coding | ENSG00000 |
| 169727872 | 169728003 | 1 | snoRNA | ENSG00000 |
| 169767185 | 169768799 | -1 | processed_pseudoge | ENSG00000 |
| 169790093 | 169790885 | 1 | processed_pseudoge | ENSG00000 |
| 169791933 | 169812064 | 1 | protein_coding | ENSG00000 |
| 169810081 | 169824931 | -1 | protein_coding | ENSG00000 |
| 169827458 | 170084131 | 1 | protein_coding | ENSG00000 |
| 170077224 | 170078294 | -1 | processed_pseudoge | ENSG00000 |
| 170119996 | 170120098 | -1 | snRNA | ENSG00000 |
| 170178145 | 170655171 | 1 | protein_coding | ENSG00000 |
| 170332085 | 170408564 | -1 | lncRNA | ENSG00000 |
| 170601662 | 170602228 | -1 | processed_pseudoge | ENSG00000 |
| 170640374 | 170695374 | -1 | lncRNA | ENSG00000 |
| 170690564 | 170690742 | 1 | misc_RNA | ENSG00000 |
| 170700368 | 170813670 | -1 | lncRNA | ENSG00000 |
| 170712451 | 170714567 | -1 | lncRNA | ENSG00000 |
| 170715337 | 170718078 | 1 | protein_coding | ENSG00000 |
| 170751805 | 170752788 | 1 | processed_pseudoge | ENSG00000 |
| 170766878 | 170778729 | 1 | lncRNA | ENSG00000 |
| 170783167 | 170783271 | 1 | misc_RNA | ENSG00000 |
| 170783786 | 170798971 | 1 | protein_coding | ENSG00000 |
| 170813213 | 170861151 | 1 | protein_coding | ENSG00000 |
| 170814686 | 170818037 | -1 | lncRNA | ENSG00000 |
| 170820066 | 170821295 | -1 | lncRNA | ENSG00000 |
| 170890393 | 170914873 | -1 | lncRNA | ENSG00000 |
| 170928464 | 170967130 | 1 | protein_coding | ENSG00000 |
| 170990823 | 171231314 | -1 | protein_coding | ENSG00000 |
| 171317405 | 171434802 | -1 | protein_coding | ENSG00000 |
| 171374931 | 171375278 | -1 | processed_pseudoge | ENSG00000 |
| 171434217 | 171485052 | 1 | protein_coding | ENSG00000 |
| 171491422 | 171491931 | 1 | processed_pseudoge | ENSG00000 |
| 171492905 | 171493095 | 1 | processed_pseudoge | ENSG00000 |
| 171517270 | 171517716 | -1 | processed_pseudoge | ENSG00000 |
| 171522247 | 171558129 | 1 | protein_coding | ENSG00000 |
| 171587093 | 171587259 | 1 | processed_pseudoge | ENSG00000 |
| 171687409 | 171748420 | 1 | protein_coding | ENSG00000 |
| 171783405 | 171999859 | -1 | protein_coding | ENSG00000 |
| 171856566 | 171856670 | -1 | snRNA | ENSG00000 |
| 171922448 | 171983686 | 1 | protein_coding | ENSG00000 |

| | | | | |
|---|---|---|---|---|
| 171999943 | 172082430 | 1 | protein_coding | ENSG00000 |
| 172084740 | 172089677 | 1 | protein_coding | ENSG00000 |
| 172099438 | 172102900 | -1 | protein_coding | ENSG00000 |
| 172103006 | 172109982 | 1 | lncRNA | ENSG00000 |
| 172137345 | 172370401 | 1 | lncRNA | ENSG00000 |
| 172234230 | 172237223 | 1 | lncRNA | ENSG00000 |
| 172323260 | 172330582 | -1 | lncRNA | ENSG00000 |
| 172423807 | 172424173 | -1 | TEC | ENSG00000 |
| 172427354 | 172506459 | 1 | protein_coding | ENSG00000 |
| 172427774 | 172428603 | -1 | lncRNA | ENSG00000 |
| 172464262 | 172466022 | -1 | lncRNA | ENSG00000 |
| 172480840 | 172556596 | -1 | lncRNA | ENSG00000 |
| 172555373 | 172608669 | 1 | protein_coding | ENSG00000 |
| 172558151 | 172558252 | -1 | misc_RNA | ENSG00000 |
| 172674212 | 172674898 | -1 | processed_pseudoge | ENSG00000 |
| 172677141 | 172736206 | -1 | lncRNA | ENSG00000 |
| 172691299 | 172691402 | 1 | snoRNA | ENSG00000 |
| 172735274 | 173052893 | 1 | protein_coding | ENSG00000 |
| 172893763 | 172894383 | -1 | processed_pseudoge | ENSG00000 |
| 173075435 | 173268015 | 1 | protein_coding | ENSG00000 |
| 173166446 | 173282036 | -1 | lncRNA | ENSG00000 |
| 173297629 | 173298512 | 1 | processed_pseudoge | ENSG00000 |
| 173338219 | 173356208 | -1 | transcribed_processe | ENSG00000 |
| 173354820 | 173368997 | 1 | protein_coding | ENSG00000 |
| 173431349 | 173431812 | -1 | processed_pseudoge | ENSG00000 |
| 173477996 | 173478400 | -1 | processed_pseudoge | ENSG00000 |
| 173485865 | 173486362 | -1 | processed_pseudoge | ENSG00000 |
| 173575110 | 173576341 | -1 | processed_pseudoge | ENSG00000 |
| 173705948 | 173706148 | -1 | processed_pseudoge | ENSG00000 |
| 173811076 | 173811392 | 1 | lncRNA | ENSG00000 |
| 173871653 | 173872465 | -1 | processed_pseudoge | ENSG00000 |
| 173880850 | 173965702 | -1 | protein_coding | ENSG00000 |
| 173968351 | 173969418 | 1 | lncRNA | ENSG00000 |
| 174011856 | 174012975 | -1 | lncRNA | ENSG00000 |
| 174025280 | 174027163 | 1 | lncRNA | ENSG00000 |
| 174043008 | 174043880 | 1 | processed_pseudoge | ENSG00000 |
| 174072447 | 174248599 | -1 | protein_coding | ENSG00000 |
| 174171891 | 174172725 | 1 | processed_pseudoge | ENSG00000 |
| 174240143 | 174240457 | -1 | misc_RNA | ENSG00000 |
| 174264878 | 174264981 | -1 | misc_RNA | ENSG00000 |
| 174299324 | 174300956 | -1 | processed_pseudoge | ENSG00000 |
| 174310015 | 174310974 | -1 | processed_pseudoge | ENSG00000 |
| 174326027 | 174330643 | 1 | lncRNA | ENSG00000 |
| 174334438 | 174337566 | -1 | TEC | ENSG00000 |
| 174334954 | 174338500 | 1 | protein_coding | ENSG00000 |
| 174348022 | 174395712 | -1 | protein_coding | ENSG00000 |
| 174395730 | 174429575 | 1 | protein_coding | ENSG00000 |
| 174431571 | 174487094 | -1 | protein_coding | ENSG00000 |
| 174487380 | 174488386 | 1 | lncRNA | ENSG00000 |
| 174501952 | 174502312 | 1 | processed_pseudoge | ENSG00000 |

| | | | | |
|---|---|---|---|---|
| 174545385 | 174546335 | -1 | TEC | ENSG0000C |
| 174547141 | 174776720 | 1 | lncRNA | ENSG0000C |
| 174557966 | 174558072 | -1 | snRNA | ENSG0000C |
| 174559572 | 174682916 | -1 | protein_coding | ENSG0000C |
| 174575227 | 174587726 | 1 | lncRNA | ENSG0000C |
| 174719908 | 174720318 | 1 | processed_pseudoge | ENSG0000C |
| 174747592 | 174787935 | -1 | protein_coding | ENSG0000C |
| 174799313 | 175005381 | -1 | protein_coding | ENSG0000C |
| 175022214 | 175022319 | 1 | snRNA | ENSG0000C |
| 175029769 | 175029877 | 1 | snRNA | ENSG0000C |
| 175033778 | 175034192 | -1 | processed_pseudoge | ENSG0000C |
| 175072250 | 175168382 | -1 | protein_coding | ENSG0000C |
| 175092401 | 175092587 | -1 | misc_RNA | ENSG0000C |
| 175167633 | 175167709 | -1 | miRNA | ENSG0000C |
| 175167904 | 175168522 | 1 | lncRNA | ENSG0000C |
| 175176258 | 175184607 | -1 | protein_coding | ENSG0000C |
| 175176493 | 175176586 | -1 | misc_RNA | ENSG0000C |
| 175184215 | 175184607 | 1 | processed_pseudoge | ENSG0000C |
| 175257250 | 175463180 | 1 | lncRNA | ENSG0000C |
| 175842887 | 175843302 | 1 | processed_pseudoge | ENSG0000C |
| 175897336 | 175904232 | 1 | lncRNA | ENSG0000C |
| 175904371 | 175905502 | -1 | lncRNA | ENSG0000C |
| 175923882 | 176002839 | -1 | protein_coding | ENSG0000C |
| 176077472 | 176083913 | -1 | protein_coding | ENSG0000C |
| 176092721 | 176095944 | 1 | protein_coding | ENSG0000C |
| 176099730 | 176101193 | 1 | protein_coding | ENSG0000C |
| 176104216 | 176109754 | 1 | protein_coding | ENSG0000C |
| 176108790 | 176119942 | 1 | protein_coding | ENSG0000C |
| 176121611 | 176137098 | -1 | lncRNA | ENSG0000C |
| 176122719 | 176124937 | 1 | protein_coding | ENSG0000C |
| 176129694 | 176132695 | 1 | protein_coding | ENSG0000C |
| 176136612 | 176173102 | 1 | protein_coding | ENSG0000C |
| 176150303 | 176150412 | 1 | miRNA | ENSG0000C |
| 176151550 | 176153226 | 1 | protein_coding | ENSG0000C |
| 176164051 | 176188958 | -1 | lncRNA | ENSG0000C |
| 176164164 | 176165716 | -1 | lncRNA | ENSG0000C |
| 176177717 | 176179008 | 1 | lncRNA | ENSG0000C |
| 176188668 | 176190907 | 1 | protein_coding | ENSG0000C |
| 176188843 | 176188901 | 1 | miRNA | ENSG0000C |
| 176189919 | 176192538 | -1 | TEC | ENSG0000C |
| 176200908 | 176201252 | -1 | processed_pseudoge | ENSG0000C |
| 176242111 | 176242993 | 1 | processed_pseudoge | ENSG0000C |
| 176269395 | 176338025 | 1 | protein_coding | ENSG0000C |
| 176338637 | 176339077 | 1 | processed_pseudoge | ENSG0000C |
| 176495255 | 176819180 | 1 | lncRNA | ENSG0000C |
| 176506245 | 176507702 | -1 | lncRNA | ENSG0000C |
| 176524879 | 176531683 | -1 | lncRNA | ENSG0000C |
| 176600980 | 176601052 | -1 | miRNA | ENSG0000C |
| 176611437 | 176612249 | -1 | lncRNA | ENSG0000C |
| 176629572 | 176637931 | -1 | lncRNA | ENSG0000C |

| | | | | |
|---|---|---|---|---|
| 176664676 | 176664766 | 1 | snRNA | ENSG00000 |
| 176723614 | 176819310 | -1 | lncRNA | ENSG00000 |
| 176804987 | 176806337 | -1 | processed_pseudoge | ENSG00000 |
| 176809426 | 176809859 | -1 | processed_pseudoge | ENSG00000 |
| 176822030 | 176823653 | 1 | TEC | ENSG00000 |
| 176830839 | 176849428 | -1 | lncRNA | ENSG00000 |
| 176929985 | 176930090 | -1 | snRNA | ENSG00000 |
| 176989418 | 177164656 | -1 | lncRNA | ENSG00000 |
| 177111036 | 177212662 | -1 | lncRNA | ENSG00000 |
| 177138699 | 177138797 | 1 | rRNA_pseudogene | ENSG00000 |
| 177177695 | 177178255 | 1 | processed_pseudoge | ENSG00000 |
| 177194983 | 177195095 | 1 | misc_RNA | ENSG00000 |
| 177197985 | 177198547 | 1 | processed_pseudoge | ENSG00000 |
| 177212563 | 177223958 | 1 | protein_coding | ENSG00000 |
| 177212726 | 177212799 | 1 | miRNA | ENSG00000 |
| 177227595 | 177392697 | -1 | protein_coding | ENSG00000 |
| 177229191 | 177229506 | -1 | processed_pseudoge | ENSG00000 |
| 177255945 | 177256010 | -1 | miRNA | ENSG00000 |
| 177264359 | 177265515 | 1 | lncRNA | ENSG00000 |
| 177283508 | 177392691 | -1 | lncRNA | ENSG00000 |
| 177300600 | 177302006 | 1 | lncRNA | ENSG00000 |
| 177306373 | 177310572 | 1 | lncRNA | ENSG00000 |
| 177313806 | 177313882 | -1 | miRNA | ENSG00000 |
| 177317715 | 177318471 | -1 | lncRNA | ENSG00000 |
| 177344443 | 177344836 | 1 | processed_pseudoge | ENSG00000 |
| 177345858 | 177345961 | -1 | snoRNA | ENSG00000 |
| 177392757 | 177559299 | 1 | protein_coding | ENSG00000 |
| 177548998 | 177552796 | -1 | protein_coding | ENSG00000 |
| 177603089 | 177618572 | 1 | lncRNA | ENSG00000 |
| 177612999 | 177618742 | -1 | protein_coding | ENSG00000 |
| 177623244 | 178072777 | -1 | protein_coding | ENSG00000 |
| 177628069 | 178108339 | -1 | protein_coding | ENSG00000 |
| 177653419 | 177723289 | 1 | lncRNA | ENSG00000 |
| 177953111 | 178011989 | -1 | lncRNA | ENSG00000 |
| 177997273 | 177998770 | -1 | processed_pseudoge | ENSG00000 |
| 178017993 | 178018465 | 1 | processed_pseudoge | ENSG00000 |
| 178038902 | 178039004 | 1 | snRNA | ENSG00000 |
| 178092503 | 178104470 | -1 | unitary_pseudogene | ENSG00000 |
| 178105637 | 178105753 | 1 | processed_pseudoge | ENSG00000 |
| 178112424 | 178139011 | 1 | protein_coding | ENSG00000 |
| 178142548 | 178142651 | 1 | snRNA | ENSG00000 |
| 178194481 | 178402893 | 1 | protein_coding | ENSG00000 |
| 178413635 | 178440243 | 1 | lncRNA | ENSG00000 |
| 178431414 | 178451512 | -1 | protein_coding | ENSG00000 |
| 178451346 | 178461409 | 1 | protein_coding | ENSG00000 |
| 178454716 | 178455428 | 1 | processed_pseudoge | ENSG00000 |
| 178463664 | 178478600 | -1 | protein_coding | ENSG00000 |
| 178480457 | 178516463 | 1 | protein_coding | ENSG00000 |
| 178521183 | 178779963 | 1 | lncRNA | ENSG00000 |
| 178525989 | 178830802 | -1 | protein_coding | ENSG00000 |

| | | | | |
|---|---|---|---|---|
| 178541125 | 178541799 | 1 | lncRNA | ENSG0000C |
| 178548884 | 178550681 | 1 | lncRNA | ENSG0000C |
| 178554561 | 178558966 | 1 | TEC | ENSG0000C |
| 178577103 | 178577622 | 1 | lncRNA | ENSG0000C |
| 178578790 | 178580906 | 1 | lncRNA | ENSG0000C |
| 178616581 | 178617123 | 1 | lncRNA | ENSG0000C |
| 178644717 | 178645179 | 1 | lncRNA | ENSG0000C |
| 178723457 | 178727046 | 1 | lncRNA | ENSG0000C |
| 178828349 | 179007807 | 1 | lncRNA | ENSG0000C |
| 178829757 | 179050086 | -1 | protein_coding | ENSG0000C |
| 178831371 | 178831449 | -1 | snRNA | ENSG0000C |
| 179001756 | 179002760 | 1 | processed_pseudoge | ENSG0000C |
| 179101678 | 179264832 | -1 | protein_coding | ENSG0000C |
| 179273831 | 179285707 | 1 | lncRNA | ENSG0000C |
| 179399728 | 179400875 | -1 | processed_pseudoge | ENSG0000C |
| 179441982 | 179861612 | -1 | protein_coding | ENSG0000C |
| 179814251 | 179814675 | 1 | processed_pseudoge | ENSG0000C |
| 179860836 | 179860908 | -1 | miRNA | ENSG0000C |
| 179934401 | 179934538 | -1 | snoRNA | ENSG0000C |
| 179934402 | 179934527 | -1 | snoRNA | ENSG0000C |
| 179944876 | 180007297 | -1 | protein_coding | ENSG0000C |
| 180473137 | 180473419 | -1 | processed_pseudoge | ENSG0000C |
| 180571712 | 180692454 | -1 | lncRNA | ENSG0000C |
| 180692104 | 180916939 | 1 | lncRNA | ENSG0000C |
| 180801227 | 180801533 | 1 | TEC | ENSG0000C |
| 180872867 | 180873414 | -1 | processed_pseudoge | ENSG0000C |
| 180938830 | 180939298 | -1 | processed_pseudoge | ENSG0000C |
| 180967248 | 181076585 | 1 | protein_coding | ENSG0000C |
| 180979427 | 180980090 | -1 | lncRNA | ENSG0000C |
| 181076051 | 181105968 | -1 | lncRNA | ENSG0000C |
| 181086076 | 181409321 | 1 | lncRNA | ENSG0000C |
| 181305593 | 181305652 | -1 | miRNA | ENSG0000C |
| 181422154 | 181425749 | -1 | lncRNA | ENSG0000C |
| 181457202 | 181538940 | 1 | protein_coding | ENSG0000C |
| 181536672 | 181680665 | -1 | protein_coding | ENSG0000C |
| 181673088 | 181680876 | -1 | protein_coding | ENSG0000C |
| 181683113 | 181685707 | 1 | lncRNA | ENSG0000C |
| 181690380 | 181693415 | -1 | lncRNA | ENSG0000C |
| 181694442 | 181694900 | 1 | processed_pseudoge | ENSG0000C |
| 181738606 | 181738715 | 1 | snRNA | ENSG0000C |
| 181887851 | 181891663 | -1 | lncRNA | ENSG0000C |
| 181891730 | 181930738 | 1 | protein_coding | ENSG0000C |
| 181954241 | 182131398 | 1 | protein_coding | ENSG0000C |
| 181961212 | 181962478 | -1 | processed_pseudoge | ENSG0000C |
| 182048822 | 182048905 | 1 | rRNA_pseudogene | ENSG0000C |
| 182140036 | 182523192 | -1 | protein_coding | ENSG0000C |
| 182278684 | 182282752 | 1 | lncRNA | ENSG0000C |
| 182314284 | 182314544 | 1 | misc_RNA | ENSG0000C |
| 182716041 | 182794464 | 1 | protein_coding | ENSG0000C |
| 182788098 | 182788435 | 1 | processed_pseudoge | ENSG0000C |

| | | | | |
|---|---|---|---|---|
| 182828124 | 182828221 | 1 | misc_RNA | ENSG00000( |
| 182833275 | 182866637 | -1 | protein_coding | ENSG00000( |
| 182873841 | 182873947 | 1 | snRNA | ENSG00000( |
| 182909115 | 183038858 | -1 | protein_coding | ENSG00000( |
| 183022859 | 183022971 | -1 | misc_RNA | ENSG00000( |
| 183064235 | 183064346 | -1 | snoRNA | ENSG00000( |
| 183071040 | 183072486 | -1 | processed_pseudoge | ENSG00000( |
| 183078559 | 183100008 | 1 | protein_coding | ENSG00000( |
| 183083405 | 183108519 | -1 | lncRNA | ENSG00000( |
| 183117513 | 183161680 | 1 | protein_coding | ENSG00000( |
| 183178806 | 183215414 | 1 | lncRNA | ENSG00000( |
| 183253652 | 183495501 | 1 | lncRNA | ENSG00000( |
| 183260642 | 183261046 | 1 | processed_pseudoge | ENSG00000( |
| 183607442 | 183608119 | -1 | processed_pseudoge | ENSG00000( |
| 183897950 | 183902147 | 1 | lncRNA | ENSG00000( |
| 183904529 | 183949112 | 1 | lncRNA | ENSG00000( |
| 183997372 | 184002264 | 1 | lncRNA | ENSG00000( |
| 184049467 | 184187573 | -1 | lncRNA | ENSG00000( |
| 184204372 | 184382869 | 1 | lncRNA | ENSG00000( |
| 184378975 | 184379044 | 1 | miRNA | ENSG00000( |
| 184593577 | 184599008 | -1 | lncRNA | ENSG00000( |
| 184598529 | 184939492 | 1 | protein_coding | ENSG00000( |
| 184902111 | 184902562 | -1 | processed_pseudoge | ENSG00000( |
| 185425063 | 185505608 | 1 | lncRNA | ENSG00000( |
| 185547135 | 185547818 | -1 | processed_pseudoge | ENSG00000( |
| 185652374 | 185800151 | -1 | lncRNA | ENSG00000( |
| 185661774 | 185661906 | 1 | snoRNA | ENSG00000( |
| 185719874 | 185740479 | -1 | lncRNA | ENSG00000( |
| 185738804 | 185833290 | 1 | protein_coding | ENSG00000( |
| 185950469 | 185953171 | -1 | lncRNA | ENSG00000( |
| 185956587 | 185957063 | -1 | processed_pseudoge | ENSG00000( |
| 186032884 | 186422432 | -1 | lncRNA | ENSG00000( |
| 186162949 | 186163245 | -1 | processed_pseudoge | ENSG00000( |
| 186261419 | 186261919 | -1 | processed_pseudoge | ENSG00000( |
| 186354570 | 186356773 | -1 | lncRNA | ENSG00000( |
| 186364598 | 186365128 | 1 | processed_pseudoge | ENSG00000( |
| 186486253 | 186509361 | 1 | protein_coding | ENSG00000( |
| 186488624 | 186489083 | -1 | processed_pseudoge | ENSG00000( |
| 186590056 | 186680901 | 1 | protein_coding | ENSG00000( |
| 186641339 | 186695287 | -1 | lncRNA | ENSG00000( |
| 186694060 | 186765959 | 1 | protein_coding | ENSG00000( |
| 186827475 | 186849208 | -1 | protein_coding | ENSG00000( |
| 187001876 | 187003377 | -1 | processed_pseudoge | ENSG00000( |
| 187003220 | 187556288 | 1 | lncRNA | ENSG00000( |
| 187122562 | 187122861 | -1 | misc_RNA | ENSG00000( |
| 187270698 | 187270804 | -1 | snRNA | ENSG00000( |
| 187341964 | 187448460 | -1 | protein_coding | ENSG00000( |
| 187415552 | 187415818 | 1 | processed_pseudoge | ENSG00000( |
| 187464230 | 187565760 | -1 | protein_coding | ENSG00000( |
| 187712816 | 188384313 | -1 | lncRNA | ENSG00000( |

| | | | | |
|---|---|---|---|---|
| 187825341 | 187826449 | 1 | processed_pseudoge | ENSG00000 |
| 188227189 | 188228382 | 1 | processed_pseudoge | ENSG00000 |
| 188276732 | 188276844 | -1 | rRNA_pseudogene | ENSG00000 |
| 188291669 | 188595931 | 1 | protein_coding | ENSG00000 |
| 188297492 | 188297588 | 1 | miRNA | ENSG00000 |
| 188598791 | 188839445 | -1 | lncRNA | ENSG00000 |
| 188733738 | 188790104 | -1 | protein_coding | ENSG00000 |
| 188974320 | 189012746 | 1 | protein_coding | ENSG00000 |
| 188978092 | 188978161 | 1 | miRNA | ENSG00000 |
| 188978093 | 188978161 | -1 | miRNA | ENSG00000 |
| 188995630 | 188995692 | 1 | miRNA | ENSG00000 |
| 189031898 | 189225312 | -1 | protein_coding | ENSG00000 |
| 189095063 | 189096158 | 1 | lncRNA | ENSG00000 |
| 189133036 | 189133111 | -1 | miRNA | ENSG00000 |
| 189311263 | 189311967 | -1 | processed_pseudoge | ENSG00000 |
| 189441446 | 189475552 | 1 | protein_coding | ENSG00000 |
| 189486480 | 189487992 | -1 | processed_pseudoge | ENSG00000 |
| 189537384 | 189540417 | -1 | lncRNA | ENSG00000 |
| 189560579 | 189583758 | -1 | protein_coding | ENSG00000 |
| 189625443 | 189626259 | -1 | unprocessed_pseudo | ENSG00000 |
| 189661385 | 189670831 | 1 | protein_coding | ENSG00000 |
| 189661452 | 189666437 | 1 | protein_coding | ENSG00000 |
| 189661519 | 189667881 | 1 | protein_coding | ENSG00000 |
| 189674290 | 189761193 | 1 | protein_coding | ENSG00000 |
| 189746660 | 189763227 | -1 | protein_coding | ENSG00000 |
| 189762704 | 189765556 | 1 | lncRNA | ENSG00000 |
| 189763859 | 189764456 | -1 | lncRNA | ENSG00000 |
| 189770267 | 189784364 | -1 | protein_coding | ENSG00000 |
| 189784085 | 189877629 | 1 | protein_coding | ENSG00000 |
| 189879609 | 190203484 | 1 | protein_coding | ENSG00000 |
| 189917254 | 189917758 | -1 | processed_pseudoge | ENSG00000 |
| 189923336 | 189924216 | 1 | processed_pseudoge | ENSG00000 |
| 189930277 | 189931010 | -1 | processed_pseudoge | ENSG00000 |
| 190055700 | 190062729 | -1 | protein_coding | ENSG00000 |
| 190189735 | 190344193 | -1 | protein_coding | ENSG00000 |
| 190343570 | 190371665 | 1 | protein_coding | ENSG00000 |
| 190408355 | 190509205 | 1 | protein_coding | ENSG00000 |
| 190454092 | 190454521 | -1 | lncRNA | ENSG00000 |
| 190504338 | 190534722 | -1 | protein_coding | ENSG00000 |
| 190534855 | 190569776 | 1 | lncRNA | ENSG00000 |
| 190553264 | 190553480 | -1 | misc_RNA | ENSG00000 |
| 190607660 | 190649840 | -1 | lncRNA | ENSG00000 |
| 190646746 | 190692766 | 1 | protein_coding | ENSG00000 |
| 190676944 | 190708716 | -1 | lncRNA | ENSG00000 |
| 190880797 | 190882059 | -1 | lncRNA | ENSG00000 |
| 190880827 | 190965552 | 1 | protein_coding | ENSG00000 |
| 190964358 | 191020960 | -1 | protein_coding | ENSG00000 |
| 190992639 | 190993567 | -1 | processed_pseudoge | ENSG00000 |
| 191017954 | 191019079 | 1 | lncRNA | ENSG00000 |
| 191021526 | 191032314 | 1 | lncRNA | ENSG00000 |

| | | | | |
|---|---|---|---|---|
| 191029576 | 191151596 | -1 | protein_coding | ENSG00000 |
| 191045458 | 191065345 | 1 | lncRNA | ENSG00000 |
| 191121950 | 191122056 | 1 | snRNA | ENSG00000 |
| 191154118 | 191156070 | -1 | TEC | ENSG00000 |
| 191174233 | 191174835 | 1 | processed_pseudoge | ENSG00000 |
| 191229165 | 191246172 | -1 | lncRNA | ENSG00000 |
| 191245185 | 191425389 | 1 | protein_coding | ENSG00000 |
| 191370938 | 191371044 | 1 | snRNA | ENSG00000 |
| 191678068 | 191696664 | 1 | protein_coding | ENSG00000 |
| 191682452 | 191682633 | 1 | misc_RNA | ENSG00000 |
| 191792797 | 191820257 | 1 | lncRNA | ENSG00000 |
| 191834310 | 191847088 | -1 | protein_coding | ENSG00000 |
| 191846539 | 192044525 | 1 | lncRNA | ENSG00000 |
| 191881182 | 191882205 | -1 | processed_pseudoge | ENSG00000 |
| 191949043 | 192194933 | -1 | protein_coding | ENSG00000 |
| 192629919 | 192645706 | 1 | lncRNA | ENSG00000 |
| 192644102 | 192645387 | -1 | lncRNA | ENSG00000 |
| 192749845 | 192776899 | 1 | lncRNA | ENSG00000 |
| 192774082 | 192774416 | 1 | processed_pseudoge | ENSG00000 |
| 192775943 | 192776073 | 1 | misc_RNA | ENSG00000 |
| 193170704 | 193172765 | 1 | lncRNA | ENSG00000 |
| 193256483 | 193257757 | 1 | processed_pseudoge | ENSG00000 |
| 193276668 | 193277614 | -1 | processed_pseudoge | ENSG00000 |
| 193730555 | 193730689 | -1 | processed_pseudoge | ENSG00000 |
| 193867722 | 193868982 | 1 | lncRNA | ENSG00000 |
| 193898367 | 193899394 | 1 | lncRNA | ENSG00000 |
| 194027919 | 194028164 | 1 | processed_pseudoge | ENSG00000 |
| 194129354 | 194130266 | -1 | processed_pseudoge | ENSG00000 |
| 194187379 | 194188309 | 1 | processed_pseudoge | ENSG00000 |
| 194344190 | 194576578 | 1 | lncRNA | ENSG00000 |
| 194454581 | 194455142 | -1 | processed_pseudoge | ENSG00000 |
| 194587925 | 194612145 | 1 | lncRNA | ENSG00000 |
| 194658376 | 194658675 | -1 | misc_RNA | ENSG00000 |
| 194664107 | 194664174 | -1 | snoRNA | ENSG00000 |
| 194730595 | 194761435 | 1 | lncRNA | ENSG00000 |
| 195003711 | 195026370 | -1 | lncRNA | ENSG00000 |
| 195448532 | 195481827 | 1 | lncRNA | ENSG00000 |
| 195451778 | 195515699 | -1 | lncRNA | ENSG00000 |
| 195514036 | 195514142 | 1 | snRNA | ENSG00000 |
| 195532945 | 195540287 | 1 | lncRNA | ENSG00000 |
| 195569628 | 195572970 | -1 | lncRNA | ENSG00000 |
| 195575977 | 195737702 | 1 | protein_coding | ENSG00000 |
| 195582794 | 195584031 | -1 | processed_pseudoge | ENSG00000 |
| 195670372 | 195670474 | -1 | snRNA | ENSG00000 |
| 195737703 | 196068837 | -1 | protein_coding | ENSG00000 |
| 195994259 | 195995680 | -1 | processed_pseudoge | ENSG00000 |
| 196063312 | 196063963 | -1 | processed_pseudoge | ENSG00000 |
| 196133583 | 196176503 | -1 | protein_coding | ENSG00000 |
| 196151263 | 196154881 | 1 | lncRNA | ENSG00000 |
| 196189099 | 196593684 | -1 | protein_coding | ENSG00000 |

| | | | | |
|---|---|---|---|---|
| 196260024 | 196264204 | 1 | lncRNA | ENSG00000 |
| 196289409 | 196289707 | 1 | misc_RNA | ENSG00000 |
| 196638422 | 196639497 | -1 | lncRNA | ENSG00000 |
| 196639554 | 196763490 | 1 | protein_coding | ENSG00000 |
| 196686148 | 196686330 | -1 | scaRNA | ENSG00000 |
| 196763035 | 196799725 | -1 | protein_coding | ENSG00000 |
| 196804415 | 196810276 | -1 | protein_coding | ENSG00000 |
| 196831975 | 196832190 | -1 | TEC | ENSG00000 |
| 196833004 | 196927796 | -1 | protein_coding | ENSG00000 |
| 196967017 | 197311173 | -1 | protein_coding | ENSG00000 |
| 197014990 | 197016015 | 1 | processed_pseudoge | ENSG00000 |
| 197028711 | 197030143 | -1 | processed_pseudoge | ENSG00000 |
| 197197991 | 197199273 | 1 | lncRNA | ENSG00000 |
| 197250858 | 197302519 | -1 | lncRNA | ENSG00000 |
| 197263111 | 197263535 | 1 | processed_pseudoge | ENSG00000 |
| 197311393 | 197312740 | 1 | lncRNA | ENSG00000 |
| 197379701 | 197380892 | -1 | processed_pseudoge | ENSG00000 |
| 197388515 | 197435091 | -1 | protein_coding | ENSG00000 |
| 197404718 | 197404854 | 1 | snoRNA | ENSG00000 |
| 197447745 | 197447851 | 1 | snRNA | ENSG00000 |
| 197453423 | 197475308 | 1 | protein_coding | ENSG00000 |
| 197486584 | 197516737 | -1 | protein_coding | ENSG00000 |
| 197500140 | 197503449 | 1 | protein_coding | ENSG00000 |
| 197500413 | 197550726 | 1 | protein_coding | ENSG00000 |
| 197515571 | 197553699 | 1 | protein_coding | ENSG00000 |
| 197533803 | 197533965 | 1 | processed_pseudoge | ENSG00000 |
| 197568224 | 197676045 | -1 | protein_coding | ENSG00000 |
| 197640441 | 197641574 | 1 | lncRNA | ENSG00000 |
| 197693106 | 197774823 | 1 | lncRNA | ENSG00000 |
| 197705369 | 197708395 | 1 | protein_coding | ENSG00000 |
| 197726879 | 197786762 | -1 | protein_coding | ENSG00000 |
| 197804593 | 198572581 | 1 | protein_coding | ENSG00000 |
| 197817423 | 197836326 | -1 | lncRNA | ENSG00000 |
| 198187802 | 198191521 | -1 | lncRNA | ENSG00000 |
| 198299276 | 198375490 | -1 | lncRNA | ENSG00000 |
| 198493242 | 198772356 | -1 | lncRNA | ENSG00000 |
| 198882573 | 199071791 | -1 | lncRNA | ENSG00000 |
| 199108584 | 199110088 | 1 | processed_pseudoge | ENSG00000 |
| 199191059 | 199191120 | 1 | snRNA | ENSG00000 |
| 199269500 | 199471266 | -1 | protein_coding | ENSG00000 |
| 199325310 | 199329362 | 1 | lncRNA | ENSG00000 |
| 199457700 | 199476935 | 1 | lncRNA | ENSG00000 |
| 199608068 | 199750341 | 1 | lncRNA | ENSG00000 |
| 199659551 | 199660729 | 1 | processed_pseudoge | ENSG00000 |
| 199760544 | 199851173 | -1 | protein_coding | ENSG00000 |
| 199867396 | 199911159 | -1 | lncRNA | ENSG00000 |
| 199872172 | 199872378 | 1 | processed_pseudoge | ENSG00000 |
| 199901081 | 199901377 | -1 | misc_RNA | ENSG00000 |
| 199911293 | 199955935 | 1 | protein_coding | ENSG00000 |
| 199928913 | 199955736 | -1 | protein_coding | ENSG00000 |

| | | | | |
|---|---|---|---|---|
| 199955317 | 200008540 | 1 | protein_coding | ENSG00000( |
| 200086870 | 200100025 | 1 | lncRNA | ENSG00000( |
| 200305881 | 200482264 | 1 | protein_coding | ENSG00000( |
| 200373175 | 200373269 | 1 | misc_RNA | ENSG00000( |
| 200390666 | 200478018 | -1 | lncRNA | ENSG00000( |
| 200488958 | 200519784 | -1 | protein_coding | ENSG00000( |
| 200510008 | 200584096 | 1 | protein_coding | ENSG00000( |
| 200586014 | 200677064 | 1 | protein_coding | ENSG00000( |
| 200678233 | 200734109 | 1 | transcribed_unproce | ENSG00000( |
| 200695723 | 200748040 | 1 | lncRNA | ENSG00000( |
| 200711535 | 200735186 | -1 | lncRNA | ENSG00000( |
| 200738608 | 200794239 | 1 | transcribed_unproce | ENSG00000( |
| 200775584 | 200775750 | 1 | snRNA | ENSG00000( |
| 200780495 | 200812170 | -1 | lncRNA | ENSG00000( |
| 200810594 | 200827338 | 1 | protein_coding | ENSG00000( |
| 200830009 | 200830116 | 1 | snRNA | ENSG00000( |
| 200844240 | 200845461 | 1 | processed_pseudoge | ENSG00000( |
| 200849038 | 200849144 | -1 | rRNA_pseudogene | ENSG00000( |
| 200853009 | 200864744 | -1 | protein_coding | ENSG00000( |
| 200863152 | 200863247 | -1 | misc_RNA | ENSG00000( |
| 200870907 | 200889303 | -1 | protein_coding | ENSG00000( |
| 200881715 | 200881821 | -1 | snRNA | ENSG00000( |
| 200889327 | 200903938 | 1 | protein_coding | ENSG00000( |
| 200899020 | 200899121 | 1 | snRNA | ENSG00000( |
| 200908977 | 200963680 | -1 | protein_coding | ENSG00000( |
| 200963263 | 201009102 | 1 | lncRNA | ENSG00000( |
| 200973718 | 201071671 | -1 | protein_coding | ENSG00000( |
| 201063089 | 201064051 | 1 | processed_pseudoge | ENSG00000( |
| 201066691 | 201067191 | 1 | processed_pseudoge | ENSG00000( |
| 201071433 | 201085750 | 1 | protein_coding | ENSG00000( |
| 201079842 | 201079944 | 1 | snRNA | ENSG00000( |
| 201097744 | 201098091 | 1 | misc_RNA | ENSG00000( |
| 201101382 | 201103136 | 1 | lncRNA | ENSG00000( |
| 201102089 | 201102957 | -1 | processed_pseudoge | ENSG00000( |
| 201116154 | 201176687 | 1 | protein_coding | ENSG00000( |
| 201137516 | 201140027 | -1 | processed_pseudoge | ENSG00000( |
| 201140278 | 201157823 | -1 | lncRNA | ENSG00000( |
| 201141904 | 201141966 | 1 | snRNA | ENSG00000( |
| 201162484 | 201162680 | -1 | processed_pseudoge | ENSG00000( |
| 201166965 | 201167546 | 1 | lncRNA | ENSG00000( |
| 201182881 | 201229406 | 1 | protein_coding | ENSG00000( |
| 201212283 | 201213081 | -1 | unprocessed_pseudc | ENSG00000( |
| 201213298 | 201214660 | -1 | unprocessed_pseudc | ENSG00000( |
| 201214653 | 201214950 | -1 | unprocessed_pseudc | ENSG00000( |
| 201233443 | 201287711 | 1 | protein_coding | ENSG00000( |
| 201288271 | 201357398 | -1 | protein_coding | ENSG00000( |
| 201377207 | 201451500 | -1 | protein_coding | ENSG00000( |
| 201387858 | 201480846 | 1 | protein_coding | ENSG00000( |
| 201410544 | 201413308 | -1 | processed_pseudoge | ENSG00000( |
| 201423513 | 201423604 | 1 | misc_RNA | ENSG00000( |

| | | | | |
|---|---|---|---|---|
| 201487421 | 201619178 | -1 | protein_coding | ENSG00000( |
| 201549631 | 201550307 | 1 | unprocessed_pseudc | ENSG00000( |
| 201550524 | 201551049 | 1 | unprocessed_pseudc | ENSG00000( |
| 201552859 | 201553612 | 1 | unprocessed_pseudc | ENSG00000( |
| 201553688 | 201554031 | 1 | unprocessed_pseudc | ENSG00000( |
| 201554096 | 201554390 | 1 | unprocessed_pseudc | ENSG00000( |
| 201554384 | 201556051 | 1 | unprocessed_pseudc | ENSG00000( |
| 201556892 | 201558265 | 1 | unprocessed_pseudc | ENSG00000( |
| 201620184 | 201643570 | -1 | protein_coding | ENSG00000( |
| 201621646 | 201623430 | -1 | processed_pseudoge | ENSG00000( |
| 201643398 | 201645990 | 1 | lncRNA | ENSG00000( |
| 201644870 | 201698694 | -1 | protein_coding | ENSG00000( |
| 201646716 | 201646820 | 1 | snRNA | ENSG00000( |
| 201700267 | 201780956 | -1 | protein_coding | ENSG00000( |
| 201762737 | 201763573 | 1 | processed_pseudoge | ENSG00000( |
| 201790461 | 201895550 | 1 | protein_coding | ENSG00000( |
| 201808681 | 201808782 | 1 | misc_RNA | ENSG00000( |
| 201824548 | 201824832 | 1 | processed_pseudoge | ENSG00000( |
| 201927439 | 201927543 | 1 | snRNA | ENSG00000( |
| 202032770 | 202033537 | -1 | lncRNA | ENSG00000( |
| 202034587 | 202038445 | 1 | protein_coding | ENSG00000( |
| 202073255 | 202205188 | 1 | protein_coding | ENSG00000( |
| 202075410 | 202117062 | -1 | lncRNA | ENSG00000( |
| 202137570 | 202149627 | -1 | lncRNA | ENSG00000( |
| 202178660 | 202179391 | -1 | lncRNA | ENSG00000( |
| 202179297 | 202179971 | 1 | processed_pseudoge | ENSG00000( |
| 202201384 | 202201886 | -1 | processed_pseudoge | ENSG00000( |
| 202206180 | 202238608 | -1 | protein_coding | ENSG00000( |
| 202231696 | 202231849 | 1 | processed_pseudoge | ENSG00000( |
| 202265736 | 202303661 | 1 | protein_coding | ENSG00000( |
| 202276431 | 202276518 | 1 | snoRNA | ENSG00000( |
| 202278108 | 202278192 | 1 | snoRNA | ENSG00000( |
| 202291332 | 202291421 | 1 | snoRNA | ENSG00000( |
| 202293051 | 202293134 | 1 | snoRNA | ENSG00000( |
| 202302122 | 202303614 | -1 | lncRNA | ENSG00000( |
| 202333439 | 202333728 | 1 | misc_RNA | ENSG00000( |
| 202336024 | 202336727 | -1 | processed_pseudoge | ENSG00000( |
| 202336739 | 202337200 | 1 | lncRNA | ENSG00000( |
| 202357076 | 202357417 | -1 | misc_RNA | ENSG00000( |
| 202374932 | 202375604 | -1 | lncRNA | ENSG00000( |
| 202376327 | 202567751 | 1 | protein_coding | ENSG00000( |
| 202505012 | 202505343 | -1 | processed_pseudoge | ENSG00000( |
| 202520401 | 202520989 | 1 | processed_pseudoge | ENSG00000( |
| 202614214 | 202614704 | 1 | unprocessed_pseudc | ENSG00000( |
| 202614853 | 202615529 | 1 | unprocessed_pseudc | ENSG00000( |
| 202615769 | 202616440 | 1 | unprocessed_pseudc | ENSG00000( |
| 202616700 | 202617212 | 1 | unprocessed_pseudc | ENSG00000( |
| 202617284 | 202617627 | 1 | unprocessed_pseudc | ENSG00000( |
| 202617695 | 202617987 | 1 | unprocessed_pseudc | ENSG00000( |
| 202617983 | 202618881 | 1 | unprocessed_pseudc | ENSG00000( |

| | | | | |
|---|---|---|---|---|
| 202619185 | 202619896 | -1 | processed_pseudoge | ENSG0000C |
| 202634969 | 202769757 | 1 | protein_coding | ENSG0000C |
| 202773150 | 202871985 | -1 | protein_coding | ENSG0000C |
| 202773720 | 202774360 | -1 | processed_pseudoge | ENSG0000C |
| 202840331 | 202841781 | -1 | processed_pseudoge | ENSG0000C |
| 202874782 | 203014798 | -1 | protein_coding | ENSG0000C |
| 202912214 | 202988263 | 1 | protein_coding | ENSG0000C |
| 202924609 | 202925904 | 1 | processed_pseudoge | ENSG0000C |
| 202995194 | 202995403 | 1 | processed_pseudoge | ENSG0000C |
| 203014879 | 203226378 | 1 | protein_coding | ENSG0000C |
| 203040108 | 203040899 | 1 | processed_pseudoge | ENSG0000C |
| 203052285 | 203052418 | -1 | snoRNA | ENSG0000C |
| 203061314 | 203062403 | -1 | processed_pseudoge | ENSG0000C |
| 203074685 | 203075142 | -1 | processed_pseudoge | ENSG0000C |
| 203190780 | 203191277 | 1 | processed_pseudoge | ENSG0000C |
| 203238940 | 203305611 | 1 | protein_coding | ENSG0000C |
| 203259604 | 203259897 | 1 | misc_RNA | ENSG0000C |
| 203315390 | 203315871 | -1 | processed_pseudoge | ENSG0000C |
| 203328239 | 203447728 | 1 | protein_coding | ENSG0000C |
| 203328459 | 203329226 | -1 | lncRNA | ENSG0000C |
| 203394345 | 203535335 | -1 | protein_coding | ENSG0000C |
| 203634577 | 203636016 | -1 | processed_pseudoge | ENSG0000C |
| 203672175 | 203672423 | -1 | processed_pseudoge | ENSG0000C |
| 203706475 | 203738912 | 1 | protein_coding | ENSG0000C |
| 203764295 | 203765593 | -1 | processed_pseudoge | ENSG0000C |
| 203772631 | 203773502 | -1 | processed_pseudoge | ENSG0000C |
| 203782037 | 203782143 | 1 | snRNA | ENSG0000C |
| 203867771 | 203873965 | 1 | protein_coding | ENSG0000C |
| 203936748 | 203961577 | 1 | protein_coding | ENSG0000C |
| 204065706 | 204067363 | 1 | processed_pseudoge | ENSG0000C |
| 204090457 | 204109849 | 1 | lncRNA | ENSG0000C |
| 204219984 | 204234985 | -1 | lncRNA | ENSG0000C |
| 204306842 | 204307364 | 1 | processed_pseudoge | ENSG0000C |
| 204379095 | 204379792 | 1 | processed_pseudoge | ENSG0000C |
| 204470076 | 204473837 | -1 | lncRNA | ENSG0000C |
| 204473834 | 204507672 | 1 | lncRNA | ENSG0000C |
| 204545475 | 205620162 | 1 | protein_coding | ENSG0000C |
| 204668863 | 204678698 | -1 | lncRNA | ENSG0000C |
| 205253304 | 205253503 | -1 | processed_pseudoge | ENSG0000C |
| 205681990 | 205798133 | 1 | protein_coding | ENSG0000C |
| 205756469 | 205764006 | -1 | lncRNA | ENSG0000C |
| 205900630 | 205900958 | 1 | misc_RNA | ENSG0000C |
| 205949091 | 205950145 | 1 | processed_pseudoge | ENSG0000C |
| 205989585 | 206000858 | 1 | lncRNA | ENSG0000C |
| 205993721 | 206086303 | -1 | protein_coding | ENSG0000C |
| 206007880 | 206007974 | -1 | misc_RNA | ENSG0000C |
| 206025593 | 206025697 | 1 | misc_RNA | ENSG0000C |
| 206040432 | 206040845 | 1 | processed_pseudoge | ENSG0000C |
| 206084605 | 206086564 | 1 | lncRNA | ENSG0000C |
| 206114817 | 206159509 | -1 | protein_coding | ENSG0000C |

| | | | | |
|---|---|---|---|---|
| 206115547 | 206122323 | 1 | lncRNA | ENSG0000C |
| 206116110 | 206116500 | -1 | processed_pseudoge | ENSG0000C |
| 206159585 | 206162928 | 1 | protein_coding | ENSG0000C |
| 206160843 | 206161024 | 1 | misc_RNA | ENSG0000C |
| 206161841 | 206161929 | 1 | snoRNA | ENSG0000C |
| 206161878 | 206161957 | 1 | snoRNA | ENSG0000C |
| 206162228 | 206162359 | 1 | snoRNA | ENSG0000C |
| 206175316 | 206218047 | -1 | protein_coding | ENSG0000C |
| 206203543 | 206265326 | 1 | lncRNA | ENSG0000C |
| 206221315 | 206221634 | 1 | misc_RNA | ENSG0000C |
| 206236236 | 206236871 | -1 | processed_pseudoge | ENSG0000C |
| 206259628 | 206259918 | -1 | processed_pseudoge | ENSG0000C |
| 206260066 | 206260214 | -1 | misc_RNA | ENSG0000C |
| 206274663 | 206314427 | 1 | protein_coding | ENSG0000C |
| 206410087 | 206410909 | -1 | processed_pseudoge | ENSG0000C |
| 206419292 | 206420238 | -1 | processed_pseudoge | ENSG0000C |
| 206443532 | 206621127 | 1 | protein_coding | ENSG0000C |
| 206606497 | 206609812 | -1 | lncRNA | ENSG0000C |
| 206640073 | 206641282 | 1 | lncRNA | ENSG0000C |
| 206642412 | 206649011 | 1 | protein_coding | ENSG0000C |
| 206651621 | 206718396 | -1 | protein_coding | ENSG0000C |
| 206686517 | 206687362 | -1 | processed_pseudoge | ENSG0000C |
| 206701065 | 206701185 | -1 | misc_RNA | ENSG0000C |
| 206737763 | 206765328 | -1 | protein_coding | ENSG0000C |
| 206765357 | 206796189 | 1 | protein_coding | ENSG0000C |
| 206783234 | 206783308 | -1 | miRNA | ENSG0000C |
| 206783234 | 206783308 | 1 | miRNA | ENSG0000C |
| 206798325 | 206926876 | 1 | lncRNA | ENSG0000C |
| 206821190 | 206822294 | -1 | lncRNA | ENSG0000C |
| 206939518 | 206969474 | 1 | protein_coding | ENSG0000C |
| 207074137 | 207167267 | -1 | protein_coding | ENSG0000C |
| 207109987 | 207110073 | -1 | miRNA | ENSG0000C |
| 207120884 | 207122044 | -1 | lncRNA | ENSG0000C |
| 207166120 | 207248668 | 1 | lncRNA | ENSG0000C |
| 207166400 | 207166498 | 1 | miRNA | ENSG0000C |
| 207173634 | 207222790 | 1 | lncRNA | ENSG0000C |
| 207186717 | 207236066 | -1 | lncRNA | ENSG0000C |
| 207217532 | 207219145 | 1 | lncRNA | ENSG0000C |
| 207226949 | 207228308 | 1 | lncRNA | ENSG0000C |
| 207235299 | 207237875 | -1 | lncRNA | ENSG0000C |
| 207239864 | 207529795 | -1 | lncRNA | ENSG0000C |
| 207260445 | 207270690 | 1 | lncRNA | ENSG0000C |
| 207269275 | 207269424 | -1 | miRNA | ENSG0000C |
| 207529737 | 207605988 | 1 | protein_coding | ENSG0000C |
| 207580631 | 207625928 | -1 | protein_coding | ENSG0000C |
| 207654781 | 207654951 | 1 | processed_pseudoge | ENSG0000C |
| 207662375 | 207667024 | 1 | lncRNA | ENSG0000C |
| 207680533 | 207680639 | -1 | snRNA | ENSG0000C |
| 207681590 | 207682033 | 1 | processed_pseudoge | ENSG0000C |
| 207687960 | 207688059 | 1 | misc_RNA | ENSG0000C |

| | | | | |
|---|---|---|---|---|
| 207711540 | 207761839 | 1 | protein_coding | ENSG00000 |
| 207753872 | 207754435 | -1 | lncRNA | ENSG00000 |
| 207754807 | 207754881 | 1 | miRNA | ENSG00000 |
| 207762598 | 207769906 | -1 | protein_coding | ENSG00000 |
| 207821288 | 208025527 | -1 | protein_coding | ENSG00000 |
| 207868582 | 207869915 | -1 | lncRNA | ENSG00000 |
| 208019638 | 208019741 | -1 | snRNA | ENSG00000 |
| 208031383 | 208031953 | -1 | processed_pseudoge | ENSG00000 |
| 208054161 | 208054258 | -1 | rRNA_pseudogene | ENSG00000 |
| 208057248 | 208057745 | -1 | processed_pseudoge | ENSG00000 |
| 208108233 | 208112511 | -1 | transcribed_unproce | ENSG00000 |
| 208121607 | 208124524 | -1 | protein_coding | ENSG00000 |
| 208128137 | 208129828 | -1 | protein_coding | ENSG00000 |
| 208135394 | 208137697 | -1 | processed_pseudoge | ENSG00000 |
| 208142573 | 208146158 | -1 | protein_coding | ENSG00000 |
| 208160740 | 208163589 | -1 | protein_coding | ENSG00000 |
| 208165343 | 208190030 | -1 | protein_coding | ENSG00000 |
| 208208135 | 208208984 | 1 | processed_pseudoge | ENSG00000 |
| 208214724 | 208222625 | -1 | processed_pseudoge | ENSG00000 |
| 208236227 | 208266074 | -1 | protein_coding | ENSG00000 |
| 208255247 | 208256181 | 1 | lncRNA | ENSG00000 |
| 208266255 | 208358746 | 1 | protein_coding | ENSG00000 |
| 208269343 | 208269957 | -1 | processed_pseudoge | ENSG00000 |
| 208359714 | 208854503 | 1 | protein_coding | ENSG00000 |
| 208360631 | 208361369 | -1 | processed_pseudoge | ENSG00000 |
| 208466238 | 208466626 | -1 | processed_pseudoge | ENSG00000 |
| 208469850 | 208540413 | 1 | lncRNA | ENSG00000 |
| 208516908 | 208517230 | 1 | processed_pseudoge | ENSG00000 |
| 208885755 | 208888143 | 1 | lncRNA | ENSG00000 |
| 208920547 | 208920664 | -1 | rRNA_pseudogene | ENSG00000 |
| 209072597 | 209074148 | -1 | processed_pseudoge | ENSG00000 |
| 209145241 | 209147687 | 1 | unprocessed_pseudc | ENSG00000 |
| 209178804 | 209182018 | 1 | processed_pseudoge | ENSG00000 |
| 209178966 | 209181123 | 1 | unprocessed_pseudc | ENSG00000 |
| 209353977 | 209387322 | -1 | lncRNA | ENSG00000 |
| 209424058 | 209734118 | 1 | protein_coding | ENSG00000 |
| 209721913 | 209722021 | -1 | rRNA_pseudogene | ENSG00000 |
| 209771993 | 209999300 | 1 | protein_coding | ENSG00000 |
| 209808804 | 209809580 | 1 | processed_pseudoge | ENSG00000 |
| 210002565 | 210022260 | 1 | protein_coding | ENSG00000 |
| 210021421 | 210171409 | -1 | protein_coding | ENSG00000 |
| 210028417 | 210029156 | 1 | lncRNA | ENSG00000 |
| 210030572 | 210064356 | 1 | lncRNA | ENSG00000 |
| 210088226 | 210089064 | 1 | processed_pseudoge | ENSG00000 |
| 210171518 | 210233976 | 1 | lncRNA | ENSG00000 |
| 210187126 | 210225447 | -1 | protein_coding | ENSG00000 |
| 210265963 | 210266074 | 1 | misc_RNA | ENSG00000 |
| 210290150 | 210315174 | -1 | protein_coding | ENSG00000 |
| 210324759 | 210469246 | 1 | lncRNA | ENSG00000 |
| 210431249 | 210477652 | -1 | protein_coding | ENSG00000 |

| | | | | |
|---|---|---|---|---|
| 210477682 | 210679107 | 1 | protein_coding | ENSG00000 |
| 210617571 | 210619876 | 1 | lncRNA | ENSG00000 |
| 211104934 | 211105732 | 1 | processed_pseudoge | ENSG00000 |
| 211298941 | 211299189 | -1 | processed_pseudoge | ENSG00000 |
| 211375717 | 212538841 | -1 | protein_coding | ENSG00000 |
| 211535653 | 211535769 | 1 | rRNA_pseudogene | ENSG00000 |
| 211777206 | 211777856 | 1 | processed_pseudoge | ENSG00000 |
| 211778273 | 211779763 | 1 | processed_pseudoge | ENSG00000 |
| 212426263 | 212426360 | -1 | miRNA | ENSG00000 |
| 212581357 | 213021545 | -1 | lncRNA | ENSG00000 |
| 212795300 | 212819264 | 1 | lncRNA | ENSG00000 |
| 212832161 | 212832945 | 1 | processed_pseudoge | ENSG00000 |
| 212911003 | 212919809 | 1 | lncRNA | ENSG00000 |
| 212926257 | 212926336 | -1 | miRNA | ENSG00000 |
| 212926257 | 212926336 | 1 | miRNA | ENSG00000 |
| 212932894 | 212937520 | -1 | lncRNA | ENSG00000 |
| 212999691 | 213152427 | -1 | protein_coding | ENSG00000 |
| 213152970 | 213153659 | 1 | lncRNA | ENSG00000 |
| 213155806 | 213167712 | -1 | lncRNA | ENSG00000 |
| 213266995 | 213276152 | -1 | lncRNA | ENSG00000 |
| 213276552 | 213284205 | -1 | lncRNA | ENSG00000 |
| 213284379 | 214410501 | 1 | protein_coding | ENSG00000 |
| 213758067 | 213758159 | 1 | miRNA | ENSG00000 |
| 214241676 | 214684246 | -1 | lncRNA | ENSG00000 |
| 214280638 | 214281452 | -1 | processed_pseudoge | ENSG00000 |
| 214411054 | 214578976 | 1 | protein_coding | ENSG00000 |
| 214510196 | 214536890 | 1 | lncRNA | ENSG00000 |
| 214697727 | 214698059 | 1 | processed_pseudoge | ENSG00000 |
| 214725646 | 214809683 | -1 | protein_coding | ENSG00000 |
| 214810181 | 214963575 | 1 | lncRNA | ENSG00000 |
| 214846947 | 214847081 | 1 | snoRNA | ENSG00000 |
| 214847128 | 214847445 | 1 | processed_pseudoge | ENSG00000 |
| 214931542 | 215138626 | -1 | protein_coding | ENSG00000 |
| 215004782 | 215085488 | 1 | lncRNA | ENSG00000 |
| 215178791 | 215310947 | -1 | lncRNA | ENSG00000 |
| 215274992 | 215277947 | 1 | lncRNA | ENSG00000 |
| 215311956 | 215349773 | 1 | protein_coding | ENSG00000 |
| 215360440 | 215436073 | -1 | protein_coding | ENSG00000 |
| 215436253 | 215436992 | 1 | lncRNA | ENSG00000 |
| 215453688 | 215486639 | 1 | lncRNA | ENSG00000 |
| 215476667 | 215480248 | 1 | lncRNA | ENSG00000 |
| 215530447 | 215545526 | -1 | lncRNA | ENSG00000 |
| 215533133 | 215713895 | 1 | lncRNA | ENSG00000 |
| 215579407 | 215582635 | -1 | lncRNA | ENSG00000 |
| 215611563 | 215848954 | -1 | lncRNA | ENSG00000 |
| 215718043 | 215719424 | 1 | lncRNA | ENSG00000 |
| 215804731 | 215804981 | -1 | processed_pseudoge | ENSG00000 |
| 215869923 | 215870518 | 1 | lncRNA | ENSG00000 |
| 215939308 | 215941719 | -1 | lncRNA | ENSG00000 |
| 215942584 | 216034096 | -1 | protein_coding | ENSG00000 |

| | | | | |
|---|---|---|---|---|
| 215996329 | 216082955 | -1 | protein_coding | ENSG00000( |
| 216081866 | 216102783 | 1 | protein_coding | ENSG00000( |
| 216107464 | 216206303 | 1 | protein_coding | ENSG00000( |
| 216156342 | 216156811 | -1 | processed_pseudoge | ENSG00000( |
| 216174896 | 216176032 | -1 | processed_pseudoge | ENSG00000( |
| 216211404 | 216213519 | 1 | TEC | ENSG00000( |
| 216217045 | 216220192 | 1 | lncRNA | ENSG00000( |
| 216257865 | 216372483 | -1 | protein_coding | ENSG00000( |
| 216259595 | 216266130 | 1 | lncRNA | ENSG00000( |
| 216303325 | 216321737 | 1 | lncRNA | ENSG00000( |
| 216351403 | 216351504 | 1 | misc_RNA | ENSG00000( |
| 216385288 | 216412696 | -1 | lncRNA | ENSG00000( |
| 216412414 | 216483053 | 1 | protein_coding | ENSG00000( |
| 216479030 | 216498761 | -1 | lncRNA | ENSG00000( |
| 216483032 | 216487196 | -1 | lncRNA | ENSG00000( |
| 216498825 | 216579180 | 1 | protein_coding | ENSG00000( |
| 216590426 | 216606938 | -1 | lncRNA | ENSG00000( |
| 216594359 | 216611206 | 1 | lncRNA | ENSG00000( |
| 216610342 | 216610959 | -1 | processed_pseudoge | ENSG00000( |
| 216632828 | 216664436 | 1 | protein_coding | ENSG00000( |
| 216672105 | 216695549 | -1 | protein_coding | ENSG00000( |
| 216694464 | 216994079 | 1 | lncRNA | ENSG00000( |
| 216785774 | 216786144 | -1 | processed_pseudoge | ENSG00000( |
| 216793489 | 216793600 | -1 | rRNA_pseudogene | ENSG00000( |
| 216799608 | 216805335 | 1 | lncRNA | ENSG00000( |
| 216805737 | 216806129 | -1 | processed_pseudoge | ENSG00000( |
| 216821077 | 216831910 | 1 | lncRNA | ENSG00000( |
| 216854389 | 216855132 | -1 | lncRNA | ENSG00000( |
| 216859458 | 216860064 | -1 | protein_coding | ENSG00000( |
| 216859948 | 216871643 | 1 | lncRNA | ENSG00000( |
| 216866332 | 216867813 | -1 | lncRNA | ENSG00000( |
| 216868012 | 216869156 | 1 | lncRNA | ENSG00000( |
| 216870441 | 216873932 | -1 | lncRNA | ENSG00000( |
| 216995906 | 216996490 | 1 | lncRNA | ENSG00000( |
| 217061067 | 217064818 | -1 | lncRNA | ENSG00000( |
| 217251556 | 217251875 | 1 | misc_RNA | ENSG00000( |
| 217261699 | 217336120 | 1 | lncRNA | ENSG00000( |
| 217283370 | 217283523 | 1 | misc_RNA | ENSG00000( |
| 217284019 | 217756593 | -1 | lncRNA | ENSG00000( |
| 217693895 | 217696763 | 1 | processed_pseudoge | ENSG00000( |
| 217760547 | 217761291 | -1 | processed_pseudoge | ENSG00000( |
| 217799588 | 218033982 | -1 | protein_coding | ENSG00000( |
| 217900513 | 217900628 | -1 | miRNA | ENSG00000( |
| 217978707 | 217993747 | 1 | lncRNA | ENSG00000( |
| 218034960 | 218090581 | 1 | protein_coding | ENSG00000( |
| 218059155 | 218065729 | -1 | transcribed_unproce | ENSG00000( |
| 218125289 | 218137251 | 1 | protein_coding | ENSG00000( |
| 218162841 | 218166962 | -1 | protein_coding | ENSG00000( |
| 218200434 | 218201388 | 1 | processed_pseudoge | ENSG00000( |
| 218217141 | 218254356 | 1 | protein_coding | ENSG00000( |

| | | | | |
|---|---|---|---|---|
| 218255319 | 218257366 | 1 | lncRNA | ENSG0000C |
| 218259496 | 218263861 | 1 | protein_coding | ENSG0000C |
| 218264123 | 218270257 | -1 | protein_coding | ENSG0000C |
| 218270392 | 218346793 | 1 | protein_coding | ENSG0000C |
| 218274197 | 218292586 | -1 | protein_coding | ENSG0000C |
| 218280125 | 218280188 | -1 | miRNA | ENSG0000C |
| 218326889 | 218357966 | -1 | lncRNA | ENSG0000C |
| 218341911 | 218341980 | 1 | miRNA | ENSG0000C |
| 218354075 | 218354572 | 1 | processed_pseudoge | ENSG0000C |
| 218356857 | 218368099 | 1 | protein_coding | ENSG0000C |
| 218366665 | 218367835 | -1 | lncRNA | ENSG0000C |
| 218382029 | 218396894 | 1 | protein_coding | ENSG0000C |
| 218398256 | 218405941 | 1 | protein_coding | ENSG0000C |
| 218398743 | 218399219 | -1 | lncRNA | ENSG0000C |
| 218402646 | 218402722 | 1 | miRNA | ENSG0000C |
| 218419121 | 218453295 | 1 | protein_coding | ENSG0000C |
| 218450251 | 218568351 | -1 | protein_coding | ENSG0000C |
| 218500729 | 218501037 | -1 | misc_RNA | ENSG0000C |
| 218568580 | 218597080 | 1 | protein_coding | ENSG0000C |
| 218589214 | 218589320 | -1 | snRNA | ENSG0000C |
| 218607855 | 218637184 | 1 | protein_coding | ENSG0000C |
| 218633256 | 218634014 | -1 | lncRNA | ENSG0000C |
| 218637916 | 218659655 | -1 | protein_coding | ENSG0000C |
| 218658764 | 218663443 | 1 | protein_coding | ENSG0000C |
| 218663892 | 218672002 | -1 | protein_coding | ENSG0000C |
| 218672069 | 218702716 | 1 | protein_coding | ENSG0000C |
| 218710835 | 218755416 | 1 | protein_coding | ENSG0000C |
| 218781749 | 218815293 | 1 | protein_coding | ENSG0000C |
| 218799729 | 218802554 | -1 | lncRNA | ENSG0000C |
| 218818690 | 218819144 | 1 | lncRNA | ENSG0000C |
| 218822383 | 218832086 | -1 | protein_coding | ENSG0000C |
| 218823090 | 218823154 | 1 | miRNA | ENSG0000C |
| 218841599 | 218842065 | 1 | processed_pseudoge | ENSG0000C |
| 218859805 | 218874233 | 1 | protein_coding | ENSG0000C |
| 218880852 | 218899581 | 1 | protein_coding | ENSG0000C |
| 218900811 | 218930636 | -1 | lncRNA | ENSG0000C |
| 218916184 | 218916287 | -1 | snRNA | ENSG0000C |
| 218944629 | 218961903 | -1 | lncRNA | ENSG0000C |
| 218953401 | 218954050 | -1 | processed_pseudoge | ENSG0000C |
| 218959666 | 218962155 | 1 | protein_coding | ENSG0000C |
| 218975393 | 218989940 | 1 | lncRNA | ENSG0000C |
| 218981087 | 218985657 | -1 | protein_coding | ENSG0000C |
| 218990189 | 218993422 | -1 | protein_coding | ENSG0000C |
| 219001645 | 219001708 | -1 | miRNA | ENSG0000C |
| 219002215 | 219015721 | 1 | lncRNA | ENSG0000C |
| 219002846 | 219041527 | -1 | protein_coding | ENSG0000C |
| 219054424 | 219060921 | -1 | protein_coding | ENSG0000C |
| 219058688 | 219058750 | -1 | miRNA | ENSG0000C |
| 219065308 | 219070022 | 1 | lncRNA | ENSG0000C |
| 219069354 | 219069809 | -1 | lncRNA | ENSG0000C |

| | | | | |
|---|---|---|---|---|
| 219075317 | 219160865 | -1 | protein_coding | ENSG00000( |
| 219075329 | 219170827 | -1 | protein_coding | ENSG00000( |
| 219102434 | 219102719 | 1 | misc_RNA | ENSG00000( |
| 219161465 | 219170095 | -1 | protein_coding | ENSG00000( |
| 219171897 | 219178106 | -1 | protein_coding | ENSG00000( |
| 219176225 | 219185475 | 1 | protein_coding | ENSG00000( |
| 219195237 | 219209651 | 1 | protein_coding | ENSG00000( |
| 219198640 | 219200034 | -1 | lncRNA | ENSG00000( |
| 219209772 | 219222738 | -1 | protein_coding | ENSG00000( |
| 219209772 | 219218994 | -1 | protein_coding | ENSG00000( |
| 219219380 | 219229717 | -1 | protein_coding | ENSG00000( |
| 219229783 | 219236679 | 1 | protein_coding | ENSG00000( |
| 219236598 | 219245478 | -1 | protein_coding | ENSG00000( |
| 219245455 | 219250337 | 1 | protein_coding | ENSG00000( |
| 219249710 | 219278170 | -1 | protein_coding | ENSG00000( |
| 219253243 | 219272197 | 1 | protein_coding | ENSG00000( |
| 219279267 | 219286900 | 1 | protein_coding | ENSG00000( |
| 219289623 | 219309648 | -1 | protein_coding | ENSG00000( |
| 219294111 | 219294200 | -1 | miRNA | ENSG00000( |
| 219299002 | 219304130 | 1 | lncRNA | ENSG00000( |
| 219327407 | 219333177 | -1 | protein_coding | ENSG00000( |
| 219358870 | 219358983 | -1 | unprocessed_pseudc | ENSG00000( |
| 219373527 | 219400022 | -1 | protein_coding | ENSG00000( |
| 219388496 | 219403633 | 1 | lncRNA | ENSG00000( |
| 219418377 | 219426734 | 1 | protein_coding | ENSG00000( |
| 219425071 | 219426184 | -1 | lncRNA | ENSG00000( |
| 219427477 | 219429319 | 1 | TEC | ENSG00000( |
| 219434843 | 219493629 | 1 | protein_coding | ENSG00000( |
| 219482073 | 219516877 | -1 | lncRNA | ENSG00000( |
| 219491867 | 219498244 | 1 | protein_coding | ENSG00000( |
| 219496125 | 219498287 | 1 | protein_coding | ENSG00000( |
| 219497611 | 219498246 | -1 | lncRNA | ENSG00000( |
| 219498867 | 219506989 | 1 | protein_coding | ENSG00000( |
| 219514170 | 219538772 | 1 | protein_coding | ENSG00000( |
| 219538948 | 219543809 | -1 | protein_coding | ENSG00000( |
| 219543663 | 219550595 | 1 | protein_coding | ENSG00000( |
| 219547211 | 219547658 | -1 | lncRNA | ENSG00000( |
| 219549073 | 219549147 | -1 | miRNA | ENSG00000( |
| 219550728 | 219571859 | -1 | protein_coding | ENSG00000( |
| 219559083 | 219559626 | 1 | lncRNA | ENSG00000( |
| 219569162 | 219575711 | 1 | protein_coding | ENSG00000( |
| 219590687 | 219590984 | -1 | misc_RNA | ENSG00000( |
| 219597857 | 219616451 | 1 | protein_coding | ENSG00000( |
| 219625059 | 219626693 | -1 | lncRNA | ENSG00000( |
| 219627394 | 219641980 | 1 | protein_coding | ENSG00000( |
| 219645090 | 219645631 | 1 | lncRNA | ENSG00000( |
| 219684030 | 219684795 | -1 | processed_pseudoge | ENSG00000( |
| 219685381 | 219737937 | 1 | lncRNA | ENSG00000( |
| 219697799 | 219698214 | 1 | processed_pseudoge | ENSG00000( |
| 219903900 | 219904923 | -1 | lncRNA | ENSG00000( |

| | | | | |
|---|---|---|---|---|
| 219904983 | 219912834 | -1 | lncRNA | ENSG0000( |
| 219906502 | 219906565 | -1 | miRNA | ENSG0000( |
| 220048822 | 220049272 | -1 | processed_pseudoge | ENSG0000( |
| 220105656 | 220450778 | 1 | lncRNA | ENSG0000( |
| 220127546 | 220212491 | 1 | lncRNA | ENSG0000( |
| 220450679 | 220741493 | 1 | lncRNA | ENSG0000( |
| 220776818 | 220855873 | 1 | lncRNA | ENSG0000( |
| 220811618 | 220853456 | -1 | lncRNA | ENSG0000( |
| 221162771 | 221163053 | 1 | processed_pseudoge | ENSG0000( |
| 221306290 | 221308329 | -1 | lncRNA | ENSG0000( |
| 221418027 | 221574202 | -1 | protein_coding | ENSG0000( |
| 221532668 | 221533008 | 1 | processed_pseudoge | ENSG0000( |
| 221572506 | 221574454 | 1 | lncRNA | ENSG0000( |
| 221592441 | 221592861 | 1 | processed_pseudoge | ENSG0000( |
| 221609324 | 221650218 | 1 | lncRNA | ENSG0000( |
| 221944223 | 221944612 | 1 | processed_pseudoge | ENSG0000( |
| 221961737 | 221963765 | 1 | processed_pseudoge | ENSG0000( |
| 222044525 | 222044994 | -1 | processed_pseudoge | ENSG0000( |
| 222199888 | 222298996 | -1 | protein_coding | ENSG0000( |
| 222298147 | 222305217 | 1 | protein_coding | ENSG0000( |
| 222317242 | 222318653 | -1 | lncRNA | ENSG0000( |
| 222318275 | 222352989 | 1 | transcribed_unproce | ENSG0000( |
| 222404985 | 222405095 | 1 | misc_RNA | ENSG0000( |
| 222424543 | 222562621 | 1 | protein_coding | ENSG0000( |
| 222452400 | 222453126 | -1 | processed_pseudoge | ENSG0000( |
| 222498291 | 222498398 | 1 | snRNA | ENSG0000( |
| 222503883 | 222504843 | -1 | processed_pseudoge | ENSG0000( |
| 222566899 | 222656092 | -1 | protein_coding | ENSG0000( |
| 222671658 | 222709930 | 1 | protein_coding | ENSG0000( |
| 222677677 | 222678289 | -1 | processed_pseudoge | ENSG0000( |
| 222698623 | 222699484 | -1 | processed_pseudoge | ENSG0000( |
| 222726457 | 222726834 | 1 | processed_pseudoge | ENSG0000( |
| 222860942 | 222944639 | 1 | protein_coding | ENSG0000( |
| 222904640 | 222905055 | -1 | processed_pseudoge | ENSG0000( |
| 222917387 | 222919363 | -1 | lncRNA | ENSG0000( |
| 222945930 | 222946388 | -1 | processed_pseudoge | ENSG0000( |
| 223051814 | 223198399 | 1 | protein_coding | ENSG0000( |
| 223248498 | 223248794 | -1 | misc_RNA | ENSG0000( |
| 223398267 | 223398546 | -1 | processed_pseudoge | ENSG0000( |
| 223498773 | 223504611 | 1 | lncRNA | ENSG0000( |
| 223505476 | 223510933 | 1 | lncRNA | ENSG0000( |
| 223596940 | 223602361 | -1 | protein_coding | ENSG0000( |
| 223704621 | 223705338 | 1 | processed_pseudoge | ENSG0000( |
| 223751686 | 223838027 | -1 | protein_coding | ENSG0000( |
| 223757258 | 223945299 | -1 | protein_coding | ENSG0000( |
| 223855716 | 223945357 | -1 | protein_coding | ENSG0000( |
| 223957463 | 223967714 | 1 | protein_coding | ENSG0000( |
| 223965701 | 223967706 | -1 | lncRNA | ENSG0000( |
| 223975045 | 224039318 | -1 | protein_coding | ENSG0000( |
| 224169664 | 224171537 | 1 | processed_pseudoge | ENSG0000( |

| | | | | |
|---|---|---|---|---|
| 224294930 | 224295753 | 1 | processed_pseudoge | ENSG00000 |
| 224378698 | 224402107 | -1 | protein_coding | ENSG00000 |
| 224464682 | 224474500 | 1 | lncRNA | ENSG00000 |
| 224470150 | 224585397 | -1 | protein_coding | ENSG00000 |
| 224499387 | 224500100 | -1 | processed_pseudoge | ENSG00000 |
| 224703764 | 224716365 | -1 | protein_coding | ENSG00000 |
| 224765090 | 225042445 | -1 | protein_coding | ENSG00000 |
| 225010461 | 225010540 | -1 | miRNA | ENSG00000 |
| 225399710 | 225654018 | 1 | protein_coding | ENSG00000 |
| 225698254 | 225698539 | 1 | processed_pseudoge | ENSG00000 |
| 225698514 | 225703654 | 1 | lncRNA | ENSG00000 |
| 225761559 | 225762112 | 1 | TEC | ENSG00000 |
| 225992981 | 225993197 | -1 | processed_pseudoge | ENSG00000 |
| 226180044 | 226185371 | -1 | lncRNA | ENSG00000 |
| 226658710 | 226658793 | -1 | miRNA | ENSG00000 |
| 226731317 | 226799759 | -1 | protein_coding | ENSG00000 |
| 226800146 | 226813741 | 1 | lncRNA | ENSG00000 |
| 226804036 | 226805061 | 1 | lncRNA | ENSG00000 |
| 226835581 | 226999215 | 1 | protein_coding | ENSG00000 |
| 226968989 | 226969122 | -1 | snoRNA | ENSG00000 |
| 227002711 | 227164453 | -1 | protein_coding | ENSG00000 |
| 227164565 | 227314792 | 1 | protein_coding | ENSG00000 |
| 227221052 | 227325711 | -1 | lncRNA | ENSG00000 |
| 227325151 | 227357836 | 1 | protein_coding | ENSG00000 |
| 227362038 | 227381995 | -1 | protein_coding | ENSG00000 |
| 227387683 | 227387949 | -1 | protein_coding | ENSG00000 |
| 227472132 | 227472187 | 1 | miRNA | ENSG00000 |
| 227472152 | 227561214 | 1 | protein_coding | ENSG00000 |
| 227587237 | 227587515 | -1 | polymorphic_pseudo | ENSG00000 |
| 227598893 | 227599255 | 1 | protein_coding | ENSG00000 |
| 227608784 | 227609101 | 1 | polymorphic_pseudo | ENSG00000 |
| 227610090 | 227648606 | -1 | protein_coding | ENSG00000 |
| 227614538 | 227614840 | -1 | protein_coding | ENSG00000 |
| 227616998 | 227618655 | 1 | protein_coding | ENSG00000 |
| 227616998 | 227617790 | -1 | lncRNA | ENSG00000 |
| 227666804 | 227667061 | 1 | protein_coding | ENSG00000 |
| 227683763 | 227718028 | -1 | protein_coding | ENSG00000 |
| 227724440 | 227724757 | -1 | protein_coding | ENSG00000 |
| 227728335 | 227728625 | 1 | protein_coding | ENSG00000 |
| 227745845 | 227746681 | 1 | protein_coding | ENSG00000 |
| 227763052 | 227763152 | -1 | rRNA_pseudogene | ENSG00000 |
| 227778510 | 227778735 | -1 | processed_pseudoge | ENSG00000 |
| 227801840 | 227804705 | 1 | processed_pseudoge | ENSG00000 |
| 227805739 | 227817564 | 1 | protein_coding | ENSG00000 |
| 227870083 | 227870644 | -1 | processed_pseudoge | ENSG00000 |
| 227871054 | 227924344 | 1 | protein_coding | ENSG00000 |
| 227887631 | 227887744 | 1 | snoRNA | ENSG00000 |
| 227979955 | 228181687 | -1 | protein_coding | ENSG00000 |
| 228169743 | 228169849 | 1 | snRNA | ENSG00000 |
| 228352120 | 228353215 | -1 | processed_pseudoge | ENSG00000 |

| | | | | |
|---|---|---|---|---|
| 228378500 | 228379773 | 1 | lncRNA | ENSG0000C |
| 228475224 | 228841689 | -1 | lncRNA | ENSG0000C |
| 228683537 | 228684580 | -1 | lncRNA | ENSG0000C |
| 228753324 | 228753786 | 1 | processed_pseudoge | ENSG0000C |
| 228850526 | 229271285 | -1 | protein_coding | ENSG0000C |
| 229003886 | 229004400 | 1 | processed_pseudoge | ENSG0000C |
| 229236340 | 229236609 | -1 | misc_RNA | ENSG0000C |
| 229357629 | 229714555 | -1 | protein_coding | ENSG0000C |
| 229407358 | 229408328 | 1 | lncRNA | ENSG0000C |
| 229606238 | 229606299 | 1 | snRNA | ENSG0000C |
| 229701794 | 229702148 | 1 | processed_pseudoge | ENSG0000C |
| 229763838 | 229923239 | -1 | protein_coding | ENSG0000C |
| 229894346 | 229894453 | 1 | snRNA | ENSG0000C |
| 229922302 | 230013119 | 1 | protein_coding | ENSG0000C |
| 229937814 | 229937882 | -1 | snRNA | ENSG0000C |
| 229942728 | 229945137 | 1 | lncRNA | ENSG0000C |
| 229973315 | 229973421 | 1 | snRNA | ENSG0000C |
| 230034982 | 230068993 | -1 | protein_coding | ENSG0000C |
| 230058229 | 230058324 | -1 | misc_RNA | ENSG0000C |
| 230121370 | 230174223 | 1 | lncRNA | ENSG0000C |
| 230138601 | 230138783 | 1 | unitary_pseudogene | ENSG0000C |
| 230167293 | 230225729 | -1 | protein_coding | ENSG0000C |
| 230203110 | 230313215 | 1 | protein_coding | ENSG0000C |
| 230327184 | 230403732 | 1 | protein_coding | ENSG0000C |
| 230410700 | 230412529 | -1 | TEC | ENSG0000C |
| 230415942 | 230545606 | 1 | protein_coding | ENSG0000C |
| 230507101 | 230507386 | 1 | misc_RNA | ENSG0000C |
| 230515023 | 230515638 | 1 | processed_pseudoge | ENSG0000C |
| 230520594 | 230580006 | -1 | lncRNA | ENSG0000C |
| 230573167 | 230573809 | -1 | processed_pseudoge | ENSG0000C |
| 230584815 | 230586715 | -1 | unitary_pseudogene | ENSG0000C |
| 230587121 | 230587227 | 1 | snRNA | ENSG0000C |
| 230690065 | 230700529 | -1 | lncRNA | ENSG0000C |
| 230712842 | 230821075 | 1 | protein_coding | ENSG0000C |
| 230725143 | 230725403 | -1 | processed_pseudoge | ENSG0000C |
| 230752141 | 230752247 | 1 | snRNA | ENSG0000C |
| 230836909 | 230837385 | 1 | processed_pseudoge | ENSG0000C |
| 230864639 | 230879248 | 1 | protein_coding | ENSG0000C |
| 230886474 | 230904693 | 1 | lncRNA | ENSG0000C |
| 230907318 | 230961066 | -1 | protein_coding | ENSG0000C |
| 230908852 | 230910102 | 1 | lncRNA | ENSG0000C |
| 230924567 | 230929919 | 1 | lncRNA | ENSG0000C |
| 230957593 | 230957950 | 1 | processed_pseudoge | ENSG0000C |
| 230984368 | 230996032 | -1 | lncRNA | ENSG0000C |
| 230990324 | 231025055 | 1 | protein_coding | ENSG0000C |
| 231037523 | 231049719 | 1 | protein_coding | ENSG0000C |
| 231052040 | 231052637 | 1 | lncRNA | ENSG0000C |
| 231056864 | 231172827 | 1 | protein_coding | ENSG0000C |
| 231108230 | 231125042 | -1 | protein_coding | ENSG0000C |
| 231133462 | 231133630 | 1 | snRNA | ENSG0000C |

| | | | | |
|---|---|---|---|---|
| 231198546 | 231374837 | 1 | protein_coding | ENSG0000C |
| 231256097 | 231256306 | 1 | processed_pseudoge | ENSG0000C |
| 231362708 | 231362793 | 1 | miRNA | ENSG0000C |
| 231388976 | 231394991 | 1 | lncRNA | ENSG0000C |
| 231395710 | 231401164 | 1 | protein_coding | ENSG0000C |
| 231433923 | 231434426 | -1 | processed_pseudoge | ENSG0000C |
| 231437101 | 231437421 | -1 | processed_pseudoge | ENSG0000C |
| 231452195 | 231453153 | 1 | lncRNA | ENSG0000C |
| 231453531 | 231483641 | -1 | protein_coding | ENSG0000C |
| 231455800 | 231455936 | -1 | snoRNA | ENSG0000C |
| 231456444 | 231456523 | -1 | snoRNA | ENSG0000C |
| 231460371 | 231460440 | -1 | snoRNA | ENSG0000C |
| 231493708 | 231495299 | -1 | processed_pseudoge | ENSG0000C |
| 231497090 | 231507256 | -1 | unprocessed_pseudc | ENSG0000C |
| 231501990 | 231502201 | -1 | snRNA | ENSG0000C |
| 231508422 | 231514366 | -1 | lncRNA | ENSG0000C |
| 231518582 | 231521067 | -1 | lncRNA | ENSG0000C |
| 231523187 | 231530445 | -1 | protein_coding | ENSG0000C |
| 231559903 | 231560364 | 1 | processed_pseudoge | ENSG0000C |
| 231585864 | 231586327 | 1 | processed_pseudoge | ENSG0000C |
| 231588959 | 231592776 | -1 | lncRNA | ENSG0000C |
| 231592864 | 231594276 | 1 | protein_coding | ENSG0000C |
| 231645998 | 231646273 | -1 | misc_RNA | ENSG0000C |
| 231706895 | 231713541 | 1 | protein_coding | ENSG0000C |
| 231711525 | 231711647 | 1 | snRNA | ENSG0000C |
| 231713314 | 231713398 | 1 | miRNA | ENSG0000C |
| 231732433 | 231786272 | -1 | protein_coding | ENSG0000C |
| 231768604 | 231768680 | 1 | pseudogene | ENSG0000C |
| 231781671 | 231809254 | 1 | protein_coding | ENSG0000C |
| 231789481 | 231789887 | -1 | processed_pseudoge | ENSG0000C |
| 231809846 | 231812352 | 1 | lncRNA | ENSG0000C |
| 231810348 | 231810844 | -1 | processed_pseudoge | ENSG0000C |
| 231892242 | 231892298 | -1 | miRNA | ENSG0000C |
| 231921809 | 231926396 | -1 | protein_coding | ENSG0000C |
| 231961245 | 232344350 | 1 | protein_coding | ENSG0000C |
| 231978488 | 232015720 | -1 | lncRNA | ENSG0000C |
| 232172653 | 232172747 | 1 | miRNA | ENSG0000C |
| 232343116 | 232343903 | 1 | processed_pseudoge | ENSG0000C |
| 232349754 | 232351309 | -1 | unprocessed_pseudc | ENSG0000C |
| 232368576 | 232368697 | 1 | unprocessed_pseudc | ENSG0000C |
| 232378724 | 232382889 | 1 | protein_coding | ENSG0000C |
| 232379635 | 232381674 | -1 | lncRNA | ENSG0000C |
| 232385750 | 232387457 | -1 | transcribed_unproce | ENSG0000C |
| 232406844 | 232410714 | 1 | protein_coding | ENSG0000C |
| 232415818 | 232419816 | -1 | unprocessed_pseudc | ENSG0000C |
| 232420661 | 232421461 | -1 | lncRNA | ENSG0000C |
| 232442034 | 232447418 | 1 | unprocessed_pseudc | ENSG0000C |
| 232456125 | 232460753 | 1 | protein_coding | ENSG0000C |
| 232479827 | 232487834 | -1 | protein_coding | ENSG0000C |
| 232520463 | 232525716 | 1 | protein_coding | ENSG0000C |

| | | | | |
|---|---|---|---|---|
| 232525993 | 232536667 | 1 | protein_coding | ENSG00000 |
| 232539692 | 232548115 | 1 | protein_coding | ENSG00000 |
| 232547970 | 232550557 | -1 | protein_coding | ENSG00000 |
| 232550474 | 232550573 | -1 | miRNA | ENSG00000 |
| 232550674 | 232583644 | 1 | protein_coding | ENSG00000 |
| 232580948 | 232611971 | -1 | lncRNA | ENSG00000 |
| 232606057 | 232682780 | 1 | protein_coding | ENSG00000 |
| 232649849 | 232650136 | 1 | misc_RNA | ENSG00000 |
| 232697299 | 232860575 | 1 | protein_coding | ENSG00000 |
| 232729523 | 232730198 | -1 | processed_pseudoge | ENSG00000 |
| 232760146 | 232767949 | -1 | lncRNA | ENSG00000 |
| 232765802 | 232776565 | -1 | protein_coding | ENSG00000 |
| 232782665 | 232782766 | 1 | snRNA | ENSG00000 |
| 232828653 | 232828751 | 1 | misc_RNA | ENSG00000 |
| 232857270 | 232878708 | 1 | protein_coding | ENSG00000 |
| 232878701 | 233013256 | -1 | protein_coding | ENSG00000 |
| 233012614 | 233015885 | 1 | lncRNA | ENSG00000 |
| 233032672 | 233035057 | 1 | protein_coding | ENSG00000 |
| 233059967 | 233207903 | 1 | protein_coding | ENSG00000 |
| 233205199 | 233205479 | 1 | misc_RNA | ENSG00000 |
| 233210051 | 233295674 | 1 | protein_coding | ENSG00000 |
| 233275727 | 233276002 | 1 | scaRNA | ENSG00000 |
| 233288676 | 233288940 | 1 | scaRNA | ENSG00000 |
| 233307816 | 233347055 | 1 | protein_coding | ENSG00000 |
| 233351132 | 233353416 | -1 | lncRNA | ENSG00000 |
| 233354507 | 233472104 | 1 | protein_coding | ENSG00000 |
| 233379843 | 233379955 | -1 | misc_RNA | ENSG00000 |
| 233475520 | 233566782 | -1 | protein_coding | ENSG00000 |
| 233545482 | 233547032 | 1 | processed_pseudoge | ENSG00000 |
| 233585439 | 233586291 | 1 | unprocessed_pseudo | ENSG00000 |
| 233603553 | 233604401 | 1 | unprocessed_pseudo | ENSG00000 |
| 233617645 | 233773310 | 1 | protein_coding | ENSG00000 |
| 233636454 | 233773305 | 1 | protein_coding | ENSG00000 |
| 233647926 | 233649026 | 1 | unprocessed_pseudo | ENSG00000 |
| 233671898 | 233773300 | 1 | protein_coding | ENSG00000 |
| 233681938 | 233773299 | 1 | protein_coding | ENSG00000 |
| 233691607 | 233773300 | 1 | protein_coding | ENSG00000 |
| 233693434 | 233708699 | -1 | lncRNA | ENSG00000 |
| 233712992 | 233773299 | 1 | protein_coding | ENSG00000 |
| 233718778 | 233773299 | 1 | protein_coding | ENSG00000 |
| 233721522 | 233722065 | -1 | processed_pseudoge | ENSG00000 |
| 233729108 | 233773299 | 1 | protein_coding | ENSG00000 |
| 233738952 | 233741957 | 1 | unprocessed_pseudo | ENSG00000 |
| 233742750 | 233744015 | -1 | transcribed_processe | ENSG00000 |
| 233747214 | 233748079 | 1 | unprocessed_pseudo | ENSG00000 |
| 233750617 | 233753656 | 1 | unprocessed_pseudo | ENSG00000 |
| 233754305 | 233755345 | -1 | transcribed_processe | ENSG00000 |
| 233760248 | 233773299 | 1 | protein_coding | ENSG00000 |
| 233775679 | 233833423 | 1 | protein_coding | ENSG00000 |
| 233833416 | 233854566 | -1 | protein_coding | ENSG00000 |

| | | | | |
|---|---|---|---|---|
| 233865437 | 233868444 | -1 | transcribed_processe | ENSG00000 |
| 233865496 | 233867359 | -1 | TEC | ENSG00000 |
| 233917373 | 234019522 | 1 | protein_coding | ENSG00000 |
| 233946146 | 233955435 | -1 | lncRNA | ENSG00000 |
| 233977872 | 233980018 | -1 | lncRNA | ENSG00000 |
| 234050679 | 234077134 | 1 | protein_coding | ENSG00000 |
| 234082259 | 234084490 | 1 | TEC | ENSG00000 |
| 234109081 | 234109481 | 1 | lncRNA | ENSG00000 |
| 234222838 | 234224514 | 1 | lncRNA | ENSG00000 |
| 234329483 | 234329838 | 1 | processed_pseudoge | ENSG00000 |
| 234351247 | 234356038 | -1 | lncRNA | ENSG00000 |
| 234438328 | 234462127 | 1 | lncRNA | ENSG00000 |
| 234444590 | 234454595 | -1 | lncRNA | ENSG00000 |
| 234455157 | 234455783 | -1 | unprocessed_pseudo | ENSG00000 |
| 234493041 | 234497081 | -1 | protein_coding | ENSG00000 |
| 234682668 | 234717764 | 1 | lncRNA | ENSG00000 |
| 234743542 | 234745606 | -1 | unprocessed_pseudo | ENSG00000 |
| 234803808 | 234913384 | -1 | lncRNA | ENSG00000 |
| 234952017 | 235055714 | 1 | protein_coding | ENSG00000 |
| 235084822 | 235088586 | -1 | lncRNA | ENSG00000 |
| 235147016 | 235147500 | -1 | processed_pseudoge | ENSG00000 |
| 235177896 | 235178905 | 1 | lncRNA | ENSG00000 |
| 235494043 | 236131800 | 1 | protein_coding | ENSG00000 |
| 235505751 | 235507566 | 1 | lncRNA | ENSG00000 |
| 235773855 | 235783387 | -1 | lncRNA | ENSG00000 |
| 235882362 | 235882493 | -1 | processed_pseudoge | ENSG00000 |
| 236037250 | 236037311 | 1 | snRNA | ENSG00000 |
| 236072295 | 236072556 | 1 | misc_RNA | ENSG00000 |
| 236153071 | 236153172 | -1 | misc_RNA | ENSG00000 |
| 236165236 | 236168386 | -1 | protein_coding | ENSG00000 |
| 236167447 | 236294927 | 1 | lncRNA | ENSG00000 |
| 236194872 | 236264409 | -1 | protein_coding | ENSG00000 |
| 236210373 | 236210548 | -1 | snRNA | ENSG00000 |
| 236324147 | 236507535 | -1 | protein_coding | ENSG00000 |
| 236367560 | 236369102 | 1 | lncRNA | ENSG00000 |
| 236391074 | 236392388 | 1 | lncRNA | ENSG00000 |
| 236407846 | 236409077 | -1 | transcribed_processe | ENSG00000 |
| 236567787 | 236582354 | 1 | protein_coding | ENSG00000 |
| 236733289 | 236754363 | 1 | lncRNA | ENSG00000 |
| 236910797 | 237085838 | -1 | lncRNA | ENSG00000 |
| 237015204 | 237015305 | -1 | snRNA | ENSG00000 |
| 237048599 | 237056167 | -1 | lncRNA | ENSG00000 |
| 237085882 | 237100474 | 1 | protein_coding | ENSG00000 |
| 237122910 | 237124097 | -1 | lncRNA | ENSG00000 |
| 237257091 | 237257676 | 1 | lncRNA | ENSG00000 |
| 237324003 | 237414375 | -1 | protein_coding | ENSG00000 |
| 237421420 | 237425276 | -1 | lncRNA | ENSG00000 |
| 237428920 | 237434822 | -1 | lncRNA | ENSG00000 |
| 237485428 | 237555322 | 1 | protein_coding | ENSG00000 |
| 237510931 | 237510988 | 1 | miRNA | ENSG00000 |

| | | | | |
|---|---|---|---|---|
| 237563165 | 237563270 | 1 | snRNA | ENSG00000 |
| 237566574 | 237567175 | 1 | protein_coding | ENSG00000 |
| 237574322 | 237601614 | -1 | protein_coding | ENSG00000 |
| 237591020 | 237595981 | 1 | lncRNA | ENSG00000 |
| 237612977 | 237626525 | 1 | lncRNA | ENSG00000 |
| 237627587 | 237813682 | 1 | protein_coding | ENSG00000 |
| 237728569 | 237731131 | -1 | lncRNA | ENSG00000 |
| 237798389 | 237842808 | 1 | protein_coding | ENSG00000 |
| 237858893 | 237912106 | 1 | protein_coding | ENSG00000 |
| 237869904 | 237869982 | 1 | snoRNA | ENSG00000 |
| 237966827 | 238042782 | 1 | protein_coding | ENSG00000 |
| 237967014 | 238099412 | 1 | protein_coding | ENSG00000 |
| 238028971 | 238029063 | -1 | snRNA | ENSG00000 |
| 238060924 | 238099413 | 1 | protein_coding | ENSG00000 |
| 238100340 | 238133287 | 1 | protein_coding | ENSG00000 |
| 238138668 | 238152947 | 1 | protein_coding | ENSG00000 |
| 238152889 | 238155994 | -1 | TEC | ENSG00000 |
| 238158970 | 238168900 | 1 | protein_coding | ENSG00000 |
| 238170402 | 238203708 | -1 | protein_coding | ENSG00000 |
| 238224552 | 238231699 | -1 | lncRNA | ENSG00000 |
| 238231684 | 238255633 | 1 | lncRNA | ENSG00000 |
| 238238267 | 238240662 | -1 | protein_coding | ENSG00000 |
| 238244044 | 238290102 | -1 | protein_coding | ENSG00000 |
| 238289127 | 238300185 | 1 | lncRNA | ENSG00000 |
| 238295392 | 238300620 | -1 | lncRNA | ENSG00000 |
| 238320441 | 238400897 | 1 | protein_coding | ENSG00000 |
| 238412642 | 238412745 | 1 | snRNA | ENSG00000 |
| 238426742 | 238452250 | 1 | protein_coding | ENSG00000 |
| 238427077 | 238427729 | -1 | lncRNA | ENSG00000 |
| 238454908 | 238457520 | 1 | unprocessed_pseudo | ENSG00000 |
| 238510690 | 238555054 | -1 | lncRNA | ENSG00000 |
| 238554541 | 238740849 | 1 | lncRNA | ENSG00000 |
| 238711224 | 238722543 | -1 | lncRNA | ENSG00000 |
| 238735263 | 238735547 | -1 | processed_pseudoge | ENSG00000 |
| 238771768 | 238784764 | 1 | lncRNA | ENSG00000 |
| 238788648 | 238789716 | -1 | lncRNA | ENSG00000 |
| 238848032 | 238910534 | 1 | protein_coding | ENSG00000 |
| 238919302 | 238926269 | -1 | lncRNA | ENSG00000 |
| 239048168 | 239401654 | -1 | protein_coding | ENSG00000 |
| 239068817 | 239068914 | -1 | miRNA | ENSG00000 |
| 239085827 | 239085926 | -1 | miRNA | ENSG00000 |
| 239114858 | 239118842 | 1 | lncRNA | ENSG00000 |
| 239194118 | 239197654 | 1 | lncRNA | ENSG00000 |
| 239305462 | 239305545 | 1 | miRNA | ENSG00000 |
| 239351724 | 239351804 | -1 | miRNA | ENSG00000 |
| 239401436 | 239402364 | 1 | lncRNA | ENSG00000 |
| 239439643 | 239531363 | 1 | lncRNA | ENSG00000 |
| 239493587 | 239496697 | -1 | lncRNA | ENSG00000 |
| 239538379 | 239544876 | 1 | lncRNA | ENSG00000 |
| 239578301 | 239586094 | 1 | lncRNA | ENSG00000 |

| | | | | |
|---|---|---|---|---|
| 239625698 | 239630839 | 1 | lncRNA | ENSG0000C |
| 239734956 | 239735538 | 1 | lncRNA | ENSG0000C |
| 239762860 | 239824764 | 1 | lncRNA | ENSG0000C |
| 239892450 | 240025345 | -1 | protein_coding | ENSG0000C |
| 239943015 | 239943094 | -1 | miRNA | ENSG0000C |
| 240028965 | 240030456 | -1 | protein_coding | ENSG0000C |
| 240045077 | 240047027 | -1 | protein_coding | ENSG0000C |
| 240079433 | 240080362 | 1 | unprocessed_pseudc | ENSG0000C |
| 240109102 | 240110052 | 1 | unitary_pseudogene | ENSG0000C |
| 240126563 | 240136807 | -1 | protein_coding | ENSG0000C |
| 240139026 | 240144562 | -1 | protein_coding | ENSG0000C |
| 240184716 | 240187878 | 1 | lncRNA | ENSG0000C |
| 240189710 | 240194408 | -1 | lncRNA | ENSG0000C |
| 240227560 | 240256035 | -1 | lncRNA | ENSG0000C |
| 240289216 | 240289443 | 1 | snoRNA | ENSG0000C |
| 240435663 | 240468076 | 1 | protein_coding | ENSG0000C |
| 240449315 | 240456714 | -1 | lncRNA | ENSG0000C |
| 240456001 | 240456089 | 1 | miRNA | ENSG0000C |
| 240479422 | 240569209 | -1 | protein_coding | ENSG0000C |
| 240558089 | 240558925 | 1 | lncRNA | ENSG0000C |
| 240560054 | 240564014 | 1 | protein_coding | ENSG0000C |
| 240565804 | 240581372 | 1 | protein_coding | ENSG0000C |
| 240582700 | 240586699 | -1 | lncRNA | ENSG0000C |
| 240586734 | 240617705 | 1 | protein_coding | ENSG0000C |
| 240605430 | 240631259 | 1 | protein_coding | ENSG0000C |
| 240672530 | 240672812 | 1 | processed_pseudoge | ENSG0000C |
| 240676418 | 240682906 | -1 | protein_coding | ENSG0000C |
| 240686334 | 240690414 | 1 | lncRNA | ENSG0000C |
| 240691851 | 240698483 | 1 | protein_coding | ENSG0000C |
| 240701204 | 240701685 | -1 | processed_pseudoge | ENSG0000C |
| 240713761 | 240821036 | -1 | protein_coding | ENSG0000C |
| 240868824 | 240880502 | 1 | protein_coding | ENSG0000C |
| 240886048 | 240896889 | -1 | protein_coding | ENSG0000C |
| 240906330 | 240993311 | 1 | protein_coding | ENSG0000C |
| 240954617 | 240967451 | -1 | lncRNA | ENSG0000C |
| 240981515 | 240986072 | -1 | lncRNA | ENSG0000C |
| 240998618 | 241095568 | 1 | protein_coding | ENSG0000C |
| 241010045 | 241010744 | -1 | lncRNA | ENSG0000C |
| 241015599 | 241064116 | -1 | lncRNA | ENSG0000C |
| 241072169 | 241102332 | -1 | protein_coding | ENSG0000C |
| 241106099 | 241150264 | -1 | protein_coding | ENSG0000C |
| 241118725 | 241118943 | 1 | processed_pseudoge | ENSG0000C |
| 241149576 | 241183652 | 1 | protein_coding | ENSG0000C |
| 241188509 | 241225377 | 1 | protein_coding | ENSG0000C |
| 241227264 | 241317061 | -1 | protein_coding | ENSG0000C |
| 241315100 | 241354027 | 1 | protein_coding | ENSG0000C |
| 241326334 | 241327039 | -1 | lncRNA | ENSG0000C |
| 241351340 | 241353104 | -1 | lncRNA | ENSG0000C |
| 241356285 | 241494841 | 1 | protein_coding | ENSG0000C |
| 241360891 | 241372749 | -1 | lncRNA | ENSG0000C |

| | | | | |
|---|---|---|---|---|
| 241477905 | 241477982 | 1 | miRNA | ENSG0000( |
| 241478316 | 241478993 | 1 | processed_pseudoge | ENSG0000( |
| 241492670 | 241509730 | -1 | protein_coding | ENSG0000( |
| 241544403 | 241558977 | -1 | lncRNA | ENSG0000( |
| 241551424 | 241574131 | 1 | protein_coding | ENSG0000( |
| 241581922 | 241582726 | 1 | lncRNA | ENSG0000( |
| 241584405 | 241637158 | -1 | protein_coding | ENSG0000( |
| 241584968 | 241585075 | 1 | rRNA_pseudogene | ENSG0000( |
| 241637213 | 241673857 | 1 | protein_coding | ENSG0000( |
| 241675747 | 241686944 | -1 | protein_coding | ENSG0000( |
| 241687085 | 241729478 | 1 | protein_coding | ENSG0000( |
| 241724615 | 241725693 | -1 | lncRNA | ENSG0000( |
| 241734602 | 241768816 | 1 | protein_coding | ENSG0000( |
| 241734715 | 241735498 | -1 | lncRNA | ENSG0000( |
| 241754793 | 241755740 | -1 | lncRNA | ENSG0000( |
| 241776822 | 241804287 | 1 | protein_coding | ENSG0000( |
| 241800916 | 241801907 | -1 | lncRNA | ENSG0000( |
| 241808312 | 241812016 | -1 | lncRNA | ENSG0000( |
| 241808825 | 241817413 | 1 | protein_coding | ENSG0000( |
| 241844380 | 241845036 | 1 | lncRNA | ENSG0000( |
| 241849884 | 241858894 | -1 | protein_coding | ENSG0000( |
| 241869600 | 241873823 | 1 | protein_coding | ENSG0000( |
| 241881363 | 242078722 | 1 | lncRNA | ENSG0000( |
| 241893985 | 241902551 | -1 | protein_coding | ENSG0000( |
| 241970683 | 241977276 | 1 | lncRNA | ENSG0000( |
| 242001209 | 242006013 | 1 | TEC | ENSG0000( |
| 242025183 | 242026176 | -1 | lncRNA | ENSG0000( |
| 242037716 | 242041501 | -1 | lncRNA | ENSG0000( |
| 242047695 | 242084233 | -1 | lncRNA | ENSG0000( |
| 242087351 | 242088457 | -1 | lncRNA | ENSG0000( |
| 242088633 | 242160153 | 1 | transcribed_unproce | ENSG0000( |
| 242119856 | 242120053 | -1 | processed_pseudoge | ENSG0000( |
| 242122287 | 242122469 | 1 | processed_pseudoge | ENSG0000( |
| 242175181 | 242175634 | 1 | processed_pseudoge | ENSG0000( |
| 11745 | 24849 | 1 | lncRNA | ENSG0000( |
| 53348 | 54346 | -1 | lncRNA | ENSG0000( |
| 109707 | 197341 | -1 | lncRNA | ENSG0000( |
| 123563 | 133500 | 1 | lncRNA | ENSG0000( |
| 196763 | 409417 | 1 | protein_coding | ENSG0000( |
| 282689 | 282792 | -1 | snRNA | ENSG0000( |
| 308714 | 309322 | -1 | processed_pseudoge | ENSG0000( |
| 363370 | 385795 | -1 | lncRNA | ENSG0000( |
| 524012 | 551657 | -1 | lncRNA | ENSG0000( |
| 536062 | 846561 | 1 | lncRNA | ENSG0000( |
| 617789 | 618314 | -1 | processed_pseudoge | ENSG0000( |
| 857124 | 858091 | 1 | lncRNA | ENSG0000( |
| 1008135 | 1012934 | 1 | lncRNA | ENSG0000( |
| 1065580 | 1065875 | -1 | misc_RNA | ENSG0000( |
| 1092658 | 1404217 | 1 | protein_coding | ENSG0000( |
| 1394173 | 1394448 | -1 | processed_pseudoge | ENSG0000( |

| | | | | |
|---|---|---|---|---|
| 1595777 | 1596245 | -1 | processed_pseudoge | ENSG00000 |
| 1730070 | 1730474 | 1 | processed_pseudoge | ENSG00000 |
| 1905651 | 1906125 | 1 | processed_pseudoge | ENSG00000 |
| 1962381 | 1987470 | -1 | lncRNA | ENSG00000 |
| 2088933 | 2089211 | -1 | misc_RNA | ENSG00000 |
| 2098813 | 3057959 | 1 | protein_coding | ENSG00000 |
| 2110409 | 2144241 | -1 | lncRNA | ENSG00000 |
| 2135757 | 2141988 | 1 | lncRNA | ENSG00000 |
| 2362258 | 2362738 | 1 | processed_pseudoge | ENSG00000 |
| 2985231 | 2985575 | -1 | processed_pseudoge | ENSG00000 |
| 3039033 | 3069242 | -1 | lncRNA | ENSG00000 |
| 3066326 | 3126613 | -1 | protein_coding | ENSG00000 |
| 3102913 | 3103015 | 1 | snoRNA | ENSG00000 |
| 3126933 | 3150879 | 1 | protein_coding | ENSG00000 |
| 3144628 | 3179727 | -1 | protein_coding | ENSG00000 |
| 3152942 | 3153435 | 1 | lncRNA | ENSG00000 |
| 3250687 | 3627296 | -1 | lncRNA | ENSG00000 |
| 3700814 | 4467273 | -1 | protein_coding | ENSG00000 |
| 3799431 | 3849834 | 1 | protein_coding | ENSG00000 |
| 3918709 | 4046722 | 1 | lncRNA | ENSG00000 |
| 3981981 | 3984295 | -1 | processed_pseudoge | ENSG00000 |
| 4269153 | 4311495 | -1 | lncRNA | ENSG00000 |
| 4303304 | 4317567 | 1 | protein_coding | ENSG00000 |
| 4414580 | 4414960 | -1 | processed_pseudoge | ENSG00000 |
| 4490891 | 4493163 | -1 | lncRNA | ENSG00000 |
| 4493345 | 4847840 | 1 | protein_coding | ENSG00000 |
| 4749192 | 4751590 | -1 | lncRNA | ENSG00000 |
| 4750333 | 4750685 | 1 | misc_RNA | ENSG00000 |
| 4814294 | 4887293 | -1 | lncRNA | ENSG00000 |
| 4896809 | 4980414 | -1 | lncRNA | ENSG00000 |
| 4979437 | 4985323 | 1 | protein_coding | ENSG00000 |
| 5068676 | 5069044 | -1 | processed_pseudoge | ENSG00000 |
| 5080212 | 5084429 | -1 | unprocessed_pseudo | ENSG00000 |
| 5086178 | 5086279 | -1 | misc_RNA | ENSG00000 |
| 5099627 | 5099861 | 1 | processed_pseudoge | ENSG00000 |
| 5122245 | 5180912 | 1 | protein_coding | ENSG00000 |
| 5156905 | 5187329 | -1 | lncRNA | ENSG00000 |
| 5187172 | 5188298 | -1 | lncRNA | ENSG00000 |
| 5187646 | 5219958 | 1 | protein_coding | ENSG00000 |
| 5250177 | 5250255 | -1 | miRNA | ENSG00000 |
| 5252926 | 5253223 | -1 | misc_RNA | ENSG00000 |
| 5255535 | 5256761 | -1 | lncRNA | ENSG00000 |
| 5962842 | 6124120 | 1 | lncRNA | ENSG00000 |
| 6148484 | 6365885 | 1 | lncRNA | ENSG00000 |
| 6208269 | 6208571 | 1 | unprocessed_pseudo | ENSG00000 |
| 6490460 | 6736750 | 1 | lncRNA | ENSG00000 |
| 6631008 | 6805479 | -1 | lncRNA | ENSG00000 |
| 6692319 | 6693088 | 1 | processed_pseudoge | ENSG00000 |
| 6770001 | 7741533 | 1 | protein_coding | ENSG00000 |
| 6773037 | 6773343 | -1 | processed_pseudoge | ENSG00000 |

| | | | | |
|---|---|---|---|---|
| 6892632 | 6894022 | -1 | lncRNA | ENSG00000 |
| 7241916 | 7250209 | 1 | lncRNA | ENSG00000 |
| 7519741 | 7560333 | -1 | lncRNA | ENSG00000 |
| 7606890 | 7790482 | -1 | lncRNA | ENSG00000 |
| 7951263 | 8611924 | -1 | lncRNA | ENSG00000 |
| 8079322 | 8125538 | 1 | lncRNA | ENSG00000 |
| 8359344 | 8366427 | -1 | lncRNA | ENSG00000 |
| 8364529 | 8364648 | -1 | snRNA | ENSG00000 |
| 8501807 | 8574668 | 1 | protein_coding | ENSG00000 |
| 8573726 | 8593124 | -1 | lncRNA | ENSG00000 |
| 8619400 | 8745040 | -1 | protein_coding | ENSG00000 |
| 8682943 | 8693701 | 1 | transcribed_unproce | ENSG00000 |
| 8733800 | 8841808 | 1 | protein_coding | ENSG00000 |
| 8750408 | 8769628 | -1 | protein_coding | ENSG00000 |
| 8775402 | 8963773 | -1 | protein_coding | ENSG00000 |
| 8931495 | 8931632 | 1 | snoRNA | ENSG00000 |
| 8931506 | 8931631 | 1 | snoRNA | ENSG00000 |
| 8980591 | 9363053 | -1 | protein_coding | ENSG00000 |
| 9014123 | 9015979 | 1 | lncRNA | ENSG00000 |
| 9100803 | 9104078 | 1 | lncRNA | ENSG00000 |
| 9192493 | 9194453 | 1 | lncRNA | ENSG00000 |
| 9216895 | 9219508 | 1 | lncRNA | ENSG00000 |
| 9249742 | 9257507 | 1 | lncRNA | ENSG00000 |
| 9292588 | 9363303 | -1 | lncRNA | ENSG00000 |
| 9348443 | 9349192 | 1 | processed_pseudoge | ENSG00000 |
| 9349689 | 9398579 | -1 | lncRNA | ENSG00000 |
| 9362971 | 9386791 | 1 | protein_coding | ENSG00000 |
| 9397615 | 9479240 | 1 | protein_coding | ENSG00000 |
| 9498361 | 9553822 | -1 | protein_coding | ENSG00000 |
| 9600748 | 9602728 | 1 | processed_pseudoge | ENSG00000 |
| 9649433 | 9702393 | 1 | protein_coding | ENSG00000 |
| 9693182 | 9721399 | -1 | lncRNA | ENSG00000 |
| 9703826 | 9729908 | 1 | protein_coding | ENSG00000 |
| 9731729 | 9748015 | 1 | protein_coding | ENSG00000 |
| 9749944 | 9788219 | 1 | protein_coding | ENSG00000 |
| 9757347 | 9769992 | -1 | protein_coding | ENSG00000 |
| 9779860 | 9793011 | -1 | protein_coding | ENSG00000 |
| 9792495 | 9807726 | 1 | protein_coding | ENSG00000 |
| 9793082 | 9835401 | 1 | protein_coding | ENSG00000 |
| 9808086 | 9855138 | 1 | protein_coding | ENSG00000 |
| 9812762 | 9813097 | -1 | lncRNA | ENSG00000 |
| 9837849 | 9844602 | -1 | protein_coding | ENSG00000 |
| 9866711 | 9880254 | -1 | protein_coding | ENSG00000 |
| 9890574 | 9894349 | 1 | protein_coding | ENSG00000 |
| 9902612 | 9916402 | 1 | protein_coding | ENSG00000 |
| 9917074 | 9933630 | 1 | protein_coding | ENSG00000 |
| 9919051 | 9919153 | -1 | snRNA | ENSG00000 |
| 9933822 | 9945413 | 1 | protein_coding | ENSG00000 |
| 9935706 | 9936258 | 1 | lncRNA | ENSG00000 |
| 9945542 | 9952408 | -1 | protein_coding | ENSG00000 |

| | | | | |
|---|---|---|---|---|
| 9947404 | 9954787 | 1 | lncRNA | ENSG0000C |
| 9958717 | 9962539 | 1 | lncRNA | ENSG0000C |
| 9962537 | 10011116 | -1 | protein_coding | ENSG0000C |
| 9986893 | 10006990 | 1 | transcribed_unproce | ENSG0000C |
| 10000647 | 10000940 | -1 | processed_pseudoge | ENSG0000C |
| 10004048 | 10011209 | -1 | lncRNA | ENSG0000C |
| 10014238 | 10026365 | -1 | transcribed_unproce | ENSG0000C |
| 10026414 | 10101930 | 1 | protein_coding | ENSG0000C |
| 10037552 | 10037655 | 1 | snRNA | ENSG0000C |
| 10057973 | 10058287 | -1 | processed_pseudoge | ENSG0000C |
| 10081317 | 10108255 | -1 | protein_coding | ENSG0000C |
| 10115675 | 10127190 | 1 | protein_coding | ENSG0000C |
| 10141008 | 10152220 | 1 | protein_coding | ENSG0000C |
| 10149986 | 10153326 | -1 | lncRNA | ENSG0000C |
| 10164919 | 10243745 | 1 | protein_coding | ENSG0000C |
| 10248023 | 10281218 | 1 | protein_coding | ENSG0000C |
| 10249372 | 10285796 | 1 | protein_coding | ENSG0000C |
| 10284419 | 10285746 | 1 | lncRNA | ENSG0000C |
| 10285666 | 10292947 | -1 | protein_coding | ENSG0000C |
| 10285754 | 10294903 | 1 | lncRNA | ENSG0000C |
| 10287413 | 10287482 | 1 | misc_RNA | ENSG0000C |
| 10293131 | 10321112 | -1 | protein_coding | ENSG0000C |
| 10324023 | 10708007 | -1 | protein_coding | ENSG0000C |
| 10330229 | 10330285 | 1 | miRNA | ENSG0000C |
| 10394489 | 10394562 | -1 | miRNA | ENSG0000C |
| 10566255 | 10570375 | -1 | lncRNA | ENSG0000C |
| 10626015 | 10629307 | -1 | lncRNA | ENSG0000C |
| 10759350 | 10764209 | -1 | lncRNA | ENSG0000C |
| 10767464 | 10771718 | 1 | lncRNA | ENSG0000C |
| 10816201 | 10940714 | 1 | protein_coding | ENSG0000C |
| 10914196 | 10977981 | -1 | lncRNA | ENSG0000C |
| 10992186 | 11039247 | 1 | protein_coding | ENSG0000C |
| 11006098 | 11019224 | -1 | lncRNA | ENSG0000C |
| 11060973 | 11091548 | 1 | lncRNA | ENSG0000C |
| 11137093 | 11263557 | 1 | protein_coding | ENSG0000C |
| 11193439 | 11225877 | -1 | lncRNA | ENSG0000C |
| 11234256 | 11234582 | -1 | processed_pseudoge | ENSG0000C |
| 11272309 | 11557665 | 1 | protein_coding | ENSG0000C |
| 11290406 | 11290663 | -1 | processed_pseudoge | ENSG0000C |
| 11488300 | 11488473 | -1 | unprocessed_pseudc | ENSG0000C |
| 11529488 | 11529755 | -1 | processed_pseudoge | ENSG0000C |
| 11556069 | 11771350 | -1 | protein_coding | ENSG0000C |
| 11607184 | 11610748 | 1 | lncRNA | ENSG0000C |
| 11611602 | 11612197 | 1 | lncRNA | ENSG0000C |
| 11790442 | 11846919 | -1 | protein_coding | ENSG0000C |
| 11859674 | 11891172 | 1 | unprocessed_pseudc | ENSG0000C |
| 11878830 | 11879129 | -1 | processed_pseudoge | ENSG0000C |
| 11900011 | 11901245 | 1 | unprocessed_pseudc | ENSG0000C |
| 11906527 | 11912313 | 1 | unprocessed_pseudc | ENSG0000C |
| 11909634 | 11910765 | -1 | unprocessed_pseudc | ENSG0000C |

| | | | | |
|---|---|---|---|---|
| 11952788 | 11953085 | -1 | misc_RNA | ENSG00000 |
| 12004388 | 12192032 | 1 | protein_coding | ENSG00000 |
| 12070195 | 12071360 | 1 | processed_pseudoge | ENSG00000 |
| 12153068 | 12158912 | -1 | protein_coding | ENSG00000 |
| 12165299 | 12165642 | -1 | processed_pseudoge | ENSG00000 |
| 12257801 | 12258460 | -1 | processed_pseudoge | ENSG00000 |
| 12287368 | 12434356 | 1 | protein_coding | ENSG00000 |
| 12328003 | 12328274 | 1 | processed_pseudoge | ENSG00000 |
| 12484432 | 12539623 | 1 | protein_coding | ENSG00000 |
| 12511095 | 12511213 | -1 | rRNA_pseudogene | ENSG00000 |
| 12514706 | 12514807 | 1 | snRNA | ENSG00000 |
| 12514934 | 12561059 | -1 | protein_coding | ENSG00000 |
| 12557087 | 12583713 | 1 | protein_coding | ENSG00000 |
| 12576961 | 12577237 | -1 | misc_RNA | ENSG00000 |
| 12583601 | 12664226 | -1 | protein_coding | ENSG00000 |
| 12673699 | 12673883 | 1 | processed_pseudoge | ENSG00000 |
| 12733528 | 12769457 | -1 | protein_coding | ENSG00000 |
| 12787393 | 12788671 | -1 | processed_pseudoge | ENSG00000 |
| 12796472 | 12871916 | 1 | protein_coding | ENSG00000 |
| 12832219 | 12832728 | -1 | lncRNA | ENSG00000 |
| 12834485 | 12841582 | -1 | protein_coding | ENSG00000 |
| 12840312 | 12840450 | -1 | snoRNA | ENSG00000 |
| 12850659 | 12850860 | 1 | transcribed_unproce | ENSG00000 |
| 12877522 | 12885211 | -1 | lncRNA | ENSG00000 |
| 12897043 | 13283281 | -1 | protein_coding | ENSG00000 |
| 13052967 | 13053122 | -1 | processed_pseudoge | ENSG00000 |
| 13316235 | 13420322 | -1 | protein_coding | ENSG00000 |
| 13460294 | 13460398 | -1 | processed_pseudoge | ENSG00000 |
| 13460424 | 13460889 | -1 | processed_pseudoge | ENSG00000 |
| 13476982 | 13480053 | -1 | lncRNA | ENSG00000 |
| 13479724 | 13506424 | 1 | protein_coding | ENSG00000 |
| 13549125 | 13638422 | 1 | protein_coding | ENSG00000 |
| 13650696 | 13746638 | 1 | lncRNA | ENSG00000 |
| 13816258 | 13880071 | -1 | protein_coding | ENSG00000 |
| 13926813 | 13927778 | 1 | unitary_pseudogene | ENSG00000 |
| 13933056 | 13937477 | 1 | transcribed_unproce | ENSG00000 |
| 13937273 | 14082811 | 1 | transcribed_processe | ENSG00000 |
| 14038647 | 14047181 | -1 | lncRNA | ENSG00000 |
| 14089107 | 14089871 | -1 | unitary_pseudogene | ENSG00000 |
| 14112077 | 14124870 | -1 | protein_coding | ENSG00000 |
| 14124940 | 14143679 | 1 | protein_coding | ENSG00000 |
| 14135217 | 14148252 | 1 | protein_coding | ENSG00000 |
| 14144637 | 14165978 | 1 | lncRNA | ENSG00000 |
| 14145147 | 14178783 | -1 | protein_coding | ENSG00000 |
| 14178817 | 14201122 | 1 | protein_coding | ENSG00000 |
| 14272373 | 14303845 | -1 | lncRNA | ENSG00000 |
| 14348451 | 14352568 | -1 | lncRNA | ENSG00000 |
| 14394648 | 14394757 | 1 | rRNA_pseudogene | ENSG00000 |
| 14402576 | 14489349 | 1 | protein_coding | ENSG00000 |
| 14489107 | 14556075 | -1 | protein_coding | ENSG00000 |

| | | | | |
|---|---|---|---|---|
| 14530982 | 14531088 | -1 | snRNA | ENSG0000C |
| 14572852 | 14574792 | -1 | processed_pseudoge | ENSG0000C |
| 14602035 | 14641443 | 1 | lncRNA | ENSG0000C |
| 14648194 | 14649432 | -1 | lncRNA | ENSG0000C |
| 14651746 | 14672659 | 1 | protein_coding | ENSG0000C |
| 14675141 | 14773036 | 1 | protein_coding | ENSG0000C |
| 14764952 | 14767440 | -1 | lncRNA | ENSG0000C |
| 14791534 | 14793504 | 1 | lncRNA | ENSG0000C |
| 14799331 | 14811414 | -1 | lncRNA | ENSG0000C |
| 14810853 | 14934571 | 1 | protein_coding | ENSG0000C |
| 14920347 | 14948424 | -1 | lncRNA | ENSG0000C |
| 14947584 | 15053600 | 1 | protein_coding | ENSG0000C |
| 15042460 | 15065317 | -1 | protein_coding | ENSG0000C |
| 15070073 | 15099163 | -1 | protein_coding | ENSG0000C |
| 15131848 | 15132208 | -1 | processed_pseudoge | ENSG0000C |
| 15141380 | 15142157 | 1 | processed_pseudoge | ENSG0000C |
| 15151833 | 15205959 | -1 | transcribed_unproce | ENSG0000C |
| 15206152 | 15252916 | 1 | protein_coding | ENSG0000C |
| 15254184 | 15264515 | 1 | lncRNA | ENSG0000C |
| 15254353 | 15341368 | -1 | protein_coding | ENSG0000C |
| 15339749 | 15339855 | 1 | snRNA | ENSG0000C |
| 15375373 | 15375645 | -1 | processed_pseudoge | ENSG0000C |
| 15381275 | 15440566 | -1 | protein_coding | ENSG0000C |
| 15427598 | 15450635 | 1 | protein_coding | ENSG0000C |
| 15433110 | 15433216 | -1 | snRNA | ENSG0000C |
| 15436171 | 15455940 | -1 | lncRNA | ENSG0000C |
| 15450133 | 15521751 | -1 | protein_coding | ENSG0000C |
| 15496239 | 15496308 | -1 | miRNA | ENSG0000C |
| 15501439 | 15504060 | 1 | lncRNA | ENSG0000C |
| 15506623 | 15506905 | -1 | misc_RNA | ENSG0000C |
| 15560699 | 15601852 | -1 | protein_coding | ENSG0000C |
| 15601341 | 15722311 | 1 | protein_coding | ENSG0000C |
| 15667236 | 15859771 | -1 | protein_coding | ENSG0000C |
| 15697298 | 15697371 | -1 | miRNA | ENSG0000C |
| 15731877 | 15732536 | 1 | TEC | ENSG0000C |
| 15732252 | 15733470 | 1 | transcribed_unproce | ENSG0000C |
| 15738515 | 15738809 | 1 | misc_RNA | ENSG0000C |
| 15860158 | 15875979 | 1 | lncRNA | ENSG0000C |
| 15873771 | 15873849 | 1 | miRNA | ENSG0000C |
| 15878047 | 15879571 | 1 | processed_pseudoge | ENSG0000C |
| 15970003 | 15971253 | 1 | lncRNA | ENSG0000C |
| 16174680 | 16231992 | 1 | protein_coding | ENSG0000C |
| 16252123 | 16253130 | -1 | lncRNA | ENSG0000C |
| 16257061 | 16264969 | -1 | protein_coding | ENSG0000C |
| 16265160 | 16350299 | 1 | protein_coding | ENSG0000C |
| 16275060 | 16276632 | -1 | processed_pseudoge | ENSG0000C |
| 16313574 | 16514026 | -1 | protein_coding | ENSG0000C |
| 16314439 | 16314987 | 1 | lncRNA | ENSG0000C |
| 16339308 | 16339871 | 1 | lncRNA | ENSG0000C |
| 16345126 | 16346440 | 1 | lncRNA | ENSG0000C |

| | | | | |
|---|---|---|---|---|
| 16361348 | 16370398 | 1 | lncRNA | ENSG0000C |
| 16444540 | 16447914 | 1 | lncRNA | ENSG0000C |
| 16524771 | 16541903 | 1 | lncRNA | ENSG0000C |
| 16524800 | 16531807 | 1 | lncRNA | ENSG0000C |
| 16586792 | 16670306 | -1 | protein_coding | ENSG0000C |
| 16687986 | 16697479 | -1 | lncRNA | ENSG0000C |
| 16692745 | 16694202 | -1 | unprocessed_pseudc | ENSG0000C |
| 16802651 | 17090604 | 1 | protein_coding | ENSG0000C |
| 16933196 | 16933260 | 1 | miRNA | ENSG0000C |
| 17042742 | 17044192 | -1 | lncRNA | ENSG0000C |
| 17141015 | 17205165 | 1 | lncRNA | ENSG0000C |
| 17157162 | 18444817 | -1 | protein_coding | ENSG0000C |
| 17333126 | 17333431 | 1 | processed_pseudoge | ENSG0000C |
| 17465150 | 17465510 | 1 | processed_pseudoge | ENSG0000C |
| 17664963 | 17665025 | 1 | snRNA | ENSG0000C |
| 17706058 | 17706233 | -1 | unprocessed_pseudc | ENSG0000C |
| 17742952 | 17745665 | 1 | lncRNA | ENSG0000C |
| 17871729 | 17872598 | -1 | processed_pseudoge | ENSG0000C |
| 17877455 | 17878169 | -1 | processed_pseudoge | ENSG0000C |
| 17990023 | 17995544 | 1 | lncRNA | ENSG0000C |
| 18013226 | 18041603 | 1 | lncRNA | ENSG0000C |
| 18345377 | 18445588 | -1 | protein_coding | ENSG0000C |
| 18408680 | 18409635 | -1 | lncRNA | ENSG0000C |
| 18445024 | 18920401 | 1 | lncRNA | ENSG0000C |
| 18538673 | 18539899 | 1 | processed_pseudoge | ENSG0000C |
| 18954943 | 18955049 | -1 | snRNA | ENSG0000C |
| 19148510 | 19535642 | 1 | protein_coding | ENSG0000C |
| 19314848 | 19314931 | -1 | miRNA | ENSG0000C |
| 19389185 | 19702795 | -1 | lncRNA | ENSG0000C |
| 19879472 | 19947025 | -1 | protein_coding | ENSG0000C |
| 19940269 | 19943310 | -1 | processed_pseudoge | ENSG0000C |
| 19947097 | 19985175 | 1 | protein_coding | ENSG0000C |
| 19979961 | 20012267 | -1 | protein_coding | ENSG0000C |
| 19996803 | 19996925 | 1 | snRNA | ENSG0000C |
| 20007852 | 20008247 | -1 | processed_pseudoge | ENSG0000C |
| 20012452 | 20012606 | 1 | processed_pseudoge | ENSG0000C |
| 20040446 | 20154404 | 1 | protein_coding | ENSG0000C |
| 20137565 | 20137641 | 1 | miRNA | ENSG0000C |
| 20160593 | 20186206 | -1 | protein_coding | ENSG0000C |
| 20174244 | 21145967 | 1 | lncRNA | ENSG0000C |
| 20268965 | 20269070 | -1 | snRNA | ENSG0000C |
| 20388249 | 20390562 | 1 | lncRNA | ENSG0000C |
| 20507644 | 20507756 | -1 | snRNA | ENSG0000C |
| 21006730 | 21226305 | -1 | lncRNA | ENSG0000C |
| 21405737 | 21406508 | 1 | processed_pseudoge | ENSG0000C |
| 21412218 | 22373321 | -1 | protein_coding | ENSG0000C |
| 21542789 | 21579959 | 1 | lncRNA | ENSG0000C |
| 21942566 | 21979828 | 1 | lncRNA | ENSG0000C |
| 22381819 | 22382929 | 1 | transcribed_processe | ENSG0000C |
| 22670201 | 22670711 | -1 | unprocessed_pseudc | ENSG0000C |

| | | | | |
|---|---|---|---|---|
| 22811390 | 22812047 | -1 | processed_pseudoge | ENSG00000( |
| 22904135 | 22905151 | 1 | processed_pseudoge | ENSG00000( |
| 22989823 | 22991582 | 1 | unprocessed_pseudc | ENSG00000( |
| 23134372 | 23134824 | 1 | processed_pseudoge | ENSG00000( |
| 23194929 | 23202703 | -1 | lncRNA | ENSG00000( |
| 23203020 | 23591794 | 1 | protein_coding | ENSG00000( |
| 23273807 | 23273910 | 1 | misc_RNA | ENSG00000( |
| 23389661 | 23389909 | -1 | unprocessed_pseudc | ENSG00000( |
| 23613661 | 23613759 | -1 | snRNA | ENSG00000( |
| 23641489 | 23641592 | -1 | snRNA | ENSG00000( |
| 23804024 | 23806905 | -1 | lncRNA | ENSG00000( |
| 23805955 | 23891640 | 1 | protein_coding | ENSG00000( |
| 23878981 | 23879498 | -1 | processed_pseudoge | ENSG00000( |
| 23891660 | 23946591 | -1 | protein_coding | ENSG00000( |
| 23916545 | 23924374 | 1 | protein_coding | ENSG00000( |
| 23945286 | 23980617 | 1 | protein_coding | ENSG00000( |
| 24096269 | 24103238 | -1 | lncRNA | ENSG00000( |
| 24096512 | 24097360 | -1 | transcribed_processe | ENSG00000( |
| 24117153 | 24495756 | -1 | protein_coding | ENSG00000( |
| 24430748 | 24431107 | 1 | processed_pseudoge | ENSG00000( |
| 24455136 | 24459434 | -1 | lncRNA | ENSG00000( |
| 24494087 | 24681711 | 1 | lncRNA | ENSG00000( |
| 24496272 | 24499580 | -1 | lncRNA | ENSG00000( |
| 24521362 | 24521435 | -1 | miRNA | ENSG00000( |
| 24555834 | 24556459 | 1 | processed_pseudoge | ENSG00000( |
| 24687919 | 25174305 | 1 | lncRNA | ENSG00000( |
| 25020166 | 25020463 | -1 | misc_RNA | ENSG00000( |
| 25025007 | 25025126 | 1 | rRNA_pseudogene | ENSG00000( |
| 25174332 | 25597932 | 1 | protein_coding | ENSG00000( |
| 25324174 | 25324659 | 1 | processed_pseudoge | ENSG00000( |
| 25352284 | 25352392 | -1 | rRNA_pseudogene | ENSG00000( |
| 25441164 | 25441502 | -1 | processed_pseudoge | ENSG00000( |
| 25555543 | 25555606 | -1 | snoRNA | ENSG00000( |
| 25597905 | 25664907 | -1 | protein_coding | ENSG00000( |
| 25618132 | 25618312 | -1 | misc_RNA | ENSG00000( |
| 25664873 | 25664939 | -1 | miRNA | ENSG00000( |
| 25686896 | 25687123 | 1 | processed_pseudoge | ENSG00000( |
| 25718944 | 25790039 | -1 | protein_coding | ENSG00000( |
| 25749488 | 25749896 | 1 | processed_pseudoge | ENSG00000( |
| 25754925 | 25755670 | 1 | processed_pseudoge | ENSG00000( |
| 25782917 | 25794534 | 1 | protein_coding | ENSG00000( |
| 25858525 | 25873748 | -1 | lncRNA | ENSG00000( |
| 26015669 | 26016369 | 1 | processed_pseudoge | ENSG00000( |
| 26216322 | 26216877 | 1 | processed_pseudoge | ENSG00000( |
| 26346681 | 26347638 | -1 | processed_pseudoge | ENSG00000( |
| 26614261 | 26622943 | -1 | lncRNA | ENSG00000( |
| 26622806 | 26717537 | 1 | protein_coding | ENSG00000( |
| 27110085 | 27369460 | -1 | protein_coding | ENSG00000( |
| 27214816 | 27215018 | -1 | processed_pseudoge | ENSG00000( |
| 27265580 | 27265683 | -1 | snRNA | ENSG00000( |

| | | | | |
|---|---|---|---|---|
| 27369568 | 27387960 | 1 | lncRNA | ENSG0000C |
| 27372721 | 27484420 | -1 | protein_coding | ENSG0000C |
| 27462506 | 27462816 | 1 | processed_pseudoge | ENSG0000C |
| 27486247 | 27486617 | 1 | processed_pseudoge | ENSG0000C |
| 27513871 | 27514032 | 1 | snRNA | ENSG0000C |
| 27525353 | 27525394 | 1 | processed_pseudoge | ENSG0000C |
| 27547757 | 27547858 | -1 | misc_RNA | ENSG0000C |
| 27624202 | 27624456 | -1 | processed_pseudoge | ENSG0000C |
| 27632976 | 27639111 | 1 | transcribed_processe | ENSG0000C |
| 27703129 | 27703568 | -1 | processed_pseudoge | ENSG0000C |
| 27712910 | 27714005 | 1 | lncRNA | ENSG0000C |
| 27715949 | 27722711 | -1 | protein_coding | ENSG0000C |
| 27797552 | 27931579 | 1 | lncRNA | ENSG0000C |
| 27830886 | 27834136 | -1 | lncRNA | ENSG0000C |
| 27916774 | 28059268 | -1 | lncRNA | ENSG0000C |
| 28241584 | 28325142 | 1 | protein_coding | ENSG0000C |
| 28315003 | 28349050 | -1 | protein_coding | ENSG0000C |
| 28348721 | 28538122 | 1 | protein_coding | ENSG0000C |
| 28349178 | 29767537 | 1 | protein_coding | ENSG0000C |
| 28427963 | 28428444 | -1 | processed_pseudoge | ENSG0000C |
| 28447725 | 28448073 | 1 | processed_pseudoge | ENSG0000C |
| 28574791 | 30010391 | 1 | protein_coding | ENSG0000C |
| 29054570 | 29290726 | -1 | lncRNA | ENSG0000C |
| 29087396 | 29088599 | 1 | processed_pseudoge | ENSG0000C |
| 29390848 | 29391218 | 1 | processed_pseudoge | ENSG0000C |
| 29526251 | 29642809 | -1 | lncRNA | ENSG0000C |
| 29797652 | 29797825 | -1 | processed_pseudoge | ENSG0000C |
| 29926811 | 29934156 | -1 | lncRNA | ENSG0000C |
| 30292548 | 30305037 | 1 | lncRNA | ENSG0000C |
| 30304320 | 30304536 | -1 | snoRNA | ENSG0000C |
| 30304321 | 30304536 | -1 | snoRNA | ENSG0000C |
| 30518833 | 30527193 | 1 | lncRNA | ENSG0000C |
| 30521664 | 30526254 | -1 | lncRNA | ENSG0000C |
| 30606502 | 30694142 | 1 | protein_coding | ENSG0000C |
| 30626423 | 30626797 | -1 | processed_pseudoge | ENSG0000C |
| 30697661 | 30699576 | 1 | lncRNA | ENSG0000C |
| 30726200 | 30894765 | -1 | protein_coding | ENSG0000C |
| 31161704 | 31161787 | -1 | miRNA | ENSG0000C |
| 31181751 | 31182650 | -1 | processed_pseudoge | ENSG0000C |
| 31228715 | 31228847 | 1 | rRNA_pseudogene | ENSG0000C |
| 31269007 | 31269406 | 1 | processed_pseudoge | ENSG0000C |
| 31452832 | 31455894 | 1 | processed_pseudoge | ENSG0000C |
| 31532638 | 31637616 | 1 | protein_coding | ENSG0000C |
| 31657890 | 32077580 | -1 | protein_coding | ENSG0000C |
| 31704058 | 31721652 | 1 | lncRNA | ENSG0000C |
| 31794470 | 31796175 | 1 | processed_pseudoge | ENSG0000C |
| 31981750 | 31991723 | 1 | protein_coding | ENSG0000C |
| 32030196 | 32030678 | 1 | processed_pseudoge | ENSG0000C |
| 32037526 | 32037653 | 1 | snoRNA | ENSG0000C |
| 32045873 | 32045979 | 1 | misc_RNA | ENSG0000C |

| | | | | |
|---|---|---|---|---|
| 32063974 | 32066611 | -1 | unprocessed_pseudc | ENSG0000C |
| 32105689 | 32168713 | 1 | protein_coding | ENSG0000C |
| 32151083 | 32151895 | -1 | unprocessed_pseudc | ENSG0000C |
| 32190747 | 32191627 | 1 | processed_pseudoge | ENSG0000C |
| 32236688 | 32238578 | -1 | lncRNA | ENSG0000C |
| 32238679 | 32370321 | 1 | protein_coding | ENSG0000C |
| 32243847 | 32244018 | -1 | processed_pseudoge | ENSG0000C |
| 32258940 | 32260230 | 1 | processed_pseudoge | ENSG0000C |
| 32391698 | 32483067 | 1 | protein_coding | ENSG0000C |
| 32481312 | 32502852 | -1 | protein_coding | ENSG0000C |
| 32506283 | 32506389 | -1 | miRNA | ENSG0000C |
| 32507955 | 32508325 | 1 | processed_pseudoge | ENSG0000C |
| 32525974 | 32570858 | -1 | protein_coding | ENSG0000C |
| 32592972 | 32593312 | -1 | processed_pseudoge | ENSG0000C |
| 32620903 | 32621902 | -1 | processed_pseudoge | ENSG0000C |
| 32635222 | 32635554 | -1 | processed_pseudoge | ENSG0000C |
| 32685145 | 32773875 | 1 | protein_coding | ENSG0000C |
| 32730635 | 32737454 | -1 | lncRNA | ENSG0000C |
| 32752910 | 32753901 | 1 | processed_pseudoge | ENSG0000C |
| 32785646 | 32786116 | 1 | processed_pseudoge | ENSG0000C |
| 32817997 | 32897824 | 1 | protein_coding | ENSG0000C |
| 32951644 | 32956349 | 1 | protein_coding | ENSG0000C |
| 32996609 | 33097202 | -1 | protein_coding | ENSG0000C |
| 33033863 | 33034810 | 1 | processed_pseudoge | ENSG0000C |
| 33077204 | 33077491 | 1 | processed_pseudoge | ENSG0000C |
| 33090421 | 33097146 | -1 | protein_coding | ENSG0000C |
| 33103008 | 33103306 | 1 | misc_RNA | ENSG0000C |
| 33113979 | 33147773 | 1 | protein_coding | ENSG0000C |
| 33144104 | 33147721 | -1 | lncRNA | ENSG0000C |
| 33150043 | 33218810 | -1 | protein_coding | ENSG0000C |
| 33190233 | 33190328 | -1 | processed_pseudoge | ENSG0000C |
| 33277025 | 33403662 | 1 | protein_coding | ENSG0000C |
| 33388336 | 33441371 | -1 | protein_coding | ENSG0000C |
| 33414150 | 33414211 | 1 | snRNA | ENSG0000C |
| 33491983 | 33492098 | -1 | rRNA_pseudogene | ENSG0000C |
| 33496245 | 33718356 | -1 | protein_coding | ENSG0000C |
| 33554636 | 33554859 | 1 | processed_pseudoge | ENSG0000C |
| 33727250 | 33727604 | -1 | processed_pseudoge | ENSG0000C |
| 33771963 | 33772295 | 1 | processed_pseudoge | ENSG0000C |
| 33793644 | 33798539 | -1 | lncRNA | ENSG0000C |
| 33798571 | 33869707 | 1 | protein_coding | ENSG0000C |
| 33956972 | 34677247 | 1 | lncRNA | ENSG0000C |
| 34152979 | 34157297 | 1 | lncRNA | ENSG0000C |
| 34203244 | 34268811 | 1 | lncRNA | ENSG0000C |
| 34524818 | 34543880 | -1 | lncRNA | ENSG0000C |
| 34692820 | 34694174 | 1 | lncRNA | ENSG0000C |
| 34872641 | 34873854 | 1 | processed_pseudoge | ENSG0000C |
| 35215705 | 35217149 | -1 | processed_pseudoge | ENSG0000C |
| 35256840 | 35256946 | -1 | snRNA | ENSG0000C |
| 35638945 | 35794496 | 1 | protein_coding | ENSG0000C |

| | | | | |
|---|---|---|---|---|
| 35650197 | 35651961 | -1 | lncRNA | ENSG0000C |
| 35744476 | 35744559 | 1 | miRNA | ENSG0000C |
| 35871866 | 35872198 | 1 | processed_pseudoge | ENSG0000C |
| 36169767 | 36170448 | -1 | processed_pseudoge | ENSG0000C |
| 36380344 | 36548007 | 1 | protein_coding | ENSG0000C |
| 36616006 | 36637457 | -1 | unprocessed_pseudc | ENSG0000C |
| 36685040 | 36685369 | 1 | misc_RNA | ENSG0000C |
| 36712422 | 36764349 | -1 | protein_coding | ENSG0000C |
| 36767117 | 36784167 | -1 | transcribed_processe | ENSG0000C |
| 36819276 | 36822498 | -1 | lncRNA | ENSG0000C |
| 36823151 | 36825158 | -1 | TEC | ENSG0000C |
| 36826820 | 36945098 | -1 | protein_coding | ENSG0000C |
| 36880184 | 36880729 | -1 | processed_pseudoge | ENSG0000C |
| 36968191 | 36968316 | -1 | snRNA | ENSG0000C |
| 36973117 | 36973672 | -1 | lncRNA | ENSG0000C |
| 36976316 | 36976840 | 1 | processed_pseudoge | ENSG0000C |
| 36985043 | 36993131 | -1 | protein_coding | ENSG0000C |
| 36993332 | 37050918 | 1 | protein_coding | ENSG0000C |
| 37016523 | 37017014 | -1 | processed_pseudoge | ENSG0000C |
| 37052626 | 37183689 | -1 | protein_coding | ENSG0000C |
| 37093133 | 37093234 | -1 | misc_RNA | ENSG0000C |
| 37134658 | 37135102 | 1 | processed_pseudoge | ENSG0000C |
| 37140327 | 37140430 | 1 | snRNA | ENSG0000C |
| 37143512 | 37143958 | -1 | processed_pseudoge | ENSG0000C |
| 37182107 | 37182734 | 1 | lncRNA | ENSG0000C |
| 37196204 | 37196666 | 1 | processed_pseudoge | ENSG0000C |
| 37216779 | 37217988 | 1 | lncRNA | ENSG0000C |
| 37241789 | 37244177 | -1 | lncRNA | ENSG0000C |
| 37243177 | 37366751 | 1 | protein_coding | ENSG0000C |
| 37275693 | 37276598 | 1 | processed_pseudoge | ENSG0000C |
| 37341649 | 37341766 | 1 | rRNA_pseudogene | ENSG0000C |
| 37381062 | 37440186 | 1 | lncRNA | ENSG0000C |
| 37452115 | 37823507 | 1 | protein_coding | ENSG0000C |
| 37573151 | 37573213 | -1 | snRNA | ENSG0000C |
| 37693655 | 37861802 | -1 | lncRNA | ENSG0000C |
| 37748110 | 37748900 | 1 | processed_pseudoge | ENSG0000C |
| 37789921 | 37790445 | -1 | processed_pseudoge | ENSG0000C |
| 37861880 | 37984469 | 1 | protein_coding | ENSG0000C |
| 37969404 | 37969480 | 1 | miRNA | ENSG0000C |
| 37988059 | 38007188 | 1 | protein_coding | ENSG0000C |
| 38007496 | 38029762 | -1 | protein_coding | ENSG0000C |
| 38014568 | 38014669 | 1 | misc_RNA | ENSG0000C |
| 38039205 | 38124025 | 1 | protein_coding | ENSG0000C |
| 38051759 | 38052395 | 1 | processed_pseudoge | ENSG0000C |
| 38103129 | 38137242 | -1 | protein_coding | ENSG0000C |
| 38125292 | 38125394 | -1 | misc_RNA | ENSG0000C |
| 38138478 | 38143022 | 1 | protein_coding | ENSG0000C |
| 38165089 | 38255484 | 1 | protein_coding | ENSG0000C |
| 38265812 | 38278757 | 1 | protein_coding | ENSG0000C |
| 38270283 | 38270767 | -1 | processed_pseudoge | ENSG0000C |

| | | | | |
|---|---|---|---|---|
| 38282294 | 38318575 | 1 | protein_coding | ENSG00000 |
| 38303773 | 38303879 | -1 | snRNA | ENSG00000 |
| 38325237 | 38329070 | -1 | unprocessed_pseudo | ENSG00000 |
| 38346760 | 38421348 | 1 | protein_coding | ENSG00000 |
| 38451027 | 38454820 | -1 | lncRNA | ENSG00000 |
| 38453890 | 38493142 | 1 | protein_coding | ENSG00000 |
| 38496127 | 38542161 | 1 | protein_coding | ENSG00000 |
| 38505480 | 38505581 | -1 | misc_RNA | ENSG00000 |
| 38515448 | 38515830 | -1 | processed_pseudoge | ENSG00000 |
| 38526802 | 38527325 | -1 | processed_pseudoge | ENSG00000 |
| 38548057 | 38649673 | -1 | protein_coding | ENSG00000 |
| 38696802 | 38816286 | -1 | protein_coding | ENSG00000 |
| 38823902 | 38825302 | 1 | lncRNA | ENSG00000 |
| 38845767 | 39051941 | -1 | protein_coding | ENSG00000 |
| 39025987 | 39026093 | 1 | snRNA | ENSG00000 |
| 39052013 | 39096671 | 1 | protein_coding | ENSG00000 |
| 39096659 | 39108363 | -1 | protein_coding | ENSG00000 |
| 39107704 | 39138903 | 1 | protein_coding | ENSG00000 |
| 39138206 | 39138266 | 1 | miRNA | ENSG00000 |
| 39141855 | 39154562 | -1 | protein_coding | ENSG00000 |
| 39148281 | 39172952 | 1 | lncRNA | ENSG00000 |
| 39152906 | 39154723 | 1 | lncRNA | ENSG00000 |
| 39161270 | 39161520 | 1 | misc_RNA | ENSG00000 |
| 39177761 | 39179586 | 1 | lncRNA | ENSG00000 |
| 39183210 | 39192596 | -1 | protein_coding | ENSG00000 |
| 39213069 | 39216585 | -1 | lncRNA | ENSG00000 |
| 39214199 | 39214672 | 1 | processed_pseudoge | ENSG00000 |
| 39258194 | 39263352 | 1 | lncRNA | ENSG00000 |
| 39263494 | 39281735 | -1 | protein_coding | ENSG00000 |
| 39285985 | 39287920 | 1 | processed_pseudoge | ENSG00000 |
| 39292556 | 39384300 | -1 | lncRNA | ENSG00000 |
| 39329709 | 39333680 | 1 | protein_coding | ENSG00000 |
| 39334979 | 39335939 | 1 | processed_pseudoge | ENSG00000 |
| 39338129 | 39338879 | -1 | processed_pseudoge | ENSG00000 |
| 39358545 | 39359922 | 1 | processed_pseudoge | ENSG00000 |
| 39383370 | 39397351 | 1 | protein_coding | ENSG00000 |
| 39406716 | 39412542 | 1 | protein_coding | ENSG00000 |
| 39408389 | 39408539 | 1 | snoRNA | ENSG00000 |
| 39411054 | 39411206 | 1 | snoRNA | ENSG00000 |
| 39420644 | 39424870 | 1 | lncRNA | ENSG00000 |
| 39425342 | 39448037 | -1 | lncRNA | ENSG00000 |
| 39467198 | 39529479 | 1 | protein_coding | ENSG00000 |
| 39494837 | 39502567 | -1 | lncRNA | ENSG00000 |
| 39643638 | 39644705 | 1 | processed_pseudoge | ENSG00000 |
| 39808914 | 40260321 | 1 | protein_coding | ENSG00000 |
| 39970125 | 39970266 | 1 | snRNA | ENSG00000 |
| 40061110 | 40309698 | -1 | lncRNA | ENSG00000 |
| 40209005 | 40209304 | 1 | misc_RNA | ENSG00000 |
| 40238608 | 40238741 | 1 | snoRNA | ENSG00000 |
| 40309707 | 40312424 | 1 | protein_coding | ENSG00000 |

| | | | | |
|---|---|---|---|---|
| 40313802 | 40453329 | -1 | lncRNA | ENSG00000 |
| 40322715 | 40323279 | -1 | processed_pseudoge | ENSG00000 |
| 40387184 | 40428744 | 1 | protein_coding | ENSG00000 |
| 40457292 | 40468587 | 1 | protein_coding | ENSG00000 |
| 40477113 | 40491053 | 1 | protein_coding | ENSG00000 |
| 40477131 | 40518736 | 1 | protein_coding | ENSG00000 |
| 40498891 | 40499003 | -1 | snRNA | ENSG00000 |
| 40524878 | 40574685 | 1 | protein_coding | ENSG00000 |
| 40586535 | 40587428 | -1 | processed_pseudoge | ENSG00000 |
| 40596207 | 40597494 | -1 | processed_pseudoge | ENSG00000 |
| 40603125 | 40605427 | -1 | lncRNA | ENSG00000 |
| 40699066 | 40699338 | -1 | processed_pseudoge | ENSG00000 |
| 40719859 | 40862626 | 1 | lncRNA | ENSG00000 |
| 40761545 | 40761799 | 1 | processed_pseudoge | ENSG00000 |
| 40869548 | 40875980 | 1 | lncRNA | ENSG00000 |
| 40970541 | 40971578 | -1 | lncRNA | ENSG00000 |
| 41162287 | 41168547 | -1 | lncRNA | ENSG00000 |
| 41189041 | 41189236 | 1 | processed_pseudoge | ENSG00000 |
| 41194741 | 41260096 | 1 | protein_coding | ENSG00000 |
| 41246599 | 41962130 | -1 | protein_coding | ENSG00000 |
| 41687868 | 41688033 | -1 | scaRNA | ENSG00000 |
| 41766615 | 41766919 | -1 | misc_RNA | ENSG00000 |
| 41859350 | 41859594 | -1 | processed_pseudoge | ENSG00000 |
| 41868998 | 41869095 | -1 | misc_RNA | ENSG00000 |
| 41899111 | 41899253 | -1 | snoRNA | ENSG00000 |
| 41955445 | 41955763 | 1 | processed_pseudoge | ENSG00000 |
| 41977008 | 41978034 | 1 | processed_pseudoge | ENSG00000 |
| 42013802 | 42225890 | 1 | protein_coding | ENSG00000 |
| 42051181 | 42051694 | -1 | processed_pseudoge | ENSG00000 |
| 42145646 | 42146371 | -1 | processed_pseudoge | ENSG00000 |
| 42221840 | 42221965 | 1 | snRNA | ENSG00000 |
| 42257825 | 42266185 | -1 | protein_coding | ENSG00000 |
| 42264989 | 42266390 | 1 | lncRNA | ENSG00000 |
| 42321005 | 42322695 | 1 | processed_pseudoge | ENSG00000 |
| 42340588 | 42342367 | 1 | processed_pseudoge | ENSG00000 |
| 42397083 | 42410610 | -1 | protein_coding | ENSG00000 |
| 42469640 | 42469879 | 1 | processed_pseudoge | ENSG00000 |
| 42489299 | 42537573 | 1 | protein_coding | ENSG00000 |
| 42506465 | 42533258 | -1 | lncRNA | ENSG00000 |
| 42547969 | 42601080 | -1 | protein_coding | ENSG00000 |
| 42581840 | 42595114 | 1 | protein_coding | ENSG00000 |
| 42600655 | 42648735 | 1 | protein_coding | ENSG00000 |
| 42601963 | 42654388 | -1 | lncRNA | ENSG00000 |
| 42653697 | 42667580 | 1 | protein_coding | ENSG00000 |
| 42665873 | 42683793 | -1 | lncRNA | ENSG00000 |
| 42676189 | 42676491 | -1 | misc_RNA | ENSG00000 |
| 42685537 | 42692544 | 1 | protein_coding | ENSG00000 |
| 42692663 | 42732922 | -1 | lncRNA | ENSG00000 |
| 42692663 | 42702824 | -1 | protein_coding | ENSG00000 |
| 42702541 | 42706776 | 1 | lncRNA | ENSG00000 |

| | | | | |
|---|---|---|---|---|
| 42705756 | 42773253 | -1 | protein_coding | ENSG0000C |
| 42705756 | 42804518 | -1 | protein_coding | ENSG0000C |
| 42719401 | 42723136 | 1 | lncRNA | ENSG0000C |
| 42732575 | 42746768 | 1 | lncRNA | ENSG0000C |
| 42770612 | 42773635 | 1 | lncRNA | ENSG0000C |
| 42784298 | 42804531 | -1 | protein_coding | ENSG0000C |
| 42785087 | 42852428 | -1 | lncRNA | ENSG0000C |
| 42804752 | 42887974 | 1 | protein_coding | ENSG0000C |
| 42809414 | 42908105 | 1 | lncRNA | ENSG0000C |
| 42809446 | 43055871 | 1 | protein_coding | ENSG0000C |
| 42809483 | 42942792 | 1 | protein_coding | ENSG0000C |
| 42856005 | 42875898 | -1 | protein_coding | ENSG0000C |
| 42889361 | 42890306 | 1 | processed_pseudoge | ENSG0000C |
| 42906142 | 42919334 | 1 | protein_coding | ENSG0000C |
| 42934252 | 42936785 | -1 | lncRNA | ENSG0000C |
| 42979267 | 43060211 | 1 | protein_coding | ENSG0000C |
| 43037121 | 43037451 | 1 | unprocessed_pseudc | ENSG0000C |
| 43079232 | 43106079 | -1 | protein_coding | ENSG0000C |
| 43087479 | 43088068 | 1 | lncRNA | ENSG0000C |
| 43088236 | 43090443 | 1 | lncRNA | ENSG0000C |
| 43286512 | 43424764 | 1 | protein_coding | ENSG0000C |
| 43314951 | 43315255 | 1 | misc_RNA | ENSG0000C |
| 43346923 | 43352152 | -1 | lncRNA | ENSG0000C |
| 43354859 | 43691594 | -1 | protein_coding | ENSG0000C |
| 43472477 | 43472596 | 1 | misc_RNA | ENSG0000C |
| 43484972 | 43485880 | -1 | processed_pseudoge | ENSG0000C |
| 43690108 | 43734371 | 1 | protein_coding | ENSG0000C |
| 43778961 | 43779982 | -1 | lncRNA | ENSG0000C |
| 43996896 | 43997443 | 1 | lncRNA | ENSG0000C |
| 43998081 | 43999149 | -1 | lncRNA | ENSG0000C |
| 44071087 | 44071186 | 1 | snRNA | ENSG0000C |
| 44114212 | 44114310 | 1 | miRNA | ENSG0000C |
| 44115818 | 44117487 | -1 | TEC | ENSG0000C |
| 44117299 | 44122365 | 1 | lncRNA | ENSG0000C |
| 44241886 | 44332098 | 1 | protein_coding | ENSG0000C |
| 44337941 | 44338552 | -1 | lncRNA | ENSG0000C |
| 44338119 | 44409451 | 1 | protein_coding | ENSG0000C |
| 44421127 | 44439946 | -1 | transcribed_unitary_ | ENSG0000C |
| 44424134 | 44429529 | 1 | lncRNA | ENSG0000C |
| 44431705 | 44477670 | -1 | protein_coding | ENSG0000C |
| 44494847 | 44510636 | -1 | protein_coding | ENSG0000C |
| 44524744 | 44525805 | -1 | processed_pseudoge | ENSG0000C |
| 44528955 | 44529391 | -1 | processed_pseudoge | ENSG0000C |
| 44555193 | 44594173 | 1 | protein_coding | ENSG0000C |
| 44557357 | 44685653 | -1 | lncRNA | ENSG0000C |
| 44578223 | 44599694 | 1 | protein_coding | ENSG0000C |
| 44579938 | 44581026 | -1 | processed_pseudoge | ENSG0000C |
| 44584888 | 44648471 | 1 | protein_coding | ENSG0000C |
| 44617128 | 44624797 | -1 | lncRNA | ENSG0000C |
| 44648732 | 44660791 | 1 | protein_coding | ENSG0000C |

| | | | | |
|---|---|---|---|---|
| 44667412 | 44669364 | 1 | lncRNA | ENSG0000( |
| 44709311 | 44710886 | -1 | processed_pseudoge | ENSG0000( |
| 44712643 | 44723831 | 1 | protein_coding | ENSG0000( |
| 44729596 | 44737083 | 1 | protein_coding | ENSG0000( |
| 44737661 | 44761619 | -1 | protein_coding | ENSG0000( |
| 44761721 | 44873376 | 1 | protein_coding | ENSG0000( |
| 44861888 | 44861981 | 1 | miRNA | ENSG0000( |
| 44861904 | 44865670 | 1 | protein_coding | ENSG0000( |
| 44874608 | 44914990 | 1 | protein_coding | ENSG0000( |
| 44899178 | 44900925 | 1 | lncRNA | ENSG0000( |
| 44915257 | 44976185 | -1 | protein_coding | ENSG0000( |
| 44937673 | 44938482 | 1 | processed_pseudoge | ENSG0000( |
| 44975241 | 45036066 | 1 | protein_coding | ENSG0000( |
| 45001548 | 45036071 | 1 | protein_coding | ENSG0000( |
| 45034474 | 45034588 | -1 | snRNA | ENSG0000( |
| 45082277 | 45146422 | -1 | protein_coding | ENSG0000( |
| 45159774 | 45160175 | -1 | processed_pseudoge | ENSG0000( |
| 45224466 | 45226287 | -1 | protein_coding | ENSG0000( |
| 45255409 | 45255620 | -1 | snoRNA | ENSG0000( |
| 45388561 | 45554726 | 1 | protein_coding | ENSG0000( |
| 45483974 | 45509545 | -1 | lncRNA | ENSG0000( |
| 45555394 | 45686341 | 1 | protein_coding | ENSG0000( |
| 45676369 | 45689200 | -1 | lncRNA | ENSG0000( |
| 45689056 | 45745409 | 1 | protein_coding | ENSG0000( |
| 45742675 | 45742970 | 1 | misc_RNA | ENSG0000( |
| 45749627 | 45750651 | 1 | TEC | ENSG0000( |
| 45755449 | 45796536 | -1 | protein_coding | ENSG0000( |
| 45823316 | 45916042 | -1 | protein_coding | ENSG0000( |
| 45842265 | 45866571 | 1 | lncRNA | ENSG0000( |
| 45883640 | 45884113 | -1 | processed_pseudoge | ENSG0000( |
| 45886504 | 45903175 | 1 | protein_coding | ENSG0000( |
| 45903187 | 45903298 | 1 | misc_RNA | ENSG0000( |
| 45917899 | 45995824 | -1 | protein_coding | ENSG0000( |
| 45940933 | 45948351 | 1 | protein_coding | ENSG0000( |
| 46017024 | 46027742 | -1 | protein_coding | ENSG0000( |
| 46023296 | 46024156 | 1 | processed_pseudoge | ENSG0000( |
| 46142358 | 46143677 | -1 | processed_pseudoge | ENSG0000( |
| 46163604 | 46266706 | 1 | protein_coding | ENSG0000( |
| 46201711 | 46208313 | -1 | protein_coding | ENSG0000( |
| 46310591 | 46311922 | -1 | processed_pseudoge | ENSG0000( |
| 46353734 | 46360928 | 1 | protein_coding | ENSG0000( |
| 46364955 | 46407059 | -1 | lncRNA | ENSG0000( |
| 46370854 | 46376206 | 1 | protein_coding | ENSG0000( |
| 46407166 | 46412997 | 1 | protein_coding | ENSG0000( |
| 46416524 | 46423591 | -1 | lncRNA | ENSG0000( |
| 46435645 | 46485234 | -1 | protein_coding | ENSG0000( |
| 46497976 | 46500950 | 1 | protein_coding | ENSG0000( |
| 46515385 | 46580099 | -1 | protein_coding | ENSG0000( |
| 46557398 | 46559694 | 1 | lncRNA | ENSG0000( |
| 46574534 | 46582457 | 1 | protein_coding | ENSG0000( |

| | | | | |
|---|---|---|---|---|
| 46581150 | 46617226 | 1 | protein_coding | ENSG00000 |
| 46612435 | 46626543 | 1 | protein_coding | ENSG00000 |
| 46647171 | 46661507 | -1 | lncRNA | ENSG00000 |
| 46650510 | 46650590 | 1 | snoRNA | ENSG00000 |
| 46668995 | 46693704 | -1 | protein_coding | ENSG00000 |
| 46694528 | 46710886 | 1 | protein_coding | ENSG00000 |
| 46712115 | 46812574 | -1 | protein_coding | ENSG00000 |
| 46712117 | 46717869 | -1 | protein_coding | ENSG00000 |
| 46719583 | 46736429 | -1 | transcribed_unitary_ | ENSG00000 |
| 46742092 | 46750891 | -1 | protein_coding | ENSG00000 |
| 46742092 | 46812558 | -1 | lncRNA | ENSG00000 |
| 46753045 | 46756857 | -1 | transcribed_unitary_ | ENSG00000 |
| 46798123 | 46798390 | 1 | processed_pseudoge | ENSG00000 |
| 46809359 | 46812486 | -1 | transcribed_unitary_ | ENSG00000 |
| 46829542 | 46834095 | -1 | transcribed_unitary_ | ENSG00000 |
| 46835111 | 46882172 | -1 | protein_coding | ENSG00000 |
| 46877721 | 46903799 | 1 | protein_coding | ENSG00000 |
| 46918807 | 46918935 | -1 | processed_pseudoge | ENSG00000 |
| 46921726 | 46982010 | -1 | protein_coding | ENSG00000 |
| 46979683 | 47009704 | 1 | protein_coding | ENSG00000 |
| 47010824 | 47013465 | 1 | transcribed_unitary_ | ENSG00000 |
| 47016429 | 47163967 | -1 | protein_coding | ENSG00000 |
| 47028855 | 47029030 | 1 | processed_pseudoge | ENSG00000 |
| 47164186 | 47246601 | 1 | lncRNA | ENSG00000 |
| 47228026 | 47283451 | -1 | protein_coding | ENSG00000 |
| 47250523 | 47250626 | -1 | snoRNA | ENSG00000 |
| 47282917 | 47346816 | 1 | protein_coding | ENSG00000 |
| 47346950 | 47349073 | 1 | TEC | ENSG00000 |
| 47379089 | 47380999 | -1 | lncRNA | ENSG00000 |
| 47381011 | 47413435 | 1 | protein_coding | ENSG00000 |
| 47413681 | 47477126 | -1 | protein_coding | ENSG00000 |
| 47469777 | 47469987 | 1 | processed_pseudoge | ENSG00000 |
| 47495640 | 47513712 | -1 | protein_coding | ENSG00000 |
| 47499841 | 47500407 | 1 | transcribed_processe | ENSG00000 |
| 47560229 | 47560524 | -1 | misc_RNA | ENSG00000 |
| 47562238 | 47580792 | -1 | protein_coding | ENSG00000 |
| 47585269 | 47782106 | -1 | protein_coding | ENSG00000 |
| 47601715 | 47604677 | -1 | lncRNA | ENSG00000 |
| 47663626 | 47663934 | 1 | processed_pseudoge | ENSG00000 |
| 47802909 | 47850195 | 1 | protein_coding | ENSG00000 |
| 47818005 | 47818128 | 1 | misc_RNA | ENSG00000 |
| 47849555 | 47849629 | 1 | miRNA | ENSG00000 |
| 47850690 | 48089272 | -1 | protein_coding | ENSG00000 |
| 47931808 | 47931914 | -1 | misc_RNA | ENSG00000 |
| 47960327 | 47961081 | -1 | processed_pseudoge | ENSG00000 |
| 48039267 | 48039490 | 1 | processed_pseudoge | ENSG00000 |
| 48094801 | 48095105 | -1 | misc_RNA | ENSG00000 |
| 48120669 | 48121215 | -1 | processed_pseudoge | ENSG00000 |
| 48157146 | 48188417 | -1 | protein_coding | ENSG00000 |
| 48161176 | 48161417 | 1 | processed_pseudoge | ENSG00000 |

| | | | | |
|---|---|---|---|---|
| 48180328 | 48180389 | 1 | snRNA | ENSG0000C |
| 48181547 | 48181726 | 1 | processed_pseudoge | ENSG0000C |
| 48196564 | 48196616 | 1 | miRNA | ENSG0000C |
| 48223347 | 48225491 | 1 | protein_coding | ENSG0000C |
| 48241100 | 48299253 | 1 | protein_coding | ENSG0000C |
| 48256350 | 48256938 | -1 | processed_pseudoge | ENSG0000C |
| 48288587 | 48288694 | 1 | misc_RNA | ENSG0000C |
| 48290722 | 48301685 | -1 | protein_coding | ENSG0000C |
| 48290793 | 48291375 | -1 | transcribed_processe | ENSG0000C |
| 48306842 | 48328341 | -1 | protein_coding | ENSG0000C |
| 48316360 | 48316459 | -1 | miRNA | ENSG0000C |
| 48372219 | 48401259 | 1 | protein_coding | ENSG0000C |
| 48379834 | 48380132 | 1 | misc_RNA | ENSG0000C |
| 48403854 | 48430086 | -1 | protein_coding | ENSG0000C |
| 48432164 | 48440456 | -1 | protein_coding | ENSG0000C |
| 48440257 | 48444208 | 1 | protein_coding | ENSG0000C |
| 48440352 | 48446656 | -1 | lncRNA | ENSG0000C |
| 48446710 | 48467645 | 1 | protein_coding | ENSG0000C |
| 48465811 | 48467645 | 1 | protein_coding | ENSG0000C |
| 48467798 | 48504826 | -1 | protein_coding | ENSG0000C |
| 48517684 | 48562015 | -1 | protein_coding | ENSG0000C |
| 48549961 | 48550021 | -1 | miRNA | ENSG0000C |
| 48561718 | 48563781 | -1 | protein_coding | ENSG0000C |
| 48564073 | 48595267 | -1 | protein_coding | ENSG0000C |
| 48578902 | 48578977 | -1 | miRNA | ENSG0000C |
| 48599002 | 48610976 | -1 | protein_coding | ENSG0000C |
| 48620759 | 48621769 | -1 | protein_coding | ENSG0000C |
| 48625723 | 48635493 | -1 | protein_coding | ENSG0000C |
| 48633636 | 48633698 | -1 | miRNA | ENSG0000C |
| 48636463 | 48662886 | -1 | protein_coding | ENSG0000C |
| 48644194 | 48644280 | -1 | miRNA | ENSG0000C |
| 48663776 | 48669174 | 1 | lncRNA | ENSG0000C |
| 48672455 | 48672733 | 1 | lncRNA | ENSG0000C |
| 48673844 | 48686364 | -1 | protein_coding | ENSG0000C |
| 48688003 | 48740353 | -1 | protein_coding | ENSG0000C |
| 48744597 | 48847874 | -1 | protein_coding | ENSG0000C |
| 48847572 | 48851981 | 1 | lncRNA | ENSG0000C |
| 48856926 | 48898904 | -1 | protein_coding | ENSG0000C |
| 48917782 | 48919385 | -1 | protein_coding | ENSG0000C |
| 48918821 | 48986382 | 1 | protein_coding | ENSG0000C |
| 48979918 | 48983985 | -1 | lncRNA | ENSG0000C |
| 48985049 | 48989988 | -1 | lncRNA | ENSG0000C |
| 48985485 | 48985963 | -1 | lncRNA | ENSG0000C |
| 48989889 | 49007153 | 1 | protein_coding | ENSG0000C |
| 48997221 | 48999442 | -1 | unitary_pseudogene | ENSG0000C |
| 49007062 | 49015953 | 1 | protein_coding | ENSG0000C |
| 49015488 | 49022293 | -1 | protein_coding | ENSG0000C |
| 49020148 | 49020234 | -1 | miRNA | ENSG0000C |
| 49020459 | 49023495 | 1 | protein_coding | ENSG0000C |
| 49020618 | 49020709 | -1 | miRNA | ENSG0000C |

| | | | | |
|---|---|---|---|---|
| 49024325 | 49029408 | -1 | protein_coding | ENSG0000C |
| 49029316 | 49029706 | 1 | lncRNA | ENSG0000C |
| 49029707 | 49094363 | -1 | protein_coding | ENSG0000C |
| 49083233 | 49083532 | 1 | misc_RNA | ENSG0000C |
| 49095932 | 49105130 | -1 | protein_coding | ENSG0000C |
| 49099854 | 49099914 | -1 | miRNA | ENSG0000C |
| 49108046 | 49120938 | -1 | protein_coding | ENSG0000C |
| 49121114 | 49133118 | -1 | protein_coding | ENSG0000C |
| 49140086 | 49160851 | -1 | unprocessed_pseudc | ENSG0000C |
| 49162535 | 49166331 | -1 | protein_coding | ENSG0000C |
| 49171598 | 49176486 | 1 | protein_coding | ENSG0000C |
| 49177634 | 49191858 | -1 | protein_coding | ENSG0000C |
| 49198428 | 49258106 | 1 | protein_coding | ENSG0000C |
| 49252129 | 49252264 | -1 | processed_pseudoge | ENSG0000C |
| 49260085 | 49261316 | 1 | lncRNA | ENSG0000C |
| 49268596 | 49277232 | -1 | protein_coding | ENSG0000C |
| 49270132 | 49270239 | 1 | misc_RNA | ENSG0000C |
| 49274120 | 49274186 | 1 | miRNA | ENSG0000C |
| 49277144 | 49340712 | -1 | protein_coding | ENSG0000C |
| 49357176 | 49358358 | -1 | polymorphic_pseudc | ENSG0000C |
| 49359145 | 49412998 | -1 | protein_coding | ENSG0000C |
| 49365145 | 49367006 | -1 | lncRNA | ENSG0000C |
| 49412206 | 49416475 | 1 | protein_coding | ENSG0000C |
| 49416775 | 49422753 | -1 | protein_coding | ENSG0000C |
| 49416777 | 49429314 | -1 | protein_coding | ENSG0000C |
| 49422946 | 49429326 | -1 | protein_coding | ENSG0000C |
| 49467115 | 49467228 | -1 | rRNA_pseudogene | ENSG0000C |
| 49468703 | 49535618 | 1 | protein_coding | ENSG0000C |
| 49549306 | 49554366 | -1 | lncRNA | ENSG0000C |
| 49554477 | 49671549 | 1 | protein_coding | ENSG0000C |
| 49640483 | 49641769 | -1 | lncRNA | ENSG0000C |
| 49674014 | 49683971 | 1 | protein_coding | ENSG0000C |
| 49683947 | 49689501 | -1 | protein_coding | ENSG0000C |
| 49684480 | 49684983 | -1 | lncRNA | ENSG0000C |
| 49689538 | 49721529 | 1 | protein_coding | ENSG0000C |
| 49716829 | 49719684 | -1 | protein_coding | ENSG0000C |
| 49716844 | 49723951 | -1 | protein_coding | ENSG0000C |
| 49724294 | 49786542 | -1 | protein_coding | ENSG0000C |
| 49758483 | 49758656 | 1 | processed_pseudoge | ENSG0000C |
| 49775285 | 49775614 | 1 | processed_pseudoge | ENSG0000C |
| 49790732 | 49799873 | -1 | protein_coding | ENSG0000C |
| 49803261 | 49805030 | 1 | protein_coding | ENSG0000C |
| 49805209 | 49813953 | -1 | protein_coding | ENSG0000C |
| 49806137 | 49806245 | -1 | miRNA | ENSG0000C |
| 49828601 | 49856574 | -1 | protein_coding | ENSG0000C |
| 49857988 | 49869935 | -1 | protein_coding | ENSG0000C |
| 49863375 | 49863660 | -1 | misc_RNA | ENSG0000C |
| 49873347 | 49877305 | -1 | unitary_pseudogene | ENSG0000C |
| 49887002 | 49903873 | -1 | protein_coding | ENSG0000C |
| 49899302 | 49903757 | 1 | lncRNA | ENSG0000C |

| | | | | |
|---|---|---|---|---|
| 49903845 | 49916937 | 1 | lncRNA | ENSG0000C |
| 49907160 | 49930173 | -1 | protein_coding | ENSG0000C |
| 49940007 | 50100045 | 1 | protein_coding | ENSG0000C |
| 50088919 | 50119021 | 1 | protein_coding | ENSG0000C |
| 50099603 | 50100988 | -1 | lncRNA | ENSG0000C |
| 50116022 | 50156085 | -1 | lncRNA | ENSG0000C |
| 50155045 | 50189075 | 1 | protein_coding | ENSG0000C |
| 50173326 | 50173419 | 1 | miRNA | ENSG0000C |
| 50191610 | 50197696 | 1 | protein_coding | ENSG0000C |
| 50205246 | 50221486 | 1 | protein_coding | ENSG0000C |
| 50226292 | 50259362 | 1 | protein_coding | ENSG0000C |
| 50227436 | 50227490 | 1 | miRNA | ENSG0000C |
| 50239618 | 50239984 | 1 | lncRNA | ENSG0000C |
| 50260303 | 50263358 | 1 | lncRNA | ENSG0000C |
| 50266641 | 50267371 | -1 | lncRNA | ENSG0000C |
| 50267558 | 50277546 | 1 | protein_coding | ENSG0000C |
| 50273236 | 50273297 | 1 | miRNA | ENSG0000C |
| 50279087 | 50288114 | 1 | protein_coding | ENSG0000C |
| 50287732 | 50292918 | -1 | protein_coding | ENSG0000C |
| 50292831 | 50299405 | -1 | protein_coding | ENSG0000C |
| 50296402 | 50299421 | -1 | protein_coding | ENSG0000C |
| 50299890 | 50312381 | -1 | protein_coding | ENSG0000C |
| 50317790 | 50322782 | -1 | protein_coding | ENSG0000C |
| 50320027 | 50328251 | -1 | protein_coding | ENSG0000C |
| 50329782 | 50340980 | -1 | protein_coding | ENSG0000C |
| 50337511 | 50338300 | 1 | lncRNA | ENSG0000C |
| 50341106 | 50345697 | 1 | lncRNA | ENSG0000C |
| 50341110 | 50345732 | -1 | protein_coding | ENSG0000C |
| 50347330 | 50350826 | -1 | protein_coding | ENSG0000C |
| 50350695 | 50358460 | 1 | protein_coding | ENSG0000C |
| 50350892 | 50367923 | 1 | protein_coding | ENSG0000C |
| 50354750 | 50359521 | -1 | protein_coding | ENSG0000C |
| 50362799 | 50504244 | -1 | protein_coding | ENSG0000C |
| 50365334 | 50368197 | 1 | lncRNA | ENSG0000C |
| 50419781 | 50419899 | 1 | rRNA_pseudogene | ENSG0000C |
| 50558025 | 50571027 | -1 | protein_coding | ENSG0000C |
| 50569152 | 50596168 | 1 | protein_coding | ENSG0000C |
| 50599879 | 50601148 | -1 | lncRNA | ENSG0000C |
| 50606490 | 50611831 | -1 | protein_coding | ENSG0000C |
| 50611520 | 50649291 | 1 | protein_coding | ENSG0000C |
| 50669989 | 50672048 | 1 | lncRNA | ENSG0000C |
| 50674927 | 51384198 | 1 | protein_coding | ENSG0000C |
| 50675080 | 50675163 | 1 | miRNA | ENSG0000C |
| 50685551 | 50685791 | -1 | processed_pseudoge | ENSG0000C |
| 50719482 | 50719951 | -1 | processed_pseudoge | ENSG0000C |
| 50786674 | 50788285 | -1 | processed_pseudoge | ENSG0000C |
| 50876712 | 50877657 | -1 | processed_pseudoge | ENSG0000C |
| 51385291 | 51389397 | 1 | protein_coding | ENSG0000C |
| 51391285 | 51397908 | 1 | protein_coding | ENSG0000C |
| 51395867 | 51500002 | -1 | protein_coding | ENSG0000C |

| | | | | |
|---|---|---|---|---|
| 51541144 | 51668667 | 1 | protein_coding | ENSG0000C |
| 51611438 | 51611562 | -1 | snRNA | ENSG0000C |
| 51662693 | 51704323 | 1 | protein_coding | ENSG0000C |
| 51668592 | 51671176 | -1 | lncRNA | ENSG0000C |
| 51694465 | 51694582 | -1 | rRNA_pseudogene | ENSG0000C |
| 51707068 | 51718613 | 1 | protein_coding | ENSG0000C |
| 51778561 | 51779241 | -1 | protein_coding | ENSG0000C |
| 51798803 | 51833389 | 1 | protein_coding | ENSG0000C |
| 51817603 | 51819635 | 1 | transcribed_unitary_ | ENSG0000C |
| 51826883 | 51830856 | 1 | protein_coding | ENSG0000C |
| 51861615 | 51863424 | 1 | protein_coding | ENSG0000C |
| 51872178 | 51872464 | 1 | misc_RNA | ENSG0000C |
| 51873596 | 51875767 | 1 | lncRNA | ENSG0000C |
| 51873721 | 51875601 | -1 | protein_coding | ENSG0000C |
| 51894876 | 51903353 | -1 | protein_coding | ENSG0000C |
| 51927071 | 51932483 | -1 | lncRNA | ENSG0000C |
| 51933429 | 51941904 | -1 | protein_coding | ENSG0000C |
| 51942345 | 51948867 | 1 | protein_coding | ENSG0000C |
| 51951849 | 51953902 | -1 | TEC | ENSG0000C |
| 51955381 | 51957499 | 1 | protein_coding | ENSG0000C |
| 51957454 | 51974016 | -1 | protein_coding | ENSG0000C |
| 51961212 | 51974031 | -1 | lncRNA | ENSG0000C |
| 51968510 | 51983409 | -1 | protein_coding | ENSG0000C |
| 51971426 | 51981199 | 1 | protein_coding | ENSG0000C |
| 51974706 | 51989183 | 1 | protein_coding | ENSG0000C |
| 51983340 | 51989197 | 1 | protein_coding | ENSG0000C |
| 51993522 | 51995895 | -1 | protein_coding | ENSG0000C |
| 52046763 | 52048657 | 1 | lncRNA | ENSG0000C |
| 52048919 | 52056571 | -1 | protein_coding | ENSG0000C |
| 52075226 | 52154690 | -1 | protein_coding | ENSG0000C |
| 52183722 | 52186834 | -1 | lncRNA | ENSG0000C |
| 52193170 | 52194785 | 1 | processed_pseudoge | ENSG0000C |
| 52198086 | 52214327 | 1 | protein_coding | ENSG0000C |
| 52217016 | 52219012 | -1 | TEC | ENSG0000C |
| 52221080 | 52226163 | -1 | protein_coding | ENSG0000C |
| 52221081 | 52231190 | -1 | protein_coding | ENSG0000C |
| 52228612 | 52239158 | -1 | protein_coding | ENSG0000C |
| 52239258 | 52241097 | 1 | lncRNA | ENSG0000C |
| 52245759 | 52250599 | 1 | protein_coding | ENSG0000C |
| 52254434 | 52288020 | -1 | protein_coding | ENSG0000C |
| 52268278 | 52268361 | -1 | miRNA | ENSG0000C |
| 52287089 | 52295257 | 1 | protein_coding | ENSG0000C |
| 52288580 | 52299067 | -1 | lncRNA | ENSG0000C |
| 52294219 | 52294308 | -1 | miRNA | ENSG0000C |
| 52316319 | 52400491 | 1 | protein_coding | ENSG0000C |
| 52373652 | 52374882 | 1 | processed_pseudoge | ENSG0000C |
| 52401008 | 52410008 | -1 | protein_coding | ENSG0000C |
| 52410660 | 52423641 | 1 | protein_coding | ENSG0000C |
| 52433035 | 52445103 | -1 | protein_coding | ENSG0000C |
| 52451100 | 52454041 | -1 | protein_coding | ENSG0000C |

| | | | | |
|---|---|---|---|---|
| 52455118 | 52493068 | 1 | protein_coding | ENSG00000 |
| 52495338 | 52524495 | 1 | protein_coding | ENSG00000 |
| 52524385 | 52535054 | -1 | protein_coding | ENSG00000 |
| 52534013 | 52579237 | 1 | protein_coding | ENSG00000 |
| 52545352 | 52685917 | -1 | protein_coding | ENSG00000 |
| 52551846 | 52551943 | -1 | snRNA | ENSG00000 |
| 52560570 | 52560707 | 1 | snRNA | ENSG00000 |
| 52681156 | 52694497 | 1 | protein_coding | ENSG00000 |
| 52688882 | 52688961 | 1 | snoRNA | ENSG00000 |
| 52688890 | 52688962 | 1 | snoRNA | ENSG00000 |
| 52689248 | 52689315 | 1 | snoRNA | ENSG00000 |
| 52690744 | 52690827 | 1 | snoRNA | ENSG00000 |
| 52691378 | 52691453 | 1 | snoRNA | ENSG00000 |
| 52692736 | 52692812 | 1 | snoRNA | ENSG00000 |
| 52694488 | 52706032 | -1 | protein_coding | ENSG00000 |
| 52704955 | 52711148 | 1 | protein_coding | ENSG00000 |
| 52708444 | 52770946 | -1 | protein_coding | ENSG00000 |
| 52732547 | 52733867 | 1 | processed_pseudoge | ENSG00000 |
| 52734735 | 52735013 | 1 | processed_pseudoge | ENSG00000 |
| 52753526 | 52754690 | 1 | processed_pseudoge | ENSG00000 |
| 52777595 | 52792068 | 1 | protein_coding | ENSG00000 |
| 52794768 | 52809009 | 1 | protein_coding | ENSG00000 |
| 52812962 | 52830688 | -1 | protein_coding | ENSG00000 |
| 52813282 | 52835729 | -1 | protein_coding | ENSG00000 |
| 52823935 | 52825314 | 1 | lncRNA | ENSG00000 |
| 52833114 | 52835019 | -1 | protein_coding | ENSG00000 |
| 52833121 | 52897562 | -1 | protein_coding | ENSG00000 |
| 52836219 | 52897548 | -1 | protein_coding | ENSG00000 |
| 52846463 | 52846552 | -1 | miRNA | ENSG00000 |
| 52848085 | 52848553 | -1 | TEC | ENSG00000 |
| 52903572 | 53046750 | -1 | protein_coding | ENSG00000 |
| 52969119 | 53099453 | -1 | protein_coding | ENSG00000 |
| 53046166 | 53048122 | 1 | TEC | ENSG00000 |
| 53064283 | 53065091 | -1 | processed_pseudoge | ENSG00000 |
| 53088483 | 53130453 | -1 | protein_coding | ENSG00000 |
| 53156009 | 53192717 | 1 | protein_coding | ENSG00000 |
| 53196820 | 53197359 | 1 | lncRNA | ENSG00000 |
| 53224712 | 53256052 | -1 | protein_coding | ENSG00000 |
| 53283429 | 53347586 | -1 | protein_coding | ENSG00000 |
| 53290640 | 53290751 | 1 | misc_RNA | ENSG00000 |
| 53328963 | 53813733 | 1 | protein_coding | ENSG00000 |
| 53333060 | 53333128 | -1 | snoRNA | ENSG00000 |
| 53387713 | 53387836 | -1 | snoRNA | ENSG00000 |
| 53474700 | 53475089 | -1 | processed_pseudoge | ENSG00000 |
| 53530322 | 53530401 | -1 | snoRNA | ENSG00000 |
| 53797764 | 53798019 | -1 | lncRNA | ENSG00000 |
| 53812335 | 53846419 | -1 | protein_coding | ENSG00000 |
| 53846568 | 53865794 | 1 | protein_coding | ENSG00000 |
| 53858994 | 53861576 | -1 | lncRNA | ENSG00000 |
| 53867066 | 53882152 | -1 | protein_coding | ENSG00000 |

| | | | | |
|---|---|---|---|---|
| 53884417 | 53891962 | -1 | protein_coding | ENSG00000( |
| 54033988 | 54058864 | 1 | lncRNA | ENSG00000( |
| 54047749 | 54048793 | -1 | processed_pseudoge | ENSG00000( |
| 54057690 | 54125377 | -1 | lncRNA | ENSG00000( |
| 54122547 | 55074557 | 1 | protein_coding | ENSG00000( |
| 54626215 | 54626646 | 1 | processed_pseudoge | ENSG00000( |
| 54632122 | 54639857 | -1 | lncRNA | ENSG00000( |
| 54874605 | 54901255 | -1 | lncRNA | ENSG00000( |
| 54918231 | 54967088 | -1 | protein_coding | ENSG00000( |
| 55166910 | 55175732 | -1 | lncRNA | ENSG00000( |
| 55178172 | 55189214 | 1 | lncRNA | ENSG00000( |
| 55226862 | 55300738 | 1 | lncRNA | ENSG00000( |
| 55335462 | 55350915 | -1 | lncRNA | ENSG00000( |
| 55360443 | 55361829 | -1 | lncRNA | ENSG00000( |
| 55465715 | 55490539 | -1 | protein_coding | ENSG00000( |
| 55487699 | 55488308 | 1 | lncRNA | ENSG00000( |
| 55493830 | 55505261 | -1 | lncRNA | ENSG00000( |
| 55508311 | 56468467 | -1 | protein_coding | ENSG00000( |
| 55610603 | 55613104 | 1 | lncRNA | ENSG00000( |
| 55657206 | 55659382 | -1 | lncRNA | ENSG00000( |
| 55852492 | 55852594 | -1 | miRNA | ENSG00000( |
| 56072104 | 56072407 | 1 | misc_RNA | ENSG00000( |
| 56185104 | 56185222 | 1 | rRNA_pseudogene | ENSG00000( |
| 56557161 | 56621837 | 1 | protein_coding | ENSG00000( |
| 56620133 | 56683237 | -1 | protein_coding | ENSG00000( |
| 56727418 | 57079329 | -1 | protein_coding | ENSG00000( |
| 56940040 | 56960854 | 1 | lncRNA | ENSG00000( |
| 57060664 | 57075432 | 1 | protein_coding | ENSG00000( |
| 57078943 | 57080101 | 1 | lncRNA | ENSG00000( |
| 57089982 | 57170306 | -1 | protein_coding | ENSG00000( |
| 57134142 | 57134723 | 1 | processed_pseudoge | ENSG00000( |
| 57197838 | 57227606 | -1 | protein_coding | ENSG00000( |
| 57227726 | 57278105 | 1 | protein_coding | ENSG00000( |
| 57268342 | 57292685 | -1 | protein_coding | ENSG00000( |
| 57293083 | 57300187 | 1 | lncRNA | ENSG00000( |
| 57293699 | 57544344 | -1 | protein_coding | ENSG00000( |
| 57493667 | 57493780 | 1 | misc_RNA | ENSG00000( |
| 57512450 | 57512555 | 1 | snRNA | ENSG00000( |
| 57519669 | 57520010 | -1 | processed_pseudoge | ENSG00000( |
| 57530345 | 57530967 | -1 | processed_pseudoge | ENSG00000( |
| 57536252 | 57536358 | -1 | snRNA | ENSG00000( |
| 57548838 | 57548945 | 1 | snRNA | ENSG00000( |
| 57555581 | 57556054 | -1 | lncRNA | ENSG00000( |
| 57556274 | 57566848 | 1 | protein_coding | ENSG00000( |
| 57571363 | 57598220 | -1 | protein_coding | ENSG00000( |
| 57597531 | 57600927 | 1 | lncRNA | ENSG00000( |
| 57619447 | 57619572 | 1 | snRNA | ENSG00000( |
| 57625454 | 57693077 | -1 | protein_coding | ENSG00000( |
| 57628810 | 57654918 | 1 | lncRNA | ENSG00000( |
| 57693205 | 57696596 | 1 | lncRNA | ENSG00000( |

| | | | | |
|---|---|---|---|---|
| 57755450 | 57930003 | 1 | protein_coding | ENSG0000( |
| 57800056 | 57801448 | -1 | processed_pseudoge | ENSG0000( |
| 57941818 | 57942304 | 1 | processed_pseudoge | ENSG0000( |
| 58008400 | 58172251 | 1 | protein_coding | ENSG0000( |
| 58162547 | 58170636 | -1 | lncRNA | ENSG0000( |
| 58174478 | 58178374 | -1 | lncRNA | ENSG0000( |
| 58180009 | 58187495 | 1 | transcribed_unitary_ | ENSG0000( |
| 58192257 | 58214697 | -1 | protein_coding | ENSG0000( |
| 58237532 | 58295693 | 1 | protein_coding | ENSG0000( |
| 58306247 | 58324695 | 1 | protein_coding | ENSG0000( |
| 58306262 | 58318574 | 1 | protein_coding | ENSG0000( |
| 58329965 | 58330118 | 1 | lncRNA | ENSG0000( |
| 58332880 | 58426127 | 1 | protein_coding | ENSG0000( |
| 58379217 | 58379525 | -1 | processed_pseudoge | ENSG0000( |
| 58427630 | 58433857 | -1 | protein_coding | ENSG0000( |
| 58428255 | 58428815 | 1 | lncRNA | ENSG0000( |
| 58490830 | 58491291 | -1 | lncRNA | ENSG0000( |
| 58492096 | 58502360 | 1 | protein_coding | ENSG0000( |
| 58505136 | 58537283 | -1 | protein_coding | ENSG0000( |
| 58511310 | 58511676 | 1 | processed_pseudoge | ENSG0000( |
| 58564117 | 58627610 | -1 | protein_coding | ENSG0000( |
| 58572744 | 58574319 | 1 | lncRNA | ENSG0000( |
| 58607080 | 58644491 | 1 | lncRNA | ENSG0000( |
| 58633946 | 58666834 | -1 | protein_coding | ENSG0000( |
| 58717365 | 59050084 | -1 | protein_coding | ENSG0000( |
| 58824437 | 59019093 | 1 | lncRNA | ENSG0000( |
| 59050164 | 59754808 | 1 | lncRNA | ENSG0000( |
| 59380024 | 59388704 | -1 | lncRNA | ENSG0000( |
| 59464330 | 59510678 | -1 | lncRNA | ENSG0000( |
| 59535863 | 59536415 | 1 | processed_pseudoge | ENSG0000( |
| 59584169 | 59584445 | 1 | misc_RNA | ENSG0000( |
| 59747277 | 61251459 | -1 | protein_coding | ENSG0000( |
| 59809269 | 59811310 | 1 | lncRNA | ENSG0000( |
| 60616822 | 60618079 | 1 | processed_pseudoge | ENSG0000( |
| 60617805 | 60617870 | -1 | miRNA | ENSG0000( |
| 60690225 | 60690719 | -1 | processed_pseudoge | ENSG0000( |
| 60730055 | 60730644 | -1 | processed_pseudoge | ENSG0000( |
| 60732144 | 60732636 | -1 | processed_pseudoge | ENSG0000( |
| 60856389 | 60856605 | -1 | snoRNA | ENSG0000( |
| 61561569 | 62297609 | 1 | protein_coding | ENSG0000( |
| 61739127 | 61739229 | -1 | misc_RNA | ENSG0000( |
| 61742455 | 61743106 | -1 | processed_pseudoge | ENSG0000( |
| 62118300 | 62118510 | -1 | snRNA | ENSG0000( |
| 62124204 | 62124608 | -1 | processed_pseudoge | ENSG0000( |
| 62221249 | 62369406 | -1 | lncRNA | ENSG0000( |
| 62318973 | 62336213 | 1 | protein_coding | ENSG0000( |
| 62369681 | 62374324 | -1 | protein_coding | ENSG0000( |
| 62398346 | 62875416 | -1 | protein_coding | ENSG0000( |
| 62779176 | 62779785 | -1 | processed_pseudoge | ENSG0000( |
| 62796124 | 62796419 | -1 | misc_RNA | ENSG0000( |

| | | | | |
|---|---|---|---|---|
| 62814382 | 62814491 | 1 | snRNA | ENSG0000C |
| 62926371 | 62926882 | 1 | processed_pseudoge | ENSG0000C |
| 62950430 | 63125062 | 1 | lncRNA | ENSG0000C |
| 63096374 | 63097711 | -1 | processed_pseudoge | ENSG0000C |
| 63203218 | 63203448 | -1 | processed_pseudoge | ENSG0000C |
| 63228315 | 63616924 | 1 | protein_coding | ENSG0000C |
| 63423596 | 63550051 | -1 | lncRNA | ENSG0000C |
| 63545440 | 63545535 | 1 | rRNA_pseudogene | ENSG0000C |
| 63652675 | 63679020 | 1 | protein_coding | ENSG0000C |
| 63741827 | 63827445 | -1 | transcribed_unitary_ | ENSG0000C |
| 63819299 | 63848636 | 1 | protein_coding | ENSG0000C |
| 63833870 | 63863868 | -1 | protein_coding | ENSG0000C |
| 63860645 | 63861819 | 1 | lncRNA | ENSG0000C |
| 63863155 | 63905838 | 1 | lncRNA | ENSG0000C |
| 63864557 | 64003462 | 1 | protein_coding | ENSG0000C |
| 63898399 | 64003453 | 1 | protein_coding | ENSG0000C |
| 63911518 | 63911772 | -1 | lncRNA | ENSG0000C |
| 64004022 | 64012148 | 1 | lncRNA | ENSG0000C |
| 64008082 | 64008692 | -1 | lncRNA | ENSG0000C |
| 64010549 | 64024010 | -1 | protein_coding | ENSG0000C |
| 64011964 | 64016246 | 1 | lncRNA | ENSG0000C |
| 64019508 | 64019925 | 1 | lncRNA | ENSG0000C |
| 64067964 | 64103131 | 1 | lncRNA | ENSG0000C |
| 64071807 | 64072019 | -1 | snoRNA | ENSG0000C |
| 64078361 | 64087363 | -1 | lncRNA | ENSG0000C |
| 64092236 | 64445476 | -1 | protein_coding | ENSG0000C |
| 64099273 | 64101122 | 1 | lncRNA | ENSG0000C |
| 64103470 | 64106056 | 1 | lncRNA | ENSG0000C |
| 64174401 | 64175147 | 1 | processed_pseudoge | ENSG0000C |
| 64187544 | 64200965 | 1 | lncRNA | ENSG0000C |
| 64445231 | 64456070 | 1 | lncRNA | ENSG0000C |
| 64511755 | 64511857 | 1 | snRNA | ENSG0000C |
| 64515654 | 64688000 | -1 | protein_coding | ENSG0000C |
| 64561322 | 64592757 | 1 | lncRNA | ENSG0000C |
| 64684909 | 65053439 | 1 | lncRNA | ENSG0000C |
| 65174916 | 65193509 | 1 | lncRNA | ENSG0000C |
| 65353525 | 66038834 | -1 | protein_coding | ENSG0000C |
| 65359268 | 65359717 | -1 | lncRNA | ENSG0000C |
| 65687301 | 65688468 | 1 | processed_pseudoge | ENSG0000C |
| 65843890 | 65844446 | 1 | processed_pseudoge | ENSG0000C |
| 65872815 | 65954558 | -1 | lncRNA | ENSG0000C |
| 65893816 | 65925297 | 1 | lncRNA | ENSG0000C |
| 66003839 | 66003940 | -1 | misc_RNA | ENSG0000C |
| 66008770 | 66009200 | -1 | processed_pseudoge | ENSG0000C |
| 66133610 | 66388116 | 1 | protein_coding | ENSG0000C |
| 66293114 | 66293220 | -1 | snRNA | ENSG0000C |
| 66323855 | 66324152 | 1 | misc_RNA | ENSG0000C |
| 66378797 | 66501263 | -1 | protein_coding | ENSG0000C |
| 66622760 | 66623102 | -1 | processed_pseudoge | ENSG0000C |
| 66637149 | 66637412 | 1 | processed_pseudoge | ENSG0000C |

| | | | | |
|---|---|---|---|---|
| 66780076 | 66824439 | -1 | lncRNA | ENSG0000( |
| 66794041 | 66796921 | 1 | lncRNA | ENSG0000( |
| 66997168 | 66998020 | -1 | lncRNA | ENSG0000( |
| 66998307 | 67011210 | 1 | protein_coding | ENSG0000( |
| 67225464 | 67225527 | 1 | miRNA | ENSG0000( |
| 67296300 | 67306359 | 1 | lncRNA | ENSG0000( |
| 67360460 | 67654614 | -1 | protein_coding | ENSG0000( |
| 67513940 | 67514320 | 1 | processed_pseudoge | ENSG0000( |
| 67654669 | 67947713 | 1 | lncRNA | ENSG0000( |
| 68004247 | 68545621 | 1 | protein_coding | ENSG0000( |
| 68145004 | 68145630 | 1 | processed_pseudoge | ENSG0000( |
| 68192398 | 68192580 | 1 | processed_pseudoge | ENSG0000( |
| 68500613 | 68505873 | 1 | lncRNA | ENSG0000( |
| 68633706 | 68634356 | -1 | processed_pseudoge | ENSG0000( |
| 68635705 | 68637024 | 1 | processed_pseudoge | ENSG0000( |
| 68731766 | 68953297 | -1 | protein_coding | ENSG0000( |
| 68882967 | 68883091 | 1 | rRNA_pseudogene | ENSG0000( |
| 68975217 | 69013684 | -1 | protein_coding | ENSG0000( |
| 69013941 | 69056622 | 1 | lncRNA | ENSG0000( |
| 69019827 | 69052339 | -1 | protein_coding | ENSG0000( |
| 69048958 | 69049035 | -1 | miRNA | ENSG0000( |
| 69054730 | 69080408 | -1 | protein_coding | ENSG0000( |
| 69084937 | 69106092 | 1 | protein_coding | ENSG0000( |
| 69106872 | 69123032 | -1 | protein_coding | ENSG0000( |
| 69168782 | 69542583 | -1 | protein_coding | ENSG0000( |
| 69471452 | 69472532 | -1 | processed_pseudoge | ENSG0000( |
| 69535664 | 69536693 | -1 | processed_pseudoge | ENSG0000( |
| 69564106 | 69571172 | -1 | lncRNA | ENSG0000( |
| 69591341 | 69593121 | 1 | lncRNA | ENSG0000( |
| 69739435 | 69968337 | 1 | protein_coding | ENSG0000( |
| 69769487 | 69769773 | -1 | misc_RNA | ENSG0000( |
| 69999550 | 70518064 | 1 | lncRNA | ENSG0000( |
| 70125070 | 70126609 | -1 | processed_pseudoge | ENSG0000( |
| 70140217 | 70140488 | 1 | processed_pseudoge | ENSG0000( |
| 70194479 | 70312638 | -1 | protein_coding | ENSG0000( |
| 70605420 | 70617605 | -1 | lncRNA | ENSG0000( |
| 70617766 | 70620325 | 1 | lncRNA | ENSG0000( |
| 70751123 | 70751350 | 1 | processed_pseudoge | ENSG0000( |
| 70753854 | 70754573 | 1 | processed_pseudoge | ENSG0000( |
| 70808350 | 70808452 | 1 | snRNA | ENSG0000( |
| 70952817 | 71583993 | -1 | protein_coding | ENSG0000( |
| 70959226 | 71754229 | -1 | protein_coding | ENSG0000( |
| 71289769 | 71305853 | 1 | lncRNA | ENSG0000( |
| 71541970 | 71542089 | -1 | miRNA | ENSG0000( |
| 71567863 | 71571112 | 1 | lncRNA | ENSG0000( |
| 71570255 | 71574457 | -1 | lncRNA | ENSG0000( |
| 71581721 | 71628558 | 1 | lncRNA | ENSG0000( |
| 71584943 | 71587409 | 1 | lncRNA | ENSG0000( |
| 71675414 | 71754773 | -1 | protein_coding | ENSG0000( |
| 71753855 | 71756496 | 1 | protein_coding | ENSG0000( |

| | | | | |
|---|---|---|---|---|
| 71771655 | 71785206 | -1 | protein_coding | ENSG00000 |
| 71785428 | 71786425 | 1 | lncRNA | ENSG00000 |
| 71827177 | 71827471 | 1 | misc_RNA | ENSG00000 |
| 71943592 | 72279503 | -1 | lncRNA | ENSG00000 |
| 72006774 | 72006975 | 1 | processed_pseudoge | ENSG00000 |
| 72061061 | 72069933 | 1 | lncRNA | ENSG00000 |
| 72080564 | 72084641 | -1 | lncRNA | ENSG00000 |
| 72085109 | 72085287 | 1 | processed_pseudoge | ENSG00000 |
| 72086888 | 72087725 | 1 | processed_pseudoge | ENSG00000 |
| 72151236 | 72207776 | 1 | lncRNA | ENSG00000 |
| 72178279 | 72178810 | 1 | lncRNA | ENSG00000 |
| 72202944 | 72209914 | -1 | lncRNA | ENSG00000 |
| 72301616 | 72319429 | -1 | lncRNA | ENSG00000 |
| 72321051 | 72324027 | -1 | lncRNA | ENSG00000 |
| 72371825 | 72446621 | -1 | protein_coding | ENSG00000 |
| 72504806 | 72550552 | -1 | lncRNA | ENSG00000 |
| 72576691 | 72576854 | 1 | snRNA | ENSG00000 |
| 72583370 | 72583535 | 1 | processed_pseudoge | ENSG00000 |
| 72691705 | 72691811 | -1 | rRNA_pseudogene | ENSG00000 |
| 72738918 | 72740525 | 1 | lncRNA | ENSG00000 |
| 72749277 | 72861914 | -1 | protein_coding | ENSG00000 |
| 72772539 | 72773471 | 1 | processed_pseudoge | ENSG00000 |
| 72876737 | 72878102 | -1 | unprocessed_pseudo | ENSG00000 |
| 72884234 | 72884679 | 1 | processed_pseudoge | ENSG00000 |
| 72888046 | 72998138 | 1 | protein_coding | ENSG00000 |
| 72926955 | 72927237 | -1 | misc_RNA | ENSG00000 |
| 72929044 | 72929593 | 1 | processed_pseudoge | ENSG00000 |
| 72996803 | 73069198 | 1 | protein_coding | ENSG00000 |
| 73027808 | 73027873 | 1 | snRNA | ENSG00000 |
| 73061659 | 73063337 | 1 | protein_coding | ENSG00000 |
| 73092148 | 73092257 | -1 | snRNA | ENSG00000 |
| 73110992 | 73111182 | 1 | snRNA | ENSG00000 |
| 73182501 | 73183291 | -1 | processed_pseudoge | ENSG00000 |
| 73241789 | 73241896 | -1 | snRNA | ENSG00000 |
| 73382431 | 73624941 | -1 | protein_coding | ENSG00000 |
| 73593940 | 73594002 | -1 | snRNA | ENSG00000 |
| 73621713 | 73626796 | 1 | lncRNA | ENSG00000 |
| 73624410 | 73627626 | 1 | TEC | ENSG00000 |
| 73648399 | 73648807 | -1 | processed_pseudoge | ENSG00000 |
| 73805754 | 73902960 | 1 | lncRNA | ENSG00000 |
| 73959642 | 73959927 | -1 | misc_RNA | ENSG00000 |
| 73999805 | 74005609 | 1 | lncRNA | ENSG00000 |
| 74101349 | 74103553 | 1 | processed_pseudoge | ENSG00000 |
| 74135959 | 74136892 | 1 | processed_pseudoge | ENSG00000 |
| 74262568 | 74521140 | -1 | protein_coding | ENSG00000 |
| 75084155 | 75085237 | 1 | processed_pseudoge | ENSG00000 |
| 75182754 | 75183048 | -1 | misc_RNA | ENSG00000 |
| 75214476 | 75214549 | 1 | miRNA | ENSG00000 |
| 75214510 | 75240770 | 1 | lncRNA | ENSG00000 |
| 75214631 | 75215636 | 1 | processed_pseudoge | ENSG00000 |

| | | | | |
|---|---|---|---|---|
| 75328549 | 75339071 | -1 | unprocessed_pseudc | ENSG00000( |
| 75347833 | 75348787 | -1 | unprocessed_pseudc | ENSG00000( |
| 75356437 | 75362268 | -1 | transcribed_unproce | ENSG00000( |
| 75370448 | 75371342 | -1 | unprocessed_pseudc | ENSG00000( |
| 75384059 | 75391810 | -1 | unprocessed_pseudc | ENSG00000( |
| 75395188 | 75395320 | 1 | processed_pseudoge | ENSG00000( |
| 75395404 | 75395536 | -1 | processed_pseudoge | ENSG00000( |
| 75415070 | 75422143 | 1 | unprocessed_pseudc | ENSG00000( |
| 75421552 | 75435110 | -1 | transcribed_unproce | ENSG00000( |
| 75435232 | 75538029 | 1 | lncRNA | ENSG00000( |
| 75446769 | 75491202 | -1 | unprocessed_pseudc | ENSG00000( |
| 75521803 | 75522012 | 1 | processed_pseudoge | ENSG00000( |
| 75540049 | 75580199 | -1 | transcribed_unproce | ENSG00000( |
| 75542149 | 75542929 | -1 | processed_pseudoge | ENSG00000( |
| 75590298 | 75590649 | -1 | processed_pseudoge | ENSG00000( |
| 75598652 | 75599673 | -1 | unprocessed_pseudc | ENSG00000( |
| 75619506 | 75623849 | -1 | unprocessed_pseudc | ENSG00000( |
| 75624275 | 75670156 | -1 | transcribed_process | ENSG00000( |
| 75624899 | 75625188 | 1 | processed_pseudoge | ENSG00000( |
| 75629993 | 75634057 | 1 | unprocessed_pseudc | ENSG00000( |
| 75630763 | 75630858 | 1 | miRNA | ENSG00000( |
| 75653561 | 75654773 | -1 | processed_pseudoge | ENSG00000( |
| 75658534 | 75659014 | 1 | processed_pseudoge | ENSG00000( |
| 75664330 | 75667220 | 1 | protein_coding | ENSG00000( |
| 75668931 | 75670185 | 1 | processed_pseudoge | ENSG00000( |
| 75672300 | 75742048 | 1 | lncRNA | ENSG00000( |
| 75678660 | 75785583 | -1 | protein_coding | ENSG00000( |
| 75738280 | 75738363 | 1 | miRNA | ENSG00000( |
| 75906695 | 77649964 | 1 | protein_coding | ENSG00000( |
| 75964980 | 75965069 | -1 | misc_RNA | ENSG00000( |
| 76434018 | 76435428 | 1 | processed_pseudoge | ENSG00000( |
| 77092057 | 77092163 | 1 | snRNA | ENSG00000( |
| 77316752 | 77317598 | -1 | processed_pseudoge | ENSG00000( |
| 77567508 | 77571881 | -1 | lncRNA | ENSG00000( |
| 77811741 | 77811844 | -1 | snRNA | ENSG00000( |
| 78038705 | 78046794 | -1 | lncRNA | ENSG00000( |
| 78087404 | 78092270 | 1 | lncRNA | ENSG00000( |
| 78115158 | 78115457 | 1 | misc_RNA | ENSG00000( |
| 78161698 | 78161992 | 1 | misc_RNA | ENSG00000( |
| 78266940 | 78294731 | 1 | lncRNA | ENSG00000( |
| 78525101 | 78525485 | 1 | processed_pseudoge | ENSG00000( |
| 78597239 | 79767998 | -1 | protein_coding | ENSG00000( |
| 78758974 | 78759374 | 1 | processed_pseudoge | ENSG00000( |
| 78938009 | 78938292 | -1 | misc_RNA | ENSG00000( |
| 79411714 | 79470827 | 1 | lncRNA | ENSG00000( |
| 79507887 | 79507969 | 1 | miRNA | ENSG00000( |
| 79957566 | 79958142 | -1 | processed_pseudoge | ENSG00000( |
| 80163327 | 80164795 | 1 | processed_pseudoge | ENSG00000( |
| 80216261 | 80217234 | 1 | processed_pseudoge | ENSG00000( |
| 80442081 | 80442305 | 1 | unprocessed_pseudc | ENSG00000( |

| | | | | |
|---|---|---|---|---|
| 80602716 | 80677898 | -1 | lncRNA | ENSG0000C |
| 80761042 | 80788963 | -1 | lncRNA | ENSG0000C |
| 80764862 | 80789388 | 1 | lncRNA | ENSG0000C |
| 80993854 | 81105182 | 1 | lncRNA | ENSG0000C |
| 81208193 | 81208914 | -1 | lncRNA | ENSG0000C |
| 81246579 | 81297345 | -1 | lncRNA | ENSG0000C |
| 81359739 | 81368191 | 1 | lncRNA | ENSG0000C |
| 81366990 | 81368021 | -1 | processed_pseudoge | ENSG0000C |
| 81489703 | 81761645 | -1 | protein_coding | ENSG0000C |
| 81509476 | 81509664 | 1 | snRNA | ENSG0000C |
| 81612238 | 81612805 | -1 | processed_pseudoge | ENSG0000C |
| 81612790 | 81613075 | 1 | processed_pseudoge | ENSG0000C |
| 81762706 | 81764756 | -1 | lncRNA | ENSG0000C |
| 81840514 | 81953534 | 1 | lncRNA | ENSG0000C |
| 81986138 | 82463675 | 1 | lncRNA | ENSG0000C |
| 82319172 | 82320279 | -1 | processed_pseudoge | ENSG0000C |
| 82806515 | 82808021 | -1 | processed_pseudoge | ENSG0000C |
| 83384445 | 83437728 | 1 | lncRNA | ENSG0000C |
| 83924157 | 83937031 | -1 | lncRNA | ENSG0000C |
| 83937045 | 83938236 | 1 | processed_pseudoge | ENSG0000C |
| 83953639 | 84051419 | 1 | lncRNA | ENSG0000C |
| 84638405 | 84881924 | -1 | lncRNA | ENSG0000C |
| 84691681 | 84692591 | -1 | processed_pseudoge | ENSG0000C |
| 84881984 | 84893379 | 1 | lncRNA | ENSG0000C |
| 84940619 | 84947061 | 1 | lncRNA | ENSG0000C |
| 84958981 | 86074429 | 1 | protein_coding | ENSG0000C |
| 85385710 | 85385792 | 1 | miRNA | ENSG0000C |
| 85799987 | 85828050 | -1 | lncRNA | ENSG0000C |
| 85992183 | 86028007 | -1 | lncRNA | ENSG0000C |
| 86012561 | 86015120 | 1 | processed_pseudoge | ENSG0000C |
| 86125216 | 86125287 | 1 | snoRNA | ENSG0000C |
| 86235321 | 86235424 | 1 | snRNA | ENSG0000C |
| 86258936 | 86267424 | -1 | lncRNA | ENSG0000C |
| 86362132 | 86362474 | 1 | misc_RNA | ENSG0000C |
| 86481943 | 86496996 | 1 | lncRNA | ENSG0000C |
| 86816740 | 86817036 | 1 | processed_pseudoge | ENSG0000C |
| 86876388 | 86991149 | -1 | protein_coding | ENSG0000C |
| 87051192 | 87052738 | -1 | processed_pseudoge | ENSG0000C |
| 87089129 | 87158363 | 1 | lncRNA | ENSG0000C |
| 87226189 | 87226277 | -1 | miRNA | ENSG0000C |
| 87227271 | 87255556 | 1 | protein_coding | ENSG0000C |
| 87259404 | 87276584 | -1 | protein_coding | ENSG0000C |
| 87323268 | 87324723 | 1 | processed_pseudoge | ENSG0000C |
| 87594838 | 87595376 | 1 | processed_pseudoge | ENSG0000C |
| 87630301 | 87631619 | -1 | processed_pseudoge | ENSG0000C |
| 87641412 | 87793629 | -1 | lncRNA | ENSG0000C |
| 87742543 | 87742649 | -1 | snRNA | ENSG0000C |
| 87910220 | 87910345 | 1 | snRNA | ENSG0000C |
| 87990696 | 87993835 | 1 | protein_coding | ENSG0000C |
| 88051944 | 88149885 | -1 | protein_coding | ENSG0000C |

| | | | | |
|---|---|---|---|---|
| 88059255 | 88144664 | 1 | protein_coding | ENSG0000C |
| 88095408 | 88095968 | -1 | processed_pseudoge | ENSG0000C |
| 88149959 | 88168729 | 1 | protein_coding | ENSG0000C |
| 88317156 | 88319022 | 1 | processed_pseudoge | ENSG0000C |
| 88338416 | 88467594 | 1 | protein_coding | ENSG0000C |
| 88431191 | 88431309 | 1 | misc_RNA | ENSG0000C |
| 88601061 | 88601402 | 1 | processed_pseudoge | ENSG0000C |
| 88948142 | 88950928 | 1 | processed_pseudoge | ENSG0000C |
| 89047442 | 89048434 | -1 | processed_pseudoge | ENSG0000C |
| 89107621 | 89482134 | 1 | protein_coding | ENSG0000C |
| 89283016 | 89314604 | -1 | lncRNA | ENSG0000C |
| 89587886 | 89588570 | -1 | processed_pseudoge | ENSG0000C |
| 89588714 | 89590097 | -1 | processed_pseudoge | ENSG0000C |
| 90030284 | 90030495 | 1 | snoRNA | ENSG0000C |
| 90184634 | 90184733 | 1 | snRNA | ENSG0000C |
| 90202316 | 90257415 | -1 | unprocessed_pseudc | ENSG0000C |
| 90261414 | 90261708 | -1 | processed_pseudoge | ENSG0000C |
| 91356755 | 91356914 | 1 | sRNA | ENSG0000C |
| 91374236 | 91513775 | -1 | unprocessed_pseudc | ENSG0000C |
| 93843764 | 93843862 | 1 | snRNA | ENSG0000C |
| 93873033 | 93980003 | -1 | protein_coding | ENSG0000C |
| 93967926 | 93968020 | 1 | misc_RNA | ENSG0000C |
| 93977029 | 93977135 | 1 | snRNA | ENSG0000C |
| 93980139 | 94055678 | 1 | protein_coding | ENSG0000C |
| 93988634 | 93988946 | -1 | processed_pseudoge | ENSG0000C |
| 94014365 | 94028597 | -1 | protein_coding | ENSG0000C |
| 94047836 | 94063389 | -1 | protein_coding | ENSG0000C |
| 94062980 | 94131832 | 1 | protein_coding | ENSG0000C |
| 94075912 | 94077175 | -1 | processed_pseudoge | ENSG0000C |
| 94506766 | 94507620 | -1 | processed_pseudoge | ENSG0000C |
| 94935760 | 94936120 | -1 | processed_pseudoge | ENSG0000C |
| 94937086 | 94938024 | -1 | processed_pseudoge | ENSG0000C |
| 94937980 | 95168351 | 1 | lncRNA | ENSG0000C |
| 95171920 | 95172377 | 1 | processed_pseudoge | ENSG0000C |
| 95654423 | 95683193 | -1 | transcribed_processe | ENSG0000C |
| 95916726 | 95919552 | 1 | lncRNA | ENSG0000C |
| 96244732 | 96245335 | -1 | processed_pseudoge | ENSG0000C |
| 96349554 | 96350939 | 1 | processed_pseudoge | ENSG0000C |
| 96359964 | 96360039 | 1 | miRNA | ENSG0000C |
| 96566357 | 96567103 | 1 | processed_pseudoge | ENSG0000C |
| 96608814 | 96608917 | -1 | snRNA | ENSG0000C |
| 96617188 | 96618236 | -1 | protein_coding | ENSG0000C |
| 96619304 | 96619831 | 1 | processed_pseudoge | ENSG0000C |
| 96753185 | 96754671 | 1 | processed_pseudoge | ENSG0000C |
| 96776945 | 96777683 | 1 | processed_pseudoge | ENSG0000C |
| 96810838 | 96814407 | -1 | lncRNA | ENSG0000C |
| 96814581 | 97752460 | 1 | protein_coding | ENSG0000C |
| 97481177 | 97481506 | -1 | processed_pseudoge | ENSG0000C |
| 97759523 | 97761532 | 1 | lncRNA | ENSG0000C |
| 97764521 | 97801229 | 1 | protein_coding | ENSG0000C |

| | | | | |
|---|---|---|---|---|
| 97800733 | 97821884 | -1 | lncRNA | ENSG0000C |
| 97822011 | 97944984 | 1 | protein_coding | ENSG0000C |
| 97836986 | 97874691 | -1 | lncRNA | ENSG0000C |
| 97941818 | 97972457 | -1 | protein_coding | ENSG0000C |
| 97973012 | 97975056 | 1 | lncRNA | ENSG0000C |
| 97986673 | 98035304 | -1 | protein_coding | ENSG0000C |
| 98053374 | 98053739 | 1 | unprocessed_pseudc | ENSG0000C |
| 98064472 | 98065396 | 1 | polymorphic_pseudc | ENSG0000C |
| 98087173 | 98088102 | 1 | protein_coding | ENSG0000C |
| 98099908 | 98148134 | -1 | lncRNA | ENSG0000C |
| 98100793 | 98105672 | 1 | transcribed_unproce | ENSG0000C |
| 98119191 | 98119731 | 1 | processed_pseudoge | ENSG0000C |
| 98130721 | 98138548 | 1 | protein_coding | ENSG0000C |
| 98147479 | 98156614 | 1 | protein_coding | ENSG0000C |
| 98166696 | 98169774 | 1 | protein_coding | ENSG0000C |
| 98197211 | 98198138 | 1 | unprocessed_pseudc | ENSG0000C |
| 98207423 | 98208341 | 1 | unprocessed_pseudc | ENSG0000C |
| 98222044 | 98222966 | 1 | unprocessed_pseudc | ENSG0000C |
| 98233651 | 98457898 | 1 | lncRNA | ENSG0000C |
| 98238352 | 98239281 | 1 | unprocessed_pseudc | ENSG0000C |
| 98263252 | 98265356 | 1 | protein_coding | ENSG0000C |
| 98282888 | 98283832 | 1 | protein_coding | ENSG0000C |
| 98306752 | 98312843 | 1 | polymorphic_pseudc | ENSG0000C |
| 98353854 | 98354819 | 1 | protein_coding | ENSG0000C |
| 98390666 | 98391631 | 1 | protein_coding | ENSG0000C |
| 98434034 | 98434847 | 1 | processed_pseudoge | ENSG0000C |
| 98463201 | 98472924 | 1 | protein_coding | ENSG0000C |
| 98497604 | 98498663 | 1 | protein_coding | ENSG0000C |
| 98497912 | 98523066 | -1 | protein_coding | ENSG0000C |
| 98521133 | 98585498 | -1 | protein_coding | ENSG0000C |
| 98522570 | 98525334 | 1 | lncRNA | ENSG0000C |
| 98525061 | 98525274 | -1 | processed_pseudoge | ENSG0000C |
| 98531978 | 98534681 | 1 | protein_coding | ENSG0000C |
| 98573061 | 98578231 | -1 | lncRNA | ENSG0000C |
| 98579446 | 98593648 | -1 | protein_coding | ENSG0000C |
| 98620065 | 98620618 | 1 | processed_pseudoge | ENSG0000C |
| 98657802 | 98660570 | 1 | processed_pseudoge | ENSG0000C |
| 98706236 | 98732757 | -1 | lncRNA | ENSG0000C |
| 98732236 | 98821201 | 1 | protein_coding | ENSG0000C |
| 98782188 | 98783193 | 1 | processed_pseudoge | ENSG0000C |
| 98795941 | 98901695 | -1 | protein_coding | ENSG0000C |
| 98804978 | 98805084 | -1 | snRNA | ENSG0000C |
| 98902424 | 99018562 | 1 | lncRNA | ENSG0000C |
| 98908459 | 98908564 | 1 | snRNA | ENSG0000C |
| 98981058 | 98983096 | 1 | lncRNA | ENSG0000C |
| 98987387 | 98987449 | 1 | snRNA | ENSG0000C |
| 99215224 | 99216266 | -1 | processed_pseudoge | ENSG0000C |
| 99326377 | 99326552 | -1 | processed_pseudoge | ENSG0000C |
| 99500347 | 99505508 | 1 | lncRNA | ENSG0000C |
| 99507187 | 99529985 | -1 | lncRNA | ENSG0000C |

| | | | | |
|---|---|---|---|---|
| 99598064 | 99637308 | 1 | lncRNA | ENSG0000C |
| 99638475 | 99799226 | 1 | protein_coding | ENSG0000C |
| 99802699 | 99806058 | -1 | lncRNA | ENSG0000C |
| 99817837 | 100181732 | 1 | protein_coding | ENSG0000C |
| 99830141 | 100114513 | -1 | protein_coding | ENSG0000C |
| 99964314 | 99964398 | -1 | miRNA | ENSG0000C |
| 100132903 | 100260264 | -1 | lncRNA | ENSG0000C |
| 100185687 | 100211766 | 1 | transcribed_unitary_ | ENSG0000C |
| 100221268 | 100223858 | -1 | unprocessed_pseudc | ENSG0000C |
| 100225374 | 100226073 | 1 | processed_pseudoge | ENSG0000C |
| 100260992 | 100325251 | 1 | protein_coding | ENSG0000C |
| 100334739 | 100361635 | 1 | protein_coding | ENSG0000C |
| 100363431 | 100401089 | -1 | protein_coding | ENSG0000C |
| 100401532 | 100456319 | 1 | protein_coding | ENSG0000C |
| 100492619 | 100577444 | 1 | protein_coding | ENSG0000C |
| 100609601 | 100695479 | 1 | protein_coding | ENSG0000C |
| 100646121 | 100646197 | 1 | snoRNA | ENSG0000C |
| 100665050 | 100665476 | -1 | processed_pseudoge | ENSG0000C |
| 100709331 | 100748964 | 1 | protein_coding | ENSG0000C |
| 100749156 | 100993515 | -1 | protein_coding | ENSG0000C |
| 100910975 | 100912175 | -1 | processed_pseudoge | ENSG0000C |
| 100952627 | 100952733 | 1 | snRNA | ENSG0000C |
| 101163094 | 101163986 | 1 | processed_pseudoge | ENSG0000C |
| 101222546 | 101320575 | -1 | protein_coding | ENSG0000C |
| 101324205 | 101513241 | -1 | protein_coding | ENSG0000C |
| 101357292 | 101359242 | 1 | processed_pseudoge | ENSG0000C |
| 101414228 | 101414633 | 1 | processed_pseudoge | ENSG0000C |
| 101457241 | 101457732 | -1 | processed_pseudoge | ENSG0000C |
| 101458530 | 101458626 | 1 | misc_RNA | ENSG0000C |
| 101521891 | 101522979 | 1 | processed_pseudoge | ENSG0000C |
| 101561868 | 101566446 | 1 | protein_coding | ENSG0000C |
| 101574180 | 101594465 | 1 | protein_coding | ENSG0000C |
| 101576489 | 101576947 | -1 | processed_pseudoge | ENSG0000C |
| 101599243 | 101599644 | -1 | processed_pseudoge | ENSG0000C |
| 101615826 | 101615928 | 1 | snRNA | ENSG0000C |
| 101628310 | 101628406 | -1 | misc_RNA | ENSG0000C |
| 101647421 | 101647522 | 1 | misc_RNA | ENSG0000C |
| 101648889 | 101677132 | -1 | protein_coding | ENSG0000C |
| 101662060 | 101662164 | -1 | misc_RNA | ENSG0000C |
| 101676475 | 101679217 | 1 | lncRNA | ENSG0000C |
| 101681091 | 101686718 | -1 | protein_coding | ENSG0000C |
| 101686827 | 101700926 | 1 | lncRNA | ENSG0000C |
| 101712472 | 101713191 | 1 | processed_pseudoge | ENSG0000C |
| 101724593 | 101770562 | 1 | protein_coding | ENSG0000C |
| 101779202 | 101828231 | 1 | protein_coding | ENSG0000C |
| 101823793 | 101824998 | -1 | lncRNA | ENSG0000C |
| 101827991 | 101861022 | 1 | protein_coding | ENSG0000C |
| 101878333 | 101925726 | 1 | lncRNA | ENSG0000C |
| 101940859 | 101997926 | 1 | lncRNA | ENSG0000C |
| 101955077 | 101955185 | 1 | misc_RNA | ENSG0000C |

| | | | | |
|---|---|---|---|---|
| 101960358 | 101997926 | 1 | lncRNA | ENSG0000C |
| 102099244 | 102479841 | 1 | protein_coding | ENSG0000C |
| 102621061 | 102673986 | -1 | lncRNA | ENSG0000C |
| 102755064 | 102755170 | -1 | snRNA | ENSG0000C |
| 103159960 | 103160124 | 1 | snRNA | ENSG0000C |
| 103240987 | 103241230 | 1 | processed_pseudoge | ENSG0000C |
| 103524033 | 103524116 | -1 | miRNA | ENSG0000C |
| 103538635 | 103540487 | -1 | processed_pseudoge | ENSG0000C |
| 103630661 | 103630912 | 1 | processed_pseudoge | ENSG0000C |
| 104063039 | 104063582 | 1 | processed_pseudoge | ENSG0000C |
| 104502700 | 104503266 | 1 | processed_pseudoge | ENSG0000C |
| 105366909 | 105576900 | 1 | protein_coding | ENSG0000C |
| 105655461 | 105869552 | -1 | protein_coding | ENSG0000C |
| 106160417 | 106224375 | 1 | lncRNA | ENSG0000C |
| 106325737 | 106326293 | -1 | processed_pseudoge | ENSG0000C |
| 106367979 | 106427977 | -1 | lncRNA | ENSG0000C |
| 106449775 | 107240671 | -1 | lncRNA | ENSG0000C |
| 106515897 | 106515998 | 1 | misc_RNA | ENSG0000C |
| 106688678 | 106688779 | 1 | misc_RNA | ENSG0000C |
| 106848671 | 106849260 | 1 | processed_pseudoge | ENSG0000C |
| 106894352 | 106895718 | -1 | unprocessed_pseudc | ENSG0000C |
| 106895729 | 106896408 | -1 | unprocessed_pseudc | ENSG0000C |
| 106896483 | 106896821 | -1 | unprocessed_pseudc | ENSG0000C |
| 106896891 | 106897671 | -1 | unprocessed_pseudc | ENSG0000C |
| 106897671 | 106898268 | -1 | unprocessed_pseudc | ENSG0000C |
| 106898889 | 106900591 | 1 | unprocessed_pseudc | ENSG0000C |
| 106900593 | 106900855 | -1 | unprocessed_pseudc | ENSG0000C |
| 106900854 | 106901081 | -1 | unprocessed_pseudc | ENSG0000C |
| 106901068 | 106901965 | 1 | unprocessed_pseudc | ENSG0000C |
| 106902173 | 106902523 | 1 | unprocessed_pseudc | ENSG0000C |
| 107007568 | 107007670 | 1 | snRNA | ENSG0000C |
| 107046026 | 107053166 | 1 | lncRNA | ENSG0000C |
| 107104911 | 107107000 | -1 | processed_pseudoge | ENSG0000C |
| 107220744 | 107348464 | 1 | lncRNA | ENSG0000C |
| 107272611 | 107421338 | -1 | lncRNA | ENSG0000C |
| 107327830 | 107329197 | -1 | processed_pseudoge | ENSG0000C |
| 107329430 | 107329962 | 1 | lncRNA | ENSG0000C |
| 107377439 | 107378635 | 1 | protein_coding | ENSG0000C |
| 107430892 | 107463912 | 1 | lncRNA | ENSG0000C |
| 107522936 | 107811339 | 1 | protein_coding | ENSG0000C |
| 107834586 | 107928907 | 1 | lncRNA | ENSG0000C |
| 107840228 | 107882250 | -1 | lncRNA | ENSG0000C |
| 108032456 | 108039916 | -1 | lncRNA | ENSG0000C |
| 108043091 | 108091862 | -1 | protein_coding | ENSG0000C |
| 108125821 | 108138610 | 1 | lncRNA | ENSG0000C |
| 108160812 | 108222435 | -1 | protein_coding | ENSG0000C |
| 108296490 | 108378285 | 1 | protein_coding | ENSG0000C |
| 108325520 | 108326814 | -1 | unprocessed_pseudc | ENSG0000C |
| 108380368 | 108529322 | -1 | protein_coding | ENSG0000C |
| 108543367 | 108543875 | -1 | processed_pseudoge | ENSG0000C |

| | | | | |
|---|---|---|---|---|
| 108549864 | 108589644 | -1 | protein_coding | ENSG0000C |
| 108574565 | 108574698 | 1 | snoRNA | ENSG0000C |
| 108574683 | 108575064 | 1 | processed_pseudoge | ENSG0000C |
| 108589705 | 108694840 | 1 | protein_coding | ENSG0000C |
| 108725440 | 108725536 | -1 | unprocessed_pseudc | ENSG0000C |
| 108743424 | 108757384 | -1 | protein_coding | ENSG0000C |
| 108822770 | 108855005 | 1 | protein_coding | ENSG0000C |
| 108907792 | 108953879 | -1 | protein_coding | ENSG0000C |
| 108958248 | 109118134 | -1 | protein_coding | ENSG0000C |
| 109018909 | 109019116 | -1 | snoRNA | ENSG0000C |
| 109101456 | 109110342 | 1 | lncRNA | ENSG0000C |
| 109118252 | 109151401 | 1 | protein_coding | ENSG0000C |
| 109176438 | 109177365 | -1 | lncRNA | ENSG0000C |
| 109178143 | 109216965 | 1 | lncRNA | ENSG0000C |
| 109225129 | 109225227 | 1 | snRNA | ENSG0000C |
| 109241507 | 109243125 | 1 | processed_pseudoge | ENSG0000C |
| 109293788 | 109316517 | -1 | protein_coding | ENSG0000C |
| 109295239 | 109295498 | 1 | misc_RNA | ENSG0000C |
| 109326141 | 109337572 | -1 | protein_coding | ENSG0000C |
| 109364047 | 109364234 | -1 | processed_pseudoge | ENSG0000C |
| 109396616 | 109397870 | 1 | processed_pseudoge | ENSG0000C |
| 109409678 | 109410084 | -1 | processed_pseudoge | ENSG0000C |
| 109409990 | 109723273 | 1 | lncRNA | ENSG0000C |
| 109471329 | 109471812 | -1 | processed_pseudoge | ENSG0000C |
| 109602828 | 109602897 | 1 | miRNA | ENSG0000C |
| 109648107 | 109810861 | 1 | lncRNA | ENSG0000C |
| 109701456 | 109702218 | -1 | processed_pseudoge | ENSG0000C |
| 109915976 | 109916940 | -1 | processed_pseudoge | ENSG0000C |
| 109977540 | 109977767 | -1 | processed_pseudoge | ENSG0000C |
| 110527482 | 110529582 | -1 | lncRNA | ENSG0000C |
| 110551845 | 110551965 | -1 | snRNA | ENSG0000C |
| 110682286 | 110683154 | -1 | processed_pseudoge | ENSG0000C |
| 110727021 | 110727121 | 1 | misc_RNA | ENSG0000C |
| 110888384 | 111071553 | -1 | lncRNA | ENSG0000C |
| 111070071 | 111275563 | 1 | protein_coding | ENSG0000C |
| 111292719 | 111665750 | 1 | protein_coding | ENSG0000C |
| 111466313 | 111497095 | -1 | lncRNA | ENSG0000C |
| 111478737 | 111478943 | 1 | processed_pseudoge | ENSG0000C |
| 111570638 | 111571054 | -1 | processed_pseudoge | ENSG0000C |
| 111592900 | 111595346 | -1 | protein_coding | ENSG0000C |
| 111633672 | 111635089 | 1 | processed_pseudoge | ENSG0000C |
| 111674676 | 111846447 | 1 | protein_coding | ENSG0000C |
| 111676736 | 111677433 | -1 | lncRNA | ENSG0000C |
| 111732497 | 111976517 | 1 | protein_coding | ENSG0000C |
| 111835723 | 111860730 | -1 | lncRNA | ENSG0000C |
| 111979010 | 111993368 | 1 | protein_coding | ENSG0000C |
| 111998739 | 112013887 | 1 | protein_coding | ENSG0000C |
| 112030931 | 112031649 | -1 | processed_pseudoge | ENSG0000C |
| 112034843 | 112081269 | 1 | protein_coding | ENSG0000C |
| 112086335 | 112131004 | 1 | protein_coding | ENSG0000C |

| | | | | |
|---|---|---|---|---|
| 112112801 | 112112898 | 1 | miRNA | ENSG0000C |
| 112120839 | 112133270 | -1 | protein_coding | ENSG0000C |
| 112133423 | 112135359 | 1 | lncRNA | ENSG0000C |
| 112140898 | 112294227 | -1 | protein_coding | ENSG0000C |
| 112185480 | 112185998 | -1 | processed_pseudoge | ENSG0000C |
| 112302478 | 112332791 | 1 | lncRNA | ENSG0000C |
| 112321140 | 112321443 | 1 | unitary_pseudogene | ENSG0000C |
| 112332347 | 112362812 | 1 | protein_coding | ENSG0000C |
| 112396647 | 112409134 | -1 | lncRNA | ENSG0000C |
| 112463966 | 112499472 | -1 | protein_coding | ENSG0000C |
| 112524187 | 112525201 | -1 | unprocessed_pseudc | ENSG0000C |
| 112525548 | 112528142 | -1 | TEC | ENSG0000C |
| 112532510 | 112562046 | -1 | protein_coding | ENSG0000C |
| 112561709 | 112585579 | 1 | protein_coding | ENSG0000C |
| 112596794 | 112649530 | -1 | protein_coding | ENSG0000C |
| 112696908 | 112697706 | -1 | unprocessed_pseudc | ENSG0000C |
| 112736447 | 112749319 | -1 | lncRNA | ENSG0000C |
| 112802478 | 112812819 | 1 | lncRNA | ENSG0000C |
| 112815709 | 112846856 | -1 | protein_coding | ENSG0000C |
| 112921205 | 112975103 | -1 | protein_coding | ENSG0000C |
| 112990447 | 112991153 | -1 | lncRNA | ENSG0000C |
| 112990984 | 113015060 | 1 | protein_coding | ENSG0000C |
| 113002444 | 113019861 | -1 | protein_coding | ENSG0000C |
| 113019468 | 113184377 | 1 | lncRNA | ENSG0000C |
| 113041390 | 113041814 | -1 | processed_pseudoge | ENSG0000C |
| 113050912 | 113063443 | -1 | lncRNA | ENSG0000C |
| 113142350 | 113167819 | -1 | lncRNA | ENSG0000C |
| 113211003 | 113287459 | 1 | protein_coding | ENSG0000C |
| 113259786 | 113266824 | -1 | lncRNA | ENSG0000C |
| 113286930 | 113441610 | -1 | protein_coding | ENSG0000C |
| 113360267 | 113361319 | 1 | processed_pseudoge | ENSG0000C |
| 113361901 | 113515151 | -1 | protein_coding | ENSG0000C |
| 113403988 | 113433992 | 1 | lncRNA | ENSG0000C |
| 113432118 | 113432200 | -1 | miRNA | ENSG0000C |
| 113442718 | 113515187 | -1 | protein_coding | ENSG0000C |
| 113532296 | 113629578 | 1 | protein_coding | ENSG0000C |
| 113588748 | 113590189 | -1 | lncRNA | ENSG0000C |
| 113594876 | 113594942 | 1 | miRNA | ENSG0000C |
| 113632704 | 113632998 | 1 | misc_RNA | ENSG0000C |
| 113648385 | 113696646 | -1 | protein_coding | ENSG0000C |
| 113716458 | 113746300 | -1 | protein_coding | ENSG0000C |
| 113740823 | 113741254 | -1 | processed_pseudoge | ENSG0000C |
| 113746872 | 113747408 | -1 | lncRNA | ENSG0000C |
| 113747033 | 113812056 | 1 | protein_coding | ENSG0000C |
| 113828182 | 113947174 | 1 | protein_coding | ENSG0000C |
| 113850237 | 113850731 | -1 | processed_pseudoge | ENSG0000C |
| 113885298 | 113886031 | 1 | processed_pseudoge | ENSG0000C |
| 113919222 | 113920200 | 1 | processed_pseudoge | ENSG0000C |
| 113947005 | 113947570 | -1 | lncRNA | ENSG0000C |
| 113947901 | 113965401 | 1 | protein_coding | ENSG0000C |

| | | | | |
|---|---|---|---|---|
| 113964137 | 114056594 | -1 | protein_coding | ENSG00000 |
| 113984037 | 113984655 | 1 | processed_pseudoge | ENSG00000 |
| 113986835 | 113992113 | 1 | lncRNA | ENSG00000 |
| 113998782 | 114016261 | 1 | lncRNA | ENSG00000 |
| 114005833 | 114088422 | 1 | protein_coding | ENSG00000 |
| 114103249 | 114103628 | -1 | processed_pseudoge | ENSG00000 |
| 114128652 | 114199407 | -1 | protein_coding | ENSG00000 |
| 114214313 | 114236204 | 1 | lncRNA | ENSG00000 |
| 114231398 | 114233021 | -1 | processed_pseudoge | ENSG00000 |
| 114234631 | 114237578 | -1 | protein_coding | ENSG00000 |
| 114276913 | 114310288 | 1 | protein_coding | ENSG00000 |
| 114314501 | 115147271 | -1 | protein_coding | ENSG00000 |
| 114314501 | 114329714 | -1 | lncRNA | ENSG00000 |
| 114316475 | 114316569 | -1 | miRNA | ENSG00000 |
| 114351771 | 114388978 | 1 | lncRNA | ENSG00000 |
| 114445521 | 114529452 | 1 | lncRNA | ENSG00000 |
| 114684580 | 114687609 | 1 | lncRNA | ENSG00000 |
| 114743445 | 114743525 | -1 | miRNA | ENSG00000 |
| 114873114 | 114876514 | 1 | lncRNA | ENSG00000 |
| 114930541 | 114931449 | 1 | processed_pseudoge | ENSG00000 |
| 115100423 | 115103061 | 1 | lncRNA | ENSG00000 |
| 115147605 | 115564389 | 1 | lncRNA | ENSG00000 |
| 115279125 | 115279860 | 1 | processed_pseudoge | ENSG00000 |
| 115413476 | 115486449 | -1 | lncRNA | ENSG00000 |
| 115418862 | 115438550 | 1 | lncRNA | ENSG00000 |
| 115623324 | 115721490 | 1 | protein_coding | ENSG00000 |
| 115658533 | 115661279 | 1 | lncRNA | ENSG00000 |
| 115802363 | 117139389 | -1 | protein_coding | ENSG00000 |
| 115837870 | 115838159 | 1 | misc_RNA | ENSG00000 |
| 116360024 | 116370090 | 1 | lncRNA | ENSG00000 |
| 116552473 | 116568264 | 1 | lncRNA | ENSG00000 |
| 116582552 | 116582830 | -1 | misc_RNA | ENSG00000 |
| 116645902 | 116646727 | -1 | processed_pseudoge | ENSG00000 |
| 116709235 | 116723581 | 1 | lncRNA | ENSG00000 |
| 116716460 | 116716624 | 1 | misc_RNA | ENSG00000 |
| 116850277 | 116850367 | -1 | miRNA | ENSG00000 |
| 116921328 | 116921479 | 1 | misc_RNA | ENSG00000 |
| 116921431 | 116932238 | 1 | lncRNA | ENSG00000 |
| 116965112 | 116965227 | -1 | snRNA | ENSG00000 |
| 117026698 | 117027039 | 1 | processed_pseudoge | ENSG00000 |
| 117544205 | 117544311 | 1 | snRNA | ENSG00000 |
| 117672154 | 117997592 | -1 | lncRNA | ENSG00000 |
| 117674342 | 117675144 | -1 | processed_pseudoge | ENSG00000 |
| 117678693 | 117691005 | -1 | lncRNA | ENSG00000 |
| 117719859 | 117794384 | 1 | lncRNA | ENSG00000 |
| 118004819 | 118810836 | -1 | lncRNA | ENSG00000 |
| 118488876 | 118491760 | 1 | lncRNA | ENSG00000 |
| 118900557 | 119146068 | -1 | protein_coding | ENSG00000 |
| 118943073 | 118948241 | 1 | lncRNA | ENSG00000 |
| 119146151 | 119160042 | 1 | protein_coding | ENSG00000 |

| | | | | |
|---|---|---|---|---|
| 119147375 | 119187807 | 1 | protein_coding | ENSG00000( |
| 119173517 | 119205143 | 1 | protein_coding | ENSG00000( |
| 119211732 | 119240946 | -1 | protein_coding | ENSG00000( |
| 119226486 | 119290666 | 1 | lncRNA | ENSG00000( |
| 119294383 | 119420714 | 1 | protein_coding | ENSG00000( |
| 119298665 | 119299012 | 1 | processed_pseudoge | ENSG00000( |
| 119314293 | 119322760 | -1 | lncRNA | ENSG00000( |
| 119341834 | 119341940 | -1 | snRNA | ENSG00000( |
| 119428949 | 119468830 | -1 | protein_coding | ENSG00000( |
| 119468963 | 119494708 | 1 | protein_coding | ENSG00000( |
| 119497678 | 119498181 | -1 | lncRNA | ENSG00000( |
| 119498547 | 119525090 | 1 | protein_coding | ENSG00000( |
| 119524293 | 119559614 | -1 | protein_coding | ENSG00000( |
| 119579212 | 119579650 | -1 | lncRNA | ENSG00000( |
| 119579268 | 119589945 | 1 | protein_coding | ENSG00000( |
| 119597875 | 119629811 | 1 | protein_coding | ENSG00000( |
| 119635526 | 119636150 | -1 | processed_pseudoge | ENSG00000( |
| 119636457 | 119665324 | -1 | protein_coding | ENSG00000( |
| 119654513 | 119677454 | -1 | protein_coding | ENSG00000( |
| 119666282 | 119670688 | 1 | lncRNA | ENSG00000( |
| 119703022 | 119767102 | 1 | protein_coding | ENSG00000( |
| 119703076 | 119807570 | 1 | protein_coding | ENSG00000( |
| 119744139 | 119750350 | -1 | lncRNA | ENSG00000( |
| 119780484 | 119818485 | 1 | protein_coding | ENSG00000( |
| 119791829 | 119792553 | 1 | processed_pseudoge | ENSG00000( |
| 119821323 | 120094417 | -1 | protein_coding | ENSG00000( |
| 120028740 | 120029868 | 1 | processed_pseudoge | ENSG00000( |
| 120041190 | 120041537 | 1 | processed_pseudoge | ENSG00000( |
| 120094895 | 120136783 | 1 | lncRNA | ENSG00000( |
| 120110498 | 120110786 | -1 | misc_RNA | ENSG00000( |
| 120121909 | 120122206 | 1 | misc_RNA | ENSG00000( |
| 120164645 | 120285094 | -1 | protein_coding | ENSG00000( |
| 120210320 | 120210421 | 1 | misc_RNA | ENSG00000( |
| 120306726 | 120307952 | 1 | processed_pseudoge | ENSG00000( |
| 120324509 | 120349354 | -1 | protein_coding | ENSG00000( |
| 120349449 | 120423826 | 1 | transcribed_unitary_ | ENSG00000( |
| 120365993 | 120368326 | -1 | lncRNA | ENSG00000( |
| 120388029 | 120389315 | 1 | processed_pseudoge | ENSG00000( |
| 120392293 | 120450993 | -1 | protein_coding | ENSG00000( |
| 120395668 | 120395729 | -1 | miRNA | ENSG00000( |
| 120448974 | 120509675 | 1 | lncRNA | ENSG00000( |
| 120484171 | 120484527 | -1 | processed_pseudoge | ENSG00000( |
| 120512850 | 120513188 | 1 | processed_pseudoge | ENSG00000( |
| 120596328 | 120602507 | 1 | protein_coding | ENSG00000( |
| 120628172 | 120682269 | -1 | protein_coding | ENSG00000( |
| 120686681 | 120742993 | -1 | protein_coding | ENSG00000( |
| 120722024 | 120722311 | 1 | unprocessed_pseudc | ENSG00000( |
| 120722458 | 120722646 | 1 | unprocessed_pseudc | ENSG00000( |
| 120742637 | 120783069 | 1 | protein_coding | ENSG00000( |
| 120805866 | 120806745 | 1 | processed_pseudoge | ENSG00000( |

| | | | | |
|---|---|---|---|---|
| 120811530 | 120908093 | -1 | lncRNA | ENSG00000 |
| 120833440 | 120836360 | -1 | lncRNA | ENSG00000 |
| 120833936 | 120871794 | 1 | lncRNA | ENSG00000 |
| 120908072 | 121424761 | 1 | protein_coding | ENSG00000 |
| 120924612 | 120925158 | -1 | processed_pseudoge | ENSG00000 |
| 121049640 | 121049715 | 1 | miRNA | ENSG00000 |
| 121099108 | 121099774 | 1 | processed_pseudoge | ENSG00000 |
| 121356991 | 121357940 | 1 | processed_pseudoge | ENSG00000 |
| 121394772 | 121431198 | -1 | lncRNA | ENSG00000 |
| 121431427 | 121546641 | -1 | protein_coding | ENSG00000 |
| 121494110 | 121494908 | 1 | processed_pseudoge | ENSG00000 |
| 121567949 | 121590622 | 1 | protein_coding | ENSG00000 |
| 121593379 | 121630295 | 1 | protein_coding | ENSG00000 |
| 121631399 | 121660927 | -1 | protein_coding | ENSG00000 |
| 121653994 | 121654296 | -1 | misc_RNA | ENSG00000 |
| 121655475 | 121655604 | -1 | snRNA | ENSG00000 |
| 121663199 | 121749767 | -1 | protein_coding | ENSG00000 |
| 121769761 | 121835079 | -1 | protein_coding | ENSG00000 |
| 121835183 | 121886526 | 1 | protein_coding | ENSG00000 |
| 121894401 | 121944188 | 1 | protein_coding | ENSG00000 |
| 121987323 | 122022247 | -1 | protein_coding | ENSG00000 |
| 122025195 | 122025304 | -1 | misc_RNA | ENSG00000 |
| 122055362 | 122121139 | 1 | protein_coding | ENSG00000 |
| 122172405 | 122174099 | -1 | processed_pseudoge | ENSG00000 |
| 122183683 | 122291629 | 1 | protein_coding | ENSG00000 |
| 122317609 | 122318741 | 1 | processed_pseudoge | ENSG00000 |
| 122325248 | 122341969 | 1 | protein_coding | ENSG00000 |
| 122359591 | 122383231 | -1 | protein_coding | ENSG00000 |
| 122384176 | 122412334 | 1 | protein_coding | ENSG00000 |
| 122411846 | 122416062 | -1 | protein_coding | ENSG00000 |
| 122416207 | 122443180 | 1 | lncRNA | ENSG00000 |
| 122416882 | 122417012 | -1 | unprocessed_pseudo | ENSG00000 |
| 122421902 | 122514945 | -1 | protein_coding | ENSG00000 |
| 122482990 | 122483229 | 1 | processed_pseudoge | ENSG00000 |
| 122515006 | 122518172 | 1 | lncRNA | ENSG00000 |
| 122527924 | 122564577 | -1 | protein_coding | ENSG00000 |
| 122564338 | 122575203 | 1 | protein_coding | ENSG00000 |
| 122577628 | 122639047 | 1 | protein_coding | ENSG00000 |
| 122660613 | 122662438 | 1 | processed_pseudoge | ENSG00000 |
| 122680839 | 122730840 | 1 | protein_coding | ENSG00000 |
| 122739999 | 122793831 | -1 | protein_coding | ENSG00000 |
| 122795069 | 122881139 | 1 | protein_coding | ENSG00000 |
| 122886941 | 122892416 | 1 | lncRNA | ENSG00000 |
| 122909082 | 123028605 | -1 | protein_coding | ENSG00000 |
| 123067025 | 123225227 | 1 | protein_coding | ENSG00000 |
| 123161794 | 123161879 | 1 | miRNA | ENSG00000 |
| 123201927 | 123274136 | 1 | protein_coding | ENSG00000 |
| 123277353 | 123277904 | 1 | lncRNA | ENSG00000 |
| 123282296 | 123449758 | -1 | protein_coding | ENSG00000 |
| 123283593 | 123283983 | 1 | lncRNA | ENSG00000 |

| | | | | |
|---|---|---|---|---|
| 123490820 | 123585053 | -1 | protein_coding | ENSG00000 |
| 123585300 | 123644568 | 1 | lncRNA | ENSG00000 |
| 123610049 | 123884331 | -1 | protein_coding | ENSG00000 |
| 123689644 | 123692407 | 1 | lncRNA | ENSG00000 |
| 123715851 | 123716399 | 1 | lncRNA | ENSG00000 |
| 123814077 | 123814207 | 1 | snoRNA | ENSG00000 |
| 123897305 | 123961408 | -1 | protein_coding | ENSG00000 |
| 123968521 | 123992178 | -1 | protein_coding | ENSG00000 |
| 124080023 | 124726325 | 1 | protein_coding | ENSG00000 |
| 124132929 | 124133025 | 1 | miRNA | ENSG00000 |
| 124151960 | 124153063 | -1 | processed_pseudoge | ENSG00000 |
| 124374332 | 124374437 | 1 | miRNA | ENSG00000 |
| 124407691 | 124407783 | 1 | snRNA | ENSG00000 |
| 124723788 | 124726325 | 1 | lncRNA | ENSG00000 |
| 124730433 | 124749273 | 1 | protein_coding | ENSG00000 |
| 124732439 | 124732516 | 1 | miRNA | ENSG00000 |
| 124733418 | 124733754 | -1 | processed_pseudoge | ENSG00000 |
| 124761948 | 124901418 | -1 | protein_coding | ENSG00000 |
| 124781155 | 124787757 | 1 | lncRNA | ENSG00000 |
| 124791119 | 124793104 | -1 | TEC | ENSG00000 |
| 124792319 | 124792562 | -1 | misc_RNA | ENSG00000 |
| 124862094 | 124863317 | -1 | processed_pseudoge | ENSG00000 |
| 124905442 | 124953819 | -1 | protein_coding | ENSG00000 |
| 124965710 | 125055997 | -1 | protein_coding | ENSG00000 |
| 125058304 | 125058438 | -1 | rRNA_pseudogene | ENSG00000 |
| 125061448 | 125063710 | 1 | lncRNA | ENSG00000 |
| 125082636 | 125212864 | -1 | protein_coding | ENSG00000 |
| 125119097 | 125119200 | -1 | snRNA | ENSG00000 |
| 125151465 | 125151552 | -1 | miRNA | ENSG00000 |
| 125225561 | 125375354 | -1 | protein_coding | ENSG00000 |
| 125310881 | 125311350 | 1 | processed_pseudoge | ENSG00000 |
| 125344085 | 125344797 | -1 | processed_pseudoge | ENSG00000 |
| 125388934 | 125388997 | 1 | snRNA | ENSG00000 |
| 125446650 | 125520202 | -1 | protein_coding | ENSG00000 |
| 125516979 | 125517086 | 1 | misc_RNA | ENSG00000 |
| 125528244 | 125528349 | -1 | misc_RNA | ENSG00000 |
| 125528858 | 125595497 | -1 | protein_coding | ENSG00000 |
| 125679573 | 125679781 | 1 | processed_pseudoge | ENSG00000 |
| 125681305 | 125681453 | 1 | processed_pseudoge | ENSG00000 |
| 125703349 | 125704312 | 1 | unprocessed_pseudc | ENSG00000 |
| 125712139 | 125713045 | 1 | unprocessed_pseudc | ENSG00000 |
| 125724539 | 125725445 | 1 | unprocessed_pseudc | ENSG00000 |
| 125734295 | 125735198 | 1 | unprocessed_pseudc | ENSG00000 |
| 125747084 | 125748098 | 1 | unprocessed_pseudc | ENSG00000 |
| 125756086 | 125756462 | 1 | processed_pseudoge | ENSG00000 |
| 125766516 | 125852936 | 1 | lncRNA | ENSG00000 |
| 125766601 | 125766818 | 1 | transcribed_unproce | ENSG00000 |
| 125774714 | 125797953 | 1 | lncRNA | ENSG00000 |
| 125787888 | 125788146 | -1 | processed_pseudoge | ENSG00000 |
| 125790404 | 125790552 | -1 | miRNA | ENSG00000 |

| | | | | |
|---|---|---|---|---|
| 125795106 | 125795885 | 1 | processed_pseudoge | ENSG00000 |
| 125799887 | 125800616 | -1 | lncRNA | ENSG00000 |
| 125816082 | 125816286 | -1 | processed_pseudoge | ENSG00000 |
| 125827238 | 125916384 | -1 | lncRNA | ENSG00000 |
| 125848223 | 125909372 | 1 | transcribed_unproce | ENSG00000 |
| 125907765 | 125916360 | -1 | lncRNA | ENSG00000 |
| 125916620 | 125930024 | 1 | transcribed_unproce | ENSG00000 |
| 125929275 | 125937039 | -1 | protein_coding | ENSG00000 |
| 125958556 | 125958817 | 1 | processed_pseudoge | ENSG00000 |
| 125969160 | 125983454 | 1 | protein_coding | ENSG00000 |
| 126006355 | 126101561 | -1 | protein_coding | ENSG00000 |
| 126056923 | 126058228 | 1 | lncRNA | ENSG00000 |
| 126083659 | 126095349 | 1 | lncRNA | ENSG00000 |
| 126103562 | 126197994 | -1 | protein_coding | ENSG00000 |
| 126103640 | 126108069 | 1 | lncRNA | ENSG00000 |
| 126160283 | 126160445 | -1 | snRNA | ENSG00000 |
| 126180012 | 126210169 | 1 | lncRNA | ENSG00000 |
| 126213204 | 126247714 | 1 | lncRNA | ENSG00000 |
| 126266747 | 126291525 | 1 | lncRNA | ENSG00000 |
| 126288123 | 126301213 | -1 | lncRNA | ENSG00000 |
| 126312385 | 126323406 | 1 | lncRNA | ENSG00000 |
| 126342635 | 126357408 | -1 | protein_coding | ENSG00000 |
| 126393032 | 126394851 | -1 | lncRNA | ENSG00000 |
| 126394909 | 126436556 | 1 | protein_coding | ENSG00000 |
| 126432796 | 126434569 | -1 | TEC | ENSG00000 |
| 126437601 | 126475891 | -1 | protein_coding | ENSG00000 |
| 126481166 | 126517773 | -1 | protein_coding | ENSG00000 |
| 126524155 | 126543291 | 1 | protein_coding | ENSG00000 |
| 126526999 | 126558965 | -1 | protein_coding | ENSG00000 |
| 126564565 | 126564678 | -1 | rRNA_pseudogene | ENSG00000 |
| 126571779 | 126608555 | -1 | protein_coding | ENSG00000 |
| 126607059 | 126655124 | -1 | protein_coding | ENSG00000 |
| 126624792 | 126626061 | 1 | processed_pseudoge | ENSG00000 |
| 126660609 | 126671939 | 1 | transcribed_processe | ENSG00000 |
| 126672106 | 126673223 | -1 | processed_pseudoge | ENSG00000 |
| 126704240 | 126960420 | 1 | protein_coding | ENSG00000 |
| 126766434 | 126767475 | -1 | processed_pseudoge | ENSG00000 |
| 126973065 | 126976426 | 1 | TEC | ENSG00000 |
| 126988594 | 127037392 | 1 | protein_coding | ENSG00000 |
| 127165506 | 127218473 | -1 | lncRNA | ENSG00000 |
| 127193131 | 127198185 | 1 | protein_coding | ENSG00000 |
| 127202372 | 127202473 | -1 | misc_RNA | ENSG00000 |
| 127221194 | 127221881 | -1 | unprocessed_pseudo | ENSG00000 |
| 127240968 | 127241074 | -1 | snRNA | ENSG00000 |
| 127274581 | 127278304 | 1 | lncRNA | ENSG00000 |
| 127283784 | 127288046 | -1 | protein_coding | ENSG00000 |
| 127322307 | 127390670 | -1 | lncRNA | ENSG00000 |
| 127480690 | 127537817 | -1 | lncRNA | ENSG00000 |
| 127489553 | 127490787 | 1 | lncRNA | ENSG00000 |
| 127537937 | 127540485 | 1 | lncRNA | ENSG00000 |

| | | | | |
|---|---|---|---|---|
| 127571232 | 127598267 | -1 | protein_coding | ENSG00000( |
| 127571232 | 127571838 | -1 | lncRNA | ENSG00000( |
| 127575266 | 127575331 | -1 | miRNA | ENSG00000( |
| 127587111 | 127587176 | -1 | miRNA | ENSG00000( |
| 127598410 | 127622436 | 1 | protein_coding | ENSG00000( |
| 127620106 | 127629049 | -1 | lncRNA | ENSG00000( |
| 127629185 | 127672802 | 1 | protein_coding | ENSG00000( |
| 127672935 | 127680926 | 1 | protein_coding | ENSG00000( |
| 127689062 | 128052190 | -1 | protein_coding | ENSG00000( |
| 127837436 | 127845499 | 1 | lncRNA | ENSG00000( |
| 127915232 | 127987671 | 1 | protein_coding | ENSG00000( |
| 127963608 | 127963724 | 1 | rRNA_pseudogene | ENSG00000( |
| 128051641 | 128071683 | 1 | protein_coding | ENSG00000( |
| 128064778 | 128153914 | -1 | protein_coding | ENSG00000( |
| 128075073 | 128075249 | -1 | snRNA | ENSG00000( |
| 128075810 | 128079056 | 1 | lncRNA | ENSG00000( |
| 128141875 | 128141977 | -1 | snRNA | ENSG00000( |
| 128153481 | 128408646 | 1 | protein_coding | ENSG00000( |
| 128181402 | 128194452 | -1 | lncRNA | ENSG00000( |
| 128462439 | 128467248 | -1 | protein_coding | ENSG00000( |
| 128463594 | 128472317 | 1 | lncRNA | ENSG00000( |
| 128479427 | 128493185 | -1 | protein_coding | ENSG00000( |
| 128489212 | 128502970 | 1 | lncRNA | ENSG00000( |
| 128538020 | 128538631 | 1 | processed_pseudoge | ENSG00000( |
| 128563316 | 128564431 | 1 | lncRNA | ENSG00000( |
| 128572000 | 128576086 | -1 | lncRNA | ENSG00000( |
| 128619969 | 128681075 | -1 | protein_coding | ENSG00000( |
| 128673681 | 128674021 | -1 | misc_RNA | ENSG00000( |
| 128674735 | 128677005 | -1 | unprocessed_pseudc | ENSG00000( |
| 128714571 | 128714705 | -1 | snoRNA | ENSG00000( |
| 128726122 | 128814796 | 1 | protein_coding | ENSG00000( |
| 128748538 | 128749489 | -1 | processed_pseudoge | ENSG00000( |
| 128764466 | 128765018 | -1 | processed_pseudoge | ENSG00000( |
| 128785147 | 128785448 | 1 | misc_RNA | ENSG00000( |
| 128798841 | 128799201 | -1 | processed_pseudoge | ENSG00000( |
| 128843564 | 128844404 | -1 | processed_pseudoge | ENSG00000( |
| 128848673 | 128849399 | -1 | processed_pseudoge | ENSG00000( |
| 128852112 | 128853008 | -1 | processed_pseudoge | ENSG00000( |
| 128859716 | 128860526 | -1 | lncRNA | ENSG00000( |
| 128860620 | 128871540 | -1 | lncRNA | ENSG00000( |
| 128869624 | 128870026 | 1 | processed_pseudoge | ENSG00000( |
| 128872349 | 128876218 | -1 | lncRNA | ENSG00000( |
| 128879596 | 128916067 | 1 | protein_coding | ENSG00000( |
| 128909866 | 129002690 | -1 | protein_coding | ENSG00000( |
| 128914833 | 128915060 | -1 | TEC | ENSG00000( |
| 129001629 | 129040742 | 1 | protein_coding | ENSG00000( |
| 129045763 | 129046921 | 1 | processed_pseudoge | ENSG00000( |
| 129060767 | 129062406 | 1 | protein_coding | ENSG00000( |
| 129087569 | 129122801 | -1 | protein_coding | ENSG00000( |
| 129087575 | 129161036 | -1 | protein_coding | ENSG00000( |

| | | | | |
|---|---|---|---|---|
| 129123439 | 129124003 | 1 | lncRNA | ENSG0000C |
| 129127415 | 129161293 | -1 | protein_coding | ENSG0000C |
| 129163606 | 129163940 | -1 | lncRNA | ENSG0000C |
| 129169484 | 129183922 | -1 | protein_coding | ENSG0000C |
| 129218093 | 129218350 | -1 | processed_pseudoge | ENSG0000C |
| 129249606 | 129277773 | 1 | protein_coding | ENSG0000C |
| 129272146 | 129272243 | 1 | miRNA | ENSG0000C |
| 129277753 | 129278782 | -1 | lncRNA | ENSG0000C |
| 129278828 | 129306186 | 1 | protein_coding | ENSG0000C |
| 129314771 | 129316286 | -1 | protein_coding | ENSG0000C |
| 129315392 | 129326225 | 1 | lncRNA | ENSG0000C |
| 129323046 | 129325426 | -1 | unprocessed_pseudc | ENSG0000C |
| 129334586 | 129335074 | 1 | processed_pseudoge | ENSG0000C |
| 129345411 | 129346164 | 1 | processed_pseudoge | ENSG0000C |
| 129381298 | 129394149 | 1 | unprocessed_pseudc | ENSG0000C |
| 129382922 | 129399655 | -1 | transcribed_unproce | ENSG0000C |
| 129397210 | 129397348 | -1 | snoRNA | ENSG0000C |
| 129401321 | 129428651 | -1 | protein_coding | ENSG0000C |
| 129430944 | 129440179 | -1 | protein_coding | ENSG0000C |
| 129440036 | 129520507 | 1 | protein_coding | ENSG0000C |
| 129528639 | 129535344 | 1 | protein_coding | ENSG0000C |
| 129543214 | 129551467 | 1 | protein_coding | ENSG0000C |
| 129555175 | 129606818 | -1 | protein_coding | ENSG0000C |
| 129591349 | 129591635 | 1 | misc_RNA | ENSG0000C |
| 129632019 | 129632492 | -1 | unprocessed_pseudc | ENSG0000C |
| 129647792 | 129893606 | -1 | protein_coding | ENSG0000C |
| 129818932 | 129819033 | 1 | misc_RNA | ENSG0000C |
| 129819777 | 129819876 | 1 | snRNA | ENSG0000C |
| 129847048 | 129847957 | -1 | lncRNA | ENSG0000C |
| 129893811 | 129918575 | 1 | lncRNA | ENSG0000C |
| 129954105 | 129969294 | -1 | lncRNA | ENSG0000C |
| 129974688 | 129977935 | 1 | protein_coding | ENSG0000C |
| 129998531 | 129998936 | 1 | processed_pseudoge | ENSG0000C |
| 130002789 | 130003007 | -1 | unprocessed_pseudc | ENSG0000C |
| 130013182 | 130013541 | -1 | processed_pseudoge | ENSG0000C |
| 130021553 | 130022520 | -1 | unprocessed_pseudc | ENSG0000C |
| 130034553 | 130035539 | -1 | unprocessed_pseudc | ENSG0000C |
| 130048143 | 130055920 | -1 | unprocessed_pseudc | ENSG0000C |
| 130081831 | 130113227 | 1 | protein_coding | ENSG0000C |
| 130089433 | 130094304 | -1 | lncRNA | ENSG0000C |
| 130099092 | 130111472 | -1 | transcribed_unproce | ENSG0000C |
| 130111669 | 130201336 | 1 | lncRNA | ENSG0000C |
| 130167790 | 130168030 | -1 | processed_pseudoge | ENSG0000C |
| 130181346 | 130188814 | 1 | lncRNA | ENSG0000C |
| 130199708 | 130199918 | 1 | processed_pseudoge | ENSG0000C |
| 130212823 | 130273806 | 1 | transcribed_unitary_ | ENSG0000C |
| 130293337 | 130293760 | 1 | processed_pseudoge | ENSG0000C |
| 130345516 | 130484844 | 1 | protein_coding | ENSG0000C |
| 130560334 | 130678155 | 1 | protein_coding | ENSG0000C |
| 130678934 | 130746829 | -1 | protein_coding | ENSG0000C |

| | | | | |
|---|---|---|---|---|
| 130811768 | 130812095 | -1 | misc_RNA | ENSG00000( |
| 130821184 | 130823914 | 1 | lncRNA | ENSG00000( |
| 130827659 | 130828376 | 1 | processed_pseudoge | ENSG00000( |
| 130850595 | 131016712 | 1 | protein_coding | ENSG00000( |
| 130868004 | 130869357 | -1 | TEC | ENSG00000( |
| 130899414 | 130929976 | -1 | lncRNA | ENSG00000( |
| 130914340 | 130914452 | -1 | misc_RNA | ENSG00000( |
| 130918226 | 130919120 | -1 | processed_pseudoge | ENSG00000( |
| 130927754 | 130928276 | -1 | processed_pseudoge | ENSG00000( |
| 131013875 | 131027649 | -1 | protein_coding | ENSG00000( |
| 131026850 | 131350465 | 1 | protein_coding | ENSG00000( |
| 131053317 | 131072339 | -1 | lncRNA | ENSG00000( |
| 131092821 | 131092927 | -1 | snRNA | ENSG00000( |
| 131325092 | 131381656 | -1 | lncRNA | ENSG00000( |
| 131361898 | 131365122 | 1 | transcribed_unitary_ | ENSG00000( |
| 131381671 | 131388830 | 1 | protein_coding | ENSG00000( |
| 131455126 | 131458598 | -1 | lncRNA | ENSG00000( |
| 131462212 | 131502983 | -1 | protein_coding | ENSG00000( |
| 131479097 | 131479233 | -1 | snoRNA | ENSG00000( |
| 131502573 | 131520880 | 1 | lncRNA | ENSG00000( |
| 131526447 | 131527179 | 1 | processed_pseudoge | ENSG00000( |
| 131533555 | 132285410 | -1 | protein_coding | ENSG00000( |
| 131537571 | 131537685 | -1 | unprocessed_pseudc | ENSG00000( |
| 131985855 | 131985931 | -1 | miRNA | ENSG00000( |
| 132175402 | 132176711 | 1 | processed_pseudoge | ENSG00000( |
| 132243528 | 132244265 | -1 | processed_pseudoge | ENSG00000( |
| 132317369 | 132368298 | 1 | protein_coding | ENSG00000( |
| 132386522 | 132389074 | 1 | processed_pseudoge | ENSG00000( |
| 132401253 | 132401796 | 1 | processed_pseudoge | ENSG00000( |
| 132417502 | 132539032 | 1 | protein_coding | ENSG00000( |
| 132558138 | 132660082 | -1 | protein_coding | ENSG00000( |
| 132558142 | 132722459 | -1 | protein_coding | ENSG00000( |
| 132597270 | 132618967 | 1 | protein_coding | ENSG00000( |
| 132645831 | 132650383 | 1 | processed_pseudoge | ENSG00000( |
| 132654446 | 132678097 | 1 | protein_coding | ENSG00000( |
| 132680609 | 132722414 | -1 | protein_coding | ENSG00000( |
| 132721750 | 132874223 | 1 | lncRNA | ENSG00000( |
| 133015004 | 133037052 | -1 | lncRNA | ENSG00000( |
| 133038391 | 133397792 | 1 | protein_coding | ENSG00000( |
| 133245603 | 133257207 | -1 | lncRNA | ENSG00000( |
| 133400056 | 133475222 | 1 | protein_coding | ENSG00000( |
| 133409955 | 133410843 | 1 | processed_pseudoge | ENSG00000( |
| 133429269 | 133455776 | -1 | lncRNA | ENSG00000( |
| 133490824 | 133491505 | 1 | processed_pseudoge | ENSG00000( |
| 133543064 | 133543466 | -1 | lncRNA | ENSG00000( |
| 133546071 | 133546272 | -1 | processed_pseudoge | ENSG00000( |
| 133551186 | 133551315 | 1 | snoRNA | ENSG00000( |
| 133573730 | 133590261 | 1 | protein_coding | ENSG00000( |
| 133598175 | 133662380 | -1 | protein_coding | ENSG00000( |
| 133661998 | 133754576 | 1 | transcribed_unproce | ENSG00000( |

| | | | | |
|---|---|---|---|---|
| 133664926 | 133665034 | 1 | snRNA | ENSG0000C |
| 133710076 | 133710167 | -1 | rRNA_pseudogene | ENSG0000C |
| 133746040 | 133796641 | 1 | protein_coding | ENSG0000C |
| 133760300 | 133762363 | 1 | processed_pseudoge | ENSG0000C |
| 133784023 | 133825772 | 1 | protein_coding | ENSG0000C |
| 133799440 | 133806139 | -1 | lncRNA | ENSG0000C |
| 133824235 | 133895882 | -1 | protein_coding | ENSG0000C |
| 133928145 | 134052184 | -1 | protein_coding | ENSG0000C |
| 134055256 | 134057648 | -1 | lncRNA | ENSG0000C |
| 134065303 | 134250744 | -1 | protein_coding | ENSG0000C |
| 134170487 | 134171085 | 1 | processed_pseudoge | ENSG0000C |
| 134313576 | 134321171 | 1 | lncRNA | ENSG0000C |
| 134347288 | 134349233 | -1 | lncRNA | ENSG0000C |
| 134351852 | 134356561 | -1 | transcribed_processe | ENSG0000C |
| 134355874 | 134375479 | -1 | protein_coding | ENSG0000C |
| 134367804 | 134367862 | -1 | miRNA | ENSG0000C |
| 134385197 | 134385494 | 1 | processed_pseudoge | ENSG0000C |
| 134437605 | 134438211 | 1 | processed_pseudoge | ENSG0000C |
| 134437827 | 134437906 | 1 | miRNA | ENSG0000C |
| 134477706 | 134486716 | -1 | protein_coding | ENSG0000C |
| 134485721 | 134575017 | 1 | protein_coding | ENSG0000C |
| 134510531 | 134511413 | -1 | processed_pseudoge | ENSG0000C |
| 134597801 | 135260467 | 1 | protein_coding | ENSG0000C |
| 134599923 | 134651636 | -1 | protein_coding | ENSG0000C |
| 134774543 | 134796493 | -1 | lncRNA | ENSG0000C |
| 134780527 | 134780627 | -1 | snRNA | ENSG0000C |
| 134783436 | 134783555 | -1 | rRNA_pseudogene | ENSG0000C |
| 135138469 | 135439888 | -1 | lncRNA | ENSG0000C |
| 135925891 | 135926167 | 1 | processed_pseudoge | ENSG0000C |
| 135959275 | 135959590 | -1 | processed_pseudoge | ENSG0000C |
| 135965728 | 136147894 | 1 | protein_coding | ENSG0000C |
| 136055184 | 136055764 | 1 | processed_pseudoge | ENSG0000C |
| 136087475 | 136087913 | -1 | lncRNA | ENSG0000C |
| 136148917 | 136197241 | -1 | protein_coding | ENSG0000C |
| 136155549 | 136155996 | 1 | processed_pseudoge | ENSG0000C |
| 136205540 | 136205915 | -1 | processed_pseudoge | ENSG0000C |
| 136250340 | 136337896 | 1 | protein_coding | ENSG0000C |
| 136336236 | 136752403 | -1 | protein_coding | ENSG0000C |
| 136430084 | 136430183 | -1 | snRNA | ENSG0000C |
| 136609050 | 136609602 | 1 | processed_pseudoge | ENSG0000C |
| 136721394 | 136721495 | -1 | snRNA | ENSG0000C |
| 136736500 | 136737229 | -1 | processed_pseudoge | ENSG0000C |
| 136752630 | 136755780 | 1 | lncRNA | ENSG0000C |
| 136778181 | 136806308 | 1 | lncRNA | ENSG0000C |
| 136808551 | 136809623 | -1 | processed_pseudoge | ENSG0000C |
| 136818647 | 136855888 | 1 | protein_coding | ENSG0000C |
| 136835345 | 136862618 | -1 | lncRNA | ENSG0000C |
| 136837338 | 136839021 | -1 | lncRNA | ENSG0000C |
| 136862208 | 136951606 | 1 | protein_coding | ENSG0000C |
| 136899076 | 136900076 | -1 | processed_pseudoge | ENSG0000C |

| | | | | |
|---|---|---|---|---|
| 136946230 | 137011085 | 1 | protein_coding | ENSG00000( |
| 136959125 | 136982196 | -1 | lncRNA | ENSG00000( |
| 137518134 | 137518255 | -1 | rRNA_pseudogene | ENSG00000( |
| 137535572 | 137536142 | -1 | processed_pseudoge | ENSG00000( |
| 137723774 | 137724543 | 1 | processed_pseudoge | ENSG00000( |
| 137764315 | 137766334 | 1 | protein_coding | ENSG00000( |
| 137771660 | 137780882 | 1 | lncRNA | ENSG00000( |
| 137791973 | 137796678 | 1 | lncRNA | ENSG00000( |
| 137880295 | 137882237 | 1 | processed_pseudoge | ENSG00000( |
| 137998735 | 138033655 | 1 | protein_coding | ENSG00000( |
| 138004649 | 138005122 | -1 | lncRNA | ENSG00000( |
| 138061990 | 138115818 | -1 | protein_coding | ENSG00000( |
| 138101478 | 138102917 | -1 | processed_pseudoge | ENSG00000( |
| 138123713 | 138132390 | -1 | protein_coding | ENSG00000( |
| 138125695 | 138125812 | -1 | unprocessed_pseudc | ENSG00000( |
| 138160988 | 138174921 | -1 | protein_coding | ENSG00000( |
| 138187248 | 138298384 | 1 | protein_coding | ENSG00000( |
| 138261437 | 138329886 | -1 | protein_coding | ENSG00000( |
| 138347648 | 138405534 | 1 | protein_coding | ENSG00000( |
| 138434586 | 138481686 | 1 | protein_coding | ENSG00000( |
| 138482065 | 138485755 | 1 | TEC | ENSG00000( |
| 138494344 | 138594538 | -1 | protein_coding | ENSG00000( |
| 138608606 | 138633376 | 1 | protein_coding | ENSG00000( |
| 138643874 | 138644694 | -1 | processed_pseudoge | ENSG00000( |
| 138652699 | 138834938 | -1 | protein_coding | ENSG00000( |
| 138777832 | 138778810 | 1 | processed_pseudoge | ENSG00000( |
| 138796851 | 138797240 | 1 | processed_pseudoge | ENSG00000( |
| 138825063 | 138826329 | 1 | processed_pseudoge | ENSG00000( |
| 138889255 | 138889957 | -1 | processed_pseudoge | ENSG00000( |
| 138935189 | 138944020 | -1 | lncRNA | ENSG00000( |
| 138944224 | 138947137 | -1 | protein_coding | ENSG00000( |
| 138947217 | 138953990 | 1 | protein_coding | ENSG00000( |
| 139003962 | 139006268 | -1 | protein_coding | ENSG00000( |
| 139005806 | 139357223 | 1 | protein_coding | ENSG00000( |
| 139019031 | 139020926 | -1 | protein_coding | ENSG00000( |
| 139042102 | 139044892 | -1 | protein_coding | ENSG00000( |
| 139081654 | 139082377 | -1 | processed_pseudoge | ENSG00000( |
| 139104185 | 139125171 | 1 | lncRNA | ENSG00000( |
| 139232992 | 139233522 | 1 | lncRNA | ENSG00000( |
| 139233414 | 139233520 | 1 | misc_RNA | ENSG00000( |
| 139316157 | 139340870 | -1 | lncRNA | ENSG00000( |
| 139349024 | 139349371 | -1 | lncRNA | ENSG00000( |
| 139355600 | 139389680 | -1 | protein_coding | ENSG00000( |
| 139388277 | 139388364 | 1 | snRNA | ENSG00000( |
| 139389761 | 139782699 | 1 | lncRNA | ENSG00000( |
| 139452884 | 139480747 | -1 | protein_coding | ENSG00000( |
| 139466430 | 139466795 | 1 | lncRNA | ENSG00000( |
| 139493809 | 139494937 | 1 | processed_pseudoge | ENSG00000( |
| 139517434 | 139539829 | -1 | protein_coding | ENSG00000( |
| 139560180 | 139678017 | -1 | protein_coding | ENSG00000( |

| | | | | |
|---|---|---|---|---|
| 139582928 | 139583593 | -1 | processed_pseudoge | ENSG00000 |
| 139584105 | 139584445 | -1 | misc_RNA | ENSG00000 |
| 139609669 | 139609939 | -1 | misc_RNA | ENSG00000 |
| 139678620 | 139682564 | -1 | lncRNA | ENSG00000 |
| 139688403 | 139689342 | -1 | lncRNA | ENSG00000 |
| 139837220 | 139859894 | 1 | lncRNA | ENSG00000 |
| 139845078 | 139845411 | -1 | processed_pseudoge | ENSG00000 |
| 139935185 | 140577397 | 1 | protein_coding | ENSG00000 |
| 140449435 | 140460351 | -1 | lncRNA | ENSG00000 |
| 140461000 | 140462825 | -1 | lncRNA | ENSG00000 |
| 140505611 | 140508789 | -1 | lncRNA | ENSG00000 |
| 140619694 | 140619837 | -1 | processed_pseudoge | ENSG00000 |
| 140678064 | 140701150 | 1 | protein_coding | ENSG00000 |
| 140865075 | 140867783 | -1 | lncRNA | ENSG00000 |
| 140902194 | 140902651 | 1 | processed_pseudoge | ENSG00000 |
| 140941830 | 140980978 | 1 | protein_coding | ENSG00000 |
| 140972744 | 140973255 | 1 | lncRNA | ENSG00000 |
| 141051347 | 141148611 | 1 | protein_coding | ENSG00000 |
| 141115124 | 141124182 | -1 | lncRNA | ENSG00000 |
| 141228726 | 141367753 | 1 | protein_coding | ENSG00000 |
| 141251004 | 141284306 | -1 | lncRNA | ENSG00000 |
| 141258418 | 141258779 | 1 | processed_pseudoge | ENSG00000 |
| 141267353 | 141367137 | -1 | lncRNA | ENSG00000 |
| 141307612 | 141307933 | -1 | processed_pseudoge | ENSG00000 |
| 141324213 | 141449792 | 1 | protein_coding | ENSG00000 |
| 141470634 | 141471902 | -1 | processed_pseudoge | ENSG00000 |
| 141487027 | 141615356 | 1 | protein_coding | ENSG00000 |
| 141525133 | 141526121 | 1 | lncRNA | ENSG00000 |
| 141600276 | 141600579 | -1 | processed_pseudoge | ENSG00000 |
| 141660536 | 141729405 | 1 | transcribed_unitary_ | ENSG00000 |
| 141709016 | 141709832 | -1 | processed_pseudoge | ENSG00000 |
| 141724425 | 141724526 | 1 | processed_pseudoge | ENSG00000 |
| 141738249 | 141747560 | 1 | protein_coding | ENSG00000 |
| 141738474 | 141807919 | 1 | protein_coding | ENSG00000 |
| 141778148 | 141818490 | 1 | protein_coding | ENSG00000 |
| 141851549 | 141861168 | 1 | lncRNA | ENSG00000 |
| 141865012 | 141865279 | -1 | processed_pseudoge | ENSG00000 |
| 141876124 | 141926549 | 1 | protein_coding | ENSG00000 |
| 141902924 | 141903030 | -1 | snRNA | ENSG00000 |
| 141918252 | 141919021 | -1 | lncRNA | ENSG00000 |
| 141936707 | 141938547 | -1 | TEC | ENSG00000 |
| 141944428 | 142149544 | -1 | protein_coding | ENSG00000 |
| 142043623 | 142044198 | -1 | processed_pseudoge | ENSG00000 |
| 142145454 | 142145555 | 1 | snRNA | ENSG00000 |
| 142157527 | 142225592 | -1 | protein_coding | ENSG00000 |
| 142306607 | 142448062 | -1 | protein_coding | ENSG00000 |
| 142407657 | 142407737 | -1 | snRNA | ENSG00000 |
| 142420205 | 142420369 | -1 | snRNA | ENSG00000 |
| 142449007 | 142578733 | -1 | protein_coding | ENSG00000 |
| 142450102 | 142452149 | 1 | processed_pseudoge | ENSG00000 |

| | | | | |
|---|---|---|---|---|
| 142465315 | 142472337 | -1 | lncRNA | ENSG0000C |
| 142580910 | 142582399 | 1 | processed_pseudoge | ENSG0000C |
| 142591677 | 142591791 | -1 | rRNA_pseudogene | ENSG0000C |
| 142596393 | 142713664 | 1 | protein_coding | ENSG0000C |
| 142654784 | 142656745 | -1 | lncRNA | ENSG0000C |
| 142673597 | 142673904 | 1 | misc_RNA | ENSG0000C |
| 142687982 | 142723881 | -1 | lncRNA | ENSG0000C |
| 142724034 | 142807888 | 1 | protein_coding | ENSG0000C |
| 142801207 | 142801268 | 1 | snRNA | ENSG0000C |
| 142815922 | 142889206 | -1 | protein_coding | ENSG0000C |
| 142827096 | 142827728 | -1 | processed_pseudoge | ENSG0000C |
| 142912116 | 142942537 | 1 | lncRNA | ENSG0000C |
| 142949164 | 142963674 | -1 | protein_coding | ENSG0000C |
| 142960372 | 142960543 | 1 | processed_pseudoge | ENSG0000C |
| 142960650 | 143001559 | 1 | lncRNA | ENSG0000C |
| 142964497 | 143060725 | 1 | protein_coding | ENSG0000C |
| 143000907 | 143001467 | -1 | lncRNA | ENSG0000C |
| 143111802 | 143112359 | 1 | lncRNA | ENSG0000C |
| 143119771 | 143124014 | 1 | protein_coding | ENSG0000C |
| 143123362 | 143131893 | 1 | lncRNA | ENSG0000C |
| 143176327 | 143177617 | 1 | processed_pseudoge | ENSG0000C |
| 143265222 | 143848485 | -1 | protein_coding | ENSG0000C |
| 143313067 | 143314429 | -1 | lncRNA | ENSG0000C |
| 143342246 | 143347071 | 1 | lncRNA | ENSG0000C |
| 143381338 | 143382161 | 1 | lncRNA | ENSG0000C |
| 143503275 | 143504273 | -1 | processed_pseudoge | ENSG0000C |
| 143517123 | 143518229 | -1 | processed_pseudoge | ENSG0000C |
| 143620528 | 143627690 | 1 | unprocessed_pseudc | ENSG0000C |
| 143855739 | 143856142 | -1 | processed_pseudoge | ENSG0000C |
| 143971798 | 144048719 | 1 | protein_coding | ENSG0000C |
| 144186905 | 144187020 | -1 | rRNA_pseudogene | ENSG0000C |
| 144442211 | 144445844 | -1 | TEC | ENSG0000C |
| 145523538 | 145524415 | 1 | processed_pseudoge | ENSG0000C |
| 145684572 | 145685092 | 1 | processed_pseudoge | ENSG0000C |
| 145824257 | 145824739 | -1 | processed_pseudoge | ENSG0000C |
| 145939912 | 145961536 | 1 | lncRNA | ENSG0000C |
| 145961755 | 145963623 | 1 | lncRNA | ENSG0000C |
| 146059585 | 146061679 | -1 | lncRNA | ENSG0000C |
| 146064042 | 146105204 | 1 | lncRNA | ENSG0000C |
| 146066344 | 146069185 | -1 | lncRNA | ENSG0000C |
| 146069440 | 146163653 | -1 | protein_coding | ENSG0000C |
| 146192335 | 146251179 | -1 | protein_coding | ENSG0000C |
| 146342996 | 146348480 | -1 | processed_pseudoge | ENSG0000C |
| 146391363 | 146416603 | -1 | transcribed_processe | ENSG0000C |
| 146391421 | 146495991 | -1 | protein_coding | ENSG0000C |
| 146504570 | 146506560 | -1 | TEC | ENSG0000C |
| 146515180 | 146544856 | -1 | protein_coding | ENSG0000C |
| 146571477 | 146571583 | 1 | snRNA | ENSG0000C |
| 146576555 | 146606216 | -1 | protein_coding | ENSG0000C |
| 146589602 | 146590325 | 1 | lncRNA | ENSG0000C |

| | | | | |
|---|---|---|---|---|
| 146909685 | 147370656 | -1 | lncRNA | ENSG00000 |
| 146921795 | 146927429 | 1 | lncRNA | ENSG00000 |
| 146996607 | 146996713 | -1 | snRNA | ENSG00000 |
| 147276768 | 147277252 | 1 | processed_pseudoge | ENSG00000 |
| 147386046 | 147406860 | -1 | protein_coding | ENSG00000 |
| 147386967 | 147387453 | 1 | lncRNA | ENSG00000 |
| 147393422 | 147510293 | 1 | protein_coding | ENSG00000 |
| 147590996 | 147591835 | -1 | processed_pseudoge | ENSG00000 |
| 147844123 | 147846657 | -1 | lncRNA | ENSG00000 |
| 147939905 | 148007127 | 1 | lncRNA | ENSG00000 |
| 148028046 | 148079637 | -1 | lncRNA | ENSG00000 |
| 148076432 | 148089134 | 1 | lncRNA | ENSG00000 |
| 148090341 | 148096921 | 1 | lncRNA | ENSG00000 |
| 148093175 | 148127233 | 1 | lncRNA | ENSG00000 |
| 148160911 | 148226606 | 1 | lncRNA | ENSG00000 |
| 148166344 | 148167298 | -1 | processed_pseudoge | ENSG00000 |
| 148192872 | 148280098 | -1 | lncRNA | ENSG00000 |
| 148279682 | 148401084 | 1 | lncRNA | ENSG00000 |
| 148368384 | 148591927 | -1 | lncRNA | ENSG00000 |
| 148695994 | 148696204 | -1 | processed_pseudoge | ENSG00000 |
| 148697784 | 148743008 | 1 | protein_coding | ENSG00000 |
| 148791102 | 148860187 | 1 | protein_coding | ENSG00000 |
| 148824782 | 148825387 | -1 | processed_pseudoge | ENSG00000 |
| 148850933 | 148960112 | -1 | lncRNA | ENSG00000 |
| 148865296 | 148897203 | 1 | protein_coding | ENSG00000 |
| 148922703 | 148956088 | 1 | lncRNA | ENSG00000 |
| 148985868 | 148987668 | -1 | processed_pseudoge | ENSG00000 |
| 148991341 | 149027668 | 1 | protein_coding | ENSG00000 |
| 149030127 | 149086554 | -1 | protein_coding | ENSG00000 |
| 149086332 | 149102823 | 1 | lncRNA | ENSG00000 |
| 149094353 | 149094462 | -1 | misc_RNA | ENSG00000 |
| 149099499 | 149099815 | -1 | processed_pseudoge | ENSG00000 |
| 149129638 | 149173732 | 1 | protein_coding | ENSG00000 |
| 149162410 | 149221829 | -1 | protein_coding | ENSG00000 |
| 149224138 | 149226334 | 1 | lncRNA | ENSG00000 |
| 149229356 | 149291800 | -1 | transcribed_unitary_ | ENSG00000 |
| 149284782 | 149333653 | 1 | lncRNA | ENSG00000 |
| 149318498 | 149334414 | -1 | protein_coding | ENSG00000 |
| 149357169 | 149357545 | -1 | processed_pseudoge | ENSG00000 |
| 149369022 | 149377692 | -1 | protein_coding | ENSG00000 |
| 149377778 | 149386583 | 1 | lncRNA | ENSG00000 |
| 149384179 | 149385800 | -1 | lncRNA | ENSG00000 |
| 149433044 | 149433274 | -1 | processed_pseudoge | ENSG00000 |
| 149462182 | 149464830 | -1 | processed_pseudoge | ENSG00000 |
| 149463779 | 149464114 | -1 | processed_pseudoge | ENSG00000 |
| 149464839 | 149465201 | -1 | processed_pseudoge | ENSG00000 |
| 149474697 | 149503394 | 1 | protein_coding | ENSG00000 |
| 149494660 | 149495995 | 1 | processed_pseudoge | ENSG00000 |
| 149517235 | 149736714 | -1 | protein_coding | ENSG00000 |
| 149525631 | 149525726 | 1 | snRNA | ENSG00000 |

| | | | | |
|---|---|---|---|---|
| 149648997 | 149650184 | -1 | lncRNA | ENSG00000( |
| 149656999 | 149661364 | 1 | lncRNA | ENSG00000( |
| 149738472 | 149752495 | -1 | protein_coding | ENSG00000( |
| 149761100 | 149968385 | -1 | protein_coding | ENSG00000( |
| 149779009 | 149779108 | -1 | snRNA | ENSG00000( |
| 149812708 | 149962139 | 1 | protein_coding | ENSG00000( |
| 149917342 | 149917449 | -1 | snRNA | ENSG00000( |
| 149938997 | 149939799 | -1 | processed_pseudoge | ENSG00000( |
| 149964904 | 150050788 | -1 | protein_coding | ENSG00000( |
| 149976755 | 149979355 | 1 | lncRNA | ENSG00000( |
| 149982181 | 149983308 | -1 | processed_pseudoge | ENSG00000( |
| 149984395 | 149984797 | 1 | processed_pseudoge | ENSG00000( |
| 149999645 | 150000690 | 1 | processed_pseudoge | ENSG00000( |
| 150039214 | 150213726 | 1 | lncRNA | ENSG00000( |
| 150050729 | 150051456 | 1 | processed_pseudoge | ENSG00000( |
| 150077494 | 150080117 | -1 | lncRNA | ENSG00000( |
| 150150538 | 150150944 | 1 | transcribed_processe | ENSG00000( |
| 150202174 | 150225190 | -1 | lncRNA | ENSG00000( |
| 150238519 | 150244232 | 1 | lncRNA | ENSG00000( |
| 150241777 | 150241860 | 1 | snRNA | ENSG00000( |
| 150265407 | 150296605 | -1 | lncRNA | ENSG00000( |
| 150339478 | 150343210 | -1 | lncRNA | ENSG00000( |
| 150408335 | 150466431 | 1 | protein_coding | ENSG00000( |
| 150541998 | 150603228 | -1 | protein_coding | ENSG00000( |
| 150546678 | 150586016 | 1 | protein_coding | ENSG00000( |
| 150602875 | 150630445 | 1 | protein_coding | ENSG00000( |
| 150611262 | 150611877 | -1 | processed_pseudoge | ENSG00000( |
| 150659885 | 150703971 | -1 | protein_coding | ENSG00000( |
| 150703564 | 150723005 | 1 | lncRNA | ENSG00000( |
| 150719997 | 150720836 | -1 | processed_pseudoge | ENSG00000( |
| 150734469 | 150738977 | -1 | lncRNA | ENSG00000( |
| 150741125 | 150763477 | -1 | protein_coding | ENSG00000( |
| 150761937 | 150873554 | 1 | lncRNA | ENSG00000( |
| 150852484 | 151080726 | 1 | lncRNA | ENSG00000( |
| 150870376 | 150905439 | -1 | protein_coding | ENSG00000( |
| 150882997 | 150941743 | -1 | protein_coding | ENSG00000( |
| 150890636 | 151038818 | 1 | lncRNA | ENSG00000( |
| 150926163 | 150972999 | -1 | protein_coding | ENSG00000( |
| 151079506 | 151079584 | 1 | misc_RNA | ENSG00000( |
| 151085697 | 151437072 | 1 | protein_coding | ENSG00000( |
| 151188099 | 151188223 | 1 | rRNA_pseudogene | ENSG00000( |
| 151197832 | 151203216 | -1 | protein_coding | ENSG00000( |
| 151212117 | 151278542 | -1 | protein_coding | ENSG00000( |
| 151269440 | 151270129 | 1 | processed_pseudoge | ENSG00000( |
| 151293317 | 151295494 | 1 | lncRNA | ENSG00000( |
| 151294086 | 151316820 | -1 | protein_coding | ENSG00000( |
| 151326312 | 151329549 | -1 | protein_coding | ENSG00000( |
| 151337380 | 151384812 | -1 | protein_coding | ENSG00000( |
| 151425384 | 151458709 | -1 | protein_coding | ENSG00000( |
| 151475804 | 151476174 | -1 | processed_pseudoge | ENSG00000( |

| | | | | |
|---|---|---|---|---|
| 151499639 | 151499806 | 1 | processed_pseudoge | ENSG0000C |
| 151565876 | 151565995 | -1 | miRNA | ENSG0000C |
| 151637174 | 151657966 | 1 | lncRNA | ENSG0000C |
| 151733916 | 151761339 | 1 | protein_coding | ENSG0000C |
| 151751443 | 151928175 | -1 | lncRNA | ENSG0000C |
| 151759747 | 151762011 | -1 | processed_pseudoge | ENSG0000C |
| 151770428 | 151784894 | 1 | transcribed_unproce | ENSG0000C |
| 151797047 | 151808249 | 1 | transcribed_unproce | ENSG0000C |
| 151814073 | 151828488 | 1 | protein_coding | ENSG0000C |
| 151873643 | 151884619 | 1 | protein_coding | ENSG0000C |
| 152118173 | 152151724 | -1 | lncRNA | ENSG0000C |
| 152243828 | 152465780 | 1 | protein_coding | ENSG0000C |
| 152245262 | 152269557 | -1 | lncRNA | ENSG0000C |
| 152339695 | 152340990 | -1 | transcribed_processe | ENSG0000C |
| 152449272 | 152449384 | 1 | misc_RNA | ENSG0000C |
| 152457759 | 152496813 | -1 | lncRNA | ENSG0000C |
| 152648146 | 152650447 | 1 | lncRNA | ENSG0000C |
| 152650519 | 152679450 | 1 | lncRNA | ENSG0000C |
| 152800434 | 152800732 | -1 | processed_pseudoge | ENSG0000C |
| 152835131 | 152841439 | 1 | protein_coding | ENSG0000C |
| 153067278 | 153067551 | 1 | processed_pseudoge | ENSG0000C |
| 153129074 | 153129640 | -1 | processed_pseudoge | ENSG0000C |
| 153156511 | 153161775 | -1 | lncRNA | ENSG0000C |
| 153162226 | 153170627 | 1 | protein_coding | ENSG0000C |
| 153199700 | 153200017 | -1 | misc_RNA | ENSG0000C |
| 153357107 | 153374186 | 1 | lncRNA | ENSG0000C |
| 153376831 | 153377562 | 1 | lncRNA | ENSG0000C |
| 153384934 | 153980186 | -1 | lncRNA | ENSG0000C |
| 153431856 | 153502697 | 1 | lncRNA | ENSG0000C |
| 153471466 | 153471571 | -1 | snRNA | ENSG0000C |
| 153607377 | 153607479 | 1 | misc_RNA | ENSG0000C |
| 153881526 | 153940836 | 1 | lncRNA | ENSG0000C |
| 154007367 | 154007502 | -1 | snoRNA | ENSG0000C |
| 154023835 | 154024298 | 1 | processed_pseudoge | ENSG0000C |
| 154024401 | 154121347 | -1 | lncRNA | ENSG0000C |
| 154121003 | 154257827 | 1 | protein_coding | ENSG0000C |
| 154272546 | 154324487 | -1 | protein_coding | ENSG0000C |
| 154329716 | 154329819 | 1 | snoRNA | ENSG0000C |
| 154334809 | 154337047 | -1 | TEC | ENSG0000C |
| 154334943 | 154430190 | -1 | protein_coding | ENSG0000C |
| 154352136 | 154354799 | -1 | processed_pseudoge | ENSG0000C |
| 154511535 | 154539309 | -1 | lncRNA | ENSG0000C |
| 154651656 | 154652423 | -1 | processed_pseudoge | ENSG0000C |
| 154657294 | 154657563 | 1 | processed_pseudoge | ENSG0000C |
| 154676588 | 154677119 | -1 | processed_pseudoge | ENSG0000C |
| 154827016 | 154861017 | -1 | lncRNA | ENSG0000C |
| 154969283 | 154970223 | -1 | lncRNA | ENSG0000C |
| 155024124 | 155183729 | 1 | protein_coding | ENSG0000C |
| 155158370 | 155183285 | -1 | lncRNA | ENSG0000C |
| 155240919 | 155258766 | 1 | lncRNA | ENSG0000C |

| | | | | |
|---|---|---|---|---|
| 155290227 | 155293683 | -1 | protein_coding | ENSG00000 |
| 155291188 | 155294176 | 1 | lncRNA | ENSG00000 |
| 155309427 | 155309807 | -1 | processed_pseudoge | ENSG00000 |
| 155375580 | 155745067 | -1 | protein_coding | ENSG00000 |
| 155375683 | 155375790 | -1 | unprocessed_pseudo | ENSG00000 |
| 155401329 | 155403399 | -1 | processed_pseudoge | ENSG00000 |
| 155431229 | 155433398 | -1 | processed_pseudoge | ENSG00000 |
| 155437538 | 155437823 | -1 | misc_RNA | ENSG00000 |
| 155449184 | 155457753 | 1 | lncRNA | ENSG00000 |
| 155485803 | 155487121 | 1 | lncRNA | ENSG00000 |
| 155657760 | 155658612 | 1 | processed_pseudoge | ENSG00000 |
| 155742142 | 155743726 | -1 | lncRNA | ENSG00000 |
| 155760617 | 155761217 | 1 | processed_pseudoge | ENSG00000 |
| 155762617 | 155806278 | -1 | protein_coding | ENSG00000 |
| 155763043 | 155854442 | -1 | protein_coding | ENSG00000 |
| 155821024 | 155854456 | -1 | protein_coding | ENSG00000 |
| 155854752 | 155863191 | 1 | lncRNA | ENSG00000 |
| 155870650 | 155944020 | 1 | protein_coding | ENSG00000 |
| 155886615 | 155890729 | 1 | lncRNA | ENSG00000 |
| 155987304 | 155988176 | 1 | processed_pseudoge | ENSG00000 |
| 156019646 | 156037647 | 1 | unitary_pseudogene | ENSG00000 |
| 156037701 | 156539138 | 1 | protein_coding | ENSG00000 |
| 156044155 | 156044948 | -1 | processed_pseudoge | ENSG00000 |
| 156111432 | 156114198 | -1 | processed_pseudoge | ENSG00000 |
| 156143570 | 156540627 | 1 | lncRNA | ENSG00000 |
| 156175852 | 156176911 | -1 | processed_pseudoge | ENSG00000 |
| 156215560 | 156227883 | -1 | lncRNA | ENSG00000 |
| 156441157 | 156446905 | -1 | lncRNA | ENSG00000 |
| 156523740 | 156524247 | -1 | lncRNA | ENSG00000 |
| 156540140 | 156555184 | -1 | protein_coding | ENSG00000 |
| 156671862 | 156674446 | -1 | lncRNA | ENSG00000 |
| 156673235 | 156706770 | 1 | protein_coding | ENSG00000 |
| 156713884 | 156714928 | -1 | processed_pseudoge | ENSG00000 |
| 156747343 | 156817062 | -1 | lncRNA | ENSG00000 |
| 156809551 | 156810732 | -1 | processed_pseudoge | ENSG00000 |
| 156825481 | 157046129 | 1 | protein_coding | ENSG00000 |
| 156977558 | 156977825 | 1 | processed_pseudoge | ENSG00000 |
| 156988117 | 156988712 | -1 | processed_pseudoge | ENSG00000 |
| 157017342 | 157017696 | 1 | misc_RNA | ENSG00000 |
| 157081667 | 157128595 | -1 | lncRNA | ENSG00000 |
| 157081738 | 157088547 | 1 | lncRNA | ENSG00000 |
| 157089634 | 157135557 | 1 | lncRNA | ENSG00000 |
| 157146508 | 157160760 | -1 | protein_coding | ENSG00000 |
| 157153548 | 157153640 | 1 | misc_RNA | ENSG00000 |
| 157162663 | 157163932 | -1 | processed_pseudoge | ENSG00000 |
| 157163452 | 157169133 | 1 | lncRNA | ENSG00000 |
| 157173918 | 157174040 | -1 | rRNA_pseudogene | ENSG00000 |
| 157174201 | 157381265 | 1 | lncRNA | ENSG00000 |
| 157230320 | 157231208 | -1 | processed_pseudoge | ENSG00000 |
| 157234580 | 157235149 | -1 | lncRNA | ENSG00000 |

| | | | | |
|---|---|---|---|---|
| 157259742 | 157533619 | -1 | protein_coding | ENSG0000C |
| 157436850 | 157443633 | 1 | protein_coding | ENSG0000C |
| 157543246 | 157677749 | 1 | protein_coding | ENSG0000C |
| 157579694 | 157587495 | -1 | lncRNA | ENSG0000C |
| 157814822 | 158088997 | -1 | lncRNA | ENSG0000C |
| 157930492 | 157930725 | -1 | misc_RNA | ENSG0000C |
| 157976561 | 157978136 | -1 | lncRNA | ENSG0000C |
| 158095954 | 158106503 | -1 | protein_coding | ENSG0000C |
| 158105855 | 158545730 | 1 | protein_coding | ENSG0000C |
| 158275809 | 158276421 | -1 | processed_pseudoge | ENSG0000C |
| 158336430 | 158336777 | -1 | processed_pseudoge | ENSG0000C |
| 158496561 | 158496934 | 1 | processed_pseudoge | ENSG0000C |
| 158545189 | 158571066 | -1 | lncRNA | ENSG0000C |
| 158571163 | 158607252 | 1 | protein_coding | ENSG0000C |
| 158609610 | 158610857 | -1 | processed_pseudoge | ENSG0000C |
| 158626656 | 158626849 | -1 | processed_pseudoge | ENSG0000C |
| 158644278 | 158692575 | 1 | protein_coding | ENSG0000C |
| 158645822 | 158672648 | -1 | protein_coding | ENSG0000C |
| 158693120 | 158693768 | -1 | lncRNA | ENSG0000C |
| 158695367 | 158695581 | 1 | lncRNA | ENSG0000C |
| 158696892 | 158732489 | -1 | protein_coding | ENSG0000C |
| 158732198 | 158829719 | 1 | protein_coding | ENSG0000C |
| 158732263 | 158784070 | 1 | lncRNA | ENSG0000C |
| 158763431 | 158763753 | 1 | processed_pseudoge | ENSG0000C |
| 158782547 | 158783124 | 1 | lncRNA | ENSG0000C |
| 158801257 | 158801935 | -1 | lncRNA | ENSG0000C |
| 158841093 | 158841960 | -1 | processed_pseudoge | ENSG0000C |
| 158869898 | 158871821 | 1 | TEC | ENSG0000C |
| 158962235 | 159897366 | 1 | protein_coding | ENSG0000C |
| 158962928 | 159266307 | 1 | protein_coding | ENSG0000C |
| 159282646 | 159282734 | 1 | miRNA | ENSG0000C |
| 159386756 | 159387440 | -1 | processed_pseudoge | ENSG0000C |
| 159711852 | 159731646 | -1 | lncRNA | ENSG0000C |
| 159721524 | 159722401 | -1 | processed_pseudoge | ENSG0000C |
| 159765387 | 159768612 | -1 | lncRNA | ENSG0000C |
| 159808416 | 159808570 | -1 | snRNA | ENSG0000C |
| 159839861 | 159897360 | 1 | protein_coding | ENSG0000C |
| 159902045 | 160225299 | -1 | lncRNA | ENSG0000C |
| 159988835 | 159996019 | 1 | protein_coding | ENSG0000C |
| 160016024 | 160031423 | 1 | lncRNA | ENSG0000C |
| 160100850 | 160102793 | 1 | processed_pseudoge | ENSG0000C |
| 160225496 | 160228213 | 1 | protein_coding | ENSG0000C |
| 160227454 | 160449829 | -1 | protein_coding | ENSG0000C |
| 160256986 | 160399880 | -1 | protein_coding | ENSG0000C |
| 160342685 | 160343015 | 1 | processed_pseudoge | ENSG0000C |
| 160399274 | 160434954 | 1 | protein_coding | ENSG0000C |
| 160404588 | 160404685 | 1 | miRNA | ENSG0000C |
| 160404745 | 160404825 | 1 | miRNA | ENSG0000C |
| 160432445 | 160485773 | -1 | protein_coding | ENSG0000C |
| 160452598 | 160453403 | 1 | processed_pseudoge | ENSG0000C |

| | | | | |
|---|---|---|---|---|
| 160472348 | 160472416 | -1 | snRNA | ENSG0000C |
| 160495007 | 160565571 | -1 | protein_coding | ENSG0000C |
| 160514907 | 160515236 | -1 | scaRNA | ENSG0000C |
| 160565447 | 160569248 | 1 | transcribed_processe | ENSG0000C |
| 160584945 | 160585787 | 1 | processed_pseudoge | ENSG0000C |
| 160677160 | 160678448 | 1 | protein_coding | ENSG0000C |
| 160696929 | 160697060 | 1 | snoRNA | ENSG0000C |
| 160753428 | 160755142 | -1 | lncRNA | ENSG0000C |
| 160755602 | 161078902 | 1 | protein_coding | ENSG0000C |
| 161083883 | 161105411 | -1 | protein_coding | ENSG0000C |
| 161104696 | 161253532 | 1 | protein_coding | ENSG0000C |
| 161222911 | 161223147 | -1 | processed_pseudoge | ENSG0000C |
| 161324913 | 161326219 | -1 | processed_pseudoge | ENSG0000C |
| 161328924 | 161330238 | -1 | processed_pseudoge | ENSG0000C |
| 161344792 | 161372880 | -1 | protein_coding | ENSG0000C |
| 161426427 | 161448242 | 1 | lncRNA | ENSG0000C |
| 161429112 | 161429613 | 1 | lncRNA | ENSG0000C |
| 161429127 | 161429597 | -1 | processed_pseudoge | ENSG0000C |
| 161496808 | 161503942 | 1 | protein_coding | ENSG0000C |
| 161600438 | 161735399 | 1 | lncRNA | ENSG0000C |
| 161816909 | 161821908 | -1 | lncRNA | ENSG0000C |
| 162486541 | 162486949 | -1 | processed_pseudoge | ENSG0000C |
| 162601627 | 162601792 | 1 | processed_pseudoge | ENSG0000C |
| 163026396 | 163232149 | 1 | lncRNA | ENSG0000C |
| 163109152 | 163361563 | -1 | lncRNA | ENSG0000C |
| 163115429 | 163116186 | 1 | processed_pseudoge | ENSG0000C |
| 163455568 | 163479500 | 1 | lncRNA | ENSG0000C |
| 163470205 | 163470266 | -1 | snRNA | ENSG0000C |
| 164001606 | 164002465 | -1 | processed_pseudoge | ENSG0000C |
| 164171471 | 164171556 | -1 | miRNA | ENSG0000C |
| 164670878 | 164686076 | -1 | lncRNA | ENSG0000C |
| 164714095 | 164831480 | -1 | lncRNA | ENSG0000C |
| 164978898 | 165078496 | -1 | protein_coding | ENSG0000C |
| 165122713 | 165122823 | -1 | misc_RNA | ENSG0000C |
| 165150006 | 165158062 | -1 | lncRNA | ENSG0000C |
| 165186720 | 165197109 | -1 | protein_coding | ENSG0000C |
| 165206929 | 165846519 | 1 | lncRNA | ENSG0000C |
| 165772904 | 165837462 | -1 | protein_coding | ENSG0000C |
| 166160021 | 166160370 | -1 | processed_pseudoge | ENSG0000C |
| 166160414 | 166160704 | -1 | processed_pseudoge | ENSG0000C |
| 166160777 | 166161119 | -1 | processed_pseudoge | ENSG0000C |
| 166161203 | 166161968 | -1 | processed_pseudoge | ENSG0000C |
| 166181313 | 166181820 | -1 | processed_pseudoge | ENSG0000C |
| 166293756 | 166294353 | 1 | processed_pseudoge | ENSG0000C |
| 166569693 | 166570763 | -1 | lncRNA | ENSG0000C |
| 166605188 | 166605455 | 1 | processed_pseudoge | ENSG0000C |
| 166670029 | 166670122 | -1 | misc_RNA | ENSG0000C |
| 166691187 | 166691712 | 1 | processed_pseudoge | ENSG0000C |
| 166721458 | 166721610 | 1 | processed_pseudoge | ENSG0000C |
| 166811009 | 166811567 | 1 | processed_pseudoge | ENSG0000C |

| | | | | |
|---|---|---|---|---|
| 166835021 | 166844956 | -1 | lncRNA | ENSG00000 |
| 166861944 | 166862110 | 1 | processed_pseudoge | ENSG00000 |
| 167065831 | 167066320 | 1 | processed_pseudoge | ENSG00000 |
| 167240287 | 167381346 | -1 | protein_coding | ENSG00000 |
| 167392796 | 167408519 | 1 | lncRNA | ENSG00000 |
| 167441789 | 167479004 | -1 | protein_coding | ENSG00000 |
| 167478684 | 167653983 | -1 | protein_coding | ENSG00000 |
| 167683298 | 167734939 | -1 | protein_coding | ENSG00000 |
| 167694355 | 167694642 | -1 | processed_pseudoge | ENSG00000 |
| 167735243 | 167825568 | 1 | protein_coding | ENSG00000 |
| 167794947 | 167811081 | -1 | lncRNA | ENSG00000 |
| 167864773 | 167924011 | 1 | transcribed_unitary_ | ENSG00000 |
| 167866500 | 167950292 | -1 | lncRNA | ENSG00000 |
| 167888414 | 167888715 | 1 | processed_pseudoge | ENSG00000 |
| 167956311 | 167956616 | -1 | processed_pseudoge | ENSG00000 |
| 167958744 | 167959082 | 1 | processed_pseudoge | ENSG00000 |
| 168008689 | 168095924 | -1 | protein_coding | ENSG00000 |
| 168094049 | 168095087 | -1 | lncRNA | ENSG00000 |
| 168249522 | 168830599 | 1 | transcribed_unitary_ | ENSG00000 |
| 168288553 | 168291228 | 1 | lncRNA | ENSG00000 |
| 168475198 | 168475388 | 1 | snRNA | ENSG00000 |
| 168551854 | 168551949 | 1 | miRNA | ENSG00000 |
| 168666559 | 168667423 | 1 | processed_pseudoge | ENSG00000 |
| 168858659 | 168859130 | -1 | processed_pseudoge | ENSG00000 |
| 168902194 | 168924590 | 1 | lncRNA | ENSG00000 |
| 168927475 | 168927919 | 1 | lncRNA | ENSG00000 |
| 168928169 | 168928598 | 1 | lncRNA | ENSG00000 |
| 169003022 | 169038416 | 1 | lncRNA | ENSG00000 |
| 169083499 | 169663775 | -1 | protein_coding | ENSG00000 |
| 169447867 | 169477052 | 1 | lncRNA | ENSG00000 |
| 169483671 | 169484080 | -1 | processed_pseudoge | ENSG00000 |
| 169613510 | 169623951 | -1 | lncRNA | ENSG00000 |
| 169709295 | 169709765 | 1 | processed_pseudoge | ENSG00000 |
| 169720187 | 169720293 | -1 | snRNA | ENSG00000 |
| 169764520 | 169765060 | -1 | lncRNA | ENSG00000 |
| 169764610 | 169765047 | -1 | misc_RNA | ENSG00000 |
| 169766921 | 169769561 | -1 | protein_coding | ENSG00000 |
| 169769649 | 169772043 | 1 | lncRNA | ENSG00000 |
| 169772831 | 169789716 | 1 | protein_coding | ENSG00000 |
| 169777192 | 169780334 | -1 | lncRNA | ENSG00000 |
| 169793428 | 169812986 | -1 | protein_coding | ENSG00000 |
| 169793495 | 169793966 | 1 | lncRNA | ENSG00000 |
| 169794962 | 169796213 | 1 | lncRNA | ENSG00000 |
| 169821922 | 169837775 | 1 | protein_coding | ENSG00000 |
| 169839172 | 169869935 | -1 | protein_coding | ENSG00000 |
| 169902980 | 169904260 | -1 | processed_pseudoge | ENSG00000 |
| 169911572 | 169939175 | 1 | protein_coding | ENSG00000 |
| 169931998 | 169933229 | 1 | processed_pseudoge | ENSG00000 |
| 169939353 | 169966734 | -1 | lncRNA | ENSG00000 |
| 169945987 | 169946754 | -1 | transcribed_processe | ENSG00000 |

| | | | | |
|---|---|---|---|---|
| 169966635 | 169998373 | 1 | protein_coding | ENSG00000 |
| 169978536 | 169985715 | -1 | lncRNA | ENSG00000 |
| 170037995 | 170085392 | 1 | protein_coding | ENSG00000 |
| 170057301 | 170057429 | 1 | snRNA | ENSG00000 |
| 170062244 | 170062951 | 1 | processed_pseudoge | ENSG00000 |
| 170086732 | 170181749 | -1 | protein_coding | ENSG00000 |
| 170149845 | 170149951 | 1 | snRNA | ENSG00000 |
| 170222424 | 170305977 | 1 | protein_coding | ENSG00000 |
| 170244782 | 170244881 | 1 | misc_RNA | ENSG00000 |
| 170345678 | 170353961 | 1 | lncRNA | ENSG00000 |
| 170357678 | 170396835 | 1 | protein_coding | ENSG00000 |
| 170410512 | 170418615 | 1 | lncRNA | ENSG00000 |
| 170418865 | 170454733 | 1 | protein_coding | ENSG00000 |
| 170418868 | 170860380 | 1 | protein_coding | ENSG00000 |
| 170423103 | 170423162 | 1 | miRNA | ENSG00000 |
| 170448639 | 170860993 | 1 | lncRNA | ENSG00000 |
| 170459548 | 170586075 | -1 | protein_coding | ENSG00000 |
| 170495627 | 170496636 | 1 | processed_pseudoge | ENSG00000 |
| 170653846 | 170654259 | -1 | processed_pseudoge | ENSG00000 |
| 170656562 | 170662123 | 1 | lncRNA | ENSG00000 |
| 170691997 | 170736061 | -1 | lncRNA | ENSG00000 |
| 170740351 | 170741365 | -1 | lncRNA | ENSG00000 |
| 170864875 | 170870208 | -1 | protein_coding | ENSG00000 |
| 170888418 | 170908644 | -1 | protein_coding | ENSG00000 |
| 170908857 | 171054973 | 1 | lncRNA | ENSG00000 |
| 170952850 | 170953897 | -1 | processed_pseudoge | ENSG00000 |
| 170994870 | 170995033 | -1 | snRNA | ENSG00000 |
| 170996347 | 171026743 | -1 | protein_coding | ENSG00000 |
| 171058414 | 171460408 | -1 | protein_coding | ENSG00000 |
| 171106664 | 171106759 | -1 | miRNA | ENSG00000 |
| 171439526 | 171439627 | 1 | misc_RNA | ENSG00000 |
| 171459248 | 171469702 | 1 | lncRNA | ENSG00000 |
| 171460612 | 171462256 | 1 | TEC | ENSG00000 |
| 171504245 | 171504351 | -1 | snRNA | ENSG00000 |
| 171534423 | 171534615 | 1 | processed_pseudoge | ENSG00000 |
| 171600404 | 171810950 | -1 | protein_coding | ENSG00000 |
| 171815586 | 171900749 | -1 | lncRNA | ENSG00000 |
| 171843349 | 171938715 | 1 | protein_coding | ENSG00000 |
| 171894436 | 171895240 | 1 | lncRNA | ENSG00000 |
| 171941607 | 171996871 | 1 | lncRNA | ENSG00000 |
| 172039578 | 172401669 | 1 | protein_coding | ENSG00000 |
| 172225428 | 172225872 | -1 | processed_pseudoge | ENSG00000 |
| 172237326 | 172237591 | 1 | misc_RNA | ENSG00000 |
| 172301654 | 172303052 | -1 | TEC | ENSG00000 |
| 172425850 | 172427089 | -1 | processed_pseudoge | ENSG00000 |
| 172443291 | 172448456 | -1 | protein_coding | ENSG00000 |
| 172505508 | 172523475 | -1 | protein_coding | ENSG00000 |
| 172522228 | 172522444 | 1 | processed_pseudoge | ENSG00000 |
| 172560888 | 172595607 | -1 | lncRNA | ENSG00000 |
| 172603440 | 172604006 | -1 | processed_pseudoge | ENSG00000 |

| | | | | |
|---|---|---|---|---|
| 172630249 | 172711218 | -1 | protein_coding | ENSG00000( |
| 172680714 | 172681224 | 1 | processed_pseudoge | ENSG00000( |
| 172703124 | 172703225 | 1 | snRNA | ENSG00000( |
| 172711151 | 172725038 | 1 | lncRNA | ENSG00000( |
| 172750682 | 172821474 | 1 | protein_coding | ENSG00000( |
| 172771277 | 172771405 | -1 | snoRNA | ENSG00000( |
| 172796429 | 172796558 | -1 | snRNA | ENSG00000( |
| 172828736 | 172829176 | -1 | processed_pseudoge | ENSG00000( |
| 172830981 | 172831229 | -1 | processed_pseudoge | ENSG00000( |
| 172889357 | 173141235 | -1 | protein_coding | ENSG00000( |
| 173153737 | 173336965 | 1 | lncRNA | ENSG00000( |
| 173396284 | 174286644 | 1 | protein_coding | ENSG00000( |
| 173644231 | 173644711 | 1 | processed_pseudoge | ENSG00000( |
| 173910498 | 173920796 | -1 | lncRNA | ENSG00000( |
| 174057641 | 174057864 | 1 | misc_RNA | ENSG00000( |
| 174302944 | 174303287 | -1 | processed_pseudoge | ENSG00000( |
| 174377244 | 174378005 | 1 | processed_pseudoge | ENSG00000( |
| 174438573 | 175810548 | 1 | protein_coding | ENSG00000( |
| 174631823 | 174632064 | 1 | misc_RNA | ENSG00000( |
| 175059361 | 175060031 | -1 | processed_pseudoge | ENSG00000( |
| 175079307 | 175115242 | -1 | lncRNA | ENSG00000( |
| 175234861 | 175271096 | -1 | lncRNA | ENSG00000( |
| 175369540 | 175369621 | 1 | miRNA | ENSG00000( |
| 175397247 | 175397348 | -1 | snRNA | ENSG00000( |
| 175473919 | 175477602 | -1 | lncRNA | ENSG00000( |
| 175609479 | 175609613 | 1 | snRNA | ENSG00000( |
| 175675087 | 175676801 | -1 | processed_pseudoge | ENSG00000( |
| 175718231 | 175718670 | -1 | processed_pseudoge | ENSG00000( |
| 175767558 | 175767661 | 1 | snRNA | ENSG00000( |
| 175773145 | 175776333 | -1 | lncRNA | ENSG00000( |
| 175938929 | 175941037 | -1 | lncRNA | ENSG00000( |
| 175977497 | 175978570 | -1 | processed_pseudoge | ENSG00000( |
| 176068042 | 176068908 | -1 | processed_pseudoge | ENSG00000( |
| 176069107 | 176069252 | 1 | processed_pseudoge | ENSG00000( |
| 176113702 | 176114537 | -1 | processed_pseudoge | ENSG00000( |
| 176415439 | 176867838 | -1 | lncRNA | ENSG00000( |
| 176515103 | 176515151 | 1 | miRNA | ENSG00000( |
| 176763233 | 176763341 | 1 | rRNA_pseudogene | ENSG00000( |
| 176814155 | 176817001 | -1 | lncRNA | ENSG00000( |
| 177010719 | 177011067 | -1 | processed_pseudoge | ENSG00000( |
| 177019340 | 177228000 | -1 | protein_coding | ENSG00000( |
| 177037405 | 177047923 | 1 | lncRNA | ENSG00000( |
| 177120691 | 177120800 | -1 | misc_RNA | ENSG00000( |
| 177126017 | 177126120 | -1 | snRNA | ENSG00000( |
| 177294442 | 177323418 | 1 | lncRNA | ENSG00000( |
| 177297161 | 177298386 | -1 | processed_pseudoge | ENSG00000( |
| 177337628 | 177349166 | -1 | lncRNA | ENSG00000( |
| 177441910 | 177767379 | 1 | lncRNA | ENSG00000( |
| 177503303 | 177503552 | -1 | misc_RNA | ENSG00000( |
| 177621382 | 177628287 | -1 | lncRNA | ENSG00000( |

| | | | | |
|---|---|---|---|---|
| 177624282 | 177624409 | -1 | snoRNA | ENSG00000 |
| 177683627 | 177691250 | 1 | lncRNA | ENSG00000 |
| 177700346 | 177701072 | -1 | processed_pseudoge | ENSG00000 |
| 177816865 | 177899224 | 1 | lncRNA | ENSG00000 |
| 177930758 | 177940299 | 1 | lncRNA | ENSG00000 |
| 177952327 | 177959817 | -1 | lncRNA | ENSG00000 |
| 178007672 | 178007779 | -1 | snRNA | ENSG00000 |
| 178028130 | 178172449 | -1 | lncRNA | ENSG00000 |
| 178164008 | 178385479 | -1 | lncRNA | ENSG00000 |
| 178272932 | 178844429 | 1 | protein_coding | ENSG00000 |
| 178419123 | 178843300 | 1 | protein_coding | ENSG00000 |
| 178419201 | 178457309 | 1 | lncRNA | ENSG00000 |
| 178429197 | 178429290 | -1 | rRNA_pseudogene | ENSG00000 |
| 178493327 | 178493810 | -1 | processed_pseudoge | ENSG00000 |
| 178526505 | 178937352 | -1 | lncRNA | ENSG00000 |
| 178960121 | 179072215 | -1 | protein_coding | ENSG00000 |
| 179097731 | 179097840 | 1 | misc_RNA | ENSG00000 |
| 179101366 | 179147973 | -1 | lncRNA | ENSG00000 |
| 179148114 | 179240093 | 1 | protein_coding | ENSG00000 |
| 179169083 | 179169210 | -1 | snoRNA | ENSG00000 |
| 179236691 | 179267002 | -1 | protein_coding | ENSG00000 |
| 179259028 | 179261273 | -1 | processed_pseudoge | ENSG00000 |
| 179323031 | 179338583 | 1 | protein_coding | ENSG00000 |
| 179340322 | 179341887 | 1 | lncRNA | ENSG00000 |
| 179347709 | 179394936 | 1 | protein_coding | ENSG00000 |
| 179396088 | 179451476 | -1 | protein_coding | ENSG00000 |
| 179396961 | 179399191 | 1 | lncRNA | ENSG00000 |
| 179405448 | 179405944 | 1 | lncRNA | ENSG00000 |
| 179452395 | 179452419 | -1 | processed_pseudoge | ENSG00000 |
| 179464346 | 179465328 | 1 | processed_pseudoge | ENSG00000 |
| 179562886 | 179588407 | 1 | protein_coding | ENSG00000 |
| 179583262 | 179583762 | -1 | lncRNA | ENSG00000 |
| 179584271 | 179584410 | -1 | lncRNA | ENSG00000 |
| 179588285 | 179604649 | -1 | protein_coding | ENSG00000 |
| 179604690 | 179627647 | 1 | protein_coding | ENSG00000 |
| 179614775 | 179614878 | -1 | snoRNA | ENSG00000 |
| 179649494 | 179650200 | -1 | processed_pseudoge | ENSG00000 |
| 179653040 | 179789401 | 1 | protein_coding | ENSG00000 |
| 179794958 | 180037053 | -1 | protein_coding | ENSG00000 |
| 179804063 | 179804366 | 1 | lncRNA | ENSG00000 |
| 179875376 | 179881609 | 1 | lncRNA | ENSG00000 |
| 179898199 | 179921895 | 1 | lncRNA | ENSG00000 |
| 179971494 | 180017083 | 1 | lncRNA | ENSG00000 |
| 180161886 | 180161977 | -1 | rRNA_pseudogene | ENSG00000 |
| 180201159 | 180201212 | 1 | processed_pseudoge | ENSG00000 |
| 180212191 | 180213108 | 1 | processed_pseudoge | ENSG00000 |
| 180323107 | 180325059 | 1 | processed_pseudoge | ENSG00000 |
| 180413008 | 180476778 | 1 | lncRNA | ENSG00000 |
| 180542701 | 180542836 | 1 | snoRNA | ENSG00000 |
| 180566718 | 180601935 | -1 | lncRNA | ENSG00000 |

| | | | | |
|---|---|---|---|---|
| 180581057 | 180581163 | 1 | snRNA | ENSG00000 |
| 180589887 | 180590047 | -1 | processed_pseudoge | ENSG00000 |
| 180602163 | 180617828 | 1 | protein_coding | ENSG00000 |
| 180602858 | 180684942 | -1 | protein_coding | ENSG00000 |
| 180614008 | 180870933 | -1 | lncRNA | ENSG00000 |
| 180680084 | 180700449 | 1 | lncRNA | ENSG00000 |
| 180707589 | 180871005 | -1 | lncRNA | ENSG00000 |
| 180736405 | 180736703 | -1 | misc_RNA | ENSG00000 |
| 180772750 | 180773914 | -1 | processed_pseudoge | ENSG00000 |
| 180820544 | 180821612 | 1 | processed_pseudoge | ENSG00000 |
| 180827977 | 180829850 | -1 | processed_pseudoge | ENSG00000 |
| 180868141 | 180982753 | 1 | protein_coding | ENSG00000 |
| 180887677 | 180888075 | -1 | processed_pseudoge | ENSG00000 |
| 180983709 | 180989774 | -1 | protein_coding | ENSG00000 |
| 180989762 | 181836880 | 1 | lncRNA | ENSG00000 |
| 181146233 | 181146463 | 1 | processed_pseudoge | ENSG00000 |
| 181231737 | 181231843 | 1 | snRNA | ENSG00000 |
| 181242345 | 181242901 | -1 | processed_pseudoge | ENSG00000 |
| 181372732 | 181442597 | -1 | lncRNA | ENSG00000 |
| 181505075 | 181505879 | -1 | processed_pseudoge | ENSG00000 |
| 181534177 | 181700611 | -1 | lncRNA | ENSG00000 |
| 181610498 | 181610729 | 1 | misc_RNA | ENSG00000 |
| 181699608 | 181699883 | 1 | misc_RNA | ENSG00000 |
| 181711925 | 181714436 | 1 | protein_coding | ENSG00000 |
| 181715185 | 181715293 | 1 | misc_RNA | ENSG00000 |
| 181813401 | 181813702 | 1 | misc_RNA | ENSG00000 |
| 181822872 | 181822990 | -1 | rRNA_pseudogene | ENSG00000 |
| 181822872 | 181822990 | -1 | rRNA | ENSG00000 |
| 181847526 | 181847885 | -1 | processed_pseudoge | ENSG00000 |
| 181866351 | 181867154 | -1 | processed_pseudoge | ENSG00000 |
| 181952343 | 182035644 | 1 | lncRNA | ENSG00000 |
| 181968459 | 181968681 | 1 | misc_RNA | ENSG00000 |
| 182049058 | 182052270 | 1 | lncRNA | ENSG00000 |
| 182121884 | 182142137 | 1 | lncRNA | ENSG00000 |
| 182149430 | 182158313 | -1 | lncRNA | ENSG00000 |
| 182365147 | 182368256 | 1 | lncRNA | ENSG00000 |
| 182446970 | 182486364 | -1 | lncRNA | ENSG00000 |
| 182460417 | 182461072 | -1 | processed_pseudoge | ENSG00000 |
| 182498187 | 182536772 | -1 | lncRNA | ENSG00000 |
| 182588093 | 182617810 | -1 | lncRNA | ENSG00000 |
| 182610674 | 182611414 | 1 | processed_pseudoge | ENSG00000 |
| 182644214 | 182655880 | 1 | lncRNA | ENSG00000 |
| 182691928 | 182692285 | 1 | processed_pseudoge | ENSG00000 |
| 182739669 | 182740848 | -1 | lncRNA | ENSG00000 |
| 182783236 | 182793394 | -1 | lncRNA | ENSG00000 |
| 182793503 | 182921629 | 1 | protein_coding | ENSG00000 |
| 182843521 | 182844619 | -1 | processed_pseudoge | ENSG00000 |
| 182921240 | 182921938 | -1 | lncRNA | ENSG00000 |
| 182938074 | 182985953 | -1 | protein_coding | ENSG00000 |
| 182950783 | 182956307 | -1 | processed_pseudoge | ENSG00000 |

| | | | | |
|---|---|---|---|---|
| 183015218 | 183116075 | -1 | protein_coding | ENSG00000 |
| 183016255 | 183017808 | 1 | lncRNA | ENSG00000 |
| 183122215 | 183163839 | -1 | protein_coding | ENSG00000 |
| 183178043 | 183428778 | -1 | protein_coding | ENSG00000 |
| 183253253 | 183298504 | 1 | protein_coding | ENSG00000 |
| 183258300 | 183258416 | -1 | rRNA_pseudogene | ENSG00000 |
| 183447608 | 183456013 | 1 | transcribed_unproce | ENSG00000 |
| 183451857 | 183451988 | 1 | snoRNA | ENSG00000 |
| 183452567 | 183452764 | 1 | snoRNA | ENSG00000 |
| 183453814 | 183453944 | 1 | snoRNA | ENSG00000 |
| 183454128 | 183454262 | 1 | snoRNA | ENSG00000 |
| 183462851 | 183463201 | -1 | processed_pseudoge | ENSG00000 |
| 183487551 | 183555706 | -1 | protein_coding | ENSG00000 |
| 183548735 | 183552326 | 1 | lncRNA | ENSG00000 |
| 183613620 | 183616742 | 1 | processed_pseudoge | ENSG00000 |
| 183635610 | 183684519 | 1 | protein_coding | ENSG00000 |
| 183697797 | 183812624 | 1 | protein_coding | ENSG00000 |
| 183806457 | 183810783 | -1 | lncRNA | ENSG00000 |
| 183815568 | 183884889 | -1 | protein_coding | ENSG00000 |
| 183815876 | 183825594 | -1 | protein_coding | ENSG00000 |
| 183829271 | 183884933 | -1 | protein_coding | ENSG00000 |
| 183884924 | 183888449 | 1 | transcribed_processe | ENSG00000 |
| 183886800 | 183886885 | 1 | miRNA | ENSG00000 |
| 183895900 | 183910936 | -1 | transcribed_unitary_ | ENSG00000 |
| 183919934 | 184017939 | -1 | protein_coding | ENSG00000 |
| 184006338 | 184011419 | 1 | lncRNA | ENSG00000 |
| 184026369 | 184027756 | -1 | processed_pseudoge | ENSG00000 |
| 184031544 | 184039369 | 1 | protein_coding | ENSG00000 |
| 184053047 | 184060673 | 1 | protein_coding | ENSG00000 |
| 184076779 | 184084219 | 1 | unprocessed_pseudc | ENSG00000 |
| 184086791 | 184086903 | -1 | misc_RNA | ENSG00000 |
| 184095118 | 184105178 | -1 | lncRNA | ENSG00000 |
| 184097064 | 184106995 | 1 | protein_coding | ENSG00000 |
| 184115352 | 184117898 | 1 | transcribed_processe | ENSG00000 |
| 184132942 | 184133561 | -1 | lncRNA | ENSG00000 |
| 184134019 | 184135238 | -1 | lncRNA | ENSG00000 |
| 184135038 | 184146127 | 1 | protein_coding | ENSG00000 |
| 184149704 | 184150435 | 1 | processed_pseudoge | ENSG00000 |
| 184155377 | 184173614 | 1 | protein_coding | ENSG00000 |
| 184174689 | 184184091 | 1 | protein_coding | ENSG00000 |
| 184186023 | 184194012 | 1 | protein_coding | ENSG00000 |
| 184230429 | 184242329 | 1 | protein_coding | ENSG00000 |
| 184241405 | 184241489 | 1 | miRNA | ENSG00000 |
| 184242301 | 184249548 | -1 | protein_coding | ENSG00000 |
| 184249650 | 184259585 | 1 | protein_coding | ENSG00000 |
| 184249695 | 184293031 | 1 | protein_coding | ENSG00000 |
| 184259213 | 184261553 | -1 | protein_coding | ENSG00000 |
| 184276011 | 184293031 | 1 | protein_coding | ENSG00000 |
| 184299198 | 184309050 | 1 | protein_coding | ENSG00000 |
| 184314495 | 184335358 | 1 | protein_coding | ENSG00000 |

| | | | | |
|---|---|---|---|---|
| 184325696 | 184325771 | 1 | snoRNA | ENSG0000C |
| 184335926 | 184348421 | 1 | protein_coding | ENSG0000C |
| 184346185 | 184361650 | -1 | protein_coding | ENSG0000C |
| 184361718 | 184368596 | 1 | protein_coding | ENSG0000C |
| 184371935 | 184381968 | -1 | protein_coding | ENSG0000C |
| 184380073 | 184390736 | 1 | protein_coding | ENSG0000C |
| 184399790 | 184457891 | 1 | lncRNA | ENSG0000C |
| 184499484 | 184500370 | -1 | unprocessed_pseudc | ENSG0000C |
| 184505113 | 184508399 | 1 | lncRNA | ENSG0000C |
| 184546714 | 184556918 | -1 | lncRNA | ENSG0000C |
| 184561785 | 184582408 | 1 | protein_coding | ENSG0000C |
| 184710364 | 184712064 | -1 | protein_coding | ENSG0000C |
| 184712245 | 184780720 | 1 | lncRNA | ENSG0000C |
| 184725013 | 184773178 | -1 | lncRNA | ENSG0000C |
| 184741937 | 184742462 | 1 | lncRNA | ENSG0000C |
| 184742818 | 184743284 | 1 | lncRNA | ENSG0000C |
| 184812143 | 185052614 | 1 | protein_coding | ENSG0000C |
| 185076838 | 185153060 | -1 | protein_coding | ENSG0000C |
| 185162871 | 185191955 | 1 | lncRNA | ENSG0000C |
| 185190624 | 185281990 | -1 | protein_coding | ENSG0000C |
| 185245460 | 185246441 | -1 | processed_pseudoge | ENSG0000C |
| 185253210 | 185253272 | -1 | miRNA | ENSG0000C |
| 185282941 | 185489094 | 1 | protein_coding | ENSG0000C |
| 185417495 | 185418778 | -1 | processed_pseudoge | ENSG0000C |
| 185476496 | 185499057 | -1 | protein_coding | ENSG0000C |
| 185499140 | 185501109 | 1 | lncRNA | ENSG0000C |
| 185506262 | 185552588 | -1 | protein_coding | ENSG0000C |
| 185582496 | 185633551 | 1 | protein_coding | ENSG0000C |
| 185606575 | 185606873 | 1 | processed_pseudoge | ENSG0000C |
| 185643739 | 185825056 | -1 | protein_coding | ENSG0000C |
| 185712528 | 185729787 | 1 | lncRNA | ENSG0000C |
| 185767847 | 185767904 | -1 | miRNA | ENSG0000C |
| 185914558 | 185938103 | -1 | protein_coding | ENSG0000C |
| 185959943 | 185980872 | 1 | transcribed_unproce | ENSG0000C |
| 186046314 | 186110318 | -1 | protein_coding | ENSG0000C |
| 186076306 | 186076591 | -1 | misc_RNA | ENSG0000C |
| 186079170 | 186080947 | 1 | lncRNA | ENSG0000C |
| 186105668 | 186362234 | -1 | protein_coding | ENSG0000C |
| 186439239 | 186457299 | 1 | lncRNA | ENSG0000C |
| 186454969 | 186493661 | -1 | lncRNA | ENSG0000C |
| 186476727 | 186478370 | 1 | lncRNA | ENSG0000C |
| 186538441 | 186546702 | -1 | protein_coding | ENSG0000C |
| 186546067 | 186570543 | -1 | protein_coding | ENSG0000C |
| 186567403 | 186585800 | 1 | protein_coding | ENSG0000C |
| 186579476 | 186772986 | -1 | lncRNA | ENSG0000C |
| 186581995 | 186597103 | 1 | protein_coding | ENSG0000C |
| 186596976 | 186597200 | 1 | processed_pseudoge | ENSG0000C |
| 186613060 | 186621318 | 1 | protein_coding | ENSG0000C |
| 186629936 | 186630211 | -1 | misc_RNA | ENSG0000C |
| 186635969 | 186653141 | 1 | protein_coding | ENSG0000C |

| | | | | |
|---|---|---|---|---|
| 186660216 | 186678240 | 1 | protein_coding | ENSG0000C |
| 186687329 | 186692506 | 1 | unprocessed_pseudc | ENSG0000C |
| 186698520 | 186708574 | 1 | unprocessed_pseudc | ENSG0000C |
| 186717348 | 186744410 | 1 | protein_coding | ENSG0000C |
| 186760693 | 186762050 | 1 | transcribed_processe | ENSG0000C |
| 186771841 | 186771947 | -1 | snRNA | ENSG0000C |
| 186772589 | 186773476 | 1 | processed_pseudoge | ENSG0000C |
| 186781780 | 186784179 | -1 | lncRNA | ENSG0000C |
| 186783205 | 186789897 | 1 | protein_coding | ENSG0000C |
| 186784796 | 186784864 | 1 | snoRNA | ENSG0000C |
| 186786323 | 186786445 | 1 | snoRNA | ENSG0000C |
| 186786672 | 186786777 | 1 | miRNA | ENSG0000C |
| 186786675 | 186786852 | 1 | snoRNA | ENSG0000C |
| 186787300 | 186787431 | 1 | snoRNA | ENSG0000C |
| 186787612 | 186787749 | 1 | snoRNA | ENSG0000C |
| 186789880 | 186807058 | -1 | protein_coding | ENSG0000C |
| 186807692 | 186818121 | 1 | lncRNA | ENSG0000C |
| 186810880 | 186825521 | -1 | lncRNA | ENSG0000C |
| 186826549 | 186827337 | 1 | unprocessed_pseudc | ENSG0000C |
| 186842704 | 186858463 | 1 | protein_coding | ENSG0000C |
| 186851886 | 186856123 | -1 | lncRNA | ENSG0000C |
| 186900198 | 186900557 | 1 | processed_pseudoge | ENSG0000C |
| 186922286 | 186922753 | -1 | processed_pseudoge | ENSG0000C |
| 186930502 | 187078553 | 1 | protein_coding | ENSG0000C |
| 187120948 | 187180908 | -1 | protein_coding | ENSG0000C |
| 187197090 | 187207629 | -1 | lncRNA | ENSG0000C |
| 187197486 | 187201462 | 1 | protein_coding | ENSG0000C |
| 187217285 | 187292022 | -1 | protein_coding | ENSG0000C |
| 187291272 | 187297933 | 1 | lncRNA | ENSG0000C |
| 187368385 | 187372076 | 1 | protein_coding | ENSG0000C |
| 187383101 | 187384849 | 1 | TEC | ENSG0000C |
| 187423315 | 187423419 | 1 | snoRNA | ENSG0000C |
| 187448845 | 187449450 | 1 | lncRNA | ENSG0000C |
| 187668912 | 187670394 | -1 | protein_coding | ENSG0000C |
| 187698259 | 187702557 | -1 | protein_coding | ENSG0000C |
| 187702313 | 187733849 | 1 | lncRNA | ENSG0000C |
| 187721377 | 187745725 | -1 | protein_coding | ENSG0000C |
| 187743686 | 187746028 | 1 | protein_coding | ENSG0000C |
| 187796187 | 187805258 | 1 | lncRNA | ENSG0000C |
| 187932764 | 187946221 | -1 | lncRNA | ENSG0000C |
| 187958775 | 187976407 | -1 | lncRNA | ENSG0000C |
| 187997422 | 188003990 | -1 | lncRNA | ENSG0000C |
| 188001700 | 188003406 | -1 | lncRNA | ENSG0000C |
| 188107072 | 188147310 | 1 | lncRNA | ENSG0000C |
| 188151206 | 188154057 | -1 | lncRNA | ENSG0000C |
| 188153284 | 188890671 | 1 | protein_coding | ENSG0000C |
| 188178543 | 188180812 | 1 | TEC | ENSG0000C |
| 188562238 | 188568666 | -1 | lncRNA | ENSG0000C |
| 188688781 | 188688866 | 1 | miRNA | ENSG0000C |
| 188941715 | 188947639 | -1 | lncRNA | ENSG0000C |

| | | | | |
|---|---|---|---|---|
| 188947214 | 189325304 | 1 | protein_coding | ENSG00000 |
| 188989382 | 188990135 | -1 | processed_pseudoge | ENSG00000 |
| 189238686 | 189240594 | -1 | lncRNA | ENSG00000 |
| 189631389 | 189897276 | 1 | protein_coding | ENSG00000 |
| 189645327 | 189645466 | -1 | processed_pseudoge | ENSG00000 |
| 189829922 | 189830009 | 1 | miRNA | ENSG00000 |
| 189956728 | 190122437 | -1 | protein_coding | ENSG00000 |
| 189969030 | 189969861 | -1 | processed_pseudoge | ENSG00000 |
| 190120964 | 190145107 | 1 | lncRNA | ENSG00000 |
| 190165417 | 190165523 | -1 | snRNA | ENSG00000 |
| 190166764 | 190167651 | 1 | processed_pseudoge | ENSG00000 |
| 190305707 | 190322446 | -1 | protein_coding | ENSG00000 |
| 190322541 | 190412143 | 1 | protein_coding | ENSG00000 |
| 190428655 | 190449876 | -1 | protein_coding | ENSG00000 |
| 190514051 | 190659750 | 1 | protein_coding | ENSG00000 |
| 190624595 | 190625169 | 1 | processed_pseudoge | ENSG00000 |
| 190642163 | 190642503 | 1 | misc_RNA | ENSG00000 |
| 190659216 | 190659750 | 1 | lncRNA | ENSG00000 |
| 190680421 | 190699314 | -1 | lncRNA | ENSG00000 |
| 190714811 | 190716399 | 1 | processed_pseudoge | ENSG00000 |
| 190830965 | 190832173 | 1 | processed_pseudoge | ENSG00000 |
| 190852737 | 190892429 | -1 | protein_coding | ENSG00000 |
| 191199241 | 191265615 | 1 | protein_coding | ENSG00000 |
| 191213291 | 191234605 | -1 | lncRNA | ENSG00000 |
| 191267168 | 191330536 | -1 | protein_coding | ENSG00000 |
| 191329085 | 191398659 | 1 | protein_coding | ENSG00000 |
| 191425493 | 191591097 | 1 | lncRNA | ENSG00000 |
| 191461163 | 191461456 | 1 | protein_coding | ENSG00000 |
| 191641135 | 191641237 | 1 | misc_RNA | ENSG00000 |
| 191952349 | 191952826 | -1 | processed_pseudoge | ENSG00000 |
| 192029880 | 192036557 | 1 | lncRNA | ENSG00000 |
| 192078950 | 192119864 | -1 | lncRNA | ENSG00000 |
| 192093122 | 192093436 | 1 | misc_RNA | ENSG00000 |
| 192139395 | 192767764 | -1 | protein_coding | ENSG00000 |
| 192238037 | 192283097 | 1 | lncRNA | ENSG00000 |
| 192515022 | 192516573 | 1 | lncRNA | ENSG00000 |
| 192516831 | 192521398 | 1 | lncRNA | ENSG00000 |
| 192534009 | 192534112 | -1 | snoRNA | ENSG00000 |
| 192729555 | 192729716 | -1 | snRNA | ENSG00000 |
| 192796815 | 192917856 | -1 | protein_coding | ENSG00000 |
| 193144464 | 193176864 | -1 | lncRNA | ENSG00000 |
| 193155456 | 193157208 | -1 | processed_pseudoge | ENSG00000 |
| 193241128 | 193277738 | 1 | protein_coding | ENSG00000 |
| 193274789 | 193378820 | -1 | protein_coding | ENSG00000 |
| 193307244 | 193314362 | 1 | lncRNA | ENSG00000 |
| 193398967 | 193593111 | -1 | protein_coding | ENSG00000 |
| 193553213 | 193555088 | 1 | lncRNA | ENSG00000 |
| 193578812 | 193593090 | -1 | transcribed_processe | ENSG00000 |
| 193593144 | 193697811 | 1 | protein_coding | ENSG00000 |
| 193618609 | 193627337 | -1 | lncRNA | ENSG00000 |

| | | | | |
|---|---|---|---|---|
| 193624189 | 193624301 | 1 | misc_RNA | ENSG00000 |
| 193740471 | 193740715 | -1 | misc_RNA | ENSG00000 |
| 193765681 | 193766003 | 1 | processed_pseudoge | ENSG00000 |
| 193770543 | 193772424 | -1 | lncRNA | ENSG00000 |
| 193771874 | 193782758 | 1 | lncRNA | ENSG00000 |
| 193842560 | 193844024 | -1 | lncRNA | ENSG00000 |
| 193957372 | 194003760 | -1 | lncRNA | ENSG00000 |
| 193980614 | 194006098 | 1 | lncRNA | ENSG00000 |
| 193993331 | 193993985 | -1 | processed_pseudoge | ENSG00000 |
| 194005259 | 194071025 | -1 | lncRNA | ENSG00000 |
| 194070103 | 194086826 | 1 | lncRNA | ENSG00000 |
| 194091561 | 194109174 | 1 | lncRNA | ENSG00000 |
| 194130616 | 194131201 | -1 | lncRNA | ENSG00000 |
| 194136148 | 194138732 | 1 | protein_coding | ENSG00000 |
| 194145674 | 194145971 | 1 | misc_RNA | ENSG00000 |
| 194202392 | 194250203 | -1 | lncRNA | ENSG00000 |
| 194247638 | 194260720 | 1 | lncRNA | ENSG00000 |
| 194276682 | 194287368 | 1 | lncRNA | ENSG00000 |
| 194296191 | 194322871 | -1 | lncRNA | ENSG00000 |
| 194300329 | 194302048 | 1 | lncRNA | ENSG00000 |
| 194339768 | 194351328 | -1 | protein_coding | ENSG00000 |
| 194355247 | 194369743 | -1 | protein_coding | ENSG00000 |
| 194394821 | 194398354 | -1 | protein_coding | ENSG00000 |
| 194402672 | 194498364 | -1 | protein_coding | ENSG00000 |
| 194471923 | 194472085 | 1 | processed_pseudoge | ENSG00000 |
| 194487140 | 194521156 | 1 | lncRNA | ENSG00000 |
| 194496317 | 194501503 | -1 | lncRNA | ENSG00000 |
| 194533503 | 194533583 | -1 | snRNA | ENSG00000 |
| 194562978 | 194564001 | -1 | processed_pseudoge | ENSG00000 |
| 194584004 | 194590260 | 1 | lncRNA | ENSG00000 |
| 194587673 | 194633689 | -1 | protein_coding | ENSG00000 |
| 194632923 | 194645401 | 1 | lncRNA | ENSG00000 |
| 194637505 | 194637664 | -1 | lncRNA | ENSG00000 |
| 194640791 | 194672463 | -1 | protein_coding | ENSG00000 |
| 194685883 | 194689037 | 1 | protein_coding | ENSG00000 |
| 194702667 | 194727675 | 1 | lncRNA | ENSG00000 |
| 194708010 | 194709395 | -1 | lncRNA | ENSG00000 |
| 194708093 | 194826008 | 1 | lncRNA | ENSG00000 |
| 194755589 | 194757422 | 1 | processed_pseudoge | ENSG00000 |
| 194765238 | 194768712 | -1 | lncRNA | ENSG00000 |
| 194827890 | 194832592 | 1 | lncRNA | ENSG00000 |
| 195068284 | 195271159 | -1 | protein_coding | ENSG00000 |
| 195094588 | 195096057 | 1 | lncRNA | ENSG00000 |
| 195134506 | 195134580 | -1 | miRNA | ENSG00000 |
| 195146728 | 195147003 | -1 | misc_RNA | ENSG00000 |
| 195147871 | 195152790 | 1 | lncRNA | ENSG00000 |
| 195214787 | 195214893 | 1 | snRNA | ENSG00000 |
| 195260632 | 195263628 | 1 | lncRNA | ENSG00000 |
| 195274745 | 195443044 | -1 | protein_coding | ENSG00000 |
| 195280723 | 195282741 | -1 | lncRNA | ENSG00000 |

| | | | | |
|---|---|---|---|---|
| 195349664 | 195350030 | 1 | processed_pseudoge | ENSG00000 |
| 195419436 | 195419546 | -1 | misc_RNA | ENSG00000 |
| 195514428 | 195543386 | -1 | protein_coding | ENSG00000 |
| 195523087 | 195523221 | -1 | snRNA | ENSG00000 |
| 195544048 | 195550581 | 1 | lncRNA | ENSG00000 |
| 195553792 | 195554043 | 1 | misc_RNA | ENSG00000 |
| 195554610 | 195555083 | 1 | processed_pseudoge | ENSG00000 |
| 195561215 | 195561580 | -1 | processed_pseudoge | ENSG00000 |
| 195568705 | 195584033 | -1 | protein_coding | ENSG00000 |
| 195614947 | 195620233 | 1 | unprocessed_pseudo | ENSG00000 |
| 195635188 | 195636301 | 1 | unprocessed_pseudo | ENSG00000 |
| 195650146 | 195651472 | -1 | processed_pseudoge | ENSG00000 |
| 195655565 | 195657927 | -1 | lncRNA | ENSG00000 |
| 195658062 | 195739964 | 1 | lncRNA | ENSG00000 |
| 195658096 | 195685904 | 1 | transcribed_unproce | ENSG00000 |
| 195699401 | 195699497 | 1 | miRNA | ENSG00000 |
| 195701618 | 195711643 | 1 | transcribed_unitary_ | ENSG00000 |
| 195720884 | 195741123 | 1 | protein_coding | ENSG00000 |
| 195746765 | 195811973 | -1 | protein_coding | ENSG00000 |
| 195836193 | 195860404 | -1 | lncRNA | ENSG00000 |
| 195843986 | 195844596 | 1 | processed_pseudoge | ENSG00000 |
| 195863364 | 195911945 | -1 | protein_coding | ENSG00000 |
| 195882329 | 195882395 | -1 | miRNA | ENSG00000 |
| 195900986 | 195903417 | 1 | lncRNA | ENSG00000 |
| 195908076 | 195913986 | 1 | lncRNA | ENSG00000 |
| 195913078 | 195954485 | -1 | transcribed_unproce | ENSG00000 |
| 195928589 | 195928775 | -1 | snRNA | ENSG00000 |
| 195937243 | 195938739 | 1 | processed_pseudoge | ENSG00000 |
| 195959748 | 195990318 | -1 | transcribed_unproce | ENSG00000 |
| 195996262 | 196027694 | 1 | lncRNA | ENSG00000 |
| 195996738 | 195998233 | 1 | processed_pseudoge | ENSG00000 |
| 196027183 | 196082096 | -1 | protein_coding | ENSG00000 |
| 196044695 | 196045254 | 1 | processed_pseudoge | ENSG00000 |
| 196072819 | 196072880 | 1 | snRNA | ENSG00000 |
| 196142525 | 196160893 | 1 | lncRNA | ENSG00000 |
| 196197452 | 196211437 | -1 | protein_coding | ENSG00000 |
| 196202349 | 196202453 | 1 | misc_RNA | ENSG00000 |
| 196211487 | 196243178 | 1 | protein_coding | ENSG00000 |
| 196214222 | 196287957 | -1 | protein_coding | ENSG00000 |
| 196248230 | 196318222 | -1 | protein_coding | ENSG00000 |
| 196250542 | 196251654 | 1 | lncRNA | ENSG00000 |
| 196291219 | 196318299 | -1 | protein_coding | ENSG00000 |
| 196298061 | 196298215 | -1 | processed_pseudoge | ENSG00000 |
| 196316082 | 196338373 | -1 | protein_coding | ENSG00000 |
| 196318330 | 196325570 | 1 | lncRNA | ENSG00000 |
| 196319342 | 196338503 | -1 | protein_coding | ENSG00000 |
| 196341609 | 196341715 | -1 | snRNA | ENSG00000 |
| 196347662 | 196432430 | -1 | protein_coding | ENSG00000 |
| 196393710 | 196393812 | -1 | snRNA | ENSG00000 |
| 196397508 | 196397778 | -1 | misc_RNA | ENSG00000 |

| | | | | |
|---|---|---|---|---|
| 196399911 | 196400207 | 1 | misc_RNA | ENSG0000C |
| 196423322 | 196423522 | 1 | processed_pseudoge | ENSG0000C |
| 196431385 | 196432530 | 1 | lncRNA | ENSG0000C |
| 196452996 | 196453055 | -1 | unprocessed_pseudc | ENSG0000C |
| 196468783 | 196503768 | -1 | protein_coding | ENSG0000C |
| 196474801 | 196475394 | 1 | lncRNA | ENSG0000C |
| 196506879 | 196515346 | -1 | protein_coding | ENSG0000C |
| 196527339 | 196528791 | -1 | unprocessed_pseudc | ENSG0000C |
| 196536532 | 196536702 | 1 | processed_pseudoge | ENSG0000C |
| 196554177 | 196568674 | -1 | protein_coding | ENSG0000C |
| 196568611 | 196589059 | 1 | protein_coding | ENSG0000C |
| 196598549 | 196609410 | -1 | lncRNA | ENSG0000C |
| 196631498 | 196632587 | -1 | lncRNA | ENSG0000C |
| 196639694 | 196662004 | 1 | protein_coding | ENSG0000C |
| 196639775 | 196736007 | 1 | protein_coding | ENSG0000C |
| 196706277 | 196712250 | -1 | protein_coding | ENSG0000C |
| 196708859 | 196708964 | -1 | snRNA | ENSG0000C |
| 196739857 | 196832647 | 1 | protein_coding | ENSG0000C |
| 196747192 | 196747324 | -1 | snRNA | ENSG0000C |
| 196784980 | 196785086 | -1 | snRNA | ENSG0000C |
| 196867856 | 196934714 | 1 | protein_coding | ENSG0000C |
| 196912646 | 196914579 | 1 | lncRNA | ENSG0000C |
| 196935402 | 196942594 | -1 | protein_coding | ENSG0000C |
| 196939877 | 196942534 | 1 | lncRNA | ENSG0000C |
| 196942674 | 196943543 | 1 | protein_coding | ENSG0000C |
| 196946356 | 196969060 | -1 | protein_coding | ENSG0000C |
| 196973709 | 196979197 | 1 | lncRNA | ENSG0000C |
| 196983890 | 196987558 | -1 | transcribed_processe | ENSG0000C |
| 196988621 | 197029817 | -1 | protein_coding | ENSG0000C |
| 196999460 | 197004744 | 1 | lncRNA | ENSG0000C |
| 197042560 | 197299330 | -1 | protein_coding | ENSG0000C |
| 197293878 | 197293948 | -1 | miRNA | ENSG0000C |
| 197298252 | 197303755 | 1 | lncRNA | ENSG0000C |
| 197333944 | 197340706 | 1 | lncRNA | ENSG0000C |
| 197445061 | 197458323 | -1 | lncRNA | ENSG0000C |
| 197450327 | 197456654 | -1 | lncRNA | ENSG0000C |
| 197458405 | 197467105 | 1 | lncRNA | ENSG0000C |
| 197505262 | 197506986 | -1 | lncRNA | ENSG0000C |
| 197509783 | 197573323 | -1 | protein_coding | ENSG0000C |
| 197578213 | 197627906 | -1 | transcribed_unproce | ENSG0000C |
| 197580344 | 197580629 | -1 | processed_pseudoge | ENSG0000C |
| 197634315 | 197635811 | 1 | processed_pseudoge | ENSG0000C |
| 197645669 | 197646017 | 1 | lncRNA | ENSG0000C |
| 197649303 | 197651482 | -1 | lncRNA | ENSG0000C |
| 197660565 | 197665757 | -1 | lncRNA | ENSG0000C |
| 197665316 | 197674863 | 1 | lncRNA | ENSG0000C |
| 197668867 | 197749727 | -1 | protein_coding | ENSG0000C |
| 197674496 | 197674576 | -1 | miRNA | ENSG0000C |
| 197737179 | 197787596 | 1 | protein_coding | ENSG0000C |
| 197789700 | 197790369 | 1 | lncRNA | ENSG0000C |

| | | | | |
|---|---|---|---|---|
| 197791226 | 197888436 | 1 | protein_coding | ENSG0000C |
| 197830685 | 197831296 | -1 | lncRNA | ENSG0000C |
| 197850400 | 197850954 | -1 | processed_pseudoge | ENSG0000C |
| 197889077 | 197960142 | -1 | protein_coding | ENSG0000C |
| 197894507 | 197894613 | 1 | snRNA | ENSG0000C |
| 197950190 | 197956610 | 1 | protein_coding | ENSG0000C |
| 197960200 | 198043720 | 1 | protein_coding | ENSG0000C |
| 197994601 | 197994709 | 1 | snRNA | ENSG0000C |
| 198027956 | 198035970 | -1 | lncRNA | ENSG0000C |
| 198038321 | 198039234 | -1 | lncRNA | ENSG0000C |
| 198053522 | 198080720 | -1 | transcribed_unproce | ENSG0000C |
| 198071311 | 198071417 | -1 | snRNA | ENSG0000C |
| 198110112 | 198111878 | 1 | transcribed_unproce | ENSG0000C |
| 198119721 | 198120778 | 1 | unprocessed_pseudc | ENSG0000C |
| 198121479 | 198123232 | -1 | unprocessed_pseudc | ENSG0000C |
| 198153287 | 198222513 | 1 | transcribed_unproce | ENSG0000C |
| 198196746 | 198196849 | 1 | snRNA | ENSG0000C |
| 198223497 | 198226310 | -1 | processed_pseudoge | ENSG0000C |
| 198228194 | 198228376 | 1 | unprocessed_pseudc | ENSG0000C |
| 49096 | 50124 | -1 | transcribed_processe | ENSG0000C |
| 53286 | 88208 | 1 | protein_coding | ENSG0000C |
| 105959 | 107114 | -1 | processed_pseudoge | ENSG0000C |
| 124501 | 202303 | 1 | protein_coding | ENSG0000C |
| 149738 | 150317 | 1 | lncRNA | ENSG0000C |
| 201409 | 205009 | -1 | unprocessed_pseudc | ENSG0000C |
| 212610 | 255985 | 1 | transcribed_unproce | ENSG0000C |
| 268982 | 269531 | -1 | processed_pseudoge | ENSG0000C |
| 269866 | 270175 | -1 | processed_pseudoge | ENSG0000C |
| 270675 | 305474 | -1 | protein_coding | ENSG0000C |
| 307335 | 337433 | -1 | lncRNA | ENSG0000C |
| 336258 | 336786 | -1 | processed_pseudoge | ENSG0000C |
| 337814 | 384868 | 1 | protein_coding | ENSG0000C |
| 338097 | 338583 | -1 | processed_pseudoge | ENSG0000C |
| 342821 | 342962 | -1 | processed_pseudoge | ENSG0000C |
| 350157 | 350252 | 1 | miRNA | ENSG0000C |
| 386174 | 454070 | 1 | lncRNA | ENSG0000C |
| 393534 | 393658 | 1 | processed_pseudoge | ENSG0000C |
| 416118 | 416537 | -1 | lncRNA | ENSG0000C |
| 425435 | 474129 | -1 | transcribed_processe | ENSG0000C |
| 425815 | 499156 | -1 | protein_coding | ENSG0000C |
| 499210 | 540200 | 1 | protein_coding | ENSG0000C |
| 573880 | 576300 | -1 | protein_coding | ENSG0000C |
| 577168 | 618076 | 1 | lncRNA | ENSG0000C |
| 625584 | 670782 | 1 | protein_coding | ENSG0000C |
| 652850 | 656213 | -1 | lncRNA | ENSG0000C |
| 661209 | 661945 | -1 | lncRNA | ENSG0000C |
| 672436 | 674330 | -1 | protein_coding | ENSG0000C |
| 673580 | 682033 | 1 | protein_coding | ENSG0000C |
| 681829 | 689441 | -1 | protein_coding | ENSG0000C |
| 705748 | 770640 | 1 | protein_coding | ENSG0000C |

| | | | | |
|---|---|---|---|---|
| 722275 | 724034 | -1 | lncRNA | ENSG00000( |
| 757022 | 757740 | -1 | lncRNA | ENSG00000( |
| 760202 | 781859 | -1 | lncRNA | ENSG00000( |
| 764487 | 765074 | -1 | lncRNA | ENSG00000( |
| 784957 | 826129 | -1 | protein_coding | ENSG00000( |
| 836512 | 837224 | -1 | lncRNA | ENSG00000( |
| 843963 | 846294 | -1 | lncRNA | ENSG00000( |
| 849276 | 932373 | -1 | protein_coding | ENSG00000( |
| 932387 | 958656 | 1 | protein_coding | ENSG00000( |
| 958887 | 986895 | -1 | protein_coding | ENSG00000( |
| 979073 | 993440 | -1 | protein_coding | ENSG00000( |
| 986997 | 1004564 | 1 | protein_coding | ENSG00000( |
| 1009936 | 1026898 | 1 | protein_coding | ENSG00000( |
| 1027678 | 1028972 | 1 | lncRNA | ENSG00000( |
| 1056250 | 1113564 | -1 | protein_coding | ENSG00000( |
| 1100016 | 1101558 | -1 | processed_pseudoge | ENSG00000( |
| 1113639 | 1132977 | 1 | lncRNA | ENSG00000( |
| 1115197 | 1153726 | -1 | transcribed_unitary_ | ENSG00000( |
| 1118884 | 1119013 | 1 | snoRNA | ENSG00000( |
| 1151372 | 1153701 | 1 | lncRNA | ENSG00000( |
| 1166932 | 1208962 | -1 | protein_coding | ENSG00000( |
| 1167778 | 1168174 | 1 | lncRNA | ENSG00000( |
| 1210120 | 1218591 | 1 | lncRNA | ENSG00000( |
| 1211448 | 1249953 | -1 | protein_coding | ENSG00000( |
| 1249300 | 1288291 | 1 | lncRNA | ENSG00000( |
| 1289887 | 1340147 | 1 | protein_coding | ENSG00000( |
| 1347208 | 1388049 | 1 | protein_coding | ENSG00000( |
| 1356581 | 1358075 | 1 | lncRNA | ENSG00000( |
| 1358479 | 1359461 | 1 | lncRNA | ENSG00000( |
| 1391552 | 1395992 | 1 | protein_coding | ENSG00000( |
| 1402932 | 1406331 | -1 | protein_coding | ENSG00000( |
| 1550284 | 1550572 | 1 | lncRNA | ENSG00000( |
| 1574062 | 1580253 | -1 | lncRNA | ENSG00000( |
| 1617915 | 1684302 | -1 | protein_coding | ENSG00000( |
| 1683420 | 1683529 | -1 | misc_RNA | ENSG00000( |
| 1692731 | 1712344 | -1 | protein_coding | ENSG00000( |
| 1712821 | 1715945 | 1 | lncRNA | ENSG00000( |
| 1712891 | 1745176 | 1 | protein_coding | ENSG00000( |
| 1715952 | 1721358 | -1 | protein_coding | ENSG00000( |
| 1793293 | 1808872 | 1 | protein_coding | ENSG00000( |
| 1811479 | 1856156 | -1 | protein_coding | ENSG00000( |
| 1871393 | 1982207 | 1 | protein_coding | ENSG00000( |
| 1974636 | 1974760 | 1 | scaRNA | ENSG00000( |
| 1982717 | 2041903 | -1 | protein_coding | ENSG00000( |
| 1986384 | 1986477 | -1 | miRNA | ENSG00000( |
| 2041993 | 2043970 | 1 | protein_coding | ENSG00000( |
| 2059512 | 2069089 | 1 | protein_coding | ENSG00000( |
| 2071918 | 2242121 | -1 | protein_coding | ENSG00000( |
| 2078998 | 2242276 | -1 | protein_coding | ENSG00000( |
| 2088718 | 2090617 | -1 | processed_pseudoge | ENSG00000( |

| | | | | |
|---|---|---|---|---|
| 2139673 | 2141058 | 1 | lncRNA | ENSG0000C |
| 2234251 | 2234499 | 1 | processed_pseudoge | ENSG0000C |
| 2247432 | 2262109 | -1 | protein_coding | ENSG0000C |
| 2250077 | 2250156 | -1 | miRNA | ENSG0000C |
| 2269582 | 2418663 | -1 | protein_coding | ENSG0000C |
| 2295584 | 2319089 | 1 | lncRNA | ENSG0000C |
| 2316197 | 2316506 | -1 | misc_RNA | ENSG0000C |
| 2324467 | 2325082 | 1 | processed_pseudoge | ENSG0000C |
| 2418974 | 2462942 | 1 | protein_coding | ENSG0000C |
| 2462220 | 2625320 | 1 | protein_coding | ENSG0000C |
| 2463797 | 2464124 | 1 | lncRNA | ENSG0000C |
| 2505081 | 2506279 | -1 | processed_pseudoge | ENSG0000C |
| 2536631 | 2732565 | 1 | protein_coding | ENSG0000C |
| 2607870 | 2608146 | -1 | unprocessed_pseudc | ENSG0000C |
| 2615124 | 2615224 | -1 | misc_RNA | ENSG0000C |
| 2741648 | 2756342 | -1 | protein_coding | ENSG0000C |
| 2793071 | 2841098 | 1 | protein_coding | ENSG0000C |
| 2843857 | 2930076 | 1 | protein_coding | ENSG0000C |
| 2859983 | 2862673 | -1 | lncRNA | ENSG0000C |
| 2930561 | 2934834 | -1 | protein_coding | ENSG0000C |
| 2934882 | 2961738 | 1 | lncRNA | ENSG0000C |
| 2937933 | 2963406 | -1 | protein_coding | ENSG0000C |
| 2963571 | 3040760 | 1 | protein_coding | ENSG0000C |
| 3041422 | 3243960 | 1 | protein_coding | ENSG0000C |
| 3044101 | 3044207 | 1 | snRNA | ENSG0000C |
| 3049094 | 3074538 | -1 | lncRNA | ENSG0000C |
| 3063472 | 3063701 | 1 | misc_RNA | ENSG0000C |
| 3069977 | 3070151 | 1 | misc_RNA | ENSG0000C |
| 3244369 | 3271738 | 1 | protein_coding | ENSG0000C |
| 3293028 | 3439913 | 1 | protein_coding | ENSG0000C |
| 3312512 | 3313058 | 1 | transcribed_processe | ENSG0000C |
| 3323977 | 3324762 | -1 | processed_pseudoge | ENSG0000C |
| 3441887 | 3449495 | 1 | protein_coding | ENSG0000C |
| 3450638 | 3453108 | 1 | lncRNA | ENSG0000C |
| 3463306 | 3501473 | 1 | protein_coding | ENSG0000C |
| 3503612 | 3532446 | -1 | protein_coding | ENSG0000C |
| 3544555 | 3548796 | 1 | lncRNA | ENSG0000C |
| 3576869 | 3590711 | 1 | lncRNA | ENSG0000C |
| 3633029 | 3633975 | 1 | lncRNA | ENSG0000C |
| 3673593 | 3677855 | -1 | lncRNA | ENSG0000C |
| 3758748 | 3763390 | -1 | lncRNA | ENSG0000C |
| 3766348 | 3768526 | 1 | protein_coding | ENSG0000C |
| 3889356 | 3890249 | -1 | unprocessed_pseudc | ENSG0000C |
| 3901566 | 3902594 | -1 | unprocessed_pseudc | ENSG0000C |
| 3910283 | 3910604 | 1 | processed_pseudoge | ENSG0000C |
| 3915183 | 3955419 | -1 | lncRNA | ENSG0000C |
| 3915187 | 3924274 | -1 | transcribed_unproce | ENSG0000C |
| 3935447 | 3942519 | 1 | unprocessed_pseudc | ENSG0000C |
| 3941760 | 3955419 | -1 | transcribed_unproce | ENSG0000C |
| 3967034 | 3967660 | -1 | processed_pseudoge | ENSG0000C |

| | | | | |
|---|---|---|---|---|
| 4048211 | 4107583 | -1 | transcribed_unproce | ENSG00000 |
| 4076711 | 4077491 | -1 | processed_pseudoge | ENSG00000 |
| 4086638 | 4086814 | 1 | processed_pseudoge | ENSG00000 |
| 4117925 | 4118284 | -1 | processed_pseudoge | ENSG00000 |
| 4126659 | 4127569 | -1 | unprocessed_pseudo | ENSG00000 |
| 4143458 | 4150421 | -1 | unprocessed_pseudo | ENSG00000 |
| 4156522 | 4157443 | -1 | unprocessed_pseudo | ENSG00000 |
| 4174319 | 4175238 | -1 | unprocessed_pseudo | ENSG00000 |
| 4188803 | 4226889 | -1 | protein_coding | ENSG00000 |
| 4235542 | 4248223 | -1 | protein_coding | ENSG00000 |
| 4267701 | 4290154 | -1 | protein_coding | ENSG00000 |
| 4290251 | 4321786 | 1 | protein_coding | ENSG00000 |
| 4321962 | 4334182 | 1 | lncRNA | ENSG00000 |
| 4348140 | 4419058 | 1 | protein_coding | ENSG00000 |
| 4415742 | 4542346 | -1 | protein_coding | ENSG00000 |
| 4476121 | 4481700 | -1 | lncRNA | ENSG00000 |
| 4507078 | 4507777 | 1 | processed_pseudoge | ENSG00000 |
| 4542131 | 4787359 | 1 | lncRNA | ENSG00000 |
| 4820405 | 4821452 | 1 | TEC | ENSG00000 |
| 4844188 | 4850827 | 1 | lncRNA | ENSG00000 |
| 4859665 | 4863936 | 1 | protein_coding | ENSG00000 |
| 4894182 | 4895173 | -1 | processed_pseudoge | ENSG00000 |
| 4920770 | 4921064 | -1 | misc_RNA | ENSG00000 |
| 5014586 | 5019458 | -1 | protein_coding | ENSG00000 |
| 5051480 | 5500994 | 1 | protein_coding | ENSG00000 |
| 5426885 | 5426984 | -1 | misc_RNA | ENSG00000 |
| 5458581 | 5458881 | -1 | misc_RNA | ENSG00000 |
| 5524569 | 5527801 | 1 | lncRNA | ENSG00000 |
| 5542772 | 5709548 | -1 | protein_coding | ENSG00000 |
| 5711201 | 5814305 | 1 | protein_coding | ENSG00000 |
| 5748084 | 5893086 | -1 | protein_coding | ENSG00000 |
| 5897373 | 6200555 | -1 | protein_coding | ENSG00000 |
| 5923275 | 5923328 | -1 | miRNA | ENSG00000 |
| 6026199 | 6200591 | -1 | protein_coding | ENSG00000 |
| 6064977 | 6070162 | -1 | protein_coding | ENSG00000 |
| 6178186 | 6184925 | 1 | lncRNA | ENSG00000 |
| 6200733 | 6239937 | 1 | transcribed_unproce | ENSG00000 |
| 6202328 | 6206649 | 1 | lncRNA | ENSG00000 |
| 6269849 | 6303265 | 1 | protein_coding | ENSG00000 |
| 6292369 | 6308636 | -1 | lncRNA | ENSG00000 |
| 6320578 | 6563600 | -1 | protein_coding | ENSG00000 |
| 6575189 | 6623362 | 1 | protein_coding | ENSG00000 |
| 6640091 | 6642745 | 1 | protein_coding | ENSG00000 |
| 6648695 | 6673965 | -1 | lncRNA | ENSG00000 |
| 6658690 | 6658981 | 1 | unprocessed_pseudo | ENSG00000 |
| 6663396 | 6676755 | 1 | protein_coding | ENSG00000 |
| 6687448 | 6690519 | -1 | lncRNA | ENSG00000 |
| 6691995 | 6692970 | -1 | TEC | ENSG00000 |
| 6693878 | 6697170 | 1 | protein_coding | ENSG00000 |
| 6697861 | 6698809 | -1 | TEC | ENSG00000 |

| | | | | |
|---|---|---|---|---|
| 6707701 | 6709865 | -1 | protein_coding | ENSG00000 |
| 6716174 | 6717664 | 1 | protein_coding | ENSG00000 |
| 6763508 | 6767660 | 1 | lncRNA | ENSG00000 |
| 6781375 | 6884170 | 1 | protein_coding | ENSG00000 |
| 6909242 | 7033118 | 1 | protein_coding | ENSG00000 |
| 6985926 | 6987036 | -1 | lncRNA | ENSG00000 |
| 6995341 | 6998958 | 1 | lncRNA | ENSG00000 |
| 6997012 | 6997328 | 1 | misc_RNA | ENSG00000 |
| 7030554 | 7046231 | -1 | lncRNA | ENSG00000 |
| 7040849 | 7043001 | -1 | protein_coding | ENSG00000 |
| 7041899 | 7057952 | 1 | protein_coding | ENSG00000 |
| 7058895 | 7068064 | -1 | protein_coding | ENSG00000 |
| 7093776 | 7103394 | -1 | lncRNA | ENSG00000 |
| 7112088 | 7112399 | -1 | misc_RNA | ENSG00000 |
| 7192538 | 7742836 | 1 | protein_coding | ENSG00000 |
| 7310450 | 7310524 | 1 | miRNA | ENSG00000 |
| 7430285 | 7434930 | -1 | protein_coding | ENSG00000 |
| 7460028 | 7460118 | 1 | miRNA | ENSG00000 |
| 7754090 | 7778928 | 1 | lncRNA | ENSG00000 |
| 7758714 | 7939926 | -1 | protein_coding | ENSG00000 |
| 7798527 | 7808051 | 1 | lncRNA | ENSG00000 |
| 7939001 | 7940296 | 1 | lncRNA | ENSG00000 |
| 7965310 | 8158832 | -1 | protein_coding | ENSG00000 |
| 8005301 | 8005381 | -1 | miRNA | ENSG00000 |
| 8022665 | 8023126 | 1 | lncRNA | ENSG00000 |
| 8066528 | 8067724 | 1 | lncRNA | ENSG00000 |
| 8174421 | 8174999 | 1 | processed_pseudoge | ENSG00000 |
| 8182072 | 8241803 | 1 | protein_coding | ENSG00000 |
| 8269754 | 8307098 | 1 | protein_coding | ENSG00000 |
| 8320105 | 8327165 | 1 | lncRNA | ENSG00000 |
| 8355090 | 8358338 | -1 | lncRNA | ENSG00000 |
| 8366282 | 8440723 | -1 | protein_coding | ENSG00000 |
| 8393015 | 8393124 | 1 | rRNA_pseudogene | ENSG00000 |
| 8436140 | 8493531 | 1 | protein_coding | ENSG00000 |
| 8453410 | 8454942 | -1 | lncRNA | ENSG00000 |
| 8482270 | 8516759 | 1 | lncRNA | ENSG00000 |
| 8558725 | 8619761 | 1 | protein_coding | ENSG00000 |
| 8592660 | 8619759 | 1 | protein_coding | ENSG00000 |
| 8745391 | 8747727 | -1 | lncRNA | ENSG00000 |
| 8846076 | 8871839 | -1 | protein_coding | ENSG00000 |
| 8858715 | 8860827 | -1 | lncRNA | ENSG00000 |
| 8949630 | 8950559 | 1 | unprocessed_pseudc | ENSG00000 |
| 8959628 | 8963981 | 1 | unprocessed_pseudc | ENSG00000 |
| 8981436 | 8982360 | 1 | unprocessed_pseudc | ENSG00000 |
| 8990455 | 8990806 | 1 | processed_pseudoge | ENSG00000 |
| 9000841 | 9001058 | 1 | unprocessed_pseudc | ENSG00000 |
| 9020857 | 9021161 | -1 | processed_pseudoge | ENSG00000 |
| 9030286 | 9031066 | 1 | processed_pseudoge | ENSG00000 |
| 9034519 | 9381269 | -1 | lncRNA | ENSG00000 |
| 9051842 | 9052051 | -1 | processed_pseudoge | ENSG00000 |

| | | | | |
|---|---|---|---|---|
| 9076949 | 9092530 | 1 | lncRNA | ENSG0000( |
| 9079023 | 9141608 | 1 | unprocessed_pseudc | ENSG0000( |
| 9153296 | 9165451 | 1 | unprocessed_pseudc | ENSG0000( |
| 9166297 | 9170270 | -1 | unprocessed_pseudc | ENSG0000( |
| 9170409 | 9176730 | 1 | protein_coding | ENSG0000( |
| 9210657 | 9212608 | 1 | protein_coding | ENSG0000( |
| 9215405 | 9217356 | 1 | protein_coding | ENSG0000( |
| 9220152 | 9221744 | 1 | protein_coding | ENSG0000( |
| 9224896 | 9226847 | 1 | protein_coding | ENSG0000( |
| 9229641 | 9231233 | 1 | unprocessed_pseudc | ENSG0000( |
| 9234385 | 9236334 | 1 | protein_coding | ENSG0000( |
| 9239130 | 9240722 | 1 | unprocessed_pseudc | ENSG0000( |
| 9243879 | 9245830 | 1 | protein_coding | ENSG0000( |
| 9248630 | 9250581 | 1 | protein_coding | ENSG0000( |
| 9253378 | 9255329 | 1 | protein_coding | ENSG0000( |
| 9258124 | 9260075 | 1 | protein_coding | ENSG0000( |
| 9262872 | 9264823 | 1 | protein_coding | ENSG0000( |
| 9267619 | 9269570 | 1 | protein_coding | ENSG0000( |
| 9272364 | 9272914 | 1 | protein_coding | ENSG0000( |
| 9325165 | 9327116 | 1 | protein_coding | ENSG0000( |
| 9329911 | 9331862 | 1 | protein_coding | ENSG0000( |
| 9334658 | 9336609 | 1 | protein_coding | ENSG0000( |
| 9339403 | 9340995 | 1 | protein_coding | ENSG0000( |
| 9344148 | 9345740 | 1 | protein_coding | ENSG0000( |
| 9348893 | 9350485 | 1 | protein_coding | ENSG0000( |
| 9353638 | 9355230 | 1 | protein_coding | ENSG0000( |
| 9358383 | 9359975 | 1 | unprocessed_pseudc | ENSG0000( |
| 9363129 | 9364721 | 1 | protein_coding | ENSG0000( |
| 9367874 | 9369070 | 1 | unprocessed_pseudc | ENSG0000( |
| 9384017 | 9388983 | -1 | unprocessed_pseudc | ENSG0000( |
| 9399204 | 9403565 | 1 | unprocessed_pseudc | ENSG0000( |
| 9411178 | 9411253 | -1 | rRNA_pseudogene | ENSG0000( |
| 9422313 | 9429702 | 1 | unprocessed_pseudc | ENSG0000( |
| 9444534 | 9450514 | 1 | protein_coding | ENSG0000( |
| 9459255 | 9460244 | 1 | unprocessed_pseudc | ENSG0000( |
| 9468974 | 9469983 | 1 | unprocessed_pseudc | ENSG0000( |
| 9483718 | 9484702 | 1 | unprocessed_pseudc | ENSG0000( |
| 9493736 | 9498066 | 1 | unprocessed_pseudc | ENSG0000( |
| 9512905 | 9513874 | 1 | unprocessed_pseudc | ENSG0000( |
| 9521910 | 9522276 | 1 | processed_pseudoge | ENSG0000( |
| 9532429 | 9532647 | 1 | unprocessed_pseudc | ENSG0000( |
| 9553614 | 9553985 | -1 | processed_pseudoge | ENSG0000( |
| 9556168 | 9556316 | -1 | miRNA | ENSG0000( |
| 9563171 | 9563950 | 1 | processed_pseudoge | ENSG0000( |
| 9567474 | 9691786 | -1 | lncRNA | ENSG0000( |
| 9583942 | 9584131 | -1 | processed_pseudoge | ENSG0000( |
| 9677308 | 9677934 | 1 | processed_pseudoge | ENSG0000( |
| 9692495 | 9702954 | 1 | unprocessed_pseudc | ENSG0000( |
| 9703754 | 9710812 | -1 | unprocessed_pseudc | ENSG0000( |
| 9711953 | 9712233 | -1 | misc_RNA | ENSG0000( |

| | | | | |
|---|---|---|---|---|
| 9733370 | 9742361 | 1 | unprocessed_pseudc | ENSG0000C |
| 9754898 | 9755778 | 1 | unprocessed_pseudc | ENSG0000C |
| 9771153 | 10054936 | -1 | protein_coding | ENSG0000C |
| 9781634 | 9784009 | 1 | protein_coding | ENSG0000C |
| 9792751 | 9792861 | -1 | rRNA_pseudogene | ENSG0000C |
| 9922814 | 9924146 | 1 | lncRNA | ENSG0000C |
| 10006482 | 10009725 | 1 | lncRNA | ENSG0000C |
| 10068089 | 10073019 | -1 | lncRNA | ENSG0000C |
| 10074339 | 10116949 | -1 | protein_coding | ENSG0000C |
| 10078611 | 10078692 | -1 | miRNA | ENSG0000C |
| 10115756 | 10115884 | -1 | rRNA_pseudogene | ENSG0000C |
| 10143367 | 10143655 | -1 | processed_pseudoge | ENSG0000C |
| 10167159 | 10167765 | -1 | processed_pseudoge | ENSG0000C |
| 10199823 | 10200672 | -1 | processed_pseudoge | ENSG0000C |
| 10238213 | 10238235 | -1 | processed_pseudoge | ENSG0000C |
| 10254926 | 10256621 | -1 | processed_pseudoge | ENSG0000C |
| 10259445 | 10260138 | -1 | processed_pseudoge | ENSG0000C |
| 10267895 | 10268231 | -1 | processed_pseudoge | ENSG0000C |
| 10284961 | 10285708 | -1 | processed_pseudoge | ENSG0000C |
| 10292195 | 10292708 | -1 | processed_pseudoge | ENSG0000C |
| 10294463 | 10295579 | -1 | processed_pseudoge | ENSG0000C |
| 10410996 | 10411644 | 1 | processed_pseudoge | ENSG0000C |
| 10439880 | 10457426 | -1 | protein_coding | ENSG0000C |
| 10456745 | 10531022 | 1 | lncRNA | ENSG0000C |
| 10478524 | 10478923 | -1 | processed_pseudoge | ENSG0000C |
| 10486395 | 10684768 | -1 | protein_coding | ENSG0000C |
| 10685003 | 10697661 | 1 | lncRNA | ENSG0000C |
| 10737558 | 10749586 | 1 | lncRNA | ENSG0000C |
| 11368827 | 11368921 | 1 | miRNA | ENSG0000C |
| 11371975 | 11372892 | 1 | processed_pseudoge | ENSG0000C |
| 11393150 | 11429564 | -1 | protein_coding | ENSG0000C |
| 11414051 | 11414160 | -1 | rRNA_pseudogene | ENSG0000C |
| 11426874 | 11467802 | 1 | lncRNA | ENSG0000C |
| 11469250 | 11478196 | 1 | lncRNA | ENSG0000C |
| 11625714 | 11823906 | 1 | lncRNA | ENSG0000C |
| 11740948 | 11769468 | -1 | lncRNA | ENSG0000C |
| 11914667 | 11919688 | 1 | lncRNA | ENSG0000C |
| 12223445 | 12251286 | 1 | lncRNA | ENSG0000C |
| 12312588 | 12312690 | 1 | snRNA | ENSG0000C |
| 12639086 | 12641009 | 1 | processed_pseudoge | ENSG0000C |
| 12640298 | 12640637 | -1 | processed_pseudoge | ENSG0000C |
| 12859101 | 12864961 | -1 | lncRNA | ENSG0000C |
| 12947574 | 12948670 | -1 | lncRNA | ENSG0000C |
| 13051342 | 13051411 | 1 | snRNA | ENSG0000C |
| 13333414 | 13338657 | 1 | transcribed_processe | ENSG0000C |
| 13361354 | 13484365 | -1 | protein_coding | ENSG0000C |
| 13491185 | 13516498 | 1 | lncRNA | ENSG0000C |
| 13526319 | 13534335 | -1 | lncRNA | ENSG0000C |
| 13540830 | 13544508 | -1 | protein_coding | ENSG0000C |
| 13546075 | 13547801 | -1 | lncRNA | ENSG0000C |

| | | | | |
|---|---|---|---|---|
| 13568738 | 13627725 | -1 | protein_coding | ENSG0000C |
| 13627865 | 13627957 | 1 | miRNA | ENSG0000C |
| 13631820 | 13632599 | -1 | processed_pseudoge | ENSG0000C |
| 13647697 | 13648400 | -1 | processed_pseudoge | ENSG0000C |
| 13654374 | 14003756 | 1 | lncRNA | ENSG0000C |
| 13777377 | 13782157 | -1 | lncRNA | ENSG0000C |
| 13977166 | 13978028 | 1 | processed_pseudoge | ENSG0000C |
| 14111968 | 14140052 | 1 | transcribed_process | ENSG0000C |
| 14134936 | 14143592 | 1 | lncRNA | ENSG0000C |
| 14164455 | 14242813 | 1 | lncRNA | ENSG0000C |
| 14359400 | 14453625 | 1 | lncRNA | ENSG0000C |
| 14383123 | 14409078 | -1 | lncRNA | ENSG0000C |
| 14390439 | 14393992 | 1 | lncRNA | ENSG0000C |
| 14470465 | 14888169 | -1 | lncRNA | ENSG0000C |
| 14505865 | 14506480 | -1 | processed_pseudoge | ENSG0000C |
| 14690710 | 14690832 | 1 | snoRNA | ENSG0000C |
| 14909961 | 15002045 | -1 | lncRNA | ENSG0000C |
| 15002674 | 15070153 | 1 | protein_coding | ENSG0000C |
| 15004942 | 15427914 | -1 | lncRNA | ENSG0000C |
| 15093767 | 15094094 | -1 | misc_RNA | ENSG0000C |
| 15339818 | 15446166 | 1 | protein_coding | ENSG0000C |
| 15358141 | 15420032 | -1 | lncRNA | ENSG0000C |
| 15469865 | 15601557 | 1 | protein_coding | ENSG0000C |
| 15492729 | 15492972 | -1 | processed_pseudoge | ENSG0000C |
| 15563698 | 15564253 | -1 | lncRNA | ENSG0000C |
| 15604381 | 15681679 | -1 | protein_coding | ENSG0000C |
| 15681662 | 15705565 | 1 | protein_coding | ENSG0000C |
| 15703065 | 15738313 | 1 | protein_coding | ENSG0000C |
| 15730962 | 15731627 | -1 | processed_pseudoge | ENSG0000C |
| 15743196 | 15743582 | 1 | processed_pseudoge | ENSG0000C |
| 15778275 | 15853232 | 1 | protein_coding | ENSG0000C |
| 15864938 | 15865940 | -1 | processed_pseudoge | ENSG0000C |
| 15935577 | 15938740 | -1 | protein_coding | ENSG0000C |
| 15960245 | 15969309 | -1 | protein_coding | ENSG0000C |
| 15963076 | 16084378 | -1 | protein_coding | ENSG0000C |
| 16114233 | 16117479 | -1 | lncRNA | ENSG0000C |
| 16160505 | 16227410 | -1 | protein_coding | ENSG0000C |
| 16178939 | 16183120 | 1 | lncRNA | ENSG0000C |
| 16226685 | 16320140 | 1 | lncRNA | ENSG0000C |
| 16256308 | 16256555 | -1 | processed_pseudoge | ENSG0000C |
| 16307456 | 16308167 | 1 | processed_pseudoge | ENSG0000C |
| 16322954 | 16326967 | 1 | lncRNA | ENSG0000C |
| 16360686 | 16397323 | -1 | transcribed_process | ENSG0000C |
| 16400430 | 16512187 | 1 | lncRNA | ENSG0000C |
| 16501541 | 16898678 | -1 | protein_coding | ENSG0000C |
| 16598718 | 16604593 | 1 | lncRNA | ENSG0000C |
| 16973275 | 17073903 | -1 | lncRNA | ENSG0000C |
| 17061878 | 17062165 | 1 | processed_pseudoge | ENSG0000C |
| 17171757 | 17186079 | -1 | lncRNA | ENSG0000C |
| 17320746 | 17320882 | -1 | snoRNA | ENSG0000C |

| | | | | |
|---|---|---|---|---|
| 17427696 | 17428274 | -1 | processed_pseudoge | ENSG00000 |
| 17460261 | 17512206 | -1 | protein_coding | ENSG00000 |
| 17515165 | 17527104 | 1 | protein_coding | ENSG00000 |
| 17552117 | 17552752 | -1 | processed_pseudoge | ENSG00000 |
| 17557892 | 17558045 | -1 | unprocessed_pseudo | ENSG00000 |
| 17577192 | 17607972 | 1 | protein_coding | ENSG00000 |
| 17586267 | 17614585 | -1 | lncRNA | ENSG00000 |
| 17614641 | 17634105 | 1 | protein_coding | ENSG00000 |
| 17629306 | 17781621 | -1 | protein_coding | ENSG00000 |
| 17800655 | 17810758 | -1 | protein_coding | ENSG00000 |
| 17810979 | 17844865 | 1 | protein_coding | ENSG00000 |
| 17841199 | 18021876 | -1 | protein_coding | ENSG00000 |
| 17911674 | 17912976 | 1 | processed_pseudoge | ENSG00000 |
| 18418662 | 18888662 | 1 | lncRNA | ENSG00000 |
| 18488062 | 18489508 | -1 | lncRNA | ENSG00000 |
| 19098199 | 19098740 | 1 | TEC | ENSG00000 |
| 19172335 | 19456994 | -1 | lncRNA | ENSG00000 |
| 19181814 | 19181856 | -1 | rRNA_pseudogene | ENSG00000 |
| 19257951 | 19258337 | -1 | TEC | ENSG00000 |
| 19455418 | 19938448 | 1 | lncRNA | ENSG00000 |
| 19747179 | 19754591 | -1 | lncRNA | ENSG00000 |
| 19813749 | 19814231 | 1 | processed_pseudoge | ENSG00000 |
| 20037560 | 20037972 | -1 | processed_pseudoge | ENSG00000 |
| 20251905 | 20620561 | 1 | protein_coding | ENSG00000 |
| 20392154 | 20394856 | 1 | lncRNA | ENSG00000 |
| 20528275 | 20528384 | 1 | miRNA | ENSG00000 |
| 20696282 | 20752907 | 1 | protein_coding | ENSG00000 |
| 20728606 | 21948772 | -1 | protein_coding | ENSG00000 |
| 20766808 | 20767372 | 1 | lncRNA | ENSG00000 |
| 21304468 | 21350673 | 1 | lncRNA | ENSG00000 |
| 21464700 | 21464758 | -1 | miRNA | ENSG00000 |
| 21520854 | 21523709 | -1 | lncRNA | ENSG00000 |
| 21582096 | 21613728 | 1 | lncRNA | ENSG00000 |
| 21656511 | 21656869 | -1 | processed_pseudoge | ENSG00000 |
| 21697450 | 21719026 | 1 | lncRNA | ENSG00000 |
| 21843341 | 21853188 | -1 | lncRNA | ENSG00000 |
| 21949015 | 22330330 | 1 | lncRNA | ENSG00000 |
| 22118983 | 22119089 | 1 | snRNA | ENSG00000 |
| 22345071 | 22516066 | -1 | protein_coding | ENSG00000 |
| 22594196 | 22594450 | 1 | processed_pseudoge | ENSG00000 |
| 22607501 | 22611063 | 1 | unprocessed_pseudo | ENSG00000 |
| 22692914 | 22819575 | 1 | polymorphic_pseudo | ENSG00000 |
| 22727375 | 22727950 | -1 | processed_pseudoge | ENSG00000 |
| 22989147 | 23193740 | 1 | lncRNA | ENSG00000 |
| 23234625 | 23285573 | -1 | lncRNA | ENSG00000 |
| 23514940 | 23515284 | -1 | processed_pseudoge | ENSG00000 |
| 23560923 | 23768652 | 1 | lncRNA | ENSG00000 |
| 23723262 | 23733579 | -1 | lncRNA | ENSG00000 |
| 23755041 | 23904089 | -1 | protein_coding | ENSG00000 |
| 23779590 | 23782560 | 1 | lncRNA | ENSG00000 |

| | | | | |
|---|---|---|---|---|
| 24517441 | 24584554 | -1 | protein_coding | ENSG0000C |
| 24520192 | 24520290 | -1 | miRNA | ENSG0000C |
| 24563193 | 24563461 | -1 | misc_RNA | ENSG0000C |
| 24658072 | 24658375 | -1 | processed_pseudoge | ENSG0000C |
| 24659856 | 24671568 | 1 | lncRNA | ENSG0000C |
| 24768049 | 24768331 | 1 | processed_pseudoge | ENSG0000C |
| 24771467 | 24772420 | 1 | processed_pseudoge | ENSG0000C |
| 24789912 | 24800842 | 1 | protein_coding | ENSG0000C |
| 24806117 | 24980204 | -1 | protein_coding | ENSG0000C |
| 24998847 | 25030946 | -1 | protein_coding | ENSG0000C |
| 25120014 | 25160449 | -1 | protein_coding | ENSG0000C |
| 25160641 | 25201440 | 1 | lncRNA | ENSG0000C |
| 25160663 | 25279204 | 1 | protein_coding | ENSG0000C |
| 25220403 | 25220913 | 1 | lncRNA | ENSG0000C |
| 25250545 | 25250639 | 1 | snRNA | ENSG0000C |
| 25312774 | 25370383 | 1 | protein_coding | ENSG0000C |
| 25377213 | 25418498 | 1 | protein_coding | ENSG0000C |
| 25472517 | 25473815 | 1 | processed_pseudoge | ENSG0000C |
| 25504997 | 25506675 | 1 | processed_pseudoge | ENSG0000C |
| 25529177 | 25619468 | 1 | lncRNA | ENSG0000C |
| 25598564 | 25598625 | -1 | snRNA | ENSG0000C |
| 25622777 | 25623601 | 1 | processed_pseudoge | ENSG0000C |
| 25648011 | 25678748 | 1 | protein_coding | ENSG0000C |
| 25678850 | 25679020 | 1 | processed_pseudoge | ENSG0000C |
| 25686202 | 25686458 | -1 | processed_pseudoge | ENSG0000C |
| 25718082 | 25719745 | -1 | unprocessed_pseudc | ENSG0000C |
| 25719755 | 25720030 | -1 | unprocessed_pseudc | ENSG0000C |
| 25720099 | 25720431 | -1 | unprocessed_pseudc | ENSG0000C |
| 25720516 | 25721056 | -1 | unprocessed_pseudc | ENSG0000C |
| 25747433 | 25863760 | -1 | protein_coding | ENSG0000C |
| 25770266 | 25773577 | 1 | lncRNA | ENSG0000C |
| 25861830 | 25929874 | 1 | protein_coding | ENSG0000C |
| 25864881 | 25869550 | -1 | lncRNA | ENSG0000C |
| 26070754 | 26104258 | 1 | lncRNA | ENSG0000C |
| 26111865 | 26113839 | 1 | processed_pseudoge | ENSG0000C |
| 26163455 | 26435131 | 1 | protein_coding | ENSG0000C |
| 26481396 | 26490484 | -1 | protein_coding | ENSG0000C |
| 26576437 | 26756223 | 1 | protein_coding | ENSG0000C |
| 26702309 | 26702388 | 1 | snoRNA | ENSG0000C |
| 26857601 | 27025381 | 1 | protein_coding | ENSG0000C |
| 26859806 | 26860632 | -1 | lncRNA | ENSG0000C |
| 26873561 | 26874349 | -1 | processed_pseudoge | ENSG0000C |
| 27133996 | 27140051 | -1 | lncRNA | ENSG0000C |
| 27207505 | 27218404 | -1 | lncRNA | ENSG0000C |
| 27217479 | 27282225 | 1 | lncRNA | ENSG0000C |
| 27223319 | 27223427 | -1 | misc_RNA | ENSG0000C |
| 27262506 | 27267254 | -1 | lncRNA | ENSG0000C |
| 27585145 | 27586144 | 1 | processed_pseudoge | ENSG0000C |
| 27917753 | 27943065 | 1 | lncRNA | ENSG0000C |
| 27964517 | 27985063 | -1 | lncRNA | ENSG0000C |

| | | | | |
|---|---|---|---|---|
| 28225969 | 28288297 | 1 | lncRNA | ENSG0000C |
| 28343862 | 28402936 | -1 | lncRNA | ENSG0000C |
| 28435449 | 28600275 | 1 | lncRNA | ENSG0000C |
| 28580991 | 28600845 | -1 | lncRNA | ENSG0000C |
| 28711198 | 28711476 | 1 | misc_RNA | ENSG0000C |
| 28819582 | 28819668 | 1 | miRNA | ENSG0000C |
| 28823244 | 28824211 | -1 | processed_pseudoge | ENSG0000C |
| 28996498 | 29017256 | 1 | lncRNA | ENSG0000C |
| 29010619 | 29011021 | -1 | processed_pseudoge | ENSG0000C |
| 29046591 | 29048765 | 1 | lncRNA | ENSG0000C |
| 29118304 | 29211696 | 1 | lncRNA | ENSG0000C |
| 29214253 | 29291869 | 1 | lncRNA | ENSG0000C |
| 29407100 | 29408012 | 1 | processed_pseudoge | ENSG0000C |
| 29465701 | 29467850 | -1 | processed_pseudoge | ENSG0000C |
| 29510635 | 29510737 | 1 | snRNA | ENSG0000C |
| 29748757 | 29750100 | 1 | processed_pseudoge | ENSG0000C |
| 29907659 | 29907791 | -1 | processed_pseudoge | ENSG0000C |
| 29962644 | 29963758 | -1 | processed_pseudoge | ENSG0000C |
| 30006951 | 30008316 | -1 | processed_pseudoge | ENSG0000C |
| 30718805 | 30721970 | -1 | lncRNA | ENSG0000C |
| 30720415 | 31146805 | 1 | protein_coding | ENSG0000C |
| 30776257 | 30793970 | -1 | lncRNA | ENSG0000C |
| 30884443 | 30885182 | -1 | unprocessed_pseudc | ENSG0000C |
| 31171013 | 31211678 | 1 | lncRNA | ENSG0000C |
| 31350284 | 31351725 | 1 | lncRNA | ENSG0000C |
| 31506422 | 31558831 | 1 | lncRNA | ENSG0000C |
| 31997376 | 32228543 | 1 | lncRNA | ENSG0000C |
| 32005076 | 32022483 | -1 | lncRNA | ENSG0000C |
| 32351038 | 32353220 | 1 | lncRNA | ENSG0000C |
| 32583855 | 32745525 | -1 | lncRNA | ENSG0000C |
| 33010948 | 33011746 | -1 | processed_pseudoge | ENSG0000C |
| 33150326 | 33208430 | 1 | lncRNA | ENSG0000C |
| 33238239 | 33238670 | -1 | processed_pseudoge | ENSG0000C |
| 33400538 | 33400679 | 1 | processed_pseudoge | ENSG0000C |
| 33433510 | 33437877 | 1 | lncRNA | ENSG0000C |
| 33467398 | 33693993 | 1 | lncRNA | ENSG0000C |
| 33775498 | 34039914 | -1 | lncRNA | ENSG0000C |
| 33850591 | 33982349 | 1 | lncRNA | ENSG0000C |
| 33968174 | 33968526 | 1 | processed_pseudoge | ENSG0000C |
| 34120090 | 34335351 | -1 | lncRNA | ENSG0000C |
| 34140598 | 34180729 | -1 | lncRNA | ENSG0000C |
| 34657606 | 34669432 | 1 | lncRNA | ENSG0000C |
| 34714270 | 34714598 | 1 | processed_pseudoge | ENSG0000C |
| 34966241 | 34966344 | -1 | snoRNA | ENSG0000C |
| 35487812 | 35489750 | 1 | processed_pseudoge | ENSG0000C |
| 35495898 | 35496003 | 1 | snRNA | ENSG0000C |
| 35948221 | 36244514 | -1 | protein_coding | ENSG0000C |
| 36065060 | 36065726 | -1 | processed_pseudoge | ENSG0000C |
| 36244116 | 36274220 | 1 | lncRNA | ENSG0000C |
| 36281622 | 36345756 | 1 | protein_coding | ENSG0000C |

| | | | | |
|---|---|---|---|---|
| 36311190 | 36392410 | -1 | lncRNA | ENSG0000C |
| 36426366 | 36426428 | -1 | miRNA | ENSG0000C |
| 36496128 | 36641939 | -1 | lncRNA | ENSG0000C |
| 36506043 | 36508869 | 1 | processed_pseudoge | ENSG0000C |
| 36902242 | 36913339 | -1 | lncRNA | ENSG0000C |
| 37001772 | 37023499 | 1 | lncRNA | ENSG0000C |
| 37073681 | 37133849 | 1 | lncRNA | ENSG0000C |
| 37241910 | 37241991 | -1 | miRNA | ENSG0000C |
| 37244743 | 37449463 | 1 | protein_coding | ENSG0000C |
| 37453925 | 37623495 | 1 | protein_coding | ENSG0000C |
| 37454698 | 37476401 | -1 | lncRNA | ENSG0000C |
| 37588087 | 37588875 | -1 | lncRNA | ENSG0000C |
| 37590800 | 37686376 | -1 | protein_coding | ENSG0000C |
| 37699895 | 37700002 | -1 | misc_RNA | ENSG0000C |
| 37821361 | 37821838 | 1 | processed_pseudoge | ENSG0000C |
| 37826660 | 37862937 | 1 | protein_coding | ENSG0000C |
| 37866561 | 37867091 | -1 | TEC | ENSG0000C |
| 37868292 | 37869978 | 1 | processed_pseudoge | ENSG0000C |
| 37891084 | 38139175 | 1 | protein_coding | ENSG0000C |
| 37960398 | 37961128 | 1 | protein_coding | ENSG0000C |
| 37995494 | 37996203 | -1 | processed_pseudoge | ENSG0000C |
| 38006784 | 38007105 | -1 | processed_pseudoge | ENSG0000C |
| 38276178 | 38279880 | -1 | lncRNA | ENSG0000C |
| 38286994 | 38287491 | 1 | lncRNA | ENSG0000C |
| 38366914 | 38385759 | 1 | lncRNA | ENSG0000C |
| 38390754 | 38391675 | 1 | TEC | ENSG0000C |
| 38420662 | 38523180 | -1 | lncRNA | ENSG0000C |
| 38509767 | 38518056 | 1 | lncRNA | ENSG0000C |
| 38564003 | 38572022 | 1 | lncRNA | ENSG0000C |
| 38602438 | 38664914 | -1 | lncRNA | ENSG0000C |
| 38618265 | 38619437 | -1 | lncRNA | ENSG0000C |
| 38664197 | 38701517 | 1 | protein_coding | ENSG0000C |
| 38758791 | 38758892 | -1 | rRNA_pseudogene | ENSG0000C |
| 38772238 | 38782990 | -1 | protein_coding | ENSG0000C |
| 38790677 | 38856817 | -1 | protein_coding | ENSG0000C |
| 38823715 | 38856817 | -1 | protein_coding | ENSG0000C |
| 38867677 | 38945739 | 1 | protein_coding | ENSG0000C |
| 38868032 | 38868127 | 1 | miRNA | ENSG0000C |
| 38966744 | 39032922 | -1 | protein_coding | ENSG0000C |
| 39045039 | 39126857 | 1 | protein_coding | ENSG0000C |
| 39066974 | 39182258 | -1 | lncRNA | ENSG0000C |
| 39133913 | 39135608 | 1 | lncRNA | ENSG0000C |
| 39182504 | 39285810 | 1 | protein_coding | ENSG0000C |
| 39287456 | 39366375 | -1 | protein_coding | ENSG0000C |
| 39297605 | 39297711 | -1 | snRNA | ENSG0000C |
| 39399149 | 39399248 | 1 | snRNA | ENSG0000C |
| 39406930 | 39451533 | 1 | protein_coding | ENSG0000C |
| 39411910 | 39411974 | 1 | miRNA | ENSG0000C |
| 39441665 | 39441764 | 1 | misc_RNA | ENSG0000C |
| 39452521 | 39458949 | -1 | protein_coding | ENSG0000C |

| | | | | |
|---|---|---|---|---|
| 39459004 | 39485109 | 1 | protein_coding | ENSG00000( |
| 39480255 | 39481905 | 1 | transcribed_processe | ENSG00000( |
| 39498755 | 39528311 | -1 | protein_coding | ENSG00000( |
| 39528019 | 39594707 | 1 | lncRNA | ENSG00000( |
| 39539395 | 39546073 | -1 | lncRNA | ENSG00000( |
| 39546336 | 39638902 | -1 | protein_coding | ENSG00000( |
| 39614465 | 39618488 | 1 | lncRNA | ENSG00000( |
| 39621012 | 39621073 | 1 | snRNA | ENSG00000( |
| 39639107 | 39666879 | 1 | lncRNA | ENSG00000( |
| 39698109 | 39782792 | 1 | protein_coding | ENSG00000( |
| 39710085 | 39710176 | -1 | misc_RNA | ENSG00000( |
| 39713842 | 39714451 | -1 | processed_pseudoge | ENSG00000( |
| 39761000 | 39761298 | 1 | misc_RNA | ENSG00000( |
| 39770081 | 39771371 | -1 | processed_pseudoge | ENSG00000( |
| 39822863 | 39977956 | -1 | protein_coding | ENSG00000( |
| 39932454 | 39933561 | 1 | processed_pseudoge | ENSG00000( |
| 39936753 | 39936855 | 1 | rRNA_pseudogene | ENSG00000( |
| 39973128 | 39973442 | -1 | processed_pseudoge | ENSG00000( |
| 39973444 | 39974338 | 1 | processed_pseudoge | ENSG00000( |
| 40020240 | 40021517 | 1 | processed_pseudoge | ENSG00000( |
| 40042917 | 40057199 | -1 | transcribed_processe | ENSG00000( |
| 40056850 | 40158252 | 1 | protein_coding | ENSG00000( |
| 40077884 | 40077982 | 1 | snRNA | ENSG00000( |
| 40082983 | 40083106 | 1 | snoRNA | ENSG00000( |
| 40142198 | 40142365 | -1 | processed_pseudoge | ENSG00000( |
| 40166675 | 40167831 | 1 | lncRNA | ENSG00000( |
| 40187170 | 40190464 | 1 | lncRNA | ENSG00000( |
| 40191053 | 40246967 | 1 | protein_coding | ENSG00000( |
| 40265472 | 40266457 | 1 | lncRNA | ENSG00000( |
| 40316485 | 40330419 | 1 | lncRNA | ENSG00000( |
| 40335333 | 40355217 | 1 | protein_coding | ENSG00000( |
| 40377452 | 40377513 | -1 | snRNA | ENSG00000( |
| 40423267 | 40630875 | -1 | protein_coding | ENSG00000( |
| 40426119 | 40427585 | 1 | lncRNA | ENSG00000( |
| 40491733 | 40492024 | 1 | processed_pseudoge | ENSG00000( |
| 40502040 | 40502119 | -1 | miRNA | ENSG00000( |
| 40743627 | 40750865 | -1 | lncRNA | ENSG00000( |
| 40749925 | 40811184 | 1 | protein_coding | ENSG00000( |
| 40786110 | 40786710 | 1 | processed_pseudoge | ENSG00000( |
| 40810027 | 41216714 | -1 | protein_coding | ENSG00000( |
| 40812779 | 40826151 | 1 | lncRNA | ENSG00000( |
| 40826655 | 40826765 | -1 | misc_RNA | ENSG00000( |
| 40990154 | 40990273 | 1 | rRNA_pseudogene | ENSG00000( |
| 41084607 | 41084713 | 1 | snRNA | ENSG00000( |
| 41113942 | 41114045 | 1 | snRNA | ENSG00000( |
| 41220074 | 41256727 | -1 | lncRNA | ENSG00000( |
| 41256413 | 41268455 | 1 | protein_coding | ENSG00000( |
| 41303237 | 41303337 | -1 | misc_RNA | ENSG00000( |
| 41359607 | 41700044 | 1 | protein_coding | ENSG00000( |
| 41389115 | 41389593 | 1 | processed_pseudoge | ENSG00000( |

| | | | | |
|---|---|---|---|---|
| 41608312 | 41612411 | -1 | lncRNA | ENSG0000C |
| 41688858 | 41692797 | -1 | lncRNA | ENSG0000C |
| 41722538 | 41723512 | -1 | unprocessed_pseudc | ENSG0000C |
| 41744082 | 41748970 | -1 | protein_coding | ENSG0000C |
| 41748293 | 41824119 | 1 | lncRNA | ENSG0000C |
| 41750345 | 41757341 | 1 | lncRNA | ENSG0000C |
| 41771945 | 41772103 | -1 | snRNA | ENSG0000C |
| 41842154 | 41843950 | -1 | processed_pseudoge | ENSG0000C |
| 41872747 | 41882955 | -1 | lncRNA | ENSG0000C |
| 41883060 | 41935075 | -1 | lncRNA | ENSG0000C |
| 41924180 | 41924380 | 1 | processed_pseudoge | ENSG0000C |
| 41935129 | 41960803 | 1 | protein_coding | ENSG0000C |
| 41981756 | 41986465 | 1 | protein_coding | ENSG0000C |
| 41988741 | 41989237 | -1 | lncRNA | ENSG0000C |
| 41990502 | 42090461 | 1 | protein_coding | ENSG0000C |
| 42029209 | 42030123 | 1 | unprocessed_pseudc | ENSG0000C |
| 42110853 | 42152878 | -1 | protein_coding | ENSG0000C |
| 42151028 | 42268110 | 1 | lncRNA | ENSG0000C |
| 42281830 | 42391651 | -1 | lncRNA | ENSG0000C |
| 42397488 | 42402487 | 1 | protein_coding | ENSG0000C |
| 42408373 | 42657105 | -1 | protein_coding | ENSG0000C |
| 42471810 | 42472380 | 1 | processed_pseudoge | ENSG0000C |
| 42561457 | 42562425 | -1 | processed_pseudoge | ENSG0000C |
| 42657496 | 42657928 | 1 | lncRNA | ENSG0000C |
| 42706107 | 42707335 | 1 | lncRNA | ENSG0000C |
| 42892396 | 42892618 | 1 | misc_RNA | ENSG0000C |
| 42893267 | 43030658 | 1 | protein_coding | ENSG0000C |
| 43133867 | 43234956 | 1 | lncRNA | ENSG0000C |
| 43340875 | 43345600 | 1 | lncRNA | ENSG0000C |
| 43388157 | 43388437 | 1 | processed_pseudoge | ENSG0000C |
| 43410041 | 43410937 | -1 | processed_pseudoge | ENSG0000C |
| 43457527 | 43492543 | -1 | lncRNA | ENSG0000C |
| 43586328 | 43588097 | 1 | processed_pseudoge | ENSG0000C |
| 43598544 | 43598844 | -1 | misc_RNA | ENSG0000C |
| 43763134 | 43972184 | 1 | lncRNA | ENSG0000C |
| 43863274 | 43863549 | 1 | misc_RNA | ENSG0000C |
| 43898962 | 43899351 | 1 | processed_pseudoge | ENSG0000C |
| 43972921 | 44025516 | -1 | lncRNA | ENSG0000C |
| 44016700 | 44022063 | 1 | lncRNA | ENSG0000C |
| 44173903 | 44448809 | -1 | protein_coding | ENSG0000C |
| 44533615 | 44533880 | 1 | processed_pseudoge | ENSG0000C |
| 44622065 | 44678556 | -1 | protein_coding | ENSG0000C |
| 44678420 | 44700928 | 1 | protein_coding | ENSG0000C |
| 44682200 | 44726588 | -1 | protein_coding | ENSG0000C |
| 44693946 | 44694386 | -1 | lncRNA | ENSG0000C |
| 44704405 | 44704965 | 1 | lncRNA | ENSG0000C |
| 44840380 | 44841402 | -1 | processed_pseudoge | ENSG0000C |
| 44944015 | 44944961 | -1 | processed_pseudoge | ENSG0000C |
| 44994426 | 44996022 | 1 | processed_pseudoge | ENSG0000C |
| 45009540 | 45051652 | 1 | lncRNA | ENSG0000C |

| | | | | |
|---|---|---|---|---|
| 45323253 | 45326028 | -1 | processed_pseudoge | ENSG00000 |
| 45414337 | 45414910 | -1 | processed_pseudoge | ENSG00000 |
| 45480030 | 45480136 | 1 | snRNA | ENSG00000 |
| 45995119 | 45995423 | 1 | misc_RNA | ENSG00000 |
| 46035769 | 46124054 | -1 | protein_coding | ENSG00000 |
| 46243548 | 46244215 | -1 | lncRNA | ENSG00000 |
| 46248427 | 46475230 | -1 | protein_coding | ENSG00000 |
| 46390255 | 46510194 | 1 | lncRNA | ENSG00000 |
| 46531237 | 46531342 | -1 | snRNA | ENSG00000 |
| 46723830 | 46724408 | -1 | processed_pseudoge | ENSG00000 |
| 46734827 | 46909245 | -1 | protein_coding | ENSG00000 |
| 46918900 | 46993581 | -1 | protein_coding | ENSG00000 |
| 46993723 | 47426447 | 1 | protein_coding | ENSG00000 |
| 47431960 | 47438959 | 1 | lncRNA | ENSG00000 |
| 47450787 | 47463702 | -1 | protein_coding | ENSG00000 |
| 47463590 | 47470477 | 1 | lncRNA | ENSG00000 |
| 47481001 | 47485210 | -1 | lncRNA | ENSG00000 |
| 47485275 | 47593486 | 1 | protein_coding | ENSG00000 |
| 47490967 | 47491471 | -1 | processed_pseudoge | ENSG00000 |
| 47556731 | 47560259 | -1 | lncRNA | ENSG00000 |
| 47593999 | 47838106 | -1 | protein_coding | ENSG00000 |
| 47652669 | 47652743 | 1 | miRNA | ENSG00000 |
| 47706372 | 47706987 | 1 | processed_pseudoge | ENSG00000 |
| 47831330 | 47897965 | 1 | lncRNA | ENSG00000 |
| 47840122 | 47844339 | -1 | lncRNA | ENSG00000 |
| 47847233 | 47914667 | -1 | protein_coding | ENSG00000 |
| 47914142 | 48040173 | 1 | protein_coding | ENSG00000 |
| 47935977 | 48016672 | -1 | protein_coding | ENSG00000 |
| 48066393 | 48134250 | -1 | protein_coding | ENSG00000 |
| 48106081 | 48106188 | -1 | snRNA | ENSG00000 |
| 48109353 | 48109410 | -1 | snRNA | ENSG00000 |
| 48135783 | 48269838 | -1 | protein_coding | ENSG00000 |
| 48153434 | 48153535 | 1 | misc_RNA | ENSG00000 |
| 48341529 | 48426201 | 1 | protein_coding | ENSG00000 |
| 48483343 | 48489526 | 1 | protein_coding | ENSG00000 |
| 48490252 | 48494389 | 1 | protein_coding | ENSG00000 |
| 48497361 | 48780322 | -1 | protein_coding | ENSG00000 |
| 48574453 | 48574520 | -1 | snRNA | ENSG00000 |
| 48805212 | 48861817 | 1 | protein_coding | ENSG00000 |
| 48852008 | 48860203 | -1 | lncRNA | ENSG00000 |
| 48885019 | 48906937 | -1 | protein_coding | ENSG00000 |
| 48932755 | 48932854 | -1 | snRNA | ENSG00000 |
| 48936582 | 48936880 | 1 | processed_pseudoge | ENSG00000 |
| 48986275 | 49062081 | 1 | protein_coding | ENSG00000 |
| 49016682 | 49017154 | -1 | processed_pseudoge | ENSG00000 |
| 49203171 | 49203726 | -1 | processed_pseudoge | ENSG00000 |
| 49212251 | 49212624 | -1 | processed_pseudoge | ENSG00000 |
| 49229573 | 49231284 | 1 | TEC | ENSG00000 |
| 49233289 | 49233965 | -1 | processed_pseudoge | ENSG00000 |
| 49238032 | 49245419 | -1 | processed_pseudoge | ENSG00000 |

| | | | | |
|---|---|---|---|---|
| 49246021 | 49246538 | 1 | processed_pseudoge | ENSG00000( |
| 49246601 | 49246915 | 1 | processed_pseudoge | ENSG00000( |
| 49486926 | 49489554 | -1 | unprocessed_pseudc | ENSG00000( |
| 49502145 | 49506022 | -1 | unprocessed_pseudc | ENSG00000( |
| 49507764 | 49508806 | -1 | processed_pseudoge | ENSG00000( |
| 49523648 | 49523857 | -1 | processed_pseudoge | ENSG00000( |
| 49524030 | 49524185 | -1 | processed_pseudoge | ENSG00000( |
| 49548564 | 49548855 | -1 | processed_pseudoge | ENSG00000( |
| 49548918 | 49549435 | -1 | processed_pseudoge | ENSG00000( |
| 49550032 | 49557400 | 1 | unprocessed_pseudc | ENSG00000( |
| 49561285 | 49561470 | -1 | processed_pseudoge | ENSG00000( |
| 49561479 | 49562149 | 1 | processed_pseudoge | ENSG00000( |
| 49579833 | 49580189 | 1 | processed_pseudoge | ENSG00000( |
| 49588772 | 49589529 | 1 | processed_pseudoge | ENSG00000( |
| 51843000 | 51916837 | 1 | protein_coding | ENSG00000( |
| 51865050 | 51865535 | -1 | processed_pseudoge | ENSG00000( |
| 51918772 | 51919381 | 1 | lncRNA | ENSG00000( |
| 51978079 | 51978372 | -1 | processed_pseudoge | ENSG00000( |
| 51993702 | 52017620 | -1 | protein_coding | ENSG00000( |
| 52020706 | 52038482 | -1 | protein_coding | ENSG00000( |
| 52044805 | 52046954 | 1 | lncRNA | ENSG00000( |
| 52051304 | 52097299 | 1 | protein_coding | ENSG00000( |
| 52252820 | 52551574 | -1 | lncRNA | ENSG00000( |
| 52440494 | 52441535 | -1 | processed_pseudoge | ENSG00000( |
| 52494849 | 52494952 | 1 | snRNA | ENSG00000( |
| 52590960 | 52659301 | -1 | protein_coding | ENSG00000( |
| 52659406 | 52661668 | 1 | lncRNA | ENSG00000( |
| 52680609 | 52692999 | 1 | lncRNA | ENSG00000( |
| 52712394 | 52866821 | 1 | lncRNA | ENSG00000( |
| 52712404 | 52720351 | 1 | lncRNA | ENSG00000( |
| 52712682 | 52712747 | 1 | miRNA | ENSG00000( |
| 52712713 | 52722377 | -1 | lncRNA | ENSG00000( |
| 52713249 | 52713370 | 1 | snoRNA | ENSG00000( |
| 52720081 | 52720831 | -1 | processed_pseudoge | ENSG00000( |
| 52722618 | 52751586 | -1 | protein_coding | ENSG00000( |
| 52748137 | 52748259 | 1 | snoRNA | ENSG00000( |
| 52786537 | 52786638 | 1 | misc_RNA | ENSG00000( |
| 52862317 | 52866835 | 1 | protein_coding | ENSG00000( |
| 52872982 | 53366066 | -1 | protein_coding | ENSG00000( |
| 52945649 | 52958087 | 1 | lncRNA | ENSG00000( |
| 53265461 | 53265567 | -1 | snRNA | ENSG00000( |
| 53377641 | 53460862 | 1 | protein_coding | ENSG00000( |
| 53377839 | 54295272 | 1 | protein_coding | ENSG00000( |
| 53459301 | 53701405 | -1 | protein_coding | ENSG00000( |
| 53496400 | 53549578 | 1 | lncRNA | ENSG00000( |
| 53575713 | 53576387 | 1 | processed_pseudoge | ENSG00000( |
| 53592956 | 53604047 | 1 | lncRNA | ENSG00000( |
| 53659208 | 53737156 | 1 | lncRNA | ENSG00000( |
| 53899871 | 53916835 | 1 | lncRNA | ENSG00000( |
| 53927499 | 53928138 | -1 | processed_pseudoge | ENSG00000( |

| | | | | |
|---|---|---|---|---|
| 53986587 | 53987058 | -1 | processed_pseudoge | ENSG00000 |
| 53997415 | 53997712 | -1 | lncRNA | ENSG00000 |
| 54009789 | 54064605 | -1 | protein_coding | ENSG00000 |
| 54059597 | 54061210 | 1 | lncRNA | ENSG00000 |
| 54065239 | 54078454 | 1 | lncRNA | ENSG00000 |
| 54076625 | 54076848 | 1 | processed_pseudoge | ENSG00000 |
| 54086926 | 54087783 | 1 | processed_pseudoge | ENSG00000 |
| 54099523 | 54102498 | 1 | protein_coding | ENSG00000 |
| 54221126 | 54221472 | 1 | processed_pseudoge | ENSG00000 |
| 54229280 | 54298245 | 1 | protein_coding | ENSG00000 |
| 54332892 | 54355954 | 1 | lncRNA | ENSG00000 |
| 54376002 | 54376752 | 1 | lncRNA | ENSG00000 |
| 54440502 | 54446723 | -1 | lncRNA | ENSG00000 |
| 54603211 | 54607131 | -1 | lncRNA | ENSG00000 |
| 54657918 | 54740715 | 1 | protein_coding | ENSG00000 |
| 54836041 | 54845470 | -1 | lncRNA | ENSG00000 |
| 54845568 | 54859241 | 1 | lncRNA | ENSG00000 |
| 54943626 | 54957832 | 1 | lncRNA | ENSG00000 |
| 54976159 | 54976371 | -1 | processed_pseudoge | ENSG00000 |
| 55031965 | 55032071 | -1 | snRNA | ENSG00000 |
| 55053060 | 55092534 | 1 | lncRNA | ENSG00000 |
| 55063738 | 55064026 | 1 | misc_RNA | ENSG00000 |
| 55078481 | 55125595 | -1 | protein_coding | ENSG00000 |
| 55157877 | 55161880 | 1 | lncRNA | ENSG00000 |
| 55215624 | 55215919 | -1 | misc_RNA | ENSG00000 |
| 55219344 | 55219973 | 1 | processed_pseudoge | ENSG00000 |
| 55294490 | 55294598 | 1 | snRNA | ENSG00000 |
| 55313011 | 55313924 | -1 | processed_pseudoge | ENSG00000 |
| 55346242 | 55373100 | 1 | protein_coding | ENSG00000 |
| 55351812 | 55352408 | -1 | processed_pseudoge | ENSG00000 |
| 55363971 | 55395847 | -1 | lncRNA | ENSG00000 |
| 55387949 | 55388271 | 1 | lncRNA | ENSG00000 |
| 55395957 | 55453397 | 1 | protein_coding | ENSG00000 |
| 55412636 | 55412738 | 1 | misc_RNA | ENSG00000 |
| 55427903 | 55546909 | -1 | protein_coding | ENSG00000 |
| 55501595 | 55501708 | -1 | misc_RNA | ENSG00000 |
| 55540502 | 55540835 | -1 | misc_RNA | ENSG00000 |
| 55547112 | 55547889 | 1 | lncRNA | ENSG00000 |
| 55556519 | 55592245 | -1 | protein_coding | ENSG00000 |
| 55595229 | 55636698 | -1 | protein_coding | ENSG00000 |
| 55798636 | 55798731 | 1 | snRNA | ENSG00000 |
| 55819819 | 55837487 | 1 | protein_coding | ENSG00000 |
| 55853648 | 55905086 | 1 | protein_coding | ENSG00000 |
| 55885595 | 55885701 | 1 | snRNA | ENSG00000 |
| 55889743 | 55903177 | -1 | lncRNA | ENSG00000 |
| 55938153 | 55947996 | -1 | lncRNA | ENSG00000 |
| 55948871 | 56033361 | 1 | protein_coding | ENSG00000 |
| 56049073 | 56328625 | 1 | protein_coding | ENSG00000 |
| 56097390 | 56097504 | 1 | rRNA_pseudogene | ENSG00000 |
| 56288485 | 56288584 | -1 | snRNA | ENSG00000 |

| | | | | |
|---|---|---|---|---|
| 56291601 | 56299873 | -1 | lncRNA | ENSG0000C |
| 56320719 | 56321334 | -1 | processed_pseudoge | ENSG0000C |
| 56338287 | 56387508 | -1 | protein_coding | ENSG0000C |
| 56356135 | 56356932 | 1 | processed_pseudoge | ENSG0000C |
| 56387625 | 56388153 | 1 | lncRNA | ENSG0000C |
| 56393362 | 56435615 | -1 | protein_coding | ENSG0000C |
| 56396312 | 56396871 | 1 | lncRNA | ENSG0000C |
| 56410516 | 56431548 | 1 | transcribed_unproce | ENSG0000C |
| 56435741 | 56464579 | 1 | protein_coding | ENSG0000C |
| 56467617 | 56503681 | 1 | protein_coding | ENSG0000C |
| 56505209 | 56525481 | 1 | protein_coding | ENSG0000C |
| 56530606 | 56603507 | 1 | protein_coding | ENSG0000C |
| 56593004 | 56595681 | 1 | transcribed_processe | ENSG0000C |
| 56647988 | 56681899 | -1 | protein_coding | ENSG0000C |
| 56662469 | 56665694 | -1 | TEC | ENSG0000C |
| 56710060 | 56710614 | 1 | processed_pseudoge | ENSG0000C |
| 56760919 | 56761485 | -1 | processed_pseudoge | ENSG0000C |
| 56794348 | 56794642 | -1 | misc_RNA | ENSG0000C |
| 56805834 | 56806131 | 1 | misc_RNA | ENSG0000C |
| 56809860 | 56821742 | -1 | protein_coding | ENSG0000C |
| 56810422 | 56810681 | 1 | misc_RNA | ENSG0000C |
| 56818218 | 56818314 | -1 | misc_RNA | ENSG0000C |
| 56907876 | 56966678 | 1 | protein_coding | ENSG0000C |
| 56960927 | 56961373 | -1 | lncRNA | ENSG0000C |
| 56963350 | 56977606 | -1 | protein_coding | ENSG0000C |
| 56977722 | 57031158 | 1 | protein_coding | ENSG0000C |
| 57002692 | 57002792 | 1 | snRNA | ENSG0000C |
| 57030773 | 57110385 | -1 | protein_coding | ENSG0000C |
| 57072683 | 57073132 | 1 | processed_pseudoge | ENSG0000C |
| 57109762 | 57207459 | 1 | lncRNA | ENSG0000C |
| 57110547 | 57111400 | 1 | lncRNA | ENSG0000C |
| 57154577 | 57156452 | 1 | lncRNA | ENSG0000C |
| 57352999 | 57353329 | -1 | processed_pseudoge | ENSG0000C |
| 57424495 | 57478341 | 1 | lncRNA | ENSG0000C |
| 57490808 | 57495844 | 1 | lncRNA | ENSG0000C |
| 57595940 | 57605872 | -1 | lncRNA | ENSG0000C |
| 57605694 | 57658800 | -1 | lncRNA | ENSG0000C |
| 57720035 | 57724655 | 1 | lncRNA | ENSG0000C |
| 57841721 | 57855271 | 1 | lncRNA | ENSG0000C |
| 58103147 | 58103434 | 1 | processed_pseudoge | ENSG0000C |
| 58117758 | 58118070 | -1 | processed_pseudoge | ENSG0000C |
| 58524515 | 58536328 | -1 | lncRNA | ENSG0000C |
| 58562160 | 58565212 | -1 | lncRNA | ENSG0000C |
| 58780626 | 58984194 | -1 | lncRNA | ENSG0000C |
| 58939288 | 58988182 | -1 | lncRNA | ENSG0000C |
| 58984215 | 59046959 | 1 | lncRNA | ENSG0000C |
| 59047020 | 59075256 | -1 | lncRNA | ENSG0000C |
| 59150924 | 59180414 | -1 | lncRNA | ENSG0000C |
| 59152834 | 59176146 | 1 | lncRNA | ENSG0000C |
| 59533786 | 59533898 | 1 | misc_RNA | ENSG0000C |

| | | | | |
|---|---|---|---|---|
| 59551142 | 59630324 | -1 | lncRNA | ENSG00000 |
| 59767816 | 59792114 | -1 | lncRNA | ENSG00000 |
| 59833175 | 59833276 | 1 | misc_RNA | ENSG00000 |
| 59834063 | 59834167 | 1 | snRNA | ENSG00000 |
| 59903404 | 59920670 | -1 | lncRNA | ENSG00000 |
| 60038269 | 60038586 | 1 | processed_pseudoge | ENSG00000 |
| 60750575 | 60796103 | 1 | lncRNA | ENSG00000 |
| 60784115 | 60784696 | 1 | processed_pseudoge | ENSG00000 |
| 60922619 | 60922684 | 1 | miRNA | ENSG00000 |
| 61143656 | 61153962 | 1 | lncRNA | ENSG00000 |
| 61201258 | 62078335 | 1 | protein_coding | ENSG00000 |
| 61211652 | 61212203 | 1 | processed_pseudoge | ENSG00000 |
| 61420246 | 61428221 | -1 | lncRNA | ENSG00000 |
| 61775449 | 61775850 | -1 | processed_pseudoge | ENSG00000 |
| 61906313 | 61906408 | 1 | misc_RNA | ENSG00000 |
| 62071752 | 62165554 | -1 | lncRNA | ENSG00000 |
| 62105660 | 62106049 | -1 | processed_pseudoge | ENSG00000 |
| 62248294 | 62248770 | 1 | processed_pseudoge | ENSG00000 |
| 62291562 | 62292337 | 1 | processed_pseudoge | ENSG00000 |
| 62489495 | 62490218 | -1 | processed_pseudoge | ENSG00000 |
| 62510469 | 62510771 | 1 | processed_pseudoge | ENSG00000 |
| 62816826 | 62818794 | 1 | processed_pseudoge | ENSG00000 |
| 63128311 | 63143881 | 1 | lncRNA | ENSG00000 |
| 63350114 | 63351472 | 1 | processed_pseudoge | ENSG00000 |
| 63465511 | 63529761 | -1 | lncRNA | ENSG00000 |
| 64275257 | 64409468 | -1 | protein_coding | ENSG00000 |
| 64397694 | 64397787 | -1 | snRNA | ENSG00000 |
| 64430909 | 64433215 | 1 | processed_pseudoge | ENSG00000 |
| 64606418 | 64606613 | 1 | unprocessed_pseudo | ENSG00000 |
| 64607703 | 64608831 | -1 | processed_pseudoge | ENSG00000 |
| 64608905 | 64609424 | 1 | unprocessed_pseudo | ENSG00000 |
| 64609454 | 64610532 | -1 | processed_pseudoge | ENSG00000 |
| 64610616 | 64610909 | -1 | unprocessed_pseudo | ENSG00000 |
| 64610980 | 64611308 | -1 | unprocessed_pseudo | ENSG00000 |
| 64611374 | 64611888 | -1 | unprocessed_pseudo | ENSG00000 |
| 64767130 | 64767897 | -1 | processed_pseudoge | ENSG00000 |
| 64836361 | 64883916 | -1 | lncRNA | ENSG00000 |
| 64914281 | 65024485 | -1 | lncRNA | ENSG00000 |
| 65034634 | 65036017 | -1 | processed_pseudoge | ENSG00000 |
| 65142703 | 65145763 | -1 | processed_pseudoge | ENSG00000 |
| 65216616 | 65217433 | 1 | processed_pseudoge | ENSG00000 |
| 65225867 | 65241814 | -1 | lncRNA | ENSG00000 |
| 65319563 | 65670495 | -1 | protein_coding | ENSG00000 |
| 65573459 | 65574328 | 1 | processed_pseudoge | ENSG00000 |
| 65669961 | 65698029 | 1 | lncRNA | ENSG00000 |
| 65702202 | 65705553 | 1 | lncRNA | ENSG00000 |
| 65807240 | 65807435 | -1 | snRNA | ENSG00000 |
| 65858761 | 65860204 | 1 | lncRNA | ENSG00000 |
| 65998846 | 66150012 | 1 | lncRNA | ENSG00000 |
| 66003281 | 66016792 | 1 | lncRNA | ENSG00000 |

| | | | | |
|---|---|---|---|---|
| 66094142 | 66094526 | 1 | processed_pseudoge | ENSG00000( |
| 66169778 | 66329377 | -1 | lncRNA | ENSG00000( |
| 66276824 | 66276928 | 1 | miRNA | ENSG00000( |
| 66431092 | 66431521 | 1 | processed_pseudoge | ENSG00000( |
| 66897262 | 66897371 | 1 | snRNA | ENSG00000( |
| 67417305 | 67468251 | -1 | lncRNA | ENSG00000( |
| 67429591 | 67429754 | -1 | snRNA | ENSG00000( |
| 67439992 | 67440118 | 1 | rRNA | ENSG00000( |
| 67446267 | 67446574 | 1 | processed_pseudoge | ENSG00000( |
| 67468762 | 67545503 | -1 | protein_coding | ENSG00000( |
| 67558727 | 67607337 | 1 | protein_coding | ENSG00000( |
| 67607856 | 67610313 | -1 | TEC | ENSG00000( |
| 67612652 | 67701155 | -1 | protein_coding | ENSG00000( |
| 67638177 | 67638359 | 1 | processed_pseudoge | ENSG00000( |
| 67701209 | 68080952 | 1 | lncRNA | ENSG00000( |
| 67716375 | 67717388 | -1 | processed_pseudoge | ENSG00000( |
| 67718996 | 67724262 | -1 | processed_pseudoge | ENSG00000( |
| 67725183 | 67726395 | -1 | processed_pseudoge | ENSG00000( |
| 67739328 | 67754360 | -1 | protein_coding | ENSG00000( |
| 67747236 | 67747386 | 1 | snoRNA | ENSG00000( |
| 67766480 | 67773854 | -1 | unprocessed_pseudc | ENSG00000( |
| 67820876 | 67884002 | -1 | protein_coding | ENSG00000( |
| 67909385 | 67964140 | -1 | protein_coding | ENSG00000( |
| 67991684 | 68015768 | -1 | transcribed_unproce | ENSG00000( |
| 68003895 | 68004001 | 1 | snRNA | ENSG00000( |
| 68038544 | 68039647 | 1 | processed_pseudoge | ENSG00000( |
| 68049706 | 68050478 | -1 | unprocessed_pseudc | ENSG00000( |
| 68053198 | 68129869 | -1 | protein_coding | ENSG00000( |
| 68061822 | 68063269 | -1 | transcribed_processe | ENSG00000( |
| 68182292 | 68212470 | 1 | transcribed_processe | ENSG00000( |
| 68184081 | 68218080 | -1 | transcribed_unproce | ENSG00000( |
| 68226653 | 68245694 | -1 | protein_coding | ENSG00000( |
| 68282461 | 68285959 | -1 | unprocessed_pseudc | ENSG00000( |
| 68293127 | 68294364 | 1 | processed_pseudoge | ENSG00000( |
| 68300325 | 68302459 | 1 | processed_pseudoge | ENSG00000( |
| 68304343 | 68304939 | -1 | processed_pseudoge | ENSG00000( |
| 68310387 | 68350089 | -1 | protein_coding | ENSG00000( |
| 68376323 | 68376505 | 1 | processed_pseudoge | ENSG00000( |
| 68447463 | 68497604 | 1 | protein_coding | ENSG00000( |
| 68509441 | 68517647 | -1 | unprocessed_pseudc | ENSG00000( |
| 68537184 | 68568527 | -1 | protein_coding | ENSG00000( |
| 68614419 | 68615013 | 1 | processed_pseudoge | ENSG00000( |
| 68615393 | 68616137 | -1 | processed_pseudoge | ENSG00000( |
| 68626847 | 68628899 | -1 | unprocessed_pseudc | ENSG00000( |
| 68646597 | 68670652 | -1 | protein_coding | ENSG00000( |
| 68704600 | 68712498 | -1 | unprocessed_pseudc | ENSG00000( |
| 68784618 | 68787474 | 1 | unprocessed_pseudc | ENSG00000( |
| 68813995 | 68814727 | 1 | unprocessed_pseudc | ENSG00000( |
| 68815994 | 68832023 | 1 | protein_coding | ENSG00000( |
| 68834213 | 68834368 | -1 | unprocessed_pseudc | ENSG00000( |

| | | | | |
|---|---|---|---|---|
| 68861332 | 68861610 | -1 | unprocessed_pseudc | ENSG0000C |
| 68877626 | 68880131 | -1 | unprocessed_pseudc | ENSG0000C |
| 68883885 | 68884440 | -1 | processed_pseudoge | ENSG0000C |
| 68900651 | 68900910 | -1 | unprocessed_pseudc | ENSG0000C |
| 68901008 | 68906983 | -1 | lncRNA | ENSG0000C |
| 68907918 | 68908034 | -1 | unprocessed_pseudc | ENSG0000C |
| 68914928 | 68924741 | -1 | unprocessed_pseudc | ENSG0000C |
| 68928463 | 68951804 | -1 | protein_coding | ENSG0000C |
| 68972999 | 68975993 | -1 | unprocessed_pseudc | ENSG0000C |
| 68985009 | 68988533 | -1 | unprocessed_pseudc | ENSG0000C |
| 68996935 | 68997709 | -1 | processed_pseudoge | ENSG0000C |
| 69004862 | 69020414 | -1 | unprocessed_pseudc | ENSG0000C |
| 69021656 | 69022340 | -1 | unprocessed_pseudc | ENSG0000C |
| 69027831 | 69044578 | 1 | unprocessed_pseudc | ENSG0000C |
| 69051363 | 69112987 | 1 | protein_coding | ENSG0000C |
| 69066395 | 69069888 | 1 | unprocessed_pseudc | ENSG0000C |
| 69125274 | 69126451 | 1 | processed_pseudoge | ENSG0000C |
| 69131351 | 69134814 | 1 | unprocessed_pseudc | ENSG0000C |
| 69144734 | 69147671 | 1 | unprocessed_pseudc | ENSG0000C |
| 69181660 | 69182372 | 1 | transcribed_unproce | ENSG0000C |
| 69182100 | 69216766 | 1 | lncRNA | ENSG0000C |
| 69199951 | 69214748 | -1 | protein_coding | ENSG0000C |
| 69215908 | 69216635 | -1 | unprocessed_pseudc | ENSG0000C |
| 69242628 | 69250815 | 1 | processed_pseudoge | ENSG0000C |
| 69280475 | 69295050 | 1 | protein_coding | ENSG0000C |
| 69306469 | 69307173 | -1 | unprocessed_pseudc | ENSG0000C |
| 69346609 | 69358177 | 1 | processed_pseudoge | ENSG0000C |
| 69387580 | 69388307 | 1 | unprocessed_pseudc | ENSG0000C |
| 69389492 | 69402081 | 1 | processed_pseudoge | ENSG0000C |
| 69406931 | 69407655 | 1 | unprocessed_pseudc | ENSG0000C |
| 69408828 | 69423164 | 1 | processed_pseudoge | ENSG0000C |
| 69437357 | 69441050 | 1 | unprocessed_pseudc | ENSG0000C |
| 69450014 | 69451613 | 1 | unprocessed_pseudc | ENSG0000C |
| 69480165 | 69526014 | -1 | protein_coding | ENSG0000C |
| 69517667 | 69518400 | -1 | unprocessed_pseudc | ENSG0000C |
| 69572391 | 69585527 | -1 | unprocessed_pseudc | ENSG0000C |
| 69588417 | 69653249 | -1 | protein_coding | ENSG0000C |
| 69589309 | 69639642 | -1 | protein_coding | ENSG0000C |
| 69721167 | 69787961 | -1 | protein_coding | ENSG0000C |
| 69791872 | 69813888 | -1 | transcribed_unitary_ | ENSG0000C |
| 69810780 | 69847786 | -1 | protein_coding | ENSG0000C |
| 69841212 | 69860145 | -1 | protein_coding | ENSG0000C |
| 69888877 | 69889406 | 1 | processed_pseudoge | ENSG0000C |
| 69931068 | 69946574 | 1 | protein_coding | ENSG0000C |
| 69955256 | 69961007 | -1 | protein_coding | ENSG0000C |
| 69995931 | 70002570 | 1 | protein_coding | ENSG0000C |
| 70028413 | 70036538 | 1 | protein_coding | ENSG0000C |
| 70050438 | 70058848 | 1 | protein_coding | ENSG0000C |
| 70067386 | 70085290 | 1 | transcribed_unitary_ | ENSG0000C |
| 70133616 | 70176799 | 1 | protein_coding | ENSG0000C |

| | | | | |
|---|---|---|---|---|
| 70196496 | 70204576 | 1 | protein_coding | ENSG00000 |
| 70226124 | 70235252 | 1 | protein_coding | ENSG00000 |
| 70242588 | 70251428 | 1 | protein_coding | ENSG00000 |
| 70334981 | 70337116 | 1 | protein_coding | ENSG00000 |
| 70360760 | 70367158 | 1 | protein_coding | ENSG00000 |
| 70370093 | 70390244 | 1 | protein_coding | ENSG00000 |
| 70397931 | 70410195 | 1 | protein_coding | ENSG00000 |
| 70430492 | 70482997 | 1 | protein_coding | ENSG00000 |
| 70518569 | 70532743 | 1 | protein_coding | ENSG00000 |
| 70592256 | 70607288 | 1 | protein_coding | ENSG00000 |
| 70628744 | 70646824 | 1 | protein_coding | ENSG00000 |
| 70637745 | 70686816 | 1 | lncRNA | ENSG00000 |
| 70655541 | 70681817 | -1 | protein_coding | ENSG00000 |
| 70688532 | 70690551 | 1 | protein_coding | ENSG00000 |
| 70701755 | 70701858 | 1 | snRNA | ENSG00000 |
| 70703018 | 70703123 | -1 | snRNA | ENSG00000 |
| 70703747 | 70704491 | -1 | lncRNA | ENSG00000 |
| 70704204 | 70808619 | 1 | protein_coding | ENSG00000 |
| 70815783 | 70839897 | -1 | protein_coding | ENSG00000 |
| 70848136 | 70848218 | -1 | snRNA | ENSG00000 |
| 70852130 | 70852240 | 1 | snRNA | ENSG00000 |
| 70902326 | 71022449 | 1 | protein_coding | ENSG00000 |
| 70992538 | 71030914 | 1 | protein_coding | ENSG00000 |
| 71062667 | 71572087 | 1 | protein_coding | ENSG00000 |
| 71300258 | 71301235 | -1 | processed_pseudoge | ENSG00000 |
| 71434405 | 71437924 | -1 | processed_pseudoge | ENSG00000 |
| 71741696 | 71804041 | -1 | protein_coding | ENSG00000 |
| 71759518 | 71759616 | -1 | rRNA_pseudogene | ENSG00000 |
| 71821305 | 71823980 | -1 | lncRNA | ENSG00000 |
| 72031804 | 72148067 | 1 | protein_coding | ENSG00000 |
| 72280969 | 72569221 | -1 | protein_coding | ENSG00000 |
| 72323028 | 72330751 | 1 | lncRNA | ENSG00000 |
| 72807267 | 72808192 | 1 | processed_pseudoge | ENSG00000 |
| 72965178 | 72965303 | 1 | snRNA | ENSG00000 |
| 73026748 | 73026873 | 1 | snRNA | ENSG00000 |
| 73052362 | 73069755 | -1 | protein_coding | ENSG00000 |
| 73073376 | 73258798 | -1 | protein_coding | ENSG00000 |
| 73098822 | 73099145 | 1 | processed_pseudoge | ENSG00000 |
| 73259209 | 73317953 | 1 | lncRNA | ENSG00000 |
| 73263960 | 73264084 | 1 | snoRNA | ENSG00000 |
| 73337233 | 73337771 | -1 | processed_pseudoge | ENSG00000 |
| 73397114 | 73421482 | 1 | protein_coding | ENSG00000 |
| 73431138 | 73456174 | 1 | protein_coding | ENSG00000 |
| 73481745 | 73504001 | 1 | protein_coding | ENSG00000 |
| 73508803 | 73534128 | 1 | lncRNA | ENSG00000 |
| 73543822 | 73544013 | 1 | processed_pseudoge | ENSG00000 |
| 73571550 | 73620631 | -1 | protein_coding | ENSG00000 |
| 73710302 | 73714527 | 1 | lncRNA | ENSG00000 |
| 73740541 | 73743716 | 1 | protein_coding | ENSG00000 |
| 73777636 | 73778060 | -1 | processed_pseudoge | ENSG00000 |

| | | | | |
|---|---|---|---|---|
| 73836640 | 73849064 | 1 | protein_coding | ENSG00000 |
| 73847866 | 73848829 | 1 | transcribed_unproce | ENSG00000 |
| 73853296 | 73854483 | 1 | protein_coding | ENSG00000 |
| 73869393 | 73871308 | 1 | protein_coding | ENSG00000 |
| 73938604 | 73939558 | -1 | processed_pseudoge | ENSG00000 |
| 73944011 | 73944324 | -1 | processed_pseudoge | ENSG00000 |
| 73980811 | 73982027 | -1 | protein_coding | ENSG00000 |
| 73986439 | 73988190 | -1 | protein_coding | ENSG00000 |
| 73995642 | 73998677 | -1 | protein_coding | ENSG00000 |
| 73997933 | 74007682 | 1 | lncRNA | ENSG00000 |
| 74011578 | 74011876 | 1 | misc_RNA | ENSG00000 |
| 74036589 | 74038807 | -1 | protein_coding | ENSG00000 |
| 74054038 | 74055399 | -1 | transcribed_unproce | ENSG00000 |
| 74076533 | 74078973 | 1 | lncRNA | ENSG00000 |
| 74085995 | 74086672 | -1 | processed_pseudoge | ENSG00000 |
| 74097040 | 74099196 | -1 | protein_coding | ENSG00000 |
| 74114174 | 74303099 | 1 | protein_coding | ENSG00000 |
| 74156511 | 74158373 | -1 | lncRNA | ENSG00000 |
| 74308470 | 74316789 | 1 | protein_coding | ENSG00000 |
| 74365145 | 74388749 | 1 | protein_coding | ENSG00000 |
| 74418917 | 74440457 | -1 | lncRNA | ENSG00000 |
| 74445136 | 74455005 | 1 | protein_coding | ENSG00000 |
| 74552584 | 74589481 | -1 | lncRNA | ENSG00000 |
| 74744759 | 74794523 | -1 | protein_coding | ENSG00000 |
| 74881174 | 74882934 | 1 | lncRNA | ENSG00000 |
| 74917822 | 74918138 | 1 | processed_pseudoge | ENSG00000 |
| 74933095 | 75050115 | 1 | protein_coding | ENSG00000 |
| 74955974 | 74970362 | -1 | lncRNA | ENSG00000 |
| 74993877 | 75034824 | -1 | lncRNA | ENSG00000 |
| 75081702 | 75084717 | -1 | lncRNA | ENSG00000 |
| 75101477 | 75101943 | 1 | transcribed_processe | ENSG00000 |
| 75194867 | 75195079 | 1 | processed_pseudoge | ENSG00000 |
| 75269068 | 75361182 | -1 | lncRNA | ENSG00000 |
| 75341279 | 75359024 | -1 | lncRNA | ENSG00000 |
| 75354076 | 75362566 | 1 | lncRNA | ENSG00000 |
| 75361207 | 75434449 | -1 | lncRNA | ENSG00000 |
| 75401195 | 75423023 | 1 | lncRNA | ENSG00000 |
| 75479037 | 75514764 | -1 | protein_coding | ENSG00000 |
| 75513946 | 75550473 | 1 | protein_coding | ENSG00000 |
| 75556048 | 75565885 | 1 | protein_coding | ENSG00000 |
| 75576496 | 75630716 | -1 | protein_coding | ENSG00000 |
| 75642782 | 75724525 | -1 | protein_coding | ENSG00000 |
| 75662120 | 75662213 | -1 | misc_RNA | ENSG00000 |
| 75724593 | 75814286 | 1 | protein_coding | ENSG00000 |
| 75822966 | 75838389 | 1 | lncRNA | ENSG00000 |
| 75829454 | 75829589 | -1 | snRNA | ENSG00000 |
| 75859864 | 75902571 | -1 | protein_coding | ENSG00000 |
| 75913660 | 75941013 | -1 | protein_coding | ENSG00000 |
| 75940950 | 75990962 | -1 | protein_coding | ENSG00000 |
| 75980790 | 76005942 | 1 | lncRNA | ENSG00000 |

| | | | | |
|---|---|---|---|---|
| 76001275 | 76007509 | -1 | protein_coding | ENSG00000( |
| 76011184 | 76112802 | 1 | protein_coding | ENSG00000( |
| 76021118 | 76023497 | -1 | protein_coding | ENSG00000( |
| 76033682 | 76041415 | -1 | protein_coding | ENSG00000( |
| 76036750 | 76037064 | -1 | processed_pseudoge | ENSG00000( |
| 76114659 | 76148444 | -1 | protein_coding | ENSG00000( |
| 76148561 | 76200666 | 1 | lncRNA | ENSG00000( |
| 76158737 | 76234536 | -1 | protein_coding | ENSG00000( |
| 76214040 | 76283783 | 1 | protein_coding | ENSG00000( |
| 76240740 | 76312098 | -1 | lncRNA | ENSG00000( |
| 76251721 | 76311129 | 1 | protein_coding | ENSG00000( |
| 76306026 | 76311599 | 1 | protein_coding | ENSG00000( |
| 76312997 | 76421868 | -1 | protein_coding | ENSG00000( |
| 76344550 | 76345590 | 1 | processed_pseudoge | ENSG00000( |
| 76356610 | 76356715 | -1 | snRNA | ENSG00000( |
| 76401251 | 76401569 | 1 | processed_pseudoge | ENSG00000( |
| 76402076 | 76402146 | 1 | snoRNA | ENSG00000( |
| 76435229 | 76783253 | 1 | protein_coding | ENSG00000( |
| 76509284 | 76509719 | -1 | processed_pseudoge | ENSG00000( |
| 76532222 | 76532327 | -1 | snRNA | ENSG00000( |
| 76573568 | 76573632 | 1 | miRNA | ENSG00000( |
| 76575551 | 76575626 | 1 | miRNA | ENSG00000( |
| 76708853 | 76802426 | -1 | lncRNA | ENSG00000( |
| 76758554 | 76801964 | -1 | lncRNA | ENSG00000( |
| 76886029 | 76886370 | -1 | processed_pseudoge | ENSG00000( |
| 76891034 | 76891445 | 1 | processed_pseudoge | ENSG00000( |
| 76894152 | 76898144 | -1 | protein_coding | ENSG00000( |
| 76949703 | 77040384 | 1 | protein_coding | ENSG00000( |
| 76958860 | 76959169 | -1 | processed_pseudoge | ENSG00000( |
| 77047155 | 77075989 | -1 | protein_coding | ENSG00000( |
| 77082403 | 77083126 | -1 | processed_pseudoge | ENSG00000( |
| 77112495 | 77113458 | -1 | processed_pseudoge | ENSG00000( |
| 77150328 | 77150437 | -1 | snRNA | ENSG00000( |
| 77157207 | 77433388 | 1 | protein_coding | ENSG00000( |
| 77216416 | 77216878 | -1 | processed_pseudoge | ENSG00000( |
| 77311397 | 77312111 | -1 | processed_pseudoge | ENSG00000( |
| 77350370 | 77351235 | -1 | processed_pseudoge | ENSG00000( |
| 77394491 | 77494286 | -1 | lncRNA | ENSG00000( |
| 77511753 | 77611834 | 1 | protein_coding | ENSG00000( |
| 77702746 | 77702803 | -1 | snoRNA | ENSG00000( |
| 77713387 | 77819615 | -1 | protein_coding | ENSG00000( |
| 77862830 | 77952785 | 1 | protein_coding | ENSG00000( |
| 77864508 | 77864804 | -1 | processed_pseudoge | ENSG00000( |
| 77958000 | 77958780 | -1 | processed_pseudoge | ENSG00000( |
| 77963940 | 77964511 | 1 | processed_pseudoge | ENSG00000( |
| 77987860 | 77988813 | -1 | processed_pseudoge | ENSG00000( |
| 78003143 | 78003439 | 1 | processed_pseudoge | ENSG00000( |
| 78008512 | 78008688 | -1 | processed_pseudoge | ENSG00000( |
| 78057323 | 78544269 | 1 | protein_coding | ENSG00000( |
| 78180866 | 78182046 | -1 | processed_pseudoge | ENSG00000( |

| | | | | |
|---|---|---|---|---|
| 78379197 | 78379514 | 1 | processed_pseudoge | ENSG00000 |
| 78551747 | 78610451 | 1 | protein_coding | ENSG00000 |
| 78632273 | 78632385 | -1 | misc_RNA | ENSG00000 |
| 78638780 | 78683185 | 1 | lncRNA | ENSG00000 |
| 78648954 | 78649242 | -1 | processed_pseudoge | ENSG00000 |
| 78669690 | 78691671 | -1 | lncRNA | ENSG00000 |
| 78768499 | 78768750 | 1 | processed_pseudoge | ENSG00000 |
| 78773654 | 78775973 | -1 | lncRNA | ENSG00000 |
| 78776342 | 78916372 | 1 | protein_coding | ENSG00000 |
| 78820752 | 78820821 | 1 | miRNA | ENSG00000 |
| 78887127 | 78939438 | -1 | protein_coding | ENSG00000 |
| 78898855 | 78899154 | 1 | misc_RNA | ENSG00000 |
| 78939485 | 79309673 | 1 | lncRNA | ENSG00000 |
| 79225694 | 79326061 | -1 | protein_coding | ENSG00000 |
| 79406361 | 79408228 | -1 | protein_coding | ENSG00000 |
| 79491802 | 79623479 | 1 | lncRNA | ENSG00000 |
| 79587302 | 79588130 | -1 | transcribed_unproce | ENSG00000 |
| 79596542 | 79597173 | 1 | lncRNA | ENSG00000 |
| 79663761 | 79696837 | 1 | lncRNA | ENSG00000 |
| 79827471 | 79877770 | 1 | lncRNA | ENSG00000 |
| 79843102 | 79843228 | 1 | snoRNA | ENSG00000 |
| 79901146 | 80125454 | -1 | protein_coding | ENSG00000 |
| 80079532 | 80081122 | 1 | processed_pseudoge | ENSG00000 |
| 80161129 | 80162011 | 1 | processed_pseudoge | ENSG00000 |
| 80182637 | 80197410 | -1 | lncRNA | ENSG00000 |
| 80183879 | 80204329 | 1 | protein_coding | ENSG00000 |
| 80266599 | 80336680 | 1 | protein_coding | ENSG00000 |
| 80335720 | 80963756 | 1 | protein_coding | ENSG00000 |
| 80386178 | 80388716 | -1 | processed_pseudoge | ENSG00000 |
| 81030708 | 81057632 | 1 | protein_coding | ENSG00000 |
| 81087370 | 81215117 | -1 | protein_coding | ENSG00000 |
| 81164922 | 81193395 | 1 | lncRNA | ENSG00000 |
| 81334303 | 81334418 | 1 | snRNA | ENSG00000 |
| 81426393 | 82044244 | -1 | protein_coding | ENSG00000 |
| 81733385 | 81734207 | -1 | processed_pseudoge | ENSG00000 |
| 81919995 | 81920387 | -1 | processed_pseudoge | ENSG00000 |
| 81928313 | 81928439 | 1 | snoRNA | ENSG00000 |
| 82010665 | 82013095 | -1 | processed_pseudoge | ENSG00000 |
| 82128535 | 82129656 | 1 | processed_pseudoge | ENSG00000 |
| 82174547 | 82174653 | 1 | snRNA | ENSG00000 |
| 82275071 | 82276600 | 1 | processed_pseudoge | ENSG00000 |
| 82284971 | 82340013 | 1 | transcribed_unitary_ | ENSG00000 |
| 82344876 | 82346842 | -1 | lncRNA | ENSG00000 |
| 82352498 | 82374503 | -1 | protein_coding | ENSG00000 |
| 82374301 | 82384027 | 1 | lncRNA | ENSG00000 |
| 82401578 | 82402456 | -1 | processed_pseudoge | ENSG00000 |
| 82402638 | 82402706 | -1 | snoRNA | ENSG00000 |
| 82422564 | 82430408 | -1 | protein_coding | ENSG00000 |
| 82430590 | 82461177 | 1 | protein_coding | ENSG00000 |
| 82483170 | 82562357 | -1 | protein_coding | ENSG00000 |

| | | | | |
|---|---|---|---|---|
| 82490823 | 82491635 | -1 | processed_pseudoge | ENSG00000 |
| 82494786 | 82496278 | -1 | transcribed_processe | ENSG00000 |
| 82571137 | 82571814 | 1 | processed_pseudoge | ENSG00000 |
| 82580117 | 82581384 | -1 | processed_pseudoge | ENSG00000 |
| 82610974 | 82621610 | -1 | lncRNA | ENSG00000 |
| 82612184 | 82615575 | 1 | lncRNA | ENSG00000 |
| 82629539 | 82798796 | -1 | protein_coding | ENSG00000 |
| 82691737 | 82692468 | 1 | processed_pseudoge | ENSG00000 |
| 82753337 | 82753430 | -1 | miRNA | ENSG00000 |
| 82775461 | 82776101 | 1 | processed_pseudoge | ENSG00000 |
| 82818509 | 82901166 | -1 | protein_coding | ENSG00000 |
| 82893009 | 82900960 | -1 | lncRNA | ENSG00000 |
| 82900684 | 82919969 | 1 | protein_coding | ENSG00000 |
| 82909973 | 83012926 | -1 | protein_coding | ENSG00000 |
| 82944738 | 82944845 | -1 | misc_RNA | ENSG00000 |
| 83003412 | 83003517 | 1 | snRNA | ENSG00000 |
| 83034447 | 83075818 | 1 | protein_coding | ENSG00000 |
| 83075957 | 83085600 | 1 | lncRNA | ENSG00000 |
| 83090048 | 83137075 | -1 | protein_coding | ENSG00000 |
| 83233512 | 83247213 | -1 | lncRNA | ENSG00000 |
| 83237303 | 83237444 | 1 | processed_pseudoge | ENSG00000 |
| 83247179 | 83248206 | 1 | processed_pseudoge | ENSG00000 |
| 83261536 | 83284914 | -1 | protein_coding | ENSG00000 |
| 83292461 | 83335153 | -1 | protein_coding | ENSG00000 |
| 83369416 | 83369523 | -1 | processed_pseudoge | ENSG00000 |
| 83377363 | 83378076 | -1 | processed_pseudoge | ENSG00000 |
| 83407343 | 83455855 | -1 | protein_coding | ENSG00000 |
| 83455932 | 83469735 | 1 | protein_coding | ENSG00000 |
| 83459517 | 83523348 | -1 | protein_coding | ENSG00000 |
| 83477524 | 83478424 | 1 | processed_pseudoge | ENSG00000 |
| 83502699 | 83503034 | 1 | processed_pseudoge | ENSG00000 |
| 83535914 | 83605875 | 1 | protein_coding | ENSG00000 |
| 83636196 | 83636290 | -1 | misc_RNA | ENSG00000 |
| 83668510 | 83731755 | -1 | lncRNA | ENSG00000 |
| 83796436 | 84299189 | -1 | lncRNA | ENSG00000 |
| 84233657 | 84233763 | 1 | snRNA | ENSG00000 |
| 84244003 | 84245488 | 1 | processed_pseudoge | ENSG00000 |
| 84371393 | 84380189 | -1 | lncRNA | ENSG00000 |
| 84491987 | 84498450 | -1 | protein_coding | ENSG00000 |
| 84544304 | 84544519 | -1 | processed_pseudoge | ENSG00000 |
| 84583127 | 84651334 | 1 | protein_coding | ENSG00000 |
| 84669610 | 84966391 | -1 | protein_coding | ENSG00000 |
| 84687725 | 84688000 | -1 | misc_RNA | ENSG00000 |
| 84796614 | 84810403 | 1 | lncRNA | ENSG00000 |
| 84886386 | 84886486 | -1 | snRNA | ENSG00000 |
| 84965534 | 85011277 | 1 | lncRNA | ENSG00000 |
| 84970180 | 84970730 | 1 | lncRNA | ENSG00000 |
| 85100496 | 85100823 | 1 | misc_RNA | ENSG00000 |
| 85246157 | 85246912 | 1 | processed_pseudoge | ENSG00000 |
| 85475150 | 86002668 | 1 | protein_coding | ENSG00000 |

| | | | | |
|---|---|---|---|---|
| 85722468 | 85722533 | 1 | miRNA | ENSG00000 |
| 85990007 | 86594625 | -1 | protein_coding | ENSG00000 |
| 86117912 | 86219926 | 1 | lncRNA | ENSG00000 |
| 86336318 | 86336614 | -1 | misc_RNA | ENSG00000 |
| 86542482 | 86542552 | -1 | miRNA | ENSG00000 |
| 86594315 | 86815171 | 1 | protein_coding | ENSG00000 |
| 86823468 | 86849384 | -1 | protein_coding | ENSG00000 |
| 86870191 | 86871404 | 1 | processed_pseudoge | ENSG00000 |
| 86876205 | 86892422 | -1 | protein_coding | ENSG00000 |
| 86876338 | 86876652 | 1 | processed_pseudoge | ENSG00000 |
| 86886472 | 86936178 | -1 | protein_coding | ENSG00000 |
| 86913266 | 86914817 | -1 | processed_pseudoge | ENSG00000 |
| 86924630 | 86936202 | -1 | lncRNA | ENSG00000 |
| 86935002 | 87141054 | 1 | protein_coding | ENSG00000 |
| 86949669 | 86950113 | -1 | processed_pseudoge | ENSG00000 |
| 87160103 | 87240314 | -1 | protein_coding | ENSG00000 |
| 87166844 | 87167734 | 1 | processed_pseudoge | ENSG00000 |
| 87207092 | 87208086 | 1 | processed_pseudoge | ENSG00000 |
| 87261931 | 87266822 | -1 | lncRNA | ENSG00000 |
| 87300495 | 87300583 | -1 | miRNA | ENSG00000 |
| 87303789 | 87322886 | -1 | protein_coding | ENSG00000 |
| 87317170 | 87345210 | -1 | lncRNA | ENSG00000 |
| 87336515 | 87391188 | -1 | protein_coding | ENSG00000 |
| 87386883 | 87387174 | -1 | misc_RNA | ENSG00000 |
| 87410644 | 87411388 | 1 | processed_pseudoge | ENSG00000 |
| 87422573 | 87459455 | 1 | protein_coding | ENSG00000 |
| 87460807 | 87462280 | -1 | lncRNA | ENSG00000 |
| 87473335 | 87531061 | -1 | protein_coding | ENSG00000 |
| 87568035 | 87733956 | -1 | lncRNA | ENSG00000 |
| 87608529 | 87616873 | 1 | protein_coding | ENSG00000 |
| 87650280 | 87664361 | 1 | protein_coding | ENSG00000 |
| 87743952 | 87744053 | -1 | misc_RNA | ENSG00000 |
| 87785920 | 87786371 | -1 | processed_pseudoge | ENSG00000 |
| 87799554 | 87812435 | 1 | protein_coding | ENSG00000 |
| 87821411 | 87846817 | 1 | protein_coding | ENSG00000 |
| 87891843 | 87894015 | 1 | processed_pseudoge | ENSG00000 |
| 87974385 | 88008477 | -1 | lncRNA | ENSG00000 |
| 87975650 | 87983426 | 1 | protein_coding | ENSG00000 |
| 88000237 | 88000401 | 1 | snRNA | ENSG00000 |
| 88007635 | 88077777 | 1 | protein_coding | ENSG00000 |
| 88090150 | 88231628 | -1 | protein_coding | ENSG00000 |
| 88163579 | 88163967 | 1 | processed_pseudoge | ENSG00000 |
| 88201703 | 88201810 | 1 | snRNA | ENSG00000 |
| 88206427 | 88206552 | -1 | snRNA | ENSG00000 |
| 88220569 | 88223837 | -1 | processed_pseudoge | ENSG00000 |
| 88226729 | 88226829 | 1 | snRNA | ENSG00000 |
| 88257620 | 88284769 | -1 | protein_coding | ENSG00000 |
| 88275205 | 88275308 | -1 | snRNA | ENSG00000 |
| 88284507 | 88331421 | 1 | lncRNA | ENSG00000 |
| 88330176 | 88330278 | -1 | misc_RNA | ENSG00000 |

| | | | | |
|---|---|---|---|---|
| 88352222 | 88358423 | -1 | lncRNA | ENSG00000( |
| 88358945 | 88361304 | -1 | lncRNA | ENSG00000( |
| 88378739 | 88443111 | 1 | protein_coding | ENSG00000( |
| 88457119 | 88506163 | 1 | protein_coding | ENSG00000( |
| 88508591 | 88510403 | -1 | processed_pseudoge | ENSG00000( |
| 88520998 | 88523776 | -1 | protein_coding | ENSG00000( |
| 88521573 | 88521789 | -1 | protein_coding | ENSG00000( |
| 88523810 | 88708450 | 1 | protein_coding | ENSG00000( |
| 88523826 | 88524983 | 1 | lncRNA | ENSG00000( |
| 88527160 | 88528170 | 1 | processed_pseudoge | ENSG00000( |
| 88583666 | 88583972 | -1 | misc_RNA | ENSG00000( |
| 88592434 | 88708541 | 1 | protein_coding | ENSG00000( |
| 88684848 | 88684954 | -1 | snRNA | ENSG00000( |
| 88695913 | 88697829 | -1 | protein_coding | ENSG00000( |
| 88709298 | 88730103 | 1 | lncRNA | ENSG00000( |
| 88710147 | 88710265 | 1 | misc_RNA | ENSG00000( |
| 88723209 | 88723316 | 1 | misc_RNA | ENSG00000( |
| 88725955 | 89111398 | -1 | protein_coding | ENSG00000( |
| 89111500 | 89114899 | 1 | protein_coding | ENSG00000( |
| 89119284 | 89119871 | 1 | lncRNA | ENSG00000( |
| 89130852 | 89130953 | 1 | snRNA | ENSG00000( |
| 89236383 | 89307800 | -1 | protein_coding | ENSG00000( |
| 89410960 | 89726752 | 1 | lncRNA | ENSG00000( |
| 89724099 | 89838315 | -1 | protein_coding | ENSG00000( |
| 89743792 | 89744305 | 1 | lncRNA | ENSG00000( |
| 89747802 | 89755853 | 1 | lncRNA | ENSG00000( |
| 89748283 | 89749154 | 1 | lncRNA | ENSG00000( |
| 89836408 | 89841978 | 1 | lncRNA | ENSG00000( |
| 89879532 | 89954629 | 1 | protein_coding | ENSG00000( |
| 89995223 | 89999409 | 1 | lncRNA | ENSG00000( |
| 90127535 | 91601913 | 1 | protein_coding | ENSG00000( |
| 90370123 | 90370427 | 1 | misc_RNA | ENSG00000( |
| 90682996 | 90683123 | 1 | processed_pseudoge | ENSG00000( |
| 90838903 | 90839037 | -1 | processed_pseudoge | ENSG00000( |
| 91108023 | 91112790 | -1 | lncRNA | ENSG00000( |
| 91319034 | 91325306 | -1 | lncRNA | ENSG00000( |
| 91603275 | 91605292 | 1 | TEC | ENSG00000( |
| 91885046 | 91885254 | -1 | processed_pseudoge | ENSG00000( |
| 91887886 | 91904335 | 1 | lncRNA | ENSG00000( |
| 92182477 | 92184052 | -1 | processed_pseudoge | ENSG00000( |
| 92183235 | 92265316 | 1 | lncRNA | ENSG00000( |
| 92260367 | 92260751 | -1 | TEC | ENSG00000( |
| 92268767 | 92277075 | -1 | lncRNA | ENSG00000( |
| 92297251 | 92304178 | -1 | lncRNA | ENSG00000( |
| 92303966 | 93810157 | 1 | protein_coding | ENSG00000( |
| 92702345 | 92702659 | 1 | processed_pseudoge | ENSG00000( |
| 92821986 | 92822918 | -1 | processed_pseudoge | ENSG00000( |
| 92833685 | 92838842 | -1 | lncRNA | ENSG00000( |
| 92884663 | 92885107 | 1 | processed_pseudoge | ENSG00000( |
| 93318623 | 93319786 | -1 | lncRNA | ENSG00000( |

| | | | | |
|---|---|---|---|---|
| 93820171 | 93820280 | 1 | rRNA_pseudogene | ENSG0000C |
| 93828753 | 93830966 | 1 | protein_coding | ENSG0000C |
| 94117792 | 94207556 | -1 | lncRNA | ENSG0000C |
| 94195940 | 94196513 | -1 | processed_pseudoge | ENSG0000C |
| 94207611 | 94291292 | 1 | protein_coding | ENSG0000C |
| 94298535 | 94342876 | -1 | protein_coding | ENSG0000C |
| 94315486 | 94343041 | 1 | lncRNA | ENSG0000C |
| 94369833 | 94370163 | 1 | processed_pseudoge | ENSG0000C |
| 94451857 | 94668227 | 1 | protein_coding | ENSG0000C |
| 94675245 | 94702570 | 1 | lncRNA | ENSG0000C |
| 94743668 | 94757681 | -1 | lncRNA | ENSG0000C |
| 94757955 | 95158448 | 1 | protein_coding | ENSG0000C |
| 95162504 | 95549206 | -1 | protein_coding | ENSG0000C |
| 95549129 | 95552457 | 1 | lncRNA | ENSG0000C |
| 95644952 | 95645293 | -1 | processed_pseudoge | ENSG0000C |
| 95840093 | 95841464 | 1 | protein_coding | ENSG0000C |
| 95868667 | 95868764 | -1 | snRNA | ENSG0000C |
| 96152297 | 96152403 | -1 | snRNA | ENSG0000C |
| 96212279 | 96213100 | 1 | processed_pseudoge | ENSG0000C |
| 96310701 | 96818864 | 1 | lncRNA | ENSG0000C |
| 96348734 | 96349038 | -1 | misc_RNA | ENSG0000C |
| 96841995 | 96857900 | 1 | lncRNA | ENSG0000C |
| 96902801 | 96903050 | 1 | processed_pseudoge | ENSG0000C |
| 97120701 | 97135059 | 1 | lncRNA | ENSG0000C |
| 97184093 | 98143476 | -1 | protein_coding | ENSG0000C |
| 97334635 | 97633823 | -1 | lncRNA | ENSG0000C |
| 97366681 | 97491899 | 1 | lncRNA | ENSG0000C |
| 97916353 | 97917401 | -1 | processed_pseudoge | ENSG0000C |
| 98025390 | 98026240 | 1 | processed_pseudoge | ENSG0000C |
| 98048555 | 98049043 | 1 | processed_pseudoge | ENSG0000C |
| 98251688 | 98261630 | -1 | lncRNA | ENSG0000C |
| 98261384 | 98443861 | 1 | protein_coding | ENSG0000C |
| 98409290 | 98409763 | 1 | processed_pseudoge | ENSG0000C |
| 98470367 | 98658611 | -1 | protein_coding | ENSG0000C |
| 98496364 | 98509935 | 1 | lncRNA | ENSG0000C |
| 98658904 | 98664550 | 1 | lncRNA | ENSG0000C |
| 98713804 | 98713971 | -1 | processed_pseudoge | ENSG0000C |
| 98740742 | 98741231 | 1 | processed_pseudoge | ENSG0000C |
| 98871684 | 98930637 | -1 | protein_coding | ENSG0000C |
| 98909537 | 98909848 | -1 | transcribed_processe | ENSG0000C |
| 98928897 | 98994994 | 1 | lncRNA | ENSG0000C |
| 98955982 | 98956767 | 1 | processed_pseudoge | ENSG0000C |
| 98961083 | 98964790 | 1 | lncRNA | ENSG0000C |
| 98966815 | 98966873 | -1 | snRNA | ENSG0000C |
| 98976800 | 98977117 | -1 | processed_pseudoge | ENSG0000C |
| 98995659 | 99062809 | 1 | protein_coding | ENSG0000C |
| 98997387 | 98997460 | 1 | miRNA | ENSG0000C |
| 99022311 | 99023131 | -1 | processed_pseudoge | ENSG0000C |
| 99067256 | 99068125 | -1 | lncRNA | ENSG0000C |
| 99070978 | 99088801 | -1 | protein_coding | ENSG0000C |

| | | | | |
|---|---|---|---|---|
| 99088805 | 99301356 | 1 | lncRNA | ENSG0000C |
| 99123657 | 99157792 | -1 | protein_coding | ENSG0000C |
| 99160514 | 99161645 | -1 | transcribed_processe | ENSG0000C |
| 99202638 | 99219537 | -1 | protein_coding | ENSG0000C |
| 99276369 | 99291003 | -1 | protein_coding | ENSG0000C |
| 99304971 | 99352760 | -1 | protein_coding | ENSG0000C |
| 99336497 | 99352746 | -1 | protein_coding | ENSG0000C |
| 99412261 | 99435737 | -1 | protein_coding | ENSG0000C |
| 99469598 | 99498698 | -1 | unprocessed_pseudc | ENSG0000C |
| 99511012 | 99542303 | 1 | protein_coding | ENSG0000C |
| 99546709 | 99564039 | -1 | protein_coding | ENSG0000C |
| 99563761 | 99623999 | 1 | protein_coding | ENSG0000C |
| 99594799 | 99625913 | -1 | lncRNA | ENSG0000C |
| 99636529 | 99654648 | -1 | protein_coding | ENSG0000C |
| 99816827 | 99870190 | 1 | protein_coding | ENSG0000C |
| 99844138 | 99844696 | 1 | TEC | ENSG0000C |
| 99878336 | 99894428 | -1 | protein_coding | ENSG0000C |
| 99896248 | 99946618 | -1 | protein_coding | ENSG0000C |
| 99942081 | 99942522 | -1 | TEC | ENSG0000C |
| 99948086 | 99950355 | -1 | protein_coding | ENSG0000C |
| 99950006 | 100195099 | 1 | lncRNA | ENSG0000C |
| 99990737 | 99991477 | -1 | processed_pseudoge | ENSG0000C |
| 100041841 | 100042104 | -1 | processed_pseudoge | ENSG0000C |
| 100185870 | 100190782 | -1 | protein_coding | ENSG0000C |
| 100190033 | 100215505 | 1 | lncRNA | ENSG0000C |
| 100395341 | 100880126 | -1 | protein_coding | ENSG0000C |
| 100421655 | 100474461 | 1 | lncRNA | ENSG0000C |
| 100660279 | 100675113 | -1 | lncRNA | ENSG0000C |
| 100778582 | 100791426 | -1 | lncRNA | ENSG0000C |
| 100812255 | 100814280 | -1 | lncRNA | ENSG0000C |
| 101023409 | 101348278 | -1 | protein_coding | ENSG0000C |
| 101240795 | 101240872 | -1 | miRNA | ENSG0000C |
| 101330302 | 101330414 | -1 | miRNA | ENSG0000C |
| 101347780 | 101348883 | 1 | lncRNA | ENSG0000C |
| 101411286 | 102074812 | 1 | protein_coding | ENSG0000C |
| 101640946 | 101649974 | -1 | lncRNA | ENSG0000C |
| 101723876 | 101723980 | 1 | snRNA | ENSG0000C |
| 101972423 | 101972798 | 1 | processed_pseudoge | ENSG0000C |
| 101976894 | 102036903 | -1 | lncRNA | ENSG0000C |
| 102251041 | 102431258 | -1 | protein_coding | ENSG0000C |
| 102348392 | 102348691 | -1 | misc_RNA | ENSG0000C |
| 102418602 | 102450010 | -1 | lncRNA | ENSG0000C |
| 102461250 | 102462152 | 1 | processed_pseudoge | ENSG0000C |
| 102500841 | 102501319 | -1 | lncRNA | ENSG0000C |
| 102501331 | 102617302 | 1 | protein_coding | ENSG0000C |
| 102630770 | 102760994 | -1 | protein_coding | ENSG0000C |
| 102662611 | 102663073 | 1 | processed_pseudoge | ENSG0000C |
| 102727274 | 102730721 | -1 | processed_pseudoge | ENSG0000C |
| 102728746 | 102730171 | -1 | processed_pseudoge | ENSG0000C |
| 102734358 | 102734612 | 1 | processed_pseudoge | ENSG0000C |

| | | | | |
|---|---|---|---|---|
| 102751401 | 102752641 | 1 | processed_pseudoge | ENSG00000( |
| 102794383 | 102868896 | -1 | protein_coding | ENSG00000( |
| 102814252 | 102819881 | 1 | lncRNA | ENSG00000( |
| 102828055 | 102844075 | 1 | lncRNA | ENSG00000( |
| 102837047 | 102837099 | -1 | snRNA | ENSG00000( |
| 102868974 | 102892807 | 1 | protein_coding | ENSG00000( |
| 102885048 | 103019719 | -1 | protein_coding | ENSG00000( |
| 102896725 | 102898237 | 1 | processed_pseudoge | ENSG00000( |
| 102961956 | 102963327 | 1 | processed_pseudoge | ENSG00000( |
| 103019868 | 103085829 | -1 | protein_coding | ENSG00000( |
| 103077592 | 103099870 | -1 | protein_coding | ENSG00000( |
| 103105349 | 103198445 | -1 | protein_coding | ENSG00000( |
| 103255822 | 103454667 | 1 | lncRNA | ENSG00000( |
| 103548745 | 103624534 | 1 | lncRNA | ENSG00000( |
| 103550586 | 103559277 | -1 | lncRNA | ENSG00000( |
| 103572089 | 103574082 | 1 | processed_pseudoge | ENSG00000( |
| 103586031 | 103719985 | -1 | protein_coding | ENSG00000( |
| 103871890 | 103878436 | 1 | lncRNA | ENSG00000( |
| 103924540 | 103924641 | 1 | snRNA | ENSG00000( |
| 103961616 | 104037543 | 1 | lncRNA | ENSG00000( |
| 104230380 | 104411035 | 1 | lncRNA | ENSG00000( |
| 104468308 | 104494901 | -1 | protein_coding | ENSG00000( |
| 104490849 | 104734546 | 1 | lncRNA | ENSG00000( |
| 104556960 | 104568877 | 1 | lncRNA | ENSG00000( |
| 104653874 | 104966793 | -1 | lncRNA | ENSG00000( |
| 104886118 | 104886955 | -1 | processed_pseudoge | ENSG00000( |
| 104900125 | 105120000 | 1 | lncRNA | ENSG00000( |
| 104974672 | 104974771 | 1 | snRNA | ENSG00000( |
| 104996900 | 104997052 | -1 | processed_pseudoge | ENSG00000( |
| 105102891 | 105104004 | -1 | processed_pseudoge | ENSG00000( |
| 105105987 | 105106094 | 1 | snoRNA | ENSG00000( |
| 105137280 | 105140619 | -1 | lncRNA | ENSG00000( |
| 105145875 | 105279816 | 1 | protein_coding | ENSG00000( |
| 105171354 | 105178063 | -1 | lncRNA | ENSG00000( |
| 105293664 | 105293946 | 1 | misc_RNA | ENSG00000( |
| 105369077 | 105474067 | -1 | protein_coding | ENSG00000( |
| 105406997 | 105407092 | 1 | snRNA | ENSG00000( |
| 105484698 | 105486080 | 1 | processed_pseudoge | ENSG00000( |
| 105526596 | 105527397 | -1 | processed_pseudoge | ENSG00000( |
| 105532475 | 105532630 | 1 | processed_pseudoge | ENSG00000( |
| 105540190 | 105552355 | -1 | lncRNA | ENSG00000( |
| 105552620 | 105708093 | 1 | protein_coding | ENSG00000( |
| 105561591 | 105570238 | 1 | lncRNA | ENSG00000( |
| 105679050 | 105680094 | -1 | processed_pseudoge | ENSG00000( |
| 105682627 | 105895986 | -1 | protein_coding | ENSG00000( |
| 105708778 | 105847728 | 1 | protein_coding | ENSG00000( |
| 105746245 | 105827172 | -1 | lncRNA | ENSG00000( |
| 105894775 | 106004027 | 1 | protein_coding | ENSG00000( |
| 105927060 | 105932722 | -1 | lncRNA | ENSG00000( |
| 106003317 | 106022478 | -1 | lncRNA | ENSG00000( |

| | | | | |
|---|---|---|---|---|
| 106041599 | 106321495 | -1 | protein_coding | ENSG00000 |
| 106315544 | 106349456 | 1 | protein_coding | ENSG00000 |
| 106357485 | 106368825 | -1 | protein_coding | ENSG00000 |
| 106433489 | 106453449 | 1 | lncRNA | ENSG00000 |
| 106525401 | 106938640 | 1 | lncRNA | ENSG00000 |
| 106836498 | 106837601 | -1 | processed_pseudoge | ENSG00000 |
| 106921802 | 107283806 | -1 | protein_coding | ENSG00000 |
| 107203349 | 107203924 | -1 | processed_pseudoge | ENSG00000 |
| 107258700 | 107301870 | 1 | lncRNA | ENSG00000 |
| 107435118 | 107435220 | 1 | snRNA | ENSG00000 |
| 107590276 | 107720234 | -1 | protein_coding | ENSG00000 |
| 107824563 | 107915047 | 1 | protein_coding | ENSG00000 |
| 107863473 | 107989679 | -1 | lncRNA | ENSG00000 |
| 107867807 | 107867910 | -1 | snRNA | ENSG00000 |
| 107931549 | 107953461 | 1 | protein_coding | ENSG00000 |
| 107936031 | 107941255 | -1 | lncRNA | ENSG00000 |
| 107989714 | 108035175 | 1 | protein_coding | ENSG00000 |
| 108047545 | 108168956 | -1 | protein_coding | ENSG00000 |
| 108167525 | 108258037 | 1 | lncRNA | ENSG00000 |
| 108407843 | 108408578 | -1 | processed_pseudoge | ENSG00000 |
| 108415220 | 108416639 | 1 | unprocessed_pseudc | ENSG00000 |
| 108417705 | 108419763 | -1 | processed_pseudoge | ENSG00000 |
| 108538190 | 108620460 | -1 | lncRNA | ENSG00000 |
| 108620566 | 108630412 | 1 | protein_coding | ENSG00000 |
| 108650585 | 108667820 | 1 | protein_coding | ENSG00000 |
| 108652150 | 108652256 | 1 | snRNA | ENSG00000 |
| 108669949 | 108678874 | 1 | lncRNA | ENSG00000 |
| 108742048 | 108763054 | -1 | protein_coding | ENSG00000 |
| 108773613 | 108774020 | 1 | processed_pseudoge | ENSG00000 |
| 108788745 | 108789779 | -1 | processed_pseudoge | ENSG00000 |
| 108808725 | 109302752 | -1 | protein_coding | ENSG00000 |
| 109303035 | 109316135 | 1 | lncRNA | ENSG00000 |
| 109346326 | 109347459 | -1 | processed_pseudoge | ENSG00000 |
| 109347475 | 109433817 | -1 | lncRNA | ENSG00000 |
| 109433772 | 109540896 | 1 | protein_coding | ENSG00000 |
| 109450772 | 109451068 | -1 | misc_RNA | ENSG00000 |
| 109488698 | 109488795 | 1 | miRNA | ENSG00000 |
| 109553243 | 109554094 | -1 | processed_pseudoge | ENSG00000 |
| 109555170 | 109555732 | 1 | processed_pseudoge | ENSG00000 |
| 109560209 | 109688719 | 1 | protein_coding | ENSG00000 |
| 109673843 | 109674124 | 1 | processed_pseudoge | ENSG00000 |
| 109688622 | 109703583 | -1 | protein_coding | ENSG00000 |
| 109692004 | 109692703 | 1 | lncRNA | ENSG00000 |
| 109709989 | 109730070 | -1 | protein_coding | ENSG00000 |
| 109713916 | 109801999 | -1 | protein_coding | ENSG00000 |
| 109740694 | 109802179 | -1 | protein_coding | ENSG00000 |
| 109815047 | 109815410 | -1 | lncRNA | ENSG00000 |
| 109815510 | 109824740 | 1 | protein_coding | ENSG00000 |
| 109827972 | 109849123 | 1 | protein_coding | ENSG00000 |
| 109848107 | 109872315 | 1 | protein_coding | ENSG00000 |

| | | | | |
|---|---|---|---|---|
| 109879070 | 109879897 | -1 | processed_pseudoge | ENSG00000 |
| 109912883 | 110013766 | 1 | protein_coding | ENSG00000 |
| 109992325 | 109992431 | 1 | snRNA | ENSG00000 |
| 110045846 | 110199199 | -1 | protein_coding | ENSG00000 |
| 110117736 | 110118070 | 1 | misc_RNA | ENSG00000 |
| 110146374 | 110147248 | 1 | processed_pseudoge | ENSG00000 |
| 110251871 | 110252099 | -1 | processed_pseudoge | ENSG00000 |
| 110264865 | 110265017 | -1 | processed_pseudoge | ENSG00000 |
| 110278185 | 110278291 | 1 | snRNA | ENSG00000 |
| 110291644 | 110293573 | 1 | processed_pseudoge | ENSG00000 |
| 110365733 | 110565285 | 1 | protein_coding | ENSG00000 |
| 110399773 | 110400511 | -1 | processed_pseudoge | ENSG00000 |
| 110415898 | 110418504 | 1 | processed_pseudoge | ENSG00000 |
| 110433109 | 110433188 | -1 | snoRNA | ENSG00000 |
| 110512488 | 110519505 | -1 | lncRNA | ENSG00000 |
| 110595513 | 110615458 | -1 | lncRNA | ENSG00000 |
| 110617423 | 110642123 | -1 | protein_coding | ENSG00000 |
| 110794403 | 110797344 | 1 | lncRNA | ENSG00000 |
| 110860582 | 110860647 | -1 | miRNA | ENSG00000 |
| 110945109 | 110945799 | -1 | processed_pseudoge | ENSG00000 |
| 111063583 | 111064227 | 1 | processed_pseudoge | ENSG00000 |
| 111331412 | 111331518 | 1 | snRNA | ENSG00000 |
| 111551854 | 111552173 | 1 | processed_pseudoge | ENSG00000 |
| 111632172 | 111648962 | 1 | lncRNA | ENSG00000 |
| 111802801 | 111839886 | 1 | lncRNA | ENSG00000 |
| 111804418 | 111809694 | -1 | lncRNA | ENSG00000 |
| 111826881 | 112072698 | -1 | lncRNA | ENSG00000 |
| 112068010 | 112068526 | 1 | processed_pseudoge | ENSG00000 |
| 112077598 | 112078400 | 1 | processed_pseudoge | ENSG00000 |
| 112116510 | 112116903 | -1 | processed_pseudoge | ENSG00000 |
| 112145454 | 112195256 | 1 | protein_coding | ENSG00000 |
| 112171215 | 112171527 | 1 | processed_pseudoge | ENSG00000 |
| 112229561 | 112231596 | -1 | lncRNA | ENSG00000 |
| 112231740 | 112273110 | 1 | protein_coding | ENSG00000 |
| 112272968 | 112273316 | -1 | lncRNA | ENSG00000 |
| 112274537 | 112285904 | -1 | protein_coding | ENSG00000 |
| 112285509 | 112442621 | 1 | protein_coding | ENSG00000 |
| 112356135 | 112370959 | 1 | transcribed_processe | ENSG00000 |
| 112455924 | 112457785 | 1 | processed_pseudoge | ENSG00000 |
| 112513516 | 112516180 | -1 | protein_coding | ENSG00000 |
| 112515385 | 112546881 | 1 | lncRNA | ENSG00000 |
| 112530371 | 112530672 | -1 | processed_pseudoge | ENSG00000 |
| 112539333 | 112636995 | -1 | protein_coding | ENSG00000 |
| 112564988 | 112565398 | 1 | processed_pseudoge | ENSG00000 |
| 112637107 | 112657592 | 1 | protein_coding | ENSG00000 |
| 112646476 | 112650051 | -1 | lncRNA | ENSG00000 |
| 112647874 | 112647941 | -1 | miRNA | ENSG00000 |
| 112648004 | 112648071 | -1 | miRNA | ENSG00000 |
| 112648183 | 112648251 | -1 | miRNA | ENSG00000 |
| 112648363 | 112648430 | -1 | miRNA | ENSG00000 |

| | | | | |
|---|---|---|---|---|
| 112648485 | 112648557 | -1 | miRNA | ENSG00000 |
| 112685055 | 112685503 | 1 | processed_pseudoge | ENSG00000 |
| 112693047 | 112706810 | -1 | lncRNA | ENSG00000 |
| 112693459 | 112693792 | -1 | transcribed_processe | ENSG00000 |
| 112738672 | 112739106 | 1 | processed_pseudoge | ENSG00000 |
| 112755816 | 112755917 | -1 | misc_RNA | ENSG00000 |
| 112788083 | 112788878 | -1 | processed_pseudoge | ENSG00000 |
| 112797050 | 112798691 | -1 | processed_pseudoge | ENSG00000 |
| 112818032 | 113384221 | 1 | protein_coding | ENSG00000 |
| 112826307 | 112827607 | 1 | processed_pseudoge | ENSG00000 |
| 112880298 | 112882330 | -1 | lncRNA | ENSG00000 |
| 112881718 | 112882132 | -1 | transcribed_processe | ENSG00000 |
| 112973272 | 113071962 | -1 | lncRNA | ENSG00000 |
| 113031287 | 113034551 | -1 | lncRNA | ENSG00000 |
| 113106863 | 113106955 | 1 | miRNA | ENSG00000 |
| 113152282 | 113152362 | 1 | miRNA | ENSG00000 |
| 113214046 | 113217170 | -1 | transcribed_processe | ENSG00000 |
| 113419840 | 113420136 | 1 | misc_RNA | ENSG00000 |
| 113420323 | 113420486 | 1 | snRNA | ENSG00000 |
| 113451032 | 113761927 | -1 | protein_coding | ENSG00000 |
| 113840050 | 113840236 | 1 | processed_pseudoge | ENSG00000 |
| 113900284 | 113979727 | -1 | protein_coding | ENSG00000 |
| 113943256 | 113944320 | 1 | lncRNA | ENSG00000 |
| 114103938 | 114104225 | 1 | processed_pseudoge | ENSG00000 |
| 114598770 | 114678225 | 1 | protein_coding | ENSG00000 |
| 114656759 | 114656854 | 1 | miRNA | ENSG00000 |
| 114713579 | 114714920 | -1 | processed_pseudoge | ENSG00000 |
| 114799331 | 114799629 | 1 | misc_RNA | ENSG00000 |
| 114827763 | 115113876 | -1 | protein_coding | ENSG00000 |
| 115142982 | 115143303 | 1 | processed_pseudoge | ENSG00000 |
| 115628812 | 115629707 | 1 | processed_pseudoge | ENSG00000 |
| 115764099 | 115764819 | -1 | processed_pseudoge | ENSG00000 |
| 115920707 | 115921978 | 1 | processed_pseudoge | ENSG00000 |
| 115922015 | 116297160 | -1 | transcribed_processe | ENSG00000 |
| 116043407 | 116044045 | -1 | processed_pseudoge | ENSG00000 |
| 116193997 | 116194751 | -1 | processed_pseudoge | ENSG00000 |
| 116298876 | 116300320 | 1 | protein_coding | ENSG00000 |
| 116299725 | 116299768 | 1 | miRNA | ENSG00000 |
| 116378919 | 116379269 | 1 | processed_pseudoge | ENSG00000 |
| 116425983 | 116430892 | -1 | processed_pseudoge | ENSG00000 |
| 116489364 | 116515619 | -1 | lncRNA | ENSG00000 |
| 116598161 | 116599611 | 1 | processed_pseudoge | ENSG00000 |
| 116714844 | 116719669 | -1 | lncRNA | ENSG00000 |
| 116720123 | 116739945 | -1 | lncRNA | ENSG00000 |
| 116751473 | 116952699 | -1 | lncRNA | ENSG00000 |
| 116754453 | 116921792 | 1 | lncRNA | ENSG00000 |
| 116839279 | 116839385 | 1 | snRNA | ENSG00000 |
| 116858093 | 116858973 | 1 | processed_pseudoge | ENSG00000 |
| 117083554 | 117085576 | -1 | protein_coding | ENSG00000 |
| 117249924 | 117250188 | 1 | unprocessed_pseudo | ENSG00000 |

| | | | | |
|---|---|---|---|---|
| 117314597 | 117360655 | -1 | lncRNA | ENSG00000 |
| 117360596 | 117412712 | 1 | lncRNA | ENSG00000 |
| 117406166 | 117691188 | 1 | lncRNA | ENSG00000 |
| 117413462 | 117414279 | 1 | processed_pseudoge | ENSG00000 |
| 117574512 | 117576174 | -1 | transcribed_processe | ENSG00000 |
| 117648832 | 117657312 | -1 | lncRNA | ENSG00000 |
| 117834145 | 117869960 | 1 | lncRNA | ENSG00000 |
| 118033618 | 118258634 | 1 | protein_coding | ENSG00000 |
| 118193405 | 118194751 | -1 | processed_pseudoge | ENSG00000 |
| 118278709 | 118285316 | 1 | lncRNA | ENSG00000 |
| 118279190 | 118279320 | 1 | snoRNA | ENSG00000 |
| 118280038 | 118353003 | -1 | protein_coding | ENSG00000 |
| 118400514 | 118400834 | -1 | processed_pseudoge | ENSG00000 |
| 118460224 | 118460301 | -1 | processed_pseudoge | ENSG00000 |
| 118467590 | 118554100 | 1 | transcribed_unproce | ENSG00000 |
| 118591773 | 118633729 | 1 | transcribed_unproce | ENSG00000 |
| 118606450 | 118606553 | 1 | snRNA | ENSG00000 |
| 118635970 | 118638782 | -1 | processed_pseudoge | ENSG00000 |
| 118640673 | 118640858 | 1 | processed_pseudoge | ENSG00000 |
| 118664087 | 118685343 | -1 | lncRNA | ENSG00000 |
| 118685392 | 118715433 | 1 | protein_coding | ENSG00000 |
| 118722823 | 118838683 | -1 | protein_coding | ENSG00000 |
| 118850688 | 119061247 | 1 | protein_coding | ENSG00000 |
| 119006032 | 119006401 | 1 | processed_pseudoge | ENSG00000 |
| 119135784 | 119187789 | 1 | protein_coding | ENSG00000 |
| 119192773 | 119212644 | -1 | transcribed_processe | ENSG00000 |
| 119212587 | 119295517 | 1 | protein_coding | ENSG00000 |
| 119296419 | 119304445 | -1 | protein_coding | ENSG00000 |
| 119317250 | 119322138 | -1 | protein_coding | ENSG00000 |
| 119334329 | 119378233 | 1 | unprocessed_pseudc | ENSG00000 |
| 119367562 | 119367646 | 1 | snRNA | ENSG00000 |
| 119391831 | 119395335 | -1 | unprocessed_pseudc | ENSG00000 |
| 119405523 | 119454638 | -1 | transcribed_unproce | ENSG00000 |
| 119409333 | 119410233 | 1 | lncRNA | ENSG00000 |
| 119434005 | 119434108 | -1 | snRNA | ENSG00000 |
| 119454791 | 119552025 | 1 | transcribed_unproce | ENSG00000 |
| 119456350 | 119457277 | 1 | lncRNA | ENSG00000 |
| 119494397 | 119628804 | -1 | protein_coding | ENSG00000 |
| 119569699 | 119569825 | 1 | snoRNA | ENSG00000 |
| 119686538 | 119693563 | 1 | lncRNA | ENSG00000 |
| 119799089 | 119804554 | -1 | lncRNA | ENSG00000 |
| 119937557 | 119963898 | 1 | transcribed_unproce | ENSG00000 |
| 119939540 | 119964293 | 1 | lncRNA | ENSG00000 |
| 119947131 | 119947232 | 1 | misc_RNA | ENSG00000 |
| 120035033 | 120040615 | -1 | lncRNA | ENSG00000 |
| 120055623 | 120066858 | -1 | protein_coding | ENSG00000 |
| 120066958 | 120564789 | 1 | lncRNA | ENSG00000 |
| 120091046 | 120092282 | -1 | processed_pseudoge | ENSG00000 |
| 120342056 | 120342647 | 1 | processed_pseudoge | ENSG00000 |
| 120639090 | 120650796 | 1 | lncRNA | ENSG00000 |

| | | | | |
|---|---|---|---|---|
| 120684919 | 120922870 | -1 | protein_coding | ENSG0000C |
| 120810469 | 120810569 | 1 | snRNA | ENSG0000C |
| 120895494 | 120897083 | -1 | processed_pseudoge | ENSG0000C |
| 121035613 | 121073021 | -1 | protein_coding | ENSG0000C |
| 121061172 | 121061478 | -1 | misc_RNA | ENSG0000C |
| 121071429 | 121080476 | 1 | lncRNA | ENSG0000C |
| 121118299 | 121118734 | 1 | processed_pseudoge | ENSG0000C |
| 121131408 | 121227466 | -1 | protein_coding | ENSG0000C |
| 121192797 | 121192903 | -1 | snRNA | ENSG0000C |
| 121328642 | 121381059 | -1 | protein_coding | ENSG0000C |
| 121329069 | 121329170 | -1 | misc_RNA | ENSG0000C |
| 121369433 | 121370345 | -1 | processed_pseudoge | ENSG0000C |
| 121451541 | 121451967 | 1 | processed_pseudoge | ENSG0000C |
| 121667946 | 121696995 | -1 | protein_coding | ENSG0000C |
| 121758930 | 121765427 | -1 | protein_coding | ENSG0000C |
| 121764585 | 121766814 | 1 | lncRNA | ENSG0000C |
| 121801317 | 121817021 | 1 | protein_coding | ENSG0000C |
| 121816444 | 121823933 | -1 | protein_coding | ENSG0000C |
| 121824329 | 121870487 | -1 | protein_coding | ENSG0000C |
| 121870583 | 121872848 | 1 | lncRNA | ENSG0000C |
| 121874481 | 121952060 | -1 | protein_coding | ENSG0000C |
| 122077840 | 122078318 | -1 | processed_pseudoge | ENSG0000C |
| 122087508 | 122087811 | -1 | misc_RNA | ENSG0000C |
| 122152333 | 122362758 | 1 | protein_coding | ENSG0000C |
| 122378966 | 122429802 | 1 | protein_coding | ENSG0000C |
| 122451470 | 122456725 | -1 | protein_coding | ENSG0000C |
| 122610108 | 122621066 | -1 | protein_coding | ENSG0000C |
| 122618983 | 122689156 | 1 | lncRNA | ENSG0000C |
| 122700445 | 122708788 | 1 | lncRNA | ENSG0000C |
| 122728532 | 122732482 | -1 | transcribed_unitary_ | ENSG0000C |
| 122732702 | 122744942 | 1 | protein_coding | ENSG0000C |
| 122815771 | 122816117 | -1 | processed_pseudoge | ENSG0000C |
| 122826708 | 122898236 | 1 | protein_coding | ENSG0000C |
| 122840343 | 122840690 | 1 | processed_pseudoge | ENSG0000C |
| 122879778 | 122885805 | -1 | lncRNA | ENSG0000C |
| 122888697 | 122922968 | -1 | protein_coding | ENSG0000C |
| 122923078 | 123319433 | 1 | protein_coding | ENSG0000C |
| 123259949 | 123260118 | -1 | processed_pseudoge | ENSG0000C |
| 123349250 | 123351221 | -1 | lncRNA | ENSG0000C |
| 123396795 | 123403760 | 1 | protein_coding | ENSG0000C |
| 123490242 | 123522106 | 1 | lncRNA | ENSG0000C |
| 123505279 | 123527525 | 1 | lncRNA | ENSG0000C |
| 123539026 | 123962698 | 1 | lncRNA | ENSG0000C |
| 123744923 | 123745673 | -1 | lncRNA | ENSG0000C |
| 123745750 | 123746226 | -1 | processed_pseudoge | ENSG0000C |
| 123861860 | 123862015 | -1 | processed_pseudoge | ENSG0000C |
| 124025295 | 124028268 | -1 | lncRNA | ENSG0000C |
| 124062462 | 124062990 | 1 | processed_pseudoge | ENSG0000C |
| 124149377 | 124252044 | -1 | lncRNA | ENSG0000C |
| 124324973 | 124339047 | -1 | lncRNA | ENSG0000C |

| | | | | |
|---|---|---|---|---|
| 124435750 | 124437120 | -1 | processed_pseudoge | ENSG00000 |
| 124462863 | 124463790 | -1 | processed_pseudoge | ENSG00000 |
| 124499940 | 124558445 | -1 | lncRNA | ENSG00000 |
| 124664048 | 124712732 | -1 | protein_coding | ENSG00000 |
| 125316399 | 125492932 | 1 | protein_coding | ENSG00000 |
| 125362819 | 125362929 | 1 | misc_RNA | ENSG00000 |
| 125507259 | 125507307 | 1 | miRNA | ENSG00000 |
| 125676718 | 126017357 | 1 | lncRNA | ENSG00000 |
| 125681177 | 125683668 | -1 | processed_pseudoge | ENSG00000 |
| 125723267 | 125724217 | -1 | processed_pseudoge | ENSG00000 |
| 126064128 | 126073370 | 1 | lncRNA | ENSG00000 |
| 126106139 | 126106923 | -1 | processed_pseudoge | ENSG00000 |
| 126431694 | 126735523 | -1 | lncRNA | ENSG00000 |
| 126563509 | 126564776 | 1 | processed_pseudoge | ENSG00000 |
| 127028107 | 127028219 | -1 | misc_RNA | ENSG00000 |
| 127043430 | 127077762 | -1 | lncRNA | ENSG00000 |
| 127094410 | 127470569 | -1 | lncRNA | ENSG00000 |
| 127228519 | 127352379 | -1 | lncRNA | ENSG00000 |
| 127493505 | 127493828 | 1 | processed_pseudoge | ENSG00000 |
| 127498327 | 127498807 | 1 | processed_pseudoge | ENSG00000 |
| 127532828 | 127623803 | -1 | lncRNA | ENSG00000 |
| 127623271 | 127726737 | 1 | protein_coding | ENSG00000 |
| 127730400 | 127774292 | 1 | protein_coding | ENSG00000 |
| 127781821 | 127840733 | 1 | protein_coding | ENSG00000 |
| 127809170 | 127809477 | 1 | processed_pseudoge | ENSG00000 |
| 127812720 | 127813200 | -1 | processed_pseudoge | ENSG00000 |
| 127840198 | 127844040 | -1 | lncRNA | ENSG00000 |
| 127880893 | 127899224 | 1 | protein_coding | ENSG00000 |
| 127891991 | 127892082 | 1 | snRNA | ENSG00000 |
| 127917732 | 127966034 | -1 | protein_coding | ENSG00000 |
| 127965306 | 128039927 | 1 | protein_coding | ENSG00000 |
| 128061286 | 128222931 | 1 | protein_coding | ENSG00000 |
| 128069014 | 128069612 | -1 | processed_pseudoge | ENSG00000 |
| 128155389 | 128156204 | 1 | processed_pseudoge | ENSG00000 |
| 128269237 | 128288829 | -1 | protein_coding | ENSG00000 |
| 128292751 | 128519398 | 1 | lncRNA | ENSG00000 |
| 128552590 | 128553416 | -1 | lncRNA | ENSG00000 |
| 128567972 | 128570531 | -1 | lncRNA | ENSG00000 |
| 128582999 | 128601407 | -1 | lncRNA | ENSG00000 |
| 128768228 | 128769948 | 1 | lncRNA | ENSG00000 |
| 128809700 | 128875224 | 1 | protein_coding | ENSG00000 |
| 128864921 | 129093600 | -1 | protein_coding | ENSG00000 |
| 128944628 | 128944956 | 1 | processed_pseudoge | ENSG00000 |
| 129041766 | 129041877 | 1 | pseudogene | ENSG00000 |
| 129093317 | 129116640 | 1 | protein_coding | ENSG00000 |
| 129133106 | 129134098 | 1 | processed_pseudoge | ENSG00000 |
| 129135926 | 129139740 | -1 | processed_pseudoge | ENSG00000 |
| 129177386 | 129179094 | 1 | processed_pseudoge | ENSG00000 |
| 129299175 | 129305022 | -1 | lncRNA | ENSG00000 |
| 129362911 | 129377504 | -1 | lncRNA | ENSG00000 |

| | | | | |
|---|---|---|---|---|
| 129612330 | 129771527 | -1 | lncRNA | ENSG00000 |
| 129771596 | 129975329 | 1 | lncRNA | ENSG00000 |
| 129903010 | 129904163 | 1 | processed_pseudoge | ENSG00000 |
| 130376229 | 130432227 | 1 | lncRNA | ENSG00000 |
| 130494673 | 130495285 | 1 | processed_pseudoge | ENSG00000 |
| 130503357 | 130504344 | -1 | processed_pseudoge | ENSG00000 |
| 130808507 | 130811933 | 1 | TEC | ENSG00000 |
| 130881027 | 130885641 | -1 | lncRNA | ENSG00000 |
| 130963925 | 130967575 | 1 | processed_pseudoge | ENSG00000 |
| 131112748 | 131113255 | 1 | processed_pseudoge | ENSG00000 |
| 131273242 | 131273348 | 1 | snRNA | ENSG00000 |
| 131379717 | 131591825 | 1 | lncRNA | ENSG00000 |
| 131540281 | 131540489 | 1 | processed_pseudoge | ENSG00000 |
| 131720260 | 131720543 | -1 | misc_RNA | ENSG00000 |
| 131727587 | 131791492 | 1 | lncRNA | ENSG00000 |
| 131768140 | 131768249 | 1 | snoRNA | ENSG00000 |
| 131768233 | 131768578 | 1 | processed_pseudoge | ENSG00000 |
| 131857343 | 131857913 | -1 | processed_pseudoge | ENSG00000 |
| 131930031 | 131975952 | -1 | lncRNA | ENSG00000 |
| 131978485 | 132345621 | -1 | lncRNA | ENSG00000 |
| 132027175 | 132028434 | -1 | lncRNA | ENSG00000 |
| 132367786 | 132368479 | -1 | processed_pseudoge | ENSG00000 |
| 132435802 | 132447235 | -1 | lncRNA | ENSG00000 |
| 132591064 | 132678503 | 1 | lncRNA | ENSG00000 |
| 132592084 | 132592536 | 1 | processed_pseudoge | ENSG00000 |
| 132836772 | 132981677 | -1 | lncRNA | ENSG00000 |
| 132985510 | 132988677 | -1 | lncRNA | ENSG00000 |
| 133075311 | 133149147 | -1 | lncRNA | ENSG00000 |
| 133124104 | 133126063 | 1 | processed_pseudoge | ENSG00000 |
| 133149294 | 133208606 | 1 | protein_coding | ENSG00000 |
| 133261022 | 133263115 | -1 | lncRNA | ENSG00000 |
| 133574566 | 133575208 | -1 | unprocessed_pseudc | ENSG00000 |
| 133871246 | 134010690 | -1 | lncRNA | ENSG00000 |
| 134026665 | 134327977 | -1 | lncRNA | ENSG00000 |
| 134057468 | 134085506 | 1 | lncRNA | ENSG00000 |
| 134196333 | 134201789 | -1 | protein_coding | ENSG00000 |
| 134327063 | 134328798 | 1 | processed_pseudoge | ENSG00000 |
| 134383467 | 134545648 | 1 | lncRNA | ENSG00000 |
| 134759399 | 134817289 | 1 | lncRNA | ENSG00000 |
| 134932889 | 134934131 | -1 | lncRNA | ENSG00000 |
| 134944275 | 134944523 | 1 | processed_pseudoge | ENSG00000 |
| 134952259 | 134952889 | -1 | processed_pseudoge | ENSG00000 |
| 135045464 | 135046850 | 1 | processed_pseudoge | ENSG00000 |
| 135067233 | 135097658 | 1 | lncRNA | ENSG00000 |
| 135112905 | 135123779 | -1 | lncRNA | ENSG00000 |
| 135315418 | 135315522 | -1 | snRNA | ENSG00000 |
| 135371177 | 135371924 | -1 | processed_pseudoge | ENSG00000 |
| 135565117 | 135567157 | -1 | processed_pseudoge | ENSG00000 |
| 135792384 | 135805904 | -1 | lncRNA | ENSG00000 |
| 135866983 | 135913680 | -1 | lncRNA | ENSG00000 |

| | | | | |
|---|---|---|---|---|
| 135995929 | 135996092 | -1 | snRNA | ENSG0000C |
| 136004509 | 136004902 | -1 | processed_pseudoge | ENSG0000C |
| 136097797 | 136099240 | -1 | lncRNA | ENSG0000C |
| 136118675 | 136395471 | -1 | lncRNA | ENSG0000C |
| 136125909 | 136138874 | 1 | lncRNA | ENSG0000C |
| 136355110 | 136356336 | -1 | processed_pseudoge | ENSG0000C |
| 136553236 | 136553739 | 1 | TEC | ENSG0000C |
| 136795919 | 137212799 | -1 | lncRNA | ENSG0000C |
| 137193756 | 137198367 | 1 | lncRNA | ENSG0000C |
| 137301001 | 137301187 | -1 | processed_pseudoge | ENSG0000C |
| 137425426 | 137425864 | -1 | processed_pseudoge | ENSG0000C |
| 137448087 | 137448501 | 1 | processed_pseudoge | ENSG0000C |
| 137518918 | 137532494 | -1 | protein_coding | ENSG0000C |
| 137545731 | 137603430 | 1 | lncRNA | ENSG0000C |
| 137601899 | 137602433 | -1 | TEC | ENSG0000C |
| 137645265 | 137655174 | -1 | lncRNA | ENSG0000C |
| 137690477 | 137734977 | -1 | lncRNA | ENSG0000C |
| 137751029 | 137755300 | 1 | lncRNA | ENSG0000C |
| 137807706 | 137816402 | 1 | lncRNA | ENSG0000C |
| 137943022 | 138051288 | 1 | lncRNA | ENSG0000C |
| 137978547 | 138131182 | -1 | lncRNA | ENSG0000C |
| 138057464 | 138178177 | 1 | lncRNA | ENSG0000C |
| 138164097 | 138242349 | -1 | protein_coding | ENSG0000C |
| 138277115 | 138281784 | -1 | lncRNA | ENSG0000C |
| 138298856 | 138312658 | -1 | lncRNA | ENSG0000C |
| 138309613 | 138663403 | 1 | lncRNA | ENSG0000C |
| 138425523 | 138436869 | 1 | lncRNA | ENSG0000C |
| 138481255 | 138482096 | -1 | processed_pseudoge | ENSG0000C |
| 138560704 | 138561941 | 1 | processed_pseudoge | ENSG0000C |
| 138664725 | 138665569 | -1 | lncRNA | ENSG0000C |
| 138773537 | 138803567 | 1 | lncRNA | ENSG0000C |
| 138819954 | 139012646 | -1 | lncRNA | ENSG0000C |
| 138923930 | 138924232 | -1 | lncRNA | ENSG0000C |
| 139015781 | 139045939 | 1 | protein_coding | ENSG0000C |
| 139028112 | 139177218 | -1 | protein_coding | ENSG0000C |
| 139055475 | 139055578 | 1 | snRNA | ENSG0000C |
| 139100377 | 139100658 | 1 | misc_RNA | ENSG0000C |
| 139114930 | 139115374 | -1 | processed_pseudoge | ENSG0000C |
| 139155153 | 139155444 | 1 | misc_RNA | ENSG0000C |
| 139185441 | 139185746 | 1 | misc_RNA | ENSG0000C |
| 139200395 | 139200513 | 1 | misc_RNA | ENSG0000C |
| 139231026 | 139231119 | -1 | snRNA | ENSG0000C |
| 139256657 | 139256760 | -1 | snRNA | ENSG0000C |
| 139266165 | 139280225 | -1 | protein_coding | ENSG0000C |
| 139266880 | 139302551 | -1 | protein_coding | ENSG0000C |
| 139301505 | 139420033 | 1 | protein_coding | ENSG0000C |
| 139318378 | 139318486 | 1 | snRNA | ENSG0000C |
| 139411927 | 139454034 | -1 | lncRNA | ENSG0000C |
| 139426299 | 139430884 | -1 | transcribed_processe | ENSG0000C |
| 139436368 | 139436494 | -1 | snoRNA | ENSG0000C |

| | | | | |
|---|---|---|---|---|
| 139453232 | 139476609 | 1 | protein_coding | ENSG00000 |
| 139495941 | 139606699 | -1 | protein_coding | ENSG00000 |
| 139546266 | 139546727 | 1 | processed_pseudoge | ENSG00000 |
| 139556799 | 139557643 | 1 | lncRNA | ENSG00000 |
| 139568573 | 139580143 | 1 | lncRNA | ENSG00000 |
| 139618136 | 139623232 | -1 | lncRNA | ENSG00000 |
| 139665768 | 139740745 | 1 | protein_coding | ENSG00000 |
| 139685127 | 139685427 | -1 | misc_RNA | ENSG00000 |
| 139698144 | 139698554 | -1 | processed_pseudoge | ENSG00000 |
| 139716753 | 140154184 | -1 | protein_coding | ENSG00000 |
| 139763080 | 139770391 | 1 | lncRNA | ENSG00000 |
| 139794125 | 139794433 | -1 | misc_RNA | ENSG00000 |
| 140128015 | 140134565 | 1 | lncRNA | ENSG00000 |
| 140257286 | 140385726 | 1 | protein_coding | ENSG00000 |
| 140283724 | 140373403 | -1 | lncRNA | ENSG00000 |
| 140314163 | 140314550 | -1 | processed_pseudoge | ENSG00000 |
| 140366240 | 140366861 | -1 | processed_pseudoge | ENSG00000 |
| 140388453 | 140427661 | -1 | protein_coding | ENSG00000 |
| 140442262 | 140498377 | -1 | protein_coding | ENSG00000 |
| 140498402 | 140501330 | 1 | TEC | ENSG00000 |
| 140524168 | 140553770 | 1 | protein_coding | ENSG00000 |
| 140559431 | 140568961 | -1 | protein_coding | ENSG00000 |
| 140577845 | 140579657 | 1 | lncRNA | ENSG00000 |
| 140577925 | 140578222 | 1 | misc_RNA | ENSG00000 |
| 140600401 | 140600575 | -1 | processed_pseudoge | ENSG00000 |
| 140620782 | 140756385 | -1 | protein_coding | ENSG00000 |
| 140641840 | 140645489 | -1 | processed_pseudoge | ENSG00000 |
| 140700835 | 140700942 | -1 | misc_RNA | ENSG00000 |
| 140712168 | 140716081 | 1 | lncRNA | ENSG00000 |
| 140756528 | 140757921 | 1 | lncRNA | ENSG00000 |
| 140779039 | 140779141 | -1 | misc_RNA | ENSG00000 |
| 140859807 | 141212877 | -1 | protein_coding | ENSG00000 |
| 141220887 | 141234697 | 1 | protein_coding | ENSG00000 |
| 141240739 | 141278789 | -1 | lncRNA | ENSG00000 |
| 141302910 | 141357096 | -1 | lncRNA | ENSG00000 |
| 141405172 | 141407586 | 1 | processed_pseudoge | ENSG00000 |
| 141430831 | 141431284 | 1 | lncRNA | ENSG00000 |
| 141569931 | 141576077 | -1 | lncRNA | ENSG00000 |
| 141636583 | 141733987 | 1 | protein_coding | ENSG00000 |
| 142023160 | 142847432 | -1 | protein_coding | ENSG00000 |
| 142286917 | 142287625 | -1 | processed_pseudoge | ENSG00000 |
| 142317238 | 142317546 | 1 | processed_pseudoge | ENSG00000 |
| 142514446 | 142518983 | 1 | lncRNA | ENSG00000 |
| 142566019 | 142660950 | 1 | lncRNA | ENSG00000 |
| 142933195 | 143184861 | -1 | lncRNA | ENSG00000 |
| 143045086 | 143045878 | 1 | processed_pseudoge | ENSG00000 |
| 143184917 | 143223874 | 1 | protein_coding | ENSG00000 |
| 143286293 | 143329858 | 1 | lncRNA | ENSG00000 |
| 143309466 | 143309566 | 1 | misc_RNA | ENSG00000 |
| 143336762 | 143474568 | 1 | protein_coding | ENSG00000 |

| | | | | |
|---|---|---|---|---|
| 143343460 | 143343535 | 1 | miRNA | ENSG00000( |
| 143349275 | 143350117 | -1 | processed_pseudoge | ENSG00000( |
| 143351515 | 143355794 | -1 | lncRNA | ENSG00000( |
| 143424753 | 143426013 | 1 | transcribed_processe | ENSG00000( |
| 143497412 | 143498126 | -1 | processed_pseudoge | ENSG00000( |
| 143513472 | 143514635 | -1 | lncRNA | ENSG00000( |
| 143513702 | 143557486 | 1 | protein_coding | ENSG00000( |
| 143555082 | 143555583 | -1 | transcribed_processe | ENSG00000( |
| 143559457 | 144343750 | 1 | lncRNA | ENSG00000( |
| 143559472 | 143561460 | 1 | transcribed_processe | ENSG00000( |
| 143577302 | 143700675 | -1 | protein_coding | ENSG00000( |
| 143760399 | 143762093 | -1 | processed_pseudoge | ENSG00000( |
| 143870864 | 143905563 | -1 | protein_coding | ENSG00000( |
| 143911514 | 143912053 | -1 | processed_pseudoge | ENSG00000( |
| 143996104 | 144019345 | -1 | protein_coding | ENSG00000( |
| 144010237 | 144012598 | 1 | lncRNA | ENSG00000( |
| 144109303 | 144140751 | -1 | protein_coding | ENSG00000( |
| 144111039 | 144870953 | -1 | lncRNA | ENSG00000( |
| 144210701 | 145098174 | -1 | protein_coding | ENSG00000( |
| 144505900 | 144509001 | 1 | lncRNA | ENSG00000( |
| 144572238 | 144572983 | 1 | processed_pseudoge | ENSG00000( |
| 144642922 | 144661357 | -1 | lncRNA | ENSG00000( |
| 144646156 | 144745271 | 1 | protein_coding | ENSG00000( |
| 144739762 | 144739941 | -1 | misc_RNA | ENSG00000( |
| 144831908 | 145098541 | -1 | protein_coding | ENSG00000( |
| 144845625 | 144847344 | 1 | processed_pseudoge | ENSG00000( |
| 144851572 | 144874787 | 1 | lncRNA | ENSG00000( |
| 144920713 | 144921164 | -1 | processed_pseudoge | ENSG00000( |
| 145035132 | 145035436 | 1 | misc_RNA | ENSG00000( |
| 145098288 | 145129524 | 1 | protein_coding | ENSG00000( |
| 145110838 | 145180589 | -1 | protein_coding | ENSG00000( |
| 145149537 | 145149649 | -1 | misc_RNA | ENSG00000( |
| 145269810 | 145270558 | -1 | processed_pseudoge | ENSG00000( |
| 145335263 | 145343559 | -1 | lncRNA | ENSG00000( |
| 145375543 | 145376253 | 1 | processed_pseudoge | ENSG00000( |
| 145376909 | 145377957 | 1 | TEC | ENSG00000( |
| 145444899 | 145469638 | -1 | lncRNA | ENSG00000( |
| 145456005 | 145456844 | 1 | processed_pseudoge | ENSG00000( |
| 145481194 | 145559176 | 1 | protein_coding | ENSG00000( |
| 145497073 | 145502971 | -1 | lncRNA | ENSG00000( |
| 145514615 | 145517118 | -1 | lncRNA | ENSG00000( |
| 145599042 | 145660033 | 1 | protein_coding | ENSG00000( |
| 145624159 | 145626035 | -1 | processed_pseudoge | ENSG00000( |
| 145642336 | 145650624 | -1 | lncRNA | ENSG00000( |
| 145680115 | 145771032 | 1 | protein_coding | ENSG00000( |
| 145757627 | 145938823 | -1 | protein_coding | ENSG00000( |
| 145830456 | 145832980 | -1 | TEC | ENSG00000( |
| 145833118 | 145839580 | 1 | lncRNA | ENSG00000( |
| 145958128 | 145958240 | -1 | misc_RNA | ENSG00000( |
| 146052604 | 146056762 | -1 | lncRNA | ENSG00000( |

| | | | | |
|---|---|---|---|---|
| 146077717 | 146121913 | -1 | lncRNA | ENSG0000C |
| 146175703 | 146200000 | 1 | protein_coding | ENSG0000C |
| 146180851 | 146181316 | 1 | TEC | ENSG0000C |
| 146214515 | 146230878 | 1 | protein_coding | ENSG0000C |
| 146231892 | 146232267 | 1 | TEC | ENSG0000C |
| 146241806 | 146243669 | 1 | lncRNA | ENSG0000C |
| 146253975 | 146521964 | -1 | protein_coding | ENSG0000C |
| 146408583 | 146408688 | 1 | miRNA | ENSG0000C |
| 146580048 | 146580210 | 1 | snRNA | ENSG0000C |
| 146638893 | 146642474 | 1 | protein_coding | ENSG0000C |
| 146706638 | 146945882 | -1 | protein_coding | ENSG0000C |
| 146934001 | 146945471 | 1 | lncRNA | ENSG0000C |
| 146957438 | 146975457 | 1 | lncRNA | ENSG0000C |
| 147005492 | 147009145 | 1 | lncRNA | ENSG0000C |
| 147025914 | 147030072 | 1 | TEC | ENSG0000C |
| 147060613 | 147159068 | 1 | lncRNA | ENSG0000C |
| 147344629 | 147344717 | -1 | miRNA | ENSG0000C |
| 147425974 | 147426351 | -1 | processed_pseudoge | ENSG0000C |
| 147458724 | 147459321 | -1 | processed_pseudoge | ENSG0000C |
| 147480917 | 147544954 | 1 | protein_coding | ENSG0000C |
| 147487971 | 147496062 | 1 | TEC | ENSG0000C |
| 147506091 | 147506765 | 1 | processed_pseudoge | ENSG0000C |
| 147567606 | 147594589 | 1 | lncRNA | ENSG0000C |
| 147609552 | 147617245 | -1 | lncRNA | ENSG0000C |
| 147617386 | 147672044 | 1 | protein_coding | ENSG0000C |
| 147637785 | 147684163 | -1 | protein_coding | ENSG0000C |
| 147732063 | 148072776 | 1 | protein_coding | ENSG0000C |
| 147745303 | 147745417 | -1 | rRNA_pseudogene | ENSG0000C |
| 147782595 | 147782668 | 1 | miRNA | ENSG0000C |
| 147808926 | 147809209 | 1 | misc_RNA | ENSG0000C |
| 147916529 | 147917386 | -1 | processed_pseudoge | ENSG0000C |
| 148078762 | 148444698 | -1 | protein_coding | ENSG0000C |
| 148146471 | 148208880 | 1 | lncRNA | ENSG0000C |
| 148445703 | 149024624 | 1 | lncRNA | ENSG0000C |
| 148526493 | 148527139 | -1 | processed_pseudoge | ENSG0000C |
| 148557947 | 148558048 | 1 | rRNA_pseudogene | ENSG0000C |
| 148693032 | 148694323 | 1 | lncRNA | ENSG0000C |
| 148703526 | 148703832 | -1 | processed_pseudoge | ENSG0000C |
| 148819351 | 148819956 | -1 | TEC | ENSG0000C |
| 148940941 | 148942415 | -1 | lncRNA | ENSG0000C |
| 148991572 | 148991633 | -1 | snRNA | ENSG0000C |
| 149027445 | 149062522 | -1 | lncRNA | ENSG0000C |
| 149147793 | 149154175 | -1 | lncRNA | ENSG0000C |
| 149154279 | 149278123 | 1 | lncRNA | ENSG0000C |
| 149351903 | 149896233 | -1 | protein_coding | ENSG0000C |
| 149377257 | 149380107 | 1 | lncRNA | ENSG0000C |
| 149666057 | 149711149 | 1 | lncRNA | ENSG0000C |
| 149789952 | 149790303 | 1 | processed_pseudoge | ENSG0000C |
| 149907560 | 149907675 | 1 | rRNA_pseudogene | ENSG0000C |
| 149934376 | 149934483 | 1 | snRNA | ENSG0000C |

| | | | | |
|---|---|---|---|---|
| 149956208 | 149958572 | 1 | lncRNA | ENSG00000( |
| 150047004 | 150051290 | -1 | lncRNA | ENSG00000( |
| 150078445 | 150257438 | 1 | protein_coding | ENSG00000( |
| 150209706 | 150209768 | 1 | snRNA | ENSG00000( |
| 150264435 | 151015727 | -1 | protein_coding | ENSG00000( |
| 150422645 | 150423642 | -1 | processed_pseudoge | ENSG00000( |
| 150579089 | 150581545 | 1 | lncRNA | ENSG00000( |
| 150582151 | 150584693 | 1 | protein_coding | ENSG00000( |
| 150720981 | 150721478 | 1 | processed_pseudoge | ENSG00000( |
| 150970671 | 150970782 | -1 | rRNA_pseudogene | ENSG00000( |
| 151027090 | 151027617 | 1 | processed_pseudoge | ENSG00000( |
| 151099624 | 151104642 | 1 | protein_coding | ENSG00000( |
| 151102057 | 151102128 | 1 | snoRNA | ENSG00000( |
| 151102751 | 151325632 | -1 | protein_coding | ENSG00000( |
| 151103827 | 151103891 | 1 | snoRNA | ENSG00000( |
| 151139991 | 151141103 | 1 | lncRNA | ENSG00000( |
| 151259503 | 151262849 | 1 | unprocessed_pseudc | ENSG00000( |
| 151277171 | 151291453 | 1 | protein_coding | ENSG00000( |
| 151279030 | 151279192 | -1 | unprocessed_pseudc | ENSG00000( |
| 151300518 | 151300613 | 1 | snRNA | ENSG00000( |
| 151333775 | 151353224 | -1 | lncRNA | ENSG00000( |
| 151407551 | 151408835 | -1 | lncRNA | ENSG00000( |
| 151409176 | 151670503 | 1 | protein_coding | ENSG00000( |
| 151576766 | 151576916 | 1 | misc_RNA | ENSG00000( |
| 151670504 | 151761007 | -1 | protein_coding | ENSG00000( |
| 151674483 | 151677893 | 1 | lncRNA | ENSG00000( |
| 151798917 | 151833097 | 1 | lncRNA | ENSG00000( |
| 151883711 | 151886239 | -1 | lncRNA | ENSG00000( |
| 151887543 | 151891263 | -1 | lncRNA | ENSG00000( |
| 151904932 | 151928710 | -1 | lncRNA | ENSG00000( |
| 151919468 | 151943933 | 1 | lncRNA | ENSG00000( |
| 151949338 | 151991925 | -1 | lncRNA | ENSG00000( |
| 151955991 | 151969381 | 1 | lncRNA | ENSG00000( |
| 151971497 | 151971613 | -1 | rRNA_pseudogene | ENSG00000( |
| 151988526 | 152045275 | 1 | lncRNA | ENSG00000( |
| 152090459 | 152104720 | 1 | lncRNA | ENSG00000( |
| 152113857 | 152116497 | -1 | lncRNA | ENSG00000( |
| 152146385 | 152178573 | -1 | lncRNA | ENSG00000( |
| 152206987 | 152225870 | -1 | lncRNA | ENSG00000( |
| 152264150 | 152264445 | -1 | misc_RNA | ENSG00000( |
| 152308283 | 152309983 | -1 | lncRNA | ENSG00000( |
| 152317902 | 152318364 | -1 | lncRNA | ENSG00000( |
| 152320544 | 152536092 | -1 | protein_coding | ENSG00000( |
| 152337655 | 152338098 | 1 | lncRNA | ENSG00000( |
| 152448275 | 152513788 | 1 | lncRNA | ENSG00000( |
| 152489327 | 152489416 | -1 | miRNA | ENSG00000( |
| 152536264 | 152539263 | 1 | lncRNA | ENSG00000( |
| 152536428 | 152536516 | 1 | miRNA | ENSG00000( |
| 152551277 | 152552364 | -1 | processed_pseudoge | ENSG00000( |
| 152613875 | 152614333 | -1 | processed_pseudoge | ENSG00000( |

| | | | | |
|---|---|---|---|---|
| 152618628 | 152680012 | -1 | protein_coding | ENSG00000( |
| 152666368 | 152670107 | 1 | lncRNA | ENSG00000( |
| 152769354 | 152779730 | -1 | protein_coding | ENSG00000( |
| 152779937 | 152918463 | 1 | protein_coding | ENSG00000( |
| 152796063 | 152796831 | -1 | processed_pseudoge | ENSG00000( |
| 152927446 | 152928207 | 1 | processed_pseudoge | ENSG00000( |
| 152934516 | 152936761 | -1 | lncRNA | ENSG00000( |
| 152936352 | 152979696 | 1 | protein_coding | ENSG00000( |
| 153034211 | 153091563 | 1 | lncRNA | ENSG00000( |
| 153152342 | 153339317 | 1 | protein_coding | ENSG00000( |
| 153265662 | 153265755 | -1 | snRNA | ENSG00000( |
| 153266909 | 153267010 | -1 | misc_RNA | ENSG00000( |
| 153307792 | 153308717 | -1 | processed_pseudoge | ENSG00000( |
| 153344649 | 153415118 | 1 | protein_coding | ENSG00000( |
| 153381729 | 153382015 | -1 | misc_RNA | ENSG00000( |
| 153466346 | 153636711 | 1 | protein_coding | ENSG00000( |
| 153633692 | 153655689 | -1 | lncRNA | ENSG00000( |
| 153640168 | 153681180 | 1 | transcribed_unproce | ENSG00000( |
| 153658572 | 153659653 | 1 | processed_pseudoge | ENSG00000( |
| 153684070 | 153705702 | 1 | protein_coding | ENSG00000( |
| 153710125 | 153760235 | -1 | protein_coding | ENSG00000( |
| 153720327 | 153727722 | 1 | lncRNA | ENSG00000( |
| 153780591 | 153789083 | -1 | protein_coding | ENSG00000( |
| 153948718 | 154300500 | 1 | lncRNA | ENSG00000( |
| 154231742 | 154491799 | -1 | protein_coding | ENSG00000( |
| 154235980 | 154237598 | 1 | lncRNA | ENSG00000( |
| 154261735 | 154262193 | 1 | lncRNA | ENSG00000( |
| 154373886 | 154374347 | 1 | processed_pseudoge | ENSG00000( |
| 154523487 | 154524807 | 1 | processed_pseudoge | ENSG00000( |
| 154535005 | 154550400 | -1 | protein_coding | ENSG00000( |
| 154543933 | 154544037 | -1 | snRNA | ENSG00000( |
| 154555208 | 154555985 | 1 | processed_pseudoge | ENSG00000( |
| 154562956 | 154571086 | 1 | protein_coding | ENSG00000( |
| 154583128 | 154590745 | -1 | protein_coding | ENSG00000( |
| 154604134 | 154612967 | -1 | protein_coding | ENSG00000( |
| 154605952 | 154606077 | -1 | unprocessed_pseudc | ENSG00000( |
| 154626945 | 154753120 | 1 | protein_coding | ENSG00000( |
| 154724586 | 154724880 | 1 | processed_pseudoge | ENSG00000( |
| 154754756 | 154781873 | -1 | lncRNA | ENSG00000( |
| 154770568 | 154770916 | 1 | processed_pseudoge | ENSG00000( |
| 154781213 | 154828813 | 1 | protein_coding | ENSG00000( |
| 154787118 | 154787278 | 1 | snRNA | ENSG00000( |
| 154790291 | 154790484 | 1 | snRNA | ENSG00000( |
| 155173716 | 155381694 | 1 | lncRNA | ENSG00000( |
| 155206529 | 155209027 | -1 | lncRNA | ENSG00000( |
| 155208636 | 155217078 | 1 | protein_coding | ENSG00000( |
| 155286162 | 155287136 | -1 | lncRNA | ENSG00000( |
| 155342658 | 155376970 | -1 | protein_coding | ENSG00000( |
| 155381042 | 155381239 | -1 | processed_pseudoge | ENSG00000( |
| 155383107 | 155384102 | -1 | processed_pseudoge | ENSG00000( |

| | | | | |
|---|---|---|---|---|
| 155385352 | 155385492 | 1 | unprocessed_pseudc | ENSG00000( |
| 155442650 | 155448610 | 1 | lncRNA | ENSG00000( |
| 155451617 | 155452405 | -1 | unprocessed_pseudc | ENSG00000( |
| 155452526 | 155453044 | 1 | unprocessed_pseudc | ENSG00000( |
| 155453049 | 155454803 | -1 | unprocessed_pseudc | ENSG00000( |
| 155455050 | 155456411 | -1 | unprocessed_pseudc | ENSG00000( |
| 155456638 | 155456981 | -1 | unprocessed_pseudc | ENSG00000( |
| 155457180 | 155457960 | -1 | unprocessed_pseudc | ENSG00000( |
| 155457961 | 155458640 | -1 | unprocessed_pseudc | ENSG00000( |
| 155458870 | 155459563 | -1 | unprocessed_pseudc | ENSG00000( |
| 155459704 | 155461237 | -1 | unprocessed_pseudc | ENSG00000( |
| 155461633 | 155462663 | -1 | unprocessed_pseudc | ENSG00000( |
| 155462873 | 155463823 | -1 | unprocessed_pseudc | ENSG00000( |
| 155496110 | 155520353 | 1 | lncRNA | ENSG00000( |
| 155666711 | 155732349 | 1 | protein_coding | ENSG00000( |
| 155734448 | 155737062 | 1 | lncRNA | ENSG00000( |
| 155758992 | 155807774 | 1 | protein_coding | ENSG00000( |
| 155829729 | 155866277 | -1 | protein_coding | ENSG00000( |
| 155854738 | 155920406 | 1 | protein_coding | ENSG00000( |
| 155924118 | 155953866 | -1 | protein_coding | ENSG00000( |
| 156006478 | 156006988 | 1 | processed_pseudoge | ENSG00000( |
| 156585979 | 156590039 | 1 | lncRNA | ENSG00000( |
| 156634494 | 156642454 | -1 | lncRNA | ENSG00000( |
| 156642580 | 156643569 | 1 | lncRNA | ENSG00000( |
| 156686664 | 156692636 | 1 | lncRNA | ENSG00000( |
| 156760454 | 156971799 | -1 | protein_coding | ENSG00000( |
| 156841359 | 156842236 | 1 | lncRNA | ENSG00000( |
| 156876435 | 156876820 | 1 | TEC | ENSG00000( |
| 156886645 | 156886862 | 1 | TEC | ENSG00000( |
| 156972631 | 156972956 | 1 | processed_pseudoge | ENSG00000( |
| 156986372 | 156986693 | -1 | ribozyme | ENSG00000( |
| 157064557 | 157064663 | 1 | misc_RNA | ENSG00000( |
| 157076125 | 157172090 | 1 | protein_coding | ENSG00000( |
| 157201562 | 157201668 | -1 | snRNA | ENSG00000( |
| 157204182 | 157366075 | 1 | protein_coding | ENSG00000( |
| 157204462 | 157204855 | -1 | processed_pseudoge | ENSG00000( |
| 157568731 | 157824256 | -1 | lncRNA | ENSG00000( |
| 157572490 | 157576154 | 1 | lncRNA | ENSG00000( |
| 157637687 | 157667044 | 1 | lncRNA | ENSG00000( |
| 157768013 | 157768113 | -1 | misc_RNA | ENSG00000( |
| 157815010 | 157815480 | 1 | processed_pseudoge | ENSG00000( |
| 157818101 | 157818572 | -1 | TEC | ENSG00000( |
| 157879902 | 157962924 | 1 | lncRNA | ENSG00000( |
| 157929071 | 158094763 | -1 | lncRNA | ENSG00000( |
| 157945817 | 157946642 | -1 | TEC | ENSG00000( |
| 157968412 | 157969615 | -1 | unprocessed_pseudc | ENSG00000( |
| 158109500 | 158110153 | -1 | TEC | ENSG00000( |
| 158111361 | 158111931 | -1 | TEC | ENSG00000( |
| 158124474 | 158173318 | -1 | protein_coding | ENSG00000( |
| 158170752 | 158202877 | 1 | lncRNA | ENSG00000( |

| | | | | |
|---|---|---|---|---|
| 158178168 | 158182806 | -1 | lncRNA | ENSG0000C |
| 158182825 | 158183393 | 1 | processed_pseudoge | ENSG0000C |
| 158199105 | 158200442 | -1 | lncRNA | ENSG0000C |
| 158201604 | 158255416 | 1 | protein_coding | ENSG0000C |
| 158262995 | 158270211 | 1 | lncRNA | ENSG0000C |
| 158270378 | 158278676 | 1 | transcribed_processe | ENSG0000C |
| 158315311 | 158653372 | 1 | protein_coding | ENSG0000C |
| 158395394 | 158395534 | 1 | snRNA | ENSG0000C |
| 158469586 | 158470702 | 1 | processed_pseudoge | ENSG0000C |
| 158490766 | 158509908 | -1 | lncRNA | ENSG0000C |
| 158666675 | 158672255 | -1 | protein_coding | ENSG0000C |
| 158672125 | 158709623 | 1 | protein_coding | ENSG0000C |
| 158700691 | 158700909 | 1 | snoRNA | ENSG0000C |
| 158709127 | 158723396 | -1 | protein_coding | ENSG0000C |
| 158769026 | 158908050 | 1 | protein_coding | ENSG0000C |
| 158794590 | 158794696 | -1 | snRNA | ENSG0000C |
| 158893134 | 159038760 | -1 | protein_coding | ENSG0000C |
| 158909373 | 158909450 | -1 | snoRNA | ENSG0000C |
| 159007119 | 159007835 | 1 | processed_pseudoge | ENSG0000C |
| 159026139 | 159026527 | 1 | processed_pseudoge | ENSG0000C |
| 159064297 | 159064398 | -1 | misc_RNA | ENSG0000C |
| 159103013 | 159360174 | 1 | protein_coding | ENSG0000C |
| 159128802 | 159128894 | -1 | miRNA | ENSG0000C |
| 159128805 | 159128891 | 1 | miRNA | ENSG0000C |
| 159375099 | 159375322 | -1 | processed_pseudoge | ENSG0000C |
| 159398533 | 159401246 | 1 | lncRNA | ENSG0000C |
| 159665833 | 159777828 | -1 | lncRNA | ENSG0000C |
| 160521829 | 160522311 | -1 | processed_pseudoge | ENSG0000C |
| 160539254 | 160586817 | 1 | lncRNA | ENSG0000C |
| 160818422 | 160818869 | 1 | processed_pseudoge | ENSG0000C |
| 161378953 | 161388417 | 1 | lncRNA | ENSG0000C |
| 161383897 | 162164004 | -1 | protein_coding | ENSG0000C |
| 162022733 | 162047567 | 1 | lncRNA | ENSG0000C |
| 162322188 | 162323907 | 1 | processed_pseudoge | ENSG0000C |
| 162521682 | 162522041 | 1 | processed_pseudoge | ENSG0000C |
| 162705359 | 162907428 | 1 | lncRNA | ENSG0000C |
| 162806370 | 162844690 | -1 | lncRNA | ENSG0000C |
| 162984795 | 162985416 | 1 | processed_pseudoge | ENSG0000C |
| 163093574 | 163093628 | -1 | miRNA | ENSG0000C |
| 163108785 | 163119965 | 1 | lncRNA | ENSG0000C |
| 163110073 | 163166890 | -1 | protein_coding | ENSG0000C |
| 163265575 | 163266583 | -1 | processed_pseudoge | ENSG0000C |
| 163323962 | 163344832 | -1 | protein_coding | ENSG0000C |
| 163343892 | 163351934 | 1 | protein_coding | ENSG0000C |
| 163382179 | 163382516 | 1 | processed_pseudoge | ENSG0000C |
| 163471095 | 163473754 | -1 | protein_coding | ENSG0000C |
| 163494442 | 163520539 | 1 | protein_coding | ENSG0000C |
| 163524298 | 164384050 | -1 | protein_coding | ENSG0000C |
| 163529771 | 163530697 | 1 | lncRNA | ENSG0000C |
| 163828800 | 163829113 | -1 | misc_RNA | ENSG0000C |

| | | | | |
|---|---|---|---|---|
| 163932526 | 163933182 | 1 | processed_pseudoge | ENSG00000 |
| 164188311 | 164190367 | 1 | processed_pseudoge | ENSG00000 |
| 164197007 | 164197711 | -1 | processed_pseudoge | ENSG00000 |
| 164597719 | 164597976 | 1 | processed_pseudoge | ENSG00000 |
| 164620732 | 164620837 | 1 | snRNA | ENSG00000 |
| 164725310 | 164726706 | 1 | processed_pseudoge | ENSG00000 |
| 164754064 | 164803795 | 1 | protein_coding | ENSG00000 |
| 164787290 | 164787396 | 1 | snRNA | ENSG00000 |
| 164877178 | 164898965 | 1 | protein_coding | ENSG00000 |
| 164915565 | 164916983 | 1 | processed_pseudoge | ENSG00000 |
| 164920618 | 164921063 | -1 | processed_pseudoge | ENSG00000 |
| 164920894 | 164931047 | 1 | lncRNA | ENSG00000 |
| 164929494 | 164929951 | -1 | processed_pseudoge | ENSG00000 |
| 164933086 | 164933506 | -1 | processed_pseudoge | ENSG00000 |
| 164938322 | 164939990 | -1 | unprocessed_pseudo | ENSG00000 |
| 164943290 | 164943937 | 1 | processed_pseudoge | ENSG00000 |
| 164948079 | 164949438 | 1 | processed_pseudoge | ENSG00000 |
| 164954446 | 164977668 | -1 | protein_coding | ENSG00000 |
| 164956948 | 164959122 | 1 | protein_coding | ENSG00000 |
| 164987265 | 164987787 | 1 | processed_pseudoge | ENSG00000 |
| 165007086 | 165007959 | -1 | lncRNA | ENSG00000 |
| 165010459 | 165011851 | 1 | processed_pseudoge | ENSG00000 |
| 165012366 | 165012845 | -1 | processed_pseudoge | ENSG00000 |
| 165016458 | 165041749 | 1 | protein_coding | ENSG00000 |
| 165016887 | 165017433 | 1 | processed_pseudoge | ENSG00000 |
| 165026261 | 165026944 | -1 | processed_pseudoge | ENSG00000 |
| 165045969 | 165046280 | 1 | processed_pseudoge | ENSG00000 |
| 165053864 | 165060554 | 1 | protein_coding | ENSG00000 |
| 165070608 | 165208549 | -1 | protein_coding | ENSG00000 |
| 165207561 | 165323156 | 1 | protein_coding | ENSG00000 |
| 165252580 | 165252661 | 1 | snRNA | ENSG00000 |
| 165277812 | 165279679 | -1 | protein_coding | ENSG00000 |
| 165327667 | 165343164 | 1 | protein_coding | ENSG00000 |
| 165361194 | 165498547 | 1 | protein_coding | ENSG00000 |
| 165386242 | 165386337 | 1 | miRNA | ENSG00000 |
| 165404067 | 165406350 | -1 | processed_pseudoge | ENSG00000 |
| 165517255 | 165517501 | -1 | processed_pseudoge | ENSG00000 |
| 165547532 | 165551060 | -1 | processed_pseudoge | ENSG00000 |
| 165647636 | 165757667 | -1 | lncRNA | ENSG00000 |
| 165664350 | 165665671 | 1 | processed_pseudoge | ENSG00000 |
| 165684639 | 165762778 | 1 | lncRNA | ENSG00000 |
| 165873237 | 166104457 | 1 | protein_coding | ENSG00000 |
| 166053187 | 166053287 | -1 | rRNA_pseudogene | ENSG00000 |
| 166312935 | 166313034 | -1 | misc_RNA | ENSG00000 |
| 166388701 | 166526119 | -1 | lncRNA | ENSG00000 |
| 166733384 | 167234796 | -1 | protein_coding | ENSG00000 |
| 166918364 | 166918478 | -1 | rRNA_pseudogene | ENSG00000 |
| 167306910 | 167308789 | -1 | lncRNA | ENSG00000 |
| 167400495 | 167400762 | -1 | misc_RNA | ENSG00000 |
| 167648430 | 167648796 | 1 | processed_pseudoge | ENSG00000 |

| | | | | |
|---|---|---|---|---|
| 167662986 | 167663302 | 1 | misc_RNA | ENSG00000 |
| 167741879 | 167742287 | -1 | processed_pseudoge | ENSG00000 |
| 167946674 | 168113331 | -1 | lncRNA | ENSG00000 |
| 168092537 | 168187736 | 1 | protein_coding | ENSG00000 |
| 168170097 | 168194861 | -1 | lncRNA | ENSG00000 |
| 168216294 | 168318804 | -1 | protein_coding | ENSG00000 |
| 168356735 | 168537786 | -1 | protein_coding | ENSG00000 |
| 168471409 | 168471539 | 1 | snoRNA | ENSG00000 |
| 168497052 | 168928457 | 1 | protein_coding | ENSG00000 |
| 168598240 | 168598757 | -1 | processed_pseudoge | ENSG00000 |
| 168647896 | 168648587 | -1 | lncRNA | ENSG00000 |
| 168686608 | 168686716 | -1 | snRNA | ENSG00000 |
| 168755816 | 168756382 | 1 | processed_pseudoge | ENSG00000 |
| 168828919 | 168832937 | -1 | lncRNA | ENSG00000 |
| 168843988 | 168844735 | 1 | TEC | ENSG00000 |
| 168863770 | 169010275 | -1 | protein_coding | ENSG00000 |
| 168885355 | 168885461 | 1 | snRNA | ENSG00000 |
| 169005249 | 169005344 | 1 | misc_RNA | ENSG00000 |
| 169010430 | 169010939 | 1 | lncRNA | ENSG00000 |
| 169094259 | 169270956 | -1 | protein_coding | ENSG00000 |
| 169166272 | 169167117 | 1 | processed_pseudoge | ENSG00000 |
| 169201794 | 169220349 | 1 | lncRNA | ENSG00000 |
| 169392857 | 169612629 | -1 | protein_coding | ENSG00000 |
| 169572328 | 169576451 | 1 | lncRNA | ENSG00000 |
| 169612633 | 169723673 | 1 | protein_coding | ENSG00000 |
| 169729470 | 169757944 | -1 | protein_coding | ENSG00000 |
| 169791221 | 169791702 | -1 | processed_pseudoge | ENSG00000 |
| 169832653 | 169833363 | 1 | processed_pseudoge | ENSG00000 |
| 169849053 | 169867346 | -1 | lncRNA | ENSG00000 |
| 169912705 | 169919663 | 1 | lncRNA | ENSG00000 |
| 169917761 | 169975904 | -1 | lncRNA | ENSG00000 |
| 169921417 | 169923442 | 1 | lncRNA | ENSG00000 |
| 169945981 | 169963337 | 1 | lncRNA | ENSG00000 |
| 169981744 | 169994488 | 1 | lncRNA | ENSG00000 |
| 169986597 | 170033031 | -1 | protein_coding | ENSG00000 |
| 170060222 | 170091699 | -1 | protein_coding | ENSG00000 |
| 170099818 | 170100104 | 1 | processed_pseudoge | ENSG00000 |
| 170141292 | 170142532 | -1 | processed_pseudoge | ENSG00000 |
| 170226535 | 170283723 | 1 | lncRNA | ENSG00000 |
| 170343089 | 170372311 | 1 | lncRNA | ENSG00000 |
| 170481695 | 170482195 | -1 | processed_pseudoge | ENSG00000 |
| 170581470 | 170605450 | -1 | transcribed_processe | ENSG00000 |
| 170742469 | 170743718 | -1 | lncRNA | ENSG00000 |
| 170814339 | 170814548 | -1 | processed_pseudoge | ENSG00000 |
| 170891467 | 170891597 | 1 | snRNA | ENSG00000 |
| 170979831 | 170994656 | 1 | lncRNA | ENSG00000 |
| 171040564 | 171059163 | 1 | lncRNA | ENSG00000 |
| 171113789 | 171114581 | 1 | processed_pseudoge | ENSG00000 |
| 171169209 | 171169942 | -1 | processed_pseudoge | ENSG00000 |
| 171186184 | 171186292 | 1 | miRNA | ENSG00000 |

| | | | | |
|---|---|---|---|---|
| 171289475 | 171298267 | -1 | lncRNA | ENSG00000 |
| 171494684 | 171525072 | -1 | lncRNA | ENSG00000 |
| 171536034 | 171638763 | -1 | lncRNA | ENSG00000 |
| 171553000 | 171564267 | -1 | lncRNA | ENSG00000 |
| 171812254 | 173041559 | 1 | protein_coding | ENSG00000 |
| 172380028 | 172380487 | -1 | processed_pseudoge | ENSG00000 |
| 173094868 | 173169652 | -1 | lncRNA | ENSG00000 |
| 173141643 | 173141938 | 1 | misc_RNA | ENSG00000 |
| 173168811 | 173323967 | 1 | protein_coding | ENSG00000 |
| 173268160 | 173268233 | 1 | miRNA | ENSG00000 |
| 173322206 | 173329694 | -1 | lncRNA | ENSG00000 |
| 173331376 | 173334432 | -1 | protein_coding | ENSG00000 |
| 173349543 | 173370721 | -1 | lncRNA | ENSG00000 |
| 173369969 | 173377532 | 1 | protein_coding | ENSG00000 |
| 173384701 | 173406380 | -1 | protein_coding | ENSG00000 |
| 173397770 | 173409559 | 1 | lncRNA | ENSG00000 |
| 173419245 | 173420137 | -1 | processed_pseudoge | ENSG00000 |
| 173445033 | 173446596 | 1 | lncRNA | ENSG00000 |
| 173446603 | 173446704 | -1 | snRNA | ENSG00000 |
| 173467833 | 173470041 | -1 | lncRNA | ENSG00000 |
| 173509633 | 173511764 | 1 | lncRNA | ENSG00000 |
| 173524969 | 173530229 | -1 | protein_coding | ENSG00000 |
| 173527270 | 173659696 | 1 | lncRNA | ENSG00000 |
| 173615936 | 173616906 | -1 | processed_pseudoge | ENSG00000 |
| 173633728 | 173634371 | -1 | processed_pseudoge | ENSG00000 |
| 173699082 | 174001488 | 1 | lncRNA | ENSG00000 |
| 173877471 | 173913252 | -1 | lncRNA | ENSG00000 |
| 173924207 | 173991024 | -1 | lncRNA | ENSG00000 |
| 174091766 | 174220398 | -1 | lncRNA | ENSG00000 |
| 174135055 | 174154637 | -1 | lncRNA | ENSG00000 |
| 174164441 | 174168913 | -1 | lncRNA | ENSG00000 |
| 174236658 | 174284264 | -1 | protein_coding | ENSG00000 |
| 174283730 | 174333380 | 1 | protein_coding | ENSG00000 |
| 174354854 | 174376445 | -1 | lncRNA | ENSG00000 |
| 174423795 | 174423864 | 1 | miRNA | ENSG00000 |
| 174490175 | 174523154 | -1 | protein_coding | ENSG00000 |
| 174522617 | 174541047 | 1 | lncRNA | ENSG00000 |
| 174523701 | 174540344 | -1 | transcribed_process | ENSG00000 |
| 174636920 | 174829247 | -1 | protein_coding | ENSG00000 |
| 174829668 | 174978180 | 1 | protein_coding | ENSG00000 |
| 174923813 | 174924038 | -1 | processed_pseudoge | ENSG00000 |
| 174977571 | 174979381 | -1 | lncRNA | ENSG00000 |
| 175017016 | 175019214 | 1 | processed_pseudoge | ENSG00000 |
| 175038080 | 175236669 | -1 | lncRNA | ENSG00000 |
| 175286605 | 175290767 | 1 | processed_pseudoge | ENSG00000 |
| 175346358 | 175403110 | -1 | lncRNA | ENSG00000 |
| 175457726 | 175473197 | -1 | lncRNA | ENSG00000 |
| 175494614 | 175496000 | 1 | processed_pseudoge | ENSG00000 |
| 175574868 | 175576140 | 1 | processed_pseudoge | ENSG00000 |
| 175632934 | 176002664 | -1 | protein_coding | ENSG00000 |

| | | | | |
|---|---|---|---|---|
| 175790307 | 175812189 | 1 | lncRNA | ENSG00000 |
| 175962061 | 175962485 | 1 | processed_pseudoge | ENSG00000 |
| 176012038 | 176014417 | -1 | processed_pseudoge | ENSG00000 |
| 176065834 | 176182818 | 1 | protein_coding | ENSG00000 |
| 176098163 | 176098285 | 1 | snoRNA | ENSG00000 |
| 176184579 | 176195585 | -1 | protein_coding | ENSG00000 |
| 176213673 | 176277571 | -1 | protein_coding | ENSG00000 |
| 176308268 | 176320458 | -1 | lncRNA | ENSG00000 |
| 176319966 | 176332245 | 1 | protein_coding | ENSG00000 |
| 176631392 | 176633231 | -1 | processed_pseudoge | ENSG00000 |
| 176669621 | 176706145 | 1 | lncRNA | ENSG00000 |
| 176683538 | 176792922 | -1 | protein_coding | ENSG00000 |
| 176844493 | 177301253 | -1 | lncRNA | ENSG00000 |
| 176866614 | 176875457 | -1 | lncRNA | ENSG00000 |
| 176875688 | 176908031 | 1 | lncRNA | ENSG00000 |
| 177181442 | 177181741 | 1 | misc_RNA | ENSG00000 |
| 177242539 | 177248773 | -1 | lncRNA | ENSG00000 |
| 177309874 | 177362936 | 1 | protein_coding | ENSG00000 |
| 177343004 | 177357032 | -1 | lncRNA | ENSG00000 |
| 177430774 | 177442437 | -1 | protein_coding | ENSG00000 |
| 177442519 | 177681381 | 1 | lncRNA | ENSG00000 |
| 177457111 | 177457228 | -1 | rRNA_pseudogene | ENSG00000 |
| 177457111 | 177457228 | -1 | rRNA | ENSG00000 |
| 177686673 | 177691068 | -1 | lncRNA | ENSG00000 |
| 177728757 | 178028971 | 1 | lncRNA | ENSG00000 |
| 177729638 | 177907936 | -1 | lncRNA | ENSG00000 |
| 178116721 | 178116861 | -1 | snRNA | ENSG00000 |
| 178286695 | 178287285 | 1 | processed_pseudoge | ENSG00000 |
| 178406695 | 178406830 | 1 | rRNA_pseudogene | ENSG00000 |
| 178685886 | 178685956 | -1 | snoRNA | ENSG00000 |
| 179069437 | 179069594 | 1 | unprocessed_pseudc | ENSG00000 |
| 179389134 | 179465305 | -1 | lncRNA | ENSG00000 |
| 179856853 | 179857581 | 1 | processed_pseudoge | ENSG00000 |
| 179923773 | 180059792 | -1 | lncRNA | ENSG00000 |
| 180095527 | 180117163 | -1 | lncRNA | ENSG00000 |
| 180573464 | 180574027 | -1 | processed_pseudoge | ENSG00000 |
| 180731164 | 180759038 | -1 | lncRNA | ENSG00000 |
| 181064089 | 181159149 | -1 | lncRNA | ENSG00000 |
| 181237358 | 181237564 | -1 | unprocessed_pseudc | ENSG00000 |
| 181260442 | 181265145 | -1 | lncRNA | ENSG00000 |
| 181522666 | 181523001 | 1 | processed_pseudoge | ENSG00000 |
| 181590255 | 181591969 | 1 | lncRNA | ENSG00000 |
| 181686182 | 181700142 | -1 | lncRNA | ENSG00000 |
| 181820019 | 181829973 | -1 | lncRNA | ENSG00000 |
| 181834865 | 181835108 | -1 | misc_RNA | ENSG00000 |
| 181874438 | 182145249 | -1 | lncRNA | ENSG00000 |
| 181905790 | 181906467 | -1 | processed_pseudoge | ENSG00000 |
| 182143987 | 182803024 | 1 | protein_coding | ENSG00000 |
| 182144852 | 182190382 | 1 | lncRNA | ENSG00000 |
| 182169293 | 182169378 | 1 | miRNA | ENSG00000 |

| | | | | |
|---|---|---|---|---|
| 182548659 | 182548779 | 1 | snRNA | ENSG00000 |
| 182631454 | 182631688 | -1 | misc_RNA | ENSG00000 |
| 182697902 | 182708042 | -1 | lncRNA | ENSG00000 |
| 182772565 | 182773931 | -1 | lncRNA | ENSG00000 |
| 182828100 | 182830597 | 1 | lncRNA | ENSG00000 |
| 182881777 | 182882203 | -1 | lncRNA | ENSG00000 |
| 182890060 | 182917936 | -1 | protein_coding | ENSG00000 |
| 183017426 | 183036100 | 1 | lncRNA | ENSG00000 |
| 183028601 | 183029720 | -1 | processed_pseudoge | ENSG00000 |
| 183037665 | 183040119 | 1 | transcribed_processe | ENSG00000 |
| 183075480 | 183076033 | -1 | lncRNA | ENSG00000 |
| 183094423 | 183095056 | -1 | lncRNA | ENSG00000 |
| 183097017 | 183099199 | -1 | lncRNA | ENSG00000 |
| 183099257 | 183320777 | 1 | protein_coding | ENSG00000 |
| 183233628 | 183240634 | -1 | lncRNA | ENSG00000 |
| 183318194 | 183320774 | -1 | protein_coding | ENSG00000 |
| 183321764 | 183322426 | -1 | protein_coding | ENSG00000 |
| 183380345 | 183381705 | 1 | processed_pseudoge | ENSG00000 |
| 183418890 | 183419805 | -1 | processed_pseudoge | ENSG00000 |
| 183444635 | 183449064 | 1 | protein_coding | ENSG00000 |
| 183483490 | 183484185 | 1 | processed_pseudoge | ENSG00000 |
| 183494884 | 183504494 | -1 | lncRNA | ENSG00000 |
| 183505058 | 183512429 | 1 | protein_coding | ENSG00000 |
| 183516894 | 183517527 | 1 | lncRNA | ENSG00000 |
| 183639635 | 183659203 | -1 | protein_coding | ENSG00000 |
| 183653572 | 183653676 | -1 | snRNA | ENSG00000 |
| 183659267 | 183713594 | 1 | protein_coding | ENSG00000 |
| 183675603 | 183675715 | -1 | snRNA | ENSG00000 |
| 183705931 | 183706031 | 1 | snRNA | ENSG00000 |
| 183725830 | 183732570 | 1 | lncRNA | ENSG00000 |
| 183762115 | 183765721 | -1 | lncRNA | ENSG00000 |
| 183797692 | 184023526 | 1 | protein_coding | ENSG00000 |
| 183987669 | 183996680 | 1 | lncRNA | ENSG00000 |
| 184014035 | 184016382 | 1 | TEC | ENSG00000 |
| 184072403 | 184073039 | 1 | processed_pseudoge | ENSG00000 |
| 184088706 | 184221230 | -1 | protein_coding | ENSG00000 |
| 184100482 | 184106038 | 1 | lncRNA | ENSG00000 |
| 184254509 | 184255330 | 1 | processed_pseudoge | ENSG00000 |
| 184278936 | 184279233 | 1 | misc_RNA | ENSG00000 |
| 184299129 | 184299640 | -1 | processed_pseudoge | ENSG00000 |
| 184334292 | 184353977 | -1 | lncRNA | ENSG00000 |
| 184365180 | 184382416 | -1 | lncRNA | ENSG00000 |
| 184387729 | 184474550 | -1 | protein_coding | ENSG00000 |
| 184431966 | 184432049 | 1 | snoRNA | ENSG00000 |
| 184471924 | 184472609 | -1 | lncRNA | ENSG00000 |
| 184474802 | 184477304 | 1 | lncRNA | ENSG00000 |
| 184484141 | 184490293 | 1 | lncRNA | ENSG00000 |
| 184503271 | 184537626 | -1 | lncRNA | ENSG00000 |
| 184559175 | 184559783 | -1 | processed_pseudoge | ENSG00000 |
| 184584093 | 184625030 | -1 | lncRNA | ENSG00000 |

| | | | | |
|---|---|---|---|---|
| 184627696 | 184649509 | -1 | protein_coding | ENSG00000 |
| 184649667 | 184694963 | 1 | protein_coding | ENSG00000 |
| 184694085 | 184734130 | -1 | protein_coding | ENSG00000 |
| 184755595 | 184826818 | -1 | protein_coding | ENSG00000 |
| 184798296 | 184799046 | -1 | transcribed_processe | ENSG00000 |
| 184813619 | 184821300 | 1 | lncRNA | ENSG00000 |
| 184826607 | 184850920 | 1 | lncRNA | ENSG00000 |
| 184844554 | 184855751 | -1 | lncRNA | ENSG00000 |
| 184851013 | 184851110 | -1 | miRNA | ENSG00000 |
| 184892858 | 184899553 | -1 | lncRNA | ENSG00000 |
| 184910889 | 184911352 | -1 | processed_pseudoge | ENSG00000 |
| 184938383 | 184938440 | -1 | miRNA | ENSG00000 |
| 184984277 | 184988127 | -1 | TEC | ENSG00000 |
| 184988997 | 185005186 | 1 | lncRNA | ENSG00000 |
| 185018841 | 185020804 | 1 | protein_coding | ENSG00000 |
| 185051081 | 185115500 | 1 | lncRNA | ENSG00000 |
| 185064981 | 185065323 | 1 | processed_pseudoge | ENSG00000 |
| 185118984 | 185127478 | -1 | lncRNA | ENSG00000 |
| 185143266 | 185150382 | 1 | protein_coding | ENSG00000 |
| 185159665 | 185209504 | -1 | protein_coding | ENSG00000 |
| 185204237 | 185370185 | 1 | protein_coding | ENSG00000 |
| 185363879 | 185395899 | -1 | protein_coding | ENSG00000 |
| 185370725 | 185390928 | 1 | lncRNA | ENSG00000 |
| 185396021 | 185400628 | 1 | protein_coding | ENSG00000 |
| 185399537 | 185425979 | -1 | protein_coding | ENSG00000 |
| 185415973 | 185416086 | -1 | misc_RNA | ENSG00000 |
| 185426249 | 185449826 | 1 | protein_coding | ENSG00000 |
| 185445182 | 185471752 | -1 | protein_coding | ENSG00000 |
| 185471461 | 185507055 | 1 | lncRNA | ENSG00000 |
| 185500660 | 185535612 | -1 | protein_coding | ENSG00000 |
| 185585444 | 185956652 | -1 | protein_coding | ENSG00000 |
| 185587909 | 185594003 | 1 | lncRNA | ENSG00000 |
| 185665885 | 185675417 | 1 | lncRNA | ENSG00000 |
| 185709873 | 185709967 | -1 | misc_RNA | ENSG00000 |
| 185756929 | 185757048 | -1 | processed_pseudoge | ENSG00000 |
| 185918140 | 185919460 | 1 | lncRNA | ENSG00000 |
| 186026456 | 186026586 | -1 | snRNA | ENSG00000 |
| 186069152 | 186088069 | 1 | protein_coding | ENSG00000 |
| 186104419 | 186172667 | 1 | protein_coding | ENSG00000 |
| 186130021 | 186130155 | 1 | snoRNA | ENSG00000 |
| 186157896 | 186158843 | 1 | processed_pseudoge | ENSG00000 |
| 186170863 | 186171257 | -1 | processed_pseudoge | ENSG00000 |
| 186189032 | 186191490 | -1 | TEC | ENSG00000 |
| 186191567 | 186213463 | 1 | protein_coding | ENSG00000 |
| 186208979 | 186258471 | 1 | protein_coding | ENSG00000 |
| 186265945 | 186288806 | 1 | protein_coding | ENSG00000 |
| 186286094 | 186500997 | -1 | lncRNA | ENSG00000 |
| 186326302 | 186336662 | 1 | lncRNA | ENSG00000 |
| 186328716 | 186329601 | -1 | processed_pseudoge | ENSG00000 |
| 186426546 | 186555328 | -1 | protein_coding | ENSG00000 |

| | | | | |
|---|---|---|---|---|
| 186501109 | 186501215 | 1 | snRNA | ENSG0000C |
| 186533655 | 186555567 | -1 | protein_coding | ENSG0000C |
| 186587794 | 186726722 | -1 | protein_coding | ENSG0000C |
| 186701579 | 186704086 | 1 | lncRNA | ENSG0000C |
| 186890969 | 186892366 | 1 | lncRNA | ENSG0000C |
| 186892552 | 186892983 | 1 | lncRNA | ENSG0000C |
| 186900540 | 186900838 | 1 | processed_pseudoge | ENSG0000C |
| 186979490 | 187027451 | -1 | lncRNA | ENSG0000C |
| 187005944 | 187060930 | 1 | lncRNA | ENSG0000C |
| 187198218 | 187201894 | -1 | lncRNA | ENSG0000C |
| 187201986 | 187209346 | 1 | lncRNA | ENSG0000C |
| 187304083 | 187309682 | -1 | lncRNA | ENSG0000C |
| 187370648 | 187371921 | 1 | lncRNA | ENSG0000C |
| 187408232 | 187415748 | 1 | lncRNA | ENSG0000C |
| 187441677 | 187444298 | -1 | processed_pseudoge | ENSG0000C |
| 187516153 | 187517208 | -1 | lncRNA | ENSG0000C |
| 187532873 | 187672755 | -1 | lncRNA | ENSG0000C |
| 187613708 | 187712329 | 1 | lncRNA | ENSG0000C |
| 187746094 | 187748199 | -1 | processed_pseudoge | ENSG0000C |
| 187942164 | 187963345 | -1 | lncRNA | ENSG0000C |
| 187970273 | 187971284 | 1 | processed_pseudoge | ENSG0000C |
| 187995771 | 188005046 | 1 | protein_coding | ENSG0000C |
| 188091273 | 188109603 | -1 | protein_coding | ENSG0000C |
| 188106558 | 188106914 | 1 | processed_pseudoge | ENSG0000C |
| 188121531 | 188121634 | -1 | snRNA | ENSG0000C |
| 188139441 | 188147743 | 1 | protein_coding | ENSG0000C |
| 188140822 | 188143379 | -1 | lncRNA | ENSG0000C |
| 188159705 | 188161157 | 1 | lncRNA | ENSG0000C |
| 188349376 | 188350024 | -1 | processed_pseudoge | ENSG0000C |
| 188387471 | 188389666 | -1 | processed_pseudoge | ENSG0000C |
| 188400736 | 188681051 | 1 | lncRNA | ENSG0000C |
| 188630924 | 188631747 | 1 | processed_pseudoge | ENSG0000C |
| 188713644 | 188715442 | 1 | processed_pseudoge | ENSG0000C |
| 188716665 | 188716725 | -1 | snRNA | ENSG0000C |
| 188738373 | 188739494 | 1 | processed_pseudoge | ENSG0000C |
| 188740507 | 188741281 | -1 | TEC | ENSG0000C |
| 188757678 | 188779343 | 1 | lncRNA | ENSG0000C |
| 188776657 | 188785511 | 1 | lncRNA | ENSG0000C |
| 188777672 | 188818522 | -1 | lncRNA | ENSG0000C |
| 188990715 | 189002075 | -1 | lncRNA | ENSG0000C |
| 189193100 | 189204877 | 1 | lncRNA | ENSG0000C |
| 189288677 | 189291336 | -1 | lncRNA | ENSG0000C |
| 189472965 | 189475192 | 1 | transcribed_processe | ENSG0000C |
| 189659605 | 189661486 | 1 | lncRNA | ENSG0000C |
| 189703881 | 189704490 | -1 | lncRNA | ENSG0000C |
| 189709539 | 189709714 | -1 | snRNA | ENSG0000C |
| 189764391 | 189940733 | -1 | lncRNA | ENSG0000C |
| 189881515 | 189884868 | 1 | lncRNA | ENSG0000C |
| 189940870 | 189963202 | 1 | protein_coding | ENSG0000C |
| 189973599 | 189973847 | -1 | processed_pseudoge | ENSG0000C |

| | | | | |
|---|---|---|---|---|
| 189979902 | 189982379 | 1 | processed_pseudoge | ENSG00000( |
| 189982523 | 189984871 | -1 | unprocessed_pseudc | ENSG00000( |
| 190015138 | 190015255 | 1 | rRNA_pseudogene | ENSG00000( |
| 190017696 | 190017814 | 1 | rRNA_pseudogene | ENSG00000( |
| 190021407 | 190022665 | -1 | unprocessed_pseudc | ENSG00000( |
| 190024351 | 190027257 | -1 | protein_coding | ENSG00000( |
| 190037916 | 190038399 | -1 | processed_pseudoge | ENSG00000( |
| 190041342 | 190043414 | 1 | processed_pseudoge | ENSG00000( |
| 190057902 | 190063005 | -1 | unprocessed_pseudc | ENSG00000( |
| 190064502 | 190067864 | 1 | lncRNA | ENSG00000( |
| 190065233 | 190065914 | 1 | unprocessed_pseudc | ENSG00000( |
| 190067935 | 190069209 | 1 | unprocessed_pseudc | ENSG00000( |
| 190071232 | 190072506 | 1 | unprocessed_pseudc | ENSG00000( |
| 190074525 | 190075799 | 1 | unprocessed_pseudc | ENSG00000( |
| 190077818 | 190079092 | 1 | unprocessed_pseudc | ENSG00000( |
| 190081118 | 190082386 | 1 | unprocessed_pseudc | ENSG00000( |
| 190084412 | 190085686 | 1 | unprocessed_pseudc | ENSG00000( |
| 190087705 | 190088979 | 1 | unprocessed_pseudc | ENSG00000( |
| 190091005 | 190092279 | 1 | unprocessed_pseudc | ENSG00000( |
| 190173774 | 190185942 | 1 | protein_coding | ENSG00000( |
| 190195727 | 190196190 | 1 | processed_pseudoge | ENSG00000( |
| 190195738 | 190195978 | -1 | processed_pseudoge | ENSG00000( |
| 58198 | 58915 | 1 | processed_pseudoge | ENSG00000( |
| 92151 | 189972 | 1 | protein_coding | ENSG00000( |
| 144087 | 144197 | -1 | misc_RNA | ENSG00000( |
| 191495 | 196334 | 1 | protein_coding | ENSG00000( |
| 195778 | 196341 | 1 | lncRNA | ENSG00000( |
| 196868 | 218153 | -1 | protein_coding | ENSG00000( |
| 213898 | 217279 | -1 | lncRNA | ENSG00000( |
| 218241 | 257082 | 1 | protein_coding | ENSG00000( |
| 218298 | 314955 | 1 | protein_coding | ENSG00000( |
| 269858 | 271516 | -1 | lncRNA | ENSG00000( |
| 271621 | 314974 | 1 | protein_coding | ENSG00000( |
| 271670 | 438291 | 1 | protein_coding | ENSG00000( |
| 288833 | 290321 | -1 | processed_pseudoge | ENSG00000( |
| 321759 | 435110 | 1 | protein_coding | ENSG00000( |
| 403731 | 404546 | -1 | processed_pseudoge | ENSG00000( |
| 418703 | 419109 | 1 | processed_pseudoge | ENSG00000( |
| 441498 | 443160 | -1 | lncRNA | ENSG00000( |
| 443175 | 471937 | 1 | protein_coding | ENSG00000( |
| 466124 | 473098 | -1 | lncRNA | ENSG00000( |
| 470456 | 524449 | -1 | protein_coding | ENSG00000( |
| 473236 | 480884 | 1 | lncRNA | ENSG00000( |
| 524109 | 527448 | 1 | lncRNA | ENSG00000( |
| 535840 | 535882 | -1 | miRNA | ENSG00000( |
| 602620 | 612210 | -1 | lncRNA | ENSG00000( |
| 612340 | 667168 | 1 | protein_coding | ENSG00000( |
| 659862 | 693352 | -1 | protein_coding | ENSG00000( |
| 675826 | 676616 | 1 | lncRNA | ENSG00000( |
| 710355 | 784729 | -1 | protein_coding | ENSG00000( |

| | | | | |
|---|---|---|---|---|
| 716808 | 766919 | -1 | protein_coding | ENSG00000 |
| 767382 | 768930 | -1 | transcribed_unproce | ENSG00000 |
| 770005 | 770113 | -1 | transcribed_unproce | ENSG00000 |
| 795606 | 850986 | -1 | protein_coding | ENSG00000 |
| 816346 | 817001 | 1 | processed_pseudoge | ENSG00000 |
| 850291 | 892801 | -1 | protein_coding | ENSG00000 |
| 892884 | 919357 | 1 | protein_coding | ENSG00000 |
| 904319 | 912587 | -1 | lncRNA | ENSG00000 |
| 946315 | 956520 | 1 | lncRNA | ENSG00000 |
| 979365 | 981300 | 1 | lncRNA | ENSG00000 |
| 987180 | 997308 | -1 | lncRNA | ENSG00000 |
| 1005039 | 1006623 | 1 | lncRNA | ENSG00000 |
| 1008802 | 1038943 | 1 | protein_coding | ENSG00000 |
| 1050384 | 1112063 | -1 | protein_coding | ENSG00000 |
| 1062896 | 1062974 | -1 | miRNA | ENSG00000 |
| 1173141 | 1178605 | -1 | lncRNA | ENSG00000 |
| 1201595 | 1225111 | 1 | protein_coding | ENSG00000 |
| 1225381 | 1246189 | 1 | protein_coding | ENSG00000 |
| 1252006 | 1255976 | 1 | lncRNA | ENSG00000 |
| 1253147 | 1295047 | -1 | protein_coding | ENSG00000 |
| 1309310 | 1309377 | -1 | miRNA | ENSG00000 |
| 1317752 | 1345099 | -1 | protein_coding | ENSG00000 |
| 1345197 | 1350786 | 1 | lncRNA | ENSG00000 |
| 1353896 | 1380107 | -1 | lncRNA | ENSG00000 |
| 1370543 | 1370945 | -1 | processed_pseudoge | ENSG00000 |
| 1383159 | 1386409 | -1 | TEC | ENSG00000 |
| 1392794 | 1445440 | -1 | protein_coding | ENSG00000 |
| 1456480 | 1523962 | -1 | protein_coding | ENSG00000 |
| 1510762 | 1510856 | -1 | miRNA | ENSG00000 |
| 1544107 | 1551710 | -1 | lncRNA | ENSG00000 |
| 1567495 | 1594620 | -1 | transcribed_unproce | ENSG00000 |
| 1594626 | 1611467 | 1 | lncRNA | ENSG00000 |
| 1598127 | 1598247 | 1 | unprocessed_pseudc | ENSG00000 |
| 1598920 | 1634005 | -1 | transcribed_unproce | ENSG00000 |
| 1614836 | 1616334 | 1 | processed_pseudoge | ENSG00000 |
| 1708785 | 1708868 | -1 | miRNA | ENSG00000 |
| 1725149 | 1728172 | 1 | lncRNA | ENSG00000 |
| 1798385 | 1801366 | -1 | protein_coding | ENSG00000 |
| 1801400 | 1816605 | 1 | protein_coding | ENSG00000 |
| 1850950 | 1851497 | 1 | lncRNA | ENSG00000 |
| 1855970 | 1856568 | -1 | lncRNA | ENSG00000 |
| 1858409 | 1859890 | -1 | TEC | ENSG00000 |
| 1877413 | 1887236 | -1 | protein_coding | ENSG00000 |
| 1883966 | 1884649 | 1 | lncRNA | ENSG00000 |
| 1887332 | 1900493 | 1 | lncRNA | ENSG00000 |
| 1931058 | 1933985 | -1 | lncRNA | ENSG00000 |
| 1931064 | 2119926 | 1 | lncRNA | ENSG00000 |
| 1963346 | 1967251 | -1 | lncRNA | ENSG00000 |
| 1968094 | 1969013 | 1 | lncRNA | ENSG00000 |
| 2184710 | 2184820 | 1 | misc_RNA | ENSG00000 |

| | | | | |
|---|---|---|---|---|
| 2261859 | 2262023 | 1 | processed_pseudoge | ENSG00000( |
| 2712591 | 2715237 | 1 | lncRNA | ENSG00000( |
| 2736662 | 2737759 | -1 | lncRNA | ENSG00000( |
| 2745845 | 2751677 | -1 | protein_coding | ENSG00000( |
| 2752131 | 2755397 | 1 | protein_coding | ENSG00000( |
| 2787451 | 2788549 | -1 | processed_pseudoge | ENSG00000( |
| 2831021 | 2835139 | -1 | lncRNA | ENSG00000( |
| 2921496 | 2967955 | 1 | lncRNA | ENSG00000( |
| 3177835 | 3181487 | 1 | lncRNA | ENSG00000( |
| 3357264 | 3536094 | -1 | lncRNA | ENSG00000( |
| 3452816 | 3461660 | 1 | lncRNA | ENSG00000( |
| 3496229 | 3504004 | -1 | lncRNA | ENSG00000( |
| 3595832 | 3601403 | 1 | protein_coding | ENSG00000( |
| 3596211 | 3600188 | -1 | lncRNA | ENSG00000( |
| 4012708 | 4013649 | -1 | lncRNA | ENSG00000( |
| 4033713 | 4041764 | -1 | lncRNA | ENSG00000( |
| 4135638 | 4147429 | 1 | lncRNA | ENSG00000( |
| 4436850 | 4440259 | -1 | lncRNA | ENSG00000( |
| 4451930 | 4866311 | -1 | lncRNA | ENSG00000( |
| 4512262 | 4516776 | 1 | lncRNA | ENSG00000( |
| 4640731 | 4648254 | 1 | lncRNA | ENSG00000( |
| 4773481 | 4774865 | 1 | lncRNA | ENSG00000( |
| 4775476 | 4805769 | -1 | lncRNA | ENSG00000( |
| 4866521 | 4874759 | 1 | lncRNA | ENSG00000( |
| 4967764 | 4970425 | -1 | lncRNA | ENSG00000( |
| 4974690 | 5034267 | -1 | lncRNA | ENSG00000( |
| 4987789 | 5070314 | 1 | lncRNA | ENSG00000( |
| 5046796 | 5049497 | -1 | lncRNA | ENSG00000( |
| 5069192 | 5078311 | -1 | lncRNA | ENSG00000( |
| 5126046 | 5126337 | 1 | misc_RNA | ENSG00000( |
| 5132101 | 5140119 | -1 | lncRNA | ENSG00000( |
| 5140330 | 5320304 | 1 | protein_coding | ENSG00000( |
| 5142138 | 5176214 | -1 | lncRNA | ENSG00000( |
| 5375723 | 5376836 | -1 | processed_pseudoge | ENSG00000( |
| 5390096 | 5422116 | -1 | lncRNA | ENSG00000( |
| 5395378 | 5395957 | -1 | processed_pseudoge | ENSG00000( |
| 5395972 | 5396335 | -1 | processed_pseudoge | ENSG00000( |
| 5396477 | 5396968 | -1 | processed_pseudoge | ENSG00000( |
| 5420664 | 5490220 | 1 | protein_coding | ENSG00000( |
| 5688805 | 5694742 | 1 | lncRNA | ENSG00000( |
| 6019029 | 6022283 | -1 | lncRNA | ENSG00000( |
| 6030035 | 6030841 | 1 | lncRNA | ENSG00000( |
| 6134303 | 6137171 | 1 | TEC | ENSG00000( |
| 6289857 | 6290624 | 1 | processed_pseudoge | ENSG00000( |
| 6310441 | 6339884 | -1 | lncRNA | ENSG00000( |
| 6371874 | 6378571 | -1 | protein_coding | ENSG00000( |
| 6378674 | 6411690 | 1 | lncRNA | ENSG00000( |
| 6448859 | 6496723 | 1 | protein_coding | ENSG00000( |
| 6532891 | 6533200 | -1 | processed_pseudoge | ENSG00000( |
| 6582136 | 6600288 | 1 | lncRNA | ENSG00000( |

| | | | | |
|---|---|---|---|---|
| 6599239 | 6633291 | -1 | protein_coding | ENSG0000C |
| 6633427 | 6674386 | 1 | protein_coding | ENSG0000C |
| 6686325 | 6707711 | -1 | lncRNA | ENSG0000C |
| 6713432 | 6757044 | 1 | protein_coding | ENSG0000C |
| 6765891 | 6833120 | 1 | lncRNA | ENSG0000C |
| 6779458 | 6779998 | 1 | lncRNA | ENSG0000C |
| 6827853 | 6827921 | -1 | miRNA | ENSG0000C |
| 6839460 | 7219291 | 1 | lncRNA | ENSG0000C |
| 6848127 | 6848462 | -1 | misc_RNA | ENSG0000C |
| 6868554 | 6888087 | 1 | lncRNA | ENSG0000C |
| 7047227 | 7047336 | 1 | rRNA_pseudogene | ENSG0000C |
| 7065091 | 7069351 | -1 | lncRNA | ENSG0000C |
| 7226156 | 7229357 | -1 | lncRNA | ENSG0000C |
| 7239781 | 7240018 | 1 | misc_RNA | ENSG0000C |
| 7272792 | 7276923 | 1 | lncRNA | ENSG0000C |
| 7287924 | 7288477 | -1 | processed_pseudoge | ENSG0000C |
| 7289891 | 7326609 | 1 | transcribed_unproce | ENSG0000C |
| 7299774 | 7306592 | -1 | unprocessed_pseudc | ENSG0000C |
| 7344579 | 7345921 | 1 | lncRNA | ENSG0000C |
| 7346986 | 7348671 | -1 | lncRNA | ENSG0000C |
| 7362979 | 7364531 | -1 | lncRNA | ENSG0000C |
| 7367929 | 7373074 | -1 | lncRNA | ENSG0000C |
| 7373115 | 7391907 | 1 | lncRNA | ENSG0000C |
| 7396138 | 7830081 | 1 | protein_coding | ENSG0000C |
| 7707802 | 7749766 | -1 | lncRNA | ENSG0000C |
| 7830378 | 7851151 | -1 | protein_coding | ENSG0000C |
| 7851186 | 7906025 | 1 | protein_coding | ENSG0000C |
| 7859159 | 7869037 | -1 | protein_coding | ENSG0000C |
| 7924292 | 7925398 | 1 | lncRNA | ENSG0000C |
| 7980146 | 7980304 | 1 | snRNA | ENSG0000C |
| 8012377 | 8014804 | 1 | lncRNA | ENSG0000C |
| 8080186 | 8080761 | -1 | TEC | ENSG0000C |
| 8143618 | 8143784 | 1 | TEC | ENSG0000C |
| 8157541 | 8157788 | -1 | unprocessed_pseudc | ENSG0000C |
| 8207019 | 8366437 | -1 | lncRNA | ENSG0000C |
| 8333481 | 8457558 | -1 | lncRNA | ENSG0000C |
| 8387638 | 8457564 | -1 | lncRNA | ENSG0000C |
| 8410789 | 8410996 | -1 | unprocessed_pseudc | ENSG0000C |
| 8444949 | 8445535 | -1 | lncRNA | ENSG0000C |
| 8450701 | 8486930 | 1 | lncRNA | ENSG0000C |
| 8455946 | 8456558 | -1 | unprocessed_pseudc | ENSG0000C |
| 8458842 | 8459183 | 1 | TEC | ENSG0000C |
| 8460925 | 8460999 | 1 | miRNA | ENSG0000C |
| 8525144 | 8561682 | 1 | lncRNA | ENSG0000C |
| 8572047 | 8572206 | 1 | TEC | ENSG0000C |
| 8614356 | 8614696 | 1 | TEC | ENSG0000C |
| 8618581 | 8619104 | -1 | processed_pseudoge | ENSG0000C |
| 8619178 | 8620310 | 1 | processed_pseudoge | ENSG0000C |
| 8659764 | 8660059 | -1 | TEC | ENSG0000C |
| 8669018 | 8669250 | -1 | TEC | ENSG0000C |

| | | | | |
|---|---|---|---|---|
| 8685581 | 8685816 | -1 | TEC | ENSG00000 |
| 8717684 | 8717883 | -1 | TEC | ENSG00000 |
| 8754869 | 8755159 | 1 | TEC | ENSG00000 |
| 8785042 | 8785468 | -1 | lncRNA | ENSG00000 |
| 8793875 | 8794197 | 1 | TEC | ENSG00000 |
| 8839700 | 8898052 | 1 | lncRNA | ENSG00000 |
| 8980404 | 8980659 | -1 | TEC | ENSG00000 |
| 9001774 | 9045940 | 1 | lncRNA | ENSG00000 |
| 9035033 | 9546075 | -1 | protein_coding | ENSG00000 |
| 9053816 | 9053895 | -1 | miRNA | ENSG00000 |
| 9363275 | 9422481 | 1 | lncRNA | ENSG00000 |
| 9498401 | 9498775 | -1 | TEC | ENSG00000 |
| 9511333 | 9518086 | 1 | lncRNA | ENSG00000 |
| 9519853 | 9523178 | -1 | lncRNA | ENSG00000 |
| 9546200 | 9550609 | 1 | lncRNA | ENSG00000 |
| 9548836 | 9548905 | 1 | snoRNA | ENSG00000 |
| 9621377 | 9658458 | -1 | lncRNA | ENSG00000 |
| 9628997 | 9712378 | -1 | protein_coding | ENSG00000 |
| 9641305 | 9903852 | -1 | lncRNA | ENSG00000 |
| 9760907 | 9761013 | 1 | rRNA_pseudogene | ENSG00000 |
| 9789295 | 9812664 | -1 | lncRNA | ENSG00000 |
| 9854371 | 9899970 | 1 | lncRNA | ENSG00000 |
| 10057502 | 10057807 | 1 | TEC | ENSG00000 |
| 10064243 | 10064476 | -1 | TEC | ENSG00000 |
| 10070531 | 10070841 | 1 | TEC | ENSG00000 |
| 10082571 | 10082772 | 1 | TEC | ENSG00000 |
| 10137138 | 10138365 | 1 | lncRNA | ENSG00000 |
| 10168783 | 10169006 | 1 | TEC | ENSG00000 |
| 10171030 | 10171214 | -1 | TEC | ENSG00000 |
| 10195121 | 10197628 | -1 | lncRNA | ENSG00000 |
| 10195187 | 10197622 | -1 | lncRNA | ENSG00000 |
| 10203600 | 10204040 | -1 | lncRNA | ENSG00000 |
| 10225507 | 10249897 | -1 | protein_coding | ENSG00000 |
| 10248325 | 10249915 | 1 | lncRNA | ENSG00000 |
| 10249929 | 10266389 | 1 | protein_coding | ENSG00000 |
| 10264597 | 10267146 | -1 | lncRNA | ENSG00000 |
| 10269489 | 10269918 | -1 | lncRNA | ENSG00000 |
| 10275875 | 10307902 | -1 | protein_coding | ENSG00000 |
| 10292041 | 10292148 | -1 | misc_RNA | ENSG00000 |
| 10337277 | 10337386 | 1 | misc_RNA | ENSG00000 |
| 10342813 | 10343131 | 1 | TEC | ENSG00000 |
| 10352701 | 10353601 | -1 | lncRNA | ENSG00000 |
| 10353695 | 10440388 | 1 | protein_coding | ENSG00000 |
| 10441290 | 10441792 | -1 | lncRNA | ENSG00000 |
| 10441524 | 10472029 | 1 | protein_coding | ENSG00000 |
| 10478037 | 10478145 | 1 | miRNA | ENSG00000 |
| 10479371 | 10482309 | 1 | lncRNA | ENSG00000 |
| 10488820 | 10489142 | 1 | processed_pseudoge | ENSG00000 |
| 10493291 | 10502728 | -1 | lncRNA | ENSG00000 |
| 10493291 | 10494094 | -1 | lncRNA | ENSG00000 |

| | | | | |
|---|---|---|---|---|
| 10504996 | 10522084 | -1 | lncRNA | ENSG0000C |
| 10541454 | 10543192 | 1 | TEC | ENSG0000C |
| 10556582 | 10556815 | 1 | TEC | ENSG0000C |
| 10564070 | 10657816 | 1 | protein_coding | ENSG0000C |
| 10627260 | 10628225 | -1 | lncRNA | ENSG0000C |
| 10652211 | 10655925 | -1 | unprocessed_pseudc | ENSG0000C |
| 10664495 | 10665113 | -1 | processed_pseudoge | ENSG0000C |
| 10675317 | 10675575 | -1 | TEC | ENSG0000C |
| 10676096 | 10676423 | -1 | TEC | ENSG0000C |
| 10679230 | 10761272 | -1 | protein_coding | ENSG0000C |
| 10761065 | 10770294 | 1 | lncRNA | ENSG0000C |
| 10775058 | 10777062 | -1 | lncRNA | ENSG0000C |
| 10971836 | 11904446 | -1 | protein_coding | ENSG0000C |
| 11027200 | 11027307 | 1 | snRNA | ENSG0000C |
| 12297399 | 12297504 | -1 | snRNA | ENSG0000C |
| 12553890 | 12574637 | -1 | lncRNA | ENSG0000C |
| 12574830 | 12804363 | 1 | lncRNA | ENSG0000C |
| 12794100 | 12794731 | -1 | processed_pseudoge | ENSG0000C |
| 12794731 | 12795522 | 1 | processed_pseudoge | ENSG0000C |
| 12914068 | 13032886 | -1 | lncRNA | ENSG0000C |
| 13144000 | 13190677 | 1 | lncRNA | ENSG0000C |
| 13250235 | 13250662 | -1 | processed_pseudoge | ENSG0000C |
| 13553654 | 13553966 | -1 | processed_pseudoge | ENSG0000C |
| 13637919 | 13638381 | -1 | processed_pseudoge | ENSG0000C |
| 13690331 | 13944543 | -1 | protein_coding | ENSG0000C |
| 13860303 | 13901036 | 1 | lncRNA | ENSG0000C |
| 14143342 | 14532128 | 1 | protein_coding | ENSG0000C |
| 14356886 | 14359285 | -1 | lncRNA | ENSG0000C |
| 14581792 | 14616180 | 1 | protein_coding | ENSG0000C |
| 14620020 | 14621313 | 1 | processed_pseudoge | ENSG0000C |
| 14639426 | 14641018 | -1 | processed_pseudoge | ENSG0000C |
| 14651941 | 14653329 | -1 | processed_pseudoge | ENSG0000C |
| 14661808 | 14664604 | -1 | lncRNA | ENSG0000C |
| 14664664 | 14699850 | 1 | protein_coding | ENSG0000C |
| 14704800 | 14871778 | -1 | protein_coding | ENSG0000C |
| 14728587 | 14734258 | 1 | lncRNA | ENSG0000C |
| 14797125 | 14798400 | -1 | processed_pseudoge | ENSG0000C |
| 14825929 | 14826012 | -1 | miRNA | ENSG0000C |
| 14872332 | 14872713 | 1 | lncRNA | ENSG0000C |
| 14874400 | 14874711 | -1 | processed_pseudoge | ENSG0000C |
| 14877667 | 14877919 | -1 | processed_pseudoge | ENSG0000C |
| 14960232 | 14961393 | 1 | processed_pseudoge | ENSG0000C |
| 14970909 | 14992961 | 1 | lncRNA | ENSG0000C |
| 15110786 | 15110920 | -1 | snoRNA | ENSG0000C |
| 15112120 | 15266541 | -1 | lncRNA | ENSG0000C |
| 15384220 | 15384853 | 1 | processed_pseudoge | ENSG0000C |
| 15425758 | 15496353 | 1 | lncRNA | ENSG0000C |
| 15450701 | 15451811 | 1 | processed_pseudoge | ENSG0000C |
| 15500180 | 15939795 | 1 | protein_coding | ENSG0000C |
| 15602189 | 15619109 | -1 | lncRNA | ENSG0000C |

| | | | | |
|---|---|---|---|---|
| 15935182 | 15935260 | 1 | miRNA | ENSG0000C |
| 16033229 | 16033356 | -1 | rRNA_pseudogene | ENSG0000C |
| 16067139 | 16180762 | -1 | protein_coding | ENSG0000C |
| 16129178 | 16141602 | 1 | lncRNA | ENSG0000C |
| 16180238 | 16185585 | 1 | lncRNA | ENSG0000C |
| 16192088 | 16192709 | -1 | processed_pseudoge | ENSG0000C |
| 16373361 | 16440081 | -1 | lncRNA | ENSG0000C |
| 16428536 | 16432436 | 1 | lncRNA | ENSG0000C |
| 16451519 | 16465800 | -1 | protein_coding | ENSG0000C |
| 16473038 | 16617058 | -1 | protein_coding | ENSG0000C |
| 16615926 | 16681905 | 1 | lncRNA | ENSG0000C |
| 16617225 | 16621276 | 1 | lncRNA | ENSG0000C |
| 16623004 | 16623095 | -1 | misc_RNA | ENSG0000C |
| 16661907 | 16936288 | -1 | protein_coding | ENSG0000C |
| 16853994 | 16854114 | 1 | rRNA_pseudogene | ENSG0000C |
| 16902294 | 16902641 | -1 | processed_pseudoge | ENSG0000C |
| 16941843 | 16941951 | -1 | snRNA | ENSG0000C |
| 17065598 | 17276843 | 1 | protein_coding | ENSG0000C |
| 17089296 | 17217047 | -1 | lncRNA | ENSG0000C |
| 17107360 | 17107789 | -1 | lncRNA | ENSG0000C |
| 17117576 | 17117686 | -1 | misc_RNA | ENSG0000C |
| 17156971 | 17157071 | -1 | rRNA_pseudogene | ENSG0000C |
| 17202274 | 17203145 | 1 | processed_pseudoge | ENSG0000C |
| 17240611 | 17240715 | 1 | snRNA | ENSG0000C |
| 17306381 | 17306960 | 1 | processed_pseudoge | ENSG0000C |
| 17345616 | 17345955 | 1 | misc_RNA | ENSG0000C |
| 17353695 | 17354624 | -1 | transcribed_processe | ENSG0000C |
| 17353910 | 17354899 | 1 | lncRNA | ENSG0000C |
| 17367588 | 17375653 | -1 | lncRNA | ENSG0000C |
| 17378906 | 17387310 | -1 | lncRNA | ENSG0000C |
| 17403889 | 17443619 | 1 | lncRNA | ENSG0000C |
| 17444010 | 17486829 | 1 | protein_coding | ENSG0000C |
| 17491325 | 17491769 | -1 | protein_coding | ENSG0000C |
| 17495658 | 17495826 | -1 | processed_pseudoge | ENSG0000C |
| 17498231 | 17498827 | 1 | protein_coding | ENSG0000C |
| 17502043 | 17502363 | -1 | processed_pseudoge | ENSG0000C |
| 17511833 | 17512238 | -1 | processed_pseudoge | ENSG0000C |
| 17518367 | 17518963 | 1 | protein_coding | ENSG0000C |
| 17521801 | 17522397 | 1 | protein_coding | ENSG0000C |
| 17525235 | 17525831 | 1 | protein_coding | ENSG0000C |
| 17528669 | 17529265 | 1 | protein_coding | ENSG0000C |
| 17585162 | 17585758 | -1 | protein_coding | ENSG0000C |
| 17587290 | 17587523 | -1 | processed_pseudoge | ENSG0000C |
| 17590364 | 17590960 | -1 | protein_coding | ENSG0000C |
| 17593798 | 17594394 | -1 | protein_coding | ENSG0000C |
| 17597232 | 17597828 | -1 | protein_coding | ENSG0000C |
| 17598432 | 17598559 | -1 | processed_pseudoge | ENSG0000C |
| 17601882 | 17602134 | -1 | processed_pseudoge | ENSG0000C |
| 17604177 | 17605377 | 1 | protein_coding | ENSG0000C |
| 17610496 | 17611583 | 1 | protein_coding | ENSG0000C |

| | | | | |
|---|---|---|---|---|
| 17614781 | 17615111 | -1 | processed_pseudoge | ENSG00000 |
| 17620533 | 17620939 | -1 | processed_pseudoge | ENSG00000 |
| 17625551 | 17625959 | -1 | processed_pseudoge | ENSG00000 |
| 17632088 | 17632684 | 1 | protein_coding | ENSG00000 |
| 17634460 | 17635053 | 1 | protein_coding | ENSG00000 |
| 17642962 | 17643228 | -1 | processed_pseudoge | ENSG00000 |
| 17647712 | 17647944 | -1 | processed_pseudoge | ENSG00000 |
| 17649046 | 17649643 | 1 | processed_pseudoge | ENSG00000 |
| 17654373 | 17654643 | -1 | processed_pseudoge | ENSG00000 |
| 17654870 | 17655847 | -1 | protein_coding | ENSG00000 |
| 17669283 | 17669518 | -1 | processed_pseudoge | ENSG00000 |
| 17670168 | 17670571 | -1 | processed_pseudoge | ENSG00000 |
| 17684584 | 17955857 | 1 | lncRNA | ENSG00000 |
| 18049232 | 18049872 | 1 | transcribed_process | ENSG00000 |
| 18236123 | 18236196 | 1 | snoRNA | ENSG00000 |
| 18547312 | 18547605 | 1 | misc_RNA | ENSG00000 |
| 18704433 | 18746173 | -1 | lncRNA | ENSG00000 |
| 18886622 | 18887095 | -1 | processed_pseudoge | ENSG00000 |
| 18893902 | 18894168 | -1 | processed_pseudoge | ENSG00000 |
| 18908866 | 18909543 | 1 | processed_pseudoge | ENSG00000 |
| 18958147 | 18958413 | -1 | misc_RNA | ENSG00000 |
| 18965861 | 19142346 | -1 | lncRNA | ENSG00000 |
| 19035197 | 19038799 | -1 | lncRNA | ENSG00000 |
| 19041009 | 19041386 | 1 | processed_pseudoge | ENSG00000 |
| 19233366 | 19234487 | 1 | processed_pseudoge | ENSG00000 |
| 19471296 | 20575873 | -1 | protein_coding | ENSG00000 |
| 20158662 | 20158956 | -1 | processed_pseudoge | ENSG00000 |
| 20304045 | 20305448 | -1 | processed_pseudoge | ENSG00000 |
| 20305565 | 20347972 | 1 | lncRNA | ENSG00000 |
| 20606710 | 20616441 | -1 | lncRNA | ENSG00000 |
| 20611806 | 20937800 | 1 | lncRNA | ENSG00000 |
| 20830166 | 20830947 | -1 | processed_pseudoge | ENSG00000 |
| 20967211 | 21032845 | -1 | lncRNA | ENSG00000 |
| 21196720 | 21196860 | 1 | processed_pseudoge | ENSG00000 |
| 21323873 | 21341375 | -1 | lncRNA | ENSG00000 |
| 21341833 | 21589372 | 1 | transcribed_unproce | ENSG00000 |
| 21481327 | 21484011 | -1 | processed_pseudoge | ENSG00000 |
| 21493436 | 21494596 | 1 | TEC | ENSG00000 |
| 21569755 | 21570045 | 1 | lncRNA | ENSG00000 |
| 21616262 | 21779522 | 1 | lncRNA | ENSG00000 |
| 21750673 | 22853344 | -1 | protein_coding | ENSG00000 |
| 21882585 | 21884310 | 1 | processed_pseudoge | ENSG00000 |
| 22139247 | 22141468 | -1 | lncRNA | ENSG00000 |
| 22142352 | 22152356 | 1 | transcribed_unproce | ENSG00000 |
| 22212476 | 22213761 | 1 | lncRNA | ENSG00000 |
| 22580225 | 22580819 | -1 | processed_pseudoge | ENSG00000 |
| 22754227 | 22766824 | 1 | lncRNA | ENSG00000 |
| 23013048 | 23014527 | 1 | lncRNA | ENSG00000 |
| 23262159 | 23263185 | -1 | processed_pseudoge | ENSG00000 |
| 23299155 | 23299262 | 1 | misc_RNA | ENSG00000 |

| | | | | |
|---|---|---|---|---|
| 23303565 | 23305143 | -1 | processed_pseudoge | ENSG00000( |
| 23329427 | 23329892 | -1 | processed_pseudoge | ENSG00000( |
| 23443586 | 23528597 | 1 | protein_coding | ENSG00000( |
| 23456468 | 23456850 | 1 | processed_pseudoge | ENSG00000( |
| 23688927 | 23689386 | 1 | processed_pseudoge | ENSG00000( |
| 23951348 | 24178263 | 1 | lncRNA | ENSG00000( |
| 23972188 | 23972294 | -1 | snoRNA | ENSG00000( |
| 23980587 | 23980938 | 1 | transcribed_process( | ENSG00000( |
| 23994502 | 23994603 | -1 | misc_RNA | ENSG00000( |
| 24135160 | 24271863 | -1 | lncRNA | ENSG00000( |
| 24170370 | 24171246 | 1 | processed_pseudoge | ENSG00000( |
| 24352309 | 24353443 | -1 | lncRNA | ENSG00000( |
| 24449351 | 24449615 | -1 | misc_RNA | ENSG00000( |
| 24478516 | 24480057 | -1 | lncRNA | ENSG00000( |
| 24487100 | 24644978 | -1 | protein_coding | ENSG00000( |
| 24554018 | 24613222 | 1 | lncRNA | ENSG00000( |
| 24578294 | 24578561 | -1 | processed_pseudoge | ENSG00000( |
| 24729379 | 24738195 | 1 | lncRNA | ENSG00000( |
| 24749374 | 24749772 | -1 | processed_pseudoge | ENSG00000( |
| 24772144 | 24850971 | -1 | lncRNA | ENSG00000( |
| 24786842 | 24787024 | -1 | processed_pseudoge | ENSG00000( |
| 24811481 | 24811548 | 1 | snoRNA | ENSG00000( |
| 24835280 | 24840583 | -1 | lncRNA | ENSG00000( |
| 24881943 | 24885461 | -1 | lncRNA | ENSG00000( |
| 24910594 | 24910695 | 1 | misc_RNA | ENSG00000( |
| 25087450 | 25190956 | -1 | lncRNA | ENSG00000( |
| 25187800 | 25324386 | 1 | lncRNA | ENSG00000( |
| 25188059 | 25188778 | -1 | lncRNA | ENSG00000( |
| 25218973 | 25298612 | -1 | lncRNA | ENSG00000( |
| 25319834 | 25321346 | 1 | lncRNA | ENSG00000( |
| 25349841 | 25350073 | 1 | TEC | ENSG00000( |
| 25404733 | 25445925 | -1 | lncRNA | ENSG00000( |
| 25425412 | 25425599 | -1 | TEC | ENSG00000( |
| 25701217 | 25701324 | -1 | snRNA | ENSG00000( |
| 25716271 | 25716715 | 1 | TEC | ENSG00000( |
| 25909503 | 25911234 | 1 | processed_pseudoge | ENSG00000( |
| 25963839 | 25990687 | 1 | lncRNA | ENSG00000( |
| 26012889 | 26013025 | 1 | snRNA | ENSG00000( |
| 26382519 | 26399819 | -1 | lncRNA | ENSG00000( |
| 26473505 | 26484711 | -1 | lncRNA | ENSG00000( |
| 26669346 | 26669736 | 1 | processed_pseudoge | ENSG00000( |
| 26669711 | 26669989 | -1 | processed_pseudoge | ENSG00000( |
| 26670334 | 26670481 | -1 | processed_pseudoge | ENSG00000( |
| 26711551 | 26749997 | -1 | lncRNA | ENSG00000( |
| 26739753 | 26741183 | -1 | processed_pseudoge | ENSG00000( |
| 26880597 | 27121150 | -1 | protein_coding | ENSG00000( |
| 27111569 | 27111674 | 1 | snRNA | ENSG00000( |
| 27217714 | 27496994 | 1 | lncRNA | ENSG00000( |
| 27406530 | 27436421 | -1 | lncRNA | ENSG00000( |
| 28213359 | 28214559 | -1 | processed_pseudoge | ENSG00000( |

| | | | | |
|---|---|---|---|---|
| 28285964 | 28287807 | -1 | lncRNA | ENSG0000C |
| 28548135 | 28809291 | -1 | lncRNA | ENSG0000C |
| 28614613 | 28654531 | 1 | lncRNA | ENSG0000C |
| 28624657 | 28624763 | 1 | snRNA | ENSG0000C |
| 28808538 | 28809255 | 1 | processed_pseudoge | ENSG0000C |
| 28809331 | 29217115 | 1 | lncRNA | ENSG0000C |
| 28897785 | 28898274 | -1 | processed_pseudoge | ENSG0000C |
| 28926816 | 28927023 | 1 | processed_pseudoge | ENSG0000C |
| 28927029 | 28927284 | 1 | processed_pseudoge | ENSG0000C |
| 28927298 | 28927459 | -1 | processed_pseudoge | ENSG0000C |
| 29001760 | 29003357 | 1 | processed_pseudoge | ENSG0000C |
| 29070496 | 29070610 | 1 | snoRNA | ENSG0000C |
| 29104880 | 29105438 | 1 | unprocessed_pseudc | ENSG0000C |
| 29105568 | 29106861 | -1 | processed_pseudoge | ENSG0000C |
| 29239922 | 29265940 | -1 | lncRNA | ENSG0000C |
| 29304305 | 29396095 | -1 | lncRNA | ENSG0000C |
| 29516005 | 29516504 | 1 | processed_pseudoge | ENSG0000C |
| 29600676 | 29600868 | 1 | processed_pseudoge | ENSG0000C |
| 29795938 | 29796268 | 1 | processed_pseudoge | ENSG0000C |
| 29848684 | 29849218 | -1 | processed_pseudoge | ENSG0000C |
| 29881709 | 29882945 | -1 | processed_pseudoge | ENSG0000C |
| 29948828 | 29949130 | 1 | misc_RNA | ENSG0000C |
| 29979669 | 29983114 | 1 | TEC | ENSG0000C |
| 30248350 | 30248893 | 1 | processed_pseudoge | ENSG0000C |
| 30365405 | 30365684 | -1 | processed_pseudoge | ENSG0000C |
| 30545177 | 30693984 | -1 | lncRNA | ENSG0000C |
| 30733557 | 30743704 | -1 | lncRNA | ENSG0000C |
| 31053565 | 31054153 | 1 | processed_pseudoge | ENSG0000C |
| 31065970 | 31066132 | -1 | processed_pseudoge | ENSG0000C |
| 31093977 | 31267610 | -1 | lncRNA | ENSG0000C |
| 31193686 | 31329146 | 1 | protein_coding | ENSG0000C |
| 31249879 | 31250987 | 1 | processed_pseudoge | ENSG0000C |
| 31400497 | 31532196 | -1 | protein_coding | ENSG0000C |
| 31505684 | 31505788 | 1 | misc_RNA | ENSG0000C |
| 31532287 | 31555053 | 1 | protein_coding | ENSG0000C |
| 31558573 | 31559875 | 1 | TEC | ENSG0000C |
| 31595565 | 31595669 | -1 | snRNA | ENSG0000C |
| 31639131 | 32110932 | 1 | protein_coding | ENSG0000C |
| 31657218 | 31665067 | 1 | lncRNA | ENSG0000C |
| 31725539 | 31725902 | -1 | transcribed_processe | ENSG0000C |
| 31738269 | 31738660 | 1 | lncRNA | ENSG0000C |
| 31741833 | 31742327 | -1 | lncRNA | ENSG0000C |
| 31743988 | 31744451 | -1 | lncRNA | ENSG0000C |
| 31747644 | 31748078 | -1 | lncRNA | ENSG0000C |
| 31754219 | 31754656 | 1 | lncRNA | ENSG0000C |
| 31820564 | 31820638 | -1 | snRNA | ENSG0000C |
| 31822561 | 31823136 | -1 | processed_pseudoge | ENSG0000C |
| 31832439 | 31832535 | 1 | snRNA | ENSG0000C |
| 31840492 | 31840971 | -1 | processed_pseudoge | ENSG0000C |
| 31847271 | 31848148 | 1 | processed_pseudoge | ENSG0000C |

| | | | | |
|---|---|---|---|---|
| 31908361 | 31909186 | 1 | processed_pseudoge | ENSG00000 |
| 31923367 | 31923532 | 1 | processed_pseudoge | ENSG00000 |
| 31936102 | 31936159 | -1 | miRNA | ENSG00000 |
| 32103445 | 32121941 | -1 | lncRNA | ENSG00000 |
| 32124716 | 32174319 | -1 | protein_coding | ENSG00000 |
| 32134738 | 32134834 | 1 | misc_RNA | ENSG00000 |
| 32174471 | 32175272 | 1 | lncRNA | ENSG00000 |
| 32221932 | 32222347 | 1 | processed_pseudoge | ENSG00000 |
| 32226994 | 32312987 | -1 | protein_coding | ENSG00000 |
| 32234766 | 32234875 | -1 | snRNA | ENSG00000 |
| 32309662 | 32309768 | 1 | snRNA | ENSG00000 |
| 32354350 | 32444740 | -1 | protein_coding | ENSG00000 |
| 32394378 | 32394475 | -1 | miRNA | ENSG00000 |
| 32522758 | 32523865 | 1 | processed_pseudoge | ENSG00000 |
| 32531633 | 32604079 | 1 | protein_coding | ENSG00000 |
| 32646564 | 32651976 | -1 | lncRNA | ENSG00000 |
| 32653321 | 32656022 | 1 | lncRNA | ENSG00000 |
| 32689070 | 32791724 | 1 | protein_coding | ENSG00000 |
| 32887990 | 32888145 | -1 | processed_pseudoge | ENSG00000 |
| 32888236 | 33298066 | -1 | lncRNA | ENSG00000 |
| 32947443 | 32962467 | 1 | lncRNA | ENSG00000 |
| 33008934 | 33026025 | 1 | lncRNA | ENSG00000 |
| 33011322 | 33017607 | -1 | lncRNA | ENSG00000 |
| 33090126 | 33090349 | -1 | processed_pseudoge | ENSG00000 |
| 33162179 | 33162807 | 1 | processed_pseudoge | ENSG00000 |
| 33229516 | 33284642 | 1 | lncRNA | ENSG00000 |
| 33424025 | 33440619 | -1 | lncRNA | ENSG00000 |
| 33440696 | 33469539 | 1 | protein_coding | ENSG00000 |
| 33502072 | 33503135 | 1 | processed_pseudoge | ENSG00000 |
| 33519616 | 33522230 | -1 | lncRNA | ENSG00000 |
| 33523535 | 33892019 | -1 | protein_coding | ENSG00000 |
| 33888056 | 33888162 | -1 | snRNA | ENSG00000 |
| 33936386 | 33938918 | 1 | protein_coding | ENSG00000 |
| 33944623 | 33984693 | -1 | protein_coding | ENSG00000 |
| 33970863 | 33972628 | 1 | processed_pseudoge | ENSG00000 |
| 33986165 | 34008104 | -1 | protein_coding | ENSG00000 |
| 33987174 | 34124528 | -1 | protein_coding | ENSG00000 |
| 33997044 | 33997724 | -1 | processed_pseudoge | ENSG00000 |
| 34019448 | 34043832 | -1 | protein_coding | ENSG00000 |
| 34105750 | 34106336 | -1 | TEC | ENSG00000 |
| 34124162 | 34124385 | -1 | TEC | ENSG00000 |
| 34158121 | 34158767 | 1 | lncRNA | ENSG00000 |
| 34164698 | 34244796 | -1 | transcribed_unproce | ENSG00000 |
| 34190056 | 34193653 | 1 | processed_pseudoge | ENSG00000 |
| 34245273 | 34288755 | 1 | lncRNA | ENSG00000 |
| 34373028 | 34373850 | 1 | processed_pseudoge | ENSG00000 |
| 34651457 | 34651888 | -1 | lncRNA | ENSG00000 |
| 34656328 | 34832612 | 1 | protein_coding | ENSG00000 |
| 34656412 | 34657250 | -1 | lncRNA | ENSG00000 |
| 34837549 | 34839278 | -1 | lncRNA | ENSG00000 |

| | | | | |
|---|---|---|---|---|
| 34838833 | 34899456 | 1 | protein_coding | ENSG0000C |
| 34860783 | 34861506 | -1 | processed_pseudoge | ENSG0000C |
| 34883057 | 34883520 | 1 | processed_pseudoge | ENSG0000C |
| 34905260 | 34918989 | -1 | protein_coding | ENSG0000C |
| 34915677 | 34925996 | 1 | protein_coding | ENSG0000C |
| 34929559 | 34958964 | 1 | protein_coding | ENSG0000C |
| 34998101 | 35048135 | -1 | protein_coding | ENSG0000C |
| 35048756 | 35230487 | -1 | protein_coding | ENSG0000C |
| 35081223 | 35081992 | 1 | processed_pseudoge | ENSG0000C |
| 35429064 | 35429277 | 1 | snoRNA | ENSG0000C |
| 35617844 | 35814611 | 1 | protein_coding | ENSG0000C |
| 35632963 | 35633024 | -1 | snRNA | ENSG0000C |
| 35678484 | 35827018 | -1 | lncRNA | ENSG0000C |
| 35852695 | 35879603 | 1 | protein_coding | ENSG0000C |
| 35895689 | 35896020 | -1 | TEC | ENSG0000C |
| 35896188 | 35896290 | -1 | unprocessed_pseudc | ENSG0000C |
| 35904288 | 35938779 | -1 | protein_coding | ENSG0000C |
| 35938822 | 35939993 | 1 | lncRNA | ENSG0000C |
| 35951006 | 36001028 | -1 | protein_coding | ENSG0000C |
| 36035021 | 36071358 | -1 | protein_coding | ENSG0000C |
| 36071791 | 36072079 | -1 | processed_pseudoge | ENSG0000C |
| 36098407 | 36151887 | -1 | protein_coding | ENSG0000C |
| 36147892 | 36147988 | -1 | miRNA | ENSG0000C |
| 36151989 | 36184319 | 1 | protein_coding | ENSG0000C |
| 36155119 | 36155222 | 1 | snRNA | ENSG0000C |
| 36192589 | 36242279 | -1 | protein_coding | ENSG0000C |
| 36221055 | 36221902 | 1 | lncRNA | ENSG0000C |
| 36248434 | 36302114 | -1 | protein_coding | ENSG0000C |
| 36343672 | 36347157 | -1 | lncRNA | ENSG0000C |
| 36485343 | 36485440 | 1 | rRNA_pseudogene | ENSG0000C |
| 36588506 | 36588661 | 1 | processed_pseudoge | ENSG0000C |
| 36606355 | 36688334 | 1 | protein_coding | ENSG0000C |
| 36636019 | 36649347 | -1 | lncRNA | ENSG0000C |
| 36666214 | 36725195 | -1 | lncRNA | ENSG0000C |
| 36667717 | 36669958 | 1 | TEC | ENSG0000C |
| 36861736 | 36867238 | -1 | lncRNA | ENSG0000C |
| 36864425 | 36876700 | -1 | lncRNA | ENSG0000C |
| 36876769 | 37066413 | 1 | protein_coding | ENSG0000C |
| 36885206 | 36886497 | 1 | processed_pseudoge | ENSG0000C |
| 37085056 | 37085852 | 1 | processed_pseudoge | ENSG0000C |
| 37106228 | 37249428 | -1 | protein_coding | ENSG0000C |
| 37130430 | 37130700 | 1 | processed_pseudoge | ENSG0000C |
| 37209364 | 37212381 | 1 | processed_pseudoge | ENSG0000C |
| 37246173 | 37246479 | -1 | misc_RNA | ENSG0000C |
| 37249026 | 37252617 | 1 | lncRNA | ENSG0000C |
| 37286449 | 37286977 | -1 | processed_pseudoge | ENSG0000C |
| 37288137 | 37371106 | -1 | protein_coding | ENSG0000C |
| 37327135 | 37327196 | 1 | snRNA | ENSG0000C |
| 37377049 | 37377971 | 1 | unprocessed_pseudc | ENSG0000C |
| 37379285 | 37753435 | 1 | protein_coding | ENSG0000C |

| | | | | |
|---|---|---|---|---|
| 37505688 | 37506857 | -1 | processed_pseudoge | ENSG00000 |
| 37619985 | 37620088 | -1 | snRNA | ENSG00000 |
| 37700331 | 37700437 | -1 | snRNA | ENSG00000 |
| 37811589 | 37953827 | 1 | lncRNA | ENSG00000 |
| 37812677 | 37840041 | -1 | protein_coding | ENSG00000 |
| 37899363 | 37920869 | -1 | lncRNA | ENSG00000 |
| 37939365 | 37947955 | -1 | lncRNA | ENSG00000 |
| 37948491 | 37951055 | 1 | lncRNA | ENSG00000 |
| 37953402 | 37966964 | 1 | lncRNA | ENSG00000 |
| 38025568 | 38184717 | 1 | lncRNA | ENSG00000 |
| 38197432 | 38198148 | 1 | processed_pseudoge | ENSG00000 |
| 38258409 | 38465480 | 1 | protein_coding | ENSG00000 |
| 38282264 | 38290986 | -1 | lncRNA | ENSG00000 |
| 38345347 | 38346551 | -1 | lncRNA | ENSG00000 |
| 38399814 | 38403471 | -1 | lncRNA | ENSG00000 |
| 38425036 | 38427376 | -1 | lncRNA | ENSG00000 |
| 38429892 | 38430160 | -1 | processed_pseudoge | ENSG00000 |
| 38460925 | 38468339 | -1 | lncRNA | ENSG00000 |
| 38474668 | 38608354 | -1 | protein_coding | ENSG00000 |
| 38556765 | 38671216 | 1 | lncRNA | ENSG00000 |
| 38557502 | 38557561 | -1 | miRNA | ENSG00000 |
| 38682070 | 38685126 | 1 | lncRNA | ENSG00000 |
| 38682581 | 38845829 | -1 | lncRNA | ENSG00000 |
| 38763158 | 38764289 | -1 | processed_pseudoge | ENSG00000 |
| 38783580 | 38792754 | 1 | lncRNA | ENSG00000 |
| 38845858 | 38945596 | 1 | protein_coding | ENSG00000 |
| 38937920 | 39074399 | -1 | protein_coding | ENSG00000 |
| 39022411 | 39022558 | 1 | unprocessed_pseudo | ENSG00000 |
| 39092658 | 39103677 | -1 | lncRNA | ENSG00000 |
| 39105252 | 39274528 | -1 | protein_coding | ENSG00000 |
| 39169210 | 39170335 | -1 | processed_pseudoge | ENSG00000 |
| 39284140 | 39424868 | -1 | protein_coding | ENSG00000 |
| 39371675 | 39462300 | -1 | protein_coding | ENSG00000 |
| 39520398 | 39524726 | 1 | lncRNA | ENSG00000 |
| 39571478 | 39572639 | -1 | processed_pseudoge | ENSG00000 |
| 39718984 | 39721513 | -1 | processed_pseudoge | ENSG00000 |
| 39888576 | 39889059 | -1 | processed_pseudoge | ENSG00000 |
| 39892138 | 40053324 | 1 | lncRNA | ENSG00000 |
| 40066651 | 40067200 | 1 | processed_pseudoge | ENSG00000 |
| 40240167 | 40267657 | 1 | lncRNA | ENSG00000 |
| 40269558 | 40269719 | 1 | snRNA | ENSG00000 |
| 40306824 | 40312905 | -1 | lncRNA | ENSG00000 |
| 40415137 | 40431491 | 1 | lncRNA | ENSG00000 |
| 40463562 | 40504563 | 1 | lncRNA | ENSG00000 |
| 40474908 | 40490430 | -1 | lncRNA | ENSG00000 |
| 40512333 | 40755963 | -1 | protein_coding | ENSG00000 |
| 40654963 | 40655094 | 1 | snoRNA | ENSG00000 |
| 40679915 | 40693735 | 1 | protein_coding | ENSG00000 |
| 40759379 | 40798374 | -1 | protein_coding | ENSG00000 |
| 40790077 | 40790204 | -1 | snoRNA | ENSG00000 |

| | | | | |
|---|---|---|---|---|
| 40825262 | 40835222 | -1 | protein_coding | ENSG0000C |
| 40832656 | 40832735 | -1 | snoRNA | ENSG0000C |
| 40841308 | 40860175 | 1 | protein_coding | ENSG0000C |
| 40909497 | 40984643 | 1 | protein_coding | ENSG0000C |
| 40911067 | 40911128 | -1 | snRNA | ENSG0000C |
| 40967505 | 40967763 | -1 | processed_pseudoge | ENSG0000C |
| 40998017 | 41071342 | -1 | protein_coding | ENSG0000C |
| 41142234 | 41261438 | -1 | protein_coding | ENSG0000C |
| 41306952 | 41510628 | -1 | protein_coding | ENSG0000C |
| 41585882 | 41587787 | -1 | processed_pseudoge | ENSG0000C |
| 41730065 | 41870425 | -1 | protein_coding | ENSG0000C |
| 41868975 | 41880210 | 1 | lncRNA | ENSG0000C |
| 41869927 | 41872241 | 1 | lncRNA | ENSG0000C |
| 41895068 | 41895599 | 1 | processed_pseudoge | ENSG0000C |
| 41904188 | 41921636 | 1 | protein_coding | ENSG0000C |
| 41925254 | 41941743 | 1 | protein_coding | ENSG0000C |
| 41951313 | 41952193 | -1 | processed_pseudoge | ENSG0000C |
| 41963868 | 41963969 | -1 | misc_RNA | ENSG0000C |
| 41967139 | 41968366 | -1 | processed_pseudoge | ENSG0000C |
| 42188266 | 42191523 | 1 | lncRNA | ENSG0000C |
| 42423439 | 42721878 | 1 | protein_coding | ENSG0000C |
| 42465400 | 42468868 | -1 | processed_pseudoge | ENSG0000C |
| 42466893 | 42468300 | -1 | processed_pseudoge | ENSG0000C |
| 42729138 | 42729519 | 1 | processed_pseudoge | ENSG0000C |
| 42756818 | 42802439 | 1 | protein_coding | ENSG0000C |
| 42799880 | 42887392 | -1 | protein_coding | ENSG0000C |
| 42806394 | 42806997 | 1 | lncRNA | ENSG0000C |
| 42813647 | 42924535 | 1 | lncRNA | ENSG0000C |
| 42892319 | 42892817 | -1 | processed_pseudoge | ENSG0000C |
| 42897262 | 42897832 | -1 | processed_pseudoge | ENSG0000C |
| 42908208 | 42908667 | -1 | processed_pseudoge | ENSG0000C |
| 42917414 | 42917984 | 1 | processed_pseudoge | ENSG0000C |
| 42972624 | 42973092 | 1 | processed_pseudoge | ENSG0000C |
| 42983636 | 42993276 | -1 | lncRNA | ENSG0000C |
| 42995134 | 42997480 | 1 | lncRNA | ENSG0000C |
| 43006733 | 43007543 | -1 | lncRNA | ENSG0000C |
| 43013060 | 43067452 | -1 | lncRNA | ENSG0000C |
| 43018429 | 43024678 | 1 | lncRNA | ENSG0000C |
| 43033716 | 43034635 | 1 | lncRNA | ENSG0000C |
| 43039081 | 43043170 | -1 | protein_coding | ENSG0000C |
| 43041575 | 43054603 | 1 | lncRNA | ENSG0000C |
| 43061395 | 43062441 | 1 | lncRNA | ENSG0000C |
| 43065176 | 43192021 | 1 | protein_coding | ENSG0000C |
| 43143362 | 43143514 | -1 | processed_pseudoge | ENSG0000C |
| 43192071 | 43280850 | 1 | protein_coding | ENSG0000C |
| 43206709 | 43207811 | 1 | processed_pseudoge | ENSG0000C |
| 43225506 | 43225813 | -1 | processed_pseudoge | ENSG0000C |
| 43231812 | 43233173 | -1 | processed_pseudoge | ENSG0000C |
| 43287470 | 43313512 | -1 | protein_coding | ENSG0000C |
| 43287601 | 43290839 | 1 | lncRNA | ENSG0000C |

| | | | | |
|---|---|---|---|---|
| 43336164 | 43348716 | 1 | transcribed_processe | ENSG00000 |
| 43376645 | 43412391 | -1 | protein_coding | ENSG00000 |
| 43444252 | 43483893 | -1 | protein_coding | ENSG00000 |
| 43483913 | 43509356 | 1 | lncRNA | ENSG00000 |
| 43486709 | 43515148 | -1 | protein_coding | ENSG00000 |
| 43495073 | 43496454 | -1 | processed_pseudoge | ENSG00000 |
| 43511058 | 43521811 | 1 | lncRNA | ENSG00000 |
| 43515274 | 43525310 | 1 | lncRNA | ENSG00000 |
| 43526267 | 43557758 | -1 | protein_coding | ENSG00000 |
| 43571594 | 43603230 | -1 | lncRNA | ENSG00000 |
| 43586367 | 43587543 | -1 | TEC | ENSG00000 |
| 43586918 | 43588223 | -1 | processed_pseudoge | ENSG00000 |
| 43602692 | 43707405 | 1 | protein_coding | ENSG00000 |
| 43666766 | 43667232 | -1 | processed_pseudoge | ENSG00000 |
| 43874367 | 43886628 | 1 | lncRNA | ENSG00000 |
| 44066624 | 44066731 | -1 | snRNA | ENSG00000 |
| 44303544 | 44389706 | -1 | protein_coding | ENSG00000 |
| 44388732 | 44413989 | 1 | lncRNA | ENSG00000 |
| 44495099 | 44658569 | -1 | lncRNA | ENSG00000 |
| 44698431 | 44700808 | -1 | lncRNA | ENSG00000 |
| 44716193 | 44716486 | 1 | misc_RNA | ENSG00000 |
| 44742420 | 44808777 | -1 | lncRNA | ENSG00000 |
| 44752949 | 44765744 | 1 | lncRNA | ENSG00000 |
| 44808947 | 44820428 | 1 | protein_coding | ENSG00000 |
| 44826076 | 44828592 | 1 | lncRNA | ENSG00000 |
| 45035199 | 45098647 | 1 | lncRNA | ENSG00000 |
| 45226286 | 45228428 | 1 | lncRNA | ENSG00000 |
| 45254948 | 45696498 | -1 | protein_coding | ENSG00000 |
| 45557919 | 45559986 | 1 | processed_pseudoge | ENSG00000 |
| 45574590 | 45576612 | 1 | processed_pseudoge | ENSG00000 |
| 45890216 | 45895970 | 1 | lncRNA | ENSG00000 |
| 50396192 | 50443248 | -1 | protein_coding | ENSG00000 |
| 50603229 | 50603511 | 1 | processed_pseudoge | ENSG00000 |
| 50633124 | 50633602 | 1 | processed_pseudoge | ENSG00000 |
| 50662859 | 50663266 | -1 | lncRNA | ENSG00000 |
| 50665899 | 50846519 | 1 | protein_coding | ENSG00000 |
| 50858760 | 50860020 | -1 | lncRNA | ENSG00000 |
| 50903608 | 50903743 | 1 | processed_pseudoge | ENSG00000 |
| 50929484 | 50934521 | -1 | lncRNA | ENSG00000 |
| 50965687 | 50967008 | -1 | lncRNA | ENSG00000 |
| 50969217 | 50970167 | -1 | lncRNA | ENSG00000 |
| 50969660 | 50970187 | 1 | lncRNA | ENSG00000 |
| 51055353 | 51055459 | -1 | snRNA | ENSG00000 |
| 51101172 | 51101643 | -1 | processed_pseudoge | ENSG00000 |
| 51167541 | 51167647 | -1 | snRNA | ENSG00000 |
| 51275253 | 51275734 | 1 | processed_pseudoge | ENSG00000 |
| 51372736 | 51383332 | -1 | lncRNA | ENSG00000 |
| 51383448 | 51394730 | 1 | protein_coding | ENSG00000 |
| 51409617 | 51410205 | -1 | processed_pseudoge | ENSG00000 |
| 51427996 | 51428427 | -1 | processed_pseudoge | ENSG00000 |

| | | | | |
|---|---|---|---|---|
| 51438792 | 51665048 | -1 | lncRNA | ENSG0000C |
| 51451158 | 51462089 | 1 | lncRNA | ENSG0000C |
| 51886688 | 51886769 | 1 | rRNA_pseudogene | ENSG0000C |
| 51930808 | 51932036 | 1 | processed_pseudoge | ENSG0000C |
| 51963992 | 51964445 | 1 | TEC | ENSG0000C |
| 52008031 | 52080987 | 1 | lncRNA | ENSG0000C |
| 52282567 | 52282859 | 1 | processed_pseudoge | ENSG0000C |
| 52579555 | 52581106 | -1 | processed_pseudoge | ENSG0000C |
| 52673186 | 52872925 | -1 | lncRNA | ENSG0000C |
| 52787916 | 52959209 | 1 | protein_coding | ENSG0000C |
| 52787916 | 52804044 | 1 | protein_coding | ENSG0000C |
| 52889666 | 52889954 | 1 | processed_pseudoge | ENSG0000C |
| 52903908 | 52904478 | 1 | processed_pseudoge | ENSG0000C |
| 52930606 | 52990278 | -1 | lncRNA | ENSG0000C |
| 52989340 | 53094779 | 1 | protein_coding | ENSG0000C |
| 53024924 | 53035884 | -1 | lncRNA | ENSG0000C |
| 53089016 | 53089468 | -1 | lncRNA | ENSG0000C |
| 53095679 | 53110063 | -1 | protein_coding | ENSG0000C |
| 53109816 | 53127673 | 1 | lncRNA | ENSG0000C |
| 53206561 | 53207176 | 1 | processed_pseudoge | ENSG0000C |
| 53297872 | 53326565 | -1 | lncRNA | ENSG0000C |
| 53413504 | 53414260 | 1 | processed_pseudoge | ENSG0000C |
| 53480626 | 53487134 | 1 | protein_coding | ENSG0000C |
| 53560633 | 53683338 | 1 | protein_coding | ENSG0000C |
| 53776048 | 53819722 | -1 | lncRNA | ENSG0000C |
| 53859166 | 53860401 | 1 | processed_pseudoge | ENSG0000C |
| 53880293 | 53881051 | -1 | lncRNA | ENSG0000C |
| 53883942 | 54310582 | -1 | protein_coding | ENSG0000C |
| 53897464 | 53899370 | 1 | lncRNA | ENSG0000C |
| 53912020 | 53912123 | -1 | snRNA | ENSG0000C |
| 53951504 | 53951599 | -1 | miRNA | ENSG0000C |
| 54048007 | 54048308 | -1 | misc_RNA | ENSG0000C |
| 54075518 | 54075583 | -1 | miRNA | ENSG0000C |
| 54286194 | 54286331 | -1 | TEC | ENSG0000C |
| 54313564 | 54415238 | 1 | lncRNA | ENSG0000C |
| 54399439 | 54400033 | 1 | processed_pseudoge | ENSG0000C |
| 54455699 | 54456377 | 1 | protein_coding | ENSG0000C |
| 54517759 | 54546586 | 1 | protein_coding | ENSG0000C |
| 54643557 | 54645987 | 1 | TEC | ENSG0000C |
| 54663327 | 54737057 | 1 | lncRNA | ENSG0000C |
| 54808210 | 54872551 | -1 | unitary_pseudogene | ENSG0000C |
| 54857272 | 54858491 | 1 | processed_pseudoge | ENSG0000C |
| 54878238 | 54878588 | 1 | processed_pseudoge | ENSG0000C |
| 54956235 | 54957846 | -1 | lncRNA | ENSG0000C |
| 54977867 | 55022671 | -1 | protein_coding | ENSG0000C |
| 55021299 | 55042844 | -1 | lncRNA | ENSG0000C |
| 55024256 | 55034570 | 1 | protein_coding | ENSG0000C |
| 55040847 | 55040945 | 1 | misc_RNA | ENSG0000C |
| 55054428 | 55055140 | 1 | lncRNA | ENSG0000C |
| 55063179 | 55066230 | 1 | lncRNA | ENSG0000C |

| | | | | |
|---|---|---|---|---|
| 55073015 | 55096553 | 1 | unprocessed_pseudc | ENSG0000C |
| 55102646 | 55110252 | 1 | protein_coding | ENSG0000C |
| 55112995 | 55173175 | -1 | protein_coding | ENSG0000C |
| 55160167 | 55167297 | 1 | protein_coding | ENSG0000C |
| 55170532 | 55170622 | -1 | miRNA | ENSG0000C |
| 55170646 | 55170742 | -1 | miRNA | ENSG0000C |
| 55172262 | 55172353 | -1 | miRNA | ENSG0000C |
| 55219580 | 55227315 | -1 | protein_coding | ENSG0000C |
| 55231152 | 55233608 | -1 | protein_coding | ENSG0000C |
| 55233934 | 55295201 | 1 | lncRNA | ENSG0000C |
| 55256055 | 55307694 | -1 | protein_coding | ENSG0000C |
| 55307989 | 55425579 | 1 | protein_coding | ENSG0000C |
| 55356308 | 55357523 | -1 | processed_pseudoge | ENSG0000C |
| 55424854 | 55534969 | -1 | protein_coding | ENSG0000C |
| 55484403 | 55485086 | 1 | TEC | ENSG0000C |
| 55508850 | 55508926 | -1 | miRNA | ENSG0000C |
| 55530156 | 55530701 | -1 | processed_pseudoge | ENSG0000C |
| 55551885 | 55552554 | -1 | processed_pseudoge | ENSG0000C |
| 55625845 | 55773194 | -1 | protein_coding | ENSG0000C |
| 55738017 | 55817157 | 1 | protein_coding | ENSG0000C |
| 55750206 | 55750401 | -1 | processed_pseudoge | ENSG0000C |
| 55777469 | 55777587 | 1 | rRNA_pseudogene | ENSG0000C |
| 55838801 | 55840025 | -1 | processed_pseudoge | ENSG0000C |
| 55851357 | 55922854 | 1 | protein_coding | ENSG0000C |
| 55935095 | 55995022 | -1 | protein_coding | ENSG0000C |
| 55936143 | 55941727 | 1 | lncRNA | ENSG0000C |
| 55944656 | 55944733 | 1 | processed_pseudoge | ENSG0000C |
| 55978248 | 56111329 | 1 | lncRNA | ENSG0000C |
| 55995167 | 56003649 | 1 | lncRNA | ENSG0000C |
| 56000518 | 56000621 | -1 | snoRNA | ENSG0000C |
| 56099680 | 56233330 | -1 | protein_coding | ENSG0000C |
| 56125671 | 56125770 | 1 | snRNA | ENSG0000C |
| 56136979 | 56137531 | 1 | processed_pseudoge | ENSG0000C |
| 56146739 | 56146846 | -1 | rRNA_pseudogene | ENSG0000C |
| 56192530 | 56193633 | 1 | processed_pseudoge | ENSG0000C |
| 56260333 | 56260446 | -1 | rRNA_pseudogene | ENSG0000C |
| 56275304 | 56276630 | -1 | processed_pseudoge | ENSG0000C |
| 56297564 | 56297689 | -1 | snRNA | ENSG0000C |
| 56313905 | 56321397 | -1 | lncRNA | ENSG0000C |
| 56381781 | 56382075 | 1 | processed_pseudoge | ENSG0000C |
| 56451627 | 56481946 | -1 | lncRNA | ENSG0000C |
| 56488486 | 56491499 | 1 | lncRNA | ENSG0000C |
| 56504635 | 56505072 | -1 | processed_pseudoge | ENSG0000C |
| 56511567 | 56606232 | -1 | protein_coding | ENSG0000C |
| 56536583 | 56537826 | -1 | lncRNA | ENSG0000C |
| 56770799 | 56772303 | 1 | lncRNA | ENSG0000C |
| 56815549 | 56896152 | 1 | protein_coding | ENSG0000C |
| 56842016 | 56862164 | -1 | lncRNA | ENSG0000C |
| 56900041 | 56910714 | -1 | lncRNA | ENSG0000C |
| 56909260 | 56925532 | 1 | protein_coding | ENSG0000C |

| | | | | |
|---|---|---|---|---|
| 56919602 | 56971675 | -1 | protein_coding | ENSG00000 |
| 56927874 | 56929573 | 1 | lncRNA | ENSG00000 |
| 56941307 | 56947152 | 1 | lncRNA | ENSG00000 |
| 56975593 | 56976224 | -1 | processed_pseudoge | ENSG00000 |
| 57081745 | 57081951 | -1 | processed_pseudoge | ENSG00000 |
| 57120796 | 57122624 | 1 | processed_pseudoge | ENSG00000 |
| 57169108 | 57169209 | -1 | misc_RNA | ENSG00000 |
| 57173948 | 57264679 | 1 | protein_coding | ENSG00000 |
| 57317989 | 57320020 | -1 | processed_pseudoge | ENSG00000 |
| 57395060 | 57534504 | 1 | lncRNA | ENSG00000 |
| 57422464 | 57424391 | 1 | processed_pseudoge | ENSG00000 |
| 57480018 | 57482811 | -1 | protein_coding | ENSG00000 |
| 57574027 | 57619816 | 1 | lncRNA | ENSG00000 |
| 57650678 | 57679842 | 1 | lncRNA | ENSG00000 |
| 57751192 | 57751717 | -1 | lncRNA | ENSG00000 |
| 57890327 | 57899162 | -1 | lncRNA | ENSG00000 |
| 58107631 | 58122375 | -1 | lncRNA | ENSG00000 |
| 58114598 | 58145996 | 1 | lncRNA | ENSG00000 |
| 58161197 | 58161752 | 1 | processed_pseudoge | ENSG00000 |
| 58453982 | 58460139 | -1 | protein_coding | ENSG00000 |
| 58491435 | 58497090 | 1 | protein_coding | ENSG00000 |
| 58503295 | 58503700 | 1 | processed_pseudoge | ENSG00000 |
| 58530043 | 58530109 | -1 | miRNA | ENSG00000 |
| 58541567 | 58558243 | -1 | lncRNA | ENSG00000 |
| 58582221 | 58859394 | 1 | protein_coding | ENSG00000 |
| 58741581 | 58817099 | -1 | lncRNA | ENSG00000 |
| 58822503 | 58823714 | 1 | processed_pseudoge | ENSG00000 |
| 58846885 | 58863414 | -1 | lncRNA | ENSG00000 |
| 58969038 | 60522120 | -1 | protein_coding | ENSG00000 |
| 59039761 | 59063503 | 1 | lncRNA | ENSG00000 |
| 59314110 | 59314800 | 1 | processed_pseudoge | ENSG00000 |
| 59528861 | 59529251 | 1 | processed_pseudoge | ENSG00000 |
| 59703606 | 59703703 | -1 | miRNA | ENSG00000 |
| 60021249 | 60033020 | 1 | lncRNA | ENSG00000 |
| 60304047 | 60304151 | -1 | snRNA | ENSG00000 |
| 60429903 | 60430276 | -1 | processed_pseudoge | ENSG00000 |
| 60459848 | 60460739 | -1 | processed_pseudoge | ENSG00000 |
| 60487713 | 60487929 | 1 | misc_RNA | ENSG00000 |
| 60487713 | 60548813 | 1 | lncRNA | ENSG00000 |
| 60488078 | 60488327 | 1 | misc_RNA | ENSG00000 |
| 60546219 | 60546349 | 1 | misc_RNA | ENSG00000 |
| 60596912 | 60700190 | -1 | protein_coding | ENSG00000 |
| 60630514 | 60630973 | 1 | processed_pseudoge | ENSG00000 |
| 60699729 | 60702010 | 1 | lncRNA | ENSG00000 |
| 60743563 | 60744970 | -1 | processed_pseudoge | ENSG00000 |
| 60751791 | 60844274 | -1 | protein_coding | ENSG00000 |
| 60866457 | 60866935 | -1 | lncRNA | ENSG00000 |
| 60873831 | 60945073 | -1 | protein_coding | ENSG00000 |
| 60891935 | 60893577 | -1 | processed_pseudoge | ENSG00000 |
| 60917490 | 60919573 | 1 | lncRNA | ENSG00000 |

| | | | | |
|---|---|---|---|---|
| 60945205 | 61153026 | 1 | protein_coding | ENSG00000( |
| 61114432 | 61116753 | -1 | TEC | ENSG00000( |
| 61145846 | 61146376 | -1 | unprocessed_pseudc | ENSG00000( |
| 61157704 | 61162468 | -1 | protein_coding | ENSG00000( |
| 61162070 | 61232040 | 1 | lncRNA | ENSG00000( |
| 61201304 | 61304811 | -1 | lncRNA | ENSG00000( |
| 61332258 | 61546172 | 1 | protein_coding | ENSG00000( |
| 61347126 | 61347716 | 1 | processed_pseudoge | ENSG00000( |
| 61375112 | 61375855 | 1 | processed_pseudoge | ENSG00000( |
| 61390668 | 61391871 | -1 | processed_pseudoge | ENSG00000( |
| 61610424 | 61793401 | 1 | lncRNA | ENSG00000( |
| 61658474 | 61698432 | -1 | lncRNA | ENSG00000( |
| 61664145 | 61664239 | -1 | snRNA | ENSG00000( |
| 61732774 | 61735715 | -1 | lncRNA | ENSG00000( |
| 61831947 | 61833024 | -1 | processed_pseudoge | ENSG00000( |
| 61878109 | 61927134 | 1 | lncRNA | ENSG00000( |
| 62146665 | 62146946 | 1 | misc_RNA | ENSG00000( |
| 62306162 | 62537249 | 1 | protein_coding | ENSG00000( |
| 62347284 | 62403939 | -1 | protein_coding | ENSG00000( |
| 62403972 | 62628582 | 1 | protein_coding | ENSG00000( |
| 62443176 | 62443282 | 1 | snRNA | ENSG00000( |
| 62512246 | 62512482 | -1 | processed_pseudoge | ENSG00000( |
| 62561859 | 62562216 | -1 | processed_pseudoge | ENSG00000( |
| 62575951 | 62576271 | -1 | processed_pseudoge | ENSG00000( |
| 62578819 | 62581446 | 1 | protein_coding | ENSG00000( |
| 62776877 | 62777263 | -1 | processed_pseudoge | ENSG00000( |
| 63022919 | 63023489 | -1 | processed_pseudoge | ENSG00000( |
| 63301523 | 63302056 | 1 | processed_pseudoge | ENSG00000( |
| 63569419 | 63571617 | 1 | lncRNA | ENSG00000( |
| 63957893 | 63981043 | -1 | lncRNA | ENSG00000( |
| 63960356 | 63962507 | -1 | protein_coding | ENSG00000( |
| 64165843 | 64372869 | 1 | protein_coding | ENSG00000( |
| 64404512 | 64486615 | -1 | unitary_pseudogene | ENSG00000( |
| 64506015 | 64612319 | 1 | protein_coding | ENSG00000( |
| 64539504 | 64539792 | 1 | misc_RNA | ENSG00000( |
| 64573569 | 64573670 | 1 | snRNA | ENSG00000( |
| 64674853 | 64675387 | 1 | processed_pseudoge | ENSG00000( |
| 64690442 | 64718190 | 1 | protein_coding | ENSG00000( |
| 64718148 | 64768691 | -1 | protein_coding | ENSG00000( |
| 64768930 | 65018750 | 1 | protein_coding | ENSG00000( |
| 65020038 | 65020551 | 1 | lncRNA | ENSG00000( |
| 65020614 | 65020989 | 1 | lncRNA | ENSG00000( |
| 65123369 | 65123474 | 1 | misc_RNA | ENSG00000( |
| 65148738 | 65481920 | -1 | protein_coding | ENSG00000( |
| 65209921 | 65210948 | 1 | processed_pseudoge | ENSG00000( |
| 65251127 | 65251784 | -1 | processed_pseudoge | ENSG00000( |
| 65415353 | 65416222 | 1 | processed_pseudoge | ENSG00000( |
| 65486444 | 65487048 | -1 | lncRNA | ENSG00000( |
| 65517766 | 65563168 | -1 | protein_coding | ENSG00000( |
| 65563236 | 65587549 | 1 | protein_coding | ENSG00000( |

| | | | | |
|---|---|---|---|---|
| 65589690 | 65625975 | -1 | protein_coding | ENSG0000C |
| 65604661 | 65604761 | -1 | snRNA | ENSG0000C |
| 65624765 | 65630891 | 1 | protein_coding | ENSG0000C |
| 65625004 | 65666233 | 1 | protein_coding | ENSG0000C |
| 65665928 | 65723035 | -1 | protein_coding | ENSG0000C |
| 65722205 | 65871725 | 1 | protein_coding | ENSG0000C |
| 65732973 | 65733654 | 1 | processed_pseudoge | ENSG0000C |
| 65924629 | 65925135 | -1 | lncRNA | ENSG0000C |
| 65926475 | 66082549 | 1 | protein_coding | ENSG0000C |
| 65961183 | 65961319 | -1 | snoRNA | ENSG0000C |
| 65965373 | 65969143 | 1 | lncRNA | ENSG0000C |
| 66085405 | 66093937 | 1 | lncRNA | ENSG0000C |
| 66139971 | 66183615 | 1 | protein_coding | ENSG0000C |
| 66144156 | 66144795 | -1 | lncRNA | ENSG0000C |
| 66205120 | 66209893 | -1 | lncRNA | ENSG0000C |
| 66298394 | 66393521 | 1 | lncRNA | ENSG0000C |
| 66350148 | 66440356 | -1 | lncRNA | ENSG0000C |
| 66507380 | 66511644 | -1 | lncRNA | ENSG0000C |
| 66572057 | 66572546 | 1 | processed_pseudoge | ENSG0000C |
| 66596361 | 67169595 | 1 | protein_coding | ENSG0000C |
| 66622868 | 66623177 | 1 | processed_pseudoge | ENSG0000C |
| 66662331 | 66662988 | 1 | lncRNA | ENSG0000C |
| 67001383 | 67003953 | -1 | lncRNA | ENSG0000C |
| 67179613 | 67196799 | -1 | protein_coding | ENSG0000C |
| 67268022 | 67270182 | 1 | lncRNA | ENSG0000C |
| 67356663 | 67366303 | -1 | lncRNA | ENSG0000C |
| 67379378 | 67806600 | 1 | lncRNA | ENSG0000C |
| 67463809 | 67475592 | -1 | lncRNA | ENSG0000C |
| 67608654 | 67609130 | 1 | processed_pseudoge | ENSG0000C |
| 67632266 | 67646789 | -1 | lncRNA | ENSG0000C |
| 67636382 | 67638942 | 1 | processed_pseudoge | ENSG0000C |
| 67690033 | 67690437 | -1 | processed_pseudoge | ENSG0000C |
| 67692812 | 67693200 | -1 | processed_pseudoge | ENSG0000C |
| 67699429 | 67902600 | -1 | lncRNA | ENSG0000C |
| 67800740 | 67890096 | -1 | lncRNA | ENSG0000C |
| 67855735 | 67855986 | 1 | processed_pseudoge | ENSG0000C |
| 68087161 | 68087494 | -1 | processed_pseudoge | ENSG0000C |
| 68159061 | 68159146 | -1 | snRNA | ENSG0000C |
| 68159218 | 68159893 | -1 | processed_pseudoge | ENSG0000C |
| 68189876 | 68198407 | 1 | lncRNA | ENSG0000C |
| 68215756 | 68301821 | 1 | protein_coding | ENSG0000C |
| 68374765 | 68375292 | 1 | lncRNA | ENSG0000C |
| 68430427 | 68434481 | -1 | lncRNA | ENSG0000C |
| 68508223 | 68565515 | -1 | lncRNA | ENSG0000C |
| 68523878 | 68530007 | 1 | lncRNA | ENSG0000C |
| 68792609 | 69007185 | -1 | lncRNA | ENSG0000C |
| 68854910 | 68855085 | -1 | processed_pseudoge | ENSG0000C |
| 68873954 | 68874087 | 1 | snoRNA | ENSG0000C |
| 68963246 | 68967845 | 1 | lncRNA | ENSG0000C |
| 68970439 | 68970752 | -1 | misc_RNA | ENSG0000C |

| | | | | |
|---|---|---|---|---|
| 68970692 | 69030165 | -1 | lncRNA | ENSG0000( |
| 69038518 | 69043821 | -1 | lncRNA | ENSG0000( |
| 69068925 | 69069200 | -1 | processed_pseudoge | ENSG0000( |
| 69093949 | 69131069 | 1 | protein_coding | ENSG0000( |
| 69113112 | 69136394 | -1 | lncRNA | ENSG0000( |
| 69160806 | 69160939 | 1 | snoRNA | ENSG0000( |
| 69160808 | 69160939 | 1 | snoRNA | ENSG0000( |
| 69167135 | 69178245 | 1 | protein_coding | ENSG0000( |
| 69186359 | 69189452 | -1 | TEC | ENSG0000( |
| 69189574 | 69210357 | 1 | protein_coding | ENSG0000( |
| 69217760 | 69230158 | 1 | protein_coding | ENSG0000( |
| 69234795 | 69277430 | 1 | protein_coding | ENSG0000( |
| 69280175 | 69332809 | -1 | protein_coding | ENSG0000( |
| 69313371 | 69313872 | -1 | processed_pseudoge | ENSG0000( |
| 69333929 | 69334249 | 1 | processed_pseudoge | ENSG0000( |
| 69349936 | 69350445 | 1 | processed_pseudoge | ENSG0000( |
| 69350984 | 69370013 | -1 | protein_coding | ENSG0000( |
| 69364743 | 69370013 | -1 | protein_coding | ENSG0000( |
| 69369293 | 69414801 | 1 | protein_coding | ENSG0000( |
| 69415065 | 69444330 | 1 | protein_coding | ENSG0000( |
| 69469883 | 69470136 | 1 | processed_pseudoge | ENSG0000( |
| 69477472 | 69502466 | -1 | lncRNA | ENSG0000( |
| 69478997 | 69479265 | -1 | misc_RNA | ENSG0000( |
| 69492292 | 69558104 | 1 | protein_coding | ENSG0000( |
| 69530613 | 69530717 | -1 | snRNA | ENSG0000( |
| 69560208 | 69594723 | 1 | protein_coding | ENSG0000( |
| 69607099 | 69624049 | 1 | transcribed_unproce | ENSG0000( |
| 69631963 | 69636399 | 1 | unprocessed_pseudc | ENSG0000( |
| 69639459 | 69679259 | -1 | transcribed_unproce | ENSG0000( |
| 69653248 | 69654334 | 1 | unprocessed_pseudc | ENSG0000( |
| 69875271 | 69920867 | 1 | unprocessed_pseudc | ENSG0000( |
| 69898867 | 69903198 | -1 | unprocessed_pseudc | ENSG0000( |
| 69985204 | 69985501 | -1 | unprocessed_pseudc | ENSG0000( |
| 70025247 | 70043113 | 1 | protein_coding | ENSG0000( |
| 70049612 | 70078522 | 1 | protein_coding | ENSG0000( |
| 70055820 | 70057416 | -1 | lncRNA | ENSG0000( |
| 70094659 | 70128434 | -1 | unprocessed_pseudc | ENSG0000( |
| 70132679 | 70132973 | 1 | unprocessed_pseudc | ENSG0000( |
| 70197255 | 70207745 | -1 | unprocessed_pseudc | ENSG0000( |
| 70219918 | 70258930 | -1 | unprocessed_pseudc | ENSG0000( |
| 70415352 | 70448015 | 1 | transcribed_unproce | ENSG0000( |
| 70449636 | 70450353 | 1 | lncRNA | ENSG0000( |
| 70462244 | 70479215 | 1 | transcribed_unproce | ENSG0000( |
| 70495100 | 70500903 | -1 | unprocessed_pseudc | ENSG0000( |
| 70516387 | 70555266 | -1 | unprocessed_pseudc | ENSG0000( |
| 70751184 | 70795914 | 1 | unprocessed_pseudc | ENSG0000( |
| 70775506 | 70778475 | -1 | processed_pseudoge | ENSG0000( |
| 70860285 | 70860579 | -1 | unprocessed_pseudc | ENSG0000( |
| 70900665 | 70918530 | 1 | protein_coding | ENSG0000( |
| 70925030 | 70953942 | 1 | protein_coding | ENSG0000( |

| | | | | |
|---|---|---|---|---|
| 70931244 | 70932840 | -1 | lncRNA | ENSG0000C |
| 70968483 | 71025114 | -1 | protein_coding | ENSG0000C |
| 71032670 | 71067689 | -1 | protein_coding | ENSG0000C |
| 71074225 | 71093193 | -1 | unprocessed_pseudc | ENSG0000C |
| 71102898 | 71128753 | -1 | unprocessed_pseudc | ENSG0000C |
| 71132993 | 71133287 | 1 | unprocessed_pseudc | ENSG0000C |
| 71197646 | 71208130 | -1 | unprocessed_pseudc | ENSG0000C |
| 71220356 | 71259238 | -1 | unprocessed_pseudc | ENSG0000C |
| 71337182 | 71446772 | -1 | lncRNA | ENSG0000C |
| 71372676 | 71374898 | -1 | lncRNA | ENSG0000C |
| 71375786 | 71385993 | 1 | transcribed_unproce | ENSG0000C |
| 71445616 | 71446569 | 1 | lncRNA | ENSG0000C |
| 71455651 | 71567820 | 1 | protein_coding | ENSG0000C |
| 71475761 | 71476316 | -1 | processed_pseudoge | ENSG0000C |
| 71537652 | 71537846 | -1 | processed_pseudoge | ENSG0000C |
| 71587288 | 71658706 | 1 | protein_coding | ENSG0000C |
| 71719275 | 71721048 | 1 | protein_coding | ENSG0000C |
| 71754378 | 71756301 | -1 | processed_pseudoge | ENSG0000C |
| 72087782 | 72108000 | -1 | lncRNA | ENSG0000C |
| 72107234 | 72209565 | 1 | protein_coding | ENSG0000C |
| 72169467 | 72169540 | 1 | miRNA | ENSG0000C |
| 72219409 | 72320646 | -1 | protein_coding | ENSG0000C |
| 72314397 | 72314688 | -1 | misc_RNA | ENSG0000C |
| 72320367 | 72368395 | 1 | protein_coding | ENSG0000C |
| 72381794 | 72382393 | 1 | processed_pseudoge | ENSG0000C |
| 72417489 | 72418335 | 1 | processed_pseudoge | ENSG0000C |
| 72439903 | 72442387 | -1 | lncRNA | ENSG0000C |
| 72439903 | 72507410 | -1 | protein_coding | ENSG0000C |
| 72574120 | 72660669 | -1 | lncRNA | ENSG0000C |
| 72687112 | 72771733 | -1 | lncRNA | ENSG0000C |
| 72725419 | 72726151 | 1 | processed_pseudoge | ENSG0000C |
| 72733833 | 72734209 | 1 | unitary_pseudogene | ENSG0000C |
| 72768702 | 72768811 | 1 | misc_RNA | ENSG0000C |
| 72794405 | 72816565 | -1 | lncRNA | ENSG0000C |
| 72816312 | 72916733 | 1 | protein_coding | ENSG0000C |
| 72878591 | 72878663 | 1 | miRNA | ENSG0000C |
| 72878868 | 72879185 | -1 | processed_pseudoge | ENSG0000C |
| 72953635 | 72954274 | -1 | lncRNA | ENSG0000C |
| 72955206 | 72955699 | -1 | lncRNA | ENSG0000C |
| 72956041 | 73090522 | 1 | protein_coding | ENSG0000C |
| 72996920 | 72997642 | 1 | processed_pseudoge | ENSG0000C |
| 73020700 | 73021279 | -1 | processed_pseudoge | ENSG0000C |
| 73120569 | 73131809 | 1 | protein_coding | ENSG0000C |
| 73132008 | 73150592 | -1 | lncRNA | ENSG0000C |
| 73152416 | 73152938 | -1 | processed_pseudoge | ENSG0000C |
| 73173193 | 73175143 | 1 | protein_coding | ENSG0000C |
| 73195314 | 73201973 | -1 | lncRNA | ENSG0000C |
| 73213930 | 73295258 | 1 | lncRNA | ENSG0000C |
| 73337906 | 73338533 | 1 | lncRNA | ENSG0000C |
| 73382384 | 73410509 | 1 | transcribed_processe | ENSG0000C |

| | | | | |
|---|---|---|---|---|
| 73444827 | 73448777 | -1 | protein_coding | ENSG0000C |
| 73446357 | 73446984 | 1 | lncRNA | ENSG0000C |
| 73451498 | 73453395 | 1 | lncRNA | ENSG0000C |
| 73454187 | 73472896 | -1 | lncRNA | ENSG0000C |
| 73497550 | 73498293 | -1 | lncRNA | ENSG0000C |
| 73498408 | 73505635 | 1 | protein_coding | ENSG0000C |
| 73508685 | 73509254 | 1 | processed_pseudoge | ENSG0000C |
| 73552190 | 73565667 | -1 | protein_coding | ENSG0000C |
| 73565443 | 73583380 | 1 | protein_coding | ENSG0000C |
| 73626158 | 73941993 | 1 | protein_coding | ENSG0000C |
| 73778039 | 73786491 | -1 | lncRNA | ENSG0000C |
| 73803296 | 73803599 | -1 | processed_pseudoge | ENSG0000C |
| 73828012 | 73828073 | -1 | snRNA | ENSG0000C |
| 73952940 | 73954014 | 1 | lncRNA | ENSG0000C |
| 74084068 | 74103216 | -1 | lncRNA | ENSG0000C |
| 74111690 | 74536976 | -1 | lncRNA | ENSG0000C |
| 74258689 | 74321255 | -1 | lncRNA | ENSG0000C |
| 74317986 | 74318285 | 1 | misc_RNA | ENSG0000C |
| 74321293 | 74341236 | 1 | lncRNA | ENSG0000C |
| 74327995 | 74334766 | -1 | lncRNA | ENSG0000C |
| 74627406 | 74641424 | -1 | protein_coding | ENSG0000C |
| 74640023 | 74722647 | 1 | protein_coding | ENSG0000C |
| 74721206 | 74767147 | -1 | protein_coding | ENSG0000C |
| 74725099 | 74725204 | -1 | snRNA | ENSG0000C |
| 74766991 | 74780113 | 1 | protein_coding | ENSG0000C |
| 74777574 | 74866966 | -1 | protein_coding | ENSG0000C |
| 74779309 | 74779413 | 1 | snRNA | ENSG0000C |
| 74798994 | 74800100 | 1 | processed_pseudoge | ENSG0000C |
| 74865893 | 74867854 | 1 | lncRNA | ENSG0000C |
| 74882656 | 74882783 | -1 | snoRNA | ENSG0000C |
| 74899115 | 74903650 | -1 | lncRNA | ENSG0000C |
| 74917726 | 75023928 | -1 | lncRNA | ENSG0000C |
| 74929880 | 74930566 | 1 | processed_pseudoge | ENSG0000C |
| 74931662 | 74931817 | 1 | processed_pseudoge | ENSG0000C |
| 74990189 | 74990637 | -1 | processed_pseudoge | ENSG0000C |
| 75006713 | 75007012 | -1 | processed_pseudoge | ENSG0000C |
| 75025346 | 75052770 | -1 | protein_coding | ENSG0000C |
| 75068275 | 75236878 | -1 | protein_coding | ENSG0000C |
| 75225424 | 75225668 | 1 | processed_pseudoge | ENSG0000C |
| 75320155 | 75336914 | -1 | lncRNA | ENSG0000C |
| 75336329 | 75362101 | 1 | protein_coding | ENSG0000C |
| 75356345 | 75512138 | -1 | protein_coding | ENSG0000C |
| 75363760 | 75364242 | 1 | lncRNA | ENSG0000C |
| 75374463 | 75374809 | 1 | processed_pseudoge | ENSG0000C |
| 75511756 | 75601144 | 1 | protein_coding | ENSG0000C |
| 75538598 | 75538658 | -1 | snRNA | ENSG0000C |
| 75598482 | 75599380 | -1 | lncRNA | ENSG0000C |
| 75608817 | 75609983 | 1 | lncRNA | ENSG0000C |
| 75611182 | 75681773 | 1 | protein_coding | ENSG0000C |
| 75674124 | 75717448 | -1 | protein_coding | ENSG0000C |

| | | | | |
|---|---|---|---|---|
| 75709495 | 75709599 | -1 | snRNA | ENSG0000C |
| 75752570 | 75753479 | -1 | processed_pseudoge | ENSG0000C |
| 75831255 | 75832764 | -1 | processed_pseudoge | ENSG0000C |
| 75910283 | 75910576 | -1 | processed_pseudoge | ENSG0000C |
| 75910565 | 75911867 | 1 | processed_pseudoge | ENSG0000C |
| 75920670 | 75921660 | -1 | processed_pseudoge | ENSG0000C |
| 76075531 | 76075920 | 1 | processed_pseudoge | ENSG0000C |
| 76078666 | 76079763 | 1 | processed_pseudoge | ENSG0000C |
| 76081946 | 76084186 | -1 | lncRNA | ENSG0000C |
| 76083383 | 76353939 | 1 | protein_coding | ENSG0000C |
| 76091708 | 76091801 | -1 | rRNA_pseudogene | ENSG0000C |
| 76170930 | 76171106 | 1 | unprocessed_pseudc | ENSG0000C |
| 76173629 | 76174183 | -1 | translated_processec | ENSG0000C |
| 76242024 | 76242404 | -1 | processed_pseudoge | ENSG0000C |
| 76280956 | 76281267 | 1 | processed_pseudoge | ENSG0000C |
| 76285542 | 76311462 | -1 | lncRNA | ENSG0000C |
| 76376675 | 76377149 | 1 | processed_pseudoge | ENSG0000C |
| 76403285 | 76708132 | 1 | protein_coding | ENSG0000C |
| 76510440 | 76510766 | 1 | processed_pseudoge | ENSG0000C |
| 76606608 | 76608970 | -1 | lncRNA | ENSG0000C |
| 76615482 | 76623413 | -1 | protein_coding | ENSG0000C |
| 76691439 | 76716215 | -1 | lncRNA | ENSG0000C |
| 76712195 | 76712327 | 1 | misc_RNA | ENSG0000C |
| 76712763 | 76712861 | 1 | misc_RNA | ENSG0000C |
| 76716126 | 76735770 | 1 | protein_coding | ENSG0000C |
| 76818933 | 76835315 | 1 | protein_coding | ENSG0000C |
| 76836900 | 76837195 | -1 | misc_RNA | ENSG0000C |
| 76850001 | 76921650 | 1 | protein_coding | ENSG0000C |
| 76948791 | 76948870 | 1 | snRNA | ENSG0000C |
| 76953045 | 76981158 | 1 | protein_coding | ENSG0000C |
| 77029251 | 77065234 | 1 | protein_coding | ENSG0000C |
| 77030902 | 77152155 | 1 | protein_coding | ENSG0000C |
| 77072072 | 77087285 | -1 | protein_coding | ENSG0000C |
| 77073881 | 77074520 | 1 | lncRNA | ENSG0000C |
| 77080434 | 77080571 | -1 | snoRNA | ENSG0000C |
| 77086688 | 77166909 | 1 | lncRNA | ENSG0000C |
| 77086732 | 77427124 | 1 | protein_coding | ENSG0000C |
| 77146568 | 77147165 | 1 | transcribed_processe | ENSG0000C |
| 77210449 | 77429807 | 1 | protein_coding | ENSG0000C |
| 77290268 | 77291800 | 1 | processed_pseudoge | ENSG0000C |
| 77425970 | 77620611 | -1 | protein_coding | ENSG0000C |
| 77545397 | 77546266 | 1 | processed_pseudoge | ENSG0000C |
| 77582376 | 77583122 | 1 | processed_pseudoge | ENSG0000C |
| 77628712 | 77639688 | -1 | protein_coding | ENSG0000C |
| 77691166 | 77868780 | -1 | protein_coding | ENSG0000C |
| 77784881 | 77786003 | 1 | processed_pseudoge | ENSG0000C |
| 77942757 | 77946438 | -1 | lncRNA | ENSG0000C |
| 77958243 | 77959092 | 1 | lncRNA | ENSG0000C |
| 78000525 | 78294755 | -1 | protein_coding | ENSG0000C |
| 78041879 | 78044138 | -1 | transcribed_processe | ENSG0000C |

| | | | | |
|---|---|---|---|---|
| 78222184 | 78222517 | 1 | processed_pseudoge | ENSG00000( |
| 78307503 | 78307914 | 1 | processed_pseudoge | ENSG00000( |
| 78314916 | 78315025 | -1 | snRNA | ENSG00000( |
| 78317036 | 78317137 | 1 | misc_RNA | ENSG00000( |
| 78342365 | 78360507 | -1 | lncRNA | ENSG00000( |
| 78360611 | 78480739 | 1 | protein_coding | ENSG00000( |
| 78485215 | 78770021 | -1 | protein_coding | ENSG00000( |
| 78644265 | 78644845 | -1 | processed_pseudoge | ENSG00000( |
| 78708734 | 78716082 | -1 | processed_pseudoge | ENSG00000( |
| 78753596 | 78753873 | 1 | processed_pseudoge | ENSG00000( |
| 78777209 | 78986087 | -1 | protein_coding | ENSG00000( |
| 78997564 | 79236038 | -1 | protein_coding | ENSG00000( |
| 79000112 | 79001124 | -1 | processed_pseudoge | ENSG00000( |
| 79069767 | 79090069 | 1 | protein_coding | ENSG00000( |
| 79111809 | 79132288 | 1 | protein_coding | ENSG00000( |
| 79170234 | 79170335 | -1 | misc_RNA | ENSG00000( |
| 79191549 | 79191747 | 1 | processed_pseudoge | ENSG00000( |
| 79220942 | 79221073 | 1 | snoRNA | ENSG00000( |
| 79229904 | 79230588 | 1 | processed_pseudoge | ENSG00000( |
| 79236131 | 79327211 | 1 | protein_coding | ENSG00000( |
| 79284132 | 79284918 | -1 | processed_pseudoge | ENSG00000( |
| 79372636 | 79514134 | -1 | protein_coding | ENSG00000( |
| 79510434 | 79510919 | 1 | processed_pseudoge | ENSG00000( |
| 79529302 | 79530093 | 1 | processed_pseudoge | ENSG00000( |
| 79534853 | 79534954 | 1 | misc_RNA | ENSG00000( |
| 79562745 | 79562846 | -1 | misc_RNA | ENSG00000( |
| 79612120 | 79686648 | 1 | protein_coding | ENSG00000( |
| 79689836 | 79800240 | 1 | protein_coding | ENSG00000( |
| 79774348 | 79816190 | -1 | lncRNA | ENSG00000( |
| 79840797 | 79845239 | 1 | lncRNA | ENSG00000( |
| 79849552 | 79849846 | -1 | processed_pseudoge | ENSG00000( |
| 79936579 | 79967626 | 1 | lncRNA | ENSG00000( |
| 79976731 | 79991265 | -1 | protein_coding | ENSG00000( |
| 79991311 | 80083287 | 1 | protein_coding | ENSG00000( |
| 80019609 | 80019920 | 1 | processed_pseudoge | ENSG00000( |
| 80033171 | 80033302 | -1 | processed_pseudoge | ENSG00000( |
| 80052374 | 80083654 | -1 | lncRNA | ENSG00000( |
| 80111651 | 80256048 | -1 | protein_coding | ENSG00000( |
| 80128361 | 80143883 | 1 | lncRNA | ENSG00000( |
| 80288449 | 80289781 | 1 | processed_pseudoge | ENSG00000( |
| 80299678 | 80302847 | -1 | processed_pseudoge | ENSG00000( |
| 80306235 | 80306368 | -1 | snoRNA | ENSG00000( |
| 80315671 | 80315826 | 1 | processed_pseudoge | ENSG00000( |
| 80319625 | 80321842 | 1 | protein_coding | ENSG00000( |
| 80331573 | 80332541 | -1 | processed_pseudoge | ENSG00000( |
| 80351021 | 80351956 | -1 | processed_pseudoge | ENSG00000( |
| 80359080 | 80360040 | -1 | processed_pseudoge | ENSG00000( |
| 80365726 | 80365832 | -1 | snRNA | ENSG00000( |
| 80408013 | 80483379 | 1 | protein_coding | ENSG00000( |
| 80411231 | 80488095 | -1 | lncRNA | ENSG00000( |

| | | | | |
|---|---|---|---|---|
| 80487969 | 80542563 | 1 | protein_coding | ENSG0000C |
| 80498534 | 80498971 | -1 | processed_pseudoge | ENSG0000C |
| 80500332 | 80501400 | 1 | processed_pseudoge | ENSG0000C |
| 80556755 | 80570280 | -1 | protein_coding | ENSG0000C |
| 80603383 | 80603644 | -1 | processed_pseudoge | ENSG0000C |
| 80608623 | 80622524 | -1 | lncRNA | ENSG0000C |
| 80626226 | 80654983 | -1 | protein_coding | ENSG0000C |
| 80630313 | 80631590 | -1 | lncRNA | ENSG0000C |
| 80650000 | 80650086 | -1 | processed_pseudoge | ENSG0000C |
| 80654652 | 80876815 | 1 | protein_coding | ENSG0000C |
| 80746339 | 80746819 | -1 | processed_pseudoge | ENSG0000C |
| 80855507 | 80855843 | -1 | processed_pseudoge | ENSG0000C |
| 80947694 | 80960907 | -1 | lncRNA | ENSG0000C |
| 80960363 | 81230162 | 1 | protein_coding | ENSG0000C |
| 80997183 | 80998047 | -1 | processed_pseudoge | ENSG0000C |
| 81113385 | 81114852 | -1 | lncRNA | ENSG0000C |
| 81201341 | 81301565 | -1 | lncRNA | ENSG0000C |
| 81233320 | 81266398 | 1 | protein_coding | ENSG0000C |
| 81242330 | 81242599 | -1 | processed_pseudoge | ENSG0000C |
| 81301587 | 81313297 | 1 | protein_coding | ENSG0000C |
| 81329996 | 81394179 | -1 | protein_coding | ENSG0000C |
| 81408517 | 81411867 | -1 | lncRNA | ENSG0000C |
| 81413021 | 81751797 | -1 | protein_coding | ENSG0000C |
| 81817531 | 81842360 | 1 | lncRNA | ENSG0000C |
| 81851601 | 81852201 | 1 | lncRNA | ENSG0000C |
| 81892490 | 81892721 | 1 | processed_pseudoge | ENSG0000C |
| 81972023 | 82276857 | 1 | protein_coding | ENSG0000C |
| 81991995 | 81992481 | 1 | lncRNA | ENSG0000C |
| 82009602 | 82010094 | -1 | processed_pseudoge | ENSG0000C |
| 82073055 | 82073702 | -1 | lncRNA | ENSG0000C |
| 82078427 | 82078724 | 1 | misc_RNA | ENSG0000C |
| 82265157 | 82265259 | -1 | processed_pseudoge | ENSG0000C |
| 82273320 | 82278396 | -1 | protein_coding | ENSG0000C |
| 82279462 | 82386977 | 1 | protein_coding | ENSG0000C |
| 82545862 | 82546310 | -1 | lncRNA | ENSG0000C |
| 82586776 | 82587411 | -1 | lncRNA | ENSG0000C |
| 82765404 | 82766333 | 1 | lncRNA | ENSG0000C |
| 82777797 | 82778586 | 1 | processed_pseudoge | ENSG0000C |
| 82824884 | 82825184 | 1 | processed_pseudoge | ENSG0000C |
| 82840155 | 82840224 | 1 | miRNA | ENSG0000C |
| 82848260 | 82859958 | -1 | lncRNA | ENSG0000C |
| 82919376 | 82921119 | -1 | lncRNA | ENSG0000C |
| 82940458 | 82940983 | -1 | lncRNA | ENSG0000C |
| 82968888 | 82977153 | 1 | processed_pseudoge | ENSG0000C |
| 83012285 | 83013109 | -1 | lncRNA | ENSG0000C |
| 83049376 | 83050964 | -1 | lncRNA | ENSG0000C |
| 83052846 | 83077863 | -1 | protein_coding | ENSG0000C |
| 83064204 | 83064337 | -1 | snoRNA | ENSG0000C |
| 83077498 | 83353787 | 1 | protein_coding | ENSG0000C |
| 83201229 | 83202141 | 1 | processed_pseudoge | ENSG0000C |

| | | | | |
|---|---|---|---|---|
| 83279921 | 83280644 | -1 | processed_pseudoge | ENSG00000 |
| 83426676 | 83427030 | 1 | processed_pseudoge | ENSG00000 |
| 83471618 | 83582303 | 1 | protein_coding | ENSG00000 |
| 83531352 | 83581320 | -1 | lncRNA | ENSG00000 |
| 83637805 | 83720855 | -1 | protein_coding | ENSG00000 |
| 83685866 | 83686018 | 1 | misc_RNA | ENSG00000 |
| 83703448 | 83703543 | 1 | snRNA | ENSG00000 |
| 83746388 | 83746989 | -1 | processed_pseudoge | ENSG00000 |
| 83803554 | 83803685 | 1 | snRNA | ENSG00000 |
| 83875166 | 83875821 | -1 | processed_pseudoge | ENSG00000 |
| 83940554 | 84384880 | -1 | protein_coding | ENSG00000 |
| 84196883 | 84196989 | 1 | snRNA | ENSG00000 |
| 84382424 | 84490765 | 1 | lncRNA | ENSG00000 |
| 84999812 | 85000315 | -1 | processed_pseudoge | ENSG00000 |
| 85112342 | 85112756 | -1 | lncRNA | ENSG00000 |
| 85190629 | 85190830 | -1 | processed_pseudoge | ENSG00000 |
| 85210060 | 85210903 | -1 | processed_pseudoge | ENSG00000 |
| 85212958 | 85213614 | -1 | lncRNA | ENSG00000 |
| 85237681 | 85238074 | -1 | processed_pseudoge | ENSG00000 |
| 85420028 | 85420451 | -1 | lncRNA | ENSG00000 |
| 85429702 | 85430184 | 1 | lncRNA | ENSG00000 |
| 85446974 | 85447758 | 1 | lncRNA | ENSG00000 |
| 85464434 | 85465310 | 1 | processed_pseudoge | ENSG00000 |
| 85549409 | 85549952 | 1 | processed_pseudoge | ENSG00000 |
| 85663232 | 85664684 | 1 | lncRNA | ENSG00000 |
| 85762559 | 85763138 | 1 | processed_pseudoge | ENSG00000 |
| 85848502 | 85849199 | 1 | lncRNA | ENSG00000 |
| 86087526 | 86087847 | 1 | processed_pseudoge | ENSG00000 |
| 86282460 | 86296463 | 1 | transcribed_unproce | ENSG00000 |
| 86305233 | 86306149 | -1 | TEC | ENSG00000 |
| 86335024 | 86335808 | 1 | lncRNA | ENSG00000 |
| 86353803 | 86368854 | 1 | lncRNA | ENSG00000 |
| 86380660 | 86381426 | -1 | lncRNA | ENSG00000 |
| 86402465 | 86402960 | 1 | processed_pseudoge | ENSG00000 |
| 86617928 | 86620962 | 1 | protein_coding | ENSG00000 |
| 86620497 | 86620575 | 1 | miRNA | ENSG00000 |
| 86663144 | 86663250 | -1 | snRNA | ENSG00000 |
| 86746818 | 86749777 | 1 | lncRNA | ENSG00000 |
| 86797685 | 86800280 | 1 | lncRNA | ENSG00000 |
| 86884231 | 86884881 | -1 | processed_pseudoge | ENSG00000 |
| 86967321 | 87137712 | 1 | lncRNA | ENSG00000 |
| 87048858 | 87240145 | -1 | lncRNA | ENSG00000 |
| 87102603 | 87102846 | -1 | misc_RNA | ENSG00000 |
| 87114879 | 87114954 | -1 | miRNA | ENSG00000 |
| 87267883 | 87391931 | 1 | protein_coding | ENSG00000 |
| 87294189 | 87294473 | -1 | misc_RNA | ENSG00000 |
| 87299687 | 87299797 | -1 | snRNA | ENSG00000 |
| 87318416 | 87412930 | -1 | protein_coding | ENSG00000 |
| 87412342 | 87499340 | 1 | lncRNA | ENSG00000 |
| 87420341 | 87420447 | -1 | snRNA | ENSG00000 |

| | | | | |
|---|---|---|---|---|
| 87662040 | 87705337 | 1 | lncRNA | ENSG0000C |
| 87665345 | 87733269 | -1 | lncRNA | ENSG0000C |
| 87767608 | 87774070 | -1 | lncRNA | ENSG0000C |
| 87863703 | 88144455 | -1 | lncRNA | ENSG0000C |
| 88189633 | 88269476 | -1 | protein_coding | ENSG0000C |
| 88268891 | 88439337 | 1 | lncRNA | ENSG0000C |
| 88274258 | 88274376 | -1 | rRNA_pseudogene | ENSG0000C |
| 88381957 | 88382725 | 1 | processed_pseudoge | ENSG0000C |
| 88382772 | 88382907 | 1 | snoRNA | ENSG0000C |
| 88382948 | 88383260 | 1 | processed_pseudoge | ENSG0000C |
| 88408982 | 88439090 | -1 | lncRNA | ENSG0000C |
| 88433892 | 88498697 | -1 | lncRNA | ENSG0000C |
| 88507546 | 88691057 | -1 | lncRNA | ENSG0000C |
| 88602780 | 88603117 | 1 | processed_pseudoge | ENSG0000C |
| 88666853 | 88666939 | -1 | miRNA | ENSG0000C |
| 88676033 | 88779088 | 1 | lncRNA | ENSG0000C |
| 88691757 | 88695041 | -1 | lncRNA | ENSG0000C |
| 88692651 | 88692859 | 1 | lncRNA | ENSG0000C |
| 88717117 | 88904257 | -1 | protein_coding | ENSG0000C |
| 88883328 | 89466398 | 1 | lncRNA | ENSG0000C |
| 89466537 | 89470039 | -1 | lncRNA | ENSG0000C |
| 89581209 | 89677701 | 1 | lncRNA | ENSG0000C |
| 89900664 | 89990883 | 1 | lncRNA | ENSG0000C |
| 90016621 | 90016720 | -1 | miRNA | ENSG0000C |
| 90153052 | 90290078 | -1 | lncRNA | ENSG0000C |
| 90353037 | 90356609 | -1 | lncRNA | ENSG0000C |
| 90388468 | 90389363 | -1 | lncRNA | ENSG0000C |
| 90392261 | 90409786 | -1 | protein_coding | ENSG0000C |
| 90410000 | 90410669 | 1 | lncRNA | ENSG0000C |
| 90458209 | 90474771 | -1 | protein_coding | ENSG0000C |
| 90471748 | 90514557 | 1 | protein_coding | ENSG0000C |
| 90515611 | 90529584 | -1 | protein_coding | ENSG0000C |
| 90529344 | 91164437 | 1 | protein_coding | ENSG0000C |
| 91030990 | 91031307 | -1 | processed_pseudoge | ENSG0000C |
| 91054834 | 91314547 | -1 | lncRNA | ENSG0000C |
| 91132303 | 91142470 | -1 | lncRNA | ENSG0000C |
| 91223419 | 91223967 | 1 | lncRNA | ENSG0000C |
| 91226475 | 91227071 | -1 | lncRNA | ENSG0000C |
| 91270727 | 91270867 | 1 | snRNA | ENSG0000C |
| 91280097 | 91281142 | -1 | lncRNA | ENSG0000C |
| 91280272 | 91281643 | 1 | lncRNA | ENSG0000C |
| 91314716 | 91314997 | -1 | misc_RNA | ENSG0000C |
| 91354478 | 91354934 | 1 | processed_pseudoge | ENSG0000C |
| 91355380 | 91356026 | 1 | lncRNA | ENSG0000C |
| 91368631 | 91383317 | -1 | protein_coding | ENSG0000C |
| 91380349 | 91612566 | 1 | lncRNA | ENSG0000C |
| 91476382 | 91477018 | 1 | processed_pseudoge | ENSG0000C |
| 91642643 | 91799234 | 1 | lncRNA | ENSG0000C |
| 91843687 | 91844529 | -1 | processed_pseudoge | ENSG0000C |
| 91844022 | 91844116 | -1 | misc_RNA | ENSG0000C |

| | | | | |
|---|---|---|---|---|
| 92082597 | 92479426 | 1 | lncRNA | ENSG00000 |
| 92410256 | 92660863 | -1 | lncRNA | ENSG00000 |
| 92654848 | 92679141 | 1 | lncRNA | ENSG00000 |
| 92675956 | 92688254 | -1 | lncRNA | ENSG00000 |
| 92823935 | 92844992 | 1 | lncRNA | ENSG00000 |
| 92840807 | 92841628 | -1 | processed_pseudoge | ENSG00000 |
| 92889387 | 92890840 | -1 | processed_pseudoge | ENSG00000 |
| 92907180 | 92939591 | -1 | lncRNA | ENSG00000 |
| 93019663 | 93068669 | -1 | lncRNA | ENSG00000 |
| 93088856 | 93091768 | 1 | lncRNA | ENSG00000 |
| 93267429 | 93268425 | 1 | processed_pseudoge | ENSG00000 |
| 93360779 | 93585649 | -1 | lncRNA | ENSG00000 |
| 93583222 | 93594611 | 1 | protein_coding | ENSG00000 |
| 93600357 | 93603598 | 1 | TEC | ENSG00000 |
| 93617725 | 94111699 | -1 | protein_coding | ENSG00000 |
| 93620696 | 93620788 | -1 | miRNA | ENSG00000 |
| 93621683 | 93675081 | 1 | lncRNA | ENSG00000 |
| 93682838 | 93683667 | -1 | processed_pseudoge | ENSG00000 |
| 93733220 | 93741600 | -1 | protein_coding | ENSG00000 |
| 93741640 | 93743500 | 1 | lncRNA | ENSG00000 |
| 93790505 | 93801320 | 1 | lncRNA | ENSG00000 |
| 93860669 | 93863825 | -1 | lncRNA | ENSG00000 |
| 94111720 | 94176029 | 1 | lncRNA | ENSG00000 |
| 94152966 | 94618597 | -1 | protein_coding | ENSG00000 |
| 94567461 | 94568909 | 1 | processed_pseudoge | ENSG00000 |
| 94568929 | 94569470 | -1 | processed_pseudoge | ENSG00000 |
| 94569540 | 94570676 | 1 | processed_pseudoge | ENSG00000 |
| 94592771 | 94594791 | 1 | processed_pseudoge | ENSG00000 |
| 94611906 | 94618122 | -1 | lncRNA | ENSG00000 |
| 94618669 | 94739436 | 1 | protein_coding | ENSG00000 |
| 94703741 | 95284575 | -1 | protein_coding | ENSG00000 |
| 94788789 | 94793599 | 1 | lncRNA | ENSG00000 |
| 94825961 | 94826694 | -1 | processed_pseudoge | ENSG00000 |
| 94877599 | 94877777 | -1 | misc_RNA | ENSG00000 |
| 94979151 | 94980893 | 1 | lncRNA | ENSG00000 |
| 95391366 | 95450454 | 1 | protein_coding | ENSG00000 |
| 95440044 | 95440687 | 1 | processed_pseudoge | ENSG00000 |
| 95461755 | 95555007 | -1 | protein_coding | ENSG00000 |
| 95521369 | 95521475 | 1 | snRNA | ENSG00000 |
| 95555101 | 95605102 | 1 | protein_coding | ENSG00000 |
| 95620087 | 95622142 | 1 | protein_coding | ENSG00000 |
| 95646754 | 95684773 | 1 | protein_coding | ENSG00000 |
| 95652181 | 95698711 | -1 | protein_coding | ENSG00000 |
| 95701249 | 95732295 | -1 | lncRNA | ENSG00000 |
| 95713522 | 95824383 | 1 | protein_coding | ENSG00000 |
| 95751319 | 95822726 | -1 | protein_coding | ENSG00000 |
| 95768999 | 95770700 | 1 | processed_pseudoge | ENSG00000 |
| 95834424 | 95835046 | -1 | processed_pseudoge | ENSG00000 |
| 95835943 | 95852721 | -1 | lncRNA | ENSG00000 |
| 95838245 | 95860133 | 1 | lncRNA | ENSG00000 |

| | | | | |
|---|---|---|---|---|
| 95849309 | 95849855 | 1 | lncRNA | ENSG0000C |
| 95861786 | 95874519 | 1 | lncRNA | ENSG0000C |
| 95885098 | 95961851 | -1 | protein_coding | ENSG0000C |
| 95962001 | 96631085 | 1 | lncRNA | ENSG0000C |
| 95964999 | 95986311 | -1 | lncRNA | ENSG0000C |
| 95973041 | 95973446 | -1 | processed_pseudoge | ENSG0000C |
| 96050115 | 96215519 | 1 | lncRNA | ENSG0000C |
| 96079138 | 96079212 | 1 | miRNA | ENSG0000C |
| 96210121 | 96210227 | -1 | snRNA | ENSG0000C |
| 96213346 | 96215075 | -1 | lncRNA | ENSG0000C |
| 96247776 | 96278751 | 1 | lncRNA | ENSG0000C |
| 96390333 | 96434143 | -1 | protein_coding | ENSG0000C |
| 96525267 | 96779595 | 1 | protein_coding | ENSG0000C |
| 96741079 | 96742698 | -1 | lncRNA | ENSG0000C |
| 96760810 | 96808100 | -1 | protein_coding | ENSG0000C |
| 96784777 | 96785999 | 1 | lncRNA | ENSG0000C |
| 96803688 | 96806105 | 1 | lncRNA | ENSG0000C |
| 96814028 | 96935809 | -1 | lncRNA | ENSG0000C |
| 96875939 | 96919716 | 1 | protein_coding | ENSG0000C |
| 96935394 | 97037513 | 1 | protein_coding | ENSG0000C |
| 96942299 | 96943184 | 1 | processed_pseudoge | ENSG0000C |
| 96962656 | 96962757 | -1 | misc_RNA | ENSG0000C |
| 97056402 | 97057358 | -1 | processed_pseudoge | ENSG0000C |
| 97089075 | 97437217 | 1 | lncRNA | ENSG0000C |
| 97091867 | 97142753 | -1 | protein_coding | ENSG0000C |
| 97160867 | 97183247 | -1 | protein_coding | ENSG0000C |
| 97175950 | 97176113 | -1 | snRNA | ENSG0000C |
| 97183827 | 97216074 | 1 | lncRNA | ENSG0000C |
| 97188090 | 97201880 | -1 | lncRNA | ENSG0000C |
| 97223371 | 97227957 | -1 | lncRNA | ENSG0000C |
| 97336858 | 97337040 | -1 | processed_pseudoge | ENSG0000C |
| 97368776 | 97370421 | 1 | processed_pseudoge | ENSG0000C |
| 97449358 | 97449741 | -1 | processed_pseudoge | ENSG0000C |
| 97504663 | 97690180 | 1 | lncRNA | ENSG0000C |
| 97677547 | 97678681 | -1 | processed_pseudoge | ENSG0000C |
| 97737894 | 97738913 | 1 | processed_pseudoge | ENSG0000C |
| 97799404 | 97882687 | -1 | lncRNA | ENSG0000C |
| 97840912 | 97923497 | 1 | lncRNA | ENSG0000C |
| 98025965 | 98026139 | 1 | processed_pseudoge | ENSG0000C |
| 98085866 | 98161352 | -1 | lncRNA | ENSG0000C |
| 98203059 | 98205193 | 1 | processed_pseudoge | ENSG0000C |
| 98213402 | 98214121 | 1 | processed_pseudoge | ENSG0000C |
| 98338744 | 98339832 | 1 | processed_pseudoge | ENSG0000C |
| 98392070 | 98393530 | -1 | processed_pseudoge | ENSG0000C |
| 98402542 | 98402732 | -1 | processed_pseudoge | ENSG0000C |
| 98409675 | 98410100 | -1 | processed_pseudoge | ENSG0000C |
| 98410235 | 98411792 | -1 | processed_pseudoge | ENSG0000C |
| 98576341 | 98577260 | 1 | processed_pseudoge | ENSG0000C |
| 98679402 | 98681367 | 1 | processed_pseudoge | ENSG0000C |
| 98768650 | 98798643 | 1 | protein_coding | ENSG0000C |

| | | | | |
|---|---|---|---|---|
| 98769618 | 98773469 | -1 | lncRNA | ENSG00000 |
| 98792861 | 98795766 | -1 | lncRNA | ENSG00000 |
| 98833382 | 98833711 | 1 | processed_pseudoge | ENSG00000 |
| 98853985 | 98928957 | -1 | protein_coding | ENSG00000 |
| 98889731 | 98889833 | 1 | snRNA | ENSG00000 |
| 98929171 | 98995013 | 1 | lncRNA | ENSG00000 |
| 98936638 | 98936747 | -1 | misc_RNA | ENSG00000 |
| 98954394 | 98954972 | 1 | processed_pseudoge | ENSG00000 |
| 99489559 | 99499367 | 1 | transcribed_unproce | ENSG00000 |
| 99522311 | 99525462 | -1 | processed_pseudoge | ENSG00000 |
| 99532628 | 99534611 | 1 | unprocessed_pseudc | ENSG00000 |
| 99549432 | 99577957 | 1 | lncRNA | ENSG00000 |
| 99594329 | 99594740 | -1 | processed_pseudoge | ENSG00000 |
| 99594880 | 99595297 | 1 | processed_pseudoge | ENSG00000 |
| 99948295 | 99949583 | -1 | processed_pseudoge | ENSG00000 |
| 99996547 | 99997932 | 1 | processed_pseudoge | ENSG00000 |
| 100021804 | 100021906 | 1 | misc_RNA | ENSG00000 |
| 100045928 | 100046374 | -1 | processed_pseudoge | ENSG00000 |
| 100046450 | 100046970 | 1 | processed_pseudoge | ENSG00000 |
| 100046977 | 100048785 | -1 | processed_pseudoge | ENSG00000 |
| 100048988 | 100050362 | -1 | processed_pseudoge | ENSG00000 |
| 100050359 | 100050650 | -1 | processed_pseudoge | ENSG00000 |
| 100050719 | 100051063 | -1 | processed_pseudoge | ENSG00000 |
| 100051133 | 100051916 | -1 | processed_pseudoge | ENSG00000 |
| 100051920 | 100052597 | -1 | processed_pseudoge | ENSG00000 |
| 100052859 | 100053539 | -1 | processed_pseudoge | ENSG00000 |
| 100053686 | 100055045 | -1 | processed_pseudoge | ENSG00000 |
| 100062275 | 100067954 | -1 | unprocessed_pseudc | ENSG00000 |
| 100153672 | 100153779 | 1 | snRNA | ENSG00000 |
| 100375700 | 100381398 | -1 | unprocessed_pseudc | ENSG00000 |
| 100388853 | 100389938 | 1 | unprocessed_pseudc | ENSG00000 |
| 100399047 | 100526912 | 1 | lncRNA | ENSG00000 |
| 100428124 | 100535262 | -1 | lncRNA | ENSG00000 |
| 100535374 | 100586741 | 1 | protein_coding | ENSG00000 |
| 100654112 | 100657970 | -1 | lncRNA | ENSG00000 |
| 100733058 | 100733386 | 1 | misc_RNA | ENSG00000 |
| 100806933 | 100903282 | -1 | protein_coding | ENSG00000 |
| 100816482 | 100816565 | -1 | miRNA | ENSG00000 |
| 101581830 | 101582128 | 1 | misc_RNA | ENSG00000 |
| 101816475 | 101816780 | -1 | unprocessed_pseudc | ENSG00000 |
| 101976296 | 101976798 | 1 | processed_pseudoge | ENSG00000 |
| 102131007 | 102131135 | -1 | rRNA_pseudogene | ENSG00000 |
| 102141893 | 102146874 | 1 | lncRNA | ENSG00000 |
| 102233986 | 102296284 | -1 | protein_coding | ENSG00000 |
| 102302504 | 102302810 | 1 | misc_RNA | ENSG00000 |
| 102371774 | 102499016 | -1 | protein_coding | ENSG00000 |
| 102500541 | 102505670 | 1 | lncRNA | ENSG00000 |
| 102581368 | 102617589 | 1 | lncRNA | ENSG00000 |
| 102604220 | 102671765 | -1 | lncRNA | ENSG00000 |
| 102605635 | 102606197 | 1 | lncRNA | ENSG00000 |

| | | | | |
|---|---|---|---|---|
| 102664610 | 102726598 | 1 | lncRNA | ENSG00000 |
| 102753981 | 103031105 | 1 | protein_coding | ENSG00000 |
| 103032376 | 103033031 | 1 | processed_pseudoge | ENSG00000 |
| 103086000 | 103120138 | -1 | protein_coding | ENSG00000 |
| 103120149 | 103212799 | 1 | protein_coding | ENSG00000 |
| 103246048 | 103253519 | -1 | lncRNA | ENSG00000 |
| 103258763 | 103278660 | 1 | protein_coding | ENSG00000 |
| 103408941 | 103412511 | -1 | lncRNA | ENSG00000 |
| 103430406 | 103526618 | -1 | transcribed_unitary_ | ENSG00000 |
| 103528434 | 103541985 | -1 | lncRNA | ENSG00000 |
| 103548855 | 103562790 | -1 | protein_coding | ENSG00000 |
| 103880129 | 103891404 | -1 | lncRNA | ENSG00000 |
| 104079847 | 104406121 | 1 | lncRNA | ENSG00000 |
| 104098874 | 104099038 | 1 | snRNA | ENSG00000 |
| 104260531 | 104260826 | -1 | misc_RNA | ENSG00000 |
| 104383298 | 105392970 | -1 | lncRNA | ENSG00000 |
| 104434772 | 104435499 | 1 | TEC | ENSG00000 |
| 104773641 | 104799772 | 1 | lncRNA | ENSG00000 |
| 104780288 | 104780389 | -1 | snRNA | ENSG00000 |
| 104917492 | 105246364 | -1 | lncRNA | ENSG00000 |
| 105099473 | 105100098 | 1 | processed_pseudoge | ENSG00000 |
| 105922994 | 105923108 | -1 | rRNA_pseudogene | ENSG00000 |
| 106415576 | 106417241 | -1 | processed_pseudoge | ENSG00000 |
| 106543066 | 106544330 | -1 | processed_pseudoge | ENSG00000 |
| 106546665 | 106546803 | 1 | snoRNA | ENSG00000 |
| 106815197 | 107011052 | -1 | lncRNA | ENSG00000 |
| 107195156 | 107195332 | 1 | processed_pseudoge | ENSG00000 |
| 107376889 | 107670937 | -1 | protein_coding | ENSG00000 |
| 107699392 | 107716841 | -1 | lncRNA | ENSG00000 |
| 107724961 | 107727117 | 1 | processed_pseudoge | ENSG00000 |
| 107734773 | 107735066 | -1 | misc_RNA | ENSG00000 |
| 107810629 | 107810924 | 1 | misc_RNA | ENSG00000 |
| 107859035 | 108382098 | -1 | protein_coding | ENSG00000 |
| 108382156 | 108418016 | 1 | lncRNA | ENSG00000 |
| 108593609 | 108593967 | 1 | processed_pseudoge | ENSG00000 |
| 108727825 | 108728260 | -1 | lncRNA | ENSG00000 |
| 108747841 | 109196841 | 1 | protein_coding | ENSG00000 |
| 108784098 | 108785552 | 1 | processed_pseudoge | ENSG00000 |
| 108818041 | 108830790 | -1 | lncRNA | ENSG00000 |
| 108891437 | 108891546 | 1 | misc_RNA | ENSG00000 |
| 108894347 | 108894628 | -1 | processed_pseudoge | ENSG00000 |
| 109014834 | 109014940 | -1 | snRNA | ENSG00000 |
| 109051315 | 109052457 | -1 | processed_pseudoge | ENSG00000 |
| 109165289 | 109187152 | -1 | lncRNA | ENSG00000 |
| 109237120 | 109326369 | -1 | lncRNA | ENSG00000 |
| 109334713 | 109409974 | -1 | protein_coding | ENSG00000 |
| 109448349 | 109449063 | -1 | processed_pseudoge | ENSG00000 |
| 109467353 | 109467952 | 1 | lncRNA | ENSG00000 |
| 109470843 | 109471586 | 1 | lncRNA | ENSG00000 |
| 109497877 | 109499108 | 1 | lncRNA | ENSG00000 |

| | | | | |
|---|---|---|---|---|
| 109588338 | 109589592 | 1 | processed_pseudoge | ENSG00000 |
| 109687802 | 109688329 | -1 | lncRNA | ENSG00000 |
| 109689366 | 109869625 | 1 | protein_coding | ENSG00000 |
| 109699500 | 109699834 | -1 | misc_RNA | ENSG00000 |
| 109840128 | 109840692 | -1 | processed_pseudoge | ENSG00000 |
| 109883182 | 109884751 | 1 | lncRNA | ENSG00000 |
| 109884610 | 109885423 | -1 | processed_pseudoge | ENSG00000 |
| 110289233 | 110738956 | -1 | protein_coding | ENSG00000 |
| 110513829 | 110513915 | -1 | miRNA | ENSG00000 |
| 110684794 | 110684927 | -1 | snoRNA | ENSG00000 |
| 110738136 | 110765161 | 1 | protein_coding | ENSG00000 |
| 110908284 | 110908713 | 1 | processed_pseudoge | ENSG00000 |
| 110948029 | 110950780 | -1 | processed_pseudoge | ENSG00000 |
| 110970951 | 111008899 | -1 | lncRNA | ENSG00000 |
| 110983563 | 110984466 | -1 | processed_pseudoge | ENSG00000 |
| 111070062 | 111078026 | 1 | protein_coding | ENSG00000 |
| 111076921 | 111092115 | -1 | lncRNA | ENSG00000 |
| 111091716 | 111130502 | 1 | protein_coding | ENSG00000 |
| 111155244 | 111156152 | -1 | processed_pseudoge | ENSG00000 |
| 111192226 | 111193042 | -1 | processed_pseudoge | ENSG00000 |
| 111223653 | 111494886 | 1 | protein_coding | ENSG00000 |
| 111265809 | 111270089 | -1 | lncRNA | ENSG00000 |
| 111277517 | 111302567 | -1 | lncRNA | ENSG00000 |
| 111496033 | 111512590 | -1 | protein_coding | ENSG00000 |
| 111510396 | 111739726 | 1 | lncRNA | ENSG00000 |
| 111572102 | 111572239 | -1 | processed_pseudoge | ENSG00000 |
| 111572236 | 111572398 | 1 | processed_pseudoge | ENSG00000 |
| 111662621 | 111997464 | -1 | protein_coding | ENSG00000 |
| 111719769 | 111720061 | -1 | misc_RNA | ENSG00000 |
| 111846050 | 111846566 | -1 | processed_pseudoge | ENSG00000 |
| 111912508 | 112017309 | 1 | lncRNA | ENSG00000 |
| 112142441 | 112419313 | -1 | protein_coding | ENSG00000 |
| 112160526 | 112164818 | 1 | lncRNA | ENSG00000 |
| 112161485 | 112161617 | 1 | snoRNA | ENSG00000 |
| 112173570 | 112175548 | 1 | lncRNA | ENSG00000 |
| 112192020 | 112210139 | 1 | lncRNA | ENSG00000 |
| 112228283 | 112257309 | 1 | lncRNA | ENSG00000 |
| 112363456 | 112363666 | -1 | processed_pseudoge | ENSG00000 |
| 112419583 | 112420978 | 1 | lncRNA | ENSG00000 |
| 112452703 | 112452989 | -1 | processed_pseudoge | ENSG00000 |
| 112546973 | 112547505 | 1 | processed_pseudoge | ENSG00000 |
| 112628436 | 112642483 | 1 | lncRNA | ENSG00000 |
| 112707498 | 112846239 | 1 | protein_coding | ENSG00000 |
| 112777630 | 112778161 | -1 | processed_pseudoge | ENSG00000 |
| 112778363 | 112778469 | 1 | snRNA | ENSG00000 |
| 112827213 | 112867582 | 1 | protein_coding | ENSG00000 |
| 112861188 | 112898371 | 1 | protein_coding | ENSG00000 |
| 112876385 | 112922289 | -1 | protein_coding | ENSG00000 |
| 112885094 | 112885745 | 1 | processed_pseudoge | ENSG00000 |
| 112891610 | 112893079 | 1 | transcribed_processe | ENSG00000 |

| | | | | |
|---|---|---|---|---|
| 112976702 | 113022195 | 1 | protein_coding | ENSG00000 |
| 113022099 | 113488823 | -1 | protein_coding | ENSG00000 |
| 113323028 | 113437174 | 1 | lncRNA | ENSG00000 |
| 113432553 | 113434989 | -1 | protein_coding | ENSG00000 |
| 113493835 | 113493960 | 1 | snRNA | ENSG00000 |
| 113513694 | 113595285 | 1 | protein_coding | ENSG00000 |
| 113738106 | 113776837 | -1 | lncRNA | ENSG00000 |
| 114055545 | 114496500 | 1 | protein_coding | ENSG00000 |
| 114253513 | 114253754 | 1 | misc_RNA | ENSG00000 |
| 114475339 | 114668410 | -1 | lncRNA | ENSG00000 |
| 114576041 | 114579970 | 1 | lncRNA | ENSG00000 |
| 115031273 | 115031894 | -1 | lncRNA | ENSG00000 |
| 115087892 | 115088475 | -1 | lncRNA | ENSG00000 |
| 115124762 | 115180546 | -1 | protein_coding | ENSG00000 |
| 115148764 | 115149644 | -1 | lncRNA | ENSG00000 |
| 115188563 | 115188932 | 1 | lncRNA | ENSG00000 |
| 115204012 | 115262877 | -1 | protein_coding | ENSG00000 |
| 115262505 | 115263448 | 1 | lncRNA | ENSG00000 |
| 115264669 | 115265464 | -1 | processed_pseudoge | ENSG00000 |
| 115267190 | 115296693 | -1 | protein_coding | ENSG00000 |
| 115289036 | 115289330 | -1 | processed_pseudoge | ENSG00000 |
| 115389643 | 115392370 | -1 | processed_pseudoge | ENSG00000 |
| 115402380 | 115402962 | 1 | processed_pseudoge | ENSG00000 |
| 115489634 | 115489931 | 1 | processed_pseudoge | ENSG00000 |
| 115493557 | 115493821 | 1 | processed_pseudoge | ENSG00000 |
| 115512077 | 115513694 | 1 | processed_pseudoge | ENSG00000 |
| 115520908 | 115544775 | -1 | protein_coding | ENSG00000 |
| 115578642 | 115626161 | -1 | protein_coding | ENSG00000 |
| 115578650 | 115602479 | -1 | protein_coding | ENSG00000 |
| 115602057 | 115623897 | 1 | lncRNA | ENSG00000 |
| 115613210 | 115632992 | -1 | protein_coding | ENSG00000 |
| 115691462 | 115692167 | -1 | lncRNA | ENSG00000 |
| 115738978 | 115756543 | 1 | lncRNA | ENSG00000 |
| 115770586 | 115770989 | 1 | processed_pseudoge | ENSG00000 |
| 115774319 | 115774502 | -1 | snRNA | ENSG00000 |
| 115804733 | 115816659 | -1 | protein_coding | ENSG00000 |
| 115828200 | 115841837 | -1 | protein_coding | ENSG00000 |
| 115841592 | 115914081 | 1 | protein_coding | ENSG00000 |
| 115956571 | 115958599 | 1 | lncRNA | ENSG00000 |
| 115962454 | 116027619 | 1 | protein_coding | ENSG00000 |
| 115974513 | 115976275 | -1 | processed_pseudoge | ENSG00000 |
| 116032324 | 116060118 | 1 | protein_coding | ENSG00000 |
| 116051917 | 116052289 | -1 | processed_pseudoge | ENSG00000 |
| 116078110 | 116078570 | -1 | lncRNA | ENSG00000 |
| 116083807 | 116085416 | -1 | lncRNA | ENSG00000 |
| 116085016 | 116412762 | 1 | protein_coding | ENSG00000 |
| 116188455 | 116188552 | -1 | snRNA | ENSG00000 |
| 116218026 | 116218295 | -1 | processed_pseudoge | ENSG00000 |
| 116302354 | 116304134 | -1 | lncRNA | ENSG00000 |
| 116443616 | 116574934 | -1 | protein_coding | ENSG00000 |

| | | | | |
|---|---|---|---|---|
| 116447547 | 116508276 | 1 | lncRNA | ENSG0000C |
| 116562562 | 116562930 | 1 | processed_pseudoge | ENSG0000C |
| 116574482 | 116611200 | 1 | lncRNA | ENSG0000C |
| 116716243 | 116716650 | 1 | processed_pseudoge | ENSG0000C |
| 116742991 | 116762209 | -1 | lncRNA | ENSG0000C |
| 116819220 | 116830370 | -1 | lncRNA | ENSG0000C |
| 117019548 | 117019679 | -1 | processed_pseudoge | ENSG0000C |
| 117031200 | 117038569 | 1 | lncRNA | ENSG0000C |
| 117301104 | 117301442 | -1 | processed_pseudoge | ENSG0000C |
| 117313085 | 117330149 | 1 | lncRNA | ENSG0000C |
| 117415509 | 117546298 | 1 | lncRNA | ENSG0000C |
| 117719240 | 117722097 | -1 | processed_pseudoge | ENSG0000C |
| 117730515 | 118266267 | 1 | lncRNA | ENSG0000C |
| 117778826 | 117779532 | -1 | processed_pseudoge | ENSG0000C |
| 118000253 | 118562199 | -1 | lncRNA | ENSG0000C |
| 118282575 | 118284782 | 1 | lncRNA | ENSG0000C |
| 118575575 | 118581324 | 1 | lncRNA | ENSG0000C |
| 118581242 | 118628178 | -1 | lncRNA | ENSG0000C |
| 118596188 | 118628092 | 1 | lncRNA | ENSG0000C |
| 118758211 | 118758274 | 1 | snRNA | ENSG0000C |
| 118760474 | 118785459 | -1 | lncRNA | ENSG0000C |
| 118836074 | 118988547 | -1 | protein_coding | ENSG0000C |
| 118973796 | 118974122 | 1 | processed_pseudoge | ENSG0000C |
| 118974586 | 118974670 | 1 | miRNA | ENSG0000C |
| 119006347 | 119070902 | -1 | lncRNA | ENSG0000C |
| 119007571 | 119007668 | 1 | snRNA | ENSG0000C |
| 119010060 | 119010542 | 1 | processed_pseudoge | ENSG0000C |
| 119011471 | 119012056 | 1 | processed_pseudoge | ENSG0000C |
| 119037772 | 119249138 | 1 | protein_coding | ENSG0000C |
| 119126782 | 119128444 | -1 | processed_pseudoge | ENSG0000C |
| 119132651 | 119132882 | -1 | processed_pseudoge | ENSG0000C |
| 119138859 | 119138965 | 1 | snRNA | ENSG0000C |
| 119154637 | 119154716 | 1 | miRNA | ENSG0000C |
| 119268692 | 119399688 | 1 | protein_coding | ENSG0000C |
| 119306343 | 119306631 | -1 | misc_RNA | ENSG0000C |
| 119445716 | 119445833 | 1 | rRNA_pseudogene | ENSG0000C |
| 119452473 | 119637199 | 1 | protein_coding | ENSG0000C |
| 119547131 | 119547606 | -1 | processed_pseudoge | ENSG0000C |
| 119555250 | 119555658 | -1 | processed_pseudoge | ENSG0000C |
| 119629559 | 119635822 | 1 | protein_coding | ENSG0000C |
| 119652523 | 119653000 | -1 | processed_pseudoge | ENSG0000C |
| 119680197 | 119681080 | 1 | processed_pseudoge | ENSG0000C |
| 120245448 | 120333502 | -1 | lncRNA | ENSG0000C |
| 120337549 | 120337653 | -1 | snRNA | ENSG0000C |
| 120345907 | 120410969 | -1 | lncRNA | ENSG0000C |
| 120464300 | 120687332 | 1 | protein_coding | ENSG0000C |
| 120658077 | 120658371 | -1 | processed_pseudoge | ENSG0000C |
| 120710698 | 120710840 | -1 | snRNA | ENSG0000C |
| 120781218 | 120790778 | 1 | lncRNA | ENSG0000C |
| 121040090 | 121044032 | 1 | processed_pseudoge | ENSG0000C |

| | | | | |
|---|---|---|---|---|
| 121059244 | 121060953 | 1 | processed_pseudoge | ENSG00000 |
| 121164687 | 121359869 | 1 | lncRNA | ENSG00000 |
| 121195980 | 121196210 | -1 | unprocessed_pseudo | ENSG00000 |
| 121363002 | 121365314 | 1 | TEC | ENSG00000 |
| 121575848 | 121576320 | -1 | processed_pseudoge | ENSG00000 |
| 121616247 | 121616807 | 1 | processed_pseudoge | ENSG00000 |
| 121671373 | 121671872 | -1 | processed_pseudoge | ENSG00000 |
| 121851882 | 121852833 | 1 | protein_coding | ENSG00000 |
| 121961975 | 122075570 | 1 | protein_coding | ENSG00000 |
| 122063195 | 122078360 | -1 | protein_coding | ENSG00000 |
| 122114598 | 122154856 | -1 | lncRNA | ENSG00000 |
| 122129546 | 122153569 | 1 | protein_coding | ENSG00000 |
| 122129622 | 122183949 | 1 | protein_coding | ENSG00000 |
| 122154496 | 122156015 | -1 | lncRNA | ENSG00000 |
| 122311297 | 122311673 | -1 | lncRNA | ENSG00000 |
| 122311354 | 122464219 | 1 | protein_coding | ENSG00000 |
| 122321291 | 122323358 | -1 | lncRNA | ENSG00000 |
| 122369762 | 122383568 | -1 | lncRNA | ENSG00000 |
| 122373512 | 122374481 | 1 | unitary_pseudogene | ENSG00000 |
| 122436497 | 122479087 | -1 | lncRNA | ENSG00000 |
| 122628952 | 122730685 | -1 | lncRNA | ENSG00000 |
| 122675795 | 122676755 | 1 | processed_pseudoge | ENSG00000 |
| 122775079 | 122834543 | 1 | protein_coding | ENSG00000 |
| 122832356 | 122834533 | 1 | lncRNA | ENSG00000 |
| 122843439 | 123029354 | 1 | protein_coding | ENSG00000 |
| 123022487 | 123022783 | -1 | misc_RNA | ENSG00000 |
| 123023250 | 123036725 | -1 | protein_coding | ENSG00000 |
| 123036271 | 123054667 | 1 | lncRNA | ENSG00000 |
| 123070782 | 123071080 | 1 | misc_RNA | ENSG00000 |
| 123087248 | 123090299 | -1 | lncRNA | ENSG00000 |
| 123089241 | 123194266 | 1 | protein_coding | ENSG00000 |
| 123236079 | 123236370 | 1 | processed_pseudoge | ENSG00000 |
| 123344885 | 123423592 | -1 | protein_coding | ENSG00000 |
| 123400922 | 123402344 | -1 | processed_pseudoge | ENSG00000 |
| 123468781 | 123469429 | -1 | processed_pseudoge | ENSG00000 |
| 123508736 | 123509187 | -1 | TEC | ENSG00000 |
| 123512099 | 123617045 | 1 | protein_coding | ENSG00000 |
| 123636110 | 123637403 | 1 | processed_pseudoge | ENSG00000 |
| 124059794 | 124438625 | -1 | lncRNA | ENSG00000 |
| 124220579 | 124221077 | -1 | processed_pseudoge | ENSG00000 |
| 124395603 | 124400655 | 1 | lncRNA | ENSG00000 |
| 124452277 | 124460552 | 1 | lncRNA | ENSG00000 |
| 124459912 | 124460549 | -1 | lncRNA | ENSG00000 |
| 124469591 | 124469901 | 1 | lncRNA | ENSG00000 |
| 124492775 | 124536348 | -1 | lncRNA | ENSG00000 |
| 124636913 | 124748807 | -1 | protein_coding | ENSG00000 |
| 124707827 | 124710737 | 1 | lncRNA | ENSG00000 |
| 124734618 | 124735175 | -1 | lncRNA | ENSG00000 |
| 124808828 | 124809233 | -1 | processed_pseudoge | ENSG00000 |
| 124808981 | 125602227 | 1 | lncRNA | ENSG00000 |

| | | | | |
|---|---|---|---|---|
| 124829472 | 124830290 | 1 | lncRNA | ENSG00000 |
| 124868972 | 124869914 | -1 | lncRNA | ENSG00000 |
| 124993408 | 124996096 | -1 | lncRNA | ENSG00000 |
| 125074665 | 125074980 | 1 | processed_pseudoge | ENSG00000 |
| 125265528 | 125265651 | 1 | processed_pseudoge | ENSG00000 |
| 125333369 | 126368755 | -1 | lncRNA | ENSG00000 |
| 125350868 | 125351131 | -1 | misc_RNA | ENSG00000 |
| 125376828 | 125377439 | -1 | lncRNA | ENSG00000 |
| 125493261 | 125602229 | 1 | lncRNA | ENSG00000 |
| 125862746 | 125863193 | -1 | processed_pseudoge | ENSG00000 |
| 125886285 | 125886753 | -1 | processed_pseudoge | ENSG00000 |
| 125966777 | 125968085 | -1 | processed_pseudoge | ENSG00000 |
| 126179565 | 126193858 | 1 | lncRNA | ENSG00000 |
| 126360132 | 126496494 | 1 | protein_coding | ENSG00000 |
| 126372477 | 126509021 | -1 | lncRNA | ENSG00000 |
| 126496279 | 126498604 | 1 | TEC | ENSG00000 |
| 126531200 | 126595362 | -1 | protein_coding | ENSG00000 |
| 126553302 | 126553402 | -1 | snRNA | ENSG00000 |
| 126600925 | 126627252 | 1 | protein_coding | ENSG00000 |
| 126628019 | 126628319 | -1 | processed_pseudoge | ENSG00000 |
| 126631705 | 126636284 | 1 | protein_coding | ENSG00000 |
| 126663337 | 126663568 | -1 | processed_pseudoge | ENSG00000 |
| 126737438 | 126737995 | -1 | processed_pseudoge | ENSG00000 |
| 126751963 | 126776486 | -1 | lncRNA | ENSG00000 |
| 126755316 | 126755417 | -1 | snRNA | ENSG00000 |
| 126776623 | 126837020 | 1 | protein_coding | ENSG00000 |
| 126867714 | 127030558 | -1 | protein_coding | ENSG00000 |
| 127042558 | 127073492 | -1 | protein_coding | ENSG00000 |
| 127073587 | 127075604 | 1 | TEC | ENSG00000 |
| 127143082 | 127143247 | -1 | processed_pseudoge | ENSG00000 |
| 127170535 | 127172266 | -1 | processed_pseudoge | ENSG00000 |
| 127176696 | 127177050 | -1 | processed_pseudoge | ENSG00000 |
| 127179756 | 127179900 | -1 | processed_pseudoge | ENSG00000 |
| 127215159 | 127229828 | -1 | lncRNA | ENSG00000 |
| 127231977 | 127290731 | -1 | lncRNA | ENSG00000 |
| 127290796 | 127465737 | 1 | protein_coding | ENSG00000 |
| 127465822 | 127466118 | 1 | processed_pseudoge | ENSG00000 |
| 127478295 | 127478737 | 1 | lncRNA | ENSG00000 |
| 127511464 | 127512842 | -1 | processed_pseudoge | ENSG00000 |
| 127517609 | 127555089 | 1 | protein_coding | ENSG00000 |
| 127588746 | 127590710 | -1 | lncRNA | ENSG00000 |
| 127649044 | 127658630 | 1 | protein_coding | ENSG00000 |
| 127651693 | 127664029 | -1 | lncRNA | ENSG00000 |
| 127703389 | 127941634 | 1 | protein_coding | ENSG00000 |
| 127784618 | 127787042 | -1 | processed_pseudoge | ENSG00000 |
| 127838486 | 127857804 | -1 | lncRNA | ENSG00000 |
| 127939152 | 128083172 | -1 | lncRNA | ENSG00000 |
| 128083766 | 128189677 | 1 | protein_coding | ENSG00000 |
| 128109486 | 128110994 | 1 | processed_pseudoge | ENSG00000 |
| 128257909 | 128659185 | -1 | protein_coding | ENSG00000 |

| | | | | |
|---|---|---|---|---|
| 128371099 | 128371212 | -1 | misc_RNA | ENSG0000C |
| 128538013 | 129033642 | 1 | protein_coding | ENSG0000C |
| 128663978 | 128742829 | 1 | lncRNA | ENSG0000C |
| 129094749 | 129114028 | 1 | protein_coding | ENSG0000C |
| 129097688 | 129097766 | 1 | miRNA | ENSG0000C |
| 129150677 | 129173257 | 1 | lncRNA | ENSG0000C |
| 129397062 | 129397147 | -1 | miRNA | ENSG0000C |
| 129424782 | 129461076 | -1 | lncRNA | ENSG0000C |
| 129460281 | 129738683 | 1 | protein_coding | ENSG0000C |
| 129500361 | 129905917 | -1 | lncRNA | ENSG0000C |
| 129748094 | 129766732 | 1 | protein_coding | ENSG0000C |
| 129904465 | 130186634 | 1 | protein_coding | ENSG0000C |
| 129974891 | 129975001 | 1 | snRNA | ENSG0000C |
| 130114409 | 130114510 | 1 | rRNA_pseudogene | ENSG0000C |
| 130140031 | 130141950 | 1 | processed_pseudoge | ENSG0000C |
| 130386621 | 130386680 | -1 | snRNA | ENSG0000C |
| 130529419 | 130529889 | 1 | processed_pseudoge | ENSG0000C |
| 130598348 | 130598779 | 1 | processed_pseudoge | ENSG0000C |
| 130655383 | 130944178 | -1 | lncRNA | ENSG0000C |
| 130989897 | 130990428 | 1 | processed_pseudoge | ENSG0000C |
| 130994253 | 130996164 | 1 | processed_pseudoge | ENSG0000C |
| 131008583 | 131008742 | 1 | processed_pseudoge | ENSG0000C |
| 131159027 | 131171735 | -1 | protein_coding | ENSG0000C |
| 131170944 | 131205428 | 1 | protein_coding | ENSG0000C |
| 131245493 | 131398447 | 1 | protein_coding | ENSG0000C |
| 131261321 | 131261607 | -1 | TEC | ENSG0000C |
| 131423921 | 131635236 | -1 | protein_coding | ENSG0000C |
| 131425891 | 131796983 | -1 | protein_coding | ENSG0000C |
| 131440031 | 131440196 | -1 | misc_RNA | ENSG0000C |
| 131641714 | 131797063 | -1 | protein_coding | ENSG0000C |
| 131659046 | 131660115 | -1 | processed_pseudoge | ENSG0000C |
| 131797415 | 131797929 | 1 | TEC | ENSG0000C |
| 131806990 | 131945698 | -1 | protein_coding | ENSG0000C |
| 131807143 | 132011637 | -1 | protein_coding | ENSG0000C |
| 131944408 | 131968220 | 1 | lncRNA | ENSG0000C |
| 131949973 | 132012243 | -1 | protein_coding | ENSG0000C |
| 132003592 | 132007022 | 1 | lncRNA | ENSG0000C |
| 132011448 | 132013199 | 1 | lncRNA | ENSG0000C |
| 132060655 | 132063204 | 1 | protein_coding | ENSG0000C |
| 132073789 | 132076170 | 1 | protein_coding | ENSG0000C |
| 132080066 | 132080133 | 1 | snoRNA | ENSG0000C |
| 132147491 | 132149559 | -1 | unprocessed_pseudc | ENSG0000C |
| 132163603 | 132163765 | -1 | unprocessed_pseudc | ENSG0000C |
| 132179234 | 132181128 | -1 | lncRNA | ENSG0000C |
| 132184876 | 132192808 | 1 | lncRNA | ENSG0000C |
| 132191838 | 132295315 | -1 | protein_coding | ENSG0000C |
| 132199456 | 132203487 | 1 | processed_pseudoge | ENSG0000C |
| 132217849 | 132217918 | -1 | miRNA | ENSG0000C |
| 132257696 | 132273454 | 1 | protein_coding | ENSG0000C |
| 132294394 | 132344190 | 1 | protein_coding | ENSG0000C |

| | | | | |
|---|---|---|---|---|
| 132311285 | 132370170 | -1 | lncRNA | ENSG0000C |
| 132365490 | 132365599 | -1 | miRNA | ENSG0000C |
| 132369752 | 132395614 | 1 | protein_coding | ENSG0000C |
| 132410636 | 132488702 | 1 | lncRNA | ENSG0000C |
| 132410832 | 132646079 | 1 | protein_coding | ENSG0000C |
| 132419416 | 132426714 | -1 | lncRNA | ENSG0000C |
| 132468147 | 132468257 | 1 | misc_RNA | ENSG0000C |
| 132468890 | 132473043 | -1 | lncRNA | ENSG0000C |
| 132481609 | 132490777 | -1 | protein_coding | ENSG0000C |
| 132541445 | 132556838 | -1 | protein_coding | ENSG0000C |
| 132556019 | 132646349 | 1 | protein_coding | ENSG0000C |
| 132630589 | 132664272 | -1 | lncRNA | ENSG0000C |
| 132656263 | 132661110 | 1 | protein_coding | ENSG0000C |
| 132673986 | 132682678 | 1 | protein_coding | ENSG0000C |
| 132688681 | 132723725 | 1 | lncRNA | ENSG0000C |
| 132692628 | 132737638 | -1 | protein_coding | ENSG0000C |
| 132747445 | 132754403 | 1 | protein_coding | ENSG0000C |
| 132750819 | 132807241 | -1 | protein_coding | ENSG0000C |
| 132813302 | 132816786 | 1 | protein_coding | ENSG0000C |
| 132817248 | 132818000 | -1 | lncRNA | ENSG0000C |
| 132822141 | 132830898 | -1 | protein_coding | ENSG0000C |
| 132848577 | 132848696 | 1 | rRNA_pseudogene | ENSG0000C |
| 132861181 | 132866884 | -1 | protein_coding | ENSG0000C |
| 132866630 | 132868847 | 1 | protein_coding | ENSG0000C |
| 132873444 | 132875046 | 1 | protein_coding | ENSG0000C |
| 132875395 | 132963634 | -1 | protein_coding | ENSG0000C |
| 132942932 | 132945050 | 1 | processed_pseudoge | ENSG0000C |
| 132945342 | 132945454 | 1 | misc_RNA | ENSG0000C |
| 132948493 | 132948718 | 1 | processed_pseudoge | ENSG0000C |
| 132963770 | 132964164 | 1 | lncRNA | ENSG0000C |
| 132987652 | 132988110 | -1 | processed_pseudoge | ENSG0000C |
| 132993099 | 132993200 | -1 | misc_RNA | ENSG0000C |
| 132996985 | 133026604 | -1 | protein_coding | ENSG0000C |
| 133003119 | 133003365 | 1 | processed_pseudoge | ENSG0000C |
| 133025847 | 133050178 | 1 | lncRNA | ENSG0000C |
| 133051065 | 133051893 | -1 | lncRNA | ENSG0000C |
| 133052013 | 133106449 | 1 | protein_coding | ENSG0000C |
| 133052769 | 133052971 | 1 | TEC | ENSG0000C |
| 133111055 | 133114475 | -1 | lncRNA | ENSG0000C |
| 133160438 | 133224413 | 1 | lncRNA | ENSG0000C |
| 133172218 | 133173053 | -1 | processed_pseudoge | ENSG0000C |
| 133196455 | 133612541 | -1 | protein_coding | ENSG0000C |
| 133243764 | 133248781 | 1 | lncRNA | ENSG0000C |
| 133256492 | 133275977 | 1 | lncRNA | ENSG0000C |
| 133387773 | 133388549 | 1 | lncRNA | ENSG0000C |
| 133427596 | 133427706 | -1 | miRNA | ENSG0000C |
| 133457805 | 133458331 | 1 | TEC | ENSG0000C |
| 133574829 | 133575298 | -1 | TEC | ENSG0000C |
| 133720495 | 133720934 | -1 | processed_pseudoge | ENSG0000C |
| 133913677 | 133917269 | 1 | lncRNA | ENSG0000C |

| | | | | |
|---|---|---|---|---|
| 133955510 | 133968674 | -1 | protein_coding | ENSG0000C |
| 133971871 | 134004975 | -1 | protein_coding | ENSG0000C |
| 134005147 | 134005562 | 1 | lncRNA | ENSG0000C |
| 134081914 | 134082040 | -1 | processed_pseudoge | ENSG0000C |
| 134114681 | 134151865 | 1 | protein_coding | ENSG0000C |
| 134148935 | 134177038 | -1 | protein_coding | ENSG0000C |
| 134167170 | 134226071 | -1 | protein_coding | ENSG0000C |
| 134194332 | 134226073 | -1 | protein_coding | ENSG0000C |
| 134205614 | 134371044 | -1 | lncRNA | ENSG0000C |
| 134225757 | 134225852 | 1 | miRNA | ENSG0000C |
| 134226410 | 134227827 | 1 | lncRNA | ENSG0000C |
| 134267421 | 134267856 | 1 | processed_pseudoge | ENSG0000C |
| 134286350 | 134371047 | -1 | protein_coding | ENSG0000C |
| 134371469 | 134392108 | 1 | protein_coding | ENSG0000C |
| 134394360 | 134395008 | -1 | TEC | ENSG0000C |
| 134399495 | 134401921 | 1 | lncRNA | ENSG0000C |
| 134402065 | 134411881 | -1 | protein_coding | ENSG0000C |
| 134422838 | 134423335 | 1 | processed_pseudoge | ENSG0000C |
| 134429051 | 134460615 | -1 | lncRNA | ENSG0000C |
| 134436711 | 134492519 | 1 | lncRNA | ENSG0000C |
| 134500178 | 134500284 | -1 | snRNA | ENSG0000C |
| 134506552 | 134509229 | 1 | lncRNA | ENSG0000C |
| 134517810 | 134518092 | 1 | misc_RNA | ENSG0000C |
| 134522613 | 134523096 | -1 | TEC | ENSG0000C |
| 134524312 | 134583230 | 1 | protein_coding | ENSG0000C |
| 134601149 | 134649271 | -1 | protein_coding | ENSG0000C |
| 134634839 | 134634945 | -1 | snRNA | ENSG0000C |
| 134648785 | 134727909 | 1 | protein_coding | ENSG0000C |
| 134701172 | 134701278 | -1 | snRNA | ENSG0000C |
| 134715483 | 134715582 | 1 | snRNA | ENSG0000C |
| 134738548 | 134752157 | 1 | protein_coding | ENSG0000C |
| 134758771 | 134855133 | 1 | protein_coding | ENSG0000C |
| 134804778 | 134805132 | -1 | processed_pseudoge | ENSG0000C |
| 134845680 | 134859735 | 1 | protein_coding | ENSG0000C |
| 134863091 | 134863433 | -1 | processed_pseudoge | ENSG0000C |
| 134873803 | 134901525 | 1 | protein_coding | ENSG0000C |
| 134904906 | 135007959 | 1 | protein_coding | ENSG0000C |
| 134905235 | 134905691 | 1 | TEC | ENSG0000C |
| 134923303 | 134924049 | -1 | processed_pseudoge | ENSG0000C |
| 134924123 | 134924644 | 1 | processed_pseudoge | ENSG0000C |
| 134924648 | 134926459 | -1 | processed_pseudoge | ENSG0000C |
| 134926660 | 134928036 | -1 | processed_pseudoge | ENSG0000C |
| 134928030 | 134928326 | -1 | processed_pseudoge | ENSG0000C |
| 134928039 | 134928112 | 1 | miRNA | ENSG0000C |
| 134928393 | 134928527 | -1 | processed_pseudoge | ENSG0000C |
| 134967907 | 135011696 | 1 | protein_coding | ENSG0000C |
| 135027734 | 135034813 | -1 | protein_coding | ENSG0000C |
| 135033280 | 135358219 | 1 | lncRNA | ENSG0000C |
| 135034521 | 135035894 | 1 | lncRNA | ENSG0000C |
| 135038831 | 135040047 | -1 | lncRNA | ENSG0000C |

| | | | | |
|---|---|---|---|---|
| 135120526 | 135139681 | 1 | lncRNA | ENSG0000C |
| 135124380 | 135131249 | -1 | lncRNA | ENSG0000C |
| 135236234 | 135248179 | -1 | lncRNA | ENSG0000C |
| 135333900 | 135399914 | -1 | protein_coding | ENSG0000C |
| 135399280 | 135401296 | 1 | lncRNA | ENSG0000C |
| 135444214 | 135447348 | -1 | protein_coding | ENSG0000C |
| 135444226 | 135452351 | -1 | protein_coding | ENSG0000C |
| 135450613 | 135458697 | 1 | lncRNA | ENSG0000C |
| 135534282 | 135535964 | -1 | protein_coding | ENSG0000C |
| 135559577 | 135634874 | 1 | lncRNA | ENSG0000C |
| 135570679 | 135579279 | -1 | protein_coding | ENSG0000C |
| 135579202 | 135889770 | 1 | protein_coding | ENSG0000C |
| 135648584 | 135653935 | -1 | lncRNA | ENSG0000C |
| 135742462 | 135746356 | -1 | unprocessed_pseudc | ENSG0000C |
| 135802985 | 135803075 | -1 | miRNA | ENSG0000C |
| 135812667 | 135827546 | 1 | lncRNA | ENSG0000C |
| 135892246 | 135895841 | -1 | protein_coding | ENSG0000C |
| 135897637 | 135900250 | 1 | lncRNA | ENSG0000C |
| 135900353 | 135918161 | -1 | lncRNA | ENSG0000C |
| 135922279 | 135954983 | -1 | protein_coding | ENSG0000C |
| 135930317 | 135951591 | 1 | transcribed_unitary_ | ENSG0000C |
| 136028988 | 136063818 | 1 | protein_coding | ENSG0000C |
| 136080470 | 136080597 | -1 | vaultRNA | ENSG0000C |
| 136080497 | 136080597 | -1 | misc_RNA | ENSG0000C |
| 136129507 | 136134890 | -1 | lncRNA | ENSG0000C |
| 136129516 | 136129781 | 1 | misc_RNA | ENSG0000C |
| 136132845 | 136188747 | 1 | protein_coding | ENSG0000C |
| 136133696 | 136133826 | 1 | misc_RNA | ENSG0000C |
| 136133912 | 136134035 | 1 | misc_RNA | ENSG0000C |
| 136134082 | 136134248 | 1 | misc_RNA | ENSG0000C |
| 136191468 | 136193162 | -1 | protein_coding | ENSG0000C |
| 136213320 | 136365545 | -1 | protein_coding | ENSG0000C |
| 136214048 | 136222159 | 1 | lncRNA | ENSG0000C |
| 136303757 | 136316100 | 1 | lncRNA | ENSG0000C |
| 136376348 | 136397042 | -1 | lncRNA | ENSG0000C |
| 136381268 | 136381511 | -1 | processed_pseudoge | ENSG0000C |
| 136428892 | 136429854 | 1 | processed_pseudoge | ENSG0000C |
| 136466645 | 136521432 | 1 | lncRNA | ENSG0000C |
| 136734830 | 136763409 | 1 | lncRNA | ENSG0000C |
| 136857628 | 136858342 | 1 | processed_pseudoge | ENSG0000C |
| 136969200 | 136969306 | -1 | rRNA_pseudogene | ENSG0000C |
| 136975298 | 137598379 | -1 | protein_coding | ENSG0000C |
| 137617500 | 137736089 | -1 | protein_coding | ENSG0000C |
| 137647572 | 137647649 | -1 | miRNA | ENSG0000C |
| 137745651 | 137754363 | -1 | protein_coding | ENSG0000C |
| 137755740 | 137760740 | -1 | transcribed_processe | ENSG0000C |
| 137761546 | 137761936 | 1 | lncRNA | ENSG0000C |
| 137801193 | 137810751 | 1 | transcribed_processe | ENSG0000C |
| 137809780 | 137889394 | -1 | lncRNA | ENSG0000C |
| 137867791 | 137887851 | 1 | protein_coding | ENSG0000C |

| | | | | |
|---|---|---|---|---|
| 137887968 | 137942747 | 1 | protein_coding | ENSG00000( |
| 137937960 | 138051961 | -1 | protein_coding | ENSG00000( |
| 138032774 | 138037266 | 1 | lncRNA | ENSG00000( |
| 138039199 | 138039979 | 1 | lncRNA | ENSG00000( |
| 138069355 | 138069458 | -1 | snRNA | ENSG00000( |
| 138083990 | 138092365 | 1 | protein_coding | ENSG00000( |
| 138115175 | 138139443 | -1 | protein_coding | ENSG00000( |
| 138116573 | 138116680 | 1 | snRNA | ENSG00000( |
| 138123011 | 138123111 | -1 | snRNA | ENSG00000( |
| 138139766 | 138178986 | -1 | protein_coding | ENSG00000( |
| 138178719 | 138187723 | 1 | protein_coding | ENSG00000( |
| 138187650 | 138213343 | -1 | protein_coding | ENSG00000( |
| 138252380 | 138274621 | -1 | protein_coding | ENSG00000( |
| 138266996 | 138267757 | -1 | processed_pseudoge | ENSG00000( |
| 138285265 | 138338355 | -1 | protein_coding | ENSG00000( |
| 138331935 | 138349729 | 1 | protein_coding | ENSG00000( |
| 138347027 | 138349641 | -1 | lncRNA | ENSG00000( |
| 138352685 | 138437028 | 1 | protein_coding | ENSG00000( |
| 138439057 | 138446969 | 1 | protein_coding | ENSG00000( |
| 138454540 | 138454732 | -1 | processed_pseudoge | ENSG00000( |
| 138465479 | 138469303 | 1 | protein_coding | ENSG00000( |
| 138473744 | 138474361 | 1 | processed_pseudoge | ENSG00000( |
| 138506095 | 138543236 | -1 | protein_coding | ENSG00000( |
| 138553756 | 138575416 | -1 | protein_coding | ENSG00000( |
| 138558970 | 138559039 | -1 | snoRNA | ENSG00000( |
| 138561043 | 138561110 | -1 | snoRNA | ENSG00000( |
| 138610967 | 138935034 | 1 | protein_coding | ENSG00000( |
| 138620329 | 138620964 | 1 | processed_pseudoge | ENSG00000( |
| 138623479 | 138623582 | 1 | snoRNA | ENSG00000( |
| 138744434 | 138753309 | -1 | lncRNA | ENSG00000( |
| 138868921 | 138875368 | -1 | protein_coding | ENSG00000( |
| 138946724 | 139293557 | -1 | protein_coding | ENSG00000( |
| 139012329 | 139012441 | 1 | rRNA | ENSG00000( |
| 139012329 | 139012441 | 1 | rRNA_pseudogene | ENSG00000( |
| 139012647 | 139051203 | 1 | lncRNA | ENSG00000( |
| 139035148 | 139035987 | -1 | processed_pseudoge | ENSG00000( |
| 139221758 | 139221837 | 1 | snRNA | ENSG00000( |
| 139273752 | 139331671 | 1 | protein_coding | ENSG00000( |
| 139274102 | 139284899 | 1 | lncRNA | ENSG00000( |
| 139276180 | 139276320 | 1 | snoRNA | ENSG00000( |
| 139278781 | 139278978 | 1 | snoRNA | ENSG00000( |
| 139293674 | 139331677 | 1 | protein_coding | ENSG00000( |
| 139299852 | 139299968 | 1 | rRNA_pseudogene | ENSG00000( |
| 139333038 | 139333365 | -1 | misc_RNA | ENSG00000( |
| 139341587 | 139369720 | 1 | protein_coding | ENSG00000( |
| 139364677 | 139369717 | -1 | lncRNA | ENSG00000( |
| 139367196 | 139384553 | -1 | protein_coding | ENSG00000( |
| 139387467 | 139390081 | -1 | protein_coding | ENSG00000( |
| 139390592 | 139395104 | -1 | protein_coding | ENSG00000( |
| 139396563 | 139404088 | -1 | protein_coding | ENSG00000( |

| | | | | |
|---|---|---|---|---|
| 139408588 | 139444491 | -1 | protein_coding | ENSG00000 |
| 139442316 | 139462707 | -1 | protein_coding | ENSG00000 |
| 139447907 | 139448017 | 1 | snRNA | ENSG00000 |
| 139448560 | 139462743 | -1 | protein_coding | ENSG00000 |
| 139470778 | 139474772 | 1 | protein_coding | ENSG00000 |
| 139475533 | 139482935 | -1 | protein_coding | ENSG00000 |
| 139492538 | 139493105 | 1 | processed_pseudoge | ENSG00000 |
| 139494256 | 139496116 | 1 | processed_pseudoge | ENSG00000 |
| 139503242 | 139511374 | 1 | lncRNA | ENSG00000 |
| 139526431 | 139628434 | 1 | protein_coding | ENSG00000 |
| 139554900 | 139555001 | -1 | misc_RNA | ENSG00000 |
| 139612002 | 139612363 | -1 | processed_pseudoge | ENSG00000 |
| 139644460 | 139645002 | -1 | lncRNA | ENSG00000 |
| 139647299 | 139683882 | 1 | protein_coding | ENSG00000 |
| 139648999 | 139649728 | -1 | lncRNA | ENSG00000 |
| 139684645 | 139745010 | 1 | lncRNA | ENSG00000 |
| 139717985 | 139718090 | -1 | snRNA | ENSG00000 |
| 139740951 | 139747406 | -1 | lncRNA | ENSG00000 |
| 139772528 | 139775406 | -1 | lncRNA | ENSG00000 |
| 139775305 | 139775472 | 1 | lncRNA | ENSG00000 |
| 139795808 | 139844466 | 1 | protein_coding | ENSG00000 |
| 139846779 | 140043299 | -1 | protein_coding | ENSG00000 |
| 139848290 | 139856389 | 1 | lncRNA | ENSG00000 |
| 140019492 | 140019592 | 1 | misc_RNA | ENSG00000 |
| 140071312 | 140109274 | -1 | lncRNA | ENSG00000 |
| 140107777 | 140125619 | 1 | protein_coding | ENSG00000 |
| 140125937 | 140129392 | 1 | protein_coding | ENSG00000 |
| 140157319 | 140173051 | 1 | lncRNA | ENSG00000 |
| 140175156 | 140282052 | 1 | protein_coding | ENSG00000 |
| 140200163 | 140203187 | 1 | lncRNA | ENSG00000 |
| 140245035 | 140303113 | -1 | protein_coding | ENSG00000 |
| 140332843 | 140346603 | -1 | protein_coding | ENSG00000 |
| 140348484 | 140357794 | -1 | lncRNA | ENSG00000 |
| 140360194 | 140375141 | 1 | protein_coding | ENSG00000 |
| 140370891 | 140401460 | -1 | lncRNA | ENSG00000 |
| 140397323 | 140397444 | -1 | snRNA | ENSG00000 |
| 140401814 | 140539856 | 1 | protein_coding | ENSG00000 |
| 140401908 | 140549569 | 1 | protein_coding | ENSG00000 |
| 140527352 | 140527428 | -1 | snoRNA | ENSG00000 |
| 140537340 | 140558252 | -1 | protein_coding | ENSG00000 |
| 140547662 | 140549576 | 1 | protein_coding | ENSG00000 |
| 140558268 | 140564781 | -1 | protein_coding | ENSG00000 |
| 140562443 | 140592123 | -1 | lncRNA | ENSG00000 |
| 140563671 | 140563751 | -1 | miRNA | ENSG00000 |
| 140564828 | 140569100 | 1 | protein_coding | ENSG00000 |
| 140569962 | 140594570 | -1 | transcribed_unitary_ | ENSG00000 |
| 140579667 | 140579792 | 1 | snoRNA | ENSG00000 |
| 140581504 | 140582335 | 1 | processed_pseudoge | ENSG00000 |
| 140609085 | 140609400 | 1 | processed_pseudoge | ENSG00000 |
| 140631728 | 140633701 | -1 | protein_coding | ENSG00000 |

| | | | | |
|---|---|---|---|---|
| 140638740 | 140647785 | -1 | protein_coding | ENSG00000 |
| 140639427 | 140645408 | 1 | protein_coding | ENSG00000 |
| 140647058 | 140662480 | 1 | protein_coding | ENSG00000 |
| 140647844 | 140647926 | 1 | miRNA | ENSG00000 |
| 140664676 | 140674124 | 1 | protein_coding | ENSG00000 |
| 140670794 | 140673576 | -1 | protein_coding | ENSG00000 |
| 140673904 | 140691537 | -1 | protein_coding | ENSG00000 |
| 140691430 | 140699305 | 1 | protein_coding | ENSG00000 |
| 140698680 | 140706686 | 1 | protein_coding | ENSG00000 |
| 140711275 | 140711373 | 1 | misc_RNA | ENSG00000 |
| 140718925 | 140719013 | 1 | misc_RNA | ENSG00000 |
| 140726158 | 140726246 | 1 | misc_RNA | ENSG00000 |
| 140732290 | 140732863 | -1 | processed_pseudoge | ENSG00000 |
| 140764110 | 140764821 | 1 | processed_pseudoge | ENSG00000 |
| 140785698 | 140828549 | -1 | lncRNA | ENSG00000 |
| 140786136 | 141012347 | 1 | protein_coding | ENSG00000 |
| 140794852 | 141012347 | 1 | protein_coding | ENSG00000 |
| 140801028 | 141012347 | 1 | protein_coding | ENSG00000 |
| 140806929 | 141012347 | 1 | protein_coding | ENSG00000 |
| 140821604 | 141012347 | 1 | protein_coding | ENSG00000 |
| 140827958 | 141012347 | 1 | protein_coding | ENSG00000 |
| 140834248 | 141012347 | 1 | protein_coding | ENSG00000 |
| 140841187 | 141012347 | 1 | protein_coding | ENSG00000 |
| 140847463 | 141012344 | 1 | protein_coding | ENSG00000 |
| 140849105 | 140849696 | -1 | lncRNA | ENSG00000 |
| 140855883 | 141012347 | 1 | protein_coding | ENSG00000 |
| 140861224 | 140863521 | 1 | transcribed_unproce | ENSG00000 |
| 140867513 | 140867959 | -1 | lncRNA | ENSG00000 |
| 140868183 | 141012344 | 1 | protein_coding | ENSG00000 |
| 140875302 | 141012347 | 1 | protein_coding | ENSG00000 |
| 140875346 | 140875922 | -1 | lncRNA | ENSG00000 |
| 140878073 | 140882642 | -1 | lncRNA | ENSG00000 |
| 140882124 | 141012347 | 1 | protein_coding | ENSG00000 |
| 140926369 | 141012344 | 1 | protein_coding | ENSG00000 |
| 140966212 | 140966490 | -1 | lncRNA | ENSG00000 |
| 140966470 | 141012347 | 1 | protein_coding | ENSG00000 |
| 141046260 | 141096402 | -1 | lncRNA | ENSG00000 |
| 141051135 | 141059344 | 1 | protein_coding | ENSG00000 |
| 141078076 | 141084481 | 1 | lncRNA | ENSG00000 |
| 141094606 | 141098703 | 1 | protein_coding | ENSG00000 |
| 141100242 | 141174391 | -1 | lncRNA | ENSG00000 |
| 141100473 | 141103827 | 1 | protein_coding | ENSG00000 |
| 141118680 | 141120765 | 1 | lncRNA | ENSG00000 |
| 141121818 | 141125623 | 1 | protein_coding | ENSG00000 |
| 141135206 | 141138615 | 1 | protein_coding | ENSG00000 |
| 141136683 | 141245380 | -1 | lncRNA | ENSG00000 |
| 141150022 | 141153287 | 1 | protein_coding | ENSG00000 |
| 141155996 | 141159061 | 1 | transcribed_unproce | ENSG00000 |
| 141168231 | 141168893 | 1 | lncRNA | ENSG00000 |
| 141172644 | 141176383 | 1 | protein_coding | ENSG00000 |

| | | | | |
|---|---|---|---|---|
| 141177790 | 141180539 | 1 | protein_coding | ENSG00000 |
| 141177849 | 141182369 | 1 | lncRNA | ENSG00000 |
| 141181399 | 141186399 | 1 | protein_coding | ENSG00000 |
| 141182409 | 141188582 | 1 | lncRNA | ENSG00000 |
| 141183401 | 141201396 | -1 | lncRNA | ENSG00000 |
| 141187127 | 141191541 | 1 | protein_coding | ENSG00000 |
| 141191599 | 141194088 | -1 | lncRNA | ENSG00000 |
| 141192353 | 141195647 | 1 | protein_coding | ENSG00000 |
| 141199610 | 141203779 | 1 | protein_coding | ENSG00000 |
| 141208697 | 141212571 | 1 | protein_coding | ENSG00000 |
| 141213919 | 141218979 | 1 | protein_coding | ENSG00000 |
| 141222932 | 141227759 | 1 | protein_coding | ENSG00000 |
| 141223372 | 141245861 | 1 | protein_coding | ENSG00000 |
| 141234333 | 141237529 | 1 | transcribed_unproce | ENSG00000 |
| 141239923 | 141243034 | 1 | transcribed_unproce | ENSG00000 |
| 141245395 | 141249365 | 1 | protein_coding | ENSG00000 |
| 141259235 | 141261364 | 1 | processed_pseudoge | ENSG00000 |
| 141260225 | 141320784 | -1 | protein_coding | ENSG00000 |
| 141302635 | 141304049 | -1 | protein_coding | ENSG00000 |
| 141320980 | 141325985 | 1 | lncRNA | ENSG00000 |
| 141326210 | 141329357 | 1 | lncRNA | ENSG00000 |
| 141330571 | 141512981 | 1 | protein_coding | ENSG00000 |
| 141338760 | 141512975 | 1 | protein_coding | ENSG00000 |
| 141343829 | 141512979 | 1 | protein_coding | ENSG00000 |
| 141350102 | 141512979 | 1 | protein_coding | ENSG00000 |
| 141350109 | 141350662 | -1 | lncRNA | ENSG00000 |
| 141355021 | 141512975 | 1 | protein_coding | ENSG00000 |
| 141360042 | 141512979 | 1 | protein_coding | ENSG00000 |
| 141364232 | 141512979 | 1 | protein_coding | ENSG00000 |
| 141370242 | 141512975 | 1 | protein_coding | ENSG00000 |
| 141373914 | 141512979 | 1 | protein_coding | ENSG00000 |
| 141382739 | 141512975 | 1 | protein_coding | ENSG00000 |
| 141387698 | 141512979 | 1 | protein_coding | ENSG00000 |
| 141390157 | 141512979 | 1 | protein_coding | ENSG00000 |
| 141397987 | 141512979 | 1 | protein_coding | ENSG00000 |
| 141402932 | 141512979 | 1 | protein_coding | ENSG00000 |
| 141408021 | 141512979 | 1 | protein_coding | ENSG00000 |
| 141412987 | 141512979 | 1 | protein_coding | ENSG00000 |
| 141417645 | 141512979 | 1 | protein_coding | ENSG00000 |
| 141421047 | 141512975 | 1 | protein_coding | ENSG00000 |
| 141426286 | 141429158 | 1 | transcribed_unitary_ | ENSG00000 |
| 141427295 | 141427752 | -1 | lncRNA | ENSG00000 |
| 141430507 | 141512975 | 1 | protein_coding | ENSG00000 |
| 141439853 | 141442449 | 1 | unprocessed_pseudo | ENSG00000 |
| 141468465 | 141471528 | 1 | lncRNA | ENSG00000 |
| 141475947 | 141512977 | 1 | protein_coding | ENSG00000 |
| 141479316 | 141479625 | 1 | misc_RNA | ENSG00000 |
| 141484997 | 141512979 | 1 | protein_coding | ENSG00000 |
| 141489121 | 141512979 | 1 | protein_coding | ENSG00000 |
| 141515016 | 141619055 | -1 | protein_coding | ENSG00000 |

| | | | | |
|---|---|---|---|---|
| 141558311 | 141565263 | 1 | lncRNA | ENSG0000C |
| 141588935 | 141589142 | 1 | processed_pseudoge | ENSG0000C |
| 141618414 | 141626481 | 1 | lncRNA | ENSG0000C |
| 141620876 | 141636849 | -1 | protein_coding | ENSG0000C |
| 141636950 | 141641064 | 1 | protein_coding | ENSG0000C |
| 141639302 | 141651418 | -1 | protein_coding | ENSG0000C |
| 141653401 | 141682230 | -1 | protein_coding | ENSG0000C |
| 141718711 | 141775187 | -1 | lncRNA | ENSG0000C |
| 141822076 | 141825488 | -1 | lncRNA | ENSG0000C |
| 141825708 | 141826325 | -1 | lncRNA | ENSG0000C |
| 141853111 | 141879246 | -1 | protein_coding | ENSG0000C |
| 141882815 | 141907939 | 1 | lncRNA | ENSG0000C |
| 141896168 | 141896689 | -1 | processed_pseudoge | ENSG0000C |
| 141923855 | 141942047 | 1 | protein_coding | ENSG0000C |
| 141943585 | 141969741 | -1 | protein_coding | ENSG0000C |
| 141952419 | 141953375 | 1 | lncRNA | ENSG0000C |
| 141958328 | 141990292 | 1 | protein_coding | ENSG0000C |
| 141970637 | 141982687 | -1 | lncRNA | ENSG0000C |
| 141991749 | 142013041 | -1 | protein_coding | ENSG0000C |
| 142036707 | 142037125 | -1 | processed_pseudoge | ENSG0000C |
| 142108779 | 142154440 | 1 | protein_coding | ENSG0000C |
| 142165767 | 142168387 | 1 | TEC | ENSG0000C |
| 142310427 | 142326455 | -1 | protein_coding | ENSG0000C |
| 142317830 | 142317920 | 1 | misc_RNA | ENSG0000C |
| 142318047 | 142318167 | 1 | misc_RNA | ENSG0000C |
| 142325293 | 142672001 | 1 | lncRNA | ENSG0000C |
| 142566700 | 142567076 | 1 | processed_pseudoge | ENSG0000C |
| 142592178 | 142698070 | -1 | protein_coding | ENSG0000C |
| 142703782 | 142705421 | 1 | lncRNA | ENSG0000C |
| 142716229 | 142761035 | 1 | lncRNA | ENSG0000C |
| 142732704 | 142737461 | -1 | lncRNA | ENSG0000C |
| 142770377 | 143229011 | 1 | protein_coding | ENSG0000C |
| 142859604 | 142868922 | -1 | lncRNA | ENSG0000C |
| 142946569 | 142947105 | -1 | TEC | ENSG0000C |
| 143192500 | 143194166 | 1 | lncRNA | ENSG0000C |
| 143277931 | 143435512 | -1 | protein_coding | ENSG0000C |
| 143332605 | 143333239 | 1 | processed_pseudoge | ENSG0000C |
| 143375628 | 143375883 | -1 | TEC | ENSG0000C |
| 143406959 | 143407420 | 1 | TEC | ENSG0000C |
| 143489855 | 143531350 | -1 | lncRNA | ENSG0000C |
| 143531331 | 143598434 | 1 | lncRNA | ENSG0000C |
| 143542514 | 143542587 | -1 | snRNA | ENSG0000C |
| 143605628 | 143828772 | 1 | lncRNA | ENSG0000C |
| 143679860 | 143679971 | 1 | miRNA | ENSG0000C |
| 143752090 | 143752704 | -1 | processed_pseudoge | ENSG0000C |
| 143812161 | 143820719 | 1 | protein_coding | ENSG0000C |
| 144140879 | 144141166 | 1 | misc_RNA | ENSG0000C |
| 144158162 | 144170714 | -1 | protein_coding | ENSG0000C |
| 144170873 | 144485686 | 1 | protein_coding | ENSG0000C |
| 144369334 | 144385310 | 1 | lncRNA | ENSG0000C |

| | | | | |
|---|---|---|---|---|
| 144439204 | 144468471 | -1 | lncRNA | ENSG00000 |
| 144439206 | 144439478 | 1 | processed_pseudoge | ENSG00000 |
| 144522140 | 144522988 | 1 | processed_pseudoge | ENSG00000 |
| 144535354 | 144535670 | -1 | misc_RNA | ENSG00000 |
| 144856890 | 145228982 | -1 | lncRNA | ENSG00000 |
| 145001853 | 145005789 | -1 | processed_pseudoge | ENSG00000 |
| 145228811 | 145230054 | 1 | processed_pseudoge | ENSG00000 |
| 145337932 | 145381670 | -1 | lncRNA | ENSG00000 |
| 145429811 | 145451108 | 1 | lncRNA | ENSG00000 |
| 145471799 | 145835369 | -1 | protein_coding | ENSG00000 |
| 145728360 | 145729349 | -1 | processed_pseudoge | ENSG00000 |
| 145858521 | 145937126 | -1 | protein_coding | ENSG00000 |
| 145936578 | 146081791 | 1 | protein_coding | ENSG00000 |
| 145997218 | 145998003 | 1 | processed_pseudoge | ENSG00000 |
| 146084313 | 146105577 | -1 | protein_coding | ENSG00000 |
| 146099406 | 146120412 | 1 | lncRNA | ENSG00000 |
| 146113034 | 146182650 | -1 | protein_coding | ENSG00000 |
| 146137755 | 146138381 | 1 | processed_pseudoge | ENSG00000 |
| 146138771 | 146139008 | 1 | processed_pseudoge | ENSG00000 |
| 146203550 | 146339251 | 1 | protein_coding | ENSG00000 |
| 146203605 | 146289223 | 1 | protein_coding | ENSG00000 |
| 146245191 | 146245253 | -1 | snRNA | ENSG00000 |
| 146307098 | 146376963 | -1 | lncRNA | ENSG00000 |
| 146338839 | 146341728 | 1 | protein_coding | ENSG00000 |
| 146375591 | 146376219 | -1 | lncRNA | ENSG00000 |
| 146400981 | 146407359 | 1 | lncRNA | ENSG00000 |
| 146447311 | 146511961 | 1 | protein_coding | ENSG00000 |
| 146510116 | 146510271 | 1 | misc_RNA | ENSG00000 |
| 146513103 | 146516190 | -1 | protein_coding | ENSG00000 |
| 146563226 | 146617004 | 1 | lncRNA | ENSG00000 |
| 146581146 | 147084784 | -1 | protein_coding | ENSG00000 |
| 146586099 | 146586200 | -1 | rRNA_pseudogene | ENSG00000 |
| 146656401 | 146656695 | -1 | misc_RNA | ENSG00000 |
| 146706381 | 146707600 | -1 | processed_pseudoge | ENSG00000 |
| 146914207 | 146919506 | -1 | lncRNA | ENSG00000 |
| 147093259 | 147097240 | 1 | lncRNA | ENSG00000 |
| 147180204 | 147234859 | -1 | lncRNA | ENSG00000 |
| 147234963 | 147387855 | 1 | protein_coding | ENSG00000 |
| 147390808 | 147510068 | -1 | protein_coding | ENSG00000 |
| 147401760 | 147401996 | 1 | lncRNA | ENSG00000 |
| 147559994 | 147662009 | 1 | lncRNA | ENSG00000 |
| 147585438 | 147782775 | -1 | protein_coding | ENSG00000 |
| 147725551 | 147728464 | 1 | lncRNA | ENSG00000 |
| 147824568 | 147831786 | -1 | protein_coding | ENSG00000 |
| 147851644 | 147852659 | -1 | processed_pseudoge | ENSG00000 |
| 147870682 | 147882191 | 1 | protein_coding | ENSG00000 |
| 147880726 | 147906538 | -1 | protein_coding | ENSG00000 |
| 147886086 | 147886878 | -1 | lncRNA | ENSG00000 |
| 147887112 | 147887704 | 1 | lncRNA | ENSG00000 |
| 147922179 | 147923421 | 1 | processed_pseudoge | ENSG00000 |

| | | | | |
|---|---|---|---|---|
| 148025683 | 148137289 | 1 | protein_coding | ENSG00000 |
| 148088125 | 148383907 | -1 | lncRNA | ENSG00000 |
| 148169733 | 148175398 | 1 | protein_coding | ENSG00000 |
| 148202794 | 148215137 | 1 | protein_coding | ENSG00000 |
| 148221360 | 148221642 | -1 | processed_pseudoge | ENSG00000 |
| 148221650 | 148243580 | 1 | protein_coding | ENSG00000 |
| 148268180 | 148286255 | 1 | protein_coding | ENSG00000 |
| 148307298 | 148308258 | 1 | processed_pseudoge | ENSG00000 |
| 148312419 | 148315922 | 1 | protein_coding | ENSG00000 |
| 148321203 | 148339852 | 1 | protein_coding | ENSG00000 |
| 148362777 | 148372152 | 1 | unprocessed_pseudo | ENSG00000 |
| 148383935 | 148442836 | 1 | protein_coding | ENSG00000 |
| 148389748 | 148390029 | 1 | misc_RNA | ENSG00000 |
| 148430159 | 148430807 | -1 | lncRNA | ENSG00000 |
| 148451032 | 148677235 | -1 | protein_coding | ENSG00000 |
| 148644687 | 148646390 | -1 | lncRNA | ENSG00000 |
| 148825245 | 148828687 | 1 | protein_coding | ENSG00000 |
| 148923639 | 149063163 | -1 | protein_coding | ENSG00000 |
| 148970340 | 148970653 | 1 | lncRNA | ENSG00000 |
| 149057554 | 149057656 | -1 | snRNA | ENSG00000 |
| 149062313 | 149062409 | -1 | miRNA | ENSG00000 |
| 149063254 | 149129578 | 1 | lncRNA | ENSG00000 |
| 149116178 | 149116480 | -1 | misc_RNA | ENSG00000 |
| 149141483 | 149260542 | 1 | protein_coding | ENSG00000 |
| 149163955 | 149276776 | -1 | lncRNA | ENSG00000 |
| 149216523 | 149276805 | -1 | lncRNA | ENSG00000 |
| 149271859 | 149343637 | 1 | protein_coding | ENSG00000 |
| 149324220 | 149345333 | -1 | lncRNA | ENSG00000 |
| 149345430 | 149354583 | 1 | protein_coding | ENSG00000 |
| 149348116 | 149357642 | -1 | lncRNA | ENSG00000 |
| 149358037 | 149369653 | 1 | protein_coding | ENSG00000 |
| 149371324 | 149404202 | -1 | protein_coding | ENSG00000 |
| 149372174 | 149375116 | 1 | lncRNA | ENSG00000 |
| 149406689 | 149432835 | 1 | lncRNA | ENSG00000 |
| 149425771 | 149428289 | -1 | lncRNA | ENSG00000 |
| 149428918 | 149429023 | 1 | miRNA | ENSG00000 |
| 149430646 | 149430733 | 1 | miRNA | ENSG00000 |
| 149492982 | 149551471 | -1 | protein_coding | ENSG00000 |
| 149494314 | 149504670 | -1 | lncRNA | ENSG00000 |
| 149545383 | 149545810 | -1 | processed_pseudoge | ENSG00000 |
| 149551947 | 149634968 | 1 | protein_coding | ENSG00000 |
| 149606637 | 149606741 | -1 | snRNA | ENSG00000 |
| 149694237 | 149696308 | 1 | lncRNA | ENSG00000 |
| 149695748 | 149695959 | -1 | snoRNA | ENSG00000 |
| 149722070 | 149722372 | 1 | misc_RNA | ENSG00000 |
| 149730298 | 149855022 | 1 | protein_coding | ENSG00000 |
| 149732825 | 149732890 | 1 | miRNA | ENSG00000 |
| 149857953 | 149944793 | -1 | protein_coding | ENSG00000 |
| 149932014 | 149932668 | -1 | processed_pseudoge | ENSG00000 |
| 149940030 | 149940131 | -1 | misc_RNA | ENSG00000 |

| | | | | |
|---|---|---|---|---|
| 149960737 | 149993455 | 1 | protein_coding | ENSG00000 |
| 149993118 | 150000654 | -1 | protein_coding | ENSG00000 |
| 150000046 | 150053142 | 1 | protein_coding | ENSG00000 |
| 150014785 | 150015154 | -1 | processed_pseudoge | ENSG00000 |
| 150021567 | 150021921 | -1 | processed_pseudoge | ENSG00000 |
| 150053291 | 150113372 | -1 | protein_coding | ENSG00000 |
| 150094302 | 150095048 | 1 | processed_pseudoge | ENSG00000 |
| 150098406 | 150098506 | 1 | misc_RNA | ENSG00000 |
| 150113839 | 150155872 | -1 | protein_coding | ENSG00000 |
| 150166778 | 150184558 | 1 | protein_coding | ENSG00000 |
| 150190062 | 150222788 | 1 | protein_coding | ENSG00000 |
| 150219491 | 150290291 | -1 | protein_coding | ENSG00000 |
| 150296343 | 150339307 | -1 | protein_coding | ENSG00000 |
| 150357629 | 150400308 | 1 | protein_coding | ENSG00000 |
| 150400041 | 150412929 | -1 | protein_coding | ENSG00000 |
| 150427904 | 150429906 | 1 | lncRNA | ENSG00000 |
| 150442635 | 150449739 | -1 | protein_coding | ENSG00000 |
| 150475531 | 150485968 | -1 | lncRNA | ENSG00000 |
| 150485818 | 150558211 | 1 | protein_coding | ENSG00000 |
| 150601080 | 150659207 | 1 | protein_coding | ENSG00000 |
| 150608428 | 150615354 | -1 | lncRNA | ENSG00000 |
| 150621007 | 150626779 | -1 | lncRNA | ENSG00000 |
| 150660874 | 150679365 | 1 | protein_coding | ENSG00000 |
| 150670658 | 150672390 | -1 | lncRNA | ENSG00000 |
| 150690792 | 150701077 | -1 | protein_coding | ENSG00000 |
| 150708440 | 150759109 | -1 | protein_coding | ENSG00000 |
| 150777946 | 150796734 | 1 | protein_coding | ENSG00000 |
| 150846523 | 150900736 | 1 | protein_coding | ENSG00000 |
| 150894392 | 150904983 | -1 | protein_coding | ENSG00000 |
| 150930645 | 150946289 | -1 | transcribed_unproce | ENSG00000 |
| 150950109 | 150952460 | -1 | unprocessed_pseudo | ENSG00000 |
| 150978419 | 150979175 | 1 | unprocessed_pseudo | ENSG00000 |
| 151020438 | 151028992 | 1 | protein_coding | ENSG00000 |
| 151029945 | 151093577 | -1 | protein_coding | ENSG00000 |
| 151100706 | 151157785 | -1 | protein_coding | ENSG00000 |
| 151158106 | 151158462 | -1 | lncRNA | ENSG00000 |
| 151173981 | 151180617 | 1 | unitary_pseudogene | ENSG00000 |
| 151181052 | 151224093 | -1 | protein_coding | ENSG00000 |
| 151212150 | 151270440 | 1 | protein_coding | ENSG00000 |
| 151276762 | 151303766 | -1 | protein_coding | ENSG00000 |
| 151314972 | 151347590 | -1 | protein_coding | ENSG00000 |
| 151319422 | 151319776 | 1 | processed_pseudoge | ENSG00000 |
| 151366299 | 151436953 | -1 | transcribed_unproce | ENSG00000 |
| 151378003 | 151378640 | -1 | processed_pseudoge | ENSG00000 |
| 151380341 | 151382381 | 1 | processed_pseudoge | ENSG00000 |
| 151437046 | 151492379 | 1 | protein_coding | ENSG00000 |
| 151477459 | 151477577 | -1 | rRNA_pseudogene | ENSG00000 |
| 151504093 | 151568944 | -1 | protein_coding | ENSG00000 |
| 151509453 | 151512769 | 1 | lncRNA | ENSG00000 |
| 151522087 | 151522148 | -1 | miRNA | ENSG00000 |

| | | | | |
|---|---|---|---|---|
| 151595264 | 151602709 | 1 | lncRNA | ENSG00000 |
| 151652275 | 151655449 | 1 | lncRNA | ENSG00000 |
| 151661096 | 151686975 | -1 | protein_coding | ENSG00000 |
| 151676945 | 151725473 | 1 | lncRNA | ENSG00000 |
| 151704289 | 151704578 | -1 | misc_RNA | ENSG00000 |
| 151724831 | 151725356 | -1 | lncRNA | ENSG00000 |
| 151742316 | 151772532 | -1 | protein_coding | ENSG00000 |
| 151753992 | 151767247 | 1 | lncRNA | ENSG00000 |
| 151765859 | 151766378 | -1 | transcribed_processe | ENSG00000 |
| 151770242 | 151771508 | -1 | lncRNA | ENSG00000 |
| 151771045 | 151812785 | 1 | protein_coding | ENSG00000 |
| 151822513 | 151924842 | -1 | protein_coding | ENSG00000 |
| 151848886 | 151850791 | 1 | processed_pseudoge | ENSG00000 |
| 151875566 | 151875692 | -1 | rRNA_pseudogene | ENSG00000 |
| 151949571 | 152270449 | 1 | lncRNA | ENSG00000 |
| 152374998 | 152726160 | 1 | lncRNA | ENSG00000 |
| 152391532 | 152433368 | -1 | protein_coding | ENSG00000 |
| 152495718 | 152496017 | -1 | processed_pseudoge | ENSG00000 |
| 152618965 | 153223543 | -1 | lncRNA | ENSG00000 |
| 153489615 | 153813869 | 1 | protein_coding | ENSG00000 |
| 153755382 | 153755680 | 1 | misc_RNA | ENSG00000 |
| 153887428 | 153898987 | -1 | lncRNA | ENSG00000 |
| 153901459 | 153903223 | 1 | lncRNA | ENSG00000 |
| 153990148 | 154038936 | -1 | protein_coding | ENSG00000 |
| 154038906 | 154220478 | 1 | protein_coding | ENSG00000 |
| 154190730 | 154420984 | 1 | protein_coding | ENSG00000 |
| 154325568 | 154445850 | -1 | lncRNA | ENSG00000 |
| 154337471 | 154338452 | -1 | processed_pseudoge | ENSG00000 |
| 154346042 | 154347834 | 1 | lncRNA | ENSG00000 |
| 154347106 | 154347247 | 1 | miRNA | ENSG00000 |
| 154349428 | 154349725 | 1 | misc_RNA | ENSG00000 |
| 154445997 | 154461053 | 1 | protein_coding | ENSG00000 |
| 154474972 | 154478227 | -1 | protein_coding | ENSG00000 |
| 154483917 | 154486150 | -1 | lncRNA | ENSG00000 |
| 154493576 | 154494743 | 1 | processed_pseudoge | ENSG00000 |
| 154559308 | 154560440 | -1 | processed_pseudoge | ENSG00000 |
| 154596012 | 154596072 | -1 | miRNA | ENSG00000 |
| 154685776 | 154685861 | 1 | miRNA | ENSG00000 |
| 154705626 | 154705727 | 1 | misc_RNA | ENSG00000 |
| 154712843 | 154817605 | 1 | protein_coding | ENSG00000 |
| 154718383 | 154718689 | -1 | misc_RNA | ENSG00000 |
| 154729231 | 154729694 | 1 | processed_pseudoge | ENSG00000 |
| 154818492 | 154859252 | -1 | protein_coding | ENSG00000 |
| 154821943 | 154822258 | 1 | misc_RNA | ENSG00000 |
| 154829458 | 154829540 | 1 | miRNA | ENSG00000 |
| 154857553 | 154876793 | 1 | protein_coding | ENSG00000 |
| 154887411 | 154938211 | -1 | protein_coding | ENSG00000 |
| 154941073 | 154969411 | 1 | protein_coding | ENSG00000 |
| 154993598 | 154994445 | 1 | processed_pseudoge | ENSG00000 |
| 155013755 | 155018141 | 1 | protein_coding | ENSG00000 |

| | | | | |
|---|---|---|---|---|
| 155087430 | 155199252 | 1 | lncRNA | ENSG00000 |
| 155111184 | 155112324 | -1 | processed_pseudoge | ENSG00000 |
| 155397291 | 155398353 | 1 | processed_pseudoge | ENSG00000 |
| 155491336 | 155493598 | 1 | processed_pseudoge | ENSG00000 |
| 155845437 | 155845528 | 1 | rRNA_pseudogene | ENSG00000 |
| 155870344 | 156767788 | 1 | protein_coding | ENSG00000 |
| 156019228 | 156039977 | -1 | lncRNA | ENSG00000 |
| 156458398 | 156458507 | -1 | unprocessed_pseudo | ENSG00000 |
| 156704058 | 156739812 | -1 | lncRNA | ENSG00000 |
| 156814130 | 156814234 | 1 | snRNA | ENSG00000 |
| 156850295 | 156852528 | 1 | protein_coding | ENSG00000 |
| 156919292 | 156963226 | -1 | protein_coding | ENSG00000 |
| 156924119 | 156924710 | 1 | processed_pseudoge | ENSG00000 |
| 157029413 | 157059119 | -1 | protein_coding | ENSG00000 |
| 157085832 | 157142869 | -1 | protein_coding | ENSG00000 |
| 157104788 | 157114900 | 1 | lncRNA | ENSG00000 |
| 157137424 | 157159019 | -1 | protein_coding | ENSG00000 |
| 157142933 | 157255191 | 1 | protein_coding | ENSG00000 |
| 157161846 | 157166264 | -1 | protein_coding | ENSG00000 |
| 157199242 | 157224380 | -1 | lncRNA | ENSG00000 |
| 157260122 | 157266625 | -1 | lncRNA | ENSG00000 |
| 157266079 | 157395595 | 1 | protein_coding | ENSG00000 |
| 157341596 | 157460100 | -1 | protein_coding | ENSG00000 |
| 157341598 | 157345677 | -1 | protein_coding | ENSG00000 |
| 157363382 | 157365501 | -1 | lncRNA | ENSG00000 |
| 157375741 | 157384950 | -1 | lncRNA | ENSG00000 |
| 157395534 | 157575775 | -1 | protein_coding | ENSG00000 |
| 157422439 | 157422873 | 1 | processed_pseudoge | ENSG00000 |
| 157460019 | 157474717 | 1 | protein_coding | ENSG00000 |
| 157565964 | 157569098 | 1 | lncRNA | ENSG00000 |
| 157579497 | 157579599 | 1 | snRNA | ENSG00000 |
| 157604707 | 157605637 | -1 | processed_pseudoge | ENSG00000 |
| 157625679 | 157671480 | -1 | protein_coding | ENSG00000 |
| 157671533 | 157680158 | 1 | protein_coding | ENSG00000 |
| 157682985 | 157690878 | -1 | transcribed_unproce | ENSG00000 |
| 157731420 | 157741449 | 1 | protein_coding | ENSG00000 |
| 157743712 | 157760709 | 1 | protein_coding | ENSG00000 |
| 157747504 | 157747707 | -1 | unprocessed_pseudo | ENSG00000 |
| 157785743 | 157859175 | -1 | protein_coding | ENSG00000 |
| 157809098 | 157809204 | 1 | snRNA | ENSG00000 |
| 157931258 | 157931563 | -1 | processed_pseudoge | ENSG00000 |
| 157976766 | 157976956 | -1 | snRNA | ENSG00000 |
| 157984658 | 157989990 | -1 | unprocessed_pseudo | ENSG00000 |
| 157986146 | 157987706 | -1 | processed_pseudoge | ENSG00000 |
| 158078099 | 158078764 | -1 | processed_pseudoge | ENSG00000 |
| 158173601 | 158176422 | -1 | lncRNA | ENSG00000 |
| 158175396 | 158200016 | 1 | lncRNA | ENSG00000 |
| 158188977 | 158189911 | -1 | processed_pseudoge | ENSG00000 |
| 158225352 | 158234376 | -1 | lncRNA | ENSG00000 |
| 158256137 | 158258436 | 1 | lncRNA | ENSG00000 |

| | | | | |
|---|---|---|---|---|
| 158275711 | 158278813 | 1 | lncRNA | ENSG0000C |
| 158320683 | 158409773 | -1 | lncRNA | ENSG0000C |
| 158424585 | 158452758 | 1 | lncRNA | ENSG0000C |
| 158464465 | 158464678 | 1 | lncRNA | ENSG0000C |
| 158485190 | 158588546 | 1 | lncRNA | ENSG0000C |
| 158695916 | 159099916 | -1 | protein_coding | ENSG0000C |
| 158983006 | 158984789 | 1 | lncRNA | ENSG0000C |
| 158985806 | 158987199 | 1 | lncRNA | ENSG0000C |
| 159100483 | 159117478 | 1 | lncRNA | ENSG0000C |
| 159106380 | 159108617 | 1 | TEC | ENSG0000C |
| 159157409 | 159210053 | -1 | protein_coding | ENSG0000C |
| 159209921 | 159248796 | 1 | lncRNA | ENSG0000C |
| 159227715 | 159245127 | 1 | lncRNA | ENSG0000C |
| 159230194 | 159230322 | 1 | snoRNA | ENSG0000C |
| 159263290 | 159286036 | 1 | protein_coding | ENSG0000C |
| 159310745 | 159805536 | 1 | lncRNA | ENSG0000C |
| 159314783 | 159330887 | -1 | protein_coding | ENSG0000C |
| 159318042 | 159318167 | 1 | snRNA | ENSG0000C |
| 159448556 | 159466291 | -1 | lncRNA | ENSG0000C |
| 159484130 | 159511687 | -1 | lncRNA | ENSG0000C |
| 159698586 | 159937766 | -1 | lncRNA | ENSG0000C |
| 159865080 | 159973012 | 1 | protein_coding | ENSG0000C |
| 159950493 | 159951498 | -1 | processed_pseudoge | ENSG0000C |
| 160008445 | 160008546 | 1 | misc_RNA | ENSG0000C |
| 160009113 | 160065543 | 1 | protein_coding | ENSG0000C |
| 160023740 | 160024045 | -1 | processed_pseudoge | ENSG0000C |
| 160061801 | 160119450 | -1 | protein_coding | ENSG0000C |
| 160187367 | 160238735 | 1 | protein_coding | ENSG0000C |
| 160195744 | 160204826 | -1 | lncRNA | ENSG0000C |
| 160247517 | 160248217 | 1 | processed_pseudoge | ENSG0000C |
| 160249106 | 160345396 | -1 | protein_coding | ENSG0000C |
| 160258253 | 160258585 | 1 | processed_pseudoge | ENSG0000C |
| 160326693 | 160326969 | 1 | processed_pseudoge | ENSG0000C |
| 160347754 | 160370641 | -1 | protein_coding | ENSG0000C |
| 160393148 | 160400054 | -1 | protein_coding | ENSG0000C |
| 160401641 | 160421711 | -1 | protein_coding | ENSG0000C |
| 160421855 | 160428739 | 1 | protein_coding | ENSG0000C |
| 160438594 | 160487426 | 1 | lncRNA | ENSG0000C |
| 160474402 | 160474483 | 1 | miRNA | ENSG0000C |
| 160485352 | 160485450 | 1 | miRNA | ENSG0000C |
| 160563120 | 160852214 | -1 | protein_coding | ENSG0000C |
| 160613873 | 160639183 | 1 | lncRNA | ENSG0000C |
| 160685351 | 160692889 | 1 | lncRNA | ENSG0000C |
| 160931778 | 160938626 | -1 | lncRNA | ENSG0000C |
| 161288429 | 161549044 | -1 | protein_coding | ENSG0000C |
| 161547063 | 161702593 | 1 | protein_coding | ENSG0000C |
| 161687347 | 161689408 | -1 | lncRNA | ENSG0000C |
| 161751448 | 161751768 | -1 | processed_pseudoge | ENSG0000C |
| 161847063 | 161899981 | 1 | protein_coding | ENSG0000C |
| 161907111 | 162001536 | -1 | lncRNA | ENSG0000C |

| | | | | |
|---|---|---|---|---|
| 162000057 | 162162977 | 1 | protein_coding | ENSG0000C |
| 162424042 | 163437326 | -1 | lncRNA | ENSG0000C |
| 162477891 | 162477993 | 1 | snRNA | ENSG0000C |
| 162761833 | 162762313 | 1 | processed_pseudoge | ENSG0000C |
| 163039291 | 163039599 | 1 | processed_pseudoge | ENSG0000C |
| 163416525 | 163416731 | 1 | unprocessed_pseudc | ENSG0000C |
| 163437569 | 163446151 | 1 | protein_coding | ENSG0000C |
| 163446526 | 163460102 | -1 | protein_coding | ENSG0000C |
| 163460203 | 163491941 | 1 | protein_coding | ENSG0000C |
| 163483065 | 163494058 | -1 | lncRNA | ENSG0000C |
| 163503114 | 163519354 | 1 | protein_coding | ENSG0000C |
| 163597352 | 163612516 | -1 | lncRNA | ENSG0000C |
| 163709210 | 163709514 | -1 | processed_pseudoge | ENSG0000C |
| 163796065 | 163796165 | -1 | snRNA | ENSG0000C |
| 163855140 | 163856642 | -1 | processed_pseudoge | ENSG0000C |
| 163887506 | 163887827 | -1 | processed_pseudoge | ENSG0000C |
| 164119098 | 164234097 | -1 | lncRNA | ENSG0000C |
| 164209631 | 164211892 | 1 | lncRNA | ENSG0000C |
| 164296696 | 165171643 | 1 | lncRNA | ENSG0000C |
| 164448161 | 164467879 | -1 | lncRNA | ENSG0000C |
| 164601002 | 164601452 | -1 | processed_pseudoge | ENSG0000C |
| 164935162 | 164935268 | -1 | snRNA | ENSG0000C |
| 165129307 | 165153014 | -1 | lncRNA | ENSG0000C |
| 165220577 | 165241756 | -1 | lncRNA | ENSG0000C |
| 165349030 | 165778689 | 1 | lncRNA | ENSG0000C |
| 165609441 | 165609679 | 1 | misc_RNA | ENSG0000C |
| 165782805 | 165783134 | -1 | processed_pseudoge | ENSG0000C |
| 165905126 | 165905608 | 1 | processed_pseudoge | ENSG0000C |
| 166028567 | 166029311 | -1 | processed_pseudoge | ENSG0000C |
| 166128498 | 166155439 | 1 | lncRNA | ENSG0000C |
| 166382305 | 166382599 | 1 | processed_pseudoge | ENSG0000C |
| 166905222 | 166926370 | -1 | lncRNA | ENSG0000C |
| 167116318 | 167119585 | -1 | lncRNA | ENSG0000C |
| 167164934 | 167168401 | -1 | lncRNA | ENSG0000C |
| 167284799 | 168264157 | 1 | protein_coding | ENSG0000C |
| 167287320 | 167294273 | -1 | lncRNA | ENSG0000C |
| 167296234 | 167303372 | -1 | lncRNA | ENSG0000C |
| 167306273 | 167309870 | -1 | lncRNA | ENSG0000C |
| 167653228 | 167660481 | -1 | lncRNA | ENSG0000C |
| 167721363 | 167729124 | 1 | lncRNA | ENSG0000C |
| 167937717 | 167953469 | -1 | lncRNA | ENSG0000C |
| 167965187 | 167967086 | -1 | lncRNA | ENSG0000C |
| 168033091 | 168033205 | -1 | snoRNA | ENSG0000C |
| 168085329 | 168187719 | -1 | lncRNA | ENSG0000C |
| 168128856 | 168130422 | 1 | TEC | ENSG0000C |
| 168229583 | 168269415 | -1 | lncRNA | ENSG0000C |
| 168291651 | 168472303 | 1 | protein_coding | ENSG0000C |
| 168486451 | 168519301 | 1 | protein_coding | ENSG0000C |
| 168529305 | 168530634 | 1 | protein_coding | ENSG0000C |
| 168548495 | 168579368 | -1 | protein_coding | ENSG0000C |

| | | | | |
|---|---|---|---|---|
| 168552277 | 168553727 | -1 | processed_pseudoge | ENSG00000 |
| 168560896 | 168560973 | -1 | miRNA | ENSG00000 |
| 168560904 | 168560965 | 1 | miRNA | ENSG00000 |
| 168616352 | 168616996 | 1 | processed_pseudoge | ENSG00000 |
| 168654513 | 168667761 | 1 | lncRNA | ENSG00000 |
| 168661733 | 169301129 | -1 | protein_coding | ENSG00000 |
| 168706567 | 168720884 | 1 | lncRNA | ENSG00000 |
| 168768146 | 168768255 | -1 | miRNA | ENSG00000 |
| 168791498 | 168791610 | 1 | misc_RNA | ENSG00000 |
| 168993000 | 168995677 | 1 | lncRNA | ENSG00000 |
| 169012148 | 169013146 | -1 | TEC | ENSG00000 |
| 169013227 | 169037998 | 1 | lncRNA | ENSG00000 |
| 169263601 | 169263694 | -1 | miRNA | ENSG00000 |
| 169552449 | 169552555 | 1 | snRNA | ENSG00000 |
| 169583636 | 169604778 | 1 | protein_coding | ENSG00000 |
| 169637268 | 170083382 | 1 | protein_coding | ENSG00000 |
| 169772966 | 169779365 | -1 | lncRNA | ENSG00000 |
| 169861303 | 169980495 | -1 | protein_coding | ENSG00000 |
| 170028488 | 170028566 | 1 | miRNA | ENSG00000 |
| 170105897 | 170109734 | 1 | protein_coding | ENSG00000 |
| 170140664 | 170141940 | -1 | processed_pseudoge | ENSG00000 |
| 170191579 | 170199141 | -1 | lncRNA | ENSG00000 |
| 170232447 | 170252575 | 1 | protein_coding | ENSG00000 |
| 170246233 | 170297818 | -1 | protein_coding | ENSG00000 |
| 170307722 | 170335508 | 1 | lncRNA | ENSG00000 |
| 170308701 | 170312716 | -1 | lncRNA | ENSG00000 |
| 170353487 | 170736632 | 1 | protein_coding | ENSG00000 |
| 170374671 | 170389634 | -1 | protein_coding | ENSG00000 |
| 170389493 | 170422844 | 1 | lncRNA | ENSG00000 |
| 170483806 | 170486407 | -1 | lncRNA | ENSG00000 |
| 170639158 | 170681437 | -1 | lncRNA | ENSG00000 |
| 170747047 | 170788650 | -1 | lncRNA | ENSG00000 |
| 170763350 | 170814047 | 1 | protein_coding | ENSG00000 |
| 170861870 | 171300015 | 1 | protein_coding | ENSG00000 |
| 170866835 | 170867144 | 1 | misc_RNA | ENSG00000 |
| 170896929 | 170904461 | 1 | processed_pseudoge | ENSG00000 |
| 171251866 | 171252982 | 1 | processed_pseudoge | ENSG00000 |
| 171305980 | 171309777 | -1 | lncRNA | ENSG00000 |
| 171309248 | 171312139 | 1 | protein_coding | ENSG00000 |
| 171345343 | 171345630 | -1 | processed_pseudoge | ENSG00000 |
| 171359117 | 171359412 | 1 | misc_RNA | ENSG00000 |
| 171365595 | 171367693 | 1 | processed_pseudoge | ENSG00000 |
| 171366118 | 171366252 | 1 | snoRNA | ENSG00000 |
| 171366299 | 171366611 | 1 | processed_pseudoge | ENSG00000 |
| 171386656 | 171386760 | -1 | miRNA | ENSG00000 |
| 171387116 | 171411137 | 1 | protein_coding | ENSG00000 |
| 171419647 | 171457626 | 1 | protein_coding | ENSG00000 |
| 171724575 | 171737519 | -1 | lncRNA | ENSG00000 |
| 171773652 | 171774739 | 1 | lncRNA | ENSG00000 |
| 171782432 | 171796126 | 1 | protein_coding | ENSG00000 |

| | | | | |
|---|---|---|---|---|
| 171861549 | 172006873 | -1 | protein_coding | ENSG00000 |
| 172042079 | 172188224 | -1 | protein_coding | ENSG00000 |
| 172194172 | 172203454 | 1 | protein_coding | ENSG00000 |
| 172209646 | 172283764 | -1 | protein_coding | ENSG00000 |
| 172252334 | 172252478 | -1 | snoRNA | ENSG00000 |
| 172254418 | 172255439 | -1 | processed_pseudoge | ENSG00000 |
| 172325000 | 172454525 | -1 | protein_coding | ENSG00000 |
| 172479274 | 172536444 | 1 | lncRNA | ENSG00000 |
| 172543403 | 172559202 | -1 | lncRNA | ENSG00000 |
| 172609094 | 172609444 | -1 | TEC | ENSG00000 |
| 172641263 | 172691540 | 1 | protein_coding | ENSG00000 |
| 172656522 | 172656713 | -1 | processed_pseudoge | ENSG00000 |
| 172662165 | 172662263 | 1 | miRNA | ENSG00000 |
| 172690454 | 172697720 | -1 | lncRNA | ENSG00000 |
| 172755457 | 172762587 | 1 | lncRNA | ENSG00000 |
| 172762521 | 172763258 | -1 | processed_pseudoge | ENSG00000 |
| 172762980 | 172782334 | 1 | lncRNA | ENSG00000 |
| 172768096 | 172771195 | -1 | protein_coding | ENSG00000 |
| 172779982 | 172780094 | -1 | misc_RNA | ENSG00000 |
| 172785262 | 172785424 | -1 | processed_pseudoge | ENSG00000 |
| 172816592 | 172819958 | 1 | lncRNA | ENSG00000 |
| 172834251 | 172952683 | 1 | protein_coding | ENSG00000 |
| 172954786 | 172959392 | -1 | lncRNA | ENSG00000 |
| 172954907 | 172957162 | 1 | lncRNA | ENSG00000 |
| 172958729 | 172969771 | 1 | protein_coding | ENSG00000 |
| 172975511 | 172976374 | 1 | processed_pseudoge | ENSG00000 |
| 172983771 | 173035445 | 1 | protein_coding | ENSG00000 |
| 172986680 | 172987198 | 1 | processed_pseudoge | ENSG00000 |
| 173020728 | 173020928 | 1 | snoRNA | ENSG00000 |
| 173052452 | 173052555 | -1 | misc_RNA | ENSG00000 |
| 173056352 | 173139284 | 1 | protein_coding | ENSG00000 |
| 173085342 | 173086210 | -1 | processed_pseudoge | ENSG00000 |
| 173144162 | 173145039 | -1 | lncRNA | ENSG00000 |
| 173144442 | 173164387 | 1 | protein_coding | ENSG00000 |
| 173225836 | 173226423 | -1 | processed_pseudoge | ENSG00000 |
| 173232109 | 173235311 | -1 | protein_coding | ENSG00000 |
| 173254190 | 173254290 | -1 | misc_RNA | ENSG00000 |
| 173292517 | 173292619 | -1 | rRNA_pseudogene | ENSG00000 |
| 173314713 | 173329503 | -1 | protein_coding | ENSG00000 |
| 173347455 | 173347536 | 1 | miRNA | ENSG00000 |
| 173362134 | 173362240 | -1 | snRNA | ENSG00000 |
| 173383941 | 173451591 | 1 | lncRNA | ENSG00000 |
| 173463484 | 173484584 | 1 | lncRNA | ENSG00000 |
| 173562478 | 173573199 | 1 | lncRNA | ENSG00000 |
| 173574938 | 173576275 | 1 | lncRNA | ENSG00000 |
| 173578275 | 173585068 | 1 | lncRNA | ENSG00000 |
| 173607145 | 173616659 | -1 | protein_coding | ENSG00000 |
| 173642519 | 173658194 | 1 | lncRNA | ENSG00000 |
| 173689459 | 173705849 | 1 | lncRNA | ENSG00000 |
| 173707614 | 173746279 | -1 | lncRNA | ENSG00000 |

| | | | | |
|---|---|---|---|---|
| 173757562 | 173773164 | 1 | lncRNA | ENSG0000( |
| 173778527 | 173809039 | -1 | lncRNA | ENSG0000( |
| 173888349 | 173961980 | 1 | protein_coding | ENSG0000( |
| 173973779 | 174006140 | 1 | protein_coding | ENSG0000( |
| 174039751 | 174040240 | -1 | processed_pseudoge | ENSG0000( |
| 174045706 | 174243501 | 1 | protein_coding | ENSG0000( |
| 174056060 | 174056585 | -1 | lncRNA | ENSG0000( |
| 174081506 | 174086618 | -1 | lncRNA | ENSG0000( |
| 174173101 | 174181814 | 1 | lncRNA | ENSG0000( |
| 174329612 | 174330503 | 1 | lncRNA | ENSG0000( |
| 174336295 | 174532457 | 1 | lncRNA | ENSG0000( |
| 174513305 | 174514307 | 1 | processed_pseudoge | ENSG0000( |
| 174561776 | 174562063 | 1 | processed_pseudoge | ENSG0000( |
| 174618164 | 174625775 | 1 | lncRNA | ENSG0000( |
| 174632932 | 174633209 | 1 | processed_pseudoge | ENSG0000( |
| 174724582 | 174730896 | 1 | protein_coding | ENSG0000( |
| 174751304 | 174850725 | 1 | lncRNA | ENSG0000( |
| 174751734 | 174751787 | 1 | miRNA | ENSG0000( |
| 174820027 | 174826324 | 1 | lncRNA | ENSG0000( |
| 174919082 | 174995513 | -1 | lncRNA | ENSG0000( |
| 174923090 | 174923945 | 1 | processed_pseudoge | ENSG0000( |
| 174926541 | 174926717 | 1 | unprocessed_pseudc | ENSG0000( |
| 174986295 | 174987050 | 1 | lncRNA | ENSG0000( |
| 175290731 | 175291252 | 1 | processed_pseudoge | ENSG0000( |
| 175440036 | 175444182 | -1 | protein_coding | ENSG0000( |
| 175471414 | 175475378 | -1 | lncRNA | ENSG0000( |
| 175477062 | 175529742 | 1 | protein_coding | ENSG0000( |
| 175483595 | 175484192 | -1 | processed_pseudoge | ENSG0000( |
| 175628188 | 175628462 | 1 | misc_RNA | ENSG0000( |
| 175658030 | 175710756 | 1 | protein_coding | ENSG0000( |
| 175703898 | 175704004 | 1 | snRNA | ENSG0000( |
| 175796310 | 175884021 | 1 | protein_coding | ENSG0000( |
| 175903874 | 175904371 | -1 | TEC | ENSG0000( |
| 175906939 | 175958538 | -1 | lncRNA | ENSG0000( |
| 175917873 | 176034680 | -1 | protein_coding | ENSG0000( |
| 175972606 | 175987666 | 1 | lncRNA | ENSG0000( |
| 176006145 | 176007387 | 1 | unprocessed_pseudc | ENSG0000( |
| 176036593 | 176042888 | 1 | lncRNA | ENSG0000( |
| 176049678 | 176062021 | -1 | lncRNA | ENSG0000( |
| 176060689 | 176116015 | 1 | protein_coding | ENSG0000( |
| 176063748 | 176126959 | 1 | protein_coding | ENSG0000( |
| 176119545 | 176156937 | 1 | transcribed_processe | ENSG0000( |
| 176124210 | 176131461 | -1 | lncRNA | ENSG0000( |
| 176143078 | 176156989 | -1 | transcribed_unproce | ENSG0000( |
| 176143085 | 176185155 | -1 | lncRNA | ENSG0000( |
| 176158610 | 176168367 | -1 | lncRNA | ENSG0000( |
| 176173345 | 176199295 | -1 | lncRNA | ENSG0000( |
| 176203314 | 176203710 | -1 | processed_pseudoge | ENSG0000( |
| 176238367 | 176345991 | 1 | protein_coding | ENSG0000( |
| 176308063 | 176309013 | 1 | processed_pseudoge | ENSG0000( |

| | | | | |
|---|---|---|---|---|
| 176346062 | 176361807 | -1 | protein_coding | ENSG0000C |
| 176347941 | 176353584 | 1 | lncRNA | ENSG0000C |
| 176354206 | 176356168 | 1 | lncRNA | ENSG0000C |
| 176365487 | 176401865 | 1 | protein_coding | ENSG0000C |
| 176367946 | 176368031 | 1 | miRNA | ENSG0000C |
| 176383938 | 176388975 | -1 | protein_coding | ENSG0000C |
| 176388751 | 176389761 | 1 | protein_coding | ENSG0000C |
| 176392455 | 176416569 | -1 | protein_coding | ENSG0000C |
| 176439090 | 176439238 | 1 | TEC | ENSG0000C |
| 176447628 | 176510074 | 1 | protein_coding | ENSG0000C |
| 176526712 | 176538025 | -1 | protein_coding | ENSG0000C |
| 176542511 | 176595974 | 1 | protein_coding | ENSG0000C |
| 176547433 | 176547726 | -1 | misc_RNA | ENSG0000C |
| 176556704 | 176556805 | 1 | misc_RNA | ENSG0000C |
| 176595802 | 176610156 | -1 | protein_coding | ENSG0000C |
| 176620082 | 176630556 | -1 | protein_coding | ENSG0000C |
| 176629439 | 176629500 | -1 | miRNA | ENSG0000C |
| 176630618 | 176646644 | 1 | protein_coding | ENSG0000C |
| 176647387 | 176659054 | 1 | protein_coding | ENSG0000C |
| 176707356 | 176726243 | -1 | lncRNA | ENSG0000C |
| 176726942 | 176739458 | -1 | lncRNA | ENSG0000C |
| 176743205 | 176743871 | 1 | lncRNA | ENSG0000C |
| 176810519 | 176880898 | 1 | protein_coding | ENSG0000C |
| 176880869 | 176899346 | -1 | protein_coding | ENSG0000C |
| 176905005 | 177022633 | -1 | protein_coding | ENSG0000C |
| 177022696 | 177081189 | 1 | protein_coding | ENSG0000C |
| 177051714 | 177052963 | 1 | lncRNA | ENSG0000C |
| 177086905 | 177098144 | 1 | protein_coding | ENSG0000C |
| 177133025 | 177300215 | 1 | protein_coding | ENSG0000C |
| 177260340 | 177263909 | 1 | lncRNA | ENSG0000C |
| 177264766 | 177265241 | 1 | processed_pseudoge | ENSG0000C |
| 177265580 | 177266588 | -1 | processed_pseudoge | ENSG0000C |
| 177301198 | 177303744 | -1 | protein_coding | ENSG0000C |
| 177301461 | 177312757 | -1 | protein_coding | ENSG0000C |
| 177303799 | 177306949 | 1 | protein_coding | ENSG0000C |
| 177331567 | 177351668 | -1 | protein_coding | ENSG0000C |
| 177346073 | 177346426 | 1 | processed_pseudoge | ENSG0000C |
| 177357924 | 177372596 | 1 | protein_coding | ENSG0000C |
| 177379235 | 177398848 | 1 | protein_coding | ENSG0000C |
| 177400109 | 177400661 | -1 | protein_coding | ENSG0000C |
| 177402140 | 177409576 | -1 | protein_coding | ENSG0000C |
| 177403204 | 177442901 | 1 | protein_coding | ENSG0000C |
| 177438503 | 177447699 | -1 | lncRNA | ENSG0000C |
| 177446445 | 177456286 | 1 | protein_coding | ENSG0000C |
| 177456608 | 177474401 | -1 | protein_coding | ENSG0000C |
| 177476631 | 177479656 | -1 | TEC | ENSG0000C |
| 177483394 | 177497606 | -1 | protein_coding | ENSG0000C |
| 177494995 | 177503647 | 1 | lncRNA | ENSG0000C |
| 177501907 | 177511274 | -1 | protein_coding | ENSG0000C |
| 177511577 | 177516961 | -1 | protein_coding | ENSG0000C |

| | | | | |
|---|---|---|---|---|
| 177519789 | 177554563 | -1 | protein_coding | ENSG00000 |
| 177554824 | 177555364 | 1 | lncRNA | ENSG00000 |
| 177592203 | 177597242 | 1 | protein_coding | ENSG00000 |
| 177600132 | 177610330 | 1 | protein_coding | ENSG00000 |
| 177611240 | 177619754 | 1 | lncRNA | ENSG00000 |
| 177619059 | 177672209 | -1 | transcribed_unproce | ENSG00000 |
| 177682294 | 177713969 | 1 | lncRNA | ENSG00000 |
| 177707981 | 177783398 | -1 | protein_coding | ENSG00000 |
| 177782197 | 177794396 | 1 | lncRNA | ENSG00000 |
| 177801204 | 177803344 | -1 | lncRNA | ENSG00000 |
| 177809407 | 177950732 | 1 | transcribed_unproce | ENSG00000 |
| 177836434 | 177837646 | -1 | unprocessed_pseudc | ENSG00000 |
| 177856262 | 177871227 | -1 | lncRNA | ENSG00000 |
| 177939622 | 177983054 | 1 | lncRNA | ENSG00000 |
| 177950335 | 177963960 | 1 | lncRNA | ENSG00000 |
| 177967004 | 177967457 | 1 | lncRNA | ENSG00000 |
| 177971316 | 177972281 | 1 | processed_pseudoge | ENSG00000 |
| 177992235 | 177996242 | -1 | protein_coding | ENSG00000 |
| 178006348 | 178086529 | 1 | lncRNA | ENSG00000 |
| 178055604 | 178056194 | 1 | transcribed_processe | ENSG00000 |
| 178113532 | 178126081 | 1 | protein_coding | ENSG00000 |
| 178130996 | 178150568 | 1 | protein_coding | ENSG00000 |
| 178149460 | 178153967 | -1 | protein_coding | ENSG00000 |
| 178165702 | 178166562 | 1 | processed_pseudoge | ENSG00000 |
| 178184503 | 178187432 | -1 | protein_coding | ENSG00000 |
| 178204533 | 178211163 | 1 | protein_coding | ENSG00000 |
| 178208471 | 178232802 | -1 | protein_coding | ENSG00000 |
| 178237618 | 178590393 | -1 | protein_coding | ENSG00000 |
| 178344843 | 178345316 | 1 | processed_pseudoge | ENSG00000 |
| 178350867 | 178352250 | 1 | lncRNA | ENSG00000 |
| 178377924 | 178378220 | 1 | misc_RNA | ENSG00000 |
| 178438681 | 178443094 | -1 | lncRNA | ENSG00000 |
| 178602664 | 178630615 | -1 | protein_coding | ENSG00000 |
| 178619728 | 178619998 | -1 | misc_RNA | ENSG00000 |
| 178690804 | 178693681 | -1 | lncRNA | ENSG00000 |
| 178694605 | 178697695 | -1 | protein_coding | ENSG00000 |
| 178694605 | 178729335 | -1 | protein_coding | ENSG00000 |
| 178711512 | 178730659 | -1 | protein_coding | ENSG00000 |
| 178764861 | 178818435 | -1 | transcribed_unproce | ENSG00000 |
| 178825382 | 178825571 | 1 | unprocessed_pseudc | ENSG00000 |
| 178840097 | 178841134 | 1 | processed_pseudoge | ENSG00000 |
| 178842028 | 178842235 | -1 | processed_pseudoge | ENSG00000 |
| 178859953 | 178888122 | 1 | protein_coding | ENSG00000 |
| 178884715 | 178884879 | 1 | snRNA | ENSG00000 |
| 178895898 | 178933212 | 1 | protein_coding | ENSG00000 |
| 178909710 | 178910609 | -1 | processed_pseudoge | ENSG00000 |
| 178922064 | 178922716 | 1 | processed_pseudoge | ENSG00000 |
| 178938677 | 178939223 | -1 | lncRNA | ENSG00000 |
| 178941191 | 178966433 | 1 | protein_coding | ENSG00000 |
| 178969390 | 178990116 | 1 | lncRNA | ENSG00000 |

| | | | | |
|---|---|---|---|---|
| 178977587 | 178996206 | -1 | protein_coding | ENSG00000 |
| 178997969 | 178998815 | -1 | processed_pseudoge | ENSG00000 |
| 179023804 | 179035064 | 1 | protein_coding | ENSG00000 |
| 179060373 | 179083977 | 1 | protein_coding | ENSG00000 |
| 179082680 | 179083194 | -1 | processed_pseudoge | ENSG00000 |
| 179110853 | 179345461 | -1 | protein_coding | ENSG00000 |
| 179377505 | 179378865 | 1 | lncRNA | ENSG00000 |
| 179377531 | 179378761 | -1 | lncRNA | ENSG00000 |
| 179385819 | 179385992 | -1 | processed_pseudoge | ENSG00000 |
| 179455415 | 179456095 | 1 | lncRNA | ENSG00000 |
| 179492076 | 179492157 | -1 | snRNA | ENSG00000 |
| 179503322 | 179515579 | -1 | lncRNA | ENSG00000 |
| 179522457 | 179525074 | 1 | unprocessed_pseudc | ENSG00000 |
| 179527224 | 179527513 | -1 | processed_pseudoge | ENSG00000 |
| 179550554 | 179610012 | 1 | protein_coding | ENSG00000 |
| 179572156 | 179572741 | 1 | processed_pseudoge | ENSG00000 |
| 179595904 | 179603741 | -1 | lncRNA | ENSG00000 |
| 179614178 | 179634784 | -1 | protein_coding | ENSG00000 |
| 179629151 | 179629410 | 1 | misc_RNA | ENSG00000 |
| 179641544 | 179645046 | -1 | lncRNA | ENSG00000 |
| 179651284 | 179652457 | 1 | unprocessed_pseudc | ENSG00000 |
| 179657762 | 179665136 | 1 | lncRNA | ENSG00000 |
| 179678560 | 179681034 | -1 | protein_coding | ENSG00000 |
| 179678628 | 179731641 | 1 | protein_coding | ENSG00000 |
| 179679032 | 179694768 | 1 | transcribed_processe | ENSG00000 |
| 179732822 | 179777283 | 1 | protein_coding | ENSG00000 |
| 179793980 | 179796647 | 1 | protein_coding | ENSG00000 |
| 179797597 | 179806952 | -1 | protein_coding | ENSG00000 |
| 179798278 | 179798346 | -1 | miRNA | ENSG00000 |
| 179806398 | 179838078 | 1 | protein_coding | ENSG00000 |
| 179835133 | 179862173 | -1 | protein_coding | ENSG00000 |
| 179845832 | 179846152 | 1 | misc_RNA | ENSG00000 |
| 179847035 | 179847130 | -1 | misc_RNA | ENSG00000 |
| 179859013 | 179861283 | 1 | lncRNA | ENSG00000 |
| 179862066 | 179907859 | -1 | protein_coding | ENSG00000 |
| 179911651 | 180072113 | -1 | protein_coding | ENSG00000 |
| 179933805 | 179934196 | -1 | processed_pseudoge | ENSG00000 |
| 179939457 | 179940132 | -1 | processed_pseudoge | ENSG00000 |
| 179963615 | 179964419 | 1 | lncRNA | ENSG00000 |
| 179974584 | 179980186 | 1 | lncRNA | ENSG00000 |
| 180015303 | 180015397 | -1 | miRNA | ENSG00000 |
| 180078357 | 180081691 | -1 | TEC | ENSG00000 |
| 180085717 | 180087038 | -1 | processed_pseudoge | ENSG00000 |
| 180100795 | 180209211 | -1 | protein_coding | ENSG00000 |
| 180226378 | 180227000 | -1 | processed_pseudoge | ENSG00000 |
| 180233143 | 180292099 | -1 | protein_coding | ENSG00000 |
| 180257680 | 180258366 | 1 | processed_pseudoge | ENSG00000 |
| 180293245 | 180295253 | 1 | lncRNA | ENSG00000 |
| 180300698 | 180353336 | -1 | protein_coding | ENSG00000 |
| 180416949 | 180443240 | 1 | lncRNA | ENSG00000 |

| | | | | |
|---|---|---|---|---|
| 180444509 | 180444612 | 1 | snRNA | ENSG00000 |
| 180494379 | 180578358 | 1 | protein_coding | ENSG00000 |
| 180534184 | 180534642 | -1 | processed_pseudoge | ENSG00000 |
| 180590105 | 180591499 | -1 | protein_coding | ENSG00000 |
| 180601506 | 180649624 | -1 | protein_coding | ENSG00000 |
| 180671892 | 180672598 | -1 | processed_pseudoge | ENSG00000 |
| 180684766 | 180686546 | 1 | lncRNA | ENSG00000 |
| 180692832 | 180693763 | -1 | unprocessed_pseudo | ENSG00000 |
| 180729586 | 180729737 | 1 | snRNA | ENSG00000 |
| 180739042 | 180740099 | -1 | protein_coding | ENSG00000 |
| 180784782 | 180815652 | -1 | protein_coding | ENSG00000 |
| 180810401 | 180811384 | 1 | TEC | ENSG00000 |
| 180826871 | 180831605 | -1 | lncRNA | ENSG00000 |
| 180830326 | 180839742 | 1 | lncRNA | ENSG00000 |
| 180833928 | 180835726 | -1 | lncRNA | ENSG00000 |
| 180847611 | 180861285 | -1 | protein_coding | ENSG00000 |
| 180899077 | 180950906 | 1 | protein_coding | ENSG00000 |
| 180909838 | 180909927 | 1 | misc_RNA | ENSG00000 |
| 180945756 | 180945907 | 1 | processed_pseudoge | ENSG00000 |
| 180967189 | 180970762 | -1 | processed_pseudoge | ENSG00000 |
| 180967848 | 180968567 | 1 | processed_pseudoge | ENSG00000 |
| 180982273 | 180982611 | 1 | processed_pseudoge | ENSG00000 |
| 180988846 | 181006727 | 1 | protein_coding | ENSG00000 |
| 181007281 | 181007384 | 1 | snRNA | ENSG00000 |
| 181040225 | 181061521 | 1 | protein_coding | ENSG00000 |
| 181043403 | 181043484 | -1 | miRNA | ENSG00000 |
| 181065086 | 181065677 | 1 | processed_pseudoge | ENSG00000 |
| 181099140 | 181100666 | -1 | processed_pseudoge | ENSG00000 |
| 181114106 | 181115674 | 1 | processed_pseudoge | ENSG00000 |
| 181123122 | 181131169 | -1 | protein_coding | ENSG00000 |
| 181147586 | 181159285 | 1 | protein_coding | ENSG00000 |
| 181178821 | 181191852 | -1 | lncRNA | ENSG00000 |
| 181191875 | 181194429 | 1 | lncRNA | ENSG00000 |
| 181193924 | 181205293 | -1 | protein_coding | ENSG00000 |
| 181195496 | 181203103 | 1 | lncRNA | ENSG00000 |
| 181205361 | 181206120 | -1 | lncRNA | ENSG00000 |
| 181207480 | 181217864 | -1 | lncRNA | ENSG00000 |
| 181222499 | 181235808 | 1 | protein_coding | ENSG00000 |
| 181222566 | 181222633 | -1 | miRNA | ENSG00000 |
| 181224646 | 181230685 | -1 | lncRNA | ENSG00000 |
| 181236897 | 181248096 | -1 | protein_coding | ENSG00000 |
| 181241814 | 181241892 | -1 | snoRNA | ENSG00000 |
| 181243312 | 181243379 | -1 | snoRNA | ENSG00000 |
| 181246507 | 181272167 | 1 | lncRNA | ENSG00000 |
| 181254417 | 181261139 | -1 | protein_coding | ENSG00000 |
| 181261169 | 181272307 | 1 | lncRNA | ENSG00000 |
| 181276848 | 181277555 | 1 | lncRNA | ENSG00000 |
| 181281375 | 181283652 | 1 | lncRNA | ENSG00000 |
| 181291673 | 181295572 | 1 | unprocessed_pseudo | ENSG00000 |
| 181301967 | 181302070 | 1 | snRNA | ENSG00000 |

| | | | | |
|---|---|---|---|---|
| 181306502 | 181324685 | 1 | lncRNA | ENSG0000C |
| 181329049 | 181331839 | -1 | processed_pseudoge | ENSG0000C |
| 181329241 | 181342213 | 1 | lncRNA | ENSG0000C |
| 181367268 | 181368262 | 1 | protein_coding | ENSG0000C |
| 181378695 | 181379038 | -1 | processed_pseudoge | ENSG0000C |
| 181471963 | 181472430 | 1 | processed_pseudoge | ENSG0000C |
| 95124 | 95454 | 1 | processed_pseudoge | ENSG0000C |
| 105919 | 106856 | -1 | unprocessed_pseudc | ENSG0000C |
| 131910 | 145083 | -1 | transcribed_processe | ENSG0000C |
| 167607 | 170631 | -1 | unprocessed_pseudc | ENSG0000C |
| 181466 | 205519 | -1 | lncRNA | ENSG0000C |
| 184339 | 186899 | 1 | lncRNA | ENSG0000C |
| 203313 | 206392 | 1 | lncRNA | ENSG0000C |
| 285443 | 292013 | -1 | lncRNA | ENSG0000C |
| 291630 | 351355 | 1 | protein_coding | ENSG0000C |
| 391752 | 411443 | 1 | protein_coding | ENSG0000C |
| 453391 | 477829 | 1 | lncRNA | ENSG0000C |
| 485154 | 693139 | -1 | protein_coding | ENSG0000C |
| 524171 | 525581 | 1 | lncRNA | ENSG0000C |
| 655939 | 656963 | -1 | protein_coding | ENSG0000C |
| 669081 | 670307 | -1 | processed_pseudoge | ENSG0000C |
| 692530 | 796781 | 1 | lncRNA | ENSG0000C |
| 708592 | 711405 | -1 | lncRNA | ENSG0000C |
| 729128 | 760936 | -1 | lncRNA | ENSG0000C |
| 761675 | 780648 | 1 | lncRNA | ENSG0000C |
| 774747 | 780214 | -1 | lncRNA | ENSG0000C |
| 905445 | 909006 | -1 | lncRNA | ENSG0000C |
| 958323 | 1101400 | -1 | lncRNA | ENSG0000C |
| 1026494 | 1027225 | -1 | lncRNA | ENSG0000C |
| 1079929 | 1104946 | 1 | lncRNA | ENSG0000C |
| 1111367 | 1114329 | 1 | lncRNA | ENSG0000C |
| 1312098 | 1314758 | 1 | protein_coding | ENSG0000C |
| 1321698 | 1324022 | -1 | lncRNA | ENSG0000C |
| 1383790 | 1385066 | -1 | lncRNA | ENSG0000C |
| 1389576 | 1395603 | 1 | protein_coding | ENSG0000C |
| 1390314 | 1390411 | -1 | miRNA | ENSG0000C |
| 1507322 | 1507598 | 1 | misc_RNA | ENSG0000C |
| 1513698 | 1515289 | -1 | processed_pseudoge | ENSG0000C |
| 1528364 | 1528911 | -1 | lncRNA | ENSG0000C |
| 1601654 | 1607354 | 1 | lncRNA | ENSG0000C |
| 1609915 | 1613897 | 1 | protein_coding | ENSG0000C |
| 1623806 | 2245605 | -1 | protein_coding | ENSG0000C |
| 2227803 | 2230261 | 1 | lncRNA | ENSG0000C |
| 2245718 | 2525976 | 1 | lncRNA | ENSG0000C |
| 2341450 | 2341711 | 1 | processed_pseudoge | ENSG0000C |
| 2353427 | 2386110 | -1 | lncRNA | ENSG0000C |
| 2437549 | 2438249 | 1 | lncRNA | ENSG0000C |
| 2503797 | 2528665 | -1 | lncRNA | ENSG0000C |
| 2561947 | 2564143 | 1 | TEC | ENSG0000C |
| 2617144 | 2640001 | 1 | lncRNA | ENSG0000C |

| | | | | |
|---|---|---|---|---|
| 2621913 | 2634619 | -1 | lncRNA | ENSG00000 |
| 2663629 | 2750922 | -1 | protein_coding | ENSG00000 |
| 2765393 | 2786952 | 1 | protein_coding | ENSG00000 |
| 2832332 | 2841959 | -1 | protein_coding | ENSG00000 |
| 2851230 | 2881407 | -1 | lncRNA | ENSG00000 |
| 2854031 | 2854107 | -1 | miRNA | ENSG00000 |
| 2887270 | 2903309 | -1 | protein_coding | ENSG00000 |
| 2910613 | 2912113 | -1 | unprocessed_pseudo | ENSG00000 |
| 2916894 | 2917733 | 1 | lncRNA | ENSG00000 |
| 2948159 | 2972165 | -1 | protein_coding | ENSG00000 |
| 2987967 | 2991173 | 1 | lncRNA | ENSG00000 |
| 2987987 | 3028869 | 1 | protein_coding | ENSG00000 |
| 2989335 | 2999604 | -1 | lncRNA | ENSG00000 |
| 3023142 | 3023772 | -1 | processed_pseudoge | ENSG00000 |
| 3033183 | 3033288 | 1 | processed_pseudoge | ENSG00000 |
| 3045384 | 3045800 | 1 | processed_pseudoge | ENSG00000 |
| 3048835 | 3049211 | 1 | unprocessed_pseudo | ENSG00000 |
| 3055849 | 3057357 | 1 | lncRNA | ENSG00000 |
| 3063991 | 3115187 | 1 | protein_coding | ENSG00000 |
| 3068045 | 3068894 | -1 | lncRNA | ENSG00000 |
| 3088686 | 3088809 | -1 | rRNA_pseudogene | ENSG00000 |
| 3118374 | 3153578 | 1 | protein_coding | ENSG00000 |
| 3138394 | 3153062 | -1 | lncRNA | ENSG00000 |
| 3153666 | 3157544 | -1 | protein_coding | ENSG00000 |
| 3177044 | 3179764 | -1 | unprocessed_pseudo | ENSG00000 |
| 3182626 | 3195784 | -1 | lncRNA | ENSG00000 |
| 3224277 | 3231730 | -1 | protein_coding | ENSG00000 |
| 3231403 | 3303373 | 1 | protein_coding | ENSG00000 |
| 3268962 | 3457022 | -1 | protein_coding | ENSG00000 |
| 3311662 | 3313650 | 1 | lncRNA | ENSG00000 |
| 3585101 | 3720081 | -1 | lncRNA | ENSG00000 |
| 3693923 | 3694075 | -1 | processed_pseudoge | ENSG00000 |
| 3722614 | 3752026 | -1 | protein_coding | ENSG00000 |
| 3751111 | 3753871 | 1 | lncRNA | ENSG00000 |
| 3754099 | 3754840 | 1 | processed_pseudoge | ENSG00000 |
| 3831933 | 3855737 | 1 | lncRNA | ENSG00000 |
| 3831936 | 3894292 | 1 | lncRNA | ENSG00000 |
| 3849373 | 3851320 | 1 | protein_coding | ENSG00000 |
| 3904920 | 3911979 | -1 | lncRNA | ENSG00000 |
| 3913922 | 3914292 | -1 | processed_pseudoge | ENSG00000 |
| 3921847 | 3922146 | 1 | processed_pseudoge | ENSG00000 |
| 3941282 | 3941801 | 1 | processed_pseudoge | ENSG00000 |
| 3978062 | 3979099 | 1 | processed_pseudoge | ENSG00000 |
| 3982673 | 3984130 | 1 | TEC | ENSG00000 |
| 4018713 | 4019202 | 1 | lncRNA | ENSG00000 |
| 4018843 | 4021215 | -1 | lncRNA | ENSG00000 |
| 4021267 | 4064983 | 1 | protein_coding | ENSG00000 |
| 4049434 | 4087344 | -1 | protein_coding | ENSG00000 |
| 4079206 | 4130951 | 1 | protein_coding | ENSG00000 |
| 4115689 | 4135597 | -1 | protein_coding | ENSG00000 |

| | | | | |
|---|---|---|---|---|
| 4135423 | 4146053 | 1 | lncRNA | ENSG0000C |
| 4136072 | 4157385 | 1 | lncRNA | ENSG0000C |
| 4185479 | 4188950 | -1 | lncRNA | ENSG0000C |
| 4189116 | 4189806 | 1 | processed_pseudoge | ENSG0000C |
| 4304942 | 4583871 | -1 | lncRNA | ENSG0000C |
| 4341756 | 4347162 | -1 | lncRNA | ENSG0000C |
| 4381211 | 4383597 | 1 | lncRNA | ENSG0000C |
| 4427963 | 4428081 | 1 | rRNA_pseudogene | ENSG0000C |
| 4449392 | 4468181 | 1 | lncRNA | ENSG0000C |
| 4491807 | 4495769 | 1 | lncRNA | ENSG0000C |
| 4529722 | 4530008 | 1 | processed_pseudoge | ENSG0000C |
| 4599287 | 4602420 | -1 | lncRNA | ENSG0000C |
| 4610612 | 4611919 | 1 | lncRNA | ENSG0000C |
| 4702463 | 4703779 | 1 | processed_pseudoge | ENSG0000C |
| 4706159 | 4955551 | 1 | protein_coding | ENSG0000C |
| 4714098 | 4724885 | -1 | lncRNA | ENSG0000C |
| 4774526 | 4775408 | -1 | lncRNA | ENSG0000C |
| 4979072 | 4979494 | 1 | processed_pseudoge | ENSG0000C |
| 4994717 | 5004063 | -1 | protein_coding | ENSG0000C |
| 5003774 | 5080946 | 1 | lncRNA | ENSG0000C |
| 5031756 | 5054423 | -1 | lncRNA | ENSG0000C |
| 5065795 | 5066982 | -1 | processed_pseudoge | ENSG0000C |
| 5084581 | 5089487 | 1 | protein_coding | ENSG0000C |
| 5102593 | 5260939 | -1 | protein_coding | ENSG0000C |
| 5148233 | 5148322 | -1 | miRNA | ENSG0000C |
| 5261044 | 5829192 | 1 | protein_coding | ENSG0000C |
| 5291030 | 5291388 | -1 | processed_pseudoge | ENSG0000C |
| 5451683 | 5458075 | -1 | lncRNA | ENSG0000C |
| 5609227 | 5610195 | -1 | processed_pseudoge | ENSG0000C |
| 5664985 | 5695272 | -1 | lncRNA | ENSG0000C |
| 5788346 | 5788699 | -1 | processed_pseudoge | ENSG0000C |
| 5817421 | 5818366 | -1 | processed_pseudoge | ENSG0000C |
| 5851506 | 5870220 | 1 | lncRNA | ENSG0000C |
| 5918564 | 5918844 | -1 | misc_RNA | ENSG0000C |
| 5972412 | 5974012 | 1 | processed_pseudoge | ENSG0000C |
| 5997999 | 6007605 | -1 | protein_coding | ENSG0000C |
| 6144085 | 6321013 | -1 | protein_coding | ENSG0000C |
| 6169304 | 6169435 | -1 | miRNA | ENSG0000C |
| 6169334 | 6169409 | 1 | miRNA | ENSG0000C |
| 6346465 | 6622771 | -1 | lncRNA | ENSG0000C |
| 6347081 | 6347381 | -1 | transcribed_processe | ENSG0000C |
| 6450356 | 6507443 | 1 | lncRNA | ENSG0000C |
| 6534697 | 6535631 | 1 | processed_pseudoge | ENSG0000C |
| 6588108 | 6654983 | 1 | protein_coding | ENSG0000C |
| 6680309 | 6683633 | -1 | lncRNA | ENSG0000C |
| 6685148 | 6686984 | 1 | TEC | ENSG0000C |
| 6692741 | 6739030 | -1 | lncRNA | ENSG0000C |
| 6710891 | 6713970 | 1 | lncRNA | ENSG0000C |
| 6725036 | 6733191 | 1 | lncRNA | ENSG0000C |
| 6758276 | 6759291 | -1 | lncRNA | ENSG0000C |

| | | | | |
|---|---|---|---|---|
| 6794915 | 6795402 | -1 | processed_pseudoge | ENSG00000 |
| 6901022 | 6918715 | -1 | lncRNA | ENSG00000 |
| 6938606 | 6938897 | -1 | misc_RNA | ENSG00000 |
| 6993191 | 6995727 | 1 | lncRNA | ENSG00000 |
| 7107597 | 7251980 | 1 | protein_coding | ENSG00000 |
| 7183083 | 7185287 | 1 | lncRNA | ENSG00000 |
| 7187582 | 7187686 | -1 | misc_RNA | ENSG00000 |
| 7232556 | 7232678 | 1 | unprocessed_pseudo | ENSG00000 |
| 7258910 | 7261993 | -1 | lncRNA | ENSG00000 |
| 7268306 | 7347446 | -1 | protein_coding | ENSG00000 |
| 7276031 | 7298872 | 1 | lncRNA | ENSG00000 |
| 7326656 | 7389742 | -1 | protein_coding | ENSG00000 |
| 7331286 | 7331676 | -1 | processed_pseudoge | ENSG00000 |
| 7338839 | 7339568 | -1 | processed_pseudoge | ENSG00000 |
| 7340854 | 7342056 | -1 | processed_pseudoge | ENSG00000 |
| 7389793 | 7418037 | 1 | protein_coding | ENSG00000 |
| 7481094 | 7482594 | -1 | processed_pseudoge | ENSG00000 |
| 7506203 | 7506549 | 1 | processed_pseudoge | ENSG00000 |
| 7516600 | 7517840 | -1 | processed_pseudoge | ENSG00000 |
| 7540451 | 7541338 | -1 | lncRNA | ENSG00000 |
| 7541617 | 7586714 | 1 | protein_coding | ENSG00000 |
| 7590198 | 7611967 | 1 | protein_coding | ENSG00000 |
| 7620422 | 7620878 | 1 | processed_pseudoge | ENSG00000 |
| 7726099 | 7881728 | 1 | protein_coding | ENSG00000 |
| 7881517 | 7910788 | -1 | protein_coding | ENSG00000 |
| 7881522 | 8064364 | -1 | protein_coding | ENSG00000 |
| 7938430 | 7939322 | -1 | processed_pseudoge | ENSG00000 |
| 7986537 | 7988192 | 1 | processed_pseudoge | ENSG00000 |
| 8013567 | 8064396 | -1 | protein_coding | ENSG00000 |
| 8015726 | 8102530 | -1 | protein_coding | ENSG00000 |
| 8073360 | 8102559 | -1 | protein_coding | ENSG00000 |
| 8102711 | 8343139 | 1 | lncRNA | ENSG00000 |
| 8158162 | 8760447 | 1 | lncRNA | ENSG00000 |
| 8341937 | 8389183 | -1 | lncRNA | ENSG00000 |
| 8411463 | 8435611 | -1 | protein_coding | ENSG00000 |
| 8435568 | 9294133 | 1 | lncRNA | ENSG00000 |
| 8653558 | 8653797 | 1 | misc_RNA | ENSG00000 |
| 9124221 | 9161777 | -1 | lncRNA | ENSG00000 |
| 9596110 | 10211608 | -1 | transcribed_unitary_ | ENSG00000 |
| 10118452 | 10119253 | 1 | processed_pseudoge | ENSG00000 |
| 10213881 | 10214362 | -1 | processed_pseudoge | ENSG00000 |
| 10222036 | 10222161 | 1 | snRNA | ENSG00000 |
| 10365593 | 10366582 | -1 | processed_pseudoge | ENSG00000 |
| 10393186 | 10419659 | -1 | protein_coding | ENSG00000 |
| 10404502 | 10407928 | 1 | lncRNA | ENSG00000 |
| 10409340 | 10416446 | 1 | lncRNA | ENSG00000 |
| 10423140 | 10426176 | -1 | lncRNA | ENSG00000 |
| 10426623 | 10429110 | 1 | lncRNA | ENSG00000 |
| 10429255 | 10435015 | -1 | lncRNA | ENSG00000 |
| 10434316 | 10456781 | 1 | lncRNA | ENSG00000 |

| | | | | |
|---|---|---|---|---|
| 10439717 | 10439794 | 1 | miRNA | ENSG0000( |
| 10459906 | 10460541 | -1 | processed_pseudoge | ENSG0000( |
| 10474500 | 10478502 | -1 | lncRNA | ENSG0000( |
| 10492223 | 10629368 | 1 | protein_coding | ENSG0000( |
| 10511036 | 10514546 | -1 | lncRNA | ENSG0000( |
| 10514282 | 10514973 | 1 | processed_pseudoge | ENSG0000( |
| 10574749 | 10575227 | -1 | processed_pseudoge | ENSG0000( |
| 10633762 | 10634695 | 1 | processed_pseudoge | ENSG0000( |
| 10659498 | 10692860 | 1 | lncRNA | ENSG0000( |
| 10671418 | 10694797 | -1 | protein_coding | ENSG0000( |
| 10674658 | 10674768 | -1 | misc_RNA | ENSG0000( |
| 10694972 | 10709782 | 1 | protein_coding | ENSG0000( |
| 10722915 | 10731129 | 1 | protein_coding | ENSG0000( |
| 10743324 | 10747663 | -1 | lncRNA | ENSG0000( |
| 10747759 | 10852753 | 1 | protein_coding | ENSG0000( |
| 10747794 | 10930423 | 1 | protein_coding | ENSG0000( |
| 10752791 | 10752926 | -1 | rRNA_pseudogene | ENSG0000( |
| 10762723 | 10838555 | -1 | protein_coding | ENSG0000( |
| 10873223 | 10881941 | -1 | protein_coding | ENSG0000( |
| 10881780 | 10884349 | 1 | lncRNA | ENSG0000( |
| 10886831 | 10979320 | 1 | protein_coding | ENSG0000( |
| 10980759 | 11044305 | -1 | protein_coding | ENSG0000( |
| 11015586 | 11015888 | 1 | processed_pseudoge | ENSG0000( |
| 11043482 | 11079154 | 1 | lncRNA | ENSG0000( |
| 11093834 | 11138733 | 1 | protein_coding | ENSG0000( |
| 11102489 | 11111725 | -1 | protein_coding | ENSG0000( |
| 11154929 | 11155150 | -1 | processed_pseudoge | ENSG0000( |
| 11173452 | 11259099 | 1 | lncRNA | ENSG0000( |
| 11183298 | 11382348 | -1 | protein_coding | ENSG0000( |
| 11291299 | 11296224 | 1 | lncRNA | ENSG0000( |
| 11414636 | 11515710 | 1 | lncRNA | ENSG0000( |
| 11417207 | 11481324 | 1 | lncRNA | ENSG0000( |
| 11503509 | 11503671 | -1 | snRNA | ENSG0000( |
| 11514192 | 11516466 | 1 | transcribed_processe | ENSG0000( |
| 11537749 | 11583524 | 1 | protein_coding | ENSG0000( |
| 11607552 | 11607981 | 1 | lncRNA | ENSG0000( |
| 11644067 | 11644343 | 1 | misc_RNA | ENSG0000( |
| 11709819 | 11709960 | 1 | snoRNA | ENSG0000( |
| 11712054 | 11807046 | -1 | protein_coding | ENSG0000( |
| 11722547 | 11731923 | 1 | lncRNA | ENSG0000( |
| 11810602 | 11811248 | 1 | lncRNA | ENSG0000( |
| 11861626 | 11862970 | -1 | processed_pseudoge | ENSG0000( |
| 11990338 | 12001286 | -1 | lncRNA | ENSG0000( |
| 12007670 | 12008856 | 1 | lncRNA | ENSG0000( |
| 12008762 | 12164999 | 1 | protein_coding | ENSG0000( |
| 12290361 | 12297194 | 1 | protein_coding | ENSG0000( |
| 12319453 | 12319592 | 1 | processed_pseudoge | ENSG0000( |
| 12406486 | 12406781 | -1 | misc_RNA | ENSG0000( |
| 12514110 | 12514724 | 1 | processed_pseudoge | ENSG0000( |
| 12582915 | 12585404 | -1 | lncRNA | ENSG0000( |

| | | | | |
|---|---|---|---|---|
| 12716805 | 13290484 | 1 | protein_coding | ENSG00000 |
| 13214056 | 13214219 | 1 | snRNA | ENSG00000 |
| 13264861 | 13295586 | -1 | lncRNA | ENSG00000 |
| 13266542 | 13328583 | -1 | protein_coding | ENSG00000 |
| 13290018 | 13290490 | -1 | lncRNA | ENSG00000 |
| 13357830 | 13487662 | -1 | protein_coding | ENSG00000 |
| 13486294 | 13486852 | 1 | lncRNA | ENSG00000 |
| 13521266 | 13522045 | 1 | processed_pseudoge | ENSG00000 |
| 13547704 | 13548013 | 1 | misc_RNA | ENSG00000 |
| 13574529 | 13615158 | 1 | protein_coding | ENSG00000 |
| 13614111 | 13615155 | -1 | lncRNA | ENSG00000 |
| 13615335 | 13632739 | 1 | protein_coding | ENSG00000 |
| 13621498 | 13711835 | -1 | protein_coding | ENSG00000 |
| 13671720 | 13678908 | 1 | lncRNA | ENSG00000 |
| 13786557 | 13814568 | -1 | protein_coding | ENSG00000 |
| 13825432 | 13826574 | 1 | lncRNA | ENSG00000 |
| 13924446 | 13980310 | 1 | protein_coding | ENSG00000 |
| 13949095 | 13949666 | 1 | processed_pseudoge | ENSG00000 |
| 14004920 | 14006901 | 1 | lncRNA | ENSG00000 |
| 14117256 | 14136918 | 1 | protein_coding | ENSG00000 |
| 14152196 | 14211186 | -1 | lncRNA | ENSG00000 |
| 14156462 | 14156523 | 1 | snRNA | ENSG00000 |
| 14210293 | 14214374 | 1 | lncRNA | ENSG00000 |
| 14229775 | 14231570 | -1 | lncRNA | ENSG00000 |
| 14280127 | 14285454 | -1 | lncRNA | ENSG00000 |
| 14391094 | 14393929 | -1 | lncRNA | ENSG00000 |
| 14394326 | 14404289 | 1 | lncRNA | ENSG00000 |
| 14431683 | 14502701 | 1 | lncRNA | ENSG00000 |
| 14597514 | 14599690 | -1 | lncRNA | ENSG00000 |
| 14646535 | 14646638 | 1 | snRNA | ENSG00000 |
| 14661829 | 15090170 | -1 | lncRNA | ENSG00000 |
| 15102946 | 15104130 | -1 | processed_pseudoge | ENSG00000 |
| 15112968 | 15113221 | 1 | misc_RNA | ENSG00000 |
| 15243923 | 15245000 | -1 | lncRNA | ENSG00000 |
| 15246069 | 15522042 | 1 | protein_coding | ENSG00000 |
| 15247815 | 15248634 | -1 | lncRNA | ENSG00000 |
| 15314920 | 15315026 | 1 | snRNA | ENSG00000 |
| 15324367 | 15324472 | -1 | snRNA | ENSG00000 |
| 15522195 | 15522300 | -1 | rRNA | ENSG00000 |
| 15522801 | 15663058 | -1 | protein_coding | ENSG00000 |
| 15555780 | 15559980 | 1 | lncRNA | ENSG00000 |
| 15934782 | 15935315 | 1 | processed_pseudoge | ENSG00000 |
| 15994944 | 15999797 | 1 | lncRNA | ENSG00000 |
| 16041381 | 16041614 | -1 | processed_pseudoge | ENSG00000 |
| 16107622 | 16108593 | 1 | processed_pseudoge | ENSG00000 |
| 16129086 | 16148248 | 1 | protein_coding | ENSG00000 |
| 16141556 | 16141624 | 1 | miRNA | ENSG00000 |
| 16160926 | 16161411 | 1 | processed_pseudoge | ENSG00000 |
| 16163377 | 16163762 | -1 | processed_pseudoge | ENSG00000 |
| 16171606 | 16172031 | 1 | processed_pseudoge | ENSG00000 |

| | | | | |
|---|---|---|---|---|
| 16205014 | 16205120 | 1 | snRNA | ENSG0000C |
| 16232231 | 16232330 | -1 | misc_RNA | ENSG0000C |
| 16238587 | 16295549 | 1 | protein_coding | ENSG0000C |
| 16259101 | 16264553 | -1 | lncRNA | ENSG0000C |
| 16299112 | 16761491 | -1 | protein_coding | ENSG0000C |
| 16761138 | 16762652 | 1 | lncRNA | ENSG0000C |
| 16764346 | 16766883 | 1 | lncRNA | ENSG0000C |
| 16901104 | 16952699 | -1 | lncRNA | ENSG0000C |
| 17015817 | 17034396 | 1 | lncRNA | ENSG0000C |
| 17092714 | 17108156 | -1 | lncRNA | ENSG0000C |
| 17102258 | 17131372 | 1 | protein_coding | ENSG0000C |
| 17224243 | 17281424 | -1 | lncRNA | ENSG0000C |
| 17281361 | 17293871 | 1 | protein_coding | ENSG0000C |
| 17381367 | 17382120 | 1 | processed_pseudoge | ENSG0000C |
| 17393505 | 17557790 | 1 | protein_coding | ENSG0000C |
| 17530899 | 17531652 | -1 | processed_pseudoge | ENSG0000C |
| 17582034 | 17582305 | 1 | processed_pseudoge | ENSG0000C |
| 17586899 | 17600191 | -1 | lncRNA | ENSG0000C |
| 17600302 | 17611715 | 1 | protein_coding | ENSG0000C |
| 17615035 | 17706834 | -1 | protein_coding | ENSG0000C |
| 17619533 | 17619634 | 1 | snRNA | ENSG0000C |
| 17706257 | 17707344 | 1 | lncRNA | ENSG0000C |
| 17721786 | 17721903 | -1 | rRNA_pseudogene | ENSG0000C |
| 17737458 | 17737556 | -1 | misc_RNA | ENSG0000C |
| 17759183 | 17987635 | -1 | protein_coding | ENSG0000C |
| 17953572 | 17953930 | 1 | unprocessed_pseudc | ENSG0000C |
| 18120440 | 18122677 | -1 | protein_coding | ENSG0000C |
| 18128311 | 18155077 | -1 | protein_coding | ENSG0000C |
| 18155329 | 18223854 | 1 | protein_coding | ENSG0000C |
| 18223860 | 18264548 | -1 | protein_coding | ENSG0000C |
| 18273165 | 18273266 | -1 | misc_RNA | ENSG0000C |
| 18304854 | 18305045 | -1 | processed_pseudoge | ENSG0000C |
| 18306973 | 18307079 | -1 | snRNA | ENSG0000C |
| 18363417 | 18365423 | -1 | processed_pseudoge | ENSG0000C |
| 18366736 | 18368644 | -1 | unprocessed_pseudc | ENSG0000C |
| 18387350 | 18468874 | 1 | protein_coding | ENSG0000C |
| 18522735 | 18674001 | 1 | lncRNA | ENSG0000C |
| 18571784 | 18571880 | 1 | miRNA | ENSG0000C |
| 19143695 | 19144160 | 1 | processed_pseudoge | ENSG0000C |
| 19290319 | 19321021 | -1 | lncRNA | ENSG0000C |
| 19323428 | 19839080 | -1 | lncRNA | ENSG0000C |
| 19348181 | 19349076 | 1 | processed_pseudoge | ENSG0000C |
| 19438283 | 19438418 | 1 | rRNA_pseudogene | ENSG0000C |
| 19612755 | 19614036 | 1 | processed_pseudoge | ENSG0000C |
| 19638040 | 19638808 | -1 | processed_pseudoge | ENSG0000C |
| 19642425 | 19642529 | -1 | snRNA | ENSG0000C |
| 19837370 | 19842197 | 1 | protein_coding | ENSG0000C |
| 19892397 | 19894214 | 1 | lncRNA | ENSG0000C |
| 20042455 | 20042940 | -1 | processed_pseudoge | ENSG0000C |
| 20099684 | 20212469 | -1 | protein_coding | ENSG0000C |

| | | | | |
|---|---|---|---|---|
| 20212087 | 20317739 | 1 | lncRNA | ENSG00000 |
| 20265720 | 20307200 | 1 | lncRNA | ENSG00000 |
| 20321461 | 20333193 | 1 | lncRNA | ENSG00000 |
| 20329922 | 20334411 | -1 | lncRNA | ENSG00000 |
| 20401879 | 20493714 | 1 | protein_coding | ENSG00000 |
| 20421583 | 20421887 | -1 | misc_RNA | ENSG00000 |
| 20437821 | 20440178 | 1 | lncRNA | ENSG00000 |
| 20452369 | 20452473 | -1 | snRNA | ENSG00000 |
| 20500175 | 20500276 | 1 | misc_RNA | ENSG00000 |
| 20534457 | 21232404 | 1 | protein_coding | ENSG00000 |
| 20722328 | 20723655 | -1 | processed_pseudoge | ENSG00000 |
| 20756103 | 20800694 | -1 | lncRNA | ENSG00000 |
| 20993474 | 20993568 | -1 | snRNA | ENSG00000 |
| 21233168 | 21271132 | -1 | lncRNA | ENSG00000 |
| 21354411 | 21355296 | -1 | lncRNA | ENSG00000 |
| 21369690 | 21383200 | -1 | lncRNA | ENSG00000 |
| 21485896 | 21521414 | -1 | lncRNA | ENSG00000 |
| 21521678 | 21523783 | -1 | lncRNA | ENSG00000 |
| 21528739 | 21595593 | -1 | lncRNA | ENSG00000 |
| 21593751 | 21598619 | 1 | protein_coding | ENSG00000 |
| 21602128 | 21602383 | 1 | processed_pseudoge | ENSG00000 |
| 21664185 | 22654455 | 1 | lncRNA | ENSG00000 |
| 21822536 | 21822737 | 1 | TEC | ENSG00000 |
| 22085544 | 22085918 | 1 | misc_RNA | ENSG00000 |
| 22133205 | 22147193 | -1 | lncRNA | ENSG00000 |
| 22213306 | 22214237 | -1 | processed_pseudoge | ENSG00000 |
| 22287244 | 22302826 | -1 | protein_coding | ENSG00000 |
| 22569566 | 22571666 | 1 | protein_coding | ENSG00000 |
| 22589137 | 22593833 | -1 | lncRNA | ENSG00000 |
| 22663507 | 22675493 | 1 | lncRNA | ENSG00000 |
| 22744395 | 23031780 | -1 | lncRNA | ENSG00000 |
| 23102680 | 23103546 | 1 | processed_pseudoge | ENSG00000 |
| 23124981 | 23125087 | -1 | snRNA | ENSG00000 |
| 23337711 | 23404132 | 1 | lncRNA | ENSG00000 |
| 23649496 | 23649774 | 1 | processed_pseudoge | ENSG00000 |
| 23854139 | 23857646 | 1 | transcribed_unproce | ENSG00000 |
| 23856698 | 23856818 | -1 | processed_pseudoge | ENSG00000 |
| 23971879 | 23972941 | 1 | processed_pseudoge | ENSG00000 |
| 24001824 | 24002433 | 1 | processed_pseudoge | ENSG00000 |
| 24126186 | 24154900 | 1 | protein_coding | ENSG00000 |
| 24166273 | 24166374 | 1 | snoRNA | ENSG00000 |
| 24171755 | 24358059 | -1 | protein_coding | ENSG00000 |
| 24356903 | 24358285 | 1 | protein_coding | ENSG00000 |
| 24365651 | 24365754 | 1 | snRNA | ENSG00000 |
| 24402908 | 24426194 | 1 | protein_coding | ENSG00000 |
| 24424565 | 24495205 | -1 | protein_coding | ENSG00000 |
| 24494867 | 24537207 | 1 | protein_coding | ENSG00000 |
| 24544104 | 24646191 | -1 | protein_coding | ENSG00000 |
| 24598012 | 24599790 | 1 | processed_pseudoge | ENSG00000 |
| 24649979 | 24666930 | -1 | protein_coding | ENSG00000 |

| | | | | |
|---|---|---|---|---|
| 24667035 | 24705065 | 1 | protein_coding | ENSG00000 |
| 24700907 | 24701793 | -1 | lncRNA | ENSG00000 |
| 24704861 | 24719998 | -1 | protein_coding | ENSG00000 |
| 24706747 | 24707151 | 1 | lncRNA | ENSG00000 |
| 24721658 | 24751960 | 1 | lncRNA | ENSG00000 |
| 24750671 | 24751625 | -1 | processed_pseudoge | ENSG00000 |
| 24774931 | 24786099 | 1 | protein_coding | ENSG00000 |
| 24797335 | 24798917 | -1 | protein_coding | ENSG00000 |
| 24797373 | 24809798 | -1 | protein_coding | ENSG00000 |
| 24804282 | 25042170 | -1 | protein_coding | ENSG00000 |
| 24947880 | 24948344 | 1 | processed_pseudoge | ENSG00000 |
| 24976419 | 24976982 | 1 | processed_pseudoge | ENSG00000 |
| 25012985 | 25015696 | 1 | lncRNA | ENSG00000 |
| 25023247 | 25024483 | -1 | processed_pseudoge | ENSG00000 |
| 25031009 | 25031287 | -1 | misc_RNA | ENSG00000 |
| 25041839 | 25056664 | 1 | lncRNA | ENSG00000 |
| 25053627 | 25057073 | -1 | lncRNA | ENSG00000 |
| 25061626 | 25452263 | -1 | transcribed_unitary_ | ENSG00000 |
| 25061796 | 25134971 | 1 | lncRNA | ENSG00000 |
| 25140003 | 25141403 | 1 | processed_pseudoge | ENSG00000 |
| 25152547 | 25152883 | 1 | processed_pseudoge | ENSG00000 |
| 25181359 | 25181745 | 1 | processed_pseudoge | ENSG00000 |
| 25218688 | 25219069 | 1 | processed_pseudoge | ENSG00000 |
| 25248263 | 25248779 | 1 | processed_pseudoge | ENSG00000 |
| 25261239 | 25261731 | 1 | processed_pseudoge | ENSG00000 |
| 25272200 | 25273751 | -1 | processed_pseudoge | ENSG00000 |
| 25279078 | 25620530 | 1 | protein_coding | ENSG00000 |
| 25287432 | 25287533 | -1 | misc_RNA | ENSG00000 |
| 25383599 | 25383702 | 1 | snRNA | ENSG00000 |
| 25632673 | 25640931 | -1 | lncRNA | ENSG00000 |
| 25652201 | 25701783 | 1 | protein_coding | ENSG00000 |
| 25678769 | 25679164 | -1 | processed_pseudoge | ENSG00000 |
| 25726132 | 25726527 | -1 | protein_coding | ENSG00000 |
| 25726777 | 25727292 | 1 | protein_coding | ENSG00000 |
| 25731782 | 25732091 | -1 | unprocessed_pseudo | ENSG00000 |
| 25732497 | 25732827 | 1 | unprocessed_pseudo | ENSG00000 |
| 25754699 | 25781199 | 1 | protein_coding | ENSG00000 |
| 25782902 | 25832052 | -1 | protein_coding | ENSG00000 |
| 25833066 | 25882286 | -1 | protein_coding | ENSG00000 |
| 25882026 | 25882395 | -1 | transcribed_processe | ENSG00000 |
| 25885419 | 25885751 | 1 | processed_pseudoge | ENSG00000 |
| 25912754 | 25930726 | -1 | protein_coding | ENSG00000 |
| 25962802 | 25991231 | 1 | protein_coding | ENSG00000 |
| 25983812 | 25999167 | -1 | lncRNA | ENSG00000 |
| 25992641 | 26001775 | 1 | lncRNA | ENSG00000 |
| 26013241 | 26013757 | 1 | lncRNA | ENSG00000 |
| 26016379 | 26016601 | 1 | unprocessed_pseudo | ENSG00000 |
| 26017085 | 26017732 | -1 | protein_coding | ENSG00000 |
| 26020490 | 26020900 | 1 | protein_coding | ENSG00000 |
| 26021679 | 26021990 | 1 | protein_coding | ENSG00000 |

| | | | | |
|---|---|---|---|---|
| 26026815 | 26027252 | -1 | protein_coding | ENSG0000C |
| 26031594 | 26032099 | -1 | protein_coding | ENSG0000C |
| 26033176 | 26033568 | -1 | protein_coding | ENSG0000C |
| 26043277 | 26043657 | -1 | protein_coding | ENSG0000C |
| 26043989 | 26044357 | 1 | unprocessed_pseudc | ENSG0000C |
| 26045411 | 26045869 | 1 | protein_coding | ENSG0000C |
| 26055787 | 26056428 | -1 | protein_coding | ENSG0000C |
| 26087281 | 26098343 | 1 | protein_coding | ENSG0000C |
| 26103876 | 26104310 | 1 | protein_coding | ENSG0000C |
| 26107419 | 26108136 | -1 | protein_coding | ENSG0000C |
| 26114873 | 26123926 | -1 | protein_coding | ENSG0000C |
| 26124145 | 26139116 | 1 | protein_coding | ENSG0000C |
| 26156354 | 26157107 | 1 | protein_coding | ENSG0000C |
| 26158146 | 26171349 | 1 | protein_coding | ENSG0000C |
| 26160765 | 26172802 | -1 | lncRNA | ENSG0000C |
| 26164072 | 26164363 | 1 | processed_pseudoge | ENSG0000C |
| 26172059 | 26184655 | 1 | protein_coding | ENSG0000C |
| 26188765 | 26189076 | -1 | protein_coding | ENSG0000C |
| 26195554 | 26196633 | 1 | unprocessed_pseudc | ENSG0000C |
| 26195595 | 26199293 | -1 | protein_coding | ENSG0000C |
| 26196840 | 26197250 | -1 | protein_coding | ENSG0000C |
| 26198851 | 26199243 | -1 | protein_coding | ENSG0000C |
| 26199520 | 26200715 | 1 | protein_coding | ENSG0000C |
| 26202156 | 26202654 | 1 | processed_pseudoge | ENSG0000C |
| 26204552 | 26206038 | 1 | protein_coding | ENSG0000C |
| 26215159 | 26216692 | -1 | protein_coding | ENSG0000C |
| 26216975 | 26217483 | 1 | protein_coding | ENSG0000C |
| 26219045 | 26219240 | 1 | unprocessed_pseudc | ENSG0000C |
| 26224199 | 26227473 | 1 | protein_coding | ENSG0000C |
| 26233122 | 26233255 | 1 | unprocessed_pseudc | ENSG0000C |
| 26234268 | 26234933 | -1 | protein_coding | ENSG0000C |
| 26240426 | 26240737 | 1 | protein_coding | ENSG0000C |
| 26246681 | 26246977 | -1 | protein_coding | ENSG0000C |
| 26250195 | 26250605 | -1 | protein_coding | ENSG0000C |
| 26251651 | 26253710 | 1 | protein_coding | ENSG0000C |
| 26269405 | 26271815 | -1 | protein_coding | ENSG0000C |
| 26272021 | 26272701 | -1 | transcribed_unproce | ENSG0000C |
| 26272976 | 26273356 | 1 | protein_coding | ENSG0000C |
| 26277609 | 26285638 | -1 | protein_coding | ENSG0000C |
| 26321876 | 26322292 | -1 | processed_pseudoge | ENSG0000C |
| 26324705 | 26331900 | -1 | lncRNA | ENSG0000C |
| 26365159 | 26378320 | 1 | protein_coding | ENSG0000C |
| 26383096 | 26394874 | 1 | protein_coding | ENSG0000C |
| 26402237 | 26415208 | 1 | protein_coding | ENSG0000C |
| 26421391 | 26432383 | 1 | transcribed_unproce | ENSG0000C |
| 26440472 | 26453415 | 1 | protein_coding | ENSG0000C |
| 26457904 | 26476621 | 1 | protein_coding | ENSG0000C |
| 26477872 | 26478533 | 1 | unprocessed_pseudc | ENSG0000C |
| 26501221 | 26510422 | 1 | protein_coding | ENSG0000C |
| 26519351 | 26519458 | -1 | snRNA | ENSG0000C |

| | | | | |
|---|---|---|---|---|
| 26521709 | 26527404 | 1 | lncRNA | ENSG0000C |
| 26538366 | 26546933 | 1 | protein_coding | ENSG0000C |
| 26569324 | 26574698 | 1 | lncRNA | ENSG0000C |
| 26596953 | 26600739 | 1 | protein_coding | ENSG0000C |
| 26602733 | 26606661 | 1 | lncRNA | ENSG0000C |
| 26611903 | 26613109 | -1 | processed_pseudoge | ENSG0000C |
| 26634383 | 26659752 | -1 | protein_coding | ENSG0000C |
| 26677003 | 26683211 | 1 | lncRNA | ENSG0000C |
| 26686241 | 26687964 | 1 | lncRNA | ENSG0000C |
| 26871484 | 26956554 | -1 | transcribed_unproce | ENSG0000C |
| 26896952 | 26898777 | 1 | processed_pseudoge | ENSG0000C |
| 26956932 | 27059749 | 1 | lncRNA | ENSG0000C |
| 27001208 | 27001648 | -1 | lncRNA | ENSG0000C |
| 27018935 | 27020280 | -1 | lncRNA | ENSG0000C |
| 27033435 | 27034310 | -1 | unitary_pseudogene | ENSG0000C |
| 27060365 | 27061249 | -1 | processed_pseudoge | ENSG0000C |
| 27083280 | 27084147 | 1 | unitary_pseudogene | ENSG0000C |
| 27109790 | 27109980 | -1 | snRNA | ENSG0000C |
| 27122657 | 27123221 | -1 | lncRNA | ENSG0000C |
| 27125897 | 27132750 | -1 | protein_coding | ENSG0000C |
| 27133042 | 27135291 | 1 | protein_coding | ENSG0000C |
| 27138588 | 27139881 | 1 | protein_coding | ENSG0000C |
| 27146418 | 27146798 | -1 | protein_coding | ENSG0000C |
| 27147129 | 27147515 | 1 | protein_coding | ENSG0000C |
| 27147626 | 27147688 | 1 | miRNA | ENSG0000C |
| 27211244 | 27211913 | 1 | processed_pseudoge | ENSG0000C |
| 27247701 | 27256624 | 1 | protein_coding | ENSG0000C |
| 27268113 | 27269229 | 1 | processed_pseudoge | ENSG0000C |
| 27285903 | 27312170 | -1 | protein_coding | ENSG0000C |
| 27324894 | 27325768 | 1 | unprocessed_pseudc | ENSG0000C |
| 27357825 | 27360221 | -1 | processed_pseudoge | ENSG0000C |
| 27374615 | 27403904 | 1 | protein_coding | ENSG0000C |
| 27404010 | 27406964 | -1 | lncRNA | ENSG0000C |
| 27407697 | 27407996 | 1 | processed_pseudoge | ENSG0000C |
| 27450743 | 27473118 | -1 | protein_coding | ENSG0000C |
| 27454568 | 27457575 | 1 | lncRNA | ENSG0000C |
| 27473194 | 27494436 | 1 | lncRNA | ENSG0000C |
| 27491422 | 27519435 | -1 | lncRNA | ENSG0000C |
| 27510501 | 27510694 | 1 | processed_pseudoge | ENSG0000C |
| 27515039 | 27515274 | -1 | processed_pseudoge | ENSG0000C |
| 27523076 | 27524339 | 1 | processed_pseudoge | ENSG0000C |
| 27560822 | 27561744 | -1 | processed_pseudoge | ENSG0000C |
| 27596412 | 27596517 | 1 | snRNA | ENSG0000C |
| 27652602 | 27652825 | -1 | processed_pseudoge | ENSG0000C |
| 27679971 | 27680555 | -1 | lncRNA | ENSG0000C |
| 27694035 | 27713352 | 1 | lncRNA | ENSG0000C |
| 27737000 | 27738494 | -1 | processed_pseudoge | ENSG0000C |
| 27759870 | 27763184 | 1 | lncRNA | ENSG0000C |
| 27780619 | 27781111 | -1 | processed_pseudoge | ENSG0000C |
| 27807075 | 27807339 | 1 | processed_pseudoge | ENSG0000C |

| | | | | |
|---|---|---|---|---|
| 27807444 | 27807931 | -1 | protein_coding | ENSG0000C |
| 27808199 | 27808701 | 1 | protein_coding | ENSG0000C |
| 27810064 | 27811300 | 1 | protein_coding | ENSG0000C |
| 27814354 | 27814740 | -1 | protein_coding | ENSG0000C |
| 27815044 | 27815424 | 1 | protein_coding | ENSG0000C |
| 27824108 | 27824480 | 1 | protein_coding | ENSG0000C |
| 27831216 | 27831527 | -1 | protein_coding | ENSG0000C |
| 27837760 | 27865798 | 1 | protein_coding | ENSG0000C |
| 27837947 | 27838339 | -1 | protein_coding | ENSG0000C |
| 27864062 | 27864401 | -1 | processed_pseudoge | ENSG0000C |
| 27865355 | 27865747 | 1 | protein_coding | ENSG0000C |
| 27866849 | 27867529 | -1 | protein_coding | ENSG0000C |
| 27871905 | 27872315 | -1 | protein_coding | ENSG0000C |
| 27873199 | 27873510 | -1 | protein_coding | ENSG0000C |
| 27890382 | 27893106 | -1 | protein_coding | ENSG0000C |
| 27892757 | 27893149 | -1 | protein_coding | ENSG0000C |
| 27893463 | 27893843 | 1 | protein_coding | ENSG0000C |
| 27897504 | 27897566 | -1 | snRNA | ENSG0000C |
| 27911185 | 27912396 | -1 | protein_coding | ENSG0000C |
| 27937465 | 27938403 | 1 | unprocessed_pseudc | ENSG0000C |
| 27957241 | 27958182 | 1 | protein_coding | ENSG0000C |
| 27965175 | 27965514 | 1 | processed_pseudoge | ENSG0000C |
| 27977150 | 27978075 | 1 | unprocessed_pseudc | ENSG0000C |
| 28010723 | 28012355 | 1 | processed_pseudoge | ENSG0000C |
| 28015122 | 28137293 | -1 | lncRNA | ENSG0000C |
| 28015568 | 28015777 | 1 | snoRNA | ENSG0000C |
| 28033947 | 28034855 | 1 | unprocessed_pseudc | ENSG0000C |
| 28046434 | 28047367 | 1 | unprocessed_pseudc | ENSG0000C |
| 28053228 | 28054165 | -1 | unprocessed_pseudc | ENSG0000C |
| 28073316 | 28074260 | 1 | unprocessed_pseudc | ENSG0000C |
| 28078792 | 28081130 | -1 | lncRNA | ENSG0000C |
| 28080568 | 28089563 | 1 | protein_coding | ENSG0000C |
| 28091154 | 28093664 | 1 | transcribed_unproce | ENSG0000C |
| 28115628 | 28116551 | 1 | unprocessed_pseudc | ENSG0000C |
| 28124609 | 28130082 | 1 | protein_coding | ENSG0000C |
| 28136849 | 28139678 | 1 | lncRNA | ENSG0000C |
| 28141883 | 28159460 | 1 | protein_coding | ENSG0000C |
| 28161769 | 28169594 | 1 | transcribed_unproce | ENSG0000C |
| 28170845 | 28172521 | 1 | TEC | ENSG0000C |
| 28176188 | 28176674 | 1 | unprocessed_pseudc | ENSG0000C |
| 28188050 | 28189432 | 1 | unprocessed_pseudc | ENSG0000C |
| 28217643 | 28218634 | -1 | processed_pseudoge | ENSG0000C |
| 28224886 | 28233482 | 1 | protein_coding | ENSG0000C |
| 28241697 | 28252269 | -1 | protein_coding | ENSG0000C |
| 28259297 | 28260958 | 1 | protein_coding | ENSG0000C |
| 28267010 | 28278224 | 1 | protein_coding | ENSG0000C |
| 28267121 | 28281580 | 1 | protein_coding | ENSG0000C |
| 28281572 | 28302549 | 1 | protein_coding | ENSG0000C |
| 28315613 | 28315883 | -1 | processed_pseudoge | ENSG0000C |
| 28319660 | 28319852 | -1 | processed_pseudoge | ENSG0000C |

| | | | | |
|---|---|---|---|---|
| 28324693 | 28356271 | -1 | protein_coding | ENSG00000( |
| 28349947 | 28369172 | 1 | protein_coding | ENSG00000( |
| 28378955 | 28399734 | -1 | protein_coding | ENSG00000( |
| 28410270 | 28410456 | 1 | snRNA | ENSG00000( |
| 28431930 | 28443502 | -1 | protein_coding | ENSG00000( |
| 28446973 | 28447807 | 1 | processed_pseudoge | ENSG00000( |
| 28455648 | 28456141 | -1 | unprocessed_pseudc | ENSG00000( |
| 28476996 | 28489455 | -1 | lncRNA | ENSG00000( |
| 28489896 | 28491661 | 1 | lncRNA | ENSG00000( |
| 28503296 | 28528215 | -1 | protein_coding | ENSG00000( |
| 28525925 | 28534952 | 1 | protein_coding | ENSG00000( |
| 28570535 | 28616212 | -1 | protein_coding | ENSG00000( |
| 28587378 | 28591747 | 1 | lncRNA | ENSG00000( |
| 28633381 | 28633594 | -1 | processed_pseudoge | ENSG00000( |
| 28634861 | 28649538 | 1 | lncRNA | ENSG00000( |
| 28648286 | 28649066 | -1 | lncRNA | ENSG00000( |
| 28732017 | 28732904 | 1 | processed_pseudoge | ENSG00000( |
| 28783633 | 28784004 | -1 | processed_pseudoge | ENSG00000( |
| 28837869 | 28839006 | -1 | lncRNA | ENSG00000( |
| 28859625 | 28864630 | -1 | lncRNA | ENSG00000( |
| 28861416 | 28862047 | 1 | processed_pseudoge | ENSG00000( |
| 28888832 | 28889906 | -1 | processed_pseudoge | ENSG00000( |
| 28896530 | 28897322 | 1 | lncRNA | ENSG00000( |
| 28903002 | 28923988 | -1 | protein_coding | ENSG00000( |
| 28915645 | 28915744 | 1 | snRNA | ENSG00000( |
| 28943877 | 28944537 | 1 | lncRNA | ENSG00000( |
| 28969130 | 28970460 | -1 | processed_pseudoge | ENSG00000( |
| 28977475 | 28977773 | 1 | misc_RNA | ENSG00000( |
| 28986203 | 28987484 | 1 | lncRNA | ENSG00000( |
| 28986800 | 28988536 | 1 | lncRNA | ENSG00000( |
| 28994785 | 29005316 | -1 | protein_coding | ENSG00000( |
| 29026680 | 29027607 | -1 | unprocessed_pseudc | ENSG00000( |
| 29036021 | 29076524 | 1 | lncRNA | ENSG00000( |
| 29044213 | 29045240 | -1 | protein_coding | ENSG00000( |
| 29071824 | 29072599 | -1 | unprocessed_pseudc | ENSG00000( |
| 29076573 | 29077168 | -1 | processed_pseudoge | ENSG00000( |
| 29086208 | 29087313 | -1 | protein_coding | ENSG00000( |
| 29099657 | 29102701 | 1 | protein_coding | ENSG00000( |
| 29108059 | 29114770 | 1 | protein_coding | ENSG00000( |
| 29124210 | 29128908 | 1 | lncRNA | ENSG00000( |
| 29137410 | 29142934 | -1 | transcribed_unproce | ENSG00000( |
| 29162475 | 29166201 | -1 | lncRNA | ENSG00000( |
| 29170907 | 29175811 | 1 | protein_coding | ENSG00000( |
| 29181510 | 29182445 | 1 | unprocessed_pseudc | ENSG00000( |
| 29212886 | 29216933 | 1 | transcribed_unproce | ENSG00000( |
| 29223973 | 29292963 | 1 | transcribed_unproce | ENSG00000( |
| 29229227 | 29230157 | -1 | unprocessed_pseudc | ENSG00000( |
| 29262703 | 29264079 | -1 | unprocessed_pseudc | ENSG00000( |
| 29268462 | 29269421 | -1 | unprocessed_pseudc | ENSG00000( |
| 29301701 | 29313017 | 1 | protein_coding | ENSG00000( |

| | | | | |
|---|---|---|---|---|
| 29329626 | 29331061 | -1 | processed_pseudoge | ENSG00000 |
| 29353749 | 29431967 | -1 | protein_coding | ENSG00000 |
| 29373423 | 29375291 | -1 | protein_coding | ENSG00000 |
| 29395631 | 29398008 | 1 | protein_coding | ENSG00000 |
| 29414928 | 29418905 | 1 | polymorphic_pseudo | ENSG00000 |
| 29425504 | 29457071 | -1 | protein_coding | ENSG00000 |
| 29439306 | 29440977 | 1 | protein_coding | ENSG00000 |
| 29450210 | 29450558 | 1 | processed_pseudoge | ENSG00000 |
| 29457181 | 29464328 | 1 | protein_coding | ENSG00000 |
| 29464596 | 29469806 | -1 | unprocessed_pseudo | ENSG00000 |
| 29475025 | 29476071 | -1 | unprocessed_pseudo | ENSG00000 |
| 29486697 | 29487956 | -1 | protein_coding | ENSG00000 |
| 29489271 | 29489676 | -1 | processed_pseudoge | ENSG00000 |
| 29497475 | 29510558 | 1 | lncRNA | ENSG00000 |
| 29528721 | 29533665 | 1 | lncRNA | ENSG00000 |
| 29537704 | 29538787 | -1 | unprocessed_pseudo | ENSG00000 |
| 29550407 | 29557721 | 1 | protein_coding | ENSG00000 |
| 29555515 | 29559732 | -1 | protein_coding | ENSG00000 |
| 29555629 | 29633976 | -1 | protein_coding | ENSG00000 |
| 29573909 | 29574836 | 1 | unprocessed_pseudo | ENSG00000 |
| 29577459 | 29577748 | 1 | processed_pseudoge | ENSG00000 |
| 29582249 | 29582332 | 1 | snoRNA | ENSG00000 |
| 29582508 | 29583025 | 1 | processed_pseudoge | ENSG00000 |
| 29585121 | 29590500 | 1 | protein_coding | ENSG00000 |
| 29636060 | 29636343 | -1 | processed_pseudoge | ENSG00000 |
| 29657002 | 29672372 | 1 | protein_coding | ENSG00000 |
| 29672392 | 29681110 | -1 | protein_coding | ENSG00000 |
| 29708125 | 29708547 | -1 | processed_pseudoge | ENSG00000 |
| 29722775 | 29738528 | 1 | protein_coding | ENSG00000 |
| 29722981 | 29723971 | -1 | unprocessed_pseudo | ENSG00000 |
| 29726601 | 29749049 | -1 | lncRNA | ENSG00000 |
| 29726669 | 29727139 | -1 | transcribed_process | ENSG00000 |
| 29741731 | 29748969 | -1 | transcribed_unproce | ENSG00000 |
| 29748289 | 29748513 | 1 | unprocessed_pseudo | ENSG00000 |
| 29750371 | 29750472 | 1 | misc_RNA | ENSG00000 |
| 29750729 | 29751148 | -1 | processed_pseudoge | ENSG00000 |
| 29751965 | 29752207 | -1 | lncRNA | ENSG00000 |
| 29752573 | 29763295 | 1 | lncRNA | ENSG00000 |
| 29763258 | 29763392 | -1 | processed_pseudoge | ENSG00000 |
| 29790954 | 29797811 | 1 | transcribed_unproce | ENSG00000 |
| 29791753 | 29792750 | -1 | unprocessed_pseudo | ENSG00000 |
| 29800415 | 29802425 | 1 | unprocessed_pseudo | ENSG00000 |
| 29803195 | 29803991 | -1 | processed_pseudoge | ENSG00000 |
| 29812565 | 29812696 | -1 | unprocessed_pseudo | ENSG00000 |
| 29826967 | 29831125 | 1 | protein_coding | ENSG00000 |
| 29827385 | 29828364 | -1 | unprocessed_pseudo | ENSG00000 |
| 29849550 | 29885615 | -1 | transcribed_unproce | ENSG00000 |
| 29871895 | 29873783 | -1 | lncRNA | ENSG00000 |
| 29887294 | 29888268 | -1 | unprocessed_pseudo | ENSG00000 |
| 29887752 | 29890482 | 1 | unprocessed_pseudo | ENSG00000 |

| | | | | |
|---|---|---|---|---|
| 29896654 | 29897786 | 1 | unprocessed_pseudc | ENSG0000C |
| 29906543 | 29906909 | 1 | unprocessed_pseudc | ENSG0000C |
| 29907783 | 29908645 | -1 | unprocessed_pseudc | ENSG0000C |
| 29925983 | 29926973 | -1 | unprocessed_pseudc | ENSG0000C |
| 29926459 | 29929232 | 1 | unprocessed_pseudc | ENSG0000C |
| 29934101 | 29934286 | 1 | unprocessed_pseudc | ENSG0000C |
| 29941260 | 29945884 | 1 | protein_coding | ENSG0000C |
| 29942075 | 29943067 | -1 | unprocessed_pseudc | ENSG0000C |
| 29956596 | 29958570 | 1 | unprocessed_pseudc | ENSG0000C |
| 29970801 | 29972464 | -1 | unprocessed_pseudc | ENSG0000C |
| 29975112 | 29978410 | 1 | lncRNA | ENSG0000C |
| 29993209 | 29993605 | 1 | unprocessed_pseudc | ENSG0000C |
| 29994437 | 29995314 | -1 | unprocessed_pseudc | ENSG0000C |
| 30001011 | 30061640 | -1 | transcribed_unitary_ | ENSG0000C |
| 30005971 | 30009956 | 1 | transcribed_unproce | ENSG0000C |
| 30006121 | 30007116 | -1 | unprocessed_pseudc | ENSG0000C |
| 30031713 | 30033003 | 1 | processed_pseudoge | ENSG0000C |
| 30035916 | 30035983 | 1 | misc_RNA | ENSG0000C |
| 30058115 | 30058190 | 1 | misc_RNA | ENSG0000C |
| 30058899 | 30064909 | 1 | protein_coding | ENSG0000C |
| 30061080 | 30061183 | 1 | misc_RNA | ENSG0000C |
| 30066709 | 30070333 | 1 | protein_coding | ENSG0000C |
| 30070266 | 30075887 | -1 | protein_coding | ENSG0000C |
| 30102897 | 30113090 | -1 | protein_coding | ENSG0000C |
| 30105240 | 30114724 | 1 | lncRNA | ENSG0000C |
| 30132805 | 30132966 | -1 | snoRNA | ENSG0000C |
| 30136124 | 30148735 | 1 | protein_coding | ENSG0000C |
| 30151943 | 30161211 | -1 | protein_coding | ENSG0000C |
| 30163206 | 30172693 | 1 | protein_coding | ENSG0000C |
| 30184455 | 30213427 | -1 | protein_coding | ENSG0000C |
| 30186798 | 30188614 | -1 | processed_pseudoge | ENSG0000C |
| 30234039 | 30326134 | -1 | lncRNA | ENSG0000C |
| 30238301 | 30242279 | 1 | unprocessed_pseudc | ENSG0000C |
| 30259584 | 30293014 | 1 | transcribed_unproce | ENSG0000C |
| 30282349 | 30286054 | -1 | TEC | ENSG0000C |
| 30286690 | 30327382 | -1 | lncRNA | ENSG0000C |
| 30326479 | 30343729 | 1 | protein_coding | ENSG0000C |
| 30328907 | 30346854 | 1 | protein_coding | ENSG0000C |
| 30345131 | 30346884 | 1 | protein_coding | ENSG0000C |
| 30351416 | 30351550 | 1 | unprocessed_pseudc | ENSG0000C |
| 30359278 | 30359911 | -1 | processed_pseudoge | ENSG0000C |
| 30363112 | 30364280 | -1 | processed_pseudoge | ENSG0000C |
| 30414715 | 30419319 | 1 | unprocessed_pseudc | ENSG0000C |
| 30451139 | 30456212 | 1 | lncRNA | ENSG0000C |
| 30466452 | 30467994 | 1 | processed_pseudoge | ENSG0000C |
| 30468882 | 30470251 | 1 | processed_pseudoge | ENSG0000C |
| 30485940 | 30486590 | 1 | processed_pseudoge | ENSG0000C |
| 30489509 | 30494194 | 1 | protein_coding | ENSG0000C |
| 30516266 | 30519217 | 1 | lncRNA | ENSG0000C |
| 30541377 | 30557174 | -1 | protein_coding | ENSG0000C |

| | | | | |
|---|---|---|---|---|
| 30556886 | 30563723 | 1 | protein_coding | ENSG0000C |
| 30571393 | 30597179 | 1 | protein_coding | ENSG0000C |
| 30584332 | 30584417 | 1 | miRNA | ENSG0000C |
| 30600413 | 30618612 | -1 | protein_coding | ENSG0000C |
| 30617840 | 30626395 | 1 | protein_coding | ENSG0000C |
| 30626842 | 30646823 | 1 | protein_coding | ENSG0000C |
| 30633632 | 30633956 | 1 | processed_pseudoge | ENSG0000C |
| 30647039 | 30653210 | 1 | protein_coding | ENSG0000C |
| 30653119 | 30673006 | -1 | protein_coding | ENSG0000C |
| 30676389 | 30687895 | -1 | protein_coding | ENSG0000C |
| 30688047 | 30691420 | -1 | protein_coding | ENSG0000C |
| 30696806 | 30697535 | -1 | processed_pseudoge | ENSG0000C |
| 30699807 | 30717447 | -1 | protein_coding | ENSG0000C |
| 30703067 | 30713184 | 1 | lncRNA | ENSG0000C |
| 30720352 | 30725426 | 1 | protein_coding | ENSG0000C |
| 30723105 | 30723877 | -1 | lncRNA | ENSG0000C |
| 30727709 | 30742732 | -1 | protein_coding | ENSG0000C |
| 30736304 | 30736414 | -1 | misc_RNA | ENSG0000C |
| 30742929 | 30743592 | 1 | lncRNA | ENSG0000C |
| 30743199 | 30744548 | -1 | protein_coding | ENSG0000C |
| 30743790 | 30792250 | 1 | lncRNA | ENSG0000C |
| 30751038 | 30751289 | 1 | misc_RNA | ENSG0000C |
| 30798654 | 30830659 | -1 | lncRNA | ENSG0000C |
| 30839526 | 30848159 | -1 | lncRNA | ENSG0000C |
| 30864250 | 30864552 | 1 | misc_RNA | ENSG0000C |
| 30866982 | 30875918 | -1 | lncRNA | ENSG0000C |
| 30876421 | 30900156 | 1 | protein_coding | ENSG0000C |
| 30890883 | 30890972 | 1 | miRNA | ENSG0000C |
| 30908207 | 30914106 | 1 | protein_coding | ENSG0000C |
| 30908242 | 30926459 | 1 | lncRNA | ENSG0000C |
| 30914205 | 30926459 | 1 | protein_coding | ENSG0000C |
| 30931353 | 30955636 | -1 | protein_coding | ENSG0000C |
| 30932618 | 30932709 | -1 | misc_RNA | ENSG0000C |
| 30934523 | 30954221 | 1 | protein_coding | ENSG0000C |
| 30945979 | 30954862 | -1 | lncRNA | ENSG0000C |
| 30961403 | 30962396 | -1 | processed_pseudoge | ENSG0000C |
| 30983718 | 30989903 | 1 | protein_coding | ENSG0000C |
| 31010474 | 31035402 | 1 | protein_coding | ENSG0000C |
| 31053450 | 31059890 | 1 | lncRNA | ENSG0000C |
| 31083010 | 31083109 | 1 | snRNA | ENSG0000C |
| 31111223 | 31112575 | -1 | protein_coding | ENSG0000C |
| 31114750 | 31140092 | 1 | protein_coding | ENSG0000C |
| 31115087 | 31120446 | -1 | protein_coding | ENSG0000C |
| 31137534 | 31139066 | -1 | protein_coding | ENSG0000C |
| 31140727 | 31140913 | 1 | processed_pseudoge | ENSG0000C |
| 31142439 | 31158238 | -1 | protein_coding | ENSG0000C |
| 31158547 | 31167159 | 1 | protein_coding | ENSG0000C |
| 31164337 | 31180731 | -1 | protein_coding | ENSG0000C |
| 31173735 | 31177899 | -1 | lncRNA | ENSG0000C |
| 31195200 | 31198037 | -1 | lncRNA | ENSG0000C |

| | | | | |
|---|---|---|---|---|
| 31197760 | 31203968 | 1 | lncRNA | ENSG0000C |
| 31200165 | 31201918 | -1 | lncRNA | ENSG0000C |
| 31222913 | 31223093 | 1 | unprocessed_pseudc | ENSG0000C |
| 31224342 | 31225058 | -1 | unprocessed_pseudc | ENSG0000C |
| 31268749 | 31272130 | -1 | protein_coding | ENSG0000C |
| 31269491 | 31357188 | -1 | protein_coding | ENSG0000C |
| 31275572 | 31278754 | 1 | processed_pseudoge | ENSG0000C |
| 31280317 | 31281519 | 1 | processed_pseudoge | ENSG0000C |
| 31287510 | 31288964 | -1 | processed_pseudoge | ENSG0000C |
| 31293908 | 31301642 | -1 | lncRNA | ENSG0000C |
| 31307815 | 31308549 | -1 | unprocessed_pseudc | ENSG0000C |
| 31355224 | 31355316 | -1 | miRNA | ENSG0000C |
| 31356647 | 31357637 | 1 | unprocessed_pseudc | ENSG0000C |
| 31366352 | 31366898 | -1 | processed_pseudoge | ENSG0000C |
| 31367057 | 31376088 | 1 | lncRNA | ENSG0000C |
| 31370134 | 31370240 | 1 | snRNA | ENSG0000C |
| 31377419 | 31378019 | 1 | processed_pseudoge | ENSG0000C |
| 31380411 | 31380839 | 1 | processed_pseudoge | ENSG0000C |
| 31382074 | 31382288 | -1 | unprocessed_pseudc | ENSG0000C |
| 31394289 | 31395495 | -1 | lncRNA | ENSG0000C |
| 31399784 | 31415315 | 1 | protein_coding | ENSG0000C |
| 31400700 | 31478903 | 1 | lncRNA | ENSG0000C |
| 31402152 | 31402250 | -1 | misc_RNA | ENSG0000C |
| 31441667 | 31446973 | 1 | lncRNA | ENSG0000C |
| 31462728 | 31463336 | -1 | unprocessed_pseudc | ENSG0000C |
| 31479973 | 31494935 | -1 | lncRNA | ENSG0000C |
| 31494881 | 31511124 | 1 | protein_coding | ENSG0000C |
| 31496689 | 31496790 | -1 | misc_RNA | ENSG0000C |
| 31515979 | 31516211 | 1 | unprocessed_pseudc | ENSG0000C |
| 31519480 | 31520291 | -1 | processed_pseudoge | ENSG0000C |
| 31528114 | 31528693 | 1 | processed_pseudoge | ENSG0000C |
| 31528962 | 31530232 | 1 | protein_coding | ENSG0000C |
| 31530219 | 31546608 | -1 | protein_coding | ENSG0000C |
| 31530219 | 31542448 | -1 | protein_coding | ENSG0000C |
| 31536374 | 31536449 | -1 | snoRNA | ENSG0000C |
| 31541101 | 31541178 | -1 | snoRNA | ENSG0000C |
| 31542304 | 31543138 | 1 | lncRNA | ENSG0000C |
| 31544462 | 31548427 | -1 | protein_coding | ENSG0000C |
| 31546870 | 31558829 | 1 | protein_coding | ENSG0000C |
| 31572054 | 31574324 | 1 | protein_coding | ENSG0000C |
| 31575565 | 31578336 | 1 | protein_coding | ENSG0000C |
| 31580525 | 31582522 | -1 | protein_coding | ENSG0000C |
| 31586124 | 31588909 | 1 | protein_coding | ENSG0000C |
| 31588895 | 31593006 | -1 | protein_coding | ENSG0000C |
| 31611083 | 31611356 | -1 | processed_pseudoge | ENSG0000C |
| 31615217 | 31617021 | 1 | protein_coding | ENSG0000C |
| 31620715 | 31637771 | 1 | protein_coding | ENSG0000C |
| 31623079 | 31623210 | 1 | snoRNA | ENSG0000C |
| 31633787 | 31633858 | 1 | miRNA | ENSG0000C |
| 31639028 | 31652705 | -1 | protein_coding | ENSG0000C |

| | | | | |
|---|---|---|---|---|
| 31652416 | 31658210 | 1 | protein_coding | ENSG0000C |
| 31658298 | 31660778 | -1 | protein_coding | ENSG0000C |
| 31658329 | 31660721 | 1 | lncRNA | ENSG0000C |
| 31661228 | 31666283 | -1 | protein_coding | ENSG0000C |
| 31663288 | 31663401 | 1 | misc_RNA | ENSG0000C |
| 31665236 | 31670343 | 1 | protein_coding | ENSG0000C |
| 31666102 | 31673546 | 1 | protein_coding | ENSG0000C |
| 31670167 | 31673776 | 1 | protein_coding | ENSG0000C |
| 31676684 | 31684040 | -1 | protein_coding | ENSG0000C |
| 31686955 | 31703356 | -1 | protein_coding | ENSG0000C |
| 31686962 | 31714072 | -1 | protein_coding | ENSG0000C |
| 31701029 | 31701091 | -1 | miRNA | ENSG0000C |
| 31706885 | 31710595 | 1 | protein_coding | ENSG0000C |
| 31706904 | 31717918 | 1 | protein_coding | ENSG0000C |
| 31711771 | 31714065 | -1 | protein_coding | ENSG0000C |
| 31715356 | 31717804 | 1 | protein_coding | ENSG0000C |
| 31718594 | 31726714 | 1 | protein_coding | ENSG0000C |
| 31718648 | 31721746 | -1 | protein_coding | ENSG0000C |
| 31727038 | 31730617 | -1 | protein_coding | ENSG0000C |
| 31730581 | 31739763 | -1 | protein_coding | ENSG0000C |
| 31739677 | 31762678 | 1 | protein_coding | ENSG0000C |
| 31740020 | 31764851 | 1 | protein_coding | ENSG0000C |
| 31756951 | 31757053 | -1 | snRNA | ENSG0000C |
| 31762996 | 31764851 | 1 | protein_coding | ENSG0000C |
| 31764310 | 31765588 | -1 | lncRNA | ENSG0000C |
| 31765590 | 31777328 | -1 | protein_coding | ENSG0000C |
| 31777518 | 31795752 | -1 | protein_coding | ENSG0000C |
| 31778817 | 31778905 | -1 | misc_RNA | ENSG0000C |
| 31797396 | 31806966 | -1 | protein_coding | ENSG0000C |
| 31809619 | 31815283 | -1 | protein_coding | ENSG0000C |
| 31815543 | 31817946 | 1 | protein_coding | ENSG0000C |
| 31827738 | 31830254 | 1 | protein_coding | ENSG0000C |
| 31828834 | 31839719 | 1 | lncRNA | ENSG0000C |
| 31834608 | 31839766 | 1 | lncRNA | ENSG0000C |
| 31835263 | 31835326 | 1 | snoRNA | ENSG0000C |
| 31837076 | 31837142 | 1 | snoRNA | ENSG0000C |
| 31857659 | 31862822 | -1 | protein_coding | ENSG0000C |
| 31863192 | 31879046 | -1 | protein_coding | ENSG0000C |
| 31877808 | 31884204 | 1 | lncRNA | ENSG0000C |
| 31879759 | 31897687 | -1 | protein_coding | ENSG0000C |
| 31897785 | 31945672 | 1 | protein_coding | ENSG0000C |
| 31899613 | 31902086 | -1 | protein_coding | ENSG0000C |
| 31927698 | 31952048 | 1 | protein_coding | ENSG0000C |
| 31934474 | 31941724 | -1 | lncRNA | ENSG0000C |
| 31945650 | 31952084 | 1 | protein_coding | ENSG0000C |
| 31952087 | 31959038 | -1 | protein_coding | ENSG0000C |
| 31956839 | 31956940 | -1 | miRNA | ENSG0000C |
| 31959117 | 31969751 | 1 | protein_coding | ENSG0000C |
| 31969810 | 31972290 | -1 | protein_coding | ENSG0000C |
| 31971091 | 31982821 | 1 | protein_coding | ENSG0000C |

| | | | | |
|---|---|---|---|---|
| 31982057 | 32002681 | 1 | protein_coding | ENSG00000( |
| 31999976 | 32003521 | -1 | lncRNA | ENSG00000( |
| 32005636 | 32008451 | 1 | transcribed_unproce | ENSG00000( |
| 32008614 | 32012472 | -1 | unprocessed_pseudc | ENSG00000( |
| 32013270 | 32013787 | 1 | unprocessed_pseudc | ENSG00000( |
| 32014795 | 32035418 | 1 | protein_coding | ENSG00000( |
| 32032713 | 32036258 | -1 | lncRNA | ENSG00000( |
| 32038327 | 32041644 | 1 | protein_coding | ENSG00000( |
| 32041153 | 32115334 | -1 | protein_coding | ENSG00000( |
| 32078508 | 32078628 | -1 | rRNA_pseudogene | ENSG00000( |
| 32098176 | 32117610 | -1 | lncRNA | ENSG00000( |
| 32108406 | 32112849 | 1 | lncRNA | ENSG00000( |
| 32115264 | 32128253 | -1 | protein_coding | ENSG00000( |
| 32128707 | 32130288 | -1 | protein_coding | ENSG00000( |
| 32148359 | 32153083 | -1 | protein_coding | ENSG00000( |
| 32150495 | 32154373 | -1 | protein_coding | ENSG00000( |
| 32152802 | 32154365 | -1 | lncRNA | ENSG00000( |
| 32153441 | 32163678 | 1 | protein_coding | ENSG00000( |
| 32153845 | 32171978 | 1 | protein_coding | ENSG00000( |
| 32164595 | 32168281 | 1 | protein_coding | ENSG00000( |
| 32168212 | 32178096 | -1 | protein_coding | ENSG00000( |
| 32170030 | 32170116 | -1 | miRNA | ENSG00000( |
| 32178405 | 32180793 | 1 | protein_coding | ENSG00000( |
| 32179816 | 32179876 | 1 | miRNA | ENSG00000( |
| 32180968 | 32184322 | -1 | protein_coding | ENSG00000( |
| 32184733 | 32190202 | -1 | protein_coding | ENSG00000( |
| 32184733 | 32185882 | -1 | lncRNA | ENSG00000( |
| 32190766 | 32195523 | -1 | protein_coding | ENSG00000( |
| 32194843 | 32224067 | -1 | protein_coding | ENSG00000( |
| 32254640 | 32407763 | 1 | lncRNA | ENSG00000( |
| 32288526 | 32371912 | -1 | protein_coding | ENSG00000( |
| 32325219 | 32326178 | 1 | processed_pseudoge | ENSG00000( |
| 32352877 | 32352983 | 1 | snRNA | ENSG00000( |
| 32393963 | 32407128 | -1 | protein_coding | ENSG00000( |
| 32439878 | 32445046 | 1 | protein_coding | ENSG00000( |
| 32459821 | 32473500 | -1 | unprocessed_pseudc | ENSG00000( |
| 32517353 | 32530287 | -1 | protein_coding | ENSG00000( |
| 32549940 | 32550090 | -1 | snRNA | ENSG00000( |
| 32552713 | 32560022 | -1 | transcribed_unproce | ENSG00000( |
| 32578769 | 32589848 | -1 | protein_coding | ENSG00000( |
| 32628179 | 32647062 | 1 | protein_coding | ENSG00000( |
| 32659467 | 32668383 | -1 | protein_coding | ENSG00000( |
| 32659880 | 32660729 | 1 | lncRNA | ENSG00000( |
| 32706124 | 32706955 | -1 | unprocessed_pseudc | ENSG00000( |
| 32718005 | 32719170 | 1 | lncRNA | ENSG00000( |
| 32730758 | 32731695 | -1 | unprocessed_pseudc | ENSG00000( |
| 32741391 | 32747198 | 1 | protein_coding | ENSG00000( |
| 32749912 | 32749979 | -1 | miRNA | ENSG00000( |
| 32756098 | 32763532 | -1 | protein_coding | ENSG00000( |
| 32812763 | 32820466 | -1 | protein_coding | ENSG00000( |

| | | | | |
|---|---|---|---|---|
| 32813767 | 32838822 | -1 | protein_coding | ENSG00000( |
| 32821833 | 32838770 | -1 | protein_coding | ENSG00000( |
| 32840717 | 32844679 | -1 | protein_coding | ENSG00000( |
| 32844078 | 32846500 | 1 | lncRNA | ENSG00000( |
| 32844136 | 32859851 | 1 | protein_coding | ENSG00000( |
| 32845209 | 32853978 | -1 | protein_coding | ENSG00000( |
| 32879171 | 32879848 | -1 | transcribed_processe | ENSG00000( |
| 32896416 | 32896490 | 1 | unprocessed_pseudo | ENSG00000( |
| 32934629 | 32941028 | -1 | protein_coding | ENSG00000( |
| 32937364 | 32953122 | -1 | protein_coding | ENSG00000( |
| 32948613 | 32969094 | -1 | protein_coding | ENSG00000( |
| 32968594 | 32981505 | 1 | protein_coding | ENSG00000( |
| 32970232 | 32970886 | 1 | lncRNA | ENSG00000( |
| 32972065 | 32972853 | -1 | lncRNA | ENSG00000( |
| 33004182 | 33009591 | -1 | protein_coding | ENSG00000( |
| 33064569 | 33080775 | -1 | protein_coding | ENSG00000( |
| 33075990 | 33089696 | 1 | protein_coding | ENSG00000( |
| 33079451 | 33079860 | 1 | processed_pseudoge | ENSG00000( |
| 33091753 | 33097295 | -1 | unprocessed_pseudo | ENSG00000( |
| 33103794 | 33107330 | -1 | unprocessed_pseudo | ENSG00000( |
| 33112451 | 33129084 | 1 | transcribed_unproce | ENSG00000( |
| 33131216 | 33143325 | -1 | unprocessed_pseudo | ENSG00000( |
| 33144783 | 33147767 | 1 | lncRNA | ENSG00000( |
| 33162681 | 33192499 | -1 | protein_coding | ENSG00000( |
| 33193588 | 33200688 | -1 | protein_coding | ENSG00000( |
| 33199601 | 33199696 | 1 | misc_RNA | ENSG00000( |
| 33200445 | 33204439 | 1 | protein_coding | ENSG00000( |
| 33204655 | 33206831 | 1 | protein_coding | ENSG00000( |
| 33207835 | 33207944 | 1 | miRNA | ENSG00000( |
| 33208500 | 33212722 | 1 | protein_coding | ENSG00000( |
| 33215705 | 33216328 | 1 | processed_pseudoge | ENSG00000( |
| 33237799 | 33239690 | 1 | processed_pseudoge | ENSG00000( |
| 33246075 | 33246856 | -1 | lncRNA | ENSG00000( |
| 33249534 | 33255167 | 1 | lncRNA | ENSG00000( |
| 33250272 | 33272047 | -1 | protein_coding | ENSG00000( |
| 33272075 | 33276511 | 1 | protein_coding | ENSG00000( |
| 33277123 | 33284832 | 1 | protein_coding | ENSG00000( |
| 33279108 | 33289247 | -1 | protein_coding | ENSG00000( |
| 33287227 | 33287289 | -1 | miRNA | ENSG00000( |
| 33289302 | 33298401 | 1 | protein_coding | ENSG00000( |
| 33290245 | 33290325 | 1 | miRNA | ENSG00000( |
| 33291654 | 33298942 | -1 | protein_coding | ENSG00000( |
| 33299694 | 33314387 | -1 | protein_coding | ENSG00000( |
| 33314406 | 33317942 | -1 | protein_coding | ENSG00000( |
| 33318558 | 33323016 | -1 | protein_coding | ENSG00000( |
| 33321386 | 33329286 | -1 | protein_coding | ENSG00000( |
| 33323625 | 33329269 | -1 | protein_coding | ENSG00000( |
| 33338978 | 33339495 | 1 | processed_pseudoge | ENSG00000( |
| 33365548 | 33366243 | -1 | processed_pseudoge | ENSG00000( |
| 33389374 | 33389475 | -1 | processed_pseudoge | ENSG00000( |

| | | | | |
|---|---|---|---|---|
| 33391823 | 33409896 | 1 | protein_coding | ENSG0000C |
| 33400059 | 33400556 | -1 | processed_pseudoge | ENSG0000C |
| 33410399 | 33416453 | 1 | protein_coding | ENSG0000C |
| 33416442 | 33418317 | -1 | protein_coding | ENSG0000C |
| 33419661 | 33453689 | 1 | protein_coding | ENSG0000C |
| 33437363 | 33454453 | -1 | lncRNA | ENSG0000C |
| 33438331 | 33438437 | 1 | miRNA | ENSG0000C |
| 33453970 | 33457544 | 1 | protein_coding | ENSG0000C |
| 33540046 | 33589026 | 1 | transcribed_unitary_ | ENSG0000C |
| 33540694 | 33541000 | 1 | misc_RNA | ENSG0000C |
| 33562880 | 33563130 | -1 | misc_RNA | ENSG0000C |
| 33572547 | 33580293 | -1 | protein_coding | ENSG0000C |
| 33586106 | 33593338 | -1 | lncRNA | ENSG0000C |
| 33620365 | 33696574 | 1 | protein_coding | ENSG0000C |
| 33694293 | 33711727 | -1 | protein_coding | ENSG0000C |
| 33698128 | 33698234 | 1 | miRNA | ENSG0000C |
| 33721662 | 33746905 | -1 | protein_coding | ENSG0000C |
| 33771202 | 33789130 | -1 | protein_coding | ENSG0000C |
| 33794673 | 33804003 | -1 | protein_coding | ENSG0000C |
| 33800142 | 33802960 | 1 | lncRNA | ENSG0000C |
| 33867506 | 33896914 | -1 | lncRNA | ENSG0000C |
| 33883170 | 33883411 | 1 | processed_pseudoge | ENSG0000C |
| 33891327 | 33927631 | 1 | lncRNA | ENSG0000C |
| 33899135 | 33899200 | 1 | miRNA | ENSG0000C |
| 33999972 | 34000051 | -1 | miRNA | ENSG0000C |
| 34018645 | 34155622 | -1 | protein_coding | ENSG0000C |
| 34189780 | 34191079 | 1 | processed_pseudoge | ENSG0000C |
| 34219439 | 34220066 | 1 | processed_pseudoge | ENSG0000C |
| 34236873 | 34246231 | 1 | protein_coding | ENSG0000C |
| 34240673 | 34240736 | 1 | miRNA | ENSG0000C |
| 34246381 | 34249108 | -1 | protein_coding | ENSG0000C |
| 34248535 | 34286768 | 1 | lncRNA | ENSG0000C |
| 34263311 | 34263673 | -1 | processed_pseudoge | ENSG0000C |
| 34279679 | 34392669 | -1 | protein_coding | ENSG0000C |
| 34284887 | 34426071 | -1 | protein_coding | ENSG0000C |
| 34417454 | 34426069 | -1 | protein_coding | ENSG0000C |
| 34466061 | 34535229 | 1 | protein_coding | ENSG0000C |
| 34537802 | 34556333 | -1 | protein_coding | ENSG0000C |
| 34576258 | 34576656 | -1 | processed_pseudoge | ENSG0000C |
| 34587288 | 34696859 | -1 | protein_coding | ENSG0000C |
| 34616538 | 34617258 | 1 | processed_pseudoge | ENSG0000C |
| 34685355 | 34685642 | -1 | misc_RNA | ENSG0000C |
| 34686602 | 34687106 | 1 | processed_pseudoge | ENSG0000C |
| 34696317 | 34697470 | 1 | lncRNA | ENSG0000C |
| 34715613 | 34715940 | 1 | processed_pseudoge | ENSG0000C |
| 34744176 | 34744673 | 1 | processed_pseudoge | ENSG0000C |
| 34757505 | 34773857 | 1 | protein_coding | ENSG0000C |
| 34792015 | 34883138 | 1 | protein_coding | ENSG0000C |
| 34821445 | 34821542 | 1 | misc_RNA | ENSG0000C |
| 34873831 | 34873927 | 1 | misc_RNA | ENSG0000C |

| | | | | |
|---|---|---|---|---|
| 34877462 | 34888071 | -1 | protein_coding | ENSG0000C |
| 34889255 | 35091406 | 1 | protein_coding | ENSG0000C |
| 35023522 | 35023831 | 1 | processed_pseudoge | ENSG0000C |
| 35041116 | 35058015 | -1 | lncRNA | ENSG0000C |
| 35070871 | 35071049 | -1 | processed_pseudoge | ENSG0000C |
| 35118071 | 35148610 | -1 | protein_coding | ENSG0000C |
| 35213956 | 35253079 | 1 | protein_coding | ENSG0000C |
| 35220370 | 35224630 | -1 | lncRNA | ENSG0000C |
| 35258909 | 35295985 | 1 | protein_coding | ENSG0000C |
| 35279177 | 35279819 | -1 | unprocessed_pseudc | ENSG0000C |
| 35297818 | 35321771 | 1 | protein_coding | ENSG0000C |
| 35342558 | 35428191 | 1 | protein_coding | ENSG0000C |
| 35443044 | 35449847 | 1 | unprocessed_pseudc | ENSG0000C |
| 35452356 | 35467103 | 1 | protein_coding | ENSG0000C |
| 35468401 | 35470785 | 1 | protein_coding | ENSG0000C |
| 35470508 | 35470579 | 1 | miRNA | ENSG0000C |
| 35473597 | 35497079 | -1 | protein_coding | ENSG0000C |
| 35497874 | 35512896 | -1 | protein_coding | ENSG0000C |
| 35539838 | 35546573 | 1 | lncRNA | ENSG0000C |
| 35555873 | 35556264 | 1 | processed_pseudoge | ENSG0000C |
| 35573585 | 35728583 | -1 | protein_coding | ENSG0000C |
| 35607628 | 35607934 | 1 | processed_pseudoge | ENSG0000C |
| 35650997 | 35652856 | 1 | lncRNA | ENSG0000C |
| 35651818 | 35651945 | 1 | snoRNA | ENSG0000C |
| 35664717 | 35664789 | -1 | miRNA | ENSG0000C |
| 35733867 | 35736947 | -1 | lncRNA | ENSG0000C |
| 35737032 | 35749079 | 1 | protein_coding | ENSG0000C |
| 35765908 | 35766927 | 1 | processed_pseudoge | ENSG0000C |
| 35776594 | 35779552 | 1 | protein_coding | ENSG0000C |
| 35781019 | 35793675 | 1 | protein_coding | ENSG0000C |
| 35794982 | 35797344 | -1 | protein_coding | ENSG0000C |
| 35797206 | 35845397 | 1 | protein_coding | ENSG0000C |
| 35832966 | 35921342 | -1 | protein_coding | ENSG0000C |
| 35943516 | 36024868 | -1 | protein_coding | ENSG0000C |
| 35989515 | 35991374 | -1 | transcribed_processe | ENSG0000C |
| 36027677 | 36111236 | 1 | protein_coding | ENSG0000C |
| 36091991 | 36092646 | -1 | processed_pseudoge | ENSG0000C |
| 36127809 | 36144524 | 1 | protein_coding | ENSG0000C |
| 36146698 | 36197205 | -1 | lncRNA | ENSG0000C |
| 36196744 | 36232790 | 1 | protein_coding | ENSG0000C |
| 36243203 | 36312229 | 1 | protein_coding | ENSG0000C |
| 36315761 | 36336888 | -1 | protein_coding | ENSG0000C |
| 36354091 | 36387800 | -1 | protein_coding | ENSG0000C |
| 36386831 | 36393462 | 1 | lncRNA | ENSG0000C |
| 36390551 | 36442854 | -1 | protein_coding | ENSG0000C |
| 36442767 | 36491143 | 1 | protein_coding | ENSG0000C |
| 36450912 | 36451201 | -1 | misc_RNA | ENSG0000C |
| 36493892 | 36547479 | -1 | protein_coding | ENSG0000C |
| 36522191 | 36522501 | 1 | misc_RNA | ENSG0000C |
| 36594353 | 36605600 | 1 | protein_coding | ENSG0000C |

| | | | | |
|---|---|---|---|---|
| 36622436 | 36622512 | -1 | miRNA | ENSG0000C |
| 36639545 | 36639672 | -1 | snRNA | ENSG0000C |
| 36672838 | 36672944 | 1 | misc_RNA | ENSG0000C |
| 36673621 | 36675126 | -1 | lncRNA | ENSG0000C |
| 36673817 | 36675270 | 1 | processed_pseudoge | ENSG0000C |
| 36676460 | 36687339 | 1 | protein_coding | ENSG0000C |
| 36677609 | 36678559 | -1 | lncRNA | ENSG0000C |
| 36697826 | 36733184 | 1 | protein_coding | ENSG0000C |
| 36737050 | 36737986 | 1 | processed_pseudoge | ENSG0000C |
| 36740775 | 36839444 | -1 | protein_coding | ENSG0000C |
| 36768485 | 36774651 | 1 | lncRNA | ENSG0000C |
| 36841430 | 36844402 | -1 | lncRNA | ENSG0000C |
| 36854827 | 36874803 | -1 | protein_coding | ENSG0000C |
| 36871870 | 36928964 | 1 | protein_coding | ENSG0000C |
| 36940071 | 36944675 | -1 | lncRNA | ENSG0000C |
| 36948263 | 36964837 | 1 | protein_coding | ENSG0000C |
| 36968141 | 36986298 | -1 | protein_coding | ENSG0000C |
| 37005646 | 37029069 | 1 | protein_coding | ENSG0000C |
| 37044860 | 37045189 | 1 | processed_pseudoge | ENSG0000C |
| 37091314 | 37091806 | 1 | processed_pseudoge | ENSG0000C |
| 37170152 | 37175428 | 1 | protein_coding | ENSG0000C |
| 37212178 | 37258155 | -1 | protein_coding | ENSG0000C |
| 37212181 | 37257637 | -1 | protein_coding | ENSG0000C |
| 37257772 | 37332970 | 1 | protein_coding | ENSG0000C |
| 37295584 | 37295742 | -1 | processed_pseudoge | ENSG0000C |
| 37301210 | 37305176 | -1 | lncRNA | ENSG0000C |
| 37353979 | 37394734 | 1 | protein_coding | ENSG0000C |
| 37361185 | 37361483 | 1 | misc_RNA | ENSG0000C |
| 37433219 | 37482827 | 1 | protein_coding | ENSG0000C |
| 37482938 | 37499893 | -1 | protein_coding | ENSG0000C |
| 37507348 | 37535616 | 1 | lncRNA | ENSG0000C |
| 37543553 | 37547280 | -1 | transcribed_processe | ENSG0000C |
| 37545145 | 37550860 | 1 | lncRNA | ENSG0000C |
| 37555365 | 37555422 | -1 | miRNA | ENSG0000C |
| 37567716 | 37571460 | 1 | TEC | ENSG0000C |
| 37630679 | 37699306 | -1 | protein_coding | ENSG0000C |
| 37815777 | 37819218 | -1 | lncRNA | ENSG0000C |
| 37819727 | 38154624 | 1 | protein_coding | ENSG0000C |
| 37832922 | 37833185 | -1 | misc_RNA | ENSG0000C |
| 37915573 | 37915742 | 1 | snRNA | ENSG0000C |
| 38002832 | 38003313 | -1 | processed_pseudoge | ENSG0000C |
| 38168451 | 38640148 | -1 | protein_coding | ENSG0000C |
| 38207274 | 38207345 | 1 | snoRNA | ENSG0000C |
| 38481692 | 38482531 | 1 | lncRNA | ENSG0000C |
| 38565950 | 38566055 | 1 | misc_RNA | ENSG0000C |
| 38587323 | 38588529 | 1 | processed_pseudoge | ENSG0000C |
| 38675925 | 38703145 | -1 | protein_coding | ENSG0000C |
| 38714051 | 38715217 | -1 | lncRNA | ENSG0000C |
| 38715311 | 39030792 | 1 | protein_coding | ENSG0000C |
| 38744086 | 38744386 | 1 | misc_RNA | ENSG0000C |

| | | | | |
|---|---|---|---|---|
| 38762905 | 38764694 | 1 | processed_pseudoge | ENSG00000( |
| 38822307 | 38822404 | -1 | snoRNA | ENSG00000( |
| 38923029 | 38953099 | -1 | lncRNA | ENSG00000( |
| 38928031 | 38928153 | 1 | processed_pseudoge | ENSG00000( |
| 39039603 | 39039985 | -1 | processed_pseudoge | ENSG00000( |
| 39048781 | 39091303 | 1 | protein_coding | ENSG00000( |
| 39104063 | 39115186 | -1 | protein_coding | ENSG00000( |
| 39110321 | 39112952 | 1 | processed_pseudoge | ENSG00000( |
| 39188971 | 39229475 | -1 | protein_coding | ENSG00000( |
| 39299001 | 39314461 | -1 | protein_coding | ENSG00000( |
| 39314698 | 39322968 | -1 | protein_coding | ENSG00000( |
| 39329990 | 39725408 | -1 | protein_coding | ENSG00000( |
| 39353747 | 39354172 | 1 | processed_pseudoge | ENSG00000( |
| 39553811 | 39554943 | 1 | processed_pseudoge | ENSG00000( |
| 39620345 | 39620464 | -1 | snRNA | ENSG00000( |
| 39792298 | 39904877 | 1 | protein_coding | ENSG00000( |
| 39818751 | 39823227 | 1 | TEC | ENSG00000( |
| 39881804 | 39900071 | -1 | lncRNA | ENSG00000( |
| 39899578 | 39934551 | -1 | protein_coding | ENSG00000( |
| 39934595 | 40000527 | 1 | transcribed_processe | ENSG00000( |
| 39958414 | 39958833 | -1 | processed_pseudoge | ENSG00000( |
| 40271566 | 40276237 | -1 | lncRNA | ENSG00000( |
| 40334775 | 40387590 | 1 | lncRNA | ENSG00000( |
| 40344344 | 40346151 | -1 | lncRNA | ENSG00000( |
| 40358744 | 40369474 | -1 | lncRNA | ENSG00000( |
| 40391591 | 40587364 | -1 | protein_coding | ENSG00000( |
| 40407853 | 40407955 | 1 | snRNA | ENSG00000( |
| 40501631 | 40502393 | -1 | lncRNA | ENSG00000( |
| 40505507 | 40523963 | -1 | lncRNA | ENSG00000( |
| 40713411 | 40715363 | -1 | lncRNA | ENSG00000( |
| 41026895 | 41039221 | -1 | protein_coding | ENSG00000( |
| 41033627 | 41097787 | -1 | protein_coding | ENSG00000( |
| 41042467 | 41044337 | 1 | protein_coding | ENSG00000( |
| 41053304 | 41064511 | 1 | protein_coding | ENSG00000( |
| 41072945 | 41099976 | 1 | protein_coding | ENSG00000( |
| 41080624 | 41101056 | -1 | lncRNA | ENSG00000( |
| 41101022 | 41140835 | 1 | transcribed_unproce | ENSG00000( |
| 41149342 | 41154337 | -1 | protein_coding | ENSG00000( |
| 41158506 | 41163186 | -1 | protein_coding | ENSG00000( |
| 41189749 | 41201149 | -1 | protein_coding | ENSG00000( |
| 41208713 | 41217947 | -1 | transcribed_unproce | ENSG00000( |
| 41228339 | 41238882 | 1 | protein_coding | ENSG00000( |
| 41239520 | 41239626 | -1 | rRNA_pseudogene | ENSG00000( |
| 41247369 | 41249649 | 1 | unprocessed_pseudc | ENSG00000( |
| 41250851 | 41251097 | 1 | processed_pseudoge | ENSG00000( |
| 41252696 | 41252858 | 1 | processed_pseudoge | ENSG00000( |
| 41267926 | 41286682 | -1 | protein_coding | ENSG00000( |
| 41269875 | 41270239 | -1 | unprocessed_pseudc | ENSG00000( |
| 41302466 | 41302572 | -1 | snRNA | ENSG00000( |
| 41308166 | 41308565 | -1 | processed_pseudoge | ENSG00000( |

| | | | | |
|---|---|---|---|---|
| 41335608 | 41350889 | 1 | protein_coding | ENSG0000C |
| 41381392 | 41381832 | -1 | TEC | ENSG0000C |
| 41405819 | 41406561 | 1 | TEC | ENSG0000C |
| 41494853 | 41548621 | -1 | lncRNA | ENSG0000C |
| 41499033 | 41519852 | 1 | lncRNA | ENSG0000C |
| 41546426 | 41602384 | 1 | protein_coding | ENSG0000C |
| 41598723 | 41598788 | 1 | miRNA | ENSG0000C |
| 41636882 | 41654246 | 1 | protein_coding | ENSG0000C |
| 41666906 | 41667649 | 1 | processed_pseudoge | ENSG0000C |
| 41683978 | 41736259 | -1 | protein_coding | ENSG0000C |
| 41720396 | 41734032 | 1 | lncRNA | ENSG0000C |
| 41736711 | 41754109 | -1 | protein_coding | ENSG0000C |
| 41764292 | 41764460 | -1 | lncRNA | ENSG0000C |
| 41770176 | 41786542 | -1 | protein_coding | ENSG0000C |
| 41780349 | 41790141 | 1 | protein_coding | ENSG0000C |
| 41780762 | 41787372 | 1 | protein_coding | ENSG0000C |
| 41787662 | 41789898 | 1 | protein_coding | ENSG0000C |
| 41789896 | 41895361 | -1 | protein_coding | ENSG0000C |
| 41791410 | 41791477 | 1 | lncRNA | ENSG0000C |
| 41832854 | 41832993 | 1 | snoRNA | ENSG0000C |
| 41868622 | 41869731 | 1 | lncRNA | ENSG0000C |
| 41905354 | 41921139 | -1 | protein_coding | ENSG0000C |
| 41917416 | 41917526 | -1 | misc_RNA | ENSG0000C |
| 41921499 | 41933046 | 1 | protein_coding | ENSG0000C |
| 41934933 | 42050357 | -1 | protein_coding | ENSG0000C |
| 42018408 | 42018514 | -1 | snRNA | ENSG0000C |
| 42030053 | 42031382 | 1 | lncRNA | ENSG0000C |
| 42050513 | 42087461 | 1 | protein_coding | ENSG0000C |
| 42092233 | 42094259 | -1 | lncRNA | ENSG0000C |
| 42101119 | 42142620 | -1 | protein_coding | ENSG0000C |
| 42155406 | 42180056 | 1 | protein_coding | ENSG0000C |
| 42155406 | 42180049 | 1 | protein_coding | ENSG0000C |
| 42155426 | 42163439 | 1 | protein_coding | ENSG0000C |
| 42183284 | 42194956 | -1 | protein_coding | ENSG0000C |
| 42206807 | 42217861 | -1 | protein_coding | ENSG0000C |
| 42224931 | 42452051 | -1 | protein_coding | ENSG0000C |
| 42499710 | 42500022 | 1 | processed_pseudoge | ENSG0000C |
| 42564062 | 42693504 | 1 | protein_coding | ENSG0000C |
| 42664162 | 42664268 | 1 | snRNA | ENSG0000C |
| 42696598 | 42722597 | -1 | protein_coding | ENSG0000C |
| 42727234 | 42727700 | 1 | processed_pseudoge | ENSG0000C |
| 42744498 | 42746103 | -1 | protein_coding | ENSG0000C |
| 42746958 | 42868560 | 1 | protein_coding | ENSG0000C |
| 42879616 | 42889925 | 1 | protein_coding | ENSG0000C |
| 42890265 | 42890821 | -1 | protein_coding | ENSG0000C |
| 42893761 | 42903072 | -1 | lncRNA | ENSG0000C |
| 42915989 | 42925835 | 1 | protein_coding | ENSG0000C |
| 42928287 | 42929457 | -1 | lncRNA | ENSG0000C |
| 42929192 | 42939294 | 1 | protein_coding | ENSG0000C |
| 42940364 | 42948360 | 1 | lncRNA | ENSG0000C |

| | | | | |
|---|---|---|---|---|
| 42956345 | 42956765 | -1 | processed_pseudoge | ENSG00000 |
| 42960754 | 42963880 | 1 | protein_coding | ENSG00000 |
| 42963865 | 42979181 | -1 | protein_coding | ENSG00000 |
| 42984553 | 43012342 | 1 | protein_coding | ENSG00000 |
| 43011143 | 43016868 | -1 | protein_coding | ENSG00000 |
| 43014103 | 43021298 | 1 | protein_coding | ENSG00000 |
| 43021623 | 43034156 | 1 | protein_coding | ENSG00000 |
| 43033897 | 43034405 | -1 | lncRNA | ENSG00000 |
| 43036196 | 43036500 | -1 | misc_RNA | ENSG00000 |
| 43037617 | 43053945 | -1 | protein_coding | ENSG00000 |
| 43040777 | 43075095 | 1 | protein_coding | ENSG00000 |
| 43054029 | 43059438 | -1 | protein_coding | ENSG00000 |
| 43074331 | 43074739 | -1 | lncRNA | ENSG00000 |
| 43076307 | 43161719 | 1 | protein_coding | ENSG00000 |
| 43171269 | 43181506 | 1 | protein_coding | ENSG00000 |
| 43182184 | 43224587 | 1 | protein_coding | ENSG00000 |
| 43213801 | 43223860 | -1 | lncRNA | ENSG00000 |
| 43225629 | 43229484 | -1 | protein_coding | ENSG00000 |
| 43243481 | 43288258 | 1 | protein_coding | ENSG00000 |
| 43295694 | 43305538 | 1 | protein_coding | ENSG00000 |
| 43299710 | 43308797 | -1 | protein_coding | ENSG00000 |
| 43307134 | 43369647 | -1 | protein_coding | ENSG00000 |
| 43310231 | 43310357 | 1 | processed_pseudoge | ENSG00000 |
| 43328134 | 43328476 | 1 | processed_pseudoge | ENSG00000 |
| 43363479 | 43364219 | 1 | processed_pseudoge | ENSG00000 |
| 43364220 | 43364294 | 1 | processed_pseudoge | ENSG00000 |
| 43370026 | 43425283 | 1 | lncRNA | ENSG00000 |
| 43403815 | 43408341 | -1 | lncRNA | ENSG00000 |
| 43427366 | 43450427 | 1 | protein_coding | ENSG00000 |
| 43434542 | 43434620 | 1 | miRNA | ENSG00000 |
| 43450352 | 43456632 | -1 | protein_coding | ENSG00000 |
| 43474186 | 43474292 | 1 | snRNA | ENSG00000 |
| 43477523 | 43506556 | 1 | protein_coding | ENSG00000 |
| 43506969 | 43510686 | -1 | protein_coding | ENSG00000 |
| 43509702 | 43562419 | 1 | protein_coding | ENSG00000 |
| 43511832 | 43516985 | -1 | protein_coding | ENSG00000 |
| 43519180 | 43519724 | -1 | lncRNA | ENSG00000 |
| 43522334 | 43576038 | -1 | protein_coding | ENSG00000 |
| 43538822 | 43539703 | -1 | processed_pseudoge | ENSG00000 |
| 43544144 | 43544270 | 1 | scaRNA | ENSG00000 |
| 43576150 | 43615660 | 1 | protein_coding | ENSG00000 |
| 43588230 | 43591362 | -1 | lncRNA | ENSG00000 |
| 43605316 | 43629264 | -1 | protein_coding | ENSG00000 |
| 43629540 | 43640952 | 1 | protein_coding | ENSG00000 |
| 43645036 | 43672600 | 1 | protein_coding | ENSG00000 |
| 43671303 | 43687791 | -1 | protein_coding | ENSG00000 |
| 43705949 | 43706751 | -1 | processed_pseudoge | ENSG00000 |
| 43722786 | 43737834 | -1 | lncRNA | ENSG00000 |
| 43770184 | 43786487 | 1 | protein_coding | ENSG00000 |
| 43770429 | 43770616 | -1 | lncRNA | ENSG00000 |

| | | | | |
|---|---|---|---|---|
| 43803193 | 43842625 | 1 | lncRNA | ENSG0000C |
| 43844878 | 43852317 | -1 | lncRNA | ENSG0000C |
| 43851757 | 43852333 | -1 | lncRNA | ENSG0000C |
| 43995723 | 44074652 | -1 | lncRNA | ENSG0000C |
| 44000580 | 44007612 | 1 | protein_coding | ENSG0000C |
| 44058792 | 44089288 | 1 | lncRNA | ENSG0000C |
| 44089242 | 44089696 | -1 | processed_pseudoge | ENSG0000C |
| 44090787 | 44095585 | -1 | lncRNA | ENSG0000C |
| 44113451 | 44127452 | -1 | protein_coding | ENSG0000C |
| 44126914 | 44155519 | 1 | protein_coding | ENSG0000C |
| 44158811 | 44184401 | 1 | protein_coding | ENSG0000C |
| 44209766 | 44210063 | 1 | misc_RNA | ENSG0000C |
| 44216926 | 44218234 | 1 | protein_coding | ENSG0000C |
| 44219505 | 44234151 | 1 | protein_coding | ENSG0000C |
| 44246166 | 44253888 | 1 | protein_coding | ENSG0000C |
| 44254096 | 44257890 | -1 | protein_coding | ENSG0000C |
| 44254206 | 44254285 | -1 | miRNA | ENSG0000C |
| 44258166 | 44265788 | -1 | protein_coding | ENSG0000C |
| 44270450 | 44307506 | 1 | protein_coding | ENSG0000C |
| 44273194 | 44378957 | 1 | protein_coding | ENSG0000C |
| 44278734 | 44297698 | -1 | protein_coding | ENSG0000C |
| 44298731 | 44313347 | -1 | protein_coding | ENSG0000C |
| 44342660 | 44377167 | 1 | protein_coding | ENSG0000C |
| 44387706 | 44450425 | 1 | protein_coding | ENSG0000C |
| 44435641 | 44435722 | 1 | miRNA | ENSG0000C |
| 44513262 | 44527448 | 1 | lncRNA | ENSG0000C |
| 44551577 | 44553281 | 1 | TEC | ENSG0000C |
| 44727921 | 44830188 | 1 | lncRNA | ENSG0000C |
| 44809317 | 45377953 | -1 | protein_coding | ENSG0000C |
| 44898711 | 44899295 | 1 | processed_pseudoge | ENSG0000C |
| 45097496 | 45097976 | 1 | unprocessed_pseudc | ENSG0000C |
| 45098021 | 45099260 | 1 | processed_pseudoge | ENSG0000C |
| 45158870 | 45159511 | 1 | processed_pseudoge | ENSG0000C |
| 45197674 | 45197770 | -1 | miRNA | ENSG0000C |
| 45328157 | 45664349 | 1 | protein_coding | ENSG0000C |
| 45421079 | 45422005 | -1 | lncRNA | ENSG0000C |
| 45573346 | 45576770 | -1 | lncRNA | ENSG0000C |
| 45880827 | 46080348 | -1 | protein_coding | ENSG0000C |
| 46018745 | 46018851 | 1 | snRNA | ENSG0000C |
| 46096004 | 46129857 | -1 | lncRNA | ENSG0000C |
| 46129989 | 46146688 | 1 | protein_coding | ENSG0000C |
| 46159185 | 46170980 | -1 | protein_coding | ENSG0000C |
| 46204729 | 46207386 | 1 | processed_pseudoge | ENSG0000C |
| 46220736 | 46491972 | -1 | protein_coding | ENSG0000C |
| 46438310 | 46438619 | 1 | processed_pseudoge | ENSG0000C |
| 46492052 | 46606162 | 1 | lncRNA | ENSG0000C |
| 46549580 | 46652830 | -1 | protein_coding | ENSG0000C |
| 46652915 | 46678190 | 1 | protein_coding | ENSG0000C |
| 46670444 | 46688165 | -1 | lncRNA | ENSG0000C |
| 46687875 | 46704319 | 1 | protein_coding | ENSG0000C |

| | | | | |
|---|---|---|---|---|
| 46704201 | 46735693 | -1 | protein_coding | ENSG00000 |
| 46746917 | 46759506 | 1 | protein_coding | ENSG00000 |
| 46758296 | 46761158 | -1 | lncRNA | ENSG00000 |
| 46793389 | 46839782 | 1 | protein_coding | ENSG00000 |
| 46852522 | 46954943 | -1 | protein_coding | ENSG00000 |
| 46903471 | 46909078 | 1 | lncRNA | ENSG00000 |
| 46997708 | 47042350 | -1 | protein_coding | ENSG00000 |
| 47231532 | 47309905 | -1 | protein_coding | ENSG00000 |
| 47368943 | 47369230 | 1 | processed_pseudoge | ENSG00000 |
| 47374561 | 47397398 | -1 | lncRNA | ENSG00000 |
| 47477243 | 47477572 | -1 | lncRNA | ENSG00000 |
| 47477789 | 47627263 | 1 | protein_coding | ENSG00000 |
| 47489067 | 47489170 | 1 | misc_RNA | ENSG00000 |
| 47562622 | 47563018 | 1 | processed_pseudoge | ENSG00000 |
| 47656436 | 47697797 | 1 | protein_coding | ENSG00000 |
| 47685864 | 47722021 | 1 | protein_coding | ENSG00000 |
| 47715283 | 47715601 | 1 | misc_RNA | ENSG00000 |
| 47729827 | 47740741 | -1 | lncRNA | ENSG00000 |
| 47752828 | 47753240 | -1 | processed_pseudoge | ENSG00000 |
| 47781982 | 47832780 | 1 | protein_coding | ENSG00000 |
| 47823390 | 47823550 | 1 | snRNA | ENSG00000 |
| 47878028 | 48068689 | -1 | protein_coding | ENSG00000 |
| 48069213 | 48111078 | -1 | lncRNA | ENSG00000 |
| 48149203 | 48150005 | -1 | processed_pseudoge | ENSG00000 |
| 48213604 | 48214794 | -1 | processed_pseudoge | ENSG00000 |
| 48701491 | 48701872 | -1 | processed_pseudoge | ENSG00000 |
| 48754649 | 48795513 | -1 | lncRNA | ENSG00000 |
| 48952070 | 48952781 | -1 | processed_pseudoge | ENSG00000 |
| 49077712 | 49078051 | -1 | processed_pseudoge | ENSG00000 |
| 49115537 | 49116115 | 1 | processed_pseudoge | ENSG00000 |
| 49273600 | 49273794 | 1 | processed_pseudoge | ENSG00000 |
| 49344800 | 49344861 | -1 | snRNA | ENSG00000 |
| 49358185 | 49360420 | -1 | processed_pseudoge | ENSG00000 |
| 49430360 | 49463253 | -1 | protein_coding | ENSG00000 |
| 49463370 | 49493107 | 1 | protein_coding | ENSG00000 |
| 49499923 | 49528078 | 1 | protein_coding | ENSG00000 |
| 49550646 | 49561907 | 1 | protein_coding | ENSG00000 |
| 49565924 | 49580166 | -1 | unitary_pseudogene | ENSG00000 |
| 49605158 | 49636884 | -1 | protein_coding | ENSG00000 |
| 49692358 | 49713590 | -1 | protein_coding | ENSG00000 |
| 49714325 | 49820503 | 1 | lncRNA | ENSG00000 |
| 49721931 | 49724007 | 1 | TEC | ENSG00000 |
| 49727384 | 49744437 | -1 | protein_coding | ENSG00000 |
| 49785660 | 49787285 | -1 | protein_coding | ENSG00000 |
| 49823712 | 49824782 | -1 | lncRNA | ENSG00000 |
| 49834257 | 49877096 | -1 | protein_coding | ENSG00000 |
| 49946021 | 49950265 | -1 | protein_coding | ENSG00000 |
| 49960249 | 49964164 | -1 | protein_coding | ENSG00000 |
| 49968677 | 49969625 | -1 | protein_coding | ENSG00000 |
| 50009138 | 50021994 | -1 | protein_coding | ENSG00000 |

| | | | | |
|---|---|---|---|---|
| 50042144 | 50049874 | -1 | protein_coding | ENSG00000 |
| 50093389 | 50181007 | 1 | lncRNA | ENSG00000 |
| 50411844 | 50415875 | 1 | lncRNA | ENSG00000 |
| 50514035 | 50521104 | 1 | lncRNA | ENSG00000 |
| 50587607 | 50637205 | -1 | lncRNA | ENSG00000 |
| 50713828 | 50772988 | 1 | protein_coding | ENSG00000 |
| 50818723 | 50847619 | 1 | protein_coding | ENSG00000 |
| 50857255 | 50857662 | -1 | processed_pseudoge | ENSG00000 |
| 50897615 | 50898225 | -1 | processed_pseudoge | ENSG00000 |
| 50912712 | 50913256 | -1 | processed_pseudoge | ENSG00000 |
| 51385282 | 51385849 | -1 | processed_pseudoge | ENSG00000 |
| 51410081 | 51410537 | -1 | processed_pseudoge | ENSG00000 |
| 51464690 | 51464765 | 1 | snoRNA | ENSG00000 |
| 51537155 | 51537543 | -1 | processed_pseudoge | ENSG00000 |
| 51599723 | 51623428 | 1 | lncRNA | ENSG00000 |
| 51615299 | 52087613 | -1 | protein_coding | ENSG00000 |
| 51975673 | 51975959 | 1 | misc_RNA | ENSG00000 |
| 52144349 | 52144434 | 1 | miRNA | ENSG00000 |
| 52146814 | 52151119 | -1 | lncRNA | ENSG00000 |
| 52148923 | 52149041 | 1 | miRNA | ENSG00000 |
| 52186375 | 52190638 | 1 | protein_coding | ENSG00000 |
| 52236681 | 52244537 | -1 | protein_coding | ENSG00000 |
| 52246460 | 52247362 | 1 | processed_pseudoge | ENSG00000 |
| 52264014 | 52284881 | -1 | protein_coding | ENSG00000 |
| 52361421 | 52407777 | 1 | protein_coding | ENSG00000 |
| 52362123 | 52529886 | 1 | protein_coding | ENSG00000 |
| 52497408 | 52577060 | -1 | protein_coding | ENSG00000 |
| 52576787 | 52643058 | 1 | lncRNA | ENSG00000 |
| 52657373 | 52657651 | 1 | processed_pseudoge | ENSG00000 |
| 52664366 | 52669155 | 1 | transcribed_unproce | ENSG00000 |
| 52671113 | 52686588 | 1 | protein_coding | ENSG00000 |
| 52687930 | 52703825 | -1 | unprocessed_pseudc | ENSG00000 |
| 52688661 | 52688998 | 1 | processed_pseudoge | ENSG00000 |
| 52739590 | 52744656 | -1 | unprocessed_pseudc | ENSG00000 |
| 52750087 | 52763475 | -1 | protein_coding | ENSG00000 |
| 52765280 | 52778347 | -1 | unprocessed_pseudc | ENSG00000 |
| 52791371 | 52803860 | -1 | protein_coding | ENSG00000 |
| 52805613 | 52813460 | -1 | unprocessed_pseudc | ENSG00000 |
| 52831655 | 52846095 | -1 | protein_coding | ENSG00000 |
| 52847910 | 52870329 | -1 | unprocessed_pseudc | ENSG00000 |
| 52873014 | 52879940 | -1 | unprocessed_pseudc | ENSG00000 |
| 52896639 | 52909698 | -1 | protein_coding | ENSG00000 |
| 52939726 | 52957521 | -1 | transcribed_unproce | ENSG00000 |
| 52970650 | 52970866 | -1 | processed_pseudoge | ENSG00000 |
| 52977948 | 52995304 | -1 | protein_coding | ENSG00000 |
| 52995620 | 52995950 | 1 | misc_RNA | ENSG00000 |
| 52995621 | 52995948 | 1 | snRNA | ENSG00000 |
| 53001279 | 53061802 | -1 | protein_coding | ENSG00000 |
| 53051991 | 53100873 | 1 | protein_coding | ENSG00000 |
| 53090961 | 53091257 | 1 | misc_RNA | ENSG00000 |

| | | | | |
|---|---|---|---|---|
| 53125644 | 53126146 | -1 | lncRNA | ENSG0000C |
| 53126961 | 53148841 | -1 | protein_coding | ENSG0000C |
| 53153795 | 53153898 | -1 | snRNA | ENSG0000C |
| 53196720 | 53197159 | -1 | processed_pseudoge | ENSG0000C |
| 53206598 | 53208697 | -1 | processed_pseudoge | ENSG0000C |
| 53219261 | 53219424 | 1 | snRNA | ENSG0000C |
| 53235621 | 53236297 | -1 | processed_pseudoge | ENSG0000C |
| 53267398 | 53349179 | -1 | protein_coding | ENSG0000C |
| 53276993 | 53277071 | 1 | miRNA | ENSG0000C |
| 53336943 | 53337346 | -1 | transcribed_processe | ENSG0000C |
| 53350158 | 53350705 | 1 | lncRNA | ENSG0000C |
| 53354715 | 53355065 | 1 | processed_pseudoge | ENSG0000C |
| 53368670 | 53369009 | 1 | processed_pseudoge | ENSG0000C |
| 53378503 | 53378861 | 1 | processed_pseudoge | ENSG0000C |
| 53381519 | 53382203 | 1 | processed_pseudoge | ENSG0000C |
| 53415294 | 53415607 | -1 | misc_RNA | ENSG0000C |
| 53418452 | 53418737 | -1 | processed_pseudoge | ENSG0000C |
| 53497341 | 53616970 | -1 | protein_coding | ENSG0000C |
| 53503109 | 53507660 | 1 | lncRNA | ENSG0000C |
| 53561289 | 53617171 | -1 | lncRNA | ENSG0000C |
| 53616471 | 53631400 | 1 | lncRNA | ENSG0000C |
| 53647916 | 53665756 | -1 | protein_coding | ENSG0000C |
| 53739266 | 53794904 | -1 | lncRNA | ENSG0000C |
| 53794497 | 53924125 | 1 | protein_coding | ENSG0000C |
| 53918974 | 53920788 | -1 | lncRNA | ENSG0000C |
| 53929982 | 54266280 | 1 | protein_coding | ENSG0000C |
| 53930022 | 53997667 | 1 | lncRNA | ENSG0000C |
| 53978549 | 54079726 | -1 | lncRNA | ENSG0000C |
| 53998890 | 54007152 | 1 | lncRNA | ENSG0000C |
| 54016479 | 54016777 | -1 | processed_pseudoge | ENSG0000C |
| 54190206 | 54190709 | 1 | lncRNA | ENSG0000C |
| 54307859 | 54390142 | 1 | protein_coding | ENSG0000C |
| 54365335 | 54369971 | -1 | lncRNA | ENSG0000C |
| 54485169 | 54485850 | -1 | processed_pseudoge | ENSG0000C |
| 54602487 | 54603150 | -1 | processed_pseudoge | ENSG0000C |
| 54624605 | 54625488 | 1 | processed_pseudoge | ENSG0000C |
| 54770583 | 54771134 | 1 | processed_pseudoge | ENSG0000C |
| 54786387 | 54786492 | 1 | snRNA | ENSG0000C |
| 54840118 | 54840855 | -1 | lncRNA | ENSG0000C |
| 54846771 | 54945099 | 1 | protein_coding | ENSG0000C |
| 54943167 | 54945099 | 1 | lncRNA | ENSG0000C |
| 55106460 | 55282620 | 1 | protein_coding | ENSG0000C |
| 55327469 | 55402493 | 1 | protein_coding | ENSG0000C |
| 55434369 | 55579214 | -1 | protein_coding | ENSG0000C |
| 55680642 | 55681021 | -1 | processed_pseudoge | ENSG0000C |
| 55753653 | 55875590 | -1 | protein_coding | ENSG0000C |
| 55939790 | 55940624 | -1 | processed_pseudoge | ENSG0000C |
| 56056590 | 56394094 | -1 | protein_coding | ENSG0000C |
| 56276529 | 56276796 | -1 | processed_pseudoge | ENSG0000C |
| 56331788 | 56332117 | -1 | lncRNA | ENSG0000C |

| | | | | |
|---|---|---|---|---|
| 56431950 | 56433213 | -1 | processed_pseudoge | ENSG00000 |
| 56457987 | 56954649 | -1 | protein_coding | ENSG00000 |
| 56843928 | 56864078 | 1 | lncRNA | ENSG00000 |
| 56871191 | 56871743 | 1 | processed_pseudoge | ENSG00000 |
| 56945730 | 56945836 | 1 | snRNA | ENSG00000 |
| 56955107 | 57027346 | 1 | protein_coding | ENSG00000 |
| 56975606 | 56976062 | -1 | processed_pseudoge | ENSG00000 |
| 57004520 | 57004799 | 1 | processed_pseudoge | ENSG00000 |
| 57029521 | 57029988 | 1 | processed_pseudoge | ENSG00000 |
| 57046532 | 57055239 | 1 | protein_coding | ENSG00000 |
| 57086844 | 57170305 | 1 | protein_coding | ENSG00000 |
| 57114894 | 57174236 | -1 | lncRNA | ENSG00000 |
| 57172326 | 57189833 | 1 | protein_coding | ENSG00000 |
| 57186992 | 57222307 | -1 | protein_coding | ENSG00000 |
| 57314805 | 57646850 | 1 | protein_coding | ENSG00000 |
| 57390132 | 57390212 | 1 | miRNA | ENSG00000 |
| 57493855 | 57497691 | 1 | lncRNA | ENSG00000 |
| 57855891 | 57856468 | -1 | lncRNA | ENSG00000 |
| 57880071 | 57884297 | -1 | lncRNA | ENSG00000 |
| 57902609 | 57903148 | -1 | lncRNA | ENSG00000 |
| 57908560 | 57913911 | -1 | lncRNA | ENSG00000 |
| 57919784 | 57930291 | -1 | transcribed_unproce | ENSG00000 |
| 57919912 | 57961382 | -1 | lncRNA | ENSG00000 |
| 57937458 | 57939015 | 1 | processed_pseudoge | ENSG00000 |
| 57946074 | 57961501 | -1 | transcribed_unproce | ENSG00000 |
| 57961438 | 58438364 | 1 | lncRNA | ENSG00000 |
| 57967687 | 57968661 | -1 | processed_pseudoge | ENSG00000 |
| 58071720 | 58073134 | -1 | processed_pseudoge | ENSG00000 |
| 58119741 | 58121029 | -1 | processed_pseudoge | ENSG00000 |
| 58386799 | 58386961 | 1 | processed_pseudoge | ENSG00000 |
| 60281444 | 60546660 | 1 | transcribed_unproce | ENSG00000 |
| 60719222 | 60720196 | -1 | processed_pseudoge | ENSG00000 |
| 60723148 | 60723898 | 1 | lncRNA | ENSG00000 |
| 60789805 | 60942327 | 1 | lncRNA | ENSG00000 |
| 60826185 | 60826533 | -1 | processed_pseudoge | ENSG00000 |
| 60873360 | 60874642 | -1 | processed_pseudoge | ENSG00000 |
| 61240898 | 61241311 | 1 | processed_pseudoge | ENSG00000 |
| 61574328 | 61574402 | 1 | processed_pseudoge | ENSG00000 |
| 61630233 | 61681049 | -1 | lncRNA | ENSG00000 |
| 61679961 | 62286225 | -1 | protein_coding | ENSG00000 |
| 61870097 | 61893882 | 1 | lncRNA | ENSG00000 |
| 62460940 | 62461503 | -1 | processed_pseudoge | ENSG00000 |
| 62547427 | 62548272 | 1 | unprocessed_pseudc | ENSG00000 |
| 62611153 | 62611327 | -1 | processed_pseudoge | ENSG00000 |
| 63193072 | 63193630 | -1 | processed_pseudoge | ENSG00000 |
| 63211446 | 63213024 | 1 | protein_coding | ENSG00000 |
| 63227485 | 63227743 | 1 | processed_pseudoge | ENSG00000 |
| 63229087 | 63230110 | -1 | processed_pseudoge | ENSG00000 |
| 63275951 | 63319983 | -1 | protein_coding | ENSG00000 |
| 63392222 | 63392606 | 1 | processed_pseudoge | ENSG00000 |

| | | | | |
|---|---|---|---|---|
| 63395007 | 63397106 | -1 | unprocessed_pseudc | ENSG00000( |
| 63440766 | 63443580 | 1 | unprocessed_pseudc | ENSG00000( |
| 63480134 | 63481876 | 1 | unprocessed_pseudc | ENSG00000( |
| 63521746 | 63583436 | 1 | protein_coding | ENSG00000( |
| 63548708 | 63549508 | -1 | processed_pseudoge | ENSG00000( |
| 63571005 | 63572408 | -1 | lncRNA | ENSG00000( |
| 63572472 | 63583587 | 1 | protein_coding | ENSG00000( |
| 63612761 | 63612903 | -1 | processed_pseudoge | ENSG00000( |
| 63615827 | 63616361 | -1 | processed_pseudoge | ENSG00000( |
| 63635820 | 63779336 | 1 | protein_coding | ENSG00000( |
| 63719980 | 65707226 | -1 | protein_coding | ENSG00000( |
| 63797189 | 63798031 | 1 | processed_pseudoge | ENSG00000( |
| 63806836 | 63822642 | 1 | lncRNA | ENSG00000( |
| 63857441 | 63857769 | 1 | processed_pseudoge | ENSG00000( |
| 64377795 | 64412779 | 1 | lncRNA | ENSG00000( |
| 64631205 | 64631682 | -1 | processed_pseudoge | ENSG00000( |
| 64728837 | 64733837 | -1 | transcribed_processe | ENSG00000( |
| 65302522 | 65304957 | 1 | processed_pseudoge | ENSG00000( |
| 65788417 | 65789287 | 1 | processed_pseudoge | ENSG00000( |
| 65836930 | 65838039 | -1 | processed_pseudoge | ENSG00000( |
| 66093431 | 66094909 | -1 | processed_pseudoge | ENSG00000( |
| 66710242 | 66710827 | -1 | processed_pseudoge | ENSG00000( |
| 66728843 | 66728904 | -1 | snRNA | ENSG00000( |
| 67210408 | 67210480 | -1 | snoRNA | ENSG00000( |
| 67456346 | 67457250 | -1 | processed_pseudoge | ENSG00000( |
| 67467231 | 67467308 | -1 | rRNA_pseudogene | ENSG00000( |
| 67546651 | 67546754 | 1 | snRNA | ENSG00000( |
| 67625356 | 67625464 | 1 | misc_RNA | ENSG00000( |
| 67878316 | 67889339 | -1 | lncRNA | ENSG00000( |
| 67884041 | 67986503 | 1 | lncRNA | ENSG00000( |
| 67888977 | 67998751 | 1 | lncRNA | ENSG00000( |
| 67933485 | 67934682 | 1 | processed_pseudoge | ENSG00000( |
| 68040290 | 68093001 | 1 | lncRNA | ENSG00000( |
| 68055351 | 68060502 | 1 | lncRNA | ENSG00000( |
| 68223667 | 68242364 | 1 | lncRNA | ENSG00000( |
| 68226972 | 68329899 | -1 | lncRNA | ENSG00000( |
| 68332019 | 68576191 | -1 | lncRNA | ENSG00000( |
| 68598204 | 68598393 | -1 | processed_pseudoge | ENSG00000( |
| 68627879 | 68635161 | -1 | lncRNA | ENSG00000( |
| 68635282 | 69389506 | 1 | protein_coding | ENSG00000( |
| 69672757 | 69867236 | -1 | protein_coding | ENSG00000( |
| 69705287 | 69706160 | -1 | processed_pseudoge | ENSG00000( |
| 69745871 | 69746851 | 1 | processed_pseudoge | ENSG00000( |
| 69866556 | 70212468 | 1 | protein_coding | ENSG00000( |
| 70098390 | 70098676 | -1 | processed_pseudoge | ENSG00000( |
| 70215061 | 70303083 | -1 | protein_coding | ENSG00000( |
| 70222758 | 70242156 | 1 | lncRNA | ENSG00000( |
| 70345919 | 70351802 | -1 | lncRNA | ENSG00000( |
| 70394881 | 70403117 | 1 | lncRNA | ENSG00000( |
| 70412828 | 70413950 | -1 | lncRNA | ENSG00000( |

| | | | | |
|---|---|---|---|---|
| 70412941 | 70561174 | 1 | protein_coding | ENSG00000 |
| 70513294 | 70513356 | 1 | snRNA | ENSG00000 |
| 70566917 | 70589569 | 1 | protein_coding | ENSG00000 |
| 70596438 | 70596980 | 1 | lncRNA | ENSG00000 |
| 70608234 | 70609334 | -1 | processed_pseudoge | ENSG00000 |
| 70663502 | 70664279 | -1 | processed_pseudoge | ENSG00000 |
| 70667776 | 70862011 | 1 | protein_coding | ENSG00000 |
| 70693839 | 70694619 | 1 | processed_pseudoge | ENSG00000 |
| 70734503 | 70734962 | 1 | processed_pseudoge | ENSG00000 |
| 70856679 | 70957060 | -1 | protein_coding | ENSG00000 |
| 71028582 | 71058476 | 1 | lncRNA | ENSG00000 |
| 71075564 | 71076375 | 1 | processed_pseudoge | ENSG00000 |
| 71126893 | 71127105 | -1 | snoRNA | ENSG00000 |
| 71165076 | 71165770 | 1 | processed_pseudoge | ENSG00000 |
| 71182612 | 71182718 | 1 | snRNA | ENSG00000 |
| 71221457 | 71328228 | -1 | lncRNA | ENSG00000 |
| 71238743 | 71240947 | 1 | lncRNA | ENSG00000 |
| 71251758 | 71252988 | 1 | processed_pseudoge | ENSG00000 |
| 71288811 | 71309059 | 1 | protein_coding | ENSG00000 |
| 71295173 | 71417436 | -1 | lncRNA | ENSG00000 |
| 71328942 | 71329806 | 1 | lncRNA | ENSG00000 |
| 71343427 | 71420745 | -1 | lncRNA | ENSG00000 |
| 71376960 | 71377031 | -1 | miRNA | ENSG00000 |
| 71386852 | 71390829 | -1 | TEC | ENSG00000 |
| 71403551 | 71403621 | -1 | miRNA | ENSG00000 |
| 71450834 | 71458874 | 1 | lncRNA | ENSG00000 |
| 71532070 | 71534715 | 1 | lncRNA | ENSG00000 |
| 71550958 | 71551643 | -1 | processed_pseudoge | ENSG00000 |
| 71584721 | 71585906 | 1 | processed_pseudoge | ENSG00000 |
| 71652474 | 71652611 | -1 | snRNA | ENSG00000 |
| 71886703 | 72403143 | 1 | protein_coding | ENSG00000 |
| 72316599 | 72317436 | -1 | processed_pseudoge | ENSG00000 |
| 72522641 | 72523238 | 1 | processed_pseudoge | ENSG00000 |
| 72598451 | 72599634 | -1 | processed_pseudoge | ENSG00000 |
| 72614386 | 72622073 | -1 | lncRNA | ENSG00000 |
| 72621792 | 73198851 | 1 | protein_coding | ENSG00000 |
| 72630495 | 72678558 | 1 | lncRNA | ENSG00000 |
| 72967687 | 72967753 | -1 | miRNA | ENSG00000 |
| 72986313 | 72987750 | 1 | processed_pseudoge | ENSG00000 |
| 73055097 | 73055857 | 1 | processed_pseudoge | ENSG00000 |
| 73130646 | 73143514 | -1 | lncRNA | ENSG00000 |
| 73157517 | 73158139 | -1 | processed_pseudoge | ENSG00000 |
| 73209083 | 73210080 | -1 | transcribed_unproce | ENSG00000 |
| 73209746 | 73263196 | -1 | protein_coding | ENSG00000 |
| 73223544 | 73225770 | -1 | protein_coding | ENSG00000 |
| 73241314 | 73310365 | -1 | protein_coding | ENSG00000 |
| 73263212 | 73301789 | 1 | lncRNA | ENSG00000 |
| 73290403 | 73291262 | 1 | processed_pseudoge | ENSG00000 |
| 73291962 | 73293277 | 1 | processed_pseudoge | ENSG00000 |
| 73322837 | 73323450 | 1 | processed_pseudoge | ENSG00000 |

| | | | | |
|---|---|---|---|---|
| 73327524 | 73329017 | 1 | processed_pseudoge | ENSG00000 |
| 73353063 | 73354276 | -1 | protein_coding | ENSG00000 |
| 73362658 | 73364171 | 1 | protein_coding | ENSG00000 |
| 73368555 | 73395133 | -1 | protein_coding | ENSG00000 |
| 73369704 | 73387717 | 1 | lncRNA | ENSG00000 |
| 73373108 | 73373263 | -1 | processed_pseudoge | ENSG00000 |
| 73391038 | 73391767 | -1 | processed_pseudoge | ENSG00000 |
| 73394828 | 73417566 | 1 | protein_coding | ENSG00000 |
| 73413515 | 73452297 | -1 | protein_coding | ENSG00000 |
| 73461578 | 73509236 | 1 | protein_coding | ENSG00000 |
| 73464073 | 73464177 | 1 | snRNA | ENSG00000 |
| 73487590 | 73487877 | -1 | misc_RNA | ENSG00000 |
| 73492025 | 73492742 | -1 | lncRNA | ENSG00000 |
| 73515750 | 73523797 | -1 | protein_coding | ENSG00000 |
| 73523618 | 73570596 | 1 | lncRNA | ENSG00000 |
| 73526744 | 73526879 | -1 | processed_pseudoge | ENSG00000 |
| 73593379 | 73653992 | -1 | protein_coding | ENSG00000 |
| 73618346 | 73618597 | 1 | processed_pseudoge | ENSG00000 |
| 73693903 | 73696131 | -1 | lncRNA | ENSG00000 |
| 73695785 | 73828316 | 1 | protein_coding | ENSG00000 |
| 73971346 | 73973115 | 1 | TEC | ENSG00000 |
| 74004520 | 74004812 | -1 | processed_pseudoge | ENSG00000 |
| 74069451 | 74690727 | 1 | lncRNA | ENSG00000 |
| 74282266 | 74282660 | 1 | processed_pseudoge | ENSG00000 |
| 74530248 | 74734319 | -1 | lncRNA | ENSG00000 |
| 74610449 | 74610715 | -1 | processed_pseudoge | ENSG00000 |
| 74642384 | 74652002 | 1 | lncRNA | ENSG00000 |
| 74849077 | 74850484 | -1 | processed_pseudoge | ENSG00000 |
| 75084326 | 75206053 | -1 | protein_coding | ENSG00000 |
| 75237675 | 75250323 | -1 | protein_coding | ENSG00000 |
| 75252924 | 75284948 | -1 | protein_coding | ENSG00000 |
| 75285014 | 75317525 | 1 | lncRNA | ENSG00000 |
| 75291859 | 75493800 | -1 | protein_coding | ENSG00000 |
| 75319101 | 75319980 | -1 | processed_pseudoge | ENSG00000 |
| 75357214 | 75399300 | 1 | lncRNA | ENSG00000 |
| 75398860 | 75399067 | -1 | snoRNA | ENSG00000 |
| 75428407 | 75428473 | 1 | miRNA | ENSG00000 |
| 75454944 | 75458900 | 1 | lncRNA | ENSG00000 |
| 75465123 | 75465570 | 1 | processed_pseudoge | ENSG00000 |
| 75473738 | 75473894 | -1 | snRNA | ENSG00000 |
| 75499705 | 75500136 | 1 | processed_pseudoge | ENSG00000 |
| 75586122 | 75586461 | 1 | processed_pseudoge | ENSG00000 |
| 75593055 | 75593145 | 1 | snRNA | ENSG00000 |
| 75601509 | 75718281 | 1 | protein_coding | ENSG00000 |
| 75610958 | 75611945 | -1 | transcribed_processe | ENSG00000 |
| 75644084 | 75644177 | 1 | snRNA | ENSG00000 |
| 75654970 | 75655299 | -1 | misc_RNA | ENSG00000 |
| 75744275 | 75744840 | 1 | processed_pseudoge | ENSG00000 |
| 75749192 | 75919537 | 1 | protein_coding | ENSG00000 |
| 75768059 | 75768158 | -1 | snRNA | ENSG00000 |

| | | | | |
|---|---|---|---|---|
| 75865239 | 75865335 | -1 | rRNA_pseudogene | ENSG0000( |
| 75921114 | 76072678 | -1 | protein_coding | ENSG0000( |
| 75928513 | 75928625 | 1 | misc_RNA | ENSG0000( |
| 76092834 | 76092940 | 1 | snRNA | ENSG0000( |
| 76446988 | 76447089 | -1 | snRNA | ENSG0000( |
| 76521640 | 76593821 | -1 | lncRNA | ENSG0000( |
| 76557653 | 76557754 | -1 | snRNA | ENSG0000( |
| 76561328 | 76562590 | 1 | lncRNA | ENSG0000( |
| 76660486 | 76662770 | 1 | processed_pseudoge | ENSG0000( |
| 76774972 | 77097788 | 1 | lncRNA | ENSG0000( |
| 77343557 | 77343654 | 1 | processed_pseudoge | ENSG0000( |
| 77461753 | 77463773 | -1 | protein_coding | ENSG0000( |
| 77496119 | 77496671 | 1 | processed_pseudoge | ENSG0000( |
| 77650274 | 77927028 | 1 | protein_coding | ENSG0000( |
| 78604467 | 78606036 | -1 | lncRNA | ENSG0000( |
| 78809715 | 78813303 | -1 | lncRNA | ENSG0000( |
| 78867551 | 78946440 | 1 | protein_coding | ENSG0000( |
| 78934419 | 79078254 | -1 | protein_coding | ENSG0000( |
| 79067692 | 79068138 | -1 | processed_pseudoge | ENSG0000( |
| 79077841 | 79081371 | 1 | lncRNA | ENSG0000( |
| 79201245 | 79234738 | -1 | protein_coding | ENSG0000( |
| 79233718 | 79236797 | 1 | lncRNA | ENSG0000( |
| 79278105 | 79278427 | -1 | processed_pseudoge | ENSG0000( |
| 79301963 | 79302066 | -1 | misc_RNA | ENSG0000( |
| 79307669 | 79313384 | -1 | lncRNA | ENSG0000( |
| 79326568 | 79327207 | -1 | processed_pseudoge | ENSG0000( |
| 79354443 | 79354713 | -1 | processed_pseudoge | ENSG0000( |
| 79406240 | 79406774 | -1 | processed_pseudoge | ENSG0000( |
| 79420172 | 79529276 | 1 | lncRNA | ENSG0000( |
| 79436908 | 79437167 | -1 | processed_pseudoge | ENSG0000( |
| 79484991 | 79537458 | -1 | protein_coding | ENSG0000( |
| 79537540 | 79539976 | 1 | lncRNA | ENSG0000( |
| 79552794 | 79553160 | 1 | processed_pseudoge | ENSG0000( |
| 79561132 | 79561602 | 1 | lncRNA | ENSG0000( |
| 79573877 | 79575530 | 1 | TEC | ENSG0000( |
| 79631329 | 79703655 | 1 | protein_coding | ENSG0000( |
| 79803582 | 79833386 | -1 | lncRNA | ENSG0000( |
| 79854684 | 79855632 | 1 | processed_pseudoge | ENSG0000( |
| 79868402 | 79870483 | 1 | lncRNA | ENSG0000( |
| 79871873 | 79874610 | 1 | lncRNA | ENSG0000( |
| 79914814 | 79947553 | -1 | protein_coding | ENSG0000( |
| 79947810 | 79980046 | 1 | lncRNA | ENSG0000( |
| 79953005 | 79954011 | -1 | processed_pseudoge | ENSG0000( |
| 79964215 | 79964519 | 1 | processed_pseudoge | ENSG0000( |
| 80003887 | 80042527 | 1 | protein_coding | ENSG0000( |
| 80046715 | 80084776 | -1 | lncRNA | ENSG0000( |
| 80064286 | 80070199 | -1 | processed_pseudoge | ENSG0000( |
| 80077729 | 80078362 | -1 | processed_pseudoge | ENSG0000( |
| 80106647 | 80346270 | 1 | protein_coding | ENSG0000( |
| 80355424 | 80356859 | 1 | lncRNA | ENSG0000( |

| | | | | |
|---|---|---|---|---|
| 80374015 | 80374569 | -1 | processed_pseudoge | ENSG00000 |
| 80440730 | 80441172 | 1 | TEC | ENSG00000 |
| 80441295 | 80466674 | 1 | lncRNA | ENSG00000 |
| 80466958 | 80469080 | 1 | lncRNA | ENSG00000 |
| 80470071 | 80470428 | -1 | processed_pseudoge | ENSG00000 |
| 80499196 | 80499392 | -1 | processed_pseudoge | ENSG00000 |
| 80555841 | 80557189 | -1 | processed_pseudoge | ENSG00000 |
| 81491439 | 81752774 | -1 | protein_coding | ENSG00000 |
| 81527102 | 81534915 | -1 | lncRNA | ENSG00000 |
| 81551686 | 81554092 | 1 | lncRNA | ENSG00000 |
| 81622200 | 81622319 | 1 | rRNA_pseudogene | ENSG00000 |
| 81724722 | 81736353 | -1 | lncRNA | ENSG00000 |
| 81764024 | 81764156 | 1 | snoRNA | ENSG00000 |
| 81764211 | 81764503 | 1 | processed_pseudoge | ENSG00000 |
| 81813286 | 81814157 | -1 | lncRNA | ENSG00000 |
| 81844602 | 82169391 | -1 | lncRNA | ENSG00000 |
| 81969453 | 81969539 | -1 | lncRNA | ENSG00000 |
| 82169986 | 82247754 | -1 | protein_coding | ENSG00000 |
| 82210338 | 82210440 | -1 | snRNA | ENSG00000 |
| 82263996 | 82264974 | -1 | processed_pseudoge | ENSG00000 |
| 82353861 | 82363657 | -1 | lncRNA | ENSG00000 |
| 82363206 | 82370828 | 1 | protein_coding | ENSG00000 |
| 82892390 | 83065841 | -1 | protein_coding | ENSG00000 |
| 82924829 | 82925122 | -1 | processed_pseudoge | ENSG00000 |
| 82932601 | 82938677 | 1 | lncRNA | ENSG00000 |
| 83067666 | 83171350 | 1 | protein_coding | ENSG00000 |
| 83161150 | 83193936 | -1 | protein_coding | ENSG00000 |
| 83193357 | 83198935 | 1 | protein_coding | ENSG00000 |
| 83210402 | 83431051 | -1 | protein_coding | ENSG00000 |
| 83392416 | 83393409 | 1 | processed_pseudoge | ENSG00000 |
| 83512534 | 83525704 | 1 | protein_coding | ENSG00000 |
| 83552880 | 83709691 | -1 | protein_coding | ENSG00000 |
| 83728055 | 83736525 | 1 | lncRNA | ENSG00000 |
| 83852312 | 83854167 | -1 | lncRNA | ENSG00000 |
| 83853266 | 83857515 | 1 | protein_coding | ENSG00000 |
| 83859656 | 83967423 | 1 | protein_coding | ENSG00000 |
| 83983728 | 84007323 | 1 | lncRNA | ENSG00000 |
| 84019204 | 84026100 | 1 | lncRNA | ENSG00000 |
| 84033772 | 84090881 | 1 | protein_coding | ENSG00000 |
| 84124241 | 84227643 | -1 | protein_coding | ENSG00000 |
| 84421028 | 84556152 | -1 | lncRNA | ENSG00000 |
| 84429584 | 84430802 | -1 | processed_pseudoge | ENSG00000 |
| 84687351 | 84764598 | -1 | protein_coding | ENSG00000 |
| 84687712 | 84709578 | 1 | lncRNA | ENSG00000 |
| 84689292 | 84689429 | -1 | processed_pseudoge | ENSG00000 |
| 84769010 | 84770219 | 1 | lncRNA | ENSG00000 |
| 84966397 | 84966625 | -1 | processed_pseudoge | ENSG00000 |
| 85155667 | 85167749 | 1 | unprocessed_pseudo | ENSG00000 |
| 85257328 | 85257694 | 1 | processed_pseudoge | ENSG00000 |
| 85286076 | 85290849 | 1 | unprocessed_pseudo | ENSG00000 |

| | | | | |
|---|---|---|---|---|
| 85287789 | 85289007 | 1 | processed_pseudoge | ENSG00000 |
| 85387219 | 85390186 | -1 | lncRNA | ENSG00000 |
| 85426657 | 85427125 | -1 | processed_pseudoge | ENSG00000 |
| 85427453 | 85427966 | -1 | processed_pseudoge | ENSG00000 |
| 85449584 | 85495791 | 1 | protein_coding | ENSG00000 |
| 85498441 | 85499058 | -1 | TEC | ENSG00000 |
| 85505496 | 85594156 | -1 | protein_coding | ENSG00000 |
| 85557978 | 85615234 | -1 | protein_coding | ENSG00000 |
| 85607785 | 85643792 | -1 | protein_coding | ENSG00000 |
| 85650491 | 85678932 | -1 | lncRNA | ENSG00000 |
| 85659892 | 85660606 | -1 | processed_pseudoge | ENSG00000 |
| 85677589 | 85677658 | -1 | snoRNA | ENSG00000 |
| 85731371 | 85732011 | 1 | processed_pseudoge | ENSG00000 |
| 85735744 | 85735921 | -1 | processed_pseudoge | ENSG00000 |
| 85751581 | 85751753 | 1 | snRNA | ENSG00000 |
| 85777639 | 85777755 | -1 | misc_RNA | ENSG00000 |
| 85868953 | 85869464 | 1 | processed_pseudoge | ENSG00000 |
| 85949690 | 85998263 | 1 | lncRNA | ENSG00000 |
| 85993021 | 85993159 | -1 | snRNA | ENSG00000 |
| 86086415 | 86087134 | -1 | processed_pseudoge | ENSG00000 |
| 86170447 | 86171018 | -1 | processed_pseudoge | ENSG00000 |
| 86214237 | 86214576 | -1 | processed_pseudoge | ENSG00000 |
| 86432244 | 86432669 | 1 | processed_pseudoge | ENSG00000 |
| 86435848 | 86436138 | 1 | misc_RNA | ENSG00000 |
| 86729708 | 86730108 | -1 | processed_pseudoge | ENSG00000 |
| 86768522 | 86768932 | -1 | processed_pseudoge | ENSG00000 |
| 86897118 | 86897421 | 1 | processed_pseudoge | ENSG00000 |
| 86897547 | 86897683 | 1 | unprocessed_pseudo | ENSG00000 |
| 86937528 | 87016679 | 1 | protein_coding | ENSG00000 |
| 86970466 | 86971193 | 1 | processed_pseudoge | ENSG00000 |
| 87000045 | 87000291 | -1 | processed_pseudoge | ENSG00000 |
| 87070156 | 87070461 | -1 | processed_pseudoge | ENSG00000 |
| 87083233 | 87083536 | 1 | misc_RNA | ENSG00000 |
| 87085498 | 87095406 | -1 | protein_coding | ENSG00000 |
| 87121693 | 87122688 | -1 | processed_pseudoge | ENSG00000 |
| 87151159 | 87155285 | -1 | lncRNA | ENSG00000 |
| 87152833 | 87265943 | 1 | protein_coding | ENSG00000 |
| 87173368 | 87174303 | -1 | processed_pseudoge | ENSG00000 |
| 87282980 | 87329278 | -1 | protein_coding | ENSG00000 |
| 87298772 | 87300447 | 1 | processed_pseudoge | ENSG00000 |
| 87322583 | 87399749 | 1 | protein_coding | ENSG00000 |
| 87344813 | 87365463 | 1 | protein_coding | ENSG00000 |
| 87356831 | 87358605 | 1 | TEC | ENSG00000 |
| 87407972 | 87464465 | 1 | protein_coding | ENSG00000 |
| 87408012 | 87511634 | 1 | protein_coding | ENSG00000 |
| 87425795 | 87426088 | -1 | processed_pseudoge | ENSG00000 |
| 87441165 | 87442146 | -1 | processed_pseudoge | ENSG00000 |
| 87444395 | 87445175 | -1 | processed_pseudoge | ENSG00000 |
| 87470623 | 87512336 | 1 | protein_coding | ENSG00000 |
| 87488445 | 87488552 | 1 | snRNA | ENSG00000 |

| | | | | |
|---|---|---|---|---|
| 87513938 | 87589987 | -1 | protein_coding | ENSG0000C |
| 87590067 | 87667453 | 1 | protein_coding | ENSG0000C |
| 87674860 | 87702233 | -1 | protein_coding | ENSG0000C |
| 87720915 | 87721018 | 1 | snoRNA | ENSG0000C |
| 87826173 | 87826285 | -1 | misc_RNA | ENSG0000C |
| 87845650 | 87845947 | 1 | misc_RNA | ENSG0000C |
| 87891424 | 87891869 | 1 | processed_pseudoge | ENSG0000C |
| 88047056 | 88047729 | -1 | lncRNA | ENSG0000C |
| 88047841 | 88066838 | 1 | protein_coding | ENSG0000C |
| 88139864 | 88166359 | -1 | protein_coding | ENSG0000C |
| 88172261 | 88175762 | 1 | TEC | ENSG0000C |
| 88177804 | 88442928 | -1 | lncRNA | ENSG0000C |
| 88265158 | 88265511 | 1 | processed_pseudoge | ENSG0000C |
| 88275882 | 88276989 | -1 | processed_pseudoge | ENSG0000C |
| 88378280 | 88385447 | 1 | lncRNA | ENSG0000C |
| 88609897 | 88963618 | -1 | protein_coding | ENSG0000C |
| 88714242 | 88714430 | -1 | snoRNA | ENSG0000C |
| 88964527 | 88966587 | 1 | TEC | ENSG0000C |
| 89001548 | 89001814 | 1 | processed_pseudoge | ENSG0000C |
| 89059628 | 89060109 | 1 | processed_pseudoge | ENSG0000C |
| 89063500 | 89063690 | -1 | snRNA | ENSG0000C |
| 89080164 | 89080667 | -1 | lncRNA | ENSG0000C |
| 89080751 | 89085160 | 1 | protein_coding | ENSG0000C |
| 89090946 | 89091242 | 1 | misc_RNA | ENSG0000C |
| 89095959 | 89118071 | -1 | protein_coding | ENSG0000C |
| 89130887 | 89145985 | -1 | lncRNA | ENSG0000C |
| 89146055 | 89165565 | 1 | protein_coding | ENSG0000C |
| 89177501 | 89231278 | -1 | protein_coding | ENSG0000C |
| 89254464 | 89315299 | -1 | protein_coding | ENSG0000C |
| 89301783 | 89303114 | 1 | processed_pseudoge | ENSG0000C |
| 89326625 | 89352722 | -1 | protein_coding | ENSG0000C |
| 89364616 | 89412273 | -1 | protein_coding | ENSG0000C |
| 89422099 | 89423048 | -1 | processed_pseudoge | ENSG0000C |
| 89433152 | 89633834 | 1 | protein_coding | ENSG0000C |
| 89478141 | 89478433 | 1 | misc_RNA | ENSG0000C |
| 89560875 | 89567044 | -1 | lncRNA | ENSG0000C |
| 89568144 | 89638749 | -1 | protein_coding | ENSG0000C |
| 89642498 | 89819794 | -1 | protein_coding | ENSG0000C |
| 89673469 | 89675783 | 1 | lncRNA | ENSG0000C |
| 89797598 | 89797877 | 1 | processed_pseudoge | ENSG0000C |
| 89814709 | 89815463 | 1 | processed_pseudoge | ENSG0000C |
| 89829894 | 89874436 | 1 | protein_coding | ENSG0000C |
| 89841329 | 89841424 | 1 | misc_RNA | ENSG0000C |
| 89876544 | 89876878 | 1 | processed_pseudoge | ENSG0000C |
| 89886156 | 89887848 | 1 | processed_pseudoge | ENSG0000C |
| 89894469 | 89921760 | 1 | protein_coding | ENSG0000C |
| 89898283 | 89898392 | -1 | misc_RNA | ENSG0000C |
| 89926528 | 90296908 | -1 | protein_coding | ENSG0000C |
| 89950116 | 89953186 | 1 | lncRNA | ENSG0000C |
| 90001222 | 90001506 | -1 | misc_RNA | ENSG0000C |

| | | | | |
|---|---|---|---|---|
| 90073051 | 90079734 | 1 | lncRNA | ENSG00000 |
| 90116700 | 90117105 | -1 | processed_pseudoge | ENSG00000 |
| 90295507 | 90363912 | 1 | lncRNA | ENSG00000 |
| 90312742 | 90312833 | 1 | miRNA | ENSG00000 |
| 90513573 | 90587072 | -1 | protein_coding | ENSG00000 |
| 91390313 | 91393000 | -1 | lncRNA | ENSG00000 |
| 91521660 | 91521737 | 1 | miRNA | ENSG00000 |
| 91557292 | 91690428 | -1 | lncRNA | ENSG00000 |
| 91726810 | 91727070 | 1 | processed_pseudoge | ENSG00000 |
| 91727159 | 91727324 | 1 | processed_pseudoge | ENSG00000 |
| 91727282 | 91727505 | -1 | processed_pseudoge | ENSG00000 |
| 91739609 | 91739897 | 1 | misc_RNA | ENSG00000 |
| 91815843 | 91816428 | -1 | processed_pseudoge | ENSG00000 |
| 92002610 | 92180156 | -1 | lncRNA | ENSG00000 |
| 92013842 | 92014734 | 1 | processed_pseudoge | ENSG00000 |
| 92119423 | 92119633 | 1 | snoRNA | ENSG00000 |
| 92387841 | 92388827 | -1 | lncRNA | ENSG00000 |
| 92631066 | 92633268 | -1 | lncRNA | ENSG00000 |
| 92723003 | 92723829 | -1 | lncRNA | ENSG00000 |
| 92790453 | 92790665 | 1 | snoRNA | ENSG00000 |
| 92887251 | 92888581 | -1 | processed_pseudoge | ENSG00000 |
| 93070387 | 93103384 | 1 | lncRNA | ENSG00000 |
| 93091976 | 93092760 | -1 | processed_pseudoge | ENSG00000 |
| 93240020 | 93419559 | -1 | protein_coding | ENSG00000 |
| 93416951 | 93677954 | 1 | lncRNA | ENSG00000 |
| 93720163 | 93745615 | 1 | lncRNA | ENSG00000 |
| 93811825 | 93813036 | 1 | TEC | ENSG00000 |
| 93879106 | 93879229 | 1 | snoRNA | ENSG00000 |
| 93886900 | 93889410 | 1 | lncRNA | ENSG00000 |
| 94163920 | 94194658 | 1 | lncRNA | ENSG00000 |
| 94344737 | 94344962 | 1 | processed_pseudoge | ENSG00000 |
| 94397664 | 94398829 | 1 | TEC | ENSG00000 |
| 94442030 | 94443809 | -1 | processed_pseudoge | ENSG00000 |
| 94446740 | 94447179 | -1 | processed_pseudoge | ENSG00000 |
| 95504182 | 95504496 | 1 | processed_pseudoge | ENSG00000 |
| 95575183 | 95577450 | -1 | lncRNA | ENSG00000 |
| 95577535 | 95609452 | 1 | protein_coding | ENSG00000 |
| 95630115 | 95632929 | -1 | processed_pseudoge | ENSG00000 |
| 95917143 | 96015391 | -1 | lncRNA | ENSG00000 |
| 95991107 | 95992349 | 1 | processed_pseudoge | ENSG00000 |
| 96015974 | 96215612 | 1 | protein_coding | ENSG00000 |
| 96199840 | 96521717 | -1 | lncRNA | ENSG00000 |
| 96282567 | 96282863 | 1 | misc_RNA | ENSG00000 |
| 96521595 | 96555276 | 1 | protein_coding | ENSG00000 |
| 96562548 | 96616636 | 1 | protein_coding | ENSG00000 |
| 96648489 | 96649069 | -1 | processed_pseudoge | ENSG00000 |
| 96649453 | 96649591 | -1 | snRNA | ENSG00000 |
| 96653274 | 96654142 | 1 | processed_pseudoge | ENSG00000 |
| 96672129 | 96672969 | -1 | processed_pseudoge | ENSG00000 |
| 96750824 | 96751728 | 1 | processed_pseudoge | ENSG00000 |

| | | | | |
|---|---|---|---|---|
| 96785137 | 96795821 | 1 | lncRNA | ENSG0000C |
| 96794125 | 96837477 | -1 | protein_coding | ENSG0000C |
| 96889315 | 96897891 | -1 | protein_coding | ENSG0000C |
| 96914699 | 96914995 | -1 | misc_RNA | ENSG0000C |
| 96924620 | 97140754 | 1 | protein_coding | ENSG0000C |
| 97142161 | 97283217 | -1 | protein_coding | ENSG0000C |
| 97283303 | 98400869 | 1 | lncRNA | ENSG0000C |
| 97710953 | 97717197 | -1 | lncRNA | ENSG0000C |
| 98024531 | 98024621 | 1 | miRNA | ENSG0000C |
| 98179499 | 98179832 | -1 | processed_pseudoge | ENSG0000C |
| 98210020 | 98214683 | -1 | lncRNA | ENSG0000C |
| 98780257 | 98780505 | -1 | processed_pseudoge | ENSG0000C |
| 98829967 | 98832508 | -1 | lncRNA | ENSG0000C |
| 98834574 | 98839458 | 1 | protein_coding | ENSG0000C |
| 98868535 | 98948006 | -1 | protein_coding | ENSG0000C |
| 99082449 | 99082761 | 1 | processed_pseudoge | ENSG0000C |
| 99124609 | 99124696 | 1 | miRNA | ENSG0000C |
| 99174744 | 99175479 | 1 | processed_pseudoge | ENSG0000C |
| 99271168 | 99350062 | -1 | protein_coding | ENSG0000C |
| 99369401 | 99394195 | -1 | protein_coding | ENSG0000C |
| 99398050 | 99425331 | -1 | protein_coding | ENSG0000C |
| 99424922 | 99432323 | 1 | lncRNA | ENSG0000C |
| 99432379 | 99521728 | -1 | protein_coding | ENSG0000C |
| 99520693 | 99589899 | 1 | protein_coding | ENSG0000C |
| 99526097 | 99526205 | 1 | misc_RNA | ENSG0000C |
| 99542387 | 99568825 | -1 | protein_coding | ENSG0000C |
| 99575712 | 99576456 | 1 | processed_pseudoge | ENSG0000C |
| 99606730 | 99615578 | 1 | protein_coding | ENSG0000C |
| 99642237 | 99642347 | -1 | misc_RNA | ENSG0000C |
| 99919910 | 99994247 | -1 | protein_coding | ENSG0000C |
| 99993934 | 100107613 | 1 | lncRNA | ENSG0000C |
| 100050372 | 100051236 | -1 | processed_pseudoge | ENSG0000C |
| 100093311 | 100093768 | -1 | processed_pseudoge | ENSG0000C |
| 100136064 | 100136663 | 1 | processed_pseudoge | ENSG0000C |
| 100177209 | 100178192 | 1 | processed_pseudoge | ENSG0000C |
| 100385015 | 100464929 | -1 | protein_coding | ENSG0000C |
| 100427118 | 100437484 | 1 | lncRNA | ENSG0000C |
| 100508194 | 100881372 | -1 | protein_coding | ENSG0000C |
| 100530276 | 100531052 | 1 | processed_pseudoge | ENSG0000C |
| 100881471 | 100882987 | 1 | lncRNA | ENSG0000C |
| 100889603 | 100890338 | 1 | processed_pseudoge | ENSG0000C |
| 101181257 | 102070083 | 1 | protein_coding | ENSG0000C |
| 101430411 | 101431553 | -1 | processed_pseudoge | ENSG0000C |
| 101452917 | 101454245 | -1 | lncRNA | ENSG0000C |
| 102078872 | 102079555 | 1 | processed_pseudoge | ENSG0000C |
| 102350271 | 102369827 | -1 | lncRNA | ENSG0000C |
| 102453367 | 102453792 | -1 | processed_pseudoge | ENSG0000C |
| 103002514 | 103003897 | 1 | processed_pseudoge | ENSG0000C |
| 103583135 | 103583268 | 1 | snoRNA | ENSG0000C |
| 103866023 | 103874687 | -1 | lncRNA | ENSG0000C |

| | | | | |
|---|---|---|---|---|
| 104017633 | 104019748 | -1 | processed_pseudoge | ENSG00000 |
| 104025540 | 104026763 | -1 | processed_pseudoge | ENSG00000 |
| 104487095 | 104528158 | -1 | lncRNA | ENSG00000 |
| 104687241 | 104687671 | 1 | processed_pseudoge | ENSG00000 |
| 104728094 | 104859919 | -1 | protein_coding | ENSG00000 |
| 104766822 | 104766926 | 1 | snRNA | ENSG00000 |
| 104831129 | 104845820 | -1 | lncRNA | ENSG00000 |
| 104860449 | 104862123 | 1 | processed_pseudoge | ENSG00000 |
| 104864464 | 104941447 | -1 | lncRNA | ENSG00000 |
| 104936616 | 105083332 | 1 | protein_coding | ENSG00000 |
| 105045562 | 105045668 | -1 | snRNA | ENSG00000 |
| 105096822 | 105137157 | -1 | protein_coding | ENSG00000 |
| 105136308 | 105169952 | 1 | lncRNA | ENSG00000 |
| 105157900 | 105180014 | -1 | protein_coding | ENSG00000 |
| 105273220 | 105454062 | -1 | protein_coding | ENSG00000 |
| 105279016 | 105281755 | 1 | lncRNA | ENSG00000 |
| 105298149 | 105298895 | 1 | processed_pseudoge | ENSG00000 |
| 105302453 | 105302823 | -1 | processed_pseudoge | ENSG00000 |
| 105403207 | 105588802 | 1 | lncRNA | ENSG00000 |
| 105612667 | 105632296 | -1 | lncRNA | ENSG00000 |
| 105666326 | 105667998 | 1 | processed_pseudoge | ENSG00000 |
| 105679378 | 105893041 | -1 | lncRNA | ENSG00000 |
| 105904373 | 105904687 | 1 | misc_RNA | ENSG00000 |
| 105993463 | 106109939 | 1 | protein_coding | ENSG00000 |
| 106045423 | 106325791 | -1 | protein_coding | ENSG00000 |
| 106100140 | 106100593 | 1 | lncRNA | ENSG00000 |
| 106271634 | 106271720 | 1 | snRNA | ENSG00000 |
| 106283413 | 106283711 | -1 | misc_RNA | ENSG00000 |
| 106304176 | 106304286 | 1 | snRNA | ENSG00000 |
| 106358566 | 106359717 | -1 | lncRNA | ENSG00000 |
| 106360808 | 106571978 | 1 | protein_coding | ENSG00000 |
| 106420706 | 106420807 | 1 | misc_RNA | ENSG00000 |
| 106449381 | 106449497 | 1 | rRNA_pseudogene | ENSG00000 |
| 106451496 | 106457248 | -1 | lncRNA | ENSG00000 |
| 106454828 | 106454929 | 1 | misc_RNA | ENSG00000 |
| 106570771 | 106629498 | -1 | protein_coding | ENSG00000 |
| 106607716 | 106607819 | 1 | snRNA | ENSG00000 |
| 106629578 | 106668417 | 1 | protein_coding | ENSG00000 |
| 106642463 | 106642928 | 1 | processed_pseudoge | ENSG00000 |
| 106695535 | 106699994 | 1 | lncRNA | ENSG00000 |
| 106705334 | 106788148 | -1 | lncRNA | ENSG00000 |
| 106738948 | 106739054 | -1 | snRNA | ENSG00000 |
| 106784125 | 106784220 | 1 | miRNA | ENSG00000 |
| 106969831 | 106975627 | -1 | protein_coding | ENSG00000 |
| 107028199 | 107051586 | 1 | protein_coding | ENSG00000 |
| 107065182 | 107115269 | -1 | protein_coding | ENSG00000 |
| 107133071 | 107133170 | 1 | snRNA | ENSG00000 |
| 107152562 | 107459564 | -1 | protein_coding | ENSG00000 |
| 107192300 | 107192573 | 1 | processed_pseudoge | ENSG00000 |
| 107229759 | 107230152 | 1 | processed_pseudoge | ENSG00000 |

| | | | | |
|---|---|---|---|---|
| 107490106 | 107661306 | 1 | protein_coding | ENSG0000( |
| 107509803 | 107511721 | -1 | lncRNA | ENSG0000( |
| 107697299 | 107700218 | -1 | TEC | ENSG0000( |
| 107702154 | 107824317 | -1 | protein_coding | ENSG0000( |
| 107867756 | 107958208 | -1 | protein_coding | ENSG0000( |
| 107930088 | 107930182 | 1 | snRNA | ENSG0000( |
| 107931800 | 107932254 | 1 | processed_pseudoge | ENSG0000( |
| 107957413 | 107959986 | 1 | lncRNA | ENSG0000( |
| 107985089 | 107985690 | -1 | processed_pseudoge | ENSG0000( |
| 107985659 | 107985773 | -1 | snoRNA | ENSG0000( |
| 107985811 | 107986031 | -1 | processed_pseudoge | ENSG0000( |
| 108004357 | 108005568 | -1 | processed_pseudoge | ENSG0000( |
| 108030249 | 108030718 | -1 | lncRNA | ENSG0000( |
| 108041409 | 108165854 | -1 | protein_coding | ENSG0000( |
| 108060818 | 108060930 | 1 | misc_RNA | ENSG0000( |
| 108123457 | 108159392 | 1 | lncRNA | ENSG0000( |
| 108166058 | 108188809 | 1 | protein_coding | ENSG0000( |
| 108178871 | 108179317 | 1 | TEC | ENSG0000( |
| 108211222 | 108261246 | -1 | protein_coding | ENSG0000( |
| 108252214 | 108252400 | 1 | processed_pseudoge | ENSG0000( |
| 108252401 | 108252507 | 1 | rRNA_pseudogene | ENSG0000( |
| 108261288 | 108269200 | 1 | lncRNA | ENSG0000( |
| 108275642 | 108276546 | 1 | lncRNA | ENSG0000( |
| 108292766 | 108292869 | 1 | snRNA | ENSG0000( |
| 108294991 | 108526796 | 1 | protein_coding | ENSG0000( |
| 108318079 | 108318404 | 1 | processed_pseudoge | ENSG0000( |
| 108359084 | 108359863 | -1 | TEC | ENSG0000( |
| 108372772 | 108372927 | -1 | processed_pseudoge | ENSG0000( |
| 108387512 | 108388080 | 1 | processed_pseudoge | ENSG0000( |
| 108392073 | 108392177 | 1 | snRNA | ENSG0000( |
| 108441880 | 108558286 | -1 | lncRNA | ENSG0000( |
| 108551018 | 108551207 | -1 | processed_pseudoge | ENSG0000( |
| 108559835 | 108684774 | 1 | protein_coding | ENSG0000( |
| 108618000 | 108618285 | -1 | processed_pseudoge | ENSG0000( |
| 108751654 | 108769942 | 1 | lncRNA | ENSG0000( |
| 108782126 | 108787053 | -1 | transcribed_processe | ENSG0000( |
| 108798929 | 108837113 | 1 | lncRNA | ENSG0000( |
| 108817680 | 108817926 | 1 | processed_pseudoge | ENSG0000( |
| 108848416 | 108974476 | 1 | protein_coding | ENSG0000( |
| 108856400 | 108856737 | -1 | processed_pseudoge | ENSG0000( |
| 108907615 | 108907873 | -1 | processed_pseudoge | ENSG0000( |
| 108922976 | 108924103 | -1 | lncRNA | ENSG0000( |
| 108986437 | 109094819 | -1 | protein_coding | ENSG0000( |
| 108998482 | 108999125 | 1 | lncRNA | ENSG0000( |
| 109059509 | 109059614 | -1 | snRNA | ENSG0000( |
| 109095110 | 109163932 | 1 | protein_coding | ENSG0000( |
| 109165831 | 109355063 | 1 | transcribed_unitary_ | ENSG0000( |
| 109288440 | 109288562 | 1 | processed_pseudoge | ENSG0000( |
| 109288571 | 109290503 | -1 | processed_pseudoge | ENSG0000( |
| 109305494 | 109305595 | 1 | misc_RNA | ENSG0000( |

| | | | | |
|---|---|---|---|---|
| 109327175 | 109327914 | -1 | processed_pseudoge | ENSG0000( |
| 109366514 | 109382467 | -1 | protein_coding | ENSG0000( |
| 109382795 | 109383666 | 1 | lncRNA | ENSG0000( |
| 109390215 | 109441171 | -1 | protein_coding | ENSG0000( |
| 109440724 | 109443919 | 1 | protein_coding | ENSG0000( |
| 109444062 | 109465968 | -1 | protein_coding | ENSG0000( |
| 109462594 | 109483237 | -1 | protein_coding | ENSG0000( |
| 109483638 | 109506800 | 1 | lncRNA | ENSG0000( |
| 109492856 | 109691217 | -1 | protein_coding | ENSG0000( |
| 109691312 | 109825428 | 1 | protein_coding | ENSG0000( |
| 109826522 | 109828785 | 1 | lncRNA | ENSG0000( |
| 109978256 | 109980718 | 1 | protein_coding | ENSG0000( |
| 110020011 | 110040241 | -1 | lncRNA | ENSG0000( |
| 110099819 | 110180004 | -1 | protein_coding | ENSG0000( |
| 110180141 | 110254275 | 1 | protein_coding | ENSG0000( |
| 110245928 | 110358272 | -1 | protein_coding | ENSG0000( |
| 110341973 | 110354563 | 1 | lncRNA | ENSG0000( |
| 110391771 | 110415575 | -1 | protein_coding | ENSG0000( |
| 110424687 | 110476641 | -1 | protein_coding | ENSG0000( |
| 110439996 | 110440303 | -1 | misc_RNA | ENSG0000( |
| 110477907 | 110479436 | 1 | lncRNA | ENSG0000( |
| 110562175 | 110562615 | -1 | processed_pseudoge | ENSG0000( |
| 110598093 | 110598705 | 1 | processed_pseudoge | ENSG0000( |
| 110609978 | 110815958 | -1 | protein_coding | ENSG0000( |
| 110645699 | 110646558 | 1 | processed_pseudoge | ENSG0000( |
| 110700562 | 110700900 | 1 | processed_pseudoge | ENSG0000( |
| 110706362 | 110706882 | -1 | processed_pseudoge | ENSG0000( |
| 110722250 | 110722348 | -1 | snRNA | ENSG0000( |
| 110848546 | 110848673 | 1 | snoRNA | ENSG0000( |
| 110856417 | 110856522 | 1 | snRNA | ENSG0000( |
| 110858239 | 110859147 | 1 | processed_pseudoge | ENSG0000( |
| 110863860 | 110864451 | 1 | processed_pseudoge | ENSG0000( |
| 110874770 | 110898879 | 1 | protein_coding | ENSG0000( |
| 110923566 | 110924107 | 1 | processed_pseudoge | ENSG0000( |
| 110958706 | 110967890 | 1 | protein_coding | ENSG0000( |
| 110982015 | 111028263 | 1 | protein_coding | ENSG0000( |
| 111046868 | 111047521 | -1 | processed_pseudoge | ENSG0000( |
| 111087503 | 111231194 | 1 | protein_coding | ENSG0000( |
| 111091213 | 111091332 | -1 | snRNA | ENSG0000( |
| 111227747 | 111259034 | -1 | lncRNA | ENSG0000( |
| 111259279 | 111445354 | 1 | protein_coding | ENSG0000( |
| 111297126 | 111298510 | 1 | lncRNA | ENSG0000( |
| 111299028 | 111483715 | -1 | protein_coding | ENSG0000( |
| 111309203 | 111313517 | 1 | lncRNA | ENSG0000( |
| 111353702 | 111354183 | 1 | processed_pseudoge | ENSG0000( |
| 111360641 | 111361607 | -1 | lncRNA | ENSG0000( |
| 111430260 | 111431781 | 1 | processed_pseudoge | ENSG0000( |
| 111483511 | 111598302 | 1 | lncRNA | ENSG0000( |
| 111494991 | 111495713 | -1 | processed_pseudoge | ENSG0000( |
| 111505307 | 111511071 | -1 | lncRNA | ENSG0000( |

| | | | | |
|---|---|---|---|---|
| 111555381 | 111606906 | -1 | protein_coding | ENSG00000( |
| 111599875 | 111602295 | 1 | lncRNA | ENSG00000( |
| 111660332 | 111873452 | -1 | protein_coding | ENSG00000( |
| 111900305 | 111909420 | -1 | lncRNA | ENSG00000( |
| 112054072 | 112070969 | 1 | protein_coding | ENSG00000( |
| 112070663 | 112087529 | -1 | protein_coding | ENSG00000( |
| 112087591 | 112102790 | 1 | protein_coding | ENSG00000( |
| 112107931 | 112254939 | -1 | protein_coding | ENSG00000( |
| 112154765 | 112166476 | 1 | lncRNA | ENSG00000( |
| 112196440 | 112196546 | -1 | snRNA | ENSG00000( |
| 112217640 | 112219570 | -1 | lncRNA | ENSG00000( |
| 112234165 | 112234758 | -1 | lncRNA | ENSG00000( |
| 112236093 | 112307009 | 1 | lncRNA | ENSG00000( |
| 112236806 | 112350094 | 1 | protein_coding | ENSG00000( |
| 112325753 | 112326749 | -1 | processed_pseudoge | ENSG00000( |
| 112347330 | 112351294 | 1 | protein_coding | ENSG00000( |
| 112355841 | 112356714 | -1 | processed_pseudoge | ENSG00000( |
| 112361437 | 112362722 | -1 | processed_pseudoge | ENSG00000( |
| 112365704 | 112367444 | -1 | processed_pseudoge | ENSG00000( |
| 112392363 | 112396364 | -1 | lncRNA | ENSG00000( |
| 112476538 | 112481636 | -1 | lncRNA | ENSG00000( |
| 112616703 | 112617880 | -1 | processed_pseudoge | ENSG00000( |
| 112825939 | 112826470 | 1 | processed_pseudoge | ENSG00000( |
| 112971493 | 112971599 | 1 | snRNA | ENSG00000( |
| 112988311 | 113140870 | 1 | lncRNA | ENSG00000( |
| 113010937 | 113011415 | 1 | processed_pseudoge | ENSG00000( |
| 113222166 | 113224072 | 1 | processed_pseudoge | ENSG00000( |
| 113357003 | 113367944 | -1 | lncRNA | ENSG00000( |
| 113428540 | 113433555 | -1 | lncRNA | ENSG00000( |
| 113531118 | 113543793 | 1 | lncRNA | ENSG00000( |
| 113581501 | 113581947 | 1 | processed_pseudoge | ENSG00000( |
| 113616927 | 113650090 | -1 | lncRNA | ENSG00000( |
| 113791829 | 113837368 | -1 | lncRNA | ENSG00000( |
| 113839279 | 113839621 | 1 | processed_pseudoge | ENSG00000( |
| 113857345 | 113863475 | 1 | protein_coding | ENSG00000( |
| 113868013 | 113873351 | -1 | lncRNA | ENSG00000( |
| 113904132 | 113921642 | 1 | lncRNA | ENSG00000( |
| 113933028 | 114011308 | -1 | protein_coding | ENSG00000( |
| 113969701 | 114471705 | 1 | lncRNA | ENSG00000( |
| 114019621 | 114020483 | -1 | processed_pseudoge | ENSG00000( |
| 114055586 | 114343045 | -1 | protein_coding | ENSG00000( |
| 114084168 | 114084792 | -1 | processed_pseudoge | ENSG00000( |
| 114220681 | 114220786 | -1 | rRNA_pseudogene | ENSG00000( |
| 114349483 | 114350648 | -1 | processed_pseudoge | ENSG00000( |
| 114477350 | 114488752 | -1 | lncRNA | ENSG00000( |
| 114523443 | 114545335 | -1 | lncRNA | ENSG00000( |
| 114822994 | 115002280 | 1 | lncRNA | ENSG00000( |
| 114866873 | 114866976 | -1 | snRNA | ENSG00000( |
| 115358498 | 115358752 | -1 | processed_pseudoge | ENSG00000( |
| 115633540 | 115643916 | -1 | lncRNA | ENSG00000( |

| | | | | |
|---|---|---|---|---|
| 115931149 | 116060891 | -1 | protein_coding | ENSG00000( |
| 116033901 | 116046753 | 1 | lncRNA | ENSG00000( |
| 116038756 | 116039495 | 1 | processed_pseudoge | ENSG00000( |
| 116100851 | 116249497 | 1 | protein_coding | ENSG00000( |
| 116118909 | 116158747 | -1 | protein_coding | ENSG00000( |
| 116137058 | 116137572 | -1 | processed_pseudoge | ENSG00000( |
| 116244187 | 116244728 | -1 | lncRNA | ENSG00000( |
| 116249964 | 116254075 | -1 | protein_coding | ENSG00000( |
| 116254173 | 116444860 | 1 | protein_coding | ENSG00000( |
| 116258493 | 116259115 | -1 | processed_pseudoge | ENSG00000( |
| 116267760 | 116279903 | -1 | protein_coding | ENSG00000( |
| 116399395 | 116463696 | 1 | protein_coding | ENSG00000( |
| 116453014 | 116453565 | -1 | processed_pseudoge | ENSG00000( |
| 116457323 | 116458669 | -1 | processed_pseudoge | ENSG00000( |
| 116460739 | 116463692 | -1 | lncRNA | ENSG00000( |
| 116461370 | 116463771 | 1 | protein_coding | ENSG00000( |
| 116492297 | 116497099 | 1 | lncRNA | ENSG00000( |
| 116494989 | 116545684 | -1 | protein_coding | ENSG00000( |
| 116511639 | 116524788 | 1 | protein_coding | ENSG00000( |
| 116529013 | 116558868 | 1 | protein_coding | ENSG00000( |
| 116571409 | 116597675 | 1 | protein_coding | ENSG00000( |
| 116616479 | 116632985 | 1 | protein_coding | ENSG00000( |
| 116635618 | 116668794 | -1 | protein_coding | ENSG00000( |
| 116681187 | 116741867 | 1 | protein_coding | ENSG00000( |
| 116752197 | 116765719 | -1 | protein_coding | ENSG00000( |
| 116792085 | 116829037 | -1 | protein_coding | ENSG00000( |
| 116877212 | 116932161 | 1 | protein_coding | ENSG00000( |
| 117048670 | 117048838 | -1 | processed_pseudoge | ENSG00000( |
| 117060682 | 117060799 | 1 | rRNA_pseudogene | ENSG00000( |
| 117262243 | 117264817 | -1 | lncRNA | ENSG00000( |
| 117265558 | 117273565 | 1 | protein_coding | ENSG00000( |
| 117288300 | 117425855 | -1 | protein_coding | ENSG00000( |
| 117299364 | 117299638 | 1 | misc_RNA | ENSG00000( |
| 117301455 | 117301758 | 1 | misc_RNA | ENSG00000( |
| 117318211 | 117573571 | -1 | protein_coding | ENSG00000( |
| 117431591 | 117432442 | 1 | processed_pseudoge | ENSG00000( |
| 117451130 | 117453525 | -1 | lncRNA | ENSG00000( |
| 117453817 | 117569858 | 1 | protein_coding | ENSG00000( |
| 117457734 | 117457838 | -1 | snRNA | ENSG00000( |
| 117560269 | 117602542 | -1 | protein_coding | ENSG00000( |
| 117633706 | 117645087 | 1 | unitary_pseudogene | ENSG00000( |
| 117675469 | 117710727 | 1 | protein_coding | ENSG00000( |
| 117907264 | 118317676 | 1 | protein_coding | ENSG00000( |
| 117998975 | 117999448 | 1 | processed_pseudoge | ENSG00000( |
| 118452469 | 118454992 | -1 | processed_pseudoge | ENSG00000( |
| 118460772 | 118710075 | -1 | protein_coding | ENSG00000( |
| 118501430 | 118502906 | 1 | processed_pseudoge | ENSG00000( |
| 118548296 | 118561716 | 1 | protein_coding | ENSG00000( |
| 118565660 | 118576840 | 1 | lncRNA | ENSG00000( |
| 118589070 | 118589634 | 1 | processed_pseudoge | ENSG00000( |

| | | | | |
|---|---|---|---|---|
| 118757518 | 118783420 | 1 | transcribed_processe | ENSG00000 |
| 118813442 | 118935162 | -1 | protein_coding | ENSG00000 |
| 118894152 | 118909171 | 1 | protein_coding | ENSG00000 |
| 118934770 | 119182745 | 1 | lncRNA | ENSG00000 |
| 118959763 | 119149387 | -1 | protein_coding | ENSG00000 |
| 119057880 | 119057974 | 1 | misc_RNA | ENSG00000 |
| 119069047 | 119069143 | -1 | miRNA | ENSG00000 |
| 119159297 | 119159476 | 1 | processed_pseudoge | ENSG00000 |
| 119177205 | 119349761 | -1 | protein_coding | ENSG00000 |
| 119269133 | 119269714 | -1 | processed_pseudoge | ENSG00000 |
| 119327281 | 119327385 | 1 | snRNA | ENSG00000 |
| 119349920 | 119391829 | 1 | lncRNA | ENSG00000 |
| 120015179 | 120015257 | 1 | miRNA | ENSG00000 |
| 120462511 | 120706789 | 1 | lncRNA | ENSG00000 |
| 120526294 | 120526400 | -1 | snRNA | ENSG00000 |
| 120686621 | 120686742 | 1 | rRNA_pseudogene | ENSG00000 |
| 120781220 | 120781512 | 1 | processed_pseudoge | ENSG00000 |
| 120840733 | 120845475 | 1 | lncRNA | ENSG00000 |
| 121079494 | 121334745 | -1 | protein_coding | ENSG00000 |
| 121354354 | 121354460 | 1 | snRNA | ENSG00000 |
| 121378797 | 121378898 | -1 | misc_RNA | ENSG00000 |
| 121381472 | 121381860 | 1 | processed_pseudoge | ENSG00000 |
| 121389324 | 121390357 | 1 | transcribed_unproce | ENSG00000 |
| 121435595 | 121449727 | 1 | protein_coding | ENSG00000 |
| 121453981 | 121454121 | -1 | snRNA | ENSG00000 |
| 121478273 | 121479908 | -1 | processed_pseudoge | ENSG00000 |
| 121542486 | 121542626 | -1 | snRNA | ENSG00000 |
| 121580332 | 121580521 | -1 | snRNA | ENSG00000 |
| 121641009 | 121641221 | -1 | processed_pseudoge | ENSG00000 |
| 121653795 | 121654690 | 1 | processed_pseudoge | ENSG00000 |
| 121679972 | 121680459 | -1 | processed_pseudoge | ENSG00000 |
| 121682925 | 121684272 | 1 | processed_pseudoge | ENSG00000 |
| 121858179 | 121858768 | -1 | processed_pseudoge | ENSG00000 |
| 122211648 | 122211811 | 1 | snRNA | ENSG00000 |
| 122399551 | 122433119 | 1 | protein_coding | ENSG00000 |
| 122436789 | 122439223 | -1 | TEC | ENSG00000 |
| 122443351 | 122471807 | -1 | protein_coding | ENSG00000 |
| 122454358 | 122454612 | 1 | processed_pseudoge | ENSG00000 |
| 122471917 | 122726373 | 1 | protein_coding | ENSG00000 |
| 122471923 | 122484161 | 1 | TEC | ENSG00000 |
| 122531123 | 122531436 | -1 | processed_pseudoge | ENSG00000 |
| 122562172 | 122621014 | -1 | lncRNA | ENSG00000 |
| 122643388 | 122644771 | 1 | lncRNA | ENSG00000 |
| 122711887 | 122718814 | -1 | lncRNA | ENSG00000 |
| 122745280 | 122745529 | 1 | misc_RNA | ENSG00000 |
| 122779716 | 122784074 | 1 | protein_coding | ENSG00000 |
| 122789049 | 122809720 | 1 | protein_coding | ENSG00000 |
| 122859678 | 122859988 | -1 | processed_pseudoge | ENSG00000 |
| 122975198 | 122996061 | -1 | lncRNA | ENSG00000 |
| 122996235 | 123072925 | 1 | protein_coding | ENSG00000 |

| | | | | |
|---|---|---|---|---|
| 123216339 | 123637093 | -1 | protein_coding | ENSG00000 |
| 123389421 | 123510247 | 1 | lncRNA | ENSG00000 |
| 123519697 | 123557670 | 1 | lncRNA | ENSG00000 |
| 123589711 | 123685323 | 1 | lncRNA | ENSG00000 |
| 123803865 | 124825640 | 1 | protein_coding | ENSG00000 |
| 123823240 | 123829285 | 1 | lncRNA | ENSG00000 |
| 124644434 | 124963165 | -1 | lncRNA | ENSG00000 |
| 124962545 | 125092633 | 1 | protein_coding | ENSG00000 |
| 125119049 | 125264407 | 1 | protein_coding | ENSG00000 |
| 125219962 | 125302078 | -1 | protein_coding | ENSG00000 |
| 125268087 | 125274828 | 1 | lncRNA | ENSG00000 |
| 125370034 | 125445193 | -1 | lncRNA | ENSG00000 |
| 125577545 | 125749190 | -1 | lncRNA | ENSG00000 |
| 125674353 | 125720218 | 1 | lncRNA | ENSG00000 |
| 125747664 | 125761269 | 1 | protein_coding | ENSG00000 |
| 125781161 | 125932034 | 1 | protein_coding | ENSG00000 |
| 125797856 | 125818858 | -1 | lncRNA | ENSG00000 |
| 125875168 | 125875498 | 1 | misc_RNA | ENSG00000 |
| 125956770 | 125980244 | 1 | protein_coding | ENSG00000 |
| 125979812 | 125979934 | -1 | rRNA_pseudogene | ENSG00000 |
| 125986479 | 126203817 | 1 | protein_coding | ENSG00000 |
| 126257921 | 126258355 | -1 | processed_pseudoge | ENSG00000 |
| 126304223 | 127213609 | -1 | transcribed_processe | ENSG00000 |
| 126340174 | 126348875 | 1 | protein_coding | ENSG00000 |
| 126484631 | 126484713 | 1 | miRNA | ENSG00000 |
| 126590287 | 126590393 | -1 | snRNA | ENSG00000 |
| 126627484 | 126628570 | -1 | processed_pseudoge | ENSG00000 |
| 126643488 | 126644390 | 1 | transcribed_processe | ENSG00000 |
| 126683036 | 126683847 | 1 | processed_pseudoge | ENSG00000 |
| 126719560 | 126721778 | 1 | lncRNA | ENSG00000 |
| 127118671 | 127199481 | 1 | protein_coding | ENSG00000 |
| 127266610 | 127288567 | 1 | protein_coding | ENSG00000 |
| 127288712 | 127343609 | -1 | protein_coding | ENSG00000 |
| 127317883 | 127317973 | -1 | rRNA_pseudogene | ENSG00000 |
| 127355756 | 127356789 | 1 | processed_pseudoge | ENSG00000 |
| 127362513 | 127363403 | 1 | processed_pseudoge | ENSG00000 |
| 127416535 | 127416952 | -1 | processed_pseudoge | ENSG00000 |
| 127435636 | 127436189 | 1 | processed_pseudoge | ENSG00000 |
| 127438406 | 127519001 | -1 | protein_coding | ENSG00000 |
| 127438406 | 127459389 | -1 | protein_coding | ENSG00000 |
| 127472794 | 127519191 | -1 | protein_coding | ENSG00000 |
| 127519455 | 127591820 | 1 | protein_coding | ENSG00000 |
| 127632958 | 127635321 | -1 | unprocessed_pseudo | ENSG00000 |
| 127654860 | 127655278 | -1 | processed_pseudoge | ENSG00000 |
| 127659424 | 127660338 | 1 | unprocessed_pseudo | ENSG00000 |
| 127659690 | 127660579 | -1 | unprocessed_pseudo | ENSG00000 |
| 127664554 | 127681555 | 1 | lncRNA | ENSG00000 |
| 127708072 | 127918631 | -1 | protein_coding | ENSG00000 |
| 127909833 | 127910052 | 1 | processed_pseudoge | ENSG00000 |
| 127968779 | 128520674 | -1 | protein_coding | ENSG00000 |

| | | | | |
|---|---|---|---|---|
| 128027886 | 128085248 | 1 | lncRNA | ENSG0000C |
| 128500527 | 128501176 | 1 | lncRNA | ENSG0000C |
| 128580113 | 128580884 | -1 | processed_pseudoge | ENSG0000C |
| 128584390 | 128584491 | 1 | misc_RNA | ENSG0000C |
| 128639188 | 128639653 | -1 | processed_pseudoge | ENSG0000C |
| 128761717 | 128761808 | -1 | misc_RNA | ENSG0000C |
| 128883141 | 129516569 | 1 | protein_coding | ENSG0000C |
| 128928995 | 128929988 | -1 | processed_pseudoge | ENSG0000C |
| 129157523 | 129159120 | -1 | processed_pseudoge | ENSG0000C |
| 129436875 | 129436980 | -1 | snRNA | ENSG0000C |
| 129439485 | 129576047 | -1 | lncRNA | ENSG0000C |
| 129479615 | 129481410 | -1 | lncRNA | ENSG0000C |
| 129576132 | 129710177 | -1 | protein_coding | ENSG0000C |
| 129756298 | 129757112 | -1 | processed_pseudoge | ENSG0000C |
| 129783769 | 129784045 | -1 | processed_pseudoge | ENSG0000C |
| 129800908 | 129802214 | 1 | processed_pseudoge | ENSG0000C |
| 129819637 | 129820170 | -1 | processed_pseudoge | ENSG0000C |
| 129831244 | 129861547 | -1 | protein_coding | ENSG0000C |
| 130013699 | 130141451 | 1 | protein_coding | ENSG0000C |
| 130133410 | 130146179 | -1 | lncRNA | ENSG0000C |
| 130144315 | 130365425 | -1 | protein_coding | ENSG0000C |
| 130365734 | 130443063 | 1 | protein_coding | ENSG0000C |
| 130573996 | 130574112 | -1 | misc_RNA | ENSG0000C |
| 130697312 | 130697778 | 1 | lncRNA | ENSG0000C |
| 130827406 | 130837135 | 1 | protein_coding | ENSG0000C |
| 130839347 | 131063322 | -1 | protein_coding | ENSG0000C |
| 131135467 | 131283535 | 1 | protein_coding | ENSG0000C |
| 131184325 | 131185486 | -1 | processed_pseudoge | ENSG0000C |
| 131469059 | 131469536 | 1 | processed_pseudoge | ENSG0000C |
| 131470832 | 131584332 | 1 | protein_coding | ENSG0000C |
| 131573966 | 131628242 | -1 | protein_coding | ENSG0000C |
| 131628442 | 131747418 | 1 | protein_coding | ENSG0000C |
| 131642818 | 131642966 | -1 | snRNA | ENSG0000C |
| 131699644 | 131701401 | -1 | protein_coding | ENSG0000C |
| 131708441 | 131711017 | -1 | protein_coding | ENSG0000C |
| 131780721 | 131781312 | -1 | processed_pseudoge | ENSG0000C |
| 131792172 | 131792231 | 1 | miRNA | ENSG0000C |
| 131808016 | 131895155 | 1 | protein_coding | ENSG0000C |
| 131819803 | 131820091 | -1 | processed_pseudoge | ENSG0000C |
| 131820334 | 131820653 | 1 | misc_RNA | ENSG0000C |
| 131825981 | 131828413 | -1 | processed_pseudoge | ENSG0000C |
| 131901963 | 131920565 | 1 | lncRNA | ENSG0000C |
| 131910191 | 131911527 | 1 | TEC | ENSG0000C |
| 131948176 | 131951372 | -1 | protein_coding | ENSG0000C |
| 131950946 | 132077393 | 1 | lncRNA | ENSG0000C |
| 132085084 | 132099279 | 1 | lncRNA | ENSG0000C |
| 132115192 | 132115263 | -1 | miRNA | ENSG0000C |
| 132130252 | 132131795 | -1 | TEC | ENSG0000C |
| 132131888 | 132169374 | 1 | lncRNA | ENSG0000C |
| 132271982 | 132273332 | 1 | processed_pseudoge | ENSG0000C |

| | | | | |
|---|---|---|---|---|
| 132296055 | 132401475 | -1 | protein_coding | ENSG00000( |
| 132445867 | 132513198 | -1 | protein_coding | ENSG00000( |
| 132518830 | 132519295 | -1 | processed_pseudoge | ENSG00000( |
| 132538290 | 132539336 | 1 | protein_coding | ENSG00000( |
| 132552693 | 132553721 | 1 | protein_coding | ENSG00000( |
| 132559024 | 132559265 | 1 | unprocessed_pseudc | ENSG00000( |
| 132570322 | 132571359 | 1 | protein_coding | ENSG00000( |
| 132588673 | 132589686 | -1 | protein_coding | ENSG00000( |
| 132594398 | 132595436 | -1 | unprocessed_pseudc | ENSG00000( |
| 132599972 | 132602740 | -1 | lncRNA | ENSG00000( |
| 132608225 | 132609302 | -1 | transcribed_unproce | ENSG00000( |
| 132617022 | 132624275 | -1 | protein_coding | ENSG00000( |
| 132644984 | 132646003 | -1 | protein_coding | ENSG00000( |
| 132674885 | 132675383 | 1 | processed_pseudoge | ENSG00000( |
| 132680849 | 132714055 | -1 | protein_coding | ENSG00000( |
| 132698783 | 132699667 | 1 | processed_pseudoge | ENSG00000( |
| 132722787 | 132734765 | -1 | protein_coding | ENSG00000( |
| 132743870 | 132763459 | -1 | protein_coding | ENSG00000( |
| 132752675 | 132753951 | 1 | lncRNA | ENSG00000( |
| 132764660 | 132765458 | 1 | processed_pseudoge | ENSG00000( |
| 132769370 | 132813339 | -1 | protein_coding | ENSG00000( |
| 132814569 | 132817564 | 1 | protein_coding | ENSG00000( |
| 132815307 | 132815379 | 1 | snoRNA | ENSG00000( |
| 132816802 | 132816877 | 1 | snoRNA | ENSG00000( |
| 132817219 | 132817348 | 1 | snoRNA | ENSG00000( |
| 132868218 | 132868859 | -1 | processed_pseudoge | ENSG00000( |
| 132891924 | 132893819 | 1 | TEC | ENSG00000( |
| 132954257 | 133107410 | 1 | lncRNA | ENSG00000( |
| 132997551 | 132998040 | -1 | processed_pseudoge | ENSG00000( |
| 133061240 | 133103128 | -1 | lncRNA | ENSG00000( |
| 133150568 | 133150816 | 1 | processed_pseudoge | ENSG00000( |
| 133240598 | 133532128 | 1 | protein_coding | ENSG00000( |
| 133435077 | 133439464 | -1 | lncRNA | ENSG00000( |
| 133452857 | 133456605 | -1 | lncRNA | ENSG00000( |
| 133502252 | 133892802 | -1 | lncRNA | ENSG00000( |
| 133510386 | 133510693 | -1 | processed_pseudoge | ENSG00000( |
| 133540784 | 133541174 | 1 | lncRNA | ENSG00000( |
| 133676729 | 133677429 | -1 | processed_pseudoge | ENSG00000( |
| 133821147 | 133872922 | 1 | lncRNA | ENSG00000( |
| 133889138 | 133895553 | 1 | protein_coding | ENSG00000( |
| 133952170 | 133990432 | 1 | protein_coding | ENSG00000( |
| 133987581 | 134052624 | -1 | protein_coding | ENSG00000( |
| 134009033 | 134009218 | 1 | processed_pseudoge | ENSG00000( |
| 134074123 | 134121433 | 1 | lncRNA | ENSG00000( |
| 134115235 | 134115736 | -1 | processed_pseudoge | ENSG00000( |
| 134133573 | 134133868 | 1 | misc_RNA | ENSG00000( |
| 134169246 | 134318112 | -1 | protein_coding | ENSG00000( |
| 134211450 | 134211527 | 1 | processed_pseudoge | ENSG00000( |
| 134257035 | 134257143 | 1 | rRNA_pseudogene | ENSG00000( |
| 134283092 | 134283193 | 1 | misc_RNA | ENSG00000( |

| | | | | |
|---|---|---|---|---|
| 134296301 | 134304297 | 1 | lncRNA | ENSG00000 |
| 134296997 | 134298695 | -1 | processed_pseudoge | ENSG00000 |
| 134343307 | 134504581 | 1 | lncRNA | ENSG00000 |
| 134345688 | 134349337 | 1 | lncRNA | ENSG00000 |
| 134393142 | 134393900 | -1 | processed_pseudoge | ENSG00000 |
| 134428239 | 134520585 | -1 | lncRNA | ENSG00000 |
| 134520163 | 134689399 | -1 | lncRNA | ENSG00000 |
| 134603564 | 134604698 | -1 | processed_pseudoge | ENSG00000 |
| 134606299 | 134609437 | -1 | lncRNA | ENSG00000 |
| 134636489 | 134642495 | 1 | lncRNA | ENSG00000 |
| 134706060 | 134707349 | -1 | lncRNA | ENSG00000 |
| 134897874 | 134898759 | 1 | processed_pseudoge | ENSG00000 |
| 134917393 | 134950115 | -1 | protein_coding | ENSG00000 |
| 134941392 | 134942450 | 1 | lncRNA | ENSG00000 |
| 134960378 | 135103056 | -1 | protein_coding | ENSG00000 |
| 134979338 | 134979432 | -1 | miRNA | ENSG00000 |
| 135055033 | 135060550 | 1 | lncRNA | ENSG00000 |
| 135181308 | 135219173 | 1 | protein_coding | ENSG00000 |
| 135195083 | 135195995 | -1 | lncRNA | ENSG00000 |
| 135239160 | 135239256 | 1 | miRNA | ENSG00000 |
| 135259996 | 135260933 | 1 | lncRNA | ENSG00000 |
| 135283532 | 135497765 | -1 | protein_coding | ENSG00000 |
| 135301568 | 135307158 | 1 | lncRNA | ENSG00000 |
| 135323399 | 135327965 | 1 | lncRNA | ENSG00000 |
| 135474504 | 135474609 | -1 | misc_RNA | ENSG00000 |
| 135497422 | 135813990 | 1 | lncRNA | ENSG00000 |
| 135619165 | 135620171 | -1 | processed_pseudoge | ENSG00000 |
| 135628787 | 135634681 | -1 | lncRNA | ENSG00000 |
| 135636086 | 135636713 | -1 | processed_pseudoge | ENSG00000 |
| 135807148 | 135808466 | -1 | lncRNA | ENSG00000 |
| 135851701 | 136195574 | 1 | protein_coding | ENSG00000 |
| 135854053 | 135880942 | -1 | lncRNA | ENSG00000 |
| 135882780 | 135888133 | -1 | lncRNA | ENSG00000 |
| 135991936 | 136225751 | -1 | lncRNA | ENSG00000 |
| 136034553 | 136034886 | -1 | transcribed_processe | ENSG00000 |
| 136206478 | 136206860 | 1 | processed_pseudoge | ENSG00000 |
| 136231024 | 136250335 | -1 | protein_coding | ENSG00000 |
| 136256627 | 136289851 | -1 | protein_coding | ENSG00000 |
| 136317961 | 136318236 | 1 | processed_pseudoge | ENSG00000 |
| 136335714 | 136336087 | -1 | lncRNA | ENSG00000 |
| 136342281 | 136550819 | -1 | protein_coding | ENSG00000 |
| 136364129 | 136365079 | 1 | transcribed_processe | ENSG00000 |
| 136419847 | 136420425 | -1 | processed_pseudoge | ENSG00000 |
| 136475862 | 136476150 | -1 | processed_pseudoge | ENSG00000 |
| 136545192 | 136545492 | 1 | misc_RNA | ENSG00000 |
| 136550661 | 136552554 | 1 | lncRNA | ENSG00000 |
| 136557046 | 136792477 | -1 | protein_coding | ENSG00000 |
| 136629066 | 136648198 | 1 | lncRNA | ENSG00000 |
| 136630243 | 136630355 | 1 | rRNA_pseudogene | ENSG00000 |
| 136784045 | 136786280 | 1 | lncRNA | ENSG00000 |

| | | | | |
|---|---|---|---|---|
| 136822564 | 136913934 | 1 | protein_coding | ENSG00000( |
| 136855698 | 136855831 | -1 | snoRNA | ENSG00000( |
| 136900233 | 136901529 | -1 | transcribed_processe | ENSG00000( |
| 136922301 | 136925660 | 1 | protein_coding | ENSG00000( |
| 136934765 | 136934866 | 1 | misc_RNA | ENSG00000( |
| 136973930 | 136974217 | -1 | processed_pseudoge | ENSG00000( |
| 136982165 | 136993234 | -1 | lncRNA | ENSG00000( |
| 136995170 | 136999504 | 1 | lncRNA | ENSG00000( |
| 136999971 | 137045180 | -1 | protein_coding | ENSG00000( |
| 137143820 | 137173648 | -1 | protein_coding | ENSG00000( |
| 137197484 | 137219449 | -1 | protein_coding | ENSG00000( |
| 137492199 | 137494394 | -1 | protein_coding | ENSG00000( |
| 137543897 | 137544372 | -1 | processed_pseudoge | ENSG00000( |
| 137657998 | 137763984 | 1 | lncRNA | ENSG00000( |
| 137693068 | 137700415 | 1 | lncRNA | ENSG00000( |
| 137705423 | 137707201 | -1 | processed_pseudoge | ENSG00000( |
| 137730170 | 137792835 | -1 | lncRNA | ENSG00000( |
| 137784374 | 137784486 | 1 | misc_RNA | ENSG00000( |
| 137823673 | 137868233 | -1 | lncRNA | ENSG00000( |
| 137867214 | 137883312 | 1 | protein_coding | ENSG00000( |
| 137900585 | 137903576 | -1 | lncRNA | ENSG00000( |
| 137943079 | 137945802 | -1 | lncRNA | ENSG00000( |
| 137945366 | 137972522 | 1 | lncRNA | ENSG00000( |
| 137995270 | 137996147 | 1 | processed_pseudoge | ENSG00000( |
| 138088505 | 138107419 | -1 | protein_coding | ENSG00000( |
| 138161939 | 138344663 | 1 | protein_coding | ENSG00000( |
| 138215986 | 138218491 | -1 | protein_coding | ENSG00000( |
| 138377905 | 138383075 | 1 | protein_coding | ENSG00000( |
| 138393373 | 138393544 | 1 | processed_pseudoge | ENSG00000( |
| 138402585 | 138403560 | -1 | processed_pseudoge | ENSG00000( |
| 138403531 | 138422197 | 1 | protein_coding | ENSG00000( |
| 138422043 | 138692571 | -1 | protein_coding | ENSG00000( |
| 138435213 | 138435294 | -1 | miRNA | ENSG00000( |
| 138464099 | 138464383 | 1 | processed_pseudoge | ENSG00000( |
| 138650226 | 138650973 | -1 | processed_pseudoge | ENSG00000( |
| 138692548 | 138697288 | 1 | lncRNA | ENSG00000( |
| 138725211 | 138773652 | -1 | protein_coding | ENSG00000( |
| 138773509 | 138793319 | 1 | protein_coding | ENSG00000( |
| 138795926 | 138904070 | 1 | protein_coding | ENSG00000( |
| 138822747 | 138823799 | 1 | processed_pseudoge | ENSG00000( |
| 138878899 | 138879407 | -1 | transcribed_processe | ENSG00000( |
| 138903493 | 138988261 | -1 | protein_coding | ENSG00000( |
| 139028745 | 139043302 | 1 | protein_coding | ENSG00000( |
| 139135080 | 139180802 | 1 | protein_coding | ENSG00000( |
| 139144204 | 139239653 | -1 | lncRNA | ENSG00000( |
| 139159157 | 139277181 | -1 | lncRNA | ENSG00000( |
| 139240061 | 139291998 | -1 | protein_coding | ENSG00000( |
| 139271362 | 139667284 | 1 | lncRNA | ENSG00000( |
| 139338018 | 139339580 | 1 | processed_pseudoge | ENSG00000( |
| 139371807 | 139374648 | -1 | protein_coding | ENSG00000( |

| | | | | |
|---|---|---|---|---|
| 139435636 | 139474658 | -1 | lncRNA | ENSG0000C |
| 139614438 | 139615204 | -1 | processed_pseudoge | ENSG0000C |
| 139659928 | 139661513 | 1 | processed_pseudoge | ENSG0000C |
| 139677639 | 139860476 | 1 | lncRNA | ENSG0000C |
| 139856104 | 139877391 | -1 | lncRNA | ENSG0000C |
| 139938864 | 139991094 | -1 | lncRNA | ENSG0000C |
| 139976352 | 140093721 | 1 | lncRNA | ENSG0000C |
| 140158591 | 140158694 | -1 | rRNA_pseudogene | ENSG0000C |
| 140205252 | 140205326 | 1 | miRNA | ENSG0000C |
| 140575788 | 140898381 | -1 | lncRNA | ENSG0000C |
| 140683814 | 140683883 | 1 | miRNA | ENSG0000C |
| 140761529 | 140762262 | 1 | processed_pseudoge | ENSG0000C |
| 140845958 | 140852924 | -1 | lncRNA | ENSG0000C |
| 140922457 | 140922910 | -1 | processed_pseudoge | ENSG0000C |
| 141019788 | 141020026 | -1 | processed_pseudoge | ENSG0000C |
| 141403240 | 141404494 | 1 | lncRNA | ENSG0000C |
| 141447011 | 141451006 | 1 | lncRNA | ENSG0000C |
| 141486141 | 141486412 | -1 | misc_RNA | ENSG0000C |
| 141635650 | 141636503 | -1 | transcribed_processe | ENSG0000C |
| 142058330 | 142088799 | -1 | protein_coding | ENSG0000C |
| 142062717 | 142063053 | 1 | processed_pseudoge | ENSG0000C |
| 142088233 | 142089219 | 1 | lncRNA | ENSG0000C |
| 142133090 | 142135151 | 1 | protein_coding | ENSG0000C |
| 142147162 | 142224685 | 1 | protein_coding | ENSG0000C |
| 142251847 | 142259632 | -1 | lncRNA | ENSG0000C |
| 142301854 | 142446266 | 1 | protein_coding | ENSG0000C |
| 142526455 | 142637889 | -1 | lncRNA | ENSG0000C |
| 142671980 | 142681267 | 1 | lncRNA | ENSG0000C |
| 142748443 | 142753759 | 1 | lncRNA | ENSG0000C |
| 142751469 | 142956698 | -1 | protein_coding | ENSG0000C |
| 142788123 | 142794086 | 1 | lncRNA | ENSG0000C |
| 142946406 | 142957945 | 1 | lncRNA | ENSG0000C |
| 142966293 | 143059682 | -1 | lncRNA | ENSG0000C |
| 143039425 | 143042324 | -1 | lncRNA | ENSG0000C |
| 143060496 | 143341058 | 1 | protein_coding | ENSG0000C |
| 143094034 | 143099645 | -1 | lncRNA | ENSG0000C |
| 143298770 | 143299554 | -1 | processed_pseudoge | ENSG0000C |
| 143327275 | 143327655 | 1 | processed_pseudoge | ENSG0000C |
| 143342246 | 143343383 | 1 | processed_pseudoge | ENSG0000C |
| 143386581 | 143387409 | 1 | processed_pseudoge | ENSG0000C |
| 143422832 | 143450695 | -1 | protein_coding | ENSG0000C |
| 143436216 | 143436710 | 1 | processed_pseudoge | ENSG0000C |
| 143449809 | 143449911 | 1 | rRNA_pseudogene | ENSG0000C |
| 143450805 | 143490616 | 1 | protein_coding | ENSG0000C |
| 143484979 | 143507327 | 1 | lncRNA | ENSG0000C |
| 143490424 | 143506406 | 1 | transcribed_processe | ENSG0000C |
| 143494812 | 143511720 | -1 | protein_coding | ENSG0000C |
| 143536845 | 143831185 | 1 | protein_coding | ENSG0000C |
| 143554325 | 143562031 | -1 | lncRNA | ENSG0000C |
| 143807092 | 143844617 | 1 | protein_coding | ENSG0000C |

| | | | | |
|---|---|---|---|---|
| 143843338 | 143863812 | 1 | protein_coding | ENSG0000C |
| 143857318 | 143938471 | 1 | protein_coding | ENSG0000C |
| 143858062 | 143858689 | -1 | processed_pseudoge | ENSG0000C |
| 143864436 | 143938343 | 1 | protein_coding | ENSG0000C |
| 143940300 | 144064599 | -1 | protein_coding | ENSG0000C |
| 144004916 | 144008262 | -1 | lncRNA | ENSG0000C |
| 144007227 | 144007464 | 1 | misc_RNA | ENSG0000C |
| 144036618 | 144037704 | 1 | processed_pseudoge | ENSG0000C |
| 144094884 | 144095573 | -1 | protein_coding | ENSG0000C |
| 144136450 | 144136775 | 1 | processed_pseudoge | ENSG0000C |
| 144150517 | 144191939 | 1 | protein_coding | ENSG0000C |
| 144200447 | 144200965 | -1 | processed_pseudoge | ENSG0000C |
| 144257034 | 144257624 | -1 | processed_pseudoge | ENSG0000C |
| 144285701 | 144853034 | 1 | protein_coding | ENSG0000C |
| 144311699 | 144333301 | -1 | lncRNA | ENSG0000C |
| 144397959 | 144398752 | 1 | processed_pseudoge | ENSG0000C |
| 144706733 | 144707047 | -1 | processed_pseudoge | ENSG0000C |
| 144708106 | 144708497 | -1 | processed_pseudoge | ENSG0000C |
| 145296545 | 145311092 | 1 | lncRNA | ENSG0000C |
| 145382535 | 145736023 | -1 | protein_coding | ENSG0000C |
| 145463238 | 145463399 | -1 | snRNA | ENSG0000C |
| 145489630 | 145492092 | -1 | processed_pseudoge | ENSG0000C |
| 145735570 | 145737218 | 1 | lncRNA | ENSG0000C |
| 145736911 | 145737173 | -1 | lncRNA | ENSG0000C |
| 145749917 | 145887430 | 1 | lncRNA | ENSG0000C |
| 145784406 | 145786559 | -1 | lncRNA | ENSG0000C |
| 145789270 | 145791973 | -1 | TEC | ENSG0000C |
| 145793502 | 145814795 | -1 | protein_coding | ENSG0000C |
| 145864245 | 145964423 | -1 | protein_coding | ENSG0000C |
| 146027646 | 146437601 | 1 | protein_coding | ENSG0000C |
| 146171315 | 146171847 | 1 | processed_pseudoge | ENSG0000C |
| 146340420 | 146340524 | 1 | rRNA_pseudogene | ENSG0000C |
| 146364931 | 146365494 | -1 | processed_pseudoge | ENSG0000C |
| 146543833 | 146554953 | 1 | protein_coding | ENSG0000C |
| 146594516 | 146598931 | -1 | lncRNA | ENSG0000C |
| 146598967 | 146815462 | 1 | protein_coding | ENSG0000C |
| 146638844 | 146639261 | -1 | processed_pseudoge | ENSG0000C |
| 146639700 | 146639769 | 1 | snRNA | ENSG0000C |
| 146802359 | 146803824 | -1 | transcribed_processe | ENSG0000C |
| 146824539 | 147204614 | -1 | lncRNA | ENSG0000C |
| 146854983 | 146863460 | 1 | lncRNA | ENSG0000C |
| 146948528 | 146951159 | -1 | lncRNA | ENSG0000C |
| 147158925 | 147180992 | 1 | lncRNA | ENSG0000C |
| 147204425 | 147390476 | 1 | protein_coding | ENSG0000C |
| 147406531 | 147406824 | 1 | processed_pseudoge | ENSG0000C |
| 147406889 | 147408281 | 1 | processed_pseudoge | ENSG0000C |
| 147508690 | 147737547 | 1 | protein_coding | ENSG0000C |
| 147660703 | 147953937 | 1 | lncRNA | ENSG0000C |
| 147741434 | 147743324 | -1 | lncRNA | ENSG0000C |
| 148001716 | 148018985 | 1 | lncRNA | ENSG0000C |

| | | | | |
|---|---|---|---|---|
| 148017422 | 148020974 | -1 | lncRNA | ENSG0000C |
| 148133809 | 148137404 | -1 | lncRNA | ENSG0000C |
| 148155120 | 148157084 | -1 | lncRNA | ENSG0000C |
| 148237585 | 148283408 | -1 | lncRNA | ENSG0000C |
| 148272304 | 148552048 | 1 | protein_coding | ENSG0000C |
| 148278672 | 148278778 | 1 | snRNA | ENSG0000C |
| 148478693 | 148480763 | 1 | processed_pseudoge | ENSG0000C |
| 148567378 | 148567654 | -1 | processed_pseudoge | ENSG0000C |
| 148647707 | 148648597 | -1 | processed_pseudoge | ENSG0000C |
| 148747030 | 149076990 | 1 | protein_coding | ENSG0000C |
| 148955628 | 148964684 | -1 | lncRNA | ENSG0000C |
| 149027700 | 149032573 | -1 | lncRNA | ENSG0000C |
| 149112009 | 149112672 | -1 | processed_pseudoge | ENSG0000C |
| 149217926 | 149245554 | 1 | lncRNA | ENSG0000C |
| 149218641 | 149411613 | 1 | protein_coding | ENSG0000C |
| 149243299 | 149257551 | -1 | lncRNA | ENSG0000C |
| 149246095 | 149255415 | 1 | lncRNA | ENSG0000C |
| 149291352 | 149291646 | 1 | misc_RNA | ENSG0000C |
| 149304107 | 149304428 | -1 | processed_pseudoge | ENSG0000C |
| 149400262 | 149401278 | 1 | protein_coding | ENSG0000C |
| 149446795 | 149485014 | -1 | protein_coding | ENSG0000C |
| 149491567 | 149492289 | -1 | processed_pseudoge | ENSG0000C |
| 149504495 | 149546043 | -1 | protein_coding | ENSG0000C |
| 149516956 | 149517017 | 1 | snRNA | ENSG0000C |
| 149561152 | 149563949 | 1 | lncRNA | ENSG0000C |
| 149566294 | 149591748 | 1 | protein_coding | ENSG0000C |
| 149576089 | 149590864 | -1 | lncRNA | ENSG0000C |
| 149591755 | 149592663 | -1 | lncRNA | ENSG0000C |
| 149592687 | 149601878 | 1 | transcribed_processe | ENSG0000C |
| 149594625 | 149594759 | 1 | snoRNA | ENSG0000C |
| 149594873 | 149648972 | -1 | protein_coding | ENSG0000C |
| 149631437 | 149631549 | 1 | misc_RNA | ENSG0000C |
| 149658153 | 149718105 | -1 | protein_coding | ENSG0000C |
| 149717621 | 149718888 | 1 | lncRNA | ENSG0000C |
| 149724315 | 149749665 | -1 | protein_coding | ENSG0000C |
| 149749443 | 149811420 | 1 | protein_coding | ENSG0000C |
| 149794763 | 149795746 | 1 | processed_pseudoge | ENSG0000C |
| 149796151 | 149799150 | -1 | lncRNA | ENSG0000C |
| 149797317 | 149826294 | -1 | lncRNA | ENSG0000C |
| 149817937 | 149896011 | -1 | protein_coding | ENSG0000C |
| 149818757 | 149864359 | -1 | protein_coding | ENSG0000C |
| 149852462 | 149853192 | 1 | lncRNA | ENSG0000C |
| 149863494 | 149919507 | 1 | lncRNA | ENSG0000C |
| 149879962 | 149881572 | 1 | processed_pseudoge | ENSG0000C |
| 149883375 | 149898102 | -1 | protein_coding | ENSG0000C |
| 149884431 | 149919508 | 1 | lncRNA | ENSG0000C |
| 149904243 | 149906418 | 1 | processed_pseudoge | ENSG0000C |
| 149916878 | 149923121 | -1 | protein_coding | ENSG0000C |
| 149934527 | 149936782 | 1 | transcribed_processe | ENSG0000C |
| 149942014 | 149949235 | 1 | protein_coding | ENSG0000C |

| | | | | |
|---|---|---|---|---|
| 149951833 | 149952239 | 1 | processed_pseudoge | ENSG00000( |
| 149963943 | 149973715 | 1 | protein_coding | ENSG00000( |
| 149977471 | 149979105 | 1 | processed_pseudoge | ENSG00000( |
| 149977922 | 149978416 | 1 | processed_pseudoge | ENSG00000( |
| 149998019 | 150005157 | -1 | transcribed_unproce | ENSG00000( |
| 150018334 | 150025532 | -1 | protein_coding | ENSG00000( |
| 150032769 | 150033566 | 1 | processed_pseudoge | ENSG00000( |
| 150040547 | 150042033 | -1 | lncRNA | ENSG00000( |
| 150042546 | 150043353 | 1 | processed_pseudoge | ENSG00000( |
| 150061053 | 150069121 | -1 | protein_coding | ENSG00000( |
| 150143044 | 150250392 | 1 | protein_coding | ENSG00000( |
| 150164584 | 150164686 | 1 | snoRNA | ENSG00000( |
| 150326623 | 150326763 | -1 | snRNA | ENSG00000( |
| 150368892 | 150405969 | 1 | protein_coding | ENSG00000( |
| 150384307 | 150385120 | -1 | processed_pseudoge | ENSG00000( |
| 150504811 | 150510300 | -1 | processed_pseudoge | ENSG00000( |
| 150599883 | 150843665 | 1 | protein_coding | ENSG00000( |
| 150600834 | 150605850 | 1 | lncRNA | ENSG00000( |
| 150624677 | 150626086 | 1 | lncRNA | ENSG00000( |
| 150650772 | 150652025 | -1 | lncRNA | ENSG00000( |
| 150827663 | 150828384 | 1 | processed_pseudoge | ENSG00000( |
| 150865549 | 151101887 | 1 | protein_coding | ENSG00000( |
| 150934968 | 150935566 | 1 | processed_pseudoge | ENSG00000( |
| 150939222 | 150939328 | 1 | snRNA | ENSG00000( |
| 151054897 | 151056057 | -1 | lncRNA | ENSG00000( |
| 151088103 | 151228447 | -1 | lncRNA | ENSG00000( |
| 151099529 | 151099933 | 1 | processed_pseudoge | ENSG00000( |
| 151196681 | 151197638 | -1 | lncRNA | ENSG00000( |
| 151201341 | 151201444 | 1 | snRNA | ENSG00000( |
| 151225550 | 151225947 | -1 | processed_pseudoge | ENSG00000( |
| 151239967 | 151358559 | 1 | protein_coding | ENSG00000( |
| 151280515 | 151280612 | -1 | snRNA | ENSG00000( |
| 151298841 | 151298933 | -1 | misc_RNA | ENSG00000( |
| 151319635 | 151319950 | 1 | misc_RNA | ENSG00000( |
| 151364115 | 151391559 | -1 | protein_coding | ENSG00000( |
| 151378000 | 151378099 | -1 | misc_RNA | ENSG00000( |
| 151404762 | 151452158 | -1 | protein_coding | ENSG00000( |
| 151411259 | 151413945 | -1 | processed_pseudoge | ENSG00000( |
| 151452258 | 151470101 | 1 | protein_coding | ENSG00000( |
| 151494017 | 151621193 | 1 | protein_coding | ENSG00000( |
| 151615773 | 151615879 | -1 | snRNA | ENSG00000( |
| 151656691 | 152129619 | 1 | protein_coding | ENSG00000( |
| 151813276 | 151814179 | -1 | lncRNA | ENSG00000( |
| 152091712 | 152094009 | -1 | lncRNA | ENSG00000( |
| 152112759 | 152118397 | 1 | lncRNA | ENSG00000( |
| 152121684 | 152637801 | -1 | protein_coding | ENSG00000( |
| 152380546 | 152381564 | 1 | lncRNA | ENSG00000( |
| 152402398 | 152404966 | 1 | lncRNA | ENSG00000( |
| 152410275 | 152410394 | 1 | rRNA_pseudogene | ENSG00000( |
| 152545930 | 152546984 | 1 | lncRNA | ENSG00000( |

| | | | | |
|---|---|---|---|---|
| 152697895 | 152724567 | 1 | protein_coding | ENSG00000 |
| 152705735 | 152706128 | 1 | processed_pseudoge | ENSG00000 |
| 152750797 | 152759765 | 1 | protein_coding | ENSG00000 |
| 152754903 | 152875991 | -1 | lncRNA | ENSG00000 |
| 152898047 | 152899280 | 1 | processed_pseudoge | ENSG00000 |
| 152959359 | 152959605 | -1 | processed_pseudoge | ENSG00000 |
| 152970519 | 152983579 | -1 | protein_coding | ENSG00000 |
| 152983331 | 152991322 | 1 | lncRNA | ENSG00000 |
| 152987362 | 153002709 | -1 | protein_coding | ENSG00000 |
| 153004459 | 153131282 | -1 | protein_coding | ENSG00000 |
| 153159220 | 153159345 | -1 | rRNA_pseudogene | ENSG00000 |
| 153231320 | 153347488 | 1 | lncRNA | ENSG00000 |
| 153282287 | 153282733 | -1 | processed_pseudoge | ENSG00000 |
| 153304595 | 153427154 | -1 | lncRNA | ENSG00000 |
| 153420438 | 153420572 | 1 | rRNA_pseudogene | ENSG00000 |
| 153665707 | 153666390 | 1 | processed_pseudoge | ENSG00000 |
| 153666466 | 153666666 | 1 | processed_pseudoge | ENSG00000 |
| 153666627 | 153667318 | 1 | processed_pseudoge | ENSG00000 |
| 153667318 | 153668106 | 1 | processed_pseudoge | ENSG00000 |
| 153668177 | 153668362 | 1 | processed_pseudoge | ENSG00000 |
| 153668589 | 153668858 | 1 | processed_pseudoge | ENSG00000 |
| 153668879 | 153669901 | 1 | processed_pseudoge | ENSG00000 |
| 153681447 | 153681553 | 1 | snRNA | ENSG00000 |
| 153938433 | 153938992 | -1 | processed_pseudoge | ENSG00000 |
| 154010496 | 154246867 | 1 | protein_coding | ENSG00000 |
| 154154496 | 154356792 | -1 | protein_coding | ENSG00000 |
| 154159828 | 154510114 | -1 | protein_coding | ENSG00000 |
| 154216931 | 154217489 | 1 | processed_pseudoge | ENSG00000 |
| 154387515 | 154510685 | -1 | protein_coding | ENSG00000 |
| 154477982 | 154479593 | 1 | lncRNA | ENSG00000 |
| 154510791 | 154522307 | 1 | lncRNA | ENSG00000 |
| 154542552 | 154542740 | 1 | processed_pseudoge | ENSG00000 |
| 154549517 | 154550242 | 1 | processed_pseudoge | ENSG00000 |
| 154576309 | 154577079 | 1 | processed_pseudoge | ENSG00000 |
| 154605616 | 154605954 | -1 | processed_pseudoge | ENSG00000 |
| 154706643 | 154707795 | 1 | processed_pseudoge | ENSG00000 |
| 154733325 | 154834244 | 1 | protein_coding | ENSG00000 |
| 154825176 | 154825279 | -1 | snRNA | ENSG00000 |
| 154832697 | 155257723 | 1 | protein_coding | ENSG00000 |
| 154845074 | 154845620 | -1 | processed_pseudoge | ENSG00000 |
| 154853360 | 154853436 | 1 | miRNA | ENSG00000 |
| 154905076 | 154905206 | -1 | snoRNA | ENSG00000 |
| 155004835 | 155005318 | -1 | unprocessed_pseudc | ENSG00000 |
| 155253139 | 155256724 | -1 | lncRNA | ENSG00000 |
| 155256134 | 155314493 | -1 | protein_coding | ENSG00000 |
| 155264013 | 155276548 | 1 | protein_coding | ENSG00000 |
| 155380511 | 155381183 | -1 | lncRNA | ENSG00000 |
| 155395368 | 155455839 | -1 | protein_coding | ENSG00000 |
| 155654963 | 155655026 | -1 | snRNA | ENSG00000 |
| 155911195 | 156397589 | -1 | lncRNA | ENSG00000 |

| | | | | |
|---|---|---|---|---|
| 155946797 | 155946879 | 1 | miRNA | ENSG00000( |
| 156350579 | 156390246 | 1 | lncRNA | ENSG00000( |
| 156378750 | 156378824 | 1 | snoRNA | ENSG00000( |
| 156493226 | 156505011 | -1 | lncRNA | ENSG00000( |
| 156662050 | 156662437 | 1 | processed_pseudoge | ENSG00000( |
| 156774217 | 156774662 | -1 | lncRNA | ENSG00000( |
| 156776020 | 157210779 | 1 | protein_coding | ENSG00000( |
| 156776360 | 156778422 | -1 | lncRNA | ENSG00000( |
| 156779678 | 156779731 | -1 | miRNA | ENSG00000( |
| 156780327 | 156780455 | -1 | lncRNA | ENSG00000( |
| 156976294 | 156977072 | 1 | processed_pseudoge | ENSG00000( |
| 157107727 | 157119706 | -1 | lncRNA | ENSG00000( |
| 157289386 | 157323601 | -1 | protein_coding | ENSG00000( |
| 157299045 | 157300126 | -1 | processed_pseudoge | ENSG00000( |
| 157323964 | 157324477 | 1 | lncRNA | ENSG00000( |
| 157365990 | 157366923 | -1 | processed_pseudoge | ENSG00000( |
| 157372556 | 157373171 | -1 | processed_pseudoge | ENSG00000( |
| 157377062 | 157380896 | -1 | lncRNA | ENSG00000( |
| 157381133 | 157678157 | 1 | protein_coding | ENSG00000( |
| 157529132 | 157529200 | 1 | miRNA | ENSG00000( |
| 157700387 | 157945077 | 1 | protein_coding | ENSG00000( |
| 157829143 | 157830573 | -1 | lncRNA | ENSG00000( |
| 157872571 | 157875210 | -1 | lncRNA | ENSG00000( |
| 157885114 | 157892786 | -1 | lncRNA | ENSG00000( |
| 157924402 | 157924676 | 1 | processed_pseudoge | ENSG00000( |
| 157934168 | 157934257 | 1 | snRNA | ENSG00000( |
| 157981863 | 158099176 | 1 | protein_coding | ENSG00000( |
| 158000109 | 158000749 | -1 | lncRNA | ENSG00000( |
| 158001107 | 158002383 | 1 | lncRNA | ENSG00000( |
| 158109519 | 158168280 | -1 | protein_coding | ENSG00000( |
| 158168350 | 158199344 | 1 | protein_coding | ENSG00000( |
| 158232236 | 158511828 | 1 | protein_coding | ENSG00000( |
| 158237336 | 158242797 | -1 | processed_pseudoge | ENSG00000( |
| 158282841 | 158428379 | 1 | lncRNA | ENSG00000( |
| 158296671 | 158303372 | -1 | lncRNA | ENSG00000( |
| 158397885 | 158398561 | -1 | lncRNA | ENSG00000( |
| 158437797 | 158438088 | 1 | misc_RNA | ENSG00000( |
| 158515605 | 158516229 | -1 | processed_pseudoge | ENSG00000( |
| 158536436 | 158635428 | 1 | protein_coding | ENSG00000( |
| 158609706 | 158621636 | -1 | processed_pseudoge | ENSG00000( |
| 158609707 | 158609790 | 1 | miRNA | ENSG00000( |
| 158636474 | 158644743 | -1 | protein_coding | ENSG00000( |
| 158650014 | 158764876 | 1 | protein_coding | ENSG00000( |
| 158725747 | 158726803 | -1 | processed_pseudoge | ENSG00000( |
| 158764661 | 158764753 | -1 | miRNA | ENSG00000( |
| 158765741 | 158819368 | -1 | protein_coding | ENSG00000( |
| 158817979 | 158822252 | 1 | lncRNA | ENSG00000( |
| 158841117 | 158857632 | -1 | transcribed_processe | ENSG00000( |
| 158853588 | 158864502 | 1 | lncRNA | ENSG00000( |
| 158869848 | 158919105 | 1 | lncRNA | ENSG00000( |

| | | | | |
|---|---|---|---|---|
| 158921271 | 158922150 | -1 | processed_pseudoge | ENSG00000( |
| 158926466 | 158926571 | -1 | snRNA | ENSG00000( |
| 158972871 | 159000187 | -1 | protein_coding | ENSG00000( |
| 158988178 | 159088114 | 1 | lncRNA | ENSG00000( |
| 159034468 | 159045152 | -1 | protein_coding | ENSG00000( |
| 159042192 | 159095823 | -1 | lncRNA | ENSG00000( |
| 159051674 | 159121495 | -1 | lncRNA | ENSG00000( |
| 159094093 | 159116254 | 1 | lncRNA | ENSG00000( |
| 159165899 | 159170007 | -1 | lncRNA | ENSG00000( |
| 159169400 | 159272108 | 1 | protein_coding | ENSG00000( |
| 159240786 | 159243329 | 1 | lncRNA | ENSG00000( |
| 159353922 | 159383339 | -1 | lncRNA | ENSG00000( |
| 159383422 | 159401147 | 1 | lncRNA | ENSG00000( |
| 159468601 | 159662627 | -1 | lncRNA | ENSG00000( |
| 159526062 | 159526527 | -1 | processed_pseudoge | ENSG00000( |
| 159555964 | 159556174 | 1 | processed_pseudoge | ENSG00000( |
| 159586955 | 159589169 | -1 | lncRNA | ENSG00000( |
| 159669069 | 159745186 | -1 | protein_coding | ENSG00000( |
| 159679119 | 159762529 | -1 | protein_coding | ENSG00000( |
| 159712801 | 159713985 | -1 | processed_pseudoge | ENSG00000( |
| 159720415 | 159720540 | -1 | snRNA | ENSG00000( |
| 159725585 | 159756319 | 1 | protein_coding | ENSG00000( |
| 159760258 | 159762332 | -1 | lncRNA | ENSG00000( |
| 159762045 | 159779112 | 1 | protein_coding | ENSG00000( |
| 159778498 | 159789703 | -1 | protein_coding | ENSG00000( |
| 159780250 | 159780381 | -1 | snoRNA | ENSG00000( |
| 159785594 | 159785733 | -1 | snoRNA | ENSG00000( |
| 159789812 | 159798436 | 1 | protein_coding | ENSG00000( |
| 159800249 | 159820704 | 1 | protein_coding | ENSG00000( |
| 159899186 | 159902510 | 1 | lncRNA | ENSG00000( |
| 159906690 | 159916530 | 1 | protein_coding | ENSG00000( |
| 159969099 | 160113507 | 1 | protein_coding | ENSG00000( |
| 160003291 | 160007664 | -1 | lncRNA | ENSG00000( |
| 160093082 | 160096212 | 1 | transcribed_processe | ENSG00000( |
| 160121815 | 160158718 | 1 | protein_coding | ENSG00000( |
| 160163539 | 160163755 | -1 | processed_pseudoge | ENSG00000( |
| 160171061 | 160277638 | -1 | protein_coding | ENSG00000( |
| 160272617 | 160276130 | 1 | lncRNA | ENSG00000( |
| 160348378 | 160452577 | 1 | protein_coding | ENSG00000( |
| 160369200 | 160386934 | -1 | lncRNA | ENSG00000( |
| 160453428 | 160520269 | -1 | transcribed_unproce | ENSG00000( |
| 160531483 | 160664259 | -1 | protein_coding | ENSG00000( |
| 160666228 | 160676523 | -1 | unprocessed_pseudo | ENSG00000( |
| 160698288 | 160700632 | 1 | lncRNA | ENSG00000( |
| 160702238 | 160753315 | 1 | protein_coding | ENSG00000( |
| 160742509 | 160767490 | -1 | lncRNA | ENSG00000( |
| 160755398 | 160757518 | -1 | unprocessed_pseudo | ENSG00000( |
| 160773988 | 160783214 | 1 | unprocessed_pseudo | ENSG00000( |
| 160872888 | 160926204 | -1 | lncRNA | ENSG00000( |
| 160905770 | 160913638 | 1 | unprocessed_pseudo | ENSG00000( |

| | | | | |
|---|---|---|---|---|
| 160926269 | 160943110 | 1 | lncRNA | ENSG0000C |
| 160931080 | 160985624 | 1 | lncRNA | ENSG0000C |
| 160990318 | 160992342 | -1 | lncRNA | ENSG0000C |
| 160991727 | 161117385 | 1 | protein_coding | ENSG0000C |
| 161129967 | 161274061 | -1 | protein_coding | ENSG0000C |
| 161347417 | 162727771 | -1 | protein_coding | ENSG0000C |
| 161353679 | 161355350 | 1 | lncRNA | ENSG0000C |
| 161435098 | 161446016 | 1 | lncRNA | ENSG0000C |
| 162568379 | 162569728 | 1 | processed_pseudoge | ENSG0000C |
| 162727132 | 163315492 | 1 | protein_coding | ENSG0000C |
| 162957258 | 162957548 | -1 | processed_pseudoge | ENSG0000C |
| 162969050 | 162975007 | -1 | lncRNA | ENSG0000C |
| 163042557 | 163054161 | -1 | lncRNA | ENSG0000C |
| 163115691 | 163137775 | -1 | lncRNA | ENSG0000C |
| 163163830 | 163192038 | -1 | lncRNA | ENSG0000C |
| 163306907 | 163324530 | -1 | lncRNA | ENSG0000C |
| 163338363 | 163346744 | -1 | lncRNA | ENSG0000C |
| 163348019 | 163353220 | -1 | lncRNA | ENSG0000C |
| 163363210 | 163363463 | 1 | processed_pseudoge | ENSG0000C |
| 163413065 | 163413960 | -1 | lncRNA | ENSG0000C |
| 163414000 | 163578592 | 1 | protein_coding | ENSG0000C |
| 163586583 | 163588165 | -1 | lncRNA | ENSG0000C |
| 163671577 | 164231610 | 1 | lncRNA | ENSG0000C |
| 163705937 | 163714306 | -1 | lncRNA | ENSG0000C |
| 163840561 | 163840852 | 1 | misc_RNA | ENSG0000C |
| 163956299 | 163960187 | -1 | lncRNA | ENSG0000C |
| 164084023 | 164085661 | -1 | lncRNA | ENSG0000C |
| 164108620 | 164109694 | -1 | lncRNA | ENSG0000C |
| 164346489 | 164348644 | -1 | lncRNA | ENSG0000C |
| 164791288 | 164827683 | -1 | lncRNA | ENSG0000C |
| 164827732 | 164832114 | 1 | lncRNA | ENSG0000C |
| 164928191 | 165077495 | 1 | lncRNA | ENSG0000C |
| 165117195 | 165117524 | -1 | processed_pseudoge | ENSG0000C |
| 165279664 | 165309605 | -1 | protein_coding | ENSG0000C |
| 165327287 | 165988078 | -1 | protein_coding | ENSG0000C |
| 165409562 | 165409676 | 1 | rRNA_pseudogene | ENSG0000C |
| 165718235 | 165718338 | -1 | snRNA | ENSG0000C |
| 165754189 | 165775780 | 1 | lncRNA | ENSG0000C |
| 165773056 | 165775221 | -1 | lncRNA | ENSG0000C |
| 165987551 | 165989615 | 1 | lncRNA | ENSG0000C |
| 166058505 | 166066028 | 1 | transcribed_processe | ENSG0000C |
| 166098111 | 166099589 | 1 | lncRNA | ENSG0000C |
| 166099665 | 166113273 | 1 | lncRNA | ENSG0000C |
| 166099853 | 166099924 | 1 | snoRNA | ENSG0000C |
| 166157656 | 166168700 | -1 | protein_coding | ENSG0000C |
| 166209403 | 166209506 | 1 | snRNA | ENSG0000C |
| 166229334 | 166229554 | 1 | unprocessed_pseudc | ENSG0000C |
| 166236969 | 166256947 | 1 | lncRNA | ENSG0000C |
| 166240290 | 166240493 | -1 | processed_pseudoge | ENSG0000C |
| 166275462 | 166277077 | -1 | lncRNA | ENSG0000C |

| | | | | |
|---|---|---|---|---|
| 166305300 | 166308448 | -1 | protein_coding | ENSG00000 |
| 166309572 | 166317470 | 1 | lncRNA | ENSG00000 |
| 166319728 | 166342576 | -1 | protein_coding | ENSG00000 |
| 166334568 | 166335471 | 1 | processed_pseudoge | ENSG00000 |
| 166342632 | 166351466 | 1 | lncRNA | ENSG00000 |
| 166364919 | 166383013 | -1 | protein_coding | ENSG00000 |
| 166383189 | 166384824 | 1 | lncRNA | ENSG00000 |
| 166388136 | 166389920 | 1 | TEC | ENSG00000 |
| 166409364 | 166906451 | -1 | protein_coding | ENSG00000 |
| 166460663 | 166465383 | -1 | lncRNA | ENSG00000 |
| 166509354 | 166509433 | -1 | miRNA | ENSG00000 |
| 166585604 | 166587594 | -1 | protein_coding | ENSG00000 |
| 166651524 | 166655610 | 1 | lncRNA | ENSG00000 |
| 166701094 | 166702839 | -1 | processed_pseudoge | ENSG00000 |
| 166858094 | 166956124 | -1 | protein_coding | ENSG00000 |
| 166903698 | 166904835 | 1 | lncRNA | ENSG00000 |
| 166929504 | 166957191 | -1 | protein_coding | ENSG00000 |
| 166969626 | 166999065 | -1 | lncRNA | ENSG00000 |
| 166995735 | 166999511 | -1 | lncRNA | ENSG00000 |
| 166997807 | 166997912 | -1 | miRNA | ENSG00000 |
| 166999317 | 167094789 | 1 | protein_coding | ENSG00000 |
| 166999405 | 167139141 | 1 | lncRNA | ENSG00000 |
| 167111807 | 167139696 | 1 | protein_coding | ENSG00000 |
| 167114957 | 167121527 | -1 | lncRNA | ENSG00000 |
| 167126223 | 167131572 | -1 | lncRNA | ENSG00000 |
| 167146414 | 167196913 | 1 | protein_coding | ENSG00000 |
| 167156271 | 167158329 | -1 | protein_coding | ENSG00000 |
| 167164565 | 167167233 | -1 | lncRNA | ENSG00000 |
| 167208631 | 167208897 | -1 | processed_pseudoge | ENSG00000 |
| 167228300 | 167237511 | 1 | lncRNA | ENSG00000 |
| 167237508 | 167240480 | -1 | lncRNA | ENSG00000 |
| 167241891 | 167245916 | 1 | lncRNA | ENSG00000 |
| 167271169 | 167316014 | 1 | protein_coding | ENSG00000 |
| 167325071 | 167359503 | 1 | protein_coding | ENSG00000 |
| 167347472 | 167348053 | 1 | processed_pseudoge | ENSG00000 |
| 167357031 | 167384510 | -1 | protein_coding | ENSG00000 |
| 167644479 | 167658518 | -1 | lncRNA | ENSG00000 |
| 167666840 | 167679270 | -1 | lncRNA | ENSG00000 |
| 167678232 | 167679523 | 1 | lncRNA | ENSG00000 |
| 167679626 | 167696290 | -1 | lncRNA | ENSG00000 |
| 167767083 | 167768568 | -1 | processed_pseudoge | ENSG00000 |
| 167784537 | 167796872 | -1 | lncRNA | ENSG00000 |
| 167795530 | 167798248 | 1 | lncRNA | ENSG00000 |
| 167796647 | 167825145 | 1 | lncRNA | ENSG00000 |
| 167822103 | 167826812 | -1 | lncRNA | ENSG00000 |
| 167826922 | 167972023 | 1 | protein_coding | ENSG00000 |
| 167975924 | 167979776 | -1 | lncRNA | ENSG00000 |
| 167992822 | 167997110 | -1 | lncRNA | ENSG00000 |
| 167996241 | 168045095 | 1 | protein_coding | ENSG00000 |
| 168053166 | 168101511 | -1 | protein_coding | ENSG00000 |

| | | | | |
|---|---|---|---|---|
| 168194358 | 168204501 | 1 | lncRNA | ENSG00000 |
| 168217032 | 168220262 | 1 | lncRNA | ENSG00000 |
| 168225279 | 168226517 | -1 | lncRNA | ENSG00000 |
| 168229732 | 168240715 | 1 | lncRNA | ENSG00000 |
| 168242938 | 168262581 | 1 | lncRNA | ENSG00000 |
| 168286905 | 168289350 | -1 | processed_pseudoge | ENSG00000 |
| 168292830 | 168319777 | -1 | protein_coding | ENSG00000 |
| 168298069 | 168302114 | -1 | lncRNA | ENSG00000 |
| 168374697 | 168375355 | 1 | lncRNA | ENSG00000 |
| 168441151 | 168673445 | 1 | protein_coding | ENSG00000 |
| 168716606 | 168723495 | 1 | lncRNA | ENSG00000 |
| 168820423 | 168820656 | 1 | TEC | ENSG00000 |
| 168920394 | 168964386 | -1 | lncRNA | ENSG00000 |
| 168999401 | 169000527 | -1 | lncRNA | ENSG00000 |
| 169033381 | 169034050 | 1 | TEC | ENSG00000 |
| 169034197 | 169035642 | -1 | lncRNA | ENSG00000 |
| 169067764 | 169069427 | 1 | lncRNA | ENSG00000 |
| 169097799 | 169102428 | 1 | lncRNA | ENSG00000 |
| 169157162 | 169162992 | -1 | lncRNA | ENSG00000 |
| 169175277 | 169182841 | -1 | lncRNA | ENSG00000 |
| 169213254 | 169245773 | 1 | lncRNA | ENSG00000 |
| 169215780 | 169254044 | -1 | protein_coding | ENSG00000 |
| 169286534 | 169289216 | -1 | lncRNA | ENSG00000 |
| 169296471 | 169297959 | -1 | processed_pseudoge | ENSG00000 |
| 169369998 | 169388385 | -1 | lncRNA | ENSG00000 |
| 169419004 | 169425156 | -1 | lncRNA | ENSG00000 |
| 169419969 | 169702047 | -1 | protein_coding | ENSG00000 |
| 169426420 | 169452475 | -1 | lncRNA | ENSG00000 |
| 169457212 | 169702067 | -1 | protein_coding | ENSG00000 |
| 169702190 | 169704856 | 1 | protein_coding | ENSG00000 |
| 169703902 | 169725566 | -1 | protein_coding | ENSG00000 |
| 169725091 | 169725854 | -1 | lncRNA | ENSG00000 |
| 169725504 | 169738992 | 1 | lncRNA | ENSG00000 |
| 169740109 | 169751587 | -1 | protein_coding | ENSG00000 |
| 169751622 | 169781600 | 1 | protein_coding | ENSG00000 |
| 169770413 | 169772042 | 1 | lncRNA | ENSG00000 |
| 169788790 | 169799549 | -1 | lncRNA | ENSG00000 |
| 169790057 | 169802873 | 1 | lncRNA | ENSG00000 |
| 169809318 | 169810102 | -1 | lncRNA | ENSG00000 |
| 170088022 | 170097234 | 1 | lncRNA | ENSG00000 |
| 170135374 | 170142926 | 1 | lncRNA | ENSG00000 |
| 170158728 | 170160471 | -1 | lncRNA | ENSG00000 |
| 170160606 | 170163142 | 1 | lncRNA | ENSG00000 |
| 170178681 | 170179660 | 1 | lncRNA | ENSG00000 |
| 170199769 | 170200258 | 1 | processed_pseudoge | ENSG00000 |
| 170254334 | 170262569 | -1 | lncRNA | ENSG00000 |
| 170266669 | 170276762 | -1 | lncRNA | ENSG00000 |
| 170272474 | 170279887 | 1 | lncRNA | ENSG00000 |
| 170282206 | 170306565 | -1 | protein_coding | ENSG00000 |
| 170290703 | 170407065 | 1 | protein_coding | ENSG00000 |

| | | | | |
|---|---|---|---|---|
| 170297876 | 170298375 | -1 | lncRNA | ENSG00000 |
| 170330761 | 170330844 | 1 | miRNA | ENSG00000 |
| 170414139 | 170419325 | 1 | lncRNA | ENSG00000 |
| 170535120 | 170553307 | -1 | protein_coding | ENSG00000 |
| 170554302 | 170572870 | 1 | protein_coding | ENSG00000 |
| 170575295 | 170584692 | -1 | protein_coding | ENSG00000 |
| 170612622 | 170638715 | 1 | lncRNA | ENSG00000 |
| 170615515 | 170738536 | 1 | lncRNA | ENSG00000 |
| 170639606 | 170640536 | 1 | unprocessed_pseudc | ENSG00000 |
| 170651012 | 170651338 | -1 | processed_pseudoge | ENSG00000 |
| 170695313 | 170696950 | 1 | lncRNA | ENSG00000 |
| 170736173 | 170737777 | -1 | lncRNA | ENSG00000 |
| 170745518 | 170745977 | 1 | processed_pseudoge | ENSG00000 |
| 12704 | 27234 | 1 | lncRNA | ENSG00000 |
| 19018 | 35489 | -1 | lncRNA | ENSG00000 |
| 37032 | 37477 | 1 | processed_pseudoge | ENSG00000 |
| 70972 | 95683 | -1 | lncRNA | ENSG00000 |
| 77038 | 81178 | 1 | lncRNA | ENSG00000 |
| 115186 | 132034 | -1 | lncRNA | ENSG00000 |
| 135853 | 149527 | -1 | lncRNA | ENSG00000 |
| 149549 | 155465 | 1 | lncRNA | ENSG00000 |
| 161765 | 164972 | -1 | lncRNA | ENSG00000 |
| 174920 | 176013 | 1 | lncRNA | ENSG00000 |
| 182935 | 194180 | -1 | lncRNA | ENSG00000 |
| 192571 | 260772 | 1 | protein_coding | ENSG00000 |
| 227554 | 229557 | -1 | TEC | ENSG00000 |
| 290170 | 294422 | 1 | protein_coding | ENSG00000 |
| 329776 | 333172 | -1 | lncRNA | ENSG00000 |
| 379359 | 382712 | 1 | lncRNA | ENSG00000 |
| 425373 | 439389 | 1 | lncRNA | ENSG00000 |
| 446281 | 448823 | 1 | lncRNA | ENSG00000 |
| 484107 | 496118 | -1 | lncRNA | ENSG00000 |
| 497258 | 520296 | -1 | protein_coding | ENSG00000 |
| 520391 | 525238 | 1 | lncRNA | ENSG00000 |
| 549197 | 727650 | -1 | protein_coding | ENSG00000 |
| 561958 | 565619 | 1 | lncRNA | ENSG00000 |
| 603185 | 608482 | 1 | lncRNA | ENSG00000 |
| 726699 | 786475 | 1 | protein_coding | ENSG00000 |
| 816615 | 896435 | 1 | protein_coding | ENSG00000 |
| 876554 | 896436 | 1 | protein_coding | ENSG00000 |
| 879790 | 886547 | -1 | lncRNA | ENSG00000 |
| 897900 | 955407 | -1 | protein_coding | ENSG00000 |
| 898778 | 975549 | -1 | protein_coding | ENSG00000 |
| 983181 | 989640 | 1 | protein_coding | ENSG00000 |
| 996986 | 1138260 | -1 | protein_coding | ENSG00000 |
| 1022933 | 1023026 | -1 | miRNA | ENSG00000 |
| 1029025 | 1043891 | 1 | lncRNA | ENSG00000 |
| 1044546 | 1059261 | 1 | protein_coding | ENSG00000 |
| 1055360 | 1059261 | -1 | lncRNA | ENSG00000 |
| 1074450 | 1078036 | 1 | lncRNA | ENSG00000 |

| | | | | |
|---|---|---|---|---|
| 1080863 | 1082178 | 1 | lncRNA | ENSG0000C |
| 1082208 | 1093815 | 1 | protein_coding | ENSG0000C |
| 1152071 | 1160759 | -1 | protein_coding | ENSG0000C |
| 1160374 | 1165607 | 1 | lncRNA | ENSG0000C |
| 1232872 | 1237326 | 1 | protein_coding | ENSG0000C |
| 1267353 | 1269267 | 1 | lncRNA | ENSG0000C |
| 1428465 | 1459470 | -1 | protein_coding | ENSG0000C |
| 1459937 | 1464008 | 1 | lncRNA | ENSG0000C |
| 1464497 | 1467522 | -1 | lncRNA | ENSG0000C |
| 1470277 | 1504389 | -1 | protein_coding | ENSG0000C |
| 1508655 | 1509427 | -1 | lncRNA | ENSG0000C |
| 1530702 | 1543043 | 1 | protein_coding | ENSG0000C |
| 1542235 | 1560821 | -1 | protein_coding | ENSG0000C |
| 1567332 | 1571005 | -1 | protein_coding | ENSG0000C |
| 1570073 | 1589626 | 1 | lncRNA | ENSG0000C |
| 1614590 | 1615325 | 1 | processed_pseudoge | ENSG0000C |
| 1620654 | 1621405 | 1 | lncRNA | ENSG0000C |
| 1688119 | 1747954 | 1 | protein_coding | ENSG0000C |
| 1691906 | 1692450 | -1 | lncRNA | ENSG0000C |
| 1692810 | 1694357 | -1 | lncRNA | ENSG0000C |
| 1738630 | 1742291 | -1 | lncRNA | ENSG0000C |
| 1815793 | 2233243 | -1 | protein_coding | ENSG0000C |
| 1815804 | 2242215 | -1 | protein_coding | ENSG0000C |
| 1838586 | 1849931 | 1 | lncRNA | ENSG0000C |
| 1844180 | 1844253 | -1 | miRNA | ENSG0000C |
| 2234222 | 2242205 | -1 | protein_coding | ENSG0000C |
| 2242222 | 2251146 | 1 | protein_coding | ENSG0000C |
| 2251770 | 2354318 | -1 | protein_coding | ENSG0000C |
| 2257515 | 2257577 | -1 | miRNA | ENSG0000C |
| 2287915 | 2288428 | 1 | processed_pseudoge | ENSG0000C |
| 2354086 | 2380745 | 1 | protein_coding | ENSG0000C |
| 2383698 | 2388652 | 1 | transcribed_unitary_ | ENSG0000C |
| 2393847 | 2394131 | -1 | processed_pseudoge | ENSG0000C |
| 2403588 | 2448484 | 1 | protein_coding | ENSG0000C |
| 2474844 | 2476894 | -1 | protein_coding | ENSG0000C |
| 2512529 | 2529177 | 1 | protein_coding | ENSG0000C |
| 2527074 | 2527145 | 1 | miRNA | ENSG0000C |
| 2537810 | 2555694 | -1 | protein_coding | ENSG0000C |
| 2558972 | 2614733 | 1 | protein_coding | ENSG0000C |
| 2631986 | 2664802 | 1 | protein_coding | ENSG0000C |
| 2679522 | 2775500 | 1 | protein_coding | ENSG0000C |
| 2728105 | 2844308 | -1 | protein_coding | ENSG0000C |
| 2835621 | 2836022 | 1 | processed_pseudoge | ENSG0000C |
| 2906141 | 3043945 | -1 | protein_coding | ENSG0000C |
| 2944035 | 2947091 | 1 | lncRNA | ENSG0000C |
| 2999094 | 2999394 | 1 | misc_RNA | ENSG0000C |
| 3083252 | 3118049 | -1 | lncRNA | ENSG0000C |
| 3140234 | 3174654 | -1 | lncRNA | ENSG0000C |
| 3196626 | 3198256 | -1 | lncRNA | ENSG0000C |
| 3264032 | 3302452 | -1 | lncRNA | ENSG0000C |

| | | | | |
|---|---|---|---|---|
| 3301252 | 4269000 | 1 | protein_coding | ENSG00000( |
| 3337889 | 3338601 | 1 | transcribed_processe | ENSG00000( |
| 3642815 | 3643784 | -1 | lncRNA | ENSG00000( |
| 4130181 | 4134837 | -1 | lncRNA | ENSG00000( |
| 4511959 | 4512033 | -1 | processed_pseudoge | ENSG00000( |
| 4642248 | 4642570 | 1 | lncRNA | ENSG00000( |
| 4682295 | 4771442 | 1 | protein_coding | ENSG00000( |
| 4725440 | 4728721 | 1 | lncRNA | ENSG00000( |
| 4775615 | 4794397 | 1 | protein_coding | ENSG00000( |
| 4788565 | 4788639 | -1 | miRNA | ENSG00000( |
| 4792983 | 4793127 | -1 | transcribed_unproce | ENSG00000( |
| 4797055 | 4883716 | -1 | protein_coding | ENSG00000( |
| 4811188 | 4811289 | -1 | misc_RNA | ENSG00000( |
| 4857738 | 4862030 | -1 | protein_coding | ENSG00000( |
| 4865432 | 4865804 | -1 | processed_pseudoge | ENSG00000( |
| 4905989 | 4959213 | -1 | protein_coding | ENSG00000( |
| 4973988 | 5040675 | 1 | transcribed_unproce | ENSG00000( |
| 4983718 | 5073221 | 1 | protein_coding | ENSG00000( |
| 5009164 | 5009328 | -1 | processed_pseudoge | ENSG00000( |
| 5031608 | 5037831 | -1 | unprocessed_pseudc | ENSG00000( |
| 5045821 | 5069487 | 1 | protein_coding | ENSG00000( |
| 5072125 | 5073223 | 1 | lncRNA | ENSG00000( |
| 5103719 | 5120245 | 1 | transcribed_unproce | ENSG00000( |
| 5121239 | 5144546 | -1 | transcribed_unproce | ENSG00000( |
| 5180061 | 5180162 | 1 | snRNA | ENSG00000( |
| 5190196 | 5233840 | 1 | protein_coding | ENSG00000( |
| 5274369 | 5306870 | 1 | protein_coding | ENSG00000( |
| 5306790 | 5425414 | -1 | protein_coding | ENSG00000( |
| 5419827 | 5420767 | 1 | lncRNA | ENSG00000( |
| 5419846 | 5423122 | 1 | lncRNA | ENSG00000( |
| 5425770 | 5426401 | 1 | lncRNA | ENSG00000( |
| 5426277 | 5428927 | -1 | lncRNA | ENSG00000( |
| 5428731 | 5429672 | 1 | lncRNA | ENSG00000( |
| 5431335 | 5513809 | -1 | protein_coding | ENSG00000( |
| 5475804 | 5479811 | 1 | lncRNA | ENSG00000( |
| 5495819 | 5495917 | -1 | miRNA | ENSG00000( |
| 5527148 | 5563784 | -1 | protein_coding | ENSG00000( |
| 5556731 | 5557245 | 1 | lncRNA | ENSG00000( |
| 5560743 | 5561055 | 1 | processed_pseudoge | ENSG00000( |
| 5592816 | 5606655 | 1 | protein_coding | ENSG00000( |
| 5620047 | 5781739 | -1 | protein_coding | ENSG00000( |
| 5662432 | 5680461 | -1 | lncRNA | ENSG00000( |
| 5711840 | 5711910 | -1 | miRNA | ENSG00000( |
| 5823160 | 5854365 | 1 | transcribed_unproce | ENSG00000( |
| 5871753 | 5872050 | -1 | misc_RNA | ENSG00000( |
| 5879827 | 5886363 | 1 | protein_coding | ENSG00000( |
| 5898725 | 5926550 | 1 | protein_coding | ENSG00000( |
| 5925550 | 5970683 | -1 | protein_coding | ENSG00000( |
| 5970925 | 6009106 | -1 | protein_coding | ENSG00000( |
| 6004283 | 6004385 | -1 | misc_RNA | ENSG00000( |

| | | | | |
|---|---|---|---|---|
| 6009255 | 6023834 | 1 | protein_coding | ENSG00000 |
| 6016877 | 6017011 | 1 | snoRNA | ENSG00000 |
| 6022247 | 6059175 | -1 | protein_coding | ENSG00000 |
| 6031376 | 6036552 | 1 | protein_coding | ENSG00000 |
| 6047721 | 6048023 | -1 | misc_RNA | ENSG00000 |
| 6057500 | 6057604 | 1 | snRNA | ENSG00000 |
| 6080704 | 6093085 | 1 | lncRNA | ENSG00000 |
| 6104884 | 6161564 | 1 | protein_coding | ENSG00000 |
| 6161776 | 6272644 | -1 | protein_coding | ENSG00000 |
| 6212638 | 6212740 | -1 | misc_RNA | ENSG00000 |
| 6329411 | 6348981 | -1 | protein_coding | ENSG00000 |
| 6329412 | 6348959 | -1 | protein_coding | ENSG00000 |
| 6365441 | 6365770 | 1 | processed_pseudoge | ENSG00000 |
| 6374527 | 6403967 | 1 | protein_coding | ENSG00000 |
| 6409126 | 6484190 | -1 | protein_coding | ENSG00000 |
| 6445953 | 6484190 | -1 | protein_coding | ENSG00000 |
| 6490793 | 6491098 | -1 | processed_pseudoge | ENSG00000 |
| 6497462 | 6551436 | -1 | protein_coding | ENSG00000 |
| 6577434 | 6589374 | 1 | protein_coding | ENSG00000 |
| 6578565 | 6588974 | -1 | lncRNA | ENSG00000 |
| 6590021 | 6608726 | 1 | protein_coding | ENSG00000 |
| 6615610 | 6624290 | 1 | protein_coding | ENSG00000 |
| 6637318 | 6658279 | 1 | protein_coding | ENSG00000 |
| 6663974 | 6708901 | 1 | lncRNA | ENSG00000 |
| 6673494 | 6676366 | 1 | unprocessed_pseudc | ENSG00000 |
| 6688433 | 6706947 | -1 | protein_coding | ENSG00000 |
| 6710128 | 6753862 | 1 | transcribed_unproce | ENSG00000 |
| 6715630 | 6720982 | -1 | unprocessed_pseudc | ENSG00000 |
| 6754109 | 6799365 | 1 | protein_coding | ENSG00000 |
| 6794134 | 6826770 | -1 | protein_coding | ENSG00000 |
| 6835420 | 6835794 | -1 | unprocessed_pseudc | ENSG00000 |
| 6855485 | 6859830 | -1 | processed_pseudoge | ENSG00000 |
| 6866658 | 6867672 | -1 | unprocessed_pseudc | ENSG00000 |
| 6879627 | 6880626 | -1 | unprocessed_pseudc | ENSG00000 |
| 6900424 | 6900623 | -1 | processed_pseudoge | ENSG00000 |
| 6923804 | 6930835 | 1 | unprocessed_pseudc | ENSG00000 |
| 6924078 | 6924513 | -1 | processed_pseudoge | ENSG00000 |
| 6931684 | 6939603 | -1 | unprocessed_pseudc | ENSG00000 |
| 6952625 | 7104482 | -1 | lncRNA | ENSG00000 |
| 7066964 | 7067045 | -1 | miRNA | ENSG00000 |
| 7078302 | 7079742 | -1 | processed_pseudoge | ENSG00000 |
| 7102573 | 7102681 | -1 | misc_RNA | ENSG00000 |
| 7156934 | 7248651 | 1 | protein_coding | ENSG00000 |
| 7177952 | 7178324 | 1 | processed_pseudoge | ENSG00000 |
| 7255154 | 7278071 | 1 | lncRNA | ENSG00000 |
| 7293508 | 7293880 | 1 | processed_pseudoge | ENSG00000 |
| 7356203 | 7535873 | -1 | protein_coding | ENSG00000 |
| 7550104 | 7552440 | -1 | lncRNA | ENSG00000 |
| 7552462 | 7566996 | -1 | lncRNA | ENSG00000 |
| 7566875 | 7608932 | 1 | protein_coding | ENSG00000 |

| | | | | |
|---|---|---|---|---|
| 7619872 | 7620543 | 1 | lncRNA | ENSG0000C |
| 7636518 | 7718607 | -1 | protein_coding | ENSG0000C |
| 7640711 | 7968020 | 1 | protein_coding | ENSG0000C |
| 7640752 | 8004053 | 1 | lncRNA | ENSG0000C |
| 7735743 | 7736851 | -1 | processed_pseudoge | ENSG0000C |
| 7738524 | 7742574 | -1 | transcribed_processe | ENSG0000C |
| 7906519 | 7906624 | 1 | snRNA | ENSG0000C |
| 7909300 | 7910449 | -1 | processed_pseudoge | ENSG0000C |
| 7949853 | 7950709 | -1 | lncRNA | ENSG0000C |
| 7958183 | 7969903 | -1 | lncRNA | ENSG0000C |
| 7968796 | 8094272 | 1 | protein_coding | ENSG0000C |
| 7980312 | 7982228 | 1 | processed_pseudoge | ENSG0000C |
| 8113184 | 8262687 | -1 | protein_coding | ENSG0000C |
| 8114025 | 8116561 | 1 | lncRNA | ENSG0000C |
| 8262233 | 8262821 | 1 | lncRNA | ENSG0000C |
| 8262264 | 8344516 | 1 | lncRNA | ENSG0000C |
| 8303741 | 8341343 | 1 | lncRNA | ENSG0000C |
| 8433609 | 8752961 | 1 | protein_coding | ENSG0000C |
| 8938787 | 8939369 | 1 | processed_pseudoge | ENSG0000C |
| 9082557 | 9189857 | -1 | lncRNA | ENSG0000C |
| 9084022 | 9084175 | 1 | processed_pseudoge | ENSG0000C |
| 9170948 | 9171273 | -1 | processed_pseudoge | ENSG0000C |
| 9614978 | 9615985 | 1 | processed_pseudoge | ENSG0000C |
| 9621764 | 9769513 | 1 | lncRNA | ENSG0000C |
| 9634270 | 9635703 | 1 | processed_pseudoge | ENSG0000C |
| 9726241 | 9728272 | 1 | processed_pseudoge | ENSG0000C |
| 9850924 | 9851386 | 1 | processed_pseudoge | ENSG0000C |
| 9869774 | 9985895 | 1 | lncRNA | ENSG0000C |
| 9981516 | 10085494 | -1 | lncRNA | ENSG0000C |
| 10207670 | 10239863 | -1 | lncRNA | ENSG0000C |
| 10222797 | 10223011 | 1 | snoRNA | ENSG0000C |
| 10449820 | 10779944 | -1 | lncRNA | ENSG0000C |
| 10451311 | 10453252 | 1 | processed_pseudoge | ENSG0000C |
| 10476581 | 10476878 | 1 | processed_pseudoge | ENSG0000C |
| 10702676 | 10707406 | 1 | lncRNA | ENSG0000C |
| 10901978 | 10902074 | 1 | misc_RNA | ENSG0000C |
| 10931943 | 10940153 | -1 | protein_coding | ENSG0000C |
| 10940423 | 10940735 | 1 | lncRNA | ENSG0000C |
| 10973336 | 11169630 | 1 | protein_coding | ENSG0000C |
| 11006399 | 11006876 | -1 | processed_pseudoge | ENSG0000C |
| 11180902 | 11520175 | 1 | lncRNA | ENSG0000C |
| 11257681 | 11258535 | -1 | processed_pseudoge | ENSG0000C |
| 11370365 | 11832198 | -1 | protein_coding | ENSG0000C |
| 11406955 | 11414078 | 1 | lncRNA | ENSG0000C |
| 11820317 | 11821294 | -1 | processed_pseudoge | ENSG0000C |
| 12211270 | 12243367 | 1 | protein_coding | ENSG0000C |
| 12330885 | 12403941 | -1 | protein_coding | ENSG0000C |
| 12439419 | 12440404 | -1 | processed_pseudoge | ENSG0000C |
| 12469621 | 12542222 | 1 | protein_coding | ENSG0000C |
| 12471352 | 12472545 | 1 | processed_pseudoge | ENSG0000C |

| | | | | |
|---|---|---|---|---|
| 12491095 | 12492004 | 1 | processed_pseudoge | ENSG00000 |
| 12506007 | 12507478 | 1 | processed_pseudoge | ENSG00000 |
| 12570125 | 12571392 | -1 | lncRNA | ENSG00000 |
| 12570577 | 12660182 | 1 | protein_coding | ENSG00000 |
| 12654179 | 12654985 | 1 | lncRNA | ENSG00000 |
| 12685637 | 12686033 | 1 | processed_pseudoge | ENSG00000 |
| 12686856 | 12690958 | 1 | protein_coding | ENSG00000 |
| 12700758 | 12700889 | -1 | snoRNA | ENSG00000 |
| 12726152 | 13751920 | 1 | lncRNA | ENSG00000 |
| 12876684 | 12876946 | 1 | misc_RNA | ENSG00000 |
| 12922545 | 12922920 | -1 | processed_pseudoge | ENSG00000 |
| 13339201 | 13365049 | -1 | lncRNA | ENSG00000 |
| 13854355 | 13862118 | -1 | lncRNA | ENSG00000 |
| 13871856 | 13872513 | 1 | processed_pseudoge | ENSG00000 |
| 13891229 | 13991425 | -1 | protein_coding | ENSG00000 |
| 14010669 | 14010763 | -1 | misc_RNA | ENSG00000 |
| 14070535 | 14071380 | -1 | processed_pseudoge | ENSG00000 |
| 14145049 | 14974777 | -1 | protein_coding | ENSG00000 |
| 14190795 | 14192157 | 1 | processed_pseudoge | ENSG00000 |
| 14814131 | 14816565 | 1 | lncRNA | ENSG00000 |
| 14985378 | 14986074 | -1 | processed_pseudoge | ENSG00000 |
| 15066207 | 15068651 | 1 | lncRNA | ENSG00000 |
| 15200317 | 15562015 | -1 | protein_coding | ENSG00000 |
| 15611212 | 15686683 | -1 | protein_coding | ENSG00000 |
| 15667947 | 15681980 | 1 | lncRNA | ENSG00000 |
| 15688378 | 15696886 | 1 | lncRNA | ENSG00000 |
| 15689075 | 15690854 | -1 | lncRNA | ENSG00000 |
| 15701435 | 15701752 | -1 | processed_pseudoge | ENSG00000 |
| 15736373 | 15796370 | -1 | lncRNA | ENSG00000 |
| 15841297 | 15842360 | -1 | lncRNA | ENSG00000 |
| 16087525 | 16421538 | -1 | protein_coding | ENSG00000 |
| 16208945 | 16209265 | 1 | processed_pseudoge | ENSG00000 |
| 16210484 | 16270604 | 1 | lncRNA | ENSG00000 |
| 16461481 | 16530580 | -1 | protein_coding | ENSG00000 |
| 16471184 | 16471373 | 1 | lncRNA | ENSG00000 |
| 16526825 | 16581568 | 1 | protein_coding | ENSG00000 |
| 16583782 | 16586757 | -1 | processed_pseudoge | ENSG00000 |
| 16593626 | 16594224 | -1 | lncRNA | ENSG00000 |
| 16599779 | 16645817 | -1 | protein_coding | ENSG00000 |
| 16603816 | 16616295 | 1 | lncRNA | ENSG00000 |
| 16646131 | 16706523 | 1 | protein_coding | ENSG00000 |
| 16695871 | 16719898 | -1 | lncRNA | ENSG00000 |
| 16753755 | 16784536 | 1 | protein_coding | ENSG00000 |
| 16791811 | 16833433 | -1 | protein_coding | ENSG00000 |
| 16849957 | 16850725 | -1 | processed_pseudoge | ENSG00000 |
| 16859412 | 16881987 | -1 | protein_coding | ENSG00000 |
| 16864267 | 16866308 | -1 | processed_pseudoge | ENSG00000 |
| 16888621 | 16888885 | 1 | processed_pseudoge | ENSG00000 |
| 16916359 | 17346152 | 1 | protein_coding | ENSG00000 |
| 16938225 | 16938557 | 1 | processed_pseudoge | ENSG00000 |

| | | | | |
|---|---|---|---|---|
| 16970320 | 17299368 | -1 | lncRNA | ENSG00000 |
| 17033089 | 17033478 | -1 | processed_pseudoge | ENSG00000 |
| 17205652 | 17205928 | 1 | misc_RNA | ENSG00000 |
| 17299295 | 17467234 | 1 | protein_coding | ENSG00000 |
| 17373201 | 17373326 | -1 | snoRNA | ENSG00000 |
| 17374867 | 17467256 | 1 | lncRNA | ENSG00000 |
| 17405479 | 17558909 | -1 | lncRNA | ENSG00000 |
| 17679549 | 17680659 | 1 | lncRNA | ENSG00000 |
| 17790761 | 17940501 | -1 | protein_coding | ENSG00000 |
| 17940503 | 17942922 | 1 | TEC | ENSG00000 |
| 17962079 | 17963029 | 1 | processed_pseudoge | ENSG00000 |
| 18000290 | 18000879 | -1 | TEC | ENSG00000 |
| 18026770 | 18027846 | -1 | protein_coding | ENSG00000 |
| 18086949 | 19002416 | 1 | protein_coding | ENSG00000 |
| 18127220 | 18127309 | -1 | miRNA | ENSG00000 |
| 18429062 | 18430738 | -1 | lncRNA | ENSG00000 |
| 18807993 | 18808279 | -1 | misc_RNA | ENSG00000 |
| 18892096 | 18899433 | -1 | lncRNA | ENSG00000 |
| 18948898 | 18949762 | -1 | processed_pseudoge | ENSG00000 |
| 19020991 | 19117636 | -1 | protein_coding | ENSG00000 |
| 19112474 | 19114271 | 1 | lncRNA | ENSG00000 |
| 19119933 | 19121916 | 1 | lncRNA | ENSG00000 |
| 19144293 | 19146253 | 1 | lncRNA | ENSG00000 |
| 19144782 | 19145421 | -1 | protein_coding | ENSG00000 |
| 19353531 | 19578606 | 1 | lncRNA | ENSG00000 |
| 19695461 | 19709037 | -1 | protein_coding | ENSG00000 |
| 19705358 | 19705436 | -1 | miRNA | ENSG00000 |
| 19719310 | 19773598 | -1 | protein_coding | ENSG00000 |
| 19813685 | 19818090 | -1 | lncRNA | ENSG00000 |
| 19918981 | 20140453 | -1 | lncRNA | ENSG00000 |
| 20002765 | 20003247 | 1 | processed_pseudoge | ENSG00000 |
| 20096137 | 20130475 | 1 | lncRNA | ENSG00000 |
| 20134655 | 20217404 | -1 | protein_coding | ENSG00000 |
| 20141916 | 20153531 | 1 | lncRNA | ENSG00000 |
| 20217577 | 20221700 | 1 | lncRNA | ENSG00000 |
| 20296637 | 20314512 | 1 | lncRNA | ENSG00000 |
| 20328299 | 20331747 | -1 | lncRNA | ENSG00000 |
| 20330702 | 20415754 | 1 | protein_coding | ENSG00000 |
| 20377331 | 20377401 | 1 | snoRNA | ENSG00000 |
| 20582072 | 20583206 | 1 | processed_pseudoge | ENSG00000 |
| 20588990 | 20590085 | 1 | processed_pseudoge | ENSG00000 |
| 20615207 | 20777038 | 1 | protein_coding | ENSG00000 |
| 20641558 | 20641948 | -1 | processed_pseudoge | ENSG00000 |
| 20782283 | 20786886 | -1 | protein_coding | ENSG00000 |
| 20827326 | 20827753 | 1 | processed_pseudoge | ENSG00000 |
| 20835376 | 21023362 | 1 | lncRNA | ENSG00000 |
| 20843627 | 20843961 | 1 | processed_pseudoge | ENSG00000 |
| 20959391 | 21028813 | -1 | lncRNA | ENSG00000 |
| 21215537 | 21215830 | -1 | misc_RNA | ENSG00000 |
| 21220213 | 21221428 | -1 | processed_pseudoge | ENSG00000 |

| | | | | |
|---|---|---|---|---|
| 21381363 | 21381506 | 1 | snRNA | ENSG0000C |
| 21400920 | 21401613 | -1 | processed_pseudoge | ENSG0000C |
| 21428043 | 21514822 | 1 | protein_coding | ENSG0000C |
| 21471058 | 21471146 | 1 | miRNA | ENSG0000C |
| 21543039 | 21901839 | 1 | protein_coding | ENSG0000C |
| 21900899 | 21945903 | -1 | protein_coding | ENSG0000C |
| 22118236 | 22357154 | -1 | protein_coding | ENSG0000C |
| 22298594 | 22298706 | 1 | rRNA_pseudogene | ENSG0000C |
| 22419444 | 22727613 | -1 | protein_coding | ENSG0000C |
| 22510616 | 22511999 | -1 | processed_pseudoge | ENSG0000C |
| 22563337 | 22573996 | 1 | lncRNA | ENSG0000C |
| 22589698 | 22602536 | 1 | lncRNA | ENSG0000C |
| 22725395 | 22727620 | -1 | lncRNA | ENSG0000C |
| 22725884 | 22732002 | 1 | protein_coding | ENSG0000C |
| 22747675 | 22748804 | -1 | processed_pseudoge | ENSG0000C |
| 22773646 | 22773993 | -1 | processed_pseudoge | ENSG0000C |
| 22812628 | 22822852 | -1 | protein_coding | ENSG0000C |
| 22822954 | 22833430 | 1 | lncRNA | ENSG0000C |
| 22854126 | 22872945 | 1 | lncRNA | ENSG0000C |
| 22856613 | 22856686 | 1 | snoRNA | ENSG0000C |
| 22863874 | 22881350 | -1 | unprocessed_pseudc | ENSG0000C |
| 22881893 | 22882390 | 1 | processed_pseudoge | ENSG0000C |
| 22889371 | 23014130 | -1 | protein_coding | ENSG0000C |
| 23100214 | 23105703 | -1 | lncRNA | ENSG0000C |
| 23105758 | 23177914 | 1 | protein_coding | ENSG0000C |
| 23129178 | 23129841 | 1 | processed_pseudoge | ENSG0000C |
| 23181841 | 23201011 | 1 | protein_coding | ENSG0000C |
| 23200131 | 23206059 | -1 | lncRNA | ENSG0000C |
| 23205787 | 23208045 | 1 | lncRNA | ENSG0000C |
| 23235967 | 23275108 | 1 | protein_coding | ENSG0000C |
| 23298739 | 23311729 | 1 | protein_coding | ENSG0000C |
| 23310209 | 23470491 | -1 | protein_coding | ENSG0000C |
| 23365618 | 23365976 | 1 | processed_pseudoge | ENSG0000C |
| 23396446 | 23396516 | 1 | snoRNA | ENSG0000C |
| 23450658 | 23450722 | 1 | snRNA | ENSG0000C |
| 23480753 | 23480959 | 1 | processed_pseudoge | ENSG0000C |
| 23490473 | 23491364 | 1 | processed_pseudoge | ENSG0000C |
| 23491792 | 23491893 | -1 | misc_RNA | ENSG0000C |
| 23504780 | 23532041 | -1 | protein_coding | ENSG0000C |
| 23562459 | 23564261 | -1 | processed_pseudoge | ENSG0000C |
| 23585062 | 23586355 | -1 | processed_pseudoge | ENSG0000C |
| 23597382 | 23644708 | 1 | protein_coding | ENSG0000C |
| 23619333 | 23622826 | 1 | processed_pseudoge | ENSG0000C |
| 23655682 | 23655861 | 1 | processed_pseudoge | ENSG0000C |
| 23680130 | 23703249 | 1 | protein_coding | ENSG0000C |
| 23680195 | 23680786 | -1 | lncRNA | ENSG0000C |
| 23710203 | 23832513 | 1 | protein_coding | ENSG0000C |
| 23721311 | 23721782 | -1 | processed_pseudoge | ENSG0000C |
| 23811144 | 23811660 | -1 | processed_pseudoge | ENSG0000C |
| 23891704 | 23897335 | -1 | lncRNA | ENSG0000C |

| | | | | |
|---|---|---|---|---|
| 24132705 | 24132837 | 1 | rRNA_pseudogene | ENSG0000C |
| 24196662 | 24255719 | -1 | lncRNA | ENSG0000C |
| 24284188 | 24291862 | 1 | protein_coding | ENSG0000C |
| 24318094 | 24445128 | -1 | lncRNA | ENSG0000C |
| 24537332 | 24537438 | 1 | snRNA | ENSG0000C |
| 24573268 | 24694193 | 1 | protein_coding | ENSG0000C |
| 24589115 | 24589407 | -1 | processed_pseudoge | ENSG0000C |
| 24674309 | 24674612 | -1 | processed_pseudoge | ENSG0000C |
| 24698355 | 24757940 | -1 | protein_coding | ENSG0000C |
| 24778444 | 24795496 | -1 | lncRNA | ENSG0000C |
| 24796540 | 24981634 | -1 | protein_coding | ENSG0000C |
| 24906135 | 24906594 | -1 | processed_pseudoge | ENSG0000C |
| 25120091 | 25125361 | -1 | protein_coding | ENSG0000C |
| 25134692 | 25180356 | -1 | protein_coding | ENSG0000C |
| 25185623 | 25188724 | 1 | processed_pseudoge | ENSG0000C |
| 25212170 | 25212965 | -1 | processed_pseudoge | ENSG0000C |
| 25224570 | 25228486 | -1 | protein_coding | ENSG0000C |
| 25254965 | 25257129 | -1 | lncRNA | ENSG0000C |
| 25264656 | 25264882 | 1 | snoRNA | ENSG0000C |
| 25265924 | 25266290 | -1 | processed_pseudoge | ENSG0000C |
| 25321302 | 25326865 | -1 | lncRNA | ENSG0000C |
| 25358235 | 25432202 | -1 | lncRNA | ENSG0000C |
| 25431980 | 25500443 | 1 | lncRNA | ENSG0000C |
| 25547762 | 25551062 | -1 | lncRNA | ENSG0000C |
| 25593304 | 25751032 | -1 | lncRNA | ENSG0000C |
| 25689580 | 25689959 | -1 | processed_pseudoge | ENSG0000C |
| 25750890 | 25756256 | 1 | lncRNA | ENSG0000C |
| 25831335 | 25831909 | -1 | lncRNA | ENSG0000C |
| 25948657 | 25949403 | -1 | lncRNA | ENSG0000C |
| 25949919 | 25949986 | -1 | miRNA | ENSG0000C |
| 26152198 | 26187137 | 1 | protein_coding | ENSG0000C |
| 26173057 | 26201529 | -1 | protein_coding | ENSG0000C |
| 26201162 | 26213607 | 1 | protein_coding | ENSG0000C |
| 26226413 | 26254709 | -1 | lncRNA | ENSG0000C |
| 26276195 | 26276606 | 1 | processed_pseudoge | ENSG0000C |
| 26291895 | 26374329 | 1 | protein_coding | ENSG0000C |
| 26369310 | 26376701 | -1 | lncRNA | ENSG0000C |
| 26398593 | 26497595 | 1 | lncRNA | ENSG0000C |
| 26533121 | 26538788 | -1 | lncRNA | ENSG0000C |
| 26538378 | 26541048 | 1 | TEC | ENSG0000C |
| 26551686 | 26558880 | 1 | lncRNA | ENSG0000C |
| 26598600 | 26617295 | -1 | lncRNA | ENSG0000C |
| 26637871 | 26647305 | 1 | lncRNA | ENSG0000C |
| 26667068 | 26995239 | -1 | protein_coding | ENSG0000C |
| 26922110 | 26922912 | 1 | processed_pseudoge | ENSG0000C |
| 26982768 | 26983367 | -1 | processed_pseudoge | ENSG0000C |
| 27023737 | 27024907 | 1 | processed_pseudoge | ENSG0000C |
| 27048169 | 27048628 | 1 | processed_pseudoge | ENSG0000C |
| 27093313 | 27095996 | -1 | protein_coding | ENSG0000C |
| 27095647 | 27100265 | 1 | lncRNA | ENSG0000C |

| | | | | |
|---|---|---|---|---|
| 27096124 | 27096248 | 1 | misc_RNA | ENSG0000C |
| 27098900 | 27099114 | 1 | misc_RNA | ENSG0000C |
| 27099778 | 27099836 | 1 | misc_RNA | ENSG0000C |
| 27099856 | 27099957 | 1 | misc_RNA | ENSG0000C |
| 27099967 | 27100111 | 1 | misc_RNA | ENSG0000C |
| 27100354 | 27102686 | -1 | protein_coding | ENSG0000C |
| 27106184 | 27152581 | -1 | protein_coding | ENSG0000C |
| 27107777 | 27134302 | 1 | lncRNA | ENSG0000C |
| 27128507 | 27130780 | -1 | protein_coding | ENSG0000C |
| 27128969 | 27152561 | -1 | lncRNA | ENSG0000C |
| 27129977 | 27155928 | 1 | lncRNA | ENSG0000C |
| 27141052 | 27143681 | -1 | protein_coding | ENSG0000C |
| 27145396 | 27150603 | -1 | protein_coding | ENSG0000C |
| 27147366 | 27152598 | -1 | lncRNA | ENSG0000C |
| 27153716 | 27157936 | -1 | protein_coding | ENSG0000C |
| 27158344 | 27158976 | 1 | lncRNA | ENSG0000C |
| 27162438 | 27175180 | -1 | protein_coding | ENSG0000C |
| 27163535 | 27180013 | -1 | protein_coding | ENSG0000C |
| 27168619 | 27171915 | 1 | lncRNA | ENSG0000C |
| 27169480 | 27169563 | -1 | miRNA | ENSG0000C |
| 27170592 | 27180261 | -1 | protein_coding | ENSG0000C |
| 27181510 | 27185223 | -1 | protein_coding | ENSG0000C |
| 27184507 | 27189298 | 1 | lncRNA | ENSG0000C |
| 27185433 | 27185530 | 1 | misc_RNA | ENSG0000C |
| 27185832 | 27186018 | 1 | misc_RNA | ENSG0000C |
| 27186166 | 27186263 | 1 | misc_RNA | ENSG0000C |
| 27186573 | 27193448 | -1 | lncRNA | ENSG0000C |
| 27186892 | 27187124 | 1 | misc_RNA | ENSG0000C |
| 27187819 | 27188021 | 1 | misc_RNA | ENSG0000C |
| 27188816 | 27188994 | 1 | misc_RNA | ENSG0000C |
| 27193503 | 27200091 | -1 | protein_coding | ENSG0000C |
| 27198575 | 27207259 | 1 | lncRNA | ENSG0000C |
| 27200465 | 27200521 | 1 | misc_RNA | ENSG0000C |
| 27201844 | 27202219 | 1 | misc_RNA | ENSG0000C |
| 27202302 | 27202638 | 1 | misc_RNA | ENSG0000C |
| 27206139 | 27206303 | 1 | misc_RNA | ENSG0000C |
| 27210801 | 27212965 | -1 | lncRNA | ENSG0000C |
| 27239243 | 27241228 | -1 | lncRNA | ENSG0000C |
| 27241429 | 27247229 | -1 | lncRNA | ENSG0000C |
| 27242700 | 27250493 | 1 | protein_coding | ENSG0000C |
| 27269357 | 27269678 | -1 | processed_pseudoge | ENSG0000C |
| 27328607 | 27329534 | -1 | processed_pseudoge | ENSG0000C |
| 27361591 | 27410358 | 1 | lncRNA | ENSG0000C |
| 27380562 | 27381206 | 1 | TEC | ENSG0000C |
| 27412164 | 27412891 | 1 | TEC | ENSG0000C |
| 27458163 | 27458849 | -1 | processed_pseudoge | ENSG0000C |
| 27461766 | 27462969 | 1 | processed_pseudoge | ENSG0000C |
| 27491682 | 27492765 | -1 | processed_pseudoge | ENSG0000C |
| 27525442 | 27662883 | -1 | protein_coding | ENSG0000C |
| 27617474 | 27627496 | 1 | lncRNA | ENSG0000C |

| | | | | |
|---|---|---|---|---|
| 27733064 | 27740395 | -1 | lncRNA | ENSG00000 |
| 27739331 | 27844564 | 1 | protein_coding | ENSG00000 |
| 27830573 | 28180795 | -1 | protein_coding | ENSG00000 |
| 27873785 | 27876546 | 1 | lncRNA | ENSG00000 |
| 28118113 | 28118219 | -1 | snRNA | ENSG00000 |
| 28180322 | 28243917 | 1 | lncRNA | ENSG00000 |
| 28279211 | 28279700 | -1 | processed_pseudoge | ENSG00000 |
| 28299321 | 28825894 | 1 | protein_coding | ENSG00000 |
| 28615097 | 28615888 | -1 | processed_pseudoge | ENSG00000 |
| 28953358 | 28958330 | -1 | protein_coding | ENSG00000 |
| 28957667 | 28959345 | 1 | lncRNA | ENSG00000 |
| 28979967 | 29013367 | 1 | lncRNA | ENSG00000 |
| 28987028 | 28988899 | 1 | lncRNA | ENSG00000 |
| 28995235 | 29195451 | -1 | protein_coding | ENSG00000 |
| 29080284 | 29082527 | 1 | lncRNA | ENSG00000 |
| 29122274 | 29128172 | 1 | lncRNA | ENSG00000 |
| 29122340 | 29514667 | 1 | protein_coding | ENSG00000 |
| 29146569 | 29514328 | 1 | protein_coding | ENSG00000 |
| 29191582 | 29194109 | -1 | lncRNA | ENSG00000 |
| 29199540 | 29208970 | -1 | lncRNA | ENSG00000 |
| 29212302 | 29213214 | 1 | transcribed_processe | ENSG00000 |
| 29514224 | 29563670 | -1 | lncRNA | ENSG00000 |
| 29551521 | 29552151 | 1 | processed_pseudoge | ENSG00000 |
| 29563835 | 29567293 | 1 | protein_coding | ENSG00000 |
| 29598795 | 29685255 | -1 | transcribed_processe | ENSG00000 |
| 29625870 | 29625983 | 1 | processed_pseudoge | ENSG00000 |
| 29650227 | 29742594 | 1 | transcribed_unproce | ENSG00000 |
| 29680734 | 29680828 | -1 | miRNA | ENSG00000 |
| 29806486 | 29917066 | 1 | protein_coding | ENSG00000 |
| 29912528 | 29912632 | 1 | snoRNA | ENSG00000 |
| 29920104 | 29990289 | -1 | protein_coding | ENSG00000 |
| 29988600 | 30027543 | 1 | lncRNA | ENSG00000 |
| 30010587 | 30026684 | -1 | protein_coding | ENSG00000 |
| 30027404 | 30130483 | 1 | protein_coding | ENSG00000 |
| 30069658 | 30075115 | -1 | lncRNA | ENSG00000 |
| 30115845 | 30115994 | -1 | sRNA | ENSG00000 |
| 30129507 | 30134714 | -1 | lncRNA | ENSG00000 |
| 30134986 | 30162762 | 1 | protein_coding | ENSG00000 |
| 30140870 | 30141417 | 1 | lncRNA | ENSG00000 |
| 30157531 | 30159534 | 1 | lncRNA | ENSG00000 |
| 30167804 | 30168004 | 1 | processed_pseudoge | ENSG00000 |
| 30176387 | 30179014 | -1 | lncRNA | ENSG00000 |
| 30284597 | 30367689 | 1 | protein_coding | ENSG00000 |
| 30289794 | 30289890 | -1 | miRNA | ENSG00000 |
| 30289794 | 30289890 | 1 | miRNA | ENSG00000 |
| 30362400 | 30412502 | 1 | protein_coding | ENSG00000 |
| 30371508 | 30372741 | 1 | processed_pseudoge | ENSG00000 |
| 30390885 | 30412375 | 1 | lncRNA | ENSG00000 |
| 30424527 | 30478784 | -1 | protein_coding | ENSG00000 |
| 30424672 | 30425412 | 1 | lncRNA | ENSG00000 |

| | | | | |
|---|---|---|---|---|
| 30496621 | 30504841 | -1 | protein_coding | ENSG00000 |
| 30497077 | 30550761 | -1 | protein_coding | ENSG00000 |
| 30516309 | 30595286 | -1 | lncRNA | ENSG00000 |
| 30523143 | 30524535 | -1 | transcribed_processe | ENSG00000 |
| 30525731 | 30533613 | 1 | lncRNA | ENSG00000 |
| 30544053 | 30544431 | 1 | processed_pseudoge | ENSG00000 |
| 30550217 | 30551569 | 1 | lncRNA | ENSG00000 |
| 30594838 | 30634033 | 1 | protein_coding | ENSG00000 |
| 30651942 | 30700129 | -1 | protein_coding | ENSG00000 |
| 30697985 | 30757602 | 1 | protein_coding | ENSG00000 |
| 30742431 | 30742818 | -1 | processed_pseudoge | ENSG00000 |
| 30752137 | 30892081 | 1 | protein_coding | ENSG00000 |
| 30771417 | 30892387 | 1 | protein_coding | ENSG00000 |
| 30796691 | 30803410 | -1 | lncRNA | ENSG00000 |
| 30852273 | 30923812 | 1 | protein_coding | ENSG00000 |
| 30911694 | 30925517 | 1 | protein_coding | ENSG00000 |
| 30938669 | 30993254 | 1 | protein_coding | ENSG00000 |
| 31052461 | 31111479 | 1 | protein_coding | ENSG00000 |
| 31119284 | 31119572 | -1 | processed_pseudoge | ENSG00000 |
| 31202831 | 31235648 | 1 | lncRNA | ENSG00000 |
| 31330711 | 31330896 | 1 | processed_pseudoge | ENSG00000 |
| 31337465 | 31340726 | -1 | protein_coding | ENSG00000 |
| 31414158 | 31421303 | -1 | lncRNA | ENSG00000 |
| 31514071 | 31658720 | 1 | protein_coding | ENSG00000 |
| 31687215 | 31708455 | 1 | protein_coding | ENSG00000 |
| 31751179 | 32428131 | -1 | protein_coding | ENSG00000 |
| 31941261 | 31941583 | 1 | processed_pseudoge | ENSG00000 |
| 32427901 | 32431804 | 1 | lncRNA | ENSG00000 |
| 32456963 | 32457758 | -1 | processed_pseudoge | ENSG00000 |
| 32471513 | 32472409 | 1 | processed_pseudoge | ENSG00000 |
| 32485338 | 32495283 | -1 | protein_coding | ENSG00000 |
| 32495426 | 32588726 | 1 | protein_coding | ENSG00000 |
| 32580949 | 32761787 | -1 | unprocessed_pseudc | ENSG00000 |
| 32728569 | 32753744 | 1 | transcribed_processe | ENSG00000 |
| 32732981 | 32733077 | 1 | miRNA | ENSG00000 |
| 32732981 | 32733077 | -1 | miRNA | ENSG00000 |
| 32758882 | 32759353 | 1 | lncRNA | ENSG00000 |
| 32760279 | 32762924 | 1 | lncRNA | ENSG00000 |
| 32788334 | 32788442 | 1 | unprocessed_pseudc | ENSG00000 |
| 32791814 | 32791946 | 1 | snoRNA | ENSG00000 |
| 32812757 | 32838570 | 1 | unprocessed_pseudc | ENSG00000 |
| 32845394 | 32846061 | 1 | lncRNA | ENSG00000 |
| 32868172 | 32894131 | -1 | protein_coding | ENSG00000 |
| 32916815 | 32943176 | -1 | transcribed_unproce | ENSG00000 |
| 32951426 | 32951656 | 1 | processed_pseudoge | ENSG00000 |
| 32957404 | 33006930 | 1 | protein_coding | ENSG00000 |
| 33001950 | 33002054 | -1 | snRNA | ENSG00000 |
| 33014114 | 33062797 | -1 | protein_coding | ENSG00000 |
| 33036212 | 33036382 | -1 | processed_pseudoge | ENSG00000 |
| 33045798 | 33046090 | -1 | misc_RNA | ENSG00000 |

| | | | | |
|---|---|---|---|---|
| 33094797 | 33109404 | -1 | protein_coding | ENSG0000C |
| 33109557 | 33877180 | 1 | protein_coding | ENSG0000C |
| 33128988 | 33129143 | -1 | processed_pseudoge | ENSG0000C |
| 33177740 | 33177838 | 1 | rRNA_pseudogene | ENSG0000C |
| 33276834 | 33277091 | -1 | processed_pseudoge | ENSG0000C |
| 33379964 | 33380886 | 1 | processed_pseudoge | ENSG0000C |
| 33725981 | 33728456 | 1 | lncRNA | ENSG0000C |
| 33730292 | 33732064 | -1 | lncRNA | ENSG0000C |
| 33793167 | 33803190 | -1 | lncRNA | ENSG0000C |
| 33868501 | 33874231 | -1 | lncRNA | ENSG0000C |
| 33904308 | 34156427 | 1 | protein_coding | ENSG0000C |
| 34124948 | 34128126 | -1 | lncRNA | ENSG0000C |
| 34209465 | 34299012 | 1 | lncRNA | ENSG0000C |
| 34297496 | 34297598 | -1 | snRNA | ENSG0000C |
| 34346512 | 34871582 | -1 | lncRNA | ENSG0000C |
| 34658218 | 34878332 | 1 | protein_coding | ENSG0000C |
| 34710757 | 34711480 | 1 | processed_pseudoge | ENSG0000C |
| 34750253 | 34752053 | -1 | processed_pseudoge | ENSG0000C |
| 34753424 | 34753719 | 1 | misc_RNA | ENSG0000C |
| 34892417 | 34892549 | 1 | snoRNA | ENSG0000C |
| 34928876 | 35038271 | -1 | protein_coding | ENSG0000C |
| 34940760 | 34940834 | -1 | miRNA | ENSG0000C |
| 35036836 | 35125012 | 1 | lncRNA | ENSG0000C |
| 35079989 | 35186041 | -1 | transcribed_unproce | ENSG0000C |
| 35169224 | 35169535 | 1 | processed_pseudoge | ENSG0000C |
| 35202430 | 35254100 | -1 | protein_coding | ENSG0000C |
| 35258381 | 35259729 | -1 | lncRNA | ENSG0000C |
| 35313838 | 35377071 | 1 | lncRNA | ENSG0000C |
| 35372349 | 35379675 | -1 | lncRNA | ENSG0000C |
| 35496021 | 35510263 | -1 | lncRNA | ENSG0000C |
| 35614650 | 35622939 | -1 | lncRNA | ENSG0000C |
| 35632659 | 35695571 | -1 | protein_coding | ENSG0000C |
| 35695214 | 35699413 | 1 | lncRNA | ENSG0000C |
| 35715032 | 35734924 | 1 | lncRNA | ENSG0000C |
| 35751856 | 35800676 | -1 | lncRNA | ENSG0000C |
| 35768122 | 35768640 | -1 | processed_pseudoge | ENSG0000C |
| 35800932 | 35907105 | 1 | protein_coding | ENSG0000C |
| 35847038 | 35847426 | -1 | processed_pseudoge | ENSG0000C |
| 35930158 | 35930264 | -1 | snRNA | ENSG0000C |
| 35933888 | 35934569 | 1 | unprocessed_pseudc | ENSG0000C |
| 35946294 | 35972587 | 1 | unprocessed_pseudc | ENSG0000C |
| 35971147 | 35971567 | -1 | processed_pseudoge | ENSG0000C |
| 35976596 | 36028174 | -1 | lncRNA | ENSG0000C |
| 36001317 | 36055431 | 1 | lncRNA | ENSG0000C |
| 36064715 | 36066679 | 1 | lncRNA | ENSG0000C |
| 36079084 | 36085736 | 1 | lncRNA | ENSG0000C |
| 36095255 | 36100660 | 1 | lncRNA | ENSG0000C |
| 36109024 | 36110872 | -1 | processed_pseudoge | ENSG0000C |
| 36146842 | 36148024 | 1 | processed_pseudoge | ENSG0000C |
| 36153254 | 36301538 | 1 | protein_coding | ENSG0000C |

| | | | | |
|---|---|---|---|---|
| 36228621 | 36228884 | -1 | processed_pseudoge | ENSG00000( |
| 36229280 | 36231107 | -1 | lncRNA | ENSG00000( |
| 36305631 | 36305904 | 1 | processed_pseudoge | ENSG00000( |
| 36319775 | 36320479 | 1 | lncRNA | ENSG00000( |
| 36324221 | 36390125 | -1 | protein_coding | ENSG00000( |
| 36337667 | 36337778 | -1 | misc_RNA | ENSG00000( |
| 36389821 | 36453791 | 1 | protein_coding | ENSG00000( |
| 36463023 | 36504513 | 1 | lncRNA | ENSG00000( |
| 36512941 | 36724549 | -1 | protein_coding | ENSG00000( |
| 36597834 | 36600120 | -1 | lncRNA | ENSG00000( |
| 36734004 | 36734535 | -1 | lncRNA | ENSG00000( |
| 36750687 | 36763081 | 1 | lncRNA | ENSG00000( |
| 36781008 | 36782789 | 1 | lncRNA | ENSG00000( |
| 36818959 | 36819690 | 1 | processed_pseudoge | ENSG00000( |
| 36838972 | 36841373 | 1 | lncRNA | ENSG00000( |
| 36854361 | 37449249 | -1 | protein_coding | ENSG00000( |
| 36919357 | 36919432 | -1 | miRNA | ENSG00000( |
| 36997796 | 37013630 | 1 | lncRNA | ENSG00000( |
| 37116779 | 37117185 | 1 | processed_pseudoge | ENSG00000( |
| 37351143 | 37351242 | 1 | snRNA | ENSG00000( |
| 37375522 | 37376023 | 1 | processed_pseudoge | ENSG00000( |
| 37419628 | 37419767 | 1 | snoRNA | ENSG00000( |
| 37435115 | 37435216 | 1 | misc_RNA | ENSG00000( |
| 37585500 | 37586329 | -1 | processed_pseudoge | ENSG00000( |
| 37683796 | 37833788 | 1 | protein_coding | ENSG00000( |
| 37683843 | 37951936 | 1 | protein_coding | ENSG00000( |
| 37796058 | 37796888 | 1 | unitary_pseudogene | ENSG00000( |
| 37821210 | 37821610 | 1 | processed_pseudoge | ENSG00000( |
| 37848597 | 37900397 | 1 | protein_coding | ENSG00000( |
| 37905932 | 38025695 | -1 | protein_coding | ENSG00000( |
| 38178222 | 38230671 | 1 | protein_coding | ENSG00000( |
| 38239580 | 38249572 | -1 | TR_C_gene | ENSG00000( |
| 38253380 | 38253429 | -1 | TR_J_gene | ENSG00000( |
| 38256337 | 38256396 | -1 | TR_J_gene | ENSG00000( |
| 38257879 | 38265678 | -1 | TR_C_gene | ENSG00000( |
| 38269491 | 38269540 | -1 | TR_J_gene | ENSG00000( |
| 38273587 | 38273647 | -1 | TR_J_gene | ENSG00000( |
| 38276259 | 38276318 | -1 | TR_J_gene | ENSG00000( |
| 38291616 | 38292078 | -1 | TR_V_gene | ENSG00000( |
| 38295763 | 38296233 | -1 | TR_V_pseudogene | ENSG00000( |
| 38299811 | 38300322 | -1 | TR_V_gene | ENSG00000( |
| 38311383 | 38311808 | 1 | processed_pseudoge | ENSG00000( |
| 38317017 | 38318861 | -1 | TR_V_gene | ENSG00000( |
| 38322446 | 38322730 | -1 | TR_V_pseudogene | ENSG00000( |
| 38326070 | 38329643 | -1 | lncRNA | ENSG00000( |
| 38330343 | 38330935 | -1 | TR_V_gene | ENSG00000( |
| 38335041 | 38335514 | -1 | TR_V_pseudogene | ENSG00000( |
| 38340700 | 38340998 | -1 | TR_V_pseudogene | ENSG00000( |
| 38341577 | 38378804 | 1 | lncRNA | ENSG00000( |
| 38345030 | 38345499 | -1 | TR_V_pseudogene | ENSG00000( |

| | | | | |
|---|---|---|---|---|
| 38349355 | 38350022 | -1 | TR_V_gene | ENSG00000 |
| 38353715 | 38354517 | -1 | TR_V_gene | ENSG00000 |
| 38358512 | 38359162 | -1 | TR_V_gene | ENSG00000 |
| 38362864 | 38363518 | -1 | TR_V_gene | ENSG00000 |
| 38367586 | 38368169 | -1 | TR_V_gene | ENSG00000 |
| 38383704 | 38631420 | -1 | protein_coding | ENSG00000 |
| 38531042 | 38531338 | -1 | misc_RNA | ENSG00000 |
| 38655205 | 38656613 | 1 | processed_pseudoge | ENSG00000 |
| 38685346 | 38687037 | -1 | processed_pseudoge | ENSG00000 |
| 38722974 | 38932394 | -1 | protein_coding | ENSG00000 |
| 38977998 | 39493095 | 1 | protein_coding | ENSG00000 |
| 38980370 | 39013551 | -1 | lncRNA | ENSG00000 |
| 39329003 | 39329134 | -1 | snoRNA | ENSG00000 |
| 39404590 | 39406579 | -1 | lncRNA | ENSG00000 |
| 39545646 | 39566239 | -1 | lncRNA | ENSG00000 |
| 39566385 | 39610320 | 1 | protein_coding | ENSG00000 |
| 39569376 | 39570198 | -1 | processed_pseudoge | ENSG00000 |
| 39609610 | 39610290 | -1 | lncRNA | ENSG00000 |
| 39609717 | 39610280 | 1 | processed_pseudoge | ENSG00000 |
| 39621253 | 39623201 | -1 | lncRNA | ENSG00000 |
| 39623565 | 39708120 | 1 | protein_coding | ENSG00000 |
| 39700341 | 39703296 | -1 | lncRNA | ENSG00000 |
| 39733430 | 39793092 | 1 | lncRNA | ENSG00000 |
| 39768341 | 39768444 | 1 | snRNA | ENSG00000 |
| 39788967 | 39789304 | -1 | processed_pseudoge | ENSG00000 |
| 39796593 | 39797911 | -1 | processed_pseudoge | ENSG00000 |
| 39816549 | 39868140 | 1 | unprocessed_pseudc | ENSG00000 |
| 39833286 | 39833793 | -1 | processed_pseudoge | ENSG00000 |
| 39833571 | 39834718 | 1 | processed_pseudoge | ENSG00000 |
| 39854494 | 39855056 | -1 | processed_pseudoge | ENSG00000 |
| 39888375 | 39888664 | 1 | misc_RNA | ENSG00000 |
| 39947522 | 39949755 | -1 | lncRNA | ENSG00000 |
| 39950121 | 40099580 | 1 | protein_coding | ENSG00000 |
| 40107288 | 40107454 | 1 | unprocessed_pseudc | ENSG00000 |
| 40115685 | 40115943 | 1 | processed_pseudoge | ENSG00000 |
| 40126027 | 40134622 | -1 | protein_coding | ENSG00000 |
| 40128121 | 40128232 | -1 | misc_RNA | ENSG00000 |
| 40134977 | 40860763 | 1 | protein_coding | ENSG00000 |
| 40151613 | 40154151 | 1 | unprocessed_pseudc | ENSG00000 |
| 40538127 | 40546928 | -1 | lncRNA | ENSG00000 |
| 40775558 | 40785217 | 1 | lncRNA | ENSG00000 |
| 40964667 | 40979939 | -1 | lncRNA | ENSG00000 |
| 41089539 | 41133507 | 1 | lncRNA | ENSG00000 |
| 41667168 | 41705834 | -1 | protein_coding | ENSG00000 |
| 41693884 | 41779388 | 1 | lncRNA | ENSG00000 |
| 41960949 | 42237870 | -1 | protein_coding | ENSG00000 |
| 42113365 | 42113620 | 1 | processed_pseudoge | ENSG00000 |
| 42661716 | 42706447 | -1 | lncRNA | ENSG00000 |
| 42794906 | 42802494 | 1 | unprocessed_pseudc | ENSG00000 |
| 42829672 | 42844438 | -1 | lncRNA | ENSG00000 |

| | | | | |
|---|---|---|---|---|
| 42901272 | 42902639 | -1 | lncRNA | ENSG0000C |
| 42908726 | 42912305 | -1 | protein_coding | ENSG0000C |
| 42909273 | 42932174 | -1 | protein_coding | ENSG0000C |
| 42916861 | 42932185 | -1 | protein_coding | ENSG0000C |
| 42918741 | 42920084 | 1 | processed_pseudoge | ENSG0000C |
| 42932376 | 42948958 | 1 | protein_coding | ENSG0000C |
| 42954135 | 43113931 | -1 | lncRNA | ENSG0000C |
| 43096732 | 43096844 | 1 | misc_RNA | ENSG0000C |
| 43112629 | 43566001 | 1 | protein_coding | ENSG0000C |
| 43117896 | 43163187 | 1 | lncRNA | ENSG0000C |
| 43150895 | 43150994 | 1 | miRNA | ENSG0000C |
| 43194421 | 43194482 | 1 | snRNA | ENSG0000C |
| 43207762 | 43207866 | -1 | snRNA | ENSG0000C |
| 43239066 | 43249268 | -1 | lncRNA | ENSG0000C |
| 43274159 | 43274582 | 1 | processed_pseudoge | ENSG0000C |
| 43508728 | 43522542 | -1 | lncRNA | ENSG0000C |
| 43582758 | 43650713 | 1 | protein_coding | ENSG0000C |
| 43608456 | 43729717 | -1 | protein_coding | ENSG0000C |
| 43758680 | 43807342 | 1 | protein_coding | ENSG0000C |
| 43767290 | 43767599 | 1 | processed_pseudoge | ENSG0000C |
| 43866558 | 43906589 | -1 | protein_coding | ENSG0000C |
| 43866558 | 43869893 | -1 | protein_coding | ENSG0000C |
| 43875894 | 43926411 | -1 | protein_coding | ENSG0000C |
| 43918697 | 43920054 | 1 | processed_pseudoge | ENSG0000C |
| 43926436 | 43956136 | 1 | protein_coding | ENSG0000C |
| 43940895 | 44019175 | -1 | lncRNA | ENSG0000C |
| 43951910 | 44019151 | -1 | lncRNA | ENSG0000C |
| 43961590 | 43973323 | -1 | transcribed_unproce | ENSG0000C |
| 43984995 | 43990207 | -1 | transcribed_unproce | ENSG0000C |
| 43997897 | 44010122 | 1 | protein_coding | ENSG0000C |
| 44004046 | 44007866 | -1 | lncRNA | ENSG0000C |
| 44011882 | 44038978 | -1 | lncRNA | ENSG0000C |
| 44013562 | 44019170 | -1 | transcribed_unproce | ENSG0000C |
| 44026951 | 44041892 | -1 | transcribed_unproce | ENSG0000C |
| 44039171 | 44042306 | 1 | lncRNA | ENSG0000C |
| 44044640 | 44069456 | 1 | protein_coding | ENSG0000C |
| 44051766 | 44051829 | 1 | miRNA | ENSG0000C |
| 44062727 | 44065567 | -1 | protein_coding | ENSG0000C |
| 44064908 | 44066079 | 1 | lncRNA | ENSG0000C |
| 44072062 | 44082530 | -1 | protein_coding | ENSG0000C |
| 44073378 | 44073433 | -1 | miRNA | ENSG0000C |
| 44104345 | 44114562 | 1 | protein_coding | ENSG0000C |
| 44110849 | 44110912 | 1 | miRNA | ENSG0000C |
| 44114681 | 44124358 | -1 | protein_coding | ENSG0000C |
| 44125752 | 44125873 | -1 | rRNA_pseudogene | ENSG0000C |
| 44138864 | 44141332 | -1 | protein_coding | ENSG0000C |
| 44143213 | 44198170 | -1 | protein_coding | ENSG0000C |
| 44200968 | 44214294 | 1 | protein_coding | ENSG0000C |
| 44217150 | 44334577 | -1 | protein_coding | ENSG0000C |
| 44367140 | 44369292 | -1 | TEC | ENSG0000C |

| | | | | |
|---|---|---|---|---|
| 44379119 | 44490658 | -1 | protein_coding | ENSG00000( |
| 44467897 | 44468304 | 1 | processed_pseudoge | ENSG00000( |
| 44503773 | 44503879 | -1 | snRNA | ENSG00000( |
| 44511025 | 44511332 | -1 | processed_pseudoge | ENSG00000( |
| 44512535 | 44541315 | -1 | protein_coding | ENSG00000( |
| 44565417 | 44575051 | -1 | protein_coding | ENSG00000( |
| 44577894 | 44582287 | -1 | protein_coding | ENSG00000( |
| 44582519 | 44583676 | 1 | lncRNA | ENSG00000( |
| 44601032 | 44601487 | 1 | processed_pseudoge | ENSG00000( |
| 44606572 | 44709066 | 1 | protein_coding | ENSG00000( |
| 44748581 | 44769881 | 1 | protein_coding | ENSG00000( |
| 44785050 | 44787340 | -1 | lncRNA | ENSG00000( |
| 44796680 | 44824564 | 1 | protein_coding | ENSG00000( |
| 44826791 | 44848083 | -1 | protein_coding | ENSG00000( |
| 44848416 | 44849568 | 1 | lncRNA | ENSG00000( |
| 44876299 | 44885530 | -1 | protein_coding | ENSG00000( |
| 44881748 | 44881800 | -1 | miRNA | ENSG00000( |
| 44884953 | 44886393 | 1 | lncRNA | ENSG00000( |
| 44919669 | 44920329 | 1 | processed_pseudoge | ENSG00000( |
| 44958999 | 44960909 | -1 | lncRNA | ENSG00000( |
| 44962662 | 44979088 | -1 | protein_coding | ENSG00000( |
| 44983023 | 44986961 | -1 | lncRNA | ENSG00000( |
| 44985378 | 44985510 | -1 | snoRNA | ENSG00000( |
| 44999475 | 45076469 | 1 | protein_coding | ENSG00000( |
| 45080437 | 45088969 | -1 | protein_coding | ENSG00000( |
| 45100100 | 45112047 | -1 | protein_coding | ENSG00000( |
| 45104349 | 45104482 | -1 | snoRNA | ENSG00000( |
| 45104906 | 45105042 | -1 | snoRNA | ENSG00000( |
| 45105968 | 45106099 | -1 | snoRNA | ENSG00000( |
| 45157791 | 45186302 | 1 | protein_coding | ENSG00000( |
| 45268681 | 45384802 | 1 | lncRNA | ENSG00000( |
| 45303620 | 45304362 | 1 | processed_pseudoge | ENSG00000( |
| 45391626 | 45392205 | -1 | processed_pseudoge | ENSG00000( |
| 45460712 | 45542182 | 1 | lncRNA | ENSG00000( |
| 45534523 | 45534681 | -1 | processed_pseudoge | ENSG00000( |
| 45574140 | 45723116 | 1 | protein_coding | ENSG00000( |
| 45723780 | 45768985 | -1 | transcribed_unproce | ENSG00000( |
| 45769105 | 45815263 | 1 | transcribed_unproce | ENSG00000( |
| 45789585 | 45789688 | 1 | snRNA | ENSG00000( |
| 45816216 | 45819016 | -1 | processed_pseudoge | ENSG00000( |
| 45816557 | 45821064 | 1 | unprocessed_pseudc | ENSG00000( |
| 45818582 | 45819097 | 1 | unprocessed_pseudc | ENSG00000( |
| 45843634 | 45843740 | 1 | snRNA | ENSG00000( |
| 45859994 | 45873082 | -1 | protein_coding | ENSG00000( |
| 45888360 | 45893660 | 1 | protein_coding | ENSG00000( |
| 45912245 | 45921874 | -1 | protein_coding | ENSG00000( |
| 45940449 | 45986472 | 1 | lncRNA | ENSG00000( |
| 45990905 | 46000898 | 1 | lncRNA | ENSG00000( |
| 45997540 | 45998071 | -1 | processed_pseudoge | ENSG00000( |
| 45999621 | 46000778 | -1 | processed_pseudoge | ENSG00000( |

| | | | | |
|---|---|---|---|---|
| 46144937 | 46145688 | 1 | processed_pseudoge | ENSG00000 |
| 46261064 | 46294469 | 1 | lncRNA | ENSG00000 |
| 46302120 | 46343621 | 1 | lncRNA | ENSG00000 |
| 46476457 | 46477940 | -1 | lncRNA | ENSG00000 |
| 46477822 | 46481578 | -1 | lncRNA | ENSG00000 |
| 46634614 | 46634946 | 1 | processed_pseudoge | ENSG00000 |
| 46673785 | 46759851 | -1 | lncRNA | ENSG00000 |
| 46781373 | 46783443 | 1 | transcribed_processe | ENSG00000 |
| 46890625 | 47049678 | -1 | lncRNA | ENSG00000 |
| 46969644 | 47027481 | 1 | lncRNA | ENSG00000 |
| 47000620 | 47079128 | -1 | lncRNA | ENSG00000 |
| 47026128 | 47026570 | -1 | processed_pseudoge | ENSG00000 |
| 47052793 | 47053674 | 1 | processed_pseudoge | ENSG00000 |
| 47252139 | 47254558 | -1 | lncRNA | ENSG00000 |
| 47275154 | 47582558 | -1 | protein_coding | ENSG00000 |
| 47608465 | 47629893 | 1 | lncRNA | ENSG00000 |
| 47655244 | 47661648 | 1 | lncRNA | ENSG00000 |
| 47740202 | 47948491 | -1 | protein_coding | ENSG00000 |
| 47761476 | 47766772 | 1 | lncRNA | ENSG00000 |
| 47795291 | 47819847 | 1 | lncRNA | ENSG00000 |
| 47956793 | 47957318 | 1 | processed_pseudoge | ENSG00000 |
| 47963288 | 47979615 | -1 | protein_coding | ENSG00000 |
| 47987148 | 48029119 | -1 | protein_coding | ENSG00000 |
| 48035511 | 48061304 | 1 | protein_coding | ENSG00000 |
| 48088628 | 48108736 | 1 | protein_coding | ENSG00000 |
| 48171458 | 48647497 | 1 | protein_coding | ENSG00000 |
| 48660125 | 48669479 | -1 | lncRNA | ENSG00000 |
| 48708327 | 48711582 | 1 | lncRNA | ENSG00000 |
| 48846426 | 48852898 | -1 | processed_pseudoge | ENSG00000 |
| 48847766 | 48851625 | 1 | TEC | ENSG00000 |
| 48902556 | 48903891 | -1 | processed_pseudoge | ENSG00000 |
| 48919765 | 48927454 | 1 | protein_coding | ENSG00000 |
| 49230137 | 49254816 | 1 | lncRNA | ENSG00000 |
| 49258493 | 49259846 | -1 | processed_pseudoge | ENSG00000 |
| 49524208 | 49587165 | 1 | lncRNA | ENSG00000 |
| 49760897 | 49762764 | -1 | TEC | ENSG00000 |
| 49773638 | 49921950 | 1 | protein_coding | ENSG00000 |
| 49850421 | 50121329 | -1 | protein_coding | ENSG00000 |
| 49942251 | 49944440 | -1 | processed_pseudoge | ENSG00000 |
| 50093279 | 50094123 | 1 | lncRNA | ENSG00000 |
| 50096036 | 50159830 | 1 | protein_coding | ENSG00000 |
| 50141540 | 50142823 | -1 | lncRNA | ENSG00000 |
| 50202001 | 50262994 | -1 | lncRNA | ENSG00000 |
| 50274790 | 50275622 | -1 | lncRNA | ENSG00000 |
| 50304068 | 50405101 | 1 | protein_coding | ENSG00000 |
| 50388489 | 50400097 | -1 | lncRNA | ENSG00000 |
| 50435380 | 50435486 | 1 | snRNA | ENSG00000 |
| 50444128 | 50542535 | -1 | protein_coding | ENSG00000 |
| 50458436 | 50565405 | -1 | protein_coding | ENSG00000 |
| 50531759 | 50543463 | 1 | lncRNA | ENSG00000 |

| | | | | |
|---|---|---|---|---|
| 50590063 | 50793462 | -1 | protein_coding | ENSG0000C |
| 50641725 | 50643868 | 1 | lncRNA | ENSG0000C |
| 50839363 | 50841992 | -1 | lncRNA | ENSG0000C |
| 50843578 | 50843732 | -1 | processed_pseudoge | ENSG0000C |
| 50866747 | 51022990 | 1 | lncRNA | ENSG0000C |
| 50935350 | 50935493 | -1 | snoRNA | ENSG0000C |
| 51016212 | 51316818 | -1 | protein_coding | ENSG0000C |
| 51259429 | 51260167 | -1 | processed_pseudoge | ENSG0000C |
| 51382284 | 51384660 | 1 | processed_pseudoge | ENSG0000C |
| 51386363 | 51387101 | -1 | processed_pseudoge | ENSG0000C |
| 51388430 | 51389114 | -1 | processed_pseudoge | ENSG0000C |
| 51471717 | 51729716 | 1 | lncRNA | ENSG0000C |
| 51600286 | 51600573 | -1 | processed_pseudoge | ENSG0000C |
| 51614251 | 51630870 | 1 | lncRNA | ENSG0000C |
| 51716657 | 51716922 | 1 | misc_RNA | ENSG0000C |
| 51773836 | 51781162 | 1 | lncRNA | ENSG0000C |
| 51849086 | 51864798 | 1 | lncRNA | ENSG0000C |
| 52165235 | 52192913 | -1 | lncRNA | ENSG0000C |
| 52269437 | 52269569 | -1 | snoRNA | ENSG0000C |
| 52493152 | 52503434 | 1 | lncRNA | ENSG0000C |
| 52891837 | 52897807 | -1 | processed_pseudoge | ENSG0000C |
| 53002420 | 53003429 | 1 | processed_pseudoge | ENSG0000C |
| 53035633 | 53036925 | 1 | protein_coding | ENSG0000C |
| 53187388 | 53188938 | 1 | processed_pseudoge | ENSG0000C |
| 53316149 | 53316740 | 1 | processed_pseudoge | ENSG0000C |
| 53366058 | 53366209 | 1 | snRNA | ENSG0000C |
| 53490148 | 53490439 | -1 | misc_RNA | ENSG0000C |
| 53514992 | 53517116 | -1 | lncRNA | ENSG0000C |
| 53559214 | 53574555 | -1 | lncRNA | ENSG0000C |
| 53655508 | 53811952 | -1 | lncRNA | ENSG0000C |
| 53776136 | 53776324 | 1 | snRNA | ENSG0000C |
| 53779783 | 53780263 | 1 | processed_pseudoge | ENSG0000C |
| 53787167 | 53787298 | 1 | processed_pseudoge | ENSG0000C |
| 53862233 | 53863339 | 1 | processed_pseudoge | ENSG0000C |
| 53926676 | 53947804 | -1 | lncRNA | ENSG0000C |
| 54185071 | 54257577 | -1 | lncRNA | ENSG0000C |
| 54201224 | 54202421 | 1 | lncRNA | ENSG0000C |
| 54330670 | 54455093 | 1 | lncRNA | ENSG0000C |
| 54419444 | 54420328 | 1 | processed_pseudoge | ENSG0000C |
| 54542325 | 54571080 | 1 | protein_coding | ENSG0000C |
| 54556970 | 54571726 | 1 | lncRNA | ENSG0000C |
| 54576052 | 54578794 | -1 | lncRNA | ENSG0000C |
| 54621025 | 54621131 | 1 | snRNA | ENSG0000C |
| 54656358 | 54656734 | 1 | processed_pseudoge | ENSG0000C |
| 54721724 | 54731047 | 1 | lncRNA | ENSG0000C |
| 54752250 | 54759974 | -1 | protein_coding | ENSG0000C |
| 54759348 | 54812727 | 1 | lncRNA | ENSG0000C |
| 54933699 | 54938184 | 1 | processed_pseudoge | ENSG0000C |
| 55019017 | 55211628 | 1 | protein_coding | ENSG0000C |
| 55179750 | 55188934 | -1 | lncRNA | ENSG0000C |

| | | | | |
|---|---|---|---|---|
| 55235965 | 55255635 | -1 | lncRNA | ENSG0000C |
| 55244247 | 55245681 | 1 | processed_pseudoge | ENSG0000C |
| 55259809 | 55260256 | -1 | processed_pseudoge | ENSG0000C |
| 55342316 | 55344958 | 1 | processed_pseudoge | ENSG0000C |
| 55365448 | 55433742 | 1 | protein_coding | ENSG0000C |
| 55436056 | 55572988 | -1 | protein_coding | ENSG0000C |
| 55573171 | 55588336 | 1 | lncRNA | ENSG0000C |
| 55592074 | 55593193 | 1 | lncRNA | ENSG0000C |
| 55593777 | 55595006 | 1 | lncRNA | ENSG0000C |
| 55638274 | 55638842 | -1 | processed_pseudoge | ENSG0000C |
| 55645620 | 55646951 | 1 | processed_pseudoge | ENSG0000C |
| 55656768 | 55678490 | -1 | unprocessed_pseudc | ENSG0000C |
| 55681074 | 55713252 | -1 | transcribed_unproce | ENSG0000C |
| 55685977 | 55686081 | 1 | snRNA | ENSG0000C |
| 55731100 | 55731403 | 1 | misc_RNA | ENSG0000C |
| 55732144 | 55732431 | 1 | processed_pseudoge | ENSG0000C |
| 55736779 | 55739605 | 1 | processed_pseudoge | ENSG0000C |
| 55741525 | 55741869 | 1 | processed_pseudoge | ENSG0000C |
| 55743073 | 55743457 | 1 | processed_pseudoge | ENSG0000C |
| 55764797 | 55773288 | 1 | unprocessed_pseudc | ENSG0000C |
| 55789368 | 55789474 | 1 | snRNA | ENSG0000C |
| 55793540 | 55862755 | -1 | protein_coding | ENSG0000C |
| 55797946 | 55798480 | -1 | processed_pseudoge | ENSG0000C |
| 55798034 | 55798876 | 1 | processed_pseudoge | ENSG0000C |
| 55810779 | 55811411 | -1 | processed_pseudoge | ENSG0000C |
| 55876738 | 55877024 | -1 | processed_pseudoge | ENSG0000C |
| 55877912 | 55878164 | 1 | processed_pseudoge | ENSG0000C |
| 55887277 | 55955239 | 1 | protein_coding | ENSG0000C |
| 55887456 | 55942530 | 1 | protein_coding | ENSG0000C |
| 55951793 | 56000181 | 1 | protein_coding | ENSG0000C |
| 55951877 | 55956500 | 1 | protein_coding | ENSG0000C |
| 56011051 | 56051604 | -1 | protein_coding | ENSG0000C |
| 56051685 | 56063989 | 1 | protein_coding | ENSG0000C |
| 56055365 | 56055502 | 1 | snoRNA | ENSG0000C |
| 56060470 | 56060602 | 1 | snoRNA | ENSG0000C |
| 56064002 | 56080670 | 1 | protein_coding | ENSG0000C |
| 56080283 | 56092996 | -1 | protein_coding | ENSG0000C |
| 56101573 | 56106476 | -1 | protein_coding | ENSG0000C |
| 56114681 | 56116417 | -1 | protein_coding | ENSG0000C |
| 56163145 | 56163533 | 1 | processed_pseudoge | ENSG0000C |
| 56175634 | 56176412 | 1 | processed_pseudoge | ENSG0000C |
| 56214979 | 56220747 | 1 | lncRNA | ENSG0000C |
| 56228634 | 56229782 | -1 | processed_pseudoge | ENSG0000C |
| 56288230 | 56291425 | 1 | processed_pseudoge | ENSG0000C |
| 56304678 | 56305526 | -1 | processed_pseudoge | ENSG0000C |
| 56322804 | 56323601 | 1 | unprocessed_pseudc | ENSG0000C |
| 56340231 | 56340337 | -1 | snRNA | ENSG0000C |
| 56360362 | 56360541 | -1 | unprocessed_pseudc | ENSG0000C |
| 56362458 | 56365270 | 1 | processed_pseudoge | ENSG0000C |
| 56380976 | 56381312 | 1 | processed_pseudoge | ENSG0000C |

| | | | | |
|---|---|---|---|---|
| 56402070 | 56402173 | -1 | snRNA | ENSG0000C |
| 56408699 | 56409210 | -1 | processed_pseudoge | ENSG0000C |
| 56421857 | 56448375 | -1 | transcribed_unproce | ENSG0000C |
| 56462536 | 56467771 | 1 | unprocessed_pseudc | ENSG0000C |
| 56477588 | 56484386 | -1 | lncRNA | ENSG0000C |
| 56490682 | 56491635 | 1 | processed_pseudoge | ENSG0000C |
| 56493124 | 56497285 | 1 | lncRNA | ENSG0000C |
| 56502284 | 56503044 | -1 | processed_pseudoge | ENSG0000C |
| 56525428 | 56526113 | -1 | processed_pseudoge | ENSG0000C |
| 56528253 | 56538676 | -1 | lncRNA | ENSG0000C |
| 56559221 | 56559958 | 1 | processed_pseudoge | ENSG0000C |
| 56567906 | 56568858 | -1 | processed_pseudoge | ENSG0000C |
| 56577848 | 56578866 | 1 | processed_pseudoge | ENSG0000C |
| 56597379 | 56598342 | 1 | processed_pseudoge | ENSG0000C |
| 56603410 | 56603553 | 1 | processed_pseudoge | ENSG0000C |
| 56615084 | 56617957 | -1 | lncRNA | ENSG0000C |
| 56631739 | 56632727 | -1 | processed_pseudoge | ENSG0000C |
| 56638943 | 56650696 | -1 | unprocessed_pseudc | ENSG0000C |
| 56659149 | 56675086 | 1 | lncRNA | ENSG0000C |
| 56685036 | 56687366 | -1 | unprocessed_pseudc | ENSG0000C |
| 56736396 | 56736590 | -1 | processed_pseudoge | ENSG0000C |
| 56805336 | 56808148 | 1 | processed_pseudoge | ENSG0000C |
| 56809214 | 56848800 | -1 | lncRNA | ENSG0000C |
| 56875385 | 56882146 | -1 | transcribed_unproce | ENSG0000C |
| 56940341 | 56944353 | -1 | unprocessed_pseudc | ENSG0000C |
| 56955785 | 56955864 | -1 | miRNA | ENSG0000C |
| 56991888 | 57009149 | 1 | unprocessed_pseudc | ENSG0000C |
| 57014285 | 57020273 | -1 | unprocessed_pseudc | ENSG0000C |
| 57060590 | 57061301 | -1 | unprocessed_pseudc | ENSG0000C |
| 57067950 | 57074664 | -1 | unprocessed_pseudc | ENSG0000C |
| 57119614 | 57139864 | -1 | protein_coding | ENSG0000C |
| 57140112 | 57153836 | 1 | lncRNA | ENSG0000C |
| 57147986 | 57150347 | -1 | unprocessed_pseudc | ENSG0000C |
| 57163168 | 57166024 | -1 | unprocessed_pseudc | ENSG0000C |
| 57167039 | 57167245 | 1 | unprocessed_pseudc | ENSG0000C |
| 57167200 | 57167880 | 1 | unprocessed_pseudc | ENSG0000C |
| 57167880 | 57168651 | 1 | unprocessed_pseudc | ENSG0000C |
| 57168725 | 57169074 | 1 | unprocessed_pseudc | ENSG0000C |
| 57169143 | 57169412 | 1 | unprocessed_pseudc | ENSG0000C |
| 57169406 | 57170771 | 1 | unprocessed_pseudc | ENSG0000C |
| 57170974 | 57172771 | 1 | unprocessed_pseudc | ENSG0000C |
| 57172772 | 57173283 | -1 | unprocessed_pseudc | ENSG0000C |
| 57173357 | 57174362 | 1 | unprocessed_pseudc | ENSG0000C |
| 57177409 | 57180052 | -1 | unprocessed_pseudc | ENSG0000C |
| 57185788 | 57186202 | 1 | unprocessed_pseudc | ENSG0000C |
| 57186412 | 57187441 | 1 | unprocessed_pseudc | ENSG0000C |
| 57187892 | 57189437 | 1 | unprocessed_pseudc | ENSG0000C |
| 57189578 | 57190226 | 1 | unprocessed_pseudc | ENSG0000C |
| 57190300 | 57190460 | 1 | unprocessed_pseudc | ENSG0000C |
| 57190461 | 57191138 | 1 | unprocessed_pseudc | ENSG0000C |

| | | | | |
|---|---|---|---|---|
| 57191141 | 57191927 | 1 | unprocessed_pseudc | ENSG00000( |
| 57191997 | 57192337 | 1 | unprocessed_pseudc | ENSG00000( |
| 57192405 | 57192698 | 1 | unprocessed_pseudc | ENSG00000( |
| 57192695 | 57194068 | 1 | unprocessed_pseudc | ENSG00000( |
| 57194257 | 57196054 | 1 | unprocessed_pseudc | ENSG00000( |
| 57196056 | 57196564 | -1 | unprocessed_pseudc | ENSG00000( |
| 57196638 | 57197772 | 1 | unprocessed_pseudc | ENSG00000( |
| 57200825 | 57203597 | -1 | unprocessed_pseudc | ENSG00000( |
| 57209865 | 57222897 | -1 | lncRNA | ENSG00000( |
| 57227001 | 57227066 | 1 | snRNA | ENSG00000( |
| 57242681 | 57245931 | 1 | lncRNA | ENSG00000( |
| 57270839 | 57275197 | 1 | unprocessed_pseudc | ENSG00000( |
| 57402181 | 57404717 | -1 | TEC | ENSG00000( |
| 57404172 | 57419535 | 1 | lncRNA | ENSG00000( |
| 57405025 | 57405090 | 1 | miRNA | ENSG00000( |
| 57422830 | 57423725 | -1 | unitary_pseudogene | ENSG00000( |
| 57424503 | 57425676 | 1 | processed_pseudoge | ENSG00000( |
| 57450177 | 57473559 | 1 | protein_coding | ENSG00000( |
| 57485166 | 57485895 | 1 | processed_pseudoge | ENSG00000( |
| 57599794 | 57610104 | 1 | unprocessed_pseudc | ENSG00000( |
| 57628479 | 57645189 | -1 | transcribed_process | ENSG00000( |
| 57629285 | 57630151 | 1 | processed_pseudoge | ENSG00000( |
| 57638725 | 57639112 | 1 | processed_pseudoge | ENSG00000( |
| 57650381 | 57651119 | -1 | unprocessed_pseudc | ENSG00000( |
| 57650521 | 57650854 | 1 | unprocessed_pseudc | ENSG00000( |
| 57652736 | 57653459 | -1 | unprocessed_pseudc | ENSG00000( |
| 57652875 | 57653208 | 1 | unprocessed_pseudc | ENSG00000( |
| 57654822 | 57655392 | 1 | unprocessed_pseudc | ENSG00000( |
| 57770619 | 57770946 | -1 | unprocessed_pseudc | ENSG00000( |
| 57770966 | 57771173 | 1 | unprocessed_pseudc | ENSG00000( |
| 57772584 | 57773199 | -1 | unprocessed_pseudc | ENSG00000( |
| 57773183 | 57773381 | 1 | unprocessed_pseudc | ENSG00000( |
| 57776972 | 57777521 | -1 | unprocessed_pseudc | ENSG00000( |
| 57817601 | 57818012 | -1 | unprocessed_pseudc | ENSG00000( |
| 57817996 | 57818197 | 1 | unprocessed_pseudc | ENSG00000( |
| 57819819 | 57820627 | 1 | unprocessed_pseudc | ENSG00000( |
| 57820032 | 57820410 | -1 | unprocessed_pseudc | ENSG00000( |
| 57822201 | 57822942 | 1 | unprocessed_pseudc | ENSG00000( |
| 57822465 | 57822801 | -1 | unprocessed_pseudc | ENSG00000( |
| 57828331 | 57837046 | 1 | unprocessed_pseudc | ENSG00000( |
| 57835309 | 57836251 | -1 | processed_pseudoge | ENSG00000( |
| 62275361 | 62275678 | -1 | processed_pseudoge | ENSG00000( |
| 63011463 | 63011569 | -1 | snRNA | ENSG00000( |
| 63044827 | 63054431 | -1 | lncRNA | ENSG00000( |
| 63057453 | 63060084 | 1 | unprocessed_pseudc | ENSG00000( |
| 63087070 | 63088261 | -1 | processed_pseudoge | ENSG00000( |
| 63113635 | 63114805 | -1 | processed_pseudoge | ENSG00000( |
| 63117501 | 63154934 | -1 | unprocessed_pseudc | ENSG00000( |
| 63175940 | 63176019 | -1 | snoRNA | ENSG00000( |
| 63209219 | 63211822 | 1 | unprocessed_pseudc | ENSG00000( |

| | | | | |
|---|---|---|---|---|
| 63233035 | 63234245 | -1 | unprocessed_pseudc | ENSG00000C |
| 63233115 | 63241645 | -1 | lncRNA | ENSG00000C |
| 63241590 | 63248321 | -1 | unprocessed_pseudc | ENSG00000C |
| 63263065 | 63263456 | 1 | processed_pseudoge | ENSG00000C |
| 63291518 | 63304056 | -1 | transcribed_unproce | ENSG00000C |
| 63326674 | 63354326 | -1 | lncRNA | ENSG00000C |
| 63342711 | 63343200 | 1 | processed_pseudoge | ENSG00000C |
| 63345840 | 63346732 | 1 | processed_pseudoge | ENSG00000C |
| 63348861 | 63351774 | 1 | lncRNA | ENSG00000C |
| 63349154 | 63350059 | -1 | processed_pseudoge | ENSG00000C |
| 63354033 | 63354365 | 1 | processed_pseudoge | ENSG00000C |
| 63354457 | 63359306 | 1 | lncRNA | ENSG00000C |
| 63367427 | 63368156 | -1 | processed_pseudoge | ENSG00000C |
| 63377329 | 63378233 | 1 | unitary_pseudogene | ENSG00000C |
| 63380466 | 63385789 | -1 | lncRNA | ENSG00000C |
| 63388808 | 63393657 | -1 | lncRNA | ENSG00000C |
| 63393745 | 63422053 | 1 | lncRNA | ENSG00000C |
| 63393749 | 63394109 | -1 | processed_pseudoge | ENSG00000C |
| 63394277 | 63395439 | 1 | lncRNA | ENSG00000C |
| 63396341 | 63399250 | -1 | lncRNA | ENSG00000C |
| 63398046 | 63398957 | 1 | processed_pseudoge | ENSG00000C |
| 63401385 | 63402271 | -1 | unitary_pseudogene | ENSG00000C |
| 63404842 | 63405400 | -1 | processed_pseudoge | ENSG00000C |
| 63449819 | 63456687 | 1 | unprocessed_pseudc | ENSG00000C |
| 63480598 | 63480989 | -1 | processed_pseudoge | ENSG00000C |
| 63495799 | 63502536 | 1 | processed_pseudoge | ENSG00000C |
| 63509303 | 63510165 | 1 | processed_pseudoge | ENSG00000C |
| 63556598 | 63562588 | 1 | unprocessed_pseudc | ENSG00000C |
| 63567743 | 63585075 | -1 | unprocessed_pseudc | ENSG00000C |
| 63621090 | 63621169 | 1 | miRNA | ENSG00000C |
| 63632608 | 63636617 | 1 | unprocessed_pseudc | ENSG00000C |
| 63699390 | 63700441 | 1 | unprocessed_pseudc | ENSG00000C |
| 63768257 | 63771047 | -1 | processed_pseudoge | ENSG00000C |
| 63789909 | 63790098 | 1 | processed_pseudoge | ENSG00000C |
| 63843314 | 63843505 | 1 | processed_pseudoge | ENSG00000C |
| 63876341 | 63877350 | 1 | processed_pseudoge | ENSG00000C |
| 63888200 | 63900794 | -1 | lncRNA | ENSG00000C |
| 63893286 | 63894641 | 1 | processed_pseudoge | ENSG00000C |
| 63900313 | 63925202 | 1 | lncRNA | ENSG00000C |
| 63908966 | 63910453 | -1 | processed_pseudoge | ENSG00000C |
| 63924787 | 63926306 | -1 | transcribed_processe | ENSG00000C |
| 63925832 | 63930920 | 1 | lncRNA | ENSG00000C |
| 63931141 | 63931968 | 1 | processed_pseudoge | ENSG00000C |
| 63934449 | 63935331 | -1 | unitary_pseudogene | ENSG00000C |
| 63937905 | 63938509 | -1 | processed_pseudoge | ENSG00000C |
| 63959844 | 63981477 | -1 | lncRNA | ENSG00000C |
| 63963431 | 63964123 | -1 | transcribed_unproce | ENSG00000C |
| 63965834 | 63966124 | -1 | unprocessed_pseudc | ENSG00000C |
| 63980385 | 63981477 | -1 | transcribed_unproce | ENSG00000C |
| 63993904 | 63994006 | 1 | unprocessed_pseudc | ENSG00000C |

| | | | | |
|---|---|---|---|---|
| 63998874 | 64016130 | 1 | protein_coding | ENSG0000C |
| 64024409 | 64030105 | -1 | lncRNA | ENSG0000C |
| 64035644 | 64037958 | -1 | lncRNA | ENSG0000C |
| 64045429 | 64114556 | 1 | lncRNA | ENSG0000C |
| 64045434 | 64085339 | 1 | protein_coding | ENSG0000C |
| 64085560 | 64086576 | 1 | processed_pseudoge | ENSG0000C |
| 64100305 | 64150721 | 1 | transcribed_unproce | ENSG0000C |
| 64104093 | 64105295 | -1 | unprocessed_pseudc | ENSG0000C |
| 64105292 | 64105576 | -1 | unprocessed_pseudc | ENSG0000C |
| 64105654 | 64105995 | -1 | unprocessed_pseudc | ENSG0000C |
| 64106065 | 64106819 | -1 | unprocessed_pseudc | ENSG0000C |
| 64106822 | 64107572 | -1 | unprocessed_pseudc | ENSG0000C |
| 64107829 | 64108486 | -1 | unprocessed_pseudc | ENSG0000C |
| 64108625 | 64110147 | -1 | unprocessed_pseudc | ENSG0000C |
| 64110547 | 64111550 | -1 | unprocessed_pseudc | ENSG0000C |
| 64111752 | 64112572 | -1 | unprocessed_pseudc | ENSG0000C |
| 64140495 | 64150058 | 1 | lncRNA | ENSG0000C |
| 64141538 | 64142401 | -1 | processed_pseudoge | ENSG0000C |
| 64149685 | 64150432 | -1 | processed_pseudoge | ENSG0000C |
| 64180006 | 64180892 | -1 | unitary_pseudogene | ENSG0000C |
| 64181710 | 64182893 | 1 | processed_pseudoge | ENSG0000C |
| 64207090 | 64220508 | 1 | protein_coding | ENSG0000C |
| 64228474 | 64266931 | 1 | protein_coding | ENSG0000C |
| 64294516 | 64300818 | -1 | lncRNA | ENSG0000C |
| 64307459 | 64356634 | 1 | protein_coding | ENSG0000C |
| 64355078 | 64356199 | 1 | unprocessed_pseudc | ENSG0000C |
| 64369241 | 64375584 | 1 | lncRNA | ENSG0000C |
| 64425623 | 64426334 | -1 | unprocessed_pseudc | ENSG0000C |
| 64433830 | 64434592 | 1 | processed_pseudoge | ENSG0000C |
| 64443802 | 64444742 | -1 | unitary_pseudogene | ENSG0000C |
| 64469185 | 64470653 | 1 | processed_pseudoge | ENSG0000C |
| 64500825 | 64501729 | 1 | processed_pseudoge | ENSG0000C |
| 64519878 | 64563075 | -1 | protein_coding | ENSG0000C |
| 64551498 | 64552066 | 1 | processed_pseudoge | ENSG0000C |
| 64566814 | 64567443 | -1 | processed_pseudoge | ENSG0000C |
| 64569428 | 64570437 | -1 | processed_pseudoge | ENSG0000C |
| 64582733 | 64583207 | 1 | processed_pseudoge | ENSG0000C |
| 64590599 | 64591133 | -1 | processed_pseudoge | ENSG0000C |
| 64612535 | 64614834 | 1 | lncRNA | ENSG0000C |
| 64651655 | 64655836 | 1 | lncRNA | ENSG0000C |
| 64666083 | 64711582 | 1 | protein_coding | ENSG0000C |
| 64678954 | 64687393 | -1 | processed_pseudoge | ENSG0000C |
| 64679064 | 64679176 | 1 | miRNA | ENSG0000C |
| 64735933 | 64736743 | -1 | processed_pseudoge | ENSG0000C |
| 64768156 | 64769101 | 1 | unprocessed_pseudc | ENSG0000C |
| 64794388 | 64833681 | 1 | protein_coding | ENSG0000C |
| 64801958 | 64804117 | -1 | lncRNA | ENSG0000C |
| 64835280 | 64836882 | -1 | processed_pseudoge | ENSG0000C |
| 64852397 | 64853354 | -1 | processed_pseudoge | ENSG0000C |
| 64862999 | 64864370 | 1 | processed_pseudoge | ENSG0000C |

| | | | | |
|---|---|---|---|---|
| 64867427 | 64870076 | -1 | lncRNA | ENSG0000C |
| 64870172 | 64930966 | 1 | protein_coding | ENSG0000C |
| 64888527 | 64890100 | -1 | lncRNA | ENSG0000C |
| 64933273 | 64933616 | 1 | processed_pseudoge | ENSG0000C |
| 64942143 | 64942926 | -1 | processed_pseudoge | ENSG0000C |
| 64947571 | 64950701 | 1 | lncRNA | ENSG0000C |
| 64971772 | 65006684 | -1 | protein_coding | ENSG0000C |
| 64990356 | 65006743 | -1 | protein_coding | ENSG0000C |
| 65023204 | 65023310 | -1 | snRNA | ENSG0000C |
| 65038354 | 65074713 | 1 | transcribed_unproce | ENSG0000C |
| 65038372 | 65038565 | 1 | unprocessed_pseudc | ENSG0000C |
| 65065999 | 65066132 | 1 | snoRNA | ENSG0000C |
| 65070538 | 65070672 | 1 | snoRNA | ENSG0000C |
| 65075023 | 65078780 | 1 | unprocessed_pseudc | ENSG0000C |
| 65084103 | 65100232 | 1 | unprocessed_pseudc | ENSG0000C |
| 65141032 | 65234216 | 1 | transcribed_unproce | ENSG0000C |
| 65235790 | 65236723 | -1 | processed_pseudoge | ENSG0000C |
| 65247608 | 65247758 | -1 | processed_pseudoge | ENSG0000C |
| 65269374 | 65306238 | 1 | lncRNA | ENSG0000C |
| 65290429 | 65292776 | 1 | unprocessed_pseudc | ENSG0000C |
| 65335174 | 65335668 | -1 | processed_pseudoge | ENSG0000C |
| 65355934 | 65357176 | 1 | processed_pseudoge | ENSG0000C |
| 65373799 | 65401135 | 1 | protein_coding | ENSG0000C |
| 65463071 | 65463438 | 1 | lncRNA | ENSG0000C |
| 65508773 | 65508944 | -1 | processed_pseudoge | ENSG0000C |
| 65525629 | 65554744 | -1 | unprocessed_pseudc | ENSG0000C |
| 65617082 | 65619153 | 1 | unprocessed_pseudc | ENSG0000C |
| 65647010 | 65770810 | -1 | transcribed_unproce | ENSG0000C |
| 65647823 | 65718688 | 1 | transcribed_unproce | ENSG0000C |
| 65751142 | 65763354 | 1 | transcribed_unproce | ENSG0000C |
| 65755526 | 65755659 | 1 | snoRNA | ENSG0000C |
| 65760052 | 65760186 | 1 | snoRNA | ENSG0000C |
| 65764535 | 65768306 | 1 | unprocessed_pseudc | ENSG0000C |
| 65773620 | 65802067 | 1 | unprocessed_pseudc | ENSG0000C |
| 65814672 | 65814775 | 1 | snRNA | ENSG0000C |
| 65840055 | 65842396 | 1 | lncRNA | ENSG0000C |
| 65859660 | 65859766 | -1 | snRNA | ENSG0000C |
| 65873074 | 65959563 | 1 | protein_coding | ENSG0000C |
| 65960684 | 65982215 | -1 | protein_coding | ENSG0000C |
| 66004017 | 66004137 | 1 | processed_pseudoge | ENSG0000C |
| 66025126 | 66031544 | -1 | processed_pseudoge | ENSG0000C |
| 66070904 | 66071318 | -1 | processed_pseudoge | ENSG0000C |
| 66075800 | 66094697 | 1 | protein_coding | ENSG0000C |
| 66087761 | 66152277 | 1 | protein_coding | ENSG0000C |
| 66114604 | 66154568 | 1 | protein_coding | ENSG0000C |
| 66119603 | 66165011 | -1 | lncRNA | ENSG0000C |
| 66205317 | 66420543 | 1 | protein_coding | ENSG0000C |
| 66344303 | 66344414 | 1 | snRNA | ENSG0000C |
| 66376044 | 66493566 | -1 | lncRNA | ENSG0000C |
| 66407288 | 66409239 | -1 | transcribed_unproce | ENSG0000C |

| | | | | |
|---|---|---|---|---|
| 66423405 | 66423893 | -1 | transcribed_processe | ENSG00000 |
| 66427949 | 66453634 | -1 | transcribed_unproce | ENSG00000 |
| 66434634 | 66435154 | -1 | processed_pseudoge | ENSG00000 |
| 66474556 | 66475749 | 1 | TEC | ENSG00000 |
| 66480394 | 66490108 | -1 | transcribed_unproce | ENSG00000 |
| 66485095 | 66486532 | -1 | transcribed_unproce | ENSG00000 |
| 66493607 | 66495758 | 1 | lncRNA | ENSG00000 |
| 66505155 | 66506223 | -1 | processed_pseudoge | ENSG00000 |
| 66511556 | 66545066 | 1 | unprocessed_pseudo | ENSG00000 |
| 66526088 | 66592397 | -1 | transcribed_unproce | ENSG00000 |
| 66553805 | 66554199 | -1 | processed_pseudoge | ENSG00000 |
| 66556216 | 66557065 | 1 | processed_pseudoge | ENSG00000 |
| 66556889 | 66557362 | -1 | processed_pseudoge | ENSG00000 |
| 66606738 | 66607107 | -1 | processed_pseudoge | ENSG00000 |
| 66628767 | 66649067 | 1 | protein_coding | ENSG00000 |
| 66628958 | 66811189 | 1 | protein_coding | ENSG00000 |
| 66654538 | 66671523 | 1 | lncRNA | ENSG00000 |
| 66682164 | 66811464 | 1 | protein_coding | ENSG00000 |
| 66739829 | 66740385 | -1 | lncRNA | ENSG00000 |
| 66748838 | 66749077 | -1 | processed_pseudoge | ENSG00000 |
| 66809993 | 66844882 | -1 | transcribed_unproce | ENSG00000 |
| 66848496 | 66858136 | 1 | unprocessed_pseudo | ENSG00000 |
| 66880708 | 66882981 | 1 | unprocessed_pseudo | ENSG00000 |
| 66891188 | 66891291 | 1 | snRNA | ENSG00000 |
| 66902857 | 66906297 | 1 | lncRNA | ENSG00000 |
| 66914581 | 66914943 | 1 | processed_pseudoge | ENSG00000 |
| 66915063 | 66915334 | -1 | misc_RNA | ENSG00000 |
| 66921225 | 66958551 | 1 | protein_coding | ENSG00000 |
| 66980120 | 66980410 | 1 | misc_RNA | ENSG00000 |
| 66987680 | 66995587 | -1 | protein_coding | ENSG00000 |
| 66995173 | 67239519 | 1 | protein_coding | ENSG00000 |
| 67089257 | 67089470 | -1 | processed_pseudoge | ENSG00000 |
| 67114322 | 67114397 | -1 | miRNA | ENSG00000 |
| 67278714 | 67286664 | 1 | protein_coding | ENSG00000 |
| 67287541 | 67302419 | -1 | transcribed_unproce | ENSG00000 |
| 67297653 | 67297754 | -1 | misc_RNA | ENSG00000 |
| 67302621 | 67362950 | 1 | transcribed_unproce | ENSG00000 |
| 67307605 | 67308356 | -1 | unprocessed_pseudo | ENSG00000 |
| 67333047 | 67334383 | 1 | lncRNA | ENSG00000 |
| 67627831 | 67628019 | 1 | unprocessed_pseudo | ENSG00000 |
| 67628022 | 67628349 | 1 | unprocessed_pseudo | ENSG00000 |
| 67691058 | 67697029 | -1 | lncRNA | ENSG00000 |
| 67769629 | 67770468 | -1 | processed_pseudoge | ENSG00000 |
| 68020235 | 68036917 | 1 | lncRNA | ENSG00000 |
| 68091223 | 68119209 | 1 | lncRNA | ENSG00000 |
| 68097737 | 68098250 | 1 | unprocessed_pseudo | ENSG00000 |
| 68149548 | 68319676 | -1 | lncRNA | ENSG00000 |
| 68241637 | 68242267 | 1 | processed_pseudoge | ENSG00000 |
| 68266319 | 68266859 | -1 | unprocessed_pseudo | ENSG00000 |
| 68640798 | 68641584 | 1 | processed_pseudoge | ENSG00000 |

| | | | | |
|---|---|---|---|---|
| 68723911 | 68724048 | -1 | rRNA_pseudogene | ENSG0000C |
| 69026433 | 69046803 | 1 | lncRNA | ENSG0000C |
| 69125270 | 69125376 | 1 | snRNA | ENSG0000C |
| 69145185 | 69174450 | -1 | lncRNA | ENSG0000C |
| 69186821 | 69430923 | -1 | lncRNA | ENSG0000C |
| 69187833 | 69189318 | -1 | lncRNA | ENSG0000C |
| 69330698 | 69331378 | -1 | unprocessed_pseudc | ENSG0000C |
| 69331835 | 69333633 | -1 | unprocessed_pseudc | ENSG0000C |
| 69401202 | 69401303 | -1 | snRNA | ENSG0000C |
| 69507510 | 69507622 | -1 | misc_RNA | ENSG0000C |
| 69594793 | 69597493 | -1 | lncRNA | ENSG0000C |
| 69598296 | 70793506 | 1 | protein_coding | ENSG0000C |
| 70837738 | 70838013 | 1 | processed_pseudoge | ENSG0000C |
| 71132144 | 71713600 | 1 | protein_coding | ENSG0000C |
| 71307672 | 71307770 | -1 | miRNA | ENSG0000C |
| 71307674 | 71307768 | 1 | miRNA | ENSG0000C |
| 71685452 | 71685747 | -1 | misc_RNA | ENSG0000C |
| 71779491 | 72447151 | -1 | protein_coding | ENSG0000C |
| 71913738 | 71913856 | -1 | rRNA_pseudogene | ENSG0000C |
| 71942259 | 71942468 | 1 | processed_pseudoge | ENSG0000C |
| 72103181 | 72104209 | 1 | unprocessed_pseudc | ENSG0000C |
| 72558744 | 72828200 | -1 | protein_coding | ENSG0000C |
| 72697903 | 72697978 | 1 | miRNA | ENSG0000C |
| 72722885 | 72723089 | 1 | processed_pseudoge | ENSG0000C |
| 72768798 | 72769191 | 1 | processed_pseudoge | ENSG0000C |
| 72822894 | 72823171 | -1 | misc_RNA | ENSG0000C |
| 72829425 | 72836701 | 1 | transcribed_unproce | ENSG0000C |
| 72841437 | 72841727 | -1 | misc_RNA | ENSG0000C |
| 72862757 | 72871581 | -1 | transcribed_unproce | ENSG0000C |
| 72879365 | 72951440 | 1 | protein_coding | ENSG0000C |
| 72924418 | 72925125 | -1 | lncRNA | ENSG0000C |
| 72947581 | 72954790 | -1 | transcribed_unproce | ENSG0000C |
| 72954797 | 72957106 | 1 | lncRNA | ENSG0000C |
| 72959485 | 72969466 | -1 | protein_coding | ENSG0000C |
| 72969696 | 73005922 | -1 | transcribed_unproce | ENSG0000C |
| 72969814 | 72971727 | 1 | unprocessed_pseudc | ENSG0000C |
| 72983360 | 72983461 | -1 | misc_RNA | ENSG0000C |
| 73005541 | 73021103 | 1 | unprocessed_pseudc | ENSG0000C |
| 73011144 | 73011245 | 1 | misc_RNA | ENSG0000C |
| 73020338 | 73029766 | -1 | protein_coding | ENSG0000C |
| 73037371 | 73049246 | 1 | unprocessed_pseudc | ENSG0000C |
| 73039085 | 73039186 | 1 | misc_RNA | ENSG0000C |
| 73039696 | 73069506 | 1 | lncRNA | ENSG0000C |
| 73048055 | 73053981 | -1 | lncRNA | ENSG0000C |
| 73048481 | 73057911 | -1 | protein_coding | ENSG0000C |
| 73065529 | 73077353 | 1 | unprocessed_pseudc | ENSG0000C |
| 73067231 | 73067332 | 1 | misc_RNA | ENSG0000C |
| 73076165 | 73086038 | -1 | protein_coding | ENSG0000C |
| 73093657 | 73105389 | 1 | unprocessed_pseudc | ENSG0000C |
| 73095357 | 73095458 | 1 | misc_RNA | ENSG0000C |

| | | | | |
|---|---|---|---|---|
| 73104201 | 73111140 | -1 | transcribed_unproce | ENSG00000 |
| 73154938 | 73207283 | 1 | transcribed_processe | ENSG00000 |
| 73187969 | 73188736 | -1 | processed_pseudoge | ENSG00000 |
| 73220624 | 73235945 | 1 | transcribed_unproce | ENSG00000 |
| 73242751 | 73280119 | -1 | transcribed_unproce | ENSG00000 |
| 73293497 | 73301161 | 1 | unprocessed_pseudo | ENSG00000 |
| 73302516 | 73308826 | -1 | protein_coding | ENSG00000 |
| 73312536 | 73328082 | -1 | protein_coding | ENSG00000 |
| 73314107 | 73315213 | 1 | processed_pseudoge | ENSG00000 |
| 73328161 | 73358637 | 1 | protein_coding | ENSG00000 |
| 73339094 | 73339200 | -1 | snRNA | ENSG00000 |
| 73403788 | 73403889 | -1 | misc_RNA | ENSG00000 |
| 73433778 | 73436120 | 1 | protein_coding | ENSG00000 |
| 73440406 | 73522293 | -1 | protein_coding | ENSG00000 |
| 73507208 | 73507274 | 1 | snRNA | ENSG00000 |
| 73536356 | 73557690 | -1 | protein_coding | ENSG00000 |
| 73567537 | 73578791 | -1 | protein_coding | ENSG00000 |
| 73593194 | 73624543 | -1 | protein_coding | ENSG00000 |
| 73609262 | 73611502 | -1 | lncRNA | ENSG00000 |
| 73667831 | 73672112 | 1 | protein_coding | ENSG00000 |
| 73680918 | 73683453 | -1 | protein_coding | ENSG00000 |
| 73683025 | 73705161 | 1 | protein_coding | ENSG00000 |
| 73699206 | 73719672 | -1 | protein_coding | ENSG00000 |
| 73711317 | 73711397 | 1 | miRNA | ENSG00000 |
| 73732569 | 73732868 | -1 | misc_RNA | ENSG00000 |
| 73735038 | 73736158 | 1 | transcribed_unitary_ | ENSG00000 |
| 73736094 | 73738867 | -1 | protein_coding | ENSG00000 |
| 73768997 | 73770270 | -1 | protein_coding | ENSG00000 |
| 73799542 | 73832693 | 1 | protein_coding | ENSG00000 |
| 73834590 | 73842516 | -1 | protein_coding | ENSG00000 |
| 73861159 | 73865890 | 1 | protein_coding | ENSG00000 |
| 73985992 | 73988767 | 1 | lncRNA | ENSG00000 |
| 74027789 | 74069907 | 1 | protein_coding | ENSG00000 |
| 74059576 | 74062284 | -1 | lncRNA | ENSG00000 |
| 74082933 | 74122525 | 1 | protein_coding | ENSG00000 |
| 74174245 | 74197101 | 1 | protein_coding | ENSG00000 |
| 74191198 | 74191294 | 1 | miRNA | ENSG00000 |
| 74199652 | 74229834 | 1 | protein_coding | ENSG00000 |
| 74231499 | 74254458 | -1 | protein_coding | ENSG00000 |
| 74280747 | 74289286 | -1 | lncRNA | ENSG00000 |
| 74289475 | 74405943 | 1 | protein_coding | ENSG00000 |
| 74453790 | 74602604 | 1 | protein_coding | ENSG00000 |
| 74487428 | 74487540 | -1 | rRNA_pseudogene | ENSG00000 |
| 74606913 | 74607299 | 1 | lncRNA | ENSG00000 |
| 74633510 | 74633884 | -1 | TEC | ENSG00000 |
| 74650231 | 74760692 | 1 | protein_coding | ENSG00000 |
| 74688864 | 74729001 | -1 | lncRNA | ENSG00000 |
| 74741457 | 74742121 | -1 | processed_pseudoge | ENSG00000 |
| 74773962 | 74789376 | 1 | protein_coding | ENSG00000 |
| 74796144 | 74851551 | -1 | protein_coding | ENSG00000 |

| | | | | |
|---|---|---|---|---|
| 74843754 | 74890610 | -1 | transcribed_unproce | ENSG00000 |
| 74894116 | 74897835 | 1 | unprocessed_pseudc | ENSG00000 |
| 74895816 | 74895917 | 1 | misc_RNA | ENSG00000 |
| 74906673 | 74913310 | -1 | unprocessed_pseudc | ENSG00000 |
| 74964705 | 75031528 | 1 | protein_coding | ENSG00000 |
| 75027122 | 75074228 | -1 | protein_coding | ENSG00000 |
| 75057877 | 75058188 | 1 | misc_RNA | ENSG00000 |
| 75092573 | 75149817 | 1 | protein_coding | ENSG00000 |
| 75156639 | 75172044 | -1 | unprocessed_pseudc | ENSG00000 |
| 75185385 | 75237696 | -1 | transcribed_unproce | ENSG00000 |
| 75203926 | 75204711 | 1 | processed_pseudoge | ENSG00000 |
| 75225433 | 75234310 | 1 | lncRNA | ENSG00000 |
| 75232928 | 75233924 | 1 | lncRNA | ENSG00000 |
| 75237293 | 75237405 | -1 | processed_pseudoge | ENSG00000 |
| 75281243 | 75287121 | 1 | protein_coding | ENSG00000 |
| 75288120 | 75299795 | -1 | transcribed_unproce | ENSG00000 |
| 75297990 | 75298091 | -1 | misc_RNA | ENSG00000 |
| 75309171 | 75315049 | 1 | protein_coding | ENSG00000 |
| 75316048 | 75327765 | -1 | transcribed_unproce | ENSG00000 |
| 75325959 | 75326060 | -1 | misc_RNA | ENSG00000 |
| 75337138 | 75343016 | 1 | protein_coding | ENSG00000 |
| 75344015 | 75359550 | -1 | translated_unproces | ENSG00000 |
| 75353885 | 75353986 | -1 | misc_RNA | ENSG00000 |
| 75359194 | 75395383 | 1 | transcribed_unproce | ENSG00000 |
| 75381638 | 75381739 | 1 | misc_RNA | ENSG00000 |
| 75391955 | 75395434 | -1 | transcribed_unproce | ENSG00000 |
| 75395063 | 75410996 | 1 | protein_coding | ENSG00000 |
| 75410322 | 75416787 | 1 | transcribed_unproce | ENSG00000 |
| 75416786 | 75486299 | -1 | protein_coding | ENSG00000 |
| 75474707 | 75486108 | -1 | transcribed_unproce | ENSG00000 |
| 75493625 | 75504304 | 1 | protein_coding | ENSG00000 |
| 75507747 | 75528148 | -1 | transcribed_unproce | ENSG00000 |
| 75513025 | 75513126 | -1 | misc_RNA | ENSG00000 |
| 75533298 | 75738962 | -1 | protein_coding | ENSG00000 |
| 75625089 | 75626970 | -1 | TEC | ENSG00000 |
| 75769533 | 75789896 | -1 | protein_coding | ENSG00000 |
| 75811665 | 75823356 | -1 | protein_coding | ENSG00000 |
| 75835663 | 75836141 | 1 | processed_pseudoge | ENSG00000 |
| 75842602 | 75888926 | 1 | protein_coding | ENSG00000 |
| 75899200 | 75986855 | 1 | protein_coding | ENSG00000 |
| 75915197 | 75915269 | 1 | miRNA | ENSG00000 |
| 75922755 | 75923443 | 1 | processed_pseudoge | ENSG00000 |
| 75943782 | 75943916 | 1 | snoRNA | ENSG00000 |
| 75986831 | 75994656 | -1 | protein_coding | ENSG00000 |
| 75996338 | 76048004 | -1 | protein_coding | ENSG00000 |
| 76043977 | 76045963 | -1 | TEC | ENSG00000 |
| 76048051 | 76067508 | 1 | protein_coding | ENSG00000 |
| 76071469 | 76074963 | -1 | processed_pseudoge | ENSG00000 |
| 76087444 | 76087547 | 1 | snRNA | ENSG00000 |
| 76090431 | 76108779 | -1 | transcribed_unproce | ENSG00000 |

| | | | | |
|---|---|---|---|---|
| 76108434 | 76132187 | -1 | transcribed_unproce | ENSG00000 |
| 76113026 | 76113400 | -1 | processed_pseudoge | ENSG00000 |
| 76173733 | 76177205 | 1 | transcribed_unproce | ENSG00000 |
| 76201896 | 76287288 | 1 | protein_coding | ENSG00000 |
| 76302673 | 76304295 | 1 | protein_coding | ENSG00000 |
| 76326799 | 76358991 | -1 | protein_coding | ENSG00000 |
| 76361768 | 76362113 | 1 | processed_pseudoge | ENSG00000 |
| 76389334 | 76409697 | -1 | protein_coding | ENSG00000 |
| 76397518 | 76442071 | 1 | protein_coding | ENSG00000 |
| 76461676 | 76505995 | 1 | protein_coding | ENSG00000 |
| 76470162 | 76471058 | -1 | processed_pseudoge | ENSG00000 |
| 76474587 | 76478856 | -1 | lncRNA | ENSG00000 |
| 76510428 | 76516521 | 1 | protein_coding | ENSG00000 |
| 76521611 | 76522074 | 1 | unprocessed_pseudc | ENSG00000 |
| 76523605 | 76523706 | 1 | misc_RNA | ENSG00000 |
| 76524515 | 76532692 | 1 | unprocessed_pseudc | ENSG00000 |
| 76531319 | 76541459 | -1 | protein_coding | ENSG00000 |
| 76549360 | 76627982 | 1 | transcribed_unproce | ENSG00000 |
| 76609986 | 76627261 | -1 | protein_coding | ENSG00000 |
| 76650401 | 76650897 | -1 | processed_pseudoge | ENSG00000 |
| 76818377 | 76903041 | -1 | lncRNA | ENSG00000 |
| 76902480 | 76919191 | 1 | lncRNA | ENSG00000 |
| 76959835 | 77043775 | 1 | lncRNA | ENSG00000 |
| 76960571 | 76978658 | 1 | lncRNA | ENSG00000 |
| 76968197 | 76969250 | -1 | processed_pseudoge | ENSG00000 |
| 76968346 | 76968518 | 1 | processed_pseudoge | ENSG00000 |
| 76972679 | 76976961 | -1 | lncRNA | ENSG00000 |
| 76978617 | 77004308 | 1 | transcribed_unproce | ENSG00000 |
| 77004662 | 77005774 | 1 | transcribed_unproce | ENSG00000 |
| 77011551 | 77015853 | 1 | transcribed_unproce | ENSG00000 |
| 77013583 | 77013684 | 1 | misc_RNA | ENSG00000 |
| 77022925 | 77032361 | -1 | unprocessed_pseudc | ENSG00000 |
| 77039944 | 77043776 | 1 | transcribed_unproce | ENSG00000 |
| 77041647 | 77041748 | 1 | misc_RNA | ENSG00000 |
| 77043721 | 77198626 | 1 | lncRNA | ENSG00000 |
| 77052785 | 77060512 | -1 | unprocessed_pseudc | ENSG00000 |
| 77071751 | 77072237 | -1 | processed_pseudoge | ENSG00000 |
| 77083787 | 77122116 | -1 | transcribed_unproce | ENSG00000 |
| 77115399 | 77116192 | -1 | processed_pseudoge | ENSG00000 |
| 77122434 | 77329533 | 1 | protein_coding | ENSG00000 |
| 77193369 | 77199848 | -1 | protein_coding | ENSG00000 |
| 77246340 | 77258123 | -1 | lncRNA | ENSG00000 |
| 77310751 | 77416349 | -1 | protein_coding | ENSG00000 |
| 77416673 | 77425444 | 1 | lncRNA | ENSG00000 |
| 77461458 | 77462139 | 1 | processed_pseudoge | ENSG00000 |
| 77487317 | 77498965 | -1 | lncRNA | ENSG00000 |
| 77537295 | 77640069 | 1 | protein_coding | ENSG00000 |
| 77657659 | 77696267 | -1 | lncRNA | ENSG00000 |
| 77696459 | 77783022 | 1 | protein_coding | ENSG00000 |
| 77793728 | 77798434 | -1 | protein_coding | ENSG00000 |

| | | | | |
|---|---|---|---|---|
| 77798792 | 77957503 | 1 | protein_coding | ENSG00000 |
| 77895880 | 77895992 | 1 | misc_RNA | ENSG00000 |
| 77990384 | 77995171 | 1 | lncRNA | ENSG00000 |
| 78017055 | 79453667 | -1 | protein_coding | ENSG00000 |
| 78134079 | 78134851 | 1 | TEC | ENSG00000 |
| 78170195 | 78170915 | 1 | TEC | ENSG00000 |
| 78347142 | 78359458 | 1 | transcribed_processe | ENSG00000 |
| 78392626 | 78448999 | -1 | lncRNA | ENSG00000 |
| 78486530 | 78487028 | 1 | processed_pseudoge | ENSG00000 |
| 78939850 | 78940895 | 1 | lncRNA | ENSG00000 |
| 79008988 | 79012277 | 1 | lncRNA | ENSG00000 |
| 79030240 | 79030345 | 1 | snRNA | ENSG00000 |
| 79124739 | 79125750 | 1 | processed_pseudoge | ENSG00000 |
| 79139829 | 79177210 | 1 | lncRNA | ENSG00000 |
| 79335780 | 79357594 | -1 | lncRNA | ENSG00000 |
| 79343266 | 79343372 | -1 | snRNA | ENSG00000 |
| 79353410 | 79354392 | 1 | processed_pseudoge | ENSG00000 |
| 79452877 | 79471208 | 1 | lncRNA | ENSG00000 |
| 79543911 | 79544853 | -1 | processed_pseudoge | ENSG00000 |
| 79654109 | 79654227 | 1 | rRNA_pseudogene | ENSG00000 |
| 79657947 | 79675015 | 1 | lncRNA | ENSG00000 |
| 79768028 | 80226181 | 1 | protein_coding | ENSG00000 |
| 79912104 | 79912208 | -1 | snRNA | ENSG00000 |
| 80096600 | 80097215 | -1 | processed_pseudoge | ENSG00000 |
| 80245926 | 80246225 | 1 | misc_RNA | ENSG00000 |
| 80246409 | 80312456 | -1 | lncRNA | ENSG00000 |
| 80312574 | 80391474 | -1 | lncRNA | ENSG00000 |
| 80318253 | 80318544 | 1 | misc_RNA | ENSG00000 |
| 80369575 | 80679277 | 1 | protein_coding | ENSG00000 |
| 80377554 | 80377835 | -1 | processed_pseudoge | ENSG00000 |
| 80458635 | 80512064 | -1 | protein_coding | ENSG00000 |
| 80662331 | 80662585 | -1 | processed_pseudoge | ENSG00000 |
| 80742538 | 80922359 | -1 | protein_coding | ENSG00000 |
| 80972516 | 80974724 | -1 | TEC | ENSG00000 |
| 81175508 | 81199115 | 1 | lncRNA | ENSG00000 |
| 81191398 | 81192009 | 1 | processed_pseudoge | ENSG00000 |
| 81335106 | 81335183 | 1 | processed_pseudoge | ENSG00000 |
| 81431731 | 81432796 | 1 | unprocessed_pseudo | ENSG00000 |
| 81463036 | 81463613 | -1 | processed_pseudoge | ENSG00000 |
| 81489204 | 81691406 | -1 | transcribed_unitary_ | ENSG00000 |
| 81610884 | 81611326 | -1 | processed_pseudoge | ENSG00000 |
| 81699010 | 81770438 | -1 | protein_coding | ENSG00000 |
| 81946444 | 82443777 | -1 | protein_coding | ENSG00000 |
| 82009177 | 82029955 | 1 | lncRNA | ENSG00000 |
| 82589848 | 82590691 | -1 | processed_pseudoge | ENSG00000 |
| 82657035 | 82657411 | 1 | processed_pseudoge | ENSG00000 |
| 82754012 | 83162930 | -1 | protein_coding | ENSG00000 |
| 83017898 | 83018011 | 1 | rRNA_pseudogene | ENSG00000 |
| 83355154 | 83362266 | 1 | lncRNA | ENSG00000 |
| 83363238 | 83649139 | -1 | protein_coding | ENSG00000 |

| | | | | |
|---|---|---|---|---|
| 83372652 | 83373096 | -1 | TEC | ENSG00000 |
| 83424880 | 83425002 | 1 | processed_pseudoge | ENSG00000 |
| 83663419 | 83664625 | 1 | processed_pseudoge | ENSG00000 |
| 83955777 | 84492724 | -1 | protein_coding | ENSG00000 |
| 84528122 | 84528830 | -1 | processed_pseudoge | ENSG00000 |
| 84532476 | 84584322 | 1 | lncRNA | ENSG00000 |
| 84847877 | 84848253 | -1 | processed_pseudoge | ENSG00000 |
| 84876554 | 84876956 | 1 | processed_pseudoge | ENSG00000 |
| 84939335 | 84940256 | -1 | TEC | ENSG00000 |
| 84939349 | 84940245 | -1 | lncRNA | ENSG00000 |
| 84983556 | 84984506 | 1 | processed_pseudoge | ENSG00000 |
| 84995553 | 85186855 | -1 | protein_coding | ENSG00000 |
| 85027828 | 85029692 | 1 | processed_pseudoge | ENSG00000 |
| 85381118 | 85381375 | -1 | processed_pseudoge | ENSG00000 |
| 85421122 | 85489293 | 1 | lncRNA | ENSG00000 |
| 85636013 | 85636344 | -1 | processed_pseudoge | ENSG00000 |
| 86216785 | 86217733 | -1 | processed_pseudoge | ENSG00000 |
| 86643914 | 86864884 | 1 | protein_coding | ENSG00000 |
| 86775081 | 86776022 | -1 | lncRNA | ENSG00000 |
| 86782357 | 86803409 | -1 | lncRNA | ENSG00000 |
| 86876906 | 87059654 | -1 | protein_coding | ENSG00000 |
| 87109539 | 87111282 | 1 | lncRNA | ENSG00000 |
| 87151422 | 87152420 | -1 | lncRNA | ENSG00000 |
| 87152361 | 87196337 | 1 | protein_coding | ENSG00000 |
| 87184849 | 87185020 | 1 | misc_RNA | ENSG00000 |
| 87196160 | 87220587 | -1 | protein_coding | ENSG00000 |
| 87218611 | 87218712 | -1 | misc_RNA | ENSG00000 |
| 87322943 | 87345528 | -1 | lncRNA | ENSG00000 |
| 87345082 | 87345233 | 1 | misc_RNA | ENSG00000 |
| 87345305 | 87345486 | 1 | misc_RNA | ENSG00000 |
| 87345681 | 87399795 | 1 | protein_coding | ENSG00000 |
| 87401697 | 87480435 | -1 | protein_coding | ENSG00000 |
| 87503633 | 87713323 | -1 | protein_coding | ENSG00000 |
| 87521461 | 87522410 | 1 | processed_pseudoge | ENSG00000 |
| 87627548 | 87832296 | 1 | protein_coding | ENSG00000 |
| 87833568 | 87876360 | -1 | protein_coding | ENSG00000 |
| 87876216 | 87909553 | 1 | protein_coding | ENSG00000 |
| 87934143 | 88202889 | 1 | protein_coding | ENSG00000 |
| 88205118 | 88226993 | -1 | protein_coding | ENSG00000 |
| 88216660 | 88219226 | 1 | lncRNA | ENSG00000 |
| 88219359 | 88304367 | 1 | lncRNA | ENSG00000 |
| 88270892 | 88306891 | -1 | protein_coding | ENSG00000 |
| 88420167 | 88756725 | 1 | lncRNA | ENSG00000 |
| 88448788 | 88449017 | -1 | processed_pseudoge | ENSG00000 |
| 88449092 | 88449231 | -1 | snoRNA | ENSG00000 |
| 88455309 | 88456191 | 1 | processed_pseudoge | ENSG00000 |
| 88512029 | 88513187 | -1 | processed_pseudoge | ENSG00000 |
| 88564793 | 88566281 | -1 | processed_pseudoge | ENSG00000 |
| 88610241 | 88610669 | -1 | processed_pseudoge | ENSG00000 |
| 88639014 | 88640416 | 1 | processed_pseudoge | ENSG00000 |

| | | | | |
|---|---|---|---|---|
| 88749470 | 88749694 | 1 | processed_pseudoge | ENSG00000 |
| 88759700 | 89338528 | 1 | protein_coding | ENSG00000 |
| 88794106 | 88795737 | -1 | protein_coding | ENSG00000 |
| 89443946 | 89496918 | 1 | lncRNA | ENSG00000 |
| 89754620 | 89754726 | -1 | snRNA | ENSG00000 |
| 89882353 | 90211635 | -1 | lncRNA | ENSG00000 |
| 90119299 | 90122890 | -1 | lncRNA | ENSG00000 |
| 90119400 | 90125600 | 1 | transcribed_unproce | ENSG00000 |
| 90154456 | 90164829 | 1 | protein_coding | ENSG00000 |
| 90167590 | 90238137 | 1 | protein_coding | ENSG00000 |
| 90245174 | 90311063 | 1 | protein_coding | ENSG00000 |
| 90251923 | 90252020 | 1 | misc_RNA | ENSG00000 |
| 90266034 | 90270216 | -1 | lncRNA | ENSG00000 |
| 90312496 | 90322592 | 1 | lncRNA | ENSG00000 |
| 90316503 | 90321308 | -1 | protein_coding | ENSG00000 |
| 90335223 | 90391455 | 1 | protein_coding | ENSG00000 |
| 90345873 | 90346218 | -1 | TEC | ENSG00000 |
| 90383721 | 90513402 | 1 | protein_coding | ENSG00000 |
| 90403434 | 90513391 | 1 | lncRNA | ENSG00000 |
| 90466424 | 91210590 | 1 | protein_coding | ENSG00000 |
| 90469787 | 90488964 | 1 | unprocessed_pseudc | ENSG00000 |
| 90590619 | 90597353 | -1 | lncRNA | ENSG00000 |
| 90631906 | 90632568 | 1 | processed_pseudoge | ENSG00000 |
| 91030149 | 91033725 | -1 | lncRNA | ENSG00000 |
| 91107374 | 91107793 | 1 | processed_pseudoge | ENSG00000 |
| 91264433 | 91271326 | 1 | protein_coding | ENSG00000 |
| 91311368 | 91515427 | 1 | lncRNA | ENSG00000 |
| 91320128 | 91321103 | -1 | processed_pseudoge | ENSG00000 |
| 91380778 | 91556848 | -1 | lncRNA | ENSG00000 |
| 91638847 | 91643512 | -1 | lncRNA | ENSG00000 |
| 91692008 | 91880702 | -1 | protein_coding | ENSG00000 |
| 91880791 | 91886490 | 1 | lncRNA | ENSG00000 |
| 91940867 | 92110673 | 1 | protein_coding | ENSG00000 |
| 92112153 | 92134803 | -1 | protein_coding | ENSG00000 |
| 92112159 | 92179511 | -1 | protein_coding | ENSG00000 |
| 92131774 | 92245924 | -1 | protein_coding | ENSG00000 |
| 92134604 | 92180725 | 1 | lncRNA | ENSG00000 |
| 92141643 | 92179531 | -1 | protein_coding | ENSG00000 |
| 92198969 | 92246166 | -1 | protein_coding | ENSG00000 |
| 92200014 | 92206857 | -1 | lncRNA | ENSG00000 |
| 92202243 | 92202334 | -1 | misc_RNA | ENSG00000 |
| 92204015 | 92204098 | -1 | miRNA | ENSG00000 |
| 92245974 | 92401383 | 1 | protein_coding | ENSG00000 |
| 92412550 | 92447813 | -1 | transcribed_unitary_ | ENSG00000 |
| 92442077 | 92442489 | 1 | processed_pseudoge | ENSG00000 |
| 92447482 | 92460075 | 1 | protein_coding | ENSG00000 |
| 92457564 | 92495601 | 1 | lncRNA | ENSG00000 |
| 92468380 | 92477986 | -1 | protein_coding | ENSG00000 |
| 92487020 | 92528520 | -1 | protein_coding | ENSG00000 |
| 92528773 | 92540481 | 1 | protein_coding | ENSG00000 |

| | | | | |
|---|---|---|---|---|
| 92540268 | 92557877 | -1 | unprocessed_pseudc | ENSG00000( |
| 92560758 | 92590393 | -1 | protein_coding | ENSG00000( |
| 92604921 | 92836594 | -1 | protein_coding | ENSG00000( |
| 92638198 | 92642316 | -1 | lncRNA | ENSG00000( |
| 92647906 | 92667083 | 1 | lncRNA | ENSG00000( |
| 92701708 | 92701814 | -1 | snRNA | ENSG00000( |
| 92836483 | 92917187 | 1 | lncRNA | ENSG00000( |
| 92971002 | 92971299 | -1 | misc_RNA | ENSG00000( |
| 93099513 | 93118023 | -1 | protein_coding | ENSG00000( |
| 93130056 | 93148385 | -1 | protein_coding | ENSG00000( |
| 93188534 | 93226469 | -1 | protein_coding | ENSG00000( |
| 93232340 | 93361123 | 1 | protein_coding | ENSG00000( |
| 93424487 | 93574730 | -1 | protein_coding | ENSG00000( |
| 93482760 | 93482855 | -1 | miRNA | ENSG00000( |
| 93483936 | 93484019 | -1 | miRNA | ENSG00000( |
| 93591573 | 93911265 | 1 | protein_coding | ENSG00000( |
| 93669826 | 93670077 | -1 | processed_pseudoge | ENSG00000( |
| 93716928 | 93717005 | 1 | miRNA | ENSG00000( |
| 93777839 | 93778514 | 1 | processed_pseudoge | ENSG00000( |
| 93844789 | 93845308 | 1 | processed_pseudoge | ENSG00000( |
| 93885396 | 93890753 | -1 | protein_coding | ENSG00000( |
| 93890913 | 93893601 | 1 | lncRNA | ENSG00000( |
| 93914987 | 93915108 | 1 | unprocessed_pseudc | ENSG00000( |
| 93921735 | 93928610 | 1 | protein_coding | ENSG00000( |
| 93954044 | 93956519 | -1 | TEC | ENSG00000( |
| 93962762 | 94004382 | -1 | protein_coding | ENSG00000( |
| 93969442 | 94011113 | 1 | lncRNA | ENSG00000( |
| 94022833 | 94066661 | 1 | lncRNA | ENSG00000( |
| 94071759 | 94077157 | -1 | lncRNA | ENSG00000( |
| 94278680 | 94395608 | -1 | lncRNA | ENSG00000( |
| 94311138 | 94347063 | 1 | lncRNA | ENSG00000( |
| 94394561 | 94431232 | 1 | protein_coding | ENSG00000( |
| 94495299 | 94495404 | 1 | snRNA | ENSG00000( |
| 94509219 | 94557019 | 1 | protein_coding | ENSG00000( |
| 94524204 | 94656572 | -1 | protein_coding | ENSG00000( |
| 94656325 | 94669695 | 1 | protein_coding | ENSG00000( |
| 94695027 | 94695819 | -1 | processed_pseudoge | ENSG00000( |
| 94712409 | 94712511 | 1 | snRNA | ENSG00000( |
| 94738652 | 94739411 | 1 | processed_pseudoge | ENSG00000( |
| 94782886 | 94783525 | 1 | processed_pseudoge | ENSG00000( |
| 94801514 | 94801812 | 1 | misc_RNA | ENSG00000( |
| 94833904 | 94834447 | 1 | processed_pseudoge | ENSG00000( |
| 94907202 | 95296415 | 1 | protein_coding | ENSG00000( |
| 95018163 | 95018397 | -1 | processed_pseudoge | ENSG00000( |
| 95018407 | 95018554 | 1 | processed_pseudoge | ENSG00000( |
| 95035731 | 95214338 | -1 | lncRNA | ENSG00000( |
| 95098227 | 95098367 | 1 | snRNA | ENSG00000( |
| 95297676 | 95324532 | -1 | protein_coding | ENSG00000( |
| 95348718 | 95348910 | -1 | processed_pseudoge | ENSG00000( |
| 95350164 | 95351229 | 1 | processed_pseudoge | ENSG00000( |

| | | | | |
|---|---|---|---|---|
| 95359872 | 95396375 | -1 | protein_coding | ENSG00000( |
| 95404862 | 95435329 | -1 | protein_coding | ENSG00000( |
| 95416108 | 95416462 | 1 | lncRNA | ENSG00000( |
| 95471835 | 95473998 | 1 | lncRNA | ENSG00000( |
| 95478444 | 95540233 | 1 | protein_coding | ENSG00000( |
| 95542145 | 95544542 | 1 | TEC | ENSG00000( |
| 95545191 | 95615132 | 1 | lncRNA | ENSG00000( |
| 95583499 | 95596516 | -1 | protein_coding | ENSG00000( |
| 95772506 | 96110322 | 1 | protein_coding | ENSG00000( |
| 96118647 | 96119995 | 1 | TEC | ENSG00000( |
| 96120220 | 96322147 | -1 | protein_coding | ENSG00000( |
| 96219662 | 96219756 | -1 | miRNA | ENSG00000( |
| 96283357 | 96283804 | 1 | processed_pseudoge | ENSG00000( |
| 96306810 | 96306874 | 1 | processed_pseudoge | ENSG00000( |
| 96330638 | 96330744 | 1 | snRNA | ENSG00000( |
| 96341140 | 96341246 | 1 | snRNA | ENSG00000( |
| 96377857 | 96377920 | 1 | snRNA | ENSG00000( |
| 96481626 | 96709891 | -1 | protein_coding | ENSG00000( |
| 96858182 | 96859618 | 1 | processed_pseudoge | ENSG00000( |
| 96940068 | 96940364 | -1 | misc_RNA | ENSG00000( |
| 96955141 | 97014088 | -1 | lncRNA | ENSG00000( |
| 96965302 | 96965481 | 1 | misc_RNA | ENSG00000( |
| 96965489 | 96965695 | 1 | misc_RNA | ENSG00000( |
| 97005553 | 97011040 | 1 | protein_coding | ENSG00000( |
| 97013925 | 97014066 | -1 | misc_RNA | ENSG00000( |
| 97020396 | 97024950 | -1 | protein_coding | ENSG00000( |
| 97116590 | 97181763 | 1 | protein_coding | ENSG00000( |
| 97135015 | 97135561 | -1 | processed_pseudoge | ENSG00000( |
| 97200708 | 97201483 | -1 | processed_pseudoge | ENSG00000( |
| 97437518 | 97437896 | 1 | processed_pseudoge | ENSG00000( |
| 97598933 | 97599260 | -1 | misc_RNA | ENSG00000( |
| 97732084 | 97740472 | 1 | protein_coding | ENSG00000( |
| 97851677 | 97872542 | -1 | protein_coding | ENSG00000( |
| 97851688 | 97972985 | -1 | lncRNA | ENSG00000( |
| 97870167 | 97870289 | 1 | processed_pseudoge | ENSG00000( |
| 97885868 | 97886074 | 1 | processed_pseudoge | ENSG00000( |
| 97898347 | 97899115 | -1 | processed_pseudoge | ENSG00000( |
| 97906429 | 97906822 | 1 | processed_pseudoge | ENSG00000( |
| 97908256 | 97908424 | 1 | processed_pseudoge | ENSG00000( |
| 97928082 | 97928300 | -1 | transcribed_unproce | ENSG00000( |
| 97938439 | 97938794 | -1 | processed_pseudoge | ENSG00000( |
| 97946987 | 97947998 | -1 | unprocessed_pseudc | ENSG00000( |
| 97963658 | 97963726 | 1 | miRNA | ENSG00000( |
| 97964405 | 97964481 | -1 | miRNA | ENSG00000( |
| 97966090 | 97967074 | -1 | unprocessed_pseudc | ENSG00000( |
| 97966377 | 97970404 | -1 | lncRNA | ENSG00000( |
| 97969005 | 97972254 | -1 | transcribed_unproce | ENSG00000( |
| 97984684 | 97991169 | -1 | protein_coding | ENSG00000( |
| 97998325 | 97998622 | 1 | misc_RNA | ENSG00000( |
| 98106862 | 98209638 | 1 | protein_coding | ENSG00000( |

| | | | | |
|---|---|---|---|---|
| 98211427 | 98212979 | 1 | protein_coding | ENSG0000C |
| 98214624 | 98252232 | -1 | protein_coding | ENSG0000C |
| 98252379 | 98310441 | 1 | protein_coding | ENSG0000C |
| 98291650 | 98401090 | -1 | protein_coding | ENSG0000C |
| 98322853 | 98323430 | 1 | lncRNA | ENSG0000C |
| 98385801 | 98386584 | -1 | processed_pseudoge | ENSG0000C |
| 98454119 | 98454617 | -1 | processed_pseudoge | ENSG0000C |
| 98478551 | 98479225 | -1 | processed_pseudoge | ENSG0000C |
| 98617285 | 98629869 | 1 | protein_coding | ENSG0000C |
| 98794718 | 98794824 | -1 | snRNA | ENSG0000C |
| 98846488 | 98870771 | -1 | protein_coding | ENSG0000C |
| 98877933 | 99050831 | 1 | protein_coding | ENSG0000C |
| 98881650 | 98881729 | 1 | miRNA | ENSG0000C |
| 98989867 | 98993778 | -1 | lncRNA | ENSG0000C |
| 98998538 | 98999930 | -1 | processed_pseudoge | ENSG0000C |
| 99027438 | 99144100 | -1 | protein_coding | ENSG0000C |
| 99032894 | 99032940 | 1 | transcribed_unproce | ENSG0000C |
| 99173574 | 99207506 | -1 | protein_coding | ENSG0000C |
| 99238829 | 99311130 | 1 | transcribed_unitary_ | ENSG0000C |
| 99252452 | 99325826 | -1 | lncRNA | ENSG0000C |
| 99325879 | 99394653 | 1 | protein_coding | ENSG0000C |
| 99325898 | 99366262 | 1 | protein_coding | ENSG0000C |
| 99374249 | 99394816 | 1 | protein_coding | ENSG0000C |
| 99392048 | 99408597 | -1 | protein_coding | ENSG0000C |
| 99408641 | 99419616 | 1 | protein_coding | ENSG0000C |
| 99416739 | 99466163 | -1 | protein_coding | ENSG0000C |
| 99419749 | 99466197 | -1 | protein_coding | ENSG0000C |
| 99438922 | 99457373 | 1 | protein_coding | ENSG0000C |
| 99442890 | 99443496 | -1 | processed_pseudoge | ENSG0000C |
| 99448475 | 99466331 | -1 | protein_coding | ENSG0000C |
| 99472890 | 99503650 | 1 | protein_coding | ENSG0000C |
| 99473877 | 99504857 | -1 | protein_coding | ENSG0000C |
| 99504651 | 99534700 | 1 | protein_coding | ENSG0000C |
| 99546300 | 99558536 | -1 | protein_coding | ENSG0000C |
| 99558406 | 99576453 | 1 | protein_coding | ENSG0000C |
| 99558695 | 99607810 | 1 | protein_coding | ENSG0000C |
| 99598267 | 99611045 | 1 | protein_coding | ENSG0000C |
| 99616946 | 99632408 | 1 | protein_coding | ENSG0000C |
| 99638242 | 99638767 | -1 | processed_pseudoge | ENSG0000C |
| 99648194 | 99679998 | -1 | protein_coding | ENSG0000C |
| 99684957 | 99735196 | -1 | protein_coding | ENSG0000C |
| 99685145 | 99700034 | -1 | unprocessed_pseudc | ENSG0000C |
| 99685226 | 99735093 | -1 | protein_coding | ENSG0000C |
| 99756960 | 99784248 | -1 | protein_coding | ENSG0000C |
| 99766543 | 99766892 | 1 | lncRNA | ENSG0000C |
| 99820018 | 99820086 | 1 | unprocessed_pseudc | ENSG0000C |
| 99828013 | 99866102 | 1 | protein_coding | ENSG0000C |
| 99869841 | 99869894 | 1 | unprocessed_pseudc | ENSG0000C |
| 99872168 | 99872258 | 1 | unprocessed_pseudc | ENSG0000C |
| 99875987 | 99877057 | -1 | protein_coding | ENSG0000C |

| | | | | |
|---|---|---|---|---|
| 99876958 | 99919600 | -1 | protein_coding | ENSG0000( |
| 99923266 | 99929620 | -1 | protein_coding | ENSG0000( |
| 99929392 | 99948620 | 1 | lncRNA | ENSG0000( |
| 99936610 | 99936711 | 1 | misc_RNA | ENSG0000( |
| 99952033 | 99952158 | 1 | snoRNA | ENSG0000( |
| 99966720 | 99976042 | -1 | protein_coding | ENSG0000( |
| 99974976 | 99979233 | 1 | lncRNA | ENSG0000( |
| 99980762 | 99987535 | 1 | transcribed_unproce | ENSG0000( |
| 99992397 | 99993050 | 1 | unprocessed_pseudo | ENSG0000( |
| 99997690 | 100011634 | 1 | lncRNA | ENSG0000( |
| 100015572 | 100041689 | 1 | protein_coding | ENSG0000( |
| 100049774 | 100065040 | 1 | protein_coding | ENSG0000( |
| 100064033 | 100082548 | -1 | protein_coding | ENSG0000( |
| 100088969 | 100092187 | 1 | protein_coding | ENSG0000( |
| 100092728 | 100101940 | -1 | protein_coding | ENSG0000( |
| 100093560 | 100093643 | -1 | miRNA | ENSG0000( |
| 100093768 | 100093847 | -1 | miRNA | ENSG0000( |
| 100093993 | 100094074 | -1 | miRNA | ENSG0000( |
| 100101549 | 100110345 | 1 | protein_coding | ENSG0000( |
| 100107070 | 100119841 | -1 | protein_coding | ENSG0000( |
| 100115214 | 100127139 | -1 | lncRNA | ENSG0000( |
| 100119634 | 100125508 | 1 | protein_coding | ENSG0000( |
| 100126785 | 100128495 | 1 | protein_coding | ENSG0000( |
| 100130964 | 100140439 | 1 | lncRNA | ENSG0000( |
| 100139629 | 100140682 | -1 | processed_pseudoge | ENSG0000( |
| 100148907 | 100155944 | 1 | protein_coding | ENSG0000( |
| 100154420 | 100158723 | -1 | protein_coding | ENSG0000( |
| 100156605 | 100156669 | -1 | miRNA | ENSG0000( |
| 100159244 | 100168617 | -1 | protein_coding | ENSG0000( |
| 100169606 | 100177381 | -1 | protein_coding | ENSG0000( |
| 100177563 | 100221488 | 1 | protein_coding | ENSG0000( |
| 100200653 | 100272232 | -1 | protein_coding | ENSG0000( |
| 100218241 | 100221490 | 1 | protein_coding | ENSG0000( |
| 100307702 | 100322196 | 1 | protein_coding | ENSG0000( |
| 100320992 | 100341908 | -1 | unprocessed_pseudo | ENSG0000( |
| 100330777 | 100330877 | -1 | misc_RNA | ENSG0000( |
| 100336079 | 100351900 | 1 | transcribed_unproce | ENSG0000( |
| 100336104 | 100367831 | 1 | lncRNA | ENSG0000( |
| 100352176 | 100367733 | 1 | protein_coding | ENSG0000( |
| 100352360 | 100353692 | 1 | transcribed_unproce | ENSG0000( |
| 100356651 | 100356721 | 1 | miRNA | ENSG0000( |
| 100367530 | 100400096 | 1 | protein_coding | ENSG0000( |
| 100388809 | 100398366 | -1 | lncRNA | ENSG0000( |
| 100400826 | 100428992 | -1 | protein_coding | ENSG0000( |
| 100428790 | 100434118 | 1 | protein_coding | ENSG0000( |
| 100435257 | 100436510 | 1 | lncRNA | ENSG0000( |
| 100435282 | 100436497 | -1 | protein_coding | ENSG0000( |
| 100436204 | 100438504 | 1 | lncRNA | ENSG0000( |
| 100456620 | 100464260 | -1 | protein_coding | ENSG0000( |
| 100463359 | 100479232 | -1 | protein_coding | ENSG0000( |

| | | | | |
|---|---|---|---|---|
| 100482221 | 100483833 | -1 | lncRNA | ENSG00000 |
| 100483927 | 100494802 | 1 | protein_coding | ENSG00000 |
| 100509416 | 100520205 | 1 | lncRNA | ENSG00000 |
| 100530362 | 100530659 | -1 | misc_RNA | ENSG00000 |
| 100539203 | 100568220 | 1 | protein_coding | ENSG00000 |
| 100570131 | 100571136 | 1 | processed_pseudoge | ENSG00000 |
| 100572228 | 100573900 | -1 | protein_coding | ENSG00000 |
| 100572232 | 100578700 | -1 | lncRNA | ENSG00000 |
| 100574011 | 100586129 | -1 | protein_coding | ENSG00000 |
| 100583982 | 100601117 | 1 | protein_coding | ENSG00000 |
| 100589402 | 100604206 | -1 | lncRNA | ENSG00000 |
| 100602363 | 100608175 | 1 | protein_coding | ENSG00000 |
| 100612102 | 100615384 | 1 | protein_coding | ENSG00000 |
| 100620416 | 100642779 | -1 | protein_coding | ENSG00000 |
| 100643097 | 100656448 | -1 | protein_coding | ENSG00000 |
| 100673567 | 100679174 | 1 | protein_coding | ENSG00000 |
| 100679507 | 100694037 | -1 | protein_coding | ENSG00000 |
| 100706121 | 100707486 | 1 | protein_coding | ENSG00000 |
| 100720468 | 100723700 | 1 | protein_coding | ENSG00000 |
| 100733595 | 100797797 | 1 | protein_coding | ENSG00000 |
| 100802565 | 100827523 | -1 | protein_coding | ENSG00000 |
| 100821027 | 100821264 | -1 | misc_RNA | ENSG00000 |
| 100826820 | 100867010 | 1 | protein_coding | ENSG00000 |
| 100837314 | 100852616 | -1 | lncRNA | ENSG00000 |
| 100867387 | 100873454 | 1 | protein_coding | ENSG00000 |
| 100868036 | 100868107 | 1 | miRNA | ENSG00000 |
| 100875103 | 100888664 | 1 | protein_coding | ENSG00000 |
| 100888721 | 100889715 | -1 | protein_coding | ENSG00000 |
| 100889994 | 100896974 | -1 | protein_coding | ENSG00000 |
| 100906291 | 100906572 | -1 | misc_RNA | ENSG00000 |
| 100928370 | 100928540 | 1 | processed_pseudoge | ENSG00000 |
| 100949534 | 100968347 | 1 | protein_coding | ENSG00000 |
| 100963828 | 100968124 | -1 | lncRNA | ENSG00000 |
| 100969623 | 101018949 | 1 | protein_coding | ENSG00000 |
| 101014320 | 101017608 | -1 | lncRNA | ENSG00000 |
| 101020072 | 101058859 | 1 | protein_coding | ENSG00000 |
| 101058299 | 101058567 | 1 | misc_RNA | ENSG00000 |
| 101085481 | 101097967 | 1 | protein_coding | ENSG00000 |
| 101127089 | 101139266 | 1 | protein_coding | ENSG00000 |
| 101154456 | 101161596 | 1 | protein_coding | ENSG00000 |
| 101159473 | 101159555 | 1 | miRNA | ENSG00000 |
| 101162509 | 101165569 | -1 | protein_coding | ENSG00000 |
| 101170496 | 101180293 | -1 | protein_coding | ENSG00000 |
| 101195007 | 101201038 | -1 | protein_coding | ENSG00000 |
| 101201809 | 101203526 | -1 | unprocessed_pseudo | ENSG00000 |
| 101203614 | 101204495 | -1 | processed_pseudoge | ENSG00000 |
| 101205977 | 101218420 | -1 | protein_coding | ENSG00000 |
| 101218165 | 101224190 | 1 | protein_coding | ENSG00000 |
| 101232092 | 101238820 | -1 | protein_coding | ENSG00000 |
| 101239458 | 101252316 | -1 | protein_coding | ENSG00000 |

| | | | | |
|---|---|---|---|---|
| 101269938 | 101270041 | -1 | snRNA | ENSG0000C |
| 101273322 | 101273715 | 1 | unprocessed_pseudc | ENSG0000C |
| 101287482 | 101289771 | -1 | unprocessed_pseudc | ENSG0000C |
| 101299578 | 101301346 | 1 | lncRNA | ENSG0000C |
| 101308270 | 101314800 | 1 | lncRNA | ENSG0000C |
| 101310914 | 101321823 | -1 | protein_coding | ENSG0000C |
| 101362875 | 101559024 | 1 | protein_coding | ENSG0000C |
| 101388868 | 101389080 | -1 | processed_pseudoge | ENSG0000C |
| 101562779 | 101569006 | -1 | lncRNA | ENSG0000C |
| 101613330 | 101629296 | -1 | protein_coding | ENSG0000C |
| 101815904 | 102283958 | 1 | protein_coding | ENSG0000C |
| 101822247 | 101824729 | -1 | lncRNA | ENSG0000C |
| 101960116 | 101961894 | 1 | lncRNA | ENSG0000C |
| 102153355 | 102154463 | -1 | lncRNA | ENSG0000C |
| 102161120 | 102161337 | -1 | pseudogene | ENSG0000C |
| 102186819 | 102187039 | -1 | pseudogene | ENSG0000C |
| 102188599 | 102188828 | -1 | pseudogene | ENSG0000C |
| 102194076 | 102194164 | -1 | snoRNA | ENSG0000C |
| 102264706 | 102264979 | -1 | processed_pseudoge | ENSG0000C |
| 102285091 | 102321711 | 1 | protein_coding | ENSG0000C |
| 102293103 | 102293187 | 1 | miRNA | ENSG0000C |
| 102327256 | 102329530 | -1 | unprocessed_pseudc | ENSG0000C |
| 102336869 | 102336970 | 1 | misc_RNA | ENSG0000C |
| 102337316 | 102339115 | 1 | unprocessed_pseudc | ENSG0000C |
| 102345746 | 102356444 | -1 | protein_coding | ENSG0000C |
| 102363872 | 102426676 | 1 | protein_coding | ENSG0000C |
| 102364162 | 102380633 | 1 | transcribed_unproce | ENSG0000C |
| 102380465 | 102382737 | -1 | unprocessed_pseudc | ENSG0000C |
| 102405742 | 102405855 | -1 | miRNA | ENSG0000C |
| 102426818 | 102434780 | -1 | lncRNA | ENSG0000C |
| 102433106 | 102456825 | 1 | protein_coding | ENSG0000C |
| 102456238 | 102464863 | -1 | protein_coding | ENSG0000C |
| 102464956 | 102473168 | 1 | protein_coding | ENSG0000C |
| 102465742 | 102465826 | 1 | miRNA | ENSG0000C |
| 102471469 | 102471531 | 1 | miRNA | ENSG0000C |
| 102473118 | 102478907 | -1 | protein_coding | ENSG0000C |
| 102482445 | 102517781 | -1 | protein_coding | ENSG0000C |
| 102537918 | 102572653 | -1 | protein_coding | ENSG0000C |
| 102538417 | 102543849 | -1 | protein_coding | ENSG0000C |
| 102541501 | 102592444 | -1 | protein_coding | ENSG0000C |
| 102551232 | 102562308 | 1 | protein_coding | ENSG0000C |
| 102562133 | 102572583 | -1 | protein_coding | ENSG0000C |
| 102573807 | 102616757 | -1 | protein_coding | ENSG0000C |
| 102579104 | 102679295 | -1 | lncRNA | ENSG0000C |
| 102582523 | 102642869 | -1 | protein_coding | ENSG0000C |
| 102637025 | 102642791 | -1 | protein_coding | ENSG0000C |
| 102637049 | 102671641 | -1 | protein_coding | ENSG0000C |
| 102650325 | 102661398 | 1 | protein_coding | ENSG0000C |
| 102665368 | 102671629 | -1 | protein_coding | ENSG0000C |
| 102681836 | 102690469 | -1 | unprocessed_pseudc | ENSG0000C |

| | | | | |
|---|---|---|---|---|
| 102699228 | 102748695 | -1 | lncRNA | ENSG0000C |
| 102748971 | 102809225 | 1 | protein_coding | ENSG0000C |
| 102755146 | 102755939 | 1 | processed_pseudoge | ENSG0000C |
| 102813230 | 103074843 | -1 | protein_coding | ENSG0000C |
| 102834605 | 102834708 | 1 | snRNA | ENSG0000C |
| 102857450 | 102857755 | 1 | misc_RNA | ENSG0000C |
| 102913000 | 102945111 | 1 | protein_coding | ENSG0000C |
| 102973522 | 102988856 | 1 | protein_coding | ENSG0000C |
| 103030104 | 103031354 | 1 | lncRNA | ENSG0000C |
| 103074881 | 103099759 | 1 | protein_coding | ENSG0000C |
| 103088664 | 103089610 | -1 | processed_pseudoge | ENSG0000C |
| 103099776 | 103149560 | -1 | protein_coding | ENSG0000C |
| 103141349 | 103141939 | -1 | processed_pseudoge | ENSG0000C |
| 103152007 | 103152325 | -1 | processed_pseudoge | ENSG0000C |
| 103161947 | 103162660 | -1 | TEC | ENSG0000C |
| 103175133 | 103280410 | -1 | transcribed_unproce | ENSG0000C |
| 103262000 | 103262311 | 1 | processed_pseudoge | ENSG0000C |
| 103297435 | 103329511 | 1 | protein_coding | ENSG0000C |
| 103312289 | 103344830 | -1 | protein_coding | ENSG0000C |
| 103344254 | 103369395 | 1 | protein_coding | ENSG0000C |
| 103348601 | 103348771 | 1 | processed_pseudoge | ENSG0000C |
| 103352730 | 103446207 | -1 | protein_coding | ENSG0000C |
| 103433461 | 103433561 | -1 | misc_RNA | ENSG0000C |
| 103434994 | 103435092 | -1 | misc_RNA | ENSG0000C |
| 103445207 | 103514007 | 1 | lncRNA | ENSG0000C |
| 103471784 | 103989658 | -1 | protein_coding | ENSG0000C |
| 103484208 | 103484539 | -1 | misc_RNA | ENSG0000C |
| 104126341 | 104208047 | -1 | protein_coding | ENSG0000C |
| 104328700 | 104907232 | 1 | protein_coding | ENSG0000C |
| 104667749 | 104669576 | 1 | processed_pseudoge | ENSG0000C |
| 104738597 | 104804107 | -1 | lncRNA | ENSG0000C |
| 104826336 | 104826796 | 1 | processed_pseudoge | ENSG0000C |
| 104894628 | 104926645 | -1 | lncRNA | ENSG0000C |
| 104911928 | 104912193 | 1 | misc_RNA | ENSG0000C |
| 104940943 | 105000713 | 1 | lncRNA | ENSG0000C |
| 104950315 | 105013044 | -1 | lncRNA | ENSG0000C |
| 105013425 | 105014321 | -1 | lncRNA | ENSG0000C |
| 105014190 | 105115019 | 1 | protein_coding | ENSG0000C |
| 105102838 | 105105483 | -1 | lncRNA | ENSG0000C |
| 105110704 | 105399308 | -1 | protein_coding | ENSG0000C |
| 105189190 | 105189814 | 1 | processed_pseudoge | ENSG0000C |
| 105204600 | 105204859 | 1 | processed_pseudoge | ENSG0000C |
| 105244652 | 105244970 | 1 | processed_pseudoge | ENSG0000C |
| 105301522 | 105302086 | 1 | processed_pseudoge | ENSG0000C |
| 105304277 | 105306642 | 1 | lncRNA | ENSG0000C |
| 105332790 | 105332896 | 1 | snRNA | ENSG0000C |
| 105439661 | 105522271 | -1 | protein_coding | ENSG0000C |
| 105530209 | 105530671 | 1 | processed_pseudoge | ENSG0000C |
| 105532169 | 105567677 | 1 | protein_coding | ENSG0000C |
| 105565120 | 105600875 | -1 | protein_coding | ENSG0000C |

| | | | | |
|---|---|---|---|---|
| 105571083 | 105573660 | 1 | lncRNA | ENSG0000C |
| 105582258 | 105583256 | -1 | processed_pseudoge | ENSG0000C |
| 105605067 | 105876604 | -1 | protein_coding | ENSG0000C |
| 105819492 | 105820101 | 1 | processed_pseudoge | ENSG0000C |
| 105876796 | 106036432 | 1 | protein_coding | ENSG0000C |
| 106035798 | 106036431 | 1 | TEC | ENSG0000C |
| 106090503 | 106112576 | -1 | protein_coding | ENSG0000C |
| 106125371 | 106126700 | 1 | processed_pseudoge | ENSG0000C |
| 106208167 | 106208267 | 1 | snRNA | ENSG0000C |
| 106248298 | 106286326 | -1 | protein_coding | ENSG0000C |
| 106285480 | 106286326 | 1 | lncRNA | ENSG0000C |
| 106315225 | 106315743 | 1 | processed_pseudoge | ENSG0000C |
| 106372251 | 106770207 | -1 | lncRNA | ENSG0000C |
| 106425278 | 106511544 | 1 | lncRNA | ENSG0000C |
| 106569876 | 106598908 | -1 | lncRNA | ENSG0000C |
| 106624072 | 106628696 | 1 | lncRNA | ENSG0000C |
| 106654360 | 106661158 | -1 | protein_coding | ENSG0000C |
| 106774955 | 106838480 | 1 | lncRNA | ENSG0000C |
| 106781600 | 106781715 | -1 | rRNA_pseudogene | ENSG0000C |
| 106865278 | 106908980 | 1 | protein_coding | ENSG0000C |
| 107044705 | 107161811 | 1 | protein_coding | ENSG0000C |
| 107066591 | 107133733 | -1 | lncRNA | ENSG0000C |
| 107168961 | 107202522 | 1 | protein_coding | ENSG0000C |
| 107192559 | 107193300 | -1 | lncRNA | ENSG0000C |
| 107201555 | 107564514 | -1 | protein_coding | ENSG0000C |
| 107450083 | 107450362 | 1 | processed_pseudoge | ENSG0000C |
| 107470018 | 107475684 | 1 | protein_coding | ENSG0000C |
| 107563484 | 107578523 | 1 | protein_coding | ENSG0000C |
| 107579557 | 107580057 | -1 | lncRNA | ENSG0000C |
| 107579977 | 107629170 | 1 | protein_coding | ENSG0000C |
| 107603363 | 107603507 | 1 | snoRNA | ENSG0000C |
| 107628553 | 107629550 | 1 | processed_pseudoge | ENSG0000C |
| 107642765 | 107643023 | -1 | processed_pseudoge | ENSG0000C |
| 107650260 | 107662204 | -1 | lncRNA | ENSG0000C |
| 107660828 | 107717809 | 1 | protein_coding | ENSG0000C |
| 107739999 | 107744581 | -1 | lncRNA | ENSG0000C |
| 107743949 | 107761667 | 1 | protein_coding | ENSG0000C |
| 107765467 | 107803225 | -1 | protein_coding | ENSG0000C |
| 107808734 | 107809568 | 1 | processed_pseudoge | ENSG0000C |
| 107891162 | 107931730 | 1 | protein_coding | ENSG0000C |
| 107923799 | 108003187 | -1 | protein_coding | ENSG0000C |
| 107942116 | 107942740 | 1 | lncRNA | ENSG0000C |
| 107999791 | 107999982 | -1 | snoRNA | ENSG0000C |
| 108023548 | 108130361 | -1 | protein_coding | ENSG0000C |
| 108147623 | 108456717 | -1 | protein_coding | ENSG0000C |
| 108470417 | 108569666 | -1 | protein_coding | ENSG0000C |
| 108510233 | 108510976 | 1 | processed_pseudoge | ENSG0000C |
| 108554543 | 108569750 | -1 | protein_coding | ENSG0000C |
| 108569867 | 108574850 | 1 | protein_coding | ENSG0000C |
| 108598352 | 108639349 | 1 | lncRNA | ENSG0000C |

| | | | | |
|---|---|---|---|---|
| 108720608 | 108721601 | -1 | processed_pseudoge | ENSG00000( |
| 108883975 | 108884587 | -1 | lncRNA | ENSG00000( |
| 108900105 | 108905226 | -1 | TEC | ENSG00000( |
| 108909453 | 108952666 | 1 | lncRNA | ENSG00000( |
| 108994031 | 108995029 | -1 | processed_pseudoge | ENSG00000( |
| 109322320 | 109326315 | -1 | lncRNA | ENSG00000( |
| 109521981 | 109597166 | -1 | lncRNA | ENSG00000( |
| 109660094 | 109660802 | 1 | processed_pseudoge | ENSG00000( |
| 109763574 | 109764357 | 1 | processed_pseudoge | ENSG00000( |
| 109959218 | 109960184 | -1 | processed_pseudoge | ENSG00000( |
| 109998434 | 109999624 | -1 | processed_pseudoge | ENSG00000( |
| 110108031 | 110534757 | -1 | lncRNA | ENSG00000( |
| 110215891 | 110216380 | 1 | unprocessed_pseudc | ENSG00000( |
| 110590082 | 110592204 | 1 | lncRNA | ENSG00000( |
| 110663051 | 111562517 | -1 | protein_coding | ENSG00000( |
| 110724159 | 110725825 | -1 | lncRNA | ENSG00000( |
| 111091006 | 111125454 | 1 | protein_coding | ENSG00000( |
| 111394875 | 111395043 | 1 | processed_pseudoge | ENSG00000( |
| 111411319 | 111411883 | 1 | processed_pseudoge | ENSG00000( |
| 111726110 | 112206411 | -1 | protein_coding | ENSG00000( |
| 111757051 | 111762216 | 1 | lncRNA | ENSG00000( |
| 111808516 | 111821773 | 1 | lncRNA | ENSG00000( |
| 111953653 | 111953765 | -1 | rRNA_pseudogene | ENSG00000( |
| 111971222 | 111971988 | -1 | processed_pseudoge | ENSG00000( |
| 112206695 | 112343934 | 1 | protein_coding | ENSG00000( |
| 112288623 | 112288952 | 1 | misc_RNA | ENSG00000( |
| 112328189 | 112409623 | -1 | lncRNA | ENSG00000( |
| 112372647 | 112373729 | 1 | unprocessed_pseudc | ENSG00000( |
| 112373733 | 112374250 | -1 | unprocessed_pseudc | ENSG00000( |
| 112374324 | 112375429 | 1 | unprocessed_pseudc | ENSG00000( |
| 112422968 | 112481017 | 1 | protein_coding | ENSG00000( |
| 112446164 | 112446375 | -1 | processed_pseudoge | ENSG00000( |
| 112480853 | 112491062 | 1 | protein_coding | ENSG00000( |
| 112520488 | 112521670 | 1 | processed_pseudoge | ENSG00000( |
| 112616440 | 112896625 | 1 | lncRNA | ENSG00000( |
| 112735166 | 112738693 | -1 | processed_pseudoge | ENSG00000( |
| 112762377 | 112790423 | -1 | protein_coding | ENSG00000( |
| 112819147 | 112939875 | -1 | protein_coding | ENSG00000( |
| 112953282 | 112995677 | -1 | lncRNA | ENSG00000( |
| 113075124 | 113077181 | -1 | TEC | ENSG00000( |
| 113078331 | 113087778 | -1 | protein_coding | ENSG00000( |
| 113100663 | 113146330 | 1 | lncRNA | ENSG00000( |
| 113116718 | 113118560 | -1 | protein_coding | ENSG00000( |
| 113415689 | 113416322 | 1 | processed_pseudoge | ENSG00000( |
| 113451072 | 113451402 | 1 | TEC | ENSG00000( |
| 113486407 | 113494870 | -1 | lncRNA | ENSG00000( |
| 113876777 | 114075920 | -1 | protein_coding | ENSG00000( |
| 114086327 | 114693772 | 1 | protein_coding | ENSG00000( |
| 114297114 | 114297429 | -1 | processed_pseudoge | ENSG00000( |
| 114414244 | 114419875 | -1 | lncRNA | ENSG00000( |

| | | | | |
|---|---|---|---|---|
| 114560961 | 114561517 | 1 | lncRNA | ENSG00000 |
| 114613787 | 114613906 | -1 | rRNA_pseudogene | ENSG00000 |
| 114653345 | 114653455 | 1 | miRNA | ENSG00000 |
| 114922154 | 115019202 | 1 | protein_coding | ENSG00000 |
| 115030564 | 115126314 | 1 | lncRNA | ENSG00000 |
| 115061537 | 115231355 | -1 | lncRNA | ENSG00000 |
| 115136475 | 115137220 | -1 | processed_pseudoge | ENSG00000 |
| 115207894 | 115207996 | -1 | misc_RNA | ENSG00000 |
| 115503367 | 115504081 | 1 | processed_pseudoge | ENSG00000 |
| 115581315 | 115581442 | -1 | snoRNA | ENSG00000 |
| 115647461 | 115676329 | -1 | lncRNA | ENSG00000 |
| 115679345 | 115682618 | -1 | lncRNA | ENSG00000 |
| 115789729 | 115791049 | 1 | lncRNA | ENSG00000 |
| 115833273 | 115833373 | -1 | misc_RNA | ENSG00000 |
| 115935148 | 116159896 | -1 | protein_coding | ENSG00000 |
| 116209234 | 116211511 | -1 | TEC | ENSG00000 |
| 116210506 | 116258783 | 1 | protein_coding | ENSG00000 |
| 116237929 | 116327896 | -1 | lncRNA | ENSG00000 |
| 116238260 | 116499465 | -1 | lncRNA | ENSG00000 |
| 116275606 | 116286734 | -1 | lncRNA | ENSG00000 |
| 116287380 | 116508541 | 1 | protein_coding | ENSG00000 |
| 116320751 | 116321031 | 1 | misc_RNA | ENSG00000 |
| 116525001 | 116561184 | 1 | protein_coding | ENSG00000 |
| 116542718 | 116551969 | -1 | lncRNA | ENSG00000 |
| 116563594 | 116663829 | -1 | lncRNA | ENSG00000 |
| 116563594 | 116571234 | -1 | lncRNA | ENSG00000 |
| 116672196 | 116798386 | 1 | protein_coding | ENSG00000 |
| 116811070 | 116922049 | 1 | protein_coding | ENSG00000 |
| 116873454 | 116876376 | -1 | processed_pseudoge | ENSG00000 |
| 116909877 | 116909964 | -1 | misc_RNA | ENSG00000 |
| 116944286 | 116944414 | -1 | rRNA_pseudogene | ENSG00000 |
| 116952446 | 116954334 | -1 | lncRNA | ENSG00000 |
| 116952698 | 116952813 | 1 | misc_RNA | ENSG00000 |
| 116953214 | 116953324 | 1 | misc_RNA | ENSG00000 |
| 116953238 | 117230103 | 1 | protein_coding | ENSG00000 |
| 116953899 | 117098806 | 1 | lncRNA | ENSG00000 |
| 116954480 | 116954679 | 1 | misc_RNA | ENSG00000 |
| 116954983 | 116955154 | 1 | misc_RNA | ENSG00000 |
| 116956431 | 116956723 | 1 | misc_RNA | ENSG00000 |
| 116959593 | 116959805 | 1 | misc_RNA | ENSG00000 |
| 116965846 | 116967472 | -1 | lncRNA | ENSG00000 |
| 116972165 | 116972911 | -1 | processed_pseudoge | ENSG00000 |
| 117020211 | 117020319 | 1 | miRNA | ENSG00000 |
| 117072072 | 117146480 | -1 | lncRNA | ENSG00000 |
| 117091678 | 117097262 | -1 | lncRNA | ENSG00000 |
| 117117912 | 117117983 | 1 | misc_RNA | ENSG00000 |
| 117184126 | 117184199 | 1 | misc_RNA | ENSG00000 |
| 117187548 | 117187740 | -1 | processed_pseudoge | ENSG00000 |
| 117190133 | 117190269 | 1 | misc_RNA | ENSG00000 |
| 117191647 | 117191761 | 1 | misc_RNA | ENSG00000 |

| | | | | |
|---|---|---|---|---|
| 117262918 | 117264888 | -1 | processed_pseudoge | ENSG0000C |
| 117263917 | 117264388 | 1 | processed_pseudoge | ENSG0000C |
| 117264393 | 117264530 | -1 | processed_pseudoge | ENSG0000C |
| 117275451 | 117323152 | -1 | protein_coding | ENSG0000C |
| 117300861 | 117322236 | 1 | lncRNA | ENSG0000C |
| 117332761 | 117350096 | -1 | lncRNA | ENSG0000C |
| 117363222 | 117428123 | -1 | protein_coding | ENSG0000C |
| 117439982 | 117440780 | -1 | processed_pseudoge | ENSG0000C |
| 117465784 | 117715971 | 1 | protein_coding | ENSG0000C |
| 117487737 | 117487929 | 1 | processed_pseudoge | ENSG0000C |
| 117560733 | 117564676 | -1 | lncRNA | ENSG0000C |
| 117586207 | 117586416 | -1 | processed_pseudoge | ENSG0000C |
| 117604791 | 117647415 | -1 | lncRNA | ENSG0000C |
| 117710651 | 117874139 | -1 | protein_coding | ENSG0000C |
| 117882859 | 117883675 | 1 | processed_pseudoge | ENSG0000C |
| 117998858 | 118004045 | -1 | lncRNA | ENSG0000C |
| 118184144 | 118204035 | 1 | protein_coding | ENSG0000C |
| 118214669 | 118496171 | 1 | protein_coding | ENSG0000C |
| 118511942 | 118513830 | 1 | lncRNA | ENSG0000C |
| 118880103 | 118881288 | -1 | processed_pseudoge | ENSG0000C |
| 118950386 | 118951259 | -1 | processed_pseudoge | ENSG0000C |
| 119178177 | 119179149 | -1 | processed_pseudoge | ENSG0000C |
| 119495024 | 119907397 | -1 | lncRNA | ENSG0000C |
| 120005976 | 120006101 | -1 | snRNA | ENSG0000C |
| 120141016 | 120227213 | -1 | lncRNA | ENSG0000C |
| 120166443 | 120186064 | 1 | lncRNA | ENSG0000C |
| 120273175 | 120750337 | 1 | protein_coding | ENSG0000C |
| 120672871 | 120672975 | 1 | snRNA | ENSG0000C |
| 120746738 | 120752514 | -1 | lncRNA | ENSG0000C |
| 120787320 | 120858402 | -1 | protein_coding | ENSG0000C |
| 120950763 | 120977216 | 1 | protein_coding | ENSG0000C |
| 120981426 | 120981553 | -1 | rRNA_pseudogene | ENSG0000C |
| 120988697 | 121297442 | 1 | protein_coding | ENSG0000C |
| 121050927 | 121051203 | 1 | processed_pseudoge | ENSG0000C |
| 121083700 | 121083813 | -1 | rRNA_pseudogene | ENSG0000C |
| 121194948 | 121195057 | -1 | snRNA | ENSG0000C |
| 121304657 | 121309842 | -1 | lncRNA | ENSG0000C |
| 121325367 | 121341104 | 1 | protein_coding | ENSG0000C |
| 121348878 | 121396364 | -1 | protein_coding | ENSG0000C |
| 121398452 | 121398767 | 1 | processed_pseudoge | ENSG0000C |
| 121419072 | 121419845 | 1 | processed_pseudoge | ENSG0000C |
| 121440834 | 121441261 | -1 | processed_pseudoge | ENSG0000C |
| 121643334 | 121686363 | -1 | lncRNA | ENSG0000C |
| 121736443 | 121736725 | 1 | misc_RNA | ENSG0000C |
| 121842368 | 121845131 | -1 | processed_pseudoge | ENSG0000C |
| 121873089 | 122062036 | 1 | protein_coding | ENSG0000C |
| 122073549 | 122144255 | -1 | protein_coding | ENSG0000C |
| 122081720 | 122081781 | 1 | snRNA | ENSG0000C |
| 122144405 | 122219277 | 1 | lncRNA | ENSG0000C |
| 122159300 | 122162243 | 1 | processed_pseudoge | ENSG0000C |

| | | | | |
|---|---|---|---|---|
| 122234531 | 122234903 | 1 | processed_pseudoge | ENSG00000( |
| 122301303 | 122310691 | -1 | protein_coding | ENSG00000( |
| 122303658 | 122310119 | 1 | lncRNA | ENSG00000( |
| 122318411 | 122886759 | -1 | protein_coding | ENSG00000( |
| 122328469 | 122440388 | 1 | lncRNA | ENSG00000( |
| 122676580 | 122676811 | 1 | processed_pseudoge | ENSG00000( |
| 122681315 | 122681662 | 1 | processed_pseudoge | ENSG00000( |
| 122697735 | 122699117 | -1 | protein_coding | ENSG00000( |
| 122701668 | 122702922 | -1 | protein_coding | ENSG00000( |
| 122849746 | 122850352 | 1 | processed_pseudoge | ENSG00000( |
| 122994704 | 122995700 | -1 | protein_coding | ENSG00000( |
| 123069249 | 123103589 | -1 | lncRNA | ENSG00000( |
| 123113531 | 123199972 | -1 | protein_coding | ENSG00000( |
| 123230120 | 123230741 | -1 | processed_pseudoge | ENSG00000( |
| 123452193 | 123535077 | -1 | protein_coding | ENSG00000( |
| 123456629 | 123459856 | 1 | lncRNA | ENSG00000( |
| 123457988 | 123458094 | -1 | snRNA | ENSG00000( |
| 123536997 | 123557904 | -1 | protein_coding | ENSG00000( |
| 123567010 | 123639481 | 1 | protein_coding | ENSG00000( |
| 123584859 | 123624957 | -1 | lncRNA | ENSG00000( |
| 123655866 | 123664290 | 1 | protein_coding | ENSG00000( |
| 123681943 | 123749003 | -1 | protein_coding | ENSG00000( |
| 123749068 | 123751166 | 1 | lncRNA | ENSG00000( |
| 123790605 | 123790711 | -1 | snRNA | ENSG00000( |
| 123814139 | 123829252 | 1 | unitary_pseudogene | ENSG00000( |
| 123828983 | 123877481 | 1 | protein_coding | ENSG00000( |
| 123925237 | 123971414 | 1 | protein_coding | ENSG00000( |
| 123932132 | 123932459 | 1 | processed_pseudoge | ENSG00000( |
| 123994622 | 124027659 | 1 | lncRNA | ENSG00000( |
| 124030921 | 124033067 | -1 | protein_coding | ENSG00000( |
| 124032126 | 124489572 | 1 | lncRNA | ENSG00000( |
| 124274671 | 124288852 | -1 | lncRNA | ENSG00000( |
| 124337380 | 124349860 | -1 | lncRNA | ENSG00000( |
| 124476371 | 124476955 | 1 | protein_coding | ENSG00000( |
| 124480524 | 124481395 | 1 | processed_pseudoge | ENSG00000( |
| 124647719 | 124647822 | 1 | snRNA | ENSG00000( |
| 124742312 | 124745017 | -1 | TEC | ENSG00000( |
| 124743885 | 124765792 | -1 | protein_coding | ENSG00000( |
| 124777292 | 124790810 | -1 | protein_coding | ENSG00000( |
| 124822386 | 124929983 | -1 | protein_coding | ENSG00000( |
| 124929873 | 125482966 | 1 | lncRNA | ENSG00000( |
| 125033453 | 125035301 | 1 | processed_pseudoge | ENSG00000( |
| 125151326 | 125153611 | -1 | lncRNA | ENSG00000( |
| 125159974 | 125160547 | -1 | processed_pseudoge | ENSG00000( |
| 125229579 | 125264291 | -1 | lncRNA | ENSG00000( |
| 125300504 | 125300878 | -1 | processed_pseudoge | ENSG00000( |
| 125344969 | 125345377 | 1 | processed_pseudoge | ENSG00000( |
| 125345825 | 125346197 | 1 | processed_pseudoge | ENSG00000( |
| 125917871 | 125933832 | 1 | lncRNA | ENSG00000( |
| 126378970 | 126424185 | 1 | lncRNA | ENSG00000( |

| | | | | |
|---|---|---|---|---|
| 126438598 | 127253294 | -1 | protein_coding | ENSG0000C |
| 126495312 | 126531336 | 1 | lncRNA | ENSG0000C |
| 126511135 | 126512879 | 1 | processed_pseudoge | ENSG0000C |
| 126533665 | 126537295 | 1 | lncRNA | ENSG0000C |
| 126868767 | 126870532 | -1 | processed_pseudoge | ENSG0000C |
| 126980967 | 126981778 | 1 | processed_pseudoge | ENSG0000C |
| 127058088 | 127058184 | -1 | miRNA | ENSG0000C |
| 127215127 | 127229927 | 1 | lncRNA | ENSG0000C |
| 127295620 | 127296203 | 1 | processed_pseudoge | ENSG0000C |
| 127346790 | 127431924 | -1 | protein_coding | ENSG0000C |
| 127349833 | 127350293 | -1 | TEC | ENSG0000C |
| 127350128 | 127351523 | 1 | lncRNA | ENSG0000C |
| 127359785 | 127364759 | -1 | TEC | ENSG0000C |
| 127476883 | 127485804 | 1 | lncRNA | ENSG0000C |
| 127580628 | 127593611 | -1 | protein_coding | ENSG0000C |
| 127588386 | 127591700 | 1 | protein_coding | ENSG0000C |
| 127591409 | 127602144 | 1 | protein_coding | ENSG0000C |
| 127610292 | 127618142 | -1 | protein_coding | ENSG0000C |
| 127644685 | 127652012 | -1 | lncRNA | ENSG0000C |
| 127652194 | 128092609 | 1 | protein_coding | ENSG0000C |
| 127857852 | 127858488 | -1 | processed_pseudoge | ENSG0000C |
| 127997597 | 128000077 | 1 | lncRNA | ENSG0000C |
| 128027071 | 128032107 | -1 | protein_coding | ENSG0000C |
| 128081861 | 128081960 | 1 | miRNA | ENSG0000C |
| 128207872 | 128207943 | 1 | miRNA | ENSG0000C |
| 128241278 | 128257629 | 1 | protein_coding | ENSG0000C |
| 128264526 | 128264889 | -1 | lncRNA | ENSG0000C |
| 128297685 | 128343908 | -1 | protein_coding | ENSG0000C |
| 128306649 | 128307678 | 1 | processed_pseudoge | ENSG0000C |
| 128344081 | 128344142 | 1 | snRNA | ENSG0000C |
| 128350325 | 128361685 | -1 | protein_coding | ENSG0000C |
| 128392277 | 128410252 | -1 | protein_coding | ENSG0000C |
| 128433422 | 128433713 | -1 | processed_pseudoge | ENSG0000C |
| 128443449 | 128443510 | -1 | snRNA | ENSG0000C |
| 128455840 | 128493859 | 1 | lncRNA | ENSG0000C |
| 128455849 | 128458418 | 1 | protein_coding | ENSG0000C |
| 128455937 | 128469197 | 1 | lncRNA | ENSG0000C |
| 128466563 | 128469171 | 1 | transcribed_unproce | ENSG0000C |
| 128476729 | 128506602 | 1 | protein_coding | ENSG0000C |
| 128483907 | 128484158 | -1 | misc_RNA | ENSG0000C |
| 128524016 | 128531069 | -1 | lncRNA | ENSG0000C |
| 128533652 | 128615970 | 1 | unprocessed_pseudc | ENSG0000C |
| 128570241 | 128570688 | -1 | processed_pseudoge | ENSG0000C |
| 128574751 | 128626473 | 1 | lncRNA | ENSG0000C |
| 128617865 | 128625905 | 1 | transcribed_unproce | ENSG0000C |
| 128627172 | 128627275 | 1 | snRNA | ENSG0000C |
| 128651185 | 128652334 | 1 | processed_pseudoge | ENSG0000C |
| 128652841 | 128653243 | 1 | processed_pseudoge | ENSG0000C |
| 128653690 | 128654019 | -1 | processed_pseudoge | ENSG0000C |
| 128653969 | 128654722 | 1 | processed_pseudoge | ENSG0000C |

| | | | | |
|---|---|---|---|---|
| 128655962 | 128658791 | -1 | processed_pseudoge | ENSG00000 |
| 128667043 | 128668156 | 1 | lncRNA | ENSG00000 |
| 128669087 | 128669892 | -1 | processed_pseudoge | ENSG00000 |
| 128672288 | 128687872 | 1 | protein_coding | ENSG00000 |
| 128690451 | 128691717 | 1 | lncRNA | ENSG00000 |
| 128693450 | 128693924 | 1 | processed_pseudoge | ENSG00000 |
| 128697439 | 128697556 | 1 | rRNA_pseudogene | ENSG00000 |
| 128697710 | 128697744 | 1 | rRNA_pseudogene | ENSG00000 |
| 128709061 | 128731743 | 1 | protein_coding | ENSG00000 |
| 128739292 | 128773400 | 1 | protein_coding | ENSG00000 |
| 128761338 | 128761598 | 1 | misc_RNA | ENSG00000 |
| 128772491 | 128775790 | -1 | protein_coding | ENSG00000 |
| 128790757 | 128822132 | 1 | protein_coding | ENSG00000 |
| 128830377 | 128859274 | 1 | protein_coding | ENSG00000 |
| 128850162 | 128862626 | -1 | lncRNA | ENSG00000 |
| 128862451 | 128910719 | -1 | protein_coding | ENSG00000 |
| 128862856 | 128865847 | 1 | protein_coding | ENSG00000 |
| 128866330 | 128872047 | 1 | protein_coding | ENSG00000 |
| 128912732 | 128914063 | -1 | unprocessed_pseudo | ENSG00000 |
| 128937457 | 128950038 | 1 | protein_coding | ENSG00000 |
| 128952527 | 128953316 | -1 | TEC | ENSG00000 |
| 128954180 | 129055173 | -1 | protein_coding | ENSG00000 |
| 128970734 | 128971029 | 1 | misc_RNA | ENSG00000 |
| 129028889 | 129030527 | 1 | processed_pseudoge | ENSG00000 |
| 129055223 | 129057239 | 1 | transcribed_process | ENSG00000 |
| 129066021 | 129066462 | -1 | processed_pseudoge | ENSG00000 |
| 129095301 | 129095811 | -1 | processed_pseudoge | ENSG00000 |
| 129096027 | 129096735 | -1 | processed_pseudoge | ENSG00000 |
| 129117513 | 129117827 | -1 | processed_pseudoge | ENSG00000 |
| 129126518 | 129130793 | 1 | protein_coding | ENSG00000 |
| 129144707 | 129169699 | 1 | protein_coding | ENSG00000 |
| 129164849 | 129164961 | 1 | misc_RNA | ENSG00000 |
| 129188633 | 129213545 | 1 | protein_coding | ENSG00000 |
| 129209775 | 129213545 | -1 | lncRNA | ENSG00000 |
| 129225023 | 129430211 | 1 | protein_coding | ENSG00000 |
| 129366827 | 129374486 | -1 | lncRNA | ENSG00000 |
| 129410113 | 129410370 | -1 | processed_pseudoge | ENSG00000 |
| 129434432 | 129488399 | 1 | protein_coding | ENSG00000 |
| 129477875 | 129478101 | 1 | processed_pseudoge | ENSG00000 |
| 129484504 | 129484656 | 1 | snRNA | ENSG00000 |
| 129502531 | 129512918 | 1 | protein_coding | ENSG00000 |
| 129604548 | 129611630 | -1 | lncRNA | ENSG00000 |
| 129611720 | 129757082 | 1 | protein_coding | ENSG00000 |
| 129756266 | 129756377 | 1 | rRNA_pseudogene | ENSG00000 |
| 129763060 | 129769202 | -1 | lncRNA | ENSG00000 |
| 129770383 | 129770492 | -1 | miRNA | ENSG00000 |
| 129774692 | 129774769 | -1 | miRNA | ENSG00000 |
| 129774905 | 129775014 | -1 | miRNA | ENSG00000 |
| 129783370 | 129785185 | -1 | lncRNA | ENSG00000 |
| 129830732 | 129952960 | -1 | protein_coding | ENSG00000 |

| | | | | |
|---|---|---|---|---|
| 129953234 | 130026989 | 1 | lncRNA | ENSG00000 |
| 129954693 | 129954794 | 1 | misc_RNA | ENSG00000 |
| 130018286 | 130051451 | -1 | protein_coding | ENSG00000 |
| 130027277 | 130027397 | 1 | rRNA_pseudogene | ENSG00000 |
| 130070534 | 130135705 | 1 | protein_coding | ENSG00000 |
| 130141707 | 130142618 | 1 | lncRNA | ENSG00000 |
| 130164713 | 130207770 | -1 | protein_coding | ENSG00000 |
| 130173718 | 130205361 | 1 | lncRNA | ENSG00000 |
| 130206344 | 130216844 | 1 | protein_coding | ENSG00000 |
| 130255902 | 130262770 | 1 | processed_pseudoge | ENSG00000 |
| 130266863 | 130289798 | 1 | protein_coding | ENSG00000 |
| 130293134 | 130324180 | 1 | protein_coding | ENSG00000 |
| 130344816 | 130368730 | 1 | protein_coding | ENSG00000 |
| 130380339 | 130388114 | 1 | protein_coding | ENSG00000 |
| 130393771 | 130442433 | -1 | protein_coding | ENSG00000 |
| 130481491 | 130484392 | -1 | lncRNA | ENSG00000 |
| 130486042 | 130491033 | -1 | lncRNA | ENSG00000 |
| 130486171 | 130506465 | 1 | protein_coding | ENSG00000 |
| 130487483 | 130487622 | 1 | misc_RNA | ENSG00000 |
| 130488304 | 130488538 | 1 | misc_RNA | ENSG00000 |
| 130489868 | 130489966 | 1 | misc_RNA | ENSG00000 |
| 130495794 | 130498427 | -1 | lncRNA | ENSG00000 |
| 130496111 | 130496204 | 1 | miRNA | ENSG00000 |
| 130506238 | 130668748 | -1 | protein_coding | ENSG00000 |
| 130507660 | 130508282 | 1 | lncRNA | ENSG00000 |
| 130538954 | 130546900 | 1 | lncRNA | ENSG00000 |
| 130602795 | 130602913 | -1 | rRNA_pseudogene | ENSG00000 |
| 130645225 | 130645594 | -1 | processed_pseudoge | ENSG00000 |
| 130668648 | 130687432 | -1 | protein_coding | ENSG00000 |
| 130668852 | 130669395 | 1 | lncRNA | ENSG00000 |
| 130731235 | 130734176 | -1 | protein_coding | ENSG00000 |
| 130790208 | 130791724 | 1 | lncRNA | ENSG00000 |
| 130791264 | 131110161 | -1 | lncRNA | ENSG00000 |
| 130822868 | 130823219 | -1 | processed_pseudoge | ENSG00000 |
| 130823205 | 130823511 | -1 | processed_pseudoge | ENSG00000 |
| 130840204 | 130843725 | -1 | processed_pseudoge | ENSG00000 |
| 130853720 | 130930682 | 1 | lncRNA | ENSG00000 |
| 130876747 | 130876810 | -1 | miRNA | ENSG00000 |
| 130876809 | 130913310 | -1 | lncRNA | ENSG00000 |
| 130877459 | 130877539 | -1 | miRNA | ENSG00000 |
| 130930209 | 130932206 | -1 | lncRNA | ENSG00000 |
| 130936464 | 130939661 | 1 | lncRNA | ENSG00000 |
| 130938963 | 131110176 | -1 | lncRNA | ENSG00000 |
| 131054886 | 131054992 | -1 | snRNA | ENSG00000 |
| 131110096 | 131496632 | 1 | protein_coding | ENSG00000 |
| 131242590 | 131242995 | 1 | processed_pseudoge | ENSG00000 |
| 131309469 | 131328312 | -1 | lncRNA | ENSG00000 |
| 131493964 | 131497694 | -1 | lncRNA | ENSG00000 |
| 131500262 | 131558217 | -1 | protein_coding | ENSG00000 |
| 131520137 | 131520503 | 1 | processed_pseudoge | ENSG00000 |

| | | | | |
|---|---|---|---|---|
| 131661952 | 131662624 | -1 | processed_pseudoge | ENSG00000 |
| 131665507 | 131665858 | 1 | processed_pseudoge | ENSG00000 |
| 131702269 | 131702653 | -1 | processed_pseudoge | ENSG00000 |
| 131753746 | 131754252 | -1 | processed_pseudoge | ENSG00000 |
| 131892616 | 131893615 | -1 | processed_pseudoge | ENSG00000 |
| 131893822 | 131894074 | 1 | processed_pseudoge | ENSG00000 |
| 131897289 | 131948953 | 1 | lncRNA | ENSG00000 |
| 132086266 | 132087992 | 1 | processed_pseudoge | ENSG00000 |
| 132123332 | 132648688 | -1 | protein_coding | ENSG00000 |
| 132264152 | 132271269 | 1 | lncRNA | ENSG00000 |
| 132352334 | 132367976 | 1 | lncRNA | ENSG00000 |
| 132648794 | 132728769 | 1 | lncRNA | ENSG00000 |
| 132753023 | 132753126 | 1 | snoRNA | ENSG00000 |
| 132758970 | 132760632 | 1 | lncRNA | ENSG00000 |
| 132784870 | 133082090 | -1 | protein_coding | ENSG00000 |
| 132830693 | 132849593 | 1 | lncRNA | ENSG00000 |
| 133034607 | 133035920 | -1 | processed_pseudoge | ENSG00000 |
| 133034860 | 133034915 | -1 | miRNA | ENSG00000 |
| 133082829 | 133082932 | 1 | snRNA | ENSG00000 |
| 133169416 | 133170514 | -1 | processed_pseudoge | ENSG00000 |
| 133253073 | 134066589 | 1 | protein_coding | ENSG00000 |
| 133290881 | 133290988 | 1 | miRNA | ENSG00000 |
| 133315009 | 133315413 | -1 | processed_pseudoge | ENSG00000 |
| 133727368 | 133727734 | -1 | processed_pseudoge | ENSG00000 |
| 133732493 | 133733595 | -1 | processed_pseudoge | ENSG00000 |
| 134127299 | 134264591 | 1 | protein_coding | ENSG00000 |
| 134284500 | 134286055 | 1 | lncRNA | ENSG00000 |
| 134289332 | 134316930 | -1 | protein_coding | ENSG00000 |
| 134346071 | 134350791 | -1 | lncRNA | ENSG00000 |
| 134368737 | 134432534 | -1 | lncRNA | ENSG00000 |
| 134442356 | 134459284 | -1 | protein_coding | ENSG00000 |
| 134527567 | 134541412 | 1 | protein_coding | ENSG00000 |
| 134549110 | 134579875 | 1 | protein_coding | ENSG00000 |
| 134646811 | 134679816 | 1 | protein_coding | ENSG00000 |
| 134684144 | 134687990 | 1 | lncRNA | ENSG00000 |
| 134734898 | 134735129 | -1 | processed_pseudoge | ENSG00000 |
| 134744252 | 134970729 | 1 | protein_coding | ENSG00000 |
| 134816543 | 134998153 | -1 | lncRNA | ENSG00000 |
| 134986508 | 135147963 | 1 | protein_coding | ENSG00000 |
| 135092363 | 135170795 | -1 | protein_coding | ENSG00000 |
| 135128444 | 135129685 | 1 | processed_pseudoge | ENSG00000 |
| 135148072 | 135166215 | 1 | protein_coding | ENSG00000 |
| 135168403 | 135169547 | 1 | lncRNA | ENSG00000 |
| 135170816 | 135172414 | 1 | lncRNA | ENSG00000 |
| 135183839 | 135211534 | -1 | protein_coding | ENSG00000 |
| 135198401 | 135209837 | 1 | lncRNA | ENSG00000 |
| 135206994 | 135207078 | -1 | miRNA | ENSG00000 |
| 135231979 | 135258492 | 1 | protein_coding | ENSG00000 |
| 135246113 | 135317752 | -1 | transcribed_unproce | ENSG00000 |
| 135361795 | 135510127 | -1 | protein_coding | ENSG00000 |

| | | | | |
|---|---|---|---|---|
| 135444461 | 135444925 | 1 | processed_pseudoge | ENSG00000( |
| 135557917 | 135648757 | 1 | protein_coding | ENSG00000( |
| 135650972 | 135651416 | 1 | processed_pseudoge | ENSG00000( |
| 135660039 | 135660647 | 1 | processed_pseudoge | ENSG00000( |
| 135662496 | 135693418 | 1 | protein_coding | ENSG00000( |
| 135665599 | 135665702 | -1 | snRNA | ENSG00000( |
| 135681237 | 135729258 | -1 | protein_coding | ENSG00000( |
| 135704537 | 135704841 | 1 | lncRNA | ENSG00000( |
| 135728348 | 135748813 | -1 | protein_coding | ENSG00000( |
| 135774521 | 135932997 | 1 | lncRNA | ENSG00000( |
| 135926760 | 135977359 | -1 | protein_coding | ENSG00000( |
| 135927274 | 135927450 | -1 | protein_coding | ENSG00000( |
| 135960703 | 135960809 | 1 | snRNA | ENSG00000( |
| 136025717 | 136084668 | 1 | lncRNA | ENSG00000( |
| 136092913 | 136437785 | 1 | lncRNA | ENSG00000( |
| 136129980 | 136130081 | -1 | misc_RNA | ENSG00000( |
| 136262558 | 136263017 | -1 | processed_pseudoge | ENSG00000( |
| 136485656 | 136488317 | 1 | processed_pseudoge | ENSG00000( |
| 136685559 | 137182107 | -1 | lncRNA | ENSG00000( |
| 136713871 | 136715188 | 1 | processed_pseudoge | ENSG00000( |
| 136868669 | 137020255 | 1 | protein_coding | ENSG00000( |
| 136903167 | 136903294 | 1 | miRNA | ENSG00000( |
| 136938280 | 136938553 | -1 | processed_pseudoge | ENSG00000( |
| 137164043 | 137164612 | 1 | processed_pseudoge | ENSG00000( |
| 137227341 | 137343774 | -1 | protein_coding | ENSG00000( |
| 137287687 | 137287762 | 1 | snoRNA | ENSG00000( |
| 137318592 | 137326953 | 1 | lncRNA | ENSG00000( |
| 137344930 | 137354483 | -1 | lncRNA | ENSG00000( |
| 137381037 | 137847092 | -1 | protein_coding | ENSG00000( |
| 137513859 | 137513933 | 1 | processed_pseudoge | ENSG00000( |
| 137721985 | 137722846 | 1 | processed_pseudoge | ENSG00000( |
| 137874979 | 138002086 | -1 | protein_coding | ENSG00000( |
| 137953348 | 137957966 | 1 | lncRNA | ENSG00000( |
| 138002324 | 138118305 | 1 | protein_coding | ENSG00000( |
| 138022458 | 138024339 | 1 | processed_pseudoge | ENSG00000( |
| 138046654 | 138047897 | 1 | processed_pseudoge | ENSG00000( |
| 138060056 | 138060293 | 1 | processed_pseudoge | ENSG00000( |
| 138091254 | 138091315 | 1 | misc_RNA | ENSG00000( |
| 138122202 | 138124595 | 1 | processed_pseudoge | ENSG00000( |
| 138123758 | 138123821 | 1 | miRNA | ENSG00000( |
| 138163440 | 138164637 | 1 | lncRNA | ENSG00000( |
| 138175045 | 138175136 | 1 | misc_RNA | ENSG00000( |
| 138187998 | 138188129 | -1 | snoRNA | ENSG00000( |
| 138298088 | 138298484 | -1 | processed_pseudoge | ENSG00000( |
| 138404195 | 138404523 | -1 | processed_pseudoge | ENSG00000( |
| 138440690 | 138442238 | 1 | processed_pseudoge | ENSG00000( |
| 138460259 | 138589996 | 1 | protein_coding | ENSG00000( |
| 138490581 | 138491361 | -1 | processed_pseudoge | ENSG00000( |
| 138594285 | 138701352 | -1 | protein_coding | ENSG00000( |
| 138625060 | 138625179 | -1 | snoRNA | ENSG00000( |

| | | | | |
|---|---|---|---|---|
| 138645671 | 138646448 | -1 | processed_pseudoge | ENSG00000( |
| 138688980 | 138689522 | 1 | processed_pseudoge | ENSG00000( |
| 138701607 | 138703062 | 1 | processed_pseudoge | ENSG00000( |
| 138706294 | 138799560 | -1 | protein_coding | ENSG00000( |
| 138797952 | 138838101 | 1 | protein_coding | ENSG00000( |
| 138831381 | 138981318 | -1 | protein_coding | ENSG00000( |
| 138876359 | 138876465 | -1 | snRNA | ENSG00000( |
| 139025706 | 139036042 | -1 | protein_coding | ENSG00000( |
| 139043515 | 139109720 | -1 | protein_coding | ENSG00000( |
| 139049456 | 139050006 | -1 | processed_pseudoge | ENSG00000( |
| 139133744 | 139191986 | 1 | protein_coding | ENSG00000( |
| 139172516 | 139174266 | -1 | processed_pseudoge | ENSG00000( |
| 139198557 | 139199142 | -1 | processed_pseudoge | ENSG00000( |
| 139227537 | 139227828 | -1 | processed_pseudoge | ENSG00000( |
| 139228221 | 139228605 | 1 | processed_pseudoge | ENSG00000( |
| 139230356 | 139308236 | 1 | protein_coding | ENSG00000( |
| 139315291 | 139315394 | -1 | snRNA | ENSG00000( |
| 139339457 | 139346328 | 1 | protein_coding | ENSG00000( |
| 139340359 | 139423457 | 1 | protein_coding | ENSG00000( |
| 139341360 | 139422599 | 1 | protein_coding | ENSG00000( |
| 139359032 | 139359566 | -1 | lncRNA | ENSG00000( |
| 139448740 | 139448843 | 1 | snRNA | ENSG00000( |
| 139452690 | 139483673 | -1 | protein_coding | ENSG00000( |
| 139502453 | 139503812 | 1 | processed_pseudoge | ENSG00000( |
| 139523685 | 139544985 | 1 | protein_coding | ENSG00000( |
| 139534240 | 139534549 | 1 | processed_pseudoge | ENSG00000( |
| 139561570 | 139777998 | -1 | protein_coding | ENSG00000( |
| 139777051 | 140020325 | 1 | protein_coding | ENSG00000( |
| 140023744 | 140063721 | -1 | protein_coding | ENSG00000( |
| 140084746 | 140176983 | -1 | protein_coding | ENSG00000( |
| 140094697 | 140094792 | -1 | misc_RNA | ENSG00000( |
| 140177184 | 140179640 | 1 | lncRNA | ENSG00000( |
| 140209563 | 140209671 | -1 | snRNA | ENSG00000( |
| 140241870 | 140242032 | 1 | snRNA | ENSG00000( |
| 140282465 | 140282920 | 1 | processed_pseudoge | ENSG00000( |
| 140292752 | 140293339 | 1 | processed_pseudoge | ENSG00000( |
| 140293693 | 140404433 | -1 | protein_coding | ENSG00000( |
| 140370441 | 140370545 | 1 | rRNA_pseudogene | ENSG00000( |
| 140386781 | 140386907 | -1 | rRNA_pseudogene | ENSG00000( |
| 140404043 | 140426250 | 1 | protein_coding | ENSG00000( |
| 140435316 | 140435787 | -1 | processed_pseudoge | ENSG00000( |
| 140453033 | 140479536 | -1 | protein_coding | ENSG00000( |
| 140518420 | 140673993 | -1 | protein_coding | ENSG00000( |
| 140609847 | 140609955 | 1 | misc_RNA | ENSG00000( |
| 140640909 | 140669476 | 1 | lncRNA | ENSG00000( |
| 140645829 | 140646126 | 1 | misc_RNA | ENSG00000( |
| 140672945 | 140696261 | 1 | protein_coding | ENSG00000( |
| 140690777 | 140722790 | 1 | protein_coding | ENSG00000( |
| 140695336 | 140697077 | -1 | lncRNA | ENSG00000( |
| 140719327 | 140924928 | -1 | protein_coding | ENSG00000( |

| | | | | |
|---|---|---|---|---|
| 140884072 | 140884178 | 1 | snRNA | ENSG0000C |
| 140934867 | 140935010 | 1 | processed_pseudoge | ENSG0000C |
| 140997952 | 140999800 | -1 | processed_pseudoge | ENSG0000C |
| 141002610 | 141015228 | -1 | protein_coding | ENSG0000C |
| 141052249 | 141052409 | 1 | snRNA | ENSG0000C |
| 141074064 | 141480380 | 1 | protein_coding | ENSG0000C |
| 141173043 | 141173216 | -1 | processed_pseudoge | ENSG0000C |
| 141351977 | 141352768 | 1 | processed_pseudoge | ENSG0000C |
| 141392155 | 141396517 | -1 | lncRNA | ENSG0000C |
| 141414383 | 141416390 | -1 | lncRNA | ENSG0000C |
| 141429711 | 141431268 | -1 | lncRNA | ENSG0000C |
| 141500079 | 141551304 | -1 | lncRNA | ENSG0000C |
| 141512698 | 141513183 | -1 | lncRNA | ENSG0000C |
| 141551278 | 141655244 | 1 | protein_coding | ENSG0000C |
| 141652381 | 141656810 | -1 | lncRNA | ENSG0000C |
| 141656728 | 141702166 | -1 | protein_coding | ENSG0000C |
| 141662922 | 141663846 | -1 | lncRNA | ENSG0000C |
| 141704003 | 141738346 | -1 | lncRNA | ENSG0000C |
| 141708353 | 141731271 | 1 | protein_coding | ENSG0000C |
| 141727984 | 141728148 | 1 | snRNA | ENSG0000C |
| 141738334 | 141787922 | 1 | protein_coding | ENSG0000C |
| 141764097 | 141765197 | 1 | protein_coding | ENSG0000C |
| 141778442 | 141780819 | 1 | protein_coding | ENSG0000C |
| 141787815 | 141788640 | 1 | processed_pseudoge | ENSG0000C |
| 141790217 | 141791367 | 1 | protein_coding | ENSG0000C |
| 141801315 | 141802059 | -1 | processed_pseudoge | ENSG0000C |
| 141802482 | 141803487 | -1 | processed_pseudoge | ENSG0000C |
| 141803701 | 141804652 | -1 | processed_pseudoge | ENSG0000C |
| 141811805 | 141812257 | 1 | processed_pseudoge | ENSG0000C |
| 141836286 | 141841487 | -1 | protein_coding | ENSG0000C |
| 141862860 | 141863737 | 1 | transcribed_unproce | ENSG0000C |
| 141887148 | 141888092 | 1 | transcribed_unproce | ENSG0000C |
| 141907813 | 142106747 | 1 | protein_coding | ENSG0000C |
| 141911217 | 141911943 | 1 | transcribed_unproce | ENSG0000C |
| 141916399 | 141920625 | 1 | protein_coding | ENSG0000C |
| 141927357 | 141947007 | -1 | protein_coding | ENSG0000C |
| 141972631 | 141973773 | -1 | protein_coding | ENSG0000C |
| 142111749 | 142222324 | 1 | protein_coding | ENSG0000C |
| 142240740 | 142247067 | -1 | transcribed_unproce | ENSG0000C |
| 142252143 | 142258058 | -1 | protein_coding | ENSG0000C |
| 142265833 | 142272235 | -1 | transcribed_unitary_ | ENSG0000C |
| 142285750 | 142288869 | 1 | lncRNA | ENSG0000C |
| 142287251 | 142291597 | -1 | transcribed_unitary_ | ENSG0000C |
| 142299177 | 142299460 | 1 | TR_V_pseudogene | ENSG0000C |
| 142300924 | 142301432 | 1 | TR_V_gene | ENSG0000C |
| 142308542 | 142309048 | 1 | TR_V_gene | ENSG0000C |
| 142313184 | 142313666 | 1 | TR_V_gene | ENSG0000C |
| 142320677 | 142321544 | 1 | TR_V_gene | ENSG0000C |
| 142328297 | 142328786 | 1 | TR_V_gene | ENSG0000C |
| 142332182 | 142332701 | 1 | TR_V_gene | ENSG0000C |

| | | | | |
|---|---|---|---|---|
| 142345421 | 142345985 | 1 | TR_V_gene | ENSG0000C |
| 142349152 | 142349664 | 1 | TR_V_gene | ENSG0000C |
| 142352819 | 142353358 | 1 | TR_V_gene | ENSG0000C |
| 142358639 | 142358917 | 1 | TR_V_pseudogene | ENSG0000C |
| 142372640 | 142372912 | 1 | TR_V_pseudogene | ENSG0000C |
| 142380806 | 142381261 | 1 | TR_V_gene | ENSG0000C |
| 142384329 | 142384841 | 1 | TR_V_gene | ENSG0000C |
| 142386378 | 142386647 | 1 | TR_V_pseudogene | ENSG0000C |
| 142389202 | 142389668 | 1 | TR_V_gene | ENSG0000C |
| 142391891 | 142392412 | 1 | TR_V_gene | ENSG0000C |
| 142399860 | 142400377 | 1 | TR_V_gene | ENSG0000C |
| 142407672 | 142408136 | 1 | TR_V_gene | ENSG0000C |
| 142415224 | 142415666 | 1 | TR_V_pseudogene | ENSG0000C |
| 142424965 | 142425465 | 1 | TR_V_gene | ENSG0000C |
| 142433895 | 142434394 | 1 | lncRNA | ENSG0000C |
| 142440883 | 142441325 | 1 | TR_V_pseudogene | ENSG0000C |
| 142450947 | 142451448 | 1 | TR_V_gene | ENSG0000C |
| 142455174 | 142455635 | 1 | TR_V_gene | ENSG0000C |
| 142462916 | 142463581 | 1 | TR_V_gene | ENSG0000C |
| 142469537 | 142470013 | 1 | TR_V_gene | ENSG0000C |
| 142474096 | 142474567 | 1 | TR_V_pseudogene | ENSG0000C |
| 142482548 | 142483019 | 1 | TR_V_gene | ENSG0000C |
| 142487863 | 142488295 | 1 | TR_V_gene | ENSG0000C |
| 142492132 | 142492673 | 1 | TR_V_gene | ENSG0000C |
| 142500028 | 142500534 | 1 | TR_V_gene | ENSG0000C |
| 142507382 | 142507810 | 1 | TR_V_gene | ENSG0000C |
| 142511626 | 142512127 | 1 | TR_V_gene | ENSG0000C |
| 142520090 | 142520556 | 1 | TR_V_gene | ENSG0000C |
| 142529290 | 142529762 | 1 | TR_V_gene | ENSG0000C |
| 142535809 | 142536292 | 1 | TR_V_gene | ENSG0000C |
| 142544212 | 142544685 | 1 | TR_V_gene | ENSG0000C |
| 142554836 | 142555318 | 1 | TR_V_gene | ENSG0000C |
| 142560423 | 142560931 | 1 | TR_V_gene | ENSG0000C |
| 142563740 | 142564245 | 1 | TR_V_gene | ENSG0000C |
| 142580917 | 142581427 | 1 | TR_V_gene | ENSG0000C |
| 142587868 | 142588359 | 1 | TR_V_gene | ENSG0000C |
| 142592928 | 142593473 | 1 | TR_V_gene | ENSG0000C |
| 142598016 | 142598469 | 1 | TR_V_gene | ENSG0000C |
| 142601628 | 142602360 | 1 | TR_V_gene | ENSG0000C |
| 142615716 | 142616415 | 1 | TR_V_gene | ENSG0000C |
| 142618849 | 142619532 | 1 | TR_V_gene | ENSG0000C |
| 142626649 | 142627399 | 1 | TR_V_gene | ENSG0000C |
| 142636924 | 142637384 | 1 | TR_V_pseudogene | ENSG0000C |
| 142641746 | 142642196 | 1 | TR_V_pseudogene | ENSG0000C |
| 142645961 | 142646467 | 1 | TR_V_gene | ENSG0000C |
| 142656701 | 142657213 | 1 | TR_V_gene | ENSG0000C |
| 142666272 | 142667718 | 1 | protein_coding | ENSG0000C |
| 142670740 | 142671244 | 1 | TR_V_gene | ENSG0000C |
| 142681415 | 142681869 | 1 | TR_V_pseudogene | ENSG0000C |
| 142695699 | 142696183 | 1 | TR_V_pseudogene | ENSG0000C |

| | | | | |
|---|---|---|---|---|
| 142711384 | 142711924 | 1 | TR_V_pseudogene | ENSG00000 |
| 142715346 | 142715861 | 1 | TR_V_gene | ENSG00000 |
| 142720660 | 142721160 | 1 | TR_V_gene | ENSG00000 |
| 142722358 | 142722764 | 1 | processed_pseudoge | ENSG00000 |
| 142725729 | 142733437 | -1 | lncRNA | ENSG00000 |
| 142740206 | 142740894 | 1 | TR_V_gene | ENSG00000 |
| 142749468 | 142753076 | 1 | protein_coding | ENSG00000 |
| 142760398 | 142774564 | 1 | protein_coding | ENSG00000 |
| 142760415 | 142763943 | 1 | unprocessed_pseudc | ENSG00000 |
| 142778642 | 142779537 | -1 | processed_pseudoge | ENSG00000 |
| 142786213 | 142786224 | 1 | TR_D_gene | ENSG00000 |
| 142786880 | 142786927 | 1 | TR_J_gene | ENSG00000 |
| 142787017 | 142787064 | 1 | TR_J_gene | ENSG00000 |
| 142787630 | 142787679 | 1 | TR_J_gene | ENSG00000 |
| 142788225 | 142788275 | 1 | TR_J_gene | ENSG00000 |
| 142788498 | 142788547 | 1 | TR_J_gene | ENSG00000 |
| 142788988 | 142789040 | 1 | TR_J_gene | ENSG00000 |
| 142791694 | 142793368 | 1 | TR_C_gene | ENSG00000 |
| 142796365 | 142796414 | 1 | TR_J_gene | ENSG00000 |
| 142796560 | 142796610 | 1 | TR_J_gene | ENSG00000 |
| 142796697 | 142796742 | 1 | TR_J_gene | ENSG00000 |
| 142796847 | 142796895 | 1 | TR_J_gene | ENSG00000 |
| 142796998 | 142797047 | 1 | TR_J_gene | ENSG00000 |
| 142797119 | 142797166 | 1 | TR_J_gene | ENSG00000 |
| 142797239 | 142797291 | 1 | TR_J_gene | ENSG00000 |
| 142797456 | 142797502 | 1 | TR_J_gene | ENSG00000 |
| 142801041 | 142802748 | 1 | TR_C_gene | ENSG00000 |
| 142812586 | 142813399 | -1 | TR_V_gene | ENSG00000 |
| 142855061 | 142871094 | 1 | protein_coding | ENSG00000 |
| 142871208 | 142885745 | -1 | protein_coding | ENSG00000 |
| 142875836 | 142892743 | 1 | lncRNA | ENSG00000 |
| 142899365 | 142900759 | -1 | lncRNA | ENSG00000 |
| 142908101 | 142933746 | -1 | protein_coding | ENSG00000 |
| 142939343 | 142940865 | 1 | protein_coding | ENSG00000 |
| 142941114 | 142962363 | -1 | protein_coding | ENSG00000 |
| 143026158 | 143027195 | -1 | protein_coding | ENSG00000 |
| 143047213 | 143047580 | -1 | unprocessed_pseudc | ENSG00000 |
| 143052320 | 143053347 | 1 | protein_coding | ENSG00000 |
| 143062330 | 143063585 | -1 | transcribed_unproce | ENSG00000 |
| 143132077 | 143139739 | 1 | protein_coding | ENSG00000 |
| 143183419 | 143184435 | 1 | protein_coding | ENSG00000 |
| 143220468 | 143222267 | -1 | lncRNA | ENSG00000 |
| 143222037 | 143223079 | 1 | protein_coding | ENSG00000 |
| 143224042 | 143224234 | -1 | TEC | ENSG00000 |
| 143244093 | 143270854 | 1 | protein_coding | ENSG00000 |
| 143255264 | 143287997 | -1 | lncRNA | ENSG00000 |
| 143279957 | 143288048 | 1 | protein_coding | ENSG00000 |
| 143288215 | 143307696 | 1 | protein_coding | ENSG00000 |
| 143290615 | 143290910 | 1 | misc_RNA | ENSG00000 |
| 143298516 | 143298814 | 1 | misc_RNA | ENSG00000 |

| | | | | |
|---|---|---|---|---|
| 143312684 | 143313061 | -1 | processed_pseudoge | ENSG00000( |
| 143316111 | 143352083 | 1 | protein_coding | ENSG00000( |
| 143353400 | 143362770 | -1 | protein_coding | ENSG00000( |
| 143363899 | 143364229 | 1 | lncRNA | ENSG00000( |
| 143379692 | 143380495 | -1 | lncRNA | ENSG00000( |
| 143381295 | 143391111 | 1 | protein_coding | ENSG00000( |
| 143382686 | 143382800 | 1 | miRNA | ENSG00000( |
| 143390289 | 143408856 | -1 | protein_coding | ENSG00000( |
| 143407813 | 143523449 | 1 | lncRNA | ENSG00000( |
| 143437034 | 143437973 | 1 | unitary_pseudogene | ENSG00000( |
| 143443453 | 143444409 | 1 | protein_coding | ENSG00000( |
| 143477873 | 143478796 | 1 | protein_coding | ENSG00000( |
| 143488462 | 143489402 | -1 | unitary_pseudogene | ENSG00000( |
| 143509256 | 143519469 | -1 | polymorphic_pseudo | ENSG00000( |
| 143532749 | 143533164 | -1 | processed_pseudoge | ENSG00000( |
| 143545186 | 143546735 | -1 | processed_pseudoge | ENSG00000( |
| 143571801 | 143574387 | 1 | protein_coding | ENSG00000( |
| 143574746 | 143574848 | 1 | snRNA | ENSG00000( |
| 143578907 | 143579072 | 1 | processed_pseudoge | ENSG00000( |
| 143598039 | 143604839 | 1 | unprocessed_pseudo | ENSG00000( |
| 143620943 | 143645675 | 1 | transcribed_unproce | ENSG00000( |
| 143620952 | 143730409 | 1 | protein_coding | ENSG00000( |
| 143639230 | 143647646 | -1 | protein_coding | ENSG00000( |
| 143754628 | 143754730 | -1 | snRNA | ENSG00000( |
| 143755089 | 143757696 | -1 | protein_coding | ENSG00000( |
| 143761790 | 143836933 | -1 | transcribed_unproce | ENSG00000( |
| 143782667 | 143784284 | 1 | processed_pseudoge | ENSG00000( |
| 143796295 | 143796720 | 1 | processed_pseudoge | ENSG00000( |
| 143800732 | 143817973 | 1 | transcribed_unproce | ENSG00000( |
| 143851375 | 143902198 | -1 | protein_coding | ENSG00000( |
| 143935166 | 143936279 | 1 | protein_coding | ENSG00000( |
| 143954844 | 143997312 | 1 | protein_coding | ENSG00000( |
| 143979121 | 143980225 | -1 | processed_pseudoge | ENSG00000( |
| 143980905 | 143981820 | 1 | unprocessed_pseudo | ENSG00000( |
| 143985142 | 143986651 | 1 | processed_pseudoge | ENSG00000( |
| 144000320 | 144008793 | 1 | protein_coding | ENSG00000( |
| 144048948 | 144058845 | 1 | protein_coding | ENSG00000( |
| 144069811 | 144075870 | 1 | protein_coding | ENSG00000( |
| 144077721 | 144077825 | 1 | unprocessed_pseudo | ENSG00000( |
| 144086278 | 144098953 | 1 | protein_coding | ENSG00000( |
| 144109514 | 144110564 | 1 | protein_coding | ENSG00000( |
| 144118461 | 144119360 | -1 | unprocessed_pseudo | ENSG00000( |
| 144123176 | 144131188 | 1 | protein_coding | ENSG00000( |
| 144142009 | 144142938 | 1 | unprocessed_pseudo | ENSG00000( |
| 144153486 | 144158905 | 1 | transcribed_unproce | ENSG00000( |
| 144176740 | 144177142 | 1 | unprocessed_pseudo | ENSG00000( |
| 144183466 | 144186053 | 1 | protein_coding | ENSG00000( |
| 144186083 | 144195655 | -1 | protein_coding | ENSG00000( |
| 144194858 | 144280547 | 1 | lncRNA | ENSG00000( |
| 144228244 | 144239605 | -1 | protein_coding | ENSG00000( |

| | | | | |
|---|---|---|---|---|
| 144250045 | 144256244 | -1 | transcribed_unproce | ENSG00000 |
| 144251264 | 144355707 | -1 | lncRNA | ENSG00000 |
| 144257663 | 144264792 | -1 | protein_coding | ENSG00000 |
| 144266701 | 144269288 | -1 | protein_coding | ENSG00000 |
| 144272445 | 144286966 | -1 | unprocessed_pseudo | ENSG00000 |
| 144294480 | 144300934 | 1 | transcribed_unproce | ENSG00000 |
| 144300395 | 144356181 | -1 | lncRNA | ENSG00000 |
| 144312464 | 144322668 | 1 | protein_coding | ENSG00000 |
| 144355288 | 144380632 | 1 | protein_coding | ENSG00000 |
| 144397240 | 144410227 | -1 | protein_coding | ENSG00000 |
| 144434826 | 144435314 | -1 | processed_pseudoge | ENSG00000 |
| 144451681 | 144451817 | 1 | snRNA | ENSG00000 |
| 144451941 | 144836395 | -1 | protein_coding | ENSG00000 |
| 144647186 | 144648575 | -1 | processed_pseudoge | ENSG00000 |
| 144849755 | 144850084 | -1 | misc_RNA | ENSG00000 |
| 145005058 | 145006135 | 1 | processed_pseudoge | ENSG00000 |
| 145009961 | 145010993 | 1 | processed_pseudoge | ENSG00000 |
| 145039999 | 145040941 | -1 | processed_pseudoge | ENSG00000 |
| 145269514 | 145270384 | 1 | processed_pseudoge | ENSG00000 |
| 145583197 | 145584817 | 1 | processed_pseudoge | ENSG00000 |
| 145675679 | 145681369 | -1 | lncRNA | ENSG00000 |
| 146050062 | 146050366 | 1 | TEC | ENSG00000 |
| 146116002 | 148420998 | 1 | protein_coding | ENSG00000 |
| 146208665 | 146208819 | -1 | misc_RNA | ENSG00000 |
| 146311720 | 146311822 | 1 | misc_RNA | ENSG00000 |
| 147080934 | 147097609 | -1 | lncRNA | ENSG00000 |
| 147146342 | 147146756 | 1 | transcribed_processe | ENSG00000 |
| 147167344 | 147167572 | 1 | processed_pseudoge | ENSG00000 |
| 147378017 | 147378121 | -1 | miRNA | ENSG00000 |
| 147671711 | 147673143 | 1 | lncRNA | ENSG00000 |
| 147704294 | 147704575 | -1 | processed_pseudoge | ENSG00000 |
| 147831770 | 147831877 | 1 | snRNA | ENSG00000 |
| 147849884 | 147849982 | 1 | rRNA_pseudogene | ENSG00000 |
| 147940007 | 147940301 | 1 | misc_RNA | ENSG00000 |
| 148389645 | 148389859 | 1 | snoRNA | ENSG00000 |
| 148438309 | 148438577 | 1 | misc_RNA | ENSG00000 |
| 148445786 | 148446299 | 1 | processed_pseudoge | ENSG00000 |
| 148473599 | 148503458 | -1 | lncRNA | ENSG00000 |
| 148543677 | 148572177 | -1 | lncRNA | ENSG00000 |
| 148580401 | 148580799 | 1 | processed_pseudoge | ENSG00000 |
| 148584818 | 148625479 | -1 | lncRNA | ENSG00000 |
| 148590766 | 148615860 | 1 | protein_coding | ENSG00000 |
| 148637179 | 148637587 | 1 | processed_pseudoge | ENSG00000 |
| 148649547 | 148650240 | 1 | processed_pseudoge | ENSG00000 |
| 148696467 | 148698664 | -1 | lncRNA | ENSG00000 |
| 148697914 | 148801110 | 1 | protein_coding | ENSG00000 |
| 148807383 | 148884321 | -1 | protein_coding | ENSG00000 |
| 148821012 | 148821073 | 1 | snRNA | ENSG00000 |
| 148876018 | 148876286 | 1 | misc_RNA | ENSG00000 |
| 148890226 | 148890513 | 1 | misc_RNA | ENSG00000 |

| | | | | |
|---|---|---|---|---|
| 148940265 | 148941228 | -1 | lncRNA | ENSG00000 |
| 148941483 | 148941638 | 1 | lncRNA | ENSG00000 |
| 148963315 | 148963410 | 1 | misc_RNA | ENSG00000 |
| 148983755 | 148983856 | 1 | misc_RNA | ENSG00000 |
| 148987136 | 148987248 | -1 | misc_RNA | ENSG00000 |
| 148987527 | 148989432 | 1 | lncRNA | ENSG00000 |
| 149003062 | 149028662 | -1 | protein_coding | ENSG00000 |
| 149033057 | 149033167 | 1 | snRNA | ENSG00000 |
| 149053961 | 149054173 | 1 | processed_pseudoge | ENSG00000 |
| 149069641 | 149090782 | -1 | protein_coding | ENSG00000 |
| 149102784 | 149126324 | -1 | protein_coding | ENSG00000 |
| 149125690 | 149125986 | -1 | misc_RNA | ENSG00000 |
| 149126416 | 149183042 | 1 | protein_coding | ENSG00000 |
| 149191043 | 149191223 | -1 | processed_pseudoge | ENSG00000 |
| 149195546 | 149226238 | 1 | protein_coding | ENSG00000 |
| 149239651 | 149255606 | 1 | protein_coding | ENSG00000 |
| 149262171 | 149297302 | 1 | protein_coding | ENSG00000 |
| 149275619 | 149277992 | -1 | unprocessed_pseudo | ENSG00000 |
| 149285281 | 149297312 | 1 | transcribed_unproce | ENSG00000 |
| 149321865 | 149322092 | -1 | processed_pseudoge | ENSG00000 |
| 149334024 | 149334857 | 1 | processed_pseudoge | ENSG00000 |
| 149398204 | 149401456 | 1 | lncRNA | ENSG00000 |
| 149422675 | 149424890 | 1 | lncRNA | ENSG00000 |
| 149431363 | 149461062 | -1 | protein_coding | ENSG00000 |
| 149472696 | 149497817 | -1 | protein_coding | ENSG00000 |
| 149547154 | 149624752 | -1 | transcribed_unproce | ENSG00000 |
| 149714781 | 149734575 | 1 | protein_coding | ENSG00000 |
| 149764182 | 149773588 | -1 | protein_coding | ENSG00000 |
| 149776042 | 149833979 | 1 | protein_coding | ENSG00000 |
| 149838375 | 149867479 | 1 | protein_coding | ENSG00000 |
| 149851572 | 149852170 | 1 | TEC | ENSG00000 |
| 149858400 | 149862492 | -1 | lncRNA | ENSG00000 |
| 149867697 | 149880610 | -1 | lncRNA | ENSG00000 |
| 149872968 | 149891204 | 1 | protein_coding | ENSG00000 |
| 149881359 | 149881580 | -1 | lncRNA | ENSG00000 |
| 149881477 | 149882105 | 1 | TEC | ENSG00000 |
| 149890739 | 149891416 | 1 | lncRNA | ENSG00000 |
| 149891191 | 149909704 | -1 | unprocessed_pseudo | ENSG00000 |
| 150000752 | 150005124 | 1 | lncRNA | ENSG00000 |
| 150033653 | 150076655 | 1 | lncRNA | ENSG00000 |
| 150047609 | 150047854 | -1 | unprocessed_pseudo | ENSG00000 |
| 150234194 | 150237608 | -1 | lncRNA | ENSG00000 |
| 150243916 | 150323725 | -1 | protein_coding | ENSG00000 |
| 150322639 | 150332721 | 1 | protein_coding | ENSG00000 |
| 150323263 | 150338156 | 1 | protein_coding | ENSG00000 |
| 150337483 | 150343346 | 1 | lncRNA | ENSG00000 |
| 150338317 | 150341662 | -1 | protein_coding | ENSG00000 |
| 150363777 | 150372590 | -1 | lncRNA | ENSG00000 |
| 150368189 | 150374044 | 1 | protein_coding | ENSG00000 |
| 150368790 | 150398630 | 1 | protein_coding | ENSG00000 |

| | | | | |
|---|---|---|---|---|
| 150379329 | 150410597 | 1 | lncRNA | ENSG0000C |
| 150379854 | 150383885 | -1 | lncRNA | ENSG0000C |
| 150400702 | 150412470 | 1 | protein_coding | ENSG0000C |
| 150433654 | 150448140 | 1 | lncRNA | ENSG0000C |
| 150450630 | 150479393 | 1 | protein_coding | ENSG0000C |
| 150484061 | 150485135 | -1 | processed_pseudoge | ENSG0000C |
| 150512559 | 150513839 | -1 | processed_pseudoge | ENSG0000C |
| 150514872 | 150521073 | 1 | protein_coding | ENSG0000C |
| 150541754 | 150542098 | -1 | processed_pseudoge | ENSG0000C |
| 150567369 | 150573953 | 1 | protein_coding | ENSG0000C |
| 150603561 | 150604644 | -1 | processed_pseudoge | ENSG0000C |
| 150614061 | 150614764 | -1 | processed_pseudoge | ENSG0000C |
| 150625375 | 150632648 | -1 | protein_coding | ENSG0000C |
| 150676833 | 150677918 | -1 | processed_pseudoge | ENSG0000C |
| 150685697 | 150693641 | 1 | protein_coding | ENSG0000C |
| 150716606 | 150724284 | 1 | protein_coding | ENSG0000C |
| 150716668 | 150743646 | 1 | protein_coding | ENSG0000C |
| 150722253 | 150750033 | 1 | protein_coding | ENSG0000C |
| 150746893 | 150747728 | -1 | processed_pseudoge | ENSG0000C |
| 150749736 | 150750094 | 1 | processed_pseudoge | ENSG0000C |
| 150791285 | 150801360 | -1 | protein_coding | ENSG0000C |
| 150800403 | 150805118 | 1 | protein_coding | ENSG0000C |
| 150824627 | 150861504 | 1 | protein_coding | ENSG0000C |
| 150944961 | 150978321 | -1 | protein_coding | ENSG0000C |
| 150991017 | 151014588 | 1 | protein_coding | ENSG0000C |
| 151012209 | 151024499 | -1 | protein_coding | ENSG0000C |
| 151028422 | 151047782 | 1 | protein_coding | ENSG0000C |
| 151028781 | 151029754 | -1 | lncRNA | ENSG0000C |
| 151048292 | 151052756 | 1 | protein_coding | ENSG0000C |
| 151053815 | 151057897 | -1 | protein_coding | ENSG0000C |
| 151057210 | 151076526 | 1 | protein_coding | ENSG0000C |
| 151074742 | 151076530 | -1 | lncRNA | ENSG0000C |
| 151076593 | 151080866 | -1 | protein_coding | ENSG0000C |
| 151081085 | 151083493 | -1 | protein_coding | ENSG0000C |
| 151085831 | 151144436 | 1 | protein_coding | ENSG0000C |
| 151148589 | 151174745 | -1 | protein_coding | ENSG0000C |
| 151175698 | 151187832 | -1 | protein_coding | ENSG0000C |
| 151190873 | 151205496 | -1 | protein_coding | ENSG0000C |
| 151207837 | 151210378 | -1 | protein_coding | ENSG0000C |
| 151207837 | 151227166 | -1 | protein_coding | ENSG0000C |
| 151211484 | 151227205 | -1 | protein_coding | ENSG0000C |
| 151232489 | 151238827 | 1 | protein_coding | ENSG0000C |
| 151238421 | 151238538 | 1 | miRNA | ENSG0000C |
| 151238764 | 151277896 | -1 | protein_coding | ENSG0000C |
| 151240399 | 151240972 | 1 | lncRNA | ENSG0000C |
| 151249325 | 151254390 | 1 | lncRNA | ENSG0000C |
| 151341699 | 151378449 | 1 | protein_coding | ENSG0000C |
| 151375909 | 151410727 | -1 | protein_coding | ENSG0000C |
| 151409161 | 151413354 | 1 | lncRNA | ENSG0000C |
| 151427683 | 151439124 | 1 | lncRNA | ENSG0000C |

| | | | | |
|---|---|---|---|---|
| 151428835 | 151440813 | -1 | protein_coding | ENSG00000 |
| 151433489 | 151433639 | -1 | miRNA | ENSG00000 |
| 151464650 | 151464938 | -1 | misc_RNA | ENSG00000 |
| 151466012 | 151520120 | -1 | protein_coding | ENSG00000 |
| 151501878 | 151503463 | 1 | processed_pseudoge | ENSG00000 |
| 151556124 | 151877125 | -1 | protein_coding | ENSG00000 |
| 151806490 | 151810820 | 1 | lncRNA | ENSG00000 |
| 151877042 | 151879223 | 1 | lncRNA | ENSG00000 |
| 151900275 | 151900374 | 1 | snRNA | ENSG00000 |
| 151916331 | 151916418 | 1 | misc_RNA | ENSG00000 |
| 151950386 | 151954985 | -1 | lncRNA | ENSG00000 |
| 151956379 | 152019929 | 1 | protein_coding | ENSG00000 |
| 152025194 | 152025472 | -1 | misc_RNA | ENSG00000 |
| 152025674 | 152122340 | 1 | protein_coding | ENSG00000 |
| 152120001 | 152121717 | -1 | lncRNA | ENSG00000 |
| 152128879 | 152129686 | 1 | processed_pseudoge | ENSG00000 |
| 152134922 | 152436005 | -1 | protein_coding | ENSG00000 |
| 152366763 | 152367015 | 1 | processed_pseudoge | ENSG00000 |
| 152367171 | 152367260 | 1 | processed_pseudoge | ENSG00000 |
| 152367519 | 152368661 | 1 | processed_pseudoge | ENSG00000 |
| 152402780 | 152402893 | 1 | misc_RNA | ENSG00000 |
| 152435695 | 152443176 | 1 | transcribed_processe | ENSG00000 |
| 152445477 | 152447150 | 1 | processed_pseudoge | ENSG00000 |
| 152463786 | 152465549 | 1 | lncRNA | ENSG00000 |
| 152590641 | 152590941 | -1 | unprocessed_pseudo | ENSG00000 |
| 152592973 | 152593091 | -1 | rRNA_pseudogene | ENSG00000 |
| 152592973 | 152593091 | -1 | rRNA | ENSG00000 |
| 152602213 | 152602892 | -1 | processed_pseudoge | ENSG00000 |
| 152609313 | 152609434 | 1 | snoRNA | ENSG00000 |
| 152644776 | 152676141 | -1 | protein_coding | ENSG00000 |
| 152749079 | 152749735 | 1 | processed_pseudoge | ENSG00000 |
| 152757662 | 152757951 | 1 | misc_RNA | ENSG00000 |
| 152759749 | 152855378 | 1 | protein_coding | ENSG00000 |
| 152925233 | 152936981 | 1 | lncRNA | ENSG00000 |
| 153144313 | 153144588 | -1 | processed_pseudoge | ENSG00000 |
| 153355365 | 153413985 | -1 | lncRNA | ENSG00000 |
| 153395484 | 153396921 | 1 | lncRNA | ENSG00000 |
| 153449429 | 153449875 | 1 | processed_pseudoge | ENSG00000 |
| 153715074 | 153748986 | -1 | lncRNA | ENSG00000 |
| 153887097 | 154894285 | 1 | protein_coding | ENSG00000 |
| 153968665 | 153968826 | -1 | processed_pseudoge | ENSG00000 |
| 154026287 | 154038604 | -1 | lncRNA | ENSG00000 |
| 154055286 | 154063325 | -1 | lncRNA | ENSG00000 |
| 154092201 | 154096043 | -1 | lncRNA | ENSG00000 |
| 154544169 | 154547285 | -1 | lncRNA | ENSG00000 |
| 154838388 | 154865483 | -1 | lncRNA | ENSG00000 |
| 154928460 | 154952188 | 1 | lncRNA | ENSG00000 |
| 154943687 | 155003124 | -1 | protein_coding | ENSG00000 |
| 154951226 | 154952188 | 1 | lncRNA | ENSG00000 |
| 154956429 | 154957107 | 1 | lncRNA | ENSG00000 |

| | | | | |
|---|---|---|---|---|
| 155003448 | 155005703 | 1 | lncRNA | ENSG0000C |
| 155067034 | 155072450 | -1 | lncRNA | ENSG0000C |
| 155070324 | 155087392 | 1 | protein_coding | ENSG0000C |
| 155074599 | 155074748 | 1 | unprocessed_pseudc | ENSG0000C |
| 155106062 | 155106400 | -1 | unprocessed_pseudc | ENSG0000C |
| 155106400 | 155106651 | 1 | unprocessed_pseudc | ENSG0000C |
| 155181881 | 155182184 | 1 | misc_RNA | ENSG0000C |
| 155196563 | 155198455 | -1 | lncRNA | ENSG0000C |
| 155204147 | 155205189 | 1 | lncRNA | ENSG0000C |
| 155213920 | 155217134 | 1 | lncRNA | ENSG0000C |
| 155222903 | 155223851 | 1 | TEC | ENSG0000C |
| 155267137 | 155270912 | 1 | lncRNA | ENSG0000C |
| 155295918 | 155297541 | -1 | lncRNA | ENSG0000C |
| 155297776 | 155310235 | 1 | protein_coding | ENSG0000C |
| 155356985 | 155367934 | -1 | lncRNA | ENSG0000C |
| 155382076 | 155396356 | 1 | lncRNA | ENSG0000C |
| 155399922 | 155401782 | 1 | lncRNA | ENSG0000C |
| 155412596 | 155421884 | 1 | lncRNA | ENSG0000C |
| 155421206 | 155457099 | -1 | lncRNA | ENSG0000C |
| 155458129 | 155464831 | 1 | protein_coding | ENSG0000C |
| 155474206 | 155555450 | -1 | protein_coding | ENSG0000C |
| 155474256 | 155644406 | -1 | lncRNA | ENSG0000C |
| 155510283 | 155518623 | 1 | lncRNA | ENSG0000C |
| 155534964 | 155536825 | 1 | lncRNA | ENSG0000C |
| 155611231 | 155645205 | -1 | lncRNA | ENSG0000C |
| 155644451 | 155781485 | 1 | protein_coding | ENSG0000C |
| 155799980 | 155812463 | -1 | protein_coding | ENSG0000C |
| 155962632 | 155966343 | 1 | lncRNA | ENSG0000C |
| 156186122 | 156186223 | -1 | misc_RNA | ENSG0000C |
| 156263547 | 156263802 | 1 | processed_pseudoge | ENSG0000C |
| 156388922 | 156640654 | -1 | lncRNA | ENSG0000C |
| 156540491 | 156541103 | 1 | lncRNA | ENSG0000C |
| 156633355 | 156633787 | 1 | processed_pseudoge | ENSG0000C |
| 156640281 | 156677130 | 1 | protein_coding | ENSG0000C |
| 156654185 | 156659582 | -1 | lncRNA | ENSG0000C |
| 156668946 | 156893216 | -1 | protein_coding | ENSG0000C |
| 156872078 | 156872218 | 1 | snRNA | ENSG0000C |
| 156944721 | 156945645 | 1 | lncRNA | ENSG0000C |
| 156949712 | 156973176 | 1 | protein_coding | ENSG0000C |
| 156994051 | 157010663 | -1 | protein_coding | ENSG0000C |
| 157006307 | 157007132 | 1 | lncRNA | ENSG0000C |
| 157010805 | 157016426 | 1 | lncRNA | ENSG0000C |
| 157057709 | 157058514 | -1 | lncRNA | ENSG0000C |
| 157087955 | 157088543 | -1 | processed_pseudoge | ENSG0000C |
| 157109980 | 157111072 | 1 | lncRNA | ENSG0000C |
| 157138916 | 157269370 | 1 | protein_coding | ENSG0000C |
| 157192368 | 157192832 | 1 | processed_pseudoge | ENSG0000C |
| 157197600 | 157198511 | -1 | processed_pseudoge | ENSG0000C |
| 157263022 | 157263229 | -1 | processed_pseudoge | ENSG0000C |
| 157281536 | 157282686 | -1 | lncRNA | ENSG0000C |

| | | | | |
|---|---|---|---|---|
| 157335381 | 157417439 | 1 | protein_coding | ENSG0000C |
| 157466231 | 157499716 | 1 | lncRNA | ENSG0000C |
| 157501322 | 157503908 | 1 | lncRNA | ENSG0000C |
| 157513294 | 157518658 | 1 | lncRNA | ENSG0000C |
| 157525783 | 157527325 | 1 | TEC | ENSG0000C |
| 157539056 | 158587823 | -1 | protein_coding | ENSG0000C |
| 157574336 | 157574422 | -1 | miRNA | ENSG0000C |
| 157614023 | 157618765 | 1 | lncRNA | ENSG0000C |
| 157739010 | 157740456 | -1 | lncRNA | ENSG0000C |
| 157854529 | 157863011 | 1 | lncRNA | ENSG0000C |
| 157868538 | 157869154 | 1 | lncRNA | ENSG0000C |
| 157987169 | 157989456 | 1 | lncRNA | ENSG0000C |
| 158027373 | 158031128 | 1 | lncRNA | ENSG0000C |
| 158532718 | 158532813 | -1 | miRNA | ENSG0000C |
| 158537495 | 158539879 | 1 | lncRNA | ENSG0000C |
| 158590629 | 158591120 | 1 | lncRNA | ENSG0000C |
| 158591616 | 158591696 | 1 | miRNA | ENSG0000C |
| 158623188 | 158623449 | -1 | processed_pseudoge | ENSG0000C |
| 158631169 | 158704804 | -1 | protein_coding | ENSG0000C |
| 158719730 | 158720161 | 1 | processed_pseudoge | ENSG0000C |
| 158730995 | 158830253 | -1 | protein_coding | ENSG0000C |
| 158856558 | 158956747 | 1 | protein_coding | ENSG0000C |
| 158957497 | 158958699 | -1 | lncRNA | ENSG0000C |
| 159006522 | 159030195 | 1 | lncRNA | ENSG0000C |
| 159028175 | 159144867 | -1 | protein_coding | ENSG0000C |
| 159107761 | 159109216 | 1 | lncRNA | ENSG0000C |
| 159231435 | 159233377 | 1 | unprocessed_pseudc | ENSG0000C |
| 64091 | 64320 | -1 | processed_pseudoge | ENSG0000C |
| 72601 | 79775 | 1 | lncRNA | ENSG0000C |
| 150584 | 150728 | -1 | processed_pseudoge | ENSG0000C |
| 155274 | 155481 | 1 | processed_pseudoge | ENSG0000C |
| 166049 | 167043 | -1 | protein_coding | ENSG0000C |
| 200385 | 200562 | -1 | unprocessed_pseudc | ENSG0000C |
| 202660 | 202897 | -1 | unprocessed_pseudc | ENSG0000C |
| 213186 | 232231 | -1 | transcribed_processe | ENSG0000C |
| 232137 | 264703 | 1 | protein_coding | ENSG0000C |
| 233119 | 233692 | 1 | lncRNA | ENSG0000C |
| 234347 | 234887 | -1 | lncRNA | ENSG0000C |
| 237045 | 237669 | 1 | lncRNA | ENSG0000C |
| 264315 | 264672 | 1 | processed_pseudoge | ENSG0000C |
| 311133 | 331026 | -1 | lncRNA | ENSG0000C |
| 375931 | 384079 | -1 | lncRNA | ENSG0000C |
| 406428 | 477967 | 1 | protein_coding | ENSG0000C |
| 450714 | 451343 | -1 | lncRNA | ENSG0000C |
| 486117 | 488512 | -1 | TEC | ENSG0000C |
| 489792 | 545781 | -1 | protein_coding | ENSG0000C |
| 614746 | 738106 | -1 | protein_coding | ENSG0000C |
| 725188 | 725877 | -1 | lncRNA | ENSG0000C |
| 737596 | 1708474 | 1 | protein_coding | ENSG0000C |
| 738548 | 740374 | 1 | lncRNA | ENSG0000C |

| | | | | |
|---|---|---|---|---|
| 891251 | 893112 | -1 | lncRNA | ENSG00000 |
| 1018757 | 1019704 | 1 | lncRNA | ENSG00000 |
| 1246789 | 1248760 | -1 | lncRNA | ENSG00000 |
| 1296034 | 1302607 | -1 | lncRNA | ENSG00000 |
| 1368642 | 1369833 | -1 | lncRNA | ENSG00000 |
| 1373685 | 1381394 | 1 | lncRNA | ENSG00000 |
| 1377867 | 1380307 | 1 | lncRNA | ENSG00000 |
| 1565509 | 1622417 | -1 | lncRNA | ENSG00000 |
| 1726273 | 1726557 | 1 | processed_pseudoge | ENSG00000 |
| 1730206 | 1731452 | 1 | processed_pseudoge | ENSG00000 |
| 1755778 | 1801711 | 1 | protein_coding | ENSG00000 |
| 1758208 | 1760447 | -1 | lncRNA | ENSG00000 |
| 1761054 | 1764508 | -1 | lncRNA | ENSG00000 |
| 1763888 | 1958627 | 1 | protein_coding | ENSG00000 |
| 1801125 | 1801192 | 1 | miRNA | ENSG00000 |
| 1810823 | 1817113 | -1 | lncRNA | ENSG00000 |
| 1817231 | 1817307 | 1 | miRNA | ENSG00000 |
| 1823926 | 1958641 | 1 | protein_coding | ENSG00000 |
| 1957921 | 1963131 | -1 | lncRNA | ENSG00000 |
| 1971153 | 1974637 | -1 | lncRNA | ENSG00000 |
| 1971397 | 1976478 | 1 | lncRNA | ENSG00000 |
| 1973677 | 2006936 | 1 | protein_coding | ENSG00000 |
| 1974818 | 1975314 | 1 | lncRNA | ENSG00000 |
| 2033508 | 2035587 | 1 | lncRNA | ENSG00000 |
| 2045046 | 2165552 | 1 | protein_coding | ENSG00000 |
| 2076589 | 2076640 | 1 | miRNA | ENSG00000 |
| 2130550 | 2134880 | -1 | lncRNA | ENSG00000 |
| 2493876 | 2512580 | -1 | lncRNA | ENSG00000 |
| 2495522 | 2499428 | 1 | lncRNA | ENSG00000 |
| 2517898 | 2556440 | -1 | lncRNA | ENSG00000 |
| 2518998 | 2522182 | 1 | lncRNA | ENSG00000 |
| 2562813 | 2623382 | -1 | lncRNA | ENSG00000 |
| 2648075 | 2728577 | -1 | lncRNA | ENSG00000 |
| 2726956 | 2926689 | 1 | lncRNA | ENSG00000 |
| 2873525 | 2877916 | -1 | lncRNA | ENSG00000 |
| 2935353 | 4994972 | -1 | protein_coding | ENSG00000 |
| 3373353 | 3375226 | -1 | lncRNA | ENSG00000 |
| 3409133 | 3426587 | 1 | lncRNA | ENSG00000 |
| 3700492 | 3700628 | 1 | rRNA_pseudogene | ENSG00000 |
| 4317388 | 4329027 | 1 | lncRNA | ENSG00000 |
| 4496392 | 4503392 | 1 | lncRNA | ENSG00000 |
| 4633065 | 4635654 | 1 | lncRNA | ENSG00000 |
| 4787332 | 4788584 | 1 | processed_pseudoge | ENSG00000 |
| 5072645 | 5072869 | -1 | processed_pseudoge | ENSG00000 |
| 5127919 | 5128233 | -1 | unprocessed_pseudc | ENSG00000 |
| 5128279 | 5128412 | -1 | snoRNA | ENSG00000 |
| 5142411 | 5142707 | -1 | misc_RNA | ENSG00000 |
| 5363371 | 5363650 | 1 | unprocessed_pseudc | ENSG00000 |
| 5455218 | 5455665 | -1 | processed_pseudoge | ENSG00000 |
| 5476548 | 5476966 | 1 | processed_pseudoge | ENSG00000 |

| | | | | |
|---|---|---|---|---|
| 5506068 | 5509126 | -1 | lncRNA | ENSG00000 |
| 5659679 | 5670140 | -1 | lncRNA | ENSG00000 |
| 5716584 | 5716804 | 1 | unprocessed_pseudo | ENSG00000 |
| 5848697 | 5849692 | 1 | processed_pseudoge | ENSG00000 |
| 5853138 | 5854977 | -1 | lncRNA | ENSG00000 |
| 5910945 | 5911236 | -1 | misc_RNA | ENSG00000 |
| 6036433 | 6036974 | -1 | lncRNA | ENSG00000 |
| 6044353 | 6257537 | -1 | lncRNA | ENSG00000 |
| 6403551 | 6407142 | -1 | lncRNA | ENSG00000 |
| 6406596 | 6648508 | 1 | protein_coding | ENSG00000 |
| 6473845 | 6474288 | -1 | processed_pseudoge | ENSG00000 |
| 6499651 | 6563409 | -1 | protein_coding | ENSG00000 |
| 6572571 | 6576219 | -1 | lncRNA | ENSG00000 |
| 6615604 | 6617198 | -1 | lncRNA | ENSG00000 |
| 6617310 | 6708224 | -1 | lncRNA | ENSG00000 |
| 6622124 | 6622220 | -1 | miRNA | ENSG00000 |
| 6708642 | 6761503 | 1 | protein_coding | ENSG00000 |
| 6745164 | 6745244 | 1 | miRNA | ENSG00000 |
| 6745168 | 6745240 | -1 | miRNA | ENSG00000 |
| 6780874 | 6789021 | 1 | lncRNA | ENSG00000 |
| 6808517 | 6835524 | -1 | protein_coding | ENSG00000 |
| 6835535 | 6885276 | 1 | lncRNA | ENSG00000 |
| 6870592 | 6877936 | -1 | protein_coding | ENSG00000 |
| 6898878 | 6899468 | -1 | processed_pseudoge | ENSG00000 |
| 6924697 | 6926076 | -1 | protein_coding | ENSG00000 |
| 6928608 | 6930473 | 1 | lncRNA | ENSG00000 |
| 6935820 | 6938306 | -1 | protein_coding | ENSG00000 |
| 6950726 | 6951599 | -1 | unprocessed_pseudo | ENSG00000 |
| 6959289 | 6960161 | -1 | unprocessed_pseudo | ENSG00000 |
| 6968141 | 6969113 | -1 | unprocessed_pseudo | ENSG00000 |
| 6977649 | 6980080 | -1 | protein_coding | ENSG00000 |
| 6987178 | 6989721 | -1 | unprocessed_pseudo | ENSG00000 |
| 6996766 | 6999195 | -1 | protein_coding | ENSG00000 |
| 7006293 | 7007389 | -1 | unprocessed_pseudo | ENSG00000 |
| 7015869 | 7018297 | -1 | protein_coding | ENSG00000 |
| 7028641 | 7029489 | -1 | unprocessed_pseudo | ENSG00000 |
| 7038571 | 7039439 | -1 | unprocessed_pseudo | ENSG00000 |
| 7055304 | 7056739 | -1 | protein_coding | ENSG00000 |
| 7082971 | 7083328 | 1 | processed_pseudoge | ENSG00000 |
| 7092102 | 7187316 | 1 | transcribed_unproce | ENSG00000 |
| 7112314 | 7113728 | -1 | processed_pseudoge | ENSG00000 |
| 7122857 | 7123704 | 1 | processed_pseudoge | ENSG00000 |
| 7143632 | 7143841 | -1 | processed_pseudoge | ENSG00000 |
| 7170550 | 7170741 | 1 | processed_pseudoge | ENSG00000 |
| 7190901 | 7191563 | -1 | processed_pseudoge | ENSG00000 |
| 7200756 | 7200857 | 1 | processed_pseudoge | ENSG00000 |
| 7221713 | 7221930 | -1 | unprocessed_pseudo | ENSG00000 |
| 7227853 | 7228073 | -1 | unprocessed_pseudo | ENSG00000 |
| 7238286 | 7238631 | -1 | processed_pseudoge | ENSG00000 |
| 7246726 | 7247571 | -1 | unprocessed_pseudo | ENSG00000 |

| | | | | |
|---|---|---|---|---|
| 7256904 | 7259914 | 1 | unprocessed_pseudc | ENSG00000( |
| 7264526 | 7267536 | 1 | unprocessed_pseudc | ENSG00000( |
| 7272148 | 7275158 | 1 | unprocessed_pseudc | ENSG00000( |
| 7279770 | 7282780 | 1 | unprocessed_pseudc | ENSG00000( |
| 7287392 | 7290402 | 1 | unprocessed_pseudc | ENSG00000( |
| 7295014 | 7298024 | 1 | unprocessed_pseudc | ENSG00000( |
| 7298744 | 7355359 | -1 | lncRNA | ENSG00000( |
| 7312846 | 7319951 | 1 | unprocessed_pseudc | ENSG00000( |
| 7332387 | 7333979 | 1 | protein_coding | ENSG00000( |
| 7337115 | 7338707 | 1 | protein_coding | ENSG00000( |
| 7341826 | 7342942 | 1 | unprocessed_pseudc | ENSG00000( |
| 7355517 | 7385558 | -1 | protein_coding | ENSG00000( |
| 7373114 | 7377500 | 1 | unprocessed_pseudc | ENSG00000( |
| 7414855 | 7416863 | -1 | protein_coding | ENSG00000( |
| 7418991 | 7420546 | 1 | processed_pseudoge | ENSG00000( |
| 7428888 | 7430348 | -1 | protein_coding | ENSG00000( |
| 7442684 | 7463674 | -1 | protein_coding | ENSG00000( |
| 7470308 | 7475082 | -1 | protein_coding | ENSG00000( |
| 7482504 | 7486400 | -1 | protein_coding | ENSG00000( |
| 7487679 | 7489593 | 1 | protein_coding | ENSG00000( |
| 7495911 | 7509311 | 1 | protein_coding | ENSG00000( |
| 7525143 | 7527297 | 1 | unprocessed_pseudc | ENSG00000( |
| 7539628 | 7542450 | 1 | protein_coding | ENSG00000( |
| 7544676 | 7545300 | 1 | unprocessed_pseudc | ENSG00000( |
| 7549053 | 7552062 | -1 | unprocessed_pseudc | ENSG00000( |
| 7556700 | 7559712 | -1 | protein_coding | ENSG00000( |
| 7564350 | 7567360 | -1 | unprocessed_pseudc | ENSG00000( |
| 7571995 | 7575006 | -1 | protein_coding | ENSG00000( |
| 7579644 | 7582653 | -1 | protein_coding | ENSG00000( |
| 7592006 | 7592916 | 1 | unprocessed_pseudc | ENSG00000( |
| 7601004 | 7601267 | 1 | processed_pseudoge | ENSG00000( |
| 7611473 | 7611690 | 1 | unprocessed_pseudc | ENSG00000( |
| 7680309 | 7680526 | -1 | unprocessed_pseudc | ENSG00000( |
| 7686423 | 7686643 | -1 | unprocessed_pseudc | ENSG00000( |
| 7696830 | 7697093 | -1 | processed_pseudoge | ENSG00000( |
| 7705182 | 7706113 | -1 | unprocessed_pseudc | ENSG00000( |
| 7715443 | 7718453 | 1 | protein_coding | ENSG00000( |
| 7723091 | 7726101 | 1 | protein_coding | ENSG00000( |
| 7730739 | 7733749 | 1 | protein_coding | ENSG00000( |
| 7738387 | 7741396 | 1 | protein_coding | ENSG00000( |
| 7746034 | 7749044 | 1 | protein_coding | ENSG00000( |
| 7753682 | 7756692 | 1 | protein_coding | ENSG00000( |
| 7761330 | 7764340 | 1 | protein_coding | ENSG00000( |
| 7768977 | 7771988 | 1 | protein_coding | ENSG00000( |
| 7775741 | 7776365 | -1 | unprocessed_pseudc | ENSG00000( |
| 7778591 | 7781413 | -1 | protein_coding | ENSG00000( |
| 7793716 | 7795870 | -1 | unprocessed_pseudc | ENSG00000( |
| 7811720 | 7815716 | -1 | protein_coding | ENSG00000( |
| 7821966 | 7823889 | -1 | protein_coding | ENSG00000( |
| 7825139 | 7829053 | 1 | protein_coding | ENSG00000( |

| | | | | |
|---|---|---|---|---|
| 7836436 | 7841242 | 1 | protein_coding | ENSG0000C |
| 7847876 | 7868867 | 1 | protein_coding | ENSG0000C |
| 7881204 | 7882664 | 1 | protein_coding | ENSG0000C |
| 7891003 | 7892555 | -1 | processed_pseudoge | ENSG0000C |
| 7894629 | 7896711 | 1 | protein_coding | ENSG0000C |
| 7926337 | 7952413 | 1 | protein_coding | ENSG0000C |
| 7934412 | 7938770 | -1 | unprocessed_pseudc | ENSG0000C |
| 7955014 | 8008755 | 1 | lncRNA | ENSG0000C |
| 7967344 | 7968544 | -1 | unprocessed_pseudc | ENSG0000C |
| 7971661 | 7973253 | -1 | protein_coding | ENSG0000C |
| 7976393 | 7977985 | -1 | protein_coding | ENSG0000C |
| 7990415 | 7997521 | -1 | unprocessed_pseudc | ENSG0000C |
| 8011782 | 8017011 | -1 | unprocessed_pseudc | ENSG0000C |
| 8019429 | 8024652 | -1 | transcribed_unproce | ENSG0000C |
| 8027077 | 8032305 | -1 | unprocessed_pseudc | ENSG0000C |
| 8039978 | 8040953 | 1 | unprocessed_pseudc | ENSG0000C |
| 8048919 | 8049267 | 1 | processed_pseudoge | ENSG0000C |
| 8059459 | 8059674 | 1 | transcribed_unproce | ENSG0000C |
| 8059572 | 8199973 | 1 | lncRNA | ENSG0000C |
| 8065599 | 8065816 | 1 | transcribed_unproce | ENSG0000C |
| 8086022 | 8086765 | -1 | processed_pseudoge | ENSG0000C |
| 8088941 | 8089089 | -1 | miRNA | ENSG0000C |
| 8095946 | 8096726 | 1 | processed_pseudoge | ENSG0000C |
| 8116745 | 8116949 | -1 | processed_pseudoge | ENSG0000C |
| 8154308 | 8216916 | 1 | transcribed_unproce | ENSG0000C |
| 8167819 | 8226614 | -1 | lncRNA | ENSG0000C |
| 8188535 | 8189195 | -1 | lncRNA | ENSG0000C |
| 8228595 | 8244865 | 1 | transcribed_unproce | ENSG0000C |
| 8236003 | 8244667 | -1 | transcribed_unproce | ENSG0000C |
| 8317736 | 8386439 | -1 | protein_coding | ENSG0000C |
| 8414204 | 8424648 | -1 | lncRNA | ENSG0000C |
| 8456909 | 8461337 | -1 | lncRNA | ENSG0000C |
| 8555738 | 8560965 | -1 | lncRNA | ENSG0000C |
| 8559894 | 8562124 | 1 | lncRNA | ENSG0000C |
| 8561386 | 8574038 | 1 | lncRNA | ENSG0000C |
| 8582352 | 8582655 | -1 | misc_RNA | ENSG0000C |
| 8665497 | 8669404 | 1 | lncRNA | ENSG0000C |
| 8701937 | 8704096 | 1 | protein_coding | ENSG0000C |
| 8714055 | 8719536 | -1 | unitary_pseudogene | ENSG0000C |
| 8723693 | 8782479 | -1 | lncRNA | ENSG0000C |
| 8783354 | 8893630 | -1 | protein_coding | ENSG0000C |
| 8895813 | 8895918 | 1 | snRNA | ENSG0000C |
| 8933721 | 8934006 | -1 | processed_pseudoge | ENSG0000C |
| 8961200 | 8979025 | 1 | transcribed_processe | ENSG0000C |
| 8961575 | 8961708 | 1 | snoRNA | ENSG0000C |
| 9002147 | 9116746 | 1 | protein_coding | ENSG0000C |
| 9048445 | 9048518 | 1 | miRNA | ENSG0000C |
| 9072455 | 9072516 | -1 | snRNA | ENSG0000C |
| 9091240 | 9091508 | 1 | misc_RNA | ENSG0000C |
| 9136255 | 9151574 | -1 | protein_coding | ENSG0000C |

| | | | | |
|---|---|---|---|---|
| 9141424 | 9145503 | 1 | lncRNA | ENSG0000C |
| 9151695 | 9425524 | 1 | lncRNA | ENSG0000C |
| 9158063 | 9158621 | 1 | lncRNA | ENSG0000C |
| 9180251 | 9182519 | -1 | lncRNA | ENSG0000C |
| 9233013 | 9233449 | 1 | processed_pseudoge | ENSG0000C |
| 9255349 | 9260295 | -1 | lncRNA | ENSG0000C |
| 9287472 | 9287578 | -1 | snRNA | ENSG0000C |
| 9351238 | 9358441 | -1 | lncRNA | ENSG0000C |
| 9367898 | 9436205 | -1 | lncRNA | ENSG0000C |
| 9371090 | 9371195 | -1 | snRNA | ENSG0000C |
| 9453698 | 9457418 | 1 | processed_pseudoge | ENSG0000C |
| 9555144 | 9556520 | -1 | lncRNA | ENSG0000C |
| 9555912 | 9782346 | 1 | protein_coding | ENSG0000C |
| 9658295 | 9658438 | -1 | unprocessed_pseudc | ENSG0000C |
| 9741672 | 9741768 | 1 | miRNA | ENSG0000C |
| 9899871 | 9906678 | -1 | lncRNA | ENSG0000C |
| 9903388 | 9903472 | -1 | miRNA | ENSG0000C |
| 9984004 | 10055041 | -1 | lncRNA | ENSG0000C |
| 10050485 | 10054254 | -1 | lncRNA | ENSG0000C |
| 10054292 | 10428891 | 1 | protein_coding | ENSG0000C |
| 10118005 | 10147423 | -1 | lncRNA | ENSG0000C |
| 10275499 | 10279737 | -1 | lncRNA | ENSG0000C |
| 10336782 | 10337677 | -1 | processed_pseudoge | ENSG0000C |
| 10433672 | 10438312 | 1 | lncRNA | ENSG0000C |
| 10474565 | 10482388 | 1 | lncRNA | ENSG0000C |
| 10474789 | 10475045 | -1 | unprocessed_pseudc | ENSG0000C |
| 10476223 | 10476329 | 1 | snRNA | ENSG0000C |
| 10476591 | 10479446 | -1 | lncRNA | ENSG0000C |
| 10482878 | 10547585 | -1 | protein_coding | ENSG0000C |
| 10486807 | 10489666 | 1 | lncRNA | ENSG0000C |
| 10525532 | 10554166 | 1 | protein_coding | ENSG0000C |
| 10606349 | 10712187 | -1 | protein_coding | ENSG0000C |
| 10666978 | 10667070 | 1 | miRNA | ENSG0000C |
| 10672628 | 10700590 | 1 | protein_coding | ENSG0000C |
| 10686569 | 10686687 | 1 | rRNA_pseudogene | ENSG0000C |
| 10723768 | 10730511 | -1 | protein_coding | ENSG0000C |
| 10725399 | 10839847 | -1 | protein_coding | ENSG0000C |
| 10729314 | 10772910 | 1 | lncRNA | ENSG0000C |
| 10764961 | 10839884 | -1 | protein_coding | ENSG0000C |
| 10825373 | 10825443 | -1 | miRNA | ENSG0000C |
| 10839964 | 10847594 | 1 | lncRNA | ENSG0000C |
| 10856085 | 10859436 | -1 | TEC | ENSG0000C |
| 10896045 | 11201833 | -1 | protein_coding | ENSG0000C |
| 11035206 | 11035302 | -1 | miRNA | ENSG0000C |
| 11062647 | 11067089 | -1 | lncRNA | ENSG0000C |
| 11104691 | 11106704 | -1 | lncRNA | ENSG0000C |
| 11107788 | 11109726 | -1 | lncRNA | ENSG0000C |
| 11123381 | 11126064 | -1 | lncRNA | ENSG0000C |
| 11136898 | 11138607 | -1 | lncRNA | ENSG0000C |
| 11175051 | 11175570 | 1 | processed_pseudoge | ENSG0000C |

| | | | | |
|---|---|---|---|---|
| 11202965 | 11203671 | 1 | lncRNA | ENSG0000C |
| 11247626 | 11267865 | -1 | lncRNA | ENSG0000C |
| 11256020 | 11256602 | 1 | processed_pseudoge | ENSG0000C |
| 11281364 | 11282501 | -1 | unprocessed_pseudc | ENSG0000C |
| 11283481 | 11285068 | -1 | lncRNA | ENSG0000C |
| 11284816 | 11328146 | 1 | protein_coding | ENSG0000C |
| 11315859 | 11325429 | -1 | lncRNA | ENSG0000C |
| 11330888 | 11332208 | 1 | protein_coding | ENSG0000C |
| 11339637 | 11368452 | 1 | transcribed_unitary_ | ENSG0000C |
| 11345748 | 11347502 | -1 | lncRNA | ENSG0000C |
| 11368402 | 11438658 | 1 | lncRNA | ENSG0000C |
| 11379395 | 11379675 | 1 | misc_RNA | ENSG0000C |
| 11392411 | 11392517 | -1 | snRNA | ENSG0000C |
| 11421476 | 11475908 | -1 | protein_coding | ENSG0000C |
| 11479203 | 11481068 | -1 | lncRNA | ENSG0000C |
| 11486894 | 11564599 | 1 | protein_coding | ENSG0000C |
| 11552488 | 11564694 | -1 | lncRNA | ENSG0000C |
| 11576257 | 11582817 | 1 | lncRNA | ENSG0000C |
| 11643404 | 11647945 | 1 | lncRNA | ENSG0000C |
| 11676959 | 11760002 | 1 | protein_coding | ENSG0000C |
| 11761256 | 11763223 | 1 | lncRNA | ENSG0000C |
| 11769639 | 11787345 | 1 | protein_coding | ENSG0000C |
| 11790005 | 11790386 | 1 | processed_pseudoge | ENSG0000C |
| 11795573 | 11839304 | 1 | protein_coding | ENSG0000C |
| 11797928 | 11802568 | -1 | lncRNA | ENSG0000C |
| 11842524 | 11869448 | -1 | protein_coding | ENSG0000C |
| 11846154 | 11846391 | -1 | lncRNA | ENSG0000C |
| 11895850 | 11896870 | -1 | unprocessed_pseudc | ENSG0000C |
| 11919900 | 11920809 | -1 | unprocessed_pseudc | ENSG0000C |
| 11928597 | 11933203 | -1 | transcribed_unproce | ENSG0000C |
| 11936033 | 11941273 | -1 | lncRNA | ENSG0000C |
| 11946461 | 11950753 | -1 | lncRNA | ENSG0000C |
| 11973937 | 11974599 | -1 | protein_coding | ENSG0000C |
| 11982321 | 11984590 | 1 | protein_coding | ENSG0000C |
| 11993174 | 11996312 | -1 | protein_coding | ENSG0000C |
| 11999087 | 11999468 | 1 | processed_pseudoge | ENSG0000C |
| 12003400 | 12004082 | -1 | transcribed_processe | ENSG0000C |
| 12006188 | 12015193 | -1 | transcribed_unproce | ENSG0000C |
| 12025244 | 12025598 | -1 | processed_pseudoge | ENSG0000C |
| 12033646 | 12034612 | -1 | unprocessed_pseudc | ENSG0000C |
| 12043506 | 12043670 | -1 | processed_pseudoge | ENSG0000C |
| 12064389 | 12071747 | -1 | protein_coding | ENSG0000C |
| 12087396 | 12087445 | -1 | rRNA_pseudogene | ENSG0000C |
| 12095176 | 12099536 | -1 | unprocessed_pseudc | ENSG0000C |
| 12104389 | 12115516 | 1 | protein_coding | ENSG0000C |
| 12115767 | 12177550 | 1 | lncRNA | ENSG0000C |
| 12128107 | 12129298 | -1 | processed_pseudoge | ENSG0000C |
| 12132417 | 12134438 | -1 | protein_coding | ENSG0000C |
| 12137168 | 12139077 | -1 | protein_coding | ENSG0000C |
| 12150895 | 12151134 | -1 | processed_pseudoge | ENSG0000C |

| | | | | |
|---|---|---|---|---|
| 12172761 | 12175771 | -1 | unprocessed_pseudo | ENSG00000 |
| 12178697 | 12182719 | 1 | unprocessed_pseudo | ENSG00000 |
| 12182096 | 12194133 | -1 | protein_coding | ENSG00000 |
| 12194467 | 12196280 | 1 | lncRNA | ENSG00000 |
| 12205759 | 12206389 | -1 | processed_pseudoge | ENSG00000 |
| 12290071 | 12296180 | -1 | unprocessed_pseudo | ENSG00000 |
| 12310962 | 12318316 | -1 | protein_coding | ENSG00000 |
| 12333644 | 12333693 | -1 | rRNA_pseudogene | ENSG00000 |
| 12341426 | 12345776 | -1 | unprocessed_pseudo | ENSG00000 |
| 12356136 | 12359391 | 1 | unprocessed_pseudo | ENSG00000 |
| 12362019 | 12388296 | 1 | lncRNA | ENSG00000 |
| 12374366 | 12375546 | -1 | processed_pseudoge | ENSG00000 |
| 12378679 | 12380265 | -1 | processed_pseudoge | ENSG00000 |
| 12393209 | 12400366 | -1 | unprocessed_pseudo | ENSG00000 |
| 12412827 | 12414373 | 1 | lncRNA | ENSG00000 |
| 12415080 | 12418090 | -1 | unprocessed_pseudo | ENSG00000 |
| 12421032 | 12425000 | 1 | unprocessed_pseudo | ENSG00000 |
| 12425614 | 12436343 | -1 | protein_coding | ENSG00000 |
| 12448013 | 12511278 | -1 | transcribed_unproce | ENSG00000 |
| 12467693 | 12665588 | -1 | lncRNA | ENSG00000 |
| 12476462 | 12477122 | 1 | lncRNA | ENSG00000 |
| 12570350 | 12571130 | -1 | processed_pseudoge | ENSG00000 |
| 12580287 | 12581063 | 1 | processed_pseudoge | ENSG00000 |
| 12601158 | 12601376 | -1 | transcribed_unproce | ENSG00000 |
| 12607291 | 12607509 | -1 | transcribed_unproce | ENSG00000 |
| 12628476 | 12629256 | -1 | processed_pseudoge | ENSG00000 |
| 12638428 | 12638602 | 1 | processed_pseudoge | ENSG00000 |
| 12659130 | 12659348 | -1 | transcribed_unproce | ENSG00000 |
| 12665278 | 12665495 | -1 | transcribed_unproce | ENSG00000 |
| 12676035 | 12676389 | -1 | processed_pseudoge | ENSG00000 |
| 12684154 | 12685153 | -1 | unprocessed_pseudo | ENSG00000 |
| 12696307 | 12697273 | -1 | unprocessed_pseudo | ENSG00000 |
| 12703066 | 12704056 | -1 | unprocessed_pseudo | ENSG00000 |
| 12719132 | 12719190 | 1 | miRNA | ENSG00000 |
| 12721906 | 12756073 | -1 | protein_coding | ENSG00000 |
| 12727232 | 12727304 | -1 | miRNA | ENSG00000 |
| 12727237 | 12727299 | 1 | miRNA | ENSG00000 |
| 12765849 | 12811478 | -1 | lncRNA | ENSG00000 |
| 12794243 | 12818291 | 1 | lncRNA | ENSG00000 |
| 12945642 | 13031503 | 1 | protein_coding | ENSG00000 |
| 12958387 | 12962200 | 1 | unprocessed_pseudo | ENSG00000 |
| 13044350 | 13044456 | 1 | snRNA | ENSG00000 |
| 13083361 | 13604610 | -1 | protein_coding | ENSG00000 |
| 13147389 | 13207125 | 1 | lncRNA | ENSG00000 |
| 13160178 | 13160279 | 1 | misc_RNA | ENSG00000 |
| 13338574 | 13342754 | 1 | lncRNA | ENSG00000 |
| 13353401 | 13353530 | -1 | processed_pseudoge | ENSG00000 |
| 13536029 | 13536135 | 1 | rRNA_pseudogene | ENSG00000 |
| 13566869 | 13568288 | 1 | protein_coding | ENSG00000 |
| 13583889 | 13584533 | -1 | processed_pseudoge | ENSG00000 |

| | | | | |
|---|---|---|---|---|
| 13629644 | 13632574 | 1 | lncRNA | ENSG0000( |
| 13695574 | 13705506 | 1 | processed_pseudoge | ENSG0000( |
| 13741809 | 13742781 | 1 | processed_pseudoge | ENSG0000( |
| 13844101 | 13849109 | -1 | lncRNA | ENSG0000( |
| 13909231 | 13909780 | -1 | processed_pseudoge | ENSG0000( |
| 14022687 | 14023422 | -1 | processed_pseudoge | ENSG0000( |
| 14084845 | 15238431 | -1 | protein_coding | ENSG0000( |
| 14161084 | 14165359 | 1 | lncRNA | ENSG0000( |
| 14309033 | 14309672 | 1 | processed_pseudoge | ENSG0000( |
| 14319455 | 14319516 | 1 | snRNA | ENSG0000( |
| 14332555 | 14332663 | 1 | misc_RNA | ENSG0000( |
| 14487776 | 14487879 | -1 | snRNA | ENSG0000( |
| 14690382 | 14690619 | 1 | processed_pseudoge | ENSG0000( |
| 14853438 | 14853510 | -1 | miRNA | ENSG0000( |
| 14879057 | 14879819 | -1 | lncRNA | ENSG0000( |
| 15325296 | 15325569 | 1 | processed_pseudoge | ENSG0000( |
| 15417215 | 15766649 | 1 | protein_coding | ENSG0000( |
| 15688914 | 15689562 | -1 | processed_pseudoge | ENSG0000( |
| 15806149 | 15807283 | -1 | processed_pseudoge | ENSG0000( |
| 15896267 | 15967950 | -1 | lncRNA | ENSG0000( |
| 15943127 | 15943532 | 1 | processed_pseudoge | ENSG0000( |
| 15973315 | 16000324 | 1 | lncRNA | ENSG0000( |
| 16107878 | 16567490 | -1 | protein_coding | ENSG0000( |
| 16275681 | 16276296 | 1 | processed_pseudoge | ENSG0000( |
| 16374618 | 16375082 | 1 | processed_pseudoge | ENSG0000( |
| 16598758 | 16915044 | -1 | lncRNA | ENSG0000( |
| 16604255 | 16664879 | 1 | lncRNA | ENSG0000( |
| 16863196 | 16863496 | -1 | misc_RNA | ENSG0000( |
| 16992181 | 17002345 | -1 | protein_coding | ENSG0000( |
| 17027238 | 17122642 | 1 | protein_coding | ENSG0000( |
| 17131181 | 17149789 | 1 | lncRNA | ENSG0000( |
| 17150301 | 17150888 | -1 | processed_pseudoge | ENSG0000( |
| 17156482 | 17224799 | 1 | protein_coding | ENSG0000( |
| 17224966 | 17246878 | -1 | protein_coding | ENSG0000( |
| 17246931 | 17302427 | 1 | protein_coding | ENSG0000( |
| 17296794 | 17413528 | -1 | protein_coding | ENSG0000( |
| 17469518 | 17471618 | 1 | unprocessed_pseudc | ENSG0000( |
| 17497088 | 17570573 | 1 | protein_coding | ENSG0000( |
| 17498647 | 17499238 | -1 | TEC | ENSG0000( |
| 17498737 | 17499157 | 1 | TEC | ENSG0000( |
| 17500012 | 17500605 | 1 | TEC | ENSG0000( |
| 17510613 | 17511300 | 1 | lncRNA | ENSG0000( |
| 17576433 | 17644071 | 1 | protein_coding | ENSG0000( |
| 17596312 | 17596378 | -1 | unprocessed_pseudc | ENSG0000( |
| 17604667 | 17610269 | -1 | lncRNA | ENSG0000( |
| 17643795 | 17800917 | -1 | protein_coding | ENSG0000( |
| 17681578 | 17681657 | -1 | miRNA | ENSG0000( |
| 17703988 | 17706187 | 1 | lncRNA | ENSG0000( |
| 17748118 | 17748230 | 1 | rRNA_pseudogene | ENSG0000( |
| 17801345 | 17861069 | 1 | lncRNA | ENSG0000( |

| | | | | |
|---|---|---|---|---|
| 17808361 | 17822183 | 1 | lncRNA | ENSG00000 |
| 17864380 | 17910365 | -1 | protein_coding | ENSG00000 |
| 17882043 | 17882661 | 1 | lncRNA | ENSG00000 |
| 17900484 | 17908011 | 1 | lncRNA | ENSG00000 |
| 17922840 | 18029944 | 1 | protein_coding | ENSG00000 |
| 18055992 | 18084998 | -1 | protein_coding | ENSG00000 |
| 18084386 | 18097644 | 1 | lncRNA | ENSG00000 |
| 18088569 | 18089687 | -1 | processed_pseudoge | ENSG00000 |
| 18104797 | 18127346 | 1 | lncRNA | ENSG00000 |
| 18170477 | 18223689 | 1 | protein_coding | ENSG00000 |
| 18200924 | 18201205 | -1 | processed_pseudoge | ENSG00000 |
| 18370607 | 18371484 | 1 | unprocessed_pseudc | ENSG00000 |
| 18383274 | 18383917 | -1 | processed_pseudoge | ENSG00000 |
| 18386311 | 18388323 | 1 | lncRNA | ENSG00000 |
| 18387108 | 18387777 | -1 | processed_pseudoge | ENSG00000 |
| 18391282 | 18401218 | 1 | protein_coding | ENSG00000 |
| 18527303 | 19084730 | -1 | protein_coding | ENSG00000 |
| 18720905 | 18734405 | 1 | lncRNA | ENSG00000 |
| 18774547 | 18774918 | -1 | processed_pseudoge | ENSG00000 |
| 18849258 | 18849707 | 1 | unprocessed_pseudc | ENSG00000 |
| 18864681 | 18865247 | -1 | TEC | ENSG00000 |
| 18979619 | 18979772 | 1 | snoRNA | ENSG00000 |
| 18989279 | 19000952 | 1 | lncRNA | ENSG00000 |
| 19001627 | 19001950 | 1 | processed_pseudoge | ENSG00000 |
| 19084992 | 19259469 | -1 | lncRNA | ENSG00000 |
| 19091703 | 19145449 | 1 | lncRNA | ENSG00000 |
| 19237737 | 19241912 | 1 | unprocessed_pseudc | ENSG00000 |
| 19246350 | 19249240 | 1 | lncRNA | ENSG00000 |
| 19254241 | 19257334 | -1 | lncRNA | ENSG00000 |
| 19277284 | 19277573 | 1 | TEC | ENSG00000 |
| 19313693 | 19396218 | 1 | protein_coding | ENSG00000 |
| 19375128 | 19375622 | -1 | TEC | ENSG00000 |
| 19404161 | 19758029 | -1 | protein_coding | ENSG00000 |
| 19678572 | 19688934 | 1 | lncRNA | ENSG00000 |
| 19817391 | 19852083 | 1 | protein_coding | ENSG00000 |
| 19901717 | 19967259 | 1 | protein_coding | ENSG00000 |
| 20079114 | 20124857 | 1 | lncRNA | ENSG00000 |
| 20113343 | 20113672 | -1 | processed_pseudoge | ENSG00000 |
| 20144855 | 20183206 | -1 | protein_coding | ENSG00000 |
| 20197381 | 20226819 | 1 | protein_coding | ENSG00000 |
| 20229779 | 20229877 | 1 | snRNA | ENSG00000 |
| 20246165 | 20303963 | -1 | protein_coding | ENSG00000 |
| 20275773 | 20290458 | 1 | lncRNA | ENSG00000 |
| 20290174 | 20290289 | 1 | rRNA_pseudogene | ENSG00000 |
| 20350210 | 20372769 | 1 | lncRNA | ENSG00000 |
| 20551201 | 20551405 | 1 | unprocessed_pseudc | ENSG00000 |
| 20655184 | 20700152 | -1 | lncRNA | ENSG00000 |
| 20934435 | 20934837 | -1 | processed_pseudoge | ENSG00000 |
| 20941428 | 20950175 | 1 | lncRNA | ENSG00000 |
| 20942863 | 21298168 | -1 | lncRNA | ENSG00000 |

| | | | | |
|---|---|---|---|---|
| 20953127 | 20968904 | 1 | lncRNA | ENSG0000C |
| 20954012 | 20968904 | 1 | lncRNA | ENSG0000C |
| 20972021 | 20995119 | 1 | lncRNA | ENSG0000C |
| 21023276 | 21030368 | 1 | lncRNA | ENSG0000C |
| 21129990 | 21130100 | -1 | TEC | ENSG0000C |
| 21167240 | 21170137 | -1 | lncRNA | ENSG0000C |
| 21297933 | 21309486 | 1 | lncRNA | ENSG0000C |
| 21690398 | 21812357 | -1 | protein_coding | ENSG0000C |
| 21797460 | 21797759 | 1 | unprocessed_pseudc | ENSG0000C |
| 21908873 | 21913690 | -1 | protein_coding | ENSG0000C |
| 21919662 | 22006585 | 1 | protein_coding | ENSG0000C |
| 22024125 | 22036897 | 1 | protein_coding | ENSG0000C |
| 22042398 | 22048809 | 1 | protein_coding | ENSG0000C |
| 22048995 | 22082527 | 1 | protein_coding | ENSG0000C |
| 22089150 | 22104911 | 1 | protein_coding | ENSG0000C |
| 22106874 | 22109419 | -1 | protein_coding | ENSG0000C |
| 22114419 | 22133384 | -1 | protein_coding | ENSG0000C |
| 22138020 | 22141951 | -1 | protein_coding | ENSG0000C |
| 22146830 | 22157084 | -1 | protein_coding | ENSG0000C |
| 22156913 | 22164479 | 1 | protein_coding | ENSG0000C |
| 22165140 | 22212326 | 1 | protein_coding | ENSG0000C |
| 22169338 | 22171710 | 1 | lncRNA | ENSG0000C |
| 22188369 | 22192578 | -1 | lncRNA | ENSG0000C |
| 22219703 | 22232101 | -1 | protein_coding | ENSG0000C |
| 22244962 | 22245043 | -1 | miRNA | ENSG0000C |
| 22245133 | 22254601 | 1 | protein_coding | ENSG0000C |
| 22254576 | 22275162 | -1 | lncRNA | ENSG0000C |
| 22275316 | 22357568 | 1 | protein_coding | ENSG0000C |
| 22367249 | 22434129 | 1 | protein_coding | ENSG0000C |
| 22389865 | 22390655 | -1 | unprocessed_pseudc | ENSG0000C |
| 22434391 | 22434497 | 1 | snRNA | ENSG0000C |
| 22440819 | 22541142 | 1 | protein_coding | ENSG0000C |
| 22481588 | 22553444 | -1 | lncRNA | ENSG0000C |
| 22492631 | 22493044 | 1 | processed_pseudoge | ENSG0000C |
| 22544986 | 22575788 | 1 | protein_coding | ENSG0000C |
| 22545560 | 22548837 | 1 | lncRNA | ENSG0000C |
| 22565997 | 22567171 | -1 | lncRNA | ENSG0000C |
| 22578279 | 22598025 | 1 | protein_coding | ENSG0000C |
| 22589274 | 22602084 | 1 | protein_coding | ENSG0000C |
| 22599599 | 22604150 | 1 | protein_coding | ENSG0000C |
| 22604632 | 22621514 | 1 | protein_coding | ENSG0000C |
| 22613908 | 22616657 | -1 | lncRNA | ENSG0000C |
| 22620418 | 22669148 | -1 | protein_coding | ENSG0000C |
| 22679013 | 22684009 | -1 | lncRNA | ENSG0000C |
| 22687659 | 22693480 | -1 | protein_coding | ENSG0000C |
| 22690150 | 22798616 | 1 | lncRNA | ENSG0000C |
| 22698384 | 22703776 | -1 | lncRNA | ENSG0000C |
| 22713251 | 23000000 | -1 | protein_coding | ENSG0000C |
| 22877972 | 22888022 | 1 | lncRNA | ENSG0000C |
| 22895434 | 22897813 | -1 | lncRNA | ENSG0000C |

| | | | | |
|---|---|---|---|---|
| 22977877 | 22978127 | 1 | misc_RNA | ENSG0000C |
| 22984596 | 23019335 | -1 | lncRNA | ENSG0000C |
| 22987417 | 23020199 | 1 | protein_coding | ENSG0000C |
| 23020133 | 23069031 | -1 | protein_coding | ENSG0000C |
| 23068229 | 23083619 | 1 | lncRNA | ENSG0000C |
| 23071377 | 23074488 | -1 | lncRNA | ENSG0000C |
| 23084355 | 23103558 | 1 | lncRNA | ENSG0000C |
| 23084403 | 23115536 | 1 | protein_coding | ENSG0000C |
| 23102590 | 23117437 | 1 | protein_coding | ENSG0000C |
| 23135588 | 23164027 | -1 | protein_coding | ENSG0000C |
| 23189279 | 23190675 | 1 | lncRNA | ENSG0000C |
| 23190452 | 23225102 | -1 | protein_coding | ENSG0000C |
| 23206918 | 23207369 | 1 | processed_pseudoge | ENSG0000C |
| 23224471 | 23230926 | 1 | lncRNA | ENSG0000C |
| 23225233 | 23230915 | -1 | lncRNA | ENSG0000C |
| 23243637 | 23262000 | 1 | protein_coding | ENSG0000C |
| 23270120 | 23296279 | 1 | protein_coding | ENSG0000C |
| 23296897 | 23425328 | -1 | protein_coding | ENSG0000C |
| 23336171 | 23366125 | 1 | lncRNA | ENSG0000C |
| 23385783 | 23457695 | -1 | protein_coding | ENSG0000C |
| 23458601 | 23484971 | 1 | lncRNA | ENSG0000C |
| 23483016 | 23483534 | -1 | processed_pseudoge | ENSG0000C |
| 23492170 | 23494897 | 1 | lncRNA | ENSG0000C |
| 23493009 | 23494198 | -1 | lncRNA | ENSG0000C |
| 23528956 | 23575463 | 1 | protein_coding | ENSG0000C |
| 23600026 | 23600138 | -1 | snRNA | ENSG0000C |
| 23634876 | 23635507 | -1 | processed_pseudoge | ENSG0000C |
| 23678697 | 23682938 | -1 | protein_coding | ENSG0000C |
| 23702451 | 23706598 | -1 | protein_coding | ENSG0000C |
| 23707141 | 23789487 | 1 | lncRNA | ENSG0000C |
| 23714066 | 23743042 | -1 | lncRNA | ENSG0000C |
| 23805933 | 23806154 | -1 | processed_pseudoge | ENSG0000C |
| 23806229 | 23806377 | -1 | snoRNA | ENSG0000C |
| 23841929 | 23854806 | -1 | protein_coding | ENSG0000C |
| 24076997 | 24077158 | -1 | snRNA | ENSG0000C |
| 24169001 | 24177080 | -1 | lncRNA | ENSG0000C |
| 24294069 | 24359014 | 1 | protein_coding | ENSG0000C |
| 24295814 | 24912073 | -1 | lncRNA | ENSG0000C |
| 24384285 | 24406013 | 1 | protein_coding | ENSG0000C |
| 24440930 | 24526970 | 1 | protein_coding | ENSG0000C |
| 24490312 | 24514856 | -1 | lncRNA | ENSG0000C |
| 24912165 | 24914717 | -1 | lncRNA | ENSG0000C |
| 24913758 | 24919098 | 1 | protein_coding | ENSG0000C |
| 24950955 | 24957110 | -1 | protein_coding | ENSG0000C |
| 24953796 | 24953867 | -1 | miRNA | ENSG0000C |
| 24956621 | 24957110 | 1 | lncRNA | ENSG0000C |
| 24992002 | 25009777 | 1 | lncRNA | ENSG0000C |
| 25000436 | 25003439 | 1 | lncRNA | ENSG0000C |
| 25141394 | 25141968 | -1 | processed_pseudoge | ENSG0000C |
| 25179113 | 25184234 | -1 | lncRNA | ENSG0000C |

| | | | | |
|---|---|---|---|---|
| 25184689 | 25418082 | 1 | protein_coding | ENSG0000C |
| 25276945 | 25277476 | -1 | processed_pseudoge | ENSG0000C |
| 25345402 | 25345474 | 1 | miRNA | ENSG0000C |
| 25419258 | 25424654 | -1 | protein_coding | ENSG0000C |
| 25425521 | 25426580 | -1 | lncRNA | ENSG0000C |
| 25427847 | 25458476 | -1 | protein_coding | ENSG0000C |
| 25459199 | 25507911 | 1 | protein_coding | ENSG0000C |
| 25593197 | 25688786 | 1 | lncRNA | ENSG0000C |
| 25776679 | 25777456 | 1 | lncRNA | ENSG0000C |
| 25820975 | 25833464 | -1 | lncRNA | ENSG0000C |
| 25834129 | 25840135 | -1 | lncRNA | ENSG0000C |
| 25841725 | 26045413 | -1 | protein_coding | ENSG0000C |
| 26136880 | 26136998 | 1 | rRNA_pseudogene | ENSG0000C |
| 26254883 | 26256628 | -1 | processed_pseudoge | ENSG0000C |
| 26291508 | 26372680 | 1 | protein_coding | ENSG0000C |
| 26375913 | 26382953 | -1 | transcribed_processe | ENSG0000C |
| 26383054 | 26505636 | 1 | protein_coding | ENSG0000C |
| 26422593 | 26423929 | 1 | lncRNA | ENSG0000C |
| 26433029 | 26434065 | -1 | processed_pseudoge | ENSG0000C |
| 26440491 | 26442595 | 1 | lncRNA | ENSG0000C |
| 26504701 | 26514092 | -1 | protein_coding | ENSG0000C |
| 26514031 | 26658178 | 1 | protein_coding | ENSG0000C |
| 26547669 | 26548463 | 1 | processed_pseudoge | ENSG0000C |
| 26748150 | 26867278 | -1 | protein_coding | ENSG0000C |
| 26771790 | 26772140 | 1 | processed_pseudoge | ENSG0000C |
| 26835817 | 26836300 | -1 | processed_pseudoge | ENSG0000C |
| 27010486 | 27010746 | 1 | unprocessed_pseudc | ENSG0000C |
| 27011121 | 27012371 | -1 | processed_pseudoge | ENSG0000C |
| 27048853 | 27048963 | -1 | miRNA | ENSG0000C |
| 27063996 | 27065278 | -1 | processed_pseudoge | ENSG0000C |
| 27086076 | 27087783 | 1 | lncRNA | ENSG0000C |
| 27171914 | 27210783 | 1 | lncRNA | ENSG0000C |
| 27235323 | 27258420 | -1 | protein_coding | ENSG0000C |
| 27284886 | 27311272 | -1 | protein_coding | ENSG0000C |
| 27311482 | 27459391 | 1 | protein_coding | ENSG0000C |
| 27433370 | 27433434 | 1 | miRNA | ENSG0000C |
| 27459756 | 27479883 | -1 | protein_coding | ENSG0000C |
| 27490781 | 27545564 | 1 | protein_coding | ENSG0000C |
| 27560274 | 27589073 | 1 | unitary_pseudogene | ENSG0000C |
| 27596917 | 27614700 | -1 | protein_coding | ENSG0000C |
| 27610601 | 27610751 | -1 | miRNA | ENSG0000C |
| 27633868 | 27676776 | 1 | protein_coding | ENSG0000C |
| 27679836 | 27679935 | 1 | snRNA | ENSG0000C |
| 27697217 | 27697526 | -1 | processed_pseudoge | ENSG0000C |
| 27701194 | 27703789 | 1 | TEC | ENSG0000C |
| 27701673 | 27701767 | -1 | miRNA | ENSG0000C |
| 27701677 | 27701759 | 1 | miRNA | ENSG0000C |
| 27732915 | 27758552 | 1 | lncRNA | ENSG0000C |
| 27733316 | 27772653 | -1 | protein_coding | ENSG0000C |
| 27771949 | 27812640 | 1 | protein_coding | ENSG0000C |

| | | | | |
|---|---|---|---|---|
| 27773645 | 27773748 | 1 | snRNA | ENSG0000C |
| 27809624 | 27838082 | -1 | protein_coding | ENSG0000C |
| 27838477 | 27840382 | 1 | processed_pseudoge | ENSG0000C |
| 27869883 | 27992673 | -1 | protein_coding | ENSG0000C |
| 27886039 | 27886116 | -1 | miRNA | ENSG0000C |
| 27903229 | 27903495 | -1 | unprocessed_pseudc | ENSG0000C |
| 27903566 | 27903715 | 1 | unprocessed_pseudc | ENSG0000C |
| 27904481 | 27910310 | 1 | lncRNA | ENSG0000C |
| 28021964 | 28083936 | -1 | protein_coding | ENSG0000C |
| 28089673 | 28191156 | 1 | protein_coding | ENSG0000C |
| 28225929 | 28227562 | 1 | processed_pseudoge | ENSG0000C |
| 28238430 | 28238905 | 1 | processed_pseudoge | ENSG0000C |
| 28250063 | 28339255 | -1 | lncRNA | ENSG0000C |
| 28299683 | 28300566 | 1 | processed_pseudoge | ENSG0000C |
| 28316986 | 28343355 | 1 | protein_coding | ENSG0000C |
| 28345590 | 28402701 | -1 | protein_coding | ENSG0000C |
| 28348287 | 28490278 | -1 | protein_coding | ENSG0000C |
| 28414499 | 28416593 | -1 | lncRNA | ENSG0000C |
| 28415524 | 28420055 | -1 | lncRNA | ENSG0000C |
| 28416448 | 28416551 | 1 | snRNA | ENSG0000C |
| 28447264 | 28455902 | 1 | lncRNA | ENSG0000C |
| 28494205 | 28574267 | 1 | protein_coding | ENSG0000C |
| 28505116 | 28505182 | -1 | miRNA | ENSG0000C |
| 28530411 | 28530519 | 1 | rRNA_pseudogene | ENSG0000C |
| 28600469 | 28755599 | 1 | protein_coding | ENSG0000C |
| 28690215 | 28702030 | -1 | lncRNA | ENSG0000C |
| 28767661 | 28890242 | -1 | protein_coding | ENSG0000C |
| 28798142 | 28802085 | 1 | lncRNA | ENSG0000C |
| 28810103 | 28810418 | 1 | processed_pseudoge | ENSG0000C |
| 28890395 | 29064764 | 1 | protein_coding | ENSG0000C |
| 28915579 | 28915864 | -1 | misc_RNA | ENSG0000C |
| 28949676 | 28955955 | 1 | lncRNA | ENSG0000C |
| 29048494 | 29048604 | -1 | rRNA_pseudogene | ENSG0000C |
| 29055935 | 29056685 | 1 | lncRNA | ENSG0000C |
| 29067278 | 29263124 | -1 | protein_coding | ENSG0000C |
| 29067279 | 29068454 | 1 | lncRNA | ENSG0000C |
| 29110573 | 29140729 | 1 | lncRNA | ENSG0000C |
| 29218905 | 29219472 | 1 | processed_pseudoge | ENSG0000C |
| 29333064 | 29350684 | -1 | protein_coding | ENSG0000C |
| 29351292 | 29360222 | 1 | lncRNA | ENSG0000C |
| 29352420 | 29353170 | -1 | lncRNA | ENSG0000C |
| 29527312 | 29530323 | 1 | lncRNA | ENSG0000C |
| 29548171 | 29564341 | 1 | lncRNA | ENSG0000C |
| 29587001 | 29588426 | 1 | TEC | ENSG0000C |
| 29632091 | 29632644 | 1 | processed_pseudoge | ENSG0000C |
| 29673922 | 29748109 | -1 | lncRNA | ENSG0000C |
| 29748309 | 29798492 | 1 | lncRNA | ENSG0000C |
| 29815004 | 29854543 | -1 | lncRNA | ENSG0000C |
| 29864644 | 29871661 | 1 | lncRNA | ENSG0000C |
| 29905735 | 29908956 | 1 | lncRNA | ENSG0000C |

| | | | | |
|---|---|---|---|---|
| 29920560 | 29953724 | 1 | lncRNA | ENSG00000 |
| 29957272 | 29957348 | -1 | miRNA | ENSG00000 |
| 30027713 | 30028900 | 1 | processed_pseudoge | ENSG00000 |
| 30043801 | 30043908 | -1 | snRNA | ENSG00000 |
| 30063003 | 30083208 | -1 | protein_coding | ENSG00000 |
| 30082758 | 30083467 | 1 | lncRNA | ENSG00000 |
| 30095408 | 30177208 | 1 | protein_coding | ENSG00000 |
| 30117384 | 30117779 | -1 | processed_pseudoge | ENSG00000 |
| 30131671 | 30144665 | -1 | protein_coding | ENSG00000 |
| 30155830 | 30156232 | -1 | lncRNA | ENSG00000 |
| 30156319 | 30183639 | 1 | protein_coding | ENSG00000 |
| 30176554 | 30180888 | -1 | lncRNA | ENSG00000 |
| 30192429 | 30201332 | 1 | lncRNA | ENSG00000 |
| 30197404 | 30198048 | 1 | lncRNA | ENSG00000 |
| 30237382 | 30240997 | 1 | processed_pseudoge | ENSG00000 |
| 30249502 | 30250092 | -1 | processed_pseudoge | ENSG00000 |
| 30279549 | 30281455 | -1 | lncRNA | ENSG00000 |
| 30331961 | 30332277 | -1 | processed_pseudoge | ENSG00000 |
| 30349866 | 30350347 | -1 | processed_pseudoge | ENSG00000 |
| 30351873 | 30353518 | 1 | transcribed_processe | ENSG00000 |
| 30382119 | 30385401 | -1 | lncRNA | ENSG00000 |
| 30384511 | 30572256 | 1 | protein_coding | ENSG00000 |
| 30552345 | 30553766 | 1 | TEC | ENSG00000 |
| 30578318 | 30658236 | -1 | protein_coding | ENSG00000 |
| 30596914 | 30597532 | 1 | lncRNA | ENSG00000 |
| 30638580 | 30646064 | 1 | protein_coding | ENSG00000 |
| 30643689 | 30643802 | 1 | snRNA | ENSG00000 |
| 30678066 | 30727846 | -1 | protein_coding | ENSG00000 |
| 30732247 | 30767006 | 1 | protein_coding | ENSG00000 |
| 30754236 | 30754539 | -1 | processed_pseudoge | ENSG00000 |
| 30774457 | 30814314 | -1 | protein_coding | ENSG00000 |
| 30831544 | 30913003 | -1 | protein_coding | ENSG00000 |
| 30921060 | 30921335 | 1 | processed_pseudoge | ENSG00000 |
| 30983459 | 30984321 | -1 | processed_pseudoge | ENSG00000 |
| 30995802 | 31033715 | -1 | protein_coding | ENSG00000 |
| 31033788 | 31175916 | 1 | protein_coding | ENSG00000 |
| 31131539 | 31131802 | -1 | processed_pseudoge | ENSG00000 |
| 31167078 | 31177167 | -1 | lncRNA | ENSG00000 |
| 31219894 | 31220984 | -1 | processed_pseudoge | ENSG00000 |
| 31275542 | 31390805 | 1 | lncRNA | ENSG00000 |
| 31339197 | 31346479 | 1 | lncRNA | ENSG00000 |
| 31497423 | 31498612 | 1 | lncRNA | ENSG00000 |
| 31536630 | 31546735 | 1 | lncRNA | ENSG00000 |
| 31598099 | 31598204 | -1 | rRNA_pseudogene | ENSG00000 |
| 31639222 | 32855666 | 1 | protein_coding | ENSG00000 |
| 31827238 | 31840709 | -1 | lncRNA | ENSG00000 |
| 32026210 | 32139495 | 1 | lncRNA | ENSG00000 |
| 32192028 | 32192153 | -1 | rRNA_pseudogene | ENSG00000 |
| 32256496 | 32256623 | 1 | rRNA_pseudogene | ENSG00000 |
| 32440704 | 32448803 | 1 | lncRNA | ENSG00000 |

| | | | | |
|---|---|---|---|---|
| 32647202 | 32647390 | 1 | protein_coding | ENSG0000C |
| 32911493 | 32911598 | -1 | snRNA | ENSG0000C |
| 32927913 | 33045445 | 1 | lncRNA | ENSG0000C |
| 33014081 | 33015026 | 1 | processed_pseudoge | ENSG0000C |
| 33015261 | 33017043 | 1 | processed_pseudoge | ENSG0000C |
| 33039523 | 33039793 | 1 | processed_pseudoge | ENSG0000C |
| 33226848 | 33227147 | -1 | processed_pseudoge | ENSG0000C |
| 33338867 | 33338966 | 1 | snRNA | ENSG0000C |
| 33360839 | 33361415 | -1 | lncRNA | ENSG0000C |
| 33370824 | 33473146 | -1 | protein_coding | ENSG0000C |
| 33436268 | 33436700 | -1 | processed_pseudoge | ENSG0000C |
| 33473386 | 33513601 | -1 | protein_coding | ENSG0000C |
| 33485182 | 33501262 | 1 | protein_coding | ENSG0000C |
| 33513475 | 33513578 | 1 | snoRNA | ENSG0000C |
| 33523154 | 33523291 | 1 | processed_pseudoge | ENSG0000C |
| 33540040 | 33540137 | 1 | snoRNA | ENSG0000C |
| 33540185 | 33540493 | 1 | processed_pseudoge | ENSG0000C |
| 33547754 | 33567128 | -1 | protein_coding | ENSG0000C |
| 33580210 | 33580508 | -1 | misc_RNA | ENSG0000C |
| 33591330 | 33600023 | -1 | protein_coding | ENSG0000C |
| 33604856 | 34039104 | 1 | lncRNA | ENSG0000C |
| 33641581 | 33641939 | -1 | processed_pseudoge | ENSG0000C |
| 33715784 | 33716070 | 1 | misc_RNA | ENSG0000C |
| 33722305 | 33723079 | -1 | processed_pseudoge | ENSG0000C |
| 33859480 | 33860223 | -1 | processed_pseudoge | ENSG0000C |
| 33969567 | 33969879 | -1 | processed_pseudoge | ENSG0000C |
| 33973701 | 34009595 | -1 | lncRNA | ENSG0000C |
| 34174886 | 34207513 | 1 | lncRNA | ENSG0000C |
| 34228439 | 34346731 | 1 | lncRNA | ENSG0000C |
| 34229294 | 34248946 | -1 | lncRNA | ENSG0000C |
| 34322992 | 34323642 | -1 | processed_pseudoge | ENSG0000C |
| 34784028 | 34865364 | 1 | lncRNA | ENSG0000C |
| 34874159 | 34874633 | 1 | processed_pseudoge | ENSG0000C |
| 35080592 | 35093509 | 1 | lncRNA | ENSG0000C |
| 35235475 | 35796550 | 1 | protein_coding | ENSG0000C |
| 35254344 | 35256605 | 1 | TEC | ENSG0000C |
| 35525176 | 35525763 | -1 | processed_pseudoge | ENSG0000C |
| 35672195 | 35707734 | -1 | lncRNA | ENSG0000C |
| 35862123 | 36191194 | 1 | lncRNA | ENSG0000C |
| 36004316 | 36095046 | -1 | lncRNA | ENSG0000C |
| 36121398 | 36121683 | -1 | processed_pseudoge | ENSG0000C |
| 36277763 | 36278886 | -1 | unprocessed_pseudc | ENSG0000C |
| 36278960 | 36279483 | 1 | unprocessed_pseudc | ENSG0000C |
| 36279491 | 36279696 | -1 | unprocessed_pseudc | ENSG0000C |
| 36308245 | 36308364 | 1 | processed_pseudoge | ENSG0000C |
| 36309584 | 36309690 | -1 | snRNA | ENSG0000C |
| 36321039 | 36321404 | -1 | processed_pseudoge | ENSG0000C |
| 36378069 | 36779209 | -1 | lncRNA | ENSG0000C |
| 36581180 | 36581591 | -1 | processed_pseudoge | ENSG0000C |
| 36764445 | 36764551 | 1 | rRNA_pseudogene | ENSG0000C |

| | | | | |
|---|---|---|---|---|
| 36784324 | 36936125 | 1 | protein_coding | ENSG00000 |
| 36795255 | 36795862 | -1 | processed_pseudoge | ENSG00000 |
| 36888482 | 36889307 | 1 | unprocessed_pseudc | ENSG00000 |
| 36987977 | 36988412 | -1 | processed_pseudoge | ENSG00000 |
| 37067441 | 37069418 | -1 | lncRNA | ENSG00000 |
| 37095150 | 37095390 | 1 | processed_pseudoge | ENSG00000 |
| 37326575 | 37331991 | -1 | lncRNA | ENSG00000 |
| 37405439 | 37406724 | -1 | lncRNA | ENSG00000 |
| 37421341 | 37554183 | -1 | lncRNA | ENSG00000 |
| 37480089 | 37480809 | 1 | lncRNA | ENSG00000 |
| 37516399 | 37521447 | -1 | lncRNA | ENSG00000 |
| 37559992 | 37567477 | 1 | lncRNA | ENSG00000 |
| 37597480 | 37599858 | -1 | lncRNA | ENSG00000 |
| 37600537 | 37625873 | -1 | lncRNA | ENSG00000 |
| 37626015 | 37674387 | -1 | lncRNA | ENSG00000 |
| 37668898 | 37669005 | 1 | snRNA | ENSG00000 |
| 37695782 | 37700019 | 1 | protein_coding | ENSG00000 |
| 37717579 | 37733756 | 1 | lncRNA | ENSG00000 |
| 37734761 | 37737426 | -1 | lncRNA | ENSG00000 |
| 37736601 | 37758422 | 1 | protein_coding | ENSG00000 |
| 37747338 | 37748004 | -1 | processed_pseudoge | ENSG00000 |
| 37762595 | 37779768 | 1 | protein_coding | ENSG00000 |
| 37784191 | 37844896 | 1 | protein_coding | ENSG00000 |
| 37843268 | 37849861 | -1 | protein_coding | ENSG00000 |
| 37858618 | 37899497 | -1 | protein_coding | ENSG00000 |
| 37858949 | 37861333 | 1 | TEC | ENSG00000 |
| 37908738 | 37909037 | 1 | misc_RNA | ENSG00000 |
| 37934281 | 37940124 | -1 | protein_coding | ENSG00000 |
| 37934340 | 37965953 | -1 | protein_coding | ENSG00000 |
| 37962990 | 37966599 | -1 | protein_coding | ENSG00000 |
| 37983070 | 37983373 | 1 | processed_pseudoge | ENSG00000 |
| 38030534 | 38060365 | 1 | protein_coding | ENSG00000 |
| 38062881 | 38063791 | 1 | lncRNA | ENSG00000 |
| 38082122 | 38099288 | 1 | lncRNA | ENSG00000 |
| 38099471 | 38099931 | 1 | lncRNA | ENSG00000 |
| 38105493 | 38144076 | 1 | protein_coding | ENSG00000 |
| 38122374 | 38122480 | 1 | snRNA | ENSG00000 |
| 38123274 | 38124651 | 1 | lncRNA | ENSG00000 |
| 38142700 | 38150992 | -1 | protein_coding | ENSG00000 |
| 38148741 | 38163772 | 1 | lncRNA | ENSG00000 |
| 38163335 | 38176730 | -1 | protein_coding | ENSG00000 |
| 38166012 | 38166118 | -1 | snRNA | ENSG00000 |
| 38176533 | 38213301 | 1 | protein_coding | ENSG00000 |
| 38201699 | 38204238 | -1 | processed_pseudoge | ENSG00000 |
| 38223957 | 38231695 | -1 | lncRNA | ENSG00000 |
| 38225218 | 38275558 | 1 | protein_coding | ENSG00000 |
| 38263130 | 38269243 | -1 | protein_coding | ENSG00000 |
| 38269704 | 38382272 | -1 | protein_coding | ENSG00000 |
| 38275888 | 38276680 | 1 | lncRNA | ENSG00000 |
| 38335981 | 38337551 | 1 | lncRNA | ENSG00000 |

| | | | | |
|---|---|---|---|---|
| 38382364 | 38383461 | 1 | lncRNA | ENSG00000( |
| 38386207 | 38409527 | 1 | protein_coding | ENSG00000( |
| 38400215 | 38468834 | -1 | protein_coding | ENSG00000( |
| 38408048 | 38408742 | -1 | lncRNA | ENSG00000( |
| 38421889 | 38426096 | 1 | lncRNA | ENSG00000( |
| 38434347 | 38435664 | -1 | transcribed_processe | ENSG00000( |
| 38510834 | 38560939 | -1 | protein_coding | ENSG00000( |
| 38543276 | 38560877 | -1 | lncRNA | ENSG00000( |
| 38600661 | 38601200 | -1 | processed_pseudoge | ENSG00000( |
| 38699274 | 38704855 | -1 | lncRNA | ENSG00000( |
| 38728186 | 38853028 | 1 | protein_coding | ENSG00000( |
| 38744020 | 38744131 | 1 | misc_RNA | ENSG00000( |
| 38799643 | 38802296 | 1 | lncRNA | ENSG00000( |
| 38844866 | 38846439 | -1 | lncRNA | ENSG00000( |
| 38901235 | 38973912 | 1 | protein_coding | ENSG00000( |
| 38970360 | 38973011 | -1 | lncRNA | ENSG00000( |
| 38974228 | 38988663 | 1 | protein_coding | ENSG00000( |
| 38988808 | 38996824 | -1 | protein_coding | ENSG00000( |
| 38996869 | 39105261 | 1 | protein_coding | ENSG00000( |
| 39018615 | 39018683 | -1 | snoRNA | ENSG00000( |
| 39106990 | 39284917 | 1 | protein_coding | ENSG00000( |
| 39157538 | 39158701 | -1 | processed_pseudoge | ENSG00000( |
| 39309909 | 39310443 | -1 | processed_pseudoge | ENSG00000( |
| 39314591 | 39417378 | 1 | transcribed_unitary_ | ENSG00000( |
| 39451045 | 39522852 | -1 | transcribed_unitary_ | ENSG00000( |
| 39522976 | 39580134 | 1 | transcribed_unitary_ | ENSG00000( |
| 39584489 | 39730065 | 1 | protein_coding | ENSG00000( |
| 39743735 | 39838289 | -1 | protein_coding | ENSG00000( |
| 39860219 | 39860525 | 1 | processed_pseudoge | ENSG00000( |
| 39867436 | 39867812 | -1 | processed_pseudoge | ENSG00000( |
| 39902275 | 39928790 | 1 | protein_coding | ENSG00000( |
| 39903775 | 39995248 | -1 | lncRNA | ENSG00000( |
| 39914229 | 39915721 | 1 | lncRNA | ENSG00000( |
| 39918076 | 39920890 | -1 | lncRNA | ENSG00000( |
| 39934614 | 40016391 | 1 | protein_coding | ENSG00000( |
| 39997869 | 39998499 | -1 | processed_pseudoge | ENSG00000( |
| 40104169 | 40107935 | 1 | lncRNA | ENSG00000( |
| 40114651 | 40127468 | 1 | lncRNA | ENSG00000( |
| 40153482 | 40155310 | 1 | protein_coding | ENSG00000( |
| 40161458 | 40172612 | 1 | lncRNA | ENSG00000( |
| 40298697 | 40353133 | -1 | lncRNA | ENSG00000( |
| 40369981 | 40402010 | 1 | lncRNA | ENSG00000( |
| 40519565 | 40520158 | -1 | lncRNA | ENSG00000( |
| 40530590 | 40897833 | -1 | protein_coding | ENSG00000( |
| 40900016 | 40901854 | -1 | lncRNA | ENSG00000( |
| 41032892 | 41032998 | 1 | snRNA | ENSG00000( |
| 41261962 | 41309473 | -1 | protein_coding | ENSG00000( |
| 41271048 | 41271143 | -1 | miRNA | ENSG00000( |
| 41271431 | 41271566 | 1 | processed_pseudoge | ENSG00000( |
| 41275115 | 41277003 | -1 | lncRNA | ENSG00000( |

| | | | | |
|---|---|---|---|---|
| 41298163 | 41298265 | 1 | snRNA | ENSG0000C |
| 41426655 | 41426727 | 1 | snoRNA | ENSG0000C |
| 41435298 | 41440419 | 1 | lncRNA | ENSG0000C |
| 41490396 | 41510980 | 1 | protein_coding | ENSG0000C |
| 41509593 | 41578421 | -1 | lncRNA | ENSG0000C |
| 41511240 | 41512521 | 1 | processed_pseudoge | ENSG0000C |
| 41529218 | 41545030 | 1 | protein_coding | ENSG0000C |
| 41540381 | 41545044 | -1 | lncRNA | ENSG0000C |
| 41566858 | 41568641 | 1 | lncRNA | ENSG0000C |
| 41577187 | 41625001 | 1 | protein_coding | ENSG0000C |
| 41609692 | 41621502 | -1 | lncRNA | ENSG0000C |
| 41645177 | 41650817 | -1 | protein_coding | ENSG0000C |
| 41653220 | 41896762 | -1 | protein_coding | ENSG0000C |
| 41660441 | 41660508 | -1 | miRNA | ENSG0000C |
| 41660444 | 41660507 | 1 | miRNA | ENSG0000C |
| 41660991 | 41665566 | 1 | lncRNA | ENSG0000C |
| 41803349 | 41803450 | 1 | misc_RNA | ENSG0000C |
| 41828165 | 41829934 | -1 | lncRNA | ENSG0000C |
| 41840059 | 41840339 | -1 | misc_RNA | ENSG0000C |
| 41929479 | 42051994 | -1 | protein_coding | ENSG0000C |
| 42043504 | 42043575 | -1 | snoRNA | ENSG0000C |
| 42101318 | 42101836 | 1 | processed_pseudoge | ENSG0000C |
| 42128774 | 42129249 | -1 | processed_pseudoge | ENSG0000C |
| 42139461 | 42139752 | -1 | lncRNA | ENSG0000C |
| 42151772 | 42152763 | -1 | lncRNA | ENSG0000C |
| 42152946 | 42171673 | 1 | protein_coding | ENSG0000C |
| 42174718 | 42207676 | -1 | protein_coding | ENSG0000C |
| 42233674 | 42271266 | -1 | lncRNA | ENSG0000C |
| 42271302 | 42332653 | 1 | protein_coding | ENSG0000C |
| 42338454 | 42371808 | 1 | protein_coding | ENSG0000C |
| 42341767 | 42342634 | -1 | processed_pseudoge | ENSG0000C |
| 42374063 | 42377229 | -1 | protein_coding | ENSG0000C |
| 42391624 | 42405937 | 1 | protein_coding | ENSG0000C |
| 42416475 | 42541926 | -1 | protein_coding | ENSG0000C |
| 42484600 | 42485019 | 1 | processed_pseudoge | ENSG0000C |
| 42537529 | 42538304 | -1 | lncRNA | ENSG0000C |
| 42541155 | 42555193 | 1 | protein_coding | ENSG0000C |
| 42697366 | 42737407 | 1 | protein_coding | ENSG0000C |
| 42705583 | 42721946 | -1 | lncRNA | ENSG0000C |
| 42725547 | 42726228 | -1 | processed_pseudoge | ENSG0000C |
| 42752620 | 42796392 | -1 | protein_coding | ENSG0000C |
| 42836674 | 42843325 | -1 | protein_coding | ENSG0000C |
| 42842989 | 42844876 | 1 | lncRNA | ENSG0000C |
| 42849637 | 42897290 | -1 | protein_coding | ENSG0000C |
| 42881236 | 42881517 | 1 | misc_RNA | ENSG0000C |
| 42896197 | 42896275 | -1 | miRNA | ENSG0000C |
| 42896946 | 43030535 | 1 | protein_coding | ENSG0000C |
| 42949388 | 42949495 | -1 | misc_RNA | ENSG0000C |
| 43018424 | 43077334 | 1 | protein_coding | ENSG0000C |
| 43034194 | 43085788 | 1 | protein_coding | ENSG0000C |

| | | | | |
|---|---|---|---|---|
| 43073474 | 43073618 | 1 | snRNA | ENSG0000C |
| 43093506 | 43123434 | 1 | protein_coding | ENSG0000C |
| 43125995 | 43126397 | -1 | processed_pseudoge | ENSG0000C |
| 43140464 | 43202855 | 1 | protein_coding | ENSG0000C |
| 43219960 | 43220809 | -1 | processed_pseudoge | ENSG0000C |
| 43246755 | 43246935 | -1 | processed_pseudoge | ENSG0000C |
| 43247191 | 43247410 | 1 | processed_pseudoge | ENSG0000C |
| 43251711 | 43251858 | 1 | processed_pseudoge | ENSG0000C |
| 43253732 | 43254089 | 1 | unprocessed_pseudc | ENSG0000C |
| 43270198 | 43271960 | -1 | processed_pseudoge | ENSG0000C |
| 43276270 | 43276823 | -1 | processed_pseudoge | ENSG0000C |
| 43284626 | 43284806 | -1 | processed_pseudoge | ENSG0000C |
| 43292483 | 43363518 | 1 | polymorphic_pseudc | ENSG0000C |
| 43303376 | 43303482 | 1 | snRNA | ENSG0000C |
| 43313948 | 43314200 | 1 | processed_pseudoge | ENSG0000C |
| 43372559 | 43372677 | 1 | processed_pseudoge | ENSG0000C |
| 43378297 | 43378510 | 1 | snoRNA | ENSG0000C |
| 43442902 | 43452912 | 1 | unprocessed_pseudc | ENSG0000C |
| 43493937 | 43494762 | 1 | processed_pseudoge | ENSG0000C |
| 43511820 | 43512033 | -1 | processed_pseudoge | ENSG0000C |
| 43513643 | 43514421 | 1 | processed_pseudoge | ENSG0000C |
| 43539137 | 43539472 | 1 | processed_pseudoge | ENSG0000C |
| 43539973 | 43540927 | 1 | unprocessed_pseudc | ENSG0000C |
| 43542076 | 43544057 | 1 | unprocessed_pseudc | ENSG0000C |
| 43672823 | 43673075 | 1 | processed_pseudoge | ENSG0000C |
| 43674270 | 43674591 | -1 | processed_pseudoge | ENSG0000C |
| 46028804 | 46028892 | -1 | misc_RNA | ENSG0000C |
| 46549089 | 46550802 | 1 | unprocessed_pseudc | ENSG0000C |
| 46579213 | 46638886 | -1 | transcribed_unproce | ENSG0000C |
| 46671839 | 46671926 | 1 | processed_pseudoge | ENSG0000C |
| 46697630 | 46698688 | 1 | unprocessed_pseudc | ENSG0000C |
| 46784800 | 46785928 | 1 | processed_pseudoge | ENSG0000C |
| 46792065 | 46793064 | 1 | processed_pseudoge | ENSG0000C |
| 46810696 | 46819098 | -1 | lncRNA | ENSG0000C |
| 46814925 | 46817964 | 1 | lncRNA | ENSG0000C |
| 46822174 | 46907309 | 1 | lncRNA | ENSG0000C |
| 46827228 | 46827894 | -1 | unprocessed_pseudc | ENSG0000C |
| 46828117 | 46828413 | -1 | unprocessed_pseudc | ENSG0000C |
| 46831050 | 46831156 | 1 | snRNA | ENSG0000C |
| 46837578 | 46838706 | -1 | processed_pseudoge | ENSG0000C |
| 46838788 | 46839205 | 1 | transcribed_processe | ENSG0000C |
| 46870902 | 46872045 | -1 | processed_pseudoge | ENSG0000C |
| 46922561 | 46928832 | 1 | lncRNA | ENSG0000C |
| 46930604 | 46951850 | -1 | lncRNA | ENSG0000C |
| 46972476 | 46974617 | -1 | processed_pseudoge | ENSG0000C |
| 47068021 | 47068323 | -1 | misc_RNA | ENSG0000C |
| 47089572 | 47198861 | -1 | unprocessed_pseudc | ENSG0000C |
| 47129262 | 47132628 | 1 | lncRNA | ENSG0000C |
| 47131781 | 47131887 | -1 | snRNA | ENSG0000C |
| 47157118 | 47157771 | 1 | processed_pseudoge | ENSG0000C |

| | | | | |
|---|---|---|---|---|
| 47164870 | 47165524 | 1 | unprocessed_pseudc | ENSG00000( |
| 47189305 | 47189560 | 1 | processed_pseudoge | ENSG00000( |
| 47190772 | 47193262 | 1 | lncRNA | ENSG00000( |
| 47194002 | 47194347 | -1 | processed_pseudoge | ENSG00000( |
| 47198032 | 47198323 | -1 | processed_pseudoge | ENSG00000( |
| 47202444 | 47202897 | -1 | IG_V_pseudogene | ENSG00000( |
| 47260878 | 47736306 | 1 | protein_coding | ENSG00000( |
| 47511034 | 47512141 | -1 | processed_pseudoge | ENSG00000( |
| 47527397 | 47528148 | 1 | lncRNA | ENSG00000( |
| 47546804 | 47547926 | -1 | unprocessed_pseudc | ENSG00000( |
| 47592147 | 47592609 | -1 | processed_pseudoge | ENSG00000( |
| 47667771 | 47667877 | 1 | snRNA | ENSG00000( |
| 47736913 | 47738164 | -1 | protein_coding | ENSG00000( |
| 47773108 | 47960183 | -1 | protein_coding | ENSG00000( |
| 47805079 | 47805179 | -1 | misc_RNA | ENSG00000( |
| 47941426 | 47943670 | 1 | lncRNA | ENSG00000( |
| 47960185 | 47978160 | 1 | protein_coding | ENSG00000( |
| 47979414 | 47979521 | 1 | snRNA | ENSG00000( |
| 48008415 | 48064708 | 1 | protein_coding | ENSG00000( |
| 48193850 | 48194627 | -1 | processed_pseudoge | ENSG00000( |
| 48294026 | 48294694 | -1 | processed_pseudoge | ENSG00000( |
| 48307927 | 48308030 | -1 | snRNA | ENSG00000( |
| 48316669 | 48316770 | 1 | rRNA | ENSG00000( |
| 48384590 | 48385049 | 1 | processed_pseudoge | ENSG00000( |
| 48428143 | 48431041 | -1 | TEC | ENSG00000( |
| 48515852 | 48518157 | -1 | lncRNA | ENSG00000( |
| 48551527 | 48556441 | -1 | lncRNA | ENSG00000( |
| 48551567 | 48698510 | 1 | lncRNA | ENSG00000( |
| 48590401 | 48594621 | 1 | lncRNA | ENSG00000( |
| 48597458 | 48621018 | -1 | lncRNA | ENSG00000( |
| 48620465 | 48623895 | 1 | lncRNA | ENSG00000( |
| 48657260 | 48658894 | -1 | lncRNA | ENSG00000( |
| 48710789 | 48735311 | -1 | protein_coding | ENSG00000( |
| 48818293 | 48824062 | 1 | lncRNA | ENSG00000( |
| 48913916 | 48914380 | 1 | processed_pseudoge | ENSG00000( |
| 48917598 | 48921740 | -1 | protein_coding | ENSG00000( |
| 48925917 | 48931288 | 1 | lncRNA | ENSG00000( |
| 49046652 | 49058536 | 1 | lncRNA | ENSG00000( |
| 49054311 | 49076090 | 1 | protein_coding | ENSG00000( |
| 49086301 | 49229174 | 1 | lncRNA | ENSG00000( |
| 49330210 | 49330547 | 1 | misc_RNA | ENSG00000( |
| 49352630 | 49352844 | -1 | processed_pseudoge | ENSG00000( |
| 49496763 | 49512180 | -1 | lncRNA | ENSG00000( |
| 49536386 | 49554414 | -1 | lncRNA | ENSG00000( |
| 49740216 | 49741321 | -1 | processed_pseudoge | ENSG00000( |
| 49817586 | 49820944 | -1 | lncRNA | ENSG00000( |
| 49906863 | 49907446 | -1 | lncRNA | ENSG00000( |
| 49909789 | 50796692 | 1 | protein_coding | ENSG00000( |
| 50381015 | 50382275 | 1 | lncRNA | ENSG00000( |
| 50490795 | 50491465 | 1 | processed_pseudoge | ENSG00000( |

| | | | | |
|---|---|---|---|---|
| 50762460 | 50762798 | -1 | processed_pseudoge | ENSG00000 |
| 50859530 | 50860062 | 1 | processed_pseudoge | ENSG00000 |
| 51015021 | 51015165 | 1 | snoRNA | ENSG00000 |
| 51257688 | 51321118 | 1 | lncRNA | ENSG00000 |
| 51319577 | 51809445 | -1 | protein_coding | ENSG00000 |
| 51435602 | 51436509 | 1 | processed_pseudoge | ENSG00000 |
| 51721670 | 51722164 | -1 | processed_pseudoge | ENSG00000 |
| 51810110 | 51810681 | -1 | lncRNA | ENSG00000 |
| 51817575 | 51899186 | -1 | protein_coding | ENSG00000 |
| 51895957 | 51896374 | -1 | lncRNA | ENSG00000 |
| 51899268 | 51949874 | 1 | lncRNA | ENSG00000 |
| 51950284 | 51950690 | 1 | lncRNA | ENSG00000 |
| 51961458 | 52022974 | 1 | lncRNA | ENSG00000 |
| 51996210 | 52008355 | -1 | lncRNA | ENSG00000 |
| 52110839 | 52460959 | -1 | protein_coding | ENSG00000 |
| 52150820 | 52154892 | 1 | lncRNA | ENSG00000 |
| 52194362 | 52199580 | 1 | lncRNA | ENSG00000 |
| 52294455 | 52301942 | 1 | lncRNA | ENSG00000 |
| 52313154 | 52313500 | -1 | processed_pseudoge | ENSG00000 |
| 52534037 | 52565430 | -1 | protein_coding | ENSG00000 |
| 52622458 | 52745843 | -1 | protein_coding | ENSG00000 |
| 52722903 | 52723141 | 1 | lncRNA | ENSG00000 |
| 52938431 | 52941117 | 1 | protein_coding | ENSG00000 |
| 52975111 | 52976116 | -1 | processed_pseudoge | ENSG00000 |
| 53162339 | 53162987 | 1 | processed_pseudoge | ENSG00000 |
| 53177571 | 53242683 | 1 | lncRNA | ENSG00000 |
| 53225716 | 53251697 | -1 | protein_coding | ENSG00000 |
| 53388701 | 53390872 | -1 | lncRNA | ENSG00000 |
| 53394110 | 53484067 | -1 | lncRNA | ENSG00000 |
| 53394215 | 53394795 | 1 | processed_pseudoge | ENSG00000 |
| 53493523 | 53524336 | -1 | lncRNA | ENSG00000 |
| 53515170 | 53515905 | 1 | lncRNA | ENSG00000 |
| 53532908 | 53533704 | 1 | processed_pseudoge | ENSG00000 |
| 53537060 | 53540300 | 1 | processed_pseudoge | ENSG00000 |
| 53712697 | 53712896 | 1 | processed_pseudoge | ENSG00000 |
| 53715543 | 53843558 | -1 | protein_coding | ENSG00000 |
| 53851795 | 53959303 | 1 | protein_coding | ENSG00000 |
| 53876150 | 53876649 | -1 | lncRNA | ENSG00000 |
| 53887008 | 53887244 | -1 | processed_pseudoge | ENSG00000 |
| 53914719 | 53914824 | 1 | snRNA | ENSG00000 |
| 53966552 | 54022456 | -1 | protein_coding | ENSG00000 |
| 53971231 | 53974210 | 1 | processed_pseudoge | ENSG00000 |
| 54042989 | 54045629 | -1 | lncRNA | ENSG00000 |
| 54046367 | 54102017 | -1 | protein_coding | ENSG00000 |
| 54073114 | 54073566 | 1 | processed_pseudoge | ENSG00000 |
| 54130582 | 54130860 | 1 | misc_RNA | ENSG00000 |
| 54135241 | 54148514 | 1 | protein_coding | ENSG00000 |
| 54152807 | 54152882 | -1 | snRNA | ENSG00000 |
| 54191582 | 54192035 | -1 | processed_pseudoge | ENSG00000 |
| 54247633 | 54247906 | 1 | processed_pseudoge | ENSG00000 |

| | | | | |
|---|---|---|---|---|
| 54330687 | 54330895 | 1 | snoRNA | ENSG0000C |
| 54354168 | 54355118 | 1 | processed_pseudoge | ENSG0000C |
| 54381698 | 54394870 | 1 | lncRNA | ENSG0000C |
| 54420468 | 54420725 | 1 | misc_RNA | ENSG0000C |
| 54457935 | 54460892 | 1 | protein_coding | ENSG0000C |
| 54467208 | 54470609 | 1 | unprocessed_pseudc | ENSG0000C |
| 54479597 | 54479963 | -1 | processed_pseudoge | ENSG0000C |
| 54509422 | 54871720 | 1 | protein_coding | ENSG0000C |
| 54522799 | 54523297 | -1 | processed_pseudoge | ENSG0000C |
| 54696398 | 54697127 | 1 | processed_pseudoge | ENSG0000C |
| 55067585 | 55081909 | -1 | lncRNA | ENSG0000C |
| 55102028 | 55542054 | 1 | protein_coding | ENSG0000C |
| 55135203 | 55142358 | -1 | lncRNA | ENSG0000C |
| 55161446 | 55164727 | 1 | lncRNA | ENSG0000C |
| 55449509 | 55455139 | -1 | processed_pseudoge | ENSG0000C |
| 55517240 | 55520977 | -1 | lncRNA | ENSG0000C |
| 55696424 | 55773407 | -1 | protein_coding | ENSG0000C |
| 55745232 | 55745342 | 1 | rRNA_pseudogene | ENSG0000C |
| 55773446 | 55826445 | 1 | protein_coding | ENSG0000C |
| 55879835 | 56014169 | 1 | protein_coding | ENSG0000C |
| 55893595 | 55895739 | 1 | lncRNA | ENSG0000C |
| 55902723 | 55902855 | -1 | snoRNA | ENSG0000C |
| 55980240 | 55980540 | -1 | misc_RNA | ENSG0000C |
| 56022019 | 56022313 | -1 | misc_RNA | ENSG0000C |
| 56050038 | 56051207 | -1 | processed_pseudoge | ENSG0000C |
| 56067295 | 56074581 | -1 | protein_coding | ENSG0000C |
| 56073835 | 56073901 | -1 | snoRNA | ENSG0000C |
| 56074592 | 56075274 | 1 | lncRNA | ENSG0000C |
| 56101199 | 56102267 | -1 | processed_pseudoge | ENSG0000C |
| 56112942 | 56113982 | -1 | protein_coding | ENSG0000C |
| 56121242 | 56121352 | -1 | snoRNA | ENSG0000C |
| 56160909 | 56211324 | -1 | protein_coding | ENSG0000C |
| 56211686 | 56218809 | 1 | protein_coding | ENSG0000C |
| 56222688 | 56223173 | 1 | lncRNA | ENSG0000C |
| 56252960 | 56253372 | -1 | processed_pseudoge | ENSG0000C |
| 56300010 | 56320776 | -1 | protein_coding | ENSG0000C |
| 56373064 | 56395365 | -1 | transcribed_unitary_ | ENSG0000C |
| 56436674 | 56446671 | -1 | protein_coding | ENSG0000C |
| 56445807 | 56556513 | 1 | lncRNA | ENSG0000C |
| 56476826 | 56477987 | -1 | processed_pseudoge | ENSG0000C |
| 56496048 | 56559823 | -1 | lncRNA | ENSG0000C |
| 56536758 | 56540497 | -1 | lncRNA | ENSG0000C |
| 56588385 | 56588787 | 1 | transcribed_processe | ENSG0000C |
| 56638894 | 56656496 | -1 | lncRNA | ENSG0000C |
| 56776283 | 56776874 | 1 | processed_pseudoge | ENSG0000C |
| 56917084 | 56917190 | 1 | snRNA | ENSG0000C |
| 56957931 | 56993867 | -1 | protein_coding | ENSG0000C |
| 57126655 | 57126762 | -1 | rRNA_pseudogene | ENSG0000C |
| 57142659 | 57244793 | 1 | lncRNA | ENSG0000C |
| 57240057 | 57240541 | 1 | processed_pseudoge | ENSG0000C |

| | | | | |
|---|---|---|---|---|
| 57261226 | 57266613 | 1 | lncRNA | ENSG00000( |
| 57279543 | 57286968 | 1 | lncRNA | ENSG00000( |
| 57289552 | 57289649 | 1 | snRNA | ENSG00000( |
| 57341021 | 57365444 | 1 | lncRNA | ENSG00000( |
| 57392546 | 57392882 | 1 | processed_pseudoge | ENSG00000( |
| 57420834 | 57423673 | 1 | lncRNA | ENSG00000( |
| 57492884 | 57592451 | 1 | lncRNA | ENSG00000( |
| 57594166 | 57805029 | -1 | lncRNA | ENSG00000( |
| 57746149 | 57750199 | 1 | lncRNA | ENSG00000( |
| 57855500 | 57984126 | 1 | lncRNA | ENSG00000( |
| 57888975 | 57934329 | -1 | lncRNA | ENSG00000( |
| 57994509 | 58204279 | 1 | protein_coding | ENSG00000( |
| 58031334 | 58032682 | -1 | lncRNA | ENSG00000( |
| 58091442 | 58106294 | -1 | lncRNA | ENSG00000( |
| 58113189 | 58113622 | -1 | processed_pseudoge | ENSG00000( |
| 58227578 | 58243984 | -1 | lncRNA | ENSG00000( |
| 58255771 | 58272137 | -1 | lncRNA | ENSG00000( |
| 58269979 | 58272450 | 1 | lncRNA | ENSG00000( |
| 58411359 | 58451501 | 1 | protein_coding | ENSG00000( |
| 58424588 | 58426685 | -1 | processed_pseudoge | ENSG00000( |
| 58490178 | 58500163 | -1 | protein_coding | ENSG00000( |
| 58503588 | 58504068 | 1 | processed_pseudoge | ENSG00000( |
| 58552924 | 58582859 | 1 | protein_coding | ENSG00000( |
| 58583508 | 58659853 | -1 | protein_coding | ENSG00000( |
| 58588420 | 58588764 | -1 | processed_pseudoge | ENSG00000( |
| 58805412 | 59119147 | -1 | protein_coding | ENSG00000( |
| 58814412 | 58814552 | -1 | snRNA | ENSG00000( |
| 58991720 | 58992938 | -1 | lncRNA | ENSG00000( |
| 59119040 | 59123478 | 1 | lncRNA | ENSG00000( |
| 59137372 | 59137502 | -1 | snoRNA | ENSG00000( |
| 59455885 | 59456010 | 1 | rRNA_pseudogene | ENSG00000( |
| 59557528 | 59557855 | -1 | processed_pseudoge | ENSG00000( |
| 59561324 | 59604774 | -1 | lncRNA | ENSG00000( |
| 59601319 | 59620227 | 1 | lncRNA | ENSG00000( |
| 59760047 | 59760263 | 1 | processed_pseudoge | ENSG00000( |
| 59821124 | 59821433 | 1 | processed_pseudoge | ENSG00000( |
| 59893856 | 59893942 | -1 | unprocessed_pseudc | ENSG00000( |
| 60046755 | 60136599 | 1 | lncRNA | ENSG00000( |
| 60141821 | 60142142 | 1 | processed_pseudoge | ENSG00000( |
| 60185412 | 60281400 | -1 | protein_coding | ENSG00000( |
| 60289928 | 60516795 | -1 | lncRNA | ENSG00000( |
| 60387262 | 60387995 | 1 | processed_pseudoge | ENSG00000( |
| 60516936 | 60623644 | 1 | protein_coding | ENSG00000( |
| 60551957 | 60553036 | -1 | lncRNA | ENSG00000( |
| 60629662 | 60653561 | -1 | lncRNA | ENSG00000( |
| 60660820 | 60664530 | 1 | lncRNA | ENSG00000( |
| 60678740 | 60868028 | 1 | protein_coding | ENSG00000( |
| 60808735 | 60809606 | -1 | lncRNA | ENSG00000( |
| 60876969 | 60904644 | 1 | lncRNA | ENSG00000( |
| 60906059 | 60906452 | 1 | processed_pseudoge | ENSG00000( |

| | | | | |
|---|---|---|---|---|
| 60910053 | 60966557 | 1 | lncRNA | ENSG00000 |
| 60937705 | 60939871 | -1 | processed_pseudoge | ENSG00000 |
| 60965802 | 60967775 | -1 | lncRNA | ENSG00000 |
| 61057158 | 61501645 | 1 | protein_coding | ENSG00000 |
| 61202350 | 61203220 | 1 | processed_pseudoge | ENSG00000 |
| 61264624 | 61292039 | -1 | lncRNA | ENSG00000 |
| 61300164 | 61301069 | -1 | lncRNA | ENSG00000 |
| 61500556 | 61714640 | -1 | protein_coding | ENSG00000 |
| 61578220 | 61579668 | 1 | processed_pseudoge | ENSG00000 |
| 61631680 | 61632012 | -1 | misc_RNA | ENSG00000 |
| 61714788 | 61714859 | 1 | miRNA | ENSG00000 |
| 61759094 | 61765969 | -1 | lncRNA | ENSG00000 |
| 61785047 | 61944180 | 1 | lncRNA | ENSG00000 |
| 61821481 | 61825252 | -1 | lncRNA | ENSG00000 |
| 61889839 | 61894206 | 1 | lncRNA | ENSG00000 |
| 61909984 | 61910802 | -1 | processed_pseudoge | ENSG00000 |
| 62110524 | 62119898 | 1 | transcribed_processe | ENSG00000 |
| 62143311 | 62144769 | -1 | processed_pseudoge | ENSG00000 |
| 62191088 | 62249879 | -1 | lncRNA | ENSG00000 |
| 62248591 | 62999652 | 1 | protein_coding | ENSG00000 |
| 62473217 | 62474269 | -1 | lncRNA | ENSG00000 |
| 62482198 | 62483811 | -1 | processed_pseudoge | ENSG00000 |
| 62658343 | 62690133 | -1 | lncRNA | ENSG00000 |
| 62855068 | 62857134 | -1 | processed_pseudoge | ENSG00000 |
| 62863082 | 62864181 | -1 | processed_pseudoge | ENSG00000 |
| 62977861 | 62984900 | 1 | lncRNA | ENSG00000 |
| 63013068 | 63014681 | 1 | lncRNA | ENSG00000 |
| 63015079 | 63038806 | -1 | protein_coding | ENSG00000 |
| 63024372 | 63025294 | 1 | lncRNA | ENSG00000 |
| 63048553 | 63086053 | -1 | protein_coding | ENSG00000 |
| 63136223 | 63138084 | -1 | unprocessed_pseudo | ENSG00000 |
| 63167725 | 63168442 | -1 | lncRNA | ENSG00000 |
| 63168553 | 63212786 | 1 | protein_coding | ENSG00000 |
| 63215981 | 63218034 | 1 | lncRNA | ENSG00000 |
| 63234634 | 63234911 | -1 | misc_RNA | ENSG00000 |
| 63384839 | 63470205 | -1 | lncRNA | ENSG00000 |
| 63409064 | 63409457 | 1 | processed_pseudoge | ENSG00000 |
| 63465831 | 63478745 | 1 | lncRNA | ENSG00000 |
| 63586000 | 63589408 | 1 | lncRNA | ENSG00000 |
| 63609354 | 63609600 | -1 | misc_RNA | ENSG00000 |
| 63651457 | 63693249 | 1 | lncRNA | ENSG00000 |
| 63687179 | 64369176 | -1 | lncRNA | ENSG00000 |
| 63703077 | 63751677 | 1 | lncRNA | ENSG00000 |
| 63769428 | 63813937 | 1 | lncRNA | ENSG00000 |
| 64331927 | 64332152 | 1 | processed_pseudoge | ENSG00000 |
| 64373151 | 64383787 | 1 | lncRNA | ENSG00000 |
| 64379149 | 64379257 | 1 | miRNA | ENSG00000 |
| 64456198 | 64462926 | -1 | lncRNA | ENSG00000 |
| 64554236 | 64554475 | -1 | processed_pseudoge | ENSG00000 |
| 64570913 | 64571332 | -1 | processed_pseudoge | ENSG00000 |

| | | | | |
|---|---|---|---|---|
| 64574306 | 64581921 | -1 | lncRNA | ENSG0000C |
| 64580365 | 64583627 | 1 | protein_coding | ENSG0000C |
| 64587763 | 64798737 | -1 | protein_coding | ENSG0000C |
| 64703774 | 64734459 | -1 | lncRNA | ENSG0000C |
| 64798804 | 64862370 | 1 | lncRNA | ENSG0000C |
| 64801236 | 64817573 | -1 | lncRNA | ENSG0000C |
| 64973488 | 64973848 | -1 | processed_pseudoge | ENSG0000C |
| 65155407 | 65155778 | 1 | processed_pseudoge | ENSG0000C |
| 65161145 | 65184210 | -1 | lncRNA | ENSG0000C |
| 65377592 | 65378111 | 1 | processed_pseudoge | ENSG0000C |
| 65526738 | 65562781 | -1 | lncRNA | ENSG0000C |
| 65591850 | 65592472 | -1 | lncRNA | ENSG0000C |
| 65592731 | 65592820 | 1 | misc_RNA | ENSG0000C |
| 65602458 | 65634217 | -1 | protein_coding | ENSG0000C |
| 65644734 | 65771261 | 1 | protein_coding | ENSG0000C |
| 65714334 | 65714778 | -1 | lncRNA | ENSG0000C |
| 65717510 | 65842322 | -1 | protein_coding | ENSG0000C |
| 65842752 | 65843331 | 1 | lncRNA | ENSG0000C |
| 66021553 | 66101245 | 1 | protein_coding | ENSG0000C |
| 66112667 | 66126632 | 1 | lncRNA | ENSG0000C |
| 66126896 | 66175485 | 1 | protein_coding | ENSG0000C |
| 66176376 | 66178464 | -1 | protein_coding | ENSG0000C |
| 66192093 | 66197315 | 1 | lncRNA | ENSG0000C |
| 66363774 | 66432370 | -1 | lncRNA | ENSG0000C |
| 66419589 | 66428977 | -1 | lncRNA | ENSG0000C |
| 66429014 | 66430733 | 1 | protein_coding | ENSG0000C |
| 66430185 | 66445324 | 1 | protein_coding | ENSG0000C |
| 66432486 | 66518524 | 1 | protein_coding | ENSG0000C |
| 66432492 | 66468907 | 1 | protein_coding | ENSG0000C |
| 66480648 | 66481486 | 1 | processed_pseudoge | ENSG0000C |
| 66493520 | 66518524 | 1 | protein_coding | ENSG0000C |
| 66501205 | 66501311 | -1 | snRNA | ENSG0000C |
| 66562175 | 66614247 | -1 | protein_coding | ENSG0000C |
| 66628487 | 66667231 | -1 | protein_coding | ENSG0000C |
| 66653469 | 66653576 | 1 | misc_RNA | ENSG0000C |
| 66667552 | 66862022 | 1 | protein_coding | ENSG0000C |
| 66767400 | 66768005 | -1 | processed_pseudoge | ENSG0000C |
| 66870749 | 66922048 | 1 | protein_coding | ENSG0000C |
| 66921684 | 66926398 | -1 | lncRNA | ENSG0000C |
| 66922467 | 66922555 | -1 | snoRNA | ENSG0000C |
| 66946501 | 66962591 | -1 | protein_coding | ENSG0000C |
| 66964099 | 67056604 | -1 | protein_coding | ENSG0000C |
| 66984135 | 66986187 | -1 | processed_pseudoge | ENSG0000C |
| 67043079 | 67083783 | -1 | protein_coding | ENSG0000C |
| 67062426 | 67196263 | 1 | protein_coding | ENSG0000C |
| 67099302 | 67099425 | 1 | rRNA_pseudogene | ENSG0000C |
| 67133554 | 67135911 | -1 | unprocessed_pseudc | ENSG0000C |
| 67173511 | 67343781 | -1 | protein_coding | ENSG0000C |
| 67313043 | 67314637 | -1 | lncRNA | ENSG0000C |
| 67343975 | 67345087 | 1 | lncRNA | ENSG0000C |

| | | | | |
|---|---|---|---|---|
| 67406515 | 67407072 | 1 | processed_pseudoge | ENSG00000 |
| 67422038 | 67746378 | -1 | protein_coding | ENSG00000 |
| 67569416 | 67570078 | 1 | processed_pseudoge | ENSG00000 |
| 67732587 | 67735665 | 1 | lncRNA | ENSG00000 |
| 67849044 | 67849338 | 1 | processed_pseudoge | ENSG00000 |
| 67952046 | 68237032 | 1 | protein_coding | ENSG00000 |
| 68082204 | 68095081 | -1 | lncRNA | ENSG00000 |
| 68176768 | 68177221 | 1 | processed_pseudoge | ENSG00000 |
| 68223318 | 68223402 | -1 | misc_RNA | ENSG00000 |
| 68293446 | 68293818 | -1 | processed_pseudoge | ENSG00000 |
| 68302246 | 68331689 | -1 | lncRNA | ENSG00000 |
| 68330955 | 68819023 | 1 | protein_coding | ENSG00000 |
| 68525517 | 68525906 | -1 | processed_pseudoge | ENSG00000 |
| 68695643 | 68695779 | 1 | rRNA_pseudogene | ENSG00000 |
| 68848737 | 68852763 | -1 | lncRNA | ENSG00000 |
| 68880139 | 69104242 | -1 | lncRNA | ENSG00000 |
| 69110513 | 69110574 | -1 | snRNA | ENSG00000 |
| 69129598 | 69130345 | 1 | processed_pseudoge | ENSG00000 |
| 69175579 | 69178189 | -1 | lncRNA | ENSG00000 |
| 69424849 | 69448282 | -1 | lncRNA | ENSG00000 |
| 69446057 | 69446260 | 1 | processed_pseudoge | ENSG00000 |
| 69466624 | 69660915 | 1 | protein_coding | ENSG00000 |
| 69667046 | 69834978 | -1 | protein_coding | ENSG00000 |
| 69700224 | 69700495 | 1 | misc_RNA | ENSG00000 |
| 69713390 | 69719722 | 1 | lncRNA | ENSG00000 |
| 69794062 | 69794734 | 1 | processed_pseudoge | ENSG00000 |
| 69834111 | 69854971 | 1 | lncRNA | ENSG00000 |
| 69851148 | 69851436 | -1 | misc_RNA | ENSG00000 |
| 69859264 | 69860540 | -1 | processed_pseudoge | ENSG00000 |
| 69922751 | 69923047 | -1 | misc_RNA | ENSG00000 |
| 69929905 | 69930022 | 1 | rRNA_pseudogene | ENSG00000 |
| 69938168 | 69938400 | 1 | TEC | ENSG00000 |
| 69942948 | 69944648 | 1 | processed_pseudoge | ENSG00000 |
| 69987415 | 69987697 | -1 | processed_pseudoge | ENSG00000 |
| 70051651 | 70071693 | -1 | protein_coding | ENSG00000 |
| 70077906 | 70078032 | -1 | snRNA | ENSG00000 |
| 70103798 | 70104179 | -1 | processed_pseudoge | ENSG00000 |
| 70109782 | 70403808 | -1 | protein_coding | ENSG00000 |
| 70273369 | 70273837 | 1 | processed_pseudoge | ENSG00000 |
| 70357347 | 70357448 | 1 | misc_RNA | ENSG00000 |
| 70424125 | 70424576 | -1 | processed_pseudoge | ENSG00000 |
| 70471112 | 70480623 | -1 | lncRNA | ENSG00000 |
| 70471134 | 70485687 | 1 | lncRNA | ENSG00000 |
| 70480192 | 70480487 | -1 | misc_RNA | ENSG00000 |
| 70506251 | 70506818 | -1 | processed_pseudoge | ENSG00000 |
| 70525053 | 70535646 | -1 | lncRNA | ENSG00000 |
| 70573218 | 70608416 | -1 | protein_coding | ENSG00000 |
| 70608577 | 70663279 | 1 | lncRNA | ENSG00000 |
| 70635318 | 70669185 | -1 | protein_coding | ENSG00000 |
| 70654578 | 70654883 | -1 | misc_RNA | ENSG00000 |

| | | | | |
|---|---|---|---|---|
| 70669339 | 70790371 | 1 | protein_coding | ENSG00000( |
| 70669369 | 71228629 | 1 | lncRNA | ENSG00000( |
| 70780914 | 70781008 | -1 | misc_RNA | ENSG00000( |
| 71155454 | 71204223 | 1 | lncRNA | ENSG00000( |
| 71197433 | 71592025 | -1 | protein_coding | ENSG00000( |
| 71448811 | 71449219 | -1 | processed_pseudoge | ENSG00000( |
| 71675300 | 71702786 | 1 | lncRNA | ENSG00000( |
| 71779605 | 71792875 | -1 | lncRNA | ENSG00000( |
| 71815494 | 71815625 | -1 | snoRNA | ENSG00000( |
| 71828167 | 72118393 | 1 | lncRNA | ENSG00000( |
| 71841549 | 71844468 | -1 | protein_coding | ENSG00000( |
| 71895991 | 71896330 | 1 | processed_pseudoge | ENSG00000( |
| 72019917 | 72075584 | -1 | protein_coding | ENSG00000( |
| 72196334 | 72202269 | 1 | lncRNA | ENSG00000( |
| 72202751 | 72251565 | -1 | unprocessed_pseudc | ENSG00000( |
| 72357728 | 72357815 | 1 | rRNA_pseudogene | ENSG00000( |
| 72429620 | 72430112 | 1 | processed_pseudoge | ENSG00000( |
| 72462311 | 72470972 | 1 | lncRNA | ENSG00000( |
| 72537225 | 72938349 | 1 | protein_coding | ENSG00000( |
| 72618722 | 72619383 | -1 | processed_pseudoge | ENSG00000( |
| 72660584 | 72661470 | -1 | unprocessed_pseudc | ENSG00000( |
| 72732045 | 72751843 | 1 | lncRNA | ENSG00000( |
| 72874859 | 72881740 | -1 | lncRNA | ENSG00000( |
| 72947150 | 72950445 | 1 | lncRNA | ENSG00000( |
| 73008864 | 73048123 | 1 | protein_coding | ENSG00000( |
| 73018945 | 73019044 | -1 | snRNA | ENSG00000( |
| 73042910 | 73043346 | -1 | lncRNA | ENSG00000( |
| 73052178 | 73063061 | 1 | lncRNA | ENSG00000( |
| 73064543 | 73124088 | -1 | protein_coding | ENSG00000( |
| 73083787 | 73085879 | -1 | lncRNA | ENSG00000( |
| 73215929 | 73216627 | -1 | processed_pseudoge | ENSG00000( |
| 73241329 | 73259502 | -1 | protein_coding | ENSG00000( |
| 73277364 | 73278046 | -1 | processed_pseudoge | ENSG00000( |
| 73290242 | 73295789 | -1 | protein_coding | ENSG00000( |
| 73294602 | 73325281 | 1 | protein_coding | ENSG00000( |
| 73297711 | 73356461 | -1 | lncRNA | ENSG00000( |
| 73326991 | 73370601 | -1 | lncRNA | ENSG00000( |
| 73358670 | 73364212 | 1 | lncRNA | ENSG00000( |
| 73420004 | 73441526 | 1 | lncRNA | ENSG00000( |
| 73420369 | 73747708 | -1 | protein_coding | ENSG00000( |
| 73511276 | 73511654 | -1 | processed_pseudoge | ENSG00000( |
| 73590574 | 73590883 | 1 | processed_pseudoge | ENSG00000( |
| 73651289 | 73651602 | 1 | processed_pseudoge | ENSG00000( |
| 73670441 | 73732897 | 1 | lncRNA | ENSG00000( |
| 73688691 | 73878854 | -1 | protein_coding | ENSG00000( |
| 73780097 | 73878910 | -1 | protein_coding | ENSG00000( |
| 73831143 | 73831947 | 1 | unprocessed_pseudc | ENSG00000( |
| 73833773 | 73834042 | -1 | misc_RNA | ENSG00000( |
| 73879046 | 73882733 | 1 | lncRNA | ENSG00000( |
| 73905357 | 73915010 | -1 | unprocessed_pseudc | ENSG00000( |

| | | | | |
|---|---|---|---|---|
| 73939169 | 73972287 | -1 | protein_coding | ENSG0000C |
| 73972437 | 73982783 | 1 | protein_coding | ENSG0000C |
| 73982125 | 73982467 | 1 | processed_pseudoge | ENSG0000C |
| 73984493 | 73984883 | 1 | processed_pseudoge | ENSG0000C |
| 73991392 | 74029079 | 1 | protein_coding | ENSG0000C |
| 74048677 | 74048784 | 1 | snRNA | ENSG0000C |
| 74052340 | 74099853 | 1 | lncRNA | ENSG0000C |
| 74055882 | 74056647 | 1 | processed_pseudoge | ENSG0000C |
| 74079537 | 74079643 | 1 | snRNA | ENSG0000C |
| 74093545 | 74208536 | 1 | lncRNA | ENSG0000C |
| 74103516 | 74106744 | 1 | lncRNA | ENSG0000C |
| 74234700 | 74321540 | -1 | protein_coding | ENSG0000C |
| 74321130 | 74488872 | 1 | protein_coding | ENSG0000C |
| 74347757 | 74350050 | -1 | lncRNA | ENSG0000C |
| 74376687 | 74376793 | 1 | misc_RNA | ENSG0000C |
| 74548543 | 74548617 | 1 | miRNA | ENSG0000C |
| 74548550 | 74548609 | -1 | miRNA | ENSG0000C |
| 74599775 | 74823313 | 1 | lncRNA | ENSG0000C |
| 74603244 | 74604244 | -1 | processed_pseudoge | ENSG0000C |
| 74609698 | 74633320 | -1 | lncRNA | ENSG0000C |
| 74705693 | 74705747 | 1 | miRNA | ENSG0000C |
| 74798784 | 74866939 | -1 | lncRNA | ENSG0000C |
| 74804287 | 74804811 | -1 | processed_pseudoge | ENSG0000C |
| 74824534 | 74855029 | 1 | protein_coding | ENSG0000C |
| 74891151 | 74896302 | -1 | lncRNA | ENSG0000C |
| 74984505 | 75034558 | 1 | protein_coding | ENSG0000C |
| 75026428 | 75029460 | -1 | lncRNA | ENSG0000C |
| 75167704 | 75176656 | -1 | lncRNA | ENSG0000C |
| 75223127 | 75324741 | -1 | lncRNA | ENSG0000C |
| 75302296 | 75302574 | 1 | processed_pseudoge | ENSG0000C |
| 75376709 | 75377014 | 1 | processed_pseudoge | ENSG0000C |
| 75407914 | 75566834 | 1 | protein_coding | ENSG0000C |
| 76162119 | 76308315 | -1 | lncRNA | ENSG0000C |
| 76265543 | 76265732 | -1 | snRNA | ENSG0000C |
| 76403998 | 76405091 | 1 | lncRNA | ENSG0000C |
| 76404058 | 76407138 | 1 | lncRNA | ENSG0000C |
| 76406654 | 76524356 | 1 | lncRNA | ENSG0000C |
| 76491200 | 76683308 | -1 | lncRNA | ENSG0000C |
| 76576551 | 76579345 | -1 | lncRNA | ENSG0000C |
| 76603240 | 76603852 | -1 | transcribed_processe | ENSG0000C |
| 76681239 | 76867281 | 1 | protein_coding | ENSG0000C |
| 76733302 | 76733476 | -1 | misc_RNA | ENSG0000C |
| 76904591 | 76913399 | 1 | lncRNA | ENSG0000C |
| 76966768 | 76966850 | -1 | miRNA | ENSG0000C |
| 76980258 | 77001044 | -1 | protein_coding | ENSG0000C |
| 77013751 | 77014028 | -1 | processed_pseudoge | ENSG0000C |
| 77399072 | 77537290 | 1 | lncRNA | ENSG0000C |
| 78148101 | 78153913 | 1 | lncRNA | ENSG0000C |
| 78268637 | 78558503 | -1 | lncRNA | ENSG0000C |
| 78398515 | 78398622 | -1 | snRNA | ENSG0000C |

| | | | | |
|---|---|---|---|---|
| 78516340 | 78605267 | 1 | protein_coding | ENSG0000C |
| 78605952 | 78609705 | 1 | lncRNA | ENSG0000C |
| 78626775 | 78665982 | -1 | lncRNA | ENSG0000C |
| 78666089 | 78719765 | 1 | protein_coding | ENSG0000C |
| 78675743 | 78805523 | -1 | protein_coding | ENSG0000C |
| 78723796 | 78724136 | -1 | processed_pseudoge | ENSG0000C |
| 78760142 | 78762431 | -1 | processed_pseudoge | ENSG0000C |
| 78805293 | 78956082 | 1 | lncRNA | ENSG0000C |
| 78835307 | 78840525 | 1 | lncRNA | ENSG0000C |
| 79259402 | 79314951 | 1 | lncRNA | ENSG0000C |
| 79520145 | 79520649 | 1 | processed_pseudoge | ENSG0000C |
| 79571153 | 79572335 | -1 | processed_pseudoge | ENSG0000C |
| 79610814 | 79666175 | 1 | protein_coding | ENSG0000C |
| 79747351 | 79749561 | 1 | lncRNA | ENSG0000C |
| 79764010 | 79767857 | -1 | protein_coding | ENSG0000C |
| 79768110 | 79802842 | 1 | lncRNA | ENSG0000C |
| 79769372 | 79871759 | -1 | lncRNA | ENSG0000C |
| 79800133 | 79800195 | -1 | snRNA | ENSG0000C |
| 79891553 | 79930892 | 1 | lncRNA | ENSG0000C |
| 79918717 | 80030289 | -1 | protein_coding | ENSG0000C |
| 79918860 | 80080775 | -1 | protein_coding | ENSG0000C |
| 79956465 | 79957381 | -1 | lncRNA | ENSG0000C |
| 80032724 | 80033300 | -1 | lncRNA | ENSG0000C |
| 80034745 | 80231232 | -1 | protein_coding | ENSG0000C |
| 80122852 | 80127670 | 1 | lncRNA | ENSG0000C |
| 80204606 | 80204902 | 1 | misc_RNA | ENSG0000C |
| 80241389 | 80241473 | 1 | miRNA | ENSG0000C |
| 80265528 | 80266155 | -1 | processed_pseudoge | ENSG0000C |
| 80265907 | 80484481 | 1 | lncRNA | ENSG0000C |
| 80300869 | 80301481 | 1 | processed_pseudoge | ENSG0000C |
| 80316939 | 80317069 | -1 | snoRNA | ENSG0000C |
| 80351132 | 80351416 | 1 | processed_pseudoge | ENSG0000C |
| 80405516 | 80405609 | 1 | snRNA | ENSG0000C |
| 80484561 | 80486699 | -1 | lncRNA | ENSG0000C |
| 80484589 | 80484683 | -1 | misc_RNA | ENSG0000C |
| 80485619 | 80526265 | 1 | protein_coding | ENSG0000C |
| 80535006 | 80539135 | -1 | lncRNA | ENSG0000C |
| 80541300 | 80543104 | 1 | lncRNA | ENSG0000C |
| 80558870 | 80559757 | 1 | processed_pseudoge | ENSG0000C |
| 80560567 | 80560625 | 1 | snRNA | ENSG0000C |
| 80568050 | 80568506 | -1 | processed_pseudoge | ENSG0000C |
| 80585585 | 80585773 | -1 | snRNA | ENSG0000C |
| 80594815 | 80595703 | -1 | processed_pseudoge | ENSG0000C |
| 80613544 | 80613826 | -1 | misc_RNA | ENSG0000C |
| 80628451 | 80874781 | -1 | protein_coding | ENSG0000C |
| 80644939 | 80645173 | -1 | processed_pseudoge | ENSG0000C |
| 80749561 | 80749693 | -1 | snRNA | ENSG0000C |
| 80776666 | 80776962 | -1 | misc_RNA | ENSG0000C |
| 80812142 | 80812581 | -1 | processed_pseudoge | ENSG0000C |
| 80894016 | 80930081 | 1 | lncRNA | ENSG0000C |

| | | | | |
|---|---|---|---|---|
| 80967810 | 81112068 | -1 | protein_coding | ENSG0000C |
| 81036964 | 81040831 | 1 | lncRNA | ENSG0000C |
| 81058523 | 81067744 | 1 | lncRNA | ENSG0000C |
| 81149107 | 81165266 | 1 | lncRNA | ENSG0000C |
| 81244156 | 81244338 | 1 | processed_pseudoge | ENSG0000C |
| 81254870 | 81255229 | 1 | processed_pseudoge | ENSG0000C |
| 81275399 | 81277570 | -1 | lncRNA | ENSG0000C |
| 81279871 | 81281446 | -1 | lncRNA | ENSG0000C |
| 81280536 | 81284777 | 1 | protein_coding | ENSG0000C |
| 81439436 | 81521818 | 1 | lncRNA | ENSG0000C |
| 81440326 | 81447439 | -1 | protein_coding | ENSG0000C |
| 81458383 | 81461579 | -1 | protein_coding | ENSG0000C |
| 81478419 | 81483236 | -1 | protein_coding | ENSG0000C |
| 81521618 | 81533275 | 1 | lncRNA | ENSG0000C |
| 81521682 | 81522232 | -1 | processed_pseudoge | ENSG0000C |
| 81524981 | 81531378 | -1 | protein_coding | ENSG0000C |
| 81605569 | 81631134 | -1 | transcribed_unproce | ENSG0000C |
| 81617220 | 81618560 | -1 | processed_pseudoge | ENSG0000C |
| 81627269 | 81627617 | 1 | processed_pseudoge | ENSG0000C |
| 81634357 | 81635662 | -1 | unprocessed_pseudc | ENSG0000C |
| 81656914 | 81686331 | -1 | protein_coding | ENSG0000C |
| 81693607 | 81695058 | -1 | protein_coding | ENSG0000C |
| 81696368 | 81698694 | 1 | TEC | ENSG0000C |
| 81701334 | 81732903 | -1 | protein_coding | ENSG0000C |
| 81732448 | 81759515 | 1 | protein_coding | ENSG0000C |
| 81791823 | 81815714 | 1 | lncRNA | ENSG0000C |
| 81799581 | 81842866 | -1 | protein_coding | ENSG0000C |
| 81807772 | 81808726 | 1 | processed_pseudoge | ENSG0000C |
| 81841952 | 81924348 | 1 | lncRNA | ENSG0000C |
| 82033533 | 82961926 | 1 | lncRNA | ENSG0000C |
| 82098451 | 82160577 | -1 | lncRNA | ENSG0000C |
| 82291624 | 82292379 | -1 | processed_pseudoge | ENSG0000C |
| 82514568 | 82677153 | -1 | lncRNA | ENSG0000C |
| 82862779 | 82958696 | -1 | lncRNA | ENSG0000C |
| 82979411 | 82979886 | 1 | processed_pseudoge | ENSG0000C |
| 83099976 | 83100846 | -1 | processed_pseudoge | ENSG0000C |
| 83200952 | 83202431 | -1 | processed_pseudoge | ENSG0000C |
| 83293925 | 83294271 | -1 | processed_pseudoge | ENSG0000C |
| 83403758 | 83408900 | 1 | lncRNA | ENSG0000C |
| 83912713 | 84142463 | 1 | lncRNA | ENSG0000C |
| 84162118 | 84162471 | 1 | processed_pseudoge | ENSG0000C |
| 84163449 | 84164564 | -1 | unprocessed_pseudc | ENSG0000C |
| 84182787 | 84921844 | 1 | protein_coding | ENSG0000C |
| 84665759 | 84665865 | -1 | snRNA | ENSG0000C |
| 84678798 | 84679842 | 1 | processed_pseudoge | ENSG0000C |
| 84828573 | 84829479 | -1 | unprocessed_pseudc | ENSG0000C |
| 84864668 | 84864907 | -1 | processed_pseudoge | ENSG0000C |
| 84948585 | 84949674 | 1 | processed_pseudoge | ENSG0000C |
| 84980073 | 84980547 | -1 | processed_pseudoge | ENSG0000C |
| 85086615 | 85087186 | 1 | processed_pseudoge | ENSG0000C |

| | | | | |
|---|---|---|---|---|
| 85107215 | 85146080 | 1 | protein_coding | ENSG0000C |
| 85172077 | 85177062 | -1 | lncRNA | ENSG0000C |
| 85177154 | 85217158 | 1 | protein_coding | ENSG0000C |
| 85177522 | 85178150 | -1 | lncRNA | ENSG0000C |
| 85214048 | 85220421 | -1 | protein_coding | ENSG0000C |
| 85220587 | 85284073 | 1 | protein_coding | ENSG0000C |
| 85222446 | 85245717 | -1 | lncRNA | ENSG0000C |
| 85246295 | 85249848 | -1 | lncRNA | ENSG0000C |
| 85327608 | 85379014 | -1 | protein_coding | ENSG0000C |
| 85373436 | 85449040 | 1 | protein_coding | ENSG0000C |
| 85440596 | 85464915 | -1 | lncRNA | ENSG0000C |
| 85463968 | 85481493 | 1 | protein_coding | ENSG0000C |
| 85495693 | 85540511 | 1 | lncRNA | ENSG0000C |
| 85617344 | 85617512 | -1 | processed_pseudoge | ENSG0000C |
| 85642777 | 85646325 | -1 | processed_pseudoge | ENSG0000C |
| 85654684 | 85658514 | -1 | processed_pseudoge | ENSG0000C |
| 85656442 | 85663497 | -1 | transcribed_processe | ENSG0000C |
| 85714922 | 85715833 | 1 | processed_pseudoge | ENSG0000C |
| 85733419 | 85737278 | 1 | processed_pseudoge | ENSG0000C |
| 85744543 | 85748373 | -1 | processed_pseudoge | ENSG0000C |
| 85762909 | 85766931 | -1 | processed_pseudoge | ENSG0000C |
| 85775106 | 85779090 | -1 | processed_pseudoge | ENSG0000C |
| 85790382 | 85794399 | -1 | processed_pseudoge | ENSG0000C |
| 85802574 | 85806435 | -1 | processed_pseudoge | ENSG0000C |
| 85814767 | 85818783 | -1 | processed_pseudoge | ENSG0000C |
| 85826865 | 85828467 | -1 | transcribed_processe | ENSG0000C |
| 85833377 | 85951083 | 1 | lncRNA | ENSG0000C |
| 85851446 | 85861562 | -1 | lncRNA | ENSG0000C |
| 85973793 | 85977154 | -1 | lncRNA | ENSG0000C |
| 85987323 | 86154225 | 1 | protein_coding | ENSG0000C |
| 86016819 | 86017126 | 1 | processed_pseudoge | ENSG0000C |
| 86043102 | 86043507 | 1 | processed_pseudoge | ENSG0000C |
| 86047109 | 86088621 | -1 | protein_coding | ENSG0000C |
| 86098965 | 86154225 | -1 | lncRNA | ENSG0000C |
| 86180418 | 86212236 | 1 | lncRNA | ENSG0000C |
| 86214063 | 86321146 | -1 | protein_coding | ENSG0000C |
| 86333274 | 86343429 | -1 | lncRNA | ENSG0000C |
| 86342547 | 86478420 | 1 | protein_coding | ENSG0000C |
| 86468257 | 86514357 | -1 | protein_coding | ENSG0000C |
| 86481754 | 86483002 | -1 | processed_pseudoge | ENSG0000C |
| 86503591 | 86505061 | 1 | processed_pseudoge | ENSG0000C |
| 86514435 | 86561498 | 1 | protein_coding | ENSG0000C |
| 86553977 | 86743675 | -1 | protein_coding | ENSG0000C |
| 86593755 | 86594589 | -1 | processed_pseudoge | ENSG0000C |
| 86657388 | 86660293 | -1 | processed_pseudoge | ENSG0000C |
| 86660525 | 86661922 | -1 | processed_pseudoge | ENSG0000C |
| 86707547 | 86765023 | 1 | lncRNA | ENSG0000C |
| 86765935 | 86818558 | 1 | lncRNA | ENSG0000C |
| 86776479 | 86776638 | -1 | processed_pseudoge | ENSG0000C |
| 86866415 | 87615219 | 1 | protein_coding | ENSG0000C |

| | | | | |
|---|---|---|---|---|
| 87540835 | 87755718 | -1 | lncRNA | ENSG0000C |
| 87575598 | 87576508 | -1 | processed_pseudoge | ENSG0000C |
| 87598462 | 87601634 | -1 | processed_pseudoge | ENSG0000C |
| 87788562 | 87789688 | -1 | processed_pseudoge | ENSG0000C |
| 87867601 | 87868607 | 1 | unprocessed_pseudc | ENSG0000C |
| 87870747 | 87874015 | -1 | protein_coding | ENSG0000C |
| 87974165 | 88019113 | 1 | lncRNA | ENSG0000C |
| 88032011 | 88328025 | -1 | protein_coding | ENSG0000C |
| 88326836 | 88887573 | 1 | lncRNA | ENSG0000C |
| 88396437 | 88396547 | -1 | rRNA_pseudogene | ENSG0000C |
| 88485110 | 88486094 | 1 | processed_pseudoge | ENSG0000C |
| 88808318 | 88808908 | -1 | lncRNA | ENSG0000C |
| 88809949 | 88813144 | -1 | lncRNA | ENSG0000C |
| 89243401 | 89243793 | -1 | processed_pseudoge | ENSG0000C |
| 89412621 | 89415071 | -1 | processed_pseudoge | ENSG0000C |
| 89585872 | 89757812 | -1 | lncRNA | ENSG0000C |
| 89757806 | 89791064 | 1 | protein_coding | ENSG0000C |
| 89823548 | 89823802 | -1 | processed_pseudoge | ENSG0000C |
| 89826593 | 89833564 | 1 | unprocessed_pseudc | ENSG0000C |
| 89900721 | 89900824 | -1 | snRNA | ENSG0000C |
| 89901849 | 89927888 | 1 | protein_coding | ENSG0000C |
| 89933336 | 90003228 | -1 | protein_coding | ENSG0000C |
| 90001405 | 90053633 | 1 | protein_coding | ENSG0000C |
| 90058608 | 90095475 | -1 | protein_coding | ENSG0000C |
| 90198389 | 90620077 | -1 | lncRNA | ENSG0000C |
| 90221341 | 90687863 | 1 | lncRNA | ENSG0000C |
| 90511128 | 90511240 | 1 | rRNA_pseudogene | ENSG0000C |
| 90592804 | 90606065 | -1 | lncRNA | ENSG0000C |
| 90621995 | 90791632 | -1 | protein_coding | ENSG0000C |
| 90646420 | 90670929 | 1 | lncRNA | ENSG0000C |
| 90701082 | 90701876 | 1 | processed_pseudoge | ENSG0000C |
| 90791741 | 90959393 | 1 | protein_coding | ENSG0000C |
| 90806474 | 90859240 | -1 | lncRNA | ENSG0000C |
| 90958471 | 90985238 | -1 | protein_coding | ENSG0000C |
| 90993802 | 91040872 | -1 | protein_coding | ENSG0000C |
| 91016588 | 91018655 | 1 | lncRNA | ENSG0000C |
| 91042690 | 91044762 | -1 | lncRNA | ENSG0000C |
| 91059318 | 91070583 | -1 | lncRNA | ENSG0000C |
| 91070196 | 91087095 | 1 | protein_coding | ENSG0000C |
| 91101832 | 91219236 | 1 | protein_coding | ENSG0000C |
| 91157285 | 91158748 | 1 | processed_pseudoge | ENSG0000C |
| 91205485 | 91205559 | 1 | miRNA | ENSG0000C |
| 91209494 | 91398155 | 1 | protein_coding | ENSG0000C |
| 91542924 | 91907619 | -1 | lncRNA | ENSG0000C |
| 91594965 | 91595241 | 1 | misc_RNA | ENSG0000C |
| 91909738 | 91910167 | -1 | processed_pseudoge | ENSG0000C |
| 91917507 | 91917918 | 1 | processed_pseudoge | ENSG0000C |
| 91954967 | 92103286 | -1 | protein_coding | ENSG0000C |
| 91975908 | 91977418 | 1 | lncRNA | ENSG0000C |
| 92144088 | 92144435 | -1 | processed_pseudoge | ENSG0000C |

| | | | | |
|---|---|---|---|---|
| 92351912 | 92352148 | -1 | processed_pseudoge | ENSG00000 |
| 92402920 | 92413681 | -1 | lncRNA | ENSG00000 |
| 92420850 | 92421798 | -1 | lncRNA | ENSG00000 |
| 92460539 | 92655576 | -1 | lncRNA | ENSG00000 |
| 92668660 | 92683450 | 1 | lncRNA | ENSG00000 |
| 92687222 | 92697826 | -1 | lncRNA | ENSG00000 |
| 92699742 | 92879502 | -1 | lncRNA | ENSG00000 |
| 92723024 | 92725111 | -1 | lncRNA | ENSG00000 |
| 92765651 | 92858297 | 1 | lncRNA | ENSG00000 |
| 92882984 | 92965645 | 1 | lncRNA | ENSG00000 |
| 92883532 | 93017673 | -1 | protein_coding | ENSG00000 |
| 92972575 | 92974738 | 1 | processed_pseudoge | ENSG00000 |
| 93029751 | 93029839 | 1 | miRNA | ENSG00000 |
| 93134095 | 93166850 | -1 | lncRNA | ENSG00000 |
| 93213302 | 93700433 | -1 | lncRNA | ENSG00000 |
| 93215925 | 93234878 | -1 | lncRNA | ENSG00000 |
| 93229659 | 93229916 | 1 | processed_pseudoge | ENSG00000 |
| 93259667 | 93298528 | 1 | lncRNA | ENSG00000 |
| 93441277 | 93441384 | 1 | rRNA_pseudogene | ENSG00000 |
| 93646066 | 93647092 | 1 | processed_pseudoge | ENSG00000 |
| 93698561 | 93731527 | 1 | protein_coding | ENSG00000 |
| 93715378 | 93716113 | -1 | lncRNA | ENSG00000 |
| 93719574 | 93721167 | 1 | lncRNA | ENSG00000 |
| 93729356 | 93741017 | -1 | protein_coding | ENSG00000 |
| 93733216 | 93734022 | 1 | lncRNA | ENSG00000 |
| 93740121 | 93740773 | 1 | TEC | ENSG00000 |
| 93741193 | 93744534 | 1 | lncRNA | ENSG00000 |
| 93754844 | 93819234 | 1 | protein_coding | ENSG00000 |
| 93834454 | 93846743 | -1 | lncRNA | ENSG00000 |
| 93849561 | 93850075 | 1 | processed_pseudoge | ENSG00000 |
| 93857807 | 93926068 | 1 | protein_coding | ENSG00000 |
| 93915734 | 93916682 | -1 | lncRNA | ENSG00000 |
| 93916022 | 93916119 | -1 | miRNA | ENSG00000 |
| 93939364 | 93942877 | -1 | processed_pseudoge | ENSG00000 |
| 93957338 | 93957691 | -1 | processed_pseudoge | ENSG00000 |
| 93974671 | 93974776 | -1 | unprocessed_pseudo | ENSG00000 |
| 94097764 | 94104322 | -1 | lncRNA | ENSG00000 |
| 94127162 | 94217303 | -1 | protein_coding | ENSG00000 |
| 94201913 | 94203052 | -1 | processed_pseudoge | ENSG00000 |
| 94249253 | 94262350 | -1 | protein_coding | ENSG00000 |
| 94352923 | 94353653 | 1 | processed_pseudoge | ENSG00000 |
| 94371960 | 94475115 | -1 | protein_coding | ENSG00000 |
| 94372170 | 94436952 | -1 | protein_coding | ENSG00000 |
| 94487689 | 94553529 | -1 | protein_coding | ENSG00000 |
| 94533628 | 94534391 | 1 | lncRNA | ENSG00000 |
| 94553668 | 94570648 | 1 | lncRNA | ENSG00000 |
| 94565036 | 94565715 | 1 | processed_pseudoge | ENSG00000 |
| 94590115 | 94590259 | 1 | processed_pseudoge | ENSG00000 |
| 94594411 | 94594601 | 1 | processed_pseudoge | ENSG00000 |
| 94637285 | 94639467 | -1 | lncRNA | ENSG00000 |

| | | | | |
|---|---|---|---|---|
| 94641074 | 94707466 | 1 | protein_coding | ENSG0000C |
| 94655911 | 94656208 | 1 | misc_RNA | ENSG0000C |
| 94688228 | 94688610 | 1 | processed_pseudoge | ENSG0000C |
| 94708952 | 94709244 | -1 | processed_pseudoge | ENSG0000C |
| 94713715 | 94713812 | -1 | misc_RNA | ENSG0000C |
| 94719703 | 94793836 | 1 | protein_coding | ENSG0000C |
| 94791643 | 94793106 | -1 | lncRNA | ENSG0000C |
| 94813311 | 94881746 | 1 | protein_coding | ENSG0000C |
| 94879770 | 94896678 | -1 | protein_coding | ENSG0000C |
| 94884609 | 94885070 | 1 | lncRNA | ENSG0000C |
| 94891239 | 94892107 | 1 | processed_pseudoge | ENSG0000C |
| 94895767 | 95116455 | 1 | protein_coding | ENSG0000C |
| 94925972 | 94949378 | -1 | protein_coding | ENSG0000C |
| 94950037 | 94951396 | 1 | lncRNA | ENSG0000C |
| 94974573 | 94974853 | -1 | processed_pseudoge | ENSG0000C |
| 95039617 | 95042209 | -1 | lncRNA | ENSG0000C |
| 95055214 | 95055320 | 1 | snRNA | ENSG0000C |
| 95066808 | 95073182 | -1 | lncRNA | ENSG0000C |
| 95071732 | 95087924 | -1 | lncRNA | ENSG0000C |
| 95072911 | 95072996 | -1 | miRNA | ENSG0000C |
| 95072914 | 95072993 | 1 | miRNA | ENSG0000C |
| 95133785 | 95156685 | 1 | protein_coding | ENSG0000C |
| 95204456 | 95811254 | 1 | lncRNA | ENSG0000C |
| 95206876 | 95216525 | -1 | lncRNA | ENSG0000C |
| 95244913 | 95269201 | -1 | protein_coding | ENSG0000C |
| 95403802 | 95403894 | 1 | processed_pseudoge | ENSG0000C |
| 95505406 | 95527524 | 1 | lncRNA | ENSG0000C |
| 95599114 | 95599219 | 1 | snRNA | ENSG0000C |
| 95947781 | 95948233 | -1 | processed_pseudoge | ENSG0000C |
| 95986108 | 95993103 | -1 | lncRNA | ENSG0000C |
| 96068729 | 96069681 | -1 | processed_pseudoge | ENSG0000C |
| 96126967 | 96127801 | -1 | processed_pseudoge | ENSG0000C |
| 96140572 | 96140944 | -1 | lncRNA | ENSG0000C |
| 96142333 | 96160806 | -1 | protein_coding | ENSG0000C |
| 96225920 | 96235634 | -1 | protein_coding | ENSG0000C |
| 96235427 | 96239149 | 1 | lncRNA | ENSG0000C |
| 96239398 | 96261610 | -1 | protein_coding | ENSG0000C |
| 96261902 | 96336995 | 1 | protein_coding | ENSG0000C |
| 96309221 | 96309322 | -1 | misc_RNA | ENSG0000C |
| 96370800 | 96387438 | -1 | lncRNA | ENSG0000C |
| 96431054 | 96431160 | -1 | snRNA | ENSG0000C |
| 96493813 | 96611790 | 1 | protein_coding | ENSG0000C |
| 96645242 | 97149654 | 1 | protein_coding | ENSG0000C |
| 97132835 | 97133379 | -1 | lncRNA | ENSG0000C |
| 97144170 | 97144723 | 1 | lncRNA | ENSG0000C |
| 97241742 | 97242204 | -1 | lncRNA | ENSG0000C |
| 97273488 | 97277928 | -1 | protein_coding | ENSG0000C |
| 97358265 | 97358478 | 1 | snoRNA | ENSG0000C |
| 97624203 | 97625079 | -1 | processed_pseudoge | ENSG0000C |
| 97643366 | 97643881 | -1 | lncRNA | ENSG0000C |

| | | | | |
|---|---|---|---|---|
| 97644184 | 97730260 | 1 | protein_coding | ENSG0000C |
| 97772313 | 97772425 | 1 | misc_RNA | ENSG0000C |
| 97775057 | 97853013 | 1 | protein_coding | ENSG0000C |
| 97788210 | 97788357 | -1 | processed_pseudoge | ENSG0000C |
| 97853021 | 97853253 | -1 | processed_pseudoge | ENSG0000C |
| 97858049 | 97858947 | -1 | processed_pseudoge | ENSG0000C |
| 97865054 | 97865485 | 1 | processed_pseudoge | ENSG0000C |
| 97868840 | 98036724 | 1 | protein_coding | ENSG0000C |
| 98024851 | 98046469 | -1 | protein_coding | ENSG0000C |
| 98041726 | 98044121 | 1 | lncRNA | ENSG0000C |
| 98042086 | 98042217 | -1 | snoRNA | ENSG0000C |
| 98048856 | 98048962 | 1 | snRNA | ENSG0000C |
| 98064522 | 98093610 | 1 | protein_coding | ENSG0000C |
| 98102344 | 98117171 | -1 | protein_coding | ENSG0000C |
| 98117293 | 98159835 | 1 | protein_coding | ENSG0000C |
| 98189826 | 98294393 | -1 | protein_coding | ENSG0000C |
| 98192205 | 98192313 | -1 | snRNA | ENSG0000C |
| 98371228 | 98942827 | -1 | protein_coding | ENSG0000C |
| 98391401 | 98391514 | -1 | TEC | ENSG0000C |
| 98412882 | 98413784 | -1 | processed_pseudoge | ENSG0000C |
| 98426958 | 98432853 | 1 | protein_coding | ENSG0000C |
| 98436669 | 98439290 | 1 | lncRNA | ENSG0000C |
| 98462944 | 98463050 | -1 | snRNA | ENSG0000C |
| 98603253 | 98606340 | 1 | lncRNA | ENSG0000C |
| 98633547 | 98633865 | 1 | processed_pseudoge | ENSG0000C |
| 98782055 | 98782628 | 1 | processed_pseudoge | ENSG0000C |
| 98904603 | 98905671 | -1 | processed_pseudoge | ENSG0000C |
| 98906260 | 98906517 | -1 | processed_pseudoge | ENSG0000C |
| 98921233 | 98921550 | -1 | misc_RNA | ENSG0000C |
| 98943595 | 98944098 | 1 | lncRNA | ENSG0000C |
| 98944403 | 98952104 | 1 | protein_coding | ENSG0000C |
| 98958277 | 99013743 | -1 | lncRNA | ENSG0000C |
| 98961428 | 98968284 | 1 | lncRNA | ENSG0000C |
| 99003407 | 99003842 | -1 | processed_pseudoge | ENSG0000C |
| 99013266 | 99877580 | 1 | protein_coding | ENSG0000C |
| 99091738 | 99094060 | -1 | lncRNA | ENSG0000C |
| 99119352 | 99120581 | 1 | lncRNA | ENSG0000C |
| 99233048 | 99234459 | -1 | processed_pseudoge | ENSG0000C |
| 99340305 | 99341761 | -1 | processed_pseudoge | ENSG0000C |
| 99346052 | 99348256 | -1 | processed_pseudoge | ENSG0000C |
| 99527826 | 99527927 | 1 | misc_RNA | ENSG0000C |
| 99536636 | 99536730 | -1 | miRNA | ENSG0000C |
| 99536786 | 99536861 | -1 | miRNA | ENSG0000C |
| 99613783 | 99614081 | -1 | misc_RNA | ENSG0000C |
| 99695957 | 99698017 | 1 | processed_pseudoge | ENSG0000C |
| 99796615 | 99799187 | -1 | lncRNA | ENSG0000C |
| 99804159 | 99804341 | -1 | processed_pseudoge | ENSG0000C |
| 99873200 | 99893707 | -1 | protein_coding | ENSG0000C |
| 99893373 | 99895121 | 1 | lncRNA | ENSG0000C |
| 99960936 | 100131268 | -1 | protein_coding | ENSG0000C |

| | | | | |
|---|---|---|---|---|
| 100004583 | 100004649 | 1 | snoRNA | ENSG0000C |
| 100023982 | 100024084 | -1 | miRNA | ENSG0000C |
| 100133351 | 100145817 | -1 | protein_coding | ENSG0000C |
| 100150623 | 100154003 | 1 | protein_coding | ENSG0000C |
| 100157906 | 100259278 | 1 | protein_coding | ENSG0000C |
| 100158868 | 100158975 | 1 | misc_RNA | ENSG0000C |
| 100165226 | 100165478 | -1 | processed_pseudoge | ENSG0000C |
| 100185138 | 100185239 | 1 | misc_RNA | ENSG0000C |
| 100257060 | 100336218 | -1 | protein_coding | ENSG0000C |
| 100337595 | 100350707 | 1 | lncRNA | ENSG0000C |
| 100380486 | 100464613 | -1 | lncRNA | ENSG0000C |
| 100382763 | 100382845 | 1 | miRNA | ENSG0000C |
| 100427559 | 100432726 | -1 | lncRNA | ENSG0000C |
| 100475667 | 100501148 | 1 | lncRNA | ENSG0000C |
| 100492528 | 100493713 | 1 | lncRNA | ENSG0000C |
| 100509752 | 100559784 | -1 | protein_coding | ENSG0000C |
| 100550724 | 100551713 | -1 | processed_pseudoge | ENSG0000C |
| 100572889 | 100663415 | -1 | protein_coding | ENSG0000C |
| 100618581 | 100619993 | 1 | lncRNA | ENSG0000C |
| 100683488 | 100683574 | 1 | snRNA | ENSG0000C |
| 100685816 | 100722809 | -1 | protein_coding | ENSG0000C |
| 100702968 | 100703024 | -1 | miRNA | ENSG0000C |
| 100739802 | 100739927 | -1 | snRNA | ENSG0000C |
| 100755099 | 100755195 | -1 | misc_RNA | ENSG0000C |
| 100776512 | 100776865 | -1 | processed_pseudoge | ENSG0000C |
| 100831728 | 100831860 | -1 | snRNA | ENSG0000C |
| 100895771 | 100896118 | 1 | processed_pseudoge | ENSG0000C |
| 100897853 | 100898136 | -1 | misc_RNA | ENSG0000C |
| 100913247 | 100914388 | -1 | lncRNA | ENSG0000C |
| 100916523 | 100953388 | -1 | protein_coding | ENSG0000C |
| 100957883 | 100958180 | 1 | misc_RNA | ENSG0000C |
| 101034868 | 101076407 | 1 | lncRNA | ENSG0000C |
| 101122145 | 101126158 | -1 | lncRNA | ENSG0000C |
| 101128987 | 101133486 | 1 | lncRNA | ENSG0000C |
| 101137959 | 101138313 | 1 | misc_RNA | ENSG0000C |
| 101139665 | 101140929 | 1 | lncRNA | ENSG0000C |
| 101166805 | 101169629 | -1 | lncRNA | ENSG0000C |
| 101177878 | 101206193 | -1 | protein_coding | ENSG0000C |
| 101208148 | 101208558 | 1 | lncRNA | ENSG0000C |
| 101223941 | 101224573 | -1 | processed_pseudoge | ENSG0000C |
| 101261541 | 101262903 | -1 | lncRNA | ENSG0000C |
| 101275425 | 101280835 | -1 | lncRNA | ENSG0000C |
| 101287445 | 101293783 | 1 | lncRNA | ENSG0000C |
| 101307367 | 101307428 | 1 | snRNA | ENSG0000C |
| 101337496 | 101337649 | 1 | processed_pseudoge | ENSG0000C |
| 101361794 | 101372707 | 1 | transcribed_processe | ENSG0000C |
| 101366417 | 101367005 | -1 | unprocessed_pseudc | ENSG0000C |
| 101387299 | 101393255 | -1 | lncRNA | ENSG0000C |
| 101397636 | 101397933 | -1 | misc_RNA | ENSG0000C |
| 101411873 | 101452452 | -1 | lncRNA | ENSG0000C |

| | | | | |
|---|---|---|---|---|
| 101461177 | 101492499 | -1 | lncRNA | ENSG0000C |
| 101492439 | 101669726 | 1 | protein_coding | ENSG0000C |
| 101528723 | 101529569 | -1 | processed_pseudoge | ENSG0000C |
| 101562008 | 101562488 | -1 | processed_pseudoge | ENSG0000C |
| 101686542 | 102124907 | -1 | protein_coding | ENSG0000C |
| 101686547 | 101689093 | 1 | lncRNA | ENSG0000C |
| 101847256 | 101847360 | -1 | snoRNA | ENSG0000C |
| 101871830 | 101872549 | -1 | processed_pseudoge | ENSG0000C |
| 102125432 | 102125515 | 1 | miRNA | ENSG0000C |
| 102154550 | 102154725 | 1 | processed_pseudoge | ENSG0000C |
| 102204502 | 102239118 | -1 | protein_coding | ENSG0000C |
| 102239394 | 102253750 | 1 | lncRNA | ENSG0000C |
| 102242040 | 102242619 | -1 | unprocessed_pseudc | ENSG0000C |
| 102252273 | 102412759 | -1 | protein_coding | ENSG0000C |
| 102256392 | 102257821 | 1 | lncRNA | ENSG0000C |
| 102417979 | 102418953 | 1 | processed_pseudoge | ENSG0000C |
| 102421197 | 102421322 | 1 | snRNA | ENSG0000C |
| 102451763 | 102452483 | 1 | processed_pseudoge | ENSG0000C |
| 102503753 | 102504137 | 1 | processed_pseudoge | ENSG0000C |
| 102528740 | 102539943 | 1 | lncRNA | ENSG0000C |
| 102533241 | 102533339 | -1 | snRNA | ENSG0000C |
| 102551589 | 102561018 | 1 | protein_coding | ENSG0000C |
| 102621509 | 102622331 | -1 | processed_pseudoge | ENSG0000C |
| 102648784 | 102655725 | -1 | protein_coding | ENSG0000C |
| 102656464 | 102687118 | 1 | lncRNA | ENSG0000C |
| 102688317 | 102688906 | -1 | processed_pseudoge | ENSG0000C |
| 102805517 | 102810039 | 1 | lncRNA | ENSG0000C |
| 102826308 | 102893864 | -1 | protein_coding | ENSG0000C |
| 102854455 | 102856075 | 1 | lncRNA | ENSG0000C |
| 102864271 | 103002424 | 1 | lncRNA | ENSG0000C |
| 102891876 | 102893608 | 1 | lncRNA | ENSG0000C |
| 102923371 | 102924255 | -1 | processed_pseudoge | ENSG0000C |
| 102950467 | 102950564 | -1 | misc_RNA | ENSG0000C |
| 103016744 | 103017589 | 1 | processed_pseudoge | ENSG0000C |
| 103020187 | 103021428 | -1 | lncRNA | ENSG0000C |
| 103021063 | 103073051 | 1 | protein_coding | ENSG0000C |
| 103085651 | 103086590 | 1 | processed_pseudoge | ENSG0000C |
| 103088796 | 103090671 | 1 | processed_pseudoge | ENSG0000C |
| 103091263 | 103091509 | 1 | processed_pseudoge | ENSG0000C |
| 103121032 | 103132966 | 1 | lncRNA | ENSG0000C |
| 103132963 | 103141475 | -1 | lncRNA | ENSG0000C |
| 103140713 | 103230305 | 1 | protein_coding | ENSG0000C |
| 103153394 | 103298772 | -1 | lncRNA | ENSG0000C |
| 103154614 | 103154689 | 1 | miRNA | ENSG0000C |
| 103228425 | 103229314 | -1 | lncRNA | ENSG0000C |
| 103298433 | 103332866 | 1 | protein_coding | ENSG0000C |
| 103319392 | 103320106 | -1 | lncRNA | ENSG0000C |
| 103371538 | 103382989 | 1 | protein_coding | ENSG0000C |
| 103383078 | 103383854 | -1 | lncRNA | ENSG0000C |
| 103384961 | 103385067 | 1 | snRNA | ENSG0000C |

| | | | | |
|---|---|---|---|---|
| 103398635 | 103415189 | -1 | protein_coding | ENSG00000( |
| 103398658 | 103501895 | -1 | protein_coding | ENSG00000( |
| 103414714 | 103443453 | 1 | protein_coding | ENSG00000( |
| 103464389 | 103468985 | 1 | lncRNA | ENSG00000( |
| 103481266 | 103481619 | -1 | lncRNA | ENSG00000( |
| 103483398 | 103501676 | -1 | lncRNA | ENSG00000( |
| 103500696 | 104256094 | 1 | protein_coding | ENSG00000( |
| 103559099 | 103559389 | -1 | processed_pseudoge | ENSG00000( |
| 103568603 | 103568978 | -1 | processed_pseudoge | ENSG00000( |
| 103768281 | 103769029 | 1 | processed_pseudoge | ENSG00000( |
| 104330324 | 104467055 | -1 | protein_coding | ENSG00000( |
| 104339087 | 104356689 | 1 | protein_coding | ENSG00000( |
| 104419512 | 104421500 | -1 | lncRNA | ENSG00000( |
| 104478539 | 104480783 | -1 | TEC | ENSG00000( |
| 104484369 | 104484465 | -1 | miRNA | ENSG00000( |
| 104489231 | 104589024 | -1 | protein_coding | ENSG00000( |
| 104566086 | 104566429 | 1 | processed_pseudoge | ENSG00000( |
| 104590733 | 105804539 | 1 | protein_coding | ENSG00000( |
| 104699479 | 104703555 | -1 | lncRNA | ENSG00000( |
| 104981164 | 104981579 | -1 | processed_pseudoge | ENSG00000( |
| 104990308 | 104990692 | -1 | processed_pseudoge | ENSG00000( |
| 105011107 | 105011568 | 1 | processed_pseudoge | ENSG00000( |
| 105142860 | 105188606 | -1 | lncRNA | ENSG00000( |
| 105194990 | 105196969 | -1 | processed_pseudoge | ENSG00000( |
| 105287830 | 105288365 | -1 | processed_pseudoge | ENSG00000( |
| 105546089 | 106060524 | -1 | lncRNA | ENSG00000( |
| 105662352 | 105685765 | -1 | lncRNA | ENSG00000( |
| 105669991 | 105679031 | 1 | lncRNA | ENSG00000( |
| 105737280 | 105737765 | -1 | processed_pseudoge | ENSG00000( |
| 105826570 | 105834038 | 1 | lncRNA | ENSG00000( |
| 105929126 | 105929238 | -1 | misc_RNA | ENSG00000( |
| 106215763 | 106217218 | 1 | unprocessed_pseudc | ENSG00000( |
| 106265435 | 106268741 | -1 | lncRNA | ENSG00000( |
| 106270144 | 106272902 | 1 | lncRNA | ENSG00000( |
| 106359476 | 106752694 | 1 | protein_coding | ENSG00000( |
| 106520474 | 106657548 | -1 | lncRNA | ENSG00000( |
| 106697427 | 106698013 | -1 | processed_pseudoge | ENSG00000( |
| 106729795 | 106729856 | -1 | snRNA | ENSG00000( |
| 106759483 | 106770244 | -1 | protein_coding | ENSG00000( |
| 106980967 | 106981571 | -1 | unprocessed_pseudc | ENSG00000( |
| 107160354 | 107166595 | 1 | lncRNA | ENSG00000( |
| 107173200 | 107173809 | -1 | processed_pseudoge | ENSG00000( |
| 107186997 | 107195938 | -1 | lncRNA | ENSG00000( |
| 107249482 | 107498055 | -1 | protein_coding | ENSG00000( |
| 107499773 | 107506416 | 1 | lncRNA | ENSG00000( |
| 107647279 | 107648032 | 1 | processed_pseudoge | ENSG00000( |
| 107857589 | 107963516 | 1 | lncRNA | ENSG00000( |
| 107884494 | 107884611 | 1 | rRNA_pseudogene | ENSG00000( |
| 107899316 | 108083642 | -1 | protein_coding | ENSG00000( |
| 107983116 | 107983965 | 1 | processed_pseudoge | ENSG00000( |

| | | | | |
|---|---|---|---|---|
| 108131100 | 108132114 | -1 | processed_pseudoge | ENSG00000( |
| 108201216 | 108435333 | -1 | protein_coding | ENSG00000( |
| 108226200 | 108227544 | 1 | lncRNA | ENSG00000( |
| 108378831 | 108379545 | -1 | processed_pseudoge | ENSG00000( |
| 108443601 | 108489196 | 1 | protein_coding | ENSG00000( |
| 108572643 | 108573174 | 1 | processed_pseudoge | ENSG00000( |
| 108581062 | 108626767 | 1 | lncRNA | ENSG00000( |
| 108606850 | 108787594 | -1 | protein_coding | ENSG00000( |
| 108652373 | 108653315 | 1 | processed_pseudoge | ENSG00000( |
| 108871128 | 109063417 | -1 | lncRNA | ENSG00000( |
| 109086621 | 109119584 | 1 | protein_coding | ENSG00000( |
| 109240919 | 109334118 | -1 | protein_coding | ENSG00000( |
| 109298598 | 109316513 | 1 | lncRNA | ENSG00000( |
| 109334324 | 109345954 | 1 | protein_coding | ENSG00000( |
| 109362461 | 109537207 | 1 | protein_coding | ENSG00000( |
| 109470131 | 109471117 | 1 | processed_pseudoge | ENSG00000( |
| 109539711 | 109565996 | 1 | protein_coding | ENSG00000( |
| 109573978 | 109691791 | -1 | protein_coding | ENSG00000( |
| 109644115 | 109648084 | 1 | lncRNA | ENSG00000( |
| 109777912 | 109782288 | 1 | processed_pseudoge | ENSG00000( |
| 109873913 | 109874018 | -1 | snRNA | ENSG00000( |
| 109899246 | 109899762 | -1 | processed_pseudoge | ENSG00000( |
| 109963636 | 109975771 | -1 | protein_coding | ENSG00000( |
| 109973943 | 109974781 | 1 | lncRNA | ENSG00000( |
| 110105076 | 110105964 | -1 | processed_pseudoge | ENSG00000( |
| 110334743 | 110349607 | -1 | lncRNA | ENSG00000( |
| 110555408 | 110556518 | 1 | processed_pseudoge | ENSG00000( |
| 110609241 | 110632394 | 1 | lncRNA | ENSG00000( |
| 110766863 | 110772759 | 1 | lncRNA | ENSG00000( |
| 110809394 | 110809894 | -1 | processed_pseudoge | ENSG00000( |
| 110899996 | 111040372 | -1 | lncRNA | ENSG00000( |
| 110933263 | 110934533 | -1 | processed_pseudoge | ENSG00000( |
| 111093263 | 111236203 | -1 | lncRNA | ENSG00000( |
| 111179193 | 111179872 | -1 | lncRNA | ENSG00000( |
| 111225585 | 111227049 | 1 | processed_pseudoge | ENSG00000( |
| 111376639 | 111757710 | 1 | lncRNA | ENSG00000( |
| 111380603 | 111380868 | 1 | processed_pseudoge | ENSG00000( |
| 111621458 | 111621797 | -1 | processed_pseudoge | ENSG00000( |
| 111699519 | 111700276 | 1 | lncRNA | ENSG00000( |
| 112148742 | 112148825 | -1 | snRNA | ENSG00000( |
| 112222928 | 113436939 | -1 | protein_coding | ENSG00000( |
| 112446575 | 112457993 | 1 | lncRNA | ENSG00000( |
| 112538889 | 112539317 | 1 | processed_pseudoge | ENSG00000( |
| 112643493 | 112643583 | 1 | miRNA | ENSG00000( |
| 113376669 | 113377227 | 1 | processed_pseudoge | ENSG00000( |
| 113432224 | 113433129 | 1 | lncRNA | ENSG00000( |
| 113438334 | 113493281 | 1 | lncRNA | ENSG00000( |
| 113600310 | 113616958 | 1 | lncRNA | ENSG00000( |
| 113950886 | 113950998 | 1 | misc_RNA | ENSG00000( |
| 114282067 | 114295839 | 1 | lncRNA | ENSG00000( |

| | | | | |
|---|---|---|---|---|
| 114318400 | 114318911 | -1 | processed_pseudoge | ENSG00000 |
| 114791653 | 114791929 | -1 | processed_pseudoge | ENSG00000 |
| 115408496 | 115809673 | -1 | protein_coding | ENSG00000 |
| 115509602 | 115511325 | 1 | lncRNA | ENSG00000 |
| 115950511 | 116325062 | -1 | lncRNA | ENSG00000 |
| 116022686 | 116022779 | -1 | rRNA_pseudogene | ENSG00000 |
| 116289622 | 116298954 | 1 | lncRNA | ENSG00000 |
| 116402543 | 116403757 | -1 | lncRNA | ENSG00000 |
| 116642130 | 116766925 | -1 | protein_coding | ENSG00000 |
| 116766505 | 116849463 | 1 | protein_coding | ENSG00000 |
| 116845935 | 116874866 | -1 | protein_coding | ENSG00000 |
| 116874424 | 116876868 | 1 | lncRNA | ENSG00000 |
| 116874728 | 116874800 | -1 | miRNA | ENSG00000 |
| 116938207 | 116944487 | 1 | protein_coding | ENSG00000 |
| 116950273 | 117176714 | 1 | protein_coding | ENSG00000 |
| 117017946 | 117018248 | 1 | misc_RNA | ENSG00000 |
| 117055818 | 117056145 | 1 | processed_pseudoge | ENSG00000 |
| 117128455 | 117130299 | 1 | lncRNA | ENSG00000 |
| 117163968 | 117164255 | 1 | misc_RNA | ENSG00000 |
| 117265272 | 117318701 | 1 | lncRNA | ENSG00000 |
| 117313528 | 117313661 | 1 | snoRNA | ENSG00000 |
| 117520713 | 117540262 | 1 | protein_coding | ENSG00000 |
| 117523271 | 117524061 | -1 | unprocessed_pseudo | ENSG00000 |
| 117794490 | 118111853 | -1 | protein_coding | ENSG00000 |
| 118189455 | 118622112 | -1 | protein_coding | ENSG00000 |
| 118282139 | 118400605 | 1 | lncRNA | ENSG00000 |
| 118620498 | 118906155 | 1 | lncRNA | ENSG00000 |
| 118761856 | 118762201 | -1 | processed_pseudoge | ENSG00000 |
| 118923557 | 118951885 | -1 | protein_coding | ENSG00000 |
| 118976513 | 118976619 | 1 | snRNA | ENSG00000 |
| 118982579 | 118982875 | -1 | processed_pseudoge | ENSG00000 |
| 118995452 | 119108455 | 1 | protein_coding | ENSG00000 |
| 119062942 | 119068782 | -1 | lncRNA | ENSG00000 |
| 119165034 | 119245673 | 1 | protein_coding | ENSG00000 |
| 119214625 | 119246848 | -1 | lncRNA | ENSG00000 |
| 119325171 | 119325265 | 1 | miRNA | ENSG00000 |
| 119350291 | 119416947 | -1 | lncRNA | ENSG00000 |
| 119388348 | 119388469 | 1 | snoRNA | ENSG00000 |
| 119416446 | 119424434 | 1 | protein_coding | ENSG00000 |
| 119419910 | 119462350 | -1 | lncRNA | ENSG00000 |
| 119480279 | 119480775 | -1 | processed_pseudoge | ENSG00000 |
| 119557086 | 119673453 | -1 | protein_coding | ENSG00000 |
| 119618162 | 119618474 | -1 | processed_pseudoge | ENSG00000 |
| 119711830 | 119712145 | -1 | misc_RNA | ENSG00000 |
| 119730774 | 119832841 | -1 | protein_coding | ENSG00000 |
| 119832875 | 119855509 | 1 | lncRNA | ENSG00000 |
| 119833976 | 119855894 | -1 | protein_coding | ENSG00000 |
| 119838736 | 119840385 | -1 | TEC | ENSG00000 |
| 119862662 | 119862956 | 1 | misc_RNA | ENSG00000 |
| 119867419 | 119874488 | -1 | lncRNA | ENSG00000 |

| | | | | |
|---|---|---|---|---|
| 119873717 | 120050918 | 1 | protein_coding | ENSG00000 |
| 119891132 | 119891229 | -1 | rRNA_pseudogene | ENSG00000 |
| 120052180 | 120056201 | 1 | lncRNA | ENSG00000 |
| 120059780 | 120373573 | 1 | protein_coding | ENSG00000 |
| 120380761 | 120445402 | -1 | protein_coding | ENSG00000 |
| 120445400 | 120542133 | 1 | protein_coding | ENSG00000 |
| 120535745 | 120813273 | -1 | protein_coding | ENSG00000 |
| 120635558 | 120638204 | -1 | processed_pseudoge | ENSG00000 |
| 120761253 | 120776832 | 1 | lncRNA | ENSG00000 |
| 120812219 | 120813359 | 1 | lncRNA | ENSG00000 |
| 120913065 | 121119754 | 1 | lncRNA | ENSG00000 |
| 121113358 | 121113919 | -1 | processed_pseudoge | ENSG00000 |
| 121380137 | 121380465 | -1 | processed_pseudoge | ENSG00000 |
| 121612116 | 121641440 | -1 | protein_coding | ENSG00000 |
| 121639293 | 121994185 | 1 | lncRNA | ENSG00000 |
| 121668934 | 121697600 | -1 | lncRNA | ENSG00000 |
| 121954640 | 122127184 | -1 | lncRNA | ENSG00000 |
| 122091480 | 122091609 | 1 | processed_pseudoge | ENSG00000 |
| 122414332 | 122428551 | -1 | lncRNA | ENSG00000 |
| 122485194 | 122733804 | -1 | lncRNA | ENSG00000 |
| 122485515 | 122489815 | -1 | lncRNA | ENSG00000 |
| 122671291 | 122671394 | 1 | snoRNA | ENSG00000 |
| 122700269 | 122703423 | -1 | lncRNA | ENSG00000 |
| 122769274 | 122770150 | 1 | processed_pseudoge | ENSG00000 |
| 122779971 | 122780830 | -1 | lncRNA | ENSG00000 |
| 122780760 | 122781071 | 1 | lncRNA | ENSG00000 |
| 122781655 | 122974510 | 1 | protein_coding | ENSG00000 |
| 122807746 | 122816517 | 1 | lncRNA | ENSG00000 |
| 122977026 | 122985629 | -1 | lncRNA | ENSG00000 |
| 123002560 | 123030751 | 1 | lncRNA | ENSG00000 |
| 123013170 | 123042302 | -1 | protein_coding | ENSG00000 |
| 123041968 | 123152153 | 1 | protein_coding | ENSG00000 |
| 123123106 | 123123682 | 1 | processed_pseudoge | ENSG00000 |
| 123157729 | 123165173 | 1 | lncRNA | ENSG00000 |
| 123178960 | 123210079 | 1 | protein_coding | ENSG00000 |
| 123180310 | 123180525 | -1 | snoRNA | ENSG00000 |
| 123181638 | 123182788 | -1 | lncRNA | ENSG00000 |
| 123192875 | 123192932 | 1 | transcribed_processe | ENSG00000 |
| 123196536 | 123202743 | -1 | lncRNA | ENSG00000 |
| 123215788 | 123215863 | -1 | miRNA | ENSG00000 |
| 123219967 | 123241377 | -1 | protein_coding | ENSG00000 |
| 123226189 | 123274284 | -1 | protein_coding | ENSG00000 |
| 123236852 | 123237062 | 1 | processed_pseudoge | ENSG00000 |
| 123248451 | 123275541 | -1 | protein_coding | ENSG00000 |
| 123262843 | 123262949 | 1 | snRNA | ENSG00000 |
| 123288355 | 123290503 | 1 | processed_pseudoge | ENSG00000 |
| 123319850 | 123416350 | -1 | protein_coding | ENSG00000 |
| 123342414 | 123342519 | 1 | snRNA | ENSG00000 |
| 123348034 | 123348130 | 1 | miRNA | ENSG00000 |
| 123348034 | 123348130 | -1 | miRNA | ENSG00000 |

| | | | | |
|---|---|---|---|---|
| 123352181 | 123352643 | 1 | processed_pseudoge | ENSG00000( |
| 123400582 | 123402095 | 1 | processed_pseudoge | ENSG00000( |
| 123416726 | 123470028 | 1 | protein_coding | ENSG00000( |
| 123454981 | 123455107 | -1 | snRNA | ENSG00000( |
| 123497889 | 123541206 | -1 | protein_coding | ENSG00000( |
| 123563070 | 123568762 | -1 | lncRNA | ENSG00000( |
| 123588759 | 123589046 | 1 | misc_RNA | ENSG00000( |
| 123614219 | 123667248 | 1 | lncRNA | ENSG00000( |
| 123645527 | 123652950 | -1 | protein_coding | ENSG00000( |
| 123680794 | 123737402 | -1 | protein_coding | ENSG00000( |
| 123768439 | 123815452 | 1 | protein_coding | ENSG00000( |
| 123851987 | 124120061 | 1 | protein_coding | ENSG00000( |
| 123924759 | 123925933 | -1 | lncRNA | ENSG00000( |
| 123984138 | 124040782 | -1 | lncRNA | ENSG00000( |
| 124036010 | 124040468 | -1 | lncRNA | ENSG00000( |
| 124044395 | 124171522 | -1 | lncRNA | ENSG00000( |
| 124141888 | 124143512 | 1 | processed_pseudoge | ENSG00000( |
| 124151923 | 124152433 | 1 | processed_pseudoge | ENSG00000( |
| 124192671 | 124247398 | -1 | lncRNA | ENSG00000( |
| 124260646 | 124260752 | 1 | snRNA | ENSG00000( |
| 124271667 | 124279580 | 1 | lncRNA | ENSG00000( |
| 124301315 | 124301804 | 1 | processed_pseudoge | ENSG00000( |
| 124306189 | 124372692 | -1 | protein_coding | ENSG00000( |
| 124450820 | 124462150 | 1 | protein_coding | ENSG00000( |
| 124462485 | 124474582 | -1 | lncRNA | ENSG00000( |
| 124474738 | 124487914 | 1 | protein_coding | ENSG00000( |
| 124488485 | 124539458 | -1 | protein_coding | ENSG00000( |
| 124488510 | 124491643 | 1 | lncRNA | ENSG00000( |
| 124508515 | 124508576 | -1 | miRNA | ENSG00000( |
| 124539101 | 124580648 | 1 | protein_coding | ENSG00000( |
| 124550790 | 124728429 | -1 | protein_coding | ENSG00000( |
| 124811042 | 124857516 | -1 | lncRNA | ENSG00000( |
| 124821978 | 124822058 | -1 | miRNA | ENSG00000( |
| 124821985 | 124822051 | 1 | miRNA | ENSG00000( |
| 124823702 | 124962531 | 1 | lncRNA | ENSG00000( |
| 124920428 | 124922076 | 1 | processed_pseudoge | ENSG00000( |
| 124936328 | 124943765 | -1 | lncRNA | ENSG00000( |
| 124940041 | 124941909 | -1 | lncRNA | ENSG00000( |
| 124973295 | 124979389 | 1 | protein_coding | ENSG00000( |
| 124996985 | 124998198 | -1 | lncRNA | ENSG00000( |
| 124998497 | 125022283 | 1 | protein_coding | ENSG00000( |
| 125024260 | 125091819 | -1 | protein_coding | ENSG00000( |
| 125040684 | 125044989 | 1 | lncRNA | ENSG00000( |
| 125091679 | 125367120 | 1 | protein_coding | ENSG00000( |
| 125270604 | 125270902 | 1 | misc_RNA | ENSG00000( |
| 125348196 | 125351416 | -1 | lncRNA | ENSG00000( |
| 125430358 | 125438403 | 1 | protein_coding | ENSG00000( |
| 125466939 | 125541373 | 1 | lncRNA | ENSG00000( |
| 125648327 | 126008930 | 1 | lncRNA | ENSG00000( |
| 125749055 | 125750241 | -1 | lncRNA | ENSG00000( |

| | | | | |
|---|---|---|---|---|
| 125859721 | 126292788 | -1 | lncRNA | ENSG0000C |
| 125900951 | 125901054 | 1 | snRNA | ENSG0000C |
| 125951861 | 125952314 | -1 | processed_pseudoge | ENSG0000C |
| 125998994 | 126001001 | -1 | lncRNA | ENSG0000C |
| 126072587 | 126074231 | 1 | processed_pseudoge | ENSG0000C |
| 126073046 | 126073532 | -1 | processed_pseudoge | ENSG0000C |
| 126174186 | 126175014 | -1 | processed_pseudoge | ENSG0000C |
| 126325454 | 126329538 | 1 | lncRNA | ENSG0000C |
| 126474189 | 126492596 | 1 | lncRNA | ENSG0000C |
| 126497385 | 126500440 | 1 | lncRNA | ENSG0000C |
| 126504763 | 126504869 | 1 | snRNA | ENSG0000C |
| 126506326 | 126522360 | 1 | lncRNA | ENSG0000C |
| 126552443 | 126558478 | -1 | protein_coding | ENSG0000C |
| 126556323 | 127419050 | 1 | lncRNA | ENSG0000C |
| 126582631 | 126583006 | -1 | processed_pseudoge | ENSG0000C |
| 126671522 | 126671654 | 1 | snRNA | ENSG0000C |
| 126766196 | 126846981 | -1 | lncRNA | ENSG0000C |
| 127072694 | 127227541 | -1 | lncRNA | ENSG0000C |
| 127079874 | 127092600 | 1 | lncRNA | ENSG0000C |
| 127086263 | 127087510 | -1 | processed_pseudoge | ENSG0000C |
| 127253213 | 127257630 | -1 | lncRNA | ENSG0000C |
| 127289817 | 127482139 | -1 | lncRNA | ENSG0000C |
| 127292233 | 127292440 | 1 | processed_pseudoge | ENSG0000C |
| 127322183 | 127420066 | 1 | protein_coding | ENSG0000C |
| 127400399 | 127402150 | 1 | lncRNA | ENSG0000C |
| 127578473 | 127636867 | 1 | lncRNA | ENSG0000C |
| 127663280 | 127670990 | -1 | lncRNA | ENSG0000C |
| 127686343 | 127738987 | -1 | lncRNA | ENSG0000C |
| 127735434 | 127742951 | 1 | protein_coding | ENSG0000C |
| 127794526 | 128187101 | 1 | lncRNA | ENSG0000C |
| 127795962 | 127796028 | 1 | miRNA | ENSG0000C |
| 127890626 | 127890720 | 1 | misc_RNA | ENSG0000C |
| 127946559 | 127948723 | -1 | TEC | ENSG0000C |
| 127960633 | 127960695 | 1 | miRNA | ENSG0000C |
| 127999131 | 127999294 | 1 | snRNA | ENSG0000C |
| 128008898 | 128008956 | 1 | miRNA | ENSG0000C |
| 128009551 | 128011938 | -1 | transcribed_processe | ENSG0000C |
| 128010382 | 128010611 | -1 | snRNA | ENSG0000C |
| 128049152 | 128049238 | 1 | miRNA | ENSG0000C |
| 128150116 | 128150188 | 1 | miRNA | ENSG0000C |
| 128220504 | 128220803 | 1 | misc_RNA | ENSG0000C |
| 128323131 | 128388636 | 1 | lncRNA | ENSG0000C |
| 128405269 | 128564679 | -1 | lncRNA | ENSG0000C |
| 128634199 | 129683770 | -1 | lncRNA | ENSG0000C |
| 129241228 | 129242469 | 1 | lncRNA | ENSG0000C |
| 129250967 | 129251138 | 1 | processed_pseudoge | ENSG0000C |
| 129290983 | 129291275 | 1 | misc_RNA | ENSG0000C |
| 129415949 | 129445852 | 1 | lncRNA | ENSG0000C |
| 129484057 | 129484142 | -1 | miRNA | ENSG0000C |
| 129685401 | 129723023 | -1 | lncRNA | ENSG0000C |

| | | | | |
|---|---|---|---|---|
| 129728744 | 129730445 | 1 | processed_pseudoge | ENSG00000 |
| 129748196 | 129786624 | -1 | protein_coding | ENSG00000 |
| 129832301 | 129844504 | 1 | lncRNA | ENSG00000 |
| 129839593 | 130017129 | -1 | protein_coding | ENSG00000 |
| 129864478 | 129865052 | 1 | processed_pseudoge | ENSG00000 |
| 129868590 | 129868716 | -1 | snoRNA | ENSG00000 |
| 129890465 | 129901758 | 1 | lncRNA | ENSG00000 |
| 129939844 | 129951056 | 1 | lncRNA | ENSG00000 |
| 129953729 | 129964634 | 1 | lncRNA | ENSG00000 |
| 130004528 | 130004601 | -1 | snRNA | ENSG00000 |
| 130008334 | 130008453 | -1 | miRNA | ENSG00000 |
| 130052104 | 130443674 | -1 | protein_coding | ENSG00000 |
| 130069541 | 130069652 | -1 | snRNA | ENSG00000 |
| 130082738 | 130084768 | -1 | lncRNA | ENSG00000 |
| 130162972 | 130163127 | 1 | snoRNA | ENSG00000 |
| 130248287 | 130251051 | 1 | lncRNA | ENSG00000 |
| 130358787 | 130362461 | 1 | lncRNA | ENSG00000 |
| 130595299 | 130655712 | -1 | lncRNA | ENSG00000 |
| 130780301 | 131040909 | -1 | protein_coding | ENSG00000 |
| 130892070 | 130892785 | -1 | lncRNA | ENSG00000 |
| 130935409 | 130949665 | 1 | lncRNA | ENSG00000 |
| 131129761 | 131144948 | -1 | lncRNA | ENSG00000 |
| 131308545 | 131317632 | 1 | lncRNA | ENSG00000 |
| 131432776 | 131432869 | -1 | snoRNA | ENSG00000 |
| 131712512 | 131712980 | 1 | processed_pseudoge | ENSG00000 |
| 131904093 | 132013642 | 1 | protein_coding | ENSG00000 |
| 132024216 | 132059382 | -1 | protein_coding | ENSG00000 |
| 132024238 | 132085655 | -1 | protein_coding | ENSG00000 |
| 132061486 | 132111159 | -1 | protein_coding | ENSG00000 |
| 132120859 | 132481095 | -1 | protein_coding | ENSG00000 |
| 132507962 | 132565246 | -1 | lncRNA | ENSG00000 |
| 132570416 | 132675592 | -1 | protein_coding | ENSG00000 |
| 132685007 | 132760712 | -1 | protein_coding | ENSG00000 |
| 132702812 | 132722283 | 1 | lncRNA | ENSG00000 |
| 132775358 | 132848807 | 1 | protein_coding | ENSG00000 |
| 132826179 | 132826903 | 1 | lncRNA | ENSG00000 |
| 132838117 | 132844298 | -1 | lncRNA | ENSG00000 |
| 132866958 | 133134903 | 1 | protein_coding | ENSG00000 |
| 133002605 | 133003075 | -1 | processed_pseudoge | ENSG00000 |
| 133036728 | 133102912 | -1 | protein_coding | ENSG00000 |
| 133046481 | 133046581 | -1 | miRNA | ENSG00000 |
| 133054959 | 133057767 | 1 | lncRNA | ENSG00000 |
| 133191039 | 133231690 | 1 | protein_coding | ENSG00000 |
| 133237171 | 133302022 | -1 | protein_coding | ENSG00000 |
| 133325997 | 133326404 | 1 | processed_pseudoge | ENSG00000 |
| 133392211 | 133392690 | -1 | processed_pseudoge | ENSG00000 |
| 133408006 | 133409107 | 1 | processed_pseudoge | ENSG00000 |
| 133454848 | 133571940 | -1 | protein_coding | ENSG00000 |
| 133515506 | 133515745 | 1 | processed_pseudoge | ENSG00000 |
| 133516184 | 133516268 | 1 | processed_pseudoge | ENSG00000 |

| | | | | |
|---|---|---|---|---|
| 133573183 | 133573861 | 1 | lncRNA | ENSG0000( |
| 133644598 | 133644725 | -1 | snoRNA | ENSG0000( |
| 133668420 | 133668701 | -1 | misc_RNA | ENSG0000( |
| 133754811 | 133755450 | 1 | processed_pseudoge | ENSG0000( |
| 133755845 | 133756646 | 1 | processed_pseudoge | ENSG0000( |
| 133771409 | 133814537 | 1 | lncRNA | ENSG0000( |
| 133775551 | 133777352 | -1 | lncRNA | ENSG0000( |
| 133786093 | 133786268 | -1 | processed_pseudoge | ENSG0000( |
| 133886496 | 133902407 | 1 | lncRNA | ENSG0000( |
| 133946677 | 133947335 | 1 | unprocessed_pseudc | ENSG0000( |
| 133962992 | 133963259 | -1 | processed_pseudoge | ENSG0000( |
| 134190383 | 134390487 | -1 | lncRNA | ENSG0000( |
| 134211865 | 134320447 | -1 | lncRNA | ENSG0000( |
| 134477788 | 134713049 | -1 | protein_coding | ENSG0000( |
| 134598071 | 134598149 | 1 | misc_RNA | ENSG0000( |
| 134598071 | 134600689 | 1 | lncRNA | ENSG0000( |
| 134598318 | 134598518 | 1 | misc_RNA | ENSG0000( |
| 134600257 | 134600336 | 1 | misc_RNA | ENSG0000( |
| 134673131 | 134673512 | 1 | processed_pseudoge | ENSG0000( |
| 134722947 | 134723949 | 1 | lncRNA | ENSG0000( |
| 134764809 | 134767276 | 1 | TEC | ENSG0000( |
| 134783694 | 134785879 | -1 | TEC | ENSG0000( |
| 134792020 | 134798272 | -1 | lncRNA | ENSG0000( |
| 134800520 | 134800607 | -1 | miRNA | ENSG0000( |
| 134804876 | 134804945 | -1 | miRNA | ENSG0000( |
| 134832747 | 134834482 | 1 | lncRNA | ENSG0000( |
| 134838069 | 134842637 | 1 | lncRNA | ENSG0000( |
| 134849935 | 134881899 | 1 | lncRNA | ENSG0000( |
| 134978827 | 134979279 | -1 | processed_pseudoge | ENSG0000( |
| 135234131 | 135299719 | 1 | lncRNA | ENSG0000( |
| 135455865 | 135456952 | -1 | lncRNA | ENSG0000( |
| 135457456 | 135656722 | 1 | protein_coding | ENSG0000( |
| 135625185 | 135625981 | -1 | processed_pseudoge | ENSG0000( |
| 135742343 | 135742495 | -1 | snRNA | ENSG0000( |
| 135859369 | 137092183 | 1 | lncRNA | ENSG0000( |
| 135977329 | 135980588 | -1 | lncRNA | ENSG0000( |
| 136237236 | 136295232 | -1 | lncRNA | ENSG0000( |
| 137105352 | 137105458 | 1 | snRNA | ENSG0000( |
| 137424904 | 137425021 | -1 | unprocessed_pseudc | ENSG0000( |
| 137524755 | 137645564 | -1 | lncRNA | ENSG0000( |
| 137696724 | 137698467 | 1 | processed_pseudoge | ENSG0000( |
| 137809444 | 138083657 | -1 | lncRNA | ENSG0000( |
| 137852204 | 138019873 | 1 | lncRNA | ENSG0000( |
| 138063268 | 138073240 | 1 | lncRNA | ENSG0000( |
| 138130023 | 138497261 | -1 | protein_coding | ENSG0000( |
| 138588235 | 138914041 | -1 | protein_coding | ENSG0000( |
| 139096305 | 139102830 | -1 | lncRNA | ENSG0000( |
| 139108430 | 139108534 | 1 | snoRNA | ENSG0000( |
| 139114193 | 139127953 | -1 | lncRNA | ENSG0000( |
| 139460062 | 139463016 | -1 | lncRNA | ENSG0000( |

| | | | | |
|---|---|---|---|---|
| 139508701 | 139508971 | -1 | unprocessed_pseudo | ENSG00000( |
| 139600838 | 139704109 | -1 | protein_coding | ENSG00000( |
| 139727725 | 140458579 | -1 | protein_coding | ENSG00000( |
| 139911238 | 139916620 | 1 | lncRNA | ENSG00000( |
| 139931172 | 139933946 | 1 | TEC | ENSG00000( |
| 140039756 | 140041037 | 1 | processed_pseudoge | ENSG00000( |
| 140094894 | 140100543 | -1 | lncRNA | ENSG00000( |
| 140505813 | 140508043 | -1 | lncRNA | ENSG00000( |
| 140511311 | 140517154 | 1 | protein_coding | ENSG00000( |
| 140520156 | 140635633 | -1 | protein_coding | ENSG00000( |
| 140572142 | 140572812 | -1 | TEC | ENSG00000( |
| 140591684 | 140593201 | 1 | TEC | ENSG00000( |
| 140636281 | 140638283 | 1 | lncRNA | ENSG00000( |
| 140657900 | 141002216 | -1 | protein_coding | ENSG00000( |
| 140732564 | 140732653 | -1 | miRNA | ENSG00000( |
| 140820599 | 140822414 | -1 | TEC | ENSG00000( |
| 140940562 | 140940663 | 1 | rRNA_pseudogene | ENSG00000( |
| 141055290 | 141060596 | 1 | lncRNA | ENSG00000( |
| 141117278 | 141195808 | 1 | protein_coding | ENSG00000( |
| 141124717 | 141130064 | -1 | lncRNA | ENSG00000( |
| 141207166 | 141308305 | -1 | protein_coding | ENSG00000( |
| 141224288 | 141226947 | -1 | TEC | ENSG00000( |
| 141252286 | 141253292 | -1 | lncRNA | ENSG00000( |
| 141254565 | 141256817 | 1 | lncRNA | ENSG00000( |
| 141278228 | 141292862 | 1 | lncRNA | ENSG00000( |
| 141326130 | 141327137 | 1 | lncRNA | ENSG00000( |
| 141340505 | 141344621 | 1 | lncRNA | ENSG00000( |
| 141353403 | 141355365 | -1 | lncRNA | ENSG00000( |
| 141356470 | 141367286 | -1 | protein_coding | ENSG00000( |
| 141389939 | 141392574 | -1 | lncRNA | ENSG00000( |
| 141391993 | 141432454 | 1 | protein_coding | ENSG00000( |
| 141433829 | 141507230 | -1 | polymorphic_pseudo | ENSG00000( |
| 141434545 | 141437954 | 1 | lncRNA | ENSG00000( |
| 141447475 | 141447549 | 1 | snoRNA | ENSG00000( |
| 141459075 | 141460031 | 1 | processed_pseudoge | ENSG00000( |
| 141507333 | 141509785 | 1 | lncRNA | ENSG00000( |
| 141514638 | 141518737 | 1 | protein_coding | ENSG00000( |
| 141581684 | 141585260 | 1 | lncRNA | ENSG00000( |
| 141786242 | 141786313 | -1 | miRNA | ENSG00000( |
| 141968091 | 141968841 | -1 | processed_pseudoge | ENSG00000( |
| 142089827 | 142091619 | 1 | lncRNA | ENSG00000( |
| 142111414 | 142185527 | -1 | lncRNA | ENSG00000( |
| 142176339 | 142176418 | 1 | miRNA | ENSG00000( |
| 142198356 | 142209003 | 1 | lncRNA | ENSG00000( |
| 142212080 | 142403182 | -1 | protein_coding | ENSG00000( |
| 142247585 | 142247866 | -1 | misc_RNA | ENSG00000( |
| 142403652 | 142407028 | 1 | lncRNA | ENSG00000( |
| 142449430 | 142545009 | 1 | protein_coding | ENSG00000( |
| 142567260 | 142577881 | -1 | transcribed_unitary_ | ENSG00000( |
| 142611049 | 142614479 | -1 | protein_coding | ENSG00000( |

| | | | | |
|---|---|---|---|---|
| 142620373 | 142621064 | 1 | lncRNA | ENSG00000 |
| 142638596 | 142640648 | 1 | lncRNA | ENSG00000 |
| 142657460 | 142681968 | -1 | protein_coding | ENSG00000 |
| 142670308 | 142682725 | 1 | protein_coding | ENSG00000 |
| 142700111 | 142705127 | 1 | protein_coding | ENSG00000 |
| 142701857 | 142727016 | -1 | lncRNA | ENSG00000 |
| 142727223 | 142736927 | 1 | protein_coding | ENSG00000 |
| 142727283 | 142727690 | -1 | lncRNA | ENSG00000 |
| 142740949 | 142742406 | -1 | protein_coding | ENSG00000 |
| 142750150 | 142752532 | -1 | protein_coding | ENSG00000 |
| 142763116 | 142766427 | 1 | lncRNA | ENSG00000 |
| 142764334 | 142778224 | -1 | protein_coding | ENSG00000 |
| 142764338 | 142769828 | -1 | protein_coding | ENSG00000 |
| 142771197 | 142777810 | -1 | protein_coding | ENSG00000 |
| 142784882 | 142786539 | -1 | protein_coding | ENSG00000 |
| 142785374 | 142812120 | 1 | lncRNA | ENSG00000 |
| 142797624 | 142806142 | -1 | unprocessed_pseudo | ENSG00000 |
| 142834138 | 142834940 | -1 | lncRNA | ENSG00000 |
| 142834247 | 142916506 | 1 | protein_coding | ENSG00000 |
| 142858354 | 142858797 | 1 | processed_pseudoge | ENSG00000 |
| 142872356 | 142879846 | -1 | protein_coding | ENSG00000 |
| 142910559 | 142917843 | -1 | protein_coding | ENSG00000 |
| 142981738 | 143018437 | -1 | lncRNA | ENSG00000 |
| 142995850 | 142996482 | 1 | processed_pseudoge | ENSG00000 |
| 143017982 | 143023832 | 1 | protein_coding | ENSG00000 |
| 143039209 | 143060684 | 1 | lncRNA | ENSG00000 |
| 143057060 | 143057816 | -1 | processed_pseudoge | ENSG00000 |
| 143065915 | 143066587 | -1 | processed_pseudoge | ENSG00000 |
| 143080457 | 143083001 | 1 | protein_coding | ENSG00000 |
| 143157914 | 143160711 | -1 | protein_coding | ENSG00000 |
| 143213218 | 143217170 | 1 | protein_coding | ENSG00000 |
| 143246821 | 143262705 | 1 | protein_coding | ENSG00000 |
| 143247110 | 143276403 | 1 | protein_coding | ENSG00000 |
| 143267433 | 143276931 | 1 | protein_coding | ENSG00000 |
| 143280161 | 143281690 | -1 | lncRNA | ENSG00000 |
| 143289676 | 143299952 | 1 | protein_coding | ENSG00000 |
| 143290399 | 143290621 | -1 | lncRNA | ENSG00000 |
| 143304384 | 143359979 | -1 | protein_coding | ENSG00000 |
| 143350172 | 143350278 | 1 | snRNA | ENSG00000 |
| 143366631 | 143368548 | -1 | lncRNA | ENSG00000 |
| 143368876 | 143384221 | 1 | protein_coding | ENSG00000 |
| 143412749 | 143417054 | 1 | lncRNA | ENSG00000 |
| 143417679 | 143419150 | 1 | lncRNA | ENSG00000 |
| 143419182 | 143430732 | -1 | protein_coding | ENSG00000 |
| 143437659 | 143541447 | -1 | protein_coding | ENSG00000 |
| 143541648 | 143545128 | 1 | lncRNA | ENSG00000 |
| 143541973 | 143549729 | -1 | lncRNA | ENSG00000 |
| 143542110 | 143542398 | 1 | misc_RNA | ENSG00000 |
| 143553207 | 143563062 | 1 | protein_coding | ENSG00000 |
| 143566192 | 143572772 | -1 | protein_coding | ENSG00000 |

| | | | | |
|---|---|---|---|---|
| 143573490 | 143577397 | 1 | lncRNA | ENSG0000C |
| 143574785 | 143578649 | -1 | protein_coding | ENSG0000C |
| 143578358 | 143583304 | 1 | TEC | ENSG0000C |
| 143579636 | 143580670 | 1 | lncRNA | ENSG0000C |
| 143579697 | 143599541 | -1 | protein_coding | ENSG0000C |
| 143597831 | 143603224 | 1 | protein_coding | ENSG0000C |
| 143603210 | 143609773 | -1 | protein_coding | ENSG0000C |
| 143612618 | 143618048 | -1 | protein_coding | ENSG0000C |
| 143632071 | 143633756 | -1 | lncRNA | ENSG0000C |
| 143636013 | 143656418 | 1 | protein_coding | ENSG0000C |
| 143684452 | 143713898 | 1 | protein_coding | ENSG0000C |
| 143696154 | 143698413 | 1 | lncRNA | ENSG0000C |
| 143706694 | 143708109 | -1 | protein_coding | ENSG0000C |
| 143709007 | 143713584 | -1 | lncRNA | ENSG0000C |
| 143716340 | 143722458 | 1 | protein_coding | ENSG0000C |
| 143718246 | 143718891 | -1 | lncRNA | ENSG0000C |
| 143723933 | 143738234 | -1 | protein_coding | ENSG0000C |
| 143733083 | 143733153 | -1 | miRNA | ENSG0000C |
| 143734140 | 143790644 | 1 | protein_coding | ENSG0000C |
| 143758153 | 143771870 | -1 | lncRNA | ENSG0000C |
| 143790920 | 143815773 | -1 | protein_coding | ENSG0000C |
| 143812957 | 143813042 | -1 | miRNA | ENSG0000C |
| 143816344 | 143829352 | -1 | protein_coding | ENSG0000C |
| 143829518 | 143832861 | 1 | lncRNA | ENSG0000C |
| 143833270 | 143834063 | -1 | lncRNA | ENSG0000C |
| 143833583 | 143840973 | -1 | protein_coding | ENSG0000C |
| 143837756 | 143837816 | -1 | miRNA | ENSG0000C |
| 143838779 | 143840256 | 1 | lncRNA | ENSG0000C |
| 143857324 | 143878464 | -1 | protein_coding | ENSG0000C |
| 143888500 | 143901588 | 1 | lncRNA | ENSG0000C |
| 143915147 | 143976734 | -1 | protein_coding | ENSG0000C |
| 143945191 | 143945279 | -1 | miRNA | ENSG0000C |
| 143977153 | 144012772 | -1 | protein_coding | ENSG0000C |
| 143990056 | 143993415 | 1 | protein_coding | ENSG0000C |
| 144012280 | 144047114 | 1 | protein_coding | ENSG0000C |
| 144049079 | 144051522 | 1 | transcribed_unitary_ | ENSG0000C |
| 144051266 | 144063965 | -1 | protein_coding | ENSG0000C |
| 144057321 | 144057380 | -1 | miRNA | ENSG0000C |
| 144078002 | 144079265 | -1 | lncRNA | ENSG0000C |
| 144078648 | 144080648 | 1 | protein_coding | ENSG0000C |
| 144079874 | 144079942 | 1 | miRNA | ENSG0000C |
| 144082590 | 144086216 | 1 | protein_coding | ENSG0000C |
| 144095039 | 144097525 | 1 | protein_coding | ENSG0000C |
| 144098633 | 144108124 | -1 | protein_coding | ENSG0000C |
| 144104461 | 144107611 | 1 | protein_coding | ENSG0000C |
| 144107726 | 144118328 | 1 | protein_coding | ENSG0000C |
| 144137774 | 144140851 | 1 | protein_coding | ENSG0000C |
| 144141214 | 144143664 | -1 | unitary_pseudogene | ENSG0000C |
| 144148016 | 144261940 | 1 | protein_coding | ENSG0000C |
| 144262045 | 144291438 | -1 | protein_coding | ENSG0000C |

| | | | | |
|---|---|---|---|---|
| 144262673 | 144262737 | -1 | miRNA | ENSG0000C |
| 144266453 | 144268481 | 1 | protein_coding | ENSG0000C |
| 144291591 | 144314720 | 1 | protein_coding | ENSG0000C |
| 144314584 | 144326910 | -1 | protein_coding | ENSG0000C |
| 144314590 | 144315138 | -1 | lncRNA | ENSG0000C |
| 144317246 | 144317315 | -1 | miRNA | ENSG0000C |
| 144330565 | 144336482 | -1 | protein_coding | ENSG0000C |
| 144352219 | 144354914 | -1 | protein_coding | ENSG0000C |
| 144353228 | 144355609 | -1 | protein_coding | ENSG0000C |
| 144354135 | 144361272 | 1 | protein_coding | ENSG0000C |
| 144355431 | 144359376 | -1 | protein_coding | ENSG0000C |
| 144373101 | 144393242 | 1 | protein_coding | ENSG0000C |
| 144393229 | 144409335 | -1 | protein_coding | ENSG0000C |
| 144394149 | 144394230 | -1 | miRNA | ENSG0000C |
| 144400086 | 144400165 | -1 | miRNA | ENSG0000C |
| 144400277 | 144400345 | -1 | miRNA | ENSG0000C |
| 144409492 | 144409976 | 1 | lncRNA | ENSG0000C |
| 144409742 | 144416895 | -1 | protein_coding | ENSG0000C |
| 144423601 | 144428563 | -1 | protein_coding | ENSG0000C |
| 144428775 | 144444440 | -1 | protein_coding | ENSG0000C |
| 144435551 | 144435619 | -1 | miRNA | ENSG0000C |
| 144437675 | 144439971 | 1 | lncRNA | ENSG0000C |
| 144449582 | 144465677 | -1 | protein_coding | ENSG0000C |
| 144463817 | 144465101 | 1 | lncRNA | ENSG0000C |
| 144466043 | 144474202 | 1 | protein_coding | ENSG0000C |
| 144473412 | 144475849 | -1 | protein_coding | ENSG0000C |
| 144477969 | 144502121 | 1 | protein_coding | ENSG0000C |
| 144495458 | 144505444 | -1 | lncRNA | ENSG0000C |
| 144502973 | 144507174 | 1 | protein_coding | ENSG0000C |
| 144509070 | 144511213 | 1 | protein_coding | ENSG0000C |
| 144511288 | 144517845 | -1 | protein_coding | ENSG0000C |
| 144512567 | 144513672 | 1 | lncRNA | ENSG0000C |
| 144517992 | 144525172 | 1 | protein_coding | ENSG0000C |
| 144522388 | 144527033 | -1 | protein_coding | ENSG0000C |
| 144525733 | 144529132 | -1 | protein_coding | ENSG0000C |
| 144529179 | 144605816 | -1 | protein_coding | ENSG0000C |
| 144584040 | 144586488 | 1 | lncRNA | ENSG0000C |
| 144683169 | 144683464 | 1 | misc_RNA | ENSG0000C |
| 144698614 | 144699185 | -1 | lncRNA | ENSG0000C |
| 144700353 | 144708517 | 1 | transcribed_unproce | ENSG0000C |
| 144713899 | 144714845 | 1 | lncRNA | ENSG0000C |
| 144720907 | 144756417 | -1 | protein_coding | ENSG0000C |
| 144773114 | 144787345 | -1 | protein_coding | ENSG0000C |
| 144785298 | 144785588 | 1 | misc_RNA | ENSG0000C |
| 144789765 | 144792587 | -1 | protein_coding | ENSG0000C |
| 144791931 | 144791991 | -1 | miRNA | ENSG0000C |
| 144798876 | 144811169 | 1 | protein_coding | ENSG0000C |
| 144827464 | 144847509 | 1 | protein_coding | ENSG0000C |
| 144841042 | 144853736 | -1 | protein_coding | ENSG0000C |
| 144854875 | 144869489 | -1 | lncRNA | ENSG0000C |

| | | | | |
|---|---|---|---|---|
| 144876497 | 144902168 | -1 | protein_coding | ENSG00000 |
| 144930358 | 144950888 | -1 | protein_coding | ENSG00000 |
| 144973589 | 145002895 | -1 | transcribed_unproce | ENSG00000 |
| 144992914 | 144994837 | 1 | lncRNA | ENSG00000 |
| 144994886 | 144995540 | 1 | processed_pseudoge | ENSG00000 |
| 145002811 | 145006046 | 1 | lncRNA | ENSG00000 |
| 145005561 | 145005906 | -1 | processed_pseudoge | ENSG00000 |
| 145047860 | 145048889 | 1 | processed_pseudoge | ENSG00000 |
| 145052465 | 145066685 | 1 | protein_coding | ENSG00000 |
| 12134 | 13783 | 1 | unprocessed_pseudo | ENSG00000 |
| 14475 | 73865 | -1 | protein_coding | ENSG00000 |
| 27657 | 30891 | 1 | lncRNA | ENSG00000 |
| 30144 | 30281 | 1 | miRNA | ENSG00000 |
| 34394 | 37269 | -1 | lncRNA | ENSG00000 |
| 52452 | 73374 | -1 | transcribed_unproce | ENSG00000 |
| 72670 | 89850 | 1 | lncRNA | ENSG00000 |
| 100804 | 110138 | -1 | lncRNA | ENSG00000 |
| 112713 | 113754 | -1 | lncRNA | ENSG00000 |
| 116231 | 118204 | -1 | protein_coding | ENSG00000 |
| 121038 | 179147 | -1 | protein_coding | ENSG00000 |
| 197065 | 209554 | -1 | lncRNA | ENSG00000 |
| 212824 | 215893 | -1 | lncRNA | ENSG00000 |
| 214854 | 465259 | 1 | protein_coding | ENSG00000 |
| 267966 | 273002 | -1 | lncRNA | ENSG00000 |
| 452492 | 492248 | -1 | lncRNA | ENSG00000 |
| 470291 | 746106 | 1 | protein_coding | ENSG00000 |
| 477750 | 478423 | 1 | transcribed_processe | ENSG00000 |
| 487774 | 495610 | 1 | lncRNA | ENSG00000 |
| 547135 | 549967 | 1 | lncRNA | ENSG00000 |
| 558826 | 558930 | 1 | snRNA | ENSG00000 |
| 659986 | 660326 | 1 | processed_pseudoge | ENSG00000 |
| 673478 | 685555 | -1 | lncRNA | ENSG00000 |
| 841690 | 969090 | 1 | protein_coding | ENSG00000 |
| 976655 | 991732 | 1 | protein_coding | ENSG00000 |
| 1001858 | 1001963 | -1 | snRNA | ENSG00000 |
| 1011085 | 1011799 | 1 | processed_pseudoge | ENSG00000 |
| 1041655 | 1042608 | -1 | lncRNA | ENSG00000 |
| 1045625 | 1048641 | 1 | lncRNA | ENSG00000 |
| 1049858 | 1057552 | 1 | protein_coding | ENSG00000 |
| 1151807 | 1152140 | -1 | processed_pseudoge | ENSG00000 |
| 1164414 | 1164867 | -1 | processed_pseudoge | ENSG00000 |
| 1328664 | 1328987 | 1 | processed_pseudoge | ENSG00000 |
| 1333837 | 1333953 | 1 | rRNA_pseudogene | ENSG00000 |
| 1540321 | 1550103 | -1 | lncRNA | ENSG00000 |
| 1977784 | 1978547 | -1 | processed_pseudoge | ENSG00000 |
| 1980290 | 2193624 | 1 | protein_coding | ENSG00000 |
| 2041900 | 2046023 | -1 | lncRNA | ENSG00000 |
| 2162763 | 2162953 | -1 | snRNA | ENSG00000 |
| 2228961 | 2229241 | -1 | misc_RNA | ENSG00000 |
| 2421597 | 2643359 | -1 | lncRNA | ENSG00000 |

| | | | | |
|---|---|---|---|---|
| 2541097 | 2541635 | 1 | lncRNA | ENSG0000C |
| 2621787 | 2660056 | 1 | protein_coding | ENSG0000C |
| 2652856 | 2654371 | -1 | lncRNA | ENSG0000C |
| 2717510 | 2730037 | 1 | protein_coding | ENSG0000C |
| 2720469 | 2844095 | -1 | protein_coding | ENSG0000C |
| 2875442 | 2876427 | -1 | processed_pseudoge | ENSG0000C |
| 2900320 | 2900804 | -1 | processed_pseudoge | ENSG0000C |
| 2907073 | 3053408 | -1 | transcribed_unproce | ENSG0000C |
| 3181342 | 3200445 | 1 | lncRNA | ENSG0000C |
| 3218297 | 3526004 | -1 | protein_coding | ENSG0000C |
| 3364329 | 3364792 | -1 | processed_pseudoge | ENSG0000C |
| 3526723 | 3691814 | 1 | lncRNA | ENSG0000C |
| 3630947 | 3648390 | 1 | lncRNA | ENSG0000C |
| 3684076 | 3689626 | -1 | lncRNA | ENSG0000C |
| 3824127 | 4348392 | -1 | protein_coding | ENSG0000C |
| 3875584 | 3878809 | 1 | lncRNA | ENSG0000C |
| 3898642 | 3901248 | 1 | lncRNA | ENSG0000C |
| 4070906 | 4071884 | -1 | lncRNA | ENSG0000C |
| 4299390 | 4306046 | 1 | lncRNA | ENSG0000C |
| 4386410 | 4386556 | -1 | snRNA | ENSG0000C |
| 4490468 | 4587469 | 1 | protein_coding | ENSG0000C |
| 4553386 | 4666674 | -1 | protein_coding | ENSG0000C |
| 4633027 | 4633756 | 1 | processed_pseudoge | ENSG0000C |
| 4662294 | 4665262 | 1 | protein_coding | ENSG0000C |
| 4676600 | 4679502 | -1 | lncRNA | ENSG0000C |
| 4679559 | 4708399 | 1 | protein_coding | ENSG0000C |
| 4709556 | 4742043 | -1 | protein_coding | ENSG0000C |
| 4741741 | 4759639 | 1 | lncRNA | ENSG0000C |
| 4768617 | 4769895 | -1 | processed_pseudoge | ENSG0000C |
| 4781475 | 4782078 | -1 | processed_pseudoge | ENSG0000C |
| 4792944 | 4885917 | 1 | protein_coding | ENSG0000C |
| 4803788 | 4804972 | -1 | processed_pseudoge | ENSG0000C |
| 4850297 | 4850375 | 1 | miRNA | ENSG0000C |
| 4850299 | 4850373 | -1 | lncRNA | ENSG0000C |
| 4944670 | 4945902 | -1 | processed_pseudoge | ENSG0000C |
| 4984390 | 5128183 | 1 | protein_coding | ENSG0000C |
| 5040945 | 5041940 | 1 | processed_pseudoge | ENSG0000C |
| 5084999 | 5086112 | -1 | processed_pseudoge | ENSG0000C |
| 5091093 | 5091604 | 1 | unprocessed_pseudc | ENSG0000C |
| 5094139 | 5095014 | 1 | unprocessed_pseudc | ENSG0000C |
| 5096666 | 5098193 | 1 | unprocessed_pseudc | ENSG0000C |
| 5098341 | 5099325 | 1 | unprocessed_pseudc | ENSG0000C |
| 5099560 | 5100229 | 1 | unprocessed_pseudc | ENSG0000C |
| 5100236 | 5101009 | 1 | unprocessed_pseudc | ENSG0000C |
| 5107937 | 5109290 | 1 | unprocessed_pseudc | ENSG0000C |
| 5109496 | 5110762 | 1 | unprocessed_pseudc | ENSG0000C |
| 5110913 | 5112849 | 1 | processed_pseudoge | ENSG0000C |
| 5113549 | 5114804 | 1 | IG_C_pseudogene | ENSG0000C |
| 5123880 | 5185647 | -1 | protein_coding | ENSG0000C |
| 5193339 | 5193558 | -1 | processed_pseudoge | ENSG0000C |

| | | | | |
|---|---|---|---|---|
| 5231419 | 5235304 | 1 | protein_coding | ENSG00000( |
| 5299864 | 5304716 | -1 | protein_coding | ENSG00000( |
| 5311444 | 5311716 | 1 | processed_pseudoge | ENSG00000( |
| 5334930 | 5339876 | -1 | protein_coding | ENSG00000( |
| 5351796 | 5352410 | -1 | lncRNA | ENSG00000( |
| 5356312 | 5356525 | -1 | processed_pseudoge | ENSG00000( |
| 5357971 | 5437925 | -1 | protein_coding | ENSG00000( |
| 5418373 | 5418976 | -1 | processed_pseudoge | ENSG00000( |
| 5450503 | 5470566 | 1 | protein_coding | ENSG00000( |
| 5457477 | 5629748 | -1 | lncRNA | ENSG00000( |
| 5510531 | 5571282 | 1 | protein_coding | ENSG00000( |
| 5584490 | 5588994 | 1 | lncRNA | ENSG00000( |
| 5629025 | 5776557 | 1 | protein_coding | ENSG00000( |
| 5719021 | 5720244 | -1 | lncRNA | ENSG00000( |
| 5765076 | 5833117 | -1 | protein_coding | ENSG00000( |
| 5855781 | 5856426 | -1 | processed_pseudoge | ENSG00000( |
| 5881596 | 6008482 | -1 | protein_coding | ENSG00000( |
| 5890889 | 5910606 | 1 | protein_coding | ENSG00000( |
| 6007826 | 6007904 | 1 | miRNA | ENSG00000( |
| 6011025 | 6015625 | -1 | protein_coding | ENSG00000( |
| 6047360 | 6066714 | 1 | lncRNA | ENSG00000( |
| 6195934 | 6197248 | 1 | processed_pseudoge | ENSG00000( |
| 6215786 | 6257983 | 1 | protein_coding | ENSG00000( |
| 6276640 | 6276974 | 1 | processed_pseudoge | ENSG00000( |
| 6278667 | 6279153 | 1 | processed_pseudoge | ENSG00000( |
| 6328375 | 6331900 | 1 | protein_coding | ENSG00000( |
| 6413151 | 6507054 | 1 | protein_coding | ENSG00000( |
| 6415145 | 6449558 | -1 | lncRNA | ENSG00000( |
| 6532464 | 6645783 | -1 | protein_coding | ENSG00000( |
| 6542094 | 6542353 | 1 | misc_RNA | ENSG00000( |
| 6568844 | 6569360 | 1 | processed_pseudoge | ENSG00000( |
| 6639139 | 6639604 | 1 | processed_pseudoge | ENSG00000( |
| 6645956 | 6670635 | 1 | lncRNA | ENSG00000( |
| 6661651 | 6661942 | 1 | misc_RNA | ENSG00000( |
| 6662424 | 6663792 | 1 | processed_pseudoge | ENSG00000( |
| 6668999 | 6670008 | -1 | processed_pseudoge | ENSG00000( |
| 6675284 | 6675614 | 1 | processed_pseudoge | ENSG00000( |
| 6704270 | 6707780 | 1 | lncRNA | ENSG00000( |
| 6716157 | 6727438 | 1 | lncRNA | ENSG00000( |
| 6720863 | 7175648 | 1 | protein_coding | ENSG00000( |
| 6734598 | 6735076 | -1 | processed_pseudoge | ENSG00000( |
| 6748703 | 6748981 | -1 | processed_pseudoge | ENSG00000( |
| 6834456 | 6835577 | 1 | processed_pseudoge | ENSG00000( |
| 6902670 | 6978859 | 1 | lncRNA | ENSG00000( |
| 7177577 | 7203458 | 1 | lncRNA | ENSG00000( |
| 7304551 | 7343556 | 1 | lncRNA | ENSG00000( |
| 7477045 | 7478320 | -1 | processed_pseudoge | ENSG00000( |
| 7597823 | 7598377 | 1 | processed_pseudoge | ENSG00000( |
| 7786105 | 7786688 | -1 | lncRNA | ENSG00000( |
| 7796500 | 7888380 | -1 | protein_coding | ENSG00000( |

| | | | | |
|---|---|---|---|---|
| 7914514 | 7961224 | -1 | lncRNA | ENSG00000( |
| 7930827 | 7935137 | -1 | lncRNA | ENSG00000( |
| 8314246 | 10612723 | -1 | protein_coding | ENSG00000( |
| 8700595 | 8701552 | -1 | lncRNA | ENSG00000( |
| 8713311 | 8713841 | 1 | processed_pseudoge | ENSG00000( |
| 8797135 | 8797194 | 1 | snRNA | ENSG00000( |
| 8858127 | 8864772 | 1 | lncRNA | ENSG00000( |
| 8936079 | 8963077 | 1 | lncRNA | ENSG00000( |
| 9090898 | 9091245 | 1 | processed_pseudoge | ENSG00000( |
| 9442060 | 9442380 | 1 | misc_RNA | ENSG00000( |
| 9799423 | 9803784 | 1 | lncRNA | ENSG00000( |
| 10532145 | 10532411 | 1 | processed_pseudoge | ENSG00000( |
| 10613202 | 10620420 | 1 | lncRNA | ENSG00000( |
| 10631750 | 10632203 | -1 | lncRNA | ENSG00000( |
| 10777357 | 10777525 | 1 | processed_pseudoge | ENSG00000( |
| 10829752 | 10830308 | -1 | processed_pseudoge | ENSG00000( |
| 10948372 | 10948481 | -1 | lncRNA | ENSG00000( |
| 11011748 | 11013096 | 1 | processed_pseudoge | ENSG00000( |
| 11275314 | 11276314 | -1 | lncRNA | ENSG00000( |
| 11313397 | 11436244 | 1 | lncRNA | ENSG00000( |
| 11618496 | 11618621 | -1 | unprocessed_pseudc | ENSG00000( |
| 11618530 | 12058227 | 1 | lncRNA | ENSG00000( |
| 12098660 | 12159148 | -1 | lncRNA | ENSG00000( |
| 12287320 | 12288238 | 1 | processed_pseudoge | ENSG00000( |
| 12300266 | 12300451 | -1 | snRNA | ENSG00000( |
| 12685439 | 12710285 | 1 | protein_coding | ENSG00000( |
| 12698554 | 12814377 | -1 | lncRNA | ENSG00000( |
| 12764761 | 12765055 | -1 | misc_RNA | ENSG00000( |
| 12775020 | 12823060 | 1 | protein_coding | ENSG00000( |
| 12885321 | 12885417 | 1 | misc_RNA | ENSG00000( |
| 12958503 | 13034326 | -1 | lncRNA | ENSG00000( |
| 12972843 | 12973438 | -1 | processed_pseudoge | ENSG00000( |
| 13021694 | 13022892 | -1 | processed_pseudoge | ENSG00000( |
| 13105704 | 13279590 | -1 | protein_coding | ENSG00000( |
| 13274510 | 13279187 | -1 | lncRNA | ENSG00000( |
| 13322806 | 13323450 | -1 | processed_pseudoge | ENSG00000( |
| 13386130 | 13836571 | 1 | lncRNA | ENSG00000( |
| 13406380 | 13488125 | -1 | lncRNA | ENSG00000( |
| 13927971 | 13945610 | 1 | lncRNA | ENSG00000( |
| 13986175 | 13987909 | 1 | processed_pseudoge | ENSG00000( |
| 14025971 | 14030115 | 1 | lncRNA | ENSG00000( |
| 14040716 | 14041861 | -1 | processed_pseudoge | ENSG00000( |
| 14068932 | 14069417 | 1 | processed_pseudoge | ENSG00000( |
| 14081843 | 14398983 | -1 | protein_coding | ENSG00000( |
| 14204248 | 14205039 | 1 | processed_pseudoge | ENSG00000( |
| 14317073 | 14358490 | 1 | lncRNA | ENSG00000( |
| 14395791 | 14472407 | 1 | lncRNA | ENSG00000( |
| 14531916 | 14532042 | -1 | processed_pseudoge | ENSG00000( |
| 14586766 | 14587326 | -1 | lncRNA | ENSG00000( |
| 14588797 | 14590065 | -1 | lncRNA | ENSG00000( |

| | | | | |
|---|---|---|---|---|
| 14593577 | 14594200 | 1 | processed_pseudoge | ENSG00000 |
| 14611071 | 14693471 | -1 | protein_coding | ENSG00000 |
| 14719724 | 14722733 | -1 | protein_coding | ENSG00000 |
| 14734666 | 14910995 | -1 | protein_coding | ENSG00000 |
| 14779930 | 14784117 | 1 | lncRNA | ENSG00000 |
| 14790635 | 14791158 | 1 | lncRNA | ENSG00000 |
| 14850429 | 14850535 | 1 | snRNA | ENSG00000 |
| 14921013 | 15146401 | -1 | transcribed_unproce | ENSG00000 |
| 14921337 | 14922334 | -1 | transcribed_processe | ENSG00000 |
| 14993312 | 15019729 | 1 | unprocessed_pseudo | ENSG00000 |
| 15055057 | 15056052 | 1 | processed_pseudoge | ENSG00000 |
| 15143289 | 15143386 | 1 | snRNA | ENSG00000 |
| 15163622 | 15307360 | -1 | protein_coding | ENSG00000 |
| 15361393 | 15362134 | 1 | processed_pseudoge | ENSG00000 |
| 15422704 | 15465953 | 1 | protein_coding | ENSG00000 |
| 15431899 | 15432006 | -1 | snRNA | ENSG00000 |
| 15464066 | 15510995 | -1 | protein_coding | ENSG00000 |
| 15524576 | 15524707 | -1 | processed_pseudoge | ENSG00000 |
| 15525782 | 15526081 | -1 | misc_RNA | ENSG00000 |
| 15527133 | 15527683 | -1 | processed_pseudoge | ENSG00000 |
| 15544188 | 15544294 | 1 | snRNA | ENSG00000 |
| 15553043 | 16061663 | 1 | protein_coding | ENSG00000 |
| 15588355 | 15588615 | 1 | processed_pseudoge | ENSG00000 |
| 15776181 | 15776286 | -1 | snRNA | ENSG00000 |
| 16203935 | 16410990 | -1 | protein_coding | ENSG00000 |
| 16409503 | 16870843 | -1 | protein_coding | ENSG00000 |
| 16473188 | 16476237 | 1 | lncRNA | ENSG00000 |
| 16625676 | 16626390 | 1 | lncRNA | ENSG00000 |
| 16726814 | 16727524 | 1 | lncRNA | ENSG00000 |
| 16775573 | 16775845 | -1 | processed_pseudoge | ENSG00000 |
| 16917511 | 17114122 | 1 | lncRNA | ENSG00000 |
| 16982723 | 16983459 | -1 | lncRNA | ENSG00000 |
| 17053901 | 17054204 | 1 | misc_RNA | ENSG00000 |
| 17096301 | 17096446 | -1 | processed_pseudoge | ENSG00000 |
| 17108304 | 17108604 | -1 | processed_pseudoge | ENSG00000 |
| 17134982 | 17503923 | 1 | protein_coding | ENSG00000 |
| 17447145 | 17448543 | -1 | processed_pseudoge | ENSG00000 |
| 17579066 | 17797124 | 1 | protein_coding | ENSG00000 |
| 17589161 | 17591318 | -1 | processed_pseudoge | ENSG00000 |
| 17906563 | 18910950 | 1 | protein_coding | ENSG00000 |
| 18360597 | 18362220 | -1 | lncRNA | ENSG00000 |
| 18573306 | 18573379 | 1 | miRNA | ENSG00000 |
| 18651147 | 18651450 | 1 | misc_RNA | ENSG00000 |
| 18717974 | 18718526 | -1 | processed_pseudoge | ENSG00000 |
| 18858906 | 19049358 | -1 | protein_coding | ENSG00000 |
| 19026892 | 19028129 | 1 | processed_pseudoge | ENSG00000 |
| 19049427 | 19051025 | 1 | protein_coding | ENSG00000 |
| 19053141 | 19102904 | -1 | protein_coding | ENSG00000 |
| 19063656 | 19063786 | -1 | scaRNA | ENSG00000 |
| 19073843 | 19073946 | 1 | snRNA | ENSG00000 |

| | | | | |
|---|---|---|---|---|
| 19089593 | 19089688 | 1 | misc_RNA | ENSG00000( |
| 19108375 | 19149290 | -1 | protein_coding | ENSG00000( |
| 19200335 | 19201046 | -1 | processed_pseudoge | ENSG00000( |
| 19230435 | 19373545 | 1 | protein_coding | ENSG00000( |
| 19291337 | 19292576 | -1 | lncRNA | ENSG00000( |
| 19371386 | 19371945 | -1 | lncRNA | ENSG00000( |
| 19375451 | 19375996 | 1 | lncRNA | ENSG00000( |
| 19375715 | 19380236 | -1 | protein_coding | ENSG00000( |
| 19383217 | 19383887 | -1 | processed_pseudoge | ENSG00000( |
| 19386927 | 19408532 | -1 | lncRNA | ENSG00000( |
| 19409009 | 19452505 | 1 | protein_coding | ENSG00000( |
| 19453209 | 19455173 | 1 | lncRNA | ENSG00000( |
| 19460926 | 19464524 | -1 | transcribed_processe | ENSG00000( |
| 19507452 | 19786928 | -1 | protein_coding | ENSG00000( |
| 19705338 | 19705655 | 1 | processed_pseudoge | ENSG00000( |
| 19789179 | 19894856 | 1 | lncRNA | ENSG00000( |
| 19895800 | 19895973 | -1 | unprocessed_pseudc | ENSG00000( |
| 19926094 | 19929937 | 1 | lncRNA | ENSG00000( |
| 19957394 | 19958147 | -1 | processed_pseudoge | ENSG00000( |
| 19967239 | 19967994 | -1 | processed_pseudoge | ENSG00000( |
| 20331446 | 20332277 | 1 | processed_pseudoge | ENSG00000( |
| 20341669 | 20622518 | -1 | protein_coding | ENSG00000( |
| 20411148 | 20411238 | -1 | miRNA | ENSG00000( |
| 20418321 | 20418427 | 1 | snRNA | ENSG00000( |
| 20502265 | 20502342 | -1 | miRNA | ENSG00000( |
| 20532552 | 20533339 | 1 | processed_pseudoge | ENSG00000( |
| 20658309 | 20995955 | 1 | protein_coding | ENSG00000( |
| 20683304 | 20684688 | -1 | lncRNA | ENSG00000( |
| 20716105 | 20716188 | 1 | miRNA | ENSG00000( |
| 20786927 | 20787055 | 1 | snoRNA | ENSG00000( |
| 20999509 | 21031640 | -1 | protein_coding | ENSG00000( |
| 21058771 | 21059323 | -1 | processed_pseudoge | ENSG00000( |
| 21077104 | 21077942 | -1 | protein_coding | ENSG00000( |
| 21106543 | 21106836 | 1 | processed_pseudoge | ENSG00000( |
| 21135598 | 21136350 | -1 | processed_pseudoge | ENSG00000( |
| 21140214 | 21142145 | -1 | protein_coding | ENSG00000( |
| 21160235 | 21160336 | -1 | misc_RNA | ENSG00000( |
| 21165637 | 21166660 | -1 | protein_coding | ENSG00000( |
| 21178589 | 21179188 | -1 | unprocessed_pseudc | ENSG00000( |
| 21186694 | 21187671 | -1 | protein_coding | ENSG00000( |
| 21190654 | 21191244 | -1 | unprocessed_pseudc | ENSG00000( |
| 21201469 | 21202205 | -1 | protein_coding | ENSG00000( |
| 21206181 | 21207143 | -1 | protein_coding | ENSG00000( |
| 21209189 | 21209779 | -1 | unprocessed_pseudc | ENSG00000( |
| 21216373 | 21217311 | -1 | protein_coding | ENSG00000( |
| 21227243 | 21228222 | -1 | protein_coding | ENSG00000( |
| 21231266 | 21231853 | -1 | unprocessed_pseudc | ENSG00000( |
| 21239002 | 21240005 | -1 | protein_coding | ENSG00000( |
| 21278050 | 21278619 | -1 | unprocessed_pseudc | ENSG00000( |
| 21304326 | 21305312 | -1 | protein_coding | ENSG00000( |

| | | | | |
|---|---|---|---|---|
| 21321300 | 21321827 | 1 | unprocessed_pseudc | ENSG0000C |
| 21329665 | 21335404 | -1 | protein_coding | ENSG0000C |
| 21349835 | 21351378 | -1 | protein_coding | ENSG0000C |
| 21367424 | 21368962 | -1 | protein_coding | ENSG0000C |
| 21384255 | 21385398 | -1 | protein_coding | ENSG0000C |
| 21394888 | 21396346 | -1 | lncRNA | ENSG0000C |
| 21398614 | 21399139 | -1 | unprocessed_pseudc | ENSG0000C |
| 21403081 | 21403651 | 1 | unprocessed_pseudc | ENSG0000C |
| 21409117 | 21410185 | 1 | protein_coding | ENSG0000C |
| 21410969 | 21559900 | -1 | lncRNA | ENSG0000C |
| 21420234 | 21420813 | 1 | unprocessed_pseudc | ENSG0000C |
| 21440439 | 21441316 | 1 | protein_coding | ENSG0000C |
| 21455484 | 21456049 | 1 | unprocessed_pseudc | ENSG0000C |
| 21480839 | 21482313 | -1 | protein_coding | ENSG0000C |
| 21512115 | 21512185 | -1 | miRNA | ENSG0000C |
| 21524307 | 21524382 | -1 | snoRNA | ENSG0000C |
| 21638285 | 21638676 | 1 | processed_pseudoge | ENSG0000C |
| 21695176 | 21696943 | 1 | processed_pseudoge | ENSG0000C |
| 21699314 | 21699596 | 1 | misc_RNA | ENSG0000C |
| 21802636 | 21937651 | 1 | protein_coding | ENSG0000C |
| 21802636 | 22029594 | 1 | protein_coding | ENSG0000C |
| 21811621 | 21812347 | -1 | processed_pseudoge | ENSG0000C |
| 21858910 | 21861926 | -1 | lncRNA | ENSG0000C |
| 21892188 | 21893857 | 1 | TEC | ENSG0000C |
| 21929457 | 21931073 | 1 | transcribed_processe | ENSG0000C |
| 21966929 | 21967751 | 1 | lncRNA | ENSG0000C |
| 21967753 | 21995301 | -1 | protein_coding | ENSG0000C |
| 21994139 | 22128103 | 1 | lncRNA | ENSG0000C |
| 21995482 | 21996013 | 1 | lncRNA | ENSG0000C |
| 22002903 | 22009305 | -1 | protein_coding | ENSG0000C |
| 22012155 | 22012536 | 1 | transcribed_processe | ENSG0000C |
| 22046758 | 22046901 | 1 | misc_RNA | ENSG0000C |
| 22117665 | 22117814 | 1 | misc_RNA | ENSG0000C |
| 22203990 | 22214672 | -1 | lncRNA | ENSG0000C |
| 22446824 | 22455740 | 1 | protein_coding | ENSG0000C |
| 22646200 | 22824213 | 1 | lncRNA | ENSG0000C |
| 22703516 | 23438700 | -1 | lncRNA | ENSG0000C |
| 22747700 | 22748234 | 1 | processed_pseudoge | ENSG0000C |
| 22767175 | 22768316 | 1 | lncRNA | ENSG0000C |
| 23291544 | 23293923 | 1 | lncRNA | ENSG0000C |
| 23500691 | 23672387 | -1 | lncRNA | ENSG0000C |
| 23632106 | 23632360 | 1 | processed_pseudoge | ENSG0000C |
| 23671788 | 23672399 | -1 | lncRNA | ENSG0000C |
| 23681894 | 23682662 | 1 | processed_pseudoge | ENSG0000C |
| 23690104 | 23826337 | -1 | protein_coding | ENSG0000C |
| 23826491 | 23828877 | 1 | lncRNA | ENSG0000C |
| 23829670 | 23849914 | 1 | lncRNA | ENSG0000C |
| 23894990 | 23898054 | -1 | lncRNA | ENSG0000C |
| 23955760 | 23956444 | 1 | unprocessed_pseudc | ENSG0000C |
| 24069731 | 24088051 | 1 | lncRNA | ENSG0000C |

| | | | | |
|---|---|---|---|---|
| 24542952 | 24545946 | -1 | protein_coding | ENSG00000( |
| 24545938 | 24592104 | 1 | lncRNA | ENSG00000( |
| 24905469 | 24905735 | 1 | ribozyme | ENSG00000( |
| 25088811 | 25089151 | -1 | misc_RNA | ENSG00000( |
| 25579657 | 25584272 | -1 | lncRNA | ENSG00000( |
| 25676389 | 25678440 | -1 | protein_coding | ENSG00000( |
| 25780032 | 25844585 | 1 | lncRNA | ENSG00000( |
| 25937912 | 25938434 | 1 | processed_pseudoge | ENSG00000( |
| 26066675 | 26118408 | 1 | lncRNA | ENSG00000( |
| 26642697 | 26644595 | -1 | TEC | ENSG00000( |
| 26746953 | 26786874 | 1 | lncRNA | ENSG00000( |
| 26796173 | 26796304 | -1 | snoRNA | ENSG00000( |
| 26801733 | 26805862 | 1 | lncRNA | ENSG00000( |
| 26840685 | 26892803 | -1 | protein_coding | ENSG00000( |
| 26880178 | 26880549 | -1 | processed_pseudoge | ENSG00000( |
| 26895678 | 26895970 | 1 | misc_RNA | ENSG00000( |
| 26903372 | 26947242 | -1 | protein_coding | ENSG00000( |
| 26947039 | 27062930 | 1 | protein_coding | ENSG00000( |
| 26955780 | 26956295 | -1 | lncRNA | ENSG00000( |
| 26993136 | 27005672 | -1 | protein_coding | ENSG00000( |
| 27102630 | 27104728 | 1 | lncRNA | ENSG00000( |
| 27109141 | 27230174 | 1 | protein_coding | ENSG00000( |
| 27198287 | 27198401 | 1 | rRNA_pseudogene | ENSG00000( |
| 27210327 | 27210626 | 1 | processed_pseudoge | ENSG00000( |
| 27245680 | 27284869 | -1 | transcribed_unproce | ENSG00000( |
| 27284654 | 27297150 | -1 | protein_coding | ENSG00000( |
| 27325209 | 27529814 | -1 | protein_coding | ENSG00000( |
| 27391734 | 27397563 | 1 | lncRNA | ENSG00000( |
| 27495857 | 27498251 | -1 | lncRNA | ENSG00000( |
| 27524314 | 27526498 | 1 | protein_coding | ENSG00000( |
| 27529085 | 27543902 | 1 | lncRNA | ENSG00000( |
| 27535640 | 27573866 | -1 | protein_coding | ENSG00000( |
| 27608382 | 27610745 | -1 | processed_pseudoge | ENSG00000( |
| 27829276 | 27844481 | 1 | lncRNA | ENSG00000( |
| 27937617 | 27944497 | -1 | lncRNA | ENSG00000( |
| 27948078 | 28670286 | -1 | protein_coding | ENSG00000( |
| 28148211 | 28149236 | 1 | processed_pseudoge | ENSG00000( |
| 28539735 | 28566733 | 1 | lncRNA | ENSG00000( |
| 28598668 | 28599292 | -1 | processed_pseudoge | ENSG00000( |
| 28863626 | 28863706 | -1 | miRNA | ENSG00000( |
| 28888879 | 28888955 | -1 | miRNA | ENSG00000( |
| 29214322 | 29229229 | 1 | lncRNA | ENSG00000( |
| 29254075 | 29254426 | -1 | processed_pseudoge | ENSG00000( |
| 29317833 | 29318952 | 1 | processed_pseudoge | ENSG00000( |
| 29636486 | 29636853 | 1 | processed_pseudoge | ENSG00000( |
| 29824742 | 29826711 | -1 | processed_pseudoge | ENSG00000( |
| 30388935 | 30575012 | -1 | lncRNA | ENSG00000( |
| 30558880 | 30559481 | -1 | processed_pseudoge | ENSG00000( |
| 30670964 | 30671966 | -1 | unprocessed_pseudc | ENSG00000( |
| 30689105 | 30690272 | 1 | processed_pseudoge | ENSG00000( |

| | | | | |
|---|---|---|---|---|
| 30773464 | 30774732 | 1 | processed_pseudoge | ENSG00000( |
| 30799471 | 30800746 | -1 | processed_pseudoge | ENSG00000( |
| 30831878 | 30832225 | -1 | processed_pseudoge | ENSG00000( |
| 30935485 | 30935977 | 1 | processed_pseudoge | ENSG00000( |
| 30988085 | 30990572 | 1 | processed_pseudoge | ENSG00000( |
| 31165618 | 31166980 | -1 | processed_pseudoge | ENSG00000( |
| 31253901 | 31254777 | 1 | processed_pseudoge | ENSG00000( |
| 31371611 | 31381490 | -1 | lncRNA | ENSG00000( |
| 31505280 | 31505943 | 1 | unprocessed_pseudc | ENSG00000( |
| 31505946 | 31506615 | 1 | unprocessed_pseudc | ENSG00000( |
| 31644576 | 31645160 | -1 | processed_pseudoge | ENSG00000( |
| 32293558 | 32293692 | 1 | rRNA_pseudogene | ENSG00000( |
| 32333457 | 32334354 | -1 | processed_pseudoge | ENSG00000( |
| 32384603 | 32454769 | 1 | protein_coding | ENSG00000( |
| 32455302 | 32526208 | -1 | protein_coding | ENSG00000( |
| 32540544 | 32552586 | -1 | protein_coding | ENSG00000( |
| 32551144 | 32568621 | 1 | protein_coding | ENSG00000( |
| 32553001 | 32573184 | -1 | protein_coding | ENSG00000( |
| 32566148 | 32567126 | -1 | processed_pseudoge | ENSG00000( |
| 32629454 | 32635669 | -1 | protein_coding | ENSG00000( |
| 32633454 | 32648685 | 1 | lncRNA | ENSG00000( |
| 32643103 | 32645026 | -1 | lncRNA | ENSG00000( |
| 32676063 | 32676520 | -1 | processed_pseudoge | ENSG00000( |
| 32727069 | 32727522 | -1 | processed_pseudoge | ENSG00000( |
| 32783540 | 32789201 | 1 | protein_coding | ENSG00000( |
| 32820261 | 32820350 | 1 | misc_RNA | ENSG00000( |
| 32836575 | 32869595 | -1 | unprocessed_pseudc | ENSG00000( |
| 32840598 | 32841762 | -1 | lncRNA | ENSG00000( |
| 32886601 | 33025130 | -1 | protein_coding | ENSG00000( |
| 32926051 | 32926450 | 1 | processed_pseudoge | ENSG00000( |
| 32945996 | 32947222 | 1 | processed_pseudoge | ENSG00000( |
| 32979560 | 32980403 | -1 | processed_pseudoge | ENSG00000( |
| 33019682 | 33020165 | -1 | processed_pseudoge | ENSG00000( |
| 33025273 | 33039907 | 1 | protein_coding | ENSG00000( |
| 33041765 | 33076674 | -1 | protein_coding | ENSG00000( |
| 33104082 | 33167356 | -1 | protein_coding | ENSG00000( |
| 33166948 | 33182865 | 1 | lncRNA | ENSG00000( |
| 33174170 | 33174295 | 1 | snRNA | ENSG00000( |
| 33218365 | 33248567 | 1 | protein_coding | ENSG00000( |
| 33247820 | 33264720 | -1 | protein_coding | ENSG00000( |
| 33264879 | 33282070 | 1 | protein_coding | ENSG00000( |
| 33290512 | 33371157 | 1 | protein_coding | ENSG00000( |
| 33352788 | 33352874 | -1 | misc_RNA | ENSG00000( |
| 33383179 | 33402682 | -1 | protein_coding | ENSG00000( |
| 33401478 | 33410553 | 1 | lncRNA | ENSG00000( |
| 33441156 | 33447596 | -1 | protein_coding | ENSG00000( |
| 33449755 | 33449941 | 1 | processed_pseudoge | ENSG00000( |
| 33461353 | 33473930 | -1 | protein_coding | ENSG00000( |
| 33467869 | 33467935 | -1 | miRNA | ENSG00000( |
| 33503655 | 33510711 | -1 | unprocessed_pseudc | ENSG00000( |

| | | | | |
|---|---|---|---|---|
| 33512959 | 33514773 | -1 | lncRNA | ENSG0000C |
| 33524394 | 33573009 | 1 | protein_coding | ENSG0000C |
| 33574361 | 33574588 | -1 | processed_pseudoge | ENSG0000C |
| 33576380 | 33577014 | 1 | processed_pseudoge | ENSG0000C |
| 33580695 | 33607874 | 1 | lncRNA | ENSG0000C |
| 33613848 | 33614009 | -1 | unprocessed_pseudc | ENSG0000C |
| 33617762 | 33618506 | 1 | TR_V_gene | ENSG0000C |
| 33624274 | 33625293 | 1 | processed_pseudoge | ENSG0000C |
| 33629121 | 33629586 | 1 | TR_V_pseudogene | ENSG0000C |
| 33634111 | 33634246 | 1 | TR_V_pseudogene | ENSG0000C |
| 33638035 | 33638494 | 1 | TR_V_pseudogene | ENSG0000C |
| 33649047 | 33649614 | 1 | TR_V_pseudogene | ENSG0000C |
| 33662166 | 33662663 | 1 | TR_V_pseudogene | ENSG0000C |
| 33673504 | 33677499 | -1 | transcribed_processe | ENSG0000C |
| 33676785 | 33688011 | 1 | lncRNA | ENSG0000C |
| 33695767 | 33696059 | 1 | TR_V_pseudogene | ENSG0000C |
| 33697459 | 33700986 | 1 | lncRNA | ENSG0000C |
| 33706679 | 33710715 | 1 | lncRNA | ENSG0000C |
| 33719690 | 33722555 | 1 | lncRNA | ENSG0000C |
| 33730345 | 33734040 | 1 | lncRNA | ENSG0000C |
| 33732975 | 33738416 | -1 | lncRNA | ENSG0000C |
| 33750466 | 33799231 | 1 | protein_coding | ENSG0000C |
| 33775183 | 33818795 | -1 | lncRNA | ENSG0000C |
| 33786221 | 33786828 | 1 | TR_V_pseudogene | ENSG0000C |
| 33817160 | 33920399 | 1 | protein_coding | ENSG0000C |
| 33830978 | 33831078 | 1 | misc_RNA | ENSG0000C |
| 33868540 | 33868665 | 1 | snRNA | ENSG0000C |
| 33921693 | 34048949 | -1 | protein_coding | ENSG0000C |
| 33934296 | 33934376 | -1 | snoRNA | ENSG0000C |
| 33952769 | 33952850 | -1 | snoRNA | ENSG0000C |
| 33968071 | 33968518 | -1 | processed_pseudoge | ENSG0000C |
| 34049966 | 34050244 | 1 | misc_RNA | ENSG0000C |
| 34084332 | 34096225 | 1 | lncRNA | ENSG0000C |
| 34086387 | 34127399 | -1 | protein_coding | ENSG0000C |
| 34109171 | 34109968 | -1 | processed_pseudoge | ENSG0000C |
| 34133157 | 34134696 | -1 | processed_pseudoge | ENSG0000C |
| 34179005 | 34252523 | 1 | protein_coding | ENSG0000C |
| 34191555 | 34191881 | 1 | processed_pseudoge | ENSG0000C |
| 34201273 | 34201515 | -1 | pseudogene | ENSG0000C |
| 34206134 | 34206241 | -1 | rRNA_pseudogene | ENSG0000C |
| 34223984 | 34224628 | -1 | processed_pseudoge | ENSG0000C |
| 34252381 | 34311371 | -1 | protein_coding | ENSG0000C |
| 34282528 | 34282670 | -1 | snRNA | ENSG0000C |
| 34318407 | 34319655 | 1 | processed_pseudoge | ENSG0000C |
| 34329506 | 34343711 | 1 | protein_coding | ENSG0000C |
| 34366666 | 34376898 | -1 | protein_coding | ENSG0000C |
| 34379019 | 34397810 | -1 | protein_coding | ENSG0000C |
| 34389943 | 34390051 | -1 | misc_RNA | ENSG0000C |
| 34398184 | 34458570 | -1 | protein_coding | ENSG0000C |
| 34457414 | 34520988 | 1 | protein_coding | ENSG0000C |

| | | | | |
|---|---|---|---|---|
| 34478622 | 34478905 | 1 | misc_RNA | ENSG0000C |
| 34521043 | 34522990 | -1 | protein_coding | ENSG0000C |
| 34521527 | 34524241 | 1 | lncRNA | ENSG0000C |
| 34546524 | 34546996 | -1 | processed_pseudoge | ENSG0000C |
| 34551432 | 34590140 | -1 | protein_coding | ENSG0000C |
| 34568012 | 34583072 | 1 | lncRNA | ENSG0000C |
| 34610486 | 34612104 | -1 | protein_coding | ENSG0000C |
| 34613545 | 34620523 | -1 | protein_coding | ENSG0000C |
| 34621379 | 34628086 | -1 | protein_coding | ENSG0000C |
| 34634722 | 34637809 | -1 | protein_coding | ENSG0000C |
| 34638133 | 34651035 | 1 | protein_coding | ENSG0000C |
| 34646645 | 34657113 | 1 | protein_coding | ENSG0000C |
| 34650702 | 34661892 | 1 | protein_coding | ENSG0000C |
| 34661880 | 34664048 | -1 | protein_coding | ENSG0000C |
| 34661903 | 34666029 | -1 | lncRNA | ENSG0000C |
| 34664163 | 34666112 | -1 | protein_coding | ENSG0000C |
| 34665607 | 34774466 | 1 | lncRNA | ENSG0000C |
| 34689570 | 34691276 | -1 | protein_coding | ENSG0000C |
| 34709005 | 34710136 | -1 | protein_coding | ENSG0000C |
| 34723053 | 34729488 | -1 | protein_coding | ENSG0000C |
| 34810020 | 34895175 | 1 | lncRNA | ENSG0000C |
| 34832295 | 34838586 | -1 | transcribed_unproce | ENSG0000C |
| 34889066 | 34895764 | -1 | protein_coding | ENSG0000C |
| 34917175 | 34918296 | 1 | processed_pseudoge | ENSG0000C |
| 34922184 | 34922921 | -1 | processed_pseudoge | ENSG0000C |
| 34957608 | 34982544 | 1 | protein_coding | ENSG0000C |
| 34985410 | 34989379 | -1 | lncRNA | ENSG0000C |
| 34989641 | 34998900 | 1 | protein_coding | ENSG0000C |
| 35001342 | 35002994 | -1 | processed_pseudoge | ENSG0000C |
| 35001913 | 35003663 | 1 | TEC | ENSG0000C |
| 35010703 | 35011285 | 1 | processed_pseudoge | ENSG0000C |
| 35012963 | 35016076 | -1 | lncRNA | ENSG0000C |
| 35023840 | 35025402 | -1 | lncRNA | ENSG0000C |
| 35038625 | 35044326 | -1 | TEC | ENSG0000C |
| 35041095 | 35045986 | 1 | protein_coding | ENSG0000C |
| 35049271 | 35049563 | 1 | misc_RNA | ENSG0000C |
| 35056064 | 35072627 | -1 | protein_coding | ENSG0000C |
| 35073835 | 35080016 | -1 | protein_coding | ENSG0000C |
| 35088688 | 35096601 | -1 | protein_coding | ENSG0000C |
| 35096378 | 35098141 | 1 | lncRNA | ENSG0000C |
| 35099776 | 35103195 | -1 | protein_coding | ENSG0000C |
| 35104112 | 35116341 | -1 | protein_coding | ENSG0000C |
| 35147410 | 35149184 | -1 | processed_pseudoge | ENSG0000C |
| 35161992 | 35405338 | 1 | protein_coding | ENSG0000C |
| 35360540 | 35362054 | -1 | TEC | ENSG0000C |
| 35406755 | 35483035 | 1 | transcribed_unproce | ENSG0000C |
| 35449748 | 35450352 | 1 | processed_pseudoge | ENSG0000C |
| 35490111 | 35561898 | 1 | protein_coding | ENSG0000C |
| 35491133 | 35491969 | -1 | lncRNA | ENSG0000C |
| 35507564 | 35507873 | 1 | processed_pseudoge | ENSG0000C |

| | | | | |
|---|---|---|---|---|
| 35561831 | 35563899 | -1 | protein_coding | ENSG00000( |
| 35592786 | 35592951 | 1 | processed_pseudoge | ENSG00000( |
| 35605262 | 35610041 | 1 | protein_coding | ENSG00000( |
| 35608094 | 35608159 | 1 | miRNA | ENSG00000( |
| 35609533 | 35646810 | -1 | protein_coding | ENSG00000( |
| 35642514 | 35643517 | 1 | lncRNA | ENSG00000( |
| 35646270 | 35647099 | 1 | lncRNA | ENSG00000( |
| 35649295 | 35650950 | -1 | protein_coding | ENSG00000( |
| 35657751 | 35658018 | -1 | lncRNA | ENSG00000( |
| 35657754 | 35658017 | -1 | ribozyme | ENSG00000( |
| 35658290 | 35661511 | 1 | protein_coding | ENSG00000( |
| 35658875 | 35675866 | -1 | protein_coding | ENSG00000( |
| 35671776 | 35672044 | 1 | misc_RNA | ENSG00000( |
| 35673918 | 35681159 | 1 | protein_coding | ENSG00000( |
| 35681992 | 35690056 | -1 | protein_coding | ENSG00000( |
| 35696948 | 35732195 | -1 | protein_coding | ENSG00000( |
| 35710676 | 35710741 | -1 | miRNA | ENSG00000( |
| 35732598 | 35736999 | 1 | protein_coding | ENSG00000( |
| 35732922 | 35732995 | 1 | miRNA | ENSG00000( |
| 35736866 | 35749228 | -1 | protein_coding | ENSG00000( |
| 35749287 | 35758585 | 1 | protein_coding | ENSG00000( |
| 35752990 | 35756613 | -1 | protein_coding | ENSG00000( |
| 35756712 | 35757940 | -1 | lncRNA | ENSG00000( |
| 35772163 | 35790432 | 1 | lncRNA | ENSG00000( |
| 35792154 | 35809732 | 1 | protein_coding | ENSG00000( |
| 35808045 | 35812272 | -1 | protein_coding | ENSG00000( |
| 35812960 | 35815354 | -1 | protein_coding | ENSG00000( |
| 35812960 | 35828732 | -1 | protein_coding | ENSG00000( |
| 35814451 | 35865518 | 1 | protein_coding | ENSG00000( |
| 35816391 | 35828747 | -1 | protein_coding | ENSG00000( |
| 35859055 | 35859980 | -1 | unitary_pseudogene | ENSG00000( |
| 35869263 | 35870601 | -1 | protein_coding | ENSG00000( |
| 35874810 | 35875194 | -1 | processed_pseudoge | ENSG00000( |
| 35906202 | 35907136 | 1 | protein_coding | ENSG00000( |
| 35909490 | 35937153 | 1 | protein_coding | ENSG00000( |
| 35922159 | 35925554 | -1 | lncRNA | ENSG00000( |
| 35943210 | 35943934 | -1 | processed_pseudoge | ENSG00000( |
| 35957108 | 35958154 | -1 | protein_coding | ENSG00000( |
| 35962195 | 36014623 | 1 | transcribed_unproce | ENSG00000( |
| 35971344 | 35972318 | -1 | processed_pseudoge | ENSG00000( |
| 35991336 | 35992285 | -1 | unprocessed_pseudc | ENSG00000( |
| 36002909 | 36003867 | -1 | polymorphic_pseudc | ENSG00000( |
| 36021692 | 36022113 | 1 | unprocessed_pseudc | ENSG00000( |
| 36036913 | 36124455 | 1 | protein_coding | ENSG00000( |
| 36072686 | 36072762 | 1 | misc_RNA | ENSG00000( |
| 36136536 | 36163913 | 1 | protein_coding | ENSG00000( |
| 36169388 | 36171334 | 1 | protein_coding | ENSG00000( |
| 36190856 | 36304781 | 1 | protein_coding | ENSG00000( |
| 36214441 | 36277056 | -1 | protein_coding | ENSG00000( |
| 36267783 | 36267922 | 1 | snRNA | ENSG00000( |

| | | | | |
|---|---|---|---|---|
| 36303499 | 36304924 | -1 | processed_pseudoge | ENSG00000 |
| 36336396 | 36487548 | -1 | protein_coding | ENSG00000 |
| 36572862 | 36677683 | 1 | protein_coding | ENSG00000 |
| 36764896 | 36830456 | 1 | lncRNA | ENSG00000 |
| 36823539 | 36823599 | -1 | miRNA | ENSG00000 |
| 36833269 | 37034268 | -1 | protein_coding | ENSG00000 |
| 36856555 | 36861375 | 1 | lncRNA | ENSG00000 |
| 36864254 | 36864308 | -1 | miRNA | ENSG00000 |
| 36893462 | 36893531 | -1 | miRNA | ENSG00000 |
| 37002697 | 37008040 | 1 | lncRNA | ENSG00000 |
| 37046835 | 37047237 | 1 | processed_pseudoge | ENSG00000 |
| 37073531 | 37075792 | -1 | lncRNA | ENSG00000 |
| 37078813 | 37079776 | -1 | lncRNA | ENSG00000 |
| 37079645 | 37090928 | 1 | lncRNA | ENSG00000 |
| 37112548 | 37120446 | -1 | lncRNA | ENSG00000 |
| 37120539 | 37358149 | 1 | protein_coding | ENSG00000 |
| 37160137 | 37160237 | 1 | misc_RNA | ENSG00000 |
| 37205112 | 37205368 | -1 | processed_pseudoge | ENSG00000 |
| 37322259 | 37322371 | 1 | misc_RNA | ENSG00000 |
| 37333857 | 37333957 | -1 | snRNA | ENSG00000 |
| 37383178 | 37384434 | -1 | lncRNA | ENSG00000 |
| 37422666 | 37436990 | 1 | protein_coding | ENSG00000 |
| 37434177 | 37434586 | 1 | processed_pseudoge | ENSG00000 |
| 37438102 | 37465450 | -1 | protein_coding | ENSG00000 |
| 37477149 | 37478317 | 1 | processed_pseudoge | ENSG00000 |
| 37485948 | 37503697 | 1 | protein_coding | ENSG00000 |
| 37490421 | 37490893 | -1 | processed_pseudoge | ENSG00000 |
| 37509150 | 37510299 | 1 | lncRNA | ENSG00000 |
| 37510892 | 37576380 | -1 | protein_coding | ENSG00000 |
| 37512547 | 37592469 | -1 | protein_coding | ENSG00000 |
| 37582646 | 37592604 | -1 | protein_coding | ENSG00000 |
| 37588413 | 38068687 | -1 | protein_coding | ENSG00000 |
| 37636688 | 37637293 | -1 | processed_pseudoge | ENSG00000 |
| 37650954 | 37746904 | 1 | protein_coding | ENSG00000 |
| 37661747 | 37662049 | -1 | misc_RNA | ENSG00000 |
| 37753803 | 37778972 | 1 | protein_coding | ENSG00000 |
| 37766978 | 37801437 | -1 | protein_coding | ENSG00000 |
| 37800554 | 37867666 | 1 | protein_coding | ENSG00000 |
| 37836044 | 37836394 | 1 | processed_pseudoge | ENSG00000 |
| 37878116 | 37879493 | 1 | processed_pseudoge | ENSG00000 |
| 37879400 | 37904353 | -1 | protein_coding | ENSG00000 |
| 37885683 | 37886390 | 1 | processed_pseudoge | ENSG00000 |
| 37915898 | 38069227 | -1 | protein_coding | ENSG00000 |
| 37936707 | 37936768 | -1 | snRNA | ENSG00000 |
| 38147428 | 38147561 | 1 | snoRNA | ENSG00000 |
| 38360427 | 38376430 | 1 | lncRNA | ENSG00000 |
| 38392702 | 38398661 | 1 | protein_coding | ENSG00000 |
| 38406528 | 38424454 | -1 | protein_coding | ENSG00000 |
| 38433696 | 38543880 | 1 | transcribed_unproce | ENSG00000 |
| 38452782 | 38454070 | -1 | processed_pseudoge | ENSG00000 |

| | | | | |
|---|---|---|---|---|
| 38478471 | 38479350 | 1 | processed_pseudoge | ENSG00000 |
| 38486859 | 38488338 | -1 | processed_pseudoge | ENSG00000 |
| 38504447 | 38505482 | -1 | processed_pseudoge | ENSG00000 |
| 38526831 | 38527032 | 1 | processed_pseudoge | ENSG00000 |
| 38527097 | 38527912 | 1 | processed_pseudoge | ENSG00000 |
| 38540567 | 38542792 | -1 | transcribed_processe | ENSG00000 |
| 38540569 | 38577207 | -1 | protein_coding | ENSG00000 |
| 38542389 | 38543215 | -1 | lncRNA | ENSG00000 |
| 38542479 | 38568422 | -1 | lncRNA | ENSG00000 |
| 38566260 | 38568211 | -1 | transcribed_unproce | ENSG00000 |
| 38571358 | 38620596 | -1 | protein_coding | ENSG00000 |
| 38620474 | 38624990 | 1 | lncRNA | ENSG00000 |
| 38643042 | 38643768 | 1 | processed_pseudoge | ENSG00000 |
| 38650215 | 38662717 | 1 | lncRNA | ENSG00000 |
| 38680006 | 38680104 | 1 | snRNA | ENSG00000 |
| 38694263 | 38720428 | -1 | protein_coding | ENSG00000 |
| 38804007 | 38851916 | 1 | lncRNA | ENSG00000 |
| 38949734 | 38998503 | 1 | lncRNA | ENSG00000 |
| 39016969 | 39039594 | 1 | unitary_pseudogene | ENSG00000 |
| 39072767 | 39288315 | -1 | protein_coding | ENSG00000 |
| 39154075 | 39155479 | 1 | processed_pseudoge | ENSG00000 |
| 39159071 | 39160319 | 1 | processed_pseudoge | ENSG00000 |
| 39173794 | 39174513 | -1 | processed_pseudoge | ENSG00000 |
| 39191100 | 39191360 | 1 | misc_RNA | ENSG00000 |
| 39355669 | 39361962 | 1 | protein_coding | ENSG00000 |
| 39359890 | 39369105 | -1 | transcribed_processe | ENSG00000 |
| 39371065 | 39376916 | 1 | lncRNA | ENSG00000 |
| 39384506 | 39405305 | -1 | lncRNA | ENSG00000 |
| 39395737 | 39397406 | -1 | lncRNA | ENSG00000 |
| 39443815 | 39508885 | -1 | transcribed_unproce | ENSG00000 |
| 39485379 | 39485865 | 1 | processed_pseudoge | ENSG00000 |
| 39631046 | 39632416 | 1 | processed_pseudoge | ENSG00000 |
| 39651901 | 39652377 | -1 | processed_pseudoge | ENSG00000 |
| 39654241 | 39655531 | -1 | processed_pseudoge | ENSG00000 |
| 39734670 | 39735628 | 1 | processed_pseudoge | ENSG00000 |
| 39739747 | 39741193 | -1 | processed_pseudoge | ENSG00000 |
| 39748514 | 39810097 | -1 | lncRNA | ENSG00000 |
| 39809562 | 39810019 | 1 | processed_pseudoge | ENSG00000 |
| 39816542 | 40106661 | -1 | transcribed_unproce | ENSG00000 |
| 39816599 | 39874562 | -1 | protein_coding | ENSG00000 |
| 39886861 | 39887144 | -1 | misc_RNA | ENSG00000 |
| 39892446 | 39892540 | 1 | snoRNA | ENSG00000 |
| 39892550 | 39892895 | 1 | processed_pseudoge | ENSG00000 |
| 39983793 | 40106611 | -1 | lncRNA | ENSG00000 |
| 39983821 | 39984401 | -1 | processed_pseudoge | ENSG00000 |
| 40105822 | 40106527 | 1 | processed_pseudoge | ENSG00000 |
| 40121962 | 40122540 | 1 | unprocessed_pseudo | ENSG00000 |
| 40130921 | 40153147 | 1 | lncRNA | ENSG00000 |
| 40131747 | 40132218 | 1 | processed_pseudoge | ENSG00000 |
| 40167392 | 40171329 | 1 | unprocessed_pseudo | ENSG00000 |

| | | | | |
|---|---|---|---|---|
| 40187792 | 40193921 | -1 | unprocessed_pseudc | ENSG0000C |
| 40223285 | 40266392 | 1 | protein_coding | ENSG0000C |
| 40223402 | 40329221 | 1 | lncRNA | ENSG0000C |
| 40233923 | 40234029 | 1 | snRNA | ENSG0000C |
| 40291275 | 40291502 | -1 | processed_pseudoge | ENSG0000C |
| 40293082 | 40293955 | 1 | unprocessed_pseudc | ENSG0000C |
| 40321299 | 40329221 | 1 | transcribed_unproce | ENSG0000C |
| 40332995 | 40333460 | -1 | unprocessed_pseudc | ENSG0000C |
| 40346144 | 40347052 | -1 | processed_pseudoge | ENSG0000C |
| 40348599 | 40349791 | 1 | processed_pseudoge | ENSG0000C |
| 40358454 | 40358682 | 1 | processed_pseudoge | ENSG0000C |
| 40481558 | 40497592 | -1 | unprocessed_pseudc | ENSG0000C |
| 40481559 | 40497591 | -1 | unprocessed_pseudc | ENSG0000C |
| 40497850 | 40566779 | 1 | transcribed_unproce | ENSG0000C |
| 40505399 | 40566281 | -1 | lncRNA | ENSG0000C |
| 40611252 | 40622155 | -1 | lncRNA | ENSG0000C |
| 40641841 | 40642117 | -1 | IG_V_pseudogene | ENSG0000C |
| 40755407 | 40767875 | -1 | unprocessed_pseudc | ENSG0000C |
| 40759600 | 40759706 | -1 | snRNA | ENSG0000C |
| 40767870 | 40806344 | 1 | lncRNA | ENSG0000C |
| 40883634 | 40883763 | 1 | processed_pseudoge | ENSG0000C |
| 40929010 | 40929092 | -1 | miRNA | ENSG0000C |
| 40992249 | 41218410 | 1 | transcribed_unproce | ENSG0000C |
| 41006999 | 41074617 | -1 | transcribed_unproce | ENSG0000C |
| 41012231 | 41014133 | 1 | TEC | ENSG0000C |
| 41061290 | 41062917 | 1 | TEC | ENSG0000C |
| 41073710 | 41076392 | 1 | lncRNA | ENSG0000C |
| 41100793 | 41119909 | -1 | lncRNA | ENSG0000C |
| 41126430 | 41128463 | -1 | protein_coding | ENSG0000C |
| 41131306 | 41199261 | -1 | protein_coding | ENSG0000C |
| 41233755 | 41233835 | 1 | miRNA | ENSG0000C |
| 41237440 | 41237557 | -1 | rRNA | ENSG0000C |
| 41269916 | 41273997 | -1 | TEC | ENSG0000C |
| 41276280 | 41282846 | 1 | lncRNA | ENSG0000C |
| 41280460 | 41280787 | 1 | transcribed_processe | ENSG0000C |
| 41292887 | 41294125 | 1 | processed_pseudoge | ENSG0000C |
| 41310816 | 41332962 | -1 | lncRNA | ENSG0000C |
| 41340823 | 41355538 | -1 | transcribed_unproce | ENSG0000C |
| 41479960 | 41480548 | 1 | processed_pseudoge | ENSG0000C |
| 41573156 | 41573453 | -1 | processed_pseudoge | ENSG0000C |
| 41573512 | 41573606 | -1 | snoRNA | ENSG0000C |
| 41578903 | 41579186 | 1 | misc_RNA | ENSG0000C |
| 41644438 | 41644756 | -1 | processed_pseudoge | ENSG0000C |
| 41644771 | 41644981 | 1 | processed_pseudoge | ENSG0000C |
| 41647825 | 41648035 | -1 | processed_pseudoge | ENSG0000C |
| 41651658 | 41654594 | -1 | lncRNA | ENSG0000C |
| 41657708 | 41699072 | -1 | lncRNA | ENSG0000C |
| 41672295 | 41672406 | -1 | misc_RNA | ENSG0000C |
| 41700480 | 41700696 | -1 | processed_pseudoge | ENSG0000C |
| 41700682 | 41701096 | 1 | processed_pseudoge | ENSG0000C |

| | | | | |
|---|---|---|---|---|
| 41760088 | 41764175 | 1 | lncRNA | ENSG0000C |
| 41890314 | 42129510 | -1 | protein_coding | ENSG0000C |
| 41974362 | 41975768 | 1 | processed_pseudoge | ENSG0000C |
| 41979352 | 41983753 | 1 | processed_pseudoge | ENSG0000C |
| 42011350 | 42011610 | 1 | misc_RNA | ENSG0000C |
| 42183659 | 42189887 | 1 | protein_coding | ENSG0000C |
| 42196333 | 42196662 | -1 | processed_pseudoge | ENSG0000C |
| 42199000 | 42204988 | 1 | lncRNA | ENSG0000C |
| 42213841 | 42213918 | -1 | processed_pseudoge | ENSG0000C |
| 42231811 | 42354454 | 1 | lncRNA | ENSG0000C |
| 42384805 | 42386136 | 1 | processed_pseudoge | ENSG0000C |
| 42408526 | 42409006 | -1 | processed_pseudoge | ENSG0000C |
| 42410830 | 42412150 | -1 | processed_pseudoge | ENSG0000C |
| 42492663 | 42493570 | 1 | processed_pseudoge | ENSG0000C |
| 42497704 | 42499134 | -1 | processed_pseudoge | ENSG0000C |
| 42566679 | 42569353 | -1 | lncRNA | ENSG0000C |
| 42568865 | 42569315 | 1 | processed_pseudoge | ENSG0000C |
| 42579414 | 42586534 | -1 | lncRNA | ENSG0000C |
| 42625436 | 42626813 | 1 | unprocessed_pseudc | ENSG0000C |
| 42636788 | 42637139 | -1 | processed_pseudoge | ENSG0000C |
| 42667506 | 42668887 | -1 | unprocessed_pseudc | ENSG0000C |
| 42707800 | 42714539 | 1 | lncRNA | ENSG0000C |
| 42725013 | 42802065 | 1 | transcribed_unproce | ENSG0000C |
| 42795392 | 42800672 | 1 | transcribed_unproce | ENSG0000C |
| 42817105 | 42823213 | -1 | unprocessed_pseudc | ENSG0000C |
| 42830337 | 42950827 | 1 | transcribed_unproce | ENSG0000C |
| 42834116 | 42838275 | 1 | transcribed_unproce | ENSG0000C |
| 42863200 | 42863306 | 1 | snRNA | ENSG0000C |
| 42921443 | 42921670 | -1 | processed_pseudoge | ENSG0000C |
| 42923277 | 42924144 | 1 | unprocessed_pseudc | ENSG0000C |
| 42950566 | 42951630 | 1 | unprocessed_pseudc | ENSG0000C |
| 42959280 | 42960066 | -1 | processed_pseudoge | ENSG0000C |
| 42972386 | 42973289 | -1 | processed_pseudoge | ENSG0000C |
| 42974855 | 43045120 | 1 | transcribed_processe | ENSG0000C |
| 43109866 | 43125905 | -1 | unprocessed_pseudc | ENSG0000C |
| 43126164 | 43139867 | 1 | transcribed_unproce | ENSG0000C |
| 43244305 | 43244508 | -1 | processed_pseudoge | ENSG0000C |
| 43244631 | 43244987 | -1 | processed_pseudoge | ENSG0000C |
| 60546920 | 60547154 | -1 | processed_pseudoge | ENSG0000C |
| 60914374 | 60920650 | 1 | protein_coding | ENSG0000C |
| 60918581 | 60927822 | -1 | transcribed_processe | ENSG0000C |
| 60929774 | 60936746 | 1 | transcribed_processe | ENSG0000C |
| 60943311 | 60964196 | -1 | lncRNA | ENSG0000C |
| 60944555 | 60944636 | -1 | processed_pseudoge | ENSG0000C |
| 61063752 | 61113954 | -1 | unprocessed_pseudc | ENSG0000C |
| 61190003 | 61196280 | 1 | protein_coding | ENSG0000C |
| 61194211 | 61203446 | -1 | lncRNA | ENSG0000C |
| 61202718 | 61203087 | -1 | transcribed_processe | ENSG0000C |
| 61205258 | 61212373 | 1 | transcribed_processe | ENSG0000C |
| 61229913 | 61231863 | -1 | lncRNA | ENSG0000C |

| | | | | |
|---|---|---|---|---|
| 61330717 | 61453030 | 1 | protein_coding | ENSG0000C |
| 61427617 | 61427877 | -1 | misc_RNA | ENSG0000C |
| 61581813 | 61582675 | 1 | unprocessed_pseudc | ENSG0000C |
| 61615835 | 61627683 | 1 | lncRNA | ENSG0000C |
| 61642383 | 61642494 | 1 | misc_RNA | ENSG0000C |
| 61659733 | 61662690 | 1 | lncRNA | ENSG0000C |
| 61665579 | 61666386 | 1 | processed_pseudoge | ENSG0000C |
| 61789143 | 61789426 | 1 | misc_RNA | ENSG0000C |
| 61854148 | 62096920 | 1 | lncRNA | ENSG0000C |
| 61861625 | 61863200 | 1 | protein_coding | ENSG0000C |
| 61861848 | 61862567 | -1 | protein_coding | ENSG0000C |
| 61865755 | 61866965 | -1 | lncRNA | ENSG0000C |
| 61868727 | 61887084 | 1 | lncRNA | ENSG0000C |
| 61883640 | 61883751 | -1 | misc_RNA | ENSG0000C |
| 61898805 | 61911022 | -1 | lncRNA | ENSG0000C |
| 61944232 | 61945136 | -1 | unprocessed_pseudc | ENSG0000C |
| 61982796 | 61986934 | 1 | TEC | ENSG0000C |
| 62266319 | 62266919 | 1 | processed_pseudoge | ENSG0000C |
| 62349941 | 62351536 | -1 | lncRNA | ENSG0000C |
| 62357834 | 62358140 | -1 | processed_pseudoge | ENSG0000C |
| 62358191 | 62358285 | -1 | snoRNA | ENSG0000C |
| 62363581 | 62363864 | 1 | misc_RNA | ENSG0000C |
| 62374128 | 62376836 | -1 | lncRNA | ENSG0000C |
| 62376161 | 62435199 | 1 | transcribed_unproce | ENSG0000C |
| 62387171 | 62394004 | -1 | lncRNA | ENSG0000C |
| 62395370 | 62395678 | -1 | unprocessed_pseudc | ENSG0000C |
| 62444256 | 62444634 | -1 | processed_pseudoge | ENSG0000C |
| 62452451 | 62522018 | 1 | lncRNA | ENSG0000C |
| 62474948 | 62476333 | -1 | unprocessed_pseudc | ENSG0000C |
| 62532337 | 62534724 | 1 | lncRNA | ENSG0000C |
| 62532397 | 62532853 | -1 | unprocessed_pseudc | ENSG0000C |
| 62559839 | 62569907 | 1 | unprocessed_pseudc | ENSG0000C |
| 62601226 | 62602672 | 1 | processed_pseudoge | ENSG0000C |
| 62606797 | 62607748 | -1 | processed_pseudoge | ENSG0000C |
| 62609725 | 62610799 | -1 | processed_pseudoge | ENSG0000C |
| 62686912 | 62688190 | 1 | processed_pseudoge | ENSG0000C |
| 62690033 | 62690534 | 1 | processed_pseudoge | ENSG0000C |
| 62710036 | 62711398 | -1 | processed_pseudoge | ENSG0000C |
| 62801461 | 62813486 | 1 | lncRNA | ENSG0000C |
| 62801534 | 62802699 | -1 | lncRNA | ENSG0000C |
| 62802320 | 62802438 | -1 | rRNA_pseudogene | ENSG0000C |
| 62837139 | 62838302 | -1 | lncRNA | ENSG0000C |
| 62838824 | 62844276 | 1 | transcribed_processe | ENSG0000C |
| 62844818 | 62845184 | 1 | processed_pseudoge | ENSG0000C |
| 62856999 | 62898095 | -1 | lncRNA | ENSG0000C |
| 62857208 | 62858484 | 1 | processed_pseudoge | ENSG0000C |
| 62860873 | 62864957 | -1 | transcribed_processe | ENSG0000C |
| 62888535 | 62889921 | 1 | unprocessed_pseudc | ENSG0000C |
| 62897368 | 62900104 | 1 | lncRNA | ENSG0000C |
| 62924911 | 62934964 | 1 | unprocessed_pseudc | ENSG0000C |

| | | | | |
|---|---|---|---|---|
| 63025597 | 63025993 | 1 | unprocessed_pseudc | ENSG0000C |
| 63055641 | 63059217 | 1 | unprocessed_pseudc | ENSG0000C |
| 63111255 | 63127100 | -1 | transcribed_unproce | ENSG0000C |
| 63127359 | 63143401 | 1 | unprocessed_pseudc | ENSG0000C |
| 63299453 | 63385117 | -1 | transcribed_unproce | ENSG0000C |
| 63317067 | 63333091 | -1 | unprocessed_pseudc | ENSG0000C |
| 63330326 | 63343660 | 1 | transcribed_unproce | ENSG0000C |
| 63581254 | 63581834 | -1 | processed_pseudoge | ENSG0000C |
| 63703065 | 63703755 | 1 | processed_pseudoge | ENSG0000C |
| 63719864 | 63720441 | 1 | processed_pseudoge | ENSG0000C |
| 63728877 | 63744910 | 1 | lncRNA | ENSG0000C |
| 63729702 | 63730173 | 1 | processed_pseudoge | ENSG0000C |
| 63761546 | 63762824 | -1 | processed_pseudoge | ENSG0000C |
| 63774870 | 63775241 | -1 | processed_pseudoge | ENSG0000C |
| 63775892 | 63776143 | -1 | processed_pseudoge | ENSG0000C |
| 63802144 | 63814159 | -1 | lncRNA | ENSG0000C |
| 63813190 | 63813307 | 1 | rRNA_pseudogene | ENSG0000C |
| 63817748 | 63818462 | -1 | unprocessed_pseudc | ENSG0000C |
| 63819574 | 63819654 | 1 | miRNA | ENSG0000C |
| 63832127 | 63859641 | -1 | unprocessed_pseudc | ENSG0000C |
| 63974762 | 64004155 | -1 | unprocessed_pseudc | ENSG0000C |
| 64082429 | 64082534 | 1 | snRNA | ENSG0000C |
| 64343044 | 64362584 | 1 | lncRNA | ENSG0000C |
| 64369394 | 64413142 | 1 | protein_coding | ENSG0000C |
| 64380194 | 64380300 | 1 | snRNA | ENSG0000C |
| 64439346 | 64440180 | 1 | processed_pseudoge | ENSG0000C |
| 64462819 | 64463196 | 1 | processed_pseudoge | ENSG0000C |
| 64465463 | 64466498 | 1 | unprocessed_pseudc | ENSG0000C |
| 64473040 | 64473826 | -1 | processed_pseudoge | ENSG0000C |
| 64486164 | 64487069 | -1 | processed_pseudoge | ENSG0000C |
| 64488600 | 64489786 | 1 | processed_pseudoge | ENSG0000C |
| 64498517 | 64498745 | 1 | processed_pseudoge | ENSG0000C |
| 64621560 | 64637586 | -1 | unprocessed_pseudc | ENSG0000C |
| 64637845 | 64652556 | 1 | transcribed_unproce | ENSG0000C |
| 64705734 | 64709311 | -1 | unprocessed_pseudc | ENSG0000C |
| 64737403 | 64737755 | -1 | processed_pseudoge | ENSG0000C |
| 64765054 | 64765535 | -1 | IG_V_pseudogene | ENSG0000C |
| 64810865 | 64811429 | -1 | processed_pseudoge | ENSG0000C |
| 64817870 | 64836341 | 1 | unprocessed_pseudc | ENSG0000C |
| 64878790 | 64889063 | -1 | unprocessed_pseudc | ENSG0000C |
| 64880369 | 64880475 | -1 | snRNA | ENSG0000C |
| 65143058 | 65153359 | 1 | transcribed_unproce | ENSG0000C |
| 65189995 | 65193610 | -1 | unprocessed_pseudc | ENSG0000C |
| 65219987 | 65230861 | -1 | transcribed_processe | ENSG0000C |
| 65222522 | 65222777 | -1 | processed_pseudoge | ENSG0000C |
| 65250543 | 65250826 | -1 | IG_V_pseudogene | ENSG0000C |
| 65282101 | 65285209 | -1 | protein_coding | ENSG0000C |
| 65287914 | 65323015 | -1 | unprocessed_pseudc | ENSG0000C |
| 65385649 | 65396372 | 1 | lncRNA | ENSG0000C |
| 65573384 | 65573495 | 1 | misc_RNA | ENSG0000C |

| | | | | |
|---|---|---|---|---|
| 65646982 | 65652679 | -1 | unprocessed_pseudc | ENSG0000C |
| 65668805 | 65734041 | 1 | protein_coding | ENSG0000C |
| 65736555 | 65738784 | 1 | protein_coding | ENSG0000C |
| 65770955 | 65771434 | 1 | IG_V_pseudogene | ENSG0000C |
| 65798726 | 65799124 | 1 | processed_pseudoge | ENSG0000C |
| 65816932 | 65817472 | -1 | processed_pseudoge | ENSG0000C |
| 65828120 | 65831696 | 1 | unprocessed_pseudc | ENSG0000C |
| 65865190 | 65877274 | -1 | transcribed_unproce | ENSG0000C |
| 65878766 | 65894790 | 1 | unprocessed_pseudc | ENSG0000C |
| 66014525 | 66020816 | -1 | lncRNA | ENSG0000C |
| 66017643 | 66017868 | -1 | transcribed_processe | ENSG0000C |
| 66022661 | 66027719 | -1 | transcribed_processe | ENSG0000C |
| 66029264 | 66030169 | 1 | processed_pseudoge | ENSG0000C |
| 66042646 | 66043342 | 1 | processed_pseudoge | ENSG0000C |
| 66047084 | 66055006 | -1 | transcribed_unproce | ENSG0000C |
| 66047107 | 66113085 | -1 | lncRNA | ENSG0000C |
| 66082350 | 66083187 | -1 | processed_pseudoge | ENSG0000C |
| 66084804 | 66085031 | 1 | processed_pseudoge | ENSG0000C |
| 66106815 | 66179474 | -1 | protein_coding | ENSG0000C |
| 66142221 | 66142327 | -1 | snRNA | ENSG0000C |
| 66182326 | 66188451 | 1 | unprocessed_pseudc | ENSG0000C |
| 66204896 | 66208833 | -1 | unprocessed_pseudc | ENSG0000C |
| 66253424 | 66254464 | -1 | unprocessed_pseudc | ENSG0000C |
| 66635754 | 66637197 | 1 | processed_pseudoge | ENSG0000C |
| 66641314 | 66642277 | -1 | processed_pseudoge | ENSG0000C |
| 66721158 | 66722760 | 1 | processed_pseudoge | ENSG0000C |
| 66724584 | 66725064 | 1 | processed_pseudoge | ENSG0000C |
| 66744561 | 66745929 | -1 | processed_pseudoge | ENSG0000C |
| 66856426 | 66932141 | 1 | protein_coding | ENSG0000C |
| 66879454 | 66879940 | -1 | processed_pseudoge | ENSG0000C |
| 66942703 | 66952344 | 1 | lncRNA | ENSG0000C |
| 66962271 | 66962348 | 1 | processed_pseudoge | ENSG0000C |
| 66976520 | 66976991 | -1 | transcribed_processe | ENSG0000C |
| 66979132 | 66988373 | 1 | lncRNA | ENSG0000C |
| 66979460 | 66979862 | 1 | unprocessed_pseudc | ENSG0000C |
| 66986304 | 66992583 | -1 | protein_coding | ENSG0000C |
| 67059560 | 67293468 | 1 | unprocessed_pseudc | ENSG0000C |
| 67175775 | 67176035 | -1 | misc_RNA | ENSG0000C |
| 67203635 | 67208072 | -1 | processed_pseudoge | ENSG0000C |
| 67211663 | 67213067 | -1 | processed_pseudoge | ENSG0000C |
| 67328207 | 67329404 | -1 | processed_pseudoge | ENSG0000C |
| 67512034 | 67515393 | -1 | lncRNA | ENSG0000C |
| 67697076 | 67697187 | 1 | misc_RNA | ENSG0000C |
| 67717411 | 67719178 | -1 | lncRNA | ENSG0000C |
| 67718236 | 67718955 | 1 | protein_coding | ENSG0000C |
| 67725690 | 67725902 | -1 | processed_pseudoge | ENSG0000C |
| 67725914 | 67726255 | 1 | protein_coding | ENSG0000C |
| 67791499 | 67791782 | -1 | misc_RNA | ENSG0000C |
| 67797087 | 67797181 | 1 | snoRNA | ENSG0000C |
| 67797221 | 67797500 | 1 | processed_pseudoge | ENSG0000C |

| | | | | |
|---|---|---|---|---|
| 67832765 | 67920552 | 1 | protein_coding | ENSG00000( |
| 67869947 | 67870053 | 1 | snRNA | ENSG00000( |
| 68228651 | 68229312 | 1 | lncRNA | ENSG00000( |
| 68232003 | 68300015 | 1 | protein_coding | ENSG00000( |
| 68250469 | 68251656 | -1 | TEC | ENSG00000( |
| 68302867 | 68305084 | 1 | protein_coding | ENSG00000( |
| 68306528 | 68330626 | 1 | lncRNA | ENSG00000( |
| 68307393 | 68308445 | 1 | lncRNA | ENSG00000( |
| 68328308 | 68531061 | 1 | protein_coding | ENSG00000( |
| 68353614 | 68357893 | -1 | lncRNA | ENSG00000( |
| 68393881 | 68406500 | -1 | lncRNA | ENSG00000( |
| 68426843 | 68429369 | 1 | lncRNA | ENSG00000( |
| 68536580 | 68540867 | -1 | protein_coding | ENSG00000( |
| 68541036 | 68644442 | 1 | lncRNA | ENSG00000( |
| 68543541 | 68546589 | -1 | lncRNA | ENSG00000( |
| 68609125 | 68611984 | 1 | processed_pseudoge | ENSG00000( |
| 68705240 | 69009176 | 1 | protein_coding | ENSG00000( |
| 68780065 | 68785566 | 1 | protein_coding | ENSG00000( |
| 68788307 | 68789670 | -1 | processed_pseudoge | ENSG00000( |
| 68822403 | 68843275 | -1 | lncRNA | ENSG00000( |
| 68935873 | 68935979 | -1 | snRNA | ENSG00000( |
| 69012529 | 69014113 | -1 | protein_coding | ENSG00000( |
| 69035563 | 69100178 | 1 | protein_coding | ENSG00000( |
| 69035747 | 69255187 | 1 | protein_coding | ENSG00000( |
| 69107926 | 69108217 | -1 | processed_pseudoge | ENSG00000( |
| 69121264 | 69274615 | 1 | protein_coding | ENSG00000( |
| 69296682 | 69311111 | -1 | lncRNA | ENSG00000( |
| 69324567 | 69392558 | 1 | protein_coding | ENSG00000( |
| 69427532 | 69672371 | -1 | protein_coding | ENSG00000( |
| 69428248 | 69428721 | -1 | lncRNA | ENSG00000( |
| 69472466 | 69494513 | 1 | lncRNA | ENSG00000( |
| 69672749 | 69673713 | 1 | lncRNA | ENSG00000( |
| 69709522 | 69760011 | -1 | protein_coding | ENSG00000( |
| 69818394 | 69820773 | -1 | lncRNA | ENSG00000( |
| 69820817 | 69906227 | 1 | protein_coding | ENSG00000( |
| 69884363 | 69884654 | -1 | misc_RNA | ENSG00000( |
| 70033921 | 70175888 | -1 | lncRNA | ENSG00000( |
| 70043848 | 70226972 | 1 | protein_coding | ENSG00000( |
| 70188571 | 70188750 | -1 | snRNA | ENSG00000( |
| 70193997 | 70258866 | -1 | lncRNA | ENSG00000( |
| 70217195 | 70217668 | 1 | processed_pseudoge | ENSG00000( |
| 70258978 | 70354873 | 1 | protein_coding | ENSG00000( |
| 70310875 | 70312120 | 1 | lncRNA | ENSG00000( |
| 70311607 | 70311701 | -1 | misc_RNA | ENSG00000( |
| 70384597 | 70414624 | -1 | protein_coding | ENSG00000( |
| 70529063 | 71446904 | -1 | protein_coding | ENSG00000( |
| 70579163 | 70579461 | -1 | misc_RNA | ENSG00000( |
| 70708047 | 70712863 | 1 | lncRNA | ENSG00000( |
| 70726040 | 70730856 | 1 | lncRNA | ENSG00000( |
| 70809975 | 70810084 | -1 | miRNA | ENSG00000( |

| | | | | |
|---|---|---|---|---|
| 70916997 | 70917501 | -1 | processed_pseudoge | ENSG00000( |
| 71589601 | 71589933 | 1 | processed_pseudoge | ENSG00000( |
| 71683366 | 71816690 | -1 | protein_coding | ENSG00000( |
| 71804283 | 71804826 | 1 | processed_pseudoge | ENSG00000( |
| 71862452 | 71910931 | -1 | protein_coding | ENSG00000( |
| 71902455 | 71902550 | -1 | misc_RNA | ENSG00000( |
| 71911510 | 71986054 | 1 | protein_coding | ENSG00000( |
| 72007881 | 72008136 | -1 | processed_pseudoge | ENSG00000( |
| 72051376 | 72072721 | -1 | protein_coding | ENSG00000( |
| 72060576 | 72081448 | 1 | lncRNA | ENSG00000( |
| 72107818 | 72108312 | -1 | processed_pseudoge | ENSG00000( |
| 72114595 | 72257193 | 1 | protein_coding | ENSG00000( |
| 72257336 | 72257783 | 1 | lncRNA | ENSG00000( |
| 72275453 | 72275973 | 1 | processed_pseudoge | ENSG00000( |
| 72299720 | 72299838 | -1 | rRNA_pseudogene | ENSG00000( |
| 72305367 | 72356721 | 1 | lncRNA | ENSG00000( |
| 72351413 | 72365235 | -1 | protein_coding | ENSG00000( |
| 72445754 | 72445969 | -1 | processed_pseudoge | ENSG00000( |
| 72470374 | 72471076 | 1 | processed_pseudoge | ENSG00000( |
| 72521608 | 72838291 | 1 | protein_coding | ENSG00000( |
| 72655832 | 72656192 | -1 | processed_pseudoge | ENSG00000( |
| 72741152 | 72741617 | -1 | processed_pseudoge | ENSG00000( |
| 72871130 | 72874135 | -1 | lncRNA | ENSG00000( |
| 72900671 | 73080442 | -1 | protein_coding | ENSG00000( |
| 73042745 | 73056528 | 1 | unprocessed_pseudc | ENSG00000( |
| 73151865 | 73170393 | 1 | protein_coding | ENSG00000( |
| 73474378 | 73476073 | -1 | processed_pseudoge | ENSG00000( |
| 73509664 | 73548331 | -1 | lncRNA | ENSG00000( |
| 73573615 | 73579498 | 1 | lncRNA | ENSG00000( |
| 73648828 | 73649839 | 1 | processed_pseudoge | ENSG00000( |
| 73750972 | 73751077 | -1 | rRNA_pseudogene | ENSG00000( |
| 73870896 | 73873297 | -1 | processed_pseudoge | ENSG00000( |
| 74371335 | 74384578 | 1 | lncRNA | ENSG00000( |
| 74485551 | 74499127 | -1 | lncRNA | ENSG00000( |
| 74497335 | 74693177 | 1 | protein_coding | ENSG00000( |
| 74630794 | 74641758 | -1 | lncRNA | ENSG00000( |
| 74722495 | 74888094 | -1 | protein_coding | ENSG00000( |
| 74799895 | 74800222 | 1 | misc_RNA | ENSG00000( |
| 74804473 | 74804773 | 1 | processed_pseudoge | ENSG00000( |
| 74857675 | 74857781 | 1 | snRNA | ENSG00000( |
| 74935176 | 74935696 | 1 | processed_pseudoge | ENSG00000( |
| 74946583 | 74952912 | -1 | protein_coding | ENSG00000( |
| 74952968 | 74996293 | 1 | lncRNA | ENSG00000( |
| 74980790 | 75028423 | -1 | protein_coding | ENSG00000( |
| 75009828 | 75016036 | -1 | lncRNA | ENSG00000( |
| 75016681 | 75016848 | -1 | misc_RNA | ENSG00000( |
| 75060573 | 75088217 | -1 | protein_coding | ENSG00000( |
| 75088514 | 75147265 | 1 | protein_coding | ENSG00000( |
| 75181520 | 75181621 | -1 | misc_RNA | ENSG00000( |
| 75198071 | 75198177 | 1 | snRNA | ENSG00000( |

| | | | | |
|---|---|---|---|---|
| 75401889 | 75402992 | -1 | processed_pseudoge | ENSG0000C |
| 75579669 | 75588548 | 1 | lncRNA | ENSG0000C |
| 75724222 | 75724575 | -1 | processed_pseudoge | ENSG0000C |
| 75890644 | 76362339 | 1 | protein_coding | ENSG0000C |
| 76286494 | 76288081 | -1 | processed_pseudoge | ENSG0000C |
| 76385526 | 76394517 | -1 | protein_coding | ENSG0000C |
| 76398699 | 76399586 | 1 | processed_pseudoge | ENSG0000C |
| 76419850 | 76651203 | 1 | protein_coding | ENSG0000C |
| 76419934 | 76420631 | 1 | processed_pseudoge | ENSG0000C |
| 76549282 | 76549771 | -1 | processed_pseudoge | ENSG0000C |
| 76611376 | 76906114 | -1 | protein_coding | ENSG0000C |
| 76691980 | 76863307 | 1 | lncRNA | ENSG0000C |
| 76779657 | 76779917 | 1 | misc_RNA | ENSG0000C |
| 76785782 | 76786190 | 1 | misc_RNA | ENSG0000C |
| 76915615 | 76919785 | -1 | lncRNA | ENSG0000C |
| 76931261 | 76932066 | -1 | processed_pseudoge | ENSG0000C |
| 76970322 | 76971028 | 1 | processed_pseudoge | ENSG0000C |
| 76992099 | 76993108 | 1 | processed_pseudoge | ENSG0000C |
| 77019655 | 77020953 | 1 | protein_coding | ENSG0000C |
| 77040416 | 77040673 | 1 | processed_pseudoge | ENSG0000C |
| 77072100 | 77073295 | 1 | processed_pseudoge | ENSG0000C |
| 77176760 | 77178180 | -1 | lncRNA | ENSG0000C |
| 77177445 | 77421537 | 1 | protein_coding | ENSG0000C |
| 77423079 | 77648322 | -1 | protein_coding | ENSG0000C |
| 77456295 | 77526697 | 1 | lncRNA | ENSG0000C |
| 77580245 | 77580628 | 1 | processed_pseudoge | ENSG0000C |
| 77635203 | 77635309 | -1 | snRNA | ENSG0000C |
| 77716097 | 78031811 | -1 | protein_coding | ENSG0000C |
| 78150532 | 78151193 | -1 | processed_pseudoge | ENSG0000C |
| 78165465 | 78165778 | 1 | misc_RNA | ENSG0000C |
| 78178263 | 78179512 | 1 | processed_pseudoge | ENSG0000C |
| 78180214 | 78180693 | -1 | processed_pseudoge | ENSG0000C |
| 78219120 | 78219357 | 1 | processed_pseudoge | ENSG0000C |
| 78236062 | 78279690 | 1 | protein_coding | ENSG0000C |
| 78297125 | 78330093 | 1 | protein_coding | ENSG0000C |
| 78589158 | 78589539 | 1 | processed_pseudoge | ENSG0000C |
| 78741720 | 78742744 | -1 | unprocessed_pseudc | ENSG0000C |
| 79036413 | 79037568 | -1 | processed_pseudoge | ENSG0000C |
| 79135374 | 79198314 | -1 | lncRNA | ENSG0000C |
| 79391304 | 79391759 | 1 | processed_pseudoge | ENSG0000C |
| 79505804 | 79567802 | -1 | lncRNA | ENSG0000C |
| 79571773 | 79726882 | 1 | protein_coding | ENSG0000C |
| 79850871 | 79851498 | 1 | processed_pseudoge | ENSG0000C |
| 79863019 | 79866875 | 1 | processed_pseudoge | ENSG0000C |
| 79874696 | 79886678 | -1 | lncRNA | ENSG0000C |
| 79889001 | 79892007 | 1 | lncRNA | ENSG0000C |
| 79913269 | 79915736 | 1 | TEC | ENSG0000C |
| 79975023 | 79990064 | 1 | lncRNA | ENSG0000C |
| 80030579 | 80034555 | 1 | lncRNA | ENSG0000C |
| 80357029 | 80360338 | -1 | processed_pseudoge | ENSG0000C |

| | | | | |
|---|---|---|---|---|
| 80563571 | 80564630 | -1 | unprocessed_pseudc | ENSG00000C |
| 80565032 | 80565962 | -1 | unprocessed_pseudc | ENSG00000C |
| 80782338 | 80782770 | -1 | processed_pseudoge | ENSG00000C |
| 80869758 | 80871295 | -1 | lncRNA | ENSG00000C |
| 81410279 | 81410630 | 1 | processed_pseudoge | ENSG00000C |
| 81488308 | 81488837 | 1 | processed_pseudoge | ENSG00000C |
| 81583683 | 81689547 | -1 | protein_coding | ENSG00000C |
| 81689713 | 81776900 | 1 | lncRNA | ENSG00000C |
| 81725243 | 81725349 | -1 | snRNA | ENSG00000C |
| 81885496 | 81885598 | 1 | rRNA_pseudogene | ENSG00000C |
| 81888918 | 81888985 | 1 | snoRNA | ENSG00000C |
| 81913437 | 81919926 | 1 | transcribed_unproce | ENSG00000C |
| 81928021 | 81931368 | -1 | lncRNA | ENSG00000C |
| 81928514 | 81934948 | 1 | protein_coding | ENSG00000C |
| 81930226 | 81977089 | -1 | lncRNA | ENSG00000C |
| 81943586 | 81950038 | 1 | protein_coding | ENSG00000C |
| 81966564 | 81972130 | 1 | unprocessed_pseudc | ENSG00000C |
| 81977614 | 82780194 | 1 | transcribed_unproce | ENSG00000C |
| 81988772 | 81995253 | 1 | protein_coding | ENSG00000C |
| 82057647 | 82063682 | -1 | unprocessed_pseudc | ENSG00000C |
| 82086155 | 82088768 | -1 | processed_pseudoge | ENSG00000C |
| 82130752 | 82131632 | 1 | processed_pseudoge | ENSG00000C |
| 82405188 | 82406239 | 1 | lncRNA | ENSG00000C |
| 82427350 | 82428025 | -1 | processed_pseudoge | ENSG00000C |
| 82431889 | 82432905 | -1 | processed_pseudoge | ENSG00000C |
| 82451656 | 82455225 | 1 | lncRNA | ENSG00000C |
| 82543718 | 82544196 | 1 | TEC | ENSG00000C |
| 82612552 | 82646584 | -1 | lncRNA | ENSG00000C |
| 82744067 | 82744792 | -1 | processed_pseudoge | ENSG00000C |
| 82979590 | 83063177 | -1 | protein_coding | ENSG00000C |
| 83063398 | 83069143 | 1 | lncRNA | ENSG00000C |
| 83200940 | 83201228 | -1 | misc_RNA | ENSG00000C |
| 83201957 | 83202065 | 1 | processed_pseudoge | ENSG00000C |
| 83219288 | 83238463 | 1 | lncRNA | ENSG00000C |
| 83242990 | 83538546 | -1 | protein_coding | ENSG00000C |
| 83272280 | 83279499 | 1 | lncRNA | ENSG00000C |
| 83275941 | 83276114 | 1 | snRNA | ENSG00000C |
| 83596444 | 83596584 | -1 | snRNA | ENSG00000C |
| 83623049 | 83644130 | 1 | protein_coding | ENSG00000C |
| 83659968 | 83707958 | -1 | protein_coding | ENSG00000C |
| 83707594 | 83713378 | 1 | lncRNA | ENSG00000C |
| 83739425 | 83829516 | -1 | protein_coding | ENSG00000C |
| 83817643 | 83837861 | 1 | lncRNA | ENSG00000C |
| 83831586 | 83868532 | 1 | lncRNA | ENSG00000C |
| 83834099 | 83921465 | -1 | protein_coding | ENSG00000C |
| 83938311 | 83956986 | -1 | protein_coding | ENSG00000C |
| 83968083 | 83980616 | -1 | protein_coding | ENSG00000C |
| 83969748 | 83969857 | -1 | miRNA | ENSG00000C |
| 83972233 | 83975777 | 1 | lncRNA | ENSG00000C |
| 83980711 | 84004074 | 1 | protein_coding | ENSG00000C |

| | | | | |
|---|---|---|---|---|
| 83992932 | 83993033 | 1 | misc_RNA | ENSG00000( |
| 84059298 | 84098984 | 1 | lncRNA | ENSG00000( |
| 84078584 | 84078857 | 1 | processed_pseudoge | ENSG00000( |
| 84275457 | 84340758 | -1 | protein_coding | ENSG00000( |
| 84278219 | 84290131 | 1 | lncRNA | ENSG00000( |
| 84316514 | 84657077 | 1 | lncRNA | ENSG00000( |
| 84604111 | 84604425 | -1 | processed_pseudoge | ENSG00000( |
| 84668551 | 85027070 | 1 | protein_coding | ENSG00000( |
| 85159682 | 85159950 | 1 | processed_pseudoge | ENSG00000( |
| 85167346 | 85194283 | 1 | lncRNA | ENSG00000( |
| 85230176 | 85233966 | -1 | lncRNA | ENSG00000( |
| 85369591 | 85444230 | -1 | lncRNA | ENSG00000( |
| 85492579 | 85494094 | 1 | processed_pseudoge | ENSG00000( |
| 85546539 | 85742029 | -1 | protein_coding | ENSG00000( |
| 85753306 | 85786575 | 1 | unprocessed_pseudc | ENSG00000( |
| 85756042 | 85765112 | -1 | lncRNA | ENSG00000( |
| 85783668 | 85805154 | -1 | lncRNA | ENSG00000( |
| 85805359 | 85849542 | 1 | transcribed_unproce | ENSG00000( |
| 85898903 | 85899716 | 1 | processed_pseudoge | ENSG00000( |
| 85934588 | 85936599 | 1 | processed_pseudoge | ENSG00000( |
| 85941146 | 86025462 | 1 | protein_coding | ENSG00000( |
| 86026146 | 86100173 | -1 | protein_coding | ENSG00000( |
| 86127507 | 86128703 | 1 | lncRNA | ENSG00000( |
| 86132622 | 86132851 | 1 | processed_pseudoge | ENSG00000( |
| 86220265 | 86259657 | -1 | protein_coding | ENSG00000( |
| 86228815 | 86228983 | 1 | processed_pseudoge | ENSG00000( |
| 86258358 | 86258646 | -1 | misc_RNA | ENSG00000( |
| 86264546 | 86283102 | -1 | protein_coding | ENSG00000( |
| 86287733 | 86354454 | -1 | protein_coding | ENSG00000( |
| 86374966 | 86375706 | -1 | processed_pseudoge | ENSG00000( |
| 86414201 | 86488607 | 1 | lncRNA | ENSG00000( |
| 86422938 | 86423128 | -1 | snRNA | ENSG00000( |
| 86632147 | 86696496 | 1 | lncRNA | ENSG00000( |
| 86669349 | 86708307 | -1 | lncRNA | ENSG00000( |
| 86944362 | 86947506 | -1 | protein_coding | ENSG00000( |
| 86948666 | 87002033 | 1 | lncRNA | ENSG00000( |
| 87008440 | 87042419 | -1 | lncRNA | ENSG00000( |
| 87011652 | 87013151 | 1 | processed_pseudoge | ENSG00000( |
| 87084161 | 87085151 | -1 | processed_pseudoge | ENSG00000( |
| 87148644 | 87159556 | 1 | lncRNA | ENSG00000( |
| 87260450 | 87260585 | 1 | snoRNA | ENSG00000( |
| 87423634 | 87423893 | 1 | processed_pseudoge | ENSG00000( |
| 87436215 | 87445795 | -1 | lncRNA | ENSG00000( |
| 87497228 | 87708634 | 1 | protein_coding | ENSG00000( |
| 87525302 | 87525475 | 1 | processed_pseudoge | ENSG00000( |
| 87553454 | 87554459 | 1 | lncRNA | ENSG00000( |
| 87726109 | 87731469 | 1 | protein_coding | ENSG00000( |
| 87741233 | 87741953 | 1 | processed_pseudoge | ENSG00000( |
| 87772453 | 87786884 | 1 | transcribed_unproce | ENSG00000( |
| 87799501 | 87821803 | -1 | unprocessed_pseudc | ENSG00000( |

| | | | | |
|---|---|---|---|---|
| 87810642 | 87837225 | -1 | transcribed_unproce | ENSG00000( |
| 87816465 | 87817970 | 1 | processed_pseudoge | ENSG00000( |
| 87825146 | 87825968 | 1 | processed_pseudoge | ENSG00000( |
| 87844154 | 87847314 | 1 | unprocessed_pseudc | ENSG00000( |
| 87848113 | 87853526 | -1 | lncRNA | ENSG00000( |
| 87857015 | 87859599 | -1 | transcribed_unproce | ENSG00000( |
| 87864096 | 87866291 | 1 | transcribed_unproce | ENSG00000( |
| 87882877 | 87888903 | 1 | protein_coding | ENSG00000( |
| 87914493 | 87923505 | 1 | transcribed_unproce | ENSG00000( |
| 87944276 | 87945023 | -1 | processed_pseudoge | ENSG00000( |
| 87956214 | 87956883 | 1 | protein_coding | ENSG00000( |
| 87966441 | 87974753 | -1 | protein_coding | ENSG00000( |
| 87998229 | 87998335 | -1 | snRNA | ENSG00000( |
| 88016300 | 88016793 | 1 | processed_pseudoge | ENSG00000( |
| 88066445 | 88076975 | -1 | lncRNA | ENSG00000( |
| 88066915 | 88068037 | 1 | processed_pseudoge | ENSG00000( |
| 88129458 | 88138446 | -1 | unprocessed_pseudc | ENSG00000( |
| 88144851 | 88145850 | -1 | processed_pseudoge | ENSG00000( |
| 88177695 | 88178578 | 1 | TEC | ENSG00000( |
| 88180673 | 88181447 | 1 | processed_pseudoge | ENSG00000( |
| 88187735 | 88190202 | -1 | processed_pseudoge | ENSG00000( |
| 88324454 | 88325339 | 1 | processed_pseudoge | ENSG00000( |
| 88384902 | 88389224 | -1 | lncRNA | ENSG00000( |
| 88388430 | 88478694 | 1 | protein_coding | ENSG00000( |
| 88391362 | 88391832 | -1 | processed_pseudoge | ENSG00000( |
| 88410529 | 88411463 | -1 | processed_pseudoge | ENSG00000( |
| 88484852 | 88485325 | -1 | processed_pseudoge | ENSG00000( |
| 88534033 | 88584274 | 1 | protein_coding | ENSG00000( |
| 88627063 | 88652180 | -1 | lncRNA | ENSG00000( |
| 88735041 | 88803716 | -1 | lncRNA | ENSG00000( |
| 88745836 | 88745905 | -1 | miRNA | ENSG00000( |
| 88970887 | 88971341 | -1 | processed_pseudoge | ENSG00000( |
| 88971638 | 88976298 | 1 | lncRNA | ENSG00000( |
| 88990863 | 89005155 | 1 | protein_coding | ENSG00000( |
| 89005771 | 89178818 | -1 | protein_coding | ENSG00000( |
| 89088604 | 89109934 | -1 | lncRNA | ENSG00000( |
| 89311195 | 89316703 | 1 | protein_coding | ENSG00000( |
| 89312225 | 89312306 | 1 | miRNA | ENSG00000( |
| 89318500 | 89359663 | 1 | protein_coding | ENSG00000( |
| 89360787 | 89498130 | -1 | protein_coding | ENSG00000( |
| 89449558 | 89450754 | 1 | processed_pseudoge | ENSG00000( |
| 89472237 | 89472692 | 1 | processed_pseudoge | ENSG00000( |
| 89563537 | 89604687 | -1 | lncRNA | ENSG00000( |
| 89605012 | 89606555 | 1 | protein_coding | ENSG00000( |
| 89639814 | 89719759 | 1 | lncRNA | ENSG00000( |
| 89797880 | 89803100 | -1 | lncRNA | ENSG00000( |
| 89799695 | 89821753 | 1 | lncRNA | ENSG00000( |
| 89825964 | 90019549 | 1 | lncRNA | ENSG00000( |
| 90020654 | 90041499 | -1 | lncRNA | ENSG00000( |
| 90023441 | 90023535 | -1 | miRNA | ENSG00000( |

| | | | | |
|---|---|---|---|---|
| 90081297 | 90082569 | -1 | processed_pseudoge | ENSG00000( |
| 90108244 | 90108518 | 1 | processed_pseudoge | ENSG00000( |
| 90158028 | 90158646 | 1 | TEC | ENSG00000( |
| 90216039 | 90216924 | 1 | unprocessed_pseudc | ENSG00000( |
| 90232185 | 90233141 | 1 | unprocessed_pseudc | ENSG00000( |
| 90300885 | 90433566 | -1 | lncRNA | ENSG00000( |
| 90462899 | 90583821 | -1 | lncRNA | ENSG00000( |
| 90609832 | 90642862 | -1 | protein_coding | ENSG00000( |
| 90742526 | 90743380 | -1 | unprocessed_pseudc | ENSG00000( |
| 90751240 | 90752230 | -1 | unprocessed_pseudc | ENSG00000( |
| 90801787 | 90898549 | 1 | protein_coding | ENSG00000( |
| 90921990 | 90922856 | 1 | lncRNA | ENSG00000( |
| 90957260 | 90965393 | -1 | lncRNA | ENSG00000( |
| 90996949 | 91001901 | 1 | lncRNA | ENSG00000( |
| 91077138 | 91163087 | -1 | lncRNA | ENSG00000( |
| 91104327 | 91213046 | -1 | lncRNA | ENSG00000( |
| 91118592 | 91182762 | 1 | lncRNA | ENSG00000( |
| 91159573 | 91166272 | 1 | lncRNA | ENSG00000( |
| 91169585 | 91170045 | -1 | processed_pseudoge | ENSG00000( |
| 91206175 | 91210299 | -1 | lncRNA | ENSG00000( |
| 91213815 | 91361913 | -1 | protein_coding | ENSG00000( |
| 91409045 | 91423832 | -1 | protein_coding | ENSG00000( |
| 91426238 | 91427144 | 1 | lncRNA | ENSG00000( |
| 91563091 | 91950228 | -1 | protein_coding | ENSG00000( |
| 91611343 | 91612492 | 1 | processed_pseudoge | ENSG00000( |
| 91636251 | 91636361 | 1 | miRNA | ENSG00000( |
| 91636264 | 91636345 | -1 | miRNA | ENSG00000( |
| 92000087 | 92115413 | -1 | protein_coding | ENSG00000( |
| 92006062 | 92006544 | 1 | processed_pseudoge | ENSG00000( |
| 92027498 | 92027824 | 1 | processed_pseudoge | ENSG00000( |
| 92104425 | 92105451 | 1 | processed_pseudoge | ENSG00000( |
| 92108858 | 92108989 | 1 | unprocessed_pseudc | ENSG00000( |
| 92109013 | 92110095 | 1 | unprocessed_pseudc | ENSG00000( |
| 92110168 | 92110514 | 1 | unprocessed_pseudc | ENSG00000( |
| 92110575 | 92110868 | 1 | unprocessed_pseudc | ENSG00000( |
| 92110865 | 92111897 | 1 | unprocessed_pseudc | ENSG00000( |
| 92132398 | 92138629 | 1 | unprocessed_pseudc | ENSG00000( |
| 92141298 | 92160114 | 1 | transcribed_unproce | ENSG00000( |
| 92150382 | 92157445 | -1 | unprocessed_pseudc | ENSG00000( |
| 92171241 | 92173494 | 1 | processed_pseudoge | ENSG00000( |
| 92177235 | 92191156 | -1 | transcribed_unproce | ENSG00000( |
| 92205356 | 92210158 | 1 | lncRNA | ENSG00000( |
| 92210207 | 92293756 | -1 | protein_coding | ENSG00000( |
| 92292458 | 92292547 | -1 | miRNA | ENSG00000( |
| 92292461 | 92292593 | -1 | snoRNA | ENSG00000( |
| 92297358 | 92325636 | -1 | protein_coding | ENSG00000( |
| 92325953 | 92620529 | 1 | protein_coding | ENSG00000( |
| 92339658 | 92341663 | -1 | processed_pseudoge | ENSG00000( |
| 92383268 | 92404696 | -1 | protein_coding | ENSG00000( |
| 92414245 | 92424461 | -1 | protein_coding | ENSG00000( |

| | | | | |
|---|---|---|---|---|
| 92456205 | 92482506 | -1 | protein_coding | ENSG00000 |
| 92493554 | 92536655 | -1 | protein_coding | ENSG00000 |
| 92527984 | 92528058 | -1 | miRNA | ENSG00000 |
| 92539053 | 92539184 | -1 | unprocessed_pseudo | ENSG00000 |
| 92600094 | 92612368 | -1 | lncRNA | ENSG00000 |
| 92613183 | 92670131 | -1 | protein_coding | ENSG00000 |
| 92676943 | 92677407 | -1 | processed_pseudoge | ENSG00000 |
| 92711363 | 92764833 | -1 | protein_coding | ENSG00000 |
| 92805431 | 92809171 | -1 | lncRNA | ENSG00000 |
| 92809388 | 92888693 | 1 | lncRNA | ENSG00000 |
| 92809568 | 92838457 | 1 | transcribed_unproce | ENSG00000 |
| 92820977 | 92821079 | 1 | snRNA | ENSG00000 |
| 92836826 | 92837668 | -1 | processed_pseudoge | ENSG00000 |
| 92840955 | 92841688 | -1 | unprocessed_pseudo | ENSG00000 |
| 92844182 | 92878038 | -1 | protein_coding | ENSG00000 |
| 92882567 | 92899608 | 1 | transcribed_unproce | ENSG00000 |
| 92893698 | 92894505 | -1 | processed_pseudoge | ENSG00000 |
| 92913156 | 92921560 | 1 | unprocessed_pseudo | ENSG00000 |
| 92947523 | 93036236 | 1 | protein_coding | ENSG00000 |
| 93058688 | 93085133 | 1 | protein_coding | ENSG00000 |
| 93091460 | 93095546 | -1 | lncRNA | ENSG00000 |
| 93096217 | 93113283 | 1 | protein_coding | ENSG00000 |
| 93121496 | 93134251 | -1 | protein_coding | ENSG00000 |
| 93147040 | 93148556 | 1 | lncRNA | ENSG00000 |
| 93184916 | 93320572 | 1 | protein_coding | ENSG00000 |
| 93318199 | 93346414 | -1 | protein_coding | ENSG00000 |
| 93375556 | 93375667 | 1 | snRNA | ENSG00000 |
| 93417855 | 93417966 | 1 | misc_RNA | ENSG00000 |
| 93430342 | 93431299 | -1 | lncRNA | ENSG00000 |
| 93431441 | 93453581 | -1 | protein_coding | ENSG00000 |
| 93451685 | 93566112 | 1 | protein_coding | ENSG00000 |
| 93576584 | 93679587 | 1 | protein_coding | ENSG00000 |
| 93594841 | 93594894 | 1 | miRNA | ENSG00000 |
| 93808347 | 93858340 | 1 | lncRNA | ENSG00000 |
| 93819357 | 93819421 | 1 | miRNA | ENSG00000 |
| 93951627 | 93955355 | -1 | protein_coding | ENSG00000 |
| 93955597 | 93959140 | 1 | lncRNA | ENSG00000 |
| 94030794 | 94109856 | 1 | protein_coding | ENSG00000 |
| 94038030 | 94038345 | 1 | processed_pseudoge | ENSG00000 |
| 94116023 | 94145693 | -1 | lncRNA | ENSG00000 |
| 94147966 | 94157219 | 1 | transcribed_unitary_ | ENSG00000 |
| 94166258 | 94200902 | 1 | lncRNA | ENSG00000 |
| 94175957 | 94176036 | 1 | miRNA | ENSG00000 |
| 94176347 | 94176433 | 1 | miRNA | ENSG00000 |
| 94176492 | 94259545 | -1 | lncRNA | ENSG00000 |
| 94178834 | 94178920 | 1 | miRNA | ENSG00000 |
| 94259298 | 94301829 | 1 | protein_coding | ENSG00000 |
| 94287699 | 94288550 | -1 | processed_pseudoge | ENSG00000 |
| 94318196 | 94328644 | -1 | protein_coding | ENSG00000 |
| 94332459 | 94363013 | 1 | lncRNA | ENSG00000 |

| | | | | |
|---|---|---|---|---|
| 94338347 | 94338733 | -1 | processed_pseudoge | ENSG00000( |
| 94347234 | 94347563 | -1 | processed_pseudoge | ENSG00000( |
| 94374569 | 94461042 | 1 | protein_coding | ENSG00000( |
| 94432631 | 94433273 | -1 | processed_pseudoge | ENSG00000( |
| 94485445 | 94511614 | -1 | lncRNA | ENSG00000( |
| 94496275 | 94512879 | 1 | unprocessed_pseudc | ENSG00000( |
| 94507738 | 94507844 | 1 | snRNA | ENSG00000( |
| 94555054 | 94603990 | 1 | lncRNA | ENSG00000( |
| 94558720 | 94593824 | -1 | protein_coding | ENSG00000( |
| 94603133 | 94640249 | -1 | protein_coding | ENSG00000( |
| 94725871 | 94726575 | -1 | lncRNA | ENSG00000( |
| 94726701 | 95087218 | 1 | protein_coding | ENSG00000( |
| 94809962 | 94810057 | 1 | miRNA | ENSG00000( |
| 94824272 | 94824773 | 1 | lncRNA | ENSG00000( |
| 94900429 | 94904754 | 1 | lncRNA | ENSG00000( |
| 94928551 | 94934946 | -1 | lncRNA | ENSG00000( |
| 94964369 | 94965624 | -1 | lncRNA | ENSG00000( |
| 95065350 | 95065446 | 1 | miRNA | ENSG00000( |
| 95085208 | 95085304 | 1 | miRNA | ENSG00000( |
| 95085445 | 95085541 | 1 | miRNA | ENSG00000( |
| 95086014 | 95086094 | -1 | miRNA | ENSG00000( |
| 95086021 | 95086088 | 1 | miRNA | ENSG00000( |
| 95099054 | 95426796 | -1 | protein_coding | ENSG00000( |
| 95113708 | 95114461 | 1 | lncRNA | ENSG00000( |
| 95123521 | 95123868 | 1 | processed_pseudoge | ENSG00000( |
| 95142449 | 95142541 | -1 | rRNA_pseudogene | ENSG00000( |
| 95292338 | 95294810 | 1 | processed_pseudoge | ENSG00000( |
| 95406990 | 95407662 | -1 | lncRNA | ENSG00000( |
| 95413267 | 95413449 | -1 | processed_pseudoge | ENSG00000( |
| 95442980 | 95517057 | -1 | protein_coding | ENSG00000( |
| 95494924 | 95495379 | 1 | lncRNA | ENSG00000( |
| 95506235 | 95507636 | 1 | lncRNA | ENSG00000( |
| 95514045 | 95514520 | 1 | lncRNA | ENSG00000( |
| 95599438 | 95606326 | -1 | lncRNA | ENSG00000( |
| 95650154 | 95715718 | 1 | lncRNA | ENSG00000( |
| 95652912 | 95653266 | 1 | processed_pseudoge | ENSG00000( |
| 95759231 | 95876049 | -1 | lncRNA | ENSG00000( |
| 95772323 | 95776282 | 1 | lncRNA | ENSG00000( |
| 95859904 | 95860016 | 1 | snRNA | ENSG00000( |
| 95871264 | 96121154 | 1 | protein_coding | ENSG00000( |
| 95905005 | 95905141 | 1 | rRNA_pseudogene | ENSG00000( |
| 95918136 | 95918704 | 1 | processed_pseudoge | ENSG00000( |
| 96019724 | 96027993 | -1 | lncRNA | ENSG00000( |
| 96065740 | 96101912 | 1 | lncRNA | ENSG00000( |
| 96103079 | 96103174 | -1 | misc_RNA | ENSG00000( |
| 96105741 | 96116411 | -1 | lncRNA | ENSG00000( |
| 96129869 | 96132214 | 1 | lncRNA | ENSG00000( |
| 96146007 | 96147856 | 1 | processed_pseudoge | ENSG00000( |
| 96235306 | 96302176 | -1 | protein_coding | ENSG00000( |
| 96235331 | 96352058 | -1 | protein_coding | ENSG00000( |

| | | | | |
|---|---|---|---|---|
| 96246462 | 96250393 | 1 | lncRNA | ENSG0000( |
| 96256216 | 96256319 | -1 | snRNA | ENSG0000( |
| 96313444 | 96383711 | -1 | protein_coding | ENSG0000( |
| 96385941 | 96418370 | -1 | protein_coding | ENSG0000( |
| 96407284 | 96407797 | 1 | processed_pseudoge | ENSG0000( |
| 96416724 | 96417377 | -1 | lncRNA | ENSG0000( |
| 96419722 | 96421129 | -1 | lncRNA | ENSG0000( |
| 96450169 | 96491336 | 1 | protein_coding | ENSG0000( |
| 96490241 | 96619843 | -1 | protein_coding | ENSG0000( |
| 96639577 | 96655317 | -1 | protein_coding | ENSG0000( |
| 96687050 | 96774318 | 1 | lncRNA | ENSG0000( |
| 96725890 | 96727469 | 1 | transcribed_processe | ENSG0000( |
| 96755865 | 96778129 | -1 | protein_coding | ENSG0000( |
| 96816269 | 96875623 | -1 | protein_coding | ENSG0000( |
| 96887373 | 97013708 | -1 | protein_coding | ENSG0000( |
| 96910076 | 96910405 | 1 | processed_pseudoge | ENSG0000( |
| 96918928 | 96919318 | 1 | processed_pseudoge | ENSG0000( |
| 96928310 | 96940253 | 1 | protein_coding | ENSG0000( |
| 96954820 | 96955461 | 1 | processed_pseudoge | ENSG0000( |
| 97029677 | 97039643 | -1 | protein_coding | ENSG0000( |
| 97067749 | 97068628 | 1 | processed_pseudoge | ENSG0000( |
| 97075671 | 97081945 | -1 | transcribed_processe | ENSG0000( |
| 97093694 | 97094638 | -1 | processed_pseudoge | ENSG0000( |
| 97121659 | 97122489 | 1 | processed_pseudoge | ENSG0000( |
| 97151894 | 97152694 | -1 | processed_pseudoge | ENSG0000( |
| 97156570 | 97221235 | -1 | unprocessed_pseudo | ENSG0000( |
| 97192923 | 97193025 | -1 | snRNA | ENSG0000( |
| 97195351 | 97197687 | -1 | lncRNA | ENSG0000( |
| 97198303 | 97199511 | -1 | processed_pseudoge | ENSG0000( |
| 97200475 | 97238745 | -1 | lncRNA | ENSG0000( |
| 97222602 | 97235017 | 1 | lncRNA | ENSG0000( |
| 97238449 | 97297314 | 1 | lncRNA | ENSG0000( |
| 97238497 | 97377287 | 1 | lncRNA | ENSG0000( |
| 97238645 | 97246184 | 1 | unprocessed_pseudo | ENSG0000( |
| 97276566 | 97295759 | 1 | unitary_pseudogene | ENSG0000( |
| 97307304 | 97378751 | 1 | protein_coding | ENSG0000( |
| 97363554 | 97363681 | -1 | miRNA | ENSG0000( |
| 97386240 | 97396112 | -1 | unprocessed_pseudo | ENSG0000( |
| 97403709 | 97411908 | -1 | lncRNA | ENSG0000( |
| 97412096 | 97496125 | 1 | protein_coding | ENSG0000( |
| 97501180 | 97601743 | 1 | protein_coding | ENSG0000( |
| 97512706 | 97517836 | -1 | lncRNA | ENSG0000( |
| 97600080 | 97633368 | -1 | protein_coding | ENSG0000( |
| 97633668 | 97673748 | 1 | protein_coding | ENSG0000( |
| 97634515 | 97636051 | -1 | lncRNA | ENSG0000( |
| 97674909 | 97697357 | -1 | protein_coding | ENSG0000( |
| 97698922 | 97700734 | 1 | processed_pseudoge | ENSG0000( |
| 97699625 | 97853080 | -1 | lncRNA | ENSG0000( |
| 97743208 | 97744935 | 1 | lncRNA | ENSG0000( |
| 97853226 | 97856717 | 1 | protein_coding | ENSG0000( |

| | | | | |
|---|---|---|---|---|
| 97904489 | 97922570 | -1 | protein_coding | ENSG00000 |
| 97915690 | 97915796 | 1 | misc_RNA | ENSG00000 |
| 97921590 | 97922181 | -1 | lncRNA | ENSG00000 |
| 97926791 | 97944856 | -1 | protein_coding | ENSG00000 |
| 97983341 | 98015943 | 1 | protein_coding | ENSG00000 |
| 97986551 | 97987656 | -1 | lncRNA | ENSG00000 |
| 98032230 | 98032335 | 1 | snRNA | ENSG00000 |
| 98056732 | 98083077 | 1 | protein_coding | ENSG00000 |
| 98069275 | 98119222 | -1 | protein_coding | ENSG00000 |
| 98115885 | 98116135 | 1 | misc_RNA | ENSG00000 |
| 98120975 | 98192637 | -1 | protein_coding | ENSG00000 |
| 98199011 | 98255649 | -1 | protein_coding | ENSG00000 |
| 98229149 | 98229217 | -1 | miRNA | ENSG00000 |
| 98234709 | 98235567 | 1 | processed_pseudoge | ENSG00000 |
| 98288109 | 98708935 | -1 | protein_coding | ENSG00000 |
| 98606833 | 98608133 | 1 | processed_pseudoge | ENSG00000 |
| 98731329 | 98796965 | -1 | protein_coding | ENSG00000 |
| 98798466 | 98807564 | -1 | lncRNA | ENSG00000 |
| 98807670 | 98850081 | 1 | protein_coding | ENSG00000 |
| 98847172 | 98872415 | -1 | lncRNA | ENSG00000 |
| 98890590 | 98891937 | 1 | processed_pseudoge | ENSG00000 |
| 98943179 | 99070792 | 1 | protein_coding | ENSG00000 |
| 98943337 | 98943775 | -1 | lncRNA | ENSG00000 |
| 99104038 | 99154192 | 1 | protein_coding | ENSG00000 |
| 99145840 | 99145966 | -1 | rRNA_pseudogene | ENSG00000 |
| 99183373 | 99184891 | 1 | lncRNA | ENSG00000 |
| 99185587 | 99185866 | -1 | misc_RNA | ENSG00000 |
| 99216425 | 99221942 | -1 | protein_coding | ENSG00000 |
| 99222064 | 99230615 | 1 | protein_coding | ENSG00000 |
| 99284022 | 99284373 | 1 | misc_RNA | ENSG00000 |
| 99305176 | 99306611 | 1 | processed_pseudoge | ENSG00000 |
| 99355337 | 99377240 | -1 | lncRNA | ENSG00000 |
| 99357827 | 99357957 | 1 | misc_RNA | ENSG00000 |
| 99359336 | 99371887 | 1 | lncRNA | ENSG00000 |
| 99368447 | 99368566 | 1 | misc_RNA | ENSG00000 |
| 99396987 | 99819921 | -1 | lncRNA | ENSG00000 |
| 99397024 | 99906605 | -1 | lncRNA | ENSG00000 |
| 99531507 | 99583301 | 1 | lncRNA | ENSG00000 |
| 99821855 | 99866891 | 1 | protein_coding | ENSG00000 |
| 99906654 | 99974534 | 1 | protein_coding | ENSG00000 |
| 99915077 | 99929896 | -1 | lncRNA | ENSG00000 |
| 99979185 | 100099000 | -1 | protein_coding | ENSG00000 |
| 99998301 | 99999069 | -1 | transcribed_processe | ENSG00000 |
| 100099243 | 100302175 | 1 | protein_coding | ENSG00000 |
| 100102559 | 100102839 | 1 | misc_RNA | ENSG00000 |
| 100116755 | 100117651 | -1 | processed_pseudoge | ENSG00000 |
| 100120435 | 100120900 | -1 | processed_pseudoge | ENSG00000 |
| 100175178 | 100176054 | 1 | transcribed_processe | ENSG00000 |
| 100302077 | 100352942 | -1 | protein_coding | ENSG00000 |
| 100427156 | 100451711 | 1 | protein_coding | ENSG00000 |

| | | | | |
|---|---|---|---|---|
| 100442271 | 100577636 | 1 | protein_coding | ENSG0000C |
| 100473149 | 100577636 | 1 | protein_coding | ENSG0000C |
| 100578079 | 100588389 | 1 | protein_coding | ENSG0000C |
| 100581079 | 100581379 | -1 | misc_RNA | ENSG0000C |
| 100731240 | 100732355 | 1 | processed_pseudoge | ENSG0000C |
| 100912320 | 100912741 | 1 | processed_pseudoge | ENSG0000C |
| 100975668 | 100976682 | 1 | processed_pseudoge | ENSG0000C |
| 101028727 | 101325135 | 1 | protein_coding | ENSG0000C |
| 101125436 | 101125921 | -1 | processed_pseudoge | ENSG0000C |
| 101322477 | 101322533 | 1 | snoRNA | ENSG0000C |
| 101334084 | 101339347 | -1 | unitary_pseudogene | ENSG0000C |
| 101356466 | 101356844 | -1 | processed_pseudoge | ENSG0000C |
| 101358932 | 101361242 | 1 | processed_pseudoge | ENSG0000C |
| 101360417 | 101383519 | -1 | protein_coding | ENSG0000C |
| 101375344 | 101376603 | -1 | processed_pseudoge | ENSG0000C |
| 101387633 | 101398618 | -1 | protein_coding | ENSG0000C |
| 101398873 | 101410660 | 1 | protein_coding | ENSG0000C |
| 101420560 | 101449664 | -1 | protein_coding | ENSG0000C |
| 101449801 | 101449933 | 1 | processed_pseudoge | ENSG0000C |
| 101468349 | 101482199 | 1 | lncRNA | ENSG0000C |
| 101473170 | 101533537 | -1 | protein_coding | ENSG0000C |
| 101501643 | 101501885 | -1 | processed_pseudoge | ENSG0000C |
| 101533853 | 101563344 | 1 | protein_coding | ENSG0000C |
| 101569353 | 101738580 | -1 | protein_coding | ENSG0000C |
| 101591604 | 101595021 | -1 | protein_coding | ENSG0000C |
| 101793454 | 101793756 | -1 | processed_pseudoge | ENSG0000C |
| 102055252 | 102055741 | -1 | processed_pseudoge | ENSG0000C |
| 102256604 | 102256710 | 1 | snRNA | ENSG0000C |
| 102444586 | 102444819 | -1 | processed_pseudoge | ENSG0000C |
| 102515204 | 102518308 | 1 | processed_pseudoge | ENSG0000C |
| 102519636 | 102657514 | 1 | lncRNA | ENSG0000C |
| 102611162 | 102612155 | -1 | processed_pseudoge | ENSG0000C |
| 102922574 | 102923244 | 1 | processed_pseudoge | ENSG0000C |
| 102995311 | 103018488 | 1 | protein_coding | ENSG0000C |
| 103065416 | 103065598 | 1 | processed_pseudoge | ENSG0000C |
| 103140523 | 103325034 | -1 | lncRNA | ENSG0000C |
| 103361952 | 103429892 | -1 | lncRNA | ENSG0000C |
| 103689473 | 103689577 | -1 | rRNA_pseudogene | ENSG0000C |
| 103999755 | 104000469 | 1 | lncRNA | ENSG0000C |
| 104080024 | 104093073 | -1 | lncRNA | ENSG0000C |
| 104094260 | 104141419 | 1 | protein_coding | ENSG0000C |
| 104270228 | 104286955 | 1 | transcribed_unitary_ | ENSG0000C |
| 104328320 | 104329396 | -1 | processed_pseudoge | ENSG0000C |
| 104504263 | 104505222 | 1 | protein_coding | ENSG0000C |
| 104519847 | 104537194 | 1 | lncRNA | ENSG0000C |
| 104526253 | 104527209 | -1 | protein_coding | ENSG0000C |
| 104535749 | 104536856 | -1 | protein_coding | ENSG0000C |
| 104569168 | 104570130 | 1 | protein_coding | ENSG0000C |
| 104590044 | 104590950 | 1 | unprocessed_pseudc | ENSG0000C |
| 104598457 | 104599413 | -1 | protein_coding | ENSG0000C |

| | | | | |
|---|---|---|---|---|
| 104604671 | 104605627 | -1 | protein_coding | ENSG00000 |
| 104617248 | 104618204 | -1 | protein_coding | ENSG00000 |
| 104629754 | 104631017 | 1 | unprocessed_pseudc | ENSG00000 |
| 104655363 | 104659055 | -1 | transcribed_unproce | ENSG00000 |
| 104694379 | 104695485 | 1 | protein_coding | ENSG00000 |
| 104722335 | 104723271 | 1 | unprocessed_pseudc | ENSG00000 |
| 104747683 | 104760120 | 1 | protein_coding | ENSG00000 |
| 104764129 | 104777764 | 1 | protein_coding | ENSG00000 |
| 104781006 | 104928155 | -1 | protein_coding | ENSG00000 |
| 104927553 | 104928892 | 1 | lncRNA | ENSG00000 |
| 104970593 | 104991818 | -1 | lncRNA | ENSG00000 |
| 105025932 | 105239221 | -1 | lncRNA | ENSG00000 |
| 105096756 | 105097061 | 1 | misc_RNA | ENSG00000 |
| 105244622 | 105439171 | 1 | protein_coding | ENSG00000 |
| 105447796 | 105552433 | 1 | protein_coding | ENSG00000 |
| 105520128 | 105526359 | 1 | processed_pseudoge | ENSG00000 |
| 105554035 | 105558304 | -1 | lncRNA | ENSG00000 |
| 105558130 | 105641118 | 1 | protein_coding | ENSG00000 |
| 105656369 | 105657989 | 1 | TEC | ENSG00000 |
| 105662457 | 105663124 | 1 | protein_coding | ENSG00000 |
| 105694541 | 105776629 | 1 | protein_coding | ENSG00000 |
| 105710290 | 105710781 | -1 | processed_pseudoge | ENSG00000 |
| 105758164 | 105760755 | 1 | processed_pseudoge | ENSG00000 |
| 105849710 | 105851425 | 1 | processed_pseudoge | ENSG00000 |
| 105993310 | 106740875 | 1 | lncRNA | ENSG00000 |
| 106061354 | 106064825 | -1 | lncRNA | ENSG00000 |
| 106584142 | 106591952 | -1 | lncRNA | ENSG00000 |
| 106600928 | 106601022 | 1 | miRNA | ENSG00000 |
| 106615078 | 106615983 | -1 | lncRNA | ENSG00000 |
| 106664754 | 106702662 | 1 | lncRNA | ENSG00000 |
| 106679977 | 106680329 | 1 | misc_RNA | ENSG00000 |
| 106722350 | 106722930 | 1 | processed_pseudoge | ENSG00000 |
| 106843749 | 106843866 | -1 | rRNA_pseudogene | ENSG00000 |
| 106863166 | 107013634 | 1 | protein_coding | ENSG00000 |
| 106974833 | 107102988 | -1 | lncRNA | ENSG00000 |
| 106978776 | 106979554 | -1 | processed_pseudoge | ENSG00000 |
| 107115784 | 107115954 | 1 | processed_pseudoge | ENSG00000 |
| 107116829 | 107117557 | 1 | processed_pseudoge | ENSG00000 |
| 107117459 | 107121705 | -1 | lncRNA | ENSG00000 |
| 107169569 | 107171148 | -1 | processed_pseudoge | ENSG00000 |
| 107283137 | 107332192 | 1 | protein_coding | ENSG00000 |
| 107420191 | 107466613 | -1 | lncRNA | ENSG00000 |
| 107426405 | 107428742 | 1 | lncRNA | ENSG00000 |
| 107458038 | 107458306 | -1 | processed_pseudoge | ENSG00000 |
| 107475036 | 107475139 | -1 | snRNA | ENSG00000 |
| 107484852 | 107490482 | -1 | protein_coding | ENSG00000 |
| 107534390 | 107535162 | 1 | processed_pseudoge | ENSG00000 |
| 107543151 | 107543616 | -1 | processed_pseudoge | ENSG00000 |
| 107663140 | 107663246 | -1 | snRNA | ENSG00000 |
| 107797744 | 107797847 | -1 | snRNA | ENSG00000 |

| | | | | |
|---|---|---|---|---|
| 107811616 | 107811914 | -1 | misc_RNA | ENSG0000C |
| 107834509 | 107834829 | -1 | processed_pseudoge | ENSG0000C |
| 107853783 | 107854258 | -1 | processed_pseudoge | ENSG0000C |
| 107918866 | 107918978 | -1 | rRNA_pseudogene | ENSG0000C |
| 107963703 | 107964045 | -1 | processed_pseudoge | ENSG0000C |
| 108040213 | 108040592 | 1 | processed_pseudoge | ENSG0000C |
| 108097432 | 108097851 | -1 | processed_pseudoge | ENSG0000C |
| 108252154 | 108254820 | -1 | lncRNA | ENSG0000C |
| 108626833 | 108627150 | 1 | processed_pseudoge | ENSG0000C |
| 108701370 | 108703092 | -1 | lncRNA | ENSG0000C |
| 108821697 | 108822004 | 1 | processed_pseudoge | ENSG0000C |
| 108836333 | 108836597 | 1 | processed_pseudoge | ENSG0000C |
| 108854588 | 108855986 | -1 | protein_coding | ENSG0000C |
| 108862266 | 108863756 | 1 | protein_coding | ENSG0000C |
| 108867517 | 108934124 | -1 | protein_coding | ENSG0000C |
| 108934400 | 108950744 | 1 | protein_coding | ENSG0000C |
| 108942569 | 109013522 | -1 | protein_coding | ENSG0000C |
| 109015135 | 109119947 | -1 | protein_coding | ENSG0000C |
| 109042386 | 109042487 | -1 | misc_RNA | ENSG0000C |
| 109046229 | 109046298 | -1 | miRNA | ENSG0000C |
| 109130293 | 109167295 | -1 | protein_coding | ENSG0000C |
| 109171975 | 109320964 | -1 | protein_coding | ENSG0000C |
| 109196201 | 109196309 | -1 | snRNA | ENSG0000C |
| 109300186 | 109301172 | 1 | unprocessed_pseudc | ENSG0000C |
| 109324685 | 109324796 | 1 | misc_RNA | ENSG0000C |
| 109375466 | 109498313 | -1 | protein_coding | ENSG0000C |
| 109511475 | 109511545 | -1 | miRNA | ENSG0000C |
| 109532830 | 109534332 | -1 | processed_pseudoge | ENSG0000C |
| 109640788 | 110172512 | 1 | protein_coding | ENSG0000C |
| 109760360 | 109772043 | -1 | lncRNA | ENSG0000C |
| 110024881 | 110025358 | -1 | processed_pseudoge | ENSG0000C |
| 110190048 | 110208189 | -1 | protein_coding | ENSG0000C |
| 110243810 | 110256507 | -1 | protein_coding | ENSG0000C |
| 110303521 | 110337884 | -1 | protein_coding | ENSG0000C |
| 110365248 | 110579880 | -1 | protein_coding | ENSG0000C |
| 110369592 | 110369698 | -1 | snRNA | ENSG0000C |
| 110599475 | 110605219 | 1 | lncRNA | ENSG0000C |
| 110668779 | 110806558 | 1 | protein_coding | ENSG0000C |
| 110727456 | 110727679 | -1 | processed_pseudoge | ENSG0000C |
| 110873263 | 111038458 | -1 | protein_coding | ENSG0000C |
| 110903337 | 110903445 | 1 | snRNA | ENSG0000C |
| 110934406 | 110934507 | 1 | misc_RNA | ENSG0000C |
| 111097325 | 111097413 | -1 | misc_RNA | ENSG0000C |
| 111139246 | 111284836 | -1 | lncRNA | ENSG0000C |
| 111271156 | 111271214 | -1 | miRNA | ENSG0000C |
| 111327483 | 111330183 | -1 | protein_coding | ENSG0000C |
| 111360692 | 111484745 | -1 | protein_coding | ENSG0000C |
| 111411644 | 111411742 | -1 | rRNA_pseudogene | ENSG0000C |
| 111525159 | 111577844 | 1 | protein_coding | ENSG0000C |
| 111549722 | 111599893 | -1 | protein_coding | ENSG0000C |

| | | | | |
|---|---|---|---|---|
| 111574192 | 111575427 | 1 | lncRNA | ENSG0000C |
| 111602831 | 111631289 | -1 | transcribed_unproce | ENSG0000C |
| 111631334 | 111654351 | 1 | protein_coding | ENSG0000C |
| 111631386 | 111670229 | 1 | protein_coding | ENSG0000C |
| 111661605 | 111670226 | 1 | protein_coding | ENSG0000C |
| 111686173 | 111795008 | -1 | protein_coding | ENSG0000C |
| 111802980 | 111803435 | 1 | processed_pseudoge | ENSG0000C |
| 111839828 | 111839934 | 1 | snRNA | ENSG0000C |
| 111896814 | 111935369 | 1 | protein_coding | ENSG0000C |
| 111932100 | 111932169 | 1 | miRNA | ENSG0000C |
| 111996635 | 111996739 | -1 | snRNA | ENSG0000C |
| 112028570 | 112039143 | -1 | lncRNA | ENSG0000C |
| 112040785 | 112175408 | -1 | protein_coding | ENSG0000C |
| 112137797 | 112137900 | 1 | snRNA | ENSG0000C |
| 112169286 | 112169397 | 1 | misc_RNA | ENSG0000C |
| 112173522 | 112173971 | -1 | processed_pseudoge | ENSG0000C |
| 112217716 | 112333667 | -1 | protein_coding | ENSG0000C |
| 112242550 | 112242839 | -1 | misc_RNA | ENSG0000C |
| 112252258 | 112252378 | 1 | rRNA_pseudogene | ENSG0000C |
| 112272596 | 112272837 | -1 | misc_RNA | ENSG0000C |
| 112280978 | 112281335 | -1 | processed_pseudoge | ENSG0000C |
| 112362714 | 112363116 | 1 | processed_pseudoge | ENSG0000C |
| 112380080 | 112472405 | 1 | protein_coding | ENSG0000C |
| 112399512 | 112487204 | -1 | lncRNA | ENSG0000C |
| 112486847 | 112669397 | 1 | protein_coding | ENSG0000C |
| 112581779 | 112582590 | -1 | unprocessed_pseudc | ENSG0000C |
| 112683926 | 112718149 | -1 | protein_coding | ENSG0000C |
| 112748902 | 112750565 | -1 | lncRNA | ENSG0000C |
| 112750760 | 112881671 | 1 | protein_coding | ENSG0000C |
| 112878920 | 112890876 | -1 | protein_coding | ENSG0000C |
| 112885158 | 112885767 | 1 | lncRNA | ENSG0000C |
| 112957722 | 113012227 | -1 | transcribed_unproce | ENSG0000C |
| 112959134 | 112962261 | -1 | transcribed_unproce | ENSG0000C |
| 112997120 | 113050043 | -1 | protein_coding | ENSG0000C |
| 113038380 | 113056759 | -1 | protein_coding | ENSG0000C |
| 113060086 | 113060507 | -1 | unprocessed_pseudc | ENSG0000C |
| 113065535 | 113065956 | -1 | unprocessed_pseudc | ENSG0000C |
| 113070983 | 113071404 | -1 | unprocessed_pseudc | ENSG0000C |
| 113076429 | 113076850 | -1 | unprocessed_pseudc | ENSG0000C |
| 113081871 | 113082292 | -1 | unprocessed_pseudc | ENSG0000C |
| 113087317 | 113087738 | -1 | unprocessed_pseudc | ENSG0000C |
| 113102117 | 113111543 | -1 | lncRNA | ENSG0000C |
| 113113042 | 113119928 | 1 | lncRNA | ENSG0000C |
| 113150976 | 113164140 | 1 | protein_coding | ENSG0000C |
| 113161006 | 113221318 | -1 | protein_coding | ENSG0000C |
| 113221544 | 113264492 | 1 | protein_coding | ENSG0000C |
| 113255835 | 113275572 | -1 | protein_coding | ENSG0000C |
| 113275343 | 113292905 | 1 | protein_coding | ENSG0000C |
| 113297093 | 113303376 | -1 | protein_coding | ENSG0000C |
| 113313222 | 113340298 | -1 | protein_coding | ENSG0000C |

| | | | | |
|---|---|---|---|---|
| 113349541 | 113371233 | 1 | protein_coding | ENSG00000 |
| 113373419 | 113376999 | -1 | protein_coding | ENSG00000 |
| 113386312 | 113401290 | -1 | protein_coding | ENSG00000 |
| 113407235 | 113410675 | -1 | protein_coding | ENSG00000 |
| 113410054 | 113429684 | 1 | protein_coding | ENSG00000 |
| 113444731 | 113597743 | 1 | protein_coding | ENSG00000 |
| 113570190 | 113590019 | -1 | lncRNA | ENSG00000 |
| 113614589 | 113617375 | -1 | TEC | ENSG00000 |
| 113619228 | 113622634 | 1 | lncRNA | ENSG00000 |
| 113650956 | 113682855 | 1 | lncRNA | ENSG00000 |
| 113758994 | 113759419 | -1 | processed_pseudoge | ENSG00000 |
| 113850687 | 113851632 | -1 | processed_pseudoge | ENSG00000 |
| 113876282 | 114056591 | 1 | protein_coding | ENSG00000 |
| 114060127 | 114078328 | -1 | protein_coding | ENSG00000 |
| 114086126 | 114099291 | -1 | protein_coding | ENSG00000 |
| 114155537 | 114312511 | 1 | protein_coding | ENSG00000 |
| 114209434 | 114209529 | 1 | miRNA | ENSG00000 |
| 114323056 | 114326475 | 1 | protein_coding | ENSG00000 |
| 114329869 | 114333256 | 1 | protein_coding | ENSG00000 |
| 114334156 | 114394405 | -1 | protein_coding | ENSG00000 |
| 114396724 | 114398503 | -1 | lncRNA | ENSG00000 |
| 114402080 | 114505473 | -1 | protein_coding | ENSG00000 |
| 114491564 | 114491741 | 1 | processed_pseudoge | ENSG00000 |
| 114572901 | 114573371 | 1 | processed_pseudoge | ENSG00000 |
| 114587769 | 114598879 | 1 | protein_coding | ENSG00000 |
| 114611206 | 114646422 | 1 | protein_coding | ENSG00000 |
| 114615455 | 114615568 | -1 | misc_RNA | ENSG00000 |
| 114656304 | 114662374 | -1 | protein_coding | ENSG00000 |
| 114666434 | 114682066 | -1 | protein_coding | ENSG00000 |
| 114695922 | 114696315 | -1 | processed_pseudoge | ENSG00000 |
| 114774737 | 114775120 | -1 | processed_pseudoge | ENSG00000 |
| 114784652 | 114806039 | -1 | protein_coding | ENSG00000 |
| 114845594 | 114848585 | 1 | processed_pseudoge | ENSG00000 |
| 114850968 | 115402644 | 1 | lncRNA | ENSG00000 |
| 114893343 | 114930595 | -1 | protein_coding | ENSG00000 |
| 115019575 | 115118257 | -1 | protein_coding | ENSG00000 |
| 115119539 | 115138409 | -1 | lncRNA | ENSG00000 |
| 115252538 | 115254251 | -1 | lncRNA | ENSG00000 |
| 115324932 | 115744330 | -1 | lncRNA | ENSG00000 |
| 115734392 | 115734491 | 1 | snRNA | ENSG00000 |
| 115888169 | 115925207 | -1 | lncRNA | ENSG00000 |
| 116012361 | 116012965 | 1 | lncRNA | ENSG00000 |
| 116153791 | 116402321 | 1 | protein_coding | ENSG00000 |
| 116285829 | 116288769 | -1 | lncRNA | ENSG00000 |
| 116288618 | 116318689 | -1 | lncRNA | ENSG00000 |
| 116398157 | 116400606 | -1 | lncRNA | ENSG00000 |
| 116425225 | 117415070 | -1 | protein_coding | ENSG00000 |
| 116504283 | 116562293 | 1 | lncRNA | ENSG00000 |
| 116567636 | 116586328 | 1 | lncRNA | ENSG00000 |
| 116687302 | 116701300 | 1 | protein_coding | ENSG00000 |

| | | | | |
|---|---|---|---|---|
| 116838144 | 116877264 | 1 | lncRNA | ENSG0000C |
| 117180628 | 117181095 | -1 | processed_pseudoge | ENSG0000C |
| 117181066 | 117181200 | -1 | snoRNA | ENSG0000C |
| 117197118 | 117197420 | 1 | misc_RNA | ENSG0000C |
| 117276985 | 117277286 | 1 | misc_RNA | ENSG0000C |
| 117462294 | 117466260 | -1 | lncRNA | ENSG0000C |
| 117643151 | 117657032 | 1 | lncRNA | ENSG0000C |
| 117685254 | 117685588 | -1 | processed_pseudoge | ENSG0000C |
| 117704157 | 118072314 | 1 | protein_coding | ENSG0000C |
| 117704175 | 117724735 | 1 | protein_coding | ENSG0000C |
| 117707996 | 117708102 | -1 | snRNA | ENSG0000C |
| 117759455 | 117879988 | 1 | lncRNA | ENSG0000C |
| 117840322 | 117897256 | -1 | lncRNA | ENSG0000C |
| 118082869 | 118083382 | -1 | processed_pseudoge | ENSG0000C |
| 118687752 | 118732772 | -1 | lncRNA | ENSG0000C |
| 118740563 | 118745312 | 1 | lncRNA | ENSG0000C |
| 118763867 | 118764345 | 1 | processed_pseudoge | ENSG0000C |
| 119153458 | 119369435 | -1 | protein_coding | ENSG0000C |
| 119495375 | 119524125 | 1 | lncRNA | ENSG0000C |
| 119935053 | 119937832 | 1 | lncRNA | ENSG0000C |
| 119973094 | 119974322 | 1 | lncRNA | ENSG0000C |
| 120244979 | 120245050 | -1 | miRNA | ENSG0000C |
| 120388869 | 120580170 | -1 | protein_coding | ENSG0000C |
| 120600811 | 120714470 | -1 | protein_coding | ENSG0000C |
| 120720673 | 120721972 | 1 | processed_pseudoge | ENSG0000C |
| 120751978 | 120793416 | -1 | protein_coding | ENSG0000C |
| 120792403 | 120799918 | 1 | protein_coding | ENSG0000C |
| 120812475 | 120812845 | -1 | processed_pseudoge | ENSG0000C |
| 120815496 | 120842951 | -1 | protein_coding | ENSG0000C |
| 120824828 | 120854385 | 1 | transcribed_unitary_ | ENSG0000C |
| 120855651 | 120894896 | -1 | protein_coding | ENSG0000C |
| 120902393 | 120929173 | -1 | protein_coding | ENSG0000C |
| 120952335 | 121050276 | -1 | protein_coding | ENSG0000C |
| 121074863 | 121177610 | 1 | protein_coding | ENSG0000C |
| 121178133 | 121223014 | -1 | protein_coding | ENSG0000C |
| 121203159 | 121203435 | 1 | misc_RNA | ENSG0000C |
| 121207794 | 121332843 | 1 | protein_coding | ENSG0000C |
| 121238117 | 121240089 | -1 | processed_pseudoge | ENSG0000C |
| 121279863 | 121282730 | -1 | lncRNA | ENSG0000C |
| 121280768 | 121285530 | -1 | lncRNA | ENSG0000C |
| 121338987 | 121370304 | -1 | protein_coding | ENSG0000C |
| 121369893 | 121463370 | 1 | lncRNA | ENSG0000C |
| 121444991 | 121500027 | -1 | transcribed_unitary_ | ENSG0000C |
| 121494519 | 121494821 | 1 | misc_RNA | ENSG0000C |
| 121519816 | 121520424 | -1 | processed_pseudoge | ENSG0000C |
| 121567057 | 121785530 | 1 | protein_coding | ENSG0000C |
| 121574891 | 121576214 | -1 | lncRNA | ENSG0000C |
| 121815674 | 121819452 | -1 | lncRNA | ENSG0000C |
| 121821928 | 122093606 | -1 | protein_coding | ENSG0000C |
| 121884636 | 121963719 | -1 | lncRNA | ENSG0000C |

| | | | | |
|---|---|---|---|---|
| 122120082 | 122120167 | -1 | miRNA | ENSG00000 |
| 122144058 | 122159779 | -1 | protein_coding | ENSG00000 |
| 122159908 | 122200088 | 1 | protein_coding | ENSG00000 |
| 122202577 | 122229626 | -1 | protein_coding | ENSG00000 |
| 122237622 | 122264840 | -1 | protein_coding | ENSG00000 |
| 122264603 | 122331337 | 1 | protein_coding | ENSG00000 |
| 122370530 | 122395703 | 1 | protein_coding | ENSG00000 |
| 122372605 | 122372825 | 1 | lncRNA | ENSG00000 |
| 122400970 | 122402906 | -1 | lncRNA | ENSG00000 |
| 122403407 | 122639685 | 1 | lncRNA | ENSG00000 |
| 122476958 | 122477926 | -1 | protein_coding | ENSG00000 |
| 122510802 | 122511743 | 1 | protein_coding | ENSG00000 |
| 122519141 | 122520082 | 1 | protein_coding | ENSG00000 |
| 122526358 | 122527296 | -1 | protein_coding | ENSG00000 |
| 122553112 | 122554214 | 1 | protein_coding | ENSG00000 |
| 122567117 | 122583384 | -1 | protein_coding | ENSG00000 |
| 122607864 | 122608800 | 1 | unprocessed_pseudo | ENSG00000 |
| 122614738 | 122615682 | 1 | protein_coding | ENSG00000 |
| 122628579 | 122629573 | -1 | protein_coding | ENSG00000 |
| 122661566 | 122662648 | 1 | protein_coding | ENSG00000 |
| 122675036 | 122676153 | 1 | protein_coding | ENSG00000 |
| 122685933 | 122688372 | 1 | processed_pseudoge | ENSG00000 |
| 122723990 | 122724925 | 1 | protein_coding | ENSG00000 |
| 122742303 | 122750783 | 1 | protein_coding | ENSG00000 |
| 122761844 | 122762233 | 1 | processed_pseudoge | ENSG00000 |
| 122788933 | 122789895 | 1 | protein_coding | ENSG00000 |
| 122798389 | 122828588 | -1 | protein_coding | ENSG00000 |
| 122800123 | 122801073 | 1 | protein_coding | ENSG00000 |
| 122837557 | 122838820 | 1 | processed_pseudoge | ENSG00000 |
| 122844556 | 122905359 | -1 | protein_coding | ENSG00000 |
| 122880213 | 122880323 | -1 | snoRNA | ENSG00000 |
| 122908056 | 122913323 | -1 | protein_coding | ENSG00000 |
| 122915566 | 122931512 | -1 | protein_coding | ENSG00000 |
| 122937623 | 122940333 | 1 | lncRNA | ENSG00000 |
| 122940733 | 122941260 | -1 | lncRNA | ENSG00000 |
| 122940833 | 123104866 | 1 | protein_coding | ENSG00000 |
| 122945988 | 122946094 | -1 | misc_RNA | ENSG00000 |
| 123029373 | 123029753 | -1 | processed_pseudoge | ENSG00000 |
| 123034527 | 123035696 | 1 | protein_coding | ENSG00000 |
| 123061845 | 123063312 | -1 | processed_pseudoge | ENSG00000 |
| 123109494 | 123115477 | -1 | lncRNA | ENSG00000 |
| 123109500 | 123268586 | -1 | protein_coding | ENSG00000 |
| 123111546 | 123111643 | -1 | miRNA | ENSG00000 |
| 123356170 | 123380324 | 1 | protein_coding | ENSG00000 |
| 123379654 | 123930152 | -1 | protein_coding | ENSG00000 |
| 123402525 | 123402603 | -1 | miRNA | ENSG00000 |
| 123485529 | 123485622 | -1 | miRNA | ENSG00000 |
| 123491884 | 123493156 | 1 | lncRNA | ENSG00000 |
| 123736437 | 123751941 | 1 | lncRNA | ENSG00000 |
| 123759220 | 123759775 | 1 | processed_pseudoge | ENSG00000 |

| | | | | |
|---|---|---|---|---|
| 123843036 | 123843686 | 1 | processed_pseudoge | ENSG0000C |
| 124001670 | 124033301 | 1 | protein_coding | ENSG0000C |
| 124006277 | 124009396 | -1 | lncRNA | ENSG0000C |
| 124010530 | 124011114 | 1 | lncRNA | ENSG0000C |
| 124031624 | 124032524 | -1 | lncRNA | ENSG0000C |
| 124257606 | 124353307 | 1 | protein_coding | ENSG0000C |
| 124259250 | 124261156 | -1 | lncRNA | ENSG0000C |
| 124262876 | 124265809 | 1 | lncRNA | ENSG0000C |
| 124353465 | 124415444 | -1 | protein_coding | ENSG0000C |
| 124451425 | 124478589 | 1 | protein_coding | ENSG0000C |
| 124481236 | 124507420 | -1 | protein_coding | ENSG0000C |
| 124514938 | 124516056 | -1 | lncRNA | ENSG0000C |
| 124517275 | 124771311 | -1 | protein_coding | ENSG0000C |
| 124598707 | 124599580 | -1 | processed_pseudoge | ENSG0000C |
| 124607159 | 124607458 | 1 | misc_RNA | ENSG0000C |
| 124658467 | 124698631 | 1 | lncRNA | ENSG0000C |
| 124692442 | 124692551 | 1 | miRNA | ENSG0000C |
| 124693710 | 124693798 | 1 | miRNA | ENSG0000C |
| 124770123 | 124772927 | 1 | lncRNA | ENSG0000C |
| 124777133 | 124814885 | 1 | protein_coding | ENSG0000C |
| 124853417 | 124857890 | 1 | protein_coding | ENSG0000C |
| 124857880 | 124861981 | -1 | protein_coding | ENSG0000C |
| 124862130 | 124877733 | 1 | protein_coding | ENSG0000C |
| 124878275 | 124948492 | -1 | protein_coding | ENSG0000C |
| 124887410 | 124887549 | -1 | snRNA | ENSG0000C |
| 124942608 | 125143506 | -1 | protein_coding | ENSG0000C |
| 124968294 | 124968840 | -1 | processed_pseudoge | ENSG0000C |
| 125146573 | 125189939 | -1 | protein_coding | ENSG0000C |
| 125150653 | 125151589 | 1 | processed_pseudoge | ENSG0000C |
| 125194649 | 125195821 | -1 | processed_pseudoge | ENSG0000C |
| 125200542 | 125234161 | 1 | protein_coding | ENSG0000C |
| 125234853 | 125241343 | -1 | protein_coding | ENSG0000C |
| 125241663 | 125257071 | 1 | lncRNA | ENSG0000C |
| 125261794 | 125367207 | 1 | protein_coding | ENSG0000C |
| 125263845 | 125264090 | -1 | processed_pseudoge | ENSG0000C |
| 125303920 | 125304026 | -1 | snRNA | ENSG0000C |
| 125372325 | 125372766 | -1 | processed_pseudoge | ENSG0000C |
| 125417305 | 125417586 | 1 | misc_RNA | ENSG0000C |
| 125426169 | 125438509 | 1 | lncRNA | ENSG0000C |
| 125437393 | 125707234 | -1 | protein_coding | ENSG0000C |
| 125512902 | 125513207 | -1 | misc_RNA | ENSG0000C |
| 125567579 | 125573023 | 1 | lncRNA | ENSG0000C |
| 125595669 | 125596628 | 1 | processed_pseudoge | ENSG0000C |
| 125743754 | 125746552 | -1 | lncRNA | ENSG0000C |
| 125747345 | 125967377 | 1 | protein_coding | ENSG0000C |
| 126057580 | 126159613 | 1 | lncRNA | ENSG0000C |
| 126270121 | 126298593 | -1 | lncRNA | ENSG0000C |
| 126326829 | 126507041 | 1 | protein_coding | ENSG0000C |
| 126408041 | 126408410 | -1 | misc_RNA | ENSG0000C |
| 126426893 | 126426970 | 1 | snoRNA | ENSG0000C |

| | | | | |
|---|---|---|---|---|
| 126516417 | 126518808 | 1 | lncRNA | ENSG0000( |
| 126518840 | 126520773 | 1 | TEC | ENSG0000( |
| 126527618 | 126530343 | 1 | lncRNA | ENSG0000( |
| 126589594 | 126613700 | -1 | lncRNA | ENSG0000( |
| 126614443 | 126701032 | 1 | protein_coding | ENSG0000( |
| 126640756 | 126641293 | 1 | lncRNA | ENSG0000( |
| 126701173 | 126704691 | -1 | lncRNA | ENSG0000( |
| 126805006 | 126838210 | 1 | protein_coding | ENSG0000( |
| 126860665 | 126885878 | 1 | protein_coding | ENSG0000( |
| 126914774 | 127223166 | 1 | protein_coding | ENSG0000( |
| 127087348 | 127122635 | -1 | protein_coding | ENSG0000( |
| 127213783 | 127221907 | -1 | lncRNA | ENSG0000( |
| 127224265 | 127393660 | 1 | protein_coding | ENSG0000( |
| 127300093 | 127300982 | -1 | processed_pseudoge | ENSG0000( |
| 127366770 | 127367397 | 1 | lncRNA | ENSG0000( |
| 127397138 | 127408424 | 1 | protein_coding | ENSG0000( |
| 127424374 | 127445372 | 1 | protein_coding | ENSG0000( |
| 127447674 | 127451406 | -1 | protein_coding | ENSG0000( |
| 127448501 | 127448630 | -1 | snoRNA | ENSG0000( |
| 127451486 | 127503501 | 1 | protein_coding | ENSG0000( |
| 127470875 | 127470987 | 1 | misc_RNA | ENSG0000( |
| 127505339 | 127578989 | -1 | protein_coding | ENSG0000( |
| 127579370 | 127696027 | 1 | protein_coding | ENSG0000( |
| 127690098 | 127690840 | -1 | lncRNA | ENSG0000( |
| 127690348 | 127724873 | -1 | protein_coding | ENSG0000( |
| 127690687 | 127690795 | -1 | miRNA | ENSG0000( |
| 127706988 | 127716002 | 1 | protein_coding | ENSG0000( |
| 127716079 | 127731590 | 1 | protein_coding | ENSG0000( |
| 127731524 | 127735313 | -1 | protein_coding | ENSG0000( |
| 127738317 | 127778710 | -1 | protein_coding | ENSG0000( |
| 127754534 | 127760560 | 1 | lncRNA | ENSG0000( |
| 127785679 | 127790792 | 1 | protein_coding | ENSG0000( |
| 127785833 | 127785923 | 1 | miRNA | ENSG0000( |
| 127785918 | 127786007 | 1 | miRNA | ENSG0000( |
| 127794597 | 127814327 | 1 | protein_coding | ENSG0000( |
| 127815012 | 127854756 | -1 | protein_coding | ENSG0000( |
| 127816066 | 127822520 | 1 | lncRNA | ENSG0000( |
| 127832635 | 127832743 | -1 | misc_RNA | ENSG0000( |
| 127841112 | 127841222 | -1 | rRNA_pseudogene | ENSG0000( |
| 127866486 | 127877675 | -1 | protein_coding | ENSG0000( |
| 127867850 | 127906022 | -1 | protein_coding | ENSG0000( |
| 127869415 | 127869495 | -1 | miRNA | ENSG0000( |
| 127885321 | 127905408 | -1 | protein_coding | ENSG0000( |
| 127907886 | 127917041 | -1 | protein_coding | ENSG0000( |
| 127920881 | 127930785 | -1 | protein_coding | ENSG0000( |
| 127934503 | 127940952 | 1 | lncRNA | ENSG0000( |
| 127935099 | 127938484 | -1 | protein_coding | ENSG0000( |
| 127940582 | 127980989 | -1 | protein_coding | ENSG0000( |
| 128039135 | 128057362 | -1 | lncRNA | ENSG0000( |
| 128061233 | 128068206 | -1 | protein_coding | ENSG0000( |

| | | | | |
|---|---|---|---|---|
| 128068201 | 128109245 | 1 | protein_coding | ENSG00000 |
| 128090969 | 128094457 | -1 | lncRNA | ENSG00000 |
| 128108581 | 128118693 | -1 | lncRNA | ENSG00000 |
| 128120693 | 128128462 | -1 | protein_coding | ENSG00000 |
| 128128529 | 128130628 | 1 | lncRNA | ENSG00000 |
| 128149071 | 128153453 | 1 | protein_coding | ENSG00000 |
| 128160265 | 128163924 | 1 | protein_coding | ENSG00000 |
| 128161251 | 128204383 | -1 | protein_coding | ENSG00000 |
| 128191655 | 128255248 | 1 | protein_coding | ENSG00000 |
| 128244721 | 128244830 | -1 | miRNA | ENSG00000 |
| 128244947 | 128245030 | -1 | miRNA | ENSG00000 |
| 128255829 | 128275995 | -1 | protein_coding | ENSG00000 |
| 128275379 | 128316123 | 1 | protein_coding | ENSG00000 |
| 128305159 | 128322447 | -1 | protein_coding | ENSG00000 |
| 128316337 | 128316909 | 1 | lncRNA | ENSG00000 |
| 128322544 | 128334072 | 1 | protein_coding | ENSG00000 |
| 128340527 | 128361470 | 1 | protein_coding | ENSG00000 |
| 128342153 | 128342770 | 1 | processed_pseudoge | ENSG00000 |
| 128371319 | 128392016 | 1 | protein_coding | ENSG00000 |
| 128391461 | 128392016 | 1 | lncRNA | ENSG00000 |
| 128392007 | 128393510 | -1 | lncRNA | ENSG00000 |
| 128392618 | 128392714 | -1 | miRNA | ENSG00000 |
| 128392621 | 128392708 | 1 | miRNA | ENSG00000 |
| 128411751 | 128437351 | 1 | protein_coding | ENSG00000 |
| 128431598 | 128432006 | 1 | lncRNA | ENSG00000 |
| 128444587 | 128446628 | 1 | processed_pseudoge | ENSG00000 |
| 128455186 | 128501292 | 1 | protein_coding | ENSG00000 |
| 128468957 | 128473012 | -1 | lncRNA | ENSG00000 |
| 128504700 | 128542288 | 1 | protein_coding | ENSG00000 |
| 128528657 | 128552410 | -1 | lncRNA | ENSG00000 |
| 128543916 | 128543977 | 1 | snRNA | ENSG00000 |
| 128552558 | 128633662 | 1 | protein_coding | ENSG00000 |
| 128595893 | 128596633 | 1 | lncRNA | ENSG00000 |
| 128630328 | 128631685 | -1 | lncRNA | ENSG00000 |
| 128633661 | 128656787 | -1 | protein_coding | ENSG00000 |
| 128650998 | 128652145 | 1 | processed_pseudoge | ENSG00000 |
| 128663134 | 128684477 | -1 | lncRNA | ENSG00000 |
| 128683424 | 128696400 | 1 | protein_coding | ENSG00000 |
| 128702503 | 128720916 | 1 | protein_coding | ENSG00000 |
| 128709780 | 128710043 | 1 | misc_RNA | ENSG00000 |
| 128720870 | 128724127 | -1 | protein_coding | ENSG00000 |
| 128724445 | 128733194 | 1 | lncRNA | ENSG00000 |
| 128729786 | 128772414 | -1 | protein_coding | ENSG00000 |
| 128745857 | 128746138 | -1 | misc_RNA | ENSG00000 |
| 128787253 | 128810430 | 1 | protein_coding | ENSG00000 |
| 128818500 | 128822676 | 1 | protein_coding | ENSG00000 |
| 128819651 | 128829794 | -1 | protein_coding | ENSG00000 |
| 128819663 | 128881931 | -1 | protein_coding | ENSG00000 |
| 128832942 | 128882494 | -1 | protein_coding | ENSG00000 |
| 128882112 | 128918039 | 1 | protein_coding | ENSG00000 |

| | | | | |
|---|---|---|---|---|
| 128920966 | 128942041 | 1 | protein_coding | ENSG00000 |
| 128941478 | 128957021 | 1 | protein_coding | ENSG00000 |
| 128945530 | 128947619 | -1 | protein_coding | ENSG00000 |
| 128947699 | 129007096 | 1 | protein_coding | ENSG00000 |
| 129004502 | 129004998 | -1 | lncRNA | ENSG00000 |
| 129007036 | 129028303 | -1 | protein_coding | ENSG00000 |
| 129036621 | 129072082 | 1 | protein_coding | ENSG00000 |
| 129080355 | 129080765 | -1 | lncRNA | ENSG00000 |
| 129081111 | 129090438 | 1 | protein_coding | ENSG00000 |
| 129094794 | 129111189 | -1 | protein_coding | ENSG00000 |
| 129097854 | 129100266 | 1 | lncRNA | ENSG00000 |
| 129110950 | 129148946 | 1 | protein_coding | ENSG00000 |
| 129170434 | 129170940 | 1 | lncRNA | ENSG00000 |
| 129175552 | 129178261 | -1 | protein_coding | ENSG00000 |
| 129175807 | 129177575 | 1 | lncRNA | ENSG00000 |
| 129176771 | 129210548 | 1 | lncRNA | ENSG00000 |
| 129258354 | 129259846 | 1 | lncRNA | ENSG00000 |
| 129259062 | 129261581 | 1 | lncRNA | ENSG00000 |
| 129282458 | 129312665 | 1 | lncRNA | ENSG00000 |
| 129321016 | 129324905 | 1 | lncRNA | ENSG00000 |
| 129328261 | 129328401 | 1 | lncRNA | ENSG00000 |
| 129332300 | 129359541 | 1 | lncRNA | ENSG00000 |
| 129370170 | 129370461 | -1 | misc_RNA | ENSG00000 |
| 129380136 | 129380470 | -1 | processed_pseudoge | ENSG00000 |
| 129430652 | 129451422 | -1 | lncRNA | ENSG00000 |
| 129438216 | 129447545 | 1 | lncRNA | ENSG00000 |
| 129476946 | 129513687 | 1 | lncRNA | ENSG00000 |
| 129480605 | 129482538 | -1 | lncRNA | ENSG00000 |
| 129503461 | 129506433 | -1 | lncRNA | ENSG00000 |
| 129575117 | 129584556 | 1 | lncRNA | ENSG00000 |
| 129608884 | 129636131 | 1 | protein_coding | ENSG00000 |
| 129612225 | 129620776 | -1 | protein_coding | ENSG00000 |
| 129634604 | 129642169 | -1 | protein_coding | ENSG00000 |
| 129640476 | 129641282 | 1 | lncRNA | ENSG00000 |
| 129665647 | 129722674 | 1 | protein_coding | ENSG00000 |
| 129712896 | 129718635 | -1 | lncRNA | ENSG00000 |
| 129738331 | 129753042 | -1 | protein_coding | ENSG00000 |
| 129791196 | 129791837 | 1 | processed_pseudoge | ENSG00000 |
| 129803157 | 129811281 | 1 | protein_coding | ENSG00000 |
| 129812942 | 129824244 | -1 | protein_coding | ENSG00000 |
| 129827290 | 129835863 | -1 | protein_coding | ENSG00000 |
| 129834698 | 129881838 | 1 | protein_coding | ENSG00000 |
| 129869605 | 129869671 | 1 | miRNA | ENSG00000 |
| 129883693 | 129884170 | -1 | processed_pseudoge | ENSG00000 |
| 129887187 | 130043194 | -1 | protein_coding | ENSG00000 |
| 129933536 | 129936541 | 1 | lncRNA | ENSG00000 |
| 130044713 | 130045081 | 1 | lncRNA | ENSG00000 |
| 130053426 | 130140169 | 1 | protein_coding | ENSG00000 |
| 130120806 | 130120912 | 1 | misc_RNA | ENSG00000 |
| 130140610 | 130144219 | -1 | lncRNA | ENSG00000 |

| | | | | |
|---|---|---|---|---|
| 130172404 | 130237303 | 1 | protein_coding | ENSG00000 |
| 130265882 | 130434123 | 1 | protein_coding | ENSG00000 |
| 130400266 | 130400565 | 1 | misc_RNA | ENSG00000 |
| 130444961 | 130501274 | 1 | protein_coding | ENSG00000 |
| 130578965 | 130638352 | 1 | protein_coding | ENSG00000 |
| 130626297 | 130626363 | 1 | miRNA | ENSG00000 |
| 130651799 | 130652383 | 1 | processed_pseudoge | ENSG00000 |
| 130664594 | 130682983 | 1 | protein_coding | ENSG00000 |
| 130693721 | 130704894 | 1 | protein_coding | ENSG00000 |
| 130713016 | 130887675 | 1 | protein_coding | ENSG00000 |
| 130724786 | 130725050 | -1 | processed_pseudoge | ENSG00000 |
| 130892707 | 130896812 | -1 | protein_coding | ENSG00000 |
| 130902438 | 130939286 | -1 | protein_coding | ENSG00000 |
| 130933851 | 130945520 | 1 | lncRNA | ENSG00000 |
| 131009174 | 131094473 | 1 | protein_coding | ENSG00000 |
| 131021168 | 131021306 | -1 | snoRNA | ENSG00000 |
| 131096476 | 131123152 | 1 | protein_coding | ENSG00000 |
| 131125586 | 131234663 | 1 | protein_coding | ENSG00000 |
| 131126990 | 131127091 | 1 | snRNA | ENSG00000 |
| 131132852 | 131167505 | -1 | lncRNA | ENSG00000 |
| 131189910 | 131194205 | -1 | lncRNA | ENSG00000 |
| 131240687 | 131241724 | 1 | processed_pseudoge | ENSG00000 |
| 131258076 | 131276510 | -1 | protein_coding | ENSG00000 |
| 131283421 | 131288278 | 1 | lncRNA | ENSG00000 |
| 131289459 | 131359022 | 1 | protein_coding | ENSG00000 |
| 131370933 | 131372514 | 1 | unprocessed_pseudo | ENSG00000 |
| 131373636 | 131500197 | 1 | protein_coding | ENSG00000 |
| 131485665 | 131485750 | 1 | snoRNA | ENSG00000 |
| 131490486 | 131490571 | 1 | snoRNA | ENSG00000 |
| 131497479 | 131500191 | -1 | lncRNA | ENSG00000 |
| 131502902 | 131523806 | 1 | protein_coding | ENSG00000 |
| 131516558 | 131522229 | -1 | lncRNA | ENSG00000 |
| 131523801 | 131531264 | -1 | protein_coding | ENSG00000 |
| 131545514 | 131558620 | 1 | protein_coding | ENSG00000 |
| 131576770 | 131740074 | -1 | protein_coding | ENSG00000 |
| 131725168 | 131725467 | -1 | misc_RNA | ENSG00000 |
| 131782570 | 131782813 | -1 | processed_pseudoge | ENSG00000 |
| 131782860 | 131782958 | -1 | snoRNA | ENSG00000 |
| 131816192 | 131820988 | 1 | lncRNA | ENSG00000 |
| 131852928 | 132079867 | -1 | protein_coding | ENSG00000 |
| 132161676 | 132244526 | 1 | protein_coding | ENSG00000 |
| 132261356 | 132354986 | -1 | protein_coding | ENSG00000 |
| 132375548 | 132406851 | -1 | protein_coding | ENSG00000 |
| 132410043 | 132573317 | 1 | protein_coding | ENSG00000 |
| 132421641 | 132421690 | 1 | snRNA | ENSG00000 |
| 132582606 | 132590252 | 1 | protein_coding | ENSG00000 |
| 132592997 | 132670401 | -1 | protein_coding | ENSG00000 |
| 132670035 | 132694953 | 1 | protein_coding | ENSG00000 |
| 132725578 | 132878777 | -1 | protein_coding | ENSG00000 |
| 132768965 | 132770212 | -1 | lncRNA | ENSG00000 |

| | | | | |
|---|---|---|---|---|
| 132878027 | 132890201 | 1 | protein_coding | ENSG00000( |
| 132891348 | 132946874 | -1 | protein_coding | ENSG00000( |
| 132944000 | 132991687 | 1 | protein_coding | ENSG00000( |
| 132945707 | 132945771 | 1 | miRNA | ENSG00000( |
| 132962511 | 132962667 | 1 | processed_pseudoge | ENSG00000( |
| 132963732 | 132963793 | 1 | snRNA | ENSG00000( |
| 133019486 | 133020874 | 1 | processed_pseudoge | ENSG00000( |
| 133030675 | 133058503 | 1 | protein_coding | ENSG00000( |
| 133051996 | 133052059 | 1 | miRNA | ENSG00000( |
| 133061978 | 133071861 | 1 | protein_coding | ENSG00000( |
| 133079900 | 133087355 | 1 | transcribed_unproce | ENSG00000( |
| 133097720 | 133149334 | -1 | protein_coding | ENSG00000( |
| 133098121 | 133163914 | -1 | protein_coding | ENSG00000( |
| 133152948 | 133163933 | -1 | protein_coding | ENSG00000( |
| 133205277 | 133209250 | -1 | protein_coding | ENSG00000( |
| 133224905 | 133228591 | -1 | unprocessed_pseudc | ENSG00000( |
| 133250401 | 133276024 | -1 | protein_coding | ENSG00000( |
| 133302470 | 133302563 | 1 | misc_RNA | ENSG00000( |
| 133317794 | 133318469 | 1 | unprocessed_pseudc | ENSG00000( |
| 133328776 | 133336188 | -1 | protein_coding | ENSG00000( |
| 133337728 | 133337824 | -1 | misc_RNA | ENSG00000( |
| 133338312 | 133348131 | -1 | protein_coding | ENSG00000( |
| 133338990 | 133339465 | 1 | processed_pseudoge | ENSG00000( |
| 133348218 | 133351426 | 1 | protein_coding | ENSG00000( |
| 133349396 | 133349470 | 1 | snoRNA | ENSG00000( |
| 133350095 | 133350168 | 1 | snoRNA | ENSG00000( |
| 133350456 | 133350528 | 1 | snoRNA | ENSG00000( |
| 133350847 | 133350912 | 1 | snoRNA | ENSG00000( |
| 133351758 | 133356676 | -1 | protein_coding | ENSG00000( |
| 133356550 | 133361158 | 1 | protein_coding | ENSG00000( |
| 133361450 | 133376166 | -1 | protein_coding | ENSG00000( |
| 133376366 | 133406096 | 1 | protein_coding | ENSG00000( |
| 133406059 | 133418096 | -1 | protein_coding | ENSG00000( |
| 133414358 | 133459402 | 1 | protein_coding | ENSG00000( |
| 133459965 | 133470848 | 1 | protein_coding | ENSG00000( |
| 133471094 | 133479127 | -1 | protein_coding | ENSG00000( |
| 133514586 | 133528612 | -1 | protein_coding | ENSG00000( |
| 133532164 | 133575519 | 1 | protein_coding | ENSG00000( |
| 133578415 | 133586197 | -1 | protein_coding | ENSG00000( |
| 133611798 | 133633099 | -1 | lncRNA | ENSG00000( |
| 133636363 | 133659329 | 1 | protein_coding | ENSG00000( |
| 133654586 | 133657313 | -1 | lncRNA | ENSG00000( |
| 133663560 | 133739955 | -1 | protein_coding | ENSG00000( |
| 133761894 | 133992604 | -1 | protein_coding | ENSG00000( |
| 134025481 | 134034666 | 1 | protein_coding | ENSG00000( |
| 134030305 | 134068535 | -1 | protein_coding | ENSG00000( |
| 134054290 | 134058805 | 1 | lncRNA | ENSG00000( |
| 134078904 | 134079444 | 1 | processed_pseudoge | ENSG00000( |
| 134132242 | 134135772 | -1 | lncRNA | ENSG00000( |
| 134135365 | 134159968 | 1 | protein_coding | ENSG00000( |

| | | | | |
|---|---|---|---|---|
| 134164439 | 134164564 | -1 | snRNA | ENSG0000C |
| 134168769 | 134169340 | 1 | lncRNA | ENSG0000C |
| 134293931 | 134295397 | -1 | lncRNA | ENSG0000C |
| 134298463 | 134300413 | 1 | TEC | ENSG0000C |
| 134317098 | 134440585 | 1 | protein_coding | ENSG0000C |
| 134379411 | 134379472 | 1 | miRNA | ENSG0000C |
| 134505472 | 134521442 | 1 | lncRNA | ENSG0000C |
| 134527182 | 134545244 | 1 | lncRNA | ENSG0000C |
| 134552738 | 134553897 | 1 | lncRNA | ENSG0000C |
| 134641774 | 134844843 | 1 | protein_coding | ENSG0000C |
| 134649385 | 134653073 | -1 | lncRNA | ENSG0000C |
| 134770651 | 134775671 | -1 | lncRNA | ENSG0000C |
| 134849298 | 134849369 | -1 | miRNA | ENSG0000C |
| 134849487 | 134849564 | -1 | miRNA | ENSG0000C |
| 134849609 | 134849682 | -1 | miRNA | ENSG0000C |
| 134850125 | 134850272 | -1 | miRNA | ENSG0000C |
| 134850277 | 134850356 | -1 | miRNA | ENSG0000C |
| 134850570 | 134850641 | -1 | miRNA | ENSG0000C |
| 134850742 | 134850807 | -1 | miRNA | ENSG0000C |
| 134868236 | 134868475 | 1 | unprocessed_pseudc | ENSG0000C |
| 134880810 | 134887523 | 1 | protein_coding | ENSG0000C |
| 134903232 | 134917912 | -1 | protein_coding | ENSG0000C |
| 134936524 | 134943207 | 1 | lncRNA | ENSG0000C |
| 134992958 | 134995854 | 1 | TEC | ENSG0000C |
| 135075422 | 135121179 | 1 | protein_coding | ENSG0000C |
| 135204722 | 135237597 | -1 | lncRNA | ENSG0000C |
| 135245696 | 135252708 | -1 | lncRNA | ENSG0000C |
| 135343249 | 135346562 | 1 | lncRNA | ENSG0000C |
| 135352673 | 135354284 | 1 | lncRNA | ENSG0000C |
| 135447446 | 135450154 | 1 | processed_pseudoge | ENSG0000C |
| 135462727 | 135480777 | -1 | lncRNA | ENSG0000C |
| 135479079 | 135488893 | 1 | protein_coding | ENSG0000C |
| 135495181 | 135501734 | -1 | protein_coding | ENSG0000C |
| 135499984 | 135504673 | 1 | protein_coding | ENSG0000C |
| 135503273 | 135506447 | -1 | lncRNA | ENSG0000C |
| 135521438 | 135526532 | 1 | protein_coding | ENSG0000C |
| 135546139 | 135549969 | 1 | protein_coding | ENSG0000C |
| 135561756 | 135566955 | 1 | protein_coding | ENSG0000C |
| 135574935 | 135587112 | 1 | lncRNA | ENSG0000C |
| 135588002 | 135591975 | 1 | transcribed_unproce | ENSG0000C |
| 135604345 | 135618952 | 1 | protein_coding | ENSG0000C |
| 135623648 | 135639540 | -1 | protein_coding | ENSG0000C |
| 135663322 | 135666422 | 1 | protein_coding | ENSG0000C |
| 135693407 | 135699528 | -1 | protein_coding | ENSG0000C |
| 135702185 | 135795508 | 1 | protein_coding | ENSG0000C |
| 135808487 | 135907546 | -1 | protein_coding | ENSG0000C |
| 135907812 | 135916126 | 1 | lncRNA | ENSG0000C |
| 135932969 | 135961373 | -1 | protein_coding | ENSG0000C |
| 136006537 | 136095289 | -1 | protein_coding | ENSG0000C |
| 136107808 | 136109424 | -1 | lncRNA | ENSG0000C |

| | | | | |
|---|---|---|---|---|
| 136114581 | 136118875 | -1 | protein_coding | ENSG00000 |
| 136122521 | 136124363 | 1 | lncRNA | ENSG00000 |
| 136196250 | 136205128 | -1 | protein_coding | ENSG00000 |
| 136206333 | 136245841 | -1 | protein_coding | ENSG00000 |
| 136249971 | 136306901 | -1 | protein_coding | ENSG00000 |
| 136249978 | 136251205 | 1 | TEC | ENSG00000 |
| 136263925 | 136267237 | 1 | lncRNA | ENSG00000 |
| 136322303 | 136327323 | -1 | lncRNA | ENSG00000 |
| 136327476 | 136359605 | 1 | protein_coding | ENSG00000 |
| 136359480 | 136363744 | -1 | protein_coding | ENSG00000 |
| 136361903 | 136373681 | -1 | protein_coding | ENSG00000 |
| 136375577 | 136400168 | -1 | protein_coding | ENSG00000 |
| 136401922 | 136410614 | -1 | protein_coding | ENSG00000 |
| 136410641 | 136423761 | 1 | protein_coding | ENSG00000 |
| 136428619 | 136439845 | -1 | protein_coding | ENSG00000 |
| 136440096 | 136483759 | -1 | protein_coding | ENSG00000 |
| 136483495 | 136486067 | 1 | lncRNA | ENSG00000 |
| 136494433 | 136546048 | -1 | protein_coding | ENSG00000 |
| 136519568 | 136519626 | -1 | miRNA | ENSG00000 |
| 136542881 | 136543835 | 1 | lncRNA | ENSG00000 |
| 136546173 | 136546259 | -1 | miRNA | ENSG00000 |
| 136546212 | 136549893 | 1 | lncRNA | ENSG00000 |
| 136612024 | 136617181 | -1 | lncRNA | ENSG00000 |
| 136648610 | 136660421 | -1 | lncRNA | ENSG00000 |
| 136658856 | 136672678 | 1 | protein_coding | ENSG00000 |
| 136670602 | 136670686 | 1 | miRNA | ENSG00000 |
| 136673143 | 136687457 | -1 | protein_coding | ENSG00000 |
| 136712572 | 136724742 | 1 | protein_coding | ENSG00000 |
| 136721366 | 136728184 | -1 | lncRNA | ENSG00000 |
| 136726104 | 136726239 | -1 | snoRNA | ENSG00000 |
| 136726105 | 136726234 | -1 | snoRNA | ENSG00000 |
| 136726747 | 136726879 | -1 | snoRNA | ENSG00000 |
| 136726748 | 136726879 | -1 | snoRNA | ENSG00000 |
| 136728953 | 136729855 | -1 | unprocessed_pseudc | ENSG00000 |
| 136738167 | 136743356 | -1 | protein_coding | ENSG00000 |
| 136738167 | 136748453 | -1 | protein_coding | ENSG00000 |
| 136744017 | 136748525 | -1 | protein_coding | ENSG00000 |
| 136746893 | 136746970 | -1 | miRNA | ENSG00000 |
| 136754386 | 136758543 | -1 | protein_coding | ENSG00000 |
| 136759634 | 136766255 | -1 | protein_coding | ENSG00000 |
| 136779939 | 136780218 | 1 | processed_pseudoge | ENSG00000 |
| 136791344 | 136793317 | 1 | protein_coding | ENSG00000 |
| 136791379 | 136800595 | 1 | protein_coding | ENSG00000 |
| 136796338 | 136807741 | 1 | protein_coding | ENSG00000 |
| 136799223 | 136810042 | 1 | lncRNA | ENSG00000 |
| 136800366 | 136829466 | 1 | lncRNA | ENSG00000 |
| 136803927 | 136808848 | -1 | lncRNA | ENSG00000 |
| 136807943 | 136841187 | 1 | protein_coding | ENSG00000 |
| 136812788 | 136815536 | -1 | processed_pseudoge | ENSG00000 |
| 136830957 | 136831023 | 1 | miRNA | ENSG00000 |

| | | | | |
|---|---|---|---|---|
| 136844415 | 136848801 | 1 | protein_coding | ENSG00000( |
| 136848724 | 136851027 | 1 | protein_coding | ENSG00000( |
| 136850943 | 136860799 | 1 | protein_coding | ENSG00000( |
| 136862119 | 136866308 | -1 | protein_coding | ENSG00000( |
| 136881912 | 136926607 | 1 | protein_coding | ENSG00000( |
| 136886733 | 136886803 | 1 | miRNA | ENSG00000( |
| 136935840 | 136937161 | 1 | processed_pseudoge | ENSG00000( |
| 136937169 | 136937988 | -1 | lncRNA | ENSG00000( |
| 136940435 | 136944738 | -1 | protein_coding | ENSG00000( |
| 136945185 | 136946975 | 1 | protein_coding | ENSG00000( |
| 136949551 | 136955497 | 1 | protein_coding | ENSG00000( |
| 136969243 | 136971990 | 1 | lncRNA | ENSG00000( |
| 136975092 | 136981742 | 1 | protein_coding | ENSG00000( |
| 136977518 | 136985435 | 1 | protein_coding | ENSG00000( |
| 136981904 | 136986410 | 1 | protein_coding | ENSG00000( |
| 136992422 | 136993984 | 1 | protein_coding | ENSG00000( |
| 136994608 | 136996568 | -1 | protein_coding | ENSG00000( |
| 137007234 | 137028922 | -1 | protein_coding | ENSG00000( |
| 137027464 | 137037957 | 1 | lncRNA | ENSG00000( |
| 137030174 | 137033010 | -1 | protein_coding | ENSG00000( |
| 137031241 | 137040436 | -1 | protein_coding | ENSG00000( |
| 137039463 | 137046179 | -1 | protein_coding | ENSG00000( |
| 137048107 | 137054061 | -1 | protein_coding | ENSG00000( |
| 137052662 | 137053375 | 1 | lncRNA | ENSG00000( |
| 137057663 | 137062461 | 1 | lncRNA | ENSG00000( |
| 137062127 | 137070557 | -1 | protein_coding | ENSG00000( |
| 137063535 | 137064581 | 1 | lncRNA | ENSG00000( |
| 137077517 | 137084539 | 1 | protein_coding | ENSG00000( |
| 137084946 | 137086817 | -1 | lncRNA | ENSG00000( |
| 137086985 | 137109183 | 1 | protein_coding | ENSG00000( |
| 137094345 | 137094430 | 1 | snoRNA | ENSG00000( |
| 137110546 | 137115177 | -1 | protein_coding | ENSG00000( |
| 137139154 | 137168756 | 1 | protein_coding | ENSG00000( |
| 137168758 | 137170051 | -1 | protein_coding | ENSG00000( |
| 137168854 | 137171984 | -1 | protein_coding | ENSG00000( |
| 137169186 | 137169270 | -1 | miRNA | ENSG00000( |
| 137170858 | 137172409 | -1 | protein_coding | ENSG00000( |
| 137174784 | 137188560 | -1 | protein_coding | ENSG00000( |
| 137188660 | 137190370 | 1 | protein_coding | ENSG00000( |
| 137191617 | 137204193 | -1 | protein_coding | ENSG00000( |
| 137204082 | 137205648 | -1 | protein_coding | ENSG00000( |
| 137205685 | 137217009 | 1 | protein_coding | ENSG00000( |
| 137217452 | 137219361 | 1 | protein_coding | ENSG00000( |
| 137220247 | 137221581 | -1 | protein_coding | ENSG00000( |
| 137224635 | 137226315 | 1 | protein_coding | ENSG00000( |
| 137227271 | 137229638 | 1 | protein_coding | ENSG00000( |
| 137230757 | 137236554 | 1 | protein_coding | ENSG00000( |
| 137241287 | 137243707 | 1 | protein_coding | ENSG00000( |
| 137243584 | 137247770 | -1 | protein_coding | ENSG00000( |
| 137250219 | 137253497 | -1 | lncRNA | ENSG00000( |

| | | | | |
|---|---|---|---|---|
| 137251261 | 137253483 | 1 | protein_coding | ENSG0000C |
| 137255327 | 137273542 | 1 | protein_coding | ENSG0000C |
| 137277726 | 137282641 | 1 | protein_coding | ENSG0000C |
| 137286112 | 137287236 | 1 | processed_pseudoge | ENSG0000C |
| 137293868 | 137295721 | -1 | lncRNA | ENSG0000C |
| 137294935 | 137300189 | 1 | lncRNA | ENSG0000C |
| 137299631 | 137302271 | -1 | protein_coding | ENSG0000C |
| 137306896 | 137423211 | -1 | protein_coding | ENSG0000C |
| 137423350 | 137434406 | 1 | protein_coding | ENSG0000C |
| 137434364 | 137441816 | -1 | protein_coding | ENSG0000C |
| 137447573 | 137459334 | -1 | protein_coding | ENSG0000C |
| 137450026 | 137450086 | -1 | miRNA | ENSG0000C |
| 137459952 | 137550402 | -1 | protein_coding | ENSG0000C |
| 137551879 | 137552555 | 1 | protein_coding | ENSG0000C |
| 137554444 | 137578925 | -1 | protein_coding | ENSG0000C |
| 137582081 | 137590512 | -1 | protein_coding | ENSG0000C |
| 137605685 | 137615360 | 1 | protein_coding | ENSG0000C |
| 137605744 | 137711018 | 1 | lncRNA | ENSG0000C |
| 137615332 | 137618906 | -1 | lncRNA | ENSG0000C |
| 137618992 | 137870016 | 1 | protein_coding | ENSG0000C |
| 137638128 | 137638408 | -1 | misc_RNA | ENSG0000C |
| 137640986 | 137641547 | -1 | processed_pseudoge | ENSG0000C |
| 137776539 | 137789235 | -1 | transcribed_process | ENSG0000C |
| 137838419 | 137838516 | 1 | miRNA | ENSG0000C |
| 137867925 | 137892570 | -1 | lncRNA | ENSG0000C |
| 137877782 | 138124624 | 1 | protein_coding | ENSG0000C |
| 138129399 | 138129526 | 1 | unprocessed_pseudc | ENSG0000C |
| 138137095 | 138143864 | 1 | unprocessed_pseudc | ENSG0000C |
| 138150075 | 138177433 | 1 | transcribed_unproce | ENSG0000C |
| 138177426 | 138179774 | -1 | unprocessed_pseudc | ENSG0000C |
| 138199933 | 138203325 | -1 | lncRNA | ENSG0000C |
| 138217068 | 138253217 | 1 | transcribed_unproce | ENSG0000C |
| 138259595 | 138259698 | 1 | snRNA | ENSG0000C |
| 138286248 | 138286430 | 1 | unprocessed_pseudc | ENSG0000C |
| 138319898 | 138320835 | 1 | unprocessed_pseudc | ENSG0000C |
| 14061 | 16544 | -1 | lncRNA | ENSG0000C |
| 44712 | 46884 | 1 | unprocessed_pseudc | ENSG0000C |
| 46892 | 74163 | -1 | protein_coding | ENSG0000C |
| 80394 | 86265 | -1 | unprocessed_pseudc | ENSG0000C |
| 134465 | 254637 | 1 | protein_coding | ENSG0000C |
| 147820 | 148751 | -1 | unprocessed_pseudc | ENSG0000C |
| 154831 | 154943 | -1 | rRNA_pseudogene | ENSG0000C |
| 274190 | 689668 | -1 | protein_coding | ENSG0000C |
| 282015 | 282125 | -1 | rRNA_pseudogene | ENSG0000C |
| 430239 | 430504 | -1 | misc_RNA | ENSG0000C |
| 437561 | 441170 | 1 | lncRNA | ENSG0000C |
| 628638 | 631255 | 1 | lncRNA | ENSG0000C |
| 641689 | 641778 | -1 | miRNA | ENSG0000C |
| 649948 | 669581 | 1 | protein_coding | ENSG0000C |
| 743992 | 744958 | 1 | lncRNA | ENSG0000C |

| | | | | |
|---|---|---|---|---|
| 806914 | 931705 | -1 | protein_coding | ENSG0000C |
| 933026 | 942743 | 1 | lncRNA | ENSG0000C |
| 971146 | 988341 | -1 | lncRNA | ENSG0000C |
| 988434 | 1019932 | 1 | protein_coding | ENSG0000C |
| 1018910 | 1025859 | -1 | protein_coding | ENSG0000C |
| 1022630 | 1045425 | 1 | lncRNA | ENSG0000C |
| 1039152 | 1049170 | -1 | protein_coding | ENSG0000C |
| 1049538 | 1132384 | 1 | protein_coding | ENSG0000C |
| 1159768 | 1289426 | 1 | lncRNA | ENSG0000C |
| 1177313 | 1737525 | -1 | protein_coding | ENSG0000C |
| 1201406 | 1208389 | -1 | lncRNA | ENSG0000C |
| 1290018 | 1293002 | -1 | lncRNA | ENSG0000C |
| 1361001 | 1361637 | -1 | lncRNA | ENSG0000C |
| 1526637 | 1556984 | 1 | lncRNA | ENSG0000C |
| 1957589 | 1960190 | 1 | lncRNA | ENSG0000C |
| 1989076 | 2015005 | -1 | lncRNA | ENSG0000C |
| 2071845 | 2081097 | 1 | lncRNA | ENSG0000C |
| 2076019 | 2076089 | -1 | miRNA | ENSG0000C |
| 2098167 | 2098269 | 1 | snRNA | ENSG0000C |
| 2150480 | 2169460 | 1 | lncRNA | ENSG0000C |
| 2169140 | 2189512 | -1 | lncRNA | ENSG0000C |
| 2199387 | 2199488 | 1 | snRNA | ENSG0000C |
| 2305083 | 2315075 | -1 | lncRNA | ENSG0000C |
| 2368561 | 2502200 | -1 | lncRNA | ENSG0000C |
| 2499721 | 2500570 | 1 | lncRNA | ENSG0000C |
| 2501600 | 2618739 | 1 | lncRNA | ENSG0000C |
| 2724819 | 2728267 | -1 | lncRNA | ENSG0000C |
| 2936021 | 2936441 | 1 | processed_pseudoge | ENSG0000C |
| 3009976 | 3013111 | 1 | lncRNA | ENSG0000C |
| 3042535 | 3047083 | -1 | lncRNA | ENSG0000C |
| 3065424 | 3066001 | -1 | lncRNA | ENSG0000C |
| 3066333 | 3137718 | 1 | protein_coding | ENSG0000C |
| 3134566 | 3145166 | 1 | lncRNA | ENSG0000C |
| 3137728 | 3172841 | -1 | protein_coding | ENSG0000C |
| 3141632 | 3167972 | 1 | lncRNA | ENSG0000C |
| 3200829 | 3277649 | -1 | lncRNA | ENSG0000C |
| 3223816 | 3227407 | 1 | lncRNA | ENSG0000C |
| 3263346 | 3267216 | 1 | lncRNA | ENSG0000C |
| 3296338 | 3298137 | 1 | lncRNA | ENSG0000C |
| 3397505 | 3400180 | 1 | lncRNA | ENSG0000C |
| 3433508 | 3502915 | -1 | lncRNA | ENSG0000C |
| 3532094 | 3556314 | -1 | lncRNA | ENSG0000C |
| 3748966 | 3763712 | 1 | lncRNA | ENSG0000C |
| 3767915 | 3768751 | 1 | lncRNA | ENSG0000C |
| 3775996 | 3785281 | -1 | protein_coding | ENSG0000C |
| 3809835 | 3816169 | 1 | lncRNA | ENSG0000C |
| 3833950 | 3834728 | 1 | lncRNA | ENSG0000C |
| 3911889 | 3935817 | 1 | lncRNA | ENSG0000C |
| 3991160 | 3991259 | 1 | miRNA | ENSG0000C |
| 4024903 | 4053018 | 1 | lncRNA | ENSG0000C |

| | | | | |
|---|---|---|---|---|
| 4051726 | 4089013 | 1 | lncRNA | ENSG00000 |
| 4201141 | 4243912 | -1 | lncRNA | ENSG00000 |
| 4384246 | 4410614 | 1 | lncRNA | ENSG00000 |
| 4415305 | 4426179 | -1 | lncRNA | ENSG00000 |
| 4437102 | 4448692 | 1 | lncRNA | ENSG00000 |
| 4497449 | 4515244 | 1 | lncRNA | ENSG00000 |
| 4514952 | 4515057 | 1 | snRNA | ENSG00000 |
| 4650158 | 4678185 | -1 | lncRNA | ENSG00000 |
| 4655427 | 4682722 | 1 | lncRNA | ENSG00000 |
| 4747846 | 4785100 | -1 | lncRNA | ENSG00000 |
| 4786629 | 4848062 | 1 | protein_coding | ENSG00000 |
| 4871901 | 4891975 | -1 | unprocessed_pseudo | ENSG00000 |
| 4962436 | 4962568 | -1 | snoRNA | ENSG00000 |
| 4963253 | 4983283 | 1 | protein_coding | ENSG00000 |
| 4987400 | 5018031 | -1 | protein_coding | ENSG00000 |
| 4995488 | 4997380 | 1 | lncRNA | ENSG00000 |
| 5004710 | 5004842 | 1 | snoRNA | ENSG00000 |
| 5035354 | 5107686 | 1 | protein_coding | ENSG00000 |
| 5093408 | 5093543 | -1 | snoRNA | ENSG00000 |
| 5122087 | 5135226 | -1 | unprocessed_pseudo | ENSG00000 |
| 5154140 | 5185187 | -1 | protein_coding | ENSG00000 |
| 5195462 | 5218949 | 1 | protein_coding | ENSG00000 |
| 5207169 | 5207742 | 1 | processed_pseudoge | ENSG00000 |
| 5215408 | 5217014 | -1 | lncRNA | ENSG00000 |
| 5234358 | 5263408 | 1 | lncRNA | ENSG00000 |
| 5266033 | 5271236 | -1 | lncRNA | ENSG00000 |
| 5275173 | 5288470 | -1 | transcribed_unproce | ENSG00000 |
| 5304431 | 5304865 | -1 | processed_pseudoge | ENSG00000 |
| 5346651 | 5347277 | 1 | TEC | ENSG00000 |
| 5364966 | 5374692 | 1 | protein_coding | ENSG00000 |
| 5393101 | 5404828 | -1 | protein_coding | ENSG00000 |
| 5412557 | 5459056 | 1 | protein_coding | ENSG00000 |
| 5498697 | 5499570 | -1 | protein_coding | ENSG00000 |
| 5510036 | 5526246 | -1 | lncRNA | ENSG00000 |
| 5524961 | 5526771 | 1 | protein_coding | ENSG00000 |
| 5524976 | 5525742 | -1 | lncRNA | ENSG00000 |
| 5549637 | 5554329 | -1 | lncRNA | ENSG00000 |
| 5563295 | 5566881 | -1 | lncRNA | ENSG00000 |
| 5594984 | 5632435 | -1 | lncRNA | ENSG00000 |
| 5608475 | 5610793 | -1 | lncRNA | ENSG00000 |
| 5616626 | 5632499 | -1 | lncRNA | ENSG00000 |
| 5624931 | 5625126 | 1 | processed_pseudoge | ENSG00000 |
| 5631463 | 5631755 | 1 | misc_RNA | ENSG00000 |
| 5638867 | 5666595 | -1 | protein_coding | ENSG00000 |
| 5684838 | 5763740 | 1 | protein_coding | ENSG00000 |
| 5712174 | 5744067 | -1 | lncRNA | ENSG00000 |
| 5765223 | 5842132 | -1 | protein_coding | ENSG00000 |
| 5776071 | 5776858 | 1 | processed_pseudoge | ENSG00000 |
| 5813985 | 5814441 | 1 | lncRNA | ENSG00000 |
| 5861616 | 5889906 | -1 | protein_coding | ENSG00000 |

| | | | | |
|---|---|---|---|---|
| 5861805 | 5862594 | -1 | processed_pseudoge | ENSG0000C |
| 5890203 | 5937594 | 1 | protein_coding | ENSG0000C |
| 5934270 | 5945900 | -1 | lncRNA | ENSG0000C |
| 5943639 | 5978187 | -1 | protein_coding | ENSG0000C |
| 6010689 | 6062370 | -1 | protein_coding | ENSG0000C |
| 6017054 | 6017194 | -1 | snoRNA | ENSG0000C |
| 6025978 | 6036427 | 1 | lncRNA | ENSG0000C |
| 6071514 | 6071918 | -1 | processed_pseudoge | ENSG0000C |
| 6089034 | 6117457 | 1 | protein_coding | ENSG0000C |
| 6144934 | 6254644 | 1 | protein_coding | ENSG0000C |
| 6149623 | 6149940 | 1 | misc_RNA | ENSG0000C |
| 6152196 | 6152277 | 1 | miRNA | ENSG0000C |
| 6152207 | 6152262 | -1 | miRNA | ENSG0000C |
| 6197612 | 6202693 | -1 | lncRNA | ENSG0000C |
| 6245731 | 6245831 | -1 | misc_RNA | ENSG0000C |
| 6271169 | 6353827 | 1 | lncRNA | ENSG0000C |
| 6293100 | 6294684 | -1 | processed_pseudoge | ENSG0000C |
| 6350316 | 6352762 | 1 | lncRNA | ENSG0000C |
| 6427143 | 6580301 | -1 | protein_coding | ENSG0000C |
| 6580419 | 6616921 | 1 | lncRNA | ENSG0000C |
| 6618716 | 6625346 | 1 | lncRNA | ENSG0000C |
| 6682513 | 6823789 | 1 | lncRNA | ENSG0000C |
| 6709530 | 6740532 | 1 | lncRNA | ENSG0000C |
| 6737418 | 6840712 | 1 | lncRNA | ENSG0000C |
| 6774307 | 6779180 | -1 | lncRNA | ENSG0000C |
| 6779549 | 6879450 | 1 | lncRNA | ENSG0000C |
| 6779608 | 6877888 | 1 | lncRNA | ENSG0000C |
| 6824275 | 7118469 | -1 | lncRNA | ENSG0000C |
| 7096216 | 7097675 | 1 | lncRNA | ENSG0000C |
| 7154120 | 7154236 | 1 | processed_pseudoge | ENSG0000C |
| 7158624 | 7411486 | -1 | protein_coding | ENSG0000C |
| 7295106 | 7295290 | -1 | processed_pseudoge | ENSG0000C |
| 7411599 | 7413467 | 1 | TEC | ENSG0000C |
| 7444340 | 7472040 | -1 | lncRNA | ENSG0000C |
| 7486703 | 7486809 | -1 | snRNA | ENSG0000C |
| 7533207 | 7536565 | 1 | lncRNA | ENSG0000C |
| 7559270 | 7666998 | -1 | protein_coding | ENSG0000C |
| 7703316 | 7749520 | 1 | protein_coding | ENSG0000C |
| 7750962 | 7787993 | -1 | protein_coding | ENSG0000C |
| 7788147 | 7807815 | 1 | protein_coding | ENSG0000C |
| 7818505 | 8016631 | 1 | protein_coding | ENSG0000C |
| 7833618 | 7833957 | -1 | processed_pseudoge | ENSG0000C |
| 8045378 | 8075198 | 1 | protein_coding | ENSG0000C |
| 8050450 | 8053484 | -1 | lncRNA | ENSG0000C |
| 8051541 | 8053084 | 1 | lncRNA | ENSG0000C |
| 8161260 | 8162192 | 1 | processed_pseudoge | ENSG0000C |
| 8259331 | 8268342 | -1 | lncRNA | ENSG0000C |
| 8298897 | 8301667 | 1 | lncRNA | ENSG0000C |
| 8400860 | 8461863 | 1 | lncRNA | ENSG0000C |
| 8472719 | 8472952 | 1 | processed_pseudoge | ENSG0000C |

| | | | | |
|---|---|---|---|---|
| 8513687 | 8515106 | -1 | processed_pseudoge | ENSG00000( |
| 8619933 | 8620608 | 1 | processed_pseudoge | ENSG00000( |
| 8656716 | 8656831 | -1 | rRNA_pseudogene | ENSG00000( |
| 8724010 | 8724288 | 1 | processed_pseudoge | ENSG00000( |
| 8897989 | 8914596 | 1 | lncRNA | ENSG00000( |
| 8970125 | 8973468 | 1 | lncRNA | ENSG00000( |
| 9275833 | 9287057 | 1 | lncRNA | ENSG00000( |
| 9292842 | 9309772 | -1 | lncRNA | ENSG00000( |
| 9721963 | 9724416 | -1 | processed_pseudoge | ENSG00000( |
| 9758783 | 9878060 | -1 | lncRNA | ENSG00000( |
| 10058722 | 10063502 | 1 | lncRNA | ENSG00000( |
| 10174230 | 10174577 | 1 | processed_pseudoge | ENSG00000( |
| 10232611 | 10232712 | 1 | misc_RNA | ENSG00000( |
| 10257169 | 10257466 | -1 | processed_pseudoge | ENSG00000( |
| 10326875 | 10337876 | -1 | lncRNA | ENSG00000( |
| 10365180 | 10366487 | -1 | lncRNA | ENSG00000( |
| 10369809 | 10462361 | -1 | lncRNA | ENSG00000( |
| 10394670 | 10395737 | 1 | TEC | ENSG00000( |
| 10400340 | 10404203 | 1 | lncRNA | ENSG00000( |
| 10776223 | 10776661 | -1 | processed_pseudoge | ENSG00000( |
| 10784437 | 10795047 | -1 | lncRNA | ENSG00000( |
| 10798397 | 11336675 | 1 | protein_coding | ENSG00000( |
| 10917866 | 10952234 | -1 | lncRNA | ENSG00000( |
| 10960148 | 10970816 | 1 | lncRNA | ENSG00000( |
| 11010164 | 11011119 | -1 | lncRNA | ENSG00000( |
| 11030334 | 11030868 | -1 | lncRNA | ENSG00000( |
| 11071541 | 11105773 | -1 | lncRNA | ENSG00000( |
| 11168922 | 11171376 | -1 | lncRNA | ENSG00000( |
| 11316833 | 11319884 | -1 | lncRNA | ENSG00000( |
| 11395808 | 11398792 | 1 | TEC | ENSG00000( |
| 11453946 | 11611754 | -1 | protein_coding | ENSG00000( |
| 11611305 | 11612227 | 1 | lncRNA | ENSG00000( |
| 11679675 | 11680514 | 1 | lncRNA | ENSG00000( |
| 11742366 | 11764070 | 1 | protein_coding | ENSG00000( |
| 11823339 | 11872277 | 1 | protein_coding | ENSG00000( |
| 11849608 | 11894700 | -1 | lncRNA | ENSG00000( |
| 11920022 | 12043170 | -1 | protein_coding | ENSG00000( |
| 11923181 | 11923288 | 1 | snRNA | ENSG00000( |
| 12068954 | 12123221 | 1 | protein_coding | ENSG00000( |
| 12116280 | 12116379 | -1 | snRNA | ENSG00000( |
| 12129637 | 12169961 | 1 | protein_coding | ENSG00000( |
| 12130760 | 12130816 | -1 | miRNA | ENSG00000( |
| 12165330 | 12196144 | -1 | protein_coding | ENSG00000( |
| 12195965 | 12250589 | 1 | protein_coding | ENSG00000( |
| 12244751 | 12247845 | -1 | lncRNA | ENSG00000( |
| 12328132 | 12328389 | -1 | misc_RNA | ENSG00000( |
| 12349482 | 12835545 | 1 | protein_coding | ENSG00000( |
| 12415928 | 12416000 | 1 | snoRNA | ENSG00000( |
| 12456483 | 12456631 | 1 | snRNA | ENSG00000( |
| 12563151 | 12567351 | 1 | lncRNA | ENSG00000( |

| | | | | |
|---|---|---|---|---|
| 12578753 | 12578823 | 1 | miRNA | ENSG0000C |
| 12653138 | 12653197 | -1 | miRNA | ENSG0000C |
| 12725254 | 12725353 | -1 | miRNA | ENSG0000C |
| 12877459 | 12965275 | 1 | lncRNA | ENSG0000C |
| 12896625 | 13099652 | -1 | protein_coding | ENSG0000C |
| 12924679 | 12924785 | -1 | rRNA_pseudogene | ENSG0000C |
| 13057902 | 13058676 | -1 | processed_pseudoge | ENSG0000C |
| 13099449 | 13138308 | 1 | protein_coding | ENSG0000C |
| 13104574 | 13104793 | -1 | processed_pseudoge | ENSG0000C |
| 13139196 | 13139533 | -1 | processed_pseudoge | ENSG0000C |
| 13142185 | 13142977 | -1 | processed_pseudoge | ENSG0000C |
| 13156684 | 13158136 | -1 | processed_pseudoge | ENSG0000C |
| 13159477 | 13159777 | -1 | processed_pseudoge | ENSG0000C |
| 13161554 | 13211104 | 1 | protein_coding | ENSG0000C |
| 13217269 | 13217375 | 1 | snRNA | ENSG0000C |
| 13221766 | 13234374 | -1 | protein_coding | ENSG0000C |
| 13277796 | 13302412 | -1 | protein_coding | ENSG0000C |
| 13308145 | 13308447 | -1 | processed_pseudoge | ENSG0000C |
| 13317428 | 13348298 | -1 | protein_coding | ENSG0000C |
| 13391347 | 13400706 | 1 | lncRNA | ENSG0000C |
| 13415202 | 13423133 | 1 | lncRNA | ENSG0000C |
| 13438484 | 13528974 | -1 | protein_coding | ENSG0000C |
| 13511206 | 13511313 | -1 | misc_RNA | ENSG0000C |
| 13528516 | 13530262 | 1 | lncRNA | ENSG0000C |
| 13586939 | 13630859 | 1 | protein_coding | ENSG0000C |
| 13591460 | 13592323 | -1 | processed_pseudoge | ENSG0000C |
| 13610047 | 13655929 | 1 | protein_coding | ENSG0000C |
| 13631143 | 13668445 | 1 | lncRNA | ENSG0000C |
| 13643706 | 14462142 | -1 | protein_coding | ENSG0000C |
| 13675646 | 13675990 | 1 | lncRNA | ENSG0000C |
| 13710415 | 13712054 | 1 | lncRNA | ENSG0000C |
| 13727796 | 13727916 | 1 | rRNA_pseudogene | ENSG0000C |
| 13729383 | 13756200 | 1 | lncRNA | ENSG0000C |
| 13870797 | 13871175 | 1 | processed_pseudoge | ENSG0000C |
| 13991815 | 14007526 | 1 | lncRNA | ENSG0000C |
| 14074284 | 14087605 | 1 | lncRNA | ENSG0000C |
| 14372001 | 14377586 | 1 | lncRNA | ENSG0000C |
| 14383200 | 14383277 | -1 | miRNA | ENSG0000C |
| 14391875 | 14392196 | 1 | misc_RNA | ENSG0000C |
| 14436576 | 14436661 | 1 | miRNA | ENSG0000C |
| 14518557 | 14774897 | -1 | protein_coding | ENSG0000C |
| 14652689 | 14661691 | 1 | lncRNA | ENSG0000C |
| 14663395 | 14663494 | -1 | rRNA_pseudogene | ENSG0000C |
| 14723171 | 14724055 | -1 | processed_pseudoge | ENSG0000C |
| 14819245 | 14838575 | -1 | protein_coding | ENSG0000C |
| 14838160 | 14847018 | 1 | protein_coding | ENSG0000C |
| 14838306 | 14871741 | 1 | protein_coding | ENSG0000C |
| 14877688 | 14878686 | -1 | lncRNA | ENSG0000C |
| 14878820 | 14904315 | 1 | protein_coding | ENSG0000C |
| 14897359 | 14954432 | -1 | protein_coding | ENSG0000C |

| | | | | |
|---|---|---|---|---|
| 14959388 | 14988050 | 1 | protein_coding | ENSG0000C |
| 14986942 | 14987515 | -1 | unprocessed_pseudc | ENSG0000C |
| 14999050 | 15000055 | -1 | unprocessed_pseudc | ENSG0000C |
| 15007861 | 15008884 | -1 | unprocessed_pseudc | ENSG0000C |
| 15015370 | 15021863 | -1 | unprocessed_pseudc | ENSG0000C |
| 15032227 | 15073852 | 1 | protein_coding | ENSG0000C |
| 15075475 | 15088776 | -1 | protein_coding | ENSG0000C |
| 15093263 | 15093611 | -1 | processed_pseudoge | ENSG0000C |
| 15095385 | 15097319 | -1 | lncRNA | ENSG0000C |
| 15097180 | 15139818 | 1 | protein_coding | ENSG0000C |
| 15102584 | 15168693 | -1 | protein_coding | ENSG0000C |
| 15154802 | 15155320 | -1 | processed_pseudoge | ENSG0000C |
| 15171036 | 15171278 | -1 | processed_pseudoge | ENSG0000C |
| 15211643 | 15371289 | -1 | protein_coding | ENSG0000C |
| 15237492 | 15241767 | 1 | lncRNA | ENSG0000C |
| 15513954 | 15719922 | -1 | protein_coding | ENSG0000C |
| 15778170 | 15860507 | -1 | protein_coding | ENSG0000C |
| 16005808 | 16005923 | 1 | snoRNA | ENSG0000C |
| 16071416 | 16071809 | -1 | processed_pseudoge | ENSG0000C |
| 16207821 | 16207927 | -1 | snRNA | ENSG0000C |
| 16276871 | 16295858 | 1 | lncRNA | ENSG0000C |
| 16436943 | 16513745 | 1 | protein_coding | ENSG0000C |
| 16476003 | 16476164 | 1 | snRNA | ENSG0000C |
| 16513734 | 16521879 | -1 | protein_coding | ENSG0000C |
| 16521714 | 16539737 | 1 | lncRNA | ENSG0000C |
| 16590611 | 16817463 | -1 | protein_coding | ENSG0000C |
| 16721352 | 16748377 | -1 | lncRNA | ENSG0000C |
| 16767235 | 16767416 | 1 | misc_RNA | ENSG0000C |
| 16817699 | 16821207 | 1 | lncRNA | ENSG0000C |
| 16823966 | 17129817 | -1 | protein_coding | ENSG0000C |
| 17137336 | 17137585 | -1 | lncRNA | ENSG0000C |
| 17137336 | 17202054 | -1 | protein_coding | ENSG0000C |
| 17214239 | 17229985 | -1 | lncRNA | ENSG0000C |
| 17228241 | 17237593 | 1 | protein_coding | ENSG0000C |
| 17233325 | 17234833 | -1 | lncRNA | ENSG0000C |
| 17315421 | 17454595 | -1 | protein_coding | ENSG0000C |
| 17386919 | 17445758 | 1 | lncRNA | ENSG0000C |
| 17403508 | 17403620 | 1 | misc_RNA | ENSG0000C |
| 17577362 | 17578212 | 1 | processed_pseudoge | ENSG0000C |
| 17589032 | 17617374 | -1 | protein_coding | ENSG0000C |
| 17602910 | 17603127 | 1 | pseudogene | ENSG0000C |
| 17641284 | 17643878 | -1 | lncRNA | ENSG0000C |
| 17644151 | 17716824 | 1 | protein_coding | ENSG0000C |
| 17695709 | 17700232 | -1 | lncRNA | ENSG0000C |
| 17752201 | 17800868 | 1 | protein_coding | ENSG0000C |
| 17809348 | 17911164 | 1 | protein_coding | ENSG0000C |
| 17845107 | 17845193 | 1 | miRNA | ENSG0000C |
| 17862887 | 17865471 | -1 | lncRNA | ENSG0000C |
| 17951839 | 18043292 | 1 | protein_coding | ENSG0000C |
| 18001786 | 18010562 | -1 | lncRNA | ENSG0000C |

| | | | | |
|---|---|---|---|---|
| 18140677 | 18543557 | 1 | protein_coding | ENSG00000 |
| 18206127 | 18261504 | -1 | lncRNA | ENSG00000 |
| 18513115 | 18545651 | -1 | lncRNA | ENSG00000 |
| 18531849 | 18533336 | -1 | lncRNA | ENSG00000 |
| 18545561 | 18659285 | -1 | protein_coding | ENSG00000 |
| 18659431 | 18681639 | 1 | protein_coding | ENSG00000 |
| 18710269 | 18748342 | 1 | lncRNA | ENSG00000 |
| 18735745 | 18738631 | 1 | processed_pseudoge | ENSG00000 |
| 18940501 | 18940953 | 1 | processed_pseudoge | ENSG00000 |
| 19048801 | 19790401 | 1 | protein_coding | ENSG00000 |
| 19051455 | 19052170 | 1 | processed_pseudoge | ENSG00000 |
| 19290223 | 19291480 | -1 | lncRNA | ENSG00000 |
| 19489101 | 19489400 | 1 | processed_pseudoge | ENSG00000 |
| 19491072 | 19491299 | -1 | processed_pseudoge | ENSG00000 |
| 19598247 | 19598463 | 1 | snoRNA | ENSG00000 |
| 19633683 | 19633812 | -1 | rRNA_pseudogene | ENSG00000 |
| 19710328 | 19728550 | -1 | lncRNA | ENSG00000 |
| 19747172 | 19747435 | -1 | processed_pseudoge | ENSG00000 |
| 19747823 | 19748621 | 1 | processed_pseudoge | ENSG00000 |
| 19792998 | 19793101 | -1 | snRNA | ENSG00000 |
| 19812372 | 19816278 | -1 | lncRNA | ENSG00000 |
| 19816239 | 20289856 | 1 | protein_coding | ENSG00000 |
| 20070805 | 20091784 | -1 | lncRNA | ENSG00000 |
| 20095125 | 20099664 | -1 | lncRNA | ENSG00000 |
| 20350049 | 20351100 | 1 | processed_pseudoge | ENSG00000 |
| 20547171 | 20547422 | 1 | processed_pseudoge | ENSG00000 |
| 20551970 | 20552046 | 1 | miRNA | ENSG00000 |
| 20779973 | 21174187 | -1 | protein_coding | ENSG00000 |
| 20804293 | 20804577 | 1 | processed_pseudoge | ENSG00000 |
| 20841375 | 20841560 | 1 | processed_pseudoge | ENSG00000 |
| 21028999 | 21030792 | 1 | processed_pseudoge | ENSG00000 |
| 21110682 | 21113388 | 1 | processed_pseudoge | ENSG00000 |
| 21125763 | 21146559 | -1 | protein_coding | ENSG00000 |
| 21174014 | 21175048 | 1 | lncRNA | ENSG00000 |
| 21217891 | 21218279 | 1 | processed_pseudoge | ENSG00000 |
| 21247752 | 21248095 | 1 | processed_pseudoge | ENSG00000 |
| 21321961 | 21322067 | 1 | snRNA | ENSG00000 |
| 21340233 | 21372950 | -1 | lncRNA | ENSG00000 |
| 21394033 | 21394592 | 1 | processed_pseudoge | ENSG00000 |
| 21452197 | 21452412 | 1 | unprocessed_pseudc | ENSG00000 |
| 21455236 | 21455515 | -1 | misc_RNA | ENSG00000 |
| 21481716 | 21481817 | 1 | misc_RNA | ENSG00000 |
| 21492658 | 21497260 | -1 | lncRNA | ENSG00000 |
| 21496562 | 21496641 | -1 | miRNA | ENSG00000 |
| 21513475 | 21526368 | -1 | protein_coding | ENSG00000 |
| 21524646 | 21743630 | 1 | protein_coding | ENSG00000 |
| 21526568 | 21526863 | 1 | lncRNA | ENSG00000 |
| 21564471 | 21564577 | -1 | snRNA | ENSG00000 |
| 21624592 | 21626286 | -1 | processed_pseudoge | ENSG00000 |
| 21661635 | 21661741 | -1 | snRNA | ENSG00000 |

| | | | | |
|---|---|---|---|---|
| 21756548 | 22003769 | -1 | protein_coding | ENSG0000( |
| 21785292 | 21785587 | 1 | misc_RNA | ENSG0000( |
| 21865335 | 21865921 | -1 | lncRNA | ENSG0000( |
| 21933143 | 21933438 | 1 | misc_RNA | ENSG0000( |
| 22162992 | 22165794 | 1 | processed_pseudoge | ENSG0000( |
| 22163092 | 22164306 | 1 | processed_pseudoge | ENSG0000( |
| 22208814 | 22210021 | -1 | protein_coding | ENSG0000( |
| 22218074 | 22221168 | 1 | lncRNA | ENSG0000( |
| 22225698 | 22226024 | 1 | processed_pseudoge | ENSG0000( |
| 22230826 | 22232973 | -1 | lncRNA | ENSG0000( |
| 22252072 | 22254589 | 1 | TEC | ENSG0000( |
| 22257786 | 22258548 | 1 | lncRNA | ENSG0000( |
| 22266991 | 22267346 | 1 | processed_pseudoge | ENSG0000( |
| 22316386 | 22320306 | 1 | protein_coding | ENSG0000( |
| 22316388 | 22329542 | 1 | protein_coding | ENSG0000( |
| 22321099 | 22331484 | 1 | protein_coding | ENSG0000( |
| 22332404 | 22332987 | 1 | lncRNA | ENSG0000( |
| 22332587 | 22332981 | -1 | lncRNA | ENSG0000( |
| 22340377 | 22340615 | -1 | lncRNA | ENSG0000( |
| 22345445 | 22454224 | 1 | protein_coding | ENSG0000( |
| 22361172 | 22413067 | 1 | lncRNA | ENSG0000( |
| 22439954 | 22479480 | -1 | lncRNA | ENSG0000( |
| 22475455 | 22478026 | -1 | protein_coding | ENSG0000( |
| 22534854 | 22714578 | -1 | protein_coding | ENSG0000( |
| 22693953 | 22694639 | -1 | TEC | ENSG0000( |
| 22787393 | 22787499 | 1 | snRNA | ENSG0000( |
| 22928024 | 23038523 | 1 | protein_coding | ENSG0000( |
| 22964926 | 23133804 | -1 | lncRNA | ENSG0000( |
| 22997255 | 22997358 | 1 | rRNA_pseudogene | ENSG0000( |
| 23095579 | 23122013 | 1 | protein_coding | ENSG0000( |
| 23136924 | 23137661 | 1 | processed_pseudoge | ENSG0000( |
| 23166985 | 23167101 | 1 | snoRNA | ENSG0000( |
| 23192312 | 23194245 | 1 | protein_coding | ENSG0000( |
| 23201916 | 23344845 | -1 | protein_coding | ENSG0000( |
| 23258211 | 23258545 | 1 | processed_pseudoge | ENSG0000( |
| 23343957 | 23345181 | 1 | lncRNA | ENSG0000( |
| 23393405 | 23393467 | 1 | miRNA | ENSG0000( |
| 23439458 | 23442390 | 1 | protein_coding | ENSG0000( |
| 23443036 | 23529801 | 1 | lncRNA | ENSG0000( |
| 23575012 | 23586224 | 1 | lncRNA | ENSG0000( |
| 23694746 | 24547848 | 1 | protein_coding | ENSG0000( |
| 24157835 | 24158781 | 1 | processed_pseudoge | ENSG0000( |
| 24255246 | 24255333 | 1 | misc_RNA | ENSG0000( |
| 24275685 | 24275781 | 1 | miRNA | ENSG0000( |
| 24410183 | 24416459 | -1 | lncRNA | ENSG0000( |
| 24481180 | 24483034 | -1 | TEC | ENSG0000( |
| 24583609 | 24723887 | -1 | protein_coding | ENSG0000( |
| 24640166 | 24640284 | 1 | rRNA_pseudogene | ENSG0000( |
| 24816331 | 24822950 | 1 | lncRNA | ENSG0000( |
| 24848614 | 24952606 | -1 | protein_coding | ENSG0000( |

| | | | | |
|---|---|---|---|---|
| 24867645 | 24868190 | 1 | processed_pseudoge | ENSG00000 |
| 24908522 | 24908812 | -1 | misc_RNA | ENSG00000 |
| 24952350 | 25161236 | 1 | lncRNA | ENSG00000 |
| 24953241 | 24953513 | 1 | lncRNA | ENSG00000 |
| 24981979 | 25062279 | -1 | protein_coding | ENSG00000 |
| 25016612 | 25026664 | 1 | protein_coding | ENSG00000 |
| 25113058 | 25161236 | 1 | lncRNA | ENSG00000 |
| 25158072 | 25176276 | -1 | lncRNA | ENSG00000 |
| 25174802 | 25602229 | 1 | protein_coding | ENSG00000 |
| 25244061 | 25244854 | -1 | unprocessed_pseudo | ENSG00000 |
| 25393557 | 25393844 | -1 | misc_RNA | ENSG00000 |
| 25647570 | 25648104 | -1 | processed_pseudoge | ENSG00000 |
| 25651712 | 25732935 | 1 | lncRNA | ENSG00000 |
| 25691537 | 25692323 | 1 | processed_pseudoge | ENSG00000 |
| 25813811 | 25813917 | -1 | rRNA_pseudogene | ENSG00000 |
| 25819363 | 25819426 | -1 | snRNA | ENSG00000 |
| 25924448 | 25933710 | -1 | lncRNA | ENSG00000 |
| 25934229 | 26212532 | 1 | protein_coding | ENSG00000 |
| 26216665 | 26304558 | 1 | protein_coding | ENSG00000 |
| 26379034 | 26387374 | -1 | unitary_pseudogene | ENSG00000 |
| 26438341 | 26567803 | 1 | protein_coding | ENSG00000 |
| 26509589 | 26509704 | 1 | rRNA_pseudogene | ENSG00000 |
| 26577676 | 26622338 | -1 | transcribed_unproce | ENSG00000 |
| 26589865 | 26594330 | 1 | transcribed_unproce | ENSG00000 |
| 26643228 | 26645016 | 1 | lncRNA | ENSG00000 |
| 26647986 | 26653454 | 1 | transcribed_unproce | ENSG00000 |
| 26664206 | 26697295 | -1 | transcribed_unitary_ | ENSG00000 |
| 26695091 | 26697625 | -1 | TEC | ENSG00000 |
| 26697701 | 26746798 | 1 | protein_coding | ENSG00000 |
| 26714622 | 26715832 | 1 | processed_pseudoge | ENSG00000 |
| 26717635 | 26718778 | 1 | lncRNA | ENSG00000 |
| 26746593 | 26861087 | -1 | protein_coding | ENSG00000 |
| 26789017 | 26789157 | 1 | snoRNA | ENSG00000 |
| 26790540 | 26790646 | -1 | snRNA | ENSG00000 |
| 26931206 | 26944418 | -1 | transcribed_unproce | ENSG00000 |
| 26991914 | 27100498 | -1 | protein_coding | ENSG00000 |
| 27088321 | 27088427 | -1 | snRNA | ENSG00000 |
| 27110112 | 27155266 | -1 | protein_coding | ENSG00000 |
| 27154824 | 27186924 | 1 | protein_coding | ENSG00000 |
| 27193358 | 27193500 | -1 | snRNA | ENSG00000 |
| 27195214 | 27242130 | -1 | protein_coding | ENSG00000 |
| 27232255 | 27232316 | -1 | snRNA | ENSG00000 |
| 27243130 | 27250804 | 1 | lncRNA | ENSG00000 |
| 27246350 | 27259455 | -1 | transcribed_processe | ENSG00000 |
| 27258762 | 27288824 | 1 | transcribed_unproce | ENSG00000 |
| 27313723 | 27316587 | 1 | unprocessed_pseudo | ENSG00000 |
| 27315120 | 27315226 | 1 | snRNA | ENSG00000 |
| 27331357 | 27341252 | -1 | unprocessed_pseudo | ENSG00000 |
| 27335501 | 27335607 | -1 | snRNA | ENSG00000 |
| 27343436 | 27344917 | 1 | lncRNA | ENSG00000 |

| | | | | |
|---|---|---|---|---|
| 27349717 | 27350290 | -1 | processed_pseudoge | ENSG0000C |
| 27378138 | 27378362 | -1 | processed_pseudoge | ENSG0000C |
| 27397121 | 27414368 | -1 | polymorphic_pseudc | ENSG0000C |
| 27504174 | 27542237 | 1 | protein_coding | ENSG0000C |
| 27564430 | 27594588 | 1 | lncRNA | ENSG0000C |
| 27672874 | 27746060 | -1 | protein_coding | ENSG0000C |
| 27744786 | 27767794 | 1 | lncRNA | ENSG0000C |
| 27775186 | 27999079 | -1 | protein_coding | ENSG0000C |
| 27783846 | 27784053 | -1 | misc_RNA | ENSG0000C |
| 27904209 | 27905162 | 1 | processed_pseudoge | ENSG0000C |
| 27999788 | 28049615 | 1 | lncRNA | ENSG0000C |
| 28004702 | 28004932 | 1 | processed_pseudoge | ENSG0000C |
| 28050993 | 28334486 | -1 | protein_coding | ENSG0000C |
| 28263703 | 28265175 | 1 | lncRNA | ENSG0000C |
| 28289258 | 28289350 | -1 | miRNA | ENSG0000C |
| 28303300 | 28307658 | 1 | lncRNA | ENSG0000C |
| 28338554 | 28340942 | -1 | processed_pseudoge | ENSG0000C |
| 28408039 | 28408251 | -1 | snoRNA | ENSG0000C |
| 28433008 | 28495813 | -1 | lncRNA | ENSG0000C |
| 28436201 | 28437078 | 1 | processed_pseudoge | ENSG0000C |
| 28442575 | 28442896 | -1 | misc_RNA | ENSG0000C |
| 28522652 | 28533066 | -1 | lncRNA | ENSG0000C |
| 28532493 | 28623112 | 1 | protein_coding | ENSG0000C |
| 28652630 | 28652755 | 1 | snRNA | ENSG0000C |
| 28658987 | 28659467 | 1 | processed_pseudoge | ENSG0000C |
| 28667919 | 28668025 | 1 | snRNA | ENSG0000C |
| 28677510 | 28682932 | 1 | protein_coding | ENSG0000C |
| 28743506 | 28941910 | 1 | lncRNA | ENSG0000C |
| 28743745 | 28744663 | 1 | lncRNA | ENSG0000C |
| 28760699 | 28760803 | -1 | snRNA | ENSG0000C |
| 28789188 | 28796113 | -1 | lncRNA | ENSG0000C |
| 28812899 | 28813139 | 1 | transcribed_processe | ENSG0000C |
| 28874393 | 28874507 | -1 | rRNA_pseudogene | ENSG0000C |
| 28898996 | 28899484 | -1 | processed_pseudoge | ENSG0000C |
| 29289061 | 29318328 | 1 | protein_coding | ENSG0000C |
| 29373027 | 29409366 | -1 | lncRNA | ENSG0000C |
| 29409402 | 29487745 | 1 | transcribed_unproce | ENSG0000C |
| 29421476 | 29422012 | -1 | unprocessed_pseudc | ENSG0000C |
| 29457338 | 29736781 | -1 | protein_coding | ENSG0000C |
| 29545004 | 29545097 | -1 | miRNA | ENSG0000C |
| 29575306 | 29575381 | 1 | snoRNA | ENSG0000C |
| 29602264 | 29602346 | -1 | miRNA | ENSG0000C |
| 29697935 | 29698174 | 1 | processed_pseudoge | ENSG0000C |
| 29711915 | 29712008 | 1 | snRNA | ENSG0000C |
| 29891032 | 29891467 | -1 | processed_pseudoge | ENSG0000C |
| 30012803 | 30115494 | -1 | protein_coding | ENSG0000C |
| 30299569 | 30299673 | -1 | snRNA | ENSG0000C |
| 30301823 | 30302072 | -1 | processed_pseudoge | ENSG0000C |
| 30302826 | 30306066 | -1 | lncRNA | ENSG0000C |
| 30309801 | 30374448 | -1 | protein_coding | ENSG0000C |

| | | | | |
|---|---|---|---|---|
| 30309803 | 30312026 | 1 | lncRNA | ENSG0000C |
| 30353706 | 30357365 | 1 | unprocessed_pseudc | ENSG0000C |
| 30364410 | 30374386 | -1 | unprocessed_pseudc | ENSG0000C |
| 30366064 | 30366299 | -1 | misc_RNA | ENSG0000C |
| 30368597 | 30368665 | -1 | miRNA | ENSG0000C |
| 30380174 | 30381062 | 1 | processed_pseudoge | ENSG0000C |
| 30383400 | 30383622 | 1 | processed_pseudoge | ENSG0000C |
| 30403197 | 30403776 | -1 | processed_pseudoge | ENSG0000C |
| 30434021 | 30461833 | 1 | protein_coding | ENSG0000C |
| 30517616 | 30518861 | -1 | processed_pseudoge | ENSG0000C |
| 30529517 | 30554625 | 1 | lncRNA | ENSG0000C |
| 30553854 | 30554483 | -1 | lncRNA | ENSG0000C |
| 30555601 | 30555904 | -1 | misc_RNA | ENSG0000C |
| 30576699 | 30580182 | 1 | lncRNA | ENSG0000C |
| 30587706 | 30591092 | 1 | lncRNA | ENSG0000C |
| 30611779 | 30629762 | -1 | protein_coding | ENSG0000C |
| 30655489 | 30717266 | 1 | transcribed_unproce | ENSG0000C |
| 30723251 | 30723740 | 1 | lncRNA | ENSG0000C |
| 30820207 | 31031937 | -1 | protein_coding | ENSG0000C |
| 30831828 | 30833387 | -1 | lncRNA | ENSG0000C |
| 30919012 | 30921235 | -1 | processed_pseudoge | ENSG0000C |
| 31033089 | 31143813 | 1 | lncRNA | ENSG0000C |
| 31187715 | 31261910 | 1 | lncRNA | ENSG0000C |
| 31206278 | 31320447 | -1 | lncRNA | ENSG0000C |
| 31259599 | 31259704 | 1 | rRNA_pseudogene | ENSG0000C |
| 31318495 | 31529814 | 1 | protein_coding | ENSG0000C |
| 31360955 | 31361215 | 1 | transcribed_processe | ENSG0000C |
| 31362799 | 31364049 | -1 | unprocessed_pseudc | ENSG0000C |
| 31602862 | 31606218 | 1 | lncRNA | ENSG0000C |
| 31608500 | 31620862 | -1 | lncRNA | ENSG0000C |
| 31628540 | 31628599 | -1 | unprocessed_pseudc | ENSG0000C |
| 31693084 | 31707388 | -1 | lncRNA | ENSG0000C |
| 31756042 | 31756136 | -1 | misc_RNA | ENSG0000C |
| 31805398 | 31928876 | -1 | protein_coding | ENSG0000C |
| 31913143 | 31913754 | 1 | processed_pseudoge | ENSG0000C |
| 31983885 | 31984554 | 1 | processed_pseudoge | ENSG0000C |
| 31992087 | 31992381 | -1 | misc_RNA | ENSG0000C |
| 32009015 | 32056425 | -1 | protein_coding | ENSG0000C |
| 32056746 | 32056839 | 1 | misc_RNA | ENSG0000C |
| 32102522 | 32103293 | -1 | processed_pseudoge | ENSG0000C |
| 32109861 | 32111666 | 1 | processed_pseudoge | ENSG0000C |
| 32111644 | 32111789 | -1 | processed_pseudoge | ENSG0000C |
| 32173684 | 32173744 | 1 | snRNA | ENSG0000C |
| 32188323 | 32188817 | -1 | processed_pseudoge | ENSG0000C |
| 32231849 | 32232245 | -1 | processed_pseudoge | ENSG0000C |
| 32266289 | 32269474 | 1 | lncRNA | ENSG0000C |
| 32267751 | 32378798 | -1 | protein_coding | ENSG0000C |
| 32281686 | 32282829 | 1 | lncRNA | ENSG0000C |
| 32346499 | 32347179 | 1 | lncRNA | ENSG0000C |
| 32347397 | 32374488 | 1 | lncRNA | ENSG0000C |

| | | | | |
|---|---|---|---|---|
| 32394558 | 32394665 | -1 | snRNA | ENSG0000( |
| 32413499 | 32446061 | -1 | lncRNA | ENSG0000( |
| 32446140 | 32882874 | 1 | protein_coding | ENSG0000( |
| 32511336 | 32511737 | -1 | processed_pseudoge | ENSG0000( |
| 32887255 | 32889311 | -1 | lncRNA | ENSG0000( |
| 32900318 | 33005792 | -1 | protein_coding | ENSG0000( |
| 32944481 | 32944999 | 1 | processed_pseudoge | ENSG0000( |
| 32958437 | 33082102 | 1 | lncRNA | ENSG0000( |
| 33023368 | 33023647 | -1 | processed_pseudoge | ENSG0000( |
| 33034655 | 33034924 | -1 | misc_RNA | ENSG0000( |
| 33035469 | 33035664 | -1 | processed_pseudoge | ENSG0000( |
| 33057615 | 33058407 | -1 | processed_pseudoge | ENSG0000( |
| 33096257 | 33116672 | -1 | lncRNA | ENSG0000( |
| 33177492 | 33336262 | -1 | protein_coding | ENSG0000( |
| 33211277 | 33213804 | 1 | lncRNA | ENSG0000( |
| 33288304 | 33288601 | -1 | misc_RNA | ENSG0000( |
| 33341655 | 33341905 | 1 | lncRNA | ENSG0000( |
| 33384891 | 33390935 | 1 | lncRNA | ENSG0000( |
| 33578931 | 33600040 | -1 | lncRNA | ENSG0000( |
| 33684755 | 33709868 | 1 | lncRNA | ENSG0000( |
| 33759713 | 33772702 | -1 | lncRNA | ENSG0000( |
| 33864313 | 33864723 | 1 | processed_pseudoge | ENSG0000( |
| 33909238 | 33917804 | 1 | lncRNA | ENSG0000( |
| 34109560 | 34815325 | -1 | protein_coding | ENSG0000( |
| 34201979 | 34202074 | -1 | misc_RNA | ENSG0000( |
| 34488583 | 34488878 | -1 | processed_pseudoge | ENSG0000( |
| 34663859 | 34664101 | -1 | processed_pseudoge | ENSG0000( |
| 34675635 | 34676041 | -1 | processed_pseudoge | ENSG0000( |
| 34815767 | 34816386 | 1 | lncRNA | ENSG0000( |
| 34875210 | 34875422 | 1 | processed_pseudoge | ENSG0000( |
| 34926775 | 34926882 | 1 | snRNA | ENSG0000( |
| 34942969 | 34943075 | 1 | snRNA | ENSG0000( |
| 34965069 | 34966259 | 1 | processed_pseudoge | ENSG0000( |
| 34969881 | 34975578 | 1 | lncRNA | ENSG0000( |
| 35008551 | 35090642 | -1 | protein_coding | ENSG0000( |
| 35079598 | 35079680 | -1 | miRNA | ENSG0000( |
| 35098006 | 35127020 | -1 | lncRNA | ENSG0000( |
| 35126791 | 35212958 | 1 | protein_coding | ENSG0000( |
| 35135443 | 35135502 | 1 | snRNA | ENSG0000( |
| 35158306 | 35158640 | 1 | processed_pseudoge | ENSG0000( |
| 35210416 | 35210750 | 1 | lncRNA | ENSG0000( |
| 35219894 | 35230598 | -1 | lncRNA | ENSG0000( |
| 35231331 | 35231394 | -1 | snRNA | ENSG0000( |
| 35247025 | 35572669 | 1 | protein_coding | ENSG0000( |
| 35250959 | 35251052 | -1 | snRNA | ENSG0000( |
| 35314552 | 35336401 | -1 | lncRNA | ENSG0000( |
| 35443411 | 35444569 | -1 | lncRNA | ENSG0000( |
| 35604485 | 35608153 | -1 | lncRNA | ENSG0000( |
| 35605341 | 35608935 | 1 | protein_coding | ENSG0000( |
| 35638249 | 35642278 | -1 | protein_coding | ENSG0000( |

| | | | | |
|---|---|---|---|---|
| 35641097 | 35647826 | 1 | lncRNA | ENSG0000C |
| 35641172 | 35641252 | -1 | miRNA | ENSG0000C |
| 35697300 | 35698037 | 1 | processed_pseudoge | ENSG0000C |
| 35778302 | 35800920 | 1 | lncRNA | ENSG0000C |
| 35896874 | 35898963 | -1 | lncRNA | ENSG0000C |
| 36089086 | 36089389 | -1 | lncRNA | ENSG0000C |
| 36432882 | 36434484 | -1 | processed_pseudoge | ENSG0000C |
| 36434710 | 36435174 | -1 | processed_pseudoge | ENSG0000C |
| 36438616 | 36442392 | 1 | lncRNA | ENSG0000C |
| 36438661 | 36438788 | 1 | snoRNA | ENSG0000C |
| 36521721 | 36524234 | -1 | processed_pseudoge | ENSG0000C |
| 36626015 | 36626138 | -1 | misc_RNA | ENSG0000C |
| 36940109 | 36943932 | 1 | lncRNA | ENSG0000C |
| 36944111 | 36944738 | -1 | processed_pseudoge | ENSG0000C |
| 36995023 | 36995585 | 1 | processed_pseudoge | ENSG0000C |
| 37125725 | 37384111 | 1 | protein_coding | ENSG0000C |
| 37135237 | 37135343 | 1 | snRNA | ENSG0000C |
| 37240887 | 37242049 | 1 | lncRNA | ENSG0000C |
| 37248118 | 37347031 | 1 | transcribed_unproce | ENSG0000C |
| 37297398 | 37297689 | 1 | misc_RNA | ENSG0000C |
| 37303000 | 37303109 | -1 | misc_RNA | ENSG0000C |
| 37337415 | 37338871 | 1 | processed_pseudoge | ENSG0000C |
| 37348616 | 37349572 | 1 | processed_pseudoge | ENSG0000C |
| 37523011 | 37544701 | 1 | unprocessed_pseudc | ENSG0000C |
| 37600796 | 37601368 | -1 | processed_pseudoge | ENSG0000C |
| 37601440 | 37602974 | -1 | protein_coding | ENSG0000C |
| 37775371 | 37784131 | -1 | lncRNA | ENSG0000C |
| 37776526 | 37858106 | -1 | protein_coding | ENSG0000C |
| 37791580 | 37791937 | 1 | lncRNA | ENSG0000C |
| 37794276 | 37796109 | -1 | processed_pseudoge | ENSG0000C |
| 37818606 | 37820982 | -1 | processed_pseudoge | ENSG0000C |
| 37857740 | 37859110 | 1 | lncRNA | ENSG0000C |
| 37883594 | 37884109 | 1 | unprocessed_pseudc | ENSG0000C |
| 37894637 | 37896666 | 1 | processed_pseudoge | ENSG0000C |
| 37949573 | 37976647 | -1 | protein_coding | ENSG0000C |
| 37976825 | 37992467 | 1 | lncRNA | ENSG0000C |
| 37994863 | 37994975 | 1 | misc_RNA | ENSG0000C |
| 38010650 | 38065088 | 1 | protein_coding | ENSG0000C |
| 38043873 | 38043975 | -1 | snRNA | ENSG0000C |
| 38068449 | 38068694 | -1 | unprocessed_pseudc | ENSG0000C |
| 38080056 | 38080877 | -1 | processed_pseudoge | ENSG0000C |
| 38091764 | 38092051 | -1 | processed_pseudoge | ENSG0000C |
| 38094334 | 38150293 | 1 | protein_coding | ENSG0000C |
| 38094368 | 38360098 | 1 | protein_coding | ENSG0000C |
| 38137337 | 38144399 | 1 | lncRNA | ENSG0000C |
| 38161570 | 38161749 | 1 | processed_pseudoge | ENSG0000C |
| 38164648 | 38212417 | 1 | transcribed_unproce | ENSG0000C |
| 38247922 | 38333155 | 1 | transcribed_unproce | ENSG0000C |
| 38250442 | 38250603 | -1 | processed_pseudoge | ENSG0000C |
| 38356380 | 38378505 | 1 | transcribed_unproce | ENSG0000C |

| | | | | |
|---|---|---|---|---|
| 38383069 | 38402916 | -1 | transcribed_unproce | ENSG00000 |
| 38403189 | 38452153 | 1 | transcribed_unproce | ENSG00000 |
| 38423625 | 38423728 | 1 | snRNA | ENSG00000 |
| 38452987 | 38455365 | -1 | processed_pseudoge | ENSG00000 |
| 38453181 | 38466176 | 1 | lncRNA | ENSG00000 |
| 38601418 | 38605609 | 1 | unprocessed_pseudc | ENSG00000 |
| 38617255 | 38617344 | 1 | processed_pseudoge | ENSG00000 |
| 38635908 | 38638082 | 1 | processed_pseudoge | ENSG00000 |
| 38640971 | 38689068 | -1 | unprocessed_pseudc | ENSG00000 |
| 38696424 | 38697902 | 1 | processed_pseudoge | ENSG00000 |
| 38713195 | 38719229 | 1 | unprocessed_pseudc | ENSG00000 |
| 38762095 | 38762491 | -1 | processed_pseudoge | ENSG00000 |
| 38783004 | 38783108 | 1 | processed_pseudoge | ENSG00000 |
| 42149310 | 42149549 | -1 | processed_pseudoge | ENSG00000 |
| 42185339 | 42185791 | 1 | IG_V_pseudogene | ENSG00000 |
| 42209990 | 42210383 | 1 | processed_pseudoge | ENSG00000 |
| 42242721 | 42276842 | 1 | transcribed_unproce | ENSG00000 |
| 42279734 | 42281819 | -1 | processed_pseudoge | ENSG00000 |
| 42331866 | 42367974 | -1 | transcribed_unproce | ENSG00000 |
| 42408168 | 42475349 | -1 | transcribed_unproce | ENSG00000 |
| 42470082 | 42470268 | -1 | transcribed_unproce | ENSG00000 |
| 42475480 | 42495337 | 1 | lncRNA | ENSG00000 |
| 42513510 | 42552822 | -1 | transcribed_processe | ENSG00000 |
| 42555440 | 42555724 | 1 | processed_pseudoge | ENSG00000 |
| 42566677 | 42567483 | 1 | processed_pseudoge | ENSG00000 |
| 42574185 | 42638570 | -1 | protein_coding | ENSG00000 |
| 42578500 | 42578727 | 1 | unprocessed_pseudc | ENSG00000 |
| 42579724 | 42579858 | 1 | unprocessed_pseudc | ENSG00000 |
| 42583241 | 42583375 | 1 | unprocessed_pseudc | ENSG00000 |
| 42605158 | 42605263 | 1 | snRNA | ENSG00000 |
| 42638314 | 42645542 | 1 | lncRNA | ENSG00000 |
| 42644445 | 42691723 | -1 | lncRNA | ENSG00000 |
| 42660728 | 42661776 | 1 | processed_pseudoge | ENSG00000 |
| 42694371 | 42714575 | -1 | unprocessed_pseudc | ENSG00000 |
| 42696012 | 42706453 | 1 | lncRNA | ENSG00000 |
| 42725857 | 42737159 | 1 | unprocessed_pseudc | ENSG00000 |
| 42747544 | 42748687 | -1 | unprocessed_pseudc | ENSG00000 |
| 42751178 | 42754297 | -1 | lncRNA | ENSG00000 |
| 42782795 | 42834937 | 1 | protein_coding | ENSG00000 |
| 42832447 | 42832550 | -1 | snRNA | ENSG00000 |
| 42871521 | 42874060 | 1 | lncRNA | ENSG00000 |
| 42979017 | 42981507 | -1 | lncRNA | ENSG00000 |
| 42997563 | 42997681 | 1 | miRNA | ENSG00000 |
| 43077064 | 43130351 | 1 | protein_coding | ENSG00000 |
| 43136824 | 43138334 | -1 | lncRNA | ENSG00000 |
| 43138445 | 43185302 | 1 | protein_coding | ENSG00000 |
| 43194535 | 43266919 | -1 | protein_coding | ENSG00000 |
| 43323665 | 43327932 | 1 | lncRNA | ENSG00000 |
| 43325833 | 43350792 | -1 | lncRNA | ENSG00000 |
| 43341786 | 43341910 | 1 | snRNA | ENSG00000 |

| | | | | |
|---|---|---|---|---|
| 43371636 | 43376335 | 1 | protein_coding | ENSG00000 |
| 43385617 | 43409166 | -1 | protein_coding | ENSG00000 |
| 43417249 | 43417438 | -1 | snoRNA | ENSG00000 |
| 43420738 | 43422100 | 1 | lncRNA | ENSG00000 |
| 43436841 | 43483181 | 1 | protein_coding | ENSG00000 |
| 43458065 | 43458129 | 1 | snRNA | ENSG00000 |
| 43523256 | 43525217 | 1 | unprocessed_pseudc | ENSG00000 |
| 43556344 | 43574618 | -1 | protein_coding | ENSG00000 |
| 43583498 | 43596625 | -1 | lncRNA | ENSG00000 |
| 43604843 | 43606251 | -1 | processed_pseudoge | ENSG00000 |
| 43606419 | 43617904 | 1 | protein_coding | ENSG00000 |
| 43628817 | 43674703 | 1 | lncRNA | ENSG00000 |
| 43630882 | 43631564 | -1 | lncRNA | ENSG00000 |
| 43630947 | 43631382 | 1 | processed_pseudoge | ENSG00000 |
| 43643860 | 43648881 | -1 | protein_coding | ENSG00000 |
| 43643872 | 43645047 | 1 | lncRNA | ENSG00000 |
| 43645942 | 43648019 | 1 | lncRNA | ENSG00000 |
| 43665668 | 43667005 | 1 | lncRNA | ENSG00000 |
| 43728217 | 43728728 | 1 | processed_pseudoge | ENSG00000 |
| 43732334 | 43732584 | 1 | processed_pseudoge | ENSG00000 |
| 43741437 | 43742350 | -1 | processed_pseudoge | ENSG00000 |
| 43777562 | 43778825 | -1 | lncRNA | ENSG00000 |
| 43778946 | 43901004 | 1 | lncRNA | ENSG00000 |
| 43789249 | 43790387 | -1 | processed_pseudoge | ENSG00000 |
| 43900874 | 43912369 | -1 | lncRNA | ENSG00000 |
| 43909244 | 43981102 | 1 | lncRNA | ENSG00000 |
| 43912444 | 43914236 | 1 | lncRNA | ENSG00000 |
| 44259602 | 44261813 | -1 | lncRNA | ENSG00000 |
| 44282489 | 44293998 | -1 | lncRNA | ENSG00000 |
| 44292569 | 44294649 | 1 | protein_coding | ENSG00000 |
| 44296805 | 44310237 | -1 | lncRNA | ENSG00000 |
| 44370165 | 44386493 | -1 | protein_coding | ENSG00000 |
| 44414649 | 44415153 | -1 | processed_pseudoge | ENSG00000 |
| 44488335 | 44488445 | -1 | processed_pseudoge | ENSG00000 |
| 44509708 | 44515421 | 1 | lncRNA | ENSG00000 |
| 44591208 | 44602104 | -1 | lncRNA | ENSG00000 |
| 44793119 | 44959709 | -1 | lncRNA | ENSG00000 |
| 44806443 | 44808223 | 1 | processed_pseudoge | ENSG00000 |
| 44899614 | 44920220 | -1 | lncRNA | ENSG00000 |
| 44911316 | 44937010 | 1 | protein_coding | ENSG00000 |
| 44937508 | 44955725 | 1 | lncRNA | ENSG00000 |
| 44958917 | 44959458 | -1 | lncRNA | ENSG00000 |
| 44959407 | 44995891 | 1 | protein_coding | ENSG00000 |
| 44970981 | 44978809 | -1 | protein_coding | ENSG00000 |
| 44997698 | 45000888 | -1 | lncRNA | ENSG00000 |
| 45000923 | 45005326 | 1 | protein_coding | ENSG00000 |
| 45002222 | 45076066 | -1 | transcribed_processe | ENSG00000 |
| 45016080 | 45147221 | -1 | lncRNA | ENSG00000 |
| 45055656 | 45056268 | 1 | processed_pseudoge | ENSG00000 |
| 45073146 | 45073541 | -1 | processed_pseudoge | ENSG00000 |

| | | | | |
|---|---|---|---|---|
| 45099487 | 45154606 | -1 | transcribed_unproce | ENSG00000( |
| 45127318 | 45148914 | 1 | unprocessed_pseudc | ENSG00000( |
| 45147394 | 45171389 | 1 | lncRNA | ENSG00000( |
| 45151445 | 45151915 | -1 | processed_pseudoge | ENSG00000( |
| 45156775 | 45176925 | 1 | unprocessed_pseudc | ENSG00000( |
| 45164014 | 45164088 | 1 | miRNA | ENSG00000( |
| 45164228 | 45181427 | -1 | lncRNA | ENSG00000( |
| 45180400 | 45180616 | 1 | unprocessed_pseudc | ENSG00000( |
| 45187292 | 45198323 | -1 | unprocessed_pseudc | ENSG00000( |
| 45188790 | 45188896 | -1 | snRNA | ENSG00000( |
| 45232026 | 45247361 | -1 | unprocessed_pseudc | ENSG00000( |
| 45255640 | 45263252 | 1 | transcribed_unitary_ | ENSG00000( |
| 45302298 | 45315608 | -1 | protein_coding | ENSG00000( |
| 45374176 | 45446119 | 1 | protein_coding | ENSG00000( |
| 45444570 | 45453121 | -1 | lncRNA | ENSG00000( |
| 45454585 | 45594906 | -1 | protein_coding | ENSG00000( |
| 45615500 | 45672780 | -1 | protein_coding | ENSG00000( |
| 45678692 | 45700532 | 1 | unprocessed_pseudc | ENSG00000( |
| 45706431 | 45727231 | -1 | transcribed_unproce | ENSG00000( |
| 45725694 | 45726038 | -1 | processed_pseudoge | ENSG00000( |
| 45727200 | 45792961 | 1 | protein_coding | ENSG00000( |
| 45812648 | 45812944 | -1 | processed_pseudoge | ENSG00000( |
| 45815428 | 45817324 | -1 | transcribed_unproce | ENSG00000( |
| 45825594 | 45853875 | -1 | protein_coding | ENSG00000( |
| 45854093 | 45972154 | -1 | transcribed_unproce | ENSG00000( |
| 45856307 | 45856422 | 1 | rRNA_pseudogene | ENSG00000( |
| 45940522 | 45940851 | 1 | processed_pseudoge | ENSG00000( |
| 45972489 | 46003742 | 1 | protein_coding | ENSG00000( |
| 45984610 | 45984810 | 1 | snoRNA | ENSG00000( |
| 46005088 | 46030714 | -1 | protein_coding | ENSG00000( |
| 46033307 | 46048180 | -1 | protein_coding | ENSG00000( |
| 46063250 | 46063715 | 1 | processed_pseudoge | ENSG00000( |
| 46109621 | 46131358 | 1 | unprocessed_pseudc | ENSG00000( |
| 46233885 | 46273557 | -1 | unprocessed_pseudc | ENSG00000( |
| 46286345 | 46330207 | 1 | protein_coding | ENSG00000( |
| 46337224 | 46358714 | 1 | unprocessed_pseudc | ENSG00000( |
| 46369113 | 46373563 | 1 | transcribed_unproce | ENSG00000( |
| 46370759 | 46382626 | 1 | lncRNA | ENSG00000( |
| 46375627 | 46391784 | 1 | protein_coding | ENSG00000( |
| 46375752 | 46537864 | 1 | lncRNA | ENSG00000( |
| 46398362 | 46453306 | 1 | lncRNA | ENSG00000( |
| 46415853 | 46416413 | 1 | processed_pseudoge | ENSG00000( |
| 46461099 | 46465958 | -1 | protein_coding | ENSG00000( |
| 46549044 | 46555530 | -1 | protein_coding | ENSG00000( |
| 46578217 | 46594173 | 1 | protein_coding | ENSG00000( |
| 46580708 | 46598150 | -1 | lncRNA | ENSG00000( |
| 46597918 | 46612154 | 1 | lncRNA | ENSG00000( |
| 46610474 | 46652025 | -1 | transcribed_unproce | ENSG00000( |
| 46632387 | 46635472 | 1 | lncRNA | ENSG00000( |
| 46634911 | 46635466 | -1 | processed_pseudoge | ENSG00000( |

| | | | | |
|---|---|---|---|---|
| 46648948 | 46649276 | 1 | misc_RNA | ENSG00000 |
| 46721002 | 46751738 | 1 | lncRNA | ENSG00000 |
| 46753605 | 46759740 | 1 | transcribed_unproce | ENSG00000 |
| 46759019 | 46786770 | -1 | transcribed_unproce | ENSG00000 |
| 46774421 | 46776180 | -1 | processed_pseudoge | ENSG00000 |
| 46786674 | 46811989 | 1 | transcribed_unproce | ENSG00000 |
| 46807458 | 46807573 | -1 | rRNA_pseudogene | ENSG00000 |
| 46816516 | 46832476 | 1 | unprocessed_pseudo | ENSG00000 |
| 46870858 | 46894562 | 1 | unprocessed_pseudo | ENSG00000 |
| 46911396 | 47002488 | 1 | protein_coding | ENSG00000 |
| 47300197 | 47313577 | 1 | protein_coding | ENSG00000 |
| 47322454 | 47327588 | 1 | protein_coding | ENSG00000 |
| 47348363 | 47357881 | 1 | protein_coding | ENSG00000 |
| 47365496 | 47384293 | -1 | protein_coding | ENSG00000 |
| 47460162 | 47484158 | -1 | protein_coding | ENSG00000 |
| 47484457 | 47484754 | -1 | processed_pseudoge | ENSG00000 |
| 47487219 | 47491693 | -1 | protein_coding | ENSG00000 |
| 47496212 | 47502195 | 1 | lncRNA | ENSG00000 |
| 47501854 | 47523638 | -1 | protein_coding | ENSG00000 |
| 47517816 | 47553514 | -1 | lncRNA | ENSG00000 |
| 47532352 | 47532467 | 1 | rRNA_pseudogene | ENSG00000 |
| 47538377 | 47551954 | -1 | transcribed_unproce | ENSG00000 |
| 47563604 | 47569697 | 1 | unprocessed_pseudo | ENSG00000 |
| 47564256 | 47566016 | 1 | processed_pseudoge | ENSG00000 |
| 47581211 | 47582321 | -1 | unprocessed_pseudo | ENSG00000 |
| 47588681 | 47619480 | -1 | lncRNA | ENSG00000 |
| 47689707 | 47730436 | 1 | unprocessed_pseudo | ENSG00000 |
| 47692399 | 47692727 | -1 | misc_RNA | ENSG00000 |
| 47706203 | 47706802 | 1 | processed_pseudoge | ENSG00000 |
| 47750864 | 47763592 | -1 | protein_coding | ENSG00000 |
| 47908064 | 47991877 | -1 | lncRNA | ENSG00000 |
| 47918739 | 47923524 | 1 | protein_coding | ENSG00000 |
| 47970043 | 47973496 | -1 | lncRNA | ENSG00000 |
| 47995322 | 47999791 | -1 | protein_coding | ENSG00000 |
| 48009873 | 48031640 | -1 | transcribed_unproce | ENSG00000 |
| 48040462 | 48040577 | 1 | rRNA_pseudogene | ENSG00000 |
| 48050282 | 48060016 | -1 | transcribed_unproce | ENSG00000 |
| 48064308 | 48119455 | -1 | unprocessed_pseudo | ENSG00000 |
| 48153088 | 48274696 | -1 | protein_coding | ENSG00000 |
| 48246525 | 48248512 | 1 | lncRNA | ENSG00000 |
| 48292699 | 48293417 | 1 | processed_pseudoge | ENSG00000 |
| 48306639 | 48439360 | 1 | protein_coding | ENSG00000 |
| 48443836 | 48445820 | -1 | TEC | ENSG00000 |
| 48446034 | 48656265 | -1 | protein_coding | ENSG00000 |
| 48510525 | 48511589 | -1 | lncRNA | ENSG00000 |
| 48546472 | 48546713 | -1 | processed_pseudoge | ENSG00000 |
| 48624004 | 48625093 | -1 | lncRNA | ENSG00000 |
| 48664199 | 48672424 | -1 | lncRNA | ENSG00000 |
| 48684876 | 48982956 | 1 | protein_coding | ENSG00000 |
| 48745545 | 48746128 | -1 | processed_pseudoge | ENSG00000 |

| | | | | |
|---|---|---|---|---|
| 48835541 | 48845945 | -1 | lncRNA | ENSG00000 |
| 48878022 | 48878649 | 1 | lncRNA | ENSG00000 |
| 48883955 | 48935213 | -1 | lncRNA | ENSG00000 |
| 48909483 | 48914232 | -1 | protein_coding | ENSG00000 |
| 48976554 | 48993046 | -1 | lncRNA | ENSG00000 |
| 48984564 | 49018897 | 1 | lncRNA | ENSG00000 |
| 48985512 | 48985587 | -1 | miRNA | ENSG00000 |
| 49014236 | 49115522 | -1 | protein_coding | ENSG00000 |
| 49121839 | 49151547 | 1 | lncRNA | ENSG00000 |
| 49131154 | 49134008 | -1 | protein_coding | ENSG00000 |
| 49154725 | 49188585 | -1 | protein_coding | ENSG00000 |
| 49296112 | 49299018 | -1 | lncRNA | ENSG00000 |
| 49299170 | 49327492 | 1 | protein_coding | ENSG00000 |
| 49364181 | 49396016 | -1 | protein_coding | ENSG00000 |
| 49419277 | 49472903 | 1 | lncRNA | ENSG00000 |
| 49454470 | 49539538 | -1 | protein_coding | ENSG00000 |
| 49551308 | 49552652 | 1 | processed_pseudoge | ENSG00000 |
| 49609095 | 49665104 | 1 | protein_coding | ENSG00000 |
| 49610310 | 49612720 | 1 | protein_coding | ENSG00000 |
| 49679651 | 49710261 | 1 | protein_coding | ENSG00000 |
| 49734641 | 49762379 | -1 | protein_coding | ENSG00000 |
| 49771841 | 49773299 | 1 | processed_pseudoge | ENSG00000 |
| 49815096 | 49815562 | 1 | processed_pseudoge | ENSG00000 |
| 49818279 | 49970203 | -1 | protein_coding | ENSG00000 |
| 49909613 | 49909942 | 1 | processed_pseudoge | ENSG00000 |
| 49942049 | 49974850 | 1 | protein_coding | ENSG00000 |
| 49942056 | 50011654 | 1 | lncRNA | ENSG00000 |
| 49979665 | 49979780 | -1 | rRNA_pseudogene | ENSG00000 |
| 49981420 | 49988899 | -1 | lncRNA | ENSG00000 |
| 49982190 | 50010499 | 1 | protein_coding | ENSG00000 |
| 50021182 | 50067803 | -1 | transcribed_unproce | ENSG00000 |
| 50025135 | 50027363 | -1 | processed_pseudoge | ENSG00000 |
| 50067888 | 50133509 | 1 | protein_coding | ENSG00000 |
| 50150603 | 50152287 | 1 | processed_pseudoge | ENSG00000 |
| 50182778 | 50279720 | -1 | protein_coding | ENSG00000 |
| 50264978 | 50266815 | 1 | processed_pseudoge | ENSG00000 |
| 50305586 | 50625163 | -1 | protein_coding | ENSG00000 |
| 50325697 | 50341585 | 1 | lncRNA | ENSG00000 |
| 50334538 | 50336123 | 1 | TEC | ENSG00000 |
| 50472814 | 50474246 | 1 | lncRNA | ENSG00000 |
| 50590652 | 50590712 | -1 | snRNA | ENSG00000 |
| 50623466 | 50641451 | 1 | lncRNA | ENSG00000 |
| 50629532 | 50631075 | -1 | processed_pseudoge | ENSG00000 |
| 50655967 | 50660472 | 1 | transcribed_processe | ENSG00000 |
| 50676743 | 50686326 | 1 | unprocessed_pseudo | ENSG00000 |
| 50695592 | 50695981 | 1 | processed_pseudoge | ENSG00000 |
| 50705380 | 50708906 | -1 | unprocessed_pseudo | ENSG00000 |
| 50726947 | 50727969 | -1 | processed_pseudoge | ENSG00000 |
| 50736078 | 50739434 | -1 | lncRNA | ENSG00000 |
| 50739318 | 50816495 | 1 | protein_coding | ENSG00000 |

| | | | | |
|---|---|---|---|---|
| 50799409 | 50885675 | -1 | protein_coding | ENSG0000C |
| 50822692 | 50824525 | -1 | lncRNA | ENSG0000C |
| 50962993 | 50963358 | -1 | processed_pseudoge | ENSG0000C |
| 50964270 | 50965220 | 1 | lncRNA | ENSG0000C |
| 50990888 | 52298423 | 1 | protein_coding | ENSG0000C |
| 51062579 | 51068553 | -1 | lncRNA | ENSG0000C |
| 51244894 | 51245806 | -1 | lncRNA | ENSG0000C |
| 51299573 | 51299655 | 1 | miRNA | ENSG0000C |
| 51302566 | 51306709 | 1 | lncRNA | ENSG0000C |
| 51319826 | 51320660 | 1 | processed_pseudoge | ENSG0000C |
| 51695486 | 51699595 | -1 | protein_coding | ENSG0000C |
| 52230398 | 52314507 | -1 | lncRNA | ENSG0000C |
| 52314281 | 52318042 | 1 | protein_coding | ENSG0000C |
| 52389112 | 52389488 | -1 | processed_pseudoge | ENSG0000C |
| 52411135 | 52413492 | 1 | processed_pseudoge | ENSG0000C |
| 52450874 | 52755507 | -1 | lncRNA | ENSG0000C |
| 52765380 | 52771700 | -1 | protein_coding | ENSG0000C |
| 52852607 | 52852703 | 1 | misc_RNA | ENSG0000C |
| 52946431 | 52955585 | -1 | lncRNA | ENSG0000C |
| 52972612 | 53030024 | -1 | lncRNA | ENSG0000C |
| 53038285 | 53038544 | 1 | processed_pseudoge | ENSG0000C |
| 53291072 | 53311065 | -1 | lncRNA | ENSG0000C |
| 53458197 | 53458288 | 1 | rRNA_pseudogene | ENSG0000C |
| 53766400 | 53766614 | 1 | processed_pseudoge | ENSG0000C |
| 53802771 | 55627942 | -1 | protein_coding | ENSG0000C |
| 54452423 | 54452511 | -1 | snRNA | ENSG0000C |
| 54486230 | 54656051 | 1 | lncRNA | ENSG0000C |
| 54599678 | 54600683 | 1 | processed_pseudoge | ENSG0000C |
| 54607874 | 54607957 | -1 | miRNA | ENSG0000C |
| 54864674 | 54869122 | -1 | lncRNA | ENSG0000C |
| 55247755 | 55597233 | -1 | lncRNA | ENSG0000C |
| 55468327 | 55469222 | 1 | lncRNA | ENSG0000C |
| 55506219 | 55513217 | 1 | lncRNA | ENSG0000C |
| 55599042 | 55600728 | 1 | protein_coding | ENSG0000C |
| 55667341 | 55668350 | -1 | processed_pseudoge | ENSG0000C |
| 55746551 | 55746908 | 1 | processed_pseudoge | ENSG0000C |
| 56357227 | 56361273 | -1 | protein_coding | ENSG0000C |
| 56361479 | 56364714 | 1 | lncRNA | ENSG0000C |
| 56595963 | 56596031 | -1 | snoRNA | ENSG0000C |
| 57304479 | 57304559 | -1 | miRNA | ENSG0000C |
| 57513157 | 57513425 | 1 | processed_pseudoge | ENSG0000C |
| 57955295 | 57957178 | 1 | processed_pseudoge | ENSG0000C |
| 57982285 | 57982993 | 1 | processed_pseudoge | ENSG0000C |
| 58106423 | 58106879 | -1 | processed_pseudoge | ENSG0000C |
| 58191517 | 58267894 | -1 | protein_coding | ENSG0000C |
| 58212541 | 58213036 | -1 | processed_pseudoge | ENSG0000C |
| 58269162 | 58289586 | 1 | protein_coding | ENSG0000C |
| 58304553 | 58305621 | 1 | lncRNA | ENSG0000C |
| 58325614 | 58327030 | -1 | lncRNA | ENSG0000C |
| 58334979 | 58370751 | 1 | protein_coding | ENSG0000C |

| | | | | |
|---|---|---|---|---|
| 58385345 | 58399220 | 1 | protein_coding | ENSG00000 |
| 58512872 | 58831435 | 1 | protein_coding | ENSG00000 |
| 58715554 | 58716257 | 1 | processed_pseudoge | ENSG00000 |
| 58869184 | 58869528 | 1 | misc_RNA | ENSG00000 |
| 58999482 | 59066396 | 1 | lncRNA | ENSG00000 |
| 59088701 | 59089039 | -1 | processed_pseudoge | ENSG00000 |
| 59136625 | 59138182 | -1 | processed_pseudoge | ENSG00000 |
| 59176643 | 59247774 | 1 | protein_coding | ENSG00000 |
| 59246130 | 59363181 | -1 | protein_coding | ENSG00000 |
| 59282147 | 59282835 | 1 | lncRNA | ENSG00000 |
| 59357007 | 59364075 | 1 | lncRNA | ENSG00000 |
| 59551404 | 59551878 | -1 | processed_pseudoge | ENSG00000 |
| 59578467 | 59650345 | 1 | lncRNA | ENSG00000 |
| 59650761 | 59736002 | -1 | protein_coding | ENSG00000 |
| 59736692 | 59756041 | -1 | protein_coding | ENSG00000 |
| 59788747 | 59906556 | -1 | protein_coding | ENSG00000 |
| 59955430 | 59960913 | -1 | lncRNA | ENSG00000 |
| 60026298 | 60733490 | -1 | protein_coding | ENSG00000 |
| 60050668 | 60060743 | 1 | lncRNA | ENSG00000 |
| 60081741 | 60081854 | 1 | misc_RNA | ENSG00000 |
| 60139912 | 60140877 | 1 | lncRNA | ENSG00000 |
| 60684505 | 60685209 | 1 | processed_pseudoge | ENSG00000 |
| 60734342 | 60741828 | 1 | lncRNA | ENSG00000 |
| 60778331 | 60794852 | 1 | protein_coding | ENSG00000 |
| 60869438 | 61001440 | -1 | protein_coding | ENSG00000 |
| 60912765 | 60912927 | -1 | snRNA | ENSG00000 |
| 61016275 | 61026420 | 1 | lncRNA | ENSG00000 |
| 61406642 | 61453381 | -1 | protein_coding | ENSG00000 |
| 61452639 | 61481956 | 1 | lncRNA | ENSG00000 |
| 61493843 | 61496499 | 1 | lncRNA | ENSG00000 |
| 61662961 | 61766766 | 1 | protein_coding | ENSG00000 |
| 61684892 | 61685388 | -1 | lncRNA | ENSG00000 |
| 61781745 | 61821246 | -1 | lncRNA | ENSG00000 |
| 61901684 | 62096944 | 1 | protein_coding | ENSG00000 |
| 62183035 | 62268844 | -1 | protein_coding | ENSG00000 |
| 62289521 | 62304033 | 1 | lncRNA | ENSG00000 |
| 62339553 | 62375166 | -1 | lncRNA | ENSG00000 |
| 62350006 | 62350297 | 1 | misc_RNA | ENSG00000 |
| 62374192 | 62670301 | 1 | protein_coding | ENSG00000 |
| 62374192 | 62672011 | 1 | lncRNA | ENSG00000 |
| 62520448 | 62672011 | 1 | protein_coding | ENSG00000 |
| 62682652 | 62805887 | -1 | lncRNA | ENSG00000 |
| 62741208 | 62741385 | -1 | processed_pseudoge | ENSG00000 |
| 62804720 | 62808479 | 1 | protein_coding | ENSG00000 |
| 62811996 | 62919900 | -1 | protein_coding | ENSG00000 |
| 63110139 | 63110246 | 1 | snRNA | ENSG00000 |
| 63123929 | 63125587 | -1 | lncRNA | ENSG00000 |
| 63133247 | 63155031 | 1 | protein_coding | ENSG00000 |
| 63167221 | 63521850 | -1 | protein_coding | ENSG00000 |
| 63222155 | 63223045 | 1 | processed_pseudoge | ENSG00000 |

| | | | | |
|---|---|---|---|---|
| 63358745 | 63359104 | 1 | processed_pseudoge | ENSG0000C |
| 63372957 | 63373048 | -1 | miRNA | ENSG0000C |
| 63427297 | 63427939 | 1 | processed_pseudoge | ENSG0000C |
| 63465229 | 63466563 | 1 | lncRNA | ENSG0000C |
| 63489461 | 63489638 | 1 | processed_pseudoge | ENSG0000C |
| 63521401 | 63625128 | 1 | protein_coding | ENSG0000C |
| 63630672 | 63655769 | 1 | lncRNA | ENSG0000C |
| 63634317 | 63634827 | -1 | processed_pseudoge | ENSG0000C |
| 63658074 | 63658521 | 1 | processed_pseudoge | ENSG0000C |
| 63664664 | 64693446 | 1 | lncRNA | ENSG0000C |
| 63902451 | 63903245 | -1 | processed_pseudoge | ENSG0000C |
| 64036345 | 64037280 | -1 | lncRNA | ENSG0000C |
| 64168959 | 64170850 | -1 | processed_pseudoge | ENSG0000C |
| 64620374 | 64620922 | -1 | processed_pseudoge | ENSG0000C |
| 64762883 | 64763172 | -1 | processed_pseudoge | ENSG0000C |
| 64825572 | 64826579 | 1 | processed_pseudoge | ENSG0000C |
| 64901136 | 65017639 | -1 | lncRNA | ENSG0000C |
| 65054460 | 65055003 | 1 | processed_pseudoge | ENSG0000C |
| 65123667 | 65141185 | 1 | lncRNA | ENSG0000C |
| 65169438 | 65169869 | -1 | processed_pseudoge | ENSG0000C |
| 65248032 | 65248792 | -1 | processed_pseudoge | ENSG0000C |
| 65271319 | 65315080 | 1 | lncRNA | ENSG0000C |
| 65570338 | 65880289 | 1 | lncRNA | ENSG0000C |
| 65912523 | 67696195 | -1 | protein_coding | ENSG0000C |
| 66079243 | 66118326 | 1 | lncRNA | ENSG0000C |
| 66837457 | 66837704 | 1 | unprocessed_pseudc | ENSG0000C |
| 66926036 | 67101551 | 1 | protein_coding | ENSG0000C |
| 67052609 | 67055028 | 1 | lncRNA | ENSG0000C |
| 67334123 | 67334882 | 1 | processed_pseudoge | ENSG0000C |
| 67403351 | 67403410 | -1 | miRNA | ENSG0000C |
| 67750284 | 67751225 | 1 | processed_pseudoge | ENSG0000C |
| 67793770 | 67794245 | 1 | processed_pseudoge | ENSG0000C |
| 67796669 | 67838188 | -1 | protein_coding | ENSG0000C |
| 67808502 | 67808608 | 1 | snRNA | ENSG0000C |
| 67825310 | 67825600 | -1 | misc_RNA | ENSG0000C |
| 67833763 | 67834081 | 1 | processed_pseudoge | ENSG0000C |
| 67849525 | 67850746 | 1 | lncRNA | ENSG0000C |
| 67858818 | 67858871 | -1 | snRNA | ENSG0000C |
| 67874332 | 67874911 | 1 | processed_pseudoge | ENSG0000C |
| 67884656 | 67918390 | 1 | protein_coding | ENSG0000C |
| 67921899 | 68075348 | -1 | protein_coding | ENSG0000C |
| 67960702 | 67961464 | -1 | processed_pseudoge | ENSG0000C |
| 68007718 | 68008017 | -1 | misc_RNA | ENSG0000C |
| 68010205 | 68010862 | -1 | processed_pseudoge | ENSG0000C |
| 68106117 | 68212017 | 1 | protein_coding | ENSG0000C |
| 68158079 | 68158408 | 1 | misc_RNA | ENSG0000C |
| 68230595 | 68232113 | -1 | protein_coding | ENSG0000C |
| 68233251 | 68242379 | -1 | lncRNA | ENSG0000C |
| 68260557 | 68261499 | 1 | processed_pseudoge | ENSG0000C |
| 68282660 | 68333049 | -1 | protein_coding | ENSG0000C |

| | | | | |
|---|---|---|---|---|
| 68331174 | 68343191 | 1 | protein_coding | ENSG0000C |
| 68341107 | 68407294 | -1 | protein_coding | ENSG0000C |
| 68414064 | 68472121 | -1 | protein_coding | ENSG0000C |
| 68424560 | 68424997 | 1 | processed_pseudoge | ENSG0000C |
| 68461407 | 68461525 | 1 | rRNA_pseudogene | ENSG0000C |
| 68477998 | 68527523 | -1 | protein_coding | ENSG0000C |
| 68499532 | 68499968 | 1 | processed_pseudoge | ENSG0000C |
| 68503152 | 68503259 | 1 | misc_RNA | ENSG0000C |
| 68544489 | 68544833 | -1 | processed_pseudoge | ENSG0000C |
| 68560337 | 68694487 | 1 | protein_coding | ENSG0000C |
| 68632371 | 68632727 | 1 | processed_pseudoge | ENSG0000C |
| 68674356 | 68675100 | 1 | processed_pseudoge | ENSG0000C |
| 68698500 | 68700794 | 1 | lncRNA | ENSG0000C |
| 68717311 | 68717795 | -1 | processed_pseudoge | ENSG0000C |
| 68721012 | 68792377 | 1 | protein_coding | ENSG0000C |
| 68735720 | 68735823 | -1 | misc_RNA | ENSG0000C |
| 68755172 | 68755238 | 1 | snoRNA | ENSG0000C |
| 68759318 | 68759414 | 1 | miRNA | ENSG0000C |
| 68827531 | 68895432 | 1 | protein_coding | ENSG0000C |
| 68846805 | 68846911 | 1 | snRNA | ENSG0000C |
| 68896920 | 68900768 | 1 | lncRNA | ENSG0000C |
| 68901286 | 68946847 | 1 | protein_coding | ENSG0000C |
| 68911653 | 68911759 | -1 | snRNA | ENSG0000C |
| 68956170 | 68985068 | 1 | protein_coding | ENSG0000C |
| 68988721 | 69043544 | 1 | protein_coding | ENSG0000C |
| 68991137 | 68991707 | 1 | processed_pseudoge | ENSG0000C |
| 69022778 | 69023866 | 1 | processed_pseudoge | ENSG0000C |
| 69088103 | 69104805 | 1 | protein_coding | ENSG0000C |
| 69098496 | 69098794 | 1 | misc_RNA | ENSG0000C |
| 69123512 | 69174412 | 1 | protein_coding | ENSG0000C |
| 69161530 | 69161906 | -1 | processed_pseudoge | ENSG0000C |
| 69180234 | 69209099 | 1 | protein_coding | ENSG0000C |
| 69215333 | 69232490 | -1 | lncRNA | ENSG0000C |
| 69220332 | 69267552 | 1 | protein_coding | ENSG0000C |
| 69265342 | 69268148 | -1 | lncRNA | ENSG0000C |
| 69269984 | 69401884 | 1 | protein_coding | ENSG0000C |
| 69300432 | 69300817 | -1 | processed_pseudoge | ENSG0000C |
| 69403903 | 69416918 | -1 | protein_coding | ENSG0000C |
| 69432972 | 69433371 | -1 | processed_pseudoge | ENSG0000C |
| 69451465 | 69507666 | 1 | protein_coding | ENSG0000C |
| 69510166 | 69544257 | 1 | lncRNA | ENSG0000C |
| 69523311 | 69523650 | -1 | processed_pseudoge | ENSG0000C |
| 69571698 | 69573422 | -1 | protein_coding | ENSG0000C |
| 69575807 | 69577154 | 1 | lncRNA | ENSG0000C |
| 69589880 | 69590355 | -1 | processed_pseudoge | ENSG0000C |
| 69590583 | 69591262 | -1 | processed_pseudoge | ENSG0000C |
| 69591406 | 69592931 | -1 | processed_pseudoge | ENSG0000C |
| 69594041 | 69594932 | -1 | processed_pseudoge | ENSG0000C |
| 69595487 | 69596436 | -1 | processed_pseudoge | ENSG0000C |
| 69630247 | 69633596 | 1 | protein_coding | ENSG0000C |

| | | | | |
|---|---|---|---|---|
| 69684899 | 69692452 | 1 | lncRNA | ENSG0000C |
| 69778962 | 69779838 | 1 | processed_pseudoge | ENSG0000C |
| 69801867 | 69964275 | 1 | protein_coding | ENSG0000C |
| 69994276 | 70019496 | -1 | lncRNA | ENSG0000C |
| 70052846 | 70112282 | 1 | protein_coding | ENSG0000C |
| 70098223 | 70132934 | -1 | protein_coding | ENSG0000C |
| 70137981 | 70146700 | -1 | protein_coding | ENSG0000C |
| 70147289 | 70170523 | -1 | protein_coding | ENSG0000C |
| 70163685 | 70164124 | 1 | processed_pseudoge | ENSG0000C |
| 70198988 | 70199239 | -1 | processed_pseudoge | ENSG0000C |
| 70202835 | 70233911 | -1 | protein_coding | ENSG0000C |
| 70247329 | 70284004 | -1 | protein_coding | ENSG0000C |
| 70298970 | 70382650 | -1 | protein_coding | ENSG0000C |
| 70389426 | 70390626 | 1 | processed_pseudoge | ENSG0000C |
| 70404145 | 70428618 | 1 | protein_coding | ENSG0000C |
| 70431936 | 70447951 | -1 | protein_coding | ENSG0000C |
| 70434829 | 70436556 | 1 | TEC | ENSG0000C |
| 70478767 | 70568450 | 1 | protein_coding | ENSG0000C |
| 70485336 | 70486325 | -1 | processed_pseudoge | ENSG0000C |
| 70597348 | 70602759 | -1 | protein_coding | ENSG0000C |
| 70672506 | 70762441 | 1 | protein_coding | ENSG0000C |
| 70771239 | 70785401 | -1 | protein_coding | ENSG0000C |
| 70794230 | 70794523 | -1 | processed_pseudoge | ENSG0000C |
| 70815948 | 70881184 | 1 | protein_coding | ENSG0000C |
| 70882280 | 70888565 | -1 | protein_coding | ENSG0000C |
| 70927055 | 71077810 | 1 | lncRNA | ENSG0000C |
| 70929925 | 70932057 | 1 | lncRNA | ENSG0000C |
| 70939036 | 70955947 | 1 | lncRNA | ENSG0000C |
| 71212570 | 71302864 | 1 | protein_coding | ENSG0000C |
| 71217224 | 71218228 | -1 | lncRNA | ENSG0000C |
| 71319259 | 71381423 | 1 | protein_coding | ENSG0000C |
| 71364243 | 71366374 | 1 | TEC | ENSG0000C |
| 71396920 | 71815947 | 1 | protein_coding | ENSG0000C |
| 71508153 | 71511873 | -1 | lncRNA | ENSG0000C |
| 71711701 | 71737824 | -1 | protein_coding | ENSG0000C |
| 71747556 | 71773520 | -1 | protein_coding | ENSG0000C |
| 71790747 | 71790800 | 1 | miRNA | ENSG0000C |
| 71816298 | 71851325 | -1 | protein_coding | ENSG0000C |
| 71844735 | 71844794 | -1 | snRNA | ENSG0000C |
| 71878356 | 71879107 | 1 | lncRNA | ENSG0000C |
| 71964395 | 72013558 | 1 | protein_coding | ENSG0000C |
| 72053294 | 72054037 | -1 | lncRNA | ENSG0000C |
| 72059034 | 72089032 | -1 | protein_coding | ENSG0000C |
| 72096032 | 72217134 | -1 | protein_coding | ENSG0000C |
| 72180858 | 72180989 | 1 | snoRNA | ENSG0000C |
| 72189941 | 72190576 | 1 | processed_pseudoge | ENSG0000C |
| 72216000 | 72235860 | 1 | protein_coding | ENSG0000C |
| 72220752 | 72220852 | 1 | misc_RNA | ENSG0000C |
| 72273924 | 72276036 | 1 | protein_coding | ENSG0000C |
| 72274915 | 72275980 | -1 | lncRNA | ENSG0000C |

| | | | | |
|---|---|---|---|---|
| 72332830 | 72355149 | -1 | protein_coding | ENSG0000( |
| 72367327 | 72626191 | -1 | protein_coding | ENSG0000( |
| 72391964 | 72392069 | 1 | snRNA | ENSG0000( |
| 72467749 | 72473304 | -1 | lncRNA | ENSG0000( |
| 72483245 | 72483812 | 1 | unprocessed_pseudc | ENSG0000( |
| 72501746 | 72502956 | 1 | lncRNA | ENSG0000( |
| 72560264 | 72560456 | 1 | processed_pseudoge | ENSG0000( |
| 72582803 | 72583086 | 1 | misc_RNA | ENSG0000( |
| 72636497 | 72636765 | -1 | processed_pseudoge | ENSG0000( |
| 72643011 | 72644650 | 1 | processed_pseudoge | ENSG0000( |
| 72654165 | 72654276 | -1 | misc_RNA | ENSG0000( |
| 72692131 | 72887694 | 1 | protein_coding | ENSG0000( |
| 72721029 | 72721100 | 1 | miRNA | ENSG0000( |
| 72730557 | 72731175 | 1 | TEC | ENSG0000( |
| 72756398 | 72757016 | 1 | TEC | ENSG0000( |
| 72766560 | 72767052 | 1 | lncRNA | ENSG0000( |
| 72893584 | 72933036 | 1 | protein_coding | ENSG0000( |
| 72917641 | 72918534 | -1 | processed_pseudoge | ENSG0000( |
| 72934762 | 72954806 | -1 | protein_coding | ENSG0000( |
| 73005833 | 73006595 | -1 | processed_pseudoge | ENSG0000( |
| 73007217 | 73096974 | -1 | protein_coding | ENSG0000( |
| 73065126 | 73065467 | 1 | processed_pseudoge | ENSG0000( |
| 73071295 | 73074008 | 1 | lncRNA | ENSG0000( |
| 73082494 | 73082595 | -1 | misc_RNA | ENSG0000( |
| 73098044 | 73101297 | -1 | lncRNA | ENSG0000( |
| 73110375 | 73131828 | 1 | protein_coding | ENSG0000( |
| 73124573 | 73125532 | -1 | lncRNA | ENSG0000( |
| 73126080 | 73126207 | -1 | snoRNA | ENSG0000( |
| 73130155 | 73169055 | -1 | protein_coding | ENSG0000( |
| 73168119 | 73244504 | 1 | protein_coding | ENSG0000( |
| 73183362 | 73248268 | -1 | protein_coding | ENSG0000( |
| 73192223 | 73192325 | -1 | misc_RNA | ENSG0000( |
| 73199826 | 73201225 | 1 | processed_pseudoge | ENSG0000( |
| 73247360 | 73276984 | 1 | lncRNA | ENSG0000( |
| 73248843 | 73252693 | -1 | protein_coding | ENSG0000( |
| 73252791 | 73254349 | 1 | lncRNA | ENSG0000( |
| 73253759 | 73358859 | -1 | protein_coding | ENSG0000( |
| 73279062 | 73279168 | 1 | snRNA | ENSG0000( |
| 73346936 | 73347048 | 1 | misc_RNA | ENSG0000( |
| 73375101 | 73414076 | -1 | protein_coding | ENSG0000( |
| 73381433 | 73383496 | 1 | lncRNA | ENSG0000( |
| 73422259 | 73422696 | -1 | processed_pseudoge | ENSG0000( |
| 73423579 | 73433561 | -1 | protein_coding | ENSG0000( |
| 73436428 | 73496024 | -1 | protein_coding | ENSG0000( |
| 73495525 | 73520070 | 1 | lncRNA | ENSG0000( |
| 73497538 | 73625953 | -1 | protein_coding | ENSG0000( |
| 73529051 | 73529157 | -1 | snRNA | ENSG0000( |
| 73537803 | 73538391 | 1 | processed_pseudoge | ENSG0000( |
| 73625996 | 73626790 | 1 | lncRNA | ENSG0000( |
| 73630556 | 73631490 | 1 | lncRNA | ENSG0000( |

| | | | | |
|---|---|---|---|---|
| 73631612 | 73641474 | -1 | protein_coding | ENSG0000C |
| 73644881 | 73663803 | -1 | protein_coding | ENSG0000C |
| 73653980 | 73675450 | 1 | lncRNA | ENSG0000C |
| 73674285 | 73698159 | -1 | protein_coding | ENSG0000C |
| 73674295 | 73730466 | -1 | lncRNA | ENSG0000C |
| 73699151 | 73730487 | -1 | lncRNA | ENSG0000C |
| 73703735 | 73713581 | -1 | lncRNA | ENSG0000C |
| 73706490 | 73706613 | 1 | rRNA_pseudogene | ENSG0000C |
| 73715843 | 73730469 | -1 | transcribed_unproce | ENSG0000C |
| 73730562 | 73737311 | 1 | transcribed_unproce | ENSG0000C |
| 73731824 | 73733638 | 1 | processed_pseudoge | ENSG0000C |
| 73740538 | 73740823 | -1 | processed_pseudoge | ENSG0000C |
| 73742995 | 73744230 | -1 | TEC | ENSG0000C |
| 73744372 | 73772161 | 1 | protein_coding | ENSG0000C |
| 73769264 | 73772862 | -1 | TEC | ENSG0000C |
| 73772276 | 73780251 | 1 | protein_coding | ENSG0000C |
| 73782047 | 73783652 | 1 | protein_coding | ENSG0000C |
| 73785582 | 73801797 | 1 | protein_coding | ENSG0000C |
| 73791218 | 73791490 | 1 | TEC | ENSG0000C |
| 73796514 | 73811651 | -1 | protein_coding | ENSG0000C |
| 73796514 | 73801399 | -1 | lncRNA | ENSG0000C |
| 73801911 | 73811798 | -1 | protein_coding | ENSG0000C |
| 73812501 | 73874591 | -1 | protein_coding | ENSG0000C |
| 73813518 | 73814737 | 1 | lncRNA | ENSG0000C |
| 73841833 | 73847115 | 1 | lncRNA | ENSG0000C |
| 73909177 | 73917496 | 1 | protein_coding | ENSG0000C |
| 73909969 | 73922777 | -1 | protein_coding | ENSG0000C |
| 73995193 | 74121363 | 1 | protein_coding | ENSG0000C |
| 74005137 | 74027915 | 1 | lncRNA | ENSG0000C |
| 74120255 | 74151063 | -1 | protein_coding | ENSG0000C |
| 74151202 | 74709963 | 1 | protein_coding | ENSG0000C |
| 74344550 | 74344805 | -1 | processed_pseudoge | ENSG0000C |
| 74371535 | 74372416 | 1 | processed_pseudoge | ENSG0000C |
| 74423435 | 74424014 | -1 | processed_pseudoge | ENSG0000C |
| 74506081 | 74530553 | -1 | lncRNA | ENSG0000C |
| 74527584 | 74527947 | 1 | processed_pseudoge | ENSG0000C |
| 74551392 | 74551493 | 1 | misc_RNA | ENSG0000C |
| 74641410 | 74641776 | -1 | processed_pseudoge | ENSG0000C |
| 74654956 | 74656131 | 1 | processed_pseudoge | ENSG0000C |
| 74744386 | 74744707 | -1 | processed_pseudoge | ENSG0000C |
| 74821610 | 74822130 | -1 | lncRNA | ENSG0000C |
| 74824927 | 75032624 | 1 | protein_coding | ENSG0000C |
| 75003055 | 75025174 | -1 | lncRNA | ENSG0000C |
| 75023475 | 75024251 | -1 | lncRNA | ENSG0000C |
| 75037836 | 75058514 | -1 | protein_coding | ENSG0000C |
| 75041876 | 75055754 | 1 | lncRNA | ENSG0000C |
| 75089248 | 75089725 | -1 | processed_pseudoge | ENSG0000C |
| 75094432 | 75109221 | -1 | protein_coding | ENSG0000C |
| 75099586 | 75182123 | 1 | protein_coding | ENSG0000C |
| 75125217 | 75125862 | 1 | processed_pseudoge | ENSG0000C |

| | | | | |
|---|---|---|---|---|
| 75210154 | 75231448 | 1 | protein_coding | ENSG0000C |
| 75233641 | 75236030 | -1 | protein_coding | ENSG0000C |
| 75243568 | 75373500 | 1 | lncRNA | ENSG0000C |
| 75269392 | 75269545 | -1 | processed_pseudoge | ENSG0000C |
| 75276376 | 75276646 | -1 | processed_pseudoge | ENSG0000C |
| 75279726 | 75401246 | -1 | lncRNA | ENSG0000C |
| 75288333 | 75288682 | -1 | transcribed_processe | ENSG0000C |
| 75382393 | 75382844 | -1 | transcribed_processe | ENSG0000C |
| 75397830 | 75401764 | -1 | protein_coding | ENSG0000C |
| 75401519 | 75408982 | 1 | lncRNA | ENSG0000C |
| 75408973 | 75409326 | -1 | lncRNA | ENSG0000C |
| 75409142 | 75411842 | 1 | lncRNA | ENSG0000C |
| 75430571 | 75431588 | -1 | lncRNA | ENSG0000C |
| 75431624 | 76560168 | 1 | protein_coding | ENSG0000C |
| 75450081 | 75451725 | 1 | TEC | ENSG0000C |
| 75552458 | 75552553 | 1 | miRNA | ENSG0000C |
| 75592644 | 75628120 | 1 | lncRNA | ENSG0000C |
| 75642476 | 75643106 | 1 | lncRNA | ENSG0000C |
| 75742740 | 75743755 | 1 | lncRNA | ENSG0000C |
| 76242022 | 76242301 | -1 | misc_RNA | ENSG0000C |
| 76260800 | 76260903 | -1 | snRNA | ENSG0000C |
| 76270206 | 76270800 | 1 | TEC | ENSG0000C |
| 76437408 | 76438775 | -1 | lncRNA | ENSG0000C |
| 76558350 | 76560994 | -1 | lncRNA | ENSG0000C |
| 76835377 | 76835502 | -1 | snoRNA | ENSG0000C |
| 76836568 | 76836861 | 1 | processed_pseudoge | ENSG0000C |
| 76869601 | 77638369 | -1 | protein_coding | ENSG0000C |
| 76888044 | 76980624 | 1 | lncRNA | ENSG0000C |
| 77147652 | 77150100 | 1 | lncRNA | ENSG0000C |
| 77313941 | 77315694 | 1 | lncRNA | ENSG0000C |
| 77354404 | 77376613 | 1 | lncRNA | ENSG0000C |
| 77433572 | 77433782 | -1 | processed_pseudoge | ENSG0000C |
| 77587049 | 77587159 | 1 | rRNA_pseudogene | ENSG0000C |
| 77730766 | 77734769 | 1 | processed_pseudoge | ENSG0000C |
| 77738246 | 77739015 | 1 | unprocessed_pseudc | ENSG0000C |
| 77776951 | 77777055 | -1 | snRNA | ENSG0000C |
| 77780337 | 77781876 | -1 | processed_pseudoge | ENSG0000C |
| 77782866 | 77793176 | 1 | lncRNA | ENSG0000C |
| 77790791 | 77926755 | -1 | protein_coding | ENSG0000C |
| 77866875 | 77869610 | 1 | lncRNA | ENSG0000C |
| 77912271 | 77912545 | -1 | misc_RNA | ENSG0000C |
| 77927372 | 77929824 | 1 | lncRNA | ENSG0000C |
| 77953495 | 77953876 | 1 | processed_pseudoge | ENSG0000C |
| 77966173 | 77966440 | 1 | misc_RNA | ENSG0000C |
| 77969251 | 78029515 | -1 | protein_coding | ENSG0000C |
| 78033760 | 78056813 | 1 | protein_coding | ENSG0000C |
| 78068623 | 78069538 | -1 | processed_pseudoge | ENSG0000C |
| 78176928 | 78178186 | 1 | lncRNA | ENSG0000C |
| 78179174 | 78675109 | 1 | lncRNA | ENSG0000C |
| 78179185 | 78551355 | 1 | lncRNA | ENSG0000C |

| | | | | |
|---|---|---|---|---|
| 78288059 | 78290565 | -1 | lncRNA | ENSG00000 |
| 78293842 | 78301003 | 1 | lncRNA | ENSG00000 |
| 78352597 | 78356100 | 1 | lncRNA | ENSG00000 |
| 78367507 | 78367623 | 1 | snoRNA | ENSG00000 |
| 78538391 | 78551719 | -1 | lncRNA | ENSG00000 |
| 78696062 | 78697022 | -1 | lncRNA | ENSG00000 |
| 78943328 | 79067895 | -1 | lncRNA | ENSG00000 |
| 79000484 | 79003803 | -1 | lncRNA | ENSG00000 |
| 79068966 | 79316528 | 1 | protein_coding | ENSG00000 |
| 79347469 | 79355334 | 1 | protein_coding | ENSG00000 |
| 79382325 | 79445624 | -1 | protein_coding | ENSG00000 |
| 79382328 | 79409274 | 1 | lncRNA | ENSG00000 |
| 79499666 | 79499984 | -1 | processed_pseudoge | ENSG00000 |
| 79504073 | 79506281 | -1 | lncRNA | ENSG00000 |
| 79507597 | 79510982 | -1 | transcribed_processe | ENSG00000 |
| 79512601 | 79516440 | 1 | protein_coding | ENSG00000 |
| 79526508 | 79526631 | -1 | processed_pseudoge | ENSG00000 |
| 79544087 | 79544356 | 1 | processed_pseudoge | ENSG00000 |
| 79555852 | 79560407 | -1 | protein_coding | ENSG00000 |
| 79582298 | 79584877 | -1 | unprocessed_pseudo | ENSG00000 |
| 79595294 | 79595660 | -1 | processed_pseudoge | ENSG00000 |
| 79610939 | 79615455 | 1 | protein_coding | ENSG00000 |
| 79628757 | 79632188 | -1 | lncRNA | ENSG00000 |
| 79660891 | 79677996 | 1 | lncRNA | ENSG00000 |
| 79661394 | 79826594 | -1 | lncRNA | ENSG00000 |
| 79663192 | 79664786 | 1 | lncRNA | ENSG00000 |
| 79681973 | 79684094 | -1 | TEC | ENSG00000 |
| 79682997 | 79685436 | 1 | processed_pseudoge | ENSG00000 |
| 79684494 | 79684983 | -1 | transcribed_processe | ENSG00000 |
| 79703227 | 79714681 | 1 | protein_coding | ENSG00000 |
| 79825902 | 79827602 | 1 | lncRNA | ENSG00000 |
| 79841358 | 79850878 | 1 | protein_coding | ENSG00000 |
| 79860573 | 79861806 | 1 | processed_pseudoge | ENSG00000 |
| 79870375 | 79873903 | -1 | unprocessed_pseudo | ENSG00000 |
| 79892148 | 79893175 | -1 | processed_pseudoge | ENSG00000 |
| 79904898 | 79951029 | 1 | lncRNA | ENSG00000 |
| 79906605 | 79907856 | 1 | transcribed_unproce | ENSG00000 |
| 79920178 | 79923119 | 1 | transcribed_unitary_ | ENSG00000 |
| 79937740 | 79982614 | -1 | protein_coding | ENSG00000 |
| 79968213 | 79973213 | 1 | lncRNA | ENSG00000 |
| 79970963 | 79973213 | -1 | TEC | ENSG00000 |
| 79981839 | 79982212 | 1 | processed_pseudoge | ENSG00000 |
| 79982910 | 79983240 | -1 | processed_pseudoge | ENSG00000 |
| 80017713 | 80017911 | -1 | unprocessed_pseudo | ENSG00000 |
| 80024737 | 80025159 | -1 | processed_pseudoge | ENSG00000 |
| 80031968 | 80032393 | -1 | processed_pseudoge | ENSG00000 |
| 80040621 | 80041044 | -1 | processed_pseudoge | ENSG00000 |
| 80046860 | 80078912 | -1 | lncRNA | ENSG00000 |
| 80053428 | 80053548 | -1 | transcribed_unproce | ENSG00000 |
| 80078646 | 80092557 | 1 | protein_coding | ENSG00000 |

| | | | | |
|---|---|---|---|---|
| 80091411 | 80091867 | -1 | TEC | ENSG0000C |
| 80125941 | 80126317 | 1 | processed_pseudoge | ENSG0000C |
| 80131682 | 80145359 | 1 | protein_coding | ENSG0000C |
| 80150889 | 80205572 | -1 | protein_coding | ENSG0000C |
| 80207372 | 80219657 | 1 | lncRNA | ENSG0000C |
| 80215851 | 80216119 | -1 | processed_pseudoge | ENSG0000C |
| 80233664 | 80245367 | 1 | unprocessed_pseudc | ENSG0000C |
| 80247219 | 80247683 | -1 | processed_pseudoge | ENSG0000C |
| 80249710 | 80254121 | 1 | TEC | ENSG0000C |
| 80271820 | 80289658 | -1 | protein_coding | ENSG0000C |
| 80297720 | 80299655 | 1 | processed_pseudoge | ENSG0000C |
| 80333771 | 80334426 | 1 | lncRNA | ENSG0000C |
| 80336105 | 80356755 | -1 | protein_coding | ENSG0000C |
| 80344745 | 80368073 | 1 | protein_coding | ENSG0000C |
| 80407829 | 80437115 | 1 | protein_coding | ENSG0000C |
| 80454166 | 80533124 | 1 | protein_coding | ENSG0000C |
| 80529597 | 80535942 | -1 | lncRNA | ENSG0000C |
| 80537902 | 80646560 | 1 | protein_coding | ENSG0000C |
| 80649797 | 80653732 | 1 | lncRNA | ENSG0000C |
| 80716483 | 80717068 | -1 | processed_pseudoge | ENSG0000C |
| 80749632 | 80751231 | -1 | processed_pseudoge | ENSG0000C |
| 80776165 | 80778088 | -1 | processed_pseudoge | ENSG0000C |
| 80940665 | 80941201 | -1 | processed_pseudoge | ENSG0000C |
| 81136540 | 81137335 | -1 | processed_pseudoge | ENSG0000C |
| 81870778 | 81875133 | -1 | lncRNA | ENSG0000C |
| 81875194 | 82987179 | 1 | protein_coding | ENSG0000C |
| 82207620 | 82210345 | -1 | lncRNA | ENSG0000C |
| 82224213 | 82229179 | -1 | lncRNA | ENSG0000C |
| 82228985 | 82232920 | -1 | lncRNA | ENSG0000C |
| 82340771 | 82340876 | 1 | snRNA | ENSG0000C |
| 83067962 | 83068071 | -1 | snRNA | ENSG0000C |
| 83311628 | 83314475 | -1 | processed_pseudoge | ENSG0000C |
| 83672406 | 83677122 | -1 | lncRNA | ENSG0000C |
| 83708841 | 83724834 | -1 | lncRNA | ENSG0000C |
| 83735506 | 83736493 | 1 | processed_pseudoge | ENSG0000C |
| 83764358 | 83764464 | 1 | snRNA | ENSG0000C |
| 83911654 | 83912922 | -1 | lncRNA | ENSG0000C |
| 84003332 | 84003479 | 1 | snRNA | ENSG0000C |
| 84081429 | 84081995 | 1 | processed_pseudoge | ENSG0000C |
| 84138420 | 84140582 | -1 | lncRNA | ENSG0000C |
| 84139509 | 84153568 | 1 | protein_coding | ENSG0000C |
| 84167228 | 84172093 | -1 | lncRNA | ENSG0000C |
| 84173801 | 84185294 | 1 | protein_coding | ENSG0000C |
| 84194537 | 84219621 | 1 | protein_coding | ENSG0000C |
| 84220495 | 84225589 | -1 | protein_coding | ENSG0000C |
| 84230666 | 84259960 | 1 | protein_coding | ENSG0000C |
| 84231520 | 84241546 | -1 | protein_coding | ENSG0000C |
| 84267747 | 84294659 | 1 | lncRNA | ENSG0000C |
| 84328586 | 84518521 | 1 | protein_coding | ENSG0000C |
| 84389170 | 84389626 | -1 | processed_pseudoge | ENSG0000C |

| | | | | |
|---|---|---|---|---|
| 84390908 | 84392045 | 1 | processed_pseudoge | ENSG00000 |
| 84454679 | 84456056 | -1 | processed_pseudoge | ENSG00000 |
| 84560443 | 84561263 | 1 | processed_pseudoge | ENSG00000 |
| 85156330 | 85157348 | 1 | processed_pseudoge | ENSG00000 |
| 85193421 | 85197958 | -1 | lncRNA | ENSG00000 |
| 85199002 | 85212800 | 1 | lncRNA | ENSG00000 |
| 85378327 | 85432775 | -1 | lncRNA | ENSG00000 |
| 85432863 | 85448961 | 1 | lncRNA | ENSG00000 |
| 85449592 | 85492001 | 1 | lncRNA | ENSG00000 |
| 85526880 | 85531115 | -1 | TEC | ENSG00000 |
| 85537379 | 85537483 | 1 | snRNA | ENSG00000 |
| 85555484 | 85557436 | -1 | TEC | ENSG00000 |
| 85577731 | 85607213 | 1 | lncRNA | ENSG00000 |
| 85599552 | 86366795 | -1 | protein_coding | ENSG00000 |
| 85613017 | 85613186 | -1 | misc_RNA | ENSG00000 |
| 85644073 | 85648066 | 1 | lncRNA | ENSG00000 |
| 85650012 | 85650260 | -1 | misc_RNA | ENSG00000 |
| 85708760 | 85709167 | 1 | processed_pseudoge | ENSG00000 |
| 86052448 | 86052566 | 1 | rRNA_pseudogene | ENSG00000 |
| 86264694 | 86264788 | -1 | miRNA | ENSG00000 |
| 86395187 | 86400943 | -1 | lncRNA | ENSG00000 |
| 86402119 | 86403201 | -1 | lncRNA | ENSG00000 |
| 86435256 | 86521792 | -1 | protein_coding | ENSG00000 |
| 86488446 | 86488549 | 1 | snRNA | ENSG00000 |
| 86521945 | 86525101 | 1 | lncRNA | ENSG00000 |
| 86630825 | 86631628 | -1 | processed_pseudoge | ENSG00000 |
| 86654547 | 86666848 | 1 | protein_coding | ENSG00000 |
| 86668449 | 86736068 | 1 | protein_coding | ENSG00000 |
| 86749754 | 86756298 | -1 | lncRNA | ENSG00000 |
| 86756601 | 86932838 | 1 | protein_coding | ENSG00000 |
| 86854696 | 86856236 | -1 | processed_pseudoge | ENSG00000 |
| 86889569 | 86889733 | 1 | snRNA | ENSG00000 |
| 86935540 | 86969481 | -1 | protein_coding | ENSG00000 |
| 86948677 | 86949875 | 1 | TEC | ENSG00000 |
| 86958599 | 86963258 | 1 | protein_coding | ENSG00000 |
| 86965345 | 86971311 | -1 | lncRNA | ENSG00000 |
| 86968432 | 86983934 | 1 | protein_coding | ENSG00000 |
| 86970237 | 86970826 | -1 | lncRNA | ENSG00000 |
| 86970741 | 87010126 | 1 | lncRNA | ENSG00000 |
| 86992253 | 87000435 | 1 | transcribed_unproce | ENSG00000 |
| 86992406 | 87010203 | 1 | lncRNA | ENSG00000 |
| 86993432 | 87024732 | 1 | lncRNA | ENSG00000 |
| 87001636 | 87009905 | 1 | transcribed_unproce | ENSG00000 |
| 87009785 | 87015782 | -1 | lncRNA | ENSG00000 |
| 87020294 | 87024730 | 1 | protein_coding | ENSG00000 |
| 87027184 | 87027477 | 1 | processed_pseudoge | ENSG00000 |
| 87041238 | 87041341 | -1 | snRNA | ENSG00000 |
| 87050202 | 87094843 | -1 | protein_coding | ENSG00000 |
| 87094161 | 87191468 | 1 | protein_coding | ENSG00000 |
| 87113954 | 87115523 | 1 | lncRNA | ENSG00000 |

| | | | | |
|---|---|---|---|---|
| 87154041 | 87154369 | -1 | misc_RNA | ENSG0000C |
| 87201647 | 87342612 | -1 | lncRNA | ENSG0000C |
| 87203717 | 87206615 | 1 | lncRNA | ENSG0000C |
| 87214136 | 87216173 | -1 | unprocessed_pseudc | ENSG0000C |
| 87225448 | 87236908 | 1 | protein_coding | ENSG0000C |
| 87341685 | 87357882 | 1 | lncRNA | ENSG0000C |
| 87357668 | 87370695 | 1 | protein_coding | ENSG0000C |
| 87376896 | 87378481 | 1 | processed_pseudoge | ENSG0000C |
| 87386685 | 87390203 | -1 | unprocessed_pseudc | ENSG0000C |
| 87396561 | 87435035 | -1 | lncRNA | ENSG0000C |
| 87408044 | 87408258 | -1 | processed_pseudoge | ENSG0000C |
| 87500337 | 87504837 | -1 | lncRNA | ENSG0000C |
| 87503881 | 87503954 | 1 | miRNA | ENSG0000C |
| 87504875 | 87553461 | 1 | protein_coding | ENSG0000C |
| 87591607 | 87606024 | -1 | lncRNA | ENSG0000C |
| 87607985 | 87659279 | 1 | lncRNA | ENSG0000C |
| 87610163 | 87660003 | -1 | lncRNA | ENSG0000C |
| 87642607 | 87642954 | 1 | processed_pseudoge | ENSG0000C |
| 87659613 | 87747705 | 1 | protein_coding | ENSG0000C |
| 87751512 | 87841343 | -1 | protein_coding | ENSG0000C |
| 87817928 | 87845612 | 1 | transcribed_unproce | ENSG0000C |
| 87832818 | 87833098 | -1 | misc_RNA | ENSG0000C |
| 87859158 | 87863437 | -1 | protein_coding | ENSG0000C |
| 87863625 | 87971930 | 1 | protein_coding | ENSG0000C |
| 87878692 | 87880427 | 1 | lncRNA | ENSG0000C |
| 87945502 | 87946024 | 1 | processed_pseudoge | ENSG0000C |
| 87994618 | 87994695 | -1 | snoRNA | ENSG0000C |
| 88048135 | 88049823 | -1 | processed_pseudoge | ENSG0000C |
| 88259129 | 88259372 | -1 | unprocessed_pseudc | ENSG0000C |
| 88273864 | 88584530 | -1 | protein_coding | ENSG0000C |
| 88585638 | 88585733 | -1 | misc_RNA | ENSG0000C |
| 88586753 | 88606976 | 1 | protein_coding | ENSG0000C |
| 88618223 | 88618934 | -1 | processed_pseudoge | ENSG0000C |
| 88664441 | 88678814 | 1 | protein_coding | ENSG0000C |
| 88699683 | 88700401 | 1 | processed_pseudoge | ENSG0000C |
| 88724544 | 88752756 | 1 | protein_coding | ENSG0000C |
| 88727485 | 88728914 | 1 | processed_pseudoge | ENSG0000C |
| 88761406 | 88778242 | 1 | protein_coding | ENSG0000C |
| 88785600 | 88787061 | -1 | processed_pseudoge | ENSG0000C |
| 88802730 | 88820546 | 1 | protein_coding | ENSG0000C |
| 88819896 | 88851844 | -1 | protein_coding | ENSG0000C |
| 88876450 | 88877544 | 1 | processed_pseudoge | ENSG0000C |
| 88879734 | 88975153 | 1 | protein_coding | ENSG0000C |
| 88932390 | 88940820 | 1 | lncRNA | ENSG0000C |
| 88935074 | 88991339 | -1 | protein_coding | ENSG0000C |
| 88990045 | 88994249 | -1 | lncRNA | ENSG0000C |
| 88990531 | 89017059 | 1 | protein_coding | ENSG0000C |
| 88992370 | 88992539 | 1 | misc_RNA | ENSG0000C |
| 89015836 | 89017059 | 1 | lncRNA | ENSG0000C |
| 89063335 | 89063411 | -1 | miRNA | ENSG0000C |

| | | | | |
|---|---|---|---|---|
| 89063336 | 89063410 | 1 | miRNA | ENSG0000C |
| 89139044 | 89139458 | 1 | processed_pseudoge | ENSG0000C |
| 89205629 | 89207317 | -1 | protein_coding | ENSG0000C |
| 89213569 | 89414557 | -1 | protein_coding | ENSG0000C |
| 89283694 | 89309271 | 1 | protein_coding | ENSG0000C |
| 89283765 | 89292125 | 1 | lncRNA | ENSG0000C |
| 89327997 | 89340971 | 1 | protein_coding | ENSG0000C |
| 89362294 | 89363487 | 1 | unprocessed_pseudo | ENSG0000C |
| 89378056 | 89385205 | 1 | protein_coding | ENSG0000C |
| 89392546 | 89406487 | 1 | protein_coding | ENSG0000C |
| 89414568 | 89420997 | 1 | protein_coding | ENSG0000C |
| 89430299 | 89556641 | -1 | protein_coding | ENSG0000C |
| 89456064 | 89468140 | 1 | lncRNA | ENSG0000C |
| 89579497 | 89645572 | -1 | protein_coding | ENSG0000C |
| 89592747 | 89592827 | -1 | miRNA | ENSG0000C |
| 89644603 | 89645223 | -1 | processed_pseudoge | ENSG0000C |
| 89645272 | 89652144 | 1 | lncRNA | ENSG0000C |
| 89646289 | 89650822 | -1 | lncRNA | ENSG0000C |
| 89667181 | 89701274 | -1 | lncRNA | ENSG0000C |
| 89694295 | 89697928 | -1 | lncRNA | ENSG0000C |
| 89701610 | 89774939 | 1 | protein_coding | ENSG0000C |
| 89786610 | 89787107 | 1 | processed_pseudoge | ENSG0000C |
| 89829483 | 89840861 | 1 | lncRNA | ENSG0000C |
| 89837612 | 89839334 | -1 | processed_pseudoge | ENSG0000C |
| 89853595 | 90225443 | 1 | lncRNA | ENSG0000C |
| 89915489 | 89957373 | -1 | lncRNA | ENSG0000C |
| 89978717 | 89979070 | 1 | processed_pseudoge | ENSG0000C |
| 90059061 | 90099083 | -1 | lncRNA | ENSG0000C |
| 90163651 | 90163960 | -1 | misc_RNA | ENSG0000C |
| 90402521 | 90628852 | -1 | lncRNA | ENSG0000C |
| 90454170 | 90502968 | 1 | lncRNA | ENSG0000C |
| 90462717 | 90467952 | 1 | lncRNA | ENSG0000C |
| 90622143 | 90623611 | -1 | lncRNA | ENSG0000C |
| 90740823 | 90858039 | -1 | protein_coding | ENSG0000C |
| 90871716 | 90908553 | 1 | protein_coding | ENSG0000C |
| 90880040 | 90880179 | -1 | misc_RNA | ENSG0000C |
| 90912096 | 90921276 | -1 | protein_coding | ENSG0000C |
| 90922731 | 90922835 | -1 | snRNA | ENSG0000C |
| 90994620 | 91006390 | 1 | lncRNA | ENSG0000C |
| 90997480 | 90997713 | -1 | lncRNA | ENSG0000C |
| 91033157 | 91062164 | 1 | lncRNA | ENSG0000C |
| 91053051 | 91055047 | 1 | processed_pseudoge | ENSG0000C |
| 91152605 | 91153349 | -1 | processed_pseudoge | ENSG0000C |
| 91153669 | 91161315 | 1 | lncRNA | ENSG0000C |
| 91163012 | 91284337 | 1 | protein_coding | ENSG0000C |
| 91409280 | 91514829 | 1 | protein_coding | ENSG0000C |
| 91464578 | 91464708 | -1 | processed_pseudoge | ENSG0000C |
| 91544852 | 91545098 | 1 | processed_pseudoge | ENSG0000C |
| 91628442 | 91633071 | -1 | protein_coding | ENSG0000C |
| 91666780 | 91667782 | -1 | processed_pseudoge | ENSG0000C |

| | | | | |
|---|---|---|---|---|
| 91765899 | 91767196 | -1 | processed_pseudoge | ENSG00000( |
| 91782835 | 91798304 | -1 | lncRNA | ENSG00000( |
| 91798426 | 91865475 | 1 | protein_coding | ENSG00000( |
| 91807179 | 91807439 | -1 | processed_pseudoge | ENSG00000( |
| 91896704 | 91896897 | 1 | unprocessed_pseudc | ENSG00000( |
| 91906584 | 91909486 | -1 | protein_coding | ENSG00000( |
| 91908131 | 91909348 | 1 | lncRNA | ENSG00000( |
| 91923770 | 92030325 | 1 | protein_coding | ENSG00000( |
| 92046692 | 92291078 | -1 | protein_coding | ENSG00000( |
| 92076242 | 92076365 | 1 | snoRNA | ENSG00000( |
| 92118664 | 92119164 | 1 | processed_pseudoge | ENSG00000( |
| 92136875 | 92138370 | 1 | processed_pseudoge | ENSG00000( |
| 92204229 | 92204306 | -1 | ribozyme | ENSG00000( |
| 92216332 | 92216793 | 1 | processed_pseudoge | ENSG00000( |
| 92272147 | 92272250 | -1 | misc_RNA | ENSG00000( |
| 92291167 | 92353964 | 1 | protein_coding | ENSG00000( |
| 92334463 | 92334564 | 1 | misc_RNA | ENSG00000( |
| 92418667 | 92420875 | 1 | processed_pseudoge | ENSG00000( |
| 92423386 | 92427620 | 1 | lncRNA | ENSG00000( |
| 92451684 | 92574093 | -1 | protein_coding | ENSG00000( |
| 92593130 | 92655395 | 1 | protein_coding | ENSG00000( |
| 92596852 | 92597287 | -1 | processed_pseudoge | ENSG00000( |
| 92659044 | 92659302 | 1 | misc_RNA | ENSG00000( |
| 92668745 | 92669743 | -1 | processed_pseudoge | ENSG00000( |
| 92689955 | 92695647 | 1 | protein_coding | ENSG00000( |
| 92710499 | 92710608 | -1 | misc_RNA | ENSG00000( |
| 92826831 | 93059493 | 1 | protein_coding | ENSG00000( |
| 92878092 | 92878289 | -1 | processed_pseudoge | ENSG00000( |
| 93059663 | 93060426 | -1 | lncRNA | ENSG00000( |
| 93060798 | 93069540 | 1 | protein_coding | ENSG00000( |
| 93063646 | 93077424 | 1 | lncRNA | ENSG00000( |
| 93071972 | 93073466 | 1 | lncRNA | ENSG00000( |
| 93073475 | 93077885 | 1 | protein_coding | ENSG00000( |
| 93106872 | 93107383 | -1 | processed_pseudoge | ENSG00000( |
| 93206857 | 93209627 | -1 | processed_pseudoge | ENSG00000( |
| 93284004 | 93284558 | -1 | processed_pseudoge | ENSG00000( |
| 93306429 | 93482334 | -1 | protein_coding | ENSG00000( |
| 93496612 | 93529092 | 1 | protein_coding | ENSG00000( |
| 93510548 | 93510680 | -1 | rRNA_pseudogene | ENSG00000( |
| 93566665 | 93604480 | 1 | protein_coding | ENSG00000( |
| 93591687 | 93601744 | -1 | protein_coding | ENSG00000( |
| 93612537 | 93666010 | 1 | protein_coding | ENSG00000( |
| 93667883 | 93702592 | -1 | protein_coding | ENSG00000( |
| 93757840 | 93806272 | 1 | protein_coding | ENSG00000( |
| 93762453 | 93771991 | 1 | lncRNA | ENSG00000( |
| 93772681 | 93788605 | -1 | lncRNA | ENSG00000( |
| 93881293 | 93881938 | -1 | processed_pseudoge | ENSG00000( |
| 93893973 | 93956062 | 1 | protein_coding | ENSG00000( |
| 93958191 | 93961540 | -1 | transcribed_processe | ENSG00000( |
| 94030812 | 94332823 | 1 | protein_coding | ENSG00000( |

| | | | | |
|---|---|---|---|---|
| 94081950 | 94108814 | -1 | lncRNA | ENSG0000C |
| 94109116 | 94121107 | 1 | lncRNA | ENSG0000C |
| 94137748 | 94139500 | 1 | processed_pseudoge | ENSG0000C |
| 94155639 | 94155745 | -1 | snRNA | ENSG0000C |
| 94225160 | 94227288 | -1 | lncRNA | ENSG0000C |
| 94278681 | 94287478 | -1 | lncRNA | ENSG0000C |
| 94314907 | 94315327 | -1 | lncRNA | ENSG0000C |
| 94333226 | 94362959 | -1 | protein_coding | ENSG0000C |
| 94402541 | 94536332 | 1 | protein_coding | ENSG0000C |
| 94501434 | 94613905 | 1 | protein_coding | ENSG0000C |
| 94577439 | 94611238 | 1 | lncRNA | ENSG0000C |
| 94646364 | 94647996 | -1 | processed_pseudoge | ENSG0000C |
| 94683729 | 94736190 | 1 | protein_coding | ENSG0000C |
| 94688154 | 94853073 | 1 | protein_coding | ENSG0000C |
| 94762681 | 94855547 | 1 | protein_coding | ENSG0000C |
| 94774156 | 94774633 | -1 | processed_pseudoge | ENSG0000C |
| 94865852 | 94873129 | -1 | processed_pseudoge | ENSG0000C |
| 94882554 | 94883343 | 1 | processed_pseudoge | ENSG0000C |
| 94938658 | 94990091 | 1 | protein_coding | ENSG0000C |
| 94940253 | 94941121 | -1 | processed_pseudoge | ENSG0000C |
| 95007126 | 95007556 | -1 | unprocessed_pseudc | ENSG0000C |
| 95009674 | 95009812 | 1 | unprocessed_pseudc | ENSG0000C |
| 95031640 | 95032048 | 1 | processed_pseudoge | ENSG0000C |
| 95036772 | 95069497 | -1 | protein_coding | ENSG0000C |
| 95109136 | 95168425 | -1 | lncRNA | ENSG0000C |
| 95173085 | 95173187 | -1 | unprocessed_pseudc | ENSG0000C |
| 95183653 | 95184217 | 1 | processed_pseudoge | ENSG0000C |
| 95194200 | 95228928 | 1 | protein_coding | ENSG0000C |
| 95228243 | 95231144 | -1 | lncRNA | ENSG0000C |
| 95237572 | 95291012 | -1 | protein_coding | ENSG0000C |
| 95311771 | 95561414 | -1 | protein_coding | ENSG0000C |
| 95594598 | 95595372 | 1 | processed_pseudoge | ENSG0000C |
| 95605941 | 95656711 | -1 | protein_coding | ENSG0000C |
| 95663396 | 95694143 | -1 | protein_coding | ENSG0000C |
| 95711779 | 95877266 | 1 | protein_coding | ENSG0000C |
| 95732976 | 96090250 | -1 | lncRNA | ENSG0000C |
| 95833508 | 95873758 | 1 | lncRNA | ENSG0000C |
| 95860548 | 95924844 | 1 | protein_coding | ENSG0000C |
| 95907603 | 96032684 | 1 | protein_coding | ENSG0000C |
| 95999778 | 96000243 | -1 | processed_pseudoge | ENSG0000C |
| 96043394 | 96060870 | 1 | protein_coding | ENSG0000C |
| 96064315 | 96064399 | -1 | miRNA | ENSG0000C |
| 96112747 | 96112851 | 1 | snRNA | ENSG0000C |
| 96129055 | 96130528 | -1 | lncRNA | ENSG0000C |
| 96129715 | 96205288 | 1 | protein_coding | ENSG0000C |
| 96189366 | 96190219 | -1 | processed_pseudoge | ENSG0000C |
| 96191702 | 96271587 | -1 | protein_coding | ENSG0000C |
| 96292685 | 96306695 | -1 | lncRNA | ENSG0000C |
| 96304409 | 96338564 | 1 | protein_coding | ENSG0000C |
| 96343221 | 96359365 | -1 | protein_coding | ENSG0000C |

| | | | | |
|---|---|---|---|---|
| 96364608 | 96513926 | -1 | protein_coding | ENSG00000 |
| 96518110 | 96587452 | -1 | protein_coding | ENSG00000 |
| 96560185 | 96560922 | 1 | processed_pseudoge | ENSG00000 |
| 96593315 | 96720514 | -1 | protein_coding | ENSG00000 |
| 96619376 | 96619483 | 1 | rRNA_pseudogene | ENSG00000 |
| 96620877 | 96621707 | -1 | processed_pseudoge | ENSG00000 |
| 96743167 | 96743273 | 1 | snRNA | ENSG00000 |
| 96750288 | 96750899 | 1 | processed_pseudoge | ENSG00000 |
| 96828669 | 96828764 | -1 | miRNA | ENSG00000 |
| 96832282 | 96995959 | 1 | protein_coding | ENSG00000 |
| 96915633 | 96915902 | -1 | processed_pseudoge | ENSG00000 |
| 96998038 | 97185959 | -1 | protein_coding | ENSG00000 |
| 97102756 | 97104355 | 1 | lncRNA | ENSG00000 |
| 97153045 | 97292637 | -1 | protein_coding | ENSG00000 |
| 97195907 | 97203721 | 1 | lncRNA | ENSG00000 |
| 97222173 | 97292673 | -1 | protein_coding | ENSG00000 |
| 97275012 | 97278097 | 1 | processed_pseudoge | ENSG00000 |
| 97297290 | 97297557 | 1 | misc_RNA | ENSG00000 |
| 97309138 | 97309958 | -1 | processed_pseudoge | ENSG00000 |
| 97319271 | 97321915 | 1 | protein_coding | ENSG00000 |
| 97332497 | 97334729 | -1 | protein_coding | ENSG00000 |
| 97334564 | 97343203 | 1 | lncRNA | ENSG00000 |
| 97356358 | 97426076 | -1 | protein_coding | ENSG00000 |
| 97393197 | 97393547 | -1 | processed_pseudoge | ENSG00000 |
| 97401115 | 97419524 | 1 | lncRNA | ENSG00000 |
| 97421195 | 97421307 | -1 | misc_RNA | ENSG00000 |
| 97426191 | 97433444 | 1 | protein_coding | ENSG00000 |
| 97435909 | 97446017 | -1 | protein_coding | ENSG00000 |
| 97446170 | 97457370 | 1 | protein_coding | ENSG00000 |
| 97458324 | 97498794 | -1 | protein_coding | ENSG00000 |
| 97498924 | 97571206 | 1 | protein_coding | ENSG00000 |
| 97572499 | 97583884 | 1 | protein_coding | ENSG00000 |
| 97584323 | 97612802 | 1 | protein_coding | ENSG00000 |
| 97584374 | 97673910 | 1 | protein_coding | ENSG00000 |
| 97589721 | 97590934 | 1 | protein_coding | ENSG00000 |
| 97614553 | 97633500 | -1 | protein_coding | ENSG00000 |
| 97640686 | 97676434 | 1 | protein_coding | ENSG00000 |
| 97677424 | 97687241 | -1 | protein_coding | ENSG00000 |
| 97713173 | 97718150 | 1 | protein_coding | ENSG00000 |
| 97737121 | 97760907 | 1 | protein_coding | ENSG00000 |
| 97766751 | 97771999 | -1 | protein_coding | ENSG00000 |
| 97828478 | 97849798 | -1 | lncRNA | ENSG00000 |
| 97849843 | 97871580 | 1 | protein_coding | ENSG00000 |
| 97865000 | 98030828 | -1 | protein_coding | ENSG00000 |
| 98076361 | 98076421 | -1 | misc_RNA | ENSG00000 |
| 98134624 | 98244897 | 1 | protein_coding | ENSG00000 |
| 98247690 | 98268194 | -1 | protein_coding | ENSG00000 |
| 98252023 | 98256575 | 1 | lncRNA | ENSG00000 |
| 98383565 | 98415182 | -1 | protein_coding | ENSG00000 |
| 98395218 | 98395307 | -1 | miRNA | ENSG00000 |

| | | | | |
|---|---|---|---|---|
| 98416198 | 98446947 | -1 | protein_coding | ENSG00000 |
| 98431292 | 98431360 | -1 | miRNA | ENSG00000 |
| 98446346 | 98453805 | 1 | lncRNA | ENSG00000 |
| 98457077 | 99235862 | -1 | protein_coding | ENSG00000 |
| 98924499 | 98924568 | -1 | miRNA | ENSG00000 |
| 99013103 | 99013952 | -1 | processed_pseudoge | ENSG00000 |
| 99047639 | 99048319 | 1 | processed_pseudoge | ENSG00000 |
| 99329356 | 99394330 | 1 | protein_coding | ENSG00000 |
| 99396870 | 99430624 | -1 | protein_coding | ENSG00000 |
| 99430919 | 99463259 | 1 | lncRNA | ENSG00000 |
| 99526350 | 99531177 | -1 | lncRNA | ENSG00000 |
| 99526948 | 99528467 | 1 | lncRNA | ENSG00000 |
| 99532942 | 99536524 | 1 | protein_coding | ENSG00000 |
| 99610522 | 99620609 | -1 | protein_coding | ENSG00000 |
| 99621055 | 99621918 | 1 | lncRNA | ENSG00000 |
| 99651595 | 99732112 | -1 | protein_coding | ENSG00000 |
| 99651595 | 99659284 | -1 | lncRNA | ENSG00000 |
| 99659509 | 99711241 | 1 | protein_coding | ENSG00000 |
| 99697407 | 99697949 | -1 | processed_pseudoge | ENSG00000 |
| 99702558 | 99756134 | 1 | protein_coding | ENSG00000 |
| 99711844 | 99732100 | -1 | protein_coding | ENSG00000 |
| 99782640 | 99852594 | 1 | protein_coding | ENSG00000 |
| 99788564 | 99789410 | -1 | processed_pseudoge | ENSG00000 |
| 99875577 | 100009947 | -1 | protein_coding | ENSG00000 |
| 99927010 | 99959050 | 1 | lncRNA | ENSG00000 |
| 100042193 | 100081869 | -1 | protein_coding | ENSG00000 |
| 100105754 | 100140151 | -1 | unitary_pseudogene | ENSG00000 |
| 100114417 | 100115223 | -1 | processed_pseudoge | ENSG00000 |
| 100143065 | 100143995 | -1 | processed_pseudoge | ENSG00000 |
| 100150094 | 100186033 | -1 | protein_coding | ENSG00000 |
| 100188300 | 100229596 | -1 | protein_coding | ENSG00000 |
| 100190036 | 100190747 | 1 | lncRNA | ENSG00000 |
| 100229629 | 100234398 | 1 | lncRNA | ENSG00000 |
| 100232298 | 100267680 | -1 | protein_coding | ENSG00000 |
| 100237156 | 100237302 | -1 | snoRNA | ENSG00000 |
| 100248271 | 100249095 | 1 | processed_pseudoge | ENSG00000 |
| 100258571 | 100258677 | 1 | snRNA | ENSG00000 |
| 100273280 | 100286680 | -1 | protein_coding | ENSG00000 |
| 100288149 | 100330264 | -1 | protein_coding | ENSG00000 |
| 100335563 | 100346390 | -1 | lncRNA | ENSG00000 |
| 100347233 | 100364826 | 1 | protein_coding | ENSG00000 |
| 100373099 | 100454043 | 1 | lncRNA | ENSG00000 |
| 100398352 | 100398446 | -1 | misc_RNA | ENSG00000 |
| 100463009 | 100483744 | 1 | protein_coding | ENSG00000 |
| 100486646 | 100519864 | -1 | protein_coding | ENSG00000 |
| 100505628 | 100529881 | -1 | protein_coding | ENSG00000 |
| 100523740 | 100530000 | -1 | protein_coding | ENSG00000 |
| 100529072 | 100559998 | 1 | protein_coding | ENSG00000 |
| 100666695 | 100667009 | -1 | misc_RNA | ENSG00000 |
| 100735603 | 100829941 | 1 | protein_coding | ENSG00000 |

| | | | | |
|---|---|---|---|---|
| 100907174 | 100907280 | -1 | misc_RNA | ENSG00000 |
| 100911103 | 100912739 | -1 | lncRNA | ENSG00000 |
| 100912963 | 100965134 | 1 | protein_coding | ENSG00000 |
| 100967688 | 100968135 | 1 | lncRNA | ENSG00000 |
| 100969458 | 100987515 | -1 | protein_coding | ENSG00000 |
| 100969518 | 100985871 | 1 | protein_coding | ENSG00000 |
| 100974985 | 100975084 | 1 | miRNA | ENSG00000 |
| 100980507 | 100985614 | -1 | lncRNA | ENSG00000 |
| 100987367 | 100994403 | 1 | protein_coding | ENSG00000 |
| 100996618 | 101007836 | 1 | protein_coding | ENSG00000 |
| 101007679 | 101032295 | -1 | protein_coding | ENSG00000 |
| 101031234 | 101041244 | 1 | protein_coding | ENSG00000 |
| 101060029 | 101061005 | -1 | lncRNA | ENSG00000 |
| 101061841 | 101068131 | 1 | protein_coding | ENSG00000 |
| 101089321 | 101131126 | -1 | lncRNA | ENSG00000 |
| 101131300 | 101137789 | 1 | protein_coding | ENSG00000 |
| 101176306 | 101194147 | 1 | lncRNA | ENSG00000 |
| 101181144 | 101181951 | -1 | processed_pseudoge | ENSG00000 |
| 101226195 | 101229794 | -1 | protein_coding | ENSG00000 |
| 101229577 | 101270148 | 1 | lncRNA | ENSG00000 |
| 101252821 | 101263550 | -1 | lncRNA | ENSG00000 |
| 101307529 | 101307718 | 1 | snRNA | ENSG00000 |
| 101311018 | 101311505 | 1 | lncRNA | ENSG00000 |
| 101322485 | 101323394 | -1 | lncRNA | ENSG00000 |
| 101354033 | 101557321 | 1 | protein_coding | ENSG00000 |
| 101364845 | 101365035 | -1 | snRNA | ENSG00000 |
| 101570560 | 101609662 | 1 | protein_coding | ENSG00000 |
| 101578882 | 101588270 | -1 | protein_coding | ENSG00000 |
| 101601417 | 101601497 | -1 | miRNA | ENSG00000 |
| 101601417 | 101601497 | 1 | miRNA | ENSG00000 |
| 101610664 | 101695295 | -1 | protein_coding | ENSG00000 |
| 101622859 | 101622965 | 1 | snRNA | ENSG00000 |
| 101694401 | 101705119 | 1 | lncRNA | ENSG00000 |
| 101701994 | 101730037 | -1 | lncRNA | ENSG00000 |
| 101770130 | 101780369 | -1 | protein_coding | ENSG00000 |
| 101781325 | 101783413 | -1 | protein_coding | ENSG00000 |
| 101784443 | 101818465 | -1 | protein_coding | ENSG00000 |
| 101818800 | 101828885 | 1 | lncRNA | ENSG00000 |
| 101825974 | 101843920 | -1 | protein_coding | ENSG00000 |
| 101845599 | 102056193 | -1 | protein_coding | ENSG00000 |
| 101912998 | 101913279 | -1 | misc_RNA | ENSG00000 |
| 101952605 | 101952704 | -1 | misc_RNA | ENSG00000 |
| 102065390 | 102068038 | 1 | protein_coding | ENSG00000 |
| 102085476 | 102085730 | -1 | processed_pseudoge | ENSG00000 |
| 102089968 | 102090062 | 1 | processed_pseudoge | ENSG00000 |
| 102107560 | 102120453 | -1 | protein_coding | ENSG00000 |
| 102132994 | 102150333 | 1 | protein_coding | ENSG00000 |
| 102152176 | 102163871 | 1 | protein_coding | ENSG00000 |
| 102226299 | 102229589 | 1 | protein_coding | ENSG00000 |
| 102230186 | 102241512 | -1 | protein_coding | ENSG00000 |

| | | | | |
|---|---|---|---|---|
| 102245532 | 102382899 | 1 | protein_coding | ENSG00000 |
| 102394110 | 102402524 | 1 | protein_coding | ENSG00000 |
| 102402617 | 102421539 | -1 | protein_coding | ENSG00000 |
| 102419189 | 102423136 | 1 | protein_coding | ENSG00000 |
| 102423245 | 102432584 | -1 | protein_coding | ENSG00000 |
| 102436512 | 102436584 | 1 | miRNA | ENSG00000 |
| 102449816 | 102461106 | 1 | lncRNA | ENSG00000 |
| 102449837 | 102451543 | -1 | protein_coding | ENSG00000 |
| 102461395 | 102477045 | 1 | protein_coding | ENSG00000 |
| 102461881 | 102502712 | -1 | protein_coding | ENSG00000 |
| 102483039 | 102483559 | 1 | lncRNA | ENSG00000 |
| 102503987 | 102633535 | 1 | protein_coding | ENSG00000 |
| 102572757 | 102573225 | -1 | processed_pseudoge | ENSG00000 |
| 102591524 | 102591630 | 1 | snRNA | ENSG00000 |
| 102642310 | 102660680 | 1 | protein_coding | ENSG00000 |
| 102642792 | 102644140 | -1 | lncRNA | ENSG00000 |
| 102673731 | 102714397 | -1 | protein_coding | ENSG00000 |
| 102714538 | 102743492 | 1 | protein_coding | ENSG00000 |
| 102743948 | 102834516 | 1 | protein_coding | ENSG00000 |
| 102803929 | 102804031 | 1 | snRNA | ENSG00000 |
| 102830531 | 102837472 | -1 | protein_coding | ENSG00000 |
| 102838011 | 102845473 | -1 | processed_pseudoge | ENSG00000 |
| 102845595 | 102845950 | 1 | processed_pseudoge | ENSG00000 |
| 102845761 | 102854513 | -1 | lncRNA | ENSG00000 |
| 102854259 | 102864961 | 1 | protein_coding | ENSG00000 |
| 102854272 | 102901899 | 1 | protein_coding | ENSG00000 |
| 102868421 | 102868519 | 1 | snRNA | ENSG00000 |
| 102869470 | 102901899 | 1 | protein_coding | ENSG00000 |
| 102887874 | 102888188 | -1 | processed_pseudoge | ENSG00000 |
| 102898649 | 102918335 | -1 | lncRNA | ENSG00000 |
| 102914585 | 102915404 | 1 | lncRNA | ENSG00000 |
| 102918294 | 103090222 | 1 | protein_coding | ENSG00000 |
| 103027336 | 103027498 | 1 | misc_RNA | ENSG00000 |
| 103088017 | 103193306 | -1 | protein_coding | ENSG00000 |
| 103125017 | 103125416 | -1 | processed_pseudoge | ENSG00000 |
| 103175554 | 103176094 | 1 | processed_pseudoge | ENSG00000 |
| 103215709 | 103216800 | 1 | processed_pseudoge | ENSG00000 |
| 103245887 | 103248016 | 1 | protein_coding | ENSG00000 |
| 103277138 | 103290346 | 1 | protein_coding | ENSG00000 |
| 103302796 | 103351144 | -1 | protein_coding | ENSG00000 |
| 103312436 | 103312568 | -1 | snRNA | ENSG00000 |
| 103367976 | 103389065 | 1 | protein_coding | ENSG00000 |
| 103389041 | 103396492 | -1 | protein_coding | ENSG00000 |
| 103394253 | 103394401 | -1 | miRNA | ENSG00000 |
| 103396626 | 103446294 | 1 | protein_coding | ENSG00000 |
| 103446786 | 103452402 | -1 | protein_coding | ENSG00000 |
| 103450196 | 103450852 | 1 | lncRNA | ENSG00000 |
| 103452846 | 103462695 | 1 | lncRNA | ENSG00000 |
| 103453387 | 103458888 | -1 | protein_coding | ENSG00000 |
| 103472804 | 103479240 | -1 | protein_coding | ENSG00000 |

| | | | | |
|---|---|---|---|---|
| 103479603 | 103517393 | -1 | lncRNA | ENSG0000C |
| 103493979 | 103592552 | 1 | protein_coding | ENSG0000C |
| 103549075 | 103550964 | 1 | lncRNA | ENSG0000C |
| 103573157 | 103583687 | -1 | lncRNA | ENSG0000C |
| 103594027 | 103855543 | -1 | protein_coding | ENSG0000C |
| 103608619 | 103610050 | 1 | lncRNA | ENSG0000C |
| 103673158 | 103673265 | 1 | misc_RNA | ENSG0000C |
| 103745966 | 103755423 | 1 | lncRNA | ENSG0000C |
| 103877374 | 103879761 | -1 | lncRNA | ENSG0000C |
| 103877569 | 103918184 | -1 | protein_coding | ENSG0000C |
| 103967140 | 104029233 | 1 | protein_coding | ENSG0000C |
| 103974515 | 103974814 | -1 | misc_RNA | ENSG0000C |
| 104031286 | 104085880 | -1 | protein_coding | ENSG0000C |
| 104048089 | 104048186 | -1 | miRNA | ENSG0000C |
| 104122058 | 104126385 | 1 | protein_coding | ENSG0000C |
| 104129888 | 104232362 | -1 | protein_coding | ENSG0000C |
| 104218789 | 104218883 | -1 | miRNA | ENSG0000C |
| 104235356 | 104267459 | 1 | protein_coding | ENSG0000C |
| 104268336 | 104268405 | -1 | miRNA | ENSG0000C |
| 104268873 | 104304950 | 1 | protein_coding | ENSG0000C |
| 104309698 | 104338465 | -1 | protein_coding | ENSG0000C |
| 104312141 | 104313881 | 1 | lncRNA | ENSG0000C |
| 104313308 | 104313479 | -1 | transcribed_unproce | ENSG0000C |
| 104323369 | 104327004 | 1 | lncRNA | ENSG0000C |
| 104351591 | 104353575 | -1 | lncRNA | ENSG0000C |
| 104353833 | 104455102 | 1 | protein_coding | ENSG0000C |
| 104474939 | 104480274 | -1 | lncRNA | ENSG0000C |
| 104566931 | 104624795 | 1 | lncRNA | ENSG0000C |
| 104641290 | 105265242 | 1 | protein_coding | ENSG0000C |
| 104664608 | 104666200 | -1 | lncRNA | ENSG0000C |
| 105534659 | 105534765 | 1 | snRNA | ENSG0000C |
| 105685383 | 105685865 | -1 | processed_pseudoge | ENSG0000C |
| 105686322 | 105687051 | 1 | processed_pseudoge | ENSG0000C |
| 105808225 | 105820349 | -1 | lncRNA | ENSG0000C |
| 105844834 | 105904771 | 1 | lncRNA | ENSG0000C |
| 105985679 | 105987727 | -1 | lncRNA | ENSG0000C |
| 106140143 | 106188722 | 1 | lncRNA | ENSG0000C |
| 106550106 | 106551121 | 1 | processed_pseudoge | ENSG0000C |
| 106573663 | 107164534 | -1 | protein_coding | ENSG0000C |
| 106648899 | 106676780 | 1 | lncRNA | ENSG0000C |
| 106970021 | 106970150 | 1 | rRNA_pseudogene | ENSG0000C |
| 107461560 | 107461669 | 1 | rRNA_pseudogene | ENSG0000C |
| 107694973 | 108197849 | -1 | lncRNA | ENSG0000C |
| 108296099 | 108296532 | 1 | processed_pseudoge | ENSG0000C |
| 108547975 | 108561815 | -1 | lncRNA | ENSG0000C |
| 108708539 | 108840686 | 1 | lncRNA | ENSG0000C |
| 108848186 | 108849561 | 1 | processed_pseudoge | ENSG0000C |
| 108940969 | 108941310 | -1 | misc_RNA | ENSG0000C |
| 109184934 | 109185001 | -1 | snRNA | ENSG0000C |
| 109239639 | 109239746 | 1 | snRNA | ENSG0000C |

| | | | | |
|---|---|---|---|---|
| 109528119 | 109529008 | 1 | processed_pseudoge | ENSG0000C |
| 109534655 | 109534990 | 1 | processed_pseudoge | ENSG0000C |
| 109661814 | 109662050 | 1 | processed_pseudoge | ENSG0000C |
| 109680442 | 109681041 | 1 | processed_pseudoge | ENSG0000C |
| 109807651 | 109808129 | 1 | processed_pseudoge | ENSG0000C |
| 109864766 | 109923553 | -1 | protein_coding | ENSG0000C |
| 109869858 | 109869992 | 1 | snRNA | ENSG0000C |
| 109940104 | 110008381 | -1 | lncRNA | ENSG0000C |
| 109996368 | 110135565 | 1 | protein_coding | ENSG0000C |
| 110139214 | 110139426 | -1 | processed_pseudoge | ENSG0000C |
| 110207605 | 110287365 | 1 | protein_coding | ENSG0000C |
| 110207850 | 110208591 | -1 | lncRNA | ENSG0000C |
| 110257064 | 110257843 | 1 | processed_pseudoge | ENSG0000C |
| 110290730 | 110304938 | -1 | protein_coding | ENSG0000C |
| 110428840 | 110496590 | -1 | lncRNA | ENSG0000C |
| 110450629 | 110451240 | 1 | processed_pseudoge | ENSG0000C |
| 110497907 | 110511533 | 1 | protein_coding | ENSG0000C |
| 110550115 | 110550854 | -1 | processed_pseudoge | ENSG0000C |
| 110567691 | 110604636 | 1 | protein_coding | ENSG0000C |
| 110642678 | 110642785 | -1 | misc_RNA | ENSG0000C |
| 110644336 | 110839468 | 1 | protein_coding | ENSG0000C |
| 110756456 | 110756723 | 1 | misc_RNA | ENSG0000C |
| 110761142 | 110761242 | 1 | rRNA_pseudogene | ENSG0000C |
| 110869743 | 110871594 | 1 | lncRNA | ENSG0000C |
| 110869868 | 110872233 | -1 | lncRNA | ENSG0000C |
| 110871795 | 110900006 | 1 | protein_coding | ENSG0000C |
| 110898090 | 110898155 | 1 | miRNA | ENSG0000C |
| 110898728 | 110919201 | -1 | protein_coding | ENSG0000C |
| 110907483 | 110908182 | 1 | lncRNA | ENSG0000C |
| 110910596 | 110912244 | 1 | lncRNA | ENSG0000C |
| 110919547 | 111013667 | 1 | protein_coding | ENSG0000C |
| 110936622 | 110937233 | -1 | processed_pseudoge | ENSG0000C |
| 110988926 | 110989013 | 1 | miRNA | ENSG0000C |
| 111077029 | 111080907 | 1 | protein_coding | ENSG0000C |
| 111104956 | 111105248 | 1 | processed_pseudoge | ENSG0000C |
| 111322778 | 111323397 | 1 | processed_pseudoge | ENSG0000C |
| 111349939 | 111352103 | -1 | lncRNA | ENSG0000C |
| 111498191 | 111498941 | -1 | processed_pseudoge | ENSG0000C |
| 111787233 | 111989909 | -1 | lncRNA | ENSG0000C |
| 112149865 | 112215377 | -1 | protein_coding | ENSG0000C |
| 112283400 | 112305038 | 1 | protein_coding | ENSG0000C |
| 112299612 | 112299690 | 1 | miRNA | ENSG0000C |
| 112299612 | 112299688 | -1 | miRNA | ENSG0000C |
| 112308181 | 112356574 | -1 | transcribed_unitary_ | ENSG0000C |
| 112374018 | 112428380 | 1 | protein_coding | ENSG0000C |
| 112395813 | 112425589 | -1 | lncRNA | ENSG0000C |
| 112424428 | 112446917 | -1 | protein_coding | ENSG0000C |
| 112446998 | 112818744 | 1 | protein_coding | ENSG0000C |
| 112548632 | 112548872 | -1 | processed_pseudoge | ENSG0000C |
| 112634170 | 112634254 | 1 | miRNA | ENSG0000C |

| | | | | |
|---|---|---|---|---|
| 112671812 | 112677819 | -1 | lncRNA | ENSG00000( |
| 112810031 | 112821838 | -1 | lncRNA | ENSG00000( |
| 112823490 | 112827726 | 1 | lncRNA | ENSG00000( |
| 112831169 | 112832454 | 1 | TEC | ENSG00000( |
| 112888735 | 112906111 | 1 | lncRNA | ENSG00000( |
| 112950247 | 113167678 | 1 | protein_coding | ENSG00000( |
| 112950646 | 112951875 | -1 | lncRNA | ENSG00000( |
| 112987476 | 112987865 | 1 | processed_pseudoge | ENSG00000( |
| 113353425 | 113353484 | 1 | snRNA | ENSG00000( |
| 113482069 | 113492054 | -1 | lncRNA | ENSG00000( |
| 113482616 | 113483113 | 1 | processed_pseudoge | ENSG00000( |
| 113550837 | 113589602 | 1 | protein_coding | ENSG00000( |
| 113588716 | 113664127 | -1 | protein_coding | ENSG00000( |
| 113679162 | 113730907 | 1 | protein_coding | ENSG00000( |
| 113710681 | 113719332 | -1 | lncRNA | ENSG00000( |
| 113751262 | 113783429 | 1 | protein_coding | ENSG00000( |
| 113777993 | 113778054 | -1 | miRNA | ENSG00000( |
| 113820459 | 113820590 | 1 | snoRNA | ENSG00000( |
| 113834725 | 113854383 | -1 | protein_coding | ENSG00000( |
| 113854661 | 113917194 | 1 | protein_coding | ENSG00000( |
| 113865811 | 113869614 | -1 | lncRNA | ENSG00000( |
| 114043866 | 114046904 | 1 | protein_coding | ENSG00000( |
| 114109856 | 114118700 | -1 | lncRNA | ENSG00000( |
| 114111414 | 114111517 | 1 | snRNA | ENSG00000( |
| 114117147 | 114117551 | 1 | processed_pseudoge | ENSG00000( |
| 114120862 | 114174232 | -1 | protein_coding | ENSG00000( |
| 114168156 | 114168280 | -1 | ribozyme | ENSG00000( |
| 114174105 | 114174179 | -1 | miRNA | ENSG00000( |
| 114179270 | 114232304 | 1 | protein_coding | ENSG00000( |
| 114239254 | 114294489 | 1 | protein_coding | ENSG00000( |
| 114246074 | 114246714 | 1 | processed_pseudoge | ENSG00000( |
| 114280725 | 114281258 | -1 | lncRNA | ENSG00000( |
| 114294824 | 114404756 | -1 | protein_coding | ENSG00000( |
| 114423770 | 114424067 | -1 | misc_RNA | ENSG00000( |
| 114424445 | 114427398 | -1 | TEC | ENSG00000( |
| 114431113 | 114768061 | -1 | protein_coding | ENSG00000( |
| 114501997 | 114502347 | -1 | processed_pseudoge | ENSG00000( |
| 114690143 | 114690634 | -1 | processed_pseudoge | ENSG00000( |
| 114764788 | 114794344 | 1 | lncRNA | ENSG00000( |
| 114809375 | 114811170 | 1 | processed_pseudoge | ENSG00000( |
| 114821744 | 114899832 | 1 | protein_coding | ENSG00000( |
| 114900722 | 114901366 | 1 | processed_pseudoge | ENSG00000( |
| 114938195 | 114977676 | 1 | protein_coding | ENSG00000( |
| 114969460 | 114969568 | -1 | snRNA | ENSG00000( |
| 114994657 | 114996593 | 1 | lncRNA | ENSG00000( |
| 115093365 | 115948999 | 1 | protein_coding | ENSG00000( |
| 115123482 | 115218819 | -1 | lncRNA | ENSG00000( |
| 115819842 | 115820487 | -1 | processed_pseudoge | ENSG00000( |
| 116056925 | 116273467 | -1 | protein_coding | ENSG00000( |
| 116274270 | 116281071 | 1 | lncRNA | ENSG00000( |

| | | | | |
|---|---|---|---|---|
| 116324448 | 116380029 | 1 | protein_coding | ENSG00000 |
| 116409256 | 116409475 | 1 | processed_pseudoge | ENSG00000 |
| 116427847 | 116477957 | 1 | protein_coding | ENSG00000 |
| 116439468 | 116440044 | -1 | processed_pseudoge | ENSG00000 |
| 116538183 | 116538276 | 1 | snRNA | ENSG00000 |
| 116545931 | 116567855 | 1 | protein_coding | ENSG00000 |
| 116577308 | 116583317 | 1 | unprocessed_pseudo | ENSG00000 |
| 116590385 | 116609175 | 1 | protein_coding | ENSG00000 |
| 116620953 | 116645143 | 1 | polymorphic_pseudo | ENSG00000 |
| 116663696 | 116670264 | -1 | protein_coding | ENSG00000 |
| 116670034 | 116672750 | 1 | lncRNA | ENSG00000 |
| 116671192 | 116850236 | -1 | protein_coding | ENSG00000 |
| 116749905 | 116751261 | 1 | processed_pseudoge | ENSG00000 |
| 116849499 | 116911788 | 1 | protein_coding | ENSG00000 |
| 116881477 | 117126586 | -1 | protein_coding | ENSG00000 |
| 117005462 | 117006416 | 1 | lncRNA | ENSG00000 |
| 117023674 | 117023851 | -1 | processed_pseudoge | ENSG00000 |
| 117128521 | 117138301 | -1 | protein_coding | ENSG00000 |
| 117144564 | 117169108 | -1 | lncRNA | ENSG00000 |
| 117167678 | 117167774 | -1 | miRNA | ENSG00000 |
| 117196168 | 117196654 | -1 | processed_pseudoge | ENSG00000 |
| 117197489 | 117210299 | 1 | protein_coding | ENSG00000 |
| 117239600 | 117241923 | -1 | lncRNA | ENSG00000 |
| 117241093 | 117279430 | 1 | protein_coding | ENSG00000 |
| 117267116 | 117268668 | -1 | lncRNA | ENSG00000 |
| 117277274 | 117375440 | -1 | protein_coding | ENSG00000 |
| 117425194 | 117490419 | 1 | lncRNA | ENSG00000 |
| 117473215 | 117545068 | -1 | lncRNA | ENSG00000 |
| 117542445 | 117549546 | 1 | protein_coding | ENSG00000 |
| 117562084 | 117572457 | -1 | lncRNA | ENSG00000 |
| 117579854 | 117593896 | 1 | lncRNA | ENSG00000 |
| 117734510 | 117753329 | 1 | lncRNA | ENSG00000 |
| 117825894 | 117831244 | -1 | lncRNA | ENSG00000 |
| 118004916 | 118046941 | -1 | protein_coding | ENSG00000 |
| 118017487 | 118045810 | 1 | lncRNA | ENSG00000 |
| 118046279 | 118210158 | 1 | lncRNA | ENSG00000 |
| 118241468 | 118267710 | 1 | lncRNA | ENSG00000 |
| 118297925 | 118342328 | -1 | protein_coding | ENSG00000 |
| 118341437 | 118398468 | 1 | lncRNA | ENSG00000 |
| 118572871 | 118573620 | 1 | processed_pseudoge | ENSG00000 |
| 118589997 | 118595648 | -1 | protein_coding | ENSG00000 |
| 118643760 | 118644198 | 1 | processed_pseudoge | ENSG00000 |
| 118674167 | 118755249 | -1 | protein_coding | ENSG00000 |
| 118692361 | 118693535 | -1 | lncRNA | ENSG00000 |
| 118784563 | 119029544 | -1 | lncRNA | ENSG00000 |
| 118872054 | 118872602 | -1 | processed_pseudoge | ENSG00000 |
| 118932674 | 118933295 | -1 | processed_pseudoge | ENSG00000 |
| 119029714 | 119033730 | 1 | protein_coding | ENSG00000 |
| 119033670 | 119080823 | -1 | protein_coding | ENSG00000 |
| 119060011 | 119060138 | -1 | snoRNA | ENSG00000 |

| | | | | |
|---|---|---|---|---|
| 119060983 | 119061111 | -1 | snoRNA | ENSG00000 |
| 119093140 | 119093341 | 1 | processed_pseudoge | ENSG00000 |
| 119104086 | 119137984 | 1 | protein_coding | ENSG00000 |
| 119140767 | 119165714 | -1 | protein_coding | ENSG00000 |
| 119167720 | 119178812 | -1 | protein_coding | ENSG00000 |
| 119207571 | 119459745 | 1 | protein_coding | ENSG00000 |
| 119208531 | 119211760 | 1 | lncRNA | ENSG00000 |
| 119212857 | 119213146 | 1 | misc_RNA | ENSG00000 |
| 119330233 | 119336182 | -1 | lncRNA | ENSG00000 |
| 119377972 | 119378043 | 1 | miRNA | ENSG00000 |
| 119499817 | 119542719 | -1 | protein_coding | ENSG00000 |
| 119562101 | 119562530 | -1 | processed_pseudoge | ENSG00000 |
| 119571802 | 119597029 | -1 | protein_coding | ENSG00000 |
| 119621068 | 119621880 | 1 | processed_pseudoge | ENSG00000 |
| 119638456 | 119639082 | 1 | processed_pseudoge | ENSG00000 |
| 119651370 | 119677819 | 1 | protein_coding | ENSG00000 |
| 119683127 | 119683823 | -1 | processed_pseudoge | ENSG00000 |
| 119726042 | 119829147 | 1 | protein_coding | ENSG00000 |
| 119768247 | 119768538 | 1 | misc_RNA | ENSG00000 |
| 119793589 | 119816642 | -1 | transcribed_processe | ENSG00000 |
| 119818866 | 119819133 | 1 | processed_pseudoge | ENSG00000 |
| 119829404 | 119892556 | -1 | protein_coding | ENSG00000 |
| 119892730 | 119944657 | 1 | protein_coding | ENSG00000 |
| 119958513 | 119958592 | 1 | miRNA | ENSG00000 |
| 119997086 | 119997709 | -1 | processed_pseudoge | ENSG00000 |
| 120069980 | 120070159 | -1 | processed_pseudoge | ENSG00000 |
| 120354701 | 120355183 | -1 | processed_pseudoge | ENSG00000 |
| 120434983 | 120440376 | 1 | unprocessed_pseudo | ENSG00000 |
| 120456954 | 120589855 | 1 | protein_coding | ENSG00000 |
| 120597949 | 120600124 | 1 | lncRNA | ENSG00000 |
| 120608580 | 120825317 | 1 | lncRNA | ENSG00000 |
| 120759898 | 120851457 | -1 | lncRNA | ENSG00000 |
| 120842521 | 120845146 | 1 | lncRNA | ENSG00000 |
| 120851305 | 120909524 | 1 | protein_coding | ENSG00000 |
| 120879256 | 120880667 | -1 | lncRNA | ENSG00000 |
| 120925547 | 120980700 | -1 | lncRNA | ENSG00000 |
| 120984966 | 120985596 | -1 | lncRNA | ENSG00000 |
| 121133090 | 121133723 | -1 | processed_pseudoge | ENSG00000 |
| 121178700 | 121185966 | 1 | lncRNA | ENSG00000 |
| 121410916 | 121411200 | 1 | misc_RNA | ENSG00000 |
| 121478334 | 121598458 | -1 | protein_coding | ENSG00000 |
| 121615425 | 121615839 | -1 | lncRNA | ENSG00000 |
| 121713607 | 121713994 | 1 | processed_pseudoge | ENSG00000 |
| 121736303 | 121739730 | 1 | lncRNA | ENSG00000 |
| 121740421 | 121928801 | -1 | protein_coding | ENSG00000 |
| 121846600 | 121846700 | 1 | misc_RNA | ENSG00000 |
| 121928312 | 121951965 | 1 | transcribed_unitary_ | ENSG00000 |
| 121949412 | 121949487 | -1 | misc_RNA | ENSG00000 |
| 121956782 | 121957098 | 1 | lncRNA | ENSG00000 |
| 121957091 | 121975217 | -1 | protein_coding | ENSG00000 |

| | | | | |
|---|---|---|---|---|
| 121965764 | 121967700 | 1 | lncRNA | ENSG0000C |
| 121989163 | 122254545 | 1 | protein_coding | ENSG0000C |
| 122017982 | 122019160 | -1 | lncRNA | ENSG0000C |
| 122145106 | 122155197 | -1 | lncRNA | ENSG0000C |
| 122271296 | 122338170 | 1 | protein_coding | ENSG0000C |
| 122345690 | 122345793 | 1 | snRNA | ENSG0000C |
| 122374696 | 122442602 | 1 | protein_coding | ENSG0000C |
| 122416965 | 122417067 | 1 | miRNA | ENSG0000C |
| 122435924 | 122461383 | -1 | lncRNA | ENSG0000C |
| 122454653 | 122457352 | 1 | protein_coding | ENSG0000C |
| 122458551 | 122514907 | 1 | protein_coding | ENSG0000C |
| 122560665 | 122643740 | 1 | protein_coding | ENSG0000C |
| 122657498 | 122664196 | 1 | unprocessed_pseudc | ENSG0000C |
| 122672850 | 122679494 | 1 | protein_coding | ENSG0000C |
| 122697709 | 122699846 | -1 | protein_coding | ENSG0000C |
| 122729429 | 122826868 | 1 | transcribed_unitary_ | ENSG0000C |
| 122832158 | 122846175 | -1 | protein_coding | ENSG0000C |
| 122832167 | 122879583 | -1 | protein_coding | ENSG0000C |
| 122849078 | 122879641 | -1 | protein_coding | ENSG0000C |
| 122879681 | 122898615 | 1 | transcribed_processe | ENSG0000C |
| 122910610 | 122913111 | 1 | protein_coding | ENSG0000C |
| 122930901 | 122954311 | -1 | protein_coding | ENSG0000C |
| 122954381 | 122997513 | 1 | protein_coding | ENSG0000C |
| 122990806 | 123008817 | -1 | protein_coding | ENSG0000C |
| 123008979 | 123058311 | 1 | protein_coding | ENSG0000C |
| 123135970 | 123139423 | 1 | protein_coding | ENSG0000C |
| 123148122 | 123150672 | 1 | protein_coding | ENSG0000C |
| 123154402 | 123170467 | 1 | protein_coding | ENSG0000C |
| 123171535 | 123171875 | -1 | processed_pseudoge | ENSG0000C |
| 123323422 | 123323586 | -1 | processed_pseudoge | ENSG0000C |
| 123356450 | 123537767 | 1 | lncRNA | ENSG0000C |
| 123425713 | 123427640 | -1 | lncRNA | ENSG0000C |
| 123560326 | 123562213 | 1 | lncRNA | ENSG0000C |
| 123574227 | 123583767 | 1 | lncRNA | ENSG0000C |
| 123666355 | 123697399 | 1 | protein_coding | ENSG0000C |
| 123706207 | 123940267 | -1 | protein_coding | ENSG0000C |
| 123776670 | 123777749 | 1 | lncRNA | ENSG0000C |
| 123913574 | 123931597 | -1 | lncRNA | ENSG0000C |
| 123943255 | 123944099 | -1 | lncRNA | ENSG0000C |
| 123991910 | 123992669 | 1 | processed_pseudoge | ENSG0000C |
| 124004304 | 124004828 | -1 | processed_pseudoge | ENSG0000C |
| 124007668 | 124093598 | -1 | protein_coding | ENSG0000C |
| 124397303 | 124418976 | -1 | protein_coding | ENSG0000C |
| 124445243 | 124450035 | -1 | protein_coding | ENSG0000C |
| 124447152 | 124450048 | 1 | lncRNA | ENSG0000C |
| 124461823 | 124617888 | 1 | protein_coding | ENSG0000C |
| 124489823 | 124490578 | -1 | processed_pseudoge | ENSG0000C |
| 124617080 | 124791727 | -1 | protein_coding | ENSG0000C |
| 124619292 | 124744378 | -1 | protein_coding | ENSG0000C |
| 124623353 | 124624079 | 1 | lncRNA | ENSG0000C |

| | | | | |
|---|---|---|---|---|
| 124703625 | 124714217 | 1 | lncRNA | ENSG00000 |
| 124748149 | 124791887 | -1 | protein_coding | ENSG00000 |
| 124766196 | 124766297 | 1 | misc_RNA | ENSG00000 |
| 124801819 | 124836667 | 1 | protein_coding | ENSG00000 |
| 124814958 | 124815062 | -1 | misc_RNA | ENSG00000 |
| 124867510 | 124868326 | 1 | processed_pseudoge | ENSG00000 |
| 124917143 | 124942881 | 1 | lncRNA | ENSG00000 |
| 124942123 | 124988189 | 1 | protein_coding | ENSG00000 |
| 124945204 | 124946432 | -1 | lncRNA | ENSG00000 |
| 124984317 | 125161170 | -1 | protein_coding | ENSG00000 |
| 124996064 | 125001491 | 1 | lncRNA | ENSG00000 |
| 125032783 | 125032870 | -1 | miRNA | ENSG00000 |
| 125162379 | 125162971 | 1 | lncRNA | ENSG00000 |
| 125166677 | 125166937 | 1 | processed_pseudoge | ENSG00000 |
| 125473686 | 125473932 | -1 | unprocessed_pseudo | ENSG00000 |
| 125574371 | 125578445 | 1 | lncRNA | ENSG00000 |
| 125576522 | 125683163 | -1 | protein_coding | ENSG00000 |
| 125666875 | 125667950 | -1 | processed_pseudoge | ENSG00000 |
| 125683229 | 125709677 | 1 | lncRNA | ENSG00000 |
| 125700436 | 125719566 | -1 | lncRNA | ENSG00000 |
| 125707416 | 125749525 | -1 | lncRNA | ENSG00000 |
| 125718771 | 125719365 | -1 | lncRNA | ENSG00000 |
| 125719515 | 125764143 | 1 | protein_coding | ENSG00000 |
| 125725634 | 125752110 | -1 | lncRNA | ENSG00000 |
| 125753580 | 125775821 | -1 | protein_coding | ENSG00000 |
| 125784980 | 125823258 | -1 | protein_coding | ENSG00000 |
| 125819740 | 125819822 | 1 | miRNA | ENSG00000 |
| 125823546 | 125853695 | 1 | protein_coding | ENSG00000 |
| 125836337 | 125896436 | -1 | protein_coding | ENSG00000 |
| 125869295 | 125869576 | 1 | misc_RNA | ENSG00000 |
| 125890566 | 125890678 | -1 | snRNA | ENSG00000 |
| 125896539 | 126009592 | 1 | protein_coding | ENSG00000 |
| 125960208 | 125960406 | -1 | processed_pseudoge | ENSG00000 |
| 125972188 | 125973126 | -1 | lncRNA | ENSG00000 |
| 126012381 | 126388455 | -1 | protein_coding | ENSG00000 |
| 126145474 | 126145582 | -1 | rRNA_pseudogene | ENSG00000 |
| 126272818 | 126273411 | 1 | processed_pseudoge | ENSG00000 |
| 126393489 | 126398789 | -1 | lncRNA | ENSG00000 |
| 126413869 | 126421879 | -1 | lncRNA | ENSG00000 |
| 126424997 | 126798708 | -1 | protein_coding | ENSG00000 |
| 126425930 | 126460957 | 1 | lncRNA | ENSG00000 |
| 126779702 | 126779774 | -1 | snoRNA | ENSG00000 |
| 126905409 | 127452517 | 1 | protein_coding | ENSG00000 |
| 126988095 | 127026507 | -1 | lncRNA | ENSG00000 |
| 127013501 | 127026475 | -1 | lncRNA | ENSG00000 |
| 127135426 | 127196591 | -1 | protein_coding | ENSG00000 |
| 127549369 | 127552639 | 1 | protein_coding | ENSG00000 |
| 127734562 | 127736996 | -1 | TEC | ENSG00000 |
| 127737185 | 127741183 | 1 | protein_coding | ENSG00000 |
| 127867881 | 127868487 | 1 | processed_pseudoge | ENSG00000 |

| | | | | |
|---|---|---|---|---|
| 127877841 | 127892941 | -1 | protein_coding | ENSG00000 |
| 127907061 | 128085855 | 1 | protein_coding | ENSG00000 |
| 127929376 | 127934517 | -1 | lncRNA | ENSG00000 |
| 127934698 | 127936167 | 1 | lncRNA | ENSG00000 |
| 128026878 | 128029444 | -1 | lncRNA | ENSG00000 |
| 128096659 | 128126423 | -1 | protein_coding | ENSG00000 |
| 128181032 | 128317726 | 1 | lncRNA | ENSG00000 |
| 128316282 | 128320214 | -1 | lncRNA | ENSG00000 |
| 128912810 | 128916429 | 1 | lncRNA | ENSG00000 |
| 128958772 | 128960062 | -1 | lncRNA | ENSG00000 |
| 128965406 | 128968740 | -1 | lncRNA | ENSG00000 |
| 129278251 | 129291722 | 1 | lncRNA | ENSG00000 |
| 129372194 | 129380753 | -1 | lncRNA | ENSG00000 |
| 129467190 | 129770983 | 1 | protein_coding | ENSG00000 |
| 129621988 | 129626664 | -1 | lncRNA | ENSG00000 |
| 129693722 | 129702117 | -1 | lncRNA | ENSG00000 |
| 129768844 | 129769435 | 1 | lncRNA | ENSG00000 |
| 129784983 | 129806971 | 1 | lncRNA | ENSG00000 |
| 129835283 | 129963841 | -1 | protein_coding | ENSG00000 |
| 129837505 | 129837794 | 1 | lncRNA | ENSG00000 |
| 129843299 | 129843374 | -1 | miRNA | ENSG00000 |
| 129845328 | 129845895 | 1 | lncRNA | ENSG00000 |
| 129857638 | 129867384 | 1 | lncRNA | ENSG00000 |
| 130020025 | 130110830 | -1 | protein_coding | ENSG00000 |
| 130104569 | 130104998 | -1 | lncRNA | ENSG00000 |
| 130106046 | 130108481 | 1 | processed_pseudoge | ENSG00000 |
| 130131770 | 130132241 | -1 | processed_pseudoge | ENSG00000 |
| 130136391 | 130184521 | 1 | protein_coding | ENSG00000 |
| 130213488 | 130483193 | 1 | lncRNA | ENSG00000 |
| 130282445 | 130286604 | -1 | unprocessed_pseudc | ENSG00000 |
| 130505106 | 130505201 | 1 | misc_RNA | ENSG00000 |
| 130525712 | 130537406 | 1 | lncRNA | ENSG00000 |
| 130962588 | 130962668 | -1 | miRNA | ENSG00000 |
| 131092391 | 131311721 | -1 | protein_coding | ENSG00000 |
| 131095218 | 131095777 | 1 | lncRNA | ENSG00000 |
| 131772398 | 131790199 | -1 | lncRNA | ENSG00000 |
| 131776386 | 131795277 | 1 | lncRNA | ENSG00000 |
| 131831910 | 131833508 | -1 | TEC | ENSG00000 |
| 131891640 | 131895297 | 1 | lncRNA | ENSG00000 |
| 131901008 | 131959834 | 1 | protein_coding | ENSG00000 |
| 131966455 | 131982013 | -1 | protein_coding | ENSG00000 |
| 131971202 | 131971533 | 1 | lncRNA | ENSG00000 |
| 131980240 | 131981337 | 1 | lncRNA | ENSG00000 |
| 131996738 | 131999846 | 1 | TEC | ENSG00000 |
| 132036336 | 132184858 | 1 | protein_coding | ENSG00000 |
| 132181225 | 132185962 | -1 | lncRNA | ENSG00000 |
| 132186948 | 132205759 | 1 | protein_coding | ENSG00000 |
| 132207492 | 132331847 | -1 | protein_coding | ENSG00000 |
| 132332154 | 132381508 | 1 | protein_coding | ENSG00000 |
| 132397168 | 132417859 | 1 | protein_coding | ENSG00000 |

| | | | | |
|---|---|---|---|---|
| 132419083 | 132420953 | 1 | lncRNA | ENSG0000C |
| 132433733 | 132441484 | -1 | lncRNA | ENSG0000C |
| 132444327 | 132449408 | 1 | lncRNA | ENSG0000C |
| 132508394 | 132518367 | -1 | lncRNA | ENSG0000C |
| 132520827 | 132522449 | -1 | lncRNA | ENSG0000C |
| 132537787 | 132783480 | 1 | protein_coding | ENSG0000C |
| 132783179 | 132786147 | -1 | protein_coding | ENSG0000C |
| 132808392 | 132942823 | -1 | protein_coding | ENSG0000C |
| 132943967 | 132965289 | -1 | lncRNA | ENSG0000C |
| 132961340 | 132962237 | -1 | lncRNA | ENSG0000C |
| 132965534 | 132976354 | 1 | lncRNA | ENSG0000C |
| 133070929 | 133131675 | 1 | protein_coding | ENSG0000C |
| 133083073 | 133088491 | -1 | lncRNA | ENSG0000C |
| 133144612 | 133145304 | -1 | processed_pseudoge | ENSG0000C |
| 133160219 | 133226412 | 1 | protein_coding | ENSG0000C |
| 133230217 | 133231558 | 1 | protein_coding | ENSG0000C |
| 133237855 | 133241928 | 1 | protein_coding | ENSG0000C |
| 133246478 | 133247891 | -1 | lncRNA | ENSG0000C |
| 133247511 | 133247620 | -1 | miRNA | ENSG0000C |
| 133257144 | 133257551 | -1 | lncRNA | ENSG0000C |
| 133262420 | 133276868 | -1 | protein_coding | ENSG0000C |
| 133278630 | 133312337 | -1 | protein_coding | ENSG0000C |
| 133295187 | 133295977 | -1 | lncRNA | ENSG0000C |
| 133308914 | 133313162 | 1 | protein_coding | ENSG0000C |
| 133309410 | 133352529 | 1 | protein_coding | ENSG0000C |
| 133324072 | 133336935 | -1 | protein_coding | ENSG0000C |
| 133344643 | 133344906 | -1 | processed_pseudoge | ENSG0000C |
| 133345754 | 133350726 | -1 | lncRNA | ENSG0000C |
| 133347368 | 133352683 | 1 | protein_coding | ENSG0000C |
| 133355158 | 133358025 | -1 | protein_coding | ENSG0000C |
| 133362485 | 133373354 | -1 | protein_coding | ENSG0000C |
| 133371556 | 133371663 | -1 | miRNA | ENSG0000C |
| 133374736 | 133374869 | -1 | lncRNA | ENSG0000C |
| 133379261 | 133391694 | 1 | protein_coding | ENSG0000C |
| 133380017 | 133420271 | 1 | protein_coding | ENSG0000C |
| 133394094 | 133422520 | 1 | protein_coding | ENSG0000C |
| 133420666 | 133424572 | -1 | protein_coding | ENSG0000C |
| 133430394 | 133431318 | -1 | unprocessed_pseudc | ENSG0000C |
| 133453928 | 133523558 | 1 | protein_coding | ENSG0000C |
| 133481238 | 133481770 | 1 | unprocessed_pseudc | ENSG0000C |
| 133520406 | 133561220 | 1 | protein_coding | ENSG0000C |
| 133526259 | 133527513 | -1 | lncRNA | ENSG0000C |
| 133541809 | 133542575 | 1 | lncRNA | ENSG0000C |
| 133553901 | 133569835 | -1 | protein_coding | ENSG0000C |
| 133565797 | 133629507 | 1 | lncRNA | ENSG0000C |
| 133575122 | 133576031 | -1 | unprocessed_pseudc | ENSG0000C |
| 133588914 | 133589413 | 1 | lncRNA | ENSG0000C |
| 133623895 | 133626795 | -1 | protein_coding | ENSG0000C |
| 133637408 | 133637892 | -1 | processed_pseudoge | ENSG0000C |
| 133640789 | 133642894 | 1 | processed_pseudoge | ENSG0000C |

| | | | | |
|---|---|---|---|---|
| 133657235 | 133660635 | -1 | unprocessed_pseudc | ENSG0000C |
| 133664429 | 133665106 | 1 | unprocessed_pseudc | ENSG0000C |
| 133667151 | 133668419 | 1 | unprocessed_pseudc | ENSG0000C |
| 133670450 | 133671718 | 1 | unprocessed_pseudc | ENSG0000C |
| 133673749 | 133675017 | 1 | unprocessed_pseudc | ENSG0000C |
| 133677059 | 133678327 | 1 | unprocessed_pseudc | ENSG0000C |
| 133680369 | 133681637 | 1 | unprocessed_pseudc | ENSG0000C |
| 133683668 | 133684936 | 1 | unprocessed_pseudc | ENSG0000C |
| 133740606 | 133741284 | 1 | unprocessed_pseudc | ENSG0000C |
| 133743332 | 133744598 | 1 | unprocessed_pseudc | ENSG0000C |
| 133746641 | 133747907 | 1 | unprocessed_pseudc | ENSG0000C |
| 133749950 | 133751216 | 1 | unprocessed_pseudc | ENSG0000C |
| 133753250 | 133754516 | 1 | unprocessed_pseudc | ENSG0000C |
| 133756559 | 133757825 | 1 | unprocessed_pseudc | ENSG0000C |
| 133759859 | 133761125 | 1 | unprocessed_pseudc | ENSG0000C |
| 133778246 | 133778699 | 1 | processed_pseudoge | ENSG0000C |
| 133778258 | 133778498 | -1 | processed_pseudoge | ENSG0000C |
| 75780 | 76143 | 1 | unprocessed_pseudc | ENSG0000C |
| 86649 | 87586 | -1 | unprocessed_pseudc | ENSG0000C |
| 112967 | 125927 | -1 | lncRNA | ENSG0000C |
| 123312 | 126123 | 1 | processed_pseudoge | ENSG0000C |
| 127204 | 139612 | -1 | lncRNA | ENSG0000C |
| 129279 | 186136 | 1 | lncRNA | ENSG0000C |
| 152999 | 153102 | -1 | snRNA | ENSG0000C |
| 167784 | 207428 | -1 | protein_coding | ENSG0000C |
| 193078 | 194575 | 1 | protein_coding | ENSG0000C |
| 196738 | 200261 | 1 | protein_coding | ENSG0000C |
| 203623 | 205470 | 1 | lncRNA | ENSG0000C |
| 207511 | 215113 | 1 | protein_coding | ENSG0000C |
| 209336 | 209406 | 1 | miRNA | ENSG0000C |
| 215030 | 236931 | -1 | protein_coding | ENSG0000C |
| 236966 | 252984 | 1 | protein_coding | ENSG0000C |
| 243099 | 243483 | -1 | processed_pseudoge | ENSG0000C |
| 278365 | 285359 | 1 | protein_coding | ENSG0000C |
| 287305 | 288987 | 1 | lncRNA | ENSG0000C |
| 289126 | 296107 | 1 | protein_coding | ENSG0000C |
| 298200 | 299526 | -1 | protein_coding | ENSG0000C |
| 303856 | 304144 | -1 | processed_pseudoge | ENSG0000C |
| 307631 | 315272 | 1 | protein_coding | ENSG0000C |
| 310139 | 311141 | -1 | lncRNA | ENSG0000C |
| 313506 | 315272 | 1 | protein_coding | ENSG0000C |
| 318640 | 325631 | 1 | lncRNA | ENSG0000C |
| 319676 | 327537 | -1 | protein_coding | ENSG0000C |
| 321991 | 322426 | 1 | lncRNA | ENSG0000C |
| 322186 | 322727 | -1 | lncRNA | ENSG0000C |
| 325703 | 326294 | -1 | lncRNA | ENSG0000C |
| 327171 | 330122 | 1 | lncRNA | ENSG0000C |
| 333192 | 333688 | 1 | lncRNA | ENSG0000C |
| 369499 | 382117 | 1 | protein_coding | ENSG0000C |
| 392614 | 404908 | 1 | protein_coding | ENSG0000C |

| | | | | |
|---|---|---|---|---|
| 405716 | 417455 | -1 | protein_coding | ENSG0000C |
| 417933 | 442011 | -1 | protein_coding | ENSG0000C |
| 440399 | 440693 | -1 | misc_RNA | ENSG0000C |
| 448268 | 491399 | 1 | protein_coding | ENSG0000C |
| 462930 | 463899 | -1 | lncRNA | ENSG0000C |
| 494512 | 507300 | -1 | protein_coding | ENSG0000C |
| 512513 | 518134 | 1 | lncRNA | ENSG0000C |
| 528907 | 529659 | 1 | lncRNA | ENSG0000C |
| 532242 | 537287 | -1 | protein_coding | ENSG0000C |
| 537527 | 554912 | 1 | protein_coding | ENSG0000C |
| 554850 | 560738 | -1 | protein_coding | ENSG0000C |
| 557595 | 560107 | 1 | lncRNA | ENSG0000C |
| 560404 | 564025 | 1 | protein_coding | ENSG0000C |
| 565660 | 568457 | -1 | lncRNA | ENSG0000C |
| 568089 | 568198 | -1 | miRNA | ENSG0000C |
| 576470 | 612222 | 1 | protein_coding | ENSG0000C |
| 612553 | 615983 | -1 | protein_coding | ENSG0000C |
| 616565 | 626078 | -1 | protein_coding | ENSG0000C |
| 626309 | 627181 | -1 | protein_coding | ENSG0000C |
| 637269 | 640706 | 1 | protein_coding | ENSG0000C |
| 644233 | 706715 | -1 | protein_coding | ENSG0000C |
| 665910 | 678391 | 1 | lncRNA | ENSG0000C |
| 694438 | 727727 | 1 | protein_coding | ENSG0000C |
| 695428 | 705028 | 1 | protein_coding | ENSG0000C |
| 708564 | 727047 | -1 | lncRNA | ENSG0000C |
| 747415 | 765012 | 1 | protein_coding | ENSG0000C |
| 767220 | 777488 | -1 | protein_coding | ENSG0000C |
| 777578 | 784297 | 1 | lncRNA | ENSG0000C |
| 779617 | 780755 | 1 | TEC | ENSG0000C |
| 781645 | 782105 | 1 | lncRNA | ENSG0000C |
| 787115 | 790113 | -1 | protein_coding | ENSG0000C |
| 790475 | 798281 | -1 | protein_coding | ENSG0000C |
| 797511 | 799185 | 1 | protein_coding | ENSG0000C |
| 799179 | 809753 | -1 | protein_coding | ENSG0000C |
| 809965 | 812880 | 1 | protein_coding | ENSG0000C |
| 811681 | 811814 | 1 | snoRNA | ENSG0000C |
| 818914 | 825573 | 1 | protein_coding | ENSG0000C |
| 823634 | 832883 | -1 | lncRNA | ENSG0000C |
| 826144 | 831991 | 1 | protein_coding | ENSG0000C |
| 832887 | 839831 | 1 | protein_coding | ENSG0000C |
| 837356 | 842529 | -1 | protein_coding | ENSG0000C |
| 842812 | 867116 | 1 | protein_coding | ENSG0000C |
| 856880 | 859795 | -1 | lncRNA | ENSG0000C |
| 867859 | 915058 | -1 | protein_coding | ENSG0000C |
| 924894 | 1012245 | 1 | protein_coding | ENSG0000C |
| 948421 | 948524 | 1 | snRNA | ENSG0000C |
| 1012823 | 1036718 | -1 | protein_coding | ENSG0000C |
| 1049880 | 1055749 | 1 | lncRNA | ENSG0000C |
| 1074875 | 1110511 | 1 | protein_coding | ENSG0000C |
| 1157953 | 1201138 | 1 | protein_coding | ENSG0000C |

| | | | | |
|---|---|---|---|---|
| 1218530 | 1220242 | 1 | lncRNA | ENSG0000C |
| 1223066 | 1262172 | 1 | protein_coding | ENSG0000C |
| 1242261 | 1249676 | -1 | lncRNA | ENSG0000C |
| 1256605 | 1256670 | 1 | miRNA | ENSG0000C |
| 1274371 | 1309654 | -1 | protein_coding | ENSG0000C |
| 1309769 | 1310707 | 1 | lncRNA | ENSG0000C |
| 1333250 | 1337068 | -1 | lncRNA | ENSG0000C |
| 1350359 | 1364286 | 1 | lncRNA | ENSG0000C |
| 1389899 | 1462689 | 1 | protein_coding | ENSG0000C |
| 1464582 | 1467632 | -1 | lncRNA | ENSG0000C |
| 1469457 | 1501247 | -1 | protein_coding | ENSG0000C |
| 1554051 | 1572271 | -1 | protein_coding | ENSG0000C |
| 1571353 | 1599184 | 1 | lncRNA | ENSG0000C |
| 1584342 | 1585283 | -1 | protein_coding | ENSG0000C |
| 1597177 | 1598294 | -1 | protein_coding | ENSG0000C |
| 1607565 | 1608463 | -1 | protein_coding | ENSG0000C |
| 1620958 | 1622138 | -1 | protein_coding | ENSG0000C |
| 1629775 | 1630707 | 1 | protein_coding | ENSG0000C |
| 1662584 | 1663343 | -1 | lncRNA | ENSG0000C |
| 1665597 | 1667856 | 1 | lncRNA | ENSG0000C |
| 1683269 | 1685629 | -1 | lncRNA | ENSG0000C |
| 1688297 | 1689056 | 1 | lncRNA | ENSG0000C |
| 1697195 | 1697755 | 1 | protein_coding | ENSG0000C |
| 1732406 | 1750595 | -1 | protein_coding | ENSG0000C |
| 1734821 | 1763954 | -1 | protein_coding | ENSG0000C |
| 1752755 | 1764573 | -1 | protein_coding | ENSG0000C |
| 1760348 | 1762486 | 1 | lncRNA | ENSG0000C |
| 1763009 | 1763749 | -1 | lncRNA | ENSG0000C |
| 1773745 | 1774903 | 1 | lncRNA | ENSG0000C |
| 1776930 | 1778388 | -1 | lncRNA | ENSG0000C |
| 1792553 | 1792870 | 1 | processed_pseudoge | ENSG0000C |
| 1802889 | 1804095 | -1 | processed_pseudoge | ENSG0000C |
| 1828307 | 1837521 | 1 | protein_coding | ENSG0000C |
| 1838981 | 1841680 | 1 | protein_coding | ENSG0000C |
| 1852970 | 1892267 | 1 | protein_coding | ENSG0000C |
| 1859464 | 1859536 | -1 | miRNA | ENSG0000C |
| 1864177 | 1866667 | 1 | lncRNA | ENSG0000C |
| 1880045 | 1880147 | 1 | miRNA | ENSG0000C |
| 1888577 | 1891895 | -1 | protein_coding | ENSG0000C |
| 1897015 | 1908656 | 1 | lncRNA | ENSG0000C |
| 1919703 | 1938706 | 1 | protein_coding | ENSG0000C |
| 1947278 | 1984522 | 1 | protein_coding | ENSG0000C |
| 1983237 | 1989920 | -1 | lncRNA | ENSG0000C |
| 1991096 | 1993670 | 1 | lncRNA | ENSG0000C |
| 1995176 | 2001470 | -1 | lncRNA | ENSG0000C |
| 1996540 | 1996610 | 1 | misc_RNA | ENSG0000C |
| 1996759 | 1996831 | -1 | miRNA | ENSG0000C |
| 1997484 | 1997552 | 1 | misc_RNA | ENSG0000C |
| 2129112 | 2141238 | -1 | protein_coding | ENSG0000C |
| 2129112 | 2158391 | -1 | protein_coding | ENSG0000C |

| | | | | |
|---|---|---|---|---|
| 2129121 | 2129964 | -1 | lncRNA | ENSG00000 |
| 2132538 | 2161209 | -1 | protein_coding | ENSG00000 |
| 2134134 | 2134209 | -1 | miRNA | ENSG00000 |
| 2140501 | 2148666 | 1 | lncRNA | ENSG00000 |
| 2159779 | 2161341 | -1 | protein_coding | ENSG00000 |
| 2163929 | 2171877 | -1 | protein_coding | ENSG00000 |
| 2173063 | 2173138 | 1 | miRNA | ENSG00000 |
| 2268495 | 2270952 | -1 | protein_coding | ENSG00000 |
| 2295645 | 2303049 | -1 | protein_coding | ENSG00000 |
| 2301997 | 2318200 | 1 | protein_coding | ENSG00000 |
| 2321718 | 2321827 | 1 | snRNA | ENSG00000 |
| 2328749 | 2377992 | -1 | lncRNA | ENSG00000 |
| 2335135 | 2347685 | -1 | transcribed_processe | ENSG00000 |
| 2372638 | 2372750 | 1 | misc_RNA | ENSG00000 |
| 2376177 | 2397419 | 1 | protein_coding | ENSG00000 |
| 2400488 | 2403878 | 1 | protein_coding | ENSG00000 |
| 2404515 | 2407908 | 1 | lncRNA | ENSG00000 |
| 2404515 | 2423045 | -1 | protein_coding | ENSG00000 |
| 2444684 | 2848991 | 1 | protein_coding | ENSG00000 |
| 2608328 | 2699994 | -1 | lncRNA | ENSG00000 |
| 2652099 | 2652562 | 1 | misc_RNA | ENSG00000 |
| 2674097 | 2674292 | 1 | misc_RNA | ENSG00000 |
| 2674625 | 2674740 | 1 | misc_RNA | ENSG00000 |
| 2697851 | 2698076 | 1 | misc_RNA | ENSG00000 |
| 2771356 | 2771577 | 1 | processed_pseudoge | ENSG00000 |
| 2840135 | 2861568 | -1 | lncRNA | ENSG00000 |
| 2870033 | 2872105 | 1 | lncRNA | ENSG00000 |
| 2871437 | 2871498 | 1 | misc_RNA | ENSG00000 |
| 2883213 | 2885773 | -1 | protein_coding | ENSG00000 |
| 2887344 | 2903575 | -1 | protein_coding | ENSG00000 |
| 2899721 | 2925246 | 1 | protein_coding | ENSG00000 |
| 2928273 | 2929420 | -1 | protein_coding | ENSG00000 |
| 2944431 | 2992377 | -1 | protein_coding | ENSG00000 |
| 2963771 | 2963893 | -1 | snoRNA | ENSG00000 |
| 2989863 | 2991344 | 1 | lncRNA | ENSG00000 |
| 3000922 | 3057613 | -1 | protein_coding | ENSG00000 |
| 3029009 | 3041260 | 1 | lncRNA | ENSG00000 |
| 3048359 | 3048521 | -1 | snRNA | ENSG00000 |
| 3057538 | 3064707 | 1 | lncRNA | ENSG00000 |
| 3084393 | 3085443 | -1 | processed_pseudoge | ENSG00000 |
| 3087107 | 3166739 | -1 | protein_coding | ENSG00000 |
| 3189546 | 3189929 | 1 | lncRNA | ENSG00000 |
| 3217944 | 3218813 | -1 | protein_coding | ENSG00000 |
| 3218332 | 3223131 | 1 | lncRNA | ENSG00000 |
| 3225030 | 3232417 | -1 | protein_coding | ENSG00000 |
| 3226061 | 3232838 | 1 | lncRNA | ENSG00000 |
| 3276045 | 3276390 | 1 | processed_pseudoge | ENSG00000 |
| 3289311 | 3289642 | -1 | processed_pseudoge | ENSG00000 |
| 3321862 | 3322202 | 1 | processed_pseudoge | ENSG00000 |
| 3335125 | 3335407 | -1 | processed_pseudoge | ENSG00000 |

| | | | | |
|---|---|---|---|---|
| 3336721 | 3340630 | 1 | lncRNA | ENSG0000C |
| 3339261 | 3379222 | -1 | protein_coding | ENSG0000C |
| 3380961 | 3408978 | 1 | transcribed_unproce | ENSG0000C |
| 3390780 | 3391752 | -1 | unprocessed_pseudc | ENSG0000C |
| 3410352 | 3422496 | -1 | unprocessed_pseudc | ENSG0000C |
| 3429758 | 3500621 | -1 | transcribed_unproce | ENSG0000C |
| 3469319 | 3531328 | 1 | lncRNA | ENSG0000C |
| 3475280 | 3477065 | -1 | lncRNA | ENSG0000C |
| 3481520 | 3581211 | -1 | lncRNA | ENSG0000C |
| 3511559 | 3520886 | 1 | lncRNA | ENSG0000C |
| 3531420 | 3531608 | 1 | processed_pseudoge | ENSG0000C |
| 3549745 | 3550519 | -1 | processed_pseudoge | ENSG0000C |
| 3559657 | 3560022 | 1 | processed_pseudoge | ENSG0000C |
| 3580927 | 3581135 | -1 | transcribed_unproce | ENSG0000C |
| 3591277 | 3591615 | -1 | processed_pseudoge | ENSG0000C |
| 3599595 | 3600706 | -1 | unprocessed_pseudc | ENSG0000C |
| 3609901 | 3637559 | 1 | transcribed_unproce | ENSG0000C |
| 3638512 | 3642316 | -1 | protein_coding | ENSG0000C |
| 3645128 | 3664416 | 1 | protein_coding | ENSG0000C |
| 3663815 | 3663911 | -1 | misc_RNA | ENSG0000C |
| 3665164 | 3665259 | -1 | misc_RNA | ENSG0000C |
| 3665587 | 3671384 | -1 | protein_coding | ENSG0000C |
| 3671083 | 3797792 | -1 | protein_coding | ENSG0000C |
| 3714772 | 3714875 | -1 | snRNA | ENSG0000C |
| 3781395 | 3781456 | 1 | snRNA | ENSG0000C |
| 3797724 | 3826371 | 1 | protein_coding | ENSG0000C |
| 3826978 | 3840959 | -1 | protein_coding | ENSG0000C |
| 3854318 | 3855509 | -1 | lncRNA | ENSG0000C |
| 3854527 | 4093210 | 1 | protein_coding | ENSG0000C |
| 3854612 | 3855399 | 1 | lncRNA | ENSG0000C |
| 3856062 | 3856141 | 1 | miRNA | ENSG0000C |
| 3892398 | 3892887 | -1 | processed_pseudoge | ENSG0000C |
| 3922567 | 3922703 | 1 | snoRNA | ENSG0000C |
| 3961277 | 3961538 | 1 | processed_pseudoge | ENSG0000C |
| 3972881 | 3974237 | -1 | processed_pseudoge | ENSG0000C |
| 3989163 | 3989463 | -1 | processed_pseudoge | ENSG0000C |
| 4018948 | 4019118 | 1 | processed_pseudoge | ENSG0000C |
| 4094707 | 4138932 | 1 | protein_coding | ENSG0000C |
| 4137116 | 4138257 | -1 | lncRNA | ENSG0000C |
| 4146112 | 4151104 | -1 | transcribed_unitary_ | ENSG0000C |
| 4179675 | 4202663 | 1 | lncRNA | ENSG0000C |
| 4210354 | 4212091 | -1 | processed_pseudoge | ENSG0000C |
| 4233288 | 4233872 | 1 | protein_coding | ENSG0000C |
| 4242056 | 4242640 | -1 | protein_coding | ENSG0000C |
| 4278705 | 4279192 | 1 | processed_pseudoge | ENSG0000C |
| 4287499 | 4288083 | -1 | protein_coding | ENSG0000C |
| 4329865 | 4330449 | 1 | protein_coding | ENSG0000C |
| 4338660 | 4339244 | -1 | protein_coding | ENSG0000C |
| 4367263 | 4368386 | -1 | protein_coding | ENSG0000C |
| 4378282 | 4379209 | 1 | unitary_pseudogene | ENSG0000C |

| | | | | |
|---|---|---|---|---|
| 4384897 | 4393702 | -1 | protein_coding | ENSG00000( |
| 4393783 | 4476361 | 1 | transcribed_unproce | ENSG00000( |
| 4418060 | 4419011 | 1 | unitary_pseudogene | ENSG00000( |
| 4431323 | 4432277 | -1 | unitary_pseudogene | ENSG00000( |
| 4449295 | 4450361 | 1 | protein_coding | ENSG00000( |
| 4482646 | 4493497 | 1 | protein_coding | ENSG00000( |
| 4514966 | 4515924 | -1 | unprocessed_pseudc | ENSG00000( |
| 4545191 | 4546144 | 1 | protein_coding | ENSG00000( |
| 4571423 | 4577820 | -1 | lncRNA | ENSG00000( |
| 4581743 | 4593340 | 1 | protein_coding | ENSG00000( |
| 4593614 | 4596574 | 1 | protein_coding | ENSG00000( |
| 4598672 | 4608231 | -1 | protein_coding | ENSG00000( |
| 4637477 | 4643060 | 1 | protein_coding | ENSG00000( |
| 4643420 | 4655488 | 1 | protein_coding | ENSG00000( |
| 4660853 | 4661763 | -1 | unitary_pseudogene | ENSG00000( |
| 4680171 | 4697854 | -1 | protein_coding | ENSG00000( |
| 4690423 | 4697831 | -1 | protein_coding | ENSG00000( |
| 4704927 | 4992429 | 1 | protein_coding | ENSG00000( |
| 4709569 | 4712421 | 1 | transcribed_unproce | ENSG00000( |
| 4718612 | 4719559 | 1 | unprocessed_pseudc | ENSG00000( |
| 4722740 | 4724171 | -1 | processed_pseudoge | ENSG00000( |
| 4736178 | 4737078 | -1 | unprocessed_pseudc | ENSG00000( |
| 4752046 | 4752944 | -1 | unprocessed_pseudc | ENSG00000( |
| 4756645 | 4756797 | 1 | snoRNA | ENSG00000( |
| 4768979 | 4769938 | -1 | protein_coding | ENSG00000( |
| 4786754 | 4787705 | 1 | unprocessed_pseudc | ENSG00000( |
| 4793778 | 4795284 | -1 | unprocessed_pseudc | ENSG00000( |
| 4803433 | 4804380 | -1 | protein_coding | ENSG00000( |
| 4821321 | 4822456 | 1 | protein_coding | ENSG00000( |
| 4830175 | 4833072 | 1 | transcribed_unproce | ENSG00000( |
| 4847489 | 4849238 | -1 | protein_coding | ENSG00000( |
| 4859610 | 4863652 | -1 | transcribed_unproce | ENSG00000( |
| 4871717 | 4878117 | 1 | transcribed_unproce | ENSG00000( |
| 4881819 | 4882883 | 1 | protein_coding | ENSG00000( |
| 4889593 | 4890596 | -1 | unprocessed_pseudc | ENSG00000( |
| 4903783 | 4909462 | 1 | protein_coding | ENSG00000( |
| 4912588 | 4919350 | -1 | protein_coding | ENSG00000( |
| 4923374 | 4924339 | -1 | protein_coding | ENSG00000( |
| 4937371 | 4938294 | -1 | unprocessed_pseudc | ENSG00000( |
| 4942831 | 4947605 | -1 | protein_coding | ENSG00000( |
| 4954772 | 4955713 | -1 | protein_coding | ENSG00000( |
| 4972943 | 4973884 | -1 | unprocessed_pseudc | ENSG00000( |
| 4994851 | 5005536 | 1 | protein_coding | ENSG00000( |
| 5015138 | 5016079 | 1 | unprocessed_pseudc | ENSG00000( |
| 5037013 | 5037951 | 1 | unprocessed_pseudc | ENSG00000( |
| 5046526 | 5047461 | 1 | protein_coding | ENSG00000( |
| 5058650 | 5059627 | -1 | protein_coding | ENSG00000( |
| 5069572 | 5070497 | 1 | polymorphic_pseudc | ENSG00000( |
| 5076130 | 5077066 | -1 | unprocessed_pseudc | ENSG00000( |
| 5092676 | 5093611 | 1 | unprocessed_pseudc | ENSG00000( |

| | | | | |
|---|---|---|---|---|
| 5104153 | 5105084 | 1 | unprocessed_pseudc | ENSG00000( |
| 5106062 | 5107530 | -1 | lncRNA | ENSG00000( |
| 5120621 | 5124513 | -1 | transcribed_unproce | ENSG00000( |
| 5128776 | 5138356 | -1 | protein_coding | ENSG00000( |
| 5143219 | 5154757 | -1 | protein_coding | ENSG00000( |
| 5170351 | 5171296 | -1 | unprocessed_pseudc | ENSG00000( |
| 5177714 | 5178664 | -1 | unprocessed_pseudc | ENSG00000( |
| 5199735 | 5200700 | -1 | protein_coding | ENSG00000( |
| 5205041 | 5207308 | 1 | lncRNA | ENSG00000( |
| 5224309 | 5225461 | -1 | TEC | ENSG00000( |
| 5224448 | 5224639 | -1 | ribozyme | ENSG00000( |
| 5225464 | 5229395 | -1 | protein_coding | ENSG00000( |
| 5232678 | 5243657 | -1 | protein_coding | ENSG00000( |
| 5241105 | 5243537 | -1 | transcribed_unproce | ENSG00000( |
| 5244554 | 5245546 | -1 | lncRNA | ENSG00000( |
| 5248079 | 5249859 | -1 | protein_coding | ENSG00000( |
| 5248270 | 5254783 | -1 | protein_coding | ENSG00000( |
| 5253188 | 5505605 | -1 | protein_coding | ENSG00000( |
| 5254392 | 5678434 | -1 | protein_coding | ENSG00000( |
| 5268345 | 5505652 | -1 | protein_coding | ENSG00000( |
| 5291761 | 5292380 | -1 | unprocessed_pseudc | ENSG00000( |
| 5301014 | 5301946 | -1 | protein_coding | ENSG00000( |
| 5303444 | 5505652 | -1 | protein_coding | ENSG00000( |
| 5314772 | 5315684 | -1 | unprocessed_pseudc | ENSG00000( |
| 5323359 | 5324297 | -1 | protein_coding | ENSG00000( |
| 5330592 | 5331360 | -1 | unprocessed_pseudc | ENSG00000( |
| 5351508 | 5352446 | 1 | protein_coding | ENSG00000( |
| 5354692 | 5355578 | 1 | processed_pseudoge | ENSG00000( |
| 5362615 | 5362849 | 1 | processed_pseudoge | ENSG00000( |
| 5383812 | 5393263 | 1 | protein_coding | ENSG00000( |
| 5402597 | 5403547 | 1 | unprocessed_pseudc | ENSG00000( |
| 5422111 | 5423206 | 1 | protein_coding | ENSG00000( |
| 5430653 | 5431602 | -1 | unprocessed_pseudc | ENSG00000( |
| 5440570 | 5441514 | -1 | protein_coding | ENSG00000( |
| 5449323 | 5456518 | 1 | protein_coding | ENSG00000( |
| 5468486 | 5469422 | -1 | unitary_pseudogene | ENSG00000( |
| 5488685 | 5489749 | 1 | protein_coding | ENSG00000( |
| 5507300 | 5509958 | -1 | protein_coding | ENSG00000( |
| 5514393 | 5516699 | -1 | protein_coding | ENSG00000( |
| 5518441 | 5524955 | -1 | unitary_pseudogene | ENSG00000( |
| 5527277 | 5528174 | -1 | unprocessed_pseudc | ENSG00000( |
| 5544489 | 5548533 | -1 | protein_coding | ENSG00000( |
| 5551662 | 5552563 | -1 | unitary_pseudogene | ENSG00000( |
| 5557096 | 5564327 | 1 | transcribed_unproce | ENSG00000( |
| 5566678 | 5567616 | -1 | unprocessed_pseudc | ENSG00000( |
| 5571247 | 5572192 | -1 | processed_pseudoge | ENSG00000( |
| 5580877 | 5581884 | 1 | protein_coding | ENSG00000( |
| 5596109 | 5612958 | 1 | protein_coding | ENSG00000( |
| 5596725 | 5644398 | 1 | protein_coding | ENSG00000( |
| 5619764 | 5644398 | 1 | protein_coding | ENSG00000( |

| | | | | |
|---|---|---|---|---|
| 5663195 | 5938619 | -1 | protein_coding | ENSG00000 |
| 5689697 | 5737089 | 1 | protein_coding | ENSG00000 |
| 5719290 | 5720212 | 1 | unprocessed_pseudo | ENSG00000 |
| 5726502 | 5727464 | 1 | unprocessed_pseudo | ENSG00000 |
| 5736448 | 5738523 | 1 | protein_coding | ENSG00000 |
| 5754243 | 5755905 | 1 | protein_coding | ENSG00000 |
| 5761153 | 5767465 | 1 | transcribed_unproce | ENSG00000 |
| 5776165 | 5783355 | -1 | protein_coding | ENSG00000 |
| 5786471 | 5791265 | -1 | protein_coding | ENSG00000 |
| 5800364 | 5801297 | 1 | unprocessed_pseudo | ENSG00000 |
| 5820314 | 5821348 | 1 | protein_coding | ENSG00000 |
| 5840928 | 5841952 | -1 | protein_coding | ENSG00000 |
| 5856674 | 5857734 | -1 | protein_coding | ENSG00000 |
| 5873960 | 5874892 | 1 | unprocessed_pseudo | ENSG00000 |
| 5880629 | 5887079 | 1 | protein_coding | ENSG00000 |
| 5893208 | 5902730 | 1 | protein_coding | ENSG00000 |
| 5903767 | 5904448 | 1 | unprocessed_pseudo | ENSG00000 |
| 5938751 | 5951347 | 1 | protein_coding | ENSG00000 |
| 5967177 | 5968494 | -1 | protein_coding | ENSG00000 |
| 5985892 | 5986985 | -1 | protein_coding | ENSG00000 |
| 5993876 | 5994995 | -1 | processed_pseudoge | ENSG00000 |
| 5999445 | 6006946 | -1 | protein_coding | ENSG00000 |
| 6016996 | 6017114 | 1 | rRNA_pseudogene | ENSG00000 |
| 6019336 | 6034338 | -1 | protein_coding | ENSG00000 |
| 6045807 | 6046747 | -1 | unprocessed_pseudo | ENSG00000 |
| 6057316 | 6058275 | -1 | unprocessed_pseudo | ENSG00000 |
| 6066873 | 6067786 | -1 | unitary_pseudogene | ENSG00000 |
| 6107684 | 6108835 | 1 | protein_coding | ENSG00000 |
| 6108135 | 6185576 | -1 | lncRNA | ENSG00000 |
| 6128610 | 6129556 | 1 | unprocessed_pseudo | ENSG00000 |
| 6151641 | 6152549 | -1 | unprocessed_pseudo | ENSG00000 |
| 6169330 | 6170408 | -1 | protein_coding | ENSG00000 |
| 6199146 | 6200259 | 1 | protein_coding | ENSG00000 |
| 6201901 | 6203253 | -1 | lncRNA | ENSG00000 |
| 6205557 | 6211135 | 1 | protein_coding | ENSG00000 |
| 6211335 | 6234711 | -1 | protein_coding | ENSG00000 |
| 6234765 | 6244479 | 1 | protein_coding | ENSG00000 |
| 6259806 | 6272127 | 1 | protein_coding | ENSG00000 |
| 6297189 | 6298606 | -1 | processed_pseudoge | ENSG00000 |
| 6318946 | 6320532 | -1 | protein_coding | ENSG00000 |
| 6319140 | 6360454 | 1 | lncRNA | ENSG00000 |
| 6362799 | 6365267 | 1 | lncRNA | ENSG00000 |
| 6390440 | 6394998 | 1 | protein_coding | ENSG00000 |
| 6395124 | 6419414 | -1 | protein_coding | ENSG00000 |
| 6431049 | 6442617 | -1 | protein_coding | ENSG00000 |
| 6448613 | 6474459 | -1 | protein_coding | ENSG00000 |
| 6474683 | 6481479 | -1 | protein_coding | ENSG00000 |
| 6481485 | 6508978 | 1 | protein_coding | ENSG00000 |
| 6481501 | 6484681 | 1 | protein_coding | ENSG00000 |
| 6488186 | 6489377 | -1 | processed_pseudoge | ENSG00000 |

| | | | | |
|---|---|---|---|---|
| 6497260 | 6593758 | 1 | protein_coding | ENSG0000C |
| 6595072 | 6603616 | -1 | protein_coding | ENSG0000C |
| 6603642 | 6604420 | -1 | lncRNA | ENSG0000C |
| 6603708 | 6610874 | 1 | protein_coding | ENSG0000C |
| 6606294 | 6612539 | -1 | protein_coding | ENSG0000C |
| 6608667 | 6610135 | -1 | lncRNA | ENSG0000C |
| 6610883 | 6616594 | -1 | lncRNA | ENSG0000C |
| 6612768 | 6619448 | -1 | protein_coding | ENSG0000C |
| 6618790 | 6619764 | 1 | lncRNA | ENSG0000C |
| 6621330 | 6655809 | -1 | protein_coding | ENSG0000C |
| 6621451 | 6622322 | 1 | lncRNA | ENSG0000C |
| 6630504 | 6658396 | 1 | lncRNA | ENSG0000C |
| 6680385 | 6683340 | -1 | protein_coding | ENSG0000C |
| 6685891 | 6686369 | -1 | processed_pseudoge | ENSG0000C |
| 6713536 | 6746439 | -1 | transcribed_unproce | ENSG0000C |
| 6748131 | 6750324 | 1 | processed_pseudoge | ENSG0000C |
| 6765626 | 6771976 | -1 | protein_coding | ENSG0000C |
| 6783020 | 6791558 | 1 | protein_coding | ENSG0000C |
| 6791736 | 6799689 | -1 | protein_coding | ENSG0000C |
| 6821942 | 6926878 | -1 | lncRNA | ENSG0000C |
| 6843664 | 6844666 | -1 | processed_pseudoge | ENSG0000C |
| 6845652 | 6846705 | 1 | protein_coding | ENSG0000C |
| 6863057 | 6874717 | 1 | protein_coding | ENSG0000C |
| 6876625 | 6877619 | 1 | protein_coding | ENSG0000C |
| 6891490 | 6892599 | -1 | protein_coding | ENSG0000C |
| 6920974 | 6922043 | 1 | protein_coding | ENSG0000C |
| 6926404 | 7001004 | 1 | protein_coding | ENSG0000C |
| 6997085 | 7020346 | -1 | protein_coding | ENSG0000C |
| 7020446 | 7071308 | 1 | protein_coding | ENSG0000C |
| 7088998 | 7091148 | 1 | protein_coding | ENSG0000C |
| 7222933 | 7230863 | 1 | lncRNA | ENSG0000C |
| 7234766 | 7234837 | 1 | miRNA | ENSG0000C |
| 7238778 | 7469043 | 1 | protein_coding | ENSG0000C |
| 7418826 | 7513644 | -1 | lncRNA | ENSG0000C |
| 7485388 | 7511377 | 1 | protein_coding | ENSG0000C |
| 7513298 | 7657127 | 1 | protein_coding | ENSG0000C |
| 7568866 | 7572046 | -1 | lncRNA | ENSG0000C |
| 7665100 | 7677222 | -1 | protein_coding | ENSG0000C |
| 7689438 | 7706421 | -1 | protein_coding | ENSG0000C |
| 7698951 | 7699393 | -1 | lncRNA | ENSG0000C |
| 7699562 | 7699988 | -1 | lncRNA | ENSG0000C |
| 7704628 | 7882947 | -1 | lncRNA | ENSG0000C |
| 7705288 | 7709517 | 1 | lncRNA | ENSG0000C |
| 7726436 | 7729684 | 1 | transcribed_unproce | ENSG0000C |
| 7745990 | 7746906 | 1 | unprocessed_pseudc | ENSG0000C |
| 7754393 | 7905955 | -1 | lncRNA | ENSG0000C |
| 7772890 | 7773814 | -1 | unprocessed_pseudc | ENSG0000C |
| 7783697 | 7784407 | 1 | processed_pseudoge | ENSG0000C |
| 7795905 | 7796973 | -1 | protein_coding | ENSG0000C |
| 7824818 | 7830840 | -1 | protein_coding | ENSG0000C |

| | | | | |
|---|---|---|---|---|
| 7848700 | 7849626 | 1 | unprocessed_pseudc | ENSG0000C |
| 7889182 | 7889279 | -1 | snRNA | ENSG0000C |
| 7924592 | 7931268 | -1 | protein_coding | ENSG0000C |
| 7937171 | 7941708 | -1 | protein_coding | ENSG0000C |
| 7957537 | 7965426 | -1 | protein_coding | ENSG0000C |
| 7970251 | 8001862 | 1 | protein_coding | ENSG0000C |
| 7977481 | 7977708 | 1 | processed_pseudoge | ENSG0000C |
| 8011278 | 8016520 | -1 | lncRNA | ENSG0000C |
| 8019244 | 8106112 | 1 | protein_coding | ENSG0000C |
| 8035446 | 8039718 | -1 | lncRNA | ENSG0000C |
| 8060038 | 8069373 | 1 | lncRNA | ENSG0000C |
| 8075573 | 8077781 | -1 | TEC | ENSG0000C |
| 8106056 | 8169055 | -1 | protein_coding | ENSG0000C |
| 8169167 | 8178903 | 1 | lncRNA | ENSG0000C |
| 8224309 | 8268716 | -1 | protein_coding | ENSG0000C |
| 8391868 | 8594289 | -1 | protein_coding | ENSG0000C |
| 8555016 | 8555146 | 1 | scaRNA | ENSG0000C |
| 8612037 | 8682694 | -1 | protein_coding | ENSG0000C |
| 8679089 | 8680913 | 1 | lncRNA | ENSG0000C |
| 8682788 | 8714759 | 1 | protein_coding | ENSG0000C |
| 8684227 | 8684356 | 1 | snoRNA | ENSG0000C |
| 8685439 | 8685569 | 1 | snoRNA | ENSG0000C |
| 8693351 | 8910951 | -1 | protein_coding | ENSG0000C |
| 8693357 | 8696607 | 1 | lncRNA | ENSG0000C |
| 8768778 | 8810231 | 1 | lncRNA | ENSG0000C |
| 8845263 | 8845358 | 1 | rRNA_pseudogene | ENSG0000C |
| 8877417 | 8877518 | -1 | misc_RNA | ENSG0000C |
| 8911139 | 8920084 | 1 | protein_coding | ENSG0000C |
| 8920076 | 8933006 | -1 | protein_coding | ENSG0000C |
| 8937579 | 8938211 | -1 | protein_coding | ENSG0000C |
| 8943380 | 8943467 | -1 | misc_RNA | ENSG0000C |
| 8947202 | 8965011 | -1 | protein_coding | ENSG0000C |
| 8964510 | 8977527 | 1 | lncRNA | ENSG0000C |
| 8980576 | 9004049 | -1 | protein_coding | ENSG0000C |
| 9004093 | 9067776 | 1 | lncRNA | ENSG0000C |
| 9019498 | 9138114 | -1 | protein_coding | ENSG0000C |
| 9089283 | 9090456 | 1 | lncRNA | ENSG0000C |
| 9090312 | 9090379 | -1 | miRNA | ENSG0000C |
| 9094377 | 9095890 | 1 | processed_pseudoge | ENSG0000C |
| 9138825 | 9265350 | -1 | protein_coding | ENSG0000C |
| 9242448 | 9245509 | -1 | lncRNA | ENSG0000C |
| 9279532 | 9279816 | -1 | misc_RNA | ENSG0000C |
| 9280654 | 9314636 | -1 | protein_coding | ENSG0000C |
| 9311284 | 9311689 | -1 | processed_pseudoge | ENSG0000C |
| 9384652 | 9448127 | 1 | protein_coding | ENSG0000C |
| 9428773 | 9428954 | 1 | snoRNA | ENSG0000C |
| 9430356 | 9433486 | -1 | lncRNA | ENSG0000C |
| 9459556 | 9460702 | -1 | lncRNA | ENSG0000C |
| 9460319 | 9528524 | 1 | protein_coding | ENSG0000C |
| 9573670 | 9593457 | 1 | protein_coding | ENSG0000C |

| | | | | |
|---|---|---|---|---|
| 9578028 | 9578131 | 1 | snoRNA | ENSG0000C |
| 9606142 | 9606609 | 1 | processed_pseudoge | ENSG0000C |
| 9631008 | 9631359 | -1 | processed_pseudoge | ENSG0000C |
| 9660438 | 9660920 | 1 | processed_pseudoge | ENSG0000C |
| 9664077 | 9752993 | 1 | protein_coding | ENSG0000C |
| 9680924 | 9681238 | 1 | misc_RNA | ENSG0000C |
| 9754770 | 9759533 | -1 | lncRNA | ENSG0000C |
| 9758268 | 9811335 | 1 | lncRNA | ENSG0000C |
| 9778667 | 10294207 | -1 | protein_coding | ENSG0000C |
| 9839143 | 9929263 | 1 | lncRNA | ENSG0000C |
| 9843815 | 9843876 | -1 | snRNA | ENSG0000C |
| 9958744 | 9959790 | -1 | processed_pseudoge | ENSG0000C |
| 9982753 | 10000290 | 1 | lncRNA | ENSG0000C |
| 10149192 | 10165109 | 1 | lncRNA | ENSG0000C |
| 10272052 | 10272259 | -1 | processed_pseudoge | ENSG0000C |
| 10302657 | 10303704 | -1 | lncRNA | ENSG0000C |
| 10305073 | 10307397 | 1 | protein_coding | ENSG0000C |
| 10308313 | 10507579 | 1 | protein_coding | ENSG0000C |
| 10399192 | 10399317 | -1 | snRNA | ENSG0000C |
| 10507894 | 10509186 | -1 | protein_coding | ENSG0000C |
| 10508270 | 10508326 | -1 | miRNA | ENSG0000C |
| 10511673 | 10541230 | -1 | protein_coding | ENSG0000C |
| 10541258 | 10599936 | 1 | lncRNA | ENSG0000C |
| 10556966 | 10611689 | -1 | protein_coding | ENSG0000C |
| 10573091 | 10693988 | -1 | protein_coding | ENSG0000C |
| 10725903 | 10726004 | 1 | misc_RNA | ENSG0000C |
| 10751246 | 10801625 | 1 | protein_coding | ENSG0000C |
| 10797050 | 10808940 | -1 | protein_coding | ENSG0000C |
| 10801467 | 10801608 | -1 | snoRNA | ENSG0000C |
| 10809204 | 10822931 | 1 | lncRNA | ENSG0000C |
| 10812074 | 10858796 | -1 | protein_coding | ENSG0000C |
| 10858179 | 10908972 | 1 | lncRNA | ENSG0000C |
| 10884158 | 10899364 | -1 | lncRNA | ENSG0000C |
| 11020883 | 11183611 | 1 | lncRNA | ENSG0000C |
| 11135482 | 11137658 | 1 | processed_pseudoge | ENSG0000C |
| 11182107 | 11182413 | -1 | processed_pseudoge | ENSG0000C |
| 11239759 | 11240280 | 1 | processed_pseudoge | ENSG0000C |
| 11243188 | 11243715 | -1 | lncRNA | ENSG0000C |
| 11270877 | 11622005 | -1 | protein_coding | ENSG0000C |
| 11351942 | 11353250 | -1 | protein_coding | ENSG0000C |
| 11352426 | 11353307 | 1 | lncRNA | ENSG0000C |
| 11570084 | 11573782 | 1 | lncRNA | ENSG0000C |
| 11656651 | 11656722 | -1 | miRNA | ENSG0000C |
| 11725332 | 11726702 | -1 | TEC | ENSG0000C |
| 11759101 | 11759433 | -1 | processed_pseudoge | ENSG0000C |
| 11781971 | 11811530 | -1 | lncRNA | ENSG0000C |
| 11783135 | 11783222 | -1 | miRNA | ENSG0000C |
| 11841423 | 11961887 | 1 | protein_coding | ENSG0000C |
| 11856490 | 11856859 | -1 | processed_pseudoge | ENSG0000C |
| 11956207 | 12009769 | -1 | protein_coding | ENSG0000C |

| | | | | |
|---|---|---|---|---|
| 12030875 | 12061785 | 1 | lncRNA | ENSG00000 |
| 12066929 | 12073014 | 1 | lncRNA | ENSG00000 |
| 12086891 | 12089441 | 1 | lncRNA | ENSG00000 |
| 12094008 | 12359144 | 1 | protein_coding | ENSG00000 |
| 12163683 | 12163767 | 1 | miRNA | ENSG00000 |
| 12261426 | 12263173 | -1 | lncRNA | ENSG00000 |
| 12303533 | 12308216 | 1 | lncRNA | ENSG00000 |
| 12377563 | 12535356 | 1 | protein_coding | ENSG00000 |
| 12538083 | 12540967 | -1 | lncRNA | ENSG00000 |
| 12619326 | 12668495 | 1 | lncRNA | ENSG00000 |
| 12674421 | 12944737 | 1 | protein_coding | ENSG00000 |
| 12822435 | 12823667 | -1 | lncRNA | ENSG00000 |
| 12848795 | 12849433 | 1 | lncRNA | ENSG00000 |
| 12904817 | 12904988 | -1 | scaRNA | ENSG00000 |
| 12921186 | 12922925 | -1 | lncRNA | ENSG00000 |
| 12961541 | 12989597 | -1 | lncRNA | ENSG00000 |
| 13001090 | 13009159 | -1 | lncRNA | ENSG00000 |
| 13009316 | 13012119 | 1 | protein_coding | ENSG00000 |
| 13054615 | 13134839 | -1 | lncRNA | ENSG00000 |
| 13152200 | 13152348 | 1 | processed_pseudoge | ENSG00000 |
| 13276652 | 13387266 | 1 | protein_coding | ENSG00000 |
| 13353208 | 13353523 | 1 | misc_RNA | ENSG00000 |
| 13388001 | 13463297 | -1 | protein_coding | ENSG00000 |
| 13463377 | 13464717 | 1 | lncRNA | ENSG00000 |
| 13478333 | 13479037 | 1 | processed_pseudoge | ENSG00000 |
| 13488612 | 13489390 | 1 | unprocessed_pseudo | ENSG00000 |
| 13492054 | 13496181 | -1 | protein_coding | ENSG00000 |
| 13610363 | 13610771 | -1 | processed_pseudoge | ENSG00000 |
| 13668668 | 13732346 | 1 | protein_coding | ENSG00000 |
| 13669327 | 13670149 | 1 | lncRNA | ENSG00000 |
| 13685499 | 13685656 | -1 | processed_pseudoge | ENSG00000 |
| 13686550 | 13686658 | -1 | misc_RNA | ENSG00000 |
| 13756027 | 13757560 | 1 | processed_pseudoge | ENSG00000 |
| 13758111 | 13758345 | -1 | processed_pseudoge | ENSG00000 |
| 13784017 | 13848002 | 1 | lncRNA | ENSG00000 |
| 13826843 | 13879499 | -1 | lncRNA | ENSG00000 |
| 13844862 | 13870616 | -1 | lncRNA | ENSG00000 |
| 13907484 | 13907590 | -1 | rRNA_pseudogene | ENSG00000 |
| 13921450 | 13924863 | -1 | lncRNA | ENSG00000 |
| 13962723 | 14268133 | 1 | protein_coding | ENSG00000 |
| 14134986 | 14135100 | 1 | rRNA_pseudogene | ENSG00000 |
| 14262846 | 14273691 | -1 | lncRNA | ENSG00000 |
| 14277922 | 14364506 | -1 | protein_coding | ENSG00000 |
| 14440787 | 14441677 | 1 | processed_pseudoge | ENSG00000 |
| 14443440 | 14500027 | -1 | protein_coding | ENSG00000 |
| 14478892 | 14478953 | 1 | snRNA | ENSG00000 |
| 14493783 | 14520344 | -1 | protein_coding | ENSG00000 |
| 14504874 | 14643635 | -1 | protein_coding | ENSG00000 |
| 14643804 | 14872044 | 1 | protein_coding | ENSG00000 |
| 14673606 | 14674555 | -1 | processed_pseudoge | ENSG00000 |

| | | | | |
|---|---|---|---|---|
| 14877440 | 14892252 | -1 | protein_coding | ENSG0000C |
| 14904997 | 15082342 | 1 | protein_coding | ENSG0000C |
| 14907518 | 14907986 | 1 | unprocessed_pseudc | ENSG0000C |
| 14966668 | 14972354 | -1 | protein_coding | ENSG0000C |
| 14987671 | 14988694 | -1 | unprocessed_pseudc | ENSG0000C |
| 15112424 | 15247208 | 1 | protein_coding | ENSG0000C |
| 15481808 | 15481893 | -1 | snoRNA | ENSG0000C |
| 15552855 | 15604169 | 1 | lncRNA | ENSG0000C |
| 15571817 | 15622447 | -1 | lncRNA | ENSG0000C |
| 15605484 | 15705376 | 1 | lncRNA | ENSG0000C |
| 15701265 | 15758898 | -1 | lncRNA | ENSG0000C |
| 15864452 | 15883653 | 1 | lncRNA | ENSG0000C |
| 15910528 | 15930951 | 1 | lncRNA | ENSG0000C |
| 15966449 | 16739591 | -1 | protein_coding | ENSG0000C |
| 15969533 | 15969621 | -1 | miRNA | ENSG0000C |
| 16023190 | 16031515 | 1 | lncRNA | ENSG0000C |
| 16484084 | 16484671 | -1 | processed_pseudoge | ENSG0000C |
| 16576851 | 16577135 | 1 | misc_RNA | ENSG0000C |
| 16613132 | 16758340 | 1 | protein_coding | ENSG0000C |
| 16777297 | 17014414 | -1 | protein_coding | ENSG0000C |
| 16842253 | 16842587 | 1 | misc_RNA | ENSG0000C |
| 16974693 | 16975013 | -1 | processed_pseudoge | ENSG0000C |
| 17013998 | 17053024 | 1 | transcribed_unproce | ENSG0000C |
| 17074388 | 17077715 | -1 | protein_coding | ENSG0000C |
| 17074654 | 17074744 | -1 | snoRNA | ENSG0000C |
| 17075779 | 17075868 | -1 | snoRNA | ENSG0000C |
| 17077730 | 17207983 | -1 | protein_coding | ENSG0000C |
| 17115652 | 17115763 | -1 | snRNA | ENSG0000C |
| 17137976 | 17138265 | -1 | processed_pseudoge | ENSG0000C |
| 17193489 | 17194308 | 1 | processed_pseudoge | ENSG0000C |
| 17208153 | 17349980 | 1 | protein_coding | ENSG0000C |
| 17228279 | 17229151 | 1 | processed_pseudoge | ENSG0000C |
| 17349053 | 17351703 | -1 | lncRNA | ENSG0000C |
| 17351800 | 17377341 | 1 | protein_coding | ENSG0000C |
| 17380649 | 17383531 | 1 | lncRNA | ENSG0000C |
| 17385859 | 17389331 | -1 | protein_coding | ENSG0000C |
| 17392498 | 17476879 | -1 | protein_coding | ENSG0000C |
| 17435672 | 17436181 | -1 | processed_pseudoge | ENSG0000C |
| 17476595 | 17492367 | 1 | lncRNA | ENSG0000C |
| 17493895 | 17544416 | -1 | protein_coding | ENSG0000C |
| 17547373 | 17647150 | 1 | protein_coding | ENSG0000C |
| 17695010 | 17697556 | 1 | lncRNA | ENSG0000C |
| 17695266 | 17696994 | -1 | lncRNA | ENSG0000C |
| 17719571 | 17722136 | 1 | protein_coding | ENSG0000C |
| 17734774 | 17856804 | 1 | protein_coding | ENSG0000C |
| 17788048 | 18013047 | -1 | protein_coding | ENSG0000C |
| 18000542 | 18022931 | -1 | lncRNA | ENSG0000C |
| 18017564 | 18042426 | -1 | protein_coding | ENSG0000C |
| 18069935 | 18106087 | -1 | protein_coding | ENSG0000C |
| 18106583 | 18106841 | 1 | processed_pseudoge | ENSG0000C |

| | | | | |
|---|---|---|---|---|
| 18112472 | 18116132 | -1 | transcribed_unproce | ENSG00000 |
| 18120955 | 18138480 | 1 | protein_coding | ENSG00000 |
| 18140186 | 18189495 | -1 | lncRNA | ENSG00000 |
| 18142341 | 18145142 | 1 | lncRNA | ENSG00000 |
| 18153277 | 18154255 | 1 | processed_pseudoge | ENSG00000 |
| 18172837 | 18174280 | 1 | protein_coding | ENSG00000 |
| 18189000 | 18189621 | 1 | processed_pseudoge | ENSG00000 |
| 18196657 | 18197337 | 1 | processed_pseudoge | ENSG00000 |
| 18209138 | 18214562 | 1 | transcribed_unproce | ENSG00000 |
| 18231355 | 18236802 | -1 | protein_coding | ENSG00000 |
| 18231423 | 18248635 | -1 | protein_coding | ENSG00000 |
| 18239223 | 18248643 | -1 | protein_coding | ENSG00000 |
| 18248389 | 18248506 | -1 | rRNA_pseudogene | ENSG00000 |
| 18261982 | 18263091 | 1 | processed_pseudoge | ENSG00000 |
| 18266260 | 18269977 | 1 | protein_coding | ENSG00000 |
| 18266393 | 18266510 | 1 | rRNA_pseudogene | ENSG00000 |
| 18278668 | 18322198 | -1 | protein_coding | ENSG00000 |
| 18322295 | 18367045 | 1 | protein_coding | ENSG00000 |
| 18387787 | 18387860 | 1 | miRNA | ENSG00000 |
| 18394560 | 18408425 | 1 | protein_coding | ENSG00000 |
| 18405609 | 18406731 | 1 | lncRNA | ENSG00000 |
| 18412318 | 18452063 | 1 | protein_coding | ENSG00000 |
| 18450112 | 18450868 | -1 | processed_pseudoge | ENSG00000 |
| 18455824 | 18479601 | 1 | protein_coding | ENSG00000 |
| 18468336 | 18526951 | -1 | protein_coding | ENSG00000 |
| 18490243 | 18490955 | -1 | processed_pseudoge | ENSG00000 |
| 18507608 | 18508820 | -1 | lncRNA | ENSG00000 |
| 18511043 | 18511475 | -1 | processed_pseudoge | ENSG00000 |
| 18529609 | 18588747 | -1 | protein_coding | ENSG00000 |
| 18538885 | 18539189 | 1 | misc_RNA | ENSG00000 |
| 18588781 | 18610255 | 1 | protein_coding | ENSG00000 |
| 18595459 | 18599683 | -1 | transcribed_processe | ENSG00000 |
| 18601882 | 18602649 | 1 | TEC | ENSG00000 |
| 18606403 | 18634791 | -1 | protein_coding | ENSG00000 |
| 18665050 | 18665548 | 1 | processed_pseudoge | ENSG00000 |
| 18693122 | 18704785 | 1 | protein_coding | ENSG00000 |
| 18704305 | 18726230 | -1 | protein_coding | ENSG00000 |
| 18706537 | 18740568 | 1 | lncRNA | ENSG00000 |
| 18727928 | 18792721 | -1 | protein_coding | ENSG00000 |
| 18761664 | 18764965 | 1 | lncRNA | ENSG00000 |
| 18836304 | 18837081 | 1 | processed_pseudoge | ENSG00000 |
| 18863558 | 18864509 | -1 | unprocessed_pseudc | ENSG00000 |
| 18870989 | 18873462 | -1 | processed_pseudoge | ENSG00000 |
| 18886933 | 18887920 | -1 | unprocessed_pseudc | ENSG00000 |
| 18910436 | 18911396 | -1 | unprocessed_pseudc | ENSG00000 |
| 18933813 | 18939507 | -1 | protein_coding | ENSG00000 |
| 18934985 | 18939721 | 1 | lncRNA | ENSG00000 |
| 18956055 | 18957015 | -1 | unprocessed_pseudc | ENSG00000 |
| 18971746 | 18972703 | -1 | unprocessed_pseudc | ENSG00000 |
| 19029697 | 19030438 | -1 | unprocessed_pseudc | ENSG00000 |

| | | | | |
|---|---|---|---|---|
| 19054455 | 19060717 | -1 | protein_coding | ENSG00000 |
| 19117099 | 19176422 | 1 | protein_coding | ENSG00000 |
| 19182030 | 19210571 | -1 | protein_coding | ENSG00000 |
| 19196775 | 19281426 | 1 | lncRNA | ENSG00000 |
| 19224063 | 19241620 | -1 | protein_coding | ENSG00000 |
| 19274540 | 19275269 | -1 | processed_pseudoge | ENSG00000 |
| 19299883 | 19308358 | 1 | lncRNA | ENSG00000 |
| 19350724 | 20121601 | 1 | protein_coding | ENSG00000 |
| 19380484 | 19385014 | 1 | lncRNA | ENSG00000 |
| 19401872 | 19401981 | -1 | rRNA_pseudogene | ENSG00000 |
| 19502672 | 19507229 | -1 | lncRNA | ENSG00000 |
| 19510890 | 19519896 | -1 | lncRNA | ENSG00000 |
| 19575310 | 19575372 | 1 | miRNA | ENSG00000 |
| 19591156 | 19591291 | -1 | snoRNA | ENSG00000 |
| 19710934 | 19712619 | -1 | lncRNA | ENSG00000 |
| 19760004 | 19760083 | -1 | miRNA | ENSG00000 |
| 19978699 | 19981337 | -1 | lncRNA | ENSG00000 |
| 20043846 | 20049303 | -1 | lncRNA | ENSG00000 |
| 20119684 | 20120632 | -1 | lncRNA | ENSG00000 |
| 20131730 | 20135465 | 1 | TEC | ENSG00000 |
| 20156155 | 20160475 | -1 | protein_coding | ENSG00000 |
| 20363685 | 20383783 | 1 | protein_coding | ENSG00000 |
| 20387558 | 20509338 | 1 | protein_coding | ENSG00000 |
| 20574186 | 20575042 | 1 | processed_pseudoge | ENSG00000 |
| 20596522 | 20596979 | -1 | lncRNA | ENSG00000 |
| 20599400 | 20659285 | 1 | protein_coding | ENSG00000 |
| 20669551 | 21575686 | 1 | protein_coding | ENSG00000 |
| 20670425 | 20671297 | -1 | lncRNA | ENSG00000 |
| 21000881 | 21000997 | -1 | rRNA_pseudogene | ENSG00000 |
| 21169748 | 21170137 | 1 | processed_pseudoge | ENSG00000 |
| 21260061 | 21262570 | -1 | lncRNA | ENSG00000 |
| 21283779 | 21284955 | 1 | processed_pseudoge | ENSG00000 |
| 21383801 | 21383909 | -1 | rRNA_pseudogene | ENSG00000 |
| 21782659 | 21843210 | 1 | lncRNA | ENSG00000 |
| 21799934 | 22283357 | 1 | protein_coding | ENSG00000 |
| 22010402 | 22170001 | -1 | lncRNA | ENSG00000 |
| 22113448 | 22117001 | 1 | lncRNA | ENSG00000 |
| 22261209 | 22262256 | -1 | lncRNA | ENSG00000 |
| 22283730 | 22338245 | -1 | lncRNA | ENSG00000 |
| 22338381 | 22379503 | 1 | protein_coding | ENSG00000 |
| 22361213 | 22362294 | -1 | lncRNA | ENSG00000 |
| 22445673 | 22492073 | -1 | lncRNA | ENSG00000 |
| 22492087 | 22510396 | 1 | lncRNA | ENSG00000 |
| 22622890 | 22625841 | -1 | protein_coding | ENSG00000 |
| 22625509 | 22813055 | 1 | protein_coding | ENSG00000 |
| 22688351 | 22688435 | 1 | rRNA_pseudogene | ENSG00000 |
| 22813799 | 22830299 | -1 | protein_coding | ENSG00000 |
| 22829380 | 22945393 | 1 | lncRNA | ENSG00000 |
| 22846922 | 22860474 | -1 | protein_coding | ENSG00000 |
| 23154683 | 23203161 | 1 | lncRNA | ENSG00000 |

| | | | | |
|---|---|---|---|---|
| 23370116 | 23377644 | -1 | processed_pseudoge | ENSG00000 |
| 23403805 | 23406103 | 1 | processed_pseudoge | ENSG00000 |
| 23419105 | 23419190 | -1 | miRNA | ENSG00000 |
| 23431677 | 23431799 | 1 | processed_pseudoge | ENSG00000 |
| 23478068 | 23480430 | 1 | processed_pseudoge | ENSG00000 |
| 23520322 | 23521202 | 1 | processed_pseudoge | ENSG00000 |
| 23730588 | 23796970 | 1 | lncRNA | ENSG00000 |
| 23761330 | 23808255 | -1 | lncRNA | ENSG00000 |
| 23849778 | 23849882 | 1 | snRNA | ENSG00000 |
| 23886135 | 23893557 | 1 | processed_pseudoge | ENSG00000 |
| 24235477 | 24262218 | -1 | lncRNA | ENSG00000 |
| 24455911 | 24456010 | 1 | misc_RNA | ENSG00000 |
| 24496970 | 25082638 | 1 | protein_coding | ENSG00000 |
| 24701275 | 24701575 | 1 | processed_pseudoge | ENSG00000 |
| 25140533 | 25141023 | -1 | processed_pseudoge | ENSG00000 |
| 25588475 | 25588795 | -1 | processed_pseudoge | ENSG00000 |
| 25630720 | 25631041 | 1 | processed_pseudoge | ENSG00000 |
| 25734757 | 25782017 | 1 | lncRNA | ENSG00000 |
| 25924188 | 25926808 | 1 | lncRNA | ENSG00000 |
| 26045987 | 26047598 | 1 | processed_pseudoge | ENSG00000 |
| 26188842 | 26663289 | 1 | protein_coding | ENSG00000 |
| 26244640 | 26269415 | -1 | lncRNA | ENSG00000 |
| 26285578 | 26331289 | -1 | lncRNA | ENSG00000 |
| 26427051 | 26427412 | 1 | processed_pseudoge | ENSG00000 |
| 26559032 | 26572263 | -1 | protein_coding | ENSG00000 |
| 26667020 | 26723427 | -1 | protein_coding | ENSG00000 |
| 26994112 | 26997087 | 1 | protein_coding | ENSG00000 |
| 27040725 | 27127809 | 1 | protein_coding | ENSG00000 |
| 27047186 | 27220113 | -1 | lncRNA | ENSG00000 |
| 27330827 | 27363234 | -1 | protein_coding | ENSG00000 |
| 27365961 | 27472790 | -1 | protein_coding | ENSG00000 |
| 27471729 | 27482433 | 1 | lncRNA | ENSG00000 |
| 27483850 | 27484128 | 1 | processed_pseudoge | ENSG00000 |
| 27494418 | 27506769 | -1 | protein_coding | ENSG00000 |
| 27506830 | 27698231 | 1 | lncRNA | ENSG00000 |
| 27514970 | 27515047 | -1 | miRNA | ENSG00000 |
| 27521605 | 27521726 | -1 | rRNA_pseudogene | ENSG00000 |
| 27581680 | 27582055 | 1 | processed_pseudoge | ENSG00000 |
| 27617626 | 27634627 | -1 | lncRNA | ENSG00000 |
| 27654893 | 27722058 | -1 | protein_coding | ENSG00000 |
| 27696312 | 27877648 | -1 | lncRNA | ENSG00000 |
| 27806443 | 27806996 | 1 | processed_pseudoge | ENSG00000 |
| 27888838 | 27891033 | -1 | processed_pseudoge | ENSG00000 |
| 27978669 | 28019575 | -1 | lncRNA | ENSG00000 |
| 28020619 | 28108156 | -1 | protein_coding | ENSG00000 |
| 28056815 | 28056910 | 1 | miRNA | ENSG00000 |
| 28108248 | 28527041 | 1 | protein_coding | ENSG00000 |
| 28144907 | 28145204 | -1 | processed_pseudoge | ENSG00000 |
| 28261194 | 28261519 | -1 | misc_RNA | ENSG00000 |
| 28287235 | 28287625 | -1 | processed_pseudoge | ENSG00000 |

| | | | | |
|---|---|---|---|---|
| 28477481 | 28477548 | -1 | miRNA | ENSG0000C |
| 28516832 | 28519341 | 1 | lncRNA | ENSG0000C |
| 28679183 | 28683469 | 1 | lncRNA | ENSG0000C |
| 28702607 | 29064322 | 1 | lncRNA | ENSG0000C |
| 28986635 | 28987498 | 1 | unprocessed_pseudc | ENSG0000C |
| 29159956 | 29266734 | 1 | lncRNA | ENSG0000C |
| 29275655 | 29276565 | 1 | processed_pseudoge | ENSG0000C |
| 29335878 | 29989328 | -1 | lncRNA | ENSG0000C |
| 29391525 | 29393844 | 1 | processed_pseudoge | ENSG0000C |
| 29445487 | 29457393 | 1 | lncRNA | ENSG0000C |
| 29482807 | 29552861 | 1 | lncRNA | ENSG0000C |
| 29594326 | 29631515 | 1 | lncRNA | ENSG0000C |
| 29668886 | 29669628 | -1 | processed_pseudoge | ENSG0000C |
| 29713909 | 29881714 | 1 | lncRNA | ENSG0000C |
| 29721399 | 29721684 | -1 | misc_RNA | ENSG0000C |
| 29725764 | 29726565 | 1 | processed_pseudoge | ENSG0000C |
| 29980113 | 29982392 | -1 | lncRNA | ENSG0000C |
| 30009730 | 30017030 | -1 | protein_coding | ENSG0000C |
| 30044053 | 30322988 | -1 | lncRNA | ENSG0000C |
| 30167163 | 30167306 | 1 | processed_pseudoge | ENSG0000C |
| 30231016 | 30235261 | 1 | protein_coding | ENSG0000C |
| 30323104 | 30338223 | 1 | protein_coding | ENSG0000C |
| 30368148 | 30368646 | 1 | processed_pseudoge | ENSG0000C |
| 30384493 | 30586872 | -1 | protein_coding | ENSG0000C |
| 30425552 | 30429268 | 1 | lncRNA | ENSG0000C |
| 30584130 | 30632583 | 1 | lncRNA | ENSG0000C |
| 30773913 | 30850559 | 1 | lncRNA | ENSG0000C |
| 30830369 | 31369810 | -1 | protein_coding | ENSG0000C |
| 31280672 | 31280987 | -1 | processed_pseudoge | ENSG0000C |
| 31305685 | 31314548 | 1 | lncRNA | ENSG0000C |
| 31369840 | 31432835 | 1 | protein_coding | ENSG0000C |
| 31432401 | 31509645 | -1 | protein_coding | ENSG0000C |
| 31509755 | 31790324 | 1 | protein_coding | ENSG0000C |
| 31618124 | 31767953 | -1 | lncRNA | ENSG0000C |
| 31784779 | 31817961 | -1 | protein_coding | ENSG0000C |
| 31812307 | 32002405 | 1 | lncRNA | ENSG0000C |
| 31812391 | 32104665 | 1 | protein_coding | ENSG0000C |
| 32035979 | 32041318 | -1 | lncRNA | ENSG0000C |
| 32052843 | 32053260 | -1 | lncRNA | ENSG0000C |
| 32064912 | 32072173 | 1 | lncRNA | ENSG0000C |
| 32081764 | 32081973 | 1 | snoRNA | ENSG0000C |
| 32085111 | 32085712 | 1 | processed_pseudoge | ENSG0000C |
| 32090904 | 32105755 | 1 | protein_coding | ENSG0000C |
| 32097143 | 32105091 | -1 | lncRNA | ENSG0000C |
| 32112049 | 32343409 | -1 | transcribed_unitary_ | ENSG0000C |
| 32132657 | 32143723 | 1 | lncRNA | ENSG0000C |
| 32387775 | 32435885 | -1 | protein_coding | ENSG0000C |
| 32435518 | 32458769 | 1 | lncRNA | ENSG0000C |
| 32435738 | 32435846 | 1 | misc_RNA | ENSG0000C |
| 32436311 | 32436447 | 1 | misc_RNA | ENSG0000C |

| | | | | |
|---|---|---|---|---|
| 32436797 | 32437030 | 1 | misc_RNA | ENSG0000C |
| 32439086 | 32439206 | 1 | misc_RNA | ENSG0000C |
| 32439716 | 32440009 | 1 | misc_RNA | ENSG0000C |
| 32440109 | 32440383 | 1 | misc_RNA | ENSG0000C |
| 32583798 | 32606264 | 1 | protein_coding | ENSG0000C |
| 32591793 | 32592771 | -1 | processed_pseudoge | ENSG0000C |
| 32602246 | 32794658 | -1 | protein_coding | ENSG0000C |
| 32758268 | 32758614 | 1 | processed_pseudoge | ENSG0000C |
| 32829927 | 32858120 | 1 | protein_coding | ENSG0000C |
| 32892820 | 32993316 | 1 | protein_coding | ENSG0000C |
| 33004250 | 33004345 | -1 | misc_RNA | ENSG0000C |
| 33015876 | 33033582 | 1 | protein_coding | ENSG0000C |
| 33039417 | 33105943 | 1 | protein_coding | ENSG0000C |
| 33075566 | 33076460 | -1 | processed_pseudoge | ENSG0000C |
| 33076149 | 33079454 | 1 | lncRNA | ENSG0000C |
| 33077188 | 33162371 | -1 | protein_coding | ENSG0000C |
| 33161657 | 33191598 | 1 | lncRNA | ENSG0000C |
| 33171381 | 33171490 | -1 | misc_RNA | ENSG0000C |
| 33190062 | 33190503 | -1 | processed_pseudoge | ENSG0000C |
| 33196547 | 33196637 | -1 | misc_RNA | ENSG0000C |
| 33237008 | 33237409 | 1 | processed_pseudoge | ENSG0000C |
| 33256672 | 33357023 | 1 | protein_coding | ENSG0000C |
| 33376083 | 33674102 | 1 | protein_coding | ENSG0000C |
| 33403216 | 33403605 | 1 | processed_pseudoge | ENSG0000C |
| 33450646 | 33450943 | 1 | processed_pseudoge | ENSG0000C |
| 33665220 | 33696701 | -1 | lncRNA | ENSG0000C |
| 33698261 | 33722467 | -1 | protein_coding | ENSG0000C |
| 33698261 | 33700801 | -1 | protein_coding | ENSG0000C |
| 33703010 | 33736491 | -1 | protein_coding | ENSG0000C |
| 33740939 | 33774543 | -1 | protein_coding | ENSG0000C |
| 33774699 | 33775670 | 1 | lncRNA | ENSG0000C |
| 33776188 | 33779495 | 1 | lncRNA | ENSG0000C |
| 33787054 | 33787795 | -1 | processed_pseudoge | ENSG0000C |
| 33804768 | 33805641 | 1 | lncRNA | ENSG0000C |
| 33810145 | 33811178 | 1 | lncRNA | ENSG0000C |
| 33813873 | 33822378 | -1 | lncRNA | ENSG0000C |
| 33858576 | 33892076 | -1 | protein_coding | ENSG0000C |
| 33880643 | 33881800 | 1 | lncRNA | ENSG0000C |
| 34049967 | 34052132 | -1 | lncRNA | ENSG0000C |
| 34051731 | 34102610 | 1 | protein_coding | ENSG0000C |
| 34105617 | 34147670 | 1 | protein_coding | ENSG0000C |
| 34150987 | 34358010 | -1 | protein_coding | ENSG0000C |
| 34189174 | 34189275 | 1 | misc_RNA | ENSG0000C |
| 34335118 | 34336003 | -1 | processed_pseudoge | ENSG0000C |
| 34404663 | 34404949 | -1 | processed_pseudoge | ENSG0000C |
| 34430880 | 34431649 | 1 | processed_pseudoge | ENSG0000C |
| 34438934 | 34472060 | 1 | protein_coding | ENSG0000C |
| 34478793 | 34513805 | -1 | protein_coding | ENSG0000C |
| 34533014 | 34557924 | -1 | lncRNA | ENSG0000C |
| 34570876 | 34573982 | 1 | lncRNA | ENSG0000C |

| | | | | |
|---|---|---|---|---|
| 34621093 | 34661057 | 1 | protein_coding | ENSG00000( |
| 34709600 | 34710161 | 1 | processed_pseudoge | ENSG00000( |
| 34742906 | 34743199 | 1 | processed_pseudoge | ENSG00000( |
| 34853094 | 34916379 | -1 | protein_coding | ENSG00000( |
| 34915829 | 35020591 | 1 | protein_coding | ENSG00000( |
| 34941837 | 34941920 | 1 | miRNA | ENSG00000( |
| 35132655 | 35138032 | -1 | lncRNA | ENSG00000( |
| 35138870 | 35232402 | 1 | protein_coding | ENSG00000( |
| 35210343 | 35214985 | -1 | lncRNA | ENSG00000( |
| 35212550 | 35214007 | 1 | lncRNA | ENSG00000( |
| 35251205 | 35420063 | -1 | protein_coding | ENSG00000( |
| 35281813 | 35286009 | 1 | lncRNA | ENSG00000( |
| 35374265 | 35374474 | 1 | processed_pseudoge | ENSG00000( |
| 35419057 | 35421002 | 1 | lncRNA | ENSG00000( |
| 35431823 | 35530300 | -1 | protein_coding | ENSG00000( |
| 35579430 | 35585310 | -1 | lncRNA | ENSG00000( |
| 35618460 | 35620865 | 1 | protein_coding | ENSG00000( |
| 35656694 | 35661339 | 1 | lncRNA | ENSG00000( |
| 35662775 | 35818007 | 1 | protein_coding | ENSG00000( |
| 35824192 | 35838523 | 1 | lncRNA | ENSG00000( |
| 35860249 | 35861498 | -1 | processed_pseudoge | ENSG00000( |
| 35894987 | 35897735 | 1 | lncRNA | ENSG00000( |
| 35915051 | 35918739 | -1 | lncRNA | ENSG00000( |
| 35943981 | 36232136 | 1 | protein_coding | ENSG00000( |
| 35967010 | 35968676 | -1 | processed_pseudoge | ENSG00000( |
| 35972428 | 35989007 | 1 | lncRNA | ENSG00000( |
| 35997226 | 35997675 | -1 | processed_pseudoge | ENSG00000( |
| 36010098 | 36010204 | 1 | miRNA | ENSG00000( |
| 36196248 | 36197901 | 1 | TEC | ENSG00000( |
| 36269284 | 36289449 | -1 | protein_coding | ENSG00000( |
| 36296288 | 36465204 | 1 | protein_coding | ENSG00000( |
| 36321158 | 36323440 | -1 | lncRNA | ENSG00000( |
| 36386521 | 36388250 | 1 | lncRNA | ENSG00000( |
| 36425447 | 36426122 | -1 | lncRNA | ENSG00000( |
| 36483769 | 36510272 | -1 | protein_coding | ENSG00000( |
| 36510361 | 36510537 | 1 | TEC | ENSG00000( |
| 36510709 | 36593156 | 1 | protein_coding | ENSG00000( |
| 36575574 | 36598279 | -1 | protein_coding | ENSG00000( |
| 36594493 | 36659290 | 1 | protein_coding | ENSG00000( |
| 36696317 | 36697867 | 1 | processed_pseudoge | ENSG00000( |
| 37702125 | 37702220 | -1 | snoRNA | ENSG00000( |
| 37725666 | 37726456 | 1 | processed_pseudoge | ENSG00000( |
| 37938601 | 37954663 | 1 | lncRNA | ENSG00000( |
| 38013510 | 38014141 | -1 | TEC | ENSG00000( |
| 38211389 | 38212799 | 1 | processed_pseudoge | ENSG00000( |
| 38498995 | 38500186 | 1 | lncRNA | ENSG00000( |
| 38618264 | 38646384 | -1 | lncRNA | ENSG00000( |
| 38646451 | 38686323 | 1 | lncRNA | ENSG00000( |
| 39024631 | 39161638 | -1 | lncRNA | ENSG00000( |
| 39161453 | 39161967 | 1 | processed_pseudoge | ENSG00000( |

| | | | | |
|---|---|---|---|---|
| 39261107 | 39261213 | 1 | snRNA | ENSG0000C |
| 39873782 | 39874388 | 1 | processed_pseudoge | ENSG0000C |
| 40083710 | 40084517 | -1 | processed_pseudoge | ENSG0000C |
| 40107244 | 40112599 | -1 | TEC | ENSG0000C |
| 40114203 | 41459773 | -1 | protein_coding | ENSG0000C |
| 40351564 | 40351672 | 1 | misc_RNA | ENSG0000C |
| 40444371 | 40513269 | -1 | transcribed_processe | ENSG0000C |
| 41122907 | 41123012 | -1 | snRNA | ENSG0000C |
| 41394595 | 41426504 | 1 | lncRNA | ENSG0000C |
| 41518895 | 41714492 | -1 | lncRNA | ENSG0000C |
| 41538434 | 41538986 | 1 | processed_pseudoge | ENSG0000C |
| 41660030 | 41660221 | 1 | processed_pseudoge | ENSG0000C |
| 41714534 | 41857733 | 1 | lncRNA | ENSG0000C |
| 41797543 | 41797964 | 1 | processed_pseudoge | ENSG0000C |
| 41855920 | 41875997 | 1 | lncRNA | ENSG0000C |
| 41876634 | 41877078 | 1 | lncRNA | ENSG0000C |
| 41993381 | 42163851 | -1 | lncRNA | ENSG0000C |
| 42183292 | 42253721 | -1 | lncRNA | ENSG0000C |
| 42939775 | 42939998 | 1 | processed_pseudoge | ENSG0000C |
| 43063637 | 43065932 | -1 | processed_pseudoge | ENSG0000C |
| 43065686 | 43066076 | -1 | processed_pseudoge | ENSG0000C |
| 43120762 | 43269393 | -1 | transcribed_processe | ENSG0000C |
| 43311963 | 43344529 | 1 | protein_coding | ENSG0000C |
| 43328748 | 43359296 | -1 | lncRNA | ENSG0000C |
| 43331261 | 43331356 | 1 | misc_RNA | ENSG0000C |
| 43358920 | 43494933 | 1 | protein_coding | ENSG0000C |
| 43378882 | 43385671 | -1 | lncRNA | ENSG0000C |
| 43390283 | 43395495 | 1 | lncRNA | ENSG0000C |
| 43435132 | 43435397 | -1 | misc_RNA | ENSG0000C |
| 43466392 | 43466871 | 1 | processed_pseudoge | ENSG0000C |
| 43522036 | 43523264 | 1 | processed_pseudoge | ENSG0000C |
| 43556436 | 43673393 | 1 | lncRNA | ENSG0000C |
| 43559656 | 43559753 | 1 | miRNA | ENSG0000C |
| 43569306 | 43570395 | 1 | lncRNA | ENSG0000C |
| 43578889 | 43840030 | 1 | lncRNA | ENSG0000C |
| 43581394 | 43581483 | 1 | miRNA | ENSG0000C |
| 43680680 | 43856617 | 1 | protein_coding | ENSG0000C |
| 43718676 | 43719143 | 1 | processed_pseudoge | ENSG0000C |
| 43718676 | 43719146 | 1 | processed_pseudoge | ENSG0000C |
| 43733759 | 43734564 | 1 | processed_pseudoge | ENSG0000C |
| 43733761 | 43734564 | 1 | processed_pseudoge | ENSG0000C |
| 43827517 | 43880726 | -1 | lncRNA | ENSG0000C |
| 43855913 | 43856404 | -1 | lncRNA | ENSG0000C |
| 43880811 | 43920274 | 1 | protein_coding | ENSG0000C |
| 43897456 | 43899636 | 1 | processed_pseudoge | ENSG0000C |
| 43909289 | 43920944 | -1 | lncRNA | ENSG0000C |
| 43921059 | 44001157 | 1 | lncRNA | ENSG0000C |
| 43925342 | 43943878 | 1 | protein_coding | ENSG0000C |
| 43943787 | 43947206 | -1 | lncRNA | ENSG0000C |
| 44047981 | 44059977 | 1 | protein_coding | ENSG0000C |

| | | | | |
|---|---|---|---|---|
| 44065925 | 44084237 | 1 | protein_coding | ENSG00000 |
| 44071462 | 44072403 | 1 | lncRNA | ENSG00000 |
| 44095673 | 44251981 | 1 | protein_coding | ENSG00000 |
| 44260440 | 44310139 | -1 | protein_coding | ENSG00000 |
| 44468464 | 44470429 | 1 | lncRNA | ENSG00000 |
| 44539778 | 44540693 | -1 | processed_pseudoge | ENSG00000 |
| 44564427 | 44620358 | 1 | protein_coding | ENSG00000 |
| 44604508 | 44605337 | -1 | lncRNA | ENSG00000 |
| 44606170 | 44608101 | -1 | lncRNA | ENSG00000 |
| 44629357 | 44629674 | 1 | processed_pseudoge | ENSG00000 |
| 44656771 | 44656920 | 1 | processed_pseudoge | ENSG00000 |
| 44694863 | 44696301 | -1 | lncRNA | ENSG00000 |
| 44719392 | 44736692 | -1 | lncRNA | ENSG00000 |
| 44726465 | 44932423 | 1 | protein_coding | ENSG00000 |
| 44885903 | 44951306 | -1 | protein_coding | ENSG00000 |
| 44973770 | 44978028 | 1 | lncRNA | ENSG00000 |
| 45008266 | 45044053 | 1 | lncRNA | ENSG00000 |
| 45095806 | 45235110 | 1 | protein_coding | ENSG00000 |
| 45215815 | 45235292 | -1 | lncRNA | ENSG00000 |
| 45240302 | 45286341 | -1 | protein_coding | ENSG00000 |
| 45253884 | 45254649 | 1 | lncRNA | ENSG00000 |
| 45355371 | 45367490 | 1 | lncRNA | ENSG00000 |
| 45371397 | 45388511 | 1 | lncRNA | ENSG00000 |
| 45387215 | 45513802 | -1 | lncRNA | ENSG00000 |
| 45397590 | 45412745 | -1 | lncRNA | ENSG00000 |
| 45399448 | 45400528 | 1 | lncRNA | ENSG00000 |
| 45486867 | 45542487 | -1 | lncRNA | ENSG00000 |
| 45514187 | 45514451 | 1 | processed_pseudoge | ENSG00000 |
| 45582525 | 45583474 | 1 | lncRNA | ENSG00000 |
| 45647689 | 45665622 | -1 | protein_coding | ENSG00000 |
| 45651529 | 45652691 | 1 | lncRNA | ENSG00000 |
| 45691704 | 45691776 | -1 | miRNA | ENSG00000 |
| 45722279 | 45748029 | 1 | lncRNA | ENSG00000 |
| 45733994 | 45735766 | -1 | lncRNA | ENSG00000 |
| 45749454 | 45752279 | -1 | lncRNA | ENSG00000 |
| 45771432 | 45772358 | 1 | lncRNA | ENSG00000 |
| 45804072 | 45813015 | 1 | protein_coding | ENSG00000 |
| 45813219 | 45825258 | 1 | lncRNA | ENSG00000 |
| 45847118 | 45883248 | 1 | protein_coding | ENSG00000 |
| 45885651 | 45906465 | 1 | protein_coding | ENSG00000 |
| 45905941 | 45906461 | -1 | lncRNA | ENSG00000 |
| 45906513 | 45907272 | -1 | protein_coding | ENSG00000 |
| 45909663 | 45918812 | -1 | protein_coding | ENSG00000 |
| 45921621 | 45929096 | 1 | protein_coding | ENSG00000 |
| 45929323 | 46121178 | -1 | protein_coding | ENSG00000 |
| 46116578 | 46117318 | 1 | lncRNA | ENSG00000 |
| 46123031 | 46177106 | 1 | lncRNA | ENSG00000 |
| 46142716 | 46143594 | 1 | processed_pseudoge | ENSG00000 |
| 46213150 | 46220438 | -1 | lncRNA | ENSG00000 |
| 46238382 | 46239267 | 1 | lncRNA | ENSG00000 |

| | | | | |
|---|---|---|---|---|
| 46256355 | 46274547 | -1 | lncRNA | ENSG00000 |
| 46277662 | 46321409 | 1 | protein_coding | ENSG00000 |
| 46332905 | 46380554 | 1 | protein_coding | ENSG00000 |
| 46376402 | 46376484 | 1 | miRNA | ENSG00000 |
| 46380756 | 46383837 | 1 | protein_coding | ENSG00000 |
| 46385098 | 46386608 | -1 | protein_coding | ENSG00000 |
| 46396414 | 46594125 | -1 | protein_coding | ENSG00000 |
| 46428653 | 46429150 | -1 | processed_pseudoge | ENSG00000 |
| 46451805 | 46451889 | -1 | miRNA | ENSG00000 |
| 46451807 | 46451887 | 1 | miRNA | ENSG00000 |
| 46572948 | 46587808 | 1 | lncRNA | ENSG00000 |
| 46602861 | 46617909 | -1 | protein_coding | ENSG00000 |
| 46617527 | 46674818 | 1 | protein_coding | ENSG00000 |
| 46677080 | 46700619 | -1 | protein_coding | ENSG00000 |
| 46701030 | 46705912 | 1 | protein_coding | ENSG00000 |
| 46719180 | 46739506 | 1 | protein_coding | ENSG00000 |
| 46743048 | 46846308 | -1 | protein_coding | ENSG00000 |
| 46753125 | 46753192 | -1 | miRNA | ENSG00000 |
| 46758766 | 46758873 | -1 | snoRNA | ENSG00000 |
| 46762389 | 46762499 | -1 | snoRNA | ENSG00000 |
| 46846412 | 46874421 | 1 | lncRNA | ENSG00000 |
| 46856717 | 46918642 | -1 | protein_coding | ENSG00000 |
| 46920880 | 46929320 | 1 | lncRNA | ENSG00000 |
| 46936689 | 47164385 | 1 | protein_coding | ENSG00000 |
| 47041027 | 47041945 | -1 | processed_pseudoge | ENSG00000 |
| 47123104 | 47130801 | -1 | lncRNA | ENSG00000 |
| 47164299 | 47177125 | -1 | protein_coding | ENSG00000 |
| 47168281 | 47169563 | -1 | lncRNA | ENSG00000 |
| 47177522 | 47186443 | -1 | protein_coding | ENSG00000 |
| 47179611 | 47179737 | -1 | miRNA | ENSG00000 |
| 47191181 | 47191542 | -1 | processed_pseudoge | ENSG00000 |
| 47214465 | 47239240 | 1 | protein_coding | ENSG00000 |
| 47220218 | 47221751 | -1 | lncRNA | ENSG00000 |
| 47239302 | 47248906 | -1 | protein_coding | ENSG00000 |
| 47248300 | 47269033 | 1 | protein_coding | ENSG00000 |
| 47269161 | 47330031 | 1 | protein_coding | ENSG00000 |
| 47270657 | 47272110 | -1 | lncRNA | ENSG00000 |
| 47323983 | 47324181 | 1 | unprocessed_pseudo | ENSG00000 |
| 47331406 | 47352702 | -1 | protein_coding | ENSG00000 |
| 47354860 | 47378576 | -1 | protein_coding | ENSG00000 |
| 47381509 | 47409369 | -1 | lncRNA | ENSG00000 |
| 47400970 | 47401042 | 1 | miRNA | ENSG00000 |
| 47407132 | 47416496 | 1 | protein_coding | ENSG00000 |
| 47418769 | 47426473 | -1 | protein_coding | ENSG00000 |
| 47437764 | 47449143 | -1 | protein_coding | ENSG00000 |
| 47465933 | 47565569 | -1 | protein_coding | ENSG00000 |
| 47513605 | 47514889 | -1 | lncRNA | ENSG00000 |
| 47557534 | 47557833 | -1 | misc_RNA | ENSG00000 |
| 47565336 | 47584562 | 1 | protein_coding | ENSG00000 |
| 47565430 | 47573461 | 1 | protein_coding | ENSG00000 |

| | | | | |
|---|---|---|---|---|
| 47572197 | 47578976 | -1 | protein_coding | ENSG00000( |
| 47576471 | 47576588 | -1 | snRNA | ENSG00000( |
| 47577725 | 47578277 | 1 | lncRNA | ENSG00000( |
| 47586693 | 47589194 | 1 | protein_coding | ENSG00000( |
| 47589667 | 47594411 | -1 | protein_coding | ENSG00000( |
| 47617315 | 47642607 | -1 | protein_coding | ENSG00000( |
| 47659591 | 47715389 | -1 | protein_coding | ENSG00000( |
| 47716494 | 47767443 | -1 | protein_coding | ENSG00000( |
| 47726894 | 47726992 | -1 | misc_RNA | ENSG00000( |
| 47778087 | 47848555 | -1 | protein_coding | ENSG00000( |
| 47796169 | 47796379 | 1 | processed_pseudoge | ENSG00000( |
| 47804296 | 47804402 | -1 | rRNA_pseudogene | ENSG00000( |
| 47820200 | 47820289 | -1 | misc_RNA | ENSG00000( |
| 47841324 | 47841682 | 1 | processed_pseudoge | ENSG00000( |
| 47905323 | 47905607 | -1 | processed_pseudoge | ENSG00000( |
| 47980558 | 48170841 | 1 | protein_coding | ENSG00000( |
| 48014406 | 48015855 | -1 | lncRNA | ENSG00000( |
| 48096782 | 48096858 | 1 | miRNA | ENSG00000( |
| 48216810 | 48217739 | 1 | protein_coding | ENSG00000( |
| 48227429 | 48228310 | 1 | unprocessed_pseudc | ENSG00000( |
| 48245056 | 48246080 | 1 | protein_coding | ENSG00000( |
| 48263861 | 48264778 | 1 | protein_coding | ENSG00000( |
| 48306223 | 48307152 | 1 | protein_coding | ENSG00000( |
| 48324920 | 48328471 | 1 | protein_coding | ENSG00000( |
| 48345351 | 48352447 | -1 | unprocessed_pseudc | ENSG00000( |
| 48365485 | 48366465 | -1 | protein_coding | ENSG00000( |
| 48420210 | 48421273 | 1 | unprocessed_pseudc | ENSG00000( |
| 48432217 | 48433147 | -1 | unprocessed_pseudc | ENSG00000( |
| 48464053 | 48464977 | -1 | unprocessed_pseudc | ENSG00000( |
| 48486363 | 48486825 | -1 | unprocessed_pseudc | ENSG00000( |
| 48488717 | 48489780 | 1 | protein_coding | ENSG00000( |
| 48492026 | 48492416 | 1 | unprocessed_pseudc | ENSG00000( |
| 48496348 | 48497280 | 1 | unprocessed_pseudc | ENSG00000( |
| 48512273 | 48513220 | 1 | unprocessed_pseudc | ENSG00000( |
| 48526064 | 48526971 | 1 | unprocessed_pseudc | ENSG00000( |
| 48579436 | 48580336 | 1 | unprocessed_pseudc | ENSG00000( |
| 48589760 | 48590687 | 1 | unprocessed_pseudc | ENSG00000( |
| 48610063 | 48611000 | 1 | unprocessed_pseudc | ENSG00000( |
| 48627546 | 48628493 | 1 | unprocessed_pseudc | ENSG00000( |
| 48880111 | 48885237 | 1 | unprocessed_pseudc | ENSG00000( |
| 48881723 | 48902181 | -1 | unprocessed_pseudc | ENSG00000( |
| 48893256 | 48898703 | 1 | unprocessed_pseudc | ENSG00000( |
| 48907103 | 48908946 | 1 | unprocessed_pseudc | ENSG00000( |
| 48945694 | 48951895 | 1 | unprocessed_pseudc | ENSG00000( |
| 48959928 | 48961810 | 1 | unprocessed_pseudc | ENSG00000( |
| 48975324 | 48983826 | -1 | protein_coding | ENSG00000( |
| 48985868 | 48991881 | -1 | unprocessed_pseudc | ENSG00000( |
| 49009959 | 49018589 | -1 | transcribed_unproce | ENSG00000( |
| 49027501 | 49038451 | 1 | protein_coding | ENSG00000( |
| 49048271 | 49050159 | -1 | processed_pseudoge | ENSG00000( |

| | | | | |
|---|---|---|---|---|
| 49053714 | 49059112 | -1 | protein_coding | ENSG00000 |
| 49067610 | 49069301 | 1 | unprocessed_pseudo | ENSG00000 |
| 49070752 | 49071000 | -1 | processed_pseudoge | ENSG00000 |
| 49081907 | 49083097 | 1 | processed_pseudoge | ENSG00000 |
| 49092404 | 49092767 | 1 | processed_pseudoge | ENSG00000 |
| 49098163 | 49101168 | -1 | processed_pseudoge | ENSG00000 |
| 49110756 | 49111220 | 1 | processed_pseudoge | ENSG00000 |
| 49117010 | 49124570 | -1 | unprocessed_pseudo | ENSG00000 |
| 49133653 | 49134800 | -1 | processed_pseudoge | ENSG00000 |
| 49145092 | 49208638 | -1 | protein_coding | ENSG00000 |
| 49305714 | 49379669 | 1 | transcribed_unproce | ENSG00000 |
| 49405091 | 49415789 | -1 | unprocessed_pseudo | ENSG00000 |
| 49406190 | 49406714 | 1 | processed_pseudoge | ENSG00000 |
| 49433480 | 49434438 | -1 | processed_pseudoge | ENSG00000 |
| 49558546 | 49810419 | 1 | transcribed_unproce | ENSG00000 |
| 49817778 | 49825553 | 1 | unprocessed_pseudo | ENSG00000 |
| 49824839 | 49826724 | -1 | unprocessed_pseudo | ENSG00000 |
| 49833140 | 49839320 | -1 | unprocessed_pseudo | ENSG00000 |
| 49843488 | 49846533 | -1 | unprocessed_pseudo | ENSG00000 |
| 49852096 | 49884421 | 1 | lncRNA | ENSG00000 |
| 49854080 | 49859402 | -1 | unprocessed_pseudo | ENSG00000 |
| 49871826 | 49874067 | -1 | unprocessed_pseudo | ENSG00000 |
| 49874945 | 49876638 | 1 | transcribed_unproce | ENSG00000 |
| 49882999 | 49884093 | 1 | transcribed_unproce | ENSG00000 |
| 49892262 | 49893777 | -1 | processed_pseudoge | ENSG00000 |
| 49898267 | 49899186 | -1 | unprocessed_pseudo | ENSG00000 |
| 49915013 | 49915644 | -1 | unprocessed_pseudo | ENSG00000 |
| 49917391 | 49918478 | -1 | unprocessed_pseudo | ENSG00000 |
| 49920312 | 49921414 | -1 | unprocessed_pseudo | ENSG00000 |
| 49923049 | 49923886 | 1 | unprocessed_pseudo | ENSG00000 |
| 49952391 | 49953419 | 1 | protein_coding | ENSG00000 |
| 49981473 | 49982535 | -1 | protein_coding | ENSG00000 |
| 50018798 | 50019695 | 1 | unprocessed_pseudo | ENSG00000 |
| 50032874 | 50033794 | 1 | polymorphic_pseudo | ENSG00000 |
| 50053410 | 50054309 | -1 | unprocessed_pseudo | ENSG00000 |
| 50100892 | 50102898 | -1 | unprocessed_pseudo | ENSG00000 |
| 50170156 | 50190441 | 1 | unprocessed_pseudo | ENSG00000 |
| 50184250 | 50208794 | 1 | lncRNA | ENSG00000 |
| 50213420 | 50223969 | 1 | lncRNA | ENSG00000 |
| 50246554 | 50247761 | -1 | unprocessed_pseudo | ENSG00000 |
| 50268161 | 50269273 | 1 | processed_pseudoge | ENSG00000 |
| 50279828 | 50298452 | -1 | transcribed_unproce | ENSG00000 |
| 50290749 | 50290933 | 1 | processed_pseudoge | ENSG00000 |
| 50298579 | 50304656 | 1 | lncRNA | ENSG00000 |
| 50307681 | 50361706 | 1 | transcribed_unproce | ENSG00000 |
| 50409042 | 50422316 | 1 | unprocessed_pseudo | ENSG00000 |
| 54591480 | 54592409 | 1 | unprocessed_pseudo | ENSG00000 |
| 54603069 | 54603998 | -1 | protein_coding | ENSG00000 |
| 54635037 | 54635955 | -1 | unprocessed_pseudo | ENSG00000 |
| 54657120 | 54657716 | -1 | unprocessed_pseudo | ENSG00000 |

| | | | | |
|---|---|---|---|---|
| 54659666 | 54660595 | 1 | unprocessed_pseudc | ENSG00000( |
| 54662580 | 54663450 | 1 | unprocessed_pseudc | ENSG00000( |
| 54667298 | 54668210 | 1 | unprocessed_pseudc | ENSG00000( |
| 54682877 | 54683821 | 1 | polymorphic_pseudc | ENSG00000( |
| 54692450 | 54693359 | 1 | unprocessed_pseudc | ENSG00000( |
| 54706832 | 54707902 | 1 | protein_coding | ENSG00000( |
| 54724847 | 54725816 | 1 | unprocessed_pseudc | ENSG00000( |
| 55262155 | 55271114 | 1 | protein_coding | ENSG00000( |
| 55267408 | 55267636 | 1 | processed_pseudoge | ENSG00000( |
| 55279447 | 55284749 | -1 | unprocessed_pseudc | ENSG00000( |
| 55279949 | 55280289 | 1 | unprocessed_pseudc | ENSG00000( |
| 55291883 | 55298270 | -1 | transcribed_unproce | ENSG00000( |
| 55302290 | 55305060 | -1 | unprocessed_pseudc | ENSG00000( |
| 55318560 | 55319465 | 1 | unprocessed_pseudc | ENSG00000( |
| 55325756 | 55326673 | 1 | unprocessed_pseudc | ENSG00000( |
| 55343151 | 55344231 | 1 | protein_coding | ENSG00000( |
| 55367884 | 55368918 | 1 | protein_coding | ENSG00000( |
| 55388529 | 55389472 | 1 | unprocessed_pseudc | ENSG00000( |
| 55411526 | 55411717 | 1 | unprocessed_pseudc | ENSG00000( |
| 55430478 | 55432370 | 1 | unprocessed_pseudc | ENSG00000( |
| 55444435 | 55445338 | 1 | unprocessed_pseudc | ENSG00000( |
| 55466771 | 55467712 | 1 | unprocessed_pseudc | ENSG00000( |
| 55475943 | 55478512 | 1 | unprocessed_pseudc | ENSG00000( |
| 55491205 | 55492148 | 1 | unprocessed_pseudc | ENSG00000( |
| 55506635 | 55511169 | 1 | transcribed_unproce | ENSG00000( |
| 55537002 | 55537924 | 1 | unprocessed_pseudc | ENSG00000( |
| 55554307 | 55555419 | 1 | protein_coding | ENSG00000( |
| 55572128 | 55573060 | 1 | protein_coding | ENSG00000( |
| 55602360 | 55607645 | -1 | protein_coding | ENSG00000( |
| 55635113 | 55640309 | 1 | protein_coding | ENSG00000( |
| 55648327 | 55652854 | 1 | protein_coding | ENSG00000( |
| 55662201 | 55666195 | 1 | protein_coding | ENSG00000( |
| 55673536 | 55674675 | 1 | unprocessed_pseudc | ENSG00000( |
| 55683239 | 55684374 | 1 | unprocessed_pseudc | ENSG00000( |
| 55684141 | 55686160 | -1 | lncRNA | ENSG00000( |
| 55714799 | 55715727 | 1 | unprocessed_pseudc | ENSG00000( |
| 55723776 | 55729621 | 1 | protein_coding | ENSG00000( |
| 55754977 | 55755957 | 1 | unprocessed_pseudc | ENSG00000( |
| 55773438 | 55774382 | 1 | protein_coding | ENSG00000( |
| 55786964 | 55787905 | 1 | unprocessed_pseudc | ENSG00000( |
| 55795556 | 55796500 | 1 | protein_coding | ENSG00000( |
| 55811367 | 55812476 | 1 | protein_coding | ENSG00000( |
| 55819607 | 55820597 | 1 | protein_coding | ENSG00000( |
| 55827219 | 55828154 | 1 | protein_coding | ENSG00000( |
| 55838752 | 55839738 | 1 | protein_coding | ENSG00000( |
| 55850347 | 55858547 | 1 | transcribed_unproce | ENSG00000( |
| 55863932 | 55881243 | 1 | unprocessed_pseudc | ENSG00000( |
| 55865709 | 55865820 | 1 | processed_pseudoge | ENSG00000( |
| 55867569 | 55874537 | -1 | unprocessed_pseudc | ENSG00000( |
| 55883297 | 55891810 | 1 | protein_coding | ENSG00000( |

| | | | | |
|---|---|---|---|---|
| 55903341 | 55904278 | -1 | unprocessed_pseudc | ENSG00000( |
| 55913650 | 55914582 | -1 | protein_coding | ENSG00000( |
| 55935456 | 55936400 | -1 | protein_coding | ENSG00000( |
| 55948120 | 55948924 | -1 | unprocessed_pseudc | ENSG00000( |
| 55957216 | 55957630 | -1 | unprocessed_pseudc | ENSG00000( |
| 55965755 | 55969945 | -1 | protein_coding | ENSG00000( |
| 55979398 | 55980103 | -1 | unprocessed_pseudc | ENSG00000( |
| 55993681 | 55994625 | -1 | protein_coding | ENSG00000( |
| 56015017 | 56015957 | -1 | unprocessed_pseudc | ENSG00000( |
| 56027654 | 56038191 | 1 | protein_coding | ENSG00000( |
| 56049785 | 56055730 | 1 | transcribed_unproce | ENSG00000( |
| 56071111 | 56072049 | 1 | unprocessed_pseudc | ENSG00000( |
| 56082801 | 56084840 | 1 | transcribed_unproce | ENSG00000( |
| 56093277 | 56094286 | 1 | protein_coding | ENSG00000( |
| 56097244 | 56097384 | -1 | unprocessed_pseudc | ENSG00000( |
| 56103687 | 56107658 | 1 | protein_coding | ENSG00000( |
| 56116639 | 56117256 | -1 | unprocessed_pseudc | ENSG00000( |
| 56122373 | 56123311 | 1 | protein_coding | ENSG00000( |
| 56132666 | 56133494 | -1 | unprocessed_pseudc | ENSG00000( |
| 56134721 | 56140201 | -1 | protein_coding | ENSG00000( |
| 56142516 | 56143387 | -1 | unprocessed_pseudc | ENSG00000( |
| 56159394 | 56160317 | -1 | protein_coding | ENSG00000( |
| 56165500 | 56165921 | 1 | unprocessed_pseudc | ENSG00000( |
| 56176618 | 56177556 | 1 | protein_coding | ENSG00000( |
| 56188379 | 56189251 | -1 | unprocessed_pseudc | ENSG00000( |
| 56205125 | 56205625 | -1 | processed_pseudoge | ENSG00000( |
| 56208985 | 56215222 | -1 | polymorphic_pseudc | ENSG00000( |
| 56231282 | 56234255 | -1 | protein_coding | ENSG00000( |
| 56252200 | 56253222 | 1 | protein_coding | ENSG00000( |
| 56274154 | 56276819 | 1 | protein_coding | ENSG00000( |
| 56288462 | 56292254 | -1 | protein_coding | ENSG00000( |
| 56296541 | 56297471 | -1 | unprocessed_pseudc | ENSG00000( |
| 56315144 | 56320639 | 1 | protein_coding | ENSG00000( |
| 56331347 | 56332488 | -1 | unprocessed_pseudc | ENSG00000( |
| 56335142 | 56336068 | 1 | unprocessed_pseudc | ENSG00000( |
| 56345945 | 56347031 | 1 | protein_coding | ENSG00000( |
| 56354291 | 56361511 | 1 | protein_coding | ENSG00000( |
| 56357328 | 56358221 | 1 | processed_pseudoge | ENSG00000( |
| 56375624 | 56376553 | 1 | protein_coding | ENSG00000( |
| 56381635 | 56382589 | 1 | unprocessed_pseudc | ENSG00000( |
| 56393729 | 56394648 | 1 | unprocessed_pseudc | ENSG00000( |
| 56412696 | 56413680 | 1 | polymorphic_pseudc | ENSG00000( |
| 56417258 | 56418232 | -1 | protein_coding | ENSG00000( |
| 56448668 | 56449592 | 1 | unprocessed_pseudc | ENSG00000( |
| 56459221 | 56473352 | -1 | protein_coding | ENSG00000( |
| 56462469 | 56463401 | -1 | protein_coding | ENSG00000( |
| 56469274 | 56473352 | -1 | protein_coding | ENSG00000( |
| 56476287 | 56484839 | -1 | transcribed_unproce | ENSG00000( |
| 56490349 | 56491395 | -1 | protein_coding | ENSG00000( |
| 56495760 | 56496412 | 1 | processed_pseudoge | ENSG00000( |

| | | | | |
|---|---|---|---|---|
| 56500300 | 56501271 | -1 | unprocessed_pseudc | ENSG0000C |
| 56509573 | 56510195 | 1 | processed_pseudoge | ENSG0000C |
| 56512235 | 56513171 | -1 | unprocessed_pseudc | ENSG0000C |
| 56526568 | 56527503 | -1 | unprocessed_pseudc | ENSG0000C |
| 56542262 | 56543281 | -1 | protein_coding | ENSG0000C |
| 56576736 | 56577787 | -1 | protein_coding | ENSG0000C |
| 56597426 | 56598362 | -1 | unprocessed_pseudc | ENSG0000C |
| 56609236 | 56614874 | -1 | protein_coding | ENSG0000C |
| 56619779 | 56620810 | -1 | unprocessed_pseudc | ENSG0000C |
| 56628932 | 56629860 | 1 | unprocessed_pseudc | ENSG0000C |
| 56633164 | 56634103 | -1 | unprocessed_pseudc | ENSG0000C |
| 56641466 | 56642471 | -1 | protein_coding | ENSG0000C |
| 56663662 | 56664687 | 1 | protein_coding | ENSG0000C |
| 56667545 | 56674664 | -1 | transcribed_unproce | ENSG0000C |
| 56690465 | 56690723 | 1 | processed_pseudoge | ENSG0000C |
| 56699095 | 56703884 | 1 | protein_coding | ENSG0000C |
| 56736316 | 56736462 | -1 | processed_pseudoge | ENSG0000C |
| 56740179 | 56741096 | 1 | unprocessed_pseudc | ENSG0000C |
| 56741223 | 56748697 | -1 | protein_coding | ENSG0000C |
| 56743873 | 56743981 | 1 | miRNA | ENSG0000C |
| 56750330 | 56750435 | -1 | unprocessed_pseudc | ENSG0000C |
| 56751023 | 56751902 | -1 | unprocessed_pseudc | ENSG0000C |
| 56775261 | 56776206 | -1 | unprocessed_pseudc | ENSG0000C |
| 56790206 | 56790965 | -1 | unprocessed_pseudc | ENSG0000C |
| 56801856 | 56820422 | -1 | transcribed_unproce | ENSG0000C |
| 56819573 | 56820516 | -1 | polymorphic_pseudc | ENSG0000C |
| 56848478 | 56878078 | 1 | lncRNA | ENSG0000C |
| 56890613 | 56927885 | 1 | unprocessed_pseudc | ENSG0000C |
| 56970850 | 56972146 | 1 | unprocessed_pseudc | ENSG0000C |
| 56988872 | 56989867 | 1 | protein_coding | ENSG0000C |
| 57018100 | 57019021 | 1 | unprocessed_pseudc | ENSG0000C |
| 57029408 | 57029722 | 1 | unprocessed_pseudc | ENSG0000C |
| 57037534 | 57038436 | 1 | unprocessed_pseudc | ENSG0000C |
| 57045091 | 57045390 | 1 | unprocessed_pseudc | ENSG0000C |
| 57055587 | 57056132 | 1 | unprocessed_pseudc | ENSG0000C |
| 57181747 | 57191717 | 1 | protein_coding | ENSG0000C |
| 57233577 | 57237314 | -1 | protein_coding | ENSG0000C |
| 57299638 | 57324952 | -1 | protein_coding | ENSG0000C |
| 57325603 | 57327958 | 1 | lncRNA | ENSG0000C |
| 57325986 | 57335892 | -1 | protein_coding | ENSG0000C |
| 57338352 | 57372396 | 1 | protein_coding | ENSG0000C |
| 57376769 | 57381150 | -1 | protein_coding | ENSG0000C |
| 57386780 | 57390650 | -1 | protein_coding | ENSG0000C |
| 57387365 | 57424059 | -1 | protein_coding | ENSG0000C |
| 57406954 | 57427580 | -1 | protein_coding | ENSG0000C |
| 57450183 | 57450299 | 1 | rRNA_pseudogene | ENSG0000C |
| 57451690 | 57452022 | 1 | misc_RNA | ENSG0000C |
| 57460528 | 57477534 | 1 | protein_coding | ENSG0000C |
| 57476493 | 57477534 | -1 | lncRNA | ENSG0000C |
| 57484534 | 57515780 | -1 | protein_coding | ENSG0000C |

| | | | | |
|---|---|---|---|---|
| 57528085 | 57528365 | -1 | misc_RNA | ENSG0000C |
| 57528464 | 57530803 | -1 | protein_coding | ENSG0000C |
| 57542641 | 57550274 | 1 | protein_coding | ENSG0000C |
| 57551656 | 57568284 | -1 | protein_coding | ENSG0000C |
| 57576292 | 57577080 | -1 | processed_pseudoge | ENSG0000C |
| 57597387 | 57614853 | 1 | protein_coding | ENSG0000C |
| 57638024 | 57652790 | 1 | lncRNA | ENSG0000C |
| 57641198 | 57641286 | 1 | miRNA | ENSG0000C |
| 57645087 | 57649944 | -1 | protein_coding | ENSG0000C |
| 57648992 | 57661865 | 1 | protein_coding | ENSG0000C |
| 57667747 | 57701182 | 1 | protein_coding | ENSG0000C |
| 57703714 | 57712323 | -1 | protein_coding | ENSG0000C |
| 57712593 | 57740973 | 1 | protein_coding | ENSG0000C |
| 57712605 | 57791586 | 1 | protein_coding | ENSG0000C |
| 57718044 | 57718530 | -1 | processed_pseudoge | ENSG0000C |
| 57741250 | 57743554 | 1 | protein_coding | ENSG0000C |
| 57742163 | 57793243 | 1 | protein_coding | ENSG0000C |
| 57743514 | 57751781 | -1 | protein_coding | ENSG0000C |
| 57753243 | 57819546 | 1 | protein_coding | ENSG0000C |
| 57861948 | 57873588 | -1 | transcribed_unproce | ENSG0000C |
| 57870322 | 57870571 | -1 | processed_pseudoge | ENSG0000C |
| 57917297 | 57918239 | -1 | unprocessed_pseudc | ENSG0000C |
| 57945598 | 57946414 | -1 | unprocessed_pseudc | ENSG0000C |
| 58005386 | 58005705 | 1 | processed_pseudoge | ENSG0000C |
| 58023881 | 58181616 | 1 | protein_coding | ENSG0000C |
| 58026706 | 58026812 | -1 | snRNA | ENSG0000C |
| 58030930 | 58031931 | 1 | protein_coding | ENSG0000C |
| 58040708 | 58041660 | -1 | unitary_pseudogene | ENSG0000C |
| 58044110 | 58060138 | -1 | lncRNA | ENSG0000C |
| 58072378 | 58075438 | -1 | processed_pseudoge | ENSG0000C |
| 58077348 | 58078603 | -1 | unprocessed_pseudc | ENSG0000C |
| 58108720 | 58109661 | -1 | unprocessed_pseudc | ENSG0000C |
| 58116742 | 58125530 | -1 | protein_coding | ENSG0000C |
| 58144495 | 58145397 | 1 | unprocessed_pseudc | ENSG0000C |
| 58171098 | 58171721 | 1 | unprocessed_pseudc | ENSG0000C |
| 58189070 | 58194053 | 1 | protein_coding | ENSG0000C |
| 58203202 | 58204181 | -1 | protein_coding | ENSG0000C |
| 58212720 | 58216084 | 1 | protein_coding | ENSG0000C |
| 58227882 | 58228918 | -1 | protein_coding | ENSG0000C |
| 58242043 | 58243097 | -1 | processed_pseudoge | ENSG0000C |
| 58266792 | 58268260 | -1 | protein_coding | ENSG0000C |
| 58291826 | 58292765 | -1 | unprocessed_pseudc | ENSG0000C |
| 58317637 | 58318375 | -1 | unprocessed_pseudc | ENSG0000C |
| 58343737 | 58344664 | -1 | unprocessed_pseudc | ENSG0000C |
| 58348989 | 58349630 | -1 | unprocessed_pseudc | ENSG0000C |
| 58354004 | 58354499 | -1 | processed_pseudoge | ENSG0000C |
| 58358124 | 58359069 | -1 | protein_coding | ENSG0000C |
| 58365836 | 58366763 | -1 | unprocessed_pseudc | ENSG0000C |
| 58387583 | 58388486 | -1 | unprocessed_pseudc | ENSG0000C |
| 58402464 | 58406874 | -1 | protein_coding | ENSG0000C |

| | | | | |
|---|---|---|---|---|
| 58421264 | 58428121 | -1 | protein_coding | ENSG00000 |
| 58439077 | 58442758 | -1 | protein_coding | ENSG00000 |
| 58491773 | 58493480 | -1 | lncRNA | ENSG00000 |
| 58497888 | 58505758 | -1 | lncRNA | ENSG00000 |
| 58507175 | 58508105 | -1 | protein_coding | ENSG00000 |
| 58526871 | 58578220 | -1 | protein_coding | ENSG00000 |
| 58579063 | 58621550 | 1 | protein_coding | ENSG00000 |
| 58579172 | 58624639 | 1 | protein_coding | ENSG00000 |
| 58611119 | 58612642 | -1 | lncRNA | ENSG00000 |
| 58622665 | 58625733 | 1 | protein_coding | ENSG00000 |
| 58627435 | 58628528 | 1 | TEC | ENSG00000 |
| 58640426 | 58731974 | -1 | protein_coding | ENSG00000 |
| 58685086 | 58691861 | 1 | processed_pseudoge | ENSG00000 |
| 58748999 | 58761053 | -1 | unprocessed_pseudo | ENSG00000 |
| 58796237 | 58796854 | -1 | processed_pseudoge | ENSG00000 |
| 58834065 | 58904215 | -1 | protein_coding | ENSG00000 |
| 58878302 | 58893460 | 1 | transcribed_unproce | ENSG00000 |
| 58905398 | 59043527 | 1 | protein_coding | ENSG00000 |
| 58917015 | 58928689 | -1 | lncRNA | ENSG00000 |
| 58933643 | 59058659 | -1 | lncRNA | ENSG00000 |
| 58967596 | 58968291 | -1 | processed_pseudoge | ENSG00000 |
| 58978787 | 58994120 | 1 | unprocessed_pseudo | ENSG00000 |
| 58995439 | 58999689 | -1 | unprocessed_pseudo | ENSG00000 |
| 59042737 | 59050971 | 1 | unprocessed_pseudo | ENSG00000 |
| 59086307 | 59094786 | 1 | protein_coding | ENSG00000 |
| 59107185 | 59127412 | 1 | protein_coding | ENSG00000 |
| 59130133 | 59143015 | -1 | lncRNA | ENSG00000 |
| 59142121 | 59142610 | -1 | processed_pseudoge | ENSG00000 |
| 59142748 | 59155039 | 1 | protein_coding | ENSG00000 |
| 59171430 | 59208588 | 1 | protein_coding | ENSG00000 |
| 59208510 | 59212927 | -1 | protein_coding | ENSG00000 |
| 59214406 | 59214699 | -1 | misc_RNA | ENSG00000 |
| 59255360 | 59259486 | 1 | unprocessed_pseudo | ENSG00000 |
| 59263389 | 59263802 | -1 | processed_pseudoge | ENSG00000 |
| 59268876 | 59284033 | -1 | lncRNA | ENSG00000 |
| 59287339 | 59289486 | -1 | processed_pseudoge | ENSG00000 |
| 59291052 | 59291333 | -1 | misc_RNA | ENSG00000 |
| 59309681 | 59310576 | -1 | unprocessed_pseudo | ENSG00000 |
| 59332757 | 59334086 | -1 | unprocessed_pseudo | ENSG00000 |
| 59358895 | 59371714 | 1 | protein_coding | ENSG00000 |
| 59391354 | 59392292 | -1 | unprocessed_pseudo | ENSG00000 |
| 59416969 | 59426412 | -1 | protein_coding | ENSG00000 |
| 59436469 | 59451380 | 1 | protein_coding | ENSG00000 |
| 59456938 | 59457991 | 1 | protein_coding | ENSG00000 |
| 59473315 | 59479361 | 1 | protein_coding | ENSG00000 |
| 59491652 | 59492581 | 1 | unprocessed_pseudo | ENSG00000 |
| 59503576 | 59504511 | 1 | protein_coding | ENSG00000 |
| 59511368 | 59520703 | 1 | protein_coding | ENSG00000 |
| 59531754 | 59532695 | 1 | unprocessed_pseudo | ENSG00000 |
| 59545602 | 59558882 | 1 | lncRNA | ENSG00000 |

| | | | | |
|---|---|---|---|---|
| 59560298 | 59566283 | -1 | lncRNA | ENSG0000C |
| 59565923 | 59639861 | 1 | lncRNA | ENSG0000C |
| 59574398 | 59615774 | -1 | protein_coding | ENSG0000C |
| 59595077 | 59595158 | -1 | miRNA | ENSG0000C |
| 59636716 | 59669037 | -1 | protein_coding | ENSG0000C |
| 59669312 | 59676041 | 1 | lncRNA | ENSG0000C |
| 59706057 | 59706383 | -1 | misc_RNA | ENSG0000C |
| 59712916 | 59713845 | -1 | protein_coding | ENSG0000C |
| 59713456 | 59805882 | 1 | protein_coding | ENSG0000C |
| 59728519 | 59729492 | 1 | unprocessed_pseudc | ENSG0000C |
| 59741325 | 59742049 | 1 | unprocessed_pseudc | ENSG0000C |
| 59748811 | 59749574 | 1 | unprocessed_pseudc | ENSG0000C |
| 59752578 | 59754975 | -1 | lncRNA | ENSG0000C |
| 59781318 | 59781722 | -1 | processed_pseudoge | ENSG0000C |
| 59806140 | 59810778 | -1 | protein_coding | ENSG0000C |
| 59829273 | 59845499 | -1 | protein_coding | ENSG0000C |
| 59852800 | 59866489 | -1 | protein_coding | ENSG0000C |
| 59878809 | 59896481 | 1 | protein_coding | ENSG0000C |
| 59898185 | 59899133 | 1 | processed_pseudoge | ENSG0000C |
| 59938432 | 59957480 | 1 | protein_coding | ENSG0000C |
| 59942879 | 59970146 | 1 | lncRNA | ENSG0000C |
| 59964033 | 59970146 | 1 | protein_coding | ENSG0000C |
| 59978020 | 60031077 | 1 | protein_coding | ENSG0000C |
| 60040409 | 60048044 | 1 | protein_coding | ENSG0000C |
| 60056587 | 60071115 | 1 | protein_coding | ENSG0000C |
| 60077350 | 60078401 | 1 | processed_pseudoge | ENSG0000C |
| 60088261 | 60098467 | 1 | protein_coding | ENSG0000C |
| 60159687 | 60160822 | -1 | lncRNA | ENSG0000C |
| 60172014 | 60184666 | -1 | protein_coding | ENSG0000C |
| 60185702 | 60317944 | 1 | protein_coding | ENSG0000C |
| 60200270 | 60243137 | -1 | protein_coding | ENSG0000C |
| 60209071 | 60209156 | -1 | miRNA | ENSG0000C |
| 60334831 | 60396596 | 1 | protein_coding | ENSG0000C |
| 60378485 | 60395951 | 1 | protein_coding | ENSG0000C |
| 60378530 | 60417756 | 1 | protein_coding | ENSG0000C |
| 60429572 | 60455214 | 1 | protein_coding | ENSG0000C |
| 60455752 | 60470760 | 1 | protein_coding | ENSG0000C |
| 60492778 | 60507430 | 1 | protein_coding | ENSG0000C |
| 60515392 | 60542721 | 1 | protein_coding | ENSG0000C |
| 60577856 | 60608418 | 1 | unprocessed_pseudc | ENSG0000C |
| 60615729 | 60687267 | 1 | transcribed_unitary_ | ENSG0000C |
| 60699585 | 60715807 | 1 | protein_coding | ENSG0000C |
| 60729304 | 60744212 | 1 | protein_coding | ENSG0000C |
| 60756867 | 60776733 | 1 | protein_coding | ENSG0000C |
| 60785333 | 60801305 | 1 | protein_coding | ENSG0000C |
| 60813932 | 60821739 | -1 | lncRNA | ENSG0000C |
| 60835996 | 60842965 | -1 | lncRNA | ENSG0000C |
| 60841806 | 60851081 | -1 | lncRNA | ENSG0000C |
| 60842113 | 60851081 | 1 | protein_coding | ENSG0000C |
| 60850933 | 60855950 | -1 | protein_coding | ENSG0000C |

| | | | | |
|---|---|---|---|---|
| 60867562 | 60875693 | 1 | protein_coding | ENSG00000( |
| 60890547 | 60906585 | -1 | protein_coding | ENSG00000( |
| 60906789 | 60909742 | 1 | lncRNA | ENSG00000( |
| 60913166 | 60914052 | -1 | lncRNA | ENSG00000( |
| 60914158 | 60923443 | 1 | protein_coding | ENSG00000( |
| 60916339 | 60925397 | -1 | lncRNA | ENSG00000( |
| 60924463 | 60937159 | 1 | protein_coding | ENSG00000( |
| 60937084 | 60952530 | -1 | protein_coding | ENSG00000( |
| 60971680 | 61020377 | 1 | protein_coding | ENSG00000( |
| 60985061 | 60985167 | -1 | snRNA | ENSG00000( |
| 61055392 | 61067597 | 1 | lncRNA | ENSG00000( |
| 61102489 | 61127852 | 1 | protein_coding | ENSG00000( |
| 61130257 | 61161615 | -1 | protein_coding | ENSG00000( |
| 61143380 | 61145464 | -1 | TEC | ENSG00000( |
| 61203307 | 61213098 | 1 | protein_coding | ENSG00000( |
| 61222216 | 61231927 | 1 | protein_coding | ENSG00000( |
| 61227168 | 61248533 | 1 | lncRNA | ENSG00000( |
| 61241042 | 61251448 | 1 | protein_coding | ENSG00000( |
| 61258286 | 61295316 | -1 | protein_coding | ENSG00000( |
| 61299451 | 61342596 | -1 | protein_coding | ENSG00000( |
| 61333210 | 61353295 | 1 | protein_coding | ENSG00000( |
| 61348745 | 61362299 | -1 | protein_coding | ENSG00000( |
| 61352244 | 61354477 | 1 | TEC | ENSG00000( |
| 61362344 | 61369509 | 1 | protein_coding | ENSG00000( |
| 61392360 | 61398863 | 1 | protein_coding | ENSG00000( |
| 61402641 | 61430031 | -1 | protein_coding | ENSG00000( |
| 61426448 | 61427325 | -1 | TEC | ENSG00000( |
| 61429220 | 61485822 | 1 | protein_coding | ENSG00000( |
| 61430042 | 61447529 | 1 | protein_coding | ENSG00000( |
| 61444149 | 61444423 | 1 | misc_RNA | ENSG00000( |
| 61481120 | 61490931 | 1 | protein_coding | ENSG00000( |
| 61496440 | 61506649 | 1 | lncRNA | ENSG00000( |
| 61508596 | 61508657 | 1 | miRNA | ENSG00000( |
| 61508749 | 61511018 | 1 | protein_coding | ENSG00000( |
| 61515313 | 61581148 | -1 | protein_coding | ENSG00000( |
| 61525708 | 61526602 | -1 | TEC | ENSG00000( |
| 61539516 | 61542259 | -1 | lncRNA | ENSG00000( |
| 61588442 | 61607550 | 1 | lncRNA | ENSG00000( |
| 61615036 | 61639449 | 1 | transcribed_processe | ENSG00000( |
| 61654665 | 61655702 | -1 | lncRNA | ENSG00000( |
| 61680391 | 61747001 | 1 | protein_coding | ENSG00000( |
| 61746493 | 61757655 | -1 | lncRNA | ENSG00000( |
| 61752642 | 61788518 | 1 | protein_coding | ENSG00000( |
| 61768501 | 61792802 | -1 | protein_coding | ENSG00000( |
| 61792495 | 61792561 | -1 | miRNA | ENSG00000( |
| 61792911 | 61797238 | 1 | protein_coding | ENSG00000( |
| 61792980 | 61867354 | 1 | protein_coding | ENSG00000( |
| 61799627 | 61829318 | -1 | protein_coding | ENSG00000( |
| 61815161 | 61815240 | -1 | miRNA | ENSG00000( |
| 61873519 | 61892051 | -1 | protein_coding | ENSG00000( |

| | | | | |
|---|---|---|---|---|
| 61878216 | 61878278 | -1 | miRNA | ENSG0000C |
| 61897301 | 61920269 | -1 | protein_coding | ENSG0000C |
| 61937894 | 61938000 | 1 | snRNA | ENSG0000C |
| 61950063 | 61965515 | 1 | protein_coding | ENSG0000C |
| 61959718 | 61967634 | -1 | protein_coding | ENSG0000C |
| 61967794 | 61969490 | 1 | TEC | ENSG0000C |
| 62011751 | 62012602 | -1 | processed_pseudoge | ENSG0000C |
| 62049863 | 62082184 | -1 | lncRNA | ENSG0000C |
| 62072882 | 62073052 | -1 | processed_pseudoge | ENSG0000C |
| 62116470 | 62123767 | -1 | lncRNA | ENSG0000C |
| 62123998 | 62153169 | 1 | protein_coding | ENSG0000C |
| 62169293 | 62169828 | 1 | processed_pseudoge | ENSG0000C |
| 62190216 | 62193539 | 1 | protein_coding | ENSG0000C |
| 62208673 | 62213943 | 1 | protein_coding | ENSG0000C |
| 62213427 | 62260549 | -1 | lncRNA | ENSG0000C |
| 62242239 | 62244812 | 1 | protein_coding | ENSG0000C |
| 62270158 | 62273160 | 1 | protein_coding | ENSG0000C |
| 62296281 | 62299075 | -1 | protein_coding | ENSG0000C |
| 62330946 | 62331813 | 1 | processed_pseudoge | ENSG0000C |
| 62336911 | 62338090 | -1 | lncRNA | ENSG0000C |
| 62337448 | 62393412 | 1 | protein_coding | ENSG0000C |
| 62371146 | 62373168 | 1 | processed_pseudoge | ENSG0000C |
| 62391516 | 62393372 | -1 | lncRNA | ENSG0000C |
| 62405103 | 62423195 | 1 | protein_coding | ENSG0000C |
| 62421845 | 62426724 | -1 | lncRNA | ENSG0000C |
| 62433542 | 62556235 | -1 | protein_coding | ENSG0000C |
| 62537312 | 62542018 | 1 | lncRNA | ENSG0000C |
| 62545999 | 62547699 | 1 | lncRNA | ENSG0000C |
| 62559596 | 62574086 | -1 | protein_coding | ENSG0000C |
| 62559603 | 62591531 | -1 | protein_coding | ENSG0000C |
| 62567011 | 62567071 | -1 | miRNA | ENSG0000C |
| 62575045 | 62591637 | -1 | protein_coding | ENSG0000C |
| 62593214 | 62601865 | -1 | protein_coding | ENSG0000C |
| 62602218 | 62612765 | -1 | protein_coding | ENSG0000C |
| 62606161 | 62606405 | 1 | lncRNA | ENSG0000C |
| 62611722 | 62615120 | 1 | protein_coding | ENSG0000C |
| 62615296 | 62622154 | -1 | protein_coding | ENSG0000C |
| 62624826 | 62646726 | -1 | protein_coding | ENSG0000C |
| 62646848 | 62653302 | -1 | protein_coding | ENSG0000C |
| 62649694 | 62665178 | -1 | protein_coding | ENSG0000C |
| 62662816 | 62665217 | -1 | protein_coding | ENSG0000C |
| 62662817 | 62672255 | -1 | protein_coding | ENSG0000C |
| 62665309 | 62668496 | 1 | protein_coding | ENSG0000C |
| 62665422 | 62665570 | 1 | snoRNA | ENSG0000C |
| 62670273 | 62673686 | 1 | protein_coding | ENSG0000C |
| 62676498 | 62679117 | -1 | protein_coding | ENSG0000C |
| 62686406 | 62689530 | -1 | protein_coding | ENSG0000C |
| 62690275 | 62709845 | -1 | protein_coding | ENSG0000C |
| 62690275 | 62727384 | -1 | protein_coding | ENSG0000C |
| 62707676 | 62709201 | 1 | protein_coding | ENSG0000C |

| | | | | |
|---|---|---|---|---|
| 62712630 | 62727457 | -1 | protein_coding | ENSG00000 |
| 62728069 | 62740293 | 1 | protein_coding | ENSG00000 |
| 62748319 | 62754160 | -1 | protein_coding | ENSG00000 |
| 62761565 | 62766710 | 1 | protein_coding | ENSG00000 |
| 62771120 | 62771606 | -1 | lncRNA | ENSG00000 |
| 62771357 | 62787342 | 1 | protein_coding | ENSG00000 |
| 62771629 | 62792006 | -1 | protein_coding | ENSG00000 |
| 62786023 | 62786785 | -1 | lncRNA | ENSG00000 |
| 62787402 | 62790400 | 1 | protein_coding | ENSG00000 |
| 62789815 | 62789885 | 1 | miRNA | ENSG00000 |
| 62792123 | 62806302 | -1 | protein_coding | ENSG00000 |
| 62792702 | 62792771 | -1 | miRNA | ENSG00000 |
| 62806860 | 62832051 | -1 | protein_coding | ENSG00000 |
| 62807682 | 62808063 | -1 | lncRNA | ENSG00000 |
| 62815966 | 62816067 | -1 | snRNA | ENSG00000 |
| 62832234 | 62834043 | 1 | lncRNA | ENSG00000 |
| 62832319 | 62832803 | -1 | protein_coding | ENSG00000 |
| 62832342 | 62841809 | -1 | protein_coding | ENSG00000 |
| 62841619 | 62841809 | -1 | snRNA | ENSG00000 |
| 62851984 | 62855953 | -1 | lncRNA | ENSG00000 |
| 62852910 | 62853035 | -1 | snoRNA | ENSG00000 |
| 62853663 | 62853732 | -1 | snoRNA | ENSG00000 |
| 62854161 | 62854285 | -1 | snoRNA | ENSG00000 |
| 62854621 | 62854695 | -1 | snoRNA | ENSG00000 |
| 62855012 | 62855083 | -1 | snoRNA | ENSG00000 |
| 62855292 | 62855366 | -1 | snoRNA | ENSG00000 |
| 62855564 | 62855632 | -1 | snoRNA | ENSG00000 |
| 62856102 | 62888875 | 1 | protein_coding | ENSG00000 |
| 62908679 | 62921807 | -1 | protein_coding | ENSG00000 |
| 62909546 | 62918361 | 1 | lncRNA | ENSG00000 |
| 62935982 | 62936234 | -1 | misc_RNA | ENSG00000 |
| 62936385 | 62984983 | -1 | protein_coding | ENSG00000 |
| 62936427 | 62959852 | -1 | transcribed_unproce | ENSG00000 |
| 62989154 | 63015841 | -1 | protein_coding | ENSG00000 |
| 63032503 | 63033103 | -1 | processed_pseudoge | ENSG00000 |
| 63046785 | 63048126 | 1 | processed_pseudoge | ENSG00000 |
| 63079940 | 63144221 | -1 | protein_coding | ENSG00000 |
| 63115880 | 63116338 | -1 | processed_pseudoge | ENSG00000 |
| 63137867 | 63369718 | 1 | protein_coding | ENSG00000 |
| 63163776 | 63229652 | -1 | protein_coding | ENSG00000 |
| 63237957 | 63239114 | -1 | unprocessed_pseudo | ENSG00000 |
| 63243085 | 63244004 | 1 | processed_pseudoge | ENSG00000 |
| 63265836 | 63266256 | -1 | unprocessed_pseudo | ENSG00000 |
| 63369785 | 63410294 | 1 | protein_coding | ENSG00000 |
| 63461404 | 63491194 | -1 | protein_coding | ENSG00000 |
| 63469376 | 63470944 | 1 | processed_pseudoge | ENSG00000 |
| 63495484 | 63500610 | 1 | lncRNA | ENSG00000 |
| 63506084 | 63516774 | 1 | protein_coding | ENSG00000 |
| 63536808 | 63546462 | 1 | protein_coding | ENSG00000 |
| 63552770 | 63563379 | -1 | protein_coding | ENSG00000 |

| | | | | |
|---|---|---|---|---|
| 63573195 | 63616883 | -1 | protein_coding | ENSG00000( |
| 63616308 | 63621671 | 1 | lncRNA | ENSG00000( |
| 63624087 | 63671921 | -1 | protein_coding | ENSG00000( |
| 63632233 | 63632720 | 1 | processed_pseudoge | ENSG00000( |
| 63637677 | 63658962 | 1 | lncRNA | ENSG00000( |
| 63681446 | 63759891 | 1 | protein_coding | ENSG00000( |
| 63698596 | 63698764 | -1 | processed_pseudoge | ENSG00000( |
| 63759892 | 63768775 | -1 | protein_coding | ENSG00000( |
| 63797784 | 63798071 | -1 | misc_RNA | ENSG00000( |
| 63813456 | 63827716 | 1 | protein_coding | ENSG00000( |
| 63834667 | 63834943 | -1 | processed_pseudoge | ENSG00000( |
| 63838928 | 63911019 | 1 | protein_coding | ENSG00000( |
| 63882587 | 63882685 | -1 | snRNA | ENSG00000( |
| 63911230 | 63917164 | -1 | protein_coding | ENSG00000( |
| 63938959 | 63957319 | 1 | protein_coding | ENSG00000( |
| 63970470 | 63970576 | 1 | snRNA | ENSG00000( |
| 63974620 | 63988346 | 1 | protein_coding | ENSG00000( |
| 63974620 | 63976543 | 1 | protein_coding | ENSG00000( |
| 63985853 | 64001811 | 1 | protein_coding | ENSG00000( |
| 63998558 | 64166113 | -1 | protein_coding | ENSG00000( |
| 64035970 | 64103653 | 1 | lncRNA | ENSG00000( |
| 64081690 | 64087175 | 1 | lncRNA | ENSG00000( |
| 64103188 | 64119173 | 1 | protein_coding | ENSG00000( |
| 64118272 | 64119210 | -1 | lncRNA | ENSG00000( |
| 64185272 | 64204543 | 1 | protein_coding | ENSG00000( |
| 64206678 | 64223886 | 1 | protein_coding | ENSG00000( |
| 64223799 | 64226254 | -1 | protein_coding | ENSG00000( |
| 64225941 | 64230686 | 1 | protein_coding | ENSG00000( |
| 64229214 | 64234352 | -1 | lncRNA | ENSG00000( |
| 64230278 | 64234286 | 1 | protein_coding | ENSG00000( |
| 64234538 | 64238793 | 1 | protein_coding | ENSG00000( |
| 64241003 | 64244134 | 1 | protein_coding | ENSG00000( |
| 64241144 | 64244134 | 1 | protein_coding | ENSG00000( |
| 64244479 | 64246943 | -1 | protein_coding | ENSG00000( |
| 64245961 | 64248218 | 1 | lncRNA | ENSG00000( |
| 64246939 | 64249494 | -1 | lncRNA | ENSG00000( |
| 64251523 | 64269150 | 1 | protein_coding | ENSG00000( |
| 64269830 | 64284704 | -1 | protein_coding | ENSG00000( |
| 64270062 | 64289500 | 1 | protein_coding | ENSG00000( |
| 64291302 | 64300031 | 1 | protein_coding | ENSG00000( |
| 64291722 | 64304769 | 1 | lncRNA | ENSG00000( |
| 64296037 | 64296138 | 1 | misc_RNA | ENSG00000( |
| 64300358 | 64304770 | 1 | protein_coding | ENSG00000( |
| 64305524 | 64316743 | 1 | protein_coding | ENSG00000( |
| 64316460 | 64318084 | -1 | protein_coding | ENSG00000( |
| 64318121 | 64321811 | 1 | protein_coding | ENSG00000( |
| 64325050 | 64329504 | -1 | lncRNA | ENSG00000( |
| 64340204 | 64357534 | 1 | protein_coding | ENSG00000( |
| 64341849 | 64341904 | -1 | miRNA | ENSG00000( |
| 64359148 | 64372215 | 1 | protein_coding | ENSG00000( |

| | | | | |
|---|---|---|---|---|
| 64368602 | 64368703 | 1 | miRNA | ENSG00000 |
| 64394342 | 64395831 | -1 | lncRNA | ENSG00000 |
| 64420311 | 64432670 | 1 | lncRNA | ENSG00000 |
| 64449074 | 64451657 | 1 | lncRNA | ENSG00000 |
| 64486136 | 64490674 | -1 | lncRNA | ENSG00000 |
| 64500846 | 64505386 | 1 | lncRNA | ENSG00000 |
| 64531044 | 64532970 | 1 | unprocessed_pseudc | ENSG00000 |
| 64555690 | 64572875 | 1 | protein_coding | ENSG00000 |
| 64590641 | 64602353 | 1 | protein_coding | ENSG00000 |
| 64606174 | 64723188 | -1 | protein_coding | ENSG00000 |
| 64646399 | 64659681 | 1 | lncRNA | ENSG00000 |
| 64726911 | 64745456 | -1 | protein_coding | ENSG00000 |
| 64746389 | 64759974 | -1 | protein_coding | ENSG00000 |
| 64764606 | 64778786 | -1 | protein_coding | ENSG00000 |
| 64778954 | 64779405 | 1 | lncRNA | ENSG00000 |
| 64784918 | 64803241 | -1 | protein_coding | ENSG00000 |
| 64803510 | 64811294 | -1 | protein_coding | ENSG00000 |
| 64823052 | 64844653 | -1 | protein_coding | ENSG00000 |
| 64851642 | 64888296 | -1 | protein_coding | ENSG00000 |
| 64881535 | 64881945 | 1 | processed_pseudoge | ENSG00000 |
| 64889560 | 64893449 | -1 | lncRNA | ENSG00000 |
| 64891137 | 64891246 | -1 | miRNA | ENSG00000 |
| 64891355 | 64891439 | -1 | miRNA | ENSG00000 |
| 64894546 | 64917211 | -1 | protein_coding | ENSG00000 |
| 64898363 | 64898437 | -1 | miRNA | ENSG00000 |
| 64902387 | 64902455 | -1 | miRNA | ENSG00000 |
| 64917553 | 64934475 | 1 | protein_coding | ENSG00000 |
| 64934471 | 64935893 | -1 | protein_coding | ENSG00000 |
| 64937517 | 64972108 | -1 | protein_coding | ENSG00000 |
| 64987945 | 64997018 | -1 | protein_coding | ENSG00000 |
| 65014160 | 65022184 | 1 | protein_coding | ENSG00000 |
| 65014182 | 65040570 | 1 | protein_coding | ENSG00000 |
| 65018505 | 65018570 | 1 | miRNA | ENSG00000 |
| 65025390 | 65026515 | -1 | processed_pseudoge | ENSG00000 |
| 65027439 | 65040572 | 1 | protein_coding | ENSG00000 |
| 65040901 | 65044828 | 1 | protein_coding | ENSG00000 |
| 65044818 | 65058553 | -1 | protein_coding | ENSG00000 |
| 65066300 | 65084164 | -1 | protein_coding | ENSG00000 |
| 65084210 | 65088398 | 1 | protein_coding | ENSG00000 |
| 65084979 | 65089375 | -1 | protein_coding | ENSG00000 |
| 65089324 | 65111862 | 1 | protein_coding | ENSG00000 |
| 65110714 | 65111695 | -1 | lncRNA | ENSG00000 |
| 65111845 | 65116396 | 1 | protein_coding | ENSG00000 |
| 65116403 | 65117701 | -1 | protein_coding | ENSG00000 |
| 65117157 | 65117458 | 1 | lncRNA | ENSG00000 |
| 65118310 | 65119111 | 1 | TEC | ENSG00000 |
| 65120630 | 65122177 | -1 | protein_coding | ENSG00000 |
| 65121780 | 65134533 | -1 | protein_coding | ENSG00000 |
| 65122183 | 65127371 | 1 | protein_coding | ENSG00000 |
| 65129916 | 65129978 | -1 | miRNA | ENSG00000 |

| | | | | |
|---|---|---|---|---|
| 65145691 | 65145972 | -1 | processed_pseudoge | ENSG0000C |
| 65147584 | 65147778 | 1 | snRNA | ENSG0000C |
| 65160194 | 65160382 | -1 | processed_pseudoge | ENSG0000C |
| 65170154 | 65173244 | 1 | protein_coding | ENSG0000C |
| 65174117 | 65176470 | -1 | transcribed_processe | ENSG0000C |
| 65177606 | 65181834 | -1 | lncRNA | ENSG0000C |
| 65180566 | 65212006 | 1 | protein_coding | ENSG0000C |
| 65213840 | 65242757 | 1 | transcribed_unitary_ | ENSG0000C |
| 65260996 | 65261748 | -1 | lncRNA | ENSG0000C |
| 65261920 | 65305589 | 1 | protein_coding | ENSG0000C |
| 65261928 | 65326543 | 1 | protein_coding | ENSG0000C |
| 65305406 | 65314244 | -1 | lncRNA | ENSG0000C |
| 65314866 | 65322417 | 1 | protein_coding | ENSG0000C |
| 65333852 | 65354262 | 1 | protein_coding | ENSG0000C |
| 65354751 | 65357613 | 1 | protein_coding | ENSG0000C |
| 65367438 | 65375299 | 1 | lncRNA | ENSG0000C |
| 65375192 | 65383701 | -1 | protein_coding | ENSG0000C |
| 65386621 | 65413525 | 1 | protein_coding | ENSG0000C |
| 65422774 | 65445540 | 1 | lncRNA | ENSG0000C |
| 65423273 | 65423392 | 1 | misc_RNA | ENSG0000C |
| 65423638 | 65423742 | 1 | misc_RNA | ENSG0000C |
| 65423960 | 65424118 | 1 | misc_RNA | ENSG0000C |
| 65444458 | 65444557 | 1 | miRNA | ENSG0000C |
| 65445541 | 65445598 | 1 | sRNA | ENSG0000C |
| 65455258 | 65466720 | 1 | transcribed_unproce | ENSG0000C |
| 65487241 | 65488714 | -1 | lncRNA | ENSG0000C |
| 65487884 | 65488896 | 1 | lncRNA | ENSG0000C |
| 65497688 | 65506516 | 1 | lncRNA | ENSG0000C |
| 65498010 | 65498405 | -1 | lncRNA | ENSG0000C |
| 65502914 | 65503008 | 1 | misc_RNA | ENSG0000C |
| 65506117 | 65506173 | 1 | sRNA | ENSG0000C |
| 65514403 | 65514923 | -1 | processed_pseudoge | ENSG0000C |
| 65525077 | 65538704 | 1 | protein_coding | ENSG0000C |
| 65538805 | 65558930 | -1 | protein_coding | ENSG0000C |
| 65561484 | 65561607 | 1 | TEC | ENSG0000C |
| 65568482 | 65570423 | -1 | lncRNA | ENSG0000C |
| 65570460 | 65573942 | 1 | protein_coding | ENSG0000C |
| 65572349 | 65574198 | 1 | protein_coding | ENSG0000C |
| 65576046 | 65592650 | 1 | protein_coding | ENSG0000C |
| 65592855 | 65595996 | -1 | protein_coding | ENSG0000C |
| 65597756 | 65615382 | -1 | protein_coding | ENSG0000C |
| 65615773 | 65637439 | 1 | protein_coding | ENSG0000C |
| 65636310 | 65636369 | 1 | miRNA | ENSG0000C |
| 65638101 | 65650918 | 1 | protein_coding | ENSG0000C |
| 65649192 | 65649253 | 1 | miRNA | ENSG0000C |
| 65653597 | 65663090 | -1 | protein_coding | ENSG0000C |
| 65663006 | 65671716 | 1 | lncRNA | ENSG0000C |
| 65695552 | 65695837 | 1 | misc_RNA | ENSG0000C |
| 65711996 | 65719604 | 1 | protein_coding | ENSG0000C |
| 65714005 | 65720947 | -1 | protein_coding | ENSG0000C |

| | | | | |
|---|---|---|---|---|
| 65726939 | 65728214 | 1 | processed_pseudoge | ENSG00000 |
| 65745729 | 65771585 | 1 | lncRNA | ENSG00000 |
| 65773898 | 65780976 | -1 | protein_coding | ENSG00000 |
| 65787063 | 65797214 | 1 | protein_coding | ENSG00000 |
| 65789051 | 65790868 | -1 | lncRNA | ENSG00000 |
| 65795946 | 65797219 | -1 | lncRNA | ENSG00000 |
| 65823022 | 65862026 | -1 | protein_coding | ENSG00000 |
| 65833834 | 65856896 | 1 | protein_coding | ENSG00000 |
| 65857126 | 65867653 | 1 | protein_coding | ENSG00000 |
| 65866441 | 65873592 | -1 | protein_coding | ENSG00000 |
| 65879837 | 65883741 | 1 | protein_coding | ENSG00000 |
| 65883741 | 65888539 | -1 | protein_coding | ENSG00000 |
| 65890673 | 65891635 | 1 | protein_coding | ENSG00000 |
| 65892049 | 65900573 | -1 | protein_coding | ENSG00000 |
| 65916810 | 65919062 | -1 | protein_coding | ENSG00000 |
| 65919274 | 65921563 | 1 | protein_coding | ENSG00000 |
| 65945445 | 65959963 | 1 | protein_coding | ENSG00000 |
| 65961728 | 65980137 | 1 | protein_coding | ENSG00000 |
| 65983679 | 65984331 | -1 | processed_pseudoge | ENSG00000 |
| 65996545 | 66002176 | -1 | protein_coding | ENSG00000 |
| 66002228 | 66004149 | 1 | protein_coding | ENSG00000 |
| 66012008 | 66013505 | 1 | protein_coding | ENSG00000 |
| 66016752 | 66026479 | -1 | protein_coding | ENSG00000 |
| 66040765 | 66049161 | -1 | protein_coding | ENSG00000 |
| 66043298 | 66044381 | 1 | lncRNA | ENSG00000 |
| 66050729 | 66069308 | 1 | protein_coding | ENSG00000 |
| 66067277 | 66069619 | -1 | lncRNA | ENSG00000 |
| 66070272 | 66244744 | 1 | protein_coding | ENSG00000 |
| 66244840 | 66246239 | -1 | lncRNA | ENSG00000 |
| 66257294 | 66267860 | 1 | protein_coding | ENSG00000 |
| 66259567 | 66261834 | -1 | lncRNA | ENSG00000 |
| 66264777 | 66265666 | -1 | lncRNA | ENSG00000 |
| 66267635 | 66268129 | -1 | lncRNA | ENSG00000 |
| 66268590 | 66277492 | 1 | protein_coding | ENSG00000 |
| 66269832 | 66278525 | -1 | lncRNA | ENSG00000 |
| 66276779 | 66277492 | -1 | lncRNA | ENSG00000 |
| 66278175 | 66285301 | 1 | protein_coding | ENSG00000 |
| 66284580 | 66289145 | -1 | protein_coding | ENSG00000 |
| 66291894 | 66296664 | 1 | protein_coding | ENSG00000 |
| 66312853 | 66319237 | 1 | lncRNA | ENSG00000 |
| 66314494 | 66317044 | -1 | protein_coding | ENSG00000 |
| 66330241 | 66336840 | -1 | protein_coding | ENSG00000 |
| 66334494 | 66339875 | 1 | lncRNA | ENSG00000 |
| 66337333 | 66345125 | -1 | protein_coding | ENSG00000 |
| 66345374 | 66347629 | -1 | protein_coding | ENSG00000 |
| 66347950 | 66364804 | 1 | lncRNA | ENSG00000 |
| 66362521 | 66372214 | -1 | protein_coding | ENSG00000 |
| 66374138 | 66374435 | -1 | misc_RNA | ENSG00000 |
| 66389609 | 66390168 | 1 | processed_pseudoge | ENSG00000 |
| 66393449 | 66393612 | 1 | snRNA | ENSG00000 |

| | | | | |
|---|---|---|---|---|
| 66409158 | 66417137 | 1 | lncRNA | ENSG0000C |
| 66421004 | 66426707 | 1 | protein_coding | ENSG0000C |
| 66432763 | 66432902 | 1 | snoRNA | ENSG0000C |
| 66435075 | 66466738 | -1 | protein_coding | ENSG0000C |
| 66454234 | 66454581 | 1 | processed_pseudoge | ENSG0000C |
| 66466327 | 66477337 | 1 | protein_coding | ENSG0000C |
| 66473490 | 66480233 | -1 | lncRNA | ENSG0000C |
| 66480013 | 66509657 | 1 | protein_coding | ENSG0000C |
| 66509079 | 66533613 | 1 | protein_coding | ENSG0000C |
| 66510606 | 66533613 | 1 | protein_coding | ENSG0000C |
| 66514306 | 66519720 | -1 | lncRNA | ENSG0000C |
| 66520637 | 66546235 | -1 | protein_coding | ENSG0000C |
| 66546395 | 66563334 | 1 | protein_coding | ENSG0000C |
| 66558866 | 66560384 | -1 | lncRNA | ENSG0000C |
| 66563464 | 66568841 | -1 | protein_coding | ENSG0000C |
| 66590176 | 66593063 | -1 | protein_coding | ENSG0000C |
| 66593153 | 66606019 | 1 | protein_coding | ENSG0000C |
| 66614964 | 66615090 | -1 | snRNA | ENSG0000C |
| 66616582 | 66627347 | 1 | protein_coding | ENSG0000C |
| 66616626 | 66646469 | 1 | protein_coding | ENSG0000C |
| 66638617 | 66668374 | 1 | protein_coding | ENSG0000C |
| 66664998 | 66677887 | -1 | protein_coding | ENSG0000C |
| 66685248 | 66729226 | -1 | protein_coding | ENSG0000C |
| 66712720 | 66713009 | -1 | misc_RNA | ENSG0000C |
| 66744451 | 66843328 | 1 | protein_coding | ENSG0000C |
| 66761575 | 66762399 | -1 | processed_pseudoge | ENSG0000C |
| 66771246 | 66771667 | -1 | processed_pseudoge | ENSG0000C |
| 66842835 | 66846552 | 1 | protein_coding | ENSG0000C |
| 66848417 | 66958386 | -1 | protein_coding | ENSG0000C |
| 66856647 | 66860475 | 1 | protein_coding | ENSG0000C |
| 66919762 | 66919823 | 1 | snRNA | ENSG0000C |
| 66934434 | 66934506 | -1 | miRNA | ENSG0000C |
| 66975277 | 66977004 | 1 | protein_coding | ENSG0000C |
| 67006778 | 67050863 | 1 | protein_coding | ENSG0000C |
| 67045643 | 67045708 | 1 | miRNA | ENSG0000C |
| 67056847 | 67072017 | 1 | protein_coding | ENSG0000C |
| 67119263 | 67258082 | 1 | protein_coding | ENSG0000C |
| 67195569 | 67197167 | -1 | transcribed_processe | ENSG0000C |
| 67252336 | 67261545 | -1 | lncRNA | ENSG0000C |
| 67266473 | 67286556 | 1 | protein_coding | ENSG0000C |
| 67288547 | 67302485 | 1 | protein_coding | ENSG0000C |
| 67303478 | 67312607 | 1 | protein_coding | ENSG0000C |
| 67316539 | 67317431 | -1 | lncRNA | ENSG0000C |
| 67317871 | 67398410 | 1 | protein_coding | ENSG0000C |
| 67350772 | 67356972 | -1 | protein_coding | ENSG0000C |
| 67351572 | 67373584 | -1 | protein_coding | ENSG0000C |
| 67353629 | 67354348 | 1 | lncRNA | ENSG0000C |
| 67362414 | 67362708 | 1 | misc_RNA | ENSG0000C |
| 67364168 | 67374177 | -1 | protein_coding | ENSG0000C |
| 67374416 | 67374932 | 1 | lncRNA | ENSG0000C |

| | | | | |
|---|---|---|---|---|
| 67395210 | 67395311 | -1 | snRNA | ENSG0000C |
| 67398183 | 67421183 | -1 | protein_coding | ENSG0000C |
| 67403915 | 67410089 | 1 | protein_coding | ENSG0000C |
| 67414968 | 67425607 | 1 | protein_coding | ENSG0000C |
| 67428460 | 67435401 | 1 | protein_coding | ENSG0000C |
| 67431367 | 67435399 | -1 | lncRNA | ENSG0000C |
| 67435510 | 67443821 | -1 | protein_coding | ENSG0000C |
| 67435510 | 67437682 | -1 | protein_coding | ENSG0000C |
| 67451301 | 67452729 | -1 | protein_coding | ENSG0000C |
| 67452406 | 67460313 | 1 | protein_coding | ENSG0000C |
| 67461710 | 67469272 | -1 | protein_coding | ENSG0000C |
| 67483026 | 67491103 | 1 | protein_coding | ENSG0000C |
| 67490245 | 67490315 | 1 | miRNA | ENSG0000C |
| 67491768 | 67506263 | -1 | protein_coding | ENSG0000C |
| 67506497 | 67508649 | -1 | protein_coding | ENSG0000C |
| 67518912 | 67524517 | -1 | protein_coding | ENSG0000C |
| 67538899 | 67540584 | 1 | unprocessed_pseudc | ENSG0000C |
| 67583595 | 67586656 | 1 | protein_coding | ENSG0000C |
| 67602880 | 67606706 | -1 | lncRNA | ENSG0000C |
| 67605521 | 67606642 | 1 | lncRNA | ENSG0000C |
| 67605653 | 67612554 | 1 | protein_coding | ENSG0000C |
| 67612651 | 67616257 | -1 | transcribed_unitary_ | ENSG0000C |
| 67627938 | 67629937 | -1 | protein_coding | ENSG0000C |
| 67631303 | 67639560 | -1 | protein_coding | ENSG0000C |
| 67642555 | 67650730 | -1 | protein_coding | ENSG0000C |
| 67651576 | 67653404 | -1 | processed_pseudoge | ENSG0000C |
| 67662162 | 67681200 | -1 | protein_coding | ENSG0000C |
| 67682772 | 67683058 | -1 | processed_pseudoge | ENSG0000C |
| 67711702 | 67716005 | -1 | unprocessed_pseudc | ENSG0000C |
| 67722483 | 67723455 | -1 | unprocessed_pseudc | ENSG0000C |
| 67735600 | 67784229 | -1 | lncRNA | ENSG0000C |
| 67735608 | 67736559 | -1 | transcribed_processe | ENSG0000C |
| 67744322 | 67744645 | 1 | processed_pseudoge | ENSG0000C |
| 67749043 | 67756968 | -1 | transcribed_unproce | ENSG0000C |
| 67785273 | 67792335 | 1 | unprocessed_pseudc | ENSG0000C |
| 67791648 | 67805336 | -1 | transcribed_unproce | ENSG0000C |
| 67812557 | 67873367 | -1 | unprocessed_pseudc | ENSG0000C |
| 67840942 | 67841049 | 1 | processed_pseudoge | ENSG0000C |
| 67886477 | 67906350 | 1 | lncRNA | ENSG0000C |
| 67895631 | 67895737 | 1 | snRNA | ENSG0000C |
| 67925651 | 67926431 | -1 | processed_pseudoge | ENSG0000C |
| 67934563 | 67955802 | -1 | transcribed_unproce | ENSG0000C |
| 67935558 | 67935926 | 1 | processed_pseudoge | ENSG0000C |
| 67965873 | 67966227 | -1 | processed_pseudoge | ENSG0000C |
| 67974286 | 67975243 | -1 | unprocessed_pseudc | ENSG0000C |
| 67991100 | 68004982 | -1 | protein_coding | ENSG0000C |
| 68008578 | 68029282 | 1 | protein_coding | ENSG0000C |
| 68024809 | 68030461 | -1 | lncRNA | ENSG0000C |
| 68030617 | 68036644 | 1 | protein_coding | ENSG0000C |
| 68032864 | 68032922 | 1 | miRNA | ENSG0000C |

| | | | | |
|---|---|---|---|---|
| 68033897 | 68033981 | 1 | miRNA | ENSG0000( |
| 68039025 | 68050895 | 1 | protein_coding | ENSG0000( |
| 68044794 | 68044957 | 1 | miRNA | ENSG0000( |
| 68050740 | 68053762 | 1 | lncRNA | ENSG0000( |
| 68052859 | 68121444 | -1 | protein_coding | ENSG0000( |
| 68121624 | 68130518 | 1 | lncRNA | ENSG0000( |
| 68154863 | 68213828 | -1 | protein_coding | ENSG0000( |
| 68261338 | 68272001 | -1 | protein_coding | ENSG0000( |
| 68272102 | 68284879 | 1 | lncRNA | ENSG0000( |
| 68312591 | 68449275 | 1 | protein_coding | ENSG0000( |
| 68460731 | 68615334 | 1 | protein_coding | ENSG0000( |
| 68488609 | 68488840 | -1 | processed_pseudoge | ENSG0000( |
| 68612899 | 68616711 | -1 | lncRNA | ENSG0000( |
| 68683779 | 68691175 | 1 | protein_coding | ENSG0000( |
| 68707440 | 68751520 | -1 | protein_coding | ENSG0000( |
| 68754620 | 68844410 | -1 | protein_coding | ENSG0000( |
| 68870664 | 68874542 | 1 | lncRNA | ENSG0000( |
| 68891276 | 68903835 | -1 | protein_coding | ENSG0000( |
| 68903842 | 68940602 | 1 | protein_coding | ENSG0000( |
| 68941503 | 68942852 | -1 | lncRNA | ENSG0000( |
| 68980021 | 68980986 | -1 | protein_coding | ENSG0000( |
| 69000765 | 69002048 | -1 | lncRNA | ENSG0000( |
| 69004394 | 69005100 | 1 | lncRNA | ENSG0000( |
| 69004395 | 69013382 | -1 | protein_coding | ENSG0000( |
| 69012283 | 69018447 | 1 | lncRNA | ENSG0000( |
| 69048932 | 69136316 | 1 | protein_coding | ENSG0000( |
| 69072915 | 69162440 | 1 | protein_coding | ENSG0000( |
| 69083176 | 69083258 | 1 | miRNA | ENSG0000( |
| 69103493 | 69109094 | 1 | lncRNA | ENSG0000( |
| 69147228 | 69171564 | -1 | lncRNA | ENSG0000( |
| 69155478 | 69159752 | 1 | protein_coding | ENSG0000( |
| 69294151 | 69367726 | 1 | protein_coding | ENSG0000( |
| 69303412 | 69303807 | -1 | processed_pseudoge | ENSG0000( |
| 69371463 | 69372512 | 1 | lncRNA | ENSG0000( |
| 69414307 | 69416832 | 1 | lncRNA | ENSG0000( |
| 69425690 | 69429621 | 1 | lncRNA | ENSG0000( |
| 69438365 | 69444743 | -1 | lncRNA | ENSG0000( |
| 69467598 | 69469705 | 1 | lncRNA | ENSG0000( |
| 69475567 | 69481545 | -1 | lncRNA | ENSG0000( |
| 69481662 | 69494952 | 1 | lncRNA | ENSG0000( |
| 69641087 | 69654474 | 1 | protein_coding | ENSG0000( |
| 69653076 | 69675416 | -1 | protein_coding | ENSG0000( |
| 69698238 | 69704022 | -1 | protein_coding | ENSG0000( |
| 69737298 | 69738017 | -1 | processed_pseudoge | ENSG0000( |
| 69771022 | 69775341 | -1 | protein_coding | ENSG0000( |
| 69809968 | 69819416 | -1 | protein_coding | ENSG0000( |
| 69909184 | 69910994 | -1 | lncRNA | ENSG0000( |
| 69921638 | 69926813 | -1 | lncRNA | ENSG0000( |
| 69985876 | 70015815 | 1 | lncRNA | ENSG0000( |
| 70014858 | 70021059 | -1 | lncRNA | ENSG0000( |

| | | | | |
|---|---|---|---|---|
| 70056230 | 70065371 | -1 | lncRNA | ENSG00000 |
| 70072434 | 70075433 | -1 | lncRNA | ENSG00000 |
| 70075363 | 70075469 | 1 | snRNA | ENSG00000 |
| 70078302 | 70189528 | 1 | protein_coding | ENSG00000 |
| 70129297 | 70149623 | 1 | lncRNA | ENSG00000 |
| 70187788 | 70188509 | -1 | lncRNA | ENSG00000 |
| 70203163 | 70207390 | 1 | protein_coding | ENSG00000 |
| 70206291 | 70207390 | -1 | lncRNA | ENSG00000 |
| 70270700 | 70385312 | 1 | protein_coding | ENSG00000 |
| 70278921 | 70279297 | -1 | processed_pseudoge | ENSG00000 |
| 70282367 | 70363368 | -1 | lncRNA | ENSG00000 |
| 70283955 | 70284070 | 1 | miRNA | ENSG00000 |
| 70319928 | 70321415 | 1 | lncRNA | ENSG00000 |
| 70324871 | 70327209 | -1 | lncRNA | ENSG00000 |
| 70358198 | 70358677 | -1 | lncRNA | ENSG00000 |
| 70372246 | 70398488 | -1 | lncRNA | ENSG00000 |
| 70398404 | 70436584 | 1 | protein_coding | ENSG00000 |
| 70467856 | 71252577 | -1 | protein_coding | ENSG00000 |
| 70477277 | 70477592 | -1 | lncRNA | ENSG00000 |
| 70603304 | 70604859 | 1 | TEC | ENSG00000 |
| 70626441 | 70635658 | 1 | lncRNA | ENSG00000 |
| 70646165 | 70647562 | 1 | lncRNA | ENSG00000 |
| 70705167 | 70705269 | -1 | misc_RNA | ENSG00000 |
| 70705605 | 70706068 | -1 | TEC | ENSG00000 |
| 70862790 | 70865115 | 1 | lncRNA | ENSG00000 |
| 70872270 | 70872368 | -1 | miRNA | ENSG00000 |
| 70886090 | 70896305 | 1 | lncRNA | ENSG00000 |
| 71382601 | 71423423 | -1 | transcribed_unitary_ | ENSG00000 |
| 71428193 | 71452868 | -1 | protein_coding | ENSG00000 |
| 71448674 | 71452157 | 1 | lncRNA | ENSG00000 |
| 71453109 | 71524107 | 1 | protein_coding | ENSG00000 |
| 71473503 | 71473568 | 1 | miRNA | ENSG00000 |
| 71506061 | 71506380 | -1 | processed_pseudoge | ENSG00000 |
| 71527267 | 71528669 | 1 | protein_coding | ENSG00000 |
| 71538025 | 71539209 | 1 | protein_coding | ENSG00000 |
| 71548418 | 71549553 | 1 | protein_coding | ENSG00000 |
| 71563389 | 71565608 | -1 | lncRNA | ENSG00000 |
| 71565563 | 71566735 | 1 | protein_coding | ENSG00000 |
| 71568680 | 71568934 | 1 | processed_pseudoge | ENSG00000 |
| 71579714 | 71603353 | -1 | protein_coding | ENSG00000 |
| 71579728 | 71579848 | -1 | processed_pseudoge | ENSG00000 |
| 71593454 | 71594382 | 1 | unprocessed_pseudo | ENSG00000 |
| 71603260 | 71608040 | 1 | transcribed_processe | ENSG00000 |
| 71620020 | 71621032 | 1 | unprocessed_pseudo | ENSG00000 |
| 71629045 | 71629421 | 1 | processed_pseudoge | ENSG00000 |
| 71639551 | 71639769 | 1 | unprocessed_pseudo | ENSG00000 |
| 71669520 | 71670297 | 1 | processed_pseudoge | ENSG00000 |
| 71673885 | 71818238 | -1 | lncRNA | ENSG00000 |
| 71690453 | 71690657 | -1 | processed_pseudoge | ENSG00000 |
| 71701268 | 71705404 | -1 | lncRNA | ENSG00000 |

| | | | | |
|---|---|---|---|---|
| 71711740 | 71781183 | 1 | transcribed_unproce | ENSG0000C |
| 71745331 | 71747051 | 1 | lncRNA | ENSG0000C |
| 71787510 | 71801237 | 1 | protein_coding | ENSG0000C |
| 71800541 | 71804640 | -1 | transcribed_unproce | ENSG0000C |
| 71814045 | 71821548 | -1 | protein_coding | ENSG0000C |
| 71833200 | 71837710 | 1 | protein_coding | ENSG0000C |
| 71845196 | 71845245 | -1 | rRNA_pseudogene | ENSG0000C |
| 71856277 | 71863658 | 1 | transcribed_unproce | ENSG0000C |
| 71865509 | 71928654 | -1 | protein_coding | ENSG0000C |
| 71878453 | 71884561 | 1 | protein_coding | ENSG0000C |
| 71893410 | 71894433 | 1 | unprocessed_pseudc | ENSG0000C |
| 71903194 | 71904081 | 1 | unprocessed_pseudc | ENSG0000C |
| 71929018 | 71997597 | 1 | protein_coding | ENSG0000C |
| 71938671 | 71939544 | -1 | processed_pseudoge | ENSG0000C |
| 71998613 | 72005715 | 1 | protein_coding | ENSG0000C |
| 72002864 | 72080693 | -1 | protein_coding | ENSG0000C |
| 72014291 | 72020910 | 1 | lncRNA | ENSG0000C |
| 72047873 | 72047951 | -1 | snRNA | ENSG0000C |
| 72072228 | 72072302 | -1 | miRNA | ENSG0000C |
| 72080331 | 72110782 | 1 | protein_coding | ENSG0000C |
| 72080777 | 72110782 | 1 | protein_coding | ENSG0000C |
| 72085895 | 72103297 | -1 | protein_coding | ENSG0000C |
| 72105924 | 72109329 | 1 | protein_coding | ENSG0000C |
| 72106378 | 72112780 | -1 | protein_coding | ENSG0000C |
| 72114869 | 72139892 | 1 | protein_coding | ENSG0000C |
| 72131282 | 72131907 | 1 | processed_pseudoge | ENSG0000C |
| 72158822 | 72159734 | 1 | unprocessed_pseudc | ENSG0000C |
| 72163322 | 72209241 | -1 | lncRNA | ENSG0000C |
| 72172455 | 72177954 | 1 | unprocessed_pseudc | ENSG0000C |
| 72189558 | 72196323 | 1 | protein_coding | ENSG0000C |
| 72216601 | 72221950 | 1 | protein_coding | ENSG0000C |
| 72223701 | 72239147 | 1 | protein_coding | ENSG0000C |
| 72239077 | 72245664 | -1 | protein_coding | ENSG0000C |
| 72249856 | 72251288 | -1 | processed_pseudoge | ENSG0000C |
| 72261731 | 72290688 | 1 | lncRNA | ENSG0000C |
| 72280151 | 72281178 | 1 | processed_pseudoge | ENSG0000C |
| 72285495 | 72434680 | -1 | protein_coding | ENSG0000C |
| 72302139 | 72302965 | 1 | lncRNA | ENSG0000C |
| 72351347 | 72353210 | 1 | lncRNA | ENSG0000C |
| 72354516 | 72363555 | 1 | lncRNA | ENSG0000C |
| 72410716 | 72412079 | 1 | lncRNA | ENSG0000C |
| 72478221 | 72479237 | 1 | unprocessed_pseudc | ENSG0000C |
| 72519986 | 72521733 | 1 | unprocessed_pseudc | ENSG0000C |
| 72551049 | 72551919 | -1 | processed_pseudoge | ENSG0000C |
| 72562044 | 72562554 | 1 | processed_pseudoge | ENSG0000C |
| 72570660 | 72573229 | 1 | lncRNA | ENSG0000C |
| 72576141 | 72674591 | -1 | protein_coding | ENSG0000C |
| 72584572 | 72587979 | 1 | lncRNA | ENSG0000C |
| 72615063 | 72615130 | -1 | miRNA | ENSG0000C |
| 72621766 | 72621827 | 1 | snRNA | ENSG0000C |

| | | | | |
|---|---|---|---|---|
| 72643237 | 72659407 | 1 | lncRNA | ENSG0000C |
| 72685069 | 72793599 | -1 | protein_coding | ENSG0000C |
| 72685075 | 72693808 | 1 | lncRNA | ENSG0000C |
| 72689650 | 72695821 | 1 | lncRNA | ENSG0000C |
| 72700474 | 72705607 | 1 | lncRNA | ENSG0000C |
| 72708186 | 72708565 | 1 | processed_pseudoge | ENSG0000C |
| 72754729 | 72793681 | -1 | protein_coding | ENSG0000C |
| 72766004 | 72766116 | 1 | misc_RNA | ENSG0000C |
| 72783530 | 72783592 | 1 | miRNA | ENSG0000C |
| 72793624 | 72814054 | -1 | lncRNA | ENSG0000C |
| 72814406 | 72843674 | 1 | protein_coding | ENSG0000C |
| 72836745 | 73142318 | -1 | protein_coding | ENSG0000C |
| 72869544 | 72869650 | -1 | snRNA | ENSG0000C |
| 72940498 | 72941112 | 1 | processed_pseudoge | ENSG0000C |
| 73157946 | 73181724 | 1 | lncRNA | ENSG0000C |
| 73214998 | 73215913 | -1 | lncRNA | ENSG0000C |
| 73218298 | 73236352 | 1 | protein_coding | ENSG0000C |
| 73238975 | 73242335 | 1 | lncRNA | ENSG0000C |
| 73248779 | 73249867 | 1 | unprocessed_pseudc | ENSG0000C |
| 73264505 | 73298617 | 1 | protein_coding | ENSG0000C |
| 73307235 | 73309361 | -1 | lncRNA | ENSG0000C |
| 73308276 | 73369388 | 1 | protein_coding | ENSG0000C |
| 73376399 | 73397474 | 1 | protein_coding | ENSG0000C |
| 73395559 | 73396436 | -1 | lncRNA | ENSG0000C |
| 73400487 | 73598189 | -1 | protein_coding | ENSG0000C |
| 73405297 | 73410682 | 1 | lncRNA | ENSG0000C |
| 73452020 | 73452344 | 1 | processed_pseudoge | ENSG0000C |
| 73510658 | 73510820 | 1 | processed_pseudoge | ENSG0000C |
| 73580253 | 73580517 | 1 | processed_pseudoge | ENSG0000C |
| 73640479 | 73641037 | 1 | processed_pseudoge | ENSG0000C |
| 73646178 | 73662819 | 1 | protein_coding | ENSG0000C |
| 73675638 | 73761137 | -1 | protein_coding | ENSG0000C |
| 73722349 | 73722694 | 1 | lncRNA | ENSG0000C |
| 73760563 | 73761070 | 1 | lncRNA | ENSG0000C |
| 73787872 | 73865133 | 1 | protein_coding | ENSG0000C |
| 73832382 | 73832690 | -1 | misc_RNA | ENSG0000C |
| 73850469 | 73851211 | -1 | processed_pseudoge | ENSG0000C |
| 73872667 | 73876901 | -1 | protein_coding | ENSG0000C |
| 73876699 | 73931124 | 1 | protein_coding | ENSG0000C |
| 73921665 | 73922196 | -1 | processed_pseudoge | ENSG0000C |
| 73950319 | 73970366 | 1 | protein_coding | ENSG0000C |
| 73963657 | 73970287 | -1 | lncRNA | ENSG0000C |
| 73974672 | 73983246 | -1 | protein_coding | ENSG0000C |
| 73983449 | 73985562 | 1 | lncRNA | ENSG0000C |
| 73991283 | 73992180 | -1 | processed_pseudoge | ENSG0000C |
| 73994972 | 73996151 | -1 | processed_pseudoge | ENSG0000C |
| 74000277 | 74009085 | -1 | protein_coding | ENSG0000C |
| 74012714 | 74171210 | -1 | protein_coding | ENSG0000C |
| 74142151 | 74142358 | 1 | processed_pseudoge | ENSG0000C |
| 74171267 | 74254703 | 1 | protein_coding | ENSG0000C |

| | | | | |
|---|---|---|---|---|
| 74198748 | 74198854 | -1 | rRNA | ENSG00000 |
| 74204869 | 74205746 | 1 | TEC | ENSG00000 |
| 74235801 | 74311640 | -1 | protein_coding | ENSG00000 |
| 74252414 | 74252552 | -1 | snoRNA | ENSG00000 |
| 74311362 | 74324705 | 1 | lncRNA | ENSG00000 |
| 74330316 | 74398433 | -1 | protein_coding | ENSG00000 |
| 74354443 | 74355720 | 1 | processed_pseudoge | ENSG00000 |
| 74397549 | 74485742 | 1 | lncRNA | ENSG00000 |
| 74398825 | 74416379 | 1 | lncRNA | ENSG00000 |
| 74399237 | 74399333 | 1 | miRNA | ENSG00000 |
| 74454841 | 74467729 | -1 | protein_coding | ENSG00000 |
| 74455348 | 74456825 | 1 | lncRNA | ENSG00000 |
| 74482250 | 74482542 | 1 | processed_pseudoge | ENSG00000 |
| 74485580 | 74486051 | -1 | processed_pseudoge | ENSG00000 |
| 74490519 | 74493724 | -1 | protein_coding | ENSG00000 |
| 74493366 | 74498533 | 1 | lncRNA | ENSG00000 |
| 74493851 | 74669117 | 1 | protein_coding | ENSG00000 |
| 74652636 | 74652911 | 1 | processed_pseudoge | ENSG00000 |
| 74685224 | 74685505 | 1 | misc_RNA | ENSG00000 |
| 74696429 | 74731385 | -1 | protein_coding | ENSG00000 |
| 74698231 | 74699658 | -1 | TEC | ENSG00000 |
| 74720268 | 74720337 | -1 | miRNA | ENSG00000 |
| 74738478 | 74738796 | -1 | processed_pseudoge | ENSG00000 |
| 74745716 | 74746114 | -1 | processed_pseudoge | ENSG00000 |
| 74748849 | 74842413 | 1 | protein_coding | ENSG00000 |
| 74807739 | 74949200 | -1 | protein_coding | ENSG00000 |
| 74830574 | 74832510 | -1 | TEC | ENSG00000 |
| 74845910 | 74846209 | 1 | misc_RNA | ENSG00000 |
| 74876286 | 74920022 | 1 | transcribed_processe | ENSG00000 |
| 74931724 | 74936937 | 1 | lncRNA | ENSG00000 |
| 74949247 | 74979033 | 1 | protein_coding | ENSG00000 |
| 74968189 | 74968292 | -1 | snRNA | ENSG00000 |
| 74988279 | 75018893 | 1 | protein_coding | ENSG00000 |
| 75069243 | 75096964 | -1 | lncRNA | ENSG00000 |
| 75071148 | 75072052 | -1 | transcribed_unitary_ | ENSG00000 |
| 75079265 | 75080131 | -1 | transcribed_processe | ENSG00000 |
| 75081753 | 75096876 | -1 | protein_coding | ENSG00000 |
| 75099172 | 75140148 | -1 | lncRNA | ENSG00000 |
| 75100563 | 75206549 | 1 | protein_coding | ENSG00000 |
| 75131138 | 75131880 | -1 | unprocessed_pseudo | ENSG00000 |
| 75206048 | 75241173 | -1 | lncRNA | ENSG00000 |
| 75228322 | 75228657 | -1 | processed_pseudoge | ENSG00000 |
| 75240774 | 75243704 | 1 | protein_coding | ENSG00000 |
| 75260122 | 75351705 | -1 | protein_coding | ENSG00000 |
| 75260127 | 75261025 | 1 | lncRNA | ENSG00000 |
| 75260129 | 75262466 | -1 | TEC | ENSG00000 |
| 75264289 | 75265170 | 1 | lncRNA | ENSG00000 |
| 75335092 | 75335186 | -1 | miRNA | ENSG00000 |
| 75399515 | 75422280 | 1 | protein_coding | ENSG00000 |
| 75400391 | 75400538 | 1 | snoRNA | ENSG00000 |

| | | | | |
|---|---|---|---|---|
| 75404421 | 75404566 | 1 | snoRNA | ENSG00000 |
| 75422394 | 75430629 | -1 | protein_coding | ENSG00000 |
| 75434640 | 75525941 | -1 | protein_coding | ENSG00000 |
| 75506937 | 75508391 | -1 | lncRNA | ENSG00000 |
| 75562056 | 75572783 | 1 | protein_coding | ENSG00000 |
| 75583196 | 75594665 | 1 | lncRNA | ENSG00000 |
| 75586918 | 75669120 | -1 | protein_coding | ENSG00000 |
| 75596144 | 75597270 | -1 | lncRNA | ENSG00000 |
| 75635883 | 75638587 | -1 | lncRNA | ENSG00000 |
| 75642600 | 75642889 | 1 | processed_pseudoge | ENSG00000 |
| 75717838 | 75732958 | 1 | protein_coding | ENSG00000 |
| 75742129 | 75742426 | 1 | misc_RNA | ENSG00000 |
| 75758455 | 75768647 | -1 | lncRNA | ENSG00000 |
| 75759512 | 75801535 | 1 | protein_coding | ENSG00000 |
| 75775904 | 75776929 | 1 | lncRNA | ENSG00000 |
| 75800877 | 75803415 | -1 | lncRNA | ENSG00000 |
| 75803431 | 75815406 | -1 | lncRNA | ENSG00000 |
| 75815210 | 76144232 | 1 | protein_coding | ENSG00000 |
| 75835215 | 75835323 | -1 | misc_RNA | ENSG00000 |
| 75837544 | 75837645 | -1 | misc_RNA | ENSG00000 |
| 75911204 | 75911628 | -1 | processed_pseudoge | ENSG00000 |
| 75914201 | 75915213 | 1 | lncRNA | ENSG00000 |
| 75934936 | 75935039 | 1 | rRNA_pseudogene | ENSG00000 |
| 75942129 | 75942435 | -1 | processed_pseudoge | ENSG00000 |
| 76137315 | 76137731 | -1 | lncRNA | ENSG00000 |
| 76186325 | 76210736 | -1 | protein_coding | ENSG00000 |
| 76190725 | 76195071 | 1 | lncRNA | ENSG00000 |
| 76210956 | 76216025 | -1 | lncRNA | ENSG00000 |
| 76349956 | 76381132 | -1 | protein_coding | ENSG00000 |
| 76381303 | 76414619 | 1 | protein_coding | ENSG00000 |
| 76404140 | 76404252 | -1 | misc_RNA | ENSG00000 |
| 76435559 | 76444687 | -1 | lncRNA | ENSG00000 |
| 76444923 | 76553025 | 1 | protein_coding | ENSG00000 |
| 76591023 | 76591211 | 1 | processed_pseudoge | ENSG00000 |
| 76607853 | 76630465 | -1 | lncRNA | ENSG00000 |
| 76625462 | 76626834 | -1 | lncRNA | ENSG00000 |
| 76654169 | 76656712 | -1 | lncRNA | ENSG00000 |
| 76657056 | 76663866 | 1 | lncRNA | ENSG00000 |
| 76657524 | 76670747 | -1 | protein_coding | ENSG00000 |
| 76675079 | 76703195 | 1 | lncRNA | ENSG00000 |
| 76694041 | 76719801 | -1 | transcribed_unproce | ENSG00000 |
| 76759916 | 76768622 | -1 | lncRNA | ENSG00000 |
| 76782251 | 76798153 | 1 | protein_coding | ENSG00000 |
| 76782581 | 76783062 | -1 | lncRNA | ENSG00000 |
| 76800364 | 76804555 | 1 | lncRNA | ENSG00000 |
| 76860867 | 77026797 | 1 | protein_coding | ENSG00000 |
| 76955417 | 76978619 | -1 | lncRNA | ENSG00000 |
| 77034398 | 77041973 | 1 | protein_coding | ENSG00000 |
| 77066961 | 77126155 | 1 | protein_coding | ENSG00000 |
| 77102840 | 77103331 | 1 | protein_coding | ENSG00000 |

| | | | | |
|---|---|---|---|---|
| 77128246 | 77215241 | 1 | protein_coding | ENSG0000C |
| 77216558 | 77301687 | -1 | protein_coding | ENSG0000C |
| 77313606 | 77314032 | 1 | processed_pseudoge | ENSG0000C |
| 77321707 | 77474635 | -1 | protein_coding | ENSG0000C |
| 77473371 | 77477030 | 1 | lncRNA | ENSG0000C |
| 77514936 | 77637794 | -1 | protein_coding | ENSG0000C |
| 77566934 | 77566994 | -1 | snRNA | ENSG0000C |
| 77589391 | 77610356 | 1 | protein_coding | ENSG0000C |
| 77659996 | 77821017 | -1 | protein_coding | ENSG0000C |
| 77691650 | 77691751 | 1 | misc_RNA | ENSG0000C |
| 77717712 | 77718411 | -1 | lncRNA | ENSG0000C |
| 77734475 | 77735026 | -1 | processed_pseudoge | ENSG0000C |
| 77738680 | 77739568 | -1 | lncRNA | ENSG0000C |
| 77813319 | 77813676 | -1 | processed_pseudoge | ENSG0000C |
| 77821109 | 77918432 | 1 | protein_coding | ENSG0000C |
| 77829654 | 77872262 | -1 | lncRNA | ENSG0000C |
| 77850604 | 77851511 | 1 | lncRNA | ENSG0000C |
| 77866412 | 77870091 | -1 | lncRNA | ENSG0000C |
| 77868722 | 77869195 | -1 | processed_pseudoge | ENSG0000C |
| 77878720 | 77994671 | -1 | protein_coding | ENSG0000C |
| 77914990 | 77915275 | -1 | processed_pseudoge | ENSG0000C |
| 78015715 | 78016495 | 1 | lncRNA | ENSG0000C |
| 78015715 | 78046191 | -1 | protein_coding | ENSG0000C |
| 78016971 | 78079865 | -1 | protein_coding | ENSG0000C |
| 78022933 | 78023721 | 1 | lncRNA | ENSG0000C |
| 78063861 | 78068351 | 1 | protein_coding | ENSG0000C |
| 78068297 | 78079862 | -1 | protein_coding | ENSG0000C |
| 78100936 | 78139653 | -1 | protein_coding | ENSG0000C |
| 78133420 | 78133522 | -1 | snRNA | ENSG0000C |
| 78139771 | 78225909 | 1 | lncRNA | ENSG0000C |
| 78171249 | 78188626 | -1 | protein_coding | ENSG0000C |
| 78188812 | 78215232 | 1 | protein_coding | ENSG0000C |
| 78215293 | 78418348 | -1 | protein_coding | ENSG0000C |
| 78324758 | 78444049 | 1 | lncRNA | ENSG0000C |
| 78384059 | 78385739 | 1 | transcribed_processe | ENSG0000C |
| 78388061 | 78392405 | 1 | lncRNA | ENSG0000C |
| 78423982 | 78429836 | -1 | lncRNA | ENSG0000C |
| 78435968 | 78574874 | -1 | protein_coding | ENSG0000C |
| 78533176 | 78558565 | 1 | lncRNA | ENSG0000C |
| 78579255 | 78579356 | -1 | snRNA | ENSG0000C |
| 78581675 | 78582156 | 1 | processed_pseudoge | ENSG0000C |
| 78652829 | 79441030 | -1 | protein_coding | ENSG0000C |
| 78749250 | 78756480 | 1 | lncRNA | ENSG0000C |
| 79092848 | 79098003 | 1 | lncRNA | ENSG0000C |
| 79191558 | 79193160 | 1 | lncRNA | ENSG0000C |
| 79402022 | 79402109 | -1 | miRNA | ENSG0000C |
| 79422169 | 79422226 | -1 | miRNA | ENSG0000C |
| 79604967 | 79612300 | -1 | lncRNA | ENSG0000C |
| 79966805 | 79968708 | 1 | TEC | ENSG0000C |
| 79987513 | 79989630 | 1 | processed_pseudoge | ENSG0000C |

| | | | | |
|---|---|---|---|---|
| 80321620 | 80327911 | -1 | lncRNA | ENSG00000 |
| 80527960 | 80528066 | 1 | snRNA | ENSG00000 |
| 80653429 | 80653988 | -1 | processed_pseudoge | ENSG00000 |
| 80751200 | 80762826 | -1 | lncRNA | ENSG00000 |
| 80957317 | 80957634 | 1 | processed_pseudoge | ENSG00000 |
| 81015849 | 81040236 | 1 | lncRNA | ENSG00000 |
| 81175201 | 81431520 | 1 | lncRNA | ENSG00000 |
| 81552226 | 81552516 | 1 | processed_pseudoge | ENSG00000 |
| 81555910 | 81556425 | -1 | unprocessed_pseudc | ENSG00000 |
| 81821272 | 82718299 | -1 | lncRNA | ENSG00000 |
| 81842435 | 81842669 | -1 | processed_pseudoge | ENSG00000 |
| 81890741 | 81890836 | -1 | miRNA | ENSG00000 |
| 81970994 | 81987813 | 1 | lncRNA | ENSG00000 |
| 82603529 | 82643445 | -1 | lncRNA | ENSG00000 |
| 82689559 | 82689768 | 1 | processed_pseudoge | ENSG00000 |
| 82729940 | 82733864 | -1 | protein_coding | ENSG00000 |
| 82781502 | 82817741 | 1 | lncRNA | ENSG00000 |
| 82805354 | 82806509 | 1 | processed_pseudoge | ENSG00000 |
| 82822936 | 82970584 | -1 | protein_coding | ENSG00000 |
| 82835509 | 82837222 | 1 | processed_pseudoge | ENSG00000 |
| 82899975 | 82958277 | 1 | protein_coding | ENSG00000 |
| 82957423 | 82958216 | 1 | processed_pseudoge | ENSG00000 |
| 82963681 | 83039115 | -1 | lncRNA | ENSG00000 |
| 82973133 | 83071923 | -1 | protein_coding | ENSG00000 |
| 83040806 | 83041403 | -1 | processed_pseudoge | ENSG00000 |
| 83041464 | 83041598 | -1 | snoRNA | ENSG00000 |
| 83072052 | 83107789 | 1 | lncRNA | ENSG00000 |
| 83072402 | 83097196 | -1 | lncRNA | ENSG00000 |
| 83083687 | 83084138 | 1 | TEC | ENSG00000 |
| 83106580 | 83106961 | -1 | processed_pseudoge | ENSG00000 |
| 83111060 | 83111442 | -1 | processed_pseudoge | ENSG00000 |
| 83132056 | 83134737 | 1 | processed_pseudoge | ENSG00000 |
| 83156988 | 83187451 | 1 | protein_coding | ENSG00000 |
| 83180144 | 83184520 | 1 | lncRNA | ENSG00000 |
| 83184491 | 83193794 | -1 | lncRNA | ENSG00000 |
| 83185521 | 83187036 | -1 | lncRNA | ENSG00000 |
| 83193712 | 83260694 | 1 | protein_coding | ENSG00000 |
| 83209399 | 83209634 | 1 | processed_pseudoge | ENSG00000 |
| 83209431 | 83213379 | -1 | lncRNA | ENSG00000 |
| 83213005 | 83213404 | 1 | processed_pseudoge | ENSG00000 |
| 83259097 | 83286407 | -1 | protein_coding | ENSG00000 |
| 83286120 | 83423516 | 1 | lncRNA | ENSG00000 |
| 83328926 | 83329246 | 1 | processed_pseudoge | ENSG00000 |
| 83386988 | 83387102 | 1 | misc_RNA | ENSG00000 |
| 83455012 | 85627922 | -1 | protein_coding | ENSG00000 |
| 83643602 | 83725390 | 1 | lncRNA | ENSG00000 |
| 83789977 | 83791578 | -1 | processed_pseudoge | ENSG00000 |
| 83814571 | 83815263 | -1 | unprocessed_pseudc | ENSG00000 |
| 84545131 | 84546846 | -1 | processed_pseudoge | ENSG00000 |
| 84639993 | 84640565 | 1 | processed_pseudoge | ENSG00000 |

| | | | | |
|---|---|---|---|---|
| 84720826 | 84800701 | 1 | lncRNA | ENSG0000C |
| 84887207 | 84887419 | 1 | misc_RNA | ENSG0000C |
| 84890744 | 84890961 | 1 | misc_RNA | ENSG0000C |
| 84936689 | 84955705 | 1 | lncRNA | ENSG0000C |
| 84997226 | 84997768 | -1 | processed_pseudoge | ENSG0000C |
| 85020785 | 85021655 | -1 | processed_pseudoge | ENSG0000C |
| 85336075 | 85336572 | -1 | processed_pseudoge | ENSG0000C |
| 85452282 | 85453698 | -1 | processed_pseudoge | ENSG0000C |
| 85509938 | 85510044 | 1 | snRNA | ENSG0000C |
| 85628573 | 85636539 | 1 | protein_coding | ENSG0000C |
| 85647967 | 85656547 | 1 | protein_coding | ENSG0000C |
| 85659708 | 85682908 | -1 | protein_coding | ENSG0000C |
| 85683848 | 85686195 | -1 | protein_coding | ENSG0000C |
| 85694224 | 85811159 | -1 | protein_coding | ENSG0000C |
| 85852557 | 85854943 | -1 | TEC | ENSG0000C |
| 85855101 | 85920021 | 1 | protein_coding | ENSG0000C |
| 85916502 | 85923682 | -1 | lncRNA | ENSG0000C |
| 85934737 | 85935663 | 1 | processed_pseudoge | ENSG0000C |
| 85957175 | 86069882 | -1 | protein_coding | ENSG0000C |
| 86153227 | 86153334 | 1 | snRNA | ENSG0000C |
| 86192787 | 86193482 | -1 | lncRNA | ENSG0000C |
| 86195132 | 86196266 | 1 | processed_pseudoge | ENSG0000C |
| 86216596 | 86216878 | 1 | misc_RNA | ENSG0000C |
| 86244753 | 86278813 | 1 | protein_coding | ENSG0000C |
| 86278333 | 86278398 | 1 | miRNA | ENSG0000C |
| 86283927 | 86284668 | 1 | processed_pseudoge | ENSG0000C |
| 86294760 | 86295576 | 1 | processed_pseudoge | ENSG0000C |
| 86302211 | 86345943 | 1 | protein_coding | ENSG0000C |
| 86324012 | 86324309 | -1 | misc_RNA | ENSG0000C |
| 86374736 | 86423109 | 1 | protein_coding | ENSG0000C |
| 86431590 | 86622867 | 1 | lncRNA | ENSG0000C |
| 86432098 | 86432575 | 1 | transcribed_processe | ENSG0000C |
| 86434924 | 86437282 | -1 | TEC | ENSG0000C |
| 86441108 | 86672636 | -1 | protein_coding | ENSG0000C |
| 86703099 | 86714092 | 1 | lncRNA | ENSG0000C |
| 86727355 | 86765467 | 1 | lncRNA | ENSG0000C |
| 86791059 | 86952910 | 1 | protein_coding | ENSG0000C |
| 86821143 | 86821728 | -1 | processed_pseudoge | ENSG0000C |
| 86832540 | 86833449 | -1 | unprocessed_pseudc | ENSG0000C |
| 86833068 | 86837615 | 1 | lncRNA | ENSG0000C |
| 86857059 | 86857947 | -1 | unprocessed_pseudc | ENSG0000C |
| 86892214 | 86925037 | -1 | lncRNA | ENSG0000C |
| 86908990 | 86912238 | -1 | TEC | ENSG0000C |
| 86945679 | 86955395 | -1 | protein_coding | ENSG0000C |
| 86955616 | 87021909 | 1 | lncRNA | ENSG0000C |
| 87024583 | 87025777 | 1 | processed_pseudoge | ENSG0000C |
| 87037844 | 87328824 | 1 | protein_coding | ENSG0000C |
| 87094734 | 87096585 | -1 | processed_pseudoge | ENSG0000C |
| 87121058 | 87121596 | 1 | TEC | ENSG0000C |
| 87258851 | 87259259 | -1 | processed_pseudoge | ENSG0000C |

| | | | | |
|---|---|---|---|---|
| 87323709 | 87324359 | -1 | processed_pseudoge | ENSG00000( |
| 87329407 | 87330052 | -1 | processed_pseudoge | ENSG00000( |
| 87480736 | 87481297 | -1 | processed_pseudoge | ENSG00000( |
| 87490144 | 87491110 | 1 | TEC | ENSG00000( |
| 87565994 | 87566421 | 1 | TEC | ENSG00000( |
| 87606783 | 87606885 | 1 | snRNA | ENSG00000( |
| 87648489 | 87648549 | -1 | snRNA | ENSG00000( |
| 87718354 | 87719411 | -1 | lncRNA | ENSG00000( |
| 87718667 | 87721233 | -1 | TEC | ENSG00000( |
| 87815035 | 87816157 | -1 | processed_pseudoge | ENSG00000( |
| 87816224 | 87816524 | 1 | processed_pseudoge | ENSG00000( |
| 87816999 | 87817567 | 1 | TEC | ENSG00000( |
| 87847259 | 88045608 | -1 | lncRNA | ENSG00000( |
| 87879473 | 87880095 | -1 | TEC | ENSG00000( |
| 87917948 | 87917999 | -1 | processed_pseudoge | ENSG00000( |
| 88050106 | 88062110 | 1 | lncRNA | ENSG00000( |
| 88061774 | 88098147 | -1 | lncRNA | ENSG00000( |
| 88098591 | 88115835 | 1 | lncRNA | ENSG00000( |
| 88099995 | 88100105 | 1 | misc_RNA | ENSG00000( |
| 88113251 | 88175443 | -1 | protein_coding | ENSG00000( |
| 88165575 | 88167691 | -1 | TEC | ENSG00000( |
| 88176502 | 88176593 | 1 | miRNA | ENSG00000( |
| 88293592 | 88337761 | -1 | protein_coding | ENSG00000( |
| 88408179 | 88409223 | -1 | processed_pseudoge | ENSG00000( |
| 88422400 | 88428176 | 1 | lncRNA | ENSG00000( |
| 88504576 | 88524054 | 1 | lncRNA | ENSG00000( |
| 88504576 | 89065945 | -1 | protein_coding | ENSG00000( |
| 88612805 | 88612911 | -1 | snRNA | ENSG00000( |
| 89177875 | 89295759 | 1 | protein_coding | ENSG00000( |
| 89293745 | 89294246 | -1 | processed_pseudoge | ENSG00000( |
| 89297717 | 89298199 | 1 | TEC | ENSG00000( |
| 89300709 | 89301477 | -1 | TEC | ENSG00000( |
| 89324353 | 89498187 | -1 | protein_coding | ENSG00000( |
| 89498748 | 89499198 | -1 | processed_pseudoge | ENSG00000( |
| 89533645 | 89534043 | -1 | TEC | ENSG00000( |
| 89545128 | 89545744 | 1 | TEC | ENSG00000( |
| 89546637 | 89589611 | -1 | lncRNA | ENSG00000( |
| 89548130 | 89548795 | -1 | TEC | ENSG00000( |
| 89556860 | 89557205 | 1 | TEC | ENSG00000( |
| 89576913 | 89577652 | 1 | TEC | ENSG00000( |
| 89639237 | 89698718 | 1 | transcribed_unproce | ENSG00000( |
| 89700764 | 89701325 | 1 | processed_pseudoge | ENSG00000( |
| 89704164 | 89704658 | 1 | TEC | ENSG00000( |
| 89710299 | 89717872 | 1 | protein_coding | ENSG00000( |
| 89723482 | 89727253 | -1 | unprocessed_pseudc | ENSG00000( |
| 89731017 | 89734782 | 1 | processed_pseudoge | ENSG00000( |
| 89752785 | 89753975 | -1 | processed_pseudoge | ENSG00000( |
| 89753982 | 89754953 | -1 | lncRNA | ENSG00000( |
| 89764884 | 89765138 | 1 | processed_pseudoge | ENSG00000( |
| 89766440 | 89768836 | -1 | unprocessed_pseudc | ENSG00000( |

| | | | | |
|---|---|---|---|---|
| 89776981 | 89782582 | 1 | transcribed_unproce | ENSG00000 |
| 89785945 | 89787838 | 1 | unprocessed_pseudo | ENSG00000 |
| 89789412 | 89789733 | -1 | processed_pseudoge | ENSG00000 |
| 89797655 | 89808575 | -1 | protein_coding | ENSG00000 |
| 89820082 | 89826922 | 1 | unprocessed_pseudo | ENSG00000 |
| 89841997 | 89850970 | 1 | unprocessed_pseudo | ENSG00000 |
| 89854953 | 89862893 | 1 | unprocessed_pseudo | ENSG00000 |
| 89863848 | 89865550 | -1 | unprocessed_pseudo | ENSG00000 |
| 89869282 | 89876017 | -1 | protein_coding | ENSG00000 |
| 89883927 | 89886213 | -1 | unprocessed_pseudo | ENSG00000 |
| 89887366 | 89889991 | 1 | unprocessed_pseudo | ENSG00000 |
| 89895799 | 89900449 | -1 | unprocessed_pseudo | ENSG00000 |
| 89908540 | 89909047 | 1 | processed_pseudoge | ENSG00000 |
| 89911111 | 89921767 | -1 | protein_coding | ENSG00000 |
| 89924064 | 89933063 | 1 | protein_coding | ENSG00000 |
| 89944035 | 89949402 | 1 | unprocessed_pseudo | ENSG00000 |
| 89954593 | 89954908 | -1 | processed_pseudoge | ENSG00000 |
| 89955146 | 89957165 | -1 | unprocessed_pseudo | ENSG00000 |
| 89960422 | 89960624 | 1 | processed_pseudoge | ENSG00000 |
| 89968502 | 89974072 | 1 | protein_coding | ENSG00000 |
| 89978540 | 89980661 | 1 | unprocessed_pseudo | ENSG00000 |
| 89981441 | 89989559 | -1 | unprocessed_pseudo | ENSG00000 |
| 89993536 | 89999741 | -1 | unprocessed_pseudo | ENSG00000 |
| 90017611 | 90024511 | -1 | unprocessed_pseudo | ENSG00000 |
| 90031106 | 90042025 | 1 | protein_coding | ENSG00000 |
| 90049945 | 90050266 | 1 | processed_pseudoge | ENSG00000 |
| 90051837 | 90053736 | -1 | unprocessed_pseudo | ENSG00000 |
| 90057605 | 90062999 | -1 | unprocessed_pseudo | ENSG00000 |
| 90071136 | 90073055 | 1 | unprocessed_pseudo | ENSG00000 |
| 90074839 | 90075096 | -1 | processed_pseudoge | ENSG00000 |
| 90085950 | 90087131 | 1 | protein_coding | ENSG00000 |
| 90098421 | 90098817 | -1 | processed_pseudoge | ENSG00000 |
| 90110638 | 90112022 | -1 | lncRNA | ENSG00000 |
| 90118391 | 90118461 | 1 | snoRNA | ENSG00000 |
| 90125658 | 90126061 | 1 | TEC | ENSG00000 |
| 90131515 | 90192894 | 1 | protein_coding | ENSG00000 |
| 90193614 | 90198120 | 1 | TEC | ENSG00000 |
| 90200429 | 90223077 | -1 | protein_coding | ENSG00000 |
| 90223153 | 90226538 | 1 | TEC | ENSG00000 |
| 90251204 | 90915052 | 1 | lncRNA | ENSG00000 |
| 90282560 | 90284172 | 1 | processed_pseudoge | ENSG00000 |
| 90555774 | 90555857 | -1 | miRNA | ENSG00000 |
| 90731110 | 90731723 | 1 | TEC | ENSG00000 |
| 90869121 | 90869202 | -1 | miRNA | ENSG00000 |
| 90972316 | 90972521 | -1 | TEC | ENSG00000 |
| 91013787 | 91015299 | 1 | TEC | ENSG00000 |
| 91093875 | 91094260 | -1 | TEC | ENSG00000 |
| 91114724 | 91115951 | -1 | processed_pseudoge | ENSG00000 |
| 91115961 | 91116555 | 1 | processed_pseudoge | ENSG00000 |
| 91157994 | 91234388 | 1 | lncRNA | ENSG00000 |

| | | | | |
|---|---|---|---|---|
| 91184934 | 91185543 | -1 | TEC | ENSG0000C |
| 91208838 | 91209590 | -1 | TEC | ENSG0000C |
| 91382909 | 91383289 | 1 | TEC | ENSG0000C |
| 91794319 | 91872086 | 1 | lncRNA | ENSG0000C |
| 91949731 | 91950034 | -1 | processed_pseudoge | ENSG0000C |
| 91992833 | 91993255 | 1 | TEC | ENSG0000C |
| 92161106 | 92161889 | 1 | processed_pseudoge | ENSG0000C |
| 92336032 | 92336496 | -1 | processed_pseudoge | ENSG0000C |
| 92352096 | 92896470 | 1 | protein_coding | ENSG0000C |
| 92366496 | 92367250 | -1 | processed_pseudoge | ENSG0000C |
| 92400191 | 92408176 | -1 | lncRNA | ENSG0000C |
| 92498152 | 92498935 | -1 | processed_pseudoge | ENSG0000C |
| 92748732 | 92766867 | -1 | lncRNA | ENSG0000C |
| 92913227 | 92915644 | -1 | lncRNA | ENSG0000C |
| 92914603 | 92931322 | 1 | transcribed_unproce | ENSG0000C |
| 92937441 | 92937666 | 1 | processed_pseudoge | ENSG0000C |
| 92965797 | 92966603 | 1 | lncRNA | ENSG0000C |
| 92969720 | 92985066 | 1 | protein_coding | ENSG0000C |
| 93052395 | 93052827 | -1 | processed_pseudoge | ENSG0000C |
| 93071529 | 93072031 | 1 | TEC | ENSG0000C |
| 93133855 | 93134164 | 1 | TEC | ENSG0000C |
| 93144174 | 93197991 | -1 | protein_coding | ENSG0000C |
| 93152075 | 93152750 | 1 | processed_pseudoge | ENSG0000C |
| 93171134 | 93171906 | 1 | TEC | ENSG0000C |
| 93206990 | 93207583 | -1 | TEC | ENSG0000C |
| 93221486 | 93221630 | -1 | processed_pseudoge | ENSG0000C |
| 93240133 | 93241168 | 1 | lncRNA | ENSG0000C |
| 93286629 | 93288903 | -1 | TEC | ENSG0000C |
| 93329971 | 93438487 | 1 | protein_coding | ENSG0000C |
| 93478472 | 93543391 | -1 | protein_coding | ENSG0000C |
| 93535468 | 93535816 | 1 | processed_pseudoge | ENSG0000C |
| 93556913 | 93557206 | 1 | misc_RNA | ENSG0000C |
| 93587703 | 93588559 | 1 | TEC | ENSG0000C |
| 93592187 | 93592669 | 1 | TEC | ENSG0000C |
| 93609760 | 93610333 | -1 | processed_pseudoge | ENSG0000C |
| 93661682 | 93730358 | 1 | protein_coding | ENSG0000C |
| 93719603 | 93719715 | 1 | misc_RNA | ENSG0000C |
| 93721513 | 93721865 | 1 | lncRNA | ENSG0000C |
| 93721542 | 93721621 | 1 | snoRNA | ENSG0000C |
| 93721797 | 93721866 | 1 | snoRNA | ENSG0000C |
| 93726654 | 93729805 | -1 | TEC | ENSG0000C |
| 93729948 | 93784391 | -1 | protein_coding | ENSG0000C |
| 93730513 | 93730646 | -1 | snoRNA | ENSG0000C |
| 93730979 | 93731099 | -1 | snoRNA | ENSG0000C |
| 93731502 | 93731574 | -1 | snoRNA | ENSG0000C |
| 93732004 | 93732133 | -1 | snoRNA | ENSG0000C |
| 93732361 | 93732499 | -1 | snoRNA | ENSG0000C |
| 93733228 | 93733300 | -1 | snoRNA | ENSG0000C |
| 93733466 | 93733597 | -1 | snoRNA | ENSG0000C |
| 93733674 | 93733764 | -1 | miRNA | ENSG0000C |

| | | | | |
|---|---|---|---|---|
| 93735111 | 93735236 | -1 | snoRNA | ENSG0000C |
| 93741591 | 93764749 | 1 | protein_coding | ENSG0000C |
| 93741664 | 93812378 | 1 | protein_coding | ENSG0000C |
| 93784227 | 93814963 | 1 | protein_coding | ENSG0000C |
| 93818232 | 93850618 | -1 | protein_coding | ENSG0000C |
| 93827622 | 93827734 | 1 | misc_RNA | ENSG0000C |
| 93991346 | 93991429 | -1 | processed_pseudoge | ENSG0000C |
| 93998643 | 93999218 | -1 | processed_pseudoge | ENSG0000C |
| 94021354 | 94114208 | 1 | protein_coding | ENSG0000C |
| 94055023 | 94055972 | -1 | processed_pseudoge | ENSG0000C |
| 94128841 | 94181968 | 1 | protein_coding | ENSG0000C |
| 94185439 | 94282203 | 1 | lncRNA | ENSG0000C |
| 94188449 | 94188997 | -1 | processed_pseudoge | ENSG0000C |
| 94305592 | 94307721 | 1 | protein_coding | ENSG0000C |
| 94377316 | 94401419 | -1 | protein_coding | ENSG0000C |
| 94415578 | 94493908 | -1 | protein_coding | ENSG0000C |
| 94466495 | 94466580 | -1 | miRNA | ENSG0000C |
| 94472908 | 94473570 | -1 | lncRNA | ENSG0000C |
| 94493979 | 94499578 | 1 | protein_coding | ENSG0000C |
| 94512461 | 94532123 | 1 | protein_coding | ENSG0000C |
| 94543840 | 94621421 | 1 | protein_coding | ENSG0000C |
| 94543840 | 94549898 | 1 | protein_coding | ENSG0000C |
| 94545330 | 94740355 | -1 | lncRNA | ENSG0000C |
| 94559018 | 94559374 | 1 | TEC | ENSG0000C |
| 94638038 | 94640833 | 1 | lncRNA | ENSG0000C |
| 94638045 | 94652884 | 1 | lncRNA | ENSG0000C |
| 94706431 | 94876748 | 1 | protein_coding | ENSG0000C |
| 94874052 | 94925521 | -1 | lncRNA | ENSG0000C |
| 94913047 | 94913766 | 1 | processed_pseudoge | ENSG0000C |
| 94962620 | 94973586 | -1 | protein_coding | ENSG0000C |
| 94973709 | 94999519 | 1 | protein_coding | ENSG0000C |
| 95011858 | 95012532 | 1 | unprocessed_pseudc | ENSG0000C |
| 95025258 | 95027596 | 1 | protein_coding | ENSG0000C |
| 95030549 | 95031245 | 1 | unprocessed_pseudc | ENSG0000C |
| 95037482 | 95039811 | 1 | transcribed_processe | ENSG0000C |
| 95040411 | 95041571 | 1 | unprocessed_pseudc | ENSG0000C |
| 95049422 | 95051338 | 1 | protein_coding | ENSG0000C |
| 95057282 | 95057984 | 1 | unprocessed_pseudc | ENSG0000C |
| 95067197 | 95071224 | 1 | protein_coding | ENSG0000C |
| 95089846 | 95132645 | 1 | protein_coding | ENSG0000C |
| 95143637 | 95145125 | 1 | processed_pseudoge | ENSG0000C |
| 95145437 | 95146004 | 1 | processed_pseudoge | ENSG0000C |
| 95150539 | 95234391 | 1 | lncRNA | ENSG0000C |
| 95165513 | 95232541 | -1 | protein_coding | ENSG0000C |
| 95203727 | 95204033 | 1 | processed_pseudoge | ENSG0000C |
| 95482406 | 95497553 | 1 | lncRNA | ENSG0000C |
| 95571040 | 95748715 | 1 | lncRNA | ENSG0000C |
| 95698086 | 95700623 | 1 | lncRNA | ENSG0000C |
| 95756959 | 95757442 | -1 | processed_pseudoge | ENSG0000C |
| 95768953 | 95790409 | -1 | protein_coding | ENSG0000C |

| | | | | |
|---|---|---|---|---|
| 95789965 | 95832693 | 1 | protein_coding | ENSG0000C |
| 95832882 | 95925315 | -1 | protein_coding | ENSG0000C |
| 95840017 | 95840124 | -1 | rRNA_pseudogene | ENSG0000C |
| 95963219 | 95963624 | 1 | processed_pseudoge | ENSG0000C |
| 95976598 | 96343195 | -1 | protein_coding | ENSG0000C |
| 96092374 | 96137827 | 1 | lncRNA | ENSG0000C |
| 96341438 | 96341526 | 1 | miRNA | ENSG0000C |
| 96352769 | 96389956 | -1 | protein_coding | ENSG0000C |
| 96389989 | 96507574 | 1 | protein_coding | ENSG0000C |
| 96447132 | 96506826 | -1 | lncRNA | ENSG0000C |
| 96474572 | 96474692 | -1 | rRNA_pseudogene | ENSG0000C |
| 96508425 | 96514748 | 1 | lncRNA | ENSG0000C |
| 96590317 | 96985810 | 1 | lncRNA | ENSG0000C |
| 96642671 | 96643225 | -1 | processed_pseudoge | ENSG0000C |
| 97000274 | 97000711 | -1 | processed_pseudoge | ENSG0000C |
| 97086160 | 97086682 | -1 | processed_pseudoge | ENSG0000C |
| 97222644 | 97259987 | -1 | lncRNA | ENSG0000C |
| 97657464 | 97657582 | 1 | rRNA_pseudogene | ENSG0000C |
| 97878475 | 97959558 | -1 | lncRNA | ENSG0000C |
| 97908253 | 97908537 | -1 | processed_pseudoge | ENSG0000C |
| 98130239 | 98131198 | -1 | processed_pseudoge | ENSG0000C |
| 98565074 | 98566827 | -1 | processed_pseudoge | ENSG0000C |
| 98676391 | 98683735 | 1 | lncRNA | ENSG0000C |
| 98938912 | 98945079 | -1 | lncRNA | ENSG0000C |
| 99020949 | 100358885 | 1 | protein_coding | ENSG0000C |
| 99120176 | 99120490 | 1 | misc_RNA | ENSG0000C |
| 100336856 | 100337625 | 1 | processed_pseudoge | ENSG0000C |
| 100641975 | 100642273 | -1 | misc_RNA | ENSG0000C |
| 100666459 | 100666962 | 1 | processed_pseudoge | ENSG0000C |
| 100684162 | 100687955 | -1 | lncRNA | ENSG0000C |
| 100687288 | 100993941 | 1 | protein_coding | ENSG0000C |
| 100839630 | 100839946 | 1 | misc_RNA | ENSG0000C |
| 101029624 | 101129813 | -1 | protein_coding | ENSG0000C |
| 101129077 | 101209591 | 1 | lncRNA | ENSG0000C |
| 101208905 | 101209450 | 1 | transcribed_processe | ENSG0000C |
| 101451564 | 101872562 | -1 | protein_coding | ENSG0000C |
| 101519820 | 101519905 | -1 | miRNA | ENSG0000C |
| 101584295 | 101595156 | 1 | transcribed_processe | ENSG0000C |
| 101765935 | 101769961 | -1 | unprocessed_pseudc | ENSG0000C |
| 101890674 | 101916522 | -1 | protein_coding | ENSG0000C |
| 101915010 | 102001062 | 1 | protein_coding | ENSG0000C |
| 102047437 | 102084554 | 1 | protein_coding | ENSG0000C |
| 102057854 | 102057960 | 1 | rRNA | ENSG0000C |
| 102107886 | 102109842 | 1 | lncRNA | ENSG0000C |
| 102109827 | 102110457 | -1 | lncRNA | ENSG0000C |
| 102110447 | 102233424 | 1 | protein_coding | ENSG0000C |
| 102229851 | 102230922 | -1 | lncRNA | ENSG0000C |
| 102295060 | 102295806 | 1 | processed_pseudoge | ENSG0000C |
| 102306112 | 102306271 | 1 | processed_pseudoge | ENSG0000C |
| 102308557 | 102308797 | -1 | processed_pseudoge | ENSG0000C |

| | | | | |
|---|---|---|---|---|
| 102313722 | 102313829 | 1 | snRNA | ENSG00000 |
| 102317450 | 102339403 | 1 | protein_coding | ENSG00000 |
| 102347211 | 102378670 | 1 | protein_coding | ENSG00000 |
| 102396332 | 102470384 | -1 | protein_coding | ENSG00000 |
| 102452919 | 102462008 | 1 | lncRNA | ENSG00000 |
| 102467255 | 102498801 | 1 | lncRNA | ENSG00000 |
| 102520508 | 102530750 | -1 | protein_coding | ENSG00000 |
| 102576832 | 102625332 | -1 | protein_coding | ENSG00000 |
| 102606916 | 102628070 | 1 | lncRNA | ENSG00000 |
| 102681310 | 102683913 | 1 | lncRNA | ENSG00000 |
| 102691487 | 102705769 | -1 | protein_coding | ENSG00000 |
| 102711796 | 102727050 | -1 | protein_coding | ENSG00000 |
| 102746968 | 102836766 | 1 | transcribed_processe | ENSG00000 |
| 102751070 | 102754943 | -1 | transcribed_unproce | ENSG00000 |
| 102766801 | 102767927 | -1 | processed_pseudoge | ENSG00000 |
| 102770502 | 102780628 | -1 | protein_coding | ENSG00000 |
| 102789919 | 102798160 | -1 | protein_coding | ENSG00000 |
| 102806948 | 102807916 | -1 | processed_pseudoge | ENSG00000 |
| 102835801 | 102843609 | -1 | protein_coding | ENSG00000 |
| 102862736 | 102874982 | -1 | protein_coding | ENSG00000 |
| 102880492 | 102880846 | 1 | processed_pseudoge | ENSG00000 |
| 102903892 | 102903953 | -1 | snRNA | ENSG00000 |
| 102942995 | 102955732 | -1 | protein_coding | ENSG00000 |
| 102963460 | 102966628 | -1 | unprocessed_pseudo | ENSG00000 |
| 103045237 | 103045712 | 1 | processed_pseudoge | ENSG00000 |
| 103050686 | 103092194 | -1 | protein_coding | ENSG00000 |
| 103050687 | 103055799 | -1 | lncRNA | ENSG00000 |
| 103109410 | 103479863 | 1 | protein_coding | ENSG00000 |
| 103252217 | 103293705 | -1 | lncRNA | ENSG00000 |
| 103402013 | 103402586 | -1 | processed_pseudoge | ENSG00000 |
| 103402588 | 103403422 | -1 | processed_pseudoge | ENSG00000 |
| 103403512 | 103404167 | -1 | processed_pseudoge | ENSG00000 |
| 103404309 | 103405834 | -1 | processed_pseudoge | ENSG00000 |
| 103406213 | 103407241 | -1 | processed_pseudoge | ENSG00000 |
| 103407441 | 103408376 | -1 | processed_pseudoge | ENSG00000 |
| 103409580 | 103409657 | -1 | processed_pseudoge | ENSG00000 |
| 103675994 | 103895271 | -1 | lncRNA | ENSG00000 |
| 103849906 | 103849980 | 1 | miRNA | ENSG00000 |
| 103907189 | 104164379 | -1 | protein_coding | ENSG00000 |
| 103945548 | 103946546 | 1 | lncRNA | ENSG00000 |
| 104036640 | 104039196 | 1 | protein_coding | ENSG00000 |
| 104252591 | 104252651 | 1 | miRNA | ENSG00000 |
| 104445868 | 104609498 | -1 | lncRNA | ENSG00000 |
| 104682383 | 104682581 | -1 | processed_pseudoge | ENSG00000 |
| 104873264 | 104873417 | 1 | unprocessed_pseudo | ENSG00000 |
| 104885718 | 104898670 | -1 | polymorphic_pseudo | ENSG00000 |
| 104901549 | 104919073 | -1 | transcribed_unproce | ENSG00000 |
| 104942866 | 104969366 | -1 | protein_coding | ENSG00000 |
| 104975356 | 104976116 | -1 | unprocessed_pseudo | ENSG00000 |
| 104994235 | 105023168 | -1 | protein_coding | ENSG00000 |

| | | | | |
|---|---|---|---|---|
| 105025443 | 105035250 | -1 | protein_coding | ENSG00000 |
| 105041326 | 105101431 | -1 | protein_coding | ENSG00000 |
| 105055567 | 105055666 | -1 | unprocessed_pseudo | ENSG00000 |
| 105063345 | 105071541 | -1 | unprocessed_pseudo | ENSG00000 |
| 105092469 | 105101431 | -1 | protein_coding | ENSG00000 |
| 105122661 | 105123339 | -1 | unprocessed_pseudo | ENSG00000 |
| 105137714 | 105531697 | -1 | protein_coding | ENSG00000 |
| 105194440 | 105194946 | -1 | unprocessed_pseudo | ENSG00000 |
| 105246880 | 105247060 | -1 | unprocessed_pseudo | ENSG00000 |
| 105352670 | 105352956 | -1 | misc_RNA | ENSG00000 |
| 105609994 | 105982092 | 1 | protein_coding | ENSG00000 |
| 105706299 | 105707937 | 1 | processed_pseudoge | ENSG00000 |
| 105824634 | 105824770 | 1 | snRNA | ENSG00000 |
| 105974826 | 105974931 | -1 | snRNA | ENSG00000 |
| 105995185 | 105999213 | 1 | lncRNA | ENSG00000 |
| 105995623 | 106022403 | -1 | protein_coding | ENSG00000 |
| 106051098 | 106077459 | -1 | protein_coding | ENSG00000 |
| 106075501 | 106098699 | 1 | protein_coding | ENSG00000 |
| 106085990 | 106101975 | -1 | lncRNA | ENSG00000 |
| 106106398 | 106107356 | -1 | processed_pseudoge | ENSG00000 |
| 106112459 | 106132116 | -1 | lncRNA | ENSG00000 |
| 106241974 | 106260146 | -1 | lncRNA | ENSG00000 |
| 106250019 | 106264905 | 1 | transcribed_unproce | ENSG00000 |
| 106310045 | 106311791 | 1 | processed_pseudoge | ENSG00000 |
| 106465985 | 106469296 | 1 | lncRNA | ENSG00000 |
| 106674012 | 107018524 | -1 | protein_coding | ENSG00000 |
| 106824025 | 106824954 | 1 | unprocessed_pseudo | ENSG00000 |
| 106826392 | 106827890 | -1 | processed_pseudoge | ENSG00000 |
| 107176286 | 107177530 | 1 | processed_pseudoge | ENSG00000 |
| 107312132 | 107316271 | -1 | lncRNA | ENSG00000 |
| 107326345 | 107457844 | -1 | protein_coding | ENSG00000 |
| 107403404 | 107404589 | -1 | processed_pseudoge | ENSG00000 |
| 107502726 | 107565746 | -1 | protein_coding | ENSG00000 |
| 107585901 | 107586903 | -1 | processed_pseudoge | ENSG00000 |
| 107591091 | 107666779 | 1 | protein_coding | ENSG00000 |
| 107600379 | 107600868 | -1 | processed_pseudoge | ENSG00000 |
| 107642887 | 107643871 | -1 | unprocessed_pseudo | ENSG00000 |
| 107707378 | 107719693 | -1 | protein_coding | ENSG00000 |
| 107736009 | 107738718 | -1 | processed_pseudoge | ENSG00000 |
| 107790991 | 107928293 | -1 | protein_coding | ENSG00000 |
| 107908420 | 107908764 | 1 | processed_pseudoge | ENSG00000 |
| 107928448 | 107963482 | 1 | protein_coding | ENSG00000 |
| 108008678 | 108009282 | -1 | lncRNA | ENSG00000 |
| 108008898 | 108107761 | 1 | protein_coding | ENSG00000 |
| 108084913 | 108085014 | 1 | misc_RNA | ENSG00000 |
| 108105074 | 108121684 | -1 | lncRNA | ENSG00000 |
| 108116695 | 108147603 | 1 | protein_coding | ENSG00000 |
| 108142458 | 108150381 | -1 | lncRNA | ENSG00000 |
| 108157215 | 108222638 | -1 | protein_coding | ENSG00000 |
| 108222484 | 108369102 | 1 | protein_coding | ENSG00000 |

| | | | | |
|---|---|---|---|---|
| 108229503 | 108229604 | -1 | misc_RNA | ENSG0000C |
| 108308519 | 108467531 | -1 | protein_coding | ENSG0000C |
| 108472112 | 108498384 | -1 | protein_coding | ENSG0000C |
| 108505435 | 108593768 | -1 | protein_coding | ENSG0000C |
| 108665069 | 108940927 | 1 | protein_coding | ENSG0000C |
| 108690289 | 108690956 | -1 | processed_pseudoge | ENSG0000C |
| 108727741 | 108730104 | -1 | processed_pseudoge | ENSG0000C |
| 108822333 | 108822650 | -1 | processed_pseudoge | ENSG0000C |
| 108957718 | 108959797 | 1 | lncRNA | ENSG0000C |
| 109002465 | 109037967 | -1 | lncRNA | ENSG0000C |
| 109005517 | 109005623 | 1 | snRNA | ENSG0000C |
| 109120878 | 109120986 | -1 | rRNA_pseudogene | ENSG0000C |
| 109263494 | 109263564 | -1 | snoRNA | ENSG0000C |
| 109355085 | 109583907 | -1 | lncRNA | ENSG0000C |
| 109422190 | 109429167 | 1 | protein_coding | ENSG0000C |
| 109486968 | 109487424 | -1 | processed_pseudoge | ENSG0000C |
| 109637468 | 109638128 | -1 | processed_pseudoge | ENSG0000C |
| 109741625 | 109823893 | -1 | lncRNA | ENSG0000C |
| 109864295 | 110296712 | -1 | protein_coding | ENSG0000C |
| 109907004 | 109907741 | -1 | processed_pseudoge | ENSG0000C |
| 109946581 | 109975996 | 1 | lncRNA | ENSG0000C |
| 109982192 | 109983043 | 1 | processed_pseudoge | ENSG0000C |
| 109985243 | 109985660 | -1 | processed_pseudoge | ENSG0000C |
| 110049528 | 110050160 | 1 | processed_pseudoge | ENSG0000C |
| 110093392 | 110171841 | 1 | protein_coding | ENSG0000C |
| 110355130 | 110407613 | 1 | lncRNA | ENSG0000C |
| 110429948 | 110464884 | 1 | protein_coding | ENSG0000C |
| 110577042 | 110713189 | -1 | protein_coding | ENSG0000C |
| 110788026 | 110788976 | -1 | processed_pseudoge | ENSG0000C |
| 110876994 | 110877177 | -1 | processed_pseudoge | ENSG0000C |
| 111013452 | 111027636 | 1 | lncRNA | ENSG0000C |
| 111040098 | 111040208 | -1 | rRNA_pseudogene | ENSG0000C |
| 111089870 | 111090368 | -1 | lncRNA | ENSG0000C |
| 111091932 | 111179641 | -1 | lncRNA | ENSG0000C |
| 111097155 | 111106001 | 1 | lncRNA | ENSG0000C |
| 111105547 | 111105895 | -1 | processed_pseudoge | ENSG0000C |
| 111107060 | 111107750 | -1 | processed_pseudoge | ENSG0000C |
| 111245726 | 111286401 | 1 | protein_coding | ENSG0000C |
| 111290787 | 111305498 | -1 | lncRNA | ENSG0000C |
| 111298555 | 111308735 | 1 | protein_coding | ENSG0000C |
| 111320379 | 111323816 | 1 | lncRNA | ENSG0000C |
| 111347757 | 111347824 | 1 | miRNA | ENSG0000C |
| 111352255 | 111455630 | -1 | protein_coding | ENSG0000C |
| 111383092 | 111383219 | 1 | snRNA | ENSG0000C |
| 111414242 | 111418186 | -1 | lncRNA | ENSG0000C |
| 111448450 | 111475397 | 1 | transcribed_unitary_ | ENSG0000C |
| 111467526 | 111512354 | -1 | protein_coding | ENSG0000C |
| 111510600 | 111513888 | 1 | lncRNA | ENSG0000C |
| 111512938 | 111513021 | 1 | miRNA | ENSG0000C |
| 111513439 | 111513515 | 1 | miRNA | ENSG0000C |

| | | | | |
|---|---|---|---|---|
| 111514043 | 111526755 | -1 | lncRNA | ENSG00000 |
| 111514785 | 111537031 | 1 | protein_coding | ENSG00000 |
| 111540280 | 111561745 | 1 | protein_coding | ENSG00000 |
| 111602449 | 111730855 | 1 | protein_coding | ENSG00000 |
| 111670956 | 111672404 | -1 | processed_pseudoge | ENSG00000 |
| 111683111 | 111683424 | 1 | misc_RNA | ENSG00000 |
| 111726908 | 111766389 | -1 | protein_coding | ENSG00000 |
| 111768668 | 111778350 | -1 | lncRNA | ENSG00000 |
| 111782195 | 111871581 | -1 | protein_coding | ENSG00000 |
| 111786286 | 111879425 | -1 | protein_coding | ENSG00000 |
| 111817214 | 111829212 | -1 | lncRNA | ENSG00000 |
| 111864254 | 111864460 | 1 | processed_pseudoge | ENSG00000 |
| 111874056 | 111881243 | -1 | protein_coding | ENSG00000 |
| 111878935 | 111885975 | 1 | protein_coding | ENSG00000 |
| 111889199 | 111889474 | -1 | processed_pseudoge | ENSG00000 |
| 111908564 | 111923722 | -1 | protein_coding | ENSG00000 |
| 111912734 | 111914093 | 1 | protein_coding | ENSG00000 |
| 111912736 | 111926871 | 1 | protein_coding | ENSG00000 |
| 111918032 | 111926871 | 1 | protein_coding | ENSG00000 |
| 111927144 | 112022653 | 1 | protein_coding | ENSG00000 |
| 111928400 | 111928508 | 1 | rRNA_pseudogene | ENSG00000 |
| 111945639 | 111946892 | -1 | processed_pseudoge | ENSG00000 |
| 112015307 | 112016124 | 1 | lncRNA | ENSG00000 |
| 112024814 | 112064390 | 1 | protein_coding | ENSG00000 |
| 112029858 | 112030367 | -1 | processed_pseudoge | ENSG00000 |
| 112032499 | 112032598 | -1 | snRNA | ENSG00000 |
| 112036627 | 112036863 | -1 | processed_pseudoge | ENSG00000 |
| 112064010 | 112074274 | -1 | protein_coding | ENSG00000 |
| 112074086 | 112085150 | 1 | protein_coding | ENSG00000 |
| 112084800 | 112086798 | -1 | protein_coding | ENSG00000 |
| 112086824 | 112120016 | 1 | protein_coding | ENSG00000 |
| 112086903 | 112193805 | 1 | protein_coding | ENSG00000 |
| 112143253 | 112164096 | -1 | protein_coding | ENSG00000 |
| 112165197 | 112172606 | -1 | lncRNA | ENSG00000 |
| 112167372 | 112172556 | 1 | protein_coding | ENSG00000 |
| 112175510 | 112224699 | 1 | protein_coding | ENSG00000 |
| 112180773 | 112183191 | 1 | processed_pseudoge | ENSG00000 |
| 112208601 | 112208898 | 1 | processed_pseudoge | ENSG00000 |
| 112218326 | 112218720 | 1 | processed_pseudoge | ENSG00000 |
| 112226367 | 112269955 | 1 | protein_coding | ENSG00000 |
| 112235371 | 112236025 | -1 | processed_pseudoge | ENSG00000 |
| 112248153 | 112260860 | -1 | protein_coding | ENSG00000 |
| 112260265 | 112261396 | 1 | lncRNA | ENSG00000 |
| 112267290 | 112268371 | -1 | processed_pseudoge | ENSG00000 |
| 112270749 | 112362534 | 1 | lncRNA | ENSG00000 |
| 112352556 | 112352662 | -1 | snRNA | ENSG00000 |
| 112393118 | 112624567 | 1 | lncRNA | ENSG00000 |
| 112461468 | 112461920 | -1 | processed_pseudoge | ENSG00000 |
| 112534220 | 112555802 | -1 | lncRNA | ENSG00000 |
| 112602354 | 112602452 | -1 | snoRNA | ENSG00000 |

| | | | | |
|---|---|---|---|---|
| 112637342 | 112751131 | 1 | lncRNA | ENSG0000C |
| 112698772 | 112721151 | -1 | lncRNA | ENSG0000C |
| 112787304 | 112795854 | 1 | lncRNA | ENSG0000C |
| 112822806 | 112835124 | -1 | lncRNA | ENSG0000C |
| 112959279 | 112963460 | -1 | lncRNA | ENSG0000C |
| 112961247 | 113278436 | 1 | protein_coding | ENSG0000C |
| 112977353 | 112977420 | 1 | snRNA | ENSG0000C |
| 113265137 | 113274032 | -1 | lncRNA | ENSG0000C |
| 113278250 | 113314482 | -1 | lncRNA | ENSG0000C |
| 113314529 | 113383544 | 1 | protein_coding | ENSG0000C |
| 113368478 | 113369117 | 1 | lncRNA | ENSG0000C |
| 113387779 | 113400416 | 1 | protein_coding | ENSG0000C |
| 113405321 | 113412117 | 1 | lncRNA | ENSG0000C |
| 113409605 | 113475691 | -1 | protein_coding | ENSG0000C |
| 113450023 | 113450088 | -1 | miRNA | ENSG0000C |
| 113687547 | 113706373 | -1 | protein_coding | ENSG0000C |
| 113711720 | 113712398 | 1 | processed_pseudoge | ENSG0000C |
| 113733187 | 113773735 | -1 | protein_coding | ENSG0000C |
| 113751116 | 113751286 | -1 | processed_pseudoge | ENSG0000C |
| 113769660 | 113769994 | 1 | processed_pseudoge | ENSG0000C |
| 113770393 | 113771691 | -1 | lncRNA | ENSG0000C |
| 113779747 | 113780500 | 1 | protein_coding | ENSG0000C |
| 113789231 | 113791366 | 1 | transcribed_processe | ENSG0000C |
| 113789242 | 113790027 | 1 | processed_pseudoge | ENSG0000C |
| 113791466 | 113791752 | 1 | processed_pseudoge | ENSG0000C |
| 113797874 | 113875570 | -1 | protein_coding | ENSG0000C |
| 113818077 | 113822458 | 1 | lncRNA | ENSG0000C |
| 113820856 | 113821885 | -1 | processed_pseudoge | ENSG0000C |
| 113859346 | 113859449 | 1 | snRNA | ENSG0000C |
| 113904677 | 113949078 | 1 | protein_coding | ENSG0000C |
| 113974881 | 113990313 | 1 | protein_coding | ENSG0000C |
| 114059711 | 114256765 | 1 | protein_coding | ENSG0000C |
| 114159791 | 114161166 | -1 | lncRNA | ENSG0000C |
| 114210616 | 114356571 | -1 | lncRNA | ENSG0000C |
| 114257787 | 114313285 | 1 | protein_coding | ENSG0000C |
| 114343052 | 114396219 | -1 | lncRNA | ENSG0000C |
| 114391443 | 114400511 | -1 | protein_coding | ENSG0000C |
| 114400030 | 114414203 | 1 | protein_coding | ENSG0000C |
| 114400682 | 114443932 | 1 | protein_coding | ENSG0000C |
| 114439386 | 114450279 | 1 | protein_coding | ENSG0000C |
| 114453064 | 114454123 | 1 | processed_pseudoge | ENSG0000C |
| 114512706 | 114513431 | -1 | processed_pseudoge | ENSG0000C |
| 114517864 | 114518371 | 1 | processed_pseudoge | ENSG0000C |
| 114521645 | 114559895 | -1 | protein_coding | ENSG0000C |
| 114527129 | 114527232 | 1 | snoRNA | ENSG0000C |
| 114570591 | 114595786 | -1 | protein_coding | ENSG0000C |
| 114583253 | 114584692 | 1 | processed_pseudoge | ENSG0000C |
| 114678527 | 114707069 | 1 | protein_coding | ENSG0000C |
| 115128218 | 115128373 | -1 | snoRNA | ENSG0000C |
| 115169218 | 115504957 | -1 | protein_coding | ENSG0000C |

| | | | | |
|---|---|---|---|---|
| 115332529 | 115333418 | -1 | lncRNA | ENSG0000( |
| 115333577 | 115340262 | 1 | lncRNA | ENSG0000( |
| 115363629 | 115377931 | 1 | lncRNA | ENSG0000( |
| 115396756 | 115397658 | 1 | lncRNA | ENSG0000( |
| 115532322 | 115532953 | -1 | lncRNA | ENSG0000( |
| 115538869 | 115539384 | -1 | processed_pseudoge | ENSG0000( |
| 115578049 | 115578243 | 1 | processed_pseudoge | ENSG0000( |
| 115582297 | 115600339 | 1 | lncRNA | ENSG0000( |
| 115627082 | 115627185 | -1 | snoRNA | ENSG0000( |
| 115628065 | 115629743 | 1 | lncRNA | ENSG0000( |
| 115638563 | 115646962 | 1 | lncRNA | ENSG0000( |
| 115659168 | 115661417 | 1 | TEC | ENSG0000( |
| 115659658 | 115843146 | 1 | lncRNA | ENSG0000( |
| 115731942 | 115734742 | -1 | TEC | ENSG0000( |
| 115734848 | 115756926 | 1 | lncRNA | ENSG0000( |
| 115753889 | 115760646 | -1 | lncRNA | ENSG0000( |
| 115920248 | 115943181 | 1 | lncRNA | ENSG0000( |
| 115950196 | 115950801 | -1 | processed_pseudoge | ENSG0000( |
| 115951105 | 115951491 | -1 | processed_pseudoge | ENSG0000( |
| 116496899 | 116500630 | -1 | lncRNA | ENSG0000( |
| 116609995 | 116610094 | -1 | TEC | ENSG0000( |
| 116639422 | 116658295 | 1 | lncRNA | ENSG0000( |
| 116748170 | 116772987 | -1 | protein_coding | ENSG0000( |
| 116773389 | 116774205 | 1 | lncRNA | ENSG0000( |
| 116773799 | 116788039 | -1 | protein_coding | ENSG0000( |
| 116789367 | 116792420 | -1 | protein_coding | ENSG0000( |
| 116813204 | 116814003 | -1 | lncRNA | ENSG0000( |
| 116820645 | 116821541 | 1 | lncRNA | ENSG0000( |
| 116820700 | 116823304 | -1 | protein_coding | ENSG0000( |
| 116829706 | 116833072 | 1 | protein_coding | ENSG0000( |
| 116835751 | 116837950 | -1 | protein_coding | ENSG0000( |
| 116836117 | 116856018 | 1 | lncRNA | ENSG0000( |
| 116843402 | 117098437 | -1 | protein_coding | ENSG0000( |
| 117015897 | 117015992 | 1 | misc_RNA | ENSG0000( |
| 117035797 | 117036051 | -1 | processed_pseudoge | ENSG0000( |
| 117098987 | 117108170 | 1 | lncRNA | ENSG0000( |
| 117117794 | 117119181 | -1 | processed_pseudoge | ENSG0000( |
| 117135528 | 117138582 | 1 | unprocessed_pseudc | ENSG0000( |
| 117143891 | 117144191 | -1 | processed_pseudoge | ENSG0000( |
| 117144284 | 117176894 | 1 | protein_coding | ENSG0000( |
| 117178736 | 117197445 | 1 | protein_coding | ENSG0000( |
| 117199370 | 117207464 | 1 | protein_coding | ENSG0000( |
| 117204337 | 117232525 | -1 | protein_coding | ENSG0000( |
| 117204967 | 117210292 | 1 | TEC | ENSG0000( |
| 117232625 | 117286454 | 1 | protein_coding | ENSG0000( |
| 117263799 | 117263952 | 1 | scaRNA | ENSG0000( |
| 117285207 | 117316259 | -1 | protein_coding | ENSG0000( |
| 117288453 | 117293571 | 1 | lncRNA | ENSG0000( |
| 117297005 | 117297328 | -1 | lncRNA | ENSG0000( |
| 117314557 | 117413266 | 1 | protein_coding | ENSG0000( |

| | | | | |
|---|---|---|---|---|
| 117316362 | 117328038 | -1 | lncRNA | ENSG0000C |
| 117336256 | 117336631 | -1 | processed_pseudoge | ENSG0000C |
| 117427772 | 117817525 | -1 | protein_coding | ENSG0000C |
| 117611101 | 117612017 | -1 | TEC | ENSG0000C |
| 117668483 | 117668858 | 1 | lncRNA | ENSG0000C |
| 117800844 | 117828698 | -1 | protein_coding | ENSG0000C |
| 117818443 | 117821992 | 1 | lncRNA | ENSG0000C |
| 117820163 | 117876667 | -1 | protein_coding | ENSG0000C |
| 117833719 | 117838942 | 1 | lncRNA | ENSG0000C |
| 117836976 | 117877486 | -1 | protein_coding | ENSG0000C |
| 117900641 | 117929459 | -1 | protein_coding | ENSG0000C |
| 117986348 | 118003037 | 1 | protein_coding | ENSG0000C |
| 118003634 | 118088299 | -1 | protein_coding | ENSG0000C |
| 118067237 | 118067539 | 1 | misc_RNA | ENSG0000C |
| 118077012 | 118121890 | 1 | protein_coding | ENSG0000C |
| 118133377 | 118152888 | -1 | protein_coding | ENSG0000C |
| 118157849 | 118176639 | -1 | protein_coding | ENSG0000C |
| 118193725 | 118225094 | -1 | protein_coding | ENSG0000C |
| 118208932 | 118209211 | -1 | processed_pseudoge | ENSG0000C |
| 118226690 | 118252365 | -1 | protein_coding | ENSG0000C |
| 118253416 | 118264536 | -1 | protein_coding | ENSG0000C |
| 118264593 | 118266817 | 1 | TEC | ENSG0000C |
| 118304545 | 118316175 | 1 | protein_coding | ENSG0000C |
| 118322789 | 118323083 | -1 | processed_pseudoge | ENSG0000C |
| 118338954 | 118342744 | -1 | protein_coding | ENSG0000C |
| 118344344 | 118355161 | 1 | protein_coding | ENSG0000C |
| 118359585 | 118399211 | 1 | protein_coding | ENSG0000C |
| 118397095 | 118401895 | -1 | lncRNA | ENSG0000C |
| 118401602 | 118482737 | 1 | protein_coding | ENSG0000C |
| 118401606 | 118431496 | 1 | protein_coding | ENSG0000C |
| 118415977 | 118416521 | 1 | lncRNA | ENSG0000C |
| 118433121 | 118435206 | -1 | lncRNA | ENSG0000C |
| 118436490 | 118526832 | 1 | protein_coding | ENSG0000C |
| 118510273 | 118531094 | -1 | lncRNA | ENSG0000C |
| 118527472 | 118531197 | 1 | protein_coding | ENSG0000C |
| 118531041 | 118547280 | 1 | protein_coding | ENSG0000C |
| 118544528 | 118572970 | -1 | protein_coding | ENSG0000C |
| 118560690 | 118561580 | 1 | processed_pseudoge | ENSG0000C |
| 118572390 | 118603033 | 1 | protein_coding | ENSG0000C |
| 118593988 | 118594093 | 1 | snRNA | ENSG0000C |
| 118606440 | 118658031 | 1 | protein_coding | ENSG0000C |
| 118636620 | 118638097 | 1 | lncRNA | ENSG0000C |
| 118644000 | 118644079 | 1 | miRNA | ENSG0000C |
| 118657316 | 118679690 | -1 | protein_coding | ENSG0000C |
| 118688033 | 118690599 | -1 | unprocessed_pseudc | ENSG0000C |
| 118700427 | 118750263 | 1 | lncRNA | ENSG0000C |
| 118702342 | 118702447 | 1 | snRNA | ENSG0000C |
| 118721104 | 118729412 | -1 | lncRNA | ENSG0000C |
| 118747763 | 118791164 | -1 | protein_coding | ENSG0000C |
| 118791254 | 118793137 | 1 | lncRNA | ENSG0000C |

| | | | | |
|---|---|---|---|---|
| 118821895 | 118822084 | 1 | processed_pseudoge | ENSG00000( |
| 118833846 | 118834556 | -1 | processed_pseudoge | ENSG00000( |
| 118836010 | 118836111 | 1 | misc_RNA | ENSG00000( |
| 118883892 | 118897787 | 1 | protein_coding | ENSG00000( |
| 118885841 | 118887742 | -1 | lncRNA | ENSG00000( |
| 118893875 | 118925608 | -1 | protein_coding | ENSG00000( |
| 118910708 | 118910787 | 1 | miRNA | ENSG00000( |
| 118925164 | 118958559 | 1 | protein_coding | ENSG00000( |
| 118931216 | 118931490 | 1 | misc_RNA | ENSG00000( |
| 118970498 | 118970595 | 1 | misc_RNA | ENSG00000( |
| 118971712 | 118981287 | 1 | protein_coding | ENSG00000( |
| 118994334 | 118994630 | 1 | misc_RNA | ENSG00000( |
| 118994560 | 118994655 | 1 | misc_RNA | ENSG00000( |
| 118994824 | 118998004 | -1 | lncRNA | ENSG00000( |
| 118998138 | 119015793 | 1 | protein_coding | ENSG00000( |
| 119003012 | 119003446 | -1 | processed_pseudoge | ENSG00000( |
| 119003742 | 119004893 | -1 | lncRNA | ENSG00000( |
| 119005727 | 119005934 | -1 | processed_pseudoge | ENSG00000( |
| 119015712 | 119018691 | -1 | protein_coding | ENSG00000( |
| 119018763 | 119025454 | 1 | protein_coding | ENSG00000( |
| 119018944 | 119019012 | 1 | miRNA | ENSG00000( |
| 119024114 | 119030906 | -1 | protein_coding | ENSG00000( |
| 119044188 | 119057227 | -1 | protein_coding | ENSG00000( |
| 119044188 | 119045493 | 1 | lncRNA | ENSG00000( |
| 119065263 | 119065677 | -1 | lncRNA | ENSG00000( |
| 119067374 | 119067698 | -1 | lncRNA | ENSG00000( |
| 119067692 | 119081978 | 1 | protein_coding | ENSG00000( |
| 119084866 | 119093549 | 1 | protein_coding | ENSG00000( |
| 119093854 | 119095467 | -1 | protein_coding | ENSG00000( |
| 119096503 | 119108331 | -1 | protein_coding | ENSG00000( |
| 119102198 | 119118544 | 1 | protein_coding | ENSG00000( |
| 119121580 | 119136059 | 1 | protein_coding | ENSG00000( |
| 119149052 | 119162653 | 1 | protein_coding | ENSG00000( |
| 119166568 | 119184016 | 1 | protein_coding | ENSG00000( |
| 119185457 | 119190223 | 1 | protein_coding | ENSG00000( |
| 119189638 | 119196769 | -1 | protein_coding | ENSG00000( |
| 119206276 | 119313926 | 1 | protein_coding | ENSG00000( |
| 119289562 | 119289677 | -1 | snRNA | ENSG00000( |
| 119308529 | 119321521 | -1 | protein_coding | ENSG00000( |
| 119312950 | 119313012 | -1 | miRNA | ENSG00000( |
| 119323293 | 119323409 | -1 | snoRNA | ENSG00000( |
| 119334527 | 119337309 | 1 | protein_coding | ENSG00000( |
| 119336249 | 119337309 | -1 | lncRNA | ENSG00000( |
| 119338942 | 119346705 | -1 | protein_coding | ENSG00000( |
| 119338942 | 119340940 | -1 | protein_coding | ENSG00000( |
| 119355215 | 119381711 | -1 | protein_coding | ENSG00000( |
| 119364359 | 119526664 | 1 | lncRNA | ENSG00000( |
| 119372706 | 119381613 | 1 | lncRNA | ENSG00000( |
| 119402792 | 119404088 | 1 | processed_pseudoge | ENSG00000( |
| 119403963 | 119404227 | -1 | processed_pseudoge | ENSG00000( |

| | | | | |
|---|---|---|---|---|
| 119417378 | 119424985 | -1 | protein_coding | ENSG00000 |
| 119417951 | 119419114 | 1 | lncRNA | ENSG00000 |
| 119444991 | 119445519 | -1 | processed_pseudoge | ENSG00000 |
| 119602875 | 119604306 | -1 | processed_pseudoge | ENSG00000 |
| 119608423 | 119659284 | 1 | lncRNA | ENSG00000 |
| 119623408 | 119729200 | -1 | protein_coding | ENSG00000 |
| 119656311 | 119656416 | 1 | snRNA | ENSG00000 |
| 119709920 | 119713925 | 1 | lncRNA | ENSG00000 |
| 119729583 | 119739623 | 1 | lncRNA | ENSG00000 |
| 119821304 | 119822129 | -1 | processed_pseudoge | ENSG00000 |
| 119832966 | 119867682 | 1 | lncRNA | ENSG00000 |
| 119892700 | 119919996 | 1 | lncRNA | ENSG00000 |
| 119976111 | 120009588 | 1 | lncRNA | ENSG00000 |
| 119987952 | 119995037 | -1 | lncRNA | ENSG00000 |
| 120008044 | 120013620 | -1 | lncRNA | ENSG00000 |
| 120026753 | 120028341 | 1 | lncRNA | ENSG00000 |
| 120111286 | 120185529 | -1 | protein_coding | ENSG00000 |
| 120168977 | 120171679 | 1 | lncRNA | ENSG00000 |
| 120211032 | 120230334 | 1 | protein_coding | ENSG00000 |
| 120236640 | 120319945 | 1 | protein_coding | ENSG00000 |
| 120249759 | 120265932 | -1 | lncRNA | ENSG00000 |
| 120325129 | 120333682 | 1 | protein_coding | ENSG00000 |
| 120336413 | 120489937 | 1 | protein_coding | ENSG00000 |
| 120365666 | 120365963 | 1 | misc_RNA | ENSG00000 |
| 120511700 | 120988904 | 1 | protein_coding | ENSG00000 |
| 120619934 | 120620267 | -1 | processed_pseudoge | ENSG00000 |
| 120867933 | 120894797 | -1 | lncRNA | ENSG00000 |
| 120936102 | 120936654 | -1 | processed_pseudoge | ENSG00000 |
| 121024072 | 121090775 | 1 | protein_coding | ENSG00000 |
| 121024125 | 121113108 | 1 | protein_coding | ENSG00000 |
| 121101173 | 121191493 | 1 | protein_coding | ENSG00000 |
| 121238304 | 121238688 | 1 | processed_pseudoge | ENSG00000 |
| 121250270 | 121250978 | -1 | processed_pseudoge | ENSG00000 |
| 121292681 | 121313410 | 1 | protein_coding | ENSG00000 |
| 121361854 | 121362865 | -1 | processed_pseudoge | ENSG00000 |
| 121426645 | 121427416 | -1 | processed_pseudoge | ENSG00000 |
| 121447331 | 121453013 | -1 | lncRNA | ENSG00000 |
| 121452314 | 121633763 | 1 | protein_coding | ENSG00000 |
| 121674261 | 121928976 | 1 | lncRNA | ENSG00000 |
| 121979571 | 122025937 | -1 | lncRNA | ENSG00000 |
| 122004355 | 122004451 | 1 | snRNA | ENSG00000 |
| 122026093 | 122027667 | 1 | lncRNA | ENSG00000 |
| 122028325 | 122556721 | -1 | lncRNA | ENSG00000 |
| 122099757 | 122099844 | -1 | miRNA | ENSG00000 |
| 122115340 | 122116215 | -1 | protein_coding | ENSG00000 |
| 122146522 | 122146593 | -1 | miRNA | ENSG00000 |
| 122152229 | 122152308 | -1 | miRNA | ENSG00000 |
| 122294938 | 122301763 | -1 | lncRNA | ENSG00000 |
| 122295190 | 122308019 | 1 | lncRNA | ENSG00000 |
| 122449210 | 122449331 | 1 | snRNA | ENSG00000 |

| | | | | |
|---|---|---|---|---|
| 122451474 | 122451599 | 1 | snRNA | ENSG00000 |
| 122579205 | 122579307 | -1 | snRNA | ENSG00000 |
| 122613444 | 122614486 | -1 | processed_pseudoge | ENSG00000 |
| 122618851 | 122619274 | 1 | processed_pseudoge | ENSG00000 |
| 122655722 | 122814473 | 1 | protein_coding | ENSG00000 |
| 122679801 | 122681942 | -1 | lncRNA | ENSG00000 |
| 122726208 | 122726311 | -1 | snoRNA | ENSG00000 |
| 122751147 | 122851617 | -1 | lncRNA | ENSG00000 |
| 122838500 | 122872643 | 1 | protein_coding | ENSG00000 |
| 122882528 | 122959798 | 1 | protein_coding | ENSG00000 |
| 122927034 | 122927173 | 1 | snRNA | ENSG00000 |
| 122960327 | 122961086 | 1 | processed_pseudoge | ENSG00000 |
| 122977570 | 122981834 | -1 | protein_coding | ENSG00000 |
| 123017638 | 123018661 | 1 | processed_pseudoge | ENSG00000 |
| 123035867 | 123036224 | 1 | processed_pseudoge | ENSG00000 |
| 123049787 | 123050353 | -1 | processed_pseudoge | ENSG00000 |
| 123051161 | 123051401 | 1 | processed_pseudoge | ENSG00000 |
| 123057489 | 123063230 | -1 | protein_coding | ENSG00000 |
| 123058077 | 123058161 | -1 | snoRNA | ENSG00000 |
| 123058909 | 123058995 | -1 | snoRNA | ENSG00000 |
| 123059335 | 123059422 | -1 | snoRNA | ENSG00000 |
| 123062655 | 123084870 | 1 | lncRNA | ENSG00000 |
| 123069872 | 123195248 | -1 | protein_coding | ENSG00000 |
| 123135923 | 123136376 | -1 | processed_pseudoge | ENSG00000 |
| 123188802 | 123228502 | 1 | lncRNA | ENSG00000 |
| 123301390 | 123301523 | 1 | snoRNA | ENSG00000 |
| 123313657 | 123315340 | -1 | lncRNA | ENSG00000 |
| 123340174 | 123340714 | -1 | processed_pseudoge | ENSG00000 |
| 123358428 | 123627774 | 1 | protein_coding | ENSG00000 |
| 123381440 | 123381512 | -1 | miRNA | ENSG00000 |
| 123597946 | 123599452 | -1 | processed_pseudoge | ENSG00000 |
| 123629187 | 123655244 | -1 | protein_coding | ENSG00000 |
| 123668499 | 123675569 | 1 | lncRNA | ENSG00000 |
| 123723914 | 123741675 | -1 | protein_coding | ENSG00000 |
| 123753488 | 123754545 | -1 | protein_coding | ENSG00000 |
| 123763927 | 123764089 | 1 | snRNA | ENSG00000 |
| 123805335 | 123806387 | -1 | protein_coding | ENSG00000 |
| 123839035 | 123841930 | -1 | transcribed_unproce | ENSG00000 |
| 123861822 | 123862521 | -1 | unprocessed_pseudc | ENSG00000 |
| 123882920 | 123885670 | -1 | protein_coding | ENSG00000 |
| 123902167 | 123909229 | 1 | protein_coding | ENSG00000 |
| 123939543 | 123940637 | 1 | protein_coding | ENSG00000 |
| 123942785 | 123943873 | -1 | protein_coding | ENSG00000 |
| 123976661 | 123977781 | -1 | protein_coding | ENSG00000 |
| 123994163 | 123995161 | -1 | protein_coding | ENSG00000 |
| 124009655 | 124010012 | 1 | unprocessed_pseudc | ENSG00000 |
| 124012997 | 124018732 | 1 | protein_coding | ENSG00000 |
| 124023013 | 124023948 | 1 | protein_coding | ENSG00000 |
| 124026798 | 124030634 | 1 | protein_coding | ENSG00000 |
| 124036013 | 124041325 | -1 | protein_coding | ENSG00000 |

| | | | | |
|---|---|---|---|---|
| 124054793 | 124055707 | 1 | unprocessed_pseudc | ENSG00000( |
| 124093582 | 124094532 | -1 | unprocessed_pseudc | ENSG00000( |
| 124104965 | 124111092 | -1 | transcribed_unproce | ENSG00000( |
| 124115404 | 124147721 | 1 | protein_coding | ENSG00000( |
| 124158421 | 124159356 | 1 | unprocessed_pseudc | ENSG00000( |
| 124183345 | 124188780 | 1 | protein_coding | ENSG00000( |
| 124207183 | 124208112 | 1 | unprocessed_pseudc | ENSG00000( |
| 124211707 | 124215828 | 1 | transcribed_unproce | ENSG00000( |
| 124224732 | 124225639 | 1 | unprocessed_pseudc | ENSG00000( |
| 124238420 | 124239486 | 1 | unprocessed_pseudc | ENSG00000( |
| 124241095 | 124254364 | 1 | protein_coding | ENSG00000( |
| 124256376 | 124266218 | 1 | protein_coding | ENSG00000( |
| 124273970 | 124274581 | 1 | processed_pseudoge | ENSG00000( |
| 124302831 | 124315099 | -1 | protein_coding | ENSG00000( |
| 124319238 | 124320288 | -1 | protein_coding | ENSG00000( |
| 124324871 | 124325789 | -1 | unprocessed_pseudc | ENSG00000( |
| 124338783 | 124339662 | -1 | unprocessed_pseudc | ENSG00000( |
| 124357560 | 124358469 | -1 | unprocessed_pseudc | ENSG00000( |
| 124365465 | 124367298 | -1 | transcribed_unproce | ENSG00000( |
| 124376877 | 124377801 | -1 | unprocessed_pseudc | ENSG00000( |
| 124382394 | 124384462 | -1 | protein_coding | ENSG00000( |
| 124395534 | 124399024 | -1 | protein_coding | ENSG00000( |
| 124404234 | 124405102 | -1 | unprocessed_pseudc | ENSG00000( |
| 124406997 | 124414792 | -1 | lncRNA | ENSG00000( |
| 124423904 | 124424893 | -1 | protein_coding | ENSG00000( |
| 124437445 | 124445696 | -1 | protein_coding | ENSG00000( |
| 124460534 | 124461485 | 1 | unprocessed_pseudc | ENSG00000( |
| 124476963 | 124482870 | -1 | transcribed_unproce | ENSG00000( |
| 124516311 | 124517242 | -1 | unprocessed_pseudc | ENSG00000( |
| 124531625 | 124531910 | 1 | unprocessed_pseudc | ENSG00000( |
| 124538500 | 124539334 | -1 | unprocessed_pseudc | ENSG00000( |
| 124539635 | 124545333 | -1 | protein_coding | ENSG00000( |
| 124566660 | 124582942 | 1 | protein_coding | ENSG00000( |
| 124601584 | 124602238 | 1 | unprocessed_pseudc | ENSG00000( |
| 124611428 | 124620356 | 1 | protein_coding | ENSG00000( |
| 124622836 | 124635926 | 1 | protein_coding | ENSG00000( |
| 124633113 | 124695707 | -1 | protein_coding | ENSG00000( |
| 124636459 | 124636592 | 1 | rRNA_pseudogene | ENSG00000( |
| 124673798 | 124697518 | 1 | protein_coding | ENSG00000( |
| 124739942 | 124747210 | 1 | protein_coding | ENSG00000( |
| 124744634 | 124746337 | -1 | lncRNA | ENSG00000( |
| 124747474 | 124752255 | -1 | protein_coding | ENSG00000( |
| 124752583 | 124762290 | -1 | protein_coding | ENSG00000( |
| 124759129 | 124766119 | 1 | lncRNA | ENSG00000( |
| 124766498 | 124800673 | -1 | protein_coding | ENSG00000( |
| 124789240 | 124792818 | -1 | lncRNA | ENSG00000( |
| 124791228 | 124791660 | 1 | processed_pseudoge | ENSG00000( |
| 124800424 | 124834487 | 1 | lncRNA | ENSG00000( |
| 124807822 | 124808269 | -1 | processed_pseudoge | ENSG00000( |
| 124865432 | 124881471 | 1 | protein_coding | ENSG00000( |

| | | | | |
|---|---|---|---|---|
| 124883691 | 124898500 | -1 | protein_coding | ENSG00000 |
| 124883691 | 124887789 | 1 | lncRNA | ENSG00000 |
| 124891304 | 124892126 | 1 | lncRNA | ENSG00000 |
| 124919205 | 124936412 | -1 | protein_coding | ENSG00000 |
| 124919244 | 124920677 | 1 | protein_coding | ENSG00000 |
| 124949205 | 124954044 | -1 | lncRNA | ENSG00000 |
| 124954121 | 125041489 | 1 | protein_coding | ENSG00000 |
| 125063067 | 125090312 | 1 | protein_coding | ENSG00000 |
| 125096545 | 125111763 | -1 | protein_coding | ENSG00000 |
| 125113161 | 125126781 | 1 | transcribed_processe | ENSG00000 |
| 125132847 | 125139587 | -1 | lncRNA | ENSG00000 |
| 125158461 | 125164609 | -1 | lncRNA | ENSG00000 |
| 125164687 | 125433389 | 1 | protein_coding | ENSG00000 |
| 125188115 | 125191404 | 1 | lncRNA | ENSG00000 |
| 125258626 | 125266798 | -1 | lncRNA | ENSG00000 |
| 125277552 | 125277657 | 1 | snRNA | ENSG00000 |
| 125442881 | 125592568 | -1 | protein_coding | ENSG00000 |
| 125495214 | 125499528 | 1 | lncRNA | ENSG00000 |
| 125569280 | 125584684 | 1 | protein_coding | ENSG00000 |
| 125574719 | 125574818 | 1 | snRNA | ENSG00000 |
| 125591712 | 125625215 | 1 | protein_coding | ENSG00000 |
| 125625665 | 125676255 | 1 | protein_coding | ENSG00000 |
| 125649746 | 125649887 | 1 | snRNA | ENSG00000 |
| 125671522 | 125680874 | -1 | protein_coding | ENSG00000 |
| 125746279 | 125749867 | 1 | protein_coding | ENSG00000 |
| 125776113 | 125778828 | -1 | protein_coding | ENSG00000 |
| 125788127 | 125791600 | 1 | protein_coding | ENSG00000 |
| 125833316 | 125840072 | 1 | protein_coding | ENSG00000 |
| 125873031 | 125874528 | -1 | lncRNA | ENSG00000 |
| 125883614 | 125900648 | 1 | protein_coding | ENSG00000 |
| 125893485 | 125903221 | -1 | protein_coding | ENSG00000 |
| 125903247 | 125938916 | -1 | lncRNA | ENSG00000 |
| 125903348 | 125943702 | 1 | protein_coding | ENSG00000 |
| 125940496 | 125941193 | -1 | lncRNA | ENSG00000 |
| 125946056 | 125958647 | 1 | protein_coding | ENSG00000 |
| 125955796 | 126063335 | -1 | protein_coding | ENSG00000 |
| 126067539 | 126068601 | 1 | processed_pseudoge | ENSG00000 |
| 126100505 | 126102413 | -1 | lncRNA | ENSG00000 |
| 126121889 | 126128081 | -1 | lncRNA | ENSG00000 |
| 126160714 | 126176035 | 1 | lncRNA | ENSG00000 |
| 126179497 | 126179698 | -1 | processed_pseudoge | ENSG00000 |
| 126202096 | 126211692 | -1 | protein_coding | ENSG00000 |
| 126208611 | 126209027 | -1 | lncRNA | ENSG00000 |
| 126211724 | 126262984 | 1 | protein_coding | ENSG00000 |
| 126214260 | 126214478 | -1 | snRNA | ENSG00000 |
| 126245035 | 126245282 | 1 | misc_RNA | ENSG00000 |
| 126262938 | 126269144 | -1 | protein_coding | ENSG00000 |
| 126269055 | 126278131 | 1 | protein_coding | ENSG00000 |
| 126278257 | 126278577 | -1 | processed_pseudoge | ENSG00000 |
| 126283065 | 126298845 | 1 | protein_coding | ENSG00000 |

| | | | | |
|---|---|---|---|---|
| 126292922 | 126294254 | -1 | lncRNA | ENSG0000C |
| 126294298 | 126304318 | -1 | lncRNA | ENSG0000C |
| 126304060 | 126350005 | 1 | protein_coding | ENSG0000C |
| 126340889 | 126355587 | -1 | lncRNA | ENSG0000C |
| 126355640 | 126440344 | 1 | protein_coding | ENSG0000C |
| 126376773 | 126376893 | -1 | processed_pseudoge | ENSG0000C |
| 126423358 | 127003460 | -1 | protein_coding | ENSG0000C |
| 126543947 | 126610948 | 1 | lncRNA | ENSG0000C |
| 126652852 | 126682104 | 1 | lncRNA | ENSG0000C |
| 126920188 | 126940659 | -1 | lncRNA | ENSG0000C |
| 126934886 | 126937150 | -1 | TEC | ENSG0000C |
| 126940746 | 126945090 | 1 | lncRNA | ENSG0000C |
| 126988458 | 126988542 | -1 | miRNA | ENSG0000C |
| 126992452 | 126995458 | 1 | lncRNA | ENSG0000C |
| 127002910 | 127006058 | 1 | lncRNA | ENSG0000C |
| 127021736 | 127172363 | 1 | lncRNA | ENSG0000C |
| 127188885 | 127223449 | 1 | lncRNA | ENSG0000C |
| 127271029 | 127337033 | 1 | lncRNA | ENSG0000C |
| 127406028 | 127406332 | -1 | misc_RNA | ENSG0000C |
| 127407851 | 127408148 | 1 | misc_RNA | ENSG0000C |
| 127657568 | 127657877 | 1 | processed_pseudoge | ENSG0000C |
| 127940913 | 127941654 | -1 | processed_pseudoge | ENSG0000C |
| 128095758 | 128183671 | 1 | lncRNA | ENSG0000C |
| 128208749 | 128241441 | 1 | lncRNA | ENSG0000C |
| 128458761 | 128587558 | -1 | protein_coding | ENSG0000C |
| 128522390 | 128522449 | 1 | miRNA | ENSG0000C |
| 128526142 | 128530389 | 1 | lncRNA | ENSG0000C |
| 128614340 | 128686922 | -1 | lncRNA | ENSG0000C |
| 128686535 | 128813267 | 1 | protein_coding | ENSG0000C |
| 128691672 | 128696023 | -1 | lncRNA | ENSG0000C |
| 128836315 | 128867373 | -1 | protein_coding | ENSG0000C |
| 128880325 | 128880668 | -1 | processed_pseudoge | ENSG0000C |
| 128891356 | 128921163 | 1 | protein_coding | ENSG0000C |
| 128899565 | 128906069 | -1 | protein_coding | ENSG0000C |
| 128934731 | 128943399 | -1 | protein_coding | ENSG0000C |
| 128965060 | 129279324 | -1 | protein_coding | ENSG0000C |
| 128993237 | 128993340 | -1 | snRNA | ENSG0000C |
| 129030459 | 129030572 | 1 | misc_RNA | ENSG0000C |
| 129202966 | 129203070 | 1 | snRNA | ENSG0000C |
| 129295286 | 129296481 | -1 | processed_pseudoge | ENSG0000C |
| 129375848 | 129452279 | 1 | protein_coding | ENSG0000C |
| 129537869 | 129538114 | 1 | processed_pseudoge | ENSG0000C |
| 129591493 | 129612110 | 1 | lncRNA | ENSG0000C |
| 129612116 | 129617269 | -1 | lncRNA | ENSG0000C |
| 129761713 | 129814690 | -1 | lncRNA | ENSG0000C |
| 129788064 | 129788190 | -1 | processed_pseudoge | ENSG0000C |
| 129815848 | 129860003 | 1 | protein_coding | ENSG0000C |
| 129863636 | 129895590 | -1 | protein_coding | ENSG0000C |
| 129899706 | 130002835 | -1 | protein_coding | ENSG0000C |
| 130002938 | 130004356 | 1 | lncRNA | ENSG0000C |

| | | | | |
|---|---|---|---|---|
| 130008586 | 130009765 | 1 | processed_pseudoge | ENSG00000 |
| 130032866 | 130033220 | -1 | processed_pseudoge | ENSG00000 |
| 130040365 | 130041071 | 1 | processed_pseudoge | ENSG00000 |
| 130068147 | 130144811 | 1 | protein_coding | ENSG00000 |
| 130082495 | 130082781 | -1 | processed_pseudoge | ENSG00000 |
| 130114251 | 130114335 | -1 | snRNA | ENSG00000 |
| 130159782 | 130210362 | 1 | protein_coding | ENSG00000 |
| 130226677 | 130314917 | -1 | protein_coding | ENSG00000 |
| 130285023 | 130285833 | -1 | processed_pseudoge | ENSG00000 |
| 130295641 | 130296946 | 1 | processed_pseudoge | ENSG00000 |
| 130310180 | 130310476 | 1 | misc_RNA | ENSG00000 |
| 130314993 | 130403657 | 1 | lncRNA | ENSG00000 |
| 130404923 | 130428609 | -1 | protein_coding | ENSG00000 |
| 130448974 | 130476641 | 1 | protein_coding | ENSG00000 |
| 130544643 | 130544856 | -1 | processed_pseudoge | ENSG00000 |
| 130546799 | 130547429 | -1 | processed_pseudoge | ENSG00000 |
| 130666735 | 130666803 | 1 | miRNA | ENSG00000 |
| 130672956 | 130717352 | 1 | lncRNA | ENSG00000 |
| 130771413 | 130785533 | -1 | lncRNA | ENSG00000 |
| 130789231 | 130791025 | -1 | processed_pseudoge | ENSG00000 |
| 130844191 | 130865561 | -1 | lncRNA | ENSG00000 |
| 130866254 | 130870247 | -1 | lncRNA | ENSG00000 |
| 130875436 | 130916509 | -1 | protein_coding | ENSG00000 |
| 131004128 | 131004429 | -1 | misc_RNA | ENSG00000 |
| 131187022 | 131189753 | 1 | lncRNA | ENSG00000 |
| 131204999 | 131215163 | 1 | lncRNA | ENSG00000 |
| 131234538 | 131251816 | 1 | lncRNA | ENSG00000 |
| 131253422 | 131350917 | 1 | lncRNA | ENSG00000 |
| 131370478 | 132336822 | 1 | protein_coding | ENSG00000 |
| 131385249 | 131385363 | 1 | snRNA | ENSG00000 |
| 131502735 | 131540867 | -1 | lncRNA | ENSG00000 |
| 131542030 | 131581145 | -1 | lncRNA | ENSG00000 |
| 131622451 | 131663583 | -1 | lncRNA | ENSG00000 |
| 131770795 | 131771638 | -1 | processed_pseudoge | ENSG00000 |
| 131877680 | 131897108 | -1 | lncRNA | ENSG00000 |
| 131981096 | 131984661 | -1 | lncRNA | ENSG00000 |
| 132284302 | 132285081 | 1 | lncRNA | ENSG00000 |
| 132403360 | 132404060 | -1 | TEC | ENSG00000 |
| 132414977 | 133532519 | -1 | protein_coding | ENSG00000 |
| 132525809 | 132525887 | 1 | snRNA | ENSG00000 |
| 132771623 | 132773371 | -1 | TEC | ENSG00000 |
| 132859245 | 132860077 | -1 | lncRNA | ENSG00000 |
| 133064697 | 133068129 | 1 | TEC | ENSG00000 |
| 133297189 | 133302634 | -1 | TEC | ENSG00000 |
| 133360029 | 133366080 | -1 | lncRNA | ENSG00000 |
| 133532415 | 133549866 | 1 | lncRNA | ENSG00000 |
| 133783671 | 133810376 | -1 | lncRNA | ENSG00000 |
| 133840631 | 133845538 | -1 | protein_coding | ENSG00000 |
| 133864877 | 133865627 | -1 | processed_pseudoge | ENSG00000 |
| 133896438 | 133956985 | -1 | protein_coding | ENSG00000 |

| | | | | |
|---|---|---|---|---|
| 133898504 | 133898581 | -1 | miRNA | ENSG0000C |
| 134008215 | 134030167 | 1 | lncRNA | ENSG0000C |
| 134032216 | 134067936 | 1 | lncRNA | ENSG0000C |
| 134035832 | 134037107 | -1 | lncRNA | ENSG0000C |
| 134069071 | 134152001 | 1 | protein_coding | ENSG0000C |
| 134123828 | 134124329 | -1 | processed_pseudoge | ENSG0000C |
| 134150119 | 134225454 | -1 | protein_coding | ENSG0000C |
| 134173095 | 134173680 | 1 | processed_pseudoge | ENSG0000C |
| 134178824 | 134186166 | 1 | lncRNA | ENSG0000C |
| 134224671 | 134247788 | 1 | protein_coding | ENSG0000C |
| 134248279 | 134253370 | -1 | protein_coding | ENSG0000C |
| 134253495 | 134265855 | 1 | protein_coding | ENSG0000C |
| 134274245 | 134319564 | 1 | protein_coding | ENSG0000C |
| 134316782 | 134317088 | -1 | processed_pseudoge | ENSG0000C |
| 134331874 | 134378341 | 1 | protein_coding | ENSG0000C |
| 134378504 | 134412242 | -1 | protein_coding | ENSG0000C |
| 134412284 | 134505665 | 1 | lncRNA | ENSG0000C |
| 134563396 | 134573056 | 1 | lncRNA | ENSG0000C |
| 134655769 | 134661206 | 1 | lncRNA | ENSG0000C |
| 134671426 | 134672024 | -1 | lncRNA | ENSG0000C |
| 134685044 | 134702004 | -1 | lncRNA | ENSG0000C |
| 134712539 | 134716250 | 1 | lncRNA | ENSG0000C |
| 134735596 | 134763810 | 1 | lncRNA | ENSG0000C |
| 134945118 | 134980574 | -1 | lncRNA | ENSG0000C |
| 134950708 | 134951568 | 1 | lncRNA | ENSG0000C |
| 134985683 | 134986799 | 1 | TEC | ENSG0000C |
| 134994982 | 135007779 | -1 | lncRNA | ENSG0000C |
| 135061781 | 135075908 | -1 | lncRNA | ENSG0000C |
| 12310 | 13501 | 1 | unprocessed_pseudc | ENSG0000C |
| 14522 | 32015 | -1 | unprocessed_pseudc | ENSG0000C |
| 36602 | 38133 | -1 | lncRNA | ENSG0000C |
| 66767 | 178455 | 1 | protein_coding | ENSG0000C |
| 106524 | 111850 | -1 | lncRNA | ENSG0000C |
| 137411 | 149169 | -1 | lncRNA | ENSG0000C |
| 164664 | 166321 | -1 | lncRNA | ENSG0000C |
| 166856 | 182399 | -1 | lncRNA | ENSG0000C |
| 190077 | 214570 | -1 | protein_coding | ENSG0000C |
| 193036 | 194130 | 1 | lncRNA | ENSG0000C |
| 203642 | 205094 | 1 | lncRNA | ENSG0000C |
| 220621 | 262873 | -1 | protein_coding | ENSG0000C |
| 253442 | 257299 | -1 | lncRNA | ENSG0000C |
| 273954 | 277123 | -1 | lncRNA | ENSG0000C |
| 280057 | 389320 | -1 | protein_coding | ENSG0000C |
| 389273 | 442642 | 1 | protein_coding | ENSG0000C |
| 459939 | 563509 | 1 | protein_coding | ENSG0000C |
| 564296 | 663779 | -1 | protein_coding | ENSG0000C |
| 585865 | 586486 | -1 | lncRNA | ENSG0000C |
| 589530 | 589589 | -1 | snRNA | ENSG0000C |
| 590633 | 591269 | 1 | lncRNA | ENSG0000C |
| 630858 | 664196 | 1 | lncRNA | ENSG0000C |

| | | | | |
|---|---|---|---|---|
| 674801 | 684127 | -1 | lncRNA | ENSG00000 |
| 752579 | 911452 | 1 | protein_coding | ENSG00000 |
| 781133 | 781258 | 1 | snRNA | ENSG00000 |
| 865288 | 867139 | -1 | lncRNA | ENSG00000 |
| 911736 | 990053 | -1 | protein_coding | ENSG00000 |
| 974133 | 991190 | -1 | lncRNA | ENSG00000 |
| 990509 | 1495933 | 1 | protein_coding | ENSG00000 |
| 1151810 | 1153059 | -1 | processed_pseudoge | ENSG00000 |
| 1380060 | 1391502 | -1 | lncRNA | ENSG00000 |
| 1385833 | 1386987 | 1 | lncRNA | ENSG00000 |
| 1500525 | 1504424 | 1 | lncRNA | ENSG00000 |
| 1529891 | 1647212 | 1 | protein_coding | ENSG00000 |
| 1565993 | 1594581 | -1 | protein_coding | ENSG00000 |
| 1660315 | 1660380 | -1 | miRNA | ENSG00000 |
| 1688574 | 1788674 | 1 | protein_coding | ENSG00000 |
| 1695830 | 1698044 | -1 | processed_pseudoge | ENSG00000 |
| 1741197 | 1747099 | -1 | lncRNA | ENSG00000 |
| 1757231 | 1758011 | -1 | processed_pseudoge | ENSG00000 |
| 1791963 | 1918666 | -1 | protein_coding | ENSG00000 |
| 1800862 | 1807486 | 1 | lncRNA | ENSG00000 |
| 1820267 | 1836753 | 1 | protein_coding | ENSG00000 |
| 1917951 | 1922867 | 1 | lncRNA | ENSG00000 |
| 1929202 | 1936574 | -1 | lncRNA | ENSG00000 |
| 1946053 | 2004535 | -1 | protein_coding | ENSG00000 |
| 1970786 | 2697950 | 1 | protein_coding | ENSG00000 |
| 2004666 | 2011392 | 1 | lncRNA | ENSG00000 |
| 2018213 | 2020532 | 1 | lncRNA | ENSG00000 |
| 2048352 | 2049463 | 1 | lncRNA | ENSG00000 |
| 2217462 | 2217920 | -1 | lncRNA | ENSG00000 |
| 2219485 | 2223479 | -1 | lncRNA | ENSG00000 |
| 2269776 | 2288937 | 1 | lncRNA | ENSG00000 |
| 2332861 | 2335702 | -1 | lncRNA | ENSG00000 |
| 2440189 | 2457392 | -1 | lncRNA | ENSG00000 |
| 2603350 | 2607440 | -1 | lncRNA | ENSG00000 |
| 2659937 | 2660969 | 1 | processed_pseudoge | ENSG00000 |
| 2668500 | 2672220 | -1 | lncRNA | ENSG00000 |
| 2676001 | 2691200 | -1 | lncRNA | ENSG00000 |
| 2695765 | 2812902 | -1 | lncRNA | ENSG00000 |
| 2726400 | 2727050 | -1 | processed_pseudoge | ENSG00000 |
| 2740064 | 2746662 | 1 | lncRNA | ENSG00000 |
| 2752597 | 2753132 | -1 | processed_pseudoge | ENSG00000 |
| 2761200 | 2775658 | -1 | transcribed_unitary_ | ENSG00000 |
| 2781904 | 2782371 | 1 | processed_pseudoge | ENSG00000 |
| 2786931 | 2794295 | -1 | transcribed_processe | ENSG00000 |
| 2794970 | 2805423 | 1 | protein_coding | ENSG00000 |
| 2796877 | 2812902 | -1 | lncRNA | ENSG00000 |
| 2797136 | 2803938 | -1 | lncRNA | ENSG00000 |
| 2812622 | 2859791 | 1 | protein_coding | ENSG00000 |
| 2825348 | 2835544 | -1 | protein_coding | ENSG00000 |
| 2849046 | 2857039 | -1 | protein_coding | ENSG00000 |

| | | | | |
|---|---|---|---|---|
| 2857681 | 2877155 | -1 | protein_coding | ENSG00000[illegible] |
| 2876258 | 2889524 | 1 | protein_coding | ENSG00000[illegible] |
| 2877223 | 2941138 | 1 | protein_coding | ENSG00000[illegible] |
| 2885819 | 2886329 | 1 | lncRNA | ENSG00000[illegible] |
| 2954240 | 2954338 | -1 | snRNA | ENSG00000[illegible] |
| 2959330 | 3040676 | 1 | protein_coding | ENSG00000[illegible] |
| 3041437 | 3044950 | 1 | lncRNA | ENSG00000[illegible] |
| 3071532 | 3072145 | -1 | processed_pseudoge | ENSG00000[illegible] |
| 3077355 | 3286564 | 1 | protein_coding | ENSG00000[illegible] |
| 3124777 | 3125063 | 1 | misc_RNA | ENSG00000[illegible] |
| 3149797 | 3151476 | 1 | lncRNA | ENSG00000[illegible] |
| 3171657 | 3174056 | 1 | lncRNA | ENSG00000[illegible] |
| 3211663 | 3211917 | -1 | processed_pseudoge | ENSG00000[illegible] |
| 3296202 | 3366122 | -1 | transcribed_unproce | ENSG00000[illegible] |
| 3300698 | 3318677 | 1 | lncRNA | ENSG00000[illegible] |
| 3318718 | 3325343 | 1 | lncRNA | ENSG00000[illegible] |
| 3367833 | 3371346 | 1 | lncRNA | ENSG00000[illegible] |
| 3381349 | 3593973 | 1 | protein_coding | ENSG00000[illegible] |
| 3462718 | 3463586 | -1 | lncRNA | ENSG00000[illegible] |
| 3468950 | 3469296 | 1 | processed_pseudoge | ENSG00000[illegible] |
| 3483714 | 3492819 | -1 | lncRNA | ENSG00000[illegible] |
| 3606633 | 3764819 | -1 | protein_coding | ENSG00000[illegible] |
| 3610307 | 3612003 | -1 | TEC | ENSG00000[illegible] |
| 3726222 | 3726327 | -1 | snRNA | ENSG00000[illegible] |
| 3752518 | 3757842 | 1 | lncRNA | ENSG00000[illegible] |
| 3791047 | 3873448 | -1 | protein_coding | ENSG00000[illegible] |
| 3839459 | 3842525 | 1 | processed_pseudoge | ENSG00000[illegible] |
| 3871579 | 3910382 | 1 | lncRNA | ENSG00000[illegible] |
| 4012902 | 4026658 | 1 | lncRNA | ENSG00000[illegible] |
| 4097451 | 4099519 | 1 | processed_pseudoge | ENSG00000[illegible] |
| 4109661 | 4118132 | -1 | transcribed_processe | ENSG00000[illegible] |
| 4247981 | 4276252 | -1 | lncRNA | ENSG00000[illegible] |
| 4273762 | 4305353 | 1 | protein_coding | ENSG00000[illegible] |
| 4273762 | 4405490 | 1 | protein_coding | ENSG00000[illegible] |
| 4307763 | 4360028 | 1 | protein_coding | ENSG00000[illegible] |
| 4323724 | 4324811 | -1 | transcribed_processe | ENSG00000[illegible] |
| 4368227 | 4379712 | -1 | protein_coding | ENSG00000[illegible] |
| 4428155 | 4445614 | -1 | protein_coding | ENSG00000[illegible] |
| 4487735 | 4538508 | -1 | protein_coding | ENSG00000[illegible] |
| 4538798 | 4560048 | 1 | protein_coding | ENSG00000[illegible] |
| 4562204 | 4615302 | 1 | protein_coding | ENSG00000[illegible] |
| 4601991 | 4603420 | -1 | processed_pseudoge | ENSG00000[illegible] |
| 4604944 | 4662643 | 1 | protein_coding | ENSG00000[illegible] |
| 4615508 | 4649051 | -1 | protein_coding | ENSG00000[illegible] |
| 4635349 | 4636140 | -1 | processed_pseudoge | ENSG00000[illegible] |
| 4645496 | 4648027 | 1 | lncRNA | ENSG00000[illegible] |
| 4649095 | 4694317 | 1 | protein_coding | ENSG00000[illegible] |
| 4649101 | 4774184 | 1 | protein_coding | ENSG00000[illegible] |
| 4700417 | 4720102 | -1 | lncRNA | ENSG00000[illegible] |
| 4720400 | 4851927 | 1 | protein_coding | ENSG00000[illegible] |

| | | | | |
|---|---|---|---|---|
| 4804499 | 4830747 | -1 | lncRNA | ENSG00000( |
| 4809176 | 4813412 | 1 | protein_coding | ENSG00000( |
| 4838955 | 4842675 | 1 | lncRNA | ENSG00000( |
| 4857283 | 4859904 | -1 | lncRNA | ENSG00000( |
| 4906393 | 4909265 | -1 | lncRNA | ENSG00000( |
| 4909867 | 5026012 | 1 | lncRNA | ENSG00000( |
| 4909905 | 4918256 | 1 | protein_coding | ENSG00000( |
| 4960113 | 4963450 | 1 | lncRNA | ENSG00000( |
| 5032388 | 5032941 | -1 | processed_pseudoge | ENSG00000( |
| 5043879 | 5046788 | 1 | protein_coding | ENSG00000( |
| 5088002 | 5092742 | 1 | lncRNA | ENSG00000( |
| 5226196 | 5244199 | -1 | lncRNA | ENSG00000( |
| 5290480 | 5383653 | 1 | lncRNA | ENSG00000( |
| 5315961 | 5319347 | 1 | lncRNA | ENSG00000( |
| 5366048 | 5367774 | -1 | lncRNA | ENSG00000( |
| 5388589 | 5406651 | 1 | lncRNA | ENSG00000( |
| 5432108 | 5521536 | 1 | protein_coding | ENSG00000( |
| 5531869 | 5946232 | -1 | protein_coding | ENSG00000( |
| 5583797 | 5584085 | 1 | processed_pseudoge | ENSG00000( |
| 5948874 | 6124770 | -1 | protein_coding | ENSG00000( |
| 6095699 | 6095970 | -1 | misc_RNA | ENSG00000( |
| 6155035 | 6160719 | -1 | lncRNA | ENSG00000( |
| 6199715 | 6238271 | 1 | protein_coding | ENSG00000( |
| 6234345 | 6234455 | -1 | misc_RNA | ENSG00000( |
| 6270168 | 6270425 | -1 | processed_pseudoge | ENSG00000( |
| 6310436 | 6328506 | 1 | protein_coding | ENSG00000( |
| 6328757 | 6342114 | -1 | protein_coding | ENSG00000( |
| 6344554 | 6344837 | 1 | misc_RNA | ENSG00000( |
| 6346843 | 6377730 | -1 | protein_coding | ENSG00000( |
| 6375045 | 6391571 | 1 | protein_coding | ENSG00000( |
| 6393879 | 6396148 | 1 | lncRNA | ENSG00000( |
| 6409637 | 6410013 | -1 | processed_pseudoge | ENSG00000( |
| 6438498 | 6438852 | 1 | processed_pseudoge | ENSG00000( |
| 6439001 | 6451567 | -1 | lncRNA | ENSG00000( |
| 6444867 | 6451718 | 1 | protein_coding | ENSG00000( |
| 6451690 | 6466517 | 1 | protein_coding | ENSG00000( |
| 6462237 | 6470987 | -1 | protein_coding | ENSG00000( |
| 6466537 | 6467135 | 1 | lncRNA | ENSG00000( |
| 6477714 | 6480207 | 1 | processed_pseudoge | ENSG00000( |
| 6491886 | 6493841 | -1 | protein_coding | ENSG00000( |
| 6493356 | 6531955 | 1 | protein_coding | ENSG00000( |
| 6510222 | 6510551 | 1 | snoRNA | ENSG00000( |
| 6510275 | 6510522 | 1 | lncRNA | ENSG00000( |
| 6532290 | 6533498 | -1 | lncRNA | ENSG00000( |
| 6534512 | 6538374 | 1 | protein_coding | ENSG00000( |
| 6537794 | 6538370 | -1 | lncRNA | ENSG00000( |
| 6538375 | 6556083 | -1 | protein_coding | ENSG00000( |
| 6543504 | 6544931 | 1 | lncRNA | ENSG00000( |
| 6556863 | 6568691 | -1 | protein_coding | ENSG00000( |
| 6556886 | 6607367 | -1 | protein_coding | ENSG00000( |

| | | | | |
|---|---|---|---|---|
| 6570082 | 6614524 | -1 | protein_coding | ENSG0000C |
| 6578622 | 6584739 | 1 | lncRNA | ENSG0000C |
| 6581474 | 6581609 | -1 | scaRNA | ENSG0000C |
| 6618835 | 6635960 | -1 | protein_coding | ENSG0000C |
| 6638075 | 6647433 | -1 | protein_coding | ENSG0000C |
| 6650301 | 6663142 | -1 | protein_coding | ENSG0000C |
| 6663260 | 6672069 | 1 | lncRNA | ENSG0000C |
| 6666477 | 6689572 | -1 | protein_coding | ENSG0000C |
| 6693792 | 6700800 | -1 | protein_coding | ENSG0000C |
| 6709452 | 6709552 | 1 | snRNA | ENSG0000C |
| 6723233 | 6723687 | 1 | lncRNA | ENSG0000C |
| 6724014 | 6731875 | 1 | protein_coding | ENSG0000C |
| 6742985 | 6743641 | 1 | lncRNA | ENSG0000C |
| 6747996 | 6767475 | -1 | protein_coding | ENSG0000C |
| 6765516 | 6770952 | 1 | protein_coding | ENSG0000C |
| 6772512 | 6778455 | 1 | protein_coding | ENSG0000C |
| 6783854 | 6784161 | -1 | misc_RNA | ENSG0000C |
| 6786858 | 6820799 | 1 | protein_coding | ENSG0000C |
| 6821624 | 6829972 | 1 | protein_coding | ENSG0000C |
| 6828407 | 6839847 | 1 | protein_coding | ENSG0000C |
| 6839954 | 6847393 | 1 | protein_coding | ENSG0000C |
| 6844793 | 6852066 | -1 | protein_coding | ENSG0000C |
| 6852128 | 6866632 | 1 | protein_coding | ENSG0000C |
| 6867119 | 6870948 | 1 | protein_coding | ENSG0000C |
| 6870935 | 6889358 | -1 | protein_coding | ENSG0000C |
| 6873389 | 6884741 | 1 | transcribed_processe | ENSG0000C |
| 6873569 | 6914241 | 1 | protein_coding | ENSG0000C |
| 6884682 | 6885786 | 1 | transcribed_unproce | ENSG0000C |
| 6899752 | 6900212 | 1 | lncRNA | ENSG0000C |
| 6913745 | 6923698 | 1 | protein_coding | ENSG0000C |
| 6924463 | 6942321 | 1 | protein_coding | ENSG0000C |
| 6942978 | 6946003 | 1 | protein_coding | ENSG0000C |
| 6943508 | 6944604 | -1 | lncRNA | ENSG0000C |
| 6943816 | 6943878 | 1 | snRNA | ENSG0000C |
| 6946468 | 6961316 | 1 | protein_coding | ENSG0000C |
| 6963246 | 6964447 | 1 | lncRNA | ENSG0000C |
| 6963699 | 6963766 | 1 | miRNA | ENSG0000C |
| 6964097 | 6964191 | 1 | miRNA | ENSG0000C |
| 6964949 | 6965382 | 1 | lncRNA | ENSG0000C |
| 6965327 | 6970753 | -1 | protein_coding | ENSG0000C |
| 6967337 | 6967606 | -1 | snoRNA | ENSG0000C |
| 6970913 | 6997428 | 1 | protein_coding | ENSG0000C |
| 6976185 | 7018477 | -1 | protein_coding | ENSG0000C |
| 6988259 | 7071032 | 1 | protein_coding | ENSG0000C |
| 7032108 | 7032398 | 1 | processed_pseudoge | ENSG0000C |
| 7041156 | 7041641 | -1 | processed_pseudoge | ENSG0000C |
| 7080214 | 7092540 | -1 | protein_coding | ENSG0000C |
| 7089587 | 7109238 | -1 | protein_coding | ENSG0000C |
| 7108052 | 7122501 | 1 | lncRNA | ENSG0000C |
| 7115736 | 7128889 | -1 | protein_coding | ENSG0000C |

| | | | | |
|---|---|---|---|---|
| 7118785 | 7118891 | 1 | snRNA | ENSG0000C |
| 7129079 | 7131198 | -1 | lncRNA | ENSG0000C |
| 7129698 | 7158945 | 1 | protein_coding | ENSG0000C |
| 7146482 | 7147817 | -1 | lncRNA | ENSG0000C |
| 7166674 | 7189069 | -1 | lncRNA | ENSG0000C |
| 7188685 | 7218574 | 1 | protein_coding | ENSG0000C |
| 7237432 | 7240986 | 1 | processed_pseudoge | ENSG0000C |
| 7304284 | 7328724 | 1 | protein_coding | ENSG0000C |
| 7346685 | 7479897 | -1 | protein_coding | ENSG0000C |
| 7438780 | 7439990 | -1 | processed_pseudoge | ENSG0000C |
| 7470813 | 7503893 | -1 | protein_coding | ENSG0000C |
| 7566369 | 7567382 | 1 | processed_pseudoge | ENSG0000C |
| 7604350 | 7605214 | -1 | processed_pseudoge | ENSG0000C |
| 7606834 | 7607331 | 1 | processed_pseudoge | ENSG0000C |
| 7619872 | 7620260 | -1 | processed_pseudoge | ENSG0000C |
| 7620542 | 7621450 | 1 | processed_pseudoge | ENSG0000C |
| 7626103 | 7626486 | 1 | processed_pseudoge | ENSG0000C |
| 7637396 | 7647086 | -1 | unprocessed_pseudc | ENSG0000C |
| 7649400 | 7665908 | -1 | protein_coding | ENSG0000C |
| 7689784 | 7695775 | -1 | protein_coding | ENSG0000C |
| 7711433 | 7717559 | 1 | protein_coding | ENSG0000C |
| 7729415 | 7751605 | -1 | protein_coding | ENSG0000C |
| 7765216 | 7774121 | 1 | protein_coding | ENSG0000C |
| 7784403 | 7784511 | -1 | misc_RNA | ENSG0000C |
| 7787794 | 7799146 | 1 | protein_coding | ENSG0000C |
| 7800361 | 7800467 | -1 | misc_RNA | ENSG0000C |
| 7812512 | 7891148 | -1 | protein_coding | ENSG0000C |
| 7838914 | 7844395 | 1 | lncRNA | ENSG0000C |
| 7850717 | 7850818 | -1 | misc_RNA | ENSG0000C |
| 7856169 | 7856471 | 1 | misc_RNA | ENSG0000C |
| 7860437 | 7860764 | 1 | processed_pseudoge | ENSG0000C |
| 7870852 | 7871370 | 1 | processed_pseudoge | ENSG0000C |
| 7890801 | 7900140 | 1 | transcribed_unproce | ENSG0000C |
| 7896608 | 7896956 | -1 | processed_pseudoge | ENSG0000C |
| 7919230 | 7936187 | -1 | protein_coding | ENSG0000C |
| 7954216 | 7972759 | 1 | lncRNA | ENSG0000C |
| 7976259 | 7976360 | 1 | misc_RNA | ENSG0000C |
| 7994770 | 7995084 | -1 | unprocessed_pseudc | ENSG0000C |
| 8014093 | 8019007 | -1 | lncRNA | ENSG0000C |
| 8015024 | 8016406 | 1 | processed_pseudoge | ENSG0000C |
| 8017201 | 8017860 | -1 | processed_pseudoge | ENSG0000C |
| 8032716 | 8055517 | 1 | protein_coding | ENSG0000C |
| 8056844 | 8066359 | -1 | protein_coding | ENSG0000C |
| 8076939 | 8097881 | 1 | protein_coding | ENSG0000C |
| 8080381 | 8082163 | -1 | TEC | ENSG0000C |
| 8082269 | 8138548 | 1 | lncRNA | ENSG0000C |
| 8123632 | 8138607 | 1 | protein_coding | ENSG0000C |
| 8133772 | 8134849 | -1 | processed_pseudoge | ENSG0000C |
| 8138140 | 8175320 | 1 | protein_coding | ENSG0000C |
| 8141168 | 8142697 | -1 | processed_pseudoge | ENSG0000C |

| | | | | |
|---|---|---|---|---|
| 8157034 | 8188537 | 1 | protein_coding | ENSG00000 |
| 8180209 | 8216151 | 1 | lncRNA | ENSG00000 |
| 8199599 | 8218911 | -1 | transcribed_unproce | ENSG00000 |
| 8205984 | 8207397 | 1 | lncRNA | ENSG00000 |
| 8217758 | 8221115 | 1 | lncRNA | ENSG00000 |
| 8221260 | 8227618 | -1 | protein_coding | ENSG00000 |
| 8227734 | 8231764 | 1 | unprocessed_pseudc | ENSG00000 |
| 8232512 | 8242948 | -1 | transcribed_unproce | ENSG00000 |
| 8235415 | 8242564 | 1 | lncRNA | ENSG00000 |
| 8243147 | 8342048 | 1 | lncRNA | ENSG00000 |
| 8254527 | 8308855 | -1 | unprocessed_pseudc | ENSG00000 |
| 8282677 | 8286518 | -1 | lncRNA | ENSG00000 |
| 8295986 | 8396803 | -1 | lncRNA | ENSG00000 |
| 8320381 | 8369555 | 1 | transcribed_process | ENSG00000 |
| 8321876 | 8329614 | 1 | lncRNA | ENSG00000 |
| 8340182 | 8340397 | 1 | processed_pseudoge | ENSG00000 |
| 8359054 | 8359834 | -1 | processed_pseudoge | ENSG00000 |
| 8390270 | 8390488 | -1 | transcribed_unproce | ENSG00000 |
| 8396423 | 8396641 | -1 | transcribed_unproce | ENSG00000 |
| 8406833 | 8407195 | -1 | processed_pseudoge | ENSG00000 |
| 8414780 | 8415805 | -1 | unprocessed_pseudc | ENSG00000 |
| 8427640 | 8428610 | -1 | unprocessed_pseudc | ENSG00000 |
| 8437403 | 8438425 | -1 | unprocessed_pseudc | ENSG00000 |
| 8455962 | 8478330 | 1 | protein_coding | ENSG00000 |
| 8460618 | 8461232 | 1 | processed_pseudoge | ENSG00000 |
| 8484746 | 8484852 | 1 | snRNA | ENSG00000 |
| 8509475 | 8522366 | 1 | protein_coding | ENSG00000 |
| 8512374 | 8512726 | -1 | processed_pseudoge | ENSG00000 |
| 8533305 | 8540905 | -1 | protein_coding | ENSG00000 |
| 8548361 | 8567613 | -1 | lncRNA | ENSG00000 |
| 8592476 | 8592585 | 1 | misc_RNA | ENSG00000 |
| 8597509 | 8597640 | 1 | scaRNA | ENSG00000 |
| 8602170 | 8612867 | -1 | protein_coding | ENSG00000 |
| 8631545 | 8632259 | -1 | TEC | ENSG00000 |
| 8634367 | 8637608 | 1 | processed_pseudoge | ENSG00000 |
| 8637346 | 8662888 | -1 | protein_coding | ENSG00000 |
| 8664011 | 8669011 | 1 | lncRNA | ENSG00000 |
| 8678773 | 8679670 | 1 | processed_pseudoge | ENSG00000 |
| 8681600 | 8783095 | 1 | protein_coding | ENSG00000 |
| 8742428 | 8743072 | 1 | processed_pseudoge | ENSG00000 |
| 8776219 | 8830947 | -1 | lncRNA | ENSG00000 |
| 8788253 | 8795789 | 1 | lncRNA | ENSG00000 |
| 8819816 | 8820713 | -1 | lncRNA | ENSG00000 |
| 8822621 | 8887001 | 1 | protein_coding | ENSG00000 |
| 8858143 | 8915276 | -1 | lncRNA | ENSG00000 |
| 8913896 | 8941467 | 1 | protein_coding | ENSG00000 |
| 8940361 | 8949761 | -1 | protein_coding | ENSG00000 |
| 8950044 | 9010760 | 1 | protein_coding | ENSG00000 |
| 8987175 | 8996566 | -1 | lncRNA | ENSG00000 |
| 9027532 | 9029089 | -1 | processed_pseudoge | ENSG00000 |

| | | | | |
|---|---|---|---|---|
| 9036344 | 9037244 | 1 | processed_pseudoge | ENSG00000 |
| 9055586 | 9065070 | -1 | lncRNA | ENSG00000 |
| 9065163 | 9068689 | 1 | lncRNA | ENSG00000 |
| 9067664 | 9116229 | -1 | protein_coding | ENSG00000 |
| 9127783 | 9128645 | 1 | processed_pseudoge | ENSG00000 |
| 9135084 | 9135591 | 1 | processed_pseudoge | ENSG00000 |
| 9148840 | 9208395 | -1 | protein_coding | ENSG00000 |
| 9155881 | 9156372 | -1 | processed_pseudoge | ENSG00000 |
| 9228533 | 9275817 | -1 | transcribed_unproce | ENSG00000 |
| 9239467 | 9239551 | -1 | miRNA | ENSG00000 |
| 9239986 | 9240331 | -1 | processed_pseudoge | ENSG00000 |
| 9240003 | 9257960 | 1 | lncRNA | ENSG00000 |
| 9240406 | 9261050 | 1 | lncRNA | ENSG00000 |
| 9277235 | 9313241 | 1 | unprocessed_pseudo | ENSG00000 |
| 9286673 | 9286822 | -1 | snoRNA | ENSG00000 |
| 9314402 | 9316173 | -1 | processed_pseudoge | ENSG00000 |
| 9347028 | 9350947 | 1 | lncRNA | ENSG00000 |
| 9347056 | 9400387 | 1 | lncRNA | ENSG00000 |
| 9348030 | 9349268 | 1 | lncRNA | ENSG00000 |
| 9398355 | 9414851 | -1 | unprocessed_pseudo | ENSG00000 |
| 9415641 | 9416718 | 1 | unprocessed_pseudo | ENSG00000 |
| 9418673 | 9448229 | -1 | unprocessed_pseudo | ENSG00000 |
| 9445058 | 9445205 | 1 | snoRNA | ENSG00000 |
| 9448295 | 9658392 | 1 | lncRNA | ENSG00000 |
| 9480608 | 9576275 | 1 | transcribed_unproce | ENSG00000 |
| 9506347 | 9508440 | -1 | processed_pseudoge | ENSG00000 |
| 9594551 | 9607916 | -1 | protein_coding | ENSG00000 |
| 9624122 | 9624228 | -1 | snRNA | ENSG00000 |
| 9641723 | 9643011 | 1 | transcribed_processe | ENSG00000 |
| 9642872 | 9657070 | 1 | lncRNA | ENSG00000 |
| 9647014 | 9648084 | -1 | lncRNA | ENSG00000 |
| 9648047 | 9658412 | 1 | transcribed_unproce | ENSG00000 |
| 9658567 | 9662085 | 1 | lncRNA | ENSG00000 |
| 9664969 | 9699553 | 1 | protein_coding | ENSG00000 |
| 9695384 | 9696227 | 1 | processed_pseudoge | ENSG00000 |
| 9704077 | 9709350 | 1 | lncRNA | ENSG00000 |
| 9715860 | 9733299 | -1 | protein_coding | ENSG00000 |
| 9752486 | 9760901 | -1 | protein_coding | ENSG00000 |
| 9817350 | 9818025 | -1 | processed_pseudoge | ENSG00000 |
| 9827481 | 9845007 | 1 | protein_coding | ENSG00000 |
| 9852984 | 9870136 | -1 | protein_coding | ENSG00000 |
| 9867027 | 9869808 | 1 | lncRNA | ENSG00000 |
| 9881489 | 9895833 | 1 | protein_coding | ENSG00000 |
| 9898673 | 9932370 | -1 | protein_coding | ENSG00000 |
| 9936504 | 9943495 | -1 | lncRNA | ENSG00000 |
| 9948137 | 9953336 | -1 | lncRNA | ENSG00000 |
| 9951316 | 9995694 | 1 | protein_coding | ENSG00000 |
| 9985642 | 10013424 | -1 | protein_coding | ENSG00000 |
| 10010627 | 10018796 | 1 | protein_coding | ENSG00000 |
| 10015240 | 10030606 | -1 | lncRNA | ENSG00000 |

| | | | | |
|---|---|---|---|---|
| 10030678 | 10066031 | 1 | protein_coding | ENSG0000C |
| 10040848 | 10045711 | 1 | unprocessed_pseudc | ENSG0000C |
| 10069554 | 10111627 | -1 | protein_coding | ENSG0000C |
| 10084186 | 10084490 | 1 | misc_RNA | ENSG0000C |
| 10111738 | 10112471 | 1 | processed_pseudoge | ENSG0000C |
| 10116777 | 10130258 | -1 | protein_coding | ENSG0000C |
| 10158301 | 10172138 | -1 | protein_coding | ENSG0000C |
| 10170542 | 10191801 | 1 | protein_coding | ENSG0000C |
| 10212458 | 10223128 | 1 | protein_coding | ENSG0000C |
| 10214161 | 10214761 | -1 | lncRNA | ENSG0000C |
| 10226058 | 10329608 | 1 | protein_coding | ENSG0000C |
| 10332861 | 10338292 | -1 | lncRNA | ENSG0000C |
| 10358464 | 10361045 | -1 | lncRNA | ENSG0000C |
| 10363638 | 10398506 | 1 | lncRNA | ENSG0000C |
| 10372353 | 10391874 | -1 | protein_coding | ENSG0000C |
| 10372353 | 10410146 | -1 | protein_coding | ENSG0000C |
| 10407382 | 10409757 | -1 | protein_coding | ENSG0000C |
| 10412312 | 10420595 | -1 | protein_coding | ENSG0000C |
| 10412312 | 10436001 | -1 | protein_coding | ENSG0000C |
| 10426854 | 10442300 | -1 | protein_coding | ENSG0000C |
| 10442264 | 10454685 | -1 | protein_coding | ENSG0000C |
| 10505602 | 10523135 | 1 | protein_coding | ENSG0000C |
| 10551266 | 10551535 | -1 | processed_pseudoge | ENSG0000C |
| 10553362 | 10572688 | 1 | lncRNA | ENSG0000C |
| 10588063 | 10599669 | -1 | transcribed_unproce | ENSG0000C |
| 10604193 | 10613609 | -1 | protein_coding | ENSG0000C |
| 10618923 | 10674318 | -1 | protein_coding | ENSG0000C |
| 10699089 | 10723323 | -1 | protein_coding | ENSG0000C |
| 10750188 | 10777446 | 1 | lncRNA | ENSG0000C |
| 10770305 | 10770605 | 1 | processed_pseudoge | ENSG0000C |
| 10801532 | 10802627 | -1 | protein_coding | ENSG0000C |
| 10806051 | 10807293 | -1 | protein_coding | ENSG0000C |
| 10809094 | 10810168 | -1 | protein_coding | ENSG0000C |
| 10824960 | 11171608 | -1 | protein_coding | ENSG0000C |
| 10825317 | 10826358 | -1 | protein_coding | ENSG0000C |
| 10831074 | 10832016 | -1 | unprocessed_pseudc | ENSG0000C |
| 10845849 | 10849475 | -1 | protein_coding | ENSG0000C |
| 10845849 | 11171600 | -1 | protein_coding | ENSG0000C |
| 10894943 | 10896952 | -1 | transcribed_processe | ENSG0000C |
| 10907926 | 10909562 | -1 | protein_coding | ENSG0000C |
| 10929236 | 10934845 | 1 | protein_coding | ENSG0000C |
| 10937406 | 11171573 | -1 | protein_coding | ENSG0000C |
| 10964425 | 10965352 | -1 | unprocessed_pseudc | ENSG0000C |
| 10985913 | 10986912 | -1 | protein_coding | ENSG0000C |
| 10996495 | 10997875 | -1 | protein_coding | ENSG0000C |
| 11021619 | 11022620 | -1 | protein_coding | ENSG0000C |
| 11030387 | 11031407 | -1 | protein_coding | ENSG0000C |
| 11048332 | 11049256 | -1 | unprocessed_pseudc | ENSG0000C |
| 11061365 | 11062294 | -1 | protein_coding | ENSG0000C |
| 11076769 | 11079171 | -1 | transcribed_unproce | ENSG0000C |

| | | | | |
|---|---|---|---|---|
| 11091287 | 11092313 | -1 | protein_coding | ENSG00000( |
| 11097179 | 11097274 | -1 | processed_pseudoge | ENSG00000( |
| 11132958 | 11134644 | -1 | protein_coding | ENSG00000( |
| 11158785 | 11159694 | -1 | unprocessed_pseudo | ENSG00000( |
| 11166090 | 11171353 | -1 | lncRNA | ENSG00000( |
| 11171194 | 11176016 | 1 | protein_coding | ENSG00000( |
| 11185993 | 11186937 | -1 | protein_coding | ENSG00000( |
| 11196075 | 11196996 | -1 | unprocessed_pseudo | ENSG00000( |
| 11212219 | 11251389 | 1 | lncRNA | ENSG00000( |
| 11265924 | 11269805 | -1 | protein_coding | ENSG00000( |
| 11307083 | 11310435 | -1 | protein_coding | ENSG00000( |
| 11351823 | 11395566 | -1 | protein_coding | ENSG00000( |
| 11391540 | 11501041 | -1 | protein_coding | ENSG00000( |
| 11399381 | 11486708 | -1 | lncRNA | ENSG00000( |
| 11481383 | 11481656 | 1 | processed_pseudoge | ENSG00000( |
| 11500499 | 11501022 | 1 | processed_pseudoge | ENSG00000( |
| 11507162 | 11507408 | -1 | processed_pseudoge | ENSG00000( |
| 11512723 | 11514136 | 1 | processed_pseudoge | ENSG00000( |
| 11516425 | 11521686 | -1 | unprocessed_pseudo | ENSG00000( |
| 11541395 | 11555838 | -1 | lncRNA | ENSG00000( |
| 11546479 | 11546540 | 1 | snRNA | ENSG00000( |
| 11548030 | 11590369 | 1 | lncRNA | ENSG00000( |
| 11649854 | 11895402 | 1 | protein_coding | ENSG00000( |
| 12049844 | 12211084 | 1 | protein_coding | ENSG00000( |
| 12111163 | 12111489 | 1 | processed_pseudoge | ENSG00000( |
| 12111952 | 12112036 | 1 | miRNA | ENSG00000( |
| 12116025 | 12267044 | -1 | protein_coding | ENSG00000( |
| 12155262 | 12155772 | 1 | processed_pseudoge | ENSG00000( |
| 12226674 | 12226777 | 1 | snRNA | ENSG00000( |
| 12248991 | 12249427 | 1 | processed_pseudoge | ENSG00000( |
| 12289227 | 12289329 | -1 | snRNA | ENSG00000( |
| 12326056 | 12350242 | -1 | protein_coding | ENSG00000( |
| 12338776 | 12339254 | 1 | processed_pseudoge | ENSG00000( |
| 12355406 | 12357067 | -1 | lncRNA | ENSG00000( |
| 12357078 | 12471233 | 1 | protein_coding | ENSG00000( |
| 12452103 | 12452481 | -1 | processed_pseudoge | ENSG00000( |
| 12473282 | 12562863 | -1 | protein_coding | ENSG00000( |
| 12485353 | 12491581 | 1 | lncRNA | ENSG00000( |
| 12568201 | 12568776 | -1 | processed_pseudoge | ENSG00000( |
| 12611827 | 12645108 | 1 | protein_coding | ENSG00000( |
| 12614359 | 12615907 | -1 | processed_pseudoge | ENSG00000( |
| 12648939 | 12649713 | 1 | lncRNA | ENSG00000( |
| 12660890 | 12696207 | -1 | protein_coding | ENSG00000( |
| 12668982 | 12685075 | 1 | lncRNA | ENSG00000( |
| 12715058 | 12722369 | 1 | protein_coding | ENSG00000( |
| 12718973 | 12719521 | 1 | lncRNA | ENSG00000( |
| 12723297 | 12724011 | -1 | lncRNA | ENSG00000( |
| 12725917 | 12829975 | 1 | protein_coding | ENSG00000( |
| 12764649 | 12764743 | 1 | miRNA | ENSG00000( |
| 12771351 | 12771705 | -1 | processed_pseudoge | ENSG00000( |

| | | | | |
|---|---|---|---|---|
| 12813316 | 12829981 | 1 | protein_coding | ENSG0000C |
| 12840804 | 12841079 | -1 | processed_pseudoge | ENSG0000C |
| 12850982 | 12851740 | 1 | processed_pseudoge | ENSG0000C |
| 12875499 | 12876107 | 1 | processed_pseudoge | ENSG0000C |
| 12890782 | 12917937 | 1 | protein_coding | ENSG0000C |
| 12915829 | 12915918 | 1 | miRNA | ENSG0000C |
| 12927726 | 12984645 | 1 | lncRNA | ENSG0000C |
| 12940775 | 12952147 | -1 | protein_coding | ENSG0000C |
| 12962300 | 12963539 | 1 | lncRNA | ENSG0000C |
| 12972053 | 12972143 | 1 | snoRNA | ENSG0000C |
| 12974870 | 13000265 | -1 | protein_coding | ENSG0000C |
| 13000420 | 13004830 | 1 | transcribed_processe | ENSG0000C |
| 13000451 | 13040679 | 1 | lncRNA | ENSG0000C |
| 13021260 | 13021606 | -1 | processed_pseudoge | ENSG0000C |
| 13044381 | 13142521 | 1 | protein_coding | ENSG0000C |
| 13083532 | 13103683 | -1 | protein_coding | ENSG0000C |
| 13196723 | 13219941 | 1 | protein_coding | ENSG0000C |
| 13238582 | 13278376 | -1 | lncRNA | ENSG0000C |
| 13371089 | 13387167 | -1 | lncRNA | ENSG0000C |
| 13437942 | 13981957 | -1 | protein_coding | ENSG0000C |
| 13440884 | 13441001 | -1 | rRNA_pseudogene | ENSG0000C |
| 13459670 | 13459776 | -1 | snRNA | ENSG0000C |
| 13526854 | 13547582 | 1 | lncRNA | ENSG0000C |
| 13540231 | 13544540 | 1 | lncRNA | ENSG0000C |
| 13582031 | 13620077 | 1 | lncRNA | ENSG0000C |
| 13683542 | 13683886 | 1 | misc_RNA | ENSG0000C |
| 14169746 | 14169877 | 1 | processed_pseudoge | ENSG0000C |
| 14192002 | 14192284 | 1 | processed_pseudoge | ENSG0000C |
| 14216590 | 14221297 | 1 | lncRNA | ENSG0000C |
| 14237091 | 14237321 | 1 | processed_pseudoge | ENSG0000C |
| 14250232 | 14250341 | -1 | snRNA | ENSG0000C |
| 14254912 | 14255226 | 1 | processed_pseudoge | ENSG0000C |
| 14356842 | 14365503 | -1 | transcribed_processe | ENSG0000C |
| 14365632 | 14502935 | 1 | protein_coding | ENSG0000C |
| 14503661 | 14567883 | -1 | protein_coding | ENSG0000C |
| 14530124 | 14543575 | 1 | lncRNA | ENSG0000C |
| 14553702 | 14553970 | -1 | misc_RNA | ENSG0000C |
| 14567393 | 14624191 | 1 | lncRNA | ENSG0000C |
| 14612632 | 14696599 | -1 | protein_coding | ENSG0000C |
| 14665655 | 14757963 | 1 | lncRNA | ENSG0000C |
| 14762504 | 14767931 | -1 | lncRNA | ENSG0000C |
| 14767999 | 14771131 | -1 | protein_coding | ENSG0000C |
| 14774383 | 14778002 | 1 | protein_coding | ENSG0000C |
| 14784582 | 14803486 | -1 | protein_coding | ENSG0000C |
| 14803666 | 14906586 | 1 | protein_coding | ENSG0000C |
| 14804650 | 14814182 | -1 | protein_coding | ENSG0000C |
| 14805108 | 14808177 | 1 | lncRNA | ENSG0000C |
| 14825569 | 14843526 | -1 | protein_coding | ENSG0000C |
| 14841092 | 14844708 | 1 | lncRNA | ENSG0000C |
| 14880864 | 14885857 | -1 | protein_coding | ENSG0000C |

| | | | | |
|---|---|---|---|---|
| 14914039 | 14938537 | -1 | protein_coding | ENSG00000 |
| 14942031 | 14961728 | -1 | protein_coding | ENSG00000 |
| 14973042 | 14981865 | 1 | protein_coding | ENSG00000 |
| 15001787 | 15006999 | 1 | lncRNA | ENSG00000 |
| 15107783 | 15348675 | -1 | protein_coding | ENSG00000 |
| 15112363 | 15114698 | -1 | lncRNA | ENSG00000 |
| 15151923 | 15155283 | 1 | lncRNA | ENSG00000 |
| 15255213 | 15255823 | -1 | processed_pseudoge | ENSG00000 |
| 15322508 | 15598331 | 1 | protein_coding | ENSG00000 |
| 15439527 | 15440401 | 1 | processed_pseudoge | ENSG00000 |
| 15620134 | 15882329 | -1 | protein_coding | ENSG00000 |
| 15671707 | 15671810 | -1 | snRNA | ENSG00000 |
| 15780068 | 15782120 | 1 | lncRNA | ENSG00000 |
| 15803164 | 15804158 | 1 | processed_pseudoge | ENSG00000 |
| 15882387 | 15903478 | 1 | protein_coding | ENSG00000 |
| 15911302 | 16037381 | 1 | protein_coding | ENSG00000 |
| 15971890 | 15972609 | -1 | processed_pseudoge | ENSG00000 |
| 16188485 | 16277685 | -1 | protein_coding | ENSG00000 |
| 16347142 | 16609259 | 1 | protein_coding | ENSG00000 |
| 16399274 | 16402406 | -1 | processed_pseudoge | ENSG00000 |
| 16420653 | 16447217 | 1 | lncRNA | ENSG00000 |
| 16513445 | 16514371 | 1 | processed_pseudoge | ENSG00000 |
| 16548372 | 16610594 | -1 | protein_coding | ENSG00000 |
| 16567411 | 16573940 | 1 | lncRNA | ENSG00000 |
| 16661766 | 16788375 | 1 | lncRNA | ENSG00000 |
| 16693800 | 16695977 | 1 | lncRNA | ENSG00000 |
| 16989126 | 16989936 | 1 | processed_pseudoge | ENSG00000 |
| 16990670 | 16992083 | 1 | processed_pseudoge | ENSG00000 |
| 17066368 | 17075631 | 1 | lncRNA | ENSG00000 |
| 17071398 | 17071505 | -1 | snRNA | ENSG00000 |
| 17083531 | 17167790 | -1 | lncRNA | ENSG00000 |
| 17121716 | 17122460 | 1 | processed_pseudoge | ENSG00000 |
| 17383352 | 17384647 | -1 | processed_pseudoge | ENSG00000 |
| 17400399 | 17400914 | 1 | processed_pseudoge | ENSG00000 |
| 17479446 | 17709867 | 1 | lncRNA | ENSG00000 |
| 17509353 | 17590049 | -1 | lncRNA | ENSG00000 |
| 17673299 | 17673394 | 1 | miRNA | ENSG00000 |
| 17710934 | 17711046 | 1 | misc_RNA | ENSG00000 |
| 18080869 | 18320107 | -1 | protein_coding | ENSG00000 |
| 18247614 | 18648416 | 1 | protein_coding | ENSG00000 |
| 18358443 | 18358967 | -1 | processed_pseudoge | ENSG00000 |
| 18424663 | 18425140 | 1 | processed_pseudoge | ENSG00000 |
| 18493051 | 18493723 | 1 | processed_pseudoge | ENSG00000 |
| 18683169 | 18738100 | -1 | protein_coding | ENSG00000 |
| 18692762 | 18694096 | 1 | processed_pseudoge | ENSG00000 |
| 18714795 | 18737878 | 1 | lncRNA | ENSG00000 |
| 18738119 | 18739188 | 1 | protein_coding | ENSG00000 |
| 19089151 | 19089888 | 1 | processed_pseudoge | ENSG00000 |
| 19107854 | 19108183 | -1 | processed_pseudoge | ENSG00000 |
| 19129752 | 19376400 | 1 | protein_coding | ENSG00000 |

| | | | | |
|---|---|---|---|---|
| 19147074 | 19154659 | 1 | lncRNA | ENSG0000( |
| 19176288 | 19176589 | -1 | misc_RNA | ENSG0000( |
| 19303480 | 19303775 | 1 | misc_RNA | ENSG0000( |
| 19404045 | 19720801 | 1 | protein_coding | ENSG0000( |
| 19413153 | 19413461 | 1 | processed_pseudoge | ENSG0000( |
| 19418927 | 19419161 | -1 | processed_pseudoge | ENSG0000( |
| 19445949 | 19446055 | -1 | snRNA | ENSG0000( |
| 19456244 | 19457648 | -1 | processed_pseudoge | ENSG0000( |
| 19508904 | 19509245 | 1 | processed_pseudoge | ENSG0000( |
| 19582317 | 19583274 | 1 | lncRNA | ENSG0000( |
| 19724435 | 19724599 | 1 | snRNA | ENSG0000( |
| 19775451 | 20012261 | 1 | lncRNA | ENSG0000( |
| 19941521 | 19943018 | 1 | processed_pseudoge | ENSG0000( |
| 20014780 | 20098868 | 1 | lncRNA | ENSG0000( |
| 20104590 | 20109893 | -1 | lncRNA | ENSG0000( |
| 20120980 | 20136163 | -1 | lncRNA | ENSG0000( |
| 20361732 | 20370262 | -1 | lncRNA | ENSG0000( |
| 20369245 | 20684381 | 1 | protein_coding | ENSG0000( |
| 20449655 | 20450083 | -1 | processed_pseudoge | ENSG0000( |
| 20551590 | 20553012 | 1 | processed_pseudoge | ENSG0000( |
| 20695355 | 20753386 | 1 | protein_coding | ENSG0000( |
| 20810702 | 20916911 | 1 | protein_coding | ENSG0000( |
| 20815674 | 21090245 | 1 | protein_coding | ENSG0000( |
| 20845065 | 20845260 | 1 | processed_pseudoge | ENSG0000( |
| 20962768 | 21092745 | 1 | protein_coding | ENSG0000( |
| 21047179 | 21176895 | 1 | protein_coding | ENSG0000( |
| 21131194 | 21239796 | 1 | protein_coding | ENSG0000( |
| 21264600 | 21419594 | -1 | protein_coding | ENSG0000( |
| 21354959 | 21379980 | 1 | protein_coding | ENSG0000( |
| 21437615 | 21471250 | 1 | protein_coding | ENSG0000( |
| 21447545 | 21448314 | -1 | processed_pseudoge | ENSG0000( |
| 21468910 | 21501669 | -1 | protein_coding | ENSG0000( |
| 21501781 | 21518408 | 1 | protein_coding | ENSG0000( |
| 21526296 | 21541249 | 1 | protein_coding | ENSG0000( |
| 21536107 | 21604847 | -1 | protein_coding | ENSG0000( |
| 21536203 | 21657798 | -1 | protein_coding | ENSG0000( |
| 21635342 | 21757857 | -1 | protein_coding | ENSG0000( |
| 21662313 | 21760032 | 1 | lncRNA | ENSG0000( |
| 21759980 | 21766705 | 1 | lncRNA | ENSG0000( |
| 21764955 | 21775600 | -1 | protein_coding | ENSG0000( |
| 21797401 | 21942529 | -1 | protein_coding | ENSG0000( |
| 21827210 | 21887957 | 1 | lncRNA | ENSG0000( |
| 21828360 | 21828564 | -1 | lncRNA | ENSG0000( |
| 22008348 | 22008880 | 1 | processed_pseudoge | ENSG0000( |
| 22046218 | 22065674 | 1 | protein_coding | ENSG0000( |
| 22063773 | 22437041 | -1 | protein_coding | ENSG0000( |
| 22104491 | 22105320 | -1 | lncRNA | ENSG0000( |
| 22116629 | 22134851 | -1 | unitary_pseudogene | ENSG0000( |
| 22195469 | 22195629 | -1 | snRNA | ENSG0000( |
| 22395088 | 22398075 | -1 | lncRNA | ENSG0000( |

| | | | | |
|---|---|---|---|---|
| 22409237 | 22618868 | 1 | lncRNA | ENSG0000C |
| 22448583 | 22544546 | -1 | protein_coding | ENSG0000C |
| 22460519 | 22463914 | -1 | lncRNA | ENSG0000C |
| 22573425 | 22575105 | 1 | processed_pseudoge | ENSG0000C |
| 22596228 | 22596551 | 1 | processed_pseudoge | ENSG0000C |
| 22609228 | 22625015 | -1 | lncRNA | ENSG0000C |
| 22625075 | 22690665 | 1 | protein_coding | ENSG0000C |
| 22699859 | 23188807 | 1 | lncRNA | ENSG0000C |
| 22722229 | 22722469 | 1 | processed_pseudoge | ENSG0000C |
| 22742582 | 22743091 | 1 | lncRNA | ENSG0000C |
| 23108458 | 23108493 | -1 | processed_pseudoge | ENSG0000C |
| 23175648 | 23191587 | 1 | lncRNA | ENSG0000C |
| 23181334 | 23251499 | -1 | lncRNA | ENSG0000C |
| 23529504 | 24562544 | -1 | protein_coding | ENSG0000C |
| 23637973 | 23638487 | 1 | lncRNA | ENSG0000C |
| 24212421 | 24212495 | 1 | miRNA | ENSG0000C |
| 24223233 | 24260056 | 1 | lncRNA | ENSG0000C |
| 24566964 | 24584168 | -1 | lncRNA | ENSG0000C |
| 24583561 | 24584549 | 1 | processed_pseudoge | ENSG0000C |
| 24642687 | 24643166 | 1 | processed_pseudoge | ENSG0000C |
| 24704503 | 24775565 | 1 | lncRNA | ENSG0000C |
| 24706788 | 24707083 | 1 | misc_RNA | ENSG0000C |
| 24810024 | 24949101 | -1 | protein_coding | ENSG0000C |
| 24830421 | 24836558 | 1 | lncRNA | ENSG0000C |
| 24902309 | 24904681 | 1 | TEC | ENSG0000C |
| 24949163 | 24960158 | 1 | lncRNA | ENSG0000C |
| 24967127 | 24967504 | -1 | processed_pseudoge | ENSG0000C |
| 24993424 | 25006403 | -1 | lncRNA | ENSG0000C |
| 25004342 | 25108334 | 1 | protein_coding | ENSG0000C |
| 25092715 | 25093900 | -1 | processed_pseudoge | ENSG0000C |
| 25096868 | 25100980 | -1 | lncRNA | ENSG0000C |
| 25103124 | 25103869 | -1 | lncRNA | ENSG0000C |
| 25108289 | 25195162 | -1 | protein_coding | ENSG0000C |
| 25195216 | 25209645 | 1 | protein_coding | ENSG0000C |
| 25205246 | 25250936 | -1 | protein_coding | ENSG0000C |
| 25210652 | 25211233 | 1 | lncRNA | ENSG0000C |
| 25225103 | 25225665 | 1 | lncRNA | ENSG0000C |
| 25385670 | 25386241 | -1 | lncRNA | ENSG0000C |
| 25404288 | 25404428 | -1 | snRNA | ENSG0000C |
| 25409293 | 25409445 | -1 | processed_pseudoge | ENSG0000C |
| 25409307 | 25648579 | -1 | protein_coding | ENSG0000C |
| 25423600 | 25424942 | 1 | TEC | ENSG0000C |
| 25435581 | 25436238 | -1 | processed_pseudoge | ENSG0000C |
| 25440875 | 25441052 | -1 | processed_pseudoge | ENSG0000C |
| 25585994 | 25592928 | -1 | lncRNA | ENSG0000C |
| 25779287 | 25779569 | -1 | misc_RNA | ENSG0000C |
| 25782706 | 25814171 | -1 | lncRNA | ENSG0000C |
| 25803298 | 25804526 | 1 | processed_pseudoge | ENSG0000C |
| 25831416 | 25834182 | -1 | lncRNA | ENSG0000C |
| 25874020 | 25874079 | -1 | miRNA | ENSG0000C |

| | | | | |
|---|---|---|---|---|
| 25882955 | 25885855 | -1 | lncRNA | ENSG00000( |
| 25936683 | 25959765 | -1 | lncRNA | ENSG00000( |
| 25959029 | 26079892 | 1 | protein_coding | ENSG00000( |
| 26120030 | 26125037 | -1 | protein_coding | ENSG00000( |
| 26121991 | 26299290 | 1 | protein_coding | ENSG00000( |
| 26125155 | 26126617 | -1 | lncRNA | ENSG00000( |
| 26139249 | 26167741 | -1 | lncRNA | ENSG00000( |
| 26211164 | 26335856 | -1 | lncRNA | ENSG00000( |
| 26229548 | 26319720 | -1 | lncRNA | ENSG00000( |
| 26273329 | 26273900 | -1 | lncRNA | ENSG00000( |
| 26301098 | 26317813 | 1 | lncRNA | ENSG00000( |
| 26318953 | 26421462 | 1 | lncRNA | ENSG00000( |
| 26335352 | 26833194 | -1 | protein_coding | ENSG00000( |
| 26335864 | 26336950 | -1 | lncRNA | ENSG00000( |
| 26373598 | 26373733 | -1 | rRNA_pseudogene | ENSG00000( |
| 26623369 | 26649479 | 1 | lncRNA | ENSG00000( |
| 26800843 | 26801302 | 1 | processed_pseudoge | ENSG00000( |
| 26905181 | 26938326 | -1 | protein_coding | ENSG00000( |
| 26938470 | 26966650 | 1 | protein_coding | ENSG00000( |
| 26971579 | 27014434 | -1 | protein_coding | ENSG00000( |
| 26971586 | 26979582 | 1 | lncRNA | ENSG00000( |
| 27022546 | 27066343 | 1 | protein_coding | ENSG00000( |
| 27037100 | 27038960 | 1 | lncRNA | ENSG00000( |
| 27081058 | 27082514 | -1 | protein_coding | ENSG00000( |
| 27093923 | 27094223 | -1 | processed_pseudoge | ENSG00000( |
| 27105151 | 27161393 | -1 | lncRNA | ENSG00000( |
| 27121910 | 27147664 | 1 | unprocessed_pseudc | ENSG00000( |
| 27243968 | 27325959 | 1 | protein_coding | ENSG00000( |
| 27332854 | 27425289 | 1 | protein_coding | ENSG00000( |
| 27389789 | 27446625 | -1 | lncRNA | ENSG00000( |
| 27466810 | 27502185 | 1 | protein_coding | ENSG00000( |
| 27496937 | 27498589 | 1 | processed_pseudoge | ENSG00000( |
| 27523431 | 27695564 | 1 | protein_coding | ENSG00000( |
| 27547050 | 27552773 | -1 | lncRNA | ENSG00000( |
| 27592461 | 27594039 | -1 | unprocessed_pseudc | ENSG00000( |
| 27608733 | 27609046 | -1 | processed_pseudoge | ENSG00000( |
| 27696388 | 27710803 | -1 | lncRNA | ENSG00000( |
| 27696447 | 27697596 | 1 | protein_coding | ENSG00000( |
| 27700066 | 27700574 | 1 | lncRNA | ENSG00000( |
| 27702338 | 27702906 | 1 | processed_pseudoge | ENSG00000( |
| 27704568 | 27708365 | 1 | lncRNA | ENSG00000( |
| 27710822 | 27756295 | 1 | protein_coding | ENSG00000( |
| 27752874 | 27752986 | 1 | misc_RNA | ENSG00000( |
| 27762738 | 27771276 | -1 | protein_coding | ENSG00000( |
| 27779821 | 27781067 | -1 | lncRNA | ENSG00000( |
| 27780048 | 27803040 | 1 | protein_coding | ENSG00000( |
| 27798641 | 27800708 | -1 | lncRNA | ENSG00000( |
| 27806082 | 27806390 | -1 | misc_RNA | ENSG00000( |
| 27824207 | 27824382 | -1 | misc_RNA | ENSG00000( |
| 27935523 | 27935749 | -1 | unprocessed_pseudc | ENSG00000( |

| | | | | |
|---|---|---|---|---|
| 27958084 | 27972733 | -1 | protein_coding | ENSG00000( |
| 27958517 | 27969813 | 1 | lncRNA | ENSG00000( |
| 28133249 | 28581511 | 1 | protein_coding | ENSG00000( |
| 28163298 | 28190738 | -1 | lncRNA | ENSG00000( |
| 28164958 | 28165662 | 1 | processed_pseudoge | ENSG00000( |
| 28185625 | 28186190 | -1 | lncRNA | ENSG00000( |
| 28236227 | 28236828 | 1 | lncRNA | ENSG00000( |
| 28319335 | 28319463 | -1 | snRNA | ENSG00000( |
| 28505394 | 28505528 | 1 | rRNA_pseudogene | ENSG00000( |
| 28564678 | 28565141 | -1 | processed_pseudoge | ENSG00000( |
| 28821975 | 28825831 | -1 | lncRNA | ENSG00000( |
| 28978584 | 28978685 | 1 | processed_pseudoge | ENSG00000( |
| 29142969 | 29158537 | -1 | lncRNA | ENSG00000( |
| 29149103 | 29340980 | 1 | protein_coding | ENSG00000( |
| 29156448 | 29156991 | -1 | lncRNA | ENSG00000( |
| 29277397 | 29277882 | -1 | lncRNA | ENSG00000( |
| 29277955 | 29317848 | -1 | lncRNA | ENSG00000( |
| 29331434 | 29331936 | -1 | lncRNA | ENSG00000( |
| 29332733 | 29333383 | -1 | lncRNA | ENSG00000( |
| 29337352 | 29381189 | -1 | protein_coding | ENSG00000( |
| 29389289 | 29487488 | 1 | lncRNA | ENSG00000( |
| 29412474 | 29497686 | -1 | protein_coding | ENSG00000( |
| 29500840 | 29784759 | -1 | protein_coding | ENSG00000( |
| 29519731 | 29529974 | 1 | lncRNA | ENSG00000( |
| 29658376 | 29658798 | -1 | processed_pseudoge | ENSG00000( |
| 30086342 | 30130590 | 1 | lncRNA | ENSG00000( |
| 30169881 | 30170091 | -1 | processed_pseudoge | ENSG00000( |
| 30200983 | 30217916 | 1 | lncRNA | ENSG00000( |
| 30230779 | 30296707 | -1 | lncRNA | ENSG00000( |
| 30306065 | 30307611 | 1 | unitary_pseudogene | ENSG00000( |
| 30306431 | 30306524 | -1 | rRNA_pseudogene | ENSG00000( |
| 30320834 | 30321617 | 1 | lncRNA | ENSG00000( |
| 30454919 | 30492302 | 1 | unprocessed_pseudo | ENSG00000( |
| 30628988 | 30695869 | -1 | protein_coding | ENSG00000( |
| 30696121 | 30700220 | 1 | lncRNA | ENSG00000( |
| 30709552 | 30754951 | -1 | protein_coding | ENSG00000( |
| 30755074 | 30780739 | 1 | lncRNA | ENSG00000( |
| 30755167 | 30802602 | 1 | lncRNA | ENSG00000( |
| 30795458 | 30879268 | 1 | lncRNA | ENSG00000( |
| 30861921 | 30862179 | 1 | processed_pseudoge | ENSG00000( |
| 30926428 | 30996602 | 1 | protein_coding | ENSG00000( |
| 30978308 | 31006010 | -1 | transcribed_unproce | ENSG00000( |
| 31012503 | 31015806 | 1 | transcribed_processe | ENSG00000( |
| 31019434 | 31073847 | -1 | lncRNA | ENSG00000( |
| 31073860 | 31104799 | 1 | protein_coding | ENSG00000( |
| 31076867 | 31077016 | -1 | snoRNA | ENSG00000( |
| 31105105 | 31106830 | -1 | unprocessed_pseudo | ENSG00000( |
| 31111652 | 31201235 | -1 | transcribed_unproce | ENSG00000( |
| 31244440 | 31244685 | 1 | processed_pseudoge | ENSG00000( |
| 31251992 | 31252597 | -1 | processed_pseudoge | ENSG00000( |

| | | | | |
|---|---|---|---|---|
| 31254361 | 31254445 | -1 | processed_pseudoge | ENSG0000C |
| 31280422 | 31280895 | -1 | lncRNA | ENSG0000C |
| 31280584 | 31327058 | -1 | protein_coding | ENSG0000C |
| 31311617 | 31312029 | 1 | processed_pseudoge | ENSG0000C |
| 31324316 | 31325829 | 1 | lncRNA | ENSG0000C |
| 31363481 | 31369301 | -1 | lncRNA | ENSG0000C |
| 31366288 | 31366366 | 1 | snRNA | ENSG0000C |
| 31382226 | 31591136 | -1 | protein_coding | ENSG0000C |
| 31396390 | 31396501 | 1 | snRNA | ENSG0000C |
| 31443792 | 31444208 | -1 | lncRNA | ENSG0000C |
| 31465062 | 31465842 | 1 | processed_pseudoge | ENSG0000C |
| 31477506 | 31477987 | 1 | processed_pseudoge | ENSG0000C |
| 31506303 | 31506404 | -1 | misc_RNA | ENSG0000C |
| 31534715 | 31535247 | -1 | processed_pseudoge | ENSG0000C |
| 31589923 | 31615666 | 1 | lncRNA | ENSG0000C |
| 31594093 | 31594206 | -1 | snRNA | ENSG0000C |
| 31597114 | 31597409 | -1 | processed_pseudoge | ENSG0000C |
| 31615771 | 31616439 | -1 | processed_pseudoge | ENSG0000C |
| 31627575 | 31652095 | -1 | lncRNA | ENSG0000C |
| 31647160 | 31673114 | 1 | protein_coding | ENSG0000C |
| 31671142 | 31729121 | -1 | protein_coding | ENSG0000C |
| 31715342 | 31715781 | -1 | processed_pseudoge | ENSG0000C |
| 31729117 | 31731204 | 1 | lncRNA | ENSG0000C |
| 31748100 | 31748240 | -1 | processed_pseudoge | ENSG0000C |
| 31754720 | 31755121 | 1 | processed_pseudoge | ENSG0000C |
| 31791185 | 31792298 | -1 | protein_coding | ENSG0000C |
| 31802958 | 31803090 | -1 | snRNA | ENSG0000C |
| 31811088 | 31811190 | 1 | snoRNA | ENSG0000C |
| 31853021 | 31853407 | -1 | processed_pseudoge | ENSG0000C |
| 31876969 | 31959316 | -1 | lncRNA | ENSG0000C |
| 31893370 | 31893702 | 1 | processed_pseudoge | ENSG0000C |
| 31897820 | 31898123 | 1 | processed_pseudoge | ENSG0000C |
| 31948334 | 31948827 | 1 | processed_pseudoge | ENSG0000C |
| 31953889 | 31955268 | 1 | processed_pseudoge | ENSG0000C |
| 31959370 | 31993107 | 1 | protein_coding | ENSG0000C |
| 32000375 | 32001222 | 1 | lncRNA | ENSG0000C |
| 32007127 | 32007435 | 1 | processed_pseudoge | ENSG0000C |
| 32072734 | 32072880 | 1 | processed_pseudoge | ENSG0000C |
| 32100485 | 32100735 | -1 | processed_pseudoge | ENSG0000C |
| 32104117 | 32107528 | -1 | lncRNA | ENSG0000C |
| 32106835 | 32383633 | 1 | protein_coding | ENSG0000C |
| 32109076 | 32109602 | 1 | lncRNA | ENSG0000C |
| 32227410 | 32228154 | -1 | processed_pseudoge | ENSG0000C |
| 32339368 | 32340724 | 1 | lncRNA | ENSG0000C |
| 32352349 | 32354144 | 1 | lncRNA | ENSG0000C |
| 32396050 | 32396147 | -1 | misc_RNA | ENSG0000C |
| 32399529 | 32646050 | 1 | protein_coding | ENSG0000C |
| 32477382 | 32477471 | -1 | snRNA | ENSG0000C |
| 32679200 | 32745650 | 1 | protein_coding | ENSG0000C |
| 32692504 | 32692848 | -1 | processed_pseudoge | ENSG0000C |

| | | | | |
|---|---|---|---|---|
| 32705403 | 32705506 | -1 | misc_RNA | ENSG0000( |
| 32725248 | 32725660 | -1 | lncRNA | ENSG0000( |
| 32726383 | 32727387 | -1 | processed_pseudoge | ENSG0000( |
| 32727490 | 32755897 | -1 | protein_coding | ENSG0000( |
| 32728169 | 32729024 | -1 | lncRNA | ENSG0000( |
| 32736930 | 32737660 | -1 | lncRNA | ENSG0000( |
| 32755510 | 32756463 | 1 | lncRNA | ENSG0000( |
| 32790745 | 32896840 | -1 | protein_coding | ENSG0000( |
| 32820142 | 32820567 | -1 | lncRNA | ENSG0000( |
| 32863118 | 32863433 | 1 | processed_pseudoge | ENSG0000( |
| 32985838 | 32987070 | 1 | processed_pseudoge | ENSG0000( |
| 32988763 | 32993754 | 1 | lncRNA | ENSG0000( |
| 33374238 | 33439819 | -1 | protein_coding | ENSG0000( |
| 33404872 | 33405896 | -1 | lncRNA | ENSG0000( |
| 33432311 | 33432853 | -1 | lncRNA | ENSG0000( |
| 33454345 | 33455452 | 1 | processed_pseudoge | ENSG0000( |
| 33576095 | 33576200 | 1 | snRNA | ENSG0000( |
| 33717853 | 33718217 | 1 | processed_pseudoge | ENSG0000( |
| 33923374 | 33923480 | -1 | snRNA | ENSG0000( |
| 33982465 | 33982566 | -1 | misc_RNA | ENSG0000( |
| 34022281 | 34046417 | -1 | lncRNA | ENSG0000( |
| 34022468 | 34029694 | 1 | protein_coding | ENSG0000( |
| 34037232 | 34058745 | 1 | lncRNA | ENSG0000( |
| 34149839 | 34162238 | 1 | lncRNA | ENSG0000( |
| 34162477 | 34163442 | 1 | unprocessed_pseudc | ENSG0000( |
| 34164773 | 34166954 | -1 | unprocessed_pseudc | ENSG0000( |
| 34190471 | 34219246 | -1 | lncRNA | ENSG0000( |
| 34205699 | 34205817 | 1 | rRNA_pseudogene | ENSG0000( |
| 34208415 | 34209675 | -1 | processed_pseudoge | ENSG0000( |
| 34249481 | 34249958 | 1 | processed_pseudoge | ENSG0000( |
| 37575587 | 37576384 | 1 | unprocessed_pseudc | ENSG0000( |
| 38078529 | 38078999 | -1 | processed_pseudoge | ENSG0000( |
| 38082046 | 38087392 | 1 | unprocessed_pseudc | ENSG0000( |
| 38150050 | 38150138 | 1 | misc_RNA | ENSG0000( |
| 38155476 | 38167631 | 1 | lncRNA | ENSG0000( |
| 38161466 | 38161582 | -1 | rRNA_pseudogene | ENSG0000( |
| 38163403 | 38163515 | -1 | rRNA_pseudogene | ENSG0000( |
| 38201566 | 38203792 | 1 | unprocessed_pseudc | ENSG0000( |
| 38205098 | 38206009 | -1 | unprocessed_pseudc | ENSG0000( |
| 38206231 | 38207192 | -1 | unprocessed_pseudc | ENSG0000( |
| 38274448 | 38274549 | -1 | misc_RNA | ENSG0000( |
| 38316762 | 38329721 | 1 | protein_coding | ENSG0000( |
| 38532895 | 38533050 | -1 | processed_pseudoge | ENSG0000( |
| 38544177 | 38589812 | -1 | lncRNA | ENSG0000( |
| 38646822 | 38907430 | -1 | protein_coding | ENSG0000( |
| 38906451 | 38909592 | 1 | lncRNA | ENSG0000( |
| 38994803 | 38996882 | -1 | unprocessed_pseudc | ENSG0000( |
| 39058028 | 39059933 | -1 | unprocessed_pseudc | ENSG0000( |
| 39087347 | 39145482 | -1 | lncRNA | ENSG0000( |
| 39170646 | 39170880 | 1 | processed_pseudoge | ENSG0000( |

| | | | | |
|---|---|---|---|---|
| 39293228 | 39443390 | -1 | protein_coding | ENSG00000 |
| 39447404 | 39448314 | 1 | lncRNA | ENSG00000 |
| 39466442 | 39467345 | -1 | processed_pseudoge | ENSG00000 |
| 39539581 | 39539851 | -1 | processed_pseudoge | ENSG00000 |
| 39550033 | 39619803 | -1 | protein_coding | ENSG00000 |
| 39611819 | 39614648 | 1 | lncRNA | ENSG00000 |
| 39626167 | 39908300 | 1 | protein_coding | ENSG00000 |
| 39755025 | 40106089 | -1 | protein_coding | ENSG00000 |
| 39819750 | 39819883 | -1 | snoRNA | ENSG00000 |
| 40068243 | 40068590 | -1 | processed_pseudoge | ENSG00000 |
| 40140926 | 40142876 | -1 | lncRNA | ENSG00000 |
| 40156113 | 40211419 | 1 | lncRNA | ENSG00000 |
| 40186009 | 40224915 | -1 | lncRNA | ENSG00000 |
| 40196744 | 40369285 | 1 | protein_coding | ENSG00000 |
| 40286289 | 40286662 | 1 | processed_pseudoge | ENSG00000 |
| 40393395 | 40570832 | 1 | protein_coding | ENSG00000 |
| 40395853 | 40443847 | -1 | lncRNA | ENSG00000 |
| 40554161 | 40554267 | -1 | snRNA | ENSG00000 |
| 40692439 | 41072415 | 1 | protein_coding | ENSG00000 |
| 40728811 | 40729897 | 1 | processed_pseudoge | ENSG00000 |
| 40978744 | 40979244 | 1 | lncRNA | ENSG00000 |
| 41188320 | 41574745 | 1 | protein_coding | ENSG00000 |
| 41409467 | 41473510 | -1 | lncRNA | ENSG00000 |
| 41658676 | 41658793 | 1 | rRNA_pseudogene | ENSG00000 |
| 41698495 | 41699182 | -1 | unprocessed_pseudo | ENSG00000 |
| 41699391 | 41700251 | -1 | unprocessed_pseudo | ENSG00000 |
| 41764144 | 41774671 | 1 | lncRNA | ENSG00000 |
| 41829898 | 41932592 | -1 | lncRNA | ENSG00000 |
| 41911256 | 41912212 | 1 | processed_pseudoge | ENSG00000 |
| 41951493 | 42024329 | -1 | lncRNA | ENSG00000 |
| 42069935 | 42070625 | -1 | processed_pseudoge | ENSG00000 |
| 42081845 | 42144874 | -1 | protein_coding | ENSG00000 |
| 42157104 | 42238349 | -1 | protein_coding | ENSG00000 |
| 42238447 | 42459715 | 1 | protein_coding | ENSG00000 |
| 42286911 | 42288560 | 1 | processed_pseudoge | ENSG00000 |
| 42312086 | 42326118 | -1 | protein_coding | ENSG00000 |
| 42323700 | 42323859 | -1 | miRNA | ENSG00000 |
| 42347061 | 42347214 | -1 | snoRNA | ENSG00000 |
| 42361267 | 42361703 | 1 | lncRNA | ENSG00000 |
| 42398550 | 42398651 | 1 | snRNA | ENSG00000 |
| 42431665 | 42433357 | -1 | processed_pseudoge | ENSG00000 |
| 42454720 | 42454821 | 1 | misc_RNA | ENSG00000 |
| 42456757 | 42590355 | -1 | protein_coding | ENSG00000 |
| 42459366 | 42466128 | 1 | lncRNA | ENSG00000 |
| 42485353 | 42498979 | -1 | lncRNA | ENSG00000 |
| 42615221 | 42646547 | -1 | lncRNA | ENSG00000 |
| 42627008 | 42627137 | -1 | processed_pseudoge | ENSG00000 |
| 42636815 | 42637111 | -1 | processed_pseudoge | ENSG00000 |
| 42646583 | 42686701 | 1 | lncRNA | ENSG00000 |
| 42678873 | 42679124 | -1 | processed_pseudoge | ENSG00000 |

| | | | | |
|---|---|---|---|---|
| 42687195 | 42717120 | 1 | lncRNA | ENSG0000C |
| 42929848 | 42930579 | -1 | processed_pseudoge | ENSG0000C |
| 42966122 | 42967353 | 1 | lncRNA | ENSG0000C |
| 42979254 | 42996486 | -1 | lncRNA | ENSG0000C |
| 43054089 | 43054386 | -1 | processed_pseudoge | ENSG0000C |
| 43155315 | 43163110 | 1 | lncRNA | ENSG0000C |
| 43353866 | 43552203 | -1 | protein_coding | ENSG0000C |
| 43560784 | 43561297 | -1 | processed_pseudoge | ENSG0000C |
| 43569769 | 43572038 | -1 | processed_pseudoge | ENSG0000C |
| 43633093 | 43633281 | 1 | processed_pseudoge | ENSG0000C |
| 43659827 | 43661101 | 1 | processed_pseudoge | ENSG0000C |
| 43718993 | 43758817 | -1 | protein_coding | ENSG0000C |
| 43736628 | 43741067 | 1 | transcribed_unproce | ENSG0000C |
| 43758944 | 43789543 | 1 | protein_coding | ENSG0000C |
| 43793723 | 43806375 | -1 | protein_coding | ENSG0000C |
| 43835967 | 44389762 | 1 | protein_coding | ENSG0000C |
| 44008620 | 44010143 | -1 | processed_pseudoge | ENSG0000C |
| 44244394 | 44263982 | -1 | lncRNA | ENSG0000C |
| 44248432 | 44249004 | -1 | processed_pseudoge | ENSG0000C |
| 44498616 | 44499158 | -1 | lncRNA | ENSG0000C |
| 44499961 | 44503596 | -1 | lncRNA | ENSG0000C |
| 44508275 | 44921848 | -1 | protein_coding | ENSG0000C |
| 44880868 | 44880969 | -1 | misc_RNA | ENSG0000C |
| 45014672 | 45051099 | -1 | protein_coding | ENSG0000C |
| 45050901 | 45103107 | 1 | lncRNA | ENSG0000C |
| 45063473 | 45065351 | -1 | processed_pseudoge | ENSG0000C |
| 45071769 | 45072843 | 1 | processed_pseudoge | ENSG0000C |
| 45117771 | 45117888 | 1 | rRNA_pseudogene | ENSG0000C |
| 45164186 | 45170809 | 1 | lncRNA | ENSG0000C |
| 45173064 | 45216041 | -1 | transcribed_processe | ENSG0000C |
| 45200817 | 45201551 | 1 | processed_pseudoge | ENSG0000C |
| 45215987 | 45440404 | 1 | protein_coding | ENSG0000C |
| 45256473 | 45256726 | 1 | lncRNA | ENSG0000C |
| 45444766 | 45445438 | 1 | processed_pseudoge | ENSG0000C |
| 45475520 | 45610620 | -1 | lncRNA | ENSG0000C |
| 45554590 | 45554659 | -1 | snRNA | ENSG0000C |
| 45718046 | 45727775 | -1 | lncRNA | ENSG0000C |
| 45729706 | 45908040 | 1 | protein_coding | ENSG0000C |
| 45789189 | 45789657 | 1 | lncRNA | ENSG0000C |
| 45874035 | 45874330 | 1 | misc_RNA | ENSG0000C |
| 45880950 | 45882266 | -1 | processed_pseudoge | ENSG0000C |
| 45919131 | 45992120 | -1 | protein_coding | ENSG0000C |
| 46004038 | 46004685 | -1 | processed_pseudoge | ENSG0000C |
| 46183063 | 46270017 | -1 | protein_coding | ENSG0000C |
| 46239106 | 46239473 | -1 | lncRNA | ENSG0000C |
| 46358188 | 46372773 | -1 | protein_coding | ENSG0000C |
| 46371463 | 46373778 | 1 | lncRNA | ENSG0000C |
| 46383652 | 46876784 | 1 | lncRNA | ENSG0000C |
| 46384233 | 46386991 | -1 | lncRNA | ENSG0000C |
| 46388856 | 46392126 | 1 | lncRNA | ENSG0000C |

| | | | | |
|---|---|---|---|---|
| 46404644 | 46407166 | 1 | TEC | ENSG0000C |
| 46537502 | 46652550 | 1 | lncRNA | ENSG0000C |
| 46592573 | 46593305 | 1 | unprocessed_pseudc | ENSG0000C |
| 46682071 | 46682713 | -1 | processed_pseudoge | ENSG0000C |
| 46764761 | 46832408 | -1 | protein_coding | ENSG0000C |
| 46970504 | 46972155 | 1 | lncRNA | ENSG0000C |
| 47075707 | 47079951 | -1 | protein_coding | ENSG0000C |
| 47079603 | 47236662 | 1 | protein_coding | ENSG0000C |
| 47135498 | 47138370 | -1 | transcribed_processe | ENSG0000C |
| 47187812 | 47187891 | 1 | miRNA | ENSG0000C |
| 47205898 | 47216456 | -1 | lncRNA | ENSG0000C |
| 47237734 | 47279021 | -1 | lncRNA | ENSG0000C |
| 47248124 | 47257539 | 1 | lncRNA | ENSG0000C |
| 47265665 | 47266108 | 1 | lncRNA | ENSG0000C |
| 47341614 | 47342089 | -1 | processed_pseudoge | ENSG0000C |
| 47346166 | 47346297 | -1 | snoRNA | ENSG0000C |
| 47353754 | 47369935 | 1 | lncRNA | ENSG0000C |
| 47364186 | 47364269 | -1 | miRNA | ENSG0000C |
| 47377638 | 47420192 | 1 | lncRNA | ENSG0000C |
| 47463087 | 47463621 | 1 | processed_pseudoge | ENSG0000C |
| 47484689 | 47485722 | 1 | processed_pseudoge | ENSG0000C |
| 47593208 | 47593440 | 1 | processed_pseudoge | ENSG0000C |
| 47661249 | 47706030 | -1 | protein_coding | ENSG0000C |
| 47699401 | 47699917 | -1 | lncRNA | ENSG0000C |
| 47706083 | 47742294 | 1 | lncRNA | ENSG0000C |
| 47709734 | 47725567 | -1 | protein_coding | ENSG0000C |
| 47728151 | 47730598 | -1 | TEC | ENSG0000C |
| 47731908 | 47732351 | 1 | lncRNA | ENSG0000C |
| 47734363 | 47771040 | -1 | protein_coding | ENSG0000C |
| 47753916 | 47782751 | 1 | protein_coding | ENSG0000C |
| 47768529 | 47769648 | 1 | lncRNA | ENSG0000C |
| 47782722 | 47833132 | -1 | protein_coding | ENSG0000C |
| 47784923 | 47786002 | 1 | lncRNA | ENSG0000C |
| 47788426 | 47788971 | 1 | lncRNA | ENSG0000C |
| 47817451 | 47817966 | -1 | lncRNA | ENSG0000C |
| 47826854 | 47837898 | 1 | lncRNA | ENSG0000C |
| 47841537 | 47943048 | -1 | protein_coding | ENSG0000C |
| 47882649 | 47901525 | 1 | lncRNA | ENSG0000C |
| 47905122 | 47906865 | 1 | lncRNA | ENSG0000C |
| 47960497 | 47963469 | -1 | TEC | ENSG0000C |
| 47963569 | 47968878 | 1 | protein_coding | ENSG0000C |
| 47972967 | 48004554 | -1 | protein_coding | ENSG0000C |
| 48005277 | 48011227 | -1 | lncRNA | ENSG0000C |
| 48011304 | 48025469 | 1 | lncRNA | ENSG0000C |
| 48019771 | 48025382 | -1 | lncRNA | ENSG0000C |
| 48039784 | 48040761 | -1 | lncRNA | ENSG0000C |
| 48042897 | 48106079 | -1 | protein_coding | ENSG0000C |
| 48054813 | 48055591 | -1 | transcribed_processe | ENSG0000C |
| 48081319 | 48081428 | 1 | snRNA | ENSG0000C |
| 48085062 | 48085421 | 1 | processed_pseudoge | ENSG0000C |

| | | | | |
|---|---|---|---|---|
| 48105139 | 48146404 | 1 | protein_coding | ENSG00000( |
| 48132797 | 48132867 | 1 | miRNA | ENSG00000( |
| 48147789 | 48181213 | -1 | protein_coding | ENSG00000( |
| 48152817 | 48153128 | 1 | lncRNA | ENSG00000( |
| 48160245 | 48160878 | -1 | processed_pseudoge | ENSG00000( |
| 48168847 | 48169659 | -1 | processed_pseudoge | ENSG00000( |
| 48171590 | 48173353 | 1 | processed_pseudoge | ENSG00000( |
| 48183644 | 48185926 | 1 | protein_coding | ENSG00000( |
| 48189349 | 48191207 | 1 | TEC | ENSG00000( |
| 48198357 | 48230275 | 1 | lncRNA | ENSG00000( |
| 48202083 | 48203387 | -1 | protein_coding | ENSG00000( |
| 48231098 | 48284210 | -1 | lncRNA | ENSG00000( |
| 48327942 | 48328472 | -1 | lncRNA | ENSG00000( |
| 48328980 | 48330279 | 1 | protein_coding | ENSG00000( |
| 48333755 | 48333901 | -1 | unprocessed_pseudc | ENSG00000( |
| 48337180 | 48351414 | -1 | protein_coding | ENSG00000( |
| 48350945 | 48442411 | 1 | lncRNA | ENSG00000( |
| 48355792 | 48356614 | -1 | unprocessed_pseudc | ENSG00000( |
| 48360920 | 48361377 | 1 | lncRNA | ENSG00000( |
| 48385417 | 48386265 | -1 | unprocessed_pseudc | ENSG00000( |
| 48394510 | 48395392 | -1 | unprocessed_pseudc | ENSG00000( |
| 48417150 | 48417337 | -1 | unprocessed_pseudc | ENSG00000( |
| 48442030 | 48442947 | -1 | unprocessed_pseudc | ENSG00000( |
| 48472665 | 48473622 | 1 | protein_coding | ENSG00000( |
| 48482498 | 48496386 | 1 | protein_coding | ENSG00000( |
| 48483287 | 48500961 | -1 | lncRNA | ENSG00000( |
| 48487946 | 48488408 | 1 | transcribed_processe | ENSG00000( |
| 48507354 | 48529897 | 1 | transcribed_processe | ENSG00000( |
| 48525632 | 48528103 | 1 | protein_coding | ENSG00000( |
| 48559882 | 48562956 | 1 | protein_coding | ENSG00000( |
| 48567684 | 48570066 | -1 | protein_coding | ENSG00000( |
| 48627879 | 48628836 | 1 | unitary_pseudogene | ENSG00000( |
| 48653401 | 48682238 | -1 | protein_coding | ENSG00000( |
| 48654382 | 48654518 | -1 | snoRNA | ENSG00000( |
| 48654444 | 48654530 | -1 | miRNA | ENSG00000( |
| 48656648 | 48656782 | -1 | snoRNA | ENSG00000( |
| 48667457 | 48667593 | -1 | snoRNA | ENSG00000( |
| 48688458 | 48716998 | -1 | protein_coding | ENSG00000( |
| 48727435 | 48765790 | 1 | protein_coding | ENSG00000( |
| 48766194 | 48789037 | -1 | protein_coding | ENSG00000( |
| 48766194 | 48767323 | 1 | lncRNA | ENSG00000( |
| 48771975 | 48772037 | -1 | miRNA | ENSG00000( |
| 48789147 | 48790535 | 1 | lncRNA | ENSG00000( |
| 48800576 | 48801086 | -1 | processed_pseudoge | ENSG00000( |
| 48809911 | 48810016 | 1 | misc_RNA | ENSG00000( |
| 48813794 | 48828941 | 1 | protein_coding | ENSG00000( |
| 48829756 | 48852842 | -1 | protein_coding | ENSG00000( |
| 48857145 | 48865870 | -1 | protein_coding | ENSG00000( |
| 48892130 | 48892228 | -1 | snRNA | ENSG00000( |
| 48903418 | 48903813 | -1 | processed_pseudoge | ENSG00000( |

| | | | | |
|---|---|---|---|---|
| 48903503 | 48957365 | -1 | protein_coding | ENSG00000( |
| 48904110 | 48931840 | 1 | protein_coding | ENSG00000( |
| 48921518 | 48926474 | -1 | protein_coding | ENSG00000( |
| 48935723 | 48957487 | -1 | protein_coding | ENSG00000( |
| 48965340 | 48971735 | -1 | protein_coding | ENSG00000( |
| 48975543 | 48975646 | 1 | snRNA | ENSG00000( |
| 48978322 | 48982620 | 1 | protein_coding | ENSG00000( |
| 48995149 | 48999375 | -1 | protein_coding | ENSG00000( |
| 48995150 | 48996334 | 1 | lncRNA | ENSG00000( |
| 48998367 | 49019235 | 1 | lncRNA | ENSG00000( |
| 49002274 | 49018807 | -1 | protein_coding | ENSG00000( |
| 49018975 | 49059774 | -1 | protein_coding | ENSG00000( |
| 49064676 | 49070025 | -1 | protein_coding | ENSG00000( |
| 49075039 | 49075802 | 1 | processed_pseudoge | ENSG00000( |
| 49086656 | 49094801 | -1 | protein_coding | ENSG00000( |
| 49090208 | 49093312 | 1 | lncRNA | ENSG00000( |
| 49096551 | 49110900 | -1 | protein_coding | ENSG00000( |
| 49127782 | 49188484 | 1 | lncRNA | ENSG00000( |
| 49127782 | 49131397 | -1 | protein_coding | ENSG00000( |
| 49132853 | 49132947 | -1 | misc_RNA | ENSG00000( |
| 49184795 | 49189080 | -1 | protein_coding | ENSG00000( |
| 49188736 | 49274603 | 1 | protein_coding | ENSG00000( |
| 49196784 | 49197062 | 1 | misc_RNA | ENSG00000( |
| 49232790 | 49264756 | -1 | lncRNA | ENSG00000( |
| 49265156 | 49273306 | -1 | lncRNA | ENSG00000( |
| 49292631 | 49324576 | -1 | lncRNA | ENSG00000( |
| 49293252 | 49298686 | 1 | protein_coding | ENSG00000( |
| 49323236 | 49331731 | 1 | protein_coding | ENSG00000( |
| 49332409 | 49337188 | -1 | protein_coding | ENSG00000( |
| 49346888 | 49357546 | 1 | protein_coding | ENSG00000( |
| 49366584 | 49527424 | 1 | protein_coding | ENSG00000( |
| 49389516 | 49390162 | 1 | processed_pseudoge | ENSG00000( |
| 49442424 | 49442652 | -1 | processed_pseudoge | ENSG00000( |
| 49536677 | 49538894 | -1 | lncRNA | ENSG00000( |
| 49539030 | 49558337 | 1 | protein_coding | ENSG00000( |
| 49556544 | 49568145 | -1 | protein_coding | ENSG00000( |
| 49568218 | 49644666 | 1 | protein_coding | ENSG00000( |
| 49576840 | 49577505 | -1 | lncRNA | ENSG00000( |
| 49582885 | 49605639 | -1 | protein_coding | ENSG00000( |
| 49593104 | 49593213 | 1 | snRNA | ENSG00000( |
| 49594091 | 49594258 | 1 | processed_pseudoge | ENSG00000( |
| 49595148 | 49595688 | 1 | processed_pseudoge | ENSG00000( |
| 49619334 | 49619612 | 1 | processed_pseudoge | ENSG00000( |
| 49636499 | 49708165 | -1 | protein_coding | ENSG00000( |
| 49707725 | 49764934 | 1 | protein_coding | ENSG00000( |
| 49709237 | 49709336 | 1 | processed_pseudoge | ENSG00000( |
| 49770960 | 49771136 | 1 | processed_pseudoge | ENSG00000( |
| 49791146 | 49828750 | -1 | protein_coding | ENSG00000( |
| 49827913 | 49841143 | 1 | lncRNA | ENSG00000( |
| 49836043 | 49843106 | -1 | protein_coding | ENSG00000( |

| | | | | |
|---|---|---|---|---|
| 49861207 | 49865802 | 1 | lncRNA | ENSG00000 |
| 49863173 | 49863503 | 1 | processed_pseudoge | ENSG00000 |
| 49866896 | 49904217 | -1 | protein_coding | ENSG00000 |
| 49900311 | 49932764 | 1 | lncRNA | ENSG00000 |
| 49908882 | 49911863 | -1 | lncRNA | ENSG00000 |
| 49950741 | 49958881 | 1 | protein_coding | ENSG00000 |
| 49951512 | 49962924 | -1 | lncRNA | ENSG00000 |
| 49954639 | 49956125 | -1 | lncRNA | ENSG00000 |
| 49961872 | 49965682 | 1 | protein_coding | ENSG00000 |
| 49967194 | 49977139 | 1 | protein_coding | ENSG00000 |
| 49976923 | 50033136 | -1 | protein_coding | ENSG00000 |
| 50057548 | 50083611 | 1 | protein_coding | ENSG00000 |
| 50085200 | 50100707 | 1 | protein_coding | ENSG00000 |
| 50103982 | 50111313 | 1 | protein_coding | ENSG00000 |
| 50112082 | 50120457 | 1 | protein_coding | ENSG00000 |
| 50112197 | 50165618 | 1 | lncRNA | ENSG00000 |
| 50129299 | 50167533 | -1 | protein_coding | ENSG00000 |
| 50175788 | 50283546 | -1 | protein_coding | ENSG00000 |
| 50185580 | 50191363 | -1 | lncRNA | ENSG00000 |
| 50217662 | 50218343 | -1 | transcribed_processe | ENSG00000 |
| 50219604 | 50229984 | 1 | lncRNA | ENSG00000 |
| 50234142 | 50234212 | -1 | miRNA | ENSG00000 |
| 50256238 | 50256344 | -1 | snRNA | ENSG00000 |
| 50286244 | 50289231 | 1 | processed_pseudoge | ENSG00000 |
| 50326230 | 50396622 | -1 | protein_coding | ENSG00000 |
| 50365652 | 50366086 | 1 | processed_pseudoge | ENSG00000 |
| 50392383 | 50480004 | 1 | protein_coding | ENSG00000 |
| 50504985 | 50748657 | 1 | protein_coding | ENSG00000 |
| 50633273 | 50633376 | 1 | snRNA | ENSG00000 |
| 50656973 | 50657078 | 1 | snRNA | ENSG00000 |
| 50743568 | 50743684 | 1 | misc_RNA | ENSG00000 |
| 50756892 | 50757433 | 1 | processed_pseudoge | ENSG00000 |
| 50763710 | 50821162 | 1 | protein_coding | ENSG00000 |
| 50806335 | 50806437 | -1 | processed_pseudoge | ENSG00000 |
| 50841498 | 50841785 | -1 | misc_RNA | ENSG00000 |
| 50842920 | 50887884 | 1 | protein_coding | ENSG00000 |
| 50847925 | 50848312 | -1 | processed_pseudoge | ENSG00000 |
| 50923472 | 50932510 | 1 | protein_coding | ENSG00000 |
| 50934942 | 50935464 | -1 | lncRNA | ENSG00000 |
| 50939927 | 50940026 | -1 | misc_RNA | ENSG00000 |
| 50949117 | 50949384 | -1 | processed_pseudoge | ENSG00000 |
| 50953922 | 50970506 | 1 | protein_coding | ENSG00000 |
| 50953924 | 50954356 | -1 | lncRNA | ENSG00000 |
| 50979401 | 51028566 | -1 | protein_coding | ENSG00000 |
| 51014134 | 51014241 | -1 | snRNA | ENSG00000 |
| 51037499 | 51037602 | 1 | snRNA | ENSG00000 |
| 51047962 | 51060424 | 1 | protein_coding | ENSG00000 |
| 51061205 | 51083664 | -1 | protein_coding | ENSG00000 |
| 51093656 | 51173135 | -1 | protein_coding | ENSG00000 |
| 51106817 | 51107144 | 1 | processed_pseudoge | ENSG00000 |

| | | | | |
|---|---|---|---|---|
| 51124628 | 51125381 | -1 | processed_pseudoge | ENSG0000C |
| 51156868 | 51156977 | -1 | misc_RNA | ENSG0000C |
| 51162901 | 51163007 | 1 | snRNA | ENSG0000C |
| 51186936 | 51217708 | -1 | protein_coding | ENSG0000C |
| 51201684 | 51202581 | -1 | lncRNA | ENSG0000C |
| 51238724 | 51271362 | 1 | protein_coding | ENSG0000C |
| 51244558 | 51270415 | -1 | protein_coding | ENSG0000C |
| 51281038 | 51324668 | -1 | protein_coding | ENSG0000C |
| 51328442 | 51346679 | -1 | protein_coding | ENSG0000C |
| 51351247 | 51392867 | -1 | protein_coding | ENSG0000C |
| 51391317 | 51515763 | 1 | protein_coding | ENSG0000C |
| 51421956 | 51424611 | -1 | lncRNA | ENSG0000C |
| 51590266 | 51812864 | 1 | protein_coding | ENSG0000C |
| 51712469 | 51713606 | -1 | processed_pseudoge | ENSG0000C |
| 51809705 | 51810600 | -1 | lncRNA | ENSG0000C |
| 51813940 | 51814926 | -1 | protein_coding | ENSG0000C |
| 51815043 | 51842106 | 1 | lncRNA | ENSG0000C |
| 51817840 | 51848766 | -1 | protein_coding | ENSG0000C |
| 51817899 | 51820150 | 1 | lncRNA | ENSG0000C |
| 51848223 | 51852729 | 1 | lncRNA | ENSG0000C |
| 51888009 | 51891727 | 1 | protein_coding | ENSG0000C |
| 51900565 | 51901336 | 1 | processed_pseudoge | ENSG0000C |
| 51906908 | 51923361 | 1 | protein_coding | ENSG0000C |
| 51951699 | 51997078 | 1 | protein_coding | ENSG0000C |
| 51987657 | 51987763 | 1 | snRNA | ENSG0000C |
| 52006946 | 52015889 | 1 | protein_coding | ENSG0000C |
| 52022832 | 52059507 | 1 | protein_coding | ENSG0000C |
| 52058459 | 52059503 | -1 | lncRNA | ENSG0000C |
| 52069246 | 52077494 | 1 | protein_coding | ENSG0000C |
| 52076841 | 52082084 | -1 | lncRNA | ENSG0000C |
| 52079696 | 52108239 | 1 | protein_coding | ENSG0000C |
| 52084744 | 52108261 | 1 | transcribed_unproce | ENSG0000C |
| 52092485 | 52104297 | -1 | lncRNA | ENSG0000C |
| 52164115 | 52223813 | 1 | lncRNA | ENSG0000C |
| 52168996 | 52192014 | -1 | protein_coding | ENSG0000C |
| 52205392 | 52213583 | -1 | lncRNA | ENSG0000C |
| 52229093 | 52229588 | -1 | processed_pseudoge | ENSG0000C |
| 52232520 | 52252186 | 1 | protein_coding | ENSG0000C |
| 52245048 | 52247448 | -1 | lncRNA | ENSG0000C |
| 52249300 | 52309163 | 1 | protein_coding | ENSG0000C |
| 52250466 | 52258867 | -1 | transcribed_unproce | ENSG0000C |
| 52263948 | 52267461 | 1 | unprocessed_pseudc | ENSG0000C |
| 52274647 | 52279156 | -1 | lncRNA | ENSG0000C |
| 52275821 | 52277736 | 1 | transcribed_unproce | ENSG0000C |
| 52285913 | 52291534 | -1 | protein_coding | ENSG0000C |
| 52298856 | 52306287 | -1 | lncRNA | ENSG0000C |
| 52306616 | 52308371 | -1 | lncRNA | ENSG0000C |
| 52314301 | 52321398 | -1 | protein_coding | ENSG0000C |
| 52341197 | 52354036 | 1 | transcribed_unproce | ENSG0000C |
| 52360006 | 52367481 | -1 | protein_coding | ENSG0000C |

| | | | | |
|---|---|---|---|---|
| 52377812 | 52385652 | -1 | protein_coding | ENSG00000 |
| 52380460 | 52402407 | 1 | lncRNA | ENSG00000 |
| 52393931 | 52406355 | -1 | protein_coding | ENSG00000 |
| 52407580 | 52428494 | 1 | lncRNA | ENSG00000 |
| 52411742 | 52421867 | -1 | unprocessed_pseudo | ENSG00000 |
| 52424070 | 52434371 | -1 | protein_coding | ENSG00000 |
| 52446651 | 52452146 | -1 | protein_coding | ENSG00000 |
| 52468516 | 52473805 | -1 | protein_coding | ENSG00000 |
| 52487176 | 52493257 | -1 | protein_coding | ENSG00000 |
| 52514575 | 52520530 | -1 | protein_coding | ENSG00000 |
| 52543909 | 52553145 | -1 | protein_coding | ENSG00000 |
| 52565782 | 52573843 | -1 | protein_coding | ENSG00000 |
| 52585589 | 52601538 | -1 | protein_coding | ENSG00000 |
| 52601467 | 52615305 | 1 | lncRNA | ENSG00000 |
| 52607570 | 52618559 | -1 | protein_coding | ENSG00000 |
| 52630339 | 52634456 | -1 | unprocessed_pseudo | ENSG00000 |
| 52644558 | 52652211 | -1 | protein_coding | ENSG00000 |
| 52674736 | 52680407 | -1 | protein_coding | ENSG00000 |
| 52689626 | 52703524 | -1 | protein_coding | ENSG00000 |
| 52692605 | 52696788 | 1 | lncRNA | ENSG00000 |
| 52715567 | 52721655 | -1 | unprocessed_pseudo | ENSG00000 |
| 52729517 | 52736142 | -1 | unprocessed_pseudo | ENSG00000 |
| 52730425 | 52732865 | 1 | processed_pseudoge | ENSG00000 |
| 52746596 | 52748308 | -1 | unprocessed_pseudo | ENSG00000 |
| 52748776 | 52750127 | 1 | processed_pseudoge | ENSG00000 |
| 52751978 | 52753821 | -1 | unprocessed_pseudo | ENSG00000 |
| 52763405 | 52763893 | 1 | processed_pseudoge | ENSG00000 |
| 52768155 | 52777345 | -1 | protein_coding | ENSG00000 |
| 52782650 | 52782839 | 1 | processed_pseudoge | ENSG00000 |
| 52789685 | 52796117 | -1 | protein_coding | ENSG00000 |
| 52806549 | 52814116 | -1 | protein_coding | ENSG00000 |
| 52821408 | 52834311 | -1 | protein_coding | ENSG00000 |
| 52837403 | 52837739 | 1 | processed_pseudoge | ENSG00000 |
| 52837804 | 52849092 | -1 | protein_coding | ENSG00000 |
| 52885429 | 52886385 | 1 | processed_pseudoge | ENSG00000 |
| 52897187 | 52949954 | -1 | protein_coding | ENSG00000 |
| 52948871 | 52952906 | 1 | protein_coding | ENSG00000 |
| 53006158 | 53042209 | 1 | protein_coding | ENSG00000 |
| 53012104 | 53013921 | 1 | lncRNA | ENSG00000 |
| 53012884 | 53054450 | -1 | lncRNA | ENSG00000 |
| 53046969 | 53064372 | 1 | protein_coding | ENSG00000 |
| 53056944 | 53057012 | 1 | miRNA | ENSG00000 |
| 53064316 | 53079404 | -1 | protein_coding | ENSG00000 |
| 53097436 | 53102345 | 1 | protein_coding | ENSG00000 |
| 53103486 | 53124535 | 1 | protein_coding | ENSG00000 |
| 53131350 | 53131985 | 1 | processed_pseudoge | ENSG00000 |
| 53138545 | 53138651 | -1 | snRNA | ENSG00000 |
| 53142053 | 53142327 | 1 | processed_pseudoge | ENSG00000 |
| 53153272 | 53154484 | 1 | processed_pseudoge | ENSG00000 |
| 53157663 | 53180909 | -1 | protein_coding | ENSG00000 |

| | | | | |
|---|---|---|---|---|
| 53159586 | 53161000 | 1 | lncRNA | ENSG0000C |
| 53180704 | 53195142 | 1 | protein_coding | ENSG0000C |
| 53191323 | 53207282 | -1 | protein_coding | ENSG0000C |
| 53210567 | 53232980 | -1 | protein_coding | ENSG0000C |
| 53241889 | 53242723 | 1 | protein_coding | ENSG0000C |
| 53251251 | 53254405 | 1 | protein_coding | ENSG0000C |
| 53268299 | 53293643 | 1 | protein_coding | ENSG0000C |
| 53295291 | 53299450 | 1 | protein_coding | ENSG0000C |
| 53298655 | 53300314 | -1 | lncRNA | ENSG0000C |
| 53299695 | 53307177 | 1 | protein_coding | ENSG0000C |
| 53307456 | 53324864 | -1 | protein_coding | ENSG0000C |
| 53326575 | 53345315 | -1 | protein_coding | ENSG0000C |
| 53380176 | 53416446 | 1 | protein_coding | ENSG0000C |
| 53423855 | 53431672 | 1 | protein_coding | ENSG0000C |
| 53441678 | 53446645 | 1 | protein_coding | ENSG0000C |
| 53441741 | 53467528 | 1 | lncRNA | ENSG0000C |
| 53452102 | 53481162 | 1 | protein_coding | ENSG0000C |
| 53464468 | 53465057 | 1 | lncRNA | ENSG0000C |
| 53464720 | 53464901 | 1 | misc_RNA | ENSG0000C |
| 53479669 | 53500063 | -1 | protein_coding | ENSG0000C |
| 53500162 | 53500936 | -1 | lncRNA | ENSG0000C |
| 53500921 | 53506431 | 1 | protein_coding | ENSG0000C |
| 53506688 | 53625979 | -1 | protein_coding | ENSG0000C |
| 53506690 | 53507638 | -1 | protein_coding | ENSG0000C |
| 53507856 | 53626410 | -1 | protein_coding | ENSG0000C |
| 53513891 | 53517608 | 1 | lncRNA | ENSG0000C |
| 53531752 | 53621218 | 1 | lncRNA | ENSG0000C |
| 53574632 | 53576103 | 1 | processed_pseudoge | ENSG0000C |
| 53632726 | 53677408 | -1 | protein_coding | ENSG0000C |
| 53632788 | 53633151 | -1 | processed_pseudoge | ENSG0000C |
| 53708517 | 53727745 | -1 | protein_coding | ENSG0000C |
| 53739611 | 53898976 | 1 | lncRNA | ENSG0000C |
| 53746337 | 53757227 | -1 | lncRNA | ENSG0000C |
| 53754358 | 53762104 | 1 | lncRNA | ENSG0000C |
| 53816185 | 53816479 | 1 | misc_RNA | ENSG0000C |
| 53935328 | 53939643 | -1 | lncRNA | ENSG0000C |
| 53938831 | 53946544 | 1 | protein_coding | ENSG0000C |
| 53954903 | 53958956 | 1 | protein_coding | ENSG0000C |
| 53962308 | 53974956 | -1 | lncRNA | ENSG0000C |
| 53963629 | 53963697 | 1 | misc_RNA | ENSG0000C |
| 53965965 | 53966061 | 1 | misc_RNA | ENSG0000C |
| 53968002 | 53968178 | 1 | misc_RNA | ENSG0000C |
| 53968351 | 53968501 | 1 | misc_RNA | ENSG0000C |
| 53973126 | 53977643 | 1 | protein_coding | ENSG0000C |
| 53981509 | 53985519 | -1 | lncRNA | ENSG0000C |
| 53985065 | 53990279 | 1 | protein_coding | ENSG0000C |
| 53985845 | 54034888 | 1 | protein_coding | ENSG0000C |
| 53990624 | 54030823 | 1 | protein_coding | ENSG0000C |
| 53991738 | 53991847 | 1 | miRNA | ENSG0000C |
| 53993810 | 53996785 | -1 | lncRNA | ENSG0000C |

| | | | | |
|---|---|---|---|---|
| 53994895 | 54003337 | 1 | protein_coding | ENSG0000C |
| 53999022 | 54000010 | -1 | lncRNA | ENSG0000C |
| 54008985 | 54012769 | 1 | protein_coding | ENSG0000C |
| 54016931 | 54056030 | 1 | protein_coding | ENSG0000C |
| 54017110 | 54035361 | 1 | lncRNA | ENSG0000C |
| 54019910 | 54022589 | 1 | lncRNA | ENSG0000C |
| 54032853 | 54035358 | 1 | protein_coding | ENSG0000C |
| 54033950 | 54034045 | 1 | miRNA | ENSG0000C |
| 54058254 | 54122235 | 1 | lncRNA | ENSG0000C |
| 54076838 | 54081903 | -1 | transcribed_unproce | ENSG0000C |
| 54081736 | 54102699 | 1 | lncRNA | ENSG0000C |
| 54085132 | 54125992 | -1 | lncRNA | ENSG0000C |
| 54099191 | 54099302 | -1 | misc_RNA | ENSG0000C |
| 54121277 | 54189008 | -1 | protein_coding | ENSG0000C |
| 54126082 | 54147485 | 1 | lncRNA | ENSG0000C |
| 54145069 | 54147225 | -1 | lncRNA | ENSG0000C |
| 54162065 | 54164452 | -1 | lncRNA | ENSG0000C |
| 54163139 | 54168595 | 1 | lncRNA | ENSG0000C |
| 54230942 | 54280133 | -1 | protein_coding | ENSG0000C |
| 54231397 | 54231476 | -1 | miRNA | ENSG0000C |
| 54255627 | 54255910 | -1 | misc_RNA | ENSG0000C |
| 54262615 | 54279063 | 1 | lncRNA | ENSG0000C |
| 54276631 | 54345083 | 1 | lncRNA | ENSG0000C |
| 54280193 | 54287088 | 1 | protein_coding | ENSG0000C |
| 54292111 | 54301015 | -1 | protein_coding | ENSG0000C |
| 54301202 | 54351846 | 1 | protein_coding | ENSG0000C |
| 54317781 | 54317887 | -1 | snRNA | ENSG0000C |
| 54337216 | 54337314 | 1 | miRNA | ENSG0000C |
| 54344610 | 54344876 | 1 | misc_RNA | ENSG0000C |
| 54353661 | 54497688 | 1 | lncRNA | ENSG0000C |
| 54353792 | 54466985 | 1 | lncRNA | ENSG0000C |
| 54362445 | 54364487 | -1 | protein_coding | ENSG0000C |
| 54369133 | 54391298 | -1 | protein_coding | ENSG0000C |
| 54395261 | 54419266 | -1 | protein_coding | ENSG0000C |
| 54428303 | 54429403 | 1 | lncRNA | ENSG0000C |
| 54455950 | 54473602 | -1 | protein_coding | ENSG0000C |
| 54497752 | 54548243 | 1 | protein_coding | ENSG0000C |
| 54543111 | 54544105 | -1 | lncRNA | ENSG0000C |
| 54549601 | 54579239 | 1 | protein_coding | ENSG0000C |
| 54575387 | 54588659 | -1 | protein_coding | ENSG0000C |
| 54608187 | 54610462 | -1 | transcribed_unitary_ | ENSG0000C |
| 54630811 | 54634895 | -1 | protein_coding | ENSG0000C |
| 54644589 | 54648493 | -1 | protein_coding | ENSG0000C |
| 54682973 | 54687434 | 1 | lncRNA | ENSG0000C |
| 54802746 | 54803589 | -1 | processed_pseudoge | ENSG0000C |
| 54830518 | 54896008 | 1 | protein_coding | ENSG0000C |
| 54948015 | 54984762 | -1 | protein_coding | ENSG0000C |
| 55009746 | 55014247 | 1 | lncRNA | ENSG0000C |
| 55019974 | 55030017 | 1 | protein_coding | ENSG0000C |
| 55113077 | 55118632 | 1 | transcribed_unitary_ | ENSG0000C |

| | | | | |
|---|---|---|---|---|
| 55122668 | 55124120 | -1 | processed_pseudoge | ENSG00000 |
| 55126406 | 55132750 | 1 | protein_coding | ENSG00000 |
| 55158847 | 55159723 | 1 | unprocessed_pseudc | ENSG00000 |
| 55193882 | 55194786 | -1 | unprocessed_pseudc | ENSG00000 |
| 55221025 | 55221975 | 1 | protein_coding | ENSG00000 |
| 55247198 | 55248302 | 1 | protein_coding | ENSG00000 |
| 55262539 | 55263431 | 1 | unprocessed_pseudc | ENSG00000 |
| 55283750 | 55284641 | 1 | unprocessed_pseudc | ENSG00000 |
| 55293988 | 55296569 | -1 | protein_coding | ENSG00000 |
| 55311835 | 55312757 | 1 | unprocessed_pseudc | ENSG00000 |
| 55314343 | 55322364 | 1 | protein_coding | ENSG00000 |
| 55330043 | 55332687 | 1 | protein_coding | ENSG00000 |
| 55342841 | 55343774 | 1 | unprocessed_pseudc | ENSG00000 |
| 55362975 | 55369279 | 1 | protein_coding | ENSG00000 |
| 55376742 | 55378797 | -1 | transcribed_unproce | ENSG00000 |
| 55385837 | 55392964 | 1 | transcribed_unproce | ENSG00000 |
| 55396340 | 55396697 | 1 | unprocessed_pseudc | ENSG00000 |
| 55400430 | 55401505 | 1 | protein_coding | ENSG00000 |
| 55411727 | 55414787 | -1 | processed_pseudoge | ENSG00000 |
| 55426254 | 55427192 | 1 | protein_coding | ENSG00000 |
| 55434734 | 55585471 | -1 | lncRNA | ENSG00000 |
| 55444069 | 55453347 | 1 | protein_coding | ENSG00000 |
| 55469200 | 55470138 | -1 | protein_coding | ENSG00000 |
| 55492378 | 55493316 | 1 | protein_coding | ENSG00000 |
| 55522593 | 55523417 | 1 | unprocessed_pseudc | ENSG00000 |
| 55549602 | 55555832 | 1 | protein_coding | ENSG00000 |
| 55572468 | 55575612 | 1 | protein_coding | ENSG00000 |
| 55608043 | 55615878 | 1 | transcribed_unproce | ENSG00000 |
| 55636860 | 55637854 | 1 | protein_coding | ENSG00000 |
| 55638912 | 55659795 | -1 | lncRNA | ENSG00000 |
| 55646195 | 55646663 | -1 | unprocessed_pseudc | ENSG00000 |
| 55654558 | 55655168 | -1 | processed_pseudoge | ENSG00000 |
| 55662337 | 55662501 | -1 | processed_pseudoge | ENSG00000 |
| 55681678 | 55684611 | 1 | protein_coding | ENSG00000 |
| 55684568 | 55716043 | -1 | protein_coding | ENSG00000 |
| 55716036 | 55724703 | 1 | protein_coding | ENSG00000 |
| 55716037 | 55720087 | 1 | protein_coding | ENSG00000 |
| 55720367 | 55724705 | 1 | protein_coding | ENSG00000 |
| 55725323 | 55729707 | -1 | protein_coding | ENSG00000 |
| 55729104 | 55730852 | 1 | lncRNA | ENSG00000 |
| 55731264 | 55731531 | -1 | misc_RNA | ENSG00000 |
| 55732366 | 55732648 | -1 | processed_pseudoge | ENSG00000 |
| 55743122 | 55757264 | 1 | protein_coding | ENSG00000 |
| 55752463 | 55817724 | -1 | protein_coding | ENSG00000 |
| 55757275 | 55827546 | -1 | protein_coding | ENSG00000 |
| 55761550 | 55762628 | -1 | lncRNA | ENSG00000 |
| 55818041 | 55821879 | 1 | protein_coding | ENSG00000 |
| 55820960 | 55830824 | -1 | protein_coding | ENSG00000 |
| 55829608 | 55836246 | 1 | transcribed_unitary_ | ENSG00000 |
| 55835433 | 55842966 | -1 | protein_coding | ENSG00000 |

| | | | | |
|---|---|---|---|---|
| 55870048 | 55872602 | -1 | processed_pseudoge | ENSG00000 |
| 55900300 | 55900618 | 1 | processed_pseudoge | ENSG00000 |
| 55901413 | 55932618 | -1 | protein_coding | ENSG00000 |
| 55927319 | 55954027 | 1 | protein_coding | ENSG00000 |
| 55929170 | 55929729 | 1 | lncRNA | ENSG00000 |
| 55954105 | 55973317 | -1 | protein_coding | ENSG00000 |
| 55966781 | 55972789 | 1 | protein_coding | ENSG00000 |
| 55966838 | 55967474 | -1 | lncRNA | ENSG00000 |
| 55973913 | 55996683 | 1 | protein_coding | ENSG00000 |
| 55980432 | 55981035 | -1 | processed_pseudoge | ENSG00000 |
| 55997180 | 56006641 | 1 | protein_coding | ENSG00000 |
| 56007659 | 56038435 | 1 | protein_coding | ENSG00000 |
| 56010091 | 56010574 | -1 | lncRNA | ENSG00000 |
| 56029649 | 56041806 | -1 | lncRNA | ENSG00000 |
| 56041351 | 56044697 | 1 | protein_coding | ENSG00000 |
| 56076799 | 56103505 | 1 | protein_coding | ENSG00000 |
| 56101331 | 56109289 | 1 | protein_coding | ENSG00000 |
| 56104537 | 56113910 | 1 | protein_coding | ENSG00000 |
| 56104614 | 56113905 | -1 | lncRNA | ENSG00000 |
| 56116590 | 56117967 | 1 | protein_coding | ENSG00000 |
| 56118250 | 56144671 | 1 | protein_coding | ENSG00000 |
| 56118260 | 56127514 | 1 | protein_coding | ENSG00000 |
| 56118968 | 56119939 | -1 | lncRNA | ENSG00000 |
| 56120033 | 56129619 | -1 | lncRNA | ENSG00000 |
| 56150796 | 56158220 | -1 | lncRNA | ENSG00000 |
| 56152256 | 56159647 | 1 | protein_coding | ENSG00000 |
| 56158161 | 56163496 | 1 | protein_coding | ENSG00000 |
| 56162359 | 56189567 | -1 | protein_coding | ENSG00000 |
| 56162359 | 56190284 | -1 | lncRNA | ENSG00000 |
| 56202179 | 56221933 | -1 | protein_coding | ENSG00000 |
| 56222015 | 56229854 | 1 | protein_coding | ENSG00000 |
| 56230049 | 56237846 | 1 | protein_coding | ENSG00000 |
| 56237807 | 56258384 | -1 | protein_coding | ENSG00000 |
| 56266858 | 56270966 | 1 | protein_coding | ENSG00000 |
| 56267793 | 56270104 | -1 | lncRNA | ENSG00000 |
| 56271699 | 56300391 | -1 | protein_coding | ENSG00000 |
| 56285916 | 56316059 | -1 | protein_coding | ENSG00000 |
| 56300076 | 56317992 | 1 | lncRNA | ENSG00000 |
| 56308868 | 56309449 | 1 | lncRNA | ENSG00000 |
| 56309842 | 56316119 | -1 | protein_coding | ENSG00000 |
| 56316223 | 56334053 | -1 | protein_coding | ENSG00000 |
| 56334174 | 56340410 | 1 | protein_coding | ENSG00000 |
| 56341597 | 56360167 | -1 | protein_coding | ENSG00000 |
| 56352387 | 56352450 | -1 | snRNA | ENSG00000 |
| 56360568 | 56362857 | -1 | protein_coding | ENSG00000 |
| 56376818 | 56377331 | -1 | processed_pseudoge | ENSG00000 |
| 56380361 | 56382711 | -1 | processed_pseudoge | ENSG00000 |
| 56411944 | 56413190 | -1 | lncRNA | ENSG00000 |
| 56413647 | 56414045 | -1 | processed_pseudoge | ENSG00000 |
| 56416363 | 56449426 | -1 | protein_coding | ENSG00000 |

| | | | | |
|---|---|---|---|---|
| 56449502 | 56469166 | -1 | protein_coding | ENSG0000C |
| 56449700 | 56488170 | -1 | protein_coding | ENSG0000C |
| 56468578 | 56479708 | 1 | protein_coding | ENSG0000C |
| 56470944 | 56488414 | -1 | protein_coding | ENSG0000C |
| 56511002 | 56512703 | 1 | processed_pseudoge | ENSG0000C |
| 56521820 | 56596193 | 1 | protein_coding | ENSG0000C |
| 56588518 | 56588621 | 1 | snRNA | ENSG0000C |
| 56595596 | 56636816 | -1 | protein_coding | ENSG0000C |
| 56638175 | 56645984 | -1 | protein_coding | ENSG0000C |
| 56645027 | 56645101 | -1 | snoRNA | ENSG0000C |
| 56663341 | 56688408 | -1 | protein_coding | ENSG0000C |
| 56670450 | 56670746 | -1 | misc_RNA | ENSG0000C |
| 56712428 | 56731628 | -1 | protein_coding | ENSG0000C |
| 56714612 | 56741535 | -1 | protein_coding | ENSG0000C |
| 56731296 | 56752374 | -1 | protein_coding | ENSG0000C |
| 56752161 | 56787790 | 1 | protein_coding | ENSG0000C |
| 56822985 | 56874268 | 1 | transcribed_processe | ENSG0000C |
| 56840973 | 56841273 | -1 | processed_pseudoge | ENSG0000C |
| 56855694 | 56855886 | 1 | unprocessed_pseudc | ENSG0000C |
| 56861375 | 56861508 | 1 | snoRNA | ENSG0000C |
| 56923133 | 56934408 | -1 | protein_coding | ENSG0000C |
| 56941174 | 56941482 | -1 | processed_pseudoge | ENSG0000C |
| 56951431 | 56959374 | -1 | protein_coding | ENSG0000C |
| 56981211 | 56985960 | -1 | unprocessed_pseudc | ENSG0000C |
| 56987734 | 56990756 | 1 | lncRNA | ENSG0000C |
| 56994492 | 56998447 | 1 | protein_coding | ENSG0000C |
| 56998836 | 57006546 | -1 | protein_coding | ENSG0000C |
| 57010000 | 57028883 | -1 | protein_coding | ENSG0000C |
| 57028517 | 57051198 | -1 | protein_coding | ENSG0000C |
| 57055643 | 57088063 | -1 | protein_coding | ENSG0000C |
| 57089043 | 57095476 | 1 | protein_coding | ENSG0000C |
| 57095408 | 57132139 | -1 | protein_coding | ENSG0000C |
| 57128483 | 57213361 | 1 | protein_coding | ENSG0000C |
| 57144620 | 57147619 | -1 | lncRNA | ENSG0000C |
| 57194504 | 57194576 | 1 | miRNA | ENSG0000C |
| 57216794 | 57226449 | 1 | protein_coding | ENSG0000C |
| 57229498 | 57230198 | -1 | lncRNA | ENSG0000C |
| 57229573 | 57234935 | 1 | protein_coding | ENSG0000C |
| 57234903 | 57240715 | -1 | protein_coding | ENSG0000C |
| 57243453 | 57251188 | -1 | protein_coding | ENSG0000C |
| 57249609 | 57296484 | -1 | protein_coding | ENSG0000C |
| 57253762 | 57431005 | -1 | protein_coding | ENSG0000C |
| 57415296 | 57415403 | 1 | snRNA | ENSG0000C |
| 57431116 | 57433935 | 1 | lncRNA | ENSG0000C |
| 57434784 | 57452062 | 1 | protein_coding | ENSG0000C |
| 57452323 | 57459280 | 1 | protein_coding | ENSG0000C |
| 57457596 | 57462818 | -1 | lncRNA | ENSG0000C |
| 57459785 | 57472268 | 1 | protein_coding | ENSG0000C |
| 57472264 | 57488814 | -1 | protein_coding | ENSG0000C |
| 57475445 | 57517569 | 1 | protein_coding | ENSG0000C |

| | | | | |
|---|---|---|---|---|
| 57512688 | 57512750 | 1 | miRNA | ENSG0000C |
| 57516588 | 57521737 | -1 | protein_coding | ENSG0000C |
| 57517712 | 57520480 | -1 | protein_coding | ENSG0000C |
| 57519163 | 57519259 | -1 | miRNA | ENSG0000C |
| 57520710 | 57530148 | 1 | protein_coding | ENSG0000C |
| 57530102 | 57547331 | -1 | protein_coding | ENSG0000C |
| 57550044 | 57586633 | 1 | protein_coding | ENSG0000C |
| 57591174 | 57603418 | 1 | protein_coding | ENSG0000C |
| 57604622 | 57609804 | 1 | protein_coding | ENSG0000C |
| 57610180 | 57617245 | 1 | protein_coding | ENSG0000C |
| 57612118 | 57619638 | -1 | lncRNA | ENSG0000C |
| 57615280 | 57626125 | 1 | protein_coding | ENSG0000C |
| 57619527 | 57626151 | 1 | protein_coding | ENSG0000C |
| 57623409 | 57633239 | -1 | protein_coding | ENSG0000C |
| 57674665 | 57675250 | 1 | processed_pseudoge | ENSG0000C |
| 57693955 | 57721557 | 1 | protein_coding | ENSG0000C |
| 57694132 | 57721510 | -1 | lncRNA | ENSG0000C |
| 57723761 | 57742157 | -1 | protein_coding | ENSG0000C |
| 57726271 | 57728356 | 1 | lncRNA | ENSG0000C |
| 57738013 | 57750219 | 1 | protein_coding | ENSG0000C |
| 57747727 | 57756013 | -1 | protein_coding | ENSG0000C |
| 57748618 | 57748682 | -1 | miRNA | ENSG0000C |
| 57755103 | 57760411 | 1 | protein_coding | ENSG0000C |
| 57762334 | 57768986 | -1 | protein_coding | ENSG0000C |
| 57768471 | 57772119 | -1 | protein_coding | ENSG0000C |
| 57771492 | 57782541 | 1 | protein_coding | ENSG0000C |
| 57772660 | 57797554 | 1 | protein_coding | ENSG0000C |
| 57782761 | 57808071 | 1 | protein_coding | ENSG0000C |
| 57797376 | 57818704 | -1 | protein_coding | ENSG0000C |
| 57803838 | 57804415 | 1 | lncRNA | ENSG0000C |
| 57806154 | 57806260 | -1 | snRNA | ENSG0000C |
| 57814494 | 57814926 | 1 | lncRNA | ENSG0000C |
| 57819927 | 57846729 | -1 | protein_coding | ENSG0000C |
| 57824609 | 57824692 | -1 | miRNA | ENSG0000C |
| 57837092 | 57842745 | 1 | lncRNA | ENSG0000C |
| 57869834 | 57896482 | -1 | lncRNA | ENSG0000C |
| 57894232 | 57896846 | 1 | lncRNA | ENSG0000C |
| 57906039 | 57959148 | 1 | protein_coding | ENSG0000C |
| 57930115 | 57936345 | -1 | lncRNA | ENSG0000C |
| 57967058 | 57968399 | 1 | processed_pseudoge | ENSG0000C |
| 57977965 | 57978277 | -1 | misc_RNA | ENSG0000C |
| 57983795 | 57984761 | -1 | processed_pseudoge | ENSG0000C |
| 58087892 | 58093362 | 1 | lncRNA | ENSG0000C |
| 58094914 | 58098013 | -1 | TEC | ENSG0000C |
| 58103907 | 58105935 | 1 | TEC | ENSG0000C |
| 58244378 | 58244865 | 1 | processed_pseudoge | ENSG0000C |
| 58394596 | 58395138 | 1 | processed_pseudoge | ENSG0000C |
| 58544124 | 58813060 | 1 | lncRNA | ENSG0000C |
| 58565959 | 58781747 | -1 | lncRNA | ENSG0000C |
| 58872149 | 58920522 | -1 | protein_coding | ENSG0000C |

| | | | | |
|---|---|---|---|---|
| 58920639 | 59064238 | 1 | lncRNA | ENSG00000( |
| 59020169 | 59020894 | 1 | processed_pseudoge | ENSG00000( |
| 59052980 | 59054184 | -1 | processed_pseudoge | ENSG00000( |
| 59129511 | 59130875 | 1 | lncRNA | ENSG00000( |
| 59322765 | 59436874 | 1 | lncRNA | ENSG00000( |
| 59366218 | 59411171 | -1 | lncRNA | ENSG00000( |
| 59369051 | 59369151 | -1 | snRNA | ENSG00000( |
| 59450673 | 59450772 | -1 | snRNA | ENSG00000( |
| 59523278 | 59524184 | -1 | lncRNA | ENSG00000( |
| 59586793 | 59586943 | -1 | snRNA | ENSG00000( |
| 59596029 | 59789855 | 1 | protein_coding | ENSG00000( |
| 59719698 | 59719944 | -1 | processed_pseudoge | ENSG00000( |
| 59812102 | 59812576 | 1 | processed_pseudoge | ENSG00000( |
| 60092864 | 60096071 | 1 | lncRNA | ENSG00000( |
| 60150187 | 60269686 | -1 | lncRNA | ENSG00000( |
| 60215994 | 60216384 | 1 | processed_pseudoge | ENSG00000( |
| 60341824 | 60341941 | -1 | misc_RNA | ENSG00000( |
| 60363822 | 60363935 | 1 | snoRNA | ENSG00000( |
| 60418983 | 60419293 | 1 | processed_pseudoge | ENSG00000( |
| 61231159 | 61232937 | -1 | processed_pseudoge | ENSG00000( |
| 61600067 | 61600843 | -1 | processed_pseudoge | ENSG00000( |
| 61668302 | 61702771 | -1 | lncRNA | ENSG00000( |
| 61708259 | 62279150 | -1 | protein_coding | ENSG00000( |
| 61775923 | 61776400 | 1 | processed_pseudoge | ENSG00000( |
| 61879318 | 61880772 | 1 | processed_pseudoge | ENSG00000( |
| 62021570 | 62022174 | -1 | processed_pseudoge | ENSG00000( |
| 62139999 | 62145643 | -1 | lncRNA | ENSG00000( |
| 62234390 | 62235515 | -1 | processed_pseudoge | ENSG00000( |
| 62260338 | 62417431 | 1 | protein_coding | ENSG00000( |
| 62260359 | 62260454 | 1 | miRNA | ENSG00000( |
| 62334091 | 62334197 | -1 | snRNA | ENSG00000( |
| 62393607 | 62393880 | -1 | misc_RNA | ENSG00000( |
| 62466817 | 62600476 | 1 | protein_coding | ENSG00000( |
| 62482349 | 62484932 | -1 | lncRNA | ENSG00000( |
| 62545582 | 62545679 | -1 | snRNA | ENSG00000( |
| 62601751 | 62603690 | -1 | lncRNA | ENSG00000( |
| 62602752 | 62622213 | 1 | lncRNA | ENSG00000( |
| 62603686 | 62603769 | 1 | miRNA | ENSG00000( |
| 62603909 | 62604399 | 1 | lncRNA | ENSG00000( |
| 62643994 | 62935150 | -1 | protein_coding | ENSG00000( |
| 62658823 | 62659225 | 1 | processed_pseudoge | ENSG00000( |
| 62755227 | 62756233 | 1 | processed_pseudoge | ENSG00000( |
| 62850738 | 62850901 | -1 | snRNA | ENSG00000( |
| 62935701 | 62935802 | 1 | misc_RNA | ENSG00000( |
| 62965325 | 62965969 | 1 | processed_pseudoge | ENSG00000( |
| 63002469 | 63002795 | 1 | processed_pseudoge | ENSG00000( |
| 63003553 | 63006902 | 1 | transcribed_processe | ENSG00000( |
| 63004338 | 63006541 | -1 | lncRNA | ENSG00000( |
| 63055275 | 63055765 | -1 | processed_pseudoge | ENSG00000( |
| 63142759 | 63150942 | -1 | protein_coding | ENSG00000( |

| | | | | |
|---|---|---|---|---|
| 63151065 | 63154006 | 1 | TEC | ENSG0000( |
| 63271404 | 63272336 | 1 | processed_pseudoge | ENSG0000( |
| 63292625 | 63360037 | -1 | lncRNA | ENSG0000( |
| 63558913 | 63668939 | -1 | protein_coding | ENSG0000( |
| 63623788 | 63795718 | -1 | lncRNA | ENSG0000( |
| 63651627 | 63651930 | 1 | processed_pseudoge | ENSG0000( |
| 63682523 | 63724935 | -1 | transcribed_unproce | ENSG0000( |
| 63760358 | 63760612 | -1 | processed_pseudoge | ENSG0000( |
| 63779842 | 63809562 | 1 | protein_coding | ENSG0000( |
| 63804739 | 63822156 | -1 | lncRNA | ENSG0000( |
| 63822021 | 63823895 | 1 | processed_pseudoge | ENSG0000( |
| 63844700 | 64162217 | 1 | protein_coding | ENSG0000( |
| 63871739 | 63872044 | -1 | processed_pseudoge | ENSG0000( |
| 63878787 | 63879474 | 1 | lncRNA | ENSG0000( |
| 63887379 | 63888598 | -1 | processed_pseudoge | ENSG0000( |
| 64032412 | 64032665 | 1 | processed_pseudoge | ENSG0000( |
| 64038562 | 64097618 | -1 | lncRNA | ENSG0000( |
| 64099415 | 64130539 | -1 | lncRNA | ENSG0000( |
| 64108763 | 64120489 | -1 | lncRNA | ENSG0000( |
| 64146388 | 64147857 | -1 | lncRNA | ENSG0000( |
| 64186316 | 64222296 | -1 | protein_coding | ENSG0000( |
| 64204793 | 64204899 | 1 | snRNA | ENSG0000( |
| 64222337 | 64397065 | 1 | transcribed_process( | ENSG0000( |
| 64228560 | 64228675 | -1 | snRNA | ENSG0000( |
| 64264762 | 64390758 | -1 | protein_coding | ENSG0000( |
| 64266413 | 64266766 | 1 | processed_pseudoge | ENSG0000( |
| 64338178 | 64338797 | -1 | processed_pseudoge | ENSG0000( |
| 64355762 | 64355865 | -1 | snoRNA | ENSG0000( |
| 64404392 | 64451125 | 1 | protein_coding | ENSG0000( |
| 64442510 | 64443330 | -1 | processed_pseudoge | ENSG0000( |
| 64451591 | 64452901 | -1 | lncRNA | ENSG0000( |
| 64452092 | 64502114 | 1 | protein_coding | ENSG0000( |
| 64507001 | 64697564 | 1 | protein_coding | ENSG0000( |
| 64507166 | 64533638 | -1 | lncRNA | ENSG0000( |
| 64575665 | 64575741 | 1 | snoRNA | ENSG0000( |
| 64599078 | 64609459 | -1 | lncRNA | ENSG0000( |
| 64622509 | 64622605 | 1 | miRNA | ENSG0000( |
| 64622509 | 64622605 | -1 | miRNA | ENSG0000( |
| 64628344 | 64629976 | 1 | lncRNA | ENSG0000( |
| 64654060 | 64655019 | -1 | lncRNA | ENSG0000( |
| 64696191 | 64696550 | -1 | TEC | ENSG0000( |
| 64709458 | 64710513 | 1 | TEC | ENSG0000( |
| 64713445 | 64759431 | -1 | protein_coding | ENSG0000( |
| 64759521 | 64779318 | 1 | lncRNA | ENSG0000( |
| 64780516 | 64881033 | 1 | protein_coding | ENSG0000( |
| 64820179 | 64825955 | -1 | lncRNA | ENSG0000( |
| 64867956 | 64868699 | -1 | processed_pseudoge | ENSG0000( |
| 64883394 | 64990646 | 1 | lncRNA | ENSG0000( |
| 64920845 | 64982524 | -1 | lncRNA | ENSG0000( |
| 64939023 | 64939117 | 1 | snRNA | ENSG0000( |

| | | | | |
|---|---|---|---|---|
| 65017468 | 65054124 | 1 | lncRNA | ENSG0000( |
| 65050626 | 65121305 | -1 | protein_coding | ENSG0000( |
| 65095749 | 65095855 | 1 | snRNA | ENSG0000( |
| 65134688 | 65135273 | 1 | processed_pseudoge | ENSG0000( |
| 65169583 | 65248355 | 1 | protein_coding | ENSG0000( |
| 65171262 | 65171917 | 1 | lncRNA | ENSG0000( |
| 65278643 | 65491430 | 1 | protein_coding | ENSG0000( |
| 65281657 | 65286728 | 1 | lncRNA | ENSG0000( |
| 65418731 | 65420027 | -1 | processed_pseudoge | ENSG0000( |
| 65466820 | 65642372 | -1 | lncRNA | ENSG0000( |
| 65556925 | 65560889 | 1 | lncRNA | ENSG0000( |
| 65589111 | 65613000 | 1 | lncRNA | ENSG0000( |
| 65622273 | 65663299 | 1 | lncRNA | ENSG0000( |
| 65645992 | 65646462 | -1 | processed_pseudoge | ENSG0000( |
| 65758020 | 65826997 | -1 | transcribed_processe | ENSG0000( |
| 65824131 | 65966295 | 1 | protein_coding | ENSG0000( |
| 65830750 | 65831050 | 1 | lncRNA | ENSG0000( |
| 65851340 | 65882167 | -1 | lncRNA | ENSG0000( |
| 65934777 | 65948479 | -1 | lncRNA | ENSG0000( |
| 66023620 | 66023726 | -1 | miRNA | ENSG0000( |
| 66027725 | 66031066 | -1 | lncRNA | ENSG0000( |
| 66037235 | 66037714 | 1 | processed_pseudoge | ENSG0000( |
| 66066221 | 66066338 | 1 | rRNA_pseudogene | ENSG0000( |
| 66116555 | 66130750 | -1 | protein_coding | ENSG0000( |
| 66123917 | 66169985 | -1 | protein_coding | ENSG0000( |
| 66130751 | 66134449 | 1 | lncRNA | ENSG0000( |
| 66135846 | 66170072 | -1 | protein_coding | ENSG0000( |
| 66189195 | 66254622 | 1 | protein_coding | ENSG0000( |
| 66234079 | 66235234 | -1 | processed_pseudoge | ENSG0000( |
| 66251082 | 66251157 | 1 | miRNA | ENSG0000( |
| 66251745 | 66257434 | -1 | lncRNA | ENSG0000( |
| 66275940 | 66276651 | -1 | processed_pseudoge | ENSG0000( |
| 66292683 | 66292987 | 1 | misc_RNA | ENSG0000( |
| 66302545 | 66347645 | 1 | protein_coding | ENSG0000( |
| 66347431 | 67069162 | -1 | protein_coding | ENSG0000( |
| 66563524 | 66563694 | 1 | processed_pseudoge | ENSG0000( |
| 66563967 | 66569194 | -1 | processed_pseudoge | ENSG0000( |
| 66950754 | 67096410 | -1 | lncRNA | ENSG0000( |
| 67065018 | 67065278 | -1 | TEC | ENSG0000( |
| 67131567 | 67132205 | -1 | processed_pseudoge | ENSG0000( |
| 67265842 | 67266966 | -1 | processed_pseudoge | ENSG0000( |
| 67269358 | 67319953 | 1 | protein_coding | ENSG0000( |
| 67329014 | 67350924 | 1 | lncRNA | ENSG0000( |
| 67351436 | 67352039 | 1 | processed_pseudoge | ENSG0000( |
| 67424272 | 67424986 | -1 | processed_pseudoge | ENSG0000( |
| 67440998 | 67442559 | -1 | lncRNA | ENSG0000( |
| 67443105 | 67590771 | 1 | lncRNA | ENSG0000( |
| 67571197 | 67589987 | -1 | lncRNA | ENSG0000( |
| 67648338 | 67665406 | 1 | protein_coding | ENSG0000( |
| 67658991 | 67670919 | -1 | lncRNA | ENSG0000( |

| | | | | |
|---|---|---|---|---|
| 67709047 | 67729475 | -1 | lncRNA | ENSG0000C |
| 67753701 | 67753817 | 1 | TEC | ENSG0000C |
| 67929235 | 67970017 | 1 | lncRNA | ENSG0000C |
| 67978107 | 67978387 | -1 | processed_pseudoge | ENSG0000C |
| 67989445 | 68234686 | 1 | lncRNA | ENSG0000C |
| 68001394 | 68001548 | -1 | processed_pseudoge | ENSG0000C |
| 68035767 | 68036853 | 1 | processed_pseudoge | ENSG0000C |
| 68143318 | 68143690 | 1 | processed_pseudoge | ENSG0000C |
| 68154768 | 68159740 | -1 | protein_coding | ENSG0000C |
| 68201349 | 68225810 | -1 | protein_coding | ENSG0000C |
| 68248242 | 68253604 | -1 | protein_coding | ENSG0000C |
| 68272443 | 68332381 | -1 | protein_coding | ENSG0000C |
| 68332888 | 68451663 | -1 | lncRNA | ENSG0000C |
| 68344664 | 68349959 | 1 | lncRNA | ENSG0000C |
| 68426331 | 68427737 | -1 | lncRNA | ENSG0000C |
| 68473741 | 68474902 | 1 | processed_pseudoge | ENSG0000C |
| 68487687 | 68487863 | 1 | processed_pseudoge | ENSG0000C |
| 68552995 | 68553882 | -1 | processed_pseudoge | ENSG0000C |
| 68575858 | 68576136 | -1 | misc_RNA | ENSG0000C |
| 68610855 | 68671901 | 1 | protein_coding | ENSG0000C |
| 68626870 | 68627187 | -1 | processed_pseudoge | ENSG0000C |
| 68627234 | 68627375 | -1 | snoRNA | ENSG0000C |
| 68642519 | 68642993 | -1 | processed_pseudoge | ENSG0000C |
| 68674371 | 68687755 | -1 | lncRNA | ENSG0000C |
| 68684595 | 68686308 | 1 | processed_pseudoge | ENSG0000C |
| 68686951 | 68745809 | 1 | protein_coding | ENSG0000C |
| 68705634 | 68707066 | 1 | processed_pseudoge | ENSG0000C |
| 68746176 | 68793964 | 1 | protein_coding | ENSG0000C |
| 68805011 | 68805479 | -1 | processed_pseudoge | ENSG0000C |
| 68808177 | 68850686 | 1 | protein_coding | ENSG0000C |
| 68828118 | 68828553 | 1 | lncRNA | ENSG0000C |
| 68841288 | 68843237 | -1 | lncRNA | ENSG0000C |
| 68841946 | 68842384 | 1 | processed_pseudoge | ENSG0000C |
| 68842197 | 68971570 | -1 | protein_coding | ENSG0000C |
| 68911921 | 68911982 | 1 | snRNA | ENSG0000C |
| 68913804 | 68914022 | 1 | processed_pseudoge | ENSG0000C |
| 68957377 | 68957677 | -1 | processed_pseudoge | ENSG0000C |
| 69212108 | 69224242 | 1 | lncRNA | ENSG0000C |
| 69239569 | 69274358 | 1 | protein_coding | ENSG0000C |
| 69273157 | 69273218 | -1 | miRNA | ENSG0000C |
| 69284397 | 69285485 | 1 | processed_pseudoge | ENSG0000C |
| 69326574 | 69331882 | -1 | lncRNA | ENSG0000C |
| 69348341 | 69354234 | 1 | protein_coding | ENSG0000C |
| 69353493 | 69354225 | -1 | lncRNA | ENSG0000C |
| 69359710 | 69390870 | 1 | protein_coding | ENSG0000C |
| 69400903 | 69401200 | 1 | misc_RNA | ENSG0000C |
| 69422656 | 69422995 | 1 | processed_pseudoge | ENSG0000C |
| 69449470 | 69460724 | 1 | lncRNA | ENSG0000C |
| 69470349 | 69579789 | 1 | protein_coding | ENSG0000C |
| 69584722 | 69584823 | -1 | miRNA | ENSG0000C |

| | | | | |
|---|---|---|---|---|
| 69584723 | 69584822 | 1 | miRNA | ENSG0000C |
| 69585426 | 69601570 | 1 | protein_coding | ENSG0000C |
| 69608564 | 69610907 | -1 | protein_coding | ENSG0000C |
| 69643360 | 69699476 | -1 | protein_coding | ENSG0000C |
| 69660034 | 69661429 | -1 | TEC | ENSG0000C |
| 69713633 | 69738574 | -1 | lncRNA | ENSG0000C |
| 69738681 | 69823204 | 1 | protein_coding | ENSG0000C |
| 69801669 | 69855363 | 1 | protein_coding | ENSG0000C |
| 69825227 | 69959097 | 1 | protein_coding | ENSG0000C |
| 69901918 | 70243379 | -1 | lncRNA | ENSG0000C |
| 69946543 | 69947081 | -1 | lncRNA | ENSG0000C |
| 70180338 | 70202004 | 1 | lncRNA | ENSG0000C |
| 70219132 | 70221862 | -1 | lncRNA | ENSG0000C |
| 70239114 | 70242430 | 1 | lncRNA | ENSG0000C |
| 70242994 | 70354993 | 1 | protein_coding | ENSG0000C |
| 70321542 | 70324274 | -1 | TEC | ENSG0000C |
| 70366290 | 70434292 | 1 | protein_coding | ENSG0000C |
| 70443784 | 70443923 | -1 | snRNA | ENSG0000C |
| 70468080 | 70543040 | 1 | lncRNA | ENSG0000C |
| 70515866 | 70637440 | -1 | protein_coding | ENSG0000C |
| 70570969 | 70571440 | 1 | lncRNA | ENSG0000C |
| 70638073 | 70920738 | -1 | protein_coding | ENSG0000C |
| 70671918 | 70672856 | 1 | processed_pseudoge | ENSG0000C |
| 70906917 | 70907018 | 1 | misc_RNA | ENSG0000C |
| 71007773 | 71032083 | -1 | lncRNA | ENSG0000C |
| 71034122 | 71104526 | -1 | lncRNA | ENSG0000C |
| 71047402 | 71118247 | -1 | lncRNA | ENSG0000C |
| 71125085 | 71441898 | -1 | protein_coding | ENSG0000C |
| 71439798 | 71586310 | 1 | protein_coding | ENSG0000C |
| 71448405 | 71448850 | 1 | lncRNA | ENSG0000C |
| 71582293 | 71594404 | -1 | lncRNA | ENSG0000C |
| 71609472 | 71667725 | -1 | protein_coding | ENSG0000C |
| 71639355 | 71639483 | 1 | snoRNA | ENSG0000C |
| 71664301 | 71680644 | 1 | protein_coding | ENSG0000C |
| 71674204 | 71698968 | 1 | protein_coding | ENSG0000C |
| 71686082 | 71705047 | 1 | protein_coding | ENSG0000C |
| 71709171 | 71710374 | -1 | lncRNA | ENSG0000C |
| 71754863 | 71800286 | 1 | protein_coding | ENSG0000C |
| 71793855 | 71799627 | -1 | lncRNA | ENSG0000C |
| 71835034 | 71835396 | -1 | processed_pseudoge | ENSG0000C |
| 71839707 | 71927248 | 1 | protein_coding | ENSG0000C |
| 71849228 | 71850428 | -1 | processed_pseudoge | ENSG0000C |
| 71938845 | 72186618 | 1 | protein_coding | ENSG0000C |
| 72046149 | 72057377 | -1 | lncRNA | ENSG0000C |
| 72087266 | 72670758 | 1 | protein_coding | ENSG0000C |
| 72249964 | 72276954 | -1 | lncRNA | ENSG0000C |
| 72431722 | 72432075 | -1 | processed_pseudoge | ENSG0000C |
| 72610678 | 72611316 | 1 | processed_pseudoge | ENSG0000C |
| 72727923 | 72728844 | -1 | lncRNA | ENSG0000C |
| 73115957 | 73143858 | -1 | lncRNA | ENSG0000C |

| | | | | |
|---|---|---|---|---|
| 73159190 | 73208317 | 1 | lncRNA | ENSG0000C |
| 73203815 | 73206835 | -1 | lncRNA | ENSG0000C |
| 73648666 | 73649039 | 1 | processed_pseudoge | ENSG0000C |
| 73679188 | 73679296 | -1 | snRNA | ENSG0000C |
| 73758657 | 73846188 | 1 | lncRNA | ENSG0000C |
| 73764279 | 73764413 | -1 | snoRNA | ENSG0000C |
| 73820315 | 74402600 | -1 | lncRNA | ENSG0000C |
| 73906940 | 73919337 | 1 | lncRNA | ENSG0000C |
| 74039024 | 74049981 | 1 | lncRNA | ENSG0000C |
| 74170445 | 74171830 | 1 | processed_pseudoge | ENSG0000C |
| 74248637 | 74283669 | -1 | lncRNA | ENSG0000C |
| 74274952 | 74275231 | 1 | processed_pseudoge | ENSG0000C |
| 74292324 | 74293096 | 1 | processed_pseudoge | ENSG0000C |
| 74537835 | 74545430 | 1 | protein_coding | ENSG0000C |
| 74538145 | 74538633 | -1 | lncRNA | ENSG0000C |
| 74656561 | 74656860 | 1 | processed_pseudoge | ENSG0000C |
| 74663774 | 74664141 | -1 | processed_pseudoge | ENSG0000C |
| 74728038 | 74728851 | -1 | processed_pseudoge | ENSG0000C |
| 75020969 | 75044793 | 1 | lncRNA | ENSG0000C |
| 75040077 | 75209868 | -1 | protein_coding | ENSG0000C |
| 75234740 | 75298508 | 1 | lncRNA | ENSG0000C |
| 75257635 | 75258634 | -1 | processed_pseudoge | ENSG0000C |
| 75275979 | 75390928 | -1 | protein_coding | ENSG0000C |
| 75305452 | 75306357 | 1 | processed_pseudoge | ENSG0000C |
| 75333798 | 75334486 | -1 | lncRNA | ENSG0000C |
| 75334670 | 75370560 | 1 | protein_coding | ENSG0000C |
| 75391089 | 75432688 | 1 | protein_coding | ENSG0000C |
| 75480753 | 75503863 | 1 | protein_coding | ENSG0000C |
| 75483454 | 75489820 | -1 | lncRNA | ENSG0000C |
| 75490863 | 75511636 | -1 | protein_coding | ENSG0000C |
| 75563202 | 75984015 | -1 | lncRNA | ENSG0000C |
| 75600047 | 75601551 | 1 | processed_pseudoge | ENSG0000C |
| 75649195 | 75691937 | 1 | lncRNA | ENSG0000C |
| 75688794 | 75689096 | -1 | processed_pseudoge | ENSG0000C |
| 75689152 | 75689304 | -1 | snoRNA | ENSG0000C |
| 75694010 | 75698816 | -1 | lncRNA | ENSG0000C |
| 75926213 | 75926467 | 1 | processed_pseudoge | ENSG0000C |
| 75957833 | 75958130 | -1 | misc_RNA | ENSG0000C |
| 75964440 | 75967062 | 1 | lncRNA | ENSG0000C |
| 75964799 | 75964995 | 1 | processed_pseudoge | ENSG0000C |
| 76025447 | 76033932 | -1 | protein_coding | ENSG0000C |
| 76030494 | 76031378 | 1 | lncRNA | ENSG0000C |
| 76032658 | 76033897 | 1 | lncRNA | ENSG0000C |
| 76036585 | 76084958 | -1 | protein_coding | ENSG0000C |
| 76057538 | 76058115 | 1 | processed_pseudoge | ENSG0000C |
| 76115711 | 76116100 | 1 | processed_pseudoge | ENSG0000C |
| 76137425 | 76138610 | 1 | processed_pseudoge | ENSG0000C |
| 76139567 | 76139671 | 1 | snRNA | ENSG0000C |
| 76259836 | 76305969 | 1 | lncRNA | ENSG0000C |
| 76315751 | 76316048 | 1 | misc_RNA | ENSG0000C |

| | | | | |
|---|---|---|---|---|
| 76344474 | 76348415 | -1 | protein_coding | ENSG0000C |
| 76351797 | 76559809 | -1 | protein_coding | ENSG0000C |
| 76562294 | 76615567 | 1 | lncRNA | ENSG0000C |
| 76589476 | 76600644 | -1 | lncRNA | ENSG0000C |
| 76599671 | 76600470 | 1 | processed_pseudoge | ENSG0000C |
| 76619386 | 76620043 | 1 | processed_pseudoge | ENSG0000C |
| 76660405 | 76660884 | 1 | processed_pseudoge | ENSG0000C |
| 76721534 | 76722286 | 1 | processed_pseudoge | ENSG0000C |
| 76757191 | 76757914 | 1 | processed_pseudoge | ENSG0000C |
| 76763588 | 76853696 | 1 | protein_coding | ENSG0000C |
| 76858709 | 76879023 | -1 | protein_coding | ENSG0000C |
| 76878193 | 76880352 | 1 | lncRNA | ENSG0000C |
| 76984079 | 76984669 | -1 | processed_pseudoge | ENSG0000C |
| 77021248 | 77065569 | -1 | protein_coding | ENSG0000C |
| 77129178 | 77163638 | 1 | lncRNA | ENSG0000C |
| 77219603 | 77317234 | -1 | lncRNA | ENSG0000C |
| 77324641 | 78213010 | 1 | protein_coding | ENSG0000C |
| 77379770 | 77390869 | 1 | lncRNA | ENSG0000C |
| 77775783 | 77783576 | 1 | lncRNA | ENSG0000C |
| 78052181 | 78091737 | -1 | lncRNA | ENSG0000C |
| 78130337 | 78131004 | -1 | processed_pseudoge | ENSG0000C |
| 78326680 | 78359746 | -1 | lncRNA | ENSG0000C |
| 78352519 | 78483030 | 1 | lncRNA | ENSG0000C |
| 78426826 | 78442952 | -1 | lncRNA | ENSG0000C |
| 78448995 | 78540675 | -1 | lncRNA | ENSG0000C |
| 78793526 | 78794114 | -1 | processed_pseudoge | ENSG0000C |
| 78794132 | 78808995 | 1 | lncRNA | ENSG0000C |
| 78863993 | 79452008 | 1 | protein_coding | ENSG0000C |
| 78960258 | 79045644 | -1 | lncRNA | ENSG0000C |
| 79341205 | 79503396 | -1 | lncRNA | ENSG0000C |
| 79419257 | 79419321 | 1 | miRNA | ENSG0000C |
| 79500119 | 79500687 | -1 | processed_pseudoge | ENSG0000C |
| 79540203 | 79550535 | 1 | lncRNA | ENSG0000C |
| 79558782 | 79559166 | 1 | processed_pseudoge | ENSG0000C |
| 79574979 | 79690964 | -1 | protein_coding | ENSG0000C |
| 79690144 | 79778451 | 1 | lncRNA | ENSG0000C |
| 79773563 | 79935460 | -1 | protein_coding | ENSG0000C |
| 79818784 | 79819465 | -1 | lncRNA | ENSG0000C |
| 79823778 | 79825496 | 1 | lncRNA | ENSG0000C |
| 79878305 | 79878419 | -1 | rRNA_pseudogene | ENSG0000C |
| 79934901 | 79942712 | 1 | lncRNA | ENSG0000C |
| 79989014 | 79989218 | -1 | processed_pseudoge | ENSG0000C |
| 80028893 | 80029631 | 1 | processed_pseudoge | ENSG0000C |
| 80068691 | 80068757 | -1 | snRNA | ENSG0000C |
| 80099537 | 80380879 | 1 | protein_coding | ENSG0000C |
| 80102899 | 80103333 | -1 | processed_pseudoge | ENSG0000C |
| 80343515 | 80343792 | -1 | misc_RNA | ENSG0000C |
| 80402178 | 80680273 | 1 | protein_coding | ENSG0000C |
| 80561017 | 80561560 | -1 | processed_pseudoge | ENSG0000C |
| 80583683 | 80593701 | -1 | lncRNA | ENSG0000C |

| | | | | |
|---|---|---|---|---|
| 80707634 | 80709474 | 1 | protein_coding | ENSG00000 |
| 80716912 | 80719671 | 1 | protein_coding | ENSG00000 |
| 80763151 | 80770717 | 1 | lncRNA | ENSG00000 |
| 80792520 | 80937925 | -1 | protein_coding | ENSG00000 |
| 80832533 | 80832629 | -1 | miRNA | ENSG00000 |
| 80860955 | 80861508 | -1 | processed_pseudoge | ENSG00000 |
| 80935736 | 80935833 | -1 | miRNA | ENSG00000 |
| 80936414 | 81261210 | 1 | protein_coding | ENSG00000 |
| 81094371 | 81125863 | -1 | lncRNA | ENSG00000 |
| 81158388 | 81158461 | 1 | miRNA | ENSG00000 |
| 81257975 | 81759553 | -1 | protein_coding | ENSG00000 |
| 81270669 | 81312422 | 1 | lncRNA | ENSG00000 |
| 81378042 | 81556637 | 1 | lncRNA | ENSG00000 |
| 81545326 | 81546450 | -1 | processed_pseudoge | ENSG00000 |
| 81868368 | 81869046 | 1 | processed_pseudoge | ENSG00000 |
| 81953719 | 81993136 | 1 | lncRNA | ENSG00000 |
| 81998632 | 81998962 | 1 | processed_pseudoge | ENSG00000 |
| 82223681 | 82358805 | -1 | protein_coding | ENSG00000 |
| 82358528 | 82479239 | 1 | protein_coding | ENSG00000 |
| 82468456 | 82468948 | 1 | processed_pseudoge | ENSG00000 |
| 82481118 | 82497560 | -1 | lncRNA | ENSG00000 |
| 82505211 | 82506188 | -1 | lncRNA | ENSG00000 |
| 82512677 | 82515817 | 1 | lncRNA | ENSG00000 |
| 82673070 | 82686632 | -1 | lncRNA | ENSG00000 |
| 82686880 | 83134870 | 1 | protein_coding | ENSG00000 |
| 83021890 | 83021992 | 1 | snRNA | ENSG00000 |
| 83151331 | 83152190 | 1 | processed_pseudoge | ENSG00000 |
| 83171590 | 83172740 | 1 | lncRNA | ENSG00000 |
| 83661009 | 83661719 | -1 | lncRNA | ENSG00000 |
| 84147304 | 84154338 | -1 | lncRNA | ENSG00000 |
| 84154434 | 84294562 | 1 | lncRNA | ENSG00000 |
| 84183324 | 84183448 | -1 | snoRNA | ENSG00000 |
| 84448625 | 84449878 | -1 | processed_pseudoge | ENSG00000 |
| 84671598 | 84672026 | 1 | processed_pseudoge | ENSG00000 |
| 84859491 | 84913615 | -1 | protein_coding | ENSG00000 |
| 84939524 | 84939668 | -1 | processed_pseudoge | ENSG00000 |
| 85014311 | 85036277 | -1 | protein_coding | ENSG00000 |
| 85036314 | 85264457 | 1 | protein_coding | ENSG00000 |
| 85280220 | 85301784 | 1 | protein_coding | ENSG00000 |
| 85318060 | 85342912 | 1 | lncRNA | ENSG00000 |
| 85567878 | 85568239 | 1 | processed_pseudoge | ENSG00000 |
| 85781892 | 85782444 | -1 | processed_pseudoge | ENSG00000 |
| 85800703 | 85836409 | -1 | protein_coding | ENSG00000 |
| 85874295 | 85882992 | 1 | protein_coding | ENSG00000 |
| 85955666 | 86838904 | -1 | protein_coding | ENSG00000 |
| 85958686 | 85960946 | 1 | lncRNA | ENSG00000 |
| 86015697 | 86016441 | 1 | processed_pseudoge | ENSG00000 |
| 86599578 | 86838998 | -1 | lncRNA | ENSG00000 |
| 87041916 | 87042423 | 1 | TEC | ENSG00000 |
| 87169985 | 87170429 | -1 | processed_pseudoge | ENSG00000 |

| | | | | |
|---|---|---|---|---|
| 87746567 | 87746683 | 1 | processed_pseudoge | ENSG0000C |
| 87750142 | 87750313 | 1 | processed_pseudoge | ENSG0000C |
| 87753414 | 87753447 | -1 | processed_pseudoge | ENSG0000C |
| 87782163 | 87784568 | -1 | transcribed_processe | ENSG0000C |
| 87795733 | 87802028 | 1 | lncRNA | ENSG0000C |
| 87816486 | 87817831 | 1 | lncRNA | ENSG0000C |
| 87820654 | 87820918 | -1 | processed_pseudoge | ENSG0000C |
| 87944885 | 87950819 | 1 | processed_pseudoge | ENSG0000C |
| 87980035 | 88034037 | -1 | protein_coding | ENSG0000C |
| 88033846 | 88050160 | 1 | protein_coding | ENSG0000C |
| 88049014 | 88142088 | -1 | protein_coding | ENSG0000C |
| 88051701 | 88051778 | 1 | rRNA_pseudogene | ENSG0000C |
| 88057876 | 88058578 | -1 | processed_pseudoge | ENSG0000C |
| 88142296 | 88199887 | 1 | protein_coding | ENSG0000C |
| 88430442 | 88430550 | 1 | misc_RNA | ENSG0000C |
| 88492793 | 88580851 | -1 | protein_coding | ENSG0000C |
| 88580531 | 88600725 | 1 | lncRNA | ENSG0000C |
| 88581639 | 88581796 | 1 | snRNA | ENSG0000C |
| 88601862 | 88602195 | 1 | lncRNA | ENSG0000C |
| 89010681 | 89309566 | -1 | lncRNA | ENSG0000C |
| 89027757 | 89028522 | 1 | processed_pseudoge | ENSG0000C |
| 89282223 | 89282285 | -1 | snRNA | ENSG0000C |
| 89319364 | 89319649 | 1 | processed_pseudoge | ENSG0000C |
| 89347235 | 89352501 | -1 | protein_coding | ENSG0000C |
| 89351015 | 89353271 | 1 | lncRNA | ENSG0000C |
| 89353798 | 89418559 | 1 | lncRNA | ENSG0000C |
| 89367807 | 89369301 | 1 | lncRNA | ENSG0000C |
| 89371820 | 89372359 | 1 | lncRNA | ENSG0000C |
| 89419718 | 89526047 | -1 | protein_coding | ENSG0000C |
| 89500093 | 89502670 | 1 | processed_pseudoge | ENSG0000C |
| 89519408 | 89526262 | -1 | protein_coding | ENSG0000C |
| 89519412 | 89524796 | -1 | protein_coding | ENSG0000C |
| 89524594 | 89548005 | 1 | lncRNA | ENSG0000C |
| 89544203 | 89544488 | -1 | TEC | ENSG0000C |
| 89561129 | 89594878 | 1 | lncRNA | ENSG0000C |
| 89588049 | 89709300 | -1 | protein_coding | ENSG0000C |
| 89708959 | 89712590 | 1 | lncRNA | ENSG0000C |
| 89712048 | 90100763 | 1 | lncRNA | ENSG0000C |
| 89726936 | 89727042 | 1 | rRNA_pseudogene | ENSG0000C |
| 89752913 | 89753831 | 1 | processed_pseudoge | ENSG0000C |
| 89754059 | 89754152 | 1 | snRNA | ENSG0000C |
| 89919437 | 89919776 | -1 | processed_pseudoge | ENSG0000C |
| 89947693 | 89949726 | 1 | lncRNA | ENSG0000C |
| 90107739 | 90112779 | 1 | lncRNA | ENSG0000C |
| 90280894 | 90300974 | 1 | lncRNA | ENSG0000C |
| 90576402 | 90889917 | 1 | lncRNA | ENSG0000C |
| 90617759 | 90808748 | -1 | lncRNA | ENSG0000C |
| 90617759 | 90622822 | -1 | lncRNA | ENSG0000C |
| 90809207 | 90810689 | 1 | lncRNA | ENSG0000C |
| 90905622 | 90955176 | -1 | protein_coding | ENSG0000C |

| | | | | |
|---|---|---|---|---|
| 90918023 | 90948669 | 1 | lncRNA | ENSG0000C |
| 90963682 | 91005026 | -1 | protein_coding | ENSG0000C |
| 91050491 | 91057983 | -1 | protein_coding | ENSG0000C |
| 91102629 | 91111494 | -1 | protein_coding | ENSG0000C |
| 91140484 | 91182824 | -1 | protein_coding | ENSG0000C |
| 91193709 | 91193942 | 1 | processed_pseudoge | ENSG0000C |
| 91327045 | 91368607 | 1 | lncRNA | ENSG0000C |
| 91423781 | 91426936 | 1 | lncRNA | ENSG0000C |
| 91632912 | 91633023 | -1 | misc_RNA | ENSG0000C |
| 91634887 | 91635644 | 1 | lncRNA | ENSG0000C |
| 91680131 | 91680873 | 1 | lncRNA | ENSG0000C |
| 91871122 | 91871230 | -1 | processed_pseudoge | ENSG0000C |
| 91876924 | 91880659 | 1 | lncRNA | ENSG0000C |
| 91901458 | 91906187 | 1 | lncRNA | ENSG0000C |
| 91984976 | 92142914 | -1 | lncRNA | ENSG0000C |
| 92008052 | 92022595 | 1 | lncRNA | ENSG0000C |
| 92026447 | 92032392 | 1 | lncRNA | ENSG0000C |
| 92140278 | 92145846 | -1 | protein_coding | ENSG0000C |
| 92145573 | 92226557 | 1 | lncRNA | ENSG0000C |
| 92168400 | 92168877 | 1 | processed_pseudoge | ENSG0000C |
| 92170353 | 92170453 | -1 | misc_RNA | ENSG0000C |
| 92247697 | 92363832 | -1 | lncRNA | ENSG0000C |
| 92420094 | 92428148 | -1 | protein_coding | ENSG0000C |
| 92421531 | 92483680 | 1 | lncRNA | ENSG0000C |
| 92421531 | 92431002 | 1 | lncRNA | ENSG0000C |
| 92466451 | 92492091 | 1 | lncRNA | ENSG0000C |
| 92530460 | 92531324 | 1 | processed_pseudoge | ENSG0000C |
| 92702843 | 92772455 | 1 | protein_coding | ENSG0000C |
| 92770637 | 92929331 | -1 | protein_coding | ENSG0000C |
| 92858079 | 92859273 | 1 | processed_pseudoge | ENSG0000C |
| 92883773 | 92884700 | 1 | processed_pseudoge | ENSG0000C |
| 92952639 | 92952745 | 1 | snRNA | ENSG0000C |
| 92999218 | 93019820 | 1 | lncRNA | ENSG0000C |
| 93003415 | 93215679 | -1 | lncRNA | ENSG0000C |
| 93021606 | 93021712 | 1 | misc_RNA | ENSG0000C |
| 93035261 | 93035744 | 1 | processed_pseudoge | ENSG0000C |
| 93083598 | 93083675 | -1 | processed_pseudoge | ENSG0000C |
| 93090480 | 93108629 | 1 | lncRNA | ENSG0000C |
| 93124063 | 93124543 | -1 | processed_pseudoge | ENSG0000C |
| 93173470 | 93182299 | 1 | lncRNA | ENSG0000C |
| 93266017 | 93266096 | 1 | snoRNA | ENSG0000C |
| 93314809 | 93315941 | 1 | transcribed_processe | ENSG0000C |
| 93316722 | 93377753 | -1 | lncRNA | ENSG0000C |
| 93377883 | 93408146 | 1 | protein_coding | ENSG0000C |
| 93405673 | 93441947 | -1 | protein_coding | ENSG0000C |
| 93460725 | 93460826 | -1 | misc_RNA | ENSG0000C |
| 93467488 | 93516214 | 1 | protein_coding | ENSG0000C |
| 93509487 | 93509768 | 1 | misc_RNA | ENSG0000C |
| 93529560 | 93529989 | -1 | processed_pseudoge | ENSG0000C |
| 93542022 | 93571768 | -1 | lncRNA | ENSG0000C |

| | | | | |
|---|---|---|---|---|
| 93569814 | 93583487 | 1 | protein_coding | ENSG00000 |
| 93640822 | 93641586 | -1 | processed_pseudoge | ENSG00000 |
| 93677375 | 93894840 | 1 | protein_coding | ENSG00000 |
| 93707791 | 93737823 | -1 | lncRNA | ENSG00000 |
| 93831076 | 93831372 | 1 | misc_RNA | ENSG00000 |
| 93836167 | 93838038 | 1 | lncRNA | ENSG00000 |
| 93894965 | 93943603 | -1 | lncRNA | ENSG00000 |
| 93945041 | 93947813 | 1 | lncRNA | ENSG00000 |
| 94008739 | 94009047 | -1 | misc_RNA | ENSG00000 |
| 94140077 | 94141337 | 1 | processed_pseudoge | ENSG00000 |
| 94148577 | 94307675 | 1 | protein_coding | ENSG00000 |
| 94167528 | 94187724 | -1 | lncRNA | ENSG00000 |
| 94260548 | 94262299 | -1 | lncRNA | ENSG00000 |
| 94272150 | 94277195 | -1 | lncRNA | ENSG00000 |
| 94277758 | 94282844 | -1 | lncRNA | ENSG00000 |
| 94306449 | 94459988 | -1 | protein_coding | ENSG00000 |
| 94361220 | 94361526 | 1 | misc_RNA | ENSG00000 |
| 94423744 | 94424969 | -1 | processed_pseudoge | ENSG00000 |
| 94460003 | 94462484 | 1 | lncRNA | ENSG00000 |
| 94491546 | 94496442 | -1 | lncRNA | ENSG00000 |
| 94515476 | 94515760 | 1 | misc_RNA | ENSG00000 |
| 94521090 | 94521869 | -1 | processed_pseudoge | ENSG00000 |
| 94548241 | 94549536 | 1 | processed_pseudoge | ENSG00000 |
| 94561789 | 94561859 | 1 | miRNA | ENSG00000 |
| 94567122 | 94650557 | -1 | protein_coding | ENSG00000 |
| 94571231 | 94571352 | -1 | miRNA | ENSG00000 |
| 94590821 | 94591093 | 1 | processed_pseudoge | ENSG00000 |
| 94834147 | 94835158 | 1 | transcribed_processe | ENSG00000 |
| 94834398 | 94834513 | 1 | miRNA | ENSG00000 |
| 94897055 | 95003748 | -1 | protein_coding | ENSG00000 |
| 95020229 | 95073628 | -1 | protein_coding | ENSG00000 |
| 95076749 | 95217482 | -1 | protein_coding | ENSG00000 |
| 95100531 | 95101045 | 1 | processed_pseudoge | ENSG00000 |
| 95125760 | 95126389 | 1 | processed_pseudoge | ENSG00000 |
| 95217746 | 95302799 | 1 | protein_coding | ENSG00000 |
| 95275770 | 95275872 | 1 | misc_RNA | ENSG00000 |
| 95286318 | 95286838 | -1 | processed_pseudoge | ENSG00000 |
| 95290692 | 95290799 | 1 | snRNA | ENSG00000 |
| 95308420 | 95308513 | 1 | miRNA | ENSG00000 |
| 95309923 | 95309984 | 1 | miRNA | ENSG00000 |
| 95311312 | 95442482 | -1 | transcribed_unitary_ | ENSG00000 |
| 95387890 | 95388558 | -1 | lncRNA | ENSG00000 |
| 95438737 | 95438843 | -1 | snRNA | ENSG00000 |
| 95467397 | 95467861 | -1 | processed_pseudoge | ENSG00000 |
| 95473520 | 95515839 | 1 | protein_coding | ENSG00000 |
| 95516560 | 95551476 | -1 | protein_coding | ENSG00000 |
| 95551582 | 95673999 | 1 | transcribed_processe | ENSG00000 |
| 95617689 | 95617786 | 1 | misc_RNA | ENSG00000 |
| 95657807 | 95791152 | -1 | protein_coding | ENSG00000 |
| 95791733 | 95858839 | -1 | lncRNA | ENSG00000 |

| | | | | |
|---|---|---|---|---|
| 95792529 | 95792635 | -1 | snRNA | ENSG00000 |
| 95803097 | 95823307 | 1 | lncRNA | ENSG00000 |
| 95858928 | 95903828 | 1 | protein_coding | ENSG00000 |
| 95867048 | 95942635 | -1 | protein_coding | ENSG00000 |
| 95885169 | 95885797 | 1 | processed_pseudoge | ENSG00000 |
| 95943331 | 95968720 | 1 | protein_coding | ENSG00000 |
| 95972662 | 95996365 | -1 | protein_coding | ENSG00000 |
| 95996521 | 96011489 | 1 | lncRNA | ENSG00000 |
| 96000753 | 96043520 | -1 | protein_coding | ENSG00000 |
| 96025323 | 96027971 | 1 | lncRNA | ENSG00000 |
| 96060024 | 96060375 | -1 | processed_pseudoge | ENSG00000 |
| 96101951 | 96102249 | -1 | misc_RNA | ENSG00000 |
| 96160692 | 96172126 | -1 | lncRNA | ENSG00000 |
| 96194375 | 96269824 | 1 | protein_coding | ENSG00000 |
| 96222797 | 96223973 | -1 | lncRNA | ENSG00000 |
| 96229563 | 96235650 | -1 | lncRNA | ENSG00000 |
| 96278261 | 96400480 | -1 | protein_coding | ENSG00000 |
| 96303375 | 96303513 | -1 | snRNA | ENSG00000 |
| 96386818 | 96387194 | -1 | processed_pseudoge | ENSG00000 |
| 96422326 | 96485442 | -1 | lncRNA | ENSG00000 |
| 96427200 | 96427486 | -1 | misc_RNA | ENSG00000 |
| 96489571 | 96875555 | 1 | protein_coding | ENSG00000 |
| 96900697 | 96902698 | -1 | processed_pseudoge | ENSG00000 |
| 96907223 | 96953777 | 1 | protein_coding | ENSG00000 |
| 96979528 | 96979629 | -1 | misc_RNA | ENSG00000 |
| 96985656 | 97185609 | 1 | lncRNA | ENSG00000 |
| 97024021 | 97127626 | 1 | lncRNA | ENSG00000 |
| 97150081 | 97154627 | -1 | TEC | ENSG00000 |
| 97155234 | 97160538 | -1 | lncRNA | ENSG00000 |
| 97272692 | 97276209 | -1 | lncRNA | ENSG00000 |
| 97430884 | 97598415 | 1 | lncRNA | ENSG00000 |
| 97491909 | 97491978 | 1 | miRNA | ENSG00000 |
| 97492462 | 97492597 | 1 | misc_RNA | ENSG00000 |
| 97493178 | 97493286 | 1 | misc_RNA | ENSG00000 |
| 97493897 | 97494003 | 1 | misc_RNA | ENSG00000 |
| 97495932 | 97496057 | 1 | misc_RNA | ENSG00000 |
| 97530656 | 97530858 | 1 | misc_RNA | ENSG00000 |
| 97532691 | 97532958 | 1 | misc_RNA | ENSG00000 |
| 97552137 | 97552228 | -1 | snoRNA | ENSG00000 |
| 97560557 | 97560721 | 1 | misc_RNA | ENSG00000 |
| 97560856 | 97561048 | 1 | misc_RNA | ENSG00000 |
| 97563257 | 97563971 | -1 | TEC | ENSG00000 |
| 97563812 | 97563911 | 1 | miRNA | ENSG00000 |
| 97564174 | 97564327 | 1 | misc_RNA | ENSG00000 |
| 97679879 | 97680047 | -1 | processed_pseudoge | ENSG00000 |
| 97713412 | 97761859 | -1 | transcribed_processe | ENSG00000 |
| 97721716 | 97721822 | 1 | snRNA | ENSG00000 |
| 97939056 | 97939121 | -1 | miRNA | ENSG00000 |
| 97995383 | 97995448 | -1 | miRNA | ENSG00000 |
| 98113014 | 98292445 | -1 | lncRNA | ENSG00000 |

| | | | | |
|---|---|---|---|---|
| 98317741 | 98318349 | 1 | processed_pseudoge | ENSG00000( |
| 98417896 | 98418038 | -1 | snRNA | ENSG00000( |
| 98420448 | 98423721 | 1 | lncRNA | ENSG00000( |
| 98453835 | 98457145 | -1 | transcribed_processe | ENSG00000( |
| 98485544 | 98503855 | -1 | lncRNA | ENSG00000( |
| 98512973 | 98516422 | -1 | lncRNA | ENSG00000( |
| 98515512 | 98550379 | 1 | protein_coding | ENSG00000( |
| 98590723 | 98591214 | -1 | processed_pseudoge | ENSG00000( |
| 98593591 | 98606379 | 1 | protein_coding | ENSG00000( |
| 98599635 | 98599883 | 1 | snoRNA | ENSG00000( |
| 98613405 | 98645113 | -1 | protein_coding | ENSG00000( |
| 98645141 | 98735433 | 1 | protein_coding | ENSG00000( |
| 98726457 | 99984654 | -1 | protein_coding | ENSG00000( |
| 98794485 | 98794964 | 1 | processed_pseudoge | ENSG00000( |
| 98931682 | 98976656 | 1 | lncRNA | ENSG00000( |
| 99083783 | 99083905 | -1 | rRNA_pseudogene | ENSG00000( |
| 99093359 | 99105011 | 1 | lncRNA | ENSG00000( |
| 99647753 | 99650114 | 1 | protein_coding | ENSG00000( |
| 99985045 | 99985619 | 1 | lncRNA | ENSG00000( |
| 100009052 | 100009838 | -1 | processed_pseudoge | ENSG00000( |
| 100026248 | 100027540 | -1 | TEC | ENSG00000( |
| 100028455 | 100142874 | -1 | protein_coding | ENSG00000( |
| 100032325 | 100034074 | -1 | TEC | ENSG00000( |
| 100143058 | 100144671 | 1 | lncRNA | ENSG00000( |
| 100156357 | 100173659 | -1 | transcribed_unproce | ENSG00000( |
| 100158502 | 100158765 | -1 | misc_RNA | ENSG00000( |
| 100173196 | 100199783 | -1 | transcribed_unproce | ENSG00000( |
| 100189884 | 100189949 | 1 | miRNA | ENSG00000( |
| 100199122 | 100241865 | 1 | protein_coding | ENSG00000( |
| 100203669 | 100267079 | -1 | protein_coding | ENSG00000( |
| 100257688 | 100257790 | -1 | misc_RNA | ENSG00000( |
| 100267140 | 100341715 | 1 | protein_coding | ENSG00000( |
| 100357074 | 100422055 | 1 | protein_coding | ENSG00000( |
| 100473708 | 100564414 | 1 | protein_coding | ENSG00000( |
| 100525186 | 100525420 | -1 | processed_pseudoge | ENSG00000( |
| 100573683 | 100628288 | 1 | protein_coding | ENSG00000( |
| 100578744 | 100580197 | 1 | processed_pseudoge | ENSG00000( |
| 100717526 | 101128641 | 1 | protein_coding | ENSG00000( |
| 100852331 | 100859262 | -1 | lncRNA | ENSG00000( |
| 101038420 | 101039094 | -1 | lncRNA | ENSG00000( |
| 101066734 | 101069232 | 1 | unprocessed_pseudc | ENSG00000( |
| 101155493 | 101210238 | -1 | protein_coding | ENSG00000( |
| 101199271 | 101199377 | -1 | snRNA | ENSG00000( |
| 101228141 | 101228876 | -1 | processed_pseudoge | ENSG00000( |
| 101280105 | 101386618 | 1 | protein_coding | ENSG00000( |
| 101393116 | 101407772 | -1 | protein_coding | ENSG00000( |
| 101408372 | 101409060 | -1 | lncRNA | ENSG00000( |
| 101411357 | 101411459 | 1 | snRNA | ENSG00000( |
| 101424441 | 101424691 | 1 | processed_pseudoge | ENSG00000( |
| 101446825 | 101446932 | 1 | misc_RNA | ENSG00000( |

| | | | | |
|---|---|---|---|---|
| 101466705 | 101466820 | 1 | snRNA | ENSG0000C |
| 101472339 | 101472456 | 1 | rRNA_pseudogene | ENSG0000C |
| 101475336 | 101486997 | 1 | protein_coding | ENSG0000C |
| 101568353 | 101686028 | 1 | protein_coding | ENSG0000C |
| 101646720 | 101659970 | -1 | lncRNA | ENSG0000C |
| 101696002 | 101696450 | -1 | lncRNA | ENSG0000C |
| 101696947 | 101744140 | 1 | protein_coding | ENSG0000C |
| 101719808 | 101719918 | 1 | misc_RNA | ENSG0000C |
| 101728648 | 101739472 | -1 | protein_coding | ENSG0000C |
| 101745499 | 101830959 | -1 | protein_coding | ENSG0000C |
| 101765406 | 101765509 | -1 | snRNA | ENSG0000C |
| 101773114 | 101773755 | -1 | transcribed_processe | ENSG0000C |
| 101779814 | 101779929 | 1 | rRNA_pseudogene | ENSG0000C |
| 101796410 | 101796502 | -1 | snRNA | ENSG0000C |
| 101836820 | 101836927 | 1 | rRNA_pseudogene | ENSG0000C |
| 101871742 | 101871801 | -1 | snRNA | ENSG0000C |
| 101873316 | 101873623 | -1 | processed_pseudoge | ENSG0000C |
| 101877580 | 102012130 | 1 | protein_coding | ENSG0000C |
| 101923410 | 101924719 | -1 | lncRNA | ENSG0000C |
| 101936216 | 101936674 | 1 | processed_pseudoge | ENSG0000C |
| 101954992 | 101962414 | -1 | lncRNA | ENSG0000C |
| 101963901 | 101964340 | 1 | processed_pseudoge | ENSG0000C |
| 102012840 | 102062149 | -1 | protein_coding | ENSG0000C |
| 102063355 | 102074820 | 1 | lncRNA | ENSG0000C |
| 102073103 | 102120120 | -1 | protein_coding | ENSG0000C |
| 102120185 | 102197520 | 1 | protein_coding | ENSG0000C |
| 102196459 | 102197833 | -1 | protein_coding | ENSG0000C |
| 102197585 | 102402596 | 1 | lncRNA | ENSG0000C |
| 102217318 | 102217613 | -1 | misc_RNA | ENSG0000C |
| 102230027 | 102230533 | -1 | processed_pseudoge | ENSG0000C |
| 102304355 | 102463491 | 1 | lncRNA | ENSG0000C |
| 102395874 | 102481744 | -1 | protein_coding | ENSG0000C |
| 102513251 | 102513655 | -1 | processed_pseudoge | ENSG0000C |
| 102809280 | 102824399 | -1 | lncRNA | ENSG0000C |
| 102836889 | 102958410 | -1 | protein_coding | ENSG0000C |
| 102953289 | 102953835 | -1 | lncRNA | ENSG0000C |
| 102957674 | 102960513 | 1 | protein_coding | ENSG0000C |
| 103013362 | 103016095 | 1 | lncRNA | ENSG0000C |
| 103079634 | 103178675 | -1 | lncRNA | ENSG0000C |
| 103151834 | 103179247 | 1 | lncRNA | ENSG0000C |
| 103237591 | 103496010 | -1 | protein_coding | ENSG0000C |
| 103503381 | 103504936 | 1 | processed_pseudoge | ENSG0000C |
| 103505411 | 103506174 | -1 | unprocessed_pseudc | ENSG0000C |
| 103518334 | 103571337 | -1 | lncRNA | ENSG0000C |
| 103547751 | 103578947 | 1 | lncRNA | ENSG0000C |
| 103587273 | 103766719 | 1 | protein_coding | ENSG0000C |
| 103592112 | 103592243 | 1 | snoRNA | ENSG0000C |
| 103654780 | 103657995 | -1 | lncRNA | ENSG0000C |
| 103668575 | 103669406 | -1 | lncRNA | ENSG0000C |
| 103746315 | 103768858 | -1 | lncRNA | ENSG0000C |

| | | | | |
|---|---|---|---|---|
| 103770453 | 103841234 | -1 | protein_coding | ENSG0000C |
| 103819610 | 103822085 | 1 | lncRNA | ENSG0000C |
| 103841451 | 103844664 | 1 | lncRNA | ENSG0000C |
| 103843749 | 103930211 | -1 | transcribed_unitary_ | ENSG0000C |
| 103930107 | 103953645 | 1 | protein_coding | ENSG0000C |
| 103930425 | 103930555 | 1 | miRNA | ENSG0000C |
| 103940763 | 103965708 | -1 | protein_coding | ENSG0000C |
| 103965822 | 103988874 | 1 | protein_coding | ENSG0000C |
| 103988984 | 104064183 | -1 | protein_coding | ENSG0000C |
| 104030779 | 104031543 | 1 | processed_pseudoge | ENSG0000C |
| 104064531 | 104106524 | 1 | protein_coding | ENSG0000C |
| 104117086 | 104138241 | -1 | protein_coding | ENSG0000C |
| 104125236 | 104125356 | 1 | rRNA_pseudogene | ENSG0000C |
| 104170395 | 104177659 | 1 | lncRNA | ENSG0000C |
| 104171600 | 104177799 | -1 | lncRNA | ENSG0000C |
| 104215779 | 104350307 | 1 | protein_coding | ENSG0000C |
| 104262314 | 104280722 | -1 | lncRNA | ENSG0000C |
| 104265309 | 104265836 | 1 | processed_pseudoge | ENSG0000C |
| 104303739 | 104305205 | 1 | protein_coding | ENSG0000C |
| 104455295 | 104762014 | 1 | protein_coding | ENSG0000C |
| 104514029 | 104514439 | 1 | processed_pseudoge | ENSG0000C |
| 104552073 | 104553447 | 1 | TEC | ENSG0000C |
| 104591633 | 104591716 | 1 | miRNA | ENSG0000C |
| 104802553 | 104958744 | -1 | protein_coding | ENSG0000C |
| 104851824 | 104851918 | 1 | misc_RNA | ENSG0000C |
| 104858838 | 104871765 | 1 | lncRNA | ENSG0000C |
| 104976749 | 104978027 | -1 | processed_pseudoge | ENSG0000C |
| 104986316 | 105074197 | 1 | protein_coding | ENSG0000C |
| 105019784 | 105107643 | -1 | protein_coding | ENSG0000C |
| 105107324 | 105169134 | 1 | protein_coding | ENSG0000C |
| 105173297 | 105236203 | -1 | protein_coding | ENSG0000C |
| 105235290 | 105396097 | 1 | protein_coding | ENSG0000C |
| 105236338 | 105243173 | 1 | lncRNA | ENSG0000C |
| 105304867 | 105327017 | -1 | lncRNA | ENSG0000C |
| 105699168 | 105703520 | -1 | lncRNA | ENSG0000C |
| 105704203 | 105744062 | 1 | lncRNA | ENSG0000C |
| 106015897 | 106017006 | 1 | processed_pseudoge | ENSG0000C |
| 106050961 | 106058254 | -1 | lncRNA | ENSG0000C |
| 106063340 | 106140033 | -1 | protein_coding | ENSG0000C |
| 106103163 | 106106165 | 1 | lncRNA | ENSG0000C |
| 106170004 | 106172437 | 1 | processed_pseudoge | ENSG0000C |
| 106237881 | 106304279 | -1 | protein_coding | ENSG0000C |
| 106245613 | 106246956 | -1 | lncRNA | ENSG0000C |
| 106250759 | 106252786 | 1 | lncRNA | ENSG0000C |
| 106282927 | 106282989 | -1 | snRNA | ENSG0000C |
| 106301929 | 106347003 | 1 | protein_coding | ENSG0000C |
| 106357748 | 106510198 | 1 | protein_coding | ENSG0000C |
| 106495958 | 106774926 | -1 | lncRNA | ENSG0000C |
| 106582907 | 106762803 | 1 | protein_coding | ENSG0000C |
| 106678583 | 106684707 | -1 | lncRNA | ENSG0000C |

| | | | | |
|---|---|---|---|---|
| 106714924 | 106733066 | -1 | lncRNA | ENSG0000C |
| 106774621 | 106889316 | 1 | protein_coding | ENSG0000C |
| 106867177 | 106871505 | -1 | lncRNA | ENSG0000C |
| 106901283 | 106901984 | -1 | processed_pseudoge | ENSG0000C |
| 106910757 | 106911109 | 1 | processed_pseudoge | ENSG0000C |
| 106954029 | 106955497 | -1 | lncRNA | ENSG0000C |
| 106955719 | 106978778 | 1 | protein_coding | ENSG0000C |
| 106977277 | 106987160 | -1 | protein_coding | ENSG0000C |
| 106991364 | 107093549 | -1 | protein_coding | ENSG0000C |
| 107093298 | 107123314 | 1 | lncRNA | ENSG0000C |
| 107219764 | 107223576 | -1 | lncRNA | ENSG0000C |
| 107271394 | 107271846 | 1 | processed_pseudoge | ENSG0000C |
| 107318421 | 107659642 | 1 | protein_coding | ENSG0000C |
| 107374747 | 107374827 | 1 | snoRNA | ENSG0000C |
| 107486511 | 107486635 | -1 | rRNA_pseudogene | ENSG0000C |
| 107610034 | 107617721 | -1 | lncRNA | ENSG0000C |
| 107650789 | 107650892 | 1 | misc_RNA | ENSG0000C |
| 107685799 | 107713162 | 1 | protein_coding | ENSG0000C |
| 107719599 | 107720133 | 1 | processed_pseudoge | ENSG0000C |
| 107732871 | 107761272 | -1 | protein_coding | ENSG0000C |
| 107736555 | 107759968 | 1 | lncRNA | ENSG0000C |
| 107774385 | 107776644 | 1 | protein_coding | ENSG0000C |
| 107809483 | 107864562 | -1 | lncRNA | ENSG0000C |
| 107880581 | 107883386 | 1 | unprocessed_pseudc | ENSG0000C |
| 107887730 | 107889886 | 1 | unprocessed_pseudc | ENSG0000C |
| 107899638 | 107899815 | 1 | unprocessed_pseudc | ENSG0000C |
| 107903270 | 107903948 | -1 | processed_pseudoge | ENSG0000C |
| 107910403 | 107912620 | -1 | transcribed_processe | ENSG0000C |
| 108129288 | 108250537 | 1 | protein_coding | ENSG0000C |
| 108281103 | 108282235 | -1 | processed_pseudoge | ENSG0000C |
| 108288044 | 108339317 | -1 | protein_coding | ENSG0000C |
| 108434130 | 108515023 | -1 | lncRNA | ENSG0000C |
| 108515277 | 108525837 | 1 | protein_coding | ENSG0000C |
| 108522214 | 108561400 | -1 | protein_coding | ENSG0000C |
| 108532074 | 108539738 | 1 | lncRNA | ENSG0000C |
| 108562582 | 108569384 | 1 | protein_coding | ENSG0000C |
| 108589851 | 108598320 | -1 | protein_coding | ENSG0000C |
| 108622277 | 108633959 | -1 | protein_coding | ENSG0000C |
| 108628668 | 108642349 | 1 | lncRNA | ENSG0000C |
| 108635810 | 108635870 | 1 | miRNA | ENSG0000C |
| 108645109 | 108731596 | -1 | protein_coding | ENSG0000C |
| 108668448 | 108668508 | 1 | snRNA | ENSG0000C |
| 108738484 | 108738604 | 1 | misc_RNA | ENSG0000C |
| 108778144 | 108778271 | -1 | snoRNA | ENSG0000C |
| 108778191 | 108857590 | -1 | protein_coding | ENSG0000C |
| 108778912 | 108779166 | 1 | TEC | ENSG0000C |
| 108833721 | 108834384 | -1 | lncRNA | ENSG0000C |
| 108836908 | 108837006 | -1 | miRNA | ENSG0000C |
| 108858932 | 108901043 | 1 | protein_coding | ENSG0000C |
| 108907741 | 109021068 | -1 | protein_coding | ENSG0000C |

| | | | | |
|---|---|---|---|---|
| 108926435 | 108926541 | -1 | snRNA | ENSG00000 |
| 109020833 | 109021240 | -1 | TEC | ENSG00000 |
| 109023089 | 109088023 | 1 | protein_coding | ENSG00000 |
| 109052350 | 109053952 | -1 | lncRNA | ENSG00000 |
| 109059986 | 109060100 | 1 | rRNA_pseudogene | ENSG00000 |
| 109088188 | 109093631 | -1 | protein_coding | ENSG00000 |
| 109097574 | 109110992 | 1 | protein_coding | ENSG00000 |
| 109111218 | 109125594 | 1 | lncRNA | ENSG00000 |
| 109116595 | 109268226 | 1 | protein_coding | ENSG00000 |
| 109277978 | 109309284 | -1 | protein_coding | ENSG00000 |
| 109347903 | 109455523 | 1 | protein_coding | ENSG00000 |
| 109354083 | 109359488 | -1 | lncRNA | ENSG00000 |
| 109445410 | 109447497 | -1 | lncRNA | ENSG00000 |
| 109448655 | 109477359 | -1 | protein_coding | ENSG00000 |
| 109477402 | 109536705 | 1 | protein_coding | ENSG00000 |
| 109553715 | 109573580 | -1 | protein_coding | ENSG00000 |
| 109567975 | 109568115 | -1 | snRNA | ENSG00000 |
| 109573255 | 109598125 | 1 | protein_coding | ENSG00000 |
| 109662863 | 109663183 | -1 | misc_RNA | ENSG00000 |
| 109714228 | 109770507 | 1 | protein_coding | ENSG00000 |
| 109734166 | 109773508 | -1 | lncRNA | ENSG00000 |
| 109783085 | 109833401 | -1 | protein_coding | ENSG00000 |
| 109833348 | 109833436 | 1 | miRNA | ENSG00000 |
| 109850945 | 109880541 | -1 | protein_coding | ENSG00000 |
| 109859767 | 109860420 | -1 | processed_pseudoge | ENSG00000 |
| 109866214 | 109866509 | -1 | misc_RNA | ENSG00000 |
| 109880676 | 109888467 | 1 | lncRNA | ENSG00000 |
| 109900264 | 109983841 | 1 | protein_coding | ENSG00000 |
| 109929802 | 109996389 | -1 | protein_coding | ENSG00000 |
| 109948389 | 109949029 | 1 | lncRNA | ENSG00000 |
| 109997419 | 109999127 | -1 | TEC | ENSG00000 |
| 109999186 | 110039763 | 1 | protein_coding | ENSG00000 |
| 110027028 | 110073634 | -1 | protein_coding | ENSG00000 |
| 110032245 | 110032803 | 1 | lncRNA | ENSG00000 |
| 110049360 | 110073686 | -1 | transcribed_unproce | ENSG00000 |
| 110056916 | 110057275 | -1 | TEC | ENSG00000 |
| 110124335 | 110218793 | 1 | protein_coding | ENSG00000 |
| 110280756 | 110351093 | 1 | protein_coding | ENSG00000 |
| 110360452 | 110360746 | -1 | misc_RNA | ENSG00000 |
| 110372900 | 110403730 | -1 | protein_coding | ENSG00000 |
| 110387463 | 110445548 | -1 | lncRNA | ENSG00000 |
| 110429959 | 110430282 | -1 | processed_pseudoge | ENSG00000 |
| 110434823 | 110450422 | -1 | protein_coding | ENSG00000 |
| 110452486 | 110469268 | -1 | protein_coding | ENSG00000 |
| 110460988 | 110461362 | 1 | processed_pseudoge | ENSG00000 |
| 110468415 | 110490385 | 1 | protein_coding | ENSG00000 |
| 110491083 | 110502111 | -1 | protein_coding | ENSG00000 |
| 110496352 | 110496421 | -1 | snoRNA | ENSG00000 |
| 110501614 | 110503441 | 1 | TEC | ENSG00000 |
| 110501655 | 110532086 | 1 | protein_coding | ENSG00000 |

| | | | | |
|---|---|---|---|---|
| 110533245 | 110583318 | -1 | protein_coding | ENSG00000 |
| 110614027 | 110649430 | 1 | protein_coding | ENSG00000 |
| 110625645 | 110625917 | -1 | misc_RNA | ENSG00000 |
| 110627841 | 110704950 | -1 | protein_coding | ENSG00000 |
| 110719680 | 110742939 | -1 | protein_coding | ENSG00000 |
| 110744549 | 110746206 | 1 | TEC | ENSG00000 |
| 110831779 | 110845963 | 1 | lncRNA | ENSG00000 |
| 110841538 | 110842387 | -1 | processed_pseudoge | ENSG00000 |
| 110846769 | 110907535 | 1 | protein_coding | ENSG00000 |
| 110910819 | 110921443 | -1 | protein_coding | ENSG00000 |
| 110934590 | 110959093 | 1 | lncRNA | ENSG00000 |
| 110951683 | 110957820 | -1 | lncRNA | ENSG00000 |
| 111034165 | 111350554 | 1 | protein_coding | ENSG00000 |
| 111304142 | 111304209 | 1 | miRNA | ENSG00000 |
| 111339527 | 111339621 | 1 | rRNA_pseudogene | ENSG00000 |
| 111360651 | 111369121 | -1 | protein_coding | ENSG00000 |
| 111369261 | 111403310 | 1 | lncRNA | ENSG00000 |
| 111381867 | 111385638 | 1 | processed_pseudoge | ENSG00000 |
| 111405948 | 111451623 | 1 | protein_coding | ENSG00000 |
| 111443485 | 111599676 | -1 | protein_coding | ENSG00000 |
| 111564821 | 111564883 | -1 | snRNA | ENSG00000 |
| 111581379 | 111581773 | 1 | processed_pseudoge | ENSG00000 |
| 111599498 | 111600256 | 1 | lncRNA | ENSG00000 |
| 111642146 | 111685956 | -1 | protein_coding | ENSG00000 |
| 111669852 | 111670362 | -1 | processed_pseudoge | ENSG00000 |
| 111686056 | 111757107 | 1 | protein_coding | ENSG00000 |
| 111753890 | 111791418 | 1 | protein_coding | ENSG00000 |
| 111766887 | 111817532 | 1 | protein_coding | ENSG00000 |
| 111799834 | 111799905 | 1 | miRNA | ENSG00000 |
| 111839764 | 111842902 | -1 | lncRNA | ENSG00000 |
| 111842228 | 111902222 | 1 | protein_coding | ENSG00000 |
| 111879338 | 111880249 | -1 | processed_pseudoge | ENSG00000 |
| 111899263 | 111901391 | 1 | unitary_pseudogene | ENSG00000 |
| 111927018 | 111929017 | 1 | unitary_pseudogene | ENSG00000 |
| 111931282 | 112013185 | -1 | protein_coding | ENSG00000 |
| 111992996 | 111993875 | 1 | processed_pseudoge | ENSG00000 |
| 112000739 | 112000985 | -1 | processed_pseudoge | ENSG00000 |
| 112013348 | 112023449 | 1 | protein_coding | ENSG00000 |
| 112018804 | 112019430 | -1 | lncRNA | ENSG00000 |
| 112026689 | 112108796 | -1 | protein_coding | ENSG00000 |
| 112037599 | 112037715 | -1 | miRNA | ENSG00000 |
| 112063909 | 112065755 | 1 | lncRNA | ENSG00000 |
| 112069104 | 112069209 | 1 | misc_RNA | ENSG00000 |
| 112125538 | 112153604 | 1 | protein_coding | ENSG00000 |
| 112149360 | 112149472 | 1 | misc_RNA | ENSG00000 |
| 112160188 | 112382439 | -1 | protein_coding | ENSG00000 |
| 112163258 | 112163321 | -1 | miRNA | ENSG00000 |
| 112256800 | 112259091 | 1 | lncRNA | ENSG00000 |
| 112267077 | 112267394 | 1 | misc_RNA | ENSG00000 |
| 112301870 | 112302663 | -1 | processed_pseudoge | ENSG00000 |

| | | | | |
|---|---|---|---|---|
| 112321759 | 112323089 | 1 | processed_pseudoge | ENSG0000C |
| 112405190 | 112418838 | -1 | protein_coding | ENSG0000C |
| 112418351 | 112509913 | 1 | protein_coding | ENSG0000C |
| 112570380 | 112898881 | 1 | protein_coding | ENSG0000C |
| 112695034 | 112695176 | -1 | miRNA | ENSG0000C |
| 112906783 | 112933222 | 1 | protein_coding | ENSG0000C |
| 112907628 | 113017751 | -1 | lncRNA | ENSG0000C |
| 112938444 | 112973251 | 1 | protein_coding | ENSG0000C |
| 112978395 | 113011723 | 1 | protein_coding | ENSG0000C |
| 113015607 | 113016098 | 1 | processed_pseudoge | ENSG0000C |
| 113054062 | 113054435 | 1 | processed_pseudoge | ENSG0000C |
| 113056709 | 113098028 | 1 | protein_coding | ENSG0000C |
| 113098819 | 113136239 | -1 | protein_coding | ENSG0000C |
| 113149858 | 113159276 | 1 | protein_coding | ENSG0000C |
| 113156763 | 113156861 | -1 | misc_RNA | ENSG0000C |
| 113157174 | 113185479 | -1 | protein_coding | ENSG0000C |
| 113159113 | 113159177 | -1 | miRNA | ENSG0000C |
| 113178423 | 113178519 | -1 | misc_RNA | ENSG0000C |
| 113185526 | 113192368 | 1 | protein_coding | ENSG0000C |
| 113185624 | 113192161 | -1 | lncRNA | ENSG0000C |
| 113195441 | 113221094 | -1 | protein_coding | ENSG0000C |
| 113221050 | 113298585 | 1 | protein_coding | ENSG0000C |
| 113249466 | 113250042 | 1 | lncRNA | ENSG0000C |
| 113291523 | 113291608 | 1 | miRNA | ENSG0000C |
| 113298759 | 113359493 | -1 | protein_coding | ENSG0000C |
| 113358566 | 113391629 | 1 | protein_coding | ENSG0000C |
| 113392445 | 113426301 | -1 | protein_coding | ENSG0000C |
| 113422380 | 113438276 | 1 | protein_coding | ENSG0000C |
| 113462033 | 113472280 | -1 | protein_coding | ENSG0000C |
| 113472003 | 113480624 | 1 | lncRNA | ENSG0000C |
| 113583886 | 113773726 | -1 | lncRNA | ENSG0000C |
| 113789542 | 113789807 | -1 | processed_pseudoge | ENSG0000C |
| 113816738 | 113966325 | -1 | protein_coding | ENSG0000C |
| 113863402 | 113871597 | 1 | lncRNA | ENSG0000C |
| 113932569 | 113937255 | 1 | lncRNA | ENSG0000C |
| 114077133 | 114079902 | 1 | lncRNA | ENSG0000C |
| 114080381 | 114113489 | 1 | lncRNA | ENSG0000C |
| 114097682 | 114098223 | 1 | processed_pseudoge | ENSG0000C |
| 114104542 | 114105931 | 1 | processed_pseudoge | ENSG0000C |
| 114238970 | 114241770 | -1 | lncRNA | ENSG0000C |
| 114353931 | 114408442 | -1 | protein_coding | ENSG0000C |
| 114408173 | 114412831 | 1 | lncRNA | ENSG0000C |
| 114504876 | 114505187 | -1 | misc_RNA | ENSG0000C |
| 114548854 | 114549852 | 1 | processed_pseudoge | ENSG0000C |
| 114588194 | 114588802 | 1 | processed_pseudoge | ENSG0000C |
| 114621761 | 114622922 | 1 | lncRNA | ENSG0000C |
| 114657949 | 114658062 | -1 | misc_RNA | ENSG0000C |
| 114670255 | 114684175 | -1 | protein_coding | ENSG0000C |
| 114682292 | 114767989 | 1 | lncRNA | ENSG0000C |
| 114733757 | 114735244 | -1 | lncRNA | ENSG0000C |

| | | | | |
|---|---|---|---|---|
| 114737704 | 114737798 | 1 | snoRNA | ENSG0000C |
| 114768674 | 114771851 | 1 | lncRNA | ENSG0000C |
| 114894632 | 114898529 | -1 | lncRNA | ENSG0000C |
| 114929757 | 114929935 | -1 | processed_pseudoge | ENSG0000C |
| 115077325 | 115080153 | 1 | lncRNA | ENSG0000C |
| 115263170 | 115270596 | 1 | lncRNA | ENSG0000C |
| 115299588 | 115300708 | -1 | lncRNA | ENSG0000C |
| 115318657 | 115320405 | -1 | lncRNA | ENSG0000C |
| 115332146 | 115332566 | -1 | lncRNA | ENSG0000C |
| 115363012 | 115364745 | 1 | lncRNA | ENSG0000C |
| 115569394 | 115581739 | 1 | lncRNA | ENSG0000C |
| 115582061 | 115639734 | -1 | lncRNA | ENSG0000C |
| 115592604 | 115592983 | -1 | processed_pseudoge | ENSG0000C |
| 115717725 | 115718013 | 1 | misc_RNA | ENSG0000C |
| 115755262 | 115799936 | 1 | lncRNA | ENSG0000C |
| 115810359 | 115886137 | -1 | lncRNA | ENSG0000C |
| 115852648 | 115852718 | -1 | snoRNA | ENSG0000C |
| 115878748 | 115885967 | 1 | lncRNA | ENSG0000C |
| 115953872 | 116277719 | -1 | protein_coding | ENSG0000C |
| 115961187 | 115962733 | 1 | lncRNA | ENSG0000C |
| 116082570 | 116082676 | 1 | snRNA | ENSG0000C |
| 116148560 | 116148654 | -1 | miRNA | ENSG0000C |
| 116169045 | 116169258 | 1 | processed_pseudoge | ENSG0000C |
| 116174502 | 116181295 | -1 | lncRNA | ENSG0000C |
| 116357578 | 116359096 | -1 | lncRNA | ENSG0000C |
| 116368764 | 116389471 | 1 | lncRNA | ENSG0000C |
| 116428252 | 116428318 | -1 | miRNA | ENSG0000C |
| 116482073 | 116521047 | 1 | lncRNA | ENSG0000C |
| 116533422 | 116536518 | 1 | lncRNA | ENSG0000C |
| 116548105 | 116576606 | 1 | protein_coding | ENSG0000C |
| 116580974 | 116583672 | 1 | lncRNA | ENSG0000C |
| 116599270 | 116603585 | 1 | lncRNA | ENSG0000C |
| 116661582 | 116698065 | 1 | lncRNA | ENSG0000C |
| 116698336 | 116703130 | 1 | lncRNA | ENSG0000C |
| 116710171 | 116738070 | -1 | protein_coding | ENSG0000C |
| 116738178 | 116853631 | 1 | protein_coding | ENSG0000C |
| 116755395 | 116755874 | -1 | processed_pseudoge | ENSG0000C |
| 116782105 | 116782208 | 1 | snRNA | ENSG0000C |
| 116801023 | 116801477 | 1 | lncRNA | ENSG0000C |
| 116856144 | 116881441 | -1 | protein_coding | ENSG0000C |
| 116908312 | 116908627 | 1 | processed_pseudoge | ENSG0000C |
| 116910950 | 117031148 | 1 | protein_coding | ENSG0000C |
| 116948738 | 116951422 | -1 | lncRNA | ENSG0000C |
| 116977442 | 116987337 | -1 | lncRNA | ENSG0000C |
| 117002463 | 117003152 | 1 | lncRNA | ENSG0000C |
| 117038923 | 117099479 | -1 | protein_coding | ENSG0000C |
| 117099481 | 117142091 | 1 | lncRNA | ENSG0000C |
| 117141988 | 117190531 | -1 | protein_coding | ENSG0000C |
| 117208142 | 117452170 | -1 | protein_coding | ENSG0000C |
| 117245551 | 117246164 | -1 | processed_pseudoge | ENSG0000C |

| | | | | |
|---|---|---|---|---|
| 117453012 | 117456986 | 1 | lncRNA | ENSG0000C |
| 117453012 | 117968990 | -1 | protein_coding | ENSG0000C |
| 117889454 | 117891008 | 1 | lncRNA | ENSG0000C |
| 118013588 | 118033130 | 1 | protein_coding | ENSG0000C |
| 118024817 | 118025518 | -1 | lncRNA | ENSG0000C |
| 118032694 | 118062430 | -1 | protein_coding | ENSG0000C |
| 118037869 | 118038081 | 1 | lncRNA | ENSG0000C |
| 118063593 | 118136026 | -1 | protein_coding | ENSG0000C |
| 118066398 | 118066725 | 1 | lncRNA | ENSG0000C |
| 118136124 | 118145584 | 1 | protein_coding | ENSG0000C |
| 118149801 | 118372907 | -1 | protein_coding | ENSG0000C |
| 118246084 | 118246962 | 1 | processed_pseudoge | ENSG0000C |
| 118375350 | 118376275 | -1 | lncRNA | ENSG0000C |
| 118376555 | 118418033 | 1 | protein_coding | ENSG0000C |
| 118428281 | 118428870 | 1 | lncRNA | ENSG0000C |
| 118430147 | 118430699 | 1 | lncRNA | ENSG0000C |
| 118645315 | 118648724 | -1 | lncRNA | ENSG0000C |
| 118737964 | 118738511 | -1 | processed_pseudoge | ENSG0000C |
| 118758217 | 118761739 | -1 | lncRNA | ENSG0000C |
| 118773031 | 118775137 | -1 | lncRNA | ENSG0000C |
| 118782926 | 118833536 | -1 | lncRNA | ENSG0000C |
| 118829681 | 118829805 | 1 | rRNA_pseudogene | ENSG0000C |
| 118849354 | 118908930 | -1 | lncRNA | ENSG0000C |
| 118888434 | 118888564 | -1 | snoRNA | ENSG0000C |
| 118981541 | 119163051 | 1 | protein_coding | ENSG0000C |
| 118988010 | 118989594 | -1 | lncRNA | ENSG0000C |
| 118993857 | 118994159 | 1 | misc_RNA | ENSG0000C |
| 119031039 | 119116961 | -1 | lncRNA | ENSG0000C |
| 119174065 | 119176484 | -1 | lncRNA | ENSG0000C |
| 119178642 | 119221131 | 1 | protein_coding | ENSG0000C |
| 119182048 | 119183259 | -1 | lncRNA | ENSG0000C |
| 119194850 | 119195394 | 1 | processed_pseudoge | ENSG0000C |
| 119225834 | 119259017 | -1 | lncRNA | ENSG0000C |
| 119283825 | 119303380 | -1 | lncRNA | ENSG0000C |
| 119334712 | 119541040 | 1 | protein_coding | ENSG0000C |
| 119361247 | 119668079 | -1 | lncRNA | ENSG0000C |
| 119593774 | 119643075 | 1 | protein_coding | ENSG0000C |
| 119631090 | 119631386 | -1 | misc_RNA | ENSG0000C |
| 119667864 | 119681624 | 1 | protein_coding | ENSG0000C |
| 119685791 | 119877320 | -1 | protein_coding | ENSG0000C |
| 119699768 | 119701011 | 1 | lncRNA | ENSG0000C |
| 119713634 | 119713724 | -1 | miRNA | ENSG0000C |
| 119959424 | 119959754 | 1 | processed_pseudoge | ENSG0000C |
| 119989869 | 120094494 | 1 | protein_coding | ENSG0000C |
| 120018900 | 120019261 | 1 | misc_RNA | ENSG0000C |
| 120057035 | 120057180 | 1 | processed_pseudoge | ENSG0000C |
| 120095099 | 120117502 | -1 | protein_coding | ENSG0000C |
| 120116907 | 120119000 | 1 | lncRNA | ENSG0000C |
| 120127202 | 120194715 | -1 | protein_coding | ENSG0000C |
| 120155434 | 120155499 | -1 | miRNA | ENSG0000C |

| | | | | |
|---|---|---|---|---|
| 120196699 | 120201235 | -1 | protein_coding | ENSG00000( |
| 120201291 | 120213231 | 1 | lncRNA | ENSG00000( |
| 120210439 | 120265771 | -1 | protein_coding | ENSG00000( |
| 120218070 | 120222668 | 1 | lncRNA | ENSG00000( |
| 120224744 | 120225421 | 1 | lncRNA | ENSG00000( |
| 120247460 | 120247791 | 1 | processed_pseudoge | ENSG00000( |
| 120282303 | 120282716 | 1 | processed_pseudoge | ENSG00000( |
| 120291763 | 120291903 | -1 | snRNA | ENSG00000( |
| 120293097 | 120293237 | -1 | snRNA | ENSG00000( |
| 120302316 | 120313249 | 1 | protein_coding | ENSG00000( |
| 120313238 | 120313341 | -1 | snRNA | ENSG00000( |
| 120322115 | 120327779 | -1 | protein_coding | ENSG00000( |
| 120341330 | 120369164 | -1 | protein_coding | ENSG00000( |
| 120369440 | 120371688 | 1 | transcribed_processe | ENSG00000( |
| 120389502 | 120395995 | 1 | lncRNA | ENSG00000( |
| 120438090 | 120440737 | 1 | protein_coding | ENSG00000( |
| 120438198 | 120460006 | 1 | protein_coding | ENSG00000( |
| 120443964 | 120446384 | -1 | protein_coding | ENSG00000( |
| 120446444 | 120463749 | 1 | protein_coding | ENSG00000( |
| 120450437 | 120450803 | -1 | processed_pseudoge | ENSG00000( |
| 120461672 | 120469748 | -1 | protein_coding | ENSG00000( |
| 120469850 | 120498493 | 1 | protein_coding | ENSG00000( |
| 120488079 | 120495946 | -1 | lncRNA | ENSG00000( |
| 120500735 | 120501090 | -1 | lncRNA | ENSG00000( |
| 120503279 | 120534434 | -1 | protein_coding | ENSG00000( |
| 120504571 | 120504659 | -1 | misc_RNA | ENSG00000( |
| 120522664 | 120523443 | -1 | processed_pseudoge | ENSG00000( |
| 120533480 | 120577588 | 1 | protein_coding | ENSG00000( |
| 120578764 | 120581402 | -1 | protein_coding | ENSG00000( |
| 120594015 | 120594548 | -1 | processed_pseudoge | ENSG00000( |
| 120628830 | 120641591 | -1 | lncRNA | ENSG00000( |
| 120640552 | 120667324 | 1 | protein_coding | ENSG00000( |
| 120687149 | 120701859 | 1 | protein_coding | ENSG00000( |
| 120697124 | 120699541 | -1 | lncRNA | ENSG00000( |
| 120703867 | 120704282 | 1 | lncRNA | ENSG00000( |
| 120709112 | 120709523 | 1 | lncRNA | ENSG00000( |
| 120710458 | 120723640 | 1 | protein_coding | ENSG00000( |
| 120721507 | 120723639 | -1 | lncRNA | ENSG00000( |
| 120723193 | 120723266 | 1 | miRNA | ENSG00000( |
| 120725774 | 120740008 | 1 | protein_coding | ENSG00000( |
| 120740470 | 120761592 | -1 | lncRNA | ENSG00000( |
| 120762510 | 120904358 | -1 | protein_coding | ENSG00000( |
| 120845110 | 120845647 | -1 | processed_pseudoge | ENSG00000( |
| 120904702 | 120907937 | 1 | lncRNA | ENSG00000( |
| 120914400 | 120915123 | 1 | processed_pseudoge | ENSG00000( |
| 120916745 | 120917236 | 1 | processed_pseudoge | ENSG00000( |
| 120941728 | 120980965 | -1 | lncRNA | ENSG00000( |
| 120978543 | 121002512 | 1 | protein_coding | ENSG00000( |
| 121000486 | 121016502 | -1 | protein_coding | ENSG00000( |
| 121019111 | 121039242 | -1 | protein_coding | ENSG00000( |

| | | | | |
|---|---|---|---|---|
| 121053732 | 121059670 | -1 | unitary_pseudogene | ENSG00000 |
| 121079842 | 121080124 | 1 | processed_pseudoge | ENSG00000 |
| 121096027 | 121096429 | -1 | TEC | ENSG00000 |
| 121107043 | 121107174 | 1 | snoRNA | ENSG00000 |
| 121107220 | 121107490 | 1 | processed_pseudoge | ENSG00000 |
| 121132819 | 121188032 | 1 | protein_coding | ENSG00000 |
| 121177919 | 121191477 | -1 | lncRNA | ENSG00000 |
| 121190868 | 121191518 | 1 | lncRNA | ENSG00000 |
| 121209857 | 121234106 | 1 | protein_coding | ENSG00000 |
| 121237675 | 121298308 | -1 | protein_coding | ENSG00000 |
| 121308245 | 121399896 | -1 | protein_coding | ENSG00000 |
| 121391962 | 121399859 | 1 | lncRNA | ENSG00000 |
| 121400041 | 121430623 | 1 | protein_coding | ENSG00000 |
| 121429096 | 121581023 | -1 | protein_coding | ENSG00000 |
| 121444273 | 121444352 | -1 | miRNA | ENSG00000 |
| 121580792 | 121593504 | 1 | lncRNA | ENSG00000 |
| 121604902 | 121605012 | 1 | snRNA | ENSG00000 |
| 121626550 | 121642677 | 1 | protein_coding | ENSG00000 |
| 121648742 | 121672631 | -1 | protein_coding | ENSG00000 |
| 121687187 | 121689380 | 1 | lncRNA | ENSG00000 |
| 121712752 | 121783001 | 1 | protein_coding | ENSG00000 |
| 121777754 | 121803403 | -1 | protein_coding | ENSG00000 |
| 121795267 | 121803906 | -1 | lncRNA | ENSG00000 |
| 121797511 | 121801972 | 1 | lncRNA | ENSG00000 |
| 121800797 | 121803403 | 1 | lncRNA | ENSG00000 |
| 121804180 | 121832584 | 1 | protein_coding | ENSG00000 |
| 121839527 | 121863596 | -1 | protein_coding | ENSG00000 |
| 121856259 | 121857059 | 1 | lncRNA | ENSG00000 |
| 121874193 | 121874337 | 1 | lncRNA | ENSG00000 |
| 121887540 | 121888643 | -1 | lncRNA | ENSG00000 |
| 121888732 | 121918297 | 1 | protein_coding | ENSG00000 |
| 121888809 | 121921470 | 1 | protein_coding | ENSG00000 |
| 121897583 | 121897826 | -1 | misc_RNA | ENSG00000 |
| 121908103 | 121908163 | 1 | snRNA | ENSG00000 |
| 121918592 | 122003927 | 1 | protein_coding | ENSG00000 |
| 122007434 | 122020063 | -1 | lncRNA | ENSG00000 |
| 122019422 | 122062044 | 1 | protein_coding | ENSG00000 |
| 122063306 | 122068616 | 1 | lncRNA | ENSG00000 |
| 122078756 | 122147344 | 1 | protein_coding | ENSG00000 |
| 122167738 | 122203471 | 1 | protein_coding | ENSG00000 |
| 122172029 | 122174221 | -1 | protein_coding | ENSG00000 |
| 122203681 | 122208952 | 1 | protein_coding | ENSG00000 |
| 122207662 | 122227534 | -1 | protein_coding | ENSG00000 |
| 122207668 | 122227456 | -1 | protein_coding | ENSG00000 |
| 122207779 | 122266423 | -1 | protein_coding | ENSG00000 |
| 122229564 | 122266494 | -1 | protein_coding | ENSG00000 |
| 122271432 | 122422632 | -1 | protein_coding | ENSG00000 |
| 122364782 | 122365632 | 1 | processed_pseudoge | ENSG00000 |
| 122395542 | 122400857 | 1 | lncRNA | ENSG00000 |
| 122471599 | 122501073 | -1 | protein_coding | ENSG00000 |

| | | | | |
|---|---|---|---|---|
| 122492113 | 122492241 | -1 | snoRNA | ENSG00000 |
| 122501187 | 122501641 | 1 | lncRNA | ENSG00000 |
| 122503454 | 122527000 | -1 | protein_coding | ENSG00000 |
| 122527246 | 122626396 | 1 | protein_coding | ENSG00000 |
| 122563735 | 122563762 | -1 | processed_pseudoge | ENSG00000 |
| 122634130 | 122634673 | 1 | lncRNA | ENSG00000 |
| 122687125 | 122715979 | 1 | lncRNA | ENSG00000 |
| 122701293 | 122703357 | -1 | protein_coding | ENSG00000 |
| 122714756 | 122716811 | -1 | protein_coding | ENSG00000 |
| 122726076 | 122730844 | -1 | protein_coding | ENSG00000 |
| 122749481 | 122749586 | -1 | processed_pseudoge | ENSG00000 |
| 122752824 | 122771064 | 1 | protein_coding | ENSG00000 |
| 122768099 | 122768200 | 1 | misc_RNA | ENSG00000 |
| 122774526 | 122827528 | 1 | protein_coding | ENSG00000 |
| 122834453 | 122862961 | 1 | protein_coding | ENSG00000 |
| 122865330 | 122896127 | -1 | protein_coding | ENSG00000 |
| 122865335 | 122867021 | 1 | lncRNA | ENSG00000 |
| 122870059 | 122888720 | -1 | TEC | ENSG00000 |
| 122920951 | 122981649 | -1 | protein_coding | ENSG00000 |
| 122974580 | 122980043 | 1 | protein_coding | ENSG00000 |
| 122975320 | 122982907 | 1 | lncRNA | ENSG00000 |
| 122980060 | 122982913 | 1 | protein_coding | ENSG00000 |
| 122983480 | 123150015 | -1 | protein_coding | ENSG00000 |
| 123010667 | 123010728 | -1 | miRNA | ENSG00000 |
| 123081384 | 123084744 | 1 | lncRNA | ENSG00000 |
| 123087256 | 123092821 | -1 | TEC | ENSG00000 |
| 123152320 | 123244014 | -1 | protein_coding | ENSG00000 |
| 123152324 | 123153377 | -1 | TEC | ENSG00000 |
| 123233436 | 123257960 | 1 | protein_coding | ENSG00000 |
| 123250112 | 123272334 | -1 | protein_coding | ENSG00000 |
| 123262060 | 123262402 | 1 | lncRNA | ENSG00000 |
| 123282916 | 123283034 | -1 | rRNA_pseudogene | ENSG00000 |
| 123289109 | 123364847 | -1 | protein_coding | ENSG00000 |
| 123348999 | 123349100 | -1 | misc_RNA | ENSG00000 |
| 123363868 | 123366113 | 1 | lncRNA | ENSG00000 |
| 123364764 | 123364843 | -1 | miRNA | ENSG00000 |
| 123383773 | 123409353 | 1 | protein_coding | ENSG00000 |
| 123410683 | 123436717 | -1 | protein_coding | ENSG00000 |
| 123441517 | 123441852 | -1 | processed_pseudoge | ENSG00000 |
| 123458088 | 123473154 | 1 | protein_coding | ENSG00000 |
| 123470054 | 123533719 | -1 | protein_coding | ENSG00000 |
| 123515275 | 123515513 | -1 | lncRNA | ENSG00000 |
| 123519390 | 123519856 | -1 | lncRNA | ENSG00000 |
| 123536409 | 123536534 | 1 | miRNA | ENSG00000 |
| 123575891 | 123585115 | -1 | lncRNA | ENSG00000 |
| 123584533 | 123598582 | 1 | protein_coding | ENSG00000 |
| 123602077 | 123620943 | 1 | protein_coding | ENSG00000 |
| 123616708 | 123616840 | 1 | snoRNA | ENSG00000 |
| 123620406 | 123633766 | -1 | protein_coding | ENSG00000 |
| 123633739 | 123662604 | 1 | protein_coding | ENSG00000 |

| | | | | |
|---|---|---|---|---|
| 123649068 | 123650485 | 1 | TEC | ENSG00000 |
| 123655528 | 123656128 | 1 | TEC | ENSG00000 |
| 123671113 | 123708399 | 1 | protein_coding | ENSG00000 |
| 123707602 | 123708386 | -1 | lncRNA | ENSG00000 |
| 123712353 | 123761755 | 1 | protein_coding | ENSG00000 |
| 123713408 | 123723169 | -1 | lncRNA | ENSG00000 |
| 123721246 | 123721639 | -1 | processed_pseudoge | ENSG00000 |
| 123754246 | 123754794 | -1 | TEC | ENSG00000 |
| 123762188 | 123936206 | 1 | protein_coding | ENSG00000 |
| 123918660 | 123972831 | -1 | protein_coding | ENSG00000 |
| 123925461 | 123934984 | -1 | lncRNA | ENSG00000 |
| 123960717 | 123961244 | -1 | lncRNA | ENSG00000 |
| 123962555 | 123962817 | -1 | lncRNA | ENSG00000 |
| 123966077 | 123966629 | -1 | lncRNA | ENSG00000 |
| 123968023 | 123968579 | -1 | lncRNA | ENSG00000 |
| 123969990 | 123970344 | -1 | lncRNA | ENSG00000 |
| 123971457 | 123971714 | -1 | lncRNA | ENSG00000 |
| 123971845 | 124015439 | 1 | protein_coding | ENSG00000 |
| 123973215 | 124088594 | 1 | lncRNA | ENSG00000 |
| 123973241 | 124316024 | 1 | protein_coding | ENSG00000 |
| 124005871 | 124006225 | -1 | processed_pseudoge | ENSG00000 |
| 124085761 | 124088598 | 1 | lncRNA | ENSG00000 |
| 124206228 | 124209018 | 1 | lncRNA | ENSG00000 |
| 124324415 | 124567589 | -1 | protein_coding | ENSG00000 |
| 124337181 | 124337242 | -1 | miRNA | ENSG00000 |
| 124383381 | 124383510 | 1 | processed_pseudoge | ENSG00000 |
| 124513222 | 124516798 | 1 | lncRNA | ENSG00000 |
| 124529722 | 124530228 | -1 | TEC | ENSG00000 |
| 124537844 | 124539677 | 1 | TEC | ENSG00000 |
| 124542503 | 124542873 | 1 | TEC | ENSG00000 |
| 124555905 | 124556432 | 1 | TEC | ENSG00000 |
| 124561869 | 124562283 | 1 | TEC | ENSG00000 |
| 124593181 | 124593691 | -1 | TEC | ENSG00000 |
| 124622290 | 124623069 | -1 | TEC | ENSG00000 |
| 124623391 | 124623987 | 1 | TEC | ENSG00000 |
| 124637999 | 124638444 | -1 | TEC | ENSG00000 |
| 124725524 | 124725934 | 1 | TEC | ENSG00000 |
| 124776856 | 124882668 | -1 | protein_coding | ENSG00000 |
| 124786783 | 124808790 | 1 | lncRNA | ENSG00000 |
| 124901512 | 124901611 | 1 | snRNA | ENSG00000 |
| 124911604 | 124917368 | -1 | protein_coding | ENSG00000 |
| 124915547 | 124915659 | 1 | miRNA | ENSG00000 |
| 124935455 | 124935895 | -1 | transcribed_processe | ENSG00000 |
| 124946825 | 124989131 | -1 | protein_coding | ENSG00000 |
| 124993645 | 125031231 | 1 | protein_coding | ENSG00000 |
| 125025434 | 125027410 | -1 | lncRNA | ENSG00000 |
| 125056359 | 125056581 | -1 | processed_pseudoge | ENSG00000 |
| 125065434 | 125143333 | 1 | protein_coding | ENSG00000 |
| 125138245 | 125141711 | 1 | TEC | ENSG00000 |
| 125150058 | 125151394 | 1 | lncRNA | ENSG00000 |

| | | | | |
|---|---|---|---|---|
| 125159331 | 125160397 | -1 | lncRNA | ENSG0000C |
| 125160525 | 125164918 | 1 | TEC | ENSG0000C |
| 125186836 | 125662377 | 1 | protein_coding | ENSG0000C |
| 125300936 | 125301917 | 1 | TEC | ENSG0000C |
| 125808668 | 125809808 | 1 | TEC | ENSG0000C |
| 125858132 | 125880724 | 1 | lncRNA | ENSG0000C |
| 125858862 | 125861119 | 1 | TEC | ENSG0000C |
| 125900421 | 125901483 | -1 | processed_pseudoge | ENSG0000C |
| 125958688 | 125983665 | -1 | lncRNA | ENSG0000C |
| 125983702 | 126043485 | 1 | lncRNA | ENSG0000C |
| 126094112 | 126177632 | 1 | lncRNA | ENSG0000C |
| 126106459 | 126107047 | -1 | TEC | ENSG0000C |
| 126130166 | 126130865 | -1 | TEC | ENSG0000C |
| 126141952 | 126142377 | 1 | TEC | ENSG0000C |
| 126165730 | 126191406 | -1 | lncRNA | ENSG0000C |
| 126191151 | 126434227 | 1 | lncRNA | ENSG0000C |
| 126400792 | 126405528 | -1 | lncRNA | ENSG0000C |
| 126438837 | 126472790 | 1 | lncRNA | ENSG0000C |
| 126458462 | 126458792 | -1 | processed_pseudoge | ENSG0000C |
| 126510910 | 126511711 | -1 | processed_pseudoge | ENSG0000C |
| 126524248 | 126524626 | 1 | processed_pseudoge | ENSG0000C |
| 126610034 | 126690318 | -1 | lncRNA | ENSG0000C |
| 126616344 | 126616554 | -1 | processed_pseudoge | ENSG0000C |
| 126628172 | 126690322 | -1 | lncRNA | ENSG0000C |
| 126675895 | 126676243 | -1 | TEC | ENSG0000C |
| 126677262 | 126677806 | 1 | TEC | ENSG0000C |
| 126688341 | 126720333 | 1 | lncRNA | ENSG0000C |
| 126723412 | 126749027 | 1 | lncRNA | ENSG0000C |
| 126729787 | 126772519 | -1 | lncRNA | ENSG0000C |
| 126864465 | 126865422 | 1 | processed_pseudoge | ENSG0000C |
| 126869010 | 126874725 | -1 | lncRNA | ENSG0000C |
| 126874804 | 126888718 | 1 | lncRNA | ENSG0000C |
| 126912034 | 126912322 | 1 | processed_pseudoge | ENSG0000C |
| 126915199 | 127060565 | -1 | lncRNA | ENSG0000C |
| 127130005 | 127207706 | 1 | lncRNA | ENSG0000C |
| 127142029 | 127146532 | -1 | lncRNA | ENSG0000C |
| 127146535 | 127151330 | 1 | lncRNA | ENSG0000C |
| 127147149 | 127150081 | 1 | lncRNA | ENSG0000C |
| 127149955 | 127161259 | -1 | lncRNA | ENSG0000C |
| 127274265 | 127284328 | -1 | lncRNA | ENSG0000C |
| 127317709 | 127319986 | -1 | unprocessed_pseudc | ENSG0000C |
| 127321355 | 127321529 | 1 | snRNA | ENSG0000C |
| 127324152 | 127340109 | -1 | lncRNA | ENSG0000C |
| 127344258 | 127344402 | -1 | processed_pseudoge | ENSG0000C |
| 127372243 | 127372669 | 1 | TEC | ENSG0000C |
| 127451346 | 127455634 | -1 | lncRNA | ENSG0000C |
| 127462507 | 127463197 | 1 | TEC | ENSG0000C |
| 127484243 | 127486675 | 1 | lncRNA | ENSG0000C |
| 127486938 | 127533242 | 1 | lncRNA | ENSG0000C |
| 127563908 | 127590930 | 1 | lncRNA | ENSG0000C |

| | | | | |
|---|---|---|---|---|
| 127598168 | 127599715 | 1 | lncRNA | ENSG0000( |
| 127619336 | 127619447 | -1 | misc_RNA | ENSG0000( |
| 127624153 | 127626267 | 1 | unprocessed_pseudc | ENSG0000( |
| 127631248 | 127636488 | -1 | lncRNA | ENSG0000( |
| 127726339 | 127821183 | -1 | lncRNA | ENSG0000( |
| 127825938 | 127826286 | -1 | TEC | ENSG0000( |
| 127876999 | 127877516 | -1 | processed_pseudoge | ENSG0000( |
| 127881616 | 127898645 | 1 | lncRNA | ENSG0000( |
| 127914707 | 127960028 | 1 | lncRNA | ENSG0000( |
| 127933689 | 127983854 | -1 | lncRNA | ENSG0000( |
| 128023788 | 128027166 | -1 | lncRNA | ENSG0000( |
| 128052122 | 128068361 | -1 | lncRNA | ENSG0000( |
| 128086621 | 128118656 | -1 | lncRNA | ENSG0000( |
| 128116730 | 128123103 | 1 | lncRNA | ENSG0000( |
| 128187225 | 128191495 | -1 | lncRNA | ENSG0000( |
| 128244506 | 128244573 | 1 | miRNA | ENSG0000( |
| 128267403 | 128707915 | 1 | protein_coding | ENSG0000( |
| 128288159 | 128289056 | 1 | TEC | ENSG0000( |
| 128294092 | 128294178 | 1 | miRNA | ENSG0000( |
| 128399978 | 128404814 | 1 | lncRNA | ENSG0000( |
| 128437778 | 128439206 | -1 | unprocessed_pseudc | ENSG0000( |
| 128793194 | 128823958 | -1 | protein_coding | ENSG0000( |
| 128813186 | 128814750 | -1 | TEC | ENSG0000( |
| 128826836 | 128827579 | 1 | lncRNA | ENSG0000( |
| 128853427 | 128984968 | 1 | protein_coding | ENSG0000( |
| 128944274 | 128945203 | -1 | unprocessed_pseudc | ENSG0000( |
| 129013336 | 129016663 | 1 | unprocessed_pseudc | ENSG0000( |
| 129071725 | 129904025 | -1 | protein_coding | ENSG0000( |
| 129109629 | 129113294 | 1 | lncRNA | ENSG0000( |
| 129170092 | 129170238 | -1 | misc_RNA | ENSG0000( |
| 129208601 | 129212662 | 1 | lncRNA | ENSG0000( |
| 129262345 | 129262824 | 1 | TEC | ENSG0000( |
| 129521303 | 129522763 | -1 | lncRNA | ENSG0000( |
| 129622929 | 129625366 | 1 | lncRNA | ENSG0000( |
| 129681427 | 129698227 | -1 | lncRNA | ENSG0000( |
| 129696599 | 129700122 | 1 | lncRNA | ENSG0000( |
| 129839928 | 129840522 | -1 | TEC | ENSG0000( |
| 129852208 | 129854944 | -1 | lncRNA | ENSG0000( |
| 130024493 | 130044956 | -1 | lncRNA | ENSG0000( |
| 130032928 | 130045057 | -1 | lncRNA | ENSG0000( |
| 130047132 | 130048376 | 1 | lncRNA | ENSG0000( |
| 130070325 | 130072685 | 1 | lncRNA | ENSG0000( |
| 130121925 | 130122974 | -1 | TEC | ENSG0000( |
| 130138693 | 130140768 | 1 | lncRNA | ENSG0000( |
| 130144315 | 130162256 | -1 | lncRNA | ENSG0000( |
| 130144954 | 130145691 | 1 | TEC | ENSG0000( |
| 130162459 | 130165740 | 1 | protein_coding | ENSG0000( |
| 130216807 | 130217157 | -1 | processed_pseudoge | ENSG0000( |
| 130249679 | 130266725 | 1 | lncRNA | ENSG0000( |
| 130337887 | 130372637 | 1 | protein_coding | ENSG0000( |

| | | | | |
|---|---|---|---|---|
| 130350996 | 130351707 | -1 | TEC | ENSG0000C |
| 130383364 | 130383890 | 1 | TEC | ENSG0000C |
| 130396137 | 130716281 | -1 | protein_coding | ENSG0000C |
| 130419535 | 130421019 | -1 | lncRNA | ENSG0000C |
| 130425004 | 130430661 | 1 | TEC | ENSG0000C |
| 130465008 | 130466315 | -1 | TEC | ENSG0000C |
| 130651342 | 130669233 | 1 | lncRNA | ENSG0000C |
| 130775694 | 130776546 | -1 | TEC | ENSG0000C |
| 130789600 | 130839266 | -1 | protein_coding | ENSG0000C |
| 130810606 | 130812438 | 1 | lncRNA | ENSG0000C |
| 130810821 | 130812622 | -1 | lncRNA | ENSG0000C |
| 130869736 | 130869837 | -1 | snRNA | ENSG0000C |
| 130872037 | 130877678 | 1 | protein_coding | ENSG0000C |
| 130906838 | 130906949 | 1 | misc_RNA | ENSG0000C |
| 130953907 | 131141469 | 1 | protein_coding | ENSG0000C |
| 130956384 | 130958031 | -1 | TEC | ENSG0000C |
| 130962175 | 130963028 | 1 | TEC | ENSG0000C |
| 130976786 | 130978812 | 1 | TEC | ENSG0000C |
| 130977562 | 130978768 | 1 | lncRNA | ENSG0000C |
| 130980667 | 130981367 | -1 | TEC | ENSG0000C |
| 130988312 | 130990028 | -1 | TEC | ENSG0000C |
| 130990138 | 130993976 | -1 | lncRNA | ENSG0000C |
| 131022769 | 131024741 | -1 | TEC | ENSG0000C |
| 131025561 | 131028060 | -1 | TEC | ENSG0000C |
| 131029110 | 131035487 | -1 | lncRNA | ENSG0000C |
| 131165011 | 131232940 | 1 | lncRNA | ENSG0000C |
| 131235511 | 131237523 | 1 | TEC | ENSG0000C |
| 131292068 | 131293176 | -1 | unprocessed_pseudc | ENSG0000C |
| 131295522 | 131298218 | 1 | TEC | ENSG0000C |
| 131296110 | 131297972 | -1 | lncRNA | ENSG0000C |
| 131299640 | 131299751 | -1 | rRNA_pseudogene | ENSG0000C |
| 131310045 | 131312811 | -1 | lncRNA | ENSG0000C |
| 131347470 | 131367555 | 1 | lncRNA | ENSG0000C |
| 131350844 | 131371721 | -1 | lncRNA | ENSG0000C |
| 131366660 | 131371059 | 1 | lncRNA | ENSG0000C |
| 131436447 | 131436960 | 1 | TEC | ENSG0000C |
| 131447337 | 131455436 | -1 | lncRNA | ENSG0000C |
| 131457018 | 131458324 | 1 | lncRNA | ENSG0000C |
| 131462853 | 131529894 | -1 | lncRNA | ENSG0000C |
| 131478844 | 131483034 | -1 | TEC | ENSG0000C |
| 131596857 | 131597115 | -1 | processed_pseudoge | ENSG0000C |
| 131620178 | 131620765 | -1 | TEC | ENSG0000C |
| 131621891 | 131623203 | -1 | lncRNA | ENSG0000C |
| 131639396 | 131661823 | -1 | lncRNA | ENSG0000C |
| 131644172 | 131645651 | 1 | lncRNA | ENSG0000C |
| 131645918 | 131646444 | 1 | TEC | ENSG0000C |
| 131661063 | 131661174 | 1 | rRNA_pseudogene | ENSG0000C |
| 131662596 | 131664704 | 1 | lncRNA | ENSG0000C |
| 131667492 | 131668246 | 1 | processed_pseudoge | ENSG0000C |
| 131711081 | 131799737 | 1 | protein_coding | ENSG0000C |

| | | | | |
|---|---|---|---|---|
| 131756966 | 131758047 | -1 | lncRNA | ENSG00000 |
| 131801315 | 131801434 | -1 | rRNA_pseudogene | ENSG00000 |
| 131815263 | 131815368 | -1 | snRNA | ENSG00000 |
| 131828393 | 131851783 | 1 | protein_coding | ENSG00000 |
| 131857420 | 131864538 | 1 | lncRNA | ENSG00000 |
| 131894622 | 131923150 | 1 | protein_coding | ENSG00000 |
| 131896500 | 131898239 | 1 | TEC | ENSG00000 |
| 131924736 | 131929351 | -1 | lncRNA | ENSG00000 |
| 131929200 | 131945896 | 1 | protein_coding | ENSG00000 |
| 131934642 | 131934928 | 1 | lncRNA | ENSG00000 |
| 131949920 | 132081102 | 1 | protein_coding | ENSG00000 |
| 132031224 | 132031359 | 1 | snoRNA | ENSG00000 |
| 132077803 | 132080460 | -1 | lncRNA | ENSG00000 |
| 132083540 | 132083779 | -1 | lncRNA | ENSG00000 |
| 132084283 | 132131639 | 1 | transcribed_unproce | ENSG00000 |
| 132100013 | 132100820 | -1 | TEC | ENSG00000 |
| 132126461 | 132126764 | -1 | processed_pseudoge | ENSG00000 |
| 132136594 | 132144319 | -1 | protein_coding | ENSG00000 |
| 132144457 | 132152473 | 1 | protein_coding | ENSG00000 |
| 132169823 | 132170043 | 1 | lncRNA | ENSG00000 |
| 132186735 | 132189695 | -1 | lncRNA | ENSG00000 |
| 132190213 | 132190997 | 1 | lncRNA | ENSG00000 |
| 132196372 | 132329589 | -1 | protein_coding | ENSG00000 |
| 132200436 | 132201287 | 1 | TEC | ENSG00000 |
| 132275391 | 132278329 | 1 | lncRNA | ENSG00000 |
| 132277349 | 132280900 | 1 | lncRNA | ENSG00000 |
| 132279256 | 132280900 | -1 | TEC | ENSG00000 |
| 132282814 | 132284606 | 1 | lncRNA | ENSG00000 |
| 132329850 | 132331575 | 1 | lncRNA | ENSG00000 |
| 132332005 | 132332432 | 1 | TEC | ENSG00000 |
| 132344477 | 132345352 | 1 | lncRNA | ENSG00000 |
| 132351855 | 132352322 | 1 | TEC | ENSG00000 |
| 132360734 | 132361198 | 1 | TEC | ENSG00000 |
| 132362810 | 132363348 | 1 | TEC | ENSG00000 |
| 132424504 | 132425208 | -1 | lncRNA | ENSG00000 |
| 132457160 | 132460024 | 1 | lncRNA | ENSG00000 |
| 132462242 | 132462856 | 1 | lncRNA | ENSG00000 |
| 132489551 | 132585188 | 1 | protein_coding | ENSG00000 |
| 132550729 | 132554947 | 1 | TEC | ENSG00000 |
| 132565071 | 132566425 | 1 | TEC | ENSG00000 |
| 132581997 | 132582061 | 1 | miRNA | ENSG00000 |
| 132593031 | 132593698 | -1 | lncRNA | ENSG00000 |
| 132603150 | 132610582 | -1 | protein_coding | ENSG00000 |
| 132618776 | 132622388 | 1 | protein_coding | ENSG00000 |
| 132621999 | 132622973 | -1 | TEC | ENSG00000 |
| 132623753 | 132687376 | -1 | protein_coding | ENSG00000 |
| 132687587 | 132704985 | 1 | protein_coding | ENSG00000 |
| 132688049 | 132714912 | 1 | lncRNA | ENSG00000 |
| 132710819 | 132722734 | 1 | protein_coding | ENSG00000 |
| 132723945 | 132724057 | -1 | rRNA_pseudogene | ENSG00000 |

| | | | | |
|---|---|---|---|---|
| 132725503 | 132761832 | -1 | protein_coding | ENSG00000( |
| 132745156 | 132745729 | 1 | processed_pseudoge | ENSG00000( |
| 132768914 | 132829078 | -1 | protein_coding | ENSG00000( |
| 132822187 | 132956304 | -1 | protein_coding | ENSG00000( |
| 132825701 | 132826184 | 1 | processed_pseudoge | ENSG00000( |
| 132835374 | 132835500 | -1 | snRNA | ENSG00000( |
| 132865855 | 132867958 | -1 | TEC | ENSG00000( |
| 132887842 | 132888583 | 1 | lncRNA | ENSG00000( |
| 132899984 | 132900368 | 1 | processed_pseudoge | ENSG00000( |
| 132911470 | 132914732 | 1 | lncRNA | ENSG00000( |
| 132918306 | 132956306 | -1 | protein_coding | ENSG00000( |
| 132972338 | 132972890 | -1 | processed_pseudoge | ENSG00000( |
| 132986365 | 133032952 | 1 | protein_coding | ENSG00000( |
| 132995190 | 132995302 | -1 | snRNA | ENSG00000( |
| 133030389 | 133037222 | -1 | lncRNA | ENSG00000( |
| 133033503 | 133033956 | 1 | processed_pseudoge | ENSG00000( |
| 133037292 | 133063304 | 1 | protein_coding | ENSG00000( |
| 133073516 | 133073983 | -1 | TEC | ENSG00000( |
| 133077762 | 133078392 | -1 | TEC | ENSG00000( |
| 133079838 | 133107544 | 1 | protein_coding | ENSG00000( |
| 133106817 | 133130473 | -1 | protein_coding | ENSG00000( |
| 133115692 | 133122340 | 1 | lncRNA | ENSG00000( |
| 133125479 | 133125952 | 1 | processed_pseudoge | ENSG00000( |
| 133130575 | 133159465 | 1 | protein_coding | ENSG00000( |
| 133130984 | 133202369 | 1 | protein_coding | ENSG00000( |
| 133181409 | 133214831 | 1 | protein_coding | ENSG00000( |
| 133217210 | 133217675 | -1 | processed_pseudoge | ENSG00000( |
| 133218312 | 133236095 | -1 | protein_coding | ENSG00000( |
| 133238335 | 133238438 | -1 | snRNA | ENSG00000( |
| 133238455 | 133238549 | 1 | misc_RNA | ENSG00000( |
| 18174010 | 18178465 | -1 | unprocessed_pseudc | ENSG00000( |
| 18195297 | 18232024 | 1 | lncRNA | ENSG00000( |
| 18206570 | 18207466 | -1 | processed_pseudoge | ENSG00000( |
| 18248913 | 18249035 | -1 | processed_pseudoge | ENSG00000( |
| 18253526 | 18270822 | -1 | unprocessed_pseudc | ENSG00000( |
| 18279320 | 18309934 | -1 | unprocessed_pseudc | ENSG00000( |
| 18297554 | 18298733 | 1 | processed_pseudoge | ENSG00000( |
| 18331933 | 18348596 | 1 | unprocessed_pseudc | ENSG00000( |
| 18408806 | 18431605 | 1 | transcribed_unproce | ENSG00000( |
| 18418526 | 18418632 | 1 | snRNA | ENSG00000( |
| 18467172 | 18485448 | -1 | unprocessed_pseudc | ENSG00000( |
| 18491904 | 18492419 | 1 | processed_pseudoge | ENSG00000( |
| 18538880 | 18551214 | 1 | lncRNA | ENSG00000( |
| 18609132 | 18609702 | 1 | processed_pseudoge | ENSG00000( |
| 18610298 | 18611675 | -1 | lncRNA | ENSG00000( |
| 18665235 | 18667221 | 1 | processed_pseudoge | ENSG00000( |
| 18672827 | 18676163 | 1 | lncRNA | ENSG00000( |
| 18680494 | 18681700 | 1 | processed_pseudoge | ENSG00000( |
| 18697003 | 18697507 | 1 | processed_pseudoge | ENSG00000( |
| 18724415 | 18724904 | -1 | processed_pseudoge | ENSG00000( |

| | | | | |
|---|---|---|---|---|
| 18726706 | 18727414 | 1 | processed_pseudoge | ENSG00000 |
| 18734080 | 18734896 | 1 | processed_pseudoge | ENSG00000 |
| 18741730 | 18742718 | -1 | unprocessed_pseudc | ENSG00000 |
| 18807279 | 18808018 | -1 | processed_pseudoge | ENSG00000 |
| 18809830 | 18810048 | 1 | processed_pseudoge | ENSG00000 |
| 18816718 | 18871967 | -1 | unprocessed_pseudc | ENSG00000 |
| 18837768 | 18838098 | -1 | processed_pseudoge | ENSG00000 |
| 18869185 | 18869291 | -1 | snRNA | ENSG00000 |
| 18880054 | 18880157 | 1 | snRNA | ENSG00000 |
| 18892260 | 18895230 | -1 | processed_pseudoge | ENSG00000 |
| 18905419 | 18935554 | 1 | lncRNA | ENSG00000 |
| 18921888 | 18944284 | -1 | lncRNA | ENSG00000 |
| 18946755 | 19062718 | -1 | transcribed_unproce | ENSG00000 |
| 19008259 | 19026802 | 1 | lncRNA | ENSG00000 |
| 19015487 | 19019336 | -1 | unprocessed_pseudc | ENSG00000 |
| 19026001 | 19032626 | -1 | unprocessed_pseudc | ENSG00000 |
| 19027837 | 19028701 | 1 | lncRNA | ENSG00000 |
| 19074038 | 19118315 | -1 | lncRNA | ENSG00000 |
| 19088487 | 19088581 | 1 | rRNA_pseudogene | ENSG00000 |
| 19126399 | 19139611 | 1 | unprocessed_pseudc | ENSG00000 |
| 19151554 | 19152307 | -1 | unprocessed_pseudc | ENSG00000 |
| 19152567 | 19171194 | -1 | unprocessed_pseudc | ENSG00000 |
| 19172773 | 19172925 | 1 | unprocessed_pseudc | ENSG00000 |
| 19173772 | 19181824 | -1 | protein_coding | ENSG00000 |
| 19182038 | 19187618 | 1 | transcribed_unproce | ENSG00000 |
| 19201637 | 19227432 | -1 | unprocessed_pseudc | ENSG00000 |
| 19262797 | 19284823 | -1 | lncRNA | ENSG00000 |
| 19290593 | 19344825 | -1 | unprocessed_pseudc | ENSG00000 |
| 19304593 | 19305625 | -1 | processed_pseudoge | ENSG00000 |
| 19345049 | 19346749 | 1 | lncRNA | ENSG00000 |
| 19349137 | 19407962 | 1 | unprocessed_pseudc | ENSG00000 |
| 19382454 | 19383072 | 1 | processed_pseudoge | ENSG00000 |
| 19408042 | 19409529 | -1 | lncRNA | ENSG00000 |
| 19411231 | 19416188 | 1 | unprocessed_pseudc | ENSG00000 |
| 19422877 | 19536762 | -1 | protein_coding | ENSG00000 |
| 19458398 | 19458718 | -1 | processed_pseudoge | ENSG00000 |
| 19552571 | 19570288 | -1 | unprocessed_pseudc | ENSG00000 |
| 19560013 | 19561269 | -1 | processed_pseudoge | ENSG00000 |
| 19561328 | 19588603 | 1 | lncRNA | ENSG00000 |
| 19563589 | 19564900 | -1 | processed_pseudoge | ENSG00000 |
| 19633659 | 19673441 | 1 | protein_coding | ENSG00000 |
| 19674624 | 19675884 | 1 | lncRNA | ENSG00000 |
| 19674752 | 19783019 | -1 | protein_coding | ENSG00000 |
| 19736975 | 19737254 | -1 | misc_RNA | ENSG00000 |
| 19818121 | 19818619 | -1 | processed_pseudoge | ENSG00000 |
| 19823482 | 19863649 | -1 | protein_coding | ENSG00000 |
| 19841827 | 19843672 | 1 | lncRNA | ENSG00000 |
| 19863858 | 19865048 | 1 | lncRNA | ENSG00000 |
| 19958670 | 20091829 | 1 | protein_coding | ENSG00000 |
| 20070250 | 20072136 | -1 | processed_pseudoge | ENSG00000 |

| | | | | |
|---|---|---|---|---|
| 20092632 | 20094144 | 1 | TEC | ENSG0000C |
| 20102701 | 20103230 | 1 | lncRNA | ENSG0000C |
| 20138255 | 20161052 | -1 | protein_coding | ENSG0000C |
| 20178978 | 20179462 | -1 | processed_pseudoge | ENSG0000C |
| 20181531 | 20181942 | 1 | lncRNA | ENSG0000C |
| 20187463 | 20192938 | -1 | protein_coding | ENSG0000C |
| 20194412 | 20196005 | 1 | TEC | ENSG0000C |
| 20221962 | 20232365 | -1 | protein_coding | ENSG0000C |
| 20403666 | 20525873 | -1 | protein_coding | ENSG0000C |
| 20433778 | 20433846 | -1 | miRNA | ENSG0000C |
| 20564708 | 20567045 | -1 | lncRNA | ENSG0000C |
| 20567069 | 20691437 | 1 | protein_coding | ENSG0000C |
| 20607268 | 20608131 | 1 | processed_pseudoge | ENSG0000C |
| 20612161 | 20612338 | 1 | snRNA | ENSG0000C |
| 20699307 | 20703718 | -1 | lncRNA | ENSG0000C |
| 20702127 | 20723098 | 1 | protein_coding | ENSG0000C |
| 20712431 | 20714146 | 1 | TEC | ENSG0000C |
| 20728731 | 20773961 | -1 | protein_coding | ENSG0000C |
| 20747035 | 20748057 | -1 | processed_pseudoge | ENSG0000C |
| 20768876 | 20769375 | -1 | lncRNA | ENSG0000C |
| 20777329 | 20903048 | -1 | protein_coding | ENSG0000C |
| 20921664 | 20923233 | -1 | processed_pseudoge | ENSG0000C |
| 20938564 | 20949599 | -1 | transcribed_processe | ENSG0000C |
| 20956179 | 20956443 | -1 | processed_pseudoge | ENSG0000C |
| 20961310 | 20962195 | 1 | processed_pseudoge | ENSG0000C |
| 20973036 | 21061586 | -1 | protein_coding | ENSG0000C |
| 21005157 | 21018122 | 1 | lncRNA | ENSG0000C |
| 21038529 | 21038667 | -1 | snRNA | ENSG0000C |
| 21094315 | 21094714 | -1 | processed_pseudoge | ENSG0000C |
| 21103003 | 21104371 | -1 | processed_pseudoge | ENSG0000C |
| 21126271 | 21126396 | -1 | misc_RNA | ENSG0000C |
| 21139886 | 21139957 | -1 | snoRNA | ENSG0000C |
| 21140514 | 21149084 | 1 | protein_coding | ENSG0000C |
| 21153595 | 21176552 | -1 | protein_coding | ENSG0000C |
| 21176658 | 21179084 | 1 | protein_coding | ENSG0000C |
| 21216555 | 21216660 | 1 | processed_pseudoge | ENSG0000C |
| 21224521 | 21235204 | 1 | lncRNA | ENSG0000C |
| 21260015 | 21295003 | 1 | transcribed_processe | ENSG0000C |
| 21287634 | 21287828 | -1 | unprocessed_pseudc | ENSG0000C |
| 21298139 | 21306373 | 1 | transcribed_unproce | ENSG0000C |
| 21302862 | 21348721 | -1 | lncRNA | ENSG0000C |
| 21319156 | 21320866 | 1 | processed_pseudoge | ENSG0000C |
| 21358373 | 21359314 | 1 | processed_pseudoge | ENSG0000C |
| 21369790 | 21369899 | -1 | rRNA_pseudogene | ENSG0000C |
| 21372573 | 21459370 | -1 | protein_coding | ENSG0000C |
| 21376977 | 21377874 | -1 | lncRNA | ENSG0000C |
| 21483834 | 21484174 | -1 | processed_pseudoge | ENSG0000C |
| 21492691 | 21604181 | -1 | protein_coding | ENSG0000C |
| 21530690 | 21531323 | 1 | processed_pseudoge | ENSG0000C |
| 21551304 | 21551410 | -1 | snRNA | ENSG0000C |

| | | | | |
|---|---|---|---|---|
| 21628413 | 21629015 | -1 | processed_pseudoge | ENSG00000 |
| 21671073 | 21704498 | 1 | protein_coding | ENSG00000 |
| 21790537 | 21790845 | 1 | misc_RNA | ENSG00000 |
| 21872763 | 21878256 | -1 | lncRNA | ENSG00000 |
| 22040972 | 22276526 | 1 | lncRNA | ENSG00000 |
| 22099163 | 22099544 | -1 | processed_pseudoge | ENSG00000 |
| 22113119 | 22113338 | -1 | processed_pseudoge | ENSG00000 |
| 22589699 | 22593871 | -1 | lncRNA | ENSG00000 |
| 22696023 | 22696574 | -1 | processed_pseudoge | ENSG00000 |
| 22704739 | 22706206 | -1 | processed_pseudoge | ENSG00000 |
| 22803145 | 22803224 | 1 | snoRNA | ENSG00000 |
| 22803296 | 22803648 | 1 | processed_pseudoge | ENSG00000 |
| 22837026 | 22837880 | -1 | processed_pseudoge | ENSG00000 |
| 22850101 | 22850561 | 1 | processed_pseudoge | ENSG00000 |
| 22859050 | 22916369 | -1 | lncRNA | ENSG00000 |
| 22897341 | 22898005 | -1 | processed_pseudoge | ENSG00000 |
| 22915651 | 22916523 | 1 | processed_pseudoge | ENSG00000 |
| 22918591 | 22924602 | 1 | lncRNA | ENSG00000 |
| 22931537 | 22931999 | 1 | lncRNA | ENSG00000 |
| 23134174 | 23134564 | 1 | processed_pseudoge | ENSG00000 |
| 23152586 | 23152686 | -1 | misc_RNA | ENSG00000 |
| 23169835 | 23170597 | -1 | lncRNA | ENSG00000 |
| 23180952 | 23325165 | 1 | protein_coding | ENSG00000 |
| 23217432 | 23217534 | -1 | snRNA | ENSG00000 |
| 23243520 | 23247184 | 1 | processed_pseudoge | ENSG00000 |
| 23267506 | 23269150 | 1 | processed_pseudoge | ENSG00000 |
| 23328826 | 23433740 | -1 | protein_coding | ENSG00000 |
| 23373423 | 23374061 | 1 | processed_pseudoge | ENSG00000 |
| 23418971 | 23428869 | 1 | lncRNA | ENSG00000 |
| 23465776 | 23487712 | 1 | lncRNA | ENSG00000 |
| 23498796 | 23502874 | 1 | lncRNA | ENSG00000 |
| 23570370 | 23676104 | 1 | protein_coding | ENSG00000 |
| 23730189 | 23889400 | -1 | protein_coding | ENSG00000 |
| 23765934 | 23766104 | -1 | processed_pseudoge | ENSG00000 |
| 23888842 | 23897263 | 1 | lncRNA | ENSG00000 |
| 23890525 | 23902655 | -1 | protein_coding | ENSG00000 |
| 23939557 | 23946782 | -1 | transcribed_unproce | ENSG00000 |
| 23954493 | 23970188 | 1 | lncRNA | ENSG00000 |
| 23979700 | 24321031 | 1 | protein_coding | ENSG00000 |
| 23979805 | 24307074 | 1 | protein_coding | ENSG00000 |
| 23979810 | 24035027 | 1 | lncRNA | ENSG00000 |
| 24122065 | 24123814 | -1 | processed_pseudoge | ENSG00000 |
| 24162416 | 24162504 | 1 | miRNA | ENSG00000 |
| 24252749 | 24254439 | -1 | lncRNA | ENSG00000 |
| 24307166 | 24322535 | 1 | protein_coding | ENSG00000 |
| 24315725 | 24321598 | -1 | lncRNA | ENSG00000 |
| 24323021 | 24323457 | 1 | processed_pseudoge | ENSG00000 |
| 24328211 | 24329072 | 1 | processed_pseudoge | ENSG00000 |
| 24331450 | 24337121 | 1 | lncRNA | ENSG00000 |
| 24400703 | 24401029 | 1 | processed_pseudoge | ENSG00000 |

| | | | | |
|---|---|---|---|---|
| 24408157 | 24597676 | 1 | transcribed_unproce | ENSG00000 |
| 24420931 | 24512778 | -1 | protein_coding | ENSG00000 |
| 24517794 | 24530461 | 1 | lncRNA | ENSG00000 |
| 24539139 | 24541720 | -1 | lncRNA | ENSG00000 |
| 24566843 | 24570728 | 1 | transcribed_unproce | ENSG00000 |
| 24566873 | 24597676 | 1 | lncRNA | ENSG00000 |
| 24611966 | 24613983 | 1 | unitary_pseudogene | ENSG00000 |
| 24680411 | 24712493 | 1 | protein_coding | ENSG00000 |
| 24703556 | 24703665 | -1 | misc_RNA | ENSG00000 |
| 24704338 | 24704788 | -1 | processed_pseudoge | ENSG00000 |
| 24723101 | 24738690 | -1 | lncRNA | ENSG00000 |
| 24742677 | 24746184 | 1 | unprocessed_pseudo | ENSG00000 |
| 24747658 | 24755201 | 1 | lncRNA | ENSG00000 |
| 24750657 | 24752171 | -1 | processed_pseudoge | ENSG00000 |
| 24764169 | 24879921 | 1 | protein_coding | ENSG00000 |
| 24882279 | 24922889 | -1 | protein_coding | ENSG00000 |
| 24924677 | 24968487 | -1 | transcribed_unproce | ENSG00000 |
| 24933006 | 24936796 | 1 | unprocessed_pseudo | ENSG00000 |
| 24943575 | 24944498 | 1 | unprocessed_pseudo | ENSG00000 |
| 24988577 | 24988925 | -1 | processed_pseudoge | ENSG00000 |
| 25017404 | 25017536 | 1 | processed_pseudoge | ENSG00000 |
| 25017766 | 25017903 | -1 | unprocessed_pseudo | ENSG00000 |
| 25095868 | 25099254 | 1 | protein_coding | ENSG00000 |
| 25141536 | 25141809 | 1 | TEC | ENSG00000 |
| 25161684 | 25172288 | -1 | protein_coding | ENSG00000 |
| 25168969 | 25190643 | 1 | lncRNA | ENSG00000 |
| 25172828 | 25180079 | -1 | lncRNA | ENSG00000 |
| 25190063 | 25190317 | -1 | TEC | ENSG00000 |
| 25193107 | 25210295 | -1 | lncRNA | ENSG00000 |
| 25206049 | 25206506 | 1 | processed_pseudoge | ENSG00000 |
| 25246201 | 25288009 | -1 | protein_coding | ENSG00000 |
| 25300124 | 25301438 | -1 | lncRNA | ENSG00000 |
| 25301556 | 25349800 | 1 | protein_coding | ENSG00000 |
| 25365799 | 25366155 | -1 | processed_pseudoge | ENSG00000 |
| 25371974 | 26025851 | 1 | protein_coding | ENSG00000 |
| 25517119 | 25517225 | 1 | snRNA | ENSG00000 |
| 25863250 | 25863667 | -1 | processed_pseudoge | ENSG00000 |
| 25944478 | 25944766 | 1 | misc_RNA | ENSG00000 |
| 25950704 | 25950821 | -1 | misc_RNA | ENSG00000 |
| 26044597 | 26051031 | -1 | protein_coding | ENSG00000 |
| 26052276 | 26053385 | -1 | lncRNA | ENSG00000 |
| 26094724 | 26154175 | -1 | unprocessed_pseudo | ENSG00000 |
| 26117548 | 26118964 | -1 | processed_pseudoge | ENSG00000 |
| 26132115 | 26222314 | -1 | protein_coding | ENSG00000 |
| 26222191 | 26222654 | 1 | lncRNA | ENSG00000 |
| 26229332 | 26229644 | -1 | processed_pseudoge | ENSG00000 |
| 26234213 | 26237129 | 1 | processed_pseudoge | ENSG00000 |
| 26254104 | 26405238 | 1 | protein_coding | ENSG00000 |
| 26359529 | 26359702 | -1 | misc_RNA | ENSG00000 |
| 26488952 | 26496120 | 1 | lncRNA | ENSG00000 |

| | | | | |
|---|---|---|---|---|
| 26557683 | 26688948 | 1 | protein_coding | ENSG00000 |
| 26576886 | 26577675 | 1 | processed_pseudoge | ENSG00000 |
| 26606544 | 26641793 | -1 | lncRNA | ENSG00000 |
| 26685640 | 26698804 | -1 | lncRNA | ENSG00000 |
| 26755200 | 26760786 | -1 | protein_coding | ENSG00000 |
| 26905362 | 26905861 | 1 | processed_pseudoge | ENSG00000 |
| 26965967 | 26991996 | -1 | lncRNA | ENSG00000 |
| 26977543 | 26977789 | 1 | processed_pseudoge | ENSG00000 |
| 27017554 | 27017909 | -1 | processed_pseudoge | ENSG00000 |
| 27066156 | 27171811 | -1 | protein_coding | ENSG00000 |
| 27077218 | 27077962 | 1 | unprocessed_pseudo | ENSG00000 |
| 27162855 | 27169135 | 1 | lncRNA | ENSG00000 |
| 27172259 | 27182998 | 1 | lncRNA | ENSG00000 |
| 27178263 | 27251288 | -1 | lncRNA | ENSG00000 |
| 27236282 | 27250711 | -1 | lncRNA | ENSG00000 |
| 27251309 | 27256691 | 1 | protein_coding | ENSG00000 |
| 27255064 | 27255135 | 1 | snoRNA | ENSG00000 |
| 27255401 | 27255526 | 1 | snoRNA | ENSG00000 |
| 27270327 | 27273690 | 1 | protein_coding | ENSG00000 |
| 27343687 | 27343746 | -1 | snRNA | ENSG00000 |
| 27373348 | 27374366 | 1 | lncRNA | ENSG00000 |
| 27402476 | 27402587 | -1 | misc_RNA | ENSG00000 |
| 27424619 | 27435823 | 1 | protein_coding | ENSG00000 |
| 27435643 | 27450591 | -1 | protein_coding | ENSG00000 |
| 27488138 | 27488232 | 1 | snRNA | ENSG00000 |
| 27545913 | 27620529 | -1 | protein_coding | ENSG00000 |
| 27620742 | 27744237 | 1 | protein_coding | ENSG00000 |
| 27667395 | 27680581 | -1 | lncRNA | ENSG00000 |
| 27696393 | 27697204 | -1 | processed_pseudoge | ENSG00000 |
| 27792483 | 27794768 | 1 | protein_coding | ENSG00000 |
| 27819376 | 27917298 | -1 | lncRNA | ENSG00000 |
| 27828763 | 27828869 | 1 | snRNA | ENSG00000 |
| 27920000 | 27926313 | 1 | protein_coding | ENSG00000 |
| 27945260 | 27945415 | 1 | processed_pseudoge | ENSG00000 |
| 27953526 | 27955370 | 1 | lncRNA | ENSG00000 |
| 27960918 | 27969315 | -1 | protein_coding | ENSG00000 |
| 27977717 | 27988693 | -1 | protein_coding | ENSG00000 |
| 27989296 | 27989584 | -1 | misc_RNA | ENSG00000 |
| 28003274 | 28100592 | -1 | protein_coding | ENSG00000 |
| 28067587 | 28068334 | 1 | processed_pseudoge | ENSG00000 |
| 28107431 | 28107877 | 1 | processed_pseudoge | ENSG00000 |
| 28136843 | 28138193 | -1 | lncRNA | ENSG00000 |
| 28138506 | 28295335 | 1 | protein_coding | ENSG00000 |
| 28214513 | 28215897 | 1 | processed_pseudoge | ENSG00000 |
| 28232501 | 28232607 | 1 | snRNA | ENSG00000 |
| 28300346 | 28495145 | -1 | protein_coding | ENSG00000 |
| 28495633 | 28497267 | 1 | TEC | ENSG00000 |
| 28598833 | 28600052 | 1 | processed_pseudoge | ENSG00000 |
| 28629794 | 28629883 | 1 | misc_RNA | ENSG00000 |
| 28659104 | 28678959 | 1 | protein_coding | ENSG00000 |

| | | | | |
|---|---|---|---|---|
| 28700064 | 28718970 | -1 | protein_coding | ENSG00000( |
| 28703702 | 28703807 | -1 | snRNA | ENSG00000( |
| 28722386 | 28724021 | -1 | processed_pseudoge | ENSG00000( |
| 28778224 | 28778448 | -1 | unprocessed_pseudc | ENSG00000( |
| 28778558 | 28778937 | 1 | processed_pseudoge | ENSG00000( |
| 28820348 | 29505947 | 1 | protein_coding | ENSG00000( |
| 29239379 | 29250554 | -1 | lncRNA | ENSG00000( |
| 29306778 | 29307785 | 1 | processed_pseudoge | ENSG00000( |
| 29339566 | 29339869 | -1 | processed_pseudoge | ENSG00000( |
| 29476515 | 29490105 | -1 | lncRNA | ENSG00000( |
| 29509414 | 29595688 | -1 | protein_coding | ENSG00000( |
| 29764371 | 29850617 | -1 | protein_coding | ENSG00000( |
| 29864372 | 29864629 | -1 | misc_RNA | ENSG00000( |
| 29865959 | 29866169 | -1 | processed_pseudoge | ENSG00000( |
| 29872613 | 29888405 | -1 | lncRNA | ENSG00000( |
| 29918647 | 29926651 | -1 | lncRNA | ENSG00000( |
| 29935771 | 29950488 | 1 | lncRNA | ENSG00000( |
| 30022021 | 30023800 | -1 | lncRNA | ENSG00000( |
| 30103178 | 30108895 | -1 | lncRNA | ENSG00000( |
| 30151886 | 30159621 | -1 | lncRNA | ENSG00000( |
| 30153744 | 30160592 | 1 | lncRNA | ENSG00000( |
| 30196443 | 30196881 | -1 | TEC | ENSG00000( |
| 30202630 | 30307551 | -1 | protein_coding | ENSG00000( |
| 30281765 | 30281869 | 1 | snRNA | ENSG00000( |
| 30296147 | 30296735 | 1 | processed_pseudoge | ENSG00000( |
| 30316360 | 30319903 | -1 | lncRNA | ENSG00000( |
| 30340267 | 30377145 | -1 | lncRNA | ENSG00000( |
| 30344672 | 30347167 | 1 | TEC | ENSG00000( |
| 30388785 | 30404967 | -1 | lncRNA | ENSG00000( |
| 30419519 | 30422237 | -1 | lncRNA | ENSG00000( |
| 30422488 | 30429758 | 1 | protein_coding | ENSG00000( |
| 30456704 | 30617597 | -1 | protein_coding | ENSG00000( |
| 30466117 | 30482312 | 1 | lncRNA | ENSG00000( |
| 30539597 | 30543461 | -1 | processed_pseudoge | ENSG00000( |
| 30541643 | 30542897 | -1 | processed_pseudoge | ENSG00000( |
| 30553670 | 30554693 | 1 | processed_pseudoge | ENSG00000( |
| 30617693 | 30660770 | 1 | protein_coding | ENSG00000( |
| 30684029 | 30684278 | -1 | misc_RNA | ENSG00000( |
| 30713478 | 30764426 | 1 | protein_coding | ENSG00000( |
| 30803206 | 30810645 | 1 | lncRNA | ENSG00000( |
| 30880912 | 30883395 | 1 | lncRNA | ENSG00000( |
| 30881933 | 30933846 | -1 | lncRNA | ENSG00000( |
| 30906271 | 30925572 | 1 | protein_coding | ENSG00000( |
| 30932656 | 30975500 | 1 | protein_coding | ENSG00000( |
| 30973289 | 30977623 | -1 | lncRNA | ENSG00000( |
| 30995162 | 30996138 | -1 | lncRNA | ENSG00000( |
| 31049715 | 31053250 | -1 | lncRNA | ENSG00000( |
| 31063306 | 31115699 | 1 | unitary_pseudogene | ENSG00000( |
| 31134973 | 31162388 | -1 | protein_coding | ENSG00000( |
| 31199975 | 31332276 | 1 | protein_coding | ENSG00000( |

| | | | | |
|---|---|---|---|---|
| 31311289 | 31312921 | 1 | processed_pseudoge | ENSG00000 |
| 31419989 | 31437999 | 1 | lncRNA | ENSG00000 |
| 31483414 | 31483699 | -1 | unprocessed_pseudo | ENSG00000 |
| 31739526 | 31803389 | 1 | protein_coding | ENSG00000 |
| 31838752 | 31846539 | -1 | lncRNA | ENSG00000 |
| 31846713 | 32299125 | 1 | protein_coding | ENSG00000 |
| 31952580 | 31953361 | 1 | transcribed_unproce | ENSG00000 |
| 31960325 | 31961946 | -1 | TEC | ENSG00000 |
| 31975167 | 31975448 | 1 | misc_RNA | ENSG00000 |
| 32025314 | 32031639 | -1 | lncRNA | ENSG00000 |
| 32303700 | 32315344 | -1 | protein_coding | ENSG00000 |
| 32315086 | 32400266 | 1 | protein_coding | ENSG00000 |
| 32384660 | 32386108 | 1 | processed_pseudoge | ENSG00000 |
| 32400723 | 32428311 | -1 | protein_coding | ENSG00000 |
| 32420390 | 32420516 | -1 | snoRNA | ENSG00000 |
| 32425358 | 32432954 | -1 | lncRNA | ENSG00000 |
| 32432417 | 32538885 | -1 | protein_coding | ENSG00000 |
| 32477274 | 32478841 | 1 | processed_pseudoge | ENSG00000 |
| 32504506 | 32509395 | -1 | lncRNA | ENSG00000 |
| 32586452 | 32778019 | 1 | protein_coding | ENSG00000 |
| 32763438 | 32763552 | -1 | misc_RNA | ENSG00000 |
| 32782874 | 32788178 | 1 | lncRNA | ENSG00000 |
| 32877431 | 32911650 | -1 | lncRNA | ENSG00000 |
| 32954057 | 32954401 | -1 | processed_pseudoge | ENSG00000 |
| 33016423 | 33066143 | 1 | protein_coding | ENSG00000 |
| 33103137 | 33350630 | -1 | protein_coding | ENSG00000 |
| 33158587 | 33164409 | -1 | lncRNA | ENSG00000 |
| 33180401 | 33281584 | 1 | lncRNA | ENSG00000 |
| 33285866 | 33290561 | 1 | lncRNA | ENSG00000 |
| 33333679 | 33335277 | -1 | lncRNA | ENSG00000 |
| 33335497 | 33349619 | 1 | lncRNA | ENSG00000 |
| 33336216 | 33336716 | -1 | lncRNA | ENSG00000 |
| 33355206 | 33676796 | -1 | lncRNA | ENSG00000 |
| 33610967 | 33611522 | 1 | lncRNA | ENSG00000 |
| 33657436 | 33659928 | 1 | lncRNA | ENSG00000 |
| 33800138 | 33802853 | 1 | lncRNA | ENSG00000 |
| 33818069 | 33966558 | 1 | protein_coding | ENSG00000 |
| 33829539 | 33829649 | -1 | snRNA | ENSG00000 |
| 33846190 | 33850825 | 1 | lncRNA | ENSG00000 |
| 34079411 | 34080134 | 1 | TEC | ENSG00000 |
| 34082429 | 34083310 | -1 | processed_pseudoge | ENSG00000 |
| 34100711 | 34100838 | -1 | snoRNA | ENSG00000 |
| 34262863 | 34263090 | 1 | unprocessed_pseudo | ENSG00000 |
| 34347986 | 34616216 | 1 | lncRNA | ENSG00000 |
| 34435450 | 34640803 | -1 | lncRNA | ENSG00000 |
| 34545929 | 34696612 | -1 | lncRNA | ENSG00000 |
| 34574204 | 34574666 | -1 | processed_pseudoge | ENSG00000 |
| 34910545 | 34910815 | -1 | misc_RNA | ENSG00000 |
| 34925834 | 34928076 | -1 | lncRNA | ENSG00000 |
| 34942287 | 35673022 | 1 | protein_coding | ENSG00000 |

| | | | | |
|---|---|---|---|---|
| 35185294 | 35186679 | 1 | processed_pseudoge | ENSG00000( |
| 35251070 | 35251829 | 1 | processed_pseudoge | ENSG00000( |
| 35473789 | 35476689 | -1 | protein_coding | ENSG00000( |
| 35494178 | 35496615 | -1 | lncRNA | ENSG00000( |
| 35697487 | 35699938 | 1 | lncRNA | ENSG00000( |
| 35768652 | 36131306 | -1 | protein_coding | ENSG00000( |
| 36168217 | 36214588 | -1 | protein_coding | ENSG00000( |
| 36168794 | 36297842 | -1 | protein_coding | ENSG00000( |
| 36222008 | 36297840 | -1 | protein_coding | ENSG00000( |
| 36267168 | 36267274 | 1 | snRNA | ENSG00000( |
| 36301638 | 36370180 | -1 | protein_coding | ENSG00000( |
| 36346431 | 36369601 | 1 | lncRNA | ENSG00000( |
| 36431520 | 36442870 | 1 | protein_coding | ENSG00000( |
| 36490078 | 36490441 | -1 | processed_pseudoge | ENSG00000( |
| 36674020 | 36697839 | 1 | protein_coding | ENSG00000( |
| 36740041 | 36740863 | 1 | processed_pseudoge | ENSG00000( |
| 36741878 | 36742465 | -1 | processed_pseudoge | ENSG00000( |
| 36789332 | 36795050 | 1 | transcribed_processe | ENSG00000( |
| 36804198 | 36805302 | -1 | processed_pseudoge | ENSG00000( |
| 36819224 | 36829104 | 1 | protein_coding | ENSG00000( |
| 36844831 | 36920765 | -1 | protein_coding | ENSG00000( |
| 36849366 | 36850046 | -1 | lncRNA | ENSG00000( |
| 36922539 | 36922908 | 1 | processed_pseudoge | ENSG00000( |
| 36926438 | 36927219 | -1 | processed_pseudoge | ENSG00000( |
| 36938400 | 36939494 | -1 | processed_pseudoge | ENSG00000( |
| 36949738 | 37000261 | -1 | protein_coding | ENSG00000( |
| 36998816 | 37009614 | 1 | protein_coding | ENSG00000( |
| 37009312 | 37059713 | -1 | protein_coding | ENSG00000( |
| 37103259 | 37105664 | -1 | protein_coding | ENSG00000( |
| 37139176 | 37139585 | -1 | unprocessed_pseudo | ENSG00000( |
| 37142967 | 37146718 | -1 | TEC | ENSG00000( |
| 37166352 | 37166658 | -1 | misc_RNA | ENSG00000( |
| 37220860 | 37221246 | -1 | processed_pseudoge | ENSG00000( |
| 37481467 | 37484769 | -1 | lncRNA | ENSG00000( |
| 37534940 | 37551536 | 1 | lncRNA | ENSG00000( |
| 37562583 | 37598844 | -1 | protein_coding | ENSG00000( |
| 37636636 | 37870425 | -1 | protein_coding | ENSG00000( |
| 37854852 | 37854970 | -1 | rRNA_pseudogene | ENSG00000( |
| 37869825 | 37874444 | 1 | lncRNA | ENSG00000( |
| 37934298 | 38066444 | 1 | lncRNA | ENSG00000( |
| 37993085 | 37994720 | -1 | processed_pseudoge | ENSG00000( |
| 38050817 | 38350259 | -1 | lncRNA | ENSG00000( |
| 38349849 | 38363619 | 1 | protein_coding | ENSG00000( |
| 38372856 | 38373935 | 1 | TEC | ENSG00000( |
| 38533343 | 38545299 | -1 | lncRNA | ENSG00000( |
| 38567715 | 38622671 | 1 | lncRNA | ENSG00000( |
| 38617419 | 38617940 | -1 | processed_pseudoge | ENSG00000( |
| 38687077 | 38887131 | 1 | protein_coding | ENSG00000( |
| 38779424 | 38779516 | 1 | snRNA | ENSG00000( |
| 38821798 | 38827570 | -1 | lncRNA | ENSG00000( |

| | | | | |
|---|---|---|---|---|
| 38902387 | 38902650 | 1 | processed_pseudoge | ENSG00000 |
| 38922727 | 38936199 | -1 | unprocessed_pseudc | ENSG00000 |
| 38965925 | 38990859 | -1 | protein_coding | ENSG00000 |
| 39009865 | 39038089 | -1 | protein_coding | ENSG00000 |
| 39038306 | 39050109 | 1 | protein_coding | ENSG00000 |
| 39053019 | 39223238 | 1 | lncRNA | ENSG00000 |
| 39169636 | 39169951 | -1 | processed_pseudoge | ENSG00000 |
| 39209116 | 39603528 | -1 | protein_coding | ENSG00000 |
| 39655627 | 39791665 | 1 | protein_coding | ENSG00000 |
| 39664034 | 39664135 | -1 | miRNA | ENSG00000 |
| 39785415 | 39785510 | -1 | misc_RNA | ENSG00000 |
| 39817741 | 39818049 | -1 | processed_pseudoge | ENSG00000 |
| 39856710 | 39857094 | -1 | processed_pseudoge | ENSG00000 |
| 39857135 | 39857227 | -1 | snoRNA | ENSG00000 |
| 40079106 | 40535807 | -1 | lncRNA | ENSG00000 |
| 40172505 | 40195361 | 1 | lncRNA | ENSG00000 |
| 40224625 | 40224862 | -1 | processed_pseudoge | ENSG00000 |
| 40226827 | 40226932 | 1 | misc_RNA | ENSG00000 |
| 40407122 | 40407440 | -1 | misc_RNA | ENSG00000 |
| 40469953 | 40666641 | -1 | protein_coding | ENSG00000 |
| 40618738 | 40621348 | 1 | processed_pseudoge | ENSG00000 |
| 40727828 | 40727875 | -1 | miRNA | ENSG00000 |
| 40729128 | 40771190 | -1 | protein_coding | ENSG00000 |
| 40730981 | 40731253 | 1 | misc_RNA | ENSG00000 |
| 40789412 | 40812460 | 1 | protein_coding | ENSG00000 |
| 40810766 | 40810861 | 1 | miRNA | ENSG00000 |
| 40812309 | 40921749 | -1 | transcribed_unproce | ENSG00000 |
| 40863599 | 40863902 | -1 | processed_pseudoge | ENSG00000 |
| 40874685 | 40877861 | -1 | lncRNA | ENSG00000 |
| 40882577 | 40921774 | -1 | transcribed_unproce | ENSG00000 |
| 40931924 | 41061440 | -1 | protein_coding | ENSG00000 |
| 40992779 | 40993331 | -1 | processed_pseudoge | ENSG00000 |
| 41061509 | 41084006 | 1 | protein_coding | ENSG00000 |
| 41101019 | 41101100 | -1 | miRNA | ENSG00000 |
| 41121876 | 41122171 | 1 | misc_RNA | ENSG00000 |
| 41127569 | 41132802 | -1 | protein_coding | ENSG00000 |
| 41132939 | 41236686 | 1 | lncRNA | ENSG00000 |
| 41170700 | 41171148 | -1 | processed_pseudoge | ENSG00000 |
| 41189834 | 41194569 | -1 | protein_coding | ENSG00000 |
| 41199832 | 41200807 | -1 | processed_pseudoge | ENSG00000 |
| 41216369 | 41263577 | -1 | protein_coding | ENSG00000 |
| 41229180 | 41229676 | -1 | lncRNA | ENSG00000 |
| 41287741 | 41288320 | -1 | processed_pseudoge | ENSG00000 |
| 41311267 | 41377030 | 1 | protein_coding | ENSG00000 |
| 41354557 | 41354659 | -1 | snRNA | ENSG00000 |
| 41384773 | 41385183 | -1 | processed_pseudoge | ENSG00000 |
| 41431248 | 41432201 | -1 | unprocessed_pseudc | ENSG00000 |
| 41439834 | 41440847 | -1 | unprocessed_pseudc | ENSG00000 |
| 41442770 | 41443666 | 1 | unprocessed_pseudc | ENSG00000 |
| 41457550 | 41470871 | 1 | protein_coding | ENSG00000 |

| | | | | |
|---|---|---|---|---|
| 41566835 | 41961120 | -1 | protein_coding | ENSG0000C |
| 41568286 | 41568395 | -1 | miRNA | ENSG0000C |
| 41807677 | 41807973 | 1 | misc_RNA | ENSG0000C |
| 41905118 | 41905224 | 1 | snRNA | ENSG0000C |
| 41925598 | 41927349 | 1 | processed_pseudoge | ENSG0000C |
| 41955808 | 41981565 | 1 | lncRNA | ENSG0000C |
| 42003430 | 42003637 | 1 | processed_pseudoge | ENSG0000C |
| 42040036 | 42256578 | 1 | protein_coding | ENSG0000C |
| 42043727 | 42044247 | -1 | lncRNA | ENSG0000C |
| 42068932 | 42071093 | -1 | processed_pseudoge | ENSG0000C |
| 42181465 | 42181921 | -1 | processed_pseudoge | ENSG0000C |
| 42255873 | 42256575 | 1 | processed_pseudoge | ENSG0000C |
| 42272152 | 42323261 | 1 | protein_coding | ENSG0000C |
| 42339188 | 42485956 | 1 | lncRNA | ENSG0000C |
| 42369259 | 42369657 | -1 | processed_pseudoge | ENSG0000C |
| 42562736 | 42608013 | 1 | protein_coding | ENSG0000C |
| 42781550 | 42791549 | 1 | protein_coding | ENSG0000C |
| 42810366 | 42814897 | -1 | lncRNA | ENSG0000C |
| 42842414 | 42842887 | -1 | lncRNA | ENSG0000C |
| 42886388 | 42992271 | -1 | protein_coding | ENSG0000C |
| 42933858 | 42934892 | 1 | processed_pseudoge | ENSG0000C |
| 43023203 | 43114213 | 1 | protein_coding | ENSG0000C |
| 43158631 | 43159466 | 1 | lncRNA | ENSG0000C |
| 43179501 | 43181193 | -1 | processed_pseudoge | ENSG0000C |
| 43213518 | 43786908 | -1 | protein_coding | ENSG0000C |
| 43394525 | 43394825 | 1 | processed_pseudoge | ENSG0000C |
| 43458465 | 43459484 | 1 | lncRNA | ENSG0000C |
| 43543918 | 43548056 | 1 | lncRNA | ENSG0000C |
| 43607313 | 43629464 | -1 | lncRNA | ENSG0000C |
| 43786781 | 43814798 | 1 | lncRNA | ENSG0000C |
| 43787591 | 43787852 | 1 | lncRNA | ENSG0000C |
| 43823909 | 43879740 | -1 | protein_coding | ENSG0000C |
| 43877715 | 43878163 | -1 | lncRNA | ENSG0000C |
| 43879284 | 43893932 | 1 | protein_coding | ENSG0000C |
| 43883800 | 44028526 | 1 | protein_coding | ENSG0000C |
| 43894725 | 43895182 | -1 | TEC | ENSG0000C |
| 43908669 | 44030461 | 1 | lncRNA | ENSG0000C |
| 43968424 | 43971199 | 1 | processed_pseudoge | ENSG0000C |
| 44094822 | 44147762 | -1 | lncRNA | ENSG0000C |
| 44110451 | 44240517 | 1 | lncRNA | ENSG0000C |
| 44143150 | 44161257 | -1 | protein_coding | ENSG0000C |
| 44146470 | 44158222 | -1 | lncRNA | ENSG0000C |
| 44196006 | 44197298 | 1 | lncRNA | ENSG0000C |
| 44196129 | 44196200 | -1 | miRNA | ENSG0000C |
| 44234118 | 44256596 | -1 | lncRNA | ENSG0000C |
| 44369365 | 44369877 | -1 | lncRNA | ENSG0000C |
| 44373665 | 44397714 | 1 | protein_coding | ENSG0000C |
| 44400250 | 44405984 | -1 | lncRNA | ENSG0000C |
| 44432143 | 44577147 | -1 | protein_coding | ENSG0000C |
| 44432144 | 44432678 | 1 | TEC | ENSG0000C |

| | | | | |
|---|---|---|---|---|
| 44437057 | 44464921 | 1 | transcribed_processe | ENSG0000C |
| 44575893 | 44580432 | 1 | lncRNA | ENSG0000C |
| 44594858 | 44596082 | 1 | processed_pseudoge | ENSG0000C |
| 44651812 | 44701016 | 1 | lncRNA | ENSG0000C |
| 44684487 | 44715393 | -1 | lncRNA | ENSG0000C |
| 44799503 | 44809630 | -1 | lncRNA | ENSG0000C |
| 44939249 | 44989471 | -1 | protein_coding | ENSG0000C |
| 44982077 | 45083125 | -1 | lncRNA | ENSG0000C |
| 44989529 | 45037669 | 1 | protein_coding | ENSG0000C |
| 45002150 | 45002447 | 1 | misc_RNA | ENSG0000C |
| 45060729 | 45061086 | 1 | misc_RNA | ENSG0000C |
| 45120510 | 45284893 | 1 | protein_coding | ENSG0000C |
| 45123093 | 45123414 | 1 | misc_RNA | ENSG0000C |
| 45124015 | 45124143 | -1 | unprocessed_pseudc | ENSG0000C |
| 45192853 | 45194717 | -1 | protein_coding | ENSG0000C |
| 45263690 | 45263915 | 1 | processed_pseudoge | ENSG0000C |
| 45333471 | 45341370 | -1 | protein_coding | ENSG0000C |
| 45336314 | 45336447 | -1 | snoRNA | ENSG0000C |
| 45337480 | 45337609 | -1 | snoRNA | ENSG0000C |
| 45340039 | 45341183 | 1 | lncRNA | ENSG0000C |
| 45341345 | 45417975 | 1 | lncRNA | ENSG0000C |
| 45350323 | 45351350 | 1 | lncRNA | ENSG0000C |
| 45390353 | 45391267 | -1 | processed_pseudoge | ENSG0000C |
| 45393316 | 45418455 | -1 | protein_coding | ENSG0000C |
| 45418162 | 45420371 | 1 | lncRNA | ENSG0000C |
| 45436121 | 45436660 | 1 | processed_pseudoge | ENSG0000C |
| 45464898 | 45536701 | 1 | protein_coding | ENSG0000C |
| 45534522 | 45615739 | -1 | protein_coding | ENSG0000C |
| 45551485 | 45551603 | -1 | rRNA_pseudogene | ENSG0000C |
| 45552863 | 45553136 | -1 | processed_pseudoge | ENSG0000C |
| 45671845 | 45672391 | 1 | processed_pseudoge | ENSG0000C |
| 45680184 | 45701184 | -1 | lncRNA | ENSG0000C |
| 45687548 | 45688469 | -1 | processed_pseudoge | ENSG0000C |
| 45702320 | 45714559 | 1 | protein_coding | ENSG0000C |
| 45732082 | 45732356 | -1 | misc_RNA | ENSG0000C |
| 45777242 | 45851753 | -1 | protein_coding | ENSG0000C |
| 45954465 | 46052759 | -1 | protein_coding | ENSG0000C |
| 46052497 | 46161379 | 1 | lncRNA | ENSG0000C |
| 46053186 | 46105033 | -1 | protein_coding | ENSG0000C |
| 46125920 | 46211871 | -1 | protein_coding | ENSG0000C |
| 46169458 | 46169748 | -1 | misc_RNA | ENSG0000C |
| 46211943 | 46277366 | 1 | protein_coding | ENSG0000C |
| 46249942 | 46250213 | 1 | misc_RNA | ENSG0000C |
| 46270077 | 46270617 | 1 | processed_pseudoge | ENSG0000C |
| 46290300 | 46298110 | 1 | lncRNA | ENSG0000C |
| 46296445 | 46297844 | -1 | lncRNA | ENSG0000C |
| 46342000 | 46438190 | -1 | protein_coding | ENSG0000C |
| 46374401 | 46374590 | -1 | snRNA | ENSG0000C |
| 46410454 | 46410557 | 1 | snRNA | ENSG0000C |
| 46437394 | 46437988 | -1 | processed_pseudoge | ENSG0000C |

| | | | | |
|---|---|---|---|---|
| 46455131 | 46515958 | -1 | lncRNA | ENSG0000( |
| 46458061 | 46459324 | -1 | transcribed_unproce | ENSG0000( |
| 46460609 | 46461121 | 1 | unprocessed_pseudo | ENSG0000( |
| 46474246 | 46493268 | 1 | lncRNA | ENSG0000( |
| 46490930 | 46491121 | 1 | processed_pseudoge | ENSG0000( |
| 46505118 | 46505436 | 1 | processed_pseudoge | ENSG0000( |
| 46553168 | 46753040 | 1 | protein_coding | ENSG0000( |
| 46717423 | 46717688 | 1 | lncRNA | ENSG0000( |
| 46771256 | 46797161 | -1 | protein_coding | ENSG0000( |
| 46831550 | 46897076 | -1 | protein_coding | ENSG0000( |
| 46852143 | 46856299 | 1 | lncRNA | ENSG0000( |
| 47307082 | 47307301 | 1 | processed_pseudoge | ENSG0000( |
| 47458941 | 47459234 | -1 | misc_RNA | ENSG0000( |
| 47745736 | 48037968 | -1 | protein_coding | ENSG0000( |
| 47763460 | 47764606 | 1 | processed_pseudoge | ENSG0000( |
| 47766640 | 47767085 | -1 | processed_pseudoge | ENSG0000( |
| 47825330 | 47835956 | -1 | lncRNA | ENSG0000( |
| 47930153 | 47932622 | -1 | lncRNA | ENSG0000( |
| 47946053 | 47956830 | 1 | lncRNA | ENSG0000( |
| 48001389 | 48002552 | 1 | lncRNA | ENSG0000( |
| 48037726 | 48047221 | 1 | protein_coding | ENSG0000( |
| 48041751 | 48042184 | -1 | lncRNA | ENSG0000( |
| 48053323 | 48095131 | -1 | protein_coding | ENSG0000( |
| 48077137 | 48079991 | 1 | lncRNA | ENSG0000( |
| 48134358 | 48134534 | -1 | processed_pseudoge | ENSG0000( |
| 48232612 | 48270357 | 1 | protein_coding | ENSG0000( |
| 48296162 | 48303661 | -1 | lncRNA | ENSG0000( |
| 48303726 | 48599436 | 1 | protein_coding | ENSG0000( |
| 48316863 | 48320475 | -1 | processed_pseudoge | ENSG0000( |
| 48328084 | 48328564 | -1 | processed_pseudoge | ENSG0000( |
| 48340797 | 48341330 | -1 | lncRNA | ENSG0000( |
| 48389567 | 48444704 | -1 | protein_coding | ENSG0000( |
| 48429269 | 48429554 | 1 | misc_RNA | ENSG0000( |
| 48488959 | 48533256 | -1 | protein_coding | ENSG0000( |
| 48531800 | 48556358 | 1 | lncRNA | ENSG0000( |
| 48532013 | 48532599 | -1 | lncRNA | ENSG0000( |
| 48570639 | 48573317 | -1 | lncRNA | ENSG0000( |
| 48576974 | 48578088 | -1 | lncRNA | ENSG0000( |
| 48653711 | 48711226 | 1 | protein_coding | ENSG0000( |
| 48771128 | 48771827 | 1 | processed_pseudoge | ENSG0000( |
| 48974967 | 48976867 | -1 | lncRNA | ENSG0000( |
| 48975912 | 49209779 | 1 | protein_coding | ENSG0000( |
| 48987915 | 48988017 | 1 | snRNA | ENSG0000( |
| 49044079 | 49046794 | -1 | processed_pseudoge | ENSG0000( |
| 49095391 | 49095492 | 1 | misc_RNA | ENSG0000( |
| 49187479 | 49187670 | -1 | processed_pseudoge | ENSG0000( |
| 49210422 | 49212294 | -1 | processed_pseudoge | ENSG0000( |
| 49220338 | 49222377 | 1 | protein_coding | ENSG0000( |
| 49232874 | 49234399 | 1 | unprocessed_pseudo | ENSG0000( |
| 49247925 | 49293485 | 1 | protein_coding | ENSG0000( |

| | | | | |
|---|---|---|---|---|
| 49308650 | 49444064 | -1 | protein_coding | ENSG0000( |
| 49444374 | 49495003 | 1 | protein_coding | ENSG0000( |
| 49488970 | 49489364 | 1 | processed_pseudoge | ENSG0000( |
| 49495610 | 49528981 | 1 | protein_coding | ENSG0000( |
| 49531946 | 49585558 | -1 | protein_coding | ENSG0000( |
| 49620314 | 49620646 | 1 | processed_pseudoge | ENSG0000( |
| 49628507 | 49633872 | 1 | protein_coding | ENSG0000( |
| 49660674 | 49691486 | -1 | protein_coding | ENSG0000( |
| 49699320 | 49792682 | -1 | protein_coding | ENSG0000( |
| 49792886 | 49793307 | 1 | lncRNA | ENSG0000( |
| 49889634 | 49889728 | -1 | misc_RNA | ENSG0000( |
| 49890591 | 49892998 | 1 | processed_pseudoge | ENSG0000( |
| 49908634 | 49908730 | -1 | misc_RNA | ENSG0000( |
| 49912702 | 49936490 | -1 | protein_coding | ENSG0000( |
| 49956670 | 50125720 | -1 | lncRNA | ENSG0000( |
| 49995888 | 50020481 | 1 | protein_coding | ENSG0000( |
| 49996415 | 49996501 | -1 | miRNA | ENSG0000( |
| 50015254 | 50020922 | 1 | protein_coding | ENSG0000( |
| 50044649 | 50044768 | 1 | misc_RNA | ENSG0000( |
| 50045143 | 50045232 | 1 | misc_RNA | ENSG0000( |
| 50048973 | 50049061 | -1 | miRNA | ENSG0000( |
| 50049119 | 50049201 | -1 | miRNA | ENSG0000( |
| 50049591 | 50049663 | 1 | misc_RNA | ENSG0000( |
| 50068107 | 50068164 | 1 | misc_RNA | ENSG0000( |
| 50075570 | 50075653 | 1 | misc_RNA | ENSG0000( |
| 50081843 | 50081978 | 1 | misc_RNA | ENSG0000( |
| 50082169 | 50906856 | 1 | lncRNA | ENSG0000( |
| 50082295 | 50082484 | 1 | misc_RNA | ENSG0000( |
| 50099331 | 50099868 | 1 | processed_pseudoge | ENSG0000( |
| 50104833 | 50105174 | 1 | misc_RNA | ENSG0000( |
| 50125816 | 50128463 | 1 | lncRNA | ENSG0000( |
| 50172089 | 50173181 | 1 | processed_pseudoge | ENSG0000( |
| 50361410 | 50361763 | -1 | processed_pseudoge | ENSG0000( |
| 50372788 | 50387158 | -1 | lncRNA | ENSG0000( |
| 50519364 | 50843939 | -1 | protein_coding | ENSG0000( |
| 50812988 | 50813106 | -1 | rRNA_pseudogene | ENSG0000( |
| 50862172 | 50910764 | -1 | lncRNA | ENSG0000( |
| 50909678 | 51024120 | 1 | protein_coding | ENSG0000( |
| 50994511 | 51080862 | -1 | transcribed_unitary_ | ENSG0000( |
| 51027747 | 51027852 | -1 | rRNA_pseudogene | ENSG0000( |
| 51082119 | 51200252 | -1 | protein_coding | ENSG0000( |
| 51161723 | 51162357 | -1 | processed_pseudoge | ENSG0000( |
| 51180178 | 51180967 | 1 | lncRNA | ENSG0000( |
| 51195881 | 51196081 | 1 | processed_pseudoge | ENSG0000( |
| 51222334 | 51284241 | 1 | protein_coding | ENSG0000( |
| 51335773 | 51364735 | 1 | protein_coding | ENSG0000( |
| 51348567 | 51348639 | -1 | miRNA | ENSG0000( |
| 51354077 | 51454264 | -1 | protein_coding | ENSG0000( |
| 51452364 | 51554678 | 1 | lncRNA | ENSG0000( |
| 51454164 | 51461315 | 1 | transcribed_processe | ENSG0000( |

| | | | | |
|---|---|---|---|---|
| 51469753 | 51470047 | 1 | misc_RNA | ENSG00000 |
| 51496020 | 51496251 | -1 | processed_pseudoge | ENSG00000 |
| 51552589 | 51552667 | 1 | miRNA | ENSG00000 |
| 51565412 | 51565509 | 1 | snRNA | ENSG00000 |
| 51584455 | 51767709 | 1 | protein_coding | ENSG00000 |
| 51586424 | 51586532 | -1 | misc_RNA | ENSG00000 |
| 51593919 | 51594215 | -1 | misc_RNA | ENSG00000 |
| 51598416 | 51599460 | 1 | processed_pseudoge | ENSG00000 |
| 51763951 | 51845159 | -1 | protein_coding | ENSG00000 |
| 51767993 | 51804162 | -1 | protein_coding | ENSG00000 |
| 51803838 | 51813832 | 1 | lncRNA | ENSG00000 |
| 51813347 | 51845177 | -1 | protein_coding | ENSG00000 |
| 51861981 | 51866232 | 1 | protein_coding | ENSG00000 |
| 51908218 | 51910621 | -1 | processed_pseudoge | ENSG00000 |
| 51921251 | 51921587 | 1 | TEC | ENSG00000 |
| 51930436 | 52012125 | -1 | protein_coding | ENSG00000 |
| 51966742 | 51967149 | 1 | processed_pseudoge | ENSG00000 |
| 52012398 | 52033600 | 1 | protein_coding | ENSG00000 |
| 52024691 | 52033600 | 1 | protein_coding | ENSG00000 |
| 52033611 | 52129092 | -1 | protein_coding | ENSG00000 |
| 52128891 | 52132723 | 1 | lncRNA | ENSG00000 |
| 52132639 | 52159861 | -1 | protein_coding | ENSG00000 |
| 52167709 | 52291557 | -1 | lncRNA | ENSG00000 |
| 52167709 | 52194465 | -1 | transcribed_unproce | ENSG00000 |
| 52167954 | 52168321 | -1 | transcribed_unproce | ENSG00000 |
| 52169987 | 52171248 | -1 | transcribed_unproce | ENSG00000 |
| 52219344 | 52334277 | -1 | transcribed_unproce | ENSG00000 |
| 52334295 | 52341896 | 1 | lncRNA | ENSG00000 |
| 52377167 | 52416373 | -1 | protein_coding | ENSG00000 |
| 52400004 | 52400099 | 1 | misc_RNA | ENSG00000 |
| 52412602 | 52450634 | -1 | protein_coding | ENSG00000 |
| 52454775 | 52455331 | -1 | lncRNA | ENSG00000 |
| 52455429 | 52476628 | 1 | protein_coding | ENSG00000 |
| 52482804 | 52489216 | -1 | lncRNA | ENSG00000 |
| 52488993 | 52587095 | 1 | transcribed_unproce | ENSG00000 |
| 52497097 | 52500195 | -1 | TEC | ENSG00000 |
| 52600042 | 52642542 | 1 | lncRNA | ENSG00000 |
| 52600119 | 52637446 | 1 | transcribed_unproce | ENSG00000 |
| 52642425 | 52643796 | 1 | protein_coding | ENSG00000 |
| 52651305 | 52652279 | -1 | lncRNA | ENSG00000 |
| 52652709 | 52700909 | 1 | protein_coding | ENSG00000 |
| 52703264 | 52739820 | -1 | protein_coding | ENSG00000 |
| 52810050 | 52810140 | 1 | miRNA | ENSG00000 |
| 52820497 | 52821018 | 1 | processed_pseudoge | ENSG00000 |
| 52842889 | 52848641 | -1 | protein_coding | ENSG00000 |
| 53028813 | 53052057 | 1 | protein_coding | ENSG00000 |
| 53099397 | 53151914 | -1 | lncRNA | ENSG00000 |
| 53193667 | 53198540 | 1 | lncRNA | ENSG00000 |
| 53199981 | 53202753 | 1 | transcribed_processe | ENSG00000 |
| 53207831 | 53801489 | 1 | lncRNA | ENSG00000 |

| | | | | |
|---|---|---|---|---|
| 53258077 | 53258372 | 1 | misc_RNA | ENSG0000C |
| 53323784 | 53331065 | -1 | lncRNA | ENSG0000C |
| 53345211 | 53410880 | 1 | lncRNA | ENSG0000C |
| 53408882 | 53409876 | -1 | unprocessed_pseudc | ENSG0000C |
| 53815419 | 53939960 | 1 | lncRNA | ENSG0000C |
| 54115783 | 54142319 | -1 | lncRNA | ENSG0000C |
| 54116476 | 54116851 | 1 | processed_pseudoge | ENSG0000C |
| 54311972 | 54312048 | -1 | miRNA | ENSG0000C |
| 54440704 | 54441315 | -1 | processed_pseudoge | ENSG0000C |
| 54572354 | 54687547 | 1 | lncRNA | ENSG0000C |
| 54816352 | 54819238 | -1 | lncRNA | ENSG0000C |
| 54938604 | 54986720 | 1 | lncRNA | ENSG0000C |
| 55062945 | 55161490 | 1 | lncRNA | ENSG0000C |
| 55128285 | 55128444 | 1 | misc_RNA | ENSG0000C |
| 55174454 | 55174548 | 1 | miRNA | ENSG0000C |
| 55535697 | 55583524 | -1 | lncRNA | ENSG0000C |
| 55999200 | 56000448 | -1 | processed_pseudoge | ENSG0000C |
| 56051034 | 56054397 | -1 | lncRNA | ENSG0000C |
| 56214373 | 56216942 | 1 | processed_pseudoge | ENSG0000C |
| 56885284 | 56885602 | -1 | misc_RNA | ENSG0000C |
| 57140918 | 57143939 | 1 | protein_coding | ENSG0000C |
| 57140918 | 57157082 | 1 | protein_coding | ENSG0000C |
| 57147488 | 57150509 | 1 | protein_coding | ENSG0000C |
| 57160632 | 57163653 | 1 | protein_coding | ENSG0000C |
| 57167197 | 57170218 | 1 | protein_coding | ENSG0000C |
| 57173453 | 57173632 | 1 | unprocessed_pseudc | ENSG0000C |
| 57204379 | 57204799 | 1 | processed_pseudoge | ENSG0000C |
| 57314739 | 57315909 | 1 | processed_pseudoge | ENSG0000C |
| 57391171 | 57391518 | -1 | processed_pseudoge | ENSG0000C |
| 57631744 | 57729311 | 1 | protein_coding | ENSG0000C |
| 57632759 | 57633575 | -1 | lncRNA | ENSG0000C |
| 57991350 | 57991464 | 1 | rRNA_pseudogene | ENSG0000C |
| 58165827 | 58209483 | 1 | lncRNA | ENSG0000C |
| 58211697 | 58233117 | -1 | lncRNA | ENSG0000C |
| 58377513 | 58377774 | 1 | unprocessed_pseudc | ENSG0000C |
| 58527655 | 58527751 | -1 | misc_RNA | ENSG0000C |
| 58528615 | 58531026 | 1 | processed_pseudoge | ENSG0000C |
| 58751047 | 58752233 | -1 | processed_pseudoge | ENSG0000C |
| 59010332 | 59010604 | 1 | processed_pseudoge | ENSG0000C |
| 59071462 | 59071803 | -1 | processed_pseudoge | ENSG0000C |
| 59121879 | 59230636 | -1 | lncRNA | ENSG0000C |
| 59262680 | 59263391 | 1 | processed_pseudoge | ENSG0000C |
| 59480070 | 59480132 | 1 | snRNA | ENSG0000C |
| 59665583 | 60163928 | -1 | protein_coding | ENSG0000C |
| 60012734 | 60044357 | 1 | lncRNA | ENSG0000C |
| 60034465 | 60034762 | -1 | misc_RNA | ENSG0000C |
| 60144648 | 60153678 | 1 | lncRNA | ENSG0000C |
| 60187738 | 60187830 | -1 | misc_RNA | ENSG0000C |
| 60214352 | 60214946 | -1 | lncRNA | ENSG0000C |
| 60274776 | 60276006 | 1 | processed_pseudoge | ENSG0000C |

| | | | | |
|---|---|---|---|---|
| 60322591 | 60322766 | 1 | unprocessed_pseudc | ENSG0000C |
| 60396457 | 60573878 | 1 | protein_coding | ENSG0000C |
| 60613608 | 60613734 | 1 | rRNA_pseudogene | ENSG0000C |
| 60618991 | 61176758 | 1 | lncRNA | ENSG0000C |
| 60636888 | 60638097 | -1 | processed_pseudoge | ENSG0000C |
| 60730176 | 60733600 | 1 | TEC | ENSG0000C |
| 60789560 | 60789661 | 1 | misc_RNA | ENSG0000C |
| 60916981 | 60945955 | 1 | lncRNA | ENSG0000C |
| 61199798 | 61199880 | -1 | miRNA | ENSG0000C |
| 61409685 | 61415522 | -1 | protein_coding | ENSG0000C |
| 61409858 | 61427849 | -1 | protein_coding | ENSG0000C |
| 61424689 | 61427946 | -1 | lncRNA | ENSG0000C |
| 61805150 | 61805676 | 1 | processed_pseudoge | ENSG0000C |
| 61927090 | 61927272 | -1 | processed_pseudoge | ENSG0000C |
| 61997136 | 62029572 | -1 | lncRNA | ENSG0000C |
| 62212577 | 62249947 | 1 | lncRNA | ENSG0000C |
| 62321305 | 62322398 | 1 | lncRNA | ENSG0000C |
| 62323657 | 62328833 | 1 | lncRNA | ENSG0000C |
| 62327988 | 62328384 | 1 | processed_pseudoge | ENSG0000C |
| 62397645 | 62397916 | 1 | unprocessed_pseudc | ENSG0000C |
| 62406058 | 62406916 | -1 | processed_pseudoge | ENSG0000C |
| 62539351 | 62574905 | -1 | lncRNA | ENSG0000C |
| 62672285 | 62796714 | -1 | lncRNA | ENSG0000C |
| 63034668 | 63097797 | 1 | lncRNA | ENSG0000C |
| 63183096 | 63328129 | -1 | lncRNA | ENSG0000C |
| 63397011 | 63513393 | 1 | lncRNA | ENSG0000C |
| 63667681 | 63738018 | -1 | lncRNA | ENSG0000C |
| 63737511 | 63742804 | 1 | transcribed_unproce | ENSG0000C |
| 63738111 | 63751083 | 1 | lncRNA | ENSG0000C |
| 63746741 | 63751080 | 1 | protein_coding | ENSG0000C |
| 63783801 | 63783904 | 1 | snRNA | ENSG0000C |
| 63811014 | 63811608 | -1 | processed_pseudoge | ENSG0000C |
| 63814653 | 63825534 | 1 | lncRNA | ENSG0000C |
| 63828840 | 63844125 | -1 | transcribed_unproce | ENSG0000C |
| 63832361 | 63833531 | -1 | protein_coding | ENSG0000C |
| 63851197 | 64076044 | -1 | lncRNA | ENSG0000C |
| 63975451 | 63976422 | -1 | processed_pseudoge | ENSG0000C |
| 64958097 | 64959396 | -1 | processed_pseudoge | ENSG0000C |
| 65309855 | 65310953 | -1 | processed_pseudoge | ENSG0000C |
| 65787932 | 65788764 | -1 | processed_pseudoge | ENSG0000C |
| 65866070 | 65922731 | 1 | lncRNA | ENSG0000C |
| 65941073 | 65973699 | 1 | lncRNA | ENSG0000C |
| 65966330 | 65966429 | -1 | miRNA | ENSG0000C |
| 66218250 | 66218324 | 1 | miRNA | ENSG0000C |
| 66268997 | 66270458 | 1 | processed_pseudoge | ENSG0000C |
| 66302834 | 67230445 | -1 | protein_coding | ENSG0000C |
| 66303871 | 66323561 | 1 | lncRNA | ENSG0000C |
| 66630715 | 66630776 | -1 | snRNA | ENSG0000C |
| 66825169 | 66915031 | 1 | lncRNA | ENSG0000C |
| 66977432 | 66985776 | 1 | lncRNA | ENSG0000C |

| | | | | |
|---|---|---|---|---|
| 66990886 | 67002007 | 1 | lncRNA | ENSG00000 |
| 67121081 | 67156624 | 1 | lncRNA | ENSG00000 |
| 67266845 | 67267706 | -1 | processed_pseudoge | ENSG00000 |
| 67326177 | 67379994 | 1 | lncRNA | ENSG00000 |
| 67577624 | 67787804 | -1 | lncRNA | ENSG00000 |
| 67790407 | 67790794 | 1 | processed_pseudoge | ENSG00000 |
| 67831876 | 67832827 | -1 | processed_pseudoge | ENSG00000 |
| 67902208 | 67903221 | 1 | unprocessed_pseudo | ENSG00000 |
| 67910986 | 67911951 | 1 | unprocessed_pseudo | ENSG00000 |
| 68025839 | 68037131 | 1 | lncRNA | ENSG00000 |
| 68289679 | 68291280 | -1 | processed_pseudoge | ENSG00000 |
| 68301605 | 68303302 | -1 | processed_pseudoge | ENSG00000 |
| 68331338 | 68331613 | -1 | processed_pseudoge | ENSG00000 |
| 68634190 | 68634486 | 1 | processed_pseudoge | ENSG00000 |
| 68654572 | 68655050 | 1 | processed_pseudoge | ENSG00000 |
| 68693804 | 68694081 | -1 | misc_RNA | ENSG00000 |
| 68861284 | 68885325 | -1 | lncRNA | ENSG00000 |
| 68878380 | 68893573 | 1 | lncRNA | ENSG00000 |
| 68985058 | 68986064 | 1 | processed_pseudoge | ENSG00000 |
| 69101832 | 69102056 | -1 | processed_pseudoge | ENSG00000 |
| 69221866 | 69322190 | 1 | lncRNA | ENSG00000 |
| 69408216 | 69408735 | -1 | processed_pseudoge | ENSG00000 |
| 69700594 | 70108493 | -1 | protein_coding | ENSG00000 |
| 69802940 | 69803041 | 1 | misc_RNA | ENSG00000 |
| 69894101 | 69895085 | -1 | processed_pseudoge | ENSG00000 |
| 70107213 | 70149092 | 1 | lncRNA | ENSG00000 |
| 70137831 | 70139431 | -1 | lncRNA | ENSG00000 |
| 70459464 | 70459570 | 1 | snRNA | ENSG00000 |
| 70564267 | 70564437 | -1 | unprocessed_pseudo | ENSG00000 |
| 70923664 | 70924455 | 1 | processed_pseudoge | ENSG00000 |
| 71015042 | 71168417 | 1 | lncRNA | ENSG00000 |
| 71296210 | 71297235 | 1 | processed_pseudoge | ENSG00000 |
| 71437966 | 71867204 | -1 | protein_coding | ENSG00000 |
| 71675067 | 71675469 | 1 | processed_pseudoge | ENSG00000 |
| 71704181 | 71704655 | 1 | processed_pseudoge | ENSG00000 |
| 71845981 | 71846317 | 1 | processed_pseudoge | ENSG00000 |
| 71913704 | 71914178 | -1 | processed_pseudoge | ENSG00000 |
| 72453902 | 72453967 | -1 | snoRNA | ENSG00000 |
| 72460132 | 72460329 | -1 | processed_pseudoge | ENSG00000 |
| 72460519 | 72460601 | -1 | snoRNA | ENSG00000 |
| 72567213 | 72567380 | 1 | misc_RNA | ENSG00000 |
| 72586680 | 72586810 | 1 | snoRNA | ENSG00000 |
| 72703693 | 72704138 | 1 | processed_pseudoge | ENSG00000 |
| 72706017 | 72706123 | -1 | snRNA | ENSG00000 |
| 72708367 | 72727629 | -1 | protein_coding | ENSG00000 |
| 72727749 | 72756198 | 1 | protein_coding | ENSG00000 |
| 72752169 | 72782096 | -1 | protein_coding | ENSG00000 |
| 72782133 | 73016461 | 1 | protein_coding | ENSG00000 |
| 72881108 | 72881214 | -1 | snRNA | ENSG00000 |
| 73036401 | 73036848 | 1 | processed_pseudoge | ENSG00000 |

| | | | | |
|---|---|---|---|---|
| 73054976 | 73077541 | 1 | protein_coding | ENSG0000C |
| 73100575 | 73100985 | -1 | processed_pseudoge | ENSG0000C |
| 73112915 | 73113018 | -1 | snRNA | ENSG0000C |
| 73191528 | 73191667 | 1 | snRNA | ENSG0000C |
| 73227147 | 73227260 | 1 | misc_RNA | ENSG0000C |
| 73399031 | 73399205 | 1 | TEC | ENSG0000C |
| 73407768 | 73408352 | -1 | processed_pseudoge | ENSG0000C |
| 73413473 | 73661891 | -1 | lncRNA | ENSG0000C |
| 73564244 | 73588070 | 1 | lncRNA | ENSG0000C |
| 73686089 | 73995056 | -1 | protein_coding | ENSG0000C |
| 73800074 | 73802203 | 1 | TEC | ENSG0000C |
| 73844683 | 73845130 | -1 | lncRNA | ENSG0000C |
| 74231457 | 74259976 | 1 | lncRNA | ENSG0000C |
| 74288070 | 74408810 | 1 | lncRNA | ENSG0000C |
| 74289100 | 74289214 | -1 | misc_RNA | ENSG0000C |
| 74377163 | 74377951 | -1 | processed_pseudoge | ENSG0000C |
| 74412957 | 74419115 | -1 | lncRNA | ENSG0000C |
| 74419158 | 74444735 | 1 | lncRNA | ENSG0000C |
| 74434538 | 74555515 | 1 | lncRNA | ENSG0000C |
| 74552503 | 74565445 | 1 | lncRNA | ENSG0000C |
| 74828553 | 74830080 | 1 | processed_pseudoge | ENSG0000C |
| 75109587 | 75109693 | 1 | snRNA | ENSG0000C |
| 75136154 | 75136940 | 1 | processed_pseudoge | ENSG0000C |
| 75237944 | 75240296 | -1 | processed_pseudoge | ENSG0000C |
| 75250480 | 75252012 | 1 | lncRNA | ENSG0000C |
| 75283503 | 75482169 | -1 | protein_coding | ENSG0000C |
| 75375511 | 75377294 | -1 | lncRNA | ENSG0000C |
| 75525219 | 75549439 | -1 | protein_coding | ENSG0000C |
| 75549480 | 75606020 | 1 | protein_coding | ENSG0000C |
| 75549773 | 75807120 | 1 | lncRNA | ENSG0000C |
| 75604700 | 75635994 | -1 | lncRNA | ENSG0000C |
| 75620434 | 75859870 | 1 | protein_coding | ENSG0000C |
| 75631271 | 75632135 | 1 | lncRNA | ENSG0000C |
| 75688556 | 75689197 | -1 | processed_pseudoge | ENSG0000C |
| 75871038 | 75883811 | 1 | lncRNA | ENSG0000C |
| 75876886 | 75881127 | 1 | lncRNA | ENSG0000C |
| 75932395 | 75935132 | -1 | lncRNA | ENSG0000C |
| 76013023 | 76014501 | -1 | lncRNA | ENSG0000C |
| 76134931 | 76135225 | 1 | misc_RNA | ENSG0000C |
| 76552807 | 76846524 | -1 | lncRNA | ENSG0000C |
| 76664144 | 76693051 | 1 | lncRNA | ENSG0000C |
| 76833839 | 76846594 | 1 | lncRNA | ENSG0000C |
| 76880175 | 76886405 | -1 | protein_coding | ENSG0000C |
| 76887551 | 76891135 | 1 | lncRNA | ENSG0000C |
| 76928066 | 76928836 | 1 | processed_pseudoge | ENSG0000C |
| 76948511 | 76958638 | 1 | protein_coding | ENSG0000C |
| 76981175 | 76981916 | -1 | processed_pseudoge | ENSG0000C |
| 76984868 | 76987943 | 1 | processed_pseudoge | ENSG0000C |
| 76990660 | 77019143 | 1 | protein_coding | ENSG0000C |
| 76992078 | 77129589 | 1 | protein_coding | ENSG0000C |

| | | | | |
|---|---|---|---|---|
| 76992598 | 77027195 | -1 | protein_coding | ENSG0000C |
| 77026767 | 77129717 | 1 | lncRNA | ENSG0000C |
| 77027944 | 77028482 | 1 | lncRNA | ENSG0000C |
| 77044657 | 77327094 | -1 | protein_coding | ENSG0000C |
| 77080511 | 77081190 | -1 | lncRNA | ENSG0000C |
| 77417821 | 77422847 | 1 | lncRNA | ENSG0000C |
| 77535674 | 77645263 | 1 | protein_coding | ENSG0000C |
| 77569175 | 77586812 | -1 | lncRNA | ENSG0000C |
| 77599755 | 77606551 | -1 | lncRNA | ENSG0000C |
| 77623372 | 77623472 | 1 | misc_RNA | ENSG0000C |
| 77659413 | 77659664 | 1 | processed_pseudoge | ENSG0000C |
| 77661376 | 77662626 | -1 | unprocessed_pseudc | ENSG0000C |
| 77697687 | 77764242 | 1 | protein_coding | ENSG0000C |
| 77698012 | 77698116 | -1 | miRNA | ENSG0000C |
| 77779723 | 77844145 | 1 | lncRNA | ENSG0000C |
| 77895481 | 77975529 | -1 | protein_coding | ENSG0000C |
| 77919689 | 78617328 | 1 | lncRNA | ENSG0000C |
| 77939016 | 77975539 | -1 | lncRNA | ENSG0000C |
| 77947497 | 77947772 | -1 | misc_RNA | ENSG0000C |
| 77979817 | 77997881 | -1 | lncRNA | ENSG0000C |
| 77997909 | 78053604 | 1 | lncRNA | ENSG0000C |
| 78275720 | 78277514 | -1 | lncRNA | ENSG0000C |
| 78285838 | 78286288 | 1 | processed_pseudoge | ENSG0000C |
| 78355227 | 78355328 | -1 | misc_RNA | ENSG0000C |
| 78422823 | 78423192 | 1 | processed_pseudoge | ENSG0000C |
| 78488182 | 78488513 | 1 | processed_pseudoge | ENSG0000C |
| 78596129 | 78599619 | 1 | lncRNA | ENSG0000C |
| 78598362 | 78603552 | -1 | protein_coding | ENSG0000C |
| 78614289 | 78659155 | -1 | protein_coding | ENSG0000C |
| 78761175 | 78761651 | 1 | processed_pseudoge | ENSG0000C |
| 78786272 | 78840051 | -1 | lncRNA | ENSG0000C |
| 78849231 | 78851208 | 1 | processed_pseudoge | ENSG0000C |
| 78908919 | 78910542 | 1 | processed_pseudoge | ENSG0000C |
| 79090783 | 79094119 | 1 | processed_pseudoge | ENSG0000C |
| 79166119 | 79166885 | 1 | processed_pseudoge | ENSG0000C |
| 79311824 | 79406477 | -1 | protein_coding | ENSG0000C |
| 79389534 | 79389631 | -1 | rRNA_pseudogene | ENSG0000C |
| 79406293 | 79427317 | 1 | lncRNA | ENSG0000C |
| 79477364 | 79481231 | -1 | lncRNA | ENSG0000C |
| 79481124 | 79556075 | 1 | protein_coding | ENSG0000C |
| 79566678 | 79571445 | 1 | lncRNA | ENSG0000C |
| 79637433 | 79667344 | 1 | lncRNA | ENSG0000C |
| 79804418 | 79805685 | 1 | lncRNA | ENSG0000C |
| 79872586 | 79918042 | 1 | lncRNA | ENSG0000C |
| 79941531 | 80074640 | -1 | lncRNA | ENSG0000C |
| 80011077 | 80028283 | 1 | lncRNA | ENSG0000C |
| 80115733 | 80134368 | -1 | lncRNA | ENSG0000C |
| 80335976 | 80341126 | -1 | protein_coding | ENSG0000C |
| 80369426 | 80369534 | -1 | snRNA | ENSG0000C |
| 80619098 | 80620067 | -1 | processed_pseudoge | ENSG0000C |

| | | | | |
|---|---|---|---|---|
| 80654755 | 80657027 | 1 | processed_pseudoge | ENSG00000( |
| 80697177 | 81017737 | -1 | lncRNA | ENSG00000( |
| 80888348 | 80888867 | 1 | processed_pseudoge | ENSG00000( |
| 81018176 | 81044691 | 1 | lncRNA | ENSG00000( |
| 81114866 | 81115295 | -1 | processed_pseudoge | ENSG00000( |
| 81225865 | 81226983 | -1 | lncRNA | ENSG00000( |
| 81250438 | 81302407 | 1 | lncRNA | ENSG00000( |
| 81265747 | 81265837 | 1 | snRNA | ENSG00000( |
| 81272891 | 81273170 | 1 | processed_pseudoge | ENSG00000( |
| 81689911 | 81691072 | 1 | transcribed_processe | ENSG00000( |
| 82778208 | 83102335 | -1 | lncRNA | ENSG00000( |
| 82823131 | 82824136 | 1 | processed_pseudoge | ENSG00000( |
| 83145540 | 83145903 | 1 | processed_pseudoge | ENSG00000( |
| 83298071 | 83298167 | -1 | snRNA | ENSG00000( |
| 83650249 | 83654789 | -1 | lncRNA | ENSG00000( |
| 83877205 | 83882393 | -1 | protein_coding | ENSG00000( |
| 83877888 | 83895211 | 1 | lncRNA | ENSG00000( |
| 83907301 | 83907781 | -1 | processed_pseudoge | ENSG00000( |
| 83996173 | 83997597 | 1 | lncRNA | ENSG00000( |
| 84020327 | 84020762 | 1 | processed_pseudoge | ENSG00000( |
| 84063106 | 84069408 | 1 | lncRNA | ENSG00000( |
| 84086802 | 84448603 | 1 | lncRNA | ENSG00000( |
| 84520103 | 84521568 | 1 | processed_pseudoge | ENSG00000( |
| 84521725 | 84523915 | 1 | processed_pseudoge | ENSG00000( |
| 84562364 | 84563236 | 1 | lncRNA | ENSG00000( |
| 84867514 | 84902424 | -1 | lncRNA | ENSG00000( |
| 85046001 | 85050096 | -1 | lncRNA | ENSG00000( |
| 85065087 | 85078920 | -1 | lncRNA | ENSG00000( |
| 85144623 | 85148121 | -1 | lncRNA | ENSG00000( |
| 85363601 | 85544570 | 1 | lncRNA | ENSG00000( |
| 85693937 | 85783203 | -1 | lncRNA | ENSG00000( |
| 85792790 | 85806683 | -1 | protein_coding | ENSG00000( |
| 85837126 | 85837532 | -1 | unprocessed_pseudo | ENSG00000( |
| 85968999 | 85969533 | 1 | processed_pseudoge | ENSG00000( |
| 86060813 | 86170890 | 1 | lncRNA | ENSG00000( |
| 86266842 | 86266926 | -1 | unprocessed_pseudo | ENSG00000( |
| 86275493 | 86277262 | 1 | processed_pseudoge | ENSG00000( |
| 86402982 | 86462734 | -1 | lncRNA | ENSG00000( |
| 86725500 | 86726053 | 1 | processed_pseudoge | ENSG00000( |
| 86909586 | 86937108 | 1 | lncRNA | ENSG00000( |
| 87219004 | 87219269 | 1 | processed_pseudoge | ENSG00000( |
| 87379472 | 87380258 | -1 | processed_pseudoge | ENSG00000( |
| 87423493 | 87458407 | 1 | lncRNA | ENSG00000( |
| 87427200 | 87447235 | -1 | lncRNA | ENSG00000( |
| 87427214 | 87674235 | -1 | lncRNA | ENSG00000( |
| 87607907 | 87609234 | -1 | lncRNA | ENSG00000( |
| 87618665 | 87618740 | -1 | miRNA | ENSG00000( |
| 87672615 | 87696272 | 1 | protein_coding | ENSG00000( |
| 87765752 | 87765881 | -1 | processed_pseudoge | ENSG00000( |
| 87801036 | 87803808 | -1 | lncRNA | ENSG00000( |

| | | | | |
|---|---|---|---|---|
| 87807634 | 87811232 | -1 | lncRNA | ENSG0000( |
| 87890172 | 87892472 | -1 | processed_pseudoge | ENSG0000( |
| 88142867 | 88236082 | 1 | lncRNA | ENSG0000( |
| 88236547 | 88236969 | -1 | processed_pseudoge | ENSG0000( |
| 88472151 | 88477267 | -1 | lncRNA | ENSG0000( |
| 88494789 | 88607814 | 1 | lncRNA | ENSG0000( |
| 88522185 | 89163186 | -1 | lncRNA | ENSG0000( |
| 88571654 | 88572312 | 1 | processed_pseudoge | ENSG0000( |
| 88626853 | 88630397 | 1 | lncRNA | ENSG0000( |
| 88978447 | 88978785 | -1 | processed_pseudoge | ENSG0000( |
| 89181324 | 89280240 | 1 | lncRNA | ENSG0000( |
| 89214847 | 89236890 | -1 | lncRNA | ENSG0000( |
| 89248818 | 89250100 | -1 | processed_pseudoge | ENSG0000( |
| 89361160 | 89363224 | -1 | processed_pseudoge | ENSG0000( |
| 89402398 | 89818453 | 1 | lncRNA | ENSG0000( |
| 89477608 | 89500509 | -1 | lncRNA | ENSG0000( |
| 89548691 | 89553804 | -1 | lncRNA | ENSG0000( |
| 89833157 | 89833941 | -1 | processed_pseudoge | ENSG0000( |
| 89837170 | 89837237 | 1 | snoRNA | ENSG0000( |
| 89991831 | 89992878 | 1 | processed_pseudoge | ENSG0000( |
| 90060247 | 90119719 | -1 | lncRNA | ENSG0000( |
| 90135712 | 90135841 | -1 | rRNA_pseudogene | ENSG0000( |
| 90170866 | 90171799 | 1 | processed_pseudoge | ENSG0000( |
| 90230384 | 90231682 | 1 | processed_pseudoge | ENSG0000( |
| 90231182 | 90231277 | 1 | miRNA | ENSG0000( |
| 90493287 | 90535080 | 1 | lncRNA | ENSG0000( |
| 90781766 | 90781869 | -1 | snRNA | ENSG0000( |
| 90890954 | 90926597 | -1 | lncRNA | ENSG0000( |
| 91023732 | 91023954 | -1 | processed_pseudoge | ENSG0000( |
| 91087254 | 91089593 | -1 | lncRNA | ENSG0000( |
| 91127613 | 91131407 | -1 | lncRNA | ENSG0000( |
| 91272081 | 91272577 | -1 | processed_pseudoge | ENSG0000( |
| 91347820 | 91354579 | 1 | lncRNA | ENSG0000( |
| 91350605 | 91350688 | 1 | miRNA | ENSG0000( |
| 91350751 | 91350821 | 1 | miRNA | ENSG0000( |
| 91350891 | 91350972 | 1 | miRNA | ENSG0000( |
| 91351065 | 91351135 | 1 | miRNA | ENSG0000( |
| 91351192 | 91351278 | 1 | miRNA | ENSG0000( |
| 91351314 | 91351391 | 1 | miRNA | ENSG0000( |
| 91398621 | 92873682 | 1 | protein_coding | ENSG0000( |
| 91485793 | 91486870 | 1 | TEC | ENSG0000( |
| 91994518 | 91996523 | 1 | lncRNA | ENSG0000( |
| 92059844 | 92059969 | 1 | snRNA | ENSG0000( |
| 92180715 | 92180885 | -1 | processed_pseudoge | ENSG0000( |
| 92239835 | 92241951 | -1 | TEC | ENSG0000( |
| 92339794 | 92379536 | -1 | lncRNA | ENSG0000( |
| 92446470 | 92449056 | 1 | TEC | ENSG0000( |
| 92484606 | 92510094 | 1 | lncRNA | ENSG0000( |
| 92490163 | 92490220 | 1 | miRNA | ENSG0000( |
| 92610646 | 92677725 | -1 | lncRNA | ENSG0000( |

| | | | | |
|---|---|---|---|---|
| 92658994 | 92659227 | -1 | TEC | ENSG0000C |
| 92701389 | 92721614 | -1 | lncRNA | ENSG0000C |
| 93032972 | 93057926 | 1 | lncRNA | ENSG0000C |
| 93150623 | 93157452 | 1 | lncRNA | ENSG0000C |
| 93226612 | 93227317 | -1 | lncRNA | ENSG0000C |
| 93226807 | 94408020 | 1 | protein_coding | ENSG0000C |
| 93309669 | 93311063 | -1 | lncRNA | ENSG0000C |
| 93394977 | 93395855 | -1 | processed_pseudoge | ENSG0000C |
| 93818424 | 93836144 | -1 | lncRNA | ENSG0000C |
| 94025470 | 94025598 | -1 | rRNA_pseudogene | ENSG0000C |
| 94154193 | 94187991 | -1 | lncRNA | ENSG0000C |
| 94436811 | 94479682 | -1 | protein_coding | ENSG0000C |
| 94574054 | 94596242 | -1 | protein_coding | ENSG0000C |
| 94601857 | 94634661 | 1 | protein_coding | ENSG0000C |
| 94651397 | 94651529 | -1 | rRNA_pseudogene | ENSG0000C |
| 94698725 | 94708475 | -1 | lncRNA | ENSG0000C |
| 94699223 | 94699502 | -1 | misc_RNA | ENSG0000C |
| 94703454 | 94803430 | 1 | lncRNA | ENSG0000C |
| 94709622 | 94712545 | -1 | protein_coding | ENSG0000C |
| 94757451 | 94759272 | -1 | processed_pseudoge | ENSG0000C |
| 94760553 | 94761015 | -1 | processed_pseudoge | ENSG0000C |
| 94960041 | 94961319 | 1 | lncRNA | ENSG0000C |
| 94960771 | 95007420 | -1 | lncRNA | ENSG0000C |
| 95019235 | 95019336 | 1 | snRNA | ENSG0000C |
| 95019835 | 95301475 | -1 | protein_coding | ENSG0000C |
| 95310830 | 95310955 | -1 | misc_RNA | ENSG0000C |
| 95336106 | 95336201 | -1 | misc_RNA | ENSG0000C |
| 95394049 | 95394454 | 1 | processed_pseudoge | ENSG0000C |
| 95433604 | 95579759 | 1 | protein_coding | ENSG0000C |
| 95479444 | 95533910 | -1 | lncRNA | ENSG0000C |
| 95578202 | 95644703 | -1 | protein_coding | ENSG0000C |
| 95648733 | 95676955 | -1 | lncRNA | ENSG0000C |
| 95677139 | 95794988 | 1 | protein_coding | ENSG0000C |
| 95692538 | 95693972 | 1 | processed_pseudoge | ENSG0000C |
| 95693983 | 95694495 | -1 | processed_pseudoge | ENSG0000C |
| 95694569 | 95695705 | 1 | processed_pseudoge | ENSG0000C |
| 95744726 | 95745765 | 1 | lncRNA | ENSG0000C |
| 95801580 | 96053482 | -1 | protein_coding | ENSG0000C |
| 95809310 | 95809607 | -1 | processed_pseudoge | ENSG0000C |
| 96090839 | 96839562 | 1 | protein_coding | ENSG0000C |
| 96169545 | 96169847 | 1 | misc_RNA | ENSG0000C |
| 96427229 | 96427292 | 1 | miRNA | ENSG0000C |
| 96573659 | 96574499 | 1 | processed_pseudoge | ENSG0000C |
| 96883842 | 96885821 | -1 | processed_pseudoge | ENSG0000C |
| 96941281 | 96984172 | -1 | transcribed_processe | ENSG0000C |
| 96948018 | 96949584 | 1 | lncRNA | ENSG0000C |
| 96949249 | 96949533 | -1 | misc_RNA | ENSG0000C |
| 96985719 | 96994730 | -1 | protein_coding | ENSG0000C |
| 97168751 | 97187578 | -1 | lncRNA | ENSG0000C |
| 97221434 | 97394120 | 1 | protein_coding | ENSG0000C |

| | | | | |
|---|---|---|---|---|
| 97278768 | 97295400 | -1 | lncRNA | ENSG0000C |
| 97362972 | 97363101 | -1 | rRNA_pseudogene | ENSG0000C |
| 97405169 | 97433994 | -1 | lncRNA | ENSG0000C |
| 97434169 | 97469128 | 1 | protein_coding | ENSG0000C |
| 97435946 | 97436168 | 1 | lncRNA | ENSG0000C |
| 97437268 | 97437630 | 1 | lncRNA | ENSG0000C |
| 97475742 | 97476459 | 1 | processed_pseudoge | ENSG0000C |
| 97675011 | 97677963 | -1 | lncRNA | ENSG0000C |
| 97725172 | 97730339 | -1 | lncRNA | ENSG0000C |
| 97901653 | 97902288 | 1 | processed_pseudoge | ENSG0000C |
| 97939356 | 97953539 | 1 | lncRNA | ENSG0000C |
| 97953658 | 98024296 | 1 | protein_coding | ENSG0000C |
| 97986950 | 97987216 | -1 | processed_pseudoge | ENSG0000C |
| 98142562 | 98455176 | 1 | protein_coding | ENSG0000C |
| 98175785 | 98177269 | -1 | protein_coding | ENSG0000C |
| 98203835 | 98204192 | -1 | misc_RNA | ENSG0000C |
| 98208524 | 98208600 | 1 | miRNA | ENSG0000C |
| 98233650 | 98234700 | 1 | lncRNA | ENSG0000C |
| 98329655 | 98333236 | -1 | lncRNA | ENSG0000C |
| 98435405 | 98435840 | -1 | lncRNA | ENSG0000C |
| 98445185 | 98577940 | -1 | protein_coding | ENSG0000C |
| 98553243 | 98553780 | 1 | processed_pseudoge | ENSG0000C |
| 98577244 | 98578830 | 1 | lncRNA | ENSG0000C |
| 98610292 | 98610876 | -1 | processed_pseudoge | ENSG0000C |
| 98629686 | 98629989 | -1 | processed_pseudoge | ENSG0000C |
| 98640814 | 98641239 | -1 | processed_pseudoge | ENSG0000C |
| 98671980 | 98672278 | 1 | misc_RNA | ENSG0000C |
| 98683801 | 98752672 | -1 | protein_coding | ENSG0000C |
| 98793429 | 99086625 | -1 | protein_coding | ENSG0000C |
| 98832084 | 98834629 | 1 | lncRNA | ENSG0000C |
| 98871742 | 98872005 | 1 | processed_pseudoge | ENSG0000C |
| 98949719 | 98950447 | -1 | processed_pseudoge | ENSG0000C |
| 99025234 | 99025345 | 1 | snRNA | ENSG0000C |
| 99087819 | 99088625 | 1 | lncRNA | ENSG0000C |
| 99173593 | 99174490 | 1 | processed_pseudoge | ENSG0000C |
| 99181223 | 99200757 | -1 | lncRNA | ENSG0000C |
| 99190579 | 99190991 | -1 | processed_pseudoge | ENSG0000C |
| 99200774 | 99386504 | 1 | protein_coding | ENSG0000C |
| 99205708 | 99206006 | 1 | misc_RNA | ENSG0000C |
| 99215372 | 99215758 | -1 | processed_pseudoge | ENSG0000C |
| 99254714 | 99261744 | -1 | protein_coding | ENSG0000C |
| 99294539 | 99307399 | -1 | protein_coding | ENSG0000C |
| 99356131 | 99356228 | 1 | miRNA | ENSG0000C |
| 99372173 | 99372686 | 1 | processed_pseudoge | ENSG0000C |
| 99404684 | 99417224 | -1 | transcribed_unproce | ENSG0000C |
| 99429211 | 99432702 | -1 | lncRNA | ENSG0000C |
| 99446311 | 99564048 | 1 | protein_coding | ENSG0000C |
| 99486962 | 99499306 | -1 | lncRNA | ENSG0000C |
| 99498737 | 99501250 | 1 | lncRNA | ENSG0000C |
| 99499727 | 99501052 | -1 | lncRNA | ENSG0000C |

| | | | | |
|---|---|---|---|---|
| 99577112 | 99580368 | 1 | lncRNA | ENSG0000C |
| 99582706 | 99583180 | 1 | processed_pseudoge | ENSG0000C |
| 99584014 | 99590508 | -1 | lncRNA | ENSG0000C |
| 99606669 | 99897134 | 1 | protein_coding | ENSG0000C |
| 99643059 | 99643149 | 1 | miRNA | ENSG0000C |
| 99690081 | 99690971 | -1 | lncRNA | ENSG0000C |
| 99726245 | 99728971 | -1 | lncRNA | ENSG0000C |
| 99783857 | 99783983 | -1 | snoRNA | ENSG0000C |
| 99835813 | 99957167 | -1 | lncRNA | ENSG0000C |
| 99962964 | 99971909 | -1 | protein_coding | ENSG0000C |
| 99973526 | 99980034 | -1 | lncRNA | ENSG0000C |
| 99981784 | 99986765 | 1 | protein_coding | ENSG0000C |
| 99994899 | 99997909 | -1 | lncRNA | ENSG0000C |
| 100027769 | 100028174 | -1 | processed_pseudoge | ENSG0000C |
| 100053074 | 100088950 | -1 | lncRNA | ENSG0000C |
| 100062296 | 100063601 | -1 | processed_pseudoge | ENSG0000C |
| 100089015 | 100530437 | 1 | protein_coding | ENSG0000C |
| 100150481 | 100151092 | -1 | processed_pseudoge | ENSG0000C |
| 100198187 | 100198469 | -1 | processed_pseudoge | ENSG0000C |
| 100338982 | 100364959 | -1 | lncRNA | ENSG0000C |
| 100464440 | 100481205 | -1 | lncRNA | ENSG0000C |
| 100530164 | 100589528 | -1 | protein_coding | ENSG0000C |
| 100535741 | 100587146 | -1 | lncRNA | ENSG0000C |
| 100537365 | 100539567 | 1 | lncRNA | ENSG0000C |
| 100539901 | 100540248 | -1 | processed_pseudoge | ENSG0000C |
| 100603927 | 100675093 | -1 | protein_coding | ENSG0000C |
| 100623875 | 100624429 | -1 | processed_pseudoge | ENSG0000C |
| 100708325 | 101059286 | 1 | lncRNA | ENSG0000C |
| 100756551 | 100757059 | -1 | processed_pseudoge | ENSG0000C |
| 100934023 | 100944553 | -1 | lncRNA | ENSG0000C |
| 101053776 | 101416508 | -1 | protein_coding | ENSG0000C |
| 101452593 | 101720856 | 1 | protein_coding | ENSG0000C |
| 101710804 | 102402457 | -1 | protein_coding | ENSG0000C |
| 101717564 | 101723106 | 1 | lncRNA | ENSG0000C |
| 101778884 | 101779048 | -1 | snRNA | ENSG0000C |
| 101801878 | 101802488 | 1 | processed_pseudoge | ENSG0000C |
| 101967642 | 101967746 | -1 | miRNA | ENSG0000C |
| 102002125 | 102003310 | 1 | processed_pseudoge | ENSG0000C |
| 102016953 | 102017069 | -1 | misc_RNA | ENSG0000C |
| 102045934 | 102046004 | -1 | miRNA | ENSG0000C |
| 102263046 | 102263202 | -1 | processed_pseudoge | ENSG0000C |
| 102292327 | 102394519 | -1 | lncRNA | ENSG0000C |
| 102299790 | 102300272 | -1 | TEC | ENSG0000C |
| 102367539 | 102373666 | 1 | lncRNA | ENSG0000C |
| 102394630 | 102395703 | 1 | lncRNA | ENSG0000C |
| 102589372 | 102589813 | -1 | lncRNA | ENSG0000C |
| 102593338 | 102593873 | -1 | lncRNA | ENSG0000C |
| 102596986 | 102679958 | 1 | protein_coding | ENSG0000C |
| 102685744 | 102694504 | -1 | protein_coding | ENSG0000C |
| 102729367 | 102759072 | -1 | protein_coding | ENSG0000C |

| | | | | |
|---|---|---|---|---|
| 102742990 | 102745224 | 1 | lncRNA | ENSG00000 |
| 102760023 | 102766653 | 1 | lncRNA | ENSG00000 |
| 102765888 | 102773811 | -1 | protein_coding | ENSG00000 |
| 102784281 | 102798976 | -1 | protein_coding | ENSG00000 |
| 102799110 | 102875994 | 1 | protein_coding | ENSG00000 |
| 102799119 | 102841533 | 1 | protein_coding | ENSG00000 |
| 102807146 | 102876001 | 1 | protein_coding | ENSG00000 |
| 102880099 | 102896033 | 1 | transcribed_unproce | ENSG00000 |
| 102888858 | 102889834 | 1 | lncRNA | ENSG00000 |
| 102903339 | 102904000 | 1 | lncRNA | ENSG00000 |
| 103043998 | 103066417 | -1 | protein_coding | ENSG00000 |
| 103343660 | 103396450 | 1 | lncRNA | ENSG00000 |
| 103420825 | 103428298 | 1 | lncRNA | ENSG00000 |
| 103444620 | 103444968 | -1 | processed_pseudoge | ENSG00000 |
| 104125930 | 104134257 | 1 | lncRNA | ENSG00000 |
| 104814327 | 104815069 | 1 | processed_pseudoge | ENSG00000 |
| 105410816 | 105414224 | 1 | lncRNA | ENSG00000 |
| 105459055 | 105505681 | -1 | lncRNA | ENSG00000 |
| 105459815 | 105460015 | 1 | misc_RNA | ENSG00000 |
| 105462302 | 105462505 | 1 | misc_RNA | ENSG00000 |
| 105465867 | 105491034 | 1 | protein_coding | ENSG00000 |
| 105690429 | 105702689 | 1 | lncRNA | ENSG00000 |
| 105702699 | 105797539 | 1 | lncRNA | ENSG00000 |
| 105897523 | 105897643 | -1 | snoRNA | ENSG00000 |
| 105950292 | 106064069 | -1 | lncRNA | ENSG00000 |
| 106155372 | 106155490 | -1 | rRNA_pseudogene | ENSG00000 |
| 106232131 | 106371082 | -1 | lncRNA | ENSG00000 |
| 106374477 | 106384315 | 1 | lncRNA | ENSG00000 |
| 106425773 | 106426122 | 1 | processed_pseudoge | ENSG00000 |
| 106489745 | 106535662 | -1 | protein_coding | ENSG00000 |
| 106491147 | 106521630 | 1 | lncRNA | ENSG00000 |
| 106506046 | 106506713 | -1 | lncRNA | ENSG00000 |
| 106541673 | 106568137 | -1 | protein_coding | ENSG00000 |
| 106568261 | 106716679 | 1 | lncRNA | ENSG00000 |
| 106653897 | 106672177 | 1 | lncRNA | ENSG00000 |
| 106663437 | 106663847 | 1 | processed_pseudoge | ENSG00000 |
| 106753247 | 106757065 | 1 | lncRNA | ENSG00000 |
| 106773623 | 106774279 | -1 | lncRNA | ENSG00000 |
| 106863816 | 106864286 | -1 | processed_pseudoge | ENSG00000 |
| 106903150 | 106904099 | -1 | lncRNA | ENSG00000 |
| 106941950 | 106942561 | -1 | TEC | ENSG00000 |
| 106955689 | 106961392 | -1 | lncRNA | ENSG00000 |
| 107163510 | 107867496 | -1 | protein_coding | ENSG00000 |
| 107320895 | 107320963 | -1 | snoRNA | ENSG00000 |
| 107465984 | 107466674 | -1 | lncRNA | ENSG00000 |
| 107531171 | 107531248 | -1 | miRNA | ENSG00000 |
| 107788342 | 107835451 | -1 | lncRNA | ENSG00000 |
| 107868823 | 107933503 | 1 | lncRNA | ENSG00000 |
| 107870383 | 107873372 | 1 | lncRNA | ENSG00000 |
| 108207439 | 108218368 | -1 | protein_coding | ENSG00000 |

| | | | | |
|---|---|---|---|---|
| 108218392 | 108234243 | 1 | protein_coding | ENSG00000( |
| 108251240 | 108308484 | 1 | protein_coding | ENSG00000( |
| 108267320 | 108267734 | 1 | lncRNA | ENSG00000( |
| 108301212 | 108301331 | -1 | rRNA_pseudogene | ENSG00000( |
| 108335354 | 108448316 | 1 | lncRNA | ENSG00000( |
| 108480226 | 108481817 | 1 | processed_pseudoge | ENSG00000( |
| 108596152 | 109208005 | 1 | protein_coding | ENSG00000( |
| 109101261 | 109101919 | -1 | lncRNA | ENSG00000( |
| 109163902 | 109201483 | -1 | lncRNA | ENSG00000( |
| 109269634 | 109278512 | 1 | lncRNA | ENSG00000( |
| 109281973 | 109297690 | 1 | lncRNA | ENSG00000( |
| 109310135 | 109311773 | 1 | lncRNA | ENSG00000( |
| 109400696 | 109401641 | 1 | lncRNA | ENSG00000( |
| 109478739 | 109731700 | -1 | lncRNA | ENSG00000( |
| 109728282 | 109730007 | 1 | lncRNA | ENSG00000( |
| 109752695 | 109786583 | -1 | protein_coding | ENSG00000( |
| 109785831 | 109808252 | 1 | lncRNA | ENSG00000( |
| 109805317 | 109805868 | 1 | lncRNA | ENSG00000( |
| 109915096 | 109915420 | 1 | misc_RNA | ENSG00000( |
| 109984858 | 109986565 | -1 | lncRNA | ENSG00000( |
| 109988097 | 110129136 | -1 | lncRNA | ENSG00000( |
| 110031101 | 110031567 | -1 | lncRNA | ENSG00000( |
| 110053285 | 110054952 | 1 | lncRNA | ENSG00000( |
| 110097851 | 110098151 | -1 | misc_RNA | ENSG00000( |
| 110123208 | 110125234 | 1 | TEC | ENSG00000( |
| 110132003 | 110136906 | -1 | lncRNA | ENSG00000( |
| 110148963 | 110307157 | -1 | protein_coding | ENSG00000( |
| 110265251 | 110266113 | -1 | TEC | ENSG00000( |
| 110305812 | 110513209 | 1 | protein_coding | ENSG00000( |
| 110340958 | 110341029 | 1 | miRNA | ENSG00000( |
| 110456396 | 110463287 | -1 | lncRNA | ENSG00000( |
| 110502575 | 110508179 | -1 | lncRNA | ENSG00000( |
| 110523066 | 110561722 | -1 | protein_coding | ENSG00000( |
| 110613082 | 110616353 | -1 | lncRNA | ENSG00000( |
| 110615460 | 110639993 | 1 | protein_coding | ENSG00000( |
| 110641412 | 110713603 | -1 | protein_coding | ENSG00000( |
| 110712736 | 110723339 | 1 | protein_coding | ENSG00000( |
| 110762674 | 110763133 | -1 | processed_pseudoge | ENSG00000( |
| 110809676 | 110813084 | -1 | lncRNA | ENSG00000( |
| 110863987 | 110870251 | -1 | lncRNA | ENSG00000( |
| 110878540 | 110915069 | -1 | protein_coding | ENSG00000( |
| 110894639 | 110899172 | -1 | lncRNA | ENSG00000( |
| 110916004 | 110917827 | 1 | lncRNA | ENSG00000( |
| 110936759 | 110939784 | 1 | processed_pseudoge | ENSG00000( |
| 110945478 | 110971603 | -1 | transcribed_unproce | ENSG00000( |
| 110965704 | 110990579 | 1 | transcribed_unproce | ENSG00000( |
| 111095838 | 111103097 | 1 | lncRNA | ENSG00000( |
| 111096586 | 111113697 | -1 | lncRNA | ENSG00000( |
| 111113812 | 111115678 | -1 | lncRNA | ENSG00000( |
| 111114559 | 111305737 | 1 | protein_coding | ENSG00000( |

| | | | | |
|---|---|---|---|---|
| 111122652 | 111144264 | 1 | lncRNA | ENSG0000C |
| 111144305 | 111144733 | -1 | lncRNA | ENSG0000C |
| 111148196 | 111153171 | -1 | lncRNA | ENSG0000C |
| 111182111 | 111182692 | -1 | unprocessed_pseudc | ENSG0000C |
| 111316184 | 111344249 | 1 | protein_coding | ENSG0000C |
| 111588201 | 111672608 | 1 | lncRNA | ENSG0000C |
| 111596009 | 111642079 | 1 | lncRNA | ENSG0000C |
| 111865136 | 111901264 | 1 | lncRNA | ENSG0000C |
| 112052078 | 112052138 | 1 | snoRNA | ENSG0000C |
| 112056845 | 112108015 | 1 | lncRNA | ENSG0000C |
| 112067149 | 112071706 | 1 | protein_coding | ENSG0000C |
| 112083549 | 112085471 | 1 | lncRNA | ENSG0000C |
| 112106095 | 112106882 | -1 | lncRNA | ENSG0000C |
| 112197350 | 112201000 | 1 | lncRNA | ENSG0000C |
| 112313467 | 112321758 | 1 | lncRNA | ENSG0000C |
| 112322567 | 112331225 | -1 | lncRNA | ENSG0000C |
| 112376355 | 112434689 | 1 | protein_coding | ENSG0000C |
| 112485011 | 112588205 | -1 | protein_coding | ENSG0000C |
| 112602828 | 112606417 | -1 | lncRNA | ENSG0000C |
| 112647044 | 112684497 | 1 | lncRNA | ENSG0000C |
| 112686769 | 112689815 | -1 | lncRNA | ENSG0000C |
| 112690329 | 112887168 | 1 | protein_coding | ENSG0000C |
| 112745449 | 112754693 | -1 | lncRNA | ENSG0000C |
| 112811566 | 112812158 | 1 | lncRNA | ENSG0000C |
| 112894378 | 113099742 | 1 | protein_coding | ENSG0000C |
| 112964835 | 112966131 | -1 | lncRNA | ENSG0000C |
| 112967484 | 112968824 | -1 | lncRNA | ENSG0000C |
| 113008778 | 113009424 | 1 | lncRNA | ENSG0000C |
| 113009671 | 113010319 | 1 | lncRNA | ENSG0000C |
| 113105788 | 113120681 | 1 | protein_coding | ENSG0000C |
| 113122799 | 113149529 | 1 | protein_coding | ENSG0000C |
| 113128155 | 113128880 | -1 | lncRNA | ENSG0000C |
| 113130416 | 113132054 | 1 | processed_pseudoge | ENSG0000C |
| 113149432 | 113154002 | -1 | lncRNA | ENSG0000C |
| 113158648 | 113172386 | 1 | protein_coding | ENSG0000C |
| 113165002 | 113165183 | -1 | lncRNA | ENSG0000C |
| 113177536 | 113208715 | -1 | protein_coding | ENSG0000C |
| 113208193 | 113267108 | 1 | protein_coding | ENSG0000C |
| 113262920 | 113262999 | 1 | miRNA | ENSG0000C |
| 113276616 | 113277734 | 1 | processed_pseudoge | ENSG0000C |
| 113297239 | 113323672 | 1 | protein_coding | ENSG0000C |
| 113324163 | 113364148 | -1 | protein_coding | ENSG0000C |
| 113339450 | 113339958 | -1 | lncRNA | ENSG0000C |
| 113351645 | 113361868 | 1 | lncRNA | ENSG0000C |
| 113399610 | 113453524 | -1 | protein_coding | ENSG0000C |
| 113455819 | 113490951 | -1 | protein_coding | ENSG0000C |
| 113468901 | 113476135 | 1 | lncRNA | ENSG0000C |
| 113478915 | 113479078 | -1 | snRNA | ENSG0000C |
| 113491021 | 113554590 | 1 | protein_coding | ENSG0000C |
| 113511747 | 113514473 | 1 | lncRNA | ENSG0000C |

| | | | | |
|---|---|---|---|---|
| 113527260 | 113530621 | 1 | lncRNA | ENSG0000C |
| 113584721 | 113641473 | 1 | protein_coding | ENSG0000C |
| 113648804 | 113658198 | -1 | protein_coding | ENSG0000C |
| 113667219 | 113737736 | 1 | protein_coding | ENSG0000C |
| 113748511 | 113751089 | -1 | TEC | ENSG0000C |
| 113759226 | 113816995 | 1 | protein_coding | ENSG0000C |
| 113815630 | 113845744 | 1 | lncRNA | ENSG0000C |
| 113820549 | 113864076 | -1 | protein_coding | ENSG0000C |
| 113864112 | 113866834 | 1 | lncRNA | ENSG0000C |
| 113877608 | 113879376 | 1 | lncRNA | ENSG0000C |
| 113878489 | 113883651 | -1 | lncRNA | ENSG0000C |
| 113883667 | 113887149 | 1 | lncRNA | ENSG0000C |
| 113888306 | 113923512 | 1 | lncRNA | ENSG0000C |
| 113926514 | 113928844 | -1 | lncRNA | ENSG0000C |
| 113953705 | 113974076 | -1 | protein_coding | ENSG0000C |
| 113977783 | 114132623 | -1 | protein_coding | ENSG0000C |
| 114107569 | 114108820 | -1 | lncRNA | ENSG0000C |
| 114154691 | 114223084 | 1 | protein_coding | ENSG0000C |
| 114175485 | 114215835 | -1 | lncRNA | ENSG0000C |
| 114234887 | 114272723 | 1 | protein_coding | ENSG0000C |
| 114244505 | 114244561 | 1 | miRNA | ENSG0000C |
| 114273828 | 114273908 | 1 | miRNA | ENSG0000C |
| 114275326 | 114277323 | -1 | lncRNA | ENSG0000C |
| 114281601 | 114305817 | 1 | protein_coding | ENSG0000C |
| 114306646 | 114307066 | 1 | processed_pseudoge | ENSG0000C |
| 114314482 | 114337626 | 1 | protein_coding | ENSG0000C |
| 114329559 | 114333948 | -1 | lncRNA | ENSG0000C |
| 114346167 | 114346637 | 1 | processed_pseudoge | ENSG0000C |
| 16057472 | 16057622 | 1 | rRNA | ENSG0000C |
| 18333726 | 18341859 | 1 | transcribed_unproce | ENSG0000C |
| 18343038 | 18343144 | 1 | snRNA | ENSG0000C |
| 18395626 | 18412264 | -1 | transcribed_unproce | ENSG0000C |
| 18418775 | 18419273 | 1 | processed_pseudoge | ENSG0000C |
| 18555677 | 18596310 | -1 | lncRNA | ENSG0000C |
| 18560047 | 18560191 | 1 | processed_pseudoge | ENSG0000C |
| 18562928 | 18568790 | 1 | transcribed_unproce | ENSG0000C |
| 18588274 | 18588315 | -1 | transcribed_processe | ENSG0000C |
| 18601045 | 18602129 | 1 | protein_coding | ENSG0000C |
| 18614388 | 18614561 | 1 | processed_pseudoge | ENSG0000C |
| 18623852 | 18626787 | -1 | processed_pseudoge | ENSG0000C |
| 18630318 | 18633634 | -1 | lncRNA | ENSG0000C |
| 18631388 | 18650966 | 1 | unprocessed_pseudc | ENSG0000C |
| 18634955 | 18637208 | -1 | processed_pseudoge | ENSG0000C |
| 18636043 | 18637421 | 1 | lncRNA | ENSG0000C |
| 18645044 | 18646221 | -1 | processed_pseudoge | ENSG0000C |
| 18645991 | 18646503 | 1 | processed_pseudoge | ENSG0000C |
| 18667249 | 18667370 | 1 | processed_pseudoge | ENSG0000C |
| 18692032 | 18692408 | 1 | processed_pseudoge | ENSG0000C |
| 18698639 | 18699978 | -1 | transcribed_processe | ENSG0000C |
| 18702281 | 18702463 | -1 | transcribed_processe | ENSG0000C |

| | | | | |
|---|---|---|---|---|
| 18709478 | 18712643 | -1 | transcribed_unproce | ENSG00000 |
| 18889084 | 18890424 | -1 | processed_pseudoge | ENSG00000 |
| 18899942 | 18909673 | -1 | unprocessed_pseudo | ENSG00000 |
| 18912805 | 18916642 | -1 | transcribed_unproce | ENSG00000 |
| 18920109 | 18934078 | 1 | lncRNA | ENSG00000 |
| 18944199 | 18945139 | 1 | lncRNA | ENSG00000 |
| 18967434 | 19003752 | 1 | protein_coding | ENSG00000 |
| 18977180 | 18980742 | -1 | lncRNA | ENSG00000 |
| 18978847 | 18978953 | 1 | snRNA | ENSG00000 |
| 19024090 | 19055551 | 1 | lncRNA | ENSG00000 |
| 19026521 | 19028189 | -1 | processed_pseudoge | ENSG00000 |
| 19062316 | 19131167 | 1 | transcribed_processe | ENSG00000 |
| 19072333 | 19075341 | 1 | transcribed_unproce | ENSG00000 |
| 19124807 | 19175530 | -1 | lncRNA | ENSG00000 |
| 19130636 | 19155089 | 1 | transcribed_unproce | ENSG00000 |
| 19131842 | 19132890 | -1 | processed_pseudoge | ENSG00000 |
| 19180291 | 19181269 | 1 | unprocessed_pseudo | ENSG00000 |
| 19199763 | 19200279 | -1 | processed_pseudoge | ENSG00000 |
| 19220796 | 19220912 | -1 | unprocessed_pseudo | ENSG00000 |
| 19228898 | 19229945 | -1 | unprocessed_pseudo | ENSG00000 |
| 19244904 | 19269380 | -1 | transcribed_unproce | ENSG00000 |
| 19244962 | 19247673 | 1 | lncRNA | ENSG00000 |
| 19267115 | 19268164 | 1 | processed_pseudoge | ENSG00000 |
| 19268853 | 19337730 | -1 | transcribed_processe | ENSG00000 |
| 19301704 | 19316914 | 1 | transcribed_unproce | ENSG00000 |
| 19324708 | 19327716 | -1 | transcribed_unproce | ENSG00000 |
| 19344578 | 19384587 | -1 | lncRNA | ENSG00000 |
| 19371822 | 19373490 | 1 | processed_pseudoge | ENSG00000 |
| 19402486 | 19434341 | -1 | protein_coding | ENSG00000 |
| 19420975 | 19425017 | 1 | lncRNA | ENSG00000 |
| 19422815 | 19422921 | -1 | snRNA | ENSG00000 |
| 19485336 | 19489148 | 1 | transcribed_processe | ENSG00000 |
| 19492281 | 19501740 | 1 | unprocessed_pseudo | ENSG00000 |
| 19499737 | 19677923 | 1 | lncRNA | ENSG00000 |
| 19511244 | 19511712 | 1 | processed_pseudoge | ENSG00000 |
| 19612191 | 19619490 | 1 | unprocessed_pseudo | ENSG00000 |
| 19629006 | 19630340 | 1 | processed_pseudoge | ENSG00000 |
| 19637225 | 19641306 | 1 | lncRNA | ENSG00000 |
| 19653623 | 19653742 | -1 | processed_pseudoge | ENSG00000 |
| 19664098 | 19683677 | -1 | unprocessed_pseudo | ENSG00000 |
| 19668542 | 19669056 | -1 | processed_pseudoge | ENSG00000 |
| 19668829 | 19670003 | 1 | processed_pseudoge | ENSG00000 |
| 19677644 | 19679016 | -1 | lncRNA | ENSG00000 |
| 19677837 | 19680104 | 1 | transcribed_processe | ENSG00000 |
| 19681420 | 19684739 | 1 | lncRNA | ENSG00000 |
| 19688261 | 19691194 | 1 | processed_pseudoge | ENSG00000 |
| 19700479 | 19700650 | -1 | unprocessed_pseudo | ENSG00000 |
| 19712904 | 19714332 | -1 | protein_coding | ENSG00000 |
| 19719015 | 19830253 | 1 | protein_coding | ENSG00000 |
| 19733355 | 19734302 | -1 | unprocessed_pseudo | ENSG00000 |

| | | | | |
|---|---|---|---|---|
| 19743571 | 19749469 | 1 | protein_coding | ENSG00000 |
| 19759868 | 19760774 | 1 | unprocessed_pseudo | ENSG00000 |
| 19773504 | 19783696 | 1 | protein_coding | ENSG00000 |
| 19796345 | 19797293 | 1 | unprocessed_pseudo | ENSG00000 |
| 19847367 | 19848263 | 1 | unprocessed_pseudo | ENSG00000 |
| 19868197 | 19869143 | -1 | polymorphic_pseudo | ENSG00000 |
| 19875142 | 19883932 | 1 | protein_coding | ENSG00000 |
| 19905917 | 19906600 | 1 | unprocessed_pseudo | ENSG00000 |
| 19908096 | 19908649 | -1 | processed_pseudoge | ENSG00000 |
| 19920582 | 19921659 | 1 | protein_coding | ENSG00000 |
| 19930917 | 19936757 | 1 | protein_coding | ENSG00000 |
| 19954490 | 19957996 | 1 | transcribed_unproce | ENSG00000 |
| 19975444 | 19976659 | 1 | protein_coding | ENSG00000 |
| 19999913 | 20004760 | 1 | polymorphic_pseudo | ENSG00000 |
| 20014143 | 20019307 | -1 | protein_coding | ENSG00000 |
| 20029399 | 20036038 | -1 | protein_coding | ENSG00000 |
| 20043916 | 20044827 | 1 | unprocessed_pseudo | ENSG00000 |
| 20060045 | 20060983 | 1 | protein_coding | ENSG00000 |
| 20090130 | 20090227 | 1 | rRNA_pseudogene | ENSG00000 |
| 20093353 | 20094263 | 1 | unprocessed_pseudo | ENSG00000 |
| 20110739 | 20122199 | 1 | protein_coding | ENSG00000 |
| 20138820 | 20145471 | 1 | protein_coding | ENSG00000 |
| 20149232 | 20149745 | 1 | processed_pseudoge | ENSG00000 |
| 20173632 | 20173813 | -1 | processed_pseudoge | ENSG00000 |
| 20179189 | 20180156 | 1 | unprocessed_pseudo | ENSG00000 |
| 20183925 | 20184298 | -1 | unprocessed_pseudo | ENSG00000 |
| 20190894 | 20201075 | 1 | protein_coding | ENSG00000 |
| 20204606 | 20210498 | 1 | transcribed_unproce | ENSG00000 |
| 20223710 | 20224702 | 1 | protein_coding | ENSG00000 |
| 20229402 | 20230346 | 1 | polymorphic_pseudo | ENSG00000 |
| 20239286 | 20244349 | 1 | protein_coding | ENSG00000 |
| 20256558 | 20305960 | -1 | protein_coding | ENSG00000 |
| 20260480 | 20264308 | -1 | lncRNA | ENSG00000 |
| 20305730 | 20306811 | 1 | lncRNA | ENSG00000 |
| 20311368 | 20333312 | -1 | protein_coding | ENSG00000 |
| 20317949 | 20319707 | 1 | lncRNA | ENSG00000 |
| 20323179 | 20323326 | -1 | snoRNA | ENSG00000 |
| 20326450 | 20326526 | -1 | snoRNA | ENSG00000 |
| 20343048 | 20343685 | -1 | lncRNA | ENSG00000 |
| 20343075 | 20343407 | -1 | ribozyme | ENSG00000 |
| 20343582 | 20357905 | 1 | protein_coding | ENSG00000 |
| 20344233 | 20346888 | -1 | lncRNA | ENSG00000 |
| 20365667 | 20413501 | -1 | protein_coding | ENSG00000 |
| 20414987 | 20415098 | 1 | rRNA_pseudogene | ENSG00000 |
| 20425852 | 20436166 | -1 | protein_coding | ENSG00000 |
| 20437982 | 20441268 | 1 | transcribed_unproce | ENSG00000 |
| 20446401 | 20455089 | -1 | protein_coding | ENSG00000 |
| 20451305 | 20451918 | 1 | lncRNA | ENSG00000 |
| 20455191 | 20457772 | 1 | protein_coding | ENSG00000 |
| 20457719 | 20461612 | -1 | protein_coding | ENSG00000 |

| | | | | |
|---|---|---|---|---|
| 20468954 | 20477094 | 1 | protein_coding | ENSG0000( |
| 20474789 | 20477089 | -1 | lncRNA | ENSG0000( |
| 20505537 | 20511169 | 1 | protein_coding | ENSG0000( |
| 20524287 | 20525422 | -1 | processed_pseudoge | ENSG0000( |
| 20528115 | 20528959 | -1 | processed_pseudoge | ENSG0000( |
| 20556093 | 20560931 | -1 | protein_coding | ENSG0000( |
| 20582892 | 20609795 | -1 | protein_coding | ENSG0000( |
| 20583559 | 20609884 | -1 | protein_coding | ENSG0000( |
| 20587644 | 20607221 | 1 | lncRNA | ENSG0000( |
| 20590193 | 20590823 | -1 | protein_coding | ENSG0000( |
| 20616644 | 20616991 | -1 | processed_pseudoge | ENSG0000( |
| 20640696 | 20641691 | -1 | protein_coding | ENSG0000( |
| 20679411 | 20679512 | 1 | misc_RNA | ENSG0000( |
| 20684177 | 20698971 | 1 | protein_coding | ENSG0000( |
| 20684560 | 20701216 | 1 | protein_coding | ENSG0000( |
| 20684587 | 20700576 | 1 | protein_coding | ENSG0000( |
| 20693480 | 20707120 | -1 | lncRNA | ENSG0000( |
| 20720742 | 20723127 | -1 | processed_pseudoge | ENSG0000( |
| 20734578 | 20735011 | 1 | unprocessed_pseudc | ENSG0000( |
| 20745887 | 20748380 | 1 | protein_coding | ENSG0000( |
| 20768404 | 20770948 | 1 | protein_coding | ENSG0000( |
| 20781268 | 20782467 | 1 | protein_coding | ENSG0000( |
| 20783888 | 20784293 | 1 | lncRNA | ENSG0000( |
| 20801228 | 20802855 | -1 | protein_coding | ENSG0000( |
| 20870278 | 20873848 | -1 | lncRNA | ENSG0000( |
| 20874305 | 20875768 | -1 | lncRNA | ENSG0000( |
| 20891399 | 20892348 | 1 | protein_coding | ENSG0000( |
| 20897985 | 20936255 | -1 | lncRNA | ENSG0000( |
| 20916766 | 20928580 | -1 | lncRNA | ENSG0000( |
| 20919361 | 20920299 | 1 | lncRNA | ENSG0000( |
| 20955487 | 20956436 | 1 | protein_coding | ENSG0000( |
| 20963579 | 20963867 | 1 | misc_RNA | ENSG0000( |
| 20983155 | 20985066 | -1 | processed_pseudoge | ENSG0000( |
| 20989770 | 20997035 | 1 | protein_coding | ENSG0000( |
| 20995837 | 21014822 | -1 | lncRNA | ENSG0000( |
| 20999255 | 21001871 | 1 | protein_coding | ENSG0000( |
| 21016763 | 21070872 | -1 | protein_coding | ENSG0000( |
| 21023314 | 21023386 | -1 | miRNA | ENSG0000( |
| 21024109 | 21036276 | 1 | protein_coding | ENSG0000( |
| 21024172 | 21029271 | 1 | lncRNA | ENSG0000( |
| 21032818 | 21034807 | -1 | protein_coding | ENSG0000( |
| 21032818 | 21033895 | 1 | lncRNA | ENSG0000( |
| 21042316 | 21044234 | 1 | protein_coding | ENSG0000( |
| 21057822 | 21058455 | 1 | protein_coding | ENSG0000( |
| 21070273 | 21090248 | 1 | protein_coding | ENSG0000( |
| 21090077 | 21104722 | -1 | protein_coding | ENSG0000( |
| 21098937 | 21103724 | 1 | protein_coding | ENSG0000( |
| 21108043 | 21116703 | 1 | lncRNA | ENSG0000( |
| 21110295 | 21113472 | -1 | processed_pseudoge | ENSG0000( |
| 21113639 | 21113745 | -1 | snRNA | ENSG0000( |

| | | | | |
|---|---|---|---|---|
| 21148370 | 21159060 | -1 | protein_coding | ENSG00000( |
| 21162513 | 21163743 | 1 | processed_pseudoge | ENSG00000( |
| 21184341 | 21184959 | -1 | processed_pseudoge | ENSG00000( |
| 21200079 | 21206900 | -1 | lncRNA | ENSG00000( |
| 21209136 | 21269494 | -1 | protein_coding | ENSG00000( |
| 21287939 | 21351301 | 1 | protein_coding | ENSG00000( |
| 21351476 | 21384019 | -1 | protein_coding | ENSG00000( |
| 21384292 | 21384920 | 1 | lncRNA | ENSG00000( |
| 21385194 | 21456126 | -1 | protein_coding | ENSG00000( |
| 21392150 | 21392253 | -1 | snoRNA | ENSG00000( |
| 21397292 | 21397401 | -1 | snoRNA | ENSG00000( |
| 21398925 | 21399282 | 1 | processed_pseudoge | ENSG00000( |
| 21419620 | 21419960 | 1 | processed_pseudoge | ENSG00000( |
| 21429508 | 21429806 | -1 | misc_RNA | ENSG00000( |
| 21459020 | 21476960 | -1 | protein_coding | ENSG00000( |
| 21476597 | 21499175 | 1 | protein_coding | ENSG00000( |
| 21498133 | 21511342 | -1 | protein_coding | ENSG00000( |
| 21521080 | 21537216 | -1 | protein_coding | ENSG00000( |
| 21521083 | 21522660 | 1 | lncRNA | ENSG00000( |
| 21563819 | 21565084 | 1 | processed_pseudoge | ENSG00000( |
| 21568520 | 21580076 | -1 | protein_coding | ENSG00000( |
| 21585290 | 21586280 | -1 | processed_pseudoge | ENSG00000( |
| 21601882 | 21602825 | -1 | unprocessed_pseudc | ENSG00000( |
| 21611924 | 21612381 | -1 | processed_pseudoge | ENSG00000( |
| 21621838 | 21622567 | 1 | TR_V_gene | ENSG00000( |
| 21633836 | 21634940 | -1 | protein_coding | ENSG00000( |
| 21642889 | 21643578 | 1 | TR_V_gene | ENSG00000( |
| 21649728 | 21650337 | -1 | processed_pseudoge | ENSG00000( |
| 21653835 | 21667642 | 1 | protein_coding | ENSG00000( |
| 21667940 | 21673818 | -1 | protein_coding | ENSG00000( |
| 21698954 | 21699151 | -1 | unprocessed_pseudc | ENSG00000( |
| 21712321 | 21712843 | 1 | TR_V_gene | ENSG00000( |
| 21714795 | 21716262 | -1 | lncRNA | ENSG00000( |
| 21723713 | 21724321 | 1 | TR_V_gene | ENSG00000( |
| 21736152 | 21736982 | 1 | TR_V_gene | ENSG00000( |
| 21749178 | 21749705 | 1 | TR_V_gene | ENSG00000( |
| 21750269 | 21751546 | 1 | processed_pseudoge | ENSG00000( |
| 21768489 | 21769080 | 1 | TR_V_gene | ENSG00000( |
| 21782993 | 21783503 | 1 | TR_V_gene | ENSG00000( |
| 21797287 | 21797886 | 1 | TR_V_gene | ENSG00000( |
| 21811502 | 21811977 | 1 | TR_V_gene | ENSG00000( |
| 21825472 | 21826075 | 1 | TR_V_gene | ENSG00000( |
| 21829539 | 21830070 | 1 | TR_V_pseudogene | ENSG00000( |
| 21841240 | 21841774 | 1 | TR_V_gene | ENSG00000( |
| 21846537 | 21847221 | 1 | TR_V_gene | ENSG00000( |
| 21852558 | 21853006 | 1 | TR_V_gene | ENSG00000( |
| 21868839 | 21869365 | 1 | TR_V_gene | ENSG00000( |
| 21887857 | 21888502 | 1 | TR_V_gene | ENSG00000( |
| 21894433 | 21895030 | 1 | TR_V_gene | ENSG00000( |
| 21903077 | 21904598 | 1 | TR_V_pseudogene | ENSG00000( |

| | | | | |
|---|---|---|---|---|
| 21918188 | 21918756 | 1 | TR_V_gene | ENSG0000C |
| 21924063 | 21924651 | 1 | TR_V_gene | ENSG0000C |
| 21941128 | 21941657 | 1 | TR_V_gene | ENSG0000C |
| 21950406 | 21950654 | 1 | TR_V_pseudogene | ENSG0000C |
| 21962819 | 21963042 | -1 | processed_pseudoge | ENSG0000C |
| 21965451 | 21966061 | 1 | TR_V_gene | ENSG0000C |
| 21978459 | 21979120 | 1 | TR_V_gene | ENSG0000C |
| 21990496 | 21990938 | 1 | TR_V_gene | ENSG0000C |
| 21997539 | 21998168 | 1 | TR_V_gene | ENSG0000C |
| 22003106 | 22003673 | 1 | TR_V_gene | ENSG0000C |
| 22007512 | 22008181 | 1 | TR_V_gene | ENSG0000C |
| 22011910 | 22014964 | 1 | processed_pseudoge | ENSG0000C |
| 22040594 | 22041153 | 1 | TR_V_gene | ENSG0000C |
| 22052514 | 22053056 | 1 | TR_V_gene | ENSG0000C |
| 22066016 | 22066373 | 1 | processed_pseudoge | ENSG0000C |
| 22070557 | 22071208 | 1 | TR_V_gene | ENSG0000C |
| 22086407 | 22086961 | 1 | TR_V_gene | ENSG0000C |
| 22096032 | 22096619 | 1 | TR_V_gene | ENSG0000C |
| 22105343 | 22105846 | 1 | TR_V_gene | ENSG0000C |
| 22112347 | 22113031 | 1 | TR_V_gene | ENSG0000C |
| 22123318 | 22124285 | 1 | TR_V_gene | ENSG0000C |
| 22132553 | 22133034 | 1 | TR_V_gene | ENSG0000C |
| 22147995 | 22148633 | 1 | TR_V_gene | ENSG0000C |
| 22155079 | 22155638 | 1 | TR_V_pseudogene | ENSG0000C |
| 22163238 | 22163870 | 1 | TR_V_gene | ENSG0000C |
| 22168429 | 22168988 | 1 | TR_V_gene | ENSG0000C |
| 22177273 | 22177864 | 1 | TR_V_pseudogene | ENSG0000C |
| 22185562 | 22186057 | 1 | TR_V_pseudogene | ENSG0000C |
| 22190158 | 22190713 | 1 | TR_V_pseudogene | ENSG0000C |
| 22197446 | 22482959 | -1 | lncRNA | ENSG0000C |
| 22202583 | 22203368 | 1 | TR_V_gene | ENSG0000C |
| 22207522 | 22208129 | 1 | TR_V_gene | ENSG0000C |
| 22221896 | 22222475 | 1 | TR_V_pseudogene | ENSG0000C |
| 22226746 | 22227254 | 1 | TR_V_gene | ENSG0000C |
| 22265749 | 22266264 | 1 | TR_V_pseudogene | ENSG0000C |
| 22271968 | 22272563 | 1 | TR_V_gene | ENSG0000C |
| 22281105 | 22281748 | 1 | TR_V_gene | ENSG0000C |
| 22304054 | 22304553 | 1 | TR_V_gene | ENSG0000C |
| 22314490 | 22314919 | 1 | TR_V_gene | ENSG0000C |
| 22320188 | 22320691 | 1 | TR_V_gene | ENSG0000C |
| 22332922 | 22334685 | -1 | processed_pseudoge | ENSG0000C |
| 22415362 | 22418657 | 1 | lncRNA | ENSG0000C |
| 22422371 | 22423042 | 1 | TR_V_gene | ENSG0000C |
| 22438547 | 22438554 | 1 | TR_D_gene | ENSG0000C |
| 22439007 | 22439015 | 1 | TR_D_gene | ENSG0000C |
| 22449113 | 22449125 | 1 | TR_D_gene | ENSG0000C |
| 22450089 | 22450139 | 1 | TR_J_gene | ENSG0000C |
| 22455249 | 22455296 | 1 | TR_J_gene | ENSG0000C |
| 22456689 | 22456742 | 1 | TR_J_gene | ENSG0000C |
| 22459098 | 22459156 | 1 | TR_J_gene | ENSG0000C |

| | | | | |
|---|---|---|---|---|
| 22462932 | 22465787 | 1 | TR_C_gene | ENSG00000 |
| 22469041 | 22469698 | -1 | TR_V_gene | ENSG00000 |
| 22475316 | 22475375 | 1 | TR_J_gene | ENSG00000 |
| 22476306 | 22476362 | 1 | TR_J_pseudogene | ENSG00000 |
| 22476553 | 22476606 | 1 | TR_J_gene | ENSG00000 |
| 22477707 | 22477769 | 1 | TR_J_gene | ENSG00000 |
| 22478872 | 22478934 | 1 | TR_J_gene | ENSG00000 |
| 22479521 | 22479582 | 1 | TR_J_gene | ENSG00000 |
| 22481697 | 22481753 | 1 | TR_J_pseudogene | ENSG00000 |
| 22482287 | 22482346 | 1 | TR_J_gene | ENSG00000 |
| 22483004 | 22483069 | 1 | TR_J_gene | ENSG00000 |
| 22486228 | 22486296 | 1 | TR_J_gene | ENSG00000 |
| 22487183 | 22487245 | 1 | TR_J_pseudogene | ENSG00000 |
| 22488593 | 22488652 | 1 | TR_J_gene | ENSG00000 |
| 22489488 | 22489543 | 1 | TR_J_gene | ENSG00000 |
| 22490491 | 22490553 | 1 | TR_J_gene | ENSG00000 |
| 22492851 | 22492907 | 1 | TR_J_gene | ENSG00000 |
| 22493403 | 22493465 | 1 | TR_J_gene | ENSG00000 |
| 22493925 | 22493990 | 1 | TR_J_gene | ENSG00000 |
| 22494821 | 22494883 | 1 | TR_J_gene | ENSG00000 |
| 22495913 | 22495966 | 1 | TR_J_gene | ENSG00000 |
| 22496887 | 22496952 | 1 | TR_J_gene | ENSG00000 |
| 22497657 | 22497718 | 1 | TR_J_gene | ENSG00000 |
| 22499689 | 22499749 | 1 | TR_J_gene | ENSG00000 |
| 22501601 | 22501663 | 1 | TR_J_gene | ENSG00000 |
| 22502231 | 22502292 | 1 | TR_J_gene | ENSG00000 |
| 22503750 | 22503814 | 1 | TR_J_gene | ENSG00000 |
| 22505110 | 22505167 | 1 | TR_J_gene | ENSG00000 |
| 22506644 | 22506702 | 1 | TR_J_gene | ENSG00000 |
| 22507666 | 22507723 | 1 | TR_J_gene | ENSG00000 |
| 22508602 | 22508658 | 1 | TR_J_gene | ENSG00000 |
| 22509341 | 22509406 | 1 | TR_J_gene | ENSG00000 |
| 22510968 | 22511024 | 1 | TR_J_gene | ENSG00000 |
| 22512852 | 22512908 | 1 | TR_J_gene | ENSG00000 |
| 22513939 | 22513998 | 1 | TR_J_gene | ENSG00000 |
| 22515623 | 22515688 | 1 | TR_J_gene | ENSG00000 |
| 22516273 | 22516331 | 1 | TR_J_gene | ENSG00000 |
| 22518446 | 22518505 | 1 | TR_J_gene | ENSG00000 |
| 22518812 | 22518871 | 1 | TR_J_gene | ENSG00000 |
| 22519969 | 22520031 | 1 | TR_J_gene | ENSG00000 |
| 22520416 | 22520478 | 1 | TR_J_gene | ENSG00000 |
| 22522040 | 22522102 | 1 | TR_J_gene | ENSG00000 |
| 22523600 | 22523654 | 1 | TR_J_gene | ENSG00000 |
| 22524325 | 22524381 | 1 | TR_J_gene | ENSG00000 |
| 22525263 | 22525322 | 1 | TR_J_gene | ENSG00000 |
| 22525650 | 22525715 | 1 | TR_J_gene | ENSG00000 |
| 22526844 | 22526906 | 1 | TR_J_gene | ENSG00000 |
| 22528527 | 22528586 | 1 | TR_J_gene | ENSG00000 |
| 22530327 | 22530378 | 1 | TR_J_gene | ENSG00000 |
| 22531076 | 22531138 | 1 | TR_J_gene | ENSG00000 |

| | | | | |
|---|---|---|---|---|
| 22531939 | 22531998 | 1 | TR_J_gene | ENSG0000C |
| 22532502 | 22532561 | 1 | TR_J_gene | ENSG0000C |
| 22533497 | 22533560 | 1 | TR_J_gene | ENSG0000C |
| 22535554 | 22535614 | 1 | TR_J_gene | ENSG0000C |
| 22536145 | 22536204 | 1 | TR_J_pseudogene | ENSG0000C |
| 22537622 | 22537680 | 1 | TR_J_gene | ENSG0000C |
| 22539073 | 22539134 | 1 | TR_J_gene | ENSG0000C |
| 22540247 | 22540306 | 1 | TR_J_gene | ENSG0000C |
| 22542199 | 22542261 | 1 | TR_J_gene | ENSG0000C |
| 22543179 | 22543240 | 1 | TR_J_gene | ENSG0000C |
| 22544071 | 22544136 | 1 | TR_J_gene | ENSG0000C |
| 22545037 | 22545098 | 1 | TR_J_gene | ENSG0000C |
| 22547506 | 22552156 | 1 | TR_C_gene | ENSG0000C |
| 22556311 | 22557062 | -1 | lncRNA | ENSG0000C |
| 22556640 | 22559037 | 1 | lncRNA | ENSG0000C |
| 22564907 | 22589224 | -1 | protein_coding | ENSG0000C |
| 22595808 | 22598946 | -1 | lncRNA | ENSG0000C |
| 22598237 | 22613215 | 1 | protein_coding | ENSG0000C |
| 22630929 | 22644352 | -1 | protein_coding | ENSG0000C |
| 22694098 | 22694535 | -1 | processed_pseudoge | ENSG0000C |
| 22701476 | 22766562 | -1 | lncRNA | ENSG0000C |
| 22702143 | 22703065 | 1 | unprocessed_pseudc | ENSG0000C |
| 22723476 | 22726572 | -1 | transcribed_unproce | ENSG0000C |
| 22756761 | 22756829 | -1 | snoRNA | ENSG0000C |
| 22766522 | 22773042 | 1 | protein_coding | ENSG0000C |
| 22773222 | 22829820 | -1 | protein_coding | ENSG0000C |
| 22829879 | 22835037 | 1 | protein_coding | ENSG0000C |
| 22836560 | 22849027 | 1 | protein_coding | ENSG0000C |
| 22852616 | 22852728 | -1 | misc_RNA | ENSG0000C |
| 22867818 | 22867917 | 1 | misc_RNA | ENSG0000C |
| 22871740 | 22881713 | 1 | protein_coding | ENSG0000C |
| 22883222 | 22887678 | 1 | protein_coding | ENSG0000C |
| 22893204 | 22919182 | -1 | protein_coding | ENSG0000C |
| 22918947 | 22926900 | 1 | lncRNA | ENSG0000C |
| 22920525 | 22929408 | -1 | protein_coding | ENSG0000C |
| 22929607 | 22956374 | 1 | lncRNA | ENSG0000C |
| 22946228 | 22957161 | -1 | protein_coding | ENSG0000C |
| 22946270 | 22982258 | -1 | protein_coding | ENSG0000C |
| 22956950 | 22957029 | -1 | miRNA | ENSG0000C |
| 22971177 | 22982551 | -1 | protein_coding | ENSG0000C |
| 22982698 | 22999078 | 1 | lncRNA | ENSG0000C |
| 22985894 | 23010166 | -1 | protein_coding | ENSG0000C |
| 23005499 | 23007123 | -1 | TEC | ENSG0000C |
| 23016543 | 23035230 | -1 | protein_coding | ENSG0000C |
| 23023083 | 23024217 | -1 | TEC | ENSG0000C |
| 23042212 | 23044060 | 1 | protein_coding | ENSG0000C |
| 23047062 | 23057538 | -1 | protein_coding | ENSG0000C |
| 23058564 | 23095614 | -1 | protein_coding | ENSG0000C |
| 23095505 | 23100456 | 1 | protein_coding | ENSG0000C |
| 23099062 | 23104989 | -1 | protein_coding | ENSG0000C |

| | | | | |
|---|---|---|---|---|
| 23117304 | 23119616 | -1 | protein_coding | ENSG00000( |
| 23125295 | 23183674 | -1 | protein_coding | ENSG00000( |
| 23156309 | 23156414 | -1 | snRNA | ENSG00000( |
| 23185316 | 23273477 | 1 | protein_coding | ENSG00000( |
| 23229961 | 23230116 | -1 | processed_pseudoge | ENSG00000( |
| 23259460 | 23259566 | -1 | snRNA | ENSG00000( |
| 23272422 | 23299447 | -1 | protein_coding | ENSG00000( |
| 23293852 | 23295798 | 1 | transcribed_unproce | ENSG00000( |
| 23295652 | 23302859 | -1 | protein_coding | ENSG00000( |
| 23298790 | 23311759 | 1 | protein_coding | ENSG00000( |
| 23306835 | 23325369 | 1 | protein_coding | ENSG00000( |
| 23321289 | 23326185 | 1 | protein_coding | ENSG00000( |
| 23346306 | 23352912 | -1 | protein_coding | ENSG00000( |
| 23356403 | 23365752 | -1 | protein_coding | ENSG00000( |
| 23356406 | 23357003 | 1 | lncRNA | ENSG00000( |
| 23372809 | 23376403 | 1 | protein_coding | ENSG00000( |
| 23376773 | 23379772 | 1 | protein_coding | ENSG00000( |
| 23381982 | 23408273 | -1 | protein_coding | ENSG00000( |
| 23388596 | 23388666 | -1 | miRNA | ENSG00000( |
| 23412740 | 23435660 | -1 | protein_coding | ENSG00000( |
| 23415339 | 23415686 | 1 | lncRNA | ENSG00000( |
| 23417987 | 23418063 | -1 | miRNA | ENSG00000( |
| 23469689 | 23509862 | 1 | protein_coding | ENSG00000( |
| 23511760 | 23560778 | 1 | lncRNA | ENSG00000( |
| 23520857 | 23556192 | -1 | protein_coding | ENSG00000( |
| 23530682 | 23532367 | 1 | lncRNA | ENSG00000( |
| 23555988 | 23560271 | 1 | protein_coding | ENSG00000( |
| 23559565 | 23568070 | -1 | protein_coding | ENSG00000( |
| 23561097 | 23568073 | 1 | lncRNA | ENSG00000( |
| 23568035 | 23578790 | -1 | protein_coding | ENSG00000( |
| 23619201 | 23620012 | 1 | lncRNA | ENSG00000( |
| 23630115 | 23645639 | 1 | protein_coding | ENSG00000( |
| 23699797 | 23729757 | -1 | lncRNA | ENSG00000( |
| 23749236 | 23751150 | 1 | processed_pseudoge | ENSG00000( |
| 23767662 | 23768998 | 1 | unprocessed_pseudc | ENSG00000( |
| 23791930 | 23792236 | 1 | misc_RNA | ENSG00000( |
| 23817281 | 23827396 | -1 | unprocessed_pseudc | ENSG00000( |
| 23875938 | 23882428 | -1 | lncRNA | ENSG00000( |
| 23922247 | 23934568 | -1 | lncRNA | ENSG00000( |
| 23938219 | 24052555 | -1 | lncRNA | ENSG00000( |
| 23953734 | 23969279 | 1 | protein_coding | ENSG00000( |
| 23969874 | 24006408 | 1 | protein_coding | ENSG00000( |
| 23969931 | 24051377 | 1 | lncRNA | ENSG00000( |
| 24007185 | 24027782 | 1 | lncRNA | ENSG00000( |
| 24036453 | 24051028 | 1 | transcribed_unproce | ENSG00000( |
| 24052009 | 24069729 | 1 | protein_coding | ENSG00000( |
| 24070837 | 24078100 | 1 | protein_coding | ENSG00000( |
| 24080107 | 24115014 | -1 | protein_coding | ENSG00000( |
| 24094053 | 24110598 | 1 | protein_coding | ENSG00000( |
| 24114195 | 24125242 | 1 | protein_coding | ENSG00000( |

| | | | | |
|---|---|---|---|---|
| 24120956 | 24131829 | 1 | protein_coding | ENSG0000C |
| 24131275 | 24132849 | 1 | protein_coding | ENSG0000C |
| 24136163 | 24138967 | 1 | protein_coding | ENSG0000C |
| 24138959 | 24141591 | -1 | protein_coding | ENSG0000C |
| 24139445 | 24140444 | 1 | lncRNA | ENSG0000C |
| 24143362 | 24147570 | -1 | protein_coding | ENSG0000C |
| 24143489 | 24143565 | -1 | miRNA | ENSG0000C |
| 24146683 | 24160660 | 1 | protein_coding | ENSG0000C |
| 24147548 | 24166452 | 1 | protein_coding | ENSG0000C |
| 24158603 | 24158722 | 1 | rRNA_pseudogene | ENSG0000C |
| 24161265 | 24166565 | 1 | protein_coding | ENSG0000C |
| 24171853 | 24180257 | 1 | protein_coding | ENSG0000C |
| 24180219 | 24188869 | -1 | protein_coding | ENSG0000C |
| 24180395 | 24190416 | -1 | protein_coding | ENSG0000C |
| 24188914 | 24193106 | 1 | lncRNA | ENSG0000C |
| 24189149 | 24195687 | -1 | protein_coding | ENSG0000C |
| 24189157 | 24213473 | -1 | protein_coding | ENSG0000C |
| 24198433 | 24199090 | 1 | lncRNA | ENSG0000C |
| 24201612 | 24202811 | -1 | lncRNA | ENSG0000C |
| 24205696 | 24208362 | 1 | protein_coding | ENSG0000C |
| 24209583 | 24213869 | -1 | protein_coding | ENSG0000C |
| 24209646 | 24215987 | -1 | lncRNA | ENSG0000C |
| 24213943 | 24216070 | -1 | protein_coding | ENSG0000C |
| 24213955 | 24232352 | -1 | protein_coding | ENSG0000C |
| 24216857 | 24232367 | -1 | protein_coding | ENSG0000C |
| 24232422 | 24239242 | 1 | protein_coding | ENSG0000C |
| 24239643 | 24242674 | -1 | protein_coding | ENSG0000C |
| 24249114 | 24264432 | -1 | protein_coding | ENSG0000C |
| 24265538 | 24271739 | -1 | protein_coding | ENSG0000C |
| 24271213 | 24298846 | 1 | lncRNA | ENSG0000C |
| 24271726 | 24279089 | 1 | lncRNA | ENSG0000C |
| 24290598 | 24299780 | -1 | protein_coding | ENSG0000C |
| 24299850 | 24309124 | 1 | protein_coding | ENSG0000C |
| 24305096 | 24311430 | -1 | protein_coding | ENSG0000C |
| 24305734 | 24312053 | 1 | protein_coding | ENSG0000C |
| 24311450 | 24318036 | 1 | protein_coding | ENSG0000C |
| 24318349 | 24335093 | -1 | protein_coding | ENSG0000C |
| 24332310 | 24337382 | -1 | lncRNA | ENSG0000C |
| 24336025 | 24340022 | -1 | protein_coding | ENSG0000C |
| 24352057 | 24352157 | -1 | snRNA | ENSG0000C |
| 24365673 | 24379604 | 1 | protein_coding | ENSG0000C |
| 24399003 | 24419283 | 1 | protein_coding | ENSG0000C |
| 24426545 | 24430954 | -1 | protein_coding | ENSG0000C |
| 24429286 | 24441843 | 1 | protein_coding | ENSG0000C |
| 24439766 | 24442905 | -1 | protein_coding | ENSG0000C |
| 24443016 | 24508688 | 1 | lncRNA | ENSG0000C |
| 24505353 | 24508265 | -1 | protein_coding | ENSG0000C |
| 24573518 | 24576250 | -1 | protein_coding | ENSG0000C |
| 24595723 | 24657774 | 1 | lncRNA | ENSG0000C |
| 24606480 | 24609699 | -1 | protein_coding | ENSG0000C |

| | | | | |
|---|---|---|---|---|
| 24630954 | 24634267 | -1 | protein_coding | ENSG00000 |
| 24809656 | 25050297 | -1 | protein_coding | ENSG00000 |
| 24818385 | 24820157 | 1 | lncRNA | ENSG00000 |
| 24861172 | 24865978 | 1 | lncRNA | ENSG00000 |
| 25124467 | 25156314 | -1 | lncRNA | ENSG00000 |
| 25267064 | 25267336 | -1 | processed_pseudoge | ENSG00000 |
| 25357764 | 25358689 | 1 | unprocessed_pseudo | ENSG00000 |
| 25431193 | 25432234 | -1 | processed_pseudoge | ENSG00000 |
| 25647304 | 26143305 | -1 | lncRNA | ENSG00000 |
| 25806648 | 25806712 | -1 | snoRNA | ENSG00000 |
| 25820366 | 25822741 | 1 | TEC | ENSG00000 |
| 25916015 | 25998805 | 1 | lncRNA | ENSG00000 |
| 26208036 | 26208424 | 1 | processed_pseudoge | ENSG00000 |
| 26443090 | 26598033 | -1 | protein_coding | ENSG00000 |
| 26592747 | 26594517 | -1 | lncRNA | ENSG00000 |
| 26598369 | 26806467 | 1 | lncRNA | ENSG00000 |
| 26599755 | 27295516 | 1 | lncRNA | ENSG00000 |
| 26775495 | 26822107 | -1 | lncRNA | ENSG00000 |
| 26809340 | 26836458 | 1 | lncRNA | ENSG00000 |
| 26831401 | 26873036 | -1 | lncRNA | ENSG00000 |
| 26873092 | 26918594 | 1 | lncRNA | ENSG00000 |
| 26908642 | 26908725 | 1 | miRNA | ENSG00000 |
| 27258717 | 27845286 | -1 | transcribed_process | ENSG00000 |
| 27612577 | 27639636 | 1 | lncRNA | ENSG00000 |
| 27633205 | 27633278 | -1 | miRNA | ENSG00000 |
| 28264390 | 28265974 | 1 | transcribed_process | ENSG00000 |
| 28318141 | 28418612 | 1 | lncRNA | ENSG00000 |
| 28489910 | 28490362 | 1 | processed_pseudoge | ENSG00000 |
| 28592223 | 28613623 | -1 | lncRNA | ENSG00000 |
| 28593779 | 28593919 | 1 | processed_pseudoge | ENSG00000 |
| 28673241 | 28673727 | -1 | processed_pseudoge | ENSG00000 |
| 28726675 | 28765292 | -1 | lncRNA | ENSG00000 |
| 28760330 | 28770277 | 1 | protein_coding | ENSG00000 |
| 28771676 | 28823817 | 1 | lncRNA | ENSG00000 |
| 28782909 | 28785798 | -1 | lncRNA | ENSG00000 |
| 28800084 | 28830204 | -1 | lncRNA | ENSG00000 |
| 28829881 | 28968400 | 1 | lncRNA | ENSG00000 |
| 28857384 | 28858701 | -1 | processed_pseudoge | ENSG00000 |
| 28875404 | 29064993 | -1 | lncRNA | ENSG00000 |
| 28901236 | 28901347 | -1 | misc_RNA | ENSG00000 |
| 28936889 | 29372406 | 1 | lncRNA | ENSG00000 |
| 28982303 | 28982410 | -1 | snRNA | ENSG00000 |
| 29210986 | 29392048 | -1 | lncRNA | ENSG00000 |
| 29390074 | 29436412 | 1 | lncRNA | ENSG00000 |
| 29415856 | 29500460 | -1 | lncRNA | ENSG00000 |
| 29476784 | 29476912 | 1 | snRNA | ENSG00000 |
| 29576479 | 30191898 | -1 | protein_coding | ENSG00000 |
| 29652809 | 29657916 | 1 | lncRNA | ENSG00000 |
| 29811916 | 29812018 | 1 | snRNA | ENSG00000 |
| 29928445 | 29932756 | 1 | lncRNA | ENSG00000 |

| | | | | |
|---|---|---|---|---|
| 29940745 | 29980212 | 1 | lncRNA | ENSG0000C |
| 29952397 | 30297043 | -1 | lncRNA | ENSG0000C |
| 30055210 | 30055954 | 1 | processed_pseudoge | ENSG0000C |
| 30370108 | 30373115 | 1 | TEC | ENSG0000C |
| 30437992 | 30573629 | -1 | lncRNA | ENSG0000C |
| 30519161 | 30519318 | 1 | misc_RNA | ENSG0000C |
| 30541776 | 30542188 | -1 | processed_pseudoge | ENSG0000C |
| 30559158 | 30620064 | 1 | protein_coding | ENSG0000C |
| 30622311 | 30735850 | 1 | protein_coding | ENSG0000C |
| 30683045 | 30683598 | -1 | processed_pseudoge | ENSG0000C |
| 30734926 | 30822702 | -1 | lncRNA | ENSG0000C |
| 30791098 | 30791489 | 1 | processed_pseudoge | ENSG0000C |
| 30819714 | 30819914 | 1 | processed_pseudoge | ENSG0000C |
| 30821419 | 30821843 | 1 | processed_pseudoge | ENSG0000C |
| 30874514 | 30895065 | 1 | protein_coding | ENSG0000C |
| 30876179 | 30889808 | -1 | lncRNA | ENSG0000C |
| 30893799 | 31026401 | -1 | protein_coding | ENSG0000C |
| 30964347 | 30964622 | 1 | processed_pseudoge | ENSG0000C |
| 31014646 | 31014742 | -1 | miRNA | ENSG0000C |
| 31018446 | 31018796 | 1 | processed_pseudoge | ENSG0000C |
| 31025106 | 31130996 | 1 | protein_coding | ENSG0000C |
| 31100112 | 31207804 | -1 | protein_coding | ENSG0000C |
| 31111569 | 31112239 | -1 | processed_pseudoge | ENSG0000C |
| 31142939 | 31143048 | 1 | snRNA | ENSG0000C |
| 31244761 | 31244861 | 1 | misc_RNA | ENSG0000C |
| 31248628 | 31302739 | 1 | lncRNA | ENSG0000C |
| 31258608 | 31258709 | -1 | misc_RNA | ENSG0000C |
| 31291788 | 31420550 | -1 | protein_coding | ENSG0000C |
| 31320224 | 31320634 | -1 | processed_pseudoge | ENSG0000C |
| 31334312 | 31457441 | -1 | protein_coding | ENSG0000C |
| 31420286 | 31452883 | 1 | lncRNA | ENSG0000C |
| 31446036 | 31457506 | -1 | protein_coding | ENSG0000C |
| 31462795 | 31463371 | -1 | processed_pseudoge | ENSG0000C |
| 31482875 | 31488039 | -1 | protein_coding | ENSG0000C |
| 31489956 | 31861224 | 1 | protein_coding | ENSG0000C |
| 31553358 | 31558498 | -1 | lncRNA | ENSG0000C |
| 31558507 | 31561334 | -1 | lncRNA | ENSG0000C |
| 31662039 | 31662143 | -1 | snRNA | ENSG0000C |
| 31686692 | 31686798 | -1 | snRNA | ENSG0000C |
| 31925801 | 31930146 | -1 | lncRNA | ENSG0000C |
| 31944853 | 31950382 | 1 | lncRNA | ENSG0000C |
| 31964550 | 31965095 | 1 | transcribed_unproce | ENSG0000C |
| 32006902 | 32018685 | 1 | lncRNA | ENSG0000C |
| 32074946 | 32076793 | -1 | lncRNA | ENSG0000C |
| 32076114 | 32159728 | 1 | protein_coding | ENSG0000C |
| 32149176 | 32158008 | -1 | lncRNA | ENSG0000C |
| 32200433 | 32201736 | -1 | lncRNA | ENSG0000C |
| 32202045 | 32202151 | 1 | snRNA | ENSG0000C |
| 32203163 | 32203269 | -1 | snRNA | ENSG0000C |
| 32241524 | 32243093 | -1 | processed_pseudoge | ENSG0000C |

| | | | | |
|---|---|---|---|---|
| 32271857 | 32272301 | -1 | processed_pseudoge | ENSG00000( |
| 32329298 | 32837684 | 1 | protein_coding | ENSG00000( |
| 32373337 | 32417974 | -1 | lncRNA | ENSG00000( |
| 32442341 | 32442639 | 1 | misc_RNA | ENSG00000( |
| 32484098 | 32484800 | -1 | unprocessed_pseudc | ENSG00000( |
| 32879246 | 32882830 | 1 | TEC | ENSG00000( |
| 32934933 | 33804176 | 1 | protein_coding | ENSG00000( |
| 33597057 | 33600989 | -1 | lncRNA | ENSG00000( |
| 33677360 | 33682886 | -1 | lncRNA | ENSG00000( |
| 33924227 | 34462774 | -1 | protein_coding | ENSG00000( |
| 34040097 | 34059300 | 1 | lncRNA | ENSG00000( |
| 34416687 | 34417602 | -1 | processed_pseudoge | ENSG00000( |
| 34432788 | 34462240 | -1 | protein_coding | ENSG00000( |
| 34485979 | 34486447 | 1 | processed_pseudoge | ENSG00000( |
| 34515938 | 34539711 | -1 | protein_coding | ENSG00000( |
| 34541044 | 34541714 | 1 | TEC | ENSG00000( |
| 34541740 | 34546329 | -1 | lncRNA | ENSG00000( |
| 34546714 | 34546877 | 1 | snRNA | ENSG00000( |
| 34556226 | 34556389 | -1 | snRNA | ENSG00000( |
| 34556774 | 34561298 | 1 | lncRNA | ENSG00000( |
| 34561093 | 34630160 | -1 | protein_coding | ENSG00000( |
| 34568689 | 34569129 | -1 | processed_pseudoge | ENSG00000( |
| 34659564 | 34659878 | -1 | misc_RNA | ENSG00000( |
| 34675891 | 34675994 | 1 | snRNA | ENSG00000( |
| 34694170 | 34694644 | 1 | processed_pseudoge | ENSG00000( |
| 34709113 | 34714823 | -1 | protein_coding | ENSG00000( |
| 34740336 | 34740833 | -1 | processed_pseudoge | ENSG00000( |
| 34752731 | 34875647 | -1 | protein_coding | ENSG00000( |
| 34845300 | 34845360 | -1 | snRNA | ENSG00000( |
| 34874343 | 34876459 | 1 | lncRNA | ENSG00000( |
| 34920858 | 34982532 | -1 | transcribed_unitary_ | ENSG00000( |
| 34920865 | 34921006 | 1 | processed_pseudoge | ENSG00000( |
| 34939324 | 34940332 | 1 | processed_pseudoge | ENSG00000( |
| 34981957 | 35029567 | 1 | protein_coding | ENSG00000( |
| 35044907 | 35113130 | 1 | protein_coding | ENSG00000( |
| 35085467 | 35122517 | -1 | protein_coding | ENSG00000( |
| 35086035 | 35088059 | 1 | TEC | ENSG00000( |
| 35121846 | 35277622 | 1 | protein_coding | ENSG00000( |
| 35122549 | 35317474 | 1 | protein_coding | ENSG00000( |
| 35144021 | 35144480 | -1 | processed_pseudoge | ENSG00000( |
| 35157904 | 35159099 | -1 | processed_pseudoge | ENSG00000( |
| 35219023 | 35219283 | -1 | processed_pseudoge | ENSG00000( |
| 35222174 | 35222746 | 1 | processed_pseudoge | ENSG00000( |
| 35235180 | 35235651 | -1 | processed_pseudoge | ENSG00000( |
| 35262462 | 35263036 | 1 | processed_pseudoge | ENSG00000( |
| 35278633 | 35317493 | 1 | protein_coding | ENSG00000( |
| 35362232 | 35363628 | -1 | lncRNA | ENSG00000( |
| 35388121 | 35389069 | -1 | processed_pseudoge | ENSG00000( |
| 35401511 | 35404749 | -1 | protein_coding | ENSG00000( |
| 35430260 | 35430631 | 1 | processed_pseudoge | ENSG00000( |

| | | | | |
|---|---|---|---|---|
| 35447003 | 35447625 | -1 | lncRNA | ENSG0000C |
| 35468272 | 35469330 | 1 | processed_pseudoge | ENSG0000C |
| 35511825 | 35513089 | -1 | processed_pseudoge | ENSG0000C |
| 35534164 | 35537054 | 1 | protein_coding | ENSG0000C |
| 35538352 | 35809304 | -1 | protein_coding | ENSG0000C |
| 35756327 | 35756463 | 1 | snoRNA | ENSG0000C |
| 35765795 | 35767110 | 1 | processed_pseudoge | ENSG0000C |
| 35819224 | 35826765 | -1 | lncRNA | ENSG0000C |
| 35819837 | 35820220 | 1 | processed_pseudoge | ENSG0000C |
| 35826338 | 35932325 | 1 | protein_coding | ENSG0000C |
| 35873857 | 35875303 | 1 | lncRNA | ENSG0000C |
| 35881001 | 35881709 | 1 | processed_pseudoge | ENSG0000C |
| 35897936 | 36068261 | 1 | lncRNA | ENSG0000C |
| 35947556 | 35950044 | -1 | lncRNA | ENSG0000C |
| 36054197 | 36055313 | -1 | processed_pseudoge | ENSG0000C |
| 36061026 | 36067190 | -1 | lncRNA | ENSG0000C |
| 36070427 | 36165288 | 1 | lncRNA | ENSG0000C |
| 36136108 | 36176468 | -1 | lncRNA | ENSG0000C |
| 36213200 | 36235608 | 1 | lncRNA | ENSG0000C |
| 36231555 | 36231774 | -1 | misc_RNA | ENSG0000C |
| 36298564 | 36320637 | -1 | protein_coding | ENSG0000C |
| 36320753 | 36656163 | 1 | lncRNA | ENSG0000C |
| 36371165 | 36372383 | 1 | transcribed_unproce | ENSG0000C |
| 36405214 | 36405274 | 1 | snRNA | ENSG0000C |
| 36473207 | 36519521 | -1 | lncRNA | ENSG0000C |
| 36473288 | 36513829 | -1 | protein_coding | ENSG0000C |
| 36516392 | 36521149 | -1 | protein_coding | ENSG0000C |
| 36519278 | 36523016 | 1 | lncRNA | ENSG0000C |
| 36533506 | 36534538 | -1 | unprocessed_pseudc | ENSG0000C |
| 36577543 | 36578005 | -1 | processed_pseudoge | ENSG0000C |
| 36580004 | 36582614 | -1 | protein_coding | ENSG0000C |
| 36593839 | 36594147 | -1 | misc_RNA | ENSG0000C |
| 36647083 | 36658801 | -1 | lncRNA | ENSG0000C |
| 36657568 | 36679715 | 1 | protein_coding | ENSG0000C |
| 36677921 | 37172606 | -1 | protein_coding | ENSG0000C |
| 36788644 | 36807163 | 1 | lncRNA | ENSG0000C |
| 36808871 | 36828729 | 1 | lncRNA | ENSG0000C |
| 36952309 | 36952391 | -1 | miRNA | ENSG0000C |
| 37097062 | 37098563 | -1 | lncRNA | ENSG0000C |
| 37171888 | 37173811 | 1 | lncRNA | ENSG0000C |
| 37187219 | 37187314 | 1 | snRNA | ENSG0000C |
| 37197913 | 37552361 | 1 | protein_coding | ENSG0000C |
| 37434469 | 37434573 | -1 | snRNA | ENSG0000C |
| 37556158 | 37567095 | -1 | lncRNA | ENSG0000C |
| 37564047 | 37579125 | 1 | lncRNA | ENSG0000C |
| 37589552 | 37596059 | -1 | protein_coding | ENSG0000C |
| 37595847 | 38041442 | 1 | protein_coding | ENSG0000C |
| 37720425 | 37720561 | 1 | snoRNA | ENSG0000C |
| 37827798 | 37827904 | -1 | snRNA | ENSG0000C |
| 37896060 | 37902372 | -1 | lncRNA | ENSG0000C |

| | | | | |
|---|---|---|---|---|
| 38003113 | 38004847 | -1 | lncRNA | ENSG00000 |
| 38034287 | 38194281 | -1 | lncRNA | ENSG00000 |
| 38190983 | 38202923 | 1 | lncRNA | ENSG00000 |
| 38207904 | 38213067 | 1 | protein_coding | ENSG00000 |
| 38254000 | 38256093 | -1 | protein_coding | ENSG00000 |
| 38256202 | 38371338 | 1 | lncRNA | ENSG00000 |
| 38605255 | 38606098 | 1 | processed_pseudoge | ENSG00000 |
| 38727275 | 38728481 | 1 | processed_pseudoge | ENSG00000 |
| 38738155 | 38948273 | -1 | lncRNA | ENSG00000 |
| 38835389 | 38835788 | 1 | processed_pseudoge | ENSG00000 |
| 38973399 | 38974085 | -1 | processed_pseudoge | ENSG00000 |
| 38987281 | 38987383 | 1 | misc_RNA | ENSG00000 |
| 39031919 | 39109646 | -1 | protein_coding | ENSG00000 |
| 39090901 | 39091101 | 1 | processed_pseudoge | ENSG00000 |
| 39103140 | 39103812 | 1 | lncRNA | ENSG00000 |
| 39108342 | 39108707 | 1 | processed_pseudoge | ENSG00000 |
| 39114223 | 39136973 | 1 | protein_coding | ENSG00000 |
| 39147811 | 39170532 | -1 | protein_coding | ENSG00000 |
| 39147833 | 39151363 | 1 | lncRNA | ENSG00000 |
| 39156742 | 39157852 | 1 | processed_pseudoge | ENSG00000 |
| 39174885 | 39175880 | -1 | lncRNA | ENSG00000 |
| 39175183 | 39183220 | 1 | protein_coding | ENSG00000 |
| 39211781 | 39212255 | 1 | processed_pseudoge | ENSG00000 |
| 39215065 | 39216335 | 1 | processed_pseudoge | ENSG00000 |
| 39230231 | 39388513 | 1 | protein_coding | ENSG00000 |
| 39265703 | 39267061 | -1 | lncRNA | ENSG00000 |
| 39370279 | 39370756 | -1 | processed_pseudoge | ENSG00000 |
| 39385404 | 39387102 | -1 | processed_pseudoge | ENSG00000 |
| 39397669 | 39432500 | -1 | protein_coding | ENSG00000 |
| 39432599 | 40390547 | 1 | lncRNA | ENSG00000 |
| 40267283 | 40348331 | -1 | lncRNA | ENSG00000 |
| 40630006 | 40905513 | 1 | lncRNA | ENSG00000 |
| 40954693 | 41149309 | 1 | lncRNA | ENSG00000 |
| 41296286 | 41298734 | 1 | processed_pseudoge | ENSG00000 |
| 41514972 | 41515350 | -1 | TEC | ENSG00000 |
| 41555662 | 41556055 | -1 | processed_pseudoge | ENSG00000 |
| 41583657 | 41604988 | -1 | lncRNA | ENSG00000 |
| 41594463 | 41594591 | 1 | snoRNA | ENSG00000 |
| 41607570 | 41904549 | 1 | protein_coding | ENSG00000 |
| 41742639 | 41742963 | 1 | processed_pseudoge | ENSG00000 |
| 42259731 | 42268586 | -1 | lncRNA | ENSG00000 |
| 42362983 | 42703655 | 1 | lncRNA | ENSG00000 |
| 42412573 | 42528888 | 1 | lncRNA | ENSG00000 |
| 42608868 | 42609577 | 1 | transcribed_processe | ENSG00000 |
| 42657763 | 42657918 | -1 | processed_pseudoge | ENSG00000 |
| 42807378 | 42808112 | -1 | processed_pseudoge | ENSG00000 |
| 42989201 | 42989483 | 1 | processed_pseudoge | ENSG00000 |
| 43274824 | 43275697 | 1 | processed_pseudoge | ENSG00000 |
| 43300263 | 43300595 | 1 | processed_pseudoge | ENSG00000 |
| 43308648 | 43314803 | 1 | lncRNA | ENSG00000 |

| | | | | |
|---|---|---|---|---|
| 43540883 | 43541826 | -1 | processed_pseudoge | ENSG00000( |
| 43592142 | 43592262 | -1 | processed_pseudoge | ENSG00000( |
| 43595136 | 43596683 | 1 | processed_pseudoge | ENSG00000( |
| 43990539 | 44386064 | -1 | lncRNA | ENSG00000( |
| 44077534 | 44077884 | -1 | transcribed_processe | ENSG00000( |
| 44120210 | 44120761 | -1 | processed_pseudoge | ENSG00000( |
| 44290997 | 44291718 | -1 | processed_pseudoge | ENSG00000( |
| 44392531 | 44393716 | -1 | lncRNA | ENSG00000( |
| 44444747 | 44480626 | -1 | lncRNA | ENSG00000( |
| 44504149 | 44507283 | -1 | protein_coding | ENSG00000( |
| 44507385 | 44567467 | 1 | lncRNA | ENSG00000( |
| 44763153 | 44782829 | -1 | lncRNA | ENSG00000( |
| 44860316 | 44866247 | -1 | processed_pseudoge | ENSG00000( |
| 44876874 | 44897077 | -1 | lncRNA | ENSG00000( |
| 44887434 | 44887741 | -1 | processed_pseudoge | ENSG00000( |
| 44888556 | 44889041 | -1 | processed_pseudoge | ENSG00000( |
| 44897275 | 44907257 | 1 | protein_coding | ENSG00000( |
| 44898908 | 44911863 | -1 | lncRNA | ENSG00000( |
| 44924319 | 45042322 | -1 | protein_coding | ENSG00000( |
| 44962190 | 45074431 | 1 | protein_coding | ENSG00000( |
| 45079849 | 45080189 | 1 | lncRNA | ENSG00000( |
| 45083045 | 45083977 | -1 | lncRNA | ENSG00000( |
| 45084107 | 45116282 | 1 | protein_coding | ENSG00000( |
| 45088243 | 45088308 | 1 | snoRNA | ENSG00000( |
| 45110883 | 45110968 | 1 | snoRNA | ENSG00000( |
| 45115599 | 45135319 | -1 | protein_coding | ENSG00000( |
| 45135930 | 45200890 | 1 | protein_coding | ENSG00000( |
| 45203190 | 45253540 | -1 | protein_coding | ENSG00000( |
| 45274359 | 45274477 | -1 | snRNA | ENSG00000( |
| 45290036 | 45290386 | 1 | processed_pseudoge | ENSG00000( |
| 45369215 | 45370839 | 1 | lncRNA | ENSG00000( |
| 45377268 | 45389286 | 1 | lncRNA | ENSG00000( |
| 45502742 | 45503140 | -1 | processed_pseudoge | ENSG00000( |
| 45567995 | 45569069 | 1 | lncRNA | ENSG00000( |
| 45706250 | 45715952 | -1 | lncRNA | ENSG00000( |
| 45891087 | 46501903 | 1 | lncRNA | ENSG00000( |
| 46651010 | 46651781 | -1 | protein_coding | ENSG00000( |
| 46840092 | 47675605 | -1 | protein_coding | ENSG00000( |
| 46960423 | 46960546 | 1 | snoRNA | ENSG00000( |
| 46996454 | 46997174 | 1 | processed_pseudoge | ENSG00000( |
| 47200575 | 47201104 | -1 | processed_pseudoge | ENSG00000( |
| 47444106 | 47444494 | -1 | processed_pseudoge | ENSG00000( |
| 47760995 | 47761104 | -1 | miRNA | ENSG00000( |
| 47764954 | 47795302 | -1 | lncRNA | ENSG00000( |
| 47801926 | 47802201 | -1 | processed_pseudoge | ENSG00000( |
| 48045481 | 48046216 | 1 | processed_pseudoge | ENSG00000( |
| 48194174 | 48228216 | 1 | lncRNA | ENSG00000( |
| 48262021 | 48265609 | -1 | lncRNA | ENSG00000( |
| 48319068 | 48319394 | -1 | processed_pseudoge | ENSG00000( |
| 48341226 | 48341332 | -1 | snRNA | ENSG00000( |

| | | | | |
|---|---|---|---|---|
| 48394815 | 48491767 | -1 | lncRNA | ENSG00000( |
| 48398310 | 48399375 | 1 | processed_pseudoge | ENSG00000( |
| 48489883 | 48490434 | 1 | processed_pseudoge | ENSG00000( |
| 48819360 | 48819593 | 1 | processed_pseudoge | ENSG00000( |
| 49057713 | 49077505 | -1 | lncRNA | ENSG00000( |
| 49373966 | 49374383 | 1 | processed_pseudoge | ENSG00000( |
| 49552633 | 49552748 | 1 | rRNA_pseudogene | ENSG00000( |
| 49570984 | 49599164 | -1 | protein_coding | ENSG00000( |
| 49577392 | 49577794 | -1 | processed_pseudoge | ENSG00000( |
| 49586580 | 49586878 | 1 | misc_RNA | ENSG00000( |
| 49592236 | 49592337 | -1 | misc_RNA | ENSG00000( |
| 49598761 | 49614672 | 1 | protein_coding | ENSG00000( |
| 49599994 | 49600572 | 1 | processed_pseudoge | ENSG00000( |
| 49601011 | 49601124 | -1 | unprocessed_pseudc | ENSG00000( |
| 49618530 | 49620626 | -1 | protein_coding | ENSG00000( |
| 49620799 | 49623481 | 1 | protein_coding | ENSG00000( |
| 49620815 | 49623480 | -1 | lncRNA | ENSG00000( |
| 49625174 | 49635230 | -1 | protein_coding | ENSG00000( |
| 49629291 | 49629890 | -1 | lncRNA | ENSG00000( |
| 49643555 | 49688422 | -1 | protein_coding | ENSG00000( |
| 49677369 | 49677893 | 1 | processed_pseudoge | ENSG00000( |
| 49681302 | 49681886 | 1 | processed_pseudoge | ENSG00000( |
| 49684544 | 49684665 | 1 | snRNA | ENSG00000( |
| 49693105 | 49753150 | 1 | protein_coding | ENSG00000( |
| 49707667 | 49708792 | -1 | lncRNA | ENSG00000( |
| 49768130 | 49786385 | 1 | protein_coding | ENSG00000( |
| 49782083 | 49852821 | -1 | protein_coding | ENSG00000( |
| 49839401 | 49839493 | -1 | misc_RNA | ENSG00000( |
| 49839725 | 49839827 | 1 | snRNA | ENSG00000( |
| 49853616 | 49853914 | -1 | misc_RNA | ENSG00000( |
| 49861176 | 49864326 | -1 | lncRNA | ENSG00000( |
| 49862550 | 49862849 | -1 | misc_RNA | ENSG00000( |
| 49863072 | 49864379 | 1 | lncRNA | ENSG00000( |
| 49893082 | 49897054 | 1 | protein_coding | ENSG00000( |
| 49901850 | 49901950 | -1 | snRNA | ENSG00000( |
| 49913493 | 49913760 | -1 | misc_RNA | ENSG00000( |
| 49927571 | 50105043 | -1 | lncRNA | ENSG00000( |
| 49966399 | 49966511 | 1 | miRNA | ENSG00000( |
| 50050368 | 50051336 | 1 | processed_pseudoge | ENSG00000( |
| 50052857 | 50053569 | 1 | processed_pseudoge | ENSG00000( |
| 50068568 | 50068873 | 1 | misc_RNA | ENSG00000( |
| 50086982 | 50087083 | 1 | misc_RNA | ENSG00000( |
| 50108632 | 50116600 | -1 | protein_coding | ENSG00000( |
| 50117130 | 50231578 | -1 | protein_coding | ENSG00000( |
| 50237563 | 50312229 | -1 | protein_coding | ENSG00000( |
| 50299855 | 50299946 | -1 | miRNA | ENSG00000( |
| 50312324 | 50335558 | 1 | protein_coding | ENSG00000( |
| 50326526 | 50327909 | -1 | lncRNA | ENSG00000( |
| 50329404 | 50416461 | -1 | protein_coding | ENSG00000( |
| 50397013 | 50398345 | 1 | lncRNA | ENSG00000( |

| | | | | |
|---|---|---|---|---|
| 50418501 | 50561126 | -1 | protein_coding | ENSG0000C |
| 50448807 | 50456742 | 1 | lncRNA | ENSG0000C |
| 50468397 | 50468498 | -1 | misc_RNA | ENSG0000C |
| 50471769 | 50472592 | 1 | processed_pseudoge | ENSG0000C |
| 50532509 | 50633068 | 1 | protein_coding | ENSG0000C |
| 50571919 | 50572140 | -1 | processed_pseudoge | ENSG0000C |
| 50632058 | 50668306 | -1 | protein_coding | ENSG0000C |
| 50637691 | 50637981 | -1 | misc_RNA | ENSG0000C |
| 50662511 | 50663178 | -1 | lncRNA | ENSG0000C |
| 50704343 | 50705511 | -1 | processed_pseudoge | ENSG0000C |
| 50719763 | 50831121 | -1 | protein_coding | ENSG0000C |
| 50723777 | 50724272 | -1 | lncRNA | ENSG0000C |
| 50821880 | 50823501 | 1 | lncRNA | ENSG0000C |
| 50831962 | 50838100 | 1 | lncRNA | ENSG0000C |
| 50844625 | 50844712 | 1 | snoRNA | ENSG0000C |
| 50848122 | 50865659 | 1 | lncRNA | ENSG0000C |
| 50857891 | 50944483 | -1 | protein_coding | ENSG0000C |
| 50872053 | 50904970 | 1 | protein_coding | ENSG0000C |
| 50912226 | 50913358 | 1 | lncRNA | ENSG0000C |
| 50956259 | 50962002 | -1 | lncRNA | ENSG0000C |
| 50975262 | 51096061 | -1 | protein_coding | ENSG0000C |
| 51022967 | 51031032 | 1 | lncRNA | ENSG0000C |
| 51088957 | 51091288 | 1 | lncRNA | ENSG0000C |
| 51240162 | 51257655 | 1 | protein_coding | ENSG0000C |
| 51244090 | 51244224 | 1 | snoRNA | ENSG0000C |
| 51253933 | 51254023 | -1 | misc_RNA | ENSG0000C |
| 51304108 | 51331918 | -1 | lncRNA | ENSG0000C |
| 51333068 | 51365557 | 1 | lncRNA | ENSG0000C |
| 51344914 | 51385603 | -1 | lncRNA | ENSG0000C |
| 51392128 | 51397135 | -1 | lncRNA | ENSG0000C |
| 51436038 | 51436963 | 1 | processed_pseudoge | ENSG0000C |
| 51454512 | 51599952 | -1 | lncRNA | ENSG0000C |
| 51489100 | 51730727 | 1 | protein_coding | ENSG0000C |
| 51522864 | 51522967 | 1 | snRNA | ENSG0000C |
| 51637348 | 51637947 | -1 | lncRNA | ENSG0000C |
| 51649516 | 51651744 | -1 | lncRNA | ENSG0000C |
| 51692752 | 51692861 | 1 | rRNA_pseudogene | ENSG0000C |
| 51724103 | 51724209 | 1 | snRNA | ENSG0000C |
| 51750560 | 51763114 | 1 | lncRNA | ENSG0000C |
| 51756996 | 51757988 | 1 | unprocessed_pseudc | ENSG0000C |
| 51763056 | 51764076 | -1 | unprocessed_pseudc | ENSG0000C |
| 51765276 | 51825422 | 1 | lncRNA | ENSG0000C |
| 51771789 | 51772086 | -1 | unprocessed_pseudc | ENSG0000C |
| 51787461 | 51787743 | 1 | unprocessed_pseudc | ENSG0000C |
| 51826195 | 51979342 | 1 | protein_coding | ENSG0000C |
| 51916145 | 51918571 | -1 | lncRNA | ENSG0000C |
| 51967003 | 51969800 | -1 | lncRNA | ENSG0000C |
| 51982520 | 51987219 | -1 | lncRNA | ENSG0000C |
| 51989514 | 52010694 | 1 | protein_coding | ENSG0000C |
| 52004803 | 52069228 | -1 | protein_coding | ENSG0000C |

| | | | | |
|---|---|---|---|---|
| 52111122 | 52129625 | -1 | lncRNA | ENSG00000 |
| 52117602 | 52117995 | 1 | processed_pseudoge | ENSG00000 |
| 52155740 | 52156179 | 1 | processed_pseudoge | ENSG00000 |
| 52267698 | 52276724 | 1 | protein_coding | ENSG00000 |
| 52286797 | 52289268 | 1 | processed_pseudoge | ENSG00000 |
| 52314305 | 52328598 | 1 | protein_coding | ENSG00000 |
| 52430596 | 52552522 | -1 | protein_coding | ENSG00000 |
| 52552836 | 52637713 | 1 | protein_coding | ENSG00000 |
| 52639915 | 52695900 | -1 | protein_coding | ENSG00000 |
| 52640839 | 52641566 | 1 | lncRNA | ENSG00000 |
| 52707178 | 52728590 | 1 | protein_coding | ENSG00000 |
| 52730166 | 52774989 | 1 | protein_coding | ENSG00000 |
| 52775193 | 52791668 | -1 | protein_coding | ENSG00000 |
| 52775237 | 52777740 | 1 | lncRNA | ENSG00000 |
| 52791756 | 52930185 | 1 | lncRNA | ENSG00000 |
| 52792875 | 52820034 | -1 | lncRNA | ENSG00000 |
| 52798904 | 52799061 | -1 | processed_pseudoge | ENSG00000 |
| 52857268 | 52952435 | -1 | protein_coding | ENSG00000 |
| 52871458 | 52872217 | 1 | processed_pseudoge | ENSG00000 |
| 52951432 | 52952864 | 1 | lncRNA | ENSG00000 |
| 53011971 | 53012267 | 1 | processed_pseudoge | ENSG00000 |
| 53023055 | 53023597 | 1 | processed_pseudoge | ENSG00000 |
| 53036745 | 53153282 | -1 | protein_coding | ENSG00000 |
| 53036755 | 53038251 | 1 | lncRNA | ENSG00000 |
| 53153354 | 53157528 | 1 | lncRNA | ENSG00000 |
| 53169010 | 53883740 | 1 | lncRNA | ENSG00000 |
| 53364146 | 53392048 | -1 | lncRNA | ENSG00000 |
| 53599832 | 53614026 | -1 | lncRNA | ENSG00000 |
| 53612724 | 53613405 | 1 | processed_pseudoge | ENSG00000 |
| 53685139 | 53850882 | -1 | lncRNA | ENSG00000 |
| 53917270 | 53917753 | -1 | processed_pseudoge | ENSG00000 |
| 53948427 | 53948484 | -1 | miRNA | ENSG00000 |
| 53949736 | 53958761 | -1 | protein_coding | ENSG00000 |
| 53954671 | 53962234 | 1 | lncRNA | ENSG00000 |
| 53991101 | 53991995 | -1 | processed_pseudoge | ENSG00000 |
| 54396849 | 54420218 | 1 | protein_coding | ENSG00000 |
| 54423561 | 54441391 | -1 | protein_coding | ENSG00000 |
| 54470709 | 54471218 | 1 | lncRNA | ENSG00000 |
| 54474484 | 54489026 | -1 | protein_coding | ENSG00000 |
| 54509812 | 54539292 | 1 | protein_coding | ENSG00000 |
| 54567097 | 54793315 | 1 | protein_coding | ENSG00000 |
| 54684753 | 54687918 | -1 | lncRNA | ENSG00000 |
| 54842008 | 54902826 | -1 | protein_coding | ENSG00000 |
| 54859215 | 54859330 | 1 | snRNA | ENSG00000 |
| 54878113 | 54878193 | -1 | miRNA | ENSG00000 |
| 54884849 | 54886213 | -1 | transcribed_process | ENSG00000 |
| 54938949 | 55027105 | -1 | protein_coding | ENSG00000 |
| 55004813 | 55005687 | -1 | processed_pseudoge | ENSG00000 |
| 55027230 | 55049489 | 1 | protein_coding | ENSG00000 |
| 55051647 | 55070194 | 1 | protein_coding | ENSG00000 |

| | | | | |
|---|---|---|---|---|
| 55124110 | 55145423 | 1 | protein_coding | ENSG0000C |
| 55148112 | 55191608 | -1 | protein_coding | ENSG0000C |
| 55196727 | 55199013 | -1 | lncRNA | ENSG0000C |
| 55229217 | 55229681 | 1 | processed_pseudoge | ENSG0000C |
| 55240622 | 55241080 | -1 | processed_pseudoge | ENSG0000C |
| 55262222 | 55272075 | -1 | lncRNA | ENSG0000C |
| 55271421 | 55361918 | 1 | protein_coding | ENSG0000C |
| 55282296 | 55282643 | -1 | processed_pseudoge | ENSG0000C |
| 55285227 | 55285558 | 1 | misc_RNA | ENSG0000C |
| 55298644 | 55299231 | 1 | processed_pseudoge | ENSG0000C |
| 55325834 | 55339501 | -1 | lncRNA | ENSG0000C |
| 55366392 | 55411858 | -1 | protein_coding | ENSG0000C |
| 55394940 | 55395233 | -1 | processed_pseudoge | ENSG0000C |
| 55413541 | 55456726 | -1 | protein_coding | ENSG0000C |
| 55413556 | 55414059 | -1 | processed_pseudoge | ENSG0000C |
| 55496786 | 55580888 | -1 | lncRNA | ENSG0000C |
| 55543401 | 55544169 | 1 | processed_pseudoge | ENSG0000C |
| 55546674 | 55548123 | 1 | unprocessed_pseudc | ENSG0000C |
| 55559072 | 55701526 | 1 | protein_coding | ENSG0000C |
| 55651534 | 55651634 | 1 | misc_RNA | ENSG0000C |
| 55748208 | 55773203 | 1 | lncRNA | ENSG0000C |
| 55766177 | 55767717 | 1 | transcribed_processe | ENSG0000C |
| 55781132 | 55796731 | -1 | lncRNA | ENSG0000C |
| 55896443 | 55962970 | 1 | lncRNA | ENSG0000C |
| 56117316 | 56117990 | -1 | lncRNA | ENSG0000C |
| 56117814 | 56301524 | 1 | protein_coding | ENSG0000C |
| 56303490 | 56310761 | -1 | lncRNA | ENSG0000C |
| 56305838 | 56306394 | 1 | lncRNA | ENSG0000C |
| 56310880 | 56427032 | 1 | lncRNA | ENSG0000C |
| 56488354 | 56650606 | 1 | protein_coding | ENSG0000C |
| 56514331 | 56551309 | 1 | lncRNA | ENSG0000C |
| 56633244 | 56648658 | -1 | lncRNA | ENSG0000C |
| 56648965 | 56730535 | -1 | lncRNA | ENSG0000C |
| 56683421 | 56683741 | -1 | processed_pseudoge | ENSG0000C |
| 56759379 | 56765757 | 1 | lncRNA | ENSG0000C |
| 56799905 | 56816693 | -1 | protein_coding | ENSG0000C |
| 56813183 | 57152177 | 1 | lncRNA | ENSG0000C |
| 56817570 | 56893710 | -1 | lncRNA | ENSG0000C |
| 56824375 | 56824656 | -1 | misc_RNA | ENSG0000C |
| 56824986 | 56825092 | 1 | snRNA | ENSG0000C |
| 56951008 | 57060715 | -1 | lncRNA | ENSG0000C |
| 56978409 | 56979621 | 1 | processed_pseudoge | ENSG0000C |
| 57040885 | 57042504 | 1 | processed_pseudoge | ENSG0000C |
| 57044646 | 57044960 | -1 | processed_pseudoge | ENSG0000C |
| 57066260 | 57112660 | -1 | lncRNA | ENSG0000C |
| 57200507 | 57268905 | -1 | protein_coding | ENSG0000C |
| 57268924 | 57298742 | 1 | protein_coding | ENSG0000C |
| 57335203 | 57344724 | -1 | lncRNA | ENSG0000C |
| 57355446 | 57359410 | 1 | lncRNA | ENSG0000C |
| 57390544 | 57415906 | 1 | protein_coding | ENSG0000C |

| | | | | |
|---|---|---|---|---|
| 57469301 | 57493867 | -1 | protein_coding | ENSG0000C |
| 57563920 | 57982194 | -1 | protein_coding | ENSG0000C |
| 57578409 | 57600404 | 1 | lncRNA | ENSG0000C |
| 57629841 | 57631162 | 1 | transcribed_processe | ENSG0000C |
| 57837354 | 57837660 | -1 | misc_RNA | ENSG0000C |
| 57993545 | 57994525 | -1 | lncRNA | ENSG0000C |
| 57999735 | 58298139 | -1 | protein_coding | ENSG0000C |
| 58100985 | 58101574 | 1 | processed_pseudoge | ENSG0000C |
| 58141279 | 58141660 | 1 | processed_pseudoge | ENSG0000C |
| 58179771 | 58180064 | 1 | misc_RNA | ENSG0000C |
| 58184576 | 58185507 | 1 | processed_pseudoge | ENSG0000C |
| 58200080 | 58235636 | 1 | protein_coding | ENSG0000C |
| 58244843 | 58272012 | 1 | protein_coding | ENSG0000C |
| 58264662 | 58269681 | 1 | lncRNA | ENSG0000C |
| 58265365 | 58298134 | -1 | lncRNA | ENSG0000C |
| 58284773 | 58285363 | 1 | processed_pseudoge | ENSG0000C |
| 58288033 | 58289158 | 1 | lncRNA | ENSG0000C |
| 58292293 | 58292388 | -1 | snRNA | ENSG0000C |
| 58298504 | 58373887 | 1 | protein_coding | ENSG0000C |
| 58370023 | 58395641 | -1 | lncRNA | ENSG0000C |
| 58395928 | 58408702 | 1 | protein_coding | ENSG0000C |
| 58398557 | 58427125 | 1 | lncRNA | ENSG0000C |
| 58408495 | 58427531 | -1 | protein_coding | ENSG0000C |
| 58427385 | 58551297 | 1 | protein_coding | ENSG0000C |
| 58441844 | 58441992 | 1 | misc_RNA | ENSG0000C |
| 58463971 | 58464136 | 1 | processed_pseudoge | ENSG0000C |
| 58521223 | 58521988 | 1 | processed_pseudoge | ENSG0000C |
| 58564146 | 58564616 | 1 | processed_pseudoge | ENSG0000C |
| 58633967 | 58648321 | 1 | protein_coding | ENSG0000C |
| 58646631 | 59184247 | 1 | lncRNA | ENSG0000C |
| 58794654 | 58795029 | 1 | processed_pseudoge | ENSG0000C |
| 58827679 | 58828176 | -1 | processed_pseudoge | ENSG0000C |
| 59135488 | 59136421 | -1 | processed_pseudoge | ENSG0000C |
| 59181593 | 59181940 | 1 | processed_pseudoge | ENSG0000C |
| 59188646 | 59371405 | 1 | protein_coding | ENSG0000C |
| 59429022 | 59465342 | -1 | protein_coding | ENSG0000C |
| 59460363 | 59484408 | -1 | protein_coding | ENSG0000C |
| 59479274 | 59525437 | -1 | lncRNA | ENSG0000C |
| 59484443 | 59505410 | 1 | protein_coding | ENSG0000C |
| 59504539 | 59576812 | -1 | protein_coding | ENSG0000C |
| 59595976 | 59870776 | -1 | protein_coding | ENSG0000C |
| 59646962 | 59647020 | -1 | miRNA | ENSG0000C |
| 59919423 | 59920339 | -1 | lncRNA | ENSG0000C |
| 59919713 | 60063559 | 1 | protein_coding | ENSG0000C |
| 59969116 | 60091783 | -1 | lncRNA | ENSG0000C |
| 60091911 | 60169133 | 1 | protein_coding | ENSG0000C |
| 60144119 | 60169856 | -1 | protein_coding | ENSG0000C |
| 60158976 | 60159324 | 1 | processed_pseudoge | ENSG0000C |
| 60179382 | 60180016 | 1 | processed_pseudoge | ENSG0000C |
| 60212462 | 60212912 | -1 | processed_pseudoge | ENSG0000C |

| | | | | |
|---|---|---|---|---|
| 60240121 | 60248765 | -1 | lncRNA | ENSG0000C |
| 60245752 | 60299087 | 1 | protein_coding | ENSG0000C |
| 60323236 | 60323907 | 1 | lncRNA | ENSG0000C |
| 60396469 | 60515543 | -1 | protein_coding | ENSG0000C |
| 60397757 | 60398281 | 1 | processed_pseudoge | ENSG0000C |
| 60398934 | 60400013 | -1 | unprocessed_pseudc | ENSG0000C |
| 60493766 | 60496885 | -1 | transcribed_processe | ENSG0000C |
| 60509146 | 60512850 | 1 | protein_coding | ENSG0000C |
| 60515119 | 60554916 | 1 | lncRNA | ENSG0000C |
| 60544400 | 60544654 | 1 | transcribed_processe | ENSG0000C |
| 60572024 | 60572311 | -1 | processed_pseudoge | ENSG0000C |
| 60602146 | 60602533 | -1 | processed_pseudoge | ENSG0000C |
| 60639216 | 60642589 | 1 | lncRNA | ENSG0000C |
| 60643421 | 60658259 | -1 | protein_coding | ENSG0000C |
| 60709539 | 60724351 | -1 | protein_coding | ENSG0000C |
| 60734742 | 60969965 | 1 | protein_coding | ENSG0000C |
| 60779978 | 60780595 | 1 | processed_pseudoge | ENSG0000C |
| 60879714 | 60982585 | 1 | lncRNA | ENSG0000C |
| 60919240 | 60920155 | -1 | processed_pseudoge | ENSG0000C |
| 60971441 | 60981170 | -1 | protein_coding | ENSG0000C |
| 60972272 | 60972466 | -1 | processed_pseudoge | ENSG0000C |
| 60974327 | 60974435 | 1 | snRNA | ENSG0000C |
| 60981114 | 61083733 | 1 | protein_coding | ENSG0000C |
| 61187559 | 61550976 | 1 | protein_coding | ENSG0000C |
| 61277370 | 61281761 | -1 | protein_coding | ENSG0000C |
| 61294909 | 61322838 | -1 | lncRNA | ENSG0000C |
| 61529128 | 61657964 | 1 | protein_coding | ENSG0000C |
| 61535385 | 61545484 | -1 | lncRNA | ENSG0000C |
| 61556273 | 61570653 | -1 | lncRNA | ENSG0000C |
| 61570405 | 61658696 | 1 | lncRNA | ENSG0000C |
| 61681041 | 61695823 | -1 | lncRNA | ENSG0000C |
| 61695513 | 61748259 | 1 | protein_coding | ENSG0000C |
| 61707564 | 61751099 | -1 | lncRNA | ENSG0000C |
| 61734138 | 61776260 | 1 | lncRNA | ENSG0000C |
| 61762420 | 61796428 | 1 | protein_coding | ENSG0000C |
| 61801713 | 61803634 | 1 | TEC | ENSG0000C |
| 61811974 | 62112550 | 1 | protein_coding | ENSG0000C |
| 61831861 | 61832369 | -1 | processed_pseudoge | ENSG0000C |
| 62069518 | 62081154 | -1 | lncRNA | ENSG0000C |
| 62103122 | 62117175 | -1 | lncRNA | ENSG0000C |
| 62114353 | 62130962 | 1 | transcribed_unitary_ | ENSG0000C |
| 62134149 | 62139973 | -1 | lncRNA | ENSG0000C |
| 62150656 | 62152258 | -1 | unprocessed_pseudc | ENSG0000C |
| 62514872 | 62515123 | -1 | processed_pseudoge | ENSG0000C |
| 62604113 | 62605717 | -1 | processed_pseudoge | ENSG0000C |
| 62699464 | 63102037 | -1 | protein_coding | ENSG0000C |
| 63122888 | 63178190 | -1 | lncRNA | ENSG0000C |
| 63123007 | 63123201 | 1 | processed_pseudoge | ENSG0000C |
| 63204114 | 63293508 | 1 | protein_coding | ENSG0000C |
| 63298126 | 63299547 | -1 | lncRNA | ENSG0000C |

| | | | | |
|---|---|---|---|---|
| 63312835 | 63318935 | -1 | protein_coding | ENSG0000C |
| 63371364 | 63543377 | -1 | protein_coding | ENSG0000C |
| 63479272 | 63479413 | 1 | scaRNA | ENSG0000C |
| 63543569 | 63544664 | 1 | lncRNA | ENSG0000C |
| 63567958 | 63568308 | 1 | processed_pseudoge | ENSG0000C |
| 63594031 | 63595291 | -1 | processed_pseudoge | ENSG0000C |
| 63597039 | 63641861 | -1 | protein_coding | ENSG0000C |
| 63598874 | 63599248 | 1 | lncRNA | ENSG0000C |
| 63601509 | 63601814 | 1 | processed_pseudoge | ENSG0000C |
| 63618043 | 63618144 | 1 | snRNA | ENSG0000C |
| 63621760 | 63621871 | 1 | misc_RNA | ENSG0000C |
| 63630678 | 63630781 | 1 | snRNA | ENSG0000C |
| 63642035 | 63665593 | 1 | lncRNA | ENSG0000C |
| 63651296 | 63651499 | -1 | snoRNA | ENSG0000C |
| 63684216 | 63728065 | -1 | protein_coding | ENSG0000C |
| 63734697 | 63735923 | 1 | processed_pseudoge | ENSG0000C |
| 63736444 | 63736505 | 1 | snRNA | ENSG0000C |
| 63762013 | 63762149 | 1 | processed_pseudoge | ENSG0000C |
| 63808546 | 63808732 | 1 | processed_pseudoge | ENSG0000C |
| 63848261 | 63848362 | -1 | misc_RNA | ENSG0000C |
| 63852983 | 64226433 | 1 | protein_coding | ENSG0000C |
| 64084232 | 64338112 | -1 | protein_coding | ENSG0000C |
| 64095024 | 64095125 | -1 | miRNA | ENSG0000C |
| 64329431 | 64338639 | -1 | lncRNA | ENSG0000C |
| 64341673 | 64387986 | -1 | transcribed_unitary_ | ENSG0000C |
| 64349677 | 64352284 | -1 | lncRNA | ENSG0000C |
| 64361392 | 64361637 | 1 | processed_pseudoge | ENSG0000C |
| 64388031 | 64463457 | 1 | protein_coding | ENSG0000C |
| 64422935 | 64448557 | -1 | lncRNA | ENSG0000C |
| 64440369 | 64442238 | -1 | lncRNA | ENSG0000C |
| 64449106 | 64505213 | -1 | protein_coding | ENSG0000C |
| 64465499 | 64474503 | 1 | protein_coding | ENSG0000C |
| 64503712 | 64533690 | 1 | protein_coding | ENSG0000C |
| 64513952 | 64540368 | -1 | lncRNA | ENSG0000C |
| 64535702 | 64535879 | -1 | processed_pseudoge | ENSG0000C |
| 64535905 | 64546173 | 1 | protein_coding | ENSG0000C |
| 64549920 | 64589381 | 1 | protein_coding | ENSG0000C |
| 64552694 | 64596536 | -1 | lncRNA | ENSG0000C |
| 64600243 | 64601644 | -1 | processed_pseudoge | ENSG0000C |
| 64704102 | 64750249 | 1 | protein_coding | ENSG0000C |
| 64746283 | 64879907 | -1 | protein_coding | ENSG0000C |
| 64785626 | 64785686 | -1 | miRNA | ENSG0000C |
| 64909488 | 64909780 | -1 | ribozyme | ENSG0000C |
| 64914361 | 64944591 | 1 | protein_coding | ENSG0000C |
| 64914485 | 65061803 | 1 | protein_coding | ENSG0000C |
| 64939152 | 64942905 | -1 | protein_coding | ENSG0000C |
| 64945814 | 64973226 | -1 | protein_coding | ENSG0000C |
| 64986895 | 65062652 | 1 | protein_coding | ENSG0000C |
| 65003325 | 65003767 | -1 | lncRNA | ENSG0000C |
| 65006174 | 65102695 | -1 | protein_coding | ENSG0000C |

| | | | | |
|---|---|---|---|---|
| 65044688 | 65044769 | 1 | miRNA | ENSG0000C |
| 65082034 | 65094212 | 1 | lncRNA | ENSG0000C |
| 65124352 | 65124542 | -1 | snRNA | ENSG0000C |
| 65197254 | 65222443 | -1 | lncRNA | ENSG0000C |
| 65234515 | 65235101 | -1 | processed_pseudoge | ENSG0000C |
| 65236480 | 65318790 | -1 | lncRNA | ENSG0000C |
| 65266638 | 65267690 | -1 | processed_pseudoge | ENSG0000C |
| 65268829 | 65269098 | 1 | processed_pseudoge | ENSG0000C |
| 65279267 | 65281166 | 1 | processed_pseudoge | ENSG0000C |
| 65335117 | 65335183 | -1 | miRNA | ENSG0000C |
| 65410592 | 65744121 | 1 | protein_coding | ENSG0000C |
| 65411170 | 65412690 | -1 | lncRNA | ENSG0000C |
| 65467807 | 65468341 | -1 | processed_pseudoge | ENSG0000C |
| 65471102 | 65471186 | 1 | miRNA | ENSG0000C |
| 65528830 | 65529574 | -1 | processed_pseudoge | ENSG0000C |
| 65879592 | 65881070 | -1 | processed_pseudoge | ENSG0000C |
| 65900932 | 65901451 | -1 | processed_pseudoge | ENSG0000C |
| 65957297 | 66004523 | -1 | lncRNA | ENSG0000C |
| 65986967 | 65987447 | -1 | lncRNA | ENSG0000C |
| 66012830 | 66013789 | -1 | processed_pseudoge | ENSG0000C |
| 66111557 | 66131716 | 1 | lncRNA | ENSG0000C |
| 66177527 | 66177635 | -1 | misc_RNA | ENSG0000C |
| 66212810 | 66509394 | -1 | lncRNA | ENSG0000C |
| 66248811 | 66278875 | 1 | lncRNA | ENSG0000C |
| 66411704 | 66498677 | 1 | protein_coding | ENSG0000C |
| 66507407 | 67181803 | 1 | protein_coding | ENSG0000C |
| 66760532 | 66761018 | 1 | processed_pseudoge | ENSG0000C |
| 66969038 | 66969816 | -1 | lncRNA | ENSG0000C |
| 67189393 | 67228550 | 1 | protein_coding | ENSG0000C |
| 67199062 | 67200327 | 1 | processed_pseudoge | ENSG0000C |
| 67241342 | 67336061 | 1 | protein_coding | ENSG0000C |
| 67283749 | 67294073 | -1 | lncRNA | ENSG0000C |
| 67294371 | 67360265 | -1 | protein_coding | ENSG0000C |
| 67340068 | 67340176 | 1 | misc_RNA | ENSG0000C |
| 67360328 | 67386516 | 1 | protein_coding | ENSG0000C |
| 67386984 | 67412167 | -1 | protein_coding | ENSG0000C |
| 67441855 | 67441930 | -1 | miRNA | ENSG0000C |
| 67447084 | 67533739 | -1 | protein_coding | ENSG0000C |
| 67533290 | 67589612 | 1 | protein_coding | ENSG0000C |
| 67581955 | 67600286 | -1 | protein_coding | ENSG0000C |
| 67610986 | 67613864 | 1 | processed_pseudoge | ENSG0000C |
| 67614929 | 67616626 | 1 | transcribed_processe | ENSG0000C |
| 67616903 | 67617084 | 1 | pseudogene | ENSG0000C |
| 67617302 | 67617415 | -1 | misc_RNA | ENSG0000C |
| 67619920 | 67651708 | 1 | protein_coding | ENSG0000C |
| 67627299 | 67639675 | -1 | lncRNA | ENSG0000C |
| 67647085 | 67674820 | -1 | protein_coding | ENSG0000C |
| 67652300 | 67652614 | -1 | processed_pseudoge | ENSG0000C |
| 67657105 | 67657211 | -1 | rRNA_pseudogene | ENSG0000C |
| 67659820 | 67690367 | -1 | protein_coding | ENSG0000C |

| | | | | |
|---|---|---|---|---|
| 67676800 | 67695793 | -1 | protein_coding | ENSG00000( |
| 67701886 | 67734451 | 1 | protein_coding | ENSG00000( |
| 67705954 | 67706252 | 1 | misc_RNA | ENSG00000( |
| 67714545 | 67715204 | 1 | processed_pseudoge | ENSG00000( |
| 67727374 | 67816590 | -1 | protein_coding | ENSG00000( |
| 67735195 | 67735350 | 1 | processed_pseudoge | ENSG00000( |
| 67751228 | 67751544 | -1 | misc_RNA | ENSG00000( |
| 67799004 | 67799609 | 1 | processed_pseudoge | ENSG00000( |
| 67819779 | 68730218 | 1 | protein_coding | ENSG00000( |
| 68113706 | 68114144 | -1 | unprocessed_pseudc | ENSG00000( |
| 68125004 | 68130196 | -1 | lncRNA | ENSG00000( |
| 68149617 | 68149908 | -1 | misc_RNA | ENSG00000( |
| 68236243 | 68236539 | -1 | misc_RNA | ENSG00000( |
| 68338728 | 68339528 | -1 | processed_pseudoge | ENSG00000( |
| 68421698 | 68422196 | -1 | processed_pseudoge | ENSG00000( |
| 68605396 | 68611732 | -1 | lncRNA | ENSG00000( |
| 68627166 | 68628445 | -1 | lncRNA | ENSG00000( |
| 68683411 | 68685565 | -1 | lncRNA | ENSG00000( |
| 68693090 | 68693583 | 1 | processed_pseudoge | ENSG00000( |
| 68693656 | 68695734 | -1 | lncRNA | ENSG00000( |
| 68731234 | 68736678 | 1 | lncRNA | ENSG00000( |
| 68735922 | 68736028 | -1 | snRNA | ENSG00000( |
| 68787660 | 68796253 | -1 | protein_coding | ENSG00000( |
| 68861762 | 68862525 | 1 | unprocessed_pseudc | ENSG00000( |
| 68867857 | 68869951 | -1 | unprocessed_pseudc | ENSG00000( |
| 68874143 | 68979440 | -1 | protein_coding | ENSG00000( |
| 68887785 | 68888084 | -1 | processed_pseudoge | ENSG00000( |
| 68936602 | 68936864 | 1 | processed_pseudoge | ENSG00000( |
| 68979682 | 68987463 | 1 | lncRNA | ENSG00000( |
| 69023150 | 69023308 | -1 | processed_pseudoge | ENSG00000( |
| 69050881 | 69153150 | -1 | protein_coding | ENSG00000( |
| 69083398 | 69085389 | 1 | processed_pseudoge | ENSG00000( |
| 69143852 | 69143953 | -1 | misc_RNA | ENSG00000( |
| 69154311 | 69154804 | 1 | lncRNA | ENSG00000( |
| 69183017 | 69214092 | -1 | lncRNA | ENSG00000( |
| 69191511 | 69244020 | 1 | protein_coding | ENSG00000( |
| 69239166 | 69260567 | -1 | lncRNA | ENSG00000( |
| 69259277 | 69357033 | 1 | protein_coding | ENSG00000( |
| 69357010 | 69359308 | 1 | TEC | ENSG00000( |
| 69380128 | 69398299 | -1 | protein_coding | ENSG00000( |
| 69398015 | 69462390 | 1 | protein_coding | ENSG00000( |
| 69479057 | 69483714 | -1 | lncRNA | ENSG00000( |
| 69484692 | 69531551 | 1 | protein_coding | ENSG00000( |
| 69561019 | 69561470 | -1 | processed_pseudoge | ENSG00000( |
| 69569799 | 69574871 | -1 | protein_coding | ENSG00000( |
| 69582916 | 69584946 | -1 | lncRNA | ENSG00000( |
| 69611596 | 69715144 | 1 | protein_coding | ENSG00000( |
| 69617122 | 69617648 | 1 | lncRNA | ENSG00000( |
| 69726900 | 69772005 | 1 | protein_coding | ENSG00000( |
| 69775416 | 69797241 | -1 | protein_coding | ENSG00000( |

| | | | | |
|---|---|---|---|---|
| 69804203 | 69804331 | -1 | snoRNA | ENSG0000C |
| 69854131 | 70032366 | 1 | protein_coding | ENSG0000C |
| 69885340 | 69886140 | -1 | processed_pseudoge | ENSG0000C |
| 70044217 | 70189070 | -1 | protein_coding | ENSG0000C |
| 70187123 | 70230187 | 1 | lncRNA | ENSG0000C |
| 70230224 | 70232426 | 1 | unprocessed_pseudc | ENSG0000C |
| 70234495 | 70235095 | 1 | processed_pseudoge | ENSG0000C |
| 70245491 | 70247801 | -1 | transcribed_processe | ENSG0000C |
| 70255629 | 70343388 | 1 | lncRNA | ENSG0000C |
| 70325081 | 70416984 | -1 | protein_coding | ENSG0000C |
| 70326064 | 70417074 | -1 | protein_coding | ENSG0000C |
| 70360892 | 70361088 | 1 | snRNA | ENSG0000C |
| 70366499 | 70417090 | -1 | protein_coding | ENSG0000C |
| 70425812 | 70547464 | -1 | lncRNA | ENSG0000C |
| 70452174 | 70459905 | 1 | protein_coding | ENSG0000C |
| 70453541 | 70483775 | -1 | transcribed_unproce | ENSG0000C |
| 70496107 | 70496210 | 1 | snRNA | ENSG0000C |
| 70522358 | 70535015 | -1 | protein_coding | ENSG0000C |
| 70556206 | 70557478 | 1 | unprocessed_pseudc | ENSG0000C |
| 70581257 | 70600690 | -1 | protein_coding | ENSG0000C |
| 70593314 | 70641204 | -1 | lncRNA | ENSG0000C |
| 70594552 | 70595770 | 1 | processed_pseudoge | ENSG0000C |
| 70608798 | 70641298 | -1 | lncRNA | ENSG0000C |
| 70610086 | 70610379 | 1 | misc_RNA | ENSG0000C |
| 70641916 | 70675366 | 1 | protein_coding | ENSG0000C |
| 70698698 | 70712153 | 1 | lncRNA | ENSG0000C |
| 70722526 | 70809534 | -1 | protein_coding | ENSG0000C |
| 70809841 | 70815994 | 1 | lncRNA | ENSG0000C |
| 70822004 | 70823984 | 1 | lncRNA | ENSG0000C |
| 70858844 | 70859429 | -1 | processed_pseudoge | ENSG0000C |
| 70906657 | 70907111 | -1 | lncRNA | ENSG0000C |
| 70907405 | 71115382 | 1 | protein_coding | ENSG0000C |
| 70945640 | 70959029 | -1 | lncRNA | ENSG0000C |
| 71031686 | 71031970 | -1 | transcribed_processe | ENSG0000C |
| 71044375 | 71044640 | -1 | transcribed_processe | ENSG0000C |
| 71085482 | 71085833 | -1 | processed_pseudoge | ENSG0000C |
| 71141125 | 71143253 | -1 | lncRNA | ENSG0000C |
| 71179053 | 71243841 | -1 | lncRNA | ENSG0000C |
| 71181456 | 71182106 | 1 | lncRNA | ENSG0000C |
| 71212633 | 71218216 | 1 | lncRNA | ENSG0000C |
| 71219100 | 71222724 | 1 | lncRNA | ENSG0000C |
| 71292729 | 71321882 | -1 | lncRNA | ENSG0000C |
| 71320449 | 71741229 | 1 | protein_coding | ENSG0000C |
| 71330342 | 71330738 | -1 | processed_pseudoge | ENSG0000C |
| 71398337 | 71398407 | 1 | snoRNA | ENSG0000C |
| 71448689 | 71448958 | 1 | misc_RNA | ENSG0000C |
| 71586269 | 71590354 | -1 | lncRNA | ENSG0000C |
| 71734927 | 71739794 | -1 | lncRNA | ENSG0000C |
| 71848606 | 71928596 | 1 | lncRNA | ENSG0000C |
| 71932429 | 72566530 | 1 | protein_coding | ENSG0000C |

| | | | | |
|---|---|---|---|---|
| 72022407 | 72022516 | -1 | misc_RNA | ENSG00000 |
| 72515407 | 72544622 | -1 | lncRNA | ENSG00000 |
| 72516820 | 72516898 | 1 | miRNA | ENSG00000 |
| 72552580 | 72595125 | -1 | lncRNA | ENSG00000 |
| 72619296 | 72894116 | -1 | protein_coding | ENSG00000 |
| 72701762 | 72704280 | 1 | lncRNA | ENSG00000 |
| 72853224 | 72853331 | -1 | misc_RNA | ENSG00000 |
| 72926377 | 72959703 | 1 | protein_coding | ENSG00000 |
| 72929612 | 72929930 | -1 | processed_pseudoge | ENSG00000 |
| 72960595 | 72961993 | 1 | processed_pseudoge | ENSG00000 |
| 72963645 | 72964008 | -1 | processed_pseudoge | ENSG00000 |
| 72969451 | 73027131 | -1 | protein_coding | ENSG00000 |
| 73040619 | 73040914 | -1 | misc_RNA | ENSG00000 |
| 73057925 | 73059415 | -1 | lncRNA | ENSG00000 |
| 73058532 | 73123899 | 1 | protein_coding | ENSG00000 |
| 73068108 | 73068618 | -1 | processed_pseudoge | ENSG00000 |
| 73136418 | 73223691 | 1 | protein_coding | ENSG00000 |
| 73189763 | 73190004 | 1 | processed_pseudoge | ENSG00000 |
| 73237497 | 73274640 | 1 | protein_coding | ENSG00000 |
| 73242651 | 73245979 | -1 | lncRNA | ENSG00000 |
| 73246818 | 73246880 | -1 | snRNA | ENSG00000 |
| 73272182 | 73274081 | -1 | lncRNA | ENSG00000 |
| 73275107 | 73458617 | -1 | protein_coding | ENSG00000 |
| 73366897 | 73463642 | -1 | lncRNA | ENSG00000 |
| 73458850 | 73466167 | -1 | lncRNA | ENSG00000 |
| 73462362 | 73477175 | 1 | lncRNA | ENSG00000 |
| 73478484 | 73558947 | -1 | protein_coding | ENSG00000 |
| 73490933 | 73493394 | 1 | protein_coding | ENSG00000 |
| 73517233 | 73520648 | -1 | lncRNA | ENSG00000 |
| 73522878 | 73530610 | 1 | lncRNA | ENSG00000 |
| 73537143 | 73543796 | 1 | protein_coding | ENSG00000 |
| 73539221 | 73539781 | -1 | transcribed_processe | ENSG00000 |
| 73555115 | 73633941 | -1 | lncRNA | ENSG00000 |
| 73567620 | 73575658 | 1 | protein_coding | ENSG00000 |
| 73571159 | 73571667 | -1 | processed_pseudoge | ENSG00000 |
| 73580694 | 73582918 | 1 | TEC | ENSG00000 |
| 73591706 | 73596496 | 1 | protein_coding | ENSG00000 |
| 73610945 | 73619888 | 1 | protein_coding | ENSG00000 |
| 73631475 | 73632019 | 1 | processed_pseudoge | ENSG00000 |
| 73644875 | 73703732 | 1 | protein_coding | ENSG00000 |
| 73659564 | 73659660 | -1 | snRNA | ENSG00000 |
| 73698103 | 73700351 | -1 | lncRNA | ENSG00000 |
| 73711783 | 73714384 | -1 | protein_coding | ENSG00000 |
| 73715122 | 73790285 | -1 | protein_coding | ENSG00000 |
| 73758747 | 73758819 | 1 | miRNA | ENSG00000 |
| 73787355 | 73803300 | 1 | lncRNA | ENSG00000 |
| 73822559 | 73830135 | -1 | lncRNA | ENSG00000 |
| 73840520 | 73840609 | -1 | processed_pseudoge | ENSG00000 |
| 73851844 | 73886827 | 1 | protein_coding | ENSG00000 |
| 73851971 | 73932278 | 1 | protein_coding | ENSG00000 |

| | | | | |
|---|---|---|---|---|
| 73858061 | 73858165 | -1 | misc_RNA | ENSG0000C |
| 73873689 | 73885555 | -1 | lncRNA | ENSG0000C |
| 73886617 | 73932511 | 1 | protein_coding | ENSG0000C |
| 73896164 | 73938114 | -1 | lncRNA | ENSG0000C |
| 73905267 | 73905636 | 1 | lncRNA | ENSG0000C |
| 73906316 | 73906427 | -1 | misc_RNA | ENSG0000C |
| 73931501 | 73950414 | -1 | protein_coding | ENSG0000C |
| 73949926 | 73963670 | 1 | protein_coding | ENSG0000C |
| 73958010 | 74019399 | -1 | protein_coding | ENSG0000C |
| 74003396 | 74003470 | 1 | processed_pseudoge | ENSG0000C |
| 74019349 | 74082863 | 1 | protein_coding | ENSG0000C |
| 74056847 | 74084492 | -1 | protein_coding | ENSG0000C |
| 74084796 | 74201235 | 1 | protein_coding | ENSG0000C |
| 74174456 | 74174753 | 1 | misc_RNA | ENSG0000C |
| 74192589 | 74192823 | 1 | transcribed_process | ENSG0000C |
| 74239449 | 74262738 | 1 | protein_coding | ENSG0000C |
| 74285269 | 74303055 | -1 | protein_coding | ENSG0000C |
| 74289127 | 74294425 | -1 | lncRNA | ENSG0000C |
| 74303069 | 74360008 | 1 | protein_coding | ENSG0000C |
| 74305425 | 74306527 | 1 | processed_pseudoge | ENSG0000C |
| 74377818 | 74378418 | -1 | processed_pseudoge | ENSG0000C |
| 74405894 | 74426210 | -1 | protein_coding | ENSG0000C |
| 74471930 | 74472360 | -1 | lncRNA | ENSG0000C |
| 74474007 | 74474864 | -1 | lncRNA | ENSG0000C |
| 74476192 | 74494177 | -1 | protein_coding | ENSG0000C |
| 74480133 | 74480204 | -1 | miRNA | ENSG0000C |
| 74482985 | 74483435 | 1 | processed_pseudoge | ENSG0000C |
| 74493756 | 74497106 | 1 | protein_coding | ENSG0000C |
| 74498183 | 74612378 | -1 | protein_coding | ENSG0000C |
| 74552181 | 74560048 | 1 | lncRNA | ENSG0000C |
| 74614693 | 74616647 | -1 | lncRNA | ENSG0000C |
| 74653437 | 74713108 | -1 | protein_coding | ENSG0000C |
| 74658049 | 74658590 | 1 | processed_pseudoge | ENSG0000C |
| 74711936 | 74712045 | 1 | snoRNA | ENSG0000C |
| 74713144 | 74738620 | 1 | protein_coding | ENSG0000C |
| 74763316 | 74859435 | 1 | protein_coding | ENSG0000C |
| 74852871 | 74871940 | -1 | protein_coding | ENSG0000C |
| 74873684 | 74874254 | 1 | processed_pseudoge | ENSG0000C |
| 74881891 | 74903743 | 1 | protein_coding | ENSG0000C |
| 74903954 | 74923396 | -1 | protein_coding | ENSG0000C |
| 74941834 | 74955626 | -1 | protein_coding | ENSG0000C |
| 75002921 | 75012366 | 1 | protein_coding | ENSG0000C |
| 75004719 | 75008481 | -1 | lncRNA | ENSG0000C |
| 75011269 | 75012851 | -1 | TEC | ENSG0000C |
| 75013764 | 75051532 | -1 | protein_coding | ENSG0000C |
| 75020761 | 75020867 | 1 | snRNA | ENSG0000C |
| 75053237 | 75069483 | -1 | protein_coding | ENSG0000C |
| 75064170 | 75079988 | 1 | protein_coding | ENSG0000C |
| 75079353 | 75127344 | -1 | protein_coding | ENSG0000C |
| 75116426 | 75116600 | -1 | processed_pseudoge | ENSG0000C |

| | | | | |
|---|---|---|---|---|
| 75127153 | 75136930 | 1 | lncRNA | ENSG0000C |
| 75131469 | 75176612 | -1 | protein_coding | ENSG0000C |
| 75176929 | 75177418 | 1 | lncRNA | ENSG0000C |
| 75202081 | 75202169 | -1 | snRNA | ENSG0000C |
| 75238616 | 75269283 | -1 | lncRNA | ENSG0000C |
| 75259411 | 75271950 | 1 | lncRNA | ENSG0000C |
| 75270700 | 75270780 | -1 | snRNA | ENSG0000C |
| 75278826 | 75282230 | 1 | protein_coding | ENSG0000C |
| 75292131 | 75292495 | -1 | processed_pseudoge | ENSG0000C |
| 75294404 | 75298109 | 1 | lncRNA | ENSG0000C |
| 75423683 | 75427741 | -1 | lncRNA | ENSG0000C |
| 75427716 | 75474111 | 1 | protein_coding | ENSG0000C |
| 75522455 | 75547004 | 1 | protein_coding | ENSG0000C |
| 75574888 | 75579588 | -1 | lncRNA | ENSG0000C |
| 75578620 | 75663214 | 1 | protein_coding | ENSG0000C |
| 75595805 | 75596206 | -1 | processed_pseudoge | ENSG0000C |
| 75604173 | 75604278 | 1 | rRNA_pseudogene | ENSG0000C |
| 75604209 | 75604261 | 1 | miRNA | ENSG0000C |
| 75633625 | 75955079 | 1 | protein_coding | ENSG0000C |
| 75649791 | 75660876 | -1 | protein_coding | ENSG0000C |
| 75814502 | 75847698 | -1 | lncRNA | ENSG0000C |
| 75838157 | 75839114 | -1 | transcribed_processe | ENSG0000C |
| 75902136 | 76084585 | 1 | protein_coding | ENSG0000C |
| 75958099 | 75982991 | -1 | protein_coding | ENSG0000C |
| 75970924 | 75971587 | 1 | lncRNA | ENSG0000C |
| 76013679 | 76014143 | 1 | processed_pseudoge | ENSG0000C |
| 76131707 | 76152627 | -1 | lncRNA | ENSG0000C |
| 76151916 | 76254342 | 1 | protein_coding | ENSG0000C |
| 76235817 | 76263474 | 1 | lncRNA | ENSG0000C |
| 76279173 | 76283103 | 1 | TEC | ENSG0000C |
| 76280943 | 76310066 | -1 | lncRNA | ENSG0000C |
| 76310712 | 76501837 | 1 | protein_coding | ENSG0000C |
| 76495363 | 76726712 | -1 | lncRNA | ENSG0000C |
| 76633714 | 76634028 | 1 | processed_pseudoge | ENSG0000C |
| 76634775 | 76635661 | -1 | transcribed_processe | ENSG0000C |
| 76690139 | 76690742 | -1 | processed_pseudoge | ENSG0000C |
| 76710658 | 76761388 | -1 | lncRNA | ENSG0000C |
| 76761468 | 76783015 | 1 | protein_coding | ENSG0000C |
| 76774284 | 76781518 | -1 | lncRNA | ENSG0000C |
| 76778952 | 76782249 | 1 | lncRNA | ENSG0000C |
| 76781733 | 76786724 | -1 | lncRNA | ENSG0000C |
| 76786168 | 76826246 | -1 | protein_coding | ENSG0000C |
| 76790267 | 76793807 | 1 | lncRNA | ENSG0000C |
| 76826372 | 76870302 | 1 | protein_coding | ENSG0000C |
| 76886377 | 76886750 | -1 | processed_pseudoge | ENSG0000C |
| 76890263 | 76890613 | 1 | misc_RNA | ENSG0000C |
| 76921840 | 76924649 | -1 | lncRNA | ENSG0000C |
| 76958098 | 77000609 | -1 | lncRNA | ENSG0000C |
| 76959638 | 76965802 | 1 | lncRNA | ENSG0000C |
| 76967518 | 76967765 | -1 | misc_RNA | ENSG0000C |

| | | | | |
|---|---|---|---|---|
| 76968683 | 76969145 | 1 | processed_pseudoge | ENSG00000 |
| 77024543 | 77028708 | -1 | protein_coding | ENSG00000 |
| 77028086 | 77031572 | 1 | lncRNA | ENSG00000 |
| 77041064 | 77069503 | 1 | lncRNA | ENSG00000 |
| 77048172 | 77086457 | -1 | lncRNA | ENSG00000 |
| 77069169 | 77076392 | -1 | lncRNA | ENSG00000 |
| 77084689 | 77085977 | 1 | lncRNA | ENSG00000 |
| 77098126 | 77117287 | 1 | protein_coding | ENSG00000 |
| 77098311 | 77225462 | 1 | protein_coding | ENSG00000 |
| 77116568 | 77259495 | 1 | protein_coding | ENSG00000 |
| 77131270 | 77142734 | -1 | protein_coding | ENSG00000 |
| 77140683 | 77142996 | 1 | lncRNA | ENSG00000 |
| 77173075 | 77173652 | -1 | processed_pseudoge | ENSG00000 |
| 77201026 | 77201140 | -1 | snoRNA | ENSG00000 |
| 77265483 | 77271312 | -1 | protein_coding | ENSG00000 |
| 77266218 | 77266290 | 1 | miRNA | ENSG00000 |
| 77274956 | 77320884 | -1 | protein_coding | ENSG00000 |
| 77320996 | 77331597 | 1 | protein_coding | ENSG00000 |
| 77335029 | 77377094 | -1 | protein_coding | ENSG00000 |
| 77359423 | 77359916 | 1 | processed_pseudoge | ENSG00000 |
| 77376689 | 77391497 | 1 | protein_coding | ENSG00000 |
| 77394021 | 77423517 | -1 | protein_coding | ENSG00000 |
| 77426675 | 77457952 | -1 | protein_coding | ENSG00000 |
| 77457870 | 77469472 | 1 | protein_coding | ENSG00000 |
| 77466284 | 77466436 | 1 | snoRNA | ENSG00000 |
| 77474394 | 77498816 | -1 | protein_coding | ENSG00000 |
| 77505997 | 77616773 | -1 | protein_coding | ENSG00000 |
| 77547566 | 77547846 | -1 | misc_RNA | ENSG00000 |
| 77652827 | 77653055 | -1 | processed_pseudoge | ENSG00000 |
| 77672404 | 77708023 | -1 | protein_coding | ENSG00000 |
| 77683202 | 77683989 | -1 | processed_pseudoge | ENSG00000 |
| 77696591 | 77697377 | 1 | processed_pseudoge | ENSG00000 |
| 77708071 | 77761104 | 1 | protein_coding | ENSG00000 |
| 77717599 | 77761207 | -1 | protein_coding | ENSG00000 |
| 77760830 | 77769742 | 1 | protein_coding | ENSG00000 |
| 77800109 | 77935014 | 1 | protein_coding | ENSG00000 |
| 77821030 | 77821142 | -1 | misc_RNA | ENSG00000 |
| 78059942 | 78060278 | -1 | processed_pseudoge | ENSG00000 |
| 78170373 | 79868290 | 1 | protein_coding | ENSG00000 |
| 78177803 | 78177921 | -1 | rRNA_pseudogene | ENSG00000 |
| 78231281 | 78231619 | -1 | processed_pseudoge | ENSG00000 |
| 78279572 | 78283376 | -1 | lncRNA | ENSG00000 |
| 78695321 | 78698122 | -1 | lncRNA | ENSG00000 |
| 78703615 | 78709934 | -1 | lncRNA | ENSG00000 |
| 78744398 | 78753878 | -1 | lncRNA | ENSG00000 |
| 79072132 | 79074767 | -1 | lncRNA | ENSG00000 |
| 79199982 | 79201632 | 1 | lncRNA | ENSG00000 |
| 79246668 | 79249023 | -1 | lncRNA | ENSG00000 |
| 79611418 | 79633311 | -1 | lncRNA | ENSG00000 |
| 79639754 | 79644431 | -1 | lncRNA | ENSG00000 |

| | | | | |
|---|---|---|---|---|
| 79661666 | 79791263 | -1 | lncRNA | ENSG0000C |
| 79893080 | 79974169 | -1 | lncRNA | ENSG0000C |
| 80197527 | 80387757 | -1 | protein_coding | ENSG0000C |
| 80211419 | 80455469 | 1 | lncRNA | ENSG0000C |
| 80476983 | 80959517 | -1 | protein_coding | ENSG0000C |
| 80822524 | 80822796 | 1 | processed_pseudoge | ENSG0000C |
| 80954989 | 81146302 | 1 | protein_coding | ENSG0000C |
| 80981988 | 80983638 | 1 | processed_pseudoge | ENSG0000C |
| 81001166 | 81001786 | -1 | unprocessed_pseudc | ENSG0000C |
| 81003325 | 81003645 | -1 | processed_pseudoge | ENSG0000C |
| 81012099 | 81170429 | -1 | lncRNA | ENSG0000C |
| 81032529 | 81033404 | 1 | processed_pseudoge | ENSG0000C |
| 81169776 | 81175400 | 1 | lncRNA | ENSG0000C |
| 81175452 | 81221377 | -1 | protein_coding | ENSG0000C |
| 81202695 | 81202834 | -1 | snoRNA | ENSG0000C |
| 81221218 | 81222460 | 1 | lncRNA | ENSG0000C |
| 81246295 | 81246563 | -1 | processed_pseudoge | ENSG0000C |
| 81259565 | 81260239 | -1 | processed_pseudoge | ENSG0000C |
| 81260656 | 81436465 | -1 | protein_coding | ENSG0000C |
| 81332961 | 81333361 | -1 | processed_pseudoge | ENSG0000C |
| 81412970 | 81413239 | 1 | processed_pseudoge | ENSG0000C |
| 81441987 | 81450371 | -1 | lncRNA | ENSG0000C |
| 81471547 | 81533853 | -1 | protein_coding | ENSG0000C |
| 81605338 | 81623086 | -1 | lncRNA | ENSG0000C |
| 81661428 | 81662874 | 1 | processed_pseudoge | ENSG0000C |
| 81727780 | 81743749 | -1 | lncRNA | ENSG0000C |
| 81741002 | 82030349 | 1 | lncRNA | ENSG0000C |
| 81777016 | 81777613 | -1 | processed_pseudoge | ENSG0000C |
| 81916231 | 81917888 | 1 | processed_pseudoge | ENSG0000C |
| 82429872 | 82430156 | 1 | misc_RNA | ENSG0000C |
| 82642605 | 82746664 | 1 | lncRNA | ENSG0000C |
| 82692704 | 82693055 | -1 | processed_pseudoge | ENSG0000C |
| 82788478 | 82796221 | -1 | lncRNA | ENSG0000C |
| 83017106 | 83018144 | -1 | lncRNA | ENSG0000C |
| 83018201 | 83039602 | -1 | lncRNA | ENSG0000C |
| 83048579 | 83048640 | 1 | snRNA | ENSG0000C |
| 83750384 | 83750501 | 1 | snRNA | ENSG0000C |
| 83805496 | 83805640 | -1 | processed_pseudoge | ENSG0000C |
| 83903437 | 83915377 | -1 | lncRNA | ENSG0000C |
| 83986776 | 84140544 | -1 | lncRNA | ENSG0000C |
| 84172841 | 84173606 | 1 | processed_pseudoge | ENSG0000C |
| 84173791 | 84175510 | 1 | processed_pseudoge | ENSG0000C |
| 84176419 | 84177088 | 1 | processed_pseudoge | ENSG0000C |
| 84739942 | 84740052 | 1 | snRNA | ENSG0000C |
| 84951274 | 84951373 | 1 | processed_pseudoge | ENSG0000C |
| 84998278 | 84999296 | -1 | processed_pseudoge | ENSG0000C |
| 85202849 | 85239474 | -1 | lncRNA | ENSG0000C |
| 85362457 | 85516934 | -1 | lncRNA | ENSG0000C |
| 85385927 | 85388765 | 1 | lncRNA | ENSG0000C |
| 85393828 | 85420550 | 1 | lncRNA | ENSG0000C |

| | | | | |
|---|---|---|---|---|
| 85518871 | 85531585 | -1 | lncRNA | ENSG00000 |
| 85528599 | 85529386 | 1 | lncRNA | ENSG00000 |
| 85530144 | 85654428 | 1 | protein_coding | ENSG00000 |
| 85911438 | 86161500 | 1 | lncRNA | ENSG00000 |
| 86006893 | 86062981 | -1 | lncRNA | ENSG00000 |
| 86283243 | 86304863 | 1 | lncRNA | ENSG00000 |
| 86332376 | 86401285 | 1 | lncRNA | ENSG00000 |
| 86905778 | 86922755 | 1 | lncRNA | ENSG00000 |
| 86936640 | 86989125 | 1 | lncRNA | ENSG00000 |
| 87251103 | 87251679 | 1 | lncRNA | ENSG00000 |
| 87323753 | 87332390 | 1 | lncRNA | ENSG00000 |
| 87353070 | 87573022 | -1 | lncRNA | ENSG00000 |
| 87568181 | 87569399 | 1 | lncRNA | ENSG00000 |
| 87634379 | 87655294 | -1 | lncRNA | ENSG00000 |
| 87710419 | 87872291 | -1 | lncRNA | ENSG00000 |
| 87810982 | 87838339 | 1 | lncRNA | ENSG00000 |
| 87837820 | 87993665 | -1 | protein_coding | ENSG00000 |
| 87952527 | 87955203 | 1 | processed_pseudoge | ENSG00000 |
| 87996345 | 87996448 | -1 | snRNA | ENSG00000 |
| 88005135 | 88014811 | 1 | protein_coding | ENSG00000 |
| 88010787 | 88015611 | -1 | lncRNA | ENSG00000 |
| 88018181 | 88036319 | -1 | lncRNA | ENSG00000 |
| 88024519 | 88098025 | 1 | lncRNA | ENSG00000 |
| 88136733 | 88159845 | -1 | lncRNA | ENSG00000 |
| 88158240 | 88160591 | -1 | lncRNA | ENSG00000 |
| 88175391 | 88180198 | 1 | lncRNA | ENSG00000 |
| 88180103 | 88326907 | -1 | protein_coding | ENSG00000 |
| 88384924 | 88470350 | 1 | protein_coding | ENSG00000 |
| 88465778 | 88555007 | -1 | protein_coding | ENSG00000 |
| 88499334 | 88515502 | 1 | lncRNA | ENSG00000 |
| 88513498 | 88513663 | 1 | snRNA | ENSG00000 |
| 88551597 | 88552493 | 1 | lncRNA | ENSG00000 |
| 88562970 | 88627596 | 1 | protein_coding | ENSG00000 |
| 88589231 | 88592408 | 1 | lncRNA | ENSG00000 |
| 88612431 | 88792752 | -1 | protein_coding | ENSG00000 |
| 88819608 | 88822151 | -1 | lncRNA | ENSG00000 |
| 88820297 | 88820432 | 1 | snRNA | ENSG00000 |
| 88824153 | 88881078 | 1 | protein_coding | ENSG00000 |
| 88840165 | 88840263 | 1 | misc_RNA | ENSG00000 |
| 89013386 | 89025807 | 1 | lncRNA | ENSG00000 |
| 89110148 | 89111223 | -1 | processed_pseudoge | ENSG00000 |
| 89124871 | 89619149 | -1 | protein_coding | ENSG00000 |
| 89156743 | 89157574 | 1 | lncRNA | ENSG00000 |
| 89290205 | 89291594 | -1 | transcribed_processe | ENSG00000 |
| 89349377 | 89366081 | 1 | lncRNA | ENSG00000 |
| 89355060 | 89356571 | 1 | lncRNA | ENSG00000 |
| 89399995 | 89409469 | 1 | lncRNA | ENSG00000 |
| 89412312 | 89954659 | -1 | lncRNA | ENSG00000 |
| 89417354 | 89419793 | 1 | lncRNA | ENSG00000 |
| 89576216 | 89577477 | 1 | lncRNA | ENSG00000 |

| | | | | |
|---|---|---|---|---|
| 89621918 | 89622207 | 1 | misc_RNA | ENSG00000 |
| 89628921 | 89642671 | -1 | lncRNA | ENSG00000 |
| 89650793 | 89650903 | -1 | misc_RNA | ENSG00000 |
| 89712511 | 89712823 | 1 | misc_RNA | ENSG00000 |
| 89736965 | 89738391 | -1 | processed_pseudoge | ENSG00000 |
| 89794669 | 89954777 | -1 | protein_coding | ENSG00000 |
| 89892128 | 89892443 | 1 | processed_pseudoge | ENSG00000 |
| 89954939 | 90044764 | 1 | protein_coding | ENSG00000 |
| 90061994 | 90185853 | 1 | protein_coding | ENSG00000 |
| 90107675 | 90107995 | 1 | processed_pseudoge | ENSG00000 |
| 90177222 | 90177330 | 1 | misc_RNA | ENSG00000 |
| 90256527 | 90275429 | 1 | protein_coding | ENSG00000 |
| 90267860 | 90331969 | -1 | protein_coding | ENSG00000 |
| 90291516 | 90294942 | 1 | lncRNA | ENSG00000 |
| 90366697 | 90367179 | -1 | processed_pseudoge | ENSG00000 |
| 90383365 | 90387973 | 1 | lncRNA | ENSG00000 |
| 90396502 | 90408268 | 1 | protein_coding | ENSG00000 |
| 90402523 | 90405235 | -1 | lncRNA | ENSG00000 |
| 90452063 | 90455127 | -1 | lncRNA | ENSG00000 |
| 90453571 | 90490763 | 1 | lncRNA | ENSG00000 |
| 90489010 | 90491637 | -1 | lncRNA | ENSG00000 |
| 90524564 | 90816479 | -1 | protein_coding | ENSG00000 |
| 90567495 | 90569976 | 1 | lncRNA | ENSG00000 |
| 90642638 | 90648894 | 1 | lncRNA | ENSG00000 |
| 90675118 | 90676294 | 1 | lncRNA | ENSG00000 |
| 90697333 | 90699374 | 1 | lncRNA | ENSG00000 |
| 90758599 | 90760061 | 1 | lncRNA | ENSG00000 |
| 90766531 | 90766989 | 1 | processed_pseudoge | ENSG00000 |
| 90820064 | 90828199 | -1 | lncRNA | ENSG00000 |
| 90847861 | 91060641 | -1 | protein_coding | ENSG00000 |
| 91060333 | 91225632 | 1 | protein_coding | ENSG00000 |
| 91091077 | 91091138 | 1 | snRNA | ENSG00000 |
| 91126425 | 91126553 | 1 | snoRNA | ENSG00000 |
| 91232532 | 91253925 | -1 | protein_coding | ENSG00000 |
| 91242759 | 91252211 | 1 | lncRNA | ENSG00000 |
| 91258299 | 91259003 | 1 | lncRNA | ENSG00000 |
| 91264903 | 91266666 | 1 | lncRNA | ENSG00000 |
| 91271323 | 91417844 | -1 | protein_coding | ENSG00000 |
| 91418266 | 91421176 | 1 | lncRNA | ENSG00000 |
| 91419449 | 91419751 | 1 | processed_pseudoge | ENSG00000 |
| 91457611 | 91510554 | -1 | protein_coding | ENSG00000 |
| 91554410 | 91554503 | 1 | unprocessed_pseudo | ENSG00000 |
| 91572444 | 91575271 | -1 | TEC | ENSG00000 |
| 91580696 | 91780707 | -1 | protein_coding | ENSG00000 |
| 91603173 | 91603433 | -1 | transcribed_process | ENSG00000 |
| 91604402 | 91605319 | -1 | transcribed_process | ENSG00000 |
| 91709111 | 91710617 | 1 | processed_pseudoge | ENSG00000 |
| 91714915 | 91715130 | -1 | snoRNA | ENSG00000 |
| 91752856 | 91759798 | -1 | lncRNA | ENSG00000 |
| 91779746 | 91867536 | -1 | protein_coding | ENSG00000 |

| | | | | |
|---|---|---|---|---|
| 91869412 | 91947987 | -1 | protein_coding | ENSG0000C |
| 91965991 | 92040896 | -1 | protein_coding | ENSG0000C |
| 92026566 | 92026887 | 1 | processed_pseudoge | ENSG0000C |
| 92044496 | 92106621 | -1 | protein_coding | ENSG0000C |
| 92116122 | 92121917 | -1 | protein_coding | ENSG0000C |
| 92121969 | 92172145 | 1 | protein_coding | ENSG0000C |
| 92253493 | 92263656 | -1 | lncRNA | ENSG0000C |
| 92265702 | 92265807 | 1 | snRNA | ENSG0000C |
| 92294771 | 92297648 | 1 | lncRNA | ENSG0000C |
| 92322581 | 92501483 | 1 | protein_coding | ENSG0000C |
| 92513781 | 92688994 | 1 | protein_coding | ENSG0000C |
| 92703807 | 92748679 | -1 | protein_coding | ENSG0000C |
| 92794305 | 92839947 | 1 | protein_coding | ENSG0000C |
| 92886352 | 92893508 | 1 | lncRNA | ENSG0000C |
| 92905697 | 92907482 | -1 | lncRNA | ENSG0000C |
| 92923150 | 92935285 | 1 | protein_coding | ENSG0000C |
| 92936914 | 93116320 | -1 | protein_coding | ENSG0000C |
| 93067452 | 93072152 | 1 | lncRNA | ENSG0000C |
| 93138066 | 93138465 | 1 | processed_pseudoge | ENSG0000C |
| 93182199 | 93184923 | -1 | protein_coding | ENSG0000C |
| 93184951 | 93187089 | 1 | protein_coding | ENSG0000C |
| 93184973 | 93218586 | 1 | protein_coding | ENSG0000C |
| 93202894 | 93207065 | -1 | protein_coding | ENSG0000C |
| 93207241 | 93229215 | 1 | protein_coding | ENSG0000C |
| 93223598 | 93224139 | -1 | processed_pseudoge | ENSG0000C |
| 93237550 | 93333092 | -1 | protein_coding | ENSG0000C |
| 93244010 | 93244323 | 1 | processed_pseudoge | ENSG0000C |
| 93333219 | 93707876 | 1 | protein_coding | ENSG0000C |
| 93334528 | 93335057 | 1 | lncRNA | ENSG0000C |
| 93347182 | 93348356 | 1 | protein_coding | ENSG0000C |
| 93397139 | 93397234 | -1 | misc_RNA | ENSG0000C |
| 93437400 | 93437510 | -1 | snRNA | ENSG0000C |
| 93637179 | 93637338 | 1 | misc_RNA | ENSG0000C |
| 93718298 | 93788485 | -1 | protein_coding | ENSG0000C |
| 93895762 | 93897759 | 1 | TEC | ENSG0000C |
| 93904730 | 93927066 | -1 | lncRNA | ENSG0000C |
| 93918894 | 93929608 | 1 | protein_coding | ENSG0000C |
| 93934166 | 93976739 | -1 | protein_coding | ENSG0000C |
| 93939497 | 93944194 | 1 | lncRNA | ENSG0000C |
| 93948226 | 93948302 | -1 | miRNA | ENSG0000C |
| 93987225 | 94008863 | 1 | protein_coding | ENSG0000C |
| 94026340 | 94048930 | 1 | protein_coding | ENSG0000C |
| 94048287 | 94081212 | -1 | protein_coding | ENSG0000C |
| 94081301 | 94103846 | 1 | protein_coding | ENSG0000C |
| 94104836 | 94116698 | 1 | protein_coding | ENSG0000C |
| 94127779 | 94130253 | -1 | protein_coding | ENSG0000C |
| 94146128 | 94279734 | 1 | protein_coding | ENSG0000C |
| 94280460 | 94293268 | -1 | protein_coding | ENSG0000C |
| 94304248 | 94323389 | -1 | protein_coding | ENSG0000C |
| 94364313 | 94366698 | -1 | polymorphic_pseudo | ENSG0000C |

| | | | | |
|---|---|---|---|---|
| 94376747 | 94390693 | -1 | protein_coding | ENSG00000 |
| 94430633 | 94464730 | 1 | lncRNA | ENSG00000 |
| 94442464 | 94452800 | -1 | protein_coding | ENSG00000 |
| 94462717 | 94479689 | -1 | protein_coding | ENSG00000 |
| 94487274 | 94517844 | -1 | protein_coding | ENSG00000 |
| 94545310 | 94547762 | 1 | lncRNA | ENSG00000 |
| 94561442 | 94569913 | 1 | protein_coding | ENSG00000 |
| 94561442 | 94593118 | 1 | protein_coding | ENSG00000 |
| 94592058 | 94624646 | 1 | protein_coding | ENSG00000 |
| 94599460 | 94607839 | 1 | lncRNA | ENSG00000 |
| 94612377 | 94624055 | 1 | protein_coding | ENSG00000 |
| 94633705 | 94634524 | 1 | processed_pseudoge | ENSG00000 |
| 94640725 | 94646994 | 1 | transcribed_unproce | ENSG00000 |
| 94674877 | 94675829 | -1 | processed_pseudoge | ENSG00000 |
| 94726288 | 94727175 | 1 | processed_pseudoge | ENSG00000 |
| 94768216 | 94770230 | -1 | protein_coding | ENSG00000 |
| 94770271 | 94774858 | 1 | lncRNA | ENSG00000 |
| 94800849 | 94801463 | -1 | processed_pseudoge | ENSG00000 |
| 94918253 | 94918734 | -1 | lncRNA | ENSG00000 |
| 94960277 | 94962976 | -1 | lncRNA | ENSG00000 |
| 95047353 | 95050957 | 1 | lncRNA | ENSG00000 |
| 95086228 | 95158010 | -1 | protein_coding | ENSG00000 |
| 95137919 | 95137986 | -1 | miRNA | ENSG00000 |
| 95157645 | 95181475 | 1 | lncRNA | ENSG00000 |
| 95181940 | 95319908 | -1 | protein_coding | ENSG00000 |
| 95185117 | 95185854 | -1 | lncRNA | ENSG00000 |
| 95199039 | 95199345 | 1 | unprocessed_pseudo | ENSG00000 |
| 95201278 | 95201439 | 1 | unprocessed_pseudo | ENSG00000 |
| 95319483 | 95335757 | 1 | lncRNA | ENSG00000 |
| 95407266 | 95475836 | -1 | protein_coding | ENSG00000 |
| 95516136 | 95517911 | 1 | lncRNA | ENSG00000 |
| 95521943 | 95534889 | -1 | lncRNA | ENSG00000 |
| 95533355 | 95533629 | -1 | scaRNA | ENSG00000 |
| 95533503 | 95544724 | 1 | protein_coding | ENSG00000 |
| 95573520 | 95582305 | 1 | lncRNA | ENSG00000 |
| 95620914 | 95643285 | 1 | lncRNA | ENSG00000 |
| 95644508 | 95645232 | 1 | lncRNA | ENSG00000 |
| 95650498 | 95679833 | 1 | lncRNA | ENSG00000 |
| 95662480 | 95662598 | 1 | misc_RNA | ENSG00000 |
| 95663256 | 95692630 | 1 | lncRNA | ENSG00000 |
| 95670441 | 95670577 | 1 | misc_RNA | ENSG00000 |
| 95686426 | 95692628 | 1 | protein_coding | ENSG00000 |
| 95709947 | 95714196 | -1 | protein_coding | ENSG00000 |
| 95711747 | 95757656 | 1 | lncRNA | ENSG00000 |
| 95828763 | 95866486 | -1 | lncRNA | ENSG00000 |
| 95876392 | 95925571 | 1 | protein_coding | ENSG00000 |
| 96019484 | 96019765 | 1 | processed_pseudoge | ENSG00000 |
| 96039324 | 96093889 | 1 | protein_coding | ENSG00000 |
| 96204679 | 96244166 | 1 | protein_coding | ENSG00000 |
| 96204844 | 96263929 | 1 | protein_coding | ENSG00000 |

| | | | | |
|---|---|---|---|---|
| 96210860 | 96214818 | 1 | lncRNA | ENSG0000C |
| 96223108 | 96223347 | 1 | processed_pseudoge | ENSG0000C |
| 96233431 | 96234101 | -1 | lncRNA | ENSG0000C |
| 96256210 | 96268967 | 1 | protein_coding | ENSG0000C |
| 96259411 | 96268624 | -1 | lncRNA | ENSG0000C |
| 96275046 | 96276723 | -1 | lncRNA | ENSG0000C |
| 96279195 | 96363341 | -1 | protein_coding | ENSG0000C |
| 96363452 | 96387288 | 1 | protein_coding | ENSG0000C |
| 96384624 | 96384815 | 1 | snRNA | ENSG0000C |
| 96392128 | 96489427 | 1 | protein_coding | ENSG0000C |
| 96423700 | 96424209 | -1 | processed_pseudoge | ENSG0000C |
| 96456576 | 96457041 | -1 | processed_pseudoge | ENSG0000C |
| 96500810 | 96502321 | -1 | lncRNA | ENSG0000C |
| 96501433 | 96567111 | 1 | protein_coding | ENSG0000C |
| 96592399 | 96605974 | 1 | lncRNA | ENSG0000C |
| 96638187 | 96638514 | -1 | misc_RNA | ENSG0000C |
| 96663791 | 96681827 | 1 | lncRNA | ENSG0000C |
| 96738021 | 96793084 | 1 | lncRNA | ENSG0000C |
| 96749695 | 96755556 | -1 | lncRNA | ENSG0000C |
| 96797382 | 96931722 | 1 | protein_coding | ENSG0000C |
| 96931367 | 96945394 | 1 | lncRNA | ENSG0000C |
| 96992864 | 96996601 | 1 | TEC | ENSG0000C |
| 97005592 | 97007624 | -1 | lncRNA | ENSG0000C |
| 97110416 | 97119247 | -1 | lncRNA | ENSG0000C |
| 97116355 | 97121501 | 1 | lncRNA | ENSG0000C |
| 97154857 | 97158736 | 1 | lncRNA | ENSG0000C |
| 97180525 | 97217778 | 1 | lncRNA | ENSG0000C |
| 97427071 | 97582146 | 1 | lncRNA | ENSG0000C |
| 97632647 | 97686658 | -1 | lncRNA | ENSG0000C |
| 97662638 | 97688600 | 1 | lncRNA | ENSG0000C |
| 97749264 | 97763989 | -1 | lncRNA | ENSG0000C |
| 97786565 | 97789256 | -1 | lncRNA | ENSG0000C |
| 97924637 | 97978124 | -1 | lncRNA | ENSG0000C |
| 98068161 | 98205143 | -1 | lncRNA | ENSG0000C |
| 98136074 | 98139242 | 1 | lncRNA | ENSG0000C |
| 98351367 | 98351664 | -1 | misc_RNA | ENSG0000C |
| 98497132 | 98514798 | 1 | lncRNA | ENSG0000C |
| 98711613 | 98717766 | 1 | protein_coding | ENSG0000C |
| 98844896 | 98845597 | 1 | lncRNA | ENSG0000C |
| 98916902 | 98917801 | -1 | TEC | ENSG0000C |
| 98925137 | 98927805 | -1 | lncRNA | ENSG0000C |
| 98972879 | 98973301 | -1 | processed_pseudoge | ENSG0000C |
| 98973314 | 98973471 | 1 | processed_pseudoge | ENSG0000C |
| 98999571 | 99003545 | 1 | lncRNA | ENSG0000C |
| 99158416 | 99159669 | 1 | lncRNA | ENSG0000C |
| 99169287 | 99272197 | -1 | protein_coding | ENSG0000C |
| 99262948 | 99264735 | 1 | lncRNA | ENSG0000C |
| 99324799 | 99332015 | -1 | lncRNA | ENSG0000C |
| 99397748 | 99480889 | -1 | protein_coding | ENSG0000C |
| 99444306 | 99444412 | 1 | snRNA | ENSG0000C |

| | | | | |
|---|---|---|---|---|
| 99481169 | 99535044 | 1 | protein_coding | ENSG0000( |
| 99500190 | 99604207 | -1 | protein_coding | ENSG0000( |
| 99512501 | 99513576 | 1 | lncRNA | ENSG0000( |
| 99583017 | 99583118 | 1 | misc_RNA | ENSG0000( |
| 99604556 | 99625740 | 1 | lncRNA | ENSG0000( |
| 99645110 | 99680569 | 1 | protein_coding | ENSG0000( |
| 99684298 | 99727301 | 1 | protein_coding | ENSG0000( |
| 99691445 | 99693648 | -1 | lncRNA | ENSG0000( |
| 99737693 | 99942060 | 1 | protein_coding | ENSG0000( |
| 99945315 | 99945477 | 1 | snRNA | ENSG0000( |
| 99964529 | 99965378 | 1 | processed_pseudoge | ENSG0000( |
| 99971449 | 100144236 | 1 | protein_coding | ENSG0000( |
| 99974336 | 99975149 | -1 | processed_pseudoge | ENSG0000( |
| 99977115 | 99978098 | -1 | lncRNA | ENSG0000( |
| 100052803 | 100074878 | -1 | lncRNA | ENSG0000( |
| 100109419 | 100109514 | -1 | miRNA | ENSG0000( |
| 100109655 | 100109753 | 1 | miRNA | ENSG0000( |
| 100143957 | 100159645 | -1 | protein_coding | ENSG0000( |
| 100207407 | 100238555 | -1 | lncRNA | ENSG0000( |
| 100238298 | 100282788 | 1 | protein_coding | ENSG0000( |
| 100263781 | 100265405 | 1 | lncRNA | ENSG0000( |
| 100277357 | 100277417 | 1 | miRNA | ENSG0000( |
| 100279959 | 100291456 | -1 | lncRNA | ENSG0000( |
| 100291116 | 100306547 | -1 | protein_coding | ENSG0000( |
| 100291117 | 100294656 | 1 | lncRNA | ENSG0000( |
| 100307859 | 100307956 | 1 | miRNA | ENSG0000( |
| 100319073 | 100319370 | -1 | misc_RNA | ENSG0000( |
| 100323339 | 100330421 | 1 | protein_coding | ENSG0000( |
| 100333790 | 100376805 | -1 | protein_coding | ENSG0000( |
| 100333790 | 100354061 | 1 | lncRNA | ENSG0000( |
| 100339832 | 100340554 | 1 | lncRNA | ENSG0000( |
| 100366628 | 100366920 | 1 | processed_pseudoge | ENSG0000( |
| 100376418 | 100530303 | 1 | protein_coding | ENSG0000( |
| 100406599 | 100408928 | 1 | lncRNA | ENSG0000( |
| 100480952 | 100481274 | 1 | transcribed_processe | ENSG0000( |
| 100487544 | 100487787 | 1 | misc_RNA | ENSG0000( |
| 100537147 | 100587413 | -1 | protein_coding | ENSG0000( |
| 100538939 | 100540409 | 1 | lncRNA | ENSG0000( |
| 100587775 | 100589326 | -1 | lncRNA | ENSG0000( |
| 100657250 | 100676069 | 1 | lncRNA | ENSG0000( |
| 100675930 | 100679884 | -1 | lncRNA | ENSG0000( |
| 100725705 | 100738224 | 1 | protein_coding | ENSG0000( |
| 100779410 | 100861031 | 1 | lncRNA | ENSG0000( |
| 100814491 | 100814574 | 1 | miRNA | ENSG0000( |
| 100816041 | 100816244 | 1 | misc_RNA | ENSG0000( |
| 100826284 | 100826370 | 1 | misc_RNA | ENSG0000( |
| 100829350 | 100829454 | 1 | misc_RNA | ENSG0000( |
| 100834432 | 100861026 | -1 | lncRNA | ENSG0000( |
| 100836269 | 100947194 | 1 | lncRNA | ENSG0000( |
| 100852390 | 100852487 | 1 | miRNA | ENSG0000( |

| | | | | |
|---|---|---|---|---|
| 100869060 | 100869148 | 1 | miRNA | ENSG00000 |
| 100874493 | 100874585 | 1 | miRNA | ENSG00000 |
| 100875033 | 100875104 | 1 | miRNA | ENSG00000 |
| 100879753 | 100903722 | -1 | protein_coding | ENSG00000 |
| 100881007 | 100881120 | 1 | miRNA | ENSG00000 |
| 100881886 | 100881978 | 1 | miRNA | ENSG00000 |
| 100882979 | 100883075 | 1 | miRNA | ENSG00000 |
| 100884483 | 100884576 | 1 | miRNA | ENSG00000 |
| 100884702 | 100884783 | 1 | miRNA | ENSG00000 |
| 100894770 | 101038859 | 1 | lncRNA | ENSG00000 |
| 100894817 | 100894913 | 1 | misc_RNA | ENSG00000 |
| 100897920 | 100897996 | 1 | snoRNA | ENSG00000 |
| 100898958 | 100899079 | 1 | misc_RNA | ENSG00000 |
| 100911139 | 100911213 | 1 | miRNA | ENSG00000 |
| 100924824 | 100924892 | 1 | snoRNA | ENSG00000 |
| 100927342 | 100927413 | 1 | snoRNA | ENSG00000 |
| 100929919 | 100929990 | 1 | snoRNA | ENSG00000 |
| 100936491 | 100936565 | 1 | snoRNA | ENSG00000 |
| 100938187 | 100938264 | 1 | snoRNA | ENSG00000 |
| 100939556 | 100939630 | 1 | snoRNA | ENSG00000 |
| 100941126 | 100941202 | 1 | snoRNA | ENSG00000 |
| 100943451 | 100943524 | 1 | snoRNA | ENSG00000 |
| 100945649 | 100945720 | 1 | snoRNA | ENSG00000 |
| 100949833 | 100949904 | 1 | snoRNA | ENSG00000 |
| 100951856 | 100951933 | 1 | snoRNA | ENSG00000 |
| 100953349 | 100953423 | 1 | snoRNA | ENSG00000 |
| 100954374 | 100954448 | 1 | snoRNA | ENSG00000 |
| 100955370 | 100955439 | 1 | snoRNA | ENSG00000 |
| 100956240 | 100956312 | 1 | snoRNA | ENSG00000 |
| 100957166 | 100957237 | 1 | snoRNA | ENSG00000 |
| 100963054 | 100963130 | 1 | snoRNA | ENSG00000 |
| 100966029 | 100966100 | 1 | snoRNA | ENSG00000 |
| 100967052 | 100967123 | 1 | snoRNA | ENSG00000 |
| 100968111 | 100968185 | 1 | snoRNA | ENSG00000 |
| 100968948 | 100969022 | 1 | snoRNA | ENSG00000 |
| 100969879 | 100969952 | 1 | snoRNA | ENSG00000 |
| 100972103 | 100972177 | 1 | snoRNA | ENSG00000 |
| 100972670 | 100972741 | 1 | snoRNA | ENSG00000 |
| 100973595 | 100973664 | 1 | snoRNA | ENSG00000 |
| 100974806 | 100974880 | 1 | snoRNA | ENSG00000 |
| 100975825 | 100975896 | 1 | snoRNA | ENSG00000 |
| 100976477 | 100976551 | 1 | snoRNA | ENSG00000 |
| 100977389 | 100977466 | 1 | snoRNA | ENSG00000 |
| 100979002 | 100979090 | 1 | snoRNA | ENSG00000 |
| 100979992 | 100980066 | 1 | snoRNA | ENSG00000 |
| 100981004 | 100981075 | 1 | snoRNA | ENSG00000 |
| 100981975 | 100982046 | 1 | snoRNA | ENSG00000 |
| 100982926 | 100982997 | 1 | snoRNA | ENSG00000 |
| 100983876 | 100983947 | 1 | snoRNA | ENSG00000 |
| 100984777 | 100984848 | 1 | snoRNA | ENSG00000 |

| | | | | |
|---|---|---|---|---|
| 100986057 | 100986128 | 1 | snoRNA | ENSG00000 |
| 100987046 | 100987117 | 1 | snoRNA | ENSG00000 |
| 100988161 | 100988230 | 1 | snoRNA | ENSG00000 |
| 100989130 | 100989201 | 1 | snoRNA | ENSG00000 |
| 100990091 | 100990160 | 1 | snoRNA | ENSG00000 |
| 100991919 | 100991990 | 1 | snoRNA | ENSG00000 |
| 100993236 | 100993310 | 1 | snoRNA | ENSG00000 |
| 100994257 | 100994330 | 1 | snoRNA | ENSG00000 |
| 100998467 | 100998541 | 1 | snoRNA | ENSG00000 |
| 101022066 | 101022132 | 1 | miRNA | ENSG00000 |
| 101023325 | 101023420 | 1 | miRNA | ENSG00000 |
| 101023794 | 101023856 | 1 | miRNA | ENSG00000 |
| 101025017 | 101025077 | 1 | miRNA | ENSG00000 |
| 101025564 | 101025651 | 1 | miRNA | ENSG00000 |
| 101025732 | 101025817 | 1 | miRNA | ENSG00000 |
| 101026020 | 101026107 | 1 | miRNA | ENSG00000 |
| 101026785 | 101026864 | 1 | miRNA | ENSG00000 |
| 101027100 | 101027183 | 1 | miRNA | ENSG00000 |
| 101029634 | 101029714 | 1 | miRNA | ENSG00000 |
| 101030052 | 101030129 | 1 | miRNA | ENSG00000 |
| 101031987 | 101032064 | 1 | miRNA | ENSG00000 |
| 101033755 | 101033836 | 1 | miRNA | ENSG00000 |
| 101039690 | 101039755 | 1 | miRNA | ENSG00000 |
| 101040069 | 101040148 | 1 | miRNA | ENSG00000 |
| 101040219 | 101040299 | 1 | miRNA | ENSG00000 |
| 101040436 | 101040535 | 1 | miRNA | ENSG00000 |
| 101040782 | 101040849 | 1 | miRNA | ENSG00000 |
| 101041363 | 101041445 | 1 | miRNA | ENSG00000 |
| 101042977 | 101043062 | 1 | miRNA | ENSG00000 |
| 101044198 | 101044283 | 1 | miRNA | ENSG00000 |
| 101045157 | 101051795 | 1 | lncRNA | ENSG00000 |
| 101045920 | 101045994 | 1 | miRNA | ENSG00000 |
| 101046455 | 101046538 | 1 | miRNA | ENSG00000 |
| 101047321 | 101047398 | 1 | miRNA | ENSG00000 |
| 101047901 | 101047979 | 1 | miRNA | ENSG00000 |
| 101048658 | 101048748 | 1 | miRNA | ENSG00000 |
| 101049550 | 101049646 | 1 | miRNA | ENSG00000 |
| 101052446 | 101052525 | 1 | miRNA | ENSG00000 |
| 101054306 | 101054381 | 1 | miRNA | ENSG00000 |
| 101054687 | 101054759 | 1 | miRNA | ENSG00000 |
| 101055258 | 101055323 | 1 | miRNA | ENSG00000 |
| 101055419 | 101055491 | 1 | miRNA | ENSG00000 |
| 101056219 | 101056300 | 1 | miRNA | ENSG00000 |
| 101059755 | 101059838 | 1 | miRNA | ENSG00000 |
| 101060573 | 101060674 | 1 | miRNA | ENSG00000 |
| 101062050 | 101062118 | 1 | miRNA | ENSG00000 |
| 101064495 | 101064578 | 1 | miRNA | ENSG00000 |
| 101065300 | 101065378 | 1 | miRNA | ENSG00000 |
| 101065447 | 101065537 | 1 | miRNA | ENSG00000 |
| 101065598 | 101065667 | 1 | miRNA | ENSG00000 |

| | | | | |
|---|---|---|---|---|
| 101065912 | 101065991 | 1 | miRNA | ENSG0000C |
| 101066724 | 101066801 | 1 | miRNA | ENSG0000C |
| 101068283 | 101072937 | 1 | lncRNA | ENSG0000C |
| 101071978 | 101077910 | -1 | lncRNA | ENSG0000C |
| 101120514 | 101121309 | -1 | lncRNA | ENSG0000C |
| 101120840 | 101123545 | 1 | lncRNA | ENSG0000C |
| 101133730 | 101151988 | -1 | lncRNA | ENSG0000C |
| 101145480 | 101145695 | -1 | snoRNA | ENSG0000C |
| 101160476 | 101172549 | 1 | lncRNA | ENSG0000C |
| 101185787 | 101197089 | -1 | lncRNA | ENSG0000C |
| 101217372 | 101218241 | -1 | processed_pseudoge | ENSG0000C |
| 101257332 | 101278318 | 1 | lncRNA | ENSG0000C |
| 101332511 | 101335497 | -1 | lncRNA | ENSG0000C |
| 101405984 | 101408258 | -1 | lncRNA | ENSG0000C |
| 101442076 | 101444315 | 1 | lncRNA | ENSG0000C |
| 101447642 | 101457117 | 1 | lncRNA | ENSG0000C |
| 101552221 | 101560431 | -1 | lncRNA | ENSG0000C |
| 101560287 | 101560422 | -1 | miRNA | ENSG0000C |
| 101561351 | 101563452 | 1 | protein_coding | ENSG0000C |
| 101628984 | 101632530 | -1 | lncRNA | ENSG0000C |
| 101634454 | 101731108 | -1 | lncRNA | ENSG0000C |
| 101677943 | 101678380 | 1 | processed_pseudoge | ENSG0000C |
| 101730437 | 101732522 | 1 | lncRNA | ENSG0000C |
| 101761798 | 101927989 | 1 | protein_coding | ENSG0000C |
| 101796555 | 101810321 | -1 | lncRNA | ENSG0000C |
| 101803227 | 101803331 | 1 | snRNA | ENSG0000C |
| 101833435 | 101839531 | -1 | lncRNA | ENSG0000C |
| 101843394 | 101844122 | -1 | processed_pseudoge | ENSG0000C |
| 101948347 | 101949425 | 1 | lncRNA | ENSG0000C |
| 101952416 | 101953063 | 1 | lncRNA | ENSG0000C |
| 101964573 | 102056443 | 1 | protein_coding | ENSG0000C |
| 101969364 | 101969601 | -1 | processed_pseudoge | ENSG0000C |
| 102036315 | 102066228 | -1 | transcribed_unitary_ | ENSG0000C |
| 102077180 | 102077472 | -1 | misc_RNA | ENSG0000C |
| 102080738 | 102139699 | -1 | protein_coding | ENSG0000C |
| 102139503 | 102224847 | 1 | protein_coding | ENSG0000C |
| 102224500 | 102305190 | -1 | protein_coding | ENSG0000C |
| 102317377 | 102342702 | 1 | protein_coding | ENSG0000C |
| 102341102 | 102362916 | -1 | protein_coding | ENSG0000C |
| 102362941 | 102502477 | 1 | protein_coding | ENSG0000C |
| 102501676 | 102501779 | 1 | snRNA | ENSG0000C |
| 102501767 | 102509799 | -1 | protein_coding | ENSG0000C |
| 102539644 | 102539726 | 1 | miRNA | ENSG0000C |
| 102545254 | 102555826 | 1 | lncRNA | ENSG0000C |
| 102582888 | 102583177 | -1 | misc_RNA | ENSG0000C |
| 102592649 | 102730561 | 1 | protein_coding | ENSG0000C |
| 102662240 | 102662708 | -1 | processed_pseudoge | ENSG0000C |
| 102695388 | 102695495 | 1 | misc_RNA | ENSG0000C |
| 102770040 | 102774791 | -1 | lncRNA | ENSG0000C |
| 102777476 | 102911500 | 1 | protein_coding | ENSG0000C |

| | | | | |
|---|---|---|---|---|
| 102865856 | 102865963 | 1 | snRNA | ENSG00000 |
| 102922656 | 102933596 | 1 | protein_coding | ENSG00000 |
| 102932380 | 103057549 | -1 | protein_coding | ENSG00000 |
| 102933574 | 102937177 | 1 | lncRNA | ENSG00000 |
| 103041469 | 103042094 | 1 | processed_pseudoge | ENSG00000 |
| 103084210 | 103090027 | 1 | protein_coding | ENSG00000 |
| 103094723 | 103098885 | 1 | lncRNA | ENSG00000 |
| 103100144 | 103110559 | 1 | protein_coding | ENSG00000 |
| 103120847 | 103123007 | -1 | lncRNA | ENSG00000 |
| 103123442 | 103137439 | 1 | protein_coding | ENSG00000 |
| 103139675 | 103141959 | 1 | transcribed_process | ENSG00000 |
| 103148597 | 103149408 | 1 | processed_pseudoge | ENSG00000 |
| 103184204 | 103185039 | 1 | processed_pseudoge | ENSG00000 |
| 103187221 | 103189028 | -1 | lncRNA | ENSG00000 |
| 103207748 | 103207789 | 1 | processed_pseudoge | ENSG00000 |
| 103208100 | 103208876 | 1 | lncRNA | ENSG00000 |
| 103241492 | 103241972 | 1 | processed_pseudoge | ENSG00000 |
| 103279223 | 103279955 | 1 | transcribed_process | ENSG00000 |
| 103282933 | 103283468 | 1 | processed_pseudoge | ENSG00000 |
| 103331674 | 103332367 | -1 | lncRNA | ENSG00000 |
| 103333544 | 103345025 | 1 | protein_coding | ENSG00000 |
| 103337849 | 103337974 | 1 | snoRNA | ENSG00000 |
| 103364239 | 103364820 | -1 | processed_pseudoge | ENSG00000 |
| 103374033 | 103374916 | -1 | processed_pseudoge | ENSG00000 |
| 103385392 | 103503831 | 1 | protein_coding | ENSG00000 |
| 103412119 | 103412761 | -1 | processed_pseudoge | ENSG00000 |
| 103519667 | 103522833 | -1 | protein_coding | ENSG00000 |
| 103525010 | 103529072 | -1 | lncRNA | ENSG00000 |
| 103529196 | 103537073 | 1 | protein_coding | ENSG00000 |
| 103550345 | 103550406 | 1 | snRNA | ENSG00000 |
| 103553421 | 103561877 | 1 | lncRNA | ENSG00000 |
| 103556544 | 103562831 | -1 | protein_coding | ENSG00000 |
| 103561896 | 103714249 | 1 | protein_coding | ENSG00000 |
| 103562962 | 103685924 | 1 | protein_coding | ENSG00000 |
| 103562962 | 103607523 | 1 | protein_coding | ENSG00000 |
| 103567405 | 103567545 | 1 | snRNA | ENSG00000 |
| 103682362 | 103684015 | -1 | lncRNA | ENSG00000 |
| 103687576 | 103688127 | 1 | lncRNA | ENSG00000 |
| 103694516 | 103695050 | -1 | lncRNA | ENSG00000 |
| 103694560 | 103695170 | 1 | lncRNA | ENSG00000 |
| 103696353 | 103697163 | 1 | lncRNA | ENSG00000 |
| 103697609 | 103715504 | -1 | protein_coding | ENSG00000 |
| 103715730 | 103733668 | 1 | protein_coding | ENSG00000 |
| 103725728 | 103725825 | -1 | misc_RNA | ENSG00000 |
| 103733195 | 103847575 | -1 | protein_coding | ENSG00000 |
| 103762803 | 103763277 | 1 | transcribed_process | ENSG00000 |
| 103797272 | 103797348 | -1 | snoRNA | ENSG00000 |
| 103847721 | 103858049 | 1 | lncRNA | ENSG00000 |
| 103854366 | 103880111 | -1 | lncRNA | ENSG00000 |
| 103873148 | 103880381 | 1 | lncRNA | ENSG00000 |

| | | | | |
|---|---|---|---|---|
| 103912288 | 103928269 | -1 | protein_coding | ENSG00000 |
| 103928456 | 104052667 | 1 | protein_coding | ENSG00000 |
| 103940426 | 103942561 | -1 | protein_coding | ENSG00000 |
| 104013642 | 104013933 | 1 | misc_RNA | ENSG00000 |
| 104085686 | 104115582 | 1 | protein_coding | ENSG00000 |
| 104117405 | 104117514 | 1 | miRNA | ENSG00000 |
| 104117418 | 104117503 | -1 | miRNA | ENSG00000 |
| 104137150 | 104137898 | -1 | lncRNA | ENSG00000 |
| 104138723 | 104180894 | 1 | protein_coding | ENSG00000 |
| 104223584 | 104288069 | 1 | lncRNA | ENSG00000 |
| 104293678 | 104294921 | 1 | lncRNA | ENSG00000 |
| 104376530 | 104378951 | -1 | lncRNA | ENSG00000 |
| 104400943 | 104401048 | -1 | snRNA | ENSG00000 |
| 104450979 | 104451433 | 1 | processed_pseudoge | ENSG00000 |
| 104474678 | 104605647 | -1 | protein_coding | ENSG00000 |
| 104579684 | 104590515 | 1 | protein_coding | ENSG00000 |
| 104589021 | 104589847 | -1 | lncRNA | ENSG00000 |
| 104653548 | 104655787 | -1 | lncRNA | ENSG00000 |
| 104661120 | 104665558 | -1 | lncRNA | ENSG00000 |
| 104677694 | 104677749 | -1 | miRNA | ENSG00000 |
| 104681146 | 104684932 | 1 | lncRNA | ENSG00000 |
| 104689606 | 104722535 | 1 | protein_coding | ENSG00000 |
| 104690091 | 104691284 | -1 | lncRNA | ENSG00000 |
| 104724186 | 104747325 | 1 | protein_coding | ENSG00000 |
| 104753147 | 104768494 | 1 | protein_coding | ENSG00000 |
| 104769349 | 104770271 | 1 | lncRNA | ENSG00000 |
| 104769349 | 104795751 | -1 | protein_coding | ENSG00000 |
| 104800596 | 104804712 | 1 | protein_coding | ENSG00000 |
| 104815449 | 104815753 | -1 | TEC | ENSG00000 |
| 104821201 | 104823718 | 1 | lncRNA | ENSG00000 |
| 104830838 | 104831171 | -1 | processed_pseudoge | ENSG00000 |
| 104835997 | 104836852 | -1 | processed_pseudoge | ENSG00000 |
| 104842663 | 104845916 | 1 | lncRNA | ENSG00000 |
| 104857792 | 104858836 | -1 | lncRNA | ENSG00000 |
| 104865268 | 104896770 | 1 | protein_coding | ENSG00000 |
| 104904342 | 104905204 | 1 | lncRNA | ENSG00000 |
| 104924713 | 104937790 | 1 | protein_coding | ENSG00000 |
| 104937244 | 104978374 | -1 | protein_coding | ENSG00000 |
| 104985775 | 105010482 | 1 | protein_coding | ENSG00000 |
| 105009573 | 105021083 | -1 | protein_coding | ENSG00000 |
| 105049395 | 105065445 | -1 | protein_coding | ENSG00000 |
| 105093609 | 105100131 | 1 | lncRNA | ENSG00000 |
| 105095045 | 105096945 | -1 | lncRNA | ENSG00000 |
| 105110347 | 105110807 | -1 | TEC | ENSG00000 |
| 105112690 | 105112863 | -1 | unprocessed_pseudc | ENSG00000 |
| 105126750 | 105136974 | -1 | lncRNA | ENSG00000 |
| 105140982 | 105168824 | -1 | protein_coding | ENSG00000 |
| 105150778 | 105150864 | -1 | miRNA | ENSG00000 |
| 105152194 | 105181194 | -1 | lncRNA | ENSG00000 |
| 105172938 | 105181323 | -1 | protein_coding | ENSG00000 |

| | | | | |
|---|---|---|---|---|
| 105209286 | 105315589 | -1 | protein_coding | ENSG0000( |
| 105248490 | 105251093 | 1 | protein_coding | ENSG0000( |
| 105300563 | 105398147 | 1 | protein_coding | ENSG0000( |
| 105370418 | 105372918 | 1 | TEC | ENSG0000( |
| 105374618 | 105374971 | 1 | processed_pseudoge | ENSG0000( |
| 105398538 | 105450106 | 1 | protein_coding | ENSG0000( |
| 105416884 | 105419739 | -1 | lncRNA | ENSG0000( |
| 105419820 | 105470729 | 1 | protein_coding | ENSG0000( |
| 105467793 | 105470617 | -1 | lncRNA | ENSG0000( |
| 105472962 | 105480170 | 1 | protein_coding | ENSG0000( |
| 105486317 | 105488947 | 1 | protein_coding | ENSG0000( |
| 105487199 | 105492267 | 1 | protein_coding | ENSG0000( |
| 105489855 | 105499575 | 1 | protein_coding | ENSG0000( |
| 105526583 | 105530202 | 1 | protein_coding | ENSG0000( |
| 105536861 | 105537239 | -1 | processed_pseudoge | ENSG0000( |
| 105546610 | 105549694 | -1 | unprocessed_pseudc | ENSG0000( |
| 105583731 | 105588395 | -1 | IG_C_gene | ENSG0000( |
| 105597691 | 105601728 | -1 | IG_C_gene | ENSG0000( |
| 105620506 | 105626066 | -1 | IG_C_gene | ENSG0000( |
| 105621116 | 105621180 | 1 | miRNA | ENSG0000( |
| 105639559 | 105644790 | -1 | IG_C_gene | ENSG0000( |
| 105640168 | 105640232 | 1 | miRNA | ENSG0000( |
| 105647924 | 105649057 | -1 | lncRNA | ENSG0000( |
| 105664633 | 105669843 | -1 | IG_C_pseudogene | ENSG0000( |
| 105672308 | 105673314 | -1 | processed_pseudoge | ENSG0000( |
| 105703995 | 105708665 | -1 | IG_C_gene | ENSG0000( |
| 105721794 | 105722527 | -1 | IG_C_pseudogene | ENSG0000( |
| 105736343 | 105743071 | -1 | IG_C_gene | ENSG0000( |
| 105764503 | 105771405 | -1 | IG_C_gene | ENSG0000( |
| 105785505 | 105785831 | -1 | processed_pseudoge | ENSG0000( |
| 105836765 | 105845678 | -1 | IG_C_gene | ENSG0000( |
| 105851705 | 105856218 | -1 | IG_C_gene | ENSG0000( |
| 105857513 | 105857572 | -1 | miRNA | ENSG0000( |
| 105858124 | 105858175 | -1 | miRNA | ENSG0000( |
| 105858165 | 105858242 | -1 | miRNA | ENSG0000( |
| 105859484 | 105859553 | -1 | miRNA | ENSG0000( |
| 105863198 | 105863258 | -1 | IG_J_gene | ENSG0000( |
| 105863416 | 105863465 | -1 | IG_J_pseudogene | ENSG0000( |
| 105863814 | 105863862 | -1 | IG_J_gene | ENSG0000( |
| 105864215 | 105864260 | -1 | IG_J_gene | ENSG0000( |
| 105864587 | 105864635 | -1 | IG_J_gene | ENSG0000( |
| 105864793 | 105864852 | -1 | IG_J_pseudogene | ENSG0000( |
| 105865199 | 105865250 | -1 | IG_J_gene | ENSG0000( |
| 105865407 | 105865458 | -1 | IG_J_gene | ENSG0000( |
| 105865551 | 105865561 | -1 | IG_D_gene | ENSG0000( |
| 105865624 | 105865678 | -1 | IG_J_pseudogene | ENSG0000( |
| 105881034 | 105881053 | -1 | IG_D_gene | ENSG0000( |
| 105881539 | 105881556 | -1 | IG_D_gene | ENSG0000( |
| 105883903 | 105883922 | -1 | IG_D_gene | ENSG0000( |
| 105884870 | 105884888 | -1 | IG_D_gene | ENSG0000( |

| | | | | |
|---|---|---|---|---|
| 105886031 | 105886061 | -1 | IG_D_gene | ENSG00000 |
| 105888551 | 105888578 | -1 | IG_D_gene | ENSG00000 |
| 105890084 | 105896577 | 1 | lncRNA | ENSG00000 |
| 105891191 | 105891207 | -1 | IG_D_gene | ENSG00000 |
| 105891699 | 105891719 | -1 | IG_D_gene | ENSG00000 |
| 105893542 | 105893561 | -1 | IG_D_gene | ENSG00000 |
| 105894508 | 105894523 | -1 | IG_D_gene | ENSG00000 |
| 105895634 | 105895670 | -1 | IG_D_gene | ENSG00000 |
| 105897957 | 105897987 | -1 | IG_D_gene | ENSG00000 |
| 105900638 | 105900654 | -1 | IG_D_gene | ENSG00000 |
| 105901142 | 105901162 | -1 | IG_D_gene | ENSG00000 |
| 105902649 | 105902671 | -1 | IG_D_gene | ENSG00000 |
| 105903616 | 105903631 | -1 | IG_D_gene | ENSG00000 |
| 105904497 | 105904527 | -1 | IG_D_gene | ENSG00000 |
| 105904681 | 105904711 | -1 | IG_D_gene | ENSG00000 |
| 105907211 | 105907241 | -1 | IG_D_gene | ENSG00000 |
| 105909907 | 105909923 | -1 | IG_D_gene | ENSG00000 |
| 105910410 | 105910427 | -1 | IG_D_gene | ENSG00000 |
| 105912257 | 105912276 | -1 | IG_D_gene | ENSG00000 |
| 105913222 | 105913237 | -1 | IG_D_gene | ENSG00000 |
| 105914359 | 105914389 | -1 | IG_D_gene | ENSG00000 |
| 105916826 | 105916856 | -1 | IG_D_gene | ENSG00000 |
| 105917979 | 105932642 | 1 | lncRNA | ENSG00000 |
| 105919502 | 105919518 | -1 | IG_D_gene | ENSG00000 |
| 105939756 | 105940253 | -1 | IG_V_gene | ENSG00000 |
| 105945210 | 105945389 | -1 | IG_V_pseudogene | ENSG00000 |
| 105969729 | 105972269 | -1 | transcribed_unproce | ENSG00000 |
| 105978520 | 105979146 | -1 | processed_pseudoge | ENSG00000 |
| 105986582 | 105987083 | -1 | IG_V_gene | ENSG00000 |
| 106001534 | 106001824 | -1 | IG_V_pseudogene | ENSG00000 |
| 106005095 | 106005574 | -1 | IG_V_gene | ENSG00000 |
| 106011922 | 106012420 | -1 | IG_V_gene | ENSG00000 |
| 106025145 | 106025630 | -1 | IG_V_gene | ENSG00000 |
| 106037902 | 106038365 | -1 | IG_V_gene | ENSG00000 |
| 106039666 | 106039764 | -1 | IG_V_pseudogene | ENSG00000 |
| 106045600 | 106045860 | -1 | IG_V_pseudogene | ENSG00000 |
| 106055549 | 106055998 | -1 | IG_V_pseudogene | ENSG00000 |
| 106062151 | 106062683 | -1 | IG_V_gene | ENSG00000 |
| 106088122 | 106088573 | -1 | IG_V_gene | ENSG00000 |
| 106107972 | 106108464 | -1 | IG_V_gene | ENSG00000 |
| 106116635 | 106117204 | -1 | IG_V_gene | ENSG00000 |
| 106120207 | 106120473 | -1 | IG_V_pseudogene | ENSG00000 |
| 106122420 | 106122709 | -1 | IG_V_pseudogene | ENSG00000 |
| 106129540 | 106130072 | -1 | IG_V_gene | ENSG00000 |
| 106142287 | 106142577 | -1 | IG_V_pseudogene | ENSG00000 |
| 106145891 | 106146131 | -1 | IG_V_pseudogene | ENSG00000 |
| 106153624 | 106154163 | -1 | IG_V_gene | ENSG00000 |
| 106163583 | 106163802 | -1 | IG_V_pseudogene | ENSG00000 |
| 106165205 | 106165730 | -1 | IG_V_gene | ENSG00000 |
| 106170749 | 106171052 | -1 | IG_V_pseudogene | ENSG00000 |

| | | | | |
|---|---|---|---|---|
| 106174342 | 106174760 | -1 | IG_V_pseudogene | ENSG00000 |
| 106179554 | 106180519 | -1 | processed_pseudoge | ENSG00000 |
| 106184901 | 106185394 | -1 | IG_V_gene | ENSG00000 |
| 106196700 | 106196990 | -1 | IG_V_pseudogene | ENSG00000 |
| 106206603 | 106208571 | 1 | lncRNA | ENSG00000 |
| 106210936 | 106211453 | -1 | IG_V_gene | ENSG00000 |
| 106235064 | 106235594 | -1 | IG_V_gene | ENSG00000 |
| 106257762 | 106258223 | -1 | IG_V_pseudogene | ENSG00000 |
| 106263362 | 106263626 | -1 | IG_V_pseudogene | ENSG00000 |
| 106264688 | 106264715 | -1 | IG_V_pseudogene | ENSG00000 |
| 106268606 | 106269140 | -1 | IG_V_gene | ENSG00000 |
| 106276548 | 106277043 | -1 | IG_V_gene | ENSG00000 |
| 106286811 | 106287657 | -1 | processed_pseudoge | ENSG00000 |
| 106287674 | 106288828 | 1 | lncRNA | ENSG00000 |
| 106289029 | 106289479 | -1 | IG_V_pseudogene | ENSG00000 |
| 106293568 | 106293808 | -1 | IG_V_pseudogene | ENSG00000 |
| 106301396 | 106301862 | -1 | IG_V_gene | ENSG00000 |
| 106309417 | 106309666 | -1 | IG_V_pseudogene | ENSG00000 |
| 106314651 | 106314975 | -1 | IG_V_pseudogene | ENSG00000 |
| 106317823 | 106318236 | -1 | IG_V_pseudogene | ENSG00000 |
| 106324254 | 106324760 | -1 | IG_V_gene | ENSG00000 |
| 106329432 | 106329684 | -1 | IG_V_pseudogene | ENSG00000 |
| 106331106 | 106331563 | -1 | IG_V_pseudogene | ENSG00000 |
| 106335082 | 106335613 | -1 | IG_V_gene | ENSG00000 |
| 106342680 | 106342980 | -1 | IG_V_pseudogene | ENSG00000 |
| 106344385 | 106344833 | -1 | IG_V_pseudogene | ENSG00000 |
| 106349283 | 106349792 | -1 | IG_V_gene | ENSG00000 |
| 106354448 | 106354704 | -1 | IG_V_pseudogene | ENSG00000 |
| 106356145 | 106356591 | -1 | IG_V_pseudogene | ENSG00000 |
| 106359793 | 106360324 | -1 | IG_V_gene | ENSG00000 |
| 106367385 | 106367664 | -1 | IG_V_pseudogene | ENSG00000 |
| 106369107 | 106369546 | -1 | IG_V_pseudogene | ENSG00000 |
| 106373663 | 106374145 | -1 | IG_V_gene | ENSG00000 |
| 106377300 | 106377733 | -1 | IG_V_pseudogene | ENSG00000 |
| 106389392 | 106389858 | -1 | IG_V_gene | ENSG00000 |
| 106392774 | 106393231 | -1 | IG_V_pseudogene | ENSG00000 |
| 106396676 | 106397114 | -1 | IG_V_pseudogene | ENSG00000 |
| 106410493 | 106411021 | -1 | IG_V_gene | ENSG00000 |
| 106418018 | 106418299 | -1 | IG_V_pseudogene | ENSG00000 |
| 106421711 | 106422218 | -1 | IG_V_gene | ENSG00000 |
| 106425359 | 106425654 | -1 | IG_V_pseudogene | ENSG00000 |
| 106436415 | 106436555 | -1 | unprocessed_pseudc | ENSG00000 |
| 106440950 | 106441007 | -1 | IG_V_pseudogene | ENSG00000 |
| 106443133 | 106443583 | -1 | IG_V_pseudogene | ENSG00000 |
| 106460092 | 106461055 | -1 | processed_pseudoge | ENSG00000 |
| 106463256 | 106463691 | -1 | IG_V_pseudogene | ENSG00000 |
| 106470264 | 106470800 | -1 | IG_V_gene | ENSG00000 |
| 106472791 | 106473000 | -1 | IG_V_pseudogene | ENSG00000 |
| 106478006 | 106478393 | -1 | IG_V_pseudogene | ENSG00000 |
| 106481424 | 106481706 | -1 | IG_V_pseudogene | ENSG00000 |

| | | | | |
|---|---|---|---|---|
| 106482435 | 106521073 | 1 | lncRNA | ENSG0000C |
| 106489345 | 106489756 | -1 | IG_V_pseudogene | ENSG0000C |
| 106494134 | 106494383 | -1 | IG_V_pseudogene | ENSG0000C |
| 106506996 | 106507491 | -1 | IG_V_gene | ENSG0000C |
| 106511117 | 106511856 | -1 | IG_V_gene | ENSG0000C |
| 106515818 | 106515854 | -1 | IG_V_pseudogene | ENSG0000C |
| 106518582 | 106519027 | -1 | IG_V_pseudogene | ENSG0000C |
| 106531141 | 106531471 | -1 | IG_V_pseudogene | ENSG0000C |
| 106537810 | 106538344 | -1 | IG_V_gene | ENSG0000C |
| 106556936 | 106557477 | -1 | IG_V_gene | ENSG0000C |
| 106564375 | 106564598 | -1 | IG_V_pseudogene | ENSG0000C |
| 106566117 | 106566555 | -1 | IG_V_pseudogene | ENSG0000C |
| 106578744 | 106579236 | -1 | IG_V_gene | ENSG0000C |
| 106583506 | 106583807 | -1 | IG_V_pseudogene | ENSG0000C |
| 106584718 | 106584976 | -1 | IG_V_pseudogene | ENSG0000C |
| 106586376 | 106586826 | -1 | IG_V_pseudogene | ENSG0000C |
| 106592676 | 106593347 | -1 | IG_V_gene | ENSG0000C |
| 106599670 | 106599924 | -1 | IG_V_pseudogene | ENSG0000C |
| 106601346 | 106601792 | -1 | IG_V_pseudogene | ENSG0000C |
| 106606101 | 106606551 | -1 | IG_V_pseudogene | ENSG0000C |
| 106609762 | 106610196 | -1 | IG_V_pseudogene | ENSG0000C |
| 106618894 | 106619173 | -1 | IG_V_pseudogene | ENSG0000C |
| 106622357 | 106622855 | -1 | IG_V_gene | ENSG0000C |
| 106627249 | 106627825 | -1 | IG_V_gene | ENSG0000C |
| 106631197 | 106631653 | -1 | IG_V_pseudogene | ENSG0000C |
| 106636672 | 106636786 | 1 | rRNA_pseudogene | ENSG0000C |
| 106637718 | 106637973 | -1 | IG_V_pseudogene | ENSG0000C |
| 106639119 | 106639657 | -1 | IG_V_gene | ENSG0000C |
| 106643142 | 106643585 | -1 | IG_V_pseudogene | ENSG0000C |
| 106650551 | 106650785 | -1 | IG_V_pseudogene | ENSG0000C |
| 106652237 | 106652681 | -1 | IG_V_pseudogene | ENSG0000C |
| 106657725 | 106658258 | -1 | IG_V_gene | ENSG0000C |
| 106666092 | 106666532 | -1 | IG_V_pseudogene | ENSG0000C |
| 106671772 | 106672073 | -1 | IG_V_pseudogene | ENSG0000C |
| 106675017 | 106675544 | -1 | IG_V_gene | ENSG0000C |
| 106680603 | 106681042 | -1 | IG_V_pseudogene | ENSG0000C |
| 106683395 | 106684350 | -1 | processed_pseudoge | ENSG0000C |
| 106686517 | 106686562 | -1 | IG_V_pseudogene | ENSG0000C |
| 106687113 | 106687200 | -1 | IG_V_pseudogene | ENSG0000C |
| 106692657 | 106692888 | -1 | IG_V_pseudogene | ENSG0000C |
| 106695102 | 106695397 | -1 | IG_V_pseudogene | ENSG0000C |
| 106703846 | 106704286 | -1 | IG_V_pseudogene | ENSG0000C |
| 106714684 | 106715181 | -1 | IG_V_gene | ENSG0000C |
| 106723574 | 106724093 | -1 | IG_V_gene | ENSG0000C |
| 106728163 | 106728615 | -1 | IG_V_pseudogene | ENSG0000C |
| 106737110 | 106737547 | -1 | IG_V_gene | ENSG0000C |
| 106750289 | 106750594 | -1 | IG_pseudogene | ENSG0000C |
| 106762092 | 106762588 | -1 | IG_V_gene | ENSG0000C |
| 106770577 | 106771020 | -1 | IG_V_gene | ENSG0000C |
| 106775157 | 106775618 | -1 | IG_V_pseudogene | ENSG0000C |

| | | | | |
|---|---|---|---|---|
| 106790691 | 106791233 | -1 | IG_V_gene | ENSG00000 |
| 106802694 | 106803233 | -1 | IG_V_gene | ENSG00000 |
| 106810442 | 106811131 | -1 | IG_V_gene | ENSG00000 |
| 106821174 | 106821414 | -1 | IG_V_pseudogene | ENSG00000 |
| 106823687 | 106824147 | -1 | IG_V_pseudogene | ENSG00000 |
| 106827869 | 106828163 | -1 | IG_V_pseudogene | ENSG00000 |
| 106831767 | 106832052 | -1 | IG_V_pseudogene | ENSG00000 |
| 106850885 | 106850999 | -1 | miRNA | ENSG00000 |
| 106851123 | 106851417 | -1 | IG_V_pseudogene | ENSG00000 |
| 106865627 | 106865895 | -1 | IG_V_pseudogene | ENSG00000 |
| 106867660 | 106868092 | -1 | IG_V_pseudogene | ENSG00000 |
| 106872783 | 106873186 | -1 | IG_V_pseudogene | ENSG00000 |
| 106874583 | 106875071 | -1 | IG_V_gene | ENSG00000 |
| 106879563 | 106879812 | -1 | IG_V_pseudogene | ENSG00000 |
| 19878555 | 19887814 | 1 | unprocessed_pseudc | ENSG00000 |
| 19883614 | 19883716 | 1 | snRNA | ENSG00000 |
| 19899334 | 19899559 | 1 | processed_pseudoge | ENSG00000 |
| 19955300 | 19955602 | -1 | unprocessed_pseudc | ENSG00000 |
| 19964666 | 19965101 | -1 | IG_V_gene | ENSG00000 |
| 19967414 | 19968352 | -1 | processed_pseudoge | ENSG00000 |
| 19972782 | 19973218 | -1 | IG_V_pseudogene | ENSG00000 |
| 19987656 | 19988117 | -1 | IG_V_gene | ENSG00000 |
| 20003840 | 20003862 | -1 | IG_D_gene | ENSG00000 |
| 20004797 | 20004815 | -1 | IG_D_gene | ENSG00000 |
| 20005905 | 20005935 | -1 | IG_D_gene | ENSG00000 |
| 20008402 | 20008432 | -1 | IG_D_gene | ENSG00000 |
| 20011153 | 20011169 | -1 | IG_D_gene | ENSG00000 |
| 20012741 | 20014208 | 1 | unprocessed_pseudc | ENSG00000 |
| 20046296 | 20047994 | -1 | processed_pseudoge | ENSG00000 |
| 20073699 | 20074334 | -1 | processed_pseudoge | ENSG00000 |
| 20095733 | 20096029 | -1 | misc_RNA | ENSG00000 |
| 20128745 | 20147953 | -1 | lncRNA | ENSG00000 |
| 20141794 | 20146430 | 1 | TEC | ENSG00000 |
| 20153776 | 20160826 | 1 | unprocessed_pseudc | ENSG00000 |
| 20228620 | 20277503 | -1 | unprocessed_pseudc | ENSG00000 |
| 20282744 | 20291586 | 1 | transcribed_unproce | ENSG00000 |
| 20301481 | 20331583 | 1 | lncRNA | ENSG00000 |
| 20344736 | 20359166 | 1 | lncRNA | ENSG00000 |
| 20379495 | 20506180 | -1 | transcribed_unproce | ENSG00000 |
| 20516252 | 20520627 | 1 | unprocessed_pseudc | ENSG00000 |
| 20531856 | 20541800 | -1 | protein_coding | ENSG00000 |
| 20545576 | 20549444 | -1 | unprocessed_pseudc | ENSG00000 |
| 20562375 | 20575696 | 1 | unprocessed_pseudc | ENSG00000 |
| 20570886 | 20571175 | 1 | misc_RNA | ENSG00000 |
| 20593494 | 20599195 | -1 | unprocessed_pseudc | ENSG00000 |
| 20626819 | 20627365 | -1 | processed_pseudoge | ENSG00000 |
| 20639464 | 20639890 | -1 | IG_V_pseudogene | ENSG00000 |
| 20642799 | 20643448 | -1 | lncRNA | ENSG00000 |
| 20657638 | 20688408 | -1 | transcribed_unproce | ENSG00000 |
| 20658900 | 20664970 | 1 | TEC | ENSG00000 |

| | | | | |
|---|---|---|---|---|
| 20729747 | 20756183 | -1 | lncRNA | ENSG0000C |
| 20738679 | 20738785 | 1 | snRNA | ENSG0000C |
| 20759311 | 20774794 | -1 | lncRNA | ENSG0000C |
| 20773084 | 20774871 | 1 | lncRNA | ENSG0000C |
| 20775464 | 20775998 | -1 | processed_pseudoge | ENSG0000C |
| 20803505 | 20826173 | -1 | transcribed_processe | ENSG0000C |
| 20813299 | 20814958 | 1 | unprocessed_pseudc | ENSG0000C |
| 20826454 | 20826640 | -1 | processed_pseudoge | ENSG0000C |
| 20832795 | 20832869 | 1 | miRNA | ENSG0000C |
| 20835372 | 20866314 | -1 | protein_coding | ENSG0000C |
| 20855251 | 20855357 | -1 | snRNA | ENSG0000C |
| 20890206 | 20890606 | -1 | processed_pseudoge | ENSG0000C |
| 20893383 | 20893980 | -1 | processed_pseudoge | ENSG0000C |
| 20901441 | 20901609 | -1 | processed_pseudoge | ENSG0000C |
| 20916686 | 20935658 | -1 | unprocessed_pseudc | ENSG0000C |
| 20940252 | 20940333 | 1 | miRNA | ENSG0000C |
| 20940380 | 21058440 | 1 | lncRNA | ENSG0000C |
| 20960675 | 20961581 | 1 | unprocessed_pseudc | ENSG0000C |
| 20969282 | 20970215 | -1 | unprocessed_pseudc | ENSG0000C |
| 21010494 | 21010516 | -1 | IG_D_gene | ENSG0000C |
| 21011451 | 21011469 | -1 | IG_D_gene | ENSG0000C |
| 21012559 | 21012589 | -1 | IG_D_gene | ENSG0000C |
| 21015048 | 21015078 | -1 | IG_D_gene | ENSG0000C |
| 21017800 | 21017816 | -1 | IG_D_gene | ENSG0000C |
| 21019388 | 21020851 | 1 | unprocessed_pseudc | ENSG0000C |
| 21053004 | 21054702 | -1 | processed_pseudoge | ENSG0000C |
| 21080812 | 21081447 | -1 | processed_pseudoge | ENSG0000C |
| 21102843 | 21103138 | -1 | misc_RNA | ENSG0000C |
| 21127698 | 21130095 | -1 | lncRNA | ENSG0000C |
| 21148334 | 21152970 | 1 | TEC | ENSG0000C |
| 21160305 | 21167357 | 1 | unprocessed_pseudc | ENSG0000C |
| 21250180 | 21260147 | -1 | unprocessed_pseudc | ENSG0000C |
| 21293653 | 21295201 | -1 | processed_pseudoge | ENSG0000C |
| 21298233 | 21325241 | -1 | lncRNA | ENSG0000C |
| 21307749 | 21307855 | 1 | snRNA | ENSG0000C |
| 21328380 | 21343881 | -1 | lncRNA | ENSG0000C |
| 21342171 | 21343958 | 1 | lncRNA | ENSG0000C |
| 21344548 | 21345085 | -1 | processed_pseudoge | ENSG0000C |
| 21368459 | 21370621 | 1 | lncRNA | ENSG0000C |
| 21372793 | 21373881 | -1 | processed_pseudoge | ENSG0000C |
| 21382383 | 21382842 | 1 | processed_pseudoge | ENSG0000C |
| 21383821 | 21384064 | 1 | processed_pseudoge | ENSG0000C |
| 21406385 | 21406459 | 1 | miRNA | ENSG0000C |
| 21408243 | 21440451 | -1 | protein_coding | ENSG0000C |
| 21428879 | 21428985 | -1 | snRNA | ENSG0000C |
| 21490354 | 21509404 | -1 | transcribed_unproce | ENSG0000C |
| 21513959 | 21514040 | 1 | miRNA | ENSG0000C |
| 21514144 | 21544394 | 1 | lncRNA | ENSG0000C |
| 21534404 | 21535346 | 1 | unprocessed_pseudc | ENSG0000C |
| 21542990 | 21544008 | -1 | unprocessed_pseudc | ENSG0000C |

| | | | | |
|---|---|---|---|---|
| 21552795 | 21653276 | 1 | protein_coding | ENSG00000 |
| 21552815 | 21557161 | 1 | lncRNA | ENSG00000 |
| 21574225 | 21575207 | -1 | unprocessed_pseudc | ENSG00000 |
| 21595684 | 21596526 | -1 | unprocessed_pseudc | ENSG00000 |
| 21609102 | 21610043 | 1 | transcribed_unproce | ENSG00000 |
| 21614118 | 21615051 | 1 | unprocessed_pseudc | ENSG00000 |
| 21637909 | 21638992 | 1 | protein_coding | ENSG00000 |
| 21651844 | 21652968 | 1 | protein_coding | ENSG00000 |
| 21683038 | 21683962 | 1 | unprocessed_pseudc | ENSG00000 |
| 21703428 | 21705627 | -1 | lncRNA | ENSG00000 |
| 21709649 | 21710270 | -1 | processed_pseudoge | ENSG00000 |
| 21717808 | 21718245 | -1 | IG_V_gene | ENSG00000 |
| 21735506 | 21735945 | -1 | IG_V_pseudogene | ENSG00000 |
| 21742364 | 21742799 | -1 | IG_V_gene | ENSG00000 |
| 21752278 | 21752710 | -1 | IG_V_pseudogene | ENSG00000 |
| 21843750 | 21843824 | 1 | miRNA | ENSG00000 |
| 21846329 | 21877703 | -1 | protein_coding | ENSG00000 |
| 21866210 | 21866316 | -1 | snRNA | ENSG00000 |
| 21911559 | 21912590 | -1 | unprocessed_pseudc | ENSG00000 |
| 21927657 | 21946641 | -1 | unprocessed_pseudc | ENSG00000 |
| 21951242 | 21951323 | 1 | miRNA | ENSG00000 |
| 21971706 | 21972645 | 1 | unprocessed_pseudc | ENSG00000 |
| 21980277 | 21981245 | -1 | unprocessed_pseudc | ENSG00000 |
| 21990074 | 22059831 | 1 | lncRNA | ENSG00000 |
| 22009547 | 22010529 | -1 | unprocessed_pseudc | ENSG00000 |
| 22015233 | 22095857 | 1 | protein_coding | ENSG00000 |
| 22030900 | 22031846 | -1 | unprocessed_pseudc | ENSG00000 |
| 22044476 | 22045400 | 1 | transcribed_unproce | ENSG00000 |
| 22056709 | 22057638 | 1 | unprocessed_pseudc | ENSG00000 |
| 22070241 | 22083221 | 1 | protein_coding | ENSG00000 |
| 22094522 | 22095472 | 1 | protein_coding | ENSG00000 |
| 22125511 | 22126434 | 1 | transcribed_unproce | ENSG00000 |
| 22145939 | 22148226 | -1 | lncRNA | ENSG00000 |
| 22152276 | 22152897 | -1 | unprocessed_pseudc | ENSG00000 |
| 22160431 | 22160868 | -1 | IG_V_gene | ENSG00000 |
| 22178107 | 22178542 | -1 | IG_V_pseudogene | ENSG00000 |
| 22184967 | 22185402 | -1 | IG_V_gene | ENSG00000 |
| 22194885 | 22195317 | -1 | IG_V_pseudogene | ENSG00000 |
| 22202392 | 22202734 | -1 | unprocessed_pseudc | ENSG00000 |
| 22225278 | 22225329 | -1 | miRNA | ENSG00000 |
| 22258138 | 22258848 | 1 | processed_pseudoge | ENSG00000 |
| 22278971 | 22282872 | 1 | unprocessed_pseudc | ENSG00000 |
| 22360639 | 22360918 | 1 | unprocessed_pseudc | ENSG00000 |
| 22371736 | 22373570 | 1 | processed_pseudoge | ENSG00000 |
| 22401208 | 22406944 | 1 | unprocessed_pseudc | ENSG00000 |
| 22427039 | 22437902 | -1 | transcribed_unproce | ENSG00000 |
| 22429200 | 22429489 | -1 | misc_RNA | ENSG00000 |
| 22451209 | 22455154 | 1 | unprocessed_pseudc | ENSG00000 |
| 22458903 | 22469230 | 1 | protein_coding | ENSG00000 |
| 22480439 | 22484840 | -1 | unprocessed_pseudc | ENSG00000 |

| | | | | |
|---|---|---|---|---|
| 22495570 | 22590815 | 1 | transcribed_unproce | ENSG0000C |
| 22606022 | 22617854 | -1 | transcribed_unproce | ENSG0000C |
| 22609141 | 22609377 | -1 | misc_RNA | ENSG0000C |
| 22664359 | 22686213 | 1 | transcribed_unproce | ENSG0000C |
| 22690454 | 22691158 | -1 | processed_pseudoge | ENSG0000C |
| 22714342 | 22717727 | 1 | unprocessed_pseudc | ENSG0000C |
| 22714939 | 22715225 | 1 | processed_pseudoge | ENSG0000C |
| 22727104 | 22744918 | 1 | transcribed_unproce | ENSG0000C |
| 22756172 | 22756729 | -1 | processed_pseudoge | ENSG0000C |
| 22757841 | 22778741 | 1 | lncRNA | ENSG0000C |
| 22770523 | 22776199 | -1 | transcribed_unproce | ENSG0000C |
| 22773063 | 22829789 | 1 | protein_coding | ENSG0000C |
| 22838641 | 22868384 | 1 | protein_coding | ENSG0000C |
| 22867052 | 22981063 | -1 | protein_coding | ENSG0000C |
| 22983192 | 23039572 | -1 | protein_coding | ENSG0000C |
| 23083094 | 23121324 | 1 | transcribed_unproce | ENSG0000C |
| 23127066 | 23136822 | -1 | protein_coding | ENSG0000C |
| 23140563 | 23144422 | -1 | unprocessed_pseudc | ENSG0000C |
| 23157366 | 23170711 | 1 | transcribed_unproce | ENSG0000C |
| 23165897 | 23166186 | 1 | misc_RNA | ENSG0000C |
| 23179869 | 23182049 | 1 | processed_pseudoge | ENSG0000C |
| 23193491 | 23193696 | 1 | processed_pseudoge | ENSG0000C |
| 23197827 | 23197920 | 1 | miRNA | ENSG0000C |
| 23225876 | 23226391 | -1 | unprocessed_pseudc | ENSG0000C |
| 23289303 | 23293667 | -1 | unprocessed_pseudc | ENSG0000C |
| 23309607 | 23314566 | 1 | transcribed_unproce | ENSG0000C |
| 23326256 | 23334127 | -1 | transcribed_unproce | ENSG0000C |
| 23337985 | 23341848 | -1 | unprocessed_pseudc | ENSG0000C |
| 23354748 | 23367231 | 1 | protein_coding | ENSG0000C |
| 23363272 | 23363561 | 1 | misc_RNA | ENSG0000C |
| 23403848 | 23407335 | 1 | lncRNA | ENSG0000C |
| 23430839 | 23435212 | 1 | unprocessed_pseudc | ENSG0000C |
| 23439038 | 23447243 | -1 | protein_coding | ENSG0000C |
| 23562062 | 23562131 | -1 | miRNA | ENSG0000C |
| 23565674 | 23630075 | 1 | protein_coding | ENSG0000C |
| 23567264 | 23569657 | -1 | lncRNA | ENSG0000C |
| 23643549 | 23647867 | -1 | protein_coding | ENSG0000C |
| 23685400 | 23687305 | -1 | protein_coding | ENSG0000C |
| 23757115 | 23856375 | 1 | lncRNA | ENSG0000C |
| 23777437 | 23777543 | -1 | snRNA | ENSG0000C |
| 23912411 | 24090821 | 1 | lncRNA | ENSG0000C |
| 24101827 | 24823365 | 1 | lncRNA | ENSG0000C |
| 24162754 | 24169948 | -1 | lncRNA | ENSG0000C |
| 24206784 | 24225854 | 1 | lncRNA | ENSG0000C |
| 24238145 | 24240192 | -1 | lncRNA | ENSG0000C |
| 24341729 | 24343464 | -1 | lncRNA | ENSG0000C |
| 24558060 | 24912924 | 1 | lncRNA | ENSG0000C |
| 24675704 | 24683393 | 1 | protein_coding | ENSG0000C |
| 24823637 | 24978723 | 1 | protein_coding | ENSG0000C |
| 24848355 | 24852253 | 1 | lncRNA | ENSG0000C |

| | | | | |
|---|---|---|---|---|
| 24908932 | 24909825 | -1 | processed_pseudoge | ENSG00000 |
| 24910874 | 24911125 | -1 | processed_pseudoge | ENSG00000 |
| 24954986 | 24977850 | 1 | protein_coding | ENSG00000 |
| 24955034 | 25000276 | 1 | protein_coding | ENSG00000 |
| 24978583 | 25420336 | 1 | lncRNA | ENSG00000 |
| 24981994 | 24982068 | 1 | snoRNA | ENSG00000 |
| 24985053 | 24988232 | 1 | TEC | ENSG00000 |
| 24985100 | 24985166 | 1 | snoRNA | ENSG00000 |
| 24986925 | 24986995 | 1 | snoRNA | ENSG00000 |
| 25027736 | 25032047 | -1 | lncRNA | ENSG00000 |
| 25041974 | 25042040 | 1 | snoRNA | ENSG00000 |
| 25051477 | 25051571 | 1 | snoRNA | ENSG00000 |
| 25054210 | 25054304 | 1 | snoRNA | ENSG00000 |
| 25056860 | 25056954 | 1 | snoRNA | ENSG00000 |
| 25059538 | 25059633 | 1 | snoRNA | ENSG00000 |
| 25062333 | 25062427 | 1 | snoRNA | ENSG00000 |
| 25065026 | 25065121 | 1 | snoRNA | ENSG00000 |
| 25067788 | 25067882 | 1 | snoRNA | ENSG00000 |
| 25070432 | 25070526 | 1 | snoRNA | ENSG00000 |
| 25073107 | 25073201 | 1 | snoRNA | ENSG00000 |
| 25074114 | 25074215 | 1 | snoRNA | ENSG00000 |
| 25075929 | 25076020 | 1 | snoRNA | ENSG00000 |
| 25077051 | 25077142 | 1 | snoRNA | ENSG00000 |
| 25079058 | 25079149 | 1 | snoRNA | ENSG00000 |
| 25080142 | 25080233 | 1 | snoRNA | ENSG00000 |
| 25081287 | 25081378 | 1 | snoRNA | ENSG00000 |
| 25082768 | 25082859 | 1 | snoRNA | ENSG00000 |
| 25083588 | 25083679 | 1 | snoRNA | ENSG00000 |
| 25085385 | 25085476 | 1 | snoRNA | ENSG00000 |
| 25086527 | 25086618 | 1 | snoRNA | ENSG00000 |
| 25087661 | 25088896 | 1 | lncRNA | ENSG00000 |
| 25087662 | 25087753 | 1 | snoRNA | ENSG00000 |
| 25088804 | 25088895 | 1 | snoRNA | ENSG00000 |
| 25089923 | 25090014 | 1 | snoRNA | ENSG00000 |
| 25091786 | 25091877 | 1 | snoRNA | ENSG00000 |
| 25091786 | 25107469 | -1 | lncRNA | ENSG00000 |
| 25091960 | 25092377 | -1 | TEC | ENSG00000 |
| 25094037 | 25094128 | 1 | snoRNA | ENSG00000 |
| 25097663 | 25097754 | 1 | snoRNA | ENSG00000 |
| 25099499 | 25099594 | 1 | snoRNA | ENSG00000 |
| 25101575 | 25101666 | 1 | snoRNA | ENSG00000 |
| 25104642 | 25104732 | 1 | snoRNA | ENSG00000 |
| 25106521 | 25106603 | 1 | snoRNA | ENSG00000 |
| 25108268 | 25108352 | 1 | snoRNA | ENSG00000 |
| 25135642 | 25138055 | 1 | TEC | ENSG00000 |
| 25159221 | 25159279 | 1 | snoRNA | ENSG00000 |
| 25169337 | 25169676 | -1 | processed_pseudoge | ENSG00000 |
| 25170723 | 25170804 | 1 | snoRNA | ENSG00000 |
| 25172635 | 25172716 | 1 | snoRNA | ENSG00000 |
| 25174927 | 25175008 | 1 | snoRNA | ENSG00000 |

| | | | | |
|---|---|---|---|---|
| 25176832 | 25176913 | 1 | snoRNA | ENSG0000( |
| 25178738 | 25178819 | 1 | snoRNA | ENSG0000( |
| 25180497 | 25180578 | 1 | snoRNA | ENSG0000( |
| 25182385 | 25182466 | 1 | snoRNA | ENSG0000( |
| 25184306 | 25184387 | 1 | snoRNA | ENSG0000( |
| 25185631 | 25185712 | 1 | snoRNA | ENSG0000( |
| 25187536 | 25187616 | 1 | snoRNA | ENSG0000( |
| 25189414 | 25189495 | 1 | snoRNA | ENSG0000( |
| 25191416 | 25191497 | 1 | snoRNA | ENSG0000( |
| 25193321 | 25193402 | 1 | snoRNA | ENSG0000( |
| 25194921 | 25195001 | 1 | snoRNA | ENSG0000( |
| 25197576 | 25197656 | 1 | snoRNA | ENSG0000( |
| 25199448 | 25199529 | 1 | snoRNA | ENSG0000( |
| 25201323 | 25201404 | 1 | snoRNA | ENSG0000( |
| 25203227 | 25203308 | 1 | snoRNA | ENSG0000( |
| 25204357 | 25204438 | 1 | snoRNA | ENSG0000( |
| 25206262 | 25206343 | 1 | snoRNA | ENSG0000( |
| 25208083 | 25208163 | 1 | snoRNA | ENSG0000( |
| 25209918 | 25209999 | 1 | snoRNA | ENSG0000( |
| 25211796 | 25211877 | 1 | snoRNA | ENSG0000( |
| 25213648 | 25213729 | 1 | snoRNA | ENSG0000( |
| 25215541 | 25215622 | 1 | snoRNA | ENSG0000( |
| 25218617 | 25218698 | 1 | snoRNA | ENSG0000( |
| 25220497 | 25220578 | 1 | snoRNA | ENSG0000( |
| 25222348 | 25222428 | 1 | snoRNA | ENSG0000( |
| 25223246 | 25223327 | 1 | snoRNA | ENSG0000( |
| 25225203 | 25225284 | 1 | snoRNA | ENSG0000( |
| 25227109 | 25227190 | 1 | snoRNA | ENSG0000( |
| 25228967 | 25229048 | 1 | snoRNA | ENSG0000( |
| 25230838 | 25230919 | 1 | snoRNA | ENSG0000( |
| 25232387 | 25232468 | 1 | snoRNA | ENSG0000( |
| 25234247 | 25234328 | 1 | snoRNA | ENSG0000( |
| 25236085 | 25236166 | 1 | snoRNA | ENSG0000( |
| 25237986 | 25238067 | 1 | snoRNA | ENSG0000( |
| 25239838 | 25239919 | 1 | snoRNA | ENSG0000( |
| 25241746 | 25241827 | 1 | snoRNA | ENSG0000( |
| 25243614 | 25243695 | 1 | snoRNA | ENSG0000( |
| 25245478 | 25245559 | 1 | snoRNA | ENSG0000( |
| 25247345 | 25247426 | 1 | snoRNA | ENSG0000( |
| 25249198 | 25249279 | 1 | snoRNA | ENSG0000( |
| 25250859 | 25250940 | 1 | snoRNA | ENSG0000( |
| 25255579 | 25257027 | 1 | TEC | ENSG0000( |
| 25264510 | 25264590 | 1 | snoRNA | ENSG0000( |
| 25266591 | 25266661 | 1 | snoRNA | ENSG0000( |
| 25268466 | 25268551 | 1 | snoRNA | ENSG0000( |
| 25269783 | 25269858 | 1 | snoRNA | ENSG0000( |
| 25278343 | 25278409 | 1 | snoRNA | ENSG0000( |
| 25333728 | 25439051 | -1 | protein_coding | ENSG0000( |
| 25345633 | 25347235 | -1 | lncRNA | ENSG0000( |
| 25456271 | 25578813 | -1 | lncRNA | ENSG0000( |

| | | | | |
|---|---|---|---|---|
| 25580969 | 25581122 | 1 | processed_pseudoge | ENSG00000 |
| 25677273 | 25865172 | -1 | protein_coding | ENSG00000 |
| 25708470 | 25710869 | -1 | lncRNA | ENSG00000 |
| 25738508 | 25738620 | 1 | rRNA_pseudogene | ENSG00000 |
| 25848747 | 25848825 | -1 | miRNA | ENSG00000 |
| 25865295 | 25877211 | 1 | lncRNA | ENSG00000 |
| 25902119 | 26053123 | 1 | lncRNA | ENSG00000 |
| 25928146 | 25928575 | -1 | processed_pseudoge | ENSG00000 |
| 26008940 | 26009074 | -1 | snoRNA | ENSG00000 |
| 26050538 | 26052276 | -1 | lncRNA | ENSG00000 |
| 26115726 | 26134001 | 1 | lncRNA | ENSG00000 |
| 26261669 | 26265918 | 1 | lncRNA | ENSG00000 |
| 26395037 | 26455217 | 1 | lncRNA | ENSG00000 |
| 26403480 | 26477840 | -1 | lncRNA | ENSG00000 |
| 26543546 | 26939539 | -1 | protein_coding | ENSG00000 |
| 26557598 | 26557937 | -1 | lncRNA | ENSG00000 |
| 26657030 | 26658763 | -1 | lncRNA | ENSG00000 |
| 26851132 | 26851610 | -1 | processed_pseudoge | ENSG00000 |
| 26866911 | 26949208 | 1 | protein_coding | ENSG00000 |
| 26964917 | 26970385 | 1 | unprocessed_pseudc | ENSG00000 |
| 26971181 | 27541984 | 1 | protein_coding | ENSG00000 |
| 27038978 | 27161319 | -1 | lncRNA | ENSG00000 |
| 27208089 | 27208635 | -1 | processed_pseudoge | ENSG00000 |
| 27280318 | 27280436 | -1 | rRNA_pseudogene | ENSG00000 |
| 27340524 | 27340804 | 1 | processed_pseudoge | ENSG00000 |
| 27362310 | 27366398 | 1 | lncRNA | ENSG00000 |
| 27418796 | 27420735 | -1 | lncRNA | ENSG00000 |
| 27422381 | 27428338 | 1 | lncRNA | ENSG00000 |
| 27483035 | 27541991 | -1 | lncRNA | ENSG00000 |
| 27621534 | 27629289 | 1 | lncRNA | ENSG00000 |
| 27662715 | 27667273 | -1 | lncRNA | ENSG00000 |
| 27684498 | 27685768 | -1 | lncRNA | ENSG00000 |
| 27754875 | 28099315 | -1 | protein_coding | ENSG00000 |
| 27775621 | 27776798 | 1 | lncRNA | ENSG00000 |
| 27895704 | 27896582 | 1 | processed_pseudoge | ENSG00000 |
| 28111040 | 28322172 | -1 | protein_coding | ENSG00000 |
| 28315651 | 28316059 | 1 | processed_pseudoge | ENSG00000 |
| 28332237 | 28336609 | 1 | unprocessed_pseudc | ENSG00000 |
| 28346490 | 28346647 | 1 | unprocessed_pseudc | ENSG00000 |
| 28349051 | 28358080 | -1 | transcribed_unproce | ENSG00000 |
| 28362796 | 28365731 | -1 | unprocessed_pseudc | ENSG00000 |
| 28378621 | 28392018 | 1 | protein_coding | ENSG00000 |
| 28387196 | 28387485 | 1 | misc_RNA | ENSG00000 |
| 28402868 | 28405033 | 1 | processed_pseudoge | ENSG00000 |
| 28416449 | 28416674 | 1 | unprocessed_pseudc | ENSG00000 |
| 28422172 | 28422373 | 1 | unprocessed_pseudc | ENSG00000 |
| 28426491 | 28426584 | 1 | miRNA | ENSG00000 |
| 28444144 | 28444264 | 1 | unprocessed_pseudc | ENSG00000 |
| 28454660 | 28457783 | -1 | unprocessed_pseudc | ENSG00000 |
| 28473322 | 28473448 | -1 | unprocessed_pseudc | ENSG00000 |

| | | | | |
|---|---|---|---|---|
| 28490752 | 28490845 | -1 | miRNA | ENSG00000( |
| 28494975 | 28495182 | -1 | unprocessed_pseudc | ENSG00000( |
| 28506625 | 28508808 | -1 | processed_pseudoge | ENSG00000( |
| 28519611 | 28533014 | -1 | protein_coding | ENSG00000( |
| 28524151 | 28524440 | -1 | misc_RNA | ENSG00000( |
| 28545912 | 28549777 | 1 | unprocessed_pseudc | ENSG00000( |
| 28553638 | 28561379 | 1 | unprocessed_pseudc | ENSG00000( |
| 28564986 | 28565152 | -1 | unprocessed_pseudc | ENSG00000( |
| 28575019 | 28579400 | -1 | unprocessed_pseudc | ENSG00000( |
| 28589492 | 28685264 | 1 | transcribed_unproce | ENSG00000( |
| 28679405 | 28682252 | -1 | unprocessed_pseudc | ENSG00000( |
| 28701954 | 28738384 | -1 | protein_coding | ENSG00000( |
| 28703554 | 28703790 | -1 | misc_RNA | ENSG00000( |
| 28719377 | 28738431 | -1 | lncRNA | ENSG00000( |
| 28737583 | 28759204 | 1 | transcribed_unproce | ENSG00000( |
| 28763440 | 28764151 | -1 | processed_pseudoge | ENSG00000( |
| 28788243 | 28792002 | 1 | transcribed_unproce | ENSG00000( |
| 28789199 | 28789486 | 1 | transcribed_process | ENSG00000( |
| 28789664 | 28859007 | 1 | transcribed_unproce | ENSG00000( |
| 28830619 | 28831182 | -1 | processed_pseudoge | ENSG00000( |
| 28841833 | 28848675 | -1 | protein_coding | ENSG00000( |
| 28857882 | 28858398 | -1 | processed_pseudoge | ENSG00000( |
| 28881667 | 28884655 | -1 | lncRNA | ENSG00000( |
| 28884483 | 29118315 | 1 | protein_coding | ENSG00000( |
| 28970196 | 28977464 | -1 | lncRNA | ENSG00000( |
| 29120254 | 29570723 | -1 | protein_coding | ENSG00000( |
| 29233673 | 29235137 | -1 | processed_pseudoge | ENSG00000( |
| 29233868 | 29233979 | 1 | snoRNA | ENSG00000( |
| 29264989 | 29269822 | -1 | protein_coding | ENSG00000( |
| 29373602 | 29376956 | -1 | lncRNA | ENSG00000( |
| 29609676 | 29611212 | -1 | lncRNA | ENSG00000( |
| 29612570 | 29614274 | 1 | processed_pseudoge | ENSG00000( |
| 29664477 | 29669482 | 1 | lncRNA | ENSG00000( |
| 29674990 | 29680957 | 1 | lncRNA | ENSG00000( |
| 29699367 | 29968865 | -1 | protein_coding | ENSG00000( |
| 29728186 | 29729531 | -1 | lncRNA | ENSG00000( |
| 29730713 | 29730985 | 1 | processed_pseudoge | ENSG00000( |
| 29822631 | 29824081 | 1 | lncRNA | ENSG00000( |
| 30005205 | 30006430 | 1 | unprocessed_pseudc | ENSG00000( |
| 30005443 | 30045848 | -1 | lncRNA | ENSG00000( |
| 30044455 | 30045296 | 1 | processed_pseudoge | ENSG00000( |
| 30052942 | 30073064 | -1 | transcribed_unproce | ENSG00000( |
| 30083053 | 30093499 | 1 | protein_coding | ENSG00000( |
| 30091662 | 30091898 | 1 | misc_RNA | ENSG00000( |
| 30098826 | 30102093 | -1 | unprocessed_pseudc | ENSG00000( |
| 30103720 | 30131757 | 1 | transcribed_unproce | ENSG00000( |
| 30112198 | 30112330 | 1 | snoRNA | ENSG00000( |
| 30135149 | 30145567 | 1 | protein_coding | ENSG00000( |
| 30143733 | 30143969 | 1 | misc_RNA | ENSG00000( |
| 30150872 | 30153409 | -1 | unprocessed_pseudc | ENSG00000( |

| | | | | |
|---|---|---|---|---|
| 30153933 | 30154110 | -1 | unprocessed_pseudc | ENSG00000( |
| 30170563 | 30179943 | -1 | lncRNA | ENSG00000( |
| 30195809 | 30217552 | 1 | lncRNA | ENSG00000( |
| 30223017 | 30225564 | 1 | processed_pseudoge | ENSG00000( |
| 30254402 | 30254508 | -1 | snRNA | ENSG00000( |
| 30282533 | 30288389 | 1 | processed_pseudoge | ENSG00000( |
| 30344307 | 30348236 | -1 | TEC | ENSG00000( |
| 30355436 | 30355540 | 1 | misc_RNA | ENSG00000( |
| 30357766 | 30393849 | -1 | protein_coding | ENSG00000( |
| 30380353 | 30386987 | -1 | transcribed_unproce | ENSG00000( |
| 30384959 | 30385091 | -1 | snoRNA | ENSG00000( |
| 30395146 | 30396368 | 1 | unprocessed_pseudc | ENSG00000( |
| 30403740 | 30414162 | -1 | protein_coding | ENSG00000( |
| 30405307 | 30405597 | -1 | misc_RNA | ENSG00000( |
| 30420283 | 30420429 | -1 | unprocessed_pseudc | ENSG00000( |
| 30470779 | 30507623 | 1 | transcribed_unproce | ENSG00000( |
| 30487963 | 30490313 | 1 | lncRNA | ENSG00000( |
| 30507377 | 30513698 | 1 | unprocessed_pseudc | ENSG00000( |
| 30516079 | 30517867 | 1 | unprocessed_pseudc | ENSG00000( |
| 30540093 | 30545969 | 1 | lncRNA | ENSG00000( |
| 30552078 | 30562501 | 1 | protein_coding | ENSG00000( |
| 30560644 | 30560934 | 1 | misc_RNA | ENSG00000( |
| 30567822 | 30571089 | -1 | unprocessed_pseudc | ENSG00000( |
| 30572738 | 30600647 | 1 | transcribed_unproce | ENSG00000( |
| 30581184 | 30581316 | 1 | snoRNA | ENSG00000( |
| 30604126 | 30614561 | 1 | protein_coding | ENSG00000( |
| 30607695 | 30608193 | -1 | lncRNA | ENSG00000( |
| 30612724 | 30612960 | 1 | misc_RNA | ENSG00000( |
| 30616958 | 30617749 | 1 | lncRNA | ENSG00000( |
| 30616998 | 30625773 | -1 | lncRNA | ENSG00000( |
| 30624494 | 30649529 | 1 | protein_coding | ENSG00000( |
| 30624548 | 30685606 | 1 | protein_coding | ENSG00000( |
| 30644633 | 30723495 | 1 | transcribed_unproce | ENSG00000( |
| 30658717 | 30772993 | 1 | lncRNA | ENSG00000( |
| 30673750 | 30673843 | 1 | misc_RNA | ENSG00000( |
| 30750406 | 30759859 | 1 | transcribed_unproce | ENSG00000( |
| 30791573 | 30801859 | 1 | unprocessed_pseudc | ENSG00000( |
| 30800160 | 30800396 | 1 | misc_RNA | ENSG00000( |
| 30815271 | 30844153 | 1 | unprocessed_pseudc | ENSG00000( |
| 30882267 | 30883231 | -1 | processed_pseudoge | ENSG00000( |
| 30890559 | 30943108 | 1 | protein_coding | ENSG00000( |
| 30899480 | 30900327 | 1 | unprocessed_pseudc | ENSG00000( |
| 30938941 | 30991628 | -1 | protein_coding | ENSG00000( |
| 30956248 | 30956350 | -1 | snRNA | ENSG00000( |
| 30992171 | 31042302 | 1 | lncRNA | ENSG00000( |
| 31001061 | 31161273 | -1 | protein_coding | ENSG00000( |
| 31065032 | 31065141 | -1 | miRNA | ENSG00000( |
| 31215622 | 31224445 | 1 | lncRNA | ENSG00000( |
| 31221999 | 31230838 | -1 | lncRNA | ENSG00000( |
| 31326835 | 31435665 | 1 | protein_coding | ENSG00000( |

| | | | | |
|---|---|---|---|---|
| 31420128 | 31420662 | 1 | processed_pseudoge | ENSG00000 |
| 31453201 | 31454139 | -1 | processed_pseudoge | ENSG00000 |
| 31475398 | 31870789 | -1 | protein_coding | ENSG00000 |
| 31765743 | 31768201 | -1 | processed_pseudoge | ENSG00000 |
| 31787494 | 31787734 | 1 | unprocessed_pseudc | ENSG00000 |
| 31923438 | 32173018 | 1 | protein_coding | ENSG00000 |
| 31928303 | 31928434 | 1 | snoRNA | ENSG00000 |
| 32175247 | 32175351 | -1 | misc_RNA | ENSG00000 |
| 32182551 | 32186475 | 1 | TEC | ENSG00000 |
| 32250302 | 32256125 | -1 | processed_pseudoge | ENSG00000 |
| 32284152 | 32284258 | 1 | snRNA | ENSG00000 |
| 32313126 | 32315654 | -1 | processed_pseudoge | ENSG00000 |
| 32359538 | 32367784 | 1 | lncRNA | ENSG00000 |
| 32384937 | 32387484 | 1 | unprocessed_pseudc | ENSG00000 |
| 32392782 | 32403292 | -1 | protein_coding | ENSG00000 |
| 32394365 | 32394613 | -1 | misc_RNA | ENSG00000 |
| 32398956 | 32435233 | -1 | lncRNA | ENSG00000 |
| 32406597 | 32434992 | -1 | transcribed_unproce | ENSG00000 |
| 32426288 | 32426420 | -1 | snoRNA | ENSG00000 |
| 32436503 | 32439767 | 1 | unprocessed_pseudc | ENSG00000 |
| 32441914 | 32455634 | -1 | protein_coding | ENSG00000 |
| 32446674 | 32446964 | -1 | misc_RNA | ENSG00000 |
| 32461666 | 32461809 | -1 | processed_pseudoge | ENSG00000 |
| 32489267 | 32491586 | -1 | unprocessed_pseudc | ENSG00000 |
| 32494003 | 32500346 | -1 | unprocessed_pseudc | ENSG00000 |
| 32519848 | 32536926 | -1 | transcribed_processe | ENSG00000 |
| 32536047 | 32587613 | 1 | lncRNA | ENSG00000 |
| 32583612 | 32584312 | 1 | lncRNA | ENSG00000 |
| 32585524 | 32615158 | -1 | lncRNA | ENSG00000 |
| 32593456 | 32607310 | 1 | protein_coding | ENSG00000 |
| 32602121 | 32602411 | 1 | misc_RNA | ENSG00000 |
| 32613733 | 32615111 | 1 | lncRNA | ENSG00000 |
| 32615144 | 32639941 | 1 | protein_coding | ENSG00000 |
| 32624106 | 32641615 | -1 | lncRNA | ENSG00000 |
| 32641676 | 32697098 | 1 | protein_coding | ENSG00000 |
| 32656084 | 32673065 | -1 | lncRNA | ENSG00000 |
| 32717270 | 32719007 | -1 | lncRNA | ENSG00000 |
| 32718004 | 32745106 | 1 | protein_coding | ENSG00000 |
| 32765545 | 33194733 | -1 | protein_coding | ENSG00000 |
| 32836983 | 32837326 | -1 | transcribed_processe | ENSG00000 |
| 32970727 | 32970791 | 1 | snoRNA | ENSG00000 |
| 33012119 | 33013068 | 1 | processed_pseudoge | ENSG00000 |
| 33105758 | 33106236 | -1 | TEC | ENSG00000 |
| 33235552 | 33247621 | -1 | transcribed_unitary_ | ENSG00000 |
| 33293568 | 33310659 | -1 | lncRNA | ENSG00000 |
| 33310945 | 33866121 | 1 | protein_coding | ENSG00000 |
| 33722492 | 33722596 | 1 | misc_RNA | ENSG00000 |
| 33850538 | 33851178 | -1 | lncRNA | ENSG00000 |
| 33851785 | 33856809 | -1 | lncRNA | ENSG00000 |
| 33858602 | 33864825 | -1 | lncRNA | ENSG00000 |

| | | | | |
|---|---|---|---|---|
| 33866227 | 34039204 | -1 | protein_coding | ENSG0000C |
| 33968720 | 34067457 | 1 | protein_coding | ENSG0000C |
| 33972059 | 33972515 | -1 | lncRNA | ENSG0000C |
| 33985406 | 33988268 | 1 | lncRNA | ENSG0000C |
| 33990686 | 33992048 | 1 | processed_pseudoge | ENSG0000C |
| 34084017 | 34101862 | -1 | protein_coding | ENSG0000C |
| 34085911 | 34086314 | -1 | processed_pseudoge | ENSG0000C |
| 34102083 | 34108686 | 1 | protein_coding | ENSG0000C |
| 34140674 | 34210096 | -1 | protein_coding | ENSG0000C |
| 34225013 | 34230156 | 1 | protein_coding | ENSG0000C |
| 34229784 | 34338060 | -1 | protein_coding | ENSG0000C |
| 34255286 | 34257832 | 1 | lncRNA | ENSG0000C |
| 34314728 | 34314828 | 1 | misc_RNA | ENSG0000C |
| 34341713 | 34343177 | -1 | protein_coding | ENSG0000C |
| 34343315 | 34357737 | 1 | protein_coding | ENSG0000C |
| 34358633 | 34367196 | -1 | protein_coding | ENSG0000C |
| 34373541 | 34374603 | -1 | processed_pseudoge | ENSG0000C |
| 34379068 | 34437466 | -1 | protein_coding | ENSG0000C |
| 34382069 | 34382150 | -1 | miRNA | ENSG0000C |
| 34394594 | 34394931 | 1 | processed_pseudoge | ENSG0000C |
| 34409625 | 34410594 | 1 | unprocessed_pseudc | ENSG0000C |
| 34415450 | 34420273 | -1 | TEC | ENSG0000C |
| 34489002 | 34490571 | -1 | unprocessed_pseudc | ENSG0000C |
| 34494420 | 34495900 | 1 | processed_pseudoge | ENSG0000C |
| 34520608 | 34521791 | 1 | unprocessed_pseudc | ENSG0000C |
| 34525207 | 34588503 | -1 | protein_coding | ENSG0000C |
| 34528290 | 34528371 | -1 | miRNA | ENSG0000C |
| 34555870 | 34556820 | 1 | unprocessed_pseudc | ENSG0000C |
| 34597458 | 34597558 | -1 | snoRNA | ENSG0000C |
| 34651135 | 34671095 | -1 | lncRNA | ENSG0000C |
| 34657736 | 34670512 | 1 | lncRNA | ENSG0000C |
| 34751032 | 34754998 | -1 | protein_coding | ENSG0000C |
| 34755084 | 34813505 | 1 | lncRNA | ENSG0000C |
| 34790230 | 34795549 | -1 | protein_coding | ENSG0000C |
| 34849137 | 34850395 | -1 | processed_pseudoge | ENSG0000C |
| 34851782 | 34969742 | -1 | protein_coding | ENSG0000C |
| 34943080 | 34943394 | 1 | processed_pseudoge | ENSG0000C |
| 34978341 | 34988287 | -1 | protein_coding | ENSG0000C |
| 34978351 | 34980084 | 1 | TEC | ENSG0000C |
| 34993646 | 35003221 | 1 | lncRNA | ENSG0000C |
| 34999902 | 35000270 | 1 | TEC | ENSG0000C |
| 35007844 | 35011191 | 1 | lncRNA | ENSG0000C |
| 35081144 | 35081513 | 1 | processed_pseudoge | ENSG0000C |
| 35084193 | 35085110 | -1 | protein_coding | ENSG0000C |
| 35092481 | 35093210 | 1 | processed_pseudoge | ENSG0000C |
| 35099022 | 35169698 | -1 | lncRNA | ENSG0000C |
| 35121573 | 35122084 | -1 | processed_pseudoge | ENSG0000C |
| 35181799 | 35238197 | 1 | transcribed_processe | ENSG0000C |
| 35217345 | 35546193 | -1 | protein_coding | ENSG0000C |
| 35282596 | 35282980 | 1 | processed_pseudoge | ENSG0000C |

| | | | | |
|---|---|---|---|---|
| 35298249 | 35299440 | -1 | processed_pseudoge | ENSG00000( |
| 35372256 | 35372364 | -1 | miRNA | ENSG00000( |
| 35396221 | 35396444 | 1 | processed_pseudoge | ENSG00000( |
| 35400691 | 35401768 | 1 | processed_pseudoge | ENSG00000( |
| 35546154 | 35860199 | 1 | lncRNA | ENSG00000( |
| 35774801 | 35774950 | 1 | unprocessed_pseudo | ENSG00000( |
| 35919892 | 36049537 | 1 | lncRNA | ENSG00000( |
| 35926856 | 35926923 | 1 | miRNA | ENSG00000( |
| 35939167 | 36252257 | -1 | lncRNA | ENSG00000( |
| 36341739 | 36344166 | 1 | lncRNA | ENSG00000( |
| 36438972 | 36471016 | -1 | lncRNA | ENSG00000( |
| 36571985 | 36572212 | 1 | processed_pseudoge | ENSG00000( |
| 36579626 | 36810248 | 1 | protein_coding | ENSG00000( |
| 36613474 | 36614198 | 1 | processed_pseudoge | ENSG00000( |
| 36617089 | 36619265 | 1 | transcribed_process | ENSG00000( |
| 36626674 | 36627775 | -1 | processed_pseudoge | ENSG00000( |
| 36641186 | 36669097 | -1 | lncRNA | ENSG00000( |
| 36798596 | 36818459 | -1 | transcribed_process | ENSG00000( |
| 36852640 | 36852859 | 1 | snoRNA | ENSG00000( |
| 36876379 | 36887035 | -1 | lncRNA | ENSG00000( |
| 36889204 | 37101299 | -1 | protein_coding | ENSG00000( |
| 36972821 | 36972901 | -1 | miRNA | ENSG00000( |
| 37050828 | 37050930 | -1 | misc_RNA | ENSG00000( |
| 37099339 | 37100173 | 1 | lncRNA | ENSG00000( |
| 37109587 | 37109984 | 1 | lncRNA | ENSG00000( |
| 37216617 | 37216898 | 1 | misc_RNA | ENSG00000( |
| 37253879 | 37254952 | 1 | processed_pseudoge | ENSG00000( |
| 37364020 | 37490864 | -1 | lncRNA | ENSG00000( |
| 37490691 | 37491103 | 1 | processed_pseudoge | ENSG00000( |
| 37921939 | 37967724 | 1 | protein_coding | ENSG00000( |
| 37980676 | 38062340 | 1 | lncRNA | ENSG00000( |
| 38068812 | 38072990 | -1 | lncRNA | ENSG00000( |
| 38139595 | 38226897 | -1 | lncRNA | ENSG00000( |
| 38252836 | 38357249 | 1 | protein_coding | ENSG00000( |
| 38454127 | 38487710 | 1 | protein_coding | ENSG00000( |
| 38458760 | 38459540 | -1 | processed_pseudoge | ENSG00000( |
| 38488103 | 38565575 | -1 | protein_coding | ENSG00000( |
| 38493308 | 38499985 | 1 | lncRNA | ENSG00000( |
| 38502853 | 38504981 | 1 | lncRNA | ENSG00000( |
| 38524070 | 38534655 | 1 | lncRNA | ENSG00000( |
| 38628005 | 39180499 | 1 | lncRNA | ENSG00000( |
| 38756552 | 38759413 | -1 | lncRNA | ENSG00000( |
| 38851306 | 38853432 | -1 | lncRNA | ENSG00000( |
| 38865322 | 39427195 | -1 | lncRNA | ENSG00000( |
| 39019193 | 39026521 | 1 | lncRNA | ENSG00000( |
| 39167676 | 39180048 | 1 | lncRNA | ENSG00000( |
| 39250684 | 39254845 | 1 | lncRNA | ENSG00000( |
| 39300418 | 39310782 | -1 | lncRNA | ENSG00000( |
| 39472703 | 39497074 | 1 | lncRNA | ENSG00000( |
| 39512335 | 39519954 | -1 | lncRNA | ENSG00000( |

| | | | | |
|---|---|---|---|---|
| 39581079 | 39599466 | 1 | protein_coding | ENSG0000( |
| 39586561 | 39587293 | 1 | lncRNA | ENSG0000( |
| 39588357 | 39588882 | -1 | lncRNA | ENSG0000( |
| 39588357 | 39782841 | -1 | protein_coding | ENSG0000( |
| 39770559 | 39801333 | -1 | lncRNA | ENSG0000( |
| 39782571 | 39785617 | 1 | lncRNA | ENSG0000( |
| 39799032 | 39920892 | -1 | protein_coding | ENSG0000( |
| 39813223 | 39815429 | -1 | lncRNA | ENSG0000( |
| 39921042 | 39925880 | 1 | lncRNA | ENSG0000( |
| 39922782 | 39922886 | -1 | misc_RNA | ENSG0000( |
| 39934115 | 40035591 | 1 | protein_coding | ENSG0000( |
| 39951375 | 39951708 | -1 | processed_pseudoge | ENSG0000( |
| 40035690 | 40039181 | -1 | protein_coding | ENSG0000( |
| 40039242 | 40076539 | 1 | lncRNA | ENSG0000( |
| 40075204 | 40078704 | -1 | lncRNA | ENSG0000( |
| 40078892 | 40079347 | 1 | lncRNA | ENSG0000( |
| 40087890 | 40108892 | -1 | protein_coding | ENSG0000( |
| 40088832 | 40089386 | 1 | lncRNA | ENSG0000( |
| 40133768 | 40135214 | -1 | processed_pseudoge | ENSG0000( |
| 40161023 | 40221136 | 1 | protein_coding | ENSG0000( |
| 40217428 | 40277487 | 1 | protein_coding | ENSG0000( |
| 40218500 | 40264890 | 1 | protein_coding | ENSG0000( |
| 40226386 | 40227069 | 1 | lncRNA | ENSG0000( |
| 40232082 | 40236109 | 1 | lncRNA | ENSG0000( |
| 40244925 | 40247366 | 1 | TEC | ENSG0000( |
| 40250664 | 40252969 | -1 | lncRNA | ENSG0000( |
| 40258501 | 40260370 | 1 | TEC | ENSG0000( |
| 40278176 | 40307935 | -1 | protein_coding | ENSG0000( |
| 40281444 | 40282586 | -1 | protein_coding | ENSG0000( |
| 40285468 | 40285909 | -1 | lncRNA | ENSG0000( |
| 40300670 | 40301820 | 1 | lncRNA | ENSG0000( |
| 40312352 | 40313289 | -1 | processed_pseudoge | ENSG0000( |
| 40312615 | 40316634 | 1 | lncRNA | ENSG0000( |
| 40323692 | 40326715 | 1 | protein_coding | ENSG0000( |
| 40331452 | 40340967 | -1 | protein_coding | ENSG0000( |
| 40331537 | 40331648 | -1 | rRNA_pseudogene | ENSG0000( |
| 40351033 | 40356434 | 1 | protein_coding | ENSG0000( |
| 40358219 | 40378621 | 1 | protein_coding | ENSG0000( |
| 40368925 | 40369640 | -1 | lncRNA | ENSG0000( |
| 40382721 | 40394246 | 1 | protein_coding | ENSG0000( |
| 40405485 | 40435947 | 1 | protein_coding | ENSG0000( |
| 40439721 | 40468236 | 1 | protein_coding | ENSG0000( |
| 40453444 | 40454639 | -1 | lncRNA | ENSG0000( |
| 40464193 | 40466726 | -1 | lncRNA | ENSG0000( |
| 40470998 | 40474571 | 1 | protein_coding | ENSG0000( |
| 40480752 | 40481315 | -1 | processed_pseudoge | ENSG0000( |
| 40488041 | 40558019 | 1 | lncRNA | ENSG0000( |
| 40528683 | 40565057 | -1 | protein_coding | ENSG0000( |
| 40529570 | 40529673 | 1 | snRNA | ENSG0000( |
| 40569299 | 40574949 | 1 | protein_coding | ENSG0000( |

| | | | | |
|---|---|---|---|---|
| 40585454 | 40585751 | -1 | misc_RNA | ENSG00000 |
| 40594020 | 40664342 | 1 | protein_coding | ENSG00000 |
| 40615665 | 40615928 | -1 | misc_RNA | ENSG00000 |
| 40642933 | 40646857 | 1 | lncRNA | ENSG00000 |
| 40686183 | 40695107 | -1 | lncRNA | ENSG00000 |
| 40694774 | 40732340 | 1 | protein_coding | ENSG00000 |
| 40735884 | 40755851 | -1 | protein_coding | ENSG00000 |
| 40760597 | 40760874 | 1 | processed_pseudoge | ENSG00000 |
| 40764068 | 40767708 | 1 | protein_coding | ENSG00000 |
| 40765161 | 40807478 | -1 | protein_coding | ENSG00000 |
| 40769980 | 40772449 | 1 | protein_coding | ENSG00000 |
| 40807086 | 40815084 | 1 | protein_coding | ENSG00000 |
| 40815445 | 40828709 | -1 | protein_coding | ENSG00000 |
| 40835808 | 40844387 | -1 | lncRNA | ENSG00000 |
| 40844018 | 40858207 | 1 | protein_coding | ENSG00000 |
| 40872214 | 40874234 | -1 | protein_coding | ENSG00000 |
| 40874433 | 40874595 | 1 | lncRNA | ENSG00000 |
| 40894450 | 40903975 | 1 | protein_coding | ENSG00000 |
| 40906811 | 40910337 | 1 | lncRNA | ENSG00000 |
| 40929340 | 40939073 | 1 | protein_coding | ENSG00000 |
| 40948736 | 40948869 | -1 | processed_pseudoge | ENSG00000 |
| 40952962 | 40956512 | 1 | protein_coding | ENSG00000 |
| 40978880 | 41116280 | -1 | protein_coding | ENSG00000 |
| 40999274 | 41004865 | 1 | lncRNA | ENSG00000 |
| 41011597 | 41013300 | 1 | lncRNA | ENSG00000 |
| 41016807 | 41027893 | 1 | lncRNA | ENSG00000 |
| 41016990 | 41018843 | -1 | lncRNA | ENSG00000 |
| 41108878 | 41109178 | 1 | processed_pseudoge | ENSG00000 |
| 41157175 | 41157342 | -1 | processed_pseudoge | ENSG00000 |
| 41163162 | 41164374 | 1 | unprocessed_pseudo | ENSG00000 |
| 41182725 | 41230743 | -1 | protein_coding | ENSG00000 |
| 41191965 | 41192265 | -1 | misc_RNA | ENSG00000 |
| 41230839 | 41281887 | 1 | protein_coding | ENSG00000 |
| 41283990 | 41309737 | 1 | lncRNA | ENSG00000 |
| 41284007 | 41380996 | 1 | protein_coding | ENSG00000 |
| 41309273 | 41332591 | -1 | protein_coding | ENSG00000 |
| 41320794 | 41381050 | 1 | protein_coding | ENSG00000 |
| 41343080 | 41344225 | 1 | lncRNA | ENSG00000 |
| 41387353 | 41402519 | -1 | protein_coding | ENSG00000 |
| 41408408 | 41483563 | 1 | protein_coding | ENSG00000 |
| 41448742 | 41448844 | -1 | snRNA | ENSG00000 |
| 41493393 | 41503551 | 1 | protein_coding | ENSG00000 |
| 41503637 | 41513887 | -1 | protein_coding | ENSG00000 |
| 41517176 | 41544269 | -1 | protein_coding | ENSG00000 |
| 41557027 | 41557435 | 1 | unprocessed_pseudo | ENSG00000 |
| 41557675 | 41583589 | 1 | protein_coding | ENSG00000 |
| 41609457 | 41630908 | -1 | lncRNA | ENSG00000 |
| 41621224 | 41773081 | 1 | protein_coding | ENSG00000 |
| 41691585 | 41691678 | 1 | miRNA | ENSG00000 |
| 41718370 | 41718672 | -1 | processed_pseudoge | ENSG00000 |

| | | | | |
|---|---|---|---|---|
| 41770756 | 41772732 | -1 | lncRNA | ENSG00000 |
| 41774434 | 41827855 | 1 | protein_coding | ENSG00000 |
| 41825099 | 41827936 | -1 | lncRNA | ENSG00000 |
| 41828092 | 41837581 | 1 | protein_coding | ENSG00000 |
| 41828095 | 41848155 | 1 | protein_coding | ENSG00000 |
| 41837775 | 41848147 | 1 | protein_coding | ENSG00000 |
| 41848146 | 41894053 | -1 | protein_coding | ENSG00000 |
| 41851913 | 41852029 | -1 | rRNA_pseudogene | ENSG00000 |
| 41866495 | 41866551 | -1 | miRNA | ENSG00000 |
| 41892777 | 41899528 | 1 | lncRNA | ENSG00000 |
| 41895933 | 41972557 | -1 | protein_coding | ENSG00000 |
| 41908204 | 41908714 | -1 | lncRNA | ENSG00000 |
| 41921417 | 41929286 | 1 | lncRNA | ENSG00000 |
| 41972763 | 41999094 | 1 | lncRNA | ENSG00000 |
| 41981582 | 42051190 | -1 | protein_coding | ENSG00000 |
| 42006132 | 42010117 | 1 | lncRNA | ENSG00000 |
| 42067009 | 42094562 | -1 | protein_coding | ENSG00000 |
| 42139034 | 42156636 | -1 | protein_coding | ENSG00000 |
| 42158701 | 42208316 | -1 | protein_coding | ENSG00000 |
| 42199570 | 42199666 | -1 | miRNA | ENSG00000 |
| 42208413 | 42226990 | 1 | lncRNA | ENSG00000 |
| 42210447 | 42273584 | -1 | protein_coding | ENSG00000 |
| 42273233 | 42356935 | 1 | protein_coding | ENSG00000 |
| 42313687 | 42314386 | 1 | processed_pseudoge | ENSG00000 |
| 42348103 | 42412317 | 1 | protein_coding | ENSG00000 |
| 42359500 | 42412318 | 1 | protein_coding | ENSG00000 |
| 42412823 | 42491141 | -1 | protein_coding | ENSG00000 |
| 42427591 | 42427697 | -1 | snRNA | ENSG00000 |
| 42491233 | 42545356 | 1 | protein_coding | ENSG00000 |
| 42497394 | 42532840 | -1 | lncRNA | ENSG00000 |
| 42497395 | 42548786 | -1 | protein_coding | ENSG00000 |
| 42537820 | 42548802 | -1 | protein_coding | ENSG00000 |
| 42548828 | 42569994 | 1 | protein_coding | ENSG00000 |
| 42567031 | 42569994 | -1 | lncRNA | ENSG00000 |
| 42571927 | 42572433 | 1 | processed_pseudoge | ENSG00000 |
| 42575606 | 42720998 | 1 | protein_coding | ENSG00000 |
| 42581156 | 42581488 | -1 | processed_pseudoge | ENSG00000 |
| 42723544 | 42737128 | -1 | protein_coding | ENSG00000 |
| 42724102 | 42724922 | 1 | lncRNA | ENSG00000 |
| 42726583 | 42727211 | 1 | lncRNA | ENSG00000 |
| 42738730 | 42920809 | -1 | protein_coding | ENSG00000 |
| 42739118 | 42743202 | 1 | lncRNA | ENSG00000 |
| 42749081 | 42749936 | 1 | lncRNA | ENSG00000 |
| 42800968 | 42802405 | -1 | processed_pseudoge | ENSG00000 |
| 42939310 | 42940358 | -1 | processed_pseudoge | ENSG00000 |
| 42942897 | 43106113 | -1 | protein_coding | ENSG00000 |
| 43106225 | 43185779 | -1 | lncRNA | ENSG00000 |
| 43109928 | 43110018 | 1 | pseudogene | ENSG00000 |
| 43115702 | 43116493 | -1 | processed_pseudoge | ENSG00000 |
| 43123279 | 43185144 | 1 | protein_coding | ENSG00000 |

| | | | | |
|---|---|---|---|---|
| 43131008 | 43131686 | 1 | processed_pseudoge | ENSG00000 |
| 43179793 | 43206481 | -1 | lncRNA | ENSG00000 |
| 43184079 | 43185141 | -1 | lncRNA | ENSG00000 |
| 43185118 | 43197177 | 1 | protein_coding | ENSG00000 |
| 43197227 | 43221125 | -1 | protein_coding | ENSG00000 |
| 43232590 | 43266928 | -1 | protein_coding | ENSG00000 |
| 43247680 | 43247781 | 1 | misc_RNA | ENSG00000 |
| 43267904 | 43268388 | -1 | processed_pseudoge | ENSG00000 |
| 43276271 | 43302255 | -1 | protein_coding | ENSG00000 |
| 43302096 | 43302772 | 1 | lncRNA | ENSG00000 |
| 43323649 | 43330582 | -1 | protein_coding | ENSG00000 |
| 43330657 | 43354555 | 1 | protein_coding | ENSG00000 |
| 43358172 | 43371025 | -1 | protein_coding | ENSG00000 |
| 43369221 | 43409771 | 1 | protein_coding | ENSG00000 |
| 43391384 | 43391727 | -1 | misc_RNA | ENSG00000 |
| 43403061 | 43510728 | -1 | protein_coding | ENSG00000 |
| 43510958 | 43531620 | 1 | protein_coding | ENSG00000 |
| 43533462 | 43590253 | -1 | protein_coding | ENSG00000 |
| 43593054 | 43604901 | 1 | protein_coding | ENSG00000 |
| 43599563 | 43618800 | -1 | protein_coding | ENSG00000 |
| 43602543 | 43602648 | 1 | snRNA | ENSG00000 |
| 43614208 | 43632283 | -1 | protein_coding | ENSG00000 |
| 43628503 | 43668118 | -1 | protein_coding | ENSG00000 |
| 43637632 | 43637738 | -1 | snRNA | ENSG00000 |
| 43642389 | 43643023 | -1 | lncRNA | ENSG00000 |
| 43649123 | 43649280 | 1 | unprocessed_pseudo | ENSG00000 |
| 43663654 | 43684339 | -1 | unprocessed_pseudo | ENSG00000 |
| 43692886 | 43699222 | 1 | protein_coding | ENSG00000 |
| 43699488 | 43718184 | -1 | unprocessed_pseudo | ENSG00000 |
| 43702363 | 43702470 | 1 | snRNA | ENSG00000 |
| 43726918 | 43747094 | -1 | transcribed_unproce | ENSG00000 |
| 43734179 | 43734285 | -1 | snRNA | ENSG00000 |
| 43746410 | 43773278 | 1 | protein_coding | ENSG00000 |
| 43772605 | 43777315 | -1 | protein_coding | ENSG00000 |
| 43772617 | 43799133 | -1 | protein_coding | ENSG00000 |
| 43777087 | 43802589 | 1 | protein_coding | ENSG00000 |
| 43793659 | 43793759 | -1 | miRNA | ENSG00000 |
| 43794162 | 43800221 | -1 | protein_coding | ENSG00000 |
| 43796142 | 43803043 | 1 | protein_coding | ENSG00000 |
| 43804492 | 43824690 | -1 | protein_coding | ENSG00000 |
| 43826980 | 43868412 | 1 | protein_coding | ENSG00000 |
| 43863893 | 43864005 | -1 | misc_RNA | ENSG00000 |
| 43870761 | 44195271 | -1 | protein_coding | ENSG00000 |
| 43875849 | 43876244 | -1 | processed_pseudoge | ENSG00000 |
| 43989061 | 43990184 | -1 | processed_pseudoge | ENSG00000 |
| 44063511 | 44064313 | 1 | processed_pseudoge | ENSG00000 |
| 44288719 | 44415758 | 1 | protein_coding | ENSG00000 |
| 44354691 | 44355692 | 1 | processed_pseudoge | ENSG00000 |
| 44387320 | 44390451 | -1 | lncRNA | ENSG00000 |
| 44402685 | 44427144 | -1 | lncRNA | ENSG00000 |

| | | | | |
|---|---|---|---|---|
| 44427622 | 44529038 | 1 | protein_coding | ENSG00000 |
| 44447577 | 44449344 | -1 | processed_pseudoge | ENSG00000 |
| 44516650 | 44517483 | -1 | lncRNA | ENSG00000 |
| 44527257 | 44537046 | -1 | lncRNA | ENSG00000 |
| 44537125 | 44563029 | 1 | protein_coding | ENSG00000 |
| 44557829 | 44559188 | 1 | lncRNA | ENSG00000 |
| 44562696 | 44663678 | -1 | protein_coding | ENSG00000 |
| 44665732 | 44711316 | -1 | protein_coding | ENSG00000 |
| 44697333 | 44697430 | -1 | misc_RNA | ENSG00000 |
| 44711487 | 44718877 | 1 | protein_coding | ENSG00000 |
| 44728988 | 44767829 | 1 | protein_coding | ENSG00000 |
| 44778196 | 44778721 | -1 | lncRNA | ENSG00000 |
| 44825747 | 44884694 | -1 | transcribed_unproce | ENSG00000 |
| 44826540 | 44827094 | 1 | lncRNA | ENSG00000 |
| 44843612 | 44843724 | 1 | misc_RNA | ENSG00000 |
| 44884616 | 44884780 | 1 | snRNA | ENSG00000 |
| 44914802 | 44930622 | -1 | unprocessed_pseudc | ENSG00000 |
| 44919376 | 44920820 | 1 | processed_pseudoge | ENSG00000 |
| 44927600 | 44927703 | -1 | snRNA | ENSG00000 |
| 44956687 | 44979229 | 1 | protein_coding | ENSG00000 |
| 44962830 | 44962933 | 1 | snRNA | ENSG00000 |
| 44970035 | 44971178 | -1 | processed_pseudoge | ENSG00000 |
| 44973558 | 44973661 | 1 | snRNA | ENSG00000 |
| 45002864 | 45003028 | -1 | snRNA | ENSG00000 |
| 45023181 | 45077185 | 1 | protein_coding | ENSG00000 |
| 45041716 | 45058707 | -1 | lncRNA | ENSG00000 |
| 45047103 | 45047215 | -1 | misc_RNA | ENSG00000 |
| 45073492 | 45074048 | -1 | lncRNA | ENSG00000 |
| 45092650 | 45114344 | -1 | protein_coding | ENSG00000 |
| 45114326 | 45118421 | 1 | protein_coding | ENSG00000 |
| 45117367 | 45129938 | -1 | protein_coding | ENSG00000 |
| 45129933 | 45165576 | 1 | protein_coding | ENSG00000 |
| 45152664 | 45167526 | -1 | lncRNA | ENSG00000 |
| 45167214 | 45201175 | -1 | protein_coding | ENSG00000 |
| 45170344 | 45199958 | 1 | lncRNA | ENSG00000 |
| 45198517 | 45199139 | -1 | lncRNA | ENSG00000 |
| 45200325 | 45200632 | -1 | lncRNA | ENSG00000 |
| 45214448 | 45215044 | -1 | TEC | ENSG00000 |
| 45214906 | 45215033 | -1 | snoRNA | ENSG00000 |
| 45215547 | 45215943 | 1 | processed_pseudoge | ENSG00000 |
| 45235930 | 45279251 | -1 | lncRNA | ENSG00000 |
| 45252234 | 45277846 | 1 | protein_coding | ENSG00000 |
| 45279423 | 45280474 | 1 | lncRNA | ENSG00000 |
| 45321077 | 45321692 | -1 | processed_pseudoge | ENSG00000 |
| 45330209 | 45332634 | -1 | processed_pseudoge | ENSG00000 |
| 45354779 | 45354885 | -1 | snRNA | ENSG00000 |
| 45361124 | 45402327 | -1 | protein_coding | ENSG00000 |
| 45378700 | 45380123 | 1 | lncRNA | ENSG00000 |
| 45402336 | 45421415 | 1 | protein_coding | ENSG00000 |
| 45430529 | 45448761 | 1 | protein_coding | ENSG00000 |

| | | | | |
|---|---|---|---|---|
| 45430652 | 45441808 | -1 | lncRNA | ENSG0000C |
| 45433050 | 45433129 | 1 | miRNA | ENSG0000C |
| 45448427 | 45513767 | 1 | lncRNA | ENSG0000C |
| 45479606 | 45522755 | -1 | protein_coding | ENSG0000C |
| 45511136 | 45586290 | 1 | transcribed_processe | ENSG0000C |
| 45537250 | 45537377 | 1 | snoRNA | ENSG0000C |
| 45558598 | 45562877 | 1 | lncRNA | ENSG0000C |
| 45566776 | 45567074 | 1 | transcribed_processe | ENSG0000C |
| 45585757 | 45586304 | -1 | lncRNA | ENSG0000C |
| 45587214 | 45615945 | 1 | protein_coding | ENSG0000C |
| 45587351 | 45676314 | 1 | protein_coding | ENSG0000C |
| 45591984 | 45592085 | 1 | misc_RNA | ENSG0000C |
| 45631148 | 45691281 | 1 | protein_coding | ENSG0000C |
| 45702640 | 45703183 | 1 | lncRNA | ENSG0000C |
| 45705078 | 45931069 | 1 | lncRNA | ENSG0000C |
| 46099193 | 46100392 | -1 | processed_pseudoge | ENSG0000C |
| 46317393 | 46351753 | 1 | lncRNA | ENSG0000C |
| 46341354 | 46341599 | 1 | processed_pseudoge | ENSG0000C |
| 46351662 | 46351746 | -1 | snoRNA | ENSG0000C |
| 46365836 | 46381043 | 1 | lncRNA | ENSG0000C |
| 46410309 | 46661103 | 1 | lncRNA | ENSG0000C |
| 46660658 | 46660760 | 1 | snRNA | ENSG0000C |
| 46693288 | 46804844 | 1 | lncRNA | ENSG0000C |
| 46720193 | 46720512 | -1 | misc_RNA | ENSG0000C |
| 46827526 | 46827958 | 1 | processed_pseudoge | ENSG0000C |
| 47133495 | 47134122 | -1 | TEC | ENSG0000C |
| 47184101 | 47774223 | 1 | protein_coding | ENSG0000C |
| 47272597 | 47272889 | 1 | misc_RNA | ENSG0000C |
| 47274183 | 47275164 | 1 | lncRNA | ENSG0000C |
| 47359430 | 47396732 | -1 | lncRNA | ENSG0000C |
| 47397501 | 47399554 | -1 | lncRNA | ENSG0000C |
| 47525169 | 47527756 | 1 | lncRNA | ENSG0000C |
| 47603880 | 47606352 | -1 | lncRNA | ENSG0000C |
| 47730144 | 47730935 | -1 | processed_pseudoge | ENSG0000C |
| 47774219 | 47846266 | -1 | lncRNA | ENSG0000C |
| 47812729 | 48050507 | 1 | lncRNA | ENSG0000C |
| 47944044 | 47949509 | 1 | lncRNA | ENSG0000C |
| 47956521 | 47988044 | -1 | lncRNA | ENSG0000C |
| 48120990 | 48142672 | 1 | protein_coding | ENSG0000C |
| 48134632 | 48178353 | -1 | protein_coding | ENSG0000C |
| 48178122 | 48203758 | 1 | protein_coding | ENSG0000C |
| 48178438 | 48304078 | 1 | protein_coding | ENSG0000C |
| 48187121 | 48191691 | -1 | lncRNA | ENSG0000C |
| 48192191 | 48201684 | -1 | lncRNA | ENSG0000C |
| 48312353 | 48331856 | -1 | lncRNA | ENSG0000C |
| 48325558 | 48325671 | -1 | misc_RNA | ENSG0000C |
| 48331011 | 48343373 | 1 | protein_coding | ENSG0000C |
| 48408306 | 48645849 | -1 | protein_coding | ENSG0000C |
| 48524308 | 48526051 | 1 | lncRNA | ENSG0000C |
| 48528980 | 48529728 | -1 | lncRNA | ENSG0000C |

| | | | | |
|---|---|---|---|---|
| 48645951 | 48652016 | 1 | lncRNA | ENSG00000 |
| 48662531 | 48663056 | -1 | lncRNA | ENSG00000 |
| 48712928 | 48811146 | -1 | protein_coding | ENSG00000 |
| 48725338 | 48725827 | 1 | processed_pseudoge | ENSG00000 |
| 48729080 | 48729844 | -1 | lncRNA | ENSG00000 |
| 48783190 | 48784121 | 1 | lncRNA | ENSG00000 |
| 48810701 | 48811909 | 1 | lncRNA | ENSG00000 |
| 48823741 | 48963919 | -1 | protein_coding | ENSG00000 |
| 48878134 | 48880173 | 1 | protein_coding | ENSG00000 |
| 48938236 | 48948031 | 1 | lncRNA | ENSG00000 |
| 48971803 | 48973361 | 1 | processed_pseudoge | ENSG00000 |
| 48988476 | 49046447 | -1 | protein_coding | ENSG00000 |
| 48995912 | 48996022 | -1 | misc_RNA | ENSG00000 |
| 49042341 | 49042643 | -1 | misc_RNA | ENSG00000 |
| 49101791 | 49103264 | 1 | lncRNA | ENSG00000 |
| 49106068 | 49155661 | -1 | protein_coding | ENSG00000 |
| 49137762 | 49137863 | -1 | misc_RNA | ENSG00000 |
| 49155656 | 49367869 | 1 | protein_coding | ENSG00000 |
| 49156329 | 49156847 | 1 | processed_pseudoge | ENSG00000 |
| 49169070 | 49169153 | -1 | miRNA | ENSG00000 |
| 49177610 | 49178741 | -1 | processed_pseudoge | ENSG00000 |
| 49198900 | 49199194 | -1 | misc_RNA | ENSG00000 |
| 49326962 | 49620929 | -1 | protein_coding | ENSG00000 |
| 49343608 | 49344254 | 1 | lncRNA | ENSG00000 |
| 49353485 | 49354034 | 1 | lncRNA | ENSG00000 |
| 49365124 | 49366685 | -1 | processed_pseudoge | ENSG00000 |
| 49423178 | 49488775 | 1 | protein_coding | ENSG00000 |
| 49621037 | 49656232 | 1 | protein_coding | ENSG00000 |
| 49794246 | 49796097 | 1 | processed_pseudoge | ENSG00000 |
| 49858238 | 50182817 | -1 | protein_coding | ENSG00000 |
| 49877299 | 49883525 | 1 | lncRNA | ENSG00000 |
| 49982443 | 49982561 | 1 | rRNA_pseudogene | ENSG00000 |
| 50182196 | 50236385 | 1 | protein_coding | ENSG00000 |
| 50221888 | 50221996 | 1 | misc_RNA | ENSG00000 |
| 50241947 | 50265965 | -1 | protein_coding | ENSG00000 |
| 50244628 | 50244931 | -1 | misc_RNA | ENSG00000 |
| 50275392 | 50355408 | -1 | protein_coding | ENSG00000 |
| 50347423 | 50347529 | -1 | snRNA | ENSG00000 |
| 50348936 | 50354861 | -1 | lncRNA | ENSG00000 |
| 50354959 | 50372202 | 1 | lncRNA | ENSG00000 |
| 50359450 | 50360194 | -1 | lncRNA | ENSG00000 |
| 50360329 | 50360410 | 1 | miRNA | ENSG00000 |
| 50417385 | 50418692 | 1 | processed_pseudoge | ENSG00000 |
| 50424380 | 50514421 | 1 | protein_coding | ENSG00000 |
| 50463381 | 50463498 | 1 | rRNA_pseudogene | ENSG00000 |
| 50466738 | 50467096 | 1 | processed_pseudoge | ENSG00000 |
| 50494018 | 50497080 | -1 | lncRNA | ENSG00000 |
| 50497195 | 50498744 | -1 | lncRNA | ENSG00000 |
| 50500562 | 50546708 | -1 | protein_coding | ENSG00000 |
| 50552473 | 50686797 | -1 | protein_coding | ENSG00000 |

| | | | | |
|---|---|---|---|---|
| 50557601 | 50560500 | 1 | lncRNA | ENSG0000( |
| 50688900 | 50689193 | 1 | misc_RNA | ENSG0000( |
| 50702266 | 50765709 | -1 | protein_coding | ENSG0000( |
| 50742702 | 50743276 | -1 | TEC | ENSG0000( |
| 50746709 | 50749829 | 1 | lncRNA | ENSG0000( |
| 50798670 | 50798951 | 1 | processed_pseudoge | ENSG0000( |
| 50839875 | 50908603 | -1 | lncRNA | ENSG0000( |
| 50908672 | 51005895 | 1 | protein_coding | ENSG0000( |
| 50944663 | 50945996 | 1 | processed_pseudoge | ENSG0000( |
| 51037488 | 51293912 | 1 | lncRNA | ENSG0000( |
| 51056598 | 51105276 | -1 | protein_coding | ENSG0000( |
| 51064609 | 51069586 | -1 | lncRNA | ENSG0000( |
| 51208057 | 51338601 | -1 | protein_coding | ENSG0000( |
| 51242190 | 51242264 | 1 | miRNA | ENSG0000( |
| 51278927 | 51280015 | -1 | lncRNA | ENSG0000( |
| 51314032 | 51314107 | -1 | miRNA | ENSG0000( |
| 51314034 | 51314109 | 1 | miRNA | ENSG0000( |
| 51315841 | 51321996 | 1 | lncRNA | ENSG0000( |
| 51341655 | 51408005 | 1 | protein_coding | ENSG0000( |
| 51351100 | 51351192 | -1 | processed_pseudoge | ENSG0000( |
| 51367377 | 51369110 | 1 | lncRNA | ENSG0000( |
| 51414097 | 51498833 | 1 | lncRNA | ENSG0000( |
| 51447711 | 51622833 | -1 | protein_coding | ENSG0000( |
| 51455268 | 51460582 | 1 | lncRNA | ENSG0000( |
| 51681492 | 51721026 | 1 | protein_coding | ENSG0000( |
| 51693216 | 51695765 | -1 | lncRNA | ENSG0000( |
| 51707995 | 51715220 | -1 | lncRNA | ENSG0000( |
| 51723011 | 51751585 | -1 | protein_coding | ENSG0000( |
| 51751597 | 51816363 | 1 | protein_coding | ENSG0000( |
| 51752485 | 51756475 | 1 | lncRNA | ENSG0000( |
| 51829628 | 51947295 | 1 | protein_coding | ENSG0000( |
| 51833134 | 51833426 | 1 | lncRNA | ENSG0000( |
| 51858070 | 51858352 | -1 | misc_RNA | ENSG0000( |
| 51859185 | 51861293 | -1 | processed_pseudoge | ENSG0000( |
| 51884556 | 51884653 | 1 | misc_RNA | ENSG0000( |
| 51887560 | 51901123 | -1 | lncRNA | ENSG0000( |
| 51907200 | 51907305 | 1 | snRNA | ENSG0000( |
| 51908902 | 51909642 | -1 | lncRNA | ENSG0000( |
| 51919073 | 51930627 | -1 | unprocessed_pseudc | ENSG0000( |
| 51938025 | 51971778 | -1 | protein_coding | ENSG0000( |
| 51952106 | 52067375 | 1 | protein_coding | ENSG0000( |
| 52010999 | 52019095 | -1 | lncRNA | ENSG0000( |
| 52017167 | 52018032 | -1 | lncRNA | ENSG0000( |
| 52033707 | 52035625 | 1 | lncRNA | ENSG0000( |
| 52056675 | 52100523 | -1 | lncRNA | ENSG0000( |
| 52082302 | 52086574 | 1 | lncRNA | ENSG0000( |
| 52095295 | 52095747 | 1 | processed_pseudoge | ENSG0000( |
| 52109263 | 52112775 | -1 | protein_coding | ENSG0000( |
| 52115105 | 52191369 | -1 | protein_coding | ENSG0000( |
| 52116574 | 52122131 | 1 | lncRNA | ENSG0000( |

| | | | | |
|---|---|---|---|---|
| 52124561 | 52140246 | 1 | lncRNA | ENSG0000C |
| 52162751 | 52162852 | 1 | misc_RNA | ENSG0000C |
| 52179999 | 52206915 | 1 | lncRNA | ENSG0000C |
| 52192322 | 52295804 | -1 | protein_coding | ENSG0000C |
| 52277117 | 52277200 | -1 | miRNA | ENSG0000C |
| 52307283 | 52529050 | -1 | protein_coding | ENSG0000C |
| 52505191 | 52505862 | 1 | processed_pseudoge | ENSG0000C |
| 52547045 | 52569883 | -1 | protein_coding | ENSG0000C |
| 52577842 | 52598709 | 1 | lncRNA | ENSG0000C |
| 52581317 | 52709817 | -1 | protein_coding | ENSG0000C |
| 52648634 | 52649866 | -1 | lncRNA | ENSG0000C |
| 52679474 | 52690077 | 1 | lncRNA | ENSG0000C |
| 52755053 | 52791078 | -1 | protein_coding | ENSG0000C |
| 52800168 | 52805972 | -1 | lncRNA | ENSG0000C |
| 52885595 | 52886470 | 1 | processed_pseudoge | ENSG0000C |
| 52937139 | 52938518 | -1 | processed_pseudoge | ENSG0000C |
| 53116365 | 53129698 | 1 | lncRNA | ENSG0000C |
| 53419865 | 53453487 | 1 | lncRNA | ENSG0000C |
| 53513741 | 53762878 | -1 | protein_coding | ENSG0000C |
| 53651977 | 53652123 | 1 | snRNA | ENSG0000C |
| 53764204 | 53764310 | 1 | snRNA | ENSG0000C |
| 53837271 | 53837560 | -1 | processed_pseudoge | ENSG0000C |
| 53910769 | 53914712 | 1 | TEC | ENSG0000C |
| 53946709 | 53984010 | -1 | lncRNA | ENSG0000C |
| 53978201 | 54633414 | 1 | protein_coding | ENSG0000C |
| 54048588 | 54050876 | -1 | processed_pseudoge | ENSG0000C |
| 54320731 | 54321565 | -1 | processed_pseudoge | ENSG0000C |
| 55056723 | 55092177 | 1 | lncRNA | ENSG0000C |
| 55171378 | 55176739 | -1 | lncRNA | ENSG0000C |
| 55180806 | 55197049 | -1 | protein_coding | ENSG0000C |
| 55202966 | 55319113 | -1 | protein_coding | ENSG0000C |
| 55288849 | 55289346 | -1 | lncRNA | ENSG0000C |
| 55317184 | 55319161 | -1 | protein_coding | ENSG0000C |
| 55318960 | 55355648 | 1 | protein_coding | ENSG0000C |
| 55340032 | 55408510 | -1 | protein_coding | ENSG0000C |
| 55343141 | 55343575 | -1 | lncRNA | ENSG0000C |
| 55346347 | 55346752 | -1 | lncRNA | ENSG0000C |
| 55355248 | 55498360 | -1 | lncRNA | ENSG0000C |
| 55372940 | 55373034 | -1 | miRNA | ENSG0000C |
| 55408495 | 55418798 | 1 | protein_coding | ENSG0000C |
| 55410525 | 55508234 | -1 | protein_coding | ENSG0000C |
| 55442635 | 55443357 | -1 | processed_pseudoge | ENSG0000C |
| 55538890 | 55588947 | -1 | protein_coding | ENSG0000C |
| 55588666 | 55589811 | 1 | lncRNA | ENSG0000C |
| 55611544 | 55743152 | -1 | protein_coding | ENSG0000C |
| 55680385 | 55681463 | 1 | lncRNA | ENSG0000C |
| 55746639 | 55746767 | 1 | processed_pseudoge | ENSG0000C |
| 55826922 | 55993746 | -1 | protein_coding | ENSG0000C |
| 56005715 | 56007176 | -1 | processed_pseudoge | ENSG0000C |
| 56029684 | 56029981 | -1 | misc_RNA | ENSG0000C |

| | | | | |
|---|---|---|---|---|
| 56071421 | 56071639 | 1 | processed_pseudoge | ENSG00000 |
| 56087280 | 56243266 | -1 | protein_coding | ENSG00000 |
| 56192353 | 56193262 | -1 | processed_pseudoge | ENSG00000 |
| 56244009 | 56445997 | 1 | protein_coding | ENSG00000 |
| 56248787 | 56249127 | 1 | lncRNA | ENSG00000 |
| 56274530 | 56274636 | -1 | snRNA | ENSG00000 |
| 56323193 | 56323719 | 1 | processed_pseudoge | ENSG00000 |
| 56394321 | 56396785 | -1 | lncRNA | ENSG00000 |
| 56408483 | 56410186 | -1 | lncRNA | ENSG00000 |
| 56421544 | 56465137 | -1 | protein_coding | ENSG00000 |
| 56447120 | 56447697 | 1 | lncRNA | ENSG00000 |
| 56542952 | 56629592 | -1 | lncRNA | ENSG00000 |
| 56630181 | 56734086 | -1 | protein_coding | ENSG00000 |
| 56634038 | 56634167 | -1 | misc_RNA | ENSG00000 |
| 56652488 | 56918571 | -1 | protein_coding | ENSG00000 |
| 56729932 | 56730611 | -1 | lncRNA | ENSG00000 |
| 56795382 | 56795522 | 1 | TEC | ENSG00000 |
| 56795460 | 56918499 | -1 | lncRNA | ENSG00000 |
| 56846340 | 56846605 | 1 | processed_pseudoge | ENSG00000 |
| 56887107 | 56888219 | 1 | lncRNA | ENSG00000 |
| 56908312 | 56908635 | -1 | lncRNA | ENSG00000 |
| 56918623 | 57299281 | 1 | protein_coding | ENSG00000 |
| 57246960 | 57247893 | -1 | processed_pseudoge | ENSG00000 |
| 57278570 | 57278673 | -1 | snoRNA | ENSG00000 |
| 57300365 | 57307769 | 1 | lncRNA | ENSG00000 |
| 57319138 | 57325039 | 1 | lncRNA | ENSG00000 |
| 57325472 | 57326447 | -1 | processed_pseudoge | ENSG00000 |
| 57375967 | 57550727 | 1 | protein_coding | ENSG00000 |
| 57405074 | 57405175 | -1 | snRNA | ENSG00000 |
| 57477484 | 57478329 | -1 | lncRNA | ENSG00000 |
| 57527187 | 57529009 | -1 | lncRNA | ENSG00000 |
| 57553955 | 57555680 | 1 | lncRNA | ENSG00000 |
| 57555951 | 57556379 | -1 | processed_pseudoge | ENSG00000 |
| 57591904 | 57685364 | 1 | protein_coding | ENSG00000 |
| 57591908 | 57714745 | 1 | protein_coding | ENSG00000 |
| 57706695 | 57782762 | 1 | protein_coding | ENSG00000 |
| 57720295 | 57720928 | 1 | lncRNA | ENSG00000 |
| 57953424 | 58497866 | -1 | protein_coding | ENSG00000 |
| 57979716 | 57980646 | -1 | processed_pseudoge | ENSG00000 |
| 57990217 | 57990636 | -1 | lncRNA | ENSG00000 |
| 58065225 | 58071043 | 1 | lncRNA | ENSG00000 |
| 58138169 | 58185911 | 1 | protein_coding | ENSG00000 |
| 58150275 | 58150563 | 1 | processed_pseudoge | ENSG00000 |
| 58150635 | 58150962 | 1 | processed_pseudoge | ENSG00000 |
| 58151033 | 58152513 | 1 | processed_pseudoge | ENSG00000 |
| 58152606 | 58154407 | 1 | processed_pseudoge | ENSG00000 |
| 58154960 | 58156061 | 1 | processed_pseudoge | ENSG00000 |
| 58200606 | 58201018 | 1 | processed_pseudoge | ENSG00000 |
| 58410569 | 58569844 | 1 | protein_coding | ENSG00000 |
| 58434890 | 58498735 | -1 | lncRNA | ENSG00000 |

| | | | | |
|---|---|---|---|---|
| 58454327 | 58454982 | 1 | lncRNA | ENSG0000C |
| 58456188 | 58456348 | 1 | lncRNA | ENSG0000C |
| 58521311 | 58523371 | 1 | lncRNA | ENSG0000C |
| 58587507 | 58591676 | 1 | lncRNA | ENSG0000C |
| 58588809 | 58749791 | -1 | protein_coding | ENSG0000C |
| 58672296 | 58672933 | 1 | processed_pseudoge | ENSG0000C |
| 58686411 | 58686617 | 1 | processed_pseudoge | ENSG0000C |
| 58691106 | 58693125 | -1 | processed_pseudoge | ENSG0000C |
| 58703852 | 58704154 | -1 | misc_RNA | ENSG0000C |
| 58732188 | 58733149 | 1 | unprocessed_pseudc | ENSG0000C |
| 58749923 | 58761004 | 1 | lncRNA | ENSG0000C |
| 58755625 | 58756247 | -1 | processed_pseudoge | ENSG0000C |
| 58760466 | 58760681 | -1 | snoRNA | ENSG0000C |
| 58768072 | 58770974 | -1 | lncRNA | ENSG0000C |
| 58771192 | 58861900 | 1 | protein_coding | ENSG0000C |
| 58815272 | 58817728 | 1 | lncRNA | ENSG0000C |
| 58856831 | 58865063 | -1 | lncRNA | ENSG0000C |
| 58865006 | 58865303 | 1 | processed_pseudoge | ENSG0000C |
| 58865175 | 59097419 | 1 | protein_coding | ENSG0000C |
| 58869659 | 58870134 | 1 | processed_pseudoge | ENSG0000C |
| 58879050 | 58933679 | -1 | protein_coding | ENSG0000C |
| 58889967 | 58894082 | 1 | lncRNA | ENSG0000C |
| 58967046 | 58967808 | -1 | processed_pseudoge | ENSG0000C |
| 59049399 | 59050385 | -1 | processed_pseudoge | ENSG0000C |
| 59105126 | 59125045 | 1 | protein_coding | ENSG0000C |
| 59115547 | 59116089 | -1 | lncRNA | ENSG0000C |
| 59121034 | 59133250 | 1 | lncRNA | ENSG0000C |
| 59132434 | 59372871 | -1 | protein_coding | ENSG0000C |
| 59171183 | 59171262 | -1 | miRNA | ENSG0000C |
| 59206843 | 59208588 | 1 | protein_coding | ENSG0000C |
| 59212648 | 59212788 | -1 | snRNA | ENSG0000C |
| 59266720 | 59271162 | -1 | lncRNA | ENSG0000C |
| 59341314 | 59341417 | -1 | snRNA | ENSG0000C |
| 59348648 | 59359889 | -1 | lncRNA | ENSG0000C |
| 59372693 | 59523555 | 1 | protein_coding | ENSG0000C |
| 59401221 | 59402004 | -1 | processed_pseudoge | ENSG0000C |
| 59406963 | 59408154 | -1 | processed_pseudoge | ENSG0000C |
| 59408905 | 59409015 | 1 | misc_RNA | ENSG0000C |
| 59525422 | 59525881 | -1 | processed_pseudoge | ENSG0000C |
| 59552615 | 59553093 | -1 | processed_pseudoge | ENSG0000C |
| 59574103 | 59574222 | -1 | rRNA_pseudogene | ENSG0000C |
| 59594875 | 59640239 | 1 | protein_coding | ENSG0000C |
| 59638062 | 59657541 | -1 | protein_coding | ENSG0000C |
| 59644298 | 59672302 | -1 | lncRNA | ENSG0000C |
| 59659146 | 59689534 | -1 | protein_coding | ENSG0000C |
| 59676757 | 59677321 | 1 | processed_pseudoge | ENSG0000C |
| 59688517 | 59689418 | 1 | lncRNA | ENSG0000C |
| 59768352 | 59769146 | 1 | processed_pseudoge | ENSG0000C |
| 59801347 | 59814749 | -1 | lncRNA | ENSG0000C |
| 59871511 | 59873495 | -1 | processed_pseudoge | ENSG0000C |

| | | | | |
|---|---|---|---|---|
| 59964942 | 59966084 | 1 | processed_pseudoge | ENSG00000( |
| 60004234 | 60061730 | 1 | protein_coding | ENSG00000( |
| 60053204 | 60054092 | -1 | processed_pseudoge | ENSG00000( |
| 60073554 | 60118320 | -1 | lncRNA | ENSG00000( |
| 60215537 | 60223272 | -1 | lncRNA | ENSG00000( |
| 60347134 | 60402883 | -1 | protein_coding | ENSG00000( |
| 60390371 | 60390682 | 1 | processed_pseudoge | ENSG00000( |
| 60404463 | 60404960 | 1 | processed_pseudoge | ENSG00000( |
| 60409234 | 60409420 | -1 | unprocessed_pseudo | ENSG00000( |
| 60419609 | 60479160 | -1 | protein_coding | ENSG00000( |
| 60479152 | 60630637 | 1 | lncRNA | ENSG00000( |
| 60488284 | 61229302 | -1 | protein_coding | ENSG00000( |
| 60529973 | 60545159 | 1 | lncRNA | ENSG00000( |
| 60558080 | 60558401 | 1 | transcribed_process | ENSG00000( |
| 60593027 | 60593460 | 1 | lncRNA | ENSG00000( |
| 60634503 | 60640915 | 1 | lncRNA | ENSG00000( |
| 60642054 | 60659791 | 1 | lncRNA | ENSG00000( |
| 60677109 | 60677272 | -1 | misc_RNA | ENSG00000( |
| 60681564 | 60687248 | 1 | lncRNA | ENSG00000( |
| 60736769 | 60736886 | 1 | rRNA_pseudogene | ENSG00000( |
| 60763593 | 60765625 | 1 | lncRNA | ENSG00000( |
| 60847757 | 60849068 | -1 | lncRNA | ENSG00000( |
| 61181249 | 61195938 | 1 | lncRNA | ENSG00000( |
| 61211913 | 61214231 | 1 | lncRNA | ENSG00000( |
| 61239906 | 61240331 | 1 | lncRNA | ENSG00000( |
| 61298791 | 61635449 | -1 | lncRNA | ENSG00000( |
| 61598514 | 61605241 | 1 | lncRNA | ENSG00000( |
| 61638928 | 61715171 | -1 | lncRNA | ENSG00000( |
| 61674767 | 61675212 | 1 | processed_pseudoge | ENSG00000( |
| 61729765 | 61731389 | 1 | lncRNA | ENSG00000( |
| 61834682 | 61935738 | 1 | lncRNA | ENSG00000( |
| 61852389 | 62060473 | -1 | protein_coding | ENSG00000( |
| 61861109 | 61894164 | 1 | lncRNA | ENSG00000( |
| 62060503 | 62062434 | 1 | lncRNA | ENSG00000( |
| 62066977 | 62070917 | 1 | protein_coding | ENSG00000( |
| 62082450 | 62083305 | -1 | processed_pseudoge | ENSG00000( |
| 62163535 | 62165285 | -1 | protein_coding | ENSG00000( |
| 62196066 | 62222911 | -1 | lncRNA | ENSG00000( |
| 62227896 | 62253235 | -1 | transcribed_unproce | ENSG00000( |
| 62230384 | 62231895 | -1 | lncRNA | ENSG00000( |
| 62244330 | 62244595 | -1 | misc_RNA | ENSG00000( |
| 62253730 | 62270680 | -1 | transcribed_unproce | ENSG00000( |
| 62255686 | 62256421 | -1 | unprocessed_pseudo | ENSG00000( |
| 62274163 | 62278365 | 1 | lncRNA | ENSG00000( |
| 62284190 | 62285006 | 1 | lncRNA | ENSG00000( |
| 62304658 | 62304734 | -1 | miRNA | ENSG00000( |
| 62343029 | 62343138 | 1 | miRNA | ENSG00000( |
| 62362029 | 62362331 | -1 | processed_pseudoge | ENSG00000( |
| 62387327 | 62388875 | -1 | lncRNA | ENSG00000( |
| 62390526 | 62844631 | 1 | protein_coding | ENSG00000( |

| | | | | |
|---|---|---|---|---|
| 62637172 | 62645291 | -1 | lncRNA | ENSG0000C |
| 62682916 | 62690448 | -1 | lncRNA | ENSG0000C |
| 62823957 | 62824041 | 1 | miRNA | ENSG0000C |
| 62827675 | 62884034 | -1 | lncRNA | ENSG0000C |
| 62895812 | 62899543 | 1 | lncRNA | ENSG0000C |
| 62953173 | 62954124 | 1 | processed_pseudoge | ENSG0000C |
| 63042632 | 63071915 | 1 | protein_coding | ENSG0000C |
| 63046034 | 63049387 | -1 | lncRNA | ENSG0000C |
| 63070025 | 63071911 | -1 | lncRNA | ENSG0000C |
| 63091085 | 63110589 | 1 | lncRNA | ENSG0000C |
| 63121833 | 63142061 | 1 | protein_coding | ENSG0000C |
| 63125872 | 63158021 | -1 | protein_coding | ENSG0000C |
| 63189560 | 63267776 | 1 | protein_coding | ENSG0000C |
| 63276018 | 63309126 | 1 | protein_coding | ENSG0000C |
| 63321378 | 63381846 | -1 | protein_coding | ENSG0000C |
| 63390136 | 63438320 | 1 | lncRNA | ENSG0000C |
| 63456294 | 63469356 | -1 | lncRNA | ENSG0000C |
| 63500737 | 63503083 | -1 | lncRNA | ENSG0000C |
| 63504511 | 63594640 | 1 | protein_coding | ENSG0000C |
| 63544247 | 63601589 | -1 | lncRNA | ENSG0000C |
| 63597353 | 63602428 | 1 | protein_coding | ENSG0000C |
| 63608618 | 63833948 | -1 | protein_coding | ENSG0000C |
| 63675146 | 63713758 | 1 | lncRNA | ENSG0000C |
| 63870930 | 63871019 | -1 | miRNA | ENSG0000C |
| 63890030 | 63890317 | 1 | unprocessed_pseudc | ENSG0000C |
| 63907036 | 64072033 | -1 | protein_coding | ENSG0000C |
| 63928274 | 63935953 | 1 | lncRNA | ENSG0000C |
| 64072565 | 64093857 | -1 | protein_coding | ENSG0000C |
| 64094123 | 64146090 | 1 | protein_coding | ENSG0000C |
| 64111458 | 64111569 | -1 | misc_RNA | ENSG0000C |
| 64151715 | 64157481 | 1 | protein_coding | ENSG0000C |
| 64155812 | 64163205 | -1 | protein_coding | ENSG0000C |
| 64165525 | 64356173 | -1 | protein_coding | ENSG0000C |
| 64181180 | 64381440 | -1 | protein_coding | ENSG0000C |
| 64342539 | 64342698 | 1 | snoRNA | ENSG0000C |
| 64364304 | 64387687 | -1 | protein_coding | ENSG0000C |
| 64387748 | 64455303 | 1 | protein_coding | ENSG0000C |
| 64445130 | 64445243 | -1 | misc_RNA | ENSG0000C |
| 64460742 | 64686068 | 1 | protein_coding | ENSG0000C |
| 64470253 | 64471364 | 1 | lncRNA | ENSG0000C |
| 64528667 | 64529671 | -1 | processed_pseudoge | ENSG0000C |
| 64592979 | 64593326 | 1 | processed_pseudoge | ENSG0000C |
| 64631109 | 64631914 | -1 | processed_pseudoge | ENSG0000C |
| 64652888 | 64652989 | 1 | misc_RNA | ENSG0000C |
| 64671263 | 64671369 | 1 | snRNA | ENSG0000C |
| 64687573 | 64703281 | -1 | protein_coding | ENSG0000C |
| 64695041 | 64695594 | 1 | lncRNA | ENSG0000C |
| 64696622 | 64696861 | 1 | lncRNA | ENSG0000C |
| 64701248 | 64719602 | 1 | lncRNA | ENSG0000C |
| 64739891 | 64775589 | -1 | protein_coding | ENSG0000C |

| | | | | |
|---|---|---|---|---|
| 64762387 | 64762515 | -1 | miRNA | ENSG0000C |
| 64815632 | 64825668 | -1 | protein_coding | ENSG0000C |
| 64841883 | 64930920 | 1 | protein_coding | ENSG0000C |
| 64841883 | 64868002 | 1 | protein_coding | ENSG0000C |
| 64878428 | 64882385 | -1 | lncRNA | ENSG0000C |
| 64911902 | 64958691 | 1 | protein_coding | ENSG0000C |
| 64943240 | 64943672 | -1 | lncRNA | ENSG0000C |
| 64950916 | 64951435 | -1 | lncRNA | ENSG0000C |
| 64963022 | 64990310 | -1 | protein_coding | ENSG0000C |
| 65001512 | 65029639 | -1 | protein_coding | ENSG0000C |
| 65026088 | 65026676 | 1 | processed_pseudoge | ENSG0000C |
| 65045387 | 65053397 | 1 | protein_coding | ENSG0000C |
| 65049188 | 65049802 | -1 | lncRNA | ENSG0000C |
| 65053337 | 65076690 | -1 | protein_coding | ENSG0000C |
| 65076816 | 65079939 | 1 | protein_coding | ENSG0000C |
| 65083042 | 65083663 | -1 | lncRNA | ENSG0000C |
| 65092770 | 65115197 | -1 | protein_coding | ENSG0000C |
| 65105534 | 65105800 | 1 | processed_pseudoge | ENSG0000C |
| 65117379 | 65133808 | -1 | protein_coding | ENSG0000C |
| 65148219 | 65185342 | -1 | protein_coding | ENSG0000C |
| 65194760 | 65211473 | -1 | protein_coding | ENSG0000C |
| 65226472 | 65226697 | -1 | TEC | ENSG0000C |
| 65229861 | 65229967 | 1 | snRNA | ENSG0000C |
| 65234460 | 65300618 | -1 | protein_coding | ENSG0000C |
| 65285461 | 65285591 | -1 | snoRNA | ENSG0000C |
| 65296051 | 65296166 | 1 | snRNA | ENSG0000C |
| 65301922 | 65302311 | 1 | processed_pseudoge | ENSG0000C |
| 65304677 | 65304792 | 1 | snRNA | ENSG0000C |
| 65327127 | 65378002 | -1 | protein_coding | ENSG0000C |
| 65381484 | 65422947 | -1 | protein_coding | ENSG0000C |
| 65442463 | 65517704 | -1 | protein_coding | ENSG0000C |
| 65458574 | 65459534 | 1 | processed_pseudoge | ENSG0000C |
| 65475318 | 65475614 | -1 | processed_pseudoge | ENSG0000C |
| 65502741 | 65502836 | 1 | misc_RNA | ENSG0000C |
| 65530418 | 65578349 | 1 | protein_coding | ENSG0000C |
| 65553081 | 65553187 | -1 | snRNA | ENSG0000C |
| 65578753 | 65611289 | -1 | protein_coding | ENSG0000C |
| 65611366 | 65660995 | 1 | protein_coding | ENSG0000C |
| 65655620 | 65656085 | -1 | lncRNA | ENSG0000C |
| 65658046 | 65792293 | -1 | protein_coding | ENSG0000C |
| 65696833 | 65696936 | -1 | snoRNA | ENSG0000C |
| 65719246 | 65719332 | -1 | miRNA | ENSG0000C |
| 65726054 | 65891989 | 1 | protein_coding | ENSG0000C |
| 65766460 | 65767398 | -1 | lncRNA | ENSG0000C |
| 65771176 | 65772139 | -1 | processed_pseudoge | ENSG0000C |
| 65816302 | 65817481 | -1 | processed_pseudoge | ENSG0000C |
| 65843484 | 65843558 | 1 | snRNA | ENSG0000C |
| 65895079 | 66253747 | -1 | protein_coding | ENSG0000C |
| 66040233 | 66040332 | 1 | miRNA | ENSG0000C |
| 66278498 | 66293357 | -1 | lncRNA | ENSG0000C |

| | | | | |
|---|---|---|---|---|
| 66293217 | 66333898 | 1 | protein_coding | ENSG0000C |
| 66314914 | 66331703 | -1 | lncRNA | ENSG0000C |
| 66336191 | 66386746 | -1 | protein_coding | ENSG0000C |
| 66347207 | 66347342 | -1 | scaRNA | ENSG0000C |
| 66348276 | 66348375 | 1 | misc_RNA | ENSG0000C |
| 66379337 | 66379915 | -1 | processed_pseudoge | ENSG0000C |
| 66386817 | 66492312 | 1 | protein_coding | ENSG0000C |
| 66414933 | 66415198 | -1 | processed_pseudoge | ENSG0000C |
| 66446455 | 66446777 | -1 | processed_pseudoge | ENSG0000C |
| 66488658 | 66492109 | -1 | lncRNA | ENSG0000C |
| 66490135 | 66497780 | -1 | protein_coding | ENSG0000C |
| 66496958 | 66497034 | -1 | miRNA | ENSG0000C |
| 66498015 | 66524532 | -1 | protein_coding | ENSG0000C |
| 66501250 | 66501318 | -1 | snoRNA | ENSG0000C |
| 66502019 | 66502089 | -1 | snoRNA | ENSG0000C |
| 66502020 | 66502091 | -1 | snoRNA | ENSG0000C |
| 66502812 | 66502910 | -1 | snoRNA | ENSG0000C |
| 66503243 | 66503314 | -1 | snoRNA | ENSG0000C |
| 66504959 | 66550130 | 1 | protein_coding | ENSG0000C |
| 66547179 | 66565979 | -1 | protein_coding | ENSG0000C |
| 66582190 | 66685794 | 1 | lncRNA | ENSG0000C |
| 66702236 | 66782849 | 1 | protein_coding | ENSG0000C |
| 66740445 | 66741151 | 1 | lncRNA | ENSG0000C |
| 66808404 | 66810560 | 1 | processed_pseudoge | ENSG0000C |
| 66860303 | 66867023 | 1 | lncRNA | ENSG0000C |
| 66919811 | 66931755 | 1 | lncRNA | ENSG0000C |
| 66931537 | 66956518 | -1 | lncRNA | ENSG0000C |
| 66984103 | 67065268 | -1 | lncRNA | ENSG0000C |
| 67000814 | 67001085 | -1 | processed_pseudoge | ENSG0000C |
| 67063763 | 67195169 | 1 | protein_coding | ENSG0000C |
| 67142734 | 67146939 | -1 | lncRNA | ENSG0000C |
| 67200667 | 67255195 | -1 | protein_coding | ENSG0000C |
| 67232446 | 67232837 | 1 | processed_pseudoge | ENSG0000C |
| 67254786 | 67502260 | 1 | protein_coding | ENSG0000C |
| 67290636 | 67521844 | -1 | lncRNA | ENSG0000C |
| 67521131 | 67530146 | 1 | protein_coding | ENSG0000C |
| 67541072 | 67542604 | -1 | lncRNA | ENSG0000C |
| 67542703 | 67807117 | 1 | protein_coding | ENSG0000C |
| 67627852 | 67629135 | 1 | processed_pseudoge | ENSG0000C |
| 67819704 | 67834582 | 1 | protein_coding | ENSG0000C |
| 67832725 | 67873866 | 1 | lncRNA | ENSG0000C |
| 67834310 | 67838879 | -1 | lncRNA | ENSG0000C |
| 67839939 | 67840045 | -1 | snRNA | ENSG0000C |
| 67974391 | 67974625 | -1 | processed_pseudoge | ENSG0000C |
| 67985059 | 67986288 | 1 | unprocessed_pseudc | ENSG0000C |
| 68054179 | 68198603 | 1 | protein_coding | ENSG0000C |
| 68184032 | 68229718 | -1 | protein_coding | ENSG0000C |
| 68190705 | 68206110 | -1 | protein_coding | ENSG0000C |
| 68206992 | 68257211 | -1 | protein_coding | ENSG0000C |
| 68254549 | 68254939 | 1 | processed_pseudoge | ENSG0000C |

| | | | | |
|---|---|---|---|---|
| 68267792 | 68277994 | -1 | lncRNA | ENSG00000 |
| 68277745 | 68295862 | 1 | protein_coding | ENSG00000 |
| 68296532 | 68432163 | -1 | protein_coding | ENSG00000 |
| 68483202 | 68487149 | -1 | lncRNA | ENSG00000 |
| 68578969 | 68727806 | 1 | protein_coding | ENSG00000 |
| 68749501 | 68750691 | 1 | processed_pseudoge | ENSG00000 |
| 68778535 | 68820897 | -1 | protein_coding | ENSG00000 |
| 68801850 | 68801925 | -1 | miRNA | ENSG00000 |
| 68818221 | 68946811 | 1 | protein_coding | ENSG00000 |
| 68833830 | 68834749 | 1 | processed_pseudoge | ENSG00000 |
| 68880663 | 68928941 | -1 | lncRNA | ENSG00000 |
| 68930504 | 69028889 | 1 | lncRNA | ENSG00000 |
| 68930525 | 69062762 | 1 | protein_coding | ENSG00000 |
| 68985795 | 68998679 | 1 | lncRNA | ENSG00000 |
| 69022230 | 69024176 | 1 | transcribed_processe | ENSG00000 |
| 69037549 | 69043565 | -1 | lncRNA | ENSG00000 |
| 69072926 | 69095820 | 1 | lncRNA | ENSG00000 |
| 69080879 | 69099987 | 1 | lncRNA | ENSG00000 |
| 69160584 | 69272217 | 1 | protein_coding | ENSG00000 |
| 69278328 | 69298795 | -1 | lncRNA | ENSG00000 |
| 69278675 | 69295322 | 1 | lncRNA | ENSG00000 |
| 69298912 | 69407780 | 1 | protein_coding | ENSG00000 |
| 69391192 | 69392149 | 1 | lncRNA | ENSG00000 |
| 69396904 | 69415029 | -1 | lncRNA | ENSG00000 |
| 69406441 | 69406552 | 1 | misc_RNA | ENSG00000 |
| 69414246 | 69448427 | 1 | protein_coding | ENSG00000 |
| 69434466 | 69435120 | -1 | processed_pseudoge | ENSG00000 |
| 69452814 | 69456205 | 1 | protein_coding | ENSG00000 |
| 69457941 | 69458149 | -1 | snoRNA | ENSG00000 |
| 69458522 | 69461806 | -1 | lncRNA | ENSG00000 |
| 69462921 | 69843120 | 1 | lncRNA | ENSG00000 |
| 69564724 | 69565790 | -1 | lncRNA | ENSG00000 |
| 69803316 | 69804046 | -1 | processed_pseudoge | ENSG00000 |
| 69820756 | 69835111 | -1 | lncRNA | ENSG00000 |
| 70047790 | 70098176 | -1 | protein_coding | ENSG00000 |
| 70079372 | 70079468 | -1 | miRNA | ENSG00000 |
| 70193236 | 70193341 | 1 | snRNA | ENSG00000 |
| 70195638 | 70198509 | -1 | lncRNA | ENSG00000 |
| 70321576 | 70326742 | 1 | lncRNA | ENSG00000 |
| 70503907 | 70505928 | -1 | lncRNA | ENSG00000 |
| 70556521 | 70562722 | 1 | lncRNA | ENSG00000 |
| 70570958 | 70586606 | -1 | lncRNA | ENSG00000 |
| 70602124 | 70602516 | 1 | processed_pseudoge | ENSG00000 |
| 70615547 | 70616567 | 1 | lncRNA | ENSG00000 |
| 70635249 | 70637314 | 1 | lncRNA | ENSG00000 |
| 70654554 | 70763558 | -1 | protein_coding | ENSG00000 |
| 70748932 | 70754177 | 1 | lncRNA | ENSG00000 |
| 70758269 | 70758856 | -1 | lncRNA | ENSG00000 |
| 70768011 | 70778868 | -1 | lncRNA | ENSG00000 |
| 70791013 | 70791486 | 1 | processed_pseudoge | ENSG00000 |

| | | | | |
|---|---|---|---|---|
| 70796597 | 70802688 | -1 | transcribed_processe | ENSG00000 |
| 70829130 | 70854157 | -1 | protein_coding | ENSG00000 |
| 70848883 | 70849770 | -1 | lncRNA | ENSG00000 |
| 70853239 | 71053657 | 1 | protein_coding | ENSG00000 |
| 70858714 | 70860162 | 1 | processed_pseudoge | ENSG00000 |
| 70881342 | 70892433 | -1 | protein_coding | ENSG00000 |
| 71063101 | 71063989 | -1 | processed_pseudoge | ENSG00000 |
| 71096952 | 71783383 | 1 | protein_coding | ENSG00000 |
| 71110244 | 71115500 | -1 | protein_coding | ENSG00000 |
| 71147650 | 71189064 | -1 | lncRNA | ENSG00000 |
| 71164770 | 71165415 | -1 | processed_pseudoge | ENSG00000 |
| 71332120 | 71352460 | 1 | lncRNA | ENSG00000 |
| 71341158 | 71341712 | -1 | processed_pseudoge | ENSG00000 |
| 71547280 | 71549832 | -1 | lncRNA | ENSG00000 |
| 71792638 | 71818259 | 1 | protein_coding | ENSG00000 |
| 71818396 | 71823384 | 1 | lncRNA | ENSG00000 |
| 71822291 | 72118577 | -1 | protein_coding | ENSG00000 |
| 71858570 | 71858686 | 1 | rRNA_pseudogene | ENSG00000 |
| 71972206 | 72040265 | 1 | lncRNA | ENSG00000 |
| 72041194 | 72041653 | -1 | processed_pseudoge | ENSG00000 |
| 72045183 | 72045367 | 1 | snRNA | ENSG00000 |
| 72114258 | 72143692 | 1 | protein_coding | ENSG00000 |
| 72134641 | 72135436 | 1 | processed_pseudoge | ENSG00000 |
| 72140504 | 72155459 | -1 | lncRNA | ENSG00000 |
| 72159806 | 72197787 | -1 | protein_coding | ENSG00000 |
| 72199029 | 72231822 | -1 | protein_coding | ENSG00000 |
| 72241181 | 72272999 | -1 | protein_coding | ENSG00000 |
| 72266746 | 72319946 | -1 | protein_coding | ENSG00000 |
| 72278867 | 72351794 | 1 | lncRNA | ENSG00000 |
| 72284727 | 72320157 | -1 | protein_coding | ENSG00000 |
| 72284727 | 72375981 | -1 | protein_coding | ENSG00000 |
| 72340919 | 72376476 | -1 | protein_coding | ENSG00000 |
| 72376113 | 72378788 | 1 | lncRNA | ENSG00000 |
| 72379215 | 72379710 | -1 | processed_pseudoge | ENSG00000 |
| 72398302 | 72408367 | 1 | protein_coding | ENSG00000 |
| 72407778 | 72475168 | -1 | lncRNA | ENSG00000 |
| 72450657 | 72451474 | 1 | processed_pseudoge | ENSG00000 |
| 72465128 | 72466262 | -1 | lncRNA | ENSG00000 |
| 72474330 | 72602987 | 1 | protein_coding | ENSG00000 |
| 72587217 | 72587313 | 1 | miRNA | ENSG00000 |
| 72589691 | 72591845 | 1 | lncRNA | ENSG00000 |
| 72604399 | 72605008 | -1 | unprocessed_pseudo | ENSG00000 |
| 72605183 | 72612326 | 1 | unprocessed_pseudo | ENSG00000 |
| 72608481 | 72636962 | -1 | lncRNA | ENSG00000 |
| 72615810 | 72618250 | 1 | lncRNA | ENSG00000 |
| 72638821 | 72640125 | 1 | unprocessed_pseudo | ENSG00000 |
| 72654738 | 72666394 | 1 | protein_coding | ENSG00000 |
| 72664577 | 72664799 | 1 | misc_RNA | ENSG00000 |
| 72675788 | 72686182 | -1 | protein_coding | ENSG00000 |
| 72682266 | 72682904 | -1 | transcribed_processe | ENSG00000 |

| | | | | |
|---|---|---|---|---|
| 72686179 | 72738475 | 1 | protein_coding | ENSG0000C |
| 72751369 | 72785846 | -1 | protein_coding | ENSG0000C |
| 72782835 | 72798199 | 1 | lncRNA | ENSG0000C |
| 72858354 | 72887128 | 1 | lncRNA | ENSG0000C |
| 72899399 | 72900260 | -1 | processed_pseudoge | ENSG0000C |
| 73051710 | 73305205 | 1 | protein_coding | ENSG0000C |
| 73143236 | 73143518 | -1 | processed_pseudoge | ENSG0000C |
| 73161779 | 73162928 | -1 | processed_pseudoge | ENSG0000C |
| 73255334 | 73256109 | -1 | lncRNA | ENSG0000C |
| 73319859 | 73368958 | -1 | protein_coding | ENSG0000C |
| 73335260 | 73342387 | 1 | lncRNA | ENSG0000C |
| 73443164 | 73560013 | 1 | protein_coding | ENSG0000C |
| 73483196 | 73483943 | -1 | processed_pseudoge | ENSG0000C |
| 73560014 | 73634134 | -1 | protein_coding | ENSG0000C |
| 73567012 | 73569294 | -1 | lncRNA | ENSG0000C |
| 73683966 | 73714514 | 1 | protein_coding | ENSG0000C |
| 73730048 | 73731711 | -1 | lncRNA | ENSG0000C |
| 73735431 | 73752747 | -1 | protein_coding | ENSG0000C |
| 73752317 | 73770613 | 1 | lncRNA | ENSG0000C |
| 73870949 | 73873540 | -1 | lncRNA | ENSG0000C |
| 73873564 | 73889214 | 1 | protein_coding | ENSG0000C |
| 73908071 | 73928248 | -1 | lncRNA | ENSG0000C |
| 73925989 | 73952137 | 1 | protein_coding | ENSG0000C |
| 73978926 | 73994622 | -1 | protein_coding | ENSG0000C |
| 73994673 | 74047827 | 1 | protein_coding | ENSG0000C |
| 74040190 | 74043332 | -1 | lncRNA | ENSG0000C |
| 74062780 | 74063785 | 1 | unprocessed_pseudc | ENSG0000C |
| 74069857 | 74082550 | -1 | protein_coding | ENSG0000C |
| 74072454 | 74072676 | -1 | misc_RNA | ENSG0000C |
| 74097106 | 74098414 | -1 | unprocessed_pseudc | ENSG0000C |
| 74100311 | 74138540 | 1 | protein_coding | ENSG0000C |
| 74125915 | 74129341 | -1 | lncRNA | ENSG0000C |
| 74152800 | 74179226 | -1 | lncRNA | ENSG0000C |
| 74162340 | 74162620 | -1 | misc_RNA | ENSG0000C |
| 74173710 | 74176872 | 1 | protein_coding | ENSG0000C |
| 74179466 | 74212267 | -1 | protein_coding | ENSG0000C |
| 74202705 | 74336472 | 1 | protein_coding | ENSG0000C |
| 74299503 | 74304527 | 1 | TEC | ENSG0000C |
| 74303005 | 74304343 | 1 | lncRNA | ENSG0000C |
| 74311516 | 74319688 | -1 | lncRNA | ENSG0000C |
| 74337759 | 74367646 | -1 | protein_coding | ENSG0000C |
| 74350768 | 74364620 | 1 | transcribed_processe | ENSG0000C |
| 74365435 | 74371211 | 1 | lncRNA | ENSG0000C |
| 74374678 | 74375511 | 1 | lncRNA | ENSG0000C |
| 74379083 | 74390535 | 1 | lncRNA | ENSG0000C |
| 74409289 | 74433958 | -1 | protein_coding | ENSG0000C |
| 74411357 | 74411432 | -1 | miRNA | ENSG0000C |
| 74429445 | 74431449 | 1 | lncRNA | ENSG0000C |
| 74445977 | 74461182 | -1 | protein_coding | ENSG0000C |
| 74461265 | 74513636 | 1 | lncRNA | ENSG0000C |

| | | | | |
|---|---|---|---|---|
| 74478070 | 74490286 | -1 | transcribed_processe | ENSG00000 |
| 74489602 | 74516959 | 1 | lncRNA | ENSG00000 |
| 74490959 | 74491028 | -1 | snoRNA | ENSG00000 |
| 74541177 | 74598131 | 1 | protein_coding | ENSG00000 |
| 74598500 | 74645414 | 1 | protein_coding | ENSG00000 |
| 74598919 | 74599397 | -1 | lncRNA | ENSG00000 |
| 74613194 | 74615596 | -1 | lncRNA | ENSG00000 |
| 74630558 | 74696292 | -1 | protein_coding | ENSG00000 |
| 74719542 | 74725536 | -1 | protein_coding | ENSG00000 |
| 74748845 | 74756607 | 1 | protein_coding | ENSG00000 |
| 74782057 | 74803198 | 1 | protein_coding | ENSG00000 |
| 74788672 | 74788757 | -1 | miRNA | ENSG00000 |
| 74812716 | 74825757 | 1 | protein_coding | ENSG00000 |
| 74816223 | 74831795 | 1 | lncRNA | ENSG00000 |
| 74826627 | 74831802 | 1 | protein_coding | ENSG00000 |
| 74836118 | 74843346 | -1 | protein_coding | ENSG00000 |
| 74840642 | 74840707 | -1 | miRNA | ENSG00000 |
| 74843730 | 74873365 | -1 | protein_coding | ENSG00000 |
| 74890005 | 74902219 | 1 | protein_coding | ENSG00000 |
| 74899992 | 74906883 | -1 | protein_coding | ENSG00000 |
| 74919791 | 74938083 | -1 | protein_coding | ENSG00000 |
| 74954416 | 74957464 | -1 | protein_coding | ENSG00000 |
| 74957219 | 75021495 | 1 | protein_coding | ENSG00000 |
| 74976240 | 74976573 | -1 | processed_pseudoge | ENSG00000 |
| 74983274 | 74983376 | -1 | misc_RNA | ENSG00000 |
| 75022980 | 75023256 | 1 | processed_pseudoge | ENSG00000 |
| 75023586 | 75117462 | 1 | protein_coding | ENSG00000 |
| 75114261 | 75114868 | 1 | processed_pseudoge | ENSG00000 |
| 75120265 | 75120562 | -1 | misc_RNA | ENSG00000 |
| 75121536 | 75121666 | -1 | scaRNA | ENSG00000 |
| 75186651 | 75186968 | -1 | processed_pseudoge | ENSG00000 |
| 75195643 | 75212169 | 1 | protein_coding | ENSG00000 |
| 75211301 | 75212167 | -1 | lncRNA | ENSG00000 |
| 75225784 | 75227185 | 1 | lncRNA | ENSG00000 |
| 75226401 | 75234834 | -1 | transcribed_unitary_ | ENSG00000 |
| 75249232 | 75249407 | -1 | unprocessed_pseudo | ENSG00000 |
| 75251743 | 75252934 | -1 | processed_pseudoge | ENSG00000 |
| 75258599 | 75270199 | 1 | protein_coding | ENSG00000 |
| 75268430 | 75268652 | 1 | misc_RNA | ENSG00000 |
| 75282835 | 75295807 | 1 | protein_coding | ENSG00000 |
| 75292706 | 75292928 | 1 | misc_RNA | ENSG00000 |
| 75299953 | 75302965 | -1 | transcribed_unproce | ENSG00000 |
| 75321397 | 75322693 | -1 | transcribed_processe | ENSG00000 |
| 75336020 | 75343224 | 1 | protein_coding | ENSG00000 |
| 75346744 | 75347161 | -1 | lncRNA | ENSG00000 |
| 75346955 | 75357115 | 1 | protein_coding | ENSG00000 |
| 75353611 | 75353685 | -1 | miRNA | ENSG00000 |
| 75355207 | 75368612 | -1 | protein_coding | ENSG00000 |
| 75368155 | 75369584 | 1 | lncRNA | ENSG00000 |
| 75369379 | 75455842 | -1 | protein_coding | ENSG00000 |

| | | | | |
|---|---|---|---|---|
| 75388267 | 75388828 | 1 | processed_pseudoge | ENSG00000 |
| 75415249 | 75416020 | 1 | processed_pseudoge | ENSG00000 |
| 75452964 | 75453947 | -1 | lncRNA | ENSG00000 |
| 75463251 | 75579315 | -1 | protein_coding | ENSG00000 |
| 75480661 | 75480836 | 1 | unprocessed_pseudc | ENSG00000 |
| 75512770 | 75513040 | -1 | processed_pseudoge | ENSG00000 |
| 75527150 | 75601205 | -1 | lncRNA | ENSG00000 |
| 75598083 | 75626469 | -1 | protein_coding | ENSG00000 |
| 75624793 | 75625690 | 1 | lncRNA | ENSG00000 |
| 75636139 | 75639239 | 1 | lncRNA | ENSG00000 |
| 75639085 | 75648706 | -1 | protein_coding | ENSG00000 |
| 75639760 | 75640976 | 1 | lncRNA | ENSG00000 |
| 75645020 | 75645442 | 1 | lncRNA | ENSG00000 |
| 75647906 | 75662301 | 1 | protein_coding | ENSG00000 |
| 75674322 | 75712848 | -1 | protein_coding | ENSG00000 |
| 75676227 | 75677162 | 1 | lncRNA | ENSG00000 |
| 75678548 | 75680752 | 1 | lncRNA | ENSG00000 |
| 75724041 | 75727688 | 1 | protein_coding | ENSG00000 |
| 75727670 | 75738623 | -1 | lncRNA | ENSG00000 |
| 75737820 | 75763322 | 1 | lncRNA | ENSG00000 |
| 75746769 | 75747263 | 1 | processed_pseudoge | ENSG00000 |
| 75759501 | 75762405 | -1 | lncRNA | ENSG00000 |
| 75762215 | 75762315 | -1 | miRNA | ENSG00000 |
| 75775553 | 75786898 | 1 | transcribed_unproce | ENSG00000 |
| 75785134 | 75785354 | 1 | misc_RNA | ENSG00000 |
| 75791470 | 75792683 | -1 | unprocessed_pseudc | ENSG00000 |
| 75843307 | 75901078 | 1 | protein_coding | ENSG00000 |
| 75895040 | 75895302 | 1 | misc_RNA | ENSG00000 |
| 75903876 | 75942511 | 1 | protein_coding | ENSG00000 |
| 75935969 | 76059795 | -1 | protein_coding | ENSG00000 |
| 75950464 | 75951144 | -1 | processed_pseudoge | ENSG00000 |
| 76059958 | 76229121 | 1 | protein_coding | ENSG00000 |
| 76174891 | 76181486 | -1 | lncRNA | ENSG00000 |
| 76215353 | 76311472 | -1 | protein_coding | ENSG00000 |
| 76230048 | 76230390 | -1 | misc_RNA | ENSG00000 |
| 76258986 | 76261690 | -1 | processed_pseudoge | ENSG00000 |
| 76263197 | 76275387 | 1 | lncRNA | ENSG00000 |
| 76336773 | 76342475 | 1 | protein_coding | ENSG00000 |
| 76339609 | 76342063 | -1 | lncRNA | ENSG00000 |
| 76343642 | 76344365 | 1 | lncRNA | ENSG00000 |
| 76347904 | 76905444 | -1 | protein_coding | ENSG00000 |
| 76472099 | 76472365 | 1 | processed_pseudoge | ENSG00000 |
| 76586647 | 76586691 | 1 | miRNA | ENSG00000 |
| 76736641 | 76736980 | 1 | misc_RNA | ENSG00000 |
| 76931738 | 76954393 | 1 | protein_coding | ENSG00000 |
| 76976915 | 76977210 | -1 | misc_RNA | ENSG00000 |
| 76979245 | 76980451 | -1 | processed_pseudoge | ENSG00000 |
| 76993359 | 77037475 | 1 | protein_coding | ENSG00000 |
| 77041404 | 77083984 | -1 | protein_coding | ENSG00000 |
| 77043680 | 77045160 | 1 | lncRNA | ENSG00000 |

| | | | | |
|---|---|---|---|---|
| 77063397 | 77064910 | -1 | TEC | ENSG00000 |
| 77067654 | 77068325 | -1 | lncRNA | ENSG00000 |
| 77100656 | 77420144 | -1 | protein_coding | ENSG00000 |
| 77204578 | 77205100 | 1 | processed_pseudoge | ENSG00000 |
| 77278465 | 77278774 | 1 | processed_pseudoge | ENSG00000 |
| 77420412 | 77485607 | 1 | protein_coding | ENSG00000 |
| 77484275 | 77485606 | -1 | TEC | ENSG00000 |
| 77525540 | 77534110 | 1 | lncRNA | ENSG00000 |
| 77568970 | 77608888 | 1 | lncRNA | ENSG00000 |
| 77613027 | 77820900 | -1 | protein_coding | ENSG00000 |
| 77641764 | 77652292 | 1 | lncRNA | ENSG00000 |
| 77660052 | 77668074 | 1 | lncRNA | ENSG00000 |
| 77784610 | 77785070 | -1 | TEC | ENSG00000 |
| 77787193 | 77788674 | 1 | lncRNA | ENSG00000 |
| 77894684 | 77904674 | 1 | transcribed_unproce | ENSG00000 |
| 77914217 | 77926846 | -1 | transcribed_unproce | ENSG00000 |
| 77916522 | 77922019 | 1 | lncRNA | ENSG00000 |
| 77916718 | 77916982 | -1 | misc_RNA | ENSG00000 |
| 77941442 | 77944582 | -1 | transcribed_unproce | ENSG00000 |
| 77954075 | 77963654 | -1 | lncRNA | ENSG00000 |
| 77976042 | 77993057 | 1 | transcribed_unproce | ENSG00000 |
| 77984036 | 78077724 | -1 | protein_coding | ENSG00000 |
| 77993405 | 77995289 | 1 | lncRNA | ENSG00000 |
| 78077808 | 78104909 | 1 | protein_coding | ENSG00000 |
| 78091172 | 78091297 | -1 | snoRNA | ENSG00000 |
| 78104606 | 78131535 | -1 | protein_coding | ENSG00000 |
| 78131498 | 78171945 | 1 | protein_coding | ENSG00000 |
| 78141243 | 78143173 | 1 | lncRNA | ENSG00000 |
| 78167468 | 78245688 | -1 | protein_coding | ENSG00000 |
| 78250502 | 78264156 | -1 | lncRNA | ENSG00000 |
| 78264086 | 78282196 | 1 | protein_coding | ENSG00000 |
| 78277835 | 78299703 | -1 | protein_coding | ENSG00000 |
| 78280950 | 78282190 | -1 | lncRNA | ENSG00000 |
| 78290527 | 78291221 | -1 | transcribed_processe | ENSG00000 |
| 78293286 | 78296049 | 1 | lncRNA | ENSG00000 |
| 78299701 | 78299924 | 1 | lncRNA | ENSG00000 |
| 78339208 | 78341757 | -1 | TEC | ENSG00000 |
| 78340353 | 78348225 | 1 | protein_coding | ENSG00000 |
| 78382573 | 78383155 | 1 | processed_pseudoge | ENSG00000 |
| 78437431 | 78501453 | 1 | protein_coding | ENSG00000 |
| 78480057 | 78481820 | -1 | lncRNA | ENSG00000 |
| 78507564 | 78537372 | 1 | protein_coding | ENSG00000 |
| 78537681 | 78538946 | 1 | TEC | ENSG00000 |
| 78540405 | 78552417 | 1 | protein_coding | ENSG00000 |
| 78565520 | 78595269 | 1 | protein_coding | ENSG00000 |
| 78589123 | 78591276 | -1 | lncRNA | ENSG00000 |
| 78593052 | 78621295 | -1 | protein_coding | ENSG00000 |
| 78599892 | 78600013 | -1 | TEC | ENSG00000 |
| 78624111 | 78727754 | -1 | protein_coding | ENSG00000 |
| 78625895 | 78628193 | 1 | lncRNA | ENSG00000 |

| | | | | |
|---|---|---|---|---|
| 78660644 | 78736684 | 1 | transcribed_unproce | ENSG00000( |
| 78661047 | 78661576 | -1 | processed_pseudoge | ENSG00000( |
| 78749833 | 78750107 | -1 | unprocessed_pseudc | ENSG00000( |
| 78752107 | 78753373 | 1 | transcribed_unproce | ENSG00000( |
| 78759206 | 78811464 | -1 | protein_coding | ENSG00000( |
| 78810487 | 78898139 | 1 | protein_coding | ENSG00000( |
| 78863247 | 78863729 | -1 | processed_pseudoge | ENSG00000( |
| 78898840 | 78898936 | -1 | snRNA | ENSG00000( |
| 78906127 | 78906809 | 1 | TEC | ENSG00000( |
| 78921058 | 78949574 | -1 | protein_coding | ENSG00000( |
| 78959947 | 79090773 | -1 | protein_coding | ENSG00000( |
| 78978889 | 78985926 | 1 | lncRNA | ENSG00000( |
| 79141939 | 79283949 | -1 | lncRNA | ENSG00000( |
| 79209788 | 79209871 | 1 | miRNA | ENSG00000( |
| 79236332 | 79237206 | 1 | processed_pseudoge | ENSG00000( |
| 79293285 | 79298235 | 1 | protein_coding | ENSG00000( |
| 79311112 | 79427432 | 1 | protein_coding | ENSG00000( |
| 79331591 | 79337277 | -1 | lncRNA | ENSG00000( |
| 79432516 | 79472290 | 1 | protein_coding | ENSG00000( |
| 79559256 | 79560304 | -1 | processed_pseudoge | ENSG00000( |
| 79668342 | 79668816 | -1 | processed_pseudoge | ENSG00000( |
| 79743311 | 79743468 | 1 | processed_pseudoge | ENSG00000( |
| 79751689 | 79752049 | -1 | processed_pseudoge | ENSG00000( |
| 79758549 | 79758762 | -1 | processed_pseudoge | ENSG00000( |
| 79817349 | 79817450 | -1 | snRNA | ENSG00000( |
| 79832466 | 79833554 | -1 | lncRNA | ENSG00000( |
| 79833585 | 79897379 | -1 | protein_coding | ENSG00000( |
| 79843547 | 79844304 | -1 | lncRNA | ENSG00000( |
| 79845150 | 79923754 | -1 | protein_coding | ENSG00000( |
| 79875920 | 79878179 | -1 | TEC | ENSG00000( |
| 79894502 | 79896695 | -1 | lncRNA | ENSG00000( |
| 79898840 | 79923702 | -1 | protein_coding | ENSG00000( |
| 79920195 | 79922455 | -1 | lncRNA | ENSG00000( |
| 79922771 | 79926993 | 1 | lncRNA | ENSG00000( |
| 79944498 | 79945578 | 1 | processed_pseudoge | ENSG00000( |
| 79960892 | 79971196 | -1 | protein_coding | ENSG00000( |
| 79984151 | 79985724 | 1 | processed_pseudoge | ENSG00000( |
| 79989905 | 79990606 | 1 | TEC | ENSG00000( |
| 80014157 | 80015707 | 1 | TEC | ENSG00000( |
| 80057425 | 80057684 | -1 | misc_RNA | ENSG00000( |
| 80059568 | 80138393 | 1 | protein_coding | ENSG00000( |
| 80152490 | 80186946 | 1 | protein_coding | ENSG00000( |
| 80165923 | 80166980 | 1 | lncRNA | ENSG00000( |
| 80195481 | 80252213 | -1 | protein_coding | ENSG00000( |
| 80263068 | 80341805 | -1 | lncRNA | ENSG00000( |
| 80344853 | 80404214 | -1 | lncRNA | ENSG00000( |
| 80404350 | 80597933 | 1 | protein_coding | ENSG00000( |
| 80433795 | 80445152 | -1 | lncRNA | ENSG00000( |
| 80440294 | 80442751 | -1 | lncRNA | ENSG00000( |
| 80554609 | 80562944 | -1 | lncRNA | ENSG00000( |

| | | | | |
|---|---|---|---|---|
| 80580029 | 80580566 | -1 | lncRNA | ENSG0000C |
| 80581103 | 80581239 | 1 | miRNA | ENSG0000C |
| 80663772 | 80663878 | 1 | snRNA | ENSG0000C |
| 80679684 | 80755621 | 1 | protein_coding | ENSG0000C |
| 80693216 | 80693707 | -1 | lncRNA | ENSG0000C |
| 80769600 | 80773135 | -1 | lncRNA | ENSG0000C |
| 80779343 | 80951776 | 1 | protein_coding | ENSG0000C |
| 80841978 | 80842073 | -1 | miRNA | ENSG0000C |
| 80896190 | 80948867 | -1 | lncRNA | ENSG0000C |
| 80896191 | 80909777 | -1 | lncRNA | ENSG0000C |
| 80946289 | 80989828 | -1 | protein_coding | ENSG0000C |
| 80990804 | 80993258 | -1 | lncRNA | ENSG0000C |
| 80997417 | 80997473 | -1 | miRNA | ENSG0000C |
| 80999593 | 80999981 | -1 | lncRNA | ENSG0000C |
| 81000923 | 81005788 | 1 | protein_coding | ENSG0000C |
| 81007033 | 81149179 | 1 | protein_coding | ENSG0000C |
| 81060953 | 81063488 | 1 | TEC | ENSG0000C |
| 81117635 | 81118340 | -1 | processed_pseudoge | ENSG0000C |
| 81159575 | 81314058 | 1 | protein_coding | ENSG0000C |
| 81303215 | 81309391 | -1 | lncRNA | ENSG0000C |
| 81309053 | 81324183 | -1 | protein_coding | ENSG0000C |
| 81315806 | 81321258 | 1 | lncRNA | ENSG0000C |
| 81324338 | 81518200 | 1 | lncRNA | ENSG0000C |
| 81331088 | 81374213 | -1 | protein_coding | ENSG0000C |
| 81335577 | 81336119 | 1 | lncRNA | ENSG0000C |
| 81385390 | 81385774 | 1 | processed_pseudoge | ENSG0000C |
| 81403026 | 81403570 | 1 | lncRNA | ENSG0000C |
| 81427448 | 81755217 | 1 | lncRNA | ENSG0000C |
| 81554003 | 81696780 | 1 | lncRNA | ENSG0000C |
| 81633426 | 82013579 | -1 | lncRNA | ENSG0000C |
| 82010212 | 82010607 | -1 | processed_pseudoge | ENSG0000C |
| 82041778 | 82046119 | -1 | protein_coding | ENSG0000C |
| 82088569 | 82097694 | 1 | lncRNA | ENSG0000C |
| 82113384 | 82119570 | 1 | lncRNA | ENSG0000C |
| 82130233 | 82262734 | -1 | protein_coding | ENSG0000C |
| 82174224 | 82174385 | -1 | snRNA | ENSG0000C |
| 82180743 | 82182700 | 1 | lncRNA | ENSG0000C |
| 82262810 | 82284930 | 1 | protein_coding | ENSG0000C |
| 82298553 | 82334609 | 1 | transcribed_unproce | ENSG0000C |
| 82339993 | 82349475 | -1 | protein_coding | ENSG0000C |
| 82355142 | 82420075 | 1 | transcribed_unproce | ENSG0000C |
| 82372196 | 82372912 | -1 | transcribed_processe | ENSG0000C |
| 82418651 | 82434235 | 1 | lncRNA | ENSG0000C |
| 82430018 | 82439153 | 1 | protein_coding | ENSG0000C |
| 82445719 | 82454837 | 1 | transcribed_unproce | ENSG0000C |
| 82459472 | 82477258 | 1 | transcribed_unproce | ENSG0000C |
| 82472203 | 82472426 | 1 | unprocessed_pseudc | ENSG0000C |
| 82472993 | 82513950 | -1 | transcribed_unproce | ENSG0000C |
| 82519590 | 82526412 | 1 | transcribed_unproce | ENSG0000C |
| 82531653 | 82534139 | -1 | unprocessed_pseudc | ENSG0000C |

| | | | | |
|---|---|---|---|---|
| 82533175 | 82540008 | -1 | lncRNA | ENSG0000C |
| 82536750 | 82540459 | -1 | protein_coding | ENSG0000C |
| 82536788 | 82573194 | -1 | protein_coding | ENSG0000C |
| 82540870 | 82562374 | 1 | lncRNA | ENSG0000C |
| 82543201 | 82648861 | -1 | protein_coding | ENSG0000C |
| 82647770 | 82692820 | 1 | lncRNA | ENSG0000C |
| 82659281 | 82709946 | -1 | protein_coding | ENSG0000C |
| 82691723 | 82692358 | -1 | processed_pseudoge | ENSG0000C |
| 82710471 | 82714029 | -1 | transcribed_unitary_ | ENSG0000C |
| 82725873 | 82738904 | -1 | transcribed_processe | ENSG0000C |
| 82738386 | 82739122 | 1 | TEC | ENSG0000C |
| 82744223 | 82750289 | 1 | lncRNA | ENSG0000C |
| 82749252 | 82750455 | -1 | lncRNA | ENSG0000C |
| 82750564 | 82757206 | 1 | lncRNA | ENSG0000C |
| 82752884 | 82757208 | 1 | lncRNA | ENSG0000C |
| 82755362 | 82806070 | -1 | protein_coding | ENSG0000C |
| 82755945 | 82756071 | 1 | scaRNA | ENSG0000C |
| 82809628 | 82836108 | 1 | protein_coding | ENSG0000C |
| 82836946 | 82986153 | -1 | protein_coding | ENSG0000C |
| 82925884 | 83103443 | 1 | lncRNA | ENSG0000C |
| 82986210 | 82991057 | 1 | protein_coding | ENSG0000C |
| 82988441 | 83011641 | -1 | protein_coding | ENSG0000C |
| 83012661 | 83061845 | 1 | lncRNA | ENSG0000C |
| 83016423 | 83067252 | -1 | protein_coding | ENSG0000C |
| 83020115 | 83020802 | 1 | lncRNA | ENSG0000C |
| 83022236 | 83024336 | -1 | lncRNA | ENSG0000C |
| 83022571 | 83090782 | 1 | lncRNA | ENSG0000C |
| 83044368 | 83044771 | 1 | processed_pseudoge | ENSG0000C |
| 83067335 | 83067415 | 1 | miRNA | ENSG0000C |
| 83095333 | 83108754 | -1 | lncRNA | ENSG0000C |
| 83107572 | 83144854 | 1 | protein_coding | ENSG0000C |
| 83112738 | 83207823 | -1 | protein_coding | ENSG0000C |
| 83113617 | 83114566 | -1 | lncRNA | ENSG0000C |
| 83179182 | 83439445 | 1 | lncRNA | ENSG0000C |
| 83255903 | 83284714 | -1 | protein_coding | ENSG0000C |
| 83447228 | 83618743 | 1 | protein_coding | ENSG0000C |
| 83654088 | 84039842 | 1 | protein_coding | ENSG0000C |
| 83717427 | 83717522 | 1 | snRNA | ENSG0000C |
| 83739784 | 83740199 | 1 | transcribed_processe | ENSG0000C |
| 83824863 | 83824953 | -1 | snRNA | ENSG0000C |
| 83962176 | 83962583 | 1 | lncRNA | ENSG0000C |
| 84053113 | 84080097 | -1 | unprocessed_pseudc | ENSG0000C |
| 84080168 | 84126604 | 1 | transcribed_unproce | ENSG0000C |
| 84166547 | 84169896 | -1 | TEC | ENSG0000C |
| 84171178 | 84173194 | -1 | lncRNA | ENSG0000C |
| 84172490 | 84182234 | 1 | protein_coding | ENSG0000C |
| 84186752 | 84197384 | 1 | unprocessed_pseudc | ENSG0000C |
| 84199311 | 84230136 | -1 | transcribed_unproce | ENSG0000C |
| 84205454 | 84205700 | -1 | misc_RNA | ENSG0000C |
| 84235773 | 84245368 | 1 | protein_coding | ENSG0000C |

| | | | | |
|---|---|---|---|---|
| 84255913 | 84256014 | 1 | unprocessed_pseudc | ENSG00000( |
| 84389729 | 84395903 | 1 | unprocessed_pseudc | ENSG00000( |
| 84394072 | 84394336 | 1 | misc_RNA | ENSG00000( |
| 84394512 | 84395514 | 1 | unprocessed_pseudc | ENSG00000( |
| 84398316 | 84411701 | -1 | transcribed_unproce | ENSG00000( |
| 84401807 | 84408005 | -1 | transcribed_unproce | ENSG00000( |
| 84412773 | 84421837 | -1 | transcribed_unproce | ENSG00000( |
| 84422618 | 84425882 | 1 | lncRNA | ENSG00000( |
| 84467901 | 84468147 | 1 | misc_RNA | ENSG00000( |
| 84491480 | 84500599 | -1 | unprocessed_pseudc | ENSG00000( |
| 84500378 | 84502381 | 1 | lncRNA | ENSG00000( |
| 84505463 | 84505495 | 1 | unprocessed_pseudc | ENSG00000( |
| 84507885 | 84516814 | -1 | transcribed_unproce | ENSG00000( |
| 84513241 | 84526949 | -1 | lncRNA | ENSG00000( |
| 84526781 | 84571216 | -1 | transcribed_unproce | ENSG00000( |
| 84570649 | 84580175 | 1 | transcribed_unproce | ENSG00000( |
| 84597809 | 84633988 | -1 | lncRNA | ENSG00000( |
| 84600986 | 84627796 | 1 | protein_coding | ENSG00000( |
| 84611689 | 84614969 | -1 | lncRNA | ENSG00000( |
| 84622015 | 84623237 | -1 | lncRNA | ENSG00000( |
| 84631451 | 84647478 | 1 | transcribed_unproce | ENSG00000( |
| 84633874 | 84634635 | 1 | transcribed_processe | ENSG00000( |
| 84639281 | 84654343 | -1 | protein_coding | ENSG00000( |
| 84642487 | 84642763 | -1 | misc_RNA | ENSG00000( |
| 84655129 | 84658563 | -1 | protein_coding | ENSG00000( |
| 84669538 | 84716460 | -1 | protein_coding | ENSG00000( |
| 84685884 | 84686946 | 1 | lncRNA | ENSG00000( |
| 84717474 | 84746837 | -1 | lncRNA | ENSG00000( |
| 84748592 | 84806445 | 1 | protein_coding | ENSG00000( |
| 84753122 | 84754502 | 1 | lncRNA | ENSG00000( |
| 84816680 | 84873482 | 1 | protein_coding | ENSG00000( |
| 84852498 | 84852615 | -1 | snoRNA | ENSG00000( |
| 84884654 | 84945798 | 1 | protein_coding | ENSG00000( |
| 84913144 | 84915705 | -1 | lncRNA | ENSG00000( |
| 84938416 | 84938522 | -1 | snRNA | ENSG00000( |
| 84939617 | 84939723 | -1 | snRNA | ENSG00000( |
| 84940504 | 84941133 | -1 | processed_pseudoge | ENSG00000( |
| 84980440 | 85139145 | 1 | protein_coding | ENSG00000( |
| 85178628 | 85178806 | -1 | processed_pseudoge | ENSG00000( |
| 85180200 | 85234795 | -1 | transcribed_unproce | ENSG00000( |
| 85181210 | 85182083 | -1 | processed_pseudoge | ENSG00000( |
| 85191438 | 85213905 | 1 | transcribed_unproce | ENSG00000( |
| 85210053 | 85210299 | -1 | misc_RNA | ENSG00000( |
| 85240472 | 85247170 | 1 | unprocessed_pseudc | ENSG00000( |
| 85255369 | 85330334 | -1 | transcribed_unproce | ENSG00000( |
| 85272527 | 85272974 | 1 | unprocessed_pseudc | ENSG00000( |
| 85380571 | 85749358 | 1 | protein_coding | ENSG00000( |
| 85380596 | 85380662 | 1 | miRNA | ENSG00000( |
| 85389823 | 85389884 | -1 | snRNA | ENSG00000( |
| 85415228 | 85415633 | 1 | transcribed_processe | ENSG00000( |

| | | | | |
|---|---|---|---|---|
| 85579046 | 85580178 | -1 | lncRNA | ENSG00000 |
| 85619623 | 85670948 | -1 | lncRNA | ENSG00000 |
| 85621264 | 85627689 | -1 | lncRNA | ENSG00000 |
| 85651522 | 85651628 | -1 | snRNA | ENSG00000 |
| 85701109 | 85702771 | 1 | lncRNA | ENSG00000 |
| 85726115 | 85727227 | -1 | lncRNA | ENSG00000 |
| 85744109 | 85750281 | -1 | lncRNA | ENSG00000 |
| 85750336 | 85752901 | -1 | lncRNA | ENSG00000 |
| 85754941 | 85756237 | -1 | lncRNA | ENSG00000 |
| 85759326 | 85794925 | -1 | protein_coding | ENSG00000 |
| 85770496 | 85770578 | -1 | miRNA | ENSG00000 |
| 85770990 | 85771163 | 1 | processed_pseudoge | ENSG00000 |
| 85821855 | 85821957 | 1 | snRNA | ENSG00000 |
| 85825635 | 85825730 | 1 | miRNA | ENSG00000 |
| 85958717 | 85958971 | -1 | processed_pseudoge | ENSG00000 |
| 86078809 | 86116747 | -1 | lncRNA | ENSG00000 |
| 86079973 | 87029052 | 1 | protein_coding | ENSG00000 |
| 86138269 | 86138379 | -1 | rRNA_pseudogene | ENSG00000 |
| 86139838 | 86140240 | -1 | processed_pseudoge | ENSG00000 |
| 86173198 | 86173270 | -1 | snoRNA | ENSG00000 |
| 86295649 | 86317173 | -1 | lncRNA | ENSG00000 |
| 86630266 | 86631136 | -1 | lncRNA | ENSG00000 |
| 86932880 | 86946331 | 1 | lncRNA | ENSG00000 |
| 86938797 | 86988426 | -1 | lncRNA | ENSG00000 |
| 87038717 | 87047303 | 1 | lncRNA | ENSG00000 |
| 87325829 | 87703852 | -1 | lncRNA | ENSG00000 |
| 87474183 | 87474289 | -1 | snRNA | ENSG00000 |
| 87576929 | 87579866 | 1 | lncRNA | ENSG00000 |
| 87638688 | 87668488 | 1 | lncRNA | ENSG00000 |
| 87859751 | 88256768 | -1 | protein_coding | ENSG00000 |
| 88143935 | 88144165 | -1 | processed_pseudoge | ENSG00000 |
| 88252730 | 88271066 | 1 | lncRNA | ENSG00000 |
| 88459477 | 88467390 | -1 | protein_coding | ENSG00000 |
| 88459501 | 88546585 | -1 | protein_coding | ENSG00000 |
| 88467453 | 88480776 | 1 | protein_coding | ENSG00000 |
| 88477871 | 88478387 | -1 | TEC | ENSG00000 |
| 88494440 | 88546675 | -1 | protein_coding | ENSG00000 |
| 88568287 | 88568562 | 1 | processed_pseudoge | ENSG00000 |
| 88585566 | 88605110 | -1 | lncRNA | ENSG00000 |
| 88608107 | 88608197 | 1 | miRNA | ENSG00000 |
| 88611825 | 88611934 | 1 | miRNA | ENSG00000 |
| 88611847 | 88611924 | -1 | miRNA | ENSG00000 |
| 88621337 | 88632281 | 1 | protein_coding | ENSG00000 |
| 88636153 | 88656483 | 1 | protein_coding | ENSG00000 |
| 88797413 | 88798734 | -1 | lncRNA | ENSG00000 |
| 88803440 | 88875354 | 1 | protein_coding | ENSG00000 |
| 88877288 | 88895626 | -1 | protein_coding | ENSG00000 |
| 88898683 | 88913381 | -1 | protein_coding | ENSG00000 |
| 88952683 | 88953460 | 1 | processed_pseudoge | ENSG00000 |
| 88960070 | 88961046 | 1 | processed_pseudoge | ENSG00000 |

| | | | | |
|---|---|---|---|---|
| 88989519 | 88990245 | -1 | processed_pseudoge | ENSG00000( |
| 89037977 | 89038286 | -1 | processed_pseudoge | ENSG00000( |
| 89041223 | 89082819 | 1 | lncRNA | ENSG00000( |
| 89087078 | 89088377 | -1 | lncRNA | ENSG00000( |
| 89087459 | 89202355 | 1 | protein_coding | ENSG00000( |
| 89120779 | 89120840 | 1 | snRNA | ENSG00000( |
| 89135547 | 89136495 | -1 | processed_pseudoge | ENSG00000( |
| 89201091 | 89201768 | -1 | processed_pseudoge | ENSG00000( |
| 89209869 | 89221614 | -1 | protein_coding | ENSG00000( |
| 89243949 | 89317261 | 1 | protein_coding | ENSG00000( |
| 89305198 | 89334861 | -1 | protein_coding | ENSG00000( |
| 89326739 | 89326810 | -1 | miRNA | ENSG00000( |
| 89335053 | 89336161 | 1 | lncRNA | ENSG00000( |
| 89358841 | 89360377 | 1 | TEC | ENSG00000( |
| 89361157 | 89362421 | -1 | lncRNA | ENSG00000( |
| 89361579 | 89398487 | 1 | lncRNA | ENSG00000( |
| 89368017 | 89368106 | 1 | miRNA | ENSG00000( |
| 89471398 | 89496589 | -1 | protein_coding | ENSG00000( |
| 89504909 | 89524049 | -1 | lncRNA | ENSG00000( |
| 89508961 | 89510416 | 1 | lncRNA | ENSG00000( |
| 89518523 | 89591956 | 1 | lncRNA | ENSG00000( |
| 89575469 | 89631056 | 1 | protein_coding | ENSG00000( |
| 89579765 | 89582297 | -1 | lncRNA | ENSG00000( |
| 89608789 | 89655467 | -1 | protein_coding | ENSG00000( |
| 89664367 | 89679427 | -1 | protein_coding | ENSG00000( |
| 89677764 | 89690783 | -1 | protein_coding | ENSG00000( |
| 89687937 | 89688243 | -1 | processed_pseudoge | ENSG00000( |
| 89690811 | 89743638 | 1 | protein_coding | ENSG00000( |
| 89704665 | 89705415 | 1 | lncRNA | ENSG00000( |
| 89714232 | 89714338 | 1 | snRNA | ENSG00000( |
| 89748661 | 89751310 | -1 | protein_coding | ENSG00000( |
| 89752296 | 89753493 | 1 | processed_pseudoge | ENSG00000( |
| 89760591 | 89778754 | 1 | protein_coding | ENSG00000( |
| 89784895 | 89815401 | -1 | protein_coding | ENSG00000( |
| 89830599 | 89894638 | -1 | protein_coding | ENSG00000( |
| 89834308 | 89912882 | -1 | protein_coding | ENSG00000( |
| 89850637 | 89850721 | -1 | miRNA | ENSG00000( |
| 89876540 | 89877285 | 1 | processed_pseudoge | ENSG00000( |
| 89883931 | 89884030 | -1 | miRNA | ENSG00000( |
| 89886485 | 89886591 | -1 | snRNA | ENSG00000( |
| 89895006 | 89912952 | -1 | protein_coding | ENSG00000( |
| 89897576 | 89897637 | 1 | snRNA | ENSG00000( |
| 89936492 | 89937005 | 1 | processed_pseudoge | ENSG00000( |
| 89950760 | 89951345 | 1 | TEC | ENSG00000( |
| 90001324 | 90082206 | 1 | protein_coding | ENSG00000( |
| 90006755 | 90006841 | 1 | miRNA | ENSG00000( |
| 90024955 | 90026003 | 1 | lncRNA | ENSG00000( |
| 90074512 | 90082207 | -1 | lncRNA | ENSG00000( |
| 90076424 | 90076486 | 1 | snRNA | ENSG00000( |
| 90083045 | 90102504 | -1 | protein_coding | ENSG00000( |

| | | | | |
|---|---|---|---|---|
| 90102649 | 90138016 | 1 | lncRNA | ENSG00000 |
| 90160604 | 90229679 | 1 | protein_coding | ENSG00000 |
| 90166233 | 90166526 | 1 | misc_RNA | ENSG00000 |
| 90206572 | 90206967 | 1 | transcribed_processe | ENSG00000 |
| 90229975 | 90234047 | -1 | protein_coding | ENSG00000 |
| 90233808 | 90245811 | 1 | protein_coding | ENSG00000 |
| 90234200 | 90265462 | 1 | protein_coding | ENSG00000 |
| 90249530 | 90265482 | 1 | protein_coding | ENSG00000 |
| 90249556 | 90272208 | 1 | protein_coding | ENSG00000 |
| 90265659 | 90278141 | 1 | protein_coding | ENSG00000 |
| 90266317 | 90314499 | 1 | protein_coding | ENSG00000 |
| 90275129 | 90275716 | 1 | transcribed_processe | ENSG00000 |
| 90281848 | 90282031 | -1 | unprocessed_pseudo | ENSG00000 |
| 90281858 | 90282270 | 1 | lncRNA | ENSG00000 |
| 90284376 | 90285580 | -1 | processed_pseudoge | ENSG00000 |
| 90291962 | 90295683 | 1 | transcribed_unproce | ENSG00000 |
| 90293939 | 90294207 | 1 | misc_RNA | ENSG00000 |
| 90347587 | 90349437 | 1 | TEC | ENSG00000 |
| 90348844 | 90349197 | -1 | processed_pseudoge | ENSG00000 |
| 90352284 | 90369146 | 1 | protein_coding | ENSG00000 |
| 90385814 | 90386063 | 1 | processed_pseudoge | ENSG00000 |
| 90388242 | 90502239 | 1 | protein_coding | ENSG00000 |
| 90393490 | 90397881 | -1 | lncRNA | ENSG00000 |
| 90529923 | 90645345 | 1 | protein_coding | ENSG00000 |
| 90595840 | 90596447 | -1 | lncRNA | ENSG00000 |
| 90604225 | 90614558 | -1 | lncRNA | ENSG00000 |
| 90620007 | 90717141 | -1 | lncRNA | ENSG00000 |
| 90630535 | 90631755 | -1 | lncRNA | ENSG00000 |
| 90634725 | 90635033 | 1 | processed_pseudoge | ENSG00000 |
| 90650631 | 90651103 | 1 | lncRNA | ENSG00000 |
| 90660207 | 90664967 | 1 | lncRNA | ENSG00000 |
| 90717346 | 90816166 | 1 | protein_coding | ENSG00000 |
| 90755243 | 90755312 | -1 | snoRNA | ENSG00000 |
| 90811528 | 90815544 | -1 | lncRNA | ENSG00000 |
| 90839641 | 90853608 | 1 | lncRNA | ENSG00000 |
| 90855935 | 90856207 | -1 | misc_RNA | ENSG00000 |
| 90868588 | 90883458 | 1 | protein_coding | ENSG00000 |
| 90883695 | 90895776 | 1 | protein_coding | ENSG00000 |
| 90902218 | 90922584 | 1 | protein_coding | ENSG00000 |
| 90920218 | 90921186 | -1 | lncRNA | ENSG00000 |
| 90929964 | 90935196 | -1 | protein_coding | ENSG00000 |
| 90930180 | 90954093 | 1 | protein_coding | ENSG00000 |
| 90952239 | 90955225 | -1 | lncRNA | ENSG00000 |
| 90954870 | 90963125 | 1 | protein_coding | ENSG00000 |
| 90966040 | 90995629 | -1 | protein_coding | ENSG00000 |
| 90966190 | 91022566 | -1 | protein_coding | ENSG00000 |
| 90966340 | 90988625 | 1 | lncRNA | ENSG00000 |
| 90977208 | 90977305 | -1 | misc_RNA | ENSG00000 |
| 90998416 | 91022603 | -1 | protein_coding | ENSG00000 |
| 91022619 | 91036611 | 1 | lncRNA | ENSG00000 |

| | | | | |
|---|---|---|---|---|
| 91022766 | 91023200 | 1 | lncRNA | ENSG0000( |
| 91029888 | 91029996 | 1 | transcribed_processe | ENSG0000( |
| 91031246 | 91031691 | -1 | unprocessed_pseudo | ENSG0000( |
| 91031712 | 91032435 | -1 | unprocessed_pseudo | ENSG0000( |
| 91094847 | 91099838 | -1 | lncRNA | ENSG0000( |
| 91099950 | 91301309 | 1 | protein_coding | ENSG0000( |
| 91303756 | 91306321 | 1 | lncRNA | ENSG0000( |
| 91408708 | 91605873 | 1 | lncRNA | ENSG0000( |
| 91532891 | 91535095 | 1 | lncRNA | ENSG0000( |
| 91643017 | 91649114 | -1 | lncRNA | ENSG0000( |
| 91676771 | 91677060 | 1 | processed_pseudoge | ENSG0000( |
| 91742841 | 91743955 | -1 | processed_pseudoge | ENSG0000( |
| 91853708 | 92172435 | 1 | protein_coding | ENSG0000( |
| 92148752 | 92331037 | -1 | lncRNA | ENSG0000( |
| 92162798 | 92172436 | -1 | lncRNA | ENSG0000( |
| 92281484 | 92282084 | 1 | processed_pseudoge | ENSG0000( |
| 92285841 | 92286678 | 1 | processed_pseudoge | ENSG0000( |
| 92393881 | 92468728 | 1 | protein_coding | ENSG0000( |
| 92417547 | 92417689 | 1 | snoRNA | ENSG0000( |
| 92420860 | 92422735 | 1 | transcribed_processe | ENSG0000( |
| 92470233 | 92471546 | -1 | lncRNA | ENSG0000( |
| 92471654 | 92501117 | 1 | lncRNA | ENSG0000( |
| 92567817 | 92574529 | -1 | lncRNA | ENSG0000( |
| 92580741 | 92600545 | 1 | lncRNA | ENSG0000( |
| 92592574 | 92596462 | -1 | lncRNA | ENSG0000( |
| 92602910 | 92620015 | 1 | lncRNA | ENSG0000( |
| 92617448 | 92809884 | -1 | protein_coding | ENSG0000( |
| 92627073 | 92627414 | 1 | processed_pseudoge | ENSG0000( |
| 92634305 | 92634665 | 1 | lncRNA | ENSG0000( |
| 92699040 | 92724277 | -1 | lncRNA | ENSG0000( |
| 92711808 | 92712110 | 1 | processed_pseudoge | ENSG0000( |
| 92713477 | 92734205 | -1 | transcribed_processe | ENSG0000( |
| 92719393 | 92719740 | 1 | processed_pseudoge | ENSG0000( |
| 92762757 | 92763750 | 1 | processed_pseudoge | ENSG0000( |
| 92779757 | 92781492 | -1 | lncRNA | ENSG0000( |
| 92795647 | 92796439 | 1 | processed_pseudoge | ENSG0000( |
| 92805770 | 92808567 | -1 | lncRNA | ENSG0000( |
| 92808451 | 92809057 | 1 | lncRNA | ENSG0000( |
| 92819540 | 92899701 | 1 | lncRNA | ENSG0000( |
| 92883413 | 92949230 | 1 | protein_coding | ENSG0000( |
| 92900189 | 93027996 | 1 | protein_coding | ENSG0000( |
| 92904399 | 92904475 | 1 | miRNA | ENSG0000( |
| 93035273 | 93089204 | -1 | protein_coding | ENSG0000( |
| 93065586 | 93066606 | -1 | transcribed_processe | ENSG0000( |
| 93177269 | 93179999 | -1 | lncRNA | ENSG0000( |
| 93230034 | 93569483 | 1 | lncRNA | ENSG0000( |
| 93257198 | 93260475 | 1 | lncRNA | ENSG0000( |
| 93301416 | 93301647 | -1 | processed_pseudoge | ENSG0000( |
| 93416850 | 93422576 | -1 | lncRNA | ENSG0000( |
| 93430659 | 93431217 | -1 | processed_pseudoge | ENSG0000( |

| | | | | |
|---|---|---|---|---|
| 93589867 | 93760886 | 1 | lncRNA | ENSG00000 |
| 93718542 | 94070898 | -1 | lncRNA | ENSG00000 |
| 93825058 | 93825159 | -1 | transcribed_processe | ENSG00000 |
| 93835490 | 93878822 | 1 | lncRNA | ENSG00000 |
| 93837611 | 93852981 | -1 | lncRNA | ENSG00000 |
| 93882082 | 93886743 | 1 | lncRNA | ENSG00000 |
| 93894431 | 93984150 | 1 | lncRNA | ENSG00000 |
| 93905405 | 94107938 | -1 | lncRNA | ENSG00000 |
| 94033658 | 94034177 | 1 | processed_pseudoge | ENSG00000 |
| 94168349 | 94168751 | 1 | processed_pseudoge | ENSG00000 |
| 94231538 | 94483952 | 1 | protein_coding | ENSG00000 |
| 94273496 | 94274005 | 1 | processed_pseudoge | ENSG00000 |
| 94293537 | 94293641 | 1 | unprocessed_pseudo | ENSG00000 |
| 94455469 | 94550572 | -1 | lncRNA | ENSG00000 |
| 94583506 | 94589259 | -1 | lncRNA | ENSG00000 |
| 94600014 | 94600821 | 1 | lncRNA | ENSG00000 |
| 94724928 | 94726519 | -1 | lncRNA | ENSG00000 |
| 94841059 | 95169864 | 1 | transcribed_processe | ENSG00000 |
| 94855586 | 94857011 | -1 | lncRNA | ENSG00000 |
| 95034108 | 95047108 | -1 | lncRNA | ENSG00000 |
| 95078481 | 95150534 | -1 | lncRNA | ENSG00000 |
| 95186634 | 95327129 | -1 | lncRNA | ENSG00000 |
| 95225326 | 95225864 | -1 | lncRNA | ENSG00000 |
| 95280753 | 95284249 | -1 | TEC | ENSG00000 |
| 95287934 | 95291643 | 1 | lncRNA | ENSG00000 |
| 95326528 | 95507865 | 1 | lncRNA | ENSG00000 |
| 95463587 | 95493843 | -1 | lncRNA | ENSG00000 |
| 95638316 | 95826499 | 1 | lncRNA | ENSG00000 |
| 95745804 | 95745994 | 1 | snRNA | ENSG00000 |
| 95990582 | 96300990 | 1 | lncRNA | ENSG00000 |
| 96045341 | 96064357 | -1 | lncRNA | ENSG00000 |
| 96080713 | 96084584 | -1 | lncRNA | ENSG00000 |
| 96110040 | 96327361 | -1 | lncRNA | ENSG00000 |
| 96112628 | 96116051 | -1 | TEC | ENSG00000 |
| 96139472 | 96140452 | -1 | lncRNA | ENSG00000 |
| 96171575 | 96174339 | 1 | lncRNA | ENSG00000 |
| 96230897 | 96231145 | 1 | lncRNA | ENSG00000 |
| 96235785 | 96237010 | 1 | lncRNA | ENSG00000 |
| 96294935 | 96298023 | -1 | TEC | ENSG00000 |
| 96325938 | 96340263 | 1 | protein_coding | ENSG00000 |
| 96333261 | 96333307 | 1 | miRNA | ENSG00000 |
| 96342953 | 96345651 | 1 | lncRNA | ENSG00000 |
| 96354237 | 96405235 | 1 | lncRNA | ENSG00000 |
| 96388214 | 96389056 | -1 | processed_pseudoge | ENSG00000 |
| 96393146 | 96405189 | -1 | lncRNA | ENSG00000 |
| 96438570 | 96444266 | 1 | lncRNA | ENSG00000 |
| 96441566 | 96442313 | -1 | processed_pseudoge | ENSG00000 |
| 96448154 | 96448493 | 1 | processed_pseudoge | ENSG00000 |
| 96448842 | 96449749 | 1 | lncRNA | ENSG00000 |
| 96486582 | 96486891 | 1 | misc_RNA | ENSG00000 |

| | | | | |
|---|---|---|---|---|
| 96496180 | 96496462 | -1 | misc_RNA | ENSG00000( |
| 96551381 | 96551517 | 1 | processed_pseudoge | ENSG00000( |
| 96756987 | 96758596 | 1 | processed_pseudoge | ENSG00000( |
| 96772005 | 96783312 | -1 | lncRNA | ENSG00000( |
| 96783389 | 96785615 | 1 | lncRNA | ENSG00000( |
| 96842069 | 96842384 | 1 | misc_RNA | ENSG00000( |
| 97215812 | 97432094 | 1 | lncRNA | ENSG00000( |
| 97272020 | 97319321 | -1 | lncRNA | ENSG00000( |
| 97339693 | 97523442 | -1 | lncRNA | ENSG00000( |
| 97353497 | 97353815 | 1 | misc_RNA | ENSG00000( |
| 97472126 | 97472252 | -1 | rRNA_pseudogene | ENSG00000( |
| 97533738 | 97541476 | -1 | TEC | ENSG00000( |
| 97540597 | 97560702 | -1 | lncRNA | ENSG00000( |
| 97557808 | 97559097 | 1 | lncRNA | ENSG00000( |
| 97560849 | 97572707 | 1 | lncRNA | ENSG00000( |
| 97572185 | 97875650 | -1 | lncRNA | ENSG00000( |
| 97876289 | 97878386 | 1 | lncRNA | ENSG00000( |
| 97945208 | 97950018 | -1 | lncRNA | ENSG00000( |
| 97960703 | 97973833 | 1 | protein_coding | ENSG00000( |
| 97982509 | 97991142 | -1 | lncRNA | ENSG00000( |
| 98003038 | 98087384 | 1 | lncRNA | ENSG00000( |
| 98026042 | 98026139 | 1 | snRNA | ENSG00000( |
| 98038571 | 98293199 | -1 | lncRNA | ENSG00000( |
| 98074667 | 98074763 | -1 | snRNA | ENSG00000( |
| 98081993 | 98103761 | -1 | lncRNA | ENSG00000( |
| 98111729 | 98131707 | -1 | lncRNA | ENSG00000( |
| 98118171 | 98118320 | 1 | processed_pseudoge | ENSG00000( |
| 98282075 | 98285907 | 1 | lncRNA | ENSG00000( |
| 98319687 | 98320237 | -1 | lncRNA | ENSG00000( |
| 98323902 | 98420998 | -1 | lncRNA | ENSG00000( |
| 98437162 | 98547728 | -1 | protein_coding | ENSG00000( |
| 98466840 | 98467412 | 1 | processed_pseudoge | ENSG00000( |
| 98646951 | 98647371 | -1 | lncRNA | ENSG00000( |
| 98648539 | 98964530 | 1 | protein_coding | ENSG00000( |
| 98660210 | 98660668 | 1 | lncRNA | ENSG00000( |
| 98784426 | 98784502 | 1 | miRNA | ENSG00000( |
| 98839292 | 98840147 | -1 | lncRNA | ENSG00000( |
| 98880659 | 98893535 | -1 | lncRNA | ENSG00000( |
| 98954149 | 99105824 | -1 | lncRNA | ENSG00000( |
| 98968229 | 99007792 | -1 | protein_coding | ENSG00000( |
| 99014526 | 99031054 | 1 | lncRNA | ENSG00000( |
| 99098217 | 99135593 | 1 | protein_coding | ENSG00000( |
| 99128832 | 99131806 | -1 | lncRNA | ENSG00000( |
| 99136323 | 99251223 | -1 | protein_coding | ENSG00000( |
| 99139317 | 99145370 | 1 | lncRNA | ENSG00000( |
| 99157231 | 99161140 | -1 | lncRNA | ENSG00000( |
| 99251362 | 99390729 | 1 | protein_coding | ENSG00000( |
| 99257632 | 99260015 | 1 | processed_pseudoge | ENSG00000( |
| 99284625 | 99285576 | -1 | processed_pseudoge | ENSG00000( |
| 99337857 | 99338481 | 1 | processed_pseudoge | ENSG00000( |

| | | | | |
|---|---|---|---|---|
| 99395179 | 99396593 | -1 | lncRNA | ENSG0000C |
| 99396613 | 99401315 | -1 | lncRNA | ENSG0000C |
| 99416584 | 99417204 | 1 | lncRNA | ENSG0000C |
| 99423769 | 99435160 | -1 | lncRNA | ENSG0000C |
| 99473454 | 99476314 | -1 | lncRNA | ENSG0000C |
| 99497812 | 99498375 | -1 | TEC | ENSG0000C |
| 99551382 | 99552117 | -1 | processed_pseudoge | ENSG0000C |
| 99565417 | 99716466 | 1 | protein_coding | ENSG0000C |
| 99681786 | 99681877 | -1 | misc_RNA | ENSG0000C |
| 99715697 | 99733561 | -1 | protein_coding | ENSG0000C |
| 99790156 | 99806927 | -1 | transcribed_unproce | ENSG0000C |
| 99791127 | 99792847 | 1 | unprocessed_pseudc | ENSG0000C |
| 99791209 | 99791410 | 1 | misc_RNA | ENSG0000C |
| 99805793 | 99916818 | 1 | lncRNA | ENSG0000C |
| 99970215 | 99974010 | 1 | lncRNA | ENSG0000C |
| 99971437 | 100342005 | -1 | protein_coding | ENSG0000C |
| 99976481 | 99980774 | 1 | lncRNA | ENSG0000C |
| 100074081 | 100074190 | -1 | rRNA_pseudogene | ENSG0000C |
| 100126106 | 100129743 | 1 | lncRNA | ENSG0000C |
| 100344047 | 100350725 | -1 | lncRNA | ENSG0000C |
| 100372936 | 100437914 | 1 | lncRNA | ENSG0000C |
| 100400395 | 100544995 | -1 | protein_coding | ENSG0000C |
| 100547765 | 100550153 | -1 | lncRNA | ENSG0000C |
| 100551820 | 100551922 | -1 | snRNA | ENSG0000C |
| 100553529 | 100558954 | -1 | processed_pseudoge | ENSG0000C |
| 100558677 | 100559798 | 1 | lncRNA | ENSG0000C |
| 100559369 | 100603230 | -1 | protein_coding | ENSG0000C |
| 100560686 | 100560792 | -1 | snRNA | ENSG0000C |
| 100602534 | 100651705 | 1 | protein_coding | ENSG0000C |
| 100638363 | 100638457 | 1 | misc_RNA | ENSG0000C |
| 100645434 | 100646655 | -1 | processed_pseudoge | ENSG0000C |
| 100716249 | 100828497 | 1 | lncRNA | ENSG0000C |
| 100782417 | 100786189 | -1 | TEC | ENSG0000C |
| 100846604 | 100846767 | -1 | processed_pseudoge | ENSG0000C |
| 100849561 | 100861756 | 1 | lncRNA | ENSG0000C |
| 100849831 | 100876836 | 1 | lncRNA | ENSG0000C |
| 100861843 | 100874369 | 1 | lncRNA | ENSG0000C |
| 100861924 | 100885329 | 1 | lncRNA | ENSG0000C |
| 100877714 | 100916626 | 1 | protein_coding | ENSG0000C |
| 100888472 | 100889106 | -1 | lncRNA | ENSG0000C |
| 100892343 | 100919391 | -1 | lncRNA | ENSG0000C |
| 100919327 | 101078257 | 1 | protein_coding | ENSG0000C |
| 101043716 | 101049456 | -1 | lncRNA | ENSG0000C |
| 101050732 | 101086066 | -1 | lncRNA | ENSG0000C |
| 101058647 | 101063904 | -1 | lncRNA | ENSG0000C |
| 101116603 | 101117546 | 1 | lncRNA | ENSG0000C |
| 101168530 | 101170821 | 1 | lncRNA | ENSG0000C |
| 101175727 | 101252048 | -1 | protein_coding | ENSG0000C |
| 101270817 | 101277500 | -1 | protein_coding | ENSG0000C |
| 101281510 | 101295282 | -1 | protein_coding | ENSG0000C |

| | | | | |
|---|---|---|---|---|
| 101295419 | 101305737 | 1 | lncRNA | ENSG00000 |
| 101297142 | 101525202 | -1 | protein_coding | ENSG00000 |
| 101299656 | 101301648 | -1 | TEC | ENSG00000 |
| 101334437 | 101337428 | 1 | lncRNA | ENSG00000 |
| 101495052 | 101495567 | -1 | lncRNA | ENSG00000 |
| 101522597 | 101524739 | -1 | lncRNA | ENSG00000 |
| 101527581 | 101528349 | -1 | lncRNA | ENSG00000 |
| 101562536 | 101562974 | 1 | processed_pseudoge | ENSG00000 |
| 101580893 | 101581556 | -1 | processed_pseudoge | ENSG00000 |
| 101602786 | 101614531 | -1 | lncRNA | ENSG00000 |
| 101621444 | 101652391 | -1 | protein_coding | ENSG00000 |
| 101648206 | 101648312 | -1 | snRNA | ENSG00000 |
| 101653596 | 101724473 | -1 | protein_coding | ENSG00000 |
| 101732201 | 101746870 | -1 | transcribed_unproce | ENSG00000 |
| 101751032 | 101794302 | 1 | transcribed_unproce | ENSG00000 |
| 101771758 | 101781989 | -1 | unprocessed_pseudo | ENSG00000 |
| 101773238 | 101773513 | -1 | misc_RNA | ENSG00000 |
| 101803509 | 101806887 | 1 | protein_coding | ENSG00000 |
| 101812202 | 101820197 | 1 | protein_coding | ENSG00000 |
| 101828741 | 101829679 | 1 | unprocessed_pseudo | ENSG00000 |
| 101842119 | 101850324 | 1 | transcribed_unproce | ENSG00000 |
| 101875964 | 101876901 | 1 | unprocessed_pseudo | ENSG00000 |
| 101887354 | 101887767 | -1 | processed_pseudoge | ENSG00000 |
| 101903154 | 101903204 | 1 | unprocessed_pseudo | ENSG00000 |
| 101922042 | 101923113 | -1 | protein_coding | ENSG00000 |
| 101926164 | 101933350 | -1 | transcribed_unproce | ENSG00000 |
| 101937591 | 101938470 | -1 | unprocessed_pseudo | ENSG00000 |
| 101954885 | 101956412 | 1 | lncRNA | ENSG00000 |
| 101959539 | 101961408 | -1 | lncRNA | ENSG00000 |
| 101960459 | 101960596 | -1 | miRNA | ENSG00000 |
| 101961603 | 101976543 | 1 | transcribed_unproce | ENSG00000 |
| 101973524 | 101973591 | 1 | miRNA | ENSG00000 |
| 101976558 | 101979093 | -1 | transcribed_unproce | ENSG00000 |
| 11555 | 14090 | 1 | transcribed_unproce | ENSG00000 |
| 14381 | 18068 | -1 | unprocessed_pseudo | ENSG00000 |
| 17052 | 17119 | -1 | miRNA | ENSG00000 |
| 17514 | 35195 | -1 | lncRNA | ENSG00000 |
| 22910 | 25123 | 1 | lncRNA | ENSG00000 |
| 29336 | 38321 | -1 | unprocessed_pseudo | ENSG00000 |
| 46407 | 53608 | -1 | protein_coding | ENSG00000 |
| 53010 | 57669 | 1 | protein_coding | ENSG00000 |
| 58059 | 76355 | -1 | protein_coding | ENSG00000 |
| 74683 | 74961 | 1 | TEC | ENSG00000 |
| 77007 | 85853 | 1 | protein_coding | ENSG00000 |
| 84271 | 138860 | -1 | protein_coding | ENSG00000 |
| 90631 | 91102 | 1 | lncRNA | ENSG00000 |
| 125737 | 127219 | 1 | lncRNA | ENSG00000 |
| 142728 | 154503 | 1 | protein_coding | ENSG00000 |
| 153892 | 166764 | 1 | protein_coding | ENSG00000 |
| 163122 | 165156 | 1 | unprocessed_pseudo | ENSG00000 |

| | | | | |
|---|---|---|---|---|
| 168679 | 169334 | 1 | unprocessed_pseudo | ENSG00000 |
| 172876 | 173710 | 1 | protein_coding | ENSG00000 |
| 176680 | 177522 | 1 | protein_coding | ENSG00000 |
| 178554 | 178656 | 1 | misc_RNA | ENSG00000 |
| 180459 | 181179 | 1 | protein_coding | ENSG00000 |
| 185748 | 186294 | -1 | lncRNA | ENSG00000 |
| 188969 | 229463 | -1 | protein_coding | ENSG00000 |
| 231134 | 231472 | 1 | processed_pseudoge | ENSG00000 |
| 234521 | 272183 | 1 | protein_coding | ENSG00000 |
| 268301 | 275980 | -1 | protein_coding | ENSG00000 |
| 268727 | 282084 | 1 | lncRNA | ENSG00000 |
| 280450 | 283010 | 1 | protein_coding | ENSG00000 |
| 283152 | 287215 | 1 | protein_coding | ENSG00000 |
| 287440 | 352723 | -1 | protein_coding | ENSG00000 |
| 366969 | 371289 | -1 | protein_coding | ENSG00000 |
| 370788 | 387113 | -1 | protein_coding | ENSG00000 |
| 382097 | 392960 | 1 | transcribed_process | ENSG00000 |
| 396725 | 410367 | 1 | protein_coding | ENSG00000 |
| 401858 | 412487 | 1 | protein_coding | ENSG00000 |
| 424243 | 424543 | -1 | lncRNA | ENSG00000 |
| 425649 | 523011 | 1 | protein_coding | ENSG00000 |
| 439297 | 439397 | -1 | misc_RNA | ENSG00000 |
| 451404 | 453461 | 1 | transcribed_unproce | ENSG00000 |
| 525155 | 527407 | -1 | lncRNA | ENSG00000 |
| 527712 | 554636 | 1 | protein_coding | ENSG00000 |
| 535316 | 535368 | 1 | miRNA | ENSG00000 |
| 543277 | 543366 | 1 | miRNA | ENSG00000 |
| 547185 | 553847 | -1 | lncRNA | ENSG00000 |
| 560394 | 565529 | 1 | protein_coding | ENSG00000 |
| 566995 | 584136 | 1 | protein_coding | ENSG00000 |
| 567005 | 569495 | 1 | protein_coding | ENSG00000 |
| 568837 | 576257 | 1 | lncRNA | ENSG00000 |
| 589357 | 629272 | 1 | protein_coding | ENSG00000 |
| 629239 | 634117 | 1 | protein_coding | ENSG00000 |
| 634427 | 636366 | -1 | protein_coding | ENSG00000 |
| 636817 | 648474 | 1 | protein_coding | ENSG00000 |
| 648473 | 649200 | -1 | lncRNA | ENSG00000 |
| 649311 | 667833 | 1 | protein_coding | ENSG00000 |
| 654611 | 656194 | -1 | lncRNA | ENSG00000 |
| 668105 | 674174 | 1 | protein_coding | ENSG00000 |
| 675666 | 678268 | 1 | protein_coding | ENSG00000 |
| 678504 | 679777 | -1 | lncRNA | ENSG00000 |
| 678645 | 679061 | 1 | TEC | ENSG00000 |
| 680224 | 682870 | 1 | protein_coding | ENSG00000 |
| 681670 | 684528 | -1 | protein_coding | ENSG00000 |
| 684622 | 690444 | -1 | protein_coding | ENSG00000 |
| 689001 | 692554 | 1 | lncRNA | ENSG00000 |
| 692498 | 705808 | -1 | protein_coding | ENSG00000 |
| 707650 | 709067 | 1 | TEC | ENSG00000 |
| 710746 | 711277 | -1 | lncRNA | ENSG00000 |

| | | | | |
|---|---|---|---|---|
| 715118 | 719655 | 1 | protein_coding | ENSG00000 |
| 720581 | 722590 | 1 | protein_coding | ENSG00000 |
| 722582 | 726954 | -1 | protein_coding | ENSG00000 |
| 726936 | 735525 | 1 | protein_coding | ENSG00000 |
| 729760 | 741329 | -1 | protein_coding | ENSG00000 |
| 760734 | 768865 | 1 | protein_coding | ENSG00000 |
| 769119 | 769443 | -1 | TEC | ENSG00000 |
| 769428 | 783370 | -1 | protein_coding | ENSG00000 |
| 770183 | 770277 | 1 | miRNA | ENSG00000 |
| 784974 | 788397 | -1 | protein_coding | ENSG00000 |
| 788046 | 800737 | 1 | protein_coding | ENSG00000 |
| 798041 | 800734 | -1 | protein_coding | ENSG00000 |
| 817489 | 862843 | 1 | lncRNA | ENSG00000 |
| 848525 | 849065 | -1 | lncRNA | ENSG00000 |
| 853634 | 981318 | -1 | protein_coding | ENSG00000 |
| 883780 | 885090 | 1 | lncRNA | ENSG00000 |
| 898967 | 905224 | -1 | lncRNA | ENSG00000 |
| 921033 | 934495 | 1 | lncRNA | ENSG00000 |
| 968375 | 969012 | -1 | lncRNA | ENSG00000 |
| 975761 | 981596 | -1 | lncRNA | ENSG00000 |
| 981770 | 986979 | 1 | protein_coding | ENSG00000 |
| 991151 | 1000926 | -1 | lncRNA | ENSG00000 |
| 1064093 | 1078731 | -1 | lncRNA | ENSG00000 |
| 1065240 | 1066502 | 1 | lncRNA | ENSG00000 |
| 1078781 | 1080142 | 1 | protein_coding | ENSG00000 |
| 1088226 | 1096244 | -1 | protein_coding | ENSG00000 |
| 1101260 | 1105461 | -1 | unprocessed_pseudo | ENSG00000 |
| 1111627 | 1113399 | 1 | lncRNA | ENSG00000 |
| 1148224 | 1148754 | -1 | lncRNA | ENSG00000 |
| 1153106 | 1221771 | 1 | protein_coding | ENSG00000 |
| 1156976 | 1157974 | -1 | lncRNA | ENSG00000 |
| 1159548 | 1160176 | -1 | lncRNA | ENSG00000 |
| 1206560 | 1207124 | -1 | lncRNA | ENSG00000 |
| 1221651 | 1225257 | -1 | protein_coding | ENSG00000 |
| 1223639 | 1224143 | 1 | lncRNA | ENSG00000 |
| 1227272 | 1230184 | -1 | protein_coding | ENSG00000 |
| 1240705 | 1242554 | 1 | protein_coding | ENSG00000 |
| 1256059 | 1259008 | 1 | protein_coding | ENSG00000 |
| 1257339 | 1258074 | 1 | lncRNA | ENSG00000 |
| 1260952 | 1263845 | -1 | transcribed_unproce | ENSG00000 |
| 1273751 | 1276522 | -1 | unprocessed_pseudo | ENSG00000 |
| 1280161 | 1280544 | -1 | unprocessed_pseudo | ENSG00000 |
| 1286351 | 1288837 | -1 | unprocessed_pseudo | ENSG00000 |
| 1294551 | 1299166 | -1 | lncRNA | ENSG00000 |
| 1305547 | 1309413 | -1 | lncRNA | ENSG00000 |
| 1308880 | 1327018 | 1 | protein_coding | ENSG00000 |
| 1317891 | 1322845 | -1 | lncRNA | ENSG00000 |
| 1329078 | 1329777 | 1 | processed_pseudoge | ENSG00000 |
| 1333601 | 1349441 | 1 | protein_coding | ENSG00000 |
| 1349240 | 1351878 | -1 | protein_coding | ENSG00000 |

| | | | | |
|---|---|---|---|---|
| 1351931 | 1364113 | 1 | protein_coding | ENSG00000 |
| 1358900 | 1361405 | -1 | lncRNA | ENSG00000 |
| 1363205 | 1414751 | -1 | protein_coding | ENSG00000 |
| 1408834 | 1412248 | 1 | lncRNA | ENSG00000 |
| 1419752 | 1420756 | -1 | protein_coding | ENSG00000 |
| 1431035 | 1433397 | 1 | protein_coding | ENSG00000 |
| 1434383 | 1444556 | -1 | protein_coding | ENSG00000 |
| 1437154 | 1439315 | 1 | TEC | ENSG00000 |
| 1444934 | 1475084 | -1 | protein_coding | ENSG00000 |
| 1445343 | 1446519 | 1 | lncRNA | ENSG00000 |
| 1451760 | 1452653 | 1 | lncRNA | ENSG00000 |
| 1467673 | 1472684 | -1 | lncRNA | ENSG00000 |
| 1477830 | 1478583 | 1 | processed_pseudoge | ENSG00000 |
| 1485886 | 1488981 | -1 | protein_coding | ENSG00000 |
| 1493344 | 1510457 | 1 | protein_coding | ENSG00000 |
| 1510427 | 1612072 | -1 | protein_coding | ENSG00000 |
| 1512979 | 1514675 | 1 | lncRNA | ENSG00000 |
| 1528688 | 1555580 | 1 | protein_coding | ENSG00000 |
| 1530715 | 1533301 | 1 | TEC | ENSG00000 |
| 1553655 | 1554130 | -1 | TEC | ENSG00000 |
| 1576716 | 1578371 | -1 | TEC | ENSG00000 |
| 1579242 | 1580308 | -1 | lncRNA | ENSG00000 |
| 1580527 | 1610328 | 1 | lncRNA | ENSG00000 |
| 1612325 | 1677908 | 1 | protein_coding | ENSG00000 |
| 1625628 | 1626160 | -1 | lncRNA | ENSG00000 |
| 1632259 | 1686715 | 1 | protein_coding | ENSG00000 |
| 1678256 | 1702280 | 1 | protein_coding | ENSG00000 |
| 1688355 | 1690536 | 1 | TEC | ENSG00000 |
| 1706183 | 1770317 | 1 | protein_coding | ENSG00000 |
| 1707252 | 1707973 | 1 | lncRNA | ENSG00000 |
| 1713527 | 1714208 | -1 | lncRNA | ENSG00000 |
| 1734985 | 1735066 | 1 | miRNA | ENSG00000 |
| 1751559 | 1752262 | -1 | lncRNA | ENSG00000 |
| 1770286 | 1771730 | -1 | protein_coding | ENSG00000 |
| 1771890 | 1773155 | -1 | protein_coding | ENSG00000 |
| 1772810 | 1781708 | 1 | protein_coding | ENSG00000 |
| 1776712 | 1793700 | -1 | protein_coding | ENSG00000 |
| 1782932 | 1789186 | 1 | protein_coding | ENSG00000 |
| 1790413 | 1794971 | -1 | protein_coding | ENSG00000 |
| 1795620 | 1827157 | -1 | protein_coding | ENSG00000 |
| 1826941 | 1840207 | 1 | protein_coding | ENSG00000 |
| 1833987 | 1884294 | -1 | protein_coding | ENSG00000 |
| 1841020 | 1843547 | -1 | lncRNA | ENSG00000 |
| 1878285 | 1884233 | -1 | lncRNA | ENSG00000 |
| 1889114 | 1890434 | 1 | lncRNA | ENSG00000 |
| 1911475 | 1918415 | -1 | protein_coding | ENSG00000 |
| 1938210 | 1943326 | -1 | protein_coding | ENSG00000 |
| 1943974 | 1957606 | -1 | protein_coding | ENSG00000 |
| 1959538 | 1961975 | 1 | protein_coding | ENSG00000 |
| 1962058 | 1964841 | -1 | protein_coding | ENSG00000 |

| | | | | |
|---|---|---|---|---|
| 1962334 | 1962466 | -1 | snoRNA | ENSG0000C |
| 1962973 | 1963106 | -1 | snoRNA | ENSG0000C |
| 1963745 | 1964095 | -1 | transcribed_processe | ENSG0000C |
| 1964959 | 1965509 | 1 | lncRNA | ENSG0000C |
| 1965184 | 1965310 | 1 | snoRNA | ENSG0000C |
| 1966823 | 1968975 | 1 | protein_coding | ENSG0000C |
| 1971655 | 1971896 | -1 | lncRNA | ENSG0000C |
| 1972053 | 1982929 | 1 | protein_coding | ENSG0000C |
| 1978917 | 1984192 | -1 | protein_coding | ENSG0000C |
| 1984193 | 1987749 | 1 | protein_coding | ENSG0000C |
| 1984877 | 1990477 | -1 | lncRNA | ENSG0000C |
| 1989660 | 1994275 | 1 | protein_coding | ENSG0000C |
| 1997654 | 1998374 | 1 | lncRNA | ENSG0000C |
| 1997654 | 2009821 | -1 | protein_coding | ENSG0000C |
| 2009926 | 2020755 | 1 | protein_coding | ENSG0000C |
| 2013181 | 2013476 | 1 | misc_RNA | ENSG0000C |
| 2025356 | 2039026 | 1 | protein_coding | ENSG0000C |
| 2039815 | 2047866 | -1 | protein_coding | ENSG0000C |
| 2047967 | 2089491 | 1 | protein_coding | ENSG0000C |
| 2088710 | 2135898 | -1 | protein_coding | ENSG0000C |
| 2090195 | 2090284 | -1 | miRNA | ENSG0000C |
| 2091436 | 2095433 | 1 | lncRNA | ENSG0000C |
| 2094830 | 2097026 | -1 | lncRNA | ENSG0000C |
| 2106669 | 2106753 | -1 | miRNA | ENSG0000C |
| 2112335 | 2113342 | 1 | lncRNA | ENSG0000C |
| 2119207 | 2120248 | 1 | lncRNA | ENSG0000C |
| 2131851 | 2131950 | 1 | misc_RNA | ENSG0000C |
| 2133119 | 2133204 | 1 | miRNA | ENSG0000C |
| 2135977 | 2136129 | -1 | miRNA | ENSG0000C |
| 2140803 | 2154165 | 1 | protein_coding | ENSG0000C |
| 2154797 | 2155358 | -1 | lncRNA | ENSG0000C |
| 2155023 | 2155105 | -1 | snoRNA | ENSG0000C |
| 2155025 | 2155104 | -1 | snoRNA | ENSG0000C |
| 2155698 | 2178129 | 1 | protein_coding | ENSG0000C |
| 2177180 | 2196605 | -1 | protein_coding | ENSG0000C |
| 2204248 | 2209453 | 1 | protein_coding | ENSG0000C |
| 2209253 | 2211950 | -1 | protein_coding | ENSG0000C |
| 2211593 | 2214840 | -1 | protein_coding | ENSG0000C |
| 2211997 | 2212863 | 1 | lncRNA | ENSG0000C |
| 2223580 | 2235742 | 1 | protein_coding | ENSG0000C |
| 2235689 | 2236913 | -1 | lncRNA | ENSG0000C |
| 2235816 | 2238711 | 1 | protein_coding | ENSG0000C |
| 2239402 | 2252300 | -1 | protein_coding | ENSG0000C |
| 2240487 | 2241818 | 1 | lncRNA | ENSG0000C |
| 2253116 | 2268397 | -1 | protein_coding | ENSG0000C |
| 2268155 | 2273418 | 1 | lncRNA | ENSG0000C |
| 2270713 | 2270772 | 1 | miRNA | ENSG0000C |
| 2271747 | 2271840 | 1 | miRNA | ENSG0000C |
| 2274620 | 2274691 | 1 | miRNA | ENSG0000C |
| 2275881 | 2340746 | -1 | protein_coding | ENSG0000C |

| | | | | |
|---|---|---|---|---|
| 2332313 | 2332732 | -1 | processed_pseudoge | ENSG00000 |
| 2339150 | 2426699 | 1 | transcribed_unitary_ | ENSG00000 |
| 2429394 | 2458854 | 1 | protein_coding | ENSG00000 |
| 2430702 | 2437103 | -1 | lncRNA | ENSG00000 |
| 2445392 | 2445457 | 1 | miRNA | ENSG00000 |
| 2452581 | 2452977 | -1 | lncRNA | ENSG00000 |
| 2456252 | 2459979 | -1 | lncRNA | ENSG00000 |
| 2460086 | 2464963 | 1 | protein_coding | ENSG00000 |
| 2463967 | 2464038 | 1 | miRNA | ENSG00000 |
| 2464950 | 2468213 | -1 | lncRNA | ENSG00000 |
| 2471297 | 2474145 | 1 | protein_coding | ENSG00000 |
| 2475051 | 2509560 | 1 | protein_coding | ENSG00000 |
| 2476558 | 2482173 | 1 | lncRNA | ENSG00000 |
| 2496032 | 2520218 | 1 | protein_coding | ENSG00000 |
| 2513952 | 2520218 | 1 | protein_coding | ENSG00000 |
| 2513965 | 2527955 | 1 | protein_coding | ENSG00000 |
| 2516658 | 2517999 | 1 | TEC | ENSG00000 |
| 2520357 | 2531422 | 1 | protein_coding | ENSG00000 |
| 2530035 | 2531417 | -1 | protein_coding | ENSG00000 |
| 2531922 | 2532005 | -1 | miRNA | ENSG00000 |
| 2537979 | 2603188 | 1 | protein_coding | ENSG00000 |
| 2554060 | 2556060 | -1 | lncRNA | ENSG00000 |
| 2554975 | 2556105 | 1 | lncRNA | ENSG00000 |
| 2561471 | 2565096 | -1 | lncRNA | ENSG00000 |
| 2569043 | 2571936 | -1 | lncRNA | ENSG00000 |
| 2571570 | 2572353 | -1 | lncRNA | ENSG00000 |
| 2575628 | 2577373 | -1 | TEC | ENSG00000 |
| 2578395 | 2579963 | 1 | TEC | ENSG00000 |
| 2592514 | 2594563 | 1 | TEC | ENSG00000 |
| 2597881 | 2599718 | -1 | lncRNA | ENSG00000 |
| 2603350 | 2630494 | 1 | transcribed_unproce | ENSG00000 |
| 2616121 | 2643296 | -1 | transcribed_unproce | ENSG00000 |
| 2620222 | 2621794 | -1 | TEC | ENSG00000 |
| 2644084 | 2645214 | -1 | lncRNA | ENSG00000 |
| 2660348 | 2682379 | -1 | lncRNA | ENSG00000 |
| 2682523 | 2709030 | 1 | protein_coding | ENSG00000 |
| 2683106 | 2684764 | 1 | TEC | ENSG00000 |
| 2712419 | 2720551 | -1 | protein_coding | ENSG00000 |
| 2737076 | 2752600 | -1 | lncRNA | ENSG00000 |
| 2752626 | 2772538 | 1 | protein_coding | ENSG00000 |
| 2771414 | 2777297 | -1 | protein_coding | ENSG00000 |
| 2777319 | 2780568 | 1 | lncRNA | ENSG00000 |
| 2783953 | 2787948 | -1 | protein_coding | ENSG00000 |
| 2796408 | 2796532 | -1 | snoRNA | ENSG00000 |
| 2798485 | 2805302 | 1 | protein_coding | ENSG00000 |
| 2817180 | 2826304 | 1 | protein_coding | ENSG00000 |
| 2822659 | 2822993 | -1 | processed_pseudoge | ENSG00000 |
| 2830169 | 2839585 | 1 | protein_coding | ENSG00000 |
| 2839568 | 2842744 | -1 | transcribed_unitary_ | ENSG00000 |
| 2852730 | 2858170 | -1 | protein_coding | ENSG00000 |

| | | | | |
|---|---|---|---|---|
| 2857898 | 2859726 | 1 | lncRNA | ENSG0000C |
| 2866325 | 2868251 | -1 | lncRNA | ENSG0000C |
| 2883213 | 2899382 | 1 | protein_coding | ENSG0000C |
| 2911937 | 2951208 | 1 | protein_coding | ENSG0000C |
| 2922662 | 2923202 | -1 | processed_pseudoge | ENSG0000C |
| 2939714 | 2954276 | -1 | lncRNA | ENSG0000C |
| 2964216 | 2968383 | 1 | protein_coding | ENSG0000C |
| 2968024 | 2980479 | -1 | protein_coding | ENSG0000C |
| 2969270 | 2973484 | 1 | protein_coding | ENSG0000C |
| 2981175 | 2981591 | -1 | lncRNA | ENSG0000C |
| 2988256 | 3002016 | 1 | protein_coding | ENSG0000C |
| 3003431 | 3005101 | -1 | lncRNA | ENSG0000C |
| 3006120 | 3007388 | 1 | lncRNA | ENSG0000C |
| 3012923 | 3014505 | 1 | protein_coding | ENSG0000C |
| 3014712 | 3020071 | -1 | protein_coding | ENSG0000C |
| 3018445 | 3022383 | 1 | protein_coding | ENSG0000C |
| 3022620 | 3024286 | -1 | protein_coding | ENSG0000C |
| 3024027 | 3027755 | 1 | protein_coding | ENSG0000C |
| 3027682 | 3036944 | -1 | protein_coding | ENSG0000C |
| 3032481 | 3039133 | 1 | lncRNA | ENSG0000C |
| 3037400 | 3059370 | -1 | lncRNA | ENSG0000C |
| 3046561 | 3060726 | 1 | protein_coding | ENSG0000C |
| 3065297 | 3082192 | 1 | protein_coding | ENSG0000C |
| 3069523 | 3069651 | 1 | snRNA | ENSG0000C |
| 3076911 | 3087111 | -1 | lncRNA | ENSG0000C |
| 3086143 | 3086300 | 1 | snRNA | ENSG0000C |
| 3088890 | 3099295 | -1 | protein_coding | ENSG0000C |
| 3106764 | 3109576 | 1 | lncRNA | ENSG0000C |
| 3110460 | 3134882 | -1 | lncRNA | ENSG0000C |
| 3112560 | 3120517 | 1 | protein_coding | ENSG0000C |
| 3129777 | 3142804 | 1 | protein_coding | ENSG0000C |
| 3144015 | 3149963 | 1 | transcribed_unitary_ | ENSG0000C |
| 3156736 | 3157483 | -1 | lncRNA | ENSG0000C |
| 3181233 | 3184018 | -1 | lncRNA | ENSG0000C |
| 3188204 | 3224779 | 1 | lncRNA | ENSG0000C |
| 3204247 | 3205188 | 1 | protein_coding | ENSG0000C |
| 3215611 | 3216543 | 1 | unprocessed_pseudc | ENSG0000C |
| 3222325 | 3236221 | -1 | protein_coding | ENSG0000C |
| 3242028 | 3256627 | -1 | protein_coding | ENSG0000C |
| 3263743 | 3267567 | 1 | lncRNA | ENSG0000C |
| 3263800 | 3301401 | 1 | protein_coding | ENSG0000C |
| 3273609 | 3275807 | 1 | TEC | ENSG0000C |
| 3292879 | 3293403 | 1 | TEC | ENSG0000C |
| 3298808 | 3305729 | -1 | protein_coding | ENSG0000C |
| 3305406 | 3318852 | 1 | protein_coding | ENSG0000C |
| 3307573 | 3308393 | -1 | lncRNA | ENSG0000C |
| 3323192 | 3323925 | -1 | processed_pseudoge | ENSG0000C |
| 3355889 | 3357306 | 1 | protein_coding | ENSG0000C |
| 3365099 | 3479550 | 1 | protein_coding | ENSG0000C |
| 3367719 | 3369246 | -1 | processed_pseudoge | ENSG0000C |

| | | | | |
|---|---|---|---|---|
| 3369957 | 3370906 | -1 | processed_pseudoge | ENSG00000 |
| 3370979 | 3372668 | -1 | protein_coding | ENSG00000 |
| 3382081 | 3401065 | -1 | protein_coding | ENSG00000 |
| 3401215 | 3409364 | 1 | protein_coding | ENSG00000 |
| 3432414 | 3443504 | -1 | protein_coding | ENSG00000 |
| 3443649 | 3486953 | 1 | protein_coding | ENSG00000 |
| 3458071 | 3515564 | 1 | lncRNA | ENSG00000 |
| 3485381 | 3485469 | -1 | miRNA | ENSG00000 |
| 3493484 | 3495489 | -1 | protein_coding | ENSG00000 |
| 3496201 | 3496768 | 1 | processed_pseudoge | ENSG00000 |
| 3500976 | 3539048 | 1 | protein_coding | ENSG00000 |
| 3533429 | 3534258 | -1 | TEC | ENSG00000 |
| 3539033 | 3577403 | -1 | protein_coding | ENSG00000 |
| 3542739 | 3545785 | 1 | lncRNA | ENSG00000 |
| 3581181 | 3611606 | -1 | protein_coding | ENSG00000 |
| 3581181 | 3583266 | 1 | lncRNA | ENSG00000 |
| 3611728 | 3680143 | 1 | protein_coding | ENSG00000 |
| 3650636 | 3651703 | -1 | lncRNA | ENSG00000 |
| 3651639 | 3717553 | -1 | protein_coding | ENSG00000 |
| 3686998 | 3687380 | 1 | lncRNA | ENSG00000 |
| 3725054 | 3880726 | -1 | protein_coding | ENSG00000 |
| 3840528 | 3841010 | 1 | processed_pseudoge | ENSG00000 |
| 3930806 | 3950586 | -1 | lncRNA | ENSG00000 |
| 3953387 | 4116442 | -1 | protein_coding | ENSG00000 |
| 4032012 | 4032936 | 1 | lncRNA | ENSG00000 |
| 4130555 | 4130820 | 1 | processed_pseudoge | ENSG00000 |
| 4180117 | 4183515 | -1 | lncRNA | ENSG00000 |
| 4189374 | 4242080 | -1 | protein_coding | ENSG00000 |
| 4243943 | 4253817 | -1 | lncRNA | ENSG00000 |
| 4257186 | 4273075 | -1 | protein_coding | ENSG00000 |
| 4314761 | 4339597 | 1 | protein_coding | ENSG00000 |
| 4324667 | 4328340 | -1 | lncRNA | ENSG00000 |
| 4331549 | 4355607 | -1 | protein_coding | ENSG00000 |
| 4335870 | 4337818 | -1 | lncRNA | ENSG00000 |
| 4340251 | 4420494 | -1 | protein_coding | ENSG00000 |
| 4346694 | 4348648 | -1 | TEC | ENSG00000 |
| 4354542 | 4425705 | -1 | protein_coding | ENSG00000 |
| 4371848 | 4383538 | 1 | protein_coding | ENSG00000 |
| 4425805 | 4456775 | 1 | protein_coding | ENSG00000 |
| 4426902 | 4427380 | 1 | lncRNA | ENSG00000 |
| 4430522 | 4431103 | 1 | lncRNA | ENSG00000 |
| 4433202 | 4433387 | -1 | processed_pseudoge | ENSG00000 |
| 4461680 | 4495763 | -1 | protein_coding | ENSG00000 |
| 4474690 | 4510347 | 1 | protein_coding | ENSG00000 |
| 4510669 | 4538828 | -1 | protein_coding | ENSG00000 |
| 4532216 | 4533670 | -1 | lncRNA | ENSG00000 |
| 4556490 | 4600714 | 1 | protein_coding | ENSG00000 |
| 4560001 | 4561662 | -1 | lncRNA | ENSG00000 |
| 4562814 | 4563128 | -1 | processed_pseudoge | ENSG00000 |
| 4608883 | 4615027 | -1 | protein_coding | ENSG00000 |

| | | | | |
|---|---|---|---|---|
| 4616493 | 4690974 | 1 | protein_coding | ENSG0000C |
| 4633591 | 4633881 | -1 | misc_RNA | ENSG0000C |
| 4634329 | 4640623 | -1 | lncRNA | ENSG0000C |
| 4658851 | 4659151 | 1 | misc_RNA | ENSG0000C |
| 4671318 | 4671390 | 1 | miRNA | ENSG0000C |
| 4693694 | 4695859 | 1 | protein_coding | ENSG0000C |
| 4696510 | 4734378 | -1 | protein_coding | ENSG0000C |
| 4730115 | 4752565 | -1 | lncRNA | ENSG0000C |
| 4734344 | 4749396 | 1 | protein_coding | ENSG0000C |
| 4748239 | 4767624 | -1 | protein_coding | ENSG0000C |
| 4756338 | 4756633 | 1 | misc_RNA | ENSG0000C |
| 4777669 | 4788521 | -1 | protein_coding | ENSG0000C |
| 4777939 | 4801423 | -1 | lncRNA | ENSG0000C |
| 4788397 | 4796491 | 1 | protein_coding | ENSG0000C |
| 4795265 | 4796532 | -1 | lncRNA | ENSG0000C |
| 4796968 | 4802950 | -1 | protein_coding | ENSG0000C |
| 4803203 | 4847288 | -1 | protein_coding | ENSG0000C |
| 4839244 | 4840334 | -1 | lncRNA | ENSG0000C |
| 4846665 | 4882401 | 1 | protein_coding | ENSG0000C |
| 4882507 | 4960741 | -1 | protein_coding | ENSG0000C |
| 4958330 | 5019157 | 1 | protein_coding | ENSG0000C |
| 5010909 | 5043999 | 1 | lncRNA | ENSG0000C |
| 5024844 | 5034141 | -1 | protein_coding | ENSG0000C |
| 5033923 | 5087379 | 1 | protein_coding | ENSG0000C |
| 5044122 | 5066110 | -1 | protein_coding | ENSG0000C |
| 5078168 | 5078605 | -1 | processed_pseudoge | ENSG0000C |
| 5084284 | 5097795 | -1 | protein_coding | ENSG0000C |
| 5098739 | 5142595 | 1 | lncRNA | ENSG0000C |
| 5103339 | 5235822 | 1 | lncRNA | ENSG0000C |
| 5110793 | 5148685 | -1 | unprocessed_pseudc | ENSG0000C |
| 5207088 | 5207439 | 1 | processed_pseudoge | ENSG0000C |
| 5215243 | 5220843 | -1 | lncRNA | ENSG0000C |
| 5239802 | 7713340 | 1 | protein_coding | ENSG0000C |
| 5260686 | 5261043 | -1 | processed_pseudoge | ENSG0000C |
| 5270174 | 5270954 | 1 | processed_pseudoge | ENSG0000C |
| 5289200 | 5289409 | -1 | processed_pseudoge | ENSG0000C |
| 5365797 | 5366688 | 1 | processed_pseudoge | ENSG0000C |
| 5596856 | 5616373 | -1 | lncRNA | ENSG0000C |
| 5632467 | 5632566 | 1 | miRNA | ENSG0000C |
| 5838131 | 5838661 | -1 | processed_pseudoge | ENSG0000C |
| 6037210 | 6038977 | 1 | TEC | ENSG0000C |
| 6056975 | 6092954 | 1 | lncRNA | ENSG0000C |
| 6380476 | 6395578 | 1 | lncRNA | ENSG0000C |
| 6573767 | 6577359 | 1 | lncRNA | ENSG0000C |
| 6703883 | 6705558 | 1 | TEC | ENSG0000C |
| 6721786 | 6721960 | 1 | processed_pseudoge | ENSG0000C |
| 6748908 | 6748969 | 1 | snRNA | ENSG0000C |
| 6873899 | 6874005 | 1 | snRNA | ENSG0000C |
| 7004007 | 7004112 | -1 | snRNA | ENSG0000C |
| 7126293 | 7126639 | -1 | processed_pseudoge | ENSG0000C |

| | | | | |
|---|---|---|---|---|
| 7301605 | 7304638 | -1 | lncRNA | ENSG00000 |
| 7431217 | 7431296 | 1 | snoRNA | ENSG00000 |
| 7614230 | 7614992 | -1 | lncRNA | ENSG00000 |
| 7726841 | 7726951 | -1 | snoRNA | ENSG00000 |
| 7878799 | 8112756 | -1 | lncRNA | ENSG00000 |
| 8276263 | 8295700 | -1 | lncRNA | ENSG00000 |
| 8298492 | 8299772 | 1 | lncRNA | ENSG00000 |
| 8309962 | 8357860 | 1 | lncRNA | ENSG00000 |
| 8369864 | 8376276 | 1 | lncRNA | ENSG00000 |
| 8508069 | 8508585 | 1 | processed_pseudoge | ENSG00000 |
| 8526549 | 8532013 | -1 | lncRNA | ENSG00000 |
| 8537605 | 8590193 | -1 | protein_coding | ENSG00000 |
| 8615855 | 8618033 | -1 | lncRNA | ENSG00000 |
| 8621683 | 8649654 | 1 | protein_coding | ENSG00000 |
| 8666762 | 8667271 | 1 | TEC | ENSG00000 |
| 8674596 | 8784575 | 1 | protein_coding | ENSG00000 |
| 8683254 | 8683551 | 1 | misc_RNA | ENSG00000 |
| 8705456 | 8705517 | -1 | snRNA | ENSG00000 |
| 8731416 | 8731559 | -1 | processed_pseudoge | ENSG00000 |
| 8780384 | 8797642 | -1 | protein_coding | ENSG00000 |
| 8788823 | 8849325 | 1 | protein_coding | ENSG00000 |
| 8847650 | 8848724 | -1 | lncRNA | ENSG00000 |
| 8848105 | 8860456 | 1 | lncRNA | ENSG00000 |
| 8852942 | 8869012 | -1 | protein_coding | ENSG00000 |
| 8853312 | 8854347 | 1 | lncRNA | ENSG00000 |
| 8869251 | 8870032 | 1 | lncRNA | ENSG00000 |
| 8881634 | 8885351 | 1 | protein_coding | ENSG00000 |
| 8892097 | 8964514 | -1 | protein_coding | ENSG00000 |
| 8962706 | 8966990 | 1 | lncRNA | ENSG00000 |
| 9068554 | 9072412 | 1 | lncRNA | ENSG00000 |
| 9091644 | 9121635 | 1 | protein_coding | ENSG00000 |
| 9104848 | 9113181 | 1 | lncRNA | ENSG00000 |
| 9105834 | 9107174 | -1 | lncRNA | ENSG00000 |
| 9156402 | 9156881 | -1 | processed_pseudoge | ENSG00000 |
| 9355588 | 9408093 | 1 | lncRNA | ENSG00000 |
| 9365540 | 9518325 | 1 | lncRNA | ENSG00000 |
| 9441294 | 9444985 | -1 | lncRNA | ENSG00000 |
| 9446010 | 9455505 | -1 | lncRNA | ENSG00000 |
| 9541970 | 9614899 | 1 | lncRNA | ENSG00000 |
| 9564390 | 9564497 | -1 | rRNA_pseudogene | ENSG00000 |
| 9600097 | 9600228 | 1 | rRNA_pseudogene | ENSG00000 |
| 9666885 | 9676843 | -1 | lncRNA | ENSG00000 |
| 9753404 | 10182754 | -1 | protein_coding | ENSG00000 |
| 9794637 | 9805571 | 1 | lncRNA | ENSG00000 |
| 9808856 | 9810726 | 1 | lncRNA | ENSG00000 |
| 10033684 | 10037297 | 1 | lncRNA | ENSG00000 |
| 10111382 | 10112885 | 1 | processed_pseudoge | ENSG00000 |
| 10214784 | 10227581 | -1 | lncRNA | ENSG00000 |
| 10221775 | 10222073 | -1 | misc_RNA | ENSG00000 |
| 10326434 | 10483638 | 1 | protein_coding | ENSG00000 |

| | | | | |
|---|---|---|---|---|
| 10351440 | 10352752 | -1 | lncRNA | ENSG00000 |
| 10445309 | 10449747 | -1 | TEC | ENSG00000 |
| 10496272 | 10496374 | 1 | snRNA | ENSG00000 |
| 10514842 | 10528202 | -1 | lncRNA | ENSG00000 |
| 10528422 | 10580632 | -1 | protein_coding | ENSG00000 |
| 10529440 | 10532082 | 1 | lncRNA | ENSG00000 |
| 10576499 | 10578183 | -1 | lncRNA | ENSG00000 |
| 10627501 | 10694930 | -1 | protein_coding | ENSG00000 |
| 10691273 | 10692973 | 1 | lncRNA | ENSG00000 |
| 10718633 | 10719762 | -1 | unprocessed_pseudo | ENSG00000 |
| 10719833 | 10720015 | 1 | unprocessed_pseudo | ENSG00000 |
| 10720015 | 10721464 | -1 | unprocessed_pseudo | ENSG00000 |
| 10721698 | 10723068 | -1 | unprocessed_pseudo | ENSG00000 |
| 10723065 | 10723355 | -1 | unprocessed_pseudo | ENSG00000 |
| 10723424 | 10723766 | -1 | unprocessed_pseudo | ENSG00000 |
| 10723836 | 10724617 | -1 | unprocessed_pseudo | ENSG00000 |
| 10724620 | 10725296 | -1 | unprocessed_pseudo | ENSG00000 |
| 10743786 | 10769351 | 1 | protein_coding | ENSG00000 |
| 10760919 | 10818794 | -1 | protein_coding | ENSG00000 |
| 10840384 | 10841544 | -1 | processed_pseudoge | ENSG00000 |
| 10864914 | 10888752 | -1 | lncRNA | ENSG00000 |
| 10866222 | 10943021 | 1 | protein_coding | ENSG00000 |
| 10928891 | 10942468 | -1 | protein_coding | ENSG00000 |
| 10933903 | 10936280 | 1 | TEC | ENSG00000 |
| 10944564 | 11182186 | 1 | protein_coding | ENSG00000 |
| 11027024 | 11027763 | 1 | processed_pseudoge | ENSG00000 |
| 11056556 | 11057034 | 1 | lncRNA | ENSG00000 |
| 11066496 | 11071102 | -1 | lncRNA | ENSG00000 |
| 11155298 | 11162818 | -1 | lncRNA | ENSG00000 |
| 11196177 | 11224969 | 1 | lncRNA | ENSG00000 |
| 11242653 | 11243552 | 1 | processed_pseudoge | ENSG00000 |
| 11249619 | 11381662 | 1 | protein_coding | ENSG00000 |
| 11254405 | 11256200 | -1 | protein_coding | ENSG00000 |
| 11267748 | 11269533 | -1 | protein_coding | ENSG00000 |
| 11273199 | 11273629 | -1 | protein_coding | ENSG00000 |
| 11274113 | 11274236 | -1 | snoRNA | ENSG00000 |
| 11275639 | 11276480 | -1 | protein_coding | ENSG00000 |
| 11280841 | 11281330 | -1 | protein_coding | ENSG00000 |
| 11290569 | 11294906 | 1 | lncRNA | ENSG00000 |
| 11306440 | 11306527 | -1 | miRNA | ENSG00000 |
| 11315067 | 11315178 | 1 | misc_RNA | ENSG00000 |
| 11341809 | 11345211 | -1 | lncRNA | ENSG00000 |
| 11348143 | 11349321 | -1 | lncRNA | ENSG00000 |
| 11359845 | 11527245 | -1 | protein_coding | ENSG00000 |
| 11380859 | 11381118 | -1 | processed_pseudoge | ENSG00000 |
| 11465260 | 11473174 | 1 | lncRNA | ENSG00000 |
| 11547722 | 11636381 | -1 | protein_coding | ENSG00000 |
| 11668455 | 11679152 | 1 | protein_coding | ENSG00000 |
| 11679080 | 11742878 | -1 | protein_coding | ENSG00000 |
| 11741910 | 11744506 | 1 | lncRNA | ENSG00000 |

| | | | | |
|---|---|---|---|---|
| 11750586 | 11797258 | -1 | protein_coding | ENSG0000C |
| 11753666 | 11753948 | -1 | misc_RNA | ENSG0000C |
| 11797468 | 11798275 | 1 | lncRNA | ENSG0000C |
| 11819829 | 11828845 | -1 | lncRNA | ENSG0000C |
| 11833850 | 11851580 | -1 | protein_coding | ENSG0000C |
| 11851649 | 11895611 | 1 | lncRNA | ENSG0000C |
| 11868128 | 11916082 | -1 | protein_coding | ENSG0000C |
| 11881075 | 11882569 | -1 | lncRNA | ENSG0000C |
| 11903923 | 11904137 | -1 | processed_pseudoge | ENSG0000C |
| 11908208 | 11908916 | 1 | lncRNA | ENSG0000C |
| 11926199 | 11926307 | 1 | unprocessed_pseudc | ENSG0000C |
| 11927259 | 11976643 | -1 | protein_coding | ENSG0000C |
| 11965210 | 11968068 | 1 | protein_coding | ENSG0000C |
| 11968508 | 11968743 | -1 | processed_pseudoge | ENSG0000C |
| 11976734 | 12574287 | 1 | protein_coding | ENSG0000C |
| 11976851 | 11977850 | -1 | lncRNA | ENSG0000C |
| 12081237 | 12081902 | 1 | processed_pseudoge | ENSG0000C |
| 12086746 | 12090302 | -1 | lncRNA | ENSG0000C |
| 12093627 | 12095307 | -1 | lncRNA | ENSG0000C |
| 12297197 | 12297323 | -1 | snoRNA | ENSG0000C |
| 12366982 | 12372582 | -1 | lncRNA | ENSG0000C |
| 12372812 | 12373899 | 1 | lncRNA | ENSG0000C |
| 12374355 | 12376032 | 1 | TEC | ENSG0000C |
| 12545482 | 12546684 | 1 | lncRNA | ENSG0000C |
| 12556353 | 12557694 | -1 | lncRNA | ENSG0000C |
| 12560756 | 12611044 | -1 | lncRNA | ENSG0000C |
| 12614451 | 12614852 | 1 | lncRNA | ENSG0000C |
| 12659799 | 12803887 | -1 | protein_coding | ENSG0000C |
| 12720321 | 12720371 | 1 | miRNA | ENSG0000C |
| 12745873 | 12757835 | 1 | lncRNA | ENSG0000C |
| 12759282 | 12761162 | 1 | lncRNA | ENSG0000C |
| 12842487 | 12842610 | -1 | snoRNA | ENSG0000C |
| 12881170 | 12881493 | 1 | processed_pseudoge | ENSG0000C |
| 12901598 | 13240416 | 1 | protein_coding | ENSG0000C |
| 13197606 | 13204907 | -1 | lncRNA | ENSG0000C |
| 13246232 | 13563388 | 1 | lncRNA | ENSG0000C |
| 13331368 | 13332583 | -1 | lncRNA | ENSG0000C |
| 13458326 | 13458698 | 1 | processed_pseudoge | ENSG0000C |
| 13728654 | 13779760 | -1 | lncRNA | ENSG0000C |
| 13920157 | 13952345 | 1 | protein_coding | ENSG0000C |
| 13922332 | 13922679 | -1 | processed_pseudoge | ENSG0000C |
| 13930677 | 13935635 | -1 | lncRNA | ENSG0000C |
| 13953155 | 13954825 | 1 | lncRNA | ENSG0000C |
| 14001320 | 14016056 | -1 | lncRNA | ENSG0000C |
| 14018880 | 14021077 | -1 | lncRNA | ENSG0000C |
| 14071321 | 14266773 | 1 | protein_coding | ENSG0000C |
| 14150833 | 14153235 | 1 | lncRNA | ENSG0000C |
| 14191820 | 14200277 | -1 | lncRNA | ENSG0000C |
| 14200357 | 14200979 | 1 | processed_pseudoge | ENSG0000C |
| 14275508 | 14275600 | 1 | misc_RNA | ENSG0000C |

| | | | | |
|---|---|---|---|---|
| 14301389 | 14331067 | 1 | lncRNA | ENSG0000C |
| 14303967 | 14304049 | 1 | miRNA | ENSG0000C |
| 14309285 | 14309371 | 1 | miRNA | ENSG0000C |
| 14363109 | 14370266 | -1 | lncRNA | ENSG0000C |
| 14407668 | 14418908 | -1 | lncRNA | ENSG0000C |
| 14435700 | 14632728 | -1 | protein_coding | ENSG0000C |
| 14601823 | 14602094 | -1 | misc_RNA | ENSG0000C |
| 14632931 | 14669236 | 1 | protein_coding | ENSG0000C |
| 14643788 | 14643849 | 1 | snRNA | ENSG0000C |
| 14672545 | 14694669 | -1 | protein_coding | ENSG0000C |
| 14695567 | 14707055 | -1 | lncRNA | ENSG0000C |
| 14708944 | 14726338 | 1 | protein_coding | ENSG0000C |
| 14727282 | 14733347 | -1 | unprocessed_pseudc | ENSG0000C |
| 14734685 | 14746177 | -1 | lncRNA | ENSG0000C |
| 14748066 | 14765413 | 1 | protein_coding | ENSG0000C |
| 14766402 | 14770075 | -1 | unprocessed_pseudc | ENSG0000C |
| 14820792 | 14824702 | -1 | transcribed_unproce | ENSG0000C |
| 14833721 | 14896157 | 1 | protein_coding | ENSG0000C |
| 14862380 | 14862749 | -1 | processed_pseudoge | ENSG0000C |
| 14901499 | 14902174 | 1 | lncRNA | ENSG0000C |
| 14901508 | 14901591 | 1 | miRNA | ENSG0000C |
| 14907717 | 14907781 | 1 | miRNA | ENSG0000C |
| 14909887 | 14911345 | -1 | lncRNA | ENSG0000C |
| 14911220 | 14911313 | 1 | miRNA | ENSG0000C |
| 14911551 | 14935708 | 1 | transcribed_unproce | ENSG0000C |
| 14915457 | 14915556 | -1 | misc_RNA | ENSG0000C |
| 14922802 | 14952060 | 1 | protein_coding | ENSG0000C |
| 14925937 | 14926003 | 1 | miRNA | ENSG0000C |
| 14930820 | 14930879 | 1 | miRNA | ENSG0000C |
| 14953012 | 14953120 | -1 | unprocessed_pseudc | ENSG0000C |
| 14974591 | 15139339 | 1 | protein_coding | ENSG0000C |
| 14988259 | 14990160 | -1 | processed_pseudoge | ENSG0000C |
| 15010321 | 15010397 | -1 | miRNA | ENSG0000C |
| 15015828 | 15016390 | -1 | lncRNA | ENSG0000C |
| 15037854 | 15056079 | -1 | protein_coding | ENSG0000C |
| 15060022 | 15094317 | -1 | protein_coding | ENSG0000C |
| 15094411 | 15109197 | 1 | transcribed_processe | ENSG0000C |
| 15104312 | 15123498 | -1 | transcribed_unproce | ENSG0000C |
| 15104723 | 15131601 | -1 | lncRNA | ENSG0000C |
| 15125242 | 15154564 | -1 | transcribed_unproce | ENSG0000C |
| 15134075 | 15134145 | -1 | miRNA | ENSG0000C |
| 15150697 | 15150796 | 1 | misc_RNA | ENSG0000C |
| 15154850 | 15155002 | -1 | miRNA | ENSG0000C |
| 15154903 | 15157020 | 1 | lncRNA | ENSG0000C |
| 15358978 | 15362674 | 1 | unprocessed_pseudc | ENSG0000C |
| 15363628 | 15381047 | -1 | protein_coding | ENSG0000C |
| 15395754 | 15413271 | 1 | protein_coding | ENSG0000C |
| 15395754 | 15515348 | 1 | protein_coding | ENSG0000C |
| 15434475 | 15625028 | 1 | protein_coding | ENSG0000C |
| 15594387 | 15643154 | -1 | protein_coding | ENSG0000C |

| | | | | |
|---|---|---|---|---|
| 15608474 | 15610563 | 1 | lncRNA | ENSG00000 |
| 15611030 | 15611095 | -1 | miRNA | ENSG00000 |
| 15643267 | 15726353 | 1 | protein_coding | ENSG00000 |
| 15643294 | 15643372 | 1 | miRNA | ENSG00000 |
| 15683290 | 15684570 | 1 | lncRNA | ENSG00000 |
| 15701237 | 15702118 | 1 | lncRNA | ENSG00000 |
| 15703135 | 15857028 | -1 | protein_coding | ENSG00000 |
| 15726674 | 15732993 | 1 | lncRNA | ENSG00000 |
| 15741151 | 15741791 | 1 | lncRNA | ENSG00000 |
| 15790975 | 15792717 | -1 | TEC | ENSG00000 |
| 15865719 | 15888625 | -1 | protein_coding | ENSG00000 |
| 15877163 | 15877266 | -1 | snRNA | ENSG00000 |
| 15885029 | 15886158 | 1 | lncRNA | ENSG00000 |
| 15931697 | 15932294 | -1 | processed_pseudoge | ENSG00000 |
| 15949577 | 16143074 | 1 | protein_coding | ENSG00000 |
| 16094192 | 16095209 | -1 | processed_pseudoge | ENSG00000 |
| 16148928 | 16223522 | -1 | protein_coding | ENSG00000 |
| 16223027 | 16224261 | 1 | lncRNA | ENSG00000 |
| 16232528 | 16294811 | 1 | protein_coding | ENSG00000 |
| 16260866 | 16261235 | -1 | processed_pseudoge | ENSG00000 |
| 16290135 | 16292242 | -1 | lncRNA | ENSG00000 |
| 16300159 | 16300242 | 1 | miRNA | ENSG00000 |
| 16306370 | 16306434 | 1 | miRNA | ENSG00000 |
| 16308542 | 16310000 | -1 | lncRNA | ENSG00000 |
| 16309879 | 16309966 | 1 | miRNA | ENSG00000 |
| 16310341 | 16334190 | 1 | unprocessed_pseudo | ENSG00000 |
| 16314112 | 16314211 | -1 | misc_RNA | ENSG00000 |
| 16317444 | 16350590 | 1 | protein_coding | ENSG00000 |
| 16324588 | 16324654 | 1 | miRNA | ENSG00000 |
| 16329305 | 16329364 | 1 | miRNA | ENSG00000 |
| 16351545 | 16351653 | -1 | unprocessed_pseudo | ENSG00000 |
| 16356224 | 16377507 | 1 | unprocessed_pseudo | ENSG00000 |
| 16368876 | 16368942 | 1 | miRNA | ENSG00000 |
| 16379055 | 16393954 | 1 | protein_coding | ENSG00000 |
| 16394912 | 16398594 | -1 | unprocessed_pseudo | ENSG00000 |
| 16499533 | 16664135 | 1 | lncRNA | ENSG00000 |
| 16541217 | 16541847 | -1 | processed_pseudoge | ENSG00000 |
| 16612378 | 16633991 | 1 | lncRNA | ENSG00000 |
| 16760905 | 16762465 | 1 | TEC | ENSG00000 |
| 16952502 | 16953468 | -1 | processed_pseudoge | ENSG00000 |
| 17082295 | 17085049 | -1 | TEC | ENSG00000 |
| 17094351 | 17094927 | -1 | unprocessed_pseudo | ENSG00000 |
| 17096202 | 17096421 | -1 | processed_pseudoge | ENSG00000 |
| 17101769 | 17470960 | -1 | protein_coding | ENSG00000 |
| 17134504 | 17138736 | 1 | lncRNA | ENSG00000 |
| 17192863 | 17194232 | -1 | TEC | ENSG00000 |
| 17227023 | 17228000 | -1 | TEC | ENSG00000 |
| 17249674 | 17251395 | -1 | TEC | ENSG00000 |
| 17405678 | 17408073 | 1 | lncRNA | ENSG00000 |
| 17445825 | 17446380 | -1 | lncRNA | ENSG00000 |

| | | | | |
|---|---|---|---|---|
| 17453470 | 17456236 | 1 | lncRNA | ENSG00000 |
| 17540331 | 17540985 | -1 | lncRNA | ENSG00000 |
| 17825252 | 17826906 | 1 | lncRNA | ENSG00000 |
| 17932720 | 17933275 | 1 | processed_pseudoge | ENSG00000 |
| 17933189 | 18151595 | -1 | lncRNA | ENSG00000 |
| 17961597 | 17962370 | 1 | TEC | ENSG00000 |
| 18002806 | 18173063 | -1 | lncRNA | ENSG00000 |
| 18170356 | 18170959 | 1 | processed_pseudoge | ENSG00000 |
| 18201501 | 18201956 | 1 | TEC | ENSG00000 |
| 18313294 | 18316984 | 1 | unprocessed_pseudo | ENSG00000 |
| 18317942 | 18336736 | -1 | protein_coding | ENSG00000 |
| 18318008 | 18344283 | -1 | lncRNA | ENSG00000 |
| 18334400 | 18352476 | -1 | transcribed_unproce | ENSG00000 |
| 18344013 | 18344079 | -1 | miRNA | ENSG00000 |
| 18357026 | 18357136 | 1 | unprocessed_pseudo | ENSG00000 |
| 18358086 | 18379331 | -1 | protein_coding | ENSG00000 |
| 18374521 | 18401940 | -1 | transcribed_unproce | ENSG00000 |
| 18379351 | 18379410 | -1 | miRNA | ENSG00000 |
| 18397937 | 18398036 | 1 | misc_RNA | ENSG00000 |
| 18402146 | 18403604 | 1 | lncRNA | ENSG00000 |
| 18402178 | 18402271 | -1 | miRNA | ENSG00000 |
| 18405698 | 18405762 | -1 | miRNA | ENSG00000 |
| 18411309 | 18411851 | -1 | lncRNA | ENSG00000 |
| 18411894 | 18411977 | -1 | miRNA | ENSG00000 |
| 18417325 | 18562211 | -1 | protein_coding | ENSG00000 |
| 18488301 | 18488365 | -1 | miRNA | ENSG00000 |
| 18494493 | 18494576 | -1 | miRNA | ENSG00000 |
| 18533418 | 18533817 | 1 | processed_pseudoge | ENSG00000 |
| 18570448 | 18571683 | -1 | lncRNA | ENSG00000 |
| 18571162 | 18598328 | 1 | transcribed_unproce | ENSG00000 |
| 18781295 | 18790383 | -1 | protein_coding | ENSG00000 |
| 18788063 | 18801519 | -1 | protein_coding | ENSG00000 |
| 18791669 | 18801572 | -1 | protein_coding | ENSG00000 |
| 18803083 | 18812181 | 1 | lncRNA | ENSG00000 |
| 18804853 | 18926454 | -1 | protein_coding | ENSG00000 |
| 18926863 | 18937043 | 1 | lncRNA | ENSG00000 |
| 18983934 | 19063942 | 1 | protein_coding | ENSG00000 |
| 19012157 | 19012263 | 1 | snRNA | ENSG00000 |
| 19062144 | 19067691 | -1 | lncRNA | ENSG00000 |
| 19065991 | 19066694 | 1 | lncRNA | ENSG00000 |
| 19067614 | 19080095 | 1 | protein_coding | ENSG00000 |
| 19086831 | 19098332 | 1 | lncRNA | ENSG00000 |
| 19111035 | 19111484 | -1 | lncRNA | ENSG00000 |
| 19113932 | 19121629 | 1 | protein_coding | ENSG00000 |
| 19114002 | 19173578 | 1 | lncRNA | ENSG00000 |
| 19119976 | 19121629 | -1 | lncRNA | ENSG00000 |
| 19167971 | 19268332 | 1 | protein_coding | ENSG00000 |
| 19180748 | 19184060 | -1 | lncRNA | ENSG00000 |
| 19285731 | 19322145 | 1 | protein_coding | ENSG00000 |
| 19315164 | 19316526 | 1 | lncRNA | ENSG00000 |

| | | | | |
|---|---|---|---|---|
| 19334011 | 19337804 | 1 | lncRNA | ENSG0000C |
| 19343647 | 19401693 | -1 | lncRNA | ENSG0000C |
| 19392213 | 19393468 | -1 | TEC | ENSG0000C |
| 19410496 | 19499113 | 1 | protein_coding | ENSG0000C |
| 19410729 | 19411662 | -1 | lncRNA | ENSG0000C |
| 19476916 | 19487899 | -1 | lncRNA | ENSG0000C |
| 19498610 | 19498747 | 1 | snRNA | ENSG0000C |
| 19501689 | 19502286 | 1 | lncRNA | ENSG0000C |
| 19501693 | 19522123 | -1 | protein_coding | ENSG0000C |
| 19503861 | 19504625 | 1 | TEC | ENSG0000C |
| 19523811 | 19553408 | 1 | protein_coding | ENSG0000C |
| 19555240 | 19706793 | 1 | protein_coding | ENSG0000C |
| 19694035 | 19694428 | -1 | lncRNA | ENSG0000C |
| 19701937 | 19718235 | -1 | protein_coding | ENSG0000C |
| 19706351 | 19715383 | 1 | lncRNA | ENSG0000C |
| 19716456 | 19858467 | 1 | protein_coding | ENSG0000C |
| 19761172 | 19766099 | -1 | lncRNA | ENSG0000C |
| 19819504 | 19819628 | -1 | processed_pseudoge | ENSG0000C |
| 19856691 | 19886167 | -1 | protein_coding | ENSG0000C |
| 20031485 | 20073917 | -1 | protein_coding | ENSG0000C |
| 20155648 | 20158391 | 1 | processed_pseudoge | ENSG0000C |
| 20208925 | 20209525 | 1 | lncRNA | ENSG0000C |
| 20238972 | 20239247 | -1 | processed_pseudoge | ENSG0000C |
| 20309572 | 20327808 | -1 | protein_coding | ENSG0000C |
| 20333052 | 20356301 | -1 | protein_coding | ENSG0000C |
| 20359175 | 20404737 | -1 | protein_coding | ENSG0000C |
| 20385957 | 20391164 | 1 | lncRNA | ENSG0000C |
| 20409534 | 20441336 | 1 | protein_coding | ENSG0000C |
| 20440266 | 20447000 | -1 | lncRNA | ENSG0000C |
| 20451461 | 20487669 | 1 | protein_coding | ENSG0000C |
| 20487702 | 20489659 | 1 | TEC | ENSG0000C |
| 20536226 | 20576427 | -1 | protein_coding | ENSG0000C |
| 20580999 | 20586640 | 1 | lncRNA | ENSG0000C |
| 20586550 | 20607107 | -1 | unprocessed_pseudc | ENSG0000C |
| 20610243 | 20797581 | 1 | protein_coding | ENSG0000C |
| 20616409 | 20616532 | -1 | unprocessed_pseudc | ENSG0000C |
| 20618205 | 20620758 | -1 | unprocessed_pseudc | ENSG0000C |
| 20623237 | 20698890 | -1 | protein_coding | ENSG0000C |
| 20702816 | 20742084 | -1 | protein_coding | ENSG0000C |
| 20711473 | 20732990 | 1 | unprocessed_pseudc | ENSG0000C |
| 20720994 | 20722281 | -1 | transcribed_unproce | ENSG0000C |
| 20743663 | 20766620 | -1 | lncRNA | ENSG0000C |
| 20780193 | 20900349 | -1 | protein_coding | ENSG0000C |
| 20784509 | 20784615 | 1 | snRNA | ENSG0000C |
| 20806429 | 20849668 | 1 | protein_coding | ENSG0000C |
| 20823443 | 20823537 | 1 | snRNA | ENSG0000C |
| 20854925 | 20900358 | -1 | protein_coding | ENSG0000C |
| 20899868 | 20925006 | 1 | protein_coding | ENSG0000C |
| 20933111 | 21159441 | -1 | protein_coding | ENSG0000C |
| 21120758 | 21130108 | 1 | lncRNA | ENSG0000C |

| | | | | |
|---|---|---|---|---|
| 21158377 | 21180616 | 1 | protein_coding | ENSG0000C |
| 21194847 | 21205977 | 1 | lncRNA | ENSG0000C |
| 21197450 | 21214510 | -1 | protein_coding | ENSG0000C |
| 21233699 | 21253850 | 1 | protein_coding | ENSG0000C |
| 21238874 | 21303083 | -1 | protein_coding | ENSG0000C |
| 21299903 | 21300506 | 1 | unitary_pseudogene | ENSG0000C |
| 21300849 | 21339037 | 1 | transcribed_unitary_ | ENSG0000C |
| 21347139 | 21352138 | 1 | unprocessed_pseudc | ENSG0000C |
| 21376648 | 21386064 | 1 | unprocessed_pseudc | ENSG0000C |
| 21401089 | 21401198 | 1 | unprocessed_pseudc | ENSG0000C |
| 21402237 | 21448567 | -1 | protein_coding | ENSG0000C |
| 21446683 | 21520444 | -1 | transcribed_unproce | ENSG0000C |
| 21506049 | 21506135 | -1 | miRNA | ENSG0000C |
| 21511635 | 21515131 | 1 | lncRNA | ENSG0000C |
| 21519830 | 21520365 | -1 | transcribed_processe | ENSG0000C |
| 21526824 | 21531141 | -1 | lncRNA | ENSG0000C |
| 21532945 | 21550373 | -1 | unprocessed_pseudc | ENSG0000C |
| 21538487 | 21538794 | 1 | processed_pseudoge | ENSG0000C |
| 21587627 | 21587864 | 1 | scaRNA | ENSG0000C |
| 21597218 | 21657473 | 1 | protein_coding | ENSG0000C |
| 21626742 | 21627569 | 1 | lncRNA | ENSG0000C |
| 21639550 | 21652608 | -1 | protein_coding | ENSG0000C |
| 21642054 | 21642160 | 1 | snRNA | ENSG0000C |
| 21642294 | 21642400 | 1 | snRNA | ENSG0000C |
| 21663971 | 21761935 | 1 | protein_coding | ENSG0000C |
| 21794095 | 21795759 | -1 | lncRNA | ENSG0000C |
| 21796106 | 21820410 | -1 | transcribed_unproce | ENSG0000C |
| 21833516 | 21833621 | 1 | unprocessed_pseudc | ENSG0000C |
| 21834569 | 21880827 | -1 | protein_coding | ENSG0000C |
| 21879338 | 21919156 | -1 | transcribed_unproce | ENSG0000C |
| 21938865 | 21944987 | 1 | transcribed_unproce | ENSG0000C |
| 21950218 | 21951708 | -1 | lncRNA | ENSG0000C |
| 21953288 | 21983660 | 1 | protein_coding | ENSG0000C |
| 21983865 | 22001110 | -1 | protein_coding | ENSG0000C |
| 21998310 | 21999656 | 1 | processed_pseudoge | ENSG0000C |
| 22007480 | 22008062 | -1 | lncRNA | ENSG0000C |
| 22007638 | 22087534 | 1 | protein_coding | ENSG0000C |
| 22083256 | 22092485 | -1 | lncRNA | ENSG0000C |
| 22092538 | 22156964 | 1 | protein_coding | ENSG0000C |
| 22165583 | 22191754 | 1 | protein_coding | ENSG0000C |
| 22170309 | 22175524 | -1 | lncRNA | ENSG0000C |
| 22206278 | 22288738 | 1 | protein_coding | ENSG0000C |
| 22286708 | 22288738 | -1 | lncRNA | ENSG0000C |
| 22297375 | 22335101 | 1 | protein_coding | ENSG0000C |
| 22331039 | 22342938 | -1 | lncRNA | ENSG0000C |
| 22345936 | 22437165 | -1 | protein_coding | ENSG0000C |
| 22365121 | 22369047 | 1 | lncRNA | ENSG0000C |
| 22374859 | 22378180 | 1 | lncRNA | ENSG0000C |
| 22382436 | 22407428 | 1 | unitary_pseudogene | ENSG0000C |
| 22418672 | 22437715 | -1 | transcribed_unproce | ENSG0000C |

| | | | | |
|---|---|---|---|---|
| 22437008 | 22492220 | 1 | transcribed_unproce | ENSG0000C |
| 22479121 | 22536521 | 1 | protein_coding | ENSG0000C |
| 22537478 | 22537584 | -1 | unprocessed_pseudc | ENSG0000C |
| 22545698 | 22576865 | 1 | transcribed_unproce | ENSG0000C |
| 22610531 | 22612196 | -1 | lncRNA | ENSG0000C |
| 22612543 | 22613483 | -1 | lncRNA | ENSG0000C |
| 22806289 | 22813680 | -1 | lncRNA | ENSG0000C |
| 22814162 | 22916338 | 1 | protein_coding | ENSG0000C |
| 23017723 | 23026285 | 1 | lncRNA | ENSG0000C |
| 23061406 | 23064173 | -1 | lncRNA | ENSG0000C |
| 23061406 | 23149270 | -1 | protein_coding | ENSG0000C |
| 23061767 | 23062232 | 1 | TEC | ENSG0000C |
| 23167577 | 23186395 | -1 | lncRNA | ENSG0000C |
| 23182745 | 23216883 | 1 | protein_coding | ENSG0000C |
| 23194878 | 23213104 | -1 | lncRNA | ENSG0000C |
| 23278231 | 23381294 | 1 | protein_coding | ENSG0000C |
| 23388493 | 23453189 | -1 | protein_coding | ENSG0000C |
| 23412380 | 23416680 | 1 | TEC | ENSG0000C |
| 23441841 | 23441921 | 1 | snoRNA | ENSG0000C |
| 23446446 | 23447282 | -1 | lncRNA | ENSG0000C |
| 23452758 | 23457606 | 1 | lncRNA | ENSG0000C |
| 23463542 | 23521995 | -1 | protein_coding | ENSG0000C |
| 23500905 | 23501630 | 1 | processed_pseudoge | ENSG0000C |
| 23522014 | 23557731 | -1 | protein_coding | ENSG0000C |
| 23537061 | 23537556 | 1 | processed_pseudoge | ENSG0000C |
| 23557721 | 23574389 | 1 | protein_coding | ENSG0000C |
| 23568673 | 23569696 | 1 | lncRNA | ENSG0000C |
| 23581014 | 23596316 | -1 | protein_coding | ENSG0000C |
| 23603160 | 23641310 | -1 | protein_coding | ENSG0000C |
| 23605841 | 23624107 | 1 | lncRNA | ENSG0000C |
| 23641466 | 23677472 | 1 | protein_coding | ENSG0000C |
| 23670011 | 23675499 | 1 | lncRNA | ENSG0000C |
| 23677656 | 23690367 | 1 | protein_coding | ENSG0000C |
| 23687049 | 23687689 | -1 | lncRNA | ENSG0000C |
| 23690310 | 23713226 | -1 | protein_coding | ENSG0000C |
| 23696665 | 23697314 | -1 | TEC | ENSG0000C |
| 23711990 | 23712793 | 1 | lncRNA | ENSG0000C |
| 23753689 | 23755939 | -1 | TEC | ENSG0000C |
| 23755026 | 23758935 | 1 | protein_coding | ENSG0000C |
| 23835983 | 24220611 | 1 | protein_coding | ENSG0000C |
| 23962951 | 23963351 | 1 | TEC | ENSG0000C |
| 24098239 | 24099971 | 1 | TEC | ENSG0000C |
| 24203116 | 24203231 | 1 | miRNA | ENSG0000C |
| 24208105 | 24208478 | 1 | TEC | ENSG0000C |
| 24236104 | 24252863 | 1 | lncRNA | ENSG0000C |
| 24255553 | 24362801 | 1 | protein_coding | ENSG0000C |
| 24333415 | 24333547 | -1 | snoRNA | ENSG0000C |
| 24400056 | 24400117 | -1 | snRNA | ENSG0000C |
| 24472798 | 24495177 | -1 | lncRNA | ENSG0000C |
| 24537693 | 24572863 | 1 | protein_coding | ENSG0000C |

| | | | | |
|---|---|---|---|---|
| 24610209 | 24827632 | 1 | protein_coding | ENSG0000C |
| 24661422 | 24671062 | -1 | lncRNA | ENSG0000C |
| 24803451 | 24819739 | -1 | lncRNA | ENSG0000C |
| 24845841 | 24911628 | 1 | protein_coding | ENSG0000C |
| 24919389 | 25015666 | -1 | protein_coding | ENSG0000C |
| 25015978 | 25021023 | 1 | processed_pseudoge | ENSG0000C |
| 25031744 | 25058109 | 1 | transcribed_unproce | ENSG0000C |
| 25066937 | 25088618 | 1 | lncRNA | ENSG0000C |
| 25069570 | 25070109 | 1 | processed_pseudoge | ENSG0000C |
| 25071490 | 25072727 | -1 | unprocessed_pseudc | ENSG0000C |
| 25085168 | 25111555 | -1 | lncRNA | ENSG0000C |
| 25093387 | 25095255 | 1 | processed_pseudoge | ENSG0000C |
| 25106569 | 25107102 | -1 | lncRNA | ENSG0000C |
| 25111731 | 25178231 | 1 | protein_coding | ENSG0000C |
| 25140577 | 25149032 | -1 | lncRNA | ENSG0000C |
| 25185253 | 25185538 | -1 | misc_RNA | ENSG0000C |
| 25215731 | 25228932 | 1 | protein_coding | ENSG0000C |
| 25236001 | 25257845 | -1 | protein_coding | ENSG0000C |
| 25238318 | 25239287 | 1 | lncRNA | ENSG0000C |
| 25257952 | 25261066 | 1 | lncRNA | ENSG0000C |
| 25419812 | 25433686 | -1 | lncRNA | ENSG0000C |
| 25454339 | 25454647 | -1 | processed_pseudoge | ENSG0000C |
| 25691959 | 26137685 | 1 | protein_coding | ENSG0000C |
| 25787987 | 26042347 | 1 | lncRNA | ENSG0000C |
| 25824710 | 25826330 | -1 | processed_pseudoge | ENSG0000C |
| 26025237 | 26025310 | 1 | miRNA | ENSG0000C |
| 26028912 | 26029025 | 1 | rRNA_pseudogene | ENSG0000C |
| 26032539 | 26032811 | -1 | processed_pseudoge | ENSG0000C |
| 26062750 | 26063097 | -1 | processed_pseudoge | ENSG0000C |
| 26302064 | 26340815 | 1 | lncRNA | ENSG0000C |
| 26354796 | 26358355 | -1 | unprocessed_pseudc | ENSG0000C |
| 26546802 | 26607396 | 1 | lncRNA | ENSG0000C |
| 26584755 | 26594813 | -1 | lncRNA | ENSG0000C |
| 26721874 | 26729126 | 1 | lncRNA | ENSG0000C |
| 27066707 | 27069165 | 1 | protein_coding | ENSG0000C |
| 27066928 | 27067858 | -1 | lncRNA | ENSG0000C |
| 27133483 | 27135179 | -1 | processed_pseudoge | ENSG0000C |
| 27158451 | 27176587 | -1 | lncRNA | ENSG0000C |
| 27203495 | 27221768 | 1 | protein_coding | ENSG0000C |
| 27213308 | 27214993 | -1 | lncRNA | ENSG0000C |
| 27224994 | 27268772 | -1 | protein_coding | ENSG0000C |
| 27268205 | 27290492 | 1 | lncRNA | ENSG0000C |
| 27313387 | 27314101 | -1 | lncRNA | ENSG0000C |
| 27313668 | 27364778 | 1 | protein_coding | ENSG0000C |
| 27402174 | 27452042 | 1 | protein_coding | ENSG0000C |
| 27447669 | 27453393 | -1 | lncRNA | ENSG0000C |
| 27459555 | 27549913 | -1 | protein_coding | ENSG0000C |
| 27550133 | 27780369 | 1 | protein_coding | ENSG0000C |
| 27643199 | 27644663 | 1 | lncRNA | ENSG0000C |
| 27678940 | 27718806 | -1 | lncRNA | ENSG0000C |

| | | | | |
|---|---|---|---|---|
| 27687182 | 27687522 | 1 | lncRNA | ENSG0000C |
| 27722284 | 27722392 | -1 | misc_RNA | ENSG0000C |
| 27787528 | 28063714 | -1 | protein_coding | ENSG0000C |
| 27863251 | 27863356 | 1 | snRNA | ENSG0000C |
| 27998295 | 27998401 | -1 | snRNA | ENSG0000C |
| 28097979 | 28211920 | -1 | protein_coding | ENSG0000C |
| 28111173 | 28111647 | -1 | processed_pseudoge | ENSG0000C |
| 28179098 | 28179223 | 1 | snoRNA | ENSG0000C |
| 28183318 | 28183430 | 1 | misc_RNA | ENSG0000C |
| 28239693 | 28240668 | -1 | processed_pseudoge | ENSG0000C |
| 28258686 | 28292173 | 1 | lncRNA | ENSG0000C |
| 28259246 | 28323849 | 1 | protein_coding | ENSG0000C |
| 28284885 | 28292064 | -1 | lncRNA | ENSG0000C |
| 28341437 | 28341543 | 1 | unprocessed_pseudc | ENSG0000C |
| 28342555 | 28363508 | -1 | protein_coding | ENSG0000C |
| 28364700 | 28365333 | 1 | processed_pseudoge | ENSG0000C |
| 28379579 | 28403879 | -1 | protein_coding | ENSG0000C |
| 28390982 | 28391051 | -1 | miRNA | ENSG0000C |
| 28413703 | 28415018 | 1 | processed_pseudoge | ENSG0000C |
| 28451644 | 28455356 | 1 | unprocessed_pseudc | ENSG0000C |
| 28454141 | 28454511 | -1 | lncRNA | ENSG0000C |
| 28456371 | 28492098 | -1 | protein_coding | ENSG0000C |
| 28456372 | 28471175 | -1 | protein_coding | ENSG0000C |
| 28474111 | 28495575 | -1 | protein_coding | ENSG0000C |
| 28494643 | 28498970 | 1 | protein_coding | ENSG0000C |
| 28499362 | 28512051 | -1 | protein_coding | ENSG0000C |
| 28532708 | 28539174 | -1 | protein_coding | ENSG0000C |
| 28548331 | 28548658 | -1 | processed_pseudoge | ENSG0000C |
| 28553709 | 28554140 | -1 | lncRNA | ENSG0000C |
| 28553915 | 28591790 | 1 | protein_coding | ENSG0000C |
| 28591943 | 28597050 | -1 | protein_coding | ENSG0000C |
| 28599241 | 28601881 | -1 | unprocessed_pseudc | ENSG0000C |
| 28602340 | 28604519 | 1 | unprocessed_pseudc | ENSG0000C |
| 28605196 | 28623625 | -1 | protein_coding | ENSG0000C |
| 28612131 | 28614884 | -1 | unprocessed_pseudc | ENSG0000C |
| 28637654 | 28658682 | 1 | protein_coding | ENSG0000C |
| 28659696 | 28740781 | -1 | unprocessed_pseudc | ENSG0000C |
| 28688558 | 28735730 | 1 | protein_coding | ENSG0000C |
| 28700294 | 28701540 | -1 | processed_pseudoge | ENSG0000C |
| 28724252 | 28724321 | 1 | miRNA | ENSG0000C |
| 28749959 | 28750595 | -1 | processed_pseudoge | ENSG0000C |
| 28751787 | 28772807 | 1 | protein_coding | ENSG0000C |
| 28773824 | 28777534 | -1 | unprocessed_pseudc | ENSG0000C |
| 28802743 | 28817828 | 1 | lncRNA | ENSG0000C |
| 28813980 | 28814673 | -1 | processed_pseudoge | ENSG0000C |
| 28820570 | 28822033 | -1 | lncRNA | ENSG0000C |
| 28822431 | 28823969 | -1 | lncRNA | ENSG0000C |
| 28823035 | 28837237 | 1 | protein_coding | ENSG0000C |
| 28830612 | 28837200 | -1 | lncRNA | ENSG0000C |
| 28842411 | 28846348 | -1 | protein_coding | ENSG0000C |

| | | | | |
|---|---|---|---|---|
| 28843919 | 28844007 | -1 | miRNA | ENSG00000 |
| 28846600 | 28874212 | 1 | protein_coding | ENSG00000 |
| 28862166 | 28863340 | -1 | lncRNA | ENSG00000 |
| 28878405 | 28904509 | 1 | protein_coding | ENSG00000 |
| 28878957 | 28879920 | -1 | lncRNA | ENSG00000 |
| 28881480 | 28881607 | -1 | snoRNA | ENSG00000 |
| 28904421 | 28936526 | -1 | protein_coding | ENSG00000 |
| 28931939 | 28939347 | 1 | protein_coding | ENSG00000 |
| 28950807 | 28967097 | 1 | protein_coding | ENSG00000 |
| 28952816 | 28966883 | -1 | lncRNA | ENSG00000 |
| 28958583 | 28958661 | 1 | miRNA | ENSG00000 |
| 28973962 | 28978824 | -1 | lncRNA | ENSG00000 |
| 28974221 | 28984548 | 1 | protein_coding | ENSG00000 |
| 28974804 | 28990775 | 1 | lncRNA | ENSG00000 |
| 28984826 | 28990783 | 1 | protein_coding | ENSG00000 |
| 28989140 | 28990778 | -1 | lncRNA | ENSG00000 |
| 29038655 | 29052726 | 1 | unprocessed_pseudo | ENSG00000 |
| 29053732 | 29108108 | -1 | transcribed_unproce | ENSG00000 |
| 29074976 | 29116718 | 1 | lncRNA | ENSG00000 |
| 29075022 | 29104162 | 1 | transcribed_unproce | ENSG00000 |
| 29113050 | 29217860 | 1 | transcribed_unproce | ENSG00000 |
| 29139379 | 29331304 | 1 | lncRNA | ENSG00000 |
| 29139661 | 29216706 | 1 | lncRNA | ENSG00000 |
| 29204633 | 29205855 | 1 | TEC | ENSG00000 |
| 29215385 | 29221040 | -1 | lncRNA | ENSG00000 |
| 29225594 | 29226348 | -1 | lncRNA | ENSG00000 |
| 29262273 | 29264479 | -1 | lncRNA | ENSG00000 |
| 29272220 | 29272772 | 1 | lncRNA | ENSG00000 |
| 29291220 | 29372110 | 1 | lncRNA | ENSG00000 |
| 29312350 | 29331277 | 1 | transcribed_unproce | ENSG00000 |
| 29350746 | 29370272 | 1 | transcribed_unproce | ENSG00000 |
| 29380242 | 29380345 | 1 | unprocessed_pseudo | ENSG00000 |
| 29381354 | 29404029 | -1 | protein_coding | ENSG00000 |
| 29425800 | 29447026 | -1 | unprocessed_pseudo | ENSG00000 |
| 29443056 | 29454964 | -1 | protein_coding | ENSG00000 |
| 29443230 | 29454651 | -1 | protein_coding | ENSG00000 |
| 29449191 | 29449625 | 1 | unprocessed_pseudo | ENSG00000 |
| 29454501 | 29458219 | 1 | protein_coding | ENSG00000 |
| 29455105 | 29464963 | 1 | lncRNA | ENSG00000 |
| 29459889 | 29464976 | 1 | protein_coding | ENSG00000 |
| 29464967 | 29467833 | -1 | transcribed_unproce | ENSG00000 |
| 29466069 | 29467578 | 1 | transcribed_unproce | ENSG00000 |
| 29475064 | 29475355 | 1 | processed_pseudoge | ENSG00000 |
| 29478879 | 29482591 | 1 | unprocessed_pseudo | ENSG00000 |
| 29483642 | 29505999 | -1 | protein_coding | ENSG00000 |
| 29527568 | 29594966 | -1 | unprocessed_pseudo | ENSG00000 |
| 29595130 | 29596137 | -1 | TEC | ENSG00000 |
| 29599179 | 29599265 | -1 | miRNA | ENSG00000 |
| 29613104 | 29613640 | -1 | unprocessed_pseudo | ENSG00000 |
| 29618785 | 29636328 | -1 | unprocessed_pseudo | ENSG00000 |

| | | | | |
|---|---|---|---|---|
| 29624517 | 29624825 | 1 | processed_pseudoge | ENSG00000 |
| 29662979 | 29670876 | 1 | protein_coding | ENSG00000 |
| 29663279 | 29698699 | 1 | protein_coding | ENSG00000 |
| 29731051 | 29731404 | -1 | misc_RNA | ENSG00000 |
| 29742463 | 29745990 | -1 | protein_coding | ENSG00000 |
| 29745247 | 29748299 | 1 | lncRNA | ENSG00000 |
| 29778256 | 29782973 | 1 | protein_coding | ENSG00000 |
| 29790719 | 29805385 | 1 | protein_coding | ENSG00000 |
| 29804430 | 29804990 | -1 | lncRNA | ENSG00000 |
| 29806106 | 29811164 | 1 | protein_coding | ENSG00000 |
| 29806496 | 29807732 | -1 | lncRNA | ENSG00000 |
| 29808636 | 29821252 | -1 | lncRNA | ENSG00000 |
| 29808679 | 29812227 | 1 | lncRNA | ENSG00000 |
| 29811382 | 29815892 | 1 | protein_coding | ENSG00000 |
| 29812261 | 29820092 | 1 | protein_coding | ENSG00000 |
| 29816152 | 29822489 | 1 | protein_coding | ENSG00000 |
| 29817239 | 29830627 | 1 | protein_coding | ENSG00000 |
| 29820394 | 29848039 | 1 | protein_coding | ENSG00000 |
| 29858357 | 29863414 | -1 | protein_coding | ENSG00000 |
| 29862760 | 29863417 | 1 | lncRNA | ENSG00000 |
| 29863593 | 29868053 | 1 | transcribed_unitary_ | ENSG00000 |
| 29871159 | 29899547 | -1 | protein_coding | ENSG00000 |
| 29900375 | 29919864 | 1 | protein_coding | ENSG00000 |
| 29905012 | 29926236 | -1 | protein_coding | ENSG00000 |
| 29913144 | 29915337 | -1 | TEC | ENSG00000 |
| 29926836 | 29928933 | 1 | lncRNA | ENSG00000 |
| 29940885 | 29973050 | 1 | protein_coding | ENSG00000 |
| 29973868 | 29992261 | 1 | protein_coding | ENSG00000 |
| 29992330 | 29996074 | -1 | protein_coding | ENSG00000 |
| 29995715 | 30005793 | 1 | protein_coding | ENSG00000 |
| 30005514 | 30023270 | -1 | protein_coding | ENSG00000 |
| 30023334 | 30027736 | 1 | protein_coding | ENSG00000 |
| 30024427 | 30052978 | -1 | protein_coding | ENSG00000 |
| 30053090 | 30070420 | 1 | protein_coding | ENSG00000 |
| 30064164 | 30070457 | 1 | protein_coding | ENSG00000 |
| 30064306 | 30064825 | -1 | lncRNA | ENSG00000 |
| 30075978 | 30085376 | 1 | protein_coding | ENSG00000 |
| 30085793 | 30091887 | -1 | protein_coding | ENSG00000 |
| 30092314 | 30096915 | -1 | protein_coding | ENSG00000 |
| 30096430 | 30104116 | 1 | lncRNA | ENSG00000 |
| 30104810 | 30113537 | -1 | protein_coding | ENSG00000 |
| 30107675 | 30110541 | 1 | lncRNA | ENSG00000 |
| 30110895 | 30111955 | 1 | lncRNA | ENSG00000 |
| 30114105 | 30123506 | -1 | protein_coding | ENSG00000 |
| 30182827 | 30189076 | 1 | protein_coding | ENSG00000 |
| 30183505 | 30184957 | -1 | lncRNA | ENSG00000 |
| 30192932 | 30194306 | -1 | protein_coding | ENSG00000 |
| 30193887 | 30197561 | 1 | protein_coding | ENSG00000 |
| 30193892 | 30204310 | 1 | lncRNA | ENSG00000 |
| 30199228 | 30204310 | 1 | protein_coding | ENSG00000 |

| | | | | |
|---|---|---|---|---|
| 30204316 | 30209071 | -1 | transcribed_unproce | ENSG00000 |
| 30205418 | 30206927 | 1 | transcribed_unproce | ENSG00000 |
| 30214409 | 30214699 | 1 | unprocessed_pseudo | ENSG00000 |
| 30218220 | 30221930 | 1 | unprocessed_pseudo | ENSG00000 |
| 30222937 | 30254510 | -1 | protein_coding | ENSG00000 |
| 30267553 | 30335374 | -1 | transcribed_unproce | ENSG00000 |
| 30336400 | 30343336 | 1 | unprocessed_pseudo | ENSG00000 |
| 30350773 | 30355308 | -1 | protein_coding | ENSG00000 |
| 30355441 | 30357104 | 1 | lncRNA | ENSG00000 |
| 30357102 | 30370494 | -1 | protein_coding | ENSG00000 |
| 30359825 | 30360336 | 1 | lncRNA | ENSG00000 |
| 30370934 | 30377991 | 1 | protein_coding | ENSG00000 |
| 30378106 | 30400108 | 1 | protein_coding | ENSG00000 |
| 30378133 | 30395991 | -1 | protein_coding | ENSG00000 |
| 30379763 | 30380110 | -1 | TEC | ENSG00000 |
| 30407414 | 30431108 | 1 | protein_coding | ENSG00000 |
| 30419625 | 30419759 | -1 | snoRNA | ENSG00000 |
| 30423615 | 30430030 | -1 | protein_coding | ENSG00000 |
| 30443631 | 30445874 | -1 | protein_coding | ENSG00000 |
| 30453997 | 30454097 | -1 | misc_RNA | ENSG00000 |
| 30471587 | 30471703 | 1 | misc_RNA | ENSG00000 |
| 30472719 | 30523185 | 1 | protein_coding | ENSG00000 |
| 30477180 | 30489353 | -1 | lncRNA | ENSG00000 |
| 30480588 | 30481346 | -1 | lncRNA | ENSG00000 |
| 30483135 | 30483195 | -1 | snRNA | ENSG00000 |
| 30498766 | 30499554 | -1 | lncRNA | ENSG00000 |
| 30503919 | 30504001 | 1 | miRNA | ENSG00000 |
| 30524004 | 30526821 | -1 | protein_coding | ENSG00000 |
| 30525923 | 30534852 | -1 | protein_coding | ENSG00000 |
| 30526918 | 30528294 | 1 | TEC | ENSG00000 |
| 30530367 | 30535347 | -1 | protein_coding | ENSG00000 |
| 30534752 | 30537149 | 1 | lncRNA | ENSG00000 |
| 30537202 | 30538231 | -1 | unprocessed_pseudo | ENSG00000 |
| 30553764 | 30558498 | -1 | protein_coding | ENSG00000 |
| 30569346 | 30572734 | -1 | protein_coding | ENSG00000 |
| 30572039 | 30583860 | 1 | lncRNA | ENSG00000 |
| 30573740 | 30585769 | -1 | protein_coding | ENSG00000 |
| 30585907 | 30608593 | 1 | lncRNA | ENSG00000 |
| 30587662 | 30587920 | -1 | processed_pseudoge | ENSG00000 |
| 30602558 | 30624012 | -1 | protein_coding | ENSG00000 |
| 30615256 | 30615679 | -1 | transcribed_processe | ENSG00000 |
| 30624285 | 30624714 | -1 | processed_pseudoge | ENSG00000 |
| 30626976 | 30627178 | 1 | processed_pseudoge | ENSG00000 |
| 30650717 | 30656440 | 1 | protein_coding | ENSG00000 |
| 30658431 | 30670810 | 1 | protein_coding | ENSG00000 |
| 30675300 | 30675402 | -1 | snRNA | ENSG00000 |
| 30697707 | 30699058 | -1 | lncRNA | ENSG00000 |
| 30698209 | 30741409 | 1 | protein_coding | ENSG00000 |
| 30701337 | 30701435 | -1 | snRNA | ENSG00000 |
| 30704067 | 30745169 | 1 | protein_coding | ENSG00000 |

| | | | | |
|---|---|---|---|---|
| 30710537 | 30710665 | 1 | snoRNA | ENSG0000( |
| 30740642 | 30745196 | 1 | protein_coding | ENSG0000( |
| 30740667 | 30751381 | 1 | lncRNA | ENSG0000( |
| 30748293 | 30761176 | 1 | protein_coding | ENSG0000( |
| 30757423 | 30762221 | -1 | protein_coding | ENSG0000( |
| 30761745 | 30776307 | 1 | protein_coding | ENSG0000( |
| 30773532 | 30776033 | -1 | TEC | ENSG0000( |
| 30778456 | 30787205 | -1 | protein_coding | ENSG0000( |
| 30816030 | 30816302 | 1 | misc_RNA | ENSG0000( |
| 30821068 | 30822110 | -1 | processed_pseudoge | ENSG0000( |
| 30821338 | 30821884 | 1 | lncRNA | ENSG0000( |
| 30833626 | 30894960 | -1 | protein_coding | ENSG0000( |
| 30875222 | 30895220 | 1 | lncRNA | ENSG0000( |
| 30875266 | 30875323 | -1 | miRNA | ENSG0000( |
| 30875766 | 30895216 | -1 | lncRNA | ENSG0000( |
| 30893903 | 30893985 | 1 | miRNA | ENSG0000( |
| 30896607 | 30903560 | 1 | protein_coding | ENSG0000( |
| 30904348 | 30910208 | -1 | unitary_pseudogene | ENSG0000( |
| 30919319 | 30923269 | -1 | lncRNA | ENSG0000( |
| 30923055 | 30948783 | 1 | protein_coding | ENSG0000( |
| 30948386 | 30956511 | 1 | lncRNA | ENSG0000( |
| 30949068 | 30956461 | 1 | protein_coding | ENSG0000( |
| 30956872 | 30957199 | -1 | lncRNA | ENSG0000( |
| 30957754 | 30984664 | 1 | protein_coding | ENSG0000( |
| 30984630 | 30988270 | -1 | TEC | ENSG0000( |
| 30985207 | 30989147 | 1 | protein_coding | ENSG0000( |
| 30989256 | 31010638 | -1 | protein_coding | ENSG0000( |
| 31032889 | 31042975 | 1 | protein_coding | ENSG0000( |
| 31043150 | 31049868 | 1 | lncRNA | ENSG0000( |
| 31056460 | 31062803 | 1 | lncRNA | ENSG0000( |
| 31060843 | 31074240 | -1 | protein_coding | ENSG0000( |
| 31065495 | 31069246 | 1 | lncRNA | ENSG0000( |
| 31074422 | 31084196 | 1 | protein_coding | ENSG0000( |
| 31083425 | 31089628 | -1 | protein_coding | ENSG0000( |
| 31083439 | 31094956 | -1 | protein_coding | ENSG0000( |
| 31090842 | 31095980 | -1 | protein_coding | ENSG0000( |
| 31093727 | 31095944 | 1 | TEC | ENSG0000( |
| 31106107 | 31112791 | 1 | protein_coding | ENSG0000( |
| 31114489 | 31131393 | 1 | protein_coding | ENSG0000( |
| 31118078 | 31118747 | 1 | lncRNA | ENSG0000( |
| 31122235 | 31124064 | 1 | lncRNA | ENSG0000( |
| 31131433 | 31135727 | -1 | protein_coding | ENSG0000( |
| 31131433 | 31131877 | 1 | lncRNA | ENSG0000( |
| 31138926 | 31150083 | -1 | protein_coding | ENSG0000( |
| 31165648 | 31165927 | 1 | processed_pseudoge | ENSG0000( |
| 31174573 | 31174822 | 1 | processed_pseudoge | ENSG0000( |
| 31180110 | 31194871 | 1 | protein_coding | ENSG0000( |
| 31182511 | 31183285 | -1 | lncRNA | ENSG0000( |
| 31196131 | 31196963 | 1 | lncRNA | ENSG0000( |
| 31201486 | 31203450 | -1 | protein_coding | ENSG0000( |

| | | | | |
|---|---|---|---|---|
| 31201885 | 31203452 | 1 | lncRNA | ENSG0000C |
| 31214091 | 31231537 | 1 | protein_coding | ENSG0000C |
| 31215962 | 31217359 | -1 | protein_coding | ENSG0000C |
| 31259967 | 31332892 | 1 | protein_coding | ENSG0000C |
| 31355134 | 31382997 | 1 | protein_coding | ENSG0000C |
| 31361278 | 31361695 | -1 | unprocessed_pseudc | ENSG0000C |
| 31393335 | 31426505 | 1 | protein_coding | ENSG0000C |
| 31402565 | 31404699 | 1 | lncRNA | ENSG0000C |
| 31427731 | 31428360 | -1 | protein_coding | ENSG0000C |
| 31428180 | 31428646 | 1 | lncRNA | ENSG0000C |
| 31432593 | 31443160 | -1 | protein_coding | ENSG0000C |
| 31449535 | 31453493 | -1 | lncRNA | ENSG0000C |
| 31456711 | 31459736 | -1 | lncRNA | ENSG0000C |
| 31458080 | 31467166 | 1 | protein_coding | ENSG0000C |
| 31470204 | 31472108 | -1 | TEC | ENSG0000C |
| 31471585 | 31477960 | 1 | protein_coding | ENSG0000C |
| 31483002 | 31490860 | 1 | protein_coding | ENSG0000C |
| 31487370 | 31488492 | -1 | lncRNA | ENSG0000C |
| 31489471 | 31509309 | -1 | protein_coding | ENSG0000C |
| 31508464 | 31509560 | 1 | lncRNA | ENSG0000C |
| 31527900 | 31528803 | 1 | protein_coding | ENSG0000C |
| 31542748 | 31553604 | -1 | lncRNA | ENSG0000C |
| 31546698 | 31547237 | -1 | lncRNA | ENSG0000C |
| 31546805 | 31547737 | 1 | unitary_pseudogene | ENSG0000C |
| 31554105 | 31554912 | 1 | unitary_pseudogene | ENSG0000C |
| 31563082 | 31567716 | -1 | transcribed_unproce | ENSG0000C |
| 31566360 | 31567000 | -1 | TEC | ENSG0000C |
| 31568386 | 31569475 | -1 | processed_pseudoge | ENSG0000C |
| 31601593 | 31603531 | -1 | processed_pseudoge | ENSG0000C |
| 31648307 | 31649222 | 1 | unprocessed_pseudc | ENSG0000C |
| 31680847 | 31682101 | 1 | processed_pseudoge | ENSG0000C |
| 31697397 | 31697607 | -1 | processed_pseudoge | ENSG0000C |
| 31699676 | 31700465 | -1 | lncRNA | ENSG0000C |
| 31700590 | 31711986 | 1 | transcribed_unproce | ENSG0000C |
| 31704728 | 31705260 | 1 | lncRNA | ENSG0000C |
| 31709113 | 31711984 | -1 | lncRNA | ENSG0000C |
| 31713229 | 31794869 | 1 | protein_coding | ENSG0000C |
| 31788202 | 31790993 | 1 | lncRNA | ENSG0000C |
| 31801585 | 31802497 | -1 | unitary_pseudogene | ENSG0000C |
| 31802947 | 31807973 | 1 | transcribed_processe | ENSG0000C |
| 31807926 | 31808844 | -1 | unitary_pseudogene | ENSG0000C |
| 31863633 | 31864876 | 1 | processed_pseudoge | ENSG0000C |
| 31870887 | 31871124 | -1 | processed_pseudoge | ENSG0000C |
| 31873807 | 31917357 | 1 | protein_coding | ENSG0000C |
| 31894542 | 31894983 | -1 | processed_pseudoge | ENSG0000C |
| 31952160 | 31952473 | 1 | processed_pseudoge | ENSG0000C |
| 31962063 | 31962381 | 1 | processed_pseudoge | ENSG0000C |
| 31963803 | 31964079 | 1 | processed_pseudoge | ENSG0000C |
| 31975803 | 32003728 | 1 | transcribed_unproce | ENSG0000C |
| 32008438 | 32084785 | -1 | transcribed_processe | ENSG0000C |

| | | | | |
|---|---|---|---|---|
| 32034853 | 32035326 | 1 | IG_V_pseudogene | ENSG0000C |
| 32051825 | 32052280 | 1 | unprocessed_pseudc | ENSG0000C |
| 32059084 | 32059372 | 1 | IG_V_pseudogene | ENSG0000C |
| 32066065 | 32066358 | 1 | IG_V_gene | ENSG0000C |
| 32103245 | 32188107 | -1 | transcribed_unproce | ENSG0000C |
| 32116700 | 32116997 | 1 | processed_pseudoge | ENSG0000C |
| 32126432 | 32132724 | 1 | lncRNA | ENSG0000C |
| 32185624 | 32186311 | 1 | TEC | ENSG0000C |
| 32188333 | 32193530 | 1 | lncRNA | ENSG0000C |
| 32189262 | 32189481 | 1 | transcribed_processe | ENSG0000C |
| 32199902 | 32202822 | 1 | unprocessed_pseudc | ENSG0000C |
| 32213380 | 32213503 | -1 | processed_pseudoge | ENSG0000C |
| 32250620 | 32254422 | -1 | lncRNA | ENSG0000C |
| 32252719 | 32255922 | 1 | protein_coding | ENSG0000C |
| 32262827 | 32265514 | -1 | lncRNA | ENSG0000C |
| 32278866 | 32279115 | 1 | processed_pseudoge | ENSG0000C |
| 32289547 | 32310555 | -1 | transcribed_processe | ENSG0000C |
| 32356981 | 32363695 | -1 | unprocessed_pseudc | ENSG0000C |
| 32379071 | 32380049 | -1 | processed_pseudoge | ENSG0000C |
| 32388135 | 32436203 | 1 | unprocessed_pseudc | ENSG0000C |
| 32439069 | 32441159 | -1 | processed_pseudoge | ENSG0000C |
| 32454612 | 32458758 | -1 | lncRNA | ENSG0000C |
| 32460266 | 32460362 | -1 | processed_pseudoge | ENSG0000C |
| 32475051 | 32478250 | -1 | unprocessed_pseudc | ENSG0000C |
| 32600299 | 32615639 | 1 | transcribed_unproce | ENSG0000C |
| 32618733 | 32622184 | 1 | unprocessed_pseudc | ENSG0000C |
| 32635673 | 32636045 | -1 | processed_pseudoge | ENSG0000C |
| 32649193 | 32663956 | 1 | lncRNA | ENSG0000C |
| 32650195 | 32650910 | -1 | transcribed_processe | ENSG0000C |
| 32663925 | 32666533 | 1 | lncRNA | ENSG0000C |
| 32673528 | 32676732 | -1 | protein_coding | ENSG0000C |
| 32675037 | 32678831 | 1 | lncRNA | ENSG0000C |
| 32715931 | 32716056 | 1 | processed_pseudoge | ENSG0000C |
| 32726615 | 32729537 | -1 | unprocessed_pseudc | ENSG0000C |
| 32735909 | 32741094 | -1 | lncRNA | ENSG0000C |
| 32739627 | 32740166 | -1 | processed_pseudoge | ENSG0000C |
| 32741314 | 32776196 | 1 | transcribed_unproce | ENSG0000C |
| 32781883 | 32784138 | 1 | TEC | ENSG0000C |
| 32786394 | 32787379 | 1 | TEC | ENSG0000C |
| 32787962 | 32788944 | 1 | TEC | ENSG0000C |
| 32809556 | 32824645 | 1 | lncRNA | ENSG0000C |
| 32811029 | 32811251 | -1 | processed_pseudoge | ENSG0000C |
| 32845964 | 32846180 | -1 | processed_pseudoge | ENSG0000C |
| 32847713 | 32848156 | 1 | IG_V_gene | ENSG0000C |
| 32869985 | 32873128 | 1 | unprocessed_pseudc | ENSG0000C |
| 32877469 | 32885501 | -1 | transcribed_unproce | ENSG0000C |
| 32882586 | 32888874 | 1 | lncRNA | ENSG0000C |
| 32903442 | 32903894 | 1 | IG_V_pseudogene | ENSG0000C |
| 32915074 | 32915536 | 1 | IG_V_pseudogene | ENSG0000C |
| 32916974 | 32946392 | 1 | unprocessed_pseudc | ENSG0000C |

| | | | | |
|---|---|---|---|---|
| 32931768 | 32932569 | -1 | processed_pseudoge | ENSG00000 |
| 32951993 | 33028037 | -1 | transcribed_processe | ENSG00000 |
| 32978461 | 32978891 | 1 | IG_V_pseudogene | ENSG00000 |
| 32995048 | 32995505 | 1 | IG_V_gene | ENSG00000 |
| 32995537 | 32996770 | 1 | lncRNA | ENSG00000 |
| 33002333 | 33002621 | 1 | IG_V_pseudogene | ENSG00000 |
| 33009175 | 33009620 | 1 | IG_V_gene | ENSG00000 |
| 33043609 | 33095431 | -1 | lncRNA | ENSG00000 |
| 33059936 | 33060098 | 1 | processed_pseudoge | ENSG00000 |
| 33094204 | 33129090 | -1 | transcribed_unproce | ENSG00000 |
| 33129316 | 33134499 | 1 | lncRNA | ENSG00000 |
| 33130235 | 33130442 | 1 | processed_pseudoge | ENSG00000 |
| 33140850 | 33143797 | 1 | unprocessed_pseudc | ENSG00000 |
| 33154352 | 33154475 | -1 | processed_pseudoge | ENSG00000 |
| 33191561 | 33195354 | -1 | lncRNA | ENSG00000 |
| 33193659 | 33196858 | 1 | protein_coding | ENSG00000 |
| 33203773 | 33206462 | -1 | lncRNA | ENSG00000 |
| 33206431 | 33330342 | -1 | lncRNA | ENSG00000 |
| 33303739 | 33306935 | 1 | protein_coding | ENSG00000 |
| 33345182 | 33345396 | 1 | processed_pseudoge | ENSG00000 |
| 33360274 | 33363478 | 1 | protein_coding | ENSG00000 |
| 33370473 | 33372915 | -1 | lncRNA | ENSG00000 |
| 33386105 | 33386349 | 1 | processed_pseudoge | ENSG00000 |
| 33459045 | 33462249 | 1 | protein_coding | ENSG00000 |
| 33484887 | 33485131 | 1 | processed_pseudoge | ENSG00000 |
| 33495943 | 33496093 | -1 | processed_pseudoge | ENSG00000 |
| 33533816 | 33534976 | -1 | lncRNA | ENSG00000 |
| 33541842 | 33545812 | -1 | lncRNA | ENSG00000 |
| 33548297 | 33554408 | -1 | lncRNA | ENSG00000 |
| 33562649 | 33570935 | -1 | transcribed_unproce | ENSG00000 |
| 33577502 | 33622547 | 1 | unprocessed_pseudc | ENSG00000 |
| 33687025 | 33707213 | -1 | transcribed_unproce | ENSG00000 |
| 33723356 | 33763773 | -1 | unprocessed_pseudc | ENSG00000 |
| 33769421 | 33769679 | 1 | processed_pseudoge | ENSG00000 |
| 33771035 | 33771248 | 1 | processed_pseudoge | ENSG00000 |
| 33802764 | 33803217 | 1 | IG_V_gene | ENSG00000 |
| 33808778 | 33810767 | 1 | processed_pseudoge | ENSG00000 |
| 33827214 | 33827661 | 1 | IG_V_gene | ENSG00000 |
| 33844784 | 33845229 | -1 | protein_coding | ENSG00000 |
| 33848476 | 33848644 | -1 | unprocessed_pseudc | ENSG00000 |
| 33851785 | 33852069 | -1 | unprocessed_pseudc | ENSG00000 |
| 33857935 | 33858864 | -1 | lncRNA | ENSG00000 |
| 33858896 | 33859352 | -1 | IG_V_pseudogene | ENSG00000 |
| 33875039 | 33875468 | -1 | unprocessed_pseudc | ENSG00000 |
| 33907419 | 33937167 | -1 | transcribed_unproce | ENSG00000 |
| 33938337 | 33938799 | -1 | IG_V_pseudogene | ENSG00000 |
| 33948278 | 33948548 | -1 | unprocessed_pseudc | ENSG00000 |
| 33949976 | 33950420 | -1 | unprocessed_pseudc | ENSG00000 |
| 33964547 | 33976346 | -1 | lncRNA | ENSG00000 |
| 33965655 | 33966091 | -1 | unprocessed_pseudc | ENSG00000 |

| | | | | |
|---|---|---|---|---|
| 33976039 | 33987630 | 1 | unprocessed_pseudc | ENSG00000( |
| 33988406 | 33991551 | -1 | unprocessed_pseudc | ENSG00000( |
| 34013394 | 34013830 | -1 | unprocessed_pseudc | ENSG00000( |
| 34015260 | 34015477 | 1 | processed_pseudoge | ENSG00000( |
| 34051140 | 34051363 | 1 | processed_pseudoge | ENSG00000( |
| 34120470 | 34120677 | 1 | processed_pseudoge | ENSG00000( |
| 34135736 | 34136792 | -1 | processed_pseudoge | ENSG00000( |
| 34139189 | 34139543 | 1 | processed_pseudoge | ENSG00000( |
| 34141256 | 34142006 | -1 | processed_pseudoge | ENSG00000( |
| 34142456 | 34142849 | 1 | processed_pseudoge | ENSG00000( |
| 34158585 | 34160036 | -1 | lncRNA | ENSG00000( |
| 34162959 | 34163110 | 1 | rRNA_pseudogene | ENSG00000( |
| 34266041 | 34268649 | -1 | processed_pseudoge | ENSG00000( |
| 34353616 | 34381110 | -1 | unprocessed_pseudc | ENSG00000( |
| 34387599 | 34389827 | -1 | lncRNA | ENSG00000( |
| 34446566 | 34448866 | 1 | unprocessed_pseudc | ENSG00000( |
| 34941977 | 34943880 | 1 | TEC | ENSG00000( |
| 34962970 | 34965270 | 1 | unprocessed_pseudc | ENSG00000( |
| 34978744 | 34979844 | -1 | lncRNA | ENSG00000( |
| 35021749 | 35024229 | 1 | lncRNA | ENSG00000( |
| 35022851 | 35024250 | -1 | lncRNA | ENSG00000( |
| 35023339 | 35023765 | 1 | processed_pseudoge | ENSG00000( |
| 35040164 | 35084313 | 1 | unprocessed_pseudc | ENSG00000( |
| 35068211 | 35085060 | 1 | unprocessed_pseudc | ENSG00000( |
| 35085896 | 35086119 | 1 | processed_pseudoge | ENSG00000( |
| 35090501 | 35104834 | 1 | unprocessed_pseudc | ENSG00000( |
| 35118238 | 35118956 | -1 | processed_pseudoge | ENSG00000( |
| 35134514 | 35135075 | 1 | processed_pseudoge | ENSG00000( |
| 35143722 | 35146129 | 1 | lncRNA | ENSG00000( |
| 35148636 | 35154366 | 1 | TEC | ENSG00000( |
| 35169692 | 35170241 | -1 | processed_pseudoge | ENSG00000( |
| 35173316 | 35174093 | 1 | processed_pseudoge | ENSG00000( |
| 35192687 | 35195661 | -1 | lncRNA | ENSG00000( |
| 35193335 | 35193734 | 1 | processed_pseudoge | ENSG00000( |
| 35195779 | 35197544 | 1 | lncRNA | ENSG00000( |
| 35207884 | 35284146 | 1 | lncRNA | ENSG00000( |
| 35207903 | 35208236 | -1 | unprocessed_pseudc | ENSG00000( |
| 35221583 | 35222048 | -1 | processed_pseudoge | ENSG00000( |
| 35232412 | 35235984 | -1 | unprocessed_pseudc | ENSG00000( |
| 35245916 | 35247534 | -1 | unprocessed_pseudc | ENSG00000( |
| 35255192 | 35255633 | -1 | processed_pseudoge | ENSG00000( |
| 35258630 | 35259629 | 1 | processed_pseudoge | ENSG00000( |
| 35335277 | 35336889 | -1 | unprocessed_pseudc | ENSG00000( |
| 35344091 | 35344559 | -1 | processed_pseudoge | ENSG00000( |
| 35347385 | 35348297 | 1 | processed_pseudoge | ENSG00000( |
| 35352482 | 35353022 | 1 | processed_pseudoge | ENSG00000( |
| 35363412 | 35390584 | 1 | lncRNA | ENSG00000( |
| 35384144 | 35385086 | -1 | unprocessed_pseudc | ENSG00000( |
| 35389203 | 35390003 | -1 | processed_pseudoge | ENSG00000( |
| 35405747 | 35407349 | -1 | unprocessed_pseudc | ENSG00000( |

| | | | | |
|---|---|---|---|---|
| 35414279 | 35414747 | -1 | processed_pseudoge | ENSG00000( |
| 35420291 | 35421209 | 1 | processed_pseudoge | ENSG00000( |
| 35425364 | 35426483 | 1 | processed_pseudoge | ENSG00000( |
| 35435303 | 35436341 | -1 | unprocessed_pseudc | ENSG00000( |
| 35446898 | 35449411 | -1 | transcribed_processe | ENSG00000( |
| 35450898 | 35452987 | 1 | lncRNA | ENSG00000( |
| 35477414 | 35480481 | -1 | transcribed_unproce | ENSG00000( |
| 35485766 | 35486225 | -1 | processed_pseudoge | ENSG00000( |
| 35489237 | 35490212 | 1 | processed_pseudoge | ENSG00000( |
| 35491174 | 35492395 | -1 | lncRNA | ENSG00000( |
| 35492324 | 35493446 | 1 | processed_pseudoge | ENSG00000( |
| 35493754 | 35496457 | -1 | lncRNA | ENSG00000( |
| 35499494 | 35522465 | 1 | lncRNA | ENSG00000( |
| 35504370 | 35506567 | -1 | lncRNA | ENSG00000( |
| 35506592 | 35507170 | -1 | transcribed_processe | ENSG00000( |
| 35528367 | 35529318 | -1 | processed_pseudoge | ENSG00000( |
| 35543279 | 35543645 | -1 | processed_pseudoge | ENSG00000( |
| 35547917 | 35548829 | 1 | processed_pseudoge | ENSG00000( |
| 35552909 | 35554001 | 1 | processed_pseudoge | ENSG00000( |
| 35574834 | 35575494 | 1 | processed_pseudoge | ENSG00000( |
| 35636687 | 35645180 | -1 | unprocessed_pseudc | ENSG00000( |
| 35640029 | 35640582 | 1 | lncRNA | ENSG00000( |
| 35722268 | 35722529 | -1 | processed_pseudoge | ENSG00000( |
| 35734393 | 35734504 | 1 | rRNA_pseudogene | ENSG00000( |
| 35734639 | 35734748 | 1 | rRNA_pseudogene | ENSG00000( |
| 35735372 | 35735471 | 1 | rRNA_pseudogene | ENSG00000( |
| 35735625 | 35735733 | 1 | rRNA_pseudogene | ENSG00000( |
| 35743268 | 35756515 | 1 | lncRNA | ENSG00000( |
| 35745989 | 35746094 | 1 | rRNA_pseudogene | ENSG00000( |
| 35746523 | 35746636 | 1 | rRNA_pseudogene | ENSG00000( |
| 35746772 | 35746894 | 1 | rRNA_pseudogene | ENSG00000( |
| 35747032 | 35747140 | 1 | rRNA_pseudogene | ENSG00000( |
| 35748520 | 35748655 | 1 | rRNA_pseudogene | ENSG00000( |
| 35750061 | 35750175 | 1 | rRNA_pseudogene | ENSG00000( |
| 35751574 | 35751684 | 1 | rRNA_pseudogene | ENSG00000( |
| 35751828 | 35751935 | 1 | rRNA_pseudogene | ENSG00000( |
| 35752826 | 35752942 | 1 | rRNA_pseudogene | ENSG00000( |
| 35753796 | 35753930 | 1 | rRNA_pseudogene | ENSG00000( |
| 35754046 | 35754182 | 1 | rRNA_pseudogene | ENSG00000( |
| 35754555 | 35754688 | 1 | rRNA_pseudogene | ENSG00000( |
| 35754923 | 35755033 | 1 | rRNA_pseudogene | ENSG00000( |
| 35755177 | 35755285 | 1 | rRNA_pseudogene | ENSG00000( |
| 35785569 | 35786887 | -1 | lncRNA | ENSG00000( |
| 35802869 | 35803280 | -1 | processed_pseudoge | ENSG00000( |
| 35836886 | 35837732 | 1 | processed_pseudoge | ENSG00000( |
| 35879036 | 35879498 | -1 | processed_pseudoge | ENSG00000( |
| 35903117 | 35904916 | 1 | TEC | ENSG00000( |
| 35912120 | 35912516 | 1 | processed_pseudoge | ENSG00000( |
| 46469341 | 46569097 | -1 | transcribed_unproce | ENSG00000( |
| 46556089 | 46556191 | -1 | snRNA | ENSG00000( |

| | | | | |
|---|---|---|---|---|
| 46571537 | 46572136 | 1 | processed_pseudoge | ENSG00000 |
| 46573763 | 46575094 | 1 | unprocessed_pseudc | ENSG00000 |
| 46578591 | 46621379 | -1 | protein_coding | ENSG00000 |
| 46622861 | 46624451 | -1 | lncRNA | ENSG00000 |
| 46626404 | 46626992 | -1 | processed_pseudoge | ENSG00000 |
| 46649126 | 46649849 | 1 | unprocessed_pseudc | ENSG00000 |
| 46656132 | 46689518 | -1 | protein_coding | ENSG00000 |
| 46660696 | 46661591 | -1 | lncRNA | ENSG00000 |
| 46673443 | 46673927 | -1 | TEC | ENSG00000 |
| 46689643 | 46698394 | 1 | protein_coding | ENSG00000 |
| 46702282 | 46790407 | -1 | protein_coding | ENSG00000 |
| 46789898 | 46792040 | 1 | processed_pseudoge | ENSG00000 |
| 46796603 | 46831180 | -1 | protein_coding | ENSG00000 |
| 46842632 | 46844618 | -1 | processed_pseudoge | ENSG00000 |
| 46850999 | 46851254 | 1 | processed_pseudoge | ENSG00000 |
| 46884362 | 46931289 | 1 | protein_coding | ENSG00000 |
| 46955362 | 46973674 | -1 | protein_coding | ENSG00000 |
| 46973989 | 46978983 | 1 | lncRNA | ENSG00000 |
| 47004513 | 47035385 | 1 | lncRNA | ENSG00000 |
| 47013957 | 47014365 | 1 | transcribed_processe | ENSG00000 |
| 47077703 | 47143945 | -1 | protein_coding | ENSG00000 |
| 47144323 | 47162749 | 1 | lncRNA | ENSG00000 |
| 47150132 | 47150566 | -1 | processed_pseudoge | ENSG00000 |
| 47154387 | 47464149 | -1 | protein_coding | ENSG00000 |
| 47196311 | 47202429 | 1 | lncRNA | ENSG00000 |
| 47218026 | 47218768 | 1 | TEC | ENSG00000 |
| 47262741 | 47263500 | 1 | processed_pseudoge | ENSG00000 |
| 47299447 | 47317814 | 1 | lncRNA | ENSG00000 |
| 47313027 | 47313121 | -1 | misc_RNA | ENSG00000 |
| 47461123 | 47701523 | 1 | protein_coding | ENSG00000 |
| 47505432 | 47505543 | 1 | rRNA_pseudogene | ENSG00000 |
| 47529190 | 47529916 | 1 | lncRNA | ENSG00000 |
| 47561307 | 47562480 | -1 | processed_pseudoge | ENSG00000 |
| 47565090 | 47566884 | -1 | processed_pseudoge | ENSG00000 |
| 47598654 | 47598996 | -1 | processed_pseudoge | ENSG00000 |
| 47724568 | 47908431 | 1 | lncRNA | ENSG00000 |
| 47760780 | 47760921 | -1 | processed_pseudoge | ENSG00000 |
| 47849314 | 47887130 | -1 | lncRNA | ENSG00000 |
| 47965620 | 47971693 | 1 | lncRNA | ENSG00000 |
| 48050588 | 48055099 | -1 | lncRNA | ENSG00000 |
| 48059671 | 48059995 | -1 | processed_pseudoge | ENSG00000 |
| 48081006 | 48156018 | -1 | protein_coding | ENSG00000 |
| 48164952 | 48167238 | -1 | lncRNA | ENSG00000 |
| 48166910 | 48247568 | -1 | protein_coding | ENSG00000 |
| 48244300 | 48363122 | 1 | protein_coding | ENSG00000 |
| 48334471 | 48334854 | -1 | processed_pseudoge | ENSG00000 |
| 48343755 | 48346058 | -1 | TEC | ENSG00000 |
| 48356364 | 48448402 | -1 | protein_coding | ENSG00000 |
| 48432927 | 48433208 | -1 | misc_RNA | ENSG00000 |
| 48447904 | 48448398 | -1 | lncRNA | ENSG00000 |

| | | | | |
|---|---|---|---|---|
| 48470953 | 48471133 | -1 | processed_pseudoge | ENSG0000C |
| 48496990 | 48498601 | 1 | processed_pseudoge | ENSG0000C |
| 48538726 | 48620148 | -1 | protein_coding | ENSG0000C |
| 48559661 | 48587403 | 1 | lncRNA | ENSG0000C |
| 48577524 | 48578369 | -1 | processed_pseudoge | ENSG0000C |
| 48600176 | 48600644 | 1 | processed_pseudoge | ENSG0000C |
| 48620319 | 48622033 | 1 | lncRNA | ENSG0000C |
| 48623422 | 48746318 | 1 | lncRNA | ENSG0000C |
| 48637143 | 48638719 | 1 | lncRNA | ENSG0000C |
| 48640473 | 48641052 | 1 | lncRNA | ENSG0000C |
| 48718083 | 48719143 | 1 | processed_pseudoge | ENSG0000C |
| 48731704 | 48731801 | -1 | snRNA | ENSG0000C |
| 48849919 | 48850015 | -1 | processed_pseudoge | ENSG0000C |
| 49066567 | 49066729 | -1 | processed_pseudoge | ENSG0000C |
| 49072543 | 49275579 | -1 | lncRNA | ENSG0000C |
| 49159508 | 49162023 | 1 | lncRNA | ENSG0000C |
| 49170552 | 49171786 | 1 | lncRNA | ENSG0000C |
| 49277917 | 49281838 | -1 | protein_coding | ENSG0000C |
| 49282021 | 49350504 | 1 | lncRNA | ENSG0000C |
| 49297143 | 49299762 | 1 | TEC | ENSG0000C |
| 49373804 | 49399431 | 1 | protein_coding | ENSG0000C |
| 49442910 | 49454109 | -1 | lncRNA | ENSG0000C |
| 49454229 | 49457269 | 1 | lncRNA | ENSG0000C |
| 49466106 | 49469875 | 1 | lncRNA | ENSG0000C |
| 49487524 | 49857919 | -1 | protein_coding | ENSG0000C |
| 49517762 | 49518601 | -1 | processed_pseudoge | ENSG0000C |
| 49708006 | 49708243 | 1 | processed_pseudoge | ENSG0000C |
| 49711273 | 49711914 | 1 | TEC | ENSG0000C |
| 49764454 | 49764688 | 1 | processed_pseudoge | ENSG0000C |
| 49847018 | 49847632 | -1 | lncRNA | ENSG0000C |
| 49900019 | 49900524 | 1 | TEC | ENSG0000C |
| 49920730 | 49924154 | -1 | lncRNA | ENSG0000C |
| 49935028 | 49935352 | 1 | processed_pseudoge | ENSG0000C |
| 49954985 | 49968730 | -1 | unitary_pseudogene | ENSG0000C |
| 50024410 | 50037088 | 1 | protein_coding | ENSG0000C |
| 50046429 | 50066224 | -1 | lncRNA | ENSG0000C |
| 50046496 | 50047798 | 1 | unprocessed_pseudc | ENSG0000C |
| 50054143 | 50054430 | 1 | misc_RNA | ENSG0000C |
| 50058150 | 50059231 | -1 | transcribed_processe | ENSG0000C |
| 50065970 | 50107272 | 1 | protein_coding | ENSG0000C |
| 50072862 | 50074986 | 1 | TEC | ENSG0000C |
| 50098264 | 50098358 | 1 | misc_RNA | ENSG0000C |
| 50100339 | 50121943 | -1 | lncRNA | ENSG0000C |
| 50120064 | 50129904 | 1 | lncRNA | ENSG0000C |
| 50151280 | 50151577 | -1 | processed_pseudoge | ENSG0000C |
| 50151642 | 50151731 | -1 | snoRNA | ENSG0000C |
| 50152918 | 50235310 | 1 | protein_coding | ENSG0000C |
| 50246137 | 50318135 | 1 | protein_coding | ENSG0000C |
| 50292616 | 50292675 | 1 | miRNA | ENSG0000C |
| 50313487 | 50368934 | -1 | protein_coding | ENSG0000C |

| | | | | |
|---|---|---|---|---|
| 50368580 | 50371480 | 1 | lncRNA | ENSG0000C |
| 50390916 | 50395133 | -1 | lncRNA | ENSG0000C |
| 50407184 | 50546347 | 1 | lncRNA | ENSG0000C |
| 50548396 | 50649249 | 1 | protein_coding | ENSG0000C |
| 50551824 | 50553940 | -1 | lncRNA | ENSG0000C |
| 50555355 | 50558719 | 1 | TEC | ENSG0000C |
| 50606076 | 50613684 | -1 | lncRNA | ENSG0000C |
| 50664903 | 50671639 | 1 | lncRNA | ENSG0000C |
| 50666300 | 50681353 | -1 | protein_coding | ENSG0000C |
| 50693588 | 50734041 | 1 | protein_coding | ENSG0000C |
| 50712844 | 50713589 | 1 | lncRNA | ENSG0000C |
| 50727417 | 50742815 | -1 | lncRNA | ENSG0000C |
| 50742050 | 50801935 | 1 | protein_coding | ENSG0000C |
| 50742305 | 50742377 | 1 | miRNA | ENSG0000C |
| 50783859 | 50803338 | -1 | lncRNA | ENSG0000C |
| 50806668 | 50807629 | 1 | lncRNA | ENSG0000C |
| 50840017 | 50901063 | 1 | lncRNA | ENSG0000C |
| 50856614 | 50879278 | -1 | lncRNA | ENSG0000C |
| 50869455 | 50869575 | -1 | rRNA_pseudogene | ENSG0000C |
| 51017544 | 51035784 | 1 | lncRNA | ENSG0000C |
| 51035118 | 51038764 | -1 | lncRNA | ENSG0000C |
| 51054728 | 51056399 | -1 | lncRNA | ENSG0000C |
| 51089466 | 51095952 | -1 | lncRNA | ENSG0000C |
| 51107571 | 51108021 | 1 | processed_pseudoge | ENSG0000C |
| 51135975 | 51151367 | -1 | protein_coding | ENSG0000C |
| 51149239 | 51149819 | 1 | lncRNA | ENSG0000C |
| 51150770 | 51442519 | 1 | lncRNA | ENSG0000C |
| 51176066 | 51178898 | 1 | TEC | ENSG0000C |
| 51197804 | 51268117 | -1 | lncRNA | ENSG0000C |
| 51244054 | 51244123 | 1 | processed_pseudoge | ENSG0000C |
| 51277852 | 51279112 | 1 | processed_pseudoge | ENSG0000C |
| 51354456 | 51525157 | -1 | lncRNA | ENSG0000C |
| 51553436 | 51647132 | 1 | protein_coding | ENSG0000C |
| 51671236 | 51671563 | -1 | misc_RNA | ENSG0000C |
| 51755325 | 51788208 | 1 | lncRNA | ENSG0000C |
| 52005487 | 52078474 | -1 | lncRNA | ENSG0000C |
| 52078622 | 52080489 | 1 | lncRNA | ENSG0000C |
| 52081568 | 52082096 | -1 | TEC | ENSG0000C |
| 52083064 | 52085109 | -1 | lncRNA | ENSG0000C |
| 52085210 | 52099120 | 1 | lncRNA | ENSG0000C |
| 52198012 | 52227956 | -1 | lncRNA | ENSG0000C |
| 52238081 | 52269574 | 1 | lncRNA | ENSG0000C |
| 52258564 | 52280638 | 1 | lncRNA | ENSG0000C |
| 52436417 | 52547802 | -1 | protein_coding | ENSG0000C |
| 52436772 | 52437474 | -1 | TEC | ENSG0000C |
| 52552084 | 52652132 | -1 | lncRNA | ENSG0000C |
| 52583694 | 52588879 | 1 | lncRNA | ENSG0000C |
| 52607006 | 52614640 | 1 | lncRNA | ENSG0000C |
| 52607762 | 52608158 | -1 | lncRNA | ENSG0000C |
| 52622115 | 52677747 | 1 | lncRNA | ENSG0000C |

| | | | | |
|---|---|---|---|---|
| 52655307 | 52656045 | 1 | processed_pseudoge | ENSG00000( |
| 52934813 | 52935635 | 1 | processed_pseudoge | ENSG00000( |
| 52942139 | 52943464 | 1 | transcribed_process | ENSG00000( |
| 53010395 | 53011286 | -1 | processed_pseudoge | ENSG00000( |
| 53035690 | 53052873 | -1 | protein_coding | ENSG00000( |
| 53055033 | 53329150 | 1 | protein_coding | ENSG00000( |
| 53168522 | 53169450 | 1 | lncRNA | ENSG00000( |
| 53227146 | 53227694 | 1 | TEC | ENSG00000( |
| 53298224 | 53299792 | 1 | transcribed_process | ENSG00000( |
| 53310797 | 53311989 | 1 | TEC | ENSG00000( |
| 53334562 | 53334664 | 1 | snoRNA | ENSG00000( |
| 53337453 | 53337571 | 1 | rRNA_pseudogene | ENSG00000( |
| 53362019 | 53363678 | -1 | unprocessed_pseudc | ENSG00000( |
| 53364982 | 53373083 | -1 | transcribed_unproce | ENSG00000( |
| 53373479 | 53385019 | 1 | lncRNA | ENSG00000( |
| 53386944 | 53389085 | 1 | lncRNA | ENSG00000( |
| 53389886 | 53390563 | 1 | TEC | ENSG00000( |
| 53391089 | 53392473 | 1 | TEC | ENSG00000( |
| 53433977 | 53491648 | 1 | protein_coding | ENSG00000( |
| 53443228 | 53443332 | -1 | snRNA | ENSG00000( |
| 53448748 | 53449590 | 1 | lncRNA | ENSG00000( |
| 53478957 | 53481550 | -1 | TEC | ENSG00000( |
| 53487607 | 53489943 | -1 | TEC | ENSG00000( |
| 53491040 | 53504411 | -1 | protein_coding | ENSG00000( |
| 53544339 | 53545689 | -1 | lncRNA | ENSG00000( |
| 53598153 | 53703938 | -1 | protein_coding | ENSG00000( |
| 53628256 | 53628816 | -1 | lncRNA | ENSG00000( |
| 53701692 | 54158512 | 1 | protein_coding | ENSG00000( |
| 53981229 | 53981912 | -1 | lncRNA | ENSG00000( |
| 53991012 | 53994454 | 1 | TEC | ENSG00000( |
| 53998313 | 53999969 | -1 | lncRNA | ENSG00000( |
| 54033997 | 54054583 | -1 | lncRNA | ENSG00000( |
| 54114697 | 54114878 | -1 | pseudogene | ENSG00000( |
| 54239693 | 54240654 | 1 | lncRNA | ENSG00000( |
| 54245544 | 54270879 | -1 | lncRNA | ENSG00000( |
| 54283304 | 54286787 | -1 | protein_coding | ENSG00000( |
| 54285604 | 54289568 | 1 | lncRNA | ENSG00000( |
| 54290965 | 54292422 | 1 | lncRNA | ENSG00000( |
| 54365996 | 54370794 | -1 | lncRNA | ENSG00000( |
| 54370265 | 54374728 | 1 | lncRNA | ENSG00000( |
| 54490281 | 54491396 | -1 | lncRNA | ENSG00000( |
| 54542808 | 54559100 | -1 | lncRNA | ENSG00000( |
| 54628963 | 54657662 | -1 | lncRNA | ENSG00000( |
| 54845189 | 54929189 | -1 | lncRNA | ENSG00000( |
| 54847316 | 54854991 | 1 | lncRNA | ENSG00000( |
| 54930865 | 54934485 | 1 | protein_coding | ENSG00000( |
| 54934913 | 54954665 | 1 | lncRNA | ENSG00000( |
| 54937786 | 54938671 | -1 | lncRNA | ENSG00000( |
| 55007982 | 55008306 | 1 | processed_pseudoge | ENSG00000( |
| 55045700 | 55049318 | -1 | lncRNA | ENSG00000( |

| | | | | |
|---|---|---|---|---|
| 55259969 | 55333588 | -1 | lncRNA | ENSG00000 |
| 55291379 | 55291675 | -1 | misc_RNA | ENSG00000 |
| 55321575 | 55322686 | -1 | lncRNA | ENSG00000 |
| 55323760 | 55330760 | 1 | protein_coding | ENSG00000 |
| 55332355 | 55332967 | 1 | lncRNA | ENSG00000 |
| 55352721 | 55353088 | -1 | processed_pseudoge | ENSG00000 |
| 55389700 | 55506691 | 1 | protein_coding | ENSG00000 |
| 55425574 | 55426130 | -1 | lncRNA | ENSG00000 |
| 55426797 | 55462297 | -1 | lncRNA | ENSG00000 |
| 55486618 | 55488777 | 1 | TEC | ENSG00000 |
| 55509072 | 55586666 | 1 | protein_coding | ENSG00000 |
| 55538200 | 55542027 | 1 | lncRNA | ENSG00000 |
| 55566684 | 55567687 | 1 | protein_coding | ENSG00000 |
| 55655604 | 55706192 | 1 | protein_coding | ENSG00000 |
| 55704119 | 55706422 | 1 | TEC | ENSG00000 |
| 55728115 | 55750111 | 1 | unprocessed_pseudo | ENSG00000 |
| 55760566 | 55793960 | 1 | transcribed_unproce | ENSG00000 |
| 55802851 | 55833337 | -1 | protein_coding | ENSG00000 |
| 55846154 | 55956031 | -1 | protein_coding | ENSG00000 |
| 55984009 | 56191118 | -1 | lncRNA | ENSG00000 |
| 56108980 | 56109970 | 1 | TEC | ENSG00000 |
| 56109537 | 56137000 | 1 | lncRNA | ENSG00000 |
| 56191489 | 56357444 | 1 | protein_coding | ENSG00000 |
| 56192614 | 56194518 | -1 | lncRNA | ENSG00000 |
| 56245520 | 56245623 | 1 | miRNA | ENSG00000 |
| 56282886 | 56299351 | -1 | lncRNA | ENSG00000 |
| 56300616 | 56302904 | 1 | lncRNA | ENSG00000 |
| 56306845 | 56308335 | 1 | TEC | ENSG00000 |
| 56351886 | 56353524 | -1 | lncRNA | ENSG00000 |
| 56361452 | 56425545 | -1 | protein_coding | ENSG00000 |
| 56409320 | 56411683 | 1 | lncRNA | ENSG00000 |
| 56429133 | 56452199 | -1 | protein_coding | ENSG00000 |
| 56451521 | 56479104 | 1 | protein_coding | ENSG00000 |
| 56465642 | 56466162 | -1 | lncRNA | ENSG00000 |
| 56466836 | 56520087 | -1 | protein_coding | ENSG00000 |
| 56565073 | 56568957 | 1 | protein_coding | ENSG00000 |
| 56589074 | 56591088 | 1 | protein_coding | ENSG00000 |
| 56608584 | 56609497 | 1 | protein_coding | ENSG00000 |
| 56609501 | 56611375 | -1 | lncRNA | ENSG00000 |
| 56617476 | 56618818 | 1 | transcribed_unproce | ENSG00000 |
| 56625475 | 56627112 | 1 | protein_coding | ENSG00000 |
| 56632659 | 56633981 | 1 | protein_coding | ENSG00000 |
| 56635739 | 56637086 | 1 | transcribed_unproce | ENSG00000 |
| 56638666 | 56640087 | 1 | protein_coding | ENSG00000 |
| 56643705 | 56644786 | 1 | transcribed_unproce | ENSG00000 |
| 56648248 | 56649514 | 1 | transcribed_unproce | ENSG00000 |
| 56648881 | 56652636 | 1 | lncRNA | ENSG00000 |
| 56651886 | 56653204 | 1 | protein_coding | ENSG00000 |
| 56657731 | 56660698 | 1 | protein_coding | ENSG00000 |
| 56666731 | 56668065 | -1 | protein_coding | ENSG00000 |

| | | | | |
|---|---|---|---|---|
| 56669814 | 56671129 | 1 | protein_coding | ENSG0000C |
| 56676132 | 56677763 | 1 | transcribed_unproce | ENSG0000C |
| 56682470 | 56684196 | 1 | protein_coding | ENSG0000C |
| 56682470 | 56687807 | 1 | lncRNA | ENSG0000C |
| 56686851 | 56688052 | 1 | processed_pseudoge | ENSG0000C |
| 56708450 | 56730010 | -1 | lncRNA | ENSG0000C |
| 56730118 | 56850286 | 1 | protein_coding | ENSG0000C |
| 56858518 | 56858601 | 1 | miRNA | ENSG0000C |
| 56865207 | 56915850 | 1 | protein_coding | ENSG0000C |
| 56904264 | 56904353 | 1 | miRNA | ENSG0000C |
| 56906534 | 56906956 | 1 | processed_pseudoge | ENSG0000C |
| 56932142 | 56944864 | 1 | protein_coding | ENSG0000C |
| 56940278 | 56941342 | 1 | lncRNA | ENSG0000C |
| 56941028 | 56941726 | 1 | lncRNA | ENSG0000C |
| 56961923 | 56983845 | 1 | protein_coding | ENSG0000C |
| 56989485 | 57083531 | 1 | protein_coding | ENSG0000C |
| 57013071 | 57014100 | -1 | processed_pseudoge | ENSG0000C |
| 57052505 | 57058497 | -1 | lncRNA | ENSG0000C |
| 57091213 | 57092303 | -1 | lncRNA | ENSG0000C |
| 57092583 | 57148369 | 1 | protein_coding | ENSG0000C |
| 57140896 | 57144586 | -1 | TEC | ENSG0000C |
| 57142652 | 57144936 | 1 | TEC | ENSG0000C |
| 57152466 | 57186116 | -1 | protein_coding | ENSG0000C |
| 57177349 | 57181390 | 1 | lncRNA | ENSG0000C |
| 57186137 | 57240469 | 1 | protein_coding | ENSG0000C |
| 57245259 | 57253635 | 1 | protein_coding | ENSG0000C |
| 57245832 | 57246396 | -1 | lncRNA | ENSG0000C |
| 57247350 | 57248492 | 1 | lncRNA | ENSG0000C |
| 57248547 | 57284672 | -1 | protein_coding | ENSG0000C |
| 57321352 | 57322117 | 1 | processed_pseudoge | ENSG0000C |
| 57332379 | 57332599 | -1 | processed_pseudoge | ENSG0000C |
| 57358783 | 57366189 | 1 | protein_coding | ENSG0000C |
| 57372477 | 57385044 | 1 | protein_coding | ENSG0000C |
| 57404767 | 57416063 | 1 | protein_coding | ENSG0000C |
| 57428187 | 57447420 | -1 | protein_coding | ENSG0000C |
| 57447425 | 57461270 | 1 | protein_coding | ENSG0000C |
| 57461055 | 57462737 | 1 | lncRNA | ENSG0000C |
| 57462660 | 57472009 | 1 | protein_coding | ENSG0000C |
| 57471922 | 57487327 | -1 | protein_coding | ENSG0000C |
| 57512181 | 57536571 | -1 | protein_coding | ENSG0000C |
| 57542643 | 57591681 | 1 | protein_coding | ENSG0000C |
| 57603137 | 57603697 | 1 | processed_pseudoge | ENSG0000C |
| 57610652 | 57665580 | 1 | protein_coding | ENSG0000C |
| 57622034 | 57631647 | -1 | lncRNA | ENSG0000C |
| 57668277 | 57689378 | 1 | protein_coding | ENSG0000C |
| 57681124 | 57701730 | -1 | lncRNA | ENSG0000C |
| 57694793 | 57731805 | 1 | protein_coding | ENSG0000C |
| 57713782 | 57714957 | 1 | processed_pseudoge | ENSG0000C |
| 57735739 | 57757244 | 1 | protein_coding | ENSG0000C |
| 57758217 | 57863053 | -1 | protein_coding | ENSG0000C |

| | | | | |
|---|---|---|---|---|
| 57759358 | 57760024 | 1 | lncRNA | ENSG0000C |
| 57772289 | 57772352 | -1 | miRNA | ENSG0000C |
| 57798186 | 57816946 | 1 | lncRNA | ENSG0000C |
| 57830536 | 57830642 | -1 | snRNA | ENSG0000C |
| 57882340 | 57971128 | -1 | protein_coding | ENSG0000C |
| 57976435 | 57988116 | 1 | protein_coding | ENSG0000C |
| 57994668 | 58000453 | -1 | protein_coding | ENSG0000C |
| 57999546 | 58021618 | 1 | protein_coding | ENSG0000C |
| 58025754 | 58046901 | 1 | protein_coding | ENSG0000C |
| 58083068 | 58083168 | -1 | misc_RNA | ENSG0000C |
| 58092053 | 58092947 | -1 | processed_pseudoge | ENSG0000C |
| 58113592 | 58129381 | -1 | protein_coding | ENSG0000C |
| 58116885 | 58119353 | 1 | lncRNA | ENSG0000C |
| 58129519 | 58163246 | 1 | lncRNA | ENSG0000C |
| 58157907 | 58198106 | -1 | protein_coding | ENSG0000C |
| 58197112 | 58217805 | 1 | lncRNA | ENSG0000C |
| 58199568 | 58199836 | -1 | misc_RNA | ENSG0000C |
| 58231157 | 58283836 | 1 | protein_coding | ENSG0000C |
| 58279997 | 58295047 | -1 | protein_coding | ENSG0000C |
| 58295080 | 58406144 | 1 | protein_coding | ENSG0000C |
| 58378052 | 58378148 | -1 | snRNA | ENSG0000C |
| 58382989 | 58383798 | -1 | lncRNA | ENSG0000C |
| 58384442 | 58384542 | -1 | snRNA | ENSG0000C |
| 58392153 | 58392807 | -1 | lncRNA | ENSG0000C |
| 58421326 | 58462470 | 1 | lncRNA | ENSG0000C |
| 58462846 | 58513628 | 1 | protein_coding | ENSG0000C |
| 58496908 | 58497014 | 1 | snRNA | ENSG0000C |
| 58515479 | 58521181 | 1 | protein_coding | ENSG0000C |
| 58519951 | 58629885 | -1 | protein_coding | ENSG0000C |
| 58522970 | 58523842 | 1 | lncRNA | ENSG0000C |
| 58548499 | 58548633 | -1 | snoRNA | ENSG0000C |
| 58559796 | 58559931 | -1 | snoRNA | ENSG0000C |
| 58559796 | 58559929 | -1 | snoRNA | ENSG0000C |
| 58597589 | 58597945 | -1 | transcribed_processe | ENSG0000C |
| 58665109 | 58684770 | -1 | protein_coding | ENSG0000C |
| 58705799 | 58706297 | -1 | TEC | ENSG0000C |
| 58707131 | 58734342 | -1 | protein_coding | ENSG0000C |
| 58733910 | 59199395 | 1 | lncRNA | ENSG0000C |
| 58744251 | 58744357 | -1 | snRNA | ENSG0000C |
| 58770524 | 58770820 | 1 | misc_RNA | ENSG0000C |
| 58776540 | 58777563 | 1 | processed_pseudoge | ENSG0000C |
| 58825891 | 58826368 | 1 | processed_pseudoge | ENSG0000C |
| 58847715 | 58880351 | -1 | lncRNA | ENSG0000C |
| 59004213 | 59004494 | 1 | misc_RNA | ENSG0000C |
| 59088837 | 59089091 | 1 | processed_pseudoge | ENSG0000C |
| 59166951 | 59167152 | -1 | processed_pseudoge | ENSG0000C |
| 59540126 | 59540266 | 1 | snRNA | ENSG0000C |
| 59655602 | 59656336 | 1 | processed_pseudoge | ENSG0000C |
| 59709993 | 59755026 | -1 | transcribed_processe | ENSG0000C |
| 59855353 | 60053973 | 1 | lncRNA | ENSG0000C |

| | | | | |
|---|---|---|---|---|
| 60024578 | 60025141 | -1 | processed_pseudoge | ENSG00000 |
| 60346478 | 60499364 | 1 | lncRNA | ENSG00000 |
| 60357214 | 60357474 | 1 | processed_pseudoge | ENSG00000 |
| 60358455 | 60359480 | -1 | unprocessed_pseudc | ENSG00000 |
| 60486819 | 60523250 | -1 | lncRNA | ENSG00000 |
| 60654235 | 60656649 | -1 | processed_pseudoge | ENSG00000 |
| 60925733 | 60928541 | -1 | unprocessed_pseudc | ENSG00000 |
| 61055399 | 61055964 | -1 | transcribed_processe | ENSG00000 |
| 61638233 | 61642577 | -1 | lncRNA | ENSG00000 |
| 61647242 | 62037035 | -1 | protein_coding | ENSG00000 |
| 61691756 | 61693750 | -1 | lncRNA | ENSG00000 |
| 61743154 | 61743476 | -1 | misc_RNA | ENSG00000 |
| 61918321 | 61940697 | 1 | lncRNA | ENSG00000 |
| 62187540 | 62187646 | -1 | snRNA | ENSG00000 |
| 62342420 | 62342557 | 1 | TEC | ENSG00000 |
| 62357371 | 62357746 | 1 | processed_pseudoge | ENSG00000 |
| 62596372 | 62598449 | 1 | lncRNA | ENSG00000 |
| 62808222 | 62808706 | -1 | processed_pseudoge | ENSG00000 |
| 63056786 | 63129654 | -1 | lncRNA | ENSG00000 |
| 63066526 | 63066990 | -1 | lncRNA | ENSG00000 |
| 63180610 | 63181071 | -1 | processed_pseudoge | ENSG00000 |
| 63314264 | 63618046 | -1 | lncRNA | ENSG00000 |
| 63350763 | 63351149 | -1 | processed_pseudoge | ENSG00000 |
| 63368844 | 63369456 | 1 | lncRNA | ENSG00000 |
| 63397707 | 63397946 | 1 | TEC | ENSG00000 |
| 63703483 | 63703665 | 1 | processed_pseudoge | ENSG00000 |
| 63730205 | 63755884 | 1 | lncRNA | ENSG00000 |
| 63935778 | 63936906 | 1 | lncRNA | ENSG00000 |
| 64155169 | 64156574 | 1 | lncRNA | ENSG00000 |
| 64242608 | 64343909 | -1 | lncRNA | ENSG00000 |
| 64260570 | 64261239 | -1 | processed_pseudoge | ENSG00000 |
| 64364781 | 64600247 | 1 | lncRNA | ENSG00000 |
| 64735925 | 64737283 | 1 | transcribed_processe | ENSG00000 |
| 64738323 | 64738564 | 1 | processed_pseudoge | ENSG00000 |
| 64943753 | 65126112 | -1 | protein_coding | ENSG00000 |
| 64981123 | 64983227 | -1 | TEC | ENSG00000 |
| 65141751 | 65176713 | -1 | lncRNA | ENSG00000 |
| 65190973 | 65234914 | -1 | lncRNA | ENSG00000 |
| 65233056 | 65432820 | 1 | lncRNA | ENSG00000 |
| 65284496 | 65576345 | -1 | lncRNA | ENSG00000 |
| 65580668 | 65646947 | 1 | lncRNA | ENSG00000 |
| 65601101 | 65601698 | -1 | lncRNA | ENSG00000 |
| 65861112 | 65863784 | -1 | lncRNA | ENSG00000 |
| 65938189 | 65938453 | -1 | processed_pseudoge | ENSG00000 |
| 66301809 | 66301940 | 1 | rRNA_pseudogene | ENSG00000 |
| 66327349 | 66335138 | 1 | lncRNA | ENSG00000 |
| 66366622 | 66404784 | 1 | protein_coding | ENSG00000 |
| 66408524 | 66412135 | 1 | lncRNA | ENSG00000 |
| 66427295 | 66493529 | 1 | protein_coding | ENSG00000 |
| 66450998 | 66451351 | -1 | processed_pseudoge | ENSG00000 |

| | | | | |
|---|---|---|---|---|
| 66469796 | 66481230 | -1 | lncRNA | ENSG0000C |
| 66469812 | 66517312 | -1 | protein_coding | ENSG0000C |
| 66508003 | 66552544 | -1 | protein_coding | ENSG0000C |
| 66509437 | 66510048 | 1 | lncRNA | ENSG0000C |
| 66549280 | 66551189 | 1 | lncRNA | ENSG0000C |
| 66550457 | 66550567 | 1 | misc_RNA | ENSG0000C |
| 66552563 | 66566251 | 1 | protein_coding | ENSG0000C |
| 66552587 | 66579135 | 1 | protein_coding | ENSG0000C |
| 66565620 | 66566001 | -1 | lncRNA | ENSG0000C |
| 66566393 | 66579137 | 1 | protein_coding | ENSG0000C |
| 66579448 | 66588275 | 1 | protein_coding | ENSG0000C |
| 66579983 | 66580716 | -1 | lncRNA | ENSG0000C |
| 66603874 | 66613892 | 1 | protein_coding | ENSG0000C |
| 66614750 | 66696707 | -1 | protein_coding | ENSG0000C |
| 66720893 | 66751609 | -1 | protein_coding | ENSG0000C |
| 66720897 | 66731785 | 1 | lncRNA | ENSG0000C |
| 66729152 | 66741416 | 1 | lncRNA | ENSG0000C |
| 66738238 | 66739722 | 1 | lncRNA | ENSG0000C |
| 66751752 | 66754740 | 1 | lncRNA | ENSG0000C |
| 66754976 | 66801620 | -1 | protein_coding | ENSG0000C |
| 66802875 | 66873256 | -1 | protein_coding | ENSG0000C |
| 66841433 | 66853448 | -1 | lncRNA | ENSG0000C |
| 66844414 | 66854153 | 1 | protein_coding | ENSG0000C |
| 66878589 | 66895754 | 1 | protein_coding | ENSG0000C |
| 66908122 | 66918917 | -1 | protein_coding | ENSG0000C |
| 66921679 | 66925536 | -1 | protein_coding | ENSG0000C |
| 66932065 | 66934423 | -1 | protein_coding | ENSG0000C |
| 66934444 | 66945096 | 1 | protein_coding | ENSG0000C |
| 66942712 | 66963256 | 1 | lncRNA | ENSG0000C |
| 66944660 | 66945096 | -1 | TEC | ENSG0000C |
| 66961245 | 66975149 | 1 | protein_coding | ENSG0000C |
| 66988589 | 67009758 | 1 | protein_coding | ENSG0000C |
| 67006672 | 67009481 | -1 | TEC | ENSG0000C |
| 67019519 | 67019817 | 1 | misc_RNA | ENSG0000C |
| 67028984 | 67101058 | 1 | protein_coding | ENSG0000C |
| 67109941 | 67148544 | 1 | protein_coding | ENSG0000C |
| 67148104 | 67150998 | -1 | protein_coding | ENSG0000C |
| 67154185 | 67159909 | -1 | protein_coding | ENSG0000C |
| 67159932 | 67164570 | 1 | protein_coding | ENSG0000C |
| 67163385 | 67165815 | 1 | protein_coding | ENSG0000C |
| 67164681 | 67169945 | 1 | protein_coding | ENSG0000C |
| 67170154 | 67175735 | 1 | protein_coding | ENSG0000C |
| 67175599 | 67184040 | -1 | protein_coding | ENSG0000C |
| 67184379 | 67190185 | -1 | protein_coding | ENSG0000C |
| 67192155 | 67198918 | 1 | protein_coding | ENSG0000C |
| 67199111 | 67204029 | 1 | protein_coding | ENSG0000C |
| 67202321 | 67202395 | -1 | miRNA | ENSG0000C |
| 67205175 | 67206849 | -1 | TEC | ENSG0000C |
| 67207139 | 67227048 | -1 | protein_coding | ENSG0000C |
| 67227103 | 67229278 | 1 | protein_coding | ENSG0000C |

| | | | | |
|---|---|---|---|---|
| 67229387 | 67247481 | -1 | protein_coding | ENSG00000 |
| 67237683 | 67272191 | 1 | protein_coding | ENSG00000 |
| 67261108 | 67263784 | 1 | lncRNA | ENSG00000 |
| 67277510 | 67289499 | 1 | protein_coding | ENSG00000 |
| 67289428 | 67326760 | -1 | protein_coding | ENSG00000 |
| 67321658 | 67321987 | -1 | misc_RNA | ENSG00000 |
| 67326798 | 67385204 | 1 | protein_coding | ENSG00000 |
| 67389809 | 67393518 | -1 | protein_coding | ENSG00000 |
| 67390691 | 67390850 | -1 | snRNA | ENSG00000 |
| 67394152 | 67416833 | -1 | protein_coding | ENSG00000 |
| 67430652 | 67437553 | 1 | protein_coding | ENSG00000 |
| 67430667 | 67431464 | -1 | lncRNA | ENSG00000 |
| 67438014 | 67481181 | -1 | protein_coding | ENSG00000 |
| 67481314 | 67505977 | 1 | lncRNA | ENSG00000 |
| 67482571 | 67483813 | -1 | protein_coding | ENSG00000 |
| 67517862 | 67528675 | -1 | lncRNA | ENSG00000 |
| 67518418 | 67546788 | 1 | protein_coding | ENSG00000 |
| 67538557 | 67540106 | -1 | lncRNA | ENSG00000 |
| 67542123 | 67542963 | -1 | lncRNA | ENSG00000 |
| 67542304 | 67542572 | -1 | lncRNA | ENSG00000 |
| 67549214 | 67563958 | -1 | lncRNA | ENSG00000 |
| 67550815 | 67552935 | -1 | TEC | ENSG00000 |
| 67561411 | 67562309 | -1 | lncRNA | ENSG00000 |
| 67562467 | 67639177 | 1 | protein_coding | ENSG00000 |
| 67614381 | 67616146 | -1 | lncRNA | ENSG00000 |
| 67644988 | 67657569 | 1 | protein_coding | ENSG00000 |
| 67657512 | 67660815 | -1 | protein_coding | ENSG00000 |
| 67660946 | 67662778 | 1 | protein_coding | ENSG00000 |
| 67662945 | 67667265 | -1 | protein_coding | ENSG00000 |
| 67666814 | 67668757 | 1 | protein_coding | ENSG00000 |
| 67674531 | 67719339 | -1 | protein_coding | ENSG00000 |
| 67723070 | 67806652 | -1 | protein_coding | ENSG00000 |
| 67738588 | 67739922 | -1 | lncRNA | ENSG00000 |
| 67806765 | 67832148 | 1 | protein_coding | ENSG00000 |
| 67828157 | 67847811 | -1 | protein_coding | ENSG00000 |
| 67842320 | 67844195 | 1 | protein_coding | ENSG00000 |
| 67846923 | 67872567 | 1 | protein_coding | ENSG00000 |
| 67855955 | 67856111 | -1 | processed_pseudoge | ENSG00000 |
| 67873052 | 67884499 | 1 | protein_coding | ENSG00000 |
| 67882461 | 67886367 | 1 | lncRNA | ENSG00000 |
| 67884885 | 67888855 | 1 | protein_coding | ENSG00000 |
| 67893254 | 67929676 | 1 | protein_coding | ENSG00000 |
| 67927640 | 67932365 | -1 | protein_coding | ENSG00000 |
| 67929614 | 67936017 | -1 | protein_coding | ENSG00000 |
| 67934506 | 67936864 | -1 | protein_coding | ENSG00000 |
| 67939750 | 67944131 | -1 | protein_coding | ENSG00000 |
| 67943474 | 67969601 | -1 | protein_coding | ENSG00000 |
| 67975663 | 67980549 | -1 | protein_coding | ENSG00000 |
| 67987394 | 68000586 | -1 | protein_coding | ENSG00000 |
| 67987746 | 68079320 | 1 | protein_coding | ENSG00000 |

| | | | | |
|---|---|---|---|---|
| 68002432 | 68004952 | -1 | unprocessed_pseudc | ENSG00000( |
| 68013436 | 68015911 | -1 | protein_coding | ENSG00000( |
| 68017710 | 68017817 | -1 | snRNA | ENSG00000( |
| 68021274 | 68023442 | -1 | protein_coding | ENSG00000( |
| 68077340 | 68122271 | 1 | lncRNA | ENSG00000( |
| 68084751 | 68229259 | 1 | protein_coding | ENSG00000( |
| 68089449 | 68089543 | -1 | misc_RNA | ENSG00000( |
| 68189287 | 68189421 | 1 | snoRNA | ENSG00000( |
| 68199795 | 68200981 | 1 | lncRNA | ENSG00000( |
| 68199841 | 68200981 | -1 | TEC | ENSG00000( |
| 68209442 | 68209778 | 1 | processed_pseudoge | ENSG00000( |
| 68212401 | 68221671 | -1 | lncRNA | ENSG00000( |
| 68224713 | 68227734 | 1 | lncRNA | ENSG00000( |
| 68225969 | 68229145 | -1 | lncRNA | ENSG00000( |
| 68229033 | 68238102 | -1 | protein_coding | ENSG00000( |
| 68233426 | 68233499 | -1 | miRNA | ENSG00000( |
| 68236845 | 68237667 | -1 | lncRNA | ENSG00000( |
| 68245304 | 68261058 | 1 | protein_coding | ENSG00000( |
| 68256162 | 68260443 | -1 | lncRNA | ENSG00000( |
| 68264516 | 68301823 | 1 | protein_coding | ENSG00000( |
| 68284223 | 68284318 | -1 | snRNA | ENSG00000( |
| 68284503 | 68310946 | -1 | protein_coding | ENSG00000( |
| 68290087 | 68292790 | -1 | TEC | ENSG00000( |
| 68310974 | 68358563 | 1 | protein_coding | ENSG00000( |
| 68316801 | 68319036 | 1 | lncRNA | ENSG00000( |
| 68329387 | 68329529 | -1 | snRNA | ENSG00000( |
| 68358327 | 68448508 | -1 | protein_coding | ENSG00000( |
| 68367325 | 68370262 | -1 | TEC | ENSG00000( |
| 68448843 | 68450002 | 1 | lncRNA | ENSG00000( |
| 68450283 | 68452318 | 1 | lncRNA | ENSG00000( |
| 68474553 | 68474664 | -1 | misc_RNA | ENSG00000( |
| 68477237 | 68477570 | -1 | processed_pseudoge | ENSG00000( |
| 68530090 | 68576072 | 1 | protein_coding | ENSG00000( |
| 68535515 | 68535658 | 1 | snRNA | ENSG00000( |
| 68573782 | 68589512 | -1 | lncRNA | ENSG00000( |
| 68591382 | 68594424 | 1 | lncRNA | ENSG00000( |
| 68630223 | 68630354 | -1 | snoRNA | ENSG00000( |
| 68630851 | 68630968 | -1 | snoRNA | ENSG00000( |
| 68636189 | 68722616 | 1 | protein_coding | ENSG00000( |
| 68644248 | 68646168 | -1 | lncRNA | ENSG00000( |
| 68651056 | 68651948 | -1 | processed_pseudoge | ENSG00000( |
| 68686399 | 68686670 | 1 | transcribed_processe | ENSG00000( |
| 68737292 | 68835541 | 1 | protein_coding | ENSG00000( |
| 68742480 | 68742588 | 1 | rRNA_pseudogene | ENSG00000( |
| 68814330 | 68823526 | -1 | lncRNA | ENSG00000( |
| 68822587 | 68823070 | 1 | processed_pseudoge | ENSG00000( |
| 68843531 | 69085182 | 1 | protein_coding | ENSG00000( |
| 68885576 | 68885677 | -1 | misc_RNA | ENSG00000( |
| 68926894 | 68926957 | -1 | snRNA | ENSG00000( |
| 68927547 | 68948261 | -1 | lncRNA | ENSG00000( |

| | | | | |
|---|---|---|---|---|
| 68933820 | 68937725 | 1 | lncRNA | ENSG0000( |
| 69007052 | 69007155 | -1 | snRNA | ENSG0000( |
| 69025860 | 69026643 | -1 | processed_pseudoge | ENSG0000( |
| 69033675 | 69036042 | 1 | transcribed_processe | ENSG0000( |
| 69105653 | 69118719 | 1 | protein_coding | ENSG0000( |
| 69118010 | 69132578 | -1 | protein_coding | ENSG0000( |
| 69118010 | 69132151 | -1 | protein_coding | ENSG0000( |
| 69131291 | 69231130 | 1 | protein_coding | ENSG0000( |
| 69175671 | 69175777 | -1 | snRNA | ENSG0000( |
| 69187129 | 69309052 | 1 | protein_coding | ENSG0000( |
| 69191123 | 69191223 | 1 | misc_RNA | ENSG0000( |
| 69240549 | 69241929 | 1 | lncRNA | ENSG0000( |
| 69299682 | 69322700 | 1 | protein_coding | ENSG0000( |
| 69311350 | 69326939 | 1 | protein_coding | ENSG0000( |
| 69320140 | 69339667 | -1 | protein_coding | ENSG0000( |
| 69326913 | 69330588 | -1 | protein_coding | ENSG0000( |
| 69328644 | 69334871 | -1 | protein_coding | ENSG0000( |
| 69335091 | 69356306 | -1 | protein_coding | ENSG0000( |
| 69335276 | 69351730 | -1 | protein_coding | ENSG0000( |
| 69337996 | 69343106 | 1 | protein_coding | ENSG0000( |
| 69343250 | 69351786 | -1 | protein_coding | ENSG0000( |
| 69355567 | 69408571 | -1 | protein_coding | ENSG0000( |
| 69424525 | 69466266 | 1 | protein_coding | ENSG0000( |
| 69443240 | 69444290 | -1 | processed_pseudoge | ENSG0000( |
| 69463844 | 69466264 | 1 | lncRNA | ENSG0000( |
| 69565094 | 69704666 | 1 | protein_coding | ENSG0000( |
| 69565808 | 69565868 | -1 | miRNA | ENSG0000( |
| 69632141 | 69632571 | 1 | lncRNA | ENSG0000( |
| 69703065 | 69704652 | 1 | lncRNA | ENSG0000( |
| 69706996 | 69726668 | -1 | protein_coding | ENSG0000( |
| 69709874 | 69710583 | 1 | lncRNA | ENSG0000( |
| 69727013 | 69742563 | 1 | lncRNA | ENSG0000( |
| 69741871 | 69754926 | -1 | protein_coding | ENSG0000( |
| 69756536 | 69757937 | -1 | processed_pseudoge | ENSG0000( |
| 69762306 | 69941741 | 1 | protein_coding | ENSG0000( |
| 69785994 | 69786113 | -1 | snoRNA | ENSG0000( |
| 69933081 | 69933180 | 1 | miRNA | ENSG0000( |
| 69950705 | 69964452 | 1 | protein_coding | ENSG0000( |
| 69975238 | 69975344 | 1 | unprocessed_pseudc | ENSG0000( |
| 69976297 | 70065948 | -1 | lncRNA | ENSG0000( |
| 69976388 | 69996188 | -1 | transcribed_unproce | ENSG0000( |
| 70010751 | 70065945 | -1 | transcribed_unproce | ENSG0000( |
| 70030346 | 70030422 | 1 | miRNA | ENSG0000( |
| 70079129 | 70079624 | -1 | processed_pseudoge | ENSG0000( |
| 70113626 | 70162537 | 1 | protein_coding | ENSG0000( |
| 70138337 | 70173483 | -1 | lncRNA | ENSG0000( |
| 70173322 | 70187361 | 1 | protein_coding | ENSG0000( |
| 70202354 | 70202459 | 1 | processed_pseudoge | ENSG0000( |
| 70219574 | 70246612 | -1 | transcribed_unproce | ENSG0000( |
| 70246778 | 70251940 | -1 | protein_coding | ENSG0000( |

| | | | | |
|---|---|---|---|---|
| 70252295 | 70289543 | -1 | protein_coding | ENSG00000[illegible] |
| 70276069 | 70276352 | 1 | misc_RNA | ENSG00000[illegible] |
| 70289663 | 70335283 | 1 | protein_coding | ENSG00000[illegible] |
| 70299194 | 70372582 | 1 | protein_coding | ENSG00000[illegible] |
| 70315442 | 70346806 | -1 | lncRNA | ENSG00000[illegible] |
| 70346861 | 70373383 | 1 | protein_coding | ENSG00000[illegible] |
| 70349085 | 70372166 | -1 | lncRNA | ENSG00000[illegible] |
| 70375977 | 70439237 | -1 | protein_coding | ENSG00000[illegible] |
| 70379457 | 70399502 | 1 | lncRNA | ENSG00000[illegible] |
| 70444674 | 70444912 | -1 | processed_pseudoge | ENSG00000[illegible] |
| 70445293 | 70445399 | -1 | snRNA | ENSG00000[illegible] |
| 70454595 | 70480274 | 1 | protein_coding | ENSG00000[illegible] |
| 70480568 | 70523558 | -1 | protein_coding | ENSG00000[illegible] |
| 70523791 | 70577670 | 1 | protein_coding | ENSG00000[illegible] |
| 70529509 | 70529588 | 1 | snoRNA | ENSG00000[illegible] |
| 70538005 | 70538098 | 1 | snoRNA | ENSG00000[illegible] |
| 70579895 | 70660682 | 1 | protein_coding | ENSG00000[illegible] |
| 70661204 | 70686053 | -1 | protein_coding | ENSG00000[illegible] |
| 70661667 | 70665836 | 1 | TEC | ENSG00000[illegible] |
| 70687439 | 70801160 | -1 | protein_coding | ENSG00000[illegible] |
| 70713987 | 70716890 | 1 | TEC | ENSG00000[illegible] |
| 70747057 | 70747926 | 1 | lncRNA | ENSG00000[illegible] |
| 70755035 | 70773251 | 1 | lncRNA | ENSG00000[illegible] |
| 70768356 | 70768785 | 1 | TEC | ENSG00000[illegible] |
| 70768967 | 70770221 | -1 | TEC | ENSG00000[illegible] |
| 70778433 | 70779145 | 1 | TEC | ENSG00000[illegible] |
| 70802083 | 70802840 | -1 | lncRNA | ENSG00000[illegible] |
| 70802084 | 71230722 | -1 | protein_coding | ENSG00000[illegible] |
| 70852698 | 70852814 | 1 | snRNA | ENSG00000[illegible] |
| 71080866 | 71093667 | 1 | lncRNA | ENSG00000[illegible] |
| 71113352 | 71114000 | 1 | lncRNA | ENSG00000[illegible] |
| 71281389 | 71289715 | -1 | protein_coding | ENSG00000[illegible] |
| 71288775 | 71309963 | 1 | lncRNA | ENSG00000[illegible] |
| 71358713 | 71390436 | 1 | protein_coding | ENSG00000[illegible] |
| 71366933 | 71426464 | -1 | lncRNA | ENSG00000[illegible] |
| 71420483 | 71421518 | 1 | unprocessed_pseudo | ENSG00000[illegible] |
| 71430262 | 71442060 | -1 | protein_coding | ENSG00000[illegible] |
| 71443055 | 71443856 | -1 | unprocessed_pseudo | ENSG00000[illegible] |
| 71447597 | 71463095 | -1 | protein_coding | ENSG00000[illegible] |
| 71447600 | 71489311 | -1 | protein_coding | ENSG00000[illegible] |
| 71462278 | 71465941 | 1 | lncRNA | ENSG00000[illegible] |
| 71464555 | 71565089 | -1 | protein_coding | ENSG00000[illegible] |
| 71513863 | 71517849 | 1 | lncRNA | ENSG00000[illegible] |
| 71525233 | 71538746 | 1 | protein_coding | ENSG00000[illegible] |
| 71539834 | 71541825 | -1 | lncRNA | ENSG00000[illegible] |
| 71563430 | 71563528 | -1 | snRNA | ENSG00000[illegible] |
| 71564952 | 71578187 | 1 | lncRNA | ENSG00000[illegible] |
| 71565660 | 71577092 | -1 | protein_coding | ENSG00000[illegible] |
| 71623708 | 71626816 | -1 | lncRNA | ENSG00000[illegible] |
| 71626161 | 71642114 | 1 | protein_coding | ENSG00000[illegible] |

| | | | | |
|---|---|---|---|---|
| 71637835 | 71724701 | -1 | protein_coding | ENSG00000 |
| 71655027 | 71664212 | 1 | lncRNA | ENSG00000 |
| 71666091 | 71666197 | 1 | snRNA | ENSG00000 |
| 71698567 | 71698701 | -1 | snoRNA | ENSG00000 |
| 71723180 | 71724230 | 1 | lncRNA | ENSG00000 |
| 71726975 | 71727992 | 1 | lncRNA | ENSG00000 |
| 71729000 | 71809201 | -1 | protein_coding | ENSG00000 |
| 71758402 | 71758487 | -1 | snoRNA | ENSG00000 |
| 71777603 | 71777834 | -1 | processed_pseudoge | ENSG00000 |
| 71833787 | 71835932 | 1 | unprocessed_pseudo | ENSG00000 |
| 71845976 | 71885268 | 1 | protein_coding | ENSG00000 |
| 71845996 | 71931199 | 1 | protein_coding | ENSG00000 |
| 71859680 | 71860676 | 1 | TEC | ENSG00000 |
| 71859680 | 71895336 | -1 | protein_coding | ENSG00000 |
| 71881553 | 71882273 | -1 | lncRNA | ENSG00000 |
| 71929538 | 71999978 | -1 | protein_coding | ENSG00000 |
| 71991151 | 71991305 | -1 | processed_pseudoge | ENSG00000 |
| 72004239 | 72004803 | -1 | unprocessed_pseudo | ENSG00000 |
| 72005037 | 72006543 | -1 | processed_pseudoge | ENSG00000 |
| 72008588 | 72027664 | 1 | protein_coding | ENSG00000 |
| 72044289 | 72094431 | -1 | protein_coding | ENSG00000 |
| 72054505 | 72061055 | 1 | protein_coding | ENSG00000 |
| 72063148 | 72077246 | 1 | protein_coding | ENSG00000 |
| 72084857 | 72087443 | 1 | lncRNA | ENSG00000 |
| 72093613 | 72112912 | 1 | protein_coding | ENSG00000 |
| 72112157 | 72176878 | -1 | protein_coding | ENSG00000 |
| 72224565 | 72225144 | -1 | lncRNA | ENSG00000 |
| 72236281 | 72665014 | -1 | lncRNA | ENSG00000 |
| 72278268 | 72278568 | 1 | processed_pseudoge | ENSG00000 |
| 72376704 | 72376760 | 1 | snRNA | ENSG00000 |
| 72425948 | 72533892 | 1 | lncRNA | ENSG00000 |
| 72665123 | 72822781 | 1 | lncRNA | ENSG00000 |
| 72673503 | 72673564 | 1 | snRNA | ENSG00000 |
| 72728616 | 72729894 | 1 | transcribed_processe | ENSG00000 |
| 72772673 | 72773272 | 1 | lncRNA | ENSG00000 |
| 72782885 | 73891871 | -1 | protein_coding | ENSG00000 |
| 72805998 | 72809872 | 1 | lncRNA | ENSG00000 |
| 72821933 | 72821992 | -1 | snRNA | ENSG00000 |
| 72973374 | 72973832 | -1 | TEC | ENSG00000 |
| 73005601 | 73006141 | -1 | TEC | ENSG00000 |
| 73014319 | 73014476 | -1 | lncRNA | ENSG00000 |
| 73015047 | 73016016 | -1 | TEC | ENSG00000 |
| 73060245 | 73062316 | 1 | lncRNA | ENSG00000 |
| 73092349 | 73099337 | 1 | lncRNA | ENSG00000 |
| 73123296 | 73137580 | 1 | lncRNA | ENSG00000 |
| 73232055 | 73233970 | 1 | lncRNA | ENSG00000 |
| 73293866 | 73294296 | -1 | TEC | ENSG00000 |
| 73386771 | 73504954 | 1 | lncRNA | ENSG00000 |
| 73543927 | 73569309 | 1 | lncRNA | ENSG00000 |
| 73794057 | 73809733 | 1 | lncRNA | ENSG00000 |

| | | | | |
|---|---|---|---|---|
| 73812560 | 73816054 | 1 | lncRNA | ENSG0000C |
| 73940639 | 73941781 | -1 | translated_processec | ENSG0000C |
| 73943078 | 74296762 | -1 | lncRNA | ENSG0000C |
| 74053365 | 74053965 | 1 | TEC | ENSG0000C |
| 74150520 | 74151299 | 1 | TEC | ENSG0000C |
| 74220929 | 74221427 | 1 | processed_pseudoge | ENSG0000C |
| 74282415 | 74287519 | 1 | lncRNA | ENSG0000C |
| 74289593 | 74291052 | -1 | lncRNA | ENSG0000C |
| 74296814 | 74306288 | 1 | protein_coding | ENSG0000C |
| 74305127 | 74335346 | -1 | lncRNA | ENSG0000C |
| 74312609 | 74313390 | -1 | processed_pseudoge | ENSG0000C |
| 74313337 | 74315634 | 1 | lncRNA | ENSG0000C |
| 74332402 | 74368240 | -1 | transcribed_unproce | ENSG0000C |
| 74367462 | 74369826 | 1 | lncRNA | ENSG0000C |
| 74377878 | 74392080 | 1 | protein_coding | ENSG0000C |
| 74393076 | 74393183 | -1 | unprocessed_pseudc | ENSG0000C |
| 74408270 | 74421953 | -1 | protein_coding | ENSG0000C |
| 74422120 | 74441937 | 1 | lncRNA | ENSG0000C |
| 74447427 | 74607144 | -1 | protein_coding | ENSG0000C |
| 74456541 | 74456611 | 1 | snRNA | ENSG0000C |
| 74463888 | 74463988 | -1 | misc_RNA | ENSG0000C |
| 74552251 | 74552746 | 1 | processed_pseudoge | ENSG0000C |
| 74612596 | 74613221 | -1 | processed_pseudoge | ENSG0000C |
| 74621399 | 74666877 | -1 | protein_coding | ENSG0000C |
| 74667506 | 74668706 | 1 | processed_pseudoge | ENSG0000C |
| 74671855 | 74700960 | -1 | protein_coding | ENSG0000C |
| 74712955 | 74774831 | -1 | protein_coding | ENSG0000C |
| 74842587 | 74843883 | 1 | unprocessed_pseudc | ENSG0000C |
| 74871362 | 75000173 | -1 | protein_coding | ENSG0000C |
| 74999024 | 75110994 | 1 | protein_coding | ENSG0000C |
| 75108601 | 75110712 | -1 | lncRNA | ENSG0000C |
| 75111860 | 75116771 | -1 | protein_coding | ENSG0000C |
| 75119558 | 75144200 | 1 | lncRNA | ENSG0000C |
| 75148494 | 75172236 | 1 | protein_coding | ENSG0000C |
| 75192465 | 75193247 | -1 | processed_pseudoge | ENSG0000C |
| 75204103 | 75207161 | -1 | protein_coding | ENSG0000C |
| 75218988 | 75226338 | 1 | protein_coding | ENSG0000C |
| 75226074 | 75228197 | 1 | transcribed_unproce | ENSG0000C |
| 75228181 | 75268053 | -1 | protein_coding | ENSG0000C |
| 75245994 | 75250077 | -1 | TEC | ENSG0000C |
| 75286712 | 75286800 | -1 | snRNA | ENSG0000C |
| 75293698 | 75433503 | -1 | protein_coding | ENSG0000C |
| 75321890 | 75325048 | -1 | lncRNA | ENSG0000C |
| 75379818 | 75381260 | -1 | lncRNA | ENSG0000C |
| 75409332 | 75409466 | -1 | snoRNA | ENSG0000C |
| 75412684 | 75464706 | -1 | protein_coding | ENSG0000C |
| 75433836 | 75436392 | 1 | lncRNA | ENSG0000C |
| 75443054 | 75465497 | -1 | protein_coding | ENSG0000C |
| 75458252 | 75460017 | -1 | lncRNA | ENSG0000C |
| 75472052 | 75495445 | -1 | protein_coding | ENSG0000C |

| | | | | |
|---|---|---|---|---|
| 75475896 | 75495407 | 1 | lncRNA | ENSG0000C |
| 75497948 | 75498289 | -1 | lncRNA | ENSG0000C |
| 75499788 | 75516280 | -1 | transcribed_unproce | ENSG0000C |
| 75528532 | 75545428 | -1 | protein_coding | ENSG0000C |
| 75528535 | 75535247 | -1 | protein_coding | ENSG0000C |
| 75536741 | 75556268 | -1 | protein_coding | ENSG0000C |
| 75541399 | 75608445 | -1 | lncRNA | ENSG0000C |
| 75556392 | 75557059 | 1 | lncRNA | ENSG0000C |
| 75566375 | 75577881 | 1 | protein_coding | ENSG0000C |
| 75572185 | 75572685 | -1 | lncRNA | ENSG0000C |
| 75596981 | 75623300 | -1 | protein_coding | ENSG0000C |
| 75627474 | 75648643 | -1 | protein_coding | ENSG0000C |
| 75646606 | 75646681 | 1 | unprocessed_pseudc | ENSG0000C |
| 75647773 | 75761872 | 1 | protein_coding | ENSG0000C |
| 75660227 | 75677009 | -1 | protein_coding | ENSG0000C |
| 75693893 | 75701461 | -1 | protein_coding | ENSG0000C |
| 75698531 | 75698816 | -1 | misc_RNA | ENSG0000C |
| 75709056 | 75709815 | -1 | processed_pseudoge | ENSG0000C |
| 75714131 | 75726490 | -1 | protein_coding | ENSG0000C |
| 75838576 | 75860869 | -1 | lncRNA | ENSG0000C |
| 75861048 | 75861157 | -1 | rRNA_pseudogene | ENSG0000C |
| 76107283 | 76147806 | -1 | lncRNA | ENSG0000C |
| 76228379 | 76262365 | -1 | lncRNA | ENSG0000C |
| 76235055 | 76235623 | -1 | processed_pseudoge | ENSG0000C |
| 76264776 | 76277303 | -1 | lncRNA | ENSG0000C |
| 76277278 | 76560757 | 1 | protein_coding | ENSG0000C |
| 76277288 | 76819624 | 1 | protein_coding | ENSG0000C |
| 76287630 | 76287739 | 1 | misc_RNA | ENSG0000C |
| 76469508 | 76469595 | -1 | snoRNA | ENSG0000C |
| 76582556 | 76583098 | 1 | lncRNA | ENSG0000C |
| 76634998 | 76658478 | 1 | lncRNA | ENSG0000C |
| 76736342 | 76928169 | 1 | lncRNA | ENSG0000C |
| 76868936 | 76869019 | 1 | miRNA | ENSG0000C |
| 76878274 | 76878936 | -1 | processed_pseudoge | ENSG0000C |
| 77104260 | 77105297 | 1 | TEC | ENSG0000C |
| 77190835 | 77202398 | 1 | protein_coding | ENSG0000C |
| 77199408 | 77213215 | 1 | protein_coding | ENSG0000C |
| 77201474 | 77249957 | -1 | lncRNA | ENSG0000C |
| 77220406 | 77221258 | 1 | unitary_pseudogene | ENSG0000C |
| 77234877 | 77299950 | 1 | lncRNA | ENSG0000C |
| 77247813 | 77435034 | -1 | protein_coding | ENSG0000C |
| 77433382 | 77446798 | 1 | lncRNA | ENSG0000C |
| 77590806 | 77610681 | 1 | lncRNA | ENSG0000C |
| 77657174 | 77660570 | 1 | TEC | ENSG0000C |
| 77722492 | 77742260 | 1 | protein_coding | ENSG0000C |
| 77741468 | 77743000 | -1 | lncRNA | ENSG0000C |
| 77752372 | 77756681 | 1 | lncRNA | ENSG0000C |
| 77788564 | 77980107 | 1 | protein_coding | ENSG0000C |
| 77800023 | 77801714 | 1 | TEC | ENSG0000C |
| 77892422 | 77892915 | 1 | processed_pseudoge | ENSG0000C |

| | | | | |
|---|---|---|---|---|
| 78014683 | 78059454 | -1 | lncRNA | ENSG00000( |
| 78022515 | 78066761 | 1 | protein_coding | ENSG00000( |
| 78048483 | 78049897 | -1 | processed_pseudoge | ENSG00000( |
| 78055824 | 78056927 | -1 | processed_pseudoge | ENSG00000( |
| 78099430 | 79212667 | 1 | protein_coding | ENSG00000( |
| 78123243 | 78124332 | 1 | lncRNA | ENSG00000( |
| 78166717 | 78168514 | -1 | TEC | ENSG00000( |
| 78234652 | 78241989 | -1 | lncRNA | ENSG00000( |
| 78273497 | 78275607 | -1 | lncRNA | ENSG00000( |
| 78355529 | 78355834 | 1 | processed_pseudoge | ENSG00000( |
| 78495926 | 78506568 | -1 | lncRNA | ENSG00000( |
| 78525189 | 78526535 | -1 | TEC | ENSG00000( |
| 78534374 | 78535648 | 1 | lncRNA | ENSG00000( |
| 78550739 | 78553296 | -1 | TEC | ENSG00000( |
| 78793584 | 78796362 | -1 | lncRNA | ENSG00000( |
| 78825281 | 78826006 | 1 | processed_pseudoge | ENSG00000( |
| 78872739 | 78873505 | 1 | TEC | ENSG00000( |
| 78890231 | 78892302 | 1 | TEC | ENSG00000( |
| 78895361 | 78899644 | 1 | lncRNA | ENSG00000( |
| 78981722 | 78981843 | -1 | TEC | ENSG00000( |
| 78994345 | 79004730 | 1 | lncRNA | ENSG00000( |
| 79016136 | 79017429 | 1 | TEC | ENSG00000( |
| 79077376 | 79078234 | -1 | lncRNA | ENSG00000( |
| 79090050 | 79101754 | 1 | lncRNA | ENSG00000( |
| 79105957 | 79110882 | 1 | lncRNA | ENSG00000( |
| 79202624 | 79206739 | -1 | lncRNA | ENSG00000( |
| 79208493 | 79211264 | -1 | lncRNA | ENSG00000( |
| 79212711 | 79229453 | -1 | lncRNA | ENSG00000( |
| 79264454 | 79264573 | 1 | rRNA_pseudogene | ENSG00000( |
| 79505603 | 79516293 | 1 | lncRNA | ENSG00000( |
| 79585843 | 79600737 | -1 | protein_coding | ENSG00000( |
| 79603572 | 79604177 | 1 | lncRNA | ENSG00000( |
| 79605802 | 79606605 | 1 | lncRNA | ENSG00000( |
| 79619469 | 79620110 | 1 | lncRNA | ENSG00000( |
| 79675685 | 79677715 | -1 | TEC | ENSG00000( |
| 79676048 | 79809716 | 1 | lncRNA | ENSG00000( |
| 79715220 | 79770651 | -1 | lncRNA | ENSG00000( |
| 79798050 | 79827150 | -1 | lncRNA | ENSG00000( |
| 80040662 | 80045745 | 1 | lncRNA | ENSG00000( |
| 80065836 | 80167577 | 1 | lncRNA | ENSG00000( |
| 80154979 | 80563354 | -1 | lncRNA | ENSG00000( |
| 80179129 | 80180682 | 1 | lncRNA | ENSG00000( |
| 80540734 | 80550811 | 1 | protein_coding | ENSG00000( |
| 80547842 | 80552514 | -1 | lncRNA | ENSG00000( |
| 80553256 | 80572808 | -1 | lncRNA | ENSG00000( |
| 80597907 | 80804598 | -1 | protein_coding | ENSG00000( |
| 80627551 | 80628379 | -1 | lncRNA | ENSG00000( |
| 80736360 | 80742305 | 1 | lncRNA | ENSG00000( |
| 80826074 | 80892682 | -1 | lncRNA | ENSG00000( |
| 80828736 | 81006894 | -1 | protein_coding | ENSG00000( |

| | | | | |
|---|---|---|---|---|
| 80966448 | 81020270 | -1 | protein_coding | ENSG0000C |
| 80982319 | 80984094 | -1 | TEC | ENSG0000C |
| 81006498 | 81033114 | 1 | protein_coding | ENSG0000C |
| 81016792 | 81035759 | -1 | lncRNA | ENSG0000C |
| 81022316 | 81096284 | -1 | protein_coding | ENSG0000C |
| 81030770 | 81031485 | 1 | lncRNA | ENSG0000C |
| 81035842 | 81047350 | 1 | protein_coding | ENSG0000C |
| 81053497 | 81077267 | -1 | protein_coding | ENSG0000C |
| 81053587 | 81096296 | -1 | protein_coding | ENSG0000C |
| 81055301 | 81056426 | 1 | lncRNA | ENSG0000C |
| 81069854 | 81076598 | 1 | lncRNA | ENSG0000C |
| 81077319 | 81078861 | 1 | lncRNA | ENSG0000C |
| 81081938 | 81096425 | -1 | protein_coding | ENSG0000C |
| 81100875 | 81220370 | -1 | polymorphic_pseudc | ENSG0000C |
| 81106533 | 81106639 | -1 | snRNA | ENSG0000C |
| 81151820 | 81153976 | 1 | lncRNA | ENSG0000C |
| 81238689 | 81291142 | 1 | protein_coding | ENSG0000C |
| 81261924 | 81264091 | -1 | TEC | ENSG0000C |
| 81310731 | 81313423 | -1 | lncRNA | ENSG0000C |
| 81314966 | 81380198 | 1 | protein_coding | ENSG0000C |
| 81385018 | 81385093 | 1 | miRNA | ENSG0000C |
| 81385463 | 81387560 | 1 | lncRNA | ENSG0000C |
| 81445170 | 81711762 | 1 | protein_coding | ENSG0000C |
| 81463892 | 81464859 | -1 | TEC | ENSG0000C |
| 81476023 | 81476519 | -1 | processed_pseudoge | ENSG0000C |
| 81517824 | 81520617 | 1 | TEC | ENSG0000C |
| 81533902 | 81533966 | 1 | miRNA | ENSG0000C |
| 81611348 | 81611408 | 1 | miRNA | ENSG0000C |
| 81617038 | 81617543 | -1 | TEC | ENSG0000C |
| 81618505 | 81620563 | 1 | TEC | ENSG0000C |
| 81630193 | 81632038 | 1 | TEC | ENSG0000C |
| 81738248 | 81767868 | 1 | lncRNA | ENSG0000C |
| 81739097 | 81962685 | 1 | protein_coding | ENSG0000C |
| 81961926 | 81962243 | 1 | misc_RNA | ENSG0000C |
| 81987948 | 81989016 | 1 | lncRNA | ENSG0000C |
| 81988855 | 82011481 | -1 | protein_coding | ENSG0000C |
| 82035004 | 82098534 | 1 | protein_coding | ENSG0000C |
| 82044336 | 82139631 | -1 | lncRNA | ENSG0000C |
| 82120638 | 82121769 | -1 | processed_pseudoge | ENSG0000C |
| 82147798 | 82170224 | -1 | protein_coding | ENSG0000C |
| 82170296 | 82175803 | 1 | lncRNA | ENSG0000C |
| 82192505 | 82192790 | -1 | misc_RNA | ENSG0000C |
| 82434828 | 82435381 | 1 | processed_pseudoge | ENSG0000C |
| 82491738 | 82575518 | -1 | lncRNA | ENSG0000C |
| 82626965 | 83800640 | 1 | protein_coding | ENSG0000C |
| 82688931 | 82689019 | 1 | miRNA | ENSG0000C |
| 82727474 | 82727694 | 1 | TEC | ENSG0000C |
| 82773319 | 82829638 | 1 | lncRNA | ENSG0000C |
| 82816831 | 82832997 | -1 | lncRNA | ENSG0000C |
| 82844455 | 82844757 | 1 | misc_RNA | ENSG0000C |

| | | | | |
|---|---|---|---|---|
| 82953230 | 82990298 | -1 | lncRNA | ENSG00000( |
| 83383007 | 83398170 | -1 | lncRNA | ENSG00000( |
| 83508346 | 83508408 | 1 | miRNA | ENSG00000( |
| 83687779 | 83717898 | -1 | lncRNA | ENSG00000( |
| 83720402 | 83772974 | -1 | lncRNA | ENSG00000( |
| 83725676 | 83726377 | -1 | lncRNA | ENSG00000( |
| 83789967 | 83807834 | -1 | lncRNA | ENSG00000( |
| 83807978 | 83819737 | 1 | protein_coding | ENSG00000( |
| 83899115 | 83951445 | 1 | protein_coding | ENSG00000( |
| 83929796 | 83931223 | -1 | lncRNA | ENSG00000( |
| 83931311 | 83966332 | 1 | protein_coding | ENSG00000( |
| 83968632 | 84002776 | 1 | protein_coding | ENSG00000( |
| 84009667 | 84042636 | -1 | protein_coding | ENSG00000( |
| 84039918 | 84040033 | -1 | rRNA_pseudogene | ENSG00000( |
| 84040790 | 84045333 | 1 | lncRNA | ENSG00000( |
| 84053761 | 84116942 | -1 | protein_coding | ENSG00000( |
| 84085979 | 84086590 | -1 | lncRNA | ENSG00000( |
| 84117051 | 84117571 | 1 | lncRNA | ENSG00000( |
| 84119293 | 84121153 | 1 | processed_pseudoge | ENSG00000( |
| 84122141 | 84145192 | -1 | protein_coding | ENSG00000( |
| 84145287 | 84178767 | 1 | protein_coding | ENSG00000( |
| 84177847 | 84187070 | -1 | protein_coding | ENSG00000( |
| 84191138 | 84197168 | 1 | protein_coding | ENSG00000( |
| 84192558 | 84197053 | -1 | lncRNA | ENSG00000( |
| 84218667 | 84239750 | -1 | protein_coding | ENSG00000( |
| 84255270 | 84261049 | -1 | lncRNA | ENSG00000( |
| 84262815 | 84262928 | -1 | rRNA_pseudogene | ENSG00000( |
| 84294846 | 84329844 | 1 | protein_coding | ENSG00000( |
| 84295498 | 84296985 | -1 | lncRNA | ENSG00000( |
| 84342464 | 84343407 | -1 | lncRNA | ENSG00000( |
| 84368527 | 84464187 | 1 | protein_coding | ENSG00000( |
| 84459259 | 84467361 | -1 | lncRNA | ENSG00000( |
| 84476355 | 84554033 | -1 | protein_coding | ENSG00000( |
| 84495599 | 84497495 | 1 | lncRNA | ENSG00000( |
| 84565596 | 84618078 | -1 | protein_coding | ENSG00000( |
| 84594393 | 84596826 | 1 | lncRNA | ENSG00000( |
| 84648511 | 84667686 | 1 | protein_coding | ENSG00000( |
| 84699978 | 84779922 | 1 | protein_coding | ENSG00000( |
| 84817182 | 84856665 | -1 | lncRNA | ENSG00000( |
| 84819985 | 84920768 | 1 | protein_coding | ENSG00000( |
| 84828263 | 84829242 | 1 | lncRNA | ENSG00000( |
| 84917377 | 84924808 | -1 | lncRNA | ENSG00000( |
| 84928353 | 84939603 | 1 | lncRNA | ENSG00000( |
| 84974181 | 85011535 | -1 | protein_coding | ENSG00000( |
| 85027751 | 85094230 | 1 | protein_coding | ENSG00000( |
| 85098358 | 85112472 | -1 | protein_coding | ENSG00000( |
| 85137150 | 85149443 | 1 | lncRNA | ENSG00000( |
| 85142696 | 85146001 | -1 | lncRNA | ENSG00000( |
| 85169525 | 85676204 | 1 | protein_coding | ENSG00000( |
| 85227203 | 85309176 | 1 | lncRNA | ENSG00000( |

| | | | | |
|---|---|---|---|---|
| 85278820 | 85279035 | -1 | processed_pseudoge | ENSG00000 |
| 85282958 | 85285963 | 1 | lncRNA | ENSG00000 |
| 85306226 | 85306325 | -1 | miRNA | ENSG00000 |
| 85345271 | 85352961 | -1 | lncRNA | ENSG00000 |
| 85489813 | 85490831 | 1 | lncRNA | ENSG00000 |
| 85580009 | 85583571 | -1 | lncRNA | ENSG00000 |
| 85592266 | 85595720 | 1 | lncRNA | ENSG00000 |
| 85659378 | 85659675 | 1 | misc_RNA | ENSG00000 |
| 85676198 | 85690073 | -1 | protein_coding | ENSG00000 |
| 85690084 | 85751129 | -1 | protein_coding | ENSG00000 |
| 85697335 | 85697868 | -1 | lncRNA | ENSG00000 |
| 85699820 | 85711186 | 1 | lncRNA | ENSG00000 |
| 85741621 | 85741700 | -1 | miRNA | ENSG00000 |
| 85743287 | 85744225 | 1 | lncRNA | ENSG00000 |
| 85771758 | 85799608 | -1 | protein_coding | ENSG00000 |
| 85781775 | 85781899 | -1 | snRNA | ENSG00000 |
| 85784382 | 85787617 | 1 | lncRNA | ENSG00000 |
| 85792415 | 85792933 | 1 | lncRNA | ENSG00000 |
| 85798633 | 85807054 | 1 | protein_coding | ENSG00000 |
| 85814425 | 85815062 | -1 | processed_pseudoge | ENSG00000 |
| 85815873 | 85816536 | -1 | processed_pseudoge | ENSG00000 |
| 85846309 | 85848138 | 1 | lncRNA | ENSG00000 |
| 85859521 | 85862012 | 1 | TEC | ENSG00000 |
| 85899162 | 85922606 | 1 | protein_coding | ENSG00000 |
| 85918347 | 85918416 | 1 | miRNA | ENSG00000 |
| 85924984 | 85948824 | 1 | lncRNA | ENSG00000 |
| 85935281 | 85936223 | -1 | lncRNA | ENSG00000 |
| 85963328 | 85985386 | -1 | lncRNA | ENSG00000 |
| 85981750 | 85984881 | 1 | lncRNA | ENSG00000 |
| 85986764 | 85995899 | -1 | lncRNA | ENSG00000 |
| 86035627 | 86036419 | 1 | TEC | ENSG00000 |
| 86081409 | 86089526 | -1 | lncRNA | ENSG00000 |
| 86158206 | 86158798 | -1 | lncRNA | ENSG00000 |
| 86192793 | 86198913 | -1 | lncRNA | ENSG00000 |
| 86195954 | 86199720 | 1 | lncRNA | ENSG00000 |
| 86221420 | 86301303 | -1 | lncRNA | ENSG00000 |
| 86275429 | 86275523 | -1 | misc_RNA | ENSG00000 |
| 86286427 | 86293389 | 1 | lncRNA | ENSG00000 |
| 86331632 | 86349688 | -1 | lncRNA | ENSG00000 |
| 86347387 | 86349611 | 1 | lncRNA | ENSG00000 |
| 86434830 | 86435884 | -1 | processed_pseudoge | ENSG00000 |
| 86474529 | 86509099 | -1 | lncRNA | ENSG00000 |
| 86510527 | 86515422 | 1 | protein_coding | ENSG00000 |
| 86520383 | 86523897 | -1 | lncRNA | ENSG00000 |
| 86525457 | 86526252 | 1 | processed_pseudoge | ENSG00000 |
| 86530178 | 86555235 | -1 | protein_coding | ENSG00000 |
| 86555320 | 86557299 | 1 | TEC | ENSG00000 |
| 86565145 | 86567761 | -1 | lncRNA | ENSG00000 |
| 86566829 | 86569728 | 1 | protein_coding | ENSG00000 |
| 86576368 | 86582160 | 1 | protein_coding | ENSG00000 |

| | | | | |
|---|---|---|---|---|
| 86595189 | 86597639 | 1 | TEC | ENSG0000C |
| 86636328 | 86637145 | -1 | lncRNA | ENSG0000C |
| 86646301 | 86668923 | 1 | lncRNA | ENSG0000C |
| 86698118 | 86698841 | 1 | lncRNA | ENSG0000C |
| 86710122 | 86742083 | 1 | lncRNA | ENSG0000C |
| 86720575 | 86722042 | -1 | lncRNA | ENSG0000C |
| 86764298 | 86771041 | 1 | lncRNA | ENSG0000C |
| 86899356 | 86961268 | 1 | lncRNA | ENSG0000C |
| 86906175 | 86906472 | -1 | processed_pseudoge | ENSG0000C |
| 87057784 | 87070105 | 1 | lncRNA | ENSG0000C |
| 87066705 | 87069752 | 1 | lncRNA | ENSG0000C |
| 87083562 | 87317420 | -1 | protein_coding | ENSG0000C |
| 87129857 | 87133107 | 1 | lncRNA | ENSG0000C |
| 87212122 | 87226430 | 1 | lncRNA | ENSG0000C |
| 87215915 | 87216452 | 1 | lncRNA | ENSG0000C |
| 87262759 | 87292438 | -1 | lncRNA | ENSG0000C |
| 87268185 | 87268337 | 1 | processed_pseudoge | ENSG0000C |
| 87296949 | 87297633 | 1 | TEC | ENSG0000C |
| 87302876 | 87334273 | -1 | protein_coding | ENSG0000C |
| 87317509 | 87318043 | 1 | lncRNA | ENSG0000C |
| 87326987 | 87392142 | -1 | protein_coding | ENSG0000C |
| 87326987 | 87327584 | 1 | lncRNA | ENSG0000C |
| 87362536 | 87367476 | 1 | lncRNA | ENSG0000C |
| 87383953 | 87404779 | 1 | protein_coding | ENSG0000C |
| 87396704 | 87400295 | -1 | lncRNA | ENSG0000C |
| 87406246 | 87493024 | -1 | protein_coding | ENSG0000C |
| 87467138 | 87467654 | 1 | unprocessed_pseudc | ENSG0000C |
| 87470370 | 87474370 | -1 | lncRNA | ENSG0000C |
| 87492555 | 87600337 | 1 | lncRNA | ENSG0000C |
| 87600526 | 87602190 | -1 | lncRNA | ENSG0000C |
| 87601835 | 87698156 | 1 | protein_coding | ENSG0000C |
| 87683945 | 87684477 | -1 | lncRNA | ENSG0000C |
| 87693537 | 87696147 | -1 | lncRNA | ENSG0000C |
| 87696485 | 87765992 | -1 | protein_coding | ENSG0000C |
| 87739771 | 87747706 | -1 | lncRNA | ENSG0000C |
| 87773304 | 87779374 | -1 | lncRNA | ENSG0000C |
| 87780134 | 87806476 | -1 | lncRNA | ENSG0000C |
| 87830023 | 87869507 | -1 | protein_coding | ENSG0000C |
| 87834592 | 87834660 | -1 | miRNA | ENSG0000C |
| 87836532 | 87837663 | 1 | lncRNA | ENSG0000C |
| 87881546 | 87936580 | -1 | protein_coding | ENSG0000C |
| 87892955 | 87893735 | 1 | processed_pseudoge | ENSG0000C |
| 87945058 | 87945202 | -1 | processed_pseudoge | ENSG0000C |
| 87949244 | 88077318 | 1 | protein_coding | ENSG0000C |
| 88079161 | 88087383 | -1 | lncRNA | ENSG0000C |
| 88087241 | 88089970 | 1 | lncRNA | ENSG0000C |
| 88088041 | 88100985 | -1 | lncRNA | ENSG0000C |
| 88177289 | 88178646 | 1 | lncRNA | ENSG0000C |
| 88177298 | 88186929 | -1 | lncRNA | ENSG0000C |
| 88186605 | 88195217 | 1 | lncRNA | ENSG0000C |

| | | | | |
|---|---|---|---|---|
| 88191486 | 88193667 | -1 | lncRNA | ENSG0000C |
| 88234785 | 88302511 | 1 | lncRNA | ENSG0000C |
| 88382989 | 88440757 | 1 | protein_coding | ENSG0000C |
| 88453280 | 88537031 | 1 | protein_coding | ENSG0000C |
| 88468918 | 88469031 | 1 | miRNA | ENSG0000C |
| 88512960 | 88531053 | -1 | lncRNA | ENSG0000C |
| 88570381 | 88631966 | 1 | protein_coding | ENSG0000C |
| 88638572 | 88640468 | 1 | protein_coding | ENSG0000C |
| 88643289 | 88651054 | -1 | protein_coding | ENSG0000C |
| 88651935 | 88663161 | -1 | protein_coding | ENSG0000C |
| 88663298 | 88687278 | 1 | lncRNA | ENSG0000C |
| 88677688 | 88686507 | -1 | protein_coding | ENSG0000C |
| 88696499 | 88706408 | -1 | protein_coding | ENSG0000C |
| 88696501 | 88698610 | 1 | lncRNA | ENSG0000C |
| 88706483 | 88715396 | 1 | protein_coding | ENSG0000C |
| 88708956 | 88710437 | -1 | lncRNA | ENSG0000C |
| 88715338 | 88785220 | -1 | protein_coding | ENSG0000C |
| 88716278 | 88716337 | -1 | miRNA | ENSG0000C |
| 88718615 | 88720459 | 1 | lncRNA | ENSG0000C |
| 88731180 | 88741425 | 1 | lncRNA | ENSG0000C |
| 88741631 | 88742367 | 1 | lncRNA | ENSG0000C |
| 88742767 | 88745748 | 1 | lncRNA | ENSG0000C |
| 88803789 | 88809258 | 1 | protein_coding | ENSG0000C |
| 88809339 | 88811937 | -1 | protein_coding | ENSG0000C |
| 88813734 | 88856970 | -1 | protein_coding | ENSG0000C |
| 88856220 | 88862686 | 1 | protein_coding | ENSG0000C |
| 88863333 | 88866660 | -1 | protein_coding | ENSG0000C |
| 88874858 | 88977207 | -1 | protein_coding | ENSG0000C |
| 88881038 | 88887136 | 1 | lncRNA | ENSG0000C |
| 88936779 | 88939651 | -1 | lncRNA | ENSG0000C |
| 88939789 | 88951524 | 1 | lncRNA | ENSG0000C |
| 88984032 | 88995369 | 1 | lncRNA | ENSG0000C |
| 89046172 | 89052966 | -1 | lncRNA | ENSG0000C |
| 89088375 | 89164121 | 1 | protein_coding | ENSG0000C |
| 89113175 | 89115279 | -1 | lncRNA | ENSG0000C |
| 89159146 | 89164245 | 1 | lncRNA | ENSG0000C |
| 89166383 | 89169147 | 1 | lncRNA | ENSG0000C |
| 89171748 | 89195492 | 1 | protein_coding | ENSG0000C |
| 89195761 | 89201651 | -1 | protein_coding | ENSG0000C |
| 89215211 | 89217653 | -1 | lncRNA | ENSG0000C |
| 89217703 | 89237071 | 1 | protein_coding | ENSG0000C |
| 89226807 | 89228692 | -1 | lncRNA | ENSG0000C |
| 89267630 | 89490561 | -1 | protein_coding | ENSG0000C |
| 89268104 | 89273044 | 1 | lncRNA | ENSG0000C |
| 89296128 | 89298317 | 1 | lncRNA | ENSG0000C |
| 89321133 | 89325110 | 1 | lncRNA | ENSG0000C |
| 89420075 | 89422226 | -1 | lncRNA | ENSG0000C |
| 89430988 | 89453678 | 1 | lncRNA | ENSG0000C |
| 89443318 | 89443424 | 1 | snRNA | ENSG0000C |
| 89458870 | 89459437 | 1 | processed_pseudoge | ENSG0000C |

| | | | | |
|---|---|---|---|---|
| 89490719 | 89557768 | 1 | protein_coding | ENSG0000C |
| 89492017 | 89504460 | -1 | lncRNA | ENSG0000C |
| 89560657 | 89566828 | 1 | protein_coding | ENSG0000C |
| 89561434 | 89561517 | 1 | snoRNA | ENSG0000C |
| 89575758 | 89597246 | 1 | protein_coding | ENSG0000C |
| 89613308 | 89638456 | 1 | protein_coding | ENSG0000C |
| 89644435 | 89657721 | -1 | protein_coding | ENSG0000C |
| 89657802 | 89671272 | 1 | protein_coding | ENSG0000C |
| 89671568 | 89685304 | 1 | lncRNA | ENSG0000C |
| 89680737 | 89696354 | 1 | protein_coding | ENSG0000C |
| 89682620 | 89686569 | -1 | lncRNA | ENSG0000C |
| 89696357 | 89701705 | -1 | protein_coding | ENSG0000C |
| 89707134 | 89720898 | -1 | protein_coding | ENSG0000C |
| 89711856 | 89718165 | 1 | lncRNA | ENSG0000C |
| 89720400 | 89740925 | 1 | protein_coding | ENSG0000C |
| 89737549 | 89816657 | -1 | protein_coding | ENSG0000C |
| 89818179 | 89871319 | 1 | protein_coding | ENSG0000C |
| 89873570 | 89913627 | 1 | protein_coding | ENSG0000C |
| 89906157 | 89918233 | 1 | lncRNA | ENSG0000C |
| 89912119 | 89920977 | 1 | protein_coding | ENSG0000C |
| 89919165 | 89936092 | 1 | protein_coding | ENSG0000C |
| 89919827 | 89922662 | -1 | lncRNA | ENSG0000C |
| 89921392 | 89938761 | 1 | protein_coding | ENSG0000C |
| 89947925 | 89968060 | 1 | protein_coding | ENSG0000C |
| 89969773 | 89972832 | -1 | protein_coding | ENSG0000C |
| 89972586 | 90002161 | 1 | transcribed_unitary_ | ENSG0000C |
| 90004865 | 90020128 | -1 | protein_coding | ENSG0000C |
| 90019629 | 90044975 | 1 | protein_coding | ENSG0000C |
| 90028908 | 90029901 | -1 | lncRNA | ENSG0000C |
| 90039761 | 90047773 | -1 | transcribed_unproce | ENSG0000C |
| 90056566 | 90092072 | -1 | protein_coding | ENSG0000C |
| 90093154 | 90096354 | 1 | transcribed_unproce | ENSG0000C |
| 90102271 | 90178344 | 1 | lncRNA | ENSG0000C |
| 90105095 | 90106316 | -1 | lncRNA | ENSG0000C |
| 90110574 | 90168225 | -1 | lncRNA | ENSG0000C |
| 90144592 | 90144695 | 1 | snRNA | ENSG0000C |
| 90173217 | 90222851 | 1 | lncRNA | ENSG0000C |
| 90174062 | 90175865 | -1 | unprocessed_pseudc | ENSG0000C |
| 64099 | 76866 | -1 | lncRNA | ENSG0000C |
| 88641 | 118578 | -1 | lncRNA | ENSG0000C |
| 90796 | 90899 | -1 | snRNA | ENSG0000C |
| 137526 | 139067 | 1 | protein_coding | ENSG0000C |
| 142789 | 181650 | -1 | protein_coding | ENSG0000C |
| 183824 | 191587 | -1 | lncRNA | ENSG0000C |
| 212389 | 386254 | -1 | protein_coding | ENSG0000C |
| 321392 | 322453 | 1 | lncRNA | ENSG0000C |
| 331205 | 333488 | 1 | lncRNA | ENSG0000C |
| 352638 | 357581 | 1 | lncRNA | ENSG0000C |
| 354737 | 356882 | 1 | lncRNA | ENSG0000C |
| 404468 | 414023 | -1 | lncRNA | ENSG0000C |

| | | | | |
|---|---|---|---|---|
| 410325 | 431062 | 1 | protein_coding | ENSG00000 |
| 439978 | 445939 | -1 | protein_coding | ENSG00000 |
| 503171 | 511347 | 1 | lncRNA | ENSG00000 |
| 508668 | 721717 | -1 | protein_coding | ENSG00000 |
| 523140 | 540576 | 1 | lncRNA | ENSG00000 |
| 552566 | 553417 | 1 | lncRNA | ENSG00000 |
| 616002 | 616595 | 1 | processed_pseudoge | ENSG00000 |
| 656794 | 657547 | -1 | processed_pseudoge | ENSG00000 |
| 732412 | 742968 | 1 | protein_coding | ENSG00000 |
| 744421 | 753999 | -1 | protein_coding | ENSG00000 |
| 752660 | 755336 | 1 | transcribed_unitary_ | ENSG00000 |
| 757097 | 783390 | -1 | protein_coding | ENSG00000 |
| 763565 | 765319 | -1 | lncRNA | ENSG00000 |
| 782353 | 792509 | 1 | protein_coding | ENSG00000 |
| 789744 | 790525 | -1 | lncRNA | ENSG00000 |
| 795306 | 795794 | 1 | lncRNA | ENSG00000 |
| 799310 | 979776 | -1 | protein_coding | ENSG00000 |
| 877902 | 880093 | 1 | lncRNA | ENSG00000 |
| 909632 | 911212 | 1 | lncRNA | ENSG00000 |
| 959034 | 959366 | 1 | processed_pseudoge | ENSG00000 |
| 997129 | 1003671 | 1 | protein_coding | ENSG00000 |
| 1003518 | 1229738 | -1 | protein_coding | ENSG00000 |
| 1022476 | 1022559 | -1 | miRNA | ENSG00000 |
| 1101007 | 1101273 | -1 | misc_RNA | ENSG00000 |
| 1155966 | 1156175 | 1 | processed_pseudoge | ENSG00000 |
| 1181893 | 1185086 | 1 | lncRNA | ENSG00000 |
| 1241939 | 1254000 | -1 | lncRNA | ENSG00000 |
| 1270559 | 1271460 | 1 | protein_coding | ENSG00000 |
| 1279662 | 1300978 | 1 | protein_coding | ENSG00000 |
| 1344275 | 1400222 | -1 | protein_coding | ENSG00000 |
| 1352070 | 1352255 | 1 | TEC | ENSG00000 |
| 1420689 | 1463162 | -1 | protein_coding | ENSG00000 |
| 1424473 | 1426484 | 1 | lncRNA | ENSG00000 |
| 1464186 | 1492686 | -1 | protein_coding | ENSG00000 |
| 1494577 | 1516742 | -1 | protein_coding | ENSG00000 |
| 1516931 | 1518096 | 1 | lncRNA | ENSG00000 |
| 1517718 | 1562792 | -1 | protein_coding | ENSG00000 |
| 1569267 | 1628886 | -1 | protein_coding | ENSG00000 |
| 1604374 | 1604672 | -1 | misc_RNA | ENSG00000 |
| 1633858 | 1645744 | -1 | protein_coding | ENSG00000 |
| 1646145 | 1649866 | -1 | protein_coding | ENSG00000 |
| 1650629 | 1684867 | -1 | protein_coding | ENSG00000 |
| 1684726 | 1685151 | 1 | lncRNA | ENSG00000 |
| 1702816 | 1710377 | -1 | protein_coding | ENSG00000 |
| 1711493 | 1717174 | -1 | lncRNA | ENSG00000 |
| 1713903 | 1713987 | -1 | miRNA | ENSG00000 |
| 1716523 | 1738599 | 1 | protein_coding | ENSG00000 |
| 1725748 | 1738585 | -1 | lncRNA | ENSG00000 |
| 1742836 | 1755268 | 1 | protein_coding | ENSG00000 |
| 1762029 | 1777565 | 1 | protein_coding | ENSG00000 |

| | | | | |
|---|---|---|---|---|
| 1779485 | 1830634 | -1 | protein_coding | ENSG00000 |
| 1829702 | 1900082 | 1 | protein_coding | ENSG00000 |
| 1858039 | 1858446 | -1 | processed_pseudoge | ENSG00000 |
| 1934677 | 2025334 | -1 | protein_coding | ENSG00000 |
| 1995614 | 2003671 | 1 | lncRNA | ENSG00000 |
| 2017716 | 2020706 | 1 | lncRNA | ENSG00000 |
| 2019952 | 2023664 | -1 | lncRNA | ENSG00000 |
| 2030110 | 2043430 | 1 | protein_coding | ENSG00000 |
| 2040953 | 2041932 | -1 | lncRNA | ENSG00000 |
| 2042022 | 2043425 | 1 | protein_coding | ENSG00000 |
| 2042900 | 2043425 | -1 | lncRNA | ENSG00000 |
| 2043475 | 2044968 | -1 | lncRNA | ENSG00000 |
| 2049908 | 2050008 | -1 | miRNA | ENSG00000 |
| 2050271 | 2050380 | -1 | miRNA | ENSG00000 |
| 2054154 | 2063241 | 1 | protein_coding | ENSG00000 |
| 2059839 | 2303785 | -1 | protein_coding | ENSG00000 |
| 2072823 | 2072956 | 1 | TEC | ENSG00000 |
| 2089681 | 2101560 | 1 | lncRNA | ENSG00000 |
| 2101542 | 2101667 | 1 | TEC | ENSG00000 |
| 2117626 | 2117772 | 1 | TEC | ENSG00000 |
| 2122342 | 2122508 | -1 | TEC | ENSG00000 |
| 2127430 | 2127904 | -1 | processed_pseudoge | ENSG00000 |
| 2199762 | 2200039 | 1 | misc_RNA | ENSG00000 |
| 2213697 | 2214414 | 1 | lncRNA | ENSG00000 |
| 2215482 | 2216015 | -1 | lncRNA | ENSG00000 |
| 2232680 | 2233610 | 1 | lncRNA | ENSG00000 |
| 2235713 | 2236086 | -1 | TEC | ENSG00000 |
| 2255719 | 2256157 | 1 | TEC | ENSG00000 |
| 2303383 | 2325260 | 1 | protein_coding | ENSG00000 |
| 2306761 | 2307715 | 1 | processed_pseudoge | ENSG00000 |
| 2322503 | 2337507 | -1 | protein_coding | ENSG00000 |
| 2329016 | 2329237 | -1 | snoRNA | ENSG00000 |
| 2330180 | 2330370 | -1 | snoRNA | ENSG00000 |
| 2337498 | 2381058 | 1 | protein_coding | ENSG00000 |
| 2366589 | 2366791 | -1 | lncRNA | ENSG00000 |
| 2375061 | 2379306 | -1 | lncRNA | ENSG00000 |
| 2384073 | 2401104 | -1 | protein_coding | ENSG00000 |
| 2384847 | 2386664 | 1 | lncRNA | ENSG00000 |
| 2405562 | 2511891 | -1 | protein_coding | ENSG00000 |
| 2424179 | 2424412 | -1 | TEC | ENSG00000 |
| 2460183 | 2460294 | -1 | TEC | ENSG00000 |
| 2557753 | 2558094 | -1 | misc_RNA | ENSG00000 |
| 2586330 | 2587866 | -1 | processed_pseudoge | ENSG00000 |
| 2593210 | 2685615 | 1 | protein_coding | ENSG00000 |
| 2613284 | 2613944 | -1 | processed_pseudoge | ENSG00000 |
| 2639297 | 2642418 | 1 | TEC | ENSG00000 |
| 2680958 | 2681236 | 1 | misc_RNA | ENSG00000 |
| 2683305 | 2685088 | -1 | lncRNA | ENSG00000 |
| 2688473 | 2688960 | 1 | lncRNA | ENSG00000 |
| 2689386 | 2712663 | -1 | protein_coding | ENSG00000 |

| | | | | |
|---|---|---|---|---|
| 2692861 | 2692919 | -1 | miRNA | ENSG0000C |
| 2712309 | 2712833 | 1 | lncRNA | ENSG0000C |
| 2720801 | 2723947 | -1 | lncRNA | ENSG0000C |
| 2724411 | 2749504 | -1 | unitary_pseudogene | ENSG0000C |
| 2748078 | 2748182 | -1 | miRNA | ENSG0000C |
| 2748078 | 2748182 | 1 | lncRNA | ENSG0000C |
| 2755705 | 3037741 | 1 | protein_coding | ENSG0000C |
| 2962248 | 2965895 | -1 | lncRNA | ENSG0000C |
| 3062669 | 3063607 | -1 | protein_coding | ENSG0000C |
| 3088484 | 3104422 | -1 | protein_coding | ENSG0000C |
| 3116427 | 3117374 | 1 | unprocessed_pseudc | ENSG0000C |
| 3126584 | 3127581 | -1 | protein_coding | ENSG0000C |
| 3134969 | 3177031 | -1 | lncRNA | ENSG0000C |
| 3153889 | 3154882 | -1 | polymorphic_pseudc | ENSG0000C |
| 3197519 | 3198448 | 1 | protein_coding | ENSG0000C |
| 3207539 | 3218896 | 1 | protein_coding | ENSG0000C |
| 3235703 | 3241614 | 1 | transcribed_unproce | ENSG0000C |
| 3265611 | 3266541 | 1 | unprocessed_pseudc | ENSG0000C |
| 3276942 | 3386317 | -1 | protein_coding | ENSG0000C |
| 3291017 | 3298360 | -1 | protein_coding | ENSG0000C |
| 3309986 | 3311446 | 1 | transcribed_unproce | ENSG0000C |
| 3311218 | 3311934 | 1 | transcribed_unproce | ENSG0000C |
| 3377527 | 3377602 | -1 | unprocessed_pseudc | ENSG0000C |
| 3385930 | 3386874 | 1 | protein_coding | ENSG0000C |
| 3397104 | 3398410 | -1 | protein_coding | ENSG0000C |
| 3411370 | 3424070 | 1 | protein_coding | ENSG0000C |
| 3432870 | 3433841 | -1 | protein_coding | ENSG0000C |
| 3439923 | 3441844 | -1 | TEC | ENSG0000C |
| 3440019 | 3513852 | -1 | protein_coding | ENSG0000C |
| 3472374 | 3503405 | 1 | protein_coding | ENSG0000C |
| 3510502 | 3557995 | -1 | protein_coding | ENSG0000C |
| 3565444 | 3609411 | -1 | protein_coding | ENSG0000C |
| 3577955 | 3578853 | -1 | lncRNA | ENSG0000C |
| 3585149 | 3636249 | -1 | protein_coding | ENSG0000C |
| 3601761 | 3602272 | -1 | lncRNA | ENSG0000C |
| 3608253 | 3636250 | -1 | protein_coding | ENSG0000C |
| 3619256 | 3619913 | -1 | processed_pseudoge | ENSG0000C |
| 3636468 | 3661542 | 1 | protein_coding | ENSG0000C |
| 3655621 | 3658092 | -1 | lncRNA | ENSG0000C |
| 3662895 | 3668679 | -1 | protein_coding | ENSG0000C |
| 3663063 | 3696194 | -1 | protein_coding | ENSG0000C |
| 3665220 | 3665702 | 1 | processed_pseudoge | ENSG0000C |
| 3668812 | 3669668 | 1 | protein_coding | ENSG0000C |
| 3672199 | 3696240 | -1 | protein_coding | ENSG0000C |
| 3714628 | 3801188 | -1 | protein_coding | ENSG0000C |
| 3721628 | 3722488 | 1 | lncRNA | ENSG0000C |
| 3723903 | 3726699 | 1 | protein_coding | ENSG0000C |
| 3729361 | 3730493 | 1 | lncRNA | ENSG0000C |
| 3802158 | 3846246 | -1 | protein_coding | ENSG0000C |
| 3860315 | 3894891 | -1 | protein_coding | ENSG0000C |

| | | | | |
|---|---|---|---|---|
| 3896592 | 3916476 | -1 | protein_coding | ENSG0000C |
| 3923870 | 3964464 | -1 | protein_coding | ENSG0000C |
| 3977103 | 3981899 | 1 | lncRNA | ENSG0000C |
| 4004445 | 4143030 | -1 | protein_coding | ENSG0000C |
| 4056756 | 4056876 | -1 | rRNA_pseudogene | ENSG0000C |
| 4143168 | 4187310 | 1 | protein_coding | ENSG0000C |
| 4163821 | 4263995 | -1 | protein_coding | ENSG0000C |
| 4163910 | 4164713 | 1 | lncRNA | ENSG0000C |
| 4186326 | 4186424 | 1 | misc_RNA | ENSG0000C |
| 4222091 | 4223828 | 1 | processed_pseudoge | ENSG0000C |
| 4267691 | 4268430 | 1 | lncRNA | ENSG0000C |
| 4269259 | 4366628 | -1 | protein_coding | ENSG0000C |
| 4301372 | 4302698 | 1 | processed_pseudoge | ENSG0000C |
| 4364164 | 4364453 | -1 | misc_RNA | ENSG0000C |
| 4433940 | 4488208 | 1 | protein_coding | ENSG0000C |
| 4481966 | 4486353 | -1 | lncRNA | ENSG0000C |
| 4497394 | 4499674 | -1 | lncRNA | ENSG0000C |
| 4498881 | 4539035 | 1 | protein_coding | ENSG0000C |
| 4538897 | 4555631 | -1 | protein_coding | ENSG0000C |
| 4556927 | 4560818 | -1 | protein_coding | ENSG0000C |
| 4565752 | 4571760 | 1 | lncRNA | ENSG0000C |
| 4572206 | 4572515 | 1 | processed_pseudoge | ENSG0000C |
| 4583999 | 4608319 | 1 | protein_coding | ENSG0000C |
| 4610449 | 4617650 | 1 | lncRNA | ENSG0000C |
| 4619749 | 4619852 | -1 | snRNA | ENSG0000C |
| 4630902 | 4642294 | -1 | protein_coding | ENSG0000C |
| 4669774 | 4704337 | -1 | protein_coding | ENSG0000C |
| 4673830 | 4696831 | 1 | lncRNA | ENSG0000C |
| 4704230 | 4705529 | 1 | lncRNA | ENSG0000C |
| 4704816 | 4705217 | -1 | processed_pseudoge | ENSG0000C |
| 4710596 | 4721499 | 1 | protein_coding | ENSG0000C |
| 4731428 | 4733607 | 1 | protein_coding | ENSG0000C |
| 4731756 | 4732371 | -1 | lncRNA | ENSG0000C |
| 4733533 | 4739928 | -1 | protein_coding | ENSG0000C |
| 4739833 | 4746119 | 1 | protein_coding | ENSG0000C |
| 4771884 | 4783213 | 1 | protein_coding | ENSG0000C |
| 4785285 | 4786433 | -1 | protein_coding | ENSG0000C |
| 4787057 | 4789162 | 1 | TEC | ENSG0000C |
| 4788964 | 4790589 | 1 | protein_coding | ENSG0000C |
| 4796144 | 4798502 | 1 | protein_coding | ENSG0000C |
| 4801159 | 4807013 | -1 | protein_coding | ENSG0000C |
| 4807152 | 4823434 | 1 | protein_coding | ENSG0000C |
| 4833340 | 4898061 | 1 | protein_coding | ENSG0000C |
| 4856757 | 4857374 | -1 | processed_pseudoge | ENSG0000C |
| 4882975 | 4883316 | 1 | misc_RNA | ENSG0000C |
| 4897771 | 4934438 | -1 | protein_coding | ENSG0000C |
| 4899418 | 4902934 | 1 | protein_coding | ENSG0000C |
| 4932277 | 4935023 | 1 | protein_coding | ENSG0000C |
| 4937130 | 4940053 | -1 | protein_coding | ENSG0000C |
| 4940008 | 4945222 | 1 | protein_coding | ENSG0000C |

| | | | | |
|---|---|---|---|---|
| 4945652 | 4949061 | -1 | protein_coding | ENSG00000 |
| 4948092 | 4957131 | 1 | protein_coding | ENSG00000 |
| 4959226 | 4967817 | -1 | protein_coding | ENSG00000 |
| 4967992 | 4987675 | -1 | protein_coding | ENSG00000 |
| 4967995 | 4968822 | 1 | lncRNA | ENSG00000 |
| 4969702 | 4969771 | -1 | miRNA | ENSG00000 |
| 4970086 | 4970150 | -1 | miRNA | ENSG00000 |
| 4972851 | 4974681 | 1 | lncRNA | ENSG00000 |
| 4986466 | 4987324 | 1 | lncRNA | ENSG00000 |
| 4987706 | 4988446 | 1 | lncRNA | ENSG00000 |
| 4988130 | 4997610 | -1 | protein_coding | ENSG00000 |
| 4997950 | 5028401 | 1 | protein_coding | ENSG00000 |
| 5019214 | 5020093 | -1 | lncRNA | ENSG00000 |
| 5032600 | 5052009 | -1 | protein_coding | ENSG00000 |
| 5060356 | 5062123 | 1 | processed_pseudoge | ENSG00000 |
| 5075678 | 5078113 | -1 | lncRNA | ENSG00000 |
| 5078467 | 5096374 | 1 | protein_coding | ENSG00000 |
| 5105541 | 5123116 | -1 | protein_coding | ENSG00000 |
| 5111468 | 5115004 | 1 | lncRNA | ENSG00000 |
| 5116438 | 5175034 | 1 | protein_coding | ENSG00000 |
| 5126740 | 5127010 | -1 | processed_pseudoge | ENSG00000 |
| 5179535 | 5191868 | -1 | protein_coding | ENSG00000 |
| 5192027 | 5248182 | 1 | lncRNA | ENSG00000 |
| 5208920 | 5234860 | -1 | protein_coding | ENSG00000 |
| 5240508 | 5241543 | -1 | lncRNA | ENSG00000 |
| 5282265 | 5386340 | 1 | protein_coding | ENSG00000 |
| 5360963 | 5419676 | -1 | protein_coding | ENSG00000 |
| 5364315 | 5371626 | -1 | lncRNA | ENSG00000 |
| 5419641 | 5432876 | 1 | protein_coding | ENSG00000 |
| 5425139 | 5432876 | -1 | lncRNA | ENSG00000 |
| 5432777 | 5448830 | -1 | protein_coding | ENSG00000 |
| 5440917 | 5468982 | -1 | protein_coding | ENSG00000 |
| 5469092 | 5470360 | 1 | lncRNA | ENSG00000 |
| 5471254 | 5486811 | -1 | protein_coding | ENSG00000 |
| 5486285 | 5490814 | 1 | protein_coding | ENSG00000 |
| 5499427 | 5619424 | -1 | protein_coding | ENSG00000 |
| 5499439 | 5501145 | -1 | protein_coding | ENSG00000 |
| 5512040 | 5512537 | -1 | processed_pseudoge | ENSG00000 |
| 5700226 | 5704056 | -1 | lncRNA | ENSG00000 |
| 5730289 | 5731085 | 1 | processed_pseudoge | ENSG00000 |
| 5772234 | 5930696 | 1 | lncRNA | ENSG00000 |
| 5772926 | 6080792 | 1 | protein_coding | ENSG00000 |
| 6057807 | 6124427 | 1 | protein_coding | ENSG00000 |
| 6163893 | 6190541 | -1 | lncRNA | ENSG00000 |
| 6207163 | 6207269 | 1 | snRNA | ENSG00000 |
| 6322413 | 6323228 | -1 | processed_pseudoge | ENSG00000 |
| 6373833 | 6375080 | -1 | lncRNA | ENSG00000 |
| 6393693 | 6435199 | -1 | protein_coding | ENSG00000 |
| 6417685 | 6418264 | 1 | processed_pseudoge | ENSG00000 |
| 6444441 | 6451469 | 1 | protein_coding | ENSG00000 |

| | | | | |
|---|---|---|---|---|
| 6451263 | 6556555 | -1 | protein_coding | ENSG00000 |
| 6521131 | 6521412 | 1 | misc_RNA | ENSG00000 |
| 6578147 | 6640711 | -1 | protein_coding | ENSG00000 |
| 6600944 | 6601051 | -1 | rRNA_pseudogene | ENSG00000 |
| 6628619 | 6640456 | -1 | protein_coding | ENSG00000 |
| 6640985 | 6644541 | 1 | protein_coding | ENSG00000 |
| 6643311 | 6651634 | -1 | protein_coding | ENSG00000 |
| 6651762 | 6693202 | 1 | protein_coding | ENSG00000 |
| 6653464 | 6655009 | -1 | lncRNA | ENSG00000 |
| 6655440 | 6655509 | -1 | miRNA | ENSG00000 |
| 6655449 | 6655502 | 1 | miRNA | ENSG00000 |
| 6657034 | 6693577 | 1 | transcribed_unproce | ENSG00000 |
| 6684713 | 6713567 | -1 | protein_coding | ENSG00000 |
| 6749685 | 6749978 | -1 | processed_pseudoge | ENSG00000 |
| 6749718 | 6750188 | 1 | processed_pseudoge | ENSG00000 |
| 6755447 | 6775647 | 1 | protein_coding | ENSG00000 |
| 6772831 | 6776116 | -1 | TEC | ENSG00000 |
| 6776215 | 6797101 | 1 | protein_coding | ENSG00000 |
| 6789133 | 6831761 | -1 | protein_coding | ENSG00000 |
| 6853861 | 6954107 | 1 | transcribed_unproce | ENSG00000 |
| 6875232 | 7012530 | -1 | lncRNA | ENSG00000 |
| 6927472 | 6928191 | -1 | processed_pseudoge | ENSG00000 |
| 6954098 | 6978960 | -1 | lncRNA | ENSG00000 |
| 6985494 | 7019414 | -1 | lncRNA | ENSG00000 |
| 6994642 | 6995189 | -1 | lncRNA | ENSG00000 |
| 6996049 | 7010754 | 1 | protein_coding | ENSG00000 |
| 7012417 | 7014532 | 1 | protein_coding | ENSG00000 |
| 7012635 | 7017520 | 1 | protein_coding | ENSG00000 |
| 7014495 | 7017525 | 1 | protein_coding | ENSG00000 |
| 7015818 | 7019659 | -1 | lncRNA | ENSG00000 |
| 7017615 | 7017701 | -1 | miRNA | ENSG00000 |
| 7017911 | 7018022 | -1 | miRNA | ENSG00000 |
| 7023050 | 7030290 | 1 | protein_coding | ENSG00000 |
| 7036015 | 7040117 | 1 | protein_coding | ENSG00000 |
| 7041630 | 7043923 | -1 | protein_coding | ENSG00000 |
| 7058934 | 7059204 | 1 | processed_pseudoge | ENSG00000 |
| 7074537 | 7080307 | -1 | protein_coding | ENSG00000 |
| 7101322 | 7115700 | -1 | protein_coding | ENSG00000 |
| 7140930 | 7141592 | 1 | processed_pseudoge | ENSG00000 |
| 7173431 | 7179564 | -1 | protein_coding | ENSG00000 |
| 7187169 | 7219841 | -1 | protein_coding | ENSG00000 |
| 7192295 | 7192448 | -1 | TEC | ENSG00000 |
| 7217125 | 7225266 | 1 | protein_coding | ENSG00000 |
| 7223297 | 7223379 | -1 | miRNA | ENSG00000 |
| 7225342 | 7234517 | -1 | protein_coding | ENSG00000 |
| 7235029 | 7239722 | -1 | protein_coding | ENSG00000 |
| 7240008 | 7242449 | -1 | protein_coding | ENSG00000 |
| 7240427 | 7244635 | -1 | protein_coding | ENSG00000 |
| 7242200 | 7243814 | 1 | TEC | ENSG00000 |
| 7243591 | 7252491 | -1 | protein_coding | ENSG00000 |

| | | | | |
|---|---|---|---|---|
| 7246829 | 7262089 | -1 | protein_coding | ENSG00000 |
| 7251416 | 7259940 | 1 | protein_coding | ENSG00000 |
| 7259903 | 7263983 | -1 | protein_coding | ENSG00000 |
| 7266055 | 7266166 | -1 | misc_RNA | ENSG00000 |
| 7281718 | 7288257 | 1 | protein_coding | ENSG00000 |
| 7282947 | 7284071 | -1 | lncRNA | ENSG00000 |
| 7288263 | 7294639 | -1 | protein_coding | ENSG00000 |
| 7306999 | 7312463 | 1 | protein_coding | ENSG00000 |
| 7311324 | 7315564 | -1 | protein_coding | ENSG00000 |
| 7312661 | 7319174 | -1 | protein_coding | ENSG00000 |
| 7315628 | 7329393 | -1 | protein_coding | ENSG00000 |
| 7330452 | 7335413 | 1 | unitary_pseudogene | ENSG00000 |
| 7336529 | 7351477 | 1 | protein_coding | ENSG00000 |
| 7351889 | 7354944 | 1 | protein_coding | ENSG00000 |
| 7352687 | 7354944 | -1 | lncRNA | ENSG00000 |
| 7355123 | 7357219 | 1 | protein_coding | ENSG00000 |
| 7380534 | 7389774 | 1 | protein_coding | ENSG00000 |
| 7389727 | 7394842 | -1 | protein_coding | ENSG00000 |
| 7389734 | 7404097 | -1 | protein_coding | ENSG00000 |
| 7402975 | 7404097 | -1 | protein_coding | ENSG00000 |
| 7404874 | 7419860 | 1 | protein_coding | ENSG00000 |
| 7420103 | 7444081 | 1 | lncRNA | ENSG00000 |
| 7420324 | 7421632 | 1 | protein_coding | ENSG00000 |
| 7425616 | 7427568 | 1 | protein_coding | ENSG00000 |
| 7428857 | 7432820 | 1 | protein_coding | ENSG00000 |
| 7428861 | 7436278 | 1 | protein_coding | ENSG00000 |
| 7435435 | 7437679 | 1 | protein_coding | ENSG00000 |
| 7436557 | 7437523 | -1 | lncRNA | ENSG00000 |
| 7438273 | 7444937 | 1 | protein_coding | ENSG00000 |
| 7439159 | 7443327 | -1 | lncRNA | ENSG00000 |
| 7439506 | 7445966 | 1 | lncRNA | ENSG00000 |
| 7445061 | 7457710 | 1 | protein_coding | ENSG00000 |
| 7459366 | 7484263 | -1 | protein_coding | ENSG00000 |
| 7481332 | 7483496 | 1 | protein_coding | ENSG00000 |
| 7484366 | 7514618 | 1 | protein_coding | ENSG00000 |
| 7537096 | 7537196 | -1 | misc_RNA | ENSG00000 |
| 7549058 | 7557890 | 1 | protein_coding | ENSG00000 |
| 7549099 | 7561601 | 1 | protein_coding | ENSG00000 |
| 7557820 | 7558245 | -1 | lncRNA | ENSG00000 |
| 7558292 | 7561608 | 1 | protein_coding | ENSG00000 |
| 7561919 | 7571969 | 1 | protein_coding | ENSG00000 |
| 7563287 | 7578715 | 1 | protein_coding | ENSG00000 |
| 7572706 | 7579006 | 1 | protein_coding | ENSG00000 |
| 7572826 | 7582024 | 1 | lncRNA | ENSG00000 |
| 7574713 | 7574847 | 1 | snoRNA | ENSG00000 |
| 7576811 | 7576952 | 1 | snoRNA | ENSG00000 |
| 7577955 | 7578091 | 1 | snoRNA | ENSG00000 |
| 7579491 | 7582111 | 1 | protein_coding | ENSG00000 |
| 7581964 | 7584086 | -1 | lncRNA | ENSG00000 |
| 7583529 | 7592789 | 1 | protein_coding | ENSG00000 |

| | | | | |
|---|---|---|---|---|
| 7588178 | 7590094 | -1 | protein_coding | ENSG00000( |
| 7591230 | 7614897 | -1 | protein_coding | ENSG00000( |
| 7613946 | 7633382 | 1 | protein_coding | ENSG00000( |
| 7626234 | 7627876 | -1 | protein_coding | ENSG00000( |
| 7646627 | 7657770 | 1 | protein_coding | ENSG00000( |
| 7661779 | 7687550 | -1 | protein_coding | ENSG00000( |
| 7686071 | 7703502 | 1 | protein_coding | ENSG00000( |
| 7705202 | 7711372 | 1 | protein_coding | ENSG00000( |
| 7717354 | 7833744 | 1 | protein_coding | ENSG00000( |
| 7754320 | 7754976 | 1 | transcribed_processe | ENSG00000( |
| 7839904 | 7854796 | 1 | protein_coding | ENSG00000( |
| 7855066 | 7856099 | 1 | protein_coding | ENSG00000( |
| 7856685 | 7885238 | -1 | protein_coding | ENSG00000( |
| 7857746 | 7862282 | 1 | protein_coding | ENSG00000( |
| 7858943 | 7866083 | 1 | TEC | ENSG00000( |
| 7884796 | 7912760 | 1 | protein_coding | ENSG00000( |
| 7906122 | 7906260 | 1 | scaRNA | ENSG00000( |
| 7913324 | 7916276 | -1 | protein_coding | ENSG00000( |
| 7921859 | 7929803 | -1 | protein_coding | ENSG00000( |
| 7925931 | 7930622 | -1 | protein_coding | ENSG00000( |
| 7930345 | 7932123 | -1 | protein_coding | ENSG00000( |
| 7932101 | 7949920 | 1 | protein_coding | ENSG00000( |
| 7954868 | 7955622 | 1 | processed_pseudoge | ENSG00000( |
| 8002594 | 8020339 | 1 | protein_coding | ENSG00000( |
| 8013327 | 8016378 | -1 | TEC | ENSG00000( |
| 8039034 | 8049134 | 1 | protein_coding | ENSG00000( |
| 8056225 | 8057621 | -1 | lncRNA | ENSG00000( |
| 8063936 | 8072533 | 1 | unprocessed_pseudc | ENSG00000( |
| 8072636 | 8087716 | -1 | protein_coding | ENSG00000( |
| 8079482 | 8081565 | 1 | lncRNA | ENSG00000( |
| 8088056 | 8088147 | 1 | miRNA | ENSG00000( |
| 8095900 | 8119047 | -1 | protein_coding | ENSG00000( |
| 8120592 | 8124106 | -1 | protein_coding | ENSG00000( |
| 8140472 | 8156506 | -1 | protein_coding | ENSG00000( |
| 8144994 | 8145071 | -1 | miRNA | ENSG00000( |
| 8150816 | 8162975 | -1 | protein_coding | ENSG00000( |
| 8159149 | 8163546 | -1 | protein_coding | ENSG00000( |
| 8172457 | 8176399 | -1 | protein_coding | ENSG00000( |
| 8173454 | 8173587 | -1 | snoRNA | ENSG00000( |
| 8176812 | 8182812 | 1 | lncRNA | ENSG00000( |
| 8186945 | 8187004 | 1 | miRNA | ENSG00000( |
| 8188345 | 8190180 | -1 | protein_coding | ENSG00000( |
| 8188933 | 8189688 | 1 | TEC | ENSG00000( |
| 8199123 | 8199437 | -1 | processed_pseudoge | ENSG00000( |
| 8204733 | 8210600 | -1 | protein_coding | ENSG00000( |
| 8220642 | 8224043 | -1 | lncRNA | ENSG00000( |
| 8224815 | 8248056 | -1 | protein_coding | ENSG00000( |
| 8247618 | 8270491 | 1 | protein_coding | ENSG00000( |
| 8277763 | 8278436 | 1 | lncRNA | ENSG00000( |
| 8287763 | 8295400 | -1 | protein_coding | ENSG00000( |

| | | | | |
|---|---|---|---|---|
| 8288654 | 8290092 | 1 | protein_coding | ENSG00000 |
| 8296505 | 8297216 | -1 | processed_pseudoge | ENSG00000 |
| 8310241 | 8322514 | 1 | protein_coding | ENSG00000 |
| 8318088 | 8318712 | -1 | lncRNA | ENSG00000 |
| 8329583 | 8329719 | 1 | snoRNA | ENSG00000 |
| 8337697 | 8338758 | 1 | lncRNA | ENSG00000 |
| 8339840 | 8346048 | 1 | protein_coding | ENSG00000 |
| 8356902 | 8376711 | -1 | protein_coding | ENSG00000 |
| 8365563 | 8381328 | 1 | lncRNA | ENSG00000 |
| 8368638 | 8383187 | -1 | protein_coding | ENSG00000 |
| 8377516 | 8383213 | -1 | protein_coding | ENSG00000 |
| 8390704 | 8397826 | -1 | protein_coding | ENSG00000 |
| 8413131 | 8490411 | 1 | protein_coding | ENSG00000 |
| 8474205 | 8630761 | -1 | protein_coding | ENSG00000 |
| 8511125 | 8511188 | 1 | snRNA | ENSG00000 |
| 8560471 | 8560962 | 1 | processed_pseudoge | ENSG00000 |
| 8561230 | 8561576 | 1 | processed_pseudoge | ENSG00000 |
| 8729935 | 8745219 | -1 | protein_coding | ENSG00000 |
| 8753106 | 8758559 | -1 | protein_coding | ENSG00000 |
| 8797110 | 8799365 | -1 | protein_coding | ENSG00000 |
| 8802723 | 8867677 | -1 | protein_coding | ENSG00000 |
| 8878911 | 8965712 | -1 | protein_coding | ENSG00000 |
| 8965896 | 8967070 | 1 | lncRNA | ENSG00000 |
| 8967523 | 8976995 | 1 | lncRNA | ENSG00000 |
| 9021510 | 9244000 | 1 | protein_coding | ENSG00000 |
| 9171068 | 9179118 | -1 | lncRNA | ENSG00000 |
| 9244666 | 9244897 | -1 | lncRNA | ENSG00000 |
| 9250471 | 9576591 | -1 | protein_coding | ENSG00000 |
| 9283174 | 9305651 | 1 | lncRNA | ENSG00000 |
| 9452197 | 9470014 | 1 | lncRNA | ENSG00000 |
| 9464742 | 9467422 | 1 | lncRNA | ENSG00000 |
| 9547298 | 9548541 | 1 | lncRNA | ENSG00000 |
| 9553323 | 9555696 | -1 | lncRNA | ENSG00000 |
| 9576627 | 9643447 | 1 | protein_coding | ENSG00000 |
| 9589902 | 9590482 | -1 | processed_pseudoge | ENSG00000 |
| 9638768 | 9645325 | -1 | lncRNA | ENSG00000 |
| 9644698 | 9729687 | 1 | protein_coding | ENSG00000 |
| 9647020 | 9647660 | -1 | lncRNA | ENSG00000 |
| 9764186 | 9805100 | 1 | lncRNA | ENSG00000 |
| 9771434 | 9791297 | -1 | protein_coding | ENSG00000 |
| 9791227 | 9808855 | -1 | lncRNA | ENSG00000 |
| 9802386 | 9822071 | -1 | protein_coding | ENSG00000 |
| 9805443 | 9811047 | 1 | lncRNA | ENSG00000 |
| 9822206 | 9892099 | 1 | protein_coding | ENSG00000 |
| 9852748 | 9853306 | -1 | processed_pseudoge | ENSG00000 |
| 9896320 | 9905271 | -1 | protein_coding | ENSG00000 |
| 9910609 | 10198551 | -1 | protein_coding | ENSG00000 |
| 10258762 | 10259232 | 1 | processed_pseudoge | ENSG00000 |
| 10291820 | 10317926 | 1 | lncRNA | ENSG00000 |
| 10300865 | 10373130 | -1 | protein_coding | ENSG00000 |

| | | | | |
|---|---|---|---|---|
| 10320392 | 10341458 | 1 | lncRNA | ENSG0000C |
| 10383132 | 10537862 | 1 | lncRNA | ENSG0000C |
| 10383144 | 10623886 | 1 | lncRNA | ENSG0000C |
| 10390322 | 10421950 | -1 | protein_coding | ENSG0000C |
| 10443290 | 10469559 | -1 | protein_coding | ENSG0000C |
| 10492307 | 10518542 | -1 | protein_coding | ENSG0000C |
| 10521148 | 10549700 | -1 | protein_coding | ENSG0000C |
| 10579040 | 10623036 | 1 | lncRNA | ENSG0000C |
| 10628526 | 10657309 | -1 | protein_coding | ENSG0000C |
| 10658542 | 10659082 | -1 | lncRNA | ENSG0000C |
| 10672474 | 10698375 | -1 | protein_coding | ENSG0000C |
| 10680734 | 10683988 | 1 | lncRNA | ENSG0000C |
| 10697594 | 10711558 | 1 | protein_coding | ENSG0000C |
| 10699015 | 10730316 | -1 | protein_coding | ENSG0000C |
| 10716002 | 10716511 | -1 | transcribed_processe | ENSG0000C |
| 10729777 | 10815164 | 1 | lncRNA | ENSG0000C |
| 10729777 | 10880895 | 1 | lncRNA | ENSG0000C |
| 10741257 | 10769089 | -1 | lncRNA | ENSG0000C |
| 10745553 | 10746543 | -1 | processed_pseudoge | ENSG0000C |
| 10792059 | 10794985 | 1 | lncRNA | ENSG0000C |
| 10794906 | 10838094 | -1 | lncRNA | ENSG0000C |
| 10794913 | 10804099 | -1 | protein_coding | ENSG0000C |
| 10821901 | 10822243 | -1 | TEC | ENSG0000C |
| 10822470 | 10838087 | -1 | protein_coding | ENSG0000C |
| 10844674 | 10880895 | 1 | lncRNA | ENSG0000C |
| 10857899 | 10858004 | -1 | snRNA | ENSG0000C |
| 10860535 | 10861462 | -1 | processed_pseudoge | ENSG0000C |
| 11157455 | 11157747 | 1 | misc_RNA | ENSG0000C |
| 11197883 | 11242062 | -1 | lncRNA | ENSG0000C |
| 11241213 | 11564063 | 1 | protein_coding | ENSG0000C |
| 11288205 | 11290811 | 1 | lncRNA | ENSG0000C |
| 11598470 | 11969748 | 1 | protein_coding | ENSG0000C |
| 11874727 | 11881670 | -1 | lncRNA | ENSG0000C |
| 11953889 | 11977594 | -1 | lncRNA | ENSG0000C |
| 11977439 | 11997475 | -1 | protein_coding | ENSG0000C |
| 11998353 | 12021508 | -1 | transcribed_processe | ENSG0000C |
| 12020829 | 12143830 | 1 | protein_coding | ENSG0000C |
| 12036496 | 12036637 | -1 | snRNA | ENSG0000C |
| 12081899 | 12081996 | 1 | miRNA | ENSG0000C |
| 12115547 | 12115885 | 1 | processed_pseudoge | ENSG0000C |
| 12201600 | 12215267 | 1 | lncRNA | ENSG0000C |
| 12390738 | 12395177 | -1 | lncRNA | ENSG0000C |
| 12549764 | 12642854 | 1 | lncRNA | ENSG0000C |
| 12596743 | 12632959 | -1 | lncRNA | ENSG0000C |
| 12665890 | 12768949 | 1 | protein_coding | ENSG0000C |
| 12671862 | 12706135 | -1 | lncRNA | ENSG0000C |
| 12760140 | 12790242 | -1 | lncRNA | ENSG0000C |
| 12789498 | 12991643 | 1 | protein_coding | ENSG0000C |
| 12917268 | 12917342 | -1 | miRNA | ENSG0000C |
| 12973033 | 12973834 | -1 | TEC | ENSG0000C |

| | | | | |
|---|---|---|---|---|
| 12982613 | 12983002 | -1 | lncRNA | ENSG0000C |
| 12990149 | 12990610 | -1 | lncRNA | ENSG0000C |
| 12991612 | 13018065 | -1 | protein_coding | ENSG0000C |
| 13095695 | 13100939 | 1 | lncRNA | ENSG0000C |
| 13299184 | 13306771 | -1 | lncRNA | ENSG0000C |
| 13494032 | 13601929 | -1 | protein_coding | ENSG0000C |
| 13543529 | 13543646 | -1 | miRNA | ENSG0000C |
| 13755574 | 14069495 | -1 | lncRNA | ENSG0000C |
| 13909231 | 13911212 | 1 | lncRNA | ENSG0000C |
| 13932720 | 13940594 | -1 | lncRNA | ENSG0000C |
| 13957748 | 13966952 | -1 | lncRNA | ENSG0000C |
| 14024514 | 14025488 | 1 | unprocessed_pseudc | ENSG0000C |
| 14069490 | 14231736 | 1 | protein_coding | ENSG0000C |
| 14177131 | 14177310 | 1 | snoRNA | ENSG0000C |
| 14235673 | 14236862 | -1 | protein_coding | ENSG0000C |
| 14295512 | 14297600 | -1 | lncRNA | ENSG0000C |
| 14301081 | 14349404 | 1 | protein_coding | ENSG0000C |
| 14303854 | 14305505 | 1 | lncRNA | ENSG0000C |
| 14327335 | 14329474 | 1 | lncRNA | ENSG0000C |
| 14374139 | 14421472 | 1 | lncRNA | ENSG0000C |
| 14705076 | 14705534 | -1 | processed_pseudoge | ENSG0000C |
| 14725729 | 14729686 | 1 | lncRNA | ENSG0000C |
| 14767583 | 14780686 | -1 | lncRNA | ENSG0000C |
| 14834557 | 14957216 | 1 | lncRNA | ENSG0000C |
| 14947741 | 14948198 | -1 | processed_pseudoge | ENSG0000C |
| 15014805 | 15052276 | 1 | lncRNA | ENSG0000C |
| 15030975 | 15031957 | 1 | lncRNA | ENSG0000C |
| 15062087 | 15104517 | -1 | lncRNA | ENSG0000C |
| 15105237 | 15106187 | -1 | lncRNA | ENSG0000C |
| 15229777 | 15265326 | -1 | protein_coding | ENSG0000C |
| 15251627 | 15251696 | -1 | miRNA | ENSG0000C |
| 15260513 | 15261841 | 1 | TEC | ENSG0000C |
| 15261015 | 15265707 | 1 | lncRNA | ENSG0000C |
| 15266848 | 15311760 | 1 | lncRNA | ENSG0000C |
| 15267461 | 15272290 | -1 | lncRNA | ENSG0000C |
| 15276562 | 15279477 | -1 | lncRNA | ENSG0000C |
| 15303811 | 15341632 | -1 | protein_coding | ENSG0000C |
| 15311345 | 15311451 | 1 | snRNA | ENSG0000C |
| 15365323 | 15365877 | 1 | TEC | ENSG0000C |
| 15379864 | 15417878 | -1 | lncRNA | ENSG0000C |
| 15408384 | 15408688 | 1 | misc_RNA | ENSG0000C |
| 15436015 | 15503608 | -1 | protein_coding | ENSG0000C |
| 15436021 | 15563561 | -1 | protein_coding | ENSG0000C |
| 15443639 | 15444191 | 1 | processed_pseudoge | ENSG0000C |
| 15502264 | 15563595 | -1 | protein_coding | ENSG0000C |
| 15506866 | 15507354 | 1 | processed_pseudoge | ENSG0000C |
| 15530773 | 15531089 | -1 | lncRNA | ENSG0000C |
| 15565483 | 15619512 | -1 | protein_coding | ENSG0000C |
| 15571491 | 15651653 | -1 | protein_coding | ENSG0000C |
| 15574019 | 15574198 | 1 | snoRNA | ENSG0000C |

| | | | | |
|---|---|---|---|---|
| 15588852 | 15589084 | 1 | unprocessed_pseudc | ENSG0000C |
| 15627960 | 15684311 | -1 | protein_coding | ENSG0000C |
| 15651590 | 15654489 | 1 | lncRNA | ENSG0000C |
| 15675476 | 15675643 | 1 | unprocessed_pseudc | ENSG0000C |
| 15699577 | 15720787 | 1 | protein_coding | ENSG0000C |
| 15699692 | 15744778 | 1 | protein_coding | ENSG0000C |
| 15704232 | 15704900 | -1 | processed_pseudoge | ENSG0000C |
| 15732247 | 15749192 | 1 | protein_coding | ENSG0000C |
| 15735023 | 15749192 | -1 | lncRNA | ENSG0000C |
| 15748703 | 15749315 | -1 | processed_pseudoge | ENSG0000C |
| 15761614 | 15763769 | -1 | lncRNA | ENSG0000C |
| 15764703 | 15765664 | 1 | unprocessed_pseudc | ENSG0000C |
| 15768717 | 15771426 | -1 | processed_pseudoge | ENSG0000C |
| 15782341 | 15782455 | 1 | rRNA_pseudogene | ENSG0000C |
| 15783288 | 15784307 | 1 | processed_pseudoge | ENSG0000C |
| 15786618 | 15787575 | 1 | processed_pseudoge | ENSG0000C |
| 15787787 | 15788205 | -1 | lncRNA | ENSG0000C |
| 15789016 | 15789705 | 1 | lncRNA | ENSG0000C |
| 15792862 | 15793045 | 1 | unprocessed_pseudc | ENSG0000C |
| 15806241 | 15847996 | -1 | lncRNA | ENSG0000C |
| 15829369 | 15908632 | 1 | transcribed_unproce | ENSG0000C |
| 15944917 | 15975746 | 1 | protein_coding | ENSG0000C |
| 15976560 | 15999717 | -1 | protein_coding | ENSG0000C |
| 15999784 | 16045015 | 1 | protein_coding | ENSG0000C |
| 16023323 | 16023653 | -1 | lncRNA | ENSG0000C |
| 16029157 | 16218185 | -1 | protein_coding | ENSG0000C |
| 16040472 | 16041273 | 1 | lncRNA | ENSG0000C |
| 16044621 | 16044961 | 1 | processed_pseudoge | ENSG0000C |
| 16084042 | 16084381 | 1 | processed_pseudoge | ENSG0000C |
| 16098704 | 16098807 | 1 | snRNA | ENSG0000C |
| 16137768 | 16137872 | -1 | snRNA | ENSG0000C |
| 16191832 | 16192151 | 1 | misc_RNA | ENSG0000C |
| 16217191 | 16351797 | 1 | protein_coding | ENSG0000C |
| 16282014 | 16282088 | 1 | miRNA | ENSG0000C |
| 16342534 | 16353656 | -1 | protein_coding | ENSG0000C |
| 16380798 | 16382745 | 1 | protein_coding | ENSG0000C |
| 16382152 | 16382669 | -1 | lncRNA | ENSG0000C |
| 16391434 | 16391882 | 1 | processed_pseudoge | ENSG0000C |
| 16414524 | 16416689 | -1 | lncRNA | ENSG0000C |
| 16415571 | 16437003 | 1 | protein_coding | ENSG0000C |
| 16427474 | 16428719 | 1 | lncRNA | ENSG0000C |
| 16438767 | 16478678 | 1 | lncRNA | ENSG0000C |
| 16439505 | 16439576 | 1 | snoRNA | ENSG0000C |
| 16440036 | 16440106 | 1 | snoRNA | ENSG0000C |
| 16440479 | 16440952 | -1 | lncRNA | ENSG0000C |
| 16441226 | 16441298 | 1 | snoRNA | ENSG0000C |
| 16441577 | 16492193 | -1 | protein_coding | ENSG0000C |
| 16525832 | 16529290 | 1 | lncRNA | ENSG0000C |
| 16546954 | 16569204 | -1 | protein_coding | ENSG0000C |
| 16557985 | 16559269 | -1 | lncRNA | ENSG0000C |

| | | | | |
|---|---|---|---|---|
| 16616848 | 16617881 | -1 | processed_pseudoge | ENSG00000( |
| 16620734 | 16653856 | -1 | protein_coding | ENSG00000( |
| 16653904 | 16654787 | 1 | lncRNA | ENSG00000( |
| 16666107 | 16667118 | -1 | processed_pseudoge | ENSG00000( |
| 16683438 | 16684227 | 1 | processed_pseudoge | ENSG00000( |
| 16689537 | 16777881 | 1 | protein_coding | ENSG00000( |
| 16690261 | 16804453 | 1 | protein_coding | ENSG00000( |
| 16691798 | 16691907 | 1 | snRNA | ENSG00000( |
| 16710059 | 16710361 | 1 | misc_RNA | ENSG00000( |
| 16745636 | 16746761 | -1 | processed_pseudoge | ENSG00000( |
| 16786489 | 16804455 | 1 | transcribed_unproce | ENSG00000( |
| 16787235 | 16788141 | -1 | processed_pseudoge | ENSG00000( |
| 16788057 | 16816540 | 1 | lncRNA | ENSG00000( |
| 16788248 | 16789234 | -1 | lncRNA | ENSG00000( |
| 16788879 | 16790501 | 1 | lncRNA | ENSG00000( |
| 16804627 | 16805707 | 1 | lncRNA | ENSG00000( |
| 16812447 | 16812651 | 1 | processed_pseudoge | ENSG00000( |
| 16817983 | 16822579 | -1 | transcribed_unproce | ENSG00000( |
| 16829999 | 16832830 | -1 | transcribed_unproce | ENSG00000( |
| 16840895 | 16845881 | -1 | transcribed_unproce | ENSG00000( |
| 16847635 | 16852777 | -1 | transcribed_unproce | ENSG00000( |
| 16853935 | 16854309 | 1 | processed_pseudoge | ENSG00000( |
| 16861146 | 16864223 | 1 | lncRNA | ENSG00000( |
| 16893198 | 16893429 | -1 | processed_pseudoge | ENSG00000( |
| 16922915 | 16934839 | -1 | transcribed_unproce | ENSG00000( |
| 16929816 | 16972118 | -1 | protein_coding | ENSG00000( |
| 16975999 | 16981353 | -1 | lncRNA | ENSG00000( |
| 16988329 | 16990177 | 1 | lncRNA | ENSG00000( |
| 17011914 | 17014990 | -1 | lncRNA | ENSG00000( |
| 17028360 | 17035057 | 1 | lncRNA | ENSG00000( |
| 17033531 | 17042305 | -1 | lncRNA | ENSG00000( |
| 17042545 | 17217679 | 1 | protein_coding | ENSG00000( |
| 17076038 | 17077752 | -1 | lncRNA | ENSG00000( |
| 17167946 | 17185554 | -1 | lncRNA | ENSG00000( |
| 17170083 | 17170189 | 1 | snRNA | ENSG00000( |
| 17178414 | 17178749 | 1 | misc_RNA | ENSG00000( |
| 17181504 | 17183257 | 1 | TEC | ENSG00000( |
| 17200995 | 17206333 | -1 | protein_coding | ENSG00000( |
| 17202649 | 17237185 | -1 | protein_coding | ENSG00000( |
| 17212212 | 17237188 | -1 | protein_coding | ENSG00000( |
| 17235433 | 17236118 | -1 | lncRNA | ENSG00000( |
| 17242010 | 17242886 | 1 | processed_pseudoge | ENSG00000( |
| 17246616 | 17281273 | -1 | protein_coding | ENSG00000( |
| 17303335 | 17347663 | 1 | protein_coding | ENSG00000( |
| 17383377 | 17384012 | -1 | processed_pseudoge | ENSG00000( |
| 17422681 | 17423704 | 1 | processed_pseudoge | ENSG00000( |
| 17456427 | 17457259 | 1 | processed_pseudoge | ENSG00000( |
| 17460444 | 17460522 | 1 | misc_RNA | ENSG00000( |
| 17476994 | 17493221 | 1 | protein_coding | ENSG00000( |
| 17494437 | 17496395 | -1 | protein_coding | ENSG00000( |

| | | | | |
|---|---|---|---|---|
| 17504005 | 17512266 | 1 | lncRNA | ENSG00000 |
| 17505563 | 17591708 | -1 | protein_coding | ENSG00000 |
| 17507351 | 17508308 | 1 | lncRNA | ENSG00000 |
| 17578631 | 17578739 | 1 | snRNA | ENSG00000 |
| 17591428 | 17610485 | 1 | lncRNA | ENSG00000 |
| 17616043 | 17617745 | 1 | TEC | ENSG00000 |
| 17674026 | 17677688 | -1 | lncRNA | ENSG00000 |
| 17675366 | 17675522 | 1 | misc_RNA | ENSG00000 |
| 17676211 | 17676371 | 1 | misc_RNA | ENSG00000 |
| 17681458 | 17811453 | 1 | protein_coding | ENSG00000 |
| 17759154 | 17770821 | -1 | lncRNA | ENSG00000 |
| 17776686 | 17779529 | -1 | lncRNA | ENSG00000 |
| 17777781 | 17779094 | 1 | lncRNA | ENSG00000 |
| 17810399 | 17837011 | -1 | protein_coding | ENSG00000 |
| 17813480 | 17813545 | -1 | miRNA | ENSG00000 |
| 17813836 | 17813931 | -1 | miRNA | ENSG00000 |
| 17843511 | 17972422 | -1 | protein_coding | ENSG00000 |
| 17858227 | 17860041 | 1 | lncRNA | ENSG00000 |
| 17972813 | 18016889 | 1 | protein_coding | ENSG00000 |
| 17977409 | 18039209 | -1 | protein_coding | ENSG00000 |
| 17981192 | 17981572 | -1 | processed_pseudoge | ENSG00000 |
| 18010643 | 18011822 | 1 | processed_pseudoge | ENSG00000 |
| 18026072 | 18026771 | -1 | TEC | ENSG00000 |
| 18039408 | 18068405 | 1 | protein_coding | ENSG00000 |
| 18087892 | 18107970 | 1 | protein_coding | ENSG00000 |
| 18107334 | 18110463 | -1 | TEC | ENSG00000 |
| 18108706 | 18179802 | 1 | protein_coding | ENSG00000 |
| 18172625 | 18184753 | -1 | lncRNA | ENSG00000 |
| 18183078 | 18209954 | 1 | protein_coding | ENSG00000 |
| 18225635 | 18244875 | 1 | protein_coding | ENSG00000 |
| 18244815 | 18258738 | -1 | protein_coding | ENSG00000 |
| 18260597 | 18266552 | 1 | protein_coding | ENSG00000 |
| 18268080 | 18268828 | 1 | lncRNA | ENSG00000 |
| 18271428 | 18315007 | -1 | protein_coding | ENSG00000 |
| 18311904 | 18312651 | 1 | processed_pseudoge | ENSG00000 |
| 18312760 | 18313533 | 1 | processed_pseudoge | ENSG00000 |
| 18315293 | 18328056 | 1 | protein_coding | ENSG00000 |
| 18327860 | 18363563 | -1 | protein_coding | ENSG00000 |
| 18340814 | 18340886 | -1 | miRNA | ENSG00000 |
| 18377778 | 18389647 | 1 | protein_coding | ENSG00000 |
| 18379855 | 18388984 | -1 | lncRNA | ENSG00000 |
| 18388871 | 18389459 | 1 | processed_pseudoge | ENSG00000 |
| 18390153 | 18390323 | -1 | unprocessed_pseudc | ENSG00000 |
| 18393480 | 18400961 | 1 | unprocessed_pseudc | ENSG00000 |
| 18411159 | 18414380 | 1 | lncRNA | ENSG00000 |
| 18415728 | 18425097 | 1 | transcribed_unproce | ENSG00000 |
| 18418195 | 18418306 | -1 | processed_pseudoge | ENSG00000 |
| 18426861 | 18431848 | 1 | transcribed_unproce | ENSG00000 |
| 18432051 | 18442895 | 1 | transcribed_unproce | ENSG00000 |
| 18450244 | 18475920 | 1 | transcribed_processe | ENSG00000 |

| | | | | |
|---|---|---|---|---|
| 18452955 | 18453062 | 1 | unprocessed_pseudc | ENSG00000( |
| 18453574 | 18453739 | -1 | processed_pseudoge | ENSG00000( |
| 18460391 | 18463055 | 1 | processed_pseudoge | ENSG00000( |
| 18476737 | 18494945 | 1 | protein_coding | ENSG00000( |
| 18497095 | 18506313 | -1 | transcribed_unproce | ENSG00000( |
| 18511221 | 18551705 | -1 | lncRNA | ENSG00000( |
| 18511262 | 18525930 | -1 | transcribed_unproce | ENSG00000( |
| 18527557 | 18528461 | 1 | processed_pseudoge | ENSG00000( |
| 18537800 | 18625617 | -1 | transcribed_unproce | ENSG00000( |
| 18572752 | 18572961 | 1 | processed_pseudoge | ENSG00000( |
| 18605054 | 18605356 | -1 | misc_RNA | ENSG00000( |
| 18635006 | 18644427 | -1 | protein_coding | ENSG00000( |
| 18638019 | 18638370 | -1 | transcribed_processe | ENSG00000( |
| 18638861 | 18661950 | -1 | lncRNA | ENSG00000( |
| 18650195 | 18651542 | 1 | processed_pseudoge | ENSG00000( |
| 18658429 | 18682228 | -1 | transcribed_unproce | ENSG00000( |
| 18667629 | 18682262 | -1 | protein_coding | ENSG00000( |
| 18677261 | 18677757 | 1 | processed_pseudoge | ENSG00000( |
| 18697998 | 18736118 | 1 | protein_coding | ENSG00000( |
| 18704640 | 18704769 | 1 | transcribed_processe | ENSG00000( |
| 18744026 | 18779349 | 1 | protein_coding | ENSG00000( |
| 18781111 | 18806714 | 1 | protein_coding | ENSG00000( |
| 18809956 | 18810940 | 1 | lncRNA | ENSG00000( |
| 18835707 | 18842364 | 1 | unprocessed_pseudc | ENSG00000( |
| 18840085 | 18931287 | 1 | protein_coding | ENSG00000( |
| 18840618 | 18840920 | 1 | misc_RNA | ENSG00000( |
| 18859354 | 18861466 | -1 | lncRNA | ENSG00000( |
| 18908279 | 18908865 | 1 | transcribed_processe | ENSG00000( |
| 18943999 | 18944073 | 1 | snoRNA | ENSG00000( |
| 18950345 | 19022595 | 1 | protein_coding | ENSG00000( |
| 18951625 | 18954149 | -1 | lncRNA | ENSG00000( |
| 18968789 | 19004764 | -1 | protein_coding | ENSG00000( |
| 18986148 | 18986334 | -1 | processed_pseudoge | ENSG00000( |
| 19020673 | 19047637 | -1 | protein_coding | ENSG00000( |
| 19030857 | 19033529 | 1 | TEC | ENSG00000( |
| 19061912 | 19062669 | 1 | snoRNA | ENSG00000( |
| 19063346 | 19064136 | -1 | snoRNA | ENSG00000( |
| 19077775 | 19087878 | 1 | lncRNA | ENSG00000( |
| 19082437 | 19084834 | 1 | processed_pseudoge | ENSG00000( |
| 19091069 | 19091825 | -1 | lncRNA | ENSG00000( |
| 19092974 | 19096837 | 1 | lncRNA | ENSG00000( |
| 19112000 | 19112636 | -1 | lncRNA | ENSG00000( |
| 19112419 | 19112636 | -1 | snoRNA | ENSG00000( |
| 19112420 | 19112636 | -1 | snoRNA | ENSG00000( |
| 19127535 | 19159187 | 1 | protein_coding | ENSG00000( |
| 19141017 | 19143689 | -1 | lncRNA | ENSG00000( |
| 19150279 | 19151286 | 1 | lncRNA | ENSG00000( |
| 19155727 | 19159111 | -1 | lncRNA | ENSG00000( |
| 19160789 | 19162274 | 1 | lncRNA | ENSG00000( |
| 19164209 | 19174436 | -1 | lncRNA | ENSG00000( |

| | | | | |
|---|---|---|---|---|
| 19166896 | 19169668 | -1 | unprocessed_pseudc | ENSG00000( |
| 19188016 | 19188714 | 1 | snoRNA | ENSG00000( |
| 19189665 | 19190245 | -1 | snoRNA | ENSG00000( |
| 19203825 | 19214045 | 1 | lncRNA | ENSG00000( |
| 19208584 | 19211359 | 1 | transcribed_unproce | ENSG00000( |
| 19215615 | 19336715 | 1 | protein_coding | ENSG00000( |
| 19219143 | 19222527 | 1 | lncRNA | ENSG00000( |
| 19296596 | 19306261 | -1 | lncRNA | ENSG00000( |
| 19334308 | 19378193 | -1 | protein_coding | ENSG00000( |
| 19344506 | 19344574 | -1 | miRNA | ENSG00000( |
| 19362760 | 19370236 | 1 | lncRNA | ENSG00000( |
| 19377721 | 19383544 | 1 | protein_coding | ENSG00000( |
| 19383442 | 19387240 | -1 | protein_coding | ENSG00000( |
| 19403283 | 19403872 | -1 | lncRNA | ENSG00000( |
| 19411125 | 19417276 | 1 | protein_coding | ENSG00000( |
| 19411786 | 19413118 | -1 | lncRNA | ENSG00000( |
| 19417769 | 19492991 | -1 | lncRNA | ENSG00000( |
| 19419035 | 19424357 | 1 | lncRNA | ENSG00000( |
| 19420279 | 19420342 | 1 | snoRNA | ENSG00000( |
| 19445970 | 19446852 | -1 | processed_pseudoge | ENSG00000( |
| 19457080 | 19458399 | 1 | lncRNA | ENSG00000( |
| 19495385 | 19579034 | 1 | protein_coding | ENSG00000( |
| 19496943 | 19497528 | 1 | processed_pseudoge | ENSG00000( |
| 19557211 | 19557911 | 1 | snoRNA | ENSG00000( |
| 19560111 | 19597922 | -1 | lncRNA | ENSG00000( |
| 19579943 | 19596058 | 1 | transcribed_unproce | ENSG00000( |
| 19582116 | 19582275 | -1 | processed_pseudoge | ENSG00000( |
| 19600860 | 19601345 | 1 | processed_pseudoge | ENSG00000( |
| 19601840 | 19602256 | 1 | processed_pseudoge | ENSG00000( |
| 19602259 | 19603208 | 1 | unprocessed_pseudc | ENSG00000( |
| 19603274 | 19603345 | -1 | unprocessed_pseudc | ENSG00000( |
| 19603797 | 19605048 | 1 | unprocessed_pseudc | ENSG00000( |
| 19605454 | 19605933 | 1 | unprocessed_pseudc | ENSG00000( |
| 19615789 | 19633825 | 1 | transcribed_unproce | ENSG00000( |
| 19648136 | 19685760 | 1 | protein_coding | ENSG00000( |
| 19649373 | 19649935 | -1 | lncRNA | ENSG00000( |
| 19649494 | 19649587 | -1 | misc_RNA | ENSG00000( |
| 19662000 | 19662090 | 1 | snoRNA | ENSG00000( |
| 19678288 | 19718979 | -1 | protein_coding | ENSG00000( |
| 19719059 | 19722428 | 1 | lncRNA | ENSG00000( |
| 19737682 | 19738542 | 1 | lncRNA | ENSG00000( |
| 19737984 | 19748943 | -1 | protein_coding | ENSG00000( |
| 19755244 | 19756207 | 1 | processed_pseudoge | ENSG00000( |
| 19770829 | 19867936 | -1 | protein_coding | ENSG00000( |
| 19840126 | 19840515 | 1 | processed_pseudoge | ENSG00000( |
| 19855747 | 19856164 | 1 | TEC | ENSG00000( |
| 19868568 | 19868689 | 1 | lncRNA | ENSG00000( |
| 19896590 | 19897287 | -1 | lncRNA | ENSG00000( |
| 19904302 | 19978343 | -1 | protein_coding | ENSG00000( |
| 19929372 | 19929737 | -1 | lncRNA | ENSG00000( |

| | | | | |
|---|---|---|---|---|
| 20008051 | 20009234 | -1 | lncRNA | ENSG0000C |
| 20009344 | 20319026 | 1 | protein_coding | ENSG0000C |
| 20056287 | 20056630 | 1 | transcribed_processe | ENSG0000C |
| 20111824 | 20112990 | 1 | transcribed_processe | ENSG0000C |
| 20126388 | 20126488 | -1 | snRNA | ENSG0000C |
| 20185082 | 20322973 | 1 | lncRNA | ENSG0000C |
| 20290257 | 20290358 | 1 | snRNA | ENSG0000C |
| 20321164 | 20403557 | 1 | transcribed_processe | ENSG0000C |
| 20323001 | 20323110 | 1 | snRNA | ENSG0000C |
| 20323058 | 20323337 | 1 | lncRNA | ENSG0000C |
| 20340743 | 20341045 | 1 | misc_RNA | ENSG0000C |
| 20375369 | 20376840 | -1 | transcribed_processe | ENSG0000C |
| 20408311 | 20434286 | 1 | transcribed_unproce | ENSG0000C |
| 20415792 | 20416697 | -1 | unprocessed_pseudo | ENSG0000C |
| 20416610 | 20447224 | 1 | lncRNA | ENSG0000C |
| 20417450 | 20417956 | 1 | processed_pseudoge | ENSG0000C |
| 20436337 | 20447249 | 1 | transcribed_unproce | ENSG0000C |
| 20449395 | 20467539 | -1 | protein_coding | ENSG0000C |
| 20481330 | 20483991 | -1 | processed_pseudoge | ENSG0000C |
| 20491323 | 20491443 | -1 | processed_pseudoge | ENSG0000C |
| 20493634 | 20494155 | -1 | processed_pseudoge | ENSG0000C |
| 20501513 | 20512357 | -1 | transcribed_unproce | ENSG0000C |
| 20507413 | 20508931 | -1 | lncRNA | ENSG0000C |
| 20512560 | 20517479 | -1 | transcribed_unproce | ENSG0000C |
| 20519312 | 20528687 | -1 | transcribed_unproce | ENSG0000C |
| 20526102 | 20526210 | 1 | processed_pseudoge | ENSG0000C |
| 20530042 | 20530881 | -1 | lncRNA | ENSG0000C |
| 20532248 | 20532881 | -1 | processed_pseudoge | ENSG0000C |
| 20543481 | 20550963 | -1 | unprocessed_pseudo | ENSG0000C |
| 20545371 | 20549952 | 1 | lncRNA | ENSG0000C |
| 20554137 | 20554380 | 1 | processed_pseudoge | ENSG0000C |
| 20554971 | 20559410 | -1 | transcribed_processe | ENSG0000C |
| 20564538 | 20564889 | 1 | processed_pseudoge | ENSG0000C |
| 20576667 | 20578748 | -1 | lncRNA | ENSG0000C |
| 20579724 | 20580911 | 1 | protein_coding | ENSG0000C |
| 20583758 | 20591180 | -1 | unprocessed_pseudo | ENSG0000C |
| 20589293 | 20590367 | 1 | processed_pseudoge | ENSG0000C |
| 20595531 | 20595708 | 1 | processed_pseudoge | ENSG0000C |
| 20609462 | 20609610 | -1 | processed_pseudoge | ENSG0000C |
| 20612667 | 20614008 | 1 | processed_pseudoge | ENSG0000C |
| 20612912 | 20633234 | -1 | lncRNA | ENSG0000C |
| 20633034 | 20633259 | -1 | transcribed_processe | ENSG0000C |
| 20647287 | 20681799 | 1 | unprocessed_pseudo | ENSG0000C |
| 20710425 | 20710533 | -1 | processed_pseudoge | ENSG0000C |
| 20716481 | 20737127 | 1 | transcribed_unproce | ENSG0000C |
| 20743333 | 20754501 | -1 | unprocessed_pseudo | ENSG0000C |
| 20771058 | 20771427 | -1 | processed_pseudoge | ENSG0000C |
| 20775642 | 20776905 | 1 | processed_pseudoge | ENSG0000C |
| 20784645 | 20785725 | 1 | processed_pseudoge | ENSG0000C |
| 20788071 | 20789584 | 1 | lncRNA | ENSG0000C |

| | | | | |
|---|---|---|---|---|
| 20789217 | 20789595 | -1 | processed_pseudoge | ENSG00000 |
| 20814620 | 20844257 | 1 | unprocessed_pseudo | ENSG00000 |
| 20836447 | 20896140 | -1 | protein_coding | ENSG00000 |
| 20855946 | 20857317 | 1 | lncRNA | ENSG00000 |
| 20868433 | 21002276 | 1 | lncRNA | ENSG00000 |
| 20894532 | 20894638 | -1 | snRNA | ENSG00000 |
| 20901939 | 20902680 | -1 | processed_pseudoge | ENSG00000 |
| 20938023 | 20938344 | -1 | processed_pseudoge | ENSG00000 |
| 20956682 | 20958404 | 1 | transcribed_unproce | ENSG00000 |
| 20963722 | 20980238 | 1 | lncRNA | ENSG00000 |
| 20981489 | 20981654 | 1 | transcribed_processe | ENSG00000 |
| 20983902 | 20984046 | 1 | processed_pseudoge | ENSG00000 |
| 20999596 | 21043760 | -1 | protein_coding | ENSG00000 |
| 20999747 | 21000323 | 1 | lncRNA | ENSG00000 |
| 21075362 | 21092351 | 1 | lncRNA | ENSG00000 |
| 21123364 | 21193265 | 1 | protein_coding | ENSG00000 |
| 21197280 | 21214624 | -1 | protein_coding | ENSG00000 |
| 21214322 | 21230035 | 1 | lncRNA | ENSG00000 |
| 21229334 | 21229628 | -1 | misc_RNA | ENSG00000 |
| 21238870 | 21253410 | -1 | protein_coding | ENSG00000 |
| 21259968 | 21260501 | -1 | unprocessed_pseudo | ENSG00000 |
| 21284672 | 21315232 | 1 | protein_coding | ENSG00000 |
| 21376357 | 21419870 | 1 | protein_coding | ENSG00000 |
| 21428263 | 21574517 | -1 | lncRNA | ENSG00000 |
| 21442805 | 21451525 | 1 | lncRNA | ENSG00000 |
| 21444833 | 21445439 | -1 | processed_pseudoge | ENSG00000 |
| 21456513 | 21460215 | -1 | lncRNA | ENSG00000 |
| 21457434 | 21458989 | 1 | lncRNA | ENSG00000 |
| 21504558 | 21505059 | 1 | processed_pseudoge | ENSG00000 |
| 21519514 | 21521269 | 1 | lncRNA | ENSG00000 |
| 21532974 | 21542786 | 1 | lncRNA | ENSG00000 |
| 21552509 | 21552711 | 1 | processed_pseudoge | ENSG00000 |
| 21573536 | 21573746 | -1 | transcribed_processe | ENSG00000 |
| 21573770 | 21574073 | 1 | processed_pseudoge | ENSG00000 |
| 21592794 | 21594180 | 1 | processed_pseudoge | ENSG00000 |
| 21614487 | 21641536 | 1 | transcribed_unproce | ENSG00000 |
| 21655843 | 21657633 | -1 | unprocessed_pseudo | ENSG00000 |
| 21678086 | 21678418 | 1 | unprocessed_pseudo | ENSG00000 |
| 21692523 | 21704612 | 1 | protein_coding | ENSG00000 |
| 21773754 | 21783465 | -1 | unprocessed_pseudo | ENSG00000 |
| 22056587 | 22056755 | -1 | processed_pseudoge | ENSG00000 |
| 22061961 | 22070538 | -1 | unprocessed_pseudo | ENSG00000 |
| 22090743 | 22205154 | 1 | transcribed_unproce | ENSG00000 |
| 22221221 | 22221605 | 1 | processed_pseudoge | ENSG00000 |
| 22233926 | 22266392 | -1 | lncRNA | ENSG00000 |
| 22266395 | 22288301 | 1 | lncRNA | ENSG00000 |
| 22277867 | 22279289 | -1 | processed_pseudoge | ENSG00000 |
| 22298691 | 22299893 | 1 | lncRNA | ENSG00000 |
| 22331597 | 22332410 | 1 | unprocessed_pseudo | ENSG00000 |
| 22406019 | 22413744 | 1 | lncRNA | ENSG00000 |

| | | | | |
|---|---|---|---|---|
| 22420022 | 22424731 | 1 | lncRNA | ENSG00000 |
| 22435390 | 22439033 | 1 | transcribed_processe | ENSG00000 |
| 22519195 | 22519504 | -1 | unprocessed_pseudo | ENSG00000 |
| 22519617 | 22520709 | 1 | unprocessed_pseudo | ENSG00000 |
| 22523111 | 22524663 | 1 | protein_coding | ENSG00000 |
| 22524563 | 22525330 | -1 | lncRNA | ENSG00000 |
| 22524738 | 22525686 | 1 | transcribed_unproce | ENSG00000 |
| 22525899 | 22526937 | 1 | unprocessed_pseudo | ENSG00000 |
| 22527324 | 22529570 | 1 | unprocessed_pseudo | ENSG00000 |
| 22528853 | 22528921 | -1 | unprocessed_pseudo | ENSG00000 |
| 22529934 | 22530609 | 1 | unprocessed_pseudo | ENSG00000 |
| 22530612 | 22531392 | 1 | unprocessed_pseudo | ENSG00000 |
| 22531463 | 22531797 | 1 | unprocessed_pseudo | ENSG00000 |
| 22531876 | 22532172 | 1 | unprocessed_pseudo | ENSG00000 |
| 22532166 | 22532522 | 1 | unprocessed_pseudo | ENSG00000 |
| 22567423 | 22568713 | -1 | unprocessed_pseudo | ENSG00000 |
| 22606388 | 22606703 | 1 | processed_pseudoge | ENSG00000 |
| 22671433 | 22706703 | 1 | transcribed_processe | ENSG00000 |
| 22676695 | 22692618 | -1 | lncRNA | ENSG00000 |
| 22693325 | 22694831 | -1 | processed_pseudoge | ENSG00000 |
| 26981694 | 26982090 | 1 | processed_pseudoge | ENSG00000 |
| 26989189 | 26999941 | -1 | lncRNA | ENSG00000 |
| 27002661 | 27003259 | -1 | processed_pseudoge | ENSG00000 |
| 27019527 | 27047934 | -1 | unprocessed_pseudo | ENSG00000 |
| 27057399 | 27058221 | 1 | processed_pseudoge | ENSG00000 |
| 27082690 | 27084036 | -1 | processed_pseudoge | ENSG00000 |
| 27089545 | 27170310 | 1 | lncRNA | ENSG00000 |
| 27237859 | 27241661 | -1 | lncRNA | ENSG00000 |
| 27274107 | 27274477 | 1 | processed_pseudoge | ENSG00000 |
| 27280482 | 27280912 | 1 | processed_pseudoge | ENSG00000 |
| 27281094 | 27281415 | 1 | processed_pseudoge | ENSG00000 |
| 27293910 | 27293996 | -1 | miRNA | ENSG00000 |
| 27294076 | 27315926 | 1 | protein_coding | ENSG00000 |
| 27333241 | 27352828 | 1 | lncRNA | ENSG00000 |
| 27351858 | 27352584 | -1 | processed_pseudoge | ENSG00000 |
| 27411438 | 27412806 | -1 | unprocessed_pseudo | ENSG00000 |
| 27417690 | 27431623 | 1 | transcribed_unproce | ENSG00000 |
| 27456470 | 27626438 | 1 | protein_coding | ENSG00000 |
| 27465112 | 27465541 | 1 | lncRNA | ENSG00000 |
| 27526386 | 27527223 | -1 | processed_pseudoge | ENSG00000 |
| 27612588 | 27613251 | -1 | processed_pseudoge | ENSG00000 |
| 27623364 | 27640777 | 1 | protein_coding | ENSG00000 |
| 27625484 | 27626438 | -1 | lncRNA | ENSG00000 |
| 27629798 | 27649560 | 1 | protein_coding | ENSG00000 |
| 27651334 | 27665712 | -1 | transcribed_unproce | ENSG00000 |
| 27666749 | 27672803 | -1 | unprocessed_pseudo | ENSG00000 |
| 27675894 | 27675997 | -1 | processed_pseudoge | ENSG00000 |
| 27677805 | 27678087 | -1 | processed_pseudoge | ENSG00000 |
| 27682476 | 27682737 | 1 | processed_pseudoge | ENSG00000 |
| 27683946 | 27684144 | -1 | processed_pseudoge | ENSG00000 |

| | | | | |
|---|---|---|---|---|
| 27707178 | 27707250 | 1 | processed_pseudoge | ENSG00000 |
| 27708111 | 27708138 | 1 | processed_pseudoge | ENSG00000 |
| 27746132 | 27754954 | 1 | unprocessed_pseudo | ENSG00000 |
| 27756766 | 27800529 | -1 | protein_coding | ENSG00000 |
| 27798806 | 27893365 | -1 | protein_coding | ENSG00000 |
| 27874645 | 27881237 | 1 | lncRNA | ENSG00000 |
| 27878314 | 27894752 | -1 | protein_coding | ENSG00000 |
| 27928924 | 27991787 | -1 | lncRNA | ENSG00000 |
| 27930177 | 27930920 | 1 | lncRNA | ENSG00000 |
| 28018770 | 28018907 | 1 | scaRNA | ENSG00000 |
| 28021668 | 28022265 | -1 | unprocessed_pseudo | ENSG00000 |
| 28022330 | 28022464 | -1 | snoRNA | ENSG00000 |
| 28041737 | 28196381 | 1 | protein_coding | ENSG00000 |
| 28061967 | 28062068 | 1 | misc_RNA | ENSG00000 |
| 28116239 | 28116406 | -1 | processed_pseudoge | ENSG00000 |
| 28204616 | 28223735 | 1 | lncRNA | ENSG00000 |
| 28219285 | 28220005 | 1 | lncRNA | ENSG00000 |
| 28226563 | 28228065 | 1 | transcribed_unproce | ENSG00000 |
| 28232590 | 28235281 | -1 | lncRNA | ENSG00000 |
| 28246454 | 28248006 | -1 | lncRNA | ENSG00000 |
| 28247444 | 28248289 | 1 | transcribed_unproce | ENSG00000 |
| 28256375 | 28256669 | -1 | processed_pseudoge | ENSG00000 |
| 28256438 | 28265551 | 1 | lncRNA | ENSG00000 |
| 28263634 | 28266369 | -1 | lncRNA | ENSG00000 |
| 28269203 | 28272365 | -1 | unprocessed_pseudo | ENSG00000 |
| 28274657 | 28275716 | -1 | unprocessed_pseudo | ENSG00000 |
| 28275986 | 28317783 | -1 | transcribed_unproce | ENSG00000 |
| 28319200 | 28328685 | 1 | protein_coding | ENSG00000 |
| 28328325 | 28335489 | -1 | protein_coding | ENSG00000 |
| 28335602 | 28347009 | 1 | protein_coding | ENSG00000 |
| 28346633 | 28357527 | -1 | protein_coding | ENSG00000 |
| 28357642 | 28363683 | 1 | protein_coding | ENSG00000 |
| 28357647 | 28381697 | 1 | lncRNA | ENSG00000 |
| 28360654 | 28360734 | 1 | miRNA | ENSG00000 |
| 28361601 | 28362859 | -1 | lncRNA | ENSG00000 |
| 28364268 | 28365244 | -1 | protein_coding | ENSG00000 |
| 28364288 | 28368012 | -1 | protein_coding | ENSG00000 |
| 28364356 | 28404049 | 1 | protein_coding | ENSG00000 |
| 28367284 | 28373091 | -1 | protein_coding | ENSG00000 |
| 28373256 | 28373562 | 1 | lncRNA | ENSG00000 |
| 28373673 | 28374301 | -1 | TEC | ENSG00000 |
| 28394642 | 28407197 | -1 | protein_coding | ENSG00000 |
| 28405240 | 28406796 | 1 | lncRNA | ENSG00000 |
| 28416922 | 28417492 | 1 | unprocessed_pseudo | ENSG00000 |
| 28444063 | 28445563 | -1 | unprocessed_pseudo | ENSG00000 |
| 28455752 | 28614197 | -1 | protein_coding | ENSG00000 |
| 28467822 | 28468406 | 1 | processed_pseudoge | ENSG00000 |
| 28473293 | 28497781 | 1 | protein_coding | ENSG00000 |
| 28506243 | 28538896 | 1 | protein_coding | ENSG00000 |
| 28546707 | 28552668 | -1 | protein_coding | ENSG00000 |

| | | | | |
|---|---|---|---|---|
| 28553383 | 28571794 | -1 | protein_coding | ENSG00000 |
| 28573115 | 28576948 | -1 | protein_coding | ENSG00000 |
| 28573117 | 28574243 | 1 | lncRNA | ENSG00000 |
| 28577565 | 28599025 | -1 | protein_coding | ENSG00000 |
| 28598790 | 28617377 | 1 | lncRNA | ENSG00000 |
| 28601827 | 28602284 | -1 | lncRNA | ENSG00000 |
| 28607963 | 28609730 | 1 | lncRNA | ENSG00000 |
| 28607964 | 28614200 | -1 | protein_coding | ENSG00000 |
| 28614446 | 28645454 | -1 | protein_coding | ENSG00000 |
| 28633206 | 28635950 | 1 | lncRNA | ENSG00000 |
| 28648346 | 28662189 | -1 | protein_coding | ENSG00000 |
| 28655557 | 28655983 | 1 | processed_pseudoge | ENSG00000 |
| 28662198 | 28702679 | 1 | protein_coding | ENSG00000 |
| 28670054 | 28672804 | -1 | lncRNA | ENSG00000 |
| 28703197 | 28711854 | -1 | protein_coding | ENSG00000 |
| 28714281 | 28718429 | -1 | protein_coding | ENSG00000 |
| 28719985 | 28724359 | 1 | protein_coding | ENSG00000 |
| 28720550 | 28720616 | 1 | snoRNA | ENSG00000 |
| 28721487 | 28722877 | -1 | lncRNA | ENSG00000 |
| 28722582 | 28722653 | 1 | snoRNA | ENSG00000 |
| 28723429 | 28723492 | 1 | snoRNA | ENSG00000 |
| 28723682 | 28723753 | 1 | snoRNA | ENSG00000 |
| 28724348 | 28727935 | -1 | protein_coding | ENSG00000 |
| 28725897 | 28743455 | 1 | protein_coding | ENSG00000 |
| 28727258 | 28727739 | 1 | lncRNA | ENSG00000 |
| 28732963 | 28743102 | -1 | lncRNA | ENSG00000 |
| 28744005 | 28750956 | 1 | protein_coding | ENSG00000 |
| 28745569 | 28747652 | -1 | lncRNA | ENSG00000 |
| 28749731 | 28750079 | -1 | lncRNA | ENSG00000 |
| 28755978 | 28855232 | -1 | protein_coding | ENSG00000 |
| 28802444 | 28802822 | 1 | processed_pseudoge | ENSG00000 |
| 28834722 | 28834822 | 1 | misc_RNA | ENSG00000 |
| 28848860 | 28849592 | 1 | unprocessed_pseudc | ENSG00000 |
| 28855010 | 28861061 | 1 | protein_coding | ENSG00000 |
| 28861072 | 28861966 | -1 | lncRNA | ENSG00000 |
| 28861369 | 28861440 | -1 | miRNA | ENSG00000 |
| 28861371 | 28861438 | 1 | miRNA | ENSG00000 |
| 28861533 | 28861618 | -1 | miRNA | ENSG00000 |
| 28861655 | 28861730 | -1 | miRNA | ENSG00000 |
| 28879335 | 28897733 | -1 | protein_coding | ENSG00000 |
| 28892469 | 28893342 | 1 | lncRNA | ENSG00000 |
| 28897738 | 28899402 | 1 | lncRNA | ENSG00000 |
| 28897781 | 28903079 | -1 | protein_coding | ENSG00000 |
| 28905250 | 28951771 | -1 | protein_coding | ENSG00000 |
| 28926275 | 28944749 | 1 | lncRNA | ENSG00000 |
| 28944796 | 28945394 | 1 | lncRNA | ENSG00000 |
| 28950513 | 29057216 | 1 | protein_coding | ENSG00000 |
| 28954901 | 29006440 | -1 | protein_coding | ENSG00000 |
| 29009799 | 29011009 | -1 | lncRNA | ENSG00000 |
| 29012865 | 29016512 | 1 | lncRNA | ENSG00000 |

| | | | | |
|---|---|---|---|---|
| 29021325 | 29021640 | -1 | processed_pseudoge | ENSG0000C |
| 29071124 | 29180412 | -1 | protein_coding | ENSG0000C |
| 29073521 | 29078857 | -1 | protein_coding | ENSG0000C |
| 29140483 | 29155489 | 1 | lncRNA | ENSG0000C |
| 29197071 | 29197785 | -1 | processed_pseudoge | ENSG0000C |
| 29203426 | 29204474 | -1 | processed_pseudoge | ENSG0000C |
| 29246863 | 29254494 | 1 | protein_coding | ENSG0000C |
| 29255839 | 29294148 | -1 | protein_coding | ENSG0000C |
| 29340482 | 29340822 | -1 | processed_pseudoge | ENSG0000C |
| 29352069 | 29404105 | 1 | lncRNA | ENSG0000C |
| 29369717 | 29390777 | -1 | lncRNA | ENSG0000C |
| 29388560 | 29388683 | 1 | snRNA | ENSG0000C |
| 29390464 | 29551904 | 1 | protein_coding | ENSG0000C |
| 29390662 | 29390730 | 1 | miRNA | ENSG0000C |
| 29424762 | 29424867 | -1 | snRNA | ENSG0000C |
| 29445300 | 29445400 | -1 | snRNA | ENSG0000C |
| 29560547 | 29707090 | 1 | lncRNA | ENSG0000C |
| 29560547 | 29567037 | -1 | protein_coding | ENSG0000C |
| 29566052 | 29573157 | 1 | protein_coding | ENSG0000C |
| 29569580 | 29570519 | 1 | lncRNA | ENSG0000C |
| 29573469 | 29594054 | -1 | protein_coding | ENSG0000C |
| 29589769 | 29614761 | 1 | protein_coding | ENSG0000C |
| 29591703 | 29592241 | -1 | lncRNA | ENSG0000C |
| 29614756 | 29622907 | -1 | protein_coding | ENSG0000C |
| 29621617 | 29622254 | -1 | lncRNA | ENSG0000C |
| 29625938 | 29930276 | -1 | protein_coding | ENSG0000C |
| 29639627 | 29640825 | 1 | lncRNA | ENSG0000C |
| 29641611 | 29641702 | -1 | snRNA | ENSG0000C |
| 29644796 | 29645847 | -1 | lncRNA | ENSG0000C |
| 29716279 | 29716685 | -1 | transcribed_processe | ENSG0000C |
| 29761103 | 29787836 | 1 | lncRNA | ENSG0000C |
| 29775747 | 29776701 | -1 | transcribed_processe | ENSG0000C |
| 29777619 | 29777744 | 1 | snoRNA | ENSG0000C |
| 29855759 | 29856332 | 1 | processed_pseudoge | ENSG0000C |
| 29863402 | 29866092 | 1 | lncRNA | ENSG0000C |
| 29883006 | 29883109 | 1 | snoRNA | ENSG0000C |
| 29929200 | 30108452 | 1 | protein_coding | ENSG0000C |
| 29972514 | 29973027 | 1 | transcribed_processe | ENSG0000C |
| 30059052 | 30059147 | -1 | misc_RNA | ENSG0000C |
| 30059054 | 30059147 | -1 | unprocessed_pseudc | ENSG0000C |
| 30059339 | 30065677 | -1 | lncRNA | ENSG0000C |
| 30068632 | 30069631 | 1 | transcribed_processe | ENSG0000C |
| 30090366 | 30117497 | -1 | lncRNA | ENSG0000C |
| 30115521 | 30186475 | 1 | protein_coding | ENSG0000C |
| 30117079 | 30117172 | 1 | miRNA | ENSG0000C |
| 30117079 | 30117172 | -1 | lncRNA | ENSG0000C |
| 30117086 | 30117160 | -1 | miRNA | ENSG0000C |
| 30122429 | 30126118 | -1 | lncRNA | ENSG0000C |
| 30144062 | 30144790 | -1 | lncRNA | ENSG0000C |
| 30194319 | 30236002 | -1 | protein_coding | ENSG0000C |

| | | | | |
|---|---|---|---|---|
| 30204318 | 30206468 | 1 | lncRNA | ENSG0000C |
| 30238741 | 30252237 | 1 | lncRNA | ENSG0000C |
| 30246757 | 30246820 | -1 | snoRNA | ENSG0000C |
| 30248203 | 30292056 | -1 | protein_coding | ENSG0000C |
| 30288071 | 30288177 | -1 | snRNA | ENSG0000C |
| 30316333 | 30334059 | -1 | protein_coding | ENSG0000C |
| 30344503 | 30344604 | -1 | misc_RNA | ENSG0000C |
| 30360463 | 30360566 | 1 | snRNA | ENSG0000C |
| 30378927 | 30469989 | 1 | protein_coding | ENSG0000C |
| 30477362 | 30527592 | 1 | protein_coding | ENSG0000C |
| 30529688 | 30542416 | 1 | unprocessed_pseudc | ENSG0000C |
| 30550493 | 30551189 | -1 | lncRNA | ENSG0000C |
| 30551193 | 30551802 | 1 | TEC | ENSG0000C |
| 30553502 | 30563493 | 1 | transcribed_unproce | ENSG0000C |
| 30557732 | 30558502 | -1 | lncRNA | ENSG0000C |
| 30564057 | 30565576 | -1 | lncRNA | ENSG0000C |
| 30567700 | 30571748 | -1 | unprocessed_pseudc | ENSG0000C |
| 30573471 | 30577000 | -1 | lncRNA | ENSG0000C |
| 30576464 | 30672789 | 1 | lncRNA | ENSG0000C |
| 30600796 | 30615980 | 1 | transcribed_unproce | ENSG0000C |
| 30617680 | 30617790 | -1 | unprocessed_pseudc | ENSG0000C |
| 30624413 | 30625494 | 1 | processed_pseudoge | ENSG0000C |
| 30629680 | 30637466 | 1 | transcribed_unproce | ENSG0000C |
| 30702504 | 30702775 | -1 | misc_RNA | ENSG0000C |
| 30709299 | 30790908 | 1 | transcribed_unproce | ENSG0000C |
| 30724982 | 30725181 | -1 | TEC | ENSG0000C |
| 30726305 | 30727564 | -1 | lncRNA | ENSG0000C |
| 30729469 | 30731202 | 1 | lncRNA | ENSG0000C |
| 30738182 | 30740275 | 1 | TEC | ENSG0000C |
| 30769388 | 30824692 | -1 | protein_coding | ENSG0000C |
| 30781493 | 30782221 | -1 | lncRNA | ENSG0000C |
| 30792372 | 30792833 | 1 | lncRNA | ENSG0000C |
| 30803654 | 30804077 | 1 | lncRNA | ENSG0000C |
| 30830901 | 30831318 | 1 | processed_pseudoge | ENSG0000C |
| 30831966 | 30895869 | 1 | protein_coding | ENSG0000C |
| 30834325 | 30863028 | -1 | lncRNA | ENSG0000C |
| 30861843 | 30861938 | 1 | snRNA | ENSG0000C |
| 30863921 | 30864940 | -1 | lncRNA | ENSG0000C |
| 30897336 | 30906238 | -1 | protein_coding | ENSG0000C |
| 30899110 | 30899651 | 1 | lncRNA | ENSG0000C |
| 30906344 | 30959322 | 1 | protein_coding | ENSG0000C |
| 30956280 | 30956961 | -1 | lncRNA | ENSG0000C |
| 30959564 | 30959850 | 1 | misc_RNA | ENSG0000C |
| 30964935 | 30965500 | -1 | lncRNA | ENSG0000C |
| 30968785 | 30970307 | 1 | lncRNA | ENSG0000C |
| 30970984 | 30999911 | 1 | protein_coding | ENSG0000C |
| 30971652 | 30973312 | 1 | lncRNA | ENSG0000C |
| 30975387 | 30975954 | 1 | processed_pseudoge | ENSG0000C |
| 30978610 | 30980250 | -1 | lncRNA | ENSG0000C |
| 31008154 | 31008319 | -1 | processed_pseudoge | ENSG0000C |

| | | | | |
|---|---|---|---|---|
| 31008497 | 31093127 | 1 | transcribed_unproce | ENSG0000C |
| 31038575 | 31059121 | -1 | transcribed_unproce | ENSG0000C |
| 31090787 | 31095450 | -1 | lncRNA | ENSG0000C |
| 31094350 | 31094425 | -1 | miRNA | ENSG0000C |
| 31094927 | 31382116 | 1 | protein_coding | ENSG0000C |
| 31133182 | 31138518 | 1 | lncRNA | ENSG0000C |
| 31272013 | 31297539 | -1 | protein_coding | ENSG0000C |
| 31303770 | 31314105 | -1 | protein_coding | ENSG0000C |
| 31305213 | 31318831 | -1 | protein_coding | ENSG0000C |
| 31317560 | 31321884 | -1 | protein_coding | ENSG0000C |
| 31345519 | 31346190 | 1 | processed_pseudoge | ENSG0000C |
| 31391675 | 31538211 | 1 | protein_coding | ENSG0000C |
| 31465543 | 31465832 | 1 | misc_RNA | ENSG0000C |
| 31518432 | 31518729 | 1 | misc_RNA | ENSG0000C |
| 31533171 | 31533938 | -1 | TEC | ENSG0000C |
| 31534883 | 31534971 | 1 | miRNA | ENSG0000C |
| 31559996 | 31560083 | 1 | miRNA | ENSG0000C |
| 31560132 | 31560573 | 1 | lncRNA | ENSG0000C |
| 31563768 | 31563894 | -1 | snRNA | ENSG0000C |
| 31571142 | 31575659 | 1 | lncRNA | ENSG0000C |
| 31575269 | 31575358 | 1 | miRNA | ENSG0000C |
| 31575411 | 31575521 | 1 | miRNA | ENSG0000C |
| 31583162 | 31637543 | 1 | lncRNA | ENSG0000C |
| 31709568 | 31709859 | 1 | processed_pseudoge | ENSG0000C |
| 31713753 | 31713853 | 1 | snRNA | ENSG0000C |
| 31759795 | 31761385 | 1 | processed_pseudoge | ENSG0000C |
| 31762440 | 31769048 | 1 | lncRNA | ENSG0000C |
| 31830731 | 31831750 | 1 | lncRNA | ENSG0000C |
| 31851871 | 31859291 | -1 | protein_coding | ENSG0000C |
| 31860904 | 31901708 | -1 | protein_coding | ENSG0000C |
| 31873926 | 31886666 | 1 | lncRNA | ENSG0000C |
| 31937007 | 32001038 | 1 | protein_coding | ENSG0000C |
| 31963805 | 31963933 | -1 | rRNA_pseudogene | ENSG0000C |
| 32003110 | 32006688 | -1 | transcribed_processe | ENSG0000C |
| 32007872 | 32053504 | 1 | protein_coding | ENSG0000C |
| 32024469 | 32024583 | -1 | processed_pseudoge | ENSG0000C |
| 32039974 | 32042828 | 1 | transcribed_processe | ENSG0000C |
| 32051030 | 32053208 | 1 | TEC | ENSG0000C |
| 32083179 | 32097065 | 1 | transcribed_processe | ENSG0000C |
| 32088160 | 32094933 | -1 | transcribed_unproce | ENSG0000C |
| 32106330 | 32114125 | -1 | transcribed_unproce | ENSG0000C |
| 32111562 | 32112590 | -1 | processed_pseudoge | ENSG0000C |
| 32127595 | 32128454 | 1 | lncRNA | ENSG0000C |
| 32141226 | 32143135 | -1 | lncRNA | ENSG0000C |
| 32142454 | 32253374 | 1 | protein_coding | ENSG0000C |
| 32150433 | 32151504 | -1 | processed_pseudoge | ENSG0000C |
| 32159311 | 32160371 | 1 | TEC | ENSG0000C |
| 32178946 | 32180074 | -1 | TEC | ENSG0000C |
| 32227890 | 32228108 | -1 | processed_pseudoge | ENSG0000C |
| 32258615 | 32265404 | -1 | lncRNA | ENSG0000C |

| | | | | |
|---|---|---|---|---|
| 32265832 | 32324659 | 1 | protein_coding | ENSG00000 |
| 32280387 | 32280953 | -1 | lncRNA | ENSG00000 |
| 32324431 | 32360700 | 1 | lncRNA | ENSG00000 |
| 32324565 | 32350023 | -1 | protein_coding | ENSG00000 |
| 32328441 | 32329395 | 1 | lncRNA | ENSG00000 |
| 32343528 | 32344124 | -1 | unprocessed_pseudo | ENSG00000 |
| 32350109 | 32350202 | 1 | miRNA | ENSG00000 |
| 32350117 | 32381886 | 1 | protein_coding | ENSG00000 |
| 32408013 | 32408309 | 1 | lncRNA | ENSG00000 |
| 32410159 | 32410746 | 1 | lncRNA | ENSG00000 |
| 32411217 | 32412420 | 1 | lncRNA | ENSG00000 |
| 32423971 | 32446038 | -1 | lncRNA | ENSG00000 |
| 32430967 | 32432841 | 1 | lncRNA | ENSG00000 |
| 32434939 | 32435028 | -1 | processed_pseudoge | ENSG00000 |
| 32444379 | 32483319 | 1 | protein_coding | ENSG00000 |
| 32463374 | 32463482 | -1 | misc_RNA | ENSG00000 |
| 32486993 | 32491253 | 1 | protein_coding | ENSG00000 |
| 32492522 | 32877177 | -1 | protein_coding | ENSG00000 |
| 32495536 | 32500910 | 1 | lncRNA | ENSG00000 |
| 32504567 | 32505644 | -1 | TEC | ENSG00000 |
| 32509954 | 32523424 | -1 | lncRNA | ENSG00000 |
| 32512869 | 32519350 | -1 | lncRNA | ENSG00000 |
| 32518322 | 32518934 | -1 | TEC | ENSG00000 |
| 32518953 | 32531492 | 1 | lncRNA | ENSG00000 |
| 32529008 | 32530049 | -1 | lncRNA | ENSG00000 |
| 32627739 | 32686484 | 1 | lncRNA | ENSG00000 |
| 32830219 | 32830329 | -1 | misc_RNA | ENSG00000 |
| 32876759 | 32881070 | 1 | lncRNA | ENSG00000 |
| 32905662 | 32906584 | 1 | lncRNA | ENSG00000 |
| 32927910 | 32945106 | 1 | protein_coding | ENSG00000 |
| 32970376 | 32997877 | 1 | protein_coding | ENSG00000 |
| 33013087 | 34174964 | -1 | protein_coding | ENSG00000 |
| 33052107 | 33052867 | -1 | lncRNA | ENSG00000 |
| 33099969 | 33100319 | -1 | TEC | ENSG00000 |
| 33111858 | 33131743 | 1 | lncRNA | ENSG00000 |
| 33529787 | 33533760 | -1 | lncRNA | ENSG00000 |
| 33534110 | 33541339 | 1 | lncRNA | ENSG00000 |
| 33565764 | 33624122 | 1 | lncRNA | ENSG00000 |
| 33627027 | 33635057 | 1 | lncRNA | ENSG00000 |
| 33680577 | 33680883 | 1 | lncRNA | ENSG00000 |
| 33688803 | 33689174 | 1 | lncRNA | ENSG00000 |
| 33692229 | 33692484 | 1 | unprocessed_pseudo | ENSG00000 |
| 33791838 | 33816761 | -1 | lncRNA | ENSG00000 |
| 33827699 | 34079100 | -1 | lncRNA | ENSG00000 |
| 33935437 | 34156000 | -1 | lncRNA | ENSG00000 |
| 33976531 | 33980851 | 1 | lncRNA | ENSG00000 |
| 34000789 | 34006772 | -1 | TEC | ENSG00000 |
| 34036681 | 34039872 | -1 | processed_pseudoge | ENSG00000 |
| 34080882 | 34082224 | -1 | lncRNA | ENSG00000 |
| 34118347 | 34118882 | -1 | lncRNA | ENSG00000 |

| | | | | |
|---|---|---|---|---|
| 34120056 | 34120167 | -1 | rRNA_pseudogene | ENSG00000 |
| 34142947 | 34147047 | 1 | lncRNA | ENSG00000 |
| 34155566 | 34165736 | 1 | lncRNA | ENSG00000 |
| 34169372 | 34195937 | -1 | lncRNA | ENSG00000 |
| 34176538 | 34178166 | 1 | lncRNA | ENSG00000 |
| 34255218 | 34257203 | 1 | protein_coding | ENSG00000 |
| 34270221 | 34272242 | 1 | protein_coding | ENSG00000 |
| 34285742 | 34288334 | 1 | protein_coding | ENSG00000 |
| 34319435 | 34321402 | 1 | protein_coding | ENSG00000 |
| 34331862 | 34332228 | -1 | processed_pseudoge | ENSG00000 |
| 34356480 | 34358610 | 1 | protein_coding | ENSG00000 |
| 34360328 | 34363233 | -1 | protein_coding | ENSG00000 |
| 34479272 | 34482148 | 1 | lncRNA | ENSG00000 |
| 34574123 | 34579369 | -1 | lncRNA | ENSG00000 |
| 34579487 | 34639318 | 1 | protein_coding | ENSG00000 |
| 34614409 | 34614809 | 1 | TEC | ENSG00000 |
| 34725509 | 34726170 | -1 | lncRNA | ENSG00000 |
| 34837871 | 34859046 | -1 | lncRNA | ENSG00000 |
| 34927859 | 34981078 | -1 | protein_coding | ENSG00000 |
| 34937306 | 34938604 | 1 | lncRNA | ENSG00000 |
| 34961540 | 34963777 | 1 | protein_coding | ENSG00000 |
| 34980512 | 35009743 | 1 | protein_coding | ENSG00000 |
| 35005990 | 35089319 | -1 | protein_coding | ENSG00000 |
| 35011349 | 35121493 | -1 | protein_coding | ENSG00000 |
| 35018660 | 35018991 | 1 | lncRNA | ENSG00000 |
| 35073831 | 35074374 | -1 | lncRNA | ENSG00000 |
| 35076737 | 35077007 | 1 | processed_pseudoge | ENSG00000 |
| 35092208 | 35121522 | -1 | protein_coding | ENSG00000 |
| 35093909 | 35093973 | 1 | snRNA | ENSG00000 |
| 35097560 | 35097657 | -1 | misc_RNA | ENSG00000 |
| 35121615 | 35130732 | 1 | protein_coding | ENSG00000 |
| 35128730 | 35142304 | -1 | protein_coding | ENSG00000 |
| 35147817 | 35189345 | 1 | protein_coding | ENSG00000 |
| 35164243 | 35167012 | -1 | lncRNA | ENSG00000 |
| 35188522 | 35191343 | 1 | lncRNA | ENSG00000 |
| 35190961 | 35191702 | 1 | transcribed_processe | ENSG00000 |
| 35192520 | 35194523 | -1 | protein_coding | ENSG00000 |
| 35231450 | 35242963 | -1 | lncRNA | ENSG00000 |
| 35243071 | 35273655 | 1 | protein_coding | ENSG00000 |
| 35313479 | 35325449 | 1 | lncRNA | ENSG00000 |
| 35344231 | 35344564 | 1 | processed_pseudoge | ENSG00000 |
| 35350305 | 35373701 | -1 | protein_coding | ENSG00000 |
| 35377416 | 35377678 | -1 | processed_pseudoge | ENSG00000 |
| 35400878 | 35403006 | -1 | lncRNA | ENSG00000 |
| 35403837 | 35404373 | -1 | lncRNA | ENSG00000 |
| 35406684 | 35409768 | -1 | lncRNA | ENSG00000 |
| 35406948 | 35407818 | -1 | transcribed_processe | ENSG00000 |
| 35409602 | 35410228 | -1 | transcribed_processe | ENSG00000 |
| 35411060 | 35433283 | -1 | protein_coding | ENSG00000 |
| 35435096 | 35448837 | -1 | protein_coding | ENSG00000 |

| | | | | |
|---|---|---|---|---|
| 35459481 | 35460550 | -1 | processed_pseudoge | ENSG00000 |
| 35464249 | 35522379 | -1 | protein_coding | ENSG00000 |
| 35470069 | 35470628 | -1 | processed_pseudoge | ENSG00000 |
| 35474904 | 35537861 | -1 | lncRNA | ENSG00000 |
| 35477994 | 35478444 | 1 | lncRNA | ENSG00000 |
| 35490009 | 35491238 | 1 | processed_pseudoge | ENSG00000 |
| 35498218 | 35499175 | 1 | processed_pseudoge | ENSG00000 |
| 35499690 | 35510270 | -1 | lncRNA | ENSG00000 |
| 35514766 | 35515186 | -1 | processed_pseudoge | ENSG00000 |
| 35522279 | 35522969 | -1 | transcribed_unproce | ENSG00000 |
| 35540039 | 35540797 | -1 | lncRNA | ENSG00000 |
| 35548125 | 35558098 | -1 | protein_coding | ENSG00000 |
| 35553205 | 35554767 | 1 | lncRNA | ENSG00000 |
| 35561539 | 35562077 | -1 | lncRNA | ENSG00000 |
| 35566517 | 35566672 | 1 | processed_pseudoge | ENSG00000 |
| 35568076 | 35574900 | 1 | lncRNA | ENSG00000 |
| 35573657 | 35573753 | 1 | snoRNA | ENSG00000 |
| 35574795 | 35578863 | -1 | protein_coding | ENSG00000 |
| 35578046 | 35726413 | 1 | protein_coding | ENSG00000 |
| 35596904 | 35597128 | -1 | processed_pseudoge | ENSG00000 |
| 35713791 | 35864615 | 1 | protein_coding | ENSG00000 |
| 35731639 | 35743521 | 1 | protein_coding | ENSG00000 |
| 35744511 | 35752878 | -1 | protein_coding | ENSG00000 |
| 35756249 | 35795707 | -1 | protein_coding | ENSG00000 |
| 35757199 | 35758325 | -1 | lncRNA | ENSG00000 |
| 35760887 | 35765079 | 1 | protein_coding | ENSG00000 |
| 35808489 | 35808897 | -1 | lncRNA | ENSG00000 |
| 35816717 | 35830293 | -1 | lncRNA | ENSG00000 |
| 35818399 | 35823713 | 1 | lncRNA | ENSG00000 |
| 35854946 | 35868891 | -1 | protein_coding | ENSG00000 |
| 35861093 | 35861566 | -1 | processed_pseudoge | ENSG00000 |
| 35868967 | 35885863 | 1 | lncRNA | ENSG00000 |
| 35871491 | 35880793 | -1 | protein_coding | ENSG00000 |
| 35889558 | 35889804 | 1 | misc_RNA | ENSG00000 |
| 35893707 | 35911651 | -1 | transcribed_processe | ENSG00000 |
| 35912635 | 35918010 | -1 | processed_pseudoge | ENSG00000 |
| 35918066 | 35930773 | -1 | protein_coding | ENSG00000 |
| 35934518 | 35943713 | -1 | protein_coding | ENSG00000 |
| 35963488 | 35965472 | -1 | transcribed_unproce | ENSG00000 |
| 35976493 | 35981497 | -1 | protein_coding | ENSG00000 |
| 35983288 | 35987004 | -1 | protein_coding | ENSG00000 |
| 35983656 | 35990270 | 1 | lncRNA | ENSG00000 |
| 35983656 | 36001621 | -1 | protein_coding | ENSG00000 |
| 35996440 | 36001553 | -1 | protein_coding | ENSG00000 |
| 36001419 | 36011618 | 1 | lncRNA | ENSG00000 |
| 36012504 | 36012891 | 1 | lncRNA | ENSG00000 |
| 36013056 | 36017972 | -1 | protein_coding | ENSG00000 |
| 36064272 | 36072032 | 1 | protein_coding | ENSG00000 |
| 36072866 | 36090134 | 1 | lncRNA | ENSG00000 |
| 36088256 | 36090169 | -1 | protein_coding | ENSG00000 |

| | | | | |
|---|---|---|---|---|
| 36103827 | 36105621 | 1 | protein_coding | ENSG0000C |
| 36110150 | 36110447 | 1 | misc_RNA | ENSG0000C |
| 36116177 | 36177510 | 1 | lncRNA | ENSG0000C |
| 36161607 | 36162926 | 1 | unprocessed_pseudc | ENSG0000C |
| 36165681 | 36176636 | -1 | protein_coding | ENSG0000C |
| 36183235 | 36196471 | 1 | lncRNA | ENSG0000C |
| 36194869 | 36196758 | -1 | protein_coding | ENSG0000C |
| 36210924 | 36212878 | 1 | protein_coding | ENSG0000C |
| 36225246 | 36226807 | -1 | lncRNA | ENSG0000C |
| 36234447 | 36234554 | -1 | unprocessed_pseudc | ENSG0000C |
| 36241701 | 36241806 | 1 | unprocessed_pseudc | ENSG0000C |
| 36253456 | 36264553 | -1 | protein_coding | ENSG0000C |
| 36274191 | 36320547 | 1 | unprocessed_pseudc | ENSG0000C |
| 36285312 | 36285413 | 1 | misc_RNA | ENSG0000C |
| 36323884 | 36334759 | 1 | protein_coding | ENSG0000C |
| 36377531 | 36388423 | -1 | protein_coding | ENSG0000C |
| 36404826 | 36404932 | -1 | snRNA | ENSG0000C |
| 36428618 | 36439566 | -1 | protein_coding | ENSG0000C |
| 36450146 | 36457535 | -1 | unprocessed_pseudc | ENSG0000C |
| 36486629 | 36499310 | 1 | protein_coding | ENSG0000C |
| 36491199 | 36491322 | -1 | rRNA_pseudogene | ENSG0000C |
| 36495633 | 36543435 | -1 | protein_coding | ENSG0000C |
| 36534987 | 36539310 | 1 | protein_coding | ENSG0000C |
| 36544912 | 36589848 | 1 | protein_coding | ENSG0000C |
| 36574462 | 36575325 | -1 | processed_pseudoge | ENSG0000C |
| 36591879 | 36600804 | 1 | protein_coding | ENSG0000C |
| 36601583 | 36608964 | 1 | protein_coding | ENSG0000C |
| 36634069 | 36634698 | 1 | lncRNA | ENSG0000C |
| 36684754 | 36685129 | 1 | lncRNA | ENSG0000C |
| 36722443 | 36726346 | 1 | lncRNA | ENSG0000C |
| 36861674 | 36936670 | -1 | lncRNA | ENSG0000C |
| 36936785 | 36944612 | 1 | protein_coding | ENSG0000C |
| 36940049 | 36943456 | -1 | lncRNA | ENSG0000C |
| 36948954 | 37056871 | 1 | protein_coding | ENSG0000C |
| 36980167 | 36980422 | 1 | lncRNA | ENSG0000C |
| 36998598 | 37000034 | 1 | lncRNA | ENSG0000C |
| 37033745 | 37033813 | 1 | miRNA | ENSG0000C |
| 37034668 | 37043252 | 1 | lncRNA | ENSG0000C |
| 37045228 | 37052890 | -1 | lncRNA | ENSG0000C |
| 37084992 | 37406836 | -1 | protein_coding | ENSG0000C |
| 37140611 | 37141424 | -1 | processed_pseudoge | ENSG0000C |
| 37143607 | 37144463 | -1 | transcribed_processe | ENSG0000C |
| 37162616 | 37163139 | 1 | processed_pseudoge | ENSG0000C |
| 37375985 | 37392708 | 1 | protein_coding | ENSG0000C |
| 37386886 | 37387926 | 1 | lncRNA | ENSG0000C |
| 37406886 | 37479725 | 1 | protein_coding | ENSG0000C |
| 37407936 | 37408594 | -1 | lncRNA | ENSG0000C |
| 37454171 | 37454931 | 1 | processed_pseudoge | ENSG0000C |
| 37489891 | 37513501 | 1 | protein_coding | ENSG0000C |
| 37514807 | 37609472 | -1 | protein_coding | ENSG0000C |

| | | | | |
|---|---|---|---|---|
| 37609739 | 37643446 | -1 | protein_coding | ENSG0000C |
| 37609739 | 37613841 | 1 | lncRNA | ENSG0000C |
| 37614931 | 37615039 | -1 | miRNA | ENSG0000C |
| 37642947 | 37684252 | 1 | lncRNA | ENSG0000C |
| 37649133 | 37650894 | 1 | lncRNA | ENSG0000C |
| 37686431 | 37745105 | -1 | protein_coding | ENSG0000C |
| 37798894 | 37802788 | -1 | lncRNA | ENSG0000C |
| 37828305 | 37884743 | -1 | transcribed_unproce | ENSG0000C |
| 37907957 | 37908063 | 1 | snRNA | ENSG0000C |
| 37924415 | 37935365 | 1 | protein_coding | ENSG0000C |
| 37940704 | 37940790 | -1 | rRNA | ENSG0000C |
| 37977972 | 37989048 | -1 | protein_coding | ENSG0000C |
| 38003976 | 38014902 | 1 | protein_coding | ENSG0000C |
| 38057693 | 38068592 | -1 | protein_coding | ENSG0000C |
| 38078262 | 38124770 | 1 | unprocessed_pseudc | ENSG0000C |
| 38089399 | 38089500 | 1 | misc_RNA | ENSG0000C |
| 38127951 | 38138862 | 1 | protein_coding | ENSG0000C |
| 38144201 | 38144287 | -1 | rRNA | ENSG0000C |
| 38181659 | 38192541 | -1 | protein_coding | ENSG0000C |
| 38195703 | 38257192 | -1 | transcribed_unproce | ENSG0000C |
| 38231043 | 38231144 | -1 | misc_RNA | ENSG0000C |
| 38297023 | 38323217 | 1 | protein_coding | ENSG0000C |
| 38325530 | 38343847 | -1 | protein_coding | ENSG0000C |
| 38351844 | 38405593 | 1 | protein_coding | ENSG0000C |
| 38419280 | 38512385 | 1 | protein_coding | ENSG0000C |
| 38450394 | 38452444 | -1 | lncRNA | ENSG0000C |
| 38530031 | 38605952 | -1 | protein_coding | ENSG0000C |
| 38601049 | 38602701 | 1 | lncRNA | ENSG0000C |
| 38671703 | 38674957 | -1 | protein_coding | ENSG0000C |
| 38702262 | 38702331 | -1 | miRNA | ENSG0000C |
| 38702452 | 38704747 | 1 | lncRNA | ENSG0000C |
| 38703480 | 38706261 | -1 | lncRNA | ENSG0000C |
| 38705273 | 38729795 | 1 | protein_coding | ENSG0000C |
| 38715328 | 38720272 | -1 | lncRNA | ENSG0000C |
| 38719691 | 38719748 | 1 | miRNA | ENSG0000C |
| 38727833 | 38728198 | -1 | TEC | ENSG0000C |
| 38730341 | 38735605 | 1 | protein_coding | ENSG0000C |
| 38731802 | 38731920 | -1 | rRNA_pseudogene | ENSG0000C |
| 38733898 | 38749817 | -1 | protein_coding | ENSG0000C |
| 38749360 | 38751457 | 1 | lncRNA | ENSG0000C |
| 38752741 | 38764225 | 1 | protein_coding | ENSG0000C |
| 38755279 | 38755400 | -1 | snRNA | ENSG0000C |
| 38765689 | 38800126 | -1 | protein_coding | ENSG0000C |
| 38800441 | 38825355 | -1 | protein_coding | ENSG0000C |
| 38825838 | 38825892 | 1 | miRNA | ENSG0000C |
| 38835088 | 38841455 | -1 | protein_coding | ENSG0000C |
| 38847860 | 38853764 | -1 | protein_coding | ENSG0000C |
| 38851524 | 38851659 | -1 | snoRNA | ENSG0000C |
| 38852863 | 38852994 | -1 | snoRNA | ENSG0000C |
| 38855314 | 38855670 | 1 | processed_pseudoge | ENSG0000C |

| | | | | |
|---|---|---|---|---|
| 38869859 | 38921770 | 1 | protein_coding | ENSG0000( |
| 38903704 | 38904600 | -1 | lncRNA | ENSG0000( |
| 38914979 | 38915042 | 1 | miRNA | ENSG0000( |
| 38918801 | 38921769 | -1 | lncRNA | ENSG0000( |
| 38925168 | 38929384 | 1 | protein_coding | ENSG0000( |
| 38936432 | 38967403 | -1 | protein_coding | ENSG0000( |
| 38996323 | 38996624 | -1 | processed_pseudoge | ENSG0000( |
| 39003248 | 39013032 | -1 | lncRNA | ENSG0000( |
| 39023394 | 39026742 | 1 | unprocessed_pseudc | ENSG0000( |
| 39026715 | 39027110 | -1 | lncRNA | ENSG0000( |
| 39027277 | 39053205 | 1 | transcribed_unproce | ENSG0000( |
| 39033926 | 39034042 | -1 | processed_pseudoge | ENSG0000( |
| 39057019 | 39113190 | 1 | transcribed_processe | ENSG0000( |
| 39063313 | 39154394 | -1 | protein_coding | ENSG0000( |
| 39086976 | 39091942 | 1 | lncRNA | ENSG0000( |
| 39156894 | 39167484 | -1 | protein_coding | ENSG0000( |
| 39165073 | 39167849 | 1 | TEC | ENSG0000( |
| 39173290 | 39177503 | 1 | lncRNA | ENSG0000( |
| 39173453 | 39197703 | -1 | protein_coding | ENSG0000( |
| 39200283 | 39204732 | 1 | protein_coding | ENSG0000( |
| 39210541 | 39225945 | -1 | protein_coding | ENSG0000( |
| 39238466 | 39250818 | 1 | transcribed_processe | ENSG0000( |
| 39252663 | 39402523 | -1 | protein_coding | ENSG0000( |
| 39401793 | 39406233 | 1 | lncRNA | ENSG0000( |
| 39404285 | 39451272 | -1 | protein_coding | ENSG0000( |
| 39461486 | 39564907 | 1 | protein_coding | ENSG0000( |
| 39546104 | 39546244 | -1 | snRNA | ENSG0000( |
| 39566915 | 39567559 | 1 | lncRNA | ENSG0000( |
| 39603536 | 39609777 | -1 | protein_coding | ENSG0000( |
| 39619613 | 39622770 | 1 | lncRNA | ENSG0000( |
| 39626740 | 39636626 | 1 | protein_coding | ENSG0000( |
| 39637090 | 39664201 | 1 | protein_coding | ENSG0000( |
| 39664187 | 39666555 | 1 | protein_coding | ENSG0000( |
| 39667981 | 39670475 | 1 | protein_coding | ENSG0000( |
| 39671122 | 39696797 | -1 | protein_coding | ENSG0000( |
| 39687914 | 39730426 | 1 | protein_coding | ENSG0000( |
| 39726495 | 39726561 | 1 | miRNA | ENSG0000( |
| 39728496 | 39730532 | -1 | protein_coding | ENSG0000( |
| 39737927 | 39747291 | 1 | protein_coding | ENSG0000( |
| 39757718 | 39864312 | -1 | protein_coding | ENSG0000( |
| 39835037 | 39836080 | -1 | processed_pseudoge | ENSG0000( |
| 39839369 | 39839524 | -1 | processed_pseudoge | ENSG0000( |
| 39868164 | 39877896 | 1 | protein_coding | ENSG0000( |
| 39904595 | 39919854 | -1 | protein_coding | ENSG0000( |
| 39921041 | 39927601 | -1 | protein_coding | ENSG0000( |
| 39927742 | 39939601 | 1 | lncRNA | ENSG0000( |
| 39941474 | 39944734 | 1 | protein_coding | ENSG0000( |
| 39953263 | 39977768 | 1 | protein_coding | ENSG0000( |
| 39980807 | 39997959 | 1 | protein_coding | ENSG0000( |
| 40012226 | 40014705 | -1 | lncRNA | ENSG0000( |

| | | | | |
|---|---|---|---|---|
| 40015361 | 40017813 | 1 | protein_coding | ENSG00000 |
| 40019097 | 40061215 | -1 | protein_coding | ENSG00000 |
| 40026332 | 40026409 | -1 | miRNA | ENSG00000 |
| 40027542 | 40027645 | -1 | snoRNA | ENSG00000 |
| 40058290 | 40093867 | 1 | protein_coding | ENSG00000 |
| 40092793 | 40100589 | -1 | protein_coding | ENSG00000 |
| 40113215 | 40115442 | -1 | TEC | ENSG00000 |
| 40121971 | 40136917 | 1 | protein_coding | ENSG00000 |
| 40140318 | 40172171 | 1 | protein_coding | ENSG00000 |
| 40161933 | 40162001 | 1 | miRNA | ENSG00000 |
| 40177010 | 40195656 | 1 | protein_coding | ENSG00000 |
| 40193597 | 40193663 | 1 | miRNA | ENSG00000 |
| 40219304 | 40284136 | 1 | protein_coding | ENSG00000 |
| 40287879 | 40304657 | 1 | protein_coding | ENSG00000 |
| 40309180 | 40357643 | 1 | protein_coding | ENSG00000 |
| 40317698 | 40318676 | -1 | lncRNA | ENSG00000 |
| 40340867 | 40343136 | -1 | lncRNA | ENSG00000 |
| 40348049 | 40350539 | -1 | TEC | ENSG00000 |
| 40360652 | 40364737 | -1 | protein_coding | ENSG00000 |
| 40360655 | 40364693 | 1 | lncRNA | ENSG00000 |
| 40367952 | 40368438 | 1 | processed_pseudoge | ENSG00000 |
| 40374089 | 40374176 | 1 | rRNA_pseudogene | ENSG00000 |
| 40380666 | 40381838 | -1 | processed_pseudoge | ENSG00000 |
| 40388525 | 40417896 | -1 | protein_coding | ENSG00000 |
| 40391232 | 40391338 | -1 | misc_RNA | ENSG00000 |
| 40439467 | 40439917 | -1 | processed_pseudoge | ENSG00000 |
| 40443450 | 40457725 | 1 | protein_coding | ENSG00000 |
| 40473554 | 40475289 | -1 | TEC | ENSG00000 |
| 40475828 | 40501623 | -1 | protein_coding | ENSG00000 |
| 40516892 | 40527002 | 1 | lncRNA | ENSG00000 |
| 40553769 | 40565472 | -1 | protein_coding | ENSG00000 |
| 40597113 | 40598585 | -1 | TEC | ENSG00000 |
| 40624962 | 40648654 | -1 | protein_coding | ENSG00000 |
| 40627356 | 40665141 | -1 | protein_coding | ENSG00000 |
| 40648300 | 40649718 | 1 | lncRNA | ENSG00000 |
| 40654665 | 40665181 | -1 | protein_coding | ENSG00000 |
| 40687038 | 40689360 | -1 | unprocessed_pseudo | ENSG00000 |
| 40697991 | 40703752 | -1 | protein_coding | ENSG00000 |
| 40717235 | 40721932 | -1 | transcribed_unproce | ENSG00000 |
| 40748021 | 40755331 | -1 | protein_coding | ENSG00000 |
| 40766238 | 40772162 | -1 | protein_coding | ENSG00000 |
| 40776808 | 40782550 | -1 | protein_coding | ENSG00000 |
| 40792196 | 40799959 | -1 | protein_coding | ENSG00000 |
| 40803744 | 40809296 | 1 | lncRNA | ENSG00000 |
| 40818117 | 40822614 | -1 | protein_coding | ENSG00000 |
| 40819106 | 40836270 | 1 | lncRNA | ENSG00000 |
| 40850800 | 40863282 | 1 | lncRNA | ENSG00000 |
| 40861303 | 40867223 | -1 | protein_coding | ENSG00000 |
| 40875889 | 40885242 | -1 | protein_coding | ENSG00000 |
| 40921430 | 40975926 | 1 | lncRNA | ENSG00000 |

| | | | | |
|---|---|---|---|---|
| 40922696 | 40937634 | -1 | protein_coding | ENSG00000 |
| 40958417 | 40966948 | -1 | protein_coding | ENSG00000 |
| 40977716 | 40987135 | -1 | protein_coding | ENSG00000 |
| 40993430 | 40994164 | -1 | protein_coding | ENSG00000 |
| 40999193 | 40999906 | -1 | protein_coding | ENSG00000 |
| 41004481 | 41004760 | -1 | unprocessed_pseudo | ENSG00000 |
| 41008521 | 41019324 | -1 | protein_coding | ENSG00000 |
| 41026026 | 41027208 | -1 | protein_coding | ENSG00000 |
| 41029697 | 41030104 | -1 | protein_coding | ENSG00000 |
| 41033884 | 41034874 | -1 | protein_coding | ENSG00000 |
| 41040541 | 41041450 | -1 | protein_coding | ENSG00000 |
| 41046541 | 41047316 | -1 | protein_coding | ENSG00000 |
| 41054498 | 41055230 | -1 | protein_coding | ENSG00000 |
| 41059240 | 41060114 | -1 | protein_coding | ENSG00000 |
| 41065116 | 41065879 | -1 | protein_coding | ENSG00000 |
| 41071946 | 41072341 | 1 | unprocessed_pseudo | ENSG00000 |
| 41084150 | 41085141 | 1 | protein_coding | ENSG00000 |
| 41090338 | 41093246 | 1 | lncRNA | ENSG00000 |
| 41096981 | 41098142 | -1 | protein_coding | ENSG00000 |
| 41101502 | 41102209 | -1 | protein_coding | ENSG00000 |
| 41105332 | 41106488 | 1 | protein_coding | ENSG00000 |
| 41117181 | 41118373 | -1 | protein_coding | ENSG00000 |
| 41123091 | 41124182 | -1 | protein_coding | ENSG00000 |
| 41139433 | 41140487 | -1 | protein_coding | ENSG00000 |
| 41148924 | 41149825 | -1 | protein_coding | ENSG00000 |
| 41159649 | 41160748 | -1 | protein_coding | ENSG00000 |
| 41167231 | 41168221 | -1 | protein_coding | ENSG00000 |
| 41177446 | 41178221 | -1 | protein_coding | ENSG00000 |
| 41184102 | 41185342 | -1 | protein_coding | ENSG00000 |
| 41186944 | 41187136 | -1 | unprocessed_pseudo | ENSG00000 |
| 41189887 | 41190639 | 1 | protein_coding | ENSG00000 |
| 41212706 | 41213237 | 1 | unprocessed_pseudo | ENSG00000 |
| 41226648 | 41227652 | 1 | protein_coding | ENSG00000 |
| 41232449 | 41233454 | 1 | protein_coding | ENSG00000 |
| 41237999 | 41239004 | 1 | protein_coding | ENSG00000 |
| 41249687 | 41250653 | 1 | protein_coding | ENSG00000 |
| 41255384 | 41256364 | 1 | protein_coding | ENSG00000 |
| 41265339 | 41266641 | 1 | protein_coding | ENSG00000 |
| 41271311 | 41271835 | 1 | unprocessed_pseudo | ENSG00000 |
| 41275659 | 41276697 | 1 | protein_coding | ENSG00000 |
| 41280604 | 41280810 | 1 | unprocessed_pseudo | ENSG00000 |
| 41301826 | 41302851 | -1 | protein_coding | ENSG00000 |
| 41307700 | 41309253 | -1 | protein_coding | ENSG00000 |
| 41314912 | 41315710 | -1 | protein_coding | ENSG00000 |
| 41323427 | 41329545 | 1 | unprocessed_pseudo | ENSG00000 |
| 41334632 | 41335733 | 1 | unprocessed_pseudo | ENSG00000 |
| 41346092 | 41350828 | -1 | protein_coding | ENSG00000 |
| 41363498 | 41369813 | -1 | protein_coding | ENSG00000 |
| 41377650 | 41382403 | -1 | protein_coding | ENSG00000 |
| 41393721 | 41397608 | -1 | protein_coding | ENSG00000 |

| | | | | |
|---|---|---|---|---|
| 41402416 | 41425049 | 1 | lncRNA | ENSG00000 |
| 41406512 | 41411775 | -1 | unprocessed_pseudo | ENSG00000 |
| 41420547 | 41424585 | -1 | protein_coding | ENSG00000 |
| 41436446 | 41440921 | -1 | protein_coding | ENSG00000 |
| 41448183 | 41454055 | -1 | unprocessed_pseudo | ENSG00000 |
| 41459811 | 41467429 | -1 | protein_coding | ENSG00000 |
| 41467956 | 41468146 | 1 | snRNA | ENSG00000 |
| 41476689 | 41481140 | -1 | protein_coding | ENSG00000 |
| 41486136 | 41492546 | -1 | protein_coding | ENSG00000 |
| 41500981 | 41505705 | -1 | protein_coding | ENSG00000 |
| 41500983 | 41502409 | 1 | lncRNA | ENSG00000 |
| 41513745 | 41522529 | -1 | protein_coding | ENSG00000 |
| 41517164 | 41517217 | -1 | miRNA | ENSG00000 |
| 41523617 | 41528308 | -1 | protein_coding | ENSG00000 |
| 41549606 | 41554495 | -1 | lncRNA | ENSG00000 |
| 41565836 | 41572059 | -1 | protein_coding | ENSG00000 |
| 41582279 | 41586895 | -1 | protein_coding | ENSG00000 |
| 41609778 | 41615899 | -1 | protein_coding | ENSG00000 |
| 41619442 | 41624842 | -1 | protein_coding | ENSG00000 |
| 41626327 | 41640199 | -1 | transcribed_unproce | ENSG00000 |
| 41665566 | 41668031 | 1 | TEC | ENSG00000 |
| 41688885 | 41692668 | 1 | protein_coding | ENSG00000 |
| 41712331 | 41715969 | 1 | protein_coding | ENSG00000 |
| 41717742 | 41734644 | -1 | protein_coding | ENSG00000 |
| 41718154 | 41718260 | 1 | rRNA_pseudogene | ENSG00000 |
| 41730127 | 41730405 | -1 | misc_RNA | ENSG00000 |
| 41754604 | 41786931 | -1 | protein_coding | ENSG00000 |
| 41801947 | 41812604 | -1 | protein_coding | ENSG00000 |
| 41812680 | 41823217 | 1 | protein_coding | ENSG00000 |
| 41825057 | 41836260 | -1 | protein_coding | ENSG00000 |
| 41835685 | 41848384 | 1 | protein_coding | ENSG00000 |
| 41848518 | 41851447 | -1 | lncRNA | ENSG00000 |
| 41848518 | 41865423 | -1 | protein_coding | ENSG00000 |
| 41866917 | 41930542 | -1 | protein_coding | ENSG00000 |
| 41867581 | 41867736 | 1 | lncRNA | ENSG00000 |
| 41930617 | 41966503 | 1 | protein_coding | ENSG00000 |
| 41966763 | 41977740 | 1 | protein_coding | ENSG00000 |
| 41976435 | 42021376 | -1 | protein_coding | ENSG00000 |
| 42011382 | 42025644 | 1 | protein_coding | ENSG00000 |
| 42025576 | 42098479 | -1 | protein_coding | ENSG00000 |
| 42038232 | 42050601 | -1 | protein_coding | ENSG00000 |
| 42101404 | 42112714 | -1 | protein_coding | ENSG00000 |
| 42113111 | 42121367 | -1 | protein_coding | ENSG00000 |
| 42119674 | 42154916 | -1 | protein_coding | ENSG00000 |
| 42122804 | 42123352 | 1 | protein_coding | ENSG00000 |
| 42124976 | 42155044 | -1 | protein_coding | ENSG00000 |
| 42154600 | 42155972 | 1 | lncRNA | ENSG00000 |
| 42156891 | 42181142 | -1 | protein_coding | ENSG00000 |
| 42184060 | 42185452 | -1 | protein_coding | ENSG00000 |
| 42188799 | 42194532 | -1 | protein_coding | ENSG00000 |

| | | | | |
|---|---|---|---|---|
| 42199168 | 42276707 | -1 | protein_coding | ENSG00000 |
| 42233832 | 42235469 | 1 | TEC | ENSG00000 |
| 42268587 | 42269807 | 1 | lncRNA | ENSG00000 |
| 42270517 | 42272683 | -1 | lncRNA | ENSG00000 |
| 42272069 | 42275571 | -1 | lncRNA | ENSG00000 |
| 42287547 | 42311943 | 1 | protein_coding | ENSG00000 |
| 42313324 | 42388568 | -1 | protein_coding | ENSG00000 |
| 42402449 | 42423256 | -1 | protein_coding | ENSG00000 |
| 42440198 | 42440259 | -1 | snRNA | ENSG00000 |
| 42458844 | 42522611 | 1 | protein_coding | ENSG00000 |
| 42494773 | 42494830 | 1 | miRNA | ENSG00000 |
| 42495867 | 42496550 | 1 | lncRNA | ENSG00000 |
| 42509784 | 42511519 | 1 | lncRNA | ENSG00000 |
| 42514188 | 42514307 | 1 | miRNA | ENSG00000 |
| 42533532 | 42534009 | 1 | processed_pseudoge | ENSG00000 |
| 42536241 | 42544449 | 1 | protein_coding | ENSG00000 |
| 42536510 | 42562062 | 1 | lncRNA | ENSG00000 |
| 42546764 | 42548706 | 1 | unprocessed_pseudo | ENSG00000 |
| 42549214 | 42555213 | 1 | protein_coding | ENSG00000 |
| 42552436 | 42554748 | -1 | lncRNA | ENSG00000 |
| 42561467 | 42566277 | 1 | protein_coding | ENSG00000 |
| 42567068 | 42573239 | 1 | protein_coding | ENSG00000 |
| 42572315 | 42577831 | -1 | protein_coding | ENSG00000 |
| 42577825 | 42578366 | 1 | lncRNA | ENSG00000 |
| 42579513 | 42610623 | -1 | protein_coding | ENSG00000 |
| 42598643 | 42599177 | 1 | TEC | ENSG00000 |
| 42609683 | 42615238 | 1 | protein_coding | ENSG00000 |
| 42647834 | 42648143 | 1 | processed_pseudoge | ENSG00000 |
| 42648178 | 42648738 | 1 | processed_pseudoge | ENSG00000 |
| 42659284 | 42667006 | 1 | protein_coding | ENSG00000 |
| 42667914 | 42676994 | -1 | protein_coding | ENSG00000 |
| 42678889 | 42683917 | -1 | protein_coding | ENSG00000 |
| 42679963 | 42682020 | 1 | lncRNA | ENSG00000 |
| 42682531 | 42699993 | 1 | protein_coding | ENSG00000 |
| 42683187 | 42699466 | -1 | lncRNA | ENSG00000 |
| 42700275 | 42745049 | -1 | protein_coding | ENSG00000 |
| 42708084 | 42708151 | -1 | miRNA | ENSG00000 |
| 42714299 | 42714643 | 1 | processed_pseudoge | ENSG00000 |
| 42751289 | 42751717 | -1 | lncRNA | ENSG00000 |
| 42753530 | 42753799 | -1 | processed_pseudoge | ENSG00000 |
| 42753914 | 42761257 | -1 | lncRNA | ENSG00000 |
| 42758447 | 42763041 | 1 | protein_coding | ENSG00000 |
| 42773449 | 42779599 | 1 | protein_coding | ENSG00000 |
| 42780610 | 42797066 | 1 | protein_coding | ENSG00000 |
| 42795147 | 42798704 | -1 | protein_coding | ENSG00000 |
| 42798800 | 42811587 | 1 | protein_coding | ENSG00000 |
| 42810134 | 42833350 | -1 | protein_coding | ENSG00000 |
| 42823880 | 42823943 | -1 | miRNA | ENSG00000 |
| 42824385 | 42843760 | 1 | protein_coding | ENSG00000 |
| 42844580 | 42850707 | 1 | protein_coding | ENSG00000 |

| | | | | |
|---|---|---|---|---|
| 42851184 | 42858130 | 1 | protein_coding | ENSG00000( |
| 42865922 | 42874369 | 1 | transcribed_unproce | ENSG00000( |
| 42874670 | 42898704 | -1 | lncRNA | ENSG00000( |
| 42891338 | 42891435 | -1 | snRNA | ENSG00000( |
| 42900797 | 42914438 | 1 | protein_coding | ENSG00000( |
| 42940575 | 42940702 | -1 | snoRNA | ENSG00000( |
| 42950526 | 42980528 | -1 | protein_coding | ENSG00000( |
| 42950526 | 42964498 | -1 | protein_coding | ENSG00000( |
| 42968088 | 42980433 | -1 | protein_coding | ENSG00000( |
| 42980565 | 42993690 | 1 | protein_coding | ENSG00000( |
| 42998273 | 43002959 | 1 | protein_coding | ENSG00000( |
| 43006725 | 43014456 | 1 | protein_coding | ENSG00000( |
| 43014607 | 43025123 | -1 | protein_coding | ENSG00000( |
| 43025231 | 43032041 | 1 | protein_coding | ENSG00000( |
| 43044295 | 43170245 | -1 | protein_coding | ENSG00000( |
| 43079302 | 43079780 | 1 | processed_pseudoge | ENSG00000( |
| 43125551 | 43153671 | 1 | lncRNA | ENSG00000( |
| 43131208 | 43131270 | 1 | transcribed_unproce | ENSG00000( |
| 43144956 | 43145255 | 1 | transcribed_processe | ENSG00000( |
| 43148368 | 43171037 | -1 | lncRNA | ENSG00000( |
| 43168170 | 43168249 | -1 | unprocessed_pseudc | ENSG00000( |
| 43169880 | 43170077 | -1 | processed_pseudoge | ENSG00000( |
| 43170481 | 43211689 | 1 | protein_coding | ENSG00000( |
| 43211835 | 43220041 | 1 | protein_coding | ENSG00000( |
| 43216941 | 43305397 | -1 | protein_coding | ENSG00000( |
| 43233787 | 43233977 | -1 | snRNA | ENSG00000( |
| 43239921 | 43240111 | -1 | snRNA | ENSG00000( |
| 43241180 | 43241370 | -1 | snRNA | ENSG00000( |
| 43245674 | 43245864 | -1 | snRNA | ENSG00000( |
| 43251829 | 43252019 | -1 | snRNA | ENSG00000( |
| 43266633 | 43266823 | -1 | snRNA | ENSG00000( |
| 43271369 | 43271559 | -1 | snRNA | ENSG00000( |
| 43273101 | 43273292 | -1 | snRNA | ENSG00000( |
| 43277978 | 43278168 | -1 | snRNA | ENSG00000( |
| 43284139 | 43284329 | -1 | snRNA | ENSG00000( |
| 43290291 | 43290481 | -1 | snRNA | ENSG00000( |
| 43296425 | 43296615 | -1 | snRNA | ENSG00000( |
| 43300041 | 43300231 | -1 | snRNA | ENSG00000( |
| 43338741 | 43389199 | -1 | lncRNA | ENSG00000( |
| 43360041 | 43361361 | -1 | TEC | ENSG00000( |
| 43387226 | 43387417 | -1 | snRNA | ENSG00000( |
| 43395993 | 43396091 | -1 | snRNA | ENSG00000( |
| 43398993 | 43401137 | 1 | protein_coding | ENSG00000( |
| 43412427 | 43412530 | 1 | snRNA | ENSG00000( |
| 43444707 | 43451200 | 1 | lncRNA | ENSG00000( |
| 43444806 | 43444885 | 1 | miRNA | ENSG00000( |
| 43445507 | 43445613 | 1 | snRNA | ENSG00000( |
| 43473585 | 43473690 | 1 | snRNA | ENSG00000( |
| 43476198 | 43476665 | 1 | processed_pseudoge | ENSG00000( |
| 43483865 | 43610338 | 1 | protein_coding | ENSG00000( |

| | | | | |
|---|---|---|---|---|
| 43527844 | 43579620 | -1 | protein_coding | ENSG00000[illegible] |
| 43607436 | 43607723 | -1 | misc_RNA | ENSG00000[illegible] |
| 43640389 | 43661922 | -1 | protein_coding | ENSG00000[illegible] |
| 43679341 | 43706709 | 1 | lncRNA | ENSG00000[illegible] |
| 43718314 | 43720436 | -1 | processed_pseudoge | ENSG00000[illegible] |
| 43753738 | 43758791 | -1 | protein_coding | ENSG00000[illegible] |
| 43766125 | 43778977 | -1 | protein_coding | ENSG00000[illegible] |
| 43780435 | 43787620 | 1 | protein_coding | ENSG00000[illegible] |
| 43782804 | 43784682 | -1 | lncRNA | ENSG00000[illegible] |
| 43800799 | 43833170 | -1 | protein_coding | ENSG00000[illegible] |
| 43847148 | 43863639 | 1 | protein_coding | ENSG00000[illegible] |
| 43875357 | 43909711 | -1 | protein_coding | ENSG00000[illegible] |
| 43914433 | 43923001 | -1 | lncRNA | ENSG00000[illegible] |
| 43917194 | 43917985 | 1 | lncRNA | ENSG00000[illegible] |
| 43922775 | 43927388 | -1 | unprocessed_pseudo | ENSG00000[illegible] |
| 43927563 | 43932622 | 1 | lncRNA | ENSG00000[illegible] |
| 43938363 | 43938959 | 1 | lncRNA | ENSG00000[illegible] |
| 43940804 | 43942468 | -1 | protein_coding | ENSG00000[illegible] |
| 43952738 | 44004469 | -1 | protein_coding | ENSG00000[illegible] |
| 44004622 | 44009068 | 1 | protein_coding | ENSG00000[illegible] |
| 44011188 | 44023946 | -1 | protein_coding | ENSG00000[illegible] |
| 44025020 | 44025223 | -1 | snoRNA | ENSG00000[illegible] |
| 44034635 | 44067619 | -1 | protein_coding | ENSG00000[illegible] |
| 44070735 | 44076344 | 1 | protein_coding | ENSG00000[illegible] |
| 44076746 | 44123702 | -1 | protein_coding | ENSG00000[illegible] |
| 44115912 | 44120595 | 1 | lncRNA | ENSG00000[illegible] |
| 44135883 | 44135989 | -1 | snRNA | ENSG00000[illegible] |
| 44141906 | 44162476 | 1 | protein_coding | ENSG00000[illegible] |
| 44170447 | 44179084 | 1 | protein_coding | ENSG00000[illegible] |
| 44175968 | 44186723 | -1 | lncRNA | ENSG00000[illegible] |
| 44186970 | 44191929 | 1 | protein_coding | ENSG00000[illegible] |
| 44191805 | 44200113 | -1 | protein_coding | ENSG00000[illegible] |
| 44198882 | 44216565 | 1 | lncRNA | ENSG00000[illegible] |
| 44205033 | 44221626 | -1 | protein_coding | ENSG00000[illegible] |
| 44207771 | 44207839 | -1 | miRNA | ENSG00000[illegible] |
| 44221401 | 44223710 | 1 | lncRNA | ENSG00000[illegible] |
| 44228760 | 44230228 | -1 | processed_pseudoge | ENSG00000[illegible] |
| 44248390 | 44268141 | -1 | protein_coding | ENSG00000[illegible] |
| 44276368 | 44281182 | -1 | lncRNA | ENSG00000[illegible] |
| 44290856 | 44291151 | -1 | misc_RNA | ENSG00000[illegible] |
| 44299574 | 44315315 | -1 | lncRNA | ENSG00000[illegible] |
| 44303965 | 44304203 | 1 | snoRNA | ENSG00000[illegible] |
| 44307088 | 44307172 | 1 | snRNA | ENSG00000[illegible] |
| 44308413 | 44318670 | 1 | protein_coding | ENSG00000[illegible] |
| 44319625 | 44324870 | -1 | protein_coding | ENSG00000[illegible] |
| 44328613 | 44331462 | 1 | lncRNA | ENSG00000[illegible] |
| 44345246 | 44353106 | 1 | protein_coding | ENSG00000[illegible] |
| 44353215 | 44363853 | -1 | protein_coding | ENSG00000[illegible] |
| 44365014 | 44365751 | -1 | processed_pseudoge | ENSG00000[illegible] |
| 44372180 | 44389649 | -1 | protein_coding | ENSG00000[illegible] |

| | | | | |
|---|---|---|---|---|
| 44395281 | 44503430 | -1 | protein_coding | ENSG00000 |
| 44409059 | 44409358 | -1 | misc_RNA | ENSG00000 |
| 44486153 | 44486815 | -1 | lncRNA | ENSG00000 |
| 44557484 | 44561262 | 1 | protein_coding | ENSG00000 |
| 44596346 | 44597344 | 1 | processed_pseudoge | ENSG00000 |
| 44645411 | 44645587 | 1 | processed_pseudoge | ENSG00000 |
| 44646364 | 44650349 | -1 | lncRNA | ENSG00000 |
| 44656404 | 44690308 | 1 | protein_coding | ENSG00000 |
| 44673069 | 44689779 | -1 | protein_coding | ENSG00000 |
| 44673689 | 44676257 | -1 | lncRNA | ENSG00000 |
| 44689858 | 44693340 | 1 | TEC | ENSG00000 |
| 44708608 | 44752264 | 1 | protein_coding | ENSG00000 |
| 44718861 | 44719130 | -1 | misc_RNA | ENSG00000 |
| 44719760 | 44720597 | -1 | processed_pseudoge | ENSG00000 |
| 44758988 | 44781846 | 1 | protein_coding | ENSG00000 |
| 44793199 | 44794474 | 1 | lncRNA | ENSG00000 |
| 44794747 | 44797783 | -1 | lncRNA | ENSG00000 |
| 44798448 | 44830816 | -1 | protein_coding | ENSG00000 |
| 44846353 | 44850476 | 1 | protein_coding | ENSG00000 |
| 44849948 | 44899445 | -1 | protein_coding | ENSG00000 |
| 44883986 | 44884274 | 1 | misc_RNA | ENSG00000 |
| 44899142 | 44905390 | 1 | protein_coding | ENSG00000 |
| 44899712 | 44905390 | 1 | protein_coding | ENSG00000 |
| 44903159 | 44916937 | -1 | protein_coding | ENSG00000 |
| 44924709 | 44947773 | -1 | protein_coding | ENSG00000 |
| 44934618 | 44934681 | -1 | miRNA | ENSG00000 |
| 44947912 | 44948939 | 1 | lncRNA | ENSG00000 |
| 44959693 | 44968303 | -1 | protein_coding | ENSG00000 |
| 44982514 | 44982772 | 1 | lncRNA | ENSG00000 |
| 45023340 | 45061109 | -1 | protein_coding | ENSG00000 |
| 45051610 | 45109016 | 1 | protein_coding | ENSG00000 |
| 45071442 | 45071543 | 1 | misc_RNA | ENSG00000 |
| 45108959 | 45133354 | -1 | protein_coding | ENSG00000 |
| 45114367 | 45114433 | -1 | miRNA | ENSG00000 |
| 45132600 | 45144181 | 1 | protein_coding | ENSG00000 |
| 45146730 | 45148470 | -1 | lncRNA | ENSG00000 |
| 45148502 | 45152101 | 1 | protein_coding | ENSG00000 |
| 45150400 | 45161510 | -1 | lncRNA | ENSG00000 |
| 45160700 | 45170040 | 1 | protein_coding | ENSG00000 |
| 45168800 | 45171584 | -1 | lncRNA | ENSG00000 |
| 45190931 | 45222222 | -1 | lncRNA | ENSG00000 |
| 45221444 | 45247319 | 1 | protein_coding | ENSG00000 |
| 45238028 | 45241734 | -1 | lncRNA | ENSG00000 |
| 45245186 | 45251129 | -1 | transcribed_unitary_ | ENSG00000 |
| 45247916 | 45269824 | 1 | lncRNA | ENSG00000 |
| 45254393 | 45262094 | -1 | protein_coding | ENSG00000 |
| 45263119 | 45317029 | -1 | protein_coding | ENSG00000 |
| 45327366 | 45327497 | -1 | rRNA_pseudogene | ENSG00000 |
| 45327366 | 45327497 | -1 | rRNA | ENSG00000 |
| 45371402 | 45372057 | -1 | lncRNA | ENSG00000 |

| | | | | |
|---|---|---|---|---|
| 45393902 | 45434421 | -1 | protein_coding | ENSG0000C |
| 45396932 | 45397477 | 1 | lncRNA | ENSG0000C |
| 45435900 | 45490749 | -1 | protein_coding | ENSG0000C |
| 45452844 | 45464065 | 1 | lncRNA | ENSG0000C |
| 45475363 | 45475435 | -1 | miRNA | ENSG0000C |
| 45506741 | 45551537 | -1 | transcribed_unproce | ENSG0000C |
| 45526466 | 45526746 | -1 | misc_RNA | ENSG0000C |
| 45531577 | 45533838 | 1 | TEC | ENSG0000C |
| 45533963 | 45534710 | 1 | lncRNA | ENSG0000C |
| 45545804 | 45563230 | 1 | lncRNA | ENSG0000C |
| 45549781 | 45558738 | -1 | lncRNA | ENSG0000C |
| 45553710 | 45559070 | -1 | transcribed_unproce | ENSG0000C |
| 45585871 | 45586929 | 1 | processed_pseudoge | ENSG0000C |
| 45586452 | 45588379 | -1 | lncRNA | ENSG0000C |
| 45592621 | 45593369 | 1 | TEC | ENSG0000C |
| 45600869 | 45602340 | -1 | processed_pseudoge | ENSG0000C |
| 45608571 | 45608918 | 1 | processed_pseudoge | ENSG0000C |
| 45620328 | 45655156 | 1 | transcribed_unitary_ | ENSG0000C |
| 45620344 | 45835826 | 1 | protein_coding | ENSG0000C |
| 45731703 | 45732977 | -1 | lncRNA | ENSG0000C |
| 45733353 | 45734669 | -1 | lncRNA | ENSG0000C |
| 45784280 | 45835828 | 1 | protein_coding | ENSG0000C |
| 45799390 | 45895680 | -1 | lncRNA | ENSG0000C |
| 45844881 | 45847067 | 1 | protein_coding | ENSG0000C |
| 45894382 | 46028334 | 1 | protein_coding | ENSG0000C |
| 45895783 | 45898798 | 1 | lncRNA | ENSG0000C |
| 45907670 | 45910779 | -1 | lncRNA | ENSG0000C |
| 45931806 | 45932083 | 1 | misc_RNA | ENSG0000C |
| 45999250 | 45999694 | 1 | protein_coding | ENSG0000C |
| 46029916 | 46225389 | -1 | protein_coding | ENSG0000C |
| 46035313 | 46035770 | 1 | lncRNA | ENSG0000C |
| 46193576 | 46196723 | 1 | lncRNA | ENSG0000C |
| 46215070 | 46215171 | -1 | misc_RNA | ENSG0000C |
| 46243606 | 46245044 | 1 | processed_pseudoge | ENSG0000C |
| 46259551 | 46260606 | -1 | processed_pseudoge | ENSG0000C |
| 46267037 | 46268694 | 1 | transcribed_unproce | ENSG0000C |
| 46274784 | 46361797 | -1 | protein_coding | ENSG0000C |
| 46292733 | 46337794 | 1 | protein_coding | ENSG0000C |
| 46319679 | 46319959 | 1 | misc_RNA | ENSG0000C |
| 46372855 | 46487141 | 1 | unprocessed_pseudc | ENSG0000C |
| 46503946 | 46509339 | 1 | unprocessed_pseudc | ENSG0000C |
| 46511511 | 46555650 | 1 | protein_coding | ENSG0000C |
| 46516702 | 46579682 | -1 | protein_coding | ENSG0000C |
| 46537534 | 46537814 | 1 | misc_RNA | ENSG0000C |
| 46558830 | 46562795 | -1 | lncRNA | ENSG0000C |
| 46590669 | 46757464 | 1 | protein_coding | ENSG0000C |
| 46721582 | 46722167 | -1 | processed_pseudoge | ENSG0000C |
| 46762506 | 46833154 | -1 | protein_coding | ENSG0000C |
| 46833201 | 46886730 | 1 | protein_coding | ENSG0000C |
| 46909742 | 46911512 | -1 | lncRNA | ENSG0000C |

| | | | | |
|---|---|---|---|---|
| 46916770 | 46923034 | -1 | lncRNA | ENSG0000C |
| 46921124 | 46921249 | 1 | snRNA | ENSG0000C |
| 46923075 | 46975524 | 1 | protein_coding | ENSG0000C |
| 46923133 | 47049932 | 1 | protein_coding | ENSG0000C |
| 46973017 | 46973100 | 1 | miRNA | ENSG0000C |
| 46978156 | 46979253 | -1 | protein_coding | ENSG0000C |
| 46978481 | 47054569 | 1 | transcribed_unproce | ENSG0000C |
| 46983287 | 47100323 | -1 | lncRNA | ENSG0000C |
| 47041825 | 47042105 | 1 | misc_RNA | ENSG0000C |
| 47117703 | 47189422 | -1 | protein_coding | ENSG0000C |
| 47159191 | 47159993 | -1 | processed_pseudoge | ENSG0000C |
| 47160398 | 47167723 | -1 | lncRNA | ENSG0000C |
| 47169826 | 47171049 | 1 | lncRNA | ENSG0000C |
| 47200446 | 47223679 | 1 | protein_coding | ENSG0000C |
| 47253846 | 47311816 | 1 | protein_coding | ENSG0000C |
| 47253897 | 47344292 | 1 | protein_coding | ENSG0000C |
| 47259358 | 47259420 | -1 | snRNA | ENSG0000C |
| 47279526 | 47280047 | -1 | lncRNA | ENSG0000C |
| 47303460 | 47323613 | -1 | lncRNA | ENSG0000C |
| 47323290 | 47441312 | 1 | protein_coding | ENSG0000C |
| 47409322 | 47423526 | -1 | lncRNA | ENSG0000C |
| 47450568 | 47492492 | -1 | transcribed_unproce | ENSG0000C |
| 47522942 | 47623276 | 1 | protein_coding | ENSG0000C |
| 47557413 | 47557514 | 1 | misc_RNA | ENSG0000C |
| 47603860 | 47649420 | -1 | lncRNA | ENSG0000C |
| 47649476 | 47685505 | 1 | protein_coding | ENSG0000C |
| 47682417 | 47682683 | -1 | lncRNA | ENSG0000C |
| 47694081 | 47712050 | 1 | protein_coding | ENSG0000C |
| 47733236 | 47746122 | 1 | protein_coding | ENSG0000C |
| 47807372 | 47821834 | -1 | protein_coding | ENSG0000C |
| 47823272 | 47831541 | -1 | protein_coding | ENSG0000C |
| 47831634 | 47837719 | 1 | protein_coding | ENSG0000C |
| 47837692 | 47841289 | -1 | protein_coding | ENSG0000C |
| 47844908 | 47855874 | -1 | protein_coding | ENSG0000C |
| 47863342 | 47865190 | 1 | lncRNA | ENSG0000C |
| 47879472 | 47882398 | -1 | lncRNA | ENSG0000C |
| 47891229 | 47895812 | -1 | lncRNA | ENSG0000C |
| 47896150 | 47928957 | 1 | protein_coding | ENSG0000C |
| 47897330 | 47941410 | -1 | lncRNA | ENSG0000C |
| 47929682 | 47933106 | -1 | lncRNA | ENSG0000C |
| 47941506 | 47949308 | 1 | protein_coding | ENSG0000C |
| 47945424 | 47981736 | 1 | lncRNA | ENSG0000C |
| 47946802 | 47948275 | -1 | lncRNA | ENSG0000C |
| 47951967 | 47957883 | -1 | protein_coding | ENSG0000C |
| 47967810 | 47981781 | 1 | protein_coding | ENSG0000C |
| 47980398 | 47996196 | -1 | lncRNA | ENSG0000C |
| 48026167 | 48038030 | -1 | protein_coding | ENSG0000C |
| 48037161 | 48037247 | -1 | miRNA | ENSG0000C |
| 48045141 | 48048073 | -1 | lncRNA | ENSG0000C |
| 48048329 | 48061545 | 1 | protein_coding | ENSG0000C |

| | | | | |
|---|---|---|---|---|
| 48060383 | 48060669 | -1 | lncRNA | ENSG00000 |
| 48066704 | 48067293 | -1 | lncRNA | ENSG00000 |
| 48070052 | 48101478 | -1 | protein_coding | ENSG00000 |
| 48088678 | 48088784 | -1 | snRNA | ENSG00000 |
| 48103357 | 48123074 | 1 | protein_coding | ENSG00000 |
| 48133442 | 48430275 | -1 | protein_coding | ENSG00000 |
| 48156427 | 48156511 | -1 | miRNA | ENSG00000 |
| 48185938 | 48204529 | 1 | lncRNA | ENSG00000 |
| 48196680 | 48196786 | -1 | snRNA | ENSG00000 |
| 48294335 | 48308762 | 1 | lncRNA | ENSG00000 |
| 48377262 | 48380370 | -1 | lncRNA | ENSG00000 |
| 48382371 | 48382586 | -1 | snoRNA | ENSG00000 |
| 48460370 | 48466040 | 1 | lncRNA | ENSG00000 |
| 48479097 | 48479198 | 1 | misc_RNA | ENSG00000 |
| 48528526 | 48531011 | -1 | protein_coding | ENSG00000 |
| 48540894 | 48544989 | -1 | protein_coding | ENSG00000 |
| 48543551 | 48551250 | 1 | lncRNA | ENSG00000 |
| 48548870 | 48604912 | -1 | protein_coding | ENSG00000 |
| 48549630 | 48606414 | 1 | lncRNA | ENSG00000 |
| 48557262 | 48560333 | 1 | lncRNA | ENSG00000 |
| 48575507 | 48578350 | -1 | protein_coding | ENSG00000 |
| 48579630 | 48582259 | -1 | lncRNA | ENSG00000 |
| 48579838 | 48579947 | -1 | miRNA | ENSG00000 |
| 48591257 | 48593961 | -1 | protein_coding | ENSG00000 |
| 48595751 | 48604992 | -1 | protein_coding | ENSG00000 |
| 48607232 | 48633572 | -1 | protein_coding | ENSG00000 |
| 48611377 | 48615292 | -1 | protein_coding | ENSG00000 |
| 48621156 | 48626358 | -1 | protein_coding | ENSG00000 |
| 48628675 | 48634932 | 1 | lncRNA | ENSG00000 |
| 48632490 | 48632559 | -1 | miRNA | ENSG00000 |
| 48635923 | 48647023 | -1 | lncRNA | ENSG00000 |
| 48646923 | 48707346 | 1 | lncRNA | ENSG00000 |
| 48690481 | 48690624 | -1 | processed_pseudoge | ENSG00000 |
| 48691116 | 48691694 | -1 | processed_pseudoge | ENSG00000 |
| 48713730 | 48713959 | 1 | processed_pseudoge | ENSG00000 |
| 48721719 | 48722518 | -1 | protein_coding | ENSG00000 |
| 48723185 | 48724758 | 1 | protein_coding | ENSG00000 |
| 48723188 | 48723314 | 1 | misc_RNA | ENSG00000 |
| 48723692 | 48723772 | 1 | misc_RNA | ENSG00000 |
| 48724408 | 48724475 | -1 | miRNA | ENSG00000 |
| 48724763 | 48729178 | -1 | protein_coding | ENSG00000 |
| 48733091 | 48733888 | 1 | lncRNA | ENSG00000 |
| 48762235 | 48817214 | -1 | protein_coding | ENSG00000 |
| 48765554 | 48765847 | -1 | misc_RNA | ENSG00000 |
| 48830988 | 48866522 | 1 | protein_coding | ENSG00000 |
| 48874860 | 48908983 | -1 | transcribed_process | ENSG00000 |
| 48880840 | 48880967 | 1 | snoRNA | ENSG00000 |
| 48881122 | 48881316 | 1 | processed_pseudoge | ENSG00000 |
| 48892765 | 48895871 | 1 | protein_coding | ENSG00000 |
| 48899131 | 48899748 | 1 | processed_pseudoge | ENSG00000 |

| | | | | |
|---|---|---|---|---|
| 48908407 | 48929056 | 1 | protein_coding | ENSG00000 |
| 48929316 | 48944842 | -1 | protein_coding | ENSG00000 |
| 48931791 | 48937100 | 1 | lncRNA | ENSG00000 |
| 48949361 | 48949520 | -1 | snRNA | ENSG00000 |
| 48956650 | 48956753 | -1 | snoRNA | ENSG00000 |
| 48958554 | 48968596 | -1 | protein_coding | ENSG00000 |
| 48995266 | 48997492 | -1 | lncRNA | ENSG00000 |
| 48997385 | 49056145 | 1 | protein_coding | ENSG00000 |
| 49004731 | 49013725 | -1 | lncRNA | ENSG00000 |
| 49017793 | 49018061 | -1 | misc_RNA | ENSG00000 |
| 49036352 | 49036455 | 1 | snRNA | ENSG00000 |
| 49037122 | 49037429 | 1 | processed_pseudoge | ENSG00000 |
| 49085511 | 49086702 | 1 | unprocessed_pseudo | ENSG00000 |
| 49132460 | 49176840 | 1 | protein_coding | ENSG00000 |
| 49148339 | 49149111 | -1 | processed_pseudoge | ENSG00000 |
| 49202791 | 49210574 | -1 | protein_coding | ENSG00000 |
| 49210411 | 49223225 | 1 | protein_coding | ENSG00000 |
| 49223362 | 49230766 | -1 | protein_coding | ENSG00000 |
| 49230853 | 49318011 | 1 | lncRNA | ENSG00000 |
| 49288346 | 49288454 | -1 | miRNA | ENSG00000 |
| 49289206 | 49362473 | -1 | protein_coding | ENSG00000 |
| 49361150 | 49369998 | 1 | lncRNA | ENSG00000 |
| 49370740 | 49476988 | 1 | lncRNA | ENSG00000 |
| 49375380 | 49380094 | 1 | lncRNA | ENSG00000 |
| 49400429 | 49400800 | -1 | processed_pseudoge | ENSG00000 |
| 49404049 | 49414905 | -1 | protein_coding | ENSG00000 |
| 49404081 | 49405197 | 1 | lncRNA | ENSG00000 |
| 49416744 | 49416850 | -1 | misc_RNA | ENSG00000 |
| 49424269 | 49424922 | -1 | processed_pseudoge | ENSG00000 |
| 49457861 | 49461749 | -1 | lncRNA | ENSG00000 |
| 49494223 | 49497800 | -1 | TEC | ENSG00000 |
| 49495293 | 49515008 | 1 | protein_coding | ENSG00000 |
| 49505657 | 49574064 | -1 | lncRNA | ENSG00000 |
| 49510817 | 49510900 | 1 | miRNA | ENSG00000 |
| 49544788 | 49577962 | -1 | lncRNA | ENSG00000 |
| 49575858 | 49583827 | 1 | protein_coding | ENSG00000 |
| 49598884 | 49678163 | -1 | protein_coding | ENSG00000 |
| 49700934 | 49709014 | -1 | protein_coding | ENSG00000 |
| 49708334 | 49720060 | 1 | lncRNA | ENSG00000 |
| 49710332 | 49789180 | -1 | protein_coding | ENSG00000 |
| 49788681 | 49835026 | 1 | protein_coding | ENSG00000 |
| 49795343 | 49795669 | 1 | processed_pseudoge | ENSG00000 |
| 49834239 | 49836454 | 1 | lncRNA | ENSG00000 |
| 49838309 | 49848017 | -1 | protein_coding | ENSG00000 |
| 49845910 | 49848837 | 1 | lncRNA | ENSG00000 |
| 49887598 | 49900893 | 1 | lncRNA | ENSG00000 |
| 49914403 | 49932918 | 1 | lncRNA | ENSG00000 |
| 49939122 | 49939819 | 1 | processed_pseudoge | ENSG00000 |
| 49966075 | 49966176 | -1 | snRNA | ENSG00000 |
| 49968970 | 49974959 | 1 | protein_coding | ENSG00000 |

| | | | | |
|---|---|---|---|---|
| 49990005 | 49995224 | -1 | protein_coding | ENSG0000C |
| 49996889 | 50016844 | 1 | lncRNA | ENSG0000C |
| 50050349 | 50055739 | -1 | lncRNA | ENSG0000C |
| 50055968 | 50090481 | 1 | protein_coding | ENSG0000C |
| 50094065 | 50094647 | 1 | lncRNA | ENSG0000C |
| 50094737 | 50112152 | 1 | protein_coding | ENSG0000C |
| 50098460 | 50098899 | -1 | lncRNA | ENSG0000C |
| 50100704 | 50101920 | -1 | lncRNA | ENSG0000C |
| 50110040 | 50129882 | -1 | protein_coding | ENSG0000C |
| 50133737 | 50150677 | -1 | protein_coding | ENSG0000C |
| 50135586 | 50146176 | 1 | lncRNA | ENSG0000C |
| 50158333 | 50161276 | -1 | lncRNA | ENSG0000C |
| 50163523 | 50165316 | -1 | lncRNA | ENSG0000C |
| 50164214 | 50175928 | 1 | protein_coding | ENSG0000C |
| 50171428 | 50181178 | -1 | transcribed_unitary_ | ENSG0000C |
| 50183289 | 50201632 | -1 | protein_coding | ENSG0000C |
| 50199876 | 50215922 | 1 | lncRNA | ENSG0000C |
| 50212933 | 50215357 | -1 | lncRNA | ENSG0000C |
| 50235845 | 50236129 | 1 | processed_pseudoge | ENSG0000C |
| 50271406 | 50281485 | 1 | protein_coding | ENSG0000C |
| 50281577 | 50287855 | -1 | lncRNA | ENSG0000C |
| 50346126 | 50363138 | 1 | protein_coding | ENSG0000C |
| 50362967 | 50365232 | -1 | TEC | ENSG0000C |
| 50367857 | 50373207 | -1 | protein_coding | ENSG0000C |
| 50373220 | 50381483 | 1 | protein_coding | ENSG0000C |
| 50375059 | 50397523 | -1 | protein_coding | ENSG0000C |
| 50386145 | 50386246 | -1 | misc_RNA | ENSG0000C |
| 50396438 | 50397888 | 1 | lncRNA | ENSG0000C |
| 50426158 | 50474845 | 1 | protein_coding | ENSG0000C |
| 50450179 | 50450725 | -1 | processed_pseudoge | ENSG0000C |
| 50464496 | 50468906 | -1 | protein_coding | ENSG0000C |
| 50475819 | 50478391 | 1 | lncRNA | ENSG0000C |
| 50478860 | 50485974 | 1 | protein_coding | ENSG0000C |
| 50503412 | 50508328 | -1 | lncRNA | ENSG0000C |
| 50508384 | 50531501 | 1 | protein_coding | ENSG0000C |
| 50525520 | 50538761 | -1 | lncRNA | ENSG0000C |
| 50532682 | 50543750 | 1 | protein_coding | ENSG0000C |
| 50543058 | 50555852 | 1 | protein_coding | ENSG0000C |
| 50556207 | 50562108 | -1 | lncRNA | ENSG0000C |
| 50560715 | 50627474 | 1 | protein_coding | ENSG0000C |
| 50563203 | 50563783 | -1 | lncRNA | ENSG0000C |
| 50627032 | 50631940 | -1 | lncRNA | ENSG0000C |
| 50634777 | 50692253 | 1 | protein_coding | ENSG0000C |
| 50693198 | 50707914 | -1 | protein_coding | ENSG0000C |
| 50693448 | 50695449 | 1 | lncRNA | ENSG0000C |
| 50696644 | 50696760 | 1 | misc_RNA | ENSG0000C |
| 50719565 | 50756219 | 1 | protein_coding | ENSG0000C |
| 50722074 | 50724224 | 1 | TEC | ENSG0000C |
| 50761029 | 50767557 | -1 | protein_coding | ENSG0000C |
| 50768650 | 50768730 | 1 | miRNA | ENSG0000C |

| | | | | |
|---|---|---|---|---|
| 50781306 | 50792493 | 1 | lncRNA | ENSG00000 |
| 50797366 | 50797651 | -1 | processed_pseudoge | ENSG00000 |
| 50834650 | 50842353 | 1 | protein_coding | ENSG00000 |
| 50840057 | 50841626 | -1 | lncRNA | ENSG00000 |
| 50847950 | 50849270 | 1 | processed_pseudoge | ENSG00000 |
| 50862223 | 50867978 | -1 | protein_coding | ENSG00000 |
| 50866679 | 50910774 | 1 | lncRNA | ENSG00000 |
| 50909637 | 50910232 | -1 | lncRNA | ENSG00000 |
| 50934989 | 50944713 | -1 | lncRNA | ENSG00000 |
| 50944104 | 50948750 | 1 | lncRNA | ENSG00000 |
| 50962174 | 51120868 | -1 | protein_coding | ENSG00000 |
| 51042524 | 51085471 | 1 | lncRNA | ENSG00000 |
| 51153536 | 51162428 | 1 | protein_coding | ENSG00000 |
| 51153590 | 51171744 | 1 | protein_coding | ENSG00000 |
| 51165435 | 51171747 | 1 | protein_coding | ENSG00000 |
| 51177425 | 51260163 | -1 | protein_coding | ENSG00000 |
| 51183191 | 51186403 | -1 | TEC | ENSG00000 |
| 51249579 | 51251748 | -1 | lncRNA | ENSG00000 |
| 51260546 | 51297936 | 1 | protein_coding | ENSG00000 |
| 51312609 | 51335165 | -1 | lncRNA | ENSG00000 |
| 51332679 | 51334090 | 1 | lncRNA | ENSG00000 |
| 51335261 | 51336240 | 1 | lncRNA | ENSG00000 |
| 51336646 | 51521401 | 1 | lncRNA | ENSG00000 |
| 51502488 | 51503233 | -1 | processed_pseudoge | ENSG00000 |
| 51630313 | 52160017 | -1 | protein_coding | ENSG00000 |
| 51761260 | 51763397 | -1 | TEC | ENSG00000 |
| 52390515 | 52535701 | 1 | lncRNA | ENSG00000 |
| 52862121 | 52899588 | 1 | lncRNA | ENSG00000 |
| 52985513 | 52987652 | 1 | lncRNA | ENSG00000 |
| 53105734 | 53106358 | 1 | processed_pseudoge | ENSG00000 |
| 53276760 | 54365634 | -1 | lncRNA | ENSG00000 |
| 53353427 | 53354134 | -1 | processed_pseudoge | ENSG00000 |
| 53434131 | 53435829 | 1 | processed_pseudoge | ENSG00000 |
| 53438556 | 53439721 | 1 | processed_pseudoge | ENSG00000 |
| 53681630 | 53682111 | -1 | processed_pseudoge | ENSG00000 |
| 53756989 | 53757870 | -1 | processed_pseudoge | ENSG00000 |
| 53822927 | 53825193 | 1 | protein_coding | ENSG00000 |
| 53952936 | 53955324 | 1 | processed_pseudoge | ENSG00000 |
| 54477796 | 54478168 | 1 | processed_pseudoge | ENSG00000 |
| 54609249 | 54670420 | -1 | lncRNA | ENSG00000 |
| 54702936 | 54703430 | 1 | processed_pseudoge | ENSG00000 |
| 54754165 | 54754649 | -1 | processed_pseudoge | ENSG00000 |
| 54770949 | 54771277 | 1 | misc_RNA | ENSG00000 |
| 54899387 | 54961956 | 1 | protein_coding | ENSG00000 |
| 54951902 | 54968785 | -1 | protein_coding | ENSG00000 |
| 54956760 | 54958426 | -1 | TEC | ENSG00000 |
| 54964474 | 54964679 | 1 | lncRNA | ENSG00000 |
| 54968727 | 55173632 | 1 | protein_coding | ENSG00000 |
| 55264960 | 55325187 | 1 | protein_coding | ENSG00000 |
| 55271504 | 55273653 | -1 | lncRNA | ENSG00000 |

| | | | | |
|---|---|---|---|---|
| 55325757 | 55327791 | -1 | lncRNA | ENSG0000C |
| 55392622 | 55421924 | -1 | protein_coding | ENSG0000C |
| 55448433 | 55511444 | -1 | protein_coding | ENSG0000C |
| 55526964 | 55561577 | -1 | transcribed_processe | ENSG0000C |
| 55550931 | 55551649 | -1 | processed_pseudoge | ENSG0000C |
| 55551810 | 55551920 | 1 | snRNA | ENSG0000C |
| 55719627 | 55732121 | -1 | protein_coding | ENSG0000C |
| 55751051 | 55842830 | 1 | protein_coding | ENSG0000C |
| 55842677 | 55872939 | -1 | lncRNA | ENSG0000C |
| 55882301 | 56511659 | 1 | protein_coding | ENSG0000C |
| 56368063 | 56368215 | 1 | processed_pseudoge | ENSG0000C |
| 56471142 | 56473647 | 1 | TEC | ENSG0000C |
| 56573040 | 56573195 | 1 | processed_pseudoge | ENSG0000C |
| 56593699 | 56595598 | 1 | protein_coding | ENSG0000C |
| 56791913 | 56838773 | -1 | protein_coding | ENSG0000C |
| 56803841 | 56804269 | -1 | lncRNA | ENSG0000C |
| 56834107 | 56869567 | 1 | protein_coding | ENSG0000C |
| 56836387 | 56914080 | -1 | protein_coding | ENSG0000C |
| 56871170 | 56871276 | 1 | snRNA | ENSG0000C |
| 56884639 | 56884819 | -1 | processed_pseudoge | ENSG0000C |
| 56888880 | 56891841 | 1 | lncRNA | ENSG0000C |
| 56891270 | 56891355 | -1 | miRNA | ENSG0000C |
| 56914186 | 56914533 | 1 | lncRNA | ENSG0000C |
| 56938199 | 56961050 | -1 | protein_coding | ENSG0000C |
| 56939932 | 56941275 | 1 | lncRNA | ENSG0000C |
| 56951664 | 56952404 | 1 | processed_pseudoge | ENSG0000C |
| 56978105 | 57006768 | 1 | protein_coding | ENSG0000C |
| 56982749 | 56985104 | 1 | lncRNA | ENSG0000C |
| 57045478 | 57046797 | 1 | transcribed_processe | ENSG0000C |
| 57071814 | 57085024 | -1 | lncRNA | ENSG0000C |
| 57085092 | 57121346 | 1 | protein_coding | ENSG0000C |
| 57092145 | 57096425 | -1 | lncRNA | ENSG0000C |
| 57196081 | 57213300 | -1 | lncRNA | ENSG0000C |
| 57253236 | 57255855 | -1 | lncRNA | ENSG0000C |
| 57255851 | 57684689 | 1 | protein_coding | ENSG0000C |
| 57277920 | 57280017 | 1 | TEC | ENSG0000C |
| 57448218 | 57450291 | 1 | TEC | ENSG0000C |
| 57522248 | 57523597 | -1 | lncRNA | ENSG0000C |
| 57531769 | 57532355 | 1 | TEC | ENSG0000C |
| 57600856 | 57608412 | -1 | lncRNA | ENSG0000C |
| 57618108 | 57618393 | 1 | misc_RNA | ENSG0000C |
| 57685529 | 57685592 | -1 | snRNA | ENSG0000C |
| 57744495 | 57745299 | -1 | protein_coding | ENSG0000C |
| 57771946 | 57834749 | -1 | lncRNA | ENSG0000C |
| 57789449 | 57789732 | 1 | misc_RNA | ENSG0000C |
| 57834781 | 57850056 | -1 | protein_coding | ENSG0000C |
| 57861243 | 57955412 | -1 | protein_coding | ENSG0000C |
| 57912333 | 57922581 | 1 | lncRNA | ENSG0000C |
| 57955965 | 57956143 | 1 | lncRNA | ENSG0000C |
| 57971547 | 57988259 | -1 | protein_coding | ENSG0000C |

| | | | | |
|---|---|---|---|---|
| 57988180 | 57988760 | 1 | TEC | ENSG0000C |
| 57989038 | 58007246 | -1 | protein_coding | ENSG0000C |
| 57989039 | 57994850 | -1 | lncRNA | ENSG0000C |
| 58000919 | 58007346 | -1 | protein_coding | ENSG0000C |
| 58006674 | 58008187 | 1 | TEC | ENSG0000C |
| 58050789 | 58060739 | -1 | lncRNA | ENSG0000C |
| 58076891 | 58083204 | -1 | lncRNA | ENSG0000C |
| 58083419 | 58095542 | 1 | protein_coding | ENSG0000C |
| 58095672 | 58101479 | 1 | TEC | ENSG0000C |
| 58148449 | 58159555 | 1 | protein_coding | ENSG0000C |
| 58157028 | 58157800 | 1 | processed_pseudoge | ENSG0000C |
| 58166982 | 58171411 | 1 | protein_coding | ENSG0000C |
| 58192726 | 58205174 | 1 | protein_coding | ENSG0000C |
| 58202352 | 58203003 | -1 | lncRNA | ENSG0000C |
| 58205437 | 58219605 | -1 | protein_coding | ENSG0000C |
| 58218548 | 58268518 | 1 | protein_coding | ENSG0000C |
| 58219375 | 58233479 | 1 | lncRNA | ENSG0000C |
| 58241164 | 58247385 | -1 | lncRNA | ENSG0000C |
| 58269855 | 58280935 | -1 | protein_coding | ENSG0000C |
| 58301228 | 58328795 | -1 | protein_coding | ENSG0000C |
| 58324472 | 58415766 | 1 | lncRNA | ENSG0000C |
| 58330884 | 58332508 | -1 | lncRNA | ENSG0000C |
| 58331232 | 58331318 | -1 | miRNA | ENSG0000C |
| 58335976 | 58336022 | -1 | miRNA | ENSG0000C |
| 58345175 | 58353093 | -1 | protein_coding | ENSG0000C |
| 58345206 | 58417595 | -1 | protein_coding | ENSG0000C |
| 58352500 | 58417595 | -1 | protein_coding | ENSG0000C |
| 58420167 | 58488408 | -1 | protein_coding | ENSG0000C |
| 58476138 | 58476950 | -1 | processed_pseudoge | ENSG0000C |
| 58489529 | 58517905 | -1 | protein_coding | ENSG0000C |
| 58519837 | 58557799 | 1 | lncRNA | ENSG0000C |
| 58520250 | 58544368 | -1 | protein_coding | ENSG0000C |
| 58556678 | 58692055 | -1 | protein_coding | ENSG0000C |
| 58631641 | 58631836 | -1 | snoRNA | ENSG0000C |
| 58660424 | 58692018 | -1 | protein_coding | ENSG0000C |
| 58666537 | 58666697 | -1 | snRNA | ENSG0000C |
| 58675286 | 58675462 | 1 | processed_pseudoge | ENSG0000C |
| 58677574 | 58677737 | -1 | snRNA | ENSG0000C |
| 58679527 | 58679690 | -1 | snRNA | ENSG0000C |
| 58692573 | 58735611 | 1 | protein_coding | ENSG0000C |
| 58747055 | 58747340 | 1 | lncRNA | ENSG0000C |
| 58755854 | 58985179 | 1 | protein_coding | ENSG0000C |
| 58785323 | 58785608 | -1 | misc_RNA | ENSG0000C |
| 58835899 | 58835994 | 1 | snRNA | ENSG0000C |
| 58897776 | 58899692 | 1 | processed_pseudoge | ENSG0000C |
| 58951185 | 58951513 | 1 | misc_RNA | ENSG0000C |
| 58965952 | 58966727 | 1 | lncRNA | ENSG0000C |
| 58982638 | 59106921 | -1 | protein_coding | ENSG0000C |
| 59059226 | 59059493 | 1 | misc_RNA | ENSG0000C |
| 59106598 | 59118267 | 1 | lncRNA | ENSG0000C |

| | | | | |
|---|---|---|---|---|
| 59109857 | 59155260 | -1 | protein_coding | ENSG0000C |
| 59129276 | 59129458 | -1 | snRNA | ENSG0000C |
| 59137758 | 59137872 | -1 | miRNA | ENSG0000C |
| 59151136 | 59151221 | -1 | miRNA | ENSG0000C |
| 59155499 | 59204705 | 1 | protein_coding | ENSG0000C |
| 59174983 | 59181787 | -1 | unprocessed_pseudc | ENSG0000C |
| 59197566 | 59215226 | 1 | protein_coding | ENSG0000C |
| 59202677 | 59203829 | -1 | lncRNA | ENSG0000C |
| 59209400 | 59215247 | 1 | protein_coding | ENSG0000C |
| 59220467 | 59275970 | 1 | protein_coding | ENSG0000C |
| 59260748 | 59260846 | 1 | misc_RNA | ENSG0000C |
| 59281130 | 59281343 | 1 | processed_pseudoge | ENSG0000C |
| 59331655 | 59401729 | 1 | protein_coding | ENSG0000C |
| 59362665 | 59363278 | 1 | TEC | ENSG0000C |
| 59366083 | 59366154 | 1 | miRNA | ENSG0000C |
| 59400488 | 59403303 | -1 | lncRNA | ENSG0000C |
| 59422088 | 59526946 | -1 | lncRNA | ENSG0000C |
| 59537359 | 59537499 | -1 | snRNA | ENSG0000C |
| 59565558 | 59608345 | 1 | protein_coding | ENSG0000C |
| 59618553 | 59619714 | -1 | lncRNA | ENSG0000C |
| 59619689 | 59696956 | 1 | protein_coding | ENSG0000C |
| 59674636 | 59707626 | -1 | protein_coding | ENSG0000C |
| 59707192 | 59842255 | 1 | protein_coding | ENSG0000C |
| 59784813 | 59785035 | 1 | lncRNA | ENSG0000C |
| 59841266 | 59841337 | 1 | miRNA | ENSG0000C |
| 59847438 | 59847544 | -1 | snRNA | ENSG0000C |
| 59859479 | 59892945 | -1 | protein_coding | ENSG0000C |
| 59885015 | 59885552 | 1 | processed_pseudoge | ENSG0000C |
| 59893046 | 59950574 | 1 | protein_coding | ENSG0000C |
| 59912197 | 59912366 | 1 | processed_pseudoge | ENSG0000C |
| 59940908 | 59973101 | 1 | protein_coding | ENSG0000C |
| 59952240 | 59964761 | -1 | protein_coding | ENSG0000C |
| 59962363 | 60018975 | -1 | lncRNA | ENSG0000C |
| 59964832 | 59996972 | 1 | lncRNA | ENSG0000C |
| 59976009 | 60002384 | -1 | transcribed_unproce | ENSG0000C |
| 60008437 | 60017498 | -1 | transcribed_unproce | ENSG0000C |
| 60019149 | 60020006 | 1 | lncRNA | ENSG0000C |
| 60021072 | 60022436 | 1 | unprocessed_pseudc | ENSG0000C |
| 60025859 | 60030746 | 1 | unprocessed_pseudc | ENSG0000C |
| 60043025 | 60043105 | -1 | miRNA | ENSG0000C |
| 60043194 | 60078931 | -1 | protein_coding | ENSG0000C |
| 60063004 | 60064782 | -1 | processed_pseudoge | ENSG0000C |
| 60079309 | 60088695 | 1 | lncRNA | ENSG0000C |
| 60083566 | 60091885 | -1 | transcribed_unitary_ | ENSG0000C |
| 60101759 | 60102919 | -1 | transcribed_processe | ENSG0000C |
| 60113793 | 60120687 | -1 | unprocessed_pseudc | ENSG0000C |
| 60122642 | 60135743 | -1 | lncRNA | ENSG0000C |
| 60124110 | 60125474 | -1 | unprocessed_pseudc | ENSG0000C |
| 60135039 | 60136271 | 1 | lncRNA | ENSG0000C |
| 60135762 | 60140081 | -1 | lncRNA | ENSG0000C |

| | | | | |
|---|---|---|---|---|
| 60149942 | 60170899 | 1 | protein_coding | ENSG0000( |
| 60177327 | 60422470 | -1 | protein_coding | ENSG0000( |
| 60231516 | 60231646 | -1 | scaRNA | ENSG0000( |
| 60245288 | 60245647 | -1 | processed_pseudoge | ENSG0000( |
| 60256720 | 60258411 | 1 | unprocessed_pseudc | ENSG0000( |
| 60315834 | 60316215 | 1 | processed_pseudoge | ENSG0000( |
| 60392429 | 60431426 | 1 | protein_coding | ENSG0000( |
| 60433678 | 60435996 | -1 | transcribed_process | ENSG0000( |
| 60443158 | 60526242 | -1 | protein_coding | ENSG0000( |
| 60513190 | 60513550 | 1 | processed_pseudoge | ENSG0000( |
| 60526293 | 60550798 | 1 | lncRNA | ENSG0000( |
| 60562010 | 60562161 | -1 | processed_pseudoge | ENSG0000( |
| 60564547 | 60586623 | 1 | lncRNA | ENSG0000( |
| 60564740 | 60565671 | -1 | processed_pseudoge | ENSG0000( |
| 60600183 | 60666280 | 1 | protein_coding | ENSG0000( |
| 60662333 | 60662437 | -1 | snRNA | ENSG0000( |
| 60677453 | 61392838 | 1 | protein_coding | ENSG0000( |
| 60696313 | 60696605 | 1 | misc_RNA | ENSG0000( |
| 60727165 | 60727482 | -1 | processed_pseudoge | ENSG0000( |
| 60748940 | 60749046 | -1 | misc_RNA | ENSG0000( |
| 60810165 | 60811462 | 1 | processed_pseudoge | ENSG0000( |
| 60851382 | 60851745 | 1 | processed_pseudoge | ENSG0000( |
| 60899891 | 60900185 | 1 | misc_RNA | ENSG0000( |
| 61034636 | 61070122 | -1 | lncRNA | ENSG0000( |
| 61069856 | 61070356 | -1 | lncRNA | ENSG0000( |
| 61069871 | 61070341 | 1 | processed_pseudoge | ENSG0000( |
| 61119095 | 61135969 | -1 | lncRNA | ENSG0000( |
| 61354763 | 61355155 | -1 | lncRNA | ENSG0000( |
| 61361668 | 61400243 | -1 | lncRNA | ENSG0000( |
| 61382785 | 61384680 | 1 | lncRNA | ENSG0000( |
| 61393456 | 61411555 | -1 | lncRNA | ENSG0000( |
| 61399843 | 61409466 | 1 | protein_coding | ENSG0000( |
| 61411751 | 61413280 | 1 | lncRNA | ENSG0000( |
| 61452404 | 61485110 | 1 | protein_coding | ENSG0000( |
| 61460224 | 61463384 | -1 | lncRNA | ENSG0000( |
| 61590421 | 61591219 | -1 | protein_coding | ENSG0000( |
| 61681266 | 61863521 | -1 | protein_coding | ENSG0000( |
| 61865367 | 61928016 | -1 | protein_coding | ENSG0000( |
| 61874084 | 61874182 | -1 | misc_RNA | ENSG0000( |
| 61942605 | 62065278 | -1 | protein_coding | ENSG0000( |
| 62003700 | 62007518 | -1 | TEC | ENSG0000( |
| 62005737 | 62006016 | -1 | misc_RNA | ENSG0000( |
| 62035944 | 62040352 | 1 | lncRNA | ENSG0000( |
| 62036833 | 62036945 | 1 | misc_RNA | ENSG0000( |
| 62122320 | 62122421 | 1 | misc_RNA | ENSG0000( |
| 62136972 | 62140639 | -1 | processed_pseudoge | ENSG0000( |
| 62231470 | 62247153 | -1 | unprocessed_pseudc | ENSG0000( |
| 62260634 | 62261936 | 1 | unprocessed_pseudc | ENSG0000( |
| 62264705 | 62275655 | -1 | transcribed_unproce | ENSG0000( |
| 62282702 | 62297026 | -1 | transcribed_unproce | ENSG0000( |

| | | | | |
|---|---|---|---|---|
| 62319884 | 62320189 | 1 | processed_pseudoge | ENSG00000 |
| 62343941 | 62416479 | 1 | protein_coding | ENSG00000 |
| 62381668 | 62382015 | 1 | processed_pseudoge | ENSG00000 |
| 62383104 | 62383166 | 1 | snRNA | ENSG00000 |
| 62397397 | 62398142 | 1 | processed_pseudoge | ENSG00000 |
| 62423867 | 62450822 | 1 | protein_coding | ENSG00000 |
| 62458658 | 62615481 | 1 | protein_coding | ENSG00000 |
| 62516321 | 62516767 | -1 | processed_pseudoge | ENSG00000 |
| 62549726 | 62552121 | 1 | lncRNA | ENSG00000 |
| 62626031 | 62626453 | 1 | lncRNA | ENSG00000 |
| 62626437 | 62627590 | -1 | lncRNA | ENSG00000 |
| 62627670 | 62693597 | 1 | protein_coding | ENSG00000 |
| 62661621 | 62661725 | -1 | misc_RNA | ENSG00000 |
| 62679403 | 62681354 | -1 | TEC | ENSG00000 |
| 62683905 | 62684010 | 1 | snRNA | ENSG00000 |
| 62699244 | 62754973 | 1 | lncRNA | ENSG00000 |
| 62701314 | 62808344 | -1 | protein_coding | ENSG00000 |
| 62789109 | 62791765 | 1 | lncRNA | ENSG00000 |
| 62808500 | 62836371 | 1 | lncRNA | ENSG00000 |
| 62841101 | 62841645 | 1 | processed_pseudoge | ENSG00000 |
| 62910021 | 62968549 | -1 | lncRNA | ENSG00000 |
| 62944215 | 62944312 | 1 | miRNA | ENSG00000 |
| 62950831 | 62951188 | -1 | transcribed_processe | ENSG00000 |
| 62978494 | 62978841 | -1 | processed_pseudoge | ENSG00000 |
| 63004672 | 63005152 | -1 | processed_pseudoge | ENSG00000 |
| 63009556 | 63427699 | 1 | protein_coding | ENSG00000 |
| 63088162 | 63089197 | -1 | processed_pseudoge | ENSG00000 |
| 63117019 | 63130947 | -1 | lncRNA | ENSG00000 |
| 63182166 | 63182817 | 1 | transcribed_processe | ENSG00000 |
| 63193930 | 63339053 | -1 | lncRNA | ENSG00000 |
| 63305262 | 63313521 | -1 | lncRNA | ENSG00000 |
| 63331984 | 63332539 | -1 | transcribed_processe | ENSG00000 |
| 63381231 | 63414430 | -1 | lncRNA | ENSG00000 |
| 63391191 | 63431089 | -1 | lncRNA | ENSG00000 |
| 63393355 | 63393586 | 1 | unprocessed_pseudo | ENSG00000 |
| 63430468 | 63432211 | -1 | lncRNA | ENSG00000 |
| 63432304 | 63446354 | -1 | protein_coding | ENSG00000 |
| 63454993 | 63455817 | 1 | lncRNA | ENSG00000 |
| 63471604 | 63472093 | 1 | processed_pseudoge | ENSG00000 |
| 63477061 | 63498380 | 1 | protein_coding | ENSG00000 |
| 63484823 | 63521848 | 1 | protein_coding | ENSG00000 |
| 63507056 | 63519806 | 1 | transcribed_unproce | ENSG00000 |
| 63523334 | 63548977 | 1 | protein_coding | ENSG00000 |
| 63550477 | 63594279 | 1 | protein_coding | ENSG00000 |
| 63573242 | 63573345 | -1 | snRNA | ENSG00000 |
| 63600882 | 63608365 | 1 | protein_coding | ENSG00000 |
| 63610772 | 63611184 | -1 | processed_pseudoge | ENSG00000 |
| 63622415 | 63696303 | 1 | protein_coding | ENSG00000 |
| 63636601 | 63637113 | 1 | transcribed_processe | ENSG00000 |
| 63682336 | 63741986 | -1 | protein_coding | ENSG00000 |

| | | | | |
|---|---|---|---|---|
| 63695888 | 63701172 | -1 | protein_coding | ENSG00000( |
| 63700847 | 63702670 | 1 | TEC | ENSG00000( |
| 63702845 | 63752097 | -1 | protein_coding | ENSG00000( |
| 63745255 | 63776351 | -1 | protein_coding | ENSG00000( |
| 63773603 | 63819317 | 1 | protein_coding | ENSG00000( |
| 63819433 | 63830012 | -1 | protein_coding | ENSG00000( |
| 63827152 | 63832026 | 1 | protein_coding | ENSG00000( |
| 63832081 | 63843065 | -1 | protein_coding | ENSG00000( |
| 63849292 | 63864379 | 1 | transcribed_unitary_ | ENSG00000( |
| 63872012 | 63873766 | -1 | protein_coding | ENSG00000( |
| 63880215 | 63881944 | -1 | protein_coding | ENSG00000( |
| 63885810 | 63886411 | 1 | unprocessed_pseudc | ENSG00000( |
| 63894909 | 63896661 | -1 | protein_coding | ENSG00000( |
| 63900643 | 63900879 | 1 | unprocessed_pseudc | ENSG00000( |
| 63909597 | 63918838 | -1 | protein_coding | ENSG00000( |
| 63917200 | 63918839 | -1 | protein_coding | ENSG00000( |
| 63917203 | 63931385 | -1 | protein_coding | ENSG00000( |
| 63928740 | 63932354 | -1 | protein_coding | ENSG00000( |
| 63938554 | 63972918 | -1 | protein_coding | ENSG00000( |
| 63972920 | 63989422 | -1 | lncRNA | ENSG00000( |
| 63995418 | 63999899 | -1 | lncRNA | ENSG00000( |
| 63998351 | 64004304 | 1 | protein_coding | ENSG00000( |
| 64002594 | 64020634 | -1 | protein_coding | ENSG00000( |
| 64039142 | 64130819 | -1 | protein_coding | ENSG00000( |
| 64067619 | 64070270 | 1 | lncRNA | ENSG00000( |
| 64145970 | 64146476 | 1 | lncRNA | ENSG00000( |
| 64146083 | 64146152 | 1 | snoRNA | ENSG00000( |
| 64146337 | 64146471 | 1 | snoRNA | ENSG00000( |
| 64146339 | 64146471 | 1 | snoRNA | ENSG00000( |
| 64147227 | 64263301 | -1 | protein_coding | ENSG00000( |
| 64308316 | 64308690 | -1 | processed_pseudoge | ENSG00000( |
| 64319415 | 64413776 | -1 | protein_coding | ENSG00000( |
| 64390158 | 64390263 | -1 | misc_RNA | ENSG00000( |
| 64425069 | 64425170 | -1 | miRNA | ENSG00000( |
| 64449037 | 64468643 | 1 | protein_coding | ENSG00000( |
| 64477785 | 64497054 | -1 | protein_coding | ENSG00000( |
| 64491523 | 64491789 | 1 | processed_pseudoge | ENSG00000( |
| 64498254 | 64508199 | -1 | protein_coding | ENSG00000( |
| 64500774 | 64500839 | -1 | miRNA | ENSG00000( |
| 64501214 | 64501313 | -1 | miRNA | ENSG00000( |
| 64506865 | 64542461 | 1 | protein_coding | ENSG00000( |
| 64542282 | 64662307 | -1 | protein_coding | ENSG00000( |
| 64572829 | 64573081 | 1 | processed_pseudoge | ENSG00000( |
| 64747264 | 64747492 | -1 | processed_pseudoge | ENSG00000( |
| 64749663 | 64781707 | -1 | lncRNA | ENSG00000( |
| 64749918 | 64758603 | -1 | transcribed_unproce | ENSG00000( |
| 64750420 | 64751311 | 1 | unprocessed_pseudc | ENSG00000( |
| 64761912 | 64762710 | -1 | unprocessed_pseudc | ENSG00000( |
| 64778842 | 64778937 | -1 | unprocessed_pseudc | ENSG00000( |
| 64779259 | 64837154 | -1 | transcribed_unproce | ENSG00000( |

| | | | | |
|---|---|---|---|---|
| 64780759 | 64780824 | 1 | miRNA | ENSG0000C |
| 64822030 | 64822102 | -1 | miRNA | ENSG0000C |
| 64825765 | 64825819 | -1 | snRNA | ENSG0000C |
| 64847386 | 64847728 | 1 | unprocessed_pseudc | ENSG0000C |
| 64854312 | 64919480 | -1 | protein_coding | ENSG0000C |
| 64871504 | 64871784 | -1 | misc_RNA | ENSG0000C |
| 64881091 | 64881181 | 1 | processed_pseudoge | ENSG0000C |
| 64892728 | 64892845 | 1 | transcribed_processe | ENSG0000C |
| 64892729 | 64910180 | 1 | lncRNA | ENSG0000C |
| 64899766 | 64900716 | 1 | lncRNA | ENSG0000C |
| 64905883 | 64906018 | -1 | transcribed_processe | ENSG0000C |
| 64927634 | 64934203 | 1 | transcribed_unproce | ENSG0000C |
| 64936485 | 64940229 | -1 | unprocessed_pseudc | ENSG0000C |
| 64940602 | 64954177 | 1 | lncRNA | ENSG0000C |
| 64940606 | 64940717 | 1 | transcribed_unproce | ENSG0000C |
| 64943262 | 64951054 | 1 | lncRNA | ENSG0000C |
| 64951498 | 64951609 | -1 | processed_pseudoge | ENSG0000C |
| 64953683 | 64956546 | 1 | transcribed_unproce | ENSG0000C |
| 64966550 | 64975576 | -1 | transcribed_unproce | ENSG0000C |
| 65009289 | 65056740 | -1 | protein_coding | ENSG0000C |
| 65051909 | 65053308 | 1 | lncRNA | ENSG0000C |
| 65100812 | 65227703 | 1 | protein_coding | ENSG0000C |
| 65185319 | 65185482 | -1 | processed_pseudoge | ENSG0000C |
| 65457541 | 65458408 | 1 | lncRNA | ENSG0000C |
| 65528563 | 65561648 | -1 | protein_coding | ENSG0000C |
| 65538102 | 65641033 | -1 | protein_coding | ENSG0000C |
| 65635537 | 66192133 | -1 | protein_coding | ENSG0000C |
| 65644308 | 65644617 | 1 | processed_pseudoge | ENSG0000C |
| 65654914 | 65655395 | 1 | processed_pseudoge | ENSG0000C |
| 66197693 | 66198238 | 1 | processed_pseudoge | ENSG0000C |
| 66212033 | 66256525 | -1 | protein_coding | ENSG0000C |
| 66267743 | 66267848 | 1 | rRNA_pseudogene | ENSG0000C |
| 66278059 | 66278664 | 1 | unprocessed_pseudc | ENSG0000C |
| 66302613 | 66810743 | 1 | protein_coding | ENSG0000C |
| 66364029 | 66364320 | 1 | misc_RNA | ENSG0000C |
| 66398069 | 66416854 | -1 | lncRNA | ENSG0000C |
| 66638271 | 66638374 | 1 | snRNA | ENSG0000C |
| 66639269 | 66639383 | 1 | rRNA_pseudogene | ENSG0000C |
| 66676372 | 66677404 | 1 | lncRNA | ENSG0000C |
| 66787072 | 66787168 | 1 | miRNA | ENSG0000C |
| 66835117 | 66885486 | 1 | protein_coding | ENSG0000C |
| 66895491 | 66895572 | 1 | rRNA_pseudogene | ENSG0000C |
| 66964707 | 67033398 | 1 | protein_coding | ENSG0000C |
| 67019934 | 67021743 | -1 | lncRNA | ENSG0000C |
| 67032409 | 67033290 | -1 | lncRNA | ENSG0000C |
| 67044554 | 67056797 | 1 | protein_coding | ENSG0000C |
| 67070444 | 67245989 | -1 | protein_coding | ENSG0000C |
| 67224303 | 67225541 | 1 | lncRNA | ENSG0000C |
| 67224728 | 67225041 | 1 | processed_pseudoge | ENSG0000C |
| 67244282 | 67273503 | 1 | lncRNA | ENSG0000C |

| | | | | |
|---|---|---|---|---|
| 67244837 | 67245806 | 1 | lncRNA | ENSG0000C |
| 67268110 | 67268228 | 1 | rRNA_pseudogene | ENSG0000C |
| 67271469 | 67271607 | 1 | snoRNA | ENSG0000C |
| 67276118 | 67276351 | -1 | processed_pseudoge | ENSG0000C |
| 67337916 | 67366605 | -1 | protein_coding | ENSG0000C |
| 67377281 | 67697261 | 1 | protein_coding | ENSG0000C |
| 67408774 | 67408881 | -1 | misc_RNA | ENSG0000C |
| 67457023 | 67457290 | -1 | misc_RNA | ENSG0000C |
| 67471489 | 67471585 | -1 | miRNA | ENSG0000C |
| 67471489 | 67471585 | 1 | miRNA | ENSG0000C |
| 67524481 | 67525422 | 1 | lncRNA | ENSG0000C |
| 67611277 | 67613822 | 1 | TEC | ENSG0000C |
| 67637774 | 67638973 | 1 | TEC | ENSG0000C |
| 67658015 | 67659465 | -1 | lncRNA | ENSG0000C |
| 67717931 | 67744531 | 1 | protein_coding | ENSG0000C |
| 67740669 | 67740799 | 1 | snoRNA | ENSG0000C |
| 67774591 | 67777324 | -1 | unprocessed_pseudc | ENSG0000C |
| 67784753 | 67785293 | 1 | processed_pseudoge | ENSG0000C |
| 67825503 | 67984378 | 1 | protein_coding | ENSG0000C |
| 67950885 | 67951746 | -1 | lncRNA | ENSG0000C |
| 67955314 | 67957291 | 1 | TEC | ENSG0000C |
| 67973934 | 67976072 | 1 | TEC | ENSG0000C |
| 67977698 | 67977972 | 1 | misc_RNA | ENSG0000C |
| 67991101 | 67996431 | -1 | protein_coding | ENSG0000C |
| 68035708 | 68046854 | 1 | protein_coding | ENSG0000C |
| 68095068 | 68095757 | 1 | processed_pseudoge | ENSG0000C |
| 68096046 | 68101496 | -1 | lncRNA | ENSG0000C |
| 68101908 | 68115518 | 1 | transcribed_unproce | ENSG0000C |
| 68125777 | 68152468 | -1 | transcribed_unproce | ENSG0000C |
| 68126666 | 68129586 | 1 | lncRNA | ENSG0000C |
| 68131462 | 68131907 | 1 | lncRNA | ENSG0000C |
| 68133201 | 68135935 | 1 | lncRNA | ENSG0000C |
| 68134675 | 68135604 | -1 | processed_pseudoge | ENSG0000C |
| 68152776 | 68159043 | -1 | unprocessed_pseudc | ENSG0000C |
| 68188547 | 68189165 | 1 | lncRNA | ENSG0000C |
| 68189884 | 68192802 | 1 | lncRNA | ENSG0000C |
| 68198921 | 68199596 | 1 | processed_pseudoge | ENSG0000C |
| 68205489 | 68207493 | 1 | lncRNA | ENSG0000C |
| 68246629 | 68247938 | -1 | lncRNA | ENSG0000C |
| 68247930 | 68257164 | 1 | protein_coding | ENSG0000C |
| 68259182 | 68422731 | 1 | protein_coding | ENSG0000C |
| 68267026 | 68291267 | -1 | protein_coding | ENSG0000C |
| 68413623 | 68524949 | 1 | lncRNA | ENSG0000C |
| 68420948 | 68457513 | -1 | protein_coding | ENSG0000C |
| 68424451 | 68424548 | -1 | miRNA | ENSG0000C |
| 68511780 | 68551319 | 1 | protein_coding | ENSG0000C |
| 68535113 | 68601367 | -1 | protein_coding | ENSG0000C |
| 68557516 | 68582577 | 1 | lncRNA | ENSG0000C |
| 68591796 | 68763882 | 1 | lncRNA | ENSG0000C |
| 68691809 | 68716064 | -1 | unitary_pseudogene | ENSG0000C |

| | | | | |
|---|---|---|---|---|
| 68793549 | 68797822 | -1 | lncRNA | ENSG0000C |
| 68867289 | 68955392 | -1 | protein_coding | ENSG0000C |
| 68944531 | 69042784 | 1 | lncRNA | ENSG0000C |
| 68974488 | 69060949 | -1 | protein_coding | ENSG0000C |
| 69078702 | 69141895 | -1 | protein_coding | ENSG0000C |
| 69094289 | 69095499 | -1 | processed_pseudoge | ENSG0000C |
| 69099542 | 69099656 | 1 | miRNA | ENSG0000C |
| 69099564 | 69099632 | -1 | miRNA | ENSG0000C |
| 69147214 | 69244846 | -1 | protein_coding | ENSG0000C |
| 69232963 | 69235012 | -1 | TEC | ENSG0000C |
| 69242230 | 69244097 | 1 | lncRNA | ENSG0000C |
| 69244311 | 69327244 | -1 | protein_coding | ENSG0000C |
| 69318733 | 69318961 | 1 | processed_pseudoge | ENSG0000C |
| 69355133 | 69355218 | -1 | snoRNA | ENSG0000C |
| 69414697 | 69553865 | 1 | protein_coding | ENSG0000C |
| 69433084 | 69435533 | 1 | TEC | ENSG0000C |
| 69477139 | 69501755 | -1 | lncRNA | ENSG0000C |
| 69577251 | 69903000 | 1 | lncRNA | ENSG0000C |
| 69707193 | 69707373 | -1 | processed_pseudoge | ENSG0000C |
| 69961568 | 69994845 | 1 | lncRNA | ENSG0000C |
| 69985986 | 69986094 | 1 | misc_RNA | ENSG0000C |
| 70017324 | 70128859 | -1 | lncRNA | ENSG0000C |
| 70051277 | 70068095 | 1 | lncRNA | ENSG0000C |
| 70053429 | 70135608 | 1 | protein_coding | ENSG0000C |
| 70166961 | 70169402 | -1 | lncRNA | ENSG0000C |
| 70168673 | 70180048 | 1 | protein_coding | ENSG0000C |
| 70241324 | 70241761 | 1 | processed_pseudoge | ENSG0000C |
| 70312831 | 70368916 | -1 | lncRNA | ENSG0000C |
| 70628917 | 70629265 | -1 | processed_pseudoge | ENSG0000C |
| 70778604 | 70779230 | -1 | lncRNA | ENSG0000C |
| 71011997 | 71013587 | 1 | processed_pseudoge | ENSG0000C |
| 71097774 | 71202203 | -1 | lncRNA | ENSG0000C |
| 71298156 | 71298218 | -1 | snRNA | ENSG0000C |
| 71596338 | 71597279 | 1 | lncRNA | ENSG0000C |
| 71620978 | 71621436 | 1 | processed_pseudoge | ENSG0000C |
| 71678899 | 71743468 | -1 | lncRNA | ENSG0000C |
| 71829870 | 71871750 | -1 | lncRNA | ENSG0000C |
| 72021851 | 72034092 | -1 | lncRNA | ENSG0000C |
| 72030291 | 72041310 | 1 | lncRNA | ENSG0000C |
| 72034107 | 72237203 | -1 | lncRNA | ENSG0000C |
| 72072333 | 72103035 | 1 | lncRNA | ENSG0000C |
| 72121020 | 72126416 | 1 | protein_coding | ENSG0000C |
| 72290091 | 72640472 | -1 | lncRNA | ENSG0000C |
| 72342692 | 72355136 | 1 | lncRNA | ENSG0000C |
| 72425939 | 72428852 | 1 | lncRNA | ENSG0000C |
| 72627231 | 72627624 | 1 | processed_pseudoge | ENSG0000C |
| 72642731 | 72644490 | 1 | lncRNA | ENSG0000C |
| 72645949 | 73092712 | -1 | protein_coding | ENSG0000C |
| 72663823 | 72664152 | -1 | misc_RNA | ENSG0000C |
| 72818836 | 72818972 | -1 | scaRNA | ENSG0000C |

| | | | | |
|---|---|---|---|---|
| 72839039 | 72839718 | 1 | lncRNA | ENSG0000C |
| 73067870 | 73068288 | -1 | processed_pseudoge | ENSG0000C |
| 73163035 | 73163359 | -1 | processed_pseudoge | ENSG0000C |
| 73164977 | 73196593 | 1 | lncRNA | ENSG0000C |
| 73165010 | 73176633 | 1 | protein_coding | ENSG0000C |
| 73192632 | 73208507 | 1 | protein_coding | ENSG0000C |
| 73202968 | 73203431 | -1 | lncRNA | ENSG0000C |
| 73207353 | 73236753 | -1 | protein_coding | ENSG0000C |
| 73232233 | 73248947 | 1 | protein_coding | ENSG0000C |
| 73243093 | 73244706 | 1 | lncRNA | ENSG0000C |
| 73248449 | 73262352 | -1 | protein_coding | ENSG0000C |
| 73283624 | 73312005 | -1 | protein_coding | ENSG0000C |
| 73334384 | 73644445 | -1 | protein_coding | ENSG0000C |
| 73513469 | 73521407 | 1 | lncRNA | ENSG0000C |
| 73641026 | 73643106 | 1 | lncRNA | ENSG0000C |
| 73662568 | 73663848 | -1 | lncRNA | ENSG0000C |
| 73737854 | 73756932 | 1 | lncRNA | ENSG0000C |
| 73749270 | 73828520 | -1 | lncRNA | ENSG0000C |
| 73786675 | 73800966 | 1 | lncRNA | ENSG0000C |
| 73831572 | 73836019 | -1 | lncRNA | ENSG0000C |
| 73894749 | 73911878 | -1 | lncRNA | ENSG0000C |
| 73912190 | 73931436 | 1 | lncRNA | ENSG0000C |
| 73926421 | 73949906 | -1 | lncRNA | ENSG0000C |
| 74043452 | 74154212 | -1 | lncRNA | ENSG0000C |
| 74203582 | 74210655 | 1 | protein_coding | ENSG0000C |
| 74209980 | 74213342 | -1 | lncRNA | ENSG0000C |
| 74213571 | 74262020 | 1 | protein_coding | ENSG0000C |
| 74256896 | 74262020 | -1 | lncRNA | ENSG0000C |
| 74274234 | 74314884 | 1 | protein_coding | ENSG0000C |
| 74307210 | 74309790 | -1 | lncRNA | ENSG0000C |
| 74326210 | 74355820 | 1 | protein_coding | ENSG0000C |
| 74356416 | 74361868 | -1 | protein_coding | ENSG0000C |
| 74367407 | 74372622 | 1 | protein_coding | ENSG0000C |
| 74417594 | 74417704 | -1 | rRNA_pseudogene | ENSG0000C |
| 74424851 | 74451653 | 1 | protein_coding | ENSG0000C |
| 74466399 | 74484794 | 1 | protein_coding | ENSG0000C |
| 74521174 | 74531475 | -1 | protein_coding | ENSG0000C |
| 74541073 | 74546115 | -1 | protein_coding | ENSG0000C |
| 74549399 | 74552737 | -1 | unprocessed_pseudc | ENSG0000C |
| 74560701 | 74567343 | -1 | polymorphic_pseudc | ENSG0000C |
| 74579365 | 74592283 | -1 | protein_coding | ENSG0000C |
| 74584679 | 74594209 | 1 | lncRNA | ENSG0000C |
| 74599840 | 74607229 | 1 | lncRNA | ENSG0000C |
| 74609885 | 74623738 | -1 | protein_coding | ENSG0000C |
| 74670077 | 74677624 | -1 | lncRNA | ENSG0000C |
| 74670578 | 74747335 | 1 | protein_coding | ENSG0000C |
| 74694311 | 74712978 | -1 | protein_coding | ENSG0000C |
| 74747319 | 74748912 | -1 | lncRNA | ENSG0000C |
| 74748613 | 74748699 | 1 | miRNA | ENSG0000C |
| 74748628 | 74769353 | 1 | protein_coding | ENSG0000C |

| | | | | |
|---|---|---|---|---|
| 74770529 | 74776367 | -1 | protein_coding | ENSG00000 |
| 74776483 | 74839779 | 1 | protein_coding | ENSG00000 |
| 74842023 | 74861504 | -1 | protein_coding | ENSG00000 |
| 74862497 | 74873031 | -1 | protein_coding | ENSG00000 |
| 74877302 | 74893781 | -1 | protein_coding | ENSG00000 |
| 74916083 | 74923256 | -1 | protein_coding | ENSG00000 |
| 74924275 | 74933912 | 1 | protein_coding | ENSG00000 |
| 74935802 | 74949992 | 1 | protein_coding | ENSG00000 |
| 74950742 | 74973166 | -1 | protein_coding | ENSG00000 |
| 74970704 | 74975728 | 1 | lncRNA | ENSG00000 |
| 74987632 | 75005800 | 1 | protein_coding | ENSG00000 |
| 75012670 | 75021261 | 1 | protein_coding | ENSG00000 |
| 75031097 | 75031186 | 1 | snRNA | ENSG00000 |
| 75032575 | 75065889 | 1 | protein_coding | ENSG00000 |
| 75038863 | 75046985 | -1 | protein_coding | ENSG00000 |
| 75041740 | 75042015 | 1 | misc_RNA | ENSG00000 |
| 75060491 | 75060620 | -1 | processed_pseudoge | ENSG00000 |
| 75074306 | 75079549 | 1 | unitary_pseudogene | ENSG00000 |
| 75087727 | 75106162 | 1 | protein_coding | ENSG00000 |
| 75089862 | 75089974 | -1 | misc_RNA | ENSG00000 |
| 75109952 | 75130272 | 1 | protein_coding | ENSG00000 |
| 75130225 | 75131757 | -1 | protein_coding | ENSG00000 |
| 75130883 | 75133043 | 1 | TEC | ENSG00000 |
| 75135248 | 75168281 | -1 | protein_coding | ENSG00000 |
| 75138416 | 75141350 | -1 | lncRNA | ENSG00000 |
| 75145261 | 75146546 | -1 | lncRNA | ENSG00000 |
| 75145670 | 75145762 | 1 | misc_RNA | ENSG00000 |
| 75165586 | 75182959 | -1 | protein_coding | ENSG00000 |
| 75205659 | 75235758 | 1 | protein_coding | ENSG00000 |
| 75236599 | 75262363 | -1 | protein_coding | ENSG00000 |
| 75261674 | 75266376 | 1 | protein_coding | ENSG00000 |
| 75266228 | 75271227 | -1 | protein_coding | ENSG00000 |
| 75271369 | 75273895 | 1 | lncRNA | ENSG00000 |
| 75272981 | 75289510 | -1 | protein_coding | ENSG00000 |
| 75318076 | 75405709 | -1 | protein_coding | ENSG00000 |
| 75338827 | 75339147 | 1 | processed_pseudoge | ENSG00000 |
| 75344405 | 75373662 | 1 | lncRNA | ENSG00000 |
| 75370947 | 75373736 | -1 | lncRNA | ENSG00000 |
| 75394123 | 75394963 | 1 | lncRNA | ENSG00000 |
| 75406069 | 75406162 | 1 | miRNA | ENSG00000 |
| 75409086 | 75409188 | -1 | snRNA | ENSG00000 |
| 75431990 | 75432087 | 1 | misc_RNA | ENSG00000 |
| 75441159 | 75500452 | 1 | protein_coding | ENSG00000 |
| 75498548 | 75498628 | 1 | miRNA | ENSG00000 |
| 75500261 | 75515537 | -1 | protein_coding | ENSG00000 |
| 75516060 | 75524735 | 1 | protein_coding | ENSG00000 |
| 75525080 | 75575208 | 1 | protein_coding | ENSG00000 |
| 75588058 | 75626849 | 1 | protein_coding | ENSG00000 |
| 75626845 | 75667189 | -1 | protein_coding | ENSG00000 |
| 75633434 | 75641404 | 1 | protein_coding | ENSG00000 |

| | | | | |
|---|---|---|---|---|
| 75646243 | 75647975 | 1 | protein_coding | ENSG0000( |
| 75667251 | 75708062 | 1 | protein_coding | ENSG0000( |
| 75679474 | 75679967 | -1 | lncRNA | ENSG0000( |
| 75683543 | 75684799 | -1 | lncRNA | ENSG0000( |
| 75721328 | 75757818 | 1 | protein_coding | ENSG0000( |
| 75751594 | 75765236 | -1 | protein_coding | ENSG0000( |
| 75776434 | 75785893 | -1 | protein_coding | ENSG0000( |
| 75784521 | 75784607 | -1 | miRNA | ENSG0000( |
| 75784806 | 75825799 | 1 | protein_coding | ENSG0000( |
| 75818815 | 75820055 | -1 | lncRNA | ENSG0000( |
| 75827225 | 75844717 | -1 | protein_coding | ENSG0000( |
| 75845699 | 75856507 | -1 | protein_coding | ENSG0000( |
| 75874164 | 75878581 | -1 | protein_coding | ENSG0000( |
| 75876372 | 75879546 | 1 | lncRNA | ENSG0000( |
| 75880335 | 75896951 | -1 | protein_coding | ENSG0000( |
| 75897060 | 75900148 | 1 | lncRNA | ENSG0000( |
| 75898643 | 75905413 | -1 | protein_coding | ENSG0000( |
| 75898645 | 75930129 | -1 | protein_coding | ENSG0000( |
| 75909574 | 75941140 | -1 | protein_coding | ENSG0000( |
| 75941507 | 75979177 | -1 | protein_coding | ENSG0000( |
| 75943832 | 75945142 | 1 | lncRNA | ENSG0000( |
| 75979231 | 76005999 | 1 | protein_coding | ENSG0000( |
| 75979240 | 76000586 | 1 | protein_coding | ENSG0000( |
| 76000906 | 76005999 | 1 | protein_coding | ENSG0000( |
| 76004502 | 76027452 | -1 | protein_coding | ENSG0000( |
| 76038775 | 76072517 | -1 | protein_coding | ENSG0000( |
| 76071961 | 76083666 | 1 | protein_coding | ENSG0000( |
| 76074781 | 76077537 | 1 | protein_coding | ENSG0000( |
| 76081016 | 76121576 | -1 | protein_coding | ENSG0000( |
| 76090308 | 76091669 | 1 | TEC | ENSG0000( |
| 76098019 | 76098076 | -1 | miRNA | ENSG0000( |
| 76136333 | 76141245 | -1 | protein_coding | ENSG0000( |
| 76138432 | 76154650 | 1 | lncRNA | ENSG0000( |
| 76142465 | 76240493 | -1 | protein_coding | ENSG0000( |
| 76202644 | 76202796 | -1 | processed_pseudoge | ENSG0000( |
| 76225751 | 76226806 | -1 | processed_pseudoge | ENSG0000( |
| 76265348 | 76271298 | 1 | protein_coding | ENSG0000( |
| 76274049 | 76307998 | -1 | protein_coding | ENSG0000( |
| 76309478 | 76384521 | -1 | protein_coding | ENSG0000( |
| 76319376 | 76319473 | 1 | misc_RNA | ENSG0000( |
| 76337316 | 76337422 | 1 | snRNA | ENSG0000( |
| 76345761 | 76346081 | 1 | processed_pseudoge | ENSG0000( |
| 76376584 | 76387860 | 1 | protein_coding | ENSG0000( |
| 76389456 | 76453152 | -1 | protein_coding | ENSG0000( |
| 76430417 | 76431124 | -1 | processed_pseudoge | ENSG0000( |
| 76453351 | 76470117 | 1 | protein_coding | ENSG0000( |
| 76470891 | 76501790 | -1 | protein_coding | ENSG0000( |
| 76527356 | 76551175 | -1 | protein_coding | ENSG0000( |
| 76527586 | 76553578 | 1 | protein_coding | ENSG0000( |
| 76545668 | 76557683 | -1 | lncRNA | ENSG0000( |

| | | | | |
|---|---|---|---|---|
| 76549951 | 76550826 | 1 | lncRNA | ENSG0000C |
| 76551352 | 76551750 | 1 | lncRNA | ENSG0000C |
| 76557764 | 76565348 | 1 | lncRNA | ENSG0000C |
| 76558791 | 76558868 | 1 | snoRNA | ENSG0000C |
| 76561109 | 76561192 | 1 | snoRNA | ENSG0000C |
| 76561634 | 76561705 | 1 | snoRNA | ENSG0000C |
| 76563710 | 76570544 | -1 | protein_coding | ENSG0000C |
| 76565377 | 76586956 | -1 | protein_coding | ENSG0000C |
| 76569792 | 76571240 | 1 | lncRNA | ENSG0000C |
| 76595584 | 76595772 | -1 | processed_pseudoge | ENSG0000C |
| 76617959 | 76619128 | -1 | lncRNA | ENSG0000C |
| 76624761 | 76643786 | -1 | protein_coding | ENSG0000C |
| 76656320 | 76656422 | 1 | snRNA | ENSG0000C |
| 76671942 | 76673658 | 1 | lncRNA | ENSG0000C |
| 76672551 | 76711016 | -1 | protein_coding | ENSG0000C |
| 76703632 | 76703727 | -1 | misc_RNA | ENSG0000C |
| 76709760 | 76710045 | 1 | lncRNA | ENSG0000C |
| 76712832 | 76726799 | -1 | protein_coding | ENSG0000C |
| 76721216 | 76721396 | -1 | misc_RNA | ENSG0000C |
| 76726830 | 76733936 | 1 | protein_coding | ENSG0000C |
| 76732978 | 76738522 | 1 | protein_coding | ENSG0000C |
| 76734115 | 76737374 | -1 | protein_coding | ENSG0000C |
| 76735865 | 76781449 | 1 | protein_coding | ENSG0000C |
| 76736450 | 76736548 | -1 | miRNA | ENSG0000C |
| 76752912 | 76753015 | 1 | snRNA | ENSG0000C |
| 76799044 | 76807102 | 1 | lncRNA | ENSG0000C |
| 76831684 | 76841099 | -1 | lncRNA | ENSG0000C |
| 76849840 | 76861836 | 1 | lncRNA | ENSG0000C |
| 76868456 | 76950393 | 1 | protein_coding | ENSG0000C |
| 76950317 | 76969156 | -1 | lncRNA | ENSG0000C |
| 76957023 | 76958222 | -1 | lncRNA | ENSG0000C |
| 76966285 | 76969530 | -1 | TEC | ENSG0000C |
| 77014880 | 77015138 | 1 | misc_RNA | ENSG0000C |
| 77086716 | 77099902 | 1 | lncRNA | ENSG0000C |
| 77088749 | 77217101 | 1 | protein_coding | ENSG0000C |
| 77089307 | 77089493 | 1 | scaRNA | ENSG0000C |
| 77089417 | 77089497 | 1 | miRNA | ENSG0000C |
| 77152561 | 77152674 | -1 | snRNA | ENSG0000C |
| 77198950 | 77199543 | -1 | processed_pseudoge | ENSG0000C |
| 77257737 | 77281897 | -1 | lncRNA | ENSG0000C |
| 77264180 | 77264776 | -1 | lncRNA | ENSG0000C |
| 77280569 | 77500596 | 1 | protein_coding | ENSG0000C |
| 77373818 | 77377236 | -1 | lncRNA | ENSG0000C |
| 77386901 | 77388546 | -1 | lncRNA | ENSG0000C |
| 77396984 | 77397054 | -1 | miRNA | ENSG0000C |
| 77442109 | 77442213 | 1 | misc_RNA | ENSG0000C |
| 77444704 | 77447167 | 1 | TEC | ENSG0000C |
| 77469068 | 77471045 | 1 | lncRNA | ENSG0000C |
| 77469162 | 77472770 | -1 | lncRNA | ENSG0000C |
| 77516887 | 77536601 | 1 | lncRNA | ENSG0000C |

| | | | | |
|---|---|---|---|---|
| 77546940 | 77563243 | 1 | lncRNA | ENSG00000( |
| 77548129 | 77550499 | 1 | lncRNA | ENSG00000( |
| 77563368 | 77568695 | -1 | lncRNA | ENSG00000( |
| 77571418 | 77572516 | 1 | TEC | ENSG00000( |
| 77590532 | 77680687 | 1 | lncRNA | ENSG00000( |
| 77722872 | 77728559 | -1 | lncRNA | ENSG00000( |
| 77877330 | 77884119 | -1 | lncRNA | ENSG00000( |
| 77920098 | 77920258 | 1 | snRNA | ENSG00000( |
| 77959240 | 78108835 | 1 | protein_coding | ENSG00000( |
| 78008421 | 78008855 | 1 | TEC | ENSG00000( |
| 78066258 | 78066761 | -1 | processed_pseudoge | ENSG00000( |
| 78066947 | 78067053 | 1 | snRNA | ENSG00000( |
| 78107398 | 78111799 | -1 | lncRNA | ENSG00000( |
| 78110458 | 78132407 | -1 | protein_coding | ENSG00000( |
| 78130770 | 78142968 | 1 | protein_coding | ENSG00000( |
| 78146353 | 78166177 | 1 | protein_coding | ENSG00000( |
| 78157403 | 78159135 | 1 | processed_pseudoge | ENSG00000( |
| 78168558 | 78173527 | 1 | protein_coding | ENSG00000( |
| 78174091 | 78187233 | -1 | protein_coding | ENSG00000( |
| 78187317 | 78207701 | 1 | protein_coding | ENSG00000( |
| 78211618 | 78211908 | -1 | processed_pseudoge | ENSG00000( |
| 78214186 | 78225636 | 1 | protein_coding | ENSG00000( |
| 78231310 | 78240987 | 1 | protein_coding | ENSG00000( |
| 78248193 | 78251440 | 1 | lncRNA | ENSG00000( |
| 78248493 | 78254416 | -1 | unitary_pseudogene | ENSG00000( |
| 78250346 | 78250932 | -1 | TEC | ENSG00000( |
| 78261349 | 78278492 | -1 | lncRNA | ENSG00000( |
| 78315729 | 78347798 | 1 | lncRNA | ENSG00000( |
| 78356778 | 78360077 | -1 | protein_coding | ENSG00000( |
| 78360453 | 78373911 | 1 | lncRNA | ENSG00000( |
| 78370852 | 78371135 | -1 | misc_RNA | ENSG00000( |
| 78378649 | 78425114 | 1 | protein_coding | ENSG00000( |
| 78399935 | 78400063 | 1 | snoRNA | ENSG00000( |
| 78423697 | 78577394 | -1 | protein_coding | ENSG00000( |
| 78484882 | 78503056 | 1 | lncRNA | ENSG00000( |
| 78522247 | 78522828 | 1 | processed_pseudoge | ENSG00000( |
| 78532813 | 78533055 | -1 | misc_RNA | ENSG00000( |
| 78605233 | 78632155 | -1 | lncRNA | ENSG00000( |
| 78674048 | 78782297 | -1 | protein_coding | ENSG00000( |
| 78691266 | 78691979 | -1 | TEC | ENSG00000( |
| 78696694 | 78696797 | -1 | snRNA | ENSG00000( |
| 78787381 | 78841441 | -1 | protein_coding | ENSG00000( |
| 78845345 | 78846686 | 1 | lncRNA | ENSG00000( |
| 78845953 | 78846868 | -1 | processed_pseudoge | ENSG00000( |
| 78852977 | 78925387 | -1 | protein_coding | ENSG00000( |
| 78855478 | 78855844 | 1 | lncRNA | ENSG00000( |
| 78870910 | 78903217 | -1 | protein_coding | ENSG00000( |
| 78897264 | 78900152 | 1 | TEC | ENSG00000( |
| 78903263 | 78905024 | 1 | lncRNA | ENSG00000( |
| 78971238 | 78979947 | -1 | protein_coding | ENSG00000( |

| | | | | |
|---|---|---|---|---|
| 78991716 | 79009867 | -1 | protein_coding | ENSG00000( |
| 79018131 | 79027673 | -1 | lncRNA | ENSG00000( |
| 79022814 | 79049788 | 1 | protein_coding | ENSG00000( |
| 79074939 | 79088599 | 1 | protein_coding | ENSG00000( |
| 79089345 | 79516148 | -1 | protein_coding | ENSG00000( |
| 79132722 | 79136402 | 1 | lncRNA | ENSG00000( |
| 79598032 | 79603921 | -1 | lncRNA | ENSG00000( |
| 79707176 | 79707249 | -1 | miRNA | ENSG00000( |
| 79707266 | 79712317 | 1 | lncRNA | ENSG00000( |
| 79730943 | 79742219 | 1 | protein_coding | ENSG00000( |
| 79778148 | 79787983 | 1 | protein_coding | ENSG00000( |
| 79792132 | 79801683 | -1 | protein_coding | ENSG00000( |
| 79800598 | 79802529 | 1 | lncRNA | ENSG00000( |
| 79819083 | 79827704 | 1 | lncRNA | ENSG00000( |
| 79833156 | 79839440 | -1 | protein_coding | ENSG00000( |
| 79848058 | 79850110 | -1 | lncRNA | ENSG00000( |
| 79915252 | 79926725 | -1 | lncRNA | ENSG00000( |
| 79919357 | 79925999 | 1 | lncRNA | ENSG00000( |
| 79932343 | 80035872 | -1 | protein_coding | ENSG00000( |
| 79952663 | 79952992 | 1 | lncRNA | ENSG00000( |
| 79991944 | 79992841 | -1 | lncRNA | ENSG00000( |
| 80023894 | 80026107 | 1 | lncRNA | ENSG00000( |
| 80036632 | 80100613 | 1 | protein_coding | ENSG00000( |
| 80098828 | 80098877 | 1 | miRNA | ENSG00000( |
| 80101556 | 80119881 | 1 | protein_coding | ENSG00000( |
| 80134369 | 80147151 | -1 | protein_coding | ENSG00000( |
| 80147250 | 80148596 | 1 | TEC | ENSG00000( |
| 80149627 | 80149798 | 1 | lncRNA | ENSG00000( |
| 80169992 | 80209331 | 1 | protein_coding | ENSG00000( |
| 80200673 | 80205949 | -1 | lncRNA | ENSG00000( |
| 80206716 | 80220923 | -1 | protein_coding | ENSG00000( |
| 80219699 | 80253500 | 1 | protein_coding | ENSG00000( |
| 80260866 | 80398786 | 1 | protein_coding | ENSG00000( |
| 80315651 | 80316633 | 1 | lncRNA | ENSG00000( |
| 80333841 | 80334366 | -1 | lncRNA | ENSG00000( |
| 80339898 | 80342058 | 1 | TEC | ENSG00000( |
| 80351828 | 80415168 | -1 | lncRNA | ENSG00000( |
| 80415165 | 80438086 | 1 | protein_coding | ENSG00000( |
| 80419418 | 80419493 | 1 | miRNA | ENSG00000( |
| 80453735 | 80454729 | -1 | lncRNA | ENSG00000( |
| 80463451 | 80465593 | -1 | TEC | ENSG00000( |
| 80466834 | 80477843 | -1 | protein_coding | ENSG00000( |
| 80515215 | 80515389 | 1 | processed_pseudoge | ENSG00000( |
| 80515657 | 80515812 | 1 | processed_pseudoge | ENSG00000( |
| 80542203 | 80542605 | 1 | processed_pseudoge | ENSG00000( |
| 80544819 | 80966371 | 1 | protein_coding | ENSG00000( |
| 80602549 | 80602926 | -1 | processed_pseudoge | ENSG00000( |
| 80801640 | 80805632 | -1 | lncRNA | ENSG00000( |
| 80824170 | 80826900 | -1 | lncRNA | ENSG00000( |
| 80940418 | 80942033 | -1 | lncRNA | ENSG00000( |

| | | | | |
|---|---|---|---|---|
| 80960766 | 80961713 | -1 | lncRNA | ENSG0000C |
| 80966239 | 80971213 | -1 | lncRNA | ENSG0000C |
| 80976454 | 80976948 | 1 | processed_pseudoge | ENSG0000C |
| 80991598 | 81009517 | 1 | protein_coding | ENSG0000C |
| 80999509 | 81000130 | -1 | lncRNA | ENSG0000C |
| 81003335 | 81005132 | 1 | TEC | ENSG0000C |
| 81017969 | 81028079 | -1 | lncRNA | ENSG0000C |
| 81023545 | 81025503 | -1 | lncRNA | ENSG0000C |
| 81029130 | 81034881 | -1 | lncRNA | ENSG0000C |
| 81035122 | 81117432 | 1 | protein_coding | ENSG0000C |
| 81110487 | 81166221 | -1 | protein_coding | ENSG0000C |
| 81125276 | 81125373 | -1 | miRNA | ENSG0000C |
| 81125877 | 81125955 | 1 | miRNA | ENSG0000C |
| 81125883 | 81125949 | -1 | miRNA | ENSG0000C |
| 81133196 | 81133308 | -1 | miRNA | ENSG0000C |
| 81135771 | 81136256 | 1 | lncRNA | ENSG0000C |
| 81165507 | 81183166 | 1 | protein_coding | ENSG0000C |
| 81189593 | 81222999 | -1 | protein_coding | ENSG0000C |
| 81197393 | 81200288 | 1 | lncRNA | ENSG0000C |
| 81228277 | 81239091 | -1 | protein_coding | ENSG0000C |
| 81228707 | 81233983 | 1 | lncRNA | ENSG0000C |
| 81239305 | 81241310 | 1 | protein_coding | ENSG0000C |
| 81244811 | 81295547 | -1 | protein_coding | ENSG0000C |
| 81251194 | 81251803 | 1 | lncRNA | ENSG0000C |
| 81262920 | 81265401 | 1 | TEC | ENSG0000C |
| 81303771 | 81309248 | -1 | lncRNA | ENSG0000C |
| 81311270 | 81330674 | -1 | lncRNA | ENSG0000C |
| 81339183 | 81345966 | -1 | lncRNA | ENSG0000C |
| 81362272 | 81374214 | 1 | lncRNA | ENSG0000C |
| 81375144 | 81385464 | -1 | lncRNA | ENSG0000C |
| 81387415 | 81387766 | 1 | lncRNA | ENSG0000C |
| 81388126 | 81390319 | -1 | lncRNA | ENSG0000C |
| 81395475 | 81466332 | 1 | protein_coding | ENSG0000C |
| 81395609 | 81397144 | -1 | TEC | ENSG0000C |
| 81400716 | 81400778 | -1 | miRNA | ENSG0000C |
| 81451104 | 81451188 | -1 | miRNA | ENSG0000C |
| 81461013 | 81461937 | -1 | lncRNA | ENSG0000C |
| 81480523 | 81481570 | -1 | lncRNA | ENSG0000C |
| 81509971 | 81523847 | -1 | protein_coding | ENSG0000C |
| 81514047 | 81527776 | 1 | lncRNA | ENSG0000C |
| 81528396 | 81537130 | 1 | protein_coding | ENSG0000C |
| 81539885 | 81553961 | -1 | protein_coding | ENSG0000C |
| 81556887 | 81648465 | -1 | protein_coding | ENSG0000C |
| 81573490 | 81573602 | 1 | misc_RNA | ENSG0000C |
| 81637171 | 81648749 | 1 | protein_coding | ENSG0000C |
| 81650459 | 81663112 | -1 | protein_coding | ENSG0000C |
| 81665036 | 81666635 | -1 | protein_coding | ENSG0000C |
| 81666737 | 81673904 | 1 | protein_coding | ENSG0000C |
| 81681174 | 81683924 | -1 | protein_coding | ENSG0000C |
| 81683326 | 81703138 | 1 | protein_coding | ENSG0000C |

| | | | | |
|---|---|---|---|---|
| 81693757 | 81693869 | 1 | miRNA | ENSG0000C |
| 81697025 | 81697714 | -1 | lncRNA | ENSG0000C |
| 81701324 | 81703300 | -1 | lncRNA | ENSG0000C |
| 81703367 | 81707517 | 1 | protein_coding | ENSG0000C |
| 81703371 | 81720539 | 1 | protein_coding | ENSG0000C |
| 81712236 | 81721016 | 1 | protein_coding | ENSG0000C |
| 81804132 | 81814008 | 1 | protein_coding | ENSG0000C |
| 81822361 | 81833302 | -1 | protein_coding | ENSG0000C |
| 81833492 | 81835050 | -1 | protein_coding | ENSG0000C |
| 81843161 | 81860624 | -1 | protein_coding | ENSG0000C |
| 81843165 | 81843958 | 1 | lncRNA | ENSG0000C |
| 81862778 | 81863102 | 1 | TEC | ENSG0000C |
| 81867721 | 81871378 | -1 | protein_coding | ENSG0000C |
| 81867721 | 81868552 | 1 | lncRNA | ENSG0000C |
| 81878425 | 81881106 | -1 | lncRNA | ENSG0000C |
| 81878667 | 81879557 | 1 | lncRNA | ENSG0000C |
| 81887835 | 81891586 | -1 | protein_coding | ENSG0000C |
| 81890790 | 81900991 | 1 | protein_coding | ENSG0000C |
| 81900745 | 81902905 | 1 | protein_coding | ENSG0000C |
| 81900965 | 81911464 | -1 | protein_coding | ENSG0000C |
| 81911939 | 81921323 | -1 | protein_coding | ENSG0000C |
| 81918270 | 81927735 | -1 | protein_coding | ENSG0000C |
| 81922899 | 81924511 | 1 | lncRNA | ENSG0000C |
| 81927829 | 81930753 | 1 | lncRNA | ENSG0000C |
| 81932384 | 81942412 | -1 | protein_coding | ENSG0000C |
| 81932398 | 81933058 | 1 | lncRNA | ENSG0000C |
| 81939645 | 81947233 | -1 | protein_coding | ENSG0000C |
| 81941869 | 81947601 | 1 | lncRNA | ENSG0000C |
| 81952507 | 81961840 | -1 | protein_coding | ENSG0000C |
| 81965632 | 81966589 | -1 | lncRNA | ENSG0000C |
| 81976807 | 82017406 | 1 | protein_coding | ENSG0000C |
| 82018702 | 82024107 | -1 | protein_coding | ENSG0000C |
| 82023305 | 82031151 | 1 | protein_coding | ENSG0000C |
| 82031678 | 82034204 | 1 | protein_coding | ENSG0000C |
| 82035136 | 82037709 | -1 | protein_coding | ENSG0000C |
| 82035403 | 82035800 | -1 | TEC | ENSG0000C |
| 82037905 | 82039380 | 1 | lncRNA | ENSG0000C |
| 82047902 | 82051831 | -1 | protein_coding | ENSG0000C |
| 82050691 | 82057470 | 1 | protein_coding | ENSG0000C |
| 82057506 | 82065887 | -1 | protein_coding | ENSG0000C |
| 82078338 | 82098294 | -1 | protein_coding | ENSG0000C |
| 82101460 | 82212830 | -1 | protein_coding | ENSG0000C |
| 82153430 | 82154815 | 1 | lncRNA | ENSG0000C |
| 82160056 | 82160452 | 1 | lncRNA | ENSG0000C |
| 82214227 | 82217352 | 1 | lncRNA | ENSG0000C |
| 82228397 | 82261129 | 1 | protein_coding | ENSG0000C |
| 82236668 | 82236728 | 1 | miRNA | ENSG0000C |
| 82239023 | 82273731 | -1 | protein_coding | ENSG0000C |
| 82244770 | 82245591 | 1 | lncRNA | ENSG0000C |
| 82249067 | 82251844 | 1 | TEC | ENSG0000C |

| | | | | |
|---|---|---|---|---|
| 82273748 | 82279645 | 1 | lncRNA | ENSG0000( |
| 82290046 | 82292814 | -1 | lncRNA | ENSG0000( |
| 82293716 | 82294910 | 1 | lncRNA | ENSG0000( |
| 82314868 | 82317608 | -1 | protein_coding | ENSG0000( |
| 82321024 | 82334074 | -1 | protein_coding | ENSG0000( |
| 82359247 | 82363775 | 1 | protein_coding | ENSG0000( |
| 82362349 | 82363196 | -1 | TEC | ENSG0000( |
| 82374230 | 82375586 | 1 | protein_coding | ENSG0000( |
| 82381110 | 82382690 | -1 | lncRNA | ENSG0000( |
| 82389210 | 82418637 | -1 | protein_coding | ENSG0000( |
| 82400703 | 82401382 | -1 | lncRNA | ENSG0000( |
| 82417226 | 82417321 | 1 | misc_RNA | ENSG0000( |
| 82418318 | 82442645 | 1 | protein_coding | ENSG0000( |
| 82425498 | 82427310 | 1 | lncRNA | ENSG0000( |
| 82442586 | 82450829 | -1 | protein_coding | ENSG0000( |
| 82450911 | 82452213 | 1 | lncRNA | ENSG0000( |
| 82454140 | 82458521 | -1 | lncRNA | ENSG0000( |
| 82458180 | 82490537 | 1 | protein_coding | ENSG0000( |
| 82462601 | 82464255 | 1 | TEC | ENSG0000( |
| 82476597 | 82477924 | -1 | lncRNA | ENSG0000( |
| 82482098 | 82483388 | 1 | lncRNA | ENSG0000( |
| 82519713 | 82644662 | 1 | protein_coding | ENSG0000( |
| 82576616 | 82577391 | -1 | processed_pseudoge | ENSG0000( |
| 82578023 | 82578616 | 1 | lncRNA | ENSG0000( |
| 82587313 | 82588411 | -1 | lncRNA | ENSG0000( |
| 82602989 | 82604178 | -1 | lncRNA | ENSG0000( |
| 82614562 | 82648553 | -1 | protein_coding | ENSG0000( |
| 82648849 | 82650657 | 1 | TEC | ENSG0000( |
| 82654973 | 82698698 | -1 | protein_coding | ENSG0000( |
| 82668233 | 82668307 | -1 | miRNA | ENSG0000( |
| 82713908 | 82716255 | -1 | lncRNA | ENSG0000( |
| 82716706 | 82730328 | 1 | protein_coding | ENSG0000( |
| 82729164 | 82734143 | -1 | lncRNA | ENSG0000( |
| 82735615 | 82751196 | 1 | protein_coding | ENSG0000( |
| 82745068 | 82745709 | 1 | lncRNA | ENSG0000( |
| 82752065 | 82945914 | 1 | protein_coding | ENSG0000( |
| 82795486 | 82796095 | -1 | lncRNA | ENSG0000( |
| 82829434 | 82840022 | -1 | protein_coding | ENSG0000( |
| 82918282 | 82918785 | 1 | lncRNA | ENSG0000( |
| 82942155 | 83051810 | -1 | protein_coding | ENSG0000( |
| 82978525 | 82981738 | 1 | lncRNA | ENSG0000( |
| 83079609 | 83095122 | 1 | protein_coding | ENSG0000( |
| 83098377 | 83098987 | 1 | lncRNA | ENSG0000( |
| 83104255 | 83106910 | 1 | lncRNA | ENSG0000( |
| 83132647 | 83136150 | -1 | TEC | ENSG0000( |
| 83135734 | 83221442 | 1 | lncRNA | ENSG0000( |
| 83220527 | 83227721 | -1 | lncRNA | ENSG0000( |
| 83226897 | 83240804 | 1 | transcribed_processe | ENSG0000( |
| 11103 | 16352 | 1 | lncRNA | ENSG0000( |
| 14195 | 16898 | -1 | lncRNA | ENSG0000( |

| | | | | |
|---|---|---|---|---|
| 45034 | 46047 | 1 | unprocessed_pseudc | ENSG00000 |
| 47390 | 49557 | -1 | protein_coding | ENSG00000 |
| 49815 | 73545 | -1 | lncRNA | ENSG00000 |
| 80140 | 88570 | -1 | unprocessed_pseudc | ENSG00000 |
| 94202 | 94330 | -1 | processed_pseudoge | ENSG00000 |
| 109065 | 122219 | 1 | transcribed_unproce | ENSG00000 |
| 112256 | 112339 | -1 | miRNA | ENSG00000 |
| 158383 | 214629 | 1 | protein_coding | ENSG00000 |
| 214520 | 268050 | -1 | protein_coding | ENSG00000 |
| 268148 | 270278 | 1 | lncRNA | ENSG00000 |
| 316737 | 500722 | -1 | protein_coding | ENSG00000 |
| 316737 | 319165 | 1 | lncRNA | ENSG00000 |
| 423744 | 424416 | -1 | lncRNA | ENSG00000 |
| 508568 | 515294 | -1 | lncRNA | ENSG00000 |
| 580380 | 582114 | 1 | protein_coding | ENSG00000 |
| 586517 | 586850 | -1 | misc_RNA | ENSG00000 |
| 596988 | 650334 | 1 | protein_coding | ENSG00000 |
| 623743 | 658340 | -1 | lncRNA | ENSG00000 |
| 650229 | 652843 | 1 | lncRNA | ENSG00000 |
| 653985 | 657259 | -1 | lncRNA | ENSG00000 |
| 657653 | 673578 | 1 | protein_coding | ENSG00000 |
| 670318 | 712662 | -1 | protein_coding | ENSG00000 |
| 706523 | 707648 | 1 | lncRNA | ENSG00000 |
| 721588 | 812546 | -1 | protein_coding | ENSG00000 |
| 735746 | 737459 | 1 | lncRNA | ENSG00000 |
| 738058 | 739662 | -1 | lncRNA | ENSG00000 |
| 738058 | 739444 | 1 | lncRNA | ENSG00000 |
| 797989 | 798135 | -1 | snRNA | ENSG00000 |
| 813274 | 813756 | 1 | lncRNA | ENSG00000 |
| 832238 | 832469 | 1 | processed_pseudoge | ENSG00000 |
| 856526 | 857045 | 1 | processed_pseudoge | ENSG00000 |
| 894435 | 907680 | -1 | lncRNA | ENSG00000 |
| 902766 | 906667 | -1 | lncRNA | ENSG00000 |
| 904871 | 912172 | 1 | protein_coding | ENSG00000 |
| 926083 | 927993 | -1 | lncRNA | ENSG00000 |
| 962592 | 976883 | -1 | lncRNA | ENSG00000 |
| 976589 | 1310018 | 1 | lncRNA | ENSG00000 |
| 1159198 | 1160404 | -1 | processed_pseudoge | ENSG00000 |
| 1176790 | 1177012 | -1 | processed_pseudoge | ENSG00000 |
| 1254383 | 1408344 | -1 | lncRNA | ENSG00000 |
| 1323794 | 1324085 | -1 | misc_RNA | ENSG00000 |
| 1357651 | 1357942 | 1 | processed_pseudoge | ENSG00000 |
| 1362445 | 1363873 | -1 | processed_pseudoge | ENSG00000 |
| 1509022 | 2049510 | 1 | lncRNA | ENSG00000 |
| 1655177 | 1779955 | 1 | lncRNA | ENSG00000 |
| 1780329 | 1782064 | 1 | lncRNA | ENSG00000 |
| 1883524 | 2489426 | -1 | lncRNA | ENSG00000 |
| 1927727 | 1929094 | -1 | processed_pseudoge | ENSG00000 |
| 2506628 | 2511580 | -1 | lncRNA | ENSG00000 |
| 2537525 | 2571509 | -1 | protein_coding | ENSG00000 |

| | | | | |
|---|---|---|---|---|
| 2555358 | 2555492 | 1 | snoRNA | ENSG0000C |
| 2561172 | 2562220 | -1 | lncRNA | ENSG0000C |
| 2568392 | 2569122 | 1 | processed_pseudoge | ENSG0000C |
| 2571557 | 2616635 | 1 | protein_coding | ENSG0000C |
| 2593674 | 2594913 | -1 | processed_pseudoge | ENSG0000C |
| 2621090 | 2621201 | 1 | misc_RNA | ENSG0000C |
| 2649761 | 2649867 | -1 | snRNA | ENSG0000C |
| 2652170 | 2655395 | -1 | transcribed_processe | ENSG0000C |
| 2655738 | 2805017 | 1 | protein_coding | ENSG0000C |
| 2688564 | 2833065 | -1 | lncRNA | ENSG0000C |
| 2778941 | 2779057 | 1 | misc_RNA | ENSG0000C |
| 2847006 | 2916003 | 1 | protein_coding | ENSG0000C |
| 2916994 | 3013315 | -1 | protein_coding | ENSG0000C |
| 2920966 | 2921685 | 1 | lncRNA | ENSG0000C |
| 2945550 | 2946890 | -1 | transcribed_processe | ENSG0000C |
| 2948238 | 2960756 | 1 | lncRNA | ENSG0000C |
| 2967018 | 2985410 | 1 | lncRNA | ENSG0000C |
| 2990652 | 2991078 | 1 | processed_pseudoge | ENSG0000C |
| 3025069 | 3025387 | -1 | processed_pseudoge | ENSG0000C |
| 3025434 | 3025566 | -1 | snoRNA | ENSG0000C |
| 3066807 | 3220108 | -1 | protein_coding | ENSG0000C |
| 3093568 | 3093629 | 1 | snRNA | ENSG0000C |
| 3190397 | 3247277 | 1 | lncRNA | ENSG0000C |
| 3246401 | 3247086 | -1 | lncRNA | ENSG0000C |
| 3247481 | 3256236 | 1 | protein_coding | ENSG0000C |
| 3255436 | 3261850 | -1 | lncRNA | ENSG0000C |
| 3261479 | 3278431 | 1 | protein_coding | ENSG0000C |
| 3284120 | 3330984 | 1 | lncRNA | ENSG0000C |
| 3347696 | 3383532 | -1 | lncRNA | ENSG0000C |
| 3377916 | 3378247 | -1 | processed_pseudoge | ENSG0000C |
| 3394889 | 3395312 | 1 | IG_C_pseudogene | ENSG0000C |
| 3406342 | 3406818 | -1 | processed_pseudoge | ENSG0000C |
| 3411608 | 3459978 | 1 | protein_coding | ENSG0000C |
| 3415145 | 3415580 | 1 | transcribed_processe | ENSG0000C |
| 3466250 | 3478978 | 1 | lncRNA | ENSG0000C |
| 3496032 | 4455307 | -1 | protein_coding | ENSG0000C |
| 3571215 | 3571511 | -1 | misc_RNA | ENSG0000C |
| 3580169 | 3580754 | 1 | processed_pseudoge | ENSG0000C |
| 3593732 | 3598363 | 1 | lncRNA | ENSG0000C |
| 3603000 | 3608336 | 1 | lncRNA | ENSG0000C |
| 3653030 | 3656282 | 1 | lncRNA | ENSG0000C |
| 3770017 | 3771430 | -1 | lncRNA | ENSG0000C |
| 3878058 | 3897069 | 1 | lncRNA | ENSG0000C |
| 3885353 | 3885432 | 1 | miRNA | ENSG0000C |
| 3894699 | 3894805 | -1 | snRNA | ENSG0000C |
| 3962353 | 4024150 | 1 | lncRNA | ENSG0000C |
| 3977649 | 3978615 | -1 | processed_pseudoge | ENSG0000C |
| 4264602 | 4295405 | 1 | lncRNA | ENSG0000C |
| 4430945 | 4431152 | 1 | processed_pseudoge | ENSG0000C |
| 4459230 | 4459458 | 1 | processed_pseudoge | ENSG0000C |

| | | | | |
|---|---|---|---|---|
| 4775054 | 5004537 | -1 | lncRNA | ENSG00000( |
| 5002759 | 5003254 | -1 | processed_pseudoge | ENSG00000( |
| 5081171 | 5098780 | 1 | lncRNA | ENSG00000( |
| 5132749 | 5133229 | -1 | processed_pseudoge | ENSG00000( |
| 5142911 | 5197503 | -1 | protein_coding | ENSG00000( |
| 5159332 | 5171026 | 1 | lncRNA | ENSG00000( |
| 5197165 | 5215497 | 1 | lncRNA | ENSG00000( |
| 5232876 | 5238526 | -1 | lncRNA | ENSG00000( |
| 5236724 | 5238598 | -1 | lncRNA | ENSG00000( |
| 5237826 | 5290608 | 1 | lncRNA | ENSG00000( |
| 5240255 | 5241167 | -1 | processed_pseudoge | ENSG00000( |
| 5289019 | 5297053 | -1 | protein_coding | ENSG00000( |
| 5310396 | 5317664 | 1 | lncRNA | ENSG00000( |
| 5381641 | 5385330 | -1 | lncRNA | ENSG00000( |
| 5392381 | 5630700 | -1 | protein_coding | ENSG00000( |
| 5463627 | 5480975 | 1 | lncRNA | ENSG00000( |
| 5567130 | 5570942 | 1 | lncRNA | ENSG00000( |
| 5721812 | 5876328 | 1 | lncRNA | ENSG00000( |
| 5840695 | 5840833 | 1 | miRNA | ENSG00000( |
| 5882072 | 5895955 | -1 | protein_coding | ENSG00000( |
| 5887487 | 5989369 | 1 | lncRNA | ENSG00000( |
| 5895745 | 6108382 | 1 | lncRNA | ENSG00000( |
| 5954706 | 6415237 | -1 | protein_coding | ENSG00000( |
| 5956101 | 5960785 | 1 | lncRNA | ENSG00000( |
| 6017723 | 6017839 | -1 | snRNA | ENSG00000( |
| 6256747 | 6260934 | 1 | lncRNA | ENSG00000( |
| 6374361 | 6374425 | -1 | miRNA | ENSG00000( |
| 6394847 | 6398020 | -1 | TEC | ENSG00000( |
| 6454962 | 6455068 | 1 | snRNA | ENSG00000( |
| 6462144 | 6463015 | -1 | transcribed_processe | ENSG00000( |
| 6493420 | 6495668 | 1 | lncRNA | ENSG00000( |
| 6511416 | 6590653 | 1 | lncRNA | ENSG00000( |
| 6557822 | 6558654 | 1 | lncRNA | ENSG00000( |
| 6601540 | 6601836 | 1 | misc_RNA | ENSG00000( |
| 6641971 | 6642478 | -1 | lncRNA | ENSG00000( |
| 6727053 | 6728981 | 1 | TEC | ENSG00000( |
| 6728821 | 6729862 | -1 | lncRNA | ENSG00000( |
| 6729718 | 6915716 | 1 | protein_coding | ENSG00000( |
| 6784614 | 6796503 | -1 | lncRNA | ENSG00000( |
| 6873399 | 6875024 | -1 | lncRNA | ENSG00000( |
| 6919496 | 6929966 | -1 | lncRNA | ENSG00000( |
| 6926944 | 6928210 | 1 | processed_pseudoge | ENSG00000( |
| 6936912 | 6937020 | -1 | snRNA | ENSG00000( |
| 6941742 | 7117797 | -1 | protein_coding | ENSG00000( |
| 6954677 | 6957419 | 1 | lncRNA | ENSG00000( |
| 7076817 | 7080123 | 1 | lncRNA | ENSG00000( |
| 7134931 | 7135819 | 1 | processed_pseudoge | ENSG00000( |
| 7231125 | 7232047 | 1 | protein_coding | ENSG00000( |
| 7301020 | 7301153 | -1 | snoRNA | ENSG00000( |
| 7455844 | 7461135 | -1 | lncRNA | ENSG00000( |

| | | | | |
|---|---|---|---|---|
| 7566782 | 8406861 | 1 | protein_coding | ENSG0000C |
| 7741310 | 7754831 | -1 | lncRNA | ENSG0000C |
| 7814118 | 7815730 | 1 | lncRNA | ENSG0000C |
| 7995570 | 7995785 | 1 | snoRNA | ENSG0000C |
| 8012257 | 8016565 | -1 | lncRNA | ENSG0000C |
| 8133203 | 8136032 | -1 | lncRNA | ENSG0000C |
| 8154558 | 8155070 | 1 | lncRNA | ENSG0000C |
| 8360820 | 8367034 | -1 | lncRNA | ENSG0000C |
| 8402847 | 8405161 | 1 | lncRNA | ENSG0000C |
| 8406761 | 8406953 | -1 | lncRNA | ENSG0000C |
| 8412129 | 8414304 | 1 | processed_pseudoge | ENSG0000C |
| 8471831 | 8472128 | 1 | misc_RNA | ENSG0000C |
| 8486997 | 8513849 | 1 | unitary_pseudogene | ENSG0000C |
| 8545635 | 8546817 | 1 | processed_pseudoge | ENSG0000C |
| 8609437 | 8639382 | 1 | protein_coding | ENSG0000C |
| 8635179 | 8636347 | -1 | lncRNA | ENSG0000C |
| 8688022 | 8688410 | -1 | processed_pseudoge | ENSG0000C |
| 8695856 | 8707621 | -1 | lncRNA | ENSG0000C |
| 8705661 | 8832778 | 1 | protein_coding | ENSG0000C |
| 8720034 | 8720132 | -1 | misc_RNA | ENSG0000C |
| 8801656 | 8802305 | 1 | lncRNA | ENSG0000C |
| 8913999 | 8919414 | 1 | lncRNA | ENSG0000C |
| 8971624 | 8974580 | 1 | lncRNA | ENSG0000C |
| 9020029 | 9020764 | 1 | processed_pseudoge | ENSG0000C |
| 9102630 | 9134345 | 1 | protein_coding | ENSG0000C |
| 9102736 | 9254346 | 1 | lncRNA | ENSG0000C |
| 9112404 | 9115877 | -1 | lncRNA | ENSG0000C |
| 9121265 | 9136645 | -1 | lncRNA | ENSG0000C |
| 9136228 | 9285985 | 1 | protein_coding | ENSG0000C |
| 9249649 | 9249746 | -1 | misc_RNA | ENSG0000C |
| 9250110 | 9250211 | 1 | misc_RNA | ENSG0000C |
| 9259388 | 9260390 | 1 | lncRNA | ENSG0000C |
| 9310522 | 9334445 | -1 | lncRNA | ENSG0000C |
| 9334767 | 9402420 | 1 | protein_coding | ENSG0000C |
| 9335269 | 9336242 | 1 | lncRNA | ENSG0000C |
| 9473421 | 9474006 | -1 | lncRNA | ENSG0000C |
| 9475009 | 9538114 | 1 | protein_coding | ENSG0000C |
| 9506242 | 9509726 | 1 | lncRNA | ENSG0000C |
| 9518119 | 9518260 | -1 | snRNA | ENSG0000C |
| 9546791 | 9615240 | -1 | protein_coding | ENSG0000C |
| 9587386 | 9590182 | 1 | lncRNA | ENSG0000C |
| 9615264 | 9619363 | 1 | lncRNA | ENSG0000C |
| 9617964 | 9618072 | -1 | snRNA | ENSG0000C |
| 9646110 | 9648502 | 1 | lncRNA | ENSG0000C |
| 9678248 | 9679493 | -1 | processed_pseudoge | ENSG0000C |
| 9708275 | 9862551 | 1 | protein_coding | ENSG0000C |
| 9734882 | 9735602 | -1 | processed_pseudoge | ENSG0000C |
| 9785108 | 9787805 | 1 | processed_pseudoge | ENSG0000C |
| 9830330 | 9830627 | -1 | misc_RNA | ENSG0000C |
| 9840340 | 9841559 | -1 | lncRNA | ENSG0000C |

| | | | | |
|---|---|---|---|---|
| 9844717 | 9844846 | -1 | rRNA_pseudogene | ENSG00000( |
| 9880580 | 9880886 | -1 | processed_pseudoge | ENSG00000( |
| 9885726 | 9889275 | 1 | protein_coding | ENSG00000( |
| 9912316 | 9912849 | -1 | lncRNA | ENSG00000( |
| 9914002 | 9960021 | 1 | protein_coding | ENSG00000( |
| 9923920 | 9924021 | 1 | rRNA_pseudogene | ENSG00000( |
| 10124588 | 10144406 | 1 | lncRNA | ENSG00000( |
| 10297299 | 10323565 | 1 | lncRNA | ENSG00000( |
| 10321380 | 10372387 | -1 | lncRNA | ENSG00000( |
| 10375103 | 10378114 | 1 | lncRNA | ENSG00000( |
| 10380423 | 10390344 | 1 | lncRNA | ENSG00000( |
| 10405133 | 10414515 | -1 | lncRNA | ENSG00000( |
| 10406558 | 10422360 | 1 | lncRNA | ENSG00000( |
| 10454635 | 10489949 | 1 | protein_coding | ENSG00000( |
| 10525905 | 10552764 | 1 | protein_coding | ENSG00000( |
| 10594590 | 10604798 | 1 | lncRNA | ENSG00000( |
| 10612304 | 10626353 | -1 | lncRNA | ENSG00000( |
| 10622123 | 10622267 | -1 | processed_pseudoge | ENSG00000( |
| 10623876 | 10624305 | -1 | processed_pseudoge | ENSG00000( |
| 10627646 | 10628846 | -1 | lncRNA | ENSG00000( |
| 10632238 | 10632396 | 1 | unprocessed_pseudc | ENSG00000( |
| 10636670 | 10637399 | 1 | processed_pseudoge | ENSG00000( |
| 10661933 | 10666887 | 1 | lncRNA | ENSG00000( |
| 10666483 | 11148762 | -1 | protein_coding | ENSG00000( |
| 10704297 | 10709599 | 1 | lncRNA | ENSG00000( |
| 10724619 | 10728539 | 1 | transcribed_processe | ENSG00000( |
| 10759584 | 10759649 | -1 | miRNA | ENSG00000( |
| 10893617 | 10908783 | 1 | lncRNA | ENSG00000( |
| 11326270 | 11328832 | 1 | unprocessed_pseudc | ENSG00000( |
| 11366913 | 11378409 | -1 | lncRNA | ENSG00000( |
| 11390033 | 11390729 | -1 | lncRNA | ENSG00000( |
| 11447756 | 11489369 | -1 | lncRNA | ENSG00000( |
| 11454478 | 11465996 | 1 | lncRNA | ENSG00000( |
| 11486205 | 11523087 | 1 | lncRNA | ENSG00000( |
| 11552366 | 11552847 | 1 | processed_pseudoge | ENSG00000( |
| 11609596 | 11610612 | 1 | protein_coding | ENSG00000( |
| 11618208 | 11618315 | 1 | processed_pseudoge | ENSG00000( |
| 11619509 | 11639699 | -1 | transcribed_unproce | ENSG00000( |
| 11620717 | 11621158 | -1 | lncRNA | ENSG00000( |
| 11644336 | 11644824 | 1 | unprocessed_pseudc | ENSG00000( |
| 11654885 | 11654941 | -1 | miRNA | ENSG00000( |
| 11666456 | 11670165 | -1 | lncRNA | ENSG00000( |
| 11689264 | 11885685 | 1 | protein_coding | ENSG00000( |
| 11745626 | 11747581 | 1 | processed_pseudoge | ENSG00000( |
| 11758977 | 11786181 | -1 | lncRNA | ENSG00000( |
| 11810406 | 11811484 | -1 | lncRNA | ENSG00000( |
| 11851413 | 11854444 | 1 | protein_coding | ENSG00000( |
| 11851414 | 11852751 | -1 | lncRNA | ENSG00000( |
| 11857155 | 11857554 | -1 | lncRNA | ENSG00000( |
| 11882622 | 11908366 | -1 | protein_coding | ENSG00000( |

| | | | | |
|---|---|---|---|---|
| 11908712 | 11909223 | 1 | lncRNA | ENSG0000C |
| 11910634 | 11914344 | -1 | lncRNA | ENSG0000C |
| 11928838 | 11929476 | 1 | processed_pseudoge | ENSG0000C |
| 11981025 | 12030877 | 1 | protein_coding | ENSG0000C |
| 11993988 | 11994938 | -1 | lncRNA | ENSG0000C |
| 12031178 | 12032181 | -1 | lncRNA | ENSG0000C |
| 12040101 | 12040300 | -1 | processed_pseudoge | ENSG0000C |
| 12050309 | 12050672 | -1 | processed_pseudoge | ENSG0000C |
| 12056676 | 12057822 | -1 | processed_pseudoge | ENSG0000C |
| 12067173 | 12068417 | -1 | processed_pseudoge | ENSG0000C |
| 12073232 | 12076654 | -1 | transcribed_processe | ENSG0000C |
| 12081540 | 12085267 | 1 | lncRNA | ENSG0000C |
| 12091674 | 12093141 | -1 | processed_pseudoge | ENSG0000C |
| 12093843 | 12129764 | 1 | protein_coding | ENSG0000C |
| 12098427 | 12098532 | 1 | snRNA | ENSG0000C |
| 12101997 | 12102470 | -1 | processed_pseudoge | ENSG0000C |
| 12192203 | 12192934 | -1 | processed_pseudoge | ENSG0000C |
| 12199345 | 12200089 | 1 | TEC | ENSG0000C |
| 12200779 | 12201979 | 1 | lncRNA | ENSG0000C |
| 12201982 | 12223539 | 1 | transcribed_processe | ENSG0000C |
| 12211378 | 12211807 | 1 | processed_pseudoge | ENSG0000C |
| 12230411 | 12231573 | -1 | lncRNA | ENSG0000C |
| 12246589 | 12247055 | 1 | processed_pseudoge | ENSG0000C |
| 12254361 | 12277595 | 1 | protein_coding | ENSG0000C |
| 12279427 | 12282739 | 1 | TEC | ENSG0000C |
| 12288308 | 12291488 | 1 | lncRNA | ENSG0000C |
| 12304091 | 12304194 | -1 | snRNA | ENSG0000C |
| 12307669 | 12344320 | 1 | protein_coding | ENSG0000C |
| 12328944 | 12377314 | -1 | protein_coding | ENSG0000C |
| 12400412 | 12400473 | 1 | snRNA | ENSG0000C |
| 12407896 | 12432238 | 1 | protein_coding | ENSG0000C |
| 12432897 | 12437635 | 1 | lncRNA | ENSG0000C |
| 12438890 | 12448205 | 1 | lncRNA | ENSG0000C |
| 12446512 | 12658134 | -1 | protein_coding | ENSG0000C |
| 12450447 | 12451042 | 1 | processed_pseudoge | ENSG0000C |
| 12658043 | 12725740 | 1 | protein_coding | ENSG0000C |
| 12661833 | 12702777 | -1 | protein_coding | ENSG0000C |
| 12670426 | 12671145 | -1 | lncRNA | ENSG0000C |
| 12738965 | 12750068 | -1 | lncRNA | ENSG0000C |
| 12739490 | 12776140 | -1 | lncRNA | ENSG0000C |
| 12760717 | 12760818 | 1 | misc_RNA | ENSG0000C |
| 12785478 | 12929643 | -1 | protein_coding | ENSG0000C |
| 12911186 | 12912023 | 1 | processed_pseudoge | ENSG0000C |
| 12912844 | 12913906 | 1 | processed_pseudoge | ENSG0000C |
| 12947133 | 12987536 | 1 | protein_coding | ENSG0000C |
| 12984694 | 12991173 | -1 | lncRNA | ENSG0000C |
| 12991362 | 13125052 | 1 | protein_coding | ENSG0000C |
| 13139780 | 13140489 | -1 | processed_pseudoge | ENSG0000C |
| 13151637 | 13152689 | -1 | processed_pseudoge | ENSG0000C |
| 13183402 | 13185617 | -1 | lncRNA | ENSG0000C |

| | | | | |
|---|---|---|---|---|
| 13203774 | 13216367 | -1 | lncRNA | ENSG00000 |
| 13217498 | 13652755 | 1 | protein_coding | ENSG00000 |
| 13234945 | 13236470 | -1 | lncRNA | ENSG00000 |
| 13239543 | 13242076 | 1 | TEC | ENSG00000 |
| 13276427 | 13277155 | -1 | lncRNA | ENSG00000 |
| 13279415 | 13280110 | -1 | TEC | ENSG00000 |
| 13362203 | 13366243 | -1 | lncRNA | ENSG00000 |
| 13419421 | 13427480 | -1 | lncRNA | ENSG00000 |
| 13459947 | 13460026 | 1 | miRNA | ENSG00000 |
| 13461020 | 13470823 | -1 | lncRNA | ENSG00000 |
| 13471012 | 13472709 | -1 | lncRNA | ENSG00000 |
| 13486462 | 13490676 | 1 | lncRNA | ENSG00000 |
| 13500641 | 13501289 | -1 | lncRNA | ENSG00000 |
| 13514520 | 13522888 | -1 | lncRNA | ENSG00000 |
| 13526078 | 13526688 | 1 | lncRNA | ENSG00000 |
| 13561399 | 13565035 | -1 | lncRNA | ENSG00000 |
| 13611114 | 13611200 | 1 | miRNA | ENSG00000 |
| 13644815 | 13645685 | -1 | lncRNA | ENSG00000 |
| 13663347 | 13726663 | -1 | protein_coding | ENSG00000 |
| 13710249 | 13710484 | -1 | TEC | ENSG00000 |
| 13715102 | 13715368 | -1 | misc_RNA | ENSG00000 |
| 13726660 | 13764558 | 1 | protein_coding | ENSG00000 |
| 13824149 | 13827323 | 1 | protein_coding | ENSG00000 |
| 13882044 | 13915707 | -1 | protein_coding | ENSG00000 |
| 13919603 | 13919923 | -1 | processed_pseudoge | ENSG00000 |
| 14010054 | 14010917 | 1 | TEC | ENSG00000 |
| 14053951 | 14054761 | 1 | processed_pseudoge | ENSG00000 |
| 14057457 | 14132490 | -1 | protein_coding | ENSG00000 |
| 14074912 | 14075741 | -1 | unprocessed_pseudo | ENSG00000 |
| 14089935 | 14091019 | 1 | lncRNA | ENSG00000 |
| 14104542 | 14105226 | 1 | lncRNA | ENSG00000 |
| 14137710 | 14139272 | 1 | processed_pseudoge | ENSG00000 |
| 14143596 | 14151139 | -1 | processed_pseudoge | ENSG00000 |
| 14153001 | 14157342 | -1 | processed_pseudoge | ENSG00000 |
| 14179097 | 14252140 | 1 | transcribed_unproce | ENSG00000 |
| 14190700 | 14190806 | 1 | snRNA | ENSG00000 |
| 14222008 | 14222331 | 1 | processed_pseudoge | ENSG00000 |
| 14225224 | 14342505 | 1 | lncRNA | ENSG00000 |
| 14253548 | 14254075 | 1 | processed_pseudoge | ENSG00000 |
| 14337423 | 14342524 | 1 | transcribed_unproce | ENSG00000 |
| 14346214 | 14346923 | -1 | processed_pseudoge | ENSG00000 |
| 14361434 | 14361943 | -1 | processed_pseudoge | ENSG00000 |
| 14377546 | 14378757 | -1 | processed_pseudoge | ENSG00000 |
| 14391913 | 14393909 | -1 | processed_pseudoge | ENSG00000 |
| 14404551 | 14430923 | -1 | lncRNA | ENSG00000 |
| 14450216 | 14450883 | -1 | processed_pseudoge | ENSG00000 |
| 14477955 | 14499278 | -1 | transcribed_processe | ENSG00000 |
| 14485806 | 14487501 | 1 | processed_pseudoge | ENSG00000 |
| 14507339 | 14543585 | -1 | protein_coding | ENSG00000 |
| 14531676 | 14531782 | -1 | snRNA | ENSG00000 |

| | | | | |
|---|---|---|---|---|
| 14570844 | 14571788 | 1 | unprocessed_pseudc | ENSG00000( |
| 14613137 | 14614184 | 1 | unprocessed_pseudc | ENSG00000( |
| 14629832 | 14632806 | -1 | processed_pseudoge | ENSG00000( |
| 14680604 | 14681106 | -1 | unprocessed_pseudc | ENSG00000( |
| 14695422 | 14706597 | 1 | unprocessed_pseudc | ENSG00000( |
| 14728272 | 14854038 | 1 | protein_coding | ENSG00000( |
| 14759188 | 14759294 | 1 | snRNA | ENSG00000( |
| 14816151 | 14825249 | -1 | lncRNA | ENSG00000( |
| 14830166 | 14830242 | 1 | miRNA | ENSG00000( |
| 14877611 | 14884052 | 1 | lncRNA | ENSG00000( |
| 14888709 | 14890086 | -1 | unprocessed_pseudc | ENSG00000( |
| 14903580 | 14915628 | 1 | lncRNA | ENSG00000( |
| 14946267 | 14974215 | 1 | lncRNA | ENSG00000( |
| 14969001 | 14970468 | -1 | lncRNA | ENSG00000( |
| 14978739 | 14979839 | 1 | lncRNA | ENSG00000( |
| 14990707 | 14993235 | -1 | TEC | ENSG00000( |
| 15003975 | 15006352 | -1 | unprocessed_pseudc | ENSG00000( |
| 15056845 | 15066598 | -1 | lncRNA | ENSG00000( |
| 15075445 | 15122767 | 1 | lncRNA | ENSG00000( |
| 15139255 | 15150980 | -1 | lncRNA | ENSG00000( |
| 15159724 | 15164467 | -1 | lncRNA | ENSG00000( |
| 15165338 | 15172007 | 1 | lncRNA | ENSG00000( |
| 15184016 | 15189760 | -1 | lncRNA | ENSG00000( |
| 15196999 | 15197775 | -1 | transcribed_processe | ENSG00000( |
| 15247429 | 15247997 | -1 | processed_pseudoge | ENSG00000( |
| 15254418 | 15271744 | 1 | unprocessed_pseudc | ENSG00000( |
| 15307663 | 15330282 | -1 | transcribed_unproce | ENSG00000( |
| 15320840 | 15320944 | -1 | snRNA | ENSG00000( |
| 20946906 | 21111813 | -1 | protein_coding | ENSG00000( |
| 21119182 | 21119661 | -1 | processed_pseudoge | ENSG00000( |
| 21120404 | 21120506 | -1 | snRNA | ENSG00000( |
| 21182503 | 21183499 | -1 | processed_pseudoge | ENSG00000( |
| 21240675 | 21241371 | -1 | lncRNA | ENSG00000( |
| 21242242 | 21525417 | 1 | protein_coding | ENSG00000( |
| 21316878 | 21317795 | 1 | lncRNA | ENSG00000( |
| 21380286 | 21451017 | -1 | lncRNA | ENSG00000( |
| 21450653 | 21450756 | -1 | snoRNA | ENSG00000( |
| 21529284 | 21600884 | -1 | protein_coding | ENSG00000( |
| 21612314 | 21633524 | 1 | protein_coding | ENSG00000( |
| 21650901 | 21704780 | -1 | protein_coding | ENSG00000( |
| 21661787 | 21662395 | -1 | lncRNA | ENSG00000( |
| 21673628 | 21682561 | 1 | lncRNA | ENSG00000( |
| 21683510 | 21683597 | 1 | miRNA | ENSG00000( |
| 21704957 | 21870957 | 1 | protein_coding | ENSG00000( |
| 21711950 | 21712119 | -1 | snoRNA | ENSG00000( |
| 21724119 | 21724229 | 1 | misc_RNA | ENSG00000( |
| 21793883 | 21794159 | 1 | misc_RNA | ENSG00000( |
| 21817624 | 21817828 | 1 | snoRNA | ENSG00000( |
| 21825487 | 21831539 | -1 | lncRNA | ENSG00000( |
| 21825698 | 21825785 | -1 | miRNA | ENSG00000( |

| | | | | |
|---|---|---|---|---|
| 21829004 | 21829088 | -1 | miRNA | ENSG0000C |
| 21854640 | 21854987 | -1 | processed_pseudoge | ENSG0000C |
| 21871446 | 21893996 | -1 | lncRNA | ENSG0000C |
| 21879801 | 21879904 | 1 | snRNA | ENSG0000C |
| 21914369 | 21914660 | -1 | processed_pseudoge | ENSG0000C |
| 21928985 | 21929094 | -1 | rRNA_pseudogene | ENSG0000C |
| 21981121 | 22051159 | 1 | lncRNA | ENSG0000C |
| 22088060 | 22098780 | 1 | lncRNA | ENSG0000C |
| 22130603 | 22130729 | -1 | snRNA | ENSG0000C |
| 22164886 | 22169878 | -1 | lncRNA | ENSG0000C |
| 22169392 | 22202528 | 1 | protein_coding | ENSG0000C |
| 22175465 | 22176662 | -1 | lncRNA | ENSG0000C |
| 22180112 | 22180218 | -1 | snRNA | ENSG0000C |
| 22200619 | 22205229 | -1 | lncRNA | ENSG0000C |
| 22213778 | 22228029 | 1 | lncRNA | ENSG0000C |
| 22347846 | 22348252 | -1 | lncRNA | ENSG0000C |
| 22413599 | 22417915 | -1 | protein_coding | ENSG0000C |
| 22418829 | 22419294 | -1 | processed_pseudoge | ENSG0000C |
| 22586465 | 22587227 | 1 | processed_pseudoge | ENSG0000C |
| 22647582 | 22647688 | 1 | snRNA | ENSG0000C |
| 22699481 | 22933764 | -1 | lncRNA | ENSG0000C |
| 22723491 | 22907721 | -1 | lncRNA | ENSG0000C |
| 22798261 | 23026488 | 1 | protein_coding | ENSG0000C |
| 22882825 | 22883357 | -1 | lncRNA | ENSG0000C |
| 22900486 | 22900995 | -1 | processed_pseudoge | ENSG0000C |
| 22933349 | 22933438 | 1 | miRNA | ENSG0000C |
| 23004564 | 23004862 | 1 | misc_RNA | ENSG0000C |
| 23024596 | 23024703 | 1 | misc_RNA | ENSG0000C |
| 23134564 | 23260467 | 1 | protein_coding | ENSG0000C |
| 23197144 | 23437961 | -1 | protein_coding | ENSG0000C |
| 23257164 | 23260354 | -1 | lncRNA | ENSG0000C |
| 23453287 | 23486603 | 1 | protein_coding | ENSG0000C |
| 23456371 | 23456467 | 1 | misc_RNA | ENSG0000C |
| 23485000 | 23485197 | 1 | processed_pseudoge | ENSG0000C |
| 23503470 | 23531822 | 1 | protein_coding | ENSG0000C |
| 23506184 | 23586506 | -1 | protein_coding | ENSG0000C |
| 23582016 | 23582121 | -1 | misc_RNA | ENSG0000C |
| 23598926 | 23662911 | -1 | protein_coding | ENSG0000C |
| 23672253 | 23672748 | -1 | processed_pseudoge | ENSG0000C |
| 23689443 | 23956222 | 1 | protein_coding | ENSG0000C |
| 23709825 | 23710287 | -1 | processed_pseudoge | ENSG0000C |
| 23957754 | 23982556 | -1 | lncRNA | ENSG0000C |
| 23992773 | 24135610 | 1 | protein_coding | ENSG0000C |
| 23994213 | 24015339 | -1 | lncRNA | ENSG0000C |
| 24076794 | 24099320 | -1 | lncRNA | ENSG0000C |
| 24113637 | 24121003 | -1 | lncRNA | ENSG0000C |
| 24124170 | 24124285 | -1 | snRNA | ENSG0000C |
| 24135468 | 24135829 | -1 | TEC | ENSG0000C |
| 24138956 | 24161603 | 1 | protein_coding | ENSG0000C |
| 24159509 | 24162211 | -1 | lncRNA | ENSG0000C |

| | | | | |
|---|---|---|---|---|
| 24162045 | 24397880 | -1 | protein_coding | ENSG00000 |
| 24170505 | 24170623 | -1 | rRNA_pseudogene | ENSG00000 |
| 24179038 | 24179295 | -1 | misc_RNA | ENSG00000 |
| 24218300 | 24218404 | 1 | snRNA | ENSG00000 |
| 24321686 | 24321735 | 1 | miRNA | ENSG00000 |
| 24364787 | 24367984 | 1 | lncRNA | ENSG00000 |
| 24402153 | 24402714 | 1 | lncRNA | ENSG00000 |
| 24426634 | 24453531 | 1 | protein_coding | ENSG00000 |
| 24429505 | 24429609 | 1 | misc_RNA | ENSG00000 |
| 24435364 | 24439638 | 1 | lncRNA | ENSG00000 |
| 24460637 | 24479961 | 1 | protein_coding | ENSG00000 |
| 24489375 | 24491080 | 1 | lncRNA | ENSG00000 |
| 24500399 | 24508133 | 1 | lncRNA | ENSG00000 |
| 24516699 | 24544154 | -1 | lncRNA | ENSG00000 |
| 24522273 | 24522919 | -1 | processed_pseudoge | ENSG00000 |
| 24534551 | 24708542 | 1 | lncRNA | ENSG00000 |
| 24685772 | 24686351 | -1 | processed_pseudoge | ENSG00000 |
| 24689914 | 24690161 | -1 | processed_pseudoge | ENSG00000 |
| 24725762 | 24986980 | 1 | lncRNA | ENSG00000 |
| 24932771 | 24987919 | -1 | lncRNA | ENSG00000 |
| 25016434 | 25017204 | -1 | processed_pseudoge | ENSG00000 |
| 25055600 | 25056760 | 1 | lncRNA | ENSG00000 |
| 25061924 | 25352190 | -1 | protein_coding | ENSG00000 |
| 25196663 | 25218241 | 1 | lncRNA | ENSG00000 |
| 25200976 | 25202837 | -1 | TEC | ENSG00000 |
| 25302126 | 25302960 | 1 | lncRNA | ENSG00000 |
| 25818234 | 25818532 | -1 | misc_RNA | ENSG00000 |
| 25953216 | 25953468 | -1 | processed_pseudoge | ENSG00000 |
| 26016253 | 26091217 | -1 | protein_coding | ENSG00000 |
| 26098839 | 26099226 | 1 | processed_pseudoge | ENSG00000 |
| 26133852 | 26193355 | 1 | protein_coding | ENSG00000 |
| 26170726 | 26171284 | -1 | processed_pseudoge | ENSG00000 |
| 26204869 | 26205904 | 1 | processed_pseudoge | ENSG00000 |
| 26225936 | 26391685 | 1 | protein_coding | ENSG00000 |
| 26255698 | 26256355 | -1 | processed_pseudoge | ENSG00000 |
| 26266009 | 26267409 | 1 | lncRNA | ENSG00000 |
| 26348347 | 26349004 | 1 | processed_pseudoge | ENSG00000 |
| 26366301 | 26366406 | -1 | snRNA | ENSG00000 |
| 26421139 | 26438041 | 1 | lncRNA | ENSG00000 |
| 26454910 | 26657401 | -1 | protein_coding | ENSG00000 |
| 26542971 | 26545791 | -1 | lncRNA | ENSG00000 |
| 26565723 | 26575626 | -1 | lncRNA | ENSG00000 |
| 26591467 | 26591535 | -1 | miRNA | ENSG00000 |
| 26599408 | 26600338 | 1 | processed_pseudoge | ENSG00000 |
| 26655737 | 27190698 | 1 | lncRNA | ENSG00000 |
| 26685954 | 26689668 | 1 | lncRNA | ENSG00000 |
| 26687621 | 26703638 | -1 | lncRNA | ENSG00000 |
| 26689316 | 26689529 | -1 | snoRNA | ENSG00000 |
| 26822341 | 26825171 | -1 | lncRNA | ENSG00000 |
| 26831520 | 26832432 | -1 | processed_pseudoge | ENSG00000 |

| | | | | |
|---|---|---|---|---|
| 26852043 | 26865771 | -1 | protein_coding | ENSG00000( |
| 26906481 | 27185308 | -1 | protein_coding | ENSG00000( |
| 26952201 | 26952558 | -1 | processed_pseudoge | ENSG00000( |
| 27225742 | 27247188 | 1 | lncRNA | ENSG00000( |
| 27335968 | 27595164 | -1 | lncRNA | ENSG00000( |
| 27343252 | 27346110 | 1 | lncRNA | ENSG00000( |
| 27355406 | 27355764 | -1 | processed_pseudoge | ENSG00000( |
| 27375310 | 27401910 | 1 | lncRNA | ENSG00000( |
| 27418884 | 27420435 | -1 | processed_pseudoge | ENSG00000( |
| 27452514 | 27453657 | -1 | processed_pseudoge | ENSG00000( |
| 27454466 | 27469673 | 1 | lncRNA | ENSG00000( |
| 27785821 | 27795637 | 1 | lncRNA | ENSG00000( |
| 27950966 | 28177130 | -1 | protein_coding | ENSG00000( |
| 27954519 | 27963687 | 1 | lncRNA | ENSG00000( |
| 28146233 | 28146703 | -1 | lncRNA | ENSG00000( |
| 28638714 | 28789580 | 1 | lncRNA | ENSG00000( |
| 28651732 | 28653208 | 1 | processed_pseudoge | ENSG00000( |
| 28824652 | 28824826 | -1 | processed_pseudoge | ENSG00000( |
| 29048620 | 29048719 | 1 | snRNA | ENSG00000( |
| 29256817 | 29257060 | -1 | processed_pseudoge | ENSG00000( |
| 29278509 | 29361963 | -1 | lncRNA | ENSG00000( |
| 29518259 | 29548241 | 1 | lncRNA | ENSG00000( |
| 29629842 | 29650440 | 1 | lncRNA | ENSG00000( |
| 30290161 | 30414073 | 1 | lncRNA | ENSG00000( |
| 30298910 | 30298960 | 1 | miRNA | ENSG00000( |
| 30713275 | 30714286 | 1 | lncRNA | ENSG00000( |
| 30780006 | 30780424 | 1 | TEC | ENSG00000( |
| 30954820 | 30980917 | -1 | lncRNA | ENSG00000( |
| 30984106 | 30984215 | 1 | snRNA | ENSG00000( |
| 30989365 | 31042815 | -1 | protein_coding | ENSG00000( |
| 31058840 | 31102522 | -1 | protein_coding | ENSG00000( |
| 31101583 | 31162789 | 1 | lncRNA | ENSG00000( |
| 31129236 | 31162856 | -1 | protein_coding | ENSG00000( |
| 31131456 | 31132494 | 1 | lncRNA | ENSG00000( |
| 31318160 | 31359246 | 1 | protein_coding | ENSG00000( |
| 31343368 | 31426988 | -1 | lncRNA | ENSG00000( |
| 31345362 | 31345467 | -1 | snRNA | ENSG00000( |
| 31376777 | 31414912 | 1 | protein_coding | ENSG00000( |
| 31382317 | 31382430 | 1 | misc_RNA | ENSG00000( |
| 31447741 | 31478702 | 1 | protein_coding | ENSG00000( |
| 31496645 | 31497195 | -1 | lncRNA | ENSG00000( |
| 31498177 | 31549008 | 1 | protein_coding | ENSG00000( |
| 31542146 | 31556948 | -1 | lncRNA | ENSG00000( |
| 31557010 | 31599021 | 1 | protein_coding | ENSG00000( |
| 31622247 | 31685836 | -1 | protein_coding | ENSG00000( |
| 31685655 | 31686823 | 1 | lncRNA | ENSG00000( |
| 31718893 | 31719208 | 1 | misc_RNA | ENSG00000( |
| 31724198 | 31726956 | 1 | transcribed_processe | ENSG00000( |
| 31726043 | 31726397 | -1 | processed_pseudoge | ENSG00000( |
| 31759562 | 31760880 | -1 | protein_coding | ENSG00000( |

| | | | | |
|---|---|---|---|---|
| 31829180 | 31953136 | -1 | protein_coding | ENSG0000C |
| 31844388 | 31845178 | 1 | lncRNA | ENSG0000C |
| 31883288 | 31883728 | 1 | processed_pseudoge | ENSG0000C |
| 31895745 | 31896410 | -1 | lncRNA | ENSG0000C |
| 31937533 | 31937639 | 1 | snRNA | ENSG0000C |
| 31942575 | 31944156 | 1 | lncRNA | ENSG0000C |
| 31962178 | 31963618 | 1 | processed_pseudoge | ENSG0000C |
| 31966960 | 31967545 | 1 | processed_pseudoge | ENSG0000C |
| 31982899 | 31986087 | 1 | TEC | ENSG0000C |
| 32018825 | 32073219 | 1 | protein_coding | ENSG0000C |
| 32018829 | 32111779 | 1 | lncRNA | ENSG0000C |
| 32031035 | 32031359 | 1 | lncRNA | ENSG0000C |
| 32075885 | 32076272 | -1 | processed_pseudoge | ENSG0000C |
| 32091295 | 32092119 | -1 | lncRNA | ENSG0000C |
| 32091874 | 32131561 | 1 | protein_coding | ENSG0000C |
| 32122400 | 32124478 | -1 | lncRNA | ENSG0000C |
| 32124877 | 32470882 | -1 | protein_coding | ENSG0000C |
| 32176050 | 32176168 | -1 | rRNA_pseudogene | ENSG0000C |
| 32185069 | 32220404 | 1 | protein_coding | ENSG0000C |
| 32196026 | 32196966 | -1 | processed_pseudoge | ENSG0000C |
| 32245946 | 32249032 | -1 | lncRNA | ENSG0000C |
| 32263135 | 32263234 | -1 | misc_RNA | ENSG0000C |
| 32287437 | 32290340 | 1 | lncRNA | ENSG0000C |
| 32412214 | 32413176 | -1 | processed_pseudoge | ENSG0000C |
| 32470288 | 32745567 | 1 | lncRNA | ENSG0000C |
| 32511663 | 32514634 | 1 | transcribed_processe | ENSG0000C |
| 32554033 | 32566634 | -1 | lncRNA | ENSG0000C |
| 32581528 | 32582828 | 1 | lncRNA | ENSG0000C |
| 32672673 | 32773023 | -1 | protein_coding | ENSG0000C |
| 32769795 | 32774413 | 1 | lncRNA | ENSG0000C |
| 32779039 | 32779286 | 1 | processed_pseudoge | ENSG0000C |
| 32833454 | 32938316 | 1 | lncRNA | ENSG0000C |
| 32937402 | 33441101 | -1 | protein_coding | ENSG0000C |
| 32954075 | 32957998 | 1 | lncRNA | ENSG0000C |
| 33552123 | 33578187 | -1 | lncRNA | ENSG0000C |
| 33578219 | 33751195 | 1 | protein_coding | ENSG0000C |
| 33851100 | 34224952 | -1 | protein_coding | ENSG0000C |
| 34001698 | 34001799 | -1 | misc_RNA | ENSG0000C |
| 34221007 | 34221603 | -1 | TEC | ENSG0000C |
| 34222965 | 34224761 | 1 | lncRNA | ENSG0000C |
| 34225930 | 34226429 | 1 | lncRNA | ENSG0000C |
| 34493290 | 34891844 | 1 | protein_coding | ENSG0000C |
| 34573987 | 34574523 | -1 | processed_pseudoge | ENSG0000C |
| 34737795 | 34767663 | -1 | lncRNA | ENSG0000C |
| 34892013 | 34903504 | -1 | lncRNA | ENSG0000C |
| 34976928 | 35143470 | 1 | protein_coding | ENSG0000C |
| 35082009 | 35082242 | 1 | TEC | ENSG0000C |
| 35144942 | 35145417 | 1 | lncRNA | ENSG0000C |
| 35241030 | 35267133 | 1 | protein_coding | ENSG0000C |
| 35251058 | 35290245 | -1 | protein_coding | ENSG0000C |

| | | | | |
|---|---|---|---|---|
| 35257040 | 35257512 | 1 | TEC | ENSG0000C |
| 35268218 | 35270238 | 1 | lncRNA | ENSG0000C |
| 35280867 | 35290201 | -1 | lncRNA | ENSG0000C |
| 35290282 | 35328390 | 1 | transcribed_unitary_ | ENSG0000C |
| 35317396 | 35317848 | 1 | lncRNA | ENSG0000C |
| 35332227 | 35345482 | -1 | protein_coding | ENSG0000C |
| 35366694 | 35377337 | -1 | protein_coding | ENSG0000C |
| 35383354 | 35384025 | -1 | processed_pseudoge | ENSG0000C |
| 35403996 | 35406528 | -1 | lncRNA | ENSG0000C |
| 35443869 | 35467088 | -1 | lncRNA | ENSG0000C |
| 35443871 | 35497940 | -1 | protein_coding | ENSG0000C |
| 35452230 | 35497991 | -1 | protein_coding | ENSG0000C |
| 35483431 | 35483667 | -1 | TEC | ENSG0000C |
| 35581117 | 35711834 | 1 | protein_coding | ENSG0000C |
| 35591737 | 35591806 | 1 | miRNA | ENSG0000C |
| 35595901 | 35597599 | 1 | lncRNA | ENSG0000C |
| 35716370 | 35717978 | 1 | lncRNA | ENSG0000C |
| 35841759 | 35842617 | 1 | processed_pseudoge | ENSG0000C |
| 35904818 | 35904926 | -1 | miRNA | ENSG0000C |
| 35934088 | 35934142 | -1 | miRNA | ENSG0000C |
| 35951803 | 35966118 | -1 | lncRNA | ENSG0000C |
| 35972151 | 35973916 | 1 | TEC | ENSG0000C |
| 35972625 | 35979286 | 1 | protein_coding | ENSG0000C |
| 35984387 | 36067576 | -1 | protein_coding | ENSG0000C |
| 36108531 | 36129385 | -1 | protein_coding | ENSG0000C |
| 36129444 | 36180557 | 1 | protein_coding | ENSG0000C |
| 36179996 | 36187448 | -1 | lncRNA | ENSG0000C |
| 36187497 | 36272157 | 1 | protein_coding | ENSG0000C |
| 36189824 | 36190272 | 1 | lncRNA | ENSG0000C |
| 36273574 | 36274599 | 1 | processed_pseudoge | ENSG0000C |
| 36278735 | 36279119 | -1 | processed_pseudoge | ENSG0000C |
| 36297714 | 36780055 | 1 | protein_coding | ENSG0000C |
| 36417055 | 36424185 | -1 | lncRNA | ENSG0000C |
| 36777647 | 36829216 | -1 | protein_coding | ENSG0000C |
| 36829106 | 37232172 | 1 | protein_coding | ENSG0000C |
| 36901946 | 36923814 | -1 | lncRNA | ENSG0000C |
| 37234119 | 37236242 | -1 | lncRNA | ENSG0000C |
| 37243040 | 37565827 | -1 | protein_coding | ENSG0000C |
| 37243776 | 37247506 | 1 | lncRNA | ENSG0000C |
| 37273706 | 37276572 | 1 | lncRNA | ENSG0000C |
| 37305295 | 37306333 | -1 | TEC | ENSG0000C |
| 37473330 | 37473977 | -1 | TEC | ENSG0000C |
| 37565281 | 37762131 | 1 | lncRNA | ENSG0000C |
| 37657135 | 37657215 | 1 | miRNA | ENSG0000C |
| 37672689 | 37673176 | 1 | processed_pseudoge | ENSG0000C |
| 37960220 | 37992413 | 1 | lncRNA | ENSG0000C |
| 38655209 | 38688033 | -1 | lncRNA | ENSG0000C |
| 39016047 | 39016344 | -1 | misc_RNA | ENSG0000C |
| 39034007 | 39034110 | 1 | snRNA | ENSG0000C |
| 39206917 | 39800322 | -1 | lncRNA | ENSG0000C |

| | | | | |
|---|---|---|---|---|
| 39314240 | 39328008 | 1 | lncRNA | ENSG0000C |
| 39334872 | 39335672 | -1 | processed_pseudoge | ENSG0000C |
| 39622123 | 39622175 | -1 | miRNA | ENSG0000C |
| 39655645 | 39665494 | 1 | lncRNA | ENSG0000C |
| 39676719 | 39676777 | -1 | miRNA | ENSG0000C |
| 39676721 | 39676779 | 1 | miRNA | ENSG0000C |
| 39678520 | 39678622 | -1 | snRNA | ENSG0000C |
| 39841174 | 39924840 | -1 | lncRNA | ENSG0000C |
| 39841709 | 39896298 | 1 | lncRNA | ENSG0000C |
| 40062903 | 40066471 | -1 | TEC | ENSG0000C |
| 40066286 | 40099233 | 1 | lncRNA | ENSG0000C |
| 40143926 | 40144504 | -1 | processed_pseudoge | ENSG0000C |
| 40245879 | 40247367 | 1 | lncRNA | ENSG0000C |
| 41341319 | 41507550 | 1 | lncRNA | ENSG0000C |
| 41465783 | 41632185 | -1 | lncRNA | ENSG0000C |
| 41789162 | 41789395 | -1 | processed_pseudoge | ENSG0000C |
| 41819053 | 41819421 | 1 | processed_pseudoge | ENSG0000C |
| 41955234 | 42087830 | 1 | protein_coding | ENSG0000C |
| 42052705 | 42053030 | 1 | lncRNA | ENSG0000C |
| 42089312 | 42089752 | 1 | processed_pseudoge | ENSG0000C |
| 42105729 | 42115538 | -1 | lncRNA | ENSG0000C |
| 42159283 | 42691422 | 1 | lncRNA | ENSG0000C |
| 42270589 | 42270715 | 1 | rRNA_pseudogene | ENSG0000C |
| 42649520 | 42657893 | -1 | lncRNA | ENSG0000C |
| 42743227 | 43115691 | -1 | protein_coding | ENSG0000C |
| 43115747 | 43129039 | 1 | lncRNA | ENSG0000C |
| 43267892 | 43277535 | -1 | protein_coding | ENSG0000C |
| 43499173 | 43499836 | 1 | TEC | ENSG0000C |
| 44039471 | 44040140 | 1 | processed_pseudoge | ENSG0000C |
| 44071583 | 44071701 | 1 | rRNA_pseudogene | ENSG0000C |
| 44214971 | 44342315 | 1 | lncRNA | ENSG0000C |
| 44280768 | 44579953 | -1 | lncRNA | ENSG0000C |
| 44320800 | 44322257 | -1 | processed_pseudoge | ENSG0000C |
| 44522366 | 44523074 | 1 | processed_pseudoge | ENSG0000C |
| 44676927 | 44679717 | -1 | lncRNA | ENSG0000C |
| 44680173 | 45068510 | 1 | protein_coding | ENSG0000C |
| 44732575 | 44733315 | 1 | TEC | ENSG0000C |
| 44970082 | 44970166 | -1 | miRNA | ENSG0000C |
| 44986109 | 44989203 | 1 | TEC | ENSG0000C |
| 45104361 | 45181382 | -1 | lncRNA | ENSG0000C |
| 45212995 | 45683686 | 1 | protein_coding | ENSG0000C |
| 45423764 | 45483218 | -1 | lncRNA | ENSG0000C |
| 45483343 | 45529855 | 1 | lncRNA | ENSG0000C |
| 45491581 | 45503824 | -1 | lncRNA | ENSG0000C |
| 45507202 | 45550183 | 1 | lncRNA | ENSG0000C |
| 45571406 | 45611540 | -1 | lncRNA | ENSG0000C |
| 45573084 | 45747215 | -1 | lncRNA | ENSG0000C |
| 45646153 | 45647937 | 1 | lncRNA | ENSG0000C |
| 45687025 | 45752520 | 1 | protein_coding | ENSG0000C |
| 45788530 | 45788670 | 1 | processed_pseudoge | ENSG0000C |

| | | | | |
|---|---|---|---|---|
| 45825675 | 45844094 | 1 | protein_coding | ENSG0000C |
| 45847609 | 45967329 | -1 | protein_coding | ENSG0000C |
| 45983536 | 46072272 | -1 | protein_coding | ENSG0000C |
| 45989653 | 45989945 | -1 | misc_RNA | ENSG0000C |
| 46028306 | 46029829 | 1 | processed_pseudoge | ENSG0000C |
| 46077004 | 46077099 | -1 | misc_RNA | ENSG0000C |
| 46080248 | 46104334 | -1 | protein_coding | ENSG0000C |
| 46104378 | 46128333 | 1 | protein_coding | ENSG0000C |
| 46121615 | 46121721 | -1 | snRNA | ENSG0000C |
| 46173553 | 46266992 | 1 | protein_coding | ENSG0000C |
| 46326809 | 46463140 | 1 | protein_coding | ENSG0000C |
| 46385707 | 46387083 | 1 | TEC | ENSG0000C |
| 46409197 | 46410645 | -1 | lncRNA | ENSG0000C |
| 46476972 | 46657220 | -1 | protein_coding | ENSG0000C |
| 46667821 | 46759257 | -1 | protein_coding | ENSG0000C |
| 46722975 | 46723223 | 1 | processed_pseudoge | ENSG0000C |
| 46756487 | 46802449 | 1 | lncRNA | ENSG0000C |
| 46786699 | 46789297 | -1 | lncRNA | ENSG0000C |
| 46803218 | 46920160 | -1 | protein_coding | ENSG0000C |
| 46852632 | 46853882 | -1 | processed_pseudoge | ENSG0000C |
| 46915021 | 46915623 | -1 | TEC | ENSG0000C |
| 46917492 | 47102243 | 1 | protein_coding | ENSG0000C |
| 46962768 | 46964644 | -1 | protein_coding | ENSG0000C |
| 46968695 | 46969912 | -1 | processed_pseudoge | ENSG0000C |
| 47022287 | 47023927 | -1 | protein_coding | ENSG0000C |
| 47028202 | 47029842 | -1 | protein_coding | ENSG0000C |
| 47032572 | 47035621 | -1 | protein_coding | ENSG0000C |
| 47076117 | 47076594 | 1 | TEC | ENSG0000C |
| 47077361 | 47091280 | -1 | lncRNA | ENSG0000C |
| 47105946 | 47108062 | 1 | lncRNA | ENSG0000C |
| 47107408 | 47150500 | -1 | protein_coding | ENSG0000C |
| 47108188 | 47176345 | -1 | protein_coding | ENSG0000C |
| 47137018 | 47137290 | 1 | processed_pseudoge | ENSG0000C |
| 47152834 | 47176364 | -1 | protein_coding | ENSG0000C |
| 47206322 | 47251603 | -1 | protein_coding | ENSG0000C |
| 47234442 | 47234548 | -1 | snRNA | ENSG0000C |
| 47263935 | 47282939 | 1 | lncRNA | ENSG0000C |
| 47285725 | 47594550 | 1 | lncRNA | ENSG0000C |
| 47380496 | 47380565 | 1 | miRNA | ENSG0000C |
| 47501172 | 47558383 | 1 | lncRNA | ENSG0000C |
| 47501172 | 47503106 | 1 | TEC | ENSG0000C |
| 47630112 | 47630848 | 1 | processed_pseudoge | ENSG0000C |
| 47703175 | 47703745 | 1 | processed_pseudoge | ENSG0000C |
| 47808957 | 47931146 | -1 | protein_coding | ENSG0000C |
| 47853233 | 47853457 | -1 | processed_pseudoge | ENSG0000C |
| 47866839 | 47867367 | 1 | TEC | ENSG0000C |
| 47878295 | 47882444 | 1 | lncRNA | ENSG0000C |
| 47912207 | 47912504 | -1 | TEC | ENSG0000C |
| 48026672 | 48410752 | -1 | protein_coding | ENSG0000C |
| 48147634 | 48150864 | -1 | TEC | ENSG0000C |

| | | | | |
|---|---|---|---|---|
| 48162039 | 48184725 | 1 | lncRNA | ENSG0000C |
| 48169784 | 48169890 | -1 | snRNA | ENSG0000C |
| 48252301 | 48252837 | 1 | TEC | ENSG0000C |
| 48475487 | 48475600 | -1 | rRNA_pseudogene | ENSG0000C |
| 48478282 | 48478491 | -1 | processed_pseudoge | ENSG0000C |
| 48478561 | 48479228 | 1 | processed_pseudoge | ENSG0000C |
| 48539031 | 48863217 | 1 | protein_coding | ENSG0000C |
| 48564795 | 48568342 | 1 | TEC | ENSG0000C |
| 48661840 | 48662635 | -1 | TEC | ENSG0000C |
| 48670600 | 48670668 | 1 | miRNA | ENSG0000C |
| 48673575 | 48688419 | -1 | lncRNA | ENSG0000C |
| 48755693 | 48756318 | 1 | TEC | ENSG0000C |
| 48826051 | 48834770 | -1 | lncRNA | ENSG0000C |
| 48919853 | 48952052 | -1 | protein_coding | ENSG0000C |
| 48962951 | 48963941 | -1 | lncRNA | ENSG0000C |
| 49013106 | 49013819 | 1 | TEC | ENSG0000C |
| 49023703 | 49048474 | 1 | lncRNA | ENSG0000C |
| 49041474 | 49461347 | -1 | protein_coding | ENSG0000C |
| 49049687 | 49049768 | -1 | miRNA | ENSG0000C |
| 49116301 | 49126479 | 1 | lncRNA | ENSG0000C |
| 49205912 | 49208781 | 1 | lncRNA | ENSG0000C |
| 49431744 | 49447420 | -1 | lncRNA | ENSG0000C |
| 49447464 | 49451827 | 1 | lncRNA | ENSG0000C |
| 49469803 | 49470223 | 1 | processed_pseudoge | ENSG0000C |
| 49477250 | 49487252 | -1 | protein_coding | ENSG0000C |
| 49481681 | 49492479 | -1 | protein_coding | ENSG0000C |
| 49484536 | 49486149 | -1 | lncRNA | ENSG0000C |
| 49487339 | 49491878 | 1 | lncRNA | ENSG0000C |
| 49487373 | 49487422 | 1 | miRNA | ENSG0000C |
| 49488453 | 49492523 | -1 | protein_coding | ENSG0000C |
| 49489245 | 49489308 | -1 | snoRNA | ENSG0000C |
| 49491283 | 49491347 | -1 | snoRNA | ENSG0000C |
| 49491664 | 49491729 | -1 | snoRNA | ENSG0000C |
| 49495147 | 49496896 | -1 | processed_pseudoge | ENSG0000C |
| 49499390 | 49501860 | 1 | lncRNA | ENSG0000C |
| 49560699 | 49599185 | 1 | protein_coding | ENSG0000C |
| 49650350 | 49651160 | 1 | processed_pseudoge | ENSG0000C |
| 49739823 | 49742063 | -1 | lncRNA | ENSG0000C |
| 49782164 | 49813953 | -1 | protein_coding | ENSG0000C |
| 49795793 | 49837623 | -1 | protein_coding | ENSG0000C |
| 49814023 | 49851059 | 1 | lncRNA | ENSG0000C |
| 49814133 | 49814276 | 1 | scaRNA | ENSG0000C |
| 49814361 | 49814443 | 1 | scaRNA | ENSG0000C |
| 49822789 | 50195147 | -1 | protein_coding | ENSG0000C |
| 49871555 | 49871666 | -1 | misc_RNA | ENSG0000C |
| 49946266 | 49946389 | -1 | rRNA_pseudogene | ENSG0000C |
| 49969721 | 49970957 | -1 | lncRNA | ENSG0000C |
| 50063795 | 50064440 | 1 | processed_pseudoge | ENSG0000C |
| 50098742 | 50099162 | 1 | processed_pseudoge | ENSG0000C |
| 50126499 | 50126563 | -1 | miRNA | ENSG0000C |

| | | | | |
|---|---|---|---|---|
| 50206998 | 50207278 | -1 | misc_RNA | ENSG00000 |
| 50227193 | 50266495 | -1 | protein_coding | ENSG00000 |
| 50256036 | 50256461 | -1 | lncRNA | ENSG00000 |
| 50266882 | 50281774 | -1 | protein_coding | ENSG00000 |
| 50282343 | 50288304 | -1 | protein_coding | ENSG00000 |
| 50302190 | 50313595 | 1 | lncRNA | ENSG00000 |
| 50318420 | 50319024 | 1 | processed_pseudoge | ENSG00000 |
| 50319002 | 50319120 | 1 | rRNA_pseudogene | ENSG00000 |
| 50375040 | 50394168 | 1 | protein_coding | ENSG00000 |
| 50457887 | 50458795 | 1 | processed_pseudoge | ENSG00000 |
| 50560087 | 50731826 | 1 | protein_coding | ENSG00000 |
| 50647315 | 50651212 | -1 | lncRNA | ENSG00000 |
| 50669510 | 50670215 | 1 | transcribed_process | ENSG00000 |
| 50795120 | 50825402 | -1 | protein_coding | ENSG00000 |
| 50814392 | 50815730 | 1 | processed_pseudoge | ENSG00000 |
| 50823577 | 50824120 | 1 | processed_pseudoge | ENSG00000 |
| 50879080 | 50954257 | 1 | protein_coding | ENSG00000 |
| 50959267 | 50959379 | -1 | misc_RNA | ENSG00000 |
| 50967991 | 50988121 | 1 | protein_coding | ENSG00000 |
| 50968019 | 51058144 | 1 | protein_coding | ENSG00000 |
| 51028394 | 51085045 | 1 | protein_coding | ENSG00000 |
| 51158282 | 51159383 | -1 | processed_pseudoge | ENSG00000 |
| 51174550 | 51218304 | -1 | protein_coding | ENSG00000 |
| 51283733 | 51283896 | -1 | snRNA | ENSG00000 |
| 51346249 | 51643939 | 1 | lncRNA | ENSG00000 |
| 51421063 | 51421674 | -1 | processed_pseudoge | ENSG00000 |
| 51433437 | 51434783 | 1 | processed_pseudoge | ENSG00000 |
| 51612639 | 51613110 | -1 | processed_pseudoge | ENSG00000 |
| 51687525 | 51688209 | -1 | processed_pseudoge | ENSG00000 |
| 51845007 | 51845628 | -1 | processed_pseudoge | ENSG00000 |
| 52223078 | 52223688 | 1 | lncRNA | ENSG00000 |
| 52283909 | 52309224 | 1 | lncRNA | ENSG00000 |
| 52336361 | 52339025 | -1 | lncRNA | ENSG00000 |
| 52340197 | 53535903 | 1 | protein_coding | ENSG00000 |
| 52409042 | 52409257 | 1 | TEC | ENSG00000 |
| 52419189 | 52419644 | -1 | unprocessed_pseudo | ENSG00000 |
| 53237101 | 53237190 | 1 | miRNA | ENSG00000 |
| 53480825 | 53581107 | -1 | lncRNA | ENSG00000 |
| 53568439 | 53629990 | 1 | lncRNA | ENSG00000 |
| 53605254 | 53605725 | -1 | processed_pseudoge | ENSG00000 |
| 54142133 | 54142577 | 1 | processed_pseudoge | ENSG00000 |
| 54151606 | 54224669 | -1 | protein_coding | ENSG00000 |
| 54222284 | 54222412 | -1 | snoRNA | ENSG00000 |
| 54268346 | 54270028 | -1 | lncRNA | ENSG00000 |
| 54269404 | 54321266 | 1 | protein_coding | ENSG00000 |
| 54324358 | 54357964 | -1 | protein_coding | ENSG00000 |
| 54357906 | 54385218 | 1 | protein_coding | ENSG00000 |
| 54406889 | 54407114 | -1 | processed_pseudoge | ENSG00000 |
| 54412763 | 54424632 | 1 | unitary_pseudogene | ENSG00000 |
| 54525434 | 54525563 | 1 | processed_pseudoge | ENSG00000 |

| | | | | |
|---|---|---|---|---|
| 54563082 | 54567303 | 1 | unprocessed_pseudc | ENSG0000C |
| 54578541 | 54578619 | 1 | unprocessed_pseudc | ENSG0000C |
| 54587757 | 54599493 | 1 | protein_coding | ENSG0000C |
| 54717860 | 54895516 | 1 | protein_coding | ENSG0000C |
| 54744750 | 54745581 | 1 | processed_pseudoge | ENSG0000C |
| 54879457 | 54880809 | 1 | lncRNA | ENSG0000C |
| 54885866 | 54898083 | -1 | lncRNA | ENSG0000C |
| 54901509 | 54959461 | -1 | protein_coding | ENSG0000C |
| 54954403 | 54954732 | 1 | processed_pseudoge | ENSG0000C |
| 55105904 | 55124306 | -1 | lncRNA | ENSG0000C |
| 55146546 | 55146645 | -1 | rRNA_pseudogene | ENSG0000C |
| 55222185 | 55664787 | -1 | protein_coding | ENSG0000C |
| 55452533 | 55482940 | 1 | lncRNA | ENSG0000C |
| 55479221 | 55479298 | 1 | miRNA | ENSG0000C |
| 55492175 | 55496392 | 1 | lncRNA | ENSG0000C |
| 55589904 | 55839184 | 1 | lncRNA | ENSG0000C |
| 55635833 | 55636310 | 1 | processed_pseudoge | ENSG0000C |
| 55776727 | 55781722 | -1 | lncRNA | ENSG0000C |
| 55817167 | 55819759 | 1 | lncRNA | ENSG0000C |
| 55863960 | 55866550 | 1 | lncRNA | ENSG0000C |
| 55881688 | 55998219 | -1 | lncRNA | ENSG0000C |
| 55893038 | 56026082 | 1 | lncRNA | ENSG0000C |
| 56003613 | 56191262 | -1 | lncRNA | ENSG0000C |
| 56026388 | 56029186 | 1 | lncRNA | ENSG0000C |
| 56031830 | 56135295 | 1 | lncRNA | ENSG0000C |
| 56061109 | 56063078 | -1 | lncRNA | ENSG0000C |
| 56079394 | 56079594 | -1 | snoRNA | ENSG0000C |
| 56083356 | 56137536 | -1 | lncRNA | ENSG0000C |
| 56137573 | 56150102 | 1 | lncRNA | ENSG0000C |
| 56597209 | 56651600 | -1 | protein_coding | ENSG0000C |
| 56651343 | 57036606 | 1 | protein_coding | ENSG0000C |
| 56753226 | 56753326 | -1 | misc_RNA | ENSG0000C |
| 56872668 | 56873970 | -1 | lncRNA | ENSG0000C |
| 56957270 | 56957469 | 1 | snoRNA | ENSG0000C |
| 57027832 | 57038845 | 1 | lncRNA | ENSG0000C |
| 57046406 | 57047692 | 1 | lncRNA | ENSG0000C |
| 57054558 | 57072119 | -1 | lncRNA | ENSG0000C |
| 57136322 | 57146998 | -1 | lncRNA | ENSG0000C |
| 57147087 | 57150410 | 1 | protein_coding | ENSG0000C |
| 57169615 | 57171194 | -1 | lncRNA | ENSG0000C |
| 57284602 | 57284708 | -1 | snRNA | ENSG0000C |
| 57352557 | 57371731 | 1 | protein_coding | ENSG0000C |
| 57427133 | 57432112 | 1 | lncRNA | ENSG0000C |
| 57435374 | 57491298 | 1 | protein_coding | ENSG0000C |
| 57544389 | 57586702 | -1 | protein_coding | ENSG0000C |
| 57588611 | 57589091 | -1 | lncRNA | ENSG0000C |
| 57600656 | 57622213 | -1 | protein_coding | ENSG0000C |
| 57630302 | 57669296 | 1 | lncRNA | ENSG0000C |
| 57639455 | 57738603 | 1 | lncRNA | ENSG0000C |
| 57646426 | 57803315 | -1 | protein_coding | ENSG0000C |

| | | | | |
|---|---|---|---|---|
| 57755394 | 57755500 | 1 | snRNA | ENSG0000C |
| 57838476 | 57838966 | 1 | processed_pseudoge | ENSG0000C |
| 57961231 | 57964434 | -1 | lncRNA | ENSG0000C |
| 58018794 | 58019061 | -1 | processed_pseudoge | ENSG0000C |
| 58044226 | 58401540 | 1 | protein_coding | ENSG0000C |
| 58189834 | 58195696 | -1 | lncRNA | ENSG0000C |
| 58371566 | 58372666 | 1 | lncRNA | ENSG0000C |
| 58398663 | 58400082 | -1 | lncRNA | ENSG0000C |
| 58416765 | 58417628 | 1 | lncRNA | ENSG0000C |
| 58445788 | 58453766 | 1 | lncRNA | ENSG0000C |
| 58451074 | 58451158 | 1 | miRNA | ENSG0000C |
| 58451080 | 58451152 | -1 | miRNA | ENSG0000C |
| 58476191 | 58477484 | 1 | lncRNA | ENSG0000C |
| 58481247 | 58629091 | -1 | protein_coding | ENSG0000C |
| 58511598 | 58512310 | -1 | lncRNA | ENSG0000C |
| 58535415 | 58538552 | 1 | lncRNA | ENSG0000C |
| 58557070 | 58566677 | 1 | lncRNA | ENSG0000C |
| 58600631 | 58600698 | 1 | snoRNA | ENSG0000C |
| 58631259 | 58632120 | 1 | processed_pseudoge | ENSG0000C |
| 58659858 | 58660524 | 1 | lncRNA | ENSG0000C |
| 58670009 | 58671877 | -1 | lncRNA | ENSG0000C |
| 58671465 | 58754477 | 1 | protein_coding | ENSG0000C |
| 58672613 | 58752730 | -1 | lncRNA | ENSG0000C |
| 58690356 | 58692153 | 1 | processed_pseudoge | ENSG0000C |
| 58752179 | 58753898 | -1 | lncRNA | ENSG0000C |
| 58754859 | 58757842 | -1 | lncRNA | ENSG0000C |
| 58805268 | 58805374 | 1 | snRNA | ENSG0000C |
| 58808468 | 58834364 | 1 | lncRNA | ENSG0000C |
| 58818883 | 58819016 | -1 | snoRNA | ENSG0000C |
| 58862600 | 58986480 | 1 | protein_coding | ENSG0000C |
| 58926197 | 58926296 | -1 | snRNA | ENSG0000C |
| 58996734 | 59069338 | 1 | transcribed_unitary_ | ENSG0000C |
| 59081911 | 59085920 | 1 | lncRNA | ENSG0000C |
| 59139866 | 59158832 | 1 | protein_coding | ENSG0000C |
| 59220158 | 59230774 | 1 | protein_coding | ENSG0000C |
| 59267035 | 59274086 | -1 | protein_coding | ENSG0000C |
| 59275156 | 59318649 | -1 | protein_coding | ENSG0000C |
| 59327823 | 59359265 | -1 | protein_coding | ENSG0000C |
| 59360700 | 59386749 | 1 | lncRNA | ENSG0000C |
| 59430939 | 59697662 | -1 | protein_coding | ENSG0000C |
| 59459072 | 59465682 | 1 | lncRNA | ENSG0000C |
| 59563709 | 59568615 | 1 | lncRNA | ENSG0000C |
| 59685820 | 59688400 | 1 | lncRNA | ENSG0000C |
| 59696459 | 59698111 | 1 | lncRNA | ENSG0000C |
| 59700523 | 59700946 | 1 | lncRNA | ENSG0000C |
| 59761558 | 59761905 | 1 | processed_pseudoge | ENSG0000C |
| 59778572 | 59778989 | 1 | processed_pseudoge | ENSG0000C |
| 59821040 | 59825261 | 1 | transcribed_processe | ENSG0000C |
| 59899948 | 59904306 | 1 | protein_coding | ENSG0000C |
| 59929226 | 59930330 | 1 | processed_pseudoge | ENSG0000C |

| | | | | |
|---|---|---|---|---|
| 59969438 | 59971396 | 1 | processed_pseudoge | ENSG0000C |
| 59972914 | 59973207 | 1 | misc_RNA | ENSG0000C |
| 60009994 | 60011143 | -1 | processed_pseudoge | ENSG0000C |
| 60010281 | 60014258 | 1 | lncRNA | ENSG0000C |
| 60016778 | 60017441 | -1 | processed_pseudoge | ENSG0000C |
| 60018625 | 60018731 | 1 | snRNA | ENSG0000C |
| 60125033 | 60161195 | 1 | lncRNA | ENSG0000C |
| 60149576 | 60150365 | 1 | processed_pseudoge | ENSG0000C |
| 60163567 | 60163707 | 1 | snRNA | ENSG0000C |
| 60196555 | 60198172 | 1 | processed_pseudoge | ENSG0000C |
| 60294136 | 60301317 | 1 | lncRNA | ENSG0000C |
| 60329190 | 60540205 | 1 | lncRNA | ENSG0000C |
| 60371062 | 60372775 | -1 | protein_coding | ENSG0000C |
| 60495868 | 60496336 | -1 | processed_pseudoge | ENSG0000C |
| 60519841 | 60520165 | 1 | processed_pseudoge | ENSG0000C |
| 60525616 | 60526156 | -1 | processed_pseudoge | ENSG0000C |
| 60611773 | 60654623 | 1 | transcribed_unproce | ENSG0000C |
| 60628963 | 60902726 | -1 | lncRNA | ENSG0000C |
| 60663115 | 60664369 | 1 | processed_pseudoge | ENSG0000C |
| 60794613 | 60794916 | -1 | processed_pseudoge | ENSG0000C |
| 61333430 | 61555779 | 1 | protein_coding | ENSG0000C |
| 61391595 | 61391703 | 1 | snRNA | ENSG0000C |
| 61404858 | 61405205 | -1 | processed_pseudoge | ENSG0000C |
| 61571342 | 61579456 | -1 | lncRNA | ENSG0000C |
| 61585746 | 61607006 | -1 | lncRNA | ENSG0000C |
| 61592375 | 61748832 | -1 | lncRNA | ENSG0000C |
| 61681653 | 61682695 | 1 | lncRNA | ENSG0000C |
| 61748082 | 61758464 | 1 | lncRNA | ENSG0000C |
| 61808067 | 61894247 | -1 | protein_coding | ENSG0000C |
| 61905255 | 62187118 | -1 | protein_coding | ENSG0000C |
| 61957273 | 61958014 | -1 | processed_pseudoge | ENSG0000C |
| 62187255 | 62310217 | 1 | protein_coding | ENSG0000C |
| 62245810 | 62247412 | 1 | lncRNA | ENSG0000C |
| 62300036 | 62300998 | -1 | lncRNA | ENSG0000C |
| 62325287 | 62391288 | 1 | protein_coding | ENSG0000C |
| 62330812 | 62330913 | 1 | misc_RNA | ENSG0000C |
| 62372894 | 62378130 | -1 | lncRNA | ENSG0000C |
| 62415678 | 62416227 | 1 | processed_pseudoge | ENSG0000C |
| 62418318 | 62418576 | 1 | processed_pseudoge | ENSG0000C |
| 62442028 | 62443126 | 1 | processed_pseudoge | ENSG0000C |
| 62523025 | 62587709 | 1 | protein_coding | ENSG0000C |
| 62525919 | 62526325 | -1 | lncRNA | ENSG0000C |
| 62551203 | 62552505 | 1 | TEC | ENSG0000C |
| 62650308 | 62650601 | -1 | misc_RNA | ENSG0000C |
| 62715541 | 62980433 | 1 | protein_coding | ENSG0000C |
| 62731226 | 62731332 | -1 | snRNA | ENSG0000C |
| 63123346 | 63320128 | -1 | protein_coding | ENSG0000C |
| 63151114 | 63151320 | -1 | lncRNA | ENSG0000C |
| 63207622 | 63208168 | 1 | lncRNA | ENSG0000C |
| 63327726 | 63367228 | -1 | protein_coding | ENSG0000C |

| | | | | |
|---|---|---|---|---|
| 63367328 | 63381629 | 1 | lncRNA | ENSG0000C |
| 63389190 | 63422483 | -1 | protein_coding | ENSG0000C |
| 63415317 | 63416469 | -1 | processed_pseudoge | ENSG0000C |
| 63476958 | 63505085 | 1 | protein_coding | ENSG0000C |
| 63485601 | 63486018 | 1 | lncRNA | ENSG0000C |
| 63496989 | 63497400 | 1 | processed_pseudoge | ENSG0000C |
| 63556160 | 63567011 | 1 | protein_coding | ENSG0000C |
| 63586989 | 63604639 | 1 | protein_coding | ENSG0000C |
| 63637259 | 63644256 | -1 | protein_coding | ENSG0000C |
| 63647579 | 63726432 | 1 | protein_coding | ENSG0000C |
| 63655197 | 63661893 | -1 | protein_coding | ENSG0000C |
| 63752935 | 63805376 | 1 | protein_coding | ENSG0000C |
| 63871692 | 63903888 | 1 | protein_coding | ENSG0000C |
| 63897174 | 63936111 | 1 | protein_coding | ENSG0000C |
| 63949301 | 63981774 | 1 | protein_coding | ENSG0000C |
| 63966406 | 63970181 | -1 | lncRNA | ENSG0000C |
| 63970029 | 64019779 | 1 | protein_coding | ENSG0000C |
| 64041555 | 64423601 | 1 | lncRNA | ENSG0000C |
| 64072158 | 64072653 | 1 | transcribed_processe | ENSG0000C |
| 64079989 | 64149088 | -1 | lncRNA | ENSG0000C |
| 64181314 | 64260055 | -1 | lncRNA | ENSG0000C |
| 64677796 | 64677993 | -1 | processed_pseudoge | ENSG0000C |
| 64872264 | 64992828 | -1 | lncRNA | ENSG0000C |
| 65105873 | 65125710 | -1 | lncRNA | ENSG0000C |
| 65186159 | 65187018 | 1 | lncRNA | ENSG0000C |
| 65423958 | 65448167 | 1 | lncRNA | ENSG0000C |
| 65606079 | 65652053 | 1 | lncRNA | ENSG0000C |
| 65750252 | 65890337 | 1 | protein_coding | ENSG0000C |
| 65865262 | 65866397 | 1 | lncRNA | ENSG0000C |
| 65917782 | 65920942 | -1 | lncRNA | ENSG0000C |
| 66068750 | 66069647 | -1 | processed_pseudoge | ENSG0000C |
| 66501083 | 66604138 | -1 | protein_coding | ENSG0000C |
| 66946116 | 66946222 | 1 | snRNA | ENSG0000C |
| 67081584 | 67081686 | 1 | miRNA | ENSG0000C |
| 67137611 | 67137751 | -1 | processed_pseudoge | ENSG0000C |
| 67375133 | 67375305 | 1 | processed_pseudoge | ENSG0000C |
| 67481791 | 67484966 | -1 | lncRNA | ENSG0000C |
| 67506587 | 67516720 | -1 | protein_coding | ENSG0000C |
| 67506589 | 67514030 | 1 | lncRNA | ENSG0000C |
| 67516546 | 67899619 | 1 | lncRNA | ENSG0000C |
| 67602692 | 67603022 | -1 | processed_pseudoge | ENSG0000C |
| 67672221 | 67767051 | -1 | lncRNA | ENSG0000C |
| 67707929 | 67708246 | -1 | processed_pseudoge | ENSG0000C |
| 67788268 | 67791035 | -1 | processed_pseudoge | ENSG0000C |
| 67808505 | 67814822 | -1 | lncRNA | ENSG0000C |
| 67830671 | 67831967 | -1 | processed_pseudoge | ENSG0000C |
| 67940949 | 67956426 | 1 | lncRNA | ENSG0000C |
| 68102087 | 68114878 | 1 | lncRNA | ENSG0000C |
| 68237100 | 68495133 | -1 | lncRNA | ENSG0000C |
| 68258080 | 68264827 | -1 | lncRNA | ENSG0000C |

| | | | | |
|---|---|---|---|---|
| 68277802 | 68278243 | -1 | processed_pseudoge | ENSG00000 |
| 68427030 | 68436918 | 1 | lncRNA | ENSG00000 |
| 68455951 | 68456877 | 1 | processed_pseudoge | ENSG00000 |
| 68527970 | 68528123 | 1 | processed_pseudoge | ENSG00000 |
| 68673688 | 68715108 | -1 | protein_coding | ENSG00000 |
| 68715209 | 69055189 | 1 | protein_coding | ENSG00000 |
| 68721082 | 68721560 | 1 | lncRNA | ENSG00000 |
| 68753179 | 68754583 | -1 | lncRNA | ENSG00000 |
| 68858934 | 68859040 | 1 | snRNA | ENSG00000 |
| 68866517 | 68867028 | 1 | processed_pseudoge | ENSG00000 |
| 68899781 | 68902255 | -1 | lncRNA | ENSG00000 |
| 68908048 | 68908599 | 1 | lncRNA | ENSG00000 |
| 68992718 | 68993375 | -1 | processed_pseudoge | ENSG00000 |
| 69212277 | 69214350 | 1 | lncRNA | ENSG00000 |
| 69240448 | 69250781 | -1 | lncRNA | ENSG00000 |
| 69398726 | 69399249 | -1 | lncRNA | ENSG00000 |
| 69400888 | 69849087 | 1 | protein_coding | ENSG00000 |
| 69469517 | 69471736 | 1 | transcribed_processe | ENSG00000 |
| 69483627 | 69618694 | -1 | lncRNA | ENSG00000 |
| 69659837 | 69663413 | 1 | TEC | ENSG00000 |
| 69704978 | 69724975 | -1 | lncRNA | ENSG00000 |
| 69831158 | 69961803 | -1 | protein_coding | ENSG00000 |
| 70003031 | 70205726 | -1 | protein_coding | ENSG00000 |
| 70205774 | 70208781 | 1 | lncRNA | ENSG00000 |
| 70289045 | 70330199 | 1 | protein_coding | ENSG00000 |
| 70335439 | 70352459 | 1 | lncRNA | ENSG00000 |
| 70335929 | 70337310 | -1 | lncRNA | ENSG00000 |
| 70380140 | 70384591 | -1 | lncRNA | ENSG00000 |
| 70412055 | 70412391 | 1 | processed_pseudoge | ENSG00000 |
| 70430231 | 70431125 | 1 | processed_pseudoge | ENSG00000 |
| 70489630 | 70491205 | 1 | TEC | ENSG00000 |
| 70497923 | 70554100 | -1 | lncRNA | ENSG00000 |
| 70512727 | 70513008 | 1 | misc_RNA | ENSG00000 |
| 70630534 | 70650744 | -1 | lncRNA | ENSG00000 |
| 71000477 | 71011490 | -1 | lncRNA | ENSG00000 |
| 71028770 | 71033056 | 1 | processed_pseudoge | ENSG00000 |
| 71137829 | 71208366 | 1 | lncRNA | ENSG00000 |
| 71215618 | 71237158 | -1 | lncRNA | ENSG00000 |
| 71225704 | 71226721 | 1 | processed_pseudoge | ENSG00000 |
| 71519962 | 71578956 | -1 | lncRNA | ENSG00000 |
| 71588537 | 71588884 | 1 | processed_pseudoge | ENSG00000 |
| 71665148 | 71665525 | 1 | processed_pseudoge | ENSG00000 |
| 71732615 | 71788128 | -1 | lncRNA | ENSG00000 |
| 71837039 | 71870788 | 1 | lncRNA | ENSG00000 |
| 71890409 | 71944577 | -1 | lncRNA | ENSG00000 |
| 71932493 | 71932796 | 1 | processed_pseudoge | ENSG00000 |
| 72536680 | 72638521 | -1 | protein_coding | ENSG00000 |
| 72618267 | 72619222 | 1 | processed_pseudoge | ENSG00000 |
| 72742314 | 72868146 | -1 | protein_coding | ENSG00000 |
| 72743756 | 72745961 | -1 | lncRNA | ENSG00000 |

| | | | | |
|---|---|---|---|---|
| 72751952 | 72752051 | 1 | rRNA_pseudogene | ENSG0000C |
| 72853321 | 72853382 | -1 | miRNA | ENSG0000C |
| 72868388 | 72881399 | 1 | lncRNA | ENSG0000C |
| 73151241 | 73264480 | -1 | lncRNA | ENSG0000C |
| 73324941 | 73349879 | 1 | lncRNA | ENSG0000C |
| 73711329 | 73715490 | -1 | lncRNA | ENSG0000C |
| 73744935 | 73745232 | 1 | misc_RNA | ENSG0000C |
| 73914405 | 74034161 | -1 | lncRNA | ENSG0000C |
| 73921939 | 73922631 | -1 | TEC | ENSG0000C |
| 73947880 | 73957974 | 1 | lncRNA | ENSG0000C |
| 74073353 | 74147843 | -1 | protein_coding | ENSG0000C |
| 74148523 | 74160531 | 1 | protein_coding | ENSG0000C |
| 74211391 | 74240555 | -1 | lncRNA | ENSG0000C |
| 74227506 | 74227803 | 1 | misc_RNA | ENSG0000C |
| 74250846 | 74291973 | -1 | protein_coding | ENSG0000C |
| 74293217 | 74298662 | 1 | lncRNA | ENSG0000C |
| 74315839 | 74359189 | 1 | protein_coding | ENSG0000C |
| 74389884 | 74390297 | -1 | processed_pseudoge | ENSG0000C |
| 74409415 | 74413209 | 1 | lncRNA | ENSG0000C |
| 74434099 | 74457944 | -1 | protein_coding | ENSG0000C |
| 74464410 | 74468707 | -1 | lncRNA | ENSG0000C |
| 74495816 | 74523454 | 1 | protein_coding | ENSG0000C |
| 74504766 | 74505248 | 1 | lncRNA | ENSG0000C |
| 74534500 | 74587212 | 1 | protein_coding | ENSG0000C |
| 74561381 | 74562100 | -1 | lncRNA | ENSG0000C |
| 74590317 | 74598885 | -1 | lncRNA | ENSG0000C |
| 74597870 | 75065671 | 1 | protein_coding | ENSG0000C |
| 75070197 | 75071091 | -1 | lncRNA | ENSG0000C |
| 75073543 | 75074205 | 1 | lncRNA | ENSG0000C |
| 75086368 | 75086514 | 1 | sRNA | ENSG0000C |
| 75103587 | 75106260 | -1 | lncRNA | ENSG0000C |
| 75112137 | 75125565 | -1 | lncRNA | ENSG0000C |
| 75195113 | 75209139 | -1 | protein_coding | ENSG0000C |
| 75210755 | 75289950 | 1 | protein_coding | ENSG0000C |
| 75387056 | 75387596 | 1 | lncRNA | ENSG0000C |
| 75407012 | 75426607 | 1 | lncRNA | ENSG0000C |
| 75409476 | 75427703 | -1 | protein_coding | ENSG0000C |
| 75432703 | 75436957 | -1 | lncRNA | ENSG0000C |
| 75455787 | 75468835 | 1 | lncRNA | ENSG0000C |
| 75472794 | 75474772 | 1 | lncRNA | ENSG0000C |
| 75506227 | 75508114 | -1 | lncRNA | ENSG0000C |
| 75642494 | 75656016 | 1 | lncRNA | ENSG0000C |
| 75696044 | 75712851 | -1 | lncRNA | ENSG0000C |
| 75844925 | 75868252 | -1 | lncRNA | ENSG0000C |
| 75927836 | 75928004 | 1 | processed_pseudoge | ENSG0000C |
| 76008118 | 76015861 | -1 | lncRNA | ENSG0000C |
| 76122968 | 76145259 | 1 | lncRNA | ENSG0000C |
| 76173304 | 76177738 | -1 | lncRNA | ENSG0000C |
| 76209422 | 76221718 | 1 | lncRNA | ENSG0000C |
| 76227990 | 76233387 | -1 | lncRNA | ENSG0000C |

| | | | | |
|---|---|---|---|---|
| 76232927 | 76252044 | 1 | lncRNA | ENSG0000C |
| 76254292 | 76255214 | -1 | lncRNA | ENSG0000C |
| 76256891 | 76258337 | 1 | lncRNA | ENSG0000C |
| 76259166 | 76260114 | 1 | lncRNA | ENSG0000C |
| 76357682 | 76495242 | -1 | protein_coding | ENSG0000C |
| 76372717 | 76378275 | 1 | lncRNA | ENSG0000C |
| 76385642 | 76389406 | -1 | TEC | ENSG0000C |
| 76491629 | 76493918 | 1 | lncRNA | ENSG0000C |
| 76495521 | 76498088 | 1 | lncRNA | ENSG0000C |
| 76528652 | 76670111 | 1 | lncRNA | ENSG0000C |
| 76615160 | 76617645 | -1 | lncRNA | ENSG0000C |
| 76622136 | 76639030 | -1 | lncRNA | ENSG0000C |
| 76686058 | 76693636 | 1 | lncRNA | ENSG0000C |
| 76703444 | 76704254 | -1 | processed_pseudoge | ENSG0000C |
| 76751305 | 76751841 | 1 | processed_pseudoge | ENSG0000C |
| 76759685 | 76759866 | -1 | processed_pseudoge | ENSG0000C |
| 76794732 | 76822295 | -1 | lncRNA | ENSG0000C |
| 76800664 | 76800774 | -1 | snRNA | ENSG0000C |
| 76805151 | 76805736 | -1 | lncRNA | ENSG0000C |
| 76822607 | 76970727 | 1 | protein_coding | ENSG0000C |
| 76834429 | 76834888 | -1 | processed_pseudoge | ENSG0000C |
| 76840838 | 76841102 | -1 | misc_RNA | ENSG0000C |
| 76884935 | 76888724 | -1 | processed_pseudoge | ENSG0000C |
| 76974690 | 76984162 | 1 | lncRNA | ENSG0000C |
| 76978827 | 77133683 | -1 | protein_coding | ENSG0000C |
| 77002849 | 77006846 | -1 | TEC | ENSG0000C |
| 77042864 | 77044739 | 1 | lncRNA | ENSG0000C |
| 77093086 | 77093446 | 1 | lncRNA | ENSG0000C |
| 77108284 | 77110537 | -1 | TEC | ENSG0000C |
| 77112602 | 77115726 | 1 | lncRNA | ENSG0000C |
| 77180106 | 77235430 | 1 | lncRNA | ENSG0000C |
| 77250549 | 77277896 | 1 | protein_coding | ENSG0000C |
| 77421619 | 77561044 | 1 | lncRNA | ENSG0000C |
| 77431563 | 77431689 | -1 | snoRNA | ENSG0000C |
| 77435253 | 77437242 | -1 | processed_pseudoge | ENSG0000C |
| 77622552 | 77624515 | 1 | lncRNA | ENSG0000C |
| 77632928 | 77633360 | -1 | processed_pseudoge | ENSG0000C |
| 77647014 | 77650973 | -1 | lncRNA | ENSG0000C |
| 77863464 | 77863567 | -1 | rRNA_pseudogene | ENSG0000C |
| 77970637 | 77993731 | -1 | lncRNA | ENSG0000C |
| 77983020 | 77986610 | -1 | lncRNA | ENSG0000C |
| 78036739 | 78036845 | 1 | snRNA | ENSG0000C |
| 78137223 | 78140887 | 1 | lncRNA | ENSG0000C |
| 78480549 | 78484867 | 1 | lncRNA | ENSG0000C |
| 78505165 | 78506410 | 1 | lncRNA | ENSG0000C |
| 78544095 | 78544188 | -1 | misc_RNA | ENSG0000C |
| 78795612 | 78796672 | -1 | lncRNA | ENSG0000C |
| 78908629 | 78918993 | -1 | lncRNA | ENSG0000C |
| 78925064 | 78927441 | -1 | lncRNA | ENSG0000C |
| 78926053 | 78928494 | 1 | lncRNA | ENSG0000C |

| | | | | |
|---|---|---|---|---|
| 78976555 | 78979074 | -1 | lncRNA | ENSG00000 |
| 78980275 | 79002677 | 1 | protein_coding | ENSG00000 |
| 79069285 | 79378287 | 1 | protein_coding | ENSG00000 |
| 79085134 | 79086435 | 1 | TEC | ENSG00000 |
| 79117207 | 79117920 | 1 | lncRNA | ENSG00000 |
| 79144752 | 79146918 | 1 | TEC | ENSG00000 |
| 79185801 | 79191559 | -1 | lncRNA | ENSG00000 |
| 79253577 | 79254856 | -1 | lncRNA | ENSG00000 |
| 79340258 | 79343972 | -1 | lncRNA | ENSG00000 |
| 79352458 | 79354567 | 1 | TEC | ENSG00000 |
| 79395856 | 79529325 | 1 | protein_coding | ENSG00000 |
| 79420836 | 79422780 | 1 | TEC | ENSG00000 |
| 79460378 | 79465843 | -1 | lncRNA | ENSG00000 |
| 79470120 | 79470940 | -1 | lncRNA | ENSG00000 |
| 79496058 | 79498348 | 1 | TEC | ENSG00000 |
| 79515798 | 79516057 | -1 | TEC | ENSG00000 |
| 79539574 | 79540840 | 1 | lncRNA | ENSG00000 |
| 79576460 | 79589010 | 1 | lncRNA | ENSG00000 |
| 79610747 | 79612303 | 1 | lncRNA | ENSG00000 |
| 79638928 | 79679745 | -1 | lncRNA | ENSG00000 |
| 79677287 | 79679358 | -1 | lncRNA | ENSG00000 |
| 79679801 | 79756623 | 1 | protein_coding | ENSG00000 |
| 79702721 | 79703651 | -1 | lncRNA | ENSG00000 |
| 79787121 | 79791432 | 1 | lncRNA | ENSG00000 |
| 79792726 | 79816529 | 1 | lncRNA | ENSG00000 |
| 79863668 | 79900184 | 1 | protein_coding | ENSG00000 |
| 79900555 | 79900903 | -1 | lncRNA | ENSG00000 |
| 79902420 | 79951657 | -1 | protein_coding | ENSG00000 |
| 79959579 | 79963178 | -1 | lncRNA | ENSG00000 |
| 79964582 | 79970822 | 1 | protein_coding | ENSG00000 |
| 79970813 | 80033949 | -1 | protein_coding | ENSG00000 |
| 80034346 | 80097088 | 1 | protein_coding | ENSG00000 |
| 80034389 | 80050651 | 1 | protein_coding | ENSG00000 |
| 80046900 | 80095482 | 1 | lncRNA | ENSG00000 |
| 80077707 | 80078259 | 1 | processed_pseudoge | ENSG00000 |
| 80109262 | 80147523 | 1 | protein_coding | ENSG00000 |
| 80147924 | 80179839 | 1 | lncRNA | ENSG00000 |
| 80157232 | 80247514 | -1 | protein_coding | ENSG00000 |
| 80161752 | 80162413 | 1 | lncRNA | ENSG00000 |
| 80183680 | 80202992 | 1 | lncRNA | ENSG00000 |
| 60951 | 71626 | -1 | transcribed_processe | ENSG00000 |
| 68403 | 69178 | 1 | TEC | ENSG00000 |
| 69167 | 69972 | 1 | TEC | ENSG00000 |
| 71778 | 72718 | 1 | lncRNA | ENSG00000 |
| 71973 | 72110 | 1 | miRNA | ENSG00000 |
| 76163 | 77686 | -1 | lncRNA | ENSG00000 |
| 94062 | 94974 | 1 | unprocessed_pseudo | ENSG00000 |
| 99175 | 106077 | 1 | transcribed_unproce | ENSG00000 |
| 107104 | 117102 | 1 | protein_coding | ENSG00000 |
| 145485 | 145812 | 1 | processed_pseudoge | ENSG00000 |

| | | | | |
|---|---|---|---|---|
| 156279 | 157215 | -1 | unprocessed_pseudc | ENSG00000( |
| 176896 | 177913 | 1 | lncRNA | ENSG00000( |
| 186373 | 195696 | -1 | transcribed_unproce | ENSG00000( |
| 193239 | 195595 | 1 | processed_pseudoge | ENSG00000( |
| 197310 | 198066 | -1 | processed_pseudoge | ENSG00000( |
| 197961 | 200775 | 1 | transcribed_processe | ENSG00000( |
| 198264 | 246544 | -1 | lncRNA | ENSG00000( |
| 200499 | 201174 | -1 | transcribed_processe | ENSG00000( |
| 223158 | 223261 | -1 | snRNA | ENSG00000( |
| 248551 | 251571 | -1 | lncRNA | ENSG00000( |
| 281040 | 291403 | -1 | protein_coding | ENSG00000( |
| 301444 | 344815 | -1 | protein_coding | ENSG00000( |
| 305573 | 306467 | 1 | lncRNA | ENSG00000( |
| 361747 | 376026 | -1 | protein_coding | ENSG00000( |
| 382725 | 387533 | 1 | lncRNA | ENSG00000( |
| 397589 | 399173 | 1 | lncRNA | ENSG00000( |
| 405445 | 409147 | -1 | protein_coding | ENSG00000( |
| 416583 | 460996 | -1 | protein_coding | ENSG00000( |
| 453134 | 453245 | 1 | rRNA_pseudogene | ENSG00000( |
| 463346 | 474983 | -1 | protein_coding | ENSG00000( |
| 489176 | 505343 | 1 | protein_coding | ENSG00000( |
| 490046 | 507833 | -1 | lncRNA | ENSG00000( |
| 507497 | 519654 | 1 | protein_coding | ENSG00000( |
| 531760 | 542092 | 1 | protein_coding | ENSG00000( |
| 544034 | 549924 | 1 | protein_coding | ENSG00000( |
| 567210 | 572228 | -1 | lncRNA | ENSG00000( |
| 571277 | 583493 | 1 | protein_coding | ENSG00000( |
| 589881 | 617159 | 1 | protein_coding | ENSG00000( |
| 590430 | 591300 | -1 | processed_pseudoge | ENSG00000( |
| 617221 | 633537 | -1 | protein_coding | ENSG00000( |
| 637105 | 637537 | -1 | lncRNA | ENSG00000( |
| 639879 | 644371 | 1 | protein_coding | ENSG00000( |
| 647526 | 663233 | -1 | protein_coding | ENSG00000( |
| 663482 | 669500 | 1 | lncRNA | ENSG00000( |
| 676392 | 683392 | 1 | protein_coding | ENSG00000( |
| 685546 | 695498 | -1 | protein_coding | ENSG00000( |
| 690353 | 691172 | 1 | processed_pseudoge | ENSG00000( |
| 708935 | 748329 | 1 | protein_coding | ENSG00000( |
| 751112 | 764318 | 1 | protein_coding | ENSG00000( |
| 781002 | 781862 | -1 | lncRNA | ENSG00000( |
| 782755 | 786092 | 1 | lncRNA | ENSG00000( |
| 797075 | 812327 | 1 | protein_coding | ENSG00000( |
| 804940 | 805001 | 1 | miRNA | ENSG00000( |
| 812488 | 821977 | -1 | protein_coding | ENSG00000( |
| 813584 | 813653 | 1 | miRNA | ENSG00000( |
| 825097 | 832018 | 1 | protein_coding | ENSG00000( |
| 840999 | 848175 | 1 | protein_coding | ENSG00000( |
| 851014 | 856247 | 1 | protein_coding | ENSG00000( |
| 859664 | 863641 | 1 | protein_coding | ENSG00000( |
| 867630 | 893218 | -1 | protein_coding | ENSG00000( |

| | | | | |
|---|---|---|---|---|
| 893484 | 893590 | 1 | snRNA | ENSG0000C |
| 896503 | 913245 | -1 | protein_coding | ENSG0000C |
| 917287 | 921005 | 1 | protein_coding | ENSG0000C |
| 925781 | 975939 | 1 | protein_coding | ENSG0000C |
| 928259 | 928817 | -1 | lncRNA | ENSG0000C |
| 956485 | 958149 | 1 | lncRNA | ENSG0000C |
| 984271 | 998438 | 1 | protein_coding | ENSG0000C |
| 999796 | 1002757 | -1 | lncRNA | ENSG0000C |
| 1000419 | 1009732 | 1 | protein_coding | ENSG0000C |
| 1009648 | 1021179 | -1 | protein_coding | ENSG0000C |
| 1010221 | 1010907 | 1 | lncRNA | ENSG0000C |
| 1021522 | 1021628 | 1 | snRNA | ENSG0000C |
| 1026586 | 1039068 | 1 | protein_coding | ENSG0000C |
| 1038727 | 1039064 | -1 | TEC | ENSG0000C |
| 1040101 | 1065572 | 1 | protein_coding | ENSG0000C |
| 1065923 | 1086628 | 1 | protein_coding | ENSG0000C |
| 1086574 | 1095380 | -1 | protein_coding | ENSG0000C |
| 1103926 | 1106791 | 1 | protein_coding | ENSG0000C |
| 1107637 | 1174268 | -1 | protein_coding | ENSG0000C |
| 1177558 | 1228435 | 1 | protein_coding | ENSG0000C |
| 1203209 | 1203842 | 1 | transcribed_processe | ENSG0000C |
| 1228287 | 1238027 | -1 | protein_coding | ENSG0000C |
| 1238179 | 1239522 | 1 | lncRNA | ENSG0000C |
| 1241746 | 1244825 | 1 | protein_coding | ENSG0000C |
| 1248553 | 1259140 | 1 | protein_coding | ENSG0000C |
| 1259384 | 1274880 | 1 | protein_coding | ENSG0000C |
| 1267471 | 1270260 | -1 | lncRNA | ENSG0000C |
| 1275530 | 1279244 | 1 | protein_coding | ENSG0000C |
| 1285873 | 1301431 | 1 | protein_coding | ENSG0000C |
| 1307401 | 1307719 | -1 | processed_pseudoge | ENSG0000C |
| 1321225 | 1322846 | 1 | lncRNA | ENSG0000C |
| 1354711 | 1378431 | 1 | protein_coding | ENSG0000C |
| 1372238 | 1374364 | -1 | TEC | ENSG0000C |
| 1376773 | 1377520 | -1 | lncRNA | ENSG0000C |
| 1383527 | 1395589 | 1 | protein_coding | ENSG0000C |
| 1385473 | 1389665 | -1 | lncRNA | ENSG0000C |
| 1392170 | 1396467 | -1 | lncRNA | ENSG0000C |
| 1397026 | 1401570 | -1 | protein_coding | ENSG0000C |
| 1407569 | 1435687 | 1 | protein_coding | ENSG0000C |
| 1438358 | 1440494 | 1 | protein_coding | ENSG0000C |
| 1440839 | 1441938 | 1 | lncRNA | ENSG0000C |
| 1446302 | 1473244 | 1 | protein_coding | ENSG0000C |
| 1457670 | 1458580 | -1 | lncRNA | ENSG0000C |
| 1461143 | 1479556 | -1 | protein_coding | ENSG0000C |
| 1481428 | 1490752 | -1 | protein_coding | ENSG0000C |
| 1491166 | 1497927 | 1 | protein_coding | ENSG0000C |
| 1505022 | 1513604 | -1 | protein_coding | ENSG0000C |
| 1508375 | 1508963 | 1 | lncRNA | ENSG0000C |
| 1524077 | 1536046 | 1 | protein_coding | ENSG0000C |
| 1554669 | 1568058 | -1 | protein_coding | ENSG0000C |

| | | | | |
|---|---|---|---|---|
| 1570575 | 1570860 | -1 | misc_RNA | ENSG0000C |
| 1573596 | 1592865 | -1 | protein_coding | ENSG0000C |
| 1576939 | 1577086 | 1 | TEC | ENSG0000C |
| 1578339 | 1605445 | -1 | protein_coding | ENSG0000C |
| 1597169 | 1605473 | -1 | protein_coding | ENSG0000C |
| 1609290 | 1652615 | -1 | protein_coding | ENSG0000C |
| 1616492 | 1616595 | -1 | snRNA | ENSG0000C |
| 1724115 | 1724395 | 1 | misc_RNA | ENSG0000C |
| 1748056 | 1748745 | 1 | lncRNA | ENSG0000C |
| 1753506 | 1780988 | 1 | protein_coding | ENSG0000C |
| 1782075 | 1812276 | -1 | protein_coding | ENSG0000C |
| 1815248 | 1848483 | -1 | protein_coding | ENSG0000C |
| 1815249 | 1815873 | 1 | lncRNA | ENSG0000C |
| 1816159 | 1816238 | -1 | miRNA | ENSG0000C |
| 1822089 | 1824542 | 1 | lncRNA | ENSG0000C |
| 1852382 | 1853622 | 1 | lncRNA | ENSG0000C |
| 1852399 | 1863579 | -1 | protein_coding | ENSG0000C |
| 1860250 | 1862019 | 1 | lncRNA | ENSG0000C |
| 1874871 | 1876169 | -1 | lncRNA | ENSG0000C |
| 1875016 | 1875992 | 1 | lncRNA | ENSG0000C |
| 1876810 | 1885547 | -1 | protein_coding | ENSG0000C |
| 1905372 | 1926013 | 1 | protein_coding | ENSG0000C |
| 1905399 | 1913447 | 1 | protein_coding | ENSG0000C |
| 1941172 | 1981338 | 1 | protein_coding | ENSG0000C |
| 1952531 | 1954586 | -1 | lncRNA | ENSG0000C |
| 1985438 | 2034881 | -1 | protein_coding | ENSG0000C |
| 1989401 | 1990370 | 1 | lncRNA | ENSG0000C |
| 2002580 | 2003581 | -1 | TEC | ENSG0000C |
| 2013728 | 2014656 | -1 | processed_pseudoge | ENSG0000C |
| 2030853 | 2031051 | -1 | transcribed_processe | ENSG0000C |
| 2037465 | 2051244 | -1 | protein_coding | ENSG0000C |
| 2071036 | 2096673 | -1 | protein_coding | ENSG0000C |
| 2096429 | 2099593 | 1 | protein_coding | ENSG0000C |
| 2100988 | 2164468 | -1 | protein_coding | ENSG0000C |
| 2163933 | 2232578 | 1 | protein_coding | ENSG0000C |
| 2212029 | 2215565 | -1 | lncRNA | ENSG0000C |
| 2230084 | 2237704 | -1 | protein_coding | ENSG0000C |
| 2234062 | 2234149 | -1 | miRNA | ENSG0000C |
| 2235829 | 2235926 | -1 | miRNA | ENSG0000C |
| 2236824 | 2248655 | 1 | protein_coding | ENSG0000C |
| 2249309 | 2252073 | 1 | protein_coding | ENSG0000C |
| 2250639 | 2250718 | 1 | miRNA | ENSG0000C |
| 2252252 | 2269759 | -1 | protein_coding | ENSG0000C |
| 2269509 | 2273490 | 1 | protein_coding | ENSG0000C |
| 2269525 | 2341172 | 1 | protein_coding | ENSG0000C |
| 2274631 | 2282175 | -1 | protein_coding | ENSG0000C |
| 2289784 | 2292024 | -1 | protein_coding | ENSG0000C |
| 2293173 | 2295781 | 1 | lncRNA | ENSG0000C |
| 2315162 | 2316875 | -1 | lncRNA | ENSG0000C |
| 2321520 | 2328611 | -1 | protein_coding | ENSG0000C |

| | | | | |
|---|---|---|---|---|
| 2328615 | 2355095 | 1 | protein_coding | ENSG00000 |
| 2360238 | 2426239 | 1 | protein_coding | ENSG00000 |
| 2425625 | 2427586 | -1 | protein_coding | ENSG00000 |
| 2427638 | 2456959 | -1 | protein_coding | ENSG00000 |
| 2434914 | 2435000 | -1 | miRNA | ENSG00000 |
| 2458935 | 2462185 | -1 | lncRNA | ENSG00000 |
| 2476122 | 2478259 | 1 | protein_coding | ENSG00000 |
| 2503047 | 2503141 | -1 | snRNA | ENSG00000 |
| 2511219 | 2702694 | -1 | protein_coding | ENSG00000 |
| 2538112 | 2538754 | 1 | processed_pseudoge | ENSG00000 |
| 2595019 | 2595291 | -1 | misc_RNA | ENSG00000 |
| 2610155 | 2611862 | -1 | lncRNA | ENSG00000 |
| 2630715 | 2630793 | 1 | miRNA | ENSG00000 |
| 2631954 | 2632849 | 1 | lncRNA | ENSG00000 |
| 2641838 | 2643853 | -1 | lncRNA | ENSG00000 |
| 2700689 | 2701266 | -1 | lncRNA | ENSG00000 |
| 2714567 | 2721372 | -1 | protein_coding | ENSG00000 |
| 2717769 | 2740048 | -1 | protein_coding | ENSG00000 |
| 2721710 | 2724517 | 1 | lncRNA | ENSG00000 |
| 2727743 | 2729327 | -1 | lncRNA | ENSG00000 |
| 2732204 | 2740028 | -1 | protein_coding | ENSG00000 |
| 2754715 | 2783282 | -1 | protein_coding | ENSG00000 |
| 2785503 | 2815807 | 1 | protein_coding | ENSG00000 |
| 2819868 | 2836735 | 1 | protein_coding | ENSG00000 |
| 2841437 | 2860471 | 1 | protein_coding | ENSG00000 |
| 2847227 | 2858716 | -1 | lncRNA | ENSG00000 |
| 2862270 | 2864428 | 1 | lncRNA | ENSG00000 |
| 2867335 | 2883445 | 1 | protein_coding | ENSG00000 |
| 2874950 | 2875592 | -1 | processed_pseudoge | ENSG00000 |
| 2896625 | 2897222 | -1 | processed_pseudoge | ENSG00000 |
| 2900898 | 2918473 | 1 | protein_coding | ENSG00000 |
| 2908371 | 2926817 | -1 | lncRNA | ENSG00000 |
| 2933218 | 2944971 | -1 | protein_coding | ENSG00000 |
| 2949035 | 2950644 | 1 | TEC | ENSG00000 |
| 2977538 | 2995179 | 1 | protein_coding | ENSG00000 |
| 2997639 | 3047635 | -1 | protein_coding | ENSG00000 |
| 3052910 | 3053724 | 1 | lncRNA | ENSG00000 |
| 3052910 | 3063107 | -1 | protein_coding | ENSG00000 |
| 3094362 | 3123999 | 1 | protein_coding | ENSG00000 |
| 3118665 | 3119304 | -1 | lncRNA | ENSG00000 |
| 3121116 | 3122128 | -1 | lncRNA | ENSG00000 |
| 3130717 | 3131398 | 1 | processed_pseudoge | ENSG00000 |
| 3136033 | 3163749 | 1 | protein_coding | ENSG00000 |
| 3141576 | 3155175 | -1 | lncRNA | ENSG00000 |
| 3172346 | 3180332 | 1 | protein_coding | ENSG00000 |
| 3185563 | 3209575 | 1 | protein_coding | ENSG00000 |
| 3224661 | 3297076 | 1 | protein_coding | ENSG00000 |
| 3296765 | 3306887 | -1 | lncRNA | ENSG00000 |
| 3314403 | 3469217 | 1 | protein_coding | ENSG00000 |
| 3355554 | 3355955 | -1 | TEC | ENSG00000 |

| | | | | |
|---|---|---|---|---|
| 3371395 | 3371602 | 1 | TEC | ENSG0000C |
| 3473986 | 3480525 | -1 | protein_coding | ENSG0000C |
| 3482804 | 3483446 | -1 | protein_coding | ENSG0000C |
| 3490821 | 3500940 | -1 | protein_coding | ENSG0000C |
| 3506273 | 3538330 | 1 | protein_coding | ENSG0000C |
| 3521248 | 3521320 | -1 | snoRNA | ENSG0000C |
| 3538261 | 3574290 | -1 | protein_coding | ENSG0000C |
| 3539154 | 3544030 | 1 | protein_coding | ENSG0000C |
| 3544199 | 3557569 | 1 | lncRNA | ENSG0000C |
| 3558015 | 3558486 | 1 | lncRNA | ENSG0000C |
| 3567278 | 3567959 | -1 | TEC | ENSG0000C |
| 3568120 | 3568273 | -1 | processed_pseudoge | ENSG0000C |
| 3572777 | 3579088 | 1 | protein_coding | ENSG0000C |
| 3585478 | 3593541 | 1 | protein_coding | ENSG0000C |
| 3594506 | 3606840 | -1 | protein_coding | ENSG0000C |
| 3607247 | 3613930 | 1 | lncRNA | ENSG0000C |
| 3610645 | 3626815 | -1 | protein_coding | ENSG0000C |
| 3630183 | 3700479 | -1 | protein_coding | ENSG0000C |
| 3672582 | 3674295 | 1 | lncRNA | ENSG0000C |
| 3708362 | 3750813 | 1 | protein_coding | ENSG0000C |
| 3750772 | 3761692 | -1 | protein_coding | ENSG0000C |
| 3753840 | 3756517 | 1 | lncRNA | ENSG0000C |
| 3754620 | 3756659 | -1 | lncRNA | ENSG0000C |
| 3762682 | 3768575 | 1 | protein_coding | ENSG0000C |
| 3769089 | 3772221 | -1 | protein_coding | ENSG0000C |
| 3775996 | 3776182 | -1 | unprocessed_pseudc | ENSG0000C |
| 3777970 | 3802129 | -1 | protein_coding | ENSG0000C |
| 3804024 | 3869038 | -1 | protein_coding | ENSG0000C |
| 3870991 | 3872107 | 1 | processed_pseudoge | ENSG0000C |
| 3879864 | 3928082 | 1 | protein_coding | ENSG0000C |
| 3920245 | 3920506 | -1 | misc_RNA | ENSG0000C |
| 3933103 | 3942416 | 1 | protein_coding | ENSG0000C |
| 3958453 | 3971123 | -1 | protein_coding | ENSG0000C |
| 3961414 | 3961512 | -1 | miRNA | ENSG0000C |
| 3976056 | 3985463 | -1 | protein_coding | ENSG0000C |
| 3982507 | 3982572 | -1 | snoRNA | ENSG0000C |
| 4007736 | 4039386 | 1 | protein_coding | ENSG0000C |
| 4043303 | 4066899 | -1 | protein_coding | ENSG0000C |
| 4061615 | 4062749 | -1 | lncRNA | ENSG0000C |
| 4090321 | 4124129 | -1 | protein_coding | ENSG0000C |
| 4127492 | 4127732 | 1 | misc_RNA | ENSG0000C |
| 4153631 | 4173054 | 1 | protein_coding | ENSG0000C |
| 4174109 | 4182566 | -1 | protein_coding | ENSG0000C |
| 4183354 | 4224814 | 1 | protein_coding | ENSG0000C |
| 4229523 | 4237528 | 1 | protein_coding | ENSG0000C |
| 4246339 | 4247358 | -1 | TEC | ENSG0000C |
| 4247080 | 4269088 | 1 | protein_coding | ENSG0000C |
| 4278601 | 4290724 | 1 | protein_coding | ENSG0000C |
| 4292232 | 4302431 | -1 | protein_coding | ENSG0000C |
| 4304598 | 4323843 | 1 | protein_coding | ENSG0000C |

| | | | | |
|---|---|---|---|---|
| 4324043 | 4342786 | -1 | protein_coding | ENSG00000 |
| 4339577 | 4343491 | 1 | lncRNA | ENSG00000 |
| 4343527 | 4360086 | 1 | protein_coding | ENSG00000 |
| 4347244 | 4354057 | -1 | lncRNA | ENSG00000 |
| 4347780 | 4349061 | -1 | unprocessed_pseudo | ENSG00000 |
| 4356637 | 4358448 | -1 | lncRNA | ENSG00000 |
| 4360370 | 4400547 | -1 | protein_coding | ENSG00000 |
| 4363789 | 4364640 | 1 | lncRNA | ENSG00000 |
| 4402640 | 4445018 | 1 | protein_coding | ENSG00000 |
| 4429689 | 4430934 | -1 | lncRNA | ENSG00000 |
| 4444999 | 4457794 | -1 | protein_coding | ENSG00000 |
| 4445978 | 4446048 | 1 | miRNA | ENSG00000 |
| 4447304 | 4448217 | 1 | lncRNA | ENSG00000 |
| 4448810 | 4450836 | 1 | TEC | ENSG00000 |
| 4454014 | 4455286 | 1 | lncRNA | ENSG00000 |
| 4457962 | 4471493 | -1 | lncRNA | ENSG00000 |
| 4472297 | 4502208 | 1 | protein_coding | ENSG00000 |
| 4502180 | 4518465 | -1 | protein_coding | ENSG00000 |
| 4522531 | 4535224 | -1 | protein_coding | ENSG00000 |
| 4528427 | 4540067 | -1 | protein_coding | ENSG00000 |
| 4536402 | 4540036 | -1 | protein_coding | ENSG00000 |
| 4542593 | 4559684 | -1 | protein_coding | ENSG00000 |
| 4621194 | 4622307 | 1 | unprocessed_pseudo | ENSG00000 |
| 4639516 | 4655568 | 1 | protein_coding | ENSG00000 |
| 4641374 | 4670370 | -1 | protein_coding | ENSG00000 |
| 4654964 | 4655524 | -1 | lncRNA | ENSG00000 |
| 4675224 | 4724673 | -1 | protein_coding | ENSG00000 |
| 4679282 | 4685948 | 1 | lncRNA | ENSG00000 |
| 4692021 | 4694340 | 1 | lncRNA | ENSG00000 |
| 4769031 | 4785077 | 1 | lncRNA | ENSG00000 |
| 4770670 | 4770779 | 1 | miRNA | ENSG00000 |
| 4785120 | 4791207 | -1 | lncRNA | ENSG00000 |
| 4791734 | 4801273 | 1 | protein_coding | ENSG00000 |
| 4791745 | 4795559 | -1 | lncRNA | ENSG00000 |
| 4815932 | 4831704 | -1 | protein_coding | ENSG00000 |
| 4838332 | 4838907 | 1 | lncRNA | ENSG00000 |
| 4838341 | 4867694 | -1 | protein_coding | ENSG00000 |
| 4890437 | 4902896 | -1 | protein_coding | ENSG00000 |
| 4903080 | 4962154 | 1 | protein_coding | ENSG00000 |
| 4932687 | 4932740 | 1 | miRNA | ENSG00000 |
| 4969113 | 5153598 | 1 | protein_coding | ENSG00000 |
| 5158495 | 5340803 | -1 | protein_coding | ENSG00000 |
| 5178119 | 5178464 | 1 | lncRNA | ENSG00000 |
| 5277734 | 5278133 | 1 | processed_pseudoge | ENSG00000 |
| 5294247 | 5294696 | 1 | lncRNA | ENSG00000 |
| 5455417 | 5456856 | 1 | protein_coding | ENSG00000 |
| 5558167 | 5578349 | -1 | protein_coding | ENSG00000 |
| 5576660 | 5576938 | -1 | processed_pseudoge | ENSG00000 |
| 5586999 | 5624046 | -1 | protein_coding | ENSG00000 |
| 5623035 | 5668478 | 1 | protein_coding | ENSG00000 |

| | | | | |
|---|---|---|---|---|
| 5674947 | 5691875 | 1 | protein_coding | ENSG00000 |
| 5678421 | 5680896 | -1 | protein_coding | ENSG00000 |
| 5680604 | 5688523 | 1 | protein_coding | ENSG00000 |
| 5691834 | 5720572 | -1 | protein_coding | ENSG00000 |
| 5720637 | 5778734 | 1 | protein_coding | ENSG00000 |
| 5782960 | 5784746 | -1 | protein_coding | ENSG00000 |
| 5784005 | 5785619 | -1 | protein_coding | ENSG00000 |
| 5784832 | 5791225 | -1 | protein_coding | ENSG00000 |
| 5823802 | 5828324 | 1 | protein_coding | ENSG00000 |
| 5830610 | 5839731 | -1 | protein_coding | ENSG00000 |
| 5842888 | 5851474 | -1 | protein_coding | ENSG00000 |
| 5847467 | 5858239 | 1 | lncRNA | ENSG00000 |
| 5865826 | 5870540 | -1 | protein_coding | ENSG00000 |
| 5866171 | 5903787 | -1 | protein_coding | ENSG00000 |
| 5891276 | 5904006 | -1 | protein_coding | ENSG00000 |
| 5899895 | 5901511 | 1 | lncRNA | ENSG00000 |
| 5904841 | 5914707 | 1 | protein_coding | ENSG00000 |
| 5904872 | 5910853 | 1 | protein_coding | ENSG00000 |
| 5911578 | 5913899 | -1 | lncRNA | ENSG00000 |
| 5912339 | 5916211 | 1 | protein_coding | ENSG00000 |
| 5916139 | 5978142 | -1 | protein_coding | ENSG00000 |
| 5978403 | 6020363 | 1 | lncRNA | ENSG00000 |
| 5993164 | 6199572 | -1 | protein_coding | ENSG00000 |
| 6067953 | 6077119 | 1 | lncRNA | ENSG00000 |
| 6072823 | 6073527 | 1 | processed_pseudoge | ENSG00000 |
| 6109322 | 6125797 | 1 | lncRNA | ENSG00000 |
| 6135247 | 6193094 | 1 | protein_coding | ENSG00000 |
| 6176256 | 6176442 | -1 | transcribed_processe | ENSG00000 |
| 6199405 | 6205121 | 1 | unprocessed_pseudo | ENSG00000 |
| 6210381 | 6279975 | -1 | protein_coding | ENSG00000 |
| 6259157 | 6259799 | 1 | lncRNA | ENSG00000 |
| 6263068 | 6263469 | -1 | TEC | ENSG00000 |
| 6306142 | 6333612 | -1 | protein_coding | ENSG00000 |
| 6343761 | 6362149 | -1 | lncRNA | ENSG00000 |
| 6361531 | 6370242 | 1 | protein_coding | ENSG00000 |
| 6372794 | 6375250 | 1 | protein_coding | ENSG00000 |
| 6375148 | 6379058 | -1 | protein_coding | ENSG00000 |
| 6379572 | 6393981 | -1 | protein_coding | ENSG00000 |
| 6386762 | 6390400 | -1 | lncRNA | ENSG00000 |
| 6389638 | 6389703 | -1 | miRNA | ENSG00000 |
| 6392921 | 6392983 | -1 | miRNA | ENSG00000 |
| 6393794 | 6412404 | 1 | transcribed_unitary_ | ENSG00000 |
| 6413348 | 6424794 | -1 | protein_coding | ENSG00000 |
| 6416410 | 6416511 | -1 | miRNA | ENSG00000 |
| 6426037 | 6433779 | -1 | protein_coding | ENSG00000 |
| 6436079 | 6465203 | -1 | protein_coding | ENSG00000 |
| 6463777 | 6467221 | 1 | protein_coding | ENSG00000 |
| 6467207 | 6482557 | -1 | protein_coding | ENSG00000 |
| 6469465 | 6470152 | 1 | lncRNA | ENSG00000 |
| 6494319 | 6502848 | -1 | protein_coding | ENSG00000 |

| | | | | |
|---|---|---|---|---|
| 6494320 | 6495025 | 1 | lncRNA | ENSG00000( |
| 6494320 | 6494805 | 1 | lncRNA | ENSG00000( |
| 6531026 | 6535924 | 1 | protein_coding | ENSG00000( |
| 6552358 | 6554494 | -1 | processed_pseudoge | ENSG00000( |
| 6562123 | 6563848 | -1 | processed_pseudoge | ENSG00000( |
| 6572956 | 6573057 | -1 | misc_RNA | ENSG00000( |
| 6583183 | 6604103 | -1 | protein_coding | ENSG00000( |
| 6593329 | 6594055 | 1 | processed_pseudoge | ENSG00000( |
| 6661253 | 6670588 | -1 | protein_coding | ENSG00000( |
| 6677704 | 6730562 | -1 | protein_coding | ENSG00000( |
| 6716386 | 6717742 | -1 | lncRNA | ENSG00000( |
| 6729914 | 6737603 | -1 | protein_coding | ENSG00000( |
| 6736712 | 6736778 | -1 | miRNA | ENSG00000( |
| 6737925 | 6751530 | 1 | protein_coding | ENSG00000( |
| 6748293 | 6751467 | -1 | lncRNA | ENSG00000( |
| 6752160 | 6767463 | -1 | protein_coding | ENSG00000( |
| 6772708 | 6857366 | 1 | protein_coding | ENSG00000( |
| 6887566 | 6940459 | 1 | protein_coding | ENSG00000( |
| 6894994 | 6896832 | -1 | TEC | ENSG00000( |
| 6898299 | 6898686 | 1 | unprocessed_pseudc | ENSG00000( |
| 6928345 | 6928629 | 1 | misc_RNA | ENSG00000( |
| 6952500 | 6997872 | -1 | transcribed_unproce | ENSG00000( |
| 6953211 | 6954904 | 1 | lncRNA | ENSG00000( |
| 7005012 | 7006074 | -1 | processed_pseudoge | ENSG00000( |
| 7011884 | 7012679 | 1 | processed_pseudoge | ENSG00000( |
| 7018969 | 7021450 | -1 | protein_coding | ENSG00000( |
| 7030578 | 7033011 | 1 | protein_coding | ENSG00000( |
| 7037748 | 7040179 | -1 | protein_coding | ENSG00000( |
| 7049321 | 7051735 | 1 | protein_coding | ENSG00000( |
| 7056209 | 7058640 | -1 | protein_coding | ENSG00000( |
| 7069703 | 7087968 | 1 | protein_coding | ENSG00000( |
| 7112255 | 7294414 | -1 | protein_coding | ENSG00000( |
| 7348943 | 7383385 | 1 | protein_coding | ENSG00000( |
| 7382834 | 7470241 | 1 | protein_coding | ENSG00000( |
| 7390522 | 7395039 | -1 | lncRNA | ENSG00000( |
| 7395113 | 7472484 | 1 | protein_coding | ENSG00000( |
| 7476875 | 7497449 | -1 | protein_coding | ENSG00000( |
| 7492976 | 7508450 | 1 | protein_coding | ENSG00000( |
| 7507052 | 7519622 | 1 | protein_coding | ENSG00000( |
| 7515292 | 7521025 | 1 | protein_coding | ENSG00000( |
| 7519916 | 7520460 | -1 | lncRNA | ENSG00000( |
| 7522624 | 7534009 | 1 | protein_coding | ENSG00000( |
| 7533595 | 7540059 | 1 | protein_coding | ENSG00000( |
| 7534004 | 7561764 | 1 | protein_coding | ENSG00000( |
| 7595902 | 7618304 | 1 | protein_coding | ENSG00000( |
| 7617439 | 7617505 | 1 | miRNA | ENSG00000( |
| 7619525 | 7629545 | -1 | protein_coding | ENSG00000( |
| 7629793 | 7631956 | 1 | protein_coding | ENSG00000( |
| 7629796 | 7643048 | 1 | protein_coding | ENSG00000( |
| 7631611 | 7633719 | -1 | protein_coding | ENSG00000( |

| | | | | |
|---|---|---|---|---|
| 7633766 | 7636990 | -1 | lncRNA | ENSG0000C |
| 7636881 | 7647873 | 1 | protein_coding | ENSG0000C |
| 7665717 | 7666476 | 1 | processed_pseudoge | ENSG0000C |
| 7669049 | 7670455 | 1 | protein_coding | ENSG0000C |
| 7676628 | 7679826 | 1 | protein_coding | ENSG0000C |
| 7678501 | 7682854 | 1 | protein_coding | ENSG0000C |
| 7680833 | 7687703 | 1 | protein_coding | ENSG0000C |
| 7688758 | 7702146 | -1 | protein_coding | ENSG0000C |
| 7728957 | 7733906 | -1 | protein_coding | ENSG0000C |
| 7739993 | 7747564 | -1 | protein_coding | ENSG0000C |
| 7750684 | 7751161 | 1 | processed_pseudoge | ENSG0000C |
| 7763149 | 7769605 | 1 | protein_coding | ENSG0000C |
| 7787549 | 7790621 | 1 | unprocessed_pseudc | ENSG0000C |
| 7813615 | 7814425 | -1 | processed_pseudoge | ENSG0000C |
| 7830233 | 7864976 | 1 | protein_coding | ENSG0000C |
| 7868719 | 7874390 | -1 | protein_coding | ENSG0000C |
| 7870561 | 7871296 | 1 | lncRNA | ENSG0000C |
| 7879445 | 7880120 | -1 | processed_pseudoge | ENSG0000C |
| 7886977 | 7887096 | -1 | rRNA_pseudogene | ENSG0000C |
| 7886977 | 7887096 | -1 | rRNA | ENSG0000C |
| 7888505 | 7902021 | 1 | protein_coding | ENSG0000C |
| 7898804 | 7903542 | -1 | lncRNA | ENSG0000C |
| 7903843 | 7914478 | 1 | protein_coding | ENSG0000C |
| 7912648 | 7913518 | -1 | lncRNA | ENSG0000C |
| 7916145 | 7919097 | 1 | protein_coding | ENSG0000C |
| 7918652 | 7919157 | -1 | lncRNA | ENSG0000C |
| 7920338 | 7923250 | 1 | protein_coding | ENSG0000C |
| 7924491 | 7926135 | -1 | protein_coding | ENSG0000C |
| 7926001 | 7926810 | 1 | lncRNA | ENSG0000C |
| 7926718 | 7943667 | -1 | protein_coding | ENSG0000C |
| 7944241 | 7945400 | 1 | lncRNA | ENSG0000C |
| 7958573 | 8005659 | -1 | protein_coding | ENSG0000C |
| 7959123 | 7960012 | 1 | lncRNA | ENSG0000C |
| 8008729 | 8016025 | 1 | lncRNA | ENSG0000C |
| 8052767 | 8062650 | 1 | protein_coding | ENSG0000C |
| 8065402 | 8149592 | -1 | protein_coding | ENSG0000C |
| 8173272 | 8175567 | 1 | unprocessed_pseudc | ENSG0000C |
| 8206736 | 8262421 | 1 | protein_coding | ENSG0000C |
| 8222153 | 8239648 | -1 | lncRNA | ENSG0000C |
| 8302127 | 8308358 | -1 | protein_coding | ENSG0000C |
| 8308283 | 8321379 | -1 | protein_coding | ENSG0000C |
| 8308768 | 8321375 | -1 | protein_coding | ENSG0000C |
| 8321158 | 8323340 | 1 | protein_coding | ENSG0000C |
| 8322584 | 8343262 | -1 | protein_coding | ENSG0000C |
| 8346517 | 8347687 | 1 | processed_pseudoge | ENSG0000C |
| 8363289 | 8374370 | 1 | protein_coding | ENSG0000C |
| 8374373 | 8390685 | -1 | lncRNA | ENSG0000C |
| 8389290 | 8389380 | -1 | miRNA | ENSG0000C |
| 8389981 | 8404434 | 1 | protein_coding | ENSG0000C |
| 8413270 | 8439017 | 1 | protein_coding | ENSG0000C |

| | | | | |
|---|---|---|---|---|
| 8423355 | 8427827 | -1 | lncRNA | ENSG0000C |
| 8444570 | 8445597 | -1 | TEC | ENSG0000C |
| 8444767 | 8489114 | 1 | protein_coding | ENSG0000C |
| 8490056 | 8502640 | -1 | protein_coding | ENSG0000C |
| 8509678 | 8514167 | -1 | protein_coding | ENSG0000C |
| 8520778 | 8577577 | -1 | protein_coding | ENSG0000C |
| 8549506 | 8550139 | 1 | lncRNA | ENSG0000C |
| 8580240 | 8610735 | -1 | protein_coding | ENSG0000C |
| 8581160 | 8582715 | 1 | lncRNA | ENSG0000C |
| 8630635 | 8677708 | 1 | protein_coding | ENSG0000C |
| 8695721 | 8696321 | -1 | TEC | ENSG0000C |
| 8697400 | 8698795 | -1 | protein_coding | ENSG0000C |
| 8721634 | 8732160 | 1 | protein_coding | ENSG0000C |
| 8728735 | 8729204 | -1 | processed_pseudoge | ENSG0000C |
| 8778665 | 8778861 | -1 | processed_pseudoge | ENSG0000C |
| 8806170 | 8832314 | -1 | protein_coding | ENSG0000C |
| 8821501 | 8824110 | 1 | lncRNA | ENSG0000C |
| 8832398 | 8843340 | 1 | protein_coding | ENSG0000C |
| 8833065 | 8833541 | 1 | lncRNA | ENSG0000C |
| 8848844 | 8981342 | -1 | protein_coding | ENSG0000C |
| 9019344 | 9019984 | 1 | processed_pseudoge | ENSG0000C |
| 9083112 | 9083398 | -1 | unprocessed_pseudc | ENSG0000C |
| 9087061 | 9095669 | 1 | protein_coding | ENSG0000C |
| 9100407 | 9107475 | -1 | protein_coding | ENSG0000C |
| 9114828 | 9115763 | -1 | protein_coding | ENSG0000C |
| 9122160 | 9122492 | -1 | unprocessed_pseudc | ENSG0000C |
| 9126012 | 9126950 | -1 | protein_coding | ENSG0000C |
| 9140397 | 9163424 | 1 | protein_coding | ENSG0000C |
| 9178979 | 9188818 | 1 | protein_coding | ENSG0000C |
| 9182369 | 9183048 | -1 | processed_pseudoge | ENSG0000C |
| 9191286 | 9192170 | 1 | unprocessed_pseudc | ENSG0000C |
| 9198354 | 9210067 | 1 | transcribed_unproce | ENSG0000C |
| 9210276 | 9219589 | -1 | protein_coding | ENSG0000C |
| 9235144 | 9236076 | 1 | unprocessed_pseudc | ENSG0000C |
| 9247344 | 9252625 | 1 | protein_coding | ENSG0000C |
| 9261015 | 9261716 | -1 | unprocessed_pseudc | ENSG0000C |
| 9265440 | 9268819 | -1 | transcribed_unproce | ENSG0000C |
| 9278448 | 9279369 | -1 | unprocessed_pseudc | ENSG0000C |
| 9291140 | 9309838 | -1 | protein_coding | ENSG0000C |
| 9291515 | 9294482 | 1 | lncRNA | ENSG0000C |
| 9323772 | 9351162 | 1 | protein_coding | ENSG0000C |
| 9324174 | 9382617 | 1 | protein_coding | ENSG0000C |
| 9348106 | 9348703 | -1 | processed_pseudoge | ENSG0000C |
| 9363020 | 9382617 | 1 | protein_coding | ENSG0000C |
| 9383581 | 9407055 | -1 | lncRNA | ENSG0000C |
| 9412461 | 9435571 | -1 | protein_coding | ENSG0000C |
| 9458119 | 9462388 | 1 | lncRNA | ENSG0000C |
| 9466355 | 9498616 | -1 | protein_coding | ENSG0000C |
| 9498678 | 9510079 | 1 | lncRNA | ENSG0000C |
| 9507820 | 9514539 | -1 | lncRNA | ENSG0000C |

| | | | | |
|---|---|---|---|---|
| 9518150 | 9519341 | 1 | processed_pseudoge | ENSG00000 |
| 9523223 | 9538645 | -1 | protein_coding | ENSG00000 |
| 9538693 | 9539734 | 1 | lncRNA | ENSG00000 |
| 9560353 | 9584533 | -1 | protein_coding | ENSG00000 |
| 9604680 | 9621236 | -1 | protein_coding | ENSG00000 |
| 9621249 | 9647203 | 1 | lncRNA | ENSG00000 |
| 9641807 | 9675100 | -1 | protein_coding | ENSG00000 |
| 9683294 | 9684073 | -1 | processed_pseudoge | ENSG00000 |
| 9689924 | 9700776 | -1 | transcribed_unproce | ENSG00000 |
| 9721903 | 9722410 | 1 | lncRNA | ENSG00000 |
| 9724008 | 9725086 | -1 | unprocessed_pseudo | ENSG00000 |
| 9730853 | 9731943 | 1 | lncRNA | ENSG00000 |
| 9751993 | 9793180 | -1 | protein_coding | ENSG00000 |
| 9756152 | 9756863 | -1 | unprocessed_pseudo | ENSG00000 |
| 9774497 | 9774960 | 1 | processed_pseudoge | ENSG00000 |
| 9793621 | 9802301 | 1 | lncRNA | ENSG00000 |
| 9810267 | 9827816 | -1 | protein_coding | ENSG00000 |
| 9819954 | 9820094 | 1 | snoRNA | ENSG00000 |
| 9820141 | 9820455 | 1 | processed_pseudoge | ENSG00000 |
| 9827892 | 9830115 | 1 | protein_coding | ENSG00000 |
| 9834079 | 9835013 | -1 | lncRNA | ENSG00000 |
| 9835257 | 9849682 | 1 | protein_coding | ENSG00000 |
| 9853718 | 9936552 | -1 | protein_coding | ENSG00000 |
| 9959561 | 10010504 | -1 | protein_coding | ENSG00000 |
| 9995997 | 9997163 | 1 | lncRNA | ENSG00000 |
| 10013249 | 10022279 | 1 | protein_coding | ENSG00000 |
| 10037803 | 10074135 | 1 | transcribed_unproce | ENSG00000 |
| 10038354 | 10038413 | 1 | miRNA | ENSG00000 |
| 10086122 | 10093252 | 1 | protein_coding | ENSG00000 |
| 10089032 | 10090377 | -1 | lncRNA | ENSG00000 |
| 10092338 | 10102796 | -1 | protein_coding | ENSG00000 |
| 10106223 | 10114780 | 1 | protein_coding | ENSG00000 |
| 10106362 | 10112012 | 1 | protein_coding | ENSG00000 |
| 10107651 | 10107735 | 1 | snoRNA | ENSG00000 |
| 10109757 | 10109835 | 1 | snoRNA | ENSG00000 |
| 10111693 | 10115372 | 1 | protein_coding | ENSG00000 |
| 10115014 | 10119918 | -1 | protein_coding | ENSG00000 |
| 10133345 | 10231286 | -1 | protein_coding | ENSG00000 |
| 10221433 | 10231331 | -1 | protein_coding | ENSG00000 |
| 10230413 | 10230485 | 1 | miRNA | ENSG00000 |
| 10251901 | 10260055 | 1 | protein_coding | ENSG00000 |
| 10252268 | 10285108 | -1 | lncRNA | ENSG00000 |
| 10259109 | 10260045 | -1 | lncRNA | ENSG00000 |
| 10271093 | 10286615 | 1 | protein_coding | ENSG00000 |
| 10285801 | 10289019 | -1 | lncRNA | ENSG00000 |
| 10286955 | 10288522 | 1 | protein_coding | ENSG00000 |
| 10289952 | 10296778 | 1 | protein_coding | ENSG00000 |
| 10304803 | 10309880 | -1 | protein_coding | ENSG00000 |
| 10305427 | 10316009 | -1 | protein_coding | ENSG00000 |
| 10310045 | 10316015 | -1 | protein_coding | ENSG00000 |

| | | | | |
|---|---|---|---|---|
| 10315471 | 10320678 | -1 | protein_coding | ENSG00000( |
| 10316212 | 10333638 | -1 | protein_coding | ENSG00000( |
| 10333436 | 10336248 | 1 | lncRNA | ENSG00000( |
| 10333776 | 10339661 | -1 | protein_coding | ENSG00000( |
| 10350529 | 10380572 | -1 | protein_coding | ENSG00000( |
| 10391133 | 10420121 | -1 | protein_coding | ENSG00000( |
| 10403458 | 10403538 | -1 | miRNA | ENSG00000( |
| 10416773 | 10469630 | 1 | protein_coding | ENSG00000( |
| 10486125 | 10503558 | -1 | protein_coding | ENSG00000( |
| 10490339 | 10491000 | -1 | lncRNA | ENSG00000( |
| 10512742 | 10517931 | -1 | protein_coding | ENSG00000( |
| 10543895 | 10553418 | 1 | protein_coding | ENSG00000( |
| 10551033 | 10551092 | -1 | snRNA | ENSG00000( |
| 10552122 | 10552204 | 1 | miRNA | ENSG00000( |
| 10553085 | 10566031 | -1 | protein_coding | ENSG00000( |
| 10566460 | 10569059 | -1 | protein_coding | ENSG00000( |
| 10572671 | 10587315 | -1 | protein_coding | ENSG00000( |
| 10598001 | 10598965 | -1 | processed_pseudoge | ENSG00000( |
| 10602457 | 10644557 | 1 | protein_coding | ENSG00000( |
| 10651862 | 10653844 | -1 | lncRNA | ENSG00000( |
| 10654261 | 10692417 | 1 | protein_coding | ENSG00000( |
| 10701430 | 10713437 | 1 | protein_coding | ENSG00000( |
| 10718079 | 10833488 | 1 | protein_coding | ENSG00000( |
| 10718404 | 10718503 | 1 | miRNA | ENSG00000( |
| 10780254 | 10780335 | 1 | miRNA | ENSG00000( |
| 10817426 | 10817496 | -1 | miRNA | ENSG00000( |
| 10828973 | 10829035 | 1 | miRNA | ENSG00000( |
| 10832067 | 10836318 | -1 | protein_coding | ENSG00000( |
| 10836575 | 10869790 | 1 | protein_coding | ENSG00000( |
| 10841839 | 10842250 | 1 | processed_pseudoge | ENSG00000( |
| 10871513 | 10923070 | 1 | protein_coding | ENSG00000( |
| 10922185 | 10928681 | -1 | protein_coding | ENSG00000( |
| 10928811 | 10933535 | 1 | protein_coding | ENSG00000( |
| 10960825 | 11079426 | 1 | protein_coding | ENSG00000( |
| 11010917 | 11016011 | -1 | lncRNA | ENSG00000( |
| 11011708 | 11012006 | 1 | misc_RNA | ENSG00000( |
| 11033727 | 11033912 | 1 | misc_RNA | ENSG00000( |
| 11089362 | 11133816 | 1 | protein_coding | ENSG00000( |
| 11113474 | 11113534 | 1 | miRNA | ENSG00000( |
| 11131520 | 11155808 | -1 | protein_coding | ENSG00000( |
| 11164267 | 11197791 | -1 | protein_coding | ENSG00000( |
| 11199295 | 11262524 | -1 | protein_coding | ENSG00000( |
| 11203628 | 11216168 | 1 | lncRNA | ENSG00000( |
| 11208195 | 11208702 | 1 | TEC | ENSG00000( |
| 11221083 | 11221573 | 1 | lncRNA | ENSG00000( |
| 11237450 | 11241943 | 1 | protein_coding | ENSG00000( |
| 11296139 | 11326996 | 1 | protein_coding | ENSG00000( |
| 11300777 | 11324441 | -1 | lncRNA | ENSG00000( |
| 11322068 | 11346270 | -1 | protein_coding | ENSG00000( |
| 11322156 | 11324195 | 1 | lncRNA | ENSG00000( |

| | | | | |
|---|---|---|---|---|
| 11342776 | 11346518 | -1 | protein_coding | ENSG0000C |
| 11344684 | 11354944 | 1 | protein_coding | ENSG0000C |
| 11355386 | 11365698 | 1 | protein_coding | ENSG0000C |
| 11368123 | 11374935 | -1 | lncRNA | ENSG0000C |
| 11374666 | 11376951 | 1 | protein_coding | ENSG0000C |
| 11377207 | 11384342 | -1 | protein_coding | ENSG0000C |
| 11384341 | 11419342 | -1 | protein_coding | ENSG0000C |
| 11408772 | 11408880 | 1 | misc_RNA | ENSG0000C |
| 11420604 | 11435782 | -1 | protein_coding | ENSG0000C |
| 11435288 | 11450968 | 1 | protein_coding | ENSG0000C |
| 11451326 | 11481046 | -1 | protein_coding | ENSG0000C |
| 11466240 | 11505698 | -1 | protein_coding | ENSG0000C |
| 11483427 | 11505839 | -1 | protein_coding | ENSG0000C |
| 11495544 | 11495622 | -1 | miRNA | ENSG0000C |
| 11505929 | 11529172 | -1 | protein_coding | ENSG0000C |
| 11516047 | 11516330 | -1 | misc_RNA | ENSG0000C |
| 11521968 | 11522498 | 1 | processed_pseudoge | ENSG0000C |
| 11523436 | 11523831 | 1 | processed_pseudoge | ENSG0000C |
| 11538767 | 11550323 | 1 | protein_coding | ENSG0000C |
| 11551147 | 11559236 | -1 | protein_coding | ENSG0000C |
| 11559374 | 11619161 | 1 | protein_coding | ENSG0000C |
| 11574660 | 11579008 | -1 | protein_coding | ENSG0000C |
| 11585246 | 11587135 | 1 | unprocessed_pseudc | ENSG0000C |
| 11639754 | 11686569 | 1 | lncRNA | ENSG0000C |
| 11648364 | 11652877 | 1 | transcribed_unproce | ENSG0000C |
| 11666069 | 11667030 | -1 | processed_pseudoge | ENSG0000C |
| 11680944 | 11686259 | 1 | transcribed_unproce | ENSG0000C |
| 11721265 | 11739009 | -1 | protein_coding | ENSG0000C |
| 11725961 | 11730783 | -1 | lncRNA | ENSG0000C |
| 11756533 | 11759933 | -1 | unprocessed_pseudc | ENSG0000C |
| 11767000 | 11784078 | 1 | protein_coding | ENSG0000C |
| 11796084 | 11798598 | -1 | lncRNA | ENSG0000C |
| 11797667 | 11809622 | 1 | protein_coding | ENSG0000C |
| 11814273 | 11835216 | 1 | protein_coding | ENSG0000C |
| 11841241 | 11842439 | -1 | lncRNA | ENSG0000C |
| 11848726 | 11883750 | 1 | protein_coding | ENSG0000C |
| 11856032 | 11858302 | 1 | TEC | ENSG0000C |
| 11887782 | 11914329 | 1 | protein_coding | ENSG0000C |
| 11925068 | 11950773 | 1 | protein_coding | ENSG0000C |
| 11925075 | 11979290 | 1 | protein_coding | ENSG0000C |
| 11939959 | 11953381 | 1 | lncRNA | ENSG0000C |
| 11965037 | 11980617 | 1 | protein_coding | ENSG0000C |
| 11987617 | 12046275 | 1 | lncRNA | ENSG0000C |
| 11987641 | 12078106 | 1 | protein_coding | ENSG0000C |
| 11996809 | 11996916 | 1 | rRNA_pseudogene | ENSG0000C |
| 12014732 | 12035741 | -1 | protein_coding | ENSG0000C |
| 12017888 | 12052967 | -1 | protein_coding | ENSG0000C |
| 12027913 | 12028021 | -1 | rRNA_pseudogene | ENSG0000C |
| 12043805 | 12052961 | -1 | protein_coding | ENSG0000C |
| 12064731 | 12081565 | 1 | protein_coding | ENSG0000C |

| | | | | |
|---|---|---|---|---|
| 12070164 | 12070271 | 1 | rRNA_pseudogene | ENSG00000 |
| 12075327 | 12113325 | 1 | protein_coding | ENSG00000 |
| 12088245 | 12088346 | -1 | misc_RNA | ENSG00000 |
| 12092263 | 12137235 | 1 | transcribed_unproce | ENSG00000 |
| 12092843 | 12140355 | -1 | protein_coding | ENSG00000 |
| 12096424 | 12096915 | 1 | processed_pseudoge | ENSG00000 |
| 12106497 | 12106614 | 1 | rRNA_pseudogene | ENSG00000 |
| 12132117 | 12156731 | -1 | protein_coding | ENSG00000 |
| 12141806 | 12143322 | -1 | transcribed_processe | ENSG00000 |
| 12142090 | 12156734 | -1 | protein_coding | ENSG00000 |
| 12163064 | 12189871 | 1 | protein_coding | ENSG00000 |
| 12195015 | 12237767 | 1 | transcribed_unproce | ENSG00000 |
| 12224686 | 12294887 | -1 | protein_coding | ENSG00000 |
| 12260126 | 12260537 | 1 | processed_pseudoge | ENSG00000 |
| 12284337 | 12285185 | 1 | processed_pseudoge | ENSG00000 |
| 12317477 | 12333720 | -1 | protein_coding | ENSG00000 |
| 12345944 | 12365905 | -1 | protein_coding | ENSG00000 |
| 12373609 | 12373731 | 1 | processed_pseudoge | ENSG00000 |
| 12379189 | 12401274 | -1 | lncRNA | ENSG00000 |
| 12379746 | 12383687 | -1 | transcribed_unproce | ENSG00000 |
| 12390016 | 12401271 | -1 | protein_coding | ENSG00000 |
| 12391949 | 12441082 | -1 | protein_coding | ENSG00000 |
| 12429706 | 12441021 | -1 | protein_coding | ENSG00000 |
| 12433103 | 12484816 | -1 | protein_coding | ENSG00000 |
| 12450935 | 12460705 | 1 | lncRNA | ENSG00000 |
| 12461184 | 12513854 | -1 | protein_coding | ENSG00000 |
| 12461559 | 12551474 | -1 | protein_coding | ENSG00000 |
| 12502675 | 12504106 | 1 | unprocessed_pseudo | ENSG00000 |
| 12504665 | 12504890 | 1 | unprocessed_pseudo | ENSG00000 |
| 12504905 | 12505976 | 1 | unprocessed_pseudo | ENSG00000 |
| 12506249 | 12508030 | 1 | unprocessed_pseudo | ENSG00000 |
| 12508129 | 12508263 | 1 | processed_pseudoge | ENSG00000 |
| 12525373 | 12551542 | -1 | protein_coding | ENSG00000 |
| 12525720 | 12580975 | -1 | protein_coding | ENSG00000 |
| 12529768 | 12532973 | -1 | lncRNA | ENSG00000 |
| 12552597 | 12553644 | 1 | lncRNA | ENSG00000 |
| 12559571 | 12561105 | 1 | processed_pseudoge | ENSG00000 |
| 12576100 | 12640098 | -1 | protein_coding | ENSG00000 |
| 12610918 | 12633840 | 1 | protein_coding | ENSG00000 |
| 12643275 | 12643919 | -1 | processed_pseudoge | ENSG00000 |
| 12643831 | 12648397 | -1 | protein_coding | ENSG00000 |
| 12646511 | 12666742 | -1 | protein_coding | ENSG00000 |
| 12664828 | 12669397 | -1 | protein_coding | ENSG00000 |
| 12666802 | 12675832 | 1 | protein_coding | ENSG00000 |
| 12668071 | 12681834 | -1 | protein_coding | ENSG00000 |
| 12668073 | 12669415 | -1 | protein_coding | ENSG00000 |
| 12670079 | 12671023 | -1 | lncRNA | ENSG00000 |
| 12675717 | 12681880 | -1 | protein_coding | ENSG00000 |
| 12682693 | 12687279 | -1 | lncRNA | ENSG00000 |
| 12687998 | 12688422 | 1 | protein_coding | ENSG00000 |

| | | | | |
|---|---|---|---|---|
| 12688053 | 12696631 | -1 | protein_coding | ENSG00000 |
| 12688922 | 12689238 | 1 | lncRNA | ENSG00000 |
| 12699194 | 12724011 | -1 | protein_coding | ENSG00000 |
| 12706449 | 12706518 | -1 | snoRNA | ENSG00000 |
| 12725517 | 12725886 | -1 | lncRNA | ENSG00000 |
| 12730640 | 12734684 | -1 | protein_coding | ENSG00000 |
| 12737139 | 12748323 | 1 | protein_coding | ENSG00000 |
| 12751702 | 12758458 | 1 | protein_coding | ENSG00000 |
| 12763003 | 12872740 | -1 | protein_coding | ENSG00000 |
| 12784539 | 12785101 | -1 | lncRNA | ENSG00000 |
| 12787128 | 12787192 | 1 | miRNA | ENSG00000 |
| 12791486 | 12793315 | 1 | protein_coding | ENSG00000 |
| 12796820 | 12801800 | -1 | protein_coding | ENSG00000 |
| 12796823 | 12801849 | 1 | lncRNA | ENSG00000 |
| 12802031 | 12813581 | 1 | protein_coding | ENSG00000 |
| 12806556 | 12815201 | 1 | protein_coding | ENSG00000 |
| 12825478 | 12835428 | -1 | protein_coding | ENSG00000 |
| 12825711 | 12832983 | 1 | lncRNA | ENSG00000 |
| 12833951 | 12874952 | 1 | protein_coding | ENSG00000 |
| 12852260 | 12852327 | 1 | miRNA | ENSG00000 |
| 12875209 | 12881466 | -1 | protein_coding | ENSG00000 |
| 12880969 | 12884088 | 1 | lncRNA | ENSG00000 |
| 12884422 | 12887199 | -1 | protein_coding | ENSG00000 |
| 12891160 | 12914207 | 1 | protein_coding | ENSG00000 |
| 12894133 | 12894880 | 1 | processed_pseudoge | ENSG00000 |
| 12898786 | 12919293 | -1 | protein_coding | ENSG00000 |
| 12920320 | 12920404 | 1 | miRNA | ENSG00000 |
| 12922479 | 12934037 | -1 | protein_coding | ENSG00000 |
| 12930522 | 12933296 | 1 | lncRNA | ENSG00000 |
| 12938578 | 12944489 | 1 | protein_coding | ENSG00000 |
| 12940484 | 12940540 | 1 | miRNA | ENSG00000 |
| 12944118 | 12944487 | -1 | lncRNA | ENSG00000 |
| 12945855 | 12953642 | 1 | protein_coding | ENSG00000 |
| 12953119 | 12957223 | -1 | protein_coding | ENSG00000 |
| 12965159 | 12974762 | 1 | protein_coding | ENSG00000 |
| 12995608 | 13098796 | 1 | protein_coding | ENSG00000 |
| 13013662 | 13014511 | -1 | lncRNA | ENSG00000 |
| 13069792 | 13071374 | -1 | lncRNA | ENSG00000 |
| 13081966 | 13082564 | 1 | TEC | ENSG00000 |
| 13099033 | 13103161 | -1 | protein_coding | ENSG00000 |
| 13104902 | 13117567 | -1 | protein_coding | ENSG00000 |
| 13116862 | 13141147 | 1 | protein_coding | ENSG00000 |
| 13131571 | 13132410 | 1 | lncRNA | ENSG00000 |
| 13139617 | 13141147 | -1 | lncRNA | ENSG00000 |
| 13144058 | 13150383 | -1 | protein_coding | ENSG00000 |
| 13150411 | 13154911 | 1 | protein_coding | ENSG00000 |
| 13152252 | 13152913 | -1 | lncRNA | ENSG00000 |
| 13153071 | 13154193 | -1 | lncRNA | ENSG00000 |
| 13157959 | 13158788 | -1 | processed_pseudoge | ENSG00000 |
| 13206442 | 13633025 | -1 | protein_coding | ENSG00000 |

| | | | | |
|---|---|---|---|---|
| 13731760 | 13763296 | 1 | protein_coding | ENSG00000( |
| 13764522 | 13774282 | 1 | protein_coding | ENSG00000( |
| 13772118 | 13774118 | -1 | lncRNA | ENSG00000( |
| 13774456 | 13778773 | 1 | protein_coding | ENSG00000( |
| 13795460 | 13832230 | 1 | protein_coding | ENSG00000( |
| 13796574 | 13796933 | -1 | lncRNA | ENSG00000( |
| 13815062 | 13815319 | -1 | misc_RNA | ENSG00000( |
| 13823880 | 13842928 | -1 | lncRNA | ENSG00000( |
| 13836287 | 13836359 | -1 | miRNA | ENSG00000( |
| 13836440 | 13836517 | -1 | miRNA | ENSG00000( |
| 13836587 | 13836659 | -1 | miRNA | ENSG00000( |
| 13862063 | 13880757 | 1 | protein_coding | ENSG00000( |
| 13874699 | 13874808 | 1 | miRNA | ENSG00000( |
| 13874875 | 13875011 | 1 | miRNA | ENSG00000( |
| 13882348 | 13906452 | -1 | protein_coding | ENSG00000( |
| 13906201 | 13930879 | 1 | protein_coding | ENSG00000( |
| 13931187 | 13953392 | -1 | protein_coding | ENSG00000( |
| 13952492 | 13961449 | 1 | protein_coding | ENSG00000( |
| 13961530 | 14007039 | -1 | protein_coding | ENSG00000( |
| 14006422 | 14006773 | 1 | lncRNA | ENSG00000( |
| 14028148 | 14031551 | 1 | protein_coding | ENSG00000( |
| 14030855 | 14031557 | -1 | lncRNA | ENSG00000( |
| 14031762 | 14053218 | 1 | protein_coding | ENSG00000( |
| 14053363 | 14062076 | -1 | protein_coding | ENSG00000( |
| 14070342 | 14071237 | 1 | processed_pseudoge | ENSG00000( |
| 14072536 | 14075062 | 1 | protein_coding | ENSG00000( |
| 14073361 | 14073479 | 1 | miRNA | ENSG00000( |
| 14081624 | 14085875 | -1 | protein_coding | ENSG00000( |
| 14087851 | 14091036 | -1 | protein_coding | ENSG00000( |
| 14091688 | 14118084 | -1 | protein_coding | ENSG00000( |
| 14119106 | 14119537 | 1 | lncRNA | ENSG00000( |
| 14119512 | 14136613 | -1 | protein_coding | ENSG00000( |
| 14137146 | 14171267 | 1 | lncRNA | ENSG00000( |
| 14147743 | 14206187 | -1 | protein_coding | ENSG00000( |
| 14213052 | 14213323 | 1 | misc_RNA | ENSG00000( |
| 14217493 | 14219605 | 1 | lncRNA | ENSG00000( |
| 14254189 | 14293205 | -1 | lncRNA | ENSG00000( |
| 14267807 | 14269377 | -1 | lncRNA | ENSG00000( |
| 14305458 | 14370196 | -1 | lncRNA | ENSG00000( |
| 14333743 | 14343916 | 1 | lncRNA | ENSG00000( |
| 14380501 | 14408725 | 1 | protein_coding | ENSG00000( |
| 14402717 | 14408723 | -1 | lncRNA | ENSG00000( |
| 14408798 | 14419383 | -1 | protein_coding | ENSG00000( |
| 14433053 | 14471867 | 1 | protein_coding | ENSG00000( |
| 14458401 | 14459366 | 1 | lncRNA | ENSG00000( |
| 14472466 | 14475354 | -1 | protein_coding | ENSG00000( |
| 14477760 | 14496149 | -1 | protein_coding | ENSG00000( |
| 14489388 | 14489609 | 1 | processed_pseudoge | ENSG00000( |
| 14514769 | 14529770 | -1 | protein_coding | ENSG00000( |
| 14517085 | 14565980 | 1 | protein_coding | ENSG00000( |

| | | | | |
|---|---|---|---|---|
| 14529543 | 14529640 | 1 | miRNA | ENSG0000( |
| 14566078 | 14572066 | -1 | protein_coding | ENSG0000( |
| 14583084 | 14611157 | 1 | protein_coding | ENSG0000( |
| 14584095 | 14584395 | 1 | misc_RNA | ENSG0000( |
| 14588357 | 14588656 | 1 | misc_RNA | ENSG0000( |
| 14619117 | 14690027 | -1 | protein_coding | ENSG0000( |
| 14621634 | 14622242 | -1 | processed_pseudoge | ENSG0000( |
| 14628599 | 14628801 | 1 | processed_pseudoge | ENSG0000( |
| 14689801 | 14733746 | 1 | protein_coding | ENSG0000( |
| 14732392 | 14778560 | -1 | protein_coding | ENSG0000( |
| 14789260 | 14835376 | -1 | protein_coding | ENSG0000( |
| 14792318 | 14792533 | 1 | processed_pseudoge | ENSG0000( |
| 14824251 | 14835285 | -1 | protein_coding | ENSG0000( |
| 14840466 | 14848922 | -1 | protein_coding | ENSG0000( |
| 14852032 | 14852944 | -1 | unprocessed_pseudc | ENSG0000( |
| 14863714 | 14867358 | -1 | transcribed_unproce | ENSG0000( |
| 14878203 | 14886132 | -1 | protein_coding | ENSG0000( |
| 14886839 | 14887115 | -1 | unprocessed_pseudc | ENSG0000( |
| 14891737 | 14897845 | -1 | transcribed_unproce | ENSG0000( |
| 14903232 | 14904138 | -1 | unprocessed_pseudc | ENSG0000( |
| 14916428 | 14917380 | 1 | unprocessed_pseudc | ENSG0000( |
| 14927052 | 14928060 | 1 | unprocessed_pseudc | ENSG0000( |
| 14941489 | 14942448 | 1 | protein_coding | ENSG0000( |
| 14950034 | 15022990 | -1 | protein_coding | ENSG0000( |
| 15010726 | 15023276 | 1 | protein_coding | ENSG0000( |
| 15049480 | 15058293 | 1 | protein_coding | ENSG0000( |
| 15082211 | 15092970 | 1 | protein_coding | ENSG0000( |
| 15107401 | 15114985 | 1 | protein_coding | ENSG0000( |
| 15114984 | 15125786 | -1 | protein_coding | ENSG0000( |
| 15124160 | 15126174 | 1 | lncRNA | ENSG0000( |
| 15141105 | 15142062 | -1 | unprocessed_pseudc | ENSG0000( |
| 15145215 | 15145318 | 1 | snRNA | ENSG0000( |
| 15159038 | 15200995 | -1 | protein_coding | ENSG0000( |
| 15179283 | 15179350 | -1 | miRNA | ENSG0000( |
| 15221015 | 15222297 | -1 | lncRNA | ENSG0000( |
| 15226919 | 15233435 | -1 | protein_coding | ENSG0000( |
| 15235519 | 15332545 | -1 | protein_coding | ENSG0000( |
| 15338948 | 15339237 | 1 | processed_pseudoge | ENSG0000( |
| 15346068 | 15348417 | -1 | lncRNA | ENSG0000( |
| 15353385 | 15379798 | -1 | protein_coding | ENSG0000( |
| 15379076 | 15381194 | 1 | lncRNA | ENSG0000( |
| 15380050 | 15419141 | -1 | protein_coding | ENSG0000( |
| 15383202 | 15383832 | -1 | TEC | ENSG0000( |
| 15398696 | 15400356 | 1 | TEC | ENSG0000( |
| 15419980 | 15449951 | -1 | protein_coding | ENSG0000( |
| 15449548 | 15449608 | 1 | miRNA | ENSG0000( |
| 15451624 | 15464544 | -1 | protein_coding | ENSG0000( |
| 15468645 | 15498956 | -1 | protein_coding | ENSG0000( |
| 15508525 | 15552317 | 1 | protein_coding | ENSG0000( |
| 15515342 | 15516836 | -1 | unprocessed_pseudc | ENSG0000( |

| | | | | |
|---|---|---|---|---|
| 15564074 | 15584709 | 1 | transcribed_unproce | ENSG00000 |
| 15611623 | 15611909 | -1 | processed_pseudoge | ENSG00000 |
| 15611657 | 15612122 | 1 | processed_pseudoge | ENSG00000 |
| 15613247 | 15614553 | -1 | lncRNA | ENSG00000 |
| 15615218 | 15630639 | 1 | protein_coding | ENSG00000 |
| 15640897 | 15662825 | 1 | protein_coding | ENSG00000 |
| 15664106 | 15669826 | 1 | unprocessed_pseudo | ENSG00000 |
| 15673018 | 15697174 | 1 | protein_coding | ENSG00000 |
| 15728020 | 15729060 | 1 | protein_coding | ENSG00000 |
| 15737985 | 15742343 | 1 | protein_coding | ENSG00000 |
| 15760241 | 15779909 | -1 | unprocessed_pseudo | ENSG00000 |
| 15760375 | 15770828 | 1 | lncRNA | ENSG00000 |
| 15787661 | 15801025 | 1 | protein_coding | ENSG00000 |
| 15804549 | 15815664 | -1 | protein_coding | ENSG00000 |
| 15821796 | 15823797 | -1 | unprocessed_pseudo | ENSG00000 |
| 15827040 | 15835853 | -1 | lncRNA | ENSG00000 |
| 15828206 | 15836328 | 1 | lncRNA | ENSG00000 |
| 15834730 | 15834804 | 1 | misc_RNA | ENSG00000 |
| 15851993 | 15864904 | -1 | transcribed_unproce | ENSG00000 |
| 15871667 | 15871852 | -1 | unprocessed_pseudo | ENSG00000 |
| 15878023 | 15898077 | -1 | protein_coding | ENSG00000 |
| 15902300 | 15903238 | -1 | processed_pseudoge | ENSG00000 |
| 15904556 | 15907234 | 1 | TEC | ENSG00000 |
| 15910582 | 15911824 | -1 | processed_pseudoge | ENSG00000 |
| 15912367 | 15934867 | -1 | protein_coding | ENSG00000 |
| 15944299 | 15950350 | 1 | protein_coding | ENSG00000 |
| 15979835 | 15999020 | -1 | unprocessed_pseudo | ENSG00000 |
| 16015287 | 16027462 | 1 | lncRNA | ENSG00000 |
| 16019719 | 16020290 | 1 | TEC | ENSG00000 |
| 16031170 | 16042142 | 1 | lncRNA | ENSG00000 |
| 16034776 | 16036537 | -1 | processed_pseudoge | ENSG00000 |
| 16037009 | 16038455 | -1 | unprocessed_pseudo | ENSG00000 |
| 16051929 | 16052689 | 1 | unprocessed_pseudo | ENSG00000 |
| 16065737 | 16066312 | 1 | lncRNA | ENSG00000 |
| 16067021 | 16103002 | 1 | protein_coding | ENSG00000 |
| 16074293 | 16077395 | -1 | TEC | ENSG00000 |
| 16103761 | 16104254 | -1 | lncRNA | ENSG00000 |
| 16111889 | 16134234 | 1 | protein_coding | ENSG00000 |
| 16123661 | 16139892 | -1 | lncRNA | ENSG00000 |
| 16134028 | 16158575 | 1 | protein_coding | ENSG00000 |
| 16161368 | 16173525 | -1 | protein_coding | ENSG00000 |
| 16185380 | 16192046 | 1 | protein_coding | ENSG00000 |
| 16186276 | 16189458 | -1 | lncRNA | ENSG00000 |
| 16187313 | 16187566 | 1 | misc_RNA | ENSG00000 |
| 16197080 | 16197738 | -1 | lncRNA | ENSG00000 |
| 16197578 | 16245907 | 1 | protein_coding | ENSG00000 |
| 16283359 | 16324514 | -1 | lncRNA | ENSG00000 |
| 16324817 | 16327874 | 1 | protein_coding | ENSG00000 |
| 16352462 | 16353182 | -1 | lncRNA | ENSG00000 |
| 16355239 | 16472085 | -1 | protein_coding | ENSG00000 |

| | | | | |
|---|---|---|---|---|
| 16356329 | 16358327 | -1 | TEC | ENSG0000( |
| 16412684 | 16413576 | 1 | processed_pseudoge | ENSG0000( |
| 16440946 | 16443584 | 1 | lncRNA | ENSG0000( |
| 16479057 | 16496192 | -1 | protein_coding | ENSG0000( |
| 16479067 | 16628204 | -1 | protein_coding | ENSG0000( |
| 16496394 | 16521352 | 1 | protein_coding | ENSG0000( |
| 16517894 | 16542437 | -1 | protein_coding | ENSG0000( |
| 16539688 | 16539984 | -1 | misc_RNA | ENSG0000( |
| 16542746 | 16544814 | 1 | lncRNA | ENSG0000( |
| 16549831 | 16572415 | -1 | protein_coding | ENSG0000( |
| 16551773 | 16552328 | 1 | lncRNA | ENSG0000( |
| 16565551 | 16566330 | 1 | lncRNA | ENSG0000( |
| 16572624 | 16575340 | 1 | lncRNA | ENSG0000( |
| 16574907 | 16629062 | -1 | protein_coding | ENSG0000( |
| 16577642 | 16660115 | -1 | protein_coding | ENSG0000( |
| 16586905 | 16587985 | -1 | lncRNA | ENSG0000( |
| 16586905 | 16589409 | 1 | TEC | ENSG0000( |
| 16610374 | 16612203 | -1 | TEC | ENSG0000( |
| 16610411 | 16636531 | 1 | lncRNA | ENSG0000( |
| 16630743 | 16643942 | 1 | lncRNA | ENSG0000( |
| 16630751 | 16660442 | -1 | protein_coding | ENSG0000( |
| 16633797 | 16635269 | -1 | lncRNA | ENSG0000( |
| 16639967 | 16640668 | -1 | lncRNA | ENSG0000( |
| 16661139 | 16690023 | 1 | protein_coding | ENSG0000( |
| 16712317 | 16712434 | 1 | processed_pseudoge | ENSG0000( |
| 16719976 | 16817963 | 1 | protein_coding | ENSG0000( |
| 16801426 | 16801714 | 1 | lncRNA | ENSG0000( |
| 16829400 | 16880353 | 1 | protein_coding | ENSG0000( |
| 16844025 | 16846473 | 1 | lncRNA | ENSG0000( |
| 16888999 | 16892606 | 1 | protein_coding | ENSG0000( |
| 16892947 | 17026815 | -1 | protein_coding | ENSG0000( |
| 16907462 | 16907723 | 1 | misc_RNA | ENSG0000( |
| 16910490 | 16910784 | -1 | misc_RNA | ENSG0000( |
| 17009189 | 17013460 | 1 | lncRNA | ENSG0000( |
| 17049729 | 17075625 | -1 | protein_coding | ENSG0000( |
| 17051994 | 17055087 | 1 | lncRNA | ENSG0000( |
| 17075781 | 17214537 | 1 | protein_coding | ENSG0000( |
| 17095418 | 17099307 | -1 | lncRNA | ENSG0000( |
| 17152588 | 17168051 | -1 | lncRNA | ENSG0000( |
| 17177511 | 17178476 | -1 | lncRNA | ENSG0000( |
| 17207138 | 17208010 | -1 | lncRNA | ENSG0000( |
| 17215346 | 17219829 | 1 | protein_coding | ENSG0000( |
| 17226213 | 17229219 | 1 | protein_coding | ENSG0000( |
| 17231883 | 17245940 | -1 | protein_coding | ENSG0000( |
| 17235940 | 17255448 | -1 | protein_coding | ENSG0000( |
| 17249171 | 17282786 | -1 | protein_coding | ENSG0000( |
| 17267376 | 17281249 | 1 | protein_coding | ENSG0000( |
| 17267418 | 17282966 | 1 | protein_coding | ENSG0000( |
| 17281645 | 17287646 | 1 | protein_coding | ENSG0000( |
| 17292131 | 17310236 | -1 | protein_coding | ENSG0000( |

| | | | | |
|---|---|---|---|---|
| 17292609 | 17306843 | 1 | protein_coding | ENSG0000C |
| 17296196 | 17296695 | -1 | lncRNA | ENSG0000C |
| 17309518 | 17323298 | 1 | protein_coding | ENSG0000C |
| 17323223 | 17334855 | -1 | protein_coding | ENSG0000C |
| 17334920 | 17342731 | 1 | protein_coding | ENSG0000C |
| 17351450 | 17377342 | -1 | protein_coding | ENSG0000C |
| 17360932 | 17362753 | 1 | lncRNA | ENSG0000C |
| 17390509 | 17394158 | -1 | protein_coding | ENSG0000C |
| 17402939 | 17405630 | -1 | protein_coding | ENSG0000C |
| 17405686 | 17419324 | 1 | lncRNA | ENSG0000C |
| 17405722 | 17433724 | 1 | protein_coding | ENSG0000C |
| 17414257 | 17422324 | -1 | lncRNA | ENSG0000C |
| 17417031 | 17418288 | -1 | processed_pseudoge | ENSG0000C |
| 17419305 | 17419774 | -1 | lncRNA | ENSG0000C |
| 17435509 | 17448668 | -1 | protein_coding | ENSG0000C |
| 17436611 | 17460804 | -1 | protein_coding | ENSG0000C |
| 17452211 | 17452969 | -1 | lncRNA | ENSG0000C |
| 17455425 | 17460954 | -1 | protein_coding | ENSG0000C |
| 17468769 | 17506168 | 1 | protein_coding | ENSG0000C |
| 17488990 | 17511889 | -1 | lncRNA | ENSG0000C |
| 17511636 | 17521288 | 1 | protein_coding | ENSG0000C |
| 17518330 | 17520565 | -1 | lncRNA | ENSG0000C |
| 17523301 | 17553839 | 1 | protein_coding | ENSG0000C |
| 17555649 | 17583162 | 1 | protein_coding | ENSG0000C |
| 17589595 | 17590071 | -1 | processed_pseudoge | ENSG0000C |
| 17601328 | 17688365 | -1 | protein_coding | ENSG0000C |
| 17613722 | 17614430 | 1 | lncRNA | ENSG0000C |
| 17622447 | 17623338 | 1 | TEC | ENSG0000C |
| 17719242 | 17734513 | 1 | protein_coding | ENSG0000C |
| 17727840 | 17734513 | -1 | lncRNA | ENSG0000C |
| 17747718 | 17788568 | 1 | protein_coding | ENSG0000C |
| 17794828 | 17813576 | 1 | protein_coding | ENSG0000C |
| 17816512 | 17821574 | -1 | protein_coding | ENSG0000C |
| 17824780 | 17848071 | -1 | protein_coding | ENSG0000C |
| 17859910 | 17864153 | 1 | protein_coding | ENSG0000C |
| 17862588 | 17862720 | 1 | snoRNA | ENSG0000C |
| 17871273 | 17871705 | -1 | TEC | ENSG0000C |
| 17871945 | 17895174 | 1 | protein_coding | ENSG0000C |
| 17933015 | 17943991 | 1 | protein_coding | ENSG0000C |
| 17951293 | 18000080 | 1 | protein_coding | ENSG0000C |
| 17972025 | 17972117 | 1 | rRNA_pseudogene | ENSG0000C |
| 18001132 | 18014102 | 1 | protein_coding | ENSG0000C |
| 18009440 | 18009881 | -1 | TEC | ENSG0000C |
| 18022403 | 18032099 | 1 | transcribed_processe | ENSG0000C |
| 18040355 | 18041083 | 1 | processed_pseudoge | ENSG0000C |
| 18048637 | 18049090 | 1 | processed_pseudoge | ENSG0000C |
| 18058995 | 18098944 | -1 | protein_coding | ENSG0000C |
| 18097793 | 18151692 | 1 | protein_coding | ENSG0000C |
| 18144522 | 18151691 | -1 | lncRNA | ENSG0000C |
| 18153158 | 18178117 | 1 | protein_coding | ENSG0000C |

| | | | | |
|---|---|---|---|---|
| 18153163 | 18170532 | 1 | protein_coding | ENSG0000( |
| 18173162 | 18178117 | 1 | protein_coding | ENSG0000( |
| 18193218 | 18196948 | 1 | protein_coding | ENSG0000( |
| 18196784 | 18204042 | -1 | protein_coding | ENSG0000( |
| 18204730 | 18220480 | 1 | lncRNA | ENSG0000( |
| 18207961 | 18255419 | -1 | protein_coding | ENSG0000( |
| 18208652 | 18248200 | -1 | protein_coding | ENSG0000( |
| 18249950 | 18255419 | -1 | protein_coding | ENSG0000( |
| 18257097 | 18274509 | -1 | protein_coding | ENSG0000( |
| 18279694 | 18281622 | -1 | protein_coding | ENSG0000( |
| 18282077 | 18282161 | 1 | miRNA | ENSG0000( |
| 18285343 | 18285495 | -1 | processed_pseudoge | ENSG0000( |
| 18306230 | 18323274 | -1 | protein_coding | ENSG0000( |
| 18333276 | 18333573 | 1 | misc_RNA | ENSG0000( |
| 18340598 | 18369950 | 1 | protein_coding | ENSG0000( |
| 18374731 | 18389176 | 1 | protein_coding | ENSG0000( |
| 18386562 | 18386634 | 1 | miRNA | ENSG0000( |
| 18391137 | 18397622 | -1 | protein_coding | ENSG0000( |
| 18418864 | 18434562 | 1 | protein_coding | ENSG0000( |
| 18434388 | 18438167 | -1 | protein_coding | ENSG0000( |
| 18441419 | 18443597 | 1 | lncRNA | ENSG0000( |
| 18442663 | 18522116 | -1 | protein_coding | ENSG0000( |
| 18448275 | 18448802 | 1 | TEC | ENSG0000( |
| 18448275 | 18448805 | -1 | TEC | ENSG0000( |
| 18531613 | 18532632 | 1 | lncRNA | ENSG0000( |
| 18531751 | 18544077 | -1 | protein_coding | ENSG0000( |
| 18532908 | 18536188 | 1 | lncRNA | ENSG0000( |
| 18557762 | 18569387 | 1 | protein_coding | ENSG0000( |
| 18557775 | 18561560 | -1 | lncRNA | ENSG0000( |
| 18568506 | 18569375 | -1 | lncRNA | ENSG0000( |
| 18571730 | 18577550 | 1 | protein_coding | ENSG0000( |
| 18572220 | 18607741 | -1 | protein_coding | ENSG0000( |
| 18588685 | 18592336 | 1 | protein_coding | ENSG0000( |
| 18607430 | 18621039 | 1 | protein_coding | ENSG0000( |
| 18631646 | 18631908 | -1 | misc_RNA | ENSG0000( |
| 18637025 | 18671721 | 1 | protein_coding | ENSG0000( |
| 18683678 | 18782333 | 1 | protein_coding | ENSG0000( |
| 18782773 | 18791305 | -1 | protein_coding | ENSG0000( |
| 18831938 | 18868236 | 1 | protein_coding | ENSG0000( |
| 18868545 | 18896727 | -1 | protein_coding | ENSG0000( |
| 18868545 | 18896158 | -1 | protein_coding | ENSG0000( |
| 18881680 | 18883376 | 1 | lncRNA | ENSG0000( |
| 18899514 | 18919387 | -1 | protein_coding | ENSG0000( |
| 18907013 | 18934397 | -1 | protein_coding | ENSG0000( |
| 18919705 | 18929189 | 1 | protein_coding | ENSG0000( |
| 18929201 | 18941261 | -1 | protein_coding | ENSG0000( |
| 18940235 | 18947452 | 1 | lncRNA | ENSG0000( |
| 18957957 | 18958258 | -1 | misc_RNA | ENSG0000( |
| 18990888 | 19034023 | -1 | protein_coding | ENSG0000( |
| 19033575 | 19060311 | 1 | protein_coding | ENSG0000( |

| | | | | |
|---|---|---|---|---|
| 19063994 | 19113030 | 1 | protein_coding | ENSG00000 |
| 19119169 | 19138513 | -1 | protein_coding | ENSG00000 |
| 19145567 | 19192131 | -1 | protein_coding | ENSG00000 |
| 19145569 | 19192158 | -1 | protein_coding | ENSG00000 |
| 19176903 | 19192591 | -1 | protein_coding | ENSG00000 |
| 19192229 | 19201869 | 1 | protein_coding | ENSG00000 |
| 19201416 | 19203424 | -1 | protein_coding | ENSG00000 |
| 19211958 | 19252233 | 1 | protein_coding | ENSG00000 |
| 19232855 | 19232947 | 1 | snRNA | ENSG00000 |
| 19254748 | 19273369 | -1 | lncRNA | ENSG00000 |
| 19254756 | 19262804 | -1 | protein_coding | ENSG00000 |
| 19255638 | 19256141 | 1 | TEC | ENSG00000 |
| 19264364 | 19273391 | -1 | protein_coding | ENSG00000 |
| 19276018 | 19320509 | -1 | protein_coding | ENSG00000 |
| 19320829 | 19358754 | 1 | protein_coding | ENSG00000 |
| 19385826 | 19508931 | 1 | protein_coding | ENSG00000 |
| 19401979 | 19402603 | -1 | lncRNA | ENSG00000 |
| 19435063 | 19435158 | 1 | miRNA | ENSG00000 |
| 19512418 | 19515548 | -1 | protein_coding | ENSG00000 |
| 19515736 | 19529054 | 1 | protein_coding | ENSG00000 |
| 19516227 | 19536076 | 1 | protein_coding | ENSG00000 |
| 19528861 | 19537581 | 1 | protein_coding | ENSG00000 |
| 19538248 | 19546659 | 1 | protein_coding | ENSG00000 |
| 19561707 | 19618693 | -1 | protein_coding | ENSG00000 |
| 19588091 | 19588415 | -1 | processed_pseudoge | ENSG00000 |
| 19606350 | 19608660 | 1 | lncRNA | ENSG00000 |
| 19623655 | 19628930 | -1 | protein_coding | ENSG00000 |
| 19629476 | 19643657 | -1 | protein_coding | ENSG00000 |
| 19645198 | 19663676 | -1 | protein_coding | ENSG00000 |
| 19668796 | 19683509 | 1 | protein_coding | ENSG00000 |
| 19699203 | 19699409 | -1 | processed_pseudoge | ENSG00000 |
| 19708965 | 19710465 | 1 | processed_pseudoge | ENSG00000 |
| 19710472 | 19733112 | -1 | protein_coding | ENSG00000 |
| 19740884 | 19750127 | 1 | unprocessed_pseudc | ENSG00000 |
| 19749637 | 19754986 | -1 | lncRNA | ENSG00000 |
| 19750618 | 19752544 | 1 | processed_pseudoge | ENSG00000 |
| 19757366 | 19776423 | -1 | lncRNA | ENSG00000 |
| 19757951 | 19774809 | 1 | lncRNA | ENSG00000 |
| 19761038 | 19821716 | -1 | lncRNA | ENSG00000 |
| 19768731 | 19768989 | 1 | processed_pseudoge | ENSG00000 |
| 19776602 | 19836073 | 1 | transcribed_unproce | ENSG00000 |
| 19785839 | 19821751 | -1 | protein_coding | ENSG00000 |
| 19788755 | 19790531 | -1 | lncRNA | ENSG00000 |
| 19856975 | 19857527 | 1 | processed_pseudoge | ENSG00000 |
| 19865882 | 19894674 | 1 | protein_coding | ENSG00000 |
| 19870723 | 19871299 | -1 | processed_pseudoge | ENSG00000 |
| 19892433 | 19895847 | -1 | lncRNA | ENSG00000 |
| 19900913 | 19963464 | 1 | protein_coding | ENSG00000 |
| 19908268 | 19909146 | -1 | unprocessed_pseudc | ENSG00000 |
| 19978190 | 19979610 | -1 | unprocessed_pseudc | ENSG00000 |

| | | | | |
|---|---|---|---|---|
| 19997058 | 20039505 | -1 | protein_coding | ENSG00000( |
| 20033444 | 20034332 | 1 | processed_pseudoge | ENSG00000( |
| 20056466 | 20056594 | -1 | unprocessed_pseudc | ENSG00000( |
| 20062755 | 20063320 | 1 | processed_pseudoge | ENSG00000( |
| 20077994 | 20127076 | 1 | protein_coding | ENSG00000( |
| 20079818 | 20080539 | -1 | processed_pseudoge | ENSG00000( |
| 20087013 | 20087576 | -1 | processed_pseudoge | ENSG00000( |
| 20122569 | 20321305 | -1 | lncRNA | ENSG00000( |
| 20125044 | 20125484 | 1 | transcribed_process | ENSG00000( |
| 20131894 | 20151908 | -1 | transcribed_unproce | ENSG00000( |
| 20136933 | 20137233 | 1 | processed_pseudoge | ENSG00000( |
| 20167214 | 20200488 | 1 | protein_coding | ENSG00000( |
| 20175013 | 20177063 | -1 | processed_pseudoge | ENSG00000( |
| 20206844 | 20206954 | -1 | unprocessed_pseudc | ENSG00000( |
| 20207192 | 20209789 | -1 | TEC | ENSG00000( |
| 20220597 | 20222186 | -1 | TEC | ENSG00000( |
| 20221869 | 20222463 | 1 | processed_pseudoge | ENSG00000( |
| 20249052 | 20250295 | 1 | unprocessed_pseudc | ENSG00000( |
| 20257720 | 20259418 | 1 | processed_pseudoge | ENSG00000( |
| 20267817 | 20268391 | 1 | processed_pseudoge | ENSG00000( |
| 20295089 | 20303794 | 1 | unprocessed_pseudc | ENSG00000( |
| 20323992 | 20324568 | -1 | processed_pseudoge | ENSG00000( |
| 20330121 | 20331517 | 1 | unprocessed_pseudc | ENSG00000( |
| 20340269 | 20424969 | -1 | transcribed_unproce | ENSG00000( |
| 20351502 | 20352079 | 1 | processed_pseudoge | ENSG00000( |
| 20389658 | 20390689 | 1 | unprocessed_pseudc | ENSG00000( |
| 20399272 | 20399354 | -1 | miRNA | ENSG00000( |
| 20416514 | 20417069 | 1 | processed_pseudoge | ENSG00000( |
| 20432552 | 20571095 | 1 | lncRNA | ENSG00000( |
| 20443215 | 20443725 | -1 | processed_pseudoge | ENSG00000( |
| 20535825 | 20565809 | -1 | protein_coding | ENSG00000( |
| 20553792 | 20661583 | -1 | lncRNA | ENSG00000( |
| 20583084 | 20583647 | -1 | unprocessed_pseudc | ENSG00000( |
| 20611559 | 20615389 | 1 | lncRNA | ENSG00000( |
| 20619939 | 20661596 | -1 | protein_coding | ENSG00000( |
| 20674923 | 20696533 | 1 | unprocessed_pseudc | ENSG00000( |
| 20691336 | 20697615 | 1 | unprocessed_pseudc | ENSG00000( |
| 20746923 | 20755250 | -1 | lncRNA | ENSG00000( |
| 20758440 | 20758952 | 1 | unprocessed_pseudc | ENSG00000( |
| 20776304 | 20809995 | 1 | protein_coding | ENSG00000( |
| 20784596 | 20785152 | -1 | processed_pseudoge | ENSG00000( |
| 20838515 | 20847417 | -1 | unprocessed_pseudc | ENSG00000( |
| 20923222 | 20950697 | 1 | protein_coding | ENSG00000( |
| 20926777 | 20928714 | -1 | TEC | ENSG00000( |
| 20961057 | 20965140 | 1 | unprocessed_pseudc | ENSG00000( |
| 20972262 | 20983210 | 1 | lncRNA | ENSG00000( |
| 21020634 | 21060050 | 1 | protein_coding | ENSG00000( |
| 21025915 | 21026041 | -1 | processed_pseudoge | ENSG00000( |
| 21063924 | 21064261 | 1 | unprocessed_pseudc | ENSG00000( |
| 21082159 | 21125270 | 1 | protein_coding | ENSG00000( |

| | | | | |
|---|---|---|---|---|
| 21113128 | 21113236 | -1 | rRNA_pseudogene | ENSG00000( |
| 21123817 | 21124188 | 1 | unprocessed_pseudc | ENSG00000( |
| 21142024 | 21196053 | 1 | protein_coding | ENSG00000( |
| 21149648 | 21150438 | -1 | processed_pseudoge | ENSG00000( |
| 21188598 | 21188953 | 1 | unprocessed_pseudc | ENSG00000( |
| 21221645 | 21222746 | -1 | unprocessed_pseudc | ENSG00000( |
| 21225098 | 21245241 | -1 | unprocessed_pseudc | ENSG00000( |
| 21233335 | 21233652 | 1 | processed_pseudoge | ENSG00000( |
| 21276023 | 21276332 | -1 | processed_pseudoge | ENSG00000( |
| 21289554 | 21289998 | -1 | unprocessed_pseudc | ENSG00000( |
| 21291160 | 21329425 | -1 | protein_coding | ENSG00000( |
| 21318161 | 21318719 | 1 | processed_pseudoge | ENSG00000( |
| 21358930 | 21379302 | 1 | protein_coding | ENSG00000( |
| 21381220 | 21381921 | -1 | processed_pseudoge | ENSG00000( |
| 21382865 | 21387177 | 1 | unprocessed_pseudc | ENSG00000( |
| 21397119 | 21427573 | 1 | protein_coding | ENSG00000( |
| 21405159 | 21491266 | 1 | protein_coding | ENSG00000( |
| 21444241 | 21463908 | -1 | lncRNA | ENSG00000( |
| 21483374 | 21503238 | 1 | lncRNA | ENSG00000( |
| 21496682 | 21556270 | 1 | protein_coding | ENSG00000( |
| 21521343 | 21521896 | -1 | processed_pseudoge | ENSG00000( |
| 21554640 | 21569237 | -1 | lncRNA | ENSG00000( |
| 21569738 | 21587322 | -1 | lncRNA | ENSG00000( |
| 21587432 | 21594628 | -1 | lncRNA | ENSG00000( |
| 21592727 | 21593041 | 1 | unprocessed_pseudc | ENSG00000( |
| 21634833 | 21635175 | -1 | processed_pseudoge | ENSG00000( |
| 21637974 | 21656300 | 1 | unprocessed_pseudc | ENSG00000( |
| 21677550 | 21678221 | -1 | processed_pseudoge | ENSG00000( |
| 21709522 | 21710191 | 1 | processed_pseudoge | ENSG00000( |
| 21719801 | 21720035 | -1 | unprocessed_pseudc | ENSG00000( |
| 21722771 | 21767579 | -1 | protein_coding | ENSG00000( |
| 21750747 | 21753438 | 1 | transcribed_processe | ENSG00000( |
| 21768772 | 21793860 | -1 | lncRNA | ENSG00000( |
| 21800480 | 21801266 | 1 | processed_pseudoge | ENSG00000( |
| 21804949 | 21852125 | -1 | protein_coding | ENSG00000( |
| 21826513 | 21827279 | 1 | processed_pseudoge | ENSG00000( |
| 21932958 | 22010949 | -1 | protein_coding | ENSG00000( |
| 21940820 | 21941533 | 1 | processed_pseudoge | ENSG00000( |
| 21965708 | 21968529 | -1 | TEC | ENSG00000( |
| 21997648 | 21998840 | 1 | processed_pseudoge | ENSG00000( |
| 22017898 | 22052370 | -1 | lncRNA | ENSG00000( |
| 22052430 | 22091480 | 1 | protein_coding | ENSG00000( |
| 22065828 | 22066398 | -1 | processed_pseudoge | ENSG00000( |
| 22128627 | 22130318 | -1 | processed_pseudoge | ENSG00000( |
| 22130325 | 22130489 | 1 | processed_pseudoge | ENSG00000( |
| 22137931 | 22138981 | 1 | processed_pseudoge | ENSG00000( |
| 22140532 | 22141117 | 1 | processed_pseudoge | ENSG00000( |
| 22174766 | 22175191 | -1 | unprocessed_pseudc | ENSG00000( |
| 22179089 | 22215801 | -1 | protein_coding | ENSG00000( |
| 22203128 | 22204003 | 1 | processed_pseudoge | ENSG00000( |

| | | | | |
|---|---|---|---|---|
| 22215634 | 22215661 | -1 | unprocessed_pseudo | ENSG00000 |
| 22243254 | 22245347 | -1 | lncRNA | ENSG00000 |
| 22259693 | 22261335 | 1 | lncRNA | ENSG00000 |
| 22286441 | 22317176 | 1 | protein_coding | ENSG00000 |
| 22298977 | 22299868 | -1 | processed_pseudoge | ENSG00000 |
| 22362589 | 22362825 | 1 | processed_pseudoge | ENSG00000 |
| 22368599 | 22368952 | -1 | processed_pseudoge | ENSG00000 |
| 22391019 | 22532485 | -1 | protein_coding | ENSG00000 |
| 22422513 | 22426045 | 1 | lncRNA | ENSG00000 |
| 22447934 | 22448777 | -1 | processed_pseudoge | ENSG00000 |
| 22455988 | 22456459 | 1 | lncRNA | ENSG00000 |
| 22457329 | 22458212 | -1 | processed_pseudoge | ENSG00000 |
| 22463922 | 22473036 | 1 | unprocessed_pseudo | ENSG00000 |
| 22501699 | 22501948 | -1 | processed_pseudoge | ENSG00000 |
| 22518414 | 22520580 | 1 | lncRNA | ENSG00000 |
| 22520995 | 22527949 | -1 | lncRNA | ENSG00000 |
| 22532626 | 22533494 | 1 | lncRNA | ENSG00000 |
| 22555261 | 22555367 | 1 | snRNA | ENSG00000 |
| 22595479 | 22596163 | -1 | unprocessed_pseudo | ENSG00000 |
| 22596257 | 22603550 | 1 | transcribed_unproce | ENSG00000 |
| 22601485 | 22601707 | 1 | misc_RNA | ENSG00000 |
| 22603269 | 22608501 | 1 | lncRNA | ENSG00000 |
| 22605739 | 22608211 | 1 | transcribed_processe | ENSG00000 |
| 22609606 | 22610601 | 1 | processed_pseudoge | ENSG00000 |
| 22615552 | 22623971 | -1 | lncRNA | ENSG00000 |
| 22620483 | 22620724 | 1 | unprocessed_pseudo | ENSG00000 |
| 22634324 | 22667671 | 1 | protein_coding | ENSG00000 |
| 22665552 | 22716991 | -1 | lncRNA | ENSG00000 |
| 22685167 | 22686732 | 1 | unprocessed_pseudo | ENSG00000 |
| 22699352 | 22699685 | 1 | processed_pseudoge | ENSG00000 |
| 22708834 | 22709314 | -1 | processed_pseudoge | ENSG00000 |
| 22742924 | 22743410 | -1 | unprocessed_pseudo | ENSG00000 |
| 22752183 | 22784151 | -1 | protein_coding | ENSG00000 |
| 22770375 | 22770622 | 1 | unprocessed_pseudo | ENSG00000 |
| 22770937 | 22771191 | 1 | unprocessed_pseudo | ENSG00000 |
| 22773853 | 22774424 | 1 | processed_pseudoge | ENSG00000 |
| 22832291 | 22858667 | 1 | protein_coding | ENSG00000 |
| 22840432 | 22840921 | -1 | processed_pseudoge | ENSG00000 |
| 22861303 | 22861594 | 1 | unprocessed_pseudo | ENSG00000 |
| 22902538 | 22914340 | -1 | lncRNA | ENSG00000 |
| 22925377 | 22925547 | 1 | unprocessed_pseudo | ENSG00000 |
| 22974883 | 23003176 | -1 | protein_coding | ENSG00000 |
| 22994968 | 22995847 | 1 | processed_pseudoge | ENSG00000 |
| 23015059 | 23031818 | -1 | lncRNA | ENSG00000 |
| 23046958 | 23063705 | -1 | lncRNA | ENSG00000 |
| 23068473 | 23071476 | -1 | lncRNA | ENSG00000 |
| 23075201 | 23100361 | 1 | lncRNA | ENSG00000 |
| 23075210 | 23147221 | 1 | protein_coding | ENSG00000 |
| 23123188 | 23124088 | -1 | processed_pseudoge | ENSG00000 |
| 23125665 | 23128543 | 1 | processed_pseudoge | ENSG00000 |

| | | | | |
|---|---|---|---|---|
| 23126569 | 23127792 | -1 | processed_pseudoge | ENSG00000( |
| 23204010 | 23204556 | 1 | processed_pseudoge | ENSG00000( |
| 23221599 | 23250390 | -1 | protein_coding | ENSG00000( |
| 23222755 | 23274221 | -1 | protein_coding | ENSG00000( |
| 23236308 | 23236851 | 1 | processed_pseudoge | ENSG00000( |
| 23255053 | 23257939 | -1 | transcribed_processe | ENSG00000( |
| 23257255 | 23274232 | -1 | lncRNA | ENSG00000( |
| 23259906 | 23274251 | -1 | lncRNA | ENSG00000( |
| 23288620 | 23289345 | 1 | unprocessed_pseudc | ENSG00000( |
| 23290849 | 23291017 | 1 | unprocessed_pseudc | ENSG00000( |
| 23304991 | 23395471 | -1 | protein_coding | ENSG00000( |
| 23323968 | 23329101 | 1 | lncRNA | ENSG00000( |
| 23380317 | 23380876 | 1 | processed_pseudoge | ENSG00000( |
| 23384530 | 23385293 | 1 | processed_pseudoge | ENSG00000( |
| 23399233 | 23416075 | -1 | lncRNA | ENSG00000( |
| 23444262 | 23445189 | 1 | unprocessed_pseudc | ENSG00000( |
| 23479762 | 23480468 | -1 | unprocessed_pseudc | ENSG00000( |
| 23492428 | 23512656 | -1 | unprocessed_pseudc | ENSG00000( |
| 23525631 | 23687220 | -1 | protein_coding | ENSG00000( |
| 23596122 | 23598159 | 1 | unprocessed_pseudc | ENSG00000( |
| 23597341 | 23597592 | 1 | processed_pseudoge | ENSG00000( |
| 23602567 | 23603106 | 1 | unprocessed_pseudc | ENSG00000( |
| 23739195 | 23758891 | -1 | protein_coding | ENSG00000( |
| 23762944 | 23833314 | 1 | transcribed_unproce | ENSG00000( |
| 23798924 | 23799477 | -1 | processed_pseudoge | ENSG00000( |
| 23817599 | 23874701 | -1 | lncRNA | ENSG00000( |
| 23827162 | 23828049 | 1 | processed_pseudoge | ENSG00000( |
| 23842549 | 23843230 | 1 | unprocessed_pseudc | ENSG00000( |
| 23897321 | 23899495 | 1 | lncRNA | ENSG00000( |
| 23914876 | 23945159 | 1 | protein_coding | ENSG00000( |
| 23919081 | 23958035 | -1 | lncRNA | ENSG00000( |
| 23927788 | 23929287 | 1 | lncRNA | ENSG00000( |
| 23935220 | 23936173 | 1 | unprocessed_pseudc | ENSG00000( |
| 24004358 | 24004507 | -1 | rRNA_pseudogene | ENSG00000( |
| 24033401 | 24077377 | 1 | lncRNA | ENSG00000( |
| 24033405 | 24129961 | 1 | protein_coding | ENSG00000( |
| 24078481 | 24080669 | 1 | TEC | ENSG00000( |
| 24098425 | 24098980 | -1 | processed_pseudoge | ENSG00000( |
| 24146701 | 24147081 | 1 | lncRNA | ENSG00000( |
| 24162370 | 24163425 | -1 | processed_pseudoge | ENSG00000( |
| 27638483 | 27646483 | -1 | lncRNA | ENSG00000( |
| 27639300 | 27639665 | -1 | TEC | ENSG00000( |
| 27681072 | 27794005 | -1 | lncRNA | ENSG00000( |
| 27757184 | 27760849 | -1 | lncRNA | ENSG00000( |
| 27793431 | 27984984 | 1 | lncRNA | ENSG00000( |
| 27802838 | 27803472 | 1 | lncRNA | ENSG00000( |
| 27803538 | 27804060 | 1 | processed_pseudoge | ENSG00000( |
| 27805423 | 27806205 | -1 | processed_pseudoge | ENSG00000( |
| 27849635 | 27889222 | -1 | lncRNA | ENSG00000( |
| 27892495 | 27898927 | -1 | unprocessed_pseudc | ENSG00000( |

| | | | | |
|---|---|---|---|---|
| 27902595 | 27984743 | 1 | transcribed_unproce | ENSG0000C |
| 27908356 | 27908554 | -1 | processed_pseudoge | ENSG0000C |
| 27918943 | 27929370 | 1 | transcribed_unproce | ENSG0000C |
| 27983032 | 27983332 | -1 | processed_pseudoge | ENSG0000C |
| 28065267 | 28125430 | 1 | lncRNA | ENSG0000C |
| 28294093 | 28313500 | -1 | lncRNA | ENSG0000C |
| 28316705 | 28386204 | -1 | lncRNA | ENSG0000C |
| 28394851 | 28396428 | -1 | lncRNA | ENSG0000C |
| 28418483 | 28429490 | -1 | lncRNA | ENSG0000C |
| 28435388 | 28727680 | -1 | lncRNA | ENSG0000C |
| 28437060 | 28535277 | -1 | lncRNA | ENSG0000C |
| 28491864 | 28632728 | -1 | lncRNA | ENSG0000C |
| 28509238 | 28518728 | 1 | lncRNA | ENSG0000C |
| 28602379 | 28648303 | -1 | lncRNA | ENSG0000C |
| 28606688 | 28615229 | 1 | lncRNA | ENSG0000C |
| 28683071 | 28727777 | -1 | transcribed_unproce | ENSG0000C |
| 28790812 | 28792871 | -1 | processed_pseudoge | ENSG0000C |
| 28869507 | 28879176 | 1 | lncRNA | ENSG0000C |
| 28883430 | 28970874 | 1 | lncRNA | ENSG0000C |
| 28892931 | 28895949 | 1 | lncRNA | ENSG0000C |
| 28949437 | 28950200 | -1 | lncRNA | ENSG0000C |
| 28960506 | 28964943 | -1 | lncRNA | ENSG0000C |
| 29001670 | 29001775 | 1 | rRNA_pseudogene | ENSG0000C |
| 29002319 | 29015160 | 1 | lncRNA | ENSG0000C |
| 29083094 | 29093031 | -1 | lncRNA | ENSG0000C |
| 29205320 | 29213151 | -1 | protein_coding | ENSG0000C |
| 29213235 | 29216604 | 1 | lncRNA | ENSG0000C |
| 29220864 | 29221666 | 1 | lncRNA | ENSG0000C |
| 29222542 | 29223137 | 1 | lncRNA | ENSG0000C |
| 29233090 | 29233385 | -1 | misc_RNA | ENSG0000C |
| 29268976 | 29298576 | 1 | lncRNA | ENSG0000C |
| 29287011 | 29525752 | -1 | lncRNA | ENSG0000C |
| 29448038 | 29448380 | 1 | lncRNA | ENSG0000C |
| 29489775 | 29526948 | -1 | lncRNA | ENSG0000C |
| 29506966 | 29509041 | -1 | TEC | ENSG0000C |
| 29526499 | 29564479 | 1 | protein_coding | ENSG0000C |
| 29606283 | 29617237 | 1 | protein_coding | ENSG0000C |
| 29665459 | 29675477 | 1 | protein_coding | ENSG0000C |
| 29698886 | 29715789 | -1 | protein_coding | ENSG0000C |
| 29775504 | 29781216 | -1 | lncRNA | ENSG0000C |
| 29811991 | 29824312 | 1 | protein_coding | ENSG0000C |
| 29838532 | 29841818 | -1 | lncRNA | ENSG0000C |
| 29901696 | 29901869 | -1 | unprocessed_pseudc | ENSG0000C |
| 29915862 | 29917830 | 1 | TEC | ENSG0000C |
| 29921182 | 29921653 | 1 | processed_pseudoge | ENSG0000C |
| 29923644 | 30016612 | 1 | protein_coding | ENSG0000C |
| 29956369 | 29956988 | -1 | processed_pseudoge | ENSG0000C |
| 30028741 | 30029916 | -1 | lncRNA | ENSG0000C |
| 30037285 | 30037411 | 1 | processed_pseudoge | ENSG0000C |
| 30037423 | 30038181 | 1 | processed_pseudoge | ENSG0000C |

| | | | | |
|---|---|---|---|---|
| 30058229 | 30060797 | 1 | lncRNA | ENSG0000( |
| 30078133 | 30085893 | 1 | lncRNA | ENSG0000( |
| 30219666 | 30228715 | -1 | lncRNA | ENSG0000( |
| 30228290 | 30713538 | 1 | protein_coding | ENSG0000( |
| 30553956 | 30558058 | 1 | TEC | ENSG0000( |
| 30665274 | 30668647 | -1 | lncRNA | ENSG0000( |
| 30850975 | 30872507 | 1 | lncRNA | ENSG0000( |
| 30907199 | 30909155 | 1 | lncRNA | ENSG0000( |
| 30946582 | 30957192 | 1 | lncRNA | ENSG0000( |
| 31100304 | 31147981 | 1 | lncRNA | ENSG0000( |
| 31149979 | 31349436 | -1 | protein_coding | ENSG0000( |
| 31167457 | 31225143 | 1 | lncRNA | ENSG0000( |
| 31348881 | 31417794 | 1 | lncRNA | ENSG0000( |
| 31364297 | 31366098 | 1 | TEC | ENSG0000( |
| 31421997 | 31427302 | 1 | lncRNA | ENSG0000( |
| 31465653 | 31466431 | 1 | lncRNA | ENSG0000( |
| 31520155 | 31554803 | 1 | lncRNA | ENSG0000( |
| 31588040 | 31595720 | -1 | lncRNA | ENSG0000( |
| 31588159 | 31593786 | 1 | lncRNA | ENSG0000( |
| 31655356 | 31655461 | -1 | rRNA_pseudogene | ENSG0000( |
| 31787148 | 31787255 | -1 | snRNA | ENSG0000( |
| 31797666 | 32072113 | -1 | lncRNA | ENSG0000( |
| 32025808 | 32073809 | 1 | lncRNA | ENSG0000( |
| 32102088 | 32105916 | -1 | lncRNA | ENSG0000( |
| 32103154 | 32106544 | 1 | lncRNA | ENSG0000( |
| 32127334 | 32184504 | -1 | lncRNA | ENSG0000( |
| 32243965 | 32244083 | -1 | rRNA_pseudogene | ENSG0000( |
| 32345594 | 32387667 | 1 | protein_coding | ENSG0000( |
| 32390050 | 32405560 | -1 | lncRNA | ENSG0000( |
| 32405543 | 32485895 | 1 | protein_coding | ENSG0000( |
| 32581190 | 32587453 | 1 | protein_coding | ENSG0000( |
| 32597006 | 32676597 | -1 | protein_coding | ENSG0000( |
| 32608337 | 32608469 | -1 | snoRNA | ENSG0000( |
| 32636078 | 32636475 | 1 | processed_pseudoge | ENSG0000( |
| 32655016 | 32655308 | -1 | misc_RNA | ENSG0000( |
| 32666298 | 32673210 | 1 | lncRNA | ENSG0000( |
| 32675848 | 32678300 | 1 | protein_coding | ENSG0000( |
| 32687089 | 32691750 | -1 | lncRNA | ENSG0000( |
| 32691961 | 32713796 | 1 | protein_coding | ENSG0000( |
| 32718298 | 32719595 | -1 | lncRNA | ENSG0000( |
| 32719753 | 32829580 | 1 | protein_coding | ENSG0000( |
| 32824142 | 32824505 | -1 | processed_pseudoge | ENSG0000( |
| 32830509 | 32869767 | -1 | protein_coding | ENSG0000( |
| 32831295 | 32833168 | 1 | lncRNA | ENSG0000( |
| 32862800 | 32863092 | -1 | misc_RNA | ENSG0000( |
| 32875925 | 32971991 | -1 | protein_coding | ENSG0000( |
| 32963499 | 32963865 | 1 | processed_pseudoge | ENSG0000( |
| 32972209 | 32978229 | 1 | protein_coding | ENSG0000( |
| 32978592 | 33064888 | -1 | protein_coding | ENSG0000( |
| 33051515 | 33051701 | 1 | processed_pseudoge | ENSG0000( |

| | | | | |
|---|---|---|---|---|
| 33067175 | 33067433 | 1 | processed_pseudoge | ENSG00000 |
| 33080899 | 33130542 | 1 | protein_coding | ENSG00000 |
| 33132090 | 33175799 | 1 | protein_coding | ENSG00000 |
| 33177603 | 33208864 | 1 | protein_coding | ENSG00000 |
| 33178056 | 33178651 | 1 | lncRNA | ENSG00000 |
| 33207129 | 33207639 | 1 | lncRNA | ENSG00000 |
| 33208664 | 33225850 | -1 | protein_coding | ENSG00000 |
| 33217817 | 33222355 | 1 | lncRNA | ENSG00000 |
| 33283978 | 33285739 | -1 | lncRNA | ENSG00000 |
| 33299934 | 33301168 | 1 | lncRNA | ENSG00000 |
| 33299934 | 33302534 | -1 | protein_coding | ENSG00000 |
| 33301279 | 33301940 | 1 | lncRNA | ENSG00000 |
| 33302857 | 33305054 | 1 | lncRNA | ENSG00000 |
| 33305036 | 33309387 | 1 | lncRNA | ENSG00000 |
| 33340686 | 33341984 | 1 | processed_pseudoge | ENSG00000 |
| 33360497 | 33361448 | -1 | processed_pseudoge | ENSG00000 |
| 33373685 | 33382686 | 1 | protein_coding | ENSG00000 |
| 33386950 | 33521791 | -1 | protein_coding | ENSG00000 |
| 33555568 | 33557470 | 1 | lncRNA | ENSG00000 |
| 33589206 | 33589688 | -1 | processed_pseudoge | ENSG00000 |
| 33621953 | 33773509 | 1 | protein_coding | ENSG00000 |
| 33795933 | 33815763 | 1 | protein_coding | ENSG00000 |
| 33906352 | 33908391 | 1 | lncRNA | ENSG00000 |
| 33927364 | 33927625 | -1 | misc_RNA | ENSG00000 |
| 33997289 | 34066283 | -1 | lncRNA | ENSG00000 |
| 34021659 | 34021895 | 1 | processed_pseudoge | ENSG00000 |
| 34022656 | 34023427 | -1 | processed_pseudoge | ENSG00000 |
| 34092561 | 34093310 | -1 | processed_pseudoge | ENSG00000 |
| 34126973 | 34127423 | 1 | processed_pseudoge | ENSG00000 |
| 34172504 | 34229515 | 1 | protein_coding | ENSG00000 |
| 34254552 | 34355571 | 1 | protein_coding | ENSG00000 |
| 34296277 | 34296592 | -1 | processed_pseudoge | ENSG00000 |
| 34348356 | 34359412 | -1 | lncRNA | ENSG00000 |
| 34359480 | 34402413 | 1 | protein_coding | ENSG00000 |
| 34396315 | 34409364 | 1 | protein_coding | ENSG00000 |
| 34404399 | 34426168 | 1 | protein_coding | ENSG00000 |
| 34421988 | 34422311 | -1 | misc_RNA | ENSG00000 |
| 34426112 | 34428273 | -1 | lncRNA | ENSG00000 |
| 34428352 | 34471251 | 1 | protein_coding | ENSG00000 |
| 34481758 | 34512304 | 1 | protein_coding | ENSG00000 |
| 34533427 | 34534084 | -1 | processed_pseudoge | ENSG00000 |
| 34542462 | 34545395 | -1 | unprocessed_pseudo | ENSG00000 |
| 34576733 | 34577701 | -1 | lncRNA | ENSG00000 |
| 34593329 | 34675699 | -1 | protein_coding | ENSG00000 |
| 34634844 | 34635717 | 1 | processed_pseudoge | ENSG00000 |
| 34637696 | 34637781 | -1 | snoRNA | ENSG00000 |
| 34638122 | 34640689 | -1 | lncRNA | ENSG00000 |
| 34645380 | 34646838 | 1 | TEC | ENSG00000 |
| 34645662 | 34646070 | 1 | processed_pseudoge | ENSG00000 |
| 34675717 | 34677581 | -1 | TEC | ENSG00000 |

| | | | | |
|---|---|---|---|---|
| 34677639 | 34686397 | 1 | protein_coding | ENSG00000( |
| 34733298 | 34733837 | -1 | lncRNA | ENSG00000( |
| 34734155 | 34745378 | 1 | protein_coding | ENSG00000( |
| 34758073 | 34773229 | -1 | protein_coding | ENSG00000( |
| 34779412 | 34780021 | -1 | processed_pseudoge | ENSG00000( |
| 34788527 | 34832869 | -1 | lncRNA | ENSG00000( |
| 34811589 | 34814345 | -1 | lncRNA | ENSG00000( |
| 34825414 | 34825996 | 1 | processed_pseudoge | ENSG00000( |
| 34837889 | 34872479 | 1 | lncRNA | ENSG00000( |
| 34849046 | 34860747 | -1 | lncRNA | ENSG00000( |
| 34866352 | 34866919 | 1 | processed_pseudoge | ENSG00000( |
| 34891605 | 34905139 | -1 | lncRNA | ENSG00000( |
| 34905315 | 34908188 | 1 | lncRNA | ENSG00000( |
| 34923299 | 34926812 | -1 | lncRNA | ENSG00000( |
| 34926903 | 34945170 | 1 | protein_coding | ENSG00000( |
| 34949038 | 34951205 | -1 | TEC | ENSG00000( |
| 34956354 | 34964229 | -1 | protein_coding | ENSG00000( |
| 34994784 | 35026471 | 1 | protein_coding | ENSG00000( |
| 34998233 | 35000170 | -1 | lncRNA | ENSG00000( |
| 35014961 | 35025335 | -1 | lncRNA | ENSG00000( |
| 35030470 | 35040449 | 1 | protein_coding | ENSG00000( |
| 35040506 | 35066571 | 1 | protein_coding | ENSG00000( |
| 35042902 | 35106307 | -1 | lncRNA | ENSG00000( |
| 35090930 | 35113664 | 1 | lncRNA | ENSG00000( |
| 35106510 | 35124297 | 1 | protein_coding | ENSG00000( |
| 35115879 | 35124324 | 1 | protein_coding | ENSG00000( |
| 35122700 | 35122764 | 1 | miRNA | ENSG00000( |
| 35124513 | 35142451 | -1 | protein_coding | ENSG00000( |
| 35137645 | 35139580 | -1 | TEC | ENSG00000( |
| 35138808 | 35143109 | 1 | protein_coding | ENSG00000( |
| 35138824 | 35151336 | 1 | lncRNA | ENSG00000( |
| 35143250 | 35154301 | 1 | protein_coding | ENSG00000( |
| 35154730 | 35169881 | 1 | protein_coding | ENSG00000( |
| 35224800 | 35228729 | -1 | protein_coding | ENSG00000( |
| 35232291 | 35233003 | -1 | unprocessed_pseudc | ENSG00000( |
| 35248330 | 35267964 | 1 | protein_coding | ENSG00000( |
| 35251440 | 35253639 | -1 | lncRNA | ENSG00000( |
| 35262846 | 35264804 | 1 | lncRNA | ENSG00000( |
| 35268962 | 35279821 | 1 | protein_coding | ENSG00000( |
| 35280716 | 35285143 | 1 | protein_coding | ENSG00000( |
| 35292125 | 35313807 | 1 | protein_coding | ENSG00000( |
| 35319261 | 35347361 | 1 | protein_coding | ENSG00000( |
| 35330843 | 35331920 | 1 | lncRNA | ENSG00000( |
| 35345513 | 35345627 | 1 | miRNA | ENSG00000( |
| 35351552 | 35353862 | 1 | protein_coding | ENSG00000( |
| 35358460 | 35360489 | 1 | protein_coding | ENSG00000( |
| 35370929 | 35372962 | 1 | protein_coding | ENSG00000( |
| 35382172 | 35384052 | 1 | processed_pseudoge | ENSG00000( |
| 35399511 | 35419385 | 1 | lncRNA | ENSG00000( |
| 35407770 | 35408493 | 1 | transcribed_unproce | ENSG00000( |

| | | | | |
|---|---|---|---|---|
| 35414881 | 35415149 | -1 | misc_RNA | ENSG0000C |
| 35424062 | 35424652 | -1 | lncRNA | ENSG0000C |
| 35432957 | 35434642 | -1 | lncRNA | ENSG0000C |
| 35443907 | 35451767 | 1 | protein_coding | ENSG0000C |
| 35487324 | 35495558 | -1 | protein_coding | ENSG0000C |
| 35497220 | 35513658 | -1 | protein_coding | ENSG0000C |
| 35523367 | 35528351 | -1 | protein_coding | ENSG0000C |
| 35533455 | 35545319 | 1 | protein_coding | ENSG0000C |
| 35540738 | 35546029 | -1 | lncRNA | ENSG0000C |
| 35545600 | 35547526 | 1 | protein_coding | ENSG0000C |
| 35550031 | 35563658 | -1 | protein_coding | ENSG0000C |
| 35557956 | 35581954 | 1 | lncRNA | ENSG0000C |
| 35586161 | 35587296 | 1 | protein_coding | ENSG0000C |
| 35596581 | 35601596 | 1 | lncRNA | ENSG0000C |
| 35608285 | 35612666 | -1 | lncRNA | ENSG0000C |
| 35612735 | 35625355 | 1 | protein_coding | ENSG0000C |
| 35629030 | 35637686 | 1 | protein_coding | ENSG0000C |
| 35641745 | 35644871 | 1 | protein_coding | ENSG0000C |
| 35648323 | 35658782 | 1 | protein_coding | ENSG0000C |
| 35666516 | 35678483 | 1 | protein_coding | ENSG0000C |
| 35667948 | 35673291 | -1 | lncRNA | ENSG0000C |
| 35683778 | 35684201 | -1 | processed_pseudoge | ENSG0000C |
| 35704558 | 35717038 | 1 | protein_coding | ENSG0000C |
| 35718019 | 35738878 | 1 | protein_coding | ENSG0000C |
| 35738801 | 35742453 | -1 | protein_coding | ENSG0000C |
| 35739252 | 35745432 | -1 | protein_coding | ENSG0000C |
| 35742464 | 35745445 | -1 | protein_coding | ENSG0000C |
| 35745600 | 35747519 | 1 | protein_coding | ENSG0000C |
| 35745678 | 35754519 | 1 | protein_coding | ENSG0000C |
| 35747057 | 35753415 | -1 | lncRNA | ENSG0000C |
| 35748361 | 35754519 | 1 | protein_coding | ENSG0000C |
| 35754566 | 35755490 | 1 | lncRNA | ENSG0000C |
| 35754566 | 35758079 | -1 | protein_coding | ENSG0000C |
| 35758143 | 35771028 | 1 | protein_coding | ENSG0000C |
| 35769144 | 35771028 | -1 | lncRNA | ENSG0000C |
| 35774532 | 35788822 | 1 | protein_coding | ENSG0000C |
| 35785697 | 35797902 | -1 | transcribed_unitary_ | ENSG0000C |
| 35799988 | 35813299 | -1 | protein_coding | ENSG0000C |
| 35825964 | 35869287 | -1 | protein_coding | ENSG0000C |
| 35855861 | 35867109 | 1 | protein_coding | ENSG0000C |
| 35867899 | 35879791 | 1 | protein_coding | ENSG0000C |
| 35881890 | 35882153 | 1 | misc_RNA | ENSG0000C |
| 35887653 | 35902303 | -1 | protein_coding | ENSG0000C |
| 35901561 | 35903855 | -1 | TEC | ENSG0000C |
| 35902529 | 35904377 | 1 | protein_coding | ENSG0000C |
| 35904401 | 35908295 | -1 | protein_coding | ENSG0000C |
| 35935358 | 35945767 | 1 | protein_coding | ENSG0000C |
| 35947115 | 35959192 | 1 | lncRNA | ENSG0000C |
| 35957121 | 35957382 | -1 | processed_pseudoge | ENSG0000C |
| 35960512 | 35964063 | 1 | lncRNA | ENSG0000C |

| | | | | |
|---|---|---|---|---|
| 35995199 | 35996315 | 1 | protein_coding | ENSG00000 |
| 36003307 | 36008813 | -1 | protein_coding | ENSG00000 |
| 36008638 | 36014235 | -1 | lncRNA | ENSG00000 |
| 36009120 | 36014239 | -1 | protein_coding | ENSG00000 |
| 36014508 | 36045972 | 1 | lncRNA | ENSG00000 |
| 36014660 | 36033343 | -1 | protein_coding | ENSG00000 |
| 36034984 | 36054739 | -1 | protein_coding | ENSG00000 |
| 36054881 | 36105108 | 1 | protein_coding | ENSG00000 |
| 36070239 | 36071070 | 1 | TEC | ENSG00000 |
| 36111151 | 36113711 | 1 | protein_coding | ENSG00000 |
| 36113709 | 36115213 | -1 | protein_coding | ENSG00000 |
| 36114289 | 36125947 | 1 | protein_coding | ENSG00000 |
| 36114362 | 36115146 | 1 | TEC | ENSG00000 |
| 36139953 | 36150353 | 1 | protein_coding | ENSG00000 |
| 36144081 | 36148348 | -1 | lncRNA | ENSG00000 |
| 36150922 | 36152449 | -1 | protein_coding | ENSG00000 |
| 36173406 | 36173853 | -1 | processed_pseudoge | ENSG00000 |
| 36182060 | 36246257 | -1 | protein_coding | ENSG00000 |
| 36203347 | 36203459 | -1 | misc_RNA | ENSG00000 |
| 36214602 | 36238774 | 1 | protein_coding | ENSG00000 |
| 36304564 | 36321274 | 1 | lncRNA | ENSG00000 |
| 36313067 | 36331770 | -1 | lncRNA | ENSG00000 |
| 36334453 | 36379201 | -1 | protein_coding | ENSG00000 |
| 36378028 | 36378832 | 1 | lncRNA | ENSG00000 |
| 36379351 | 36380912 | -1 | lncRNA | ENSG00000 |
| 36383120 | 36418644 | -1 | protein_coding | ENSG00000 |
| 36421536 | 36422534 | 1 | processed_pseudoge | ENSG00000 |
| 36445119 | 36489902 | -1 | protein_coding | ENSG00000 |
| 36489649 | 36491040 | 1 | lncRNA | ENSG00000 |
| 36496916 | 36498159 | -1 | processed_pseudoge | ENSG00000 |
| 36510687 | 36528271 | -1 | protein_coding | ENSG00000 |
| 36528318 | 36535235 | 1 | lncRNA | ENSG00000 |
| 36534774 | 36605276 | -1 | protein_coding | ENSG00000 |
| 36573070 | 36594708 | 1 | lncRNA | ENSG00000 |
| 36604817 | 36634114 | 1 | protein_coding | ENSG00000 |
| 36636618 | 36666853 | -1 | protein_coding | ENSG00000 |
| 36668102 | 36669404 | 1 | lncRNA | ENSG00000 |
| 36682636 | 36687449 | -1 | lncRNA | ENSG00000 |
| 36687612 | 36727701 | 1 | protein_coding | ENSG00000 |
| 36714383 | 36772825 | -1 | protein_coding | ENSG00000 |
| 36773112 | 36777078 | 1 | lncRNA | ENSG00000 |
| 36773712 | 36775908 | -1 | lncRNA | ENSG00000 |
| 36797502 | 36831596 | 1 | lncRNA | ENSG00000 |
| 36817428 | 36850787 | -1 | protein_coding | ENSG00000 |
| 36850361 | 36913029 | 1 | protein_coding | ENSG00000 |
| 36858227 | 36858329 | -1 | misc_RNA | ENSG00000 |
| 36888124 | 36916291 | -1 | protein_coding | ENSG00000 |
| 36901742 | 36902117 | 1 | processed_pseudoge | ENSG00000 |
| 36916329 | 36998700 | 1 | protein_coding | ENSG00000 |
| 37007857 | 37130368 | 1 | protein_coding | ENSG00000 |

| | | | | |
|---|---|---|---|---|
| 37024886 | 37025254 | -1 | unprocessed_pseudc | ENSG00000C |
| 37075113 | 37078605 | -1 | lncRNA | ENSG00000C |
| 37091341 | 37092564 | 1 | lncRNA | ENSG00000C |
| 37093019 | 37210512 | -1 | protein_coding | ENSG00000C |
| 37106734 | 37172741 | -1 | protein_coding | ENSG00000C |
| 37128679 | 37141640 | -1 | lncRNA | ENSG00000C |
| 37181579 | 37218153 | -1 | protein_coding | ENSG00000C |
| 37217926 | 37248738 | 1 | protein_coding | ENSG00000C |
| 37235507 | 37304395 | -1 | lncRNA | ENSG00000C |
| 37251885 | 37265535 | 1 | lncRNA | ENSG00000C |
| 37304451 | 37309641 | 1 | lncRNA | ENSG00000C |
| 37312837 | 37369365 | 1 | protein_coding | ENSG00000C |
| 37314868 | 37315620 | -1 | lncRNA | ENSG00000C |
| 37337236 | 37337743 | 1 | lncRNA | ENSG00000C |
| 37371061 | 37393066 | 1 | protein_coding | ENSG00000C |
| 37401404 | 37403846 | -1 | TEC | ENSG00000C |
| 37411155 | 37469275 | -1 | protein_coding | ENSG00000C |
| 37467585 | 37488652 | 1 | protein_coding | ENSG00000C |
| 37497159 | 37507122 | -1 | lncRNA | ENSG00000C |
| 37503906 | 37504465 | -1 | lncRNA | ENSG00000C |
| 37506939 | 37548762 | 1 | protein_coding | ENSG00000C |
| 37545470 | 37549171 | -1 | lncRNA | ENSG00000C |
| 37548914 | 37587348 | 1 | lncRNA | ENSG00000C |
| 37551406 | 37614179 | 1 | protein_coding | ENSG00000C |
| 37554782 | 37594792 | -1 | protein_coding | ENSG00000C |
| 37613749 | 37692337 | -1 | protein_coding | ENSG00000C |
| 37667751 | 37692315 | -1 | protein_coding | ENSG00000C |
| 37689865 | 37719188 | -1 | protein_coding | ENSG00000C |
| 37693280 | 37693564 | -1 | processed_pseudoge | ENSG00000C |
| 37696371 | 37719761 | -1 | protein_coding | ENSG00000C |
| 37728586 | 37730643 | -1 | lncRNA | ENSG00000C |
| 37728596 | 37730733 | 1 | lncRNA | ENSG00000C |
| 37735833 | 37817300 | -1 | protein_coding | ENSG00000C |
| 37779686 | 37792865 | 1 | transcribed_processe | ENSG00000C |
| 37817359 | 37832160 | 1 | transcribed_unproce | ENSG00000C |
| 37823722 | 37855215 | -1 | transcribed_unproce | ENSG00000C |
| 37884823 | 37906677 | -1 | protein_coding | ENSG00000C |
| 37907208 | 38208369 | 1 | protein_coding | ENSG00000C |
| 37963853 | 37964790 | 1 | lncRNA | ENSG00000C |
| 38108500 | 38109533 | -1 | lncRNA | ENSG00000C |
| 38184376 | 38186265 | 1 | lncRNA | ENSG00000C |
| 38199836 | 38200934 | 1 | lncRNA | ENSG00000C |
| 38200234 | 38200532 | 1 | misc_RNA | ENSG00000C |
| 38211006 | 38229714 | -1 | protein_coding | ENSG00000C |
| 38244035 | 38292615 | 1 | protein_coding | ENSG00000C |
| 38251237 | 38256532 | -1 | protein_coding | ENSG00000C |
| 38289151 | 38304910 | 1 | protein_coding | ENSG00000C |
| 38290117 | 38292423 | -1 | lncRNA | ENSG00000C |
| 38294544 | 38294636 | 1 | misc_RNA | ENSG00000C |
| 38303558 | 38317273 | -1 | protein_coding | ENSG00000C |

| | | | | |
|---|---|---|---|---|
| 38304164 | 38305006 | 1 | protein_coding | ENSG00000( |
| 38319845 | 38332076 | 1 | protein_coding | ENSG00000( |
| 38335775 | 38370943 | 1 | protein_coding | ENSG00000( |
| 38361795 | 38362484 | -1 | lncRNA | ENSG00000( |
| 38374536 | 38383824 | 1 | protein_coding | ENSG00000( |
| 38384267 | 38388082 | -1 | protein_coding | ENSG00000( |
| 38385522 | 38386759 | 1 | lncRNA | ENSG00000( |
| 38388421 | 38399587 | 1 | protein_coding | ENSG00000( |
| 38403135 | 38409088 | 1 | protein_coding | ENSG00000( |
| 38409051 | 38426305 | -1 | protein_coding | ENSG00000( |
| 38433699 | 38587564 | 1 | protein_coding | ENSG00000( |
| 38526954 | 38528555 | -1 | lncRNA | ENSG00000( |
| 38535517 | 38537128 | -1 | lncRNA | ENSG00000( |
| 38587641 | 38618882 | -1 | protein_coding | ENSG00000( |
| 38596346 | 38601694 | 1 | lncRNA | ENSG00000( |
| 38619082 | 38636955 | 1 | protein_coding | ENSG00000( |
| 38647649 | 38731589 | 1 | protein_coding | ENSG00000( |
| 38683873 | 38693606 | -1 | lncRNA | ENSG00000( |
| 38730187 | 38769904 | -1 | protein_coding | ENSG00000( |
| 38738284 | 38739863 | 1 | lncRNA | ENSG00000( |
| 38770971 | 38773492 | -1 | protein_coding | ENSG00000( |
| 38789200 | 38791754 | 1 | protein_coding | ENSG00000( |
| 38797002 | 38797109 | -1 | snRNA | ENSG00000( |
| 38801671 | 38812945 | -1 | protein_coding | ENSG00000( |
| 38815422 | 38831841 | -1 | protein_coding | ENSG00000( |
| 38817471 | 38840178 | -1 | protein_coding | ENSG00000( |
| 38820166 | 38823223 | 1 | lncRNA | ENSG00000( |
| 38836388 | 38852347 | -1 | protein_coding | ENSG00000( |
| 38844729 | 38845499 | -1 | lncRNA | ENSG00000( |
| 38867830 | 38878275 | -1 | protein_coding | ENSG00000( |
| 38870159 | 38873763 | 1 | lncRNA | ENSG00000( |
| 38878555 | 38899862 | -1 | protein_coding | ENSG00000( |
| 38899700 | 38908893 | 1 | protein_coding | ENSG00000( |
| 38908980 | 38912158 | -1 | protein_coding | ENSG00000( |
| 38915266 | 38930896 | -1 | protein_coding | ENSG00000( |
| 38915404 | 38949855 | -1 | protein_coding | ENSG00000( |
| 38930548 | 38933162 | 1 | protein_coding | ENSG00000( |
| 38935297 | 38938632 | -1 | lncRNA | ENSG00000( |
| 38941401 | 38975742 | -1 | protein_coding | ENSG00000( |
| 38975161 | 39014527 | 1 | lncRNA | ENSG00000( |
| 38990714 | 39032785 | -1 | protein_coding | ENSG00000( |
| 39030436 | 39031323 | 1 | lncRNA | ENSG00000( |
| 39083913 | 39111493 | 1 | protein_coding | ENSG00000( |
| 39125770 | 39182816 | 1 | protein_coding | ENSG00000( |
| 39134882 | 39136463 | -1 | lncRNA | ENSG00000( |
| 39196964 | 39201884 | 1 | protein_coding | ENSG00000( |
| 39202831 | 39204266 | -1 | protein_coding | ENSG00000( |
| 39236433 | 39237293 | -1 | unprocessed_pseudc | ENSG00000( |
| 39243553 | 39245129 | -1 | protein_coding | ENSG00000( |
| 39246314 | 39248856 | -1 | polymorphic_pseudc | ENSG00000( |

| | | | | |
|---|---|---|---|---|
| 39254916 | 39255269 | -1 | processed_pseudoge | ENSG00000 |
| 39264382 | 39265817 | 1 | unprocessed_pseudo | ENSG00000 |
| 39268514 | 39270092 | 1 | protein_coding | ENSG00000 |
| 39296407 | 39298673 | 1 | protein_coding | ENSG00000 |
| 39306568 | 39315336 | -1 | protein_coding | ENSG00000 |
| 39314651 | 39320858 | -1 | lncRNA | ENSG00000 |
| 39328353 | 39342372 | -1 | protein_coding | ENSG00000 |
| 39341773 | 39341945 | -1 | lncRNA | ENSG00000 |
| 39342396 | 39385710 | 1 | protein_coding | ENSG00000 |
| 39369153 | 39369448 | 1 | misc_RNA | ENSG00000 |
| 39385629 | 39391154 | -1 | protein_coding | ENSG00000 |
| 39391303 | 39400641 | 1 | protein_coding | ENSG00000 |
| 39406847 | 39409412 | 1 | protein_coding | ENSG00000 |
| 39409623 | 39409678 | -1 | miRNA | ENSG00000 |
| 39412669 | 39428415 | 1 | protein_coding | ENSG00000 |
| 39433137 | 39435949 | -1 | protein_coding | ENSG00000 |
| 39436156 | 39476670 | 1 | protein_coding | ENSG00000 |
| 39480412 | 39493785 | 1 | protein_coding | ENSG00000 |
| 39498895 | 39508481 | 1 | protein_coding | ENSG00000 |
| 39515113 | 39520686 | 1 | protein_coding | ENSG00000 |
| 39530987 | 39532852 | -1 | protein_coding | ENSG00000 |
| 39532412 | 39534422 | 1 | lncRNA | ENSG00000 |
| 39534463 | 39535021 | 1 | processed_pseudoge | ENSG00000 |
| 39538707 | 39540161 | -1 | protein_coding | ENSG00000 |
| 39553034 | 39555356 | 1 | unprocessed_pseudo | ENSG00000 |
| 39584042 | 39586489 | 1 | unprocessed_pseudo | ENSG00000 |
| 39588955 | 39589125 | 1 | processed_pseudoge | ENSG00000 |
| 39590981 | 39591151 | 1 | processed_pseudoge | ENSG00000 |
| 39597832 | 39598002 | 1 | processed_pseudoge | ENSG00000 |
| 39602501 | 39607474 | 1 | protein_coding | ENSG00000 |
| 39631300 | 39631446 | -1 | processed_pseudoge | ENSG00000 |
| 39638685 | 39642401 | 1 | transcribed_unproce | ENSG00000 |
| 39644502 | 39644612 | 1 | processed_pseudoge | ENSG00000 |
| 39655913 | 39660647 | 1 | protein_coding | ENSG00000 |
| 39679374 | 39686373 | 1 | transcribed_unproce | ENSG00000 |
| 39685244 | 39694577 | 1 | lncRNA | ENSG00000 |
| 39693925 | 39696258 | 1 | transcribed_unproce | ENSG00000 |
| 39697904 | 39698073 | 1 | processed_pseudoge | ENSG00000 |
| 39704481 | 39709444 | 1 | protein_coding | ENSG00000 |
| 39726030 | 39726775 | -1 | unprocessed_pseudo | ENSG00000 |
| 39731255 | 39738029 | -1 | protein_coding | ENSG00000 |
| 39776595 | 39786167 | 1 | protein_coding | ENSG00000 |
| 39812972 | 39813555 | -1 | lncRNA | ENSG00000 |
| 39825350 | 39834201 | -1 | protein_coding | ENSG00000 |
| 39829716 | 39829802 | -1 | miRNA | ENSG00000 |
| 39834458 | 39846379 | -1 | protein_coding | ENSG00000 |
| 39863323 | 39934626 | -1 | protein_coding | ENSG00000 |
| 39943239 | 39943680 | -1 | processed_pseudoge | ENSG00000 |
| 39971165 | 39981764 | 1 | protein_coding | ENSG00000 |
| 39984134 | 40021038 | 1 | protein_coding | ENSG00000 |

| | | | | |
|---|---|---|---|---|
| 40023384 | 40025502 | -1 | processed_pseudoge | ENSG00000 |
| 40028260 | 40056231 | -1 | protein_coding | ENSG00000 |
| 40048757 | 40090925 | -1 | lncRNA | ENSG00000 |
| 40069152 | 40090938 | -1 | protein_coding | ENSG00000 |
| 40090754 | 40094406 | 1 | lncRNA | ENSG00000 |
| 40121311 | 40122168 | -1 | unitary_pseudogene | ENSG00000 |
| 40191426 | 40215575 | 1 | protein_coding | ENSG00000 |
| 40216058 | 40218399 | -1 | protein_coding | ENSG00000 |
| 40222208 | 40226697 | -1 | protein_coding | ENSG00000 |
| 40230317 | 40285536 | -1 | protein_coding | ENSG00000 |
| 40273489 | 40275479 | -1 | lncRNA | ENSG00000 |
| 40282543 | 40282641 | -1 | miRNA | ENSG00000 |
| 40299478 | 40299584 | -1 | snRNA | ENSG00000 |
| 40319536 | 40348527 | -1 | protein_coding | ENSG00000 |
| 40339059 | 40339164 | -1 | misc_RNA | ENSG00000 |
| 40342329 | 40342441 | 1 | misc_RNA | ENSG00000 |
| 40348456 | 40380439 | 1 | protein_coding | ENSG00000 |
| 40355842 | 40360606 | 1 | TEC | ENSG00000 |
| 40369846 | 40369907 | 1 | miRNA | ENSG00000 |
| 40379271 | 40390181 | -1 | protein_coding | ENSG00000 |
| 40393768 | 40413366 | -1 | protein_coding | ENSG00000 |
| 40421589 | 40425992 | -1 | protein_coding | ENSG00000 |
| 40426115 | 40426702 | 1 | lncRNA | ENSG00000 |
| 40440844 | 40444335 | -1 | protein_coding | ENSG00000 |
| 40443436 | 40444087 | 1 | lncRNA | ENSG00000 |
| 40447765 | 40465764 | -1 | protein_coding | ENSG00000 |
| 40450225 | 40450480 | 1 | misc_RNA | ENSG00000 |
| 40466241 | 40576464 | 1 | protein_coding | ENSG00000 |
| 40576853 | 40591399 | 1 | protein_coding | ENSG00000 |
| 40592883 | 40629818 | 1 | protein_coding | ENSG00000 |
| 40665905 | 40690972 | -1 | protein_coding | ENSG00000 |
| 40691530 | 40718207 | -1 | protein_coding | ENSG00000 |
| 40708055 | 40708163 | -1 | snRNA | ENSG00000 |
| 40717112 | 40740860 | 1 | protein_coding | ENSG00000 |
| 40740856 | 40751553 | -1 | protein_coding | ENSG00000 |
| 40750637 | 40765389 | 1 | protein_coding | ENSG00000 |
| 40771648 | 40796943 | 1 | protein_coding | ENSG00000 |
| 40771648 | 40777490 | 1 | protein_coding | ENSG00000 |
| 40778216 | 40796942 | 1 | protein_coding | ENSG00000 |
| 40778242 | 40808418 | 1 | protein_coding | ENSG00000 |
| 40779780 | 40796943 | -1 | lncRNA | ENSG00000 |
| 40798996 | 40808434 | 1 | protein_coding | ENSG00000 |
| 40801297 | 40898282 | 1 | protein_coding | ENSG00000 |
| 40808474 | 40812100 | -1 | transcribed_unitary_ | ENSG00000 |
| 40818414 | 40826772 | -1 | transcribed_processe | ENSG00000 |
| 40831221 | 40837210 | -1 | lncRNA | ENSG00000 |
| 40840159 | 40842039 | 1 | TEC | ENSG00000 |
| 40843541 | 40850447 | -1 | protein_coding | ENSG00000 |
| 40875439 | 40882752 | -1 | protein_coding | ENSG00000 |
| 40890826 | 40912357 | 1 | translated_unproces | ENSG00000 |

| | | | | |
|---|---|---|---|---|
| 40908480 | 40910685 | -1 | unprocessed_pseudc | ENSG0000C |
| 40924219 | 40950660 | 1 | transcribed_unproce | ENSG0000C |
| 40946347 | 40947450 | 1 | lncRNA | ENSG0000C |
| 40991282 | 41018398 | 1 | protein_coding | ENSG0000C |
| 41024268 | 41027720 | -1 | unprocessed_pseudc | ENSG0000C |
| 41050449 | 41062444 | -1 | unprocessed_pseudc | ENSG0000C |
| 41073804 | 41074107 | -1 | processed_pseudoge | ENSG0000C |
| 41088451 | 41096195 | 1 | protein_coding | ENSG0000C |
| 41114432 | 41128381 | 1 | protein_coding | ENSG0000C |
| 41134722 | 41137308 | 1 | unprocessed_pseudc | ENSG0000C |
| 41169861 | 41170166 | 1 | processed_pseudoge | ENSG0000C |
| 41190927 | 41191229 | -1 | misc_RNA | ENSG0000C |
| 41193210 | 41207539 | 1 | protein_coding | ENSG0000C |
| 41219223 | 41261766 | 1 | protein_coding | ENSG0000C |
| 41221426 | 41222051 | -1 | lncRNA | ENSG0000C |
| 41262496 | 41307787 | 1 | protein_coding | ENSG0000C |
| 41286424 | 41286693 | -1 | misc_RNA | ENSG0000C |
| 41301587 | 41353922 | -1 | protein_coding | ENSG0000C |
| 41307397 | 41310412 | -1 | lncRNA | ENSG0000C |
| 41310172 | 41324873 | 1 | protein_coding | ENSG0000C |
| 41350853 | 41425001 | 1 | protein_coding | ENSG0000C |
| 41350911 | 41384083 | 1 | protein_coding | ENSG0000C |
| 41354417 | 41364165 | -1 | protein_coding | ENSG0000C |
| 41373971 | 41374419 | 1 | lncRNA | ENSG0000C |
| 41386374 | 41397359 | -1 | protein_coding | ENSG0000C |
| 41397808 | 41425002 | 1 | protein_coding | ENSG0000C |
| 41399372 | 41400365 | 1 | lncRNA | ENSG0000C |
| 41425359 | 41426237 | 1 | lncRNA | ENSG0000C |
| 41425359 | 41428730 | -1 | protein_coding | ENSG0000C |
| 41431318 | 41440717 | -1 | protein_coding | ENSG0000C |
| 41442102 | 41442392 | -1 | misc_RNA | ENSG0000C |
| 41443156 | 41444765 | 1 | protein_coding | ENSG0000C |
| 41449520 | 41501223 | -1 | lncRNA | ENSG0000C |
| 41506152 | 41506898 | 1 | processed_pseudoge | ENSG0000C |
| 41521043 | 41521989 | -1 | processed_pseudoge | ENSG0000C |
| 41531206 | 41532174 | 1 | lncRNA | ENSG0000C |
| 41535183 | 41536904 | 1 | lncRNA | ENSG0000C |
| 41545192 | 41555462 | 1 | lncRNA | ENSG0000C |
| 41549518 | 41586844 | 1 | protein_coding | ENSG0000C |
| 41551568 | 41552274 | -1 | TEC | ENSG0000C |
| 41599735 | 41605984 | 1 | transcribed_unproce | ENSG0000C |
| 41605341 | 41605671 | -1 | processed_pseudoge | ENSG0000C |
| 41618971 | 41627074 | -1 | protein_coding | ENSG0000C |
| 41638569 | 41638899 | -1 | processed_pseudoge | ENSG0000C |
| 41639659 | 41640602 | -1 | processed_pseudoge | ENSG0000C |
| 41652676 | 41652963 | 1 | processed_pseudoge | ENSG0000C |
| 41673303 | 41706976 | -1 | protein_coding | ENSG0000C |
| 41708585 | 41729798 | 1 | protein_coding | ENSG0000C |
| 41708657 | 41756737 | 1 | protein_coding | ENSG0000C |
| 41750977 | 41772211 | 1 | protein_coding | ENSG0000C |

| | | | | |
|---|---|---|---|---|
| 41757898 | 41786893 | -1 | lncRNA | ENSG00000 |
| 41796587 | 41811554 | 1 | protein_coding | ENSG00000 |
| 41829692 | 41830938 | 1 | processed_pseudoge | ENSG00000 |
| 41833686 | 41835950 | -1 | lncRNA | ENSG00000 |
| 41837074 | 41844697 | -1 | protein_coding | ENSG00000 |
| 41844743 | 41852333 | 1 | protein_coding | ENSG00000 |
| 41860255 | 41872925 | 1 | protein_coding | ENSG00000 |
| 41869627 | 41869698 | 1 | miRNA | ENSG00000 |
| 41877120 | 41881372 | 1 | protein_coding | ENSG00000 |
| 41883161 | 41930150 | 1 | protein_coding | ENSG00000 |
| 41907705 | 41928516 | -1 | protein_coding | ENSG00000 |
| 41956681 | 41959321 | -1 | protein_coding | ENSG00000 |
| 41956879 | 41994232 | -1 | protein_coding | ENSG00000 |
| 41966582 | 41997497 | -1 | protein_coding | ENSG00000 |
| 41998321 | 42069498 | -1 | protein_coding | ENSG00000 |
| 42068477 | 42081552 | 1 | protein_coding | ENSG00000 |
| 42086110 | 42196585 | -1 | protein_coding | ENSG00000 |
| 42132555 | 42137099 | 1 | lncRNA | ENSG00000 |
| 42133445 | 42133513 | -1 | miRNA | ENSG00000 |
| 42152569 | 42157523 | 1 | lncRNA | ENSG00000 |
| 42198598 | 42220140 | -1 | protein_coding | ENSG00000 |
| 42220312 | 42228201 | 1 | protein_coding | ENSG00000 |
| 42230190 | 42242625 | -1 | protein_coding | ENSG00000 |
| 42234583 | 42255132 | -1 | protein_coding | ENSG00000 |
| 42242775 | 42244973 | 1 | TEC | ENSG00000 |
| 42247569 | 42255128 | -1 | protein_coding | ENSG00000 |
| 42268537 | 42295797 | 1 | protein_coding | ENSG00000 |
| 42297033 | 42303546 | -1 | protein_coding | ENSG00000 |
| 42302098 | 42310821 | 1 | protein_coding | ENSG00000 |
| 42313309 | 42325064 | 1 | protein_coding | ENSG00000 |
| 42325609 | 42378769 | 1 | protein_coding | ENSG00000 |
| 42351131 | 42351205 | 1 | miRNA | ENSG00000 |
| 42387019 | 42390297 | -1 | protein_coding | ENSG00000 |
| 42397128 | 42652355 | 1 | lncRNA | ENSG00000 |
| 42401514 | 42427388 | -1 | protein_coding | ENSG00000 |
| 42424384 | 42425071 | -1 | lncRNA | ENSG00000 |
| 42428278 | 42442946 | -1 | protein_coding | ENSG00000 |
| 42472875 | 42473009 | 1 | snRNA | ENSG00000 |
| 42485076 | 42485714 | -1 | processed_pseudoge | ENSG00000 |
| 42507304 | 42561234 | -1 | protein_coding | ENSG00000 |
| 42560433 | 42561750 | -1 | unprocessed_pseudo | ENSG00000 |
| 42580243 | 42595055 | -1 | protein_coding | ENSG00000 |
| 42620212 | 42633034 | -1 | unprocessed_pseudo | ENSG00000 |
| 42654200 | 42669076 | 1 | unprocessed_pseudo | ENSG00000 |
| 42670005 | 42670816 | 1 | processed_pseudoge | ENSG00000 |
| 42721638 | 42740481 | -1 | protein_coding | ENSG00000 |
| 42752686 | 42855691 | -1 | protein_coding | ENSG00000 |
| 42781206 | 42786595 | -1 | unprocessed_pseudo | ENSG00000 |
| 42821863 | 42826878 | 1 | lncRNA | ENSG00000 |
| 42839368 | 42855718 | -1 | transcribed_unproce | ENSG00000 |

| | | | | |
|---|---|---|---|---|
| 42866464 | 42879822 | -1 | protein_coding | ENSG00000 |
| 42902079 | 42919563 | -1 | protein_coding | ENSG00000 |
| 42924132 | 42937207 | -1 | protein_coding | ENSG00000 |
| 42949503 | 42950387 | -1 | unprocessed_pseudo | ENSG00000 |
| 42977463 | 43171515 | 1 | lncRNA | ENSG00000 |
| 43007656 | 43026474 | -1 | protein_coding | ENSG00000 |
| 43035869 | 43041248 | -1 | unprocessed_pseudo | ENSG00000 |
| 43064209 | 43083045 | -1 | protein_coding | ENSG00000 |
| 43096456 | 43097360 | -1 | unprocessed_pseudo | ENSG00000 |
| 43153279 | 43160862 | 1 | lncRNA | ENSG00000 |
| 43166256 | 43186536 | -1 | protein_coding | ENSG00000 |
| 43192702 | 43207299 | -1 | protein_coding | ENSG00000 |
| 43211791 | 43248646 | -1 | transcribed_processe | ENSG00000 |
| 43211791 | 43269530 | -1 | protein_coding | ENSG00000 |
| 43278933 | 43284318 | -1 | unprocessed_pseudo | ENSG00000 |
| 43303171 | 43304523 | -1 | unprocessed_pseudo | ENSG00000 |
| 43329295 | 43331430 | -1 | lncRNA | ENSG00000 |
| 43353686 | 43363172 | 1 | protein_coding | ENSG00000 |
| 43360685 | 43368970 | -1 | lncRNA | ENSG00000 |
| 43372742 | 43379123 | -1 | unprocessed_pseudo | ENSG00000 |
| 43401496 | 43418597 | 1 | protein_coding | ENSG00000 |
| 43460787 | 43465608 | -1 | protein_coding | ENSG00000 |
| 43474954 | 43504935 | -1 | protein_coding | ENSG00000 |
| 43506719 | 43527230 | -1 | protein_coding | ENSG00000 |
| 43525497 | 43536130 | 1 | protein_coding | ENSG00000 |
| 43543040 | 43580473 | -1 | protein_coding | ENSG00000 |
| 43553445 | 43555494 | 1 | lncRNA | ENSG00000 |
| 43554692 | 43593036 | -1 | protein_coding | ENSG00000 |
| 43574638 | 43575618 | 1 | lncRNA | ENSG00000 |
| 43575801 | 43583964 | 1 | protein_coding | ENSG00000 |
| 43584369 | 43596135 | -1 | protein_coding | ENSG00000 |
| 43596392 | 43601157 | 1 | protein_coding | ENSG00000 |
| 43596617 | 43614498 | 1 | protein_coding | ENSG00000 |
| 43607224 | 43619629 | -1 | protein_coding | ENSG00000 |
| 43622368 | 43639850 | -1 | protein_coding | ENSG00000 |
| 43646095 | 43670547 | -1 | protein_coding | ENSG00000 |
| 43660500 | 43660776 | -1 | misc_RNA | ENSG00000 |
| 43690002 | 43700508 | 1 | transcribed_unproce | ENSG00000 |
| 43716076 | 43720021 | 1 | protein_coding | ENSG00000 |
| 43727983 | 43754962 | -1 | protein_coding | ENSG00000 |
| 43766533 | 43781257 | -1 | protein_coding | ENSG00000 |
| 43785874 | 43827206 | -1 | protein_coding | ENSG00000 |
| 43794309 | 43795658 | -1 | lncRNA | ENSG00000 |
| 43827321 | 43852017 | 1 | protein_coding | ENSG00000 |
| 43872365 | 43901385 | -1 | protein_coding | ENSG00000 |
| 43891233 | 43895411 | 1 | TEC | ENSG00000 |
| 43891804 | 43901805 | -1 | lncRNA | ENSG00000 |
| 43897211 | 43898533 | -1 | lncRNA | ENSG00000 |
| 43901909 | 43927767 | 1 | lncRNA | ENSG00000 |
| 43912624 | 43935282 | -1 | protein_coding | ENSG00000 |

| | | | | |
|---|---|---|---|---|
| 43951223 | 43967709 | 1 | protein_coding | ENSG00000 |
| 43967862 | 43998326 | 1 | protein_coding | ENSG00000 |
| 43976815 | 43977448 | 1 | lncRNA | ENSG00000 |
| 43978376 | 43978663 | 1 | lncRNA | ENSG00000 |
| 43996896 | 44003298 | -1 | lncRNA | ENSG00000 |
| 44002957 | 44013924 | 1 | protein_coding | ENSG00000 |
| 44023118 | 44025262 | -1 | lncRNA | ENSG00000 |
| 44025342 | 44033110 | 1 | protein_coding | ENSG00000 |
| 44025354 | 44087318 | 1 | protein_coding | ENSG00000 |
| 44051367 | 44067999 | 1 | protein_coding | ENSG00000 |
| 44072159 | 44089613 | 1 | protein_coding | ENSG00000 |
| 44085213 | 44085303 | -1 | snRNA | ENSG00000 |
| 44094339 | 44109886 | 1 | protein_coding | ENSG00000 |
| 44103007 | 44113183 | -1 | lncRNA | ENSG00000 |
| 44112181 | 44134822 | 1 | protein_coding | ENSG00000 |
| 44141554 | 44160313 | 1 | protein_coding | ENSG00000 |
| 44162527 | 44164955 | -1 | lncRNA | ENSG00000 |
| 44165073 | 44178381 | 1 | protein_coding | ENSG00000 |
| 44207547 | 44237268 | 1 | protein_coding | ENSG00000 |
| 44228729 | 44305046 | -1 | protein_coding | ENSG00000 |
| 44246478 | 44248374 | -1 | TEC | ENSG00000 |
| 44259880 | 44275317 | 1 | protein_coding | ENSG00000 |
| 44290081 | 44290911 | 1 | processed_pseudoge | ENSG00000 |
| 44297104 | 44297355 | 1 | processed_pseudoge | ENSG00000 |
| 44326555 | 44367217 | -1 | protein_coding | ENSG00000 |
| 44329695 | 44401608 | -1 | protein_coding | ENSG00000 |
| 44382298 | 44401608 | -1 | protein_coding | ENSG00000 |
| 44392439 | 44448452 | -1 | lncRNA | ENSG00000 |
| 44417519 | 44448578 | -1 | protein_coding | ENSG00000 |
| 44467641 | 44473227 | 1 | unprocessed_pseudc | ENSG00000 |
| 44474428 | 44500524 | -1 | protein_coding | ENSG00000 |
| 44501677 | 44529788 | -1 | protein_coding | ENSG00000 |
| 44536263 | 44536613 | -1 | lncRNA | ENSG00000 |
| 44537058 | 44620821 | -1 | transcribed_unproce | ENSG00000 |
| 44570304 | 44570564 | -1 | unprocessed_pseudc | ENSG00000 |
| 44613563 | 44636781 | 1 | protein_coding | ENSG00000 |
| 44631573 | 44725217 | -1 | lncRNA | ENSG00000 |
| 44643798 | 44666162 | 1 | protein_coding | ENSG00000 |
| 44653686 | 44653732 | -1 | miRNA | ENSG00000 |
| 44662278 | 44684359 | 1 | protein_coding | ENSG00000 |
| 44664131 | 44666158 | 1 | TEC | ENSG00000 |
| 44699151 | 44710718 | 1 | protein_coding | ENSG00000 |
| 44725984 | 44726058 | -1 | snoRNA | ENSG00000 |
| 44747705 | 44760044 | 1 | protein_coding | ENSG00000 |
| 44758657 | 44758721 | 1 | miRNA | ENSG00000 |
| 44764950 | 44765100 | 1 | snoRNA | ENSG00000 |
| 44765148 | 44765464 | 1 | processed_pseudoge | ENSG00000 |
| 44777869 | 44800652 | 1 | protein_coding | ENSG00000 |
| 44803701 | 44804010 | 1 | lncRNA | ENSG00000 |
| 44809059 | 44821421 | 1 | protein_coding | ENSG00000 |

| | | | | |
|---|---|---|---|---|
| 44846175 | 44889223 | 1 | protein_coding | ENSG0000C |
| 44882027 | 44890876 | -1 | lncRNA | ENSG0000C |
| 44890569 | 44903689 | 1 | protein_coding | ENSG0000C |
| 44905791 | 44909393 | 1 | protein_coding | ENSG0000C |
| 44909375 | 44914968 | 1 | TEC | ENSG0000C |
| 44914247 | 44919349 | 1 | protein_coding | ENSG0000C |
| 44926804 | 44931386 | 1 | transcribed_unproce | ENSG0000C |
| 44942237 | 44945496 | 1 | protein_coding | ENSG0000C |
| 44942238 | 44949565 | 1 | protein_coding | ENSG0000C |
| 44945971 | 44949566 | 1 | protein_coding | ENSG0000C |
| 44950044 | 44954007 | -1 | lncRNA | ENSG0000C |
| 44954585 | 44993341 | 1 | protein_coding | ENSG0000C |
| 45001449 | 45038198 | 1 | protein_coding | ENSG0000C |
| 45039045 | 45070956 | 1 | protein_coding | ENSG0000C |
| 45047458 | 45047561 | 1 | snRNA | ENSG0000C |
| 45071500 | 45076587 | -1 | protein_coding | ENSG0000C |
| 45076510 | 45092635 | -1 | lncRNA | ENSG0000C |
| 45079195 | 45091518 | 1 | protein_coding | ENSG0000C |
| 45079288 | 45305284 | 1 | protein_coding | ENSG0000C |
| 45091396 | 45148077 | 1 | protein_coding | ENSG0000C |
| 45128569 | 45129030 | 1 | processed_pseudoge | ENSG0000C |
| 45135993 | 45136977 | -1 | lncRNA | ENSG0000C |
| 45149750 | 45158737 | -1 | protein_coding | ENSG0000C |
| 45162928 | 45178237 | -1 | protein_coding | ENSG0000C |
| 45178745 | 45216933 | 1 | protein_coding | ENSG0000C |
| 45212621 | 45245431 | -1 | protein_coding | ENSG0000C |
| 45306413 | 45322875 | -1 | protein_coding | ENSG0000C |
| 45332698 | 45333124 | -1 | processed_pseudoge | ENSG0000C |
| 45333434 | 45351520 | 1 | protein_coding | ENSG0000C |
| 45349837 | 45370918 | -1 | protein_coding | ENSG0000C |
| 45379638 | 45406349 | -1 | protein_coding | ENSG0000C |
| 45406209 | 45410766 | 1 | protein_coding | ENSG0000C |
| 45407333 | 45478828 | -1 | protein_coding | ENSG0000C |
| 45436654 | 45436704 | 1 | miRNA | ENSG0000C |
| 45467995 | 45475179 | 1 | protein_coding | ENSG0000C |
| 45485294 | 45497055 | -1 | protein_coding | ENSG0000C |
| 45488777 | 45502510 | 1 | protein_coding | ENSG0000C |
| 45506579 | 45526983 | 1 | protein_coding | ENSG0000C |
| 45527427 | 45602212 | -1 | protein_coding | ENSG0000C |
| 45589764 | 45602212 | -1 | protein_coding | ENSG0000C |
| 45606994 | 45645629 | -1 | protein_coding | ENSG0000C |
| 45638994 | 45639087 | -1 | miRNA | ENSG0000C |
| 45641494 | 45642840 | 1 | lncRNA | ENSG0000C |
| 45651510 | 45651805 | 1 | misc_RNA | ENSG0000C |
| 45668221 | 45683722 | 1 | protein_coding | ENSG0000C |
| 45674928 | 45675024 | 1 | miRNA | ENSG0000C |
| 45674932 | 45675008 | -1 | miRNA | ENSG0000C |
| 45687454 | 45692569 | -1 | protein_coding | ENSG0000C |
| 45692666 | 45703989 | 1 | protein_coding | ENSG0000C |
| 45710629 | 45730896 | -1 | protein_coding | ENSG0000C |

| | | | | |
|---|---|---|---|---|
| 45714387 | 45717381 | -1 | TEC | ENSG0000C |
| 45733251 | 45764534 | 1 | protein_coding | ENSG0000C |
| 45764785 | 45769806 | 1 | lncRNA | ENSG0000C |
| 45764785 | 45769252 | -1 | protein_coding | ENSG0000C |
| 45767796 | 45772504 | 1 | lncRNA | ENSG0000C |
| 45769717 | 45782552 | -1 | protein_coding | ENSG0000C |
| 45779437 | 45785973 | -1 | protein_coding | ENSG0000C |
| 45782947 | 45792845 | -1 | protein_coding | ENSG0000C |
| 45795713 | 45815308 | -1 | protein_coding | ENSG0000C |
| 45815410 | 45863194 | -1 | protein_coding | ENSG0000C |
| 45830164 | 45831108 | 1 | lncRNA | ENSG0000C |
| 45863989 | 45873797 | 1 | protein_coding | ENSG0000C |
| 45883608 | 45886141 | -1 | protein_coding | ENSG0000C |
| 45890023 | 45902613 | -1 | protein_coding | ENSG0000C |
| 45913214 | 45914778 | -1 | protein_coding | ENSG0000C |
| 45933734 | 45973865 | -1 | protein_coding | ENSG0000C |
| 45995461 | 46021318 | 1 | protein_coding | ENSG0000C |
| 46018932 | 46019049 | 1 | miRNA | ENSG0000C |
| 46019153 | 46023053 | -1 | protein_coding | ENSG0000C |
| 46027648 | 46031667 | 1 | lncRNA | ENSG0000C |
| 46039748 | 46077118 | -1 | protein_coding | ENSG0000C |
| 46101122 | 46196218 | 1 | lncRNA | ENSG0000C |
| 46111079 | 46113863 | -1 | processed_pseudoge | ENSG0000C |
| 46120067 | 46124688 | -1 | protein_coding | ENSG0000C |
| 46136746 | 46137547 | -1 | processed_pseudoge | ENSG0000C |
| 46143106 | 46161299 | 1 | protein_coding | ENSG0000C |
| 46163893 | 46180647 | -1 | lncRNA | ENSG0000C |
| 46189029 | 46203160 | -1 | lncRNA | ENSG0000C |
| 46196238 | 46197323 | 1 | unprocessed_pseudc | ENSG0000C |
| 46201552 | 46214838 | 1 | lncRNA | ENSG0000C |
| 46219423 | 46219920 | 1 | processed_pseudoge | ENSG0000C |
| 46228964 | 46230388 | -1 | lncRNA | ENSG0000C |
| 46229742 | 46231243 | 1 | protein_coding | ENSG0000C |
| 46249394 | 46250163 | -1 | unprocessed_pseudc | ENSG0000C |
| 46260420 | 46260858 | -1 | unprocessed_pseudc | ENSG0000C |
| 46297046 | 46343433 | 1 | protein_coding | ENSG0000C |
| 46306470 | 46306577 | -1 | snRNA | ENSG0000C |
| 46320197 | 46340004 | -1 | lncRNA | ENSG0000C |
| 46347087 | 46392981 | 1 | protein_coding | ENSG0000C |
| 46382492 | 46383169 | -1 | lncRNA | ENSG0000C |
| 46390515 | 46390852 | -1 | lncRNA | ENSG0000C |
| 46410329 | 46413564 | -1 | protein_coding | ENSG0000C |
| 46424697 | 46428951 | -1 | protein_coding | ENSG0000C |
| 46466491 | 46471563 | -1 | protein_coding | ENSG0000C |
| 46480796 | 46601200 | -1 | protein_coding | ENSG0000C |
| 46486906 | 46495879 | -1 | protein_coding | ENSG0000C |
| 46494508 | 46496502 | 1 | lncRNA | ENSG0000C |
| 46533669 | 46534351 | 1 | unprocessed_pseudc | ENSG0000C |
| 46547056 | 46600861 | -1 | lncRNA | ENSG0000C |
| 46558641 | 46559642 | 1 | processed_pseudoge | ENSG0000C |

| | | | | |
|---|---|---|---|---|
| 46601074 | 46610782 | 1 | protein_coding | ENSG0000C |
| 46609277 | 46610779 | -1 | lncRNA | ENSG0000C |
| 46620468 | 46625089 | -1 | protein_coding | ENSG0000C |
| 46634076 | 46634685 | -1 | protein_coding | ENSG0000C |
| 46636810 | 46639161 | -1 | TEC | ENSG0000C |
| 46647551 | 46661182 | -1 | protein_coding | ENSG0000C |
| 46660364 | 46677447 | 1 | lncRNA | ENSG0000C |
| 46674275 | 46717127 | -1 | protein_coding | ENSG0000C |
| 46688797 | 46689068 | -1 | misc_RNA | ENSG0000C |
| 46709293 | 46709345 | -1 | miRNA | ENSG0000C |
| 46719511 | 46746994 | -1 | protein_coding | ENSG0000C |
| 46728603 | 46732700 | 1 | lncRNA | ENSG0000C |
| 46736312 | 46736419 | 1 | misc_RNA | ENSG0000C |
| 46746046 | 46776988 | 1 | protein_coding | ENSG0000C |
| 46774883 | 46788594 | -1 | protein_coding | ENSG0000C |
| 46787815 | 46789043 | 1 | lncRNA | ENSG0000C |
| 46809583 | 46810182 | 1 | processed_pseudoge | ENSG0000C |
| 46809585 | 46810183 | 1 | processed_pseudoge | ENSG0000C |
| 46826895 | 46828440 | -1 | processed_pseudoge | ENSG0000C |
| 46838136 | 46850992 | -1 | protein_coding | ENSG0000C |
| 46860997 | 47005077 | 1 | protein_coding | ENSG0000C |
| 46946535 | 46949156 | 1 | TEC | ENSG0000C |
| 47019837 | 47045775 | 1 | protein_coding | ENSG0000C |
| 47045909 | 47048624 | -1 | protein_coding | ENSG0000C |
| 47064187 | 47113776 | -1 | protein_coding | ENSG0000C |
| 47113274 | 47210636 | 1 | protein_coding | ENSG0000C |
| 47220822 | 47232766 | -1 | protein_coding | ENSG0000C |
| 47226942 | 47227021 | 1 | miRNA | ENSG0000C |
| 47226944 | 47227019 | -1 | miRNA | ENSG0000C |
| 47255850 | 47256477 | -1 | lncRNA | ENSG0000C |
| 47255980 | 47273701 | 1 | protein_coding | ENSG0000C |
| 47274885 | 47275723 | 1 | protein_coding | ENSG0000C |
| 47290023 | 47322066 | 1 | protein_coding | ENSG0000C |
| 47332175 | 47347329 | 1 | protein_coding | ENSG0000C |
| 47349315 | 47382704 | 1 | protein_coding | ENSG0000C |
| 47403124 | 47419527 | -1 | protein_coding | ENSG0000C |
| 47428017 | 47471893 | -1 | protein_coding | ENSG0000C |
| 47469910 | 47478550 | 1 | lncRNA | ENSG0000C |
| 47475150 | 47484265 | -1 | protein_coding | ENSG0000C |
| 47484282 | 47501597 | 1 | lncRNA | ENSG0000C |
| 47487637 | 47515091 | -1 | protein_coding | ENSG0000C |
| 47487638 | 47489519 | -1 | TEC | ENSG0000C |
| 47520685 | 47555856 | -1 | protein_coding | ENSG0000C |
| 47581023 | 47581321 | 1 | misc_RNA | ENSG0000C |
| 47607524 | 47733098 | 1 | lncRNA | ENSG0000C |
| 47607873 | 47608454 | -1 | lncRNA | ENSG0000C |
| 47608196 | 47703277 | 1 | protein_coding | ENSG0000C |
| 47615699 | 47626395 | -1 | lncRNA | ENSG0000C |
| 47661657 | 47663895 | -1 | TEC | ENSG0000C |
| 47713422 | 47743134 | 1 | protein_coding | ENSG0000C |

| | | | | |
|---|---|---|---|---|
| 47745546 | 47757058 | 1 | protein_coding | ENSG0000C |
| 47755853 | 47755962 | 1 | snoRNA | ENSG0000C |
| 47757036 | 47768840 | -1 | lncRNA | ENSG0000C |
| 47778585 | 47784686 | 1 | protein_coding | ENSG0000C |
| 47794467 | 47794890 | -1 | processed_pseudoge | ENSG0000C |
| 47794650 | 47794920 | 1 | processed_pseudoge | ENSG0000C |
| 47801243 | 47819051 | -1 | protein_coding | ENSG0000C |
| 47819779 | 47843330 | 1 | protein_coding | ENSG0000C |
| 47860113 | 47861512 | 1 | unprocessed_pseudc | ENSG0000C |
| 47863409 | 47865459 | -1 | lncRNA | ENSG0000C |
| 47870467 | 47886315 | -1 | protein_coding | ENSG0000C |
| 47968046 | 47992170 | -1 | protein_coding | ENSG0000C |
| 47994632 | 48025154 | 1 | protein_coding | ENSG0000C |
| 48029383 | 48044079 | -1 | protein_coding | ENSG0000C |
| 48047843 | 48110817 | -1 | protein_coding | ENSG0000C |
| 48061371 | 48064945 | 1 | lncRNA | ENSG0000C |
| 48115445 | 48170603 | -1 | protein_coding | ENSG0000C |
| 48118432 | 48127706 | 1 | lncRNA | ENSG0000C |
| 48145908 | 48147413 | 1 | lncRNA | ENSG0000C |
| 48170680 | 48197620 | 1 | protein_coding | ENSG0000C |
| 48180770 | 48255946 | -1 | protein_coding | ENSG0000C |
| 48198107 | 48199720 | -1 | TEC | ENSG0000C |
| 48204071 | 48214751 | 1 | lncRNA | ENSG0000C |
| 48255675 | 48258199 | 1 | lncRNA | ENSG0000C |
| 48262900 | 48271283 | -1 | lncRNA | ENSG0000C |
| 48270081 | 48287608 | 1 | protein_coding | ENSG0000C |
| 48296457 | 48321894 | -1 | protein_coding | ENSG0000C |
| 48321509 | 48330553 | 1 | protein_coding | ENSG0000C |
| 48332356 | 48364059 | -1 | protein_coding | ENSG0000C |
| 48364367 | 48376377 | 1 | protein_coding | ENSG0000C |
| 48382575 | 48391551 | -1 | protein_coding | ENSG0000C |
| 48394875 | 48444931 | 1 | protein_coding | ENSG0000C |
| 48445841 | 48457022 | 1 | protein_coding | ENSG0000C |
| 48455509 | 48466980 | 1 | protein_coding | ENSG0000C |
| 48465348 | 48469693 | -1 | lncRNA | ENSG0000C |
| 48465837 | 48472431 | 1 | protein_coding | ENSG0000C |
| 48469208 | 48482314 | 1 | protein_coding | ENSG0000C |
| 48485271 | 48513935 | -1 | protein_coding | ENSG0000C |
| 48547637 | 48597238 | -1 | lncRNA | ENSG0000C |
| 48552075 | 48599425 | 1 | protein_coding | ENSG0000C |
| 48600810 | 48614854 | -1 | protein_coding | ENSG0000C |
| 48606742 | 48607714 | 1 | protein_coding | ENSG0000C |
| 48615328 | 48619184 | -1 | protein_coding | ENSG0000C |
| 48619272 | 48624132 | -1 | lncRNA | ENSG0000C |
| 48619291 | 48630717 | 1 | protein_coding | ENSG0000C |
| 48630030 | 48637379 | -1 | protein_coding | ENSG0000C |
| 48637946 | 48646187 | -1 | protein_coding | ENSG0000C |
| 48638071 | 48682245 | 1 | transcribed_unitary_ | ENSG0000C |
| 48661407 | 48673081 | -1 | protein_coding | ENSG0000C |
| 48695971 | 48705951 | 1 | protein_coding | ENSG0000C |

| | | | | |
|---|---|---|---|---|
| 48712725 | 48719725 | -1 | protein_coding | ENSG00000( |
| 48720585 | 48740610 | -1 | protein_coding | ENSG00000( |
| 48740852 | 48746909 | -1 | protein_coding | ENSG00000( |
| 48748011 | 48755390 | -1 | protein_coding | ENSG00000( |
| 48755524 | 48758333 | 1 | protein_coding | ENSG00000( |
| 48793152 | 48793258 | -1 | snRNA | ENSG00000( |
| 48795062 | 48811029 | -1 | protein_coding | ENSG00000( |
| 48807321 | 48810717 | 1 | lncRNA | ENSG00000( |
| 48813018 | 48836510 | -1 | protein_coding | ENSG00000( |
| 48837097 | 48868617 | -1 | protein_coding | ENSG00000( |
| 48872421 | 48876058 | 1 | protein_coding | ENSG00000( |
| 48880967 | 48898744 | -1 | protein_coding | ENSG00000( |
| 48900312 | 48923372 | 1 | protein_coding | ENSG00000( |
| 48910930 | 48918891 | -1 | lncRNA | ENSG00000( |
| 48926171 | 48926444 | 1 | lncRNA | ENSG00000( |
| 48933699 | 48944969 | 1 | protein_coding | ENSG00000( |
| 48954815 | 48961798 | 1 | protein_coding | ENSG00000( |
| 48963975 | 48965158 | 1 | lncRNA | ENSG00000( |
| 48965309 | 48966879 | 1 | protein_coding | ENSG00000( |
| 48968130 | 48993310 | -1 | protein_coding | ENSG00000( |
| 48993562 | 49015970 | 1 | protein_coding | ENSG00000( |
| 49009906 | 49009972 | 1 | miRNA | ENSG00000( |
| 49015980 | 49017090 | -1 | protein_coding | ENSG00000( |
| 49017496 | 49020523 | -1 | protein_coding | ENSG00000( |
| 49022869 | 49024333 | -1 | protein_coding | ENSG00000( |
| 49022953 | 49036895 | -1 | protein_coding | ENSG00000( |
| 49025461 | 49026020 | -1 | unprocessed_pseudc | ENSG00000( |
| 49029726 | 49030341 | 1 | unprocessed_pseudc | ENSG00000( |
| 49031912 | 49033238 | 1 | protein_coding | ENSG00000( |
| 49035610 | 49036895 | -1 | protein_coding | ENSG00000( |
| 49038479 | 49039038 | -1 | unprocessed_pseudc | ENSG00000( |
| 49042174 | 49042731 | 1 | unprocessed_pseudc | ENSG00000( |
| 49043848 | 49045311 | 1 | protein_coding | ENSG00000( |
| 49047638 | 49049106 | -1 | protein_coding | ENSG00000( |
| 49050217 | 49050846 | -1 | unprocessed_pseudc | ENSG00000( |
| 49054275 | 49058860 | -1 | protein_coding | ENSG00000( |
| 49055793 | 49065076 | -1 | protein_coding | ENSG00000( |
| 49056088 | 49058761 | -1 | transcribed_unproce | ENSG00000( |
| 49060613 | 49061132 | 1 | lncRNA | ENSG00000( |
| 49061066 | 49065054 | -1 | protein_coding | ENSG00000( |
| 49064259 | 49064856 | 1 | lncRNA | ENSG00000( |
| 49067418 | 49072941 | -1 | protein_coding | ENSG00000( |
| 49081332 | 49081630 | 1 | misc_RNA | ENSG00000( |
| 49085419 | 49108605 | 1 | protein_coding | ENSG00000( |
| 49114324 | 49118460 | 1 | protein_coding | ENSG00000( |
| 49118402 | 49119143 | -1 | protein_coding | ENSG00000( |
| 49119544 | 49151026 | 1 | protein_coding | ENSG00000( |
| 49151198 | 49155396 | -1 | protein_coding | ENSG00000( |
| 49157741 | 49211836 | 1 | protein_coding | ENSG00000( |
| 49255472 | 49256102 | -1 | processed_pseudoge | ENSG00000( |

| | | | | |
|---|---|---|---|---|
| 49275119 | 49275420 | 1 | processed_pseudoge | ENSG00000 |
| 49289638 | 49325215 | -1 | protein_coding | ENSG00000 |
| 49308797 | 49308868 | -1 | miRNA | ENSG00000 |
| 49331525 | 49340303 | -1 | lncRNA | ENSG00000 |
| 49335171 | 49343335 | 1 | protein_coding | ENSG00000 |
| 49340595 | 49362457 | -1 | protein_coding | ENSG00000 |
| 49356152 | 49357769 | 1 | lncRNA | ENSG00000 |
| 49361783 | 49375116 | 1 | protein_coding | ENSG00000 |
| 49368558 | 49388091 | -1 | lncRNA | ENSG00000 |
| 49388219 | 49417990 | 1 | protein_coding | ENSG00000 |
| 49422419 | 49423441 | -1 | protein_coding | ENSG00000 |
| 49423749 | 49428818 | 1 | protein_coding | ENSG00000 |
| 49429401 | 49442360 | -1 | protein_coding | ENSG00000 |
| 49446298 | 49453497 | -1 | protein_coding | ENSG00000 |
| 49453225 | 49471050 | 1 | protein_coding | ENSG00000 |
| 49462752 | 49486231 | 1 | protein_coding | ENSG00000 |
| 49474207 | 49486231 | 1 | protein_coding | ENSG00000 |
| 49474586 | 49487637 | 1 | lncRNA | ENSG00000 |
| 49487554 | 49492308 | 1 | protein_coding | ENSG00000 |
| 49489965 | 49490048 | 1 | snoRNA | ENSG00000 |
| 49490615 | 49490699 | 1 | snoRNA | ENSG00000 |
| 49490904 | 49490974 | 1 | snoRNA | ENSG00000 |
| 49491175 | 49491260 | 1 | snoRNA | ENSG00000 |
| 49496365 | 49499708 | 1 | protein_coding | ENSG00000 |
| 49497720 | 49497806 | 1 | snoRNA | ENSG00000 |
| 49500785 | 49500868 | -1 | miRNA | ENSG00000 |
| 49502432 | 49502639 | -1 | processed_pseudoge | ENSG00000 |
| 49506816 | 49526428 | 1 | protein_coding | ENSG00000 |
| 49528003 | 49546962 | 1 | protein_coding | ENSG00000 |
| 49555468 | 49590262 | -1 | protein_coding | ENSG00000 |
| 49566672 | 49566772 | 1 | misc_RNA | ENSG00000 |
| 49580646 | 49591004 | 1 | protein_coding | ENSG00000 |
| 49591182 | 49626439 | 1 | protein_coding | ENSG00000 |
| 49625994 | 49626439 | -1 | lncRNA | ENSG00000 |
| 49635292 | 49640143 | -1 | protein_coding | ENSG00000 |
| 49642209 | 49658642 | 1 | protein_coding | ENSG00000 |
| 49659569 | 49665875 | -1 | protein_coding | ENSG00000 |
| 49665142 | 49673916 | 1 | protein_coding | ENSG00000 |
| 49675786 | 49689029 | 1 | protein_coding | ENSG00000 |
| 49682117 | 49682195 | 1 | miRNA | ENSG00000 |
| 49688664 | 49690575 | 1 | protein_coding | ENSG00000 |
| 49688853 | 49690573 | -1 | lncRNA | ENSG00000 |
| 49690898 | 49713731 | 1 | protein_coding | ENSG00000 |
| 49719498 | 49720081 | -1 | processed_pseudoge | ENSG00000 |
| 49739753 | 49763306 | -1 | protein_coding | ENSG00000 |
| 49764524 | 49764630 | -1 | snRNA | ENSG00000 |
| 49766968 | 49807113 | 1 | protein_coding | ENSG00000 |
| 49791866 | 49791934 | 1 | miRNA | ENSG00000 |
| 49806866 | 49817376 | -1 | protein_coding | ENSG00000 |
| 49808933 | 49809738 | 1 | lncRNA | ENSG00000 |

| | | | | |
|---|---|---|---|---|
| 49818279 | 49840383 | 1 | protein_coding | ENSG00000 |
| 49832018 | 49832099 | 1 | miRNA | ENSG00000 |
| 49838639 | 49851676 | -1 | lncRNA | ENSG00000 |
| 49850735 | 49860744 | 1 | protein_coding | ENSG00000 |
| 49852887 | 49854967 | -1 | lncRNA | ENSG00000 |
| 49854591 | 49854651 | 1 | miRNA | ENSG00000 |
| 49856970 | 49859289 | -1 | lncRNA | ENSG00000 |
| 49859882 | 49878351 | -1 | protein_coding | ENSG00000 |
| 49869033 | 49878459 | -1 | protein_coding | ENSG00000 |
| 49877425 | 49888750 | 1 | protein_coding | ENSG00000 |
| 49888175 | 49888230 | 1 | miRNA | ENSG00000 |
| 49889654 | 49929539 | -1 | protein_coding | ENSG00000 |
| 49906825 | 49929763 | -1 | protein_coding | ENSG00000 |
| 49909501 | 49958391 | -1 | protein_coding | ENSG00000 |
| 49928702 | 49933935 | 1 | protein_coding | ENSG00000 |
| 49933064 | 49933137 | 1 | miRNA | ENSG00000 |
| 49948985 | 49961172 | -1 | protein_coding | ENSG00000 |
| 49949316 | 49949527 | -1 | snoRNA | ENSG00000 |
| 49969673 | 49975814 | 1 | transcribed_unproce | ENSG00000 |
| 49976468 | 50025946 | -1 | protein_coding | ENSG00000 |
| 50019019 | 50019320 | 1 | misc_RNA | ENSG00000 |
| 50025714 | 50053414 | 1 | protein_coding | ENSG00000 |
| 50043196 | 50051062 | -1 | lncRNA | ENSG00000 |
| 50050589 | 50066793 | 1 | lncRNA | ENSG00000 |
| 50058882 | 50060232 | -1 | processed_pseudoge | ENSG00000 |
| 50075123 | 50075902 | 1 | lncRNA | ENSG00000 |
| 50136859 | 50148230 | -1 | lncRNA | ENSG00000 |
| 50152548 | 50163195 | -1 | protein_coding | ENSG00000 |
| 50188186 | 50310542 | 1 | protein_coding | ENSG00000 |
| 50258443 | 50258722 | 1 | misc_RNA | ENSG00000 |
| 50310022 | 50310539 | -1 | lncRNA | ENSG00000 |
| 50311937 | 50333515 | -1 | protein_coding | ENSG00000 |
| 50329653 | 50383388 | 1 | protein_coding | ENSG00000 |
| 50333796 | 50344767 | -1 | transcribed_unproce | ENSG00000 |
| 50358477 | 50365830 | -1 | protein_coding | ENSG00000 |
| 50365880 | 50369331 | -1 | lncRNA | ENSG00000 |
| 50384204 | 50418018 | 1 | protein_coding | ENSG00000 |
| 50415799 | 50428409 | 1 | protein_coding | ENSG00000 |
| 50418938 | 50431314 | 1 | protein_coding | ENSG00000 |
| 50432892 | 50466321 | 1 | protein_coding | ENSG00000 |
| 50466785 | 50476848 | -1 | protein_coding | ENSG00000 |
| 50476400 | 50490870 | 1 | protein_coding | ENSG00000 |
| 50480119 | 50483351 | -1 | lncRNA | ENSG00000 |
| 50486810 | 50487638 | -1 | lncRNA | ENSG00000 |
| 50505998 | 50511353 | -1 | protein_coding | ENSG00000 |
| 50511600 | 50514690 | -1 | protein_coding | ENSG00000 |
| 50516892 | 50568435 | -1 | protein_coding | ENSG00000 |
| 50555370 | 50557969 | -1 | lncRNA | ENSG00000 |
| 50621307 | 50639881 | -1 | protein_coding | ENSG00000 |
| 50649445 | 50659310 | 1 | protein_coding | ENSG00000 |

| | | | | |
|---|---|---|---|---|
| 50661827 | 50719450 | -1 | protein_coding | ENSG00000 |
| 50723364 | 50725718 | 1 | protein_coding | ENSG00000 |
| 50758382 | 50759192 | 1 | unprocessed_pseudo | ENSG00000 |
| 50770464 | 50771732 | 1 | protein_coding | ENSG00000 |
| 50776141 | 50793142 | -1 | protein_coding | ENSG00000 |
| 50790415 | 50795219 | 1 | protein_coding | ENSG00000 |
| 50792685 | 50793584 | -1 | lncRNA | ENSG00000 |
| 50797704 | 50804929 | -1 | protein_coding | ENSG00000 |
| 50799032 | 50799122 | -1 | snoRNA | ENSG00000 |
| 50799442 | 50799532 | -1 | snoRNA | ENSG00000 |
| 50802328 | 50802418 | -1 | snoRNA | ENSG00000 |
| 50808145 | 50812707 | -1 | unitary_pseudogene | ENSG00000 |
| 50808874 | 50818880 | 1 | transcribed_unitary_ | ENSG00000 |
| 50819146 | 50823787 | -1 | protein_coding | ENSG00000 |
| 50825289 | 50837213 | -1 | protein_coding | ENSG00000 |
| 50830530 | 50851089 | 1 | lncRNA | ENSG00000 |
| 50854915 | 50860764 | 1 | protein_coding | ENSG00000 |
| 50861568 | 50880567 | 1 | protein_coding | ENSG00000 |
| 50882096 | 50896398 | -1 | transcribed_unproce | ENSG00000 |
| 50906352 | 50910738 | -1 | protein_coding | ENSG00000 |
| 50927079 | 50927571 | -1 | processed_pseudoge | ENSG00000 |
| 50943303 | 50953093 | -1 | protein_coding | ENSG00000 |
| 50950185 | 50963649 | 1 | lncRNA | ENSG00000 |
| 50958631 | 50969673 | -1 | protein_coding | ENSG00000 |
| 50962197 | 50962781 | -1 | TEC | ENSG00000 |
| 50968193 | 51012129 | 1 | lncRNA | ENSG00000 |
| 50976468 | 50984099 | -1 | protein_coding | ENSG00000 |
| 50996007 | 51002711 | -1 | protein_coding | ENSG00000 |
| 50996018 | 51009581 | -1 | protein_coding | ENSG00000 |
| 51002508 | 51009634 | -1 | protein_coding | ENSG00000 |
| 51012739 | 51020175 | -1 | protein_coding | ENSG00000 |
| 51014374 | 51014734 | 1 | lncRNA | ENSG00000 |
| 51022216 | 51028039 | -1 | protein_coding | ENSG00000 |
| 51029092 | 51035230 | -1 | protein_coding | ENSG00000 |
| 51030076 | 51034335 | 1 | lncRNA | ENSG00000 |
| 51043806 | 51052077 | 1 | lncRNA | ENSG00000 |
| 51055626 | 51065114 | -1 | protein_coding | ENSG00000 |
| 51077495 | 51084245 | -1 | protein_coding | ENSG00000 |
| 51097606 | 51108409 | -1 | protein_coding | ENSG00000 |
| 51119987 | 51122965 | 1 | unprocessed_pseudo | ENSG00000 |
| 51124908 | 51136651 | 1 | protein_coding | ENSG00000 |
| 51137970 | 51138634 | 1 | unprocessed_pseudo | ENSG00000 |
| 51142299 | 51153526 | 1 | protein_coding | ENSG00000 |
| 51143826 | 51144196 | 1 | transcribed_unproce | ENSG00000 |
| 51152923 | 51181966 | -1 | lncRNA | ENSG00000 |
| 51155288 | 51155624 | -1 | processed_pseudoge | ENSG00000 |
| 51167328 | 51173524 | 1 | transcribed_unproce | ENSG00000 |
| 51184905 | 51190809 | 1 | unprocessed_pseudo | ENSG00000 |
| 51194118 | 51195540 | -1 | processed_pseudoge | ENSG00000 |
| 51198855 | 51201449 | 1 | unprocessed_pseudo | ENSG00000 |

| | | | | |
|---|---|---|---|---|
| 51206929 | 51207009 | -1 | miRNA | ENSG0000C |
| 51211080 | 51212088 | 1 | transcribed_unproce | ENSG0000C |
| 51225064 | 51243860 | 1 | protein_coding | ENSG0000C |
| 51246348 | 51269330 | 1 | protein_coding | ENSG0000C |
| 51251231 | 51271179 | -1 | lncRNA | ENSG0000C |
| 51271266 | 51281234 | 1 | lncRNA | ENSG0000C |
| 51280862 | 51283022 | 1 | transcribed_unproce | ENSG0000C |
| 51311848 | 51330354 | 1 | protein_coding | ENSG0000C |
| 51331536 | 51342124 | -1 | protein_coding | ENSG0000C |
| 51340695 | 51344117 | 1 | lncRNA | ENSG0000C |
| 51345169 | 51353293 | 1 | lncRNA | ENSG0000C |
| 51345169 | 51366418 | -1 | protein_coding | ENSG0000C |
| 51350373 | 51368099 | -1 | protein_coding | ENSG0000C |
| 51361712 | 51365544 | -1 | lncRNA | ENSG0000C |
| 51367098 | 51369003 | -1 | protein_coding | ENSG0000C |
| 51371606 | 51372701 | -1 | protein_coding | ENSG0000C |
| 51379909 | 51387960 | -1 | protein_coding | ENSG0000C |
| 51388289 | 51390591 | -1 | protein_coding | ENSG0000C |
| 51394488 | 51403650 | 1 | lncRNA | ENSG0000C |
| 51396755 | 51403526 | -1 | processed_pseudoge | ENSG0000C |
| 51410020 | 51417803 | -1 | protein_coding | ENSG0000C |
| 51414298 | 51414965 | 1 | lncRNA | ENSG0000C |
| 51415724 | 51417425 | 1 | lncRNA | ENSG0000C |
| 51426347 | 51426668 | -1 | unprocessed_pseudc | ENSG0000C |
| 51442379 | 51442650 | 1 | processed_pseudoge | ENSG0000C |
| 51450847 | 51458456 | -1 | protein_coding | ENSG0000C |
| 51468360 | 51468636 | -1 | unprocessed_pseudc | ENSG0000C |
| 51478622 | 51490952 | 1 | protein_coding | ENSG0000C |
| 51491227 | 51501800 | -1 | protein_coding | ENSG0000C |
| 51505059 | 51505362 | -1 | unprocessed_pseudc | ENSG0000C |
| 51508573 | 51509311 | 1 | unprocessed_pseudc | ENSG0000C |
| 51517819 | 51531856 | -1 | protein_coding | ENSG0000C |
| 51519731 | 51520260 | 1 | lncRNA | ENSG0000C |
| 51560122 | 51560590 | -1 | unprocessed_pseudc | ENSG0000C |
| 51563797 | 51564910 | -1 | unprocessed_pseudc | ENSG0000C |
| 51571283 | 51592510 | 1 | protein_coding | ENSG0000C |
| 51594561 | 51606150 | 1 | transcribed_processe | ENSG0000C |
| 51599289 | 51600470 | 1 | lncRNA | ENSG0000C |
| 51611927 | 51645545 | -1 | protein_coding | ENSG0000C |
| 51612091 | 51646889 | -1 | protein_coding | ENSG0000C |
| 51621136 | 51621701 | 1 | processed_pseudoge | ENSG0000C |
| 51639478 | 51639931 | -1 | lncRNA | ENSG0000C |
| 51642553 | 51646801 | -1 | protein_coding | ENSG0000C |
| 51648862 | 51649610 | 1 | processed_pseudoge | ENSG0000C |
| 51685363 | 51693456 | -1 | lncRNA | ENSG0000C |
| 51689128 | 51712387 | 1 | protein_coding | ENSG0000C |
| 51692612 | 51692681 | 1 | miRNA | ENSG0000C |
| 51692786 | 51692864 | 1 | miRNA | ENSG0000C |
| 51693254 | 51693339 | 1 | miRNA | ENSG0000C |
| 51713112 | 51723994 | -1 | protein_coding | ENSG0000C |

| | | | | |
|---|---|---|---|---|
| 51745172 | 51804110 | -1 | protein_coding | ENSG0000C |
| 51752026 | 51770531 | 1 | protein_coding | ENSG0000C |
| 51771311 | 51773169 | -1 | lncRNA | ENSG0000C |
| 51795157 | 51826190 | 1 | protein_coding | ENSG0000C |
| 51804816 | 51890950 | -1 | protein_coding | ENSG0000C |
| 51839771 | 51840945 | -1 | lncRNA | ENSG0000C |
| 51842878 | 51843315 | -1 | transcribed_unproce | ENSG0000C |
| 51860242 | 51863236 | -1 | transcribed_unproce | ENSG0000C |
| 51888025 | 51900463 | 1 | lncRNA | ENSG0000C |
| 51889235 | 51905040 | -1 | protein_coding | ENSG0000C |
| 51914675 | 51917703 | -1 | unprocessed_pseudc | ENSG0000C |
| 51927147 | 51948759 | 1 | protein_coding | ENSG0000C |
| 51949134 | 51981367 | 1 | lncRNA | ENSG0000C |
| 51964340 | 51986856 | -1 | protein_coding | ENSG0000C |
| 51991332 | 52008230 | -1 | protein_coding | ENSG0000C |
| 52012765 | 52030240 | -1 | protein_coding | ENSG0000C |
| 52031378 | 52095738 | -1 | protein_coding | ENSG0000C |
| 52049007 | 52049754 | 1 | lncRNA | ENSG0000C |
| 52058490 | 52063703 | -1 | lncRNA | ENSG0000C |
| 52064466 | 52095765 | -1 | protein_coding | ENSG0000C |
| 52110318 | 52110726 | -1 | processed_pseudoge | ENSG0000C |
| 52113091 | 52139938 | -1 | protein_coding | ENSG0000C |
| 52130548 | 52171533 | -1 | lncRNA | ENSG0000C |
| 52142626 | 52144156 | 1 | lncRNA | ENSG0000C |
| 52143043 | 52143336 | 1 | processed_pseudoge | ENSG0000C |
| 52153864 | 52171643 | -1 | protein_coding | ENSG0000C |
| 52171158 | 52177002 | 1 | lncRNA | ENSG0000C |
| 52190048 | 52229518 | 1 | protein_coding | ENSG0000C |
| 52222020 | 52222098 | 1 | miRNA | ENSG0000C |
| 52222225 | 52231294 | -1 | lncRNA | ENSG0000C |
| 52241765 | 52242440 | -1 | processed_pseudoge | ENSG0000C |
| 52259076 | 52259187 | -1 | misc_RNA | ENSG0000C |
| 52269587 | 52296046 | 1 | protein_coding | ENSG0000C |
| 52281797 | 52281893 | 1 | miRNA | ENSG0000C |
| 52284780 | 52297920 | -1 | lncRNA | ENSG0000C |
| 52297169 | 52325922 | 1 | protein_coding | ENSG0000C |
| 52300693 | 52345229 | -1 | lncRNA | ENSG0000C |
| 52336243 | 52367778 | 1 | protein_coding | ENSG0000C |
| 52369917 | 52385795 | 1 | protein_coding | ENSG0000C |
| 52388842 | 52397783 | -1 | lncRNA | ENSG0000C |
| 52392659 | 52392755 | 1 | unprocessed_pseudc | ENSG0000C |
| 52397849 | 52418412 | 1 | protein_coding | ENSG0000C |
| 52425863 | 52426202 | -1 | processed_pseudoge | ENSG0000C |
| 52429187 | 52452315 | 1 | protein_coding | ENSG0000C |
| 52453553 | 52516882 | 1 | protein_coding | ENSG0000C |
| 52453580 | 52458836 | 1 | lncRNA | ENSG0000C |
| 52473336 | 52475239 | -1 | unprocessed_pseudc | ENSG0000C |
| 52511282 | 52512342 | -1 | lncRNA | ENSG0000C |
| 52527652 | 52564464 | 1 | protein_coding | ENSG0000C |
| 52555457 | 52587174 | 1 | protein_coding | ENSG0000C |

| | | | | |
|---|---|---|---|---|
| 52564076 | 52564231 | 1 | transcribed_unproce | ENSG0000C |
| 52575883 | 52585561 | -1 | lncRNA | ENSG0000C |
| 52588505 | 52597345 | 1 | transcribed_unproce | ENSG0000C |
| 52594060 | 52690496 | -1 | protein_coding | ENSG0000C |
| 52597699 | 52598887 | -1 | lncRNA | ENSG0000C |
| 52650437 | 52653284 | -1 | lncRNA | ENSG0000C |
| 52702813 | 52735073 | -1 | protein_coding | ENSG0000C |
| 52749942 | 52786807 | -1 | protein_coding | ENSG0000C |
| 52797408 | 52857600 | -1 | protein_coding | ENSG0000C |
| 52809824 | 52810708 | 1 | processed_pseudoge | ENSG0000C |
| 52838010 | 52890375 | -1 | protein_coding | ENSG0000C |
| 52861807 | 52864151 | -1 | unprocessed_pseudc | ENSG0000C |
| 52863790 | 52897693 | -1 | protein_coding | ENSG0000C |
| 52865369 | 52865429 | -1 | unprocessed_pseudc | ENSG0000C |
| 52904417 | 52923481 | -1 | protein_coding | ENSG0000C |
| 52923382 | 52924075 | 1 | lncRNA | ENSG0000C |
| 52927135 | 52942601 | -1 | transcribed_processe | ENSG0000C |
| 52928475 | 52962823 | -1 | protein_coding | ENSG0000C |
| 52949379 | 52962911 | -1 | protein_coding | ENSG0000C |
| 52956041 | 52981654 | -1 | lncRNA | ENSG0000C |
| 52966896 | 53037898 | -1 | transcribed_processe | ENSG0000C |
| 53004040 | 53005150 | 1 | processed_pseudoge | ENSG0000C |
| 53007512 | 53013180 | 1 | lncRNA | ENSG0000C |
| 53013921 | 53016122 | 1 | protein_coding | ENSG0000C |
| 53044740 | 53051680 | 1 | protein_coding | ENSG0000C |
| 53066606 | 53103436 | -1 | protein_coding | ENSG0000C |
| 53107879 | 53133077 | -1 | protein_coding | ENSG0000C |
| 53124072 | 53159075 | -1 | protein_coding | ENSG0000C |
| 53159213 | 53193386 | -1 | protein_coding | ENSG0000C |
| 53162428 | 53163563 | 1 | lncRNA | ENSG0000C |
| 53197111 | 53214522 | 1 | transcribed_unproce | ENSG0000C |
| 53223973 | 53224879 | -1 | processed_pseudoge | ENSG0000C |
| 53235381 | 53254898 | -1 | protein_coding | ENSG0000C |
| 53240675 | 53241913 | 1 | lncRNA | ENSG0000C |
| 53258292 | 53261837 | 1 | protein_coding | ENSG0000C |
| 53266676 | 53267723 | -1 | protein_coding | ENSG0000C |
| 53267870 | 53270932 | -1 | lncRNA | ENSG0000C |
| 53281559 | 53284944 | 1 | transcribed_unproce | ENSG0000C |
| 53289601 | 53291426 | -1 | transcribed_processe | ENSG0000C |
| 53299051 | 53308644 | -1 | transcribed_unproce | ENSG0000C |
| 53315137 | 53316043 | 1 | unitary_pseudogene | ENSG0000C |
| 53333749 | 53356906 | 1 | protein_coding | ENSG0000C |
| 53365693 | 53392217 | 1 | protein_coding | ENSG0000C |
| 53386233 | 53386388 | 1 | transcribed_processe | ENSG0000C |
| 53389793 | 53430413 | 1 | protein_coding | ENSG0000C |
| 53413190 | 53413345 | 1 | processed_pseudoge | ENSG0000C |
| 53426019 | 53426499 | -1 | processed_pseudoge | ENSG0000C |
| 53431984 | 53461862 | 1 | protein_coding | ENSG0000C |
| 53431984 | 53444670 | 1 | transcribed_processe | ENSG0000C |
| 53434661 | 53473936 | -1 | lncRNA | ENSG0000C |

| | | | | |
|---|---|---|---|---|
| 53445001 | 53458261 | 1 | protein_coding | ENSG00000 |
| 53467733 | 53496255 | 1 | protein_coding | ENSG00000 |
| 53479350 | 53480091 | 1 | processed_pseudoge | ENSG00000 |
| 53495887 | 53496037 | 1 | transcribed_processe | ENSG00000 |
| 53503392 | 53512687 | 1 | transcribed_processe | ENSG00000 |
| 53512092 | 53512616 | -1 | unprocessed_pseudo | ENSG00000 |
| 53519527 | 53580269 | 1 | protein_coding | ENSG00000 |
| 53599628 | 53608451 | -1 | transcribed_unproce | ENSG00000 |
| 53616802 | 53617254 | 1 | processed_pseudoge | ENSG00000 |
| 53629713 | 53630041 | 1 | misc_RNA | ENSG00000 |
| 53632056 | 53637009 | 1 | protein_coding | ENSG00000 |
| 53653685 | 53653791 | 1 | snRNA | ENSG00000 |
| 53657400 | 53658462 | -1 | unprocessed_pseudo | ENSG00000 |
| 53666679 | 53666762 | 1 | miRNA | ENSG00000 |
| 53669157 | 53669254 | 1 | miRNA | ENSG00000 |
| 53671968 | 53672040 | 1 | miRNA | ENSG00000 |
| 53674197 | 53674320 | 1 | miRNA | ENSG00000 |
| 53675711 | 53675797 | 1 | miRNA | ENSG00000 |
| 53679003 | 53679085 | 1 | miRNA | ENSG00000 |
| 53679940 | 53680023 | 1 | miRNA | ENSG00000 |
| 53682159 | 53682245 | 1 | miRNA | ENSG00000 |
| 53685009 | 53685091 | 1 | miRNA | ENSG00000 |
| 53686469 | 53686555 | 1 | miRNA | ENSG00000 |
| 53688481 | 53688567 | 1 | miRNA | ENSG00000 |
| 53690881 | 53690965 | 1 | miRNA | ENSG00000 |
| 53694393 | 53694475 | 1 | miRNA | ENSG00000 |
| 53695213 | 53695293 | 1 | miRNA | ENSG00000 |
| 53697533 | 53697617 | 1 | miRNA | ENSG00000 |
| 53698385 | 53698471 | 1 | miRNA | ENSG00000 |
| 53700015 | 53700101 | 1 | miRNA | ENSG00000 |
| 53701227 | 53701287 | 1 | miRNA | ENSG00000 |
| 53702737 | 53702819 | 1 | miRNA | ENSG00000 |
| 53706252 | 53706336 | 1 | miRNA | ENSG00000 |
| 53707453 | 53707539 | 1 | miRNA | ENSG00000 |
| 53708735 | 53708835 | 1 | miRNA | ENSG00000 |
| 53711002 | 53711088 | 1 | miRNA | ENSG00000 |
| 53712268 | 53712354 | 1 | miRNA | ENSG00000 |
| 53713347 | 53713434 | 1 | miRNA | ENSG00000 |
| 53716594 | 53716680 | 1 | miRNA | ENSG00000 |
| 53719064 | 53719165 | -1 | snRNA | ENSG00000 |
| 53720096 | 53720182 | 1 | miRNA | ENSG00000 |
| 53721076 | 53721142 | 1 | miRNA | ENSG00000 |
| 53722166 | 53722255 | 1 | miRNA | ENSG00000 |
| 53725442 | 53725526 | 1 | miRNA | ENSG00000 |
| 53726922 | 53726986 | 1 | miRNA | ENSG00000 |
| 53729838 | 53729925 | 1 | miRNA | ENSG00000 |
| 53731006 | 53731090 | 1 | miRNA | ENSG00000 |
| 53732981 | 53733083 | -1 | snRNA | ENSG00000 |
| 53734877 | 53734963 | 1 | miRNA | ENSG00000 |
| 53736845 | 53736934 | 1 | miRNA | ENSG00000 |

| | | | | |
|---|---|---|---|---|
| 53739333 | 53739419 | 1 | miRNA | ENSG00000 |
| 53741313 | 53741407 | 1 | miRNA | ENSG00000 |
| 53742512 | 53742599 | 1 | miRNA | ENSG00000 |
| 53745004 | 53745105 | -1 | snRNA | ENSG00000 |
| 53748636 | 53748722 | 1 | miRNA | ENSG00000 |
| 53749851 | 53749951 | -1 | snRNA | ENSG00000 |
| 53751211 | 53751297 | 1 | miRNA | ENSG00000 |
| 53752397 | 53752481 | 1 | miRNA | ENSG00000 |
| 53754018 | 53754102 | 1 | miRNA | ENSG00000 |
| 53756741 | 53756830 | 1 | miRNA | ENSG00000 |
| 53758232 | 53758318 | 1 | miRNA | ENSG00000 |
| 53760087 | 53760188 | -1 | snRNA | ENSG00000 |
| 53761133 | 53761222 | 1 | miRNA | ENSG00000 |
| 53761961 | 53764199 | 1 | lncRNA | ENSG00000 |
| 53762344 | 53762430 | 1 | miRNA | ENSG00000 |
| 53766534 | 53766632 | -1 | snRNA | ENSG00000 |
| 53767971 | 53768553 | -1 | processed_pseudoge | ENSG00000 |
| 53772506 | 53773280 | -1 | processed_pseudoge | ENSG00000 |
| 53775599 | 53775882 | 1 | processed_pseudoge | ENSG00000 |
| 53787597 | 53788169 | -1 | lncRNA | ENSG00000 |
| 53787675 | 53787741 | 1 | miRNA | ENSG00000 |
| 53787677 | 53787742 | -1 | miRNA | ENSG00000 |
| 53787890 | 53787956 | 1 | miRNA | ENSG00000 |
| 53788705 | 53788773 | 1 | miRNA | ENSG00000 |
| 53788782 | 53789168 | 1 | lncRNA | ENSG00000 |
| 53793603 | 53824394 | -1 | protein_coding | ENSG00000 |
| 53854581 | 53869107 | -1 | lncRNA | ENSG00000 |
| 53864763 | 53876435 | 1 | protein_coding | ENSG00000 |
| 53874626 | 53876049 | -1 | lncRNA | ENSG00000 |
| 53879190 | 53907652 | 1 | protein_coding | ENSG00000 |
| 53909335 | 53943941 | 1 | protein_coding | ENSG00000 |
| 53963040 | 53990215 | 1 | protein_coding | ENSG00000 |
| 53982307 | 53982397 | 1 | miRNA | ENSG00000 |
| 53992288 | 54012669 | 1 | protein_coding | ENSG00000 |
| 54031412 | 54031504 | 1 | unprocessed_pseudc | ENSG00000 |
| 54040825 | 54063953 | -1 | protein_coding | ENSG00000 |
| 54069895 | 54081365 | -1 | protein_coding | ENSG00000 |
| 54094668 | 54102692 | -1 | protein_coding | ENSG00000 |
| 54102728 | 54109257 | 1 | protein_coding | ENSG00000 |
| 54107013 | 54115675 | -1 | protein_coding | ENSG00000 |
| 54115410 | 54131719 | 1 | protein_coding | ENSG00000 |
| 54119511 | 54125343 | -1 | lncRNA | ENSG00000 |
| 54137749 | 54155681 | 1 | protein_coding | ENSG00000 |
| 54155161 | 54159721 | -1 | protein_coding | ENSG00000 |
| 54160108 | 54173250 | -1 | protein_coding | ENSG00000 |
| 54173412 | 54189882 | -1 | protein_coding | ENSG00000 |
| 54189938 | 54194536 | 1 | protein_coding | ENSG00000 |
| 54199454 | 54199971 | -1 | lncRNA | ENSG00000 |
| 54200809 | 54249003 | 1 | protein_coding | ENSG00000 |
| 54209092 | 54209418 | -1 | unprocessed_pseudc | ENSG00000 |

| | | | | |
|---|---|---|---|---|
| 54216278 | 54223506 | -1 | protein_coding | ENSG00000 |
| 54224523 | 54224881 | 1 | lncRNA | ENSG00000 |
| 54229036 | 54230213 | 1 | processed_pseudoge | ENSG00000 |
| 54236592 | 54242791 | -1 | protein_coding | ENSG00000 |
| 54246101 | 54247322 | -1 | unprocessed_pseudo | ENSG00000 |
| 54249431 | 54257301 | -1 | protein_coding | ENSG00000 |
| 54263951 | 54264053 | -1 | snRNA | ENSG00000 |
| 54268005 | 54269045 | 1 | processed_pseudoge | ENSG00000 |
| 54273821 | 54281184 | -1 | protein_coding | ENSG00000 |
| 54282109 | 54282180 | 1 | miRNA | ENSG00000 |
| 54294049 | 54294366 | -1 | unprocessed_pseudo | ENSG00000 |
| 54296194 | 54296252 | -1 | unprocessed_pseudo | ENSG00000 |
| 54298105 | 54298814 | -1 | unprocessed_pseudo | ENSG00000 |
| 54301758 | 54306026 | -1 | unprocessed_pseudo | ENSG00000 |
| 54307070 | 54313166 | -1 | protein_coding | ENSG00000 |
| 54308915 | 54337168 | -1 | lncRNA | ENSG00000 |
| 54320204 | 54321212 | -1 | transcribed_unproce | ENSG00000 |
| 54328272 | 54331434 | -1 | transcribed_unproce | ENSG00000 |
| 54333185 | 54339162 | -1 | protein_coding | ENSG00000 |
| 54351384 | 54370558 | -1 | protein_coding | ENSG00000 |
| 54379489 | 54379960 | 1 | processed_pseudoge | ENSG00000 |
| 54415219 | 54436904 | 1 | protein_coding | ENSG00000 |
| 54430654 | 54434698 | -1 | lncRNA | ENSG00000 |
| 54434183 | 54434665 | 1 | lncRNA | ENSG00000 |
| 54438010 | 54438346 | -1 | lncRNA | ENSG00000 |
| 54438665 | 54439544 | -1 | lncRNA | ENSG00000 |
| 54444807 | 54449045 | -1 | lncRNA | ENSG00000 |
| 54446830 | 54447047 | 1 | processed_pseudoge | ENSG00000 |
| 54448887 | 54462037 | 1 | protein_coding | ENSG00000 |
| 54461732 | 54463778 | -1 | protein_coding | ENSG00000 |
| 54465026 | 54473296 | -1 | protein_coding | ENSG00000 |
| 54497879 | 54510687 | 1 | protein_coding | ENSG00000 |
| 54521723 | 54522031 | 1 | unprocessed_pseudo | ENSG00000 |
| 54532692 | 54545771 | 1 | transcribed_unproce | ENSG00000 |
| 54551970 | 54552256 | 1 | unprocessed_pseudo | ENSG00000 |
| 54572920 | 54590287 | 1 | protein_coding | ENSG00000 |
| 54589441 | 54590287 | 1 | lncRNA | ENSG00000 |
| 54593582 | 54602381 | 1 | protein_coding | ENSG00000 |
| 54607870 | 54608376 | 1 | unprocessed_pseudo | ENSG00000 |
| 54617158 | 54637528 | 1 | protein_coding | ENSG00000 |
| 54635722 | 54638892 | -1 | lncRNA | ENSG00000 |
| 54643889 | 54670359 | 1 | protein_coding | ENSG00000 |
| 54645309 | 54645383 | 1 | miRNA | ENSG00000 |
| 54647944 | 54648853 | 1 | transcribed_processe | ENSG00000 |
| 54652085 | 54652230 | 1 | unprocessed_pseudo | ENSG00000 |
| 54671231 | 54672591 | 1 | unprocessed_pseudo | ENSG00000 |
| 54696604 | 54700905 | 1 | unprocessed_pseudo | ENSG00000 |
| 54703952 | 54704235 | 1 | unprocessed_pseudo | ENSG00000 |
| 54707916 | 54713453 | 1 | transcribed_processe | ENSG00000 |
| 54724479 | 54736536 | 1 | protein_coding | ENSG00000 |

| | | | | |
|---|---|---|---|---|
| 54724496 | 54798285 | 1 | lncRNA | ENSG00000 |
| 54738513 | 54753052 | 1 | protein_coding | ENSG00000 |
| 54755023 | 54767371 | 1 | unprocessed_pseudo | ENSG00000 |
| 54769793 | 54784322 | 1 | protein_coding | ENSG00000 |
| 54786355 | 54790318 | 1 | transcribed_unproce | ENSG00000 |
| 54803535 | 54814517 | 1 | protein_coding | ENSG00000 |
| 54816468 | 54830778 | 1 | protein_coding | ENSG00000 |
| 54832676 | 54848569 | 1 | polymorphic_pseudo | ENSG00000 |
| 54850443 | 54867207 | 1 | protein_coding | ENSG00000 |
| 54868939 | 54869045 | -1 | snRNA | ENSG00000 |
| 54874235 | 54891420 | 1 | protein_coding | ENSG00000 |
| 54890673 | 54891420 | 1 | lncRNA | ENSG00000 |
| 54906148 | 54916140 | 1 | protein_coding | ENSG00000 |
| 54923509 | 54966312 | -1 | protein_coding | ENSG00000 |
| 54948864 | 54948993 | 1 | processed_pseudoge | ENSG00000 |
| 54953130 | 55001142 | 1 | protein_coding | ENSG00000 |
| 54973817 | 54974417 | 1 | processed_pseudoge | ENSG00000 |
| 55006193 | 55048086 | 1 | lncRNA | ENSG00000 |
| 55013705 | 55038264 | -1 | protein_coding | ENSG00000 |
| 55027593 | 55053928 | -1 | lncRNA | ENSG00000 |
| 55039108 | 55071291 | -1 | protein_coding | ENSG00000 |
| 55072020 | 55087923 | 1 | protein_coding | ENSG00000 |
| 55090914 | 55117637 | -1 | protein_coding | ENSG00000 |
| 55116420 | 55123169 | 1 | lncRNA | ENSG00000 |
| 55119250 | 55120383 | -1 | processed_pseudoge | ENSG00000 |
| 55123225 | 55123292 | -1 | miRNA | ENSG00000 |
| 55132698 | 55149206 | -1 | protein_coding | ENSG00000 |
| 55151767 | 55157773 | -1 | protein_coding | ENSG00000 |
| 55154757 | 55160671 | -1 | protein_coding | ENSG00000 |
| 55158661 | 55166722 | -1 | protein_coding | ENSG00000 |
| 55158939 | 55177540 | 1 | lncRNA | ENSG00000 |
| 55167171 | 55167274 | 1 | snoRNA | ENSG00000 |
| 55171196 | 55180289 | -1 | protein_coding | ENSG00000 |
| 55181247 | 55209506 | -1 | protein_coding | ENSG00000 |
| 55214287 | 55217281 | -1 | lncRNA | ENSG00000 |
| 55216660 | 55221616 | 1 | lncRNA | ENSG00000 |
| 55226639 | 55229264 | -1 | protein_coding | ENSG00000 |
| 55227219 | 55230279 | -1 | lncRNA | ENSG00000 |
| 55229779 | 55259017 | -1 | protein_coding | ENSG00000 |
| 55230885 | 55230952 | -1 | miRNA | ENSG00000 |
| 55239912 | 55239976 | -1 | miRNA | ENSG00000 |
| 55245186 | 55245250 | -1 | miRNA | ENSG00000 |
| 55262223 | 55280381 | -1 | protein_coding | ENSG00000 |
| 55282072 | 55312562 | 1 | protein_coding | ENSG00000 |
| 55312029 | 55312495 | -1 | lncRNA | ENSG00000 |
| 55312801 | 55334048 | -1 | protein_coding | ENSG00000 |
| 55338945 | 55339283 | -1 | lncRNA | ENSG00000 |
| 55339853 | 55348121 | 1 | protein_coding | ENSG00000 |
| 55349306 | 55354719 | -1 | protein_coding | ENSG00000 |
| 55352301 | 55359516 | -1 | protein_coding | ENSG00000 |

| | | | | |
|---|---|---|---|---|
| 55354908 | 55363260 | -1 | protein_coding | ENSG00000( |
| 55364382 | 55370463 | -1 | protein_coding | ENSG00000( |
| 55376826 | 55378246 | 1 | protein_coding | ENSG00000( |
| 55377579 | 55378125 | -1 | lncRNA | ENSG00000( |
| 55379244 | 55384292 | -1 | protein_coding | ENSG00000( |
| 55385345 | 55403250 | 1 | protein_coding | ENSG00000( |
| 55388181 | 55388242 | 1 | miRNA | ENSG00000( |
| 55399745 | 55407788 | -1 | protein_coding | ENSG00000( |
| 55428740 | 55442863 | -1 | protein_coding | ENSG00000( |
| 55452985 | 55462343 | -1 | protein_coding | ENSG00000( |
| 55463002 | 55464707 | -1 | protein_coding | ENSG00000( |
| 55467438 | 55467934 | -1 | TEC | ENSG00000( |
| 55475983 | 55476482 | -1 | lncRNA | ENSG00000( |
| 55476617 | 55484487 | 1 | protein_coding | ENSG00000( |
| 55485188 | 55487568 | 1 | protein_coding | ENSG00000( |
| 55488404 | 55519099 | 1 | protein_coding | ENSG00000( |
| 55529733 | 55537089 | -1 | protein_coding | ENSG00000( |
| 55540656 | 55545543 | -1 | protein_coding | ENSG00000( |
| 55576774 | 55580848 | -1 | protein_coding | ENSG00000( |
| 55591376 | 55601970 | -1 | protein_coding | ENSG00000( |
| 55600022 | 55603138 | 1 | protein_coding | ENSG00000( |
| 55605405 | 55617269 | 1 | protein_coding | ENSG00000( |
| 55612490 | 55613097 | -1 | lncRNA | ENSG00000( |
| 55620742 | 55624601 | -1 | protein_coding | ENSG00000( |
| 55635016 | 55643470 | 1 | protein_coding | ENSG00000( |
| 55635459 | 55645623 | 1 | protein_coding | ENSG00000( |
| 55641062 | 55653161 | 1 | protein_coding | ENSG00000( |
| 55654146 | 55674715 | 1 | protein_coding | ENSG00000( |
| 55661901 | 55674715 | -1 | lncRNA | ENSG00000( |
| 55670632 | 55672069 | 1 | lncRNA | ENSG00000( |
| 55675226 | 55709858 | 1 | protein_coding | ENSG00000( |
| 55694118 | 55694558 | -1 | lncRNA | ENSG00000( |
| 55705978 | 55706172 | -1 | processed_pseudoge | ENSG00000( |
| 55707550 | 55707855 | 1 | processed_pseudoge | ENSG00000( |
| 55708438 | 55738402 | -1 | protein_coding | ENSG00000( |
| 55737285 | 55737600 | 1 | misc_RNA | ENSG00000( |
| 55759014 | 55763175 | 1 | protein_coding | ENSG00000( |
| 55769141 | 55773179 | 1 | protein_coding | ENSG00000( |
| 55777553 | 55778900 | 1 | processed_pseudoge | ENSG00000( |
| 55785397 | 55836800 | -1 | protein_coding | ENSG00000( |
| 55836540 | 55881855 | 1 | protein_coding | ENSG00000( |
| 55891699 | 55932336 | -1 | protein_coding | ENSG00000( |
| 55947832 | 55988629 | 1 | protein_coding | ENSG00000( |
| 55999726 | 56061810 | 1 | protein_coding | ENSG00000( |
| 56066684 | 56078801 | 1 | lncRNA | ENSG00000( |
| 56087366 | 56121295 | -1 | protein_coding | ENSG00000( |
| 56105416 | 56105510 | 1 | misc_RNA | ENSG00000( |
| 56127236 | 56127466 | -1 | TEC | ENSG00000( |
| 56132599 | 56160893 | 1 | protein_coding | ENSG00000( |
| 56149817 | 56150490 | 1 | TEC | ENSG00000( |

| | | | | |
|---|---|---|---|---|
| 56176008 | 56185775 | 1 | protein_coding | ENSG00000 |
| 56189570 | 56197920 | -1 | protein_coding | ENSG00000 |
| 56202301 | 56209452 | 1 | protein_coding | ENSG00000 |
| 56217168 | 56217777 | -1 | processed_pseudoge | ENSG00000 |
| 56219670 | 56368383 | -1 | protein_coding | ENSG00000 |
| 56244043 | 56247482 | 1 | unprocessed_pseudo | ENSG00000 |
| 56272748 | 56310454 | 1 | protein_coding | ENSG00000 |
| 56311928 | 56312486 | -1 | lncRNA | ENSG00000 |
| 56314703 | 56365316 | 1 | lncRNA | ENSG00000 |
| 56316524 | 56316636 | -1 | misc_RNA | ENSG00000 |
| 56317177 | 56318008 | -1 | processed_pseudoge | ENSG00000 |
| 56319790 | 56322310 | -1 | processed_pseudoge | ENSG00000 |
| 56338288 | 56338648 | -1 | processed_pseudoge | ENSG00000 |
| 56342432 | 56343419 | -1 | transcribed_processe | ENSG00000 |
| 56347081 | 56348183 | -1 | lncRNA | ENSG00000 |
| 56354493 | 56368053 | -1 | lncRNA | ENSG00000 |
| 56368099 | 56379828 | 1 | transcribed_unproce | ENSG00000 |
| 56375846 | 56393545 | -1 | protein_coding | ENSG00000 |
| 56376704 | 56377284 | -1 | lncRNA | ENSG00000 |
| 56377037 | 56377792 | -1 | transcribed_processe | ENSG00000 |
| 56380230 | 56383924 | 1 | lncRNA | ENSG00000 |
| 56386226 | 56386690 | 1 | processed_pseudoge | ENSG00000 |
| 56387412 | 56388424 | 1 | lncRNA | ENSG00000 |
| 56393656 | 56399172 | 1 | lncRNA | ENSG00000 |
| 56397966 | 56436035 | 1 | protein_coding | ENSG00000 |
| 56439325 | 56478065 | -1 | protein_coding | ENSG00000 |
| 56477250 | 56504362 | 1 | lncRNA | ENSG00000 |
| 56480397 | 56483977 | -1 | TEC | ENSG00000 |
| 56485923 | 56494280 | -1 | unprocessed_pseudo | ENSG00000 |
| 56507850 | 56530221 | 1 | protein_coding | ENSG00000 |
| 56535299 | 56535763 | 1 | lncRNA | ENSG00000 |
| 56536156 | 56538575 | -1 | lncRNA | ENSG00000 |
| 56538948 | 56556808 | 1 | protein_coding | ENSG00000 |
| 56545566 | 56567411 | -1 | lncRNA | ENSG00000 |
| 56567468 | 56588911 | 1 | protein_coding | ENSG00000 |
| 56589495 | 56590281 | -1 | unprocessed_pseudo | ENSG00000 |
| 56595264 | 56626481 | 1 | protein_coding | ENSG00000 |
| 56595313 | 56799602 | 1 | lncRNA | ENSG00000 |
| 56643159 | 56657247 | 1 | protein_coding | ENSG00000 |
| 56661980 | 56671783 | -1 | protein_coding | ENSG00000 |
| 56669941 | 56823395 | 1 | lncRNA | ENSG00000 |
| 56760933 | 56761215 | 1 | transcribed_unproce | ENSG00000 |
| 56774547 | 56840729 | -1 | protein_coding | ENSG00000 |
| 56793048 | 56793249 | -1 | processed_pseudoge | ENSG00000 |
| 56810077 | 56840728 | -1 | protein_coding | ENSG00000 |
| 56840866 | 56848556 | 1 | lncRNA | ENSG00000 |
| 56848068 | 56848204 | 1 | misc_RNA | ENSG00000 |
| 56922018 | 56922213 | 1 | processed_pseudoge | ENSG00000 |
| 57065334 | 57066139 | 1 | processed_pseudoge | ENSG00000 |
| 57111120 | 57112027 | -1 | unprocessed_pseudo | ENSG00000 |

| | | | | |
|---|---|---|---|---|
| 57119138 | 57131926 | 1 | protein_coding | ENSG0000C |
| 57134096 | 57145202 | -1 | protein_coding | ENSG0000C |
| 57150513 | 57150978 | 1 | processed_pseudoge | ENSG0000C |
| 57151587 | 57151860 | -1 | processed_pseudoge | ENSG0000C |
| 57154021 | 57167443 | -1 | protein_coding | ENSG0000C |
| 57175233 | 57177921 | 1 | processed_pseudoge | ENSG0000C |
| 57191500 | 57222846 | 1 | protein_coding | ENSG0000C |
| 57230802 | 57235548 | 1 | protein_coding | ENSG0000C |
| 57240632 | 57262728 | 1 | protein_coding | ENSG0000C |
| 57261354 | 57262738 | 1 | lncRNA | ENSG0000C |
| 57267211 | 57280334 | -1 | lncRNA | ENSG0000C |
| 57280051 | 57294069 | 1 | protein_coding | ENSG0000C |
| 57293083 | 57294469 | -1 | TEC | ENSG0000C |
| 57304305 | 57308562 | 1 | lncRNA | ENSG0000C |
| 57317208 | 57317562 | -1 | processed_pseudoge | ENSG0000C |
| 57320472 | 57330770 | 1 | protein_coding | ENSG0000C |
| 57350848 | 57352012 | -1 | lncRNA | ENSG0000C |
| 57351307 | 57359898 | 1 | protein_coding | ENSG0000C |
| 57363477 | 57379565 | 1 | protein_coding | ENSG0000C |
| 57363551 | 57365405 | 1 | protein_coding | ENSG0000C |
| 57363566 | 57398584 | 1 | protein_coding | ENSG0000C |
| 57389850 | 57402992 | 1 | protein_coding | ENSG0000C |
| 57389884 | 57419994 | 1 | protein_coding | ENSG0000C |
| 57394183 | 57438457 | 1 | protein_coding | ENSG0000C |
| 57411163 | 57421939 | 1 | protein_coding | ENSG0000C |
| 57435325 | 57447101 | 1 | protein_coding | ENSG0000C |
| 57437970 | 57477536 | -1 | protein_coding | ENSG0000C |
| 57449689 | 57453011 | 1 | lncRNA | ENSG0000C |
| 57454790 | 57457142 | -1 | protein_coding | ENSG0000C |
| 57459912 | 57460120 | 1 | unprocessed_pseudc | ENSG0000C |
| 57466663 | 57477570 | -1 | protein_coding | ENSG0000C |
| 57477649 | 57482996 | 1 | lncRNA | ENSG0000C |
| 57487711 | 57496097 | 1 | protein_coding | ENSG0000C |
| 57487718 | 57517559 | 1 | protein_coding | ENSG0000C |
| 57499915 | 57518404 | 1 | protein_coding | ENSG0000C |
| 57527325 | 57557542 | 1 | protein_coding | ENSG0000C |
| 57535257 | 57559863 | -1 | protein_coding | ENSG0000C |
| 57571566 | 57578911 | -1 | protein_coding | ENSG0000C |
| 57578456 | 57593890 | 1 | protein_coding | ENSG0000C |
| 57599885 | 57612722 | 1 | protein_coding | ENSG0000C |
| 57614233 | 57624724 | 1 | protein_coding | ENSG0000C |
| 57625032 | 57626950 | 1 | lncRNA | ENSG0000C |
| 57630395 | 57644041 | 1 | protein_coding | ENSG0000C |
| 57664280 | 57665039 | 1 | processed_pseudoge | ENSG0000C |
| 57668935 | 57679152 | 1 | protein_coding | ENSG0000C |
| 57681969 | 57717301 | 1 | protein_coding | ENSG0000C |
| 57682045 | 57753575 | 1 | protein_coding | ENSG0000C |
| 57697367 | 57709194 | -1 | protein_coding | ENSG0000C |
| 57719751 | 57727624 | -1 | protein_coding | ENSG0000C |
| 57746815 | 57758148 | 1 | protein_coding | ENSG0000C |

| | | | | |
|---|---|---|---|---|
| 57769655 | 57819939 | 1 | protein_coding | ENSG0000C |
| 57803841 | 57814913 | -1 | protein_coding | ENSG0000C |
| 57819719 | 57858941 | 1 | protein_coding | ENSG0000C |
| 57819721 | 57846238 | 1 | protein_coding | ENSG0000C |
| 57826461 | 57826778 | -1 | processed_pseudoge | ENSG0000C |
| 57848731 | 57889074 | -1 | protein_coding | ENSG0000C |
| 57849859 | 57865117 | 1 | protein_coding | ENSG0000C |
| 57867038 | 57868172 | -1 | lncRNA | ENSG0000C |
| 57867885 | 57868834 | 1 | transcribed_processe | ENSG0000C |
| 57876765 | 57916591 | -1 | protein_coding | ENSG0000C |
| 57900296 | 57916610 | -1 | protein_coding | ENSG0000C |
| 57921884 | 57935393 | -1 | protein_coding | ENSG0000C |
| 57940833 | 57947706 | -1 | protein_coding | ENSG0000C |
| 57953158 | 57953759 | -1 | processed_pseudoge | ENSG0000C |
| 57958437 | 57974534 | -1 | protein_coding | ENSG0000C |
| 57977053 | 58003349 | -1 | protein_coding | ENSG0000C |
| 58002061 | 58011232 | 1 | lncRNA | ENSG0000C |
| 58012589 | 58025926 | -1 | unitary_pseudogene | ENSG0000C |
| 58016695 | 58025589 | -1 | lncRNA | ENSG0000C |
| 58034025 | 58054631 | 1 | protein_coding | ENSG0000C |
| 58044592 | 58046769 | 1 | processed_pseudoge | ENSG0000C |
| 58049046 | 58049346 | -1 | processed_pseudoge | ENSG0000C |
| 58059239 | 58086310 | 1 | protein_coding | ENSG0000C |
| 58069970 | 58070288 | 1 | misc_RNA | ENSG0000C |
| 58083838 | 58118427 | -1 | protein_coding | ENSG0000C |
| 58126248 | 58155110 | -1 | protein_coding | ENSG0000C |
| 58136713 | 58137054 | 1 | processed_pseudoge | ENSG0000C |
| 58155864 | 58177576 | 1 | lncRNA | ENSG0000C |
| 58166166 | 58167145 | 1 | processed_pseudoge | ENSG0000C |
| 58183029 | 58213562 | 1 | protein_coding | ENSG0000C |
| 58216180 | 58216561 | 1 | processed_pseudoge | ENSG0000C |
| 58224506 | 58225469 | 1 | processed_pseudoge | ENSG0000C |
| 58228594 | 58277495 | 1 | protein_coding | ENSG0000C |
| 58228914 | 58315183 | 1 | protein_coding | ENSG0000C |
| 58257270 | 58278808 | -1 | lncRNA | ENSG0000C |
| 58266635 | 58267685 | 1 | lncRNA | ENSG0000C |
| 58278955 | 58302791 | 1 | protein_coding | ENSG0000C |
| 58278966 | 58315197 | 1 | lncRNA | ENSG0000C |
| 58305319 | 58315663 | 1 | protein_coding | ENSG0000C |
| 58309727 | 58327248 | -1 | lncRNA | ENSG0000C |
| 58319277 | 58320489 | 1 | processed_pseudoge | ENSG0000C |
| 58326994 | 58342332 | 1 | protein_coding | ENSG0000C |
| 58334688 | 58334751 | 1 | miRNA | ENSG0000C |
| 58345178 | 58353492 | -1 | protein_coding | ENSG0000C |
| 58346854 | 58362751 | -1 | lncRNA | ENSG0000C |
| 58347718 | 58355455 | 1 | lncRNA | ENSG0000C |
| 58350795 | 58351720 | 1 | TEC | ENSG0000C |
| 58354357 | 58362848 | -1 | protein_coding | ENSG0000C |
| 58357999 | 58359603 | 1 | lncRNA | ENSG0000C |
| 58362583 | 58367593 | 1 | lncRNA | ENSG0000C |

| | | | | |
|---|---|---|---|---|
| 58363438 | 58363562 | 1 | rRNA_pseudogene | ENSG00000( |
| 58367618 | 58381030 | -1 | protein_coding | ENSG00000( |
| 58386400 | 58394806 | 1 | protein_coding | ENSG00000( |
| 58386770 | 58386858 | -1 | miRNA | ENSG00000( |
| 58396090 | 58397079 | 1 | protein_coding | ENSG00000( |
| 58400221 | 58400679 | 1 | lncRNA | ENSG00000( |
| 58401504 | 58418327 | 1 | protein_coding | ENSG00000( |
| 58404238 | 58408484 | -1 | lncRNA | ENSG00000( |
| 58418560 | 58419310 | 1 | TEC | ENSG00000( |
| 58428632 | 58431148 | -1 | lncRNA | ENSG00000( |
| 58432814 | 58440153 | -1 | protein_coding | ENSG00000( |
| 58440448 | 58445849 | 1 | lncRNA | ENSG00000( |
| 58451611 | 58457833 | 1 | protein_coding | ENSG00000( |
| 58467045 | 58475436 | 1 | protein_coding | ENSG00000( |
| 58474017 | 58481230 | 1 | protein_coding | ENSG00000( |
| 58475355 | 58475763 | -1 | lncRNA | ENSG00000( |
| 58479512 | 58512413 | -1 | protein_coding | ENSG00000( |
| 58483749 | 58483843 | -1 | snRNA | ENSG00000( |
| 58490797 | 58491075 | 1 | misc_RNA | ENSG00000( |
| 58500505 | 58501137 | 1 | lncRNA | ENSG00000( |
| 58513530 | 58538911 | -1 | protein_coding | ENSG00000( |
| 58543299 | 58543578 | -1 | misc_RNA | ENSG00000( |
| 58544064 | 58550722 | 1 | protein_coding | ENSG00000( |
| 58550285 | 58550376 | 1 | miRNA | ENSG00000( |
| 58551566 | 58555105 | -1 | protein_coding | ENSG00000( |
| 58555712 | 58558954 | -1 | protein_coding | ENSG00000( |
| 58559129 | 58574797 | 1 | lncRNA | ENSG00000( |
| 58561931 | 58573575 | -1 | protein_coding | ENSG00000( |
| 58573503 | 58605223 | 1 | transcribed_processe | ENSG00000( |
| 58575445 | 58601316 | 1 | lncRNA | ENSG00000( |
| 58586158 | 58586356 | 1 | transcribed_processe | ENSG00000( |
| 58593134 | 58599355 | -1 | lncRNA | ENSG00000( |
| 87250 | 97094 | 1 | protein_coding | ENSG00000( |
| 142590 | 145751 | 1 | protein_coding | ENSG00000( |
| 157454 | 159163 | 1 | protein_coding | ENSG00000( |
| 187853 | 189711 | -1 | protein_coding | ENSG00000( |
| 227258 | 229886 | 1 | protein_coding | ENSG00000( |
| 257724 | 261096 | 1 | protein_coding | ENSG00000( |
| 267186 | 268857 | 1 | lncRNA | ENSG00000( |
| 270863 | 290778 | -1 | protein_coding | ENSG00000( |
| 297570 | 300321 | 1 | protein_coding | ENSG00000( |
| 311731 | 313237 | 1 | lncRNA | ENSG00000( |
| 316860 | 348490 | -1 | lncRNA | ENSG00000( |
| 325595 | 330224 | 1 | protein_coding | ENSG00000( |
| 346782 | 359660 | 1 | protein_coding | ENSG00000( |
| 362835 | 397559 | 1 | protein_coding | ENSG00000( |
| 407498 | 430966 | 1 | protein_coding | ENSG00000( |
| 435480 | 462543 | -1 | protein_coding | ENSG00000( |
| 450663 | 450762 | 1 | misc_RNA | ENSG00000( |
| 472498 | 543835 | -1 | protein_coding | ENSG00000( |

| | | | | |
|---|---|---|---|---|
| 603797 | 610398 | -1 | protein_coding | ENSG00000 |
| 646615 | 653370 | -1 | protein_coding | ENSG00000 |
| 646626 | 675800 | -1 | protein_coding | ENSG00000 |
| 661596 | 675802 | -1 | protein_coding | ENSG00000 |
| 760080 | 776015 | -1 | protein_coding | ENSG00000 |
| 833715 | 857463 | 1 | protein_coding | ENSG00000 |
| 839447 | 839977 | -1 | processed_pseudoge | ENSG00000 |
| 869900 | 916334 | -1 | protein_coding | ENSG00000 |
| 958452 | 1002311 | -1 | protein_coding | ENSG00000 |
| 1023874 | 1118467 | -1 | lncRNA | ENSG00000 |
| 1113240 | 1189415 | 1 | protein_coding | ENSG00000 |
| 1160584 | 1161697 | 1 | processed_pseudoge | ENSG00000 |
| 1180561 | 1185415 | -1 | protein_coding | ENSG00000 |
| 1186092 | 1207036 | 1 | lncRNA | ENSG00000 |
| 1203454 | 1208279 | 1 | protein_coding | ENSG00000 |
| 1226056 | 1296421 | 1 | protein_coding | ENSG00000 |
| 1266280 | 1309328 | 1 | protein_coding | ENSG00000 |
| 1309909 | 1329139 | -1 | protein_coding | ENSG00000 |
| 1317571 | 1393096 | -1 | protein_coding | ENSG00000 |
| 1320710 | 1324367 | 1 | TEC | ENSG00000 |
| 1325405 | 1378735 | 1 | lncRNA | ENSG00000 |
| 1361622 | 1362585 | 1 | lncRNA | ENSG00000 |
| 1368978 | 1393172 | -1 | protein_coding | ENSG00000 |
| 1392900 | 1392961 | -1 | miRNA | ENSG00000 |
| 1442162 | 1473842 | -1 | protein_coding | ENSG00000 |
| 1470741 | 1491587 | -1 | protein_coding | ENSG00000 |
| 1516759 | 1557653 | -1 | unprocessed_pseudo | ENSG00000 |
| 1529056 | 1529159 | -1 | snRNA | ENSG00000 |
| 1534251 | 1557705 | -1 | protein_coding | ENSG00000 |
| 1540144 | 1620009 | -1 | protein_coding | ENSG00000 |
| 1563521 | 1620061 | -1 | protein_coding | ENSG00000 |
| 1591392 | 1591848 | 1 | processed_pseudoge | ENSG00000 |
| 1629152 | 1657779 | -1 | protein_coding | ENSG00000 |
| 1633508 | 1648472 | 1 | lncRNA | ENSG00000 |
| 1665283 | 1708200 | -1 | unprocessed_pseudo | ENSG00000 |
| 1666360 | 1666640 | 1 | misc_RNA | ENSG00000 |
| 1676662 | 1676907 | -1 | unprocessed_pseudo | ENSG00000 |
| 1722045 | 1722537 | -1 | unprocessed_pseudo | ENSG00000 |
| 1729038 | 1817765 | -1 | lncRNA | ENSG00000 |
| 1756756 | 1758940 | -1 | processed_pseudoge | ENSG00000 |
| 1767957 | 1779995 | -1 | transcribed_unitary_ | ENSG00000 |
| 1888128 | 1894374 | -1 | lncRNA | ENSG00000 |
| 1894167 | 1940592 | 1 | protein_coding | ENSG00000 |
| 1947246 | 2030028 | 1 | lncRNA | ENSG00000 |
| 1978757 | 1994285 | -1 | protein_coding | ENSG00000 |
| 2075555 | 2075822 | 1 | processed_pseudoge | ENSG00000 |
| 2101827 | 2177038 | 1 | protein_coding | ENSG00000 |
| 2107900 | 2109991 | -1 | lncRNA | ENSG00000 |
| 2194074 | 2201329 | 1 | lncRNA | ENSG00000 |
| 2206683 | 2207397 | -1 | lncRNA | ENSG00000 |

| | | | | |
|---|---|---|---|---|
| 2207217 | 2213151 | 1 | lncRNA | ENSG00000 |
| 2207328 | 2341059 | 1 | protein_coding | ENSG00000 |
| 2296001 | 2341079 | 1 | protein_coding | ENSG00000 |
| 2380901 | 2432753 | 1 | protein_coding | ENSG00000 |
| 2453010 | 2453117 | -1 | misc_RNA | ENSG00000 |
| 2461634 | 2470853 | -1 | protein_coding | ENSG00000 |
| 2467212 | 2508907 | -1 | protein_coding | ENSG00000 |
| 2481817 | 2524702 | -1 | protein_coding | ENSG00000 |
| 2536607 | 2641784 | 1 | protein_coding | ENSG00000 |
| 2652593 | 2658393 | 1 | protein_coding | ENSG00000 |
| 2652777 | 2652842 | 1 | miRNA | ENSG00000 |
| 2654212 | 2654286 | 1 | snoRNA | ENSG00000 |
| 2655067 | 2655198 | 1 | snoRNA | ENSG00000 |
| 2656097 | 2656182 | 1 | snoRNA | ENSG00000 |
| 2656624 | 2656694 | 1 | snoRNA | ENSG00000 |
| 2656939 | 2657010 | 1 | snoRNA | ENSG00000 |
| 2658395 | 2664219 | -1 | protein_coding | ENSG00000 |
| 2664352 | 2665874 | 1 | lncRNA | ENSG00000 |
| 2692878 | 2760108 | 1 | protein_coding | ENSG00000 |
| 2780614 | 2781041 | 1 | processed_pseudoge | ENSG00000 |
| 2794074 | 2800627 | -1 | protein_coding | ENSG00000 |
| 2814968 | 2818066 | 1 | protein_coding | ENSG00000 |
| 2814987 | 2815833 | 1 | protein_coding | ENSG00000 |
| 2816302 | 2820284 | 1 | protein_coding | ENSG00000 |
| 2835314 | 2841190 | -1 | protein_coding | ENSG00000 |
| 2840703 | 2866732 | 1 | protein_coding | ENSG00000 |
| 2864184 | 3039076 | 1 | protein_coding | ENSG00000 |
| 3043622 | 3045747 | 1 | protein_coding | ENSG00000 |
| 3046052 | 3048250 | 1 | protein_coding | ENSG00000 |
| 3071620 | 3072517 | 1 | protein_coding | ENSG00000 |
| 3082556 | 3084724 | -1 | protein_coding | ENSG00000 |
| 3094171 | 3094509 | 1 | misc_RNA | ENSG00000 |
| 3106913 | 3150867 | 1 | lncRNA | ENSG00000 |
| 3107573 | 3160196 | -1 | protein_coding | ENSG00000 |
| 3146519 | 3159865 | -1 | protein_coding | ENSG00000 |
| 3162617 | 3173592 | -1 | protein_coding | ENSG00000 |
| 3190350 | 3204685 | -1 | protein_coding | ENSG00000 |
| 3208868 | 3223870 | 1 | protein_coding | ENSG00000 |
| 3227417 | 3239559 | -1 | protein_coding | ENSG00000 |
| 3239705 | 3245382 | 1 | lncRNA | ENSG00000 |
| 3249305 | 3407625 | -1 | protein_coding | ENSG00000 |
| 3360036 | 3360142 | 1 | snRNA | ENSG00000 |
| 3361880 | 3362324 | -1 | processed_pseudoge | ENSG00000 |
| 3384229 | 3384440 | 1 | snoRNA | ENSG00000 |
| 3465177 | 3465444 | -1 | processed_pseudoge | ENSG00000 |
| 3471018 | 3651118 | 1 | protein_coding | ENSG00000 |
| 3489529 | 3491005 | -1 | processed_pseudoge | ENSG00000 |
| 3659292 | 3663399 | -1 | protein_coding | ENSG00000 |
| 3667965 | 3682246 | -1 | protein_coding | ENSG00000 |
| 3686970 | 3707128 | -1 | protein_coding | ENSG00000 |

| | | | | |
|---|---|---|---|---|
| 3732685 | 3753111 | 1 | protein_coding | ENSG0000C |
| 3753508 | 3768387 | -1 | protein_coding | ENSG0000C |
| 3777504 | 3781448 | -1 | protein_coding | ENSG0000C |
| 3783851 | 3786740 | -1 | protein_coding | ENSG0000C |
| 3786772 | 3806121 | 1 | protein_coding | ENSG0000C |
| 3808357 | 3812434 | 1 | lncRNA | ENSG0000C |
| 3811384 | 3823882 | 1 | lncRNA | ENSG0000C |
| 3820524 | 3828838 | 1 | protein_coding | ENSG0000C |
| 3846799 | 3876123 | 1 | protein_coding | ENSG0000C |
| 3888239 | 3888868 | -1 | lncRNA | ENSG0000C |
| 3888839 | 3929882 | 1 | protein_coding | ENSG0000C |
| 3917494 | 3917571 | 1 | miRNA | ENSG0000C |
| 3917502 | 3917563 | -1 | miRNA | ENSG0000C |
| 3921279 | 3923400 | -1 | lncRNA | ENSG0000C |
| 3927309 | 4015558 | -1 | protein_coding | ENSG0000C |
| 4023917 | 4024444 | 1 | processed_pseudoge | ENSG0000C |
| 4059633 | 4060113 | -1 | processed_pseudoge | ENSG0000C |
| 4070152 | 4075165 | -1 | lncRNA | ENSG0000C |
| 4120980 | 4187747 | 1 | protein_coding | ENSG0000C |
| 4192932 | 4195953 | 1 | lncRNA | ENSG0000C |
| 4220630 | 4249287 | -1 | protein_coding | ENSG0000C |
| 4422186 | 4431732 | -1 | lncRNA | ENSG0000C |
| 4475638 | 4525200 | 1 | lncRNA | ENSG0000C |
| 4590993 | 4591300 | -1 | processed_pseudoge | ENSG0000C |
| 4627758 | 4685148 | -1 | transcribed_processe | ENSG0000C |
| 4686350 | 4701590 | 1 | protein_coding | ENSG0000C |
| 4721909 | 4728460 | 1 | protein_coding | ENSG0000C |
| 4731279 | 4740668 | -1 | lncRNA | ENSG0000C |
| 4735669 | 4736673 | -1 | processed_pseudoge | ENSG0000C |
| 4761300 | 4761696 | 1 | lncRNA | ENSG0000C |
| 4780023 | 4823608 | -1 | protein_coding | ENSG0000C |
| 4852356 | 5010293 | -1 | protein_coding | ENSG0000C |
| 4868984 | 4869118 | 1 | snoRNA | ENSG0000C |
| 5013520 | 5013724 | -1 | processed_pseudoge | ENSG0000C |
| 5045761 | 5046276 | -1 | lncRNA | ENSG0000C |
| 5049857 | 5050321 | -1 | lncRNA | ENSG0000C |
| 5061037 | 5061340 | -1 | lncRNA | ENSG0000C |
| 5066082 | 5068154 | -1 | lncRNA | ENSG0000C |
| 5068232 | 5113103 | -1 | protein_coding | ENSG0000C |
| 5098387 | 5098476 | -1 | rRNA_pseudogene | ENSG0000C |
| 5113993 | 5114083 | -1 | misc_RNA | ENSG0000C |
| 5114953 | 5126626 | -1 | protein_coding | ENSG0000C |
| 5121417 | 5121539 | -1 | snoRNA | ENSG0000C |
| 5126879 | 5197887 | 1 | protein_coding | ENSG0000C |
| 5292388 | 5292831 | -1 | processed_pseudoge | ENSG0000C |
| 5302040 | 5314369 | -1 | protein_coding | ENSG0000C |
| 5318268 | 5503189 | 1 | lncRNA | ENSG0000C |
| 5346006 | 5346160 | -1 | rRNA_pseudogene | ENSG0000C |
| 5407564 | 5407876 | 1 | lncRNA | ENSG0000C |
| 5426892 | 5471094 | -1 | lncRNA | ENSG0000C |

| | | | | |
|---|---|---|---|---|
| 5445838 | 5475483 | 1 | lncRNA | ENSG0000C |
| 5482736 | 5526709 | -1 | lncRNA | ENSG0000C |
| 5507875 | 5510002 | -1 | lncRNA | ENSG0000C |
| 5532878 | 5533335 | 1 | processed_pseudoge | ENSG0000C |
| 5533628 | 5535889 | 1 | lncRNA | ENSG0000C |
| 5544399 | 5611006 | -1 | protein_coding | ENSG0000C |
| 5548840 | 5549481 | 1 | processed_pseudoge | ENSG0000C |
| 5750393 | 5864395 | 1 | protein_coding | ENSG0000C |
| 5890055 | 5890212 | 1 | snRNA | ENSG0000C |
| 5911510 | 5925353 | 1 | protein_coding | ENSG0000C |
| 5928085 | 5929318 | -1 | processed_pseudoge | ENSG0000C |
| 5937228 | 5950558 | -1 | protein_coding | ENSG0000C |
| 5950652 | 5998977 | 1 | protein_coding | ENSG0000C |
| 5950949 | 6039856 | 1 | protein_coding | ENSG0000C |
| 5964538 | 5964650 | 1 | misc_RNA | ENSG0000C |
| 5965180 | 5965511 | 1 | processed_pseudoge | ENSG0000C |
| 5990943 | 6005821 | -1 | lncRNA | ENSG0000C |
| 6000418 | 6000941 | 1 | lncRNA | ENSG0000C |
| 6001733 | 6002030 | 1 | misc_RNA | ENSG0000C |
| 6006093 | 6040053 | 1 | protein_coding | ENSG0000C |
| 6040546 | 6054060 | -1 | protein_coding | ENSG0000C |
| 6065966 | 6067897 | -1 | lncRNA | ENSG0000C |
| 6074845 | 6123544 | -1 | protein_coding | ENSG0000C |
| 6200989 | 6202191 | -1 | processed_pseudoge | ENSG0000C |
| 6213699 | 6215381 | 1 | processed_pseudoge | ENSG0000C |
| 6219199 | 6219443 | 1 | processed_pseudoge | ENSG0000C |
| 6220330 | 6220759 | 1 | processed_pseudoge | ENSG0000C |
| 6446723 | 6528459 | 1 | lncRNA | ENSG0000C |
| 6731080 | 6736326 | -1 | lncRNA | ENSG0000C |
| 6767686 | 6780246 | 1 | protein_coding | ENSG0000C |
| 7069614 | 7146656 | 1 | lncRNA | ENSG0000C |
| 7146467 | 7254202 | -1 | lncRNA | ENSG0000C |
| 7255053 | 7258323 | -1 | lncRNA | ENSG0000C |
| 7302056 | 7307432 | -1 | lncRNA | ENSG0000C |
| 7347451 | 7367933 | 1 | lncRNA | ENSG0000C |
| 7371608 | 7371692 | 1 | miRNA | ENSG0000C |
| 7633562 | 7633851 | -1 | misc_RNA | ENSG0000C |
| 7831798 | 7832322 | -1 | processed_pseudoge | ENSG0000C |
| 7867327 | 7867614 | 1 | processed_pseudoge | ENSG0000C |
| 7882985 | 7940458 | -1 | protein_coding | ENSG0000C |
| 7977346 | 8019805 | -1 | protein_coding | ENSG0000C |
| 8019180 | 8043512 | 1 | lncRNA | ENSG0000C |
| 8077251 | 8968360 | 1 | protein_coding | ENSG0000C |
| 8097824 | 8099774 | -1 | processed_pseudoge | ENSG0000C |
| 8248704 | 8256918 | 1 | lncRNA | ENSG0000C |
| 8831186 | 8831393 | 1 | snoRNA | ENSG0000C |
| 8929895 | 8930038 | -1 | processed_pseudoge | ENSG0000C |
| 8934984 | 8935481 | -1 | processed_pseudoge | ENSG0000C |
| 8998849 | 8999100 | 1 | misc_RNA | ENSG0000C |
| 9068763 | 9481242 | 1 | protein_coding | ENSG0000C |

| | | | | |
|---|---|---|---|---|
| 9505180 | 9514998 | -1 | lncRNA | ENSG0000C |
| 9514358 | 9530524 | 1 | protein_coding | ENSG0000C |
| 9537389 | 9839041 | -1 | protein_coding | ENSG0000C |
| 9562941 | 9571257 | 1 | lncRNA | ENSG0000C |
| 9575608 | 9577689 | 1 | lncRNA | ENSG0000C |
| 9835556 | 9873900 | 1 | lncRNA | ENSG0000C |
| 9986088 | 10007116 | 1 | lncRNA | ENSG0000C |
| 9986126 | 10058303 | 1 | protein_coding | ENSG0000C |
| 10006381 | 10368776 | -1 | lncRNA | ENSG0000C |
| 10025917 | 10026168 | 1 | misc_RNA | ENSG0000C |
| 10172522 | 10200824 | 1 | lncRNA | ENSG0000C |
| 10172701 | 10186740 | 1 | lncRNA | ENSG0000C |
| 10218830 | 10307418 | 1 | protein_coding | ENSG0000C |
| 10317889 | 10318753 | -1 | lncRNA | ENSG0000C |
| 10334419 | 10334698 | 1 | processed_pseudoge | ENSG0000C |
| 10349157 | 10349628 | -1 | processed_pseudoge | ENSG0000C |
| 10385779 | 10387806 | 1 | processed_pseudoge | ENSG0000C |
| 10401009 | 10434222 | -1 | protein_coding | ENSG0000C |
| 10413520 | 10431922 | -1 | protein_coding | ENSG0000C |
| 10420546 | 10420737 | -1 | protein_coding | ENSG0000C |
| 10435305 | 10636829 | 1 | protein_coding | ENSG0000C |
| 10612861 | 10614229 | -1 | processed_pseudoge | ENSG0000C |
| 10637684 | 10673999 | -1 | protein_coding | ENSG0000C |
| 10649636 | 10649695 | -1 | miRNA | ENSG0000C |
| 10672695 | 10994924 | 1 | lncRNA | ENSG0000C |
| 10696949 | 10754033 | 1 | lncRNA | ENSG0000C |
| 10753278 | 10765286 | -1 | lncRNA | ENSG0000C |
| 10875333 | 10909279 | -1 | lncRNA | ENSG0000C |
| 10888582 | 10889088 | -1 | processed_pseudoge | ENSG0000C |
| 10996293 | 11029455 | 1 | lncRNA | ENSG0000C |
| 11001304 | 11001763 | 1 | lncRNA | ENSG0000C |
| 11234170 | 11301525 | -1 | lncRNA | ENSG0000C |
| 11345383 | 11345855 | -1 | processed_pseudoge | ENSG0000C |
| 11536406 | 11541647 | -1 | lncRNA | ENSG0000C |
| 11566484 | 11581335 | -1 | lncRNA | ENSG0000C |
| 11608436 | 11609048 | -1 | processed_pseudoge | ENSG0000C |
| 11621051 | 11621477 | 1 | processed_pseudoge | ENSG0000C |
| 11685144 | 11687323 | -1 | lncRNA | ENSG0000C |
| 11800463 | 11878429 | -1 | lncRNA | ENSG0000C |
| 11818821 | 11819091 | -1 | misc_RNA | ENSG0000C |
| 11838355 | 11838574 | 1 | processed_pseudoge | ENSG0000C |
| 11890723 | 11926609 | 1 | protein_coding | ENSG0000C |
| 11909404 | 11918677 | -1 | lncRNA | ENSG0000C |
| 12243308 | 12318221 | -1 | lncRNA | ENSG0000C |
| 12305613 | 12316485 | -1 | lncRNA | ENSG0000C |
| 12380056 | 12381255 | 1 | processed_pseudoge | ENSG0000C |
| 12865202 | 12952519 | -1 | lncRNA | ENSG0000C |
| 12934877 | 12937097 | -1 | lncRNA | ENSG0000C |
| 12936262 | 12940902 | 1 | lncRNA | ENSG0000C |
| 12950288 | 13008417 | 1 | lncRNA | ENSG0000C |

| | | | | |
|---|---|---|---|---|
| 13008972 | 13169103 | 1 | protein_coding | ENSG00000 |
| 13073667 | 13074312 | -1 | processed_pseudoge | ENSG00000 |
| 13221274 | 13300651 | 1 | protein_coding | ENSG00000 |
| 13237801 | 13239674 | -1 | lncRNA | ENSG00000 |
| 13244064 | 13245369 | -1 | lncRNA | ENSG00000 |
| 13368291 | 13368703 | -1 | lncRNA | ENSG00000 |
| 13389392 | 13638932 | -1 | protein_coding | ENSG00000 |
| 13392677 | 13393674 | 1 | processed_pseudoge | ENSG00000 |
| 13714322 | 13784886 | -1 | protein_coding | ENSG00000 |
| 13785007 | 13821580 | 1 | protein_coding | ENSG00000 |
| 13849247 | 13996443 | -1 | protein_coding | ENSG00000 |
| 13973316 | 13973424 | -1 | snRNA | ENSG00000 |
| 13995369 | 16053197 | 1 | protein_coding | ENSG00000 |
| 14003508 | 14003699 | 1 | processed_pseudoge | ENSG00000 |
| 14127400 | 14128336 | 1 | processed_pseudoge | ENSG00000 |
| 14223041 | 14223325 | 1 | misc_RNA | ENSG00000 |
| 14322985 | 14337614 | -1 | protein_coding | ENSG00000 |
| 14352320 | 14352425 | 1 | snRNA | ENSG00000 |
| 14522081 | 14523314 | -1 | lncRNA | ENSG00000 |
| 14547184 | 14547623 | -1 | processed_pseudoge | ENSG00000 |
| 14554384 | 14636524 | 1 | lncRNA | ENSG00000 |
| 14757563 | 14758056 | 1 | processed_pseudoge | ENSG00000 |
| 14884250 | 14929528 | -1 | lncRNA | ENSG00000 |
| 14933843 | 14935363 | -1 | lncRNA | ENSG00000 |
| 15021553 | 15021659 | -1 | snRNA | ENSG00000 |
| 15022852 | 15022958 | 1 | snRNA | ENSG00000 |
| 15280764 | 15280873 | -1 | rRNA_pseudogene | ENSG00000 |
| 15552157 | 15552885 | -1 | lncRNA | ENSG00000 |
| 15619494 | 15619819 | -1 | processed_pseudoge | ENSG00000 |
| 15892355 | 15985876 | -1 | lncRNA | ENSG00000 |
| 16177933 | 16259603 | -1 | lncRNA | ENSG00000 |
| 16223750 | 16224535 | -1 | processed_pseudoge | ENSG00000 |
| 16272104 | 16573434 | -1 | protein_coding | ENSG00000 |
| 16576068 | 16579615 | 1 | lncRNA | ENSG00000 |
| 16580258 | 16580622 | 1 | lncRNA | ENSG00000 |
| 16586333 | 16586693 | -1 | processed_pseudoge | ENSG00000 |
| 16670641 | 16670742 | -1 | misc_RNA | ENSG00000 |
| 16684108 | 16684404 | -1 | processed_pseudoge | ENSG00000 |
| 16714844 | 16730948 | -1 | lncRNA | ENSG00000 |
| 16729961 | 16742564 | 1 | protein_coding | ENSG00000 |
| 16748358 | 16770062 | 1 | protein_coding | ENSG00000 |
| 16857962 | 16864148 | -1 | lncRNA | ENSG00000 |
| 16982831 | 16983033 | -1 | snoRNA | ENSG00000 |
| 17187510 | 17187616 | 1 | snRNA | ENSG00000 |
| 17188134 | 17188344 | 1 | processed_pseudoge | ENSG00000 |
| 17209060 | 17209220 | 1 | snRNA | ENSG00000 |
| 17216779 | 17221913 | -1 | lncRNA | ENSG00000 |
| 17222851 | 17226146 | -1 | lncRNA | ENSG00000 |
| 17226107 | 17484578 | 1 | protein_coding | ENSG00000 |
| 17479085 | 17479431 | -1 | processed_pseudoge | ENSG00000 |

| | | | | |
|---|---|---|---|---|
| 17493905 | 17569220 | -1 | protein_coding | ENSG00000( |
| 17516843 | 17517307 | 1 | processed_pseudoge | ENSG00000( |
| 17565501 | 17565851 | -1 | lncRNA | ENSG00000( |
| 17570075 | 17609919 | 1 | protein_coding | ENSG00000( |
| 17576206 | 17576510 | -1 | misc_RNA | ENSG00000( |
| 17599255 | 17599807 | -1 | processed_pseudoge | ENSG00000( |
| 17613678 | 17682295 | -1 | protein_coding | ENSG00000( |
| 17693672 | 17735871 | 1 | protein_coding | ENSG00000( |
| 17887363 | 17888160 | -1 | lncRNA | ENSG00000( |
| 17903762 | 17903861 | 1 | snRNA | ENSG00000( |
| 17941597 | 17968980 | -1 | protein_coding | ENSG00000( |
| 17956979 | 18059188 | -1 | protein_coding | ENSG00000( |
| 17962710 | 17962946 | -1 | snoRNA | ENSG00000( |
| 17969018 | 17991122 | 1 | protein_coding | ENSG00000( |
| 18011955 | 18012284 | 1 | processed_pseudoge | ENSG00000( |
| 18059493 | 18071008 | 1 | lncRNA | ENSG00000( |
| 18090116 | 18090715 | -1 | processed_pseudoge | ENSG00000( |
| 18095571 | 18095632 | -1 | snRNA | ENSG00000( |
| 18113467 | 18113579 | -1 | misc_RNA | ENSG00000( |
| 18137863 | 18143169 | 1 | protein_coding | ENSG00000( |
| 18138118 | 18188387 | 1 | protein_coding | ENSG00000( |
| 18220113 | 18220512 | 1 | processed_pseudoge | ENSG00000( |
| 18258554 | 18258851 | -1 | misc_RNA | ENSG00000( |
| 18265690 | 18265879 | -1 | snRNA | ENSG00000( |
| 18270630 | 18271069 | -1 | lncRNA | ENSG00000( |
| 18288283 | 18316996 | 1 | protein_coding | ENSG00000( |
| 18289343 | 18359300 | -1 | lncRNA | ENSG00000( |
| 18295143 | 18295451 | -1 | misc_RNA | ENSG00000( |
| 18313765 | 18315111 | -1 | lncRNA | ENSG00000( |
| 18379049 | 18381484 | 1 | lncRNA | ENSG00000( |
| 18383367 | 18467281 | -1 | protein_coding | ENSG00000( |
| 18420160 | 18421454 | -1 | processed_pseudoge | ENSG00000( |
| 18433842 | 18433944 | 1 | rRNA_pseudogene | ENSG00000( |
| 18467389 | 18484646 | 1 | protein_coding | ENSG00000( |
| 18470615 | 18470691 | 1 | miRNA | ENSG00000( |
| 18485265 | 18486052 | 1 | processed_pseudoge | ENSG00000( |
| 18486540 | 18497225 | -1 | protein_coding | ENSG00000( |
| 18504633 | 18505066 | 1 | processed_pseudoge | ENSG00000( |
| 18507520 | 18561415 | 1 | protein_coding | ENSG00000( |
| 18567347 | 18569563 | 1 | protein_coding | ENSG00000( |
| 18567453 | 18744216 | 1 | protein_coding | ENSG00000( |
| 18587893 | 18766644 | 1 | protein_coding | ENSG00000( |
| 18592330 | 18592619 | 1 | misc_RNA | ENSG00000( |
| 18611047 | 18611167 | -1 | snRNA | ENSG00000( |
| 18642782 | 18643634 | -1 | processed_pseudoge | ENSG00000( |
| 18674395 | 18698709 | -1 | lncRNA | ENSG00000( |
| 18782424 | 18783690 | -1 | processed_pseudoge | ENSG00000( |
| 18786065 | 18794579 | -1 | lncRNA | ENSG00000( |
| 18794529 | 18796067 | 1 | lncRNA | ENSG00000( |
| 18809728 | 18830153 | -1 | lncRNA | ENSG00000( |

| | | | | |
|---|---|---|---|---|
| 18813783 | 18814378 | 1 | protein_coding | ENSG0000C |
| 19000709 | 19056796 | -1 | lncRNA | ENSG0000C |
| 19208652 | 19212164 | -1 | lncRNA | ENSG0000C |
| 19212642 | 19722926 | 1 | protein_coding | ENSG0000C |
| 19212852 | 19213477 | -1 | lncRNA | ENSG0000C |
| 19242302 | 19284596 | -1 | lncRNA | ENSG0000C |
| 19693209 | 19697576 | -1 | lncRNA | ENSG0000C |
| 19756390 | 19758037 | -1 | lncRNA | ENSG0000C |
| 19757606 | 20002459 | 1 | protein_coding | ENSG0000C |
| 19823443 | 19823943 | -1 | processed_pseudoge | ENSG0000C |
| 20017310 | 20033655 | 1 | protein_coding | ENSG0000C |
| 20034368 | 20056046 | -1 | protein_coding | ENSG0000C |
| 20052514 | 20360698 | 1 | protein_coding | ENSG0000C |
| 20094401 | 20095684 | -1 | lncRNA | ENSG0000C |
| 20214299 | 20215262 | -1 | lncRNA | ENSG0000C |
| 20253303 | 20253838 | -1 | processed_pseudoge | ENSG0000C |
| 20258407 | 20267809 | -1 | lncRNA | ENSG0000C |
| 20316755 | 20316994 | 1 | misc_RNA | ENSG0000C |
| 20363691 | 20365657 | 1 | TEC | ENSG0000C |
| 20368104 | 20370949 | 1 | protein_coding | ENSG0000C |
| 20389530 | 20712644 | -1 | protein_coding | ENSG0000C |
| 20659710 | 20659964 | 1 | processed_pseudoge | ENSG0000C |
| 20721187 | 20721879 | 1 | processed_pseudoge | ENSG0000C |
| 20738433 | 20738731 | -1 | misc_RNA | ENSG0000C |
| 20854121 | 20854642 | -1 | processed_pseudoge | ENSG0000C |
| 20970448 | 20972562 | 1 | lncRNA | ENSG0000C |
| 21085576 | 21106514 | -1 | lncRNA | ENSG0000C |
| 21089696 | 21092539 | 1 | lncRNA | ENSG0000C |
| 21114723 | 21115197 | 1 | processed_pseudoge | ENSG0000C |
| 21125983 | 21246622 | 1 | protein_coding | ENSG0000C |
| 21138956 | 21139074 | -1 | rRNA_pseudogene | ENSG0000C |
| 21148741 | 21162890 | -1 | lncRNA | ENSG0000C |
| 21154023 | 21218289 | -1 | lncRNA | ENSG0000C |
| 21166206 | 21166596 | -1 | processed_pseudoge | ENSG0000C |
| 21265854 | 21266465 | 1 | unprocessed_pseudc | ENSG0000C |
| 21302731 | 21303704 | -1 | lncRNA | ENSG0000C |
| 21303331 | 21389825 | 1 | protein_coding | ENSG0000C |
| 21395365 | 21397526 | -1 | protein_coding | ENSG0000C |
| 21397740 | 21400391 | 1 | lncRNA | ENSG0000C |
| 21424758 | 21425041 | -1 | misc_RNA | ENSG0000C |
| 21488244 | 21488625 | -1 | processed_pseudoge | ENSG0000C |
| 21499261 | 21502934 | -1 | lncRNA | ENSG0000C |
| 21501395 | 21501968 | 1 | processed_pseudoge | ENSG0000C |
| 21511017 | 21514064 | -1 | protein_coding | ENSG0000C |
| 21511447 | 21512309 | 1 | lncRNA | ENSG0000C |
| 21530122 | 21535665 | 1 | lncRNA | ENSG0000C |
| 21569976 | 21666621 | 1 | lncRNA | ENSG0000C |
| 21610787 | 21703585 | -1 | lncRNA | ENSG0000C |
| 21614978 | 21615435 | 1 | misc_RNA | ENSG0000C |
| 21705659 | 21718486 | 1 | protein_coding | ENSG0000C |

| | | | | |
|---|---|---|---|---|
| 21742303 | 21743077 | -1 | processed_pseudoge | ENSG00000 |
| 21755270 | 21755350 | 1 | processed_pseudoge | ENSG00000 |
| 21947647 | 21970783 | 1 | lncRNA | ENSG00000 |
| 22054074 | 22074654 | 1 | lncRNA | ENSG00000 |
| 22220554 | 22223283 | 1 | lncRNA | ENSG00000 |
| 22263065 | 22284029 | -1 | lncRNA | ENSG00000 |
| 22272052 | 22273454 | -1 | processed_pseudoge | ENSG00000 |
| 22343693 | 22343930 | -1 | processed_pseudoge | ENSG00000 |
| 22371216 | 22471557 | -1 | lncRNA | ENSG00000 |
| 22547671 | 22578642 | -1 | lncRNA | ENSG00000 |
| 22581005 | 22585455 | -1 | protein_coding | ENSG00000 |
| 22587522 | 22607517 | -1 | lncRNA | ENSG00000 |
| 22668480 | 22685344 | -1 | lncRNA | ENSG00000 |
| 22684958 | 22746991 | 1 | lncRNA | ENSG00000 |
| 22732802 | 22734089 | -1 | processed_pseudoge | ENSG00000 |
| 22885670 | 22886068 | -1 | processed_pseudoge | ENSG00000 |
| 22960534 | 22966063 | -1 | processed_pseudoge | ENSG00000 |
| 23010209 | 23030785 | -1 | lncRNA | ENSG00000 |
| 23030863 | 23039241 | 1 | lncRNA | ENSG00000 |
| 23035386 | 23036812 | 1 | protein_coding | ENSG00000 |
| 23045633 | 23049741 | -1 | protein_coding | ENSG00000 |
| 23071547 | 23072722 | -1 | TEC | ENSG00000 |
| 23079360 | 23086324 | -1 | protein_coding | ENSG00000 |
| 23125068 | 23132635 | -1 | lncRNA | ENSG00000 |
| 23148086 | 23149939 | 1 | TEC | ENSG00000 |
| 23160857 | 23160971 | -1 | rRNA_pseudogene | ENSG00000 |
| 23179272 | 23190419 | 1 | lncRNA | ENSG00000 |
| 23320958 | 23325352 | -1 | lncRNA | ENSG00000 |
| 23346941 | 23351486 | -1 | lncRNA | ENSG00000 |
| 23350791 | 23354771 | 1 | protein_coding | ENSG00000 |
| 23356594 | 23358116 | -1 | lncRNA | ENSG00000 |
| 23362182 | 23373062 | 1 | protein_coding | ENSG00000 |
| 23374519 | 23421519 | -1 | protein_coding | ENSG00000 |
| 23380896 | 23381012 | -1 | rRNA_pseudogene | ENSG00000 |
| 23439685 | 23444930 | 1 | protein_coding | ENSG00000 |
| 23450403 | 23452876 | -1 | protein_coding | ENSG00000 |
| 23452623 | 23461207 | 1 | lncRNA | ENSG00000 |
| 23457584 | 23457685 | -1 | misc_RNA | ENSG00000 |
| 23475700 | 23479798 | 1 | unprocessed_pseudc | ENSG00000 |
| 23481645 | 23519054 | -1 | lncRNA | ENSG00000 |
| 23489416 | 23489973 | -1 | lncRNA | ENSG00000 |
| 23491101 | 23496010 | 1 | protein_coding | ENSG00000 |
| 23519146 | 23542018 | 1 | transcribed_unproce | ENSG00000 |
| 23547764 | 23550652 | -1 | unprocessed_pseudc | ENSG00000 |
| 23564732 | 23568484 | -1 | protein_coding | ENSG00000 |
| 23583645 | 23586510 | 1 | unprocessed_pseudc | ENSG00000 |
| 23602410 | 23605876 | -1 | protein_coding | ENSG00000 |
| 23626706 | 23638473 | -1 | protein_coding | ENSG00000 |
| 23631826 | 23632316 | -1 | lncRNA | ENSG00000 |
| 23655225 | 23656390 | -1 | lncRNA | ENSG00000 |

| | | | | |
|---|---|---|---|---|
| 23656151 | 23674888 | 1 | lncRNA | ENSG0000C |
| 23685640 | 23689040 | -1 | protein_coding | ENSG0000C |
| 23711320 | 23711406 | -1 | unprocessed_pseudc | ENSG0000C |
| 23722589 | 23745524 | 1 | lncRNA | ENSG0000C |
| 23747553 | 23751268 | -1 | protein_coding | ENSG0000C |
| 23772785 | 23776169 | -1 | unprocessed_pseudc | ENSG0000C |
| 23798092 | 23805983 | 1 | lncRNA | ENSG0000C |
| 23823769 | 23826729 | -1 | protein_coding | ENSG0000C |
| 23875934 | 23879748 | -1 | protein_coding | ENSG0000C |
| 23918538 | 23928954 | -1 | transcribed_unproce | ENSG0000C |
| 23980585 | 23980734 | -1 | unprocessed_pseudc | ENSG0000C |
| 23985050 | 23988779 | -1 | protein_coding | ENSG0000C |
| 23989292 | 23990454 | 1 | unprocessed_pseudc | ENSG0000C |
| 24063255 | 24226013 | 1 | lncRNA | ENSG0000C |
| 24142590 | 24144386 | -1 | lncRNA | ENSG0000C |
| 24237255 | 24237398 | 1 | snRNA | ENSG0000C |
| 24297585 | 24318086 | -1 | lncRNA | ENSG0000C |
| 24445391 | 24446314 | 1 | processed_pseudoge | ENSG0000C |
| 24469629 | 24666616 | 1 | protein_coding | ENSG0000C |
| 24491472 | 24502345 | -1 | lncRNA | ENSG0000C |
| 24679547 | 24680991 | 1 | lncRNA | ENSG0000C |
| 24931840 | 24932983 | 1 | lncRNA | ENSG0000C |
| 24949269 | 24959928 | 1 | protein_coding | ENSG0000C |
| 24962925 | 24992751 | -1 | protein_coding | ENSG0000C |
| 24974171 | 24974280 | 1 | snRNA | ENSG0000C |
| 24992312 | 24993322 | 1 | lncRNA | ENSG0000C |
| 25006230 | 25058980 | -1 | protein_coding | ENSG0000C |
| 25062962 | 25063591 | -1 | lncRNA | ENSG0000C |
| 25070885 | 25082365 | -1 | protein_coding | ENSG0000C |
| 25099105 | 25099928 | 1 | processed_pseudoge | ENSG0000C |
| 25139652 | 25149372 | -1 | lncRNA | ENSG0000C |
| 25161167 | 25162061 | -1 | processed_pseudoge | ENSG0000C |
| 25184658 | 25187849 | -1 | lncRNA | ENSG0000C |
| 25195693 | 25228075 | 1 | protein_coding | ENSG0000C |
| 25214698 | 25214798 | 1 | misc_RNA | ENSG0000C |
| 25226448 | 25226735 | -1 | TEC | ENSG0000C |
| 25229150 | 25231933 | 1 | lncRNA | ENSG0000C |
| 25239007 | 25245229 | -1 | lncRNA | ENSG0000C |
| 25248085 | 25298012 | 1 | protein_coding | ENSG0000C |
| 25251008 | 25251304 | -1 | lncRNA | ENSG0000C |
| 25284915 | 25285588 | -1 | lncRNA | ENSG0000C |
| 25294742 | 25390835 | -1 | protein_coding | ENSG0000C |
| 25368275 | 25368766 | -1 | processed_pseudoge | ENSG0000C |
| 25407673 | 25452628 | 1 | protein_coding | ENSG0000C |
| 25452697 | 25585531 | -1 | protein_coding | ENSG0000C |
| 25611153 | 25611223 | 1 | snRNA | ENSG0000C |
| 25612935 | 25624014 | -1 | protein_coding | ENSG0000C |
| 25624045 | 25689032 | 1 | lncRNA | ENSG0000C |
| 25654595 | 25654881 | -1 | misc_RNA | ENSG0000C |
| 25669857 | 25670386 | -1 | processed_pseudoge | ENSG0000C |

| | | | | |
|---|---|---|---|---|
| 25673195 | 25696853 | -1 | protein_coding | ENSG00000 |
| 25680781 | 25681246 | 1 | lncRNA | ENSG00000 |
| 25697003 | 25752776 | 1 | lncRNA | ENSG00000 |
| 25734264 | 25735093 | 1 | unprocessed_pseudo | ENSG00000 |
| 25762384 | 25868225 | -1 | lncRNA | ENSG00000 |
| 25783553 | 25783919 | 1 | processed_pseudoge | ENSG00000 |
| 25835985 | 25836832 | -1 | unprocessed_pseudo | ENSG00000 |
| 25845497 | 25845862 | 1 | processed_pseudoge | ENSG00000 |
| 25919499 | 25919681 | -1 | unprocessed_pseudo | ENSG00000 |
| 25955795 | 25969288 | 1 | lncRNA | ENSG00000 |
| 25985210 | 26086917 | 1 | lncRNA | ENSG00000 |
| 26008791 | 26010531 | -1 | lncRNA | ENSG00000 |
| 26009792 | 26011243 | 1 | lncRNA | ENSG00000 |
| 26018832 | 26020684 | 1 | unprocessed_pseudo | ENSG00000 |
| 26071988 | 26072351 | -1 | processed_pseudoge | ENSG00000 |
| 26103416 | 26114028 | -1 | transcribed_unproce | ENSG00000 |
| 26132887 | 26134198 | -1 | lncRNA | ENSG00000 |
| 26135983 | 26188191 | 1 | lncRNA | ENSG00000 |
| 26167817 | 26251546 | -1 | lncRNA | ENSG00000 |
| 26188408 | 26196891 | 1 | lncRNA | ENSG00000 |
| 26208186 | 26208278 | -1 | miRNA | ENSG00000 |
| 28563850 | 28602843 | -1 | transcribed_unproce | ENSG00000 |
| 29080390 | 29102761 | 1 | unprocessed_pseudo | ENSG00000 |
| 29297095 | 29297246 | -1 | rRNA | ENSG00000 |
| 29318824 | 29319579 | 1 | unprocessed_pseudo | ENSG00000 |
| 29321310 | 29321666 | -1 | unprocessed_pseudo | ENSG00000 |
| 29324092 | 29325049 | 1 | unprocessed_pseudo | ENSG00000 |
| 29410348 | 29411600 | -1 | unprocessed_pseudo | ENSG00000 |
| 29415264 | 29416911 | -1 | unprocessed_pseudo | ENSG00000 |
| 29427400 | 29427883 | -1 | processed_pseudoge | ENSG00000 |
| 29430723 | 29432913 | 1 | processed_pseudoge | ENSG00000 |
| 29448517 | 29449306 | -1 | unprocessed_pseudo | ENSG00000 |
| 29480147 | 29497179 | -1 | unprocessed_pseudo | ENSG00000 |
| 29548661 | 29559705 | 1 | lncRNA | ENSG00000 |
| 29575060 | 29575171 | 1 | misc_RNA | ENSG00000 |
| 29641778 | 29641976 | 1 | processed_pseudoge | ENSG00000 |
| 29741510 | 29741661 | -1 | rRNA | ENSG00000 |
| 29746181 | 29748090 | 1 | processed_pseudoge | ENSG00000 |
| 29874059 | 29874177 | 1 | rRNA | ENSG00000 |
| 29878636 | 29879363 | -1 | unprocessed_pseudo | ENSG00000 |
| 30214765 | 30214976 | -1 | processed_pseudoge | ENSG00000 |
| 30278906 | 30289956 | -1 | lncRNA | ENSG00000 |
| 30281837 | 30282954 | 1 | lncRNA | ENSG00000 |
| 30294550 | 30312480 | 1 | lncRNA | ENSG00000 |
| 30309015 | 30309126 | -1 | misc_RNA | ENSG00000 |
| 30323367 | 30362276 | -1 | lncRNA | ENSG00000 |
| 30377372 | 30399257 | 1 | unprocessed_pseudo | ENSG00000 |
| 30403123 | 30403384 | -1 | processed_pseudoge | ENSG00000 |
| 30484925 | 30485076 | -1 | rRNA | ENSG00000 |
| 30545601 | 30551470 | 1 | unprocessed_pseudo | ENSG00000 |

| | | | | |
|---|---|---|---|---|
| 30656033 | 30723932 | -1 | transcribed_unproce | ENSG0000C |
| 30713240 | 30713346 | -1 | snRNA | ENSG0000C |
| 30816156 | 30816274 | -1 | rRNA | ENSG0000C |
| 31257664 | 31259632 | 1 | protein_coding | ENSG0000C |
| 31274170 | 31274867 | 1 | processed_pseudoge | ENSG0000C |
| 31285317 | 31286835 | -1 | processed_pseudoge | ENSG0000C |
| 31303212 | 31308585 | -1 | protein_coding | ENSG0000C |
| 31311303 | 31311962 | -1 | processed_pseudoge | ENSG0000C |
| 31341371 | 31344471 | -1 | processed_pseudoge | ENSG0000C |
| 31356870 | 31356978 | 1 | rRNA_pseudogene | ENSG0000C |
| 31360790 | 31360930 | 1 | processed_pseudoge | ENSG0000C |
| 31368601 | 31373923 | 1 | protein_coding | ENSG0000C |
| 31376319 | 31376446 | 1 | snoRNA | ENSG0000C |
| 31377164 | 31390590 | -1 | protein_coding | ENSG0000C |
| 31404845 | 31412838 | -1 | protein_coding | ENSG0000C |
| 31421436 | 31429180 | -1 | transcribed_unproce | ENSG0000C |
| 31440632 | 31450257 | 1 | protein_coding | ENSG0000C |
| 31465506 | 31476757 | -1 | protein_coding | ENSG0000C |
| 31475288 | 31484895 | 1 | protein_coding | ENSG0000C |
| 31485778 | 31487574 | 1 | lncRNA | ENSG0000C |
| 31514410 | 31577923 | 1 | protein_coding | ENSG0000C |
| 31535263 | 31535626 | 1 | lncRNA | ENSG0000C |
| 31547412 | 31548081 | 1 | protein_coding | ENSG0000C |
| 31563166 | 31564076 | 1 | lncRNA | ENSG0000C |
| 31567707 | 31573263 | -1 | lncRNA | ENSG0000C |
| 31580890 | 31581214 | 1 | lncRNA | ENSG0000C |
| 31581089 | 31581192 | 1 | snRNA | ENSG0000C |
| 31605283 | 31606515 | 1 | protein_coding | ENSG0000C |
| 31607186 | 31607240 | 1 | miRNA | ENSG0000C |
| 31637912 | 31645006 | 1 | protein_coding | ENSG0000C |
| 31664452 | 31723989 | -1 | protein_coding | ENSG0000C |
| 31686216 | 31716825 | 1 | lncRNA | ENSG0000C |
| 31721507 | 31723409 | 1 | lncRNA | ENSG0000C |
| 31739271 | 31801805 | 1 | protein_coding | ENSG0000C |
| 31808503 | 31808749 | -1 | processed_pseudoge | ENSG0000C |
| 31819308 | 31834689 | 1 | protein_coding | ENSG0000C |
| 31844303 | 31845604 | -1 | protein_coding | ENSG0000C |
| 31847637 | 31870747 | -1 | protein_coding | ENSG0000C |
| 31870702 | 31944963 | 1 | protein_coding | ENSG0000C |
| 31901254 | 31901426 | 1 | snRNA | ENSG0000C |
| 31944337 | 31952046 | -1 | protein_coding | ENSG0000C |
| 31968151 | 32003387 | 1 | protein_coding | ENSG0000C |
| 31970181 | 31970831 | 1 | lncRNA | ENSG0000C |
| 32005671 | 32031591 | -1 | lncRNA | ENSG0000C |
| 32008351 | 32008472 | 1 | rRNA_pseudogene | ENSG0000C |
| 32010438 | 32032180 | 1 | protein_coding | ENSG0000C |
| 32050588 | 32050705 | 1 | rRNA_pseudogene | ENSG0000C |
| 32052188 | 32101856 | 1 | protein_coding | ENSG0000C |
| 32109714 | 32167258 | 1 | protein_coding | ENSG0000C |
| 32116171 | 32116629 | 1 | lncRNA | ENSG0000C |

| | | | | |
|---|---|---|---|---|
| 32170390 | 32170790 | -1 | processed_pseudoge | ENSG00000 |
| 32186477 | 32190527 | -1 | transcribed_processe | ENSG00000 |
| 32192504 | 32207743 | -1 | protein_coding | ENSG00000 |
| 32207880 | 32238658 | 1 | protein_coding | ENSG00000 |
| 32237795 | 32237847 | 1 | miRNA | ENSG00000 |
| 32275117 | 32277510 | -1 | lncRNA | ENSG00000 |
| 32277651 | 32335011 | 1 | protein_coding | ENSG00000 |
| 32355053 | 32355734 | 1 | lncRNA | ENSG00000 |
| 32358330 | 32439319 | 1 | protein_coding | ENSG00000 |
| 32443059 | 32585074 | -1 | protein_coding | ENSG00000 |
| 32449755 | 32453607 | 1 | lncRNA | ENSG00000 |
| 32485149 | 32487019 | 1 | lncRNA | ENSG00000 |
| 32509959 | 32520285 | 1 | lncRNA | ENSG00000 |
| 32561093 | 32573888 | 1 | lncRNA | ENSG00000 |
| 32564992 | 32608893 | 1 | lncRNA | ENSG00000 |
| 32581963 | 32593900 | 1 | lncRNA | ENSG00000 |
| 32631625 | 32673941 | -1 | protein_coding | ENSG00000 |
| 32632183 | 32743567 | -1 | protein_coding | ENSG00000 |
| 32690180 | 32690815 | -1 | processed_pseudoge | ENSG00000 |
| 32702691 | 32743997 | -1 | protein_coding | ENSG00000 |
| 32762385 | 32809356 | 1 | protein_coding | ENSG00000 |
| 32819954 | 32850405 | 1 | protein_coding | ENSG00000 |
| 32843128 | 32854257 | -1 | lncRNA | ENSG00000 |
| 32856621 | 32858751 | 1 | lncRNA | ENSG00000 |
| 32858923 | 32961609 | 1 | protein_coding | ENSG00000 |
| 32983773 | 33004433 | -1 | protein_coding | ENSG00000 |
| 33007704 | 33023703 | 1 | protein_coding | ENSG00000 |
| 33031648 | 33044047 | 1 | protein_coding | ENSG00000 |
| 33047517 | 33048152 | -1 | processed_pseudoge | ENSG00000 |
| 33055424 | 33073628 | 1 | protein_coding | ENSG00000 |
| 33079644 | 33111751 | 1 | protein_coding | ENSG00000 |
| 33161768 | 33181412 | 1 | protein_coding | ENSG00000 |
| 33187391 | 33188292 | 1 | processed_pseudoge | ENSG00000 |
| 33193585 | 33210462 | 1 | transcribed_unproce | ENSG00000 |
| 33214920 | 33217067 | -1 | lncRNA | ENSG00000 |
| 33217310 | 33227806 | 1 | protein_coding | ENSG00000 |
| 33235995 | 33243311 | 1 | protein_coding | ENSG00000 |
| 33272262 | 33272657 | -1 | processed_pseudoge | ENSG00000 |
| 33273480 | 33309871 | 1 | protein_coding | ENSG00000 |
| 33316869 | 33329969 | 1 | unitary_pseudogene | ENSG00000 |
| 33347720 | 33354444 | 1 | transcribed_unitary_ | ENSG00000 |
| 33358839 | 33401561 | -1 | protein_coding | ENSG00000 |
| 33407955 | 33443892 | -1 | protein_coding | ENSG00000 |
| 33490075 | 33650036 | 1 | protein_coding | ENSG00000 |
| 33655701 | 33656423 | -1 | lncRNA | ENSG00000 |
| 33657087 | 33674463 | -1 | protein_coding | ENSG00000 |
| 33662327 | 33665619 | 1 | protein_coding | ENSG00000 |
| 33666943 | 33668525 | 1 | protein_coding | ENSG00000 |
| 33674517 | 33675380 | 1 | lncRNA | ENSG00000 |
| 33675477 | 33686385 | -1 | protein_coding | ENSG00000 |

| | | | | |
|---|---|---|---|---|
| 33702758 | 33720319 | -1 | protein_coding | ENSG00000( |
| 33728931 | 33731828 | -1 | lncRNA | ENSG00000( |
| 33731657 | 33792269 | 1 | protein_coding | ENSG00000( |
| 33752167 | 33752543 | -1 | processed_pseudoge | ENSG00000( |
| 33787373 | 33811109 | -1 | lncRNA | ENSG00000( |
| 33811348 | 33854366 | 1 | protein_coding | ENSG00000( |
| 33912323 | 33913336 | -1 | processed_pseudoge | ENSG00000( |
| 33965162 | 33965576 | 1 | processed_pseudoge | ENSG00000( |
| 33980679 | 33994357 | -1 | lncRNA | ENSG00000( |
| 33993646 | 34108308 | 1 | protein_coding | ENSG00000( |
| 34014969 | 34017749 | -1 | lncRNA | ENSG00000( |
| 34049119 | 34049190 | 1 | miRNA | ENSG00000( |
| 34088309 | 34112243 | -1 | protein_coding | ENSG00000( |
| 34122470 | 34123290 | 1 | processed_pseudoge | ENSG00000( |
| 34132270 | 34136485 | 1 | lncRNA | ENSG00000( |
| 34194569 | 34269344 | 1 | protein_coding | ENSG00000( |
| 34213495 | 34215892 | -1 | processed_pseudoge | ENSG00000( |
| 34234840 | 34281173 | -1 | lncRNA | ENSG00000( |
| 34245953 | 34246537 | -1 | processed_pseudoge | ENSG00000( |
| 34280268 | 34311802 | -1 | protein_coding | ENSG00000( |
| 34281632 | 34286466 | -1 | lncRNA | ENSG00000( |
| 34347671 | 34347785 | -1 | processed_pseudoge | ENSG00000( |
| 34363241 | 34540748 | 1 | protein_coding | ENSG00000( |
| 34384165 | 34384265 | -1 | misc_RNA | ENSG00000( |
| 34402149 | 34402696 | -1 | processed_pseudoge | ENSG00000( |
| 34441592 | 34442025 | -1 | lncRNA | ENSG00000( |
| 34450930 | 34454552 | 1 | lncRNA | ENSG00000( |
| 34466325 | 34466418 | 1 | miRNA | ENSG00000( |
| 34475924 | 34476474 | -1 | processed_pseudoge | ENSG00000( |
| 34476205 | 34476787 | 1 | lncRNA | ENSG00000( |
| 34516409 | 34540958 | 1 | protein_coding | ENSG00000( |
| 34526510 | 34526606 | -1 | misc_RNA | ENSG00000( |
| 34546854 | 34560345 | 1 | protein_coding | ENSG00000( |
| 34560542 | 34698790 | -1 | protein_coding | ENSG00000( |
| 34688688 | 34688978 | 1 | processed_pseudoge | ENSG00000( |
| 34689097 | 34825651 | -1 | protein_coding | ENSG00000( |
| 34704339 | 34713439 | 1 | protein_coding | ENSG00000( |
| 34741401 | 34807447 | 1 | lncRNA | ENSG00000( |
| 34833575 | 34834462 | -1 | processed_pseudoge | ENSG00000( |
| 34844720 | 34872856 | -1 | protein_coding | ENSG00000( |
| 34872146 | 34927962 | 1 | protein_coding | ENSG00000( |
| 34928432 | 34956027 | -1 | protein_coding | ENSG00000( |
| 34975403 | 35002437 | 1 | protein_coding | ENSG00000( |
| 34990376 | 34990497 | 1 | miRNA | ENSG00000( |
| 34990400 | 34990472 | -1 | miRNA | ENSG00000( |
| 35002404 | 35092807 | -1 | protein_coding | ENSG00000( |
| 35030317 | 35030420 | -1 | snRNA | ENSG00000( |
| 35115364 | 35147336 | -1 | protein_coding | ENSG00000( |
| 35117136 | 35117207 | -1 | snoRNA | ENSG00000( |
| 35172072 | 35216240 | 1 | protein_coding | ENSG00000( |

| | | | | |
|---|---|---|---|---|
| 35174355 | 35174919 | -1 | lncRNA | ENSG0000C |
| 35195306 | 35195423 | 1 | rRNA_pseudogene | ENSG0000C |
| 35201745 | 35278131 | -1 | protein_coding | ENSG0000C |
| 35218009 | 35218185 | 1 | processed_pseudoge | ENSG0000C |
| 35226690 | 35276998 | 1 | protein_coding | ENSG0000C |
| 35262727 | 35264187 | -1 | TEC | ENSG0000C |
| 35267885 | 35280043 | -1 | protein_coding | ENSG0000C |
| 35278911 | 35284985 | -1 | protein_coding | ENSG0000C |
| 35285251 | 35285756 | 1 | lncRNA | ENSG0000C |
| 35285731 | 35292425 | -1 | protein_coding | ENSG0000C |
| 35302566 | 35412031 | -1 | protein_coding | ENSG0000C |
| 35433029 | 35435450 | 1 | protein_coding | ENSG0000C |
| 35433347 | 35454746 | -1 | protein_coding | ENSG0000C |
| 35453954 | 35454097 | -1 | miRNA | ENSG0000C |
| 35455164 | 35519280 | 1 | protein_coding | ENSG0000C |
| 35476203 | 35490982 | -1 | lncRNA | ENSG0000C |
| 35520852 | 35528255 | -1 | transcribed_unproce | ENSG0000C |
| 35523186 | 35529652 | -1 | protein_coding | ENSG0000C |
| 35542021 | 35557634 | 1 | protein_coding | ENSG0000C |
| 35544430 | 35544747 | -1 | processed_pseudoge | ENSG0000C |
| 35558737 | 35607562 | -1 | transcribed_unitary_ | ENSG0000C |
| 35588324 | 35588607 | 1 | processed_pseudoge | ENSG0000C |
| 35615829 | 35621094 | 1 | protein_coding | ENSG0000C |
| 35626031 | 35664956 | -1 | protein_coding | ENSG0000C |
| 35632340 | 35674544 | -1 | protein_coding | ENSG0000C |
| 35643143 | 35643462 | 1 | misc_RNA | ENSG0000C |
| 35647328 | 35647430 | -1 | snRNA | ENSG0000C |
| 35648925 | 35664956 | -1 | protein_coding | ENSG0000C |
| 35668052 | 35699355 | -1 | protein_coding | ENSG0000C |
| 35699272 | 35700984 | 1 | protein_coding | ENSG0000C |
| 35701347 | 35742312 | -1 | protein_coding | ENSG0000C |
| 35752814 | 35753719 | 1 | processed_pseudoge | ENSG0000C |
| 35769785 | 35769867 | -1 | snRNA | ENSG0000C |
| 35771974 | 35950370 | 1 | protein_coding | ENSG0000C |
| 35850674 | 35850915 | -1 | processed_pseudoge | ENSG0000C |
| 35853252 | 35853364 | -1 | misc_RNA | ENSG0000C |
| 35878476 | 35878733 | 1 | processed_pseudoge | ENSG0000C |
| 35887659 | 35887801 | 1 | snRNA | ENSG0000C |
| 35928570 | 35928675 | -1 | snRNA | ENSG0000C |
| 35953617 | 35959472 | -1 | protein_coding | ENSG0000C |
| 35954564 | 36030700 | 1 | protein_coding | ENSG0000C |
| 36045618 | 36051018 | -1 | lncRNA | ENSG0000C |
| 36051491 | 36085703 | 1 | lncRNA | ENSG0000C |
| 36064243 | 36064563 | -1 | lncRNA | ENSG0000C |
| 36076018 | 36076601 | -1 | processed_pseudoge | ENSG0000C |
| 36086252 | 36088192 | -1 | lncRNA | ENSG0000C |
| 36091504 | 36232799 | 1 | protein_coding | ENSG0000C |
| 36147334 | 36155760 | -1 | lncRNA | ENSG0000C |
| 36233851 | 36234297 | 1 | lncRNA | ENSG0000C |
| 36236459 | 36270918 | 1 | protein_coding | ENSG0000C |

| | | | | |
|---|---|---|---|---|
| 36306336 | 36528637 | 1 | protein_coding | ENSG0000C |
| 36507702 | 36573391 | -1 | lncRNA | ENSG0000C |
| 36541497 | 36551447 | 1 | protein_coding | ENSG0000C |
| 36561237 | 36561530 | -1 | misc_RNA | ENSG0000C |
| 36573488 | 36593950 | 1 | protein_coding | ENSG0000C |
| 36574553 | 36612384 | 1 | protein_coding | ENSG0000C |
| 36605779 | 36612557 | 1 | protein_coding | ENSG0000C |
| 36612318 | 36646196 | -1 | protein_coding | ENSG0000C |
| 36620403 | 36621423 | 1 | unprocessed_pseudc | ENSG0000C |
| 36651766 | 36746090 | -1 | protein_coding | ENSG0000C |
| 36668559 | 36668670 | -1 | misc_RNA | ENSG0000C |
| 36751791 | 36773818 | -1 | protein_coding | ENSG0000C |
| 36777447 | 36863538 | -1 | protein_coding | ENSG0000C |
| 36870099 | 36870402 | 1 | misc_RNA | ENSG0000C |
| 36876121 | 36894235 | 1 | protein_coding | ENSG0000C |
| 36889773 | 36951901 | -1 | protein_coding | ENSG0000C |
| 36996349 | 37095997 | -1 | protein_coding | ENSG0000C |
| 37021743 | 37021984 | -1 | processed_pseudoge | ENSG0000C |
| 37049254 | 37049707 | 1 | processed_pseudoge | ENSG0000C |
| 37095785 | 37097178 | 1 | lncRNA | ENSG0000C |
| 37101226 | 37179588 | -1 | protein_coding | ENSG0000C |
| 37165774 | 37166312 | -1 | processed_pseudoge | ENSG0000C |
| 37178410 | 37241619 | 1 | protein_coding | ENSG0000C |
| 37251082 | 37261835 | -1 | protein_coding | ENSG0000C |
| 37289638 | 37317260 | 1 | protein_coding | ENSG0000C |
| 37338886 | 37344109 | -1 | lncRNA | ENSG0000C |
| 37344685 | 37406050 | 1 | protein_coding | ENSG0000C |
| 37444733 | 37445534 | -1 | processed_pseudoge | ENSG0000C |
| 37492472 | 37527931 | -1 | protein_coding | ENSG0000C |
| 37521206 | 37523690 | 1 | protein_coding | ENSG0000C |
| 37526642 | 37527060 | -1 | lncRNA | ENSG0000C |
| 37530039 | 37530543 | 1 | processed_pseudoge | ENSG0000C |
| 37571103 | 37591493 | 1 | lncRNA | ENSG0000C |
| 37602852 | 37603185 | -1 | processed_pseudoge | ENSG0000C |
| 37619298 | 37623119 | -1 | lncRNA | ENSG0000C |
| 37676889 | 37683234 | 1 | lncRNA | ENSG0000C |
| 37693955 | 37872129 | 1 | protein_coding | ENSG0000C |
| 37903111 | 37945350 | 1 | protein_coding | ENSG0000C |
| 37975156 | 37975481 | -1 | misc_RNA | ENSG0000C |
| 37983007 | 38033461 | -1 | protein_coding | ENSG0000C |
| 38033725 | 38127780 | 1 | protein_coding | ENSG0000C |
| 38098414 | 38098700 | -1 | misc_RNA | ENSG0000C |
| 38103524 | 38104961 | -1 | lncRNA | ENSG0000C |
| 38127385 | 38166578 | -1 | protein_coding | ENSG0000C |
| 38210503 | 38260772 | -1 | protein_coding | ENSG0000C |
| 38233251 | 38233799 | -1 | lncRNA | ENSG0000C |
| 38260149 | 38304332 | 1 | lncRNA | ENSG0000C |
| 38260175 | 38293892 | 1 | lncRNA | ENSG0000C |
| 38270778 | 38274020 | 1 | lncRNA | ENSG0000C |
| 38288331 | 38326536 | -1 | lncRNA | ENSG0000C |

| | | | | |
|---|---|---|---|---|
| 38304123 | 38337505 | 1 | protein_coding | ENSG00000 |
| 38346482 | 38377013 | 1 | protein_coding | ENSG00000 |
| 38378825 | 38383179 | 1 | lncRNA | ENSG00000 |
| 38404893 | 38416797 | -1 | lncRNA | ENSG00000 |
| 38418483 | 38419202 | -1 | lncRNA | ENSG00000 |
| 38419638 | 38435409 | -1 | lncRNA | ENSG00000 |
| 38421647 | 38421774 | -1 | snoRNA | ENSG00000 |
| 38425083 | 38425354 | -1 | snoRNA | ENSG00000 |
| 38427309 | 38427442 | -1 | snoRNA | ENSG00000 |
| 38429670 | 38429803 | -1 | snoRNA | ENSG00000 |
| 38433865 | 38433998 | -1 | snoRNA | ENSG00000 |
| 38442036 | 38442169 | 1 | snoRNA | ENSG00000 |
| 38446343 | 38450940 | 1 | lncRNA | ENSG00000 |
| 38448084 | 38448215 | 1 | snoRNA | ENSG00000 |
| 38449370 | 38449503 | 1 | snoRNA | ENSG00000 |
| 38472816 | 38578859 | 1 | protein_coding | ENSG00000 |
| 38516563 | 38516632 | 1 | miRNA | ENSG00000 |
| 38537758 | 38538458 | 1 | processed_pseudoge | ENSG00000 |
| 38581195 | 38588463 | 1 | protein_coding | ENSG00000 |
| 38601934 | 38651035 | 1 | protein_coding | ENSG00000 |
| 38614913 | 38615476 | -1 | processed_pseudoge | ENSG00000 |
| 38711821 | 38712103 | -1 | misc_RNA | ENSG00000 |
| 38724486 | 38729372 | 1 | protein_coding | ENSG00000 |
| 38748460 | 38772520 | 1 | protein_coding | ENSG00000 |
| 38761528 | 38761842 | 1 | misc_RNA | ENSG00000 |
| 38805697 | 38923024 | 1 | protein_coding | ENSG00000 |
| 38872770 | 38873074 | 1 | misc_RNA | ENSG00000 |
| 38926312 | 38953106 | 1 | protein_coding | ENSG00000 |
| 38955910 | 38956547 | 1 | lncRNA | ENSG00000 |
| 38961925 | 38962111 | -1 | lncRNA | ENSG00000 |
| 38962299 | 39039723 | 1 | protein_coding | ENSG00000 |
| 38962472 | 38962993 | 1 | lncRNA | ENSG00000 |
| 38977711 | 38978422 | -1 | processed_pseudoge | ENSG00000 |
| 39213777 | 39224748 | -1 | lncRNA | ENSG00000 |
| 39231115 | 39233836 | -1 | lncRNA | ENSG00000 |
| 39329014 | 39329924 | -1 | processed_pseudoge | ENSG00000 |
| 39349094 | 39349392 | -1 | misc_RNA | ENSG00000 |
| 39655325 | 39663552 | -1 | lncRNA | ENSG00000 |
| 39757202 | 39815097 | 1 | lncRNA | ENSG00000 |
| 39932922 | 39933239 | -1 | processed_pseudoge | ENSG00000 |
| 40004216 | 40011323 | 1 | lncRNA | ENSG00000 |
| 40031585 | 40043889 | 1 | lncRNA | ENSG00000 |
| 40121041 | 40126357 | 1 | lncRNA | ENSG00000 |
| 40685848 | 40689236 | -1 | protein_coding | ENSG00000 |
| 40696499 | 40698616 | -1 | lncRNA | ENSG00000 |
| 40717257 | 40758636 | 1 | lncRNA | ENSG00000 |
| 40854119 | 40854229 | -1 | rRNA_pseudogene | ENSG00000 |
| 41024205 | 41024394 | 1 | snRNA | ENSG00000 |
| 41028822 | 41124487 | 1 | protein_coding | ENSG00000 |
| 41098019 | 41138003 | -1 | lncRNA | ENSG00000 |

| | | | | |
|---|---|---|---|---|
| 41136960 | 41196801 | 1 | protein_coding | ENSG0000C |
| 41169023 | 41169078 | 1 | miRNA | ENSG0000C |
| 41178448 | 41317672 | -1 | protein_coding | ENSG0000C |
| 41196647 | 41196938 | -1 | transcribed_processe | ENSG0000C |
| 41196691 | 41197157 | 1 | transcribed_processe | ENSG0000C |
| 41290280 | 41290577 | -1 | misc_RNA | ENSG0000C |
| 41331123 | 41331660 | 1 | processed_pseudoge | ENSG0000C |
| 41340920 | 41360582 | 1 | protein_coding | ENSG0000C |
| 41359962 | 41366818 | -1 | protein_coding | ENSG0000C |
| 41382383 | 41383107 | 1 | processed_pseudoge | ENSG0000C |
| 41402083 | 41618384 | -1 | protein_coding | ENSG0000C |
| 41459354 | 41461944 | -1 | processed_pseudoge | ENSG0000C |
| 41485571 | 41486225 | -1 | lncRNA | ENSG0000C |
| 41547422 | 41547922 | 1 | processed_pseudoge | ENSG0000C |
| 41684986 | 41685091 | -1 | snRNA | ENSG0000C |
| 41712479 | 41712600 | -1 | snoRNA | ENSG0000C |
| 41991140 | 42063969 | 1 | lncRNA | ENSG0000C |
| 41997131 | 41999397 | -1 | processed_pseudoge | ENSG0000C |
| 42014034 | 42015478 | -1 | lncRNA | ENSG0000C |
| 42072752 | 43189970 | -1 | protein_coding | ENSG0000C |
| 42606694 | 42607559 | 1 | processed_pseudoge | ENSG0000C |
| 42685404 | 42688562 | 1 | lncRNA | ENSG0000C |
| 42968743 | 42971492 | 1 | lncRNA | ENSG0000C |
| 43063326 | 43063609 | 1 | misc_RNA | ENSG0000C |
| 43189998 | 43202599 | 1 | lncRNA | ENSG0000C |
| 43222206 | 43222509 | 1 | misc_RNA | ENSG0000C |
| 43230760 | 43231260 | 1 | processed_pseudoge | ENSG0000C |
| 43304555 | 43304679 | 1 | scaRNA | ENSG0000C |
| 43388642 | 43389041 | -1 | processed_pseudoge | ENSG0000C |
| 43405473 | 43405578 | 1 | snRNA | ENSG0000C |
| 43423196 | 43423538 | -1 | processed_pseudoge | ENSG0000C |
| 43451159 | 43451260 | -1 | misc_RNA | ENSG0000C |
| 43457893 | 43466046 | 1 | protein_coding | ENSG0000C |
| 43458054 | 43541881 | 1 | protein_coding | ENSG0000C |
| 43473013 | 43473119 | 1 | snRNA | ENSG0000C |
| 43489442 | 43550950 | 1 | protein_coding | ENSG0000C |
| 43523261 | 43523572 | -1 | processed_pseudoge | ENSG0000C |
| 43540171 | 43569498 | 1 | protein_coding | ENSG0000C |
| 43549389 | 43550949 | -1 | lncRNA | ENSG0000C |
| 43558968 | 43588237 | 1 | protein_coding | ENSG0000C |
| 43590009 | 43590596 | -1 | lncRNA | ENSG0000C |
| 43590937 | 43647299 | 1 | protein_coding | ENSG0000C |
| 43652523 | 43653291 | 1 | processed_pseudoge | ENSG0000C |
| 43667019 | 43716495 | 1 | protein_coding | ENSG0000C |
| 43726164 | 43727002 | -1 | protein_coding | ENSG0000C |
| 43796437 | 43796543 | -1 | snRNA | ENSG0000C |
| 43894720 | 43895468 | 1 | lncRNA | ENSG0000C |
| 43914852 | 44069616 | 1 | protein_coding | ENSG0000C |
| 44007248 | 44007537 | 1 | misc_RNA | ENSG0000C |
| 44111695 | 44187578 | -1 | protein_coding | ENSG0000C |

| | | | | |
|---|---|---|---|---|
| 44181111 | 44182712 | 1 | lncRNA | ENSG0000C |
| 44195939 | 44210771 | -1 | protein_coding | ENSG0000C |
| 44210907 | 44226027 | 1 | lncRNA | ENSG0000C |
| 44247099 | 44280917 | 1 | protein_coding | ENSG0000C |
| 44302840 | 44311202 | -1 | protein_coding | ENSG0000C |
| 44336986 | 44351242 | 1 | protein_coding | ENSG0000C |
| 44344839 | 44344933 | 1 | misc_RNA | ENSG0000C |
| 44347552 | 44355185 | -1 | lncRNA | ENSG0000C |
| 44355700 | 44434596 | 1 | protein_coding | ENSG0000C |
| 44372746 | 44395706 | -1 | lncRNA | ENSG0000C |
| 44389624 | 44391537 | -1 | lncRNA | ENSG0000C |
| 44408120 | 44408203 | 1 | miRNA | ENSG0000C |
| 44436172 | 44465349 | -1 | lncRNA | ENSG0000C |
| 44466564 | 44466842 | -1 | processed_pseudoge | ENSG0000C |
| 44475874 | 44494603 | 1 | protein_coding | ENSG0000C |
| 44496221 | 44522085 | -1 | protein_coding | ENSG0000C |
| 44531785 | 44624247 | 1 | protein_coding | ENSG0000C |
| 44619522 | 44652233 | -1 | protein_coding | ENSG0000C |
| 44656451 | 44663498 | -1 | lncRNA | ENSG0000C |
| 44656451 | 44696096 | -1 | lncRNA | ENSG0000C |
| 44694892 | 44746021 | -1 | lncRNA | ENSG0000C |
| 44714844 | 44728509 | 1 | protein_coding | ENSG0000C |
| 44745865 | 44752313 | 1 | protein_coding | ENSG0000C |
| 44746642 | 44747201 | 1 | lncRNA | ENSG0000C |
| 44751808 | 44810338 | -1 | protein_coding | ENSG0000C |
| 44841925 | 44842539 | -1 | unprocessed_pseudc | ENSG0000C |
| 44864435 | 44885604 | -1 | lncRNA | ENSG0000C |
| 44885702 | 44908532 | 1 | protein_coding | ENSG0000C |
| 44910062 | 44959035 | 1 | protein_coding | ENSG0000C |
| 44942130 | 44960397 | -1 | protein_coding | ENSG0000C |
| 44963794 | 44966402 | -1 | lncRNA | ENSG0000C |
| 44966479 | 45080021 | 1 | protein_coding | ENSG0000C |
| 45004348 | 45004454 | 1 | misc_RNA | ENSG0000C |
| 45091214 | 45101127 | -1 | protein_coding | ENSG0000C |
| 45109452 | 45115172 | -1 | protein_coding | ENSG0000C |
| 45123425 | 45124465 | -1 | protein_coding | ENSG0000C |
| 45137706 | 45138976 | -1 | transcribed_unproce | ENSG0000C |
| 45174902 | 45176544 | 1 | protein_coding | ENSG0000C |
| 45207033 | 45209768 | 1 | protein_coding | ENSG0000C |
| 45221373 | 45224458 | 1 | protein_coding | ENSG0000C |
| 45229817 | 45230191 | 1 | unprocessed_pseudc | ENSG0000C |
| 45230868 | 45233331 | 1 | transcribed_unproce | ENSG0000C |
| 45252239 | 45254564 | -1 | protein_coding | ENSG0000C |
| 45293445 | 45308529 | -1 | protein_coding | ENSG0000C |
| 45306840 | 45317824 | 1 | protein_coding | ENSG0000C |
| 45325288 | 45348424 | -1 | protein_coding | ENSG0000C |
| 45345115 | 45345823 | -1 | lncRNA | ENSG0000C |
| 45361937 | 45376798 | 1 | protein_coding | ENSG0000C |
| 45363200 | 45410610 | 1 | protein_coding | ENSG0000C |
| 45372557 | 45407889 | -1 | protein_coding | ENSG0000C |

| | | | | |
|---|---|---|---|---|
| 45388347 | 45388705 | -1 | processed_pseudoge | ENSG00000 |
| 45406057 | 45410610 | 1 | protein_coding | ENSG00000 |
| 45416084 | 45456934 | 1 | protein_coding | ENSG00000 |
| 45425510 | 45425573 | 1 | miRNA | ENSG00000 |
| 45435272 | 45448325 | -1 | lncRNA | ENSG00000 |
| 45469753 | 45481532 | 1 | protein_coding | ENSG00000 |
| 45487613 | 45490248 | 1 | lncRNA | ENSG00000 |
| 45490818 | 45491671 | -1 | processed_pseudoge | ENSG00000 |
| 45512461 | 45515622 | -1 | protein_coding | ENSG00000 |
| 45534196 | 45539495 | -1 | protein_coding | ENSG00000 |
| 45534196 | 45547662 | -1 | protein_coding | ENSG00000 |
| 45536796 | 45537014 | -1 | processed_pseudoge | ENSG00000 |
| 45539983 | 45540227 | -1 | processed_pseudoge | ENSG00000 |
| 45540626 | 45547752 | -1 | protein_coding | ENSG00000 |
| 45551153 | 45579326 | -1 | protein_coding | ENSG00000 |
| 45559758 | 45559876 | -1 | rRNA_pseudogene | ENSG00000 |
| 45567990 | 45568807 | -1 | processed_pseudoge | ENSG00000 |
| 45607939 | 45631284 | -1 | protein_coding | ENSG00000 |
| 45629739 | 45631196 | 1 | protein_coding | ENSG00000 |
| 45640486 | 45641370 | -1 | processed_pseudoge | ENSG00000 |
| 45648563 | 45670270 | -1 | protein_coding | ENSG00000 |
| 45684653 | 45705019 | -1 | protein_coding | ENSG00000 |
| 45702038 | 45708817 | 1 | protein_coding | ENSG00000 |
| 45705102 | 45705180 | -1 | miRNA | ENSG00000 |
| 45722347 | 45725830 | 1 | protein_coding | ENSG00000 |
| 45745737 | 45746287 | -1 | processed_pseudoge | ENSG00000 |
| 45747944 | 45791932 | -1 | protein_coding | ENSG00000 |
| 45749920 | 45750074 | 1 | processed_pseudoge | ENSG00000 |
| 45778698 | 45778823 | 1 | snRNA | ENSG00000 |
| 45791954 | 45811427 | 1 | protein_coding | ENSG00000 |
| 45812576 | 45816957 | 1 | protein_coding | ENSG00000 |
| 45823214 | 45833745 | -1 | protein_coding | ENSG00000 |
| 45833799 | 45843276 | 1 | protein_coding | ENSG00000 |
| 45841721 | 45857405 | -1 | protein_coding | ENSG00000 |
| 45847630 | 45847940 | -1 | misc_RNA | ENSG00000 |
| 45857614 | 45879122 | 1 | protein_coding | ENSG00000 |
| 45881227 | 45885266 | 1 | protein_coding | ENSG00000 |
| 45886489 | 45887635 | -1 | protein_coding | ENSG00000 |
| 45888625 | 45891287 | -1 | protein_coding | ENSG00000 |
| 45890144 | 45898820 | 1 | protein_coding | ENSG00000 |
| 45892694 | 45893419 | -1 | lncRNA | ENSG00000 |
| 45898621 | 45912155 | -1 | protein_coding | ENSG00000 |
| 45926518 | 45935055 | -1 | lncRNA | ENSG00000 |
| 45934683 | 45948023 | 1 | protein_coding | ENSG00000 |
| 45948660 | 45972203 | -1 | protein_coding | ENSG00000 |
| 45975249 | 45976075 | -1 | processed_pseudoge | ENSG00000 |
| 46008908 | 46016561 | 1 | protein_coding | ENSG00000 |
| 46013500 | 46022073 | -1 | lncRNA | ENSG00000 |
| 46021690 | 46060152 | 1 | protein_coding | ENSG00000 |
| 46060991 | 46089962 | -1 | protein_coding | ENSG00000 |

| | | | | |
|---|---|---|---|---|
| 46099518 | 46100176 | 1 | processed_pseudoge | ENSG00000 |
| 46118278 | 46129863 | 1 | protein_coding | ENSG00000 |
| 46168404 | 46171237 | -1 | lncRNA | ENSG00000 |
| 46173739 | 46308498 | -1 | protein_coding | ENSG00000 |
| 46345980 | 46364458 | -1 | protein_coding | ENSG00000 |
| 46364551 | 46397994 | 1 | lncRNA | ENSG00000 |
| 46366050 | 46432985 | -1 | protein_coding | ENSG00000 |
| 46414228 | 46459276 | -1 | transcribed_unitary_ | ENSG00000 |
| 46455033 | 46496731 | -1 | lncRNA | ENSG00000 |
| 46463719 | 46477232 | -1 | transcribed_processe | ENSG00000 |
| 46484461 | 46492640 | 1 | unprocessed_pseudo | ENSG00000 |
| 46499630 | 46513559 | -1 | protein_coding | ENSG00000 |
| 46540946 | 46550654 | -1 | protein_coding | ENSG00000 |
| 46557823 | 46684467 | -1 | protein_coding | ENSG00000 |
| 46681676 | 46682375 | -1 | lncRNA | ENSG00000 |
| 46684365 | 46689444 | -1 | protein_coding | ENSG00000 |
| 46689659 | 46690289 | 1 | lncRNA | ENSG00000 |
| 46709649 | 46736347 | 1 | protein_coding | ENSG00000 |
| 46859333 | 46859672 | 1 | misc_RNA | ENSG00000 |
| 46863563 | 46864104 | -1 | processed_pseudoge | ENSG00000 |
| 46894624 | 47188844 | 1 | protein_coding | ENSG00000 |
| 46901143 | 46901726 | -1 | lncRNA | ENSG00000 |
| 46997596 | 46998488 | 1 | processed_pseudoge | ENSG00000 |
| 47071865 | 47072236 | 1 | processed_pseudoge | ENSG00000 |
| 47160838 | 47161700 | -1 | processed_pseudoge | ENSG00000 |
| 47166967 | 47167058 | 1 | miRNA | ENSG00000 |
| 47209214 | 47356889 | -1 | protein_coding | ENSG00000 |
| 47318502 | 47320754 | 1 | lncRNA | ENSG00000 |
| 47348137 | 47349142 | -1 | lncRNA | ENSG00000 |
| 47352561 | 47354633 | 1 | lncRNA | ENSG00000 |
| 47391925 | 47412327 | -1 | lncRNA | ENSG00000 |
| 47478214 | 47479010 | -1 | processed_pseudoge | ENSG00000 |
| 47484278 | 47484381 | -1 | snRNA | ENSG00000 |
| 47492978 | 47493085 | -1 | snRNA | ENSG00000 |
| 47501887 | 47656877 | 1 | protein_coding | ENSG00000 |
| 47630266 | 47631438 | -1 | processed_pseudoge | ENSG00000 |
| 47656348 | 47786616 | -1 | protein_coding | ENSG00000 |
| 47677901 | 47686297 | 1 | lncRNA | ENSG00000 |
| 47826969 | 47827044 | -1 | snRNA | ENSG00000 |
| 47859788 | 47860672 | 1 | processed_pseudoge | ENSG00000 |
| 47873191 | 47873273 | 1 | rRNA_pseudogene | ENSG00000 |
| 47904348 | 47904423 | 1 | snRNA | ENSG00000 |
| 47950797 | 47954808 | 1 | lncRNA | ENSG00000 |
| 47958022 | 47958603 | 1 | lncRNA | ENSG00000 |
| 47974137 | 47977584 | 1 | lncRNA | ENSG00000 |
| 47979102 | 47990127 | -1 | lncRNA | ENSG00000 |
| 47983404 | 47984313 | 1 | lncRNA | ENSG00000 |
| 48007875 | 48022237 | -1 | lncRNA | ENSG00000 |
| 48025245 | 48060271 | 1 | lncRNA | ENSG00000 |
| 48073869 | 48074188 | 1 | lncRNA | ENSG00000 |

| | | | | |
|---|---|---|---|---|
| 48120269 | 48136844 | 1 | lncRNA | ENSG00000( |
| 48264969 | 48275392 | -1 | lncRNA | ENSG00000( |
| 48324812 | 48367976 | -1 | lncRNA | ENSG00000( |
| 48359884 | 48370636 | 1 | lncRNA | ENSG00000( |
| 48383323 | 48384895 | -1 | lncRNA | ENSG00000( |
| 48406777 | 48410716 | 1 | lncRNA | ENSG00000( |
| 48471308 | 48472414 | 1 | lncRNA | ENSG00000( |
| 48476999 | 48477553 | 1 | lncRNA | ENSG00000( |
| 48494234 | 48494309 | -1 | snRNA | ENSG00000( |
| 48624252 | 48827999 | -1 | protein_coding | ENSG00000( |
| 48657073 | 48659119 | 1 | lncRNA | ENSG00000( |
| 48821688 | 48849458 | -1 | lncRNA | ENSG00000( |
| 48921711 | 49036693 | 1 | protein_coding | ENSG00000( |
| 49038357 | 49038602 | -1 | processed_pseudoge | ENSG00000( |
| 49040463 | 49046168 | -1 | lncRNA | ENSG00000( |
| 49046246 | 49096960 | 1 | protein_coding | ENSG00000( |
| 49113339 | 49188367 | -1 | protein_coding | ENSG00000( |
| 49134480 | 49135012 | 1 | processed_pseudoge | ENSG00000( |
| 49219295 | 49244077 | 1 | protein_coding | ENSG00000( |
| 49237946 | 49278426 | -1 | protein_coding | ENSG00000( |
| 49278178 | 49299600 | 1 | lncRNA | ENSG00000( |
| 49278427 | 49278624 | 1 | misc_RNA | ENSG00000( |
| 49278940 | 49279028 | 1 | snoRNA | ENSG00000( |
| 49279116 | 49279208 | 1 | misc_RNA | ENSG00000( |
| 49280319 | 49280409 | 1 | snoRNA | ENSG00000( |
| 49280485 | 49280571 | 1 | misc_RNA | ENSG00000( |
| 49280683 | 49280772 | 1 | snoRNA | ENSG00000( |
| 49293394 | 49484297 | -1 | protein_coding | ENSG00000( |
| 49318337 | 49441654 | 1 | transcribed_unproce | ENSG00000( |
| 49503874 | 49568137 | -1 | protein_coding | ENSG00000( |
| 49632945 | 49713878 | -1 | protein_coding | ENSG00000( |
| 49721949 | 49722247 | 1 | misc_RNA | ENSG00000( |
| 49729591 | 49729842 | -1 | processed_pseudoge | ENSG00000( |
| 49794811 | 49794914 | 1 | snRNA | ENSG00000( |
| 49812713 | 49892242 | 1 | protein_coding | ENSG00000( |
| 49829125 | 49831085 | 1 | processed_pseudoge | ENSG00000( |
| 49903391 | 49915529 | -1 | protein_coding | ENSG00000( |
| 49920597 | 49920694 | -1 | misc_RNA | ENSG00000( |
| 49932795 | 49933078 | 1 | misc_RNA | ENSG00000( |
| 49936336 | 49953885 | 1 | protein_coding | ENSG00000( |
| 49956745 | 49958032 | 1 | processed_pseudoge | ENSG00000( |
| 49972106 | 49972212 | 1 | snRNA | ENSG00000( |
| 49982980 | 49988886 | 1 | protein_coding | ENSG00000( |
| 50040716 | 50041504 | -1 | lncRNA | ENSG00000( |
| 50081124 | 50115959 | -1 | protein_coding | ENSG00000( |
| 50081124 | 50153637 | -1 | protein_coding | ENSG00000( |
| 50118254 | 50153734 | -1 | protein_coding | ENSG00000( |
| 50162765 | 50166216 | -1 | lncRNA | ENSG00000( |
| 50166362 | 50171742 | 1 | lncRNA | ENSG00000( |
| 50171809 | 50176676 | 1 | lncRNA | ENSG00000( |

| | | | | |
|---|---|---|---|---|
| 50184598 | 50191498 | -1 | lncRNA | ENSG0000C |
| 50190830 | 50192668 | 1 | protein_coding | ENSG0000C |
| 50247643 | 50283250 | 1 | lncRNA | ENSG0000C |
| 50292709 | 50315488 | 1 | lncRNA | ENSG0000C |
| 50310711 | 50321342 | -1 | lncRNA | ENSG0000C |
| 50429190 | 50429483 | -1 | misc_RNA | ENSG0000C |
| 50479767 | 50479991 | 1 | processed_pseudoge | ENSG0000C |
| 50510321 | 50585241 | 1 | protein_coding | ENSG0000C |
| 50557682 | 50557998 | -1 | misc_RNA | ENSG0000C |
| 50563009 | 50563110 | 1 | misc_RNA | ENSG0000C |
| 50570975 | 50578041 | -1 | lncRNA | ENSG0000C |
| 50585786 | 50585879 | 1 | miRNA | ENSG0000C |
| 50586108 | 50691542 | -1 | protein_coding | ENSG0000C |
| 50614636 | 50614785 | -1 | miRNA | ENSG0000C |
| 50620763 | 50621077 | 1 | processed_pseudoge | ENSG0000C |
| 50645471 | 50662275 | 1 | lncRNA | ENSG0000C |
| 50731580 | 50756795 | 1 | protein_coding | ENSG0000C |
| 50794894 | 50882676 | 1 | protein_coding | ENSG0000C |
| 50840615 | 50840749 | -1 | processed_pseudoge | ENSG0000C |
| 50888916 | 50931437 | -1 | protein_coding | ENSG0000C |
| 50913731 | 50914411 | 1 | processed_pseudoge | ENSG0000C |
| 50930984 | 50945148 | 1 | lncRNA | ENSG0000C |
| 50934867 | 50958555 | -1 | protein_coding | ENSG0000C |
| 50958818 | 50963929 | 1 | protein_coding | ENSG0000C |
| 50999370 | 51010019 | 1 | lncRNA | ENSG0000C |
| 51003656 | 51023107 | -1 | protein_coding | ENSG0000C |
| 51098138 | 51098845 | 1 | TEC | ENSG0000C |
| 51130788 | 51131667 | -1 | processed_pseudoge | ENSG0000C |
| 51386957 | 51562831 | -1 | protein_coding | ENSG0000C |
| 51452905 | 51452977 | -1 | miRNA | ENSG0000C |
| 51596514 | 51768390 | -1 | protein_coding | ENSG0000C |
| 51734533 | 51735067 | 1 | processed_pseudoge | ENSG0000C |
| 51782331 | 51802521 | -1 | protein_coding | ENSG0000C |
| 51831797 | 51862912 | -1 | lncRNA | ENSG0000C |
| 51902291 | 51902395 | 1 | snRNA | ENSG0000C |
| 51904969 | 51905030 | 1 | snRNA | ENSG0000C |
| 52051663 | 52204308 | -1 | protein_coding | ENSG0000C |
| 52072685 | 52096480 | 1 | lncRNA | ENSG0000C |
| 52110344 | 52111049 | -1 | processed_pseudoge | ENSG0000C |
| 52192159 | 52670578 | 1 | lncRNA | ENSG0000C |
| 52451464 | 52455619 | -1 | lncRNA | ENSG0000C |
| 52482801 | 52482991 | -1 | misc_RNA | ENSG0000C |
| 52487922 | 52500591 | -1 | lncRNA | ENSG0000C |
| 52511252 | 52511568 | 1 | processed_pseudoge | ENSG0000C |
| 52671735 | 52699472 | 1 | lncRNA | ENSG0000C |
| 52840936 | 52843775 | -1 | lncRNA | ENSG0000C |
| 52858338 | 52862855 | -1 | lncRNA | ENSG0000C |
| 52890162 | 52890386 | 1 | processed_pseudoge | ENSG0000C |
| 52972358 | 53495330 | 1 | protein_coding | ENSG0000C |
| 53097652 | 53097984 | 1 | misc_RNA | ENSG0000C |

| | | | | |
|---|---|---|---|---|
| 53137293 | 53141169 | -1 | lncRNA | ENSG0000C |
| 53168643 | 53179550 | -1 | lncRNA | ENSG0000C |
| 53196229 | 53208705 | -1 | lncRNA | ENSG0000C |
| 53255979 | 53487274 | -1 | lncRNA | ENSG0000C |
| 53397661 | 53412910 | -1 | lncRNA | ENSG0000C |
| 53421199 | 53421656 | 1 | processed_pseudoge | ENSG0000C |
| 53453058 | 53504314 | -1 | lncRNA | ENSG0000C |
| 53544615 | 53544725 | 1 | misc_RNA | ENSG0000C |
| 53552770 | 53575863 | 1 | lncRNA | ENSG0000C |
| 53567065 | 53609907 | -1 | protein_coding | ENSG0000C |
| 53608354 | 53634590 | 1 | lncRNA | ENSG0000C |
| 53668697 | 53668758 | -1 | snRNA | ENSG0000C |
| 53738728 | 53741797 | 1 | lncRNA | ENSG0000C |
| 53799823 | 53827297 | -1 | lncRNA | ENSG0000C |
| 53875252 | 53875557 | -1 | processed_pseudoge | ENSG0000C |
| 53936777 | 54070594 | -1 | protein_coding | ENSG0000C |
| 53940160 | 53942508 | 1 | lncRNA | ENSG0000C |
| 54068408 | 54068485 | -1 | miRNA | ENSG0000C |
| 54153446 | 54173986 | -1 | protein_coding | ENSG0000C |
| 54178414 | 54183129 | 1 | lncRNA | ENSG0000C |
| 54208087 | 54228052 | 1 | protein_coding | ENSG0000C |
| 54232758 | 54269542 | 1 | lncRNA | ENSG0000C |
| 54475593 | 54651169 | 1 | protein_coding | ENSG0000C |
| 54859688 | 54859812 | -1 | snRNA | ENSG0000C |
| 55074644 | 55075140 | 1 | processed_pseudoge | ENSG0000C |
| 55408898 | 55684915 | 1 | lncRNA | ENSG0000C |
| 55420336 | 55427197 | -1 | lncRNA | ENSG0000C |
| 55504906 | 55505215 | -1 | lncRNA | ENSG0000C |
| 55607852 | 55744938 | -1 | lncRNA | ENSG0000C |
| 55775176 | 55781231 | 1 | lncRNA | ENSG0000C |
| 55997357 | 56005519 | -1 | protein_coding | ENSG0000C |
| 56205052 | 56205173 | -1 | rRNA_pseudogene | ENSG0000C |
| 56248732 | 56249815 | 1 | protein_coding | ENSG0000C |
| 56267872 | 56269231 | 1 | processed_pseudoge | ENSG0000C |
| 56295967 | 56299160 | -1 | lncRNA | ENSG0000C |
| 56358974 | 56368663 | 1 | protein_coding | ENSG0000C |
| 56369389 | 56392337 | -1 | protein_coding | ENSG0000C |
| 56392371 | 56406362 | 1 | protein_coding | ENSG0000C |
| 56412112 | 56460387 | 1 | protein_coding | ENSG0000C |
| 56459428 | 56459583 | -1 | processed_pseudoge | ENSG0000C |
| 56468585 | 56519449 | 1 | protein_coding | ENSG0000C |
| 56491492 | 56525925 | -1 | protein_coding | ENSG0000C |
| 56517187 | 56526152 | 1 | protein_coding | ENSG0000C |
| 56522252 | 56523499 | 1 | TEC | ENSG0000C |
| 56533246 | 56536520 | 1 | protein_coding | ENSG0000C |
| 56550605 | 56555579 | 1 | lncRNA | ENSG0000C |
| 56577563 | 56598621 | 1 | lncRNA | ENSG0000C |
| 56601701 | 56602113 | 1 | processed_pseudoge | ENSG0000C |
| 56629306 | 56639283 | 1 | protein_coding | ENSG0000C |
| 56667430 | 56667531 | 1 | snRNA | ENSG0000C |

| | | | | |
|---|---|---|---|---|
| 56688401 | 56688695 | 1 | misc_RNA | ENSG0000C |
| 56700837 | 56701514 | -1 | processed_pseudoge | ENSG0000C |
| 56730397 | 56731460 | 1 | lncRNA | ENSG0000C |
| 56785981 | 56786087 | -1 | snRNA | ENSG0000C |
| 57105741 | 57107128 | 1 | lncRNA | ENSG0000C |
| 57110640 | 57122745 | 1 | lncRNA | ENSG0000C |
| 57116566 | 57119543 | -1 | lncRNA | ENSG0000C |
| 57161822 | 57163775 | 1 | TEC | ENSG0000C |
| 57168753 | 57266641 | -1 | protein_coding | ENSG0000C |
| 57214872 | 57215866 | 1 | lncRNA | ENSG0000C |
| 57266797 | 57282998 | 1 | lncRNA | ENSG0000C |
| 57311854 | 57311958 | 1 | misc_RNA | ENSG0000C |
| 57321502 | 57321591 | -1 | miRNA | ENSG0000C |
| 57328546 | 57328696 | 1 | processed_pseudoge | ENSG0000C |
| 57329803 | 57343994 | 1 | protein_coding | ENSG0000C |
| 57351223 | 57379211 | 1 | protein_coding | ENSG0000C |
| 57357426 | 57358369 | -1 | processed_pseudoge | ENSG0000C |
| 57358447 | 57359498 | -1 | protein_coding | ENSG0000C |
| 57377140 | 57393062 | -1 | lncRNA | ENSG0000C |
| 57391396 | 57409333 | 1 | protein_coding | ENSG0000C |
| 57442742 | 57458298 | 1 | lncRNA | ENSG0000C |
| 57488392 | 57489027 | -1 | processed_pseudoge | ENSG0000C |
| 57495966 | 57525652 | -1 | protein_coding | ENSG0000C |
| 57561080 | 57568121 | 1 | protein_coding | ENSG0000C |
| 57599695 | 57601200 | -1 | lncRNA | ENSG0000C |
| 57603846 | 57620576 | -1 | protein_coding | ENSG0000C |
| 57648392 | 57711536 | -1 | protein_coding | ENSG0000C |
| 57710156 | 57712780 | 1 | lncRNA | ENSG0000C |
| 57895394 | 57895444 | 1 | miRNA | ENSG0000C |
| 57957126 | 57960176 | -1 | lncRNA | ENSG0000C |
| 58003904 | 58004648 | 1 | lncRNA | ENSG0000C |
| 58069054 | 58072070 | -1 | lncRNA | ENSG0000C |
| 58097267 | 58097591 | 1 | processed_pseudoge | ENSG0000C |
| 58150902 | 58161150 | 1 | protein_coding | ENSG0000C |
| 58218495 | 58228653 | -1 | protein_coding | ENSG0000C |
| 58228940 | 58309451 | -1 | transcribed_unproce | ENSG0000C |
| 58309715 | 58367507 | 1 | protein_coding | ENSG0000C |
| 58389122 | 58451101 | 1 | protein_coding | ENSG0000C |
| 58459101 | 58515399 | -1 | protein_coding | ENSG0000C |
| 58515379 | 58619888 | 1 | lncRNA | ENSG0000C |
| 58594417 | 58603973 | 1 | lncRNA | ENSG0000C |
| 58634772 | 58635738 | 1 | lncRNA | ENSG0000C |
| 58651253 | 58679526 | 1 | protein_coding | ENSG0000C |
| 58651434 | 58715410 | 1 | protein_coding | ENSG0000C |
| 58689131 | 58719238 | 1 | protein_coding | ENSG0000C |
| 58710795 | 58711633 | 1 | lncRNA | ENSG0000C |
| 58740532 | 58783821 | -1 | unprocessed_pseudc | ENSG0000C |
| 58754351 | 58756838 | 1 | lncRNA | ENSG0000C |
| 58817132 | 58817725 | -1 | lncRNA | ENSG0000C |
| 58817615 | 58817694 | -1 | miRNA | ENSG0000C |

| | | | | |
|---|---|---|---|---|
| 58818226 | 58818313 | -1 | miRNA | ENSG0000( |
| 58818919 | 58850903 | -1 | lncRNA | ENSG0000( |
| 58833809 | 58836529 | -1 | lncRNA | ENSG0000( |
| 58839718 | 58911192 | 1 | protein_coding | ENSG0000( |
| 58842030 | 58842132 | 1 | misc_RNA | ENSG0000( |
| 58842433 | 58842546 | 1 | misc_RNA | ENSG0000( |
| 58850530 | 58850641 | 1 | misc_RNA | ENSG0000( |
| 58850783 | 58850903 | 1 | misc_RNA | ENSG0000( |
| 58863528 | 58888809 | -1 | lncRNA | ENSG0000( |
| 58876592 | 58876981 | -1 | lncRNA | ENSG0000( |
| 58916000 | 58981169 | -1 | lncRNA | ENSG0000( |
| 58948089 | 58948267 | -1 | unprocessed_pseudc | ENSG0000( |
| 58981208 | 58995133 | 1 | protein_coding | ENSG0000( |
| 58995185 | 59007254 | -1 | protein_coding | ENSG0000( |
| 59019254 | 59026654 | 1 | protein_coding | ENSG0000( |
| 59025475 | 59032345 | -1 | protein_coding | ENSG0000( |
| 59033145 | 59042809 | -1 | protein_coding | ENSG0000( |
| 59113224 | 59113631 | 1 | processed_pseudoge | ENSG0000( |
| 59123381 | 59259113 | 1 | protein_coding | ENSG0000( |
| 59300443 | 59325992 | 1 | protein_coding | ENSG0000( |
| 59352195 | 59357774 | -1 | lncRNA | ENSG0000( |
| 59360927 | 59364773 | -1 | lncRNA | ENSG0000( |
| 59395496 | 59397109 | -1 | lncRNA | ENSG0000( |
| 59460619 | 59460837 | -1 | processed_pseudoge | ENSG0000( |
| 59467755 | 59469853 | -1 | lncRNA | ENSG0000( |
| 59515191 | 59516039 | 1 | lncRNA | ENSG0000( |
| 59577509 | 59847711 | 1 | protein_coding | ENSG0000( |
| 59624904 | 59628295 | -1 | lncRNA | ENSG0000( |
| 59852667 | 59852728 | -1 | snRNA | ENSG0000( |
| 59863564 | 59933655 | -1 | protein_coding | ENSG0000( |
| 59933764 | 59948680 | 1 | protein_coding | ENSG0000( |
| 59936663 | 59940305 | -1 | protein_coding | ENSG0000( |
| 59958423 | 60034011 | 1 | protein_coding | ENSG0000( |
| 60055853 | 60072956 | 1 | lncRNA | ENSG0000( |
| 60083360 | 60083872 | -1 | lncRNA | ENSG0000( |
| 60087826 | 60653784 | 1 | lncRNA | ENSG0000( |
| 60308474 | 60308567 | 1 | miRNA | ENSG0000( |
| 60414396 | 60414872 | -1 | processed_pseudoge | ENSG0000( |
| 60478111 | 60478181 | 1 | miRNA | ENSG0000( |
| 60564562 | 60564625 | 1 | miRNA | ENSG0000( |
| 60755001 | 60767279 | 1 | lncRNA | ENSG0000( |
| 60812004 | 60814211 | -1 | lncRNA | ENSG0000( |
| 61077224 | 61077816 | -1 | lncRNA | ENSG0000( |
| 61079049 | 61083750 | 1 | lncRNA | ENSG0000( |
| 61252261 | 61940617 | 1 | protein_coding | ENSG0000( |
| 61267525 | 61271827 | -1 | lncRNA | ENSG0000( |
| 61369879 | 61370855 | 1 | lncRNA | ENSG0000( |
| 61432868 | 61437630 | -1 | lncRNA | ENSG0000( |
| 61717506 | 61719748 | -1 | lncRNA | ENSG0000( |
| 61738219 | 61755056 | -1 | lncRNA | ENSG0000( |

| | | | | |
|---|---|---|---|---|
| 61953469 | 62065810 | -1 | protein_coding | ENSG0000C |
| 61953546 | 61953662 | -1 | miRNA | ENSG0000C |
| 62037429 | 62037956 | -1 | processed_pseudoge | ENSG0000C |
| 62064802 | 62064885 | 1 | miRNA | ENSG0000C |
| 62066830 | 62068437 | 1 | lncRNA | ENSG0000C |
| 62122461 | 62135374 | 1 | protein_coding | ENSG0000C |
| 62136733 | 62143440 | -1 | protein_coding | ENSG0000C |
| 62143769 | 62182514 | 1 | protein_coding | ENSG0000C |
| 62183029 | 62203568 | 1 | protein_coding | ENSG0000C |
| 62214960 | 62220278 | -1 | protein_coding | ENSG0000C |
| 62231922 | 62296213 | 1 | protein_coding | ENSG0000C |
| 62302093 | 62308862 | 1 | protein_coding | ENSG0000C |
| 62305432 | 62306325 | -1 | lncRNA | ENSG0000C |
| 62307955 | 62367312 | -1 | protein_coding | ENSG0000C |
| 62332487 | 62332557 | -1 | miRNA | ENSG0000C |
| 62352995 | 62356480 | 1 | lncRNA | ENSG0000C |
| 62386303 | 62386970 | -1 | lncRNA | ENSG0000C |
| 62387103 | 62388520 | 1 | protein_coding | ENSG0000C |
| 62388632 | 62407285 | -1 | protein_coding | ENSG0000C |
| 62402236 | 62405935 | -1 | lncRNA | ENSG0000C |
| 62410237 | 62427539 | -1 | protein_coding | ENSG0000C |
| 62427827 | 62447677 | 1 | lncRNA | ENSG0000C |
| 62463497 | 62475995 | -1 | protein_coding | ENSG0000C |
| 62477870 | 62478594 | 1 | lncRNA | ENSG0000C |
| 62513909 | 62516096 | -1 | lncRNA | ENSG0000C |
| 62544343 | 62551526 | -1 | lncRNA | ENSG0000C |
| 62550453 | 62570764 | 1 | lncRNA | ENSG0000C |
| 62554306 | 62554376 | 1 | miRNA | ENSG0000C |
| 62564912 | 62565013 | 1 | miRNA | ENSG0000C |
| 62572671 | 62573384 | 1 | processed_pseudoge | ENSG0000C |
| 62575590 | 62577507 | 1 | lncRNA | ENSG0000C |
| 62596732 | 62603355 | -1 | lncRNA | ENSG0000C |
| 62633681 | 62635504 | -1 | lncRNA | ENSG0000C |
| 62633772 | 62635862 | 1 | lncRNA | ENSG0000C |
| 62640719 | 62643304 | -1 | lncRNA | ENSG0000C |
| 62642503 | 62685785 | 1 | protein_coding | ENSG0000C |
| 62648961 | 62650767 | -1 | lncRNA | ENSG0000C |
| 62651272 | 62652186 | -1 | lncRNA | ENSG0000C |
| 62663019 | 62666724 | -1 | lncRNA | ENSG0000C |
| 62695024 | 62700480 | -1 | lncRNA | ENSG0000C |
| 62708836 | 62762771 | 1 | protein_coding | ENSG0000C |
| 62732566 | 62735347 | -1 | lncRNA | ENSG0000C |
| 62774121 | 62776996 | -1 | lncRNA | ENSG0000C |
| 62796473 | 62801729 | 1 | protein_coding | ENSG0000C |
| 62800627 | 62805587 | -1 | lncRNA | ENSG0000C |
| 62804835 | 62814000 | 1 | protein_coding | ENSG0000C |
| 62816244 | 62841159 | 1 | protein_coding | ENSG0000C |
| 62841005 | 62861822 | -1 | protein_coding | ENSG0000C |
| 62845664 | 62845912 | 1 | processed_pseudoge | ENSG0000C |
| 62872250 | 62873076 | -1 | processed_pseudoge | ENSG0000C |

| | | | | |
|---|---|---|---|---|
| 62877738 | 62937952 | -1 | protein_coding | ENSG00000 |
| 62928621 | 62929297 | -1 | lncRNA | ENSG00000 |
| 62938147 | 62948475 | 1 | protein_coding | ENSG00000 |
| 62952707 | 62969585 | 1 | protein_coding | ENSG00000 |
| 63005927 | 63007035 | -1 | protein_coding | ENSG00000 |
| 63009383 | 63085071 | 1 | lncRNA | ENSG00000 |
| 63034217 | 63037028 | -1 | lncRNA | ENSG00000 |
| 63038011 | 63053863 | 1 | lncRNA | ENSG00000 |
| 63090806 | 63102631 | -1 | lncRNA | ENSG00000 |
| 63102142 | 63102259 | 1 | misc_RNA | ENSG00000 |
| 63102205 | 63104386 | 1 | lncRNA | ENSG00000 |
| 63116218 | 63117205 | 1 | lncRNA | ENSG00000 |
| 63127495 | 63129459 | -1 | lncRNA | ENSG00000 |
| 63144311 | 63152087 | -1 | lncRNA | ENSG00000 |
| 63166797 | 63180903 | -1 | lncRNA | ENSG00000 |
| 63178500 | 63178586 | 1 | miRNA | ENSG00000 |
| 63195429 | 63216139 | -1 | protein_coding | ENSG00000 |
| 63218041 | 63218502 | 1 | lncRNA | ENSG00000 |
| 63235883 | 63240495 | 1 | protein_coding | ENSG00000 |
| 63238779 | 63238842 | 1 | miRNA | ENSG00000 |
| 63240784 | 63272694 | -1 | protein_coding | ENSG00000 |
| 63253978 | 63261615 | 1 | lncRNA | ENSG00000 |
| 63272785 | 63289790 | 1 | protein_coding | ENSG00000 |
| 63286808 | 63286866 | 1 | miRNA | ENSG00000 |
| 63293186 | 63334851 | 1 | protein_coding | ENSG00000 |
| 63337339 | 63337449 | 1 | snRNA | ENSG00000 |
| 63343223 | 63378401 | -1 | protein_coding | ENSG00000 |
| 63359988 | 63371177 | 1 | lncRNA | ENSG00000 |
| 63400210 | 63472677 | -1 | protein_coding | ENSG00000 |
| 63422269 | 63424555 | 1 | lncRNA | ENSG00000 |
| 63448051 | 63449329 | 1 | lncRNA | ENSG00000 |
| 63488014 | 63499185 | -1 | protein_coding | ENSG00000 |
| 63502287 | 63505111 | 1 | lncRNA | ENSG00000 |
| 63520765 | 63522206 | 1 | protein_coding | ENSG00000 |
| 63528001 | 63537376 | -1 | protein_coding | ENSG00000 |
| 63540810 | 63547504 | -1 | protein_coding | ENSG00000 |
| 63543411 | 63543738 | 1 | lncRNA | ENSG00000 |
| 63547891 | 63556695 | 1 | protein_coding | ENSG00000 |
| 63558086 | 63574239 | -1 | protein_coding | ENSG00000 |
| 63587602 | 63627101 | -1 | protein_coding | ENSG00000 |
| 63627227 | 63628824 | 1 | lncRNA | ENSG00000 |
| 63639705 | 63657682 | -1 | protein_coding | ENSG00000 |
| 63657810 | 63696253 | 1 | protein_coding | ENSG00000 |
| 63659300 | 63698684 | 1 | protein_coding | ENSG00000 |
| 63696652 | 63698684 | 1 | protein_coding | ENSG00000 |
| 63698642 | 63708025 | -1 | protein_coding | ENSG00000 |
| 63707465 | 63736142 | 1 | protein_coding | ENSG00000 |
| 63708864 | 63739103 | 1 | protein_coding | ENSG00000 |
| 63730072 | 63730377 | -1 | lncRNA | ENSG00000 |
| 63736283 | 63739103 | 1 | protein_coding | ENSG00000 |

| | | | | |
|---|---|---|---|---|
| 63738270 | 63740398 | 1 | protein_coding | ENSG00000 |
| 63739776 | 63744050 | 1 | protein_coding | ENSG00000 |
| 63743666 | 63832038 | -1 | protein_coding | ENSG00000 |
| 63744689 | 63745958 | 1 | lncRNA | ENSG00000 |
| 63808076 | 63816521 | 1 | lncRNA | ENSG00000 |
| 63843436 | 63843915 | -1 | TEC | ENSG00000 |
| 63861212 | 63864293 | -1 | lncRNA | ENSG00000 |
| 63861498 | 63862988 | 1 | protein_coding | ENSG00000 |
| 63865228 | 63891545 | 1 | protein_coding | ENSG00000 |
| 63895126 | 63936031 | 1 | protein_coding | ENSG00000 |
| 63908955 | 63909119 | 1 | snRNA | ENSG00000 |
| 63919449 | 63919520 | 1 | miRNA | ENSG00000 |
| 63919505 | 63919576 | 1 | miRNA | ENSG00000 |
| 63919561 | 63919632 | 1 | miRNA | ENSG00000 |
| 63919756 | 63919827 | 1 | miRNA | ENSG00000 |
| 63919868 | 63919939 | 1 | miRNA | ENSG00000 |
| 63939829 | 63956416 | -1 | protein_coding | ENSG00000 |
| 63941465 | 63941544 | -1 | miRNA | ENSG00000 |
| 63942631 | 63942726 | -1 | miRNA | ENSG00000 |
| 63953384 | 63956985 | 1 | lncRNA | ENSG00000 |
| 63956704 | 63969930 | -1 | protein_coding | ENSG00000 |
| 63974116 | 63980008 | -1 | protein_coding | ENSG00000 |
| 63981132 | 64033100 | 1 | protein_coding | ENSG00000 |
| 64034344 | 64039962 | 1 | protein_coding | ENSG00000 |
| 64047582 | 64049639 | -1 | protein_coding | ENSG00000 |
| 64049836 | 64072347 | 1 | protein_coding | ENSG00000 |
| 64056803 | 64057084 | 1 | lncRNA | ENSG00000 |
| 64073181 | 64079988 | -1 | protein_coding | ENSG00000 |
| 64076955 | 64077010 | -1 | miRNA | ENSG00000 |
| 64080082 | 64100643 | 1 | protein_coding | ENSG00000 |
| 64083380 | 64084359 | -1 | protein_coding | ENSG00000 |
| 64097208 | 64101162 | -1 | lncRNA | ENSG00000 |
| 64102394 | 64242253 | 1 | protein_coding | ENSG00000 |
| 64105820 | 64107171 | -1 | protein_coding | ENSG00000 |
| 64255695 | 64287821 | 1 | protein_coding | ENSG00000 |
| 64278306 | 64278561 | 1 | processed_pseudoge | ENSG00000 |
| 64290187 | 64292998 | -1 | processed_pseudoge | ENSG00000 |
| 64290385 | 64313132 | 1 | lncRNA | ENSG00000 |
| 64327418 | 64327972 | -1 | processed_pseudoge | ENSG00000 |
| 5011799 | 5017145 | 1 | protein_coding | ENSG00000 |
| 5022493 | 5040666 | 1 | protein_coding | ENSG00000 |
| 5073458 | 5087867 | 1 | lncRNA | ENSG00000 |
| 5079294 | 5128413 | -1 | protein_coding | ENSG00000 |
| 5116343 | 5133805 | -1 | protein_coding | ENSG00000 |
| 5130871 | 5154658 | -1 | protein_coding | ENSG00000 |
| 5155499 | 5165472 | -1 | protein_coding | ENSG00000 |
| 5232668 | 5243833 | -1 | lncRNA | ENSG00000 |
| 5499151 | 5502542 | -1 | lncRNA | ENSG00000 |
| 5553637 | 5614880 | 1 | lncRNA | ENSG00000 |
| 5597390 | 5597490 | 1 | misc_RNA | ENSG00000 |

| | | | | |
|---|---|---|---|---|
| 5703182 | 5705637 | -1 | TEC | ENSG00000 |
| 5705345 | 5707160 | 1 | lncRNA | ENSG00000 |
| 5707004 | 5709456 | -1 | lncRNA | ENSG00000 |
| 5972924 | 5973383 | -1 | protein_coding | ENSG00000 |
| 6008604 | 6008810 | 1 | processed_pseudoge | ENSG00000 |
| 6034690 | 6036093 | 1 | TEC | ENSG00000 |
| 6060340 | 6076305 | 1 | lncRNA | ENSG00000 |
| 6081193 | 6082585 | 1 | lncRNA | ENSG00000 |
| 6084364 | 6091407 | -1 | lncRNA | ENSG00000 |
| 6111131 | 6123778 | 1 | protein_coding | ENSG00000 |
| 6215291 | 6217681 | -1 | lncRNA | ENSG00000 |
| 6223480 | 6223580 | -1 | misc_RNA | ENSG00000 |
| 6228966 | 6267317 | -1 | lncRNA | ENSG00000 |
| 6272135 | 6276532 | -1 | unprocessed_pseudc | ENSG00000 |
| 6309161 | 6312948 | 1 | unprocessed_pseudc | ENSG00000 |
| 6318434 | 6360415 | 1 | lncRNA | ENSG00000 |
| 6365955 | 6366055 | 1 | misc_RNA | ENSG00000 |
| 6444869 | 6468040 | -1 | protein_coding | ENSG00000 |
| 6484623 | 6499261 | -1 | protein_coding | ENSG00000 |
| 6507017 | 6507389 | -1 | processed_pseudoge | ENSG00000 |
| 6520344 | 6520670 | -1 | TEC | ENSG00000 |
| 6550749 | 6553955 | -1 | lncRNA | ENSG00000 |
| 6560714 | 6564489 | 1 | protein_coding | ENSG00000 |
| 6630182 | 6670695 | -1 | lncRNA | ENSG00000 |
| 6667304 | 6670667 | -1 | lncRNA | ENSG00000 |
| 6676178 | 6679962 | -1 | unprocessed_pseudc | ENSG00000 |
| 6712596 | 6716361 | 1 | unprocessed_pseudc | ENSG00000 |
| 6721812 | 6725209 | 1 | lncRNA | ENSG00000 |
| 6789592 | 6812297 | 1 | lncRNA | ENSG00000 |
| 6858539 | 6897263 | 1 | lncRNA | ENSG00000 |
| 6859171 | 6859256 | 1 | miRNA | ENSG00000 |
| 6897291 | 6899280 | 1 | processed_pseudoge | ENSG00000 |
| 6904884 | 6906692 | 1 | lncRNA | ENSG00000 |
| 6986450 | 6997765 | -1 | lncRNA | ENSG00000 |
| 7003248 | 7007022 | -1 | unprocessed_pseudc | ENSG00000 |
| 7020599 | 7025166 | 1 | lncRNA | ENSG00000 |
| 7048891 | 7087229 | 1 | lncRNA | ENSG00000 |
| 7092616 | 7092716 | 1 | misc_RNA | ENSG00000 |
| 7129103 | 7130287 | 1 | processed_pseudoge | ENSG00000 |
| 7135334 | 7137789 | -1 | TEC | ENSG00000 |
| 7385058 | 7388799 | -1 | unprocessed_pseudc | ENSG00000 |
| 7421442 | 7425839 | -1 | unprocessed_pseudc | ENSG00000 |
| 7430659 | 7469007 | 1 | lncRNA | ENSG00000 |
| 7474394 | 7474494 | 1 | misc_RNA | ENSG00000 |
| 7626344 | 7627528 | -1 | processed_pseudoge | ENSG00000 |
| 7663911 | 7664011 | -1 | misc_RNA | ENSG00000 |
| 7669397 | 7681742 | -1 | lncRNA | ENSG00000 |
| 7744962 | 7777853 | 1 | protein_coding | ENSG00000 |
| 7768884 | 7770591 | -1 | protein_coding | ENSG00000 |
| 7784482 | 7793954 | -1 | protein_coding | ENSG00000 |

| | | | | |
|---|---|---|---|---|
| 7816675 | 7829926 | -1 | protein_coding | ENSG00000( |
| 7826098 | 7826225 | -1 | snoRNA | ENSG00000( |
| 8101251 | 8103706 | -1 | TEC | ENSG00000( |
| 8197620 | 8227646 | 1 | lncRNA | ENSG00000( |
| 8205315 | 8205406 | 1 | miRNA | ENSG00000( |
| 8205851 | 8205940 | 1 | misc_RNA | ENSG00000( |
| 8208473 | 8208652 | 1 | miRNA | ENSG00000( |
| 8208844 | 8208904 | 1 | miRNA | ENSG00000( |
| 8210384 | 8211306 | -1 | lncRNA | ENSG00000( |
| 8212572 | 8212724 | 1 | rRNA | ENSG00000( |
| 8235354 | 8235832 | 1 | TEC | ENSG00000( |
| 8237649 | 8237979 | 1 | TEC | ENSG00000( |
| 8238859 | 8239092 | 1 | TEC | ENSG00000( |
| 8240736 | 8241047 | 1 | TEC | ENSG00000( |
| 8249505 | 8249596 | 1 | miRNA | ENSG00000( |
| 8250060 | 8250149 | 1 | misc_RNA | ENSG00000( |
| 8254592 | 8255514 | -1 | lncRNA | ENSG00000( |
| 8256781 | 8256933 | 1 | rRNA | ENSG00000( |
| 8380643 | 8454792 | 1 | lncRNA | ENSG00000( |
| 8388362 | 8388453 | 1 | miRNA | ENSG00000( |
| 8388898 | 8388987 | 1 | misc_RNA | ENSG00000( |
| 8393419 | 8394341 | -1 | lncRNA | ENSG00000( |
| 8395607 | 8395759 | 1 | rRNA | ENSG00000( |
| 8417757 | 8418235 | 1 | TEC | ENSG00000( |
| 8420675 | 8421003 | 1 | TEC | ENSG00000( |
| 8421883 | 8422116 | 1 | TEC | ENSG00000( |
| 8423760 | 8424071 | 1 | TEC | ENSG00000( |
| 8432530 | 8432621 | 1 | miRNA | ENSG00000( |
| 8433085 | 8433174 | 1 | misc_RNA | ENSG00000( |
| 8437629 | 8438551 | -1 | lncRNA | ENSG00000( |
| 8439823 | 8439975 | 1 | rRNA | ENSG00000( |
| 8462320 | 8462798 | 1 | TEC | ENSG00000( |
| 8465254 | 8465586 | 1 | TEC | ENSG00000( |
| 8466466 | 8466699 | 1 | TEC | ENSG00000( |
| 8468320 | 8468631 | 1 | TEC | ENSG00000( |
| 8603547 | 8603984 | 1 | processed_pseudoge | ENSG00000( |
| 8680538 | 8680934 | -1 | processed_pseudoge | ENSG00000( |
| 8701461 | 8701562 | 1 | processed_pseudoge | ENSG00000( |
| 8759077 | 8761335 | 1 | lncRNA | ENSG00000( |
| 8762386 | 8836565 | -1 | lncRNA | ENSG00000( |
| 8801362 | 8801880 | -1 | processed_pseudoge | ENSG00000( |
| 8845566 | 8846646 | 1 | processed_pseudoge | ENSG00000( |
| 8857260 | 8880976 | 1 | unprocessed_pseudc | ENSG00000( |
| 8986999 | 8987178 | 1 | miRNA | ENSG00000( |
| 8987370 | 8987430 | 1 | miRNA | ENSG00000( |
| 8996496 | 9018670 | -1 | lncRNA | ENSG00000( |
| 9026821 | 9027329 | -1 | processed_pseudoge | ENSG00000( |
| 9053122 | 9059060 | 1 | unprocessed_pseudc | ENSG00000( |
| 9068361 | 9129752 | -1 | transcribed_unproce | ENSG00000( |
| 9082601 | 9083101 | -1 | processed_pseudoge | ENSG00000( |

| | | | | |
|---|---|---|---|---|
| 9088188 | 9088391 | 1 | processed_pseudoge | ENSG0000C |
| 9325013 | 9368775 | 1 | lncRNA | ENSG0000C |
| 9364467 | 9365225 | -1 | processed_pseudoge | ENSG0000C |
| 9369285 | 9376645 | -1 | unprocessed_pseudc | ENSG0000C |
| 9529934 | 9559268 | 1 | transcribed_processe | ENSG0000C |
| 9580024 | 9580415 | -1 | processed_pseudoge | ENSG0000C |
| 9634244 | 9679906 | 1 | lncRNA | ENSG0000C |
| 9646825 | 9647000 | 1 | snRNA | ENSG0000C |
| 9781910 | 9838011 | -1 | transcribed_processe | ENSG0000C |
| 9810501 | 9810848 | 1 | processed_pseudoge | ENSG0000C |
| 9902344 | 9902627 | -1 | misc_RNA | ENSG0000C |
| 9907916 | 9908010 | 1 | snoRNA | ENSG0000C |
| 9913117 | 9914846 | 1 | lncRNA | ENSG0000C |
| 9975017 | 10119309 | -1 | transcribed_processe | ENSG0000C |
| 10028361 | 10029855 | -1 | TEC | ENSG0000C |
| 10122273 | 10129029 | 1 | lncRNA | ENSG0000C |
| 10136419 | 10137004 | 1 | unprocessed_pseudc | ENSG0000C |
| 10328411 | 10342737 | -1 | lncRNA | ENSG0000C |
| 10330732 | 10331537 | 1 | processed_pseudoge | ENSG0000C |
| 10357400 | 10358620 | 1 | processed_pseudoge | ENSG0000C |
| 10397644 | 10397778 | 1 | unprocessed_pseudc | ENSG0000C |
| 10413477 | 10516431 | 1 | transcribed_unproce | ENSG0000C |
| 10482738 | 10605716 | 1 | lncRNA | ENSG0000C |
| 10521553 | 10606140 | 1 | protein_coding | ENSG0000C |
| 10576292 | 10576587 | 1 | processed_pseudoge | ENSG0000C |
| 10612455 | 10613379 | -1 | unprocessed_pseudc | ENSG0000C |
| 10640028 | 10640338 | -1 | unprocessed_pseudc | ENSG0000C |
| 10649400 | 10649835 | -1 | IG_V_gene | ENSG0000C |
| 12999676 | 13016692 | -1 | lncRNA | ENSG0000C |
| 13038160 | 13067033 | 1 | transcribed_unproce | ENSG0000C |
| 13047583 | 13047689 | 1 | snRNA | ENSG0000C |
| 13095305 | 13113790 | -1 | unprocessed_pseudc | ENSG0000C |
| 13120244 | 13120807 | 1 | processed_pseudoge | ENSG0000C |
| 13305152 | 13314944 | 1 | lncRNA | ENSG0000C |
| 13349265 | 13350648 | 1 | unprocessed_pseudc | ENSG0000C |
| 13384249 | 13427773 | -1 | unprocessed_pseudc | ENSG0000C |
| 13406384 | 13406460 | -1 | miRNA | ENSG0000C |
| 13443373 | 13489176 | -1 | unprocessed_pseudc | ENSG0000C |
| 13476371 | 13477263 | 1 | processed_pseudoge | ENSG0000C |
| 13516107 | 13574999 | 1 | lncRNA | ENSG0000C |
| 13525599 | 13528579 | 1 | processed_pseudoge | ENSG0000C |
| 13544210 | 13545185 | -1 | unprocessed_pseudc | ENSG0000C |
| 13581060 | 13582004 | -1 | unprocessed_pseudc | ENSG0000C |
| 13609858 | 13641585 | 1 | protein_coding | ENSG0000C |
| 13621577 | 13621683 | 1 | snRNA | ENSG0000C |
| 13644775 | 13644850 | -1 | miRNA | ENSG0000C |
| 13654073 | 13654356 | 1 | processed_pseudoge | ENSG0000C |
| 13665868 | 13667546 | -1 | processed_pseudoge | ENSG0000C |
| 13676022 | 13677116 | 1 | processed_pseudoge | ENSG0000C |
| 13679300 | 13681138 | -1 | TEC | ENSG0000C |

| | | | | |
|---|---|---|---|---|
| 13704853 | 13705518 | 1 | processed_pseudoge | ENSG00000 |
| 13724189 | 13724274 | 1 | miRNA | ENSG00000 |
| 13762338 | 13764332 | 1 | processed_pseudoge | ENSG00000 |
| 13769932 | 13771740 | 1 | lncRNA | ENSG00000 |
| 13776086 | 13777266 | 1 | processed_pseudoge | ENSG00000 |
| 13792635 | 13793141 | 1 | processed_pseudoge | ENSG00000 |
| 13824406 | 13825635 | -1 | processed_pseudoge | ENSG00000 |
| 13826696 | 13827627 | 1 | processed_pseudoge | ENSG00000 |
| 13839118 | 13839409 | 1 | processed_pseudoge | ENSG00000 |
| 13843133 | 13848364 | -1 | transcribed_unproce | ENSG00000 |
| 13905960 | 13906488 | -1 | processed_pseudoge | ENSG00000 |
| 13909574 | 13980437 | -1 | transcribed_unproce | ENSG00000 |
| 13936993 | 13937325 | -1 | processed_pseudoge | ENSG00000 |
| 13968489 | 13968595 | -1 | snRNA | ENSG00000 |
| 14000927 | 14005279 | 1 | unprocessed_pseudo | ENSG00000 |
| 14007134 | 14014880 | 1 | unprocessed_pseudo | ENSG00000 |
| 14019060 | 14020725 | -1 | processed_pseudoge | ENSG00000 |
| 14027419 | 14144468 | 1 | lncRNA | ENSG00000 |
| 14064325 | 14069087 | 1 | unprocessed_pseudo | ENSG00000 |
| 14070871 | 14070986 | 1 | rRNA_pseudogene | ENSG00000 |
| 14075950 | 14076038 | -1 | misc_RNA | ENSG00000 |
| 14108813 | 14210891 | -1 | protein_coding | ENSG00000 |
| 14143581 | 14144158 | 1 | processed_pseudoge | ENSG00000 |
| 14216130 | 14228372 | 1 | protein_coding | ENSG00000 |
| 14236206 | 14362754 | 1 | transcribed_unproce | ENSG00000 |
| 14371115 | 14383484 | -1 | protein_coding | ENSG00000 |
| 14485228 | 14658821 | -1 | protein_coding | ENSG00000 |
| 14582202 | 14598303 | 1 | lncRNA | ENSG00000 |
| 14728396 | 14753925 | -1 | lncRNA | ENSG00000 |
| 14757588 | 14758352 | 1 | processed_pseudoge | ENSG00000 |
| 14761710 | 14763090 | -1 | lncRNA | ENSG00000 |
| 14774301 | 14775262 | 1 | processed_pseudoge | ENSG00000 |
| 14818843 | 15014430 | -1 | lncRNA | ENSG00000 |
| 14819699 | 14918552 | -1 | lncRNA | ENSG00000 |
| 14829154 | 14829989 | -1 | processed_pseudoge | ENSG00000 |
| 14918534 | 14947096 | 1 | lncRNA | ENSG00000 |
| 14961235 | 15065936 | -1 | protein_coding | ENSG00000 |
| 14961309 | 14964233 | 1 | lncRNA | ENSG00000 |
| 14971470 | 14992854 | 1 | lncRNA | ENSG00000 |
| 15050189 | 15052379 | 1 | TEC | ENSG00000 |
| 15067070 | 15067837 | 1 | lncRNA | ENSG00000 |
| 15346652 | 15346738 | 1 | TEC | ENSG00000 |
| 15368966 | 15627342 | -1 | lncRNA | ENSG00000 |
| 15490530 | 15490767 | -1 | processed_pseudoge | ENSG00000 |
| 15493932 | 15496124 | -1 | lncRNA | ENSG00000 |
| 15614283 | 15614389 | 1 | snRNA | ENSG00000 |
| 15694300 | 15696581 | -1 | processed_pseudoge | ENSG00000 |
| 15730025 | 15880069 | 1 | protein_coding | ENSG00000 |
| 15744004 | 15745080 | -1 | processed_pseudoge | ENSG00000 |
| 15928296 | 16645467 | 1 | lncRNA | ENSG00000 |

| | | | | |
|---|---|---|---|---|
| 16035413 | 16035509 | -1 | snRNA | ENSG00000 |
| 16094415 | 16095372 | -1 | processed_pseudoge | ENSG00000 |
| 16121310 | 16121644 | -1 | processed_pseudoge | ENSG00000 |
| 16284696 | 16284768 | -1 | snoRNA | ENSG00000 |
| 16291079 | 16310382 | -1 | lncRNA | ENSG00000 |
| 16417139 | 16419080 | -1 | lncRNA | ENSG00000 |
| 16505555 | 16507916 | -1 | lncRNA | ENSG00000 |
| 16525919 | 16527019 | 1 | TEC | ENSG00000 |
| 16539089 | 16539169 | 1 | miRNA | ENSG00000 |
| 16539828 | 16539911 | 1 | miRNA | ENSG00000 |
| 16574718 | 16582637 | -1 | lncRNA | ENSG00000 |
| 16590237 | 16590325 | 1 | miRNA | ENSG00000 |
| 16718998 | 16719157 | 1 | snRNA | ENSG00000 |
| 16754519 | 16815688 | -1 | lncRNA | ENSG00000 |
| 16862875 | 16873691 | -1 | lncRNA | ENSG00000 |
| 17210469 | 17211347 | -1 | processed_pseudoge | ENSG00000 |
| 17296219 | 17296596 | 1 | processed_pseudoge | ENSG00000 |
| 17431547 | 17431647 | 1 | snRNA | ENSG00000 |
| 17438821 | 17450104 | 1 | lncRNA | ENSG00000 |
| 17500679 | 17500824 | -1 | processed_pseudoge | ENSG00000 |
| 17506453 | 17506728 | 1 | misc_RNA | ENSG00000 |
| 17513043 | 17593579 | 1 | protein_coding | ENSG00000 |
| 17518526 | 17518993 | -1 | processed_pseudoge | ENSG00000 |
| 17527140 | 17527247 | -1 | misc_RNA | ENSG00000 |
| 17576798 | 17576906 | 1 | misc_RNA | ENSG00000 |
| 17593653 | 17612947 | -1 | protein_coding | ENSG00000 |
| 17611744 | 17633199 | 1 | lncRNA | ENSG00000 |
| 17659276 | 17660384 | -1 | lncRNA | ENSG00000 |
| 17705408 | 17707455 | 1 | TEC | ENSG00000 |
| 17763315 | 17792523 | -1 | lncRNA | ENSG00000 |
| 17788974 | 17819386 | -1 | protein_coding | ENSG00000 |
| 17793488 | 17810845 | 1 | lncRNA | ENSG00000 |
| 17835016 | 17885608 | -1 | lncRNA | ENSG00000 |
| 17894193 | 17894485 | -1 | processed_pseudoge | ENSG00000 |
| 17901263 | 18267373 | 1 | protein_coding | ENSG00000 |
| 18022370 | 18114904 | -1 | lncRNA | ENSG00000 |
| 18269116 | 18485879 | -1 | protein_coding | ENSG00000 |
| 18477358 | 18486599 | -1 | lncRNA | ENSG00000 |
| 18561265 | 18760320 | -1 | lncRNA | ENSG00000 |
| 18614431 | 18650360 | 1 | lncRNA | ENSG00000 |
| 18686090 | 18686164 | -1 | miRNA | ENSG00000 |
| 18857779 | 18858276 | 1 | processed_pseudoge | ENSG00000 |
| 18917213 | 18917429 | 1 | TEC | ENSG00000 |
| 18917934 | 18935859 | -1 | lncRNA | ENSG00000 |
| 18953264 | 18967058 | -1 | lncRNA | ENSG00000 |
| 19046310 | 19047684 | -1 | lncRNA | ENSG00000 |
| 19130523 | 19134423 | 1 | lncRNA | ENSG00000 |
| 19244543 | 19246392 | 1 | processed_pseudoge | ENSG00000 |
| 19301613 | 19303739 | 1 | lncRNA | ENSG00000 |
| 19345148 | 19345312 | 1 | snRNA | ENSG00000 |

| | | | | |
|---|---|---|---|---|
| 19593841 | 19594108 | 1 | processed_pseudoge | ENSG00000 |
| 19620695 | 19621545 | 1 | processed_pseudoge | ENSG00000 |
| 19739709 | 19740150 | 1 | processed_pseudoge | ENSG00000 |
| 19759359 | 19760128 | -1 | processed_pseudoge | ENSG00000 |
| 19893113 | 19900043 | -1 | lncRNA | ENSG00000 |
| 20256752 | 20258820 | -1 | lncRNA | ENSG00000 |
| 20355748 | 20355852 | -1 | snRNA | ENSG00000 |
| 20356653 | 20356896 | 1 | misc_RNA | ENSG00000 |
| 20388334 | 20388845 | 1 | processed_pseudoge | ENSG00000 |
| 20424949 | 20426206 | -1 | processed_pseudoge | ENSG00000 |
| 20430443 | 20430762 | -1 | processed_pseudoge | ENSG00000 |
| 20597953 | 20598163 | 1 | processed_pseudoge | ENSG00000 |
| 20651450 | 20803816 | -1 | lncRNA | ENSG00000 |
| 20828127 | 20828622 | -1 | processed_pseudoge | ENSG00000 |
| 20998409 | 21543329 | 1 | protein_coding | ENSG00000 |
| 21223295 | 21226829 | -1 | lncRNA | ENSG00000 |
| 21566763 | 21615437 | 1 | lncRNA | ENSG00000 |
| 21655038 | 21686329 | -1 | lncRNA | ENSG00000 |
| 21723293 | 21737319 | -1 | lncRNA | ENSG00000 |
| 21746973 | 21797415 | 1 | lncRNA | ENSG00000 |
| 21933315 | 21975681 | -1 | lncRNA | ENSG00000 |
| 22008756 | 22098518 | -1 | lncRNA | ENSG00000 |
| 22098616 | 22116605 | 1 | lncRNA | ENSG00000 |
| 22134698 | 22136083 | -1 | processed_pseudoge | ENSG00000 |
| 22153950 | 22154299 | -1 | processed_pseudoge | ENSG00000 |
| 22205192 | 22205332 | 1 | snRNA | ENSG00000 |
| 22209920 | 22420819 | 1 | lncRNA | ENSG00000 |
| 22436290 | 22438349 | -1 | processed_pseudoge | ENSG00000 |
| 22500604 | 22520058 | -1 | lncRNA | ENSG00000 |
| 22882582 | 22884000 | 1 | processed_pseudoge | ENSG00000 |
| 23065491 | 23131206 | 1 | lncRNA | ENSG00000 |
| 23079284 | 23079392 | 1 | miRNA | ENSG00000 |
| 23090028 | 23091831 | -1 | processed_pseudoge | ENSG00000 |
| 23101223 | 23103074 | 1 | processed_pseudoge | ENSG00000 |
| 23106928 | 23119903 | -1 | lncRNA | ENSG00000 |
| 23281736 | 23281909 | -1 | snRNA | ENSG00000 |
| 23352929 | 23423778 | -1 | lncRNA | ENSG00000 |
| 23390258 | 23390996 | 1 | processed_pseudoge | ENSG00000 |
| 23407658 | 23409128 | -1 | processed_pseudoge | ENSG00000 |
| 23432181 | 23432282 | -1 | misc_RNA | ENSG00000 |
| 23477641 | 23490724 | 1 | lncRNA | ENSG00000 |
| 23704260 | 23709345 | 1 | lncRNA | ENSG00000 |
| 23857081 | 23889150 | 1 | lncRNA | ENSG00000 |
| 23888798 | 23890541 | -1 | lncRNA | ENSG00000 |
| 23960569 | 23968747 | -1 | lncRNA | ENSG00000 |
| 23982645 | 24114352 | -1 | lncRNA | ENSG00000 |
| 24043257 | 24050286 | 1 | lncRNA | ENSG00000 |
| 24154871 | 24192924 | -1 | lncRNA | ENSG00000 |
| 24304546 | 24321377 | -1 | lncRNA | ENSG00000 |
| 24306939 | 24547942 | 1 | lncRNA | ENSG00000 |

| | | | | |
|---|---|---|---|---|
| 24840550 | 25057746 | 1 | lncRNA | ENSG00000( |
| 24886676 | 24902756 | -1 | lncRNA | ENSG00000( |
| 24938431 | 24992817 | -1 | lncRNA | ENSG00000( |
| 25031005 | 25031529 | -1 | processed_pseudoge | ENSG00000( |
| 25055535 | 25069906 | 1 | lncRNA | ENSG00000( |
| 25095022 | 25103670 | 1 | lncRNA | ENSG00000( |
| 25101132 | 25101415 | 1 | misc_RNA | ENSG00000( |
| 25127469 | 25135247 | 1 | lncRNA | ENSG00000( |
| 25169431 | 25361868 | -1 | lncRNA | ENSG00000( |
| 25202207 | 25202315 | 1 | rRNA_pseudogene | ENSG00000( |
| 25361821 | 25381756 | 1 | transcribed_processe | ENSG00000( |
| 25371827 | 25372043 | 1 | TEC | ENSG00000( |
| 25385388 | 25431701 | -1 | lncRNA | ENSG00000( |
| 25515349 | 25518865 | -1 | lncRNA | ENSG00000( |
| 25561909 | 25575168 | 1 | lncRNA | ENSG00000( |
| 25573980 | 25574044 | 1 | miRNA | ENSG00000( |
| 25582770 | 25583326 | -1 | lncRNA | ENSG00000( |
| 25585656 | 25607517 | -1 | protein_coding | ENSG00000( |
| 25639258 | 25717562 | 1 | protein_coding | ENSG00000( |
| 25643489 | 25644216 | 1 | processed_pseudoge | ENSG00000( |
| 25646113 | 25654671 | -1 | lncRNA | ENSG00000( |
| 25692180 | 25692554 | 1 | processed_pseudoge | ENSG00000( |
| 25716503 | 25735673 | -1 | protein_coding | ENSG00000( |
| 25734570 | 25772460 | 1 | protein_coding | ENSG00000( |
| 25762938 | 25763333 | -1 | processed_pseudoge | ENSG00000( |
| 25880550 | 26171128 | -1 | protein_coding | ENSG00000( |
| 25928754 | 25952614 | 1 | lncRNA | ENSG00000( |
| 25943044 | 25943145 | -1 | snRNA | ENSG00000( |
| 26158091 | 26175824 | 1 | lncRNA | ENSG00000( |
| 26170871 | 26217381 | 1 | lncRNA | ENSG00000( |
| 26190707 | 26190813 | 1 | snRNA | ENSG00000( |
| 26317390 | 26320773 | -1 | lncRNA | ENSG00000( |
| 26349780 | 26350634 | -1 | lncRNA | ENSG00000( |
| 26378552 | 26471698 | 1 | lncRNA | ENSG00000( |
| 26393635 | 26569252 | 1 | lncRNA | ENSG00000( |
| 26459903 | 26460214 | -1 | lncRNA | ENSG00000( |
| 26466209 | 26573286 | -1 | protein_coding | ENSG00000( |
| 26835755 | 26845409 | -1 | protein_coding | ENSG00000( |
| 26889376 | 26939742 | 1 | lncRNA | ENSG00000( |
| 26917922 | 26967088 | -1 | protein_coding | ENSG00000( |
| 26953961 | 26954043 | 1 | miRNA | ENSG00000( |
| 27143344 | 27143949 | -1 | processed_pseudoge | ENSG00000( |
| 27358885 | 27448579 | -1 | lncRNA | ENSG00000( |
| 27361121 | 27395787 | 1 | lncRNA | ENSG00000( |
| 27367014 | 27367947 | -1 | processed_pseudoge | ENSG00000( |
| 27420380 | 27421023 | -1 | processed_pseudoge | ENSG00000( |
| 27492118 | 27492261 | 1 | processed_pseudoge | ENSG00000( |
| 27638613 | 27675024 | 1 | lncRNA | ENSG00000( |
| 27722379 | 27751233 | 1 | lncRNA | ENSG00000( |
| 27954922 | 27985295 | 1 | lncRNA | ENSG00000( |

| | | | | |
|---|---|---|---|---|
| 28013363 | 28023233 | 1 | lncRNA | ENSG00000( |
| 28048404 | 28137611 | 1 | lncRNA | ENSG00000( |
| 28116094 | 28228667 | -1 | lncRNA | ENSG00000( |
| 28328738 | 28674848 | -1 | lncRNA | ENSG00000( |
| 28539318 | 28540355 | 1 | lncRNA | ENSG00000( |
| 28743208 | 28743291 | 1 | snRNA | ENSG00000( |
| 28872191 | 28885371 | -1 | protein_coding | ENSG00000( |
| 28887280 | 28889272 | -1 | processed_pseudoge | ENSG00000( |
| 28901779 | 28903697 | -1 | processed_pseudoge | ENSG00000( |
| 28928144 | 28992956 | -1 | protein_coding | ENSG00000( |
| 28997613 | 28998033 | -1 | processed_pseudoge | ENSG00000( |
| 29004384 | 29019360 | -1 | protein_coding | ENSG00000( |
| 29024255 | 29024890 | -1 | lncRNA | ENSG00000( |
| 29024629 | 29054488 | 1 | protein_coding | ENSG00000( |
| 29055805 | 29073797 | -1 | protein_coding | ENSG00000( |
| 29058073 | 29060095 | -1 | lncRNA | ENSG00000( |
| 29063446 | 29064368 | 1 | lncRNA | ENSG00000( |
| 29077471 | 29175889 | 1 | protein_coding | ENSG00000( |
| 29127701 | 29128188 | -1 | processed_pseudoge | ENSG00000( |
| 29139283 | 29139384 | -1 | misc_RNA | ENSG00000( |
| 29180425 | 29180639 | -1 | snoRNA | ENSG00000( |
| 29182027 | 29187795 | -1 | lncRNA | ENSG00000( |
| 29193480 | 29288205 | 1 | lncRNA | ENSG00000( |
| 29194071 | 29630751 | 1 | protein_coding | ENSG00000( |
| 29222321 | 29223257 | 1 | processed_pseudoge | ENSG00000( |
| 29351634 | 29361894 | 1 | lncRNA | ENSG00000( |
| 29359002 | 29359453 | 1 | lncRNA | ENSG00000( |
| 29370019 | 29376339 | -1 | lncRNA | ENSG00000( |
| 29370497 | 29373709 | 1 | lncRNA | ENSG00000( |
| 29496047 | 29500386 | 1 | lncRNA | ENSG00000( |
| 29536933 | 29940033 | -1 | protein_coding | ENSG00000( |
| 29657406 | 29657831 | 1 | lncRNA | ENSG00000( |
| 29748175 | 29764002 | 1 | lncRNA | ENSG00000( |
| 30089717 | 30097836 | 1 | lncRNA | ENSG00000( |
| 30165565 | 30166805 | -1 | protein_coding | ENSG00000( |
| 30209151 | 30211783 | -1 | lncRNA | ENSG00000( |
| 30214006 | 30216097 | -1 | protein_coding | ENSG00000( |
| 30263380 | 30263881 | -1 | processed_pseudoge | ENSG00000( |
| 30281309 | 30282950 | -1 | protein_coding | ENSG00000( |
| 30289145 | 30289514 | -1 | protein_coding | ENSG00000( |
| 30319124 | 30320315 | -1 | protein_coding | ENSG00000( |
| 30337013 | 30337694 | -1 | protein_coding | ENSG00000( |
| 30348399 | 30348609 | -1 | protein_coding | ENSG00000( |
| 30356515 | 30356885 | -1 | unprocessed_pseudc | ENSG00000( |
| 30371390 | 30372271 | -1 | protein_coding | ENSG00000( |
| 30375294 | 30375378 | -1 | miRNA | ENSG00000( |
| 30396031 | 30396822 | 1 | protein_coding | ENSG00000( |
| 30425265 | 30425968 | -1 | protein_coding | ENSG00000( |
| 30430230 | 30431026 | 1 | protein_coding | ENSG00000( |
| 30436466 | 30436964 | -1 | unprocessed_pseudc | ENSG00000( |

| | | | | |
|---|---|---|---|---|
| 30440275 | 30440945 | 1 | protein_coding | ENSG00000 |
| 30479706 | 30480367 | -1 | protein_coding | ENSG00000 |
| 30487043 | 30487436 | -1 | protein_coding | ENSG00000 |
| 30491462 | 30492013 | -1 | protein_coding | ENSG00000 |
| 30496824 | 30497165 | -1 | protein_coding | ENSG00000 |
| 30501657 | 30502141 | -1 | protein_coding | ENSG00000 |
| 30510307 | 30510497 | -1 | unprocessed_pseudo | ENSG00000 |
| 30514613 | 30514799 | -1 | unprocessed_pseudo | ENSG00000 |
| 30526546 | 30526701 | -1 | unprocessed_pseudo | ENSG00000 |
| 30537278 | 30537381 | -1 | unprocessed_pseudo | ENSG00000 |
| 30541535 | 30541864 | -1 | protein_coding | ENSG00000 |
| 30560875 | 30561314 | -1 | protein_coding | ENSG00000 |
| 30590105 | 30590397 | -1 | protein_coding | ENSG00000 |
| 30592440 | 30593075 | 1 | protein_coding | ENSG00000 |
| 30598590 | 30598902 | -1 | protein_coding | ENSG00000 |
| 30601087 | 30601382 | 1 | protein_coding | ENSG00000 |
| 30613431 | 30613930 | -1 | protein_coding | ENSG00000 |
| 30616425 | 30616699 | 1 | protein_coding | ENSG00000 |
| 30620627 | 30620850 | 1 | protein_coding | ENSG00000 |
| 30635206 | 30635619 | 1 | protein_coding | ENSG00000 |
| 30642864 | 30643136 | 1 | protein_coding | ENSG00000 |
| 30718525 | 30718777 | -1 | protein_coding | ENSG00000 |
| 30741780 | 30742595 | -1 | unprocessed_pseudo | ENSG00000 |
| 30746794 | 30747259 | -1 | protein_coding | ENSG00000 |
| 30755015 | 30755428 | -1 | protein_coding | ENSG00000 |
| 30762682 | 30762882 | -1 | unprocessed_pseudo | ENSG00000 |
| 30802242 | 30802427 | -1 | unprocessed_pseudo | ENSG00000 |
| 30806932 | 30807119 | -1 | unprocessed_pseudo | ENSG00000 |
| 30812697 | 30813274 | -1 | protein_coding | ENSG00000 |
| 30829039 | 30829759 | -1 | protein_coding | ENSG00000 |
| 30880644 | 30881580 | -1 | protein_coding | ENSG00000 |
| 31038159 | 31038476 | -1 | protein_coding | ENSG00000 |
| 31060600 | 31063168 | -1 | processed_pseudoge | ENSG00000 |
| 31118416 | 31559977 | -1 | protein_coding | ENSG00000 |
| 31347736 | 31359157 | 1 | lncRNA | ENSG00000 |
| 31452466 | 31453223 | 1 | processed_pseudoge | ENSG00000 |
| 31559245 | 31560487 | 1 | lncRNA | ENSG00000 |
| 31627127 | 31628600 | -1 | processed_pseudoge | ENSG00000 |
| 31653593 | 31659500 | -1 | lncRNA | ENSG00000 |
| 31659622 | 31668931 | 1 | protein_coding | ENSG00000 |
| 31664306 | 31664482 | 1 | snoRNA | ENSG00000 |
| 31666728 | 31667247 | -1 | lncRNA | ENSG00000 |
| 31671000 | 31732118 | -1 | protein_coding | ENSG00000 |
| 31706555 | 31706864 | 1 | processed_pseudoge | ENSG00000 |
| 31735732 | 31736407 | 1 | lncRNA | ENSG00000 |
| 31840341 | 31840858 | 1 | processed_pseudoge | ENSG00000 |
| 31873020 | 32044633 | 1 | protein_coding | ENSG00000 |
| 32020966 | 32021782 | -1 | lncRNA | ENSG00000 |
| 32080316 | 32197813 | -1 | lncRNA | ENSG00000 |
| 32259804 | 32261585 | -1 | lncRNA | ENSG00000 |

| | | | | |
|---|---|---|---|---|
| 32268228 | 32279049 | -1 | protein_coding | ENSG00000 |
| 32277863 | 32280988 | 1 | lncRNA | ENSG00000 |
| 32291813 | 32314784 | 1 | protein_coding | ENSG00000 |
| 32306464 | 32308737 | -1 | lncRNA | ENSG00000 |
| 32311018 | 32393012 | -1 | protein_coding | ENSG00000 |
| 32377187 | 32377322 | -1 | snoRNA | ENSG00000 |
| 32393130 | 32393960 | 1 | lncRNA | ENSG00000 |
| 32412006 | 32515397 | 1 | protein_coding | ENSG00000 |
| 32496812 | 32497311 | 1 | processed_pseudoge | ENSG00000 |
| 32538299 | 32538434 | 1 | snoRNA | ENSG00000 |
| 32563075 | 32563210 | 1 | rRNA_pseudogene | ENSG00000 |
| 32572238 | 32587373 | -1 | protein_coding | ENSG00000 |
| 32578822 | 32612281 | -1 | protein_coding | ENSG00000 |
| 32592079 | 32612603 | -1 | protein_coding | ENSG00000 |
| 32621084 | 32622083 | 1 | unprocessed_pseudc | ENSG00000 |
| 32624416 | 32625096 | -1 | processed_pseudoge | ENSG00000 |
| 32628759 | 32728048 | -1 | protein_coding | ENSG00000 |
| 32728097 | 32747065 | 1 | lncRNA | ENSG00000 |
| 32733899 | 32771792 | -1 | protein_coding | ENSG00000 |
| 32772100 | 32955437 | 1 | lncRNA | ENSG00000 |
| 32790673 | 32791504 | -1 | TEC | ENSG00000 |
| 32790673 | 32813743 | -1 | protein_coding | ENSG00000 |
| 32841496 | 32841811 | -1 | processed_pseudoge | ENSG00000 |
| 32841861 | 32841995 | -1 | snoRNA | ENSG00000 |
| 32844367 | 32849934 | -1 | lncRNA | ENSG00000 |
| 32913649 | 33071104 | -1 | lncRNA | ENSG00000 |
| 32958888 | 32960566 | 1 | lncRNA | ENSG00000 |
| 33025935 | 33029196 | 1 | protein_coding | ENSG00000 |
| 33057829 | 33064983 | 1 | lncRNA | ENSG00000 |
| 33070141 | 33072413 | 1 | protein_coding | ENSG00000 |
| 33110867 | 33123725 | -1 | lncRNA | ENSG00000 |
| 33152614 | 33159110 | 1 | lncRNA | ENSG00000 |
| 33164809 | 33170649 | -1 | lncRNA | ENSG00000 |
| 33229901 | 33297160 | 1 | protein_coding | ENSG00000 |
| 33229901 | 33265675 | 1 | protein_coding | ENSG00000 |
| 33263873 | 33266260 | -1 | lncRNA | ENSG00000 |
| 33266367 | 33310187 | 1 | protein_coding | ENSG00000 |
| 33324429 | 33359864 | 1 | protein_coding | ENSG00000 |
| 33334571 | 33335096 | 1 | processed_pseudoge | ENSG00000 |
| 33402896 | 33479348 | 1 | protein_coding | ENSG00000 |
| 33432485 | 33479974 | -1 | protein_coding | ENSG00000 |
| 33481580 | 33482195 | -1 | processed_pseudoge | ENSG00000 |
| 33482499 | 33484258 | -1 | lncRNA | ENSG00000 |
| 33485530 | 33491720 | -1 | protein_coding | ENSG00000 |
| 33503931 | 33543491 | -1 | protein_coding | ENSG00000 |
| 33518610 | 33519108 | -1 | processed_pseudoge | ENSG00000 |
| 33543038 | 33577514 | 1 | protein_coding | ENSG00000 |
| 33550662 | 33550728 | 1 | miRNA | ENSG00000 |
| 33559542 | 33588706 | -1 | protein_coding | ENSG00000 |
| 33584687 | 33931607 | -1 | protein_coding | ENSG00000 |

| | | | | |
|---|---|---|---|---|
| 33589341 | 33643926 | -1 | protein_coding | ENSG0000( |
| 33642400 | 33899861 | 1 | protein_coding | ENSG0000( |
| 33903453 | 33915814 | -1 | protein_coding | ENSG0000( |
| 33915534 | 33977691 | 1 | lncRNA | ENSG0000( |
| 33918995 | 33919338 | -1 | misc_RNA | ENSG0000( |
| 33967101 | 33968573 | -1 | lncRNA | ENSG0000( |
| 34073224 | 34143034 | 1 | protein_coding | ENSG0000( |
| 34073578 | 34106260 | 1 | protein_coding | ENSG0000( |
| 34073592 | 34360033 | 1 | lncRNA | ENSG0000( |
| 34135432 | 34136071 | -1 | processed_pseudoge | ENSG0000( |
| 34157724 | 34190244 | 1 | lncRNA | ENSG0000( |
| 34205055 | 34325034 | 1 | lncRNA | ENSG0000( |
| 34353099 | 34375364 | -1 | lncRNA | ENSG0000( |
| 34364024 | 34371389 | 1 | protein_coding | ENSG0000( |
| 34375480 | 34416961 | 1 | protein_coding | ENSG0000( |
| 34400112 | 34401072 | -1 | protein_coding | ENSG0000( |
| 34412200 | 34412587 | 1 | lncRNA | ENSG0000( |
| 34418715 | 34423966 | -1 | protein_coding | ENSG0000( |
| 34425508 | 34426017 | 1 | lncRNA | ENSG0000( |
| 34446688 | 34512210 | -1 | protein_coding | ENSG0000( |
| 34456110 | 34456237 | -1 | snoRNA | ENSG0000( |
| 34513142 | 34615113 | -1 | protein_coding | ENSG0000( |
| 34669389 | 34718227 | 1 | protein_coding | ENSG0000( |
| 34723807 | 34737204 | -1 | lncRNA | ENSG0000( |
| 34745757 | 34784886 | 1 | lncRNA | ENSG0000( |
| 34787801 | 36004667 | -1 | protein_coding | ENSG0000( |
| 34836286 | 34884882 | 1 | lncRNA | ENSG0000( |
| 35136638 | 35139222 | 1 | lncRNA | ENSG0000( |
| 35472095 | 35472432 | -1 | processed_pseudoge | ENSG0000( |
| 35599732 | 35600022 | -1 | processed_pseudoge | ENSG0000( |
| 35713139 | 35732942 | -1 | lncRNA | ENSG0000( |
| 35720715 | 35720808 | 1 | miRNA | ENSG0000( |
| 35724747 | 35725100 | -1 | processed_pseudoge | ENSG0000( |
| 35887195 | 35887807 | 1 | processed_pseudoge | ENSG0000( |
| 36005338 | 36007838 | 1 | lncRNA | ENSG0000( |
| 36016079 | 36016546 | -1 | processed_pseudoge | ENSG0000( |
| 36034541 | 36079389 | -1 | protein_coding | ENSG0000( |
| 36050214 | 36051377 | 1 | processed_pseudoge | ENSG0000( |
| 36060432 | 36064408 | 1 | lncRNA | ENSG0000( |
| 36066545 | 36066652 | 1 | snRNA | ENSG0000( |
| 36069642 | 36126640 | -1 | lncRNA | ENSG0000( |
| 36069941 | 36073166 | 1 | protein_coding | ENSG0000( |
| 36082859 | 36090414 | 1 | lncRNA | ENSG0000( |
| 36104881 | 36109690 | 1 | lncRNA | ENSG0000( |
| 36130489 | 36131376 | 1 | processed_pseudoge | ENSG0000( |
| 36131767 | 36175815 | -1 | lncRNA | ENSG0000( |
| 36132450 | 36133032 | 1 | processed_pseudoge | ENSG0000( |
| 36135079 | 36146562 | 1 | protein_coding | ENSG0000( |
| 36156782 | 36294274 | 1 | protein_coding | ENSG0000( |
| 36168970 | 36170180 | -1 | processed_pseudoge | ENSG0000( |

| | | | | |
|---|---|---|---|---|
| 36295173 | 36295702 | -1 | processed_pseudoge | ENSG00000 |
| 36319792 | 36320670 | -1 | lncRNA | ENSG00000 |
| 36320189 | 36386148 | 1 | protein_coding | ENSG00000 |
| 36360630 | 36362040 | 1 | lncRNA | ENSG00000 |
| 36385392 | 36419015 | 1 | protein_coding | ENSG00000 |
| 36388878 | 36389112 | -1 | processed_pseudoge | ENSG00000 |
| 36430360 | 36481070 | 1 | lncRNA | ENSG00000 |
| 36445731 | 36532408 | 1 | lncRNA | ENSG00000 |
| 36460621 | 36576569 | -1 | protein_coding | ENSG00000 |
| 36485867 | 36487760 | 1 | TEC | ENSG00000 |
| 36485983 | 36487411 | -1 | processed_pseudoge | ENSG00000 |
| 36526280 | 36542855 | -1 | lncRNA | ENSG00000 |
| 36580172 | 36581229 | 1 | lncRNA | ENSG00000 |
| 36632681 | 36637033 | -1 | lncRNA | ENSG00000 |
| 36698773 | 36701564 | -1 | lncRNA | ENSG00000 |
| 36699115 | 36749917 | 1 | protein_coding | ENSG00000 |
| 36750888 | 36990236 | -1 | protein_coding | ENSG00000 |
| 36803984 | 36806284 | -1 | lncRNA | ENSG00000 |
| 36851911 | 36852028 | 1 | rRNA_pseudogene | ENSG00000 |
| 36943267 | 36944125 | 1 | processed_pseudoge | ENSG00000 |
| 36966461 | 36975164 | 1 | lncRNA | ENSG00000 |
| 36986739 | 36986851 | -1 | misc_RNA | ENSG00000 |
| 36994643 | 36995075 | 1 | processed_pseudoge | ENSG00000 |
| 37006150 | 37019662 | 1 | protein_coding | ENSG00000 |
| 37045530 | 37045636 | 1 | snRNA | ENSG00000 |
| 37059170 | 37073170 | -1 | protein_coding | ENSG00000 |
| 37073226 | 37203112 | 1 | protein_coding | ENSG00000 |
| 37100814 | 37101343 | 1 | lncRNA | ENSG00000 |
| 37187666 | 37193926 | -1 | lncRNA | ENSG00000 |
| 37208503 | 37221736 | 1 | lncRNA | ENSG00000 |
| 37215605 | 37215903 | 1 | misc_RNA | ENSG00000 |
| 37221419 | 37237744 | 1 | lncRNA | ENSG00000 |
| 37223420 | 37267919 | -1 | protein_coding | ENSG00000 |
| 37267784 | 37268497 | 1 | lncRNA | ENSG00000 |
| 37337382 | 37366858 | -1 | lncRNA | ENSG00000 |
| 37365477 | 37365932 | -1 | lncRNA | ENSG00000 |
| 37365573 | 37526358 | 1 | protein_coding | ENSG00000 |
| 37593977 | 37661496 | 1 | lncRNA | ENSG00000 |
| 37607373 | 38121345 | -1 | protein_coding | ENSG00000 |
| 37717102 | 37719569 | -1 | lncRNA | ENSG00000 |
| 37951425 | 38121360 | -1 | lncRNA | ENSG00000 |
| 38006544 | 38010618 | -1 | lncRNA | ENSG00000 |
| 38121451 | 38156511 | 1 | lncRNA | ENSG00000 |
| 38155549 | 38307357 | 1 | protein_coding | ENSG00000 |
| 38204141 | 38206080 | -1 | transcribed_processe | ENSG00000 |
| 38206156 | 38208644 | 1 | lncRNA | ENSG00000 |
| 38237217 | 38238201 | -1 | lncRNA | ENSG00000 |
| 38238227 | 38238664 | -1 | processed_pseudoge | ENSG00000 |
| 38323635 | 38333421 | -1 | lncRNA | ENSG00000 |
| 38380027 | 38661780 | -1 | protein_coding | ENSG00000 |

| | | | | |
|---|---|---|---|---|
| 38502445 | 38502621 | 1 | processed_pseudoge | ENSG00000 |
| 38503562 | 38543336 | 1 | lncRNA | ENSG00000 |
| 38733890 | 38747460 | -1 | lncRNA | ENSG00000 |
| 38805183 | 38824955 | 1 | protein_coding | ENSG00000 |
| 38812878 | 38956467 | -1 | lncRNA | ENSG00000 |
| 38846247 | 38848644 | 1 | lncRNA | ENSG00000 |
| 38857539 | 38858861 | 1 | lncRNA | ENSG00000 |
| 38888772 | 38903905 | 1 | lncRNA | ENSG00000 |
| 38894785 | 38894867 | 1 | snoRNA | ENSG00000 |
| 38894917 | 38895252 | 1 | processed_pseudoge | ENSG00000 |
| 38974429 | 38977774 | -1 | lncRNA | ENSG00000 |
| 38988569 | 39062894 | -1 | lncRNA | ENSG00000 |
| 39023000 | 39023835 | -1 | TEC | ENSG00000 |
| 39028536 | 39029128 | -1 | lncRNA | ENSG00000 |
| 39127568 | 39128040 | 1 | processed_pseudoge | ENSG00000 |
| 39171130 | 39172106 | -1 | processed_pseudoge | ENSG00000 |
| 39171462 | 39171560 | -1 | misc_RNA | ENSG00000 |
| 39174769 | 39183488 | -1 | protein_coding | ENSG00000 |
| 39184176 | 39321559 | -1 | protein_coding | ENSG00000 |
| 39184469 | 39184899 | 1 | lncRNA | ENSG00000 |
| 39216624 | 39217506 | 1 | processed_pseudoge | ENSG00000 |
| 39217093 | 39219805 | -1 | lncRNA | ENSG00000 |
| 39235386 | 39236020 | -1 | processed_pseudoge | ENSG00000 |
| 39313935 | 39314962 | 1 | lncRNA | ENSG00000 |
| 39315707 | 39323218 | 1 | lncRNA | ENSG00000 |
| 39342315 | 39349647 | -1 | protein_coding | ENSG00000 |
| 39344537 | 39344628 | 1 | misc_RNA | ENSG00000 |
| 39373763 | 39377066 | -1 | processed_pseudoge | ENSG00000 |
| 39377698 | 39428528 | 1 | protein_coding | ENSG00000 |
| 39380332 | 39515506 | 1 | protein_coding | ENSG00000 |
| 39405844 | 39445805 | -1 | protein_coding | ENSG00000 |
| 39445855 | 39515506 | 1 | protein_coding | ENSG00000 |
| 39447010 | 39447069 | 1 | miRNA | ENSG00000 |
| 39488327 | 39488760 | 1 | processed_pseudoge | ENSG00000 |
| 39491544 | 39491898 | -1 | processed_pseudoge | ENSG00000 |
| 39525583 | 39529855 | 1 | processed_pseudoge | ENSG00000 |
| 39556442 | 39673137 | 1 | protein_coding | ENSG00000 |
| 39597147 | 39612910 | -1 | lncRNA | ENSG00000 |
| 39630271 | 39726085 | 1 | lncRNA | ENSG00000 |
| 39727755 | 39730680 | 1 | lncRNA | ENSG00000 |
| 39745407 | 39802096 | 1 | protein_coding | ENSG00000 |
| 39867438 | 39929397 | 1 | protein_coding | ENSG00000 |
| 40010999 | 40847158 | -1 | protein_coding | ENSG00000 |
| 40212352 | 40212431 | -1 | miRNA | ENSG00000 |
| 40383083 | 40385358 | 1 | lncRNA | ENSG00000 |
| 40513144 | 40513279 | 1 | snoRNA | ENSG00000 |
| 40615378 | 40630767 | -1 | lncRNA | ENSG00000 |
| 40736318 | 40754888 | 1 | lncRNA | ENSG00000 |
| 40863994 | 40864473 | 1 | processed_pseudoge | ENSG00000 |
| 41007244 | 41037362 | -1 | lncRNA | ENSG00000 |

| | | | | |
|---|---|---|---|---|
| 41141493 | 41148198 | -1 | lncRNA | ENSG0000C |
| 41167557 | 41167629 | 1 | miRNA | ENSG0000C |
| 41167801 | 41282530 | 1 | protein_coding | ENSG0000C |
| 41175231 | 41186788 | -1 | lncRNA | ENSG0000C |
| 41180097 | 41180626 | 1 | lncRNA | ENSG0000C |
| 41304212 | 41357727 | 1 | protein_coding | ENSG0000C |
| 41361999 | 41409393 | 1 | protein_coding | ENSG0000C |
| 41420304 | 41459214 | 1 | protein_coding | ENSG0000C |
| 41441056 | 41445708 | -1 | lncRNA | ENSG0000C |
| 41464305 | 41531116 | -1 | protein_coding | ENSG0000C |
| 41539206 | 41539326 | 1 | snoRNA | ENSG0000C |
| 41559125 | 41562958 | -1 | lncRNA | ENSG0000C |
| 41576135 | 41581319 | -1 | lncRNA | ENSG0000C |
| 41580475 | 41582887 | 1 | lncRNA | ENSG0000C |
| 41679181 | 41697336 | 1 | lncRNA | ENSG0000C |
| 41711520 | 41715775 | -1 | lncRNA | ENSG0000C |
| 41715806 | 41717975 | 1 | lncRNA | ENSG0000C |
| 41739369 | 41767089 | -1 | protein_coding | ENSG0000C |
| 41739373 | 41741308 | 1 | lncRNA | ENSG0000C |
| 41746772 | 41746841 | -1 | miRNA | ENSG0000C |
| 41798225 | 41879482 | -1 | protein_coding | ENSG0000C |
| 41870633 | 41872054 | 1 | lncRNA | ENSG0000C |
| 41874756 | 41877613 | -1 | lncRNA | ENSG0000C |
| 41882205 | 41882300 | 1 | snoRNA | ENSG0000C |
| 41885112 | 41954018 | -1 | protein_coding | ENSG0000C |
| 41986831 | 42010387 | -1 | protein_coding | ENSG0000C |
| 42009194 | 42055130 | 1 | lncRNA | ENSG0000C |
| 42062959 | 42143453 | 1 | protein_coding | ENSG0000C |
| 42102134 | 42108534 | -1 | lncRNA | ENSG0000C |
| 42199689 | 42297244 | 1 | protein_coding | ENSG0000C |
| 42221173 | 42221286 | 1 | rRNA_pseudogene | ENSG0000C |
| 42311667 | 42315651 | -1 | protein_coding | ENSG0000C |
| 42346357 | 42350997 | -1 | protein_coding | ENSG0000C |
| 42362282 | 42366535 | -1 | protein_coding | ENSG0000C |
| 42371837 | 42396091 | -1 | protein_coding | ENSG0000C |
| 42403447 | 42447684 | 1 | protein_coding | ENSG0000C |
| 42417497 | 42417593 | -1 | snRNA | ENSG0000C |
| 42472486 | 42496246 | -1 | protein_coding | ENSG0000C |
| 42496008 | 42581440 | 1 | protein_coding | ENSG0000C |
| 42496539 | 42497443 | 1 | lncRNA | ENSG0000C |
| 42508624 | 42509661 | 1 | lncRNA | ENSG0000C |
| 42560374 | 42561934 | -1 | lncRNA | ENSG0000C |
| 42599280 | 42615058 | -1 | lncRNA | ENSG0000C |
| 42648271 | 42651244 | -1 | lncRNA | ENSG0000C |
| 42653621 | 42775509 | 1 | protein_coding | ENSG0000C |
| 42733594 | 42741758 | -1 | lncRNA | ENSG0000C |
| 42777819 | 42779994 | -1 | lncRNA | ENSG0000C |
| 42781074 | 42782229 | -1 | lncRNA | ENSG0000C |
| 42831040 | 42836477 | -1 | lncRNA | ENSG0000C |
| 42843094 | 42879568 | -1 | protein_coding | ENSG0000C |

| | | | | |
|---|---|---|---|---|
| 42879644 | 42913304 | 1 | protein_coding | ENSG0000( |
| 42916803 | 42925646 | -1 | lncRNA | ENSG0000( |
| 42950928 | 42951014 | -1 | miRNA | ENSG0000( |
| 42964639 | 42965363 | -1 | lncRNA | ENSG0000( |
| 42974510 | 43033931 | 1 | protein_coding | ENSG0000( |
| 43053191 | 43076943 | -1 | protein_coding | ENSG0000( |
| 43092956 | 43107587 | -1 | protein_coding | ENSG0000( |
| 43107942 | 43168646 | 1 | lncRNA | ENSG0000( |
| 43115334 | 43115709 | -1 | processed_pseudoge | ENSG0000( |
| 43140523 | 43141092 | -1 | lncRNA | ENSG0000( |
| 43159066 | 43162277 | -1 | lncRNA | ENSG0000( |
| 43169008 | 43172805 | 1 | protein_coding | ENSG0000( |
| 43322417 | 43332039 | -1 | lncRNA | ENSG0000( |
| 43358147 | 43362349 | -1 | lncRNA | ENSG0000( |
| 43363332 | 43366566 | 1 | lncRNA | ENSG0000( |
| 43414483 | 43427131 | -1 | protein_coding | ENSG0000( |
| 43446601 | 43453902 | 1 | lncRNA | ENSG0000( |
| 43461960 | 43479534 | -1 | lncRNA | ENSG0000( |
| 43465309 | 43467298 | 1 | lncRNA | ENSG0000( |
| 43529186 | 43659488 | -1 | protein_coding | ENSG0000( |
| 43551229 | 43551600 | 1 | processed_pseudoge | ENSG0000( |
| 43565182 | 43565648 | 1 | protein_coding | ENSG0000( |
| 43609887 | 43609989 | -1 | miRNA | ENSG0000( |
| 43659560 | 43696079 | 1 | protein_coding | ENSG0000( |
| 43719094 | 43762307 | 1 | protein_coding | ENSG0000( |
| 43748365 | 43748468 | -1 | pseudogene | ENSG0000( |
| 43768320 | 43771666 | 1 | lncRNA | ENSG0000( |
| 43772511 | 43776445 | -1 | protein_coding | ENSG0000( |
| 43783123 | 43783573 | 1 | processed_pseudoge | ENSG0000( |
| 43789513 | 43805293 | 1 | protein_coding | ENSG0000( |
| 43805758 | 43812567 | -1 | lncRNA | ENSG0000( |
| 43855890 | 43856342 | -1 | processed_pseudoge | ENSG0000( |
| 43865223 | 43987592 | 1 | protein_coding | ENSG0000( |
| 43919795 | 43919901 | -1 | snRNA | ENSG0000( |
| 43996322 | 43996420 | -1 | snRNA | ENSG0000( |
| 44012309 | 44106552 | 1 | protein_coding | ENSG0000( |
| 44046347 | 44047041 | -1 | processed_pseudoge | ENSG0000( |
| 44107373 | 44131181 | 1 | protein_coding | ENSG0000( |
| 44133610 | 44210114 | 1 | protein_coding | ENSG0000( |
| 44158740 | 44160076 | -1 | lncRNA | ENSG0000( |
| 44172169 | 44194338 | -1 | lncRNA | ENSG0000( |
| 44175489 | 44176453 | 1 | lncRNA | ENSG0000( |
| 44200602 | 44207399 | -1 | lncRNA | ENSG0000( |
| 44217014 | 44240966 | -1 | protein_coding | ENSG0000( |
| 44241847 | 44242081 | 1 | lncRNA | ENSG0000( |
| 44244545 | 44244993 | 1 | lncRNA | ENSG0000( |
| 44246339 | 44262216 | -1 | protein_coding | ENSG0000( |
| 44250813 | 44251520 | 1 | lncRNA | ENSG0000( |
| 44285838 | 44298648 | 1 | protein_coding | ENSG0000( |
| 44300051 | 44327376 | 1 | protein_coding | ENSG0000( |

| | | | | |
|---|---|---|---|---|
| 44328944 | 44339402 | -1 | protein_coding | ENSG0000C |
| 44328944 | 44330221 | -1 | lncRNA | ENSG0000C |
| 44331234 | 44335851 | 1 | lncRNA | ENSG0000C |
| 44339376 | 44340470 | 1 | lncRNA | ENSG0000C |
| 44347767 | 44348295 | -1 | lncRNA | ENSG0000C |
| 44350163 | 44443081 | 1 | protein_coding | ENSG0000C |
| 44414588 | 44425272 | -1 | lncRNA | ENSG0000C |
| 44437121 | 44437175 | 1 | snoRNA | ENSG0000C |
| 44439035 | 44439110 | 1 | snoRNA | ENSG0000C |
| 44450986 | 44455284 | -1 | lncRNA | ENSG0000C |
| 44455510 | 44462196 | 1 | protein_coding | ENSG0000C |
| 44469929 | 44472516 | -1 | processed_pseudoge | ENSG0000C |
| 44472895 | 44473739 | 1 | processed_pseudoge | ENSG0000C |
| 44473723 | 44475097 | -1 | processed_pseudoge | ENSG0000C |
| 44477850 | 44478493 | 1 | lncRNA | ENSG0000C |
| 44485577 | 44490288 | 1 | lncRNA | ENSG0000C |
| 44494874 | 44495519 | 1 | lncRNA | ENSG0000C |
| 44497893 | 44711572 | -1 | protein_coding | ENSG0000C |
| 44506807 | 44516575 | 1 | lncRNA | ENSG0000C |
| 44517216 | 44525952 | 1 | lncRNA | ENSG0000C |
| 44530172 | 44533145 | 1 | lncRNA | ENSG0000C |
| 44538981 | 44540195 | -1 | protein_coding | ENSG0000C |
| 44550357 | 44551505 | -1 | protein_coding | ENSG0000C |
| 44557790 | 44558795 | -1 | protein_coding | ENSG0000C |
| 44573724 | 44638284 | 1 | protein_coding | ENSG0000C |
| 44579455 | 44580604 | -1 | protein_coding | ENSG0000C |
| 44591268 | 44592505 | -1 | protein_coding | ENSG0000C |
| 44600597 | 44602174 | 1 | protein_coding | ENSG0000C |
| 44612079 | 44612954 | 1 | protein_coding | ENSG0000C |
| 44627093 | 44628378 | 1 | protein_coding | ENSG0000C |
| 44637356 | 44638455 | 1 | protein_coding | ENSG0000C |
| 44646414 | 44647650 | 1 | protein_coding | ENSG0000C |
| 44654213 | 44654659 | -1 | protein_coding | ENSG0000C |
| 44657932 | 44658341 | 1 | protein_coding | ENSG0000C |
| 44666189 | 44666927 | -1 | protein_coding | ENSG0000C |
| 44675868 | 44678086 | -1 | processed_pseudoge | ENSG0000C |
| 44681576 | 44682163 | -1 | protein_coding | ENSG0000C |
| 44686358 | 44686629 | 1 | processed_pseudoge | ENSG0000C |
| 44697172 | 44698044 | 1 | protein_coding | ENSG0000C |
| 44702221 | 44702791 | 1 | unprocessed_pseudc | ENSG0000C |
| 44768580 | 44801826 | -1 | protein_coding | ENSG0000C |
| 44802577 | 44804717 | 1 | lncRNA | ENSG0000C |
| 44805617 | 44818779 | -1 | protein_coding | ENSG0000C |
| 44849585 | 44873903 | -1 | protein_coding | ENSG0000C |
| 44885953 | 44931989 | -1 | protein_coding | ENSG0000C |
| 44921051 | 44929678 | 1 | lncRNA | ENSG0000C |
| 44929653 | 44930112 | -1 | lncRNA | ENSG0000C |
| 44932814 | 44939937 | -1 | lncRNA | ENSG0000C |
| 44936303 | 44936954 | 1 | lncRNA | ENSG0000C |
| 44940012 | 44976989 | 1 | protein_coding | ENSG0000C |

| | | | | |
|---|---|---|---|---|
| 44978832 | 44979274 | 1 | lncRNA | ENSG00000 |
| 44989864 | 44994086 | -1 | lncRNA | ENSG00000 |
| 44994362 | 44995185 | -1 | lncRNA | ENSG00000 |
| 44999208 | 45004727 | -1 | lncRNA | ENSG00000 |
| 45070952 | 45074165 | -1 | transcribed_processe | ENSG00000 |
| 45073853 | 45226560 | 1 | protein_coding | ENSG00000 |
| 45100487 | 45101094 | -1 | lncRNA | ENSG00000 |
| 45234308 | 45264548 | 1 | lncRNA | ENSG00000 |
| 45263928 | 45287898 | -1 | protein_coding | ENSG00000 |
| 45288050 | 45297806 | 1 | lncRNA | ENSG00000 |
| 45299929 | 45302964 | 1 | lncRNA | ENSG00000 |
| 45300245 | 45305257 | -1 | lncRNA | ENSG00000 |
| 45336707 | 45338665 | 1 | lncRNA | ENSG00000 |
| 45338590 | 45341990 | -1 | lncRNA | ENSG00000 |
| 45376191 | 45376382 | -1 | processed_pseudoge | ENSG00000 |
| 45378201 | 45379635 | -1 | lncRNA | ENSG00000 |
| 45403809 | 45404369 | -1 | lncRNA | ENSG00000 |
| 45405165 | 45513720 | 1 | protein_coding | ENSG00000 |
| 45407386 | 45410065 | -1 | lncRNA | ENSG00000 |
| 45419716 | 45425070 | -1 | lncRNA | ENSG00000 |
| 45478266 | 45478326 | 1 | miRNA | ENSG00000 |
| 45493572 | 45573365 | -1 | protein_coding | ENSG00000 |
| 45593654 | 45603088 | 1 | lncRNA | ENSG00000 |
| 45596364 | 45625675 | -1 | lncRNA | ENSG00000 |
| 45607386 | 45609194 | 1 | lncRNA | ENSG00000 |
| 45643694 | 45942454 | 1 | protein_coding | ENSG00000 |
| 45759804 | 45763758 | -1 | lncRNA | ENSG00000 |
| 45827961 | 45837987 | -1 | lncRNA | ENSG00000 |
| 45861830 | 45888638 | -1 | lncRNA | ENSG00000 |
| 45870854 | 45873345 | 1 | lncRNA | ENSG00000 |
| 45914296 | 45919483 | 1 | lncRNA | ENSG00000 |
| 45974489 | 45974953 | 1 | lncRNA | ENSG00000 |
| 45981737 | 46005050 | 1 | protein_coding | ENSG00000 |
| 46037052 | 46039807 | 1 | lncRNA | ENSG00000 |
| 46052596 | 46053105 | 1 | lncRNA | ENSG00000 |
| 46056516 | 46057567 | 1 | lncRNA | ENSG00000 |
| 46072085 | 46072248 | -1 | processed_pseudoge | ENSG00000 |
| 46093264 | 46097530 | -1 | lncRNA | ENSG00000 |
| 46098097 | 46132849 | 1 | protein_coding | ENSG00000 |
| 46136262 | 46155579 | -1 | protein_coding | ENSG00000 |
| 46151614 | 46152647 | 1 | lncRNA | ENSG00000 |
| 46161148 | 46184476 | -1 | protein_coding | ENSG00000 |
| 46185079 | 46188941 | 1 | lncRNA | ENSG00000 |
| 46188141 | 46228824 | -1 | protein_coding | ENSG00000 |
| 46220269 | 46225364 | 1 | lncRNA | ENSG00000 |
| 46229196 | 46259390 | 1 | lncRNA | ENSG00000 |
| 46235133 | 46286297 | -1 | protein_coding | ENSG00000 |
| 46246890 | 46247682 | 1 | lncRNA | ENSG00000 |
| 46251549 | 46254133 | -1 | lncRNA | ENSG00000 |
| 46286342 | 46297751 | 1 | protein_coding | ENSG00000 |

| | | | | |
|---|---|---|---|---|
| 46300181 | 46323875 | -1 | protein_coding | ENSG00000 |
| 46324141 | 46445769 | 1 | protein_coding | ENSG00000 |
| 46326288 | 46326491 | 1 | protein_coding | ENSG00000 |
| 46420553 | 46421034 | -1 | processed_pseudoge | ENSG00000 |
| 46458891 | 46570015 | 1 | protein_coding | ENSG00000 |
| 46462471 | 46469306 | 1 | lncRNA | ENSG00000 |
| 46525618 | 46525722 | 1 | snRNA | ENSG00000 |
| 46598604 | 46605208 | -1 | protein_coding | ENSG00000 |
| 46635595 | 46665124 | 1 | protein_coding | ENSG00000 |
| 46653558 | 46654022 | -1 | processed_pseudoge | ENSG00000 |
| 46690764 | 46691226 | 1 | processed_pseudoge | ENSG00000 |
| 10736171 | 10736283 | -1 | snRNA | ENSG00000 |
| 10939388 | 10961338 | -1 | unprocessed_pseudo | ENSG00000 |
| 11066418 | 11068174 | 1 | lncRNA | ENSG00000 |
| 11124337 | 11125705 | 1 | processed_pseudoge | ENSG00000 |
| 11249809 | 11249959 | -1 | rRNA | ENSG00000 |
| 11474744 | 11479643 | 1 | unprocessed_pseudo | ENSG00000 |
| 11827523 | 11910358 | 1 | transcribed_unproce | ENSG00000 |
| 12602466 | 12626642 | 1 | unprocessed_pseudo | ENSG00000 |
| 15273855 | 15273961 | 1 | snRNA | ENSG00000 |
| 15282557 | 15288670 | -1 | lncRNA | ENSG00000 |
| 15290718 | 15297196 | 1 | unprocessed_pseudo | ENSG00000 |
| 15298378 | 15304556 | -1 | lncRNA | ENSG00000 |
| 15326037 | 15343065 | -1 | unprocessed_pseudo | ENSG00000 |
| 15349467 | 15350043 | 1 | processed_pseudoge | ENSG00000 |
| 15489990 | 15492564 | 1 | unprocessed_pseudo | ENSG00000 |
| 15528159 | 15529139 | 1 | protein_coding | ENSG00000 |
| 15541436 | 15578006 | 1 | unprocessed_pseudo | ENSG00000 |
| 15550904 | 15553838 | -1 | processed_pseudoge | ENSG00000 |
| 15557577 | 15560694 | -1 | lncRNA | ENSG00000 |
| 15562017 | 15564276 | -1 | processed_pseudoge | ENSG00000 |
| 15572089 | 15573265 | -1 | processed_pseudoge | ENSG00000 |
| 15588361 | 15588478 | 1 | unprocessed_pseudo | ENSG00000 |
| 15600908 | 15604882 | -1 | lncRNA | ENSG00000 |
| 15611759 | 15613096 | -1 | processed_pseudoge | ENSG00000 |
| 15615402 | 15615578 | -1 | unprocessed_pseudo | ENSG00000 |
| 15622601 | 15632051 | -1 | unprocessed_pseudo | ENSG00000 |
| 15635180 | 15644330 | -1 | unprocessed_pseudo | ENSG00000 |
| 15690026 | 15721631 | 1 | protein_coding | ENSG00000 |
| 15699361 | 15703403 | -1 | lncRNA | ENSG00000 |
| 15701459 | 15701565 | 1 | snRNA | ENSG00000 |
| 15721486 | 15722608 | 1 | processed_pseudoge | ENSG00000 |
| 15746630 | 15778297 | 1 | lncRNA | ENSG00000 |
| 15749156 | 15750825 | -1 | processed_pseudoge | ENSG00000 |
| 15784959 | 15829984 | 1 | lncRNA | ENSG00000 |
| 15794901 | 15797909 | 1 | transcribed_unproce | ENSG00000 |
| 15805263 | 15820884 | -1 | transcribed_unproce | ENSG00000 |
| 15823197 | 15823890 | 1 | lncRNA | ENSG00000 |
| 15826566 | 15827187 | 1 | transcribed_processe | ENSG00000 |
| 15852990 | 15893942 | 1 | unprocessed_pseudo | ENSG00000 |

| | | | | |
|---|---|---|---|---|
| 15854195 | 15855243 | -1 | processed_pseudoge | ENSG00000 |
| 15901791 | 15901911 | 1 | unprocessed_pseudc | ENSG00000 |
| 15914721 | 15915800 | -1 | lncRNA | ENSG00000 |
| 16141767 | 16141872 | 1 | snRNA | ENSG00000 |
| 16387694 | 16390887 | 1 | unprocessed_pseudc | ENSG00000 |
| 16405330 | 16405417 | 1 | processed_pseudoge | ENSG00000 |
| 16423943 | 16426031 | 1 | unprocessed_pseudc | ENSG00000 |
| 16428947 | 16476870 | -1 | unprocessed_pseudc | ENSG00000 |
| 16486985 | 16488322 | 1 | processed_pseudoge | ENSG00000 |
| 16503304 | 16509475 | 1 | unprocessed_pseudc | ENSG00000 |
| 16521059 | 16527237 | 1 | lncRNA | ENSG00000 |
| 16572027 | 16574637 | 1 | unprocessed_pseudc | ENSG00000 |
| 16575363 | 16582707 | -1 | transcribed_unproce | ENSG00000 |
| 16590751 | 16592810 | -1 | protein_coding | ENSG00000 |
| 16592895 | 16600110 | 1 | lncRNA | ENSG00000 |
| 16595089 | 16595485 | -1 | processed_pseudoge | ENSG00000 |
| 16601887 | 16698742 | 1 | transcribed_unproce | ENSG00000 |
| 16601911 | 16615111 | 1 | lncRNA | ENSG00000 |
| 16616170 | 16617114 | 1 | lncRNA | ENSG00000 |
| 16645695 | 16646581 | -1 | unprocessed_pseudc | ENSG00000 |
| 16653892 | 16659628 | -1 | transcribed_unproce | ENSG00000 |
| 16654066 | 16675540 | -1 | lncRNA | ENSG00000 |
| 16671090 | 16679493 | -1 | transcribed_unproce | ENSG00000 |
| 16676016 | 16694648 | -1 | lncRNA | ENSG00000 |
| 16690103 | 16704477 | -1 | transcribed_unproce | ENSG00000 |
| 16725072 | 16747761 | 1 | lncRNA | ENSG00000 |
| 16746628 | 16748645 | -1 | lncRNA | ENSG00000 |
| 16783412 | 16821699 | -1 | protein_coding | ENSG00000 |
| 16827474 | 16829335 | 1 | transcribed_unproce | ENSG00000 |
| 16837522 | 16837666 | 1 | processed_pseudoge | ENSG00000 |
| 16844399 | 16846357 | 1 | unprocessed_pseudc | ENSG00000 |
| 16855804 | 16856162 | 1 | processed_pseudoge | ENSG00000 |
| 16857619 | 16860718 | -1 | processed_pseudoge | ENSG00000 |
| 16869478 | 16871126 | 1 | lncRNA | ENSG00000 |
| 16876218 | 16876421 | -1 | processed_pseudoge | ENSG00000 |
| 16876552 | 16876902 | -1 | processed_pseudoge | ENSG00000 |
| 16897445 | 16897546 | 1 | processed_pseudoge | ENSG00000 |
| 16904219 | 16904696 | -1 | IG_V_pseudogene | ENSG00000 |
| 16914235 | 16914981 | -1 | IG_V_pseudogene | ENSG00000 |
| 16921443 | 16922173 | -1 | IG_V_pseudogene | ENSG00000 |
| 16925952 | 16926444 | -1 | IG_V_pseudogene | ENSG00000 |
| 16933521 | 16934003 | -1 | IG_V_pseudogene | ENSG00000 |
| 16953895 | 16956808 | 1 | unprocessed_pseudc | ENSG00000 |
| 16961936 | 17008222 | -1 | protein_coding | ENSG00000 |
| 17012404 | 17012932 | 1 | processed_pseudoge | ENSG00000 |
| 17021398 | 17022570 | -1 | processed_pseudoge | ENSG00000 |
| 17036570 | 17060825 | 1 | transcribed_unproce | ENSG00000 |
| 17067821 | 17070675 | 1 | lncRNA | ENSG00000 |
| 17080701 | 17081456 | 1 | lncRNA | ENSG00000 |
| 17084954 | 17115694 | 1 | protein_coding | ENSG00000 |

| | | | | |
|---|---|---|---|---|
| 17093612 | 17093939 | -1 | processed_pseudoge | ENSG00000 |
| 17116297 | 17121367 | -1 | protein_coding | ENSG00000 |
| 17121560 | 17132104 | 1 | lncRNA | ENSG00000 |
| 17137511 | 17165287 | -1 | protein_coding | ENSG00000 |
| 17159384 | 17165445 | 1 | lncRNA | ENSG00000 |
| 17178790 | 17258235 | -1 | protein_coding | ENSG00000 |
| 17192936 | 17193308 | 1 | processed_pseudoge | ENSG00000 |
| 17213516 | 17214155 | 1 | processed_pseudoge | ENSG00000 |
| 17256859 | 17266733 | -1 | lncRNA | ENSG00000 |
| 17329034 | 17329232 | 1 | TEC | ENSG00000 |
| 17352881 | 17353177 | -1 | misc_RNA | ENSG00000 |
| 17359949 | 17558149 | 1 | protein_coding | ENSG00000 |
| 17418697 | 17419828 | 1 | processed_pseudoge | ENSG00000 |
| 17424388 | 17424775 | -1 | processed_pseudoge | ENSG00000 |
| 17518676 | 17519029 | -1 | processed_pseudoge | ENSG00000 |
| 17521514 | 17570078 | -1 | lncRNA | ENSG00000 |
| 17563450 | 17590995 | 1 | protein_coding | ENSG00000 |
| 17580157 | 17589192 | -1 | lncRNA | ENSG00000 |
| 17592136 | 17628749 | -1 | protein_coding | ENSG00000 |
| 17628855 | 17730855 | 1 | protein_coding | ENSG00000 |
| 17734138 | 17774770 | -1 | protein_coding | ENSG00000 |
| 17764180 | 17764259 | -1 | miRNA | ENSG00000 |
| 17777322 | 17779481 | 1 | lncRNA | ENSG00000 |
| 17787649 | 18024561 | -1 | protein_coding | ENSG00000 |
| 17787652 | 17811497 | -1 | lncRNA | ENSG00000 |
| 17800722 | 17802035 | 1 | lncRNA | ENSG00000 |
| 17980868 | 17980961 | -1 | miRNA | ENSG00000 |
| 17983205 | 17983619 | -1 | lncRNA | ENSG00000 |
| 18004270 | 18007308 | -1 | lncRNA | ENSG00000 |
| 18019214 | 18020200 | 1 | processed_pseudoge | ENSG00000 |
| 18029385 | 18037968 | 1 | lncRNA | ENSG00000 |
| 18076527 | 18078884 | -1 | lncRNA | ENSG00000 |
| 18077920 | 18131138 | 1 | protein_coding | ENSG00000 |
| 18081466 | 18081675 | -1 | processed_pseudoge | ENSG00000 |
| 18101612 | 18101888 | 1 | unprocessed_pseudc | ENSG00000 |
| 18102457 | 18103649 | -1 | processed_pseudoge | ENSG00000 |
| 18110331 | 18146554 | 1 | protein_coding | ENSG00000 |
| 18110759 | 18131154 | -1 | lncRNA | ENSG00000 |
| 18150170 | 18177397 | 1 | protein_coding | ENSG00000 |
| 18178038 | 18205915 | -1 | lncRNA | ENSG00000 |
| 18340145 | 18345899 | 1 | unprocessed_pseudc | ENSG00000 |
| 18361820 | 18391105 | -1 | lncRNA | ENSG00000 |
| 18374106 | 18375689 | 1 | lncRNA | ENSG00000 |
| 18400407 | 18404206 | -1 | unprocessed_pseudc | ENSG00000 |
| 18422244 | 18500594 | 1 | lncRNA | ENSG00000 |
| 18486999 | 18491247 | -1 | lncRNA | ENSG00000 |
| 18501794 | 18512187 | -1 | lncRNA | ENSG00000 |
| 18516344 | 18518161 | -1 | protein_coding | ENSG00000 |
| 18524620 | 18524935 | 1 | misc_RNA | ENSG00000 |
| 18527157 | 18527626 | 1 | processed_pseudoge | ENSG00000 |

| | | | | |
|---|---|---|---|---|
| 18527802 | 18530573 | 1 | protein_coding | ENSG00000 |
| 18533646 | 18577968 | -1 | transcribed_unproce | ENSG00000 |
| 18605355 | 18605497 | 1 | scaRNA | ENSG00000 |
| 18605802 | 18606102 | -1 | misc_RNA | ENSG00000 |
| 18605815 | 18611919 | -1 | protein_coding | ENSG00000 |
| 18615581 | 18615900 | 1 | misc_RNA | ENSG00000 |
| 18623339 | 18625289 | -1 | unprocessed_pseudo | ENSG00000 |
| 18633984 | 18634682 | 1 | protein_coding | ENSG00000 |
| 18636445 | 18638405 | -1 | unprocessed_pseudo | ENSG00000 |
| 18650023 | 18653617 | -1 | processed_pseudoge | ENSG00000 |
| 18715815 | 18719341 | 1 | processed_pseudoge | ENSG00000 |
| 18733914 | 18757906 | 1 | lncRNA | ENSG00000 |
| 18773665 | 18843399 | -1 | transcribed_unproce | ENSG00000 |
| 18794414 | 18794804 | -1 | transcribed_processe | ENSG00000 |
| 18833694 | 18834438 | 1 | processed_pseudoge | ENSG00000 |
| 18846811 | 18861049 | 1 | transcribed_processe | ENSG00000 |
| 18855621 | 18858640 | -1 | unprocessed_pseudo | ENSG00000 |
| 18873543 | 18894407 | -1 | lncRNA | ENSG00000 |
| 18883502 | 18884682 | -1 | lncRNA | ENSG00000 |
| 18906028 | 18914238 | 1 | protein_coding | ENSG00000 |
| 18906238 | 18947732 | 1 | protein_coding | ENSG00000 |
| 18912777 | 18936553 | -1 | protein_coding | ENSG00000 |
| 18936411 | 18947741 | 1 | lncRNA | ENSG00000 |
| 18951934 | 18959512 | -1 | lncRNA | ENSG00000 |
| 18970514 | 18994628 | 1 | transcribed_unproce | ENSG00000 |
| 18970525 | 19031242 | 1 | lncRNA | ENSG00000 |
| 18980695 | 18981325 | -1 | unprocessed_pseudo | ENSG00000 |
| 18997138 | 18997595 | -1 | processed_pseudoge | ENSG00000 |
| 19014353 | 19014506 | 1 | transcribed_unproce | ENSG00000 |
| 19018043 | 19018916 | -1 | processed_pseudoge | ENSG00000 |
| 19023056 | 19023550 | 1 | lncRNA | ENSG00000 |
| 19029524 | 19031242 | 1 | polymorphic_pseudo | ENSG00000 |
| 19031564 | 19034564 | -1 | unitary_pseudogene | ENSG00000 |
| 19036282 | 19122454 | -1 | protein_coding | ENSG00000 |
| 19045256 | 19045367 | 1 | misc_RNA | ENSG00000 |
| 19046162 | 19048375 | -1 | lncRNA | ENSG00000 |
| 19055801 | 19056512 | 1 | processed_pseudoge | ENSG00000 |
| 19061041 | 19061843 | -1 | lncRNA | ENSG00000 |
| 19121529 | 19124503 | 1 | lncRNA | ENSG00000 |
| 19124309 | 19128449 | -1 | lncRNA | ENSG00000 |
| 19124879 | 19125646 | 1 | unitary_pseudogene | ENSG00000 |
| 19130279 | 19144684 | -1 | protein_coding | ENSG00000 |
| 19131308 | 19132622 | 1 | protein_coding | ENSG00000 |
| 19148576 | 19150283 | -1 | protein_coding | ENSG00000 |
| 19171395 | 19172839 | 1 | lncRNA | ENSG00000 |
| 19175581 | 19178739 | -1 | protein_coding | ENSG00000 |
| 19179473 | 19291719 | -1 | protein_coding | ENSG00000 |
| 19196402 | 19200083 | 1 | lncRNA | ENSG00000 |
| 19238443 | 19238739 | -1 | processed_pseudoge | ENSG00000 |
| 19249873 | 19249966 | 1 | snoRNA | ENSG00000 |

| | | | | |
|---|---|---|---|---|
| 19257520 | 19258806 | 1 | processed_pseudoge | ENSG0000C |
| 19291969 | 19328901 | 1 | lncRNA | ENSG0000C |
| 19330698 | 19447450 | -1 | protein_coding | ENSG0000C |
| 19351368 | 19448232 | -1 | protein_coding | ENSG0000C |
| 19365766 | 19366044 | 1 | misc_RNA | ENSG0000C |
| 19431902 | 19436075 | 1 | protein_coding | ENSG0000C |
| 19447893 | 19450105 | 1 | lncRNA | ENSG0000C |
| 19449911 | 19479202 | -1 | protein_coding | ENSG0000C |
| 19454179 | 19454605 | 1 | lncRNA | ENSG0000C |
| 19456503 | 19456962 | 1 | lncRNA | ENSG0000C |
| 19479457 | 19520612 | 1 | protein_coding | ENSG0000C |
| 19520572 | 19522769 | -1 | lncRNA | ENSG0000C |
| 19523024 | 19527545 | -1 | protein_coding | ENSG0000C |
| 19565203 | 19566839 | -1 | lncRNA | ENSG0000C |
| 19630750 | 19633574 | -1 | processed_pseudoge | ENSG0000C |
| 19667023 | 19667555 | 1 | lncRNA | ENSG0000C |
| 19714503 | 19724224 | 1 | protein_coding | ENSG0000C |
| 19717220 | 19724772 | 1 | protein_coding | ENSG0000C |
| 19722945 | 19724771 | 1 | protein_coding | ENSG0000C |
| 19756703 | 19783593 | 1 | protein_coding | ENSG0000C |
| 19783223 | 19854939 | -1 | protein_coding | ENSG0000C |
| 19792294 | 19793094 | 1 | processed_pseudoge | ENSG0000C |
| 19846138 | 19854896 | -1 | protein_coding | ENSG0000C |
| 19875517 | 19941820 | -1 | protein_coding | ENSG0000C |
| 19887289 | 19887970 | 1 | processed_pseudoge | ENSG0000C |
| 19941607 | 19969975 | 1 | protein_coding | ENSG0000C |
| 19963753 | 19963834 | 1 | miRNA | ENSG0000C |
| 19969896 | 20016823 | -1 | protein_coding | ENSG0000C |
| 20017014 | 20067164 | 1 | protein_coding | ENSG0000C |
| 20033139 | 20033220 | 1 | miRNA | ENSG0000C |
| 20058030 | 20070569 | -1 | lncRNA | ENSG0000C |
| 20064552 | 20065705 | -1 | lncRNA | ENSG0000C |
| 20070716 | 20070965 | 1 | misc_RNA | ENSG0000C |
| 20080232 | 20111877 | 1 | protein_coding | ENSG0000C |
| 20085746 | 20085833 | 1 | miRNA | ENSG0000C |
| 20086058 | 20086142 | 1 | miRNA | ENSG0000C |
| 20110821 | 20111875 | -1 | lncRNA | ENSG0000C |
| 20111875 | 20117392 | -1 | protein_coding | ENSG0000C |
| 20114686 | 20114751 | -1 | miRNA | ENSG0000C |
| 20115938 | 20127355 | 1 | protein_coding | ENSG0000C |
| 20126402 | 20126526 | 1 | snoRNA | ENSG0000C |
| 20129456 | 20148007 | 1 | protein_coding | ENSG0000C |
| 20148416 | 20151055 | -1 | protein_coding | ENSG0000C |
| 20198729 | 20204918 | -1 | lncRNA | ENSG0000C |
| 20206397 | 20208524 | 1 | lncRNA | ENSG0000C |
| 20241415 | 20283246 | -1 | protein_coding | ENSG0000C |
| 20249134 | 20249211 | -1 | miRNA | ENSG0000C |
| 20299760 | 20313216 | 1 | unprocessed_pseudc | ENSG0000C |
| 20314238 | 20320080 | -1 | protein_coding | ENSG0000C |
| 20318119 | 20318749 | -1 | lncRNA | ENSG0000C |

| | | | | |
|---|---|---|---|---|
| 20320739 | 20321203 | 1 | lncRNA | ENSG0000C |
| 20338805 | 20354372 | 1 | lncRNA | ENSG0000C |
| 20343203 | 20344365 | -1 | lncRNA | ENSG0000C |
| 20348758 | 20354387 | -1 | lncRNA | ENSG0000C |
| 20350578 | 20390758 | -1 | protein_coding | ENSG0000C |
| 20394115 | 20408461 | 1 | protein_coding | ENSG0000C |
| 20416797 | 20416899 | -1 | snRNA | ENSG0000C |
| 20424815 | 20437826 | -1 | protein_coding | ENSG0000C |
| 20429241 | 20446620 | -1 | protein_coding | ENSG0000C |
| 20441519 | 20495844 | -1 | protein_coding | ENSG0000C |
| 20450122 | 20451824 | 1 | processed_pseudoge | ENSG0000C |
| 20475203 | 20475311 | -1 | misc_RNA | ENSG0000C |
| 20481158 | 20481268 | 1 | misc_RNA | ENSG0000C |
| 20481935 | 20482213 | -1 | misc_RNA | ENSG0000C |
| 20482815 | 20484105 | -1 | processed_pseudoge | ENSG0000C |
| 20495913 | 20587632 | 1 | protein_coding | ENSG0000C |
| 20522070 | 20523870 | -1 | lncRNA | ENSG0000C |
| 20587496 | 20592218 | -1 | unprocessed_pseudc | ENSG0000C |
| 20602595 | 20656914 | -1 | transcribed_unproce | ENSG0000C |
| 20632619 | 20632751 | 1 | unprocessed_pseudc | ENSG0000C |
| 20652730 | 20654750 | 1 | processed_pseudoge | ENSG0000C |
| 20667836 | 20670984 | -1 | processed_pseudoge | ENSG0000C |
| 20689929 | 20698938 | 1 | transcribed_unproce | ENSG0000C |
| 20697962 | 20698934 | -1 | unprocessed_pseudc | ENSG0000C |
| 20701114 | 20704606 | 1 | transcribed_unproce | ENSG0000C |
| 20707691 | 20859417 | -1 | protein_coding | ENSG0000C |
| 20774113 | 20787720 | 1 | protein_coding | ENSG0000C |
| 20859007 | 20891214 | 1 | protein_coding | ENSG0000C |
| 20889206 | 20891214 | -1 | lncRNA | ENSG0000C |
| 20917407 | 20953747 | 1 | protein_coding | ENSG0000C |
| 20957092 | 20964679 | 1 | lncRNA | ENSG0000C |
| 20965108 | 20981360 | 1 | protein_coding | ENSG0000C |
| 20979462 | 20998121 | 1 | lncRNA | ENSG0000C |
| 20981361 | 20981755 | -1 | lncRNA | ENSG0000C |
| 20982269 | 20999032 | 1 | protein_coding | ENSG0000C |
| 20999104 | 21002196 | -1 | protein_coding | ENSG0000C |
| 21001886 | 21010342 | 1 | lncRNA | ENSG0000C |
| 21002895 | 21009453 | -1 | transcribed_unproce | ENSG0000C |
| 21008193 | 21014292 | -1 | lncRNA | ENSG0000C |
| 21009808 | 21028830 | 1 | protein_coding | ENSG0000C |
| 21011384 | 21014217 | -1 | transcribed_unproce | ENSG0000C |
| 21028718 | 21032840 | -1 | protein_coding | ENSG0000C |
| 21031354 | 21044117 | 1 | lncRNA | ENSG0000C |
| 21034176 | 21034272 | -1 | miRNA | ENSG0000C |
| 21035243 | 21045017 | -1 | transcribed_unproce | ENSG0000C |
| 21045960 | 21064168 | 1 | protein_coding | ENSG0000C |
| 21065462 | 21070097 | 1 | unprocessed_pseudc | ENSG0000C |
| 21071082 | 21081018 | -1 | unprocessed_pseudc | ENSG0000C |
| 21103016 | 21122285 | 1 | transcribed_unproce | ENSG0000C |
| 21114607 | 21124451 | -1 | lncRNA | ENSG0000C |

| | | | | |
|---|---|---|---|---|
| 21125660 | 21128063 | -1 | processed_pseudoge | ENSG00000( |
| 21141624 | 21142371 | -1 | processed_pseudoge | ENSG00000( |
| 21167758 | 21192156 | 1 | lncRNA | ENSG00000( |
| 21177892 | 21179875 | -1 | lncRNA | ENSG00000( |
| 21193360 | 21203755 | -1 | transcribed_unproce | ENSG00000( |
| 21207973 | 21227637 | -1 | protein_coding | ENSG00000( |
| 21228760 | 21229138 | -1 | unprocessed_pseudc | ENSG00000( |
| 21268161 | 21268903 | 1 | processed_pseudoge | ENSG00000( |
| 21282881 | 21284161 | 1 | processed_pseudoge | ENSG00000( |
| 21290760 | 21294586 | -1 | unprocessed_pseudc | ENSG00000( |
| 21300990 | 21325042 | -1 | lncRNA | ENSG00000( |
| 21312920 | 21314904 | 1 | lncRNA | ENSG00000( |
| 21341595 | 21345258 | 1 | processed_pseudoge | ENSG00000( |
| 21354563 | 21358067 | 1 | lncRNA | ENSG00000( |
| 21360601 | 21361299 | -1 | protein_coding | ENSG00000( |
| 21370064 | 21371945 | 1 | processed_pseudoge | ENSG00000( |
| 21379382 | 21379701 | -1 | misc_RNA | ENSG00000( |
| 21383374 | 21389478 | 1 | protein_coding | ENSG00000( |
| 21389191 | 21389491 | 1 | misc_RNA | ENSG00000( |
| 21389796 | 21389938 | -1 | scaRNA | ENSG00000( |
| 21396273 | 21396590 | -1 | misc_RNA | ENSG00000( |
| 21417371 | 21451463 | 1 | protein_coding | ENSG00000( |
| 21463963 | 21464279 | 1 | misc_RNA | ENSG00000( |
| 21466423 | 21471269 | 1 | protein_coding | ENSG00000( |
| 21473000 | 21517533 | -1 | transcribed_unitary_ | ENSG00000( |
| 21544897 | 21545039 | 1 | scaRNA | ENSG00000( |
| 21545344 | 21545644 | -1 | misc_RNA | ENSG00000( |
| 21545357 | 21551461 | -1 | protein_coding | ENSG00000( |
| 21549447 | 21624034 | 1 | protein_coding | ENSG00000( |
| 21555138 | 21555457 | 1 | misc_RNA | ENSG00000( |
| 21628089 | 21630064 | -1 | protein_coding | ENSG00000( |
| 21632716 | 21637329 | 1 | protein_coding | ENSG00000( |
| 21640844 | 21641284 | 1 | lncRNA | ENSG00000( |
| 21642302 | 21644299 | 1 | protein_coding | ENSG00000( |
| 21652270 | 21670237 | 1 | lncRNA | ENSG00000( |
| 21652981 | 21653058 | 1 | miRNA | ENSG00000( |
| 21653304 | 21653385 | 1 | miRNA | ENSG00000( |
| 21657811 | 21661021 | 1 | transcribed_unitary_ | ENSG00000( |
| 21661242 | 21661295 | -1 | unitary_pseudogene | ENSG00000( |
| 21661476 | 21661593 | -1 | unitary_pseudogene | ENSG00000( |
| 21661859 | 21661929 | -1 | unitary_pseudogene | ENSG00000( |
| 21661934 | 21662363 | 1 | lncRNA | ENSG00000( |
| 21666009 | 21700015 | 1 | protein_coding | ENSG00000( |
| 21697536 | 21735794 | -1 | protein_coding | ENSG00000( |
| 21712761 | 21714013 | 1 | lncRNA | ENSG00000( |
| 21722895 | 21723182 | -1 | misc_RNA | ENSG00000( |
| 21737448 | 21744108 | -1 | lncRNA | ENSG00000( |
| 21754500 | 21867680 | -1 | protein_coding | ENSG00000( |
| 21792434 | 21792524 | 1 | rRNA_pseudogene | ENSG00000( |
| 21885282 | 21885673 | -1 | processed_pseudoge | ENSG00000( |

| | | | | |
|---|---|---|---|---|
| 21919425 | 21952848 | -1 | protein_coding | ENSG0000C |
| 21938269 | 21977632 | 1 | lncRNA | ENSG0000C |
| 21957025 | 21982816 | -1 | protein_coding | ENSG0000C |
| 21991099 | 22043934 | -1 | transcribed_unproce | ENSG0000C |
| 22026076 | 22026593 | 1 | IG_V_pseudogene | ENSG0000C |
| 22030934 | 22031472 | 1 | IG_V_gene | ENSG0000C |
| 22032745 | 22033194 | 1 | IG_V_pseudogene | ENSG0000C |
| 22036344 | 22040065 | 1 | lncRNA | ENSG0000C |
| 22038643 | 22039126 | 1 | unprocessed_pseudc | ENSG0000C |
| 22038643 | 22215270 | 1 | IG_V_gene | ENSG0000C |
| 22043618 | 22043942 | 1 | IG_V_pseudogene | ENSG0000C |
| 22046430 | 22046591 | 1 | unprocessed_pseudc | ENSG0000C |
| 22058433 | 22058892 | 1 | IG_V_pseudogene | ENSG0000C |
| 22061077 | 22061538 | 1 | IG_V_pseudogene | ENSG0000C |
| 22070225 | 22070707 | 1 | IG_V_pseudogene | ENSG0000C |
| 22075366 | 22075666 | 1 | IG_V_pseudogene | ENSG0000C |
| 22079770 | 22080263 | 1 | IG_V_pseudogene | ENSG0000C |
| 22086561 | 22087110 | 1 | IG_V_pseudogene | ENSG0000C |
| 22087924 | 22088085 | 1 | unprocessed_pseudc | ENSG0000C |
| 22098700 | 22099212 | 1 | IG_V_gene | ENSG0000C |
| 22103219 | 22104857 | 1 | unprocessed_pseudc | ENSG0000C |
| 22114844 | 22118005 | -1 | processed_pseudoge | ENSG0000C |
| 22155582 | 22155942 | 1 | unprocessed_pseudc | ENSG0000C |
| 22157382 | 22157631 | 1 | unprocessed_pseudc | ENSG0000C |
| 22162199 | 22162681 | 1 | IG_V_gene | ENSG0000C |
| 22168289 | 22168634 | 1 | unprocessed_pseudc | ENSG0000C |
| 22170983 | 22171556 | -1 | processed_pseudoge | ENSG0000C |
| 22179228 | 22179477 | 1 | IG_V_pseudogene | ENSG0000C |
| 22182582 | 22182885 | 1 | IG_V_pseudogene | ENSG0000C |
| 22185941 | 22186619 | 1 | processed_pseudoge | ENSG0000C |
| 22195799 | 22196276 | 1 | IG_V_gene | ENSG0000C |
| 22198560 | 22199011 | 1 | IG_V_pseudogene | ENSG0000C |
| 22201663 | 22202161 | 1 | IG_V_gene | ENSG0000C |
| 22219647 | 22220133 | 1 | IG_V_pseudogene | ENSG0000C |
| 22223187 | 22224566 | -1 | processed_pseudoge | ENSG0000C |
| 22228719 | 22230469 | -1 | processed_pseudoge | ENSG0000C |
| 22244780 | 22245515 | 1 | protein_coding | ENSG0000C |
| 22264601 | 22273020 | 1 | lncRNA | ENSG0000C |
| 22283928 | 22287220 | -1 | lncRNA | ENSG0000C |
| 22293733 | 22294794 | 1 | lncRNA | ENSG0000C |
| 22298124 | 22329122 | 1 | lncRNA | ENSG0000C |
| 22303224 | 22310401 | 1 | transcribed_unproce | ENSG0000C |
| 22318727 | 22319226 | 1 | IG_V_gene | ENSG0000C |
| 22322472 | 22322969 | 1 | IG_V_gene | ENSG0000C |
| 22327300 | 22327814 | 1 | IG_V_gene | ENSG0000C |
| 22339573 | 22341214 | 1 | unprocessed_pseudc | ENSG0000C |
| 22343187 | 22343732 | 1 | IG_V_gene | ENSG0000C |
| 22347807 | 22348164 | 1 | unprocessed_pseudc | ENSG0000C |
| 22352940 | 22353433 | 1 | IG_V_gene | ENSG0000C |
| 22357739 | 22358260 | 1 | IG_V_gene | ENSG0000C |

| | | | | |
|---|---|---|---|---|
| 22366210 | 22367770 | 1 | processed_pseudoge | ENSG0000C |
| 22369614 | 22370087 | 1 | IG_V_gene | ENSG0000C |
| 22370829 | 22371120 | 1 | unprocessed_pseudc | ENSG0000C |
| 22375986 | 22376505 | 1 | IG_V_gene | ENSG0000C |
| 22380766 | 22381347 | 1 | IG_V_gene | ENSG0000C |
| 22395018 | 22395489 | 1 | IG_V_gene | ENSG0000C |
| 22397032 | 22397524 | 1 | IG_V_pseudogene | ENSG0000C |
| 22398849 | 22398999 | 1 | IG_V_pseudogene | ENSG0000C |
| 22404207 | 22404721 | 1 | IG_V_pseudogene | ENSG0000C |
| 22409766 | 22410282 | 1 | IG_V_gene | ENSG0000C |
| 22420326 | 22421310 | 1 | processed_pseudoge | ENSG0000C |
| 22425821 | 22426314 | 1 | IG_V_pseudogene | ENSG0000C |
| 22427540 | 22428035 | 1 | IG_V_gene | ENSG0000C |
| 22431958 | 22432465 | 1 | IG_V_gene | ENSG0000C |
| 22446686 | 22448336 | 1 | processed_pseudoge | ENSG0000C |
| 22450202 | 22450652 | 1 | IG_V_pseudogene | ENSG0000C |
| 22465659 | 22465986 | 1 | unprocessed_pseudc | ENSG0000C |
| 22484421 | 22508742 | -1 | protein_coding | ENSG0000C |
| 22513736 | 22520270 | -1 | protein_coding | ENSG0000C |
| 22547701 | 22559361 | -1 | protein_coding | ENSG0000C |
| 22559343 | 22566602 | 1 | lncRNA | ENSG0000C |
| 22580014 | 22580296 | 1 | IG_V_pseudogene | ENSG0000C |
| 22588155 | 22588688 | 1 | IG_V_gene | ENSG0000C |
| 22594528 | 22595056 | 1 | IG_V_gene | ENSG0000C |
| 22604445 | 22605165 | 1 | IG_V_pseudogene | ENSG0000C |
| 22618572 | 22619163 | 1 | IG_V_pseudogene | ENSG0000C |
| 22630065 | 22636153 | 1 | unprocessed_pseudc | ENSG0000C |
| 22630546 | 22633849 | -1 | lncRNA | ENSG0000C |
| 22639177 | 22644473 | -1 | transcribed_unproce | ENSG0000C |
| 22644475 | 22647903 | 1 | protein_coding | ENSG0000C |
| 22661299 | 22661542 | 1 | IG_V_pseudogene | ENSG0000C |
| 22664473 | 22664907 | 1 | IG_V_pseudogene | ENSG0000C |
| 22668288 | 22668806 | 1 | IG_V_gene | ENSG0000C |
| 22673152 | 22673602 | 1 | IG_V_pseudogene | ENSG0000C |
| 22679097 | 22679345 | 1 | IG_V_pseudogene | ENSG0000C |
| 22686726 | 22687271 | 1 | IG_V_gene | ENSG0000C |
| 22689831 | 22689978 | 1 | unprocessed_pseudc | ENSG0000C |
| 22692778 | 22693312 | -1 | lncRNA | ENSG0000C |
| 22694439 | 22694952 | 1 | IG_V_pseudogene | ENSG0000C |
| 22697789 | 22698407 | 1 | IG_V_gene | ENSG0000C |
| 22700760 | 22700970 | 1 | IG_V_pseudogene | ENSG0000C |
| 22704265 | 22704822 | 1 | IG_V_gene | ENSG0000C |
| 22711689 | 22713203 | 1 | IG_V_gene | ENSG0000C |
| 22715290 | 22715419 | 1 | IG_V_pseudogene | ENSG0000C |
| 22720623 | 22721145 | 1 | IG_V_gene | ENSG0000C |
| 22730725 | 22731090 | -1 | processed_pseudoge | ENSG0000C |
| 22734607 | 22735089 | 1 | IG_V_gene | ENSG0000C |
| 22738944 | 22739187 | 1 | IG_V_pseudogene | ENSG0000C |
| 22747383 | 22747921 | 1 | IG_V_gene | ENSG0000C |
| 22755554 | 22755797 | 1 | IG_V_pseudogene | ENSG0000C |

| | | | | |
|---|---|---|---|---|
| 22758700 | 22759218 | 1 | IG_V_gene | ENSG00000 |
| 22762294 | 22762516 | 1 | IG_V_pseudogene | ENSG00000 |
| 22771824 | 22772582 | 1 | IG_V_gene | ENSG00000 |
| 22774529 | 22774691 | 1 | unprocessed_pseudo | ENSG00000 |
| 22785643 | 22787268 | -1 | processed_pseudoge | ENSG00000 |
| 22792491 | 22793007 | 1 | IG_V_gene | ENSG00000 |
| 22803076 | 22803360 | 1 | unprocessed_pseudo | ENSG00000 |
| 22811747 | 22812281 | 1 | IG_V_gene | ENSG00000 |
| 22814671 | 22814926 | 1 | unprocessed_pseudo | ENSG00000 |
| 22819010 | 22819756 | 1 | IG_V_gene | ENSG00000 |
| 22822658 | 22823289 | 1 | IG_V_gene | ENSG00000 |
| 22822776 | 22822871 | 1 | miRNA | ENSG00000 |
| 22834865 | 22835117 | 1 | unprocessed_pseudo | ENSG00000 |
| 22838602 | 22838872 | 1 | IG_V_pseudogene | ENSG00000 |
| 22850375 | 22850624 | 1 | IG_V_pseudogene | ENSG00000 |
| 22856762 | 22857038 | 1 | IG_V_pseudogene | ENSG00000 |
| 22858025 | 22858268 | 1 | IG_V_pseudogene | ENSG00000 |
| 22871507 | 22872035 | 1 | IG_V_gene | ENSG00000 |
| 22873212 | 22873440 | 1 | IG_V_pseudogene | ENSG00000 |
| 22880706 | 22881396 | 1 | IG_V_gene | ENSG00000 |
| 22886267 | 22886379 | 1 | miRNA | ENSG00000 |
| 22887780 | 22896111 | 1 | protein_coding | ENSG00000 |
| 22893692 | 22893818 | 1 | IG_J_gene | ENSG00000 |
| 22895375 | 22895834 | 1 | IG_C_gene | ENSG00000 |
| 22899481 | 22899655 | 1 | IG_J_gene | ENSG00000 |
| 22900976 | 22901437 | 1 | IG_C_gene | ENSG00000 |
| 22904850 | 22905025 | 1 | IG_J_gene | ENSG00000 |
| 22906342 | 22906803 | 1 | IG_C_gene | ENSG00000 |
| 22910574 | 22910606 | 1 | IG_J_gene | ENSG00000 |
| 22910828 | 22911075 | 1 | IG_C_pseudogene | ENSG00000 |
| 22914237 | 22914308 | 1 | IG_J_gene | ENSG00000 |
| 22915635 | 22915927 | 1 | IG_C_pseudogene | ENSG00000 |
| 22918132 | 22918201 | 1 | IG_J_gene | ENSG00000 |
| 22919535 | 22919851 | 1 | IG_C_pseudogene | ENSG00000 |
| 22921390 | 22921435 | 1 | IG_J_gene | ENSG00000 |
| 22922594 | 22923034 | 1 | IG_C_gene | ENSG00000 |
| 22988263 | 22988581 | 1 | processed_pseudoge | ENSG00000 |
| 22990690 | 23008356 | 1 | unprocessed_pseudo | ENSG00000 |
| 23059415 | 23145021 | -1 | protein_coding | ENSG00000 |
| 23070519 | 23125032 | 1 | protein_coding | ENSG00000 |
| 23132730 | 23133005 | -1 | misc_RNA | ENSG00000 |
| 23137805 | 23137864 | -1 | snRNA | ENSG00000 |
| 23145326 | 23164350 | 1 | protein_coding | ENSG00000 |
| 23179704 | 23318037 | 1 | protein_coding | ENSG00000 |
| 23179918 | 23180172 | -1 | processed_pseudoge | ENSG00000 |
| 23261782 | 23262071 | -1 | misc_RNA | ENSG00000 |
| 23262767 | 23265005 | 1 | processed_pseudoge | ENSG00000 |
| 23322543 | 23322927 | -1 | processed_pseudoge | ENSG00000 |
| 23326616 | 23328493 | 1 | lncRNA | ENSG00000 |
| 23359603 | 23387731 | -1 | transcribed_unproce | ENSG00000 |

| | | | | |
|---|---|---|---|---|
| 23390606 | 23402726 | -1 | transcribed_unproce | ENSG00000 |
| 23392553 | 23394792 | 1 | unprocessed_pseudo | ENSG00000 |
| 23433564 | 23435071 | 1 | lncRNA | ENSG00000 |
| 23446598 | 23460833 | 1 | transcribed_processe | ENSG00000 |
| 23462086 | 23486980 | -1 | lncRNA | ENSG00000 |
| 23510154 | 23512662 | 1 | lncRNA | ENSG00000 |
| 23512494 | 23513602 | -1 | lncRNA | ENSG00000 |
| 23536881 | 23547797 | 1 | lncRNA | ENSG00000 |
| 23567064 | 23573507 | 1 | lncRNA | ENSG00000 |
| 23573125 | 23580302 | -1 | protein_coding | ENSG00000 |
| 23580880 | 23583859 | -1 | lncRNA | ENSG00000 |
| 23607041 | 23608407 | 1 | unprocessed_pseudo | ENSG00000 |
| 23608452 | 23632321 | -1 | protein_coding | ENSG00000 |
| 23610329 | 23610538 | 1 | processed_pseudoge | ENSG00000 |
| 23630618 | 23638941 | -1 | lncRNA | ENSG00000 |
| 23638487 | 23717356 | -1 | lncRNA | ENSG00000 |
| 23638492 | 23640762 | -1 | transcribed_unproce | ENSG00000 |
| 23649213 | 23653334 | -1 | unprocessed_pseudo | ENSG00000 |
| 23658094 | 23717356 | -1 | transcribed_unproce | ENSG00000 |
| 23688136 | 23699176 | 1 | protein_coding | ENSG00000 |
| 23738682 | 23751112 | -1 | protein_coding | ENSG00000 |
| 23752743 | 23754425 | -1 | protein_coding | ENSG00000 |
| 23763021 | 23765861 | 1 | protein_coding | ENSG00000 |
| 23765834 | 23768443 | -1 | protein_coding | ENSG00000 |
| 23768226 | 23784316 | 1 | protein_coding | ENSG00000 |
| 23786931 | 23838008 | 1 | protein_coding | ENSG00000 |
| 23834503 | 23839128 | -1 | protein_coding | ENSG00000 |
| 23856427 | 23857039 | -1 | lncRNA | ENSG00000 |
| 23856703 | 23886309 | 1 | protein_coding | ENSG00000 |
| 23862188 | 23895223 | 1 | protein_coding | ENSG00000 |
| 23865248 | 23873277 | -1 | lncRNA | ENSG00000 |
| 23879493 | 23879790 | -1 | misc_RNA | ENSG00000 |
| 23894383 | 23895227 | 1 | protein_coding | ENSG00000 |
| 23894426 | 23898930 | -1 | lncRNA | ENSG00000 |
| 23901432 | 23907068 | -1 | lncRNA | ENSG00000 |
| 23926900 | 23929574 | 1 | processed_pseudoge | ENSG00000 |
| 23939998 | 23942798 | 1 | unprocessed_pseudo | ENSG00000 |
| 23949918 | 23954042 | 1 | unprocessed_pseudo | ENSG00000 |
| 23957414 | 23961186 | -1 | protein_coding | ENSG00000 |
| 23963780 | 23964374 | 1 | processed_pseudoge | ENSG00000 |
| 23966888 | 23972556 | 1 | protein_coding | ENSG00000 |
| 23969211 | 23969873 | 1 | lncRNA | ENSG00000 |
| 23971365 | 23979828 | -1 | protein_coding | ENSG00000 |
| 23971378 | 23980469 | -1 | protein_coding | ENSG00000 |
| 23976872 | 23991421 | -1 | transcribed_unproce | ENSG00000 |
| 23976904 | 23977585 | -1 | unprocessed_pseudo | ENSG00000 |
| 23980058 | 23983915 | 1 | polymorphic_pseudo | ENSG00000 |
| 23998401 | 24005453 | -1 | protein_coding | ENSG00000 |
| 24006305 | 24008258 | -1 | unprocessed_pseudo | ENSG00000 |
| 24011192 | 24178628 | 1 | protein_coding | ENSG00000 |

| | | | | |
|---|---|---|---|---|
| 24101689 | 24103354 | 1 | lncRNA | ENSG0000C |
| 24181487 | 24189106 | 1 | protein_coding | ENSG0000C |
| 24219654 | 24245142 | -1 | protein_coding | ENSG0000C |
| 24248074 | 24249466 | -1 | unprocessed_pseudc | ENSG0000C |
| 24251828 | 24265525 | 1 | transcribed_unproce | ENSG0000C |
| 24259929 | 24263764 | -1 | unprocessed_pseudc | ENSG0000C |
| 24269346 | 24270134 | -1 | processed_pseudoge | ENSG0000C |
| 24270817 | 24417739 | 1 | protein_coding | ENSG0000C |
| 24270898 | 24442356 | 1 | protein_coding | ENSG0000C |
| 24417879 | 24442357 | 1 | protein_coding | ENSG0000C |
| 24429206 | 24495074 | -1 | lncRNA | ENSG0000C |
| 24494107 | 24528390 | 1 | protein_coding | ENSG0000C |
| 24516508 | 24518386 | 1 | lncRNA | ENSG0000C |
| 24540423 | 24555935 | -1 | protein_coding | ENSG0000C |
| 24555958 | 24582052 | 1 | protein_coding | ENSG0000C |
| 24556007 | 24629005 | 1 | protein_coding | ENSG0000C |
| 24585620 | 24593208 | -1 | protein_coding | ENSG0000C |
| 24594811 | 24629005 | 1 | protein_coding | ENSG0000C |
| 24598054 | 24599398 | 1 | lncRNA | ENSG0000C |
| 24632915 | 24650012 | 1 | lncRNA | ENSG0000C |
| 24633272 | 24633443 | 1 | transcribed_unproce | ENSG0000C |
| 24644791 | 24653356 | 1 | transcribed_unproce | ENSG0000C |
| 24645171 | 24651657 | -1 | lncRNA | ENSG0000C |
| 24657481 | 24658765 | -1 | processed_pseudoge | ENSG0000C |
| 24686168 | 24686679 | 1 | processed_pseudoge | ENSG0000C |
| 24686923 | 24688087 | 1 | processed_pseudoge | ENSG0000C |
| 24691122 | 24696003 | -1 | lncRNA | ENSG0000C |
| 24703348 | 24703503 | 1 | processed_pseudoge | ENSG0000C |
| 24712292 | 24712520 | 1 | processed_pseudoge | ENSG0000C |
| 24719034 | 24774720 | -1 | protein_coding | ENSG0000C |
| 24806169 | 24927578 | 1 | protein_coding | ENSG0000C |
| 24901147 | 24901214 | -1 | snoRNA | ENSG0000C |
| 24934954 | 24946695 | -1 | protein_coding | ENSG0000C |
| 24952730 | 25197448 | 1 | protein_coding | ENSG0000C |
| 25009685 | 25015072 | -1 | lncRNA | ENSG0000C |
| 25022594 | 25024652 | -1 | lncRNA | ENSG0000C |
| 25048967 | 25051966 | -1 | lncRNA | ENSG0000C |
| 25052122 | 25065241 | -1 | lncRNA | ENSG0000C |
| 25102433 | 25112692 | -1 | lncRNA | ENSG0000C |
| 25179309 | 25179554 | -1 | processed_pseudoge | ENSG0000C |
| 25199858 | 25207359 | 1 | protein_coding | ENSG0000C |
| 25212085 | 25213826 | 1 | lncRNA | ENSG0000C |
| 25212564 | 25231870 | 1 | protein_coding | ENSG0000C |
| 25251523 | 25252602 | -1 | lncRNA | ENSG0000C |
| 25279529 | 25282674 | -1 | lncRNA | ENSG0000C |
| 25282613 | 25320086 | 1 | lncRNA | ENSG0000C |
| 25282943 | 25300791 | 1 | transcribed_unproce | ENSG0000C |
| 25318256 | 25320080 | 1 | unprocessed_pseudc | ENSG0000C |
| 25327252 | 25339975 | 1 | unprocessed_pseudc | ENSG0000C |
| 25349543 | 25350322 | -1 | processed_pseudoge | ENSG0000C |

| | | | | |
|---|---|---|---|---|
| 25351418 | 25405377 | -1 | protein_coding | ENSG00000( |
| 25434324 | 25435070 | -1 | lncRNA | ENSG00000( |
| 25436312 | 25436915 | 1 | lncRNA | ENSG00000( |
| 25437306 | 25437823 | -1 | IG_V_pseudogene | ENSG00000( |
| 25448105 | 25520854 | 1 | transcribed_unproce | ENSG00000( |
| 25455646 | 25455711 | 1 | miRNA | ENSG00000( |
| 25476218 | 25479971 | 1 | lncRNA | ENSG00000( |
| 25561117 | 25564719 | -1 | lncRNA | ENSG00000( |
| 25564675 | 25729294 | 1 | protein_coding | ENSG00000( |
| 25580241 | 25581382 | -1 | lncRNA | ENSG00000( |
| 25647261 | 25649232 | 1 | processed_pseudoge | ENSG00000( |
| 25715041 | 25715169 | -1 | rRNA_pseudogene | ENSG00000( |
| 25742144 | 26031041 | 1 | protein_coding | ENSG00000( |
| 25756318 | 25756669 | -1 | lncRNA | ENSG00000( |
| 25876854 | 25903247 | -1 | lncRNA | ENSG00000( |
| 25959074 | 25983526 | -1 | lncRNA | ENSG00000( |
| 25963983 | 25964273 | -1 | misc_RNA | ENSG00000( |
| 26006389 | 26006532 | -1 | processed_pseudoge | ENSG00000( |
| 26161834 | 26164303 | 1 | lncRNA | ENSG00000( |
| 26161905 | 26169341 | -1 | lncRNA | ENSG00000( |
| 26169462 | 26383597 | 1 | protein_coding | ENSG00000( |
| 26241331 | 26254092 | -1 | lncRNA | ENSG00000( |
| 26390667 | 26390776 | -1 | rRNA_pseudogene | ENSG00000( |
| 26422071 | 26423193 | -1 | processed_pseudoge | ENSG00000( |
| 26429260 | 26445015 | 1 | protein_coding | ENSG00000( |
| 26443423 | 26483837 | -1 | protein_coding | ENSG00000( |
| 26483877 | 26494658 | 1 | protein_coding | ENSG00000( |
| 26491225 | 26512505 | -1 | protein_coding | ENSG00000( |
| 26512537 | 26514568 | 1 | lncRNA | ENSG00000( |
| 26521996 | 26596717 | -1 | protein_coding | ENSG00000( |
| 26555212 | 26555323 | -1 | miRNA | ENSG00000( |
| 26560526 | 26561088 | 1 | processed_pseudoge | ENSG00000( |
| 26597028 | 26611453 | 1 | lncRNA | ENSG00000( |
| 26599278 | 26618027 | -1 | protein_coding | ENSG00000( |
| 26621963 | 26630669 | 1 | protein_coding | ENSG00000( |
| 26644889 | 26645336 | -1 | processed_pseudoge | ENSG00000( |
| 26646411 | 26676475 | 1 | lncRNA | ENSG00000( |
| 26657521 | 26657655 | 1 | misc_RNA | ENSG00000( |
| 26667693 | 26672654 | -1 | lncRNA | ENSG00000( |
| 26671402 | 26671550 | 1 | misc_RNA | ENSG00000( |
| 26672164 | 26672248 | 1 | misc_RNA | ENSG00000( |
| 26672747 | 26780893 | 1 | lncRNA | ENSG00000( |
| 26673254 | 26673630 | 1 | misc_RNA | ENSG00000( |
| 26858634 | 26968763 | 1 | lncRNA | ENSG00000( |
| 26886909 | 26887548 | -1 | processed_pseudoge | ENSG00000( |
| 26903292 | 26920935 | 1 | lncRNA | ENSG00000( |
| 27019369 | 27024550 | 1 | lncRNA | ENSG00000( |
| 27035887 | 27035991 | 1 | snRNA | ENSG00000( |
| 27040197 | 27041494 | 1 | lncRNA | ENSG00000( |
| 27044001 | 27045315 | 1 | lncRNA | ENSG00000( |

| | | | | |
|---|---|---|---|---|
| 27048144 | 27061247 | -1 | lncRNA | ENSG00000 |
| 27102766 | 27107019 | 1 | lncRNA | ENSG00000 |
| 27143478 | 27159878 | 1 | lncRNA | ENSG00000 |
| 27204211 | 27205126 | 1 | lncRNA | ENSG00000 |
| 27221349 | 27225134 | -1 | lncRNA | ENSG00000 |
| 27276240 | 27286349 | 1 | lncRNA | ENSG00000 |
| 27307483 | 27318538 | -1 | lncRNA | ENSG00000 |
| 27307777 | 27318501 | 1 | lncRNA | ENSG00000 |
| 27331634 | 27379265 | 1 | lncRNA | ENSG00000 |
| 27414462 | 27419713 | -1 | lncRNA | ENSG00000 |
| 27436292 | 27441774 | 1 | lncRNA | ENSG00000 |
| 27457247 | 27460060 | -1 | lncRNA | ENSG00000 |
| 27563839 | 27573573 | -1 | lncRNA | ENSG00000 |
| 27676559 | 27714970 | -1 | lncRNA | ENSG00000 |
| 27707580 | 27709931 | 1 | lncRNA | ENSG00000 |
| 27716480 | 27721677 | 1 | lncRNA | ENSG00000 |
| 27748277 | 27801756 | -1 | protein_coding | ENSG00000 |
| 27851669 | 27920134 | -1 | protein_coding | ENSG00000 |
| 27919376 | 28008581 | 1 | lncRNA | ENSG00000 |
| 27919384 | 27919545 | 1 | misc_RNA | ENSG00000 |
| 27920525 | 27920612 | -1 | miRNA | ENSG00000 |
| 27920526 | 27920611 | 1 | miRNA | ENSG00000 |
| 27922022 | 27922178 | 1 | misc_RNA | ENSG00000 |
| 27978014 | 28679865 | -1 | protein_coding | ENSG00000 |
| 27994474 | 27994654 | 1 | misc_RNA | ENSG00000 |
| 28002484 | 28002595 | 1 | misc_RNA | ENSG00000 |
| 28056153 | 28056453 | -1 | misc_RNA | ENSG00000 |
| 28232756 | 28232823 | -1 | snoRNA | ENSG00000 |
| 28303754 | 28303850 | 1 | misc_RNA | ENSG00000 |
| 28459869 | 28459948 | 1 | miRNA | ENSG00000 |
| 28642979 | 28643270 | -1 | misc_RNA | ENSG00000 |
| 28687743 | 28742422 | -1 | protein_coding | ENSG00000 |
| 28742039 | 28757515 | 1 | protein_coding | ENSG00000 |
| 28772674 | 28789301 | 1 | protein_coding | ENSG00000 |
| 28794555 | 28800597 | -1 | protein_coding | ENSG00000 |
| 28800683 | 28848559 | 1 | lncRNA | ENSG00000 |
| 28814914 | 28815662 | 1 | lncRNA | ENSG00000 |
| 28835893 | 28836137 | -1 | processed_pseudoge | ENSG00000 |
| 28883572 | 29057488 | 1 | protein_coding | ENSG00000 |
| 28992721 | 29018620 | 1 | lncRNA | ENSG00000 |
| 29024999 | 29031476 | -1 | lncRNA | ENSG00000 |
| 29058672 | 29061844 | -1 | protein_coding | ENSG00000 |
| 29073078 | 29168333 | 1 | protein_coding | ENSG00000 |
| 29099041 | 29111683 | -1 | lncRNA | ENSG00000 |
| 29134014 | 29134120 | 1 | snRNA | ENSG00000 |
| 29191697 | 29191808 | -1 | snRNA | ENSG00000 |
| 29205896 | 29259597 | 1 | protein_coding | ENSG00000 |
| 29259872 | 29268209 | -1 | protein_coding | ENSG00000 |
| 29260889 | 29262037 | 1 | lncRNA | ENSG00000 |
| 29268009 | 29300525 | 1 | protein_coding | ENSG00000 |

| | | | | |
|---|---|---|---|---|
| 29306582 | 29312785 | 1 | protein_coding | ENSG00000 |
| 29312933 | 29319679 | -1 | protein_coding | ENSG00000 |
| 29316744 | 29317220 | -1 | lncRNA | ENSG00000 |
| 29327680 | 29423179 | -1 | protein_coding | ENSG00000 |
| 29333158 | 29333267 | -1 | miRNA | ENSG00000 |
| 29333163 | 29333258 | -1 | snoRNA | ENSG00000 |
| 29387909 | 29417757 | 1 | transcribed_unproce | ENSG00000 |
| 29419155 | 29423193 | -1 | unprocessed_pseudo | ENSG00000 |
| 29432944 | 29435005 | 1 | unprocessed_pseudo | ENSG00000 |
| 29436534 | 29478868 | -1 | lncRNA | ENSG00000 |
| 29438583 | 29442455 | 1 | protein_coding | ENSG00000 |
| 29480218 | 29491390 | 1 | protein_coding | ENSG00000 |
| 29480220 | 29481113 | -1 | lncRNA | ENSG00000 |
| 29505879 | 29555216 | -1 | protein_coding | ENSG00000 |
| 29513889 | 29535390 | -1 | lncRNA | ENSG00000 |
| 29554808 | 29581327 | -1 | protein_coding | ENSG00000 |
| 29603556 | 29698598 | 1 | protein_coding | ENSG00000 |
| 29608900 | 29609501 | 1 | processed_pseudoge | ENSG00000 |
| 29705256 | 29719859 | -1 | lncRNA | ENSG00000 |
| 29711820 | 29719714 | -1 | lncRNA | ENSG00000 |
| 29720003 | 29731833 | 1 | protein_coding | ENSG00000 |
| 29730956 | 29767011 | -1 | protein_coding | ENSG00000 |
| 29758758 | 29758859 | 1 | misc_RNA | ENSG00000 |
| 29767091 | 29768096 | -1 | lncRNA | ENSG00000 |
| 29767369 | 29770413 | 1 | protein_coding | ENSG00000 |
| 29788609 | 29838304 | -1 | protein_coding | ENSG00000 |
| 29883155 | 30030866 | 1 | protein_coding | ENSG00000 |
| 29899587 | 29899693 | -1 | snRNA | ENSG00000 |
| 29978950 | 30028236 | -1 | lncRNA | ENSG00000 |
| 30007049 | 30007113 | 1 | miRNA | ENSG00000 |
| 30008742 | 30080480 | -1 | lncRNA | ENSG00000 |
| 30046276 | 30047193 | -1 | processed_pseudoge | ENSG00000 |
| 30080174 | 30177075 | 1 | protein_coding | ENSG00000 |
| 30105094 | 30105408 | -1 | processed_pseudoge | ENSG00000 |
| 30136673 | 30137681 | -1 | processed_pseudoge | ENSG00000 |
| 30182818 | 30207195 | -1 | lncRNA | ENSG00000 |
| 30239194 | 30240538 | 1 | lncRNA | ENSG00000 |
| 30240453 | 30246759 | -1 | protein_coding | ENSG00000 |
| 30246205 | 30246998 | 1 | lncRNA | ENSG00000 |
| 30262829 | 30266851 | -1 | protein_coding | ENSG00000 |
| 30275215 | 30276951 | -1 | TEC | ENSG00000 |
| 30285117 | 30289627 | -1 | protein_coding | ENSG00000 |
| 30285238 | 30299482 | -1 | protein_coding | ENSG00000 |
| 30291990 | 30327046 | -1 | protein_coding | ENSG00000 |
| 30331988 | 30356919 | -1 | protein_coding | ENSG00000 |
| 30356635 | 30378658 | 1 | protein_coding | ENSG00000 |
| 30368811 | 30421771 | -1 | protein_coding | ENSG00000 |
| 30396941 | 30425303 | 1 | protein_coding | ENSG00000 |
| 30409255 | 30428990 | 1 | protein_coding | ENSG00000 |
| 30420512 | 30420912 | 1 | lncRNA | ENSG00000 |

| | | | | |
|---|---|---|---|---|
| 30421045 | 30421148 | 1 | snRNA | ENSG0000C |
| 30421206 | 30421536 | -1 | lncRNA | ENSG0000C |
| 30425623 | 30429054 | 1 | protein_coding | ENSG0000C |
| 30435544 | 30436247 | 1 | lncRNA | ENSG0000C |
| 30439837 | 30440318 | 1 | processed_pseudoge | ENSG0000C |
| 30447959 | 30472049 | -1 | protein_coding | ENSG0000C |
| 30475364 | 30492804 | 1 | lncRNA | ENSG0000C |
| 30481311 | 30481911 | -1 | processed_pseudoge | ENSG0000C |
| 30488902 | 30505711 | -1 | protein_coding | ENSG0000C |
| 30522799 | 30546682 | -1 | protein_coding | ENSG0000C |
| 30542536 | 30544305 | 1 | processed_pseudoge | ENSG0000C |
| 30554635 | 30574665 | -1 | protein_coding | ENSG0000C |
| 30576625 | 30607083 | -1 | protein_coding | ENSG0000C |
| 30598309 | 30606687 | 1 | lncRNA | ENSG0000C |
| 30607003 | 30627271 | 1 | protein_coding | ENSG0000C |
| 30635781 | 30669016 | 1 | protein_coding | ENSG0000C |
| 30652051 | 30667890 | -1 | protein_coding | ENSG0000C |
| 30653877 | 30654814 | -1 | unprocessed_pseudc | ENSG0000C |
| 30664961 | 30674134 | 1 | unprocessed_pseudc | ENSG0000C |
| 30693782 | 30907824 | 1 | protein_coding | ENSG0000C |
| 30731557 | 30731641 | 1 | miRNA | ENSG0000C |
| 30902219 | 30902781 | 1 | processed_pseudoge | ENSG0000C |
| 30922308 | 30932449 | 1 | lncRNA | ENSG0000C |
| 30925130 | 30968774 | -1 | protein_coding | ENSG0000C |
| 30969245 | 30979395 | 1 | lncRNA | ENSG0000C |
| 30971005 | 30971149 | 1 | misc_RNA | ENSG0000C |
| 30971296 | 30971382 | 1 | misc_RNA | ENSG0000C |
| 30972857 | 30973093 | 1 | misc_RNA | ENSG0000C |
| 30973417 | 30973597 | 1 | misc_RNA | ENSG0000C |
| 30976515 | 30978848 | -1 | lncRNA | ENSG0000C |
| 30977516 | 30977858 | -1 | lncRNA | ENSG0000C |
| 31051630 | 31068327 | 1 | lncRNA | ENSG0000C |
| 31059989 | 31060285 | 1 | misc_RNA | ENSG0000C |
| 31064105 | 31104757 | 1 | protein_coding | ENSG0000C |
| 31082156 | 31083565 | -1 | lncRNA | ENSG0000C |
| 31104772 | 31120069 | -1 | protein_coding | ENSG0000C |
| 31113017 | 31113396 | 1 | processed_pseudoge | ENSG0000C |
| 31122731 | 31134697 | 1 | protein_coding | ENSG0000C |
| 31134807 | 31140508 | -1 | protein_coding | ENSG0000C |
| 31155966 | 31156647 | -1 | processed_pseudoge | ENSG0000C |
| 31160062 | 31160119 | -1 | miRNA | ENSG0000C |
| 31160182 | 31207019 | 1 | protein_coding | ENSG0000C |
| 31205264 | 31205616 | -1 | lncRNA | ENSG0000C |
| 31212239 | 31280080 | 1 | protein_coding | ENSG0000C |
| 31222745 | 31222853 | -1 | snRNA | ENSG0000C |
| 31230071 | 31230172 | -1 | misc_RNA | ENSG0000C |
| 31281594 | 31292534 | -1 | protein_coding | ENSG0000C |
| 31292499 | 31338021 | 1 | lncRNA | ENSG0000C |
| 31305404 | 31305512 | 1 | rRNA_pseudogene | ENSG0000C |
| 31325804 | 31346346 | -1 | protein_coding | ENSG0000C |

| | | | | |
|---|---|---|---|---|
| 31346777 | 31348719 | 1 | lncRNA | ENSG0000C |
| 31388733 | 31388837 | -1 | snRNA | ENSG0000C |
| 31399604 | 31530634 | 1 | protein_coding | ENSG0000C |
| 31436977 | 31496108 | -1 | protein_coding | ENSG0000C |
| 31452881 | 31464304 | 1 | lncRNA | ENSG0000C |
| 31479383 | 31479489 | -1 | snRNA | ENSG0000C |
| 31488688 | 31618588 | 1 | protein_coding | ENSG0000C |
| 31490150 | 31491022 | 1 | lncRNA | ENSG0000C |
| 31521199 | 31521592 | -1 | processed_pseudoge | ENSG0000C |
| 31538352 | 31538589 | -1 | processed_pseudoge | ENSG0000C |
| 31559483 | 31559939 | 1 | processed_pseudoge | ENSG0000C |
| 31618491 | 31662432 | -1 | protein_coding | ENSG0000C |
| 31621467 | 31621531 | -1 | miRNA | ENSG0000C |
| 31676256 | 31750140 | -1 | protein_coding | ENSG0000C |
| 31753867 | 31908033 | 1 | protein_coding | ENSG0000C |
| 31754899 | 31956243 | 1 | protein_coding | ENSG0000C |
| 31757202 | 31757498 | 1 | misc_RNA | ENSG0000C |
| 31816379 | 31817491 | -1 | processed_pseudoge | ENSG0000C |
| 31837238 | 31837298 | 1 | snRNA | ENSG0000C |
| 31890344 | 31891434 | -1 | processed_pseudoge | ENSG0000C |
| 31926214 | 31945518 | -1 | lncRNA | ENSG0000C |
| 31944522 | 31957603 | 1 | protein_coding | ENSG0000C |
| 31963357 | 31970400 | 1 | lncRNA | ENSG0000C |
| 31970823 | 32037995 | 1 | lncRNA | ENSG0000C |
| 31981147 | 31981427 | -1 | misc_RNA | ENSG0000C |
| 32039490 | 32039896 | 1 | processed_pseudoge | ENSG0000C |
| 32043261 | 32113029 | 1 | protein_coding | ENSG0000C |
| 32121977 | 32133469 | 1 | transcribed_unproce | ENSG0000C |
| 32129689 | 32130612 | 1 | transcribed_unproce | ENSG0000C |
| 32134028 | 32134133 | 1 | unprocessed_pseudc | ENSG0000C |
| 32141267 | 32143272 | 1 | lncRNA | ENSG0000C |
| 32149006 | 32159322 | -1 | protein_coding | ENSG0000C |
| 32159454 | 32160392 | -1 | lncRNA | ENSG0000C |
| 32188400 | 32188487 | -1 | misc_RNA | ENSG0000C |
| 32190435 | 32205001 | -1 | protein_coding | ENSG0000C |
| 32199919 | 32200234 | 1 | IG_C_pseudogene | ENSG0000C |
| 32205115 | 32278382 | 1 | lncRNA | ENSG0000C |
| 32218476 | 32255341 | -1 | protein_coding | ENSG0000C |
| 32269381 | 32273110 | 1 | transcribed_processe | ENSG0000C |
| 32284683 | 32285131 | 1 | lncRNA | ENSG0000C |
| 32327064 | 32343105 | -1 | lncRNA | ENSG0000C |
| 32354885 | 32361161 | 1 | protein_coding | ENSG0000C |
| 32356676 | 32356988 | -1 | IG_C_pseudogene | ENSG0000C |
| 32359886 | 32382106 | -1 | lncRNA | ENSG0000C |
| 32362972 | 32363059 | 1 | misc_RNA | ENSG0000C |
| 32368497 | 32368567 | 1 | misc_RNA | ENSG0000C |
| 32376664 | 32384343 | 1 | lncRNA | ENSG0000C |
| 32376682 | 32377135 | 1 | IG_V_pseudogene | ENSG0000C |
| 32383786 | 32385631 | 1 | lncRNA | ENSG0000C |
| 32386668 | 32386868 | 1 | unprocessed_pseudc | ENSG0000C |

| | | | | |
|---|---|---|---|---|
| 32387582 | 32412248 | -1 | protein_coding | ENSG00000 |
| 32413847 | 32464484 | -1 | protein_coding | ENSG00000 |
| 32474676 | 32498829 | 1 | protein_coding | ENSG00000 |
| 32512552 | 33058372 | -1 | protein_coding | ENSG00000 |
| 32583300 | 32584204 | 1 | lncRNA | ENSG00000 |
| 32635102 | 32635198 | -1 | rRNA_pseudogene | ENSG00000 |
| 32783177 | 32784982 | 1 | lncRNA | ENSG00000 |
| 32801701 | 32863043 | 1 | protein_coding | ENSG00000 |
| 33014980 | 33015461 | 1 | processed_pseudoge | ENSG00000 |
| 33104330 | 33145861 | 1 | lncRNA | ENSG00000 |
| 33105383 | 33106059 | -1 | lncRNA | ENSG00000 |
| 33137140 | 33137338 | 1 | processed_pseudoge | ENSG00000 |
| 33162226 | 33922841 | -1 | protein_coding | ENSG00000 |
| 33164063 | 33166439 | 1 | lncRNA | ENSG00000 |
| 33320865 | 33322813 | 1 | lncRNA | ENSG00000 |
| 33436582 | 33436669 | -1 | miRNA | ENSG00000 |
| 33704786 | 33704920 | -1 | snoRNA | ENSG00000 |
| 33704786 | 33704922 | -1 | snoRNA | ENSG00000 |
| 33723832 | 33724912 | -1 | lncRNA | ENSG00000 |
| 33725007 | 33750843 | 1 | lncRNA | ENSG00000 |
| 33922422 | 33922766 | 1 | lncRNA | ENSG00000 |
| 34017422 | 34218796 | 1 | lncRNA | ENSG00000 |
| 34191194 | 34210564 | -1 | lncRNA | ENSG00000 |
| 34589085 | 34591278 | -1 | processed_pseudoge | ENSG00000 |
| 34641179 | 34651862 | 1 | lncRNA | ENSG00000 |
| 34703126 | 34704885 | -1 | lncRNA | ENSG00000 |
| 34711908 | 34724919 | -1 | lncRNA | ENSG00000 |
| 34756676 | 35002862 | -1 | lncRNA | ENSG00000 |
| 35066136 | 35087387 | 1 | protein_coding | ENSG00000 |
| 35119824 | 35231056 | -1 | transcribed_processe | ENSG00000 |
| 35122661 | 35127419 | 1 | lncRNA | ENSG00000 |
| 35164304 | 35165347 | 1 | processed_pseudoge | ENSG00000 |
| 35249772 | 35249833 | 1 | snRNA | ENSG00000 |
| 35257452 | 35295807 | 1 | protein_coding | ENSG00000 |
| 35298838 | 35299541 | -1 | lncRNA | ENSG00000 |
| 35299275 | 35347992 | 1 | protein_coding | ENSG00000 |
| 35335640 | 35335758 | 1 | miRNA | ENSG00000 |
| 35336721 | 35336799 | -1 | miRNA | ENSG00000 |
| 35372174 | 35372621 | 1 | lncRNA | ENSG00000 |
| 35376422 | 35377261 | 1 | lncRNA | ENSG00000 |
| 35380361 | 35394207 | 1 | protein_coding | ENSG00000 |
| 35400134 | 35425431 | 1 | protein_coding | ENSG00000 |
| 35501861 | 35503586 | -1 | processed_pseudoge | ENSG00000 |
| 35526932 | 35527415 | 1 | lncRNA | ENSG00000 |
| 35540831 | 35553999 | 1 | protein_coding | ENSG00000 |
| 35606764 | 35637951 | -1 | protein_coding | ENSG00000 |
| 35626988 | 35635134 | -1 | lncRNA | ENSG00000 |
| 35648446 | 35668404 | 1 | protein_coding | ENSG00000 |
| 35685940 | 35689373 | 1 | lncRNA | ENSG00000 |
| 35703803 | 35704176 | 1 | processed_pseudoge | ENSG00000 |

| | | | | |
|---|---|---|---|---|
| 35717872 | 35729483 | 1 | protein_coding | ENSG00000( |
| 35738736 | 36028824 | -1 | protein_coding | ENSG00000( |
| 35897270 | 35898404 | -1 | processed_pseudoge | ENSG00000( |
| 35992321 | 36000469 | 1 | lncRNA | ENSG00000( |
| 36066533 | 36070110 | 1 | lncRNA | ENSG00000( |
| 36071000 | 36085573 | 1 | lncRNA | ENSG00000( |
| 36091148 | 36093352 | 1 | lncRNA | ENSG00000( |
| 36137430 | 36139350 | -1 | TEC | ENSG00000( |
| 36140330 | 36166177 | -1 | protein_coding | ENSG00000( |
| 36172422 | 36172779 | 1 | processed_pseudoge | ENSG00000( |
| 36172934 | 36173948 | 1 | processed_pseudoge | ENSG00000( |
| 36174261 | 36174822 | 1 | processed_pseudoge | ENSG00000( |
| 36175167 | 36175854 | 1 | processed_pseudoge | ENSG00000( |
| 36175857 | 36176526 | 1 | processed_pseudoge | ENSG00000( |
| 36178554 | 36179385 | 1 | processed_pseudoge | ENSG00000( |
| 36189124 | 36204840 | -1 | protein_coding | ENSG00000( |
| 36226209 | 36239954 | -1 | protein_coding | ENSG00000( |
| 36253010 | 36267530 | 1 | protein_coding | ENSG00000( |
| 36281277 | 36388067 | -1 | protein_coding | ENSG00000( |
| 36286847 | 36286907 | -1 | miRNA | ENSG00000( |
| 36335421 | 36343530 | 1 | lncRNA | ENSG00000( |
| 36388626 | 36396517 | 1 | lncRNA | ENSG00000( |
| 36421273 | 36421644 | -1 | processed_pseudoge | ENSG00000( |
| 36440880 | 36440970 | -1 | misc_RNA | ENSG00000( |
| 36445395 | 36454944 | -1 | lncRNA | ENSG00000( |
| 36467046 | 36481640 | -1 | protein_coding | ENSG00000( |
| 36487190 | 36507101 | -1 | protein_coding | ENSG00000( |
| 36510855 | 36529184 | -1 | protein_coding | ENSG00000( |
| 36536656 | 36537604 | -1 | unprocessed_pseudc | ENSG00000( |
| 36552165 | 36552700 | 1 | lncRNA | ENSG00000( |
| 36560870 | 36562915 | -1 | lncRNA | ENSG00000( |
| 36563921 | 36703558 | -1 | protein_coding | ENSG00000( |
| 36703918 | 36721472 | 1 | lncRNA | ENSG00000( |
| 36754086 | 36755731 | -1 | lncRNA | ENSG00000( |
| 36758202 | 36776256 | -1 | protein_coding | ENSG00000( |
| 36766478 | 36769022 | 1 | lncRNA | ENSG00000( |
| 36800684 | 36819479 | -1 | protein_coding | ENSG00000( |
| 36816326 | 36819736 | 1 | lncRNA | ENSG00000( |
| 36847372 | 36870441 | -1 | lncRNA | ENSG00000( |
| 36860988 | 36878017 | 1 | protein_coding | ENSG00000( |
| 36913628 | 36940439 | 1 | protein_coding | ENSG00000( |
| 36950529 | 36953723 | -1 | unprocessed_pseudc | ENSG00000( |
| 36965248 | 36968172 | 1 | lncRNA | ENSG00000( |
| 36991120 | 37007851 | -1 | protein_coding | ENSG00000( |
| 37010859 | 37020183 | -1 | protein_coding | ENSG00000( |
| 37013225 | 37013319 | 1 | misc_RNA | ENSG00000( |
| 37019635 | 37029822 | 1 | protein_coding | ENSG00000( |
| 37033124 | 37035513 | 1 | lncRNA | ENSG00000( |
| 37051736 | 37063390 | 1 | protein_coding | ENSG00000( |
| 37055033 | 37055347 | -1 | misc_RNA | ENSG00000( |

| | | | | |
|---|---|---|---|---|
| 37065436 | 37109713 | -1 | protein_coding | ENSG00000 |
| 37080068 | 37082847 | 1 | lncRNA | ENSG00000 |
| 37125843 | 37175054 | -1 | protein_coding | ENSG00000 |
| 37166757 | 37182850 | 1 | lncRNA | ENSG00000 |
| 37180166 | 37199385 | -1 | protein_coding | ENSG00000 |
| 37204237 | 37212477 | -1 | protein_coding | ENSG00000 |
| 37215738 | 37215891 | -1 | processed_pseudoge | ENSG00000 |
| 37225270 | 37244448 | -1 | protein_coding | ENSG00000 |
| 37282027 | 37315341 | 1 | protein_coding | ENSG00000 |
| 37339583 | 37427445 | -1 | lncRNA | ENSG00000 |
| 37352190 | 37355002 | -1 | lncRNA | ENSG00000 |
| 37367960 | 37427479 | -1 | protein_coding | ENSG00000 |
| 37371684 | 37372858 | 1 | lncRNA | ENSG00000 |
| 37469063 | 37486393 | -1 | protein_coding | ENSG00000 |
| 37490362 | 37519542 | -1 | protein_coding | ENSG00000 |
| 37550026 | 37551735 | 1 | lncRNA | ENSG00000 |
| 37560480 | 37569405 | 1 | protein_coding | ENSG00000 |
| 37570248 | 37582616 | -1 | protein_coding | ENSG00000 |
| 37578669 | 37578922 | 1 | misc_RNA | ENSG00000 |
| 37608725 | 37633564 | 1 | protein_coding | ENSG00000 |
| 37634654 | 37656119 | 1 | protein_coding | ENSG00000 |
| 37639693 | 37666932 | 1 | protein_coding | ENSG00000 |
| 37641832 | 37658377 | -1 | lncRNA | ENSG00000 |
| 37658723 | 37666932 | 1 | protein_coding | ENSG00000 |
| 37669464 | 37669739 | -1 | misc_RNA | ENSG00000 |
| 37675636 | 37679802 | 1 | protein_coding | ENSG00000 |
| 37681673 | 37693476 | 1 | protein_coding | ENSG00000 |
| 37686414 | 37726503 | 1 | protein_coding | ENSG00000 |
| 37697048 | 37776556 | 1 | protein_coding | ENSG00000 |
| 37805093 | 37807436 | 1 | protein_coding | ENSG00000 |
| 37807905 | 37817176 | 1 | protein_coding | ENSG00000 |
| 37823382 | 37825485 | 1 | protein_coding | ENSG00000 |
| 37830855 | 37849327 | -1 | protein_coding | ENSG00000 |
| 37844272 | 37844371 | -1 | miRNA | ENSG00000 |
| 37847678 | 37847774 | -1 | miRNA | ENSG00000 |
| 37848868 | 37889407 | 1 | protein_coding | ENSG00000 |
| 37876148 | 37895563 | 1 | lncRNA | ENSG00000 |
| 37891347 | 37891448 | 1 | snRNA | ENSG00000 |
| 37905657 | 37942822 | 1 | protein_coding | ENSG00000 |
| 37943050 | 37953669 | -1 | protein_coding | ENSG00000 |
| 37943050 | 37944898 | 1 | TEC | ENSG00000 |
| 37950965 | 37951778 | 1 | lncRNA | ENSG00000 |
| 37952607 | 38041915 | 1 | protein_coding | ENSG00000 |
| 37967563 | 37967624 | 1 | miRNA | ENSG00000 |
| 37970686 | 37987422 | -1 | protein_coding | ENSG00000 |
| 37988794 | 37988853 | 1 | miRNA | ENSG00000 |
| 38032165 | 38034228 | -1 | lncRNA | ENSG00000 |
| 38043230 | 38043920 | 1 | lncRNA | ENSG00000 |
| 38056311 | 38075701 | 1 | protein_coding | ENSG00000 |
| 38057180 | 38073940 | -1 | lncRNA | ENSG00000 |

| | | | | |
|---|---|---|---|---|
| 38078134 | 38084093 | -1 | protein_coding | ENSG00000 |
| 38084900 | 38110684 | -1 | protein_coding | ENSG00000 |
| 38090127 | 38091559 | 1 | lncRNA | ENSG00000 |
| 38111495 | 38214778 | -1 | protein_coding | ENSG00000 |
| 38130216 | 38150612 | 1 | lncRNA | ENSG00000 |
| 38200767 | 38216507 | 1 | protein_coding | ENSG00000 |
| 38219291 | 38273010 | -1 | protein_coding | ENSG00000 |
| 38231320 | 38232248 | 1 | lncRNA | ENSG00000 |
| 38231720 | 38231993 | 1 | misc_RNA | ENSG00000 |
| 38281028 | 38281289 | -1 | misc_RNA | ENSG00000 |
| 38290691 | 38318084 | -1 | protein_coding | ENSG00000 |
| 38291918 | 38398522 | -1 | protein_coding | ENSG00000 |
| 38335762 | 38398929 | -1 | transcribed_unproce | ENSG00000 |
| 38387918 | 38388857 | -1 | lncRNA | ENSG00000 |
| 38416565 | 38423933 | -1 | lncRNA | ENSG00000 |
| 38424288 | 38427336 | 1 | lncRNA | ENSG00000 |
| 38426327 | 38455199 | -1 | protein_coding | ENSG00000 |
| 38434307 | 38436686 | 1 | lncRNA | ENSG00000 |
| 38468078 | 38483447 | 1 | protein_coding | ENSG00000 |
| 38483438 | 38507660 | -1 | protein_coding | ENSG00000 |
| 38518948 | 38570204 | -1 | protein_coding | ENSG00000 |
| 38574931 | 38575431 | -1 | processed_pseudoge | ENSG00000 |
| 38578120 | 38656629 | -1 | protein_coding | ENSG00000 |
| 38656636 | 38673854 | 1 | protein_coding | ENSG00000 |
| 38666508 | 38668750 | -1 | lncRNA | ENSG00000 |
| 38667585 | 38681847 | -1 | lncRNA | ENSG00000 |
| 38675876 | 38677800 | 1 | lncRNA | ENSG00000 |
| 38681957 | 38685421 | 1 | protein_coding | ENSG00000 |
| 38685543 | 38701556 | -1 | protein_coding | ENSG00000 |
| 38705742 | 38738299 | 1 | protein_coding | ENSG00000 |
| 38722743 | 38723505 | -1 | processed_pseudoge | ENSG00000 |
| 38734725 | 38738765 | 1 | lncRNA | ENSG00000 |
| 38734725 | 38794143 | -1 | protein_coding | ENSG00000 |
| 38734730 | 38738990 | 1 | lncRNA | ENSG00000 |
| 38736610 | 38736792 | -1 | lncRNA | ENSG00000 |
| 38739003 | 38749041 | 1 | lncRNA | ENSG00000 |
| 38742625 | 38743115 | 1 | lncRNA | ENSG00000 |
| 38743495 | 38743910 | 1 | lncRNA | ENSG00000 |
| 38778508 | 38794198 | -1 | protein_coding | ENSG00000 |
| 38818452 | 38844028 | -1 | protein_coding | ENSG00000 |
| 38861422 | 38872249 | -1 | protein_coding | ENSG00000 |
| 38886837 | 38903794 | 1 | lncRNA | ENSG00000 |
| 38921227 | 38924708 | 1 | lncRNA | ENSG00000 |
| 38952741 | 38992778 | 1 | protein_coding | ENSG00000 |
| 38982347 | 38992804 | 1 | protein_coding | ENSG00000 |
| 38991559 | 38998209 | -1 | lncRNA | ENSG00000 |
| 39014257 | 39020352 | 1 | protein_coding | ENSG00000 |
| 39014363 | 39033276 | 1 | protein_coding | ENSG00000 |
| 39021113 | 39033277 | 1 | protein_coding | ENSG00000 |
| 39040604 | 39055972 | 1 | protein_coding | ENSG00000 |

| | | | | |
|---|---|---|---|---|
| 39070414 | 39070671 | 1 | unprocessed_pseudc | ENSG0000C |
| 39077067 | 39087743 | 1 | protein_coding | ENSG0000C |
| 39091586 | 39092855 | 1 | transcribed_unproce | ENSG0000C |
| 39097224 | 39104067 | 1 | protein_coding | ENSG0000C |
| 39120167 | 39152680 | -1 | protein_coding | ENSG0000C |
| 39120293 | 39120566 | -1 | processed_pseudoge | ENSG0000C |
| 39133090 | 39136760 | 1 | lncRNA | ENSG0000C |
| 39155525 | 39155945 | 1 | processed_pseudoge | ENSG0000C |
| 39223359 | 39244982 | -1 | protein_coding | ENSG0000C |
| 39242188 | 39279897 | 1 | lncRNA | ENSG0000C |
| 39292988 | 39295589 | 1 | lncRNA | ENSG0000C |
| 39312882 | 39320389 | -1 | protein_coding | ENSG0000C |
| 39313819 | 39313911 | -1 | snoRNA | ENSG0000C |
| 39315213 | 39315307 | -1 | snoRNA | ENSG0000C |
| 39319050 | 39319113 | -1 | snoRNA | ENSG0000C |
| 39349925 | 39385575 | 1 | protein_coding | ENSG0000C |
| 39379610 | 39380015 | 1 | TEC | ENSG0000C |
| 39399778 | 39437060 | 1 | protein_coding | ENSG0000C |
| 39457012 | 39492194 | 1 | protein_coding | ENSG0000C |
| 39475807 | 39476822 | -1 | lncRNA | ENSG0000C |
| 39499432 | 39518132 | 1 | protein_coding | ENSG0000C |
| 39504231 | 39504443 | 1 | protein_coding | ENSG0000C |
| 39519695 | 39522685 | 1 | protein_coding | ENSG0000C |
| 39529093 | 39532761 | -1 | protein_coding | ENSG0000C |
| 39570753 | 39689735 | 1 | protein_coding | ENSG0000C |
| 39743044 | 39893864 | -1 | protein_coding | ENSG0000C |
| 39750127 | 39751293 | 1 | processed_pseudoge | ENSG0000C |
| 39874255 | 39874515 | -1 | misc_RNA | ENSG0000C |
| 39875289 | 39876108 | 1 | processed_pseudoge | ENSG0000C |
| 39901084 | 39973721 | 1 | protein_coding | ENSG0000C |
| 39960397 | 39964718 | 1 | protein_coding | ENSG0000C |
| 39994954 | 40043534 | 1 | protein_coding | ENSG0000C |
| 40043068 | 40044530 | -1 | lncRNA | ENSG0000C |
| 40044817 | 40335808 | 1 | protein_coding | ENSG0000C |
| 40106325 | 40107066 | -1 | processed_pseudoge | ENSG0000C |
| 40346500 | 40390463 | 1 | protein_coding | ENSG0000C |
| 40346529 | 40410054 | 1 | protein_coding | ENSG0000C |
| 40370591 | 40410289 | 1 | protein_coding | ENSG0000C |
| 40372664 | 40388290 | -1 | lncRNA | ENSG0000C |
| 40410281 | 40636719 | -1 | protein_coding | ENSG0000C |
| 40415003 | 40415445 | 1 | lncRNA | ENSG0000C |
| 40433136 | 40433641 | 1 | processed_pseudoge | ENSG0000C |
| 40521800 | 40526707 | 1 | lncRNA | ENSG0000C |
| 40569286 | 40569895 | 1 | processed_pseudoge | ENSG0000C |
| 40586700 | 40588003 | -1 | processed_pseudoge | ENSG0000C |
| 40673484 | 40674451 | -1 | processed_pseudoge | ENSG0000C |
| 40678750 | 40682814 | 1 | protein_coding | ENSG0000C |
| 40769630 | 40819399 | -1 | protein_coding | ENSG0000C |
| 40788777 | 40789267 | -1 | processed_pseudoge | ENSG0000C |
| 40798039 | 40798285 | 1 | processed_pseudoge | ENSG0000C |

| | | | | |
|---|---|---|---|---|
| 40813883 | 40813958 | -1 | miRNA | ENSG0000( |
| 40824535 | 40856639 | -1 | protein_coding | ENSG0000( |
| 40857077 | 40932815 | 1 | protein_coding | ENSG0000( |
| 40859549 | 40862113 | -1 | protein_coding | ENSG0000( |
| 40917111 | 40917381 | 1 | misc_RNA | ENSG0000( |
| 40917743 | 40917846 | 1 | snRNA | ENSG0000( |
| 40951347 | 40973309 | 1 | protein_coding | ENSG0000( |
| 41045497 | 41045591 | 1 | misc_RNA | ENSG0000( |
| 41064052 | 41064633 | 1 | processed_pseudoge | ENSG0000( |
| 41065554 | 41065646 | -1 | misc_RNA | ENSG0000( |
| 41074180 | 41075239 | -1 | processed_pseudoge | ENSG0000( |
| 41092513 | 41092566 | 1 | miRNA | ENSG0000( |
| 41092592 | 41180077 | 1 | protein_coding | ENSG0000( |
| 41168217 | 41168316 | 1 | snRNA | ENSG0000( |
| 41169190 | 41183144 | -1 | lncRNA | ENSG0000( |
| 41174591 | 41197456 | -1 | lncRNA | ENSG0000( |
| 41189727 | 41190188 | -1 | processed_pseudoge | ENSG0000( |
| 41205282 | 41231271 | 1 | protein_coding | ENSG0000( |
| 41207592 | 41228500 | -1 | lncRNA | ENSG0000( |
| 41229510 | 41240931 | -1 | protein_coding | ENSG0000( |
| 41245611 | 41286251 | -1 | protein_coding | ENSG0000( |
| 41252992 | 41253050 | -1 | miRNA | ENSG0000( |
| 41301525 | 41360147 | 1 | protein_coding | ENSG0000( |
| 41367333 | 41399326 | 1 | protein_coding | ENSG0000( |
| 41377177 | 41377283 | -1 | snRNA | ENSG0000( |
| 41430934 | 41431375 | 1 | lncRNA | ENSG0000( |
| 41433494 | 41446801 | -1 | protein_coding | ENSG0000( |
| 41459717 | 41468692 | -1 | protein_coding | ENSG0000( |
| 41469117 | 41528989 | 1 | protein_coding | ENSG0000( |
| 41525799 | 41544606 | -1 | protein_coding | ENSG0000( |
| 41551710 | 41560905 | -1 | lncRNA | ENSG0000( |
| 41561010 | 41577741 | 1 | protein_coding | ENSG0000( |
| 41576900 | 41589871 | -1 | protein_coding | ENSG0000( |
| 41598028 | 41621043 | -1 | protein_coding | ENSG0000( |
| 41621119 | 41664048 | 1 | protein_coding | ENSG0000( |
| 41648089 | 41648190 | -1 | misc_RNA | ENSG0000( |
| 41673933 | 41690504 | -1 | protein_coding | ENSG0000( |
| 41676745 | 41677125 | -1 | processed_pseudoge | ENSG0000( |
| 41679711 | 41679817 | 1 | snRNA | ENSG0000( |
| 41688877 | 41698136 | 1 | unitary_pseudogene | ENSG0000( |
| 41699503 | 41799456 | 1 | protein_coding | ENSG0000( |
| 41709225 | 41709489 | -1 | processed_pseudoge | ENSG0000( |
| 41722043 | 41722131 | -1 | snRNA | ENSG0000( |
| 41747312 | 41747413 | 1 | misc_RNA | ENSG0000( |
| 41800679 | 41826299 | 1 | protein_coding | ENSG0000( |
| 41831215 | 41834665 | -1 | lncRNA | ENSG0000( |
| 41833079 | 41907307 | 1 | protein_coding | ENSG0000( |
| 41900944 | 41901012 | 1 | miRNA | ENSG0000( |
| 41909554 | 41914667 | -1 | protein_coding | ENSG0000( |
| 41922023 | 41926818 | -1 | protein_coding | ENSG0000( |

| | | | | |
|---|---|---|---|---|
| 41923222 | 41923297 | -1 | miRNA | ENSG0000C |
| 41938737 | 41947152 | -1 | protein_coding | ENSG0000C |
| 41952174 | 41958933 | 1 | protein_coding | ENSG0000C |
| 41969475 | 41998221 | 1 | protein_coding | ENSG0000C |
| 41981304 | 41982217 | -1 | lncRNA | ENSG0000C |
| 41998725 | 42058456 | 1 | protein_coding | ENSG0000C |
| 42000446 | 42001991 | -1 | transcribed_processe | ENSG0000C |
| 42058334 | 42070842 | -1 | protein_coding | ENSG0000C |
| 42070949 | 42071488 | 1 | lncRNA | ENSG0000C |
| 42074248 | 42079438 | 1 | protein_coding | ENSG0000C |
| 42076058 | 42076161 | -1 | snoRNA | ENSG0000C |
| 42079691 | 42084284 | 1 | protein_coding | ENSG0000C |
| 42085526 | 42090884 | -1 | protein_coding | ENSG0000C |
| 42089630 | 42090028 | -1 | lncRNA | ENSG0000C |
| 42090931 | 42137742 | 1 | lncRNA | ENSG0000C |
| 42107765 | 42108953 | -1 | processed_pseudoge | ENSG0000C |
| 42126499 | 42130881 | -1 | protein_coding | ENSG0000C |
| 42132543 | 42132998 | 1 | processed_pseudoge | ENSG0000C |
| 42136433 | 42139927 | -1 | lncRNA | ENSG0000C |
| 42138060 | 42139726 | 1 | lncRNA | ENSG0000C |
| 42140203 | 42149455 | -1 | polymorphic_pseudc | ENSG0000C |
| 42149886 | 42155001 | -1 | unprocessed_pseudc | ENSG0000C |
| 42160013 | 42343616 | -1 | protein_coding | ENSG0000C |
| 42269703 | 42279534 | 1 | lncRNA | ENSG0000C |
| 42364390 | 42369284 | -1 | lncRNA | ENSG0000C |
| 42380410 | 42432395 | -1 | protein_coding | ENSG0000C |
| 42438023 | 42446195 | 1 | lncRNA | ENSG0000C |
| 42500568 | 42512560 | 1 | transcribed_unitary_ | ENSG0000C |
| 42508344 | 42519796 | -1 | protein_coding | ENSG0000C |
| 42553617 | 42574382 | 1 | protein_coding | ENSG0000C |
| 42555223 | 42582038 | -1 | transcribed_unproce | ENSG0000C |
| 42565048 | 42565330 | -1 | misc_RNA | ENSG0000C |
| 42569147 | 42569250 | -1 | snRNA | ENSG0000C |
| 42583721 | 42614962 | -1 | protein_coding | ENSG0000C |
| 42615244 | 42615393 | 1 | snRNA | ENSG0000C |
| 42615244 | 42615907 | 1 | lncRNA | ENSG0000C |
| 42617840 | 42649392 | -1 | protein_coding | ENSG0000C |
| 42639803 | 42640601 | -1 | protein_coding | ENSG0000C |
| 42692121 | 42721298 | -1 | protein_coding | ENSG0000C |
| 42762860 | 42762959 | -1 | misc_RNA | ENSG0000C |
| 42776406 | 42777296 | -1 | processed_pseudoge | ENSG0000C |
| 42784462 | 42787100 | 1 | unprocessed_pseudc | ENSG0000C |
| 42791814 | 42794313 | -1 | processed_pseudoge | ENSG0000C |
| 42796502 | 42858106 | -1 | protein_coding | ENSG0000C |
| 42835412 | 43015149 | -1 | protein_coding | ENSG0000C |
| 42853403 | 42853799 | -1 | processed_pseudoge | ENSG0000C |
| 43038585 | 43052366 | 1 | lncRNA | ENSG0000C |
| 43039516 | 43089419 | -1 | protein_coding | ENSG0000C |
| 43092167 | 43092524 | 1 | processed_pseudoge | ENSG0000C |
| 43110750 | 43129712 | 1 | protein_coding | ENSG0000C |

| | | | | |
|---|---|---|---|---|
| 43132209 | 43143398 | -1 | protein_coding | ENSG0000C |
| 43151547 | 43163242 | 1 | protein_coding | ENSG0000C |
| 43166622 | 43187134 | -1 | protein_coding | ENSG0000C |
| 43197280 | 43343372 | -1 | protein_coding | ENSG0000C |
| 43212674 | 43213661 | 1 | lncRNA | ENSG0000C |
| 43232141 | 43239010 | 1 | lncRNA | ENSG0000C |
| 43275955 | 43283830 | 1 | lncRNA | ENSG0000C |
| 43400331 | 43409681 | 1 | lncRNA | ENSG0000C |
| 43411196 | 43507848 | 1 | protein_coding | ENSG0000C |
| 43507824 | 43513727 | -1 | lncRNA | ENSG0000C |
| 43516107 | 43560063 | 1 | lncRNA | ENSG0000C |
| 43528744 | 43812337 | -1 | protein_coding | ENSG0000C |
| 43759199 | 43759910 | -1 | processed_pseudoge | ENSG0000C |
| 43802544 | 43802772 | 1 | processed_pseudoge | ENSG0000C |
| 43812456 | 43817394 | 1 | lncRNA | ENSG0000C |
| 43824509 | 43862513 | -1 | protein_coding | ENSG0000C |
| 43879678 | 43892013 | -1 | protein_coding | ENSG0000C |
| 43923792 | 43964488 | 1 | protein_coding | ENSG0000C |
| 43955442 | 44010531 | 1 | protein_coding | ENSG0000C |
| 43957027 | 43957348 | 1 | processed_pseudoge | ENSG0000C |
| 43999211 | 44172949 | 1 | protein_coding | ENSG0000C |
| 44102779 | 44116232 | 1 | lncRNA | ENSG0000C |
| 44139365 | 44153626 | 1 | lncRNA | ENSG0000C |
| 44172956 | 44219533 | 1 | protein_coding | ENSG0000C |
| 44183031 | 44187739 | -1 | TEC | ENSG0000C |
| 44210442 | 44229512 | -1 | lncRNA | ENSG0000C |
| 44243665 | 44312951 | -1 | protein_coding | ENSG0000C |
| 44365551 | 44366284 | 1 | lncRNA | ENSG0000C |
| 44413485 | 44414814 | -1 | processed_pseudoge | ENSG0000C |
| 44443327 | 44444788 | 1 | lncRNA | ENSG0000C |
| 44483495 | 44483717 | -1 | processed_pseudoge | ENSG0000C |
| 44486295 | 44486914 | -1 | TEC | ENSG0000C |
| 44492583 | 44498233 | -1 | protein_coding | ENSG0000C |
| 44566965 | 44568247 | 1 | processed_pseudoge | ENSG0000C |
| 44569295 | 44572453 | 1 | lncRNA | ENSG0000C |
| 44606939 | 44625419 | -1 | lncRNA | ENSG0000C |
| 44651835 | 44652379 | 1 | processed_pseudoge | ENSG0000C |
| 44668547 | 44737681 | 1 | protein_coding | ENSG0000C |
| 44702233 | 44862706 | 1 | protein_coding | ENSG0000C |
| 44752558 | 44862788 | 1 | protein_coding | ENSG0000C |
| 44881162 | 45010005 | -1 | protein_coding | ENSG0000C |
| 44901547 | 44902471 | 1 | transcribed_processe | ENSG0000C |
| 45000565 | 45003619 | -1 | lncRNA | ENSG0000C |
| 45133020 | 45163781 | -1 | lncRNA | ENSG0000C |
| 45155539 | 45156011 | -1 | lncRNA | ENSG0000C |
| 45163925 | 45188017 | 1 | protein_coding | ENSG0000C |
| 45178990 | 45179310 | 1 | lncRNA | ENSG0000C |
| 45190338 | 45240769 | -1 | protein_coding | ENSG0000C |
| 45200954 | 45201019 | -1 | miRNA | ENSG0000C |
| 45262273 | 45263585 | -1 | lncRNA | ENSG0000C |

| | | | | |
|---|---|---|---|---|
| 45275599 | 45276085 | 1 | processed_pseudoge | ENSG00000( |
| 45284949 | 45295874 | 1 | protein_coding | ENSG00000( |
| 45308968 | 45341955 | 1 | protein_coding | ENSG00000( |
| 45344063 | 45413619 | -1 | protein_coding | ENSG00000( |
| 45413693 | 45432509 | 1 | protein_coding | ENSG00000( |
| 45435864 | 45448743 | -1 | lncRNA | ENSG00000( |
| 45502238 | 45601135 | 1 | protein_coding | ENSG00000( |
| 45604432 | 45605621 | -1 | lncRNA | ENSG00000( |
| 45624531 | 45624632 | -1 | snRNA | ENSG00000( |
| 45657019 | 45680130 | 1 | lncRNA | ENSG00000( |
| 45671798 | 45845307 | 1 | protein_coding | ENSG00000( |
| 45760524 | 45760598 | 1 | miRNA | ENSG00000( |
| 45792418 | 45793636 | -1 | processed_pseudoge | ENSG00000( |
| 45875932 | 45879247 | 1 | lncRNA | ENSG00000( |
| 45875999 | 45891438 | -1 | lncRNA | ENSG00000( |
| 45920366 | 45977162 | -1 | protein_coding | ENSG00000( |
| 46006616 | 46010777 | 1 | TEC | ENSG00000( |
| 46013606 | 46015498 | 1 | lncRNA | ENSG00000( |
| 46039907 | 46044853 | -1 | lncRNA | ENSG00000( |
| 46049478 | 46054144 | -1 | lncRNA | ENSG00000( |
| 46053093 | 46053560 | 1 | lncRNA | ENSG00000( |
| 46053705 | 46057210 | 1 | lncRNA | ENSG00000( |
| 46053869 | 46113928 | 1 | lncRNA | ENSG00000( |
| 46055740 | 46058160 | -1 | lncRNA | ENSG00000( |
| 46067356 | 46069891 | -1 | lncRNA | ENSG00000( |
| 46091044 | 46091126 | 1 | miRNA | ENSG00000( |
| 46112749 | 46112822 | 1 | miRNA | ENSG00000( |
| 46113566 | 46113657 | 1 | miRNA | ENSG00000( |
| 46113686 | 46113768 | 1 | miRNA | ENSG00000( |
| 46150521 | 46243756 | 1 | protein_coding | ENSG00000( |
| 46163303 | 46165347 | 1 | lncRNA | ENSG00000( |
| 46244011 | 46250311 | -1 | protein_coding | ENSG00000( |
| 46255663 | 46263343 | -1 | protein_coding | ENSG00000( |
| 46267961 | 46294008 | 1 | protein_coding | ENSG00000( |
| 46295143 | 46296660 | -1 | lncRNA | ENSG00000( |
| 46296870 | 46330810 | 1 | protein_coding | ENSG00000( |
| 46330875 | 46357340 | 1 | protein_coding | ENSG00000( |
| 46360834 | 46537170 | -1 | protein_coding | ENSG00000( |
| 46481329 | 46481760 | -1 | processed_pseudoge | ENSG00000( |
| 46541495 | 46548196 | 1 | lncRNA | ENSG00000( |
| 46576012 | 46679790 | 1 | protein_coding | ENSG00000( |
| 46684410 | 46738252 | -1 | protein_coding | ENSG00000( |
| 46761894 | 46762563 | -1 | lncRNA | ENSG00000( |
| 46762617 | 47175699 | 1 | protein_coding | ENSG00000( |
| 46914092 | 46916042 | -1 | lncRNA | ENSG00000( |
| 47115838 | 47117217 | -1 | TEC | ENSG00000( |
| 47259366 | 47261007 | 1 | TEC | ENSG00000( |
| 47345569 | 47373541 | 1 | lncRNA | ENSG00000( |
| 47461299 | 47487111 | -1 | lncRNA | ENSG00000( |
| 47612231 | 47631569 | -1 | lncRNA | ENSG00000( |

| | | | | |
|---|---|---|---|---|
| 47630827 | 48023004 | 1 | lncRNA | ENSG0000C |
| 47662559 | 47663603 | -1 | processed_pseudoge | ENSG0000C |
| 47916728 | 47926264 | 1 | lncRNA | ENSG0000C |
| 48044710 | 48048549 | -1 | lncRNA | ENSG0000C |
| 48052646 | 48058275 | 1 | lncRNA | ENSG0000C |
| 48139461 | 48141001 | -1 | lncRNA | ENSG0000C |
| 48141542 | 48143229 | 1 | lncRNA | ENSG0000C |
| 48254903 | 48259116 | -1 | lncRNA | ENSG0000C |
| 48274364 | 48274415 | 1 | miRNA | ENSG0000C |
| 48403462 | 48415571 | 1 | lncRNA | ENSG0000C |
| 48448890 | 48451217 | 1 | lncRNA | ENSG0000C |
| 48474135 | 48489324 | -1 | lncRNA | ENSG0000C |
| 48489460 | 48850912 | 1 | protein_coding | ENSG0000C |
| 48538900 | 48547387 | -1 | lncRNA | ENSG0000C |
| 48780295 | 48780353 | 1 | miRNA | ENSG0000C |
| 48866770 | 48898386 | 1 | lncRNA | ENSG0000C |
| 49047484 | 49052549 | -1 | lncRNA | ENSG0000C |
| 49166421 | 49188062 | -1 | lncRNA | ENSG0000C |
| 49213911 | 49214247 | -1 | processed_pseudoge | ENSG0000C |
| 49264201 | 49266080 | 1 | lncRNA | ENSG0000C |
| 49372924 | 49377119 | 1 | lncRNA | ENSG0000C |
| 49414524 | 49657542 | -1 | lncRNA | ENSG0000C |
| 49500568 | 49501585 | 1 | lncRNA | ENSG0000C |
| 49534062 | 49534926 | 1 | lncRNA | ENSG0000C |
| 49543393 | 49543466 | -1 | miRNA | ENSG0000C |
| 49548681 | 49556473 | 1 | lncRNA | ENSG0000C |
| 49572264 | 49575426 | -1 | lncRNA | ENSG0000C |
| 49612657 | 49615716 | -1 | lncRNA | ENSG0000C |
| 49662312 | 49664968 | -1 | lncRNA | ENSG0000C |
| 49718053 | 49724506 | -1 | lncRNA | ENSG0000C |
| 49738677 | 49739003 | 1 | misc_RNA | ENSG0000C |
| 49757627 | 49758980 | -1 | unprocessed_pseudc | ENSG0000C |
| 49773283 | 49827512 | -1 | protein_coding | ENSG0000C |
| 49805452 | 49807208 | 1 | lncRNA | ENSG0000C |
| 49817248 | 49817712 | 1 | processed_pseudoge | ENSG0000C |
| 49832449 | 49838901 | -1 | lncRNA | ENSG0000C |
| 49845929 | 49846090 | 1 | processed_pseudoge | ENSG0000C |
| 49846456 | 49846720 | 1 | misc_RNA | ENSG0000C |
| 49851425 | 49853364 | 1 | lncRNA | ENSG0000C |
| 49853844 | 49890080 | 1 | protein_coding | ENSG0000C |
| 49900229 | 49918438 | -1 | protein_coding | ENSG0000C |
| 49902228 | 49904576 | 1 | lncRNA | ENSG0000C |
| 49918167 | 49927540 | 1 | protein_coding | ENSG0000C |
| 49933198 | 49934074 | -1 | lncRNA | ENSG0000C |
| 49960768 | 49964072 | 1 | protein_coding | ENSG0000C |
| 49962866 | 49962939 | 1 | miRNA | ENSG0000C |
| 49994513 | 50012659 | -1 | protein_coding | ENSG0000C |
| 50015123 | 50056935 | -1 | protein_coding | ENSG0000C |
| 50059391 | 50085902 | -1 | protein_coding | ENSG0000C |
| 50089879 | 50161690 | 1 | protein_coding | ENSG0000C |

| | | | | |
|---|---|---|---|---|
| 50092745 | 50096437 | 1 | lncRNA | ENSG0000C |
| 50170731 | 50180295 | 1 | protein_coding | ENSG0000C |
| 50185915 | 50199598 | 1 | protein_coding | ENSG0000C |
| 50191724 | 50192402 | -1 | lncRNA | ENSG0000C |
| 50199090 | 50200837 | -1 | lncRNA | ENSG0000C |
| 50200979 | 50217616 | 1 | protein_coding | ENSG0000C |
| 50205585 | 50206062 | -1 | lncRNA | ENSG0000C |
| 50208461 | 50209542 | -1 | lncRNA | ENSG0000C |
| 50217689 | 50245023 | -1 | protein_coding | ENSG0000C |
| 50245183 | 50251405 | -1 | protein_coding | ENSG0000C |
| 50245450 | 50261716 | -1 | protein_coding | ENSG0000C |
| 50263713 | 50270767 | -1 | protein_coding | ENSG0000C |
| 50274979 | 50307646 | -1 | protein_coding | ENSG0000C |
| 50309030 | 50327012 | -1 | protein_coding | ENSG0000C |
| 50314631 | 50316008 | 1 | lncRNA | ENSG0000C |
| 50316035 | 50317025 | 1 | lncRNA | ENSG0000C |
| 50343304 | 50445090 | 1 | protein_coding | ENSG0000C |
| 50445000 | 50475035 | -1 | protein_coding | ENSG0000C |
| 50481543 | 50486440 | 1 | protein_coding | ENSG0000C |
| 50486784 | 50490648 | 1 | protein_coding | ENSG0000C |
| 50502949 | 50507702 | -1 | protein_coding | ENSG0000C |
| 50508224 | 50524780 | 1 | protein_coding | ENSG0000C |
| 50523568 | 50526461 | -1 | protein_coding | ENSG0000C |
| 50523926 | 50524780 | 1 | lncRNA | ENSG0000C |
| 50525752 | 50530032 | -1 | protein_coding | ENSG0000C |
| 50529710 | 50532580 | -1 | protein_coding | ENSG0000C |
| 50541108 | 50543011 | 1 | lncRNA | ENSG0000C |
| 50542305 | 50542906 | -1 | lncRNA | ENSG0000C |
| 50545899 | 50551023 | 1 | protein_coding | ENSG0000C |
| 50551112 | 50562919 | -1 | protein_coding | ENSG0000C |
| 50568861 | 50578465 | -1 | protein_coding | ENSG0000C |
| 50568869 | 50582965 | -1 | protein_coding | ENSG0000C |
| 50578949 | 50601455 | -1 | protein_coding | ENSG0000C |
| 50583026 | 50595634 | 1 | lncRNA | ENSG0000C |
| 50597152 | 50597599 | 1 | lncRNA | ENSG0000C |
| 50600793 | 50613981 | 1 | protein_coding | ENSG0000C |
| 50622754 | 50628173 | -1 | protein_coding | ENSG0000C |
| 50669804 | 50669901 | 1 | misc_RNA | ENSG0000C |
| 50672714 | 50674174 | 1 | lncRNA | ENSG0000C |
| 50674415 | 50733298 | 1 | protein_coding | ENSG0000C |
| 50691260 | 50691363 | -1 | snRNA | ENSG0000C |
| 50735825 | 50738139 | -1 | lncRNA | ENSG0000C |
| 50738196 | 50745339 | 1 | protein_coding | ENSG0000C |
| 50740593 | 50743520 | -1 | lncRNA | ENSG0000C |
| 50754675 | 50755434 | -1 | processed_pseudoge | ENSG0000C |
| 50756948 | 50801309 | 1 | transcribed_unproce | ENSG0000C |
| 50767501 | 50783663 | -1 | protein_coding | ENSG0000C |

0283350
0283640
0284971
0283158
0288360
0288519
0288240
0288498
0288126
0288183
0288462
0288505
0288397
0288329
0288152
0288293
0288195
0288453
0288283
0288351
0288474
0288441
0288185
0288318
0288278
0288191
0288366
0288239
0288488
0288516
0288496
0288210
0288348
0288324
0288289
0288425
0288182
0288433
0281766
0280864
0281822
0280505
0280775
0288175
0288457
0288244
0288157
0288304
0288481

0288424
0288201
0288196
0288134
0288522
0288204
0288247
0288372
0288288
0288129
0288472
0288417
0288502
0288311
0288449
0288499
0288159
0288431
0283595
0287911
0283264
0283243
0283539
0283287
0283252
0283658
0288269
0288213
0288186
0282162
0281934
0282116
0282175
0282817
0282584
0282663
0279195
0282338
0282549
0282119
0282791
0282437
0282559
0282566
0282258
0282562
0282530
0282468
0282341
0281987

)282095
)282298
)282741
)282212
)282289
)282424
)282216
)282288
)282583
)282125
)281449
)281442
)283344
)281324
)280475
)206783
)281874
)207477
)281818
)281051
)281234
)280689
)281168
)280962
)281671
)281525
)288277
)288406
)288275
)288260
)288430
)288297
)288274
)288243
)288388
)288138
)288512
)288124
)288352
)288369
)284403
)288383
)288427
)288281
)288116
)288217
)288463
)281819
)263150
)262851

0261897
0262784
0262611
0280712
0262755
0281773
0280617
0263328
0281820
0262796
0262315
0281038
0280577
0288494
0288135
0288322
0288356
0288117
0288319
0288521
0288394
0288228
0288342
0288315
0288272
0261958
0262191
0281683
0262268
0262647
0281777
0281313
0281279
0262535
0281107
0281839
0288359
0288263
0288305
0285380
0285001
0285019
0285196
0285308
0285028
0285455
0285477
0288404
0282467
0284255

0284278
0284123
0284767
0285070
0285310
0285682
0284817
0284953
0286298
0285383
0285149
0285233
0284850
0285479
0285355
0285291
0285266
0285324
0285107
0284775
0285924
0285983
0284942
0284759
0285101
0285335
0285134
0280963
0281230
0281067
0281552
0280680
0287371
0281499
0287648
0280937
0285111
0284831
0285274
0284911
0284904
0285965
0285289
0285282
0284807
0286673
0284898
0285036
0284765
0285221

0285493
0285241
0284796
0284818
0285321
0284916
0285377
0285298
0284889
0285146
0285152
0284845
0285306
0285048
0285007
0285248
0285125
0287341
0285465
0285502
0284802
0285309
0285140
0285302
0285296
0285200
0285425
0285246
0284752
0284915
0285348
0285193
0287436
0286762
0284868
0285326
0285199
0285203
0285456
0284814
0285511
0285379
0285386
0285485
0285512
0285344
0285435
0284795
0284841
0285449

0284998
0285234
0285050
0285426
0285153
0285166
0285285
0285168
0284853
0285158
0284877
0285311
0285183
0285319
0284984
0280774
0281677
0281320
0281403
0280548
0281123
0280932
0281221
0281526
0281233
0281601
0281009
0280568
0280797
0283272
0283276
0283668
0283153
0283508
0283127
0283593
0283700
0283581
0283181
0285063
0283449
0283363
0284383
0284496
0284262
0283767
0284210
0283884
0283955
0284025

0283878
0284181
0283820
0283949
0283660
0283223
0280584
0280820
0281879
0280632
0281430
0281309
0281393
0281022
0281699
0280858
0280889
0281243
0281907
0283984
0281541
0280627
0281024
0280951
0281245
0280706
0281244
0280479
0281165
0283781
0284009
0284483
0283771
0283859
0283843
0284598
0283714
0284090
0286184
0284228
0283895
0283903
0284222
0284115
0283946
0283868
0284515
0284832
0285481
0285371

0285222
0284750
0285415
0285209
0284885
0285893
0285312
0284941
0285226
0284936
0281316
0281877
0280948
0281092
0280876
0281679
0280806
0280926
0281805
0285164
0285500
0285429
0285143
0281211
0283530
0283651
0283163
0283034
0281626
0280781
0281707
0281894
0281456
0280661
0280795
0281709
0280907
0281306
0281102
0280872
0281448
0281661
0281665
0280610
0284275
0280516
0281886
0280701
0284058
0284367

0287606
0283068
0282867
0282877
0287304
0283085
0288307
0288264
0288485
0288292
0288215
0288174
0288371
0288504
0288145
0288325
0288456
0288301
0288279
0288169
0288399
0288409
0288511
0284126
0284068
0281226
0283081
0281459
0280549
0281810
0280908
0280949
0281391
0281540
0281917
0280751
0284271
0281557
0288248
0288411
0288479
0288445
0288114
0288358
0288299
0288379
0285387
0285206
0284900
0285092

0285139
0284938
0284838
0285121
0285501
0285182
0285489
0285503
0285046
0285031
0284852
0288198
0288378
0288343
0288233
0288267
0288163
0288420
0288166
0288401
0283114
0283062
0288121
0288332
0288475
0288443
0288421
0288402
0288435
0288500
0252303
0281771
0284833
0285086
0285436
0281256
0281813
0281770
0281086
0281660
0285951
0288334
0288508
0288257
0288407
0288132
0288365
0288125
0288362
0288314

0288150
0288179
0288178
0288455
0288497
0288212
0288408
0288390
0288227
0288276
0288341
0282825
0283080
0282900
0282959
0283002
0282923
0281277
0281029
0283318
0281705
0280975
0281913
0280582
0281697
0280831
0280495
0284577
0281500
0281860
0280682
0281163
0280967
0280616
0281614
0280669
0281089
0280652
0281388
0281857
0280928
0280873
0281623
0280792
0281384
0282956
0282739
0282510
0282585
0284345

0284109
0284017
0283960
0284477
0287604
0288400
0288454
0288447
0288503
0288146
0288256
0288291
0288151
0288490
0288298
0288209
0288510
0288327
0288384
0288458
0288376
0288450
0288514
0288333
0288120
0282873
0280651
0281178
0283035
0281865
0285194
0285145
0285099
0285395
0284910
0284843
0285003
0285009
0286814
0285170
0285903
0284096
0283847
0284406
0284369
0284045
0283812
0284103
0284582
0283852

0284201
0284189
0283985
0283979
0284165
0283883
0284075
0284323
0284390
0284153
0283302
0280477
0280531
0282172
0282050
0282699
0282448
0282801
0282163
0282310
0283113
0282758
0282248
0282823
0282694
0282603
0282251
0282447
0282282
0281978
0282200
0282244
0282402
0282210
0282762
0282093
0282671
0282062
0282622
0282000
0282666
0282025
0282635
0281933
0282743
0282571
0282811
0282149
0282157
0282646

0282394
0282387
0282120
0282182
0281509
0281252
0281175
0283864
0284712
0281616
0281084
0280895
0281878
0283024
0282866
0283092
0283059
0284792
0283055
0283120
0282931
0283067
0288229
0288222
0288340
0288189
0288375
0288429
0288133
0288270
0288137
0288280
0288112
0288480
0288202
0288350
0288382
0288265
0288337
0288113
0288194
0288501
0288164
0288190
0288313
0288377
0288345
0288282
0288518
0288465

0288171
0288142
0288469
0288193
0288165
0288413
0288221
0288483
0288432
0288286
0288347
0288211
0288419
0288442
0288176
0288354
0288423
0283465
0288478
0288461
0288241
0288214
0288344
0288339
0288173
0288440
0288128
0288517
0288405
0288188
0288364
0288448
0285384
0284955
0285874
0285232
0286287
0287491
0285301
0284774
0285482
0284758
0285428
0284923
0285272
0285047
0285112
0285442
0285451
0285049

0284918
0285207
0285088
0284888
0285243
0285339
0285060
0285389
0287013
0288130
0288115
0288216
0288353
0288238
0288513
0288495
0288123
0288464
0288246
0288468
0288207
0288355
0288170
0288336
0288284
0288392
0288236
0288506
0288515
0288259
0288147
0288385
0288416
0288391
0288410
0288285
0288230
0288180
0288438
0288482
0288192
0288444
0288387
0288491
0288395
0288203
0288451
0288308
0288346
0288254

0288326
0288168
0288386
0288493
0288287
0288144
0288294
0288290
0288446
0288158
0288489
0288303
0288184
0288249
0282928
0282982
0282986
0282844
0282976
0282869
0282902
0283082
0283011
0282992
0282853
0282832
0283077
0282947
0282846
0282895
0283121
0283100
0282905
0283099
0283104
0282835
0283018
0282930
0282954
0286702
0288149
0288219
0288199
0288415
0288452
0288331
0288160
0288250
0288317
0288141

0288320
0288486
0288237
0288428
0288367
0288361
0283802
0284315
0284358
0284579
0284494
0283939
0283896
0284549
0284124
0284260
0284556
0283808
0284254
0284069
0284506
0284445
0283718
0283777
0284207
0283780
0283887
0284501
0283912
0284099
0283918
0284026
0284437
0284274
0284400
0288295
0288368
0288412
0288226
0288466
0288328
0288273
0288477
0288251
0288363
0288197
0288220
0288118
0288140
0288374

0288167
0288310
0288205
0288268
0288296
0288143
0288154
0288223
0288155
0288338
0288509
0288261
0288426
0288231
0288224
0288492
0288414
0288122
0288470
0285443
0285487
0284816
0285468
0285054
0285341
0284982
0285299
0285252
0285052
0285495
0285172
0284766
0285123
0285277
0285393
0285351
0284965
0284866
0285391
0284945
0285242
0284939
0288507
0288403
0288335
0285262
0285267
0285400
0284905
0284980

0284880
0285083
0286726
0285327
0285156
0284881
0285136
0285227
0285318
0284863
0284789
0285475
0284983
0284913
0285176
0284924
0284835
0285439
0285113
0285251
0284960
0285014
0285129
0284771
0285418
0285381
0285466
0284760
0285211
0284777
0285474
0285832
0285767
0284870
0285678
0284979
0285015
0285617
0285034
0285507
0285376
0285504
0285433
0284958
0285120
0285494
0284781
0284806
0284851
0284799

0285074
0285795
0284947
0285452
0285316
0284926
0284975
0284991
0285124
0284778
0284894
0285249
0284782
0285490
0285349
0285431
0284893
0284754
0285365
0285087
0287890
0287681
0285469
0285225
0285132
0284973
0284967
0285261
0285173
0284811
0285217
0285109
0287572
0285457
0285369
0285157
0285072
0285275
0284798
0285464
0285295
0284856
0285084
0285340
0285032
0285067
0285118
0285284
0285195
0284784

0284927
0285026
0285276
0284974
0285459
0284801
0285247
0285056
0285403
0284764
0284875
0285187
0285438
0285147
0285126
0285364
0285141
0285392
0285229
0284756
0284813
0284793
0285250
0285320
0285787
0285995
0284859
0285161
0285372
0285033
0284854
0284878
0285359
0285236
0285484
0285061
0284864
0285423
0285185
0285127
0284919
0285343
0285002
0284887
0285167
0285514
0285368
0285264
0284860
0285414

0285022
0284943
0285290
0285098
0287874
0284857
0285260
0285345
0285450
0285119
0285027
0284929
0284972
0285962
0285432
0285411
0284937
0285356
0288309
0284978
0285066
0285181
0284907
0285286
0285097
0285497
0284990
0285038
0285397
0285186
0284920
0284909
0284992
0285037
0284751
0284865
0284995
0284822
0285419
0285315
0284785
0285360
0285270
0284855
0285506
0285297
0284812
0284872
0284861
0285265

0288242
0284761
0283997
0284382
0284588
0284290
0283738
0283892
0284218
0288373
0284282
0284161
0284487
0284316
0284493
0283826
0283855
0284559
0284548
0283934
0284330
0284454
0287573
0282878
0281764
0286783
0281518
0281876
0276777
0278036
0275336
0273501
0273681
0278364
0276595
0275985
0278121
0282156
0275615
0273495
0274650
0277495
0275565
0276446
0278545
0282764
0282178
0282271
0282372
0276274

0282534
0282233
0274429
0282690
0276516
0282770
0283079
0283007
0282971
0281118
0284476
0280931
0280808
0281056
0281563
0280899
0281680
0281166
0285029
0285510
0285375
0285463
0285422
0284903
0284935
0281427
0276025
0278253
0280497
0281367
0278379
0276451
0273953
0276780
0273793
0281801
0276565
0278210
0278489
0275920
0277068
0277291
0281645
0281733
0281783
0281240
0282076
0282441
0282343
0278855

0275660
0273707
0274454
0276130
0274538
0273562
0273525
0282438
0278191
0282355
0275205
0282367
0276230
0282464
0276536
0273831
0275873
0282005
0273859
0282810
0274780
0276561
0273572
0274473
0276825
0282712
0282598
0280759
0274980
0277518
0278165
0281176
0280585
0281533
0281191
0278620
0278302
0281111
0281261
0281829
0281633
0277389
0281618
0284890
0288058
0285180
0285421
0285516
0285305
0285023

0285065
0284997
0281385
0276186
0277281
0281524
0278466
0272877
0262435
0262927
0282676
0262607
0262226
0263137
0262016
0262193
0262349
0262145
0262819
0262776
0262082
0262201
0273326
0272712
0273086
0263247
0272805
0272718
0272751
0273457
0272803
0261984
0273144
0273431
0273092
0263028
0263097
0263158
0262525
0263083
0262612
0262293
0262829
0261936
0262111
0261853
0262531
0262986
0262159
0262036

0262592
0274327
0277316
0273713
0277823
0277238
0282174
0282269
0277454
0285332
0282363
0277254
0275679
0276541
0276217
0276167
0282612
0278307
0282256
0282293
0278111
0282079
0282537
0274699
0282675
0282546
0281930
0273505
0281977
0282539
0275624
0274839
0282673
0262139
0262217
0261867
0263114
0275428
0282391
0282388
0282329
0276371
0282551
0282176
0278208
0278711
0276226
0275517
0275051
0282480

0282766
0281935
0275035
0281954
0285137
0285478
0286166
0286560
0287228
0282647
0282706
0282737
0275865
0282003
0282037
0282611
0282352
0282782
0282677
0282536
0282715
0278853
0252379
0281988
0282664
0282107
0282311
0280633
0274338
0281924
0276238
0282153
0274581
0282241
0276776
0276342
0274126
0281161
0282751
0282219
0276173
0276192
0277016
0274497
0282430
0282094
0282360
0282474
0282633
0281947

)277633
)282184
)282819
)278801
)282657
)282797
)282541
)282615
)282065
)282471
)282778
)282342
)282409
)282455
)282672
)282323
)282724
)282674
)282640
)282396
)281984
)282135
)282592
)282487
)282346
)282274
)282232
)282818
)282736
)282042
)282102
)281940
)282373
)281939
)282132
)282495
)282010
)282769
)282227
)282754
)282578
)277059
)282714
)282745
)282197
)282626
)282039
)282550
)282108
)282213

0282223
0282678
0277318
0282066
0282139
0281944
0282211
0282644
0282401
0282322
0281964
0282366
0282286
0282569
0282796
0282290
0282631
0282653
0282561
0282659
0282716
0281996
0281928
0282476
0282281
0282609
0282127
0282687
0281962
0282284
0282601
0282710
0281926
0282344
0282239
0282452
0282267
0282726
0282702
0282777
0282136
0282356
0282404
0282761
0281983
0282642
0281959
0278082
0282595
0282150

0282552
0282526
0276491
0282123
0282104
0281986
0281973
0282425
0282576
0282579
0282194
0282533
0282292
0282783
0281971
0282180
0282151
0282802
0282553
0281956
0282776
0282131
0282196
0282398
0282708
0282627
0275316
0282465
0282138
0277574
0282294
0282312
0282188
0282593
0281923
0282383
0282616
0282504
0282794
0282337
0282691
0282586
0273928
0282082
0282451
0282414
0282099
0282560
0274236
0282821

0282018
0282045
0282439
0282117
0282652
0282016
0282161
0282808
0282055
0282350
0282482
0282421
0275148
0282822
0282399
0282453
0282679
0278042
0273539
0282305
0282186
0276060
0281966
0282473
0276708
0280805
0281811
0276781
0274958
0280802
0281141
0278268
0275947
0281847
0275619
0278787
0280562
0277222
0281292
0276416
0281396
0277018
0277032
0276881
0274710
0274089
0281432
0281032
0273533
0278693

)281498
)280668
)277276
)273761
)276880
)275214
)276879
)278326
)278767
)277405
)274821
)277377
)287764
)285281
)284819
)284769
)285008
)285273
)288232
)281804
)274240
)281836
)275809
)281507
)281263
)281208
)281803
)274439
)281153
)281718
)278699
)273624
)281688
)281604
)281622
)280555
)278776
)278178
)274846
)280463
)280618
)276856
)280500
)275009
)280476
)278070
)274040
)280461
)281012
)280679

0274700
0277278
0274801
0282126
0276455
0275855
0276896
0274122
0276143
0282731
0282516
0282581
0282587
0273635
0273969
0281946
0274192
0273651
0276767
0282058
0276485
0282531
0282090
0276005
0280807
0281725
0278247
0281737
0278508
0277046
0281173
0280490
0281254
0281754
0277848
0282280
0277844
0281422
0281537
0281846
0278429
0276377
0280700
0274404
0281828
0280787
0281356
0276062
0282700
0207390

0282077
0273686
0278211
0282195
0273845
0273636
0277361
0282545
0276802
0282105
0282544
0282006
0282475
0282190
0278310
0273839
0282479
0282779
0282038
0282073
0282224
0282060
0282696
0282169
0282660
0282368
0282118
0282023
0282004
0277090
0278003
0282145
0282444
0282422
0282567
0282417
0278186
0275257
0282540
0282072
0282648
0282198
0282655
0276495
0273564
0281925
0276132
0282130
0282503
0277401

0272101
0278119
0276120
0275394
0273660
0276253
0277419
0275698
0275917
0276367
0278118
0276439
0275648
0277167
0274734
0277980
0276152
0276787
0277086
0277262
0274965
0275746
0273538
0274542
0278470
0276847
0278115
0275568
0277614
0276886
0281976
0274042
0282377
0275049
0274166
0282084
0273673
0282814
0274320
0276228
0282457
0282514
0282183
0282209
0282596
0278229
0282755
0277445
0283102
0282193

0282362
0288148
0282654
0282324
0282509
0282767
0284076
0282328
0282400
0282316
0282249
0282009
0282722
0282729
0282088
0286139
0282554
0282670
0282275
0282207
0282689
0282177
0282701
0282773
0282547
0281960
0282109
0282068
0283589
0282629
0282168
0285828
0281931
0282081
0282555
0282046
0282410
0274829
0263947
0275618
0251964
0276870
0282427
0278308
0275283
0282086
0282091
0278494
0275779
0275498

0282717
0278440
0281994
0282788
0281965
0277625
0276709
0273905
0274779
0282247
0273808
0282705
0275543
0282074
0278823
0277140
0282314
0275911
0282255
0282638
0276480
0282287
0282167
0276914
0251771
0282412
0282445
0278183
0281929
0275331
0282574
0278087
0282597
0282332
0282522
0277242
0273807
0277983
0282528
0282744
0281975
0282713
0281948
0262243
0261972
0262788
0262579
0260873
0262354
0274636

0282703
0282228
0278760
0282011
0282725
0274334
0276726
0282450
0282590
0282374
0274684
0277424
0274336
0282650
0282469
0282069
0276333
0283022
0283015
0284106
0282830
0287711
0282838
0282845
0283056
0283070
0282937
0283112
0283049
0285178
0285337
0284810
0285346
0285350
0285460
0285307
0274296
0282201
0275338
0274018
0282378
0282485
0282214
0278006
0282775
0278567
0277943
0274169
0282763
0282389

0282750
0282229
0277796
0282604
0282512
0278632
0282319
0277768
0275313
0282747
0282768
0282403
0275861
0283825
0285039
0282235
0277660
0282518
0277730
0272670
0282365
0262334
0282030
0263267
0282634
0282720
0278550
0273968
0276336
0274145
0274442
0275246
0282800
0276021
0276838
0282307
0263186
0263341
0262591
0262282
0261986
0263012
0262045
0262748
0262993
0262603
0262040
0263022
0283454
0281837

0276836
0281898
0277111
0274332
0278357
0281364
0280810
0275748
0278742
0278553
0281642
0281815
0275867
0274754
0281569
0280735
0276155
0281139
0277055
0280786
0281284
0281578
0278194
0278232
0281283
0281144
0275775
0281487
0281336
0281922
0281797
0275621
0275949
0275574
0281693
0278174
0277626
0278759
0277033
0281945
0275031
0282013
0282357
0275886
0282336
0282426
0280563
0277649
0278741
0273728

0281251
0281590
0280938
0274474
0282128
0282637
0275780
0282563
0276907
0281982
0276358
0282325
0282331
0275587
0282727
0282491
0282002
0282614
0282351
0282238
0282171
0282456
0282523
0282361
0276191
0282067
0277106
0277956
0282620
0282165
0275602
0278458
0282279
0282799
0276222
0282723
0281927
0282032
0276276
0277221
0274171
0276262
0282733
0282052
0277641
0276799
0284137
0284571
0284495
0284449

0284238
0284212
0283753
0283741
0284320
0284492
0284166
0284566
0284422
0284444
0284576
0284456
0284350
0283850
0284257
0284527
0284285
0284205
0284014
0283924
0284021
0284312
0281300
0278501
0276016
0275991
0282649
0282753
0282302
0264450
0263243
0263057
0262487
0263309
0262164
0262845
0263101
0263296
0262616
0262068
0261862
0263242
0262347
0262808
0261852
0262731
0262862
0263091
0262376
0266286

0261968
0262520
0262121
0263229
0263236
0263090
0262749
0267208
0262512
0267728
0267635
0263103
0262218
0262214
0262573
0262896
0262594
0263263
0262946
0262221
0263115
0262542
0278843
0282384
0276833
0282202
0282083
0276986
0282115
0282498
0274233
0274262
0282124
0282486
0276432
0277042
0282525
0275095
0277236
0282521
0282446
0275528
0282588
0282071
0276114
0278167
0282721
0274221
0275824
0278781

0282348
0275347
0274939
0282231
0277336
0276125
0282370
0281967
0281992
0282144
0282411
0278320
0275760
0277974
0278515
0282087
0282656
0278574
0277395
0278372
0275600
0275099
0282707
0275397
0274853
0282570
0282028
0282237
0281998
0274577
0282454
0276072
0282636
0282805
0282043
0282166
0275176
0282532
0282695
0282489
0278145
0282806
0277104
0282790
0282015
0275932
0274047
0277594
0276194
0282632

0282630
0277891
0275153
0276349
0282641
0276153
0277936
0276469
0274229
0273780
0282205
0273608
0282217
0277303
0282494
0275851
0277005
0274768
0278779
0278644
0282749
0275903
0277546
0277292
0276761
0276913
0282257
0275193
0282291
0282385
0284899
0285208
0282443
0282085
0262132
0262634
0262892
0282110
0282433
0262104
0282429
0279612
0263322
0262102
0282220
0282112
0262280
0263310
0274273
0281942

0281995
0282606
0282330
0282752
0281949
0282841
0283107
0282658
0273770
0282765
0277015
0282266
0282460
0283837
0284355
0283800
0284086
0283702
0283832
0288389
0284451
0283953
0283715
0283872
0283942
0284104
0283962
0284504
0283723
0284105
0283961
0283954
0284307
0284384
0283746
0284563
0283794
0284511
0283716
0284089
0284039
0283860
0284580
0284417
0284457
0283719
0288381
0284150
0284053
0284590

0284472
0284405
0283915
0283778
0283708
0284248
0284145
0284102
0283869
0283731
0283779
0283975
0284581
0284348
0284097
0284136
0284249
0284100
0284119
0283986
0283729
0283870
0284213
0284545
0284245
0284389
0284296
0284593
0284371
0284423
0284333
0283725
0284347
0284397
0284460
0283827
0283846
0283951
0283964
0283810
0284339
0284127
0284441
0284241
0284313
0288316
0284007
0284340
0284093
0283882

0284192
0284279
0276806
0278327
0275815
0276625
0278833
0277850
0275717
0275353
0278809
0275671
0275136
0277442
0277570
0276306
0275442
0274696
0277484
0274851
0274782
0273688
0274955
0276379
0278325
0276974
0274826
0276930
0277317
0273522
0278755
0277742
0275699
0275288
0275938
0278726
0273929
0283747
0283916
0284051
0283875
0283910
0284132
0284497
0284514
0283919
0284365
0284426
0283727
0284295

0284211
0284326
0284141
0283823
0284322
0283996
0284088
0284530
0283974
0284509
0284177
0284408
0284063
0284409
0284183
0284030
0284304
0284510
0283763
0284401
0283713
0284562
0284111
0288357
0283937
0284466
0284524
0283750
0284113
0283806
0284164
0284374
0283966
0283790
0284533
0284169
0283902
0284301
0288487
0284016
0283968
0283784
0283889
0288206
0284120
0284101
0284470
0284133
0283732
0284044

0284478
0283722
0284022
0284004
0277028
0273887
0277375
0278495
0277691
0275317
0275786
0276885
0275629
0278063
0275970
0276450
0277552
0277885
0276011
0275937
0278306
0275795
0276820
0277835
0275732
0274465
0278116
0275500
0276327
0276882
0274597
0276985
0278025
0277392
0274108
0278329
0277356
0275751
0277750
0278368
0276465
0278710
0277567
0276858
0275521
0274788
0274480
0275452
0277725
0274610

0275237
0276423
0274714
0274722
0273752
0275269
0277334
0275433
0273947
0278339
0273510
0277560
0276779
0278079
0273526
0275416
0278140
0273738
0275637
0273881
0277940
0275289
0277596
0278369
0274707
0274563
0275059
0275080
0275659
0278345
0276044
0277078
0278474
0276552
0274339
0276808
0274861
0273502
0276218
0274912
0276310
0278051
0277362
0273775
0276395
0278850
0278190
0277070
0273755
0274724

0276258
0275407
0275946
0276676
0277376
0278805
0277176
0274189
0276329
0277345
0276424
0277551
0277240
0276324
0277963
0274556
0274830
0273955
0273794
0278786
0273575
0274948
0274807
0276004
0275021
0277894
0278432
0273666
0274394
0275737
0276731
0278481
0277163
0276279
0275522
0278691
0277540
0275037
0276177
0278086
0275306
0278403
0274623
0273670
0277456
0273528
0276084
0274402
0274476
0277833

0275255
0278606
0275486
0275351
0275083
0274545
0278360
0278288
0278490
0276139
0275546
0275887
0277076
0274146
0276209
0278656
0274771
0276307
0273918
0273883
0274763
0275623
0278614
0278821
0275593
0278430
0277175
0274957
0278361
0277642
0275513
0274438
0275914
0275436
0277657
0277185
0278207
0275422
0275848
0274920
0276254
0277271
0275062
0275008
0275247
0274692
0277729
0278201
0274283
0276686

)275599
)278304
)277982
)278230
)276512
)274254
)275253
)277251
)274971
)278271
)278427
)275731
)275626
)284033
)284225
)283742
)284217
)284560
)284528
)284303
)284061
)283925
)284208
)284462
)283748
)284314
)284480
)283901
)284236
)284232
)284551
)284006
)284206
)284589
)284264
)284046
)284302
)283833
)283983
)284318
)284555
)284013
)284342
)284244
)284381
)274639
)276459
)275225
)278248
)275245

0275456
0278856
0276154
0276357
0278411
0273546
0276994
0273922
0276086
0275735
0278692
0278277
0273498
0275608
0275596
0275047
0273603
0275511
0275782
0274271
0277267
0276433
0278152
0273661
0276518
0274193
0276534
0274676
0277650
0273517
0273735
0277921
0274009
0274842
0276328
0277554
0274942
0275276
0274163
0278074
0273518
0274406
0275262
0277153
0275112
0274130
0276247
0277620
0274518
0278731

0278414
0274243
0276013
0277616
0274945
0276501
0274324
0276001
0274082
0276243
0274786
0276590
0276970
0275750
0276035
0277964
0277272
0274947
0278442
0275168
0276099
0274120
0275172
0276425
0273578
0276212
0277355
0276498
0276392
0277193
0275421
0278758
0274708
0275330
0276604
0273511
0274359
0274073
0277154
0277114
0277808
0277789
0276935
0278622
0274646
0277816
0275116
0275584
0277414
0274323

0277366
0276146
0276175
0274914
0276537
0278409
0276458
0278312
0273633
0274084
0276848
0274000
0274935
0274669
0274101
0278555
0273740
0278795
0278723
0274952
0277286
0278738
0277477
0275822
0277786
0275082
0276211
0278149
0275577
0275384
0275644
0276220
0276323
0275117
0274176
0276774
0276260
0278322
0274796
0274626
0273884
0278355
0276798
0276163
0275196
0276912
0276979
0275434
0274143
0274739

0278120
0278707
0275304
0278415
0278362
0274174
0275796
0275633
0276826
0276066
0277178
0278533
0276766
0276263
0274651
0274941
0276628
0274905
0278519
0274078
0277359
0275583
0274412
0274921
0274498
0273911
0273506
0277179
0274638
0278670
0276341
0276363
0276145
0276982
0273642
0275086
0274894
0276082
0275935
0274873
0277025
0274672
0275323
0276875
0276468
0275921
0277709
0277629
0276804
0275843

0276065
0273944
0276159
0274889
0274703
0278365
0276296
0276421
0275979
0277457
0277996
0277733
0275165
0278081
0274533
0276739
0278296
0277824
0277776
0275399
0274566
0275474
0274953
0275123
0275551
0275724
0273833
0277953
0273943
0278426
0277667
0277923
0273896
0274005
0276572
0275658
0273753
0276387
0275838
0273519
0273916
0277071
0278789
0274050
0274504
0274887
0275806
0277088
0275605
0278161

0277707
0274616
0278516
0274384
0273592
0274129
0274950
0275019
0276720
0275539
0273991
0275889
0278137
0277751
0274110
0276887
0278827
0274305
0274495
0273622
0275819
0276033
0278634
0277398
0277807
0278031
0276042
0276447
0276976
0278729
0277924
0278169
0274926
0273880
0274609
0275545
0276634
0275566
0276339
0275564
0273535
0278173
0277060
0278316
0274418
0277607
0273875
0278378
0277722
0276022

0274144
0277615
0278257
0278363
0274194
0278712
0277079
0277177
0274311
0275752
0273726
0274513
0278046
0275404
0274185
0278154
0276452
0276906
0278279
0276526
0275462
0275960
0274511
0278300
0278157
0274053
0275483
0278682
0275931
0278284
0275321
0280030
0275050
0280428
0274449
0274106
0278091
0279318
0277976
0279974
0278780
0262860
0279389
0262113
0276999
0280066
0279280
0280218
0274746
0279112

0280271
0279614
0279566
0282735
0282746
0282019
0274875
0279577
0275641
0279661
0280210
0276219
0279350
0279902
0279731
0279552
0279984
0280726
0276944
0274857
0281382
0280540
0278565
0281556
0281224
0280470
0281505
0274131
0281659
0280545
0281424
0280501
0278788
0275962
0276355
0275736
0276061
0276504
0275885
0274041
0277600
0276959
0273722
0275118
0278605
0276287
0278270
0273655
0274587
0277190

0274345
0274148
0277080
0278437
0275299
0240042
0275463
0275841
0277961
0275951
0278402
0274113
0275893
0276466
0274785
0277092
0276053
0276551
0275650
0275137
0274045
0275797
0275470
0276681
0273574
0278444
0277335
0274968
0275918
0276983
0275290
0275440
0275525
0278826
0277134
0275274
0275730
0275984
0274881
0274517
0273864
0276196
0274410
0276503
0278503
0276041
0274232
0274036
0273931
0277181

0273938
0274580
0275156
0274571
0273992
0277439
0274413
0276684
0282283
0278624
0277804
0277571
0274619
0282092
0281870
0280892
0281104
0276429
0281225
0281274
0273494
0273529
0281734
0281197
0273936
0280609
0281740
0275125
0277131
0280645
0275571
0275531
0262767
0276295
0262994
0262469
0262826
0263130
0263275
0269823
0263163
0262710
0262299
0261992
0262025
0281216
0280946
0276816
0281597
0281229

0274109
0281711
0276299
0273865
0281573
0274107
0277853
0281073
0276420
0281121
0252239
0278021
0276370
0262734
0261905
0263231
0263037
0262446
0262666
0263290
0261893
0263324
0262785
0274870
0278689
0275244
0281395
0276821
0278377
0273827
0283119
0263014
0262375
0262418
0262237
0262210
0280891
0280534
0280994
0277580
0276725
0276111
0275199
0280906
0282934
0283124
0284647
0282958
0283009
0283106

0282829
0283012
0282941
0283105
0282854
0282899
0282990
0283060
0285916
0273990
0273757
0274574
0281416
0280708
0275974
0280579
0280460
0283185
0283425
0283292
0283348
0283661
0283410
0283370
0283543
0283624
0283216
0283231
0283139
0283565
0283150
0283650
0202250
0283387
0283488
0282984
0282856
0285104
0282862
0282880
0282918
0283066
0283014
0282824
0282857
0283116
0282999
0282972
0283019
0284539

0284338
0284468
0284319
0283911
0284432
0284110
0284199
0285317
0285390
0285279
0284786
0285069
0284944
0288467
0284996
0275617
0278659
0273508
0281489
0275754
0277113
0281658
0275102
0280815
0262517
0262640
0262802
0280641
0280529
0277044
0277339
0280695
0276876
0275018
0280663
0281151
0281914
0281651
0262510
0262752
0261847
0262704
0262476
0263141
0263140
0280072
0262938
0262795
0263160
0262148

0263314
0262911
0262473
0281775
0280541
0274679
0281753
0281289
0281484
0281317
0280972
0280882
0280542
0280681
0280973
0272901
0272765
0273616
0272813
0262303
0263181
0262055
0262024
0262453
0262536
0262156
0262679
0281145
0274854
0278293
0281361
0281536
0281074
0281565
0274575
0281502
0280941
0281471
0273737
0280754
0280825
0281326
0277421
0275499
0275013
0281296
0281055
0281345
0280866
0281493

0281031
0280984
0280782
0277390
0277564
0274665
0281401
0281093
0275990
0280839
0281076
0281149
0280986
0281374
0274657
0281077
0273579
0273607
0275365
0267954
0275837
0274437
0281354
0275744
0281287
0281180
0277187
0276701
0281335
0281582
0281900
0281065
0280601
0278695
0281265
0275758
0281400
0275003
0281722
0277897
0281276
0281748
0280929
0278401
0276636
0277507
0277656
0281310
0280940
0281670

0272835
0273397
0280750
0280613
0273194
0272532
0281417
0272000
0280821
0274112
0277365
0281542
0280811
0278846
0275211
0280556
0280880
0278088
0273785
0276461
0281013
0281750
0280714
0280705
0281908
0281757
0281004
0280467
0280905
0281387
0284131
0284297
0281436
0281897
0283032
0283089
0282969
0282837
0282884
0283026
0282919
0282966
0283006
0282979
0282874
0283008
0282926
0282892
0283284
0283173

0281053
0281266
0283373
0283628
0283681
0274891
0280770
0281723
0280830
0280733
0280591
0280691
0280980
0281782
0278476
0279215
0277354
0281095
0281668
0280917
0280944
0275326
0277909
0278373
0276582
0281806
0280538
0281362
0281213
0281099
0281268
0280943
0281519
0281078
0281759
0275777
0280768
0280572
0280569
0280491
0276734
0281486
0276126
0281082
0281624
0277926
0281648
0276977
0277949
0280686

0281553
0274392
0281848
0274362
0281020
0281232
0277603
0283135
0283134
0283315
0283510
0283129
0283687
0283641
0283192
0283447
0283260
0283168
0283391
0283254
0283323
0283671
0283482
0283448
0283643
0283395
0283357
0281402
0283649
0283169
0283305
0286304
0284963
0285005
0284897
0285055
0285342
0285271
0285058
0285434
0285011
0279643
0282035
0266488
0264064
0265773
0266576
0266695
0266381
0262411

0261846
0262466
0281188
0281562
0281564
0282953
0281334
0276583
0277585
0281157
0277101
0281763
0280611
0280749
0281360
0275544
0280552
0281798
0280629
0280993
0280909
0274843
0273822
0284871
0284804
0286574
0281246
0280982
0281785
0280597
0280996
0281363
0280813
0281743
0281094
0281794
0281237
0278114
0278468
0288306
0288418
0288439
0288127
0288161
0288396
0288323
0288312
0288471
0288200
0288131

0288476
0281250
0281738
0281821
0280865
0280482
0281603
0280521
0277753
0278303
0285394
0284925
0285293
0285044
0285255
0280506
0280631
0280512
0280912
0275501
0275164
0281583
0281654
0281915
0281418
0281630
0273984
0285406
0285461
0286706
0281901
0280748
0281060
0281687
0275430
0276613
0284901
0284805
0284869
0285949
0281617
0274706
0281062
0281503
0281577
0276938
0281496
0278747
0267639
0267237

0283244
0283630
0283397
0283437
0283655
0283308
0283282
0283184
0285202
0283250
0283570
0284475
0284002
0283830
0283898
0284502
0283893
0284156
0284507
0284066
0284174
0283834
0284298
0283917
0284227
0284467
0283841
0283957
0262505
0262635
0263238
0281758
0280838
0281559
0281470
0263270
0275272
0278061
0274058
0281547
0277132
0281154
0280740
0280626
0275190
0280720
0281174
0281890
0280578
0280794

0281704
0281127
0281612
0278216
0281510
0280504
0280525
0281308
0280621
0282962
0281916
0276052
0280716
0281830
0281122
0281482
0281800
0281580
0280590
0277908
0281543
0281015
0275145
0280930
0280732
0173876
0277584
0275999
0276616
0274972
0281409
0281610
0280513
0280853
0274208
0281787
0281377
0283109
0282938
0283030
0282891
0282967
0283046
0283090
0282868
0273954
0278019
0281703
0280638
0277429

0273772
0285263
0281380
0280983
0275581
0275349
0285240
0276068
0276910
0280518
0281650
0281425
0278562
0281321
0281210
0280840
0280619
0274803
0281742
0262513
0262922
0277995
0275151
0274882
0279969
0278461
0277273
0277868
0275282
0275346
0279830
0279247
0276463
0276618
0274671
0274567
0278096
0275420
0273814
0277139
0280514
0275045
0274434
0278613
0279234
0280235
0277773
0285499
0278839
0273656

0275311
0277036
0273847
0276909
0275825
0275536
0277606
0276794
0273715
0273909
0274675
0274741
0277219
0273935
0273799
0275192
0277680
0275396
0276297
0273873
0274506
0274138
0275300
0281464
0275495
0281081
0281048
0281513
0274681
0280635
0274811
0281736
0281511
0280841
0276996
0275079
0280559
0278666
0280647
0281644
0280971
0281351
0274382
0281019
0281521
0276920
0276482
0281036
0276134
0281609

281861
280666
280978
280688
281690
277091
277069
281919
281198
283963
284036
284420
283989
284287
284434
284230
284151
284072
283730
278697
273605
282623
273741
277783
281421
281625
281410
275368
276187
281833
280868
281305
281249
280592
280883
275555
275333
281761
280722
281184
280737
281567
277364
280814
280658
281054
281270
275122
281841
273993

0280612
0281853
0280784
0280765
0280557
0280898
0275798
0277263
0272603
0272793
0272857
0272613
0236614
0234411
0234495
0264025
0224642
0241408
0225351
0236070
0226463
0233054
0234305
0226271
0224229
0229040
0225550
0226799
0230754
0237690
0236927
0242289
0232289
0230505
0229181
0229125
0237421
0229136
0234954
0232974
0224753
0235788
0227576
0226896
0232569
0226898
0229469
0225305
0265634
0224053

0224319
0236971
0234096
0227858
0206507
0236225
0260538
0237554
0235714
0276051
0277480
0278443
0228490
0229215
0231904
0232988
0235697
0224538
0231287
0233964
0227366
0234831
0235144
0235809
0236203
0236598
0223798
0228524
0235372
0273601
0276608
0235443
0275424
0233314
0236967
0224542
0235722
0265674
0224496
0237703
0235259
0228881
0236798
0236729
0249434
0235727
0230308
0241779
0232232
0229149

0223560
0236727
0225382
0206412
0227222
0230341
0228854
0236100
0234012
0231860
0224156
0223654
0229767
0231350
0237108
0228147
0231179
0266548
0230616
0233965
0225228
0237889
0265849
0228978
0231314
0235915
0236315
0238126
0233345
0266231
0264619
0227861
0236912
0238203
0231116
0264393
0224979
0226704
0235941
0232804
0234728
0263864
0266300
0228691
0235336
0232045
0232266
0229341
0266653
0238056

0236649
0240389
0236873
0223767
0231268
0225987
0237052
0238196
0236672
0234216
0266435
0235135
0226127
0227384
0264998
0230463
0237577
0226343
0233192
0230675
0243612
0225967
0230669
0233089
0242711
0227816
0234642
0237334
0242092
0242361
0235844
0236693
0231580
0234260
0227577
0226071
0231895
0217281
0281185
0257770
0248496
0275478
0249032
0249814
0230574
0237006
0227966
0226714
0235184
0230004

0237071
0263937
0237634
0233505
0263419
0227380
0239940
0223687
0227626
0234101
0274063
0225838
0226657
0233180
0234804
0232178
0232272
0224925
0231676
0226610
0234092
0233950
0225150
0237366
0226844
0231279
0225291
0229045
0233046
0251394
0235966
0249030
0230780
0224234
0235132
0227228
0224444
0233141
0232302
0232652
0230674
0237698
0229467
0228913
0232632
0226796
0225248
0266672
0230181
0229185

)223771
)237834
)226858
)235507
)137403
)233935
)206446
)241880
)251581
)224253
)263876
)228912
)230238
)225346
)260055
)233462
)229995
)230413
)234681
)224118
)226152
)224320
)226893
)227027
)233241
)231998
)227956
)232331
)223352
)223708
)232011
)237395
)236579
)234242
)235340
)229653
)224006
)228220
)232141
)276049
)274071
)224859
)276034
)234058
)227171
)226402
)238027
)265495
)228001
)237192

0227147
0230230
0228199
0241701
0223840
0243492
0228894
0224994
0241863
0227334
0234417
0232095
0233466
0223933
0223412
0227703
0233904
0223827
0235986
0228186
0236149
0230995
0227420
0223752
0225453
0233641
0233561
0231247
0235773
0235342
0224587
0236766
0227739
0232280
0266119
0237155
0263728
0224641
0236006
0231522
0265903
0224360
0137332
0236895
0285224
0236178
0224246
0266178
0232308
0223454

0234137
0234063
0238234
0265921
0229432
0235487
0137197
0224544
0226993
0234114
0224379
0233911
0223364
0232758
0228583
0233841
0223532
0233486
0228461
0231433
0225939
0230016
0264185
0223971
0232659
0225145
0233051
0264692
0234858
0237105
0264846
0234218
0236415
0230211
0226923
0225073
0263359
0265607
0232256
0227587
0234530
0238130
0204490
0204487
0226182
0237808
0227650
0237727
0225164
0263879

0229524
0224290
0235360
0235516
0236011
0266759
0228875
0241713
0231325
0235676
0240008
0226603
0224708
0230060
0236183
0226634
0230685
0235222
0254405
0265409
0229176
0238017
0227144
0226589
0264955
0236826
0236251
0237724
0224501
0226710
0264202
0263362
0227315
0231479
0236759
0223557
0235017
0237900
0227884
0239754
0231044
0225737
0225682
0234947
0228267
0236587
0233151
0233323
0265816
0234539

)230907
)229071
)206256
)254778
)242038
)226467
)228405
)234729
)237344
)233490
)235396
)234280
)231275
)263869
)225834
)224770
)227993
)227526
)265223
)231679
)264909
)232156
)264259
)206240
)257473
)206237
)231677
)243252
)228845
)232026
)224793
)224305
)241386
)228582
)235715
)235540
)243067
)232367
)236093
)228785
)234154
)243719
)234507
)234673
)232962
)224103
)230708
)228579
)223990
)230816

0225132
0227936
0237196
0227801
0227322
0266781
0226614
0228712
0226788
0230949
0237536
0224096
0225590
0235650
0226936
0204221
0237335
0224841
0112493
0229253
0227046
0226575
0232639
0230511
0235460
0225604
0224158
0263491
0235405
0239675
0235178
0225048
0229328
0275606
0237608
0229179
0225736
0226192
0229866
0227448
0236771
0232945
0228483
0229441
0235273
0224990
0233168
0234351
0232469
0237777

0228197
0242022
0227446
0249103
0230742
0234347
0229412
0232984
0233989
0229889
0228515
0223769
0232356
0224446
0225274
0237725
0231968
0237112
0225955
0232189
0265926
0227961
0229680
0229210
0234623
0223852
0238053
0235220
0233617
0230017
0240845
0248230
0235458
0264218
0228102
0223606
0235317
0261814
0227030
0225763
0235346
0231430
0235536
0226517
0235657
0227669
0229293
0228891
0230357
0231941

0232708
0233239
0225114
0234849
0226713
0234313
0228458
0231719
0225618
0227025
0234074
0235764
0275415
0273764
0236949
0274401
0227720
0235022
0233573
0232510
0265138
0227458
0227472
0235960
0226060
0225109
0241053
0225659
0241166
0224705
0232839
0243009
0230509
0235561
0232473
0229651
0236369
0231157
0227652
0229252
0233315
0229470
0223887
0232169
0227804
0233813
0229061
0237156
0237012
0233418

0234000
0236843
0233091
0225589
0229681
0235067
0206379
0263950
0227231
0264417
0224923
0227596
0231733
0265255
0236492
0230456
0226384
0285192
0230985
0225454
0266841
0232251
0235747
0237962
0234461
0229501
0263467
0225543
0233439
0237123
0232127
0232007
0206355
0233890
0229094
0231450
0234079
0228469
0228299
0236374
0229430
0229103
0231710
0263833
0224378
0231118
0229535
0231975
0230624
0266419

264591
236019
234668
236196
223919
228849
236925
223465
237103
232683
234836
231370
266004
227761
227567
228435
227665
233210
264353
230700
258589
244672
226404
231488
243003
259101
237482
234443
224393
228859
227317
226248
230293
266027
263896
227074
227856
230048
096171
266113
231502
234258
234475
231555
229300
264086
265707
234343
228263
227333

0236903
0231543
0234852
0233692
0241534
0206268
0223493
0236765
0226257
0244207
0232565
0231759
0269167
0248352
0249480
0236625
0225512
0232414
0231608
0266615
0168468
0224200
0225141
0228116
0226892
0230209
0265351
0226228
0229741
0228987
0237591
0235577
0234470
0233240
0235391
0265983
0236884
0236418
0233209
0237455
0240684
0236629
0234331
0231596
0223793
0228813
0243496
0223481
0230034
0206232

)240118
)224212
)227435
)237871
)226264
)241394
)230678
)235744
)229685
)237710
)226879
)223894
)232802
)225429
)223696
)230134
)230930
)228333
)266410
)229802
)225312
)228520
)229394
)224052
)233772
)228425
)223367
)235155
)226916
)224782
)237825
)236490
)227780
)206206
)229342
)273083
)273040
)273200
)273286
)237462
)264777
)231041
)226462
)265326
)233035
)240111
)235589
)232781
)228652
)277403

0233019
0227529
0231319
0225441
0233636
0231960
0228846
0226347
0230136
0231919
0225188
0231320
0236918
0249415
0227178
0234100
0225927
0243441
0224487
0233481
0248182
0237258
0232397
0224395
0226365
0230957
0228377
0228840
0228370
0225426
0237811
0223441
0224654
0237051
0225034
0224241
0265663
0224146
0227044
0228130
0230885
0232099
0226303
0237508
0229573
0226378
0242039
0250154
0228900
0264495

233023
224736
226379
261134
233465
229461
235680
224801
235498
231197
223463
226727
225926
226940
226563
232816
226354
231834
226109
230915
230745
227647
224362
231745
235394
225086
223439
224108
278049
277415
235176
277085
236560
237733
225130
237050
263589
237046
235025
233599
231002
226175
240924
231928
239618
230660
226437
239865
234095
236422

0231659
0229187
0233700
0227754
0224554
0236632
0235801
0232143
0223766
0231129
0231737
0226111
0235658
0223851
0233164
0233049
0229998
0228867
0226494
0228575
0224508
0232575
0236271
0230128
0263671
0228762
0229170
0231854
0266263
0231444
0234078
0233149
0285089
0223494
0230072
0265323
0223590
0231474
0237894
0263367
0231624
0231094
0237114
0234605
0225714
0223533
0224941
0230336
0257215
0226422

0238148
0229823
0237022
0224608
0229265
0237448
0234980
0228954
0233604
0263701
0224131
0226732
0228179
0235233
0266747
0231836
0231372
0229216
0236273
0238238
0225859
0265116
0265325
0231647
0226850
0227565
0231408
0228321
0223448
0231048
0236979
0226618
0266847
0228760
0231974
0228177
0233169
0223932
0264915
0232960
0258543
0240433
0228883
0224552
0239741
0258635
0236210
0225993
0228090
0235452

0233076
0226651
0230961
0223957
0203463
0204364
0223747
0204359
0236221
0263418
0229488
0168452
0243897
0227642
0228907
0229058
0232005
0236697
0232339
0206245
0233160
0266660
0229903
0225412
0226260
0234164
0223359
0265059
0224230
0266081
0227826
0264640
0233853
0265074
0238248
0237120
0229099
0223844
0263871
0229074
0225890
0225824
0238226
0242138
0237145
0226139
0228684
0231823
0229493
0206235

0236443
0231591
0239836
0226173
0236488
0225459
0239329
0243189
0234704
0236657
0231558
0228163
0229295
0233972
0235528
0226591
0233584
0234372
0233043
0223699
0231321
0265159
0227402
0228357
0235107
0236910
0230012
0231037
0223618
0235668
0234966
0234904
0232017
0236835
0233948
0263598
0237313
0225567
0228977
0274733
0228635
0238070
0233687
0226931
0230855
0228698
0196231
0220379
0233182
0224633

0112459
0232435
0112461
0204692
0204690
0249571
0206472
0204689
0206471
0234856
0227668
0206470
0223376
0227886
0233857
0228993
0232049
0206468
0206466
0223853
0224293
0265013
0231190
0206467
0230888
0232641
0223858
0236926
0234487
0249663
0228491
0243430
0228202
0264646
0233298
0229194
0224564
0259930
0233982
0233592
0237216
0232122
0234364
0225120
0247700
0237098
0236265
0232649
0227715
0228921

0236746
0235600
0234599
0227389
0236113
0232691
0223892
0226586
0228765
0278347
0276640
0236808
0276594
0237403
0230332
0223531
0225038
0263517
0236399
0229381
0235905
0231641
0234574
0230887
0225201
0234057
0228581
0233564
0236342
0235291
0229861
0231257
0227655
0224120
0231377
0225060
0234809
0229575
0237095
0233883
0229684
0224740
0266523
0224862
0265633
0227422
0223680
0234370
0285515
0234032

0237814
0264847
0257232
0228448
0227506
0227132
0225629
0263474
0224105
0233734
0237165
0227246
0233504
0224180
0224472
0235068
0258002
0230983
0223577
0226408
0225691
0232126
0226050
0227656
0231309
0228713
0227429
0263679
0238289
0229706
0236547
0228662
0229496
0263625
0266755
0234082
0230900
0235125
0226275
0223952
0236237
0234514
0225211
0235985
0225748
0265754
0234651
0226215
0226531
0225390

0232312
0265738
0224398
0241132
0237495
0236063
0240957
0236902
0235769
0237459
0228250
0225635
0226417
0227314
0266442
0264951
0234951
0235765
0223757
0224264
0264887
0225998
0227129
0229077
0238134
0233649
0226560
0234196
0223661
0243570
0229363
0232616
0224313
0226033
0224639
0229856
0224140
0230354
0227746
0225540
0235134
0206258
0266726
0228628
0223666
0227122
0231618
0240592
0235758
0225452

0230514
0236353
0234243
0223355
0237881
0225237
0264557
0230602
0225845
0230726
0228260
0232441
0264893
0235074
0263971
0231021
0228284
0231286
0232351
0242044
0228025
0226550
0234510
0206301
0228254
0241910
0237599
0226201
0234365
0240508
0230705
0233498
0226691
0242386
0239463
0236227
0237348
0230141
0168384
0230763
0231288
0230878
0234249
0223372
0235708
0235712
0266464
0224399
0232357
0231115

0237723
0227750
0238064
0224455
0226225
0236802
0236222
0235692
0228736
0206208
0237056
0231617
0224874
0205677
0231602
0056678
0273323
0272907
0272629
0273122
0229714
0227559
0215641
0265795
0206526
0235541
0264866
0225368
0243286
0169260
0237043
0206525
0278714
0235195
0226795
0206524
0206523
0206522
0225184
0233506
0234746
0218297
0235401
0224392
0251547
0240130
0206517
0220550
0206516
0224044

0227859
0206515
0229439
0232902
0226687
0224894
0231973
0206511
0206513
0235630
0229737
0266132
0206519
0206512
0234528
0137345
0206510
0228371
0206509
0224921
0206508
0239784
0251197
0236619
0265105
0230722
0206444
0237905
0260638
0233113
0224355
0206506
0225023
0232253
0230419
0227296
0229552
0236441
0229411
0220391
0224526
0229428
0206505
0235952
0228029
0232345
0206504
0234140
0234970
0235814

237884
243336
232516
278511
278397
206502
275133
206501
206500
137397
229223
265602
172524
137394
137384
137313
228227
244695
235923
244721
234893
206495
242726
236688
227100
236122
237034
235223
223693
229038
206493
236934
206492
206491
206490
206489
203624
206488
235757
206487
206486
206485
206484
238255
206481
227899
183311
206480
265955
206478

0264688
0224133
0215523
0225888
0264415
0223405
0215522
0221974
0285213
0206476
0206465
0264602
0229284
0230541
0206463
0229199
0227974
0266375
0206461
0206458
0206460
0206459
0237545
0206457
0206455
0206454
0224744
0220890
0223868
0206452
0206450
0228904
0223927
0225954
0225571
0236851
0266886
0232276
0231773
0237559
0183214
0265896
0265852
0206449
0225317
0232572
0206447
0215425
0266295
0266646

0238071
0206445
0206440
0173503
0206439
0206437
0206433
0206430
0229189
0206428
0206427
0264153
0096155
0206409
0203623
0224835
0206408
0265278
0206406
0239285
0206404
0206403
0243804
0206402
0206401
0206396
0206398
0206395
0206394
0235569
0264556
0227686
0172850
0206383
0215328
0212866
0206380
0265732
0265156
0184494
0206338
0206378
0206376
0231869
0206372
0206366
0235250
0241253
0206357
0206353

0206346
0206342
0272295
0206340
0230075
0229353
0266035
0206331
0206329
0254875
0244444
0206324
0183574
0206320
0206315
0206314
0206312
0206310
0235171
0265920
0228738
0229597
0206308
0226035
0266884
0196101
0264389
0227442
0266689
0206306
0206305
0206302
0233744
0240110
0229710
0230150
0224197
0225103
0196610
0206299
0206298
0206296
0243594
0206297
0226914
0233053
0241674
0242685
0215077
0228104

0206292
0206291
0215048
0233194
0227286
0228164
0234834
0224929
0238100
0206290
0206289
0265071
0206288
0112474
0206287
0231030
0235684
0231343
0206286
0096150
0206285
0206284
0206283
0206282
0206281
0206280
0206279
0230193
0232376
0227168
0204197
0228099
0239756
0248563
0248655
0233826
0229773
0231746
0232997
0237010
0229006
0266870
0233566
0228990
0266191
0237898
0241124
0233831
0233650
0227639

0278450
0229100
0231592
0226832
0235632
0233618
0233119
0231670
0226000
0229069
0225592
0230089
0230874
0230752
0226932
0232146
0234195
0227843
0227137
0250364
0225247
0250294
0223898
0235441
0229408
0237369
0237544
0237284
0231000
0225432
0229383
0234992
0235666
0236561
0234669
0229898
0229698
0236881
0230545
0242511
0249123
0226267
0229942
0229529
0228706
0260221
0227326
0229499
0233095
0230504

0236642
0226736
0237218
0224693
0229364
0223860
0236033
0229959
0236706
0223980
0231067
0234786
0235380
0237812
0235305
0229034
0196299
0231385
0235309
0223996
0275472
0274187
0233795
0276383
0237829
0230467
0224168
0225732
0263942
0233931
0229346
0224145
0234046
0224954
0240833
0233350
0242227
0224619
0229929
0239927
0234799
0235877
0230441
0231785
0230422
0237865
0137321
0230254
0232523
0226882

0229202
0225989
0238104
0223775
0234549
0224889
0237100
0226171
0236428
0231135
0236522
0232421
0230143
0263610
0235030
0263912
0238216
0237890
0229126
0237093
0266626
0234674
0237582
0224056
0231281
0229075
0206435
0228964
0225029
0227672
0225403
0226834
0233161
0266784
0232244
0227405
0226027
0231225
0266250
0226768
0230389
0265509
0227772
0235596
0231623
0226841
0235439
0254899
0263358
0263442

0228245
0234920
0236346
0230279
0230108
0238114
0230791
0223833
0228926
0235588
0231825
0263994
0233348
0235754
0226103
0227853
0228605
0266716
0224774
0239497
0111971
0230475
0241822
0258880
0226187
0235302
0231003
0235925
0228128
0223639
0237333
0263638
0255419
0237918
0234433
0231945
0264778
0234846
0232180
0224143
0224766
0235696
0225773
0242335
0233801
0228896
0234798
0236250
0233312
0232577

0239882
0234328
0248989
0228454
0238170
0198457
0236236
0235956
0227600
0239974
0228892
0227277
0237405
0224952
0234508
0234876
0232106
0237295
0265048
0224507
0224242
0234794
0232705
0235228
0264704
0226227
0265838
0227357
0264960
0233697
0265248
0235324
0235116
0227099
0238101
0266860
0228080
0232062
0231939
0230414
0243245
0233317
0232343
0234968
0231526
0226165
0239457
0232326
0231631
0233140

0243958
0224748
0223939
0228498
0241296
0243215
0235307
0225788
0232957
0236177
0226826
0224098
0236898
0226513
0168379
0223447
0232541
0225696
0234251
0231573
0236014
0227794
0229396
0226538
0233450
0232733
0225553
0226492
0227460
0236515
0281993
0282155
0282226
0282075
0281767
0281349
0281788
0281429
0282813
0282789
0275723
0282463
0282063
0281979
0282026
0262327
0262671
0262599
0261994
0263042

0262656
0262771
0282185
0281951
0282333
0282645
0282557
0282662
0282461
0282781
0282309
0282575
0282147
0282379
0276184
0282349
0281950
0282254
0276464
0282364
0282580
0282607
0278563
0282809
0276785
0282170
0282264
0282303
0282423
0282568
0282624
0282014
0282803
0281970
0282225
0282285
0282719
0282505
0282543
0282353
0282506
0281952
0282236
0282240
0282203
0282529
0282148
0282204
0282618
0282711

0282760
0282007
0282436
0282326
0277110
0282756
0282466
0282459
0282709
0282577
0282470
0282704
0282098
0282134
0282179
0282748
0282610
0282040
0282054
0281943
0282407
0282340
0282242
0282208
0282354
0282605
0282252
0282497
0282415
0282483
0282771
0282621
0282064
0282462
0282036
0282449
0282730
0282017
0281963
0281972
0281968
0282047
0282234
0282812
0282270
0282628
0274247
0282295
0282667
0282565

0282049
0282492
0282253
0282078
0281989
0282500
0282273
0282215
0281957
0281981
0282488
0282272
0282589
0282371
0282380
0282820
0282154
0282392
0276849
0282297
0275482
0276971
0282313
0274348
0275821
0276615
0273914
0282335
0277378
0274360
0282507
0284921
0285440
0285110
0284896
0285496
0284949
0285071
0284891
0288370
0288422
0284836
0285059
0284749
0288181
0288136
0285362
0285105
0285408
0284912

0284827
0285013
0285154
0284933
0285328
0288119
0278728
0277205
0274906
0282044
0281999
0282734
0281953
0281997
0282141
0282152
0282001
0282395
0282061
0282027
0282106
0278101
0282114
0282113
0282472
0282261
0282428
0282524
0282619
0278062
0282146
0280677
0277802
0281561
0274195
0276378
0275444
0282496
0273641
0274405
0277006
0275794
0274729
0277530
0274540
0280471
0281040
0281150
0281407
0281862

0280960
0280939
0282684
0277845
0278718
0277781
0282613
0278488
0282189
0273594
0278016
0276657
0278243
0276340
0274352
0277126
0278749
0282538
0274205
0282617
0273889
0282685
0282519
0274287
0274081
0282347
0282665
0282103
0282245
0282681
0278032
0278812
0282669
0282683
0282693
0274488
0274161
0282643
0282594
0282573
0275181
0275575
0282181
0282477
0276275
0282315
0282029
0282515
0282020
0282318

)282786
)278220
)282493
)282481
)285363
)285114
)274726
)282101
)282405
)282334
)282053
)273645
)282260
)282191
)274137
)282484
)282263
)281936
)282397
)281980
)282435
)282661
)282230
)275594
)282698
)282490
)282250
)278548
)276286
)282680
)282382
)263127
)280745
)276497
)274039
)278608
)277409
)277517
)274806
)276374
)278688
)282008
)276539
)282376
)276714
)282259
)282406
)282159
)276559
)282345

0282932
0282942
0280741
0276438
0280593
0276895
0280532
0280935
0280844
0277800
0273850
0280484
0280769
0280877
0281050
0278704
0277400
0274847
0277428
0276256
0278198
0276932
0273496
0277630
0278793
0274202
0278384
0273748
0276197
0274917
0273937
0273730
0276711
0278294
0278457
0271254
0275405
0275987
0277475
0268674
0276345
0273532
0276351
0275661
0277856
0275063
0278673
0277890
0277666
0276760

0277927
0275249
0274792
0276872
0274175
0273843
0275869
0278510
0273554
0278782
0277761
0277836
0278633
0278066
0273739
0276700
0276312
0275757
0278573
0276017
0278817
0277196
0278625
0277374

[illegible]210049
[illegible]211459
[illegible]210077
[illegible]210082
[illegible]209082
[illegible]198888
[illegible]210100
[illegible]210107
[illegible]210112
[illegible]198763
[illegible]210117
[illegible]210127
[illegible]210135
[illegible]210140
[illegible]210144
[illegible]198804
[illegible]210151

0210154
0198712
0210156
0228253
0198899
0198938
0210164
0198840
0210174
0212907
0198886
0210176
0210184
0210191
0198786
0198695
0210194
0198727
0210195
0210196
0228572
0182378
0178605
0226179
0167393
0281849
0275287
0280767
0234958
0229232
0185960
0237531
0225661
0205755
0198223
0265658
0223274
0185291
0169100
0236871
0236017
0169093
0182162
0197976
0196433
0223511
0234622
0169084
0223571
0214717

0277120
0223773
0230542
0002586
0124343
0056998
0235483
0006756
0229851
0157399
0205667
0062096
0239228
0230249
0228459
0101825
0213487
0212214
0207435
0183943
0207332
0207320
0236188
0182888
0285756
0286431
0225722
0236513
0235039
0226313
0238193
0201660
0236120
0146938
0252291
0265284
0237730
0169059
0130021
0237490
0228550
0101846
0264268
0279682
0182583
0274923
0006757
0285679
0207628
0177504

0205642
0287171
0011201
0225574
0183304
0231619
0229012
0285896
0225271
0177138
0200852
0219870
0228543
0232111
0282306
0282501
0282358
0282668
0282299
0282697
0282222
0281955
0281941
0282300
0101849
0101850
0146950
0215319
0230326
0206844
0234469
0047644
0225076
0202469
0073464
0227042
0101871
0207202
0234129
0004961
0207151
0047648
0278218
0275194
0125363
0263652
0232392
0282840
0223418
0005302

0169933
0206792
0206795
0223487
0231211
0273800
0271814
0101911
0229083
0230691
0196664
0233338
0101916
0205542
0187268
0261030
0235385
0231216
0123594
0235592
0226985
0197582
0198759
0233357
0283493
0199622
0176896
0123595
0196459
0046651
0046653
0233535
0212663
0046647
0233247
0101958
0231066
0181544
0130150
0235929
0236929
0102048
0165192
0165195
0165197
0087842
0102010
0130234
0285602
0225833

0147003
0237539
0186312
0169239
0279413
0200620
0281371
0169249
0182287
0200566
0226536
0126010
0238178
0182798
0215313
0235071
0239333
0047230
0265144
0169906
0169895
0238709
0086712
0230180
0102054
0222736
0223498
0199410
0169891
0201467
0214322
0214321
0238764
0188158
0265465
0235834
0230020
0226153
0047634
0227873
0131831
0276467
0225066
0225882
0177324
0102098
0235211
0207025
0008086
0236187

0102104
0086717
0237221
0196564
0224224
0237836
0044446
0266710
0286724
0173698
0131828
0180815
0202144
0275800
0147010
0173681
0206663
0184368
0264566
0173674
0201882
0201592
0225037
0177189
0283631
0283400
0252978
0283380
0283546
0206716
0149970
0231913
0185915
0091482
0012174
0275681
0230797
0206639
0271286
0102172
0102174
0224204
0233067
0175809
0231846
0224396
0201095
0231011
0237019
0184735

)174028
)228610
)165186
)226774
)123131
)123130
)233785
)202272
)130066
)184831
)238103
)165182
)174010
)130741
)201407
)234230
)005889
)223611
)223731
)225071
)067992
)214124
)102230
)226222
)236836
)236487
)101868
)251869
)232472
)004848
)214111
)240439
)176812
)229310
)228519
)233403
)176774
)232446
)232030
)227511
)176746
)232644
)188408
)237171
)228464
)231388
)230265
)259849
)236160
)216938

0236683
0252491
0242021
0228933
0233807
0252486
0235642
0224960
0223391
0238247
0189186
0177689
0229731
0226199
0232834
0228345
0233243
0224589
0226372
0277569
0223742
0169306
0226436
0201666
0237994
0201356
0227393
0264090
0227592
0270794
0099399
0198798
0120289
0214107
0276940
0169297
0120280
0178556
0232368
0198814
0229331
0241886
0243055
0277876
0157625
0231542
0235512
0132446
0198947
0252903

0285900
0222922
0236285
0263600
0221348
0236828
0226781
0215310
0287700
0233928
0185448
0185638
0233571
0236072
0271136
0147027
0225384
0273704
0189132
0229347
0252411
0230227
0233663
0271088
0233880
0189023
0271519
0165164
0206733
0229384
0226484
0233921
0286475
0249467
0275314
0235510
0223723
0231069
0198173
0243048
0214071
0233810
0130962
0250349
0241607
0219186
0234355
0174678
0147036
0047597

0285944
0165168
0165169
0206983
0259977
0224339
0187516
0229674
0147041
0221466
0101955
0226679
0156313
0252050
0036473
0227988
0232370
0229735
0233007
0156298
0238606
0235806
0287729
0165175
0238123
0207122
0224610
0235304
0236747
0234191
0263730
0264618
0228232
0270759
0261435
0263972
0214047
0285709
0223486
0183337
0236393
0214045
0206677
0235685
0238730
0182220
0287098
0238205
0185753
0180182

234636
270791
214031
237661
238222
232801
189145
216866
287549
237931
124486
223714
199564
238426
226641
215301
264573
188937
225238
147044
233033
212207
217624
242559
171659
171657
229822
237875
251807
233103
228847
212560
228780
206896
231772
283737
287223
102055
270280
224294
237206
285899
231750
189221
069535
124479
236276
214019
183690
198414

0214018
0200702
0224054
0215035
0069509
0233543
0229168
0214016
0242065
0189037
0147050
0147113
0229491
0237782
0229563
0214282
0204915
0270069
0207870
0207725
0269902
0231566
0212347
0215286
0234613
0224701
0213100
0214009
0278838
0236751
0229145
0286306
0231235
0235587
0147121
0224072
0251192
0199226
0230844
0277721
0270499
0147119
0065923
0213997
0226048
0215284
0102218
0204904
0102221
0130988

0206685
0201659
0147123
0182872
0237351
0130985
0224975
0102225
0102226
0237623
0266158
0203402
0147117
0221459
0232009
0226556
0147124
0277605
0196741
0235636
0078061
0008056
0283743
0102265
0263858
0126759
0126767
0126756
0267064
0187893
0224656
0229968
0188459
0197779
0224401
0147118
0221994
0171489
0277541
0287757
0171483
0171478
0230241
0233139
0201517
0225198
0238032
0165583
0234780
0252856

204368
231568
229662
126752
232813
241207
237345
204648
234391
227302
230100
270747
165584
225055
234448
268009
269791
231603
231489
235350
226971
017483
068438
224292
102312
147155
286268
276823
068354
204620
102317
228343
279528
220125
101940
231725
015285
101945
232828
206723
171433
233585
207367
102145
094631
187682
102109
102103
126768
102100

0102096
0068308
0279155
0223309
0102057
0068400
0068323
0147144
0243279
0288053
0196998
0206936
0068394
0265033
0280116
0269313
0279422
0102007
0012211
0102003
0237341
0102001
0230216
0101997
0049768
0286181
0049769
0270012
0286031
0215274
0224659
0274274
0275113
0236737
0237671
0227488
0236362
0216649
0215269
0224902
0205777
0189064
0213856
0227391
0068985
0101951
0234390
0273820
0220132
0171365

0201341
0207758
0207768
0207785
0208015
0211538
0239057
0207970
0272080
0181977
0202495
0147081
0147082
0274588
0158352
0222303
0226881
0130385
0238190
0227493
0230926
0230317
0237926
0229151
0287767
0122824
0226530
0187690
0196368
0234766
0224109
0283093
0223591
0189369
0179222
0236576
0252377
0228827
0232951
0237528
0232562
0182776
0279750
0187243
0221705
0154545
0221475
0287215
0179028
0276474

0275335
0155622
0225957
0276897
0224556
0250084
0278358
0278160
0275643
0204379
0223395
0204382
0277289
0227058
0231593
0228771
0185319
0197185
0234792
0157965
0187754
0228354
0222608
0225508
0237275
0241476
0223958
0225397
0229826
0268447
0269281
0226867
0204363
0229885
0171405
0196395
0224781
0171402
0179304
0286118
0268350
0184194
0270497
0184205
0234019
0232593
0235224
0231340
0227329
0126012

0276575
0277474
0235262
0124313
0226299
0207408
0236571
0234413
0072501
0278204
0158423
0072506
0233250
0235416
0199422
0086758
0271886
0208012
0224735
0226110
0229037
0235916
0201618
0172943
0184083
0224949
0196632
0234566
0207104
0158526
0102302
0130119
0215197
0102313
0102316
0221716
0275387
0067445
0221750
0158571
0169188
0158578
0278283
0234466
0238269
0234068
0278319
0276929
0182518
0186678

229760
231947
238047
256045
158639
204279
232765
266328
283334
187601
247746
277516
200635
235136
189299
229594
169164
083750
227486
214745
102349
234135
188021
230105
204272
237748
204271
224799
237708
186787
226310
147059
286977
231765
165591
231831
198455
215174
227427
198205
228004
235124
236852
235170
228182
234570
277499
235437
230823
186767

0233661
0131089
0231729
0236813
0222532
0232794
0226010
0184675
0287370
0244006
0198881
0102043
0213754
0283349
0230889
0227261
0226502
0225814
0236982
0126970
0237971
0225569
0231857
0237963
0102053
0228158
0227534
0001497
0227942
0147065
0231809
0276689
0233833
0230039
0237311
0217835
0274536
0284567
0155659
0215168
0237294
0089472
0231356
0226820
0235892
0131080
0222667
0226280
0233710
0169083

234201
260118
079482
228328
252145
228125
206646
225925
181704
206747
236735
234463
130052
232267
213740
090776
181191
172186
215162
231706
130054
228160
158813
285776
272351
212434
233229
211991
286077
147160
189401
226579
235382
089289
220925
203588
184210
242657
204195
200473
186912
120500
147127
120509
090889
231053
201377
130055
284391
082458

0200431
0231651
0120498
0230187
0277492
0280704
0213747
0165349
0230629
0229324
0212605
0227536
0147164
0184481
0204165
0285171
0147168
0184634
0196338
0228427
0169562
0147130
0147140
0147166
0224208
0213706
0147133
0275975
0229601
0213018
0199751
0243468
0215120
0147162
0147174
0186810
0237265
0283599
0230934
0281852
0215115
0226870
0237182
0237717
0234442
0215113
0224617
0229030
0283178
0204131

0201392
0196933
0242732
0225471
0102309
0239577
0186871
0198034
0238926
0125931
0285547
0147099
0222810
0067177
0225858
0231944
0273788
0213698
0201271
0184911
0269911
0268994
0225396
0269502
0283594
0275520
0226725
0184388
0186288
0231963
0204118
0186462
0230311
0237014
0225664
0207486
0131264
0233162
0269904
0237475
0204116
0230146
0270641
0229807
0274655
0277577
0278039
0182707
0230590
0225470

0274430
0276784
0228906
0271199
0233984
0203397
0234969
0223650
0232928
0280375
0230986
0224430
0231549
0204113
0228492
0271533
0202566
0212027
0283534
0265727
0278421
0273974
0275254
0275625
0207820
0199168
0277922
0187969
0226515
0213269
0239745
0230665
0213695
0147100
0231206
0131263
0230673
0199885
0050030
0271589
0131269
0094841
0229661
0199601
0232186
0102383
0233254
0237089
0224775
0215105

0186675
0233063
0226753
0226854
0232332
0102390
0237252
0201826
0274398
0213684
0198934
0223259
0235447
0280870
0283242
0207995
0201447
0234404
0196468
0085224
0237106
0102158
0200906
0263410
0131174
0165240
0102144
0248503
0226784
0187325
0223592
0173198
0224755
0235061
0179300
0271433
0147145
0227645
0283622
0078589
0232168
0147138
0271005
0225025
0078596
0270268
0122145
0278530
0174016
0230247

0228051
0228612
0165288
0225715
0239008
0212623
0073905
0212546
0284618
0198157
0131171
0231875
0224523
0288059
0233446
0206826
0228041
0213653
0202183
0236365
0232735
0279437
0196767
0236870
0215102
0183035
0072133
0221494
0165259
0233484
0237211
0229547
0230781
0155008
0184788
0283446
0276391
0235461
0147180
0124429
0283435
0223707
0276892
0188419
0199051
0199567
0234039
0225536
0223574
0126733

)215093
)223772
)277717
)231478
)271290
)236190
)271457
)200422
)230478
)102271
)237506
)282914
)271205
)270446
)271056
)230706
)230956
)252016
)271693
)147183
)271157
)232576
)270745
)224065
)225012
)153779
)227581
)226081
)274022
)227434
)229667
)207515
)206062
)234161
)236413
)174740
)230275
)102290
)222440
)234702
)229132
)215089
)230543
)225912
)285445
)237873
)234130
)234322
)237381
)228301

)186310
)179083
)270826
)212495
)202410
)227521
)232293
)231001
)221187
)271094
)223467
)201567
)215085
)236185
)226307
)235425
)270583
)232842
)242998
)203262
)286523
)232069
)271209
)223260
)229693
)147202
)207426
)204086
)214628
)277900
)265260
)236256
)233887
)237752
)234889
)236243
)271322
)213704
)226837
)281566
)270308
)215070
)229946
)252296
)236988
)230592
)165194
)000005
)000003
)102359

)102362
)223819
)238116
)275320
)101811
)202231
)007952
)239345
)233680
)179031
)182489
)182574
)174225
)188917
)225839
)126950
)102384
)231643
)102385
)102387
)202024
)270813
)183911
)126953
)010671
)241343
)257529
)102393
)126945
)196440
)234557
)237682
)126947
)204072
)198960
)102401
)228275
)261101
)207291
)286000
)184867
)223753
)212584
)229090
)178146
)126952
)166432
)235196
)252810
)235976

0226119
0234901
0184905
0204071
0286794
0184515
0251525
0268235
0269405
0273746
0284800
0269437
0231590
0215029
0158164
0176318
0196970
0125962
0271147
0270050
0227211
0198932
0286327
0252294
0230320
0158301
0286237
0198908
0223546
0232282
0229954
0237300
0237628
0228316
0224471
0226056
0234886
0227649
0229794
0239407
0102128
0234050
0286846
0224632
0133169
0147206
0226241
0233071
0102409
0180964

0224692
0204065
0133134
0182916
0238049
0185222
0166681
0281091
0234405
0172476
0287619
0133142
0279894
0196507
0224031
0172465
0180284
0123562
0231154
0188828
0179363
0123560
0123570
0222419
0231728
0231787
0276298
0234345
0158427
0230312
0227067
0123569
0101812
0235946
0269226
0269743
0166707
0123575
0123576
0189108
0232665
0223733
0268629
0181819
0206864
0123572
0123561
0230788
0157502
0256709

0202407
0224816
0227657
0277673
0147231
0263515
0133135
0133138
0133131
0147223
0165376
0231144
0089682
0198088
0213569
0199832
0080572
0204053
0215007
0214720
0227610
0147234
0147224
0157514
0170935
0080561
0236064
0207363
0225728
0277883
0170925
0101842
0101843
0101844
0197565
0189372
0188153
0207146
0261409
0260548
0280322
0133124
0237863
0201443
0275110
0215000
0101890
0101888
0233874
0176076

0068366
0237319
0252069
0225381
0157600
0208013
0265584
0101935
0208883
0224142
0133136
0243978
0287685
0225366
0228042
0101938
0077264
0226112
0234349
0077274
0077279
0251954
0223904
0260802
0229939
0225031
0236189
0101901
0201420
0229487
0072315
0204025
0271421
0271464
0271400
0187823
0182508
0200436
0213652
0126016
0283775
0243011
0286072
0252930
0286767
0235329
0220541
0241743
0270524
0201674

0207220
0240327
0212611
0230502
0271686
0226547
0147246
0224664
0208839
0212100
0222321
0276158
0221710
0222715
0224773
0199001
0270413
0123496
0222122
0238223
0236080
0130224
0271907
0212241
0175718
0235656
0102021
0274374
0271021
0271826
0252441
0102024
0270431
0227881
0235244
0234785
0228532
0229335
0266828
0231371
0234558
0180772
0207033
0268104
0204019
0236077
0237860
0230159
0224217
0206752

0213526
0238066
0224314
0279587
0003096
0200291
0237458
0234524
0270619
0131725
0221463
0147251
0206862
0230399
0131724
0239182
0250995
0174460
0203650
0224341
0231270
0175556
0277535
0250423
0227864
0230170
0270625
0101856
0214992
0230392
0226661
0227814
0077713
0237903
0207175
0266420
0252147
0224281
0005022
0018610
0077721
0186416
0125354
0211578
0279601
0187808
0198918
0206622
0286366
0125351

0125352
0125356
0186471
0101882
0282933
0234493
0282850
0258545
0203989
0101883
0228139
0233382
0131721
0227541
0237757
0177485
0125355
0232042
0101892
0005893
0251707
0158290
0232119
0171155
0278317
0181511
0237828
0236446
0271346
0278646
0226685
0270678
0226929
0270493
0224089
0271154
0226600
0271041
0230347
0270397
0228517
0271502
0226023
0271211
0237957
0270736
0230594
0270771
0236126
0270646

0242362
0270456
0236371
0270251
0233433
0182890
0287379
0286163
0234176
0265456
0213498
0260585
0237891
0280142
0287776
0232496
0212321
0224236
0125675
0179101
0229733
0228466
0125676
0232149
0230376
0252178
0226902
0264933
0199705
0101966
0237331
0252627
0237025
0232412
0101972
0237426
0252693
0234564
0183918
0282419
0232054
0226532
0009694
0236880
0232599
0282815
0213489
0280223
0237208
0198354

0230916
0231501
0232177
0274901
0198889
0226414
0183631
0230538
0280275
0274631
0260683
0235274
0224548
0225689
0123165
0236042
0235334
0235239
0235735
0226802
0218728
0223995
0234592
0226339
0201518
0102038
0252170
0122126
0171388
0287024
0122121
0122122
0240143
0275070
0188706
0235189
0238040
0156697
0085185
0102034
0156709
0286650
0134594
0056277
0202304
0225046
0102078
0147262
0134597
0274775

0221930
0287160
0271677
0165675
0238188
0228659
0147256
0241198
0237650
0276205
0147255
0185903
0229082
0287806
0242610
0234288
0230973
0228450
0171054
0237225
0286060
0232787
0235730
0227183
0213468
0241418
0228089
0235190
0200587
0134602
0165694
0228819
0123728
0232160
0076770
0206900
0171004
0235849
0227064
0134588
0225949
0207100
0183434
0202199
0076716
0206765
0147257
0227008
0233260
0286096

0232068
0200572
0224908
0283638
0284499
0284538
0284107
0284043
0283931
0284157
0232664
0203952
0214961
0156531
0165704
0226107
0216001
0199132
0207755
0207784
0223749
0208005
0284231
0227060
0170965
0235813
0236509
0237432
0199920
0156504
0156500
0280249
0101928
0207081
0236091
0231929
0184785
0212747
0134590
0286964
0203950
0228372
0234228
0196972
0224107
0223809
0232582
0169551
0236491
0228922

0269833
0186376
0238210
0274024
0283644
0284994
0173275
0229015
0178947
0201440
0225769
0273988
0271158
0225235
0165359
0237833
0224539
0277486
0198022
0274601
0268940
0233861
0238200
0269096
0228836
0278289
0271449
0273696
0278085
0270946
0269586
0181433
0230162
0169446
0198689
0219930
0231513
0239080
0022267
0129680
0230079
0156920
0102239
0102241
0102243
0216060
0199424
0252320
0233093
0102245

0129675
0212510
0271339
0231164
0225125
0202279
0232611
0225400
0147274
0206979
0234062
0232379
0237307
0165370
0214748
0237584
0224765
0156925
0283692
0239579
0229562
0222114
0252163
0183981
0214930
0285989
0129682
0232183
0207800
0226031
0235975
0173954
0232549
0101981
0101977
0238485
0101974
0221375
0252719
0207633
0203933
0233145
0214653
0241248
0230707
0231110
0224942
0275268
0225758
0134595

0230668
0203930
0184258
0221081
0229269
0227234
0182195
0207041
0214925
0277215
0198573
0213423
0227779
0198021
0203926
0223438
0196406
0165509
0155495
0288098
0236205
0225469
0250625
0046774
0275508
0201000
0189326
0202473
0199878
0227303
0226064
0235539
0230130
0189252
0179542
0213924
0268988
0225970
0235729
0276380
0224630
0235598
0215054
0201912
0233548
0234712
0203923
0231093
0286085
0270709

)279585
)185985
)252219
)216075
)216005
)215943
)216098
)216064
)216056
)201594
)234748
)226388
)231686
)235699
)224440
)225660
)271660
)284377
)216171
)207871
)207873
)207984
)207731
)207969
)207589
)252583
)212013
)212014
)208000
)207641
)207868
)287017
)207866
)207867
)274579
)228855
)227505
)271175
)202051
)287796
)227083
)268066
)274086
)102081
)236337
)176988
)201285
)219507
)233467
)176970

0237741
0155966
0223516
0222313
0242186
0287585
0010404
0241489
0238039
0176289
0241769
0197620
0283697
0267978
0268738
0269556
0229322
0185247
0228833
0223377
0231694
0235254
0171116
0123584
0224732
0229829
0280752
0230899
0156009
0197021
0283463
0235703
0183171
0252454
0234825
0013619
0171100
0063601
0102181
0228728
0230153
0222796
0029993
0230508
0270916
0212595
0265789
0234696
0102195
0269993

0228717
0278724
0287918
0160131
0270052
0214915
0166049
0130032
0147378
0183862
0229967
0147381
0102287
0284363
0283751
0228965
0242520
0266560
0124260
0231937
0011677
0252205
0207957
0211583
0207818
0223940
0287394
0268089
0269475
0221867
0268902
0183305
0198930
0213401
0242599
0231447
0268606
0268916
0197172
0286051
0147400
0147383
0147394
0286939
0198883
0234200
0229979
0183837
0235961
0268883

0224931
0198681
0235219
0273877
0231600
0278803
0225110
0063587
0232249
0214897
0265170
0189420
0183479
0213397
0224963
0182492
0067842
0262919
0260081
0217889
0130829
0237793
0244335
0130822
0130821
0185825
0101986
0232725
0198753
0184343
0067829
0180879
0067840
0184844
0198910
0284987
0273769
0126895
0089820
0102030
0102032
0172534
0235802
0177854
0265176
0283694
0184216
0284286
0169057
0102076

0277008
0268221
0274962
0166160
0277009
0269433
0278057
0007350
0231830
0196924
0285018
0102119
0147403
0280195
0207165
0013563
0102125
0197180
0071553
0203879
0071859
0276350
0130827
0196976
0102178
0126903
0264484
0071889
0261773
0160211
0269335
0279245
0273041
0268651
0232155
0184033
0273773
0213338
0272681
0275882
0126890
0226141
0237619
0160219
0130826
0206948
0284450
0206693
0130830
0229237

0203870
0185010
0274183
0277203
0221533
0224069
0237859
0233688
0165775
0288258
0182712
0214827
0185515
0235991
0155959
0224978
0155961
0155962
0225008
0228986
0230491
0224216
0230578
0277858
0274791
0221190
0277150
0221603
0277745
0224533
0225393
0185973
0168939
0237801
0237040
0124333
0228410
0223484
0124334
0270726
0185203
0182484
0227159
0251841
0184895
0237659
0232195
0286130
0129824
0227289

0229163
0278847
0067646
0233070
0223600
0231535
0234652
0216777
0176679
0232927
0225653
0286050
0218410
0229518
0229308
0252900
0225878
0277930
0099715
0252468
0231341
0235001
0214207
0278585
0235175
0285470
0237195
0250204
0168757
0234830
0238235
0275352
0234583
0235094
0223744
0229643
0129816
0212856
0237563
0147753
0185700
0235895
0232620
0224953
0274365
0233634
0099721
0237701
0092377
0273906

0232583
0099725
0252633
0251996
0252155
0252472
0275280
0238073
0223422
0236435
0236690
0234795
0237048
0278854
0215603
0273863
0244246
0229745
0241200
0239893
0227204
0225809
0215601
0224634
0223955
0234179
0226529
0234850
0227830
0216824
0275866
0286217
0233699
0232419
0180910
0228207
0173357
0273731
0230025
0231988
0232235
0224408
0238154
0232924
0197038
0275310
0232808
0233803
0225560
0229549

0228383
0286120
0188656
0237802
0286173
0228927
0225516
0258992
0274196
0238074
0230066
0270073
0234803
0236424
0224336
0236786
0231436
0234110
0226223
0232617
0183385
0147761
0228890
0212855
0129845
0224075
0228379
0229208
0274445
0231874
0187657
0239225
0234950
0236718
0243643
0225685
0278478
0251766
0252289
0231411
0226061
0237447
0225840
0251705
0273858
0273693
0271365
0271309
0270570
0274231

0278212
0274502
0274162
0258567
0259154
0259029
0258991
0270455
0183704
0270242
0275828
0225433
0224567
0271375
0234385
0215583
0235649
0206159
0236477
0232226
0243980
0232730
0224060
0225117
0227447
0114374
0226863
0233774
0286009
0228411
0067048
0234059
0224035
0235462
0223915
0183878
0215414
0154620
0241859
0129864
0129862
0235451
0223517
0165246
0226555
0228787
0236131
0227949
0224518
0234620

0227166
0252689
0252173
0252323
0286201
0228518
0271595
0224989
0283076
0228578
0230663
0252315
0252012
0273966
0238191
0226353
0275578
0226449
0225896
0224166
0229129
0250868
0223555
0235521
0252513
0224964
0226011
0225740
0129873
0229138
0224485
0226116
0237558
0233378
0237616
0224033
0182415
0230412
0226611
0225895
0233120
0252166
0232744
0244646
0229465
0231311
0236599
0232899
0229302
0228945

0233522
0278391
0251953
0252667
0249726
0282909
0226362
0225716
0229406
0227494
0252209
0249634
0251510
0238067
0233546
0230377
0250951
0172468
0230458
0131007
0240438
0235479
0249606
0242153
0169953
0236429
0176728
0249501
0228193
0248573
0230904
0238135
0224151
0251970
0238088
0217179
0217896
0185275
0252766
0286265
0274282
0234529
0215580
0131002
0267793
0260197
0012817
0288049
0274837
0237997

232522
228764
229236
277438
198692
286247
225624
280969
224482
216844
229551
226504
254488
237427
232634
226918
183146
223637
236615
288057
237269
169849
169811
242393
242875
234414
228257
184991
271123
226042
237023
229416
230029
277519
286187
229725
244395
251618
242389
227444
226092
237968
231159
232475
229553
228850
232029
169807
236951
131548

0169800
0229159
0225615
0229234
0215560
0229940
0243040
0259247
0237902
0226941
0235059
0131538
0169789
0234131
0224873
0230814
0228571
0230476
0273589
0224657
0224917
0230727
0228240
0276829
0215537
0226906
0183753
0244000
0232764
0233126
0227251
0188120
0205944
0205936
0251925
0286744
0225189
0230073
0215540
0236647
0225466
0224169
0223407
0226873
0227915
0169763
0236620
0224866
0232424
0252681

0232910
0232845
0237302
0231540
0223362
0231026
0224571
0226369
0280961
0227871
0251917
0231423
0228465
0232585
0234081
0172352
0231375
0237467
0229250
0232205
0235981
0223698
0242854
0230977
0244231
0172342
0284071
0239533
0274234
0225491
0224827
0233740
0248792
0279274
0234399
0227439
0224210
0232003
0235412
0183795
0240566
0232976
0226270
0227837
0187191
0205916
0278602
0223978
0278197
0230819

0185894
0228296
0236379
0274899
0223641
0227633
0279115
0228786
0233652
0233944
0223636
0172297
0278242
0284380
0172294
0240450
0239304
0233619
0197092
0235583
0237823
0229709
0232064
0225609
0172288
0270535
0225287
0225256
0223856
0235004
0252426
0231141
0232614
0225326
0223406
0229343
0232695
0235511
0232914
0223655
0230854
0252625
0227635
0237546
0172283
0227867
0231716
0234744
0233156
0234888

0215507
0235014
0225876
0251796
0229238
0252948
0233843
0188399
0277146
0215506
0224240
0227629
0237917
0231514
0235857
0223972
0227232
0278267
0243485
0284332
0237613
0268020
0240361
0186092
0238009
0239945
0233750
0268903
0269981
0239906
0241860
0222623
0241599
0279928
0279457
0273874
0228463
0286448
0236679
0236601
0237094
0269732
0284733
0233653
0250575
0278757
0230021
0235146
0225972
0225630

0237973
0278791
0229344
0240409
0248527
0198744
0268663
0284662
0229376
0228327
0223181
0229905
0237491
0285268
0230092
0177757
0228794
0225880
0230368
0234711
0283040
0272438
0230699
0241180
0223764
0187634
0188976
0187961
0187583
0187642
0272512
0188290
0187608
0231702
0224969
0188157
0242590
0217801
0273443
0237330
0131591
0285812
0223823
0207730
0207607
0198976
0272141
0205231
0162571
0186891

0186827
0078808
0176022
0184163
0260179
0160087
0230415
0162572
0131584
0278073
0169972
0127054
0283712
0240731
0224051
0169962
0107404
0284372
0162576
0175756
0223663
0221978
0224870
0242485
0264293
0272455
0235098
0225905
0205116
0225285
0179403
0215915
0160072
0284740
0197785
0205090
0160075
0215014
0274481
0228594
0286989
0272106
0197530
0189409
0248333
0272004
0189339
0269737
0215914
0008128

0268575
0227775
0215790
0008130
0078369
0231050
0169885
0178821
0142609
0233542
0187730
0226969
0067606
0271806
0182873
0162585
0234396
0157933
0287356
0116151
0272161
0269896
0272420
0157916
0157911
0149527
0224387
0229393
0157881
0197921
0272449
0238164
0157873
0225931
0228037
0157870
0142606
0237058
0215912
0285945
0283259
0233234
0287396
0231630
0284745
0169717
0177133
0142611
0283572
0226286

0279839
0272235
0287828
0286518
0130762
0272088
0162591
0207776
0238260
0158109
0116213
0078900
0227589
0235131
0227372
0276189
0162592
0235169
0130764
0266075
0272153
0116198
0169598
0198912
0236423
0233304
0226374
0284668
0284703
0229280
0235054
0284694
0227169
0232596
0196581
0287586
0284739
0284674
0231510
0284616
0284692
0284666
0260972
0283356
0236948
0264341
0264101
0131697
0069424
0116254

0285629
0116251
0226944
0158286
0116237
0225077
0173673
0158292
0097021
0271746
0069812
0187017
0265392
0231868
0215788
0171680
0162408
0253022
0229519
0173662
0204859
0162413
0116273
0041988
0007923
0228750
0284744
0237436
0171735
0227950
0237365
0207056
0237402
0237728
0225126
0270171
0270035
0269978
0049245
0269925
0049246
0236266
0049247
0049249
0270330
0116288
0200344
0284747
0284716
0116285

0238290
0232848
0229305
0227634
0200975
0233645
0251977
0162426
0252254
0142599
0232912
0221083
0270282
0234619
0224315
0222511
0226545
0228423
0229716
0074800
0274258
0230679
0201725
0238249
0232208
0232663
0131686
0265141
0197241
0142583
0252404
0180758
0228526
0284357
0234546
0049239
0171621
0284652
0284693
0252956
0235263
0171612
0231181
0188807
0171608
0179840
0233268
0231789
0171603
0271264

0280113
0178585
0223989
0285701
0162441
0228150
0173614
0202415
0283611
0265521
0241326
0162444
0130939
0224340
0233623
0201746
0054523
0264881
0199562
0264501
0284735
0284642
0142657
0271989
0251503
0175279
0241563
0160049
0203469
0142655
0287727
0243267
0130940
0272078
0225158
0175262
0215785
0120948
0277726
0009724
0271895
0116649
0171824
0226849
0230337
0198793
0225602
0253086
0171819
0207451

0238173
0120942
0238199
0284646
0224338
0204624
0285833
0235643
0284708
0116661
0132879
0116663
0116670
0162490
0177674
0215910
0177553
0177000
0011021
0276470
0278852
0175206
0275915
0120937
0224904
0285646
0199347
0201801
0116685
0083444
0287384
0116688
0270914
0116691
0202461
0276869
0243539
0120949
0201135
0227875
0285604
0283789
0028137
0263676
0225196
0252969
0048707
0239149
0229484
0162496

0221340
0276830
0272482
0204518
0226166
0188984
0157330
0116726
0116721
0228338
0239810
0231103
0179172
0120952
0243073
0187545
0204510
0207434
0234064
0232423
0274178
0274764
0277058
0229571
0277862
0236179
0275774
0229978
0233336
0280267
0179412
0204505
0279169
0279804
0250894
0270601
0231219
0207511
0182330
0237700
0204501
0228229
0204481
0204480
0204479
0204478
0162494
0259961
0215909
0162493

0222952
0237445
0253085
0116731
0189337
0252151
0231606
0234593
0235829
0287756
0175147
0171729
0204464
0215720
0278480
0142621
0233485
0236045
0228140
0142634
0162438
0142615
0215704
0132906
0116138
0251866
0272510
0116771
0237301
0235084
0252417
0236500
0197312
0215695
0271742
0116786
0237938
0162461
0224459
0162460
0162458
0224321
0233954
0178715
0179743
0065526
0270620
0116809
0234607
0231353

0275503
0224966
0183888
0233078
0173641
0186510
0232456
0184908
0185519
0142627
0227959
0224621
0234166
0142632
0237276
0288398
0132881
0233929
0037637
0055070
0226457
0187144
0157191
0226029
0080947
0206652
0261135
0285853
0224174
0233421
0277344
0225710
0219481
0271732
0215908
0283773
0186301
0282143
0207513
0280114
0236698
0268869
0279151
0282740
0058453
0227684
0207389
0186715
0223643
0230239

0263811
0228549
0277234
0238142
0235241
0207005
0272426
0117122
0286898
0226526
0159363
0117118
0117115
0204362
0282843
0142623
0142619
0266634
0159339
0276747
0179051
0227751
0074964
0236648
0117148
0284653
0261781
0117154
0280222
0230035
0225387
0179023
0235282
0225478
0009709
0179002
0159423
0265606
0221662
0169991
0272084
0127481
0230424
0127463
0053372
0286064
0211454
0162482
0270728
0053371

0200403
0040487
0077549
0240490
0173436
0201609
0235185
0158747
0270136
0226396
0158748
0162542
0178828
0235434
0169914
0188784
0242688
0188257
0127472
0117215
0158786
0227066
0187980
0225986
0162543
0228105
0284710
0158816
0226664
0236963
0226487
0284641
0284743
0162545
0090432
0215520
0183114
0158825
0158828
0284005
0117242
0244038
0117245
0235432
0189410
0233069
0127483
0075151
0251914
0221808

0233072
0236073
0235112
0287192
0117298
0236936
0231105
0236216
0228823
0236009
0227001
0187952
0186543
0142794
0176378
0162551
0233431
0076864
0090686
0231978
0187942
0142798
0232037
0283234
0219073
0285959
0273610
0201919
0142789
0278124
0201273
0228397
0218510
0070831
0285752
0285794
0230068
0271428
0162552
0285873
0266564
0234397
0279625
0184677
0237200
0070886
0276835
0173372
0159189
0173369

0133216
0265422
0225952
0264014
0215906
0227868
0004487
0240553
0263793
0266802
0169641
0229010
0206935
0252578
0179546
0215381
0261326
0284726
0125944
0125945
0249087
0201405
0271420
0204219
0232482
0088280
0007968
0235052
0117318
0285802
0197880
0229239
0142676
0236810
0011007
0057757
0011009
0117308
0117305
0179163
0188822
0229106
0232557
0264926
0230228
0189266
0188529
0215905
0225315
0142661

0230703
0142677
0185436
0230023
0232298
0158055
0001460
0001461
0286061
0117602
0264443
0184454
0227312
0239106
0133226
0284699
0169504
0238482
0020633
0278034
0229162
0261025
0264371
0285823
0233755
0233419
0231953
0117614
0284602
0284657
0117616
0272432
0261349
0187010
0224183
0259984
0183726
0188672
0238084
0204178
0157978
0225643
0117643
0233478
0162430
0255054
0228172
0117640
0272478
0127423

0236528
0182749
0207302
0117632
0229786
0223624
0283938
0158006
0207237
0252190
0252691
0158008
0158014
0284309
0158022
0175087
0197245
0282872
0236782
0142684
0142675
0188782
0236155
0278572
0130695
0270733
0142669
0158062
0169442
0176092
0225854
0176083
0131914
0117682
0223583
0225891
0198830
0224129
0117676
0238705
0243905
0252802
0252429
0229247
0238316
0235069
0260063
0117713
0060642
0241188

0287810
0204160
0175793
0142751
0226698
0198746
0090273
0131910
0175707
0172746
0224064
0225294
0253368
0158246
0090020
0229985
0214973
0225159
0225990
0224311
0142784
0252098
0243659
0186501
0206888
0241547
0142765
0142733
0142748
0174950
0231207
0181773
0158195
0241169
0237429
0126705
0000938
0235912
0233975
0126709
0225886
0206767
0226662
0207095
0287244
0009780
0231344
0230833
0117758
0269971

0270031
0223062
0286433
0117751
0252947
0130775
0117748
0130768
0227050
0158156
0158161
0238231
0240750
0228589
0228943
0214812
0253005
0169403
0126698
0130770
0270605
0271398
0130766
0130772
0204138
0206779
0229820
0221216
0180198
0242125
0274266
0200087
0231468
0279443
0180098
0197989
0221539
0278274
0273544
0280498
0274582
0120656
0188060
0229388
0274978
0270103
0162419
0252777
0198492
0116329

)233427
)199756
)159023
)225616
)206704
)277958
)253304
)225750
)116350
)277916
)116353
)237934
)060656
)230523
)225011
)228176
)284676
)270927
)233399
)233372
)231251
)236335
)231949
)162510
)186056
)287510
)162511
)264773
)229607
)223382
)222784
)235143
)162512
)266210
)134644
)237329
)232768
)215900
)225142
)084628
)060688
)229447
)121766
)229044
)121769
)168528
)215899
)206981
)260386
)265961

0284543
0229167
0142910
0121764
0162517
0235790
0264078
0084636
0121753
0266580
0254545
0134668
0203620
0269967
0184007
0228634
0284702
0121774
0203325
0121775
0266203
0276493
0025800
0250135
0084652
0160050
0224066
0160051
0222046
0160055
0084623
0220785
0183615
0182866
0277617
0252450
0116478
0175130
0233775
0162526
0225828
0160058
0224333
0270850
0229259
0263529
0254553
0273274
0160062
0176261

0200591
0162521
0162520
0224409
0162522
0134684
0116497
0160097
0121905
0121900
0287691
0116514
0278966
0217644
0239670
0236065
0004455
0276645
0142920
0278997
0116525
0279179
0284721
0160094
0270115
0184389
0225313
0134686
0284154
0222112
0233246
0234512
0121903
0235907
0121904
0227000
0176256
0231163
0201148
0142698
0270241
0287703
0207941
0163866
0189280
0189433
0255811
0188910
0187513
0230163

)116544
)284773
)241014
)271741
)243749
)163867
)197056
)233940
)228348
)284640
)116560
)201868
)146463
)240374
)201542
)229994
)227409
)278702
)142687
)236274
)020129
)239636
)116819
)126067
)142686
)092853
)232335
)134698
)092847
)271164
)286899
)271914
)126070
)271554
)232862
)092850
)116863
)171812
)054116
)116871
)264592
)054118
)231849
)214193
)142694
)286379
)196182
)181817
)222821
)116885

0201448
0116898
0119535
0230214
0284705
0284720
0234481
0163873
0284650
0223944
0264698
0207328
0252368
0230451
0233621
0163874
0283724
0163875
0263675
0163877
0232273
0237749
0163879
0134697
0284748
0169218
0116922
0134690
0183317
0200421
0214141
0227416
0185090
0233728
0196449
0197982
0188786
0235673
0204084
0230955
0252448
0222282
0277168
0183431
0212541
0183386
0183520
0185668
0224592
0223589

0265596
0284623
0286552
0287987
0237290
0275350
0200796
0238063
0215895
0284632
0116954
0273637
0228436
0214114
0274944
0131233
0158315
0199963
0207466
0235549
0174574
0168653
0127603
0206654
0222378
0270040
0287422
0226438
0217897
0183682
0237624
0243970
0090621
0228060
0201457
0163909
0225903
0116981
0116983
0084072
0225333
0252413
0116985
0198754
0285905
0261798
0229213
0284719
0284678
0043514

0202222
0116990
0236546
0207356
0168389
0228477
0131236
0131238
0226745
0117000
0207508
0188800
0259943
0084073
0231296
0049089
0227311
0084070
0213172
0284677
0187801
0260920
0279667
0187815
0238287
0164002
0227278
0238186
0117010
0286838
0236505
0233910
0117016
0287743
0237899
0272145
0066136
0198974
0207962
0117013
0264582
0179862
0229528
0228940
0286668
0233118
0171793
0281207
0171790
0010803

215893
287400
235358
213987
286640
204060
229901
252563
127129
127124
230881
230638
284895
287587
227538
228776
044012
197273
198815
227527
177181
235002
066185
127125
186409
230116
236876
200254
236180
171960
285728
234917
065978
164007
117385
164008
283580
228452
274386
177868
164010
228192
164011
233708
237090
225099
117394
284138
227533
283973

0207256
0283136
0277513
0186973
0117395
0284410
0243710
0199240
0179178
0253313
0066056
0117400
0287113
0234694
0117399
0066322
0283836
0159479
0229431
0198198
0229372
0274975
0178922
0229348
0142949
0284989
0066135
0236200
0126091
0229444
0283414
0233674
0283477
0224751
0117407
0237950
0117408
0285649
0132768
0117410
0117411
0159214
0196517
0230615
0227163
0271647
0239560
0171872
0233514
0283039

0226804
0200171
0238037
0227994
0271329
0178028
0117419
0233602
0199385
0187147
0263381
0126106
0198520
0199377
0200169
0142945
0225721
0142937
0264294
0200913
0202031
0281859
0207421
0234093
0142959
0173846
0188396
0222009
0117425
0202444
0226499
0070785
0253039
0229815
0232369
0268949
0126107
0126088
0162415
0234315
0281912
0186603
0132781
0288208
0132773
0070759
0236872
0280670
0132763
0117450

)234379
)117448
)281133
)280836
)132780
)281112
)281825
)159588
)159592
)225447
)234329
)159596
)197429
)230896
)236111
)086015
)232467
)117461
)278139
)226957
)233114
)227857
)117472
)250719
)085998
)171357
)085999
)132128
)173660
)117481
)225779
)117480
)232022
)224863
)197587
)282881
)269956
)162456
)079277
)142961
)123472
)233647
)186118
)228237
)159658
)236476
)142973
)154198
)238145
)187048

0259832
0261196
0261593
0261290
0186377
0186160
0225154
0225506
0162365
0225514
0224805
0162366
0162367
0226252
0123473
0162368
0225762
0186790
0237424
0186564
0271355
0229554
0227883
0227947
0269113
0223814
0225028
0226133
0231413
0237180
0242267
0279096
0223720
0117834
0272491
0279214
0132122
0224986
0206700
0235105
0223175
0232514
0186094
0287661
0162373
0286597
0229846
0230114
0225623
0237478

0215887
0279324
0233595
0162374
0230828
0237337
0233407
0284696
0284645
0234080
0229316
0225671
0142700
0284700
0185104
0225767
0207194
0230585
0237130
0123080
0265538
0275816
0204006
0200262
0213478
0203356
0123091
0233406
0236434
0085831
0261664
0085832
0232027
0206595
0238140
0227070
0252032
0227742
0117859
0236973
0232846
0212624
0078618
0266993
0272175
0283899
0230604
0169213
0200839
0229032

264613
117862
285839
198841
223390
228369
134717
157077
241745
228407
223429
224255
236004
237663
213735
154222
272100
287078
224680
085840
134748
134744
272371
236290
116157
182183
162377
233664
239640
162378
252018
228929
206627
203995
242391
121310
230953
231866
116171
264834
266188
265815
237279
236360
226147
174348
232993
227644
162383
235563

0277397
0157184
0236723
0162384
0259818
0162385
0226754
0157193
0234578
0228838
0232762
0225675
0285954
0226938
0230138
0236253
0143006
0174332
0239007
0058804
0201003
0229687
0058799
0211452
0081870
0116212
0219102
0203985
0116209
0280378
0116205
0283749
0225183
0279049
0256407
0157211
0280425
0215883
0116221
0157216
0198711
0225632
0232245
0287582
0235741
0237173
0227453
0162390
0162391
0230728

0184313
0271723
0243725
0162396
0006555
0237453
0162398
0116133
0287724
0242396
0233203
0284601
0143001
0233271
0162399
0169174
0162402
0231090
0231095
0265822
0234810
0231900
0232198
0199831
0252162
0272051
0233079
0230250
0227104
0235612
0260971
0284686
0229871
0223956
0162407
0236776
0224980
0225475
0162409
0187889
0229913
0157131
0021852
0236341
0173406
0227935
0236888
0226759
0229887
0223416

0235038
0230546
0233216
0162600
0286918
0270209
0264128
0184292
0185839
0162601
0283445
0177606
0234807
0272226
0231740
0232453
0237352
0228121
0241042
0235215
0224609
0270457
0172456
0266150
0226883
0134709
0134716
0265535
0162598
0229332
0238242
0226476
0231252
0162599
0237928
0237853
0284808
0287224
0223920
0162604
0271200
0132849
0200575
0212360
0236646
0242860
0263908
0283690
0234204
0234088

0240563
0237227
0200174
0132854
0201153
0162607
0116641
0270549
0132855
0278967
0213703
0235545
0125703
0234318
0237163
0229537
0286455
0227485
0229225
0203605
0224209
0252259
0228734
0223683
0230798
0187140
0275836
0252784
0236674
0088035
0142856
0286429
0203965
0264271
0251720
0116652
0276884
0278287
0277565
0275823
0277689
0274199
0079739
0244256
0185483
0177452
0228056
0207190
0200508
0223949

0284928
0177414
0238653
0158966
0264470
0162437
0162434
0234784
0233877
0226891
0185031
0272506
0212257
0265996
0199135
0235804
0234496
0162433
0231622
0116675
0229294
0234144
0222624
0213625
0116678
0237852
0224570
0285079
0239319
0184588
0227466
0223152
0118473
0264720
0248458
0152760
0172410
0152763
0231080
0198160
0116704
0203963
0252433
0275678
0162594
0253074
0252116
0228523
0081985
0142864

0223263
0235200
0207504
0238139
0242482
0232048
0116717
0172380
0238778
0232284
0235055
0162595
0228431
0116729
0234383
0221203
0229133
0225242
0237622
0229440
0116745
0024526
0233589
0234264
0225167
0285041
0285407
0285473
0223883
0235446
0229639
0287453
0033122
0270615
0287283
0240692
0229359
0237919
0066557
0244389
0116754
0228988
0118454
0197568
0226088
0116761
0278636
0271992
0233020
0237033

)229051
)226208
)050628
)235782
)269933
)235079
)132485
)207721
)229956
)226324
)172260
)231985
)271618
)228853
)233994
)286863
)227207
)225087
)212366
)227016
)285808
)280317
)229636
)251825
)233973
)223479
)200601
)285778
)252005
)162620
)254685
)259030
)116783
)237324
)233894
)162621
)178965
)234497
)272864
)116791
)162623
)229943
)261213
)224127
)162624
)224149
)206999
)137968
)224493
)252338

0213579
0230863
0117054
0181227
0137955
0206620
0207241
0201487
0057468
0154007
0225605
0230027
0184005
0223905
0272855
0226415
0199921
0237552
0117069
0276081
0230498
0142892
0226084
0228187
0287870
0154027
0233099
0251767
0287647
0036549
0251785
0243437
0077254
0232672
0180488
0222849
0219201
0235613
0232015
0235927
0162614
0162613
0162616
0137960
0273338
0285928
0213561
0251958
0213560
0202263

)233333
)237413
)122420
)227556
)212308
)137959
)137965
)238015
)235400
)162618
)230787
)282898
)235011
)285462
)232676
)285409
)235756
)260322
)277670
)236407
)225598
)213559
)227062
)234108
)224326
)234184
)227937
)229752
)235089
)285179
)223026
)117114
)224494
)227288
)227960
)234953
)242598
)236676
)234683
)215875
)233290
)230817
)236268
)285782
)231364
)233008
)237076
)137941
)233061
)229486

0285201
0271576
0142875
0213540
0226970
0199959
0203943
0249237
0240520
0137976
0285361
0117133
0174021
0285851
0122432
0117151
0284882
0285374
0180869
0232622
0117155
0285325
0171517
0153898
0162643
0055732
0266110
0280099
0097096
0162642
0223254
0142867
0223653
0153904
0199459
0273264
0282057
0272691
0142871
0199934
0230285
0117174
0171502
0201620
0229505
0285210
0122417
0278281
0231349
0137975

0016490
0016602
0224466
0236915
0153923
0261737
0097033
0230721
0284846
0183291
0153936
0267734
0267561
0235251
0225568
0267272
0284240
0143013
0227290
0199318
0237505
0279778
0270507
0286758
0239504
0230053
0274321
0237959
0065243
0207234
0241975
0137947
0235308
0137944
0213516
0117226
0202385
0117228
0234518
0162645
0286802
0213512
0162654
0284734
0284637
0154451
0237568
0238081
0233235
0183347

0215874
0225492
0286548
0197147
0231999
0230735
0171488
0271949
0226394
0231613
0272931
0171492
0272672
0252797
0228175
0162664
0287015
0287406
0212459
0287372
0229201
0199666
0143032
0287076
0233593
0285532
0225446
0272094
0232882
0122482
0232393
0162669
0206817
0215873
0097046
0241318
0229067
0203914
0069702
0239794
0137948
0172031
0233228
0230667
0189195
0224678
0233401
0273487
0226773
0203910

0233125
0174842
0122484
0242764
0162676
0067208
0223787
0229992
0201317
0225505
0234202
0223896
0122406
0154511
0206680
0207523
0207022
0229052
0252121
0239710
0287797
0252752
0222664
0251837
0143033
0117500
0122483
0237003
0225297
0223745
0229567
0117505
0229635
0207379
0137942
0212601
0137936
0224093
0211575
0260464
0067334
0023909
0223490
0234099
0235794
0232752
0229842
0250890
0236689
0226960

0233129
0198691
0233482
0137962
0264963
0226835
0178596
0236098
0232918
0286692
0117528
0117525
0223675
0224081
0232857
0263526
0143036
0117519
0235501
0172339
0230427
0152078
0271092
0231992
0206679
0226026
0122481
0228852
0287919
0228504
0230434
0237954
0233907
0228971
0271252
0237435
0200800
0231987
0225923
0234422
0228502
0241992
0234226
0137970
0223229
0117569
0188641
0232878
0232542
0228057

226128
230718
235777
270911
225206
259946
284247
284202
285922
233557
226053
280040
162627
117598
232825
227034
117600
233983
224445
099260
223656
156869
201491
202254
231996
162688
228084
117620
283761
212248
202259
156875
241073
156876
226952
122435
122477
137992
271415
225169
285530
230287
224616
137996
207750
215871
223906
079335
234332
228086

0181656
0213421
0230402
0285525
0224685
0162692
0162694
0273204
0162695
0230946
0117543
0235795
0233184
0252765
0271578
0225938
0170989
0228399
0173810
0231671
0226715
0252530
0183298
0118733
0252717
0271277
0162699
0233359
0230864
0060718
0234257
0230759
0224613
0222069
0185946
0240038
0236085
0243480
0237763
0234441
0174876
0227408
0187733
0225191
0215869
0228347
0285981
0225036
0230768
0227850

0237897
0232952
0233047
0237480
0230932
0270342
0270976
0233449
0232551
0237460
0198890
0162631
0224550
0134215
0275455
0230489
0226822
0085491
0260879
0280186
0196427
0238122
0241361
0243967
0238118
0224698
0186086
0283354
0227832
0226483
0162636
0162639
0285923
0134186
0143107
0232971
0116266
0228076
0162641
0203897
0121957
0121940
0085433
0237349
0244716
0228665
0228054
0197780
0273382
0215717

0278249
0270066
0251484
0179902
0116299
0031698
0143126
0134222
0221986
0134243
0143106
0143028
0162650
0174151
0181754
0156097
0254942
0065135
0206832
0225113
0284443
0134183
0116337
0228703
0235045
0168765
0213366
0134184
0260246
0134201
0241720
0134202
0198758
0224927
0228420
0235005
0261055
0184371
0168710
0143093
0258634
0156150
0258673
0186150
0197106
0227091
0235526
0224965
0116396
0282852

0200536
0227963
0162775
0224699
0168679
0273373
0134248
0143125
0270380
0282887
0283999
0240194
0235407
0143105
0177301
0259834
0177272
0199710
0237415
0232791
0143119
0261654
0232811
0121931
0273010
0225672
0252760
0156171
0134255
0272982
0273221
0162777
0064886
0236040
0203878
0134216
0229283
0232240
0173947
0234020
0236012
0227179
0085465
0260948
0233337
0243960
0116455
0199890
0116459
0143110

0121933
0200360
0282608
0116473
0231346
0215867
0201028
0197852
0227811
0284830
0064703
0171385
0232558
0237556
0231437
0231246
0234388
0238761
0143079
0283296
0134245
0273483
0007341
0116489
0235299
0252750
0155363
0225075
0155366
0271810
0155367
0261595
0184599
0234790
0215866
0224167
0228040
0155380
0229020
0226419
0287807
0238198
0198799
0237278
0233839
0236887
0081026
0232499
0238256
0116793

0232450
0081019
0231128
0134242
0226167
0188761
0134262
0118655
0235527
0163349
0116774
0270780
0282048
0134207
0232895
0273698
0271419
0197323
0226984
0225650
0201114
0236480
0116752
0175984
0116748
0242769
0213281
0009307
0201900
0052723
0227970
0198765
0134200
0134198
0233730
0285698
0228035
0134259
0287103
0226973
0274468
0229101
0232185
0270631
0239984
0207502
0173218
0118729
0177551
0214204

0228127
0271143
0163393
0237993
0173212
0235933
0221040
0287217
0163399
0203865
0212385
0233154
0230381
0200547
0224950
0286276
0116815
0177173
0264419
0143061
0211543
0203864
0231072
0223489
0116824
0225079
0272715
0134247
0252510
0134256
0236137
0116830
0215930
0134253
0228453
0134258
0277141
0235202
0271427
0198162
0279513
0226755
0236866
0183508
0213262
0212266
0270719
0228217
0196505
0065183

0155761
0287980
0222209
0226126
0229911
0092607
0226172
0239216
0116874
0224238
0223575
0236804
0231365
0227056
0227712
0232650
0194297
0116882
0230921
0203859
0232780
0203858
0235595
0187481
0229570
0203857
0226443
0249798
0239873
0203855
0240244
0260941
0198857
0143067
0092621
0134240
0134193
0215864
0227205
0226446
0134249
0134250
0273406
0207149
0265808
0274642
0277313
0223804
0273727
0270231

0270392
0275131
0273694
0275933
0286106
0273136
0287979
0275538
0263353
0281741
0226067
0277610
0285215
0275585
0273516
0275296
0198019
0234998
0233029
0223345
0227193
0213244
0268993
0188610
0171943
0275129
0230806
0227082
0228826
0233432
0272583
0231752
0224857
0276118
0277095
0277702
0252830
0264145
0274927
0242943
0223495
0235887
0203825
0179571
0226028
0202167
0207349
0252826
0206737
0232721

0223779
0242352
0270962
0263590
0252105
0233430
0237503
0265531
0234571
0273213
0230186
0268074
0263513
0270872
0276674
0275075
0275767
0232527
0276442
0175658
0276756
0271567
0271439
0207205
0237343
0233586
0266338
0227212
0229002
0275291
0278099
0277918
0279782
0277199
0224363
0278273
0222854
0273825
0287607
0196369
0271644
0215784
0283752
0272419
0277678
0281571
0223612
0256374
0271291
0207501

0276216
0236140
0162825
0234367
0270384
0286620
0270722
0287374
0278182
0275064
0272150
0117262
0174827
0117281
0265491
0186141
0186364
0131788
0278549
0198483
0143127
0244619
0131779
0265241
0211451
0234222
0280778
0271601
0272031
0278431
0121851
0265972
0168509
0287190
0201558
0233396
0271425
0264343
0276509
0201789
0271121
0277406
0270339
0273768
0287978
0274428
0277655
0276975
0274408
0274450

)268043
)244371
)234225
)227242
)225603
)201142
)230832
)286172
)283342
)131791
)180867
)237188
)131781
)226015
)236806
)131778
)278811
)213226
)226653
)207209
)272443
)272480
)116128
)162836
)277762
)227139
)265107
)274415
)234190
)223728
)121634
)188092
)215859
)273059
)274372
)206791
)206585
)229828
)263956
)225080
)198658
)232265
)227733
)238765
)224335
)227700
)199879
)234283
)228626
)274020

0238107
0224481
0235988
0252744
0283676
0202408
0272824
0207340
0252135
0278596
0238825
0231551
0201183
0225871
0285062
0280649
0236334
0274210
0270629
0286019
0252515
0177144
0275140
0271546
0178104
0252656
0222788
0254913
0255148
0254539
0269713
0274019
0201699
0274265
0215861
0272755
0273591
0274423
0286219
0286185
0271383
0263464
0277147
0284964
0275557
0206828
0284842
0234232
0275229
0226500

150337
203814
233030
183598
270882
203811
203812
220323
261716
272196
203852
270276
264207
184678
184260
184270
178096
159164
143368
014914
264522
285184
223945
136631
023902
285554
266187
143401
222222
276110
118298
117362
118292
159208
266472
117360
163125
143374
283933
143369
228126
237781
143382
264553
203804
277452
274963
143384
253047
143420

0276103
0143457
0143452
0206931
0163131
0236713
0143387
0224690
0143437
0200175
0224800
0283324
0229357
0143379
0143418
0259357
0231073
0143412
0143409
0143363
0200759
0163141
0143443
0197622
0213190
0143458
0233717
0261168
0143434
0163154
0163156
0163155
0163157
0163159
0143398
0159352
0232671
0143373
0143393
0265753
0273481
0143390
0224645
0237976
0143416
0159377
0143442
0238711
0143375
0252840

0143367
0232536
0207606
0223861
0143376
0250734
0206635
0159409
0227045
0178796
0206980
0269621
0236940
0143436
0143450
0249602
0232937
0268288
0182134
0203288
0269489
0213171
0143365
0225556
0201134
0234614
0196407
0159445
0285651
0286581
0223810
0197747
0229021
0240667
0163191
0232678
0285818
0182898
0159450
0226716
0233792
0215853
0237975
0236427
0197915
0143631
0143520
0183586
0143536
0186207

0169509
0185966
0163202
0244057
0187238
0185962
0226947
0285753
0187223
0187180
0159455
0187173
0187170
0198854
0229713
0233819
0203786
0240386
0186226
0172155
0197084
0196734
0186844
0235942
0285946
0163206
0163207
0224308
0283227
0184148
0169474
0163209
0252920
0234262
0169469
0163216
0241794
0196805
0203785
0244094
0229035
0159516
0229699
0203784
0203783
0230779
0203782
0159527
0163218
0207321

0163220
0163221
0237008
0143546
0224784
0184330
0203781
0197364
0143556
0263841
0238279
0197956
0196420
0196154
0188015
0196754
0285867
0188643
0189334
0189171
0271853
0160678
0272030
0160679
0143553
0143621
0169418
0274940
0242565
0235015
0199565
0143624
0243613
0233222
0143554
0207039
0143614
0231827
0223599
0236327
0284738
0198837
0160741
0143570
0285779
0276584
0282386
0273026
0143578
0143543

0285641
0272654
0143545
0177954
0143552
0231416
0200220
0231837
0263987
0143549
0264384
0215938
0163263
0143612
0143569
0201129
0143575
0252682
0212292
0222457
0278694
0143595
0143515
0238365
0237920
0226855
0228013
0160712
0228264
0169291
0273110
0163239
0160714
0229780
0286391
0160716
0233875
0160710
0287064
0143603
0163344
0270361
0163346
0268154
0163348
0271380
0160691
0173207
0264349
0160688

0163352
0160685
0163354
0163357
0232093
0143537
0243364
0251246
0143590
0202027
0169242
0169241
0179085
0223452
0163463
0273088
0163462
0185499
0231064
0284586
0169231
0173171
0236263
0160766
0236675
0177628
0160767
0116521
0176444
0143630
0143627
0160752
0225855
0160753
0116539
0283701
0207134
0227773
0207144
0237872
0235919
0271267
0225082
0223503
0232519
0125459
0246203
0287839
0163374
0132676

0227673
0203761
0116580
0234937
0132718
0283442
0143622
0132680
0207475
0281394
0280466
0173080
0116584
0277817
0273002
0224276
0285677
0163479
0160803
0116586
0132698
0254726
0160789
0196189
0252236
0160785
0260238
0160783
0242252
0160781
0198952
0163472
0198715
0189030
0163468
0163467
0132677
0237390
0125462
0207933
0116604
0206651
0260460
0183856
0187862
0163382
0160818
0272971
0132702
0272068

0132692
0272405
0229953
0132688
0237588
0285570
0143320
0223356
0143319
0143303
0143314
0143321
0143294
0287624
0027869
0198400
0027644
0187800
0160838
0132694
0211581
0266160
0237842
0224520
0253831
0117036
0235700
0229961
0284592
0228239
0271736
0286073
0286151
0286005
0143297
0163518
0236731
0233712
0160856
0227217
0231700
0232366
0132704
0163534
0073754
0227425
0236957
0183853
0226520
0288460

0231666
0227295
0158473
0229914
0176320
0158477
0230942
0158481
0158485
0158488
0186306
0180708
0203758
0236257
0173285
0198965
0236656
0198703
0227445
0231434
0197532
0186440
0279111
0198967
0163554
0231237
0196171
0203757
0180437
0226949
0197403
0180433
0188340
0229724
0180409
0163563
0229849
0163564
0163565
0163568
0233473
0162706
0222552
0225670
0213088
0215846
0248642
0179639
0196266
0228560

0248442
0251686
0249197
0249730
0196184
0224943
0184155
0231100
0271890
0132703
0158731
0276632
0223603
0132693
0235226
0158716
0181036
0158714
0256029
0188004
0212161
0243284
0272668
0213085
0266458
0287040
0279430
0158710
0085552
0162723
0224259
0273933
0177807
0237409
0143315
0225279
0162728
0162729
0018625
0132681
0143318
0227741
0162734
0132716
0258465
0228606
0224261
0162735
0122218
0235082

0202078
0162736
0171786
0201608
0162738
0162739
0234425
0066294
0117090
0235101
0228863
0117091
0026751
0275801
0274562
0122224
0122223
0226003
0179914
0213080
0232188
0198358
0233691
0158764
0158769
0270149
0232879
0215845
0158773
0186517
0143217
0228917
0162755
0143256
0158793
0158796
0238934
0143222
0224985
0143258
0143224
0158850
0158859
0158864
0158869
0158874
0158882
0284035
0143257
0248485

0158887
0143252
0188931
0236484
0283696
0288093
0206921
0283360
0215840
0277882
0283317
0143226
0273112
0173110
0224203
0203747
0224515
0244682
0225217
0231586
0162747
0072694
0234211
0213075
0199595
0132185
0162746
0241347
0081721
0226889
0118217
0229808
0162745
0227818
0198929
0285636
0266144
0227094
0252262
0207729
0254706
0171722
0239887
0198574
0227243
0152332
0227941
0259788
0117143
0272574

0162733
0243173
0132196
0185860
0230739
0227667
0117152
0143248
0232892
0225755
0232995
0237756
0143228
0225122
0200327
0212527
0228289
0212538
0225719
0199849
0213070
0185630
0207082
0269887
0276359
0233693
0271917
0201987
0225487
0201270
0162761
0224702
0238022
0143171
0237463
0230748
0162763
0237783
0236206
0143198
0143149
0206990
0230175
0225272
0215838
0273365
0143183
0224358
0230659
0143179

0236364
0283936
0271527
0226292
0207341
0188859
0203307
0230898
0215952
0229588
0215835
0225325
0230231
0234142
0238087
0215834
0234984
0232341
0232148
0232223
0143157
0152382
0225171
0143195
0143194
0200036
0227907
0143167
0198842
0213068
0231605
0272205
0143190
0272033
0198821
0287218
0233411
0273160
0227776
0143162
0231029
0241666
0198771
0197965
0143199
0250762
0232194
0143158
0143164
0221545

0224837
0143147
0143155
0237049
0237131
0213064
0214262
0206880
0143178
0227722
0207974
0228697
0283255
0215833
0143185
0227777
0143184
0229757
0143196
0285622
0235736
0225826
0229612
0287831
0203601
0213063
0287282
0252987
0237707
0143153
0143156
0235575
0117475
0117477
0117479
0234604
0213062
0198734
0174175
0000460
0188404
0007908
0230704
0171806
0000457
0239494
0288139
0075945
0243051
0235378

232959
224896
203740
283340
263390
233985
238051
224286
226967
231407
120370
225545
235303
116132
271811
231424
117501
226552
007933
221390
283598
117507
094963
225243
010932
206692
230547
076258
285665
268062
239428
234333
117523
283683
034971
233328
230364
117533
232261
271459
236741
253060
010165
213060
224600
197959
233540
230630
283844
283152

0208024
0287336
0206684
0135845
0180999
0094975
0251943
0117560
0229785
0279061
0224228
0227177
0224000
0120337
0230849
0226375
0117586
0232751
0231615
0203739
0117592
0162753
0238272
0285777
0183831
0232113
0076321
0225591
0200674
0120334
0252357
0117593
0234741
0270084
0185278
0117601
0252231
0135870
0200755
0236535
0224977
0225713
0227373
0152061
0237317
0203737
0223828
0231378
0237249
0223525

235869
229531
252552
287697
116161
120333
206659
181101
120332
235750
230687
233946
260990
116147
235628
224718
227740
230777
143207
252906
228686
236021
273629
232463
253025
227815
231020
225904
116183
237514
286754
152092
202609
198797
224968
227579
236720
120341
254154
242193
276563
224687
075391
270575
213058
188585
264809
285910
240021
213057

0201347
0273384
0266417
0273062
0116191
0228196
0201791
0116194
0116199
0232750
0186283
0143322
0234041
0231905
0212338
0201619
0231564
0225711
0223794
0228191
0057252
0162779
0224620
0236539
0261250
0116218
0251875
0227141
0261060
0162782
0243062
0143340
0229407
0231966
0169905
0272906
0143337
0264916
0261831
0260360
0135837
0235145
0223450
0261817
0116260
0121454
0230124
0235060
0265435
0206905

0236719
0143324
0283575
0225857
0135835
0243155
0232586
0251520
0270711
0135823
0153029
0162783
0237292
0225359
0179452
0198216
0273573
0252977
0287452
0272880
0222397
0179930
0224810
0225982
0228918
0232309
0232096
0206764
0228664
0231684
0135821
0203730
0203729
0121446
0270443
0233537
0135828
0276839
0143333
0261504
0135824
0227181
0271269
0135838
0287808
0135829
0157060
0224040
0285609
0199837

226570
176855
207472
135862
224468
058085
157064
287929
233562
286372
232860
116698
116701
286966
162704
143344
173627
270994
227554
198756
198860
286655
199840
207080
231791
244568
271387
116667
230470
232036
285847
286378
116406
135842
252222
260976
201312
121481
232792
121486
116668
231084
213051
201421
206640
233583
252612
238061
279838
116679

0273004
0261024
0252407
0270920
0279401
0228309
0207108
0201596
0261729
0143341
0224691
0288078
0116690
0047410
0202025
0157181
0262180
0229739
0116703
0273198
0228238
0233196
0073756
0273129
0116711
0230426
0271558
0225462
0274702
0236401
0223847
0285894
0150732
0222240
0261182
0224278
0225006
0235083
0231022
0237283
0238054
0226486
0271036
0226196
0252553
0232212
0238270
0287472
0230987
0162670

0225811
0237457
0241505
0231175
0236025
0238108
0285638
0228020
0261642
0271187
0233882
0228215
0223344
0150681
0285280
0226723
0236792
0253148
0236069
0090104
0127074
0231767
0283795
0228687
0226814
0116741
0223075
0234473
0116750
0238754
0116747
0023572
0134371
0221680
0162630
0276735
0232077
0286285
0227240
0226640
0237011
0237213
0233057
0231714
0251813
0228167
0285718
0287364
0227421
0162687

0265986
0000971
0116785
0244414
0285986
0134365
0080910
0273844
0134389
0143278
0066279
0235748
0177888
0237861
0134376
0221879
0230260
0213047
0207139
0213046
0235582
0233040
0224901
0203724
0143355
0151414
0232824
0287989
0151418
0223881
0081237
0261573
0236278
0229989
0207975
0207759
0227747
0225172
0233410
0235492
0286541
0231718
0228190
0236468
0231984
0228530
0200139
0229747
0202491
0202329

0116833
0252860
0229220
0233827
0203721
0273093
0213045
0230623
0162702
0282849
0118193
0118197
0260088
0118200
0231333
0170128
0163362
0233217
0201032
0116852
0229191
0234132
0081248
0232237
0116857
0163395
0229821
0081277
0118194
0286600
0159166
0282221
0159173
0227048
0174307
0224818
0159176
0224536
0223396
0134369
0236390
0231871
0264802
0235121
0200942
0221028
0198700
0277681
0198892
0223774

)163431
)134375
)206637
)238003
)176393
)234678
)275207
)163435
)249007
)232296
)235811
)212512
)227109
)170075
)143862
)143851
)243323
)133067
)077152
)077157
)229120
)272262
)232626
)236439
)253042
)143858
)226862
)225620
)117139
)235449
)226148
)260021
)228288
)184774
)214796
)234996
)183155
)117153
)230280
)199471
)159346
)159348
)234761
)231547
)163444
)143847
)122180
)163485
)234775
)224671

0133055
0133048
0133063
0236523
0231507
0233791
0159388
0202300
0122176
0229652
0271588
0188783
0188770
0058668
0236035
0221643
0122188
0058673
0257315
0227417
0223505
0182004
0236430
0232917
0237379
0286383
0176754
0286572
0143842
0143845
0261065
0230550
0143839
0225522
0170498
0174567
0143850
0231691
0219133
0280924
0226330
0237848
0158615
0133056
0198625
0236779
0200408
0240710
0170382
0240219

0228153
0252650
0251861
0287197
0163531
0229657
0184144
0174529
0117222
0271580
0236108
0133059
0213041
0133069
0225063
0163545
0162873
0235363
0226235
0186007
0281406
0199059
0224717
0117266
0253097
0174514
0206762
0158711
0236942
0158715
0275213
0069275
0201944
0117280
0285521
0133065
0286619
0162877
0174502
0227687
0276600
0196188
0285417
0263961
0252692
0198049
0226780
0287432
0225483
0236889

196550
266028
229509
263528
278465
162888
261000
266094
234981
279946
143486
237605
143479
162889
224114
199349
136634
142224
162891
162892
162894
271680
226945
162896
162897
182795
123836
180667
123843
123838
275392
226565
224462
243636
232621
237074
196352
283044
117322
203710
236911
197721
244703
226289
274059
285239
117335
234219
203709
284214

0284203
0174059
0285719
0284237
0076356
0287220
0226843
0286198
0261453
0259815
0227887
0232812
0236950
0278684
0287046
0232537
0228081
0224540
0225712
0227940
0231648
0230937
0284485
0224260
0287902
0008118
0196878
0283876
0227591
0123689
0117594
0236136
0009790
0162757
0117595
0117597
0143469
0287343
0203706
0082497
0228110
0233455
0054392
0287354
0200972
0287157
0270936
0143473
0279333
0284299

0224668
0283952
0284376
0234233
0234004
0226986
0117625
0082512
0287033
0153363
0236809
0238137
0198570
0223649
0170385
0261314
0228792
0227764
0117650
0224763
0231057
0226868
0123684
0241395
0212205
0229258
0233626
0143493
0212187
0143476
0220749
0264358
0252879
0229983
0226251
0066027
0230063
0234915
0225338
0201544
0229832
0065600
0287445
0117691
0224535
0286213
0228067
0260805
0288007
0162772

0235862
0162771
0226036
0237980
0123685
0117697
0203705
0236905
0185523
0198468
0235182
0236317
0162769
0282718
0143494
0174606
0136643
0228646
0225233
0228255
0230461
0281664
0117707
0274895
0272167
0283518
0143499
0228470
0152104
0213036
0215819
0117724
0250536
0224584
0185221
0223869
0287008
0082482
0232809
0282265
0136636
0042781
0202498
0229242
0233620
0236292
0226013
0196482
0092978
0162814

0234070
0228247
0231814
0230714
0232100
0286775
0201493
0067533
0229016
0223375
0232480
0092969
0281453
0228208
0212610
0226693
0225561
0228063
0143353
0225920
0223842
0277007
0287676
0228536
0238232
0215817
0196660
0252086
0221673
0136628
0162813
0067704
0207590
0207624
0232134
0118873
0281696
0222370
0270585
0270708
0218283
0213033
0230318
0232883
0226927
0260505
0235163
0234863
0226226
0231529

0116141
0243872
0233012
0162817
0117791
0186205
0286231
0221571
0276642
0272823
0238078
0257551
0136630
0226211
0234754
0232436
0286398
0227585
0286421
0143507
0227925
0200033
0238042
0228437
0232679
0236230
0227613
0226643
0234419
0276997
0222399
0267305
0143512
0143498
0225265
0154305
0229399
0272750
0186063
0162819
0197520
0287684
0228106
0278610
0154309
0234523
0287338
0187554
0236846
0226601

0143502
0251789
0178395
0203697
0227002
0212398
0162909
0143514
0227621
0188460
0233771
0272645
0212157
0185495
0278467
0229930
0252484
0143756
0236773
0143753
0201898
0232628
0237101
0143748
0221406
0229742
0206887
0143771
0162923
0266618
0143786
0229991
0233384
0234247
0286174
0229400
0225334
0286719
0185842
0143815
0234476
0282418
0154380
0227496
0226349
0223306
0143742
0234112
0143819
0242861

0196187
0243709
0255835
0143811
0284519
0248322
0143768
0213032
0143751
0272562
0163041
0225518
0270598
0182827
0234478
0223570
0185155
0183814
0236994
0275406
0143799
0270287
0229965
0223282
0177770
0287627
0203685
0143772
0228382
0228548
0238183
0287259
0143801
0163050
0228525
0143776
0233706
0287532
0228625
0228729
0236636
0287525
0234277
0232864
0242757
0224834
0237193
0278180
0237469
0235827

0227711
0287205
0202264
0181450
0225934
0215812
0143740
0081692
0230005
0286389
0185888
0143816
0264483
0233003
0274886
0154342
0276255
0143761
0284163
0143793
0162910
0224566
0287895
0280157
0227625
0143774
0198835
0203684
0181873
0162913
0154358
0269934
0270110
0286773
0287315
0270104
0154370
0270094
0284311
0231563
0162931
0168148
0230841
0181218
0196890
0266174
0181201
0279306
0168159
0215811

0202056
0251823
0199352
0201588
0199337
0200381
0199396
0200624
0202521
0200343
0201321
0199910
0199334
0199270
0202526
0201355
0201925
0202257
0200370
0212237
0183929
0206878
0234975
0116574
0271475
0271399
0282111
0229840
0236975
0233920
0177800
0177788
0168118
0213029
0237481
0154429
0252506
0252051
0230331
0143632
0069248
0226920
0223635
0135776
0199672
0201492
0229367
0135801
0135763
0213028

0225984

0227043

0228058

0143641

0227006

0224407

0177614

0282564

0232964

0235817

0135775

0286859

0135744

0244137

0135773

0202063

0225656

0119280

0223393

0288037

0287395

0240502

0143643

0173409

0182118

0283799

0227934

0229595

0119283

0235710

0143633

0116906

0222407

0116903

0010072

0287450

0135766

0287856

0223982

0233461

0116918

0270106

0229228

0162946

0222986

0286071

0274121

0226758

0235152

0236372

0224436
0242794
0116991
0238382
0225919
0236497
0212916
0206835
0286774
0232175
0135778
0135749
0231940
0251508
0143674
0252501
0287423
0135750
0265744
0286109
0225486
0286666
0287921
0183780
0236101
0233332
0231272
0236244
0232166
0264377
0235605
0236358
0273367
0231663
0168275
0059588
0231768
0286210
0287633
0230404
0212144
0168264
0228830
0224939
0272362
0228044
0241475
0230628
0282097
0227630

0279261
0286263
0201638
0239690
0237520
0238005
0258082
0280587
0233018
0273416
0173726
0207181
0188739
0054267
0263439
0236863
0152904
0285053
0227962
0230026
0285177
0284770
0229795
0162885
0238236
0235638
0231199
0226663
0231582
0232714
0235761
0168243
0270710
0227236
0143669
0235674
0229463
0222831
0206803
0237845
0229291
0116962
0252822
0235371
0286142
0077585
0086619
0222650
0237922
0186197

0244457
0116977
0223776
0273058
0230325
0119285
0077522
0116984
0226498
0237991
0244020
0288099
0198626
0252396
0266262
0252290
0237250
0116996
0243781
0283174
0283166
0283377
0270188
0232154
0225723
0227860
0230019
0234464
0232989
0252371
0215808
0231877
0215149
0224783
0287177
0227854
0287589
0259776
0133019
0233355
0234601
0231979
0215805
0287102
0217327
0155816
0228818
0202041
0223694
0233735

0252317
0233519
0180875
0224359
0226014
0231440
0228844
0251750
0234872
0199241
0230015
0226919
0223963
0182901
0215802
0233615
0238085
0265831
0224348
0286496
0287516
0287738
0091483
0117009
0054277
0203668
0287513
0162843
0229022
0174371
0235990
0196289
0213026
0197769
0224625
0277099
0180287
0272865
0252084
0224525
0226750
0235467
0213690
0270859
0215800
0270818
0277704
0230199
0234116
0231512

0278455
0253326
0214837
0212230
0143702
0227230
0232085
0054282
0224727
0265201
0117020
0236031
0226766
0232184
0228939
0226828
0179456
0279774
0229960
0286486
0244066
0237759
0287531
0173728
0240963
0225401
0035687
0179397
0227735
0121644
0232059
0284188
0282317
0287601
0203667
0153187
0273175
0201758
0252073
0272195
0203666
0252282
0251754
0223353
0201170
0162849
0232192
0232793
0238224
0231612

185420
235096
226876
225222
202184
230184
207326
227728
242042
230813
229112
162851
162852
225300
228955
260698
235021
143653
228879
223519
260855
227953
235386
252011
153207
135747
197472
236775
277462
215796
228982
188295
286747
221953
226164
196418
224014
215795
259865
227671
283881
252516
232347
214144
235818
197617
202079
162714
286216
287116

0162711
0177535
0203664
0169224
0196242
0284824
0236817
0177489
0177476
0283369
0198452
0235749
0230411
0197437
0169214
0239395
0241128
0153230
0221888
0228336
0196772
0232908
0230576
0197591
0162722
0238243
0177462
0177275
0226191
0224830
0279263
0187080
0237492
0224227
0199442
0232215
0197454
0200085
0203663
0200982
0198128
0197067
0196071
0177233
0162727
0198601
0228198
0171180
0177212
0177201

0177186
0177174
0196944
0198104
0175143
0227152
0196240
0196539
0229255
0203661
0271934
0188558
0224521
0227102
0182783
0183310
0184022
0201602
0271701
0279301
0177151
0187701
0286015
0189181
0229703
0276520
0242529
0259823
0229663
0175137
0264500
0171161
0171163
0227237
0185220
0200495
0233084
0184731
0227061
0035115
0143727
0189292
0228643
0235779
0277997
0233684
0236856
0226277
0223985
0284452

)225942
)233633
)151353
)233296
)280360
)233970
)227713
)272342
)231173
)223751
)237667
)228799
)234796
)235688
)172554
)235403
)236665
)233553
)231482
)115705
)228613
)283766
)203635
)130508
)186487
)284600
)232057
)225619
)234929
)287583
)237720
)227364
)228391
)234423
)236760
)226649
)032389
)224885
)171853
)225234
)271868
)182551
)235078
)242282
)286905
)171865
)234171
)287126
)171863
)118004

0237370
0188765
0151360
0232718
0224661
0214866
0237401
0229550
0231821
0231532
0252238
0207192
0228585
0232835
0224128
0230090
0176887
0242540
0236106
0237266
0232044
0276733
0234275
0227007
0226488
0231610
0236172
0286851
0283394
0205837
0225964
0134326
0134321
0271947
0228203
0151692
0223884
0223145
0272002
0213774
0229727
0221255
0287119
0230515
0235576
0232979
0229405
0226506
0235665
0286857

0236790
0229740
0236008
0285591
0231083
0225642
0235092
0115738
0134313
0143797
0230886
0223640
0242136
0261104
0244310
0240980
0151693
0213772
0119185
0119203
0134330
0151694
0239300
0271855
0134308
0207086
0240687
0244260
0200034
0240960
0243491
0115750
0269973
0287305
0134317
0188525
0260476
0260077
0172059
0271787
0205795
0233502
0212558
0171848
0206647
0265418
0264030
0285872
0206898
0115756

)115758
)206633
)257135
)115761
)234818
)243819
)277446
)236162
)143882
)239053
)272275
)143870
)238962
)271952
)232056
)270488
)162975
)226080
)145063
)150873
)162976
)134318
)207267
)230154
)226961
)230952
)231403
)224177
)285569
)169016
)196208
)201610
)264010
)251718
)285626
)202248
)239899
)169006
)230044
)276411
)134324
)228496
)203643
)230790
)265056
)224184
)265172
)238503
)264089
)207183

223360
286427
261117
071575
264370
225649
285876
229370
230448
227718
251859
237261
228505
162981
287291
235127
151779
229087
234022
079785
224194
231031
237326
202160
214657
238371
233718
223850
134323
243541
226764
236289
236989
271315
251704
226041
228876
231266
285354
232444
224400
237633
279663
197872
223536
224604
237670
222842
214843
236348

0214842
0163032
0163029
0178295
0151379
0170745
0212455
0287881
0287849
0260331
0240857
0250741
0185013
0207170
0224626
0287284
0236204
0266738
0143867
0234597
0237992
0235911
0213403
0228784
0183891
0271991
0118965
0227210
0132031
0068697
0223734
0234378
0213729
0266059
0200763
0115884
0235411
0238735
0055917
0232800
0212171
0143878
0228950
0200829
0227047
0279307
0118960
0269976
0231948
0233416

0143869
0270100
0118961
0236436
0261012
0084674
0280390
0287956
0218819
0271629
0233005
0231204
0270654
0235537
0228538
0229621
0264192
0228999
0223691
0234189
0206882
0234207
0233587
0222616
0232451
0223530
0233714
0223634
0235497
0119771
0224361
0279526
0283031
0119778
0224464
0238111
0173960
0234946
0243847
0205639
0163026
0222940
0119782
0115128
0219626
0115129
0176732
0276087
0232642
0186453

0223754
0198399
0242628
0251805
0232963
0233747
0286829
0084676
0202430
0206732
0184924
0138092
0138031
0271936
0115137
0237953
0202479
0224165
0286645
0207069
0084710
0276653
0229593
0115138
0230452
0119772
0221445
0235072
0138101
0224220
0201160
0143970
0187536
0217643
0218682
0217718
0084731
0237629
0268412
0084733
0199872
0235440
0231655
0214465
0227133
0237951
0157833
0084754
0138029
0286707

0173567
0138018
0157856
0115155
0173557
0157884
0225378
0225024
0171303
0213699
0115163
0145835
0157851
0230286
0084764
0205500
0119777
0084693
0231636
0272056
0229122
0272148
0228474
0138080
0138030
0138028
0143994
0138073
0186143
0163792
0236099
0138074
0138085
0084774
0115194
0163793
0138100
0163794
0115204
0115207
0234945
0234072
0115211
0115234
0163795
0115241
0213453
0115216
0157992
0138002

0235267
0200003
0115226
0084734
0221843
0243943
0259080
0176714
0198522
0206731
0119760
0163798
0205334
0243147
0171174
0158019
0270846
0265321
0270528
0277035
0223522
0225991
0229951
0075426
0270640
0270210
0227938
0163803
0242915
0230730
0222232
0226833
0213639
0163806
0273233
0171103
0163811
0264994
0265145
0222224
0265706
0189350
0200283
0179270
0229224
0115295
0171094
0286963
0242699
0233862

0230737
0197644
0225943
0119801
0207187
0231712
0213626
0287658
0235997
0280154
0172954
0278908
0162949
0230118
0285984
0158089
0234579
0214711
0201671
0013016
0213620
0158125
0277893
0228563
0270422
0223647
0237007
0242272
0273165
0162959
0162961
0271228
0021574
0272716
0152683
0229816
0201113
0091106
0119820
0272754
0115760
0230046
0276334
0276517
0207653
0274159
0279897
0279544
0018699
0265057

0230876
0236854
0206951
0049323
0223951
0285577
0212204
0252246
0283591
0152689
0237133
0119812
0286415
0238038
0187999
0228262
0239649
0235064
0237790
0285754
0272027
0232153
0226994
0264352
0237320
0196085
0265301
0207255
0229013
0234587
0233752
0260025
0150938
0273090
0171055
0270677
0205221
0115808
0279519
0252756
0008869
0152133
0055332
0224891
0138068
0233159
0218739
0115816
0272054
0003509

0115825
0232028
0207364
0285925
0115828
0253078
0287316
0236213
0163171
0236572
0225402
0237803
0115841
0235848
0138061
0232973
0280276
0229160
0199603
0273006
0227292
0227770
0213594
0119787
0231367
0287313
0235009
0213553
0271443
0143889
0235586
0143891
0232518
0115875
0152147
0227238
0232114
0163214
0223922
0225284
0188010
0207295
0214694
0265905
0269210
0115904
0229692
0206985
0225900
0202309

0205111
0011566
0252473
0273035
0231312
0152154
0138050
0227028
0235653
0183023
0287255
0285898
0287468
0233128
0237442
0287145
0236090
0236533
0207218
0214691
0223245
0234362
0205086
0226398
0162878
0224875
0143924
0115944
0221300
0171126
0232526
0236913
0057935
0273913
0215263
0226523
0200550
0162881
0162882
0226491
0232202
0224739
0231826
0233978
0207087
0230587
0286796
0287387
0152518
0279873

0234936
0115970
0252804
0264071
0207411
0152527
0223658
0229695
0252490
0138036
0138075
0143921
0138095
0200924
0264047
0219391
0252599
0224637
0273106
0138032
0285542
0226972
0138079
0138078
0143919
0259439
0278957
0274520
0277311
0276229
0277809
0276048
0274547
0276670
0236502
0138083
0225156
0231848
0170577
0231156
0205054
0231054
0286728
0286519
0068784
0239396
0171132
0228481
0233829
0232696

0231336
0233335
0278764
0116016
0228100
0187600
0284608
0241791
0284701
0250565
0253515
0119729
0250116
0151665
0119878
0279254
0222005
0171150
0239332
0226548
0228925
0180398
0068724
0272814
0233845
0239605
0273269
0225187
0143933
0234690
0226087
0236824
0119888
0222685
0207923
0277396
0095002
0184261
0235760
0230979
0116062
0224443
0224058
0283502
0138081
0235369
0236549
0233230
0230773
0226473

0275423
0201010
0170802
0251889
0272663
0162869
0202227
0243244
0068781
0242441
0279956
0239995
0138039
0232285
0282890
0214602
0170820
0212036
0282998
0206915
0282828
0234253
0212580
0179915
0285548
0230327
0283058
0216191
0238165
0286412
0231918
0287867
0224344
0223710
0230840
0226366
0287344
0236538
0202195
0225444
0225842
0236837
0204993
0233083
0264975
0228033
0232604
0232668
0251942
0223897

115239
207456
143942
068912
265452
119737
068878
170634
235937
233266
241114
272156
252934
178021
177994
115306
228108
237887
234943
238018
214595
231334
285519
252507
115310
200086
162994
203327
227799
143947
085760
162997
240401
206964
115355
233594
238619
238493
239189
163001
275052
212175
272606
138035
115380
202344
272180
226702
207548
207798

0211520
0229805
0271894
0233251
0055813
0199395
0231570
0271115
0285755
0270569
0212168
0271615
0028116
0287875
0115392
0273063
0233426
0225226
0233723
0231043
0285673
0222030
0271955
0233891
0231815
0270447
0200101
0252726
0286604
0285611
0228590
0287640
0200807
0266078
0119866
0233953
0242158
0271134
0213495
0252718
0271707
0115421
0228414
0241524
0237024
0162924
0201076
0267520
0237522
0232713

0162927
0222251
0162928
0162929
0274769
0212978
0237651
0271889
0173209
0115464
0273302
0271146
0234624
0206937
0270820
0082898
0285857
0271243
0226884
0206973
0213486
0227361
0229831
0170264
0230702
0277498
0115484
0236498
0286360
0173163
0278523
0229839
0242735
0271657
0170340
0266097
0239958
0270335
0228541
0241625
0186889
0233702
0229503
0226622
0232144
0230595
0115504
0252436
0286524
0226605

0237162
0231609
0115507
0270437
0286480
0232403
0242412
0143951
0283636
0233716
0279485
0014641
0228305
0229920
0234488
0228872
0229221
0169764
0251775
0143952
0228079
0197329
0225889
0230923
0270354
0238012
0238201
0264297
0283222
0236148
0223935
0237217
0119862
0223863
0260101
0119844
0283204
0252414
0233694
0179833
0226756
0227791
0275588
0199964
0287123
0234572
0237638
0244534
0227394
0115902

0252892
0232613
0011523
0138069
0272025
0273763
0172974
0138071
0198369
0232693
0204929
0234255
0237979
0223739
0243029
0251900
0235725
0281920
0214533
0280257
0227293
0225815
0264525
0232164
0226819
0143995
0230749
0232046
0237179
0224071
0230355
0230525
0232688
0223859
0235885
0236780
0236605
0143971
0224173
0235495
0286002
0237013
0235968
0197223
0284932
0243667
0236259
0273398
0115946
0221823

)273275
)273064
)119865
)203395
)115956
)229462
)204923
)169621
)214525
)169618
)163219
)228329
)237576
)163217
)283586
)183607
)169605
)169604
)286347
)266649
)251850
)199460
)271597
)198380
)252250
)169599
)115977
)244236
)207016
)232228
)229948
)196975
)231024
)087338
)275381
)270723
)124380
)059728
)244617
)179818
)263669
)169564
)233060
)225392
)231327
)115998
)116001
)221238
)231414
)116005

0143977
0035141
0233849
0235289
0229229
0163235
0224606
0075340
0286979
0235035
0183733
0228034
0152672
0231386
0116031
0237751
0277764
0116035
0116039
0258881
0239322
0226186
0241159
0144031
0144043
0228384
0236469
0205847
0234485
0234896
0124357
0272735
0124370
0288065
0124383
0233870
0124374
0075292
0281195
0200779
0135636
0233971
0003137
0144036
0228568
0222536
0212378
0116096
0135638
0278060

0144040
0135631
0272702
0214513
0135632
0135617
0135624
0163013
0135625
0252988
0287435
0285068
0214484
0116127
0230002
0284902
0144035
0163016
0204872
0144034
0273245
0280037
0144048
0187833
0287250
0124356
0286244
0271536
0163017
0114956
0237883
0201876
0235499
0187605
0279181
0257800
0217702
0163170
0225439
0114978
0279201
0279070
0065911
0264324
0188687
0229798
0252214
0234632
0225658
0204843

0237737
0159239
0234521
0284308
0286623
0233155
0005448
0114993
0115274
0274049
0239779
0115275
0286883
0204822
0135637
0115282
0179528
0272183
0257702
0115289
0115297
0144045
0115307
0115317
0115318
0115325
0159374
0233872
0108442
0135622
0287687
0224646
0286739
0272711
0159399
0204792
0236209
0230836
0115350
0115353
0263909
0270571
0236167
0115363
0237293
0238410
0231172
0230477
0221638
0115364

0226247
0005436
0270696
0271452
0270462
0286045
0270996
0287474
0233107
0287172
0287749
0228209
0271356
0200488
0251947
0287026
0287130
0286202
0176204
0200028
0287025
0234653
0286932
0227088
0230968
0237077
0233605
0229494
0214429
0270470
0234877
0286260
0202423
0200048
0143954
0172023
0115386
0204787
0172016
0224879
0066032
0286227
0229385
0277171
0230237
0252031
0266436
0276917
0224731
0223427

0162951
0237031
0230975
0234429
0231722
0235519
0200890
0233444
0242118
0231781
0229560
0252897
0201311
0237498
0226860
0226700
0225045
0224695
0227789
0286211
0168129
0223977
0232548
0224627
0199295
0228225
0228902
0182814
0228272
0163541
0115423
0259056
0235847
0186854
0276362
0213400
0034510
0213399
0287625
0176407
0260840
0235463
0225787
0152284
0231134
0200701
0230299
0246575
0152291
0239143

0270532
0042445
0115459
0271014
0202382
0278590
0273196
0042493
0286011
0152292
0207207
0244399
0199936
0229648
0286532
0168906
0115486
0118640
0276281
0168899
0168894
0168890
0168883
0168887
0168878
0115523
0203363
0199687
0168874
0278783
0229498
0252321
0115525
0232504
0272564
0287628
0068654
0132300
0208772
0132305
0200241
0265420
0273080
0132313
0068615
0202537
0251974
0115548
0115561
0249884

)200563
)228363
)239305
)153561
)153563
)172116
)233673
)213605
)187627
)231351
)226114
)183281
)285793
)231259
)266931
)287931
)230395
)204745
)264793
)233671
)281477
)271097
)287763
)233999
)224881
)280721
)222041
)284879
)273445
)214354
)265507
)225933
)234231
)125551
)185304
)229604
)225428
)233205
)275597
)288437
)222724
)287670
)172086
)115593
)265003
)163586
)144115
)232508
)283262
)212133

0252395
0224291
0214336
0172073
0225420
0172071
0234028
0287686
0153574
0230006
0240040
0283270
0265510
0211592
0211593
0211594
0211595
0211596
0211597
0211598
0211599
0197794
0225136
0253435
0243466
0239855
0243063
0240671
0241755
0253278
0241351
0243290
0253497
0253265
0244437
0240864
0240382
0254157
0253732
0239951
0211611
0253578
0253625
0241294
0253202
0254098
0277852
0244575
0244116
0253998

0243238
0253158
0253870
0242076
0253860
0253461
0253487
0239862
0253592
0242371
0273962
0283427
0283196
0276861
0283132
0283214
0283659
0251039
0251546
0254009
0250036
0253127
0232747
0253501
0239975
0253191
0253999
0239571
0243264
0242534
0224607
0211623
0254097
0241566
0254345
0253365
0225523
0211625
0253765
0254220
0211626
0242766
0241244
0224041
0254292
0276566
0278857
0211632
0253906
0211633

0242580
0239819
0228325
0275490
0270999
0265897
0281904
0230964
0261600
0278131
0143429
0222835
0184303
0230572
0271627
0223703
0232531
0286698
0236969
0235235
0214330
0237085
0227497
0214329
0226481
0234898
0235147
0223816
0286156
0261522
0259916
0237383
0227265
0213355
0234837
0223564
0232502
0235571
0229089
0212140
0278631
0259848
0163060
0237055
0232128
0233158
0231331
0232594
0233105
0224585

)233447
)235186
)277591
)233850
)172005
)231062
)287362
)144029
)144026
)275111
)275768
)233757
)155066
)115041
)235129
)115042
)248821
)233275
)227120
)204705
)277747
)234903
)144010
)238162
)223917
)237308
)224719
)226185
)272913
)229689
)236431
)273305
)233037
)168992
)232717
)274028
)230083
)144015
)236026
)271003
)270193
)204699
)275497
)235959
)236750
)235584
)237510
)232931
)174501
)275961

0231584
0186281
0274286
0188886
0158050
0228873
0084090
0204685
0135956
0144021
0144028
0230747
0198885
0121152
0163121
0235480
0196843
0114982
0249715
0114988
0158158
0264157
0273265
0168763
0163126
0213337
0168758
0168754
0286654
0274206
0252845
0275094
0273634
0248134
0144188
0230343
0236847
0270748
0271402
0204670
0188383
0144199
0279791
0275655
0135976
0277701
0237837
0271351
0276050
0271015

0270252
0271569
0270450
0278537
0270187
0230606
0278766
0196912
0235833
0286036
0228970
0135940
0115073
0228486
0201806
0115085
0075568
0235734
0238719
0168658
0231635
0222000
0144191
0227987
0040933
0183513
0115446
0071073
0236564
0226791
0201070
0196872
0238830
0135951
0231822
0273045
0241962
0144182
0273155
0158411
0185414
0185674
0144214
0115514
0158417
0135945
0273306
0144218
0230393
0232084

0170500
0269383
0270699
0115526
0238029
0204640
0204637
0115539
0227259
0234988
0287832
0226034
0234169
0228488
0286737
0170485
0232034
0230140
0223947
0071082
0204634
0271532
0272902
0158435
0163162
0212283
0264857
0175874
0226058
0196460
0223826
0237549
0232556
0286047
0286101
0071054
0281162
0115590
0287771
0115594
0226925
0115598
0205716
0115602
0115604
0236785
0115607
0264764
0236525
0234389

180251
115616
135953
170417
277527
224095
227680
229209
227623
286290
236596
221849
231723
232597
236141
256637
273515
228968
233955
227157
225765
287793
235597
234781
231626
236109
233639
230690
225669
228528
279904
207249
234177
213301
269707
198914
229743
287135
224509
135972
231851
226508
135973
272861
135966
235319
272994
135974
115641
238273

0287036
0287308
0071051
0235522
0234162
0119147
0115652
0233339
0273595
0213295
0224061
0230353
0233939
0227513
0229682
0235486
0240935
0153165
0237666
0254126
0232001
0231505
0214199
0144057
0238250
0227294
0286218
0229360
0226905
0229457
0225588
0237880
0224568
0286769
0232991
0230651
0196862
0233494
0227270
0237745
0235325
0115665
0231221
0229890
0225328
0236877
0196228
0227597
0198203
0261209

0237223
0271070
0198075
0271284
0135968
0214184
0169756
0228763
0233648
0235644
0153201
0163006
0227470
0135960
0277628
0259863
0172985
0264934
0228551
0265965
0186522
0237916
0230696
0198142
0230603
0231099
0233307
0233707
0282033
0015568
0256977
0284337
0186148
0283684
0240069
0225108
0204588
0235721
0229730
0283283
0265682
0144063
0264979
0144061
0233370
0175701
0261760
0282304
0282785
0263881

283167
226991
230650
175772
226065
230596
273471
184115
257207
256671
227574
231583
183054
231536
216867
169679
235881
153093
236330
172965
204581
153094
230499
227992
271590
229118
270190
226928
266139
279191
223973
285016
280878
240350
224959
271635
225744
153107
237583
227842
228615
266063
230958
236307
153208
240183
225496
232277
286904
153214

0287141
0144152
0278962
0144161
0286545
0188177
0235258
0236555
0169629
0283698
0114999
0125630
0207383
0125611
0243389
0228251
0231747
0237753
0144136
0144130
0169607
0115008
0287937
0125538
0280228
0180152
0125571
0227368
0136688
0236124
0232090
0136694
0136696
0136695
0136697
0201805
0136689
0125637
0234174
0189223
0125618
0234997
0231292
0272563
0136682
0184492
0234148
0231943
0225398
0226516

0287165
0221055
0146556
0236397
0279267
0240356
0144134
0144158
0175509
0233479
0227359
0202427
0115084
0265429
0244063
0228857
0270019
0115091
0279957
0234199
0243179
0212182
0286513
0235242
0231926
0222923
0175497
0231538
0235717
0286776
0235026
0230391
0287451
0271667
0224869
0226341
0235871
0230158
0230950
0228898
0239185
0227291
0286370
0238207
0088205
0236255
0125631
0125633
0224967
0235066

0272895
0279227
0243510
0125629
0226856
0258910
0163064
0019169
0259094
0144119
0264833
0115107
0229867
0186132
0155368
0171227
0080293
0231013
0163075
0144120
0229326
0088179
0233741
0115109
0224789
0202046
0236878
0227788
0233286
0243104
0234065
0223549
0226479
0144118
0235840
0163083
0280409
0237614
0232140
0201006
0229774
0074047
0271709
0201584
0234455
0236145
0115112
0232723
0074054
0265451

0264229
0222467
0236859
0230993
0279721
0213239
0155438
0211460
0270798
0224655
0286481
0287871
0237856
0232740
0287854
0226708
0235640
0227738
0279874
0223118
0228400
0155052
0238046
0286384
0270815
0252674
0206726
0286206
0235491
0224410
0226813
0286515
0235774
0236655
0213236
0136732
0235128
0206963
0237524
0238788
0260634
0286971
0260163
0136717
0286400
0237630
0186684
0226789
0163161
0286145

0169967
0200250
0236682
0287742
0115718
0264075
0163166
0231731
0202429
0169994
0272789
0072163
0144230
0136709
0173349
0201470
0200390
0274010
0144231
0202532
0244563
0273950
0144233
0272667
0286873
0204399
0136715
0234044
0200432
0136731
0229173
0232101
0229758
0185631
0238180
0136720
0252742
0238379
0227653
0237532
0238277
0204460
0233428
0229536
0286957
0214100
0237574
0234559
0225449
0222014

0284619
0286026
0250207
0217289
0231147
0180178
0286058
0231240
0248313
0196604
0228471
0224087
0252223
0223760
0152076
0136699
0152082
0152086
0227632
0230756
0213225
0234398
0236595
0237722
0231954
0236211
0230496
0232990
0249997
0283940
0284171
0230548
0234934
0248136
0233550
0226915
0271435
0224352
0136710
0136718
0072135
0178162
0230646
0239402
0196834
0251973
0225819
0274711
0152093
0225341

184761
183292
232408
136698
277668
222038
251757
214081
282944
240253
223349
173302
229797
178171
232503
233221
136002
281159
281516
284479
152102
115762
230450
225448
236956
285819
248796
188219
212199
231431
236516
284604
227817
284706
224425
233087
283303
233285
227275
235080
231777
284003
283865
227661
236590
234480
237910
234939
234576
226831

0276460
0223631
0284659
0214077
0217950
0232742
0273073
0173272
0075886
0284149
0152117
0163040
0224679
0286208
0204434
0251956
0224559
0270930
0287151
0288031
0197927
0213222
0287071
0186825
0274353
0229203
0232760
0244337
0235615
0236485
0287414
0226886
0271179
0225594
0227745
0163046
0239108
0202174
0221288
0233786
0230803
0018607
0272769
0286833
0230992
0252705
0183840
0232349
0200718
0150551

0230065
0176771
0286068
0228721
0233729
0279512
0226953
0252688
0222068
0232474
0223430
0200708
0286697
0152127
0263813
0222921
0234060
0152128
0287463
0224043
0234682
0228098
0153086
0263783
0082258
0176601
0115839
0208308
0121988
0235651
0048991
0201308
0207654
0144224
0115850
0200664
0226806
0076003
0214070
0115866
0231890
0237262
0121966
0227347
0279024
0230037
0251976
0229568
0144229
0212425

0222394
0150540
0224231
0233045
0234645
0237939
0232915
0228043
0230569
0144228
0241772
0260059
0237772
0144227
0222826
0229104
0232992
0287147
0286778
0226939
0229131
0206901
0235232
0228374
0223554
0240977
0224028
0227149
0231567
0223921
0168702
0277306
0283506
0236440
0238065
0201892
0252015
0233329
0244125
0228134
0229781
0115919
0270390
0075884
0224288
0226839
0224852
0232772
0236300
0227557

0225476
0284167
0231766
0257284
0228655
0283994
0231758
0257277
0257226
0236356
0200287
0232377
0121964
0169554
0283118
0279166
0238057
0275372
0273537
0277444
0287115
0232606
0226674
0234940
0276706
0235352
0268580
0235435
0214064
0225107
0286225
0226218
0234119
0233842
0279876
0251905
0230559
0227134
0198526
0271583
0231040
0227902
0287847
0207225
0207161
0202074
0253083
0229143
0121989
0212389

0223911
0115947
0204406
0281469
0281772
0212181
0224007
0232238
0135999
0222126
0223962
0231079
0228848
0226340
0168280
0286167
0150556
0228236
0286335
0224337
0187123
0223604
0168288
0231969
0207270
0162947
0286081
0230645
0231420
0115963
0236049
0237220
0232359
0224048
0228064
0222031
0236885
0213201
0184898
0123609
0123610
0283569
0264684
0242113
0080345
0183091
0162980
0288066
0283228
0182389

)225214
)115145
)234272
)286234
)199884
)157827
)213197
)287085
)270776
)286207
)196504
)177917
)234932
)226213
)214025
)224612
)226338
)227708
)214024
)242575
)227400
)200377
)177519
)236917
)144278
)230251
)224675
)238004
)223290
)287900
)162989
)228837
)230991
)223197
)232982
)229701
)227682
)233709
)233039
)233248
)229935
)206718
)286679
)226383
)235548
)282440
)153234
)115159
)283436
)238069

0287048
0286889
0286530
0213194
0136542
0222404
0136541
0230562
0115165
0123612
0234584
0115170
0007001
0270557
0251980
0271320
0228937
0212410
0231158
0237327
0227480
0153237
0234890
0243723
0144283
0227331
0204380
0228586
0231675
0163331
0286621
0243792
0251491
0188668
0222300
0115183
0213189
0202029
0226878
0275141
0196151
0123636
0223642
0270893
0225369
0207117
0224152
0226266
0230783
0136536

0241399
0248672
0287091
0054219
0227055
0153246
0115221
0285155
0285373
0153250
0283734
0279844
0244372
0232337
0224467
0136560
0235724
0227403
0225813
0115233
0232320
0271924
0136535
0224076
0251621
0144290
0280105
0226121
0238570
0226946
0288091
0235716
0234732
0197635
0230918
0231445
0224674
0233397
0115263
0236841
0078098
0115267
0115271
0184611
0212312
0237750
0230282
0200902
0182263
0237844

0226847
0229641
0115290
0277998
0082438
0224331
0206869
0236283
0199508
0223318
0169507
0153253
0136531
0233204
0288089
0178662
0115339
0235192
0229195
0232411
0123607
0224490
0236107
0144285
0238813
0222376
0169432
0136546
0283657
0163092
0225182
0254552
0228222
0238357
0214211
0235335
0172318
0198648
0238259
0241074
0172292
0265694
0199348
0227617
0163072
0152253
0152254
0073734
0073737
0235321

)235995
)081479
)251569
)163093
)239474
)138399
)229827
)138398
)154479
)226072
)144362
)213160
)206961
)231550
)214369
)138385
)138382
)144357
)227176
)252807
)071909
)231898
)214821
)213981
)275414
)234350
)222033
)204335
)128692
)239467
)222509
)204334
)128683
)235934
)286611
)286115
)115806
)198586
)123600
)224553
)115827
)228507
)234061
)233762
)071967
)233934
)077380
)115840
)252779
)128708

)172878
)144355
)115844
)236651
)232555
)288048
)286557
)278924
)091409
)226963
)232788
)225205
)152256
)199866
)230104
)228016
)239041
)091428
)232292
)091436
)238133
)231735
)236391
)144354
)213148
)236832
)226131
)237016
)237617
)270460
)234415
)172845
)271151
)273258
)260868
)235414
)138430
)270799
)243770
)201088
)227180
)237804
)231453
)280414
)217236
)138433
)144306
)163328
)226853
)179997

0279884
0236449
0206965
0115935
0237798
0235655
0138435
0128656
0212167
0199827
0227187
0115966
0277948
0215973
0229750
0154518
0200121
0233131
0229066
0223976
0235047
0229779
0144320
0174279
0128714
0170178
0128713
0128710
0237380
0128709
0175879
0128652
0207744
0170166
0224189
0272729
0226363
0128645
0283880
0279160
0218175
0230384
0128654
0237260
0224577
0229434
0236231
0283203
0272551
0163364

0252027
0230552
0237053
0223777
0279205
0227098
0206866
0236501
0229337
0251746
0226273
0201102
0213069
0170144
0284252
0116044
0225808
0265396
0222043
0213963
0271825
0271996
0278418
0280374
0237178
0238295
0018510
0196659
0237655
0197557
0128655
0284741
0229941
0236664
0213962
0229744
0206788
0249909
0227241
0155636
0223096
0079156
0223960
0180228
0204311
0238082
0079150
0116095
0237298
0155657

0270956
0270277
0279598
0271011
0270574
0271141
0271401
0267784
0287149
0163492
0238542
0231541
0187231
0237477
0237964
0144331
0213126
0221240
0280636
0202216
0163510
0270289
0225258
0281131
0279559
0226564
0236667
0170035
0236153
0238171
0234663
0266705
0226681
0115232
0188452
0162992
0234595
0225570
0223873
0221023
0260742
0138434
0150722
0213958
0222418
0115252
0287536
0242121
0077232
0232430

0212404
0162998
0207178
0061676
0202141
0221498
0231203
0162999
0224643
0163002
0272800
0287621
0213120
0177855
0286879
0234172
0286807
0286152
0286980
0266808
0283839
0170396
0237824
0286797
0225406
0231646
0212581
0226747
0188738
0287129
0224019
0237877
0226410
0232568
0259915
0227692
0065548
0224323
0138448
0227227
0144369
0163012
0226553
0224063
0222845
0199626
0064989
0231739
0003436
0231689

0235616
0213115
0200650
0144366
0207951
0223523
0174325
0168542
0221502
0283238
0284180
0204262
0228073
0264725
0213601
0115368
0233996
0287418
0138449
0276828
0138381
0286053
0286165
0151687
0128694
0253559
0273240
0128699
0064933
0187699
0223824
0204253
0213110
0138379
0198130
0151689
0151690
0272979
0189362
0233654
0251935
0284052
0138386
0228509
0235852
0115419
0115415
0229023
0230686
0231858

0138378
0288064
0207402
0280083
0230611
0227542
0128641
0252130
0173559
0274845
0225884
0168497
0233766
0213946
0144339
0229395
0232227
0227418
0226242
0278406
0286358
0232658
0235218
0224219
0286868
0260142
0224670
0215043
0236695
0225539
0225394
0286539
0276123
0252517
0230173
0235056
0224099
0271893
0202206
0223466
0234919
0196950
0213107
0201813
0118997
0203435
0227878
0081320
0272211
0138411

0229056
0276200
0282836
0144395
0252923
0119041
0187944
0280176
0197121
0065413
0225636
0224442
0231621
0236977
0230482
0225979
0213104
0115524
0202434
0206836
0115520
0144381
0115541
0270757
0115540
0231699
0162944
0287858
0222017
0247626
0152430
0115896
0286020
0237035
0236653
0225421
0231557
0230407
0238698
0119042
0257045
0225953
0238217
0213938
0226124
0232732
0227004
0240302
0178074
0162971

0162972
0287027
0196141
0201649
0287299
0155729
0163535
0138356
0244301
0284084
0237166
0243478
0201737
0230408
0082153
0207116
0224346
0223092
0013441
0252759
0240344
0201499
0196290
0252916
0115942
0183308
0155744
0225695
0226658
0119013
0252148
0273975
0232719
0224672
0003402
0232133
0226312
0238829
0235579
0234431
0003400
0227348
0225796
0235318
0064012
0155749
0115993
0082146
0213090
0202008

0155754
0234842
0224299
0227736
0224590
0234731
0249177
0227841
0155755
0241790
0287133
0082126
0222972
0003393
0230799
0138395
0202137
0231124
0212184
0273209
0155760
0182329
0222035
0287041
0231903
0227890
0223825
0116030
0224463
0055044
0212534
0212309
0271852
0238317
0286223
0264275
0226261
0272966
0242696
0273456
0204217
0270915
0236047
0223619
0233888
0235433
0237542
0233134
0241221
0227454

0233311
0138439
0163596
0240761
0233579
0138442
0138380
0232479
0270664
0144426
0232994
0202059
0228513
0225124
0204196
0119004
0264041
0238285
0138443
0256458
0173166
0233459
0233031
0178562
0224791
0213925
0206970
0163599
0163600
0235951
0234714
0237271
0236681
0225304
0236634
0237843
0116117
0287524
0223890
0118257
0225216
0201875
0213739
0225610
0114933
0252485
0207386
0237041
0227946
0023228

0231955
0237580
0114942
0277502
0281780
0207047
0207406
0183671
0279220
0222496
0159186
0224070
0252716
0204186
0180105
0234491
0114948
0228577
0231653
0235118
0232125
0213083
0207319
0138400
0118246
0283469
0263468
0229321
0260171
0144410
0118263
0253008
0237892
0229647
0277590
0224342
0234902
0286568
0225916
0240440
0223725
0225064
0221628
0118260
0144401
0223433
0224137
0199251
0213082
0200764

0163249
0272851
0284079
0163251
0178385
0225111
0212246
0185078
0251941
0224839
0229150
0118231
0163254
0224529
0182187
0168582
0188674
0233869
0228111
0138413
0231908
0115020
0228380
0144407
0213081
0227430
0231896
0232409
0287662
0202164
0235864
0232799
0240286
0229670
0286249
0078018
0222317
0144406
0237194
0197713
0144445
0272807
0229127
0231063
0279317
0115361
0199203
0168530
0234281
0115365

021826
280837
271074
231017
178568
199585
232631
236007
221782
273118
234308
231254
223373
283637
266354
286965
030419
270659
236193
272519
196096
144451
265734
197585
213917
174453
224257
177261
138376
229267
207274
230076
144452
227769
286836
237571
138363
115414
230695
225166
228618
234938
237525
227981
235770
230838
216721
227824
226276
118242

0115425
0163449
0079246
0232970
0236478
0279348
0260804
0144583
0231092
0233581
0207303
0235042
0138375
0232485
0241520
0197756
0224391
0225794
0235444
0115457
0115461
0236886
0237930
0222832
0229352
0234588
0286601
0231597
0118245
0223874
0237479
0227021
0236295
0259855
0287498
0251982
0233143
0278004
0231672
0224414
0231035
0079308
0275458
0223923
0188282
0229754
0180871
0163464
0224159
0163466

0261338
0179921
0127837
0127838
0135926
0284173
0237281
0278524
0230580
0158428
0225062
0018280
0144579
0273361
0199121
0127831
0135913
0222714
0144580
0207393
0115556
0273466
0115568
0074582
0163481
0163482
0135912
0135929
0286154
0272555
0115592
0283639
0232625
0115596
0135925
0228135
0200029
0235024
0224928
0171450
0236445
0163497
0163499
0198973
0224090
0181378
0163501
0264755
0286606
0272644

0187736
0280537
0240317
0213901
0115649
0144567
0158552
0287529
0284820
0115657
0198925
0163516
0163521
0115661
0127824
0243910
0135924
0054356
0207647
0230432
0182698
0232789
0123992
0229525
0175084
0234638
0280323
0072195
0227432
0286143
0286095
0268603
0144591
0072182
0123989
0188760
0268896
0265252
0124006
0269068
0123999
0222678
0144589
0228973
0114923
0267919
0231802
0227308
0223903
0228909

0286199
0266518
0225911
0239498
0235337
0236451
0224819
0286272
0232089
0286170
0116106
0226726
0272944
0230406
0234446
0233868
0226666
0226331
0135903
0163081
0267034
0237732
0200822
0163082
0228670
0202016
0234285
0116120
0124003
0233751
0203531
0235225
0123983
0224121
0234193
0278083
0152049
0243125
0233299
0231189
0235610
0171951
0226539
0152056
0286239
0085449
0135900
0228802
0135919
0224826

0228681
0124019
0274629
0036257
0228446
0283428
0135905
0263828
0144460
0228206
0273301
0279015
0230621
0235070
0263363
0169047
0272622
0261379
0144468
0212391
0081052
0169031
0236432
0168958
0168955
0284629
0284092
0173744
0285843
0284643
0284622
0042304
0284704
0284631
0236116
0284667
0135917
0284725
0284718
0284635
0252346
0226718
0229172
0115009
0183514
0123977
0272237
0153820
0200281
0203387

0286772
0232023
0228226
0237322
0153823
0226297
0253006
0187957
0287791
0238782
0235993
0153827
0222344
0153832
0252333
0231534
0206725
0163053
0199400
0225963
0283451
0135899
0079263
0185404
0279964
0067066
0243650
0250011
0235419
0183022
0283202
0199791
0226125
0135932
0237758
0201044
0270891
0135916
0232520
0135898
0230385
0283164
0235013
0238062
0173699
0204128
0261829
0173692
0135914
0201574

0135931
0237039
0263641
0224376
0156966
0172799
0230047
0233538
0115053
0206885
0207280
0202400
0283312
0283491
0223198
0181798
0286942
0171596
0231494
0228884
0286399
0177673
0239202
0187514
0277986
0284378
0156973
0281072
0144524
0237641
0261096
0260622
0222246
0163273
0144535
0227033
0207626
0251485
0230122
0275157
0163283
0224516
0244280
0163286
0204121
0237087
0223788
0163295
0171551
0237412

0135902
0196811
0221944
0284407
0135930
0237126
0115468
0244384
0204120
0213055
0241409
0115474
0252452
0199781
0182600
0066248
0222001
0115488
0168918
0263941
0085978
0252010
0251791
0130561
0259793
0077044
0201013
0085982
0271639
0228949
0227846
0242366
0242515
0234143
0241119
0244122
0167165
0224814
0240224
0244474
0233445
0243135
0248114
0227802
0228445
0248705
0178836
0241635
0185038
0123485

0224287
0279809
0144481
0237581
0286059
0072080
0279218
0226442
0261298
0237784
0288040
0228162
0231682
0232690
0188042
0280744
0278593
0235726
0130147
0226542
0231032
0235293
0157985
0235529
0222007
0228499
0238812
0264676
0207049
0168505
0233611
0182177
0200745
0132321
0270540
0232893
0231546
0144476
0232328
0227252
0202341
0124835
0198612
0224844
0224132
0163359
0222022
0222032
0115648
0277842

0222449
0071677
0124839
0234949
0227107
0124831
0286864
0177483
0132329
0263723
0184182
0258984
0251971
0132330
0144488
0168427
0279484
0178752
0132323
0186235
0225057
0144485
0132326
0283635
0232306
0204104
0202099
0065802
0229915
0274346
0225493
0234279
0286160
0224252
0286748
0226992
0233608
0227744
0068024
0266109
0264810
0286307
0287405
0265215
0264292
0222020
0286525
0288082
0286188
0196758

0227459
0259889
0220256
0130414
0264279
0182083
0178586
0231278
0213048
0172428
0178602
0286325
0286183
0227479
0276366
0063660
0218416
0207611
0144504
0287049
0188542
0142327
0260942
0142330
0178623
0276949
0185176
0219159
0184945
0274977
0130294
0172482
0172478
0226321
0233392
0223991
0162804
0229996
0225521
0122085
0115687
0235861
0115685
0146205
0115677
0168385
0286588
0223374
0006607
0288080

0263752
0224232
0115694
0234235
0176720
0273113
0176946
0207186
0168397
0168393
0168395
0235351
0180902
0215692
0234793
0154252
0215023
0224272
0204099
0235151
0188389
0188011
0233806
0216921
0237940
0280119
0226423
0287971
0232002
0261186
0220804
0224160
0244528
0271153
0223587
0224918
0224318
0287140
0134121
0252017
0231660
0234661
0287505
0224957
0238075
0224239
0235158
0244169
0134115
0232563

)184423
)214074
)214073
)225044
)223040
)144619
)227588
)286363
)230398
)225310
)237990
)091181
)253049
)072756
)113851
)271870
)223727
)144455
)175928
)287720
)229241
)286579
)170364
)234325
)231249
)150995
)235947
)275981
)235978
)235831
)134107
)230182
)224082
)207283
)237566
)134108
)233912
)268509
)134109
)265180
)241227
)230944
)229642
)226022
)232958
)189229
)226258
)223719
)196277
)236152

0237665
0288030
0236202
0270207
0227110
0228351
0231401
0253081
0071282
0224884
0125046
0215160
0182533
0180914
0070950
0281863
0199815
0196220
0224808
0287623
0228723
0227929
0235830
0254485
0214041
0206573
0134077
0168137
0156959
0235338
0163719
0287878
0144550
0156983
0114026
0134072
0171148
0241553
0250151
0214021
0269886
0156990
0187288
0171135
0163701
0163702
0212327
0163703
0269894
0163704

0230082
0269982
0125037
0180385
0232901
0206567
0186162
0144554
0201182
0229217
0163705
0254999
0134086
0287086
0134070
0157014
0272410
0231177
0157017
0240288
0276678
0157020
0157087
0264534
0216135
0236999
0224771
0226567
0230599
0132164
0286962
0157103
0232287
0287964
0196639
0285906
0213964
0197548
0226621
0235240
0270531
0144560
0271716
0272483
0144559
0230342
0224263
0225275
0231897
0251038

0263988
0157152
0226642
0157150
0233026
0236062
0132170
0225026
0154743
0200114
0251774
0225526
0075975
0274514
0132155
0233252
0088726
0213943
0144712
0272263
0144713
0207496
0250939
0232746
0144711
0231362
0132182
0238138
0223398
0244502
0163517
0163520
0224514
0154764
0233121
0275340
0180438
0287059
0224752
0163528
0170876
0268279
0228242
0154767
0170860
0255021
0251576
0199609
0131389
0144596

0207163
0234123
0283392
0235629
0154781
0131379
0230172
0285969
0260902
0154783
0225733
0177463
0131368
0131381
0228874
0227578
0230524
0131375
0224660
0131370
0201162
0226427
0206562
0144597
0206926
0249786
0206561
0263573
0288077
0241064
0131373
0169814
0206560
0264354
0279465
0270409
0263740
0287042
0207815
0224728
0286689
0131386
0285654
0154813
0154814
0270739
0131378
0271964
0272498
0272529

0287377
0288050
0233570
0269391
0092345
0229271
0224039
0154822
0283885
0226441
0285731
0131374
0231780
0227309
0271841
0233133
0286352
0213383
0214286
0226238
0274840
0182568
0272477
0228956
0236570
0251761
0183960
0271898
0287069
0163576
0232953
0144566
0183977
0201545
0230697
0229468
0114166
0266745
0129810
0231304
0199577
0261734
0206807
0282987
0236380
0151789
0225542
0223351
0132967
0234723

0271235
0226183
0231915
0215016
0233153
0182247
0206728
0228171
0252562
0212269
0223791
0170142
0231395
0197885
0174748
0174738
0224074
0242109
0151090
0229756
0224822
0228791
0272554
0265028
0233544
0237838
0244404
0201713
0077092
0213877
0201962
0229421
0272166
0077097
0275227
0283971
0238225
0151092
0213872
0213871
0151093
0230891
0234158
0237603
0230465
0225386
0179796
0163491
0224237
0207251

0287348
0033867
0227523
0230807
0201033
0232167
0238912
0223968
0271943
0232766
0272282
0163508
0225548
0229243
0235493
0187118
0163512
0206559
0283563
0234048
0230480
0144642
0235904
0215004
0226216
0203506
0227795
0235593
0287774
0199927
0281710
0227260
0227549
0163513
0224618
0261468
0144644
0265376
0213854
0222983
0230111
0227339
0163527
0144645
0232490
0223965
0197385
0235670
0201701
0252398

0251590
0152642
0236732
0237433
0261572
0170293
0270377
0234737
0153551
0091317
0276147
0213849
0144635
0226955
0231552
0237676
0182973
0251224
0213842
0227337
0206557
0183813
0170266
0223899
0235424
0188167
0275090
0170275
0272149
0173705
0230119
0153558
0153560
0252700
0202175
0163539
0238008
0228112
0227707
0271643
0170248
0226320
0287981
0236452
0227498
0287641
0235534
0228767
0212442
0172995

0230830
0207625
0226489
0236158
0144681
0231382
0201315
0163673
0230067
0279727
0281100
0168016
0234073
0210181
0272334
0236165
0178567
0076242
0224858
0093167
0206645
0225558
0199594
0224080
0271993
0231449
0271653
0270194
0144674
0230409
0222208
0198590
0144668
0239105
0235257
0235482
0236411
0144677
0199075
0136059
0187091
0200579
0008226
0231668
0060971
0201965
0172936
0172939
0172940
0237982

0144671
0199514
0230123
0093217
0229589
0114739
0157036
0207512
0230683
0232439
0183873
0185313
0231243
0168356
0206708
0114742
0114745
0168026
0284158
0144655
0284669
0283849
0214305
0287620
0168334
0287995
0215034
0287958
0168329
0270367
0287780
0179934
0228168
0238267
0223668
0144659
0168028
0206760
0202363
0286996
0286781
0168314
0285885
0230757
0170011
0201570
0280739
0265327
0202517
0114784

)223797
)230274
)168032
)188846
)177873
)177842
)199906
)172888
)234067
)226845
)226302
)229072
)231873
)234287
)287415
)233096
)233919
)275256
)168036
)168038
)252409
)222942
)223513
)251986
)212145
)234270
)235339
)182606
)229339
)234562
)222872
)187094
)281160
)231280
)229572
)157093
)235787
)114812
)232354
)093183
)008324
)114857
)230084
)114853
)287629
)240203
)157119
)285539
)010282
)230970

0244607
0280571
0287417
0173811
0225611
0181061
0235288
0144648
0273328
0273291
0240747
0180432
0236459
0182983
0206552
0144649
0232375
0144647
0227245
0285766
0163788
0241939
0234617
0160746
0251811
0223916
0011198
0235886
0271192
0272121
0252980
0207954
0280435
0261786
0173769
0271937
0179152
0225873
0283036
0185219
0178917
0214825
0277509
0196345
0236869
0144792
0214820
0186448
0233509
0169981

0272077
0235805
0196653
0186446
0163807
0163808
0284498
0169964
0163810
0235845
0163812
0227123
0075914
0163815
0252410
0163814
0224094
0249992
0202268
0011376
0232455
0144791
0230530
0211456
0244357
0279017
0163817
0163818
0285788
0236930
0173585
0201635
0163820
0172215
0173578
0226933
0229515
0183625
0163823
0236736
0121807
0223552
0160791
0121797
0283646
0012223
0163825
0163827
0268324
0241186

0283877
0283473
0286570
0251967
0178038
0181585
0206549
0283706
0261603
0188086
0284672
0286651
0238013
0226074
0178055
0160808
0160801
0273682
0160799
0160796
0236409
0181555
0275940
0227398
0088727
0239128
0114648
0280173
0260236
0076201
0114650
0276925
0163832
0271161
0244732
0114646
0173473
0214773
0235695
0132153
0274420
0221585
0047849
0199591
0224895
0270285
0265078
0236768
0164045
0235627

0253056
0231884
0265483
0164047
0164048
0229759
0199476
0172113
0228638
0229453
0252466
0164049
0277483
0164050
0164051
0232112
0244380
0164053
0213689
0164054
0114268
0278799
0145040
0114270
0284251
0010256
0183396
0225697
0284364
0008300
0284575
0228350
0271973
0213672
0068745
0114302
0224424
0178537
0221883
0177479
0235236
0223343
0273211
0178467
0285416
0178252
0178149
0199032
0178057
0207605

)178035
)272434
)198218
)241461
)172053
)284488
)172046
)172037
)270441
)177352
)185909
)236980
)173421
)270538
)225399
)188315
)199546
)264633
)114316
)233276
)067560
)235908
)145022
)145020
)283189
)145029
)201301
)173402
)226913
)164061
)235120
)164062
)173531
)259970
)164068
)176020
)173540
)176095
)239576
)244730
)187492
)185614
)182179
)283726
)183763
)164076
)264706
)234667
)164078
)230698

0228008
0164077
0004534
0003756
0281691
0235016
0001617
0207922
0114349
0188338
0114353
0275334
0213600
0230454
0232352
0012171
0283848
0179564
0214706
0186792
0243477
0114378
0068001
0114383
0068028
0281358
0235058
0004838
0114388
0114395
0272104
0126062
0007402
0271858
0202322
0088543
0114735
0280382
0114737
0114738
0273356
0088538
0272543
0228337
0213587
0235278
0240959
0145050
0259956
0145041

0164080
0221015
0164081
0288063
0201595
0164082
0214686
0285749
0224792
0229972
0184345
0241789
0235455
0214681
0173389
0287707
0114767
0041880
0280422
0180929
0090097
0272762
0114779
0248487
0114786
0243989
0162244
0285685
0164086
0164087
0286419
0242849
0023330
0280003
0239732
0173366
0247596
0243224
0164088
0164091
0199150
0168237
0242797
0207926
0114841
0239557
0163930
0010318
0010319
0114854

0010322
0010327
0168268
0168273
0163939
0252768
0221518
0163938
0280904
0252787
0212493
0238862
0222345
0212452
0016864
0114902
0114904
0275956
0270941
0271137
0055957
0162267
0055955
0243696
0239799
0272573
0248592
0213533
0275789
0279144
0163935
0272305
0280417
0242142
0163933
0163932
0286531
0163931
0272886
0274967
0157388
0207109
0212608
0227224
0251987
0271916
0016391
0056736
0271976
0113812

0113811
0288029
0242983
0286353
0157445
0229806
0265992
0243715
0144771
0239991
0243572
0240708
0242568
0242775
0114251
0244586
0242317
0187672
0240777
0281708
0264013
0200379
0200238
0180376
0163946
0163947
0240198
0186451
0272202
0144730
0229863
0163666
0157500
0239388
0286952
0174844
0212392
0206918
0239831
0237186
0202428
0206815
0287502
0174840
0168374
0272146
0210841
0174839
0239801
0241933

0163681
0270772
0213509
0136068
0244161
0283148
0283511
0163687
0163686
0163684
0255154
0273493
0168297
0275607
0168291
0272182
0272360
0168301
0168306
0236722
0168309
0243384
0244383
0198643
0163689
0242428
0243295
0243903
0242545
0241804
0277592
0189283
0287649
0240097
0281426
0232407
0225673
0244183
0212211
0144724
0252420
0226360
0252184
0237456
0241472
0114405
0153266
0163618
0225662
0244632

252449
226082
244342
242841
240359
163630
241359
252301
188817
224479
163632
163634
240549
285399
285258
163635
280620
239653
271843
163636
243410
272181
241111
200222
189196
163637
241572
241101
239926
226017
244564
212340
163638
241158
241684
243149
151276
270059
244226
223409
272610
240175
202071
203647
144741
206759
241587
144749
271174
214560

0288039
0285738
0287873
0163376
0283298
0241884
0172340
0242510
0241316
0183662
0214552
0240568
0286967
0244640
0241506
0163377
0252828
0163378
0244513
0144747
0265355
0144744
0144746
0163380
0114541
0239263
0240265
0240842
0241667
0187098
0241665
0240405
0271034
0242728
0242120
0287487
0287669
0240156
0243873
0206939
0114861
0285708
0244203
0221264
0274387
0242094
0277855
0270562
0163412
0170837

0163421
0287131
0239250
0241163
0244130
0273461
0276471
0239350
0242609
0243083
0272774
0242618
0287882
0260670
0163602
0244345
0199469
0239568
0222838
0287595
0144736
0241537
0240135
0271266
0172986
0276890
0242960
0163605
0238959
0255423
0252651
0223247
0182921
0252937
0121440
0239119
0239677
0278934
0242364
0242741
0278554
0239454
0223738
0213452
0113805
0241022
0242638
0266780
0287398
0213300

0228868
0243674
0179799
0230655
0244699
0240665
0242953
0238278
0244026
0272690
0239959
0243794
0272710
0244674
0239383
0244222
0172971
0243422
0282912
0282511
0283894
0241532
0241622
0172969
0236138
0242516
0227124
0266396
0185008
0222883
0240809
0199520
0213450
0287567
0206891
0244461
0240573
0243957
0222574
0240241
0239226
0169855
0240752
0240964
0288105
0265193
0243877
0271389
0239699
0242586

0286329
0242828
0242781
0243694
0242145
0241593
0287053
0244281
0114480
0222389
0241319
0240404
0242190
0242009
0239440
0241367
0241095
0288004
0286567
0242195
0287158
0242641
0241776
0242339
0287273
0175161
0264084
0241648
0239519
0243915
0201410
0251773
0239767
0222934
0241328
0277746
0206538
0241293
0281392
0264119
0083937
0064835
0239829
0240902
0240255
0239572
0200703
0221059
0179097
0163320

0175105
0241535
0179021
0242159
0283434
0207316
0242365
0242705
0241861
0044524
0285780
0240951
0240309
0212598
0222490
0189002
0271024
0283367
0283544
0251727
0184500
0201339
0200366
0169379
0239614
0178750
0178700
0178694
0242457
0178660
0242911
0234629
0239589
0244048
0244681
0286516
0271671
0243547
0278103
0241815
0199286
0269028
0218754
0241018
0176243
0286447
0080224
0251172
0261292
0113966

0282527
0080200
0279658
0170854
0285614
0183185
0250750
0213439
0196578
0249225
0251090
0248365
0231192
0236032
0233412
0249321
0249975
0248507
0251088
0187900
0230301
0197938
0232535
0196098
0206536
0244653
0232382
0231861
0080822
0285635
0248839
0250562
0154165
0286602
0080819
0240827
0244153
0239445
0064225
0249274
0057019
0206712
0239462
0207331
0240476
0275108
0241305
0240572
0243296
0243089

| |
|---|
| 244464 |
| 144810 |
| 273374 |
| 184220 |
| 168386 |
| 264897 |
| 287378 |
| 235156 |
| 277000 |
| 243974 |
| 036054 |
| 114021 |
| 154174 |
| 206535 |
| 181458 |
| 144820 |
| 252989 |
| 243495 |
| 114354 |
| 154175 |
| 213431 |
| 201642 |
| 242573 |
| 081148 |
| 138468 |
| 241738 |
| 232830 |
| 282978 |
| 201554 |
| 175841 |
| 174173 |
| 081154 |
| 242299 |
| 243396 |
| 252348 |
| 212319 |
| 201511 |
| 066422 |
| 201121 |
| 256628 |
| 114391 |
| 286699 |
| 244119 |
| 182504 |
| 144815 |
| 249474 |
| 144802 |
| 287682 |
| 214407 |
| 202177 |

0241280
0170044
0241754
0201065
0201922
0236691
0265328
0240107
0243195
0214405
0243635
0170017
0114423
0287421
0244652
0286854
0242759
0200610
0200361
0239997
0241787
0239632
0242214
0242836
0249580
0243658
0240034
0250630
0239586
0240796
0252626
0286956
0239455
0243701
0239828
0241218
0272597
0138483
0273125
0114439
0240423
0241469
0279277
0196776
0271856
0114446
0114455
0243188
0144821
0241634

0163507
0202379
0276763
0198919
0241777
0163515
0163519
0138472
0114487
0221633
0239314
0241224
0261364
0214381
0252889
0239288
0163530
0277635
0121570
0241257
0244561
0230618
0228980
0244196
0265956
0242029
0271249
0214380
0243945
0240895
0221206
0244722
0201426
0242242
0177707
0153283
0239311
0244142
0240787
0177494
0244720
0240891
0240766
0144824
0286492
0144827
0144834
0239439
0176040
0114529

0207940
0174500
0261488
0172139
0244144
0239482
0283669
0091972
0243081
0186265
0242525
0279349
0144848
0138459
0091986
0242308
0240893
0242770
0206531
0163606
0272844
0163607
0163608
0240057
0243483
0241219
0243795
0144857
0288079
0206530
0240751
0285943
0243849
0275670
0163611
0072858
0239453
0265253
0241529
0176542
0121579
0239280
0242659
0114573
0178075
0240776
0241889
0240375
0273394
0184307

0163617
0270880
0287805
0285836
0151576
0240912
0151577
0241490
0240487
0174255
0181847
0181722
0259976
0284134
0241560
0242290
0241295
0264623
0239946
0240002
0242767
0242880
0244157
0287795
0288074
0172020
0241596
0185565
0243359
0240922
0241397
0241156
0198406
0243197
0278072
0265433
0274732
0242385
0200372
0243014
0206889
0239268
0240393
0241213
0242816
0243276
0286660
0144847
0239877
0163424

0251012
0114638
0121578
0240254
0031081
0242829
0241155
0207310
0176142
0163389
0272662
0113845
0121594
0272967
0144843
0144837
0242052
0121577
0138495
0286584
0183833
0285585
0239994
0144852
0240471
0082701
0239835
0241546
0242622
0244308
0244139
0175697
0206721
0240882
0163428
0240661
0282950
0242613
0163430
0284121
0286735
0240774
0244441
0065518
0113924
0144840
0240452
0243257
0153767
0213371

0286827
0243593
0287366
0145087
0270782
0264477
0243813
0242103
0287022
0051341
0242445
0186103
0163833
0180353
0243544
0222057
0173230
0173226
0145088
0163406
0145103
0202388
0114013
0287977
0036828
0240236
0121552
0160124
0114023
0196981
0272758
0242531
0114030
0275348
0287207
0138496
0163840
0173200
0243918
0173193
0169087
0138463
0273033
0082684
0065485
0284421
0121542
0273454
0173175
0272678

206527
239523
065534
250174
273123
207002
175455
065371
160145
266383
240622
276656
252751
260391
114491
265981
242199
082781
244286
280042
276626
243986
173702
173706
252642
288022
221955
199327
264986
163848
229048
243508
252953
114520
201800
252965
144909
243072
242001
234458
243429
228963
239978
179170
239432
284624
284660
272840
239804
221737

0242229
0287617
0241346
0241288
0241278
0248787
0171084
0189366
0241439
0114547
0114544
0248607
0250012
0144908
0250218
0272020
0246022
0287232
0250934
0251448
0287440
0163884
0249833
0163885
0279328
0070476
0159650
0180767
0180697
0201968
0288436
0197763
0248557
0198284
0248850
0159685
0249558
0280053
0114554
0287784
0214324
0212556
0243016
0201288
0286919
0239620
0244215
0239921
0240562
0282975

0163870
0282860
0275067
0276083
0073111
0285600
0114631
0114626
0074416
0287143
0187715
0201210
0058262
0175792
0222627
0239608
0212344
0132394
0285619
0179407
0242049
0179348
0244300
0239405
0286806
0198685
0163902
0277250
0242551
0207130
0075785
0239503
0242992
0244232
0239483
0244065
0243022
0242062
0261159
0231305
0250796
0287110
0177646
0114656
0279507
0114654
0249505
0169704
0172780
0261796

)273174
)240682
)273437
)169714
)241673
)181789
)278658
)286729
)183624
)184897
)206417
)248916
)270492
)270773
)244932
)251474
)207088
)172771
)129071
)163913
)163914
)178804
)004399
)239437
)248659
)172765
)202412
)283509
)203644
)271270
)250643
)170893
)248459
)284731
)249869
)180770
)251178
)251610
)251287
)248243
)253540
)249846
)248618
)288111
)250580
)228252
)249253
)172752
)206384
)196455

222783
285631
232829
017260
284095
250592
252894
249098
251579
034533
114670
250129
201550
250608
246082
198585
261167
114686
249020
248468
248530
196353
251447
265859
240854
242899
014257
240890
213264
138246
240303
274810
129048
251614
081307
113971
248724
249725
144868
251011
170819
249820
249993
214301
272832
250983
201827
091527
163781
242337

0252641
0201394
0091513
0244062
0144867
0285908
0154917
0174640
0243832
0163785
0231845
0240006
0260633
0214289
0114019
0277723
0248377
0214288
0284561
0129055
0182923
0249691
0154928
0174611
0286982
0253087
0201413
0240086
0242222
0241905
0073711
0227267
0273455
0174579
0241438
0244740
0114054
0118007
0200571
0244101
0252914
0240695
0261758
0286915
0242968
0168917
0239213
0273486
0158092
0214283

0174564
0249407
0201325
0243886
0242070
0168875
0239513
0261146
0241478
0066405
0272609
0158163
0240668
0118017
0286396
0138231
0114098
0181322
0158186
0158220
0280399
0114107
0158234
0174715
0051382
0239207
0242321
0241429
0242244
0244578
0183770
0206262
0206260
0175110
0184814
0233701
0251137
0232416
0281473
0276304
0286988
0272656
0184432
0283418
0248932
0114113
0248790
0178631
0114115
0163864

0214280
0201600
0244692
0261763
0250543
0251471
0250161
0158258
0249290
0251058
0250433
0249305
0155890
0261826
0224085
0114120
0248773
0175093
0251270
0155893
0287155
0241400
0249417
0249540
0177311
0215606
0155903
0250170
0249986
0242104
0241899
0241526
0114125
0285558
0114124
0286822
0228398
0069849
0200389
0244124
0279147
0114126
0233597
0206604
0175066
0114127
0251804
0202125
0175054
0242479

0244327
0242390
0252745
0120756
0239641
0199319
0287045
0144935
0251787
0163710
0240950
0243818
0188582
0241592
0241570
0163714
0268129
0243733
0175040
0241679
0244171
0181804
0240888
0240012
0244493
0243033
0243759
0248374
0235110
0181744
0222778
0280397
0244024
0241695
0239797
0244358
0240095
0261051
0243415
0240032
0152952
0114698
0243596
0241358
0163746
0279320
0188313
0199812
0231213
0241457

0243620
0242671
0207156
0239412
0174963
0241202
0152977
0240804
0239661
0239922
0285798
0241131
0285660
0241792
0244650
0205412
0243347
0242440
0285557
0241993
0144891
0153002
0244089
0240521
0163751
0287198
0241627
0163754
0071794
0239718
0201031
0240511
0163755
0047457
0286714
0240216
0244468
0163762
0243822
0169908
0240541
0243885
0240246
0241929
0282502
0239408
0169903
0244503
0018408
0252581

0241985
0241313
0114744
0206199
0251854
0082996
0252172
0239351
0070087
0242791
0224831
0182000
0241947
0243944
0240477
0243321
0241143
0241048
0244541
0278234
0243550
0286501
0196428
0120742
0144895
0198843
0271711
0163645
0240137
0244668
0287688
0181788
0244265
0239265
0214237
0260234
0243273
0163646
0276055
0144893
0199994
0174946
0174944
0239570
0286273
0138271
0181631
0169313
0152580
0242810

0242119
0264330
0241151
0197953
0242908
0242097
0240602
0250271
0114771
0198829
0239941
0152601
0229619
0221962
0201217
0243305
0244545
0287706
0242561
0169860
0241810
0241671
0244268
0181467
0265813
0241220
0241912
0238755
0237787
0207323
0200162
0243486
0201398
0240068
0243069
0114790
0174953
0271922
0279305
0174948
0240048
0243861
0243154
0242151
0240601
0243969
0243953
0196549
0240666
0241336

0240045
0242790
0239443
0114805
0248772
0243518
0242152
0272096
0239508
0242925
0239212
0214919
0241307
0174928
0284952
0169359
0286585
0163655
0288032
0240489
0174930
0169282
0242017
0242537
0287916
0227709
0240596
0242370
0272990
0114850
0243926
0163659
0174912
0240875
0230457
0197980
0241879
0239706
0222499
0243629
0241544
0241135
0163660
0201778
0244515
0241770
0222675
0243176
0270981
0286361

0197415
0163661
0174899
0286892
0242536
0251751
0239716
0168779
0174891
0241723
0277143
0243314
0243150
0178053
0243025
0243165
0168827
0079257
0272087
0272440
0118849
0118855
0240207
0243675
0271778
0272247
0232374
0279311
0283154
0214216
0263634
0244249
0286913
0243116
0241211
0252763
0151967
0244040
0168811
0242107
0184100
0180044
0248710
0068885
0243687
0113810
0207779
0198987
0213186
0244009

0238427
0186432
0238741
0229320
0242285
0179674
0207084
0260572
0163590
0169255
0169251
0271052
0240138
0243838
0196542
0240567
0269888
0234851
0182447
0288087
0240354
0241874
0243044
0241168
0241369
0244571
0240792
0238924
0214210
0221251
0241636
0241767
0090402
0200052
0239205
0121871
0244128
0114200
0242178
0240084
0240824
0242456
0242326
0244429
0244321
0274149
0251759
0224962
0240458
0231516

0240033
0239227
0220343
0169064
0241669
0114204
0174776
0114209
0241120
0163536
0287319
0244227
0244706
0239238
0242036
0241203
0173905
0286994
0206120
0241882
0222357
0207717
0226765
0231688
0242268
0272797
0273177
0241383
0085276
0241479
0213178
0242795
0244381
0206777
0270141
0277925
0184378
0269889
0085274
0269984
0171757
0270135
0270096
0188306
0114248
0240128
0187033
0244039
0239219
0244193

)008952
)240373
)173890
)201342
)242583
)173889
)199536
)163558
)200118
)239628
)136603
)242578
)013297
)285218
)283474
)285051
)013293
)241416
)213174
)244738
)286695
)239739
)163584
)163577
)286856
)240704
)199488
)163581
)154310
)207963
)222506
)240497
)279673
)207114
)234594
)075651
)234717
)186329
)235943
)287899
)075420
)237483
)243398
)279862
)236686
)121853
)121858
)224611
)223387
)224426

0144959
0231384
0252082
0287012
0114346
0200355
0201458
0227440
0237501
0144962
0237473
0169760
0237645
0228213
0252982
0232601
0213169
0177694
0253020
0230497
0230292
0226779
0264974
0199792
0288104
0201648
0236241
0214192
0201452
0225552
0260217
0230362
0236357
0271413
0238026
0223715
0283333
0252302
0228308
0231888
0177565
0231310
0201343
0200882
0203645
0236893
0226782
0228221
0252028
0236385

200288
228561
277241
231574
225790
270321
199858
286003
223930
197584
275163
223941
252336
271131
237978
172667
200616
229102
121879
201957
171121
240429
121864
260743
171109
114450
242539
272699
239255
181260
136518
272910
272660
136522
136521
239096
243977
058056
114757
270178
243799
244302
287645
201822
282327
213158
239880
241696
201810
239774

199986
241946
163728
284862
145075
243187
285336
242811
242068
241473
239516
114416
213157
205981
242808
242140
206932
237806
241231
235294
239381
276074
276690
181449
278026
266317
285546
199839
239805
213155
242512
252257
286453
240478
287605
242012
241098
244199
244247
287941
228766
238837
243378
260377
240063
058063
270554
273261
043093
244346

0078070
0243368
0078081
0053524
0176597
0202502
0240024
0201229
0253092
0199363
0251730
0271682
0172578
0242522
0243584
0114796
0163872
0233885
0283765
0180834
0175193
0234371
0263932
0233441
0114770
0223882
0223529
0186090
0178084
0233576
0207351
0238020
0186038
0205955
0273181
0272721
0145191
0228205
0161202
0161203
0161204
0145198
0221120
0214160
0284753
0284917
0163888
0145194
0175166
0114867

212158
175182
114859
163882
090534
090539
224153
283625
227509
230215
182580
177383
272970
229433
273403
272922
156931
187068
223358
113790
229704
264614
073803
229638
163900
286086
163898
163904
236489
073792
163915
265470
136527
171658
244405
277118
234197
058866
234548
224406
226859
213139
113838
090520
197099
283149
283390
145192
275696
090512

)113905
)225175
)226724
)113889
)226652
)207505
)230951
)263826
)156976
)238942
)200418
)283958
)221420
)200320
)263776
)163918
)231724
)232233
)231982
)181092
)226482
)223803
)233565
)073849
)163923
)198491
)175077
)127241
)283175
)136514
)280270
)239093
)228952
)157005
)198471
)228804
)113916
)285938
)224049
)236864
)224187
)225058
)236412
)234238
)270959
)145012
)279891
)224563
)207651
)234076

0188001
0236182
0230115
0073282
0227971
0216058
0090530
0230077
0225764
0212489
0238077
0163347
0113946
0198398
0196083
0231047
0223117
0273370
0228730
0232610
0214147
0205835
0188729
0233308
0188958
0152492
0223812
0253548
0201860
0237653
0232353
0287178
0222583
0114279
0231383
0230126
0226709
0238902
0200184
0180611
0232130
0236229
0127252
0187527
0236508
0127249
0225473
0236297
0198836
0224855

0207370
0243991
0223986
0286624
0287206
0229155
0214146
0238043
0224403
0230102
0276407
0229964
0232874
0114315
0242201
0225742
0238097
0228271
0214145
0286317
0178772
0172061
0178732
0133657
0271483
0233058
0244675
0252053
0232873
0231770
0145014
0229334
0272707
0041802
0185112
0233987
0287073
0237222
0234805
0230401
0238031
0173950
0233303
0265333
0244314
0230266
0206600
0287005
0114331
0229325

0227433
0207368
0184203
0252620
0223711
0265882
0236844
0236425
0189058
0224769
0282206
0271662
0229178
0242086
0215837
0207650
0283426
0176945
0145113
0223783
0235836
0061938
0276489
0286004
0224614
0260261
0239122
0231443
0185485
0286168
0231464
0072274
0228413
0252174
0224652
0163958
0222335
0163959
0161217
0272741
0228028
0213123
0237418
0273331
0235897
0145107
0212146
0163960
0206644
0241868

0243339
0226345
0225822
0225240
0163961
0273013
0214097
0213358
0225770
0185798
0174013
0286661
0232065
0174004
0163964
0174007
0201441
0180370
0272359
0206892
0119231
0230732
0114503
0225578
0270170
0119227
0286017
0233487
0163975
0228109
0075711
0265850
0227375
0286870
0235126
0229912
0237167
0272989
0161267
0214135
0213519
0249626
0273308
0286909
0236833
0287286
0145016
0284146
0122068
0273375

0186001
0234136
0234742
0114473
0199306
0182899
0185621
0201622
0286755
0232832
0226435
0212297
0232783
0213113
0236246
0236438
0222792
0230304
0226008
0248302
0272602
0250238
0250312
0275426
0250892
0198155
0251188
0251689
0186777
0286705
0251535
0131127
0248564
0250321
0207642
0281016
0249025
0272885
0251595
0182903
0174227
0273238
0283183
0133256
0242686
0272927
0169020
0215375
0169026
0185619

248416
272588
249592
233799
168993
260262
286205
178950
127419
145214
145217
127415
127418
250259
178222
251639
251652
215367
212458
227189
159674
273179
280927
159692
196810
090316
163945
254094
253399
288380
235608
272783
244459
174137
207009
163950
270195
013810
168936
068078
168924
109685
249784
185049
284587
243449
185818
130997
214367
249006

0250623
0249047
0123933
0284276
0159733
0251148
0239247
0249077
0206113
0063978
0248399
0251229
0125386
0248155
0206738
0168884
0087266
0087274
0287099
0109736
0249673
0087269
0125388
0197386
0199335
0251075
0278778
0274693
0188981
0159788
0248840
0248843
0109758
0286900
0175920
0163956
0270090
0216560
0250681
0250632
0250986
0184160
0251166
0234386
0249522
0253917
0271977
0251271
0251669
0250476

0284727
0243894
0250940
0248669
0198787
0250381
0250710
0249844
0163982
0132406
0145220
0168826
0248516
0168824
0168818
0248221
0237170
0247708
0280310
0273396
0163132
0251587
0200761
0170891
0152953
0252266
0201496
0082929
0173040
0072840
0072832
0181215
0263631
0152969
0284684
0287786
0249896
0251408
0109501
0286176
0074211
0013288
0179010
0251580
0284847
0170846
0246526
0279859
0163993
0282742

0178988
0186222
0287331
0170871
0132405
0287104
0187904
0202392
0245748
0173013
0173011
0109519
0245468
0200867
0184985
0264658
0178597
0266690
0272620
0196526
0287164
0228919
0163995
0207807
0273267
0251460
0250471
0125089
0170801
0249145
0251615
0087008
0202054
0155275
0251186
0205959
0155269
0109625
0250915
0215612
0258507
0197468
0249272
0250804
0249347
0284636
0271057
0251313
0288075
0250342

0287972
0249767
0163612
0250268
0229924
0231396
0233136
0227551
0232399
0249482
0223569
0250231
0249104
0250844
0248920
0250745
0249811
0248933
0250913
0232264
0230430
0229579
0227140
0235780
0231051
0231637
0251694
0228856
0205946
0219492
0188438
0252307
0232773
0186146
0248364
0284689
0250884
0251624
0249866
0249443
0284648
0242034
0221275
0244004
0287117
0249799
0250942
0186234
0251087
0203436

0249563
0250425
0109667
0169676
0252002
0249219
0250413
0261490
0071127
0264931
0223086
0250393
0251296
0248777
0271544
0250040
0250573
0250074
0250505
0251278
0251338
0250613
0178163
0287154
0248419
0109684
0249334
0283083
0207716
0250896
0002587
0222182
0287778
0251152
0249631
0248300
0248262
0251210
0212282
0249995
0249780
0250371
0250098
0283271
0205940
0157869
0286297
0281202
0109705
0246095

0038219
0266240
0249148
0242209
0250634
0250497
0224569
0248698
0251679
0249988
0287360
0251412
0248425
0248360
0251252
0202449
0247624
0137449
0249252
0199420
0163145
0251379
0048342
0249930
0273133
0118564
0237765
0109743
0214846
0248188
0004468
0251644
0137440
0137441
0007062
0286280
0169762
0249234
0263327
0242358
0248851
0286888
0249506
0248138
0169744
0287081
0249998
0251212
0250819
0206780

215278
151552
249581
249488
283219
002549
249502
118579
047662
163257
109805
178177
235413
286046
251048
279918
248238
251816
279379
248515
249441
241612
250611
145147
248228
207732
163138
185774
272995
250243
283281
286318
250092
242768
248343
280650
250039
239001
152990
244538
251516
249948
237350
248837
249547
251220
250137
251292
109819
249453

0109606
0207697
0243005
0249256
0261121
0251638
0248444
0109610
0181982
0153012
0109618
0281501
0038210
0271172
0278135
0168228
0053900
0249320
0248608
0248545
0238632
0228154
0157765
0242640
0249277
0250050
0248133
0251353
0248654
0091490
0250541
0250317
0251009
0283314
0248452
0168214
0163394
0109680
0200999
0109689
0240005
0248340
0251080
0205830
0249699
0222206
0251325
0250858
0287389
0251410

)286321
)250038
)250064
)286141
)243548
)283275
)250927
)250190
)276507
)249228
)248176
)248338
)251373
)251113
)283666
)249542
)249564
)244031
)248281
)286596
)169851
)249678
)250624
)251182
)251434
)249882
)251129
)286212
)251329
)286784
)250076
)287968
)249452
)249766
)248417
)288321
)250954
)250723
)239983
)248685
)287320
)250597
)275250
)238694
)250416
)206665
)047365
)248466
)247193
)197057

0251438

0283207

0248215

0251588

0287416

0261761

0247810

0264319

0174145

0154274

0288073

0248936

0181826

0207075

0239532

0169299

0279386

0249863

0065882

0250254

0251332

0249649

0287659

0251642

0248227

0280015

0249534

0249667

0251635

0231160

0271958

0109787

0222230

0174123

0174125

0174130

0197712

0207944

0121895

0109790

0249207

0249685

0157796

0035928

0206675

0222592

0134962

0264621

0238797

0163682

0121897
0224097
0109814
0249348
0287262
0163683
0286349
0252796
0255458
0078140
0252975
0249019
0243260
0180610
0121892
0271278
0252970
0250568
0231707
0249064
0205794
0078177
0200455
0201863
0248977
0260296
0286089
0168421
0250338
0249241
0174343
0239010
0163694
0250893
0242197
0263642
0287182
0179299
0248780
0163697
0250906
0207195
0201736
0199790
0207198
0251173
0154277
0207481
0064042
0239627

0287762
0249216
0249081
0109132
0250467
0249122
0200338
0249745
0245870
0249771
0249887
0109133
0182308
0272862
0014824
0249212
0188848
0285454
0250781
0178343
0124406
0240167
0248283
0260519
0251517
0252595
0215203
0248939
0250657
0277096
0213851
0248143
0251501
0239464
0286891
0239679
0124399
0250863
0251350
0183783
0251159
0177752
0151806
0163281
0273369
0272936
0249729
0248987
0275959
0248744

0248320
0224560
0200720
0222257
0163285
0260878
0151834
0249330
0252243
0249936
0170516
0109158
0163288
0260918
0169019
0282904
0286294
0145246
0242431
0248254
0145244
0277388
0242262
0282917
0259959
0170448
0163293
0198515
0074966
0200269
0283499
0135605
0202014
0109171
0145248
0182223
0075539
0251722
0109180
0248256
0145247
0252913
0251334
0109182
0225455
0249828
0249079
0280043
0250016
0237961

0268736
0248946
0248583
0251056
0250769
0248532
0269848
0268967
0268391
0232471
0251630
0251539
0250753
0249337
0109184
0250473
0272576
0243929
0188993
0163069
0251040
0163071
0287999
0251286
0199924
0109189
0248866
0260120
0250302
0226950
0264585
0286161
0212588
0248375
0226887
0212490
0199901
0128045
0184178
0248115
0207385
0145216
0282278
0072201
0250930
0173966
0248494
0249341
0249706
0250815

)229585
)272650
)109220
)269506
)287534
)251105
)250147
)180613
)242670
)134853
)248184
)248518
)251264
)250456
)157404
)249727
)251256
)248317
)244422
)252815
)250646
)244034
)128052
)287060
)239545
)250812
)206839
)250464
)128039
)251111
)249700
)273257
)134851
)239040
)134852
)238585
)223305
)272969
)163440
)109255
)251923
)250821
)090989
)202358
)286599
)287382
)174799
)109265
)200741
)252489

0286093
0250324
0157426
0242083
0269921
0128059
0270147
0283156
0128050
0174780
0196503
0249693
0248590
0171476
0280285
0241250
0227040
0240545
0265894
0128040
0278226
0251799
0084093
0269949
0084092
0047315
0251703
0163453
0250384
0245067
0287369
0250192
0243920
0248505
0251459
0251049
0250333
0249831
0285998
0248660
0249105
0250340
0250078
0251339
0249382
0251266
0250375
0286097
0249111
0199780

0249392
0249892
0201676
0201775
0287174
0283043
0249066
0248237
0265829
0240152
0150471
0241261
0205682
0226439
0202169
0248692
0225405
0241344
0250249
0248847
0248336
0180673
0251055
0231442
0250775
0205678
0252104
0250768
0251430
0251385
0249893
0250169
0248259
0249045
0248328
0244669
0286664
0250125
0248447
0248998
0249351
0285711
0145242
0243199
0250846
0248479
0222765
0251527
0249413
0272304

0236562
0287383
0221563
0244099
0252890
0250075
0206629
0274759
0250030
0145241
0035720
0279464
0033178
0270257
0248049
0250656
0270228
0270292
0109163
0202374
0232398
0153802
0187054
0229009
0207428
0227725
0283131
0198092
0215127
0250426
0250026
0185873
0250642
0248639
0249531
0248122
0083896
0162840
0087128
0250566
0197888
0249472
0251101
0251074
0196620
0248635
0250376
0251236
0109181
0251424

0249235
0251529
0248547
0248613
0272626
0249985
0198277
0135220
0251498
0249735
0250277
0251685
0249890
0250919
0171234
0248763
0251284
0250100
0251427
0196472
0250696
0213759
0268803
0249763
0135226
0251691
0250612
0268209
0215110
0248824
0249956
0250828
0249686
0156096
0248886
0251177
0173610
0271271
0173597
0236959
0284695
0109193
0251489
0126545
0135222
0126549
0205649
0126550
0234124
0187533

0109205
0181617
0171209
0145309
0109208
0171201
0171199
0171195
0187689
0178522
0132464
0286848
0132465
0132467
0207448
0207058
0272986
0018189
0132463
0283313
0207007
0173542
0156136
0080493
0249170
0270098
0145321
0222236
0248567
0056291
0156140
0250877
0270665
0252955
0210839
0163626
0132466
0248641
0250220
0221639
0249976
0163631
0081051
0079557
0250436
0249970
0169435
0228277
0169429
0249051

0124875
0250550
0109272
0163739
0249920
0250339
0163737
0163736
0163735
0287037
0244194
0163734
0248848
0288019
0251017
0081041
0163738
0269559
0182585
0124882
0250532
0109321
0249942
0174808
0287375
0250356
0169116
0249717
0248165
0260265
0250315
0250560
0251185
0251454
0251383
0248646
0250735
0163743
0174796
0174792
0138769
0138757
0201644
0138768
0229717
0222644
0156194
0138744
0198301
0245928

0138755
0156219
0169245
0169248
0225846
0138750
0286074
0138760
0189157
0287401
0272414
0118804
0163749
0249858
0212368
0240426
0199857
0138771
0249278
0207307
0263445
0283682
0233860
0224218
0270244
0242727
0186212
0138758
0234036
0118816
0214062
0271676
0201641
0138764
0270669
0250006
0248831
0249036
0156234
0221711
0138767
0169288
0250214
0248926
0248956
0248640
0248128
0249072
0138759
0249565

248117
138772
200998
251442
251019
287632
239793
260278
138756
266270
163291
242175
249307
156269
196475
250334
249646
251399
248408
251321
212620
163297
236530
244002
248719
152784
138675
197826
232327
152785
138669
251059
207065
138670
248282
248448
252762
248543
250826
202485
234568
284516
273156
138668
272677
250325
202440
152795
145293
249242

0214988
0227304
0249960
0248113
0231782
0286490
0145284
0270480
0207746
0270842
0138674
0251022
0168152
0189308
0199530
0251874
0138663
0286035
0145287
0250057
0250677
0248401
0173085
0173083
0274154
0251647
0163312
0163319
0163322
0213608
0239525
0138678
0201633
0249171
0250546
0200108
0214980
0248749
0163623
0258013
0163624
0163625
0239466
0251260
0252062
0180769
0286189
0201901
0248725
0138639

0266421
0109339
0250062
0252757
0266515
0163629
0145283
0243312
0163633
0250202
0285458
0251411
0284968
0172493
0235043
0145332
0248196
0248180
0250572
0263981
0170509
0255723
0198189
0240966
0249262
0170502
0272856
0152583
0249001
0152591
0152592
0200024
0249973
0029559
0152595
0183199
0286618
0118785
0202000
0118762
0118777
0241853
0201707
0221440
0251285
0199326
0163644
0200469
0246375
0207480

0287181
0286978
0138642
0138646
0249934
0145337
0255072
0287542
0285122
0249755
0252322
0138641
0207524
0177432
0248019
0278151
0276003
0138640
0180346
0270720
0212226
0185477
0251095
0145335
0274238
0277695
0276542
0247775
0138722
0287239
0184305
0252087
0251401
0187653
0248984
0249049
0279013
0249891
0249052
0248817
0287392
0280056
0249152
0248511
0152208
0251449
0249239
0248627
0248750
0250908

0252342
0172238
0246541
0250897
0163104
0163106
0287552
0241103
0163110
0249951
0249599
0138696
0182168
0271474
0242936
0163114
0200622
0201744
0250655
0248510
0201640
0287841
0236764
0253170
0163116
0254044
0251620
0250293
0242697
0214904
0214559
0138698
0242318
0168785
0251523
0260641
0249764
0227118
0151247
0249055
0263923
0214886
0287512
0252548
0251437
0164024
0265213
0250496
0272777
0197894

0246090
0198099
0249065
0172955
0187758
0196616
0248144
0196344
0250300
0138813
0145331
0138823
0248676
0248713
0070190
0279913
0109270
0164031
0279098
0164032
0245322
0250403
0248820
0145358
0249710
0164035
0286150
0250223
0251219
0251636
0138814
0273882
0221265
0254531
0153064
0286124
0199529
0251149
0251309
0138821
0241144
0248161
0251572
0260651
0109320
0109323
0241981
0230069
0248971
0224207

0251288
0109332
0286242
0246560
0252739
0145354
0164037
0250651
0249283
0164038
0164039
0138778
0248740
0251577
0250920
0251571
0169836
0286291
0252460
0249614
0251170
0168772
0245384
0250670
0248242
0244470
0248373
0251906
0276992
0251473
0252136
0251259
0168769
0251586
0243383
0138777
0200917
0249264
0216425
0250922
0250522
0236699
0249885
0248778
0138785
0138780
0251175
0168743
0250740
0249635

0145348
0164022
0250298
0249613
0286147
0248915
0155011
0213522
0251081
0252470
0138801
0164023
0245293
0222691
0155016
0249604
0138796
0138795
0232021
0239939
0278589
0250485
0234492
0109475
0198856
0206601
0286136
0164089
0249257
0250185
0188517
0246774
0249465
0247950
0138802
0243227
0207988
0225094
0249965
0005059
0248785
0138794
0273447
0123739
0285330
0205403
0272795
0109534
0180245
0183423

0249681
0138798
0207260
0170522
0263940
0248200
0166530
0250594
0212160
0250304
0138792
0251464
0251046
0199894
0250511
0250103
0164093
0249519
0215961
0251483
0251312
0200963
0241722
0248656
0250855
0248432
0249815
0250556
0250307
0240459
0174749
0243280
0260526
0138660
0272567
0145365
0073331
0251441
0248697
0178403
0249509
0248439
0138658
0250966
0174720
0249532
0199169
0199145
0207927
0199102

0284463
0220924
0250046
0248633
0241002
0202536
0241741
0250500
0145362
0234841
0251126
0228814
0248152
0249373
0283193
0275810
0196656
0239279
0206820
0145349
0250229
0180801
0249304
0248716
0174607
0207931
0250215
0240637
0138653
0251308
0250693
0248488
0250363
0249259
0250442
0250627
0270394
0284253
0250272
0248139
0250791
0213493
0287845
0287520
0287290
0286637
0201752
0271053
0174599
0223605

224932
228358
236922
237684
213492
286269
229565
164100
251463
269893
275994
164099
230671
273716
154608
260404
223225
248213
251155
281731
145388
150961
172403
249002
172399
178636
145390
164096
145384
250412
201186
249244
283050
260091
223208
245958
250950
138735
221245
286241
250772
250183
250392
200553
253825
164109
250938
250219
225665
249409

0138738
0212359
0251609
0173376
0223006
0250344
0271302
0050730
0252183
0186867
0207148
0213480
0224062
0164111
0164112
0226757
0123737
0145386
0138686
0261501
0138741
0237868
0241037
0138688
0164113
0109471
0138684
0227145
0287692
0224786
0181004
0244083
0138685
0240775
0273007
0170917
0145375
0250322
0287951
0164056
0249125
0251526
0249464
0226655
0228489
0250458
0286700
0250484
0249837
0286734

0248506
0248565
0261083
0151458
0196159
0201297
0283550
0250149
0251436
0249555
0251649
0251005
0286251
0223620
0199862
0250945
0248491
0287101
0248209
0249795
0287021
0164066
0151475
0164070
0282855
0244021
0261668
0142731
0251821
0164073
0164074
0138709
0241651
0248394
0164040
0251432
0273077
0248187
0248802
0280711
0077684
0151466
0275647
0281369
0151470
0251345
0248958
0251195
0250193
0287748

0246876
0249618
0234125
0251326
0248877
0249018
0280059
0250503
0287924
0250213
0206782
0250102
0248715
0241392
0251676
0252014
0249416
0184139
0251555
0251598
0249463
0251051
0248369
0251398
0249038
0251488
0249513
0250241
0248735
0138650
0286467
0250191
0251388
0251199
0286253
0254535
0248191
0251291
0249000
0249847
0251228
0241411
0250144
0248434
0248456
0283432
0250834
0248104
0250865
0248330

0207322
0251492
0261166
0251567
0250665
0249311
0279749
0248869
0250341
0250813
0251440
0213448
0189184
0251632
0280352
0250126
0250777
0287441
0250034
0287144
0248307
0250033
0151012
0249091
0248397
0251372
0249381
0248305
0248795
0250977
0250501
0250195
0250577
0151014
0109381
0252503
0240723
0179967
0276396
0265892
0207384
0252362
0206722
0137463
0109390
0164134
0200520
0273247
0248863
0239005

0172007
0145391
0249302
0272717
0286896
0261268
0085871
0201533
0178458
0196782
0286320
0252233
0250698
0153130
0196951
0249754
0241923
0153132
0205301
0280262
0179387
0109424
0287449
0242102
0248621
0109436
0249661
0207155
0248335
0273472
0222613
0170153
0109445
0248747
0248810
0250189
0272632
0250141
0164136
0109452
0242134
0250835
0251248
0249806
0250326
0251300
0170185
0250969
0200049
0109458

0265623
0228981
0286771
0243175
0250540
0245112
0153147
0248924
0251600
0236296
0183090
0250345
0197465
0249741
0250361
0286055
0170180
0285783
0285713
0261129
0250504
0248890
0164161
0273621
0164162
0249193
0250406
0251452
0222594
0164163
0164164
0238713
0240992
0272727
0251333
0278943
0286420
0248745
0170365
0250582
0250902
0151611
0249245
0248356
0237136
0151612
0279481
0251687
0201314
0260303

0248809
0164167
0279845
0250673
0280250
0251010
0120519
0278438
0207182
0151615
0137473
0248764
0248346
0260090
0279376
0286371
0221369
0241281
0249004
0151617
0280219
0251254
0251688
0251298
0164168
0164169
0071205
0252951
0264274
0240014
0270387
0151623
0250354
0287292
0251418
0201846
0250357
0225176
0279016
0249317
0239168
0249752
0250252
0248210
0234828
0287565
0250905
0251482
0202331
0207121

0250704
0286711
0170390
0238721
0198589
0248521
0249690
0181541
0183432
0252451
0235606
0145425
0201264
0109686
0208797
0270681
0249012
0189099
0250350
0252544
0270265
0251611
0164142
0253019
0059691
0251455
0249184
0286761
0249708
0249818
0248991
0286066
0251249
0253077
0286720
0245954
0287504
0250706
0251377
0244544
0270302
0270883
0109670
0270751
0287555
0264678
0268471
0283792
0243417
0240669

0170006
0248571
0169989
0164144
0250980
0231841
0287642
0137460
0249022
0109654
0252030
0201786
0213406
0121211
0243883
0121210
0287216
0250771
0248208
0137462
0145428
0249309
0145423
0280241
0197410
0278981
0280005
0249200
0250609
0171566
0200350
0248611
0171564
0171560
0171557
0287770
0121207
0249074
0249041
0249692
0151962
0222582
0252604
0250910
0250538
0185149
0249924
0164114
0250205
0250398

0271606
0287730
0249247
0249770
0248748
0248548
0249127
0249698
0248420
0249386
0251223
0248907
0251407
0251244
0164116
0260244
0061918
0256394
0151790
0256043
0249013
0251511
0251283
0249479
0248912
0145431
0248629
0280284
0278880
0250622
0252350
0252524
0109738
0199859
0120251
0249819
0286133
0234111
0249275
0200521
0249627
0279703
0286862
0287226
0279994
0250475
0279139
0280140
0164125
0248429

0287822
0251073
0250604
0164124
0286558
0251429
0171509
0199634
0248387
0249901
0205208
0171503
0271817
0171497
0052795
0206703
0164123
0274535
0248988
0249367
0206978
0109756
0264105
0283379
0248552
0250180
0250488
0250997
0249425
0242635
0249568
0168843
0249419
0251253
0248396
0248431
0287072
0251123
0283468
0250027
0145414
0250547
0164128
0164129
0227283
0151005
0198498
0145416
0273449
0253001

0177803
0250746
0248546
0251162
0223037
0248162
0248771
0201050
0248329
0250227
0244146
0285563
0221891
0270960
0248645
0121089
0236941
0183439
0250486
0248229
0250887
0250712
0240674
0176979
0276623
0250430
0249617
0250374
0170088
0109466
0200974
0229894
0052802
0109472
0207559
0251596
0248287
0248327
0287424
0250725
0249500
0038295
0252959
0206711
0249675
0196104
0212373
0250150
0240772
0248913

0222721
0270336
0248601
0109511
0287188
0137628
0181381
0207171
0129116
0229226
0249623
0206613
0214676
0249609
0279384
0145439
0201176
0201818
0251445
0154447
0214671
0251171
0137601
0286302
0109572
0056050
0229204
0248625
0287660
0288003
0248319
0287298
0251200
0249955
0198948
0109576
0248822
0248739
0250266
0251061
0250082
0181359
0250968
0249943
0251961
0286580
0248872
0249525
0248755
0278502

0248339
0287210
0249297
0251176
0174473
0248637
0245213
0241652
0109586
0221296
0248774
0164104
0272870
0164105
0164106
0288025
0242882
0286226
0252898
0261646
0286039
0164107
0237125
0234801
0213370
0250708
0250043
0251216
0248174
0250957
0251213
0164117
0164118
0249875
0265846
0164120
0251584
0250596
0145451
0168594
0250200
0250431
0249411
0286494
0249284
0248551
0249945
0251174
0248574
0150625

0249106
0248249
0250372
0150627
0201516
0150628
0164122
0248980
0129128
0248480
0248388
0150630
0287544
0249458
0250263
0222859
0249008
0109674
0249084
0038002
0250131
0285574
0201931
0251416
0231171
0251504
0253000
0241877
0201727
0212191
0250839
0250993
0248921
0287083
0287775
0251025
0249883
0248197
0248770
0249460
0225356
0286860
0287948
0251336
0251742
0177822
0251433
0218336
0248266
0221227

0252343
0251714
0248816
0248694
0286119
0272646
0129187
0286821
0250636
0230219
0273180
0272784
0251359
0151718
0251128
0177300
0185758
0250292
0248376
0168564
0213434
0232648
0168556
0272744
0182552
0252157
0168538
0201433
0251739
0287349
0283383
0173320
0248206
0283355
0250726
0164303
0287433
0240159
0244512
0227765
0180712
0249096
0168310
0251878
0270426
0271646
0286292
0271538
0249056
0254233

0164305
0164306
0151725
0151726
0249138
0251139
0286256
0251230
0266698
0249173
0251147
0263458
0279460
0248517
0187821
0250754
0249520
0287183
0151729
0164323
0109762
0109771
0250410
0186352
0109775
0207231
0205129
0168491
0249679
0154553
0154556
0233110
0235902
0207497
0285745
0235149
0222727
0164342
0109794
0253013
0250327
0251008
0279943
0145476
0164344
0088926
0251165
0287667
0213332
0272297

)212387
)168412
)083857
)287169
)250829
)272218
)249539
)249742
)250971
)287571
)249747
)250658
)250042
)250620
)251643
)251490
)250590
)249642
)249162
)286641
)213331
)179059
)179046
)241011
)201085
)184108
)247130
)248370
)242169
)248880
)249378
)250523
)275826
)252275
)180015
)278974
)249269
)251619
)250626
)250024
)251310
)249877
)205100
)250739
)272566
)202215
)245685
)250666
)109536
)249462

271307
251297
199572
201145
224807
205097
250053
214273
249003
281591
277162
281720
281652
281058
280799
258834
280757
281627
280457
260596
261914
282994
250020
153404
199540
185028
261434
164366
260774
073578
286001
248925
249915
286169
250848
063438
214278
250645
221990
180104
188242
066230
225138
250385
264233
249650
112877
171368
271781
206077

0286094
0249908
0285522
0188818
0247872
0028310
0071539
0287600
0248949
0250173
0215246
0272347
0145506
0113504
0263834
0249201
0174358
0164363
0287486
0164362
0263670
0049656
0286388
0250584
0248951
0279908
0142319
0153395
0273742
0251532
0185986
0271119
0248126
0188002
0232615
0263746
0260066
0171421
0145494
0249966
0250417
0279222
0113430
0249116
0249326
0288108
0248994
0248597
0249731
0201026

275733
281560
259757
170561
186493
249541
248137
248736
249808
248118
249830
250716
170549
259603
248962
249994
250921
260763
248973
231291
251371
249521
248311
249941
251506
250481
215231
251426
250529
223007
250579
145536
250866
251568
286753
248729
250808
251476
164151
286986
261037
259861
278913
250730
250490
133398
287031
215218
271230
250056

0037474
0145545
0248677
0112941
0251365
0272071
0283420
0250974
0200243
0250060
0212258
0287790
0286254
0237611
0285689
0248775
0251250
0205976
0251678
0250668
0249638
0249865
0223597
0078295
0250761
0215217
0124275
0124279
0251168
0199773
0287947
0280338
0280251
0249486
0288002
0245729
0249159
0251549
0272049
0247516
0250037
0279075
0284000
0249782
0279097
0279264
0250389
0248765
0279002
0280373

0279125
0280438
0278877
0260981
0280185
0250198
0279635
0250619
0112902
0266415
0248537
0279698
0251370
0250001
0250786
0239112
0248525
0169777
0249781
0222054
0286556
0250814
0279011
0279755
0278874
0280016
0249807
0280449
0280368
0260515
0271998
0272417
0150756
0248968
0150753
0271980
0271715
0164237
0252341
0252671
0280369
0259802
0145495
0250600
0145491
0277073
0246016
0244361
0249396
0249166

0249160
0280017
0279450
0164236
0250106
0251196
0251548
0279979
0278865
0112977
0272324
0250576
0169862
0207312
0212305
0248783
0248131
0251112
0249053
0248457
0248145
0242608
0248462
0243744
0039139
0251423
0038382
0287581
0145569
0248791
0250526
0250182
0261360
0154124
0154122
0286970
0249485
0264792
0272057
0249928
0250662
0250247
0286441
0202269
0249332
0248834
0286377
0248699
0183580
0250250

0216077
0222312
0183654
0250981
0250448
0248403
0248150
0249737
0173545
0154153
0246214
0250415
0201370
0145555
0253093
0243538
0201436
0176788
0215196
0271892
0206814
0251990
0250816
0252908
0185296
0201715
0223361
0248223
0249199
0250822
0248455
0249662
0268799
0250667
0284373
0248205
0249620
0284439
0284283
0284234
0284042
0284465
0250088
0283967
0283988
0284356
0248422
0249282
0283740
0249156

0271296
0250351
0250386
0283776
0250782
0248471
0248160
0248542
0248337
0269466
0274832
0250715
0249937
0240785
0212278
0241552
0248693
0249191
0249927
0249285
0276764
0249174
0251487
0242654
0249557
0145526
0248766
0214132
0249854
0249490
0251629
0248308
0286751
0248286
0249359
0183666
0233974
0279107
0272130
0253766
0154162
0213430
0253104
0168967
0287471
0249206
0286961
0249404
0250319
0206912

250332
249289
164256
248591
249607
248874
252083
234865
200600
286134
226432
250694
277455
287199
040731
251294
249562
286454
250099
286238
248156
223004
248908
250524
199911
251273
245662
285042
251654
260761
279671
248605
279531
201016
279869
251593
286625
222560
249099
286204
230571
248426
250703
251033
249616
113100
207261
250337
248490
250079

0249785
0250453
0286432
0252867
0270495
0259786
0271035
0162685
0226400
0277878
0249226
0252601
0248687
0270430
0286182
0248391
0276098
0248418
0250984
0249744
0233787
0223136
0280305
0250587
0250118
0287176
0287940
0241668
0253940
0254138
0113361
0250482
0113360
0206682
0082213
0279923
0199765
0133401
0248624
0248878
0249114
0248813
0248378
0250164
0249688
0252373
0224282
0251887
0241739
0243914

0214108
0249774
0266243
0250764
0113384
0200065
0272086
0230829
0150712
0199731
0207052
0056097
0207956
0249142
0113387
0250885
0286741
0113389
0251062
0250697
0248279
0249102
0251281
0250672
0240376
0251443
0249572
0113407
0249360
0250583
0151388
0201623
0182631
0164175
0250914
0242110
0273294
0250767
0082196
0278900
0279995
0271771
0215158
0215156
0286543
0249380
0271874
0039560
0250234
0272323

205838
250140
241829
113456
113460
168724
113492
113494
250886
201368
152582
238441
248969
168685
279737
248349
152611
249748
145626
168671
251591
164187
207756
145604
201223
152620
245711
164188
287634
222178
242292
079215
274441
250155
279400
272103
285967
164190
249850
244073
197603
271378
228212
242493
286193
270558
113569
251880
250455
082068

0248216
0222743
0206743
0248587
0168621
0248539
0250032
0251518
0248533
0248461
0276332
0164318
0248730
0249071
0248572
0249491
0248234
0251257
0113594
0244968
0265304
0251189
0249740
0248577
0250629
0145623
0164327
0249089
0286809
0082074
0251215
0113600
0153071
0271334
0251515
0250492
0248651
0250048
0249668
0250585
0199361
0287597
0283286
0285552
0285616
0113638
0199552
0171522
0132356
0212567

0145592
0212296
0132357
0112936
0253098
0248120
0171495
0039537
0182836
0251478
0083720
0286164
0248668
0240281
0205765
0151876
0239694
0199487
0249266
0260786
0112964
0248873
0250860
0270513
0198865
0250722
0272234
0286271
0249092
0251163
0251656
0249516
0250711
0287263
0286656
0271788
0177738
0251131
0272382
0177721
0215068
0272144
0172262
0278487
0177453
0249779
0249888
0247763
0112972
0261604

188850
151882
151881
249492
172244
249855
248554
248240
172239
248092
279557
249286
112992
224993
249276
212561
070193
248464
249203
250418
263556
251141
248779
112996
272335
248148
287540
164588
250422
250017
250122
170571
271455
213956
271752
151883
249405
270830
251601
250360
251573
248918
207060
278734
206606
240627
259663
016082
249338
248891

0288035
0251125
0251873
0249270
0279470
0248789
0244621
0250827
0248898
0213949
0152684
0251237
0241809
0249899
0164171
0286048
0272123
0164172
0247796
0213940
0285831
0240052
0134363
0164258
0250447
0217791
0272416
0185305
0287087
0222960
0207627
0241230
0265421
0279638
0249069
0251597
0169271
0178996
0280159
0287367
0232517
0226810
0249843
0249016
0164283
0240535
0113088
0207229
0251376
0248029

249454
145649
164287
164294
198983
207728
251856
234602
152669
251307
067248
039123
271410
067113
279883
265135
250853
249410
177058
152670
251368
199322
248854
164509
134352
262211
264281
249236
227908
238326
164512
253032
243592
223003
251435
212265
248781
210678
249697
250787
248727
285997
226221
225940
234553
228650
095015
237705
225230
155542

0155545
0271828
0235635
0233943
0248652
0250480
0207293
0062194
0248586
0250961
0250308
0169067
0249436
0287709
0249198
0249584
0248132
0286853
0248271
0145632
0175857
0271257
0265699
0250127
0152932
0248475
0240412
0248733
0113448
0247345
0276214
0251306
0202601
0248935
0202017
0213896
0214244
0273701
0152931
0275634
0276233
0035499
0251303
0287434
0250148
0164181
0272308
0049167
0215032
0233847

0164182
0279985
0276945
0188725
0251279
0249279
0130449
0251682
0250461
0213891
0178722
0248529
0212215
0251575
0249878
0248199
0251983
0068796
0086189
0086200
0199279
0268942
0241991
0240436
0186105
0217416
0249579
0248185
0287179
0248285
0178394
0164197
0236068
0186479
0242547
0201869
0224296
0145642
0153006
0153015
0272370
0272354
0207439
0049192
0248288
0250732
0205644
0250081
0123219
0113593

0113595
0207352
0253251
0113597
0197860
0123213
0251648
0269961
0112851
0252904
0286314
0285999
0153914
0253744
0248846
0251391
0286062
0250313
0234838
0069020
0248803
0249057
0229666
0134061
0251206
0286647
0249364
0250978
0239870
0249736
0249238
0250669
0249721
0250421
0249894
0273860
0253801
0252108
0213864
0253787
0145675
0251044
0248884
0248359
0249588
0249335
0250289
0212249
0250066
0276309

0249352
0250237
0249183
0145740
0248664
0220986
0280894
0134057
0280187
0153044
0134056
0134058
0183323
0213830
0215006
0271045
0085231
0273841
0152942
0152939
0244717
0249295
0266477
0197822
0212260
0183474
0248477
0250138
0253203
0248769
0198237
0251158
0249230
0205572
0205571
0285151
0179978
0253492
0254701
0253816
0226259
0269983
0250687
0253333
0196302
0253366
0288349
0254335
0172058
0172062

0285204
0249437
0145736
0230847
0251634
0253900
0215630
0235558
0250387
0253985
0169040
0249981
0145734
0253536
0247911
0131844
0164326
0278824
0285804
0131711
0264099
0113048
0244748
0049883
0244061
0251108
0261269
0178175
0248371
0251613
0226544
0223497
0206770
0249085
0083312
0263593
0242602
0271926
0272081
0157107
0251467
0248930
0157111
0251599
0250747
0164325
0287090
0249743
0251543
0249149

0251493
0247993
0251604
0251324
0272525
0145741
0255883
0164331
0164338
0214944
0249293
0184084
0238913
0259968
0248474
0248673
0248942
0244326
0249343
0250446
0171617
0049860
0164347
0207336
0164346
0198780
0199645
0250071
0271714
0212363
0287701
0249856
0196390
0239517
0182383
0249157
0176928
0145700
0251663
0247372
0113161
0113163
0271815
0242198
0122008
0251724
0248881
0272040
0189045
0152359

207333
251078
251419
249301
248684
228367
249777
251342
248127
122012
252833
251668
254893
249014
251107
250348
251235
145703
250574
249713
164220
225407
277188
278237
181104
164251
264391
171643
252684
145708
164252
285000
132846
250615
238961
250802
284762
251566
113231
251400
164253
244019
244363
171540
171530
213763
253572
287162
132842
253558

0249372
0251605
0200331
0206714
0245556
0085365
0145685
0248909
0254170
0254310
0113273
0132837
0214890
0132840
0145692
0201955
0249829
0206592
0250330
0152409
0213757
0152413
0213755
0243160
0207391
0277818
0164329
0164309
0250258
0286818
0238254
0250888
0177034
0113296
0251050
0249778
0249825
0164300
0251675
0249438
0184188
0251828
0244076
0164299
0227087
0248569
0220157
0206774
0039319
0249042

0152380
0242706
0240003
0189127
0251366
0251221
0228716
0249655
0271043
0113318
0248967
0244630
0251450
0113319
0251001
0249772
0247572
0131730
0248794
0131732
0172497
0286721
0145687
0287938
0249483
0214857
0152348
0236447
0251495
0248192
0265684
0251374
0186468
0205464
0248393
0248870
0249100
0240411
0248105
0283359
0281327
0248112
0249857
0248400
0249664
0271862
0174695
0238835
0152422
0243385

0251552
0251395
0038427
0249835
0145681
0253073
0206744
0242858
0202157
0250013
0164176
0252861
0250320
0214853
0249349
0248674
0250253
0249792
0248170
0248667
0249023
0249842
0243859
0251066
0250874
0241965
0249153
0250359
0205449
0280009
0248195
0248363
0250124
0251585
0127184
0284078
0207013
0250544
0248701
0242477
0249061
0233828
0251951
0265919
0145715
0241243
0207452
0134480
0285190
0200462

0249362
0248708
0287783
0287862
0164180
0247828
0207129
0240083
0206958
0250306
0250156
0271904
0245526
0250705
0284447
0245864
0250377
0250555
0081189
0248309
0250831
0250437
0214942
0264342
0248555
0253321
0260871
0153140
0255647
0176055
0113356
0176018
0164199
0249514
0248323
0285618
0212930
0251093
0199643
0234292
0286638
0277001
0240388
0241059
0113369
0281357
0251356
0286121
0248826
0276426

0250049
0249776
0249169
0249984
0248588
0248950
0249958
0248528
0251361
0286509
0213730
0237187
0175745
0278905
0113391
0284336
0287180
0249353
0248483
0272406
0287447
0251023
0286577
0185261
0251544
0254132
0251340
0232578
0270133
0133302
0175471
0249175
0243806
0276514
0249545
0153347
0225165
0198677
0212259
0164291
0178015
0175449
0145757
0250240
0164292
0173221
0251348
0251000
0251409
0236882

0272021
0250362
0118985
0251314
0250955
0213716
0250551
0207578
0206997
0249746
0250158
0175426
0153113
0249180
0164307
0248734
0272109
0247121
0164308
0113441
0227836
0200884
0213701
0251513
0145721
0058729
0206698
0286828
0248758
0251054
0249761
0251389
0249977
0250331
0248931
0251193
0286953
0248202
0249101
0248901
0249135
0238000
0176183
0250221
0250830
0249990
0249448
0251102
0251553
0174136

0246763
0279232
0250162
0153922
0222610
0248489
0200351
0241597
0249444
0206356
0227762
0249515
0248236
0248928
0250391
0249963
0199378
0270232
0270230
0271687
0270906
0270350
0271480
0270388
0270307
0270225
0271207
0274455
0207077
0275374
0273957
0249787
0247877
0174132
0250806
0207269
0113532
0221263
0277506
0251261
0249495
0199786
0248261
0173930
0222987
0205359
0249017
0250958
0250682
0247402

0286338
0145730
0175749
0145723
0145725
0248203
0181751
0250567
0226926
0248757
0112874
0283462
0251026
0201910
0239808
0251574
0278958
0253776
0252881
0253584
0232159
0201790
0251204
0250145
0252337
0251027
0250273
0184349
0249959
0248827
0239708
0251732
0145743
0286503
0244245
0272523
0151422
0249921
0250383
0207404
0250441
0206593
0176857
0286882
0249476
0198961
0248867
0251367
0248876
0248440

0214794
0271849
0112893
0202512
0249068
0254106
0253224
0186952
0221436
0207177
0164209
0270779
0248966
0248428
0251627
0145777
0253613
0134987
0250728
0214784
0152495
0249318
0248268
0164211
0246859
0249619
0251099
0134986
0253057
0249439
0250095
0129595
0224032
0238363
0250882
0251187
0251076
0248350
0278921
0248430
0251014
0250358
0134982
0250045
0212370
0258864
0153037
0129625
0249947
0212643

0172795
0171444
0232633
0212122
0251741
0047188
0286839
0080709
0222706
0246316
0251542
0251132
0250242
0152503
0250472
0249791
0164219
0271797
0243304
0164221
0250197
0249026
0248722
0249944
0249433
0250444
0145780
0251201
0243414
0249249
0134970
0249021
0248709
0251118
0222598
0129596
0145782
0177879
0287148
0172901
0248295
0268223
0185641
0272265
0271918
0145781
0212457
0232916
0250015
0092421

)248445
)239528
)249167
)197575
)251311
)249150
)250650
)250949
)243758
)285809
)248663
)251477
)249797
)249582
)250891
)250427
)249551
)286745
)249128
)238904
)249426
)169570
)197744
)283498
)249494
)223179
)250678
)250928
)172869
)248885
)248979
)206786
)264536
)145779
)243333
)202092
)133835
)213663
)249919
)164334
)250895
)234259
)251293
)251975
)261036
)184838
)270741
)222609
)248927
)251135

0248853
0229855
0250847
0279860
0244088
0213661
0249600
0181867
0151304
0113083
0247311
0164185
0250803
0249610
0272139
0064692
0249621
0249916
0287794
0250328
0251538
0240058
0205302
0260686
0064652
0263432
0168938
0249996
0276047
0223652
0061455
0213655
0168944
0250539
0249400
0279772
0151292
0235275
0253807
0250862
0251421
0248600
0251214
0250438
0248296
0168916
0249112
0251456
0242814
0260192

248443
249261
288008
250194
250721
248752
222107
250402
286111
249950
249904
241888
248107
155324
250602
279118
164904
202082
164902
230929
196900
213630
249768
251072
252185
113368
173926
164241
279446
214743
250080
271585
250956
283897
286615
145794
249577
271766
250859
164244
282925
205279
248799
230561
251032
250603
245937
064651
250329
138829

0207290
0113396
0248634
0066583
0264563
0286274
0265691
0249421
0145808
0251680
0186367
0198108
0221562
0199455
0248610
0252514
0249312
0249418
0287390
0244192
0250405
0248955
0169567
0186687
0158985
0279370
0158987
0273217
0275939
0217128
0286676
0279584
0239642
0281938
0234758
0164398
0223548
0231585
0164399
0164400
0253067
0227616
0235775
0224015
0237714
0072682
0224431
0283678
0131435
0197208

0233006
0263597
0197375
0197536
0283782
0238160
0202533
0234290
0125347
0113525
0113522
0223442
0169194
0113520
0230612
0131437
0205089
0164402
0198944
0272203
0164403
0201274
0164404
0164405
0164406
0072364
0235253
0202417
0225364
0272023
0229738
0199677
0155329
0248648
0287054
0286408
0170606
0279691
0251616
0248245
0240611
0053108
0249478
0250244
0250409
0221287
0280104
0279197
0251243
0249073

0113583
0213585
0271737
0271128
0081059
0113558
0272772
0113575
0273345
0266751
0270177
0239615
0006837
0119048
0279469
0248559
0237190
0243648
0250564
0250994
0207222
0251169
0240250
0280420
0043143
0152700
0207459
0113615
0201298
0222266
0164615
0145833
0241179
0181904
0270892
0113621
0132570
0279799
0244921
0249119
0248923
0247627
0198868
0263963
0249192
0152705
0069011
0224186
0277619
0249082

0248753
0248482
0249647
0113648
0270021
0251380
0255833
0249639
0181965
0250167
0145824
0145832
0251045
0223539
0265924
0250378
0145839
0254239
0253927
0145826
0164616
0120708
0270123
0278815
0164621
0277859
0113658
0275646
0277524
0274934
0271824
0069018
0248211
0250947
0279240
0248351
0213568
0249803
0250284
0248616
0222285
0152377
0146021
0216009
0177733
0287067
0270697
0226306
0250159
0120729

0078795
0031003
0246323
0250260
0222924
0061492
0112981
0212460
0199880
0112983
0112984
0094880
0146013
0249851
0158402
0120709
0249971
0120733
0132563
0271196
0120738
0241458
0120705
0113013
0222937
0206989
0044115
0232174
0238745
0253404
0146006
0120725
0201532
0285956
0249593
0242683
0252533
0280987
0281398
0252213
0200959
0015479
0199545
0253015
0120727
0272742
0170482
0170476
0228672
0170469

0170464
0279686
0200378
0249751
0283288
0184584
0249689
0249215
0286403
0131508
0207020
0275506
0272255
0171604
0250635
0249526
0200756
0249131
0251387
0271792
0146005
0158458
0250692
0207317
0245146
0185129
0182700
0254363
0120306
0250069
0113068
0113070
0253965
0113073
0249637
0222790
0131503
0254996
0200051
0213523
0243056
0113108
0283155
0283717
0176087
0283602
0200235
0251453
0240518
0170458

0131495
0113119
0113141
0284325
0120314
0256453
0170445
0112855
0146007
0199990
0202111
0202515
0250515
0248106
0279726
0204970
0204969
0255408
0204967
0204965
0081842
0204963
0204962
0204961
0278907
0250120
0249504
0278946
0249158
0251664
0278915
0278901
0239389
0248383
0279028
0243232
0279047
0171815
0279558
0112852
0272154
0113205
0272108
0081818
0113209
0280029
0113211
0255622
0280336
0113212

0120322
0279472
0272674
0279068
0278936
0177839
0279375
0120324
0197479
0120328
0187372
0120327
0279983
0146001
0262096
0113248
0277795
0178913
0120329
0288095
0272070
0204956
0081853
0254245
0254221
0280026
0262576
0253910
0253485
0262209
0253731
0253537
0253953
0253767
0276547
0261934
0253305
0253846
0254122
0253873
0248449
0279855
0253159
0276545
0273557
0240184
0242020
0242419
0240764
0131504

)246422
)240974
)228737
)171720
)164620
)197948
)120318
)286736
)287527
)278925
)156453
)287726
)226040
)081791
)113555
)271871
)013561
)254099
)113552
)250347
)131507
)280047
)187678
)277968
)274721
)231185
)225147
)113578
)261757
)236714
)287203
)145819
)226272
)285366
)230789
)113580
)231579
)279679
)279130
)249429
)251205
)253023
)249881
)266478
)225869
)158497
)239390
)145817
)183775
)249605

285605
250972
213488
201423
251031
249229
215325
250842
248125
186314
249363
204928
156463
248493
173261
251556
133706
239602
239317
275740
091009
238374
250025
091010
271724
248842
113649
277568
173250
250407
156475
280646
241291
248568
249553
286468
250343
169302
113657
272239
280780
176049
287630
164266
249518
164265
178776
248362
251320
250346

0133710
0247199
0196800
0178172
0249503
0248109
0214510
0249834
0145879
0204909
0283653
0145868
0274799
0251330
0164270
0253297
0169252
0169247
0272411
0251842
0207714
0250072
0200161
0173210
0248647
0253406
0157510
0285736
0164284
0253618
0145882
0127743
0253865
0249669
0275871
0284182
0276365
0113712
0230551
0241112
0183111
0222213
0287323
0221043
0242928
0155846
0199047
0132915
0251583
0200334

0155850
0164296
0113716
0248696
0241907
0182578
0214485
0199409
0113721
0113722
0011083
0070808
0183876
0070814
0019582
0286331
0164587
0254333
0070614
0171992
0253852
0286657
0164591
0250309
0086589
0132912
0256235
0237693
0145908
0197083
0274235
0277866
0211445
0145901
0197043
0254298
0285420
0198624
0196743
0186334
0186335
0270904
0253897
0253472
0271494
0123643
0200227
0086570
0271795
0276622

0288081
0260581
0113140
0249035
0222102
0272112
0177556
0253921
0213433
0275765
0145907
0145888
0270978
0222308
0254226
0286749
0132911
0242080
0249484
0155511
0242976
0251183
0261382
0055147
0037749
0164574
0245275
0253119
0287695
0221430
0241963
0164576
0113196
0254293
0253886
0253146
0264760
0221552
0274472
0155506
0272197
0241187
0170271
0277039
0263361
0155508
0082516
0082515
0253261
0226650

0287963
0253646
0271477
0270442
0200275
0170624
0286690
0253370
0254163
0199468
0231989
0145850
0270560
0113249
0135077
0254246
0155868
0113263
0170613
0253980
0253653
0055163
0285868
0172568
0251405
0248544
0135074
0253155
0172548
0253519
0252068
0254336
0039600
0187658
0253798
0113272
0155858
0253630
0113282
0206819
0244331
0222626
0253743
0213414
0253422
0253792
0253673
0254047
0253449
0253424

0253134
0276778
0254350
0272085
0254135
0164330
0253811
0253456
0245812
0279204
0145860
0253256
0253315
0252458
0164332
0249738
0113302
0221601
0253298
0283413
0253311
0170214
0248626
0199398
0113312
0223908
0170234
0170231
0247699
0253772
0135083
0254373
0270612
0145861
0221886
0164609
0164611
0253522
0265237
0283733
0118322
0253687
0254391
0253417
0145864
0145863
0253403
0118990
0022355
0280776

0113327
0254186
0201474
0254351
0249449
0250114
0113328
0170584
0072571
0251018
0038274
0285590
0224012
0251998
0273569
0271679
0253331
0253538
0241956
0253236
0253600
0206845
0285549
0254066
0253693
0252794
0253512
0253946
0254252
0254171
0229119
0254130
0253713
0253469
0145934
0254365
0254297
0253527
0254187
0253357
0253660
0253947
0253065
0253925
0279739
0253978
0113645
0113643
0188573
0120137

)253861
)199035
)283612
)233913
)254192
)184347
)254042
)207739
)202345
)248965
)279752
)248222
)207619
)206614
)040275
)134516
)253269
)204767
)263831
)168269
)253219
)249601
)234511
)043462
)235172
)250274
)182132
)145936
)253647
)253858
)253591
)253348
)094755
)204764
)244501
)253966
)253213
)275038
)164438
)213393
)265740
)254256
)206909
)254012
)264249
)181163
)156427
)285914
)253110
)185662

0072803
0072786
0214360
0168246
0212529
0253218
0174705
0287814
0254299
0280318
0214357
0253683
0265160
0253445
0253295
0213386
0253736
0120129
0206741
0270243
0253628
0113719
0204758
0254295
0037241
0253785
0113732
0254328
0212402
0207210
0164463
0253439
0253172
0113734
0254200
0183072
0201277
0252169
0113739
0274994
0199219
0253968
0253141
0253768
0254164
0253955
0145919
0253959
0253428
0253686

0287003
0254211
0113742
0185056
0253109
0170091
0253244
0253447
0254011
0249722
0249306
0213376
0249031
0250256
0248929
0120149
0251670
0266890
0251144
0204754
0251122
0248293
0249403
0248355
0184845
0286350
0164466
0270839
0222111
0113749
0200648
0145920
0280068
0250820
0051596
0250801
0272494
0250992
0248469
0182230
0285476
0251623
0251458
0283559
0248596
0287579
0283235
0270989
0170085
0251667

0122203
0250909
0251414
0175414
0221464
0048162
0146066
0175416
0280139
0113194
0146083
0074276
0243959
0207420
0169258
0074317
0266329
0175766
0048140
0248484
0251446
0248859
0113763
0160883
0087206
0113761
0251666
0160867
0165671
0286634
0240729
0270706
0169228
0213347
0169230
0169223
0239393
0169220
0131183
0196570
0131187
0198055
0246334
0131188
0113758
0279821
0196923
0248996
0146094
0183258

0146067
0272459
0184840
0027847
0247679
0246596
0249849
0170074
0249109
0248943
0170089
0214351
0249186
0250101
0249684
0280207
0249129
0175325
0204677
0218227
0145911
0145916
0145912
0251211
0244234
0197451
0175309
0050767
0271681
0253698
0242341
0245688
0113240
0252464
0286713
0285891
0285978
0169131
0250420
0274813
0253725
0278441
0178338
0206624
0198939
0213328
0253869
0272431
0178187
0254035

0113262
0254158
0234284
0177932
0253144
0087116
0253979
0253163
0253617
0250674
0274716
0253652
0251545
0253520
0176783
0249140
0244945
0169045
0273549
0204661
0228259
0250999
0204659
0127022
0225051
0161021
0213316
0161013
0221394
0161011
0161010
0222448
0252202
0245317
0197226
0113269
0242979
0249876
0249412
0285865
0198995
0278965
0253480
0146090
0229721
0050748
0253908
0248367
0131459
0250509

0200555
0113300
0250749
0161055
0037280
0251352
0248717
0236791
0199892
0174339
0131446
0280161
0278970
0245060
0286128
0196670
0113303
0222852
0243664
0249287
0248761
0231228
0168903
0206686
0165810
0277912
0241870
0214295
0248560
0185372
0182613
0248473
0250222
0146054
0250900
0248514
0286644
0146063
0264732
0247049
0204628
0272296
0264549
0233937
0183718
0248275
0286310
0248103
0274525
0222533

0238035
0274553
0250765
0230178
0251035
0213285
0271530
0220212
0217929
0218577
0263667
0285578
0205653
0287265
0112679
0137265
0286364
0112685
0230433
0188996
0219712
0271727
0272463
0287293
0271911
0272485
0276687
0286785
0270937
0176515
0286324
0164379
0281809
0261730
0137273
0275859
0243439
0218027
0272279
0280916
0054598
0112699
0285603
0250903
0236086
0286512
0272465
0287903
0280423
0228718

)164385
)145949
)124535
)021355
)230438
)266750
)170542
)232999
)228365
)124570
)244041
)124588
)224846
)271361
)274855
)232654
)217707
)229282
)137275
)272277
)222248
)137274
)228170
)137267
)216819
)230269
)137285
)180822
)137266
)272320
)228793
)270477
)168994
)270504
)219992
)238158
)233068
)145945
)260604
)218472
)217135
)217566
)216657
)279926
)272248
)230648
)112739
)145975
)185689
)198721

0254821
0234817
0237716
0216307
0285424
0231811
0287761
0201185
0285694
0260239
0270695
0260673
0205444
0217385
0153046
0284823
0236336
0217030
0124787
0272142
0271978
0220685
0219607
0214113
0265083
0145982
0220446
0269985
0218574
0270174
0217239
0216360
0233064
0239472
0215057
0124785
0124491
0263572
0283616
0216863
0241216
0285717
0218868
0112799
0261211
0279368
0226281
0287497
0288046
0224532

0219986
0223342
0240936
0203498
0124782
0225092
0201483
0287002
0285771
0124783
0238221
0164304
0219375
0220472
0230570
0124784
0233503
0219993
0234115
0261189
0096696
0168566
0236512
0153162
0239264
0259040
0217746
0219294
0188428
0265818
0124802
0232234
0285216
0234763
0124786
0285219
0276019
0225775
0181355
0216368
0229765
0221583
0216781
0137203
0285278
0229950
0228478
0285759
0183674
0229401

0263705
0238076
0231056
0111846
0237685
0216359
0217181
0205318
0285763
0137434
0201048
0111845
0111843
0272097
0137210
0272162
0222383
0111837
0124827
0235051
0153157
0197977
0274541
0230314
0224531
0244476
0271245
0247925
0111859
0287920
0242753
0233656
0202313
0218020
0205269
0271897
0277473
0207419
0111863
0287593
0234427
0215124
0225102
0229896
0095951
0078401
0218893
0223321
0212802
0234015

0112137
0206702
0215022
0145979
0272379
0145990
0237786
0218265
0202351
0124523
0261071
0225921
0010017
0286316
0050393
0272209
0180537
0220868
0233981
0112149
0285639
0238987
0286281
0230631
0226673
0237346
0234540
0286277
0229646
0206960
0234261
0216754
0242989
0271888
0008083
0235488
0201367
0201519
0277411
0047579
0287626
0214027
0232947
0274212
0217585
0007944
0263712
0218073
0217078
0216364

0251831
0207193
0137198
0282024
0124788
0229931
0272341
0287347
0286631
0287359
0230873
0287559
0112183
0219314
0112186
0184612
0218359
0286885
0137414
0124789
0206881
0272269
0201415
0201442
0137177
0220920
0187566
0137364
0165097
0124795
0199715
0278102
0200681
0217414
0218976
0137393
0283408
0207775
0217379
0231662
0228412
0218643
0201523
0214012
0217314
0200957
0172201
0287138
0229700
0172197

0227803
0287614
0286590
0287483
0112242
0240869
0224707
0222431
0201683
0145996
0225832
0233848
0252046
0287404
0227089
0286368
0280989
0231754
0283480
0124766
0220771
0272168
0280443
0222515
0260455
0219404
0172179
0112273
0228772
0286954
0233358
0218476
0207394
0235743
0219453
0278128
0218512
0220748
0219582
0152954
0251830
0146038
0146049
0202050
0124532
0112293
0112294
0137261
0218186
0111802

0112304
0272345
0112308
0272402
0224164
0218806
0112312
0260286
0282804
0111913
0217083
0214975
0286633
0220517
0244618
0229313
0223623
0168405
0285801
0219682
0262400
0217805
0219681
0271108
0216718
0213972
0079691
0207286
0207490
0286933
0079689
0217408
0164508
0146047
0178762
0216436
0146039
0124568
0124564
0242387
0277410
0112337
0112343
0272558
0272462
0272810
0224447
0124610
0275714
0278637

0278705
0286522
0278463
0276410
0234816
0287080
0187837
0010704
0197061
0187475
0180596
0180573
0168298
0158373
0283064
0217159
0274290
0277157
0216331
0282988
0197409
0196866
0277224
0217275
0276966
0273802
0277075
0277720
0274750
0218281
0124575
0274618
0275663
0277775
0275713
0273983
0218690
0278588
0158406
0220875
0287050
0186470
0124508
0026950
0124549
0111801
0112763
0284688
0124557
0199289

0228223
0182952
0284607
0146109
0275846
0271071
0181315
0285571
0261584
0241549
0243307
0224843
0272312
0287966
0220581
0216762
0219770
0222800
0272468
0124635
0196787
0276180
0197903
0274997
0265565
0174572
0112812
0270935
0158553
0220758
0204789
0124613
0271755
0237154
0096654
0285849
0286652
0285703
0278332
0270666
0218347
0219738
0206671
0216676
0287674
0281706
0216915
0287252
0218069
0217862

0185130
0196747
0278828
0276368
0273703
0197238
0273542
0233822
0275221
0217646
0276903
0184357
0275379
0275126
0197153
0278677
0274641
0238610
0168131
0168126
0124657
0214374
0216629
0217539
0269293
0199851
0217315
0187763
0182477
0220721
0272009
0197279
0219891
0219392
0196812
0261839
0198315
0226314
0280107
0216901
0218016
0176933
0137185
0187626
0189134
0197062
0276302
0137338
0273712
0270326

0235109
0189298
0158691
0251877
0187987
0231162
0219262
0287804
0286819
0198704
0224586
0232040
0246350
0271440
0287279
0235570
0237425
0235559
0225173
0225595
0213916
0233366
0224157
0204713
0212240
0204709
0228666
0263426
0227214
0244349
0197935
0223677
0277661
0204704
0236909
0228364
0204703
0204702
0204701
0227206
0203492
0284656
0204700
0224233
0230598
0197171
0213911
0204697
0242524
0204695

)230056
)243729
)112462
)280236
)251608
)204694
)206474
)277881
)204688
)225797
)230164
)204687
)213900
)229274
)224582
)229281
)237988
)213886
)204681
)232173
)227609
)201330
)231301
)204657
)235238
)204655
)204644
)233916
)204642
)225864
)214922
)239257
)273340
)227758
)199290
)235821
)272236
)285761
)270896
)181126
)176998
)261548
)213880
)237042
)204632
)229142
)285799
)281831
)230521
)206341

0231130
0233677
0235963
0227262
0230795
0228078
0206503
0227766
0235290
0229390
0204625
0238024
0224312
0204623
0204622
0237669
0232757
0278104
0275856
0066379
0278773
0204619
0204618
0204616
0231226
0252228
0204614
0204613
0204610
0234127
0233892
0270604
0236475
0243753
0280128
0231074
0204599
0248167
0241370
0224372
0224486
0236405
0226577
0286301
0229068
0224936
0236603
0204592
0235781
0204590

0204576
0204574
0216101
0204569
0204568
0137343
0228415
0204564
0204560
0146112
0137404
0230449
0137337
0224328
0196230
0272540
0137312
0201988
0272273
0137331
0228022
0263608
0214894
0237923
0202241
0237775
0204580
0284370
0213780
0288473
0137411
0196260
0252761
0168631
0233529
0275906
0204544
0261272
0228789
0222895
0204542
0204540
0204539
0204538
0238211
0204536
0137310
0204531
0204528
0272501

206344
271821
255726
255899
204525
234745
214892
227939
231402
256166
229836
277402
271581
228432
285647
201658
230994
223702
225851
272221
204520
206337
199332
230174
233902
286940
204516
201680
256851
219797
225499
204511
254870
198563
201785
265236
234006
213760
204498
226979
232810
227507
204482
204475
230622
204472
204469
200816
274494
204463

0204444
0204439
0227198
0204438
0201207
0204435
0263020
0240053
0204428
0204427
0204422
0266776
0204424
0250641
0255552
0244355
0204420
0204421
0213722
0213719
0204410
0255152
0252743
0228727
0235663
0204396
0204394
0201555
0204392
0204390
0204389
0204388
0285565
0204387
0201823
0201754
0204386
0204385
0237080
0204371
0166278
0204366
0244255
0281756
0243649
0204356
0284446
0204351
0204348
0204344

)244731
)233627
)204338
)248290
)250535
)224389
)229776
)231852
)168477
)252512
)284829
)286974
)213676
)204315
)204314
)285085
)284954
)221988
)258388
)241404
)204310
)284469
)204308
)277264
)204305
)204304
)273333
)213654
)204301
)225914
)204296
)237285
)223335
)204290
)204287
)196301
)198502
)251916
)229391
)196126
)196735
)179344
)223534
)235040
)232080
)226030
)237541
)263649
)232629
)241106

)250264
)204267
)204264
)204261
)240065
)168394
)234515
)235301
)242574
)248993
)204257
)204256
)223837
)263756
)204252
)231389
)223865
)224796
)231461
)228688
)224557
)237398
)230313
)204248
)204231
)202441
)112473
)204228
)199036
)204227
)225463
)223457
)272217
)232940
)223501
)231500
)235863
)227057
)284517
)204220
)284064
)237441
)231925
)236104
)204209
)285064
)286920
)229596
)228285
)225644

0237649
0204194
0112511
0112514
0197283
0274259
0284256
0213588
0204188
0242014
0276746
0030110
0197251
0096433
0137288
0266509
0161896
0161904
0096395
0287089
0249346
0271362
0233183
0276824
0221697
0124493
0271231
0214810
0137309
0276404
0186577
0225339
0220583
0272325
0270800
0124614
0124507
0124664
0196114
0196821
0216636
0265123
0220643
0272288
0186328
0217130
0124562
0065060
0206717
0252106

0064995
0064999
0187762
0286550
0273870
0124678
0146197
0272374
0065029
0217004
0023892
0112033
0227154
0112039
0198755
0276712
0007866
0112041
0228559
0219023
0096060
0218749
0287458
0212579
0265527
0232909
0157343
0220734
0196748
0204140
0137392
0197753
0096063
0112053
0271304
0112062
0237719
0156711
0246982
0096070
0180316
0189325
0010030
0224666
0179165
0112078
0240733
0112079
0239964
0112081

0263894
0238554
0251864
0281450
0213500
0124762
0285244
0255587
0220349
0124772
0285888
0287891
0137168
0198663
0232598
0164530
0137409
0146192
0226976
0216412
0137193
0172738
0286105
0065491
0275550
0286672
0112130
0239953
0137200
0198937
0204110
0229559
0227920
0263926
0279942
0112139
0225945
0156639
0243313
0200597
0218521
0183826
0200706
0226533
0199938
0217227
0124767
0236355
0124721
0244297

0219273
0212586
0231150
0220076
0220556
0112164
0112167
0217165
0164626
0124780
0095981
0164627
0218986
0231276
0252990
0146122
0281969
0235033
0124615
0220586
0180211
0227131
0204091
0204092
0226070
0156564
0252767
0236075
0226454
0237947
0124602
0124596
0112212
0124701
0001167
0287678
0161912
0161911
0095970
0112195
0184106
0188056
0212176
0225690
0274256
0278745
0124731
0218809
0206745
0213461

096264
279284
280371
234753
226917
137166
266494
112559
230913
112561
231102
096088
269387
137218
124593
278224
214736
164663
268745
206977
227516
124641
207371
112578
112576
206875
223946
137413
261068
188112
287363
048545
214732
112599
048544
124496
237107
024048
206848
112619
213435
124659
112624
146223
221821
288010
171611
287825
137161
231113

0181524
0124713
0124587
0112640
0124733
0124702
0124541
0272223
0240625
0044090
0137171
0112651
0272170
0112655
0112658
0112659
0245261
0112667
0146216
0137204
0146215
0171467
0219222
0220614
0179157
0226558
0287055
0287266
0124574
0276770
0171462
0207076
0137221
0204052
0171453
0137207
0271754
0124571
0219470
0252218
0170734
0203362
0172432
0124688
0172426
0096080
0218107
0236961
0112715
0272114

)283573
)283215
)223469
)237686
)181577
)231881
)183239
)287562
)180992
)137216
)137225
)265439
)262179
)112759
)096384
)157593
)284427
)146232
)178233
)272442
)146221
)124608
)249481
)096401
)266619
)237530
)279076
)286417
)196284
)270900
)276156
)277769
)219384
)207769
)124813
)271857
)261080
)112782
)212468
)231769
)001561
)112796
)229349
)172348
)226541
)236466
)146233
)153291
)242973
)180113

)146070
)230062
)236164
)112818
)069122
)225730
)153292
)146072
)271328
)287485
)270761
)198087
)199762
)216813
)164393
)153294
)252711
)226594
)216616
)124818
)200340
)244694
)286811
)226068
)216835
)278736
)287137
)271162
)218337
)274867
)217631
)252457
)277043
)146085
)031691
)203972
)197261
)216998
)112077
)124490
)235122
)279127
)096006
)170950
)226707
)124812
)214643
)177684
)214642
)203970

0180872
0226733
0286405
0233470
0230472
0008197
0008196
0060303
0216913
0230204
0270306
0232702
0212532
0214641
0228689
0170927
0240641
0207604
0225613
0199080
0112115
0112116
0218363
0112118
0170915
0096093
0065308
0225791
0270382
0216775
0096092
0220377
0271227
0237674
0244067
0217228
0243955
0223622
0182793
0230375
0230845
0174156
0243236
0288014
0170899
0202198
0283293
0112144
0112146
0242865

)261745
)137270
)253026
)219074
)218713
)206908
)219163
)012660
)264056
)217527
)271367
)219863
)218049
)218261
)271338
)253010
)271373
)001084
)249379
)231683
)235899
)124743
)227885
)137269
)271218
)146147
)236740
)235050
)236996
)218143
)287745
)137251
)228231
)213335
)216347
)220773
)220635
)251946
)224984
)168143
)261116
)137252
)187871
)146151
)220725
)112175
)214604
)124749
)219095
)227602

0220666
0151914
0231441
0216854
0200224
0151917
0219790
0218980
0276576
0168116
0112200
0226803
0112208
0112210
0146143
0212017
0266579
0272541
0286723
0271761
0272316
0239650
0283352
0218189
0215190
0225096
0214563
0216687
0214561
0275046
0283453
0277172
0275773
0287598
0274844
0276290
0220030
0255633
0250686
0112232
0287679
0216859
0217067
0262566
0217477
0198225
0219940
0218520
0146166
0218617

0218274
0218048
0225121
0112245
0213312
0266680
0285976
0262651
0213309
0118482
0188107
0218813
0236345
0219653
0232120
0217811
0243828
0214558
0220483
0233859
0218890
0270521
0238949
0212229
0271111
0251965
0201015
0206672
0227706
0286680
0287557
0270509
0285838
0223504
0286564
0226497
0288088
0218834
0230910
0135298
0168216
0219085
0219881
0082293
0216365
0112280
0253809
0287300
0237643
0224349

0082269
0238386
0154079
0271967
0232389
0218870
0112305
0262803
0217767
0112309
0287939
0219669
0221345
0218350
0207180
0232295
0287380
0276127
0119900
0269966
0230597
0233237
0199094
0279289
0207827
0223765
0285963
0219575
0218014
0202069
0079841
0218732
0217483
0181514
0285844
0185760
0233844
0266180
0232422
0220515
0229154
0234025
0251360
0243501
0256980
0135314
0229852
0239626
0234882
0223389

0226840
0203909
0203908
0203907
0231332
0235174
0224221
0080007
0164430
0135297
0207023
0271986
0223821
0156508
0229862
0223967
0119899
0218459
0231652
0156535
0279790
0217268
0223786
0217178
0272243
0218483
0224583
0219736
0111799
0112695
0112697
0225793
0118407
0203489
0237174
0221332
0263533
0238156
0217680
0201291
0219722
0220730
0252137
0112701
0217488
0252498
0200091
0285401
0196586
0252156

0223169
0112706
0200040
0222652
0238697
0236635
0272445
0280511
0231762
0271945
0276064
0135312
0219253
0269964
0230309
0229495
0146243
0146247
0217512
0286340
0118418
0270362
0220154
0201464
0286042
0217786
0218029
0262048
0231533
0218748
0135338
0287811
0261970
0272137
0279659
0198478
0235357
0220918
0285586
0233835
0118402
0287816
0218582
0220311
0112742
0288071
0218418
0217377
0083123
0272129

0219547
0279022
0233967
0260645
0219361
0181705
0216352
0112773
0226089
0277797
0212336
0232031
0206886
0219702
0224995
0226453
0260574
0005700
0223044
0220537
0286875
0146242
0118420
0220091
0227215
0083097
0013375
0013392
0065833
0219951
0146250
0065609
0237874
0287705
0203877
0065615
0228679
0230852
0135324
0135315
0231776
0226309
0112837
0228290
0219604
0287294
0216523
0220291
0216902
0217060

0213211
0234155
0216439
0214460
0135318
0280232
0135317
0271793
0135316
0203875
0220563
0275072
0217334
0198738
0222686
0200427
0218766
0288021
0202103
0220069
0217272
0220831
0219500
0243124
0218561
0219190
0217769
0220240
0168830
0214457
0217169
0217776
0252697
0135346
0215477
0272008
0188994
0218313
0164411
0216990
0111850
0203872
0279616
0272514
0213204
0216324
0218793
0218698
0164414
0206715

0146282
0135336
0135334
0238628
0207131
0240997
0271114
0286278
0118434
0118432
0279565
0234426
0220131
0220267
0286871
0111880
0222145
0280277
0220130
0220960
0223001
0271931
0146278
0264017
0154548
0288009
0146281
0146276
0111886
0220418
0198833
0025039
0218965
0135299
0239316
0237027
0083099
0112159
0228124
0220871
0217653
0118412
0199630
0218300
0219240
0135355
0252891
0112182
0226455
0222078

0287789
0218730
0260271
0266760
0135341
0224605
0263734
0224944
0262095
0219087
0220340
0240961
0219867
0285961
0220553
0200492
0261038
0288101
0220908
0221455
0219387
0235099
0213187
0135333
0287683
0288085
0280451
0252249
0229600
0231143
0220370
0279403
0270484
0216853
0219627
0261366
0172469
0219549
0287578
0219941
0172461
0233797
0242560
0014123
0112214
0213183
0207309
0219139
0217331
0216518

0224384

0112218

0123545

0263479

0186231

0146263

0271860

0226207

0238367

0220505

0236920

0187472

0283010

0184486

0112234

0271042

0265226

0232699

0146267

0132423

0132424

0228506

0123552

0279170

0274603

0112237

0219755

0112238

0199303

0152034

0229315

0219257

0275716

0213177

0216378

0112246

0228082

0112249

0220695

0260000

0270987

0164418

0218475

0287397

0218536

0287499

0219757

0217120

0202283

0288084

0216817
0214428
0287616
0218173
0085382
0252944
0227535
0271099
0203809
0187772
0212147
0112276
0203808
0132429
0085377
0231628
0216639
0219902
0286084
0237567
0219088
0284999
0200198
0057657
0057663
0269919
0273744
0244710
0252444
0233941
0112297
0207499
0202386
0282408
0200314
0130347
0200295
0130348
0216977
0229654
0235142
0202285
0207577
0272398
0130349
0178409
0252125
0164494
0276620
0220237

0112320
0234206
0280135
0146285
0025796
0238420
0232778
0272476
0224987
0253090
0231112
0225093
0271734
0081087
0238490
0225174
0112333
0279398
0112335
0233511
0212525
0286562
0230248
0206974
0135537
0238055
0279498
0238079
0227072
0200799
0287044
0271608
0118689
0228976
0203801
0219565
0286511
0231559
0118690
0218872
0228834
0230290
0080546
0271730
0199944
0183137
0203799
0233908
0219700
0201023

0218632
0135535
0260273
0185250
0135587
0135596
0112365
0223537
0155085
0112367
0232311
0146360
0286691
0112290
0168438
0053328
0287268
0203797
0004809
0241162
0260188
0219559
0270934
0155111
0213150
0217041
0219150
0200522
0212587
0199360
0213149
0271607
0123505
0219329
0155115
0197498
0218803
0112394
0200926
0231046
0173214
0271789
0009413
0272356
0220392
0229276
0218676
0231889
0220506
0286616

0056972
0255389
0010810
0227012
0112761
0074935
0203778
0112769
0237234
0207044
0270661
0271208
0226440
0281613
0216663
0251258
0219773
0215807
0217770
0286914
0232299
0225580
0271498
0201386
0287728
0218490
0216904
0223811
0232316
0236347
0218208
0230943
0231912
0219758
0277443
0227502
0175967
0196591
0228624
0216723
0249853
0216471
0212333
0218089
0232395
0226079
0287097
0251882
0219619
0228777

0111816
0287933
0186743
0178425
0123500
0218428
0236326
0187189
0111817
0233558
0189241
0285446
0217241
0218153
0244158
0188820
0234117
0173626
0178033
0164451
0111832
0111834
0153975
0196911
0183807
0173612
0185002
0217089
0200248
0287253
0170162
0047936
0251814
0222713
0282218
0219703
0226181
0164465
0207461
0047932
0218233
0153989
0196376
0230202
0216809
0111860
0169075
0198523
0286339
0217330

0225549
0111877
0111875
0253194
0111879
0222649
0207982
0220139
0111885
0216316
0200732
0287100
0265725
0286540
0206857
0200310
0216710
0287992
0146350
0206598
0207403
0220447
0272714
0152661
0199458
0219784
0201379
0222659
0220326
0218180
0219135
0217139
0220184
0199932
0025156
0279453
0111897
0275339
0135549
0279114
0277408
0287258
0272472
0287818
0240606
0164434
0172594
0232362
0285652
0146352

0186439
0235535
0285691
0285941
0188580
0237321
0236548
0146373
0111907
0111906
0286299
0226409
0237742
0224506
0135547
0111912
0232131
0222445
0111911
0251920
0066651
0213133
0286215
0203760
0207632
0201613
0275953
0217325
0217404
0287731
0146374
0118518
0093144
0251768
0213131
0220744
0220522
0220694
0255330
0189367
0214338
0184530
0219284
0218187
0260212
0219699
0281248
0172673
0274493
0152894

0227945
0224733
0213130
0206982
0220110
0202200
0196569
0220884
0219642
0252554
0226149
0233351
0146376
0218689
0219666
0218725
0219302
0203756
0198945
0227678
0164483
0164484
0202438
0229923
0256162
0079819
0118507
0218857
0219776
0118520
0112282
0154269
0252560
0180658
0236761
0216917
0266807
0197594
0219532
0200895
0237115
0236673
0280058
0118523
0227220
0236166
0265669
0280155
0228495
0223912

0079931
0079950
0225343
0237110
0146385
0235998
0146383
0135569
0237449
0286663
0179073
0146378
0146399
0218194
0112299
0218991
0093134
0112303
0234484
0271488
0146409
0112306
0206754
0221500
0200534
0220557
0279960
0231023
0213122
0286438
0220113
0112319
0223542
0234567
0227954
0238059
0272428
0218213
0223586
0118526
0028839
0146411
0278744
0287413
0216753
0266875
0118515
0276703
0200058
0201309

0286887
0220378
0236700
0287974
0213118
0231971
0232310
0218772
0229722
0227723
0236389
0213117
0118514
0240056
0112339
0283409
0232876
0118513
0236703
0207689
0224374
0135541
0234084
0287094
0201012
0231028
0226540
0233534
0217482
0236378
0171408
0286313
0288054
0237596
0213111
0277973
0146410
0029363
0216519
0260418
0135525
0220660
0216613
0218424
0271765
0272189
0197442
0234263
0212242
0286646

112357
201807
213109
182747
207247
230350
225391
235399
016402
164485
027697
177468
213108
230533
283265
220412
234956
207300
237499
118503
287393
226004
229922
219463
112378
112379
254440
262543
218499
215878
051620
135540
266555
274594
216802
225177
279968
024862
203734
225415
220600
135597
146386
112406
231329
272446
164440
226571
218565
164442

0238099
0219806
0205695
0231426
0225148
0287820
0236013
0252107
0263514
0285875
0264390
0217684
0234147
0216548
0275138
0286452
0259828
0222764
0218351
0135577
0270983
0236822
0203733
0009844
0237494
0112414
0236366
0287084
0233138
0010818
0237851
0232618
0229017
0227192
0146416
0225752
0217495
0216642
0217648
0270655
0189007
0218896
0223203
0034693
0278206
0229036
0001036
0112419
0235740
0257065

135521
280148
270890
118491
118495
283122
276680
217231
169976
219297
135604
217027
216475
152818
225311
219409
217195
220739
285598
112425
201119
217612
270638
270828
235652
288056
280184
118496
146414
152822
187103
222971
216811
118508
237468
118492
273719
207431
228283
233452
227748
272397
196634
164506
273997
220494
203727
227681
226249
287976

0234675
0230205
0287473
0224658
0111961
0207345
0215223
0217824
0231368
0111962
0227660
0236591
0219487
0228408
0055208
0225924
0283608
0241391
0216265
0177688
0178199
0219553
0131013
0252244
0235168
0055211
0281021
0233330
0220848
0202343
0186625
0207292
0131023
0278899
0120253
0120265
0219433
0231760
0285889
0285991
0120256
0273132
0268592
0217733
0164520
0223701
0216906
0203722
0216621
0131015

0215094
0111981
0219298
0231120
0218358
0155918
0216444
0235972
0213091
0131019
0198729
0238594
0201628
0009765
0220598
0219622
0120278
0232891
0224029
0226599
0213089
0120254
0218996
0202119
0232290
0223598
0219747
0231883
0238616
0213087
0131016
0252431
0252487
0223039
0181472
0252615
0155906
0219395
0146476
0120262
0207422
0091831
0233823
0287914
0284615
0131018
0234577
0226193
0202044
0233464

0120279
0218792
0146469
0224893
0217372
0220745
0112029
0227627
0112031
0091844
0201939
0213121
0218426
0237312
0222806
0217160
0242246
0218776
0217044
0220733
0220514
0216713
0199246
0217085
0112038
0074706
0288520
0237593
0153721
0286553
0287260
0275419
0220181
0219146
0231316
0205625
0213079
0200594
0146426
0218757
0264814
0238963
0274451
0235381
0029639
0171217
0232529
0074771
0238789
0287092

0221456
0285797
0212295
0227360
0220347
0271265
0049618
0271551
0271899
0270487
0218596
0233044
0215712
0217783
0271040
0213078
0218631
0286482
0175048
0266617
0130340
0234361
0229502
0236324
0220867
0252658
0078269
0285642
0233496
0122335
0272047
0130338
0188451
0274023
0287591
0238019
0243373
0219188
0146433
0218226
0278571
0146425
0164674
0219249
0265558
0092820
0233893
0243775
0285159
0203711

0213076
0223191
0130363
0271913
0164691
0226032
0285492
0224478
0233682
0164694
0235086
0231178
0234777
0286533
0216480
0276413
0237927
0112096
0285441
0220305
0251988
0146457
0285427
0120437
0120438
0207392
0206910
0112110
0146453
0236823
0130368
0197081
0268257
0213073
0175003
0216516
0112499
0230234
0146477
0287656
0213071
0198670
0243831
0224477
0122194
0287558
0276960
0274903
0224371
0220913

0233342
0231863
0272841
0085511
0026652
0185345
0286498
0286805
0227913
0112530
0217557
0285610
0225437
0285553
0225683
0281692
0285564
0285726
0217514
0270419
0112531
0228692
0235538
0285858
0266128
0286573
0260422
0230627
0226739
0287550
0227455
0287877
0217878
0112539
0112541
0199523
0206681
0236627
0223942
0281832
0216624
0287824
0231297
0276643
0164458
0201292
0280991
0233365
0213536
0280850

0176381
0287189
0198818
0233231
0286760
0060762
0261420
0280057
0071242
0232082
0222958
0235272
0287690
0235008
0249141
0231654
0026297
0227598
0285730
0265828
0213066
0272980
0112486
0284825
0284914
0166984
0120436
0286674
0216966
0280707
0231720
0228648
0112494
0120440
0217447
0203690
0287297
0272549
0287594
0203688
0213065
0146521
0269155
0285212
0198221
0130396
0235994
0229921
0125337
0153303

0234768
0287056
0237987
0228361
0234647
0230711
0164488
0224417
0260771
0112562
0235815
0280665
0225879
0285917
0279810
0229720
0234519
0286826
0223485
0261039
0226445
0186340
0233085
0224999
0226194
0285887
0285733
0272848
0184465
0185127
0130024
0232640
0227704
0184786
0130023
0266896
0229214
0231690
0236173
0287823
0288162
0287407
0232197
0273100
0218716
0230960
0271820
0227508
0198719
0112584

0271234
0266245
0261003
0008018
0112592
0071994
0287411
0230423
0217874
0273640
0234986
0225532
0219814
0283061
0282572
0273644
0232325
0242611
0287883
0242474
0240859
0261795
0239715
0240093
0177706
0249852
0248767
0287342
0249574
0286710
0286764
0234471
0197461
0223855
0188191
0229380
0237181
0164818
0164828
0239857
0273151
0105963
0240230
0073067
0146540
0199023
0225146
0164849
0257607
0224079

0226291
0164850
0178381
0229043
0164853
0233082
0164877
0225981
0273230
0164880
0231927
0198517
0164855
0157778
0230487
0230444
0231476
0225968
0234977
0205971
0236081
0002822
0286192
0176349
0265089
0122687
0106268
0106266
0284047
0271098
0106263
0230914
0271163
0136213
0275572
0106003
0264357
0106009
0106012
0136295
0174945
0146535
0231357
0198286
0237286
0201794
0228334
0217455
0231892
0236708

0146555
0224593
0236510
0283991
0261809
0286874
0164916
0287665
0242802
0264474
0287541
0157927
0206895
0218823
0213935
0136297
0196204
0272968
0270822
0275328
0146587
0273313
0272595
0159904
0202039
0157954
0164638
0182095
0241269
0188365
0272953
0234432
0273084
0155034
0230733
0207973
0075624
0272719
0228974
0075618
0011275
0237738
0283419
0235944
0277265
0122543
0122674
0155026
0122512
0201990

0106305
0207217
0086232
0157999
0240718
0252929
0231704
0106346
0008256
0202273
0178397
0286075
0274319
0136238
0164535
0136240
0231359
0215045
0136247
0232581
0146576
0236609
0205903
0228010
0198580
0164631
0187953
0277327
0169402
0146574
0273252
0155070
0205897
0250561
0243531
0226943
0241539
0242731
0287860
0266287
0239696
0201218
0106392
0244167
0230825
0236712
0215018
0272732
0272894
0164654

0272745
0106399
0219545
0283549
0270314
0234718
0201747
0214998
0234141
0233108
0106415
0233264
0003147
0227719
0244239
0229970
0272328
0122584
0236748
0278254
0271526
0278446
0233876
0234710
0271077
0197320
0270611
0235431
0287409
0286639
0212422
0283117
0229091
0234356
0236414
0201774
0189043
0271185
0106443
0230370
0230333
0226827
0005108
0230435
0235920
0106460
0146530
0274970
0226690
0236048

0203523
0224667
0225606
0006747
0229233
0271253
0122644
0199470
0229618
0222974
0230879
0237713
0224330
0213875
0006468
0252374
0223445
0136267
0233167
0226323
0225816
0227489
0187546
0106511
0237070
0229108
0225974
0235828
0286376
0229379
0214960
0224683
0229688
0171243
0272537
0205858
0224280
0272361
0106524
0287799
0136261
0235837
0106537
0106541
0270593
0173467
0232400
0227965
0106546
0223867

0237773
0232841
0278569
0283321
0199473
0236318
0236039
0228675
0071189
0279048
0236195
0280340
0229937
0048052
0221393
0225000
0222164
0226522
0232492
0122691
0232821
0236536
0229533
0146618
0223838
0105849
0265932
0173452
0237921
0243004
0213860
0267055
0183742
0228598
0233834
0226097
0271133
0105855
0200753
0179460
0231525
0004846
0223866
0164651
0219451
0232790
0234839
0285592
0243633
0225308

)195024
)229949
)105866
)221783
)105877
)164649
)136237
)251993
)105889
)233476
)232759
)232949
)179428
)136244
)228803
)234513
)196683
)286703
)228649
)221740
)226329
)226279
)122591
)230658
)122550
)230042
)136243
)285926
)226816
)136235
)156928
)136231
)232627
)212264
)252590
)236654
)232818
)200847
)164548
)224785
)232553
)169193
)227436
)228953
)188732
)234286
)196335
)234800
)231328
)227930

0252041
0228944
0122585
0287523
0206877
0105926
0271096
0271330
0105928
0287093
0070882
0234999
0172115
0153790
0234406
0213798
0105954
0285716
0202233
0234922
0233824
0260951
0285960
0227386
0223561
0239622
0287096
0286725
0270933
0199085
0050344
0122566
0122565
0285860
0232383
0086300
0225792
0214870
0122548
0280255
0233760
0286507
0222004
0005020
0213787
0226059
0214869
0213786
0105991
0233429

0276528
0276771
0274864
0274396
0277694
0105996
0105997
0253552
0197576
0273433
0254369
0106004
0106006
0272801
0122592
0270182
0078399
0257184
0253187
0283745
0253293
0005073
0240990
0273961
0278334
0276496
0253508
0277966
0278592
0278020
0106031
0243766
0276609
0278708
0277469
0277553
0286323
0253308
0253405
0106038
0213783
0236569
0224322
0279655
0279429
0233830
0213781
0235574
0106049
0286478

0229893
0106052
0153814
0287489
0206623
0234336
0174495
0146592
0225286
0255690
0228421
0272568
0233517
0106066
0229452
0285412
0285162
0106069
0285081
0235669
0237065
0223813
0270196
0176532
0225264
0223437
0227855
0212024
0122574
0238906
0136193
0227014
0106080
0106086
0231519
0283618
0286847
0180354
0230751
0251660
0225907
0227017
0180233
0283237
0207771
0281593
0235859
0281404
0106100
0272638

0006625
0281039
0196295
0244480
0264520
0226874
0263683
0106105
0106113
0241644
0223419
0254959
0106125
0229263
0250424
0240583
0106128
0078549
0232887
0286487
0271027
0164600
0231418
0180347
0106341
0154678
0236460
0284839
0226468
0226421
0106355
0105778
0229358
0237004
0207573
0283461
0273014
0281332
0250756
0251999
0231952
0272905
0170852
0205763
0224573
0122642
0252821
0122643
0229054
0241420

)164610
)122507
)228015
)222741
)224946
)238090
)173862
)227481
)236494
)233219
)164619
)286300
)236212
)202431
)197085
)187258
)235379
)230356
)242653
)251802
)173852
)221669
)287249
)189212
)236592
)164532
)226063
)287651
)286526
)235464
)287032
)122557
)271122
)227544
)228878
)232736
)122545
)274328
)200446
)273580
)226102
)270889
)287893
)232930
)285573
)205745
)261184
)228815
)271117
)122547

0228770
0229679
0234457
0260070
0164542
0201432
0011426
0237400
0136250
0230539
0272984
0229424
0235728
0225739
0230831
0155849
0221325
0224101
0224706
0222869
0213744
0200113
0201566
0178503
0187037
0086289
0283444
0225314
0086288
0106483
0010270
0227191
0211687
0211688
0211689
0211690
0211691
0211692
0211693
0231202
0211694
0231916
0211695
0225992
0272908
0211696
0249978
0226212
0281103
0228668

0211697
0211698
0211699
0233306
0211701
0078053
0241756
0227404
0164556
0006715
0106536
0233854
0206947
0224122
0227172
0241127
0236015
0231951
0106540
0287584
0006451
0228554
0188185
0252480
0270797
0226220
0237251
0231198
0233225
0175741
0242855
0259826
0065883
0273444
0270577
0168303
0199273
0175600
0248711
0203446
0227658
0232458
0224017
0122641
0224116
0106571
0213721
0238284
0229161
0231539

0261019
0136197
0256646
0106588
0234983
0106591
0232006
0201933
0002746
0181211
0264069
0252983
0223258
0228680
0213717
0231638
0164543
0106603
0106605
0229497
0270617
0062582
0106608
0237972
0078967
0214783
0273432
0241057
0239556
0136206
0228434
0285596
0272655
0228903
0235314
0136279
0284067
0164708
0239775
0122678
0283969
0106624
0284412
0106628
0252848
0106631
0106633
0106636
0058404
0279853

0015676
0146677
0200456
0225443
0015520
0136271
0158604
0287574
0271586
0105953
0122515
0228596
0196262
0105968
0234183
0146676
0284261
0272768
0230160
0260997
0136286
0232956
0277184
0136280
0136274
0136270
0206838
0201772
0200656
0122679
0286738
0275295
0234359
0225537
0226999
0164742
0214765
0272556
0212450
0235955
0226838
0251378
0202350
0283247
0146678
0146674
0237471
0229628
0233952
0221971

0224547
0229459
0237760
0240355
0230680
0224519
0233539
0242948
0232072
0223829
0229192
0242552
0214754
0286315
0136205
0236078
0221845
0158683
0146666
0136275
0225507
0136273
0164744
0164746
0183696
0179869
0228173
0285536
0225705
0279578
0229165
0218305
0234686
0270763
0286995
0279104
0188730
0042813
0223893
0287521
0164500
0228005
0231681
0235620
0185811
0285165
0200815
0132436
0132437
0226122

)106070
)286658
)236046
)228085
)228204
)201133
)106078
)231490
)227233
)228897
)227080
)285741
)229478
)229403
)242650
)286987
)287039
)233960
)253028
)286908
)234304
)223970
)221900
)227344
)271194
)199629
)222154
)234105
)228627
)205628
)222355
)271306
)232418
)235454
)238131
)287953
)280920
)231427
)213673
)170419
)224223
)228735
)200471
)232612
)286404
)132432
)234707
)218586
)146648
)224057

0280890
0237210
0230564
0230936
0132434
0154978
0223475
0233977
0231394
0234844
0235207
0231317
0176826
0252857
0275853
0224971
0237799
0275875
0273720
0226278
0206729
0154997
0235738
0225270
0229762
0225018
0227499
0249773
0178665
0146729
0239789
0146733
0146731
0206603
0207168
0129103
0164776
0106153
0185290
0230191
0233028
0285670
0227491
0223559
0224155
0203462
0202296
0231299
0225371
0224370

0223330
0232944
0237268
0278205
0225488
0226485
0261275
0231504
0223740
0233288
0224947
0213650
0234716
0213067
0271047
0287019
0229909
0228303
0286477
0277174
0282879
0277253
0279072
0233437
0225244
0264426
0189166
0214668
0229508
0275061
0185177
0286436
0235095
0283431
0284474
0283547
0231537
0283230
0283481
0225295
0228440
0283311
0224880
0224723
0230764
0237309
0236692
0284291
0284098
0283626

0228851
0284558
0284221
0227584
0230531
0284572
0234671
0169325
0234089
0238431
0287517
0274299
0280225
0260653
0266168
0227182
0235368
0182111
0270957
0233601
0243981
0270749
0229627
0230796
0237639
0233962
0236261
0234085
0227015
0232161
0230271
0250923
0224484
0236907
0250618
0224653
0223836
0237236
0231232
0231484
0271696
0233918
0207443
0227148
0233454
0233782
0234312
0231523
0199231
0237572

0226075
0244550
0240651
0236638
0185037
0226587
0227298
0232165
0230000
0227545
0227923
0229301
0236284
0230361
0230600
0232817
0227305
0223889
0234387
0268181
0234467
0227426
0224368
0197990
0230132
0164740
0228645
0214660
0223566
0265214
0226401
0226411
0271466
0227397
0236574
0225451
0236299
0235720
0226581
0226342
0227910
0287588
0213645
0233173
0231264
0284534
0228660
0230639
0176232
0278577

0241149
0228564
0286397
0285544
0214652
0223558
0224458
0248704
0248315
0230052
0233385
0227035
0234234
0234700
0230252
0235646
0229881
0236976
0213643
0235227
0213644
0223614
0197123
0287699
0234444
0227986
0287985
0276650
0232727
0223845
0228014
0228653
0173041
0223974
0235349
0213642
0224669
0276475
0286456
0287317
0196247
0271550
0275667
0224172
0277206
0197008
0287580
0234338
0182722
0213640

0287869
0198039
0189316
0223476
0270948
0286342
0152926
0213462
0201165
0234585
0239985
0207344
0207062
0227113
0226002
0164669
0237026
0229064
0282381
0277544
0231556
0232906
0146757
0238124
0226767
0235421
0230386
0272693
0273024
0228409
0206634
0206785
0275833
0224316
0200670
0236529
0201560
0196715
0169919
0224365
0229886
0205596
0126522
0249319
0241258
0234185
0169902
0252126
0179406
0251451

0231234
0230189
0236928
0279785
0244510
0244657
0237310
0179342
0234500
0229180
0275400
0232491
0177418
0225573
0243335
0284461
0226824
0154710
0272831
0271064
0230583
0232546
0230295
0212568
0273142
0179131
0274135
0106609
0275878
0126524
0198874
0233383
0266525
0230358
0067601
0201565
0106610
0228019
0273448
0226392
0237466
0233423
0223948
0225209
0236531
0232975
0226829
0237754
0225959
0228429

0222428
0233689
0200189
0286466
0225718
0236839
0230654
0229031
0252423
0200857
0234215
0158321
0236978
0185274
0265878
0283278
0199940
0183166
0201014
0235581
0228769
0277149
0264494
0270694
0270555
0266569
0225648
0274656
0179994
0196313
0272843
0106133
0285886
0155428
0174353
0205584
0276178
0229018
0201282
0273520
0273897
0273927
0285702
0275976
0286228
0277125
0274984
0262461
0174384
0199913

0274570
0233369
0233523
0182487
0214544
0205578
0130305
0146755
0225005
0077800
0206709
0202035
0188763
0009954
0252713
0106635
0106638
0009950
0274080
0176428
0176410
0071462
0106089
0265724
0241709
0225969
0106077
0165215
0189143
0165171
0175877
0261467
0049540
0232415
0106683
0106682
0207741
0086730
0049541
0287815
0106665
0006704
0238391
0273069
0279005
0263001
0232729
0262648
0158517
0196275

0277072
0123965
0239069
0184616
0274070
0274523
0277127
0174428
0165178
0277053
0276592
0277675
0263081
0275121
0286038
0276840
0267828
0286137
0273598
0202021
0286014
0278416
0199870
0205583
0275930
0146722
0178809
0223705
0272391
0242073
0170092
0127957
0200874
0127946
0279996
0006606
0106178
0219039
0005486
0127948
0284334
0235090
0201643
0189077
0127952
0280388
0146701
0230882
0251798
0227038

0224273
0213549
0228829
0177679
0106211
0170027
0213548
0146700
0188372
0091073
0233980
0250614
0243566
0250778
0238560
0230305
0185040
0205485
0146707
0214243
0285666
0231183
0265479
0283398
0231087
0236280
0225703
0186704
0242435
0241350
0201885
0186645
0233448
0201959
0259628
0205482
0225726
0214439
0213542
0135205
0127951
0250990
0186088
0273341
0237620
0287519
0127947
0214293
0187257
0135211

006576
222432
232756
187391
281008
280958
231322
285892
226230
251276
226978
212545
229110
281120
286855
212482
222024
234456
226285
200786
287488
127955
206818
230853
244392
234223
232667
240347
135218
223550
214415
229436
075223
279326
237896
226671
223514
224134
232968
233491
230196
019991
153956
223770
213530
228711
186472
202281
235503
170381

0285398
0230617
0230244
0075213
0227982
0235139
0235243
0232605
0280325
0232019
0229251
0153993
0224059
0230614
0225128
0227785
0236610
0198822
0231255
0233073
0164659
0261462
0224046
0135164
0278236
0135185
0200397
0182165
0273623
0276185
0005469
0005471
0085563
0231956
0105784
0075303
0006634
0008277
0075142
0254003
0228113
0127954
0233420
0225701
0207094
0270812
0235639
0228928
0231763
0237709

0227979
0182348
0164645
0227863
0239075
0227646
0238358
0235436
0164647
0157214
0105792
0207488
0234459
0225498
0283267
0105793
0280440
0157224
0273299
0058091
0251154
0223969
0223584
0286411
0226254
0157240
0243144
0225843
0235450
0223665
0127989
0287672
0127914
0001630
0285772
0285953
0188693
0240720
0001631
0243107
0200769
0221520
0001629
0230927
0270453
0157259
0244055
0242950
0127980
0127993

0278819
0234545
0105810
0287932
0286742
0206763
0237819
0266794
0205413
0177409
0188175
0004766
0004948
0208014
0207656
0127928
0237729
0265423
0225898
0213483
0105825
0234695
0278388
0127920
0278959
0105829
0236861
0236453
0236938
0285090
0285964
0164692
0222870
0127995
0127990
0242265
0232385
0239030
0214342
0230682
0222220
0219807
0158528
0271159
0231153
0236197
0201607
0005421
0237160
0228751

0105852
0105854
0273138
0233942
0005981
0279525
0231170
0004799
0158560
0279067
0004864
0208025
0237551
0275834
0207045
0207115
0252872
0127922
0227284
0244318
0231764
0274409
0277124
0006377
0274709
0105880
0196636
0224551
0224475
0226046
0199475
0006128
0070669
0284707
0226744
0231812
0237818
0231859
0232032
0284627
0232097
0238228
0266318
0266668
0183444
0285725
0243554
0135175
0277946
0164715

0180535
0205356
0164713
0006453
0272950
0214389
0213455
0223402
0106236
0207204
0166448
0196367
0266019
0286305
0238109
0198742
0284840
0185467
0002079
0284523
0284292
0241685
0130429
0106244
0106245
0106246
0248919
0160917
0228335
0241468
0198556
0160908
0196652
0221909
0197343
0272647
0244219
0197037
0224448
0106258
0160870
0282277
0282301
0160868
0273407
0261511
0021461
0260445
0260524
0244623

)146833
)176402
)237640
)199711
)222966
)160862
)286921
)214313
)235713
)286923
)106261
)166529
)166526
)168090
)166508
)207547
)207757
)208036
)221838
)106290
)242798
)166997
)214309
)235077
)214142
)188186
)146826
)284518
)197093
)213420
)066923
)239521
)213413
)214300
)078319
)201913
)242294
)272752
)121716
)235333
)284012
)085514
)287631
)078487
)146834
)241357
)160813
)240211
)185955
)166925

0286938
0166924
0225807
0242101
0106351
0184414
0205307
0274272
0077454
0106336
0224729
0106333
0106330
0106327
0077080
0172354
0146830
0172336
0130427
0146839
0196411
0263672
0146828
0236305
0087077
0283821
0087087
0176125
0087085
0239825
0235926
0169894
0274993
0205277
0227053
0169876
0222636
0169871
0106366
0106367
0264425
0128564
0167011
0106384
0205267
0213394
0106397
0106400
0106404
0214253

0207454
0278683
0214252
0232301
0232445
0128581
0160963
0233683
0233123
0106436
0257923
0259294
0272219
0261535
0280404
0280004
0279724
0252824
0278586
0160999
0264675
0170409
0200552
0228546
0260097
0128563
0239969
0239486
0221510
0239480
0160991
0160993
0161036
0284176
0284596
0005075
0170667
0168255
0284981
0270249
0205238
0285437
0105808
0279168
0205236
0267368
0267645
0173678
0228049
0233297

0286830
0222011
0213385
0161040
0252643
0238324
0128606
0230257
0236226
0170632
0233025
0161048
0224415
0236848
0279482
0170629
0237632
0105819
0105821
0161057
0235354
0170615
0251852
0252047
0234715
0189056
0222386
0164815
0187416
0197258
0226869
0237606
0225329
0243352
0237513
0228393
0239569
0005483
0272918
0135250
0270764
0271482
0213361
0244490
0242154
0201179
0091127
0223886
0135249
0185055

0272604
0231167
0146776
0226624
0128536
0280439
0008282
0242562
0201796
0105835
0273320
0243438
0243797
0286076
0267052
0286013
0253276
0228742
0251978
0105851
0005249
0243193
0105856
0272072
0164597
0228341
0172209
0105865
0272854
0075790
0238832
0250474
0225935
0233705
0091137
0241764
0105879
0091138
0229626
0091140
0091136
0273055
0238297
0091128
0091129
0135241
0231006
0177683
0128590
0225647

0270425
0205174
0279043
0229603
0227968
0230941
0234273
0230192
0271368
0237064
0232199
0226965
0278424
0287186
0184903
0228540
0173114
0243621
0227948
0128512
0287011
0225572
0222346
0234358
0198839
0202406
0226851
0230999
0232922
0236232
0006652
0180019
0181016
0182965
0223646
0225358
0146802
0164603
0234520
0279288
0164604
0225457
0214194
0270997
0278894
0287592
0154415
0128573
0236580
0224595

0282859
0202334
0272230
0135272
0225535
0233607
0226962
0199764
0244565
0202377
0242072
0240973
0243345
0199224
0105967
0279086
0135269
0243243
0237813
0237870
0105971
0213167
0105974
0235427
0231210
0243220
0105976
0198898
0235945
0252115
0222150
0227199
0275359
0277583
0004866
0214188
0273596
0273571
0276100
0274606
0228368
0213050
0283923
0226367
0287829
0274344
0275054
0227532
0278526
0278482

0230785
0224042
0230605
0105989
0238202
0286390
0154438
0214684
0001626
0237974
0232661
0234001
0083622
0077063
0224136
0234826
0128534
0106013
0287827
0229982
0230520
0270516
0225546
0271739
0233417
0233969
0184408
0252023
0231295
0106025
0071243
0202225
0106034
0234927
0212628
0207090
0287554
0002745
0196937
0213310
0227371
0234985
0227743
0252704
0233272
0106278
0008311
0252003
0234418
0226636

0213302
0128610
0230316
0081803
0240499
0225795
0235459
0188050
0235631
0227573
0128519
0226680
0081800
0223595
0164675
0232524
0207338
0128609
0146809
0230442
0170807
0106299
0272686
0201104
0228211
0106302
0106304
0225583
0241345
0234224
0242593
0243574
0241324
0230268
0213296
0207214
0279419
0170775
0275356
0128513
0224897
0186676
0224899
0205898
0219445
0213291
0234071
0227847
0197462
0241921

| |
|---|
| 0179603 |
| 0287702 |
| 0227249 |
| 0287568 |
| 0224981 |
| 0230820 |
| 0207692 |
| 0236340 |
| 0236105 |
| 0048405 |
| 0279265 |
| 0224138 |
| 0280347 |
| 0226770 |
| 0179562 |
| 0004059 |
| 0106328 |
| 0106331 |
| 0240790 |
| 0197157 |
| 0240579 |
| 0279078 |
| 0128594 |
| 0207588 |
| 0207705 |
| 0174697 |
| 0272915 |
| 0106344 |
| 0242261 |
| 0238750 |
| 0224940 |
| 0106348 |
| 0228700 |
| 0238590 |
| 0273184 |
| 0135245 |
| 0240758 |
| 0272601 |
| 0165055 |
| 0276988 |
| 0273270 |
| 0280828 |
| 0213280 |
| 0242588 |
| 0281896 |
| 0223189 |
| 0243302 |
| 0230715 |
| 0229413 |
| 0243679 |

0281490
0271553
0241493
0205085
0271344
0239722
0201041
0252866
0135248
0128595
0244218
0128617
0128596
0128591
0242902
0135253
0128524
0272899
0224163
0128604
0275106
0064419
0242241
0244556
0230359
0242162
0241102
0241573
0240416
0230626
0158457
0200629
0128602
0243230
0158467
0286722
0238072
0128578
0239919
0199846
0240204
0273329
0106459
0212238
0286380
0207990
0199158
0207691
0242078
0186591

0244036
0199516
0091732
0201109
0128607
0229196
0146842
0240571
0165120
0270992
0158516
0128510
0158525
0091704
0106477
0259920
0272701
0106484
0277269
0278055
0274500
0270823
0199043
0158623
0270953
0286214
0239021
0226045
0213265
0287547
0266265
0271522
0285106
0229858
0240680
0232716
0233559
0284032
0226380
0283797
0271204
0273319
0231721
0199627
0128585
0233287
0236753
0273489
0128567
0224545

0213261
0236238
0236395
0236281
0233263
0236386
0224865
0229532
0221866
0225144
0223436
0183470
0201009
0225881
0106554
0227197
0283041
0284135
0272393
0237821
0131558
0276137
0235429
0235967
0226205
0155530
0231098
0205060
0229177
0273297
0085662
0198074
0227471
0172331
0224375
0230101
0122786
0286458
0146856
0122783
0223500
0146859
0272941
0287733
0105875
0231794
0277138
0146857
0276067
0080802

227073
155561
271414
223718
243317
212303
164707
273219
189320
224746
105887
267697
199700
230649
232053
206705
236503
226066
234352
231244
181072
207597
223657
224469
105894
202023
231114
228031
157680
279483
230383
182158
237243
122787
234639
231923
231931
222514
233951
284028
225559
252820
201465
213238
213237
214815
122779
234995
157703
252188

)234658
)243099
)223721
)105929
)214128
)122778
)199646
)146858
)105939
)229677
)105948
)239254
)271537
)228360
)271404
)157741
)206843
)164898
)146963
)269955
)273391
)252332
)188883
)253183
)236279
)254270
)064393
)059377
)059378
)006459
)251728
)260231
)199971
)199283
)270512
)241661
)157800
)223113
)202472
)146955
)103200
)133606
)146966
)201354
)285904
)240173
)133597
)090266
)240889
)157764

0271932
0271611
0242703
0090263
0223212
0261115
0261778
0261624
0285841
0204990
0261629
0261570
0261797
0006530
0244701
0257093
0270157
0228775
0214102
0212153
0106028
0127362
0127364
0270923
0127366
0241926
0224354
0239648
0240677
0165076
0240031
0237621
0257335
0244292
0258083
0258227
0257138
0257743
0240268
0258223
0186163
0241881
0277083
0242736
0226660
0237702
0211710
0211734
0211706
0211707

0211745
0283063
0282939
0243889
0242771
0211713
0211714
0239944
0211715
0211716
0211717
0211720
0244661
0229769
0241657
0211739
0211721
0253409
0230099
0211724
0227550
0211725
0253188
0211727
0211728
0253534
0253291
0211731
0278030
0276405
0275791
0276597
0274752
0276953
0275158
0275743
0276819
0275243
0277880
0276557
0211746
0211747
0251578
0240578
0211749
0211750
0270672
0282499
0239992
0249912

)241911
)211752
)211753
)244273
)286831
)232869
)204983
)275896
)250591
)243661
)282431
)282320
)282420
)282133
)281958
)282173
)282780
)211751
)211764
)211765
)211766
)211767
)211768
)211769
)211770
)211771
)211772
)237254
)106123
)165125
)224970
)286569
)127412
)165131
)197993
)179468
)239293
)225781
)179420
)159763
)236398
)268170
)221937
)279223
)197448
)231840
)178826
)106144
)239419
)240322

0231531
0188037
0159784
0272619
0232533
0159840
0278449
0146904
0229153
0234066
0185899
0221855
0236853
0176510
0229977
0229877
0271079
0252037
0230556
0223459
0283537
0170379
0283528
0252523
0271321
0253882
0241136
0232145
0159860
0198420
0221910
0213215
0270729
0273377
0242280
0221813
0221836
0221933
0239967
0221858
0221989
0239981
0221938
0273234
0183122
0240621
0225932
0213214
0244198
0212807

170356
284644
243896
244693
204959
228960
244479
221970
050327
106410
242978
221507
196511
243746
200673
236343
214035
225918
226592
231839
230746
280144
174469
274127
274118
236795
226857
232391
221442
230190
270634
207318
251936
242266
199370
243374
274133
283504
283648
213210
287636
170279
213209
270200
273314
055130
106462
251712
275586
239468

286180
286171
252316
202354
201098
281189
155660
202528
234565
197362
204947
240877
197024
239719
170265
170260
204946
271664
244560
231397
215311
228151
261842
196453
181220
133624
133619
181444
197558
106479
280149
273419
204934
171130
273293
279536
273011
224016
260555
241449
276538
286912
106526
188707
127399
261305
106538
240449
214022
196456

0284048
0239377
0284691
0242258
0171115
0271582
0213205
0179144
0243853
0133574
0271568
0232361
0133561
0270990
0106560
0213203
0281887
0196329
0177590
0241134
0106565
0002933
0002726
0055118
0164867
0181652
0197150
0243433
0213199
0164885
0164889
0244151
0164896
0164897
0133612
0164900
0146926
0278685
0285480
0285292
0033050
0033100
0284191
0082014
0272661
0243018
0013374
0187260
0243836
0241456

0127377
0265810
0241959
0106615
0240132
0106617
0242048
0239911
0212219
0222941
0227562
0106648
0276213
0178234
0229591
0213188
0055609
0270405
0204894
0241007
0253088
0241735
0020219
0261455
0231139
0285912
0199404
0225416
0212590
0196584
0214003
0244328
0133627
0286565
0224104
0234722
0285524
0233489
0287744
0130226
0226509
0270114
0203335
0233363
0286461
0236408
0214106
0157212
0272760
0273183

0273344
0220575
0157219
0283339
0274637
0228806
0201066
0217825
0228521
0287865
0278998
0225612
0273117
0186480
0204960
0218672
0260426
0288051
0236544
0164778
0146910
0283128
0227365
0287610
0216895
0184863
0164690
0204876
0207061
0270584
0182648
0279418
0234450
0105982
0224903
0105983
0206938
0230033
0146909
0130675
0235029
0243479
0236102
0229365
0269446
0009335
0225224
0236018
0224629
0229660

0105993
0234210
0223872
0233191
0280266
0155093
0207960
0222012
0231515
0233038
0272839
0285821
0231980
0207637
0225365
0232715
0265598
0224324
0146918
0228030
0117868
0126870
0286407
0231419
0106018
0287184
0229435
0253620
0253896
0254193
0253612
0176269
0255464
0253166
0223508
0172748
0272812
0272240
0273402
0254104
0249868
0182366
0147364
0272293
0279852
0180190
0104714
0254207
0198010
0282375

0282759
0282692
0254160
0275427
0260721
0282160
0281961
0253267
0283576
0254265
0182372
0282021
0253982
0283239
0283858
0287464
0207826
0104728
0285957
0253764
0253696
0176595
0270988
0253805
0036448
0273593
0287970
0253444
0282686
0282142
0253855
0277526
0254319
0253853
0287740
0183117
0253162
0285861
0222546
0286934
0288110
0285977
0254244
0253262
0273876
0206661
0242079
0278256
0271248
0240486

253510
254007
275461
253571
286969
222589
253189
253880
246089
147316
253550
091879
288076
271743
249898
276580
155189
266038
283386
287747
275591
245857
164825
250752
164822
254625
164821
223629
233238
233531
206047
215378
240247
232039
239839
234178
206042
164816
255273
284620
234120
228744
254683
230106
214268
231930
284614
284728
255025
177306

0230045
0233132
0249005
0223885
0215373
0233295
0215374
0206034
0230549
0236125
0235778
0215372
0215371
0177257
0255128
0177243
0164871
0177023
0187082
0186599
0198129
0285659
0255251
0285927
0248944
0285975
0234749
0285687
0285765
0233176
0254796
0284617
0284655
0284698
0254889
0254715
0285814
0285913
0285620
0285937
0285720
0285657
0285607
0285950
0285862
0255378
0254776
0186572
0186562
0186579

)176782
)178287
)176797
)254543
)171711
)215356
)229907
)225725
)255211
)237038
)225327
)205989
)233115
)215354
)254229
)227013
)253881
)284603
)284606
)284661
)255459
)221305
)244427
)254311
)249188
)253893
)268955
)173295
)253981
)275342
)253505
)254153
)253343
)253774
)253130
)263616
)286910
)253958
)284717
)254367
)147324
)200713
)270966
)233609
)207244
)104626
)263407
)239078
)197334
)173281

0254340
0248538
0279949
0253252
0254091
0253887
0207415
0253735
0254237
0201815
0249050
0272267
0173273
0253697
0207701
0253230
0284321
0286622
0260093
0175806
0285675
0285606
0249848
0261451
0253641
0285237
0207128
0253678
0253649
0272505
0184647
0183638
0263762
0171060
0212433
0171056
0258724
0248896
0254093
0283210
0253695
0280294
0171044
0207600
0254839
0269918
0255310
0254936
0254556
0231965

0270076
0236827
0239315
0249316
0280273
0104643
0246477
0177710
0154316
0255020
0184608
0242483
0199368
0154319
0284957
0136573
0269954
0170983
0286985
0136574
0255394
0154328
0227203
0079459
0255046
0164733
0269899
0255144
0254948
0206014
0287755
0287798
0205884
0205883
0205882
0255174
0254507
0284610
0255016
0254817
0254700
0233050
0252029
0255544
0215343
0255052
0254923
0226430
0223443
0254866

0205879
0249889
0186523
0255495
0254527
0237215
0232948
0252535
0254623
0215339
0227888
0254423
0255556
0242296
0270074
0251402
0250794
0145002
0255549
0283674
0270154
0242607
0255122
0284746
0284663
0244289
0255253
0284724
0284613
0254581
0177400
0254917
0254724
0264512
0154359
0266206
0283523
0254813
0255494
0250305
0251468
0206996
0164741
0287134
0200630
0253932
0253257
0252179
0164743
0254491

0286798
0255457
0255413
0254753
0254425
0255437
0185053
0254575
0214366
0252264
0206950
0252035
0255365
0199127
0255076
0249368
0104723
0248423
0250483
0287476
0226836
0253648
0038945
0253951
0253934
0253496
0253184
0264092
0078579
0155970
0249258
0244113
0104219
0198791
0155975
0003987
0214607
0003989
0279271
0280453
0279903
0279932
0104213
0277466
0287303
0129422
0265520
0253168
0212280
0253944

)253671
)104760
)253215
)254054
)078674
)104763
)245281
)253384
)286542
)171428
)254015
)253937
)244593
)285624
)271654
)156006
)156011
)187229
)244018
)283402
)278886
)201157
)253335
)270255
)253557
)254242
)250316
)254201
)253280
)279919
)104611
)279858
)147408
)253270
)104613
)175445
)253775
)242709
)036565
)147416
)222267
)061337
)253733
)274467
)253164
)253178
)253300
)253866
)254260
)254092

)253147
)254026
)253199
)254040
)280215
)286346
)254334
)168546
)273484
)147443
)130227
)158806
)158815
)158856
)158863
)275074
)168453
)168476
)168481
)168484
)168487
)177725
)285881
)168490
)208037
)168495
)254064
)197181
)104635
)241026
)201761
)120910
)251034
)254272
)120896
)287467
)254230
)120913
)248235
)241852
)158941
)253200
)147439
)261026
)179388
)253125
)287812
)134020
)248738
)253142

0240116
0245025
0008853
0120889
0246130
0253616
0284948
0284956
0173535
0173530
0253930
0104689
0243265
0246582
0250714
0147457
0104679
0134013
0253837
0197217
0253390
0253483
0287166
0253986
0147454
0252067
0253488
0167034
0180053
0253471
0254002
0271118
0207027
0159167
0207201
0253891
0042980
0253535
0134028
0069206
0253643
0272163
0104722
0277586
0283327
0272157
0253832
0288055
0253670
0287185

0147459
0284858
0274988
0147437
0253476
0104756
0184661
0253100
0253342
0285890
0254178
0221818
0199620
0249213
0221914
0228451
0104765
0253430
0251203
0254362
0240694
0092964
0253208
0120907
0240519
0253813
0275115
0215288
0221616
0253899
0253903
0253888
0015592
0104228
0120899
0273836
0120903
0120915
0234770
0120885
0284280
0168077
0222862
0253560
0279302
0265075
0284351
0253875
0147419
0171320

0202242
0168078
0253176
0168079
0265847
0253290
0253615
0253397
0189233
0134014
0250516
0253907
0253690
0239198
0168081
0186918
0214050
0286804
0254370
0207361
0253567
0104290
0265251
0202411
0012232
0246339
0104299
0254034
0242958
0147421
0273710
0259196
0200719
0259366
0197892
0259607
0254129
0253770
0120875
0285601
0260253
0253632
0253182
0280147
0243844
0251191
0253490
0248964
0254086
0253604

253279
264788
254013
239175
133872
272256
104660
241511
177669
271869
104671
253708
285669
272375
248159
254316
287590
253346
254152
127589
254109
157110
279041
197265
253112
253457
254172
104687
104691
271081
104695
133863
253204
253793
172733
165392
253579
253961
229211
253377
254095
253275
287204
222468
157168
286648
253974
252261
200246
254049

0286131
0212407
0247134
0254346
0254090
0253393
0253993
0252735
0272338
0172728
0254061
0129696
0198042
0239039
0253232
0252505
0253995
0133874
0240189
0133878
0253642
0253684
0263794
0253569
0240738
0188512
0254194
0254302
0253108
0254033
0215264
0254344
0243181
0287152
0156687
0279099
0232024
0286693
0253452
0253524
0254309
0253504
0254384
0253685
0253610
0206871
0254320
0253363
0244708
0199985

0215262
0274489
0219491
0243503
0254038
0253650
0253123
0253746
0253161
0253388
0253414
0254111
0254290
0254306
0253344
0223215
0183779
0253181
0183154
0147475
0233170
0147471
0020181
0104221
0156675
0280064
0241032
0169154
0285880
0188778
0270865
0187840
0260949
0286677
0272128
0129691
0253437
0240015
0147465
0253356
0175324
0253739
0156735
0248585
0285632
0085788
0147535
0147548
0255487
0254898

0272092
0165046
0077782
0272159
0255201
0239218
0196166
0253361
0253570
0254321
0147526
0207258
0253586
0253829
0169499
0253645
0169495
0169490
0168615
0207199
0197140
0243095
0253160
0196115
0197475
0254067
0168619
0104755
0253332
0253233
0131203
0253939
0253838
0254287
0188676
0253790
0253381
0253665
0176907
0254143
0253802
0253354
0272479
0165061
0254383
0206867
0104332
0263372
0239281
0253509

0206852
0238936
0253396
0147533
0253133
0253943
0147536
0253174
0287535
0158669
0264578
0165066
0029534
0274705
0283450
0253389
0207101
0260588
0241834
0083168
0238966
0244235
0253135
0271938
0261449
0070718
0104368
0253408
0104365
0070501
0240395
0104371
0078668
0168575
0254131
0254165
0176209
0147432
0255101
0255361
0147434
0131931
0286837
0120925
0242719
0284062
0168172
0202514
0254673
0168522

200731
185900
254198
165102
253611
253884
255166
253312
253186
253606
234713
253707
188877
238509
255497
254342
201329
250637
254069
253319
253418
254145
253748
253195
253845
253198
276685
253425
248498
254070
251470
228984
250857
286484
286554
253314
253828
282321
252594
254118
254156
253803
253745
253782
215179
222099
248347
285992
200986
188873

0248531
0254348
0255366
0253502
0253817
0215177
0164808
0253330
0269924
0251354
0254065
0207369
0221869
0253729
0207450
0287959
0104738
0222522
0169139
0253843
0253654
0252710
0277418
0224594
0279881
0253140
0253688
0253608
0233858
0253702
0253455
0253892
0034239
0286878
0233109
0019549
0287788
0286701
0168333
0253474
0199640
0254268
0253380
0253206
0253810
0253484
0271148
0147481
0253849
0254392

0254358
0253340
0201316
0253664
0147485
0254025
0254225
0272076
0168300
0253475
0228801
0272024
0253844
0254136
0147488
0254314
0253551
0253924
0240534
0196711
0023287
0271555
0183729
0248911
0250494
0254687
0082556
0260484
0253369
0274295
0237807
0254204
0240919
0206144
0253668
0047249
0147509
0272457
0244159
0200528
0187735
0253667
0260955
0120992
0254274
0276353
0137547
0251835
0270980
0253840

0199212
0276605
0286471
0244107
0164736
0274310
0270275
0104237
0226098
0253114
0286348
0206579
0253976
0253857
0248522
0254357
0167904
0222955
0137574
0254087
0254325
0199405
0240905
0241997
0236814
0008988
0238650
0253603
0248578
0172680
0221093
0181690
0170791
0272343
0254216
0170786
0253542
0181195
0254254
0253541
0246430
0253139
0241431
0254055
0270328
0206975
0104331
0222205
0253301
0251142

)253614
)215117
)202265
)253871
)243050
)287301
)253821
)253322
)254139
)205293
)253376
)169122
)253523
)253116
)242595
)288041
)253281
)286578
)215114
)250031
)167910
)254103
)137575
)035681
)242970
)198846
)201231
)254048
)167912
)206853
)201763
)254150
)253260
)253879
)253413
)253723
)278428
)254775
)254954
)178538
)251396
)226301
)104388
)255321
)228862
)255289
)171316
)254432
)287975
)271377

0254777
0255397
0254802
0177182
0213881
0254222
0253711
0198363
0254285
0222898
0264408
0251127
0254119
0253834
0254120
0253544
0253789
0237810
0285971
0185942
0254050
0253975
0287105
0253517
0253431
0274956
0240915
0137563
0253121
0137561
0253555
0270673
0185728
0261542
0241964
0253205
0215096
0253894
0253583
0252358
0287844
0253554
0253762
0253734
0253231
0254377
0207816
0287615
0253750
0253800

254102
180828
172817
254330
287998
254006
253493
239261
280725
213873
254081
272010
200714
104442
066855
272155
205268
272192
147570
253190
147573
147571
253138
287127
246145
179041
285655
285791
147576
253623
169085
206949
185697
175073
201365
104205
213005
178460
245910
254341
261787
178125
253223
121022
104218
252637
250869
066777
287214
271966

0241961
0165078
0254253
0255130
0255531
0046889
0255206
0254051
0252210
0253964
0248801
0165084
0236191
0239184
0254337
0253658
0238808
0254332
0253373
0253479
0253216
0137573
0137571
0222889
0254557
0271113
0246528
0277740
0255107
0239580
0223293
0280393
0247570
0253721
0147596
0212473
0253173
0140396
0213003
0207036
0241204
0253143
0253967
0275128
0241105
0287230
0067167
0246366
0147592
0243532

0221947
0285579
0223220
0254031
0104313
0253124
0254277
0253287
0200191
0235531
0178860
0241942
0104321
0253317
0250979
0252509
0212989
0254347
0182674
0253226
0278193
0253726
0253784
0260838
0147601
0222488
0272254
0253636
0164764
0237061
0253564
0274443
0254213
0147604
0121039
0250295
0253235
0253339
0253302
0040341
0254273
0253383
0254105
0254538
0258677
0104343
0253273
0272469
0286113
0251633

0154582
0175606
0244295
0253833
0154589
0212348
0254288
0240047
0212485
0253983
0253115
0104369
0104381
0253596
0207417
0266502
0283505
0254349
0253341
0254080
0283188
0253706
0253777
0137558
0254043
0121005
0254238
0253712
0249395
0253229
0254023
0164749
0287352
0222231
0270866
0271167
0254300
0253661
0253681
0254305
0091656
0276673
0253416
0266712
0164751
0253952
0254366
0254001
0254266
0252935

0171033
0260398
0286675
0104427
0104432
0254352
0253919
0285744
0261618
0253659
0253391
0240540
0104435
0287052
0164683
0272138
0249328
0238884
0285758
0147586
0276418
0272518
0272264
0076554
0254205
0241550
0265588
0241746
0253238
0213791
0206649
0253237
0252884
0251867
0277604
0205189
0254162
0260317
0188856
0252057
0253760
0223327
0254181
0243424
0164684
0254331
0212429
0243951
0253516
0286763

0076641
0254060
0254177
0253214
0254014
0253677
0272425
0254027
0164687
0253859
0147588
0205186
0170323
0253374
0237264
0197416
0251521
0254151
0240427
0205184
0133731
0253598
0280193
0104231
0164695
0253334
0104497
0231942
0254689
0254394
0254123
0206228
0253503
0253836
0253581
0254189
0253423
0274116
0253898
0254202
0254195
0253362
0184672
0206701
0253137
0248978
0254212
0203413
0253655
0253283

0133739
0260493
0133740
0254208
0176731
0185015
0258256
0257962
0133742
0164879
0253549
0104267
0254380
0253680
0270971
0273567
0205176
0276006
0275928
0276540
0271381
0276998
0274343
0270416
0277436
0277422
0253154
0285768
0287927
0147614
0227753
0243761
0147613
0253675
0253699
0164893
0254231
0123124
0176623
0250569
0254088
0085719
0170289
0253292
0205333
0233915
0254115
0253778
0253568
0176571

0253500
0253401
0251563
0254376
0273978
0176566
0253171
0156103
0253553
0251904
0250962
0271156
0286984
0253621
0248231
0251136
0104312
0253349
0275103
0207359
0164823
0104320
0104325
0104327
0254180
0253394
0199354
0253799
0180694
0246792
0254182
0123119
0254251
0253250
0155099
0253358
0254219
0253738
0155100
0214954
0253525
0266194
0147606
0253901
0200151
0271202
0229182
0079102
0253576
0233778

253118
253682
253749
253634
253177
288005
248858
254020
253577
253197
205133
251347
277063
279847
246662
253960
268080
254089
222416
253781
188343
271971
253854
183808
253722
279331
253848
164953
254057
254387
164951
264448
284288
251234
240509
275386
253585
079112
249510
164949
244217
197275
265817
164944
254315
253704
253175
254190
254283
261437

0104413
0274701
0241003
0253429
0199701
0156162
0254307
0164941
0175305
0272509
0243979
0156170
0164938
0253878
0253528
0287322
0200525
0245080
0254248
0284147
0283522
0175895
0253773
0253351
0156172
0253945
0253872
0202112
0254079
0248762
0183663
0271349
0270131
0156466
0156467
0254224
0156469
0156471
0199732
0253105
0202095
0169439
0104324
0272249
0270077
0260640
0180543
0207215
0213750
0287654

)147649
)202399
)104341
)270682
)270861
)253639
)243504
)132561
)156482
)245970
)207067
)252755
)177459
)132541
)104356
)104361
)252558
)104375
)278914
)253468
)156486
)272321
)207378
)253911
)254112
)240870
)253415
)254312
)223281
)271930
)164920
)253948
)229625
)253180
)132549
)253562
)253532
)236583
)237704
)225017
)206699
)207804
)216069
)243254
)235683
)253539
)253912
)164919
)287952
)132554

0212414
0221753
0156509
0147669
0104450
0202310
0253299
0199667
0034677
0253217
0253824
0265599
0253842
0253666
0254364
0186106
0251013
0174226
0253740
0252764
0070756
0284570
0221527
0202001
0242399
0222795
0212994
0275041
0260368
0164924
0242315
0248599
0253395
0253355
0202360
0253737
0261087
0120963
0271882
0253942
0253679
0287479
0253282
0239115
0271560
0228224
0254063
0253644
0239211
0254084

0254024
0083307
0253991
0253153
0104490
0253629
0238372
0282989
0266756
0254492
0048392
0246263
0253328
0104517
0272037
0253923
0221387
0271612
0240724
0253633
0252593
0155087
0253382
0155090
0283959
0250684
0253669
0155096
0253385
0253320
0253263
0244052
0201216
0253663
0254236
0155097
0253795
0253556
0253977
0250929
0236939
0164929
0247081
0265657
0253851
0164930
0286337
0164932
0266289
0207399

0164933
0285982
0164934
0261670
0271830
0253477
0176406
0255105
0254644
0239344
0147647
0164935
0179577
0279138
0208032
0147650
0253454
0169946
0283157
0243939
0253997
0240271
0254041
0253318
0243171
0251003
0253420
0254141
0243601
0253526
0201627
0213708
0254021
0254615
0164830
0253582
0253676
0251892
0174429
0275905
0287289
0254146
0287208
0154188
0286766
0248838
0287949
0200806
0147655
0253228

0230948
0104408
0253754
0243287
0104412
0253657
0253949
0164841
0253580
0253796
0174417
0120526
0250267
0120533
0205038
0254185
0147654
0147642
0248050
0255099
0283555
0241385
0164794
0255402
0240590
0253956
0253107
0253122
0253717
0253994
0253877
0254241
0253103
0253489
0254124
0253434
0255033
0253156
0287942
0222146
0164796
0286946
0230169
0238399
0227514
0287819
0286937
0253207
0206719
0254339

0253499
0253756
0104447
0227170
0249917
0199450
0287826
0253672
0147677
0147679
0164754
0253327
0283956
0205002
0164756
0244033
0253271
0253622
0240151
0286308
0252852
0164758
0239917
0182197
0177570
0225885
0281641
0242206
0164761
0207334
0254247
0184374
0254278
0147676
0253972
0276479
0286282
0206776
0136999
0253398
0254037
0136960
0253495
0201896
0064313
0286362
0136982
0279347
0244642
0245330

0155792
0202356
0254343
0187955
0172172
0172167
0172164
0250290
0248318
0272502
0253619
0271219
0239872
0170961
0248690
0254209
0254303
0253814
0255364
0253819
0255325
0238901
0263443
0235165
0272384
0272043
0178764
0253372
0259631
0253607
0136986
0156787
0253463
0287657
0147689
0221461
0253258
0253410
0204949
0266324
0189376
0259305
0243498
0165156
0222607
0253336
0156802
0252297
0207704
0283172

0254388
0254294
0156795
0278374
0156804
0254113
0251840
0253286
0175946
0104537
0176853
0214814
0254000
0181171
0254249
0253868
0250727
0253885
0214803
0206724
0253227
0237586
0164983
0183665
0245149
0170881
0147687
0253106
0284087
0147684
0170873
0255491
0263735
0283609
0249816
0250428
0255313
0286955
0180938
0253513
0104549
0164961
0253167
0156831
0242170
0254431
0173334
0253111
0286122
0253470

0245164
0206695
0253651
0254227
0250563
0253841
0248720
0244791
0253427
0253220
0207138
0253530
0168672
0253438
0253543
0212451
0253573
0254166
0282961
0224722
0287781
0246228
0271509
0212993
0280997
0286266
0286010
0249375
0136997
0249859
0283710
0278324
0280055
0221771
0207110
0283200
0278275
0222501
0221176
0221261
0201782
0286034
0254275
0229140
0287785
0253225
0222755
0253926
0263763
0285108

0224110
0147697
0254317
0153310
0243402
0200075
0254263
0253720
0287195
0253043
0264653
0153317
0200304
0280543
0212342
0287201
0286535
0253656
0155897
0253259
0253992
0253407
0253507
0252158
0254255
0132294
0253117
0258417
0132297
0184156
0253521
0129295
0165071
0287095
0129292
0270137
0223697
0042832
0240237
0155926
0276961
0287736
0104415
0104419
0243537
0253638
0225259
0008513
0283176
0283197

0261220
0212273
0276193
0253970
0253916
0253561
0253165
0254313
0253593
0277576
0271240
0288067
0253627
0066827
0276140
0248492
0277732
0278454
0271048
0272456
0278898
0279518
0259820
0207582
0199153
0254028
0253433
0251218
0244413
0254083
0254372
0131773
0253915
0199652
0254101
0286927
0285683
0206678
0274572
0285817
0254076
0253288
0254361
0253988
0147724
0169436
0253432
0251744
0253783
0254258

275078
169427
167632
253574
250733
274064
282164
259891
104472
123908
279766
280326
280303
169398
254324
279786
252864
253266
105339
253210
022567
280035
253307
254019
254197
254291
253595
261655
204882
244998
184489
226807
271959
238854
253605
287325
226490
287677
221768
254183
261710
287332
265247
254008
171045
253602
261693
181790
232722
198576

0261044
0277332
0234616
0167653
0160886
0253741
0130193
0253806
0126233
0197353
0253196
0284505
0283992
0180155
0167656
0253715
0283686
0253728
0104499
0253421
0160882
0179142
0247317
0253971
0160932
0177335
0250115
0243328
0261667
0176956
0277494
0181638
0264668
0250571
0253716
0185730
0272172
0184428
0212221
0254389
0158106
0253931
0254338
0182759
0014164
0287576
0254859
0275558
0104518
0204839

0255050
0147813
0279605
0254741
0104529
0179886
0104524
0104522
0254812
0183309
0181135
0181097
0255181
0254574
0181085
0254548
0180921
0265660
0203499
0254973
0180900
0284224
0179950
0287222
0255343
0185189
0284142
0285599
0261150
0288096
0178209
0207574
0178685
0178719
0186583
0204791
0178814
0277888
0255224
0178896
0284200
0197858
0179091
0179526
0179632
0179698
0235173
0227473
0179832
0261236

0284054
0260428
0185122
0185000
0254690
0284229
0261678
0261587
0271698
0185803
0182325
0173137
0071894
0284310
0284139
0283770
0272115
0147804
0160948
0160949
0276598
0232600
0187954
0254578
0167702
0160973
0160972
0255182
0167701
0167700
0160957
0265393
0160959
0254402
0213563
0147799
0255456
0274772
0254533
0255085
0263612
0198169
0196378
0244307
0161016
0283764
0197363
0147789
0170619
0286681

)196150
)170631
)196922
)255164
)254618
)255559
)255530
)255000
)182307
)236875
)181404
)227518
)283921
)218839
)284986
)277631
)227917
)231808
)170122
)172785
)287480
)183784
)107099
)235880
)227155
)107104
)235330
)228115
)226403
)202172
)229875
)227914
)137090
)064218
)252991
)228783
)286684
)281769
)173253
)224836
)236594
)233575
)202383
)287465
)224092
)080503
)236199
)222973
)264615
)236404

0231052
0147852
0286670
0168263
0080608
0236496
0229184
0227835
0236511
0080298
0215297
0232104
0237359
0226669
0107249
0236724
0237009
0230001
0228322
0200941
0106688
0106686
0225893
0205808
0273061
0106993
0147853
0287759
0270639
0220105
0120158
0233186
0199065
0228165
0220937
0096968
0234534
0182347
0227730
0228166
0224083
0235917
0228097
0237711
0236254
0230225
0236567
0254017
0120210
0237553

0120211
0107014
0214195
0107018
0272866
0231967
0107020
0214190
0120217
0286162
0197646
0231509
0107036
0225408
0099219
0223621
0183354
0120215
0263575
0137040
0228739
0237274
0137033
0228132
0233846
0170777
0147854
0233367
0178445
0264530
0237158
0206147
0236924
0266863
0229057
0231381
0232946
0229611
0225489
0107077
0231108
0256968
0230581
0224972
0285688
0288042
0230207
0178654
0273056
0137038

0231902
0226376
0153707
0235389
0212695
0239102
0225706
0234021
0212829
0265735
0230920
0234360
0226717
0232035
0227138
0230589
0270372
0234146
0230365
0286474
0275130
0285784
0224935
0231491
0222581
0107165
0235448
0243115
0153714
0222658
0285637
0214121
0234586
0107186
0234740
0223672
0226197
0270547
0205636
0229268
0235533
0223568
0213954
0147862
0234982
0225472
0287708
0237137
0273399
0272871

0229029
0175893
0147869
0164946
0286500
0283647
0199814
0232000
0214110
0215237
0234297
0251960
0155158
0215236
0164975
0251834
0164985
0278754
0240613
0213362
0207433
0164989
0231376
0207360
0205549
0173068
0231756
0230694
0234779
0233551
0237153
0285169
0241152
0226470
0237083
0044459
0224488
0107295
0228810
0178031
0227167
0264638
0252960
0236857
0155875
0236680
0155876
0147874
0251733
0252943

0201118
0147872
0232176
0137145
0232978
0272842
0273226
0137154
0234853
0285911
0177076
0260912
0231909
0155886
0232158
0286685
0276667
0261402
0270712
0277711
0237434
0171843
0263790
0222202
0264941
0237088
0188352
0227071
0207609
0202189
0188921
0230208
0171855
0225027
0275980
0177047
0199635
0137080
0232281
0236637
0226597
0214042
0186803
0223684
0147885
0234829
0232138
0228083
0224416
0147873

0226393
0198642
0120235
0233816
0188379
0261696
0231195
0235108
0120242
0171889
0237691
0197919
0238271
0184995
0199177
0264379
0227443
0229835
0244230
0099810
0264545
0229298
0265194
0279670
0264801
0224854
0147889
0240498
0266446
0147883
0215221
0274055
0277653
0236921
0176399
0234840
0284418
0236739
0224549
0287696
0283982
0227607
0233906
0226626
0107105
0287843
0231460
0237414
0277608
0287139

0205442
0269957
0223088
0222693
0287318
0198680
0236306
0270683
0276759
0279256
0254261
0252580
0253400
0120159
0232466
0240306
0137055
0096872
0234676
0184434
0254396
0120156
0223162
0232867
0283945
0120160
0120162
0237734
0287355
0147896
0285103
0147894
0215441
0260390
0260412
0174482
0229060
0287551
0224955
0215966
0215939
0287038
0230276
0277646
0229834
0230097
0280683
0232060
0239792
0215210

0235626
0226246
0233278
0214020
0228557
0223421
0232203
0260720
0226998
0229080
0235148
0252313
0225347
0122729
0107201
0197579
0235453
0165264
0232303
0122728
0223440
0230516
0238001
0230867
0188133
0272113
0236796
0231193
0137074
0223807
0237437
0236184
0225693
0086061
0122692
0086062
0233554
0252224
0122711
0107262
0086065
0086102
0200544
0165269
0286322
0165272
0270542
0165271
0275651
0226823

228072
230453
225123
234373
226562
204982
205274
231991
183938
234122
229063
215206
240215
237984
281128
231165
260947
287063
227301
287495
233776
010438
235481
223394
107341
200834
251748
137073
238300
238886
226801
222259
228352
198876
224802
239392
165006
227383
279336
252164
233668
186638
222426
229207
164978
164976
164972
207032
164970
122735

0251697
0168913
0230729
0232983
0122756
0237159
0164967
0137100
0205143
0147955
0213930
0258728
0137070
0213927
0261215
0187186
0230074
0172724
0137077
0205108
0287368
0257198
0187791
0178723
0215199
0122733
0281491
0137094
0273597
0279489
0233148
0286782
0287868
0279608
0174038
0241568
0165280
0221829
0165282
0234181
0165283
0005238
0215198
0198722
0279083
0179766
0244213
0198853
0287287
0235619

0215187
0227852
0107140
0263892
0137101
0231393
0227933
0137078
0269900
0277027
0159884
0137135
0243136
0107159
0198467
0137076
0284195
0107175
0273945
0070610
0107185
0215183
0228843
0227388
0159899
0137098
0137133
0285645
0137103
0204930
0236083
0168828
0233659
0196196
0235387
0287986
0225639
0278889
0228307
0213866
0179443
0243641
0236110
0122707
0276730
0122694
0185972
0122705
0159921
0222760

0215283
0137075
0165304
0287514
0266255
0196092
0258885
0264922
0265368
0250850
0125385
0232920
0260100
0281649
0233242
0147905
0200502
0228292
0206784
0252723
0224648
0137106
0236156
0168795
0187988
0137054
0230188
0234160
0147912
0256966
0175768
0255872
0233111
0070601
0201287
0165275
0107371
0122741
0232454
0226055
0122696
0213839
0107338
0251745
0200026
0223716
0137124
0137142
0283162
0225254

)215165
)229511
)237487
)243085
)176007
)273036
)283486
)272934
)272904
)225345
)180071
)204860
)237557
)284612
)252725
)283329
)228467
)287838
)234680
)106714
)226625
)271454
)243695
)278825
)204849
)277024
)215112
)224038
)231140
)198416
)229707
)269692
)273994
)234720
)226108
)232239
)278175
)229273
)204837
)283886
)277774
)252724
)232798
)284116
)184961
)237846
)230804
)228136
)236966
)283211

0283246
0183148
0240240
0212441
0234812
0231390
0223839
0224681
0227069
0232433
0274157
0278122
0274067
0184906
0276500
0276266
0225380
0276963
0278331
0275390
0276868
0275377
0276291
0277778
0280077
0279456
0225655
0276462
0273514
0215126
0276029
0277049
0279561
0275450
0278647
0274680
0276502
0276832
0275160
0277340
0276094
0276562
0288225
0278077
0278054
0276412
0275239
0278028
0274688
0278001

0275297
0154529
0237207
0270909
0273717
0185775
0239986
0277260
0276277
0275649
0268234
0277213
0234861
0227353
0227924
0237357
0212952
0229311
0204816
0230245
0216829
0233424
0287276
0204818
0224537
0236816
0286793
0238529
0273900
0275144
0230635
0229079
0213816
0276128
0277737
0286506
0287800
0286916
0277598
0276581
0275628
0274516
0276422
0278485
0283541
0276040
0277412
0276149
0274583
0275676

0283378
0276522
0277109
0231248
0238933
0204814
0231454
0273940
0231527
0204805
0170152
0235659
0223379
0238749
0229697
0276845
0284471
0231995
0231212
0236450
0252133
0275230
0204802
0227449
0233651
0226429
0224603
0232827
0204801
0226007
0212951
0278213
0234217
0236825
0234451
0241571
0234276
0235256
0238113
0287168
0202474
0229422
0184523
0237451
0234665
0238245
0227582
0219693
0170161
0273509

0244081
0198312
0182021
0176115
0237238
0186466
0228522
0233555
0234042
0268951
0224267
0225360
0228656
0226020
0239684
0225411
0207277
0232815
0266017
0215548
0204790
0275068
0285580
0172014
0238551
0224185
0229156
0224106
0224762
0235832
0235310
0277869
0181997
0176134
0225883
0235932
0156755
0271225
0197550
0223602
0206804
0287988
0224599
0236252
0267085
0204780
0204779
0204778
0287329
0276723

0280286
0147996
0184659
0204776
0237904
0235893
0231701
0279955
0279688
0274893
0273668
0275026
0278068
0278720
0274628
0286945
0229146
0276257
0276203
0275030
0284611
0284658
0233765
0274181
0276894
0277297
0273849
0276457
0274349
0277490
0277350
0276039
0274355
0276348
0274820
0275969
0276386
0274852
0277575
0273999
0276204
0276898
0275904
0274026
0286079
0278763
0275493
0274098
0278010
0277618

0260691
0278551
0226904
0196873
0279706
0187559
0234394
0225337
0154330
0224958
0233178
0229019
0181778
0226337
0234506
0224025
0107242
0187866
0236998
0236733
0207000
0165059
0165060
0285130
0270236
0119139
0278910
0135063
0107282
0243888
0229312
0261447
0188647
0225626
0204711
0241174
0204706
0165072
0222465
0268364
0236801
0198887
0274421
0239180
0119138
0083067
0272232
0232086
0223966
0207935

0224437
0229814
0135048
0271166
0107362
0200922
0155621
0236060
0204669
0286823
0224820
0119125
0235523
0233933
0252428
0225434
0107372
0230003
0225947
0165091
0236862
0229753
0236849
0165092
0228024
0135046
0221844
0286840
0232590
0236701
0212454
0223368
0227809
0224825
0198963
0286944
0119121
0202217
0270954
0252758
0224297
0135045
0203321
0156017
0229587
0278040
0106733
0134996
0200041
0212316

0235707
0236376
0234644
0099139
0231137
0135002
0234618
0187210
0233814
0219149
0106772
0225937
0274326
0277320
0286332
0230335
0236604
0229857
0204612
0232851
0233526
0232998
0197969
0156049
0231373
0228248
0201711
0156052
0233457
0222452
0233947
0228781
0234819
0148019
0135069
0270627
0237257
0226234
0260995
0186940
0233086
0106829
0270691
0238002
0232494
0260564
0279715
0280366
0230945
0236521

0232981
0225901
0234637
0226798
0228375
0237072
0196781
0233926
0207206
0252167
0200969
0240632
0230846
0189357
0267559
0186788
0237770
0228430
0214929
0231649
0230360
0235377
0225085
0228046
0228123
0232749
0278988
0286612
0233309
0165105
0286451
0251776
0273806
0229109
0172159
0237529
0251752
0199483
0148057
0135018
0254473
0165113
0226877
0231616
0165115
0165118
0165119
0284179
0235298
0178966

0202523
0227463
0270166
0197506
0233262
0285987
0228095
0148053
0226590
0285556
0287930
0285634
0227412
0135049
0234424
0229454
0230303
0165121
0213711
0231741
0135040
0135052
0287333
0233713
0187753
0270688
0223012
0135070
0083223
0214908
0235819
0222293
0232211
0285686
0180447
0226237
0269994
0231007
0224887
0204446
0212421
0228373
0286842
0196730
0219928
0236709
0135047
0237747
0188029
0276478

0237743
0232743
0277357
0234575
0233884
0214888
0234460
0177992
0230246
0238110
0283205
0156345
0271923
0217716
0287750
0225385
0177910
0276535
0279154
0204429
0229288
0226327
0287769
0106723
0275017
0227638
0231804
0130045
0228189
0286414
0265873
0225856
0228658
0213694
0148082
0224945
0123975
0265112
0187742
0187764
0231799
0225460
0285907
0130222
0237372
0236507
0226206
0237626
0227555
0263967

0234910
0278382
0279503
0185074
0225072
0231107
0229613
0165023
0236316
0270432
0165025
0287212
0228216
0260454
0237422
0230537
0229694
0235641
0230147
0273381
0148090
0165030
0233081
0169071
0228595
0266855
0283351
0090054
0227399
0237306
0275756
0232179
0228069
0238288
0240791
0227321
0234537
0225511
0236717
0224809
0276717
0236115
0196305
0281156
0239183
0198000
0188312
0229530
0106809
0127083

0106819
0106823
0264158
0225551
0285650
0127080
0213669
0185963
0288062
0288271
0187984
0252772
0226721
0275318
0127081
0226668
0232843
0188101
0127084
0157303
0223446
0165233
0131669
0203364
0165238
0204352
0200570
0201074
0237385
0188938
0048828
0197724
0283146
0231990
0265347
0131668
0235601
0158079
0227666
0286834
0286203
0269929
0199165
0199072
0230262
0199133
0175787
0237679
0130950
0232063

0224815
0282886
0148110
0278486
0286850
0224245
0202445
0231806
0130957
0165140
0288023
0148120
0252153
0230815
0236095
0224764
0285807
0274115
0207563
0207864
0207617
0284459
0158169
0229065
0223849
0251884
0230221
0271384
0213761
0185920
0271314
0271155
0271659
0229345
0228142
0227358
0175611
0268926
0252847
0182150
0199202
0237631
0225194
0237212
0201938
0275465
0287187
0224546
0130948
0285269

0232283
0207276
0130958
0165244
0238251
0286835
0287302
0130956
0081377
0158122
0224848
0233820
0081386
0196597
0196312
0269907
0227257
0188152
0236745
0136943
0197214
0228376
0228038
0213612
0277103
0159712
0252721
0242375
0188801
0203279
0235494
0254876
0255036
0254483
0254633
0197816
0221269
0228957
0286375
0196116
0136842
0228174
0136925
0136937
0231521
0136936
0214417
0236130
0260677
0178919

0136932
0201830
0287070
0136929
0136938
0236896
0212521
0095380
0106785
0274011
0106789
0095383
0278412
0224001
0136928
0229897
0165138
0285706
0119514
0267026
0229338
0204291
0270412
0106799
0252942
0227269
0240235
0119523
0106803
0222337
0255815
0271086
0277828
0287955
0277465
0237461
0255145
0234860
0119508
0136874
0254571
0023318
0234709
0119509
0239908
0238172
0270995
0231697
0136891
0066697

0251349
0241697
0170681
0200966
0237999
0230161
0229541
0148123
0238143
0253041
0227411
0236763
0230328
0136881
0234502
0136897
0136870
0136872
0270982
0225376
0165152
0233055
0155827
0198785
0188386
0230302
0230418
0238882
0286243
0235901
0204250
0232242
0234269
0155833
0229830
0225564
0226566
0253031
0285483
0270332
0136824
0242111
0203396
0186881
0283001
0148136
0204246
0186943
0237443
0277556

0276119
0136839
0273423
0235248
0179055
0188712
0136783
0165028
0165029
0226334
0230013
0286536
0201583
0070214
0106701
0229419
0228317
0106692
0279609
0186051
0095209
0228366
0232486
0228948
0234323
0225002
0229297
0277889
0237062
0234229
0200131
0228143
0212276
0148143
0230782
0231562
0223615
0225531
0226047
0226535
0119318
0226825
0287233
0213558
0199905
0136826
0236005
0227350
0202308
0222558

0244104
0231293
0233727
0222459
0236783
0213557
0237360
0230030
0169253
0231678
0231810
0213556
0148156
0187003
0070061
0119328
0119326
0106771
0199331
0207698
0260230
0095203
0200106
0237124
0207329
0070159
0263861
0213539
0157654
0232939
0228053
0188959
0136810
0204193
0165124
0206658
0232896
0030304
0223918
0198121
0206923
0207401
0212409
0227531
0274263
0171133
0136813
0223238
0173258
0106853

0224644
0204173
0059769
0244115
0242616
0165181
0224067
0206907
0148154
0266315
0222356
0259953
0106868
0222327
0200261
0225513
0119314
0242256
0199525
0243911
0230081
0213529
0119471
0230185
0165185
0250068
0148153
0226609
0148158
0119457
0271631
0228623
0237906
0285447
0136866
0285115
0284763
0284962
0285076
0284867
0285370
0225684
0231528
0136867
0119321
0136868
0176386
0136875
0165188
0148225

0119411
0119431
0148218
0148229
0157653
0138835
0237073
0282897
0233817
0227482
0214669
0232905
0157657
0106927
0136883
0196739
0207726
0229314
0228278
0106948
0235119
0095397
0271214
0235797
0136888
0157693
0201749
0230054
0230601
0230284
0274919
0181634
0229817
0173077
0106952
0041982
0236461
0234692
0228714
0274755
0204148
0228707
0182752
0226604
0244757
0256040
0148219
0229105
0230894
0119401

0285820
0230734
0207268
0201140
0222413
0233516
0233569
0235133
0285082
0136869
0201444
0284977
0231901
0234782
0225050
0235201
0231166
0078725
0260970
0261432
0225960
0207814
0136861
0106780
0238181
0119402
0214654
0270917
0095261
0226752
0119403
0056558
0106804
0119397
0119396
0243738
0148180
0214651
0239593
0235865
0148175
0227355
0204136
0240299
0213467
0136848
0229854
0261534
0175764
0237548

266583
119421
185681
106852
119446
148187
095303
269970
233616
234156
136834
197233
239590
171505
171501
171496
228914
165202
280094
173679
171481
226783
136939
171459
232387
148215
136940
165204
233425
056586
212447
186130
171448
261094
286718
011454
222351
213216
188394
225078
236901
165209
283941
148204
119522
207991
274932
237336
230826
229115

)234881
)106689
)236668
)231227
)234921
)119408
)235204
)227200
)136930
)180264
)136931
)231149
)148200
)231744
)241246
)224020
)207595
)207737
)236643
)185585
)136918
)136942
)136950
)136935
)199313
)173611
)177736
)119414
)232630
)235332
)136933
)044574
)239705
)165219
)214593
)200554
)227845
)274510
)287782
)119487
)243845
)227068
)214761
)229582
)167081
)232413
)228392
)196814
)253079
)252985

0233721
0279137
0226078
0224842
0136944
0238010
0286428
0169155
0177125
0136828
0136859
0228487
0136895
0229369
0271833
0136856
0196152
0197958
0201302
0148356
0200788
0136830
0136854
0279571
0187024
0283874
0160401
0167094
0160404
0095370
0286196
0136807
0283863
0284547
0136877
0106991
0225032
0222421
0222455
0106992
0257524
0263979
0160408
0136840
0167103
0227218
0136908
0167106
0230848
0171169

0148339
0230536
0234771
0148334
0232850
0148346
0171159
0148337
0106976
0207581
0264823
0167110
0175854
0167112
0272696
0167113
0167114
0223551
0167118
0272593
0207955
0284185
0283335
0167123
0273186
0228510
0136811
0225951
0119392
0228395
0238406
0197694
0280474
0280758
0119333
0227759
0234705
0119335
0160447
0264438
0160446
0223478
0160445
0203286
0107021
0167136
0198917
0286112
0171097
0136802

0175287
0251184
0175283
0095319
0267834
0148341
0148343
0287234
0167130
0095321
0234055
0119383
0268707
0188483
0268050
0204055
0235007
0241084
0224307
0179082
0268615
0233901
0242281
0236146
0230676
0226355
0204054
0261334
0286078
0227619
0148335
0179058
0148331
0234789
0167157
0236024
0148344
0233909
0136816
0136827
0136819
0136878
0276124
0224988
0187239
0230684
0270755
0148358
0223188
0196979

0107130
0148357
0264169
0130707
0107164
0277713
0232172
0130711
0130713
0097007
0224797
0188710
0130720
0236658
0050555
0252582
0126878
0126883
0252622
0236986
0246851
0226987
0126882
0286547
0160539
0285287
0130723
0235284
0231587
0176868
0130714
0230289
0130717
0283526
0107263
0240853
0231018
0212395
0236687
0160563
0196358
0107290
0125482
0188523
0252521
0125492
0125485
0125484
0165695
0224992

0165698
0165699
0165702
0263816
0234907
0238558
0196205
0148308
0273932
0170835
0170827
0160271
0285245
0148288
0171102
0119440
0175164
0201843
0204031
0148296
0201451
0148297
0230064
0148303
0206611
0200831
0199744
0252542
0148290
0148291
0148248
0198870
0148300
0160323
0160325
0160326
0187616
0197859
0196990
0261018
0123454
0225756
0123453
0160293
0235106
0169925
0235138
0271525
0273249
0196363

)221676
)273473
)223729
)279929
)186350
)263897
)223795
)228877
)227150
)130635
)204011
)286502
)264744
)265872
)265848
)264163
)264136
)266827
)266514
)232355
)160339
)085265
)236403
)279935
)130558
)227958
)233936
)178243
)235572
)224682
)225361
)196422
)160345
)122140
)226706
)160349
)122136
)122133
)237339
)224045
)236543
)204007
)148386
)165643
)107147
)130559
)238058
)130560
)148411
)260193

0238227
0275329
0107187
0165661
0260220
0279813
0283769
0262075
0160360
0213221
0187796
0165684
0165689
0165688
0148384
0148396
0196366
0148400
0263403
0227512
0265181
0237886
0279141
0228401
0172889
0199161
0169692
0165716
0233016
0280496
0276161
0274998
0281808
0274356
0187922
0204003
0267206
0278756
0204001
0177984
0224662
0244187
0272896
0213213
0273066
0272679
0228544
0196642
0213212
0265806

0232434
0054148
0177943
0107223
0127191
0266507
0213590
0260190
0159069
0176919
0184925
0198454
0107317
0284341
0214402
0148362
0169583
0107331
0180539
0180549
0279073
0107281
0054179
0236394
0229257
0186193
0231864
0197355
0268996
0177239
0199411
0176978
0176884
0184709
0261793
0263697
0185863
0176248
0176101
0176058
0187713
0188566
0284976
0212864
0197191
0233198
0198569
0188229
0188163
0275549

0197768
0188986
0198113
0270259
0260996
0288234
0198435
0187609
0188747
0188833
0165802
0284317
0130653
0182154
0148399
0165724
0197070
0283411
0203993
0181090
0274841
0233998
0255585
0207693
0203987
0148408
0276388
0229926
0159247
0278130
0237419
0233013
0223256
0250210
0286255
0260370
0237297
0261456
0232420
0015171
0276662
0212331
0151240
0201861
0239822
0233021
0225140
0263511
0180525
0231601

0107929
0229869
0205740
0107937
0148377
0232656
0067064
0047056
0229205
0185736
0287043
0282390
0233052
0205696
0287653
0234962
0226694
0278069
0252998
0227912
0231326
0212156
0234556
0234170
0237500
0235281
0286858
0231743
0234182
0287560
0273001
0067057
0278419
0107959
0237399
0226762
0236968
0227338
0287428
0286610
0233321
0236892
0229672
0228353
0067082
0287175
0230573
0226005
0274809
0227101

0236990
0233117
0224382
0230096
0287246
0286687
0207124
0231298
0225269
0287023
0165568
0151631
0238840
0187134
0151632
0224251
0239148
0196139
0239142
0225418
0264006
0198610
0228291
0287848
0231039
0224034
0215267
0223987
0280450
0178473
0178462
0173848
0178372
0205488
0178363
0256462
0287566
0287235
0242147
0231483
0228951
0228685
0240577
0196372
0108021
0226647
0057608
0270427
0272764
0134461

0240180
0134452
0232807
0134470
0134460
0251922
0229664
0214015
0134453
0170525
0201581
0263628
0283634
0213994
0238366
0215244
0226705
0212743
0065675
0237943
0225948
0285743
0223784
0285988
0281186
0238266
0285845
0287277
0223581
0270811
0198879
0235141
0278982
0232591
0207453
0230121
0123243
0151655
0151657
0165629
0165632
0233990
0107485
0197308
0232638
0225053
0232170
0226861
0226990
0271251

0213771
0236420
0212505
0270234
0234752
0223808
0230014
0287120
0235838
0228636
0224788
0229919
0200849
0272436
0286439
0234306
0270111
0279575
0287309
0228997
0225383
0048740
0229240
0231496
0228027
0229206
0237986
0230322
0181800
0280086
0148429
0271360
0271046
0134463
0148426
0225778
0151461
0251957
0181192
0272507
0065665
0265653
0165609
0151465
0228302
0264036
0183049
0252438
0252118
0285994

0263584
0283699
0221331
0285520
0151468
0212499
0225251
0123240
0235763
0234175
0228330
0203414
0237540
0065328
0272055
0165623
0107537
0226209
0086475
0286514
0233256
0165626
0202508
0227175
0165630
0227474
0282246
0239665
0151474
0273474
0225112
0199407
0234091
0215227
0229751
0235410
0285671
0266321
0278100
0221371
0065809
0236495
0201766
0237032
0185267
0284024
0187522
0272853
0152455
0152457

0197889
0235623
0228547
0182531
0237470
0152463
0176244
0228312
0176236
0152464
0152465
0206448
0228181
0148468
0232739
0077943
0148481
0252537
0237913
0201260
0226140
0165983
0223156
0165985
0287925
0148484
0225213
0275001
0286958
0107611
0273153
0107614
0229124
0026025
0234961
0148488
0204832
0251803
0236582
0165996
0280126
0260589
0136738
0229190
0148483
0260314
0207938
0235637
0148482
0226083

0165995
0235020
0240291
0225527
0241058
0165997
0234244
0225509
0234813
0204740
0226255
0227734
0239539
0236960
0200545
0252001
0233968
0283456
0230818
0252832
0285852
0120594
0238246
0287613
0228339
0270727
0265372
0078114
0235940
0230112
0228753
0228219
0204683
0231920
0228860
0232853
0207264
0230109
0231553
0278650
0273566
0199222
0204682
0222071
0180592
0078403
0286881
0207347
0223984
0201390

0136770
0252634
0279623
0201364
0232497
0236777
0223601
0260205
0227462
0286832
0279631
0261671
0233318
0148444
0269897
0168283
0272516
0271981
0272366
0077327
0233451
0286810
0285254
0150867
0279819
0206842
0165309
0286924
0223131
0148450
0229932
0252049
0168267
0179133
0227537
0224215
0264574
0165312
0287124
0286442
0120549
0223928
0277563
0207930
0286951
0279029
0107863
0199535
0285750
0099256

0228667
0240294
0285859
0273107
0151023
0185875
0231422
0233642
0151025
0274242
0222543
0227465
0280809
0271563
0223019
0251711
0226304
0095777
0136750
0284951
0077420
0199733
0236894
0233261
0234296
0231976
0227932
0279212
0148459
0234788
0235843
0136754
0223027
0206605
0283709
0107890
0199855
0136758
0120539
0252639
0107897
0251839
0262412
0230445
0238021
0278830
0200369
0215409
0207135
0260151

)213770
)237956
)182077
)099246
)285737
)150051
)230500
)169126
)251810
)226016
)233472
)226395
)150054
)286413
)275036
)282996
)237746
)222666
)237128
)224439
)222705
)254635
)095787
)252401
)231911
)207483
)095739
)232624
)236716
)201001
)235824
)271220
)199402
)229605
)120563
)287402
)224597
)274985
)197321
)207612
)212411
)216035
)225976
)222092
)237936
)165757
)200887
)230125
)259994
)107951

0285824
0250833
0241621
0239625
0284277
0214434
0235679
0224473
0107968
0213778
0287420
0228800
0239744
0230298
0285630
0151033
0234814
0230720
0183621
0272914
0237135
0287564
0272381
0237036
0252479
0148516
0230397
0285781
0223834
0286694
0231855
0237797
0222412
0165322
0231148
0230240
0239731
0170759
0206660
0234335
0235113
0234811
0252482
0217094
0233190
0227253
0120616
0226842
0233825
0229327

200484
286409
216937
231245
273038
150093
228816
229656
234163
265319
229878
235962
233387
099250
238258
244356
273012
287278
226412
203565
261683
226407
223470
148498
199200
234167
229235
215184
226386
232525
223047
222909
213724
283000
108094
266228
230534
095794
253054
233343
269952
233200
253070
108100
200097
271335
287945
273312
177291
177283

0287528
0284265
0226646
0280719
0227313
0227475
0236626
0228694
0237002
0274458
0229644
0207271
0238176
0233533
0225355
0148513
0206796
0234918
0235687
0244402
0252621
0237970
0236313
0224920
0225276
0256892
0226578
0198105
0260137
0225192
0228828
0236514
0151963
0235197
0175395
0226447
0206840
0189180
0252132
0275675
0226086
0235964
0075407
0283930
0272983
0225965
0224761
0226113
0234941
0099251

0120555
0276805
0222314
0228882
0203496
0215151
0277133
0225638
0233867
0227264
0234541
0227679
0233832
0229485
0237592
0271650
0274167
0228755
0215146
0182632
0279239
0185904
0234420
0270762
0233837
0196693
0277479
0272319
0272387
0251783
0285884
0233515
0232109
0231009
0234864
0230425
0270552
0259869
0165733
0252416
0234944
0229630
0263795
0165731
0273008
0169826
0198915
0224265
0285712
0221468

0150201
0169813
0221400
0230555
0243660
0252532
0236114
0196793
0287852
0232004
0198298
0223910
0237389
0238263
0169740
0226245
0230565
0228426
0271173
0224162
0223746
0234973
0226808
0226790
0236769
0233395
0227029
0237590
0229116
0277288
0287901
0107562
0214089
0270767
0287173
0234580
0224812
0227395
0234504
0187783
0223462
0273363
0107551
0165507
0165511
0165512
0226937
0277757
0270258
0228462

0232554
0235690
0243349
0236442
0230501
0263476
0227683
0275312
0278531
0207071
0234993
0224402
0256574
0012779
0231964
0165406
0172671
0230869
0237840
0228702
0172661
0270434
0231122
0188234
0239883
0239152
0264217
0265354
0265733
0266412
0263639
0280012
0264204
0263482
0274209
0279058
0273225
0279458
0264230
0285402
0285294
0213412
0204174
0204175
0204176
0231187
0229227
0165874
0272430
0229927

0274186
0285662
0280913
0265366
0215097
0204177
0238405
0243289
0150175
0204179
0266524
0263761
0265203
0265763
0265190
0270369
0189090
0224919
0204172
0254929
0252877
0251079
0265630
0215065
0266217
0264404
0189014
0276544
0226964
0277758
0276850
0264717
0273760
0276430
0265018
0252149
0270025
0278561
0170324
0285786
0225482
0107643
0279822
0128805
0248682
0231906
0251413
0236800
0128815
0228754

0286993
0228403
0241577
0165383
0233665
0226576
0264800
0165633
0234736
0172538
0204161
0236208
0177354
0165606
0235939
0225830
0271237
0070748
0187714
0178645
0197444
0226389
0229870
0227345
0230166
0204152
0178440
0222108
0285803
0204149
0235618
0226631
0099290
0233011
0188611
0225137
0198964
0286401
0279863
0225303
0238523
0226200
0231588
0231345
0232706
0226168
0230011
0213667
0287221
0204147

0148584
0229711
0236944
0261368
0185532
0223502
0231132
0207813
0235279
0213659
0177613
0236671
0107984
0213657
0227972
0231131
0165471
0201196
0285958
0227121
0231399
0226296
0252161
0233805
0150275
0252252
0234173
0224697
0221594
0236958
0236744
0223800
0228048
0249860
0236993
0270541
0122952
0287016
0238707
0264747
0235810
0223841
0270192
0232115
0151151
0180221
0122873
0228527
0261076
0072401

0108064
0122870
0183055
0252076
0237949
0226888
0226557
0165443
0148541
0236556
0287969
0235469
0235140
0165449
0227877
0108091
0235931
0151150
0232682
0202190
0282121
0122872
0254271
0170312
0072422
0223107
0227244
0196932
0237233
0288052
0183346
0272592
0233643
0150347
0182010
0234756
0288011
0240940
0138311
0285837
0285551
0238280
0234489
0181915
0122877
0199446
0224301
0148572
0171988
0234369

)270421
)221063
)235816
)272767
)224412
)165476
)286373
)232075
)270352
)228566
)213609
)226426
)235489
)233900
)270648
)216740
)224714
)270754
)282906
)227173
)233724
)235356
)228065
)183230
)273360
)277056
)198739
)225299
)226448
)276866
)213606
)225922
)108176
)212520
)239942
)237141
)272892
)252113
)219932
)096717
)148634
)226318
)266467
)236375
)138347
)222371
)179774
)233590
)234102
)108187

0096746
0204130
0138346
0234806
0199638
0122912
0226493
0207438
0214881
0138336
0213025
0227583
0260400
0270494
0060339
0195401
0283551
0221184
0165730
0200218
0285871
0107625
0206855
0165732
0198954
0214875
0229001
0122862
0274008
0122958
0233163
0156502
0229261
0156510
0231748
0156515
0229197
0075073
0232734
0099282
0287306
0233197
0122859
0236154
0230755
0235803
0235524
0231152
0226794
0171224

0235645
0230469
0197467
0224222
0099284
0042286
0156521
0079332
0229097
0232142
0180817
0148734
0172731
0224725
0148730
0156574
0280401
0107719
0230661
0180644
0138316
0166220
0231366
0166224
0166228
0285300
0271409
0259267
0107731
0237512
0198246
0279406
0107736
0223817
0214688
0107738
0274824
0197746
0238446
0230526
0122863
0272988
0107742
0138303
0200294
0226701
0166295
0200170
0168209
0269926

0148719
0107745
0202513
0282915
0276531
0226163
0230609
0272536
0231471
0238215
0201881
0156026
0266719
0280166
0279502
0272627
0138315
0215086
0138308
0213700
0122884
0232646
0237768
0201047
0272630
0166321
0272599
0221164
0122882
0138286
0213551
0199201
0227382
0236756
0182180
0227540
0156042
0200356
0207473
0138279
0233144
0229659
0166343
0107758
0221817
0166348
0207327
0237139
0268584
0272791

0177791
0166317
0271848
0172650
0242288
0271816
0272140
0252072
0242338
0250959
0235316
0270808
0279088
0176986
0279689
0196968
0172586
0214655
0280238
0272916
0272589
0166507
0148660
0224195
0229990
0122861
0222047
0035403
0225761
0185009
0156110
0236900
0214629
0213731
0232342
0226253
0206756
0271710
0214626
0213525
0272748
0156650
0234149
0227186
0188716
0285810
0237882
0079393
0156671
0232437

0165637
0165644
0226051
0227271
0233313
0270087
0223428
0236797
0165655
0237149
0273248
0272692
0236842
0148655
0279814
0207583
0286715
0268659
0228280
0265486
0201954
0278992
0230575
0269256
0252888
0224948
0156113
0236467
0225497
0224500
0225652
0228092
0199592
0213514
0277981
0199664
0213513
0228748
0151208
0204049
0243446
0233871
0234612
0274686
0148606
0138326
0231957
0283096
0282863
0230417

0288090
0229543
0230229
0201393
0282952
0228683
0224596
0229569
0108175
0108179
0165424
0235426
0259021
0241317
0235705
0253626
0271985
0232965
0185303
0219430
0225827
0122852
0226676
0244733
0225484
0272489
0280355
0278616
0224886
0188199
0272447
0228570
0214695
0204437
0214584
0283913
0283101
0242600
0133661
0273372
0280259
0235924
0225100
0229616
0224760
0236217
0227515
0230091
0282432
0133678

0279399
0230171
0189129
0122359
0237523
0234319
0234382
0234743
0279359
0151224
0232950
0235858
0170788
0133665
0122378
0108219
0226659
0178217
0231082
0233353
0233703
0231060
0227209
0226466
0287358
0185737
0285739
0229458
0225738
0207275
0200774
0234931
0233945
0286876
0223565
0212324
0233258
0200789
0212769
0271933
0165678
0285972
0188373
0148600
0204033
0148604
0148602
0229404
0107771
0233189

)225423
)225928
)178429
)288218
)237267
)287057
)223993
)224504
)230962
)279910
)223064
)280134
)234942
)182771
)278333
)270002
)252292
)236606
)200487
)199104
)287475
)286359
)062650
)212332
)227896
)230507
)122375
)122367
)272631
)107779
)270874
)200176
)173269
)281974
)173267
)272734
)148671
)273413
)271880
)240089
)151303
)272508
)261011
)229969
)188100
)271573
)200253
)148672
)122376
)237280

0273946
0223482
0287077
0285257
0184923
0224914
0214562
0237740
0236417
0228055
0231569
0286445
0283672
0107789
0223761
0225913
0196566
0234192
0198682
0138138
0223820
0243782
0227268
0171862
0224745
0213613
0200891
0227905
0275015
0184719
0201548
0204022
0237205
0182333
0271408
0204021
0226358
0204020
0235150
0173239
0152766
0234256
0138134
0180139
0107796
0286116
0026103
0275693
0261438
0265375

0283664
0233292
0138135
0107798
0119922
0232110
0119917
0224289
0204010
0185745
0152778
0152779
0234452
0152782
0198997
0249962
0232936
0225836
0235100
0240996
0138182
0283578
0232229
0270670
0280560
0226159
0228938
0287150
0222451
0236373
0224750
0238291
0283053
0148680
0148688
0222305
0148677
0201604
0225519
0273124
0224851
0220585
0234043
0286981
0180628
0165338
0271385
0223876
0119938
0213449

0228759
0228701
0107854
0180581
0273497
0174721
0272817
0095564
0107864
0252993
0214549
0213484
0221042
0105988
0199881
0198060
0202297
0232709
0286795
0119912
0138160
0184111
0264313
0236493
0152804
0201412
0138190
0225118
0226425
0187553
0285846
0279607
0095596
0232805
0237417
0213432
0138119
0138180
0212396
0186188
0138207
0095464
0148690
0108231
0280660
0280485
0227995
0176273
0180764
0138193

0232913
0231575
0234724
0206631
0228553
0268894
0273450
0173145
0108239
0119969
0244332
0235746
0108242
0276490
0165841
0230338
0235199
0225354
0138109
0233377
0228460
0260863
0237885
0138115
0231829
0274691
0225533
0173124
0234026
0107438
0095637
0214362
0059573
0119977
0138185
0226688
0240527
0270099
0188649
0227805
0107443
0266407
0200728
0287009
0177853
0229417
0095585
0229418
0107447
0197430

)095587
)077147
)235677
)155629
)202047
)227032
)207287
)236552
)207976
)196233
)232875
)187122
)234855
)269891
)286683
)213390
)231398
)277308
)237169
)165879
)181274
)225850
)052749
)231508
)231970
)200737
)171314
)171311
)171307
)155229
)165886
)165887
)241935
)249967
)203942
)171160
)155252
)119986
)155254
)155256
)120057
)227356
)155265
)095713
)278752
)166024
)138131
)230928
)119943
)284415

0107521
0264610
0287261
0172987
0274056
0236818
0234109
0119946
0120053
0224934
0257582
0228778
0119919
0155287
0260475
0285932
0229278
0198018
0233690
0119929
0014919
0023839
0227351
0107554
0227695
0120054
0283232
0232230
0270623
0107566
0213341
0236308
0227492
0095485
0212464
0230224
0207362
0196072
0107593
0231188
0099194
0235823
0212325
0075290
0075826
0255339
0166136
0166135
0274660
0075891

0222072
0272572
0119906
0273476
0055950
0095539
0207551
0236662
0107815
0107816
0186862
0107819
0273162
0107821
0236311
0107807
0237579
0270973
0138136
0227128
0229649
0222238
0225208
0230967
0166167
0222414
0166171
0166169
0283558
0263436
0107829
0222051
0286489
0224817
0107831
0107833
0198408
0226009
0120049
0120029
0274742
0222430
0166189
0224302
0229629
0198728
0148840
0166197
0119915
0107859

0107862
0077150
0059915
0107872
0107874
0202569
0269609
0120055
0138111
0138107
0273262
0107882
0228657
0207029
0171206
0272933
0138175
0156398
0166272
0252994
0148795
0213061
0236937
0282772
0166275
0270316
0283337
0214435
0235266
0286575
0272912
0148842
0278498
0076685
0237827
0213277
0229256
0235376
0148798
0156374
0212413
0148835
0173915
0283867
0148843
0138172
0273485
0234699
0185933
0183128

235470
107954
226431
287419
107957
273108
222503
280693
260461
107960
065613
266754
065618
283788
156384
197748
208033
148834
266852
065621
148841
270075
287779
228261
231233
120051
225768
237761
156395
226387
207068
233477
213260
229466
287732
286609
229775
215002
108018
287047
200626
200079
229981
234544
203434
223381
224498
222436
252574
200217

0232728
0237242
0270333
0270294
0224907
0108039
0272160
0203876
0148700
0226734
0119950
0228417
0214067
0119953
0273143
0237642
0138166
0233518
0108055
0223302
0203867
0252036
0199364
0232470
0203497
0150593
0284482
0214413
0278601
0270589
0108061
0234118
0221214
0150594
0213247
0237618
0227851
0230809
0287231
0119927
0119913
0276857
0283688
0243316
0197142
0232934
0023041
0151532
0234017
0264763

233340
285676
260917
279315
225292
148737
233547
227560
238380
286289
230018
148702
197893
165806
234393
148735
283656
212589
198924
196865
287857
043591
285540
253066
226582
165813
273904
284442
095627
165816
230165
275024
169129
242912
279796
099204
229668
228169
228484
215760
151553
226428
165832
252611
236799
107518
285582
270965
151892
285587

0182645
0205433
0203837
0227096
0200935
0175535
0232091
0187021
0266200
0165863
0232767
0165868
0228195
0188316
0187164
0287655
0235742
0148704
0234474
0266782
0236171
0186795
0225936
0165646
0277879
0165650
0258114
0229847
0170370
0235198
0288172
0234952
0263041
0107560
0231104
0177640
0238276
0165669
0232139
0234309
0119973
0225155
0151893
0277687
0229272
0236058
0213574
0188613
0107581
0207468

0222588
0271343
0119979
0183605
0165672
0198873
0228485
0242853
0236426
0265719
0148908
0271353
0151923
0234569
0227437
0151929
0232823
0198825
0242818
0270300
0282413
0197771
0107651
0265370
0180230
0223455
0220842
0234522
0203805
0177234
0274461
0227165
0287936
0120008
0271670
0227307
0278484
0236762
0227143
0222979
0066468
0223432
0223540
0224250
0107669
0201884
0226864
0273767
0276742
0107672

0273891
0138162
0285715
0285973
0138152
0202245
0107679
0265442
0285955
0254636
0166033
0187908
0286114
0286135
0183559
0176584
0138161
0286088
0213185
0255624
0203795
0119965
0179988
0095574
0196177
0188620
0188816
0154473
0227586
0270902
0230131
0225152
0286344
0234542
0154478
0121898
0231138
0234677
0287768
0213438
0232334
0182022
0065154
0229544
0271434
0107902
0229455
0258539
0189319
0278831

0233334
0203791
0199466
0165660
0207494
0214807
0249456
0019995
0226899
0175029
0273599
0264572
0282787
0214298
0233135
0237675
0175018
0214297
0228021
0224023
0280561
0234134
0107938
0236991
0154485
0188690
0265092
0107949
0089876
0277371
0222629
0203780
0234016
0233409
0148848
0222740
0213170
0287609
0235180
0154493
0287326
0199321
0150760
0223528
0232935
0188916
0214285
0279725
0186766
0270778

0180745
0132334
0232259
0227076
0287427
0148773
0280953
0234640
0224190
0232985
0287361
0225424
0285615
0170430
0287466
0237224
0227374
0283141
0108001
0275327
0266676
0275005
0287997
0237489
0277218
0233122
0233319
0108010
0236303
0276375
0222997
0285529
0264803
0176769
0230098
0189275
0286295
0280446
0283095
0175470
0176171
0277959
0273521
0279982
0188385
0235010
0151640
0165752
0148814
0171813

0231705
0234311
0180066
0226900
0229081
0068383
0148826
0171811
0232903
0224758
0240707
0197177
0256925
0235264
0171798
0171794
0151650
0166917
0284219
0273980
0151651
0130640
0235245
0198546
0283496
0130643
0230306
0226699
0165828
0148803
0127884
0265395
0274685
0148832
0254536
0148824
0203772
0273327
0214279
0273336
0130649
0278518
0286789
0171772
0288107
0233685
0287327
0225899
0213543
0233435

230468
279626
280337
274599
276964
277274
275610
276164
237635
278664
278761
278790
276046
276904
278641
213147
282875
253826
224777
254468
270921
230724
255229
222225
177951
188076
177947
254559
177963
283920
142082
185627
277290
174885
255026
142102
206013
255533
185201
254910
185885
251661
142089
255089
270105
270030
255328
270972
182272
184363

0185187
0185101
0243562
0174915
0255237
0023191
0288033
0254739
0174775
0161328
0185522
0254815
0099849
0247095
0199038
0070047
0185507
0099834
0070031
0069696
0177030
0255158
0177106
0177042
0269915
0177156
0177225
0255284
0279672
0255142
0184524
0177542
0274897
0177595
0177600
0199785
0177666
0255108
0177685
0177697
0177700
0214063
0250397
0177830
0183020
0222561
0184956
0254872
0198788
0215182

286275
117983
255177
275669
078902
255153
287935
285985
174672
285581
182208
184545
233930
205869
205867
196224
241598
185940
231487
205866
205865
227306
205864
244242
250644
117984
235027
229512
287484
230834
230980
230080
149043
130598
130592
264493
249086
284594
283787
229671
130595
214026
226416
232987
130600
275266
284010
274866
167244
284779

0240801
0129965
0207805
0099869
0254647
0180176
0265258
0183734
0110665
0064201
0207308
0238184
0236264
0199550
0110651
0184281
0230483
0070985
0053918
0269821
0275666
0276105
0276494
0276015
0236041
0229414
0237941
0274740
0129757
0254827
0110628
0181649
0205531
0207008
0183562
0110619
0247473
0201616
0285644
0236710
0021762
0234791
0182170
0236301
0184350
0224513
0236987
0276951
0232390
0273813

0254592
0005801
0223756
0189398
0166492
0284605
0254757
0287906
0255367
0287312
0255522
0237226
0223688
0284639
0230851
0238161
0182048
0167311
0129744
0201279
0200201
0129749
0110713
0251934
0238304
0148985
0177105
0228661
0167323
0229368
0284525
0230593
0206976
0255232
0255141
0224555
0240385
0167325
0255276
0231548
0254480
0255387
0284018
0284306
0236175
0283873
0284546
0284438
0221996
0175800

)132109
)225101
)237272
)171999
)181963
)196778
)226616
)197790
)171987
)226288
)232268
)167333
)197428
)180785
)180723
)167332
)197674
)167346
)272634
)272664
)271602
)272559
)273051
)201980
)280021
)176951
)231070
)279270
)176925
)197984
)176922
)176904
)198217
)176900
)197023
)176895
)176893
)278870
)227023
)205497
)205496
)225003
)176798
)224300
)230261
)205495
)176787
)273085
)237354
)226157

0203560
0236897
0205494
0171944
0182070
0176752
0176748
0176742
0224091
0285498
0221031
0244734
0223609
0229988
0260629
0213934
0284931
0196565
0239920
0213931
0229093
0183251
0167355
0223735
0279012
0236359
0176239
0230087
0236248
0184698
0184321
0167360
0248424
0167359
0187918
0230484
0181609
0175520
0175518
0224295
0249633
0181616
0248553
0231049
0233646
0229534
0187747
0121236
0258588
0258659

)132256
)132274
)232381
)181109
)181023
)181074
)181017
)181009
)181001
)236965
)180988
)205409
)183269
)233563
)180974
)277932
)277514
)184478
)188691
)183313
)254831
)183389
)202147
)180934
)255481
)262980
)234895
)180919
)254444
)180913
)180909
)255307
)175485
)267940
)180878
)051009
)132259
)110148
)270435
)170955
)282556
)254443
)166311
)166313
)110169
)110171
)132254
)283977
)132286
)254595

179532
132275
254400
166333
166337
254641
285338
166340
255680
166341
255390
255410
158042
241678
254838
254781
188124
279486
184933
283415
254712
166363
170790
170782
166368
178358
149054
149050
158077
170748
255179
221703
170743
251364
183801
166387
254864
166394
183378
275612
276668
271758
254707
176716
227892
254951
166408
241008
183303
182334

0170688
0252769
0279000
0170683
0182261
0175390
0254642
0255420
0166402
0254921
0248332
0279391
0166405
0246820
0166407
0130413
0252778
0166436
0254900
0166441
0200983
0212607
0166444
0254665
0255159
0252905
0202276
0166452
0176029
0176009
0223080
0175348
0254860
0175352
0253973
0175356
0286935
0264984
0213538
0184014
0255097
0273725
0166471
0254884
0205339
0201998
0254397
0268403
0166478
0166483

0238387
0243964
0255463
0239470
0133789
0201564
0245522
0246273
0133812
0255476
0252568
0254765
0286561
0254865
0254719
0254554
0148926
0133805
0221574
0255823
0283813
0110315
0177112
0133800
0072952
0206858
0198730
0110321
0238622
0246308
0236287
0247271
0250041
0254401
0254957
0270897
0255123
0255260
0110328
0254598
0255351
0255462
0266645
0280288
0255309
0286959
0278529
0170242
0255492
0050165

0254486
0254991
0255400
0133816
0275373
0254680
0254983
0197702
0254847
0286521
0187079
0254688
0255067
0252329
0203258
0251381
0254670
0189431
0255558
0255018
0133794
0222162
0148925
0287548
0254583
0255401
0152266
0228901
0197601
0254791
0240454
0200685
0255075
0255088
0255411
0254927
0254930
0201856
0254438
0262655
0212365
0254418
0133818
0255074
0129083
0251991
0256206
0129084
0152270
0255040

186104
175868
253179
110680
250640
188487
253072
254946
254789
254695
254645
286960
254661
110693
284094
254878
213785
241943
110696
166689
199883
244398
184669
110700
272034
201403
011405
201586
240808
213779
070081
197149
285545
188211
260196
187486
006071
255035
287898
006611
188162
254586
255335
129152
129159
129158
255448
129167
166788
255254

166787
179826
255470
285562
254857
179817
255138
254546
189332
148965
255071
134339
200336
212789
173432
201695
110756
110768
264603
134333
256006
166796
256734
166800
074319
256464
256588
256361
151116
274589
247595
256282
279837
179119
257043
151117
179057
254966
110786
286998
276789
254529
255511
254541
255536
170255
255244
254738
254720
255424

0183695
0177054
0129170
0255308
0129173
0254544
0255160
0166833
0255270
0200687
0255043
0254622
0265210
0207407
0270607
0264309
0254542
0254453
0254894
0279266
0109851
0109854
0185238
0255291
0286382
0165970
0165973
0254906
0201059
0254816
0255167
0225477
0252816
0287753
0171714
0287962
0254820
0255372
0254768
0091664
0254540
0255323
0255357
0183161
0148935
0222427
0198168
0246225
0255359
0255418

0271491
0255293
0275868
0254564
0254465
0240881
0255193
0254861
0252519
0277803
0254594
0207252
0187398
0255489
0255368
0240298
0255454
0254456
0254754
0255086
0134343
0287227
0254819
0255505
0169550
0148942
0176971
0129151
0254560
0109881
0205213
0254862
0242527
0148943
0245573
0278483
0212289
0240036
0254934
0176697
0255496
0177447
0224411
0255094
0121621
0207874
0169519
0271332
0222385
0238149

0273912
0254606
0248990
0249867
0272882
0254526
0255450
0254530
0255281
0255227
0255029
0255117
0255047
0254734
0243505
0240554
0261340
0182255
0254532
0255288
0131808
0152219
0242353
0066382
0255480
0254489
0287373
0170959
0255370
0255525
0170946
0148950
0109911
0228061
0007372
0281880
0285283
0254584
0254836
0254627
0212551
0255490
0049449
0255252
0227160
0255375
0184937
0183242
0278822
0273677

0278045
0276530
0273908
0277119
0149100
0270903
0186714
0244535
0135378
0060749
0200615
0121690
0176148
0213713
0280798
0176102
0247151
0206808
0241950
0200485
0241255
0110422
0110427
0254537
0255207
0255202
0284969
0205177
0085063
0110429
0254508
0255272
0255161
0255169
0254619
0255133
0135363
0184566
0286626
0135387
0135372
0166016
0201867
0254708
0220204
0216708
0121691
0135374
0255271
0255427

135373
271369
270491
149089
110435
251862
255521
026508
255443
251194
110436
255004
215380
255542
149090
254669
179431
254686
166326
285705
186082
285740
254919
179241
254672
255256
241228
263389
280321
110442
135362
254566
255186
255060
175104
280331
166349
175097
166352
254498
251838
242834
254516
279004
254784
255477
255175
254562
285751
240975

)201591
)270588
)255347
)279675
)148948
)206806
)254579
)252922
)255132
)255388
)242729
)255563
)255171
)254668
)255279
)254822
)255300
)255109
)254914
)271028
)254725
)251557
)166181
)254907
)252652
)052841
)255340
)254577
)252355
)254463
)213697
)283217
)211568
)235661
)283341
)199077
)149084
)283393
)243721
)254512
)283483
)246250
)283375
)166199
)213693
)244926
)254409
)187479
)244953
)205126

0110455
0255165
0151348
0052850
0255451
0255106
0085117
0254693
0255092
0244267
0275725
0255079
0255032
0157570
0175274
0254654
0287984
0019485
0254664
0019505
0254497
0254427
0255267
0255041
0286418
0254651
0254746
0255519
0254514
0175264
0255091
0276639
0254519
0255426
0255226
0205106
0181830
0255447
0121671
0121653
0255498
0234776
0121680
0165905
0135365
0254653
0255314
0254871
0255269
0254639

)255007
)157613
)149091
)264102
)110492
)180720
)110497
)244313
)265014
)283497
)285658
)180423
)175224
)175220
)175213
)180210
)175216
)263540
)252427
)212135
)247675
)134569
)286450
)149179
)271350
)255520
)149182
)270060
)165912
)275208
)243802
)134574
)256897
)134575
)025434
)110514
)256746
)283338
)134571
)066336
)255197
)264583
)165915
)165916
)165917
)149187
)270072
)265559
)213619
)110536

0123444
0200376
0231880
0196666
0172247
0109919
0165923
0109920
0200090
0030066
0271543
0252941
0252874
0271557
0254780
0149177
0254879
0263693
0175619
0255514
0172208
0176567
0176555
0176547
0197161
0176540
0182565
0184789
0254925
0255215
0237388
0255113
0255304
0254832
0255534
0213607
0255297
0254674
0255053
0255551
0254728
0254920
0255527
0249910
0254517
0220948
0254764
0205044
0182053
0288434

0214891
0276637
0254801
0229361
0255452
0254412
0255338
0254993
0254567
0086205
0255532
0123447
0254496
0226268
0205035
0254487
0255268
0219061
0265973
0288177
0255550
0255214
0255190
0254800
0255111
0254408
0254472
0254637
0254940
0254624
0258817
0221954
0283605
0260811
0260595
0255199
0255442
0287538
0286983
0236919
0255500
0255042
0255001
0254518
0254840
0214883
0237610
0185926
0255312
0255560

0254769
0254464
0227547
0225997
0255349
0221840
0255078
0150244
0255543
0255283
0255110
0166007
0254828
0254663
0254723
0181961
0181958
0254891
0255077
0255493
0255196
0181950
0255294
0181943
0254576
0233301
0181939
0279514
0172188
0181927
0174982
0181903
0254841
0188439
0254804
0254457
0186886
0181837
0279761
0255019
0186113
0279395
0186119
0205030
0205029
0186124
0255204
0255499
0254547
0124900

0227806
0187612
0167825
0255416
0254807
0174970
0214880
0149133
0182365
0181785
0184741
0181780
0225538
0172154
0254524
0181767
0255140
0181761
0254848
0167822
0254947
0181752
0255217
0174957
0230213
0213604
0254658
0181718
0172489
0181698
0181693
0255461
0280314
0254722
0255134
0150261
0172487
0242208
0172199
0254903
0255466
0272987
0279961
0230416
0284732
0150269
0174937
0254752
0181371
0254411

0254490
0254660
0254603
0185701
0255223
0254834
0166693
0255012
0255172
0197866
0254795
0172464
0172459
0232511
0238041
0174914
0254953
0181325
0172457
0274770
0286704
0254825
0181296
0255485
0205025
0241356
0255433
0274150
0181282
0181273
0255083
0273228
0283307
0272676
0273255
0183908
0134817
0149115
0254662
0149136
0109991
0156575
0186652
0254979
0134802
0252070
0222998
0186907
0255301
0149150

0265566
0134809
0214872
0156587
0240371
0149131
0254602
0208009
0166793
0172409
0156599
0156603
0213593
0254462
0213592
0211450
0254732
0233436
0198561
0255303
0255266
0180714
0254749
0254601
0186509
0200817
0279051
0236541
0255146
0254853
0255131
0272685
0172377
0186508
0272569
0186513
0197887
0280204
0180475
0254589
0172772
0272900
0255218
0255431
0255096
0255333
0197786
0233499
0273408
0172769

0172365
0172362
0285895
0255299
0198283
0110031
0186660
0255073
0269570
0242689
0280010
0149124
0197254
0254926
0255265
0156689
0254717
0166840
0255523
0255240
0254877
0255189
0254786
0254399
0255151
0189057
0245571
0255381
0166801
0110042
0197629
0263999
0225802
0254799
0214797
0254704
0263944
0235678
0255386
0176495
0198261
0172324
0172320
0166884
0254466
0204989
0176200
0172742
0254770
0287264

)255008
)255139
)110048
)264559
)166889
)255355
)223223
)172289
)166900
)254403
)254743
)255538
)254477
)234964
)166902
)134812
)134827
)285231
)270929
)284873
)214788
)285010
)255393
)149507
)149516
)255331
)149534
)254952
)110077
)110079
)214787
)275344
)166926
)166927
)166928
)166930
)156738
)071203
)204979
)283601
)181995
)166959
)214782
)166961
)172689
)255845
)255959
)256813
)110104
)183134

0149506
0110107
0257052
0256944
0110108
0256196
0006118
0110446
0013725
0207153
0256733
0110448
0167987
0279549
0229859
0229183
0256220
0256713
0167992
0167986
0149476
0162144
0279246
0149483
0187049
0149532
0279632
0256591
0167985
0240823
0162148
0255931
0266006
0204950
0011347
0279878
0256443
0250230
0243742
0255947
0134780
0124915
0124920
0134825
0284108
0168496
0134824
0149485
0284416
0221968

0277892
0167994
0200898
0167995
0167996
0279491
0244176
0255553
0254424
0285656
0149503
0236607
0168515
0124939
0254404
0124935
0110484
0197745
0255213
0255118
0162174
0254454
0255126
0149021
0255446
0124942
0250659
0257058
0254772
0255508
0276102
0149016
0149480
0149499
0254964
0149489
0149541
0089597
0185085
0255432
0278615
0162194
0214756
0206597
0204922
0162191
0177363
0168000
0234857
0162188

0214753
0162222
0185670
0168002
0267811
0162227
0168569
0269176
0185475
0284535
0162231
0274066
0162236
0269463
0252361
0256690
0283268
0133316
0222328
0255717
0277194
0277846
0278527
0274544
0275996
0276788
0275043
0168003
0168539
0257002
0241082
0197901
0285388
0149452
0256041
0253547
0197658
0239924
0184999
0196600
0256863
0256847
0256181
0149742
0168004
0275598
0287412
0133317
0133321
0133328

)176485
)203520
)184743
)270269
)256789
)133318
)271100
)188070
)264519
)168005
)256280
)072518
)202089
)167771
)110583
)207200
)256100
)176340
)167770
)133315
)256824
)256481
)126500
)256341
)168439
)149781
)149743
)149761
)256116
)110011
)173511
)173486
)286264
)173457
)256940
)257086
)149782
)002330
)173264
)182450
)257069
)207024
)219435
)173153
)173113
)126432
)236935
)168071
)278359
)162302

0221273
0231680
0231492
0181908
0286816
0232500
0237363
0168065
0197891
0110076
0237410
0068831
0068976
0168066
0269038
0168067
0133895
0171219
0110047
0230835
0229719
0283926
0284155
0110046
0274986
0274314
0068971
0149735
0168070
0168062
0213465
0273003
0278851
0270775
0110025
0168061
0168060
0146670
0162300
0187066
0149823
0254501
0149809
0174276
0255173
0278952
0149806
0162298
0149792
0284489

0254455
0222477
0255058
0204710
0255200
0254614
0014216
0197847
0287821
0014138
0285816
0287917
0149798
0133884
0173825
0255478
0162241
0126391
0245532
0278144
0278050
0277599
0283791
0275948
0173727
0279243
0286756
0251562
0270117
0278217
0274072
0270526
0142186
0168056
0279093
0260233
0173465
0176973
0173442
0173338
0173327
0197136
0266041
0213445
0265874
0173039
0285533
0239356
0172977
0172922

)214659
)255557
)254470
)172818
)255120
)255404
)172757
)172803
)172732
)172638
)172543
)172500
)175602
)175592
)175573
)175550
)175513
)175467
)254867
)175376
)175334
)175315
)175294
)175229
)286459
)087365
)255038
)175115
)255320
)174996
)254461
)254855
)254762
)174903
)245156
)254452
)174871
)174851
)179292
)254458
)174807
)174791
)254756
)174744
)174684
)255468
)174669
)276918
)254736
)202317

0254510
0174576
0201733
0174547
0254596
0174516
0255517
0254986
0256349
0174483
0286688
0174165
0248746
0250105
0174080
0182791
0173992
0199325
0239306
0248643
0173933
0173914
0173898
0239553
0173715
0213409
0254984
0173653
0173599
0173621
0239099
0266423
0173237
0173227
0283326
0173156
0173120
0179038
0287934
0173020
0172932
0172830
0287851
0172613
0175482
0256514
0274251
0201684
0175505
0275484

253024
172531
175463
172508
175634
255949
172725
213402
175514
175544
172663
110711
276769
110697
167797
167791
283257
084207
184224
255119
167792
231793
167799
167800
132744
227834
132746
239559
184795
235855
254447
285537
254883
255147
227620
160172
273819
254792
251637
206587
243672
254610
254850
255230
255554
110057
006534
255306
110717
284293

0284435
0110719
0275022
0255031
0110721
0255236
0110066
0171067
0286369
0162337
0110075
0254856
0260808
0069482
0132749
0110090
0250508
0197345
0132740
0255741
0172938
0261625
0261276
0172935
0256508
0162341
0287725
0265539
0260895
0261070
0284713
0172927
0213275
0260877
0259854
0255980
0255606
0261347
0255774
0285094
0110092
0149716
0162344
0256441
0075388
0186895
0260348
0260679
0255191
0254605

0248844
0254417
0202070
0131620
0255143
0254902
0168040
0254721
0131626
0255329
0254604
0221333
0254484
0255539
0254495
0246889
0085733
0162105
0227726
0279459
0226627
0236262
0207196
0280089
0171671
0263744
0286708
0172900
0172893
0254682
0172890
0278349
0254924
0244411
0241233
0254997
0286948
0204572
0248903
0204571
0249387
0184055
0255562
0255261
0255415
0284625
0239481
0248671
0254805
0254972

0255319
0285933
0158483
0254978
0214534
0184276
0252902
0255157
0254469
0225805
0228915
0254593
0137522
0255296
0137496
0137497
0251143
0207012
0263742
0184154
0284922
0149357
0284844
0110200
0110203
0257075
0256518
0204971
0255860
0110195
0165457
0165458
0165462
0228286
0285864
0213365
0162129
0255843
0256739
0255672
0256403
0237408
0238048
0276109
0274664
0227467
0186642
0256633
0272036
0251919

0255808
0186635
0256007
0286555
0245148
0244134
0214530
0202522
0265064
0285693
0168010
0137478
0206638
0234751
0256568
0215841
0175591
0260401
0256347
0171631
0257038
0110237
0054967
0256928
0054965
0256448
0255786
0256148
0231726
0270323
0021300
0175582
0255928
0256034
0175581
0199975
0256681
0181924
0175575
0256898
0187726
0255847
0175567
0287425
0256189
0256723
0175564
0168014
0256098
0214517

0284736
0280269
0149380
0206913
0246211
0165434
0212961
0254631
0255440
0264402
0175538
0254928
0255444
0254974
0175536
0254837
0077514
0254500
0223202
0054938
0279353
0265902
0241097
0227615
0166439
0166435
0278879
0242999
0241170
0286943
0118363
0200152
0162139
0284722
0234611
0213333
0171561
0255395
0137491
0254475
0255136
0255460
0261594
0137486
0254429
0279117
0254963
0199090
0149273
0206941

0207445
0149243
0158555
0254460
0149257
0255326
0171533
0255434
0254630
0255280
0166391
0263993
0247867
0062282
0254826
0254814
0255507
0198382
0200256
0199362
0255255
0255081
0223013
0254915
0255421
0085741
0254933
0255362
0137492
0179240
0201756
0255135
0158636
0254755
0255363
0255479
0254810
0236304
0137507
0254975
0204529
0254632
0182704
0255100
0261578
0078124
0254988
0198488
0149260
0254550

0137474
0178795
0224829
0149269
0268635
0074201
0238880
0178301
0048649
0206816
0254985
0227376
0255409
0219529
0087884
0254459
0254691
0255449
0241782
0149262
0255115
0254829
0151364
0259112
0254675
0151365
0151366
0159063
0252494
0246174
0188997
0118369
0033327
0254420
0288393
0254649
0251323
0137513
0255084
0252033
0254697
0149256
0254563
0255345
0254885
0211997
0266570
0255209
0279295
0254471

0254434
0200146
0255014
0255178
0254437
0285568
0287912
0254747
0255316
0245832
0270701
0264110
0254511
0255246
0227097
0182103
0255382
0255063
0137509
0254441
0165490
0242279
0254698
0137502
0254522
0207221
0246067
0255503
0279900
0255010
0254965
0255051
0165494
0254676
0247137
0269939
0137494
0255541
0254551
0241020
0137500
0255234
0255003
0274004
0150672
0255311
0254629
0250257
0254713
0240174

0254787
0274006
0276179
0255555
0255396
0213305
0254684
0254897
0206583
0171204
0171202
0137504
0179071
0137501
0279742
0150676
0255005
0254479
0073921
0200877
0254699
0254502
0276669
0074266
0283744
0254783
0255222
0149196
0239856
0149201
0254733
0254727
0278989
0151376
0254731
0255250
0150687
0271318
0197882
0269895
0255203
0255471
0280339
0174804
0246523
0254748
0166575
0254843
0279836
0213287

0270510
0254582
0255391
0279701
0278929
0223015
0283412
0255183
0279341
0254650
0255022
0279793
0285835
0280307
0254794
0255516
0255102
0255241
0206690
0123892
0279733
0266581
0109861
0249489
0288018
0255082
0168959
0207113
0077498
0197692
0279297
0279603
0086991
0255295
0279248
0279697
0255429
0279045
0279056
0280201
0134612
0255385
0279163
0214414
0255486
0255540
0234376
0254971
0255170
0204456

0254751
0255162
0255285
0168930
0250946
0166013
0204455
0254558
0189253
0255235
0254617
0254785
0255055
0223417
0233802
0254818
0278837
0255360
0254655
0204450
0255305
0237706
0225581
0255184
0204449
0254803
0254916
0223709
0255011
0254888
0255009
0254436
0286665
0207299
0279299
0077616
0280385
0110172
0280367
0261645
0214391
0266703
0280430
0221586
0280085
0279209
0279438
0254677
0255070
0255187

0280124
0278980
0280093
0255332
0254961
0278859
0242248
0235547
0165323
0249255
0254705
0244244
0255506
0282916
0255127
0255287
0254874
0134640
0242807
0278892
0279304
0180773
0255445
0280379
0279269
0233737
0255233
0279684
0165325
0166002
0234106
0240202
0279454
0278953
0255515
0166004
0199875
0254911
0275146
0273885
0279696
0166012
0207112
0206799
0202314
0206834
0207304
0239195
0207145
0284458

0210825
0182919
0284057
0042429
0214376
0206911
0255995
0256752
0181333
0227593
0110218
0250519
0256745
0183560
0123901
0020922
0221230
0255893
0168876
0257057
0134627
0196371
0255929
0280167
0233536
0255666
0166025
0256469
0213368
0150316
0186280
0255653
0235268
0274486
0256779
0257012
0255855
0277984
0263465
0149218
0270689
0270578
0245552
0149212
0282834
0250390
0285842
0255605
0273600
0077458

0166037
0087053
0222578
0242993
0184384
0285921
0266192
0149231
0183340
0255679
0200411
0256684
0254587
0255376
0255205
0270753
0255039
0199315
0255502
0254555
0255380
0254939
0254599
0254830
0149972
0223269
0255547
0242165
0255059
0248027
0165895
0200047
0082175
0282728
0255504
0137672
0263885
0254506
0254534
0187151
0110318
0137691
0274097
0260008
0277459
0137693
0254422
0243777
0270868
0271600

0212466
0023445
0110330
0152558
0255337
0255482
0137673
0137674
0256916
0281655
0137675
0118113
0255282
0225678
0256254
0166670
0196611
0256035
0149968
0262406
0255469
0238562
0137745
0276521
0239861
0137692
0260966
0187240
0285878
0254895
0254931
0255417
0255350
0254565
0253245
0254824
0254987
0264200
0170962
0255548
0170967
0280773
0256422
0270449
0254569
0204403
0235505
0196954
0284715
0137757

0137752
0204397
0288255
0254750
0255221
0255430
0255501
0254767
0255336
0278640
0152578
0255403
0222663
0252081
0285813
0170903
0182359
0149313
0254433
0287846
0254580
0286345
0254811
0255065
0287802
0152402
0274584
0213252
0255353
0261098
0152404
0225215
0137760
0270969
0110675
0255483
0254758
0170290
0254702
0110660
0240652
0179331
0288012
0166266
0200855
0255467
0075239
0285696
0149308
0149311

0206967
0166323
0178202
0110723
0178105
0240698
0254730
0255379
0203334
0255528
0201243
0200613
0264997
0255028
0185742
0254890
0254482
0254659
0137710
0254798
0287245
0224776
0270423
0255210
0149289
0254416
0137714
0137727
0233497
0255149
0287556
0200168
0271390
0271584
0287028
0239354
0270202
0150750
0196167
0214290
0288097
0264032
0110777
0253099
0255428
0255093
0137707
0286028
0207811
0207562

0254980
0183644
0204381
0170145
0235286
0223324
0137713
0254990
0086848
0258529
0254450
0254949
0255561
0137720
0176343
0109846
0170276
0254445
0149300
0150764
0238965
0255286
0255334
0150768
0230911
0238444
0271025
0150773
0150776
0150779
0204370
0255292
0150782
0254638
0150783
0197580
0214264
0255231
0243930
0150787
0243250
0188771
0268472
0213234
0250303
0206772
0254968
0213233
0255484
0212397

0285769
0287006
0254626
0288070
0247416
0149294
0238998
0227487
0255129
0149292
0270179
0170209
0256757
0149295
0265140
0166682
0256091
0086827
0243353
0257070
0256603
0228607
0256167
0255870
0257101
0048028
0256452
0255710
0201687
0149305
0166736
0109906
0286463
0256947
0166741
0256195
0180425
0076053
0255663
0076043
0256533
0255982
0270204
0095110
0238724
0137634
0256674
0204361
0252870
0182985

0287897
0256972
0255580
0256315
0256281
0257087
0255903
0255689
0239153
0255599
0260254
0279771
0256717
0279818
0288020
0246100
0257067
0244259
0239600
0236437
0278945
0237937
0137656
0226645
0109917
0110243
0236267
0285513
0110244
0110245
0118137
0235910
0160584
0200537
0234268
0224077
0225236
0254851
0254678
0168092
0149577
0149591
0160613
0280143
0167257
0252992
0186318
0278768
0276505
0110274

0250699
0271412
0177103
0279586
0270403
0137731
0254844
0255245
0254528
0137726
0137747
0110324
0255274
0272075
0137648
0177098
0149575
0160593
0255163
0160588
0149573
0280032
0198851
0254992
0167286
0160654
0110344
0254873
0285827
0167283
0269944
0255384
0118058
0255435
0172425
0149582
0118096
0243431
0095139
0207185
0019144
0255176
0284246
0118094
0255239
0255422
0207462
0287238
0110367
0278376

0254478
0254621
0201535
0160683
0245869
0186174
0264211
0110375
0239726
0222529
0176302
0280502
0283691
0255121
0186166
0240970
0254428
0254909
0118181
0196655
0283704
0137700
0149428
0255114
0271751
0272186
0160695
0256269
0188486
0172269
0172375
0172273
0172350
0160703
0172367
0248712
0110395
0222249
0076706
0284148
0252119
0173456
0245385
0235718
0223953
0036672
0245248
0254740
0254590
0254892

0154096
0263873
0255377
0230716
0254561
0110400
0199217
0255247
0254854
0254406
0287545
0288047
0286992
0254863
0255216
0286731
0137699
0176984
0184232
0137709
0259541
0181264
0196914
0274303
0149403
0255006
0250493
0254735
0154114
0285509
0109927
0254515
0244400
0109929
0232460
0271022
0246790
0137642
0286044
0286023
0252556
0286497
0255248
0207971
0259571
0198975
0207994
0255015
0255219
0252776

0253035
0223265
0254572
0254935
0154127
0286341
0239079
0285909
0109943
0109944
0199709
0254944
0188909
0213184
0243669
0244641
0233541
0109971
0200879
0207118
0202252
0288061
0166250
0255342
0254710
0200496
0254823
0254407
0023171
0265357
0254449
0166257
0254467
0166261
0221931
0200197
0196099
0254613
0254430
0204300
0181518
0171014
0181499
0196248
0198674
0226461
0254737
0236981
0234560
0182634

0213182
0186268
0213181
0110002
0196403
0197309
0239426
0255425
0181214
0276316
0197849
0255298
0255116
0196341
0279116
0254976
0255225
0255030
0227737
0229254
0284680
0284609
0255048
0285561
0280090
0197125
0255510
0229386
0254646
0255315
0254937
0170953
0196119
0255341
0154143
0154144
0110013
0200278
0064199
0154146
0255045
0019102
0149564
0250073
0120458
0279342
0254509
0245498
0254941
0154134

0154133
0254568
0254943
0165478
0221932
0285825
0149548
0134955
0150433
0187686
0255475
0254880
0165495
0259290
0254932
0222844
0149557
0255537
0149547
0222297
0134910
0149554
0252255
0134940
0171053
0196844
0236027
0237353
0260209
0198331
0110060
0255027
0109832
0254790
0283703
0064309
0254759
0261257
0287666
0254833
0254771
0165526
0254694
0197798
0222067
0240098
0182934
0110074
0243607
0150455

0254905
0255062
0110063
0280832
0110080
0278215
0149571
0257271
0254607
0254938
0279358
0254960
0266215
0255317
0218109
0255087
0272642
0273409
0222774
0223315
0254591
0254612
0273415
0272575
0134954
0276176
0254588
0245008
0151702
0254703
0151704
0255369
0120457
0174370
0120471
0134909
0212597
0206847
0199260
0255259
0043039
0240121
0281386
0281097
0287426
0255188
0151715
0170322
0170325
0233220

0255057
0255262
0255535
0084234
0244451
0274197
0149418
0196323
0255358
0255220
0272412
0175773
0134917
0166106
0255155
0236616
0274999
0175728
0288013
0255352
0254842
0255455
0120451
0242673
0236129
0227125
0237612
0231698
0182667
0252351
0237654
0285980
0224795
0271624
0238117
0224700
0238262
0279090
0183715
0283489
0279497
0255371
0279711
0279189
0254896
0287068
0255258
0166118
0213153
0080854

0284020
0255406
0204241
0254648
0166086
0254481
0151503
0254616
0255348
0151502
0151500
0151498
0166105
0254442
0149328
0109956
0255545
0254573
0286374
0254989
0285640
0255344
0251226
0287251
0255327
0279046
0285735
0255512
0256263
0226210
0249054
0120645
0256948
0249695
0256694
0256540
0111181
0255671
0256577
0010379
0255746
0261799
0073614
0120647
0139044
0171840
0256020
0238370
0255825
0177406

0256672
0060237
0221439
0285704
0002016
0250132
0082805
0275811
0249028
0267285
0249628
0111186
0171823
0266043
0006831
0271500
0285627
0243663
0151062
0287698
0166159
0256706
0235049
0151065
0151067
0203593
0256837
0256257
0278255
0256025
0256721
0285555
0285734
0256769
0203648
0256271
0246627
0256150
0283587
0274659
0256902
0255669
0224438
0256030
0004478
0258325
0258092
0111203
0053702
0283297

0111206
0171792
0078246
0278356
0252996
0197905
0250899
0234589
0011105
0278469
0256197
0275850
0227081
0250770
0287712
0236908
0257048
0111218
0256691
0243507
0287243
0130038
0280202
0222338
0283138
0111224
0118976
0256862
0256969
0256356
0242444
0255920
0118971
0285901
0078237
0213970
0118972
0111241
0047621
0111247
0010219
0256748
0272921
0111254
0256799
0256381
0139180
0255639
0255474
0130035

)256988
)151079
)255608
)286866
)287835
)256654
)111262
)284513
)255996
)130037
)285523
)256115
)256417
)256146
)256218
)255983
)185652
)047617
)255973
)110799
)240533
)255775
)010278
)202318
)256103
)008323
)067182
)265388
)111319
)111321
)256433
)213942
)224906
)215039
)139193
)139192
)139190
)276718
)256913
)111639
)010292
)239002
)276232
)255966
)111640
)269968
)010295
)245667
)111641
)285238

0111642
0247853
0251898
0184574
0111644
0111653
0219410
0126746
0139200
0252186
0270068
0111652
0269892
0089693
0159335
0089692
0244532
0010610
0250510
0110811
0111664
0111665
0111667
0111669
0111671
0240370
0010626
0248593
0275703
0111674
0111676
0111678
0272173
0238923
0111679
0257084
0207713
0207708
0271969
0215021
0238795
0126749
0111684
0182326
0239701
0257078
0159403
0139178
0205885
0139194

0200345
0256967
0139182
0285770
0255572
0139197
0256480
0215009
0177675
0255836
0177575
0235868
0256331
0255977
0284327
0284591
0284522
0285142
0111701
0184344
0187569
0198178
0205857
0201549
0111704
0199912
0173262
0287713
0201663
0278459
0241828
0255885
0176654
0255356
0059804
0288043
0206636
0240739
0282080
0255964
0255581
0065970
0171860
0089818
0279865
0284697
0111729
0235602
0284393
0256171

0196946
0226711
0244050
0276417
0275367
0171847
0254016
0164845
0215241
0286895
0256589
0286504
0226091
0256552
0256101
0277952
0242661
0284687
0284673
0256136
0238152
0214487
0177586
0205846
0243830
0201780
0166527
0246115
0166523
0255801
0238943
0252727
0111732
0280345
0255967
0197614
0255829
0256004
0166532
0244131
0256661
0249790
0256904
0166535
0282022
0111752
0003056
0139187
0257105
0213049

0255776
0214851
0245105
0175899
0256937
0256720
0126838
0256817
0256069
0283429
0231503
0237248
0256427
0111788
0212440
0255753
0286911
0260423
0287577
0256673
0278635
0214826
0212432
0284634
0214776
0256975
0111796
0212345
0233719
0272917
0256442
0256594
0257027
0069493
0213443
0256582
0184293
0110848
0256912
0150045
0110852
0255882
0256797
0188393
0225231
0231560
0172322
0165682
0256660
0256803

0197992
0256626
0150048
0223042
0255734
0172243
0173391
0165685
0139112
0255958
0134539
0256288
0256155
0245648
0213809
0255819
0183542
0205810
0255641
0205809
0134545
0180574
0257016
0256039
0256667
0111196
0060140
0060138
0256888
0235930
0121377
0121314
0121381
0231887
0121318
0256651
0111215
0275778
0256682
0212128
0134551
0212127
0212125
0212126
0255837
0212124
0256436
0256019
0226761
0256274

0255374
0256400
0256188
0256981
0256712
0256537
0186136
0256657
0275119
0197870
0230657
0251655
0121335
0255790
0237303
0278654
0277705
0270863
0274551
0256237
0251747
0247157
0139083
0121380
0198134
0283475
0070018
0242405
0199551
0270857
0207010
0111261
0243141
0205791
0165714
0271370
0111266
0255670
0240424
0111269
0271215
0275963
0183150
0257004
0111276
0275560
0256658
0178878
0207983
0270296

0213782
0242329
0241352
0234498
0013588
0283759
0247498
0111291
0256011
0221611
0013583
0183935
0255621
0243129
0084444
0111305
0134531
0286869
0180861
0273079
0201476
0201909
0256306
0277935
0287928
0200475
0256346
0227760
0256084
0255763
0251908
0255749
0244573
0171681
0121316
0255649
0200830
0256751
0070019
0214772
0261324
0197837
0246705
0084463
0182993
0179256
0286107
0111339
0256339
0111341

0139055
0111348
0139053
0255727
0134533
0256650
0255660
0256283
0151490
0243916
0151491
0200105
0255565
0256627
0023734
0023697
0256134
0188991
0008394
0256238
0286303
0256450
0048540
0256564
0283240
0255871
0256211
0213235
0287455
0212358
0285724
0242991
0256896
0256756
0256389
0256994
0283665
0206662
0111404
0139144
0255892
0255960
0256138
0139151
0255993
0255648
0177938
0227525
0256844
0052126

0253284
0240993
0243854
0139154
0255909
0271462
0200247
0245205
0240328
0287074
0200885
0255910
0256849
0256287
0287075
0256499
0256879
0172572
0131982
0256663
0139155
0111700
0257046
0256157
0205754
0257062
0134538
0084453
0121351
0121350
0256021
0004700
0111711
0134548
0111713
0285854
0111716
0256615
0255644
0121361
0069431
0257022
0275567
0255627
0111726
0111728
0274624
0283582
0212172
0256714

250166
111731
256973
256371
256774
257023
139163
256995
241451
256923
256353
256321
115934
134532
256473
216192
256120
197503
256852
244662
255745
240481
060982
256482
278969
255921
213736
226397
118308
258546
258449
276842
118307
205707
133703
274987
275197
278743
201439
241846
152936
279173
256079
255988
257009
222950
256686
255725
256440
283603

0256256
0246695
0123094
0123095
0123096
0256894
0286752
0256234
0255750
0278095
0255858
0255968
0123104
0256185
0212549
0234428
0256640
0064102
0111790
0064115
0247903
0152944
0275764
0214700
0271418
0256226
0256625
0211455
0029153
0245311
0165935
0257037
0110841
0248100
0256056
0256557
0256377
0174236
0276261
0230519
0256504
0061794
0252585
0205693
0256747
0087448
0256512
0201612
0274075
0256378

0087494
0257042
0123106
0247934
0255951
0278733
0273989
0252237
0202187
0244712
0256513
0278687
0257258
0064763
0275278
0275476
0257176
0274315
0273680
0087502
0257599
0187950
0133687
0257456
0242986
0286566
0276431
0257262
0257756
0284682
0251781
0257932
0258202
0133704
0285870
0110888
0246331
0235884
0285517
0258118
0110900
0226472
0256984
0245614
0013573
0212533
0270926
0177359
0270766
0243517

0270395
0275097
0139146
0256176
0177340
0256232
0283256
0170456
0200388
0275769
0203437
0256159
0201228
0225349
0255867
0252390
0231788
0233381
0287388
0139160
0151743
0276058
0276900
0256843
0223722
0188375
0252421
0252204
0256372
0255760
0256465
0244436
0242314
0256987
0174718
0276136
0257094
0257852
0271357
0257530
0151746
0277342
0258134
0274964
0276115
0207176
0139132
0253063
0087470
0257643

201563
276148
257511
139131
274105
275854
286950
057294
274976
244743
257316
257435
110975
259937
273853
256070
212475
270807
207026
201624
245482
139133
256538
255628
256986
250788
255652
272435
258794
256614
258368
257173
258278
274293
287442
199843
201708
254381
257292
258360
253016
175548
257645
258119
139117
257718
257586
257237
258144
270718

0139116
0286576
0255991
0270273
0173208
0285732
0180116
0151229
0199571
0229014
0260943
0223914
0225342
0188906
0229899
0205592
0258167
0207492
0018236
0257680
0274682
0165966
0257228
0252364
0258239
0257858
0257784
0257239
0257128
0286591
0271379
0151233
0015153
0134283
0257674
0139168
0278520
0252917
0274943
0199886
0257376
0207142
0139174
0257225
0258068
0257510
0257687
0257405
0257164
0240759

0257114
0271272
0257849
0257373
0257927
0258331
0173157
0241251
0257813
0271508
0257907
0129317
0257896
0198001
0151239
0139173
0258212
0257947
0257864
0275286
0286671
0184613
0201788
0185610
0257319
0257331
0257572
0199556
0287386
0134297
0257738
0177119
0278351
0257368
0257657
0283545
0273015
0189079
0278475
0265093
0270429
0139218
0239397
0111371
0274591
0134294
0258096
0257261
0257496
0275481

0278896
0272369
0272963
0271642
0139209
0274723
0139211
0179715
0258352
0263838
0247774
0257925
0258181
0276454
0258116
0199566
0257924
0264906
0257906
0258369
0257807
0257905
0005175
0276390
0257433
0111405
0280054
0274902
0079337
0211584
0277173
0061273
0268069
0276691
0274737
0257488
0111424
0205537
0278385
0279840
0134291
0139219
0258203
0257985
0257955
0276814
0079387
0240399
0222635
0226138

0152556
0275770
0177981
0274124
0258051
0257885
0257848
0177875
0279875
0269514
0172640
0258234
0275228
0187166
0258273
0167528
0257735
0257763
0273765
0258024
0257792
0273434
0226413
0139223
0177627
0258121
0240443
0197376
0284723
0198678
0167531
0224709
0139620
0221491
0281842
0206612
0207313
0129315
0257987
0174233
0257653
0264201
0257660
0271547
0207467
0167535
0174243
0172602
0199394
0239617

272822
139537
134285
134287
169884
200303
125084
181418
258283
257913
181929
167548
167550
277672
139549
257346
139636
258017
123416
200309
167552
167553
274017
258101
258232
258334
135406
135451
186897
178401
123352
257954
257464
287537
135519
187778
110844
257964
135436
199237
258284
257243
258104
161791
139644
257570
257730
167566
258057
186666

0257253
0242041
0135472
0257771
0258135
0167580
0257588
0257378
0161798
0086159
0161800
0110881
0066117
0167588
0178449
0272368
0139624
0050405
0257298
0257531
0257256
0221604
0212496
0278842
0185958
0244266
0161813
0066084
0207136
0200183
0200428
0271018
0123268
0271490
0243075
0186452
0271621
0185432
0274156
0199740
0271596
0214511
0274797
0110911
0272028
0199903
0050426
0110925
0135457
0244540

0257246
0200953
0199824
0184271
0278126
0183283
0170545
0110934
0139610
0139629
0050438
0271065
0196876
0257851
0260122
0284730
0260473
0261308
0261586
0259887
0167612
0258021
0139567
0135503
0206992
0161835
0123358
0259884
0123395
0257663
0284791
0257542
0257989
0258279
0167767
0257137
0257649
0135480
0257671
0170442
0135477
0284883
0257830
0285102
0205426
0287051
0257829
0170523
0274928
0135443

0161849
0258253
0161850
0257500
0257844
0170454
0185479
0170465
0205420
0186081
0139648
0170484
0170486
0257495
0186049
0258120
0172867
0167768
0189182
0257700
0257402
0258036
0257616
0257343
0257675
0257125
0258071
0185069
0257404
0186442
0170477
0185640
0257389
0170423
0241146
0170421
0111057
0063046
0257475
0257337
0111077
0278108
0167778
0167779
0167780
0258074
0238669
0258016
0257790
0139631

0257808
0139651
0139626
0172819
0283536
0182544
0135476
0123349
0257605
0139637
0094914
0170374
0185591
0135409
0205352
0257379
0197111
0282977
0273658
0139625
0270175
0139546
0267281
0139574
0170653
0257550
0285692
0275488
0135390
0213470
0012822
0286069
0260492
0260030
0253053
0249641
0123364
0123407
0228630
0275589
0277994
0274817
0277129
0123388
0251151
0180818
0273049
0197757
0207924
0250133

0180806
0250451
0037965
0198353
0273046
0260597
0172789
0207571
0248265
0250654
0249388
0250432
0202146
0123415
0250742
0260470
0257534
0248576
0094916
0283803
0241785
0257596
0258344
0135486
0123405
0111481
0207381
0199122
0264028
0258137
0258086
0139572
0161642
0161638
0257477
0170627
0123338
0257824
0123360
0135447
0257780
0135413
0161634
0257634
0270858
0172551
0135426
0257870
0123307
0196534

0257350
0170605
0224622
0227423
0179919
0197706
0213451
0205331
0188324
0230307
0205330
0205329
0257757
0187857
0203408
0233606
0257414
0205328
0179899
0185821
0258763
0179695
0184954
0205327
0234670
0179626
0179615
0272937
0175398
0258921
0272837
0258907
0270458
0170439
0135424
0258311
0135441
0135437
0135404
0258056
0258045
0257384
0135414
0205323
0257390
0257509
0123353
0135392
0182796
0123342

0257966
0257569
0170473
0065357
0273973
0185664
0123374
0258554
0111540
0237493
0139531
0123411
0273890
0257449
0197728
0065361
0257411
0170515
0257553
0229117
0139641
0135482
0258345
0258317
0257809
0196465
0092841
0139613
0258199
0181852
0139579
0139540
0139645
0135469
0258260
0062485
0144785
0257303
0257740
0257727
0135473
0110944
0170581
0252206
0175336
0274823
0259099
0276272
0274569
0111602

0135517
0285528
0176422
0135423
0257576
0076067
0201579
0076108
0110955
0207031
0110958
0241217
0196531
0285625
0198056
0025423
0258679
0273536
0258816
0212383
0170426
0278400
0139547
0278399
0287715
0166856
0166860
0166863
0166866
0166881
0166886
0166888
0123384
0259125
0221365
0182379
0276727
0182199
0185633
0185482
0258830
0179912
0201198
0258001
0175189
0139269
0287200
0111087
0123329
0166986

0284152
0175197
0285133
0208028
0166987
0175203
0155980
0166908
0178498
0240771
0224713
0287908
0135502
0135454
0242990
0135506
0257342
0135439
0255737
0135452
0135446
0283621
0139266
0111012
0037897
0123427
0257921
0123297
0135407
0270039
0206749
0269903
0175215
0207789
0257953
0245651
0273805
0166896
0257698
0257159
0222210
0257668
0257541
0279134
0279411
0241423
0258133
0258231
0257259
0139263

0257443
0244390
0257288
0286351
0286279
0258214
0212599
0251931
0257230
0202471
0118596
0257146
0237176
0257294
0287014
0258184
0252607
0251822
0257865
0257308
0258336
0286402
0198673
0242922
0257528
0213363
0257880
0258316
0135655
0284360
0200814
0275881
0061987
0257568
0202034
0221949
0257354
0199179
0275180
0111110
0241941
0213352
0200296
0252660
0139239
0257779
0250517
0257235
0257664
0166148

0279444
0257332
0258117
0177990
0249753
0257005
0255583
0240027
0118600
0255850
0255642
0196935
0242087
0256571
0256399
0255780
0255886
0287618
0255817
0255629
0174206
0212298
0243024
0251788
0185306
0256192
0256293
0238440
0184575
0255566
0277895
0183735
0153179
0256199
0223294
0256670
0207546
0266016
0225195
0256314
0280320
0280181
0135677
0215159
0111490
0286682
0243020
0255693
0248995
0221564

0198671
0156076
0207099
0255700
0174106
0276853
0174099
0250280
0215208
0250748
0255866
0256268
0256915
0213344
0241749
0149948
0277945
0197301
0256083
0276878
0256259
0213343
0251857
0139233
0228144
0239335
0155957
0090376
0225422
0284011
0256072
0256777
0222744
0127311
0155974
0256285
0256420
0257083
0279310
0255976
0237766
0111530
0286563
0256037
0256355
0256172
0203585
0256077
0127334
0235872

0255970
0279505
0255772
0256835
0255733
0244432
0236946
0256708
0111537
0111536
0127318
0111554
0251301
0256273
0273824
0256075
0256917
0240087
0274459
0127314
0234040
0206650
0234925
0247363
0241825
0111581
0233560
0175782
0256678
0135679
0256325
0257181
0256664
0135678
0252770
0257289
0257336
0258140
0111605
0283476
0257818
0274979
0090382
0257764
0127337
0266185
0241765
0257507
0166225
0264405

283677
166226
198812
127325
279551
247131
127328
258052
166268
257815
257241
257139
257613
287132
111596
279530
135643
222405
258168
127329
277247
153233
257302
207387
257265
257454
258053
127324
139292
277223
257761
133858
212461
173451
258064
139291
257515
080371
257410
257991
121749
257802
139287
258115
072657
236333
257287
257723
258235
258294

0258123
0257682
0243164
0252415
0257750
0201809
0251138
0257587
0258332
0258320
0257183
0257113
0257364
0253719
0257386
0257323
0188646
0257998
0257434
0166006
0254451
0258112
0180881
0257823
0273987
0173401
0180481
0139278
0257497
0111615
0258077
0257777
0286259
0258245
0251893
0258088
0258091
0243420
0257329
0198923
0139289
0257453
0257839
0187109
0257941
0258304
0257872
0252858
0257219
0223273

0179941
0091039
0257526
0287030
0213272
0271142
0243071
0241045
0258090
0186908
0175183
0257910
0257199
0165891
0286476
0257677
0067798
0257835
0258066
0258225
0257220
0258220
0258084
0257879
0257165
0257564
0257948
0067715
0257191
0257894
0221788
0257604
0257474
0257956
0177425
0258048
0058272
0277130
0258044
0201942
0257557
0271496
0239473
0238842
0165899
0230291
0222880
0139304
0257786
0257429

0111046
0111049
0257741
0111052
0207763
0258215
0208022
0111058
0258026
0265227
0139220
0257467
0258162
0258375
0257609
0257747
0258193
0133773
0127720
0257875
0258170
0257837
0257431
0286043
0179104
0252220
0213270
0258338
0257124
0285191
0257729
0221148
0258358
0257296
0072041
0258073
0231738
0133640
0180318
0258815
0257855
0258206
0198774
0133636
0182050
0258178
0257897
0258185
0280049
0242850

0258205
0257648
0258173
0258128
0257912
0257940
0258179
0240721
0165805
0133641
0198707
0252747
0257752
0139324
0199245
0049130
0281333
0252850
0278011
0246363
0257629
0238302
0257820
0139318
0274021
0286608
0271327
0271259
0139323
0226982
0259075
0257594
0270344
0279939
0258302
0070961
0271614
0258216
0252207
0257480
0252823
0258290
0257725
0257194
0258183
0286021
0288102
0257995
0257787
0197651

0196243
0083782
0139330
0139329
0011465
0258148
0258100
0286049
0274647
0277738
0258125
0258224
0257893
0257863
0257242
0286907
0287714
0133639
0245904
0239201
0199895
0277851
0205057
0266923
0257127
0205056
0258262
0187510
0102189
0258012
0257195
0202534
0257345
0257322
0207365
0270506
0256393
0257746
0257210
0258171
0201502
0257512
0257252
0173598
0177889
0200011
0198015
0243015
0258271
0246985

)120833
)186076
)169372
)258274
)264978
)258303
)257283
)271382
)223126
)257360
)136040
)258035
)287237
)258172
)258365
)173588
)244391
)213250
)278916
)257400
)244091
)258357
)236349
)266099
)057704
)283818
)257847
)216306
)283998
)184752
)120798
)180263
)257121
)237774
)028203
)202470
)257666
)212535
)199172
)265917
)257943
)258313
)222544
)241556
)111142
)136014
)257150
)212448
)074527
)258343

0199506
0258292
0139343
0165972
0285030
0139344
0084110
0257878
0111144
0257715
0258251
0277062
0283422
0111145
0258177
0287454
0059758
0201435
0257169
0258272
0252827
0188596
0258846
0139350
0202368
0258131
0257470
0279190
0286434
0257920
0255794
0221479
0273942
0274230
0275231
0273570
0275971
0274819
0251844
0278802
0276398
0279798
0207586
0277081
0257157
0257501
0206899
0265861
0263890
0258312

0257580
0201296
0286056
0227825
0245017
0257167
0120802
0225185
0075415
0212443
0166130
0120868
0185046
0258204
0257458
0202324
0258039
0180219
0287675
0214203
0279148
0111647
0280088
0257696
0238105
0266610
0257489
0221476
0075089
0166153
0206790
0136021
0179520
0012504
0271177
0139354
0257803
0151572
0257325
0258007
0258033
0256870
0207414
0241744
0120800
0120805
0257543
0222890
0239413
0206929

202249
252366
166211
196091
257514
274560
111666
199933
139351
111670
222255
258230
201168
222932
223046
252863
258153
136048
257202
241917
258308
258288
120860
257222
075188
185480
183395
281344
264554
258142
283052
017427
257254
258169
171759
282964
139352
288092
257703
257860
179088
257415
257994
286197
257762
136011
212594
257737
257766
257681

0111696
0257754
0257327
0214198
0166598
0284503
0204954
0139372
0120820
0216285
0111727
0120837
0199415
0257193
0286258
0198431
0257732
0213442
0255150
0171310
0258111
0279176
0264295
0136052
0222579
0286410
0257758
0151131
0136010
0136051
0136044
0235162
0286903
0257642
0286520
0257859
0257773
0257890
0074590
0257438
0257611
0136026
0258355
0277715
0238609
0166046
0013503
0257545
0111783
0257711

0257918
0111785
0287957
0257119
0240631
0260329
0151135
0120832
0008405
0257548
0286678
0257444
0151136
0200897
0222302
0257579
0222160
0136045
0240441
0110851
0258136
0187855
0257398
0257951
0257392
0257426
0258072
0257129
0075035
0257281
0174600
0247213
0198855
0075856
0287202
0136003
0183160
0110876
0257221
0284388
0110880
0238457
0200060
0273499
0084112
0280110
0274598
0207622
0110887
0166111

0201324
0279630
0135093
0256262
0212138
0189046
0076248
0256139
0076555
0139445
0174527
0256560
0255655
0110906
0151148
0139428
0200274
0110921
0200794
0139438
0255650
0111199
0263510
0139433
0256351
0241413
0249094
0139437
0139436
0277299
0280426
0076513
0174456
0277595
0286220
0279360
0122970
0174437
0276956
0196510
0258210
0258011
0111229
0111231
0241680
0204856
0111237
0202335
0278993
0151164

0196850
0204852
0263537
0122986
0186298
0279925
0257268
0241286
0173093
0111245
0185847
0258240
0111249
0273700
0253080
0198324
0257595
0257539
0111252
0204842
0272215
0238168
0258099
0089234
0258359
0111271
0257767
0111275
0274697
0234608
0089022
0244060
0229186
0226469
0198270
0258373
0257624
0089248
0274227
0111300
0266370
0258323
0200688
0135148
0200135
0173064
0283793
0257494
0201428
0213152

0257658
0089009
0179295
0089169
0283507
0089127
0257452
0111331
0111335
0258159
0242461
0135144
0111344
0186710
0199899
0123064
0276908
0201208
0139405
0257286
0166578
0186815
0277566
0274822
0089060
0151176
0135094
0139410
0089116
0257935
0249550
0257231
0122965
0287229
0257359
0257603
0257997
0257300
0257915
0257476
0089225
0255399
0222982
0255830
0258244
0258254
0199220
0135111
0257817
0287982

)252459
)257517
)283459
)274879
)257958
)257519
)271579
)276308
)276972
)257726
)258108
)257407
)241656
)240205
)257781
)257683
)200112
)286991
)123066
)258034
)200665
)207967
)257979
)258337
)286354
)258346
)264037
)258018
)196668
)258102
)275898
)258249
)257883
)257654
)111412
)135119
)239374
)201382
)275467
)135116
)240100
)174989
)274554
)257279
)277840
)088992
)258285
)135108
)089250
)257465

0255686
0171435
0256071
0111445
0277873
0176871
0276292
0176834
0274859
0089220
0135090
0240342
0270482
0111707
0275759
0275409
0255714
0241590
0256750
0255618
0255814
0199809
0256149
0201042
0139767
0286233
0244112
0257095
0256884
0152137
0256609
0213144
0256311
0281196
0183273
0248636
0224982
0252886
0111725
0122966
0255692
0283768
0226729
0135127
0278250
0255575
0111737
0277283
0089154
0266704

)089157
)255857
)089159
)286067
)275936
)240925
)123009
)202538
)200795
)089163
)252659
)170890
)135097
)239881
)256695
)111775
)111780
)170855
)257218
)219355
)111786
)088986
)248008
)278344
)110871
)222601
)241030
)022840
)167272
)242137
)256008
)157782
)110917
)256364
)277423
)276188
)175970
)256569
)284143
)122971
)255946
)157837
)213137
)286493
)231313
)244585
)241388
)135100
)157895
)135114

0283542
0256963
0279001
0201945
0223538
0089041
0286248
0274029
0135124
0110931
0089053
0258435
0170633
0089094
0284402
0256742
0252393
0276045
0139714
0286586
0188735
0139725
0212694
0272849
0274292
0139718
0158104
0256811
0278084
0287493
0110801
0256950
0283389
0239082
0158023
0255856
0110987
0256546
0175727
0158113
0204671
0176383
0284934
0184047
0256861
0139719
0130779
0214760
0257097
0033030

)252192
)275265
)111011
)184445
)255972
)274191
)256249
)182782
)255398
)196917
)256044
)139726
)206914
)130783
)130787
)139722
)256152
)280138
)150967
)111325
)256028
)182196
)090975
)265526
)251497
)280381
)051825
)280120
)130921
)111328
)269980
)199845
)139697
)207189
)256092
)284425
)183955
)150977
)270445
)184209
)188026
)274427
)278112
)266655
)247373
)086598
)111364
)206897
)111361
)111358

0279953
0278861
0168778
0255839
0185344
0286092
0239215
0280300
0197653
0119242
0250091
0270130
0270048
0269997
0269938
0270061
0270095
0179195
0274874
0178882
0256827
0275389
0256596
0196498
0275967
0255965
0214650
0279527
0279071
0280354
0279121
0279334
0280097
0278973
0280444
0279087
0279931
0073060
0287242
0252008
0150991
0265345
0241129
0150990
0184992
0280634
0256093
0081760
0279233
0256814

0287240
0280117
0139364
0279462
0280125
0286209
0280415
0271184
0249267
0275212
0284848
0280364
0280398
0280272
0286386
0255595
0256494
0214043
0256125
0255853
0256310
0256732
0255900
0255944
0278866
0279171
0255998
0189238
0256128
0255923
0249873
0256286
0257021
0249345
0286016
0256001
0286791
0278266
0287339
0256292
0256496
0253089
0256362
0256502
0280024
0287365
0279952
0287112
0286922
0287622

255945
199584
257035
256022
286662
279475
255757
256597
256193
256971
256643
287311
256551
256659
276487
265061
181234
279180
265635
257025
256630
139370
279500
274695
151948
255838
256581
151952
249196
277217
256699
280051
256085
286246
256137
286394
280196
251536
261650
214039
256298
256906
279193
276122
250208
280405
111432
242963
259862
125207

279343
279478
060709
256064
279905
279146
256343
279650
111450
256250
256299
222438
132341
206850
111452
279940
279976
280200
256204
279581
280002
256151
279128
279993
256810
204603
280217
226356
279746
248703
212251
286462
230294
287792
256484
280319
275232
287136
256209
279388
255704
279455
256258
256424
287639
279153
212154
249926
243499
061936

0255933
0200516
0200483
0198598
0256955
0177169
0279283
0255992
0177192
0273568
0183495
0208892
0257000
0274670
0185684
0279640
0256804
0185163
0184967
0276693
0256576
0256312
0182870
0279480
0256943
0256542
0278872
0277011
0255916
0279113
0286008
0279704
0279023
0280150
0274373
0256783
0256875
0112787
0280287
0279700
0277052
0277186
0204583
0187848
0280311
0177084
0176894
0256632
0247077
0201469

0176915
0256108
0090615
0072609
0232888
0252079
0278949
0236617
0255703
0250790
0196458
0256563
0198393
0252269
0246394
0255807
0198040
0279466
0279817
0196387
0214029
0271963
0240991
0256223
0256825
0090612
0255919
0227059
0239190
0238443
0279924
0279516
0279081
0284679
0280208
0279124
0279730
0279231
0268486
0276183
0215604
0270496
0231238
0233905
0229788
0235876
0231914
0226361
0228797
0230931

0204718
0227784
0244155
0227568
0225016
0206192
0242756
0223024
0206787
0235500
0226250
0231358
0226057
0232685
0229375
0187721
0226619
0287861
0252287
0224778
0234894
0229147
0278337
0198033
0121388
0227879
0232243
0237636
0215349
0236834
0224976
0231184
0229791
0237164
0132958
0223625
0232894
0215572
0225316
0238132
0196199
0226352
0121390
0275014
0233469
0132950
0223576
0275964
0121741
0237672

0279951
0236076
0121743
0226633
0260131
0165474
0279065
0121742
0165475
0263978
0277020
0032742
0238286
0222726
0278291
0172458
0279135
0150456
0233524
0275485
0132953
0236704
0233780
0234363
0213621
0150457
0233851
0201821
0180172
0225585
0275689
0252128
0150459
0165480
0173141
0271250
0233405
0235438
0277077
0233325
0224429
0215571
0235076
0222747
0180776
0236953
0226908
0165487
0230022
0207518

0226525
0102678
0244197
0226722
0276476
0237175
0226118
0287357
0232187
0225777
0253094
0237952
0226507
0231650
0262619
0230535
0234685
0262198
0283075
0233440
0207157
0229483
0102683
0252952
0235205
0236803
0151835
0232163
0229558
0232977
0227893
0127863
0027001
0232102
0205861
0205863
0196593
0287530
0273167
0182957
0228741
0225674
0252695
0227213
0240654
0240868
0277040
0234627
0261498
0233545

0205822
0102699
0288103
0287376
0244615
0243008
0285228
0075673
0207474
0225404
0285806
0231417
0287887
0226695
0132972
0151849
0253771
0273628
0251299
0232858
0269099
0276958
0151846
0279695
0165566
0238169
0234056
0279550
0225105
0235315
0139505
0260509
0139496
0234152
0132932
0199767
0180803
0242381
0223158
0180730
0231983
0233963
0234706
0127870
0277368
0271559
0227882
0132964
0278413
0285929

0132970
0234031
0237001
0285935
0132975
0230256
0283579
0223782
0218198
0152484
0278496
0232162
0230641
0275294
0234772
0122026
0207500
0207051
0122035
0252247
0229609
0201242
0122034
0122033
0252499
0139517
0186184
0273552
0226703
0169840
0247381
0207330
0139515
0180389
0260704
0165556
0183463
0242360
0122025
0271026
0235115
0261485
0152520
0232587
0200840
0102755
0279393
0235472
0273964
0132963

0139508
0202237
0233588
0277227
0236339
0132938
0236758
0223460
0270680
0179141
0139514
0122042
0278172
0236258
0224329
0224405
0122043
0266505
0224511
0232117
0232643
0279499
0102781
0252928
0215049
0236463
0238121
0279149
0285773
0225039
0236444
0189403
0285840
0213411
0232489
0224117
0132952
0275788
0132965
0237879
0236094
0224743
0102802
0175664
0238185
0230403
0286286
0231894
0120694
0187676

0229427
0278305
0277319
0133105
0287904
0073910
0229715
0279314
0278732
0237637
0189167
0139618
0215515
0139597
0212293
0270008
0244754
0229800
0281026
0083642
0207325
0277151
0226968
0227604
0133116
0133121
0230300
0236581
0287274
0227911
0285621
0277386
0230490
0234535
0232986
0286779
0133119
0252397
0276672
0280431
0227254
0199196
0278477
0271850
0225179
0271901
0231910
0273886
0261728
0172915

232795
234835
180660
287650
236036
133083
120669
250709
242715
207203
133104
120664
133101
223383
180440
235306
223543
215486
271337
133111
120693
236711
228825
232378
226161
120697
120699
102710
180138
274549
279511
202395
230283
230390
275226
133110
133107
199276
287477
274317
228036
223685
120686
279025
236354
229437
234701
150893
252795
225350

0225221
0235550
0133115
0120685
0188811
0273507
0214823
0183722
0133103
0264171
0200526
0224351
0270444
0212553
0215483
0230710
0271429
0207458
0252812
0150907
0229456
0211491
0102738
0277794
0102743
0207652
0168852
0176268
0287837
0239827
0120690
0229473
0120688
0264226
0244264
0165572
0278390
0215482
0120696
0225133
0120662
0277662
0231404
0172766
0223280
0214222
0205240
0234655
0215480
0102760

0102763
0284584
0241406
0206962
0229696
0278338
0226093
0102780
0231530
0237263
0270809
0234213
0023516
0283554
0233259
0120659
0179813
0271216
0229546
0133106
0238025
0120675
0229928
0276569
0120658
0232540
0238189
0233821
0287842
0287029
0270522
0151773
0274001
0179630
0281883
0280060
0233725
0179611
0226519
0227258
0139656
0235285
0276854
0276319
0230731
0277831
0151778
0237361
0102804
0280357

0225727
0278156
0230793
0237585
0276527
0235097
0083635
0223732
0133114
0265802
0223341
0188342
0271818
0277358
0180332
0270381
0133112
0253051
0199477
0273149
0170919
0228886
0214455
0174032
0251015
0229303
0136152
0165837
0200709
0232786
0228869
0235366
0237876
0174015
0274357
0215475
0123200
0235903
0080618
0136167
0240767
0173988
0252854
0251106
0287863
0261097
0102445
0223336
0252385
0241353

0231817
0285576
0227019
0277228
0205105
0230243
0136141
0278462
0139684
0102468
0224517
0234590
0244521
0136143
0234145
0229423
0228573
0260388
0287456
0227848
0136159
0276968
0136146
0229111
0224510
0136156
0231473
0139687
0238086
0177197
0277684
0139679
0277233
0136161
0287599
0274929
0233456
0233610
0152207
0225131
0275202
0102531
0201662
0223724
0199788
0235957
0231665
0102539
0274351
0102543

0102547
0136169
0236577
0136147
0136144
0176882
0152213
0123179
0102753
0276436
0222148
0181358
0200064
0123178
0231607
0204977
0264864
0198553
0277913
0277066
0208006
0283785
0274878
0273612
0277752
0275559
0176124
0276089
0214359
0273541
0274270
0232150
0223676
0286191
0186047
0200711
0233672
0136104
0123201
0222920
0226792
0214878
0274652
0271242
0150510
0253309
0266072
0102786
0236778
0224892

0243900
0229906
0266611
0252141
0139668
0206920
0242893
0224451
0285444
0102796
0231856
0281106
0123171
0232872
0279677
0123191
0236044
0253710
0253797
0197168
0273523
0136098
0217576
0243406
0285160
0285323
0272281
0271564
0136114
0199605
0136100
0278238
0136108
0235660
0244471
0280296
0273784
0250299
0139675
0273723
0165416
0136110
0211579
0150276
0136099
0102837
0237092
0279325
0225510
0287722

0241613
0287247
0273919
0274316
0261517
0234787
0271652
0283126
0136149
0287460
0286537
0277047
0278630
0275071
0264387
0274090
0228611
0286709
0228319
0201665
0204919
0229665
0204918
0227151
0234278
0271140
0230009
0234421
0225352
0118946
0278722
0202422
0277448
0232954
0276208
0222733
0214335
0223717
0214281
0232647
0288015
0271190
0239003
0139734
0227528
0265745
0223815
0202151
0227336
0238213

0277549
0083544
0223076
0225249
0236608
0279657
0206854
0227510
0266663
0280165
0197991
0279797
0234033
0270856
0229578
0230142
0227611
0229307
0232105
0276586
0271465
0286469
0228669
0273550
0227564
0287996
0231061
0274052
0285764
0237378
0202478
0213232
0287128
0219926
0285566
0227674
0237849
0214269
0214266
0236565
0234767
0286395
0263581
0263561
0235737
0184226
0234527
0238500
0228842
0225263

233840
285588
214263
230040
287876
271287
227969
231667
226537
286820
234765
229910
236433
230405
229824
243671
281778
280580
232380
225501
235221
224640
150361
207495
232314
230223
288330
202433
276841
226554
226846
225801
276644
236534
237816
233460
232881
212377
227952
251715
278113
199282
226617
206922
204899
136122
083520
083535
199381
232041

0102554
0236972
0200267
0201253
0206697
0279754
0229708
0224853
0228295
0118922
0280348
0279463
0235532
0286330
0206617
0232701
0177596
0226240
0281167
0236678
0225944
0206812
0226665
0214249
0223880
0136111
0225203
0188243
0118939
0261553
0261105
0136153
0228444
0230902
0178734
0223458
0261206
0224933
0243274
0285572
0287018
0284809
0178695
0278727
0118903
0102794
0233873
0228002
0102805
0283208

0005812
0236051
0274898
0005810
0229521
0285714
0136155
0287270
0224347
0206800
0227922
0285746
0139737
0266325
0225579
0136160
0234377
0233379
0243035
0236133
0229249
0235625
0271386
0200686
0229865
0224686
0271776
0152192
0152193
0237109
0225427
0215357
0230584
0236277
0226670
0139746
0252496
0227354
0232132
0102471
0227676
0285902
0229011
0229175
0284196
0281721
0285684
0136158
0202398
0227862

0226683
0286746
0231222
0229246
0270415
0260094
0229309
0222486
0237148
0214182
0286385
0237099
0270400
0222791
0286446
0178235
0285680
0215037
0286284
0224188
0285742
0288016
0228597
0232447
0233349
0288038
0287438
0226370
0286906
0226317
0285834
0184564
0278580
0228241
0232252
0283371
0277914
0287880
0231879
0233528
0228074
0228473
0285699
0232636
0228824
0272515
0266052
0165300
0277640
0223404

0272046
0232204
0231019
0225289
0285811
0229443
0286422
0283130
0261666
0233211
0234660
0232225
0227247
0225530
0271395
0226037
0236176
0234227
0200733
0215418
0261446
0252696
0235879
0224419
0283783
0234384
0251753
0231674
0227983
0234625
0229557
0214178
0215417
0284536
0283815
0284204
0283762
0284375
0283705
0179399
0282864
0282997
0252508
0229287
0282865
0232885
0283097
0236240
0263741
0287159

0282993
0235984
0232849
0288001
0278177
0183098
0286843
0235784
0224394
0212559
0236520
0080166
0088451
0152749
0222129
0238230
0274168
0227640
0125285
0235768
0237924
0260962
0287635
0252335
0125257
0223298
0200211
0271492
0134873
0223392
0134874
0247400
0102580
0229622
0227714
0232373
0276809
0102595
0226977
0185352
0242614
0283487
0233273
0235137
0234267
0272329
0222472
0165621
0233124
0139793

0286334
0202290
0286416
0125249
0276573
0276704
0271625
0226134
0286544
0233291
0270725
0065150
0232217
0152767
0139797
0222969
0263399
0269189
0269599
0231194
0102572
0271437
0224418
0217686
0233549
0233662
0243366
0088386
0088387
0229918
0275919
0231981
0207298
0260992
0230871
0228889
0234005
0134882
0201793
0218502
0125245
0169508
0207719
0228808
0238241
0285448
0125304
0280734
0280710
0203441

234168
234239
287746
125246
263615
227659
234303
201245
286757
139800
288060
043355
260738
225999
274605
236554
175198
228501
270632
287330
234650
134864
224356
280169
234354
125247
237082
233009
214051
229599
102452
198542
102466
276012
201155
215399
265164
229975
201690
264482
236860
243319
279702
234445
272143
261057
276957
134900
139780
175820

0231633
0286312
0151287
0134901
0270181
0134897
0134899
0250878
0272542
0278445
0125255
0285789
0234551
0230474
0287734
0225823
0286929
0232307
0275968
0275502
0182346
0280455
0226620
0252550
0287923
0222682
0286780
0233532
0174418
0125266
0284966
0274204
0134884
0272274
0230156
0227590
0287546
0278698
0174408
0277863
0279157
0285643
0204442
0201847
0276740
0221650
0227248
0286343
0274718
0174405

0139826
0102524
0277246
0223177
0283384
0232087
0041515
0229938
0236242
0275216
0223617
0236053
0229792
0285534
0234854
0185950
0275741
0275248
0201161
0274331
0275830
0276269
0231428
0265885
0279965
0287575
0187498
0279237
0134871
0276765
0224821
0232814
0139832
0275880
0213995
0134905
0153487
0225239
0259831
0255874
0088448
0229152
0277767
0227105
0275989
0225760
0225870
0286824
0235875
0102606

0233644
0227352
0285856
0274614
0153495
0204398
0215881
0226903
0200072
0224243
0182968
0260385
0229520
0260102
0260343
0223626
0153498
0126216
0277159
0197595
0274922
0068650
0232684
0277545
0126217
0267868
0235280
0269356
0269376
0057593
0126218
0231882
0230371
0283828
0126231
0269125
0126226
0139842
0277942
0232662
0185896
0139835
0276434
0225083
0153531
0150401
0233613
0202347
0150403
0276916

0276248
0198176
0186009
0185974
0279770
0184497
0233695
0183087
0272695
0237699
0226921
0274766
0229373
0260910
0283199
0185989
0232487
0283361
0283347
0130177
0264539
0265450
0285672
0169062
0226158
0198824
0229723
0260615
0283274
0215398
0206906
0257171
0257356
0257959
0257721
0258367
0257825
0257115
0257672
0258233
0257504
0257644
0257175
0258364
0257731
0258076
0257558
0239200
0257635
0257224

0277529
0277367
0278143
0257853
0257891
0258265
0222036
0275563
0212612
0258314
0274369
0225210
0286614
0277128
0277710
0278594
0196143
0271632
0277252
0258781
0285990
0257310
0277156
0244306
0228294
0287515
0274827
0278301
0187537
0276888
0212516
0257162
0258027
0273639
0274649
0278184
0257432
0273558
0258324
0257493
0257751
0257846
0259069
0257884
0257395
0258080
0257142
0258453
0176294
0258438

0182652
0176312
0176299
0258564
0228304
0176290
0165762
0258541
0259016
0176281
0155249
0258822
0169488
0196383
0169484
0176253
0258899
0176246
0252999
0258641
0176230
0184394
0259074
0259095
0258468
0284742
0196832
0258625
0176219
0258806
0176198
0136319
0258768
0258459
0100814
0287735
0222489
0238344
0259001
0277209
0129484
0254846
0129566
0200225
0185271
0259162
0092094
0258515
0100823
0165782

0198805
0258908
0182545
0258456
0258863
0188655
0259060
0173464
0258573
0258436
0239327
0181803
0199461
0214274
0258818
0259171
0258451
0259144
0258963
0181562
0181552
0169413
0258810
0129538
0259130
0258642
0169397
0258918
0258772
0136315
0169385
0243817
0258599
0165792
0258471
0165794
0165795
0284122
0179636
0258604
0206150
0255472
0165799
0173431
0165801
0165804
0232070
0178107
0258722
0200102

169327
259102
258906
258441
092199
092200
092201
260830
100888
199436
200785
241035
258870
274475
129472
092203
165819
165821
257096
256737
169208
256081
273586
256385
255569
255582
256553
255664
221977
276240
256350
211776
248358
211777
211778
211779
240954
211780
211781
211782
211783
211784
256474
211785
211786
211787
211788
211789
211790
256379

0211791
0211792
0211793
0258482
0258557
0211794
0211795
0211796
0211797
0211798
0211799
0258705
0211800
0211801
0276428
0211802
0211803
0211804
0211805
0211806
0211807
0211808
0211809
0258835
0211810
0259092
0249048
0258905
0258812
0251002
0211812
0211813
0211814
0211815
0259068
0211816
0211817
0211818
0211819
0211820
0256221
0259042
0211821
0223997
0237235
0228985
0211825
0211826
0211827
0211828

0211829
0256590
0211831
0249446
0211832
0211833
0211834
0211835
0250688
0211836
0211837
0211838
0248366
0211839
0211840
0211841
0211842
0211843
0211844
0211845
0211846
0211847
0211848
0211849
0211850
0211851
0278661
0276699
0211854
0211855
0211856
0211857
0211858
0211859
0211860
0211861
0211862
0211863
0211864
0211865
0211866
0211867
0211868
0211869
0211870
0211871
0211872
0211873
0211875
0211876

0211877
0211878
0211879
0211880
0211881
0211882
0211883
0211884
0211885
0211886
0211887
0211888
0277734
0259054
0259003
0129562
0275552
0100439
0255804
0242061
0258458
0235213
0258823
0212302
0155463
0155465
0172590
0157227
0212335
0199716
0197324
0139890
0100461
0237054
0100462
0257285
0092036
0259132
0284118
0129474
0258457
0100802
0280129
0100804
0279656
0222028
0139880
0100813
0179933
0283654

0092067
0092068
0202229
0215277
0240131
0252347
0215271
0270433
0235194
0129473
0258643
0100836
0092096
0100842
0259018
0166090
0166091
0197616
0199157
0092054
0258444
0215991
0129460
0157306
0136367
0273618
0259431
0213983
0258727
0092051
0274002
0100867
0258464
0259043
0270618
0222931
0258887
0285727
0259334
0215256
0157326
0187630
0285467
0286931
0225766
0186648
0100884
0129535
0100889
0100897

0259371
0139914
0092010
0100908
0259321
0100911
0283856
0092098
0259529
0199804
0213928
0100918
0196497
0259522
0287765
0100926
0254692
0276698
0278784
0139908
0254505
0260669
0213920
0255526
0129559
0100938
0092330
0092295
0100949
0288044
0286825
0157379
0196943
0136305
0213906
0213903
0129467
0258973
0129465
0283290
0100968
0205978
0139899
0100441
0100445
0258744
0092009
0100448
0258657
0100450

100453
168952
288100
287481
258098
257179
258275
257976
257986
212270
279521
285633
257503
139910
262119
257842
258081
257845
257185
286479
257612
265165
258932
258548
264657
197358
257869
240179
258175
257662
258282
257126
176165
186960
257056
257748
258038
258065
257523
207164
258028
252065
257522
258107
286040
212518
184304
257120
252686
287142

0257904
0248975
0240201
0279032
0257636
0278587
0258350
0092140
0092108
0258648
0258558
0243855
0240513
0213867
0100473
0258525
0196792
0258886
0207952
0258968
0100478
0092148
0258640
0207440
0202402
0257831
0199291
0129493
0283016
0203546
0250365
0129480
0239323
0214943
0151413
0258196
0257155
0223087
0207412
0257472
0258474
0275002
0258386
0258655
0100852
0285608
0286527
0201654
0202337
0258619

0259086
0151320
0258580
0278746
0259045
0280281
0151322
0287153
0287777
0129521
0258897
0223593
0165389
0237450
0129518
0279423
0259002
0206596
0206588
0259135
0129515
0240463
0275978
0253003
0241582
0165410
0243824
0198604
0251726
0258738
0258704
0258493
0226677
0100883
0151327
0092020
0279434
0100890
0258790
0241052
0259090
0282625
0282308
0256243
0240392
0100902
0258860
0240023
0100906
0259022

0259017
0259105
0258783
0168348
0174373
0253059
0257957
0258938
0258300
0100916
0257272
0257307
0258342
0257614
0257720
0257826
0257585
0259104
0258844
0252312
0151332
0283098
0188831
0238540
0257520
0229415
0136352
0253563
0258467
0243730
0136327
0201395
0258661
0198807
0183032
0258690
0258601
0266327
0258827
0258708
0252746
0151338
0207046
0259087
0258414
0129514
0139865
0200385
0201431
0259091

0258696
0259048
0258649
0139874
0176435
0259072
0271468
0258948
0259070
0258680
0270713
0199285
0100934
0259121
0258651
0258591
00092208
0182400
0285830
0241984
0259083
0100941
0258671
0270503
0150527
0258940
0271424
0259100
0165355
0258526
0286123
0286099
0251363
0258385
0279972
0258901
0258636
0251858
0165379
0258536
0259149
0258394
0258850
0258505
0258622
0258934
0259047
0258406
0259051
0286464

0258828
0258387
0258894
0258969
0259019
0259137
0259148
0259126
0258487
0189139
0258747
0258998
0259000
0258633
0258509
0259046
0179476
0258949
0179454
0198718
0278147
0249163
0185246
0212615
0239043
0100442
0187790
0129534
0199739
0258608
0260046
0257900
0258746
0258845
0258616
0258700
0165496
0139915
0251735
0258392
0244053
0214903
0263945
0259129
0259117
0258639
0286595
0287492
0270359
0207366

0258751
0258750
0242417
0258381
0258868
0258401
0252424
0213741
0244270
0276168
0252919
0165501
0258568
0278009
0165502
0168282
0258377
0165506
0258450
0100479
0259155
0258462
0221114
0197776
0258400
0165516
0165525
0277634
0252474
0278771
0282885
0274012
0278002
0165527
0251929
0274704
0214900
0278038
0270788
0243038
0201358
0251792
0100483
0100485
0087299
0264712
0125375
0259071
0100490
0258857

0012983
0259113
0201573
0273675
0198513
0258577
0151748
0243103
0269906
0258491
0100503
0270062
0258843
0285747
0252945
0258687
0100504
0131969
0258745
0258711
0100505
0287990
0259055
0139921
0201376
0201820
0258955
0258479
0258942
0258808
0258893
0258537
0139926
0252400
0277050
0273888
0251756
0207004
0259111
0258423
0258550
0258535
0258849
0259157
0186469
0258928
0259007
0258854
0087302
0087303

0258694
0258981
0258440
0168229
0261208
0125384
0087301
0180998
0197930
0258757
0100519
0198252
0100522
0259049
0285664
0258698
0258802
0073712
0271075
0288045
0258503
0258993
0100523
0258985
0258731
0237356
0286730
0225680
0226132
0235269
0231922
0266431
0125378
0287156
0224004
0100526
0100528
0258753
0197045
0100532
0020577
0285774
0131979
0252019
0265432
0258872
0198554
0233924
0180008
0168175

131981
126787
258050
279392
271180
258413
178974
239686
278818
258469
258455
126775
259318
182521
239199
186615
213641
271484
126777
200742
258784
177350
258791
259868
277763
139946
259483
275569
259719
070269
258803
258428
259133
242995
258703
165588
248550
258776
243801
200818
286257
243388
258758
259136
258592
070367
053770
286355
259008
139977

0100557
0151812
0259039
0258856
0252837
0259969
0139971
0258836
0243023
0243700
0258519
0131966
0100567
0258682
0257621
0180189
0279636
0252782
0032219
0258658
0196860
0258378
0100575
0100578
0252198
0258650
0258900
0239510
0165617
0258583
0213598
0258734
0258685
0258996
0100592
0181619
0126790
0258782
0050130
0151838
0139970
0263629
0261120
0131951
0258553
0126773
0100612
0258896
0258563
0242135

0254718
0100614
0258556
0179008
0258427
0259169
0258692
0184302
0258670
0258764
0244756
0241354
0258952
0126778
0100625
0020426
0258445
0258892
0258485
0126814
0258656
0206870
0139974
0027075
0182107
0247287
0258989
0258926
0250548
0232774
0258777
0100644
0258667
0258964
0023608
0279633
0139973
0258956
0258842
0258903
0186369
0259142
0270852
0258638
0258877
0140015
0259093
0258995
0126785
0258943

)179600
)154001
)252800
)261242
)205767
)270878
)140006
)258800
)258645
)202490
)206822
)207172
)274015
)200693
)126821
)258778
)252749
)258411
)270912
)202182
)054654
)140009
)221537
)214770
)234911
)284664
)271511
)100714
)258824
)272909
)089775
)179841
)126804
)259116
)259160
)126803
)165807
)259076
)259010
)126822
)070182
)284269
)252497
)258289
)125954
)176153
)139998
)257365
)272158
)125952

0266531
0259118
0222985
0255002
0258878
0258760
0258613
0214759
0259078
0266740
0033170
0276116
0180662
0207781
0258980
0258629
0258960
0258847
0277651
0224861
0258502
0200860
0258561
0287833
0196553
0171723
0258796
0258490
0172717
0258988
0072415
0286319
0100554
0199755
0134001
0100558
0265993
0198133
0054690
0100564
0259502
0259648
0281228
0201529
0081181
0286861
0100568
0258626
0272253
0258466

0072042
0139988
0241542
0258618
0072121
0271697
0239820
0258759
0182185
0274666
0258837
0244677
0243546
0240210
0258477
0287385
0259038
0258623
0242941
0287167
0287913
0207089
0185650
0270975
0258565
0072110
0258967
0258531
0259062
0242293
0139990
0259165
0206768
0276063
0258957
0081177
0258520
0100626
0280073
0100632
0029364
0286424
0175985
0244237
0267909
0287310
0100647
0273797
0100650
0100652

0221060
0198732
0242071
0100678
0258422
0274330
0270610
0235812
0259033
0133983
0258644
0222640
0213463
0257759
0139985
0259158
0252263
0134007
0274397
0133997
0259115
0258951
0245466
0240837
0133985
0258689
0006432
0259153
0275630
0258431
0274818
0100731
0287643
0258930
0258966
0258571
0269927
0286423
0259907
0258517
0269945
0259146
0197555
0259079
0207444
0242330
0285612
0285518
0266869
0182732

0200298
0286333
0273830
0258871
0205683
0285936
0206751
0119599
0176043
0259015
0239544
0165861
0241487
0258813
0119707
0258448
0080815
0259109
0100767
0258376
0252839
0258944
0133961
0285006
0284930
0251393
0187105
0170468
0258443
0258695
0184227
0258408
0258603
0119673
0258524
0279026
0177465
0205669
0258972
0119661
0251907
0258660
0176903
0156030
0264741
0259065
0258586
0270576
0140043
0258653

0223300
0273711
0119725
0258891
0270140
0199472
0156050
0119723
0187097
0279461
0119636
0119711
0205659
0239910
0258513
0119614
0119688
0258559
0133980
0258737
0177776
0183379
0270000
0259005
0119655
0283944
0258769
0165898
0119681
0258425
0258976
0119682
0258439
0222604
0119616
0119596
0119608
0258587
0119689
0198208
0119630
0119718
0258646
0279594
0119684
0206924
0119640
0119703
0119638
0258978

0259138
0170348
0273565
0252013
0258740
0258820
0283519
0170345
0119660
0259687
0259319
0140044
0156127
0224721
0119686
0242951
0201096
0280494
0119685
0133935
0259103
0242488
0119650
0119699
0258876
0258578
0283629
0089916
0258454
0280078
0258402
0119715
0259124
0271423
0242952
0258566
0258569
0071246
0259081
0258610
0258301
0013523
0286437
0100565
0241081
0223174
0259058
0285966
0258602
0266553

0243225
0119669
0273729
0246548
0258473
0258819
0287004
0198894
0259164
0165548
0177108
0269883
0258552
0201384
0165553
0221754
0009830
0100577
0100580
0258510
0100583
0165555
0151445
0100591
0212371
0100593
0100596
0240233
0258668
0100601
0239272
0258917
0119705
0100603
0197734
0063761
0199440
0258421
0021645
0199454
0239438
0258723
0258719
0258874
0258478
0258829
0258419
0258662
0259106
0287118

0258637
0258416
0211448
0258766
0100629
0258452
0165409
0258915
0271705
0271656
0284959
0259167
0286686
0165417
0221303
0273783
0205579
0270343
0140022
0242163
0258605
0258675
0071537
0258977
0258841
0258718
0259035
0241891
0258501
0276782
0258743
0259032
0258683
0258600
0258426
0252369
0238561
0258549
0258532
0285826
0258762
0258923
0259012
0251895
0258506
0271712
0259044
0258902
0258814
0259107

0205562
0258945
0185070
0258733
0258912
0286582
0258994
0258804
0259061
0258496
0258958
0258859
0259028
0258770
0258807
0259077
0054983
0259096
0252783
0140030
0258407
0258826
0258867
0258975
0259998
0258538
0100433
0042317
0070778
0258983
0222990
0258789
0100722
0278576
0165521
0274492
0200653
0165533
0207342
0283627
0258990
0053254
0277801
0259151
0258752
0258699
0258380
0259053
0258920
0259073

0201027
0258792
0252655
0200312
0259050
0140025
0230997
0042088
0152315
0258544
0201351
0100764
0119720
0287437
0242571
0275198
0198668
0258424
0258678
0233208
0259789
0165914
0258935
0259163
0258437
0258716
0287663
0213315
0258884
0100784
0133943
0252644
0221102
0119714
0258875
0260810
0286607
0015133
0258798
0258446
0100796
0278342
0279286
0133962
0268295
0258774
0258962
0275154
0260711
0165929

0140092
0100815
0259020
0066427
0183648
0165934
0258481
0201097
0286630
0140090
0100599
0100600
0066455
0258742
0258499
0100604
0100605
0258730
0258595
0165943
0153485
0259066
0170270
0012963
0240096
0011114
0241499
0133958
0278396
0187581
0202306
0252720
0277826
0175785
0279593
0258584
0140067
0100628
0258987
0265768
0175699
0089723
0089737
0165948
0165949
0119632
0119698
0140093
0170099
0258597

197249
256357
186910
170054
165953
287668
100665
188488
273259
288028
196136
258805
187483
259140
239269
133937
285205
240914
274379
258933
258866
100697
264607
235706
165959
259143
258615
287111
258630
176438
258572
247092
252481
182512
258390
258927
270038
187621
275033
259084
277468
213231
100721
257275
258516
250366
271358
227051
168398
258691

0258412
0258788
0259036
0100739
0258793
0258729
0066739
0100744
0222276
0140057
0259059
0240051
0260806
0090060
0258702
0223299
0287350
0258979
0287772
0100749
0225163
0280070
0288034
0259026
0258393
0259110
0285584
0246084
0197176
0287501
0258869
0258379
0246223
0259097
0258511
0241757
0258795
0176605
0237342
0278990
0259119
0232573
0234648
0287991
0229402
0127152
0235785
0258383
0183576
0272439

0090061
0205476
0258749
0200506
0247970
0182218
0036530
0258672
0066629
0199836
0271198
0196405
0242888
0258693
0258560
0265154
0199082
0168350
0258982
0100811
0258904
0283840
0259052
0197119
0258504
0198984
0243976
0140107
0140105
0258666
0258521
0258581
0176473
0258620
0258500
0253075
0183092
0259031
0258576
0196273
0258717
0185559
0214548
0266461
0275161
0274884
0276919
0258663
0258399
0211574

0207989
0199151
0283159
0254656
0208001
0207569
0207608
0272458
0207942
0225746
0276225
0275662
0275134
0199005
0202191
0212384
0201700
0201672
0272474
0200215
0200632
0200367
0201950
0199575
0200823
0201839
0200832
0199798
0200150
0201263
0199390
0201240
0200279
0200608
0202270
0201247
0199593
0201557
0199914
0201569
0202142
0199942
0201500
0201036
0201710
0202048
0272344
0202293
0200406
0201899

0200612
0200413
0200636
0200480
0201689
0201318
0200089
0222095
0222185
0199088
0199109
0207749
0198982
0221745
0199069
0211582
0207761
0207762
0194717
0221036
0212040
0207743
0283279
0283561
0207934
0283556
0283588
0215957
0221525
0221614
0258861
0199020
0207754
0202560
0216099
0207587
0207646
0207742
0283170
0207993
0276352
0208027
0208004
0207978
0207961
0199015
0216179
0199107
0199012
0199025

0199092
0207959
0223403
0230805
0288302
0259004
0286551
0277754
0288252
0288245
0258681
0258497
0258460
0259023
0259082
0259166
0258498
0283857
0197406
0258919
0258404
0241494
0258512
0078304
0259088
0207208
0256705
0259161
0271780
0272444
0197102
0259013
0258959
0240457
0080824
0140153
0080823
0022976
0100865
0196663
0212330
0156381
0266015
0259230
0239224
0089902
0230988
0202459
0259508
0131323

0206969
0166126
0198752
0259515
0213176
0283071
0259444
0205436
0259717
0185215
0259374
0259535
0259525
0251533
0270705
0278071
0242837
0270938
0240624
0259775
0100664
0272533
0259428
0243904
0075413
0244691
0166165
0260285
0166166
0252469
0258851
0166170
0126214
0256500
0256053
0201184
0246451
0270108
0269910
0269940
0269958
0126215
0100711
0253096
0088808
0270591
0202275
0258735
0258534
0287346

0156411
0156414
0227729
0242894
0166183
0207568
0283165
0258748
0066735
0258913
0288459
0258598
0222761
0213159
0258986
0184601
0259037
0258736
0260792
0265291
0256050
0203485
0258858
0185100
0184990
0258430
0142208
0179627
0279140
0258701
0239365
0196183
0258811
0258593
0099814
0259067
0166428
0185567
0140104
0170779
0183484
0257556
0286636
0280244
0258154
0287459
0184916
0273778
0257622
0183828

0185024
0184887
0179364
0279495
0242085
0226174
0251602
0182979
0257270
0182809
0213145
0257341
0185347
0184986
0230157
0234402
0211890
0211891
0211892
0274172
0211893
0277030
0253364
0253755
0213140
0211895
0253692
0211896
0211897
0270371
0211898
0211899
0265612
0264024
0263413
0264781
0211900
0237111
0242472
0240041
0242887
0237547
0211904
0211905
0236597
0227335
0211907
0225825
0211909
0227196

0211911
0211912
0244620
0237020
0211914
0211915
0227800
0211917
0211918
0227108
0211920
0211921
0232543
0211923
0211924
0211925
0237197
0228131
0211928
0233655
0211930
0211931
0226777
0236170
0211933
0254279
0271968
0225200
0211934
0253780
0211935
0276775
0282122
0211937
0253387
0254053
0253763
0211938
0282639
0282651
0211941
0254215
0254174
0211942
0253412
0253709
0211943
0253458
0211944
0283948

0254046
0273894
0211945
0253883
0271201
0211946
0211947
0253957
0253345
0254045
0211949
0211950
0238275
0276210
0253441
0253367
0211951
0253462
0253482
0254326
0211952
0253149
0254289
0270550
0253491
0253587
0231475
0283607
0270474
0211955
0254203
0253440
0211956
0253325
0211957
0253240
0253359
0211958
0253989
0211959
0253467
0228757
0253294
0278473
0228966
0254228
0232216
0283195
0283562
0283464

0270816
0253465
0253895
0211961
0211962
0253808
0229092
0253862
0211964
0211965
0253386
0253241
0211966
0254167
0253941
0253545
0211967
0253714
0275475
0254395
0253131
0253759
0211968
0224373
0253742
0199480
0254329
0211970
0253132
0253747
0253936
0223648
0253209
0253169
0211972
0253274
0223506
0253820
0254036
0253635
0254052
0253703
0211973
0211974
0282600
0281990
0283571
0280411
0274576
0254056

)225698
)211976
)224650
)254134
)254176
)253247
)253310
)265929
)211978
)253674
)275747
)283906
)211979
)253303
)215567
)201241
)258463
)274347
)188403
)258706
)259337
)259490
)270961
)271317
)282520
)282599
)271336
)277988
)259769
)258397
)239471
)258410
)278935
)258780
)258628
)259156
)287122
)258654
)180229
)258488
)277322
)258797
)181984
)278497
)258916
)258707
)270685
)278626
)258590
)279628

)260409
)238478
)259383
)259443
)258420
)274499
)259089
)259214
)265322
)230031
)238489
)261048
)261554
)258799
)270831
)266545
)258710
)258855
)258494
)270824
)270451
)282089
)282268
)270185
)259698
)259025
)258771
)243059
)260986
)279595
)258684
)258883
)237161
)247765
)207289
)280881
)280703
)258415
)268531
)258712
)258767
)258848
)277243
)278522
)275174
)281347
)277841
)287345
)284834
)284788

0280709
0284988
0279639
0259324
0285491
0285116
0182974
0279408
0279445
0281113
0281087
0281179
0281550
0278263
0280655
0265793
0233917
0200790
0258585
0258997
0264902
0283524
0285405
0284500
0258721
0285472
0283680
0285135
0284892
0274102
0183706
0259435
0260739
0223877
0270505
0271288
0259261
0270356
0271507
0221641
0259098
0258732
0274835
0273679
0274279
0185182
0274568
0275363
0277865
0281909

0276550
0277561
0277515
0276141
0277435
0283273
0277867
0275325
0277755
0274253
0259425
0170113
0140157
0273749
0275835
0276172
0273976
0277505
0175676
0273981
0274099
0278541
0276941
0278552
0259414
0274471
0273756
0260399
0261739
0278222
0286598
0261418
0174450
0265673
0179455
0260978
0254585
0182636
0286973
0206616
0260232
0259905
0260551
0260780
0260959
0287888
0286110
0185823
0128739
0280118

0241061
0242088
0273173
0214265
0224078
0276314
0279192
0276610
0239014
0257647
0274640
0207063
0207001
0207014
0275529
0207191
0207442
0207133
0207093
0206727
0200661
0206609
0207197
0207137
0206621
0207174
0207263
0206656
0206688
0207460
0261069
0278715
0277785
0275127
0207375
0287482
0279735
0207279
0252326
0251815
0251896
0278123
0207245
0252277
0279050
0212428
0270814
0201831
0199712
0199970

200680
200503
200812
278089
200726
199782
201943
200486
199453
273835
199960
201679
200757
201482
200163
199968
201969
199833
201326
201331
200398
199489
275524
201300
200801
199704
200987
202188
200949
200593
199311
201992
202499
200638
201907
200564
272460
200478
201143
202373
202261
280234
212380
276844
212528
201634
239169
114062
261529
235731

0223628
0206190
0274307
0222123
0266517
0259011
0206187
0258965
0212604
0258931
0259150
0286884
0235160
0287280
0166206
0278840
0235518
0242793
0186297
0259122
0182256
0228740
0214254
0200326
0258624
0261426
0258970
0259152
0259168
0258594
0287922
0258853
0104044
0232394
0228992
0128731
0256338
0261401
0261561
0237850
0261497
0153684
0276891
0261524
0276928
0274532
0266039
0284628
0260159
0261365

263964
274814
260053
183629
276955
261041
227717
260444
261581
206149
254398
188626
274632
261480
248334
260844
232431
256951
261377
271616
261649
270301
287894
034053
259277
104059
259299
252868
185115
259690
259814
259300
287334
256802
104067
259644
234664
259523
259749
259647
260174
229389
179938
273818
261792
178081
207432
261247
277637
260897

0261747
0259906
0225930
0259993
0206972
0270955
0279417
0238783
0166664
0263070
0238519
0274966
0186399
0274424
0271333
0215302
0270055
0283345
0260784
0260693
0178115
0277031
0259890
0260128
0207430
0261794
0270016
0277467
0269930
0247728
0285077
0284906
0285035
0187951
0206846
0284783
0103832
0278696
0259845
0260382
0198690
0261628
0166912
0212526
0259720
0134160
0207702
0259448
0259772
0169926

0259179
0259696
0169918
0259358
0277500
0175344
0206849
0201084
0279912
0271078
0207257
0260211
0260642
0261491
0249931
0275776
0215304
0261279
0206987
0261708
0206127
0274076
0270789
0261375
0254912
0223509
0261064
0276724
0262728
0232653
0277464
0244952
0198826
0285948
0166922
0241818
0259721
0166923
0248905
0259392
0212415
0259686
0279958
0215296
0259446
0198838
0200008
0276702
0259408
0259287

169857
184984
278472
287840
259505
134153
243094
182405
134152
128463
140199
259468
207091
182117
184507
176454
259904
175265
221649
260639
261081
279092
259917
261559
278582
215252
221065
259892
252425
261304
287973
159248
250007
159251
259751
021776
231409
198146
279364
259181
280396
259587
259665
255192
259397
259336
259592
259516
134146
259710

0259585
0265102
0259688
0259411
0248079
0259217
0259245
0265098
0259639
0259395
0259737
0270500
0186073
0261635
0261315
0260582
0261191
0223518
0212511
0259280
0134138
0277202
0206676
0259460
0274444
0274216
0270304
0259434
0243122
0166069
0259225
0236914
0259380
0166068
0171262
0259192
0172575
0286786
0259326
0259598
0175779
0259731
0274281
0259345
0259278
0259423
0175746
0259447
0259269
0259390

0137801
0276107
0278621
0150667
0259580
0261136
0166073
0259432
0246863
0200305
0128829
0259389
0140319
0248508
0259584
0275636
0104081
0259409
0259239
0156970
0137843
0259288
0273786
0260648
0279537
0176753
0279409
0137841
0230778
0273855
0259307
0244705
0259198
0259330
0188549
0252714
0233041
0140323
0259368
0128944
0128928
0140320
0259364
0259211
0169105
0244251
0259536
0128891
0223313
0166133

0203573
0137812
0265814
0259254
0245849
0051180
0137824
0271236
0137880
0104129
0188277
0166140
0166143
0261183
0166145
0104140
0274798
0104142
0251161
0128917
0259405
0128965
0128908
0259617
0259463
0259521
0280036
0259547
0283133
0157021
0178997
0240847
0187446
0247556
0285920
0104147
0137804
0259396
0137806
0137815
0200407
0137825
0062524
0103932
0259838
0092445
0260926
0174197
0207766
0270702

0260814
0137802
0250379
0243789
0168970
0243708
0137877
0200293
0264850
0174171
0103966
0278493
0259883
0246740
0188089
0257797
0159337
0168907
0166887
0207712
0261002
0103978
0214013
0260571
0258461
0092529
0103994
0201077
0092531
0261684
0285942
0180979
0137814
0261822
0260490
0159433
0260282
0140326
0278769
0274403
0128881
0246283
0286877
0260799
0254793
0159459
0285080
0281269
0205871
0137842

0260928
0285117
0261687
0166946
0166947
0104055
0202211
0261679
0159495
0260406
0168806
0168803
0140265
0137822
0243871
0067369
0166963
0168781
0237289
0242866
0222398
0284772
0166762
0206991
0275601
0224677
0249839
0223572
0166763
0201136
0205771
0206589
0167004
0128886
0262560
0140264
0221792
0184716
0242028
0140259
0092470
0200391
0171877
0227973
0185607
0224055
0166734
0250130
0259499
0259595

0137770
0259335
0259563
0179523
0104131
0259659
0104133
0229474
0206957
0166710
0185880
0276121
0259479
0259187
0238845
0212424
0259742
0259231
0252117
0167014
0252475
0259304
0252288
0212550
0140263
0259418
0252171
0259352
0140279
0140274
0140254
0137857
0259539
0138606
0259519
0259932
0260035
0279014
0261709
0259338
0259520
0137860
0285597
0235390
0259433
0238941
0171766
0275672
0171763
0166920

0259342
0284386
0259354
0104154
0179362
0207516
0259681
0259427
0275709
0104164
0260170
0200419
0137767
0273972
0259200
0259743
0287020
0259593
0238819
0285934
0287704
0238456
0259255
0223308
0259509
0284540
0137872
0222343
0259554
0259221
0259360
0259740
0259588
0213862
0259572
0259754
0259730
0287439
0188467
0104177
0233932
0074803
0259235
0259385
0259488
0202542
0128951
0166147
0286947
0274654

259705
273925
103995
259216
259469
259670
259700
185634
255302
259602
259500
138593
207105
265942
259656
166200
200120
156958
259467
264210
259545
243338
166262
275580
276593
259531
140285
104047
259388
104043
259188
201086
140284
200605
140287
241175
104064
271819
285410
244879
259715
284284
259465
138592
252376
242737
259618
259298
170236
092439

0259684
0243027
0138600
0279665
0259773
0241130
0273674
0081014
0259378
0259240
0183578
0259204
0137869
0266593
0259701
0283614
0278098
0259306
0186417
0270378
0259194
0259678
0104093
0259668
0104112
0259241
0259296
0140280
0128872
0259377
0138594
0274954
0277677
0259632
0207484
0259201
0272337
0259185
0259556
0166477
0069956
0259438
0274528
0259398
0259712
0259178
0242327
0137875
0069966
0259327

0259709
0206995
0259577
0128833
0221052
0197535
0231169
0128989
0260618
0047346
0259935
0287500
0169856
0259203
0214896
0259612
0259237
0287596
0166415
0252066
0206641
0259301
0280362
0259669
0137766
0259619
0259550
0260937
0279145
0137876
0069974
0276533
0225973
0069943
0260916
0274765
0277548
0261771
0283891
0261652
0256061
0260036
0171016
0285805
0166450
0259180
0259369
0069869
0230183
0239703

261333
181827
261072
151575
275417
199784
261174
259941
261823
138587
277245
260392
137871
276696
285253
274667
279683
285331
225625
276524
275343
140262
260689
239035
247982
260172
260870
128849
252984
261489
261265
276772
240874
263155
137878
255529
261219
128918
259477
274719
259285
103569
259651
259626
259600
259379
259671
240934
166035
259293

0277229
0277790
0259476
0259250
0137845
0270986
0182625
0282100
0199730
0278513
0286897
0259173
0200318
0245975
0128923
0259402
0259353
0259738
0157450
0242747
0137776
0225798
0259393
0241640
0157456
0277144
0259732
0157483
0253030
0171989
0200070
0259771
0199512
0259735
0157470
0259750
0259674
0200252
0243780
0240238
0201704
0140297
0140307
0227161
0140299
0259657
0259238
0234797
0286495
0259601

0259573
0171956
0259558
0259223
0287915
0182718
0240163
0242667
0273721
0128915
0245534
0069667
0259274
0259513
0273958
0286457
0287508
0277413
0259482
0212625
0259591
0273792
0259481
0259575
0276473
0259616
0259530
0259675
0271195
0259284
0259564
0129003
0285863
0259251
0198535
0259466
0205502
0259697
0166104
0259878
0276713
0260062
0260375
0261296
0260957
0277919
0275451
0259557
0274340
0171914

0259458
0259756
0211137
0259370
0259727
0259394
0140416
0259498
0259627
0259672
0103642
0185088
0166128
0138613
0074410
0259459
0259268
0276651
0140455
0259248
0197361
0103657
0259589
0199156
0275785
0035664
0259351
0166797
0028528
0201071
0157734
0166794
0169118
0259316
0252774
0166803
0103671
0206781
0180357
0259682
0248415
0243403
0239465
0207223
0207162
0180304
0265967
0264937
0259635
0166831

0221033
0140451
0249240
0241839
0287494
0166839
0275332
0274383
0090487
0103707
0270218
0186198
0277351
0103710
0234438
0278737
0246922
0259258
0090470
0166855
0138615
0279268
0201709
0138617
0206903
0199568
0259346
0200156
0174498
0103742
0074603
0270690
0259232
0252217
0074696
0207449
0138614
0074621
0275638
0174485
0238311
0264737
0103769
0261544
0259924
0249271
0283249
0157890
0263512
0261318

0166938
0260773
0075131
0252712
0223271
0240821
0169032
0261102
0243404
0261351
0174446
0266589
0174444
0199574
0280554
0202529
0199673
0200623
0174442
0188501
0259471
0137834
0274995
0270490
0277152
0259437
0259289
0259347
0259246
0166949
0259202
0103591
0242634
0103599
0259673
0189227
0270964
0137764
0259512
0188779
0259410
0245719
0206625
0261702
0260109
0033800
0260007
0129007
0128973
0260602

)260657
)169018
)137809
)286770
)103647
)271232
)140350
)265195
)258484
)259286
)285919
)137808
)255346
)288068
)273851
)259265
)212766
)259222
)138604
)261634
)259504
)137819
)259645
)259426
)207395
)137807
)259191
)137818
)207119
)259215
)245750
)259457
)259496
)259309
)140332
)207965
)200216
)259252
)259503
)259473
)287741
)280639
)259227
)281264
)280515
)137831
)259532
)274297
)259624
)259452

235420
166173
259744
137821
259470
129028
242276
187720
225362
259964
259781
260586
212664
278408
278570
260037
066933
200873
261632
260576
222094
166192
243568
260173
175318
067225
137817
273025
261460
140488
260729
213614
260339
243007
187806
261423
271072
261187
166233
283798
260534
278313
260144
259783
260672
259909
215186
274350
175202
261281

0140463
0159322
0260898
0285729
0259259
0067141
0259691
0259440
0259528
0138622
0259650
0183324
0259186
0156642
0281183
0103855
0276807
0205363
0260469
0260624
0167139
0261801
0129038
0067221
0140464
0287543
0260096
0159289
0278604
0214702
0167178
0248540
0261543
0278348
0129009
0137868
0140481
0280309
0261384
0277749
0140459
0260266
0261821
0274937
0277718
0138623
0277391
0286696
0138629
0247240

260103
261775
212279
179361
179335
275527
260919
179151
140465
140505
103653
264386
140506
261606
213578
140474
275411
140497
178802
178761
178741
178718
198794
261813
199580
260483
138621
260152
203411
252722
240366
167173
261779
287926
260660
260104
261205
167195
277295
140478
275992
260357
259790
140365
275645
140398
284343
140400
260274
169375

241890
261714
276744
169410
260165
259931
260269
169371
260235
260206
177971
275454
274515
173548
173546
260892
203392
182950
246877
260288
260815
261043
284385
187812
241807
261820
140367
266308
167196
169752
260685
169758
259422
140374
196274
259581
287503
159556
259514
270036
140386
261232
266449
201510
117906
243365
259565
140368
140391
259652

0278991
0269951
0173517
0259722
0230459
0140382
0279033
0259362
0259420
0169783
0259666
0259281
0280359
0259213
0260139
0260776
0261244
0242066
0214646
0261403
0261143
0167202
0259792
0183476
0199633
0136425
0166411
0260988
0103740
0259708
0140403
0140395
0272418
0259562
0259322
0277482
0279694
0166426
0259413
0136381
0259474
0188266
0279373
0041357
0169684
0261762
0080644
0279421
0117971
0259555

261303
213307
273512
238166
136378
185787
243711
252061
279235
103811
058335
177699
259234
207695
259419
235711
166557
287408
169330
259493
259208
260674
213527
259229
252995
259548
136371
261229
259332
279394
286813
180953
278600
259642
259770
140379
271364
279719
279705
275460
086666
103876
259967
259417
259361
259495
172379
258010
287813
259175

0270919
0284059
0222139
0136379
0271983
0287108
0103888
0208003
0259546
0259649
0117899
0260619
0265855
0277782
0140406
0156206
0279834
0259461
0172349
0271725
0172345
0286488
0259343
0188869
0273920
0284709
0275995
0259543
0259594
0259692
0259622
0183496
0259518
0287872
0140598
0212170
0285974
0188659
0274376
0278662
0259429
0237550
0278202
0197978
0275695
0276710
0278603
0255769
0277741
0259637

284803
182774
260836
278013
214575
259462
103723
259707
228141
259315
279758
259442
286817
250988
252690
186628
277864
156232
103942
259805
169612
169609
260608
064726
273747
259767
260351
259308
263643
286872
136404
166503
260579
259986
169594
140600
156218
212374
261665
200444
274128
259609
259404
280038
259694
259511
259726
225151
278422
184206

0259633
0259683
0244056
0259570
0235370
0272887
0259538
0259774
0278785
0197627
0259551
0259302
0230373
0259244
0189136
0259728
0275120
0176371
0256278
0276278
0176700
0254779
0177082
0277578
0197696
0140612
0259654
0285667
0166716
0259415
0136383
0200991
0156222
0286505
0207037
0212388
0271044
0073417
0259270
0229212
0259449
0259295
0277582
0188388
0218052
0259317
0170776
0284160
0251891
0259630

0259375
0259367
0259276
0202081
0259453
0259544
0259407
0259762
0259416
0183655
0221634
0259195
0252931
0264406
0259608
0260477
0273540
0252964
0261077
0200206
0260125
0259734
0259636
0259620
0259761
0259560
0207150
0259527
0287391
0140538
0259183
0260305
0259494
0173867
0181991
0279945
0140543
0259569
0249487
0221630
0207703
0283484
0181026
0172183
0259676
0157766
0140511
0140545
0259350
0259489

0261549
0260275
0260123
0261407
0140526
0239151
0259699
0259948
0140522
0140525
0140521
0275101
0261441
0279708
0287717
0255571
0284329
0140519
0259218
0261478
0259713
0140534
0259615
0166813
0166819
0166821
0241362
0140527
0259561
0206590
0166823
0259441
0188095
0166825
0157823
0250021
0264966
0259677
0264796
0212420
0242498
0252645
0259273
0280377
0140548
0265871
0273691
0259291
0276376
0182054

)259685
)185033
)240470
)225193
)185043
)183208
)284626
)213471
)261147
)182768
)275674
)228998
)270322
)275345
)271396
)214433
)275803
)279980
)238244
)196391
)259262
)140575
)259177
)140577
)259212
)259314
)259736
)285560
)259746
)278370
)245479
)197299
)200677
)287061
)259704
)241869
)140564
)182511
)196547
)259661
)184508
)140553
)258384
)166965
)198901
)284946
)258725
)200142
)184056
)214432

)278514
)258542
)275929
)258665
)285661
)185518
)285883
)258551
)258954
)258765
)258540
)259057
)176463
)260661
)258761
)258726
)258925
)140557
)239197
)258739
)259170
)183643
)258647
)258676
)260337
)271763
)185442
)259064
)275965
)272298
)253954
)258741
)240484
)258562
)260361
)258527
)258922
)274215
)272888
)279765
)173575
)284324
)182175
)258611
)285663
)257060
)258888
)258971
)258909
)258787

0258631
0258754
0258627
0258476
0259123
0258831
0258785
0259724
0258895
0258570
0140563
0258632
0287328
0287950
0285866
0259331
0286917
0277654
0270017
0258433
0258773
0248441
0273923
0280304
0285700
0259134
0258489
0275016
0222076
0275443
0259702
0273771
0247809
0280048
0277135
0259359
0278146
0259763
0280291
0185551
0283888
0275322
0259275
0259614
0259542
0259312
0250825
0242295
0259534
0222617

0274079
0259340
0214424
0259282
0185594
0223120
0259485
0259403
0259664
0242863
0251729
0279091
0259582
0287267
0287321
0251209
0259870
0287000
0140450
0259478
0258483
0283372
0259199
0222361
0259611
0259320
0259266
0259041
0274312
0277987
0283597
0259507
0259424
0140443
0278022
0264480
0287191
0259621
0259475
0183571
0278090
0182253
0261054
0103852
0261616
0259921
0168904
0259706
0259228
0259257

)259590
)259760
)274294
)259341
)286893
)279846
)241562
)068305
)201724
)183060
)182397
)259655
)241588
)259363
)254744
)140470
)259219
)252957
)259356
)189419
)259430
)154227
)270127
)251819
)259205
)259540
)140471
)200095
)183475
)212306
)259381
)259579
)279588
)259586
)232386
)272808
)259604
)272639
)184254
)278456
)259583
)154237
)259376
)259755
)287892
)259365
)259182
)131873
)131871
)131876

259172
140479
279970
259764
274139
277210
282793
259464
259695
286733
184277
252614
185418
259658
259660
259236
240927
184140
182854
232459
214344
257109
259454
259646
177693
183909
275771
248893
259553
284557
185596
275449
248472
233614
234769
278739
260803
231439
226942
161980
161981
007384
278995
103152
103148
269482
228779
130656
206177
206178

0225323
0188536
0206172
0207243
0086506
0268836
0007392
0206168
0167930
0076344
0286901
0242173
0185615
0103126
0086504
0129925
0236829
0103202
0242612
0276984
0090565
0201034
0230428
0277142
0103326
0266124
0266235
0261691
0161992
0007541
0257108
0282907
0197562
0127578
0130731
0172366
0228201
0161996
0262528
0140983
0103269
0260394
0279441
0103266
0161999
0127580
0261659
0127585
0279255
0259840

0103260
0103254
0162004
0103253
0103245
0102854
0279136
0162006
0207579
0007376
0127586
0127588
0287855
0275438
0103227
0260022
0260316
0260439
0276931
0260807
0005513
0260496
0261713
0261720
0162009
0184471
0260702
0260532
0273551
0196557
0260403
0259910
0261294
0116176
0277010
0197253
0172236
0095917
0260182
0196364
0261564
0260991
0260284
0260710
0274751
0103275
0261505
0219274
0007516
0007520

0090581
0260425
0059145
0260132
0174109
0284395
0197599
0279930
0103249
0261641
0260051
0261430
0219027
0251692
0100726
0187535
0260646
0131634
0280062
0280231
0280327
0260954
0260989
0007545
0226890
0261732
0206053
0278987
0138834
0275092
0261399
0265820
0261207
0103024
0074071
0197774
0162032
0095906
0099769
0063854
0180185
0162039
0260541
0281219
0259947
0162040
0198736
0140986
0140990
0140988

206811
207405
255513
255198
273587
179580
277602
183751
196408
127554
261790
127561
260107
167962
183971
265386
065054
065057
103197
008710
221656
259933
261123
284008
261240
260447
200059
265867
264397
167964
260260
206630
281010
131653
167971
167965
182685
184207
261532
167967
259780
167968
167969
261663
205937
260778
266643
284346
264004
167972

243445
238098
162063
285970
283180
260095
260874
162062
274805
259895
162068
162065
260293
260272
185883
259784
279520
162066
205923
266232
140992
261613
261288
269937
261140
260436
279568
280402
279162
261093
215154
205918
279057
260176
260565
167977
279901
172382
205913
167978
103363
276791
103355
221719
215148
007038
262154
162078
172460
005001

0263325
0263280
0162076
0059122
0262959
0262482
0131650
0127564
0162073
0274367
0262152
0262362
0272079
0213937
0184697
0006327
0103145
0131652
0162069
0205890
0261971
0008516
0008517
0252561
0262370
0200204
0130182
0263011
0263072
0122386
0085644
0228146
0261889
0262521
0262668
0168124
0203581
0010539
0103313
0281005
0006194
0279330
0279031
0140993
0162086
0262899
0262316
0168158
0285329
0262118

0263177
0232196
0140987
0103343
0167981
0122390
0263212
0273776
0215131
0240311
0103351
0278942
0167984
0262312
0188827
0261938
0213918
0263235
0126602
0276754
0005339
0262516
0262185
0162104
0263159
0262471
0263105
0185739
0262468
0090447
0126603
0262686
0217930
0262712
0103426
0280063
0262246
0168140
0103423
0277170
0277440
0275515
0153406
0103415
0089486
0278434
0205832
0261789
0261612
0153443

102858
263955
261442
278700
276641
168101
168096
266994
166246
103199
276096
140623
285952
267795
267077
067836
140632
275056
118900
118898
103184
267072
103174
033011
153446
260882
118894
267070
285567
273963
260431
261811
078328
260817
242278
267182
261007
260338
273648
270440
280000
257180
286635
260835
278975
259963
252080
200869
212189
240419

0287340
0283584
0261198
0252138
0260289
0260003
0261319
0260058
0259819
0260379
0261066
0232258
0287522
0067365
0279357
0183044
0243954
0238417
0260764
0184857
0140650
0260350
0260276
0153048
0259939
0261481
0283516
0187555
0260979
0261392
0182831
0263244
0260349
0220793
0261617
0260071
0261397
0261075
0283003
0200468
0252927
0260362
0183454
0283221
0287830
0261810
0261647
0285543
0266439
0166669

0260756
0279662
0199482
0260468
0213853
0256013
0260310
0153060
0263279
0263241
0262107
0261904
0262322
0262259
0263196
0262158
0262381
0103274
0166676
0262488
0262151
0179583
0182108
0280153
0038532
0229917
0274038
0262020
0287121
0263033
0263179
0175643
0185338
0178279
0178257
0212228
0122304
0175646
0287429
0221801
0199668
0263080
0262703
0188897
0262636
0262999
0189067
0184602
0153066
0262420

0122299
0274505
0277369
0262117
0171490
0263307
0103342
0261560
0260318
0261216
0277458
0234719
0048462
0260224
0048471
0260488
0241641
0175604
0261293
0238685
0261394
0260723
0278979
0260601
0259876
0259899
0274834
0103381
0264733
0260378
0261158
0199474
0241932
0237515
0262116
0262801
0262011
0262235
0262267
0175595
0242307
0262732
0263234
0262097
0261856
0186260
0276564
0262529
0262470
0201075

0262454
0207639
0199130
0260009
0263257
0140694
0242430
0103429
0238964
0069764
0261523
0224712
0254609
0261695
0254852
0255037
0255277
0103512
0270734
0257264
0284305
0263918
0258354
0265537
0183458
0207425
0183426
0275259
0278265
0273804
0179889
0261819
0238728
0275910
0157045
0085721
0260735
0188599
0270580
0250251
0283346
0207294
0264115
0257391
0277770
0183793
0156968
0261130
0166780
0166783

0257769
0284184
0072864
0283736
0262380
0280206
0133392
0263335
0263065
0280429
0133393
0206778
0262171
0215003
0103222
0241067
0091262
0262332
0103226
0263029
0262848
0257381
0264722
0257366
0265373
0244257
0206706
0183889
0278221
0277698
0277920
0227827
0276311
0214967
0261181
0260126
0261362
0287905
0279729
0261748
0279490
0260675
0260405
0103489
0261448
0279410
0279415
0279620
0286849
0275927

0285850
0260945
0261555
0228071
0259929
0280283
0260242
0260933
0278994
0276484
0214940
0285628
0205746
0273613
0277647
0233024
0254681
0275391
0206927
0257563
0266291
0265776
0257527
0266454
0185164
0274025
0277014
0262974
0260550
0256340
0134419
0260342
0170540
0260017
0157106
0260352
0170537
0207167
0261465
0261357
0167186
0260430
0278389
0205730
0261427
0261759
0103528
0286790
0261210
0259871

0285869
0259925
0280265
0103534
0260681
0260592
0222750
0260934
0006007
0279747
0103540
0103544
0276571
0103550
0261312
0174628
0261195
0261939
0167191
0180269
0277211
0263253
0227624
0169347
0169344
0169340
0263237
0183549
0262995
0183747
0280306
0066813
0260201
0260762
0005187
0275315
0274730
0166743
0066654
0283399
0285303
0260510
0196678
0200893
0005189
0275908
0188215
0102897
0158486
0263331

0011638
0262983
0103310
0175311
0103316
0283421
0189149
0257403
0158482
0276548
0169246
0180747
0265462
0287809
0258186
0273956
0257639
0271609
0252798
0197006
0261596
0140749
0207248
0207042
0155719
0260306
0248124
0276721
0185864
0185710
0257408
0274460
0140740
0155714
0260395
0275445
0185716
0260277
0175267
0183921
0287739
0103319
0261113
0058600
0286808
0140743
0260635
0260790
0230872
0257122

0237296
0243716
0277041
0257838
0260973
0260905
0283213
0122254
0261090
0260566
0103404
0280400
0285613
0166828
0260741
0168447
0168434
0280039
0212593
0259782
0260136
0103365
0261528
0103356
0260247
0103353
0260751
0004779
0083093
0261723
0166847
0260482
0166851
0261266
0134398
0278912
0261583
0279756
0166869
0166501
0279276
0279202
0274466
0279554
0260264
0006116
0201541
0239081
0260072
0122257

0090905
0224310
0261669
0158865
0140750
0260714
0262587
0262155
0271105
0263326
0260448
0262700
0275494
0205629
0260034
0266166
0103375
0155592
0259955
0274925
0261636
0260690
0182601
0285882
0260255
0265005
0199929
0230330
0237140
0231876
0228480
0286712
0236481
0260580
0234186
0261482
0261557
0261045
0155666
0259940
0169189
0245888
0274092
0077238
0103522
0259954
0077235
0047578
0261736
0261329

0273553
0201371
0169181
0212382
0207266
0169180
0260452
0200652
0200138
0250982
0283662
0188322
0246465
0277191
0198156
0271623
0205609
0275429
0261089
0275654
0270118
0261832
0233232
0188603
0184730
0197272
0176046
0271495
0275441
0176476
0197165
0278665
0278725
0196502
0285020
0255524
0261419
0184110
0259982
0278340
0270424
0196993
0278528
0251417
0240634
0260796
0275807
0168488
0260570
0178952

0283845
0178188
0261766
0196296
0260442
0252461
0177548
0177455
0176953
0260853
0284541
0260367
0169682
0261067
0213658
0261552
0196796
0260908
0103472
0284671
0284649
0284685
0260517
0279106
0259807
0273582
0260953
0277999
0198106
0260413
0271699
0278078
0254206
0254634
0183336
0261740
0260335
0181625
0260280
0213648
0258150
0279228
0257691
0257506
0169203
0205534
0279583
0266758
0260727
0260133

0260514
0197471
0103485
0222375
0185905
0260719
0174992
0079616
0275857
0103495
0259952
0238045
0280607
0167371
0280893
0280789
0281348
0013364
0103502
0278713
0214725
0174938
0174939
0174943
0279789
0247735
0149932
0149930
0149929
0169592
0149927
0167194
0149926
0285043
0149925
0274904
0149923
0149922
0090238
0250616
0102886
0261367
0275371
0102882
0102879
0261416
0169627
0132207
0213599
0261052

0258130
0278887
0261444
0261203
0198064
0183604
0273724
0169217
0260219
0169221
0274653
0180209
0180035
0180096
0280137
0179965
0199787
0179958
0179918
0202476
0222701
0005844
0261346
0260487
0251706
0261332
0266305
0169957
0261459
0278922
0169955
0235560
0260494
0169951
0229809
0239791
0197162
0260167
0261588
0156853
0260678
0261680
0260113
0156858
0156860
0252074
0261840
0080603
0222287
0282034

0206755
0281991
0260899
0156873
0196118
0103549
0280211
0102870
0274848
0260082
0274678
0099385
0260083
0265991
0262721
0211591
0150281
0230447
0260852
0099364
0261487
0175938
0275263
0099381
0279196
0099377
0099365
0103496
0260911
0232748
0167394
0261124
0167395
0151006
0255439
0167397
0280160
0103507
0103510
0262766
0278133
0052344
0261385
0178226
0263343
0262366
0089280
0260304
0260060
0103490

0261359
0177238
0169900
0169896
0140678
0261245
0156886
0260757
0156885
0277543
0176723
0261474
0260267
0140691
0280132
0140682
0140675
0260740
0140688
0260625
0169877
0261475
0261648
0213547
0260883
0261741
0278885
0261614
0260010
0237185
0260631
0260472
0260568
0131797
0276867
0261731
0197302
0259810
0260722
0261457
0180663
0261284
0259874
0185947
0259950
0261289
0197476
0260218
0260628
0261727

261704
223931
260048
270472
230267
260584
286473
279997
260847
260649
260540
260344
260575
205456
260402
259822
261127
260516
260327
260866
260307
261541
260662
261233
259966
262187
261111
261569
261263
260311
183632
260974
261391
260864
260845
261108
260644
278950
279780
279795
260158
261719
259934
259303
260047
214617
214614
261834
270864
277304

260610
261259
259852
233732
260921
259997
271130
260141
260900
261599
261682
259800
261536
260414
260419
205457
260827
261507
275034
282973
261509
261009
283110
278848
282927
262090
260626
261200
261466
261405
259882
260518
282924
261580
260308
270467
261153
271178
259680
270924
271691
260312
270318
260525
260781
271620
261607
259842
205452
270401

| |
|---|
| 0198555 |
| 0261217 |
| 0259990 |
| 0283065 |
| 0261197 |
| 0262561 |
| 0261440 |
| 0260087 |
| 0259987 |
| 0260207 |
| 0256642 |
| 0200434 |
| 0279800 |
| 0284209 |
| 0288300 |
| 0286968 |
| 0279165 |
| 0287353 |
| 0260854 |
| 0179755 |
| 0261566 |
| 0262885 |
| 0216671 |
| 0278138 |
| 0261053 |
| 0260809 |
| 0259979 |
| 0260724 |
| 0261800 |
| 0280180 |
| 0261461 |
| 0261608 |
| 0260073 |
| 0260590 |
| 0261398 |
| 0260958 |
| 0261274 |
| 0260967 |
| 0260984 |
| 0259897 |
| 0261010 |
| 0261752 |
| 0260846 |
| 0261486 |
| 0260427 |
| 0260480 |
| 0259841 |
| 0260291 |
| 0261689 |
| 0260598 |

260812
260680
261299
278681
214581
287163
261711
260153
259836
261350
260341
261445
287448
260857
269622
282946
259996
260994
261046
260134
260746
259791
260522
251915
222207
251924
252624
261122
252094
276599
277004
252182
274228
253055
252647
252268
251760
251717
252806
252673
252587
252936
260611
261620
261515
261782
279666
261084
261239
199448

0261356
0260251
0171241
0261512
0259856
0260909
0069329
0261131
0280376
0091651
0140795
0260782
0155330
0260889
0261725
0166123
0069345
0261173
0259821
0259983
0171208
0260281
0239840
0129636
0260744
0279509
0260012
0261369
0222767
0102893
0222268
0262038
0259866
0261336
0259918
0261231
0260497
0261325
0260450
0288026
0261017
0140798
0261538
0121270
0102910
0240793
0280067
0196470
0275909
0261802

0260688
0260347
0102921
0261267
0243609
0260033
0259912
0260086
0260052
0272545
0260726
0222170
0260184
0260067
0287469
0261421
0261815
0102924
0279249
0280189
0166152
0262950
0263110
0261623
0102935
0260089
0260024
0280131
0259978
0275155
0279842
0261751
0244551
0283365
0205423
0259843
0261835
0273829
0260298
0155393
0279356
0202124
0260381
0287444
0260031
0252526
0121274
0121281
0274969
0166164

0261393
0261092
0260573
0140807
0260029
0278909
0205414
0260249
0167208
0167207
0270120
0261644
0083799
0264947
0260616
0261703
0261241
0260042
0199771
0261637
0260605
0260620
0287718
0261282
0103449
0261238
0285367
0280344
0287256
0261047
0260818
0260850
0224578
0223168
0260057
0261190
0261470
0279779
0260268
0261749
0260975
0260478
0260887
0103460
0279427
0249231
0285800
0260963
0260393
0261261

)234337
)260326
)260258
)260939
)277639
)177200
)261291
)280227
)261056
)279589
)238645
)202193
)259962
)260078
)261804
)262714
)279741
)280027
)103479
)252569
)259926
)279344
)279722
)166971
)287909
)103494
)275191
)140718
)261630
)280392
)260194
)261049
)280454
)259759
)259517
)177508
)283689
)277559
)261803
)287885
)286845
)283304
)283258
)245694
)286357
)176842
)259711
)259725
)260147
)285775

0259283
0239555
0283423
0159387
0263207
0240760
0087245
0274508
0260135
0279030
0087253
0261997
0256812
0103546
0278928
0260765
0228695
0198848
0159398
0246379
0279523
0260041
0087258
0261439
0265281
0287866
0260198
0279764
0272372
0159461
0260621
0167005
0087263
0274031
0125124
0102891
0087250
0125148
0260823
0260549
0169715
0205364
0255986
0205362
0205361
0205360
0259923
0169688
0198417
0125144

205358
275691
187193
259827
261429
261302
102900
207649
070915
283471
243988
051108
261270
261114
087237
140853
259922
260145
260148
140848
279019
279803
172775
285979
159579
102931
276663
260038
102934
260965
271456
102962
006210
102970
005194
088682
260345
102978
125170
135736
159618
260828
205336
261633
182885
260467
159625
260438
140854
140859

)276166
)274816
)187185
)206833
)070729
)159648
)166188
)103005
)102996
)207173
)261151
)070761
)260545
)260927
)070770
)260867
)240863
)103021
)103023
)181938
)212379
)261078
)252271
)276131
)260186
)103034
)200556
)103037
)125107
)276259
)207493
)281910
)206952
)260865
)103042
)279889
)125166
)245768
)200424
)244003
)260451
)242423
)261638
)275808
)244043
)261144
)200062
)270222
)260240
)261807

0261278
0261436
0261590
0278499
0261310
0259844
0261178
0224631
0286653
0150394
0260600
0201289
0260115
0207441
0280261
0237901
0231589
0260332
0261502
0269986
0261199
0260658
0241808
0261743
0279225
0259864
0261014
0261653
0260715
0261028
0260668
0259846
0260187
0260068
0140937
0278893
0259847
0260834
0260364
0261742
0261818
0260004
0260695
0261234
0201999
0286928
0179776
0246898
0166546
0260146

261656
260851
166548
260755
261519
275745
217555
254788
277978
089505
140932
260650
140931
183723
135720
260465
287965
260558
246777
249961
159593
258122
168748
172840
166589
166592
166595
172831
265408
280416
172828
172824
280334
265918
067955
125149
237172
102871
135722
265690
102878
140939
196123
179044
205250
102890
207948
280163
125122
168701

0135723
0135740
0262691
0196155
0168676
0201201
0159708
0159713
0239194
0159714
0176387
0261320
0159720
0270049
0159723
0276075
0039523
0261396
0259945
0260894
0261386
0280214
0259804
0102974
0237718
0159753
0102977
0102981
0124074
0159761
0141098
0141084
0270165
0102904
0102901
0168286
0102898
0262141
0038358
0263126
0188038
0159792
0141086
0261884
0205220
0213398
0124067
0141096
0167261
0167264

0262962
0263201
0202336
0182810
0261864
0072736
0201850
0212445
0262514
0279621
0240435
0260891
0263276
0262160
0103067
0284259
0261469
0103066
0260441
0103064
0252026
0103061
0279649
0132600
0259797
0222177
0103056
0279693
0287760
0274698
0199263
0241334
0184939
0201164
0260084
0275383
0252040
0252640
0062038
0260577
0260860
0261395
0039068
0200558
0260798
0260459
0103047
0199979
0275504
0260999

0266801
0202497
0271129
0244378
0103044
0168802
0286140
0141076
0207083
0168807
0200164
0265113
0260914
0132612
0213380
0258429
0272617
0260371
0259900
0132603
0157315
0132604
0103018
0261527
0260108
0102908
0223109
0274093
0260772
0181019
0261602
0262136
0141101
0260290
0198373
0252443
0208017
0157322
0277003
0196696
0226232
0255185
0239118
0241183
0090857
0247228
0157335
0261145
0261556
0223496

)090861
)265648
)157349
)260537
)261777
)168872
)285710
)157350
)260111
)240043
)200153
)157353
)103051
)189091
)221514
)221066
)157368
)132613
)279569
)103043
)279412
)260156
)214353
)279815
)279122
)279123
)262890
)157423
)221725
)259833
)259798
)180917
)261260
)172137
)261348
)262120
)260734
)262008
)167377
)261611
)247324
)157429
)260520
)140835
)260015
)252339
)260886
)198650
)260593
)140832

0040199
0260185
0199301
0263666
0261513
0260612
0166747
0223224
0283057
0262140
0224470
0182149
0279592
0102984
0261765
0277481
0242176
0263311
0263232
0102967
0140830
0257017
0261701
0260252
0140829
0118557
0261673
0261008
0261774
0283564
0260664
0259768
0238731
0261764
0279206
0140836
0259209
0251868
0279757
0280092
0272330
0279591
0271009
0260880
0284675
0261227
0279120
0258779
0260848
0283457

0259817
0284670
0261404
0279420
0279100
0260471
0261248
0275236
0103035
0259972
0239763
0260884
0214331
0261079
0196436
0276822
0140839
0261170
0090863
0251794
0207525
0240861
0270945
0168411
0262904
0168404
0103089
0261458
0103091
0186187
0247033
0166816
0261058
0184517
0213318
0168928
0168925
0240338
0050820
0280152
0252101
0153774
0259999
0261783
0252122
0261717
0274220
0166822
0261476
0183196

0203472
0273971
0262583
0260092
0135702
0205084
0259992
0276408
0034713
0276523
0065457
0065427
0287194
0166848
0284484
0282757
0240199
0261424
0283755
0261313
0199299
0260223
0260983
0244485
0286841
0152910
0287694
0252814
0252022
0273659
0250514
0259995
0266426
0260797
0279856
0103111
0205078
0261063
0226348
0260922
0140873
0260701
0261154
0278862
0140876
0261707
0285538
0171724
0278926
0261330

0261540
0166509
0260731
0261776
0186153
0276007
0279991
0260969
0287106
0260014
0260733
0279887
0261837
0279129
0286894
0243101
0280115
0280190
0260816
0279732
0261537
0280274
0260694
0260479
0260511
0277954
0285918
0261722
0222244
0261472
0178573
0277214
0278058
0275040
0279378
0260876
0261390
0261082
0260237
0260706
0259867
0261376
0168589
0260183
0260737
0166446
0260064
0260594
0260896
0286221

0103121
0278985
0166451
0260213
0284512
0261061
0166454
0166455
0260643
0261141
0261838
0245059
0140905
0166473
0252608
0286759
0135697
0280182
0260495
0261609
0284473
0272923
0153815
0279021
0260229
0279476
0277255
0275109
0279681
0279294
0279841
0261218
0197943
0260682
0276519
0184860
0086696
0261235
0260161
0135698
0261029
0251888
0259873
0261285
0140945
0277387
0280044
0260862
0286528
0263360

260832
261410
263785
261103
260788
259914
260228
230989
103150
260932
140961
103154
166558
199350
285792
140943
274677
260018
260410
103160
154099
103168
140955
250685
168418
260858
201361
103175
260530
260504
064270
261286
140950
261243
103187
261471
135686
103194
285848
103196
260859
279622
260340
153786
135709
153789
278214
275088
131149
287787

0270313
0179219
0266307
0287946
0278716
0270124
0260417
0263968
0131153
0154102
0275393
0287125
0283416
0269898
0131148
0252311
0270184
0270159
0131143
0270698
0214259
0286510
0279907
0140968
0274134
0285163
0268804
0285040
0269667
0285012
0279604
0261177
0268078
0268505
0269186
0268754
0199949
0268532
0168367
0269826
0268473
0268388
0103241
0270020
0243697
0103248
0280278
0260944
0176692
0176678

0279142
0260387
0261161
0274478
0261175
0260026
0287010
0287161
0260795
0232190
0261363
0260456
0285527
0261095
0261651
0261697
0243705
0280419
0131152
0270006
0103264
0270082
0269901
0140941
0261592
0140948
0276393
0276337
0260750
0269935
0154118
0277504
0226180
0104731
0260671
0260498
0260177
0103257
0278598
0260466
0174990
0270385
0260443
0172530
0261193
0260166
0205037
0260162
0261327
0260420

0261816
0261273
0225614
0179588
0263456
0261744
0158545
0124391
0051523
0167508
0260630
0185669
0158717
0259813
0174177
0278341
0103335
0284585
0260121
0224888
0260617
0182376
0167513
0198931
0141012
0167515
0205022
0129993
0259881
0261226
0205018
0259989
0256982
0176715
0261546
0180422
0205015
0129910
0259803
0259877
0170100
0260659
0167522
0268218
0260279
0261253
0261692
0287351
0252887
0270832

0197912
0261118
0167526
0200084
0178773
0015413
0131165
0167523
0275734
0185324
0260259
0158792
0075399
0261373
0158805
0187741
0204991
0141002
0267048
0258839
0198211
0259006
0258947
0140995
0177946
0223959
0003249
0141013
0221819
0222019
0126856
0261812
0260528
0260507
0261172
0222359
0260923
0277349
0280279
0280136
0277613
0268320
0272636
0273172
0181031
0262920
0262061
0262294
0262558
0241525

0187624
0183688
0263015
0141252
0263300
0262905
0262708
0244097
0167695
0179409
0231784
0167699
0262228
0171861
0262434
0277491
0167693
0262133
0262003
0263017
0177370
0159842
0264429
0278794
0275997
0235361
0262213
0205899
0184811
0108953
0280408
0167193
0262777
0197879
0132376
0236618
0174238
0167703
0243704
0074660
0167705
0174231
0277597
0185561
0186594
0283824
0167716
0262791
0167711
0132386

0186532
0132383
0108958
0185924
0262445
0228133
0262869
0108963
0287553
0262664
0262533
0263050
0267200
0267195
0177374
0070366
0280333
0262810
0279040
0279562
0279340
0262402
0265777
0236838
0236457
0225084
0278941
0280242
0167720
0262333
0167721
0275084
0212163
0141258
0274758
0263345
0070444
0262456
0127804
0279872
0280280
0243049
0262953
0007168
0262480
0279432
0239884
0272770
0272911
0132361

0273606
0262050
0277200
0261963
0221200
0272920
0132359
0262884
0262628
0184166
0142163
0183024
0261848
0262085
0172150
0172146
0255095
0262106
0221882
0180090
0180068
0285760
0267129
0180042
0180016
0159961
0127780
0280268
0141255
0108381
0167723
0196689
0263338
0262304
0261916
0197417
0262248
0040531
0262903
0213977
0257950
0240898
0127774
0083454
0083457
0262692
0177602
0262358
0074356
0004660

0108405
0074370
0263312
0074755
0252456
0167740
0185722
0263165
0251812
0263219
0274363
0132388
0261868
0263882
0182557
0262823
0229782
0183018
0132382
0167741
0262921
0262519
0188176
0261863
0222429
0161905
0141456
0235085
0244184
0213939
0141480
0161920
0261898
0161921
0141497
0142484
0182853
0280254
0182327
0142507
0262165
0129219
0141503
0230201
0264113
0108556
0205710
0185245
0108528
0108523

0108518
0108515
0091640
0108509
0262429
0274300
0278517
0234203
0262678
0262227
0196388
0129250
0227495
0132517
0262495
0262693
0180787
0167840
0234327
0129204
0262855
0180626
0261879
0161929
0263164
0029725
0108559
0263220
0129197
0263272
0108561
0005100
0262099
0072849
0167842
0091592
0286190
0282980
0286356
0262961
0284837
0285471
0179314
0262231
0206618
0262609
0285593
0129221
0262818
0129195

0091622
0274860
0198920
0200914
0282936
0129235
0108590
0256806
0261996
0264468
0283301
0274114
0141485
0262990
0244283
0132530
0279296
0177294
0167858
0262943
0215067
0262503
0263316
0267047
0262089
0108839
0219200
0161939
0258315
0267532
0284112
0284027
0161940
0174327
0174326
0262067
0132514
0161944
0213876
0141505
0132535
0279812
0072778
0199053
0004975
0040633
0170296
0262526
0279641
0175826

0262302
0170291
0181885
0202251
0181856
0263342
0006047
0132507
0132522
0261915
0215041
0224647
0072818
0213859
0263171
0182896
0174292
0187838
0262481
0205544
0169992
0262880
0181323
0184560
0283439
0286007
0181284
0262624
0161958
0263301
0272884
0170175
0174282
0259224
0181222
0201489
0239697
0248871
0276384
0161955
0161956
0277957
0161960
0264772
0209582
0238917
0277985
0129226
0233223
0129255

0129194
0129245
0129214
0141504
0129244
0141510
0141499
0108947
0183914
0240480
0132510
0167874
0183011
0182224
0280046
0170004
0252835
0179859
0170049
0262730
0170043
0170037
0262492
0132518
0279174
0179593
0263427
0275011
0179477
0214999
0264005
0179148
0179111
0179094
0284117
0263620
0220205
0179029
0200463
0266824
0283160
0196544
0279152
0271002
0178999
0178977
0178971
0178921
0269947
0125434

0108961
0271029
0198844
0226871
0212206
0266279
0184650
0184619
0265749
0263809
0161970
0189051
0166579
0133026
0252363
0270606
0244604
0161973
0183318
0185156
0276231
0141506
0266389
0265975
0065320
0262966
0273816
0170310
0263708
0225751
0262815
0263051
0271851
0166596
0241654
0265349
0154914
0262296
0282882
0184544
0278944
0214978
0283025
0065325
0265593
0109047
0007237
0242692
0273388
0006788

0264067
0272736
0272975
0133020
0264424
0109061
0125414
0214970
0109063
0261433
0133028
0263388
0170222
0187824
0264176
0263400
0285220
0264016
0240813
0266114
0284876
0263429
0279542
0233670
0263508
0207242
0244753
0243346
0285541
0188803
0264727
0007174
0263684
0266368
0154957
0241185
0065559
0252707
0266297
0263648
0265400
0286617
0179136
0287462
0141052
0227274
0265489
0006740
0265503
0280168

0277621
0263707
0006744
0273098
0234634
0153976
0221698
0236088
0265494
0233852
0266744
0141028
0006695
0252305
0223510
0262786
0125430
0266709
0266378
0230647
0230897
0226130
0205325
0238212
0226156
0232058
0259944
0287307
0265163
0109099
0265110
0279660
0265445
0286792
0230971
0237377
0125409
0200437
0279567
0266667
0239888
0239704
0259024
0234324
0175106
0223878
0266538
0241322
0251537
0252129

0223544
0221926
0266261
0108452
0187607
0255104
0233966
0214946
0233002
0267227
0233193
0227255
0235430
0251829
0227018
0179277
0265519
0276855
0233090
0237057
0227790
0170425
0214941
0011295
0275413
0141027
0227782
0243686
0233979
0252383
0199674
0239221
0108474
0221355
0166582
0170315
0265401
0265095
0264739
0187688
0239203
0175061
0277108
0277370
0266651
0277512
0181350
0286743
0141040
0266803

0235554
0197566
0264765
0264359
0231458
0170160
0266302
0252446
0276088
0226478
0188933
0266129
0272815
0264673
0288235
0264729
0264892
0226145
0227300
0131885
0205312
0229749
0283033
0265853
0128438
0240505
0230709
0230969
0260328
0287114
0287910
0133030
0225442
0263624
0206859
0243370
0278864
0179598
0264187
0154803
0266498
0226359
0141030
0205309
0215030
0227158
0230865
0201741
0141026
0108551

0286430
0133027
0265511
0206730
0264666
0279200
0223979
0273948
0274836
0108557
0237328
0226746
0264167
0072310
0274111
0207839
0175662
0197815
0171962
0171953
0230273
0232344
0280198
0141034
0108591
0279428
0091536
0266677
0091542
0131899
0177731
0177427
0260647
0177302
0228000
0237382
0176994
0176974
0283772
0214860
0264177
0267441
0275206
0204471
0220161
0205266
0266162
0186831
0214856
0229526

0227919
0267492
0230662
0171916
0167494
0273018
0233327
0227689
0154874
0227077
0276532
0189375
0262262
0287058
0174977
0249459
0240445
0224126
0108448
0265693
0171931
0171928
0265478
0250111
0141127
0274747
0264273
0235672
0212618
0154025
0196893
0188522
0263045
0154016
0279825
0265185
0262074
0230197
0265746
0263394
0228157
0262202
0277947
0281000
0189152
0262319
0266664
0197665
0262292
0266311

0261987
0263934
0264940
0236022
0263206
0072134
0281856
0235397
0108641
0221540
0286799
0166484
0166482
0287196
0128482
0265126
0235979
0227078
0283675
0189343
0231477
0142494
0228331
0266079
0262769
0228983
0263107
0266179
0271900
0227225
0242752
0228725
0224254
0231625
0072210
0264932
0275982
0252349
0180638
0262681
0264785
0108602
0230607
0083290
0231096
0279251
0276406
0270091
0108599
0266126

)261033
)128487
)225681
)225014
)212186
)263494
)252971
)154898
)276577
)266839
)278701
)214832
)189423
)237911
)287490
)287751
)230528
)170298
)229586
)263976
)230088
)214822
)266925
)231645
)205215
)266042
)260907
)265916
)235203
)267075
)230339
)263946
)235323
)264422
)214819
)231258
)188013
)232889
)226096
)235546
)227685
)266364
)236504
)264874
)226521
)187870
)218454
)213671
)226549
)264215

264662
226981
205212
266369
233098
252483
265790
265099
244456
264660
266009
263729
124422
263986
236819
109016
178307
235530
263815
274180
266563
034152
184185
212719
287671
265881
265043
264023
244582
265265
266466
266673
266050
264245
263375
260157
264617
274748
260458
283566
265233
265019
263563
263725
263609
264956
266691
178130
266806
276399

0261020
0266885
0283879
0266529
0256618
0243655
0264168
0263914
0240103
0241225
0263483
0264316
0284300
0284328
0284263
0275480
0264930
0264970
0264262
0264431
0263433
0264734
0264172
0266736
0264837
0265646
0285822
0265246
0239223
0213664
0265453
0263583
0109046
0266313
0265683
0266356
0266433
0141068
0265801
0263368
0266853
0266728
0266872
0168961
0265788
0265041
0266771
0264240
0264689
0263604

0265833
0266416
0266786
0007171
0266202
0266527
0232859
0260019
0286589
0251818
0266190
0202389
0087095
0252283
0240873
0287721
0267644
0237575
0267259
0266830
0265060
0263788
0264486
0260777
0266306
0264029
0265480
0109084
0109083
0109079
0004142
0244045
0258924
0284532
0265618
0274529
0273171
0004139
0109072
0277450
0279532
0076351
0265254
0265541
0263870
0258472
0263266
0007216
0109101
0109103

0087111
0109107
0265168
0076382
0227543
0264608
0265287
0167524
0007202
0264044
0132581
0240494
0109111
0265205
0167525
0109113
0198242
0238423
0264577
0238578
0238649
0238597
0160606
0160602
0267729
0265073
0076604
0265474
0265840
0173065
0243053
0238516
0264138
0132591
0264066
0284565
0283364
0283819
0284085
0132589
0264304
0266642
0167536
0109118
0265845
0265908
0179761
0063015
0263613
0238007

)263781
)196535
)221995
)263709
)264050
)178082
)108255
)108256
)239256
)266111
)264808
)222363
)160551
)284162
)253064
)202205
)264031
)168792
)167543
)264290
)108262
)198720
)264647
)167549
)264007
)141298
)263370
)222858
)265625
)240531
)263657
)265713
)252657
)240074
)263477
)239129
)176927
)264435
)207011
)267482
)265289
)266876
)265394
)126653
)283935
)266919
)284399
)265739
)266987
)108576

0264125
0266120
0252112
0108578
0222679
0182271
0207132
0201255
0108582
0108587
0264958
0263860
0279431
0266733
0266775
0259928
0231870
0231421
0214719
0248121
0273609
0264943
0250462
0241631
0264538
0279337
0265443
0263603
0280069
0176390
0265791
0266490
0276250
0242439
0176208
0265334
0212190
0263531
0172171
0275185
0184060
0230113
0266274
0265743
0286194
0181481
0264456
0264743
0266340
0264148

0266865
0265798
0264107
0265444
0196712
0266371
0126861
0185862
0265118
0126860
0263535
0131242
0239595
0264862
0278977
0284424
0207614
0278546
0221038
0228768
0265976
0283978
0266877
0266448
0202026
0265863
0263567
0265046
0172301
0108651
0263990
0178691
0253058
0264300
0185158
0264497
0266777
0278867
0264164
0266379
0263369
0265574
0277511
0214708
0126858
0265293
0280033
0280020
0266282
0287506

0141314
0263674
0265794
0108666
0265139
0263749
0283774
0010244
0274341
0278668
0266385
0279762
0264083
0264373
0108671
0202100
0176749
0176658
0266718
0280245
0265222
0263717
0279781
0266599
0265337
0264458
0201178
0226377
0236377
0006042
0141316
0108684
0264598
0280349
0266535
0265544
0265697
0265125
0265115
0264643
0265689
0283381
0283417
0279668
0265356
0263435
0279674
0226049
0263485
0265775

0252370
0264791
0263571
0261156
0264990
0108691
0108688
0172156
0108700
0265614
0181374
0108702
0225582
0197322
0181291
0278860
0264174
0264622
0132141
0285559
0198783
0005156
0092871
0267618
0273687
0267457
0243423
0185379
0253084
0252328
0073598
0073536
0141161
0267782
0266981
0267744
0164729
0266947
0166750
0267364
0267271
0172716
0267315
0267711
0267547
0267745
0267648
0267102
0172123
0154760

267312
205045
267554
286065
267349
267046
267369
267074
255987
286030
267035
236320
267359
267625
242660
267321
207297
108733
006125
267592
270647
270885
270765
271447
271392
270806
270829
270871
270894
270379
271268
270240
271503
271394
270977
271013
278023
275722
273736
275152
276409
275431
275688
275718
275944
278690
274736
275385
277089
277632

0275302
0276014
0276241
0282738
0274808
0274767
0276085
0276070
0278097
0276755
0278395
0274933
0261499
0265889
0260287
0274226
0277029
0275954
0274658
0273611
0274164
0278259
0277161
0278311
0274756
0278535
0278619
0275720
0276508
0275613
0277268
0273706
0276707
0275700
0277911
0277589
0276326
0277579
0278638
0278540
0274308
0275029
0276810
0278505
0276054
0276234
0273965
0275839
0276023
0275066

0278053
0277688
0274620
0277501
0274244
0275410
0275173
0276715
0277341
0273513
0277488
0274512
0274419
0278299
0274615
0275006
0278599
0274663
0274611
0274487
0274284
0278845
0277399
0274211
0275832
0276170
0277363
0274996
0273604
0275238
0277969
0275532
0275023
0275665
0273627
0279119
0277972
0276036
0277258
0277182
0277791
0274607
0276293
0273559
0274054
0275489
0125691
0252699
0199293
0244086

0002834
0263466
0277057
0265784
0263874
0204952
0265428
0266588
0277426
0266013
0214553
0266048
0263818
0161381
0286443
0141748
0280177
0266101
0067191
0108298
0141750
0265460
0108306
0266469
0125686
0167258
0222777
0273576
0171532
0214546
0131771
0131748
0173991
0141744
0161395
0141736
0265178
0141741
0141738
0161405
0264663
0226117
0186075
0073605
0172057
0264968
0204913
0167914
0108344
0265799

0108342
0008838
0283721
0238793
0126351
0126368
0279199
0188895
0108349
0284144
0108352
0274621
0171475
0094804
0131759
0287644
0265666
0278918
0183153
0266208
0264655
0252618
0278346
0131747
0222881
0240125
0141753
0279806
0131746
0266088
0126353
0279775
0073584
0264058
0278834
0213424
0224244
0167916
0229028
0204897
0186393
0171446
0173908
0264488
0186395
0167920
0265359
0187242
0171431
0234477

)108244
)196859
)204889
)212899
)212900
)226776
)212901
)221852
)204887
)221880
)188581
)212725
)214518
)212724
)213417
)237230
)240871
)287602
)204880
)241241
)212722
)212721
)213416
)198090
)198271
)171396
)196156
)244537
)198443
)251439
)240542
)248807
)239886
)204873
)187272
)241595
)198083
)212659
)229351
)180386
)237183
)212658
)212657
)186860
)233014
)108516
)006059
)131738
)131737
)094796

0234859
0225438
0108417
0171360
0236473
0108759
0223125
0197079
0126337
0171401
0229732
0171346
0283796
0171345
0226629
0171403
0186847
0186832
0128422
0214514
0278954
0173812
0184502
0173805
0201920
0239542
0173801
0141696
0141756
0141698
0161594
0259623
0178502
0131473
0274630
0204815
0173786
0168259
0168256
0187595
0267221
0108771
0108773
0267261
0260325
0108774
0267658
0089558
0161610
0167925

0173757
0280295
0267758
0236194
0278829
0126561
0168610
0177469
0238704
0033627
0264314
0267222
0267632
0283929
0267185
0108784
0266929
0108785
0108786
0266962
0068120
0108788
0131470
0287710
0141699
0280183
0131462
0267189
0267132
0037042
0068137
0184451
0267042
0108797
0267765
0108799
0275273
0239671
0273650
0214578
0197291
0131477
0131475
0126562
0183978
0176563
0126581
0278447
0131467
0131480

0131471
0260105
0213373
0252039
0131482
0212149
0108825
0266967
0267060
0198863
0131469
0068079
0108828
0108830
0012048
0240828
0198496
0285057
0267681
0267002
0267595
0267340
0188554
0184988
0236383
0274585
0275616
0273709
0274432
0278048
0276596
0277903
0278774
0274862
0274062
0274452
0275219
0278591
0188825
0279602
0277084
0252882
0175906
0251763
0267151
0284344
0252279
0252729
0237888
0067596

0175832
0278109
0005102
0267440
0267253
0167941
0108861
0231256
0267604
0161647
0161649
0108852
0282199
0267496
0267166
0267420
0261514
0108849
0131096
0161653
0091947
0221044
0161654
0141349
0108840
0267638
0212446
0125319
0161664
0267080
0168591
0087152
0267394
0108312
0275107
0260793
0267691
0004939
0285486
0239702
0267750
0221496
0252628
0108309
0013306
0267211
0030582
0161682
0267634
0005961

0186566
0240589
0283045
0180340
0271222
0267384
0266979
0180336
0180329
0267160
0279879
0161692
0264251
0240305
0073670
0267505
0267405
0182963
0131097
0108883
0264540
0167131
0214447
0131095
0186185
0278223
0267334
0131094
0267788
0172992
0136448
0200888
0161714
0277249
0181513
0276728
0186834
0224505
0168517
0267288
0267121
0184922
0233175
0233483
0267278
0184361
0006062
0285757
0199953
0267446

0159314
0267344
0225190
0236234
0284055
0214425
0278770
0266918
0131484
0267198
0267246
0266504
0264070
0285668
0280022
0263503
0204652
0204650
0263715
0265547
0265964
0120088
0264589
0185294
0186868
0279685
0262881
0274883
0256762
0120071
0262372
0214401
0238723
0262500
0262539
0261575
0228696
0176681
0265411
0260075
0266497
0238083
0185829
0265315
0232300
0073969
0213326
0108379
0158955
0262031

0261886
0210709
0108433
0262633
0264999
0179673
0263142
0262879
0277198
0004897
0239291
0262265
0261872
0198336
0259207
0259753
0238419
0276790
0263293
0178852
0253347
0228782
0141279
0206734
0263766
0108424
0264558
0198933
0073861
0006025
0159111
0141294
0141295
0189120
0264243
0266821
0264920
0167182
0234494
0266601
0108439
0263798
0264019
0167183
0108465
0264701
0005243
0207947
0263412
0082641

0266341
0278765
0108468
0206954
0002919
0141293
0221739
0263787
0207306
0235300
0266222
0200538
0264451
0206805
0120094
0173917
0230148
0120093
0233101
0239552
0182742
0257178
0284038
0120075
0108511
0260027
0120068
0170689
0242207
0210741
0272763
0244649
0244559
0242067
0241788
0159182
0229637
0273750
0277542
0263602
0159184
0242407
0170703
0244514
0136436
0248278
0212565
0248895
0159199
0270781

159202
159210
230532
200903
238804
159224
250838
159217
251461
255851
252636
251550
248883
167080
240995
167083
108798
173868
177369
273500
198740
248714
262039
250948
186244
167085
250186
207127
229944
249497
279036
064300
249906
277478
250310
182575
121067
121073
250751
121104
136504
235183
262837
176358
204584
248954
254039
249176
199492
108813

0064195
0248172
0246640
0005884
0276851
0005882
0275025
0250282
0167100
0108819
0236472
0253730
0261959
0108823
0188662
0108821
0249406
0263176
0249838
0167105
0251179
0015532
0279792
0108826
0154920
0108829
0206824
0253102
0167107
0262759
0136457
0275897
0136444
0249451
0136449
0250286
0049283
0006282
0250107
0006283
0250976
0251239
0108846
0154945
0262967
0201340
0108848
0278963
0167117
0278672

0285829
0248702
0173714
0261976
0243521
0141232
0229980
0262006
0247011
0251665
0008294
0249870
0239672
0011052
0243678
0011258
0279089
0264895
0011260
0249383
0249982
0249658
0267452
0187504
0154975
0279743
0263317
0226364
0227011
0262902
0285939
0263252
0262271
0262518
0262384
0233380
0141200
0263189
0261965
0288036
0263199
0263125
0251809
0141198
0166260
0279059
0275710
0166263
0108924
0263096

0262079
0108960
0261873
0236319
0262135
0200107
0166292
0141179
0262052
0153930
0261882
0279606
0243058
0183691
0214226
0283209
0153933
0121060
0212469
0262408
0262112
0284542
0274213
0121058
0262298
0262623
0121064
0263120
0261192
0263004
0121057
0263089
0285785
0286732
0153944
0279555
0279281
0266100
0280028
0263499
0242889
0238447
0166329
0265542
0222976
0181610
0180891
0264914
0278642
0136451

280233
266086
264112
136450
279069
285930
266290
264364
279207
141194
229590
255713
121053
278627
011143
167419
287337
286788
005381
005379
265148
265206
284353
264399
213246
285897
108375
176160
266648
108389
264672
108387
121101
212195
266826
199426
266561
206917
200997
108384
274198
175175
276601
251865
241462
275680
263938
108395
263558
224738

0182628
0252212
0211514
0207996
0068489
0266537
0265303
0265415
0167447
0153982
0207235
0271063
0175155
0278863
0263857
0266002
0265313
0200889
0108406
0273702
0141367
0141378
0062716
0267637
0284190
0201524
0108423
0270264
0108443
0241913
0267318
0189050
0267104
0267302
0266992
0238283
0287664
0275860
0274378
0264049
0068097
0267526
0267416
0261040
0280852
0267295
0267248
0277532
0286435
0267095

167434
170832
252577
241157
275562
271626
141371
213228
062725
270553
259349
267678
267772
266943
170836
200013
141376
239932
224011
252534
267083
254114
264322
267207
267449
227694
286997
273982
267131
266934
267280
121068
187013
121075
267137
253506
136492
108506
202361
108510
279133
242398
285879
200842
207123
266066
266365
266647
188755
263887

0242123
0172421
0239809
0251981
0239527
0087995
0146872
0239246
0264546
0277463
0274565
0011028
0207382
0279713
0199697
0265702
0173838
0287457
0265000
0264907
0283538
0207552
0265262
0270324
0272164
0170921
0272383
0264513
0264529
0233635
0263501
0271919
0226797
0263644
0271749
0265282
0008283
0265971
0228979
0159640
0264813
0224353
0173826
0136485
0200560
0136463
0270587
0198909
0263883
0266173

)136490
)279369
)125695
)108588
)198231
)108592
)087191
)108604
)240280
)213218
)136487
)259533
)136488
)259333
)204414
)259384
)285947
)007312
)007314
)263489
)264954
)224383
)108622
)178607
)286629
)266402
)199753
)281311
)277887
)136478
)256358
)261371
)239823
)265031
)271605
)256525
)256826
)108654
)284564
)284368
)258890
)108854
)271779
)270714
)215769
)266820
)265298
)263647
)265982
)214176

0278581
0284172
0252242
0272400
0176809
0264840
0272134
0272220
0265218
0266644
0271974
0264057
0266155
0280299
0272346
0265912
0272006
0232457
0214174
0120063
0266598
0108370
0265883
0263520
0168646
0266076
0154240
0271587
0270870
0266707
0091583
0252653
0284242
0154229
0244044
0264630
0252685
0251698
0229330
0207943
0075429
0251764
0075461
0264491
0265664
0108878
0198265
0264421
0213180
0285877

0266473
0200619
0207410
0264093
0197170
0154217
0201547
0244610
0263690
0207688
0264754
0279917
0279361
0288109
0130935
0200394
0266858
0213179
0171634
0266717
0279573
0279880
0265052
0186665
0182481
0266117
0265055
0237854
0267023
0278730
0274561
0277476
0267352
0267708
0278740
0267731
0267472
0274712
0265100
0196704
0141337
0108932
0267009
0070540
0207561
0108946
0108950
0267461
0267659
0283376

0267250
0141338
0231749
0154258
0154262
0267361
0265331
0283385
0154263
0278873
0285931
0154265
0267203
0252352
0108984
0278972
0267653
0227517
0267184
0237560
0199801
0230258
0267603
0153822
0267365
0123700
0267170
0267109
0267471
0271239
0271101
0260785
0238759
0225818
0228118
0286387
0283517
0228639
0256124
0234899
0226101
0125398
0227036
0264026
0263680
0243514
0263893
0133195
0200783
0252274

0264196
0266501
0265469
0264860
0180616
0166685
0277728
0133193
0141219
0265010
0187959
0179604
0069188
0264985
0264750
0265935
0263574
0177338
0234721
0265927
0266765
0288006
0285648
0266357
0172809
0246731
0141540
0264272
0171595
0266106
0196169
0204347
0257008
0200021
0170412
0167851
0178789
0167850
0236770
0284690
0204345
0182352
0261222
0186407
0264659
0172794
0186074
0266036
0284186
0109062

0109065
0109066
0161509
0161513
0172782
0182040
0183034
0182938
0167861
0263586
0109089
0167862
0252545
0180901
0167863
0239607
0270993
0232724
0170190
0206713
0125449
0125458
0279035
0189159
0265800
0263786
0252042
0188612
0125450
0125447
0125445
0125457
0263843
0125454
0177885
0228007
0265987
0264853
0265342
0264511
0223217
0202332
0177728
0284595
0177303
0182173
0073350
0266714
0108469
0204323

0259120
0161526
0264829
0264270
0132470
0108479
0132475
0283853
0132478
0266980
0092929
0132471
0132481
0267801
0141569
0267342
0204316
0267426
0188878
0161533
0267615
0261408
0257949
0250506
0167880
0167881
0186919
0182687
0182473
0279382
0275360
0129654
0267128
0141576
0267376
0228218
0185262
0129646
0161542
0251779
0207169
0267518
0176170
0175931
0242931
0129673
0129667
0161544
0214140
0267546

0267543
0272386
0163597
0274091
0199961
0278261
0284526
0070731
0267078
0267084
0286750
0070526
0199735
0261335
0182534
0212418
0277382
0070495
0274051
0181038
0267168
0161547
0092931
0283805
0200257
0267065
0287836
0267535
0167889
0267568
0267521
0280122
0276202
0234912
0129657
0275143
0284250
0222808
0262870
0263718
0265121
0184640
0266998
0287257
0264060
0200651
0279801
0267263
0267016
0267665

0267506
0286534
0267466
0279572
0285535
0267790
0204283
0238898
0078687
0279600
0267463
0238658
0204282
0141524
0167895
0187997
0267679
0108639
0167900
0183077
0267624
0089685
0204278
0283914
0267676
0279886
0204277
0267737
0184557
0266970
0243870
0087157
0200063
0187775
0267432
0267770
0243426
0267123
0108669
0279570
0252391
0055483
0287539
0267355
0035862
0267601
0178404
0279339
0267491
0108679

)171302
)265096
)173918
)167280
)167281
)266711
)263278
)266665
)267719
)182156
)173894
)141570
)262768
)262772
)141582
)262343
)262585
)262188
)167291
)275516
)262172
)261978
)141519
)265561
)171298
)141543
)279259
)275479
)141527
)262580
)181523
)181045
)173821
)262979
)276863
)280248
)263069
)173818
)264961
)260369
)279273
)171246
)262000
)271340
)213128
)141564
)232083
)262833
)287403
)262313

0262662
0261924
0240270
0176108
0263218
0280351
0262873
0262098
0226137
0175866
0181409
0207736
0211563
0283604
0221025
0263167
0225180
0141577
0262115
0167302
0260005
0224877
0157637
0276101
0279187
0185168
0185332
0287431
0263154
0262223
0274833
0262877
0266074
0279692
0266392
0266189
0263271
0278200
0184009
0229848
0186765
0185504
0182446
0207021
0182612
0185527
0204237
0185298
0214087
0185359

0277784
0275902
0262049
0262814
0262660
0183048
0215644
0225663
0182676
0185624
0262831
0278876
0141522
0262413
0263731
0263859
0183684
0141552
0183979
0185813
0187531
0197063
0264769
0265688
0183010
0263585
0185105
0235296
0185269
0264735
0169696
0169689
0169683
0169750
0169738
0279372
0264569
0169733
0169727
0169718
0169710
0176155
0265678
0266654
0264548
0141526
0284574
0141551
0275888
0280407

0287737
0265692
0260563
0173762
0141574
0182459
0278964
0181408
0260011
0181396
0264812
0201239
0169660
0279066
0178927
0287193
0265458
0141562
0279744
0266445
0266236
0141568
0265136
0262652
0262147
0261845
0141580
0279613
0141542
0266107
0263063
0141560
0263321
0167363
0262410
0141556
0263098
0141579
0262663
0175711
0262339
0176845
0274370
0261888
0279143
0262898
0262094
0232938
0262352
0263305

)262181
)173213
)282965
)262081
)274446
)263006
)277521
)101557
)079134
)263884
)158270
)264514
)265477
)264265
)177143
)222174
)079101
)176912
)266456
)263727
)176890
)132199
)265490
)176105
)266171
)264635
)264339
)206687
)273355
)266575
)264179
)265671
)265179
)141433
)259837
)272461
)263551
)263505
)266251
)132204
)199197
)263396
)265417
)266602
)266450
)264080
)263745
)264189
)286975
)101574

0238575
0266783
0264150
0080986
0235422
0202224
0200875
0266405
0101596
0266049
0207034
0132205
0101577
0272625
0263606
0265907
0266397
0264186
0277447
0252258
0101605
0252353
0265399
0272688
0101608
0264235
0118680
0266578
0259256
0240364
0265243
0243720
0177426
0263460
0266835
0170579
0241223
0266373
0177337
0262001
0266401
0266513
0263724
0275518
0207278
0263878
0250933
0261520
0264246
0265545

0266268
0264775
0265514
0215368
0231824
0266153
0285575
0265091
0264575
0263753
0265188
0198081
0264254
0266288
0082397
0265316
0264000
0261738
0272380
0206432
0264449
0266846
0154655
0265487
0200637
0264707
0283532
0279885
0199272
0235552
0266065
0265944
0265174
0243591
0261627
0279479
0266441
0088756
0287834
0263797
0265933
0264127
0251897
0101680
0265069
0264475
0263716
0206422
0212626
0286126

0173482
0266767
0264596
0199856
0286627
0286649
0273368
0266149
0265413
0263970
0266613
0242985
0229442
0225212
0206418
0266708
0263935
0265962
0168502
0252319
0272788
0287719
0286765
0242058
0178127
0265257
0263847
0266053
0101745
0200883
0207092
0273284
0264964
0128791
0287509
0273335
0017797
0266805
0251994
0154845
0266541
0263627
0212572
0285594
0264644
0168461
0266891
0266127
0242651
0285653

0252680
0265564
0168454
0272799
0101558
0223138
0265554
0287065
0264876
0266102
0286499
0260913
0287563
0154856
0134265
0265728
0266604
0264503
0264311
0264072
0266470
0265038
0260779
0154864
0264843
0264714
0274838
0263952
0263682
0267051
0273119
0267564
0267425
0267252
0267371
0236396
0267794
0257513
0267143
0267773
0283492
0260759
0141404
0267333
0286812
0267455
0255112
0267165
0272703
0154889

)273141
)267079
)267661
)141401
)266955
)267480
)266995
)267292
)212712
)267533
)256616
)267722
)267478
)181626
)200827
)267162
)267116
)279285
)267643
)267733
)267076
)267247
)267663
)176194
)280302
)267069
)199702
)176014
)141385
)251937
)141391
)267108
)267199
)134278
)267515
)128789
)101624
)267249
)266961
)260302
)201466
)175354
)267570
)267336
)085415
)266969
)101639
)267656
)267032
)267286

0267239
0168675
0267177
0279020
0286926
0279275
0267393
0267690
0266146
0267694
0267529
0267702
0267366
0267136
0272746
0267503
0263527
0266954
0177150
0279962
0242707
0101654
0176136
0185231
0243779
0279352
0270469
0175322
0267756
0267150
0267356
0267704
0267324
0175319
0186481
0212329
0266145
0283294
0274214
0265787
0277743
0265437
0187589
0265296
0286593
0266605
0265766
0266242
0183206
0200132

0265340
0266693
0264570
0273732
0276318
0180777
0200645
0265737
0265499
0265786
0264061
0263618
0266554
0264301
0263821
0279042
0266522
0278923
0263635
0286815
0264880
0287723
0286293
0265015
0265631
0266818
0215512
0222087
0067900
0244527
0251886
0263748
0265984
0141449
0265948
0265751
0221139
0141446
0167088
0158201
0264188
0265656
0221493
0101752
0252677
0200750
0242971
0199977
0265142
0283927

0284453
0243303
0264149
0252325
0264289
0222520
0265758
0263588
0221363
0266010
0141448
0264012
0201852
0266283
0264825
0273348
0212710
0270855
0239490
0206827
0265943
0266850
0101773
0273232
0265939
0284331
0242182
0223023
0134508
0134490
0266495
0101782
0199357
0283084
0141452
0141458
0199366
0154065
0267752
0053747
0267301
0265752
0168234
0264745
0265204
0264924
0206863
0279332
0154040
0265750

0141447
0199874
0273524
0206766
0212051
0264365
0273321
0154059
0199636
0265094
0134489
0264695
0287611
0266489
0266181
0265485
0266520
0266809
0266573
0264345
0229248
0265552
0198795
0285595
0279439
0264434
0242216
0264193
0141380
0243925
0154611
0188985
0266227
0141384
0263829
0266237
0264911
0207160
0263862
0134504
0277534
0275805
0276221
0263846
0260372
0263677
0265369
0275900
0263382
0266184

0171885
0154080
0266549
0265374
0264151
0264790
0244329
0263611
0265097
0263405
0286465
0286426
0170558
0227279
0274578
0265994
0266296
0271238
0251719
0266479
0265980
0264587
0287803
0286605
0283218
0266196
0279734
0265670
0251702
0134762
0134755
0265888
0134765
0263698
0134760
0266729
0207019
0175065
0207240
0134757
0266521
0046604
0264859
0118271
0118276
0259985
0200472
0265158
0263772
0141437

0153339
0263924
0276486
0276282
0222320
0263823
0265273
0266105
0280380
0101695
0263917
0274275
0214917
0265008
0134758
0263393
0141441
0199373
0141434
0265279
0263904
0222395
0264982
0215492
0285095
0260389
0287994
0263688
0197705
0228835
0243355
0263765
0166960
0263450
0267642
0141431
0101746
0199204
0280406
0267746
0278052
0134769
0267585
0268873
0278464
0166974
0280096
0274400
0186812
0186814

0279989
0274184
0268573
0257267
0274918
0172466
0186496
0239683
0275512
0267583
0267140
0153391
0279074
0141429
0283514
0266965
0267397
0267797
0207797
0265641
0267627
0278986
0141428
0141425
0141424
0134759
0260552
0075643
0274849
0267404
0267456
0134775
0286890
0134779
0150477
0267039
0267651
0101489
0267707
0267202
0279678
0279543
0285940
0266530
0243516
0286716
0268566
0222704
0253038
0267374

0283458
0175886
0283615
0286559
0283645
0266148
0212354
0267413
0283503
0279034
0261715
0267664
0286844
0286328
0267313
0267652
0225253
0078142
0276174
0267088
0287007
0267586
0253040
0286976
0152214
0287209
0132872
0278933
0267716
0202060
0285550
0267337
0267573
0277899
0267414
0152217
0280365
0265957
0279319
0267101
0132874
0267097
0266988
0285790
0267354
0287943
0267193
0266968
0141469
0267558

0197046
0152223
0152229
0222179
0267293
0252034
0152234
0152240
0251939
0152242
0141622
0279829
0285993
0167210
0101638
0240983
0270112
0274776
0078043
0267587
0279354
0167216
0183791
0275553
0278674
0274744
0206181
0280212
0266957
0267724
0167220
0267228
0267800
0134049
0215474
0201825
0287673
0267761
0266582
0261307
0279367
0267156
0267187
0175387
0267541
0279707
0269365
0279366
0184828
0278891

286583
206944
280197
200872
277907
266984
134030
278983
279629
266276
267762
279250
267764
101665
266905
280335
264269
141627
263849
265128
266696
263895
287518
264971
177576
215472
263916
265496
284239
265681
202093
206602
271982
265182
264464
101670
267444
267674
167315
266997
267322
251992
252139
167306
201668
200527
267196
267511
267591
283343

0240663
0172361
0277310
0141644
0154832
0267695
0274930
0251997
0154839
0239780
0141639
0287381
0267713
0134042
0267233
0239689
0082212
0200752
0141642
0267699
0141646
0267717
0176624
0207154
0227115
0266201
0266003
0265402
0265712
0215457
0286283
0285695
0266335
0187323
0280098
0273762
0263872
0260433
0263438
0242945
0264296
0134046
0207233
0277324
0101751
0174448
0166845
0264350
0283613
0264564

0267428
0288484
0178690
0041353
0267524
0267311
0267112
0166510
0267647
0267013
0252437
0196628
0267028
0264571
0267402
0267284
0242320
0267325
0267172
0287944
0260930
0267327
0206129
0287248
0267057
0267399
0201816
0267712
0267732
0091164
0091157
0206865
0267146
0212539
0267225
0286777
0258609
0267789
0228075
0267726
0202240
0177511
0277837
0119547
0066926
0278703
0134440
0267040
0267787
0081923

0202159
0231544
0267209
0235028
0049759
0267743
0267504
0267396
0267675
0267391
0284440
0207778
0276403
0198796
0274354
0267257
0267579
0252284
0239659
0275418
0267226
0172175
0267476
0266946
0267705
0267501
0201598
0267593
0199713
0074657
0251870
0224367
0267215
0166562
0134443
0134438
0166569
0074695
0267677
0183287
0267000
0267061
0267269
0267774
0273664
0224161
0267486
0267669
0141682
0267066

0267497
0239398
0256704
0267462
0267742
0252555
0267686
0242060
0202468
0267620
0267401
0285681
0166603
0267400
0266915
0267687
0267382
0267098
0267153
0267513
0101542
0206769
0243256
0267316
0267279
0267175
0267038
0260440
0176641
0197563
0267597
0134444
0266900
0267560
0141655
0199867
0267341
0241088
0267487
0266920
0141664
0278017
0279236
0243549
0081913
0206746
0171791
0267766
0283125
0119537

0267390
0119541
0267430
0206075
0269989
0214318
0166634
0197641
0206073
0206072
0057149
0166396
0197632
0242550
0221887
0283667
0166401
0267134
0237314
0179676
0266952
0275448
0286800
0263863
0264131
0264232
0265217
0081138
0263720
0263622
0264685
0071991
0238680
0263354
0264054
0265936
0260578
0171451
0263424
0265533
0264040
0263594
0265472
0264104
0265555
0265240
0264634
0264869
0287907
0264699

260747
265425
263355
264697
166479
150636
276092
263637
207072
264444
264472
273584
264236
264705
285748
278532
206052
264845
265643
279221
266840
150637
176225
287646
170677
266258
266304
266278
265698
214255
278971
286642
239748
263417
265781
265639
287693
265484
263733
260676
264388
265946
265352
260457
265995
271313
141668
264315
166342
265579

)252598
)263750
)263958
)263711
)261780
)265380
)241866
)263655
)278902
)287314
)141665
)075336
)260569
)243856
)166347
)286529
)206043
)235297
)259779
)187773
)287689
)133313
)276934
)150656
)264340
)264247
)215421
)278607
)273669
)283452
)278911
)285915
)180011
)179981
)276993
)264116
)206026
)266774
)287281
)286603
)287295
)287993
)261194
)266460
)275827
)264108
)263547
)266743
)264260
)264595

0266014
0265717
0264212
0266614
0101493
0263982
0280205
0265778
0275763
0266256
0286090
0266312
0276397
0265934
0265836
0265626
0264278
0252097
0278107
0130856
0244229
0275355
0265261
0266844
0197971
0279811
0287680
0275178
0279433
0278330
0265844
0166573
0264015
0199392
0264736
0264693
0271702
0286727
0252260
0265843
0264334
0199728
0266213
0265644
0265378
0201723
0266273
0266586
0265101
0287758

0263146
0256463
0166377
0279416
0267655
0279474
0286245
0267628
0267015
0279077
0131196
0279637
0286186
0275186
0279981
0279397
0286296
0267287
0267409
0178412
0274828
0060069
0266901
0266924
0267780
0178342
0278075
0122490
0287879
0226742
0141759
0267127
0101546
0261126
0266909
0101544
0267270
0178184
0278000
0267251
0282458
0282542
0282798
0282807
0283801
0282591
0282137
0267310
0176695
0267792

0266971
0282535
0281379
0282059
0282416
0282051
0282508
0282339
0222329
0282393
0141934
0105556
0267124
0105549
0286667
0267443
0183186
0129946
0252539
0181781
0099866
0266933
0141933
0099804
0197540
0267751
0172270
0099822
0267036
0099821
0261204
0070388
0070423
0267666
0070404
0185198
0213726
0099864
0099812
0272473
0267530
0011304
0284361
0129951
0263414
0172232
0196415
0197561
0197766
0175221

0207507
0198858
0116014
0116017
0267159
0275162
0065268
0266990
0116032
0182087
0274177
0207357
0064666
0279753
0064687
0180448
0099817
0167468
0064932
0118046
0267736
0099625
0267778
0099624
0167470
0099622
0267493
0228300
0099617
0240846
0267372
0160953
0279488
0267755
0115286
0280486
0248015
0130005
0071626
0115268
0268798
0115266
0267317
0119559
0115257
0115255
0185761
0267092
0185988
0181588

0239367
0071655
0279009
0267059
0127540
0071564
0252933
0275642
0267073
0205922
0130270
0079313
0267007
0284216
0267244
0267125
0129911
0267141
0261526
0267232
0129968
0227500
0213638
0133275
0180846
0133243
0267283
0279653
0275468
0267180
0099875
0172081
0099840
0065000
0104885
0267122
0104886
0221411
0284129
0104897
0104899
0284272
0167476
0104904
0273734
0188305
0220008
0288156
0286880
0130332

0005206
0178297
0099800
0176619
0283728
0267201
0099860
0252962
0176533
0105694
0263974
0267412
0277904
0267214
0267749
0276097
0176490
0267001
0286460
0261342
0141873
0104969
0172009
0172006
0186300
0267063
0287766
0172000
0267398
0266938
0171970
0253392
0175691
0279452
0104953
0065717
0267469
0104964
0088256
0267139
0267688
0266944
0060558
0267551
0125910
0125912
0161082
0267448
0141905
0280103

0279599
0095932
0284638
0129932
0105325
0252408
0161091
0183397
0267436
0267231
0279590
0266959
0064961
0179855
0006638
0226800
0105298
0186111
0267304
0105289
0011132
0267205
0267138
0183617
0173976
0267148
0007264
0105278
0266989
0167654
0266627
0077009
0167657
0283928
0167658
0206775
0105229
0178951
0268670
0126934
0278007
0060566
0077463
0089847
0105246
0280247
0105248
0105251
0167664
0105255

0178078
0269425
0008382
0269807
0267938
0269318
0141985
0267980
0167670
0267255
0167671
0266437
0267030
0280239
0267769
0267011
0167674
0167676
0214456
0267385
0171236
0167680
0267408
0185361
0074842
0268565
0142002
0205790
0287884
0176840
0207630
0268536
0141965
0269604
0127666
0267484
0105355
0205784
0276043
0263409
0127663
0105426
0267550
0239524
0274447
0105428
0223573
0233270
0130254
0160633

0130255
0174917
0167733
0196365
0174898
0212123
0267157
0141994
0171119
0156413
0171124
0267709
0130383
0267740
0174886
0266941
0267314
0187650
0267571
0105519
0031823
0266983
0087903
0267262
0267589
0267299
0130377
0267415
0284490
0130382
0267563
0280394
0167769
0269802
0125656
0125652
0125650
0125651
0269444
0274390
0273758
0214347
0088247
0284159
0181240
0125648
0130545
0205744
0275234
0104833

)268191
)268203
)125657
)268742
)269546
)199223
)125726
)268785
)125735
)125730
)276980
)125734
)283950
)125733
)269680
)125731
)141968
)174837
)280091
)269385
)276568
)268758
)268845
)269761
)283679
)196589
)237247
)205718
)230522
)182315
)130544
)171105
)263264
)268861
)267852
)104880
)104883
)198723
)267952
)198816
)269371
)090674
)268614
)032444
)076826
)273657
)076924
)229833
)268400
)174788

)268204
)076944
)269228
)104918
)183019
)269711
)181029
)104921
)182566
)090659
)242636
)104938
)268297
)269763
)142459
)183248
)268120
)269153
)285674
)199900
)171017
)214248
)076984
)268149
)260001
)260500
)104976
)178531
)269139
)104980
)286138
)066044
)269813
)267939
)131142
)142449
)269651
)090661
)271717
)167775
)167774
)267855
)233927
)186994
)268884
)167772
)269386
)265390
)185236
)099785

0268931
0279827
0099783
0133246
0133250
0142347
0268618
0142303
0267986
0268480
0279426
0181786
0181733
0269244
0268922
0167785
0269300
0170948
0269400
0181143
0267332
0267154
0170929
0170923
0161807
0275787
0170920
0130803
0188000
0267418
0267775
0187847
0174667
0188314
0237521
0180926
0225980
0226480
0196110
0267510
0188321
0270011
0270614
0188629
0283108
0174652
0287275
0198028
0267273
0287960

)260112
)130818
)278611
)197961
)171469
)267106
)171466
)239830
)224689
)277587
)267490
)267612
)196605
)267370
)267741
)266950
)127452
)200237
)267119
)198258
)267289
)127445
)105088
)080573
)267650
)080511
)167798
)266204
)130813
)267387
)130812
)243207
)130810
)209645
)238531
)244165
)130811
)130816
)267534
)264266
)105364
)266978
)267105
)090339
)267607
)105371
)105376
)220201
)167807
)267673

0267303
0161847
0274425
0076662
0105397
0105401
0284268
0065989
0079999
0267197
0180739
0130734
0238364
0221410
0129347
0129355
0129354
0271010
0129353
0267100
0129351
0213339
0079805
0207972
0265879
0207752
0275640
0099203
0214212
0267544
0142453
0130733
0142444
0127616
0266936
0276757
0278643
0130164
0284553
0161888
0197256
0130158
0267082
0279807
0273733
0130173
0130167
0267174
0105514
0267576

0105518
0183401
0105520
0267277
0173928
0187266
0205517
0202357
0198003
0130175
0196361
0267477
0161914
0274713
0130159
0244080
0213304
0213303
0130176
0130165
0198551
0102575
0249840
0197332
0267500
0214223
0267343
0197933
0266922
0220949
0197044
0267646
0177599
0171295
0267089
0171291
0278897
0198429
0196757
0267179
0267274
0197054
0219665
0286098
0252060
0197647
0257355
0212497
0257446
0223547

252546
286132
199444
214189
132010
237273
212576
213297
242615
257591
196646
234773
197857
213293
234750
188868
198342
231361
268744
248406
196466
268870
180855
269755
234848
242852
196826
236483
232579
230321
234003
235686
249709
269693
269560
230310
213290
188033
173875
178464
269242
104774
269590
123154
285589
105583
285583
095059
268945
283980

0132004
0267791
0105576
0209702
0267004
0123144
0198356
0039987
0095066
0267212
0263800
0171223
0167815
0267062
0284491
0104889
0132026
0267424
0105613
0276181
0105612
0267735
0105610
0105607
0240616
0161860
0266721
0179115
0266975
0179218
0274417
0267458
0179262
0179271
0179284
0008441
0267417
0267610
0279044
0104903
0104907
0160877
0266897
0267512
0104915
0160888
0287296
0267598
0213253
0141837

0104957
0037757
0267633
0104979
0132003
0267582
0266770
0267519
0284387
0207808
0207980
0187556
0207613
0207585
0132016
0132024
0132000
0132017
0132005
0275091
0171136
0267670
0104998
0187867
0228887
0141854
0284081
0188032
0141858
0072062
0267783
0105011
0267169
0072071
0240803
0287254
0267723
0267406
0266913
0267147
0123146
0267379
0123136
0123143
0267474
0160951
0123159
0224543
0132002
0099797

0283822
0099795
0187912
0243488
0243066
0131355
0269378
0275432
0160961
0127507
0127530
0256210
0188269
0127515
0269552
0172148
0185385
0269678
0268834
0269787
0198367
0176923
0127529
0105143
0160994
0105141
0094661
0105137
0105135
0268564
0267961
0252661
0074181
0275711
0269635
0105131
0141867
0270802
0273218
0105127
0268189
0011243
0279203
0279716
0011451
0269782
0105122
0161031
0171954
0269765

0269516
0267135
0225067
0266951
0186526
0186529
0267241
0186204
0171942
0171936
0267594
0268673
0172519
0186723
0267235
0267308
0214049
0273782
0267453
0273330
0186115
0267218
0279717
0267056
0171903
0176231
0267392
0205396
0280412
0167459
0267609
0277766
0267450
0267517
0167460
0279198
0267373
0167461
0269243
0196684
0141977
0105058
0267033
0275901
0287961
0072958
0267275
0127528
0277825
0127527

0280332
0240418
0267703
0269058
0141979
0105072
0085872
0240106
0269399
0127526
0268309
0269085
0279529
0105085
0268790
0269578
0279748
0279977
0268087
0269427
0214046
0269044
0267904
0072954
0269419
0188039
0269799
0127511
0268743
0127533
0160111
0264250
0263595
0268985
0131351
0268289
0099331
0269066
0268056
0269836
0269480
0053501
0099330
0160113
0269095
0130307
0105393
0269307
0160117
0127220

)130312
)269815
)130311
)074855
)130299
)130300
)269350
)269720
)130303
)282851
)141971
)269481
)254503
)269053
)188051
)269035
)269736
)171773
)130304
)269439
)130313
)269161
)167483
)130309
)243771
)130477
)269752
)279300
)130479
)268112
)130475
)179913
)248099
)105639
)105640
)207166
)279617
)105641
)007080
)105642
)252591
)105643
)279172
)273654
)268623
)243455
)096996
)099308
)269145
)268173

0105647
0216490
0254858
0105649
0268650
0285188
0105650
0284797
0130518
0130522
0267959
0241464
0130520
0239821
0130517
0130513
0264175
0175489
0130511
0105655
0268199
0105656
0279262
0280121
0268938
0105701
0269191
0105700
0268030
0268983
0221983
0006016
0006015
0105696
0263490
0167487
0105662
0105664
0005007
0223802
0130283
0269694
0105669
0268193
0105671
0051128
0269019
0241172
0064607
0105676

0181035
0064545
0213999
0064489
0254901
0064490
0184162
0130287
0252973
0267629
0187664
0280282
0213996
0105705
0129933
0167491
0267234
0207821
0178093
0186010
0258674
0250067
0160161
0105717
0267178
0259242
0064547
0089639
0105726
0181896
0271283
0270402
0105708
0275540
0286080
0271499
0266904
0287533
0267581
0267693
0267419
0081665
0267481
0270325
0256771
0270270
0267565
0184635
0273791
0274800

0197124
0268070
0274033
0269154
0213988
0240673
0274612
0267383
0224864
0267220
0270716
0256229
0267641
0267771
0279039
0280079
0271524
0268620
0213985
0269055
0269274
0271638
0276294
0231205
0271704
0268461
0238517
0268174
0269043
0268335
0237440
0269110
0268272
0268560
0188171
0280171
0269779
0268416
0269885
0160229
0269040
0269373
0105750
0279404
0269012
0268326
0118620
0271598
0268598
0160352

201035
268357
196705
240522
268995
268892
269839
242291
268747
268521
182141
268322
172687
268731
213976
196268
269237
268119
268658
197013
268705
268081
268555
268240
274503
271182
268278
268433
268535
268117
197020
269845
268184
269136
198521
268438
160321
269615
279377
271095
269504
197134
269742
269505
268116
268140
269575
269345
196109
268079

282732
267872
268597
196350
270191
268685
236312
197360
269796
268830
270947
268717
269138
270317
286157
260599
269364
207397
271149
269332
240713
268823
269025
269316
268038
268589
229676
268981
198153
241282
270475
268789
213973
279966
270518
269509
268696
271655
268541
268296
269458
269067
271661
261558
261615
269543
267886
183850
270479
268105

0267920
0269471
0196081
0283201
0268469
0269837
0269107
0284428
0268794
0268148
0167232
0267934
0269431
0268992
0269416
0268908
0275201
0268100
0197372
0268455
0240231
0269320
0196172
0233836
0269662
0267924
0225178
0277493
0287399
0213967
0269289
0269397
0269421
0201966
0268362
0213096
0279425
0268058
0276030
0268442
0267696
0279882
0261824
0261770
0267575
0281468
0267264
0267323
0266977
0267389

0267623
0267768
0267630
0266906
0267509
0266928
0267320
0267238
0260725
0283403
0267243
0266893
0267423
0267537
0266976
0283269
0267799
0267617
0267339
0267683
0267027
0267240
0252272
0267014
0288017
0169021
0267498
0276251
0276631
0241604
0287272
0264515
0267574
0266248
0279387
0187135
0105171
0166289
0131943
0286867
0105173
0286538
0267433
0278895
0267258
0105176
0242100
0267006
0267692
0266910

0287859
0288072
0267223
0198597
0280061
0267760
0267081
0261316
0267378
0284430
0121297
0267777
0267662
0278875
0267636
0267599
0267407
0261400
0267465
0252780
0201894
0267489
0267779
0267204
0267133
0286041
0200738
0168813
0267213
0178904
0105185
0105186
0201388
0242325
0242436
0267557
0186326
0267475
0213965
0267567
0173809
0267245
0021488
0267555
0201967
0121289
0239988
0131944
0131941
0267611

0267725
0076650
0166359
0130881
0267054
0273420
0130876
0267714
0270212
0267580
0245848
0267727
0267296
0267130
0243864
0266899
0153879
0124299
0266985
0232429
0124302
0153885
0277013
0240626
0287526
0239253
0249987
0186008
0270849
0257103
0166398
0257074
0267219
0105220
0266953
0126249
0264317
0267024
0126261
0142279
0239210
0248830
0268751
0205209
0271049
0252230
0268683
0279971
0269811
0279329

0089335
0274104
0197841
0153896
0270281
0267767
0261754
0271666
0269086
0271109
0270624
0271171
0269037
0270876
0168661
0280106
0180884
0089351
0269303
0271032
0105711
0105707
0227392
0179066
0285526
0089356
0284065
0153902
0279619
0266964
0221857
0221946
0089327
0177558
0262497
0105699
0271366
0268947
0105698
0105697
0105695
0012124
0269553
0284034
0126266
0185897
0126251
0268222
0205786
0283479

0264400
0233214
0232680
0126262
0188508
0161249
0189001
0105679
0236144
0105677
0105675
0283907
0283758
0224910
0267626
0249115
0126254
0105672
0126267
0105668
0226510
0267190
0011590
0272333
0126246
0267120
0161265
0205155
0188223
0267439
0267796
0267328
0004776
0167595
0267049
0004777
0225872
0250799
0161270
0126259
0105290
0243968
0167604
0280194
0126264
0011600
0126243
0267786
0267113
0267388

0205138
0181392
0248101
0239382
0267698
0105270
0161277
0075702
0280023
0105261
0105258
0105254
0279504
0126247
0270760
0161281
0267005
0196357
0200179
0167635
0267053
0232677
0142065
0266973
0267363
0181007
0267142
0186017
0267309
0267091
0254004
0228629
0186020
0233527
0161298
0197808
0276071
0225975
0189042
0267041
0267260
0267353
0267254
0197863
0251247
0206669
0185869
0243297
0198453
0197050

275547
267345
267672
267360
196967
267437
245680
188283
267605
226686
286449
181666
276846
267682
189164
279936
196437
171827
266916
267115
188227
180458
267470
171817
180479
120784
196381
267552
267319
198182
267152
287496
189144
267422
267640
225868
171804
105738
268051
268764
269653
268499
275132
011332
167642
167641
267748
286037
200209
167645

0167644
0099337
0099338
0266963
0099341
0179168
0267090
0188766
0130244
0171777
0196218
0269445
0268055
0104814
0267291
0178982
0130402
0267375
0182472
0267892
0205076
0178934
0207296
0171747
0104823
0268083
0268756
0104824
0269688
0187994
0269050
0068903
0104825
0262484
0104835
0269547
0128626
0269486
0269190
0287271
0161243
0267992
0183760
0130669
0269172
0188505
0179751
0268510
0197110
0272395

213926
272311
183709
182393
128011
268262
130755
269246
179134
241983
006712
063322
128016
266559
090924
105193
196235
105197
090932
186838
176401
269792
269584
176396
269588
269188
244253
243581
239320
105198
269519
268153
268243
249861
226025
269460
268088
240320
006659
278813
105205
213921
276445
105204
277759
105202
275395
187534
013275
187187

0269069
0128000
0269749
0197782
0269296
0230086
0130758
0174521
0105219
0105221
0205041
0207631
0206674
0160392
0207281
0206925
0105223
0279759
0275652
0160396
0105227
0197019
0277453
0167565
0268366
0090013
0277510
0160460
0160410
0090006
0105245
0123815
0223284
0086544
0188493
0077312
0268975
0261857
0167578
0171570
0282951
0269858
0268797
0233622
0237118
0269843
0279108
0255974
0198077
0130612

)198251
)256612
)269353
)197408
)213908
)268101
)271299
)197838
)197446
)268529
)224476
)264423
)167600
)167601
)269652
)105323
)239868
)105329
)286177
)142039
)255730
)142046
)123810
)277744
)077348
)248098
)268987
)268475
)177191
)105341
)277136
)204978
)267107
)268034
)269800
)268355
)270164
)268027
)007129
)280014
)239736
)269266
)105352
)268997
)269354
)268582
)007306
)105388
)267881
)086548

0268833
0170956
0269119
0278239
0273111
0142025
0105372
0276926
0105369
0076928
0268041
0105404
0285505
0105409
0105737
0105732
0028277
0254887
0266226
0259436
0160570
0167625
0105723
0268643
0279539
0105722
0079432
0079462
0188368
0167619
0105429
0277533
0105427
0213904
0079435
0268605
0189377
0253048
0269374
0079385
0230558
0124469
0233681
0231561
0243167
0221826
0124467
0238092
0225877
0248257

0231924
0170848
0221878
0227349
0282943
0243130
0236932
0242221
0230529
0227606
0204941
0243137
0241104
0183668
0236123
0230681
0231412
0204936
0272396
0204933
0131126
0124466
0176531
0105755
0176472
0073050
0269177
0268361
0269583
0234465
0167378
0124444
0226763
0131116
0105767
0011422
0239948
0283525
0124449
0105771
0104783
0159871
0268601
0167637
0176222
0278917
0267058
0286613
0267191
0124459

0159905
0204920
0277806
0278492
0266921
0159882
0267144
0159885
0267022
0178386
0186026
0253021
0267680
0186019
0256294
0263002
0287035
0167380
0131115
0159917
0279103
0159915
0270679
0267590
0062370
0267173
0267508
0267188
0278318
0176761
0167384
0273777
0267614
0230666
0224366
0216588
0266903
0073008
0283673
0186567
0279095
0213892
0252200
0069399
0284258
0253027
0236807
0142273
0285128
0187244

0130202
0267282
0130204
0130203
0280087
0130208
0214855
0267467
0224916
0234906
0267114
0104853
0104856
0104859
0207003
0170684
0267348
0142252
0007047
0104866
0267346
0267044
0179846
0007255
0189114
0283632
0104879
0242675
0104892
0104884
0104881
0117877
0012061
0275726
0125740
0125744
0213889
0125753
0125741
0177464
0125746
0284544
0267757
0241226
0010310
0207773
0283212
0125743
0011478
0177051

0279407
0237452
0259605
0177045
0267395
0104936
0268434
0185800
0104941
0125755
0269148
0170608
0170604
0176182
0188425
0104967
0104983
0211580
0008438
0268401
0204869
0267922
0268053
0188624
0259339
0204866
0269729
0268621
0268879
0268460
0269637
0269745
0188293
0277748
0268238
0124440
0199796
0269124
0011485
0268810
0269151
0169515
0277531
0182013
0230510
0204851
0204850
0278797
0268423
0249210

0160014
0269292
0160013
0167414
0279161
0197380
0245598
0105287
0243560
0211513
0090372
0269487
0222614
0181027
0105281
0275719
0275862
0269576
0269014
0042753
0160007
0279948
0130751
0130748
0130749
0142230
0105327
0265134
0284411
0286669
0105321
0257704
0197405
0134830
0134815
0105419
0118160
0287896
0118162
0268061
0105402
0279861
0118156
0276588
0268746
0277383
0063169
0269806
0278999
0024422

0105373
0221803
0269656
0178980
0213035
0271150
0178928
0105392
0259009
0259108
0105398
0188334
0169393
0105507
0105499
0269420
0105486
0269534
0269321
0185453
0105483
0280353
0268583
0268001
0268186
0178150
0105479
0142227
0161558
0105467
0105438
0105464
0105447
0182324
0268530
0268465
0105443
0142235
0287603
0088002
0105523
0177202
0063177
0268093
0063176
0105516
0063180
0232871
0142233
0176920

0176909
0105538
0182264
0174951
0105550
0206758
0105552
0286024
0087076
0105559
0087074
0104804
0104805
0235191
0260366
0104808
0087088
0267898
0087086
0104812
0183207
0273898
0104826
0268655
0104827
0267335
0267019
0267099
0104818
0267631
0267685
0268669
0189052
0213030
0269706
0196337
0283663
0283251
0268108
0225950
0268287
0104848
0265675
0104852
0104863
0221916
0177380
0130528
0130529
0269009

235555
063127
265407
197813
104894
074219
268157
104901
268686
161609
142538
261949
104888
104872
161618
269469
090554
273189
142541
201675
199631
202503
200259
142534
200530
207782
268681
104870
142552
142546
207073
126460
126464
268636
126458
126461
126456
126453
126457
274306
224420
268677
169169
243829
126467
206599
196961
276270
010361
269194

0104973
0284114
0268006
0104960
0268047
0265954
0269352
0039650
0204673
0104946
0284394
0104951
0213024
0269179
0169136
0283842
0161640
0221125
0161643
0105053
0276729
0142528
0269091
0204666
0269540
0267890
0287001
0161652
0105357
0276002
0267815
0131398
0131408
0131401
0131400
0269392
0062822
0142539
0269404
0086967
0142530
0161671
0268854
0268518
0161677
0204653
0131409
0268231
0213023
0235034

0161681
0105472
0269565
0142511
0261341
0142513
0268375
0167747
0221381
0221241
0220988
0284711
0180279
0167748
0174562
0267968
0142515
0167751
0197588
0167749
0228674
0167754
0269495
0167755
0279676
0267879
0169035
0129455
0269741
0213022
0129451
0268739
0167757
0186474
0268906
0268390
0167759
0129437
0142544
0268581
0129450
0267990
0168995
0267895
0269072
0269681
0171101
0268336
0268652
0269253

0278617
0268849
0105383
0179213
0268595
0268318
0269021
0142549
0186806
0267905
0267984
0105379
0269403
0268520
0160318
0105374
0105370
0262874
0268889
0268144
0142512
0254760
0255441
0268711
0269181
0105366
0268957
0213822
0254521
0268839
0269580
0105492
0268777
0269130
0268267
0105497
0269388
0267808
0105501
0268500
0244071
0273837
0254415
0243469
0269959
0182310
0207550
0198972
0208008
0105509

0171051
0171049
0286236
0187474
0161551
0268316
0269483
0283236
0268095
0198093
0283285
0176024
0269235
0256683
0197619
0142556
0256087
0275055
0260160
0197608
0270248
0204611
0267827
0268458
0243680
0196267
0267927
0105568
0274380
0268015
0277562
0207265
0196214
0208002
0269102
0198464
0269535
0167554
0221923
0269834
0277977
0167555
0269776
0198633
0258405
0288253
0278543
0269349
0198482
0167562

0242255
0268886
0123870
0167766
0268970
0269825
0213020
0189190
0198538
0255875
0204604
0268225
0182986
0283695
0213793
0269646
0213801
0221874
0180257
0269082
0242779
0268282
0267943
0269526
0268964
0170949
0170954
0197937
0197497
0268842
0269001
0267809
0197928
0269288
0196131
0228567
0269758
0189348
0163098
0269118
0174677
0213799
0203326
0241069
0196417
0239912
0249435
0286261
0241015
0287608

0160336
0198346
0250731
0241434
0213777
0274892
0130844
0268864
0269608
0242668
0204595
0200393
0270347
0207645
0207644
0221017
0207869
0207599
0207616
0207810
0283540
0207615
0207788
0221421
0207594
0207580
0207825
0207711
0283455
0207706
0207722
0207862
0207629
0207738
0283490
0283289
0207734
0207981
0207549
0251843
0207735
0207837
0207799
0207925
0211532
0207987
0207803
0252734
0283330
0207946

0207699
0207838
0207861
0252025
0207634
0252191
0283685
0207992
0207979
0207767
0221548
0252063
0207620
0269842
0284362
0252979
0237589
0229000
0238150
0269877
0199031
0284568
0199095
0199143
0269564
0142405
0228323
0179820
0232220
0126583
0105605
0142408
0284583
0130433
0276206
0189068
0248385
0170909
0170906
0105619
0105618
0237017
0088038
0105617
0167608
0125505
0170892
0223660
0170889
0224579

0204577
0269524
0235081
0244482
0249189
0105609
0212314
0240296
0131042
0264703
0240197
0276758
0251431
0234436
0187116
0275210
0249654
0250507
0239961
0167613
0237955
0167614
0227407
0268496
0269873
0267838
0226696
0235681
0167615
0275183
0167617
0167618
0213016
0104970
0235181
0239998
0269271
0104974
0238094
0104972
0224730
0186818
0277636
0242856
0237323
0225370
0186152
0228790
0170858
0242019

0215765
0243772
0242473
0125498
0264257
0189013
0167633
0221957
0240403
0199308
0186431
0268734
0189430
0167634
0275012
0022556
0243494
0267265
0088053
0267149
0160439
0131037
0125503
0287170
0267507
0275863
0105048
0129991
0267110
0167646
0267577
0239137
0129990
0080031
0286230
0267649
0180089
0276570
0105063
0275519
0278328
0278264
0133265
0160469
0268729
0180061
0276831
0133247
0160471
0267706

0180043
0095752
0160472
0269275
0233493
0108107
0284029
0108106
0187902
0063241
0283567
0279063
0269859
0197483
0090971
0179954
0187550
0231274
0218891
0179943
0171443
0261221
0276488
0179922
0213015
0171425
0173581
0063244
0267523
0267096
0063245
0267117
0267689
0268593
0185792
0222524
0223638
0229292
0255238
0179873
0160505
0173572
0179709
0171487
0267522
0142409
0223060
0279959
0167685
0279794

0197487
0197213
0204532
0242411
0131848
0267908
0267710
0267298
0267549
0200646
0213014
0266907
0267145
0266932
0267429
0267606
0240225
0018869
0267776
0267459
0285996
0267781
0267192
0267454
0198440
0198046
0166770
0279405
0276578
0196263
0267224
0267421
0196867
0269696
0197016
0268847
0197951
0269793
0268182
0127903
0286125
0268067
0269699
0268407
0198300
0268654
0274777
0269786
0268483
0274219

0131864
0141946
0270293
0270790
0258873
0268379
0083844
0105146
0204524
0268713
0267871
0197714
0279541
0268205
0268723
0178229
0268678
0131845
0152433
0256060
0268133
0188785
0269533
0268533
0186272
0186230
0268163
0276449
0178201
0268545
0197128
0268266
0105136
0268107
0152439
0121406
0251369
0083817
0171649
0183647
0213762
0268392
0121417
0269097
0180532
0204519
0269026
0179909
0083814
0152443

0083828
0178935
0268750
0269343
0269304
0204514
0198466
0269867
0270804
0269476
0173480
0196724
0152454
0243234
0177025
0166704
0176593
0235974
0286949
0152467
0268660
0270837
0176293
0243642
0121413
0181894
0268293
0286632
0269236
0171606
0213013
0268866
0198131
0283515
0268516
0268201
0278129
0267216
0142396
0283103
0269794
0182318
0277588
0121410
0268230
0268895
0279611
0174586
0268049
0269054

252211
152475
083845
266640
269855
268307
171574
232098
268543
268912
131849
269473
249471
083812
083838
269106
083807
252334
265272
273901
119574
264910
130726
275924
130724
130725
267858
099326
213753
286104
269032
269600
178591
125788
088782
185982
125903
186458
272874
196476
247315
286158
225377
177732
125841
101255
125826
125875
206797
101266

0125878
0271303
0270299
0215397
0101276
0125898
0101278
0101280
0101282
0286787
0125818
0215388
0125895
0274269
0215595
0244588
0101298
0125775
0274322
0280162
0234684
0229728
0088832
0284436
0088833
0196209
0242324
0252103
0125900
0260861
0101307
0271523
0089012
0237914
0223750
0242348
0236550
0234282
0286288
0271461
0232528
0276649
0198053
0233896
0101327
0227663
0125834
0286108
0287213
0287561

0226644
0286022
0125780
0166948
0207305
0125835
0256566
0088876
0149488
0101361
0284481
0221116
0271798
0212498
0229686
0226572
0101365
0228293
0088881
0226217
0088882
0241690
0258713
0198326
0132635
0215305
0132670
0125787
0125901
0101405
0101200
0263905
0235958
0185019
0215251
0088899
0198171
0125877
0088836
0277287
0088854
0201294
0226632
0201346
0225116
0088812
0230568
0125861
0149451
0088827

0132622
0101220
0101222
0125817
0101224
0275491
0274949
0125843
0088888
0229539
0125779
0199024
0283320
0275582
0101236
0226608
0215063
0205300
0088826
0230176
0171873
0236387
0238282
0228210
0226948
0171867
0171864
0180259
0231071
0278595
0101265
0089057
0252096
0234573
0277087
0276768
0278816
0277425
0089063
0212536
0252367
0132646
0212517
0101290
0225022
0101292
0266908
0200313
0276688
0226995

0230563
0205181
0225640
0225831
0235820
0125772
0233578
0171984
0202380
0089199
0215589
0089195
0125885
0286235
0207380
0231961
0278719
0275632
0263755
0088766
0125872
0278192
0101311
0233402
0234241
0271565
0276915
0229876
0228482
0125845
0232271
0228888
0238102
0261249
0224711
0278159
0240584
0215586
0277315
0101323
0125827
0229766
0182621
0226262
0225479
0201348
0228438
0276385
0275222
0101333

0225988
0125869
0101349
0286740
0232738
0286470
0243961
0132623
0227906
0276525
0230506
0232448
0132639
0286936
0237005
0214835
0230750
0125863
0285508
0285723
0149346
0235036
0101384
0273745
0270792
0224961
0234900
0232900
0217809
0125899
0270777
0230990
0235544
0282478
0287754
0271397
0237256
0261411
0228422
0222281
0271298
0132640
0236526
0235292
0230437
0235698
0233048
0269931
0286087
0234265

0172296
0271426
0101230
0226263
0225956
0278153
0089123
0236811
0089048
0101247
0101251
0212247
0172264
0230935
0234187
0239923
0125848
0199570
0287062
0227563
0227927
0213950
0235914
0237832
0199803
0201180
0222500
0225181
0224274
0286546
0287410
0225622
0089177
0273998
0275728
0226229
0200494
0235208
0228809
0125870
0125879
0236985
0212165
0251955
0220540
0238833
0286571
0287241
0125851
0232241

0125864
0232333
0277373
0125868
0202260
0230495
0125844
0125888
0233625
0212555
0089006
0125850
0212232
0125871
0218902
0229262
0227343
0252597
0207282
0232838
0149474
0232943
0266720
0252635
0287954
0125846
0230010
0199719
0237259
0237282
0089091
0236474
0252422
0132664
0265137
0225419
0089050
0214612
0101310
0232388
0284776
0125821
0243702
0221806
0232136
0233993
0233324
0179935
0273148
0149443

0132631
0274863
0287430
0185052
0235996
0179447
0232675
0268628
0132669
0236992
0173418
0101343
0089101
0241054
0225417
0229500
0243810
0264879
0280229
0173404
0188559
0213384
0270793
0278525
0225803
0234104
0225127
0283405
0235065
0088970
0199509
0228604
0232712
0214535
0226875
0275457
0088930
0125816
0225280
0223128
0226405
0278041
0227693
0125820
0258197
0233492
0279082
0280350
0278410
0125813

0233015
0227063
0238034
0234435
0234862
0225321
0225170
0270838
0204684
0259974
0125798
0237396
0230400
0283072
0215553
0213882
0228935
0230492
0234646
0132671
0178726
0279447
0125810
0233746
0279164
0201527
0230387
0277901
0234832
0132661
0232645
0125812
0125814
0201728
0125823
0125831
0273838
0200208
0226344
0260202
0276638
0125815
0204663
0204662
0101435
0204658
0173335
0101439
0270001
0225056

0286117
0101441
0232595
0286270
0170373
0277975
0230908
0170369
0170367
0228476
0236168
0149435
0167390
0230133
0236151
0200231
0232200
0234069
0101463
0228539
0231015
0274173
0077984
0101474
0207355
0286444
0154930
0275358
0100987
0225069
0230725
0225344
0286472
0197586
0202414
0279322
0277938
0274414
0100994
0274507
0276952
0100997
0227379
0101003
0101004
0251883
0170191
0213742
0242236
0236542

130684
278383
226465
230772
175170
275048
250090
274973
228593
226203
125804
231081
287569
204556
277508
240108
230440
287211
227195
238129
284419
282826
282870
283568
283069
282949
282894
282935
282842
283123
282968
282911
282995
287889
283592
282921
283291
283005
283433
283020
204555
205611
286483
287288
200325
231934
149531
238151
275877
283443

0277112
0274385
0275305
0215547
0215546
0224628
0215545
0237929
0236121
0252649
0212717
0131068
0212134
0180483
0204548
0204547
0180424
0180383
0088320
0233354
0101294
0274364
0101898
0235313
0230613
0278276
0201770
0125968
0264395
0131055
0171552
0236559
0281376
0088325
0230801
0101306
0179772
0149599
0131044
0199497
0088356
0260903
0278012
0226239
0252623
0101331
0212571
0101336
0101337
0275576

0224452
0235217
0126003
0101346
0284125
0287850
0101350
0277692
0171456
0197183
0236772
0286311
0277301
0233293
0204393
0228156
0198547
0285382
0175730
0149600
0088305
0101367
0260536
0260257
0215529
0167098
0078898
0167104
0214245
0186190
0186191
0131050
0237662
0183566
0224876
0131059
0198183
0233106
0125999
0233146
0125997
0101391
0101400
0078699
0275223
0125967
0149609
0182584
0271803
0101412

101417
229188
131061
230837
230753
101421
214200
228958
285230
125970
228386
264616
125977
225246
287853
101440
214185
250917
236456
101444
272945
234087
078747
201498
234933
236388
231795
207997
181741
277558
125971
206582
101460
101464
231034
198646
078804
286803
225336
131067
131069
100983
078814
207635
283441
100991
202150
088298
201151
101000

0278367
0200301
0126005
0229230
0125966
0279253
0261582
0242372
0235214
0125998
0101019
0204183
0125965
0221763
0126001
0230155
0242507
0125975
0125991
0224497
0088340
0227401
0061656
0214078
0272897
0222460
0252549
0244462
0244005
0125995
0131051
0225357
0283527
0025293
0234274
0199676
0219608
0201221
0207053
0171222
0149646
0260032
0234139
0277235
0231470
0237063
0088367
0232406
0278035
0131043

0080845
0232907
0101335
0273986
0118707
0259399
0101084
0101082
0227688
0101079
0202255
0149636
0149639
0242509
0101342
0101347
0080839
0231241
0226144
0269846
0101353
0236159
0118705
0118702
0101363
0286240
0197122
0213979
0166619
0053438
0276603
0235044
0274469
0228234
0225759
0204117
0132792
0132821
0199683
0101407
0101413
0264767
0277550
0198959
0149633
0277829
0285144
0285347
0285100
0206249

0101425
0129988
0275285
0224635
0275401
0196756
0277034
0235408
0225091
0201512
0200354
0201811
0174365
0274309
0199266
0170471
0275453
0231307
0182035
0124143
0213959
0277965
0101438
0101442
0199691
0101445
0240474
0101447
0274825
0261431
0101452
0277581
0223864
0230324
0287983
0224357
0275337
0286083
0229976
0234878
0237767
0233415
0223410
0204103
0229771
0286972
0238908
0222612
0198900
0226648

0124181
0276169
0174306
0272407
0242748
0263989
0235102
0132793
0183798
0229398
0124177
0235088
0273951
0232493
0252386
0212224
0225458
0223651
0228959
0196090
0229892
0229042
0227599
0223190
0233508
0240639
0226038
0252193
0237555
0201021
0233711
0206998
0124193
0288000
0201372
0185513
0234271
0277611
0226143
0101049
0282876
0101052
0215467
0101057
0124196
0206801
0233277
0124191
0241229
0149596

0287570
0132823
0223891
0124194
0197296
0101074
0201363
0226812
0101076
0229005
0233376
0266151
0168746
0226243
0124120
0132824
0168734
0196839
0283440
0132832
0244558
0064205
0124249
0276223
0101098
0277708
0286100
0166913
0101104
0025772
0227477
0101109
0252021
0124134
0175121
0168703
0283142
0124102
0124233
0124157
0233352
0243995
0124107
0124159
0124232
0124145
0275894
0204070
0254806
0124251

0232880
0244274
0124155
0273555
0277022
0101443
0237464
0237068
0101446
0243543
0249139
0232135
0229900
0101448
0158901
0201923
0230860
0180205
0180305
0235508
0180083
0182931
0168634
0266071
0149651
0226188
0124116
0168630
0221046
0101457
0175063
0101470
0124104
0101473
0277051
0132801
0168612
0149634
0124257
0064601
0271984
0100979
0285796
0100982
0198026
0235525
0100985
0204044
0124140
0124160

0213820
0101017
0229957
0149654
0080189
0273828
0062598
0215452
0283757
0225849
0184617
0198185
0149635
0158296
0273893
0172315
0271784
0197496
0222874
0235922
0064655
0236028
0226769
0239783
0215450
0264901
0101040
0267882
0273451
0231119
0224565
0234075
0202186
0201742
0124151
0229853
0196562
0255438
0238950
0215448
0251770
0238785
0234967
0228503
0287443
0237423
0229522
0286179
0287335
0276923

0287107
0288083
0286063
0235621
0236874
0274261
0281365
0281460
0238452
0124126
0286159
0278231
0124198
0230758
0227431
0124207
0124214
0226284
0124228
0124201
0177410
0277830
0209042
0274760
0222365
0277967
0212304
0158445
0283529
0124212
0158470
0276031
0231878
0252540
0197818
0237788
0158480
0199490
0269103
0124226
0229222
0206710
0124216
0231265
0244687
0124208
0240849
0237595
0234698
0231742

277449
172216
224397
203999
233077
244376
231715
196396
239742
201501
232043
208018
042062
221091
231302
234693
124171
124243
230043
101126
228663
259456
000419
124217
232358
026559
279835
228820
101096
266761
054793
215444
101115
227964
252467
252654
020256
287511
233347
234948
225785
275958
232465
234821
232294
286250
232286
230636
182463
199218

0286865
0223492
0227705
0271774
0231703
0224530
0259723
0206959
0197670
0171940
0235415
0238468
0287886
0225563
0241721
0064787
0236352
0265595
0019186
0286587
0101132
0286229
0101134
0252089
0185834
0235166
0224008
0275923
0287478
0286006
0054803
0222118
0124089
0235594
0230843
0124098
0087586
0101138
0087589
0228601
0022277
0124091
0124103
0280387
0213714
0287079
0225657
0214541
0087510
0252091

0241821
0237000
0231604
0207158
0231078
0203266
0288086
0280220
0101144
0235032
0226308
0252064
0266666
0231005
0054796
0101146
0228995
0256222
0218018
0132819
0286052
0124097
0124092
0124253
0227297
0124256
0124225
0278709
0263453
0203971
0275852
0225708
0270411
0124237
0124227
0124224
0124209
0124164
0198768
0231290
0254620
0268941
0124222
0254995
0215440
0254419
0237121
0270279
0268649
0284040

0216031
0235590
0268333
0087460
0276859
0276722
0274491
0275069
0225806
0270951
0285091
0284985
0101158
0101160
0101162
0124172
0101166
0232925
0124203
0124205
0225129
0230352
0286309
0233686
0287752
0227282
0087495
0238194
0238777
0196074
0196227
0132825
0124215
0176659
0276093
0228340
0207802
0229690
0266140
0265617
0276627
0261035
0229755
0229606
0179242
0286232
0276786
0277270
0179253
0225106

0130699
0284193
0235018
0265306
0283078
0149657
0101182
0184402
0101181
0101180
0130703
0130706
0226332
0130702
0284074
0228812
0273619
0171858
0149679
0275437
0130701
0233017
0130700
0272050
0260542
0174403
0174407
0199017
0284508
0226094
0264490
0273812
0232121
0229882
0167046
0101187
0277496
0276317
0232803
0237687
0101188
0223669
0228705
0101189
0229873
0060491
0092758
0101190
0233838
0236527

0101191
0273759
0101193
0101194
0125533
0272259
0125514
0237119
0231133
0274915
0225978
0287324
0260416
0286628
0231977
0207598
0149658
0273821
0101197
0266463
0101198
0275620
0101199
0266104
0101203
0222439
0101204
0203900
0075043
0280936
0226390
0101210
0230226
0125534
0101213
0125508
0275812
0125531
0130589
0101216
0232442
0197457
0258366
0026036
0243509
0101246
0197114
0273154
0274501
0203896

0273047
0125520
0130584
0229299
0231208
0279610
0268858
0183260
0101150
0101152
0199805
0283206
0216141
0283513
0216195
0275842
0198276
0284433
0207554
0280213
0196700
0130590
0101161
0196421
0203883
0171703
0237371
0171700
0274034
0125510
0171695
0286999
0196132
0125522
0203880
0278171
0232852
0149656
0274727
0279493
0277117
0279687
0280071
0276612
0275464
0280433
0279669
0279094
0274333
0277777

)279186
)279784
)279064
)274559
)280013
)279769
)280191
)280095
)280179
)275993
)288187
)274790
)275496
)280019
)279709
)278903
)276902
)274276
)275895
)280346
)279226
)278927
)276076
)280145
)278878
)279788
)279728
)280164
)279998
)278955
)284550
)280172
)279751
)280018
)279477
)278884
)277067
)274046
)280330
)279303
)279647
)279895
)276077
)274484
)279177
)276546
)277991
)273590
)277277
)278961

0276289
0273614
0279967
0278996
0275950
0275664
0275708
0277437
0280800
0278233
0286155
0286252
0286057
0286012
0274060
0277671
0281383
0277739
0280441
0277379
0274868
0280614
0275215
0286267
0286146
0286054
0286178
0275692
0278775
0281181
0278189
0286149
0286032
0286091
0286148
0279990
0279615
0279167
0279579
0286033
0279213
0280243
0278931
0264462
0264063
0279501
0279414
0280436
0188681
0279720

279381
278932
279718
279208
279864
279211
286103
275631
280081
279783
264002
252199
279321
270533
280372
279851
279062
277693
275945
275592
278106
187172
273840
274391
278678
273872
276556
277282
229306
224309
207097
168122
237202
280108
185390
175302
266211
226930
219280
228159
228184
179381
218549
166351
207476
266299
278381
227874
214319
279773

0273692
0278618
0229231
0224922
0173231
0232797
0224860
0215562
0234538
0228314
0230965
0215559
0203616
0223287
0183249
0233442
0275170
0224905
0249493
0201812
0277572
0188992
0240755
0185272
0243064
0155304
0155307
0223662
0232884
0228600
0226751
0225502
0235609
0281903
0226406
0235277
0180530
0229047
0231201
0279390
0236471
0280082
0229425
0224524
0224247
0212564
0226298
0155313
0234159
0215386

0252273
0214976
0224602
0201025
0270071
0270139
0287066
0280432
0207638
0199030
0269950
0207863
0239023
0237735
0232886
0225735
0278181
0252462
0232560
0226580
0266195
0154639
0232260
0200754
0199962
0154640
0280594
0226956
0279648
0240770
0154642
0244676
0231755
0215369
0154645
0227330
0154646
0228708
0224141
0230233
0265841
0198618
0279156
0232193
0226204
0235965
0229289
0226818
0231620
0212609

0230198
0271486
0229336
0215353
0233480
0233236
0251851
0252963
0233676
0234439
0233206
0276738
0224924
0215351
0154654
0226771
0234730
0237527
0238265
0233997
0227075
0233215
0184856
0234340
0231136
0202339
0226043
0223488
0234034
0225906
0230972
0275469
0219368
0231058
0286222
0223078
0228592
0223822
0237569
0199698
0227716
0288094
0287612
0231986
0235564
0287801
0226996
0224018
0224832
0237484

0226983
0227090
0230379
0237444
0244278
0232512
0274662
0223870
0222042
0238627
0213885
0279534
0185433
0229962
0234883
0283904
0260583
0154719
0154721
0237731
0287109
0227054
0154723
0154727
0235514
0142192
0233783
0251972
0224541
0273492
0206802
0286902
0229025
0232692
0197934
0273115
0166265
0154734
0223563
0154736
0266133
0215326
0231236
0236332
0233300
0217026
0219592
0234052
0225298
0234083

0178457
0232079
0236532
0232855
0226935
0283300
0156239
0215005
0215317
0198862
0176054
0156253
0273254
0156256
0156261
0231125
0286018
0156265
0232687
0201984
0212479
0224649
0215533
0156273
0236056
0248476
0273017
0232118
0228817
0234293
0171189
0273464
0174680
0259981
0156282
0227342
0156284
0225267
0188694
0232263
0197683
0206107
0186980
0250973
0182816
0265007
0198390
0240432
0186971
0233640

0186970
0184351
0186965
0244025
0186967
0186977
0237841
0235312
0218125
0236612
0186925
0244362
0206106
0212938
0186930
0186924
0184724
0244624
0206105
0184032
0206104
0231068
0236400
0187026
0187005
0238257
0233036
0227840
0183640
0274749
0182591
0206102
0228941
0156299
0286643
0237325
0237594
0230870
0234509
0142168
0238390
0273271
0156304
0229046
0273091
0234107
0142149
0237138
0230323
0261610

159055
227256
170262
232623
142207
200792
256073
166979
229007
252045
252950
242220
265590
159079
228433
238220
159082
238197
159086
205930
279690
205929
224427
207098
232360
227757
228961
205927
232539
184221
226527
226433
229086
249624
159110
223799
243646
142166
226501
159128
142188
215088
231355
177692
159131
233956
159140
284448
159147
249209

0205758
0205726
0241837
0237945
0244294
0273102
0243927
0198743
0272657
0224598
0227456
0214955
0225555
0159197
0205670
0222018
0273104
0243627
0272958
0180509
0221398
0159200
0159212
0230978
0234380
0159216
0286153
0234703
0223671
0231300
0230794
0211590
0229761
0234008
0231106
0214914
0185917
0189089
0236677
0200213
0230212
0159228
0236119
0233393
0226054
0236830
0214889
0159231
0142197
0228149

0214867
0273199
0159256
0228107
0159259
0224421
0230479
0233818
0159261
0279365
0223741
0286517
0286852
0231324
0224269
0159263
0159267
0237646
0199806
0270652
0224790
0207416
0215734
0183145
0212136
0185808
0182670
0270116
0228677
0230366
0263969
0242553
0157538
0272948
0287637
0273210
0157540
0286717
0157542
0233213
0184029
0223608
0198054
0157551
0270835
0233316
0226012
0231123
0231231
0157554

0231480
0287470
0223806
0157557
0205622
0229986
0287082
0229925
0272015
0227721
0232837
0235888
0288266
0237609
0228861
0235701
0276873
0183527
0185658
0272991
0232608
0237373
0229623
0255568
0238141
0205581
0252915
0227406
0182093
0285815
0157578
0185437
0275523
0235808
0228349
0235012
0183778
0184809
0225330
0231713
0183067
0183036
0171587
0263973
0235123
0207147
0233756
0288069
0230859
0286425

0226496
0263681
0182240
0280109
0224388
0183844
0183486
0157601
0228318
0184012
0207503
0232806
0223400
0286027
0227702
0236384
0232401
0183421
0236883
0275166
0141956
0236545
0227698
0251778
0157617
0173276
0237232
0177398
0184385
0160179
0223262
0160180
0160181
0160182
0160183
0160185
0252619
0160188
0160190
0239930
0231867
0235772
0225431
0235023
0160191
0225731
0283051
0233754
0225218
0160193

0160194
0233056
0264580
0224100
0160199
0160200
0160201
0286549
0232777
0236663
0228120
0160202
0237864
0237989
0225637
0142178
0188660
0185186
0223975
0160207
0214326
0234289
0278433
0160208
0160209
0281420
0287507
0160213
0234030
0160214
0215458
0226543
0160216
0199598
0252606
0160218
0213440
0241945
0160221
0225331
0285413
0237604
0232124
0160223
0278158
0275799
0142182
0232010
0160224
0141959

0160226
0241728
0184441
0232969
0260256
0142185
0230061
0266692
0264452
0229356
0160233
0224747
0223431
0227999
0274225
0228709
0277352
0175894
0235890
0182912
0287088
0215455
0205445
0212935
0215454
0241123
0188155
0272804
0187766
0221837
0221859
0243489
0212933
0205439
0221864
0229880
0187175
0276647
0189169
0236382
0184787
0236519
0184900
0183255
0160255
0227039
0273027
0183250
0272825
0160256

0276529
0234880
0261706
0275874
0235374
0197381
0267857
0182586
0186866
0223768
0286262
0184274
0229382
0237664
0228930
0228355
0273796
0182871
0224574
0183535
0275167
0173638
0233922
0286082
0286594
0183570
0275139
0205424
0286886
0276633
0280604
0274248
0142156
0224413
0228235
0226115
0233767
0227438
0142173
0160282
0237338
0160284
0228404
0160285
0223901
0215424
0160294
0228137
0239415
0182362

0160298
0160299
0286224
0225043
0160305
0223692
0202239
0160307
0160310
0230982
0212932
0277248
0283047
0279973
0226444
0276871
0288262
0286406
0283023
0276138
0280341
0236235
0279442
0184624
0236097
0236831
0130538
0237299
0235759
0233408
0215268
0213727
0230643
0230471
0231565
0226474
0224435
0234381
0198062
0236666
0212216
0241838
0225255
0235992
0206195
0271127
0232775
0272872
0271672
0223875

215270
229286
233866
278188
225293
226160
229658
227367
280263
235969
273362
233995
239435
198445
287285
240122
100181
283633
267338
249288
249263
189295
259271
273244
241717
288024
235343
172967
229027
234121
283535
237835
220248
237689
228122
237358
225480
230481
253691
253460
254264
253481
285045
215568
270226
276400
237438
273203
273442
177663

0235328
0183307
0235478
0069998
0185837
0093072
0236325
0203416
0241832
0231004
0239547
0099954
0229492
0232305
0228045
0286195
0182902
0236754
0131100
0099968
0015475
0264757
0269220
0243156
0093100
0286990
0207780
0235445
0234913
0237446
0235295
0225335
0215193
0236890
0235617
0225225
0183785
0280007
0184979
0187979
0276160
0274044
0276874
0275060
0277870
0277017
0278044
0274252
0275319
0277690

0278558
0274602
0274625
0222630
0275793
0275362
0278110
0286175
0273907
0273643
0278008
0182824
0197421
0239989
0234764
0161103
0215544
0276095
0279987
0183628
0283809
0100033
0284294
0280418
0237517
0273032
0283583
0271275
0283366
0270393
0273164
0286025
0241527
0070413
0206739
0273311
0237347
0281530
0223461
0272682
0231086
0100056
0206203
0063515
0260924
0100075
0070371
0286367
0230194
0251940

233471
287146
100084
242259
244296
185608
185065
070010
273300
273212
093009
287652
184113
281548
230485
225007
184702
284874
203618
184058
185838
235776
215012
184470
232926
093010
284031
099889
183597
208023
236540
268292
277950
128191
284140
284464
243762
099899
278278
099901
264346
099904
234409
249923
236499
040608
221039
161132
128185
273343

0273139
0188280
0235578
0287446
0161133
0185252
0207343
0244486
0277971
0099910
0215493
0255156
0201916
0242876
0236670
0099917
0236003
0250261
0223553
0276427
0238125
0229107
0217261
0235062
0226287
0241973
0099937
0099940
0272600
0099942
0237476
0183773
0285314
0272829
0099949
0184436
0230513
0284060
0161149
0099957
0284130
0099960
0226872
0207575
0206145
0187905
0249680
0284665
0169668
0197210

0239511
0224688
0215498
0234503
0283972
0133475
0283145
0226885
0229266
0169662
0206142
0237407
0232771
0279920
0286102
0226534
0273846
0274600
0200057
0252020
0275840
0169635
0276800
0206140
0183506
0252143
0222352
0183246
0185651
0275293
0161179
0161180
0273342
0128228
0207751
0212102
0283871
0284630
0284651
0284654
0284621
0272954
0100023
0100027
0286365
0244671
0286127
0100030
0200985
0225544

100034
224086
100038
197549
253239
211637
254355
278657
249333
211642
253794
228161
253874
254075
254161
253242
253752
253823
275361
211638
225741
263366
236118
233720
211639
227710
224465
254308
253637
236323
211640
253126
211641
253935
228050
234726
169575
279278
274422
234630
286129
272779
211643
211644
211645
225724
223350
232515
211647
211648

0237428
0211649
0232603
0211650
0211651
0211652
0254175
0254073
0253818
0211653
0224144
0253889
0211654
0211655
0230821
0253631
0226595
0275004
0169548
0185686
0220891
0253120
0211656
0211657
0253920
0253329
0215456
0286941
0183169
0100121
0253338
0253451
0211658
0253913
0253779
0211659
0232350
0223999
0253822
0211660
0253546
0211661
0211662
0254240
0211663
0278667
0211664
0253152
0211665
0253786

0211666
0253590
0211667
0231392
0275772
0211668
0274381
0211669
0226477
0211670
0278196
0284049
0237127
0254077
0253448
0253234
0226420
0211672
0253963
0211673
0264824
0254709
0211674
0211675
0211676
0211677
0211678
0211679
0211680
0254029
0211681
0254030
0211682
0222037
0211684
0211685
0237472
0223679
0100218
0128266
0283438
0275635
0100228
0186716
0236794
0240160
0230701
0214526
0236611
0215478

0133519
0248409
0234928
0227755
0178248
0287864
0225413
0280623
0224277
0128322
0272733
0278655
0189269
0275609
0211683
0228315
0234353
0244723
0272578
0159496
0187792
0128218
0169314
0250479
0099953
0099956
0099958
0272973
0133460
0251357
0279467
0244425
0240972
0218537
0273295
0225282
0206090
0231271
0133433
0228039
0099974
0272787
0099977
0285762
0224205
0250470
0099984
0276950
0235689
0099991

0232545
0099994
0099998
0230712
0128262
0225098
0215464
0100014
0258555
0128271
0178803
0100024
0228923
0138867
0100028
0286070
0178026
0100031
0230947
0215481
0284233
0284128
0284070
0224124
0224806
0224334
0282012
0237601
0182351
0184571
0167037
0199783
0206069
0197077
0279085
0279548
0279006
0279110
0203280
0271181
0100053
0236641
0244752
0279298
0233577
0285968
0285690
0206066
0230637
0231466

0100068
0272942
0272798
0271138
0100058
0277872
0272977
0234884
0100077
0237387
0224003
0222585
0133454
0278960
0231933
0232464
0223056
0232050
0236464
0229770
0100095
0224192
0252267
0230203
0128203
0100099
0100104
0100109
0261188
0128294
0221760
0213707
0286326
0100122
0196431
0226912
0225783
0278404
0206028
0277941
0275942
0244625
0276991
0223704
0231369
0235271
0279440
0200443
0279914
0279848

0236858
0280025
0231405
0238195
0233521
0233574
0223726
0226741
0230051
0280445
0287034
0227880
0224027
0227838
0231495
0226169
0169184
0180957
0235954
0277460
0264073
0283225
0276150
0100154
0274342
0276675
0239249
0201209
0199797
0276162
0266503
0183765
0100209
0159873
0100219
0226471
0272858
0224256
0183579
0235786
0177993
0100249
0183762
0226772
0200871
0251952
0186998
0100263
0237015
0182944

0185340
0100276
0273216
0100280
0280968
0239127
0285697
0242156
0278292
0225465
0128250
0100285
0282816
0100296
0234208
0184117
0186575
0233010
0232396
0239446
0100314
0100319
0200976
0287967
0184076
0100325
0100330
0207455
0279159
0275818
0227117
0227201
0176635
0226573
0225971
0225676
0232530
0128342
0268812
0099985
0279699
0239282
0248751
0099992
0099995
0187860
0099999
0100003
0249590
0273350

222915
272689
242114
273428
230137
100012
181123
214161
133488
214491
225774
128242
100029
223831
185339
100036
167065
250318
288153
184792
264661
237977
235989
133422
253352
276664
276057
276965
275307
269987
269972
278920
240186
183963
273387
198832
235573
185133
100078
214125
264141
138942
254835
182541
199695
202019
100100
228839
212542
100105

0213888
0206615
0185721
0184708
0240591
0199248
0198089
0287817
0233733
0229169
0214093
0241878
0284082
0183530
0100150
0285404
0242251
0236132
0252909
0227813
0128254
0128245
0227519
0230866
0240647
0232346
0100170
0234479
0237439
0232707
0239674
0205856
0230120
0275656
0128253
0254127
0242082
0100191
0214076
0273325
0234626
0128276
0271093
0205853
0274617
0274170
0230736
0243519
0241954
0232218

0100220
0184459
0100225
0185666
0236054
0251890
0229673
0100234
0228194
0231253
0228622
0233632
0133424
0232073
0230741
0266012
0253007
0281295
0232081
0224973
0273082
0236052
0224404
0235485
0280356
0234891
0237316
0286592
0175329
0233080
0273068
0238153
0238584
0100281
0273176
0100284
0266320
0274280
0282602
0282041
0100292
0100297
0233388
0273044
0100302
0198125
0268818
0221963
0228587
0226336

0128313
0100320
0237406
0279652
0279927
0279217
0279714
0279805
0128284
0233764
0231576
0237948
0225557
0237129
0229088
0100336
0128335
0100342
0100345
0278420
0287269
0223695
0237668
0252225
0228719
0100348
0100350
0100353
0188078
0229971
0261675
0166862
0234688
0287773
0100360
0286485
0100362
0234979
0183822
0100365
0100368
0232254
0215403
0185264
0128311
0252412
0128309
0287236
0100379
0201078

0187045
0231467
0100385
0235237
0133466
0278195
0226233
0128340
0100055
0243902
0237862
0166897
0272694
0100060
0100065
0225867
0128283
0100079
0273866
0100083
0100092
0285304
0233360
0241360
0241333
0100097
0273899
0100101
0100106
0189060
0100116
0128310
0100124
0284197
0207696
0100129
0279738
0207227
0100139
0128346
0278948
0272582
0100142
0284289
0100146
0284266
0222044
0273214
0100151
0233739

0100156
0128298
0272720
0184381
0279080
0185022
0198792
0235246
0241693
0277385
0213923
0283900
0244627
0229955
0286144
0228620
0168135
0286440
0100196
0100201
0100206
0224279
0184949
0100211
0230912
0228274
0287219
0100216
0100221
0100226
0229598
0244491
0100242
0230149
0273096
0225450
0272669
0273076
0100246
0221890
0183741
0288106
0235209
0128383
0179750
0249310
0244509
0284554
0243811
0128394

259421
239713
225720
100298
100307
226024
279833
227009
100311
284633
224794
100316
209480
209482
263764
100321
280216
100324
128268
227188
100335
285025
128272
187051
100346
176177
235647
238875
226085
100351
225528
133477
261202
100354
228599
239900
284431
100359
229999
196588
272834
227413
232564
172912
230071
231907
128285
100372
230769
233603

)266594
)100380
)196236
)172404
)273979
)200683
)100387
)222698
)237214
)199515
)213857
)284015
)100393
)252859
)232754
)231993
)215771
)100395
)235513
)100399
)100401
)274552
)100403
)167074
)199865
)273424
)183864
)100410
)100412
)100413
)286491
)172346
)100417
)100418
)196419
)200332
)100138
)285531
)207457
)184208
)167077
)231261
)252603
)206813
)100147
)184068
)198911
)207932
)234965
)159958

0283829
0100162
0205704
0100167
0224883
0183066
0215347
0198951
0286659
0177096
0238498
0183172
0184983
0270083
0237037
0213790
0100197
0227370
0232710
0281538
0205702
0226450
0100207
0182057
0229891
0235568
0230107
0172250
0189306
0183569
0182841
0202058
0251913
0100227
0276027
0270022
0100243
0249222
0128274
0200448
0234009
0229608
0274717
0242247
0100266
0226989
0230319
0100271
0234735
0100290

100294
100300
100304
159307
236272
234892
203527
236117
186732
286801
223843
186976
233157
180150
261251
130540
100341
100344
100347
224856
188677
279175
280434
138964
279933
280011
138944
220702
236079
232655
228807
279642
188636
215333
187012
234300
231801
186654
248405
241484
056487
223730
230922
226328
273243
093000
273353
100364
221598
273287

0213752
0100373
0100376
0077935
0128408
0232363
0077942
0238120
0251985
0280383
0130638
0264160
0235347
0279954
0235091
0188064
0280424
0273145
0231711
0182257
0273289
0241990
0197182
0231010
0235159
0266533
0283990
0284175
0284520
0186951
0280384
0205643
0130943
0075234
0277232
0075218
0100416
0075275
0230634
0234869
0075240
0100422
0260708
0054611
0280080
0278881
0280312
0224715
0218357
0205634

0224271
0232897
0279712
0235154
0278890
0238192
0229275
0285707
0266508
0285855
0233179
0287225
0219438
0281732
0266887
0205632
0286381
0285623
0226142
0280199
0285722
0188511
0280224
0279597
0264139
0212939
0213279
0235111
0286393
0226954
0223142
0214727
0100425
0279345
0236867
0260613
0229409
0277771
0279494
0100426
0182858
0273192
0184164
0278869
0198355
0276753
0188263
0138892
0100427
0073146

0280395
0073150
0170638
0273188
0273253
0073169
0272836
0273137
0128159
0100429
0188130
0185386
0196576
0205593
0227484
0279182
0100239
0100241
0128165
0100253
0100258
0025770
0284194
0272821
0025708
0177989
0273272
0272666
0130487
0217442
0205560
0254413
0100288
0205559
0272940
0008735
0100299
0212569
0286508
0251322
0206841
0225929
0100312
0254499
0213683
0184319
0079974
