## Supplemental file 2 for "Evolutionary context can clarify teleosts gene names"

| gene | all extra copies | total number of extra copies genes |
|---|---|---|
| a1cf | a1cf | 1 |
| aak1a | aak1a,aak1b | 2 |
| aak1b | aak1a,aak1b | 2 |
| abca1a | abca1a,abca1b | 2 |
| abca1b | abca1a,abca1b | 2 |
| abca3b | abca3b | 1 |
| abca4a | abca4b,abca4a | 2 |
| abca4b | abca4b,abca4a | 2 |
| abcb11a | abcb11a,abcb11b | 2 |
| abcb11b | abcb11a,abcb11b | 2 |
| abcb6a | abcb6a,abcb6b | 2 |
| abcb6b | abcb6a,abcb6b | 2 |
| abcc6a | abcc6a | 1 |
| abcc8b | abcc8,abcc8b,abcc8 | 3 |
| abcd3a | abcd3a,abcd3a,abcd3b,al | 4 |
| abcd3a | abcd3a,abcd3a,abcd3b,al | 4 |
| abcd3a | abcd3a,abcd3a,abcd3b,al | 4 |
| abcd3b | abcd3a,abcd3a,abcd3b,al | 4 |
| abcf2a | abcf2a,abcf2b | 2 |
| abcf2b | abcf2a,abcf2b | 2 |
| abcg2a | abcg2b,abcg2b,abcg2d,ak | 5 |
| abcg2b | abcg2b,abcg2b,abcg2d,ak | 5 |
| abcg2b | abcg2b,abcg2b,abcg2d,ak | 5 |
| abcg2c | abcg2b,abcg2b,abcg2d,ak | 5 |
| abcg2d | abcg2b,abcg2b,abcg2d,ak | 5 |
| abcg4a | abcg4a,abcg4b | 2 |
| abcg4b | abcg4a,abcg4b | 2 |
| abhd10a | abhd10a | 1 |
| abhd14a | abhd14b,abhd14a | 2 |
| abhd14b | abhd14b,abhd14a | 2 |
| abhd15a | abhd15a | 1 |
| abhd16a | abhd16a | 1 |
| abhd17aa | abhd17c,abhd17ab,abhd1 | 6 |
| abhd17ab | abhd17c,abhd17ab,abhd1 | 6 |
| abhd17ab | abhd17c,abhd17ab,abhd1 | 6 |
| abhd17b | abhd17c,abhd17ab,abhd1 | 6 |
| abhd17c | abhd17c,abhd17ab,abhd1 | 6 |
| abhd17c | abhd17c,abhd17ab,abhd1 | 6 |
| abhd2a | abhd2b,abhd2a,abhd2b | 3 |
| abhd2b | abhd2b,abhd2a,abhd2b | 3 |
| abhd2b | abhd2b,abhd2a,abhd2b | 3 |
| abhd5b | abhd5b | 1 |
| abhd6a | abhd6b,abhd6a,abhd6b | 3 |
| abhd6b | abhd6b,abhd6a,abhd6b | 3 |
| abhd6b | abhd6b,abhd6a,abhd6b | 3 |
| abhd8a | abhd8a,abhd8b | 2 |
| abhd8b | abhd8a,abhd8b | 2 |
| abi1a | abi1b,abi1a | 2 |
| abi1b | abi1b,abi1a | 2 |

| | | |
|---|---|---|
| abi2a | abi2a,abi2b,abi2a | 3 |
| abi2a | abi2a,abi2b,abi2a | 3 |
| abi2b | abi2a,abi2b,abi2a | 3 |
| abi3a | abi3bpb,abi3bpb,abi3a,al | 6 |
| abi3b | abi3bpb,abi3bpb,abi3a,al | 6 |
| abi3bpa | abi3bpb,abi3bpb,abi3a,al | 6 |
| abi3bpb | abi3bpb,abi3bpb,abi3a,al | 6 |
| abi3bpb | abi3bpb,abi3bpb,abi3a,al | 6 |
| abi3bpb | abi3bpb,abi3bpb,abi3a,al | 6 |
| ablim1a | ablim1b,ablim1b,ablim1b | 4 |
| ablim1b | ablim1b,ablim1b,ablim1b | 4 |
| ablim1b | ablim1b,ablim1b,ablim1b | 4 |
| ablim1b | ablim1b,ablim1b,ablim1b | 4 |
| abtb2a | abtb2a,abtb2b | 2 |
| abtb2b | abtb2a,abtb2b | 2 |
| acap3a | acap3b,acap3a | 2 |
| acap3b | acap3b,acap3a | 2 |
| acbd5a | acbd5a | 1 |
| acin1a | acin1a,acin1b | 2 |
| acin1b | acin1a,acin1b | 2 |
| ackr3a | ackr3b,ackr3b,ackr3a | 3 |
| ackr3b | ackr3b,ackr3b,ackr3a | 3 |
| ackr3b | ackr3b,ackr3b,ackr3a | 3 |
| ackr4a | ackr4a,ackr4b | 2 |
| ackr4b | ackr4a,ackr4b | 2 |
| acot11a | acot11a,acot11b | 2 |
| acot11b | acot11a,acot11b | 2 |
| acp5a | acp5b,acp5b,acp5a,acp5a | 5 |
| acp5a | acp5b,acp5b,acp5a,acp5a | 5 |
| acp5b | acp5b,acp5b,acp5a,acp5a | 5 |
| acp5b | acp5b,acp5b,acp5a,acp5a | 5 |
| acp5b | acp5b,acp5b,acp5a,acp5a | 5 |
| acsl1a | acsl1a,acsl1a,acsl1b | 3 |
| acsl1a | acsl1a,acsl1a,acsl1b | 3 |
| acsl1b | acsl1a,acsl1a,acsl1b | 3 |
| acsl3a | acsl3b,acsl3a | 2 |
| acsl3b | acsl3b,acsl3a | 2 |
| acsl4a | acsl4b,acsl4a | 2 |
| acsl4b | acsl4b,acsl4a | 2 |
| acta1a | acta1a,acta1b | 2 |
| acta1b | acta1a,acta1b | 2 |
| actc1a | actc1a,actc1b,actc1a,actc | 5 |
| actc1a | actc1a,actc1b,actc1a,actc | 5 |
| actc1a | actc1a,actc1b,actc1a,actc | 5 |
| actc1b | actc1a,actc1b,actc1a,actc | 5 |
| actc1c | actc1a,actc1b,actc1a,actc | 5 |
| actl6a | actl6a,actl6a,actl6b | 3 |
| actl6a | actl6a,actl6a,actl6b | 3 |
| actl6b | actl6a,actl6a,actl6b | 3 |
| actn2b | actn2b,actn2b | 2 |

| | | |
|---|---|---|
| actn2b | actn2b,actn2b | 2 |
| actn3a | actn3a,actn3b,actn3a | 3 |
| actn3a | actn3a,actn3b,actn3a | 3 |
| actn3b | actn3a,actn3b,actn3a | 3 |
| actr2a | actr2a,actr2b | 2 |
| actr2b | actr2a,actr2b | 2 |
| actr3b | actr3,actr3b | 2 |
| acvr1ba | acvr1bb,acvr1ba | 2 |
| acvr1bb | acvr1bb,acvr1ba | 2 |
| acvr2aa | acvr2bb,acvr2ab,acvr2aa, | 4 |
| acvr2ab | acvr2bb,acvr2ab,acvr2aa, | 4 |
| acvr2ba | acvr2bb,acvr2ab,acvr2aa, | 4 |
| acvr2bb | acvr2bb,acvr2ab,acvr2aa, | 4 |
| ada2a | ada2b,ada2a | 2 |
| ada2b | ada2b,ada2a | 2 |
| adam10a | adam10a,adam10b | 2 |
| adam10b | adam10a,adam10b | 2 |
| adam17a | adam17a,adam17b | 2 |
| adam17b | adam17a,adam17b | 2 |
| adam19a | adam19a,adam19b | 2 |
| adam19b | adam19a,adam19b | 2 |
| adam23a | adam23a | 1 |
| adam8a | adam8b,adam8a | 2 |
| adam8b | adam8b,adam8a | 2 |
| adamts15a | adamts15a,adamts15a,ac | 4 |
| adamts15a | adamts15a,adamts15a,ac | 4 |
| adamts15b | adamts15a,adamts15a,ac | 4 |
| adamts15b | adamts15a,adamts15a,ac | 4 |
| adamts8a | adamts8a | 1 |
| adarb1a | adarb1b,adarb1a | 2 |
| adarb1b | adarb1b,adarb1a | 2 |
| adcy1a | adcy1b,adcy1a | 2 |
| adcy1b | adcy1b,adcy1a | 2 |
| adcy2a | adcy2a,adcy2b | 2 |
| adcy2b | adcy2a,adcy2b | 2 |
| adcy3a | adcy3a,adcy3b | 2 |
| adcy3b | adcy3a,adcy3b | 2 |
| adcy6a | adcy6b,adcy6a,adcy6a,ac | 4 |
| adcy6a | adcy6b,adcy6a,adcy6a,ac | 4 |
| adcy6b | adcy6b,adcy6a,adcy6a,ac | 4 |
| adcy6b | adcy6b,adcy6a,adcy6a,ac | 4 |
| adcyap1a | adcyap1b,adcyap1a | 2 |
| adcyap1b | adcyap1b,adcyap1a | 2 |
| adcyap1r1a | adcyap1r1a,adcyap1r1a,a | 3 |
| adcyap1r1a | adcyap1r1a,adcyap1r1a,a | 3 |
| adcyap1r1a | adcyap1r1a,adcyap1r1a,a | 3 |
| add3a | add3b,add3b,add3a | 3 |
| add3b | add3b,add3b,add3a | 3 |
| add3b | add3b,add3b,add3a | 3 |
| adgra1a | adgra1a | 1 |

| | | |
|---|---|---|
| adgrb1a | adgrb1a,adgrb1b | 2 |
| adgrb1b | adgrb1a,adgrb1b | 2 |
| adgre5a | adgre5a | 1 |
| adgrf3a | adgrf3a,adgrf3b | 2 |
| adgrf3b | adgrf3a,adgrf3b | 2 |
| adgrg2a | adgrg2a | 1 |
| adgrg4a | adgrg4b,adgrg4a | 2 |
| adgrg4b | adgrg4b,adgrg4a | 2 |
| adgrl1a | adgrl1a | 1 |
| adgrl2a | adgrl2a | 1 |
| adh8a | adh8b,adh8a,adh8b,adh8 | 4 |
| adh8a | adh8b,adh8a,adh8b,adh8 | 4 |
| adh8b | adh8b,adh8a,adh8b,adh8 | 4 |
| adh8b | adh8b,adh8a,adh8b,adh8 | 4 |
| adipor1a | adipor1a,adipor1b,adipor | 3 |
| adipor1a | adipor1a,adipor1b,adipor | 3 |
| adipor1b | adipor1a,adipor1b,adipor | 3 |
| adm2a | adm2a,adm2a,adm2b | 3 |
| adm2a | adm2a,adm2a,adm2b | 3 |
| adm2b | adm2a,adm2a,adm2b | 3 |
| adnp2a | adnp2a,adnp2b | 2 |
| adnp2b | adnp2a,adnp2b | 2 |
| adora1a | adora1b,adora1a | 2 |
| adora1b | adora1b,adora1a | 2 |
| adora2aa | adora2aa,adora2b,adora2 | 4 |
| adora2aa | adora2aa,adora2b,adora2 | 4 |
| adora2ab | adora2aa,adora2b,adora2 | 4 |
| adora2b | adora2aa,adora2b,adora2 | 4 |
| adora4a | adora4a | 1 |
| adra1aa | adra1bb,adra1d,adra1aa, | 6 |
| adra1ab | adra1bb,adra1d,adra1aa, | 6 |
| adra1ba | adra1bb,adra1d,adra1aa, | 6 |
| adra1bb | adra1bb,adra1d,adra1aa, | 6 |
| adra1bb | adra1bb,adra1d,adra1aa, | 6 |
| adra1d | adra1bb,adra1d,adra1aa, | 6 |
| adra2a | adra2c,adra2c,adra2db,a | 10 |
| adra2a | adra2c,adra2c,adra2db,a | 10 |
| adra2a | adra2c,adra2c,adra2db,a | 10 |
| adra2b | adra2c,adra2c,adra2db,a | 10 |
| adra2c | adra2c,adra2c,adra2db,a | 10 |
| adra2c | adra2c,adra2c,adra2db,a | 10 |
| adra2c | adra2c,adra2c,adra2db,a | 10 |
| adra2da | adra2c,adra2c,adra2db,a | 10 |
| adra2db | adra2c,adra2c,adra2db,a | 10 |
| adra2db | adra2c,adra2c,adra2db,a | 10 |
| adrb2a | adrb2b,adrb2a,adrb2b | 3 |
| adrb2b | adrb2b,adrb2a,adrb2b | 3 |
| adrb2b | adrb2b,adrb2a,adrb2b | 3 |
| adrb3a | adrb3a,adrb3b | 2 |
| adrb3b | adrb3a,adrb3b | 2 |

| | | |
|---|---|---|
| afap1l1a | afap1l1a,afap1l1a,afap1l | 4 |
| afap1l1a | afap1l1a,afap1l1a,afap1l | 4 |
| afap1l1a | afap1l1a,afap1l1a,afap1l | 4 |
| afap1l1b | afap1l1a,afap1l1a,afap1l | 4 |
| agfg1a | agfg1b,agfg1a | 2 |
| agfg1b | agfg1b,agfg1a | 2 |
| ago3a | ago3a,ago3b | 2 |
| ago3b | ago3a,ago3b | 2 |
| agtr1a | agtr1a,agtr1b | 2 |
| agtr1b | agtr1a,agtr1b | 2 |
| ahr1a | ahr1a,ahr1b | 2 |
| ahr1b | ahr1a,ahr1b | 2 |
| ahsa1a | ahsa1b,ahsa1a | 2 |
| ahsa1b | ahsa1b,ahsa1a | 2 |
| aimp1a | aimp1a,aimp1b | 2 |
| aimp1b | aimp1a,aimp1b | 2 |
| ak7a | ak7b,ak7a | 2 |
| ak7b | ak7b,ak7a | 2 |
| akap12a | akap12b,akap12a,akap12 | 3 |
| akap12b | akap12b,akap12a,akap12 | 3 |
| akap12b | akap12b,akap12a,akap12 | 3 |
| akap17a | akap17a | 1 |
| akap1a | akap1a,akap1b | 2 |
| akap1b | akap1a,akap1b | 2 |
| akr1a1a | akr1a1a,akr1a1b,akr1a1a | 4 |
| akr1a1a | akr1a1a,akr1a1b,akr1a1a | 4 |
| akr1a1b | akr1a1a,akr1a1b,akr1a1a | 4 |
| akr1a1b | akr1a1a,akr1a1b,akr1a1a | 4 |
| akt3a | akt3b,akt3a | 2 |
| akt3b | akt3b,akt3a | 2 |
| aldh3a2a | aldh3a2a,aldh3a2b | 2 |
| aldh3a2b | aldh3a2a,aldh3a2b | 2 |
| aldh9a1b | aldh9a1b | 1 |
| alkal2a | alkal2b,alkal2a | 2 |
| alkal2b | alkal2b,alkal2a | 2 |
| alox5a | alox5ap,alox5a,alox5ap,a | 4 |
| alox5a | alox5ap,alox5a,alox5ap,a | 4 |
| alox5ap | alox5ap,alox5a,alox5ap,a | 4 |
| alox5ap | alox5ap,alox5a,alox5ap,a | 4 |
| alpk3a | alpk3b,alpk3a | 2 |
| alpk3b | alpk3b,alpk3a | 2 |
| als2a | als2b,als2a | 2 |
| als2b | als2b,als2a | 2 |
| alx4a | alx4a,alx4b | 2 |
| alx4b | alx4a,alx4b | 2 |
| ambra1a | ambra1a,ambra1b | 2 |
| ambra1b | ambra1a,ambra1b | 2 |
| amotl2a | amotl2b,amotl2a | 2 |
| amotl2b | amotl2b,amotl2a | 2 |
| ampd2a | ampd2b,ampd2a | 2 |

| | | |
|---|---|---|
| ampd2b | ampd2b,ampd2a | 2 |
| ampd3a | ampd3b,ampd3a,ampd3b | 3 |
| ampd3b | ampd3b,ampd3a,ampd3b | 3 |
| ampd3b | ampd3b,ampd3a,ampd3b | 3 |
| amy2a | amy2a | 1 |
| angpt2a | angpt2a,angpt2b | 2 |
| angpt2b | angpt2a,angpt2b | 2 |
| angptl1a | angptl1b,angptl1a | 2 |
| angptl1b | angptl1b,angptl1a | 2 |
| angptl2a | angptl2a,angptl2b | 2 |
| angptl2b | angptl2a,angptl2b | 2 |
| ank1a | ank1a,ank1a,ank1b | 3 |
| ank1a | ank1a,ank1a,ank1b | 3 |
| ank1b | ank1a,ank1a,ank1b | 3 |
| ank2a | ank2a,ank2b | 2 |
| ank2b | ank2a,ank2b | 2 |
| ank3a | ank3b,ank3b,ank3a | 3 |
| ank3b | ank3b,ank3b,ank3a | 3 |
| ank3b | ank3b,ank3b,ank3a | 3 |
| ankdd1a | ankdd1b,ankdd1a | 2 |
| ankdd1b | ankdd1b,ankdd1a | 2 |
| ankef1a | ankef1b,ankef1a | 2 |
| ankef1b | ankef1b,ankef1a | 2 |
| ankib1a | ankib1b,ankib1a | 2 |
| ankib1b | ankib1b,ankib1a | 2 |
| ankmy2a | ankmy2a,ankmy2b | 2 |
| ankmy2b | ankmy2a,ankmy2b | 2 |
| ankrd10a | ankrd10a,ankrd10b | 2 |
| ankrd10b | ankrd10a,ankrd10b | 2 |
| ankrd13a | ankrd13a,ankrd13c,ankrc | 4 |
| ankrd13b | ankrd13a,ankrd13c,ankrc | 4 |
| ankrd13c | ankrd13a,ankrd13c,ankrc | 4 |
| ankrd13d | ankrd13a,ankrd13c,ankrc | 4 |
| ankrd1a | ankrd1a,ankrd1a,ankrd1b | 4 |
| ankrd1a | ankrd1a,ankrd1a,ankrd1b | 4 |
| ankrd1a | ankrd1a,ankrd1a,ankrd1b | 4 |
| ankrd1b | ankrd1a,ankrd1a,ankrd1b | 4 |
| ankrd28b | ankrd28b | 1 |
| ankrd33aa | ankrd33bb,ankrd33ab,an | 4 |
| ankrd33ab | ankrd33bb,ankrd33ab,an | 4 |
| ankrd33ba | ankrd33bb,ankrd33ab,an | 4 |
| ankrd33bb | ankrd33bb,ankrd33ab,an | 4 |
| ankrd34ba | ankrd34bb,ankrd34ba | 2 |
| ankrd34bb | ankrd34bb,ankrd34ba | 2 |
| ankrd46a | ankrd46b,ankrd46a | 2 |
| ankrd46b | ankrd46b,ankrd46a | 2 |
| ankrd52a | ankrd52a,ankrd52a | 2 |
| ankrd52a | ankrd52a,ankrd52a | 2 |
| ankrd6a | ankrd6a,ankrd6b | 2 |
| ankrd6b | ankrd6a,ankrd6b | 2 |

| | | |
|---|---|---|
| anks1aa | anks1b,anks1ab,anks1aa | 3 |
| anks1ab | anks1b,anks1ab,anks1aa | 3 |
| anks1b | anks1b,anks1ab,anks1aa | 3 |
| anks4b | anks4b | 1 |
| ano10a | ano10a,ano10b | 2 |
| ano10b | ano10a,ano10b | 2 |
| ano2a | ano2b,ano2a | 2 |
| ano2b | ano2b,ano2a | 2 |
| ano5a | ano5b,ano5a | 2 |
| ano5b | ano5b,ano5a | 2 |
| ano8a | ano8a,ano8b | 2 |
| ano8b | ano8a,ano8b | 2 |
| ano9a | ano9b,ano9a,ano9a | 3 |
| ano9a | ano9b,ano9a,ano9a | 3 |
| ano9b | ano9b,ano9a,ano9a | 3 |
| anos1a | anos1a,anos1b | 2 |
| anos1b | anos1a,anos1b | 2 |
| anp32a | anp32b,anp32a,anp32e | 3 |
| anp32b | anp32b,anp32a,anp32e | 3 |
| anp32e | anp32b,anp32a,anp32e | 3 |
| antxr1a | antxr1a,antxr1b,antxr1c,a | 4 |
| antxr1b | antxr1a,antxr1b,antxr1c,a | 4 |
| antxr1c | antxr1a,antxr1b,antxr1c,a | 4 |
| antxr1d | antxr1a,antxr1b,antxr1c,a | 4 |
| antxr2a | antxr2b,antxr2a,antxr2a,a | 4 |
| antxr2a | antxr2b,antxr2a,antxr2a,a | 4 |
| antxr2b | antxr2b,antxr2a,antxr2a,a | 4 |
| antxr2b | antxr2b,antxr2a,antxr2a,a | 4 |
| anxa11a | anxa11b,anxa11a | 2 |
| anxa11b | anxa11b,anxa11a | 2 |
| anxa1a | anxa1a,anxa1d,anxa1c,ar | 4 |
| anxa1b | anxa1a,anxa1d,anxa1c,ar | 4 |
| anxa1c | anxa1a,anxa1d,anxa1c,ar | 4 |
| anxa1d | anxa1a,anxa1d,anxa1c,ar | 4 |
| anxa2a | anxa2b,anxa2b,anxa2b,ai | 4 |
| anxa2b | anxa2b,anxa2b,anxa2b,ai | 4 |
| anxa2b | anxa2b,anxa2b,anxa2b,ai | 4 |
| anxa2b | anxa2b,anxa2b,anxa2b,ai | 4 |
| anxa3a | anxa3b,anxa3a,anxa3a,ar | 4 |
| anxa3a | anxa3b,anxa3a,anxa3a,ar | 4 |
| anxa3b | anxa3b,anxa3a,anxa3a,ar | 4 |
| anxa3b | anxa3b,anxa3a,anxa3a,ar | 4 |
| anxa5a | anxa5b,anxa5a | 2 |
| anxa5b | anxa5b,anxa5a | 2 |
| ap1ar | ap1ar | 1 |
| ap1s3a | ap1s3b,ap1s3a | 2 |
| ap1s3b | ap1s3b,ap1s3a | 2 |
| ap2m1a | ap2m1a,ap2m1b | 2 |
| ap2m1b | ap2m1a,ap2m1b | 2 |
| ap3b1a | ap3b1a | 1 |

| | | |
|---|---|---|
| apba1a | apba1a,apba1b | 2 |
| apba1b | apba1a,apba1b | 2 |
| apba2a | apba2b,apba2a | 2 |
| apba2b | apba2b,apba2a | 2 |
| apbb1ip | apbb1,apbb1ip | 2 |
| apbb2b | apbb2b | 1 |
| aph1b | aph1b,aph1b | 2 |
| aph1b | aph1b,aph1b | 2 |
| apoa1a | apoa1a,apoa1a,apoa1b | 3 |
| apoa1a | apoa1a,apoa1a,apoa1b | 3 |
| apoa1b | apoa1a,apoa1a,apoa1b | 3 |
| apoa4a | apoa4a | 1 |
| apobec2a | apobec2b,apobec2a,apol | 3 |
| apobec2b | apobec2b,apobec2a,apol | 3 |
| apobec2b | apobec2b,apobec2a,apol | 3 |
| aqp10a | aqp10a,aqp10a,aqp10b | 3 |
| aqp10a | aqp10a,aqp10a,aqp10b | 3 |
| aqp10b | aqp10a,aqp10a,aqp10b | 3 |
| aqp3a | aqp3a,aqp3b | 2 |
| aqp3b | aqp3a,aqp3b | 2 |
| aqp8b | aqp8b | 1 |
| aqp9b | aqp9b | 1 |
| arap1a | arap1a | 1 |
| arcn1a | arcn1a,arcn1b | 2 |
| arcn1b | arcn1a,arcn1b | 2 |
| arf2a | arf2a,arf2b,arf2b | 3 |
| arf2b | arf2a,arf2b,arf2b | 3 |
| arf2b | arf2a,arf2b,arf2b | 3 |
| arf3a | arf3b,arf3b,arf3a | 3 |
| arf3b | arf3b,arf3b,arf3a | 3 |
| arf3b | arf3b,arf3b,arf3a | 3 |
| arf4a | arf4b,arf4a,arf4a | 3 |
| arf4a | arf4b,arf4a,arf4a | 3 |
| arf4b | arf4b,arf4a,arf4a | 3 |
| arf6a | arf6b,arf6a | 2 |
| arf6b | arf6b,arf6a | 2 |
| arfip2a | arfip2b,arfip2b,arfip2a | 3 |
| arfip2b | arfip2b,arfip2b,arfip2a | 3 |
| arfip2b | arfip2b,arfip2b,arfip2a | 3 |
| arglu1a | arglu1b,arglu1a | 2 |
| arglu1b | arglu1b,arglu1a | 2 |
| arhgap11a | arhgap11a | 1 |
| arhgap12a | arhgap12b,arhgap12a,arl | 3 |
| arhgap12b | arhgap12b,arhgap12a,arl | 3 |
| arhgap12b | arhgap12b,arhgap12a,arl | 3 |
| arhgap17a | arhgap17a,arhgap17b,arl | 3 |
| arhgap17a | arhgap17a,arhgap17b,arl | 3 |
| arhgap17b | arhgap17a,arhgap17b,arl | 3 |
| arhgap20b | arhgap20b,arhgap20 | 2 |
| arhgap21a | arhgap21b,arhgap21a | 2 |

| | | |
|---|---|---|
| arhgap21b | arhgap21b,arhgap21a | 2 |
| arhgap23a | arhgap23a,arhgap23b | 2 |
| arhgap23b | arhgap23a,arhgap23b | 2 |
| arhgap29a | arhgap29b,arhgap29a | 2 |
| arhgap29b | arhgap29b,arhgap29a | 2 |
| arhgap32a | arhgap32a,arhgap32b | 2 |
| arhgap32b | arhgap32a,arhgap32b | 2 |
| arhgap35a | arhgap35b,arhgap35b,arł | 3 |
| arhgap35b | arhgap35b,arhgap35b,arł | 3 |
| arhgap35b | arhgap35b,arhgap35b,arł | 3 |
| arhgap42a | arhgap42a,arhgap42b,arł | 3 |
| arhgap42a | arhgap42a,arhgap42b,arł | 3 |
| arhgap42b | arhgap42a,arhgap42b,arł | 3 |
| arhgap45a | arhgap45a,arhgap45b,arł | 3 |
| arhgap45b | arhgap45a,arhgap45b,arł | 3 |
| arhgap45b | arhgap45a,arhgap45b,arł | 3 |
| arhgap4a | arhgap4b,arhgap4a | 2 |
| arhgap4b | arhgap4b,arhgap4a | 2 |
| arhgef12a | arhgef12b,arhgef12a | 2 |
| arhgef12b | arhgef12b,arhgef12a | 2 |
| arhgef18a | arhgef18a,arhgef18b | 2 |
| arhgef18b | arhgef18a,arhgef18b | 2 |
| arhgef1a | arhgef1b,arhgef1a | 2 |
| arhgef1b | arhgef1b,arhgef1a | 2 |
| arhgef25b | arhgef25b | 1 |
| arhgef28a | arhgef28a,arhgef28b | 2 |
| arhgef28b | arhgef28a,arhgef28b | 2 |
| arhgef7a | arhgef7a,arhgef7b,arhgef | 3 |
| arhgef7a | arhgef7a,arhgef7b,arhgef | 3 |
| arhgef7b | arhgef7a,arhgef7b,arhgef | 3 |
| arhgef9a | arhgef9b,arhgef9a,arhgef | 3 |
| arhgef9b | arhgef9b,arhgef9a,arhgef | 3 |
| arhgef9b | arhgef9b,arhgef9a,arhgef | 3 |
| arid1aa | arid1aa,arid1ab,arid1b | 3 |
| arid1ab | arid1aa,arid1ab,arid1b | 3 |
| arid1b | arid1aa,arid1ab,arid1b | 3 |
| arid3a | arid3c,arid3c,arid3a,arid3 | 4 |
| arid3b | arid3c,arid3c,arid3a,arid3 | 4 |
| arid3c | arid3c,arid3c,arid3a,arid3 | 4 |
| arid3c | arid3c,arid3c,arid3a,arid3 | 4 |
| arid4a | arid4a | 1 |
| arid5a | arid5a,arid5b | 2 |
| arid5b | arid5a,arid5b | 2 |
| arl13a | arl13b,arl13a | 2 |
| arl13b | arl13b,arl13a | 2 |
| arl14ep | arl14,arl14ep | 2 |
| arl15a | arl15b,arl15a,arl15a,arl15 | 4 |
| arl15a | arl15b,arl15a,arl15a,arl15 | 4 |
| arl15b | arl15b,arl15a,arl15a,arl15 | 4 |
| arl15b | arl15b,arl15a,arl15a,arl15 | 4 |

| | | |
|---|---|---|
| arl2bp | arl2bp,arl2,arl2,arl2bp | 4 |
| arl2bp | arl2bp,arl2,arl2,arl2bp | 4 |
| arl3a | arl3b,arl3b,arl3a | 3 |
| arl3b | arl3b,arl3b,arl3a | 3 |
| arl3b | arl3b,arl3b,arl3a | 3 |
| arl4aa | arl4d,arl4ab,arl4cb,arl4cb | 8 |
| arl4ab | arl4d,arl4ab,arl4cb,arl4cb | 8 |
| arl4ab | arl4d,arl4ab,arl4cb,arl4cb | 8 |
| arl4ca | arl4d,arl4ab,arl4cb,arl4cb | 8 |
| arl4cb | arl4d,arl4ab,arl4cb,arl4cb | 8 |
| arl4cb | arl4d,arl4ab,arl4cb,arl4cb | 8 |
| arl4d | arl4d,arl4ab,arl4cb,arl4cb | 8 |
| arl4d | arl4d,arl4ab,arl4cb,arl4cb | 8 |
| arl5a | arl5a,arl5c,arl5a | 3 |
| arl5a | arl5a,arl5c,arl5a | 3 |
| arl5c | arl5a,arl5c,arl5a | 3 |
| arl6ip5a | arl6ip5a,arl6ip5a,arl6ip5b | 3 |
| arl6ip5a | arl6ip5a,arl6ip5a,arl6ip5b | 3 |
| arl6ip5b | arl6ip5a,arl6ip5a,arl6ip5b | 3 |
| arl8a | arl8ba,arl8,arl8bb,arl8a | 4 |
| arl8ba | arl8ba,arl8,arl8bb,arl8a | 4 |
| arl8bb | arl8ba,arl8,arl8bb,arl8a | 4 |
| arntl1a | arntl1b,arntl1b,arntl1a | 3 |
| arntl1b | arntl1b,arntl1b,arntl1a | 3 |
| arntl1b | arntl1b,arntl1b,arntl1a | 3 |
| arpc1a | arpc1a,arpc1b | 2 |
| arpc1b | arpc1a,arpc1b | 2 |
| arpc5a | arpc5a,arpc5b | 2 |
| arpc5b | arpc5a,arpc5b | 2 |
| arpp19a | arpp19a,arpp19a,arpp19b | 3 |
| arpp19a | arpp19a,arpp19a,arpp19b | 3 |
| arpp19b | arpp19a,arpp19a,arpp19b | 3 |
| arr3a | arr3a,arr3a,arr3b | 3 |
| arr3a | arr3a,arr3a,arr3b | 3 |
| arr3b | arr3a,arr3a,arr3b | 3 |
| arrb2a | arrb2b,arrb2a | 2 |
| arrb2b | arrb2b,arrb2a | 2 |
| arrdc1a | arrdc1a,arrdc1b | 2 |
| arrdc1b | arrdc1a,arrdc1b | 2 |
| arrdc3a | arrdc3a,arrdc3b | 2 |
| arrdc3b | arrdc3a,arrdc3b | 2 |
| as3mt | as3mt | 1 |
| asah1a | asah1a,asah1b,asah1b,as | 4 |
| asah1a | asah1a,asah1b,asah1b,as | 4 |
| asah1b | asah1a,asah1b,asah1b,as | 4 |
| asah1b | asah1a,asah1b,asah1b,as | 4 |
| asap1a | asap1a,asap1b | 2 |
| asap1b | asap1a,asap1b | 2 |
| asap2a | asap2a,asap2b | 2 |
| asap2b | asap2a,asap2b | 2 |

| | | |
|---|---|---|
| asb12a | asb12b,asb12b,asb12a,as | 5 |
| asb12a | asb12b,asb12b,asb12a,as | 5 |
| asb12b | asb12b,asb12b,asb12a,as | 5 |
| asb12b | asb12b,asb12b,asb12a,as | 5 |
| asb12b | asb12b,asb12b,asb12a,as | 5 |
| asb13b | asb13b | 1 |
| asb14a | asb14b,asb14a,asb14b | 3 |
| asb14b | asb14b,asb14a,asb14b | 3 |
| asb14b | asb14b,asb14a,asb14b | 3 |
| asb15a | asb15b,asb15a | 2 |
| asb15b | asb15b,asb15a | 2 |
| asb2b | asb2b | 1 |
| asb5a | asb5a,asb5b,asb5a | 3 |
| asb5a | asb5a,asb5b,asb5a | 3 |
| asb5b | asb5a,asb5b,asb5a | 3 |
| ascl1a | ascl1a,ascl1b,ascl1b | 3 |
| ascl1b | ascl1a,ascl1b,ascl1b | 3 |
| ascl1b | ascl1a,ascl1b,ascl1b | 3 |
| asf1ba | asf1bb,asf1ba | 2 |
| asf1bb | asf1bb,asf1ba | 2 |
| asic1a | asic1a,asic1b,asic1c | 3 |
| asic1b | asic1a,asic1b,asic1c | 3 |
| asic1c | asic1a,asic1b,asic1c | 3 |
| asic4a | asic4b,asic4a | 2 |
| asic4b | asic4b,asic4a | 2 |
| asip2b | asip2b | 1 |
| aste1a | aste1a | 1 |
| atad1a | atad1a,atad1b | 2 |
| atad1b | atad1a,atad1b | 2 |
| atad2b | atad2,atad2b | 2 |
| atad5a | atad5b,atad5b,atad5a,ata | 4 |
| atad5b | atad5b,atad5b,atad5a,ata | 4 |
| atad5b | atad5b,atad5b,atad5a,ata | 4 |
| atad5b | atad5b,atad5b,atad5a,ata | 4 |
| atf4a | atf4b,atf4b,atf4b,atf4a | 4 |
| atf4b | atf4b,atf4b,atf4b,atf4a | 4 |
| atf4b | atf4b,atf4b,atf4b,atf4a | 4 |
| atf4b | atf4b,atf4b,atf4b,atf4a | 4 |
| atf5a | atf5a,atf5b | 2 |
| atf5b | atf5a,atf5b | 2 |
| atf7a | atf7b,atf7b,atf7ip,atf7b,a | 5 |
| atf7b | atf7b,atf7b,atf7ip,atf7b,a | 5 |
| atf7b | atf7b,atf7b,atf7ip,atf7b,a | 5 |
| atf7b | atf7b,atf7b,atf7ip,atf7b,a | 5 |
| atf7ip | atf7b,atf7b,atf7ip,atf7b,a | 5 |
| atg2b | atg2b | 1 |
| atg4a | atg4da,atg4db,atg4b,atg4 | 5 |
| atg4b | atg4da,atg4db,atg4b,atg4 | 5 |
| atg4c | atg4da,atg4db,atg4b,atg4 | 5 |
| atg4da | atg4da,atg4db,atg4b,atg4 | 5 |

| | | |
|---|---|---|
| atg4db | atg4da,atg4db,atg4b,atg4 | 5 |
| atg9a | atg9a,atg9b | 2 |
| atg9b | atg9a,atg9b | 2 |
| atoh1a | atoh1b,atoh1a,atoh1c | 3 |
| atoh1b | atoh1b,atoh1a,atoh1c | 3 |
| atoh1c | atoh1b,atoh1a,atoh1c | 3 |
| atp10a | atp10a,atp10a,atp10d,atp | 4 |
| atp10a | atp10a,atp10a,atp10d,atp | 4 |
| atp10b | atp10a,atp10a,atp10d,atp | 4 |
| atp10d | atp10a,atp10a,atp10d,atp | 4 |
| atp11a | atp11c,atp11a,atp11c | 3 |
| atp11c | atp11c,atp11a,atp11c | 3 |
| atp11c | atp11c,atp11a,atp11c | 3 |
| atp1a1b | atp1a1b,atp1a1b | 2 |
| atp1a1b | atp1a1b,atp1a1b | 2 |
| atp1a2a | atp1a2a,atp1a2a,atp1a2a | 3 |
| atp1a2a | atp1a2a,atp1a2a,atp1a2a | 3 |
| atp1a2a | atp1a2a,atp1a2a,atp1a2a | 3 |
| atp1a3a | atp1a3b,atp1a3a | 2 |
| atp1a3b | atp1a3b,atp1a3a | 2 |
| atp1b1a | atp1b1b,atp1b1a | 2 |
| atp1b1b | atp1b1b,atp1b1a | 2 |
| atp1b2a | atp1b2b,atp1b2b,atp1b2 | 3 |
| atp1b2b | atp1b2b,atp1b2b,atp1b2 | 3 |
| atp1b2b | atp1b2b,atp1b2b,atp1b2 | 3 |
| atp1b3a | atp1b3a,atp1b3b | 2 |
| atp1b3b | atp1b3a,atp1b3b | 2 |
| atp2a2a | atp2a2a,atp2a2b | 2 |
| atp2a2b | atp2a2a,atp2a2b | 2 |
| atp2b1a | atp2b1a,atp2b1b | 2 |
| atp2b1b | atp2b1a,atp2b1b | 2 |
| atp2b3a | atp2b3b,atp2b3a,atp2b3b | 3 |
| atp2b3b | atp2b3b,atp2b3a,atp2b3b | 3 |
| atp2b3b | atp2b3b,atp2b3a,atp2b3b | 3 |
| atp5f1b | atp5f1c,atp5f1e,atp5f1b, | 4 |
| atp5f1c | atp5f1c,atp5f1e,atp5f1b, | 4 |
| atp5f1d | atp5f1c,atp5f1e,atp5f1b, | 4 |
| atp5f1e | atp5f1c,atp5f1e,atp5f1b, | 4 |
| atp5if1a | atp5if1b,atp5if1b,atp5if1 | 3 |
| atp5if1b | atp5if1b,atp5if1b,atp5if1 | 3 |
| atp5if1b | atp5if1b,atp5if1b,atp5if1 | 3 |
| atp5mc3a | atp5mc3b,atp5mc3a | 2 |
| atp5mc3b | atp5mc3b,atp5mc3a | 2 |
| atp5md | atp5meb,atp5pf,atp5md, | 9 |
| atp5mea | atp5meb,atp5pf,atp5md, | 9 |
| atp5meb | atp5meb,atp5pf,atp5md, | 9 |
| atp5meb | atp5meb,atp5pf,atp5md, | 9 |
| atp5mf | atp5meb,atp5pf,atp5md, | 9 |
| atp5pb | atp5meb,atp5pf,atp5md, | 9 |
| atp5pd | atp5meb,atp5pf,atp5md, | 9 |

| | | |
|---|---|---|
| atp5pf | atp5meb,atp5pf,atp5md, | 9 |
| atp5po | atp5meb,atp5pf,atp5md, | 9 |
| atp6ap1a | atp6ap1a,atp6ap1b | 2 |
| atp6ap1b | atp6ap1a,atp6ap1b | 2 |
| atp6v0a1a | atp6v0a1a,atp6v0a1b | 2 |
| atp6v0a1b | atp6v0a1a,atp6v0a1b | 2 |
| atp6v0a2a | atp6v0a2b,atp6v0a2b,atp | 3 |
| atp6v0a2b | atp6v0a2b,atp6v0a2b,atp | 3 |
| atp6v0a2b | atp6v0a2b,atp6v0a2b,atp | 3 |
| atp6v0b | atp6v0b,atp6v0ca,atp6v0 | 3 |
| atp6v0ca | atp6v0b,atp6v0ca,atp6v0 | 3 |
| atp6v0cb | atp6v0b,atp6v0ca,atp6v0 | 3 |
| atp6v1aa | atp6v1ba,atp6v1ab,atp6v | 9 |
| atp6v1aa | atp6v1ba,atp6v1ab,atp6v | 9 |
| atp6v1ab | atp6v1ba,atp6v1ab,atp6v | 9 |
| atp6v1ab | atp6v1ba,atp6v1ab,atp6v | 9 |
| atp6v1ba | atp6v1ba,atp6v1ab,atp6v | 9 |
| atp6v1ba | atp6v1ba,atp6v1ab,atp6v | 9 |
| atp6v1c1a | atp6v1c1b,atp6v1c1a,atp | 3 |
| atp6v1c1b | atp6v1c1b,atp6v1c1a,atp | 3 |
| atp6v1c1b | atp6v1c1b,atp6v1c1a,atp | 3 |
| atp6v1d | atp6v1ba,atp6v1ab,atp6v | 9 |
| atp6v1e1a | atp6v1e1b,atp6v1e1a | 2 |
| atp6v1e1b | atp6v1e1b,atp6v1e1a | 2 |
| atp6v1f | atp6v1ba,atp6v1ab,atp6v | 9 |
| atp6v1h | atp6v1ba,atp6v1ab,atp6v | 9 |
| atp7a | atp7b,atp7a | 2 |
| atp7b | atp7b,atp7a | 2 |
| atp8b5a | atp8b5a | 1 |
| atp9b | atp9b | 1 |
| atrnl1a | atrnl1b,atrnl1a | 2 |
| atrnl1b | atrnl1b,atrnl1a | 2 |
| atxn1a | atxn1a,atxn1b,atxn1a | 3 |
| atxn1a | atxn1a,atxn1b,atxn1a | 3 |
| atxn1b | atxn1a,atxn1b,atxn1a | 3 |
| atxn7l2a | atxn7l2a,atxn7l2b,atxn7l | 3 |
| atxn7l2a | atxn7l2a,atxn7l2b,atxn7l | 3 |
| atxn7l2b | atxn7l2a,atxn7l2b,atxn7l | 3 |
| auts2a | auts2a,auts2b | 2 |
| auts2b | auts2a,auts2b | 2 |
| avpr1aa | avpr1ab,avpr1aa | 2 |
| avpr1ab | avpr1ab,avpr1aa | 2 |
| avpr2aa | avpr2ab,avpr2aa | 2 |
| avpr2ab | avpr2ab,avpr2aa | 2 |
| azin1a | azin1b,azin1a,azin1b | 3 |
| azin1b | azin1b,azin1a,azin1b | 3 |
| azin1b | azin1b,azin1a,azin1b | 3 |
| b2m | b2m | 1 |
| b3galt1a | b3galt1a | 1 |
| b3gat1a | b3gat1a,b3gat1b,b3gat1a | 3 |

| | | |
|---|---|---|
| b3gat1a | b3gat1a,b3gat1b,b3gat1a | 3 |
| b3gat1b | b3gat1a,b3gat1b,b3gat1a | 3 |
| b3gnt2a | b3gnt2a,b3gnt2b | 2 |
| b3gnt2b | b3gnt2a,b3gnt2b | 2 |
| b3gnt5a | b3gnt5b,b3gnt5a | 2 |
| b3gnt5b | b3gnt5b,b3gnt5a | 2 |
| b4galnt1a | b4galnt1a,b4galnt1b,b4ga | 3 |
| b4galnt1a | b4galnt1a,b4galnt1b,b4ga | 3 |
| b4galnt1b | b4galnt1a,b4galnt1b,b4ga | 3 |
| b4galnt3a | b4galnt3a,b4galnt3b | 2 |
| b4galnt3b | b4galnt3a,b4galnt3b | 2 |
| b4galnt4a | b4galnt4a,b4galnt4b,b4ga | 3 |
| b4galnt4a | b4galnt4a,b4galnt4b,b4ga | 3 |
| b4galnt4b | b4galnt4a,b4galnt4b,b4ga | 3 |
| bach1a | bach1b,bach1a | 2 |
| bach1b | bach1b,bach1a | 2 |
| bach2a | bach2a,bach2b | 2 |
| bach2b | bach2a,bach2b | 2 |
| bahcc1a | bahcc1a,bahcc1b | 2 |
| bahcc1b | bahcc1a,bahcc1b | 2 |
| baiap2a | baiap2b,baiap2b,baiap2a | 5 |
| baiap2b | baiap2b,baiap2b,baiap2a | 5 |
| baiap2b | baiap2b,baiap2b,baiap2a | 5 |
| baiap2b | baiap2b,baiap2b,baiap2a | 5 |
| baiap2b | baiap2b,baiap2b,baiap2a | 5 |
| baiap2l1a | baiap2l1b,baiap2l1a | 2 |
| baiap2l1b | baiap2l1b,baiap2l1a | 2 |
| baiap2l2a | baiap2l2a,baiap2l2b | 2 |
| baiap2l2b | baiap2l2a,baiap2l2b | 2 |
| barhl1a | barhl1b,barhl1a,barhl1a,b | 5 |
| barhl1a | barhl1b,barhl1a,barhl1a,b | 5 |
| barhl1a | barhl1b,barhl1a,barhl1a,b | 5 |
| barhl1b | barhl1b,barhl1a,barhl1a,b | 5 |
| barhl1b | barhl1b,barhl1a,barhl1a,b | 5 |
| baz1a | baz1b,baz1a,baz1a,baz1b | 4 |
| baz1a | baz1b,baz1a,baz1a,baz1b | 4 |
| baz1b | baz1b,baz1a,baz1a,baz1b | 4 |
| baz1b | baz1b,baz1a,baz1a,baz1b | 4 |
| baz2a | baz2ba,baz2a | 2 |
| baz2ba | baz2ba,baz2a | 2 |
| bcdin3d | bcdin3d | 1 |
| bcl11aa | bcl11ab,bcl11aa,bcl11ba | 3 |
| bcl11ab | bcl11ab,bcl11aa,bcl11ba | 3 |
| bcl11ba | bcl11ab,bcl11aa,bcl11ba | 3 |
| bcl2a | bcl2b,bcl2a | 2 |
| bcl2b | bcl2b,bcl2a | 2 |
| bcl6aa | bcl6aa,bcl6aa,bcl6aa,bcl6 | 6 |
| bcl6aa | bcl6aa,bcl6aa,bcl6aa,bcl6 | 6 |
| bcl6aa | bcl6aa,bcl6aa,bcl6aa,bcl6 | 6 |
| bcl6aa | bcl6aa,bcl6aa,bcl6aa,bcl6 | 6 |

| | | |
|---|---|---|
| bcl6ab | bcl6aa,bcl6aa,bcl6aa,bcl6 | 6 |
| bcl6b | bcl6aa,bcl6aa,bcl6aa,bcl6 | 6 |
| bcl7a | bcl7bb,bcl7a,bcl7a,bcl7ba | 5 |
| bcl7a | bcl7bb,bcl7a,bcl7a,bcl7ba | 5 |
| bcl7ba | bcl7bb,bcl7a,bcl7a,bcl7ba | 5 |
| bcl7bb | bcl7bb,bcl7a,bcl7a,bcl7ba | 5 |
| bcl7bb | bcl7bb,bcl7a,bcl7a,bcl7ba | 5 |
| bco2a | bco2a,bco2b | 2 |
| bco2b | bco2a,bco2b | 2 |
| bicc1a | bicc1b,bicc1a | 2 |
| bicc1b | bicc1b,bicc1a | 2 |
| bicd1a | bicd1a | 1 |
| bin1a | bin1b,bin1a | 2 |
| bin1b | bin1b,bin1a | 2 |
| bin2a | bin2b,bin2a | 2 |
| bin2b | bin2b,bin2a | 2 |
| birc5a | birc5b,birc5a,birc5b | 3 |
| birc5b | birc5b,birc5a,birc5b | 3 |
| birc5b | birc5b,birc5a,birc5b | 3 |
| bmi1a | bmi1b,bmi1a | 2 |
| bmi1b | bmi1b,bmi1a | 2 |
| bmp1a | bmp1a,bmp1b | 2 |
| bmp1b | bmp1a,bmp1b | 2 |
| bmp2a | bmp2b,bmp2k,bmp2k,bm | 5 |
| bmp2b | bmp2b,bmp2k,bmp2k,bm | 5 |
| bmp2b | bmp2b,bmp2k,bmp2k,bm | 5 |
| bmp2k | bmp2b,bmp2k,bmp2k,bm | 5 |
| bmp2k | bmp2b,bmp2k,bmp2k,bm | 5 |
| bmp7a | bmp7a,bmp7b | 2 |
| bmp7b | bmp7a,bmp7b | 2 |
| bmp8a | bmp8a | 1 |
| bmpr1aa | bmpr1aa,bmpr1ba,bmpr1 | 5 |
| bmpr1aa | bmpr1aa,bmpr1ba,bmpr1 | 5 |
| bmpr1ab | bmpr1aa,bmpr1ba,bmpr1 | 5 |
| bmpr1ba | bmpr1aa,bmpr1ba,bmpr1 | 5 |
| bmpr1bb | bmpr1aa,bmpr1ba,bmpr1 | 5 |
| bmpr2a | bmpr2a,bmpr2b | 2 |
| bmpr2b | bmpr2a,bmpr2b | 2 |
| bnip1a | bnip1a,bnip1b | 2 |
| bnip1b | bnip1a,bnip1b | 2 |
| brd1a | brd1a,brd1a,brd1b | 3 |
| brd1a | brd1a,brd1a,brd1b | 3 |
| brd1b | brd1a,brd1a,brd1b | 3 |
| brd2a | brd2b,brd2a | 2 |
| brd2b | brd2b,brd2a | 2 |
| brd3a | brd3b,brd3a | 2 |
| brd3b | brd3b,brd3a | 2 |
| brf1a | brf1b,brf1a,brf1b | 3 |
| brf1b | brf1b,brf1a,brf1b | 3 |
| brf1b | brf1b,brf1a,brf1b | 3 |

| | | |
|---|---|---|
| bri3bp | bri3,bri3bp | 2 |
| brinp3b | brinp3b | 1 |
| brpf3a | brpf3a,brpf3b,brpf3a | 3 |
| brpf3a | brpf3a,brpf3b,brpf3a | 3 |
| brpf3b | brpf3a,brpf3b,brpf3a | 3 |
| brsk2b | brsk2b | 1 |
| btbd10a | btbd10b,btbd10a | 2 |
| btbd10b | btbd10b,btbd10a | 2 |
| btbd11a | btbd11a,btbd11b | 2 |
| btbd11b | btbd11a,btbd11b | 2 |
| btbd17a | btbd17b,btbd17b,btbd17 | 3 |
| btbd17b | btbd17b,btbd17b,btbd17 | 3 |
| btbd17b | btbd17b,btbd17b,btbd17 | 3 |
| btbd2a | btbd2a,btbd2b,btbd2a | 3 |
| btbd2a | btbd2a,btbd2b,btbd2a | 3 |
| btbd2b | btbd2a,btbd2b,btbd2a | 3 |
| btbd3a | btbd3a,btbd3b | 2 |
| btbd3b | btbd3a,btbd3b | 2 |
| btbd6a | btbd6b,btbd6a,btbd6b | 3 |
| btbd6b | btbd6b,btbd6a,btbd6b | 3 |
| btbd6b | btbd6b,btbd6a,btbd6b | 3 |
| bub1ba | bub1bb,bub1,bub1ba | 3 |
| bub1bb | bub1bb,bub1,bub1ba | 3 |
| bzw1a | bzw1b,bzw1a | 2 |
| bzw1b | bzw1b,bzw1a | 2 |
| c1d | c1qc,c1qa,c1qbp,c1d,c1r, | 9 |
| c1galt1a | c1galt1a,c1galt1b,c1galt1 | 4 |
| c1galt1a | c1galt1a,c1galt1b,c1galt1 | 4 |
| c1galt1b | c1galt1a,c1galt1b,c1galt1 | 4 |
| c1galt1b | c1galt1a,c1galt1b,c1galt1 | 4 |
| c1qa | c1qc,c1qa,c1qbp,c1d,c1r, | 9 |
| c1qa | c1qc,c1qa,c1qbp,c1d,c1r, | 9 |
| c1qb | c1qc,c1qa,c1qbp,c1d,c1r, | 9 |
| c1qbp | c1qc,c1qa,c1qbp,c1d,c1r, | 9 |
| c1qc | c1qc,c1qa,c1qbp,c1d,c1r, | 9 |
| c1qc | c1qc,c1qa,c1qbp,c1d,c1r, | 9 |
| c1ql3a | c1ql3b,c1ql3b,c1ql3a | 3 |
| c1ql3b | c1ql3b,c1ql3b,c1ql3a | 3 |
| c1ql3b | c1ql3b,c1ql3b,c1ql3a | 3 |
| c1ql4a | c1ql4b,c1ql4a | 2 |
| c1ql4b | c1ql4b,c1ql4a | 2 |
| c1qtnf6a | c1qtnf6a,c1qtnf6b | 2 |
| c1qtnf6b | c1qtnf6a,c1qtnf6b | 2 |
| c1r | c1qc,c1qa,c1qbp,c1d,c1r, | 9 |
| c1s | c1qc,c1qa,c1qbp,c1d,c1r, | 9 |
| c2cd4a | c2cd4a | 1 |
| c4b | c4,c4b | 2 |
| c7a | c7b,c7a,c7b | 3 |
| c7b | c7b,c7a,c7b | 3 |
| c7b | c7b,c7a,c7b | 3 |

| | | |
|---|---|---|
| c8a | c8b,c8a,c8g | 3 |
| c8b | c8b,c8a,c8g | 3 |
| c8g | c8b,c8a,c8g | 3 |
| ca10a | ca10b,ca10a | 2 |
| ca10b | ca10b,ca10a | 2 |
| ca15a | ca15b,ca15a,ca15c | 3 |
| ca15b | ca15b,ca15a,ca15c | 3 |
| ca15c | ca15b,ca15a,ca15c | 3 |
| ca16b | ca16b | 1 |
| ca4a | ca4a,ca4b,ca4c | 3 |
| ca4b | ca4a,ca4b,ca4c | 3 |
| ca4c | ca4a,ca4b,ca4c | 3 |
| ca5a | ca5a,ca5a | 2 |
| ca5a | ca5a,ca5a | 2 |
| cables2a | cables2b,cables2a | 2 |
| cables2b | cables2b,cables2a | 2 |
| cabp1a | cabp1a,cabp1b | 2 |
| cabp1b | cabp1a,cabp1b | 2 |
| cabp2a | cabp2a,cabp2b | 2 |
| cabp2b | cabp2a,cabp2b | 2 |
| cabp5a | cabp5b,cabp5b,cabp5a | 3 |
| cabp5b | cabp5b,cabp5b,cabp5a | 3 |
| cabp5b | cabp5b,cabp5b,cabp5a | 3 |
| cabp7b | cabp7b | 1 |
| cacna1aa | cacna1bb,cacna1fb,cacna | 24 |
| cacna1ab | cacna1bb,cacna1fb,cacna | 24 |
| cacna1ba | cacna1bb,cacna1fb,cacna | 24 |
| cacna1bb | cacna1bb,cacna1fb,cacna | 24 |
| cacna1bb | cacna1bb,cacna1fb,cacna | 24 |
| cacna1c | cacna1bb,cacna1fb,cacna | 24 |
| cacna1da | cacna1bb,cacna1fb,cacna | 24 |
| cacna1db | cacna1bb,cacna1fb,cacna | 24 |
| cacna1db | cacna1bb,cacna1fb,cacna | 24 |
| cacna1db | cacna1bb,cacna1fb,cacna | 24 |
| cacna1db | cacna1bb,cacna1fb,cacna | 24 |
| cacna1ea | cacna1bb,cacna1fb,cacna | 24 |
| cacna1eb | cacna1bb,cacna1fb,cacna | 24 |
| cacna1fa | cacna1bb,cacna1fb,cacna | 24 |
| cacna1fa | cacna1bb,cacna1fb,cacna | 24 |
| cacna1fb | cacna1bb,cacna1fb,cacna | 24 |
| cacna1fb | cacna1bb,cacna1fb,cacna | 24 |
| cacna1g | cacna1bb,cacna1fb,cacna | 24 |
| cacna1ha | cacna1bb,cacna1fb,cacna | 24 |
| cacna1hb | cacna1bb,cacna1fb,cacna | 24 |
| cacna1ia | cacna1bb,cacna1fb,cacna | 24 |
| cacna1ib | cacna1bb,cacna1fb,cacna | 24 |
| cacna1sa | cacna1bb,cacna1fb,cacna | 24 |
| cacna1sb | cacna1bb,cacna1fb,cacna | 24 |
| cacna2d1a | cacna2d1a | 1 |
| cacna2d2a | cacna2d2a,cacna2d2b | 2 |

| | | |
|---|---|---|
| cacna2d2b | cacna2d2a,cacna2d2b | 2 |
| cacna2d4a | cacna2d4a,cacna2d4a,ca | 3 |
| cacna2d4a | cacna2d4a,cacna2d4a,ca | 3 |
| cacna2d4b | cacna2d4a,cacna2d4a,ca | 3 |
| cacnb2a | cacnb2b,cacnb2a | 2 |
| cacnb2b | cacnb2b,cacnb2a | 2 |
| cacnb3a | cacnb3a,cacnb3a,cacnb3 | 4 |
| cacnb3a | cacnb3a,cacnb3a,cacnb3 | 4 |
| cacnb3a | cacnb3a,cacnb3a,cacnb3 | 4 |
| cacnb3b | cacnb3a,cacnb3a,cacnb3 | 4 |
| cacnb4a | cacnb4b,cacnb4a | 2 |
| cacnb4b | cacnb4b,cacnb4a | 2 |
| cacng1a | cacng1b,cacng1a | 2 |
| cacng1b | cacng1b,cacng1a | 2 |
| cacng2a | cacng2b,cacng2a | 2 |
| cacng2b | cacng2b,cacng2a | 2 |
| cacng3a | cacng3a,cacng3b | 2 |
| cacng3b | cacng3a,cacng3b | 2 |
| cacng4a | cacng4b,cacng4a | 2 |
| cacng4b | cacng4b,cacng4a | 2 |
| cacng5a | cacng5b,cacng5a | 2 |
| cacng5b | cacng5b,cacng5a | 2 |
| cacng6a | cacng6b,cacng6a | 2 |
| cacng6b | cacng6b,cacng6a | 2 |
| cacng7a | cacng7b,cacng7a | 2 |
| cacng7b | cacng7b,cacng7a | 2 |
| cacng8a | cacng8b,cacng8a | 2 |
| cacng8b | cacng8b,cacng8a | 2 |
| cadm1a | cadm1b,cadm1a | 2 |
| cadm1b | cadm1b,cadm1a | 2 |
| cadm2a | cadm2a,cadm2a,cadm2b | 3 |
| cadm2a | cadm2a,cadm2a,cadm2b | 3 |
| cadm2b | cadm2a,cadm2a,cadm2b | 3 |
| calb2a | calb2a,calb2b,calb2a | 3 |
| calb2a | calb2a,calb2b,calb2a | 3 |
| calb2b | calb2a,calb2b,calb2a | 3 |
| calcoco1a | calcoco1b,calcoco1a | 2 |
| calcoco1b | calcoco1b,calcoco1a | 2 |
| cald1a | cald1a,cald1a,cald1b | 3 |
| cald1a | cald1a,cald1a,cald1b | 3 |
| cald1b | cald1a,cald1a,cald1b | 3 |
| calm1a | calm1a | 1 |
| calm2a | calm2a,calm2b,calm2a | 3 |
| calm2a | calm2a,calm2b,calm2a | 3 |
| calm2b | calm2a,calm2b,calm2a | 3 |
| calm3a | calm3a,calm3a,calm3b | 3 |
| calm3a | calm3a,calm3a,calm3b | 3 |
| calm3b | calm3a,calm3a,calm3b | 3 |
| calml4a | calml4a,calml4b | 2 |
| calml4b | calml4a,calml4b | 2 |

| | | |
|---|---|---|
| calr3a | calr3b,calr3a | 2 |
| calr3b | calr3b,calr3a | 2 |
| camk1a | camk1da,camk1a,camk1c | 8 |
| camk1a | camk1da,camk1a,camk1c | 8 |
| camk1b | camk1da,camk1a,camk1c | 8 |
| camk1da | camk1da,camk1a,camk1c | 8 |
| camk1da | camk1da,camk1a,camk1c | 8 |
| camk1db | camk1da,camk1a,camk1c | 8 |
| camk1ga | camk1da,camk1a,camk1c | 8 |
| camk1gb | camk1da,camk1a,camk1c | 8 |
| camk2a | camk2a | 1 |
| camk2n1a | camk2n1a | 1 |
| camkk1a | camkk1b,camkk1a,camkk | 4 |
| camkk1a | camkk1b,camkk1a,camkk | 4 |
| camkk1b | camkk1b,camkk1a,camkk | 4 |
| camkk1b | camkk1b,camkk1a,camkk | 4 |
| camsap1a | camsap1b,camsap1a | 2 |
| camsap1b | camsap1b,camsap1a | 2 |
| camsap2a | camsap2a,camsap2b,cam | 3 |
| camsap2a | camsap2a,camsap2b,cam | 3 |
| camsap2b | camsap2a,camsap2b,cam | 3 |
| camta1a | camta1a,camta1b,camta1 | 4 |
| camta1a | camta1a,camta1b,camta1 | 4 |
| camta1b | camta1a,camta1b,camta1 | 4 |
| cant1a | cant1a,cant1b | 2 |
| cant1b | cant1a,cant1b | 2 |
| capn1a | capn1b,capn1a,capn1b | 3 |
| capn1b | capn1b,capn1a,capn1b | 3 |
| capn1b | capn1b,capn1a,capn1b | 3 |
| capn2a | capn2a,capn2b | 2 |
| capn2b | capn2a,capn2b | 2 |
| capn3a | capn3b,capn3a,capn3b | 3 |
| capn3b | capn3b,capn3a,capn3b | 3 |
| capn3b | capn3b,capn3a,capn3b | 3 |
| capn5a | capn5a,capn5b | 2 |
| capn5b | capn5a,capn5b | 2 |
| capns1a | capns1a,capns1b | 2 |
| capns1b | capns1a,capns1b | 2 |
| caprin1a | caprin1a,caprin1a,caprin1 | 3 |
| caprin1a | caprin1a,caprin1a,caprin1 | 3 |
| caprin1b | caprin1a,caprin1a,caprin1 | 3 |
| capza1a | capza1b,capza1b,capza1a | 4 |
| capza1b | capza1b,capza1b,capza1a | 4 |
| capza1b | capza1b,capza1b,capza1a | 4 |
| capza1b | capza1b,capza1b,capza1a | 4 |
| casp3a | casp3a,casp3a,casp3b | 3 |
| casp3a | casp3a,casp3a,casp3b | 3 |
| casp3b | casp3a,casp3a,casp3b | 3 |
| casp6a | casp6a,casp6a | 2 |
| casp6a | casp6a,casp6a | 2 |

| | | |
|---|---|---|
| casq1a | casq1a,casq1a,casq1b | 3 |
| casq1a | casq1a,casq1a,casq1b | 3 |
| casq1b | casq1a,casq1a,casq1b | 3 |
| cavin1a | cavin1a,cavin1b | 2 |
| cavin1b | cavin1a,cavin1b | 2 |
| cavin2a | cavin2a,cavin2b,cavin2a | 3 |
| cavin2a | cavin2a,cavin2b,cavin2a | 3 |
| cavin2b | cavin2a,cavin2b,cavin2a | 3 |
| cavin4a | cavin4a,cavin4b | 2 |
| cavin4b | cavin4a,cavin4b | 2 |
| cbln2a | cbln2a,cbln2b | 2 |
| cbln2b | cbln2a,cbln2b | 2 |
| cbx1a | cbx1a,cbx1a,cbx1b | 3 |
| cbx1a | cbx1a,cbx1a,cbx1b | 3 |
| cbx1b | cbx1a,cbx1a,cbx1b | 3 |
| cbx3a | cbx3b,cbx3a | 2 |
| cbx3b | cbx3b,cbx3a | 2 |
| cbx6a | cbx6b,cbx6a | 2 |
| cbx6b | cbx6b,cbx6a | 2 |
| cbx7a | cbx7a,cbx7b | 2 |
| cbx7b | cbx7a,cbx7b | 2 |
| cbx8a | cbx8b,cbx8b,cbx8a,cbx8b | 4 |
| cbx8b | cbx8b,cbx8b,cbx8a,cbx8b | 4 |
| cbx8b | cbx8b,cbx8b,cbx8a,cbx8b | 4 |
| cbx8b | cbx8b,cbx8b,cbx8a,cbx8b | 4 |
| cc2d1b | cc2d1b | 1 |
| cc2d2a | cc2d2a | 1 |
| ccdc102a | ccdc102a | 1 |
| ccdc106a | ccdc106a,ccdc106b | 2 |
| ccdc106b | ccdc106a,ccdc106b | 2 |
| ccdc127a | ccdc127a,ccdc127b | 2 |
| ccdc127b | ccdc127a,ccdc127b | 2 |
| ccdc136b | ccdc136b,ccdc136b | 2 |
| ccdc136b | ccdc136b,ccdc136b | 2 |
| ccdc149a | ccdc149b,ccdc149b,ccdc1 | 3 |
| ccdc149b | ccdc149b,ccdc149b,ccdc1 | 3 |
| ccdc149b | ccdc149b,ccdc149b,ccdc1 | 3 |
| ccdc28a | ccdc28b,ccdc28a,ccdc28a | 3 |
| ccdc28a | ccdc28b,ccdc28a,ccdc28a | 3 |
| ccdc28b | ccdc28b,ccdc28a,ccdc28a | 3 |
| ccdc3a | ccdc3b,ccdc3a,ccdc3a,ccc | 4 |
| ccdc3a | ccdc3b,ccdc3a,ccdc3a,ccc | 4 |
| ccdc3b | ccdc3b,ccdc3a,ccdc3a,ccc | 4 |
| ccdc3b | ccdc3b,ccdc3a,ccdc3a,ccc | 4 |
| ccdc6a | ccdc6a,ccdc6b,ccdc6a | 3 |
| ccdc6a | ccdc6a,ccdc6b,ccdc6a | 3 |
| ccdc6b | ccdc6a,ccdc6b,ccdc6a | 3 |
| ccdc85a | ccdc85a,ccdc85b,ccdc85c | 4 |
| ccdc85b | ccdc85a,ccdc85b,ccdc85c | 4 |
| ccdc85ca | ccdc85a,ccdc85b,ccdc85c | 4 |

| | | |
|---|---|---|
| ccdc85cb | ccdc85a,ccdc85b,ccdc85c | 4 |
| ccdc88aa | ccdc88aa,ccdc88aa,ccdc8 | 4 |
| ccdc88aa | ccdc88aa,ccdc88aa,ccdc8 | 4 |
| ccdc88b | ccdc88aa,ccdc88aa,ccdc8 | 4 |
| ccdc88c | ccdc88aa,ccdc88aa,ccdc8 | 4 |
| ccdc90b | ccdc90b,ccdc90b | 2 |
| ccdc90b | ccdc90b,ccdc90b | 2 |
| ccdc9b | ccdc9,ccdc9b | 2 |
| ccl19b | ccl19b | 1 |
| ccl20b | ccl20b,ccl20b | 2 |
| ccl20b | ccl20b,ccl20b | 2 |
| ccl25a | ccl25a,ccl25b | 2 |
| ccl25b | ccl25a,ccl25b | 2 |
| ccl27a | ccl27a,ccl27b | 2 |
| ccl27b | ccl27a,ccl27b | 2 |
| ccn2a | ccn2a,ccn2b,ccn2a | 3 |
| ccn2a | ccn2a,ccn2b,ccn2a | 3 |
| ccn2b | ccn2a,ccn2b,ccn2a | 3 |
| ccn4a | ccn4b,ccn4a | 2 |
| ccn4b | ccn4b,ccn4a | 2 |
| ccnd2a | ccnd2b,ccnd2a | 2 |
| ccnd2b | ccnd2b,ccnd2a | 2 |
| ccnl1a | ccnl1a,ccnl1b,ccnl1a | 3 |
| ccnl1a | ccnl1a,ccnl1b,ccnl1a | 3 |
| ccnl1b | ccnl1a,ccnl1b,ccnl1a | 3 |
| ccnt2a | ccnt2a,ccnt2b | 2 |
| ccnt2b | ccnt2a,ccnt2b | 2 |
| ccr12a | ccr12a | 1 |
| ccr6a | ccr6b,ccr6a | 2 |
| ccr6b | ccr6b,ccr6a | 2 |
| ccr9a | ccr9a,ccr9b | 2 |
| ccr9b | ccr9a,ccr9b | 2 |
| ccser2a | ccser2b,ccser2a | 2 |
| ccser2b | ccser2b,ccser2a | 2 |
| cct6a | cct6a | 1 |
| cd248a | cd248a,cd248a,cd248b | 3 |
| cd248a | cd248a,cd248a,cd248b | 3 |
| cd248b | cd248a,cd248a,cd248b | 3 |
| cd2ap | cd2ap | 1 |
| cd3eap | cd3eap | 1 |
| cd44a | cd44a,cd44b | 2 |
| cd44b | cd44a,cd44b | 2 |
| cd74a | cd74a,cd74b | 2 |
| cd74b | cd74a,cd74b | 2 |
| cd79a | cd79b,cd79a | 2 |
| cd79b | cd79b,cd79a | 2 |
| cd81a | cd81a,cd81b | 2 |
| cd81b | cd81a,cd81b | 2 |
| cd82a | cd82a,cd82b | 2 |
| cd82b | cd82a,cd82b | 2 |

| | | |
|---|---|---|
| cd8a | cd8b,cd8a | 2 |
| cd8b | cd8b,cd8a | 2 |
| cd9a | cd9b,cd9a | 2 |
| cd9b | cd9b,cd9a | 2 |
| cdc14aa | cdc14aa,cdc14b,cdc14ab | 3 |
| cdc14ab | cdc14aa,cdc14b,cdc14ab | 3 |
| cdc14b | cdc14aa,cdc14b,cdc14ab | 3 |
| cdc25b | cdc25d,cdc25b | 2 |
| cdc25d | cdc25d,cdc25b | 2 |
| cdc34a | cdc34b,cdc34a | 2 |
| cdc34b | cdc34b,cdc34a | 2 |
| cdc42bpaa | cdc42,cdc42bpaa,cdc42b | 4 |
| cdc42bpab | cdc42,cdc42bpaa,cdc42b | 4 |
| cdc42bpb | cdc42,cdc42bpaa,cdc42b | 4 |
| cdc42ep1a | cdc42ep1b,cdc42ep1a | 2 |
| cdc42ep1b | cdc42ep1b,cdc42ep1a | 2 |
| cdc42ep4a | cdc42ep4a,cdc42ep4a,cd | 3 |
| cdc42ep4a | cdc42ep4a,cdc42ep4a,cd | 3 |
| cdc42ep4b | cdc42ep4a,cdc42ep4a,cd | 3 |
| cdca7a | cdca7a,cdca7b | 2 |
| cdca7b | cdca7a,cdca7b | 2 |
| cdcp1a | cdcp1a | 1 |
| cdh10a | cdh10a | 1 |
| cdh12a | cdh12a | 1 |
| cdh18a | cdh18a | 1 |
| cdh24a | cdh24b,cdh24a | 2 |
| cdh24b | cdh24b,cdh24a | 2 |
| cdh7a | cdh7a,cdh7b | 2 |
| cdh7b | cdh7a,cdh7b | 2 |
| cdhr1a | cdhr1a | 1 |
| cdhr5b | cdhr5b | 1 |
| cdk11b | cdk11b | 1 |
| cdk5r1a | cdk5r1a,cdk5r1b | 2 |
| cdk5r1b | cdk5r1a,cdk5r1b | 2 |
| cdk5r2a | cdk5r2a,cdk5r2b | 2 |
| cdk5r2b | cdk5r2a,cdk5r2b | 2 |
| cdkn1a | cdkn1d,cdkn1ba,cdkn1ca | 6 |
| cdkn1ba | cdkn1d,cdkn1ba,cdkn1ca | 6 |
| cdkn1bb | cdkn1d,cdkn1ba,cdkn1ca | 6 |
| cdkn1ca | cdkn1d,cdkn1ba,cdkn1ca | 6 |
| cdkn1cb | cdkn1d,cdkn1ba,cdkn1ca | 6 |
| cdkn1d | cdkn1d,cdkn1ba,cdkn1ca | 6 |
| cdkn2aip | cdkn2d,cdkn2aip,cdkn2c | 3 |
| cdkn2c | cdkn2d,cdkn2aip,cdkn2c | 3 |
| cdkn2d | cdkn2d,cdkn2aip,cdkn2c | 3 |
| cdr2a | cdr2a | 1 |
| cdx1a | cdx1a,cdx1b | 2 |
| cdx1b | cdx1a,cdx1b | 2 |
| celf3a | celf3b,celf3a | 2 |
| celf3b | celf3b,celf3a | 2 |

| | | |
|---|---|---|
| celf5a | celf5a,celf5b,celf5a | 3 |
| celf5a | celf5a,celf5b,celf5a | 3 |
| celf5b | celf5a,celf5b,celf5a | 3 |
| celsr1a | celsr1a,celsr1b | 2 |
| celsr1b | celsr1a,celsr1b | 2 |
| cep170aa | cep170ab,cep170ab,cep1 | 4 |
| cep170ab | cep170ab,cep170ab,cep1 | 4 |
| cep170ab | cep170ab,cep170ab,cep1 | 4 |
| cep170b | cep170ab,cep170ab,cep1 | 4 |
| cept1a | cept1a,cept1b | 2 |
| cept1b | cept1a,cept1b | 2 |
| cers2a | cers2b,cers2a | 2 |
| cers2b | cers2b,cers2a | 2 |
| cers3a | cers3a,cers3b | 2 |
| cers3b | cers3a,cers3b | 2 |
| cers4a | cers4b,cers4a | 2 |
| cers4b | cers4b,cers4a | 2 |
| cert1a | cert1a,cert1a,cert1b | 3 |
| cert1a | cert1a,cert1a,cert1b | 3 |
| cert1b | cert1a,cert1a,cert1b | 3 |
| ch25h | ch25h | 1 |
| chaf1a | chaf1a,chaf1b,chaf1a | 3 |
| chaf1a | chaf1a,chaf1b,chaf1a | 3 |
| chaf1b | chaf1a,chaf1b,chaf1a | 3 |
| chchd3a | chchd3a,chchd3b | 2 |
| chchd3b | chchd3a,chchd3b | 2 |
| chchd4a | chchd4a,chchd4b,chchd4 | 3 |
| chchd4a | chchd4a,chchd4b,chchd4 | 3 |
| chchd4b | chchd4a,chchd4b,chchd4 | 3 |
| chchd6a | chchd6b,chchd6a | 2 |
| chchd6b | chchd6b,chchd6a | 2 |
| chd4a | chd4a,chd4b,chd4a | 3 |
| chd4a | chd4a,chd4b,chd4a | 3 |
| chd4b | chd4a,chd4b,chd4a | 3 |
| chl1a | chl1b,chl1a | 2 |
| chl1b | chl1b,chl1a | 2 |
| chmp1a | chmp1b,chmp1b,chmp1a | 3 |
| chmp1b | chmp1b,chmp1b,chmp1a | 3 |
| chmp1b | chmp1b,chmp1b,chmp1a | 3 |
| chmp2a | chmp2ba,chmp2ba,chmp | 5 |
| chmp2ba | chmp2ba,chmp2ba,chmp | 5 |
| chmp2ba | chmp2ba,chmp2ba,chmp | 5 |
| chmp2ba | chmp2ba,chmp2ba,chmp | 5 |
| chmp2bb | chmp2ba,chmp2ba,chmp | 5 |
| chmp4ba | chmp4c,chmp4ba,chmp4 | 4 |
| chmp4bb | chmp4c,chmp4ba,chmp4 | 4 |
| chmp4c | chmp4c,chmp4ba,chmp4 | 4 |
| chmp4c | chmp4c,chmp4ba,chmp4 | 4 |
| chmp5a | chmp5a,chmp5b | 2 |
| chmp5b | chmp5a,chmp5b | 2 |

| | | |
|---|---|---|
| chmp6a | chmp6a,chmp6b | 2 |
| chmp6b | chmp6a,chmp6b | 2 |
| chordc1a | chordc1b,chordc1a | 2 |
| chordc1b | chordc1b,chordc1a | 2 |
| chrm1a | chrm1a | 1 |
| chrm2a | chrm2b,chrm2a,chrm2b | 3 |
| chrm2b | chrm2b,chrm2a,chrm2b | 3 |
| chrm2b | chrm2b,chrm2a,chrm2b | 3 |
| chrm3a | chrm3b,chrm3a | 2 |
| chrm3b | chrm3b,chrm3a | 2 |
| chrm4a | chrm4a | 1 |
| chrm5a | chrm5a,chrm5b | 2 |
| chrm5b | chrm5a,chrm5b | 2 |
| chrna10a | chrna10a | 1 |
| chrna2a | chrna2a,chrna2b | 2 |
| chrna2b | chrna2a,chrna2b | 2 |
| chrna4a | chrna4a,chrna4b | 2 |
| chrna4b | chrna4a,chrna4b | 2 |
| chrnb3a | chrnb3a,chrnb3a,chrnb3l | 3 |
| chrnb3a | chrnb3a,chrnb3a,chrnb3l | 3 |
| chrnb3b | chrnb3a,chrnb3a,chrnb3l | 3 |
| chrnb5a | chrnb5a,chrnb5a,chrnb5l | 3 |
| chrnb5a | chrnb5a,chrnb5a,chrnb5l | 3 |
| chrnb5b | chrnb5a,chrnb5a,chrnb5l | 3 |
| chst12a | chst12b,chst12b,chst12a | 3 |
| chst12b | chst12b,chst12b,chst12a | 3 |
| chst12b | chst12b,chst12b,chst12a | 3 |
| chst2a | chst2a,chst2b | 2 |
| chst2b | chst2a,chst2b | 2 |
| chst3a | chst3b,chst3a | 2 |
| chst3b | chst3b,chst3a | 2 |
| ciao2a | ciao2b,ciao2b,ciao2a | 3 |
| ciao2b | ciao2b,ciao2b,ciao2a | 3 |
| ciao2b | ciao2b,ciao2b,ciao2a | 3 |
| cip2a | cip2a | 1 |
| cited4a | cited4b,cited4b,cited4a | 3 |
| cited4b | cited4b,cited4b,cited4a | 3 |
| cited4b | cited4b,cited4b,cited4a | 3 |
| ciz1a | ciz1a,ciz1b | 2 |
| ciz1b | ciz1a,ciz1b | 2 |
| ckmt2a | ckmt2a,ckmt2b,ckmt2b,c | 4 |
| ckmt2a | ckmt2a,ckmt2b,ckmt2b,c | 4 |
| ckmt2b | ckmt2a,ckmt2b,ckmt2b,c | 4 |
| ckmt2b | ckmt2a,ckmt2b,ckmt2b,c | 4 |
| cks1b | cks1b | 1 |
| clasp1a | clasp1a | 1 |
| clcn1a | clcn1b,clcn1a | 2 |
| clcn1b | clcn1b,clcn1a | 2 |
| clcn2a | clcn2b,clcn2a,clcn2c | 3 |
| clcn2b | clcn2b,clcn2a,clcn2c | 3 |

| | | |
|---|---|---|
| clcn2c | clcn2b,clcn2a,clcn2c | 3 |
| clcn5a | clcn5b,clcn5a | 2 |
| clcn5b | clcn5b,clcn5a | 2 |
| cldn10a | cldn10a,cldn10a,cldn10b | 3 |
| cldn10a | cldn10a,cldn10a,cldn10b | 3 |
| cldn10b | cldn10a,cldn10a,cldn10b | 3 |
| cldn11a | cldn11a,cldn11b | 2 |
| cldn11b | cldn11a,cldn11b | 2 |
| cldn15a | cldn15b,cldn15a | 2 |
| cldn15b | cldn15b,cldn15a | 2 |
| cldn23a | cldn23a,cldn23b | 2 |
| cldn23b | cldn23a,cldn23b | 2 |
| cldn5a | cldn5a,cldn5b | 2 |
| cldn5b | cldn5a,cldn5b | 2 |
| cldn7a | cldn7b,cldn7a,cldn7a,cldr | 5 |
| cldn7a | cldn7b,cldn7a,cldn7a,cldr | 5 |
| cldn7a | cldn7b,cldn7a,cldn7a,cldr | 5 |
| cldn7b | cldn7b,cldn7a,cldn7a,cldr | 5 |
| cldn7b | cldn7b,cldn7a,cldn7a,cldr | 5 |
| cldnd1a | cldnd1b,cldnd1a | 2 |
| cldnd1b | cldnd1b,cldnd1a | 2 |
| clec11a | clec11a | 1 |
| clec14a | clec14a | 1 |
| clec16a | clec16a | 1 |
| clec19a | clec19a,clec19a | 2 |
| clec19a | clec19a,clec19a | 2 |
| clec3ba | clec3ba,clec3bb | 2 |
| clec3bb | clec3ba,clec3bb | 2 |
| clic5a | clic5a,clic5a,clic5b | 3 |
| clic5a | clic5a,clic5a,clic5b | 3 |
| clic5b | clic5a,clic5a,clic5b | 3 |
| clint1a | clint1a,clint1b | 2 |
| clint1b | clint1a,clint1b | 2 |
| clip1a | clip1a | 1 |
| clk2a | clk2b,clk2a | 2 |
| clk2b | clk2b,clk2a | 2 |
| clk4a | clk4a,clk4b | 2 |
| clk4b | clk4a,clk4b | 2 |
| cln6a | cln6a,cln6b | 2 |
| cln6b | cln6a,cln6b | 2 |
| clns1a | clns1a | 1 |
| cmtm8a | cmtm8b,cmtm8b,cmtm8 | 3 |
| cmtm8b | cmtm8b,cmtm8b,cmtm8 | 3 |
| cmtm8b | cmtm8b,cmtm8b,cmtm8 | 3 |
| cnga1a | cnga1a,cnga1b | 2 |
| cnga1b | cnga1a,cnga1b | 2 |
| cnga2a | cnga2b,cnga2a | 2 |
| cnga2b | cnga2b,cnga2a | 2 |
| cnga3a | cnga3b,cnga3a | 2 |
| cnga3b | cnga3b,cnga3a | 2 |

| | | |
|---|---|---|
| cngb1a | cngb1a | 1 |
| cnksr2a | cnksr2b,cnksr2a | 2 |
| cnksr2b | cnksr2b,cnksr2a | 2 |
| cnn1a | cnn1a,cnn1b | 2 |
| cnn1b | cnn1a,cnn1b | 2 |
| cnn3a | cnn3b,cnn3a | 2 |
| cnn3b | cnn3b,cnn3a | 2 |
| cnnm2a | cnnm2b,cnnm2a | 2 |
| cnnm2b | cnnm2b,cnnm2a | 2 |
| cnnm4a | cnnm4b,cnnm4a | 2 |
| cnnm4b | cnnm4b,cnnm4a | 2 |
| cnot3a | cnot3a,cnot3b | 2 |
| cnot3b | cnot3a,cnot3b | 2 |
| cnot4a | cnot4b,cnot4a | 2 |
| cnot4b | cnot4b,cnot4a | 2 |
| cnot6a | cnot6a,cnot6b,cnot6a | 3 |
| cnot6a | cnot6a,cnot6b,cnot6a | 3 |
| cnot6b | cnot6a,cnot6b,cnot6a | 3 |
| cnrip1a | cnrip1a,cnrip1b | 2 |
| cnrip1b | cnrip1a,cnrip1b | 2 |
| cntn1a | cntn1b,cntn1a | 2 |
| cntn1b | cntn1b,cntn1a | 2 |
| cntn3b | cntn3b | 1 |
| cntnap2a | cntnap2b,cntnap2b,cntna | 4 |
| cntnap2b | cntnap2b,cntnap2b,cntna | 4 |
| cntnap2b | cntnap2b,cntnap2b,cntna | 4 |
| cntnap2b | cntnap2b,cntnap2b,cntna | 4 |
| cntnap5a | cntnap5a,cntnap5b | 2 |
| cntnap5b | cntnap5a,cntnap5b | 2 |
| coa3a | coa3b,coa3a | 2 |
| coa3b | coa3b,coa3a | 2 |
| cobll1a | cobll1a,cobll1b | 2 |
| cobll1b | cobll1a,cobll1b | 2 |
| col10a1a | col10a1a,col10a1b | 2 |
| col10a1b | col10a1a,col10a1b | 2 |
| col11a1a | col11a1a,col11a1b,col11a | 3 |
| col11a1a | col11a1a,col11a1b,col11a | 3 |
| col11a1b | col11a1a,col11a1b,col11a | 3 |
| col12a1a | col12a1a,col12a1b | 2 |
| col12a1b | col12a1a,col12a1b | 2 |
| col14a1a | col14a1a,col14a1b | 2 |
| col14a1b | col14a1a,col14a1b | 2 |
| col15a1b | col15a1b | 1 |
| col17a1a | col17a1a,col17a1b | 2 |
| col17a1b | col17a1a,col17a1b | 2 |
| col18a1a | col18a1b,col18a1a | 2 |
| col18a1b | col18a1b,col18a1a | 2 |
| col1a1a | col1a1a,col1a1a,col1a1b | 3 |
| col1a1a | col1a1a,col1a1a,col1a1b | 3 |
| col1a1b | col1a1a,col1a1a,col1a1b | 3 |

| | | |
|---|---|---|
| col27a1b | col27a1b | 1 |
| col28a1a | col28a1b,col28a1a | 2 |
| col28a1b | col28a1b,col28a1a | 2 |
| col28a2a | col28a2b,col28a2a | 2 |
| col28a2b | col28a2b,col28a2a | 2 |
| col2a1a | col2a1a,col2a1a,col2a1b | 3 |
| col2a1a | col2a1a,col2a1a,col2a1b | 3 |
| col2a1b | col2a1a,col2a1a,col2a1b | 3 |
| col5a2a | col5a2b,col5a2a | 2 |
| col5a2b | col5a2b,col5a2a | 2 |
| col5a3a | col5a3b,col5a3a | 2 |
| col5a3b | col5a3b,col5a3a | 2 |
| col6a4a | col6a4a | 1 |
| col8a1a | col8a1a,col8a1a,col8a1b | 3 |
| col8a1a | col8a1a,col8a1a,col8a1b | 3 |
| col8a1b | col8a1a,col8a1a,col8a1b | 3 |
| col9a1a | col9a1b,col9a1a,col9a1b | 3 |
| col9a1b | col9a1b,col9a1a,col9a1b | 3 |
| col9a1b | col9a1b,col9a1a,col9a1b | 3 |
| cops7a | cops7a,cops7a | 2 |
| cops7a | cops7a,cops7a | 2 |
| coq10b | coq10b | 1 |
| coq8aa | coq8ab,coq8b,coq8ab,co | 4 |
| coq8ab | coq8ab,coq8b,coq8ab,co | 4 |
| coq8ab | coq8ab,coq8b,coq8ab,co | 4 |
| coq8b | coq8ab,coq8b,coq8ab,co | 4 |
| coro1a | coro1a,coro1cb,coro1a,c | 6 |
| coro1a | coro1a,coro1cb,coro1a,c | 6 |
| coro1b | coro1a,coro1cb,coro1a,c | 6 |
| coro1ca | coro1a,coro1cb,coro1a,c | 6 |
| coro1cb | coro1a,coro1cb,coro1a,c | 6 |
| coro1cb | coro1a,coro1cb,coro1a,c | 6 |
| coro2a | coro2a,coro2ba,coro2bb | 3 |
| coro2ba | coro2a,coro2ba,coro2bb | 3 |
| coro2bb | coro2a,coro2ba,coro2bb | 3 |
| cox5aa | cox5ab,cox5aa | 2 |
| cox5ab | cox5ab,cox5aa | 2 |
| cox6c | cox6c | 1 |
| cox7a2a | cox7a2a | 1 |
| cox7b | cox7c,cox7b | 2 |
| cox7c | cox7c,cox7b | 2 |
| cox8a | cox8a,cox8b | 2 |
| cox8b | cox8a,cox8b | 2 |
| cpeb1a | cpeb1b,cpeb1a | 2 |
| cpeb1b | cpeb1b,cpeb1a | 2 |
| cpeb4a | cpeb4a,cpeb4a,cpeb4b | 3 |
| cpeb4a | cpeb4a,cpeb4a,cpeb4b | 3 |
| cpeb4b | cpeb4a,cpeb4a,cpeb4b | 3 |
| cplx3a | cplx3a,cplx3a,cplx3b,cplx | 4 |
| cplx3a | cplx3a,cplx3a,cplx3b,cplx | 4 |

| | | |
|---|---|---|
| cplx3a | cplx3a,cplx3a,cplx3b,cplx | 4 |
| cplx3b | cplx3a,cplx3a,cplx3b,cplx | 4 |
| cplx4a | cplx4c,cplx4b,cplx4a | 3 |
| cplx4b | cplx4c,cplx4b,cplx4a | 3 |
| cplx4c | cplx4c,cplx4b,cplx4a | 3 |
| cpne4a | cpne4a,cpne4b | 2 |
| cpne4b | cpne4a,cpne4b | 2 |
| cpne5a | cpne5a,cpne5a,cpne5b | 3 |
| cpne5a | cpne5a,cpne5a,cpne5b | 3 |
| cpne5b | cpne5a,cpne5a,cpne5b | 3 |
| cpt1aa | cpt1cb,cpt1aa,cpt1b,cpt1 | 4 |
| cpt1ab | cpt1cb,cpt1aa,cpt1b,cpt1 | 4 |
| cpt1b | cpt1cb,cpt1aa,cpt1b,cpt1 | 4 |
| cpt1cb | cpt1cb,cpt1aa,cpt1b,cpt1 | 4 |
| cpxm1a | cpxm1a,cpxm1b | 2 |
| cpxm1b | cpxm1a,cpxm1b | 2 |
| crabp1a | crabp1b,crabp1a | 2 |
| crabp1b | crabp1b,crabp1a | 2 |
| crabp2a | crabp2a,crabp2b | 2 |
| crabp2b | crabp2a,crabp2b | 2 |
| cracr2ab | cracr2ab,cracr2b | 2 |
| cracr2b | cracr2ab,cracr2b | 2 |
| crb2a | crb2b,crb2a | 2 |
| crb2b | crb2b,crb2a | 2 |
| crb3a | crb3b,crb3a | 2 |
| crb3b | crb3b,crb3a | 2 |
| creb1a | creb1a,creb1b | 2 |
| creb1b | creb1a,creb1b | 2 |
| creb3l3a | creb3l3a,creb3l3b,creb3l | 3 |
| creb3l3a | creb3l3a,creb3l3b,creb3l | 3 |
| creb3l3b | creb3l3a,creb3l3b,creb3l | 3 |
| creb5a | creb5b,creb5a | 2 |
| creb5b | creb5b,creb5a | 2 |
| creld1a | creld1b,creld1a | 2 |
| creld1b | creld1b,creld1a | 2 |
| crispld1a | crispld1b,crispld1a | 2 |
| crispld1b | crispld1b,crispld1a | 2 |
| crlf1a | crlf1a,crlf1b | 2 |
| crlf1b | crlf1a,crlf1b | 2 |
| crtac1a | crtac1a,crtac1b | 2 |
| crtac1b | crtac1a,crtac1b | 2 |
| crtc1a | crtc1a,crtc1a,crtc1b | 3 |
| crtc1a | crtc1a,crtc1a,crtc1b | 3 |
| crtc1b | crtc1a,crtc1a,crtc1b | 3 |
| cry1a | cry1a,cry1b | 2 |
| cry1b | cry1a,cry1b | 2 |
| cry3a | cry3a,cry3a,cry3b | 3 |
| cry3a | cry3a,cry3a,cry3b | 3 |
| cry3b | cry3a,cry3a,cry3b | 3 |
| cryba1a | cryba1a,cryba1b | 2 |

| | | |
|---|---|---|
| cryba1b | cryba1a,cryba1b | 2 |
| cryba2a | cryba2a,cryba2b | 2 |
| cryba2b | cryba2a,cryba2b | 2 |
| crybg1a | crybg1b,crybg1a | 2 |
| crybg1b | crybg1b,crybg1a | 2 |
| crygm2a | crygm2f,crygm2c,crygm2 | 6 |
| crygm2b | crygm2f,crygm2c,crygm2 | 6 |
| crygm2c | crygm2f,crygm2c,crygm2 | 6 |
| crygm2e | crygm2f,crygm2c,crygm2 | 6 |
| crygm2f | crygm2f,crygm2c,crygm2 | 6 |
| crygm2f | crygm2f,crygm2c,crygm2 | 6 |
| csdc2a | csdc2a,csdc2a | 2 |
| csdc2a | csdc2a,csdc2a | 2 |
| csf1a | csf1b,csf1a,csf1ra,csf1rb | 4 |
| csf1b | csf1b,csf1a,csf1ra,csf1rb | 4 |
| csf1ra | csf1b,csf1a,csf1ra,csf1rb | 4 |
| csf1rb | csf1b,csf1a,csf1ra,csf1rb | 4 |
| csf2rb | csf2rb,csf2rb,csf2rb | 3 |
| csf2rb | csf2rb,csf2rb,csf2rb | 3 |
| csf2rb | csf2rb,csf2rb,csf2rb | 3 |
| csf3a | csf3a,csf3r,csf3b | 3 |
| csf3b | csf3a,csf3r,csf3b | 3 |
| csf3r | csf3a,csf3r,csf3b | 3 |
| csgalnact1a | csgalnact1b,csgalnact1a | 2 |
| csgalnact1b | csgalnact1b,csgalnact1a | 2 |
| csmd1a | csmd1a | 1 |
| csmd3a | csmd3a,csmd3a | 2 |
| csmd3a | csmd3a,csmd3a | 2 |
| csnk1da | csnk1e,csnk1da,csnk1e,cs | 4 |
| csnk1db | csnk1e,csnk1da,csnk1e,cs | 4 |
| csnk1e | csnk1e,csnk1da,csnk1e,cs | 4 |
| csnk1e | csnk1e,csnk1da,csnk1e,cs | 4 |
| csnk1g2a | csnk1g2b,csnk1g2a,csnk1 | 3 |
| csnk1g2b | csnk1g2b,csnk1g2a,csnk1 | 3 |
| csnk1g2b | csnk1g2b,csnk1g2a,csnk1 | 3 |
| csnk2a2a | csnk2a2a,csnk2a2b | 2 |
| csnk2a2b | csnk2a2a,csnk2a2b | 2 |
| csnk2b | csnk2b | 1 |
| cspg5a | cspg5a,cspg5b | 2 |
| cspg5b | cspg5a,cspg5b | 2 |
| cspp1a | cspp1b,cspp1a | 2 |
| cspp1b | cspp1b,cspp1a | 2 |
| csrnp1a | csrnp1a,csrnp1b | 2 |
| csrnp1b | csrnp1a,csrnp1b | 2 |
| csrp1a | csrp1b,csrp1a | 2 |
| csrp1b | csrp1b,csrp1a | 2 |
| ctbp2a | ctbp2a | 1 |
| ctdnep1a | ctdnep1a,ctdnep1a,ctdne | 3 |
| ctdnep1a | ctdnep1a,ctdnep1a,ctdne | 3 |
| ctdnep1b | ctdnep1a,ctdnep1a,ctdne | 3 |

| | | |
|---|---|---|
| ctdspl2a | ctdspl2b,ctdspl2a | 2 |
| ctdspl2b | ctdspl2b,ctdspl2a | 2 |
| cthrc1a | cthrc1b,cthrc1a | 2 |
| cthrc1b | cthrc1b,cthrc1a | 2 |
| ctnnd2a | ctnnd2a,ctnnd2b,ctnnd2a | 3 |
| ctnnd2a | ctnnd2a,ctnnd2b,ctnnd2a | 3 |
| ctnnd2b | ctnnd2a,ctnnd2b,ctnnd2a | 3 |
| ctps1a | ctps1b,ctps1b,ctps1a | 3 |
| ctps1b | ctps1b,ctps1b,ctps1a | 3 |
| ctps1b | ctps1b,ctps1b,ctps1a | 3 |
| cuedc1a | cuedc1a,cuedc1b | 2 |
| cuedc1b | cuedc1a,cuedc1b | 2 |
| cul1a | cul1a,cul1a,cul1a,cul1b | 4 |
| cul1a | cul1a,cul1a,cul1a,cul1b | 4 |
| cul1a | cul1a,cul1a,cul1a,cul1b | 4 |
| cul1b | cul1a,cul1a,cul1a,cul1b | 4 |
| cul3a | cul3a,cul3b | 2 |
| cul3b | cul3a,cul3b | 2 |
| cul4a | cul4a,cul4b | 2 |
| cul4b | cul4a,cul4b | 2 |
| cul5a | cul5a,cul5a,cul5a,cul5b | 4 |
| cul5a | cul5a,cul5a,cul5a,cul5b | 4 |
| cul5a | cul5a,cul5a,cul5a,cul5b | 4 |
| cul5b | cul5a,cul5a,cul5a,cul5b | 4 |
| cux1a | cux1a,cux1b | 2 |
| cux1b | cux1a,cux1b | 2 |
| cux2b | cux2b | 1 |
| cxcl12a | cxcl12b,cxcl12a,cxcl12b | 3 |
| cxcl12b | cxcl12b,cxcl12a,cxcl12b | 3 |
| cxcl12b | cxcl12b,cxcl12a,cxcl12b | 3 |
| cxcl18b | cxcl18b | 1 |
| cxcl8a | cxcl8a | 1 |
| cxcr4a | cxcr4a,cxcr4b | 2 |
| cxcr4b | cxcr4a,cxcr4b | 2 |
| cxl34c | cxl34c | 1 |
| cxxc1a | cxxc1b,cxxc1a,cxxc1b | 3 |
| cxxc1b | cxxc1b,cxxc1a,cxxc1b | 3 |
| cxxc1b | cxxc1b,cxxc1a,cxxc1b | 3 |
| cxxc5a | cxxc5b,cxxc5a | 2 |
| cxxc5b | cxxc5b,cxxc5a | 2 |
| cyb561a3a | cyb561a3a,cyb561a3b | 2 |
| cyb561a3b | cyb561a3a,cyb561a3b | 2 |
| cyb5a | cyb5b,cyb5a | 2 |
| cyb5b | cyb5b,cyb5a | 2 |
| cyp19a1a | cyp19a1a,cyp19a1a,cyp1 | 3 |
| cyp19a1a | cyp19a1a,cyp19a1a,cyp1 | 3 |
| cyp19a1b | cyp19a1a,cyp19a1a,cyp1 | 3 |
| cyp1a | cyp1a | 1 |
| cyth1a | cyth1a,cyth1a,cyth1b | 3 |
| cyth1a | cyth1a,cyth1a,cyth1b | 3 |

| | | |
|---|---|---|
| cyth1b | cyth1a,cyth1a,cyth1b | 3 |
| cyth3a | cyth3a | 1 |
| cyth4a | cyth4a,cyth4b | 2 |
| cyth4b | cyth4a,cyth4b | 2 |
| d2hgdh | d2hgdh | 1 |
| daam1a | daam1a,daam1b | 2 |
| daam1b | daam1a,daam1b | 2 |
| dab1a | dab1b,dab1a | 2 |
| dab1b | dab1b,dab1a | 2 |
| dab2ipa | dab2ipb,dab2,dab2ipb,da | 4 |
| dab2ipb | dab2ipb,dab2,dab2ipb,da | 4 |
| dab2ipb | dab2ipb,dab2,dab2ipb,da | 4 |
| dact3a | dact3a | 1 |
| dap1b | dap1b,dap1b | 2 |
| dap1b | dap1b,dap1b | 2 |
| dapk2a | dapk2a,dapk2a,dapk2b | 3 |
| dapk2a | dapk2a,dapk2a,dapk2b | 3 |
| dapk2b | dapk2a,dapk2a,dapk2b | 3 |
| dbf4b | dbf4b,dbf4 | 2 |
| dbx1a | dbx1b,dbx1a,dbx1b | 3 |
| dbx1b | dbx1b,dbx1a,dbx1b | 3 |
| dbx1b | dbx1b,dbx1a,dbx1b | 3 |
| dcdc2b | dcdc2b | 1 |
| dchs1a | dchs1b,dchs1a | 2 |
| dchs1b | dchs1b,dchs1a | 2 |
| dclk1a | dclk1a,dclk1b | 2 |
| dclk1b | dclk1a,dclk1b | 2 |
| dclk2a | dclk2a,dclk2b | 2 |
| dclk2b | dclk2a,dclk2b | 2 |
| dclre1a | dclre1c,dclre1c,dclre1c,d | 5 |
| dclre1b | dclre1c,dclre1c,dclre1c,d | 5 |
| dclre1c | dclre1c,dclre1c,dclre1c,d | 5 |
| dclre1c | dclre1c,dclre1c,dclre1c,d | 5 |
| dclre1c | dclre1c,dclre1c,dclre1c,d | 5 |
| dcp1a | dcp1b,dcp1a | 2 |
| dcp1b | dcp1b,dcp1a | 2 |
| dctn1a | dctn1a,dctn1b | 2 |
| dctn1b | dctn1a,dctn1b | 2 |
| dcun1d2a | dcun1d2a,dcun1d2b,dcu | 3 |
| dcun1d2a | dcun1d2a,dcun1d2b,dcu | 3 |
| dcun1d2b | dcun1d2a,dcun1d2b,dcu | 3 |
| ddhd1a | ddhd1b,ddhd1b,ddhd1a, | 4 |
| ddhd1b | ddhd1b,ddhd1b,ddhd1a, | 4 |
| ddhd1b | ddhd1b,ddhd1b,ddhd1a, | 4 |
| ddhd1b | ddhd1b,ddhd1b,ddhd1a, | 4 |
| ddr2a | ddr2b,ddr2a | 2 |
| ddr2b | ddr2b,ddr2a | 2 |
| ddx39aa | ddx39ab,ddx39aa,ddx39a | 4 |
| ddx39ab | ddx39ab,ddx39aa,ddx39a | 4 |
| ddx39ab | ddx39ab,ddx39aa,ddx39a | 4 |

| | | |
|---|---|---|
| ddx39b | ddx39ab,ddx39aa,ddx39a | 4 |
| ddx3xa | ddx3xb,ddx3xa | 2 |
| ddx3xb | ddx3xb,ddx3xa | 2 |
| def6a | def6c,def6a,def6b | 3 |
| def6b | def6c,def6a,def6b | 3 |
| def6c | def6c,def6a,def6b | 3 |
| dennd1a | dennd1a,dennd1b | 2 |
| dennd1b | dennd1a,dennd1b | 2 |
| dennd2c | dennd2c,dennd2da,denn | 3 |
| dennd2da | dennd2c,dennd2da,denn | 3 |
| dennd2db | dennd2c,dennd2da,denn | 3 |
| dennd3a | dennd3b,dennd3a | 2 |
| dennd3b | dennd3b,dennd3a | 2 |
| dennd4a | dennd4a,dennd4c,dennd | 4 |
| dennd4a | dennd4a,dennd4c,dennd | 4 |
| dennd4b | dennd4a,dennd4c,dennd | 4 |
| dennd4c | dennd4a,dennd4c,dennd | 4 |
| dennd5a | dennd5b,dennd5a | 2 |
| dennd5b | dennd5b,dennd5a | 2 |
| dennd6aa | dennd6b,dennd6aa | 2 |
| dennd6b | dennd6b,dennd6aa | 2 |
| depdc1a | depdc1a,depdc1a | 2 |
| depdc1a | depdc1a,depdc1a | 2 |
| depdc7a | depdc7a,depdc7b | 2 |
| depdc7b | depdc7a,depdc7b | 2 |
| desi1a | desi1b,desi1b,desi1a,desi | 4 |
| desi1b | desi1b,desi1b,desi1a,desi | 4 |
| desi1b | desi1b,desi1b,desi1a,desi | 4 |
| desi1b | desi1b,desi1b,desi1a,desi | 4 |
| dgat1a | dgat1b,dgat1a | 2 |
| dgat1b | dgat1b,dgat1a | 2 |
| dhrs11a | dhrs11a,dhrs11a,dhrs11b | 3 |
| dhrs11a | dhrs11a,dhrs11a,dhrs11b | 3 |
| dhrs11b | dhrs11a,dhrs11a,dhrs11b | 3 |
| dhrs3a | dhrs3a,dhrs3b,dhrs3a | 3 |
| dhrs3a | dhrs3a,dhrs3b,dhrs3a | 3 |
| dhrs3b | dhrs3a,dhrs3b,dhrs3a | 3 |
| dhrs7b | dhrs7b,dhrs7ca,dhrs7cb,c | 4 |
| dhrs7ca | dhrs7b,dhrs7ca,dhrs7cb,c | 4 |
| dhrs7cb | dhrs7b,dhrs7ca,dhrs7cb,c | 4 |
| dhx32a | dhx32a,dhx32b | 2 |
| dhx32b | dhx32a,dhx32b | 2 |
| dio3b | dio3b | 1 |
| dip2a | dip2ca,dip2ba,dip2ba,dip | 8 |
| dip2ba | dip2ca,dip2ba,dip2ba,dip | 8 |
| dip2ba | dip2ca,dip2ba,dip2ba,dip | 8 |
| dip2ba | dip2ca,dip2ba,dip2ba,dip | 8 |
| dip2bb | dip2ca,dip2ba,dip2ba,dip | 8 |
| dip2ca | dip2ca,dip2ba,dip2ba,dip | 8 |
| dip2ca | dip2ca,dip2ba,dip2ba,dip | 8 |

| | | |
|---|---|---|
| dip2cb | dip2ca,dip2ba,dip2ba,dip | 8 |
| dipk1aa | dipk1aa,dipk1ab,dipk1b,c | 4 |
| dipk1ab | dipk1aa,dipk1ab,dipk1b,c | 4 |
| dipk1b | dipk1aa,dipk1ab,dipk1b,c | 4 |
| dipk1c | dipk1aa,dipk1ab,dipk1b,c | 4 |
| dipk2aa | dipk2aa,dipk2b,dipk2ab | 3 |
| dipk2ab | dipk2aa,dipk2b,dipk2ab | 3 |
| dipk2b | dipk2aa,dipk2b,dipk2ab | 3 |
| diras1a | diras1b,diras1a | 2 |
| diras1b | diras1b,diras1a | 2 |
| dixdc1a | dixdc1a,dixdc1b | 2 |
| dixdc1b | dixdc1a,dixdc1b | 2 |
| dkk1a | dkk1b,dkk1b,dkk1a | 3 |
| dkk1b | dkk1b,dkk1b,dkk1a | 3 |
| dkk1b | dkk1b,dkk1b,dkk1a | 3 |
| dkk3a | dkk3a,dkk3b | 2 |
| dkk3b | dkk3a,dkk3b | 2 |
| dlg4a | dlg4a,dlg4b,dlg4b | 3 |
| dlg4b | dlg4a,dlg4b,dlg4b | 3 |
| dlg4b | dlg4a,dlg4b,dlg4b | 3 |
| dlg5a | dlg5a | 1 |
| dlgap1a | dlgap1a,dlgap1b | 2 |
| dlgap1b | dlgap1a,dlgap1b | 2 |
| dlgap2a | dlgap2a,dlgap2b,dlgap2b | 3 |
| dlgap2b | dlgap2a,dlgap2b,dlgap2b | 3 |
| dlgap2b | dlgap2a,dlgap2b,dlgap2b | 3 |
| dlgap4a | dlgap4b,dlgap4a | 2 |
| dlgap4b | dlgap4b,dlgap4a | 2 |
| dlx1a | dlx1a | 1 |
| dlx2a | dlx2b,dlx2b,dlx2a | 3 |
| dlx2b | dlx2b,dlx2b,dlx2a | 3 |
| dlx2b | dlx2b,dlx2b,dlx2a | 3 |
| dlx3b | dlx3b | 1 |
| dlx4a | dlx4a,dlx4b | 2 |
| dlx4b | dlx4a,dlx4b | 2 |
| dlx5a | dlx5a | 1 |
| dlx6a | dlx6a | 1 |
| dmbx1a | dmbx1a,dmbx1b | 2 |
| dmbx1b | dmbx1a,dmbx1b | 2 |
| dmrt2a | dmrt2a,dmrt2b,dmrt2b,d | 5 |
| dmrt2a | dmrt2a,dmrt2b,dmrt2b,d | 5 |
| dmrt2b | dmrt2a,dmrt2b,dmrt2b,d | 5 |
| dmrt2b | dmrt2a,dmrt2b,dmrt2b,d | 5 |
| dmrt2b | dmrt2a,dmrt2b,dmrt2b,d | 5 |
| dmrt3a | dmrt3a,dmrt3a | 2 |
| dmrt3a | dmrt3a,dmrt3a | 2 |
| dnai2a | dnai2a,dnai2b | 2 |
| dnai2b | dnai2a,dnai2b | 2 |
| dnaja2a | dnaja2a,dnaja2b | 2 |
| dnaja2b | dnaja2a,dnaja2b | 2 |

| | | |
|---|---|---|
| dnaja3a | dnaja3a,dnaja3a,dnaja3b | 3 |
| dnaja3a | dnaja3a,dnaja3a,dnaja3b | 3 |
| dnaja3b | dnaja3a,dnaja3a,dnaja3b | 3 |
| dnajb12a | dnajb12b,dnajb12a | 2 |
| dnajb12b | dnajb12b,dnajb12a | 2 |
| dnajb1a | dnajb1b,dnajb1a | 2 |
| dnajb1b | dnajb1b,dnajb1a | 2 |
| dnajb6a | dnajb6b,dnajb6a,dnajb6 | 3 |
| dnajb6b | dnajb6b,dnajb6a,dnajb6 | 3 |
| dnajb6b | dnajb6b,dnajb6a,dnajb6 | 3 |
| dnajb9a | dnajb9b,dnajb9a | 2 |
| dnajb9b | dnajb9b,dnajb9a | 2 |
| dnajc11a | dnajc11a,dnajc11b,dnajc | 3 |
| dnajc11a | dnajc11a,dnajc11b,dnajc | 3 |
| dnajc11b | dnajc11a,dnajc11b,dnajc | 3 |
| dnajc30b | dnajc30b | 1 |
| dnajc3a | dnajc3a,dnajc3b,dnajc3a | 3 |
| dnajc3a | dnajc3a,dnajc3b,dnajc3a | 3 |
| dnajc3b | dnajc3a,dnajc3b,dnajc3a | 3 |
| dnajc5aa | dnajc5gb,dnajc5aa,dnajc | 6 |
| dnajc5ab | dnajc5gb,dnajc5aa,dnajc | 6 |
| dnajc5b | dnajc5gb,dnajc5aa,dnajc | 6 |
| dnajc5ga | dnajc5gb,dnajc5aa,dnajc | 6 |
| dnajc5gb | dnajc5gb,dnajc5aa,dnajc | 6 |
| dnajc5gb | dnajc5gb,dnajc5aa,dnajc | 6 |
| dnal4a | dnal4b,dnal4b,dnal4a,dn | 5 |
| dnal4a | dnal4b,dnal4b,dnal4a,dn | 5 |
| dnal4b | dnal4b,dnal4b,dnal4a,dn | 5 |
| dnal4b | dnal4b,dnal4b,dnal4a,dn | 5 |
| dnal4b | dnal4b,dnal4b,dnal4a,dn | 5 |
| dnase2b | dnase2b,dnase2 | 2 |
| dnm1a | dnm1a,dnm1b | 2 |
| dnm1b | dnm1a,dnm1b | 2 |
| dnm2a | dnm2b,dnm2a | 2 |
| dnm2b | dnm2b,dnm2a | 2 |
| dnm3a | dnm3a | 1 |
| dnmt3aa | dnmt3ba,dnmt3ab,dnmt | 3 |
| dnmt3ab | dnmt3ba,dnmt3ab,dnmt | 3 |
| dnmt3ba | dnmt3ba,dnmt3ab,dnmt | 3 |
| doc2a | doc2a,doc2a,doc2b,doc2 | 4 |
| doc2a | doc2a,doc2a,doc2b,doc2 | 4 |
| doc2b | doc2a,doc2a,doc2b,doc2 | 4 |
| doc2d | doc2a,doc2a,doc2b,doc2 | 4 |
| dock4b | dock4b | 1 |
| dock9b | dock9b | 1 |
| dok1a | dok1a,dok1b | 2 |
| dok1b | dok1a,dok1b | 2 |
| dop1a | dop1a,dop1a,dop1b,dop | 4 |
| dop1a | dop1a,dop1a,dop1b,dop | 4 |
| dop1a | dop1a,dop1a,dop1b,dop | 4 |

| | | |
|---|---|---|
| dop1b | dop1a,dop1a,dop1b,dop[illegible] | 4 |
| dpp6a | dpp6b,dpp6a | 2 |
| dpp6b | dpp6b,dpp6a | 2 |
| dpysl2b | dpysl2b,dpysl2b | 2 |
| dpysl2b | dpysl2b,dpysl2b | 2 |
| dpysl5a | dpysl5b,dpysl5a | 2 |
| dpysl5b | dpysl5b,dpysl5a | 2 |
| dram2a | dram2a,dram2b | 2 |
| dram2b | dram2a,dram2b | 2 |
| drd1a | drd1a,drd1b,drd1a | 3 |
| drd1a | drd1a,drd1b,drd1a | 3 |
| drd1b | drd1a,drd1b,drd1a | 3 |
| drd2a | drd2b,drd2b,drd2a | 3 |
| drd2b | drd2b,drd2b,drd2a | 3 |
| drd2b | drd2b,drd2b,drd2a | 3 |
| drd4a | drd4b,drd4a | 2 |
| drd4b | drd4b,drd4a | 2 |
| drd6b | drd6b | 1 |
| dre-let-7b | dre-let-7b,dre-let-7j,dre-l | 6 |
| dre-let-7e | dre-let-7b,dre-let-7j,dre-l | 6 |
| dre-let-7f | dre-let-7b,dre-let-7j,dre-l | 6 |
| dre-let-7h | dre-let-7b,dre-let-7j,dre-l | 6 |
| dre-let-7i | dre-let-7b,dre-let-7j,dre-l | 6 |
| dre-let-7j | dre-let-7b,dre-let-7j,dre-l | 6 |
| dre-mir-10 | dre-mir-101b,dre-mir-10[illegible] | 2 |
| dre-mir-10 | dre-mir-101b,dre-mir-10[illegible] | 2 |
| dre-mir-10[illegible] | dre-mir-107a,dre-mir-10[illegible] | 2 |
| dre-mir-10[illegible] | dre-mir-107a,dre-mir-10[illegible] | 2 |
| dre-mir-10[illegible] | dre-mir-10c,dre-mir-10d, | 3 |
| dre-mir-10[illegible] | dre-mir-10c,dre-mir-10d, | 3 |
| dre-mir-10[illegible] | dre-mir-10c,dre-mir-10d, | 3 |
| dre-mir-12[illegible] | dre-mir-125c | 1 |
| dre-mir-12[illegible] | dre-mir-126a,dre-mir-12[illegible] | 2 |
| dre-mir-12[illegible] | dre-mir-126a,dre-mir-12[illegible] | 2 |
| dre-mir-13[illegible] | dre-mir-130b,dre-mir-13[illegible] | 2 |
| dre-mir-13[illegible] | dre-mir-130b,dre-mir-13[illegible] | 2 |
| dre-mir-13[illegible] | dre-mir-133c,dre-mir-13[illegible] | 2 |
| dre-mir-13[illegible] | dre-mir-133c,dre-mir-13[illegible] | 2 |
| dre-mir-13[illegible] | dre-mir-135b,dre-mir-13[illegible] | 2 |
| dre-mir-13[illegible] | dre-mir-135b,dre-mir-13[illegible] | 2 |
| dre-mir-14[illegible] | dre-mir-142a,dre-mir-14[illegible] | 2 |
| dre-mir-14[illegible] | dre-mir-142a,dre-mir-14[illegible] | 2 |
| dre-mir-14[illegible] | dre-mir-146a,dre-mir-14[illegible] | 2 |
| dre-mir-14[illegible] | dre-mir-146a,dre-mir-14[illegible] | 2 |
| dre-mir-15[illegible] | dre-mir-153b,dre-mir-15[illegible] | 3 |
| dre-mir-15[illegible] | dre-mir-153b,dre-mir-15[illegible] | 3 |
| dre-mir-15[illegible] | dre-mir-153b,dre-mir-15[illegible] | 3 |
| dre-mir-15[illegible] | dre-mir-15b,dre-mir-15c | 2 |
| dre-mir-15[illegible] | dre-mir-15b,dre-mir-15c | 2 |
| dre-mir-16[illegible] | dre-mir-16b,dre-mir-16a | 2 |

| | | |
|---|---|---|
| dre-mir-16 | dre-mir-16b,dre-mir-16a | 2 |
| dre-mir-18 | dre-mir-181c | 1 |
| dre-mir-18 | dre-mir-18a,dre-mir-18b, | 3 |
| dre-mir-18 | dre-mir-18a,dre-mir-18b, | 3 |
| dre-mir-18 | dre-mir-18a,dre-mir-18b, | 3 |
| dre-mir-19 | dre-mir-190b,dre-mir-19 | 2 |
| dre-mir-19 | dre-mir-190b,dre-mir-19 | 2 |
| dre-mir-19 | dre-mir-193b | 1 |
| dre-mir-19 | dre-mir-194a,dre-mir-19 | 2 |
| dre-mir-19 | dre-mir-194a,dre-mir-19 | 2 |
| dre-mir-19 | dre-mir-19b,dre-mir-19a, | 4 |
| dre-mir-19 | dre-mir-19b,dre-mir-19a, | 4 |
| dre-mir-19 | dre-mir-19b,dre-mir-19a, | 4 |
| dre-mir-19 | dre-mir-19b,dre-mir-19a, | 4 |
| dre-mir-20 | dre-mir-200c,dre-mir-20 | 3 |
| dre-mir-20 | dre-mir-200c,dre-mir-20 | 3 |
| dre-mir-20 | dre-mir-200c,dre-mir-20 | 3 |
| dre-mir-20 | dre-mir-203a,dre-mir-20 | 2 |
| dre-mir-20 | dre-mir-203a,dre-mir-20 | 2 |
| dre-mir-20 | dre-mir-20a,dre-mir-20b | 2 |
| dre-mir-20 | dre-mir-20a,dre-mir-20b | 2 |
| dre-mir-21 | dre-mir-216a,dre-mir-21 | 2 |
| dre-mir-21 | dre-mir-216a,dre-mir-21 | 2 |
| dre-mir-21 | dre-mir-218b | 1 |
| dre-mir-22 | dre-mir-222a | 1 |
| dre-mir-22 | dre-mir-22b,dre-mir-22a | 2 |
| dre-mir-22 | dre-mir-22b,dre-mir-22a | 2 |
| dre-mir-23 | dre-mir-23b | 1 |
| dre-mir-26 | dre-mir-26b | 1 |
| dre-mir-27 | dre-mir-27a,dre-mir-27c, | 5 |
| dre-mir-27 | dre-mir-27a,dre-mir-27c, | 5 |
| dre-mir-27 | dre-mir-27a,dre-mir-27c, | 5 |
| dre-mir-27 | dre-mir-27a,dre-mir-27c, | 5 |
| dre-mir-27 | dre-mir-27a,dre-mir-27c, | 5 |
| dre-mir-30 | dre-mir-301b,dre-mir-30 | 3 |
| dre-mir-30 | dre-mir-301b,dre-mir-30 | 3 |
| dre-mir-30 | dre-mir-301b,dre-mir-30 | 3 |
| dre-mir-30 | dre-mir-30c,dre-mir-30d, | 4 |
| dre-mir-30 | dre-mir-30c,dre-mir-30d, | 4 |
| dre-mir-30 | dre-mir-30c,dre-mir-30d, | 4 |
| dre-mir-30 | dre-mir-30c,dre-mir-30d, | 4 |
| dre-mir-34 | dre-mir-34c,dre-mir-34b, | 3 |
| dre-mir-34 | dre-mir-34c,dre-mir-34b, | 3 |
| dre-mir-34 | dre-mir-34c,dre-mir-34b, | 3 |
| dre-mir-42 | dre-mir-429b,dre-mir-42 | 2 |
| dre-mir-42 | dre-mir-429b,dre-mir-42 | 2 |
| dre-mir-45 | dre-mir-454b,dre-mir-45 | 2 |
| dre-mir-45 | dre-mir-454b,dre-mir-45 | 2 |
| dre-mir-45 | dre-mir-457a,dre-mir-45 | 2 |
| dre-mir-45 | dre-mir-457a,dre-mir-45 | 2 |

| | | |
|---|---|---|
| dre-mir-7b | dre-mir-7b | 1 |
| dre-mir-92[illegible] | dre-mir-92b | 1 |
| dtnbp1a | dtnbp1a,dtnbp1b | 2 |
| dtnbp1b | dtnbp1a,dtnbp1b | 2 |
| dtx4a | dtx4b,dtx4a | 2 |
| dtx4b | dtx4b,dtx4a | 2 |
| dusp13a | dusp13a | 1 |
| dusp19a | dusp19b,dusp19a | 2 |
| dusp19b | dusp19b,dusp19a | 2 |
| dusp22a | dusp22b,dusp22a | 2 |
| dusp22b | dusp22b,dusp22a | 2 |
| dusp23a | dusp23a,dusp23b,dusp2[illegible] | 3 |
| dusp23a | dusp23a,dusp23b,dusp2[illegible] | 3 |
| dusp23b | dusp23a,dusp23b,dusp2[illegible] | 3 |
| dusp3a | dusp3a,dusp3b,dusp3a | 3 |
| dusp3a | dusp3a,dusp3b,dusp3a | 3 |
| dusp3b | dusp3a,dusp3b,dusp3a | 3 |
| dusp8a | dusp8a | 1 |
| dvl1a | dvl1a,dvl1b,dvl1a | 3 |
| dvl1a | dvl1a,dvl1b,dvl1a | 3 |
| dvl1b | dvl1a,dvl1b,dvl1a | 3 |
| dvl3a | dvl3a,dvl3b | 2 |
| dvl3b | dvl3a,dvl3b | 2 |
| dync1i2a | dync1i2b,dync1i2a | 2 |
| dync1i2b | dync1i2b,dync1i2a | 2 |
| dynll2a | dynll2a,dynll2a,dynll2b | 3 |
| dynll2a | dynll2a,dynll2a,dynll2b | 3 |
| dynll2b | dynll2a,dynll2a,dynll2b | 3 |
| dyrk1aa | dyrk1aa,dyrk1ab,dyrk1b | 3 |
| dyrk1ab | dyrk1aa,dyrk1ab,dyrk1b | 3 |
| dyrk1b | dyrk1aa,dyrk1ab,dyrk1b | 3 |
| ebf1b | ebf1b | 1 |
| ebf3a | ebf3a,ebf3a,ebf3b | 3 |
| ebf3a | ebf3a,ebf3a,ebf3b | 3 |
| ebf3b | ebf3a,ebf3a,ebf3b | 3 |
| ece2a | ece2a,ece2b | 2 |
| ece2b | ece2a,ece2b | 2 |
| ecm1a | ecm1b,ecm1a | 2 |
| ecm1b | ecm1b,ecm1a | 2 |
| ecrg4a | ecrg4b,ecrg4a | 2 |
| ecrg4b | ecrg4b,ecrg4a | 2 |
| edil3a | edil3a,edil3b | 2 |
| edil3b | edil3a,edil3b | 2 |
| edn3b | edn3b | 1 |
| eef1a1a | eef1a1b,eef1a1a | 2 |
| eef1a1b | eef1a1b,eef1a1a | 2 |
| eef1da | eef1da,eef1g,eef1db | 3 |
| eef1db | eef1da,eef1g,eef1db | 3 |
| eef1g | eef1da,eef1g,eef1db | 3 |
| eef2b | eef2b,eef2b,eef2kmt,eef[illegible] | 4 |

| | | |
|---|---|---|
| eef2b | eef2b,eef2b,eef2kmt,eef2 | 4 |
| eef2k | eef2b,eef2b,eef2kmt,eef2 | 4 |
| eef2kmt | eef2b,eef2b,eef2kmt,eef2 | 4 |
| efemp2a | efemp2a,efemp2b | 2 |
| efemp2b | efemp2a,efemp2b | 2 |
| efna1a | efna1b,efna1a | 2 |
| efna1b | efna1b,efna1a | 2 |
| efna2a | efna2a,efna2b | 2 |
| efna2b | efna2a,efna2b | 2 |
| efna3a | efna3b,efna3a | 2 |
| efna3b | efna3b,efna3a | 2 |
| efna5a | efna5b,efna5a | 2 |
| efna5b | efna5b,efna5a | 2 |
| efnb2a | efnb2b,efnb2a | 2 |
| efnb2b | efnb2b,efnb2a | 2 |
| efnb3a | efnb3b,efnb3a | 2 |
| efnb3b | efnb3b,efnb3a | 2 |
| efr3a | efr3bb,efr3a,efr3a,efr3ba | 5 |
| efr3a | efr3bb,efr3a,efr3a,efr3ba | 5 |
| efr3ba | efr3bb,efr3a,efr3a,efr3ba | 5 |
| efr3bb | efr3bb,efr3a,efr3a,efr3ba | 5 |
| efr3bb | efr3bb,efr3a,efr3a,efr3ba | 5 |
| egln1a | egln1b,egln1b,egln1a | 3 |
| egln1b | egln1b,egln1b,egln1a | 3 |
| egln1b | egln1b,egln1b,egln1a | 3 |
| egr2a | egr2b,egr2a | 2 |
| egr2b | egr2b,egr2a | 2 |
| ehbp1l1a | ehbp1l1a,ehbp1l1b | 2 |
| ehbp1l1b | ehbp1l1a,ehbp1l1b | 2 |
| ehd1a | ehd1b,ehd1b,ehd1b,ehd1 | 4 |
| ehd1b | ehd1b,ehd1b,ehd1b,ehd1 | 4 |
| ehd1b | ehd1b,ehd1b,ehd1b,ehd1 | 4 |
| ehd1b | ehd1b,ehd1b,ehd1b,ehd1 | 4 |
| ehd2a | ehd2a,ehd2b | 2 |
| ehd2b | ehd2a,ehd2b | 2 |
| ehmt1a | ehmt1b,ehmt1a,ehmt1b | 3 |
| ehmt1b | ehmt1b,ehmt1a,ehmt1b | 3 |
| ehmt1b | ehmt1b,ehmt1a,ehmt1b | 3 |
| eif1ad | eif1ad,eif1axb,eif1axa,eif | 7 |
| eif1ad | eif1ad,eif1axb,eif1axa,eif | 7 |
| eif1axa | eif1ad,eif1axb,eif1axa,eif | 7 |
| eif1axa | eif1ad,eif1axb,eif1axa,eif | 7 |
| eif1axb | eif1ad,eif1axb,eif1axa,eif | 7 |
| eif1axb | eif1ad,eif1axb,eif1axa,eif | 7 |
| eif1b | eif1ad,eif1axb,eif1axa,eif | 7 |
| eif2a | eif2d,eif2a | 2 |
| eif2d | eif2d,eif2a | 2 |
| eif2s1a | eif2s1a,eif2s1b | 2 |
| eif2s1b | eif2s1a,eif2s1b | 2 |
| eif3ba | eif3eb,eif3f,eif3bb,eif3d,e | 16 |

| | | |
|---|---|---|
| eif3bb | eif3eb,eif3f,eif3bb,eif3d,e | 16 |
| eif3c | eif3eb,eif3f,eif3bb,eif3d,e | 16 |
| eif3d | eif3eb,eif3f,eif3bb,eif3d,e | 16 |
| eif3ea | eif3eb,eif3f,eif3bb,eif3d,e | 16 |
| eif3eb | eif3eb,eif3f,eif3bb,eif3d,e | 16 |
| eif3eb | eif3eb,eif3f,eif3bb,eif3d,e | 16 |
| eif3f | eif3eb,eif3f,eif3bb,eif3d,e | 16 |
| eif3g | eif3eb,eif3f,eif3bb,eif3d,e | 16 |
| eif3ha | eif3eb,eif3f,eif3bb,eif3d,e | 16 |
| eif3hb | eif3eb,eif3f,eif3bb,eif3d,e | 16 |
| eif3i | eif3eb,eif3f,eif3bb,eif3d,e | 16 |
| eif3ja | eif3eb,eif3f,eif3bb,eif3d,e | 16 |
| eif3jb | eif3eb,eif3f,eif3bb,eif3d,e | 16 |
| eif3k | eif3eb,eif3f,eif3bb,eif3d,e | 16 |
| eif3m | eif3eb,eif3f,eif3bb,eif3d,e | 16 |
| eif3s6ip | eif3s6ip | 1 |
| eif4a1a | eif4a1b,eif4a1b,eif4a1a | 3 |
| eif4a1b | eif4a1b,eif4a1b,eif4a1a | 3 |
| eif4a1b | eif4a1b,eif4a1b,eif4a1a | 3 |
| eif4ba | eif4eb,eif4ea,eif4ba,eif4b | 9 |
| eif4ba | eif4eb,eif4ea,eif4ba,eif4b | 9 |
| eif4bb | eif4eb,eif4ea,eif4ba,eif4b | 9 |
| eif4bb | eif4eb,eif4ea,eif4ba,eif4b | 9 |
| eif4e1b | eif4e1c,eif4e1b,eif4e1c | 3 |
| eif4e1c | eif4e1c,eif4e1b,eif4e1c | 3 |
| eif4e1c | eif4e1c,eif4e1b,eif4e1c | 3 |
| eif4ea | eif4eb,eif4ea,eif4ba,eif4b | 9 |
| eif4ea | eif4eb,eif4ea,eif4ba,eif4b | 9 |
| eif4eb | eif4eb,eif4ea,eif4ba,eif4b | 9 |
| eif4eb | eif4eb,eif4ea,eif4ba,eif4b | 9 |
| eif4g1a | eif4g1a | 1 |
| eif4g2a | eif4g2b,eif4g2a,eif4g2b | 3 |
| eif4g2b | eif4g2b,eif4g2a,eif4g2b | 3 |
| eif4g2b | eif4g2b,eif4g2a,eif4g2b | 3 |
| eif4g3a | eif4g3b,eif4g3a,eif4g3b | 3 |
| eif4g3b | eif4g3b,eif4g3a,eif4g3b | 3 |
| eif4g3b | eif4g3b,eif4g3a,eif4g3b | 3 |
| eif4h | eif4eb,eif4ea,eif4ba,eif4b | 9 |
| eif5a | eif5b,eif5,eif5a | 3 |
| eif5b | eif5b,eif5,eif5a | 3 |
| elavl1a | elavl1a,elavl1b | 2 |
| elavl1b | elavl1a,elavl1b | 2 |
| elf2a | elf2b,elf2b,elf2a | 3 |
| elf2b | elf2b,elf2b,elf2a | 3 |
| elf2b | elf2b,elf2b,elf2a | 3 |
| elfn1b | elfn1b | 1 |
| elfn2a | elfn2a | 1 |
| elmsan1a | elmsan1b,elmsan1a | 2 |
| elmsan1b | elmsan1b,elmsan1a | 2 |
| elovl1a | elovl1a,elovl1b | 2 |

| | | |
|---|---|---|
| elovl1b | elovl1a,elovl1b | 2 |
| elovl4a | elovl4a,elovl4b | 2 |
| elovl4b | elovl4a,elovl4b | 2 |
| elovl7a | elovl7a,elovl7b,elovl7a | 3 |
| elovl7a | elovl7a,elovl7b,elovl7a | 3 |
| elovl7b | elovl7a,elovl7b,elovl7a | 3 |
| elovl8a | elovl8a,elovl8b | 2 |
| elovl8b | elovl8a,elovl8b | 2 |
| emilin1a | emilin1b,emilin1a | 2 |
| emilin1b | emilin1b,emilin1a | 2 |
| emilin2a | emilin2a,emilin2b | 2 |
| emilin2b | emilin2a,emilin2b | 2 |
| emilin3a | emilin3a | 1 |
| emp3b | emp3b | 1 |
| en1a | en1b,en1a | 2 |
| en1b | en1b,en1a | 2 |
| en2a | en2a,en2b,en2a | 3 |
| en2a | en2a,en2b,en2a | 3 |
| en2b | en2a,en2b,en2a | 3 |
| eno1a | eno1a,eno1b,eno1a | 3 |
| eno1a | eno1a,eno1b,eno1a | 3 |
| eno1b | eno1a,eno1b,eno1a | 3 |
| entpd2b | entpd2b | 1 |
| entpd5a | entpd5a,entpd5b | 2 |
| entpd5b | entpd5a,entpd5b | 2 |
| ep300a | ep300b,ep300a | 2 |
| ep300b | ep300b,ep300a | 2 |
| epas1a | epas1b,epas1b,epas1a,ep | 4 |
| epas1b | epas1b,epas1b,epas1a,ep | 4 |
| epas1b | epas1b,epas1b,epas1a,ep | 4 |
| epas1b | epas1b,epas1b,epas1a,ep | 4 |
| epb41a | epb41b,epb41a | 2 |
| epb41b | epb41b,epb41a | 2 |
| epb41l3a | epb41l3b,epb41l3a | 2 |
| epb41l3b | epb41l3b,epb41l3a | 2 |
| epb41l4a | epb41l4a,epb41l4b | 2 |
| epb41l4b | epb41l4a,epb41l4b | 2 |
| epc1a | epc1b,epc1a | 2 |
| epc1b | epc1b,epc1a | 2 |
| epha2a | epha2a,epha2a,epha2b | 3 |
| epha2a | epha2a,epha2a,epha2b | 3 |
| epha2b | epha2a,epha2a,epha2b | 3 |
| epha4a | epha4a,epha4b | 2 |
| epha4b | epha4a,epha4b | 2 |
| ephb2a | ephb2a,ephb2b,ephb2a | 3 |
| ephb2a | ephb2a,ephb2b,ephb2a | 3 |
| ephb2b | ephb2a,ephb2b,ephb2a | 3 |
| ephb3a | ephb3a | 1 |
| ephb4a | ephb4a,ephb4b | 2 |
| ephb4b | ephb4a,ephb4b | 2 |

| | | |
|---|---|---|
| epm2a | epm2a | 1 |
| epn3a | epn3a,epn3b,epn3a | 3 |
| epn3a | epn3a,epn3b,epn3a | 3 |
| epn3b | epn3a,epn3b,epn3a | 3 |
| eps15l1a | eps15l1a | 1 |
| eps8a | eps8a,eps8b | 2 |
| eps8b | eps8a,eps8b | 2 |
| eps8l1a | eps8l1a,eps8l1a,eps8l1b | 3 |
| eps8l1a | eps8l1a,eps8l1a,eps8l1b | 3 |
| eps8l1b | eps8l1a,eps8l1a,eps8l1b | 3 |
| eps8l3a | eps8l3b,eps8l3a | 2 |
| eps8l3b | eps8l3b,eps8l3a | 2 |
| erap1a | erap1b,erap1a | 2 |
| erap1b | erap1b,erap1a | 2 |
| erbb3a | erbb3b,erbb3a,erbb3b | 3 |
| erbb3b | erbb3b,erbb3a,erbb3b | 3 |
| erbb3b | erbb3b,erbb3a,erbb3b | 3 |
| erbb4a | erbb4a,erbb4b | 2 |
| erbb4b | erbb4a,erbb4b | 2 |
| erc1a | erc1a,erc1b,erc1a | 3 |
| erc1a | erc1a,erc1b,erc1a | 3 |
| erc1b | erc1a,erc1b,erc1a | 3 |
| ero1a | ero1b,ero1a | 2 |
| ero1b | ero1b,ero1a | 2 |
| errfi1a | errfi1a | 1 |
| esr2a | esr2b,esr2a | 2 |
| esr2b | esr2b,esr2a | 2 |
| esyt1a | esyt1b,esyt1a | 2 |
| esyt1b | esyt1b,esyt1a | 2 |
| esyt2a | esyt2a,esyt2b | 2 |
| esyt2b | esyt2a,esyt2b | 2 |
| etf1a | etf1b,etf1b,etf1a | 3 |
| etf1b | etf1b,etf1b,etf1a | 3 |
| etf1b | etf1b,etf1b,etf1a | 3 |
| etv5a | etv5b,etv5a,etv5a,etv5b, | 5 |
| etv5a | etv5b,etv5a,etv5a,etv5b, | 5 |
| etv5a | etv5b,etv5a,etv5a,etv5b, | 5 |
| etv5b | etv5b,etv5a,etv5a,etv5b, | 5 |
| etv5b | etv5b,etv5a,etv5a,etv5b, | 5 |
| eva1a | eva1c,eva1ba,eva1a,eva1 | 4 |
| eva1ba | eva1c,eva1ba,eva1a,eva1 | 4 |
| eva1bb | eva1c,eva1ba,eva1a,eva1 | 4 |
| eva1c | eva1c,eva1ba,eva1a,eva1 | 4 |
| evi5a | evi5a,evi5b | 2 |
| evi5b | evi5a,evi5b | 2 |
| ewsr1a | ewsr1a,ewsr1b,ewsr1a,e | 4 |
| ewsr1a | ewsr1a,ewsr1b,ewsr1a,e | 4 |
| ewsr1b | ewsr1a,ewsr1b,ewsr1a,e | 4 |
| ewsr1b | ewsr1a,ewsr1b,ewsr1a,e | 4 |
| exoc3l2a | exoc3l2b,exoc3l2a,exoc3 | 3 |

| | | |
|---|---|---|
| exoc3l2b | exoc3l2b,exoc3l2a,exoc3 | 3 |
| exoc3l2b | exoc3l2b,exoc3l2a,exoc3 | 3 |
| exoc6b | exoc6b,exoc6b,exoc6 | 3 |
| exoc6b | exoc6b,exoc6b,exoc6 | 3 |
| ext1a | ext1a,ext1b,ext1c | 3 |
| ext1b | ext1a,ext1b,ext1c | 3 |
| ext1c | ext1a,ext1b,ext1c | 3 |
| f13a1b | f13a1b | 1 |
| f13b | f13b | 1 |
| f2r | f2r,f2 | 2 |
| f3a | f3a,f3a,f3b,f3a | 4 |
| f3a | f3a,f3a,f3b,f3a | 4 |
| f3a | f3a,f3a,f3b,f3a | 4 |
| f3b | f3a,f3a,f3b,f3a | 4 |
| f7i | f7,f7i | 2 |
| f9a | f9b,f9a,f9b,f9a | 4 |
| f9a | f9b,f9a,f9b,f9a | 4 |
| f9b | f9b,f9a,f9b,f9a | 4 |
| f9b | f9b,f9a,f9b,f9a | 4 |
| fa2h | fa2h | 1 |
| faah2a | faah2a,faah2b,faah2a,faa | 4 |
| faah2a | faah2a,faah2b,faah2a,faa | 4 |
| faah2b | faah2a,faah2b,faah2a,faa | 4 |
| faah2b | faah2a,faah2b,faah2a,faa | 4 |
| fabp10a | fabp10a,fabp10b | 2 |
| fabp10b | fabp10a,fabp10b | 2 |
| fabp11a | fabp11b,fabp11a | 2 |
| fabp11b | fabp11b,fabp11a | 2 |
| fabp1a | fabp1a | 1 |
| fabp7a | fabp7a,fabp7b | 2 |
| fabp7b | fabp7a,fabp7b | 2 |
| fahd2a | fahd2a | 1 |
| faim2a | faim2a,faim2b | 2 |
| faim2b | faim2a,faim2b | 2 |
| fam102aa | fam102ab,fam102ba,fam | 5 |
| fam102ab | fam102ab,fam102ba,fam | 5 |
| fam102ab | fam102ab,fam102ba,fam | 5 |
| fam102ba | fam102ab,fam102ba,fam | 5 |
| fam102bb | fam102ab,fam102ba,fam | 5 |
| fam107b | fam107b,fam107b,fam10 | 3 |
| fam107b | fam107b,fam107b,fam10 | 3 |
| fam107b | fam107b,fam107b,fam10 | 3 |
| fam110a | fam110b,fam110a,fam11 | 3 |
| fam110b | fam110b,fam110a,fam11 | 3 |
| fam110c | fam110b,fam110a,fam11 | 3 |
| fam117aa | fam117ab,fam117aa,fam | 5 |
| fam117ab | fam117ab,fam117aa,fam | 5 |
| fam117ab | fam117ab,fam117aa,fam | 5 |
| fam117ba | fam117ab,fam117aa,fam | 5 |
| fam117bb | fam117ab,fam117aa,fam | 5 |

| | | |
|---|---|---|
| fam118b | fam118b | 1 |
| fam120a | fam120a,fam120b,fam12 | 3 |
| fam120b | fam120a,fam120b,fam12 | 3 |
| fam120c | fam120a,fam120b,fam12 | 3 |
| fam122b | fam122b,fam122b | 2 |
| fam122b | fam122b,fam122b | 2 |
| fam124b | fam124b | 1 |
| fam126a | fam126a | 1 |
| fam131a | fam131a,fam131bb,fam1 | 4 |
| fam131ba | fam131a,fam131bb,fam1 | 4 |
| fam131bb | fam131a,fam131bb,fam1 | 4 |
| fam131c | fam131a,fam131bb,fam1 | 4 |
| fam133b | fam133b | 1 |
| fam135a | fam135a | 1 |
| fam136a | fam136a | 1 |
| fam13a | fam13a,fam13b | 2 |
| fam13b | fam13a,fam13b | 2 |
| fam149a | fam149a | 1 |
| fam151a | fam151a,fam151b | 2 |
| fam151b | fam151a,fam151b | 2 |
| fam155a | fam155b,fam155a,fam15 | 3 |
| fam155b | fam155b,fam155a,fam15 | 3 |
| fam155b | fam155b,fam155a,fam15 | 3 |
| fam160a1a | fam160a1a,fam160a1a,fa | 3 |
| fam160a1a | fam160a1a,fam160a1a,fa | 3 |
| fam160a1b | fam160a1a,fam160a1a,fa | 3 |
| fam161a | fam161a,fam161b | 2 |
| fam161b | fam161a,fam161b | 2 |
| fam162a | fam162a | 1 |
| fam163ba | fam163ba | 1 |
| fam166b | fam166b,fam166c | 2 |
| fam166c | fam166b,fam166c | 2 |
| fam167aa | fam167aa,fam167b,fam1 | 5 |
| fam167aa | fam167aa,fam167b,fam1 | 5 |
| fam167ab | fam167aa,fam167b,fam1 | 5 |
| fam167b | fam167aa,fam167b,fam1 | 5 |
| fam167b | fam167aa,fam167b,fam1 | 5 |
| fam168a | fam168b,fam168b,fam16 | 4 |
| fam168b | fam168b,fam168b,fam16 | 4 |
| fam168b | fam168b,fam168b,fam16 | 4 |
| fam168b | fam168b,fam168b,fam16 | 4 |
| fam169aa | fam169aa,fam169aa,fam | 4 |
| fam169aa | fam169aa,fam169aa,fam | 4 |
| fam169ab | fam169aa,fam169aa,fam | 4 |
| fam169b | fam169aa,fam169aa,fam | 4 |
| fam171a2a | fam171a2a,fam171a2b | 2 |
| fam171a2b | fam171a2a,fam171a2b | 2 |
| fam172a | fam172a | 1 |
| fam174b | fam174b | 1 |
| fam183a | fam183a,fam183a,fam18 | 3 |

| | | |
|---|---|---|
| fam183a | fam183a,fam183a,fam18 | 3 |
| fam183a | fam183a,fam183a,fam18 | 3 |
| fam184a | fam184b,fam184a | 2 |
| fam184b | fam184b,fam184a | 2 |
| fam185a | fam185a | 1 |
| fam192a | fam192a | 1 |
| fam193a | fam193a,fam193b | 2 |
| fam193b | fam193a,fam193b | 2 |
| fam199x | fam199x | 1 |
| fam204a | fam204a,fam204a | 2 |
| fam204a | fam204a,fam204a | 2 |
| fam207a | fam207a | 1 |
| fam20a | fam20ca,fam20a,fam20cl | 4 |
| fam20b | fam20ca,fam20a,fam20cl | 4 |
| fam20ca | fam20ca,fam20a,fam20cl | 4 |
| fam20cb | fam20ca,fam20a,fam20cl | 4 |
| fam210aa | fam210b,fam210ab,fam2 | 3 |
| fam210ab | fam210b,fam210ab,fam2 | 3 |
| fam210b | fam210b,fam210ab,fam2 | 3 |
| fam214a | fam214a,fam214b,fam21 | 3 |
| fam214a | fam214a,fam214b,fam21 | 3 |
| fam214b | fam214a,fam214b,fam21 | 3 |
| fam217b | fam217b | 1 |
| fam219aa | fam219ab,fam219b,fam2 | 3 |
| fam219ab | fam219ab,fam219b,fam2 | 3 |
| fam219b | fam219ab,fam219b,fam2 | 3 |
| fam221a | fam221a | 1 |
| fam222a | fam222a,fam222ba,fam2 | 3 |
| fam222ba | fam222a,fam222ba,fam2 | 3 |
| fam222bb | fam222a,fam222ba,fam2 | 3 |
| fam228a | fam228a,fam228a | 2 |
| fam228a | fam228a,fam228a | 2 |
| fam234a | fam234b,fam234a | 2 |
| fam234b | fam234b,fam234a | 2 |
| fam241a | fam241a,fam241a | 2 |
| fam241a | fam241a,fam241a | 2 |
| fam32a | fam32a,fam32a | 2 |
| fam32a | fam32a,fam32a | 2 |
| fam3a | fam3a,fam3c,fam3a | 3 |
| fam3a | fam3a,fam3c,fam3a | 3 |
| fam3c | fam3a,fam3c,fam3a | 3 |
| fam43a | fam43a,fam43b | 2 |
| fam43b | fam43a,fam43b | 2 |
| fam49a | fam49ba,fam49a,fam49b | 3 |
| fam49ba | fam49ba,fam49a,fam49b | 3 |
| fam49bb | fam49ba,fam49a,fam49b | 3 |
| fam50a | fam50a | 1 |
| fam53b | fam53b | 1 |
| fam76b | fam76b | 1 |
| fam78aa | fam78ba,fam78aa,fam78 | 4 |

| | | |
|---|---|---|
| fam78ab | fam78ba,fam78aa,fam78 | 4 |
| fam78ba | fam78ba,fam78aa,fam78 | 4 |
| fam78bb | fam78ba,fam78aa,fam78 | 4 |
| fam81b | fam81b | 1 |
| fam83b | fam83fa,fam83fb,fam83c | 8 |
| fam83c | fam83fa,fam83fb,fam83c | 8 |
| fam83d | fam83fa,fam83fb,fam83c | 8 |
| fam83e | fam83fa,fam83fb,fam83c | 8 |
| fam83fa | fam83fa,fam83fb,fam83c | 8 |
| fam83fb | fam83fa,fam83fb,fam83c | 8 |
| fam83ha | fam83fa,fam83fb,fam83c | 8 |
| fam83hb | fam83fa,fam83fb,fam83c | 8 |
| fam89a | fam89b,fam89a | 2 |
| fam89b | fam89b,fam89a | 2 |
| fam8a1a | fam8a1b,fam8a1a | 2 |
| fam8a1b | fam8a1b,fam8a1a | 2 |
| fam98a | fam98a,fam98b,fam98a,f | 5 |
| fam98a | fam98a,fam98b,fam98a,f | 5 |
| fam98a | fam98a,fam98b,fam98a,f | 5 |
| fam98b | fam98a,fam98b,fam98a,f | 5 |
| fam98b | fam98a,fam98b,fam98a,f | 5 |
| fat1a | fat1a,fat1a,fat1b | 3 |
| fat1a | fat1a,fat1a,fat1b | 3 |
| fat1b | fat1a,fat1a,fat1b | 3 |
| fat3a | fat3b,fat3a | 2 |
| fat3b | fat3b,fat3a | 2 |
| fbn2a | fbn2a,fbn2b | 2 |
| fbn2b | fbn2a,fbn2b | 2 |
| fbp1a | fbp1b,fbp1b,fbp1a | 3 |
| fbp1b | fbp1b,fbp1b,fbp1a | 3 |
| fbp1b | fbp1b,fbp1b,fbp1a | 3 |
| fbxl14a | fbxl14a,fbxl14b,fbxl14a | 3 |
| fbxl14a | fbxl14a,fbxl14b,fbxl14a | 3 |
| fbxl14b | fbxl14a,fbxl14b,fbxl14a | 3 |
| fbxl3a | fbxl3a,fbxl3b,fbxl3a | 3 |
| fbxl3a | fbxl3a,fbxl3b,fbxl3a | 3 |
| fbxl3b | fbxl3a,fbxl3b,fbxl3a | 3 |
| fbxo11a | fbxo11a,fbxo11a,fbxo11a | 3 |
| fbxo11a | fbxo11a,fbxo11a,fbxo11a | 3 |
| fbxo11a | fbxo11a,fbxo11a,fbxo11a | 3 |
| fbxo30a | fbxo30a,fbxo30b | 2 |
| fbxo30b | fbxo30a,fbxo30b | 2 |
| fbxo36a | fbxo36a,fbxo36b,fbxo36a | 3 |
| fbxo36a | fbxo36a,fbxo36b,fbxo36a | 3 |
| fbxo36b | fbxo36a,fbxo36b,fbxo36a | 3 |
| fbxw11a | fbxw11b,fbxw11b,fbxw1 | 3 |
| fbxw11b | fbxw11b,fbxw11b,fbxw1 | 3 |
| fbxw11b | fbxw11b,fbxw11b,fbxw1 | 3 |
| fcer1g | fcer1g | 1 |
| fdx1b | fdx1b,fdx1,fdx1b,fdx1 | 4 |

| | | |
|---|---|---|
| fdx1b | fdx1b,fdx1,fdx1b,fdx1 | 4 |
| fem1a | fem1a,fem1c,fem1b | 3 |
| fem1b | fem1a,fem1c,fem1b | 3 |
| fem1c | fem1a,fem1c,fem1b | 3 |
| fermt3b | fermt3b | 1 |
| fgd4a | fgd4a,fgd4b,fgd4a | 3 |
| fgd4a | fgd4a,fgd4b,fgd4a | 3 |
| fgd4b | fgd4a,fgd4b,fgd4a | 3 |
| fgd5a | fgd5b,fgd5a | 2 |
| fgd5b | fgd5b,fgd5a | 2 |
| fgf10a | fgf10a,fgf10b,fgf10a | 3 |
| fgf10a | fgf10a,fgf10b,fgf10a | 3 |
| fgf10b | fgf10a,fgf10b,fgf10a | 3 |
| fgf11a | fgf11a,fgf11a,fgf11a,fgf1[illegible] | 4 |
| fgf11a | fgf11a,fgf11a,fgf11a,fgf1[illegible] | 4 |
| fgf11a | fgf11a,fgf11a,fgf11a,fgf1[illegible] | 4 |
| fgf11b | fgf11a,fgf11a,fgf11a,fgf1[illegible] | 4 |
| fgf12a | fgf12a,fgf12b | 2 |
| fgf12b | fgf12a,fgf12b | 2 |
| fgf13a | fgf13b,fgf13a | 2 |
| fgf13b | fgf13b,fgf13a | 2 |
| fgf18a | fgf18b,fgf18b,fgf18a | 3 |
| fgf18b | fgf18b,fgf18b,fgf18a | 3 |
| fgf18b | fgf18b,fgf18b,fgf18a | 3 |
| fgf1a | fgf1a,fgf1a,fgf1a,fgf1b | 4 |
| fgf1a | fgf1a,fgf1a,fgf1a,fgf1b | 4 |
| fgf1a | fgf1a,fgf1a,fgf1a,fgf1b | 4 |
| fgf1b | fgf1a,fgf1a,fgf1a,fgf1b | 4 |
| fgf20a | fgf20a,fgf20a,fgf20b | 3 |
| fgf20a | fgf20a,fgf20a,fgf20b | 3 |
| fgf20b | fgf20a,fgf20a,fgf20b | 3 |
| fgf6a | fgf6b,fgf6a | 2 |
| fgf6b | fgf6b,fgf6a | 2 |
| fgf8a | fgf8b,fgf8a | 2 |
| fgf8b | fgf8b,fgf8a | 2 |
| fgfbp1b | fgfbp1b,fgfbp1b | 2 |
| fgfbp1b | fgfbp1b,fgfbp1b | 2 |
| fgfbp2a | fgfbp2b,fgfbp2b,fgfbp2b, | 4 |
| fgfbp2b | fgfbp2b,fgfbp2b,fgfbp2b, | 4 |
| fgfbp2b | fgfbp2b,fgfbp2b,fgfbp2b, | 4 |
| fgfbp2b | fgfbp2b,fgfbp2b,fgfbp2b, | 4 |
| fgfr1a | fgfr1a,fgfr1b,fgfr1op | 3 |
| fgfr1b | fgfr1a,fgfr1b,fgfr1op | 3 |
| fgfr1op | fgfr1a,fgfr1b,fgfr1op | 3 |
| fgfrl1a | fgfrl1b,fgfrl1a,fgfrl1b | 3 |
| fgfrl1b | fgfrl1b,fgfrl1a,fgfrl1b | 3 |
| fgfrl1b | fgfrl1b,fgfrl1a,fgfrl1b | 3 |
| fgl2a | fgl2a,fgl2b | 2 |
| fgl2b | fgl2a,fgl2b | 2 |
| fhl1a | fhl1a,fhl1b,fhl1a | 3 |

| | | |
|---|---|---|
| fhl1a | fhl1a,fhl1b,fhl1a | 3 |
| fhl1b | fhl1a,fhl1b,fhl1a | 3 |
| fhl2a | fhl2b,fhl2a | 2 |
| fhl2b | fhl2b,fhl2a | 2 |
| fhl3a | fhl3a,fhl3a,fhl3b | 3 |
| fhl3a | fhl3a,fhl3a,fhl3b | 3 |
| fhl3b | fhl3a,fhl3a,fhl3b | 3 |
| fhod3a | fhod3b,fhod3a | 2 |
| fhod3b | fhod3b,fhod3a | 2 |
| filip1a | filip1a,filip1b | 2 |
| filip1b | filip1a,filip1b | 2 |
| fip1l1a | fip1l1b,fip1l1a | 2 |
| fip1l1b | fip1l1b,fip1l1a | 2 |
| fkbp10a | fkbp10b,fkbp10a | 2 |
| fkbp10b | fkbp10b,fkbp10a | 2 |
| fkbp1aa | fkbp1ab,fkbp1ab,fkbp1aa | 5 |
| fkbp1ab | fkbp1ab,fkbp1ab,fkbp1aa | 5 |
| fkbp1ab | fkbp1ab,fkbp1ab,fkbp1aa | 5 |
| fkbp1ab | fkbp1ab,fkbp1ab,fkbp1aa | 5 |
| fkbp1b | fkbp1ab,fkbp1ab,fkbp1aa | 5 |
| fli1a | fli1b,fli1a | 2 |
| fli1b | fli1b,fli1a | 2 |
| flot1a | flot1b,flot1a | 2 |
| flot1b | flot1b,flot1a | 2 |
| flot2a | flot2a,flot2b | 2 |
| flot2b | flot2a,flot2b | 2 |
| flrt1b | flrt1b | 1 |
| flvcr2a | flvcr2a,flvcr2b | 2 |
| flvcr2b | flvcr2a,flvcr2b | 2 |
| fmn2a | fmn2b,fmn2a | 2 |
| fmn2b | fmn2b,fmn2a | 2 |
| fmnl1a | fmnl1a | 1 |
| fmnl2a | fmnl2b,fmnl2a | 2 |
| fmnl2b | fmnl2b,fmnl2a | 2 |
| fn1a | fn1a,fn1a,fn1b,fn1a | 4 |
| fn1a | fn1a,fn1a,fn1b,fn1a | 4 |
| fn1a | fn1a,fn1a,fn1b,fn1a | 4 |
| fn1b | fn1a,fn1a,fn1b,fn1a | 4 |
| fn3krp | fn3krp | 1 |
| fnbp1a | fnbp1a,fnbp1b,fnbp1b | 3 |
| fnbp1b | fnbp1a,fnbp1b,fnbp1b | 3 |
| fnbp1b | fnbp1a,fnbp1b,fnbp1b | 3 |
| fndc3a | fndc3ba,fndc3a,fndc3bb | 3 |
| fndc3ba | fndc3ba,fndc3a,fndc3bb | 3 |
| fndc3bb | fndc3ba,fndc3a,fndc3bb | 3 |
| fndc4a | fndc4b,fndc4a | 2 |
| fndc4b | fndc4b,fndc4a | 2 |
| fndc5a | fndc5a,fndc5b | 2 |
| fndc5b | fndc5a,fndc5b | 2 |
| fndc7a | fndc7a,fndc7b | 2 |

| | | |
|---|---|---|
| fndc7b | fndc7a,fndc7b | 2 |
| fosl1a | fosl1a,fosl1b | 2 |
| fosl1b | fosl1a,fosl1b | 2 |
| foxb1a | foxb1b,foxb1b,foxb1b,fo› | 4 |
| foxb1b | foxb1b,foxb1b,foxb1b,fo› | 4 |
| foxb1b | foxb1b,foxb1b,foxb1b,fo› | 4 |
| foxb1b | foxb1b,foxb1b,foxb1b,fo› | 4 |
| foxc1a | foxc1a,foxc1b | 2 |
| foxc1b | foxc1a,foxc1b | 2 |
| foxf2a | foxf2a,foxf2b | 2 |
| foxf2b | foxf2a,foxf2b | 2 |
| foxg1a | foxg1c,foxg1b,foxg1d,fox | 5 |
| foxg1b | foxg1c,foxg1b,foxg1d,fox | 5 |
| foxg1c | foxg1c,foxg1b,foxg1d,fox | 5 |
| foxg1c | foxg1c,foxg1b,foxg1d,fox | 5 |
| foxg1d | foxg1c,foxg1b,foxg1d,fox | 5 |
| foxi3a | foxi3b,foxi3b,foxi3b,foxi3 | 4 |
| foxi3b | foxi3b,foxi3b,foxi3b,foxi3 | 4 |
| foxi3b | foxi3b,foxi3b,foxi3b,foxi3 | 4 |
| foxi3b | foxi3b,foxi3b,foxi3b,foxi3 | 4 |
| foxj1a | foxj1a,foxj1b | 2 |
| foxj1b | foxj1a,foxj1b | 2 |
| foxl2a | foxl2a,foxl2b,foxl2a | 3 |
| foxl2a | foxl2a,foxl2b,foxl2a | 3 |
| foxl2b | foxl2a,foxl2b,foxl2a | 3 |
| foxn2b | foxn2b,foxn2b | 2 |
| foxn2b | foxn2b,foxn2b | 2 |
| foxo1a | foxo1b,foxo1a | 2 |
| foxo1b | foxo1b,foxo1a | 2 |
| foxo3a | foxo3a,foxo3b | 2 |
| foxo3b | foxo3a,foxo3b | 2 |
| foxo6a | foxo6a,foxo6b | 2 |
| foxo6b | foxo6a,foxo6b | 2 |
| foxp1a | foxp1b,foxp1a | 2 |
| foxp1b | foxp1b,foxp1a | 2 |
| foxp3a | foxp3b,foxp3a | 2 |
| foxp3b | foxp3b,foxp3a | 2 |
| foxq1a | foxq1a,foxq1b | 2 |
| foxq1b | foxq1a,foxq1b | 2 |
| frem1a | frem1a,frem1b | 2 |
| frem1b | frem1a,frem1b | 2 |
| frem2a | frem2a,frem2b | 2 |
| frem2b | frem2a,frem2b | 2 |
| frmd4ba | frmd4ba,frmd4ba,frmd4l | 3 |
| frmd4ba | frmd4ba,frmd4ba,frmd4l | 3 |
| frmd4bb | frmd4ba,frmd4ba,frmd4l | 3 |
| frmpd1a | frmpd1a,frmpd1b | 2 |
| frmpd1b | frmpd1a,frmpd1b | 2 |
| frrs1a | frrs1a,frrs1b | 2 |
| frrs1b | frrs1a,frrs1b | 2 |

| | | |
|---|---|---|
| frs2a | frs2a,frs2b | 2 |
| frs2b | frs2a,frs2b | 2 |
| fscn1a | fscn1b,fscn1b,fscn1a | 3 |
| fscn1b | fscn1b,fscn1b,fscn1a | 3 |
| fscn1b | fscn1b,fscn1b,fscn1a | 3 |
| fscn2a | fscn2a,fscn2b | 2 |
| fscn2b | fscn2a,fscn2b | 2 |
| fstl1a | fstl1a,fstl1b | 2 |
| fstl1b | fstl1a,fstl1b | 2 |
| fth1a | fth1a,fth1b | 2 |
| fth1b | fth1a,fth1b | 2 |
| ftr39p | ftr39p | 1 |
| ftr52p | ftr52p | 1 |
| fut8a | fut8a,fut8b | 2 |
| fut8b | fut8a,fut8b | 2 |
| fut9a | fut9b,fut9d,fut9a | 3 |
| fut9b | fut9b,fut9d,fut9a | 3 |
| fut9d | fut9b,fut9d,fut9a | 3 |
| fyco1a | fyco1b,fyco1a | 2 |
| fyco1b | fyco1b,fyco1a | 2 |
| fzd3a | fzd3b,fzd3a | 2 |
| fzd3b | fzd3b,fzd3a | 2 |
| fzd7a | fzd7a,fzd7b,fzd7a | 3 |
| fzd7a | fzd7a,fzd7b,fzd7a | 3 |
| fzd7b | fzd7a,fzd7b,fzd7a | 3 |
| fzd8a | fzd8b,fzd8b,fzd8a | 3 |
| fzd8b | fzd8b,fzd8b,fzd8a | 3 |
| fzd8b | fzd8b,fzd8b,fzd8a | 3 |
| fzd9a | fzd9b,fzd9a,fzd9b | 3 |
| fzd9b | fzd9b,fzd9a,fzd9b | 3 |
| fzd9b | fzd9b,fzd9a,fzd9b | 3 |
| fzr1a | fzr1b,fzr1a | 2 |
| fzr1b | fzr1b,fzr1a | 2 |
| g6pcb | g6pd,g6pcb,g6pd | 3 |
| g6pd | g6pd,g6pcb,g6pd | 3 |
| g6pd | g6pd,g6pcb,g6pd | 3 |
| gabbr1a | gabbr1a,gabbr1b | 2 |
| gabbr1b | gabbr1a,gabbr1b | 2 |
| gabpb2a | gabpb2b,gabpb2b,gabpb2 | 3 |
| gabpb2b | gabpb2b,gabpb2b,gabpb2 | 3 |
| gabpb2b | gabpb2b,gabpb2b,gabpb2 | 3 |
| gabra2a | gabra2a | 1 |
| gabra6a | gabra6a,gabra6b | 2 |
| gabra6b | gabra6a,gabra6b | 2 |
| gabrb2a | gabrb2a | 1 |
| gabrr2a | gabrr2a,gabrr2b | 2 |
| gabrr2b | gabrr2a,gabrr2b | 2 |
| gabrr3a | gabrr3a,gabrr3b | 2 |
| gabrr3b | gabrr3a,gabrr3b | 2 |
| gad1a | gad1b,gad1a | 2 |

| | | |
|---|---|---|
| gad1b | gad1b,gad1a | 2 |
| gadd45aa | gadd45ab,gadd45ga,gadc | 7 |
| gadd45ab | gadd45ab,gadd45ga,gadc | 7 |
| gadd45ab | gadd45ab,gadd45ga,gadc | 7 |
| gadd45ba | gadd45ab,gadd45ga,gadc | 7 |
| gadd45bb | gadd45ab,gadd45ga,gadc | 7 |
| gadd45ga | gadd45ab,gadd45ga,gadc | 7 |
| gadd45ga | gadd45ab,gadd45ga,gadc | 7 |
| gal3st1a | gal3st1a,gal3st1a,gal3st1 | 3 |
| gal3st1a | gal3st1a,gal3st1a,gal3st1 | 3 |
| gal3st1b | gal3st1a,gal3st1a,gal3st1 | 3 |
| galnt18a | galnt18b,galnt18a | 2 |
| galnt18b | galnt18b,galnt18a | 2 |
| galr1a | galr1b,galr1a | 2 |
| galr1b | galr1b,galr1a | 2 |
| galr2a | galr2a,galr2b,galr2a | 3 |
| galr2a | galr2a,galr2b,galr2a | 3 |
| galr2b | galr2a,galr2b,galr2a | 3 |
| gas1a | gas1b,gas1a | 2 |
| gas1b | gas1b,gas1a | 2 |
| gas2a | gas2b,gas2a | 2 |
| gas2b | gas2b,gas2a | 2 |
| gas7a | gas7a,gas7a | 2 |
| gas7a | gas7a,gas7a | 2 |
| gask1a | gask1b,gask1a | 2 |
| gask1b | gask1b,gask1a | 2 |
| gata1a | gata1a,gata1b,gata1a | 3 |
| gata1a | gata1a,gata1b,gata1a | 3 |
| gata1b | gata1a,gata1b,gata1a | 3 |
| gata2a | gata2b,gata2a | 2 |
| gata2b | gata2b,gata2a | 2 |
| gatad2ab | gatad2b,gatad2ab | 2 |
| gatad2b | gatad2b,gatad2ab | 2 |
| gatd3a | gatd3a | 1 |
| gbe1a | gbe1a,gbe1a,gbe1b | 3 |
| gbe1a | gbe1a,gbe1a,gbe1b | 3 |
| gbe1b | gbe1a,gbe1a,gbe1b | 3 |
| gcnt4a | gcnt4a,gcnt4a,gcnt4b | 3 |
| gcnt4a | gcnt4a,gcnt4a,gcnt4b | 3 |
| gcnt4b | gcnt4a,gcnt4a,gcnt4b | 3 |
| gdf10a | gdf10b,gdf10a | 2 |
| gdf10b | gdf10b,gdf10a | 2 |
| gdf6a | gdf6a,gdf6b | 2 |
| gdf6b | gdf6a,gdf6b | 2 |
| gdpd3a | gdpd3a,gdpd3b | 2 |
| gdpd3b | gdpd3a,gdpd3b | 2 |
| gdpd4a | gdpd4a,gdpd4b | 2 |
| gdpd4b | gdpd4a,gdpd4b | 2 |
| gdpd5a | gdpd5a,gdpd5b | 2 |
| gdpd5b | gdpd5a,gdpd5b | 2 |

| | | |
|---|---|---|
| gfi1aa | gfi1ab,gfi1aa,gfi1ab,gfi1b | 4 |
| gfi1ab | gfi1ab,gfi1aa,gfi1ab,gfi1b | 4 |
| gfi1ab | gfi1ab,gfi1aa,gfi1ab,gfi1b | 4 |
| gfi1b | gfi1ab,gfi1aa,gfi1ab,gfi1b | 4 |
| gfra1a | gfra1b,gfra1a | 2 |
| gfra1b | gfra1b,gfra1a | 2 |
| gfra2a | gfra2a,gfra2b | 2 |
| gfra2b | gfra2a,gfra2b | 2 |
| gfra4a | gfra4a,gfra4a,gfra4b | 3 |
| gfra4a | gfra4a,gfra4a,gfra4b | 3 |
| gfra4b | gfra4a,gfra4a,gfra4b | 3 |
| gga3a | gga3b,gga3a | 2 |
| gga3b | gga3b,gga3a | 2 |
| ggt1a | ggt1a,ggt1b | 2 |
| ggt1b | ggt1a,ggt1b | 2 |
| ggt5a | ggt5a | 1 |
| gid8a | gid8a,gid8b | 2 |
| gid8b | gid8a,gid8b | 2 |
| gig2d | gig2k,gig2j,gig2i,gig2h,gig | 19 |
| gig2d | gig2k,gig2j,gig2i,gig2h,gig | 19 |
| gig2e | gig2k,gig2j,gig2i,gig2h,gig | 19 |
| gig2e | gig2k,gig2j,gig2i,gig2h,gig | 19 |
| gig2f | gig2k,gig2j,gig2i,gig2h,gig | 19 |
| gig2f | gig2k,gig2j,gig2i,gig2h,gig | 19 |
| gig2g | gig2k,gig2j,gig2i,gig2h,gig | 19 |
| gig2g | gig2k,gig2j,gig2i,gig2h,gig | 19 |
| gig2h | gig2k,gig2j,gig2i,gig2h,gig | 19 |
| gig2h | gig2k,gig2j,gig2i,gig2h,gig | 19 |
| gig2i | gig2k,gig2j,gig2i,gig2h,gig | 19 |
| gig2i | gig2k,gig2j,gig2i,gig2h,gig | 19 |
| gig2j | gig2k,gig2j,gig2i,gig2h,gig | 19 |
| gig2j | gig2k,gig2j,gig2i,gig2h,gig | 19 |
| gig2k | gig2k,gig2j,gig2i,gig2h,gig | 19 |
| gig2k | gig2k,gig2j,gig2i,gig2h,gig | 19 |
| gig2o | gig2k,gig2j,gig2i,gig2h,gig | 19 |
| gig2p | gig2k,gig2j,gig2i,gig2h,gig | 19 |
| gig2q | gig2k,gig2j,gig2i,gig2h,gig | 19 |
| gigyf1a | gigyf1a,gigyf1b | 2 |
| gigyf1b | gigyf1a,gigyf1b | 2 |
| git2a | git2b,git2a,git2b | 3 |
| git2b | git2b,git2a,git2b | 3 |
| git2b | git2b,git2a,git2b | 3 |
| gja5a | gja5b,gja5a | 2 |
| gja5b | gja5b,gja5a | 2 |
| gja8a | gja8b,gja8a | 2 |
| gja8b | gja8b,gja8a | 2 |
| gjd1a | gjd1a,gjd1a | 2 |
| gjd1a | gjd1a,gjd1a | 2 |
| gjd2b | gjd2b,gjd2b,gjd2b | 3 |
| gjd2b | gjd2b,gjd2b,gjd2b | 3 |

| | | |
|---|---|---|
| gjd2b | gjd2b,gjd2b,gjd2b | 3 |
| glcci1a | glcci1a | 1 |
| glg1a | glg1b,glg1a | 2 |
| glg1b | glg1b,glg1a | 2 |
| gli2a | gli2a,gli2b | 2 |
| gli2b | gli2a,gli2b | 2 |
| glipr1a | glipr1a,glipr1b | 2 |
| glipr1b | glipr1a,glipr1b | 2 |
| glis1a | glis1a,glis1b | 2 |
| glis1b | glis1a,glis1b | 2 |
| glis2a | glis2b,glis2a | 2 |
| glis2b | glis2b,glis2a | 2 |
| glra4a | glra4a,glra4b,glra4b,glra4 | 4 |
| glra4a | glra4a,glra4b,glra4b,glra4 | 4 |
| glra4b | glra4a,glra4b,glra4b,glra4 | 4 |
| glra4b | glra4a,glra4b,glra4b,glra4 | 4 |
| gls2a | gls2a,gls2b | 2 |
| gls2b | gls2a,gls2b | 2 |
| glud1a | glud1a,glud1b,glud1a | 3 |
| glud1a | glud1a,glud1b,glud1a | 3 |
| glud1b | glud1a,glud1b,glud1a | 3 |
| gm2a | gm2a | 1 |
| gna11a | gna11a,gna11b,gna11a | 3 |
| gna11a | gna11a,gna11b,gna11a | 3 |
| gna11b | gna11a,gna11b,gna11a | 3 |
| gna12a | gna12a | 1 |
| gna13a | gna13b,gna13a | 2 |
| gna13b | gna13b,gna13a | 2 |
| gna14a | gna14,gna14a | 2 |
| gnai2a | gnai2b,gnai2a | 2 |
| gnai2b | gnai2b,gnai2a | 2 |
| gnao1a | gnao1b,gnao1b,gnao1a | 3 |
| gnao1b | gnao1b,gnao1b,gnao1a | 3 |
| gnao1b | gnao1b,gnao1b,gnao1a | 3 |
| gnb1a | gnb1b,gnb1a | 2 |
| gnb1b | gnb1b,gnb1a | 2 |
| gnb3a | gnb3a,gnb3a,gnb3b | 3 |
| gnb3a | gnb3a,gnb3a,gnb3b | 3 |
| gnb3b | gnb3a,gnb3a,gnb3b | 3 |
| gnb4b | gnb4b | 1 |
| gnb5a | gnb5b,gnb5a | 2 |
| gnb5b | gnb5b,gnb5a | 2 |
| gng12a | gng12a | 1 |
| gng13a | gng13a,gng13a,gng13b | 3 |
| gng13a | gng13a,gng13a,gng13b | 3 |
| gng13b | gng13a,gng13a,gng13b | 3 |
| gngt2a | gngt2a,gngt2a,gngt2b | 3 |
| gngt2a | gngt2a,gngt2a,gngt2b | 3 |
| gngt2b | gngt2a,gngt2a,gngt2b | 3 |
| golga7ba | golga7ba,golga7,golga7bl | 3 |

| | | |
|---|---|---|
| golga7bb | golga7ba,golga7,golga7bl | 3 |
| golim4a | golim4a,golim4a,golim4b | 4 |
| golim4a | golim4a,golim4a,golim4b | 4 |
| golim4a | golim4a,golim4a,golim4b | 4 |
| golim4b | golim4a,golim4a,golim4b | 4 |
| golt1a | golt1ba,golt1a,golt1bb | 3 |
| golt1ba | golt1ba,golt1a,golt1bb | 3 |
| golt1bb | golt1ba,golt1a,golt1bb | 3 |
| gorasp1a | gorasp1a,gorasp1b | 2 |
| gorasp1b | gorasp1a,gorasp1b | 2 |
| got2a | got2a,got2b,got2a | 3 |
| got2a | got2a,got2b,got2a | 3 |
| got2b | got2a,got2b,got2a | 3 |
| gp1bb | gp1bb | 1 |
| gpa33a | gpa33a,gpa33b | 2 |
| gpa33b | gpa33a,gpa33b | 2 |
| gpc1a | gpc1a,gpc1b,gpc1a | 3 |
| gpc1a | gpc1a,gpc1b,gpc1a | 3 |
| gpc1b | gpc1a,gpc1b,gpc1a | 3 |
| gpc5a | gpc5b,gpc5a,gpc5c,gpc5b | 4 |
| gpc5b | gpc5b,gpc5a,gpc5c,gpc5b | 4 |
| gpc5b | gpc5b,gpc5a,gpc5c,gpc5b | 4 |
| gpc5c | gpc5b,gpc5a,gpc5c,gpc5b | 4 |
| gpc6a | gpc6a,gpc6b | 2 |
| gpc6b | gpc6a,gpc6b | 2 |
| gpd1a | gpd1a,gpd1c,gpd1b | 3 |
| gpd1b | gpd1a,gpd1c,gpd1b | 3 |
| gpd1c | gpd1a,gpd1c,gpd1b | 3 |
| gpm6aa | gpm6aa,gpm6ba,gpm6ba | 6 |
| gpm6aa | gpm6aa,gpm6ba,gpm6ba | 6 |
| gpm6ab | gpm6aa,gpm6ba,gpm6ba | 6 |
| gpm6ba | gpm6aa,gpm6ba,gpm6ba | 6 |
| gpm6ba | gpm6aa,gpm6ba,gpm6ba | 6 |
| gpm6bb | gpm6aa,gpm6ba,gpm6ba | 6 |
| gpr132a | gpr132a,gpr132b | 2 |
| gpr132b | gpr132a,gpr132b | 2 |
| gpr137ba | gpr137bb,gpr137ba,gpr1: | 4 |
| gpr137bb | gpr137bb,gpr137ba,gpr1: | 4 |
| gpr137c | gpr137bb,gpr137ba,gpr1: | 4 |
| gpr155a | gpr155b,gpr155a | 2 |
| gpr155b | gpr155b,gpr155a | 2 |
| gpr158a | gpr158b,gpr158a | 2 |
| gpr158b | gpr158b,gpr158a | 2 |
| gpr183a | gpr183b,gpr183a | 2 |
| gpr183b | gpr183b,gpr183a | 2 |
| gpr185a | gpr185b,gpr185a | 2 |
| gpr185b | gpr185b,gpr185a | 2 |
| gpr22a | gpr22a,gpr22b | 2 |
| gpr22b | gpr22a,gpr22b | 2 |
| gpr34b | gpr34b | 1 |

| | | |
|---|---|---|
| gpr37a | gpr37b,gpr37a | 2 |
| gpr37b | gpr37b,gpr37a | 2 |
| gpr37l1a | gpr37l1b,gpr37l1b,gpr37l | 3 |
| gpr37l1b | gpr37l1b,gpr37l1b,gpr37l | 3 |
| gpr37l1b | gpr37l1b,gpr37l1b,gpr37l | 3 |
| gpr55a | gpr55a | 1 |
| gpr78a | gpr78a,gpr78b | 2 |
| gpr78b | gpr78a,gpr78b | 2 |
| gprc5ba | gprc5bb,gprc5ba,gprc5c | 3 |
| gprc5bb | gprc5bb,gprc5ba,gprc5c | 3 |
| gprc5c | gprc5bb,gprc5ba,gprc5c | 3 |
| gprc6a | gprc6a | 1 |
| gpsm1a | gpsm1a,gpsm1b | 2 |
| gpsm1b | gpsm1a,gpsm1b | 2 |
| gpx1a | gpx1a,gpx1b,gpx1a | 3 |
| gpx1a | gpx1a,gpx1b,gpx1a | 3 |
| gpx1b | gpx1a,gpx1b,gpx1a | 3 |
| gpx4a | gpx4b,gpx4a | 2 |
| gpx4b | gpx4b,gpx4a | 2 |
| gramd1a | gramd1ba,gramd1a,gram | 4 |
| gramd1ba | gramd1ba,gramd1a,gram | 4 |
| gramd1bb | gramd1ba,gramd1a,gram | 4 |
| gramd1c | gramd1ba,gramd1a,gram | 4 |
| gramd2aa | gramd2aa | 1 |
| gramd4a | gramd4a,gramd4b | 2 |
| gramd4b | gramd4a,gramd4b | 2 |
| grap2a | grap2a,grap2b | 2 |
| grap2b | grap2a,grap2b | 2 |
| grb10a | grb10b,grb10a | 2 |
| grb10b | grb10b,grb10a | 2 |
| grb2a | grb2a,grb2a,grb2b,grb2a | 4 |
| grb2a | grb2a,grb2a,grb2b,grb2a | 4 |
| grb2a | grb2a,grb2a,grb2b,grb2a | 4 |
| grb2b | grb2a,grb2a,grb2b,grb2a | 4 |
| grem1a | grem1b,grem1a,grem1a | 3 |
| grem1a | grem1b,grem1a,grem1a | 3 |
| grem1b | grem1b,grem1a,grem1a | 3 |
| grem2a | grem2b,grem2a | 2 |
| grem2b | grem2b,grem2a | 2 |
| grhl2a | grhl2a,grhl2b | 2 |
| grhl2b | grhl2a,grhl2b | 2 |
| gria1a | gria1a,gria1a,gria1a,gria1 | 6 |
| gria1a | gria1a,gria1a,gria1a,gria1 | 6 |
| gria1a | gria1a,gria1a,gria1a,gria1 | 6 |
| gria1a | gria1a,gria1a,gria1a,gria1 | 6 |
| gria1b | gria1a,gria1a,gria1a,gria1 | 6 |
| gria1b | gria1a,gria1a,gria1a,gria1 | 6 |
| gria2a | gria2a,gria2a,gria2a,gria2 | 4 |
| gria2a | gria2a,gria2a,gria2a,gria2 | 4 |
| gria2a | gria2a,gria2a,gria2a,gria2 | 4 |

| | | |
|---|---|---|
| gria2b | gria2a,gria2a,gria2a,gria2 | 4 |
| gria3a | gria3b,gria3a,gria3b | 3 |
| gria3b | gria3b,gria3a,gria3b | 3 |
| gria3b | gria3b,gria3a,gria3b | 3 |
| gria4a | gria4b,gria4b,gria4a,gria4 | 4 |
| gria4b | gria4b,gria4b,gria4a,gria4 | 4 |
| gria4b | gria4b,gria4b,gria4a,gria4 | 4 |
| gria4b | gria4b,gria4b,gria4a,gria4 | 4 |
| grid1a | grid1b,grid1a | 2 |
| grid1b | grid1b,grid1a | 2 |
| grid2ipa | grid2ipb,grid2,grid2ipa,gr | 5 |
| grid2ipb | grid2ipb,grid2,grid2ipa,gr | 5 |
| grid2ipb | grid2ipb,grid2,grid2ipa,gr | 5 |
| grik1a | grik1a,grik1b | 2 |
| grik1b | grik1a,grik1b | 2 |
| grin1a | grin1b,grin1a | 2 |
| grin1b | grin1b,grin1a | 2 |
| grin2aa | grin2ab,grin2bb,grin2ca,g | 7 |
| grin2ab | grin2ab,grin2bb,grin2ca,g | 7 |
| grin2bb | grin2ab,grin2bb,grin2ca,g | 7 |
| grin2ca | grin2ab,grin2bb,grin2ca,g | 7 |
| grin2cb | grin2ab,grin2bb,grin2ca,g | 7 |
| grin2da | grin2ab,grin2bb,grin2ca,g | 7 |
| grin2db | grin2ab,grin2bb,grin2ca,g | 7 |
| grin3a | grin3ba,grin3a,grin3bb | 3 |
| grin3ba | grin3ba,grin3a,grin3bb | 3 |
| grin3bb | grin3ba,grin3a,grin3bb | 3 |
| grip2a | grip2a,grip2b | 2 |
| grip2b | grip2a,grip2b | 2 |
| grk1a | grk1a,grk1b | 2 |
| grk1b | grk1a,grk1b | 2 |
| grk7a | grk7a,grk7b | 2 |
| grk7b | grk7a,grk7b | 2 |
| grm1a | grm1b,grm1b,grm1a | 3 |
| grm1b | grm1b,grm1b,grm1a | 3 |
| grm1b | grm1b,grm1b,grm1a | 3 |
| grm2a | grm2b,grm2a,grm2b | 3 |
| grm2b | grm2b,grm2a,grm2b | 3 |
| grm2b | grm2b,grm2a,grm2b | 3 |
| grm5a | grm5a,grm5b | 2 |
| grm5b | grm5a,grm5b | 2 |
| grm6a | grm6b,grm6a,grm6b | 3 |
| grm6b | grm6b,grm6a,grm6b | 3 |
| grm6b | grm6b,grm6a,grm6b | 3 |
| grm8a | grm8a,grm8b | 2 |
| grm8b | grm8a,grm8b | 2 |
| grtp1a | grtp1a,grtp1b | 2 |
| grtp1b | grtp1a,grtp1b | 2 |
| grxcr1a | grxcr1b,grxcr1a | 2 |
| grxcr1b | grxcr1b,grxcr1a | 2 |

| | | |
|---|---|---|
| gsg1l2a | gsg1l2a,gsg1l2b | 2 |
| gsg1l2b | gsg1l2a,gsg1l2b | 2 |
| gsk3aa | gsk3bb,gsk3ba,gsk3ab,gsl | 4 |
| gsk3ab | gsk3bb,gsk3ba,gsk3ab,gsl | 4 |
| gsk3ba | gsk3bb,gsk3ba,gsk3ab,gsl | 4 |
| gsk3bb | gsk3bb,gsk3ba,gsk3ab,gsl | 4 |
| gstt1a | gstt1b,gstt1a,gstt1a,gstt1 | 4 |
| gstt1a | gstt1b,gstt1a,gstt1a,gstt1 | 4 |
| gstt1b | gstt1b,gstt1a,gstt1a,gstt1 | 4 |
| gstt1b | gstt1b,gstt1a,gstt1a,gstt1 | 4 |
| gtf2b | gtf2b | 1 |
| gtf2f2a | gtf2f2a,gtf2f2b | 2 |
| gtf2f2b | gtf2f2a,gtf2f2b | 2 |
| gtf3aa | gtf3ab,gtf3aa,gtf3ab | 3 |
| gtf3ab | gtf3ab,gtf3aa,gtf3ab | 3 |
| gtf3ab | gtf3ab,gtf3aa,gtf3ab | 3 |
| gtpbp2b | gtpbp2b,gtpbp2b | 2 |
| gtpbp2b | gtpbp2b,gtpbp2b | 2 |
| guca1a | guca1d,guca1b,guca1g,gu | 9 |
| guca1b | guca1d,guca1b,guca1g,gu | 9 |
| guca1b | guca1d,guca1b,guca1g,gu | 9 |
| guca1c | guca1d,guca1b,guca1g,gu | 9 |
| guca1d | guca1d,guca1b,guca1g,gu | 9 |
| guca1d | guca1d,guca1b,guca1g,gu | 9 |
| guca1e | guca1d,guca1b,guca1g,gu | 9 |
| guca1g | guca1d,guca1b,guca1g,gu | 9 |
| guca1g | guca1d,guca1b,guca1g,gu | 9 |
| gucy2c | gucy2c,gucy2d,gucy2c,gu | 6 |
| gucy2c | gucy2c,gucy2d,gucy2c,gu | 6 |
| gucy2d | gucy2c,gucy2d,gucy2c,gu | 6 |
| gucy2d | gucy2c,gucy2d,gucy2c,gu | 6 |
| gucy2f | gucy2c,gucy2d,gucy2c,gu | 6 |
| gucy2g | gucy2c,gucy2d,gucy2c,gu | 6 |
| guk1a | guk1a,guk1b | 2 |
| guk1b | guk1a,guk1b | 2 |
| gulp1a | gulp1b,gulp1a | 2 |
| gulp1b | gulp1b,gulp1a | 2 |
| gxylt1b | gxylt1b | 1 |
| gyg1a | gyg1a,gyg1b | 2 |
| gyg1b | gyg1a,gyg1b | 2 |
| h1fx | h1m,h1fx | 2 |
| h1m | h1m,h1fx | 2 |
| h2afva | h2afva,h2afx,h2afvb,h2af | 4 |
| h2afvb | h2afva,h2afx,h2afvb,h2af | 4 |
| h2afx | h2afva,h2afx,h2afvb,h2af | 4 |
| h2afy | h2afva,h2afx,h2afvb,h2af | 4 |
| h3f3a | h3f3a,h3f3c | 2 |
| h3f3c | h3f3a,h3f3c | 2 |
| h6pd | h6pd,h6pd | 2 |
| h6pd | h6pd,h6pd | 2 |

| | | |
|---|---|---|
| hapln1a | hapln1a,hapln1b | 2 |
| hapln1b | hapln1a,hapln1b | 2 |
| hcfc1a | hcfc1a,hcfc1b,hcfc1a | 3 |
| hcfc1a | hcfc1a,hcfc1b,hcfc1a | 3 |
| hcfc1b | hcfc1a,hcfc1b,hcfc1a | 3 |
| hcn2b | hcn2b | 1 |
| hdac7a | hdac7b,hdac7a | 2 |
| hdac7b | hdac7b,hdac7a | 2 |
| hdac9b | hdac9b | 1 |
| heatr5a | heatr5b,heatr5a | 2 |
| heatr5b | heatr5b,heatr5a | 2 |
| hecw1b | hecw1b,hecw1b,hecw1b | 3 |
| hecw1b | hecw1b,hecw1b,hecw1b | 3 |
| hecw1b | hecw1b,hecw1b,hecw1b | 3 |
| hecw2a | hecw2b,hecw2a | 2 |
| hecw2b | hecw2b,hecw2a | 2 |
| hephl1a | hephl1b,hephl1b,hephl1a | 3 |
| hephl1b | hephl1b,hephl1b,hephl1a | 3 |
| hephl1b | hephl1b,hephl1b,hephl1a | 3 |
| her8a | her8a | 1 |
| hiat1a | hiat1a,hiat1b | 2 |
| hiat1b | hiat1a,hiat1b | 2 |
| hid1a | hid1a,hid1b | 2 |
| hid1b | hid1a,hid1b | 2 |
| hif1aa | hif1an,hif1aa,hif1ab | 3 |
| hif1ab | hif1an,hif1aa,hif1ab | 3 |
| hif1an | hif1an,hif1aa,hif1ab | 3 |
| higd1a | higd1a | 1 |
| higd2a | higd2a | 1 |
| hip1ra | hip1,hip1rb,hip1ra,hip1 | 4 |
| hip1rb | hip1,hip1rb,hip1ra,hip1 | 4 |
| hipk1a | hipk1a | 1 |
| hipk3a | hipk3b,hipk3a | 2 |
| hipk3b | hipk3b,hipk3a | 2 |
| hist1h2ba | hist1h2ba | 1 |
| hist2h3c | hist2h3c | 1 |
| hivep2a | hivep2b,hivep2a | 2 |
| hivep2b | hivep2b,hivep2a | 2 |
| hivep3a | hivep3b,hivep3a | 2 |
| hivep3b | hivep3b,hivep3a | 2 |
| hmbox1a | hmbox1a | 1 |
| hmg20a | hmg20b,hmg20b,hmg20a | 3 |
| hmg20b | hmg20b,hmg20b,hmg20a | 3 |
| hmg20b | hmg20b,hmg20b,hmg20a | 3 |
| hmga1a | hmga1b,hmga1a | 2 |
| hmga1b | hmga1b,hmga1a | 2 |
| hmgb1a | hmgb1a,hmgb1a,hmgb1b | 3 |
| hmgb1a | hmgb1a,hmgb1a,hmgb1b | 3 |
| hmgb1b | hmgb1a,hmgb1a,hmgb1b | 3 |
| hmgb2a | hmgb2a,hmgb2b | 2 |

| | | |
|---|---|---|
| hmgb2b | hmgb2a,hmgb2b | 2 |
| hmgb3a | hmgb3a,hmgb3b | 2 |
| hmgb3b | hmgb3a,hmgb3b | 2 |
| hmgxb4a | hmgxb4a | 1 |
| hmox1a | hmox1a | 1 |
| hmox2a | hmox2a,hmox2b,hmox2b | 4 |
| hmox2a | hmox2a,hmox2b,hmox2b | 4 |
| hmox2b | hmox2a,hmox2b,hmox2b | 4 |
| hmox2b | hmox2a,hmox2b,hmox2b | 4 |
| hmx3a | hmx3a,hmx3b,hmx3a | 3 |
| hmx3a | hmx3a,hmx3b,hmx3a | 3 |
| hmx3b | hmx3a,hmx3b,hmx3a | 3 |
| hnf1a | hnf1ba,hnf1a,hnf1ba,hnf | 4 |
| hnf1ba | hnf1ba,hnf1a,hnf1ba,hnf | 4 |
| hnf1ba | hnf1ba,hnf1a,hnf1ba,hnf | 4 |
| hnf1bb | hnf1ba,hnf1a,hnf1ba,hnf | 4 |
| hnf4a | hnf4a,hnf4b,hnf4a,hnf4g | 4 |
| hnf4a | hnf4a,hnf4b,hnf4a,hnf4g | 4 |
| hnf4b | hnf4a,hnf4b,hnf4a,hnf4g | 4 |
| hnf4g | hnf4a,hnf4b,hnf4a,hnf4g | 4 |
| hnrnpa0a | hnrnpa0b,hnrnpa0a,hnrn | 3 |
| hnrnpa0b | hnrnpa0b,hnrnpa0a,hnrn | 3 |
| hnrnpa0b | hnrnpa0b,hnrnpa0a,hnrn | 3 |
| hnrnpa1a | hnrnpa1a,hnrnpa1b | 2 |
| hnrnpa1b | hnrnpa1a,hnrnpa1b | 2 |
| homer1b | homer1b | 1 |
| homer3a | homer3a,homer3b | 2 |
| homer3b | homer3a,homer3b | 2 |
| hoxa10b | hoxa10b | 1 |
| hoxa11a | hoxa11b,hoxa11a | 2 |
| hoxa11b | hoxa11b,hoxa11a | 2 |
| hoxa13a | hoxa13b,hoxa13a | 2 |
| hoxa13b | hoxa13b,hoxa13a | 2 |
| hoxa1a | hoxa1a | 1 |
| hoxa2b | hoxa2b | 1 |
| hoxa4a | hoxa4a | 1 |
| hoxa5a | hoxa5a | 1 |
| hoxa9a | hoxa9b,hoxa9a | 2 |
| hoxa9b | hoxa9b,hoxa9a | 2 |
| hoxb10a | hoxb10a | 1 |
| hoxb13a | hoxb13a | 1 |
| hoxb1a | hoxb1b,hoxb1a,hoxb1b | 3 |
| hoxb1b | hoxb1b,hoxb1a,hoxb1b | 3 |
| hoxb1b | hoxb1b,hoxb1a,hoxb1b | 3 |
| hoxb2a | hoxb2a | 1 |
| hoxb3a | hoxb3a | 1 |
| hoxb4a | hoxb4a | 1 |
| hoxb5a | hoxb5b,hoxb5a,hoxb5b | 3 |
| hoxb5b | hoxb5b,hoxb5a,hoxb5b | 3 |
| hoxb5b | hoxb5b,hoxb5a,hoxb5b | 3 |

| | | |
|---|---|---|
| hoxb6a | hoxb6b,hoxb6a,hoxb6b | 3 |
| hoxb6b | hoxb6b,hoxb6a,hoxb6b | 3 |
| hoxb6b | hoxb6b,hoxb6a,hoxb6b | 3 |
| hoxb7a | hoxb7a | 1 |
| hoxb8a | hoxb8b,hoxb8a,hoxb8b | 3 |
| hoxb8b | hoxb8b,hoxb8a,hoxb8b | 3 |
| hoxb8b | hoxb8b,hoxb8a,hoxb8b | 3 |
| hoxb9a | hoxb9a | 1 |
| hoxc10a | hoxc10a | 1 |
| hoxc11a | hoxc11b,hoxc11a | 2 |
| hoxc11b | hoxc11b,hoxc11a | 2 |
| hoxc12a | hoxc12b,hoxc12a | 2 |
| hoxc12b | hoxc12b,hoxc12a | 2 |
| hoxc13a | hoxc13b,hoxc13a | 2 |
| hoxc13b | hoxc13b,hoxc13a | 2 |
| hoxc1a | hoxc1a | 1 |
| hoxc3a | hoxc3a | 1 |
| hoxc4a | hoxc4a | 1 |
| hoxc5a | hoxc5a | 1 |
| hoxc6a | hoxc6b,hoxc6a | 2 |
| hoxc6b | hoxc6b,hoxc6a | 2 |
| hoxc8a | hoxc8a | 1 |
| hoxc9a | hoxc9a | 1 |
| hoxd10a | hoxd10a | 1 |
| hoxd11a | hoxd11a | 1 |
| hoxd12a | hoxd12a | 1 |
| hoxd13a | hoxd13a | 1 |
| hoxd3a | hoxd3a | 1 |
| hoxd4a | hoxd4a | 1 |
| hoxd9a | hoxd9a | 1 |
| hrh2a | hrh2b,hrh2b,hrh2a | 3 |
| hrh2b | hrh2b,hrh2b,hrh2a | 3 |
| hrh2b | hrh2b,hrh2b,hrh2a | 3 |
| hs2st1a | hs2st1a,hs2st1b | 2 |
| hs2st1b | hs2st1a,hs2st1b | 2 |
| hs3st3b1a | hs3st3b1b,hs3st3b1a,hs3 | 4 |
| hs3st3b1a | hs3st3b1b,hs3st3b1a,hs3 | 4 |
| hs3st3b1b | hs3st3b1b,hs3st3b1a,hs3 | 4 |
| hs3st3b1b | hs3st3b1b,hs3st3b1a,hs3 | 4 |
| hs6st1a | hs6st1b,hs6st1a,hs6st1b | 3 |
| hs6st1b | hs6st1b,hs6st1a,hs6st1b | 3 |
| hs6st1b | hs6st1b,hs6st1a,hs6st1b | 3 |
| hs6st3a | hs6st3a,hs6st3b | 2 |
| hs6st3b | hs6st3a,hs6st3b | 2 |
| hsbp1a | hsbp1a,hsbp1b | 2 |
| hsbp1b | hsbp1a,hsbp1b | 2 |
| hsd17b12a | hsd17b12b,hsd17b12a | 2 |
| hsd17b12b | hsd17b12b,hsd17b12a | 2 |
| hsf2bp | hsf2bp,hsf2 | 2 |
| hsh2d | hsh2d,hsh2d | 2 |

| | | |
|---|---|---|
| hsh2d | hsh2d,hsh2d | 2 |
| hspa12a | hspa12b,hspa12a | 2 |
| hspa12b | hspa12b,hspa12a | 2 |
| hspa4a | hspa4b,hspa4b,hspa4b,hs | 4 |
| hspa4b | hspa4b,hspa4b,hspa4b,hs | 4 |
| hspa4b | hspa4b,hspa4b,hspa4b,hs | 4 |
| hspa4b | hspa4b,hspa4b,hspa4b,hs | 4 |
| htr1aa | htr1d,htr1ab,htr1fa,htr1f | 9 |
| htr1ab | htr1d,htr1ab,htr1fa,htr1f | 9 |
| htr1ab | htr1d,htr1ab,htr1fa,htr1f | 9 |
| htr1b | htr1d,htr1ab,htr1fa,htr1f | 9 |
| htr1d | htr1d,htr1ab,htr1fa,htr1f | 9 |
| htr1d | htr1d,htr1ab,htr1fa,htr1f | 9 |
| htr1fa | htr1d,htr1ab,htr1fa,htr1f | 9 |
| htr1fa | htr1d,htr1ab,htr1fa,htr1f | 9 |
| htr1fb | htr1d,htr1ab,htr1fa,htr1f | 9 |
| htr2aa | htr2ab,htr2aa,htr2b | 3 |
| htr2ab | htr2ab,htr2aa,htr2b | 3 |
| htr2b | htr2ab,htr2aa,htr2b | 3 |
| htr3a | htr3b,htr3a | 2 |
| htr3b | htr3b,htr3a | 2 |
| htr5aa | htr5aa,htr5ab | 2 |
| htr5ab | htr5aa,htr5ab | 2 |
| htr7a | htr7a,htr7a,htr7c,htr7b,h | 5 |
| htr7a | htr7a,htr7a,htr7c,htr7b,h | 5 |
| htr7a | htr7a,htr7a,htr7c,htr7b,h | 5 |
| htr7b | htr7a,htr7a,htr7c,htr7b,h | 5 |
| htr7c | htr7a,htr7a,htr7c,htr7b,h | 5 |
| htra1a | htra1a,htra1b,htra1a | 3 |
| htra1a | htra1a,htra1b,htra1a | 3 |
| htra1b | htra1a,htra1b,htra1a | 3 |
| htra3a | htra3a | 1 |
| hyal2a | hyal2a,hyal2b | 2 |
| hyal2b | hyal2a,hyal2b | 2 |
| id2a | id2a,id2b | 2 |
| id2b | id2a,id2b | 2 |
| idh3a | idh3b,idh3b,idh3g,idh3a | 4 |
| idh3b | idh3b,idh3b,idh3g,idh3a | 4 |
| idh3b | idh3b,idh3b,idh3g,idh3a | 4 |
| idh3g | idh3b,idh3b,idh3g,idh3a | 4 |
| ier2a | ier2a,ier2b | 2 |
| ier2b | ier2a,ier2b | 2 |
| iffo1a | iffo1b,iffo1a | 2 |
| iffo1b | iffo1b,iffo1a | 2 |
| iffo2a | iffo2a,iffo2b,iffo2a | 3 |
| iffo2a | iffo2a,iffo2b,iffo2a | 3 |
| iffo2b | iffo2a,iffo2b,iffo2a | 3 |
| ifng1r | ifng1,ifng1r | 2 |
| igf1ra | igf1ra,igf1ra,igf1rb,igf1,ig | 6 |
| igf1ra | igf1ra,igf1ra,igf1rb,igf1,ig | 6 |

| | | |
|---|---|---|
| igf1ra | igf1ra,igf1ra,igf1rb,igf1,ig | 6 |
| igf1rb | igf1ra,igf1ra,igf1rb,igf1,ig | 6 |
| igf1rb | igf1ra,igf1ra,igf1rb,igf1,ig | 6 |
| igf2a | igf2b,igf2a,igf2r,igf2b | 4 |
| igf2b | igf2b,igf2a,igf2r,igf2b | 4 |
| igf2b | igf2b,igf2a,igf2r,igf2b | 4 |
| igf2bp2a | igf2bp2b,igf2bp2b,igf2bp | 4 |
| igf2bp2b | igf2bp2b,igf2bp2b,igf2bp | 4 |
| igf2bp2b | igf2bp2b,igf2bp2b,igf2bp | 4 |
| igf2bp2b | igf2bp2b,igf2bp2b,igf2bp | 4 |
| igf2r | igf2b,igf2a,igf2r,igf2b | 4 |
| igfbp1a | igfbp1b,igfbp1a | 2 |
| igfbp1b | igfbp1b,igfbp1a | 2 |
| igfbp2a | igfbp2a,igfbp2a,igfbp2b | 3 |
| igfbp2a | igfbp2a,igfbp2a,igfbp2b | 3 |
| igfbp2b | igfbp2a,igfbp2a,igfbp2b | 3 |
| igfbp5a | igfbp5a,igfbp5a,igfbp5b | 3 |
| igfbp5a | igfbp5a,igfbp5a,igfbp5b | 3 |
| igfbp5b | igfbp5a,igfbp5a,igfbp5b | 3 |
| igfbp6a | igfbp6a,igfbp6b | 2 |
| igfbp6b | igfbp6a,igfbp6b | 2 |
| igsf21a | igsf21a,igsf21b | 2 |
| igsf21b | igsf21a,igsf21b | 2 |
| igsf5a | igsf5a,igsf5b | 2 |
| igsf5b | igsf5a,igsf5b | 2 |
| igsf9a | igsf9b,igsf9a,igsf9ba,igsf9 | 4 |
| igsf9b | igsf9b,igsf9a,igsf9ba,igsf9 | 4 |
| igsf9ba | igsf9b,igsf9a,igsf9ba,igsf9 | 4 |
| igsf9bb | igsf9b,igsf9a,igsf9ba,igsf9 | 4 |
| il10ra | il10rb,il10,il10ra | 3 |
| il10rb | il10rb,il10,il10ra | 3 |
| il11a | il11ra,il11a,il11b | 3 |
| il11b | il11ra,il11a,il11b | 3 |
| il11ra | il11ra,il11a,il11b | 3 |
| il12a | il12ba,il12a,il12bb | 3 |
| il12ba | il12ba,il12a,il12bb | 3 |
| il12bb | il12ba,il12a,il12bb | 3 |
| il15ra | il15ra,il15,il15ra | 3 |
| il15ra | il15ra,il15,il15ra | 3 |
| il17c | il17c,il17d,il17d,il17rc,il1 | 6 |
| il17c | il17c,il17d,il17d,il17rc,il1 | 6 |
| il17d | il17c,il17d,il17d,il17rc,il1 | 6 |
| il17d | il17c,il17d,il17d,il17rc,il1 | 6 |
| il17ra1a | il17ra1a,il17ra1b | 2 |
| il17ra1b | il17ra1a,il17ra1b | 2 |
| il17rc | il17c,il17d,il17d,il17rc,il1 | 6 |
| il17rd | il17c,il17d,il17d,il17rc,il1 | 6 |
| il1b | il1b,il1fma | 2 |
| il1fma | il1b,il1fma | 2 |
| il1rapl1a | il1rapl1a,il1rapl1a,il1rapl | 3 |

| | | |
|---|---|---|
| il1rapl1a | il1rapl1a,il1rapl1a,il1rapl: | 3 |
| il1rapl1b | il1rapl1a,il1rapl1a,il1rapl: | 3 |
| il20ra | il20ra | 1 |
| il23r | il23r | 1 |
| il2rb | il2rga,il2rb,il2rga,il2rgb | 4 |
| il2rga | il2rga,il2rb,il2rga,il2rgb | 4 |
| il2rga | il2rga,il2rb,il2rga,il2rgb | 4 |
| il2rgb | il2rga,il2rb,il2rga,il2rgb | 4 |
| il6r | il6st,il6r,il6 | 3 |
| il6st | il6st,il6r,il6 | 3 |
| il7r | il7r | 1 |
| ildr1a | ildr1a,ildr1b | 2 |
| ildr1b | ildr1a,ildr1b | 2 |
| ilf3a | ilf3a,ilf3b,ilf3a | 3 |
| ilf3a | ilf3a,ilf3b,ilf3a | 3 |
| ilf3b | ilf3a,ilf3b,ilf3a | 3 |
| impdh1a | impdh1a,impdh1a,impdh | 4 |
| impdh1a | impdh1a,impdh1a,impdh | 4 |
| impdh1a | impdh1a,impdh1a,impdh | 4 |
| impdh1b | impdh1a,impdh1a,impdh | 4 |
| impg1a | impg1b,impg1a | 2 |
| impg1b | impg1b,impg1a | 2 |
| impg2a | impg2a,impg2b,impg2b | 3 |
| impg2b | impg2a,impg2b,impg2b | 3 |
| impg2b | impg2a,impg2b,impg2b | 3 |
| ing5a | ing5b,ing5a | 2 |
| ing5b | ing5b,ing5a | 2 |
| inka1a | inka1b,inka1a | 2 |
| inka1b | inka1b,inka1a | 2 |
| ino80b | ino80db,ino80b,ino80da, | 7 |
| ino80c | ino80db,ino80b,ino80da, | 7 |
| ino80da | ino80db,ino80b,ino80da, | 7 |
| ino80db | ino80db,ino80b,ino80da, | 7 |
| ino80e | ino80db,ino80b,ino80da, | 7 |
| ino80e | ino80db,ino80b,ino80da, | 7 |
| inpp4aa | inpp4b,inpp4ab,inpp4aa | 3 |
| inpp4ab | inpp4b,inpp4ab,inpp4aa | 3 |
| inpp4b | inpp4b,inpp4ab,inpp4aa | 3 |
| inpp5b | inpp5d,inpp5e,inpp5jb,in | 9 |
| inpp5d | inpp5d,inpp5e,inpp5jb,in | 9 |
| inpp5d | inpp5d,inpp5e,inpp5jb,in | 9 |
| inpp5e | inpp5d,inpp5e,inpp5jb,in | 9 |
| inpp5f | inpp5d,inpp5e,inpp5jb,in | 9 |
| inpp5ja | inpp5d,inpp5e,inpp5jb,in | 9 |
| inpp5jb | inpp5d,inpp5e,inpp5jb,in | 9 |
| inpp5ka | inpp5d,inpp5e,inpp5jb,in | 9 |
| inpp5kb | inpp5d,inpp5e,inpp5jb,in | 9 |
| inppl1a | inppl1b,inppl1a | 2 |
| inppl1b | inppl1b,inppl1a | 2 |
| insl5a | insl5a,insl5b,insl5a | 3 |

| | | |
|---|---|---|
| insl5a | insl5a,insl5b,insl5a | 3 |
| insl5b | insl5a,insl5b,insl5a | 3 |
| insm1a | insm1b,insm1a | 2 |
| insm1b | insm1b,insm1a | 2 |
| ip6k2a | ip6k2b,ip6k2a | 2 |
| ip6k2b | ip6k2b,ip6k2a | 2 |
| iqsec1b | iqsec1b | 1 |
| iqsec2a | iqsec2a,iqsec2a,iqsec2b | 3 |
| iqsec2a | iqsec2a,iqsec2a,iqsec2b | 3 |
| iqsec2b | iqsec2a,iqsec2a,iqsec2b | 3 |
| iqsec3a | iqsec3a,iqsec3a,iqsec3b | 3 |
| iqsec3a | iqsec3a,iqsec3a,iqsec3b | 3 |
| iqsec3b | iqsec3a,iqsec3a,iqsec3b | 3 |
| irf1a | irf1a,irf1b | 2 |
| irf1b | irf1a,irf1b | 2 |
| irf2a | irf2a,irf2 | 2 |
| irf2bp2a | irf2bp2b,irf2bp2a | 2 |
| irf2bp2b | irf2bp2b,irf2bp2a | 2 |
| irf4a | irf4b,irf4a,irf4b | 3 |
| irf4b | irf4b,irf4a,irf4b | 3 |
| irf4b | irf4b,irf4a,irf4b | 3 |
| irs2a | irs2a,irs2b | 2 |
| irs2b | irs2a,irs2b | 2 |
| irs4a | irs4a | 1 |
| irx1a | irx1a,irx1b | 2 |
| irx1b | irx1a,irx1b | 2 |
| irx2a | irx2a | 1 |
| irx3a | irx3a,irx3b | 2 |
| irx3b | irx3a,irx3b | 2 |
| irx4a | irx4a,irx4b | 2 |
| irx4b | irx4a,irx4b | 2 |
| irx5a | irx5a,irx5b | 2 |
| irx5b | irx5a,irx5b | 2 |
| irx6a | irx6a | 1 |
| isl2a | isl2b,isl2a | 2 |
| isl2b | isl2b,isl2a | 2 |
| ism2a | ism2a,ism2b | 2 |
| ism2b | ism2a,ism2b | 2 |
| itga11a | itga11a,itga11a,itga11b | 3 |
| itga11a | itga11a,itga11a,itga11b | 3 |
| itga11b | itga11a,itga11a,itga11b | 3 |
| itga2b | itga2b | 1 |
| itga3a | itga3a,itga3b | 2 |
| itga3b | itga3a,itga3b | 2 |
| itga6a | itga6b,itga6b,itga6a | 3 |
| itga6b | itga6b,itga6b,itga6a | 3 |
| itga6b | itga6b,itga6b,itga6a | 3 |
| itgb1a | itgb1b,itgb1a | 2 |
| itgb1b | itgb1b,itgb1a | 2 |
| itgb3a | itgb3a,itgb3a,itgb3b | 3 |

| | | |
|---|---|---|
| itgb3a | itgb3a,itgb3a,itgb3b | 3 |
| itgb3b | itgb3a,itgb3a,itgb3b | 3 |
| itih3a | itih3a,itih3b | 2 |
| itih3b | itih3a,itih3b | 2 |
| itm2ba | itm2cb,itm2ba,itm2bb,itr | 4 |
| itm2bb | itm2cb,itm2ba,itm2bb,itr | 4 |
| itm2ca | itm2cb,itm2ba,itm2bb,itr | 4 |
| itm2cb | itm2cb,itm2ba,itm2bb,itr | 4 |
| itpk1a | itpk1b,itpk1a | 2 |
| itpk1b | itpk1b,itpk1a | 2 |
| itpr1a | itpr1a,itpr1a,itpr1b | 3 |
| itpr1a | itpr1a,itpr1a,itpr1b | 3 |
| itpr1b | itpr1a,itpr1a,itpr1b | 3 |
| itsn2a | itsn2b,itsn2b,itsn2a | 3 |
| itsn2b | itsn2b,itsn2b,itsn2a | 3 |
| itsn2b | itsn2b,itsn2b,itsn2a | 3 |
| ivns1abpa | ivns1abpb,ivns1abpa | 2 |
| ivns1abpb | ivns1abpb,ivns1abpa | 2 |
| jag1a | jag1a,jag1b | 2 |
| jag1b | jag1a,jag1b | 2 |
| jag2a | jag2b,jag2a,jag2b | 3 |
| jag2b | jag2b,jag2a,jag2b | 3 |
| jag2b | jag2b,jag2a,jag2b | 3 |
| jagn1a | jagn1a,jagn1a,jagn1b | 3 |
| jagn1a | jagn1a,jagn1a,jagn1b | 3 |
| jagn1b | jagn1a,jagn1a,jagn1b | 3 |
| jak2a | jak2b,jak2a | 2 |
| jak2b | jak2b,jak2a | 2 |
| jam2a | jam2a,jam2b | 2 |
| jam2b | jam2a,jam2b | 2 |
| jam3a | jam3a,jam3b | 2 |
| jam3b | jam3a,jam3b | 2 |
| jarid2a | jarid2a,jarid2b | 2 |
| jarid2b | jarid2a,jarid2b | 2 |
| jazf1a | jazf1b,jazf1a | 2 |
| jazf1b | jazf1b,jazf1a | 2 |
| jdp2a | jdp2a,jdp2b | 2 |
| jdp2b | jdp2a,jdp2b | 2 |
| jmjd1cb | jmjd1cb | 1 |
| jph1a | jph1b,jph1a | 2 |
| jph1b | jph1b,jph1a | 2 |
| jpt1a | jpt1b,jpt1a | 2 |
| jpt1b | jpt1b,jpt1a | 2 |
| kank1b | kank1b | 1 |
| kansl1a | kansl1b,kansl1a | 2 |
| kansl1b | kansl1b,kansl1a | 2 |
| kat2a | kat2a,kat2b | 2 |
| kat2b | kat2a,kat2b | 2 |
| kat5a | kat5a,kat5b | 2 |
| kat5b | kat5a,kat5b | 2 |

| | | |
|---|---|---|
| kat6a | kat6a,kat6b | 2 |
| kat6b | kat6a,kat6b | 2 |
| kat7a | kat7a,kat7a,kat7b | 3 |
| kat7a | kat7a,kat7a,kat7b | 3 |
| kat7b | kat7a,kat7a,kat7b | 3 |
| kcna1a | kcna1b,kcna1a | 2 |
| kcna1b | kcna1b,kcna1a | 2 |
| kcna2a | kcna2a,kcna2a,kcna2b | 3 |
| kcna2a | kcna2a,kcna2a,kcna2b | 3 |
| kcna2b | kcna2a,kcna2a,kcna2b | 3 |
| kcna6a | kcna6a | 1 |
| kcnab1a | kcnab1b,kcnab1a | 2 |
| kcnab1b | kcnab1b,kcnab1a | 2 |
| kcnab2a | kcnab2b,kcnab2a,kcnab2 | 3 |
| kcnab2b | kcnab2b,kcnab2a,kcnab2 | 3 |
| kcnab2b | kcnab2b,kcnab2a,kcnab2 | 3 |
| kcnc1a | kcnc1b,kcnc1a,kcnc1b | 3 |
| kcnc1b | kcnc1b,kcnc1a,kcnc1b | 3 |
| kcnc1b | kcnc1b,kcnc1a,kcnc1b | 3 |
| kcnc3a | kcnc3a,kcnc3b | 2 |
| kcnc3b | kcnc3a,kcnc3b | 2 |
| kcnf1a | kcnf1b,kcnf1a | 2 |
| kcnf1b | kcnf1b,kcnf1a | 2 |
| kcng4a | kcng4b,kcng4a | 2 |
| kcng4b | kcng4b,kcng4a | 2 |
| kcnh1a | kcnh1a,kcnh1b | 2 |
| kcnh1b | kcnh1a,kcnh1b | 2 |
| kcnh2a | kcnh2a | 1 |
| kcnh4a | kcnh4a,kcnh4b | 2 |
| kcnh4b | kcnh4a,kcnh4b | 2 |
| kcnh5a | kcnh5a,kcnh5b,kcnh5a | 3 |
| kcnh5a | kcnh5a,kcnh5b,kcnh5a | 3 |
| kcnh5b | kcnh5a,kcnh5b,kcnh5a | 3 |
| kcnh6a | kcnh6a,kcnh6b | 2 |
| kcnh6b | kcnh6a,kcnh6b | 2 |
| kcnip1b | kcnip1b,kcnip1b | 2 |
| kcnip1b | kcnip1b,kcnip1b | 2 |
| kcnip3a | kcnip3b,kcnip3a,kcnip3b | 3 |
| kcnip3b | kcnip3b,kcnip3a,kcnip3b | 3 |
| kcnip3b | kcnip3b,kcnip3a,kcnip3b | 3 |
| kcnj10a | kcnj10a | 1 |
| kcnj12a | kcnj12b,kcnj12a | 2 |
| kcnj12b | kcnj12b,kcnj12a | 2 |
| kcnj19a | kcnj19a,kcnj19b | 2 |
| kcnj19b | kcnj19a,kcnj19b | 2 |
| kcnj1b | kcnj1b | 1 |
| kcnj2a | kcnj2a,kcnj2b,kcnj2a | 3 |
| kcnj2a | kcnj2a,kcnj2b,kcnj2a | 3 |
| kcnj2b | kcnj2a,kcnj2b,kcnj2a | 3 |
| kcnj3a | kcnj3b,kcnj3a | 2 |

| | | |
|---|---|---|
| kcnj3b | kcnj3b,kcnj3a | 2 |
| kcnk10a | kcnk10b,kcnk10a | 2 |
| kcnk10b | kcnk10b,kcnk10a | 2 |
| kcnk13a | kcnk13a,kcnk13b | 2 |
| kcnk13b | kcnk13a,kcnk13b | 2 |
| kcnk1a | kcnk1a,kcnk1b | 2 |
| kcnk1b | kcnk1a,kcnk1b | 2 |
| kcnk2a | kcnk2a,kcnk2b | 2 |
| kcnk2b | kcnk2a,kcnk2b | 2 |
| kcnk3a | kcnk3b,kcnk3a | 2 |
| kcnk3b | kcnk3b,kcnk3a | 2 |
| kcnk4a | kcnk4a | 1 |
| kcnk5a | kcnk5a,kcnk5a,kcnk5b | 3 |
| kcnk5a | kcnk5a,kcnk5a,kcnk5b | 3 |
| kcnk5b | kcnk5a,kcnk5a,kcnk5b | 3 |
| kcnma1a | kcnma1a | 1 |
| kcnn1a | kcnn1a,kcnn1b | 2 |
| kcnn1b | kcnn1a,kcnn1b | 2 |
| kcnq2a | kcnq2b,kcnq2a | 2 |
| kcnq2b | kcnq2b,kcnq2a | 2 |
| kcnq5a | kcnq5b,kcnq5a | 2 |
| kcnq5b | kcnq5b,kcnq5a | 2 |
| kcns3a | kcns3a,kcns3b | 2 |
| kcns3b | kcns3a,kcns3b | 2 |
| kcnv2a | kcnv2b,kcnv2b,kcnv2b,kc | 4 |
| kcnv2b | kcnv2b,kcnv2b,kcnv2b,kc | 4 |
| kcnv2b | kcnv2b,kcnv2b,kcnv2b,kc | 4 |
| kcnv2b | kcnv2b,kcnv2b,kcnv2b,kc | 4 |
| kctd12b | kctd12b,kctd12b | 2 |
| kctd12b | kctd12b,kctd12b | 2 |
| kctd15a | kctd15a | 1 |
| kctd16a | kctd16a,kctd16a,kctd16a | 4 |
| kctd16a | kctd16a,kctd16a,kctd16a | 4 |
| kctd16a | kctd16a,kctd16a,kctd16a | 4 |
| kctd16b | kctd16a,kctd16a,kctd16a | 4 |
| kctd5a | kctd5a | 1 |
| kctd6a | kctd6b,kctd6a,kctd6b | 3 |
| kctd6b | kctd6b,kctd6a,kctd6b | 3 |
| kctd6b | kctd6b,kctd6a,kctd6b | 3 |
| kctd9a | kctd9a,kctd9b | 2 |
| kctd9b | kctd9a,kctd9b | 2 |
| kdelr2a | kdelr2a,kdelr2b | 2 |
| kdelr2b | kdelr2a,kdelr2b | 2 |
| kdf1a | kdf1a,kdf1b | 2 |
| kdf1b | kdf1a,kdf1b | 2 |
| kdm1a | kdm1a | 1 |
| kdm2aa | kdm2aa,kdm2ba,kdm2bb | 4 |
| kdm2ab | kdm2aa,kdm2ba,kdm2bb | 4 |
| kdm2ba | kdm2aa,kdm2ba,kdm2bb | 4 |
| kdm2bb | kdm2aa,kdm2ba,kdm2bb | 4 |

| | | |
|---|---|---|
| kdm3b | kdm3b | 1 |
| kdm4aa | kdm4c,kdm4c,kdm4c,kdr | 6 |
| kdm4ab | kdm4c,kdm4c,kdm4c,kdr | 6 |
| kdm4b | kdm4c,kdm4c,kdm4c,kdr | 6 |
| kdm4c | kdm4c,kdm4c,kdm4c,kdr | 6 |
| kdm4c | kdm4c,kdm4c,kdm4c,kdr | 6 |
| kdm4c | kdm4c,kdm4c,kdm4c,kdr | 6 |
| kdm5a | kdm5ba,kdm5a,kdm5bb, | 5 |
| kdm5ba | kdm5ba,kdm5a,kdm5bb, | 5 |
| kdm5ba | kdm5ba,kdm5a,kdm5bb, | 5 |
| kdm5bb | kdm5ba,kdm5a,kdm5bb, | 5 |
| kdm5c | kdm5ba,kdm5a,kdm5bb, | 5 |
| kdm6a | kdm6ba,kdm6a,kdm6bb | 3 |
| kdm6ba | kdm6ba,kdm6a,kdm6bb | 3 |
| kdm6bb | kdm6ba,kdm6a,kdm6bb | 3 |
| kdm7aa | kdm7ab,kdm7aa,kdm7ab | 3 |
| kdm7ab | kdm7ab,kdm7aa,kdm7ab | 3 |
| kdm7ab | kdm7ab,kdm7aa,kdm7ab | 3 |
| keap1a | keap1a,keap1b | 2 |
| keap1b | keap1a,keap1b | 2 |
| khdrbs1a | khdrbs1a,khdrbs1b | 2 |
| khdrbs1b | khdrbs1a,khdrbs1b | 2 |
| kidins220a | kidins220a,kidins220b | 2 |
| kidins220b | kidins220a,kidins220b | 2 |
| kif13a | kif13a,kif13ba,kif13bb | 3 |
| kif13ba | kif13a,kif13ba,kif13bb | 3 |
| kif13bb | kif13a,kif13ba,kif13bb | 3 |
| kif16ba | kif16ba,kif16bb | 2 |
| kif16bb | kif16ba,kif16bb | 2 |
| kif18a | kif18a | 1 |
| kif1aa | kif1bp,kif1ab,kif1aa,kif1b | 6 |
| kif1ab | kif1bp,kif1ab,kif1aa,kif1b | 6 |
| kif1b | kif1bp,kif1ab,kif1aa,kif1b | 6 |
| kif1bp | kif1bp,kif1ab,kif1aa,kif1b | 6 |
| kif1bp | kif1bp,kif1ab,kif1aa,kif1b | 6 |
| kif1c | kif1bp,kif1ab,kif1aa,kif1b | 6 |
| kif20a | kif20bb,kif20ba,kif20a | 3 |
| kif20ba | kif20bb,kif20ba,kif20a | 3 |
| kif20bb | kif20bb,kif20ba,kif20a | 3 |
| kif21a | kif21b,kif21a | 2 |
| kif21b | kif21b,kif21a | 2 |
| kif26aa | kif26ba,kif26ab,kif26aa,k | 4 |
| kif26ab | kif26ba,kif26ab,kif26aa,k | 4 |
| kif26ba | kif26ba,kif26ab,kif26aa,k | 4 |
| kif26bb | kif26ba,kif26ab,kif26aa,k | 4 |
| kif2c | kif2c | 1 |
| kif3a | kif3a,kif3a,kif3a,kif3cb,kif | 6 |
| kif3a | kif3a,kif3a,kif3a,kif3cb,kif | 6 |
| kif3a | kif3a,kif3a,kif3a,kif3cb,kif | 6 |
| kif3b | kif3a,kif3a,kif3a,kif3cb,kif | 6 |

| | | |
|---|---|---|
| kif3ca | kif3a,kif3a,kif3a,kif3cb,kif | 6 |
| kif3cb | kif3a,kif3a,kif3a,kif3cb,kif | 6 |
| kif5aa | kif5ba,kif5ab,kif5aa,kif5c, | 5 |
| kif5ab | kif5ba,kif5ab,kif5aa,kif5c, | 5 |
| kif5ba | kif5ba,kif5ab,kif5aa,kif5c, | 5 |
| kif5ba | kif5ba,kif5ab,kif5aa,kif5c, | 5 |
| kif5c | kif5ba,kif5ab,kif5aa,kif5c, | 5 |
| kifap3a | kifap3b,kifap3a | 2 |
| kifap3b | kifap3b,kifap3a | 2 |
| kirrel1a | kirrel1b,kirrel1a | 2 |
| kirrel1b | kirrel1b,kirrel1a | 2 |
| kirrel3a | kirrel3a | 1 |
| kiss1ra | kiss1ra,kiss1rb,kiss1 | 3 |
| kiss1rb | kiss1ra,kiss1rb,kiss1 | 3 |
| klc1a | klc1a | 1 |
| klf11a | klf11b,klf11a | 2 |
| klf11b | klf11b,klf11a | 2 |
| klf12a | klf12a,klf12a,klf12b | 3 |
| klf12a | klf12a,klf12a,klf12b | 3 |
| klf12b | klf12a,klf12a,klf12b | 3 |
| klf2a | klf2a,klf2b,klf2a | 3 |
| klf2a | klf2a,klf2b,klf2a | 3 |
| klf2b | klf2a,klf2b,klf2a | 3 |
| klf5b | klf5b | 1 |
| klf6a | klf6b,klf6a | 2 |
| klf6b | klf6b,klf6a | 2 |
| klf7a | klf7a,klf7b | 2 |
| klf7b | klf7a,klf7b | 2 |
| klhdc7a | klhdc7a | 1 |
| klhdc8a | klhdc8a,klhdc8b | 2 |
| klhdc8b | klhdc8a,klhdc8b | 2 |
| klhl10a | klhl10a | 1 |
| klhl24a | klhl24a,klhl24a,klhl24b | 3 |
| klhl24a | klhl24a,klhl24a,klhl24b | 3 |
| klhl24b | klhl24a,klhl24a,klhl24b | 3 |
| klhl38a | klhl38a,klhl38b,klhl38a | 3 |
| klhl38a | klhl38a,klhl38b,klhl38a | 3 |
| klhl38b | klhl38a,klhl38b,klhl38a | 3 |
| klhl40a | klhl40a,klhl40b | 2 |
| klhl40b | klhl40a,klhl40b | 2 |
| klhl41a | klhl41b,klhl41a | 2 |
| klhl41b | klhl41b,klhl41a | 2 |
| kmt2a | kmt2cb,kmt2e,kmt2a,km | 7 |
| kmt2ba | kmt2cb,kmt2e,kmt2a,km | 7 |
| kmt2bb | kmt2cb,kmt2e,kmt2a,km | 7 |
| kmt2ca | kmt2cb,kmt2e,kmt2a,km | 7 |
| kmt2cb | kmt2cb,kmt2e,kmt2a,km | 7 |
| kmt2d | kmt2cb,kmt2e,kmt2a,km | 7 |
| kmt2e | kmt2cb,kmt2e,kmt2a,km | 7 |
| kmt5aa | kmt5b,kmt5ab,kmt5ab,kı | 7 |

| | | |
|---|---|---|
| kmt5ab | kmt5b,kmt5ab,kmt5ab,kı | 7 |
| kmt5ab | kmt5b,kmt5ab,kmt5ab,kı | 7 |
| kmt5ab | kmt5b,kmt5ab,kmt5ab,kı | 7 |
| kmt5b | kmt5b,kmt5ab,kmt5ab,kı | 7 |
| kmt5b | kmt5b,kmt5ab,kmt5ab,kı | 7 |
| kmt5c | kmt5b,kmt5ab,kmt5ab,kı | 7 |
| krt1-19d | krt1-19d | 1 |
| krt18b | krt18b,krt18b | 2 |
| krt18b | krt18b,krt18b | 2 |
| krtt1c19e | krtt1c19e | 1 |
| ksr1a | ksr1a,ksr1b | 2 |
| ksr1b | ksr1a,ksr1b | 2 |
| l1cama | l1cama,l1camb,l1cama | 3 |
| l1cama | l1cama,l1camb,l1cama | 3 |
| l1camb | l1cama,l1camb,l1cama | 3 |
| l2hgdh | l2hgdh | 1 |
| l3hypdh | l3hypdh | 1 |
| l3mbtl1a | l3mbtl1b,l3mbtl1,l3mbtl1 | 3 |
| l3mbtl1b | l3mbtl1b,l3mbtl1,l3mbtl1 | 3 |
| lactbl1a | lactbl1a,lactbl1a,lactbl1b, | 4 |
| lactbl1a | lactbl1a,lactbl1a,lactbl1b, | 4 |
| lactbl1a | lactbl1a,lactbl1a,lactbl1b, | 4 |
| lactbl1b | lactbl1a,lactbl1a,lactbl1b, | 4 |
| lamb1a | lamb1b,lamb1b,lamb1a | 3 |
| lamb1b | lamb1b,lamb1b,lamb1a | 3 |
| lamb1b | lamb1b,lamb1b,lamb1a | 3 |
| lamp1a | lamp1a,lamp1b | 2 |
| lamp1b | lamp1a,lamp1b | 2 |
| laptm4a | laptm4b,laptm4b,laptm4 | 3 |
| laptm4b | laptm4b,laptm4b,laptm4 | 3 |
| laptm4b | laptm4b,laptm4b,laptm4 | 3 |
| larp1b | larp1b,larp1 | 2 |
| larp4aa | larp4ab,larp4ab,larp4aa,l | 4 |
| larp4ab | larp4ab,larp4ab,larp4aa,l | 4 |
| larp4ab | larp4ab,larp4ab,larp4aa,l | 4 |
| larp4b | larp4ab,larp4ab,larp4aa,l | 4 |
| larp6a | larp6b,larp6a | 2 |
| larp6b | larp6b,larp6a | 2 |
| lbx1a | lbx1b,lbx1a | 2 |
| lbx1b | lbx1b,lbx1a | 2 |
| lcp2a | lcp2b,lcp2a | 2 |
| lcp2b | lcp2b,lcp2a | 2 |
| ldb1a | ldb1b,ldb1a | 2 |
| ldb1b | ldb1b,ldb1a | 2 |
| ldb2a | ldb2b,ldb2b,ldb2b,ldb2a | 4 |
| ldb2b | ldb2b,ldb2b,ldb2b,ldb2a | 4 |
| ldb2b | ldb2b,ldb2b,ldb2b,ldb2a | 4 |
| ldb2b | ldb2b,ldb2b,ldb2b,ldb2a | 4 |
| ldb3a | ldb3a,ldb3b,ldb3a | 3 |
| ldb3a | ldb3a,ldb3b,ldb3a | 3 |

| | | |
|---|---|---|
| ldb3b | ldb3a,ldb3b,ldb3a | 3 |
| ldlrad4a | ldlrad4b,ldlrad4a | 2 |
| ldlrad4b | ldlrad4b,ldlrad4a | 2 |
| ldlrap1a | ldlrap1b,ldlrap1a | 2 |
| ldlrap1b | ldlrap1b,ldlrap1a | 2 |
| lgals2a | lgals2a,lgals2b | 2 |
| lgals2b | lgals2a,lgals2b | 2 |
| lgals3a | lgals3bpa,lgals3bpb,lgals3 | 4 |
| lgals3b | lgals3bpa,lgals3bpb,lgals3 | 4 |
| lgals3bpa | lgals3bpa,lgals3bpb,lgals3 | 4 |
| lgals3bpb | lgals3bpa,lgals3bpb,lgals3 | 4 |
| lgals8a | lgals8b,lgals8a | 2 |
| lgals8b | lgals8b,lgals8a | 2 |
| lgi1a | lgi1b,lgi1a | 2 |
| lgi1b | lgi1b,lgi1a | 2 |
| lgi2a | lgi2b,lgi2b,lgi2a | 3 |
| lgi2b | lgi2b,lgi2b,lgi2a | 3 |
| lgi2b | lgi2b,lgi2b,lgi2a | 3 |
| lhfpl2a | lhfpl2b,lhfpl2a | 2 |
| lhfpl2b | lhfpl2b,lhfpl2a | 2 |
| lhfpl4a | lhfpl4a,lhfpl4b | 2 |
| lhfpl4b | lhfpl4a,lhfpl4b | 2 |
| lhfpl5a | lhfpl5b,lhfpl5b,lhfpl5a | 3 |
| lhfpl5b | lhfpl5b,lhfpl5b,lhfpl5a | 3 |
| lhfpl5b | lhfpl5b,lhfpl5b,lhfpl5a | 3 |
| lhx1a | lhx1b,lhx1b,lhx1a | 3 |
| lhx1b | lhx1b,lhx1b,lhx1a | 3 |
| lhx1b | lhx1b,lhx1b,lhx1a | 3 |
| lhx2a | lhx2b,lhx2a | 2 |
| lhx2b | lhx2b,lhx2a | 2 |
| lhx8a | lhx8a,lhx8b | 2 |
| lhx8b | lhx8a,lhx8b | 2 |
| lima1a | lima1a | 1 |
| limch1a | limch1a,limch1b | 2 |
| limch1b | limch1a,limch1b | 2 |
| limk1a | limk1a,limk1b | 2 |
| limk1b | limk1a,limk1b | 2 |
| lin28a | lin28a,lin28b | 2 |
| lin28b | lin28a,lin28b | 2 |
| lin7a | lin7b,lin7a,lin7c | 3 |
| lin7b | lin7b,lin7a,lin7c | 3 |
| lin7c | lin7b,lin7a,lin7c | 3 |
| lingo1a | lingo1b,lingo1a | 2 |
| lingo1b | lingo1b,lingo1a | 2 |
| lingo2a | lingo2b,lingo2b,lingo2a | 3 |
| lingo2b | lingo2b,lingo2b,lingo2a | 3 |
| lingo2b | lingo2b,lingo2b,lingo2a | 3 |
| lingo3a | lingo3a | 1 |
| lingo4a | lingo4b,lingo4a | 2 |
| lingo4b | lingo4b,lingo4a | 2 |

| | | |
|---|---|---|
| lmbrd2a | lmbrd2a,lmbrd2b | 2 |
| lmbrd2b | lmbrd2a,lmbrd2b | 2 |
| lmf2a | lmf2b,lmf2b,lmf2a | 3 |
| lmf2b | lmf2b,lmf2b,lmf2a | 3 |
| lmf2b | lmf2b,lmf2b,lmf2a | 3 |
| lmo4a | lmo4a,lmo4a,lmo4b,lmo4 | 5 |
| lmo4a | lmo4a,lmo4a,lmo4b,lmo4 | 5 |
| lmo4a | lmo4a,lmo4a,lmo4b,lmo4 | 5 |
| lmo4b | lmo4a,lmo4a,lmo4b,lmo4 | 5 |
| lmo4b | lmo4a,lmo4a,lmo4b,lmo4 | 5 |
| lmo7a | lmo7b,lmo7a | 2 |
| lmo7b | lmo7b,lmo7a | 2 |
| lmod1a | lmod1a,lmod1b | 2 |
| lmod1b | lmod1a,lmod1b | 2 |
| lmod2a | lmod2a,lmod2a,lmod2b,l | 4 |
| lmod2a | lmod2a,lmod2a,lmod2b,l | 4 |
| lmod2a | lmod2a,lmod2a,lmod2b,l | 4 |
| lmod2b | lmod2a,lmod2a,lmod2b,l | 4 |
| lmx1a | lmx1ba,lmx1bb,lmx1a | 3 |
| lmx1ba | lmx1ba,lmx1bb,lmx1a | 3 |
| lmx1bb | lmx1ba,lmx1bb,lmx1a | 3 |
| lnx2a | lnx2a,lnx2b,lnx2a | 3 |
| lnx2a | lnx2a,lnx2b,lnx2a | 3 |
| lnx2b | lnx2a,lnx2b,lnx2a | 3 |
| loxhd1a | loxhd1b,loxhd1a | 2 |
| loxhd1b | loxhd1b,loxhd1a | 2 |
| loxl2a | loxl2b,loxl2a | 2 |
| loxl2b | loxl2b,loxl2a | 2 |
| loxl3a | loxl3b,loxl3a | 2 |
| loxl3b | loxl3b,loxl3a | 2 |
| loxl5a | loxl5b,loxl5a | 2 |
| loxl5b | loxl5b,loxl5a | 2 |
| lpar2a | lpar2b,lpar2a | 2 |
| lpar2b | lpar2b,lpar2a | 2 |
| lpar5a | lpar5a,lpar5b | 2 |
| lpar5b | lpar5a,lpar5b | 2 |
| lpar6a | lpar6a,lpar6b | 2 |
| lpar6b | lpar6a,lpar6b | 2 |
| lrfn2b | lrfn2b | 1 |
| lrfn4a | lrfn4b,lrfn4a | 2 |
| lrfn4b | lrfn4b,lrfn4a | 2 |
| lrfn5a | lrfn5a,lrfn5b | 2 |
| lrfn5b | lrfn5a,lrfn5b | 2 |
| lrit1a | lrit1b,lrit1a | 2 |
| lrit1b | lrit1b,lrit1a | 2 |
| lrit3a | lrit3a,lrit3b,lrit3a | 3 |
| lrit3a | lrit3a,lrit3b,lrit3a | 3 |
| lrit3b | lrit3a,lrit3b,lrit3a | 3 |
| lrp1aa | lrp1bb,lrp1aa,lrp1ba,lrp1 | 5 |
| lrp1ab | lrp1bb,lrp1aa,lrp1ba,lrp1 | 5 |

| | | |
|---|---|---|
| lrp1ba | lrp1bb,lrp1aa,lrp1ba,lrp1 | 5 |
| lrp1ba | lrp1bb,lrp1aa,lrp1ba,lrp1 | 5 |
| lrp1bb | lrp1bb,lrp1aa,lrp1ba,lrp1 | 5 |
| lrp2a | lrp2a,lrp2b,lrp2bp | 3 |
| lrp2b | lrp2a,lrp2b,lrp2bp | 3 |
| lrp2bp | lrp2a,lrp2b,lrp2bp | 3 |
| lrrc14b | lrrc14b | 1 |
| lrrc18a | lrrc18b,lrrc18a | 2 |
| lrrc18b | lrrc18b,lrrc18a | 2 |
| lrrc30a | lrrc30a | 1 |
| lrrc38a | lrrc38b,lrrc38a | 2 |
| lrrc38b | lrrc38b,lrrc38a | 2 |
| lrrc3b | lrrc3cb,lrrc3,lrrc3ca,lrrc3l | 4 |
| lrrc3ca | lrrc3cb,lrrc3,lrrc3ca,lrrc3l | 4 |
| lrrc3cb | lrrc3cb,lrrc3,lrrc3ca,lrrc3l | 4 |
| lrrc4ba | lrrc4ba,lrrc4ba,lrrc4cb,lrr | 5 |
| lrrc4ba | lrrc4ba,lrrc4ba,lrrc4cb,lrr | 5 |
| lrrc4bb | lrrc4ba,lrrc4ba,lrrc4cb,lrr | 5 |
| lrrc4ca | lrrc4ba,lrrc4ba,lrrc4cb,lrr | 5 |
| lrrc4cb | lrrc4ba,lrrc4ba,lrrc4cb,lrr | 5 |
| lrrc58a | lrrc58a,lrrc58b | 2 |
| lrrc58b | lrrc58a,lrrc58b | 2 |
| lrrc74a | lrrc74b,lrrc74a | 2 |
| lrrc74b | lrrc74b,lrrc74a | 2 |
| lrrc75ba | lrrc75ba,lrrc75bb | 2 |
| lrrc75bb | lrrc75ba,lrrc75bb | 2 |
| lrrc8aa | lrrc8da,lrrc8ab,lrrc8db,lrr | 5 |
| lrrc8ab | lrrc8da,lrrc8ab,lrrc8db,lrr | 5 |
| lrrc8c | lrrc8da,lrrc8ab,lrrc8db,lrr | 5 |
| lrrc8da | lrrc8da,lrrc8ab,lrrc8db,lrr | 5 |
| lrrc8db | lrrc8da,lrrc8ab,lrrc8db,lrr | 5 |
| lrrfip1a | lrrfip1b,lrrfip1a | 2 |
| lrrfip1b | lrrfip1b,lrrfip1a | 2 |
| lrrn3a | lrrn3a,lrrn3b,lrrn3a | 3 |
| lrrn3a | lrrn3a,lrrn3b,lrrn3a | 3 |
| lrrn3b | lrrn3a,lrrn3b,lrrn3a | 3 |
| lrtm2a | lrtm2a,lrtm2b | 2 |
| lrtm2b | lrtm2a,lrtm2b | 2 |
| lsm12a | lsm12a,lsm12a,lsm12b | 3 |
| lsm12a | lsm12a,lsm12a,lsm12b | 3 |
| lsm12b | lsm12a,lsm12a,lsm12b | 3 |
| lsm14aa | lsm14ab,lsm14b,lsm14aa | 3 |
| lsm14ab | lsm14ab,lsm14b,lsm14aa | 3 |
| lsm14b | lsm14ab,lsm14b,lsm14aa | 3 |
| lta4h | lta4h,lta4h | 2 |
| lta4h | lta4h,lta4h | 2 |
| ltb4r | ltb4r | 1 |
| ltb4r2a | ltb4r2b,ltb4r2a | 2 |
| ltb4r2b | ltb4r2b,ltb4r2a | 2 |
| ltc4s | ltc4s | 1 |

| | | |
|---|---|---|
| ly6pge | ly6pge | 1 |
| lypd6b | lypd6,lypd6b | 2 |
| lyrm5a | lyrm5a,lyrm5b | 2 |
| lyrm5b | lyrm5a,lyrm5b | 2 |
| lyve1a | lyve1a,lyve1b | 2 |
| lyve1b | lyve1a,lyve1b | 2 |
| lzts2a | lzts2a,lzts2b,lzts2a | 3 |
| lzts2a | lzts2a,lzts2b,lzts2a | 3 |
| lzts2b | lzts2a,lzts2b,lzts2a | 3 |
| lzts3a | lzts3a,lzts3b | 2 |
| lzts3b | lzts3a,lzts3b | 2 |
| m1ap | m1ap | 1 |
| m6pr | m6pr | 1 |
| macf1a | macf1b,macf1a | 2 |
| macf1b | macf1b,macf1a | 2 |
| maco1a | maco1b,maco1a | 2 |
| maco1b | maco1b,maco1a | 2 |
| mad2l1bp | mad2l1,mad2l1bp | 2 |
| magi1a | magi1a,magi1b | 2 |
| magi1b | magi1a,magi1b | 2 |
| magi2a | magi2a,magi2a,magi2b | 3 |
| magi2a | magi2a,magi2a,magi2b | 3 |
| magi2b | magi2a,magi2a,magi2b | 3 |
| magi3b | magi3b | 1 |
| mamdc2a | mamdc2a,mamdc2b | 2 |
| mamdc2b | mamdc2a,mamdc2b | 2 |
| man1b1a | man1b1b,man1b1b,man1 | 4 |
| man1b1b | man1b1b,man1b1b,man1 | 4 |
| man1b1b | man1b1b,man1b1b,man1 | 4 |
| man1b1b | man1b1b,man1b1b,man1 | 4 |
| map1aa | map1sa,map1ab,map1b,m | 8 |
| map1ab | map1sa,map1ab,map1b,m | 8 |
| map1ab | map1sa,map1ab,map1b,m | 8 |
| map1b | map1sa,map1ab,map1b,m | 8 |
| map1b | map1sa,map1ab,map1b,m | 8 |
| map1lc3a | map1lc3a,map1lc3c,map | 3 |
| map1lc3b | map1lc3a,map1lc3c,map | 3 |
| map1lc3c | map1lc3a,map1lc3c,map | 3 |
| map1sa | map1sa,map1ab,map1b,m | 8 |
| map1sa | map1sa,map1ab,map1b,m | 8 |
| map1sb | map1sa,map1ab,map1b,m | 8 |
| map2k2a | map2k2b,map2k2a | 2 |
| map2k2b | map2k2b,map2k2a | 2 |
| map2k4a | map2k4a,map2k4b | 2 |
| map2k4b | map2k4a,map2k4b | 2 |
| map3k14a | map3k14a | 1 |
| map4k3a | map4k3b,map4k3a,map4 | 3 |
| map4k3b | map4k3b,map4k3a,map4 | 3 |
| map4k3b | map4k3b,map4k3a,map4 | 3 |
| map6a | map6a,map6b | 2 |

| | | |
|---|---|---|
| map6b | map6a,map6b | 2 |
| map7a | map7a | 1 |
| map7d1a | map7d1a,map7d1b | 2 |
| map7d1b | map7d1a,map7d1b | 2 |
| map7d2a | map7d2a,map7d2b | 2 |
| map7d2b | map7d2a,map7d2b | 2 |
| mapk12a | mapk12b,mapk12a | 2 |
| mapk12b | mapk12b,mapk12a | 2 |
| mapk14a | mapk14a,mapk14a,mapk | 3 |
| mapk14a | mapk14a,mapk14a,mapk | 3 |
| mapk14b | mapk14a,mapk14a,mapk | 3 |
| mapk8a | mapk8b,mapk8a | 2 |
| mapk8b | mapk8b,mapk8a | 2 |
| mapk8ip1a | mapk8ip1a | 1 |
| mapkapk2a | mapkapk2a,mapkapk2b,r | 3 |
| mapkapk2a | mapkapk2a,mapkapk2b,r | 3 |
| mapkapk2b | mapkapk2a,mapkapk2b,r | 3 |
| mapre1a | mapre1a,mapre1b | 2 |
| mapre1b | mapre1a,mapre1b | 2 |
| mapre3a | mapre3b,mapre3a | 2 |
| mapre3b | mapre3b,mapre3a | 2 |
| marcksl1a | marcksl1a,marcksl1b | 2 |
| marcksl1b | marcksl1a,marcksl1b | 2 |
| mark2a | mark2b,mark2a | 2 |
| mark2b | mark2b,mark2a | 2 |
| mark3a | mark3a,mark3b | 2 |
| mark3b | mark3a,mark3b | 2 |
| mark4a | mark4a,mark4a | 2 |
| mark4a | mark4a,mark4a | 2 |
| marveld2a | marveld2b,marveld2a | 2 |
| marveld2b | marveld2b,marveld2a | 2 |
| mast1a | mast1a,mast1b | 2 |
| mast1b | mast1a,mast1b | 2 |
| mast3a | mast3a,mast3b | 2 |
| mast3b | mast3a,mast3b | 2 |
| mat1a | mat1a,mat1a | 2 |
| mat1a | mat1a,mat1a | 2 |
| mat2aa | mat2ab,mat2aa,mat2b | 3 |
| mat2ab | mat2ab,mat2aa,mat2b | 3 |
| mat2b | mat2ab,mat2aa,mat2b | 3 |
| matn3a | matn3b,matn3a | 2 |
| matn3b | matn3b,matn3a | 2 |
| mbd1a | mbd1b,mbd1a,mbd1b | 3 |
| mbd1b | mbd1b,mbd1a,mbd1b | 3 |
| mbd1b | mbd1b,mbd1a,mbd1b | 3 |
| mbd3a | mbd3b,mbd3a,mbd3b | 3 |
| mbd3b | mbd3b,mbd3a,mbd3b | 3 |
| mbd3b | mbd3b,mbd3a,mbd3b | 3 |
| mboat2a | mboat2a,mboat2b | 2 |
| mboat2b | mboat2a,mboat2b | 2 |

| | | |
|---|---|---|
| mc1r | mc1r | 1 |
| mc2r | mc2r | 1 |
| mc3r | mc3r,mc3r | 2 |
| mc3r | mc3r,mc3r | 2 |
| mc4r | mc4r | 1 |
| mc5ra | mc5rb,mc5ra | 2 |
| mc5rb | mc5rb,mc5ra | 2 |
| mcf2a | mcf2b,mcf2b,mcf2a | 3 |
| mcf2b | mcf2b,mcf2b,mcf2a | 3 |
| mcf2b | mcf2b,mcf2b,mcf2a | 3 |
| mchr1a | mchr1b,mchr1a | 2 |
| mchr1b | mchr1b,mchr1a | 2 |
| mcl1a | mcl1b,mcl1a | 2 |
| mcl1b | mcl1b,mcl1a | 2 |
| mcm3ap | mcm3ap,mcm3 | 2 |
| mcoln1a | mcoln1a,mcoln1b | 2 |
| mcoln1b | mcoln1a,mcoln1b | 2 |
| mcoln3a | mcoln3a | 1 |
| mctp1a | mctp1a,mctp1b | 2 |
| mctp1b | mctp1a,mctp1b | 2 |
| mctp2a | mctp2b,mctp2a | 2 |
| mctp2b | mctp2b,mctp2a | 2 |
| mdga2a | mdga2a | 1 |
| mdh1aa | mdh1b,mdh1aa,mdh1ab | 3 |
| mdh1ab | mdh1b,mdh1aa,mdh1ab | 3 |
| mdh1b | mdh1b,mdh1aa,mdh1ab | 3 |
| med13a | med13a,med13a,med13l | 3 |
| med13a | med13a,med13a,med13l | 3 |
| med13b | med13a,med13a,med13l | 3 |
| med19a | med19a,med19b,med19a | 3 |
| med19a | med19a,med19b,med19a | 3 |
| med19b | med19a,med19b,med19a | 3 |
| mef2aa | mef2ab,mef2cb,mef2ab,r | 7 |
| mef2ab | mef2ab,mef2cb,mef2ab,r | 7 |
| mef2ab | mef2ab,mef2cb,mef2ab,r | 7 |
| mef2b | mef2ab,mef2cb,mef2ab,r | 7 |
| mef2ca | mef2ab,mef2cb,mef2ab,r | 7 |
| mef2cb | mef2ab,mef2cb,mef2ab,r | 7 |
| mef2d | mef2ab,mef2cb,mef2ab,r | 7 |
| megf6a | megf6b,megf6a | 2 |
| megf6b | megf6b,megf6a | 2 |
| meis1a | meis1a,meis1b | 2 |
| meis1b | meis1a,meis1b | 2 |
| meis2a | meis2a,meis2b | 2 |
| meis2b | meis2a,meis2b | 2 |
| meox2a | meox2b,meox2a,meox2b | 3 |
| meox2b | meox2b,meox2a,meox2b | 3 |
| meox2b | meox2b,meox2a,meox2b | 3 |
| mep1b | mep1b,mep1b | 2 |
| mep1b | mep1b,mep1b | 2 |

| | | |
|---|---|---|
| metap1d | metap1,metap1d,metap1 | 4 |
| metap1d | metap1,metap1d,metap1 | 4 |
| metap2a | metap2a,metap2b | 2 |
| metap2b | metap2a,metap2b | 2 |
| mettl11b | mettl11b | 1 |
| mettl21a | mettl21a | 1 |
| mettl2a | mettl2a,mettl2a | 2 |
| mettl2a | mettl2a,mettl2a | 2 |
| mettl7a | mettl7a,mettl7a,mettl7a | 3 |
| mettl7a | mettl7a,mettl7a,mettl7a | 3 |
| mettl7a | mettl7a,mettl7a,mettl7a | 3 |
| mex3b | mex3b | 1 |
| mfge8a | mfge8b,mfge8a,mfge8b | 3 |
| mfge8b | mfge8b,mfge8a,mfge8b | 3 |
| mfge8b | mfge8b,mfge8a,mfge8b | 3 |
| mfn1b | mfn1b | 1 |
| mfsd12a | mfsd12a | 1 |
| mfsd13a | mfsd13a,mfsd13a | 2 |
| mfsd13a | mfsd13a,mfsd13a | 2 |
| mfsd2aa | mfsd2ab,mfsd2aa,mfsd2a | 4 |
| mfsd2ab | mfsd2ab,mfsd2aa,mfsd2a | 4 |
| mfsd2ab | mfsd2ab,mfsd2aa,mfsd2a | 4 |
| mfsd2b | mfsd2ab,mfsd2aa,mfsd2a | 4 |
| mfsd4aa | mfsd4aa,mfsd4b,mfsd4al | 3 |
| mfsd4ab | mfsd4aa,mfsd4b,mfsd4al | 3 |
| mfsd4b | mfsd4aa,mfsd4b,mfsd4al | 3 |
| mfsd6a | mfsd6b,mfsd6a | 2 |
| mfsd6b | mfsd6b,mfsd6a | 2 |
| mgat1a | mgat1a,mgat1b | 2 |
| mgat1b | mgat1a,mgat1b | 2 |
| mgat3a | mgat3a,mgat3b | 2 |
| mgat3b | mgat3a,mgat3b | 2 |
| mgat4a | mgat4c,mgat4a,mgat4b,r | 4 |
| mgat4b | mgat4c,mgat4a,mgat4b,r | 4 |
| mgat4c | mgat4c,mgat4a,mgat4b,r | 4 |
| mgat4c | mgat4c,mgat4a,mgat4b,r | 4 |
| mgrn1a | mgrn1b,mgrn1a | 2 |
| mgrn1b | mgrn1b,mgrn1a | 2 |
| mgst3a | mgst3b,mgst3a | 2 |
| mgst3b | mgst3b,mgst3a | 2 |
| mhc1uba | mhc1zka,mhc1zea,mhc1z | 11 |
| mhc1uka | mhc1zka,mhc1zea,mhc1z | 11 |
| mhc1uma | mhc1zka,mhc1zea,mhc1z | 11 |
| mhc1zaa | mhc1zka,mhc1zea,mhc1z | 11 |
| mhc1zba | mhc1zka,mhc1zea,mhc1z | 11 |
| mhc1zca | mhc1zka,mhc1zea,mhc1z | 11 |
| mhc1zda | mhc1zka,mhc1zea,mhc1z | 11 |
| mhc1zea | mhc1zka,mhc1zea,mhc1z | 11 |
| mhc1zfa | mhc1zka,mhc1zea,mhc1z | 11 |
| mhc1zja | mhc1zka,mhc1zea,mhc1z | 11 |

| | | |
|---|---|---|
| mhc1zka | mhc1zka,mhc1zea,mhc1z | 11 |
| mhc2a | mhc2b,mhc2dab,mhc2a,r | 4 |
| mhc2b | mhc2b,mhc2dab,mhc2a,r | 4 |
| mhc2dab | mhc2b,mhc2dab,mhc2a,r | 4 |
| mhc2dbb | mhc2b,mhc2dab,mhc2a,r | 4 |
| mical2a | mical2a,mical2b | 2 |
| mical2b | mical2a,mical2b | 2 |
| mical3a | mical3b,mical3a,mical3a | 3 |
| mical3a | mical3b,mical3a,mical3a | 3 |
| mical3b | mical3b,mical3a,mical3a | 3 |
| micall1a | micall1a | 1 |
| micall2a | micall2b,micall2a | 2 |
| micall2b | micall2b,micall2a | 2 |
| micu3a | micu3a,micu3b | 2 |
| micu3b | micu3a,micu3b | 2 |
| mid1ip1a | mid1ip1a,mid1ip1b | 2 |
| mid1ip1b | mid1ip1a,mid1ip1b | 2 |
| mier1a | mier1b,mier1a | 2 |
| mier1b | mier1b,mier1a | 2 |
| mier3a | mier3b,mier3b,mier3a | 3 |
| mier3b | mier3b,mier3b,mier3a | 3 |
| mier3b | mier3b,mier3b,mier3a | 3 |
| mif4gda | mif4gda,mif4gda,mif4gdb | 4 |
| mif4gda | mif4gda,mif4gda,mif4gdb | 4 |
| mif4gda | mif4gda,mif4gda,mif4gdb | 4 |
| mif4gdb | mif4gda,mif4gda,mif4gdb | 4 |
| mindy4b | mindy4b | 1 |
| minpp1a | minpp1a,minpp1a,minpp | 4 |
| minpp1a | minpp1a,minpp1a,minpp | 4 |
| minpp1a | minpp1a,minpp1a,minpp | 4 |
| minpp1b | minpp1a,minpp1a,minpp | 4 |
| mir196b | mir196b,mir196c,mir196 | 3 |
| mir196c | mir196b,mir196c,mir196 | 3 |
| mir196d | mir196b,mir196c,mir196 | 3 |
| mis18a | mis18a | 1 |
| mknk2a | mknk2a,mknk2b | 2 |
| mknk2b | mknk2a,mknk2b | 2 |
| mllt1a | mllt1b,mllt1a | 2 |
| mllt1b | mllt1b,mllt1a | 2 |
| mmd2a | mmd2b,mmd2b,mmd2a | 3 |
| mmd2b | mmd2b,mmd2b,mmd2a | 3 |
| mmd2b | mmd2b,mmd2b,mmd2a | 3 |
| mmp11a | mmp11a,mmp11b | 2 |
| mmp11b | mmp11a,mmp11b | 2 |
| mmp13a | mmp13a,mmp13a,mmp1 | 3 |
| mmp13a | mmp13a,mmp13a,mmp1 | 3 |
| mmp13b | mmp13a,mmp13a,mmp1 | 3 |
| mmp14a | mmp14b,mmp14b,mmp1 | 3 |
| mmp14b | mmp14b,mmp14b,mmp1 | 3 |
| mmp14b | mmp14b,mmp14b,mmp1 | 3 |

| | | |
|---|---|---|
| mmp15a | mmp15b,mmp15a | 2 |
| mmp15b | mmp15b,mmp15a | 2 |
| mmp16b | mmp16b | 1 |
| mmp17a | mmp17b,mmp17a | 2 |
| mmp17b | mmp17b,mmp17a | 2 |
| mmp20a | mmp20a,mmp20b | 2 |
| mmp20b | mmp20a,mmp20b | 2 |
| mmp23bb | mmp23bb | 1 |
| mmp25a | mmp25a,mmp25b | 2 |
| mmp25b | mmp25a,mmp25b | 2 |
| mmrn2a | mmrn2b,mmrn2a | 2 |
| mmrn2b | mmrn2b,mmrn2a | 2 |
| mn1a | mn1b,mn1a | 2 |
| mn1b | mn1b,mn1a | 2 |
| mnx2a | mnx2b,mnx2a | 2 |
| mnx2b | mnx2b,mnx2a | 2 |
| mob1a | mob1a,mob1a,mob1ba,n | 5 |
| mob1a | mob1a,mob1a,mob1ba,n | 5 |
| mob1a | mob1a,mob1a,mob1ba,n | 5 |
| mob1ba | mob1a,mob1a,mob1ba,n | 5 |
| mob1bb | mob1a,mob1a,mob1ba,n | 5 |
| mob2a | mob2a,mob2b,mob2a | 3 |
| mob2a | mob2a,mob2b,mob2a | 3 |
| mob2b | mob2a,mob2b,mob2a | 3 |
| mob3a | mob3a,mob3c | 2 |
| mob3c | mob3a,mob3c | 2 |
| mogat3a | mogat3b,mogat3a | 2 |
| mogat3b | mogat3b,mogat3a | 2 |
| mon1a | mon1bb,mon1ba,mon1b | 4 |
| mon1ba | mon1bb,mon1ba,mon1b | 4 |
| mon1bb | mon1bb,mon1ba,mon1b | 4 |
| mon1bb | mon1bb,mon1ba,mon1b | 4 |
| morc3a | morc3b,morc3a | 2 |
| morc3b | morc3b,morc3a | 2 |
| mov10a | mov10a | 1 |
| mpdu1a | mpdu1a,mpdu1a,mpdu1l | 3 |
| mpdu1a | mpdu1a,mpdu1a,mpdu1l | 3 |
| mpdu1b | mpdu1a,mpdu1a,mpdu1l | 3 |
| mpp2a | mpp2a,mpp2b | 2 |
| mpp2b | mpp2a,mpp2b | 2 |
| mpp3a | mpp3a,mpp3b | 2 |
| mpp3b | mpp3a,mpp3b | 2 |
| mpp4a | mpp4a | 1 |
| mpp5a | mpp5a,mpp5b | 2 |
| mpp5b | mpp5a,mpp5b | 2 |
| mpp6a | mpp6a,mpp6b | 2 |
| mpp6b | mpp6a,mpp6b | 2 |
| mpp7a | mpp7b,mpp7a | 2 |
| mpp7b | mpp7b,mpp7a | 2 |
| mpped2a | mpped2,mpped2a,mppe | 3 |

| | | |
|---|---|---|
| mpzl2b | mpzl2b,mpzl2b | 2 |
| mpzl2b | mpzl2b,mpzl2b | 2 |
| mrap2a | mrap2b,mrap2a | 2 |
| mrap2b | mrap2b,mrap2a | 2 |
| mrc1a | mrc1b,mrc1a | 2 |
| mrc1b | mrc1b,mrc1a | 2 |
| mre11a | mre11a | 1 |
| mrm3a | mrm3a,mrm3a,mrm3b | 3 |
| mrm3a | mrm3a,mrm3a,mrm3b | 3 |
| mrm3b | mrm3a,mrm3a,mrm3b | 3 |
| mrps18a | mrps18c,mrps18a,mrps1[illegible] | 4 |
| mrps18b | mrps18c,mrps18a,mrps1[illegible] | 4 |
| mrps18c | mrps18c,mrps18a,mrps1[illegible] | 4 |
| mrps18c | mrps18c,mrps18a,mrps1[illegible] | 4 |
| msi2a | msi2a,msi2a,msi2b | 3 |
| msi2a | msi2a,msi2a,msi2b | 3 |
| msi2b | msi2a,msi2a,msi2b | 3 |
| msl1a | msl1a,msl1b | 2 |
| msl1b | msl1a,msl1b | 2 |
| msl2a | msl2a,msl2b | 2 |
| msl2b | msl2a,msl2b | 2 |
| msrb1a | msrb1b,msrb1a,msrb1b,[illegible] | 4 |
| msrb1b | msrb1b,msrb1a,msrb1b,[illegible] | 4 |
| msrb1b | msrb1b,msrb1a,msrb1b,[illegible] | 4 |
| msrb1b | msrb1b,msrb1a,msrb1b,[illegible] | 4 |
| mst1ra | mst1rb,mst1ra,mst1,mst[illegible] | 4 |
| mst1rb | mst1rb,mst1ra,mst1,mst[illegible] | 4 |
| mst1rb | mst1rb,mst1ra,mst1,mst[illegible] | 4 |
| msx1a | msx1b,msx1a | 2 |
| msx1b | msx1b,msx1a | 2 |
| msx2a | msx2a,msx2a,msx2b | 3 |
| msx2a | msx2a,msx2a,msx2b | 3 |
| msx2b | msx2a,msx2a,msx2b | 3 |
| mthfd1a | mthfd1b,mthfd1a | 2 |
| mthfd1b | mthfd1b,mthfd1a | 2 |
| mtmr1a | mtmr1a,mtmr1b | 2 |
| mtmr1b | mtmr1a,mtmr1b | 2 |
| mtmr7a | mtmr7a,mtmr7b,mtmr7[illegible] | 4 |
| mtmr7a | mtmr7a,mtmr7b,mtmr7[illegible] | 4 |
| mtmr7b | mtmr7a,mtmr7b,mtmr7[illegible] | 4 |
| mtmr7b | mtmr7a,mtmr7b,mtmr7[illegible] | 4 |
| mtnr1aa | mtnr1aa,mtnr1c,mtnr1a[illegible] | 7 |
| mtnr1aa | mtnr1aa,mtnr1c,mtnr1a[illegible] | 7 |
| mtnr1ab | mtnr1aa,mtnr1c,mtnr1a[illegible] | 7 |
| mtnr1ba | mtnr1aa,mtnr1c,mtnr1a[illegible] | 7 |
| mtnr1bb | mtnr1aa,mtnr1c,mtnr1a[illegible] | 7 |
| mtnr1c | mtnr1aa,mtnr1c,mtnr1a[illegible] | 7 |
| mtnr1c | mtnr1aa,mtnr1c,mtnr1a[illegible] | 7 |
| mtus1a | mtus1a,mtus1b,mtus1a | 3 |
| mtus1a | mtus1a,mtus1b,mtus1a | 3 |

| | | |
|---|---|---|
| mtus1b | mtus1a,mtus1b,mtus1a | 3 |
| mtus2a | mtus2a | 1 |
| mtx1a | mtx1a,mtx1b | 2 |
| mtx1b | mtx1a,mtx1b | 2 |
| muc13a | muc13a,muc13b | 2 |
| muc13b | muc13a,muc13b | 2 |
| muc5f | muc5f | 1 |
| mul1a | mul1a,mul1b | 2 |
| mul1b | mul1a,mul1b | 2 |
| mustn1a | mustn1b,mustn1a,mustn | 3 |
| mustn1b | mustn1b,mustn1a,mustn | 3 |
| mustn1b | mustn1b,mustn1a,mustn | 3 |
| mvb12a | mvb12a,mvb12ba,mvb12 | 3 |
| mvb12ba | mvb12a,mvb12ba,mvb12 | 3 |
| mvb12bb | mvb12a,mvb12ba,mvb12 | 3 |
| mxra5a | mxra5a,mxra5b | 2 |
| mxra5b | mxra5a,mxra5b | 2 |
| mxra8a | mxra8a,mxra8b | 2 |
| mxra8b | mxra8a,mxra8b | 2 |
| mybbp1a | mybbp1a | 1 |
| mybl2a | mybl2b,mybl2a | 2 |
| mybl2b | mybl2b,mybl2a | 2 |
| mybpc2a | mybpc2a,mybpc2b | 2 |
| mybpc2b | mybpc2a,mybpc2b | 2 |
| myct1a | myct1a,myct1b,myct1a | 3 |
| myct1a | myct1a,myct1b,myct1a | 3 |
| myct1b | myct1a,myct1b,myct1a | 3 |
| myh11a | myh11b,myh11a | 2 |
| myh11b | myh11b,myh11a | 2 |
| myh7ba | myh7,myh7ba,myh7bb | 3 |
| myh7bb | myh7,myh7ba,myh7bb | 3 |
| myh9a | myh9b,myh9a | 2 |
| myh9b | myh9b,myh9a | 2 |
| myl2a | myl2a,myl2b | 2 |
| myl2b | myl2a,myl2b | 2 |
| myl9a | myl9b,myl9a | 2 |
| myl9b | myl9b,myl9a | 2 |
| mylk4a | mylk4a,mylk4b | 2 |
| mylk4b | mylk4a,mylk4b | 2 |
| myo15aa | myo15ab,myo15aa,myo1 | 3 |
| myo15ab | myo15ab,myo15aa,myo1 | 3 |
| myo15b | myo15ab,myo15aa,myo1 | 3 |
| myo18aa | myo18aa,myo18ab | 2 |
| myo18ab | myo18aa,myo18ab | 2 |
| myo1b | myo1eb,myo1ha,myo1hb | 10 |
| myo1ca | myo1eb,myo1ha,myo1hb | 10 |
| myo1cb | myo1eb,myo1ha,myo1hb | 10 |
| myo1ea | myo1eb,myo1ha,myo1hb | 10 |
| myo1eb | myo1eb,myo1ha,myo1hb | 10 |
| myo1eb | myo1eb,myo1ha,myo1hb | 10 |

| | | |
|---|---|---|
| myo1f | myo1eb,myo1ha,myo1hk | 10 |
| myo1g | myo1eb,myo1ha,myo1hk | 10 |
| myo1ha | myo1eb,myo1ha,myo1hk | 10 |
| myo1hb | myo1eb,myo1ha,myo1hk | 10 |
| myo3a | myo3b,myo3a | 2 |
| myo3b | myo3b,myo3a | 2 |
| myo5aa | myo5aa,myo5aa,myo5b,r | 5 |
| myo5aa | myo5aa,myo5aa,myo5b,r | 5 |
| myo5ab | myo5aa,myo5aa,myo5b,r | 5 |
| myo5b | myo5aa,myo5aa,myo5b,r | 5 |
| myo5c | myo5aa,myo5aa,myo5b,r | 5 |
| myo6a | myo6b,myo6a | 2 |
| myo6b | myo6b,myo6a | 2 |
| myo7aa | myo7bb,myo7aa,myo7ak | 3 |
| myo7ab | myo7bb,myo7aa,myo7ak | 3 |
| myo7bb | myo7bb,myo7aa,myo7ak | 3 |
| myo9aa | myo9b,myo9aa,myo9ab | 3 |
| myo9ab | myo9b,myo9aa,myo9ab | 3 |
| myo9b | myo9b,myo9aa,myo9ab | 3 |
| myom1a | myom1a,myom1b | 2 |
| myom1b | myom1a,myom1b | 2 |
| myom2a | myom2a | 1 |
| myoz1a | myoz1a,myoz1b,myoz1a | 3 |
| myoz1a | myoz1a,myoz1b,myoz1a | 3 |
| myoz1b | myoz1a,myoz1b,myoz1a | 3 |
| myoz2a | myoz2b,myoz2a | 2 |
| myoz2b | myoz2b,myoz2a | 2 |
| myoz3a | myoz3a | 1 |
| myt1a | myt1a,myt1b,myt1a | 3 |
| myt1a | myt1a,myt1b,myt1a | 3 |
| myt1b | myt1a,myt1b,myt1a | 3 |
| mzt2b | mzt2b | 1 |
| naa15a | naa15b,naa15a | 2 |
| naa15b | naa15b,naa15a | 2 |
| nab1a | nab1b,nab1a | 2 |
| nab1b | nab1b,nab1a | 2 |
| nabp1a | nabp1a,nabp1b,nabp1a | 3 |
| nabp1a | nabp1a,nabp1b,nabp1a | 3 |
| nabp1b | nabp1a,nabp1b,nabp1a | 3 |
| nacc1a | nacc1a,nacc1b | 2 |
| nacc1b | nacc1a,nacc1b | 2 |
| nap1l4a | nap1l4a,nap1l4b,nap1l4a | 3 |
| nap1l4a | nap1l4a,nap1l4b,nap1l4a | 3 |
| nap1l4b | nap1l4a,nap1l4b,nap1l4a | 3 |
| nav1b | nav1b | 1 |
| nav2a | nav2a,nav2b | 2 |
| nav2b | nav2a,nav2b | 2 |
| nbr1a | nbr1b,nbr1a | 2 |
| nbr1b | nbr1b,nbr1a | 2 |
| ncam1a | ncam1b,ncam1a | 2 |

| | | |
|---|---|---|
| ncam1b | ncam1b,ncam1a | 2 |
| nceh1a | nceh1a | 1 |
| nck1a | nck1a,nck1b | 2 |
| nck1b | nck1a,nck1b | 2 |
| nck2a | nck2b,nck2a | 2 |
| nck2b | nck2b,nck2a | 2 |
| ncs1a | ncs1a,ncs1b | 2 |
| ncs1b | ncs1a,ncs1b | 2 |
| ndel1a | ndel1b,ndel1b,ndel1a | 3 |
| ndel1b | ndel1b,ndel1b,ndel1a | 3 |
| ndel1b | ndel1b,ndel1b,ndel1a | 3 |
| ndrg1a | ndrg1b,ndrg1a | 2 |
| ndrg1b | ndrg1b,ndrg1a | 2 |
| ndrg3a | ndrg3b,ndrg3a,ndrg3b | 3 |
| ndrg3b | ndrg3b,ndrg3a,ndrg3b | 3 |
| ndrg3b | ndrg3b,ndrg3a,ndrg3b | 3 |
| ndst1a | ndst1a,ndst1b | 2 |
| ndst1b | ndst1a,ndst1b | 2 |
| ndst2a | ndst2b,ndst2a | 2 |
| ndst2b | ndst2b,ndst2a | 2 |
| ndufa4l2a | ndufa4l2b,ndufa4l2a | 2 |
| ndufa4l2b | ndufa4l2b,ndufa4l2a | 2 |
| ndufa9a | ndufa9b,ndufa9a | 2 |
| ndufa9b | ndufa9b,ndufa9a | 2 |
| ndufab1a | ndufab1a,ndufab1a,nduf[illegible] | 3 |
| ndufab1a | ndufab1a,ndufab1a,nduf[illegible] | 3 |
| ndufab1b | ndufab1a,ndufab1a,nduf[illegible] | 3 |
| ndufs8a | ndufs8a,ndufs8b | 2 |
| ndufs8b | ndufs8a,ndufs8b | 2 |
| nectin1a | nectin1a,nectin1b,nectin[illegible] | 3 |
| nectin1a | nectin1a,nectin1b,nectin[illegible] | 3 |
| nectin1b | nectin1a,nectin1b,nectin[illegible] | 3 |
| nectin3b | nectin3b | 1 |
| nectin4a | nectin4b,nectin4a | 2 |
| nectin4b | nectin4b,nectin4a | 2 |
| nedd4a | nedd4a | 1 |
| nell2a | nell2b,nell2a | 2 |
| nell2b | nell2b,nell2a | 2 |
| neo1a | neo1a,neo1b,neo1b | 3 |
| neo1b | neo1a,neo1b,neo1b | 3 |
| neo1b | neo1a,neo1b,neo1b | 3 |
| neto2b | neto2b,neto2b | 2 |
| neto2b | neto2b,neto2b | 2 |
| neurl1aa | neurl1ab,neurl1aa,neurl1 | 4 |
| neurl1ab | neurl1ab,neurl1aa,neurl1 | 4 |
| neurl1ab | neurl1ab,neurl1aa,neurl1 | 4 |
| neurl1b | neurl1ab,neurl1aa,neurl1 | 4 |
| neurod6a | neurod6b,neurod6a | 2 |
| neurod6b | neurod6b,neurod6a | 2 |
| nf1a | nf1b,nf1a | 2 |

| | | |
|---|---|---|
| nf1b | nf1b,nf1a | 2 |
| nf2a | nf2a,nf2b | 2 |
| nf2b | nf2a,nf2b | 2 |
| nfat5a | nfat5b,nfat5a | 2 |
| nfat5b | nfat5b,nfat5a | 2 |
| nfatc2a | nfatc2ip,nfatc2b,nfatc2a | 3 |
| nfatc2b | nfatc2ip,nfatc2b,nfatc2a | 3 |
| nfatc2ip | nfatc2ip,nfatc2b,nfatc2a | 3 |
| nfatc3a | nfatc3a,nfatc3b | 2 |
| nfatc3b | nfatc3a,nfatc3b | 2 |
| nfe2l1a | nfe2l1b,nfe2l1a,nfe2l1a,r | 4 |
| nfe2l1a | nfe2l1b,nfe2l1a,nfe2l1a,r | 4 |
| nfe2l1b | nfe2l1b,nfe2l1a,nfe2l1a,r | 4 |
| nfe2l1b | nfe2l1b,nfe2l1a,nfe2l1a,r | 4 |
| nfe2l2a | nfe2l2b,nfe2l2a | 2 |
| nfe2l2b | nfe2l2b,nfe2l2a | 2 |
| nhsl1a | nhsl1a,nhsl1b | 2 |
| nhsl1b | nhsl1a,nhsl1b | 2 |
| niban1a | niban1b,niban1a | 2 |
| niban1b | niban1b,niban1a | 2 |
| niban2a | niban2b,niban2b,niban2a | 3 |
| niban2b | niban2b,niban2b,niban2a | 3 |
| niban2b | niban2b,niban2b,niban2a | 3 |
| nid1a | nid1b,nid1a | 2 |
| nid1b | nid1b,nid1a | 2 |
| nid2a | nid2a,nid2b | 2 |
| nid2b | nid2a,nid2b | 2 |
| nim1k | nim1k,nim1k | 2 |
| nim1k | nim1k,nim1k | 2 |
| nipsnap3a | nipsnap3a | 1 |
| nitr10a | nitr10a | 1 |
| nitr11a | nitr11a | 1 |
| nitr14a | nitr14b,nitr14a | 2 |
| nitr14b | nitr14b,nitr14a | 2 |
| nitr1b | nitr1b,nitr1b,nitr1f,nitr1i, | 7 |
| nitr1b | nitr1b,nitr1b,nitr1f,nitr1i, | 7 |
| nitr1b | nitr1b,nitr1b,nitr1f,nitr1i, | 7 |
| nitr1f | nitr1b,nitr1b,nitr1f,nitr1i, | 7 |
| nitr1i | nitr1b,nitr1b,nitr1f,nitr1i, | 7 |
| nitr1k | nitr1b,nitr1b,nitr1f,nitr1i, | 7 |
| nitr1m | nitr1b,nitr1b,nitr1f,nitr1i, | 7 |
| nitr2a | nitr2a,nitr2b,nitr2a,nitr2t | 4 |
| nitr2a | nitr2a,nitr2b,nitr2a,nitr2t | 4 |
| nitr2b | nitr2a,nitr2b,nitr2a,nitr2t | 4 |
| nitr2b | nitr2a,nitr2b,nitr2a,nitr2t | 4 |
| nitr3a | nitr3a,nitr3c,nitr3b,nitr3c | 5 |
| nitr3a | nitr3a,nitr3c,nitr3b,nitr3c | 5 |
| nitr3b | nitr3a,nitr3c,nitr3b,nitr3c | 5 |
| nitr3c | nitr3a,nitr3c,nitr3b,nitr3c | 5 |
| nitr3c | nitr3a,nitr3c,nitr3b,nitr3c | 5 |

| | | |
|---|---|---|
| nitr4a | nitr4a | 1 |
| nitr6a | nitr6b,nitr6b,nitr6a,nitr6l | 4 |
| nitr6b | nitr6b,nitr6b,nitr6a,nitr6l | 4 |
| nitr6b | nitr6b,nitr6b,nitr6a,nitr6l | 4 |
| nitr6b | nitr6b,nitr6b,nitr6a,nitr6l | 4 |
| nitr7a | nitr7a,nitr7b | 2 |
| nitr7b | nitr7a,nitr7b | 2 |
| nkd2a | nkd2a | 1 |
| nkx2.2a | nkx2.2a,nkx2.2b | 2 |
| nkx2.2b | nkx2.2a,nkx2.2b | 2 |
| nkx2.4a | nkx2.4a,nkx2.4b | 2 |
| nkx2.4b | nkx2.4a,nkx2.4b | 2 |
| nlgn2a | nlgn2b,nlgn2a,nlgn2b | 3 |
| nlgn2b | nlgn2b,nlgn2a,nlgn2b | 3 |
| nlgn2b | nlgn2b,nlgn2a,nlgn2b | 3 |
| nlgn3a | nlgn3b,nlgn3a | 2 |
| nlgn3b | nlgn3b,nlgn3a | 2 |
| nlgn4xa | nlgn4xa,nlgn4xb | 2 |
| nlgn4xb | nlgn4xa,nlgn4xb | 2 |
| nme2a | nme2a | 1 |
| nmnat1-rb | nmnat1-rbp7a | 1 |
| nmt1a | nmt1a,nmt1b | 2 |
| nmt1b | nmt1a,nmt1b | 2 |
| nmur1a | nmur1b,nmur1a | 2 |
| nmur1b | nmur1b,nmur1a | 2 |
| nos1apa | nos1apb,nos1,nos1apa,n | 4 |
| nos1apa | nos1apb,nos1,nos1apa,n | 4 |
| nos1apb | nos1apb,nos1,nos1apa,n | 4 |
| nos2a | nos2a,nos2b | 2 |
| nos2b | nos2a,nos2b | 2 |
| notch1a | notch1b,notch1a | 2 |
| notch1b | notch1b,notch1a | 2 |
| notum1a | notum1a,notum1b | 2 |
| notum1b | notum1a,notum1b | 2 |
| noxo1a | noxo1a,noxo1b | 2 |
| noxo1b | noxo1a,noxo1b | 2 |
| npas4a | npas4a | 1 |
| npbwr2a | npbwr2a,npbwr2a | 2 |
| npbwr2a | npbwr2a,npbwr2a | 2 |
| npdc1a | npdc1b,npdc1a | 2 |
| npdc1b | npdc1b,npdc1a | 2 |
| npffr2b | npffr2b | 1 |
| npm1a | npm1a,npm1a,npm1b | 3 |
| npm1a | npm1a,npm1a,npm1b | 3 |
| npm1b | npm1a,npm1a,npm1b | 3 |
| npm2a | npm2a,npm2b | 2 |
| npm2b | npm2a,npm2b | 2 |
| npr1a | npr1b,npr1a | 2 |
| npr1b | npr1b,npr1a | 2 |
| nptx2a | nptx2a,nptx2b | 2 |

| | | |
|---|---|---|
| nptx2b | nptx2a,nptx2b | 2 |
| npy1r | npy1r,npy1r,npy1r,npy1r | 4 |
| npy1r | npy1r,npy1r,npy1r,npy1r | 4 |
| npy1r | npy1r,npy1r,npy1r,npy1r | 4 |
| npy1r | npy1r,npy1r,npy1r,npy1r | 4 |
| npy2r | npy2r,npy2r | 2 |
| npy2r | npy2r,npy2r | 2 |
| npy4r | npy4r,npy4r | 2 |
| npy4r | npy4r,npy4r | 2 |
| npy7r | npy7r | 1 |
| npy8ar | npy8br,npy8ar | 2 |
| npy8br | npy8br,npy8ar | 2 |
| nr0b2a | nr0b2a,nr0b2b | 2 |
| nr0b2b | nr0b2a,nr0b2b | 2 |
| nr1d2a | nr1d2a,nr1d2b | 2 |
| nr1d2b | nr1d2a,nr1d2b | 2 |
| nr1d4a | nr1d4b,nr1d4a | 2 |
| nr1d4b | nr1d4b,nr1d4a | 2 |
| nr2c2ap | nr2c2,nr2c2ap | 2 |
| nr2f1a | nr2f1a,nr2f1b | 2 |
| nr2f1b | nr2f1a,nr2f1b | 2 |
| nr2f6a | nr2f6a,nr2f6b | 2 |
| nr2f6b | nr2f6a,nr2f6b | 2 |
| nr4a2a | nr4a2b,nr4a2a | 2 |
| nr4a2b | nr4a2b,nr4a2a | 2 |
| nr5a1a | nr5a1a,nr5a1b | 2 |
| nr5a1b | nr5a1a,nr5a1b | 2 |
| nr6a1a | nr6a1a,nr6a1a,nr6a1b | 3 |
| nr6a1a | nr6a1a,nr6a1a,nr6a1b | 3 |
| nr6a1b | nr6a1a,nr6a1a,nr6a1b | 3 |
| nrbf2b | nrbf2b | 1 |
| nrbp2a | nrbp2a,nrbp2b | 2 |
| nrbp2b | nrbp2a,nrbp2b | 2 |
| nrd1a | nrd1a,nrd1b | 2 |
| nrd1b | nrd1a,nrd1b | 2 |
| nrg2a | nrg2b,nrg2a | 2 |
| nrg2b | nrg2b,nrg2a | 2 |
| nrg3b | nrg3b | 1 |
| nrip1a | nrip1b,nrip1a | 2 |
| nrip1b | nrip1b,nrip1a | 2 |
| nrn1a | nrn1b,nrn1a | 2 |
| nrn1b | nrn1b,nrn1a | 2 |
| nrp1a | nrp1b,nrp1a | 2 |
| nrp1b | nrp1b,nrp1a | 2 |
| nrp2a | nrp2a,nrp2b | 2 |
| nrp2b | nrp2a,nrp2b | 2 |
| nrxn1a | nrxn1a,nrxn1b,nrxn1a,nr: | 4 |
| nrxn1a | nrxn1a,nrxn1b,nrxn1a,nr: | 4 |
| nrxn1b | nrxn1a,nrxn1b,nrxn1a,nr: | 4 |
| nrxn1b | nrxn1a,nrxn1b,nrxn1a,nr: | 4 |

| | | |
|---|---|---|
| nrxn2a | nrxn2b,nrxn2a | 2 |
| nrxn2b | nrxn2b,nrxn2a | 2 |
| nrxn3a | nrxn3a,nrxn3b | 2 |
| nrxn3b | nrxn3a,nrxn3b | 2 |
| nsd1a | nsd1a,nsd1b | 2 |
| nsd1b | nsd1a,nsd1b | 2 |
| nsfl1c | nsfl1c | 1 |
| nsmce4a | nsmce4a | 1 |
| nt5c1aa | nt5c1aa,nt5c1ba,nt5c1ab | 5 |
| nt5c1aa | nt5c1aa,nt5c1ba,nt5c1ab | 5 |
| nt5c1ab | nt5c1aa,nt5c1ba,nt5c1ab | 5 |
| nt5c1ba | nt5c1aa,nt5c1ba,nt5c1ab | 5 |
| nt5c1bb | nt5c1aa,nt5c1ba,nt5c1ab | 5 |
| nt5c2a | nt5c2b,nt5c2a | 2 |
| nt5c2b | nt5c2b,nt5c2a | 2 |
| nt5c3a | nt5c3a | 1 |
| nt5e | nt5e | 1 |
| ntn1a | ntn1a,ntn1b,ntn1a | 3 |
| ntn1a | ntn1a,ntn1b,ntn1a | 3 |
| ntn1b | ntn1a,ntn1b,ntn1a | 3 |
| ntng1a | ntng1a | 1 |
| ntng2a | ntng2a,ntng2b | 2 |
| ntng2b | ntng2a,ntng2b | 2 |
| ntrk2a | ntrk2b,ntrk2a | 2 |
| ntrk2b | ntrk2b,ntrk2a | 2 |
| ntrk3a | ntrk3b,ntrk3a | 2 |
| ntrk3b | ntrk3b,ntrk3a | 2 |
| nuak1a | nuak1a,nuak1b | 2 |
| nuak1b | nuak1a,nuak1b | 2 |
| nucb2a | nucb2a,nucb2a,nucb2b | 3 |
| nucb2a | nucb2a,nucb2a,nucb2b | 3 |
| nucb2b | nucb2a,nucb2a,nucb2b | 3 |
| nucks1a | nucks1a,nucks1b | 2 |
| nucks1b | nucks1a,nucks1b | 2 |
| nudt3a | nudt3b,nudt3a | 2 |
| nudt3b | nudt3b,nudt3a | 2 |
| nudt4a | nudt4a,nudt4b,nudt4a | 3 |
| nudt4a | nudt4a,nudt4b,nudt4a | 3 |
| nudt4b | nudt4a,nudt4b,nudt4a | 3 |
| nupr1a | nupr1b,nupr1b,nupr1a | 3 |
| nupr1b | nupr1b,nupr1b,nupr1a | 3 |
| nupr1b | nupr1b,nupr1b,nupr1a | 3 |
| nxph2a | nxph2a,nxph2b | 2 |
| nxph2b | nxph2a,nxph2b | 2 |
| nyap2a | nyap2b,nyap2a | 2 |
| nyap2b | nyap2b,nyap2a | 2 |
| oaz1a | oaz1b,oaz1a | 2 |
| oaz1b | oaz1b,oaz1a | 2 |
| oaz2a | oaz2b,oaz2a | 2 |
| oaz2b | oaz2b,oaz2a | 2 |

| | | |
|---|---|---|
| obsl1a | obsl1a,obsl1b | 2 |
| obsl1b | obsl1a,obsl1b | 2 |
| odf2a | odf2b,odf2a | 2 |
| odf2b | odf2b,odf2a | 2 |
| odf3b | odf3b,odf3b | 2 |
| odf3b | odf3b,odf3b | 2 |
| odf3l2a | odf3l2b,odf3l2a,odf3l2b | 3 |
| odf3l2b | odf3l2b,odf3l2a,odf3l2b | 3 |
| odf3l2b | odf3l2b,odf3l2a,odf3l2b | 3 |
| olfm1a | olfm1a,olfm1b | 2 |
| olfm1b | olfm1a,olfm1b | 2 |
| olfm2a | olfm2b,olfm2a | 2 |
| olfm2b | olfm2b,olfm2a | 2 |
| olfm3a | olfm3a,olfm3b,olfm3a | 3 |
| olfm3a | olfm3a,olfm3b,olfm3a | 3 |
| olfm3b | olfm3a,olfm3b,olfm3a | 3 |
| olfml2a | olfml2bb,olfml2a,olfml2b | 3 |
| olfml2ba | olfml2bb,olfml2a,olfml2b | 3 |
| olfml2bb | olfml2bb,olfml2a,olfml2b | 3 |
| olfml3a | olfml3b,olfml3a | 2 |
| olfml3b | olfml3b,olfml3a | 2 |
| onecut3a | onecut3b,onecut3a | 2 |
| onecut3b | onecut3b,onecut3a | 2 |
| opn4a | opn4a,opn4xb,opn4xa,op | 5 |
| opn4a | opn4a,opn4xb,opn4xa,op | 5 |
| opn4b | opn4a,opn4xb,opn4xa,op | 5 |
| opn4xa | opn4a,opn4xb,opn4xa,op | 5 |
| opn4xb | opn4a,opn4xb,opn4xa,op | 5 |
| opn6a | opn6a,opn6b | 2 |
| opn6b | opn6a,opn6b | 2 |
| opn7a | opn7c,opn7d,opn7a,opn7 | 4 |
| opn7b | opn7c,opn7d,opn7a,opn7 | 4 |
| opn7c | opn7c,opn7d,opn7a,opn7 | 4 |
| opn7d | opn7c,opn7d,opn7a,opn7 | 4 |
| opn8a | opn8c,opn8a,opn8b | 3 |
| opn8b | opn8c,opn8a,opn8b | 3 |
| opn8c | opn8c,opn8a,opn8b | 3 |
| oprd1a | oprd1b,oprd1a | 2 |
| oprd1b | oprd1b,oprd1a | 2 |
| orai1a | orai1b,orai1a,orai1b | 3 |
| orai1b | orai1b,orai1a,orai1b | 3 |
| orai1b | orai1b,orai1a,orai1b | 3 |
| osbpl10b | osbpl10b | 1 |
| osbpl1a | osbpl1a | 1 |
| osbpl2a | osbpl2b,osbpl2b,osbpl2a | 3 |
| osbpl2b | osbpl2b,osbpl2b,osbpl2a | 3 |
| osbpl2b | osbpl2b,osbpl2b,osbpl2a | 3 |
| osbpl3a | osbpl3b,osbpl3a | 2 |
| osbpl3b | osbpl3b,osbpl3a | 2 |
| oscp1a | oscp1a,oscp1a | 2 |

| | | |
|---|---|---|
| oscp1a | oscp1a,oscp1a | 2 |
| otol1a | otol1a,otol1b | 2 |
| otol1b | otol1a,otol1b | 2 |
| otub1a | otub1b,otub1a | 2 |
| otub1b | otub1b,otub1a | 2 |
| otud5a | otud5b,otud5a,otud5b | 3 |
| otud5b | otud5b,otud5a,otud5b | 3 |
| otud5b | otud5b,otud5a,otud5b | 3 |
| otud6b | otud6b | 1 |
| otud7b | otud7b | 1 |
| otx2a | otx2a,otx2b | 2 |
| otx2b | otx2a,otx2b | 2 |
| ovol1a | ovol1a,ovol1b | 2 |
| ovol1b | ovol1a,ovol1b | 2 |
| oxct1a | oxct1a,oxct1b | 2 |
| oxct1b | oxct1a,oxct1b | 2 |
| oxgr1b | oxgr1b | 1 |
| oxr1a | oxr1a,oxr1b | 2 |
| oxr1b | oxr1a,oxr1b | 2 |
| oxsr1a | oxsr1a,oxsr1b | 2 |
| oxsr1b | oxsr1a,oxsr1b | 2 |
| p2rx3a | p2rx3a,p2rx3b,p2rx3a | 3 |
| p2rx3a | p2rx3a,p2rx3b,p2rx3a | 3 |
| p2rx3b | p2rx3a,p2rx3b,p2rx3a | 3 |
| p2rx4a | p2rx4a,p2rx4b,p2rx4a | 3 |
| p2rx4a | p2rx4a,p2rx4b,p2rx4a | 3 |
| p2rx4b | p2rx4a,p2rx4b,p2rx4a | 3 |
| p4ha1a | p4ha1a,p4ha1b,p4ha1a,p | 4 |
| p4ha1a | p4ha1a,p4ha1b,p4ha1a,p | 4 |
| p4ha1b | p4ha1a,p4ha1b,p4ha1a,p | 4 |
| p4ha1b | p4ha1a,p4ha1b,p4ha1a,p | 4 |
| p4hb | p4hb,p4hb,p4htm | 3 |
| p4hb | p4hb,p4hb,p4htm | 3 |
| p4htm | p4hb,p4hb,p4htm | 3 |
| pa2g4a | pa2g4b,pa2g4a,pa2g4b | 3 |
| pa2g4b | pa2g4b,pa2g4a,pa2g4b | 3 |
| pa2g4b | pa2g4b,pa2g4a,pa2g4b | 3 |
| pabpc1a | pabpc1a,pabpc1b | 2 |
| pabpc1b | pabpc1a,pabpc1b | 2 |
| pacs1a | pacs1a,pacs1a | 2 |
| pacs1a | pacs1a,pacs1a | 2 |
| pacsin1a | pacsin1b,pacsin1a | 2 |
| pacsin1b | pacsin1b,pacsin1a | 2 |
| pafah1b1a | pafah1b1a,pafah1b1b | 2 |
| pafah1b1b | pafah1b1a,pafah1b1b | 2 |
| paip2b | paip2b | 1 |
| pak2a | pak2a,pak2b | 2 |
| pak2b | pak2a,pak2b | 2 |
| pak6a | pak6a,pak6b | 2 |
| pak6b | pak6a,pak6b | 2 |

| | | |
|---|---|---|
| pald1a | pald1a,pald1b | 2 |
| pald1b | pald1a,pald1b | 2 |
| palm1a | palm1a,palm1b,palm1a | 3 |
| palm1a | palm1a,palm1b,palm1a | 3 |
| palm1b | palm1a,palm1b,palm1a | 3 |
| pank1a | pank1a,pank1a,pank1b,p | 4 |
| pank1a | pank1a,pank1a,pank1b,p | 4 |
| pank1a | pank1a,pank1a,pank1b,p | 4 |
| pank1b | pank1a,pank1a,pank1b,p | 4 |
| panx1a | panx1b,panx1b,panx1a | 3 |
| panx1b | panx1b,panx1b,panx1a | 3 |
| panx1b | panx1b,panx1b,panx1a | 3 |
| papss2a | papss2a,papss2a,papss2b | 4 |
| papss2a | papss2a,papss2a,papss2b | 4 |
| papss2a | papss2a,papss2a,papss2b | 4 |
| papss2b | papss2a,papss2a,papss2b | 4 |
| paqr3a | paqr3b,paqr3a,paqr3a,pa | 4 |
| paqr3a | paqr3b,paqr3a,paqr3a,pa | 4 |
| paqr3b | paqr3b,paqr3a,paqr3a,pa | 4 |
| paqr3b | paqr3b,paqr3a,paqr3a,pa | 4 |
| paqr4a | paqr4a,paqr4a,paqr4b | 3 |
| paqr4a | paqr4a,paqr4a,paqr4b | 3 |
| paqr4b | paqr4a,paqr4a,paqr4b | 3 |
| paqr5a | paqr5a,paqr5b,paqr5a | 3 |
| paqr5a | paqr5a,paqr5b,paqr5a | 3 |
| paqr5b | paqr5a,paqr5b,paqr5a | 3 |
| paqr7a | paqr7b,paqr7a | 2 |
| paqr7b | paqr7b,paqr7a | 2 |
| pard3aa | pard3ba,pard3ba,pard3al | 5 |
| pard3ab | pard3ba,pard3ba,pard3al | 5 |
| pard3ba | pard3ba,pard3ba,pard3al | 5 |
| pard3ba | pard3ba,pard3ba,pard3al | 5 |
| pard3bb | pard3ba,pard3ba,pard3al | 5 |
| pard6a | pard6a,pard6ga,pard6a,p | 5 |
| pard6a | pard6a,pard6ga,pard6a,p | 5 |
| pard6b | pard6a,pard6ga,pard6a,p | 5 |
| pard6ga | pard6a,pard6ga,pard6a,p | 5 |
| pard6gb | pard6a,pard6ga,pard6a,p | 5 |
| parp12a | parp12b,parp12b,parp12 | 3 |
| parp12b | parp12b,parp12b,parp12 | 3 |
| parp12b | parp12b,parp12b,parp12 | 3 |
| parp6a | parp6a,parp6b | 2 |
| parp6b | parp6a,parp6b | 2 |
| pax1a | pax1a,pax1b | 2 |
| pax1b | pax1a,pax1b | 2 |
| pax2a | pax2b,pax2a | 2 |
| pax2b | pax2b,pax2a | 2 |
| pax3a | pax3a,pax3b | 2 |
| pax3b | pax3a,pax3b | 2 |
| pax6a | pax6b,pax6a | 2 |

| | | |
|---|---|---|
| pax6b | pax6b,pax6a | 2 |
| pax7a | pax7a,pax7a,pax7b | 3 |
| pax7a | pax7a,pax7a,pax7b | 3 |
| pax7b | pax7a,pax7a,pax7b | 3 |
| pbx1a | pbx1a,pbx1b | 2 |
| pbx1b | pbx1a,pbx1b | 2 |
| pbx3a | pbx3b,pbx3a,pbx3b | 3 |
| pbx3b | pbx3b,pbx3a,pbx3b | 3 |
| pbx3b | pbx3b,pbx3a,pbx3b | 3 |
| pbxip1a | pbxip1b,pbxip1a | 2 |
| pbxip1b | pbxip1b,pbxip1a | 2 |
| pcdh10a | pcdh10a,pcdh10b | 2 |
| pcdh10b | pcdh10a,pcdh10b | 2 |
| pcdh15a | pcdh15b,pcdh15a | 2 |
| pcdh15b | pcdh15b,pcdh15a | 2 |
| pcdh18a | pcdh18b,pcdh18a,pcdh1 | 3 |
| pcdh18b | pcdh18b,pcdh18a,pcdh1 | 3 |
| pcdh18b | pcdh18b,pcdh18a,pcdh1 | 3 |
| pcdh1a | pcdh1a,pcdh1b | 2 |
| pcdh1b | pcdh1a,pcdh1b | 2 |
| pcdh2ac | pcdh2ac | 1 |
| pcdh7a | pcdh7a,pcdh7b | 2 |
| pcdh7b | pcdh7a,pcdh7b | 2 |
| pcgf5a | pcgf5a,pcgf5a,pcgf5b,pcg | 4 |
| pcgf5a | pcgf5a,pcgf5a,pcgf5b,pcg | 4 |
| pcgf5a | pcgf5a,pcgf5a,pcgf5b,pcg | 4 |
| pcgf5b | pcgf5a,pcgf5a,pcgf5b,pcg | 4 |
| pcmtd2a | pcmtd2a,pcmtd2a | 2 |
| pcmtd2a | pcmtd2a,pcmtd2a | 2 |
| pcolce2a | pcolce2a,pcolce2b | 2 |
| pcolce2b | pcolce2a,pcolce2b | 2 |
| pcp4a | pcp4a | 1 |
| pcsk5a | pcsk5a,pcsk5b | 2 |
| pcsk5b | pcsk5a,pcsk5b | 2 |
| pcyt1aa | pcyt1aa,pcyt1ab,pcyt1bb | 4 |
| pcyt1ab | pcyt1aa,pcyt1ab,pcyt1bb | 4 |
| pcyt1ba | pcyt1aa,pcyt1ab,pcyt1bb | 4 |
| pcyt1bb | pcyt1aa,pcyt1ab,pcyt1bb | 4 |
| pdap1a | pdap1a,pdap1b,pdap1a | 3 |
| pdap1a | pdap1a,pdap1b,pdap1a | 3 |
| pdap1b | pdap1a,pdap1b,pdap1a | 3 |
| pdcd10a | pdcd10a,pdcd10b,pdcd1 | 3 |
| pdcd10a | pdcd10a,pdcd10b,pdcd1 | 3 |
| pdcd10b | pdcd10a,pdcd10b,pdcd1 | 3 |
| pdcd4a | pdcd4b,pdcd4b,pdcd4a,p | 4 |
| pdcd4b | pdcd4b,pdcd4b,pdcd4a,p | 4 |
| pdcd4b | pdcd4b,pdcd4b,pdcd4a,p | 4 |
| pdcd4b | pdcd4b,pdcd4b,pdcd4a,p | 4 |
| pdcd6ip | pdcd6ip,pdcd6 | 2 |
| pde10a | pde10a | 1 |

| | | |
|---|---|---|
| pde11a | pde11a | 1 |
| pde1a | pde1a | 1 |
| pde3a | pde3a,pde3b | 2 |
| pde3b | pde3a,pde3b | 2 |
| pde4a | pde4ca,pde4ba,pde4bb,p | 8 |
| pde4ba | pde4ca,pde4ba,pde4bb,p | 8 |
| pde4ba | pde4ca,pde4ba,pde4bb,p | 8 |
| pde4bb | pde4ca,pde4ba,pde4bb,p | 8 |
| pde4ca | pde4ca,pde4ba,pde4bb,p | 8 |
| pde4ca | pde4ca,pde4ba,pde4bb,p | 8 |
| pde4cb | pde4ca,pde4ba,pde4bb,p | 8 |
| pde4d | pde4ca,pde4ba,pde4bb,p | 8 |
| pde5aa | pde5aa,pde5ab | 2 |
| pde5ab | pde5aa,pde5ab | 2 |
| pde6a | pde6d,pde6d,pde6ga,pde | 9 |
| pde6b | pde6d,pde6d,pde6ga,pde | 9 |
| pde6c | pde6d,pde6d,pde6ga,pde | 9 |
| pde6d | pde6d,pde6d,pde6ga,pde | 9 |
| pde6d | pde6d,pde6d,pde6ga,pde | 9 |
| pde6d | pde6d,pde6d,pde6ga,pde | 9 |
| pde6ga | pde6d,pde6d,pde6ga,pde | 9 |
| pde6gb | pde6d,pde6d,pde6ga,pde | 9 |
| pde6ha | pde6d,pde6d,pde6ga,pde | 9 |
| pde7a | pde7a | 1 |
| pde8a | pde8a,pde8b | 2 |
| pde8b | pde8a,pde8b | 2 |
| pde9a | pde9a | 1 |
| pdha1a | pdha1b,pdha1a,pdha1a,p | 4 |
| pdha1a | pdha1b,pdha1a,pdha1a,p | 4 |
| pdha1b | pdha1b,pdha1a,pdha1a,p | 4 |
| pdha1b | pdha1b,pdha1a,pdha1a,p | 4 |
| pdk2a | pdk2a,pdk2b | 2 |
| pdk2b | pdk2a,pdk2b | 2 |
| pdk3a | pdk3b,pdk3a | 2 |
| pdk3b | pdk3b,pdk3a | 2 |
| pdlim3a | pdlim3b,pdlim3a,pdlim3a | 4 |
| pdlim3a | pdlim3b,pdlim3a,pdlim3a | 4 |
| pdlim3b | pdlim3b,pdlim3a,pdlim3a | 4 |
| pdlim3b | pdlim3b,pdlim3a,pdlim3a | 4 |
| pdlim5a | pdlim5a,pdlim5b | 2 |
| pdlim5b | pdlim5a,pdlim5b | 2 |
| pdpk1a | pdpk1a,pdpk1b | 2 |
| pdpk1b | pdpk1a,pdpk1b | 2 |
| pds5a | pds5a,pds5a,pds5b | 3 |
| pds5a | pds5a,pds5a,pds5b | 3 |
| pds5b | pds5a,pds5a,pds5b | 3 |
| pdzd3a | pdzd3b,pdzd3a | 2 |
| pdzd3b | pdzd3b,pdzd3a | 2 |
| pdzd7a | pdzd7a,pdzd7a | 2 |
| pdzd7a | pdzd7a,pdzd7a | 2 |

| | | |
|---|---|---|
| pdzrn3a | pdzrn3b,pdzrn3a | 2 |
| pdzrn3b | pdzrn3b,pdzrn3a | 2 |
| peli1a | peli1b,peli1a,peli1b | 3 |
| peli1b | peli1b,peli1a,peli1b | 3 |
| peli1b | peli1b,peli1a,peli1b | 3 |
| per1a | per1b,per1a | 2 |
| per1b | per1b,per1a | 2 |
| pex11a | pex11a,pex11g,pex11a,p | 4 |
| pex11a | pex11a,pex11g,pex11a,p | 4 |
| pex11b | pex11a,pex11g,pex11a,p | 4 |
| pex11g | pex11a,pex11g,pex11a,p | 4 |
| pfkfb2a | pfkfb2b,pfkfb2a | 2 |
| pfkfb2b | pfkfb2b,pfkfb2a | 2 |
| pfkfb4a | pfkfb4b,pfkfb4a | 2 |
| pfkfb4b | pfkfb4b,pfkfb4a | 2 |
| pgam1a | pgam1b,pgam1a | 2 |
| pgam1b | pgam1b,pgam1a | 2 |
| pggt1b | pggt1b | 1 |
| phactr3a | phactr3b,phactr3a | 2 |
| phactr3b | phactr3b,phactr3a | 2 |
| phactr4a | phactr4b,phactr4a | 2 |
| phactr4b | phactr4b,phactr4a | 2 |
| phb2a | phb2b,phb2b,phb2a,phb | 4 |
| phb2b | phb2b,phb2b,phb2a,phb | 4 |
| phb2b | phb2b,phb2b,phb2a,phb | 4 |
| phb2b | phb2b,phb2b,phb2a,phb | 4 |
| phc2a | phc2a,phc2b,phc2a | 3 |
| phc2a | phc2a,phc2b,phc2a | 3 |
| phc2b | phc2a,phc2b,phc2a | 3 |
| phf12a | phf12a,phf12b | 2 |
| phf12b | phf12a,phf12b | 2 |
| phf20a | phf20b,phf20b,phf20a,ph | 4 |
| phf20b | phf20b,phf20b,phf20a,ph | 4 |
| phf20b | phf20b,phf20b,phf20a,ph | 4 |
| phf20b | phf20b,phf20b,phf20a,ph | 4 |
| phf21aa | phf21aa,phf21ab | 2 |
| phf21ab | phf21aa,phf21ab | 2 |
| phf23a | phf23a,phf23a,phf23b | 3 |
| phf23a | phf23a,phf23a,phf23b | 3 |
| phf23b | phf23a,phf23a,phf23b | 3 |
| phf5a | phf5a,phf5a | 2 |
| phf5a | phf5a,phf5a | 2 |
| phka1b | phka1b | 1 |
| phkg1a | phkg1a,phkg1a,phkg1b | 3 |
| phkg1a | phkg1a,phkg1a,phkg1b | 3 |
| phkg1b | phkg1a,phkg1a,phkg1b | 3 |
| phldb1a | phldb1a,phldb1b | 2 |
| phldb1b | phldb1a,phldb1b | 2 |
| phldb2a | phldb2a,phldb2b | 2 |
| phldb2b | phldb2a,phldb2b | 2 |

| | | |
|---|---|---|
| phox2a | phox2ba,phox2bb,phox2a | 3 |
| phox2ba | phox2ba,phox2bb,phox2a | 3 |
| phox2bb | phox2ba,phox2bb,phox2a | 3 |
| pi15a | pi15b,pi15a | 2 |
| pi15b | pi15b,pi15a | 2 |
| pi4k2a | pi4k2b,pi4k2a | 2 |
| pi4k2b | pi4k2b,pi4k2a | 2 |
| pi4kaa | pi4kb,pi4kab,pi4kaa,pi4kb | 4 |
| pi4kab | pi4kb,pi4kab,pi4kaa,pi4kb | 4 |
| pi4kb | pi4kb,pi4kab,pi4kaa,pi4kb | 4 |
| pi4kb | pi4kb,pi4kab,pi4kaa,pi4kb | 4 |
| pias1a | pias1a,pias1b | 2 |
| pias1b | pias1a,pias1b | 2 |
| pias4a | pias4b,pias4a | 2 |
| pias4b | pias4b,pias4a | 2 |
| pik3c2a | pik3c2b,pik3c2a,pik3c2g | 3 |
| pik3c2b | pik3c2b,pik3c2a,pik3c2g | 3 |
| pik3c2g | pik3c2b,pik3c2a,pik3c2g | 3 |
| pik3ca | pik3cg,pik3ca,pik3cb,pik3 | 4 |
| pik3cb | pik3cg,pik3ca,pik3cb,pik3 | 4 |
| pik3cd | pik3cg,pik3ca,pik3cb,pik3 | 4 |
| pik3cg | pik3cg,pik3ca,pik3cb,pik3 | 4 |
| pik3r3a | pik3r3b,pik3r3b,pik3r3a,p | 4 |
| pik3r3b | pik3r3b,pik3r3b,pik3r3a,p | 4 |
| pik3r3b | pik3r3b,pik3r3b,pik3r3a,p | 4 |
| pik3r3b | pik3r3b,pik3r3b,pik3r3a,p | 4 |
| pik3r6a | pik3r6b,pik3r6a | 2 |
| pik3r6b | pik3r6b,pik3r6a | 2 |
| pip4k2aa | pip4k2ab,pip4k2cb,pip4k | 4 |
| pip4k2ab | pip4k2ab,pip4k2cb,pip4k | 4 |
| pip4k2ca | pip4k2ab,pip4k2cb,pip4k | 4 |
| pip4k2cb | pip4k2ab,pip4k2cb,pip4k | 4 |
| pip4p1a | pip4p1a,pip4p1a,pip4p1b | 4 |
| pip4p1a | pip4p1a,pip4p1a,pip4p1b | 4 |
| pip4p1a | pip4p1a,pip4p1a,pip4p1b | 4 |
| pip4p1b | pip4p1a,pip4p1a,pip4p1b | 4 |
| pip5k1aa | pip5k1cb,pip5k1bb,pip5k | 6 |
| pip5k1ab | pip5k1cb,pip5k1bb,pip5k | 6 |
| pip5k1ba | pip5k1cb,pip5k1bb,pip5k | 6 |
| pip5k1bb | pip5k1cb,pip5k1bb,pip5k | 6 |
| pip5k1ca | pip5k1cb,pip5k1bb,pip5k | 6 |
| pip5k1cb | pip5k1cb,pip5k1bb,pip5k | 6 |
| pitpnc1b | pitpnc1b | 1 |
| pkd1a | pkd1a,pkd1b | 2 |
| pkd1b | pkd1a,pkd1b | 2 |
| pkd1l2a | pkd1l2a,pkd1l2b,pkd1l2b | 3 |
| pkd1l2b | pkd1l2a,pkd1l2b,pkd1l2b | 3 |
| pkd1l2b | pkd1l2a,pkd1l2b,pkd1l2b | 3 |
| pkn1a | pkn1a,pkn1b | 2 |
| pkn1b | pkn1a,pkn1b | 2 |

| | | |
|---|---|---|
| pkp1a | pkp1b,pkp1a | 2 |
| pkp1b | pkp1b,pkp1a | 2 |
| pkp3a | pkp3b,pkp3a | 2 |
| pkp3b | pkp3b,pkp3a | 2 |
| pla1a | pla1a,pla1a | 2 |
| pla1a | pla1a,pla1a | 2 |
| pla2g12a | pla2g12a,pla2g12b | 2 |
| pla2g12b | pla2g12a,pla2g12b | 2 |
| pla2g1b | pla2g1b | 1 |
| pla2g4aa | pla2g4aa,pla2g4ab | 2 |
| pla2g4ab | pla2g4aa,pla2g4ab | 2 |
| plcd1a | plcd1b,plcd1a | 2 |
| plcd1b | plcd1b,plcd1a | 2 |
| plcd3a | plcd3a,plcd3b | 2 |
| plcd3b | plcd3a,plcd3b | 2 |
| plcd4a | plcd4b,plcd4a | 2 |
| plcd4b | plcd4b,plcd4a | 2 |
| plch2a | plch2b,plch2a,plch2b | 3 |
| plch2b | plch2b,plch2a,plch2b | 3 |
| plch2b | plch2b,plch2a,plch2b | 3 |
| pld1a | pld1a,pld1b | 2 |
| pld1b | pld1a,pld1b | 2 |
| plekha1a | plekha1b,plekha1a,plekh | 4 |
| plekha1a | plekha1b,plekha1a,plekh | 4 |
| plekha1b | plekha1b,plekha1a,plekh | 4 |
| plekha1b | plekha1b,plekha1a,plekh | 4 |
| plekha7a | plekha7b,plekha7b,plekh | 3 |
| plekha7b | plekha7b,plekha7b,plekh | 3 |
| plekha7b | plekha7b,plekha7b,plekh | 3 |
| plekhg5a | plekhg5a,plekhg5b | 2 |
| plekhg5b | plekhg5a,plekhg5b | 2 |
| plekho1a | plekho1b,plekho1a | 2 |
| plekho1b | plekho1b,plekho1a | 2 |
| plk2a | plk2b,plk2b,plk2a | 3 |
| plk2b | plk2b,plk2b,plk2a | 3 |
| plk2b | plk2b,plk2b,plk2a | 3 |
| plod1a | plod1a | 1 |
| plp1a | plp1b,plp1a | 2 |
| plp1b | plp1b,plp1a | 2 |
| plpp1a | plpp1a | 1 |
| plpp2a | plpp2b,plpp2a,plpp2b | 3 |
| plpp2b | plpp2b,plpp2a,plpp2b | 3 |
| plpp2b | plpp2b,plpp2a,plpp2b | 3 |
| plpp7a | plpp7a | 1 |
| plppr2a | plppr2a,plppr2b | 2 |
| plppr2b | plppr2a,plppr2b | 2 |
| plppr3a | plppr3a,plppr3b | 2 |
| plppr3b | plppr3a,plppr3b | 2 |
| plppr4a | plppr4a,plppr4b | 2 |
| plppr4b | plppr4a,plppr4b | 2 |

| | | |
|---|---|---|
| plppr5a | plppr5a,plppr5b | 2 |
| plppr5b | plppr5a,plppr5b | 2 |
| plscr3a | plscr3a,plscr3b | 2 |
| plscr3b | plscr3a,plscr3b | 2 |
| plxna1a | plxna1a | 1 |
| plxnb1a | plxnb1b,plxnb1a | 2 |
| plxnb1b | plxnb1b,plxnb1a | 2 |
| plxnb2a | plxnb2a,plxnb2a,plxnb2b | 3 |
| plxnb2a | plxnb2a,plxnb2a,plxnb2b | 3 |
| plxnb2b | plxnb2a,plxnb2a,plxnb2b | 3 |
| pmp22a | pmp22a,pmp22a,pmp22l | 3 |
| pmp22a | pmp22a,pmp22a,pmp22l | 3 |
| pmp22b | pmp22a,pmp22a,pmp22l | 3 |
| pnp4a | pnp4a,pnp4b | 2 |
| pnp4b | pnp4a,pnp4b | 2 |
| pnp5a | pnp5b,pnp5a | 2 |
| pnp5b | pnp5b,pnp5a | 2 |
| pnpla7a | pnpla7b,pnpla7a | 2 |
| pnpla7b | pnpla7b,pnpla7a | 2 |
| poc1a | poc1b,poc1a,poc1b | 3 |
| poc1b | poc1b,poc1a,poc1b | 3 |
| poc1b | poc1b,poc1a,poc1b | 3 |
| pof1b | pof1b | 1 |
| polr1a | polr1b,polr1e,polr1e,polr | 7 |
| polr1b | polr1b,polr1e,polr1e,polr | 7 |
| polr1b | polr1b,polr1e,polr1e,polr | 7 |
| polr1c | polr1b,polr1e,polr1e,polr | 7 |
| polr1d | polr1b,polr1e,polr1e,polr | 7 |
| polr1e | polr1b,polr1e,polr1e,polr | 7 |
| polr1e | polr1b,polr1e,polr1e,polr | 7 |
| polr2a | polr2i,polr2c,polr2c,polr2 | 16 |
| polr2b | polr2i,polr2c,polr2c,polr2 | 16 |
| polr2c | polr2i,polr2c,polr2c,polr2 | 16 |
| polr2c | polr2i,polr2c,polr2c,polr2 | 16 |
| polr2c | polr2i,polr2c,polr2c,polr2 | 16 |
| polr2d | polr2i,polr2c,polr2c,polr2 | 16 |
| polr2eb | polr2i,polr2c,polr2c,polr2 | 16 |
| polr2f | polr2i,polr2c,polr2c,polr2 | 16 |
| polr2f | polr2i,polr2c,polr2c,polr2 | 16 |
| polr2h | polr2i,polr2c,polr2c,polr2 | 16 |
| polr2i | polr2i,polr2c,polr2c,polr2 | 16 |
| polr2i | polr2i,polr2c,polr2c,polr2 | 16 |
| polr2j | polr2i,polr2c,polr2c,polr2 | 16 |
| polr2k | polr2i,polr2c,polr2c,polr2 | 16 |
| polr2m | polr2i,polr2c,polr2c,polr2 | 16 |
| polr2m | polr2i,polr2c,polr2c,polr2 | 16 |
| polr3a | polr3b,polr3f,polr3d,polr | 11 |
| polr3b | polr3b,polr3f,polr3d,polr | 11 |
| polr3b | polr3b,polr3f,polr3d,polr | 11 |
| polr3c | polr3b,polr3f,polr3d,polr | 11 |

| | | |
|---|---|---|
| polr3d | polr3b,polr3f,polr3d,polr. | 11 |
| polr3e | polr3b,polr3f,polr3d,polr. | 11 |
| polr3f | polr3b,polr3f,polr3d,polr. | 11 |
| polr3f | polr3b,polr3f,polr3d,polr. | 11 |
| polr3g | polr3b,polr3f,polr3d,polr. | 11 |
| polr3h | polr3b,polr3f,polr3d,polr. | 11 |
| polr3k | polr3b,polr3f,polr3d,polr. | 11 |
| pou2f1b | pou2f1b | 1 |
| pou2f2a | pou2f2a,pou2f2a | 2 |
| pou2f2a | pou2f2a,pou2f2a | 2 |
| pou3f2a | pou3f2b,pou3f2a,pou3f2 | 3 |
| pou3f2b | pou3f2b,pou3f2a,pou3f2 | 3 |
| pou3f2b | pou3f2b,pou3f2a,pou3f2 | 3 |
| pou3f3a | pou3f3b,pou3f3a | 2 |
| pou3f3b | pou3f3b,pou3f3a | 2 |
| ppa1a | ppa1a,ppa1b | 2 |
| ppa1b | ppa1a,ppa1b | 2 |
| ppap2d | ppap2d,ppap2d | 2 |
| ppap2d | ppap2d,ppap2d | 2 |
| ppargc1a | ppargc1a,ppargc1b | 2 |
| ppargc1b | ppargc1a,ppargc1b | 2 |
| ppfibp1a | ppfibp1b,ppfibp1b,ppfibp | 5 |
| ppfibp1a | ppfibp1b,ppfibp1b,ppfibp | 5 |
| ppfibp1b | ppfibp1b,ppfibp1b,ppfibp | 5 |
| ppfibp1b | ppfibp1b,ppfibp1b,ppfibp | 5 |
| ppfibp1b | ppfibp1b,ppfibp1b,ppfibp | 5 |
| ppfibp2a | ppfibp2a,ppfibp2b | 2 |
| ppfibp2b | ppfibp2a,ppfibp2b | 2 |
| ppip5k1a | ppip5k1a,ppip5k1b | 2 |
| ppip5k1b | ppip5k1a,ppip5k1b | 2 |
| ppm1aa | ppm1db,ppm1f,ppm1db, | 18 |
| ppm1ab | ppm1db,ppm1f,ppm1db, | 18 |
| ppm1ba | ppm1db,ppm1f,ppm1db, | 18 |
| ppm1bb | ppm1db,ppm1f,ppm1db, | 18 |
| ppm1da | ppm1db,ppm1f,ppm1db, | 18 |
| ppm1db | ppm1db,ppm1f,ppm1db, | 18 |
| ppm1db | ppm1db,ppm1f,ppm1db, | 18 |
| ppm1db | ppm1db,ppm1f,ppm1db, | 18 |
| ppm1db | ppm1db,ppm1f,ppm1db, | 18 |
| ppm1e | ppm1db,ppm1f,ppm1db, | 18 |
| ppm1f | ppm1db,ppm1f,ppm1db, | 18 |
| ppm1f | ppm1db,ppm1f,ppm1db, | 18 |
| ppm1g | ppm1db,ppm1f,ppm1db, | 18 |
| ppm1h | ppm1db,ppm1f,ppm1db, | 18 |
| ppm1j | ppm1db,ppm1f,ppm1db, | 18 |
| ppm1k | ppm1db,ppm1f,ppm1db, | 18 |
| ppm1na | ppm1db,ppm1f,ppm1db, | 18 |
| ppm1nb | ppm1db,ppm1f,ppm1db, | 18 |
| ppp1caa | ppp1caa,ppp1caa,ppp1ca | 4 |
| ppp1caa | ppp1caa,ppp1caa,ppp1ca | 4 |

| | | |
|---|---|---|
| ppp1cab | ppp1caa,ppp1caa,ppp1ca | 4 |
| ppp1cb | ppp1caa,ppp1caa,ppp1ca | 4 |
| ppp1r12a | ppp1r12a,ppp1r12c | 2 |
| ppp1r12c | ppp1r12a,ppp1r12c | 2 |
| ppp1r13ba | ppp1r13ba,ppp1r13bb | 2 |
| ppp1r13bb | ppp1r13ba,ppp1r13bb | 2 |
| ppp1r14aa | ppp1r14ab,ppp1r14aa,pp | 6 |
| ppp1r14ab | ppp1r14ab,ppp1r14aa,pp | 6 |
| ppp1r14ab | ppp1r14ab,ppp1r14aa,pp | 6 |
| ppp1r14ba | ppp1r14ab,ppp1r14aa,pp | 6 |
| ppp1r14bb | ppp1r14ab,ppp1r14aa,pp | 6 |
| ppp1r14c | ppp1r14ab,ppp1r14aa,pp | 6 |
| ppp1r15a | ppp1r15b,ppp1r15a,ppp1 | 3 |
| ppp1r15b | ppp1r15b,ppp1r15a,ppp1 | 3 |
| ppp1r15b | ppp1r15b,ppp1r15a,ppp1 | 3 |
| ppp1r16a | ppp1r16b,ppp1r16a | 2 |
| ppp1r16b | ppp1r16b,ppp1r16a | 2 |
| ppp1r1b | ppp1r1c,ppp1r1b | 2 |
| ppp1r1c | ppp1r1c,ppp1r1b | 2 |
| ppp1r27b | ppp1r27b,ppp1r27b | 2 |
| ppp1r27b | ppp1r27b,ppp1r27b | 2 |
| ppp1r3aa | ppp1r3ca,ppp1r3ca,ppp1 | 11 |
| ppp1r3ab | ppp1r3ca,ppp1r3ca,ppp1 | 11 |
| ppp1r3b | ppp1r3ca,ppp1r3ca,ppp1 | 11 |
| ppp1r3b | ppp1r3ca,ppp1r3ca,ppp1 | 11 |
| ppp1r3b | ppp1r3ca,ppp1r3ca,ppp1 | 11 |
| ppp1r3ca | ppp1r3ca,ppp1r3ca,ppp1 | 11 |
| ppp1r3ca | ppp1r3ca,ppp1r3ca,ppp1 | 11 |
| ppp1r3ca | ppp1r3ca,ppp1r3ca,ppp1 | 11 |
| ppp1r3cb | ppp1r3ca,ppp1r3ca,ppp1 | 11 |
| ppp1r3da | ppp1r3ca,ppp1r3ca,ppp1 | 11 |
| ppp1r3db | ppp1r3ca,ppp1r3ca,ppp1 | 11 |
| ppp1r8a | ppp1r8b,ppp1r8b,ppp1r8 | 3 |
| ppp1r8b | ppp1r8b,ppp1r8b,ppp1r8 | 3 |
| ppp1r8b | ppp1r8b,ppp1r8b,ppp1r8 | 3 |
| ppp1r9a | ppp1r9ba,ppp1r9bb,ppp1 | 3 |
| ppp1r9ba | ppp1r9ba,ppp1r9bb,ppp1 | 3 |
| ppp1r9bb | ppp1r9ba,ppp1r9bb,ppp1 | 3 |
| ppp2ca | ppp2cb,ppp2cb,ppp2ca | 3 |
| ppp2cb | ppp2cb,ppp2cb,ppp2ca | 3 |
| ppp2cb | ppp2cb,ppp2cb,ppp2ca | 3 |
| ppp2r1ba | ppp2r1ba,ppp2r1bb | 2 |
| ppp2r1bb | ppp2r1ba,ppp2r1bb | 2 |
| ppp2r2aa | ppp2r2ab,ppp2r2ca,ppp2 | 8 |
| ppp2r2ab | ppp2r2ab,ppp2r2ca,ppp2 | 8 |
| ppp2r2ab | ppp2r2ab,ppp2r2ca,ppp2 | 8 |
| ppp2r2bb | ppp2r2ab,ppp2r2ca,ppp2 | 8 |
| ppp2r2ca | ppp2r2ab,ppp2r2ca,ppp2 | 8 |
| ppp2r2ca | ppp2r2ab,ppp2r2ca,ppp2 | 8 |
| ppp2r2cb | ppp2r2ab,ppp2r2ca,ppp2 | 8 |

| | | |
|---|---|---|
| ppp2r2d | ppp2r2ab,ppp2r2ca,ppp2 | 8 |
| ppp2r3a | ppp2r3a,ppp2r3b,ppp2r3 | 3 |
| ppp2r3b | ppp2r3a,ppp2r3b,ppp2r3 | 3 |
| ppp2r3c | ppp2r3a,ppp2r3b,ppp2r3 | 3 |
| ppp2r5a | ppp2r5b,ppp2r5d,ppp2r5 | 7 |
| ppp2r5b | ppp2r5b,ppp2r5d,ppp2r5 | 7 |
| ppp2r5ca | ppp2r5b,ppp2r5d,ppp2r5 | 7 |
| ppp2r5cb | ppp2r5b,ppp2r5d,ppp2r5 | 7 |
| ppp2r5d | ppp2r5b,ppp2r5d,ppp2r5 | 7 |
| ppp2r5ea | ppp2r5b,ppp2r5d,ppp2r5 | 7 |
| ppp2r5eb | ppp2r5b,ppp2r5d,ppp2r5 | 7 |
| ppp3ca | ppp3ccb,ppp3cca,ppp3cc | 6 |
| ppp3cb | ppp3ccb,ppp3cca,ppp3cc | 6 |
| ppp3cca | ppp3ccb,ppp3cca,ppp3cc | 6 |
| ppp3cca | ppp3ccb,ppp3cca,ppp3cc | 6 |
| ppp3ccb | ppp3ccb,ppp3cca,ppp3cc | 6 |
| ppp3ccb | ppp3ccb,ppp3cca,ppp3cc | 6 |
| ppp3r1a | ppp3r1b,ppp3r1a | 2 |
| ppp3r1b | ppp3r1b,ppp3r1a | 2 |
| ppp4ca | ppp4ca,ppp4cb | 2 |
| ppp4cb | ppp4ca,ppp4cb | 2 |
| ppp4r2a | ppp4r2b,ppp4r2a | 2 |
| ppp4r2b | ppp4r2b,ppp4r2a | 2 |
| ppp4r3b | ppp4r3b | 1 |
| ppp5c | ppp5c | 1 |
| ppp6c | ppp6c | 1 |
| ppp6r2a | ppp6r2a,ppp6r2b | 2 |
| ppp6r2b | ppp6r2a,ppp6r2b | 2 |
| pptc7a | pptc7a,pptc7b | 2 |
| pptc7b | pptc7a,pptc7b | 2 |
| prc1a | prc1b,prc1a | 2 |
| prc1b | prc1b,prc1a | 2 |
| prdm12b | prdm12b | 1 |
| prdm1a | prdm1a,prdm1b | 2 |
| prdm1b | prdm1a,prdm1b | 2 |
| prdm2a | prdm2a,prdm2b | 2 |
| prdm2b | prdm2a,prdm2b | 2 |
| prdm8b | prdm8b,prdm8b,prdm8 | 3 |
| prdm8b | prdm8b,prdm8b,prdm8 | 3 |
| prelid1a | prelid1a,prelid1b | 2 |
| prelid1b | prelid1a,prelid1b | 2 |
| prelid3a | prelid3b,prelid3a | 2 |
| prelid3b | prelid3b,prelid3a | 2 |
| prg4a | prg4a,prg4b | 2 |
| prg4b | prg4a,prg4b | 2 |
| prickle1a | prickle1b,prickle1a | 2 |
| prickle1b | prickle1b,prickle1a | 2 |
| prickle2a | prickle2a,prickle2b | 2 |
| prickle2b | prickle2a,prickle2b | 2 |
| prkab1a | prkab1b,prkab1a | 2 |

| | | |
|---|---|---|
| prkab1b | prkab1b,prkab1a | 2 |
| prkag2a | prkag2b,prkag2a | 2 |
| prkag2b | prkag2b,prkag2a | 2 |
| prkag3a | prkag3a,prkag3b | 2 |
| prkag3b | prkag3a,prkag3b | 2 |
| prkar1aa | prkar1b,prkar1b,prkar1a | 4 |
| prkar1ab | prkar1b,prkar1b,prkar1a | 4 |
| prkar1b | prkar1b,prkar1b,prkar1a | 4 |
| prkar1b | prkar1b,prkar1b,prkar1a | 4 |
| prkar2aa | prkar2aa,prkar2ab | 2 |
| prkar2ab | prkar2aa,prkar2ab | 2 |
| prkg1a | prkg1b,prkg1a | 2 |
| prkg1b | prkg1b,prkg1a | 2 |
| prlh2r | prlh2r,prlh2,prlh2r,prlh2 | 4 |
| prlh2r | prlh2r,prlh2,prlh2r,prlh2 | 4 |
| prlhr2a | prlhr2a,prlhr2b | 2 |
| prlhr2b | prlhr2a,prlhr2b | 2 |
| prmt8b | prmt8b | 1 |
| prokr1a | prokr1a,prokr1b | 2 |
| prokr1b | prokr1a,prokr1b | 2 |
| prom1a | prom1b,prom1b,prom1b | 4 |
| prom1b | prom1b,prom1b,prom1b | 4 |
| prom1b | prom1b,prom1b,prom1b | 4 |
| prom1b | prom1b,prom1b,prom1b | 4 |
| prox1a | prox1a | 1 |
| prpf38a | prpf38a,prpf38b,prpf38a | 3 |
| prpf38a | prpf38a,prpf38b,prpf38a | 3 |
| prpf38b | prpf38a,prpf38b,prpf38a | 3 |
| prpf40a | prpf40a | 1 |
| prpf4ba | prpf4,prpf4,prpf4ba,prpf | 5 |
| prpf4bb | prpf4,prpf4,prpf4ba,prpf | 5 |
| prph2a | prph2b,prph2a,prph2b | 3 |
| prph2b | prph2b,prph2a,prph2b | 3 |
| prph2b | prph2b,prph2a,prph2b | 3 |
| prps1a | prps1b,prps1a,prps1b | 3 |
| prps1b | prps1b,prps1a,prps1b | 3 |
| prps1b | prps1b,prps1a,prps1b | 3 |
| prr12a | prr12b,prr12a | 2 |
| prr12b | prr12b,prr12a | 2 |
| prr5a | prr5a | 1 |
| prrc2a | prrc2c,prrc2a,prrc2c | 3 |
| prrc2c | prrc2c,prrc2a,prrc2c | 3 |
| prrc2c | prrc2c,prrc2a,prrc2c | 3 |
| prrx1a | prrx1a,prrx1b | 2 |
| prrx1b | prrx1a,prrx1b | 2 |
| prxl2b | prxl2c,prxl2b | 2 |
| prxl2c | prxl2c,prxl2b | 2 |
| psip1a | psip1a,psip1b | 2 |
| psip1b | psip1a,psip1b | 2 |
| psma6a | psma6a,psma6b | 2 |

| | | |
|---|---|---|
| psma6b | psma6a,psma6b | 2 |
| psmb11a | psmb11a,psmb11a,psmb | 3 |
| psmb11a | psmb11a,psmb11a,psmb | 3 |
| psmb11b | psmb11a,psmb11a,psmb | 3 |
| psmb13a | psmb13a | 1 |
| psmb8a | psmb8a | 1 |
| psmb9a | psmb9a | 1 |
| psmc1a | psmc1a,psmc1a,psmc1b | 3 |
| psmc1a | psmc1a,psmc1a,psmc1b | 3 |
| psmc1b | psmc1a,psmc1a,psmc1b | 3 |
| psmc3ip | psmc3ip,psmc3 | 2 |
| psmd11a | psmd11a,psmd11b | 2 |
| psmd11b | psmd11a,psmd11b | 2 |
| psmd4a | psmd4b,psmd4a | 2 |
| psmd4b | psmd4b,psmd4a | 2 |
| psme4a | psme4b,psme4b,psme4b | 4 |
| psme4b | psme4b,psme4b,psme4b | 4 |
| psme4b | psme4b,psme4b,psme4b | 4 |
| psme4b | psme4b,psme4b,psme4b | 4 |
| pstpip1a | pstpip1b,pstpip1b,pstpip | 3 |
| pstpip1b | pstpip1b,pstpip1b,pstpip | 3 |
| pstpip1b | pstpip1b,pstpip1b,pstpip | 3 |
| ptbp1a | ptbp1b,ptbp1a,ptbp1b | 3 |
| ptbp1b | ptbp1b,ptbp1a,ptbp1b | 3 |
| ptbp1b | ptbp1b,ptbp1a,ptbp1b | 3 |
| ptbp2a | ptbp2b,ptbp2a | 2 |
| ptbp2b | ptbp2b,ptbp2a | 2 |
| ptdss1a | ptdss1a,ptdss1b | 2 |
| ptdss1b | ptdss1a,ptdss1b | 2 |
| ptf1a | ptf1a | 1 |
| ptger1a | ptger1c,ptger1b,ptger1c, | 4 |
| ptger1b | ptger1c,ptger1b,ptger1c, | 4 |
| ptger1c | ptger1c,ptger1b,ptger1c, | 4 |
| ptger1c | ptger1c,ptger1b,ptger1c, | 4 |
| ptger2a | ptger2b,ptger2a,ptger2b | 3 |
| ptger2b | ptger2b,ptger2a,ptger2b | 3 |
| ptger2b | ptger2b,ptger2a,ptger2b | 3 |
| ptger4a | ptger4b,ptger4c,ptger4b, | 4 |
| ptger4b | ptger4b,ptger4c,ptger4b, | 4 |
| ptger4b | ptger4b,ptger4c,ptger4b, | 4 |
| ptger4c | ptger4b,ptger4c,ptger4b, | 4 |
| ptges3a | ptges3a,ptges3b,ptges3a | 3 |
| ptges3a | ptges3a,ptges3b,ptges3a | 3 |
| ptges3b | ptges3a,ptges3b,ptges3a | 3 |
| ptgs2a | ptgs2a,ptgs2b | 2 |
| ptgs2b | ptgs2a,ptgs2b | 2 |
| pth1a | pth1b,pth1ra,pth1b,pth1 | 5 |
| pth1b | pth1b,pth1ra,pth1b,pth1 | 5 |
| pth1b | pth1b,pth1ra,pth1b,pth1 | 5 |
| pth1ra | pth1b,pth1ra,pth1b,pth1 | 5 |

| | | |
|---|---|---|
| pth1rb | pth1b,pth1ra,pth1b,pth1 | 5 |
| pth2r | pth2r,pth2r,pth2 | 3 |
| pth2r | pth2r,pth2r,pth2 | 3 |
| ptk2aa | ptk2ab,ptk2ba,ptk2aa,ptl | 4 |
| ptk2ab | ptk2ab,ptk2ba,ptk2aa,ptl | 4 |
| ptk2ba | ptk2ab,ptk2ba,ptk2aa,ptl | 4 |
| ptk2bb | ptk2ab,ptk2ba,ptk2aa,ptl | 4 |
| ptk6a | ptk6a,ptk6b | 2 |
| ptk6b | ptk6a,ptk6b | 2 |
| ptk7a | ptk7a | 1 |
| ptp4a2a | ptp4a2a,ptp4a2b | 2 |
| ptp4a2b | ptp4a2a,ptp4a2b | 2 |
| ptp4a3a | ptp4a3b,ptp4a3b,ptp4a3 | 3 |
| ptp4a3b | ptp4a3b,ptp4a3b,ptp4a3 | 3 |
| ptp4a3b | ptp4a3b,ptp4a3b,ptp4a3 | 3 |
| ptpdc1a | ptpdc1a,ptpdc1b | 2 |
| ptpdc1b | ptpdc1a,ptpdc1b | 2 |
| ptpn11a | ptpn11b,ptpn11a,ptpn11 | 3 |
| ptpn11b | ptpn11b,ptpn11a,ptpn11 | 3 |
| ptpn11b | ptpn11b,ptpn11a,ptpn11 | 3 |
| ptpn23a | ptpn23a,ptpn23b | 2 |
| ptpn23b | ptpn23a,ptpn23b | 2 |
| ptpn2a | ptpn2a,ptpn2b | 2 |
| ptpn2b | ptpn2a,ptpn2b | 2 |
| ptpn4a | ptpn4a,ptpn4a,ptpn4b | 3 |
| ptpn4a | ptpn4a,ptpn4a,ptpn4b | 3 |
| ptpn4b | ptpn4a,ptpn4a,ptpn4b | 3 |
| ptpn9a | ptpn9b,ptpn9a | 2 |
| ptpn9b | ptpn9b,ptpn9a | 2 |
| ptprz1a | ptprz1b,ptprz1a | 2 |
| ptprz1b | ptprz1b,ptprz1a | 2 |
| pttg1ipa | pttg1ipa,pttg1ipb,pttg1 | 3 |
| pttg1ipb | pttg1ipa,pttg1ipb,pttg1 | 3 |
| ptx3a | ptx3b,ptx3a | 2 |
| ptx3b | ptx3b,ptx3a | 2 |
| puf60a | puf60a,puf60b | 2 |
| puf60b | puf60a,puf60b | 2 |
| pwp2h | pwp2h | 1 |
| pwwp2a | pwwp2b,pwwp2a | 2 |
| pwwp2b | pwwp2b,pwwp2a | 2 |
| pxdc1a | pxdc1a,pxdc1b | 2 |
| pxdc1b | pxdc1a,pxdc1b | 2 |
| pycr1a | pycr1a,pycr1b | 2 |
| pycr1b | pycr1a,pycr1b | 2 |
| rab11a | rab11a,rab11bb,rab11a,r | 4 |
| rab11a | rab11a,rab11bb,rab11a,r | 4 |
| rab11ba | rab11a,rab11bb,rab11a,r | 4 |
| rab11bb | rab11a,rab11bb,rab11a,r | 4 |
| rab11fip1a | rab11fip1a,rab11fip1b | 2 |
| rab11fip1b | rab11fip1a,rab11fip1b | 2 |

| | | |
|---|---|---|
| rab11fip4a | rab11fip4b,rab11fip4b,ra | 4 |
| rab11fip4b | rab11fip4b,rab11fip4b,ra | 4 |
| rab11fip4b | rab11fip4b,rab11fip4b,ra | 4 |
| rab11fip4b | rab11fip4b,rab11fip4b,ra | 4 |
| rab11fip5a | rab11fip5a | 1 |
| rab18a | rab18a,rab18b,rab18a | 3 |
| rab18a | rab18a,rab18b,rab18a | 3 |
| rab18b | rab18a,rab18b,rab18a | 3 |
| rab1aa | rab1bb,rab1aa,rab1ba,ra | 5 |
| rab1ab | rab1bb,rab1aa,rab1ba,ra | 5 |
| rab1ba | rab1bb,rab1aa,rab1ba,ra | 5 |
| rab1bb | rab1bb,rab1aa,rab1ba,ra | 5 |
| rab1bb | rab1bb,rab1aa,rab1ba,ra | 5 |
| rab22a | rab22a,rab22a | 2 |
| rab22a | rab22a,rab22a | 2 |
| rab25a | rab25b,rab25a | 2 |
| rab25b | rab25b,rab25a | 2 |
| rab27a | rab27a,rab27b | 2 |
| rab27b | rab27a,rab27b | 2 |
| rab2a | rab2a | 1 |
| rab32a | rab32b,rab32a | 2 |
| rab32b | rab32b,rab32a | 2 |
| rab33a | rab33a,rab33a,rab33ba | 3 |
| rab33a | rab33a,rab33a,rab33ba | 3 |
| rab33ba | rab33a,rab33a,rab33ba | 3 |
| rab34a | rab34a,rab34a,rab34b | 3 |
| rab34a | rab34a,rab34a,rab34b | 3 |
| rab34b | rab34a,rab34a,rab34b | 3 |
| rab35b | rab35b | 1 |
| rab38b | rab38b,rab38c | 2 |
| rab38c | rab38b,rab38c | 2 |
| rab39ba | rab39bb,rab39ba | 2 |
| rab39bb | rab39bb,rab39ba | 2 |
| rab3aa | rab3aa,rab3c,rab3da,rab3 | 9 |
| rab3aa | rab3aa,rab3c,rab3da,rab3 | 9 |
| rab3ab | rab3aa,rab3c,rab3da,rab3 | 9 |
| rab3b | rab3aa,rab3c,rab3da,rab3 | 9 |
| rab3c | rab3aa,rab3c,rab3da,rab3 | 9 |
| rab3c | rab3aa,rab3c,rab3da,rab3 | 9 |
| rab3da | rab3aa,rab3c,rab3da,rab3 | 9 |
| rab3db | rab3aa,rab3c,rab3da,rab3 | 9 |
| rab3ip | rab3aa,rab3c,rab3da,rab3 | 9 |
| rab40b | rab40b,rab40c,rab40b | 3 |
| rab40b | rab40b,rab40c,rab40b | 3 |
| rab40c | rab40b,rab40c,rab40b | 3 |
| rab42a | rab42b,rab42b,rab42a | 3 |
| rab42b | rab42b,rab42b,rab42a | 3 |
| rab42b | rab42b,rab42b,rab42a | 3 |
| rab4a | rab4a,rab4b | 2 |
| rab4b | rab4a,rab4b | 2 |

| | | |
|---|---|---|
| rab5aa | rab5b,rab5c,rab5b,rab5al | 6 |
| rab5ab | rab5b,rab5c,rab5b,rab5al | 6 |
| rab5b | rab5b,rab5c,rab5b,rab5al | 6 |
| rab5b | rab5b,rab5c,rab5b,rab5al | 6 |
| rab5c | rab5b,rab5c,rab5b,rab5al | 6 |
| rab5if | rab5b,rab5c,rab5b,rab5al | 6 |
| rab6a | rab6bb,rab6ba,rab6a | 3 |
| rab6ba | rab6bb,rab6ba,rab6a | 3 |
| rab6bb | rab6bb,rab6ba,rab6a | 3 |
| rab7a | rab7a,rab7a,rab7b,rab7a | 4 |
| rab7a | rab7a,rab7a,rab7b,rab7a | 4 |
| rab7a | rab7a,rab7a,rab7b,rab7a | 4 |
| rab7b | rab7a,rab7a,rab7b,rab7a | 4 |
| rab8a | rab8b,rab8b,rab8a,rab8b | 4 |
| rab8b | rab8b,rab8b,rab8a,rab8b | 4 |
| rab8b | rab8b,rab8b,rab8a,rab8b | 4 |
| rab8b | rab8b,rab8b,rab8a,rab8b | 4 |
| rab9a | rab9a,rab9b | 2 |
| rab9b | rab9a,rab9b | 2 |
| rabl6a | rabl6a,rabl6b | 2 |
| rabl6b | rabl6a,rabl6b | 2 |
| rac1a | rac1b,rac1a | 2 |
| rac1b | rac1b,rac1a | 2 |
| rac3a | rac3a,rac3b | 2 |
| rac3b | rac3a,rac3b | 2 |
| rad21a | rad21a,rad21b | 2 |
| rad21b | rad21a,rad21b | 2 |
| rad23aa | rad23aa,rad23b,rad23ab | 3 |
| rad23ab | rad23aa,rad23b,rad23ab | 3 |
| rad23b | rad23aa,rad23b,rad23ab | 3 |
| rad51b | rad51,rad51d,rad51c,rad. | 5 |
| rad51c | rad51,rad51d,rad51c,rad. | 5 |
| rad51d | rad51,rad51d,rad51c,rad. | 5 |
| rad54b | rad54b | 1 |
| rad9a | rad9a,rad9b,rad9a | 3 |
| rad9a | rad9a,rad9b,rad9a | 3 |
| rad9b | rad9a,rad9b,rad9a | 3 |
| raf1a | raf1b,raf1a,raf1a | 3 |
| raf1a | raf1b,raf1a,raf1a | 3 |
| raf1b | raf1b,raf1a,raf1a | 3 |
| ranbp3a | ranbp3b,ranbp3a | 2 |
| ranbp3b | ranbp3b,ranbp3a | 2 |
| rangap1a | rangap1b,rangap1a,ranga | 3 |
| rangap1b | rangap1b,rangap1a,ranga | 3 |
| rangap1b | rangap1b,rangap1a,ranga | 3 |
| rap1aa | rap1ab,rap1gap,rap1b,ra | 6 |
| rap1ab | rap1ab,rap1gap,rap1b,ra | 6 |
| rap1ab | rap1ab,rap1gap,rap1b,ra | 6 |
| rap1b | rap1ab,rap1gap,rap1b,ra | 6 |
| rap1gap | rap1ab,rap1gap,rap1b,ra | 6 |

| | | |
|---|---|---|
| rap1gap | rap1ab,rap1gap,rap1b,ra| | 6 |
| rap1gap2a | rap1gap2a,rap1gap2b | 2 |
| rap1gap2b | rap1gap2a,rap1gap2b | 2 |
| rap2aa | rap2ab,rap2aa,rap2c,rap2 | 4 |
| rap2ab | rap2ab,rap2aa,rap2c,rap2 | 4 |
| rap2b | rap2ab,rap2aa,rap2c,rap2 | 4 |
| rap2c | rap2ab,rap2aa,rap2c,rap2 | 4 |
| rapgef1a | rapgef1a,rapgef1b | 2 |
| rapgef1b | rapgef1a,rapgef1b | 2 |
| rapgef5a | rapgef5b,rapgef5a | 2 |
| rapgef5b | rapgef5b,rapgef5a | 2 |
| raph1a | raph1b,raph1b,raph1a | 3 |
| raph1b | raph1b,raph1b,raph1a | 3 |
| raph1b | raph1b,raph1b,raph1a | 3 |
| rasa1a | rasa1a,rasa1b | 2 |
| rasa1b | rasa1a,rasa1b | 2 |
| rasal1b | rasal1b | 1 |
| rasgef1ba | rasgef1bb,rasgef1ba,rasg | 4 |
| rasgef1ba | rasgef1bb,rasgef1ba,rasg | 4 |
| rasgef1bb | rasgef1bb,rasgef1ba,rasg | 4 |
| rasgef1bb | rasgef1bb,rasgef1ba,rasg | 4 |
| rasgrf2a | rasgrf2b,rasgrf2b,rasgrf2a | 3 |
| rasgrf2b | rasgrf2b,rasgrf2b,rasgrf2a | 3 |
| rasgrf2b | rasgrf2b,rasgrf2b,rasgrf2a | 3 |
| rasl10a | rasl10a | 1 |
| rasl11a | rasl11a,rasl11b | 2 |
| rasl11b | rasl11a,rasl11b | 2 |
| rassf2a | rassf2b,rassf2a,rassf2a,ra | 4 |
| rassf2a | rassf2b,rassf2a,rassf2a,ra | 4 |
| rassf2b | rassf2b,rassf2a,rassf2a,ra | 4 |
| rassf2b | rassf2b,rassf2a,rassf2a,ra | 4 |
| rassf4a | rassf4a | 1 |
| rassf7a | rassf7b,rassf7a | 2 |
| rassf7b | rassf7b,rassf7a | 2 |
| rassf8a | rassf8b,rassf8a | 2 |
| rassf8b | rassf8b,rassf8a | 2 |
| rbfox3a | rbfox3b,rbfox3a | 2 |
| rbfox3b | rbfox3b,rbfox3a | 2 |
| rbm12b | rbm12,rbm12b | 2 |
| rbm14a | rbm14a,rbm14b,rbm14a | 3 |
| rbm14a | rbm14a,rbm14b,rbm14a | 3 |
| rbm14b | rbm14a,rbm14b,rbm14a | 3 |
| rbm15b | rbm15b,rbm15,rbm15,rb | 4 |
| rbm15b | rbm15b,rbm15,rbm15,rb | 4 |
| rbm24a | rbm24b,rbm24b,rbm24a | 3 |
| rbm24b | rbm24b,rbm24b,rbm24a | 3 |
| rbm24b | rbm24b,rbm24b,rbm24a | 3 |
| rbm25a | rbm25a,rbm25b | 2 |
| rbm25b | rbm25a,rbm25b | 2 |
| rbm33a | rbm33b,rbm33a | 2 |

| | | |
|---|---|---|
| rbm33b | rbm33b,rbm33a | 2 |
| rbm39a | rbm39b,rbm39a | 2 |
| rbm39b | rbm39b,rbm39a | 2 |
| rbm8a | rbm8a | 1 |
| rbms1a | rbms1b,rbms1a | 2 |
| rbms1b | rbms1b,rbms1a | 2 |
| rbms2a | rbms2b,rbms2a,rbms2b | 3 |
| rbms2b | rbms2b,rbms2a,rbms2b | 3 |
| rbms2b | rbms2b,rbms2a,rbms2b | 3 |
| rbp2a | rbp2a,rbp2b,rbp2b,rbp2a | 4 |
| rbp2a | rbp2a,rbp2b,rbp2b,rbp2a | 4 |
| rbp2b | rbp2a,rbp2b,rbp2b,rbp2a | 4 |
| rbp2b | rbp2a,rbp2b,rbp2b,rbp2a | 4 |
| rbp7a | rbp7b,rbp7b,rbp7a | 3 |
| rbp7b | rbp7b,rbp7b,rbp7a | 3 |
| rbp7b | rbp7b,rbp7b,rbp7a | 3 |
| rbpms2a | rbpms2b,rbpms2a | 2 |
| rbpms2b | rbpms2b,rbpms2a | 2 |
| rc3h1a | rc3h1a,rc3h1a,rc3h1b,rc3 | 4 |
| rc3h1a | rc3h1a,rc3h1a,rc3h1b,rc3 | 4 |
| rc3h1a | rc3h1a,rc3h1a,rc3h1b,rc3 | 4 |
| rc3h1b | rc3h1a,rc3h1a,rc3h1b,rc3 | 4 |
| rcan1a | rcan1a,rcan1a | 2 |
| rcan1a | rcan1a,rcan1a | 2 |
| rce1a | rce1a,rce1b | 2 |
| rce1b | rce1a,rce1b | 2 |
| rdh10a | rdh10a,rdh10b,rdh10a | 3 |
| rdh10a | rdh10a,rdh10b,rdh10a | 3 |
| rdh10b | rdh10a,rdh10b,rdh10a | 3 |
| rdh14a | rdh14a,rdh14b | 2 |
| rdh14b | rdh14a,rdh14b | 2 |
| rdh8a | rdh8b,rdh8a | 2 |
| rdh8b | rdh8b,rdh8a | 2 |
| rec8a | rec8a,rec8b | 2 |
| rec8b | rec8a,rec8b | 2 |
| reep3a | reep3b,reep3a | 2 |
| reep3b | reep3b,reep3a | 2 |
| rex1bd | rex1bd | 1 |
| rftn1a | rftn1a | 1 |
| rfx1a | rfx1b,rfx1a | 2 |
| rfx1b | rfx1b,rfx1a | 2 |
| rfx7a | rfx7a,rfx7b | 2 |
| rfx7b | rfx7a,rfx7b | 2 |
| rgl3a | rgl3a | 1 |
| rgs12a | rgs12b,rgs12a | 2 |
| rgs12b | rgs12b,rgs12a | 2 |
| rgs14a | rgs14a | 1 |
| rgs3a | rgs3b,rgs3a | 2 |
| rgs3b | rgs3b,rgs3a | 2 |
| rgs5a | rgs5a,rgs5b,rgs5a | 3 |

| | | |
|---|---|---|
| rgs5a | rgs5a,rgs5b,rgs5a | 3 |
| rgs5b | rgs5a,rgs5b,rgs5a | 3 |
| rgs7a | rgs7a,rgs7bpb,rgs7bpa,rg | 6 |
| rgs7a | rgs7a,rgs7bpb,rgs7bpa,rg | 6 |
| rgs7b | rgs7a,rgs7bpb,rgs7bpa,rg | 6 |
| rgs7bpa | rgs7a,rgs7bpb,rgs7bpa,rg | 6 |
| rgs7bpb | rgs7a,rgs7bpb,rgs7bpa,rg | 6 |
| rgs7bpb | rgs7a,rgs7bpb,rgs7bpa,rg | 6 |
| rgs9a | rgs9b,rgs9a,rgs9bp,rgs9b | 4 |
| rgs9b | rgs9b,rgs9a,rgs9bp,rgs9b | 4 |
| rgs9b | rgs9b,rgs9a,rgs9bp,rgs9b | 4 |
| rgs9bp | rgs9b,rgs9a,rgs9bp,rgs9b | 4 |
| rhbdf1a | rhbdf1b,rhbdf1a,rhbdf1b | 3 |
| rhbdf1b | rhbdf1b,rhbdf1a,rhbdf1b | 3 |
| rhbdf1b | rhbdf1b,rhbdf1a,rhbdf1b | 3 |
| rhobtb2a | rhobtb2a,rhobtb2b | 2 |
| rhobtb2b | rhobtb2a,rhobtb2b | 2 |
| rhot1a | rhot1b,rhot1b,rhot1a,rhc | 4 |
| rhot1b | rhot1b,rhot1b,rhot1a,rhc | 4 |
| rhot1b | rhot1b,rhot1b,rhot1a,rhc | 4 |
| rhot1b | rhot1b,rhot1b,rhot1a,rhc | 4 |
| ric8a | ric8b,ric8b,ric8a | 3 |
| ric8b | ric8b,ric8b,ric8a | 3 |
| ric8b | ric8b,ric8b,ric8a | 3 |
| rims1a | rims1b,rims1b,rims1a | 3 |
| rims1b | rims1b,rims1b,rims1a | 3 |
| rims1b | rims1b,rims1b,rims1a | 3 |
| rims2a | rims2a,rims2b | 2 |
| rims2b | rims2a,rims2b | 2 |
| rin1a | rin1a | 1 |
| rlbp1a | rlbp1b,rlbp1a,rlbp1b | 3 |
| rlbp1b | rlbp1b,rlbp1a,rlbp1b | 3 |
| rlbp1b | rlbp1b,rlbp1a,rlbp1b | 3 |
| rln3a | rln3b,rln3a | 2 |
| rln3b | rln3b,rln3a | 2 |
| rmnd5b | rmnd5b | 1 |
| rn7sk | rn7sk | 1 |
| rnaseh2a | rnaseh2b,rnaseh2b,rnase | 5 |
| rnaseh2b | rnaseh2b,rnaseh2b,rnase | 5 |
| rnaseh2b | rnaseh2b,rnaseh2b,rnase | 5 |
| rnaseh2b | rnaseh2b,rnaseh2b,rnase | 5 |
| rnaseh2c | rnaseh2b,rnaseh2b,rnase | 5 |
| rnd1a | rnd1b,rnd1a,rnd1b | 3 |
| rnd1b | rnd1b,rnd1a,rnd1b | 3 |
| rnd1b | rnd1b,rnd1a,rnd1b | 3 |
| rnd3a | rnd3b,rnd3a | 2 |
| rnd3b | rnd3b,rnd3a | 2 |
| rnf113a | rnf113a | 1 |
| rnf11a | rnf11a,rnf11b | 2 |
| rnf11b | rnf11a,rnf11b | 2 |

| | | |
|---|---|---|
| rnf128a | rnf128a | 1 |
| rnf144aa | rnf144b,rnf144aa,rnf144a | 3 |
| rnf144ab | rnf144b,rnf144aa,rnf144a | 3 |
| rnf144b | rnf144b,rnf144aa,rnf144a | 3 |
| rnf145a | rnf145a,rnf145b | 2 |
| rnf145b | rnf145a,rnf145b | 2 |
| rnf150a | rnf150a,rnf150b | 2 |
| rnf150b | rnf150a,rnf150b | 2 |
| rnf165a | rnf165b,rnf165a | 2 |
| rnf165b | rnf165b,rnf165a | 2 |
| rnf19a | rnf19a,rnf19b | 2 |
| rnf19b | rnf19a,rnf19b | 2 |
| rnf207a | rnf207b,rnf207a,rnf207b | 3 |
| rnf207b | rnf207b,rnf207a,rnf207b | 3 |
| rnf207b | rnf207b,rnf207a,rnf207b | 3 |
| rnf212b | rnf212b,rnf212 | 2 |
| rnf213a | rnf213a,rnf213b | 2 |
| rnf213b | rnf213a,rnf213b | 2 |
| rnf220a | rnf220a,rnf220b | 2 |
| rnf220b | rnf220a,rnf220b | 2 |
| rnf34a | rnf34a,rnf34b | 2 |
| rnf34b | rnf34a,rnf34b | 2 |
| rock2a | rock2a,rock2b,rock2b | 3 |
| rock2b | rock2a,rock2b,rock2b | 3 |
| rock2b | rock2a,rock2b,rock2b | 3 |
| rom1a | rom1a,rom1b | 2 |
| rom1b | rom1a,rom1b | 2 |
| rp1l1a | rp1l1a,rp1l1b | 2 |
| rp1l1b | rp1l1a,rp1l1b | 2 |
| rpe65a | rpe65b,rpe65c,rpe65a,rp | 5 |
| rpe65b | rpe65b,rpe65c,rpe65a,rp | 5 |
| rpe65b | rpe65b,rpe65c,rpe65a,rp | 5 |
| rpe65c | rpe65b,rpe65c,rpe65a,rp | 5 |
| rpe65c | rpe65b,rpe65c,rpe65a,rp | 5 |
| rph3aa | rph3ab,rph3aa | 2 |
| rph3ab | rph3ab,rph3aa | 2 |
| rpl10a | rpl10,rpl10,rpl10a,rpl10a | 5 |
| rpl10a | rpl10,rpl10,rpl10a,rpl10a | 5 |
| rpl13a | rpl13a,rpl13,rpl13a | 3 |
| rpl13a | rpl13a,rpl13,rpl13a | 3 |
| rpl18a | rpl18a,rpl18 | 2 |
| rpl23a | rpl23,rpl23a | 2 |
| rpl35a | rpl35a,rpl35 | 2 |
| rpl36a | rpl36a,rpl36a,rpl36 | 3 |
| rpl36a | rpl36a,rpl36a,rpl36 | 3 |
| rpl5a | rpl5b,rpl5a,rpl5b | 3 |
| rpl5b | rpl5b,rpl5a,rpl5b | 3 |
| rpl5b | rpl5b,rpl5a,rpl5b | 3 |
| rpl7a | rpl7,rpl7a | 2 |
| rpp25a | rpp25b,rpp25a | 2 |

| | | |
|---|---|---|
| rpp25b | rpp25b,rpp25a | 2 |
| rprd1a | rprd1a,rprd1b | 2 |
| rprd1b | rprd1a,rprd1b | 2 |
| rprd2a | rprd2b,rprd2a | 2 |
| rprd2b | rprd2b,rprd2a | 2 |
| rps15a | rps15,rps15a,rps15 | 3 |
| rps27a | rps27a,rps27a | 2 |
| rps27a | rps27a,rps27a | 2 |
| rps3a | rps3a,rps3a,rps3a,rps3 | 4 |
| rps3a | rps3a,rps3a,rps3a,rps3 | 4 |
| rps3a | rps3a,rps3a,rps3a,rps3 | 4 |
| rps4x | rps4x | 1 |
| rps6ka3a | rps6ka3b,rps6ka3a,rps6k | 4 |
| rps6ka3a | rps6ka3b,rps6ka3a,rps6k | 4 |
| rps6ka3b | rps6ka3b,rps6ka3a,rps6k | 4 |
| rps6ka3b | rps6ka3b,rps6ka3a,rps6k | 4 |
| rps6kb1a | rps6kb1b,rps6kb1a,rps6k | 3 |
| rps6kb1b | rps6kb1b,rps6kb1a,rps6k | 3 |
| rps6kb1b | rps6kb1b,rps6kb1a,rps6k | 3 |
| rps8a | rps8a,rps8b | 2 |
| rps8b | rps8a,rps8b | 2 |
| rrbp1a | rrbp1b,rrbp1a,rrbp1b | 3 |
| rrbp1b | rrbp1b,rrbp1a,rrbp1b | 3 |
| rrbp1b | rrbp1b,rrbp1a,rrbp1b | 3 |
| rreb1a | rreb1b,rreb1a | 2 |
| rreb1b | rreb1b,rreb1a | 2 |
| rrm2b | rrm2b,rrm2b,rrm2,rrm2 | 4 |
| rrm2b | rrm2b,rrm2b,rrm2,rrm2 | 4 |
| rrp7a | rrp7a | 1 |
| rs1a | rs1a | 1 |
| rsph10b | rsph10b,rsph10b | 2 |
| rsph10b | rsph10b,rsph10b | 2 |
| rsph4a | rsph4a | 1 |
| rtkn2a | rtkn2a,rtkn2b | 2 |
| rtkn2b | rtkn2a,rtkn2b | 2 |
| rtn1a | rtn1a,rtn1b | 2 |
| rtn1b | rtn1a,rtn1b | 2 |
| rtn2a | rtn2a,rtn2b | 2 |
| rtn2b | rtn2a,rtn2b | 2 |
| rtn4a | rtn4a,rtn4r,rtn4a,rtn4b | 4 |
| rtn4a | rtn4a,rtn4r,rtn4a,rtn4b | 4 |
| rtn4b | rtn4a,rtn4r,rtn4a,rtn4b | 4 |
| rtn4r | rtn4a,rtn4r,rtn4a,rtn4b | 4 |
| rtn4rl1a | rtn4rl1a,rtn4rl1b | 2 |
| rtn4rl1b | rtn4rl1a,rtn4rl1b | 2 |
| rtn4rl2a | rtn4rl2a,rtn4rl2b | 2 |
| rtn4rl2b | rtn4rl2a,rtn4rl2b | 2 |
| rundc3aa | rundc3ab,rundc3aa,rund | 4 |
| rundc3ab | rundc3ab,rundc3aa,rund | 4 |
| rundc3ab | rundc3ab,rundc3aa,rund | 4 |

| | | |
|---|---|---|
| rundc3b | rundc3ab,rundc3aa,rund[illegible] | 4 |
| runx2a | runx2a,runx2b | 2 |
| runx2b | runx2a,runx2b | 2 |
| rwdd2b | rwdd2b | 1 |
| rxfp2a | rxfp2a,rxfp2a,rxfp2b | 3 |
| rxfp2a | rxfp2a,rxfp2a,rxfp2b | 3 |
| rxfp2b | rxfp2a,rxfp2a,rxfp2b | 3 |
| rxfp3.2b | rxfp3.2b,rxfp3.2b | 2 |
| rxfp3.2b | rxfp3.2b,rxfp3.2b | 2 |
| rxfp3.3b | rxfp3.3b | 1 |
| ryr1a | ryr1a,ryr1b | 2 |
| ryr1b | ryr1a,ryr1b | 2 |
| ryr2b | ryr2b | 1 |
| s100a10a | s100a10b,s100a10a | 2 |
| s100a10b | s100a10b,s100a10a | 2 |
| s100b | s100t,s100w,s100u,s100[illegible] | 6 |
| s100s | s100t,s100w,s100u,s100[illegible] | 6 |
| s100t | s100t,s100w,s100u,s100[illegible] | 6 |
| s100u | s100t,s100w,s100u,s100[illegible] | 6 |
| s100w | s100t,s100w,s100u,s100[illegible] | 6 |
| s100z | s100t,s100w,s100u,s100[illegible] | 6 |
| s1pr3a | s1pr3a | 1 |
| s1pr5a | s1pr5a,s1pr5b | 2 |
| s1pr5b | s1pr5a,s1pr5b | 2 |
| sall1a | sall1a,sall1a,sall1b | 3 |
| sall1a | sall1a,sall1a,sall1b | 3 |
| sall1b | sall1a,sall1a,sall1b | 3 |
| sall3a | sall3a,sall3b | 2 |
| sall3b | sall3a,sall3b | 2 |
| samd10a | samd10a,samd10a,samd[illegible] | 3 |
| samd10a | samd10a,samd10a,samd[illegible] | 3 |
| samd10b | samd10a,samd10a,samd[illegible] | 3 |
| samd1a | samd1b,samd1a | 2 |
| samd1b | samd1b,samd1a | 2 |
| samd4a | samd4a | 1 |
| samsn1a | samsn1b,samsn1a | 2 |
| samsn1b | samsn1b,samsn1a | 2 |
| sap130a | sap130b,sap130a | 2 |
| sap130b | sap130b,sap130a | 2 |
| sap30bp | sap30bp | 1 |
| sar1ab | sar1ab,sar1b | 2 |
| sar1b | sar1ab,sar1b | 2 |
| sash1a | sash1b,sash1a | 2 |
| sash1b | sash1b,sash1a | 2 |
| sat1b | sat1b,sat1b | 2 |
| sat1b | sat1b,sat1b | 2 |
| sat2a | sat2a,sat2b,sat2a | 3 |
| sat2a | sat2a,sat2b,sat2a | 3 |
| sat2b | sat2a,sat2b,sat2a | 3 |
| satb1a | satb1b,satb1a | 2 |

| | | |
|---|---|---|
| satb1b | satb1b,satb1a | 2 |
| sbno2a | sbno2a | 1 |
| sc5d | sc5d | 1 |
| scaf4a | scaf4a,scaf4b | 2 |
| scaf4b | scaf4a,scaf4b | 2 |
| scamp5a | scamp5b,scamp5a | 2 |
| scamp5b | scamp5b,scamp5a | 2 |
| scarb2a | scarb2a,scarb2c | 2 |
| scarb2c | scarb2a,scarb2c | 2 |
| scg2b | scg2b | 1 |
| scn12aa | scn12aa | 1 |
| scn1ba | scn1ba,scn1ba,scn1bb | 3 |
| scn1ba | scn1ba,scn1ba,scn1bb | 3 |
| scn1bb | scn1ba,scn1ba,scn1bb | 3 |
| scn2b | scn2b | 1 |
| scn3b | scn3b | 1 |
| scn4aa | scn4aa,scn4bb,scn4ab,sc | 6 |
| scn4aa | scn4aa,scn4bb,scn4ab,sc | 6 |
| scn4ab | scn4aa,scn4bb,scn4ab,sc | 6 |
| scn4ba | scn4aa,scn4bb,scn4ab,sc | 6 |
| scn4bb | scn4aa,scn4bb,scn4ab,sc | 6 |
| scn4bb | scn4aa,scn4bb,scn4ab,sc | 6 |
| scn8aa | scn8ab,scn8ab,scn8aa | 3 |
| scn8ab | scn8ab,scn8ab,scn8aa | 3 |
| scn8ab | scn8ab,scn8ab,scn8aa | 3 |
| scp2a | scp2a,scp2b | 2 |
| scp2b | scp2a,scp2b | 2 |
| scrt1a | scrt1b,scrt1b,scrt1a | 3 |
| scrt1b | scrt1b,scrt1b,scrt1a | 3 |
| scrt1b | scrt1b,scrt1b,scrt1a | 3 |
| sdk1a | sdk1b,sdk1a | 2 |
| sdk1b | sdk1b,sdk1a | 2 |
| sdk2a | sdk2a,sdk2b | 2 |
| sdk2b | sdk2a,sdk2b | 2 |
| sdr16c5a | sdr16c5a,sdr16c5b | 2 |
| sdr16c5b | sdr16c5a,sdr16c5b | 2 |
| sec11a | sec11a,sec11a | 2 |
| sec11a | sec11a,sec11a | 2 |
| sec16b | sec16b | 1 |
| sec22a | sec22ba,sec22bb,sec22a, | 4 |
| sec22ba | sec22ba,sec22bb,sec22a, | 4 |
| sec22bb | sec22ba,sec22bb,sec22a, | 4 |
| sec22c | sec22ba,sec22bb,sec22a, | 4 |
| sec23a | sec23b,sec23ip,sec23a | 3 |
| sec23b | sec23b,sec23ip,sec23a | 3 |
| sec23ip | sec23b,sec23ip,sec23a | 3 |
| sec24a | sec24d,sec24c,sec24b,se | 4 |
| sec24b | sec24d,sec24c,sec24b,se | 4 |
| sec24c | sec24d,sec24c,sec24b,se | 4 |
| sec24d | sec24d,sec24c,sec24b,se | 4 |

| | | |
|---|---|---|
| sec31a | sec31a,sec31a,sec31a,sec | 4 |
| sec31a | sec31a,sec31a,sec31a,sec | 4 |
| sec31a | sec31a,sec31a,sec31a,sec | 4 |
| sec31b | sec31a,sec31a,sec31a,sec | 4 |
| sec61b | sec61g,sec61b,sec61g | 3 |
| sec61g | sec61g,sec61b,sec61g | 3 |
| sec61g | sec61g,sec61b,sec61g | 3 |
| selenot1a | selenot1b,selenot1a,sele | 3 |
| selenot1b | selenot1b,selenot1a,sele | 3 |
| selenot1b | selenot1b,selenot1a,sele | 3 |
| selenou1a | selenou1a,selenou1b,sele | 3 |
| selenou1a | selenou1a,selenou1b,sele | 3 |
| selenou1b | selenou1a,selenou1b,sele | 3 |
| selenow2a | selenow2b,selenow2a,se | 4 |
| selenow2a | selenow2b,selenow2a,se | 4 |
| selenow2b | selenow2b,selenow2a,se | 4 |
| selenow2b | selenow2b,selenow2a,se | 4 |
| sema3aa | sema3ab,sema3e,sema3a | 15 |
| sema3aa | sema3ab,sema3e,sema3a | 15 |
| sema3ab | sema3ab,sema3e,sema3a | 15 |
| sema3ab | sema3ab,sema3e,sema3a | 15 |
| sema3b | sema3ab,sema3e,sema3a | 15 |
| sema3b | sema3ab,sema3e,sema3a | 15 |
| sema3c | sema3ab,sema3e,sema3a | 15 |
| sema3d | sema3ab,sema3e,sema3a | 15 |
| sema3e | sema3ab,sema3e,sema3a | 15 |
| sema3e | sema3ab,sema3e,sema3a | 15 |
| sema3fa | sema3ab,sema3e,sema3a | 15 |
| sema3fb | sema3ab,sema3e,sema3a | 15 |
| sema3ga | sema3ab,sema3e,sema3a | 15 |
| sema3gb | sema3ab,sema3e,sema3a | 15 |
| sema3h | sema3ab,sema3e,sema3a | 15 |
| sema4aa | sema4e,sema4c,sema4bb | 8 |
| sema4ab | sema4e,sema4c,sema4bb | 8 |
| sema4ba | sema4e,sema4c,sema4bb | 8 |
| sema4bb | sema4e,sema4c,sema4bb | 8 |
| sema4c | sema4e,sema4c,sema4bb | 8 |
| sema4e | sema4e,sema4c,sema4bb | 8 |
| sema4ga | sema4e,sema4c,sema4bb | 8 |
| sema4gb | sema4e,sema4c,sema4bb | 8 |
| sema5a | sema5bb,sema5ba,sema5 | 3 |
| sema5ba | sema5bb,sema5ba,sema5 | 3 |
| sema5bb | sema5bb,sema5ba,sema5 | 3 |
| sema6a | sema6a,sema6ba,sema6b | 5 |
| sema6ba | sema6a,sema6ba,sema6b | 5 |
| sema6bb | sema6a,sema6ba,sema6b | 5 |
| sema6d | sema6a,sema6ba,sema6b | 5 |
| sema6e | sema6a,sema6ba,sema6b | 5 |
| sema7a | sema7a | 1 |
| senp3a | senp3a,senp3a,senp3b | 3 |

| | | |
|---|---|---|
| senp3a | senp3a,senp3a,senp3b | 3 |
| senp3b | senp3a,senp3a,senp3b | 3 |
| senp6a | senp6b,senp6a | 2 |
| senp6b | senp6b,senp6a | 2 |
| senp7a | senp7a,senp7a,senp7b | 3 |
| senp7a | senp7a,senp7a,senp7b | 3 |
| senp7b | senp7a,senp7a,senp7b | 3 |
| sept4a | sept4a | 1 |
| sept5a | sept5a,sept5b | 2 |
| sept5b | sept5a,sept5b | 2 |
| sept7a | sept7b,sept7a | 2 |
| sept7b | sept7b,sept7a | 2 |
| sept8a | sept8b,sept8a | 2 |
| sept8b | sept8b,sept8a | 2 |
| sept9a | sept9b,sept9a,sept9b | 3 |
| sept9b | sept9b,sept9a,sept9b | 3 |
| sept9b | sept9b,sept9a,sept9b | 3 |
| serbp1a | serbp1b,serbp1a | 2 |
| serbp1b | serbp1b,serbp1a | 2 |
| serpina10a | serpina10a | 1 |
| serpinf2a | serpinf2b,serpinf2a | 2 |
| serpinf2b | serpinf2b,serpinf2a | 2 |
| serpinh1a | serpinh1a,serpinh1b | 2 |
| serpinh1b | serpinh1a,serpinh1b | 2 |
| sertad2a | sertad2a,sertad2b | 2 |
| sertad2b | sertad2a,sertad2b | 2 |
| setd1a | setd1ba,setd1a,setd1bb,s | 4 |
| setd1ba | setd1ba,setd1a,setd1bb,s | 4 |
| setd1ba | setd1ba,setd1a,setd1bb,s | 4 |
| setd1bb | setd1ba,setd1a,setd1bb,s | 4 |
| setdb1a | setdb1b,setdb1a | 2 |
| setdb1b | setdb1b,setdb1a | 2 |
| sez6a | sez6b,sez6a,sez6b | 3 |
| sez6b | sez6b,sez6a,sez6b | 3 |
| sez6b | sez6b,sez6a,sez6b | 3 |
| sfrp1a | sfrp1b,sfrp1a,sfrp1b | 3 |
| sfrp1b | sfrp1b,sfrp1a,sfrp1b | 3 |
| sfrp1b | sfrp1b,sfrp1a,sfrp1b | 3 |
| sft2d2a | sft2d2b,sft2d2b,sft2d2a | 3 |
| sft2d2b | sft2d2b,sft2d2b,sft2d2a | 3 |
| sft2d2b | sft2d2b,sft2d2b,sft2d2a | 3 |
| sfxn5a | sfxn5a,sfxn5b | 2 |
| sfxn5b | sfxn5a,sfxn5b | 2 |
| sgip1a | sgip1a,sgip1b,sgip1a | 3 |
| sgip1a | sgip1a,sgip1b,sgip1a | 3 |
| sgip1b | sgip1a,sgip1b,sgip1a | 3 |
| sgk2a | sgk2b,sgk2a | 2 |
| sgk2b | sgk2b,sgk2a | 2 |
| sgms2a | sgms2a,sgms2b | 2 |
| sgms2b | sgms2a,sgms2b | 2 |

| | | |
|---|---|---|
| sgsm1a | sgsm1a,sgsm1b | 2 |
| sgsm1b | sgsm1a,sgsm1b | 2 |
| sh2d1aa | sh2d1ab,sh2d1aa,sh2d1a | 3 |
| sh2d1ab | sh2d1ab,sh2d1aa,sh2d1a | 3 |
| sh2d1ab | sh2d1ab,sh2d1aa,sh2d1a | 3 |
| sh2d3ca | sh2d3cb,sh2d3cb,sh2d3c | 3 |
| sh2d3cb | sh2d3cb,sh2d3cb,sh2d3c | 3 |
| sh2d3cb | sh2d3cb,sh2d3cb,sh2d3c | 3 |
| sh2d4a | sh2d4a,sh2d4bb,sh2d4ba | 3 |
| sh2d4ba | sh2d4a,sh2d4bb,sh2d4ba | 3 |
| sh2d4bb | sh2d4a,sh2d4bb,sh2d4ba | 3 |
| sh3bgr | sh3bgr | 1 |
| sh3bp4a | sh3bp4,sh3bp4,sh3bp4a | 3 |
| sh3bp5a | sh3bp5b,sh3bp5a | 2 |
| sh3bp5b | sh3bp5b,sh3bp5a | 2 |
| sh3gl1a | sh3gl1b,sh3gl1a,sh3gl1b | 3 |
| sh3gl1b | sh3gl1b,sh3gl1a,sh3gl1b | 3 |
| sh3gl1b | sh3gl1b,sh3gl1a,sh3gl1b | 3 |
| sh3gl2a | sh3gl2a,sh3gl2b | 2 |
| sh3gl2b | sh3gl2a,sh3gl2b | 2 |
| sh3gl3a | sh3gl3a,sh3gl3b | 2 |
| sh3gl3b | sh3gl3a,sh3gl3b | 2 |
| sh3glb1a | sh3glb1a,sh3glb1b,sh3glb | 3 |
| sh3glb1a | sh3glb1a,sh3glb1b,sh3glb | 3 |
| sh3glb1b | sh3glb1a,sh3glb1b,sh3glb | 3 |
| sh3glb2a | sh3glb2b,sh3glb2b,sh3glb | 3 |
| sh3glb2b | sh3glb2b,sh3glb2b,sh3glb | 3 |
| sh3glb2b | sh3glb2b,sh3glb2b,sh3glb | 3 |
| sh3pxd2aa | sh3pxd2aa,sh3pxd2ab,sh | 3 |
| sh3pxd2ab | sh3pxd2aa,sh3pxd2ab,sh | 3 |
| sh3pxd2b | sh3pxd2aa,sh3pxd2ab,sh | 3 |
| shank2a | shank2a,shank2b | 2 |
| shank2b | shank2a,shank2b | 2 |
| shank3a | shank3b,shank3a | 2 |
| shank3b | shank3b,shank3a | 2 |
| shisa2a | shisa2a,shisa2a,shisa2b | 3 |
| shisa2a | shisa2a,shisa2a,shisa2b | 3 |
| shisa2b | shisa2a,shisa2a,shisa2b | 3 |
| shisa7a | shisa7a,shisa7b | 2 |
| shisa7b | shisa7a,shisa7b | 2 |
| shisa8b | shisa8b | 1 |
| shisa9a | shisa9a,shisa9b,shisa9a | 3 |
| shisa9a | shisa9a,shisa9b,shisa9a | 3 |
| shisa9b | shisa9a,shisa9b,shisa9a | 3 |
| shisal1a | shisal1a,shisal1b | 2 |
| shisal1b | shisal1a,shisal1b | 2 |
| shroom2a | shroom2a | 1 |
| shrprbck1r | shrprbck1r | 1 |
| sik2a | sik2a,sik2a,sik2b | 3 |
| sik2a | sik2a,sik2a,sik2b | 3 |

| | | |
|---|---|---|
| sik2b | sik2a,sik2a,sik2b | 3 |
| sim1a | sim1b,sim1a | 2 |
| sim1b | sim1b,sim1a | 2 |
| sin3aa | sin3ab,sin3b,sin3aa | 3 |
| sin3ab | sin3ab,sin3b,sin3aa | 3 |
| sin3b | sin3ab,sin3b,sin3aa | 3 |
| six1a | six1a,six1b | 2 |
| six1b | six1a,six1b | 2 |
| six2a | six2a,six2b,six2a | 3 |
| six2a | six2a,six2b,six2a | 3 |
| six2b | six2a,six2b,six2a | 3 |
| six3a | six3a,six3b,six3a | 3 |
| six3a | six3a,six3b,six3a | 3 |
| six3b | six3a,six3b,six3a | 3 |
| six4a | six4a,six4b | 2 |
| six4b | six4a,six4b | 2 |
| six6a | six6a,six6b | 2 |
| six6b | six6a,six6b | 2 |
| skor1a | skor1b,skor1a,skor1b | 3 |
| skor1b | skor1b,skor1a,skor1b | 3 |
| skor1b | skor1b,skor1a,skor1b | 3 |
| sla1a | sla1a | 1 |
| slain1a | slain1a,slain1a,slain1a,sla | 5 |
| slain1a | slain1a,slain1a,slain1a,sla | 5 |
| slain1a | slain1a,slain1a,slain1a,sla | 5 |
| slain1a | slain1a,slain1a,slain1a,sla | 5 |
| slain1b | slain1a,slain1a,slain1a,sla | 5 |
| slc12a5a | slc12a5a,slc12a5b | 2 |
| slc12a5b | slc12a5a,slc12a5b | 2 |
| slc12a7a | slc12a7a,slc12a7b | 2 |
| slc12a7b | slc12a7a,slc12a7b | 2 |
| slc13a5a | slc13a5b,slc13a5a | 2 |
| slc13a5b | slc13a5b,slc13a5a | 2 |
| slc15a1a | slc15a1b,slc15a1a | 2 |
| slc15a1b | slc15a1b,slc15a1a | 2 |
| slc16a12a | slc16a12b,slc16a12a | 2 |
| slc16a12b | slc16a12b,slc16a12a | 2 |
| slc16a1a | slc16a1a,slc16a1b,slc16a | 4 |
| slc16a1a | slc16a1a,slc16a1b,slc16a | 4 |
| slc16a1b | slc16a1a,slc16a1b,slc16a | 4 |
| slc16a1b | slc16a1a,slc16a1b,slc16a | 4 |
| slc16a5a | slc16a5a,slc16a5b,slc16a | 3 |
| slc16a5a | slc16a5a,slc16a5b,slc16a | 3 |
| slc16a5b | slc16a5a,slc16a5b,slc16a | 3 |
| slc16a6a | slc16a6a,slc16a6b | 2 |
| slc16a6b | slc16a6a,slc16a6b | 2 |
| slc16a9a | slc16a9a,slc16a9b,slc16a | 3 |
| slc16a9a | slc16a9a,slc16a9b,slc16a | 3 |
| slc16a9b | slc16a9a,slc16a9b,slc16a | 3 |
| slc17a6a | slc17a6b,slc17a6a | 2 |

| | | |
|---|---|---|
| slc17a6b | slc17a6b,slc17a6a | 2 |
| slc17a7a | slc17a7a,slc17a7a,slc17a7 | 3 |
| slc17a7a | slc17a7a,slc17a7a,slc17a7 | 3 |
| slc17a7b | slc17a7a,slc17a7a,slc17a7 | 3 |
| slc17a9a | slc17a9a,slc17a9b | 2 |
| slc17a9b | slc17a9a,slc17a9b | 2 |
| slc18a3a | slc18a3b,slc18a3a | 2 |
| slc18a3b | slc18a3b,slc18a3a | 2 |
| slc19a3a | slc19a3a,slc19a3b | 2 |
| slc19a3b | slc19a3a,slc19a3b | 2 |
| slc1a2a | slc1a2a,slc1a2b | 2 |
| slc1a2b | slc1a2a,slc1a2b | 2 |
| slc1a3a | slc1a3a,slc1a3b | 2 |
| slc1a3b | slc1a3a,slc1a3b | 2 |
| slc1a7a | slc1a7b,slc1a7a,slc1a7b | 3 |
| slc1a7b | slc1a7b,slc1a7a,slc1a7b | 3 |
| slc1a7b | slc1a7b,slc1a7a,slc1a7b | 3 |
| slc1a8a | slc1a8b,slc1a8a | 2 |
| slc1a8b | slc1a8b,slc1a8a | 2 |
| slc20a1a | slc20a1a,slc20a1a,slc20a1 | 3 |
| slc20a1a | slc20a1a,slc20a1a,slc20a1 | 3 |
| slc20a1b | slc20a1a,slc20a1a,slc20a1 | 3 |
| slc22a13a | slc22a13a,slc22a13b | 2 |
| slc22a13b | slc22a13a,slc22a13b | 2 |
| slc22a7a | slc22a7a | 1 |
| slc24a4a | slc24a4a,slc24a4b | 2 |
| slc24a4b | slc24a4a,slc24a4b | 2 |
| slc24a6a | slc24a6a | 1 |
| slc25a15a | slc25a15b,slc25a15b,slc2 | 3 |
| slc25a15b | slc25a15b,slc25a15b,slc2 | 3 |
| slc25a15b | slc25a15b,slc25a15b,slc2 | 3 |
| slc25a1a | slc25a1b,slc25a1a | 2 |
| slc25a1b | slc25a1b,slc25a1a | 2 |
| slc25a22a | slc25a22b,slc25a22a | 2 |
| slc25a22b | slc25a22b,slc25a22a | 2 |
| slc25a23a | slc25a23a,slc25a23b | 2 |
| slc25a23b | slc25a23a,slc25a23b | 2 |
| slc25a25a | slc25a25b,slc25a25b,slc2 | 3 |
| slc25a25b | slc25a25b,slc25a25b,slc2 | 3 |
| slc25a25b | slc25a25b,slc25a25b,slc2 | 3 |
| slc25a32a | slc25a32b,slc25a32a | 2 |
| slc25a32b | slc25a32b,slc25a32a | 2 |
| slc25a36a | slc25a36a,slc25a36a,slc25 | 3 |
| slc25a36a | slc25a36a,slc25a36a,slc25 | 3 |
| slc25a36b | slc25a36a,slc25a36a,slc25 | 3 |
| slc25a38a | slc25a38a,slc25a38b | 2 |
| slc25a38b | slc25a38a,slc25a38b | 2 |
| slc25a3a | slc25a3a,slc25a3b | 2 |
| slc25a3b | slc25a3a,slc25a3b | 2 |
| slc25a44a | slc25a44b,slc25a44a | 2 |

| | | |
|---|---|---|
| slc25a44b | slc25a44b,slc25a44a | 2 |
| slc25a47a | slc25a47b,slc25a47a | 2 |
| slc25a47b | slc25a47b,slc25a47a | 2 |
| slc25a51a | slc25a51a,slc25a51b | 2 |
| slc25a51b | slc25a51a,slc25a51b | 2 |
| slc25a55a | slc25a55a,slc25a55a,slc2 | 3 |
| slc25a55a | slc25a55a,slc25a55a,slc2 | 3 |
| slc25a55b | slc25a55a,slc25a55a,slc2 | 3 |
| slc27a1a | slc27a1a,slc27a1b,slc27a | 4 |
| slc27a1a | slc27a1a,slc27a1b,slc27a | 4 |
| slc27a1b | slc27a1a,slc27a1b,slc27a | 4 |
| slc27a1b | slc27a1a,slc27a1b,slc27a | 4 |
| slc27a2a | slc27a2a | 1 |
| slc29a1a | slc29a1b,slc29a1a | 2 |
| slc29a1b | slc29a1b,slc29a1a | 2 |
| slc2a11a | slc2a11b,slc2a11b,slc2a1 | 3 |
| slc2a11b | slc2a11b,slc2a11b,slc2a1 | 3 |
| slc2a11b | slc2a11b,slc2a11b,slc2a1 | 3 |
| slc2a13b | slc2a13b | 1 |
| slc2a15a | slc2a15b,slc2a15a | 2 |
| slc2a15b | slc2a15b,slc2a15a | 2 |
| slc2a1a | slc2a1a,slc2a1b,slc2a1a | 3 |
| slc2a1a | slc2a1a,slc2a1b,slc2a1a | 3 |
| slc2a1b | slc2a1a,slc2a1b,slc2a1a | 3 |
| slc2a3a | slc2a3b,slc2a3a | 2 |
| slc2a3b | slc2a3b,slc2a3a | 2 |
| slc30a1a | slc30a1b,slc30a1a | 2 |
| slc30a1b | slc30a1b,slc30a1a | 2 |
| slc34a1a | slc34a1a,slc34a1b | 2 |
| slc34a1b | slc34a1a,slc34a1b | 2 |
| slc34a2a | slc34a2a,slc34a2b | 2 |
| slc34a2b | slc34a2a,slc34a2b | 2 |
| slc35a3a | slc35a3b,slc35a3a | 2 |
| slc35a3b | slc35a3b,slc35a3a | 2 |
| slc35d1a | slc35d1b,slc35d1a | 2 |
| slc35d1b | slc35d1b,slc35d1a | 2 |
| slc35f3a | slc35f3a,slc35f3b | 2 |
| slc35f3b | slc35f3a,slc35f3b | 2 |
| slc35g2a | slc35g2a,slc35g2b | 2 |
| slc35g2b | slc35g2a,slc35g2b | 2 |
| slc37a4a | slc37a4b,slc37a4b,slc37a | 3 |
| slc37a4b | slc37a4b,slc37a4b,slc37a | 3 |
| slc37a4b | slc37a4b,slc37a4b,slc37a | 3 |
| slc38a3a | slc38a3a,slc38a3b,slc38a | 3 |
| slc38a3a | slc38a3a,slc38a3b,slc38a | 3 |
| slc38a3b | slc38a3a,slc38a3b,slc38a | 3 |
| slc38a5a | slc38a5b,slc38a5a | 2 |
| slc38a5b | slc38a5b,slc38a5a | 2 |
| slc38a8a | slc38a8b,slc38a8a | 2 |
| slc38a8b | slc38a8b,slc38a8a | 2 |

| | | |
|---|---|---|
| slc3a2a | slc3a2a,slc3a2a,slc3a2b | 3 |
| slc3a2a | slc3a2a,slc3a2a,slc3a2b | 3 |
| slc3a2b | slc3a2a,slc3a2a,slc3a2b | 3 |
| slc41a2a | slc41a2b,slc41a2a | 2 |
| slc41a2b | slc41a2b,slc41a2a | 2 |
| slc43a1a | slc43a1a,slc43a1b,slc43a | 3 |
| slc43a1a | slc43a1a,slc43a1b,slc43a | 3 |
| slc43a1b | slc43a1a,slc43a1b,slc43a | 3 |
| slc43a2a | slc43a2b,slc43a2b,slc43a | 3 |
| slc43a2b | slc43a2b,slc43a2b,slc43a | 3 |
| slc43a2b | slc43a2b,slc43a2b,slc43a | 3 |
| slc43a3a | slc43a3a,slc43a3b | 2 |
| slc43a3b | slc43a3a,slc43a3b | 2 |
| slc44a1a | slc44a1a,slc44a1b | 2 |
| slc44a1b | slc44a1a,slc44a1b | 2 |
| slc44a5a | slc44a5a,slc44a5b | 2 |
| slc44a5b | slc44a5a,slc44a5b | 2 |
| slc45a4b | slc45a4b | 1 |
| slc48a1a | slc48a1b,slc48a1a | 2 |
| slc48a1b | slc48a1b,slc48a1a | 2 |
| slc4a10a | slc4a10b,slc4a10a | 2 |
| slc4a10b | slc4a10b,slc4a10a | 2 |
| slc4a1a | slc4a1a,slc4a1b,slc4a1ap | 3 |
| slc4a1ap | slc4a1a,slc4a1b,slc4a1ap | 3 |
| slc4a1b | slc4a1a,slc4a1b,slc4a1ap | 3 |
| slc4a2a | slc4a2a,slc4a2b | 2 |
| slc4a2b | slc4a2a,slc4a2b | 2 |
| slc4a4a | slc4a4a,slc4a4b | 2 |
| slc4a4b | slc4a4a,slc4a4b | 2 |
| slc51a | slc51a | 1 |
| slc5a3a | slc5a3a,slc5a3b | 2 |
| slc5a3b | slc5a3a,slc5a3b | 2 |
| slc5a6a | slc5a6b,slc5a6b,slc5a6a | 3 |
| slc5a6b | slc5a6b,slc5a6b,slc5a6a | 3 |
| slc5a6b | slc5a6b,slc5a6b,slc5a6a | 3 |
| slc5a7a | slc5a7a | 1 |
| slc6a11a | slc6a11a,slc6a11b | 2 |
| slc6a11b | slc6a11a,slc6a11b | 2 |
| slc6a16a | slc6a16a,slc6a16b | 2 |
| slc6a16b | slc6a16a,slc6a16b | 2 |
| slc6a19b | slc6a19b | 1 |
| slc6a1a | slc6a1b,slc6a1a | 2 |
| slc6a1b | slc6a1b,slc6a1a | 2 |
| slc6a4a | slc6a4b,slc6a4a | 2 |
| slc6a4b | slc6a4b,slc6a4a | 2 |
| slc6a6a | slc6a6a,slc6a6b | 2 |
| slc6a6b | slc6a6a,slc6a6b | 2 |
| slc7a10a | slc7a10a,slc7a10b | 2 |
| slc7a10b | slc7a10a,slc7a10b | 2 |
| slc7a14a | slc7a14a,slc7a14b | 2 |

| | | |
|---|---|---|
| slc7a14b | slc7a14a,slc7a14b | 2 |
| slc7a1a | slc7a1a,slc7a1a | 2 |
| slc7a1a | slc7a1a,slc7a1a | 2 |
| slc7a3a | slc7a3b,slc7a3a | 2 |
| slc7a3b | slc7a3b,slc7a3a | 2 |
| slc7a6os | slc7a6os,slc7a6 | 2 |
| slc7a8a | slc7a8b,slc7a8b,slc7a8a | 3 |
| slc7a8b | slc7a8b,slc7a8b,slc7a8a | 3 |
| slc7a8b | slc7a8b,slc7a8b,slc7a8a | 3 |
| slc8a1a | slc8a1b,slc8a1a,slc8a1b | 3 |
| slc8a1b | slc8a1b,slc8a1a,slc8a1b | 3 |
| slc8a1b | slc8a1b,slc8a1a,slc8a1b | 3 |
| slc8a2a | slc8a2a,slc8a2a,slc8a2b | 3 |
| slc8a2a | slc8a2a,slc8a2a,slc8a2b | 3 |
| slc8a2b | slc8a2a,slc8a2a,slc8a2b | 3 |
| slc8a4a | slc8a4b,slc8a4a | 2 |
| slc8a4b | slc8a4b,slc8a4a | 2 |
| slc9a1b | slc9a1b | 1 |
| slc9a3r1a | slc9a3r1a,slc9a3r1b,slc9a | 3 |
| slc9a3r1a | slc9a3r1a,slc9a3r1b,slc9a | 3 |
| slc9a3r1b | slc9a3r1a,slc9a3r1b,slc9a | 3 |
| slc9a6a | slc9a6a,slc9a6b,slc9a6a | 3 |
| slc9a6a | slc9a6a,slc9a6b,slc9a6a | 3 |
| slc9a6b | slc9a6a,slc9a6b,slc9a6a | 3 |
| slco5a1a | slco5a1a,slco5a1a | 2 |
| slco5a1a | slco5a1a,slco5a1a | 2 |
| slit1a | slit1a,slit1b | 2 |
| slit1b | slit1a,slit1b | 2 |
| slitrk3a | slitrk3b,slitrk3a | 2 |
| slitrk3b | slitrk3b,slitrk3a | 2 |
| slitrk5a | slitrk5b,slitrk5a | 2 |
| slitrk5b | slitrk5b,slitrk5a | 2 |
| slx1b | slx1b | 1 |
| slx4ip | slx4ip,slx4,slx4 | 3 |
| smad3a | smad3a,smad3a,smad3b | 3 |
| smad3a | smad3a,smad3a,smad3b | 3 |
| smad3b | smad3a,smad3a,smad3b | 3 |
| smad4a | smad4a | 1 |
| smad6a | smad6a,smad6a,smad6b | 3 |
| smad6a | smad6a,smad6a,smad6b | 3 |
| smad6b | smad6a,smad6a,smad6b | 3 |
| smarca4a | smarca4a | 1 |
| smarcad1a | smarcad1a,smarcad1a,sn | 3 |
| smarcad1a | smarcad1a,smarcad1a,sn | 3 |
| smarcad1b | smarcad1a,smarcad1a,sn | 3 |
| smarcb1a | smarcb1b,smarcb1a,sma | 3 |
| smarcb1b | smarcb1b,smarcb1a,sma | 3 |
| smarcb1b | smarcb1b,smarcb1a,sma | 3 |
| smarcc1a | smarcc1a,smarcc1b | 2 |
| smarcc1b | smarcc1a,smarcc1b | 2 |

| | | |
|---|---|---|
| smarcd3a | smarcd3a,smarcd3b | 2 |
| smarcd3b | smarcd3a,smarcd3b | 2 |
| smc1a | smc1a,smc1b | 2 |
| smc1b | smc1a,smc1b | 2 |
| smcr8a | smcr8b,smcr8a | 2 |
| smcr8b | smcr8b,smcr8a | 2 |
| smdt1a | smdt1b,smdt1a | 2 |
| smdt1b | smdt1b,smdt1a | 2 |
| smpd2a | smpd2b,smpd2a | 2 |
| smpd2b | smpd2b,smpd2a | 2 |
| smpdl3a | smpdl3b,smpdl3a | 2 |
| smpdl3b | smpdl3b,smpdl3a | 2 |
| smu1a | smu1b,smu1a | 2 |
| smu1b | smu1b,smu1a | 2 |
| smyd1a | smyd1a,smyd1b | 2 |
| smyd1b | smyd1a,smyd1b | 2 |
| smyd2a | smyd2a,smyd2b | 2 |
| smyd2b | smyd2a,smyd2b | 2 |
| snai1a | snai1a,snai1b | 2 |
| snai1b | snai1a,snai1b | 2 |
| snap25a | snap25b,snap25a | 2 |
| snap25b | snap25b,snap25a | 2 |
| snap91a | snap91b,snap91a | 2 |
| snap91b | snap91b,snap91a | 2 |
| snapc1a | snapc1b,snapc1a | 2 |
| snapc1b | snapc1b,snapc1a | 2 |
| snu13a | snu13a,snu13b | 2 |
| snu13b | snu13a,snu13b | 2 |
| snx10a | snx10b,snx10a | 2 |
| snx10b | snx10b,snx10a | 2 |
| snx18a | snx18b,snx18a,snx18a,sn | 4 |
| snx18a | snx18b,snx18a,snx18a,sn | 4 |
| snx18b | snx18b,snx18a,snx18a,sn | 4 |
| snx18b | snx18b,snx18a,snx18a,sn | 4 |
| snx19a | snx19a,snx19b | 2 |
| snx19b | snx19a,snx19b | 2 |
| snx1a | snx1b,snx1a | 2 |
| snx1b | snx1b,snx1a | 2 |
| snx27a | snx27b,snx27a | 2 |
| snx27b | snx27b,snx27a | 2 |
| snx8a | snx8b,snx8a | 2 |
| snx8b | snx8b,snx8a | 2 |
| snx9a | snx9b,snx9a,snx9b | 3 |
| snx9b | snx9b,snx9a,snx9b | 3 |
| snx9b | snx9b,snx9a,snx9b | 3 |
| socs1a | socs1b,socs1b,socs1a | 3 |
| socs1b | socs1b,socs1b,socs1a | 3 |
| socs1b | socs1b,socs1b,socs1a | 3 |
| socs3a | socs3a,socs3b | 2 |
| socs3b | socs3a,socs3b | 2 |

| | | |
|---|---|---|
| socs5a | socs5b,socs5b,socs5a,soc | 4 |
| socs5b | socs5b,socs5b,socs5a,soc | 4 |
| socs5b | socs5b,socs5b,socs5a,soc | 4 |
| socs5b | socs5b,socs5b,socs5a,soc | 4 |
| socs6a | socs6b,socs6a | 2 |
| socs6b | socs6b,socs6a | 2 |
| sod3a | sod3b,sod3a | 2 |
| sod3b | sod3b,sod3a | 2 |
| soga3a | soga3a,soga3a,soga3b | 3 |
| soga3a | soga3a,soga3a,soga3b | 3 |
| soga3b | soga3a,soga3a,soga3b | 3 |
| sorbs2a | sorbs2a,sorbs2b | 2 |
| sorbs2b | sorbs2a,sorbs2b | 2 |
| sorcs3b | sorcs3b | 1 |
| sort1a | sort1a,sort1b,sort1a | 3 |
| sort1a | sort1a,sort1b,sort1a | 3 |
| sort1b | sort1a,sort1b,sort1a | 3 |
| sostdc1a | sostdc1b,sostdc1a,sostdc | 3 |
| sostdc1b | sostdc1b,sostdc1a,sostdc | 3 |
| sostdc1b | sostdc1b,sostdc1a,sostdc | 3 |
| sox11a | sox11a,sox11b | 2 |
| sox11b | sox11a,sox11b | 2 |
| sox19a | sox19a,sox19a,sox19b | 3 |
| sox19a | sox19a,sox19a,sox19b | 3 |
| sox19b | sox19a,sox19a,sox19b | 3 |
| sox1a | sox1a,sox1a,sox1b,sox1a | 4 |
| sox1a | sox1a,sox1a,sox1b,sox1a | 4 |
| sox1a | sox1a,sox1a,sox1b,sox1a | 4 |
| sox1b | sox1a,sox1a,sox1b,sox1a | 4 |
| sox21a | sox21a,sox21a,sox21b | 3 |
| sox21a | sox21a,sox21a,sox21b | 3 |
| sox21b | sox21a,sox21a,sox21b | 3 |
| sox4a | sox4b,sox4a,sox4a | 3 |
| sox4a | sox4b,sox4a,sox4a | 3 |
| sox4b | sox4b,sox4a,sox4a | 3 |
| sox8a | sox8b,sox8a | 2 |
| sox8b | sox8b,sox8a | 2 |
| sox9a | sox9b,sox9a | 2 |
| sox9b | sox9b,sox9a | 2 |
| sp3a | sp3b,sp3a | 2 |
| sp3b | sp3b,sp3a | 2 |
| sp5a | sp5a | 1 |
| sp8a | sp8b,sp8a | 2 |
| sp8b | sp8b,sp8a | 2 |
| spag1a | spag1b,spag1a | 2 |
| spag1b | spag1b,spag1a | 2 |
| spag9a | spag9a | 1 |
| spi1a | spi1b,spi1a | 2 |
| spi1b | spi1b,spi1a | 2 |
| spint1a | spint1a,spint1b | 2 |

| | | |
|---|---|---|
| spint1b | spint1a,spint1b | 2 |
| spire1a | spire1a,spire1b,spire1a | 3 |
| spire1a | spire1a,spire1b,spire1a | 3 |
| spire1b | spire1a,spire1b,spire1a | 3 |
| spon1a | spon1b,spon1a,spon1b | 3 |
| spon1b | spon1b,spon1a,spon1b | 3 |
| spon1b | spon1b,spon1a,spon1b | 3 |
| spon2a | spon2a,spon2b,spon2a | 3 |
| spon2a | spon2a,spon2b,spon2a | 3 |
| spon2b | spon2a,spon2b,spon2a | 3 |
| spred2a | spred2a,spred2b | 2 |
| spred2b | spred2a,spred2b | 2 |
| sprydd7a | sprydd7b,sprydd7b,sprydd7a | 4 |
| sprydd7b | sprydd7b,sprydd7b,sprydd7a | 4 |
| sprydd7b | sprydd7b,sprydd7b,sprydd7a | 4 |
| sprydd7b | sprydd7b,sprydd7b,sprydd7a | 4 |
| spsb3a | spsb3a,spsb3b | 2 |
| spsb3b | spsb3a,spsb3b | 2 |
| spsb4a | spsb4a,spsb4a,spsb4b | 3 |
| spsb4a | spsb4a,spsb4a,spsb4b | 3 |
| spsb4b | spsb4a,spsb4a,spsb4b | 3 |
| sptbn4a | sptbn4a,sptbn4b,sptbn4k | 3 |
| sptbn4b | sptbn4a,sptbn4b,sptbn4k | 3 |
| sptbn4b | sptbn4a,sptbn4b,sptbn4k | 3 |
| sptlc2a | sptlc2a,sptlc2b | 2 |
| sptlc2b | sptlc2a,sptlc2b | 2 |
| srd5a2a | srd5a2a,srd5a2b | 2 |
| srd5a2b | srd5a2a,srd5a2b | 2 |
| srgap1a | srgap1b,srgap1a | 2 |
| srgap1b | srgap1b,srgap1a | 2 |
| srpk1a | srpk1a,srpk1a,srpk1b | 3 |
| srpk1a | srpk1a,srpk1a,srpk1b | 3 |
| srpk1b | srpk1a,srpk1a,srpk1b | 3 |
| srsf10b | srsf10b | 1 |
| srsf1a | srsf1a,srsf1a,srsf1b | 3 |
| srsf1a | srsf1a,srsf1a,srsf1b | 3 |
| srsf1b | srsf1a,srsf1a,srsf1b | 3 |
| srsf2a | srsf2b,srsf2a | 2 |
| srsf2b | srsf2b,srsf2a | 2 |
| srsf3a | srsf3a,srsf3a,srsf3b | 3 |
| srsf3a | srsf3a,srsf3a,srsf3b | 3 |
| srsf3b | srsf3a,srsf3a,srsf3b | 3 |
| srsf5a | srsf5a,srsf5b | 2 |
| srsf5b | srsf5a,srsf5b | 2 |
| srsf6a | srsf6b,srsf6a | 2 |
| srsf6b | srsf6b,srsf6a | 2 |
| srsf7a | srsf7a,srsf7a,srsf7b | 3 |
| srsf7a | srsf7a,srsf7a,srsf7b | 3 |
| srsf7b | srsf7a,srsf7a,srsf7b | 3 |
| ssbp3a | ssbp3a,ssbp3b | 2 |

| | | |
|---|---|---|
| ssbp3b | ssbp3a,ssbp3b | 2 |
| ssh1a | ssh1a,ssh1b | 2 |
| ssh1b | ssh1a,ssh1b | 2 |
| ssh2a | ssh2a,ssh2b | 2 |
| ssh2b | ssh2a,ssh2b | 2 |
| ssrp1a | ssrp1a,ssrp1b,ssrp1a | 3 |
| ssrp1a | ssrp1a,ssrp1b,ssrp1a | 3 |
| ssrp1b | ssrp1a,ssrp1b,ssrp1a | 3 |
| sstr1a | sstr1b,sstr1a | 2 |
| sstr1b | sstr1b,sstr1a | 2 |
| sstr2a | sstr2b,sstr2a | 2 |
| sstr2b | sstr2b,sstr2a | 2 |
| ssx2ipa | ssx2ipa,ssx2ipb | 2 |
| ssx2ipb | ssx2ipa,ssx2ipb | 2 |
| st14a | st14a,st14b | 2 |
| st14b | st14a,st14b | 2 |
| st3gal3a | st3gal3b,st3gal3a | 2 |
| st3gal3b | st3gal3b,st3gal3a | 2 |
| st6gal2a | st6gal2b,st6gal2a | 2 |
| st6gal2b | st6gal2b,st6gal2a | 2 |
| st6galnac5a | st6galnac5b,st6galnac5a, | 3 |
| st6galnac5l | st6galnac5b,st6galnac5a, | 3 |
| st6galnac5l | st6galnac5b,st6galnac5a, | 3 |
| stag1a | stag1a,stag1b | 2 |
| stag1b | stag1a,stag1b | 2 |
| stag2a | stag2b,stag2a,stag2b | 3 |
| stag2b | stag2b,stag2a,stag2b | 3 |
| stag2b | stag2b,stag2a,stag2b | 3 |
| stap2a | stap2a,stap2b | 2 |
| stap2b | stap2a,stap2b | 2 |
| stard13a | stard13a,stard13b | 2 |
| stard13b | stard13a,stard13b | 2 |
| stat1a | stat1b,stat1a | 2 |
| stat1b | stat1b,stat1a | 2 |
| stat5a | stat5a,stat5a,stat5b | 3 |
| stat5a | stat5a,stat5a,stat5b | 3 |
| stat5b | stat5a,stat5a,stat5b | 3 |
| stc2a | stc2a,stc2a,stc2b | 3 |
| stc2a | stc2a,stc2a,stc2b | 3 |
| stc2b | stc2a,stc2a,stc2b | 3 |
| stim1a | stim1a,stim1b | 2 |
| stim1b | stim1a,stim1b | 2 |
| stim2a | stim2a,stim2b | 2 |
| stim2b | stim2a,stim2b | 2 |
| stk11ip | stk11ip,stk11,stk11ip | 3 |
| stk11ip | stk11ip,stk11,stk11ip | 3 |
| stk17a | stk17b,stk17b,stk17a | 3 |
| stk17b | stk17b,stk17b,stk17a | 3 |
| stk17b | stk17b,stk17b,stk17a | 3 |
| stk24a | stk24b,stk24a | 2 |

| | | |
|---|---|---|
| stk24b | stk24b,stk24a | 2 |
| stk25a | stk25b,stk25a | 2 |
| stk25b | stk25b,stk25a | 2 |
| stk32a | stk32a | 1 |
| stk38a | stk38a,stk38b | 2 |
| stk38b | stk38a,stk38b | 2 |
| stmn1a | stmn1b,stmn1a | 2 |
| stmn1b | stmn1b,stmn1a | 2 |
| stmn2a | stmn2a,stmn2b | 2 |
| stmn2b | stmn2a,stmn2b | 2 |
| stoml3a | stoml3b,stoml3a | 2 |
| stoml3b | stoml3b,stoml3a | 2 |
| stox2a | stox2a,stox2b | 2 |
| stox2b | stox2a,stox2b | 2 |
| stt3a | stt3a,stt3b | 2 |
| stt3b | stt3a,stt3b | 2 |
| stx11a | stx11a | 1 |
| stx1a | stx1a,stx1b | 2 |
| stx1b | stx1a,stx1b | 2 |
| stx2a | stx2b,stx2a | 2 |
| stx2b | stx2b,stx2a | 2 |
| stx3a | stx3a | 1 |
| stx5a | stx5a,stx5a | 2 |
| stx5a | stx5a,stx5a | 2 |
| stxbp1a | stxbp1b,stxbp1b,stxbp1a | 3 |
| stxbp1b | stxbp1b,stxbp1b,stxbp1a | 3 |
| stxbp1b | stxbp1b,stxbp1b,stxbp1a | 3 |
| stxbp5a | stxbp5b,stxbp5a | 2 |
| stxbp5b | stxbp5b,stxbp5a | 2 |
| styk1a | styk1b,styk1a | 2 |
| styk1b | styk1b,styk1a | 2 |
| sub1a | sub1b,sub1b,sub1a | 3 |
| sub1b | sub1b,sub1b,sub1a | 3 |
| sub1b | sub1b,sub1b,sub1a | 3 |
| sulf2a | sulf2a,sulf2b,sulf2a,sulf2l | 4 |
| sulf2a | sulf2a,sulf2b,sulf2a,sulf2l | 4 |
| sulf2b | sulf2a,sulf2b,sulf2a,sulf2l | 4 |
| sulf2b | sulf2a,sulf2b,sulf2a,sulf2l | 4 |
| sumo2a | sumo2b,sumo2a | 2 |
| sumo2b | sumo2b,sumo2a | 2 |
| sumo3a | sumo3a,sumo3b | 2 |
| sumo3b | sumo3a,sumo3b | 2 |
| supt16h | supt16h | 1 |
| supt3h | supt3h | 1 |
| supt5h | supt5h,supt5h | 2 |
| supt5h | supt5h,supt5h | 2 |
| supt6h | supt6h | 1 |
| suv39h1a | suv39h1a,suv39h1b | 2 |
| suv39h1b | suv39h1a,suv39h1b | 2 |
| suz12a | suz12b,suz12b,suz12a,su: | 4 |

| | | |
|---|---|---|
| suz12b | suz12b,suz12b,suz12a,su: | 4 |
| suz12b | suz12b,suz12b,suz12a,su: | 4 |
| suz12b | suz12b,suz12b,suz12a,su: | 4 |
| sv2a | sv2ca,sv2ba,sv2a,sv2,sv2 | 5 |
| sv2ba | sv2ca,sv2ba,sv2a,sv2,sv2 | 5 |
| sv2bb | sv2ca,sv2ba,sv2a,sv2,sv2 | 5 |
| sv2ca | sv2ca,sv2ba,sv2a,sv2,sv2 | 5 |
| swap70a | swap70b,swap70b,swap7 | 4 |
| swap70b | swap70b,swap70b,swap7 | 4 |
| swap70b | swap70b,swap70b,swap7 | 4 |
| swap70b | swap70b,swap70b,swap7 | 4 |
| syn2a | syn2b,syn2a | 2 |
| syn2b | syn2b,syn2a | 2 |
| syne1a | syne1a,syne1b,syne1a | 3 |
| syne1a | syne1a,syne1b,syne1a | 3 |
| syne1b | syne1a,syne1b,syne1a | 3 |
| syne2b | syne2b | 1 |
| syngap1a | syngap1b,syngap1a | 2 |
| syngap1b | syngap1b,syngap1a | 2 |
| syngr1a | syngr1b,syngr1a | 2 |
| syngr1b | syngr1b,syngr1a | 2 |
| syngr2a | syngr2a,syngr2b | 2 |
| syngr2b | syngr2a,syngr2b | 2 |
| syngr3a | syngr3b,syngr3a | 2 |
| syngr3b | syngr3b,syngr3a | 2 |
| synj2bp | synj2bp | 1 |
| synpo2b | synpo2b | 1 |
| sypl2a | sypl2a,sypl2b,sypl2a | 3 |
| sypl2a | sypl2a,sypl2b,sypl2a | 3 |
| sypl2b | sypl2a,sypl2b,sypl2a | 3 |
| syt11a | syt11b,syt11a | 2 |
| syt11b | syt11b,syt11a | 2 |
| syt14a | syt14a,syt14b | 2 |
| syt14b | syt14a,syt14b | 2 |
| syt1a | syt1b,syt1a,syt1b | 3 |
| syt1b | syt1b,syt1a,syt1b | 3 |
| syt1b | syt1b,syt1a,syt1b | 3 |
| syt2a | syt2a | 1 |
| syt5a | syt5a,syt5b | 2 |
| syt5b | syt5a,syt5b | 2 |
| syt6a | syt6b,syt6b,syt6a | 3 |
| syt6b | syt6b,syt6b,syt6a | 3 |
| syt6b | syt6b,syt6b,syt6a | 3 |
| syt7a | syt7b,syt7a | 2 |
| syt7b | syt7b,syt7a | 2 |
| syt9a | syt9a,syt9b | 2 |
| syt9b | syt9a,syt9b | 2 |
| sytl2a | sytl2a,sytl2b | 2 |
| sytl2b | sytl2a,sytl2b | 2 |
| taar10a | taar10c,taar10a,taar10,ta | 12 |

| | | |
|---|---|---|
| taar10a | taar10c,taar10a,taar10,ta | 12 |
| taar10a | taar10c,taar10a,taar10,ta | 12 |
| taar10b | taar10c,taar10a,taar10,ta | 12 |
| taar10c | taar10c,taar10a,taar10,ta | 12 |
| taar10c | taar10c,taar10a,taar10,ta | 12 |
| taar10c | taar10c,taar10a,taar10,ta | 12 |
| taar10c | taar10c,taar10a,taar10,ta | 12 |
| taar10d | taar10c,taar10a,taar10,ta | 12 |
| taar12a | taar12j,taar12e,taar12g,t | 22 |
| taar12b | taar12j,taar12e,taar12g,t | 22 |
| taar12b | taar12j,taar12e,taar12g,t | 22 |
| taar12b | taar12j,taar12e,taar12g,t | 22 |
| taar12c | taar12j,taar12e,taar12g,t | 22 |
| taar12c | taar12j,taar12e,taar12g,t | 22 |
| taar12c | taar12j,taar12e,taar12g,t | 22 |
| taar12e | taar12j,taar12e,taar12g,t | 22 |
| taar12e | taar12j,taar12e,taar12g,t | 22 |
| taar12e | taar12j,taar12e,taar12g,t | 22 |
| taar12f | taar12j,taar12e,taar12g,t | 22 |
| taar12g | taar12j,taar12e,taar12g,t | 22 |
| taar12g | taar12j,taar12e,taar12g,t | 22 |
| taar12g | taar12j,taar12e,taar12g,t | 22 |
| taar12h | taar12j,taar12e,taar12g,t | 22 |
| taar12i | taar12j,taar12e,taar12g,t | 22 |
| taar12i | taar12j,taar12e,taar12g,t | 22 |
| taar12i | taar12j,taar12e,taar12g,t | 22 |
| taar12j | taar12j,taar12e,taar12g,t | 22 |
| taar12j | taar12j,taar12e,taar12g,t | 22 |
| taar12j | taar12j,taar12e,taar12g,t | 22 |
| taar12m | taar12j,taar12e,taar12g,t | 22 |
| taar13a | taar13e,taar13c,taar13e, | 9 |
| taar13b | taar13e,taar13c,taar13e, | 9 |
| taar13c | taar13e,taar13c,taar13e, | 9 |
| taar13c | taar13e,taar13c,taar13e, | 9 |
| taar13c | taar13e,taar13c,taar13e, | 9 |
| taar13d | taar13e,taar13c,taar13e, | 9 |
| taar13e | taar13e,taar13c,taar13e, | 9 |
| taar13e | taar13e,taar13c,taar13e, | 9 |
| taar13e | taar13e,taar13c,taar13e, | 9 |
| taar14a | taar14e,taar14f,taar14d,t | 13 |
| taar14b | taar14e,taar14f,taar14d,t | 13 |
| taar14d | taar14e,taar14f,taar14d,t | 13 |
| taar14d | taar14e,taar14f,taar14d,t | 13 |
| taar14e | taar14e,taar14f,taar14d,t | 13 |
| taar14e | taar14e,taar14f,taar14d,t | 13 |
| taar14f | taar14e,taar14f,taar14d,t | 13 |
| taar14f | taar14e,taar14f,taar14d,t | 13 |
| taar14g | taar14e,taar14f,taar14d,t | 13 |
| taar14h | taar14e,taar14f,taar14d,t | 13 |
| taar14i | taar14e,taar14f,taar14d,t | 13 |

| | | |
|---|---|---|
| taar14j | taar14e,taar14f,taar14d,t | 13 |
| taar14j | taar14e,taar14f,taar14d,t | 13 |
| taar16a | taar16b,taar16a,taar16f,t | 4 |
| taar16b | taar16b,taar16a,taar16f,t | 4 |
| taar16f | taar16b,taar16a,taar16f,t | 4 |
| taar16g | taar16b,taar16a,taar16f,t | 4 |
| taar17a | taar17a | 1 |
| taar18a | taar18a,taar18g,taar18d, | 10 |
| taar18b | taar18a,taar18g,taar18d, | 10 |
| taar18c | taar18a,taar18g,taar18d, | 10 |
| taar18d | taar18a,taar18g,taar18d, | 10 |
| taar18e | taar18a,taar18g,taar18d, | 10 |
| taar18f | taar18a,taar18g,taar18d, | 10 |
| taar18g | taar18a,taar18g,taar18d, | 10 |
| taar18h | taar18a,taar18g,taar18d, | 10 |
| taar18i | taar18a,taar18g,taar18d, | 10 |
| taar18j | taar18a,taar18g,taar18d, | 10 |
| taar19a | taar19k,taar19n,taar19t,t | 20 |
| taar19b | taar19k,taar19n,taar19t,t | 20 |
| taar19c | taar19k,taar19n,taar19t,t | 20 |
| taar19d | taar19k,taar19n,taar19t,t | 20 |
| taar19e | taar19k,taar19n,taar19t,t | 20 |
| taar19g | taar19k,taar19n,taar19t,t | 20 |
| taar19h | taar19k,taar19n,taar19t,t | 20 |
| taar19i | taar19k,taar19n,taar19t,t | 20 |
| taar19j | taar19k,taar19n,taar19t,t | 20 |
| taar19k | taar19k,taar19n,taar19t,t | 20 |
| taar19m | taar19k,taar19n,taar19t,t | 20 |
| taar19n | taar19k,taar19n,taar19t,t | 20 |
| taar19n | taar19k,taar19n,taar19t,t | 20 |
| taar19o | taar19k,taar19n,taar19t,t | 20 |
| taar19o | taar19k,taar19n,taar19t,t | 20 |
| taar19p | taar19k,taar19n,taar19t,t | 20 |
| taar19q | taar19k,taar19n,taar19t,t | 20 |
| taar19s | taar19k,taar19n,taar19t,t | 20 |
| taar19t | taar19k,taar19n,taar19t,t | 20 |
| taar19u | taar19k,taar19n,taar19t,t | 20 |
| taar1b | taar1b,taar1b,taar1b | 3 |
| taar1b | taar1b,taar1b,taar1b | 3 |
| taar1b | taar1b,taar1b,taar1b | 3 |
| taar20a | taar20z,taar20i,taar20a,t | 20 |
| taar20b | taar20z,taar20i,taar20a,t | 20 |
| taar20c | taar20z,taar20i,taar20a,t | 20 |
| taar20f | taar20z,taar20i,taar20a,t | 20 |
| taar20g | taar20z,taar20i,taar20a,t | 20 |
| taar20h | taar20z,taar20i,taar20a,t | 20 |
| taar20i | taar20z,taar20i,taar20a,t | 20 |
| taar20i | taar20z,taar20i,taar20a,t | 20 |
| taar20j | taar20z,taar20i,taar20a,t | 20 |
| taar20n | taar20z,taar20i,taar20a,t | 20 |

| | | |
|---|---|---|
| taar20o | taar20z,taar20i,taar20a,t | 20 |
| taar20o | taar20z,taar20i,taar20a,t | 20 |
| taar20p | taar20z,taar20i,taar20a,t | 20 |
| taar20q | taar20z,taar20i,taar20a,t | 20 |
| taar20r | taar20z,taar20i,taar20a,t | 20 |
| taar20t | taar20z,taar20i,taar20a,t | 20 |
| taar20w | taar20z,taar20i,taar20a,t | 20 |
| taar20x | taar20z,taar20i,taar20a,t | 20 |
| taar20z | taar20z,taar20i,taar20a,t | 20 |
| taar20z | taar20z,taar20i,taar20a,t | 20 |
| tac3a | tac3a,tac3b,tac3b,tac3a | 4 |
| tac3a | tac3a,tac3b,tac3b,tac3a | 4 |
| tac3b | tac3a,tac3b,tac3b,tac3a | 4 |
| tac3b | tac3a,tac3b,tac3b,tac3a | 4 |
| tacr1a | tacr1a,tacr1a,tacr1b | 3 |
| tacr1a | tacr1a,tacr1a,tacr1b | 3 |
| tacr1b | tacr1a,tacr1a,tacr1b | 3 |
| tacr3a | tacr3a | 1 |
| tada2a | tada2a,tada2b,tada2a | 3 |
| tada2a | tada2a,tada2b,tada2a | 3 |
| tada2b | tada2a,tada2b,tada2a | 3 |
| taf1a | taf1,taf1b,taf1a | 3 |
| taf1b | taf1,taf1b,taf1a | 3 |
| taf4a | taf4a,taf4b | 2 |
| taf4b | taf4a,taf4b | 2 |
| tafa1a | tafa1a | 1 |
| tafa4a | tafa4b,tafa4a | 2 |
| tafa4b | tafa4b,tafa4a | 2 |
| tafa5a | tafa5a,tafa5b | 2 |
| tafa5b | tafa5a,tafa5b | 2 |
| tagln3a | tagln3b,tagln3a,tagln3b | 3 |
| tagln3b | tagln3b,tagln3a,tagln3b | 3 |
| tagln3b | tagln3b,tagln3a,tagln3b | 3 |
| tanc1a | tanc1a,tanc1b | 2 |
| tanc1b | tanc1a,tanc1b | 2 |
| tanc2a | tanc2a | 1 |
| taok1a | taok1a,taok1b | 2 |
| taok1b | taok1a,taok1b | 2 |
| taok2a | taok2a,taok2b | 2 |
| taok2b | taok2a,taok2b | 2 |
| taok3a | taok3a,taok3b | 2 |
| taok3b | taok3a,taok3b | 2 |
| tap2a | tap2t,tap2a | 2 |
| tap2t | tap2t,tap2a | 2 |
| tapt1a | tapt1b,tapt1b,tapt1b,tap | 4 |
| tapt1b | tapt1b,tapt1b,tapt1b,tap | 4 |
| tapt1b | tapt1b,tapt1b,tapt1b,tap | 4 |
| tapt1b | tapt1b,tapt1b,tapt1b,tap | 4 |
| tax1bp1a | tax1bp1b,tax1bp1a | 2 |
| tax1bp1b | tax1bp1b,tax1bp1a | 2 |

| | | |
|---|---|---|
| tbc1d10aa | tbc1d10b,tbc1d10b,tbc1c | 5 |
| tbc1d10ab | tbc1d10b,tbc1d10b,tbc1c | 5 |
| tbc1d10b | tbc1d10b,tbc1d10b,tbc1c | 5 |
| tbc1d10b | tbc1d10b,tbc1d10b,tbc1c | 5 |
| tbc1d10c | tbc1d10b,tbc1d10b,tbc1c | 5 |
| tbc1d12a | tbc1d12a,tbc1d12b | 2 |
| tbc1d12b | tbc1d12a,tbc1d12b | 2 |
| tbc1d22a | tbc1d22a,tbc1d22b | 2 |
| tbc1d22b | tbc1d22a,tbc1d22b | 2 |
| tbc1d2b | tbc1d2,tbc1d2b | 2 |
| tbl1x | tbl1x | 1 |
| tbl1xr1a | tbl1xr1b,tbl1xr1a | 2 |
| tbl1xr1b | tbl1xr1b,tbl1xr1a | 2 |
| tbr1a | tbr1b,tbr1a | 2 |
| tbr1b | tbr1b,tbr1a | 2 |
| tbx2a | tbx2b,tbx2a,tbx2a,tbx2b | 4 |
| tbx2a | tbx2b,tbx2a,tbx2a,tbx2b | 4 |
| tbx2b | tbx2b,tbx2a,tbx2a,tbx2b | 4 |
| tbx2b | tbx2b,tbx2a,tbx2a,tbx2b | 4 |
| tbx3a | tbx3b,tbx3a | 2 |
| tbx3b | tbx3b,tbx3a | 2 |
| tbx5a | tbx5b,tbx5a | 2 |
| tbx5b | tbx5b,tbx5a | 2 |
| tbxa2r | tbxa2r | 1 |
| tcerg1a | tcerg1a | 1 |
| tcf3a | tcf3b,tcf3a,tcf3b | 3 |
| tcf3b | tcf3b,tcf3a,tcf3b | 3 |
| tcf3b | tcf3b,tcf3a,tcf3b | 3 |
| tcf7l1a | tcf7l1b,tcf7l1a | 2 |
| tcf7l1b | tcf7l1b,tcf7l1a | 2 |
| tcirg1a | tcirg1a,tcirg1b | 2 |
| tcirg1b | tcirg1a,tcirg1b | 2 |
| tdo2a | tdo2a,tdo2b,tdo2a | 3 |
| tdo2a | tdo2a,tdo2b,tdo2a | 3 |
| tdo2b | tdo2a,tdo2b,tdo2a | 3 |
| tdp2a | tdp2b,tdp2a,tdp2b | 3 |
| tdp2b | tdp2b,tdp2a,tdp2b | 3 |
| tdp2b | tdp2b,tdp2a,tdp2b | 3 |
| tdrd7a | tdrd7a,tdrd7b | 2 |
| tdrd7b | tdrd7a,tdrd7b | 2 |
| tead1a | tead1b,tead1a | 2 |
| tead1b | tead1b,tead1a | 2 |
| tead3a | tead3b,tead3a | 2 |
| tead3b | tead3b,tead3a | 2 |
| tecpr1a | tecpr1b,tecpr1a | 2 |
| tecpr1b | tecpr1b,tecpr1a | 2 |
| tecrl2a | tecrl2b,tecrl2a | 2 |
| tecrl2b | tecrl2b,tecrl2a | 2 |
| tent4a | tent4b,tent4a | 2 |
| tent4b | tent4b,tent4a | 2 |

| | | |
|---|---|---|
| tent5ab | tent5d,tent5d,tent5c,ten | 7 |
| tent5ba | tent5d,tent5d,tent5c,ten | 7 |
| tent5bb | tent5d,tent5d,tent5c,ten | 7 |
| tent5c | tent5d,tent5d,tent5c,ten | 7 |
| tent5d | tent5d,tent5d,tent5c,ten | 7 |
| tent5d | tent5d,tent5d,tent5c,ten | 7 |
| tent5d | tent5d,tent5d,tent5c,ten | 7 |
| terf2ip | terf2ip | 1 |
| tex264a | tex264a | 1 |
| tfap2a | tfap2c,tfap2e,tfap2b,tfap | 5 |
| tfap2b | tfap2c,tfap2e,tfap2b,tfap | 5 |
| tfap2c | tfap2c,tfap2e,tfap2b,tfap | 5 |
| tfap2d | tfap2c,tfap2e,tfap2b,tfap | 5 |
| tfap2e | tfap2c,tfap2e,tfap2b,tfap | 5 |
| tfb1m | tfb1m | 1 |
| tfb2m | tfb2m | 1 |
| tfdp1a | tfdp1a,tfdp1b,tfdp1a | 3 |
| tfdp1a | tfdp1a,tfdp1b,tfdp1a | 3 |
| tfdp1b | tfdp1a,tfdp1b,tfdp1a | 3 |
| tfe3a | tfe3b,tfe3a,tfe3b | 3 |
| tfe3b | tfe3b,tfe3a,tfe3b | 3 |
| tfe3b | tfe3b,tfe3a,tfe3b | 3 |
| tfr1a | tfr1b,tfr1a,tfr1b | 3 |
| tfr1b | tfr1b,tfr1a,tfr1b | 3 |
| tfr1b | tfr1b,tfr1a,tfr1b | 3 |
| tgfb1a | tgfb1a,tgfb1b | 2 |
| tgfb1b | tgfb1a,tgfb1b | 2 |
| tgfbr1a | tgfbr1a,tgfbr1b | 2 |
| tgfbr1b | tgfbr1a,tgfbr1b | 2 |
| tgfbr2a | tgfbr2b,tgfbr2a | 2 |
| tgfbr2b | tgfbr2b,tgfbr2a | 2 |
| tgm2a | tgm2b,tgm2a | 2 |
| tgm2b | tgm2b,tgm2a | 2 |
| thap12a | thap12a,thap12b | 2 |
| thap12b | thap12a,thap12b | 2 |
| thbs1a | thbs1a,thbs1b | 2 |
| thbs1b | thbs1a,thbs1b | 2 |
| thbs2a | thbs2b,thbs2a | 2 |
| thbs2b | thbs2b,thbs2a | 2 |
| thbs3a | thbs3a,thbs3b | 2 |
| thbs3b | thbs3a,thbs3b | 2 |
| thbs4a | thbs4a,thbs4b | 2 |
| thbs4b | thbs4a,thbs4b | 2 |
| thrap3a | thrap3a,thrap3b | 2 |
| thrap3b | thrap3a,thrap3b | 2 |
| thsd7aa | thsd7ba,thsd7bb,thsd7ab | 4 |
| thsd7ab | thsd7ba,thsd7bb,thsd7ab | 4 |
| thsd7ba | thsd7ba,thsd7bb,thsd7ab | 4 |
| thsd7bb | thsd7ba,thsd7bb,thsd7ab | 4 |
| tiam1a | tiam1a,tiam1b | 2 |

| | | |
|---|---|---|
| tiam1b | tiam1a,tiam1b | 2 |
| tiam2a | tiam2a,tiam2b | 2 |
| tiam2b | tiam2a,tiam2b | 2 |
| timm10b | timm10,timm10b | 2 |
| timm17a | timm17a,timm17b,timm1 | 3 |
| timm17a | timm17a,timm17b,timm1 | 3 |
| timm17b | timm17a,timm17b,timm1 | 3 |
| timm23a | timm23a,timm23b | 2 |
| timm23b | timm23a,timm23b | 2 |
| timm8a | timm8a,timm8b,timm8a | 3 |
| timm8a | timm8a,timm8b,timm8a | 3 |
| timm8b | timm8a,timm8b,timm8a | 3 |
| timp2a | timp2a,timp2b,timp2a | 3 |
| timp2a | timp2a,timp2b,timp2a | 3 |
| timp2b | timp2a,timp2b,timp2a | 3 |
| tjp1a | tjp1a,tjp1b | 2 |
| tjp1b | tjp1a,tjp1b | 2 |
| tjp2a | tjp2b,tjp2a | 2 |
| tjp2b | tjp2b,tjp2a | 2 |
| tlcd3a | tlcd3ba,tlcd3a,tlcd3bb | 3 |
| tlcd3ba | tlcd3ba,tlcd3a,tlcd3bb | 3 |
| tlcd3bb | tlcd3ba,tlcd3a,tlcd3bb | 3 |
| tlcd4a | tlcd4b,tlcd4a | 2 |
| tlcd4b | tlcd4b,tlcd4a | 2 |
| tlcd5a | tlcd5b,tlcd5b,tlcd5a | 3 |
| tlcd5b | tlcd5b,tlcd5b,tlcd5a | 3 |
| tlcd5b | tlcd5b,tlcd5b,tlcd5a | 3 |
| tle2a | tle2c,tle2a,tle2b | 3 |
| tle2b | tle2c,tle2a,tle2b | 3 |
| tle2c | tle2c,tle2a,tle2b | 3 |
| tle3a | tle3a,tle3b,tle3a | 3 |
| tle3a | tle3a,tle3b,tle3a | 3 |
| tle3b | tle3a,tle3b,tle3a | 3 |
| tlk1a | tlk1b,tlk1a | 2 |
| tlk1b | tlk1b,tlk1a | 2 |
| tln2a | tln2a,tln2a,tln2b | 3 |
| tln2a | tln2a,tln2a,tln2b | 3 |
| tln2b | tln2a,tln2a,tln2b | 3 |
| tlr4ba | tlr4ba,tlr4bb,tlr4ba,tlr4bb | 4 |
| tlr4ba | tlr4ba,tlr4bb,tlr4ba,tlr4bb | 4 |
| tlr4bb | tlr4ba,tlr4bb,tlr4ba,tlr4bb | 4 |
| tlr4bb | tlr4ba,tlr4bb,tlr4ba,tlr4bb | 4 |
| tlr5a | tlr5b,tlr5a | 2 |
| tlr5b | tlr5b,tlr5a | 2 |
| tlr8a | tlr8a,tlr8b | 2 |
| tlr8b | tlr8a,tlr8b | 2 |
| tlx3b | tlx3b,tlx3b | 2 |
| tlx3b | tlx3b,tlx3b | 2 |
| tmbim1a | tmbim1a,tmbim1b | 2 |
| tmbim1b | tmbim1a,tmbim1b | 2 |

| | | |
|---|---|---|
| tmc2a | tmc2a,tmc2b,tmc2a | 3 |
| tmc2a | tmc2a,tmc2b,tmc2a | 3 |
| tmc2b | tmc2a,tmc2b,tmc2a | 3 |
| tmc6a | tmc6a,tmc6b | 2 |
| tmc6b | tmc6a,tmc6b | 2 |
| tmcc1b | tmcc1b,tmcc1b | 2 |
| tmcc1b | tmcc1b,tmcc1b | 2 |
| tmed1a | tmed1b,tmed1a | 2 |
| tmed1b | tmed1b,tmed1a | 2 |
| tmeff1a | tmeff1a,tmeff1b | 2 |
| tmeff1b | tmeff1a,tmeff1b | 2 |
| tmeff2a | tmeff2b,tmeff2a | 2 |
| tmeff2b | tmeff2b,tmeff2a | 2 |
| tmem106a | tmem106a,tmem106bb,t | 6 |
| tmem106a | tmem106a,tmem106bb,t | 6 |
| tmem106b | tmem106a,tmem106bb,t | 6 |
| tmem106b | tmem106a,tmem106bb,t | 6 |
| tmem106b | tmem106a,tmem106bb,t | 6 |
| tmem106c | tmem106a,tmem106bb,t | 6 |
| tmem119a | tmem119b,tmem119a,tn | 3 |
| tmem119b | tmem119b,tmem119a,tn | 3 |
| tmem119b | tmem119b,tmem119a,tn | 3 |
| tmem120a | tmem120b,tmem120a,tn | 3 |
| tmem120b | tmem120b,tmem120a,tn | 3 |
| tmem120b | tmem120b,tmem120a,tn | 3 |
| tmem121a | tmem121b,tmem121aa,t | 3 |
| tmem121a | tmem121b,tmem121aa,t | 3 |
| tmem121b | tmem121b,tmem121aa,t | 3 |
| tmem125b | tmem125b | 1 |
| tmem126a | tmem126a | 1 |
| tmem132a | tmem132a,tmem132e | 2 |
| tmem132e | tmem132a,tmem132e | 2 |
| tmem144a | tmem144a,tmem144a,tn | 5 |
| tmem144a | tmem144a,tmem144a,tn | 5 |
| tmem144a | tmem144a,tmem144a,tn | 5 |
| tmem144a | tmem144a,tmem144a,tn | 5 |
| tmem144b | tmem144a,tmem144a,tn | 5 |
| tmem14ca | tmem14ca,tmem14cb | 2 |
| tmem14cb | tmem14ca,tmem14cb | 2 |
| tmem150a | tmem150ab,tmem150aa, | 4 |
| tmem150a | tmem150ab,tmem150aa, | 4 |
| tmem150a | tmem150ab,tmem150aa, | 4 |
| tmem150c | tmem150ab,tmem150aa, | 4 |
| tmem151a | tmem151bb,tmem151ba | 3 |
| tmem151b | tmem151bb,tmem151ba | 3 |
| tmem151b | tmem151bb,tmem151ba | 3 |
| tmem161a | tmem161b,tmem161a | 2 |
| tmem161b | tmem161b,tmem161a | 2 |
| tmem163a | tmem163a,tmem163a,tn | 3 |
| tmem163a | tmem163a,tmem163a,tn | 3 |

| | | |
|---|---|---|
| tmem163b | tmem163a,tmem163a,tm | 3 |
| tmem167a | tmem167a,tmem167b | 2 |
| tmem167b | tmem167a,tmem167b | 2 |
| tmem168a | tmem168a,tmem168b | 2 |
| tmem168b | tmem168a,tmem168b | 2 |
| tmem169a | tmem169b,tmem169a | 2 |
| tmem169b | tmem169b,tmem169a | 2 |
| tmem170a | tmem170a,tmem170a,tm | 3 |
| tmem170a | tmem170a,tmem170a,tm | 3 |
| tmem170b | tmem170a,tmem170a,tm | 3 |
| tmem176l. | tmem176l.3b,tmem176l. | 2 |
| tmem176l. | tmem176l.3b,tmem176l. | 2 |
| tmem178b | tmem178b,tmem178 | 2 |
| tmem179a | tmem179aa,tmem179b | 2 |
| tmem179b | tmem179aa,tmem179b | 2 |
| tmem182a | tmem182b,tmem182a | 2 |
| tmem182b | tmem182b,tmem182a | 2 |
| tmem183a | tmem183a | 1 |
| tmem184a | tmem184ba,tmem184c,t | 6 |
| tmem184b | tmem184ba,tmem184c,t | 6 |
| tmem184b | tmem184ba,tmem184c,t | 6 |
| tmem184b | tmem184ba,tmem184c,t | 6 |
| tmem184c | tmem184ba,tmem184c,t | 6 |
| tmem184c | tmem184ba,tmem184c,t | 6 |
| tmem196b | tmem196b | 1 |
| tmem198a | tmem198b,tmem198b,tm | 3 |
| tmem198b | tmem198b,tmem198b,tm | 3 |
| tmem198b | tmem198b,tmem198b,tm | 3 |
| tmem200a | tmem200b,tmem200a | 2 |
| tmem200b | tmem200b,tmem200a | 2 |
| tmem222a | tmem222a,tmem222b | 2 |
| tmem222b | tmem222a,tmem222b | 2 |
| tmem229b | tmem229b | 1 |
| tmem230a | tmem230a,tmem230b,tm | 3 |
| tmem230a | tmem230a,tmem230b,tm | 3 |
| tmem230b | tmem230a,tmem230b,tm | 3 |
| tmem235b | tmem235b | 1 |
| tmem237a | tmem237b,tmem237a | 2 |
| tmem237b | tmem237b,tmem237a | 2 |
| tmem238a | tmem238a | 1 |
| tmem240a | tmem240a,tmem240b,tm | 3 |
| tmem240a | tmem240a,tmem240b,tm | 3 |
| tmem240b | tmem240a,tmem240b,tm | 3 |
| tmem243a | tmem243a,tmem243b,tm | 5 |
| tmem243a | tmem243a,tmem243b,tm | 5 |
| tmem243b | tmem243a,tmem243b,tm | 5 |
| tmem243b | tmem243a,tmem243b,tm | 5 |
| tmem243b | tmem243a,tmem243b,tm | 5 |
| tmem255a | tmem255a | 1 |
| tmem26a | tmem26b,tmem26a | 2 |

| | | |
|---|---|---|
| tmem26b | tmem26b,tmem26a | 2 |
| tmem30aa | tmem30c,tmem30c,tmer | 5 |
| tmem30ab | tmem30c,tmem30c,tmer | 5 |
| tmem30b | tmem30c,tmem30c,tmer | 5 |
| tmem30c | tmem30c,tmem30c,tmer | 5 |
| tmem30c | tmem30c,tmem30c,tmer | 5 |
| tmem38a | tmem38a | 1 |
| tmem39a | tmem39a,tmem39b | 2 |
| tmem39b | tmem39a,tmem39b | 2 |
| tmem41aa | tmem41ab,tmem41ab,tn | 5 |
| tmem41ab | tmem41ab,tmem41ab,tn | 5 |
| tmem41ab | tmem41ab,tmem41ab,tn | 5 |
| tmem41ab | tmem41ab,tmem41ab,tn | 5 |
| tmem41b | tmem41ab,tmem41ab,tn | 5 |
| tmem42a | tmem42a,tmem42b | 2 |
| tmem42b | tmem42a,tmem42b | 2 |
| tmem45a | tmem45a,tmem45b | 2 |
| tmem45b | tmem45a,tmem45b | 2 |
| tmem50a | tmem50a | 1 |
| tmem51a | tmem51a,tmem51a,tmer | 3 |
| tmem51a | tmem51a,tmem51a,tmer | 3 |
| tmem51b | tmem51a,tmem51a,tmer | 3 |
| tmem54a | tmem54a,tmem54b,tmer | 4 |
| tmem54a | tmem54a,tmem54b,tmer | 4 |
| tmem54b | tmem54a,tmem54b,tmer | 4 |
| tmem54b | tmem54a,tmem54b,tmer | 4 |
| tmem63a | tmem63c,tmem63a,tmer | 5 |
| tmem63ba | tmem63c,tmem63a,tmer | 5 |
| tmem63bb | tmem63c,tmem63a,tmer | 5 |
| tmem63c | tmem63c,tmem63a,tmer | 5 |
| tmem63c | tmem63c,tmem63a,tmer | 5 |
| tmem74b | tmem74b | 1 |
| tmem79a | tmem79b,tmem79a | 2 |
| tmem79b | tmem79b,tmem79a | 2 |
| tmem86a | tmem86b,tmem86a | 2 |
| tmem86b | tmem86b,tmem86a | 2 |
| tmem88a | tmem88b,tmem88a | 2 |
| tmem88b | tmem88b,tmem88a | 2 |
| tmem8a | tmem8a | 1 |
| tmem9b | tmem9b,tmem9 | 2 |
| tmprss13a | tmprss13b,tmprss13a | 2 |
| tmprss13b | tmprss13b,tmprss13a | 2 |
| tmprss3a | tmprss3b,tmprss3a | 2 |
| tmprss3b | tmprss3b,tmprss3a | 2 |
| tmprss4a | tmprss4b,tmprss4a | 2 |
| tmprss4b | tmprss4b,tmprss4a | 2 |
| tmsb4x | tmsb4x | 1 |
| tmtc2a | tmtc2b,tmtc2a,tmtc2a,tn | 4 |
| tmtc2a | tmtc2b,tmtc2a,tmtc2a,tn | 4 |
| tmtc2b | tmtc2b,tmtc2a,tmtc2a,tn | 4 |

| | | |
|---|---|---|
| tmtc2b | tmtc2b,tmtc2a,tmtc2a,tn | 4 |
| tmtops2a | tmtops2a,tmtops2b | 2 |
| tmtops2b | tmtops2a,tmtops2b | 2 |
| tmtops3a | tmtops3b,tmtops3a | 2 |
| tmtops3b | tmtops3b,tmtops3a | 2 |
| tmx2a | tmx2a,tmx2b,tmx2a | 3 |
| tmx2a | tmx2a,tmx2b,tmx2a | 3 |
| tmx2b | tmx2a,tmx2b,tmx2a | 3 |
| tmx3a | tmx3a,tmx3b | 2 |
| tmx3b | tmx3a,tmx3b | 2 |
| tnfaip2a | tnfaip2b,tnfaip2a | 2 |
| tnfaip2b | tnfaip2b,tnfaip2a | 2 |
| tnfaip8l2a | tnfaip8l2b,tnfaip8l2a | 2 |
| tnfaip8l2b | tnfaip8l2b,tnfaip8l2a | 2 |
| tnfrsf11a | tnfrsf11b,tnfrsf11a | 2 |
| tnfrsf11b | tnfrsf11b,tnfrsf11a | 2 |
| tnfrsf1a | tnfrsf1b,tnfrsf1a | 2 |
| tnfrsf1b | tnfrsf1b,tnfrsf1a | 2 |
| tnfrsf9a | tnfrsf9a,tnfrsf9b,tnfrsf9a | 3 |
| tnfrsf9a | tnfrsf9a,tnfrsf9b,tnfrsf9a | 3 |
| tnfrsf9b | tnfrsf9a,tnfrsf9b,tnfrsf9a | 3 |
| tnfsf13b | tnfsf13,tnfsf13b | 2 |
| tnk2a | tnk2b,tnk2a | 2 |
| tnk2b | tnk2b,tnk2a | 2 |
| tnnc1a | tnnc1a,tnnc1b | 2 |
| tnnc1b | tnnc1a,tnnc1b | 2 |
| tnni1a | tnni1d,tnni1d,tnni1d,tnni | 6 |
| tnni1b | tnni1d,tnni1d,tnni1d,tnni | 6 |
| tnni1c | tnni1d,tnni1d,tnni1d,tnni | 6 |
| tnni1d | tnni1d,tnni1d,tnni1d,tnni | 6 |
| tnni1d | tnni1d,tnni1d,tnni1d,tnni | 6 |
| tnni1d | tnni1d,tnni1d,tnni1d,tnni | 6 |
| tnni3k | tnni3k | 1 |
| tnni4a | tnni4a | 1 |
| tnnt2a | tnnt2e,tnnt2e,tnnt2c,tnn | 6 |
| tnnt2b | tnnt2e,tnnt2e,tnnt2c,tnn | 6 |
| tnnt2c | tnnt2e,tnnt2e,tnnt2c,tnn | 6 |
| tnnt2d | tnnt2e,tnnt2e,tnnt2c,tnn | 6 |
| tnnt2e | tnnt2e,tnnt2e,tnnt2c,tnn | 6 |
| tnnt2e | tnnt2e,tnnt2e,tnnt2c,tnn | 6 |
| tnnt3a | tnnt3b,tnnt3a | 2 |
| tnnt3b | tnnt3b,tnnt3a | 2 |
| tnrc6a | tnrc6a,tnrc6b | 2 |
| tnrc6b | tnrc6a,tnrc6b | 2 |
| tns1a | tns1a,tns1b | 2 |
| tns1b | tns1a,tns1b | 2 |
| tns2a | tns2b,tns2a | 2 |
| tns2b | tns2b,tns2a | 2 |
| tob1a | tob1a,tob1b | 2 |
| tob1b | tob1a,tob1b | 2 |

| | | |
|---|---|---|
| tomm20a | tomm20b,tomm20a,tom | 3 |
| tomm20a | tomm20b,tomm20a,tom | 3 |
| tomm20b | tomm20b,tomm20a,tom | 3 |
| tomm70a | tomm70a,tomm70a | 2 |
| tomm70a | tomm70a,tomm70a | 2 |
| top1mt | top1,top1mt,top1mt | 3 |
| top1mt | top1,top1mt,top1mt | 3 |
| top2a | top2a,top2b | 2 |
| top2b | top2a,top2b | 2 |
| top3a | top3a,top3b | 2 |
| top3b | top3a,top3b | 2 |
| tor2a | tor2a,tor2a | 2 |
| tor2a | tor2a,tor2a | 2 |
| tor3a | tor3a | 1 |
| tor4aa | tor4aa,tor4ab,tor4ab,tor4 | 6 |
| tor4aa | tor4aa,tor4ab,tor4ab,tor4 | 6 |
| tor4ab | tor4aa,tor4ab,tor4ab,tor4 | 6 |
| tor4ab | tor4aa,tor4ab,tor4ab,tor4 | 6 |
| tor4ab | tor4aa,tor4ab,tor4ab,tor4 | 6 |
| tor4ab | tor4aa,tor4ab,tor4ab,tor4 | 6 |
| tox4a | tox4a,tox4a,tox4b | 3 |
| tox4a | tox4a,tox4a,tox4b | 3 |
| tox4b | tox4a,tox4a,tox4b | 3 |
| tp53bp2a | tp53bp2a,tp53bp2b,tp53 | 4 |
| tp53bp2a | tp53bp2a,tp53bp2b,tp53 | 4 |
| tp53bp2b | tp53bp2a,tp53bp2b,tp53 | 4 |
| tp53bp2b | tp53bp2a,tp53bp2b,tp53 | 4 |
| tp53i11a | tp53i11a,tp53i11b | 2 |
| tp53i11b | tp53i11a,tp53i11b | 2 |
| tp53rk | tp53,tp53,tp53rk | 3 |
| tpd52l2a | tpd52l2a,tpd52l2b | 2 |
| tpd52l2b | tpd52l2a,tpd52l2b | 2 |
| tph1a | tph1a,tph1b,tph1b,tph1a | 4 |
| tph1a | tph1a,tph1b,tph1b,tph1a | 4 |
| tph1b | tph1a,tph1b,tph1b,tph1a | 4 |
| tph1b | tph1a,tph1b,tph1b,tph1a | 4 |
| tpi1a | tpi1b,tpi1b,tpi1a | 3 |
| tpi1b | tpi1b,tpi1b,tpi1a | 3 |
| tpi1b | tpi1b,tpi1b,tpi1a | 3 |
| tpm4a | tpm4b,tpm4a | 2 |
| tpm4b | tpm4b,tpm4a | 2 |
| tra2a | tra2b,tra2a | 2 |
| tra2b | tra2b,tra2a | 2 |
| trabd2a | trabd2a,trabd2b | 2 |
| trabd2b | trabd2a,trabd2b | 2 |
| traf2a | traf2a,traf2b | 2 |
| traf2b | traf2a,traf2b | 2 |
| traf4a | traf4a,traf4a,traf4b | 3 |
| traf4a | traf4a,traf4a,traf4b | 3 |
| traf4b | traf4a,traf4a,traf4b | 3 |

| | | |
|---|---|---|
| trak1a | trak1a | 1 |
| trappc6b | trappc6b | 1 |
| trarg1a | trarg1a,trarg1b | 2 |
| trarg1b | trarg1a,trarg1b | 2 |
| trim2a | trim2b,trim2a | 2 |
| trim2b | trim2b,trim2a | 2 |
| trim3a | trim3b,trim3a | 2 |
| trim3b | trim3b,trim3a | 2 |
| trim46a | trim46a,trim46b | 2 |
| trim46b | trim46a,trim46b | 2 |
| trim55a | trim55a,trim55b | 2 |
| trim55b | trim55a,trim55b | 2 |
| trim63a | trim63a,trim63a,trim63a, | 4 |
| trim63a | trim63a,trim63a,trim63a, | 4 |
| trim63a | trim63a,trim63a,trim63a, | 4 |
| trim63b | trim63a,trim63a,trim63a, | 4 |
| trim8a | trim8b,trim8b,trim8a | 3 |
| trim8b | trim8b,trim8b,trim8a | 3 |
| trim8b | trim8b,trim8b,trim8a | 3 |
| trip10a | trip10b,trip10a | 2 |
| trip10b | trip10b,trip10a | 2 |
| trmt10a | trmt10c,trmt10b,trmt10a | 3 |
| trmt10b | trmt10c,trmt10b,trmt10a | 3 |
| trmt10c | trmt10c,trmt10b,trmt10a | 3 |
| trmt2a | trmt2a,trmt2a,trmt2b | 3 |
| trmt2a | trmt2a,trmt2a,trmt2b | 3 |
| trmt2b | trmt2a,trmt2a,trmt2b | 3 |
| trmt61a | trmt61a,trmt61b | 2 |
| trmt61b | trmt61a,trmt61b | 2 |
| trmt9b | trmt9b | 1 |
| trnau1apa | trnau1apb,trnau1apa | 2 |
| trnau1apb | trnau1apb,trnau1apa | 2 |
| trpa1a | trpa1a,trpa1b | 2 |
| trpa1b | trpa1a,trpa1b | 2 |
| trpc2a | trpc2a,trpc2b | 2 |
| trpc2b | trpc2a,trpc2b | 2 |
| trpc4a | trpc4a,trpc4apa,trpc4b,tr | 4 |
| trpc4apa | trpc4a,trpc4apa,trpc4b,tr | 4 |
| trpc4apb | trpc4a,trpc4apa,trpc4b,tr | 4 |
| trpc4b | trpc4a,trpc4apa,trpc4b,tr | 4 |
| trpc5a | trpc5b,trpc5a,trpc5b | 3 |
| trpc5b | trpc5b,trpc5a,trpc5b | 3 |
| trpc5b | trpc5b,trpc5a,trpc5b | 3 |
| trpc6a | trpc6a,trpc6b,trpc6a | 3 |
| trpc6a | trpc6a,trpc6b,trpc6a | 3 |
| trpc6b | trpc6a,trpc6b,trpc6a | 3 |
| trpc7a | trpc7a,trpc7a,trpc7b | 3 |
| trpc7a | trpc7a,trpc7a,trpc7b | 3 |
| trpc7b | trpc7a,trpc7a,trpc7b | 3 |
| trpm1a | trpm1a,trpm1a,trpm1a,tr | 4 |

| | | |
|---|---|---|
| trpm1a | trpm1a,trpm1a,trpm1a,tı | 4 |
| trpm1a | trpm1a,trpm1a,trpm1a,tı | 4 |
| trpm1b | trpm1a,trpm1a,trpm1a,tı | 4 |
| trpm4a | trpm4a | 1 |
| tsc1a | tsc1a,tsc1a,tsc1a,tsc1b | 4 |
| tsc1a | tsc1a,tsc1a,tsc1a,tsc1b | 4 |
| tsc1a | tsc1a,tsc1a,tsc1a,tsc1b | 4 |
| tsc1b | tsc1a,tsc1a,tsc1a,tsc1b | 4 |
| tsg101a | tsg101b,tsg101a | 2 |
| tsg101b | tsg101b,tsg101a | 2 |
| tshz3a | tshz3b,tshz3a | 2 |
| tshz3b | tshz3b,tshz3a | 2 |
| tspan13a | tspan13a,tspan13b | 2 |
| tspan13b | tspan13a,tspan13b | 2 |
| tspan18a | tspan18a,tspan18b | 2 |
| tspan18b | tspan18a,tspan18b | 2 |
| tspan2a | tspan2a,tspan2a,tspan2b | 3 |
| tspan2a | tspan2a,tspan2a,tspan2b | 3 |
| tspan2b | tspan2a,tspan2a,tspan2b | 3 |
| tspan33a | tspan33a,tspan33b | 2 |
| tspan33b | tspan33a,tspan33b | 2 |
| tspan3a | tspan3a,tspan3a,tspan3b | 5 |
| tspan3a | tspan3a,tspan3a,tspan3b | 5 |
| tspan3a | tspan3a,tspan3a,tspan3b | 5 |
| tspan3b | tspan3a,tspan3a,tspan3b | 5 |
| tspan3b | tspan3a,tspan3a,tspan3b | 5 |
| tspan4a | tspan4b,tspan4b,tspan4a | 3 |
| tspan4b | tspan4b,tspan4b,tspan4a | 3 |
| tspan4b | tspan4b,tspan4b,tspan4a | 3 |
| tspan5a | tspan5b,tspan5a | 2 |
| tspan5b | tspan5b,tspan5a | 2 |
| tspan7b | tspan7,tspan7b | 2 |
| tspan9a | tspan9b,tspan9b,tspan9b | 4 |
| tspan9b | tspan9b,tspan9b,tspan9b | 4 |
| tspan9b | tspan9b,tspan9b,tspan9b | 4 |
| tspan9b | tspan9b,tspan9b,tspan9b | 4 |
| ttbk1a | ttbk1a,ttbk1b | 2 |
| ttbk1b | ttbk1a,ttbk1b | 2 |
| ttbk2a | ttbk2b,ttbk2a,ttbk2b | 3 |
| ttbk2b | ttbk2b,ttbk2a,ttbk2b | 3 |
| ttbk2b | ttbk2b,ttbk2a,ttbk2b | 3 |
| ttc21b | ttc21b | 1 |
| ttc39a | ttc39c,ttc39a,ttc39c,ttc3 | 4 |
| ttc39b | ttc39c,ttc39a,ttc39c,ttc3 | 4 |
| ttc39c | ttc39c,ttc39a,ttc39c,ttc3 | 4 |
| ttc39c | ttc39c,ttc39a,ttc39c,ttc3 | 4 |
| ttc7a | ttc7a,ttc7b | 2 |
| ttc7b | ttc7a,ttc7b | 2 |
| ttc9b | ttc9c,ttc9b | 2 |
| ttc9c | ttc9c,ttc9b | 2 |

| | | |
|---|---|---|
| ttyh3a | ttyh3b,ttyh3a | 2 |
| ttyh3b | ttyh3b,ttyh3a | 2 |
| tuba1a | tuba1c,tuba1a,tuba1b | 3 |
| tuba1b | tuba1c,tuba1a,tuba1b | 3 |
| tuba1c | tuba1c,tuba1a,tuba1b | 3 |
| tubb2b | tubb2,tubb2b | 2 |
| tubb4b | tubb4b | 1 |
| tuft1a | tuft1a,tuft1a,tuft1b | 3 |
| tuft1a | tuft1a,tuft1a,tuft1b | 3 |
| tuft1b | tuft1a,tuft1a,tuft1b | 3 |
| tulp1a | tulp1b,tulp1a | 2 |
| tulp1b | tulp1b,tulp1a | 2 |
| tulp4a | tulp4a,tulp4b | 2 |
| tulp4b | tulp4a,tulp4b | 2 |
| tusc2a | tusc2a,tusc2b | 2 |
| tusc2b | tusc2a,tusc2b | 2 |
| tvp23b | tvp23b | 1 |
| twf1a | twf1b,twf1a | 2 |
| twf1b | twf1b,twf1a | 2 |
| twf2a | twf2b,twf2a,twf2b | 3 |
| twf2b | twf2b,twf2a,twf2b | 3 |
| twf2b | twf2b,twf2a,twf2b | 3 |
| twist1a | twist1b,twist1a | 2 |
| twist1b | twist1b,twist1a | 2 |
| twsg1a | twsg1a | 1 |
| txnl4a | txnl4b,txnl4a | 2 |
| txnl4b | txnl4b,txnl4a | 2 |
| tyrp1a | tyrp1b,tyrp1a | 2 |
| tyrp1b | tyrp1b,tyrp1a | 2 |
| u2af2a | u2af2a,u2af2b | 2 |
| u2af2b | u2af2a,u2af2b | 2 |
| u2surp | u2surp | 1 |
| ubald1a | ubald1b,ubald1a | 2 |
| ubald1b | ubald1b,ubald1a | 2 |
| ubap2a | ubap2b,ubap2b,ubap2a | 3 |
| ubap2b | ubap2b,ubap2b,ubap2a | 3 |
| ubap2b | ubap2b,ubap2b,ubap2a | 3 |
| ubash3ba | ubash3ba,ubash3bb | 2 |
| ubash3bb | ubash3ba,ubash3bb | 2 |
| ube2a | ube2s,ube2kb,ube2f,ube[illegible] | 18 |
| ube2b | ube2s,ube2kb,ube2f,ube[illegible] | 18 |
| ube2c | ube2s,ube2kb,ube2f,ube[illegible] | 18 |
| ube2c | ube2s,ube2kb,ube2f,ube[illegible] | 18 |
| ube2d1a | ube2d1a,ube2d1b,ube2d | 3 |
| ube2d1a | ube2d1a,ube2d1b,ube2d | 3 |
| ube2d1b | ube2d1a,ube2d1b,ube2d | 3 |
| ube2f | ube2s,ube2kb,ube2f,ube[illegible] | 18 |
| ube2g1a | ube2g1a,ube2g1a,ube2g[illegible] | 3 |
| ube2g1a | ube2g1a,ube2g1a,ube2g[illegible] | 3 |
| ube2g1b | ube2g1a,ube2g1a,ube2g[illegible] | 3 |

| | | |
|---|---|---|
| ube2h | ube2s,ube2kb,ube2f,ube2 | 18 |
| ube2ia | ube2s,ube2kb,ube2f,ube2 | 18 |
| ube2ib | ube2s,ube2kb,ube2f,ube2 | 18 |
| ube2ka | ube2s,ube2kb,ube2f,ube2 | 18 |
| ube2kb | ube2s,ube2kb,ube2f,ube2 | 18 |
| ube2kb | ube2s,ube2kb,ube2f,ube2 | 18 |
| ube2l3a | ube2l3a,ube2l3b,ube2l3a | 3 |
| ube2l3a | ube2l3a,ube2l3b,ube2l3a | 3 |
| ube2l3b | ube2l3a,ube2l3b,ube2l3a | 3 |
| ube2na | ube2s,ube2kb,ube2f,ube2 | 18 |
| ube2nb | ube2s,ube2kb,ube2f,ube2 | 18 |
| ube2s | ube2s,ube2kb,ube2f,ube2 | 18 |
| ube2s | ube2s,ube2kb,ube2f,ube2 | 18 |
| ube2t | ube2s,ube2kb,ube2f,ube2 | 18 |
| ube2w | ube2s,ube2kb,ube2f,ube2 | 18 |
| ube2z | ube2s,ube2kb,ube2f,ube2 | 18 |
| ube3a | ube3c,ube3b,ube3a,ube3 | 5 |
| ube3b | ube3c,ube3b,ube3a,ube3 | 5 |
| ube3c | ube3c,ube3b,ube3a,ube3 | 5 |
| ube3c | ube3c,ube3b,ube3a,ube3 | 5 |
| ube3d | ube3c,ube3b,ube3a,ube3 | 5 |
| ube4a | ube4a,ube4b | 2 |
| ube4b | ube4a,ube4b | 2 |
| ubl3a | ubl3a,ubl3a,ubl3b | 3 |
| ubl3a | ubl3a,ubl3a,ubl3b | 3 |
| ubl3b | ubl3a,ubl3a,ubl3b | 3 |
| ubl7a | ubl7b,ubl7a | 2 |
| ubl7b | ubl7b,ubl7a | 2 |
| ubn2a | ubn2b,ubn2a | 2 |
| ubn2b | ubn2b,ubn2a | 2 |
| ubtd1a | ubtd1b,ubtd1a | 2 |
| ubtd1b | ubtd1b,ubtd1a | 2 |
| ubxn2a | ubxn2a | 1 |
| uck2a | uck2a,uck2b,uck2a | 3 |
| uck2a | uck2a,uck2b,uck2a | 3 |
| uck2b | uck2a,uck2b,uck2a | 3 |
| uckl1a | uckl1a,uckl1b | 2 |
| uckl1b | uckl1a,uckl1b | 2 |
| ugp2a | ugp2b,ugp2a,ugp2b | 3 |
| ugp2b | ugp2b,ugp2a,ugp2b | 3 |
| ugp2b | ugp2b,ugp2a,ugp2b | 3 |
| ugt1b6p | ugt1b6p | 1 |
| ulk1a | ulk1a,ulk1b | 2 |
| ulk1b | ulk1a,ulk1b | 2 |
| unc119a | unc119a,unc119b | 2 |
| unc119b | unc119a,unc119b | 2 |
| unc13ba | unc13ba,unc13d | 2 |
| unc13d | unc13ba,unc13d | 2 |
| unc45a | unc45b,unc45a | 2 |
| unc45b | unc45b,unc45a | 2 |

| | | |
|---|---|---|
| unc5a | unc5db,unc5c,unc5b,unc! | 5 |
| unc5b | unc5db,unc5c,unc5b,unc! | 5 |
| unc5c | unc5db,unc5c,unc5b,unc! | 5 |
| unc5da | unc5db,unc5c,unc5b,unc! | 5 |
| unc5db | unc5db,unc5c,unc5b,unc! | 5 |
| unc93a | unc93a | 1 |
| upf3a | upf3b,upf3a,upf3a,upf3b | 4 |
| upf3a | upf3b,upf3a,upf3a,upf3b | 4 |
| upf3b | upf3b,upf3a,upf3a,upf3b | 4 |
| upf3b | upf3b,upf3a,upf3a,upf3b | 4 |
| upk1a | upk1a | 1 |
| upk3b | upk3b | 1 |
| uqcrc2a | uqcrc2a,uqcrc2b | 2 |
| uqcrc2b | uqcrc2a,uqcrc2b | 2 |
| ush1c | ush1c,ush1ga,ush1gb,ush | 4 |
| ush1c | ush1c,ush1ga,ush1gb,ush | 4 |
| ush1ga | ush1c,ush1ga,ush1gb,ush | 4 |
| ush1gb | ush1c,ush1ga,ush1gb,ush | 4 |
| ush2a | ush2a | 1 |
| usp12a | usp12b,usp12a | 2 |
| usp12b | usp12b,usp12a | 2 |
| usp2a | usp2b,usp2a | 2 |
| usp2b | usp2b,usp2a | 2 |
| usp43a | usp43a,usp43a,usp43b,us | 4 |
| usp43a | usp43a,usp43a,usp43b,us | 4 |
| usp43a | usp43a,usp43a,usp43b,us | 4 |
| usp43b | usp43a,usp43a,usp43b,us | 4 |
| usp53b | usp53b | 1 |
| usp54a | usp54a,usp54b,usp54a | 3 |
| usp54a | usp54a,usp54b,usp54a | 3 |
| usp54b | usp54a,usp54b,usp54a | 3 |
| uts2a | uts2d,uts2a,uts2b,uts2d,u | 5 |
| uts2a | uts2d,uts2a,uts2b,uts2d,u | 5 |
| uts2b | uts2d,uts2a,uts2b,uts2d,u | 5 |
| uts2d | uts2d,uts2a,uts2b,uts2d,u | 5 |
| uts2d | uts2d,uts2a,uts2b,uts2d,u | 5 |
| vav3a | vav3b,vav3b,vav3a | 3 |
| vav3b | vav3b,vav3b,vav3a | 3 |
| vav3b | vav3b,vav3b,vav3a | 3 |
| vcam1a | vcam1b,vcam1a,vcam1b | 3 |
| vcam1b | vcam1b,vcam1a,vcam1b | 3 |
| vcam1b | vcam1b,vcam1a,vcam1b | 3 |
| vezf1a | vezf1a,vezf1b | 2 |
| vezf1b | vezf1a,vezf1b | 2 |
| vgll2a | vgll2b,vgll2a | 2 |
| vgll2b | vgll2b,vgll2a | 2 |
| vgll4a | vgll4b,vgll4a | 2 |
| vgll4b | vgll4b,vgll4a | 2 |
| vipr1a | vipr1a,vipr1b | 2 |
| vipr1b | vipr1a,vipr1b | 2 |

| | | |
|---|---|---|
| vmo1a | vmo1a,vmo1b | 2 |
| vmo1b | vmo1a,vmo1b | 2 |
| vps13a | vps13c,vps13c,vps13a,vp | 4 |
| vps13c | vps13c,vps13c,vps13a,vp | 4 |
| vps13c | vps13c,vps13c,vps13a,vp | 4 |
| vps13d | vps13c,vps13c,vps13a,vp | 4 |
| vps26a | vps26a,vps26b,vps26c,vp | 5 |
| vps26a | vps26a,vps26b,vps26c,vp | 5 |
| vps26b | vps26a,vps26b,vps26c,vp | 5 |
| vps26b | vps26a,vps26b,vps26c,vp | 5 |
| vps26c | vps26a,vps26b,vps26c,vp | 5 |
| vps33a | vps33a,vps33a,vps33b | 3 |
| vps33a | vps33a,vps33a,vps33b | 3 |
| vps33b | vps33a,vps33a,vps33b | 3 |
| vps37a | vps37a,vps37b,vps37a,vp | 5 |
| vps37a | vps37a,vps37b,vps37a,vp | 5 |
| vps37b | vps37a,vps37b,vps37a,vp | 5 |
| vps37b | vps37a,vps37b,vps37a,vp | 5 |
| vps37c | vps37a,vps37b,vps37a,vp | 5 |
| vps4a | vps4b,vps4a | 2 |
| vps4b | vps4b,vps4a | 2 |
| vps72a | vps72b,vps72a | 2 |
| vps72b | vps72b,vps72a | 2 |
| vsig8a | vsig8a,vsig8b,vsig8a | 3 |
| vsig8a | vsig8a,vsig8b,vsig8a | 3 |
| vsig8b | vsig8a,vsig8b,vsig8a | 3 |
| vsnl1a | vsnl1b,vsnl1a | 2 |
| vsnl1b | vsnl1b,vsnl1a | 2 |
| vstm4a | vstm4b,vstm4a | 2 |
| vstm4b | vstm4b,vstm4a | 2 |
| vti1a | vti1a,vti1b | 2 |
| vti1b | vti1a,vti1b | 2 |
| vwa3a | vwa3a | 1 |
| wasf3a | wasf3b,wasf3a,wasf3b | 3 |
| wasf3b | wasf3b,wasf3a,wasf3b | 3 |
| wasf3b | wasf3b,wasf3a,wasf3b | 3 |
| washc2c | washc2c | 1 |
| wdr20a | wdr20a,wdr20b | 2 |
| wdr20b | wdr20a,wdr20b | 2 |
| wdr26a | wdr26a,wdr26b | 2 |
| wdr26b | wdr26a,wdr26b | 2 |
| wdr45b | wdr45,wdr45b | 2 |
| wdr47a | wdr47a,wdr47b | 2 |
| wdr47b | wdr47a,wdr47b | 2 |
| wdr48a | wdr48a,wdr48b,wdr48a | 3 |
| wdr48a | wdr48a,wdr48b,wdr48a | 3 |
| wdr48b | wdr48a,wdr48b,wdr48a | 3 |
| wdr83os | wdr83,wdr83os,wdr83 | 3 |
| wfikkn2a | wfikkn2a,wfikkn2b | 2 |
| wfikkn2b | wfikkn2a,wfikkn2b | 2 |

| | | |
|---|---|---|
| wfs1a | wfs1a,wfs1b | 2 |
| wfs1b | wfs1a,wfs1b | 2 |
| wipf1a | wipf1b,wipf1a | 2 |
| wipf1b | wipf1b,wipf1a | 2 |
| wipf2a | wipf2a,wipf2b | 2 |
| wipf2b | wipf2a,wipf2b | 2 |
| wnk1a | wnk1b,wnk1a | 2 |
| wnk1b | wnk1b,wnk1a | 2 |
| wnk4b | wnk4b | 1 |
| wnt10a | wnt10a,wnt10b | 2 |
| wnt10b | wnt10a,wnt10b | 2 |
| wnt11r | wnt11,wnt11,wnt11r | 3 |
| wnt2ba | wnt2,wnt2bb,wnt2ba,wn | 5 |
| wnt2bb | wnt2,wnt2bb,wnt2ba,wn | 5 |
| wnt2bb | wnt2,wnt2bb,wnt2ba,wn | 5 |
| wnt3a | wnt3,wnt3a,wnt3 | 3 |
| wnt4b | wnt4b,wnt4,wnt4b | 3 |
| wnt4b | wnt4b,wnt4,wnt4b | 3 |
| wnt5a | wnt5b,wnt5a | 2 |
| wnt5b | wnt5b,wnt5a | 2 |
| wnt6a | wnt6a,wnt6b | 2 |
| wnt6b | wnt6a,wnt6b | 2 |
| wnt7aa | wnt7ba,wnt7aa,wnt7ab,v | 4 |
| wnt7ab | wnt7ba,wnt7aa,wnt7ab,v | 4 |
| wnt7ba | wnt7ba,wnt7aa,wnt7ab,v | 4 |
| wnt7bb | wnt7ba,wnt7aa,wnt7ab,v | 4 |
| wnt8a | wnt8b,wnt8a,wnt8a | 3 |
| wnt8a | wnt8b,wnt8a,wnt8a | 3 |
| wnt8b | wnt8b,wnt8a,wnt8a | 3 |
| wnt9a | wnt9a,wnt9b | 2 |
| wnt9b | wnt9a,wnt9b | 2 |
| wscd1b | wscd1b | 1 |
| wt1a | wt1b,wt1a | 2 |
| wt1b | wt1b,wt1a | 2 |
| xirp2a | xirp2a,xirp2b | 2 |
| xirp2b | xirp2a,xirp2b | 2 |
| xkr5a | xkr5b,xkr5a | 2 |
| xkr5b | xkr5b,xkr5a | 2 |
| xkr6a | xkr6a,xkr6b | 2 |
| xkr6b | xkr6a,xkr6b | 2 |
| xpo1a | xpo1a,xpo1b | 2 |
| xpo1b | xpo1a,xpo1b | 2 |
| xpr1a | xpr1b,xpr1a | 2 |
| xpr1b | xpr1b,xpr1a | 2 |
| yif1a | yif1a,yif1b | 2 |
| yif1b | yif1a,yif1b | 2 |
| yme1l1a | yme1l1b,yme1l1a | 2 |
| yme1l1b | yme1l1b,yme1l1a | 2 |
| ypel2a | ypel2b,ypel2a,ypel2b | 3 |
| ypel2b | ypel2b,ypel2a,ypel2b | 3 |

| | | |
|---|---|---|
| ypel2b | ypel2b,ypel2a,ypel2b | 3 |
| yy1a | yy1a,yy1b | 2 |
| yy1b | yy1a,yy1b | 2 |
| zbtb16a | zbtb16b,zbtb16a | 2 |
| zbtb16b | zbtb16b,zbtb16a | 2 |
| zbtb22a | zbtb22a,zbtb22b | 2 |
| zbtb22b | zbtb22a,zbtb22b | 2 |
| zbtb2a | zbtb2b,zbtb2a,zbtb2b | 3 |
| zbtb2b | zbtb2b,zbtb2a,zbtb2b | 3 |
| zbtb2b | zbtb2b,zbtb2a,zbtb2b | 3 |
| zbtb47a | zbtb47a,zbtb47b | 2 |
| zbtb47b | zbtb47a,zbtb47b | 2 |
| zbtb7a | zbtb7b,zbtb7c,zbtb7a | 3 |
| zbtb7b | zbtb7b,zbtb7c,zbtb7a | 3 |
| zbtb7c | zbtb7b,zbtb7c,zbtb7a | 3 |
| zbtb8a | zbtb8os,zbtb8a,zbtb8b | 3 |
| zbtb8b | zbtb8os,zbtb8a,zbtb8b | 3 |
| zbtb8os | zbtb8os,zbtb8a,zbtb8b | 3 |
| zc2hc1a | zc2hc1a,zc2hc1c,zc2hc1a | 3 |
| zc2hc1a | zc2hc1a,zc2hc1c,zc2hc1a | 3 |
| zc2hc1c | zc2hc1a,zc2hc1c,zc2hc1a | 3 |
| zc3h11a | zc3h11a | 1 |
| zc3h12a | zc3h12b,zc3h12b,zc3h12 | 3 |
| zc3h12b | zc3h12b,zc3h12b,zc3h12 | 3 |
| zc3h12b | zc3h12b,zc3h12b,zc3h12 | 3 |
| zc3h7a | zc3h7ba,zc3h7a,zc3h7bb | 3 |
| zc3h7ba | zc3h7ba,zc3h7a,zc3h7bb | 3 |
| zc3h7bb | zc3h7ba,zc3h7a,zc3h7bb | 3 |
| zdhhc12a | zdhhc12b,zdhhc12a | 2 |
| zdhhc12b | zdhhc12b,zdhhc12a | 2 |
| zdhhc15a | zdhhc15a,zdhhc15a,zdhh | 3 |
| zdhhc15a | zdhhc15a,zdhhc15a,zdhh | 3 |
| zdhhc15b | zdhhc15a,zdhhc15a,zdhh | 3 |
| zdhhc16a | zdhhc16b,zdhhc16a | 2 |
| zdhhc16b | zdhhc16b,zdhhc16a | 2 |
| zdhhc18a | zdhhc18a,zdhhc18b | 2 |
| zdhhc18b | zdhhc18a,zdhhc18b | 2 |
| zdhhc20a | zdhhc20a,zdhhc20b | 2 |
| zdhhc20b | zdhhc20a,zdhhc20b | 2 |
| zdhhc23a | zdhhc23b,zdhhc23a,zdhh | 3 |
| zdhhc23b | zdhhc23b,zdhhc23a,zdhh | 3 |
| zdhhc23b | zdhhc23b,zdhhc23a,zdhh | 3 |
| zdhhc3a | zdhhc3a,zdhhc3b | 2 |
| zdhhc3b | zdhhc3a,zdhhc3b | 2 |
| zdhhc5a | zdhhc5a,zdhhc5b | 2 |
| zdhhc5b | zdhhc5a,zdhhc5b | 2 |
| zdhhc8a | zdhhc8b,zdhhc8a | 2 |
| zdhhc8b | zdhhc8b,zdhhc8a | 2 |
| zeb1a | zeb1b,zeb1a,zeb1b | 3 |
| zeb1b | zeb1b,zeb1a,zeb1b | 3 |

| | | |
|---|---|---|
| zeb1b | zeb1b,zeb1a,zeb1b | 3 |
| zeb2a | zeb2a,zeb2a,zeb2b,zeb2 | 4 |
| zeb2a | zeb2a,zeb2a,zeb2b,zeb2 | 4 |
| zeb2a | zeb2a,zeb2a,zeb2b,zeb2 | 4 |
| zeb2b | zeb2a,zeb2a,zeb2b,zeb2 | 4 |
| zfand2a | zfand2a | 1 |
| zfand5a | zfand5a,zfand5b | 2 |
| zfand5b | zfand5a,zfand5b | 2 |
| zfp36l1a | zfp36l1a,zfp36l1b | 2 |
| zfp36l1b | zfp36l1a,zfp36l1b | 2 |
| zfpm2a | zfpm2a,zfpm2b | 2 |
| zfpm2b | zfpm2a,zfpm2b | 2 |
| zfyve9a | zfyve9a,zfyve9b | 2 |
| zfyve9b | zfyve9a,zfyve9b | 2 |
| zhx2a | zhx2a,zhx2b | 2 |
| zhx2b | zhx2a,zhx2b | 2 |
| zic2a | zic2b,zic2b,zic2a | 3 |
| zic2b | zic2b,zic2b,zic2a | 3 |
| zic2b | zic2b,zic2b,zic2a | 3 |
| zmat4a | zmat4a,zmat4a,zmat4b | 3 |
| zmat4a | zmat4a,zmat4a,zmat4b | 3 |
| zmat4b | zmat4a,zmat4a,zmat4b | 3 |
| zmiz1a | zmiz1b,zmiz1a | 2 |
| zmiz1b | zmiz1b,zmiz1a | 2 |
| znf106a | znf106a,znf106b | 2 |
| znf106b | znf106a,znf106b | 2 |
| znf143a | znf143b,znf143b,znf143a | 4 |
| znf143b | znf143b,znf143b,znf143a | 4 |
| znf143b | znf143b,znf143b,znf143a | 4 |
| znf143b | znf143b,znf143b,znf143a | 4 |
| znf207a | znf207b,znf207b,znf207a | 4 |
| znf207b | znf207b,znf207b,znf207a | 4 |
| znf207b | znf207b,znf207b,znf207a | 4 |
| znf207b | znf207b,znf207b,znf207a | 4 |
| znf280d | znf280d | 1 |
| znf281a | znf281a,znf281b | 2 |
| znf281b | znf281a,znf281b | 2 |
| znf292a | znf292a,znf292b | 2 |
| znf292b | znf292a,znf292b | 2 |
| znf319b | znf319b | 1 |
| znf362a | znf362a,znf362b | 2 |
| znf362b | znf362a,znf362b | 2 |
| znf385a | znf385c,znf385b,znf385d | 4 |
| znf385b | znf385c,znf385b,znf385d | 4 |
| znf385c | znf385c,znf385b,znf385d | 4 |
| znf385d | znf385c,znf385b,znf385d | 4 |
| znf395a | znf395a,znf395b | 2 |
| znf395b | znf395a,znf395b | 2 |
| znf513a | znf513b,znf513a | 2 |
| znf513b | znf513b,znf513a | 2 |

| | | |
|---|---|---|
| znf609a | znf609b,znf609a | 2 |
| znf609b | znf609b,znf609a | 2 |
| znf644a | znf644a,znf644b | 2 |
| znf644b | znf644a,znf644b | 2 |
| znf687a | znf687a,znf687b | 2 |
| znf687b | znf687a,znf687b | 2 |
| znf710a | znf710b,znf710a | 2 |
| znf710b | znf710b,znf710a | 2 |
| znf740a | znf740a,znf740a,znf740b | 3 |
| znf740a | znf740a,znf740a,znf740b | 3 |
| znf740b | znf740a,znf740a,znf740b | 3 |
| znf800a | znf800b,znf800a | 2 |
| znf800b | znf800b,znf800a | 2 |
| znf804a | znf804a,znf804b | 2 |
| znf804b | znf804a,znf804b | 2 |
| znfl1b | znfl1,znfl1b,znfl1i,znfl1g,z | 7 |
| znfl1c | znfl1,znfl1b,znfl1i,znfl1g,z | 7 |
| znfl1g | znfl1,znfl1b,znfl1i,znfl1g,z | 7 |
| znfl1h | znfl1,znfl1b,znfl1i,znfl1g,z | 7 |
| znfl1i | znfl1,znfl1b,znfl1i,znfl1g,z | 7 |
| znfl1k | znfl1,znfl1b,znfl1i,znfl1g,z | 7 |
| znfl2a | znfl2a | 1 |
| znrf2a | znrf2b,znrf2a | 2 |
| znrf2b | znrf2b,znrf2a | 2 |
| zp3b | zp3c,zp3b,zp3e,zp3 | 4 |
| zp3c | zp3c,zp3b,zp3e,zp3 | 4 |
| zp3e | zp3c,zp3b,zp3e,zp3 | 4 |
| zpld1a | zpld1a,zpld1b | 2 |
| zpld1b | zpld1a,zpld1b | 2 |
| zranb1a | zranb1b,zranb1a,zranb1b | 3 |
| zranb1b | zranb1b,zranb1a,zranb1b | 3 |
| zranb1b | zranb1b,zranb1a,zranb1b | 3 |
