## Supplemental file 3 for "Evolutionary context can clarify teleosts gene names"

| gene | all duplicates | total number of duplicates |
|---|---|---|
| abcg2a | abcg2a,abcg2c | 2 |
| abcg2c | abcg2a,abcg2c | 2 |
| abhd17aa | abhd17b,abhd17c,abhd17 | 3 |
| abhd17b | abhd17b,abhd17c,abhd17 | 3 |
| abhd17c | abhd17b,abhd17c,abhd17 | 3 |
| abi3a | abi3a,abi3bpb | 2 |
| abi3bpb | abi3a,abi3bpb | 2 |
| acp5a | acp5a,acp5b | 2 |
| acp5b | acp5a,acp5b | 2 |
| actc1a | actc1a,actc1c | 2 |
| actc1c | actc1a,actc1c | 2 |
| actl6a | actl6b,actl6a | 2 |
| actl6b | actl6b,actl6a | 2 |
| actr3b | actr3b,actr3 | 2 |
| acvr2aa | acvr2bb,acvr2aa | 2 |
| acvr2bb | acvr2bb,acvr2aa | 2 |
| adam19a | adam19a,adam19b | 2 |
| adam19b | adam19a,adam19b | 2 |
| adamts15b | adamts15b,adamts15b | 2 |
| adamts15b | adamts15b,adamts15b | 2 |
| adora2aa | adora2aa,adora2b | 2 |
| adora2b | adora2aa,adora2b | 2 |
| adra1aa | adra1aa,adra1d,adra1bb | 3 |
| adra1bb | adra1aa,adra1d,adra1bb | 3 |
| adra1d | adra1aa,adra1d,adra1bb | 3 |
| adra2a | adra2b,adra2c,adra2a,adr | 4 |
| adra2b | adra2b,adra2c,adra2a,adr | 4 |
| adra2c | adra2b,adra2c,adra2a,adr | 4 |
| adra2da | adra2b,adra2c,adra2a,adr | 4 |
| akr1a1a | akr1a1a,akr1a1b | 2 |
| akr1a1b | akr1a1a,akr1a1b | 2 |
| alox5a | alox5ap,alox5a | 2 |
| alox5ap | alox5ap,alox5a | 2 |
| angpt2a | angpt2b,angpt2a | 2 |
| angpt2b | angpt2b,angpt2a | 2 |
| ankdd1a | ankdd1b,ankdd1a | 2 |
| ankdd1b | ankdd1b,ankdd1a | 2 |
| ankrd13a | ankrd13d,ankrd13a,ankrd | 3 |
| ankrd13b | ankrd13d,ankrd13a,ankrd | 3 |
| ankrd13d | ankrd13d,ankrd13a,ankrd | 3 |
| ankrd33ab | ankrd33ab,ankrd33bb | 2 |
| ankrd33bb | ankrd33ab,ankrd33bb | 2 |
| ano10a | ano10a,ano10b | 2 |
| ano10b | ano10a,ano10b | 2 |
| anos1a | anos1b,anos1a | 2 |
| anos1b | anos1b,anos1a | 2 |
| anp32a | anp32a,anp32b,anp32e | 3 |
| anp32b | anp32a,anp32b,anp32e | 3 |
| anp32e | anp32a,anp32b,anp32e | 3 |

| | | |
|---|---|---|
| antxr1a | antxr1a,antxr1d | 2 |
| antxr1d | antxr1a,antxr1d | 2 |
| apbb1ip | apbb1,apbb1ip | 2 |
| arid1aa | arid1b,arid1aa | 2 |
| arid1b | arid1b,arid1aa | 2 |
| arid3a | arid3c,arid3b,arid3a | 3 |
| arid3b | arid3c,arid3b,arid3a | 3 |
| arid3c | arid3c,arid3b,arid3a | 3 |
| arl13a | arl13a,arl13b | 2 |
| arl13b | arl13a,arl13b | 2 |
| arl2bp | arl2bp,arl2 | 2 |
| arl4aa | arl4aa,arl4cb,arl4d | 3 |
| arl4cb | arl4aa,arl4cb,arl4d | 3 |
| arl4d | arl4aa,arl4cb,arl4d | 3 |
| arl8a | arl8a,arl8bb,arl8 | 3 |
| arl8bb | arl8a,arl8bb,arl8 | 3 |
| arpc1a | arpc1b,arpc1a | 2 |
| arpc1b | arpc1b,arpc1a | 2 |
| ascl1b | ascl1b,ascl1b | 2 |
| ascl1b | ascl1b,ascl1b | 2 |
| atad1a | atad1a,atad1b | 2 |
| atad1b | atad1a,atad1b | 2 |
| atf7a | atf7a,atf7ip | 2 |
| atf7ip | atf7a,atf7ip | 2 |
| atg4a | atg4db,atg4a,atg4c,atg4b | 4 |
| atg4b | atg4db,atg4a,atg4c,atg4b | 4 |
| atg4c | atg4db,atg4a,atg4c,atg4b | 4 |
| atg4db | atg4db,atg4a,atg4c,atg4b | 4 |
| atg9a | atg9b,atg9a | 2 |
| atg9b | atg9b,atg9a | 2 |
| atoh1a | atoh1a,atoh1c,atoh1b | 3 |
| atoh1b | atoh1a,atoh1c,atoh1b | 3 |
| atoh1c | atoh1a,atoh1c,atoh1b | 3 |
| atp10a | atp10d,atp10b,atp10a | 3 |
| atp10b | atp10d,atp10b,atp10a | 3 |
| atp10d | atp10d,atp10b,atp10a | 3 |
| atp11a | atp11c,atp11a | 2 |
| atp11c | atp11c,atp11a | 2 |
| atp5f1b | atp5f1d,atp5f1c,atp5f1b,a | 4 |
| atp5f1c | atp5f1d,atp5f1c,atp5f1b,a | 4 |
| atp5f1d | atp5f1d,atp5f1c,atp5f1b,a | 4 |
| atp5f1e | atp5f1d,atp5f1c,atp5f1b,a | 4 |
| atp5md | atp5meb,atp5pb,atp5md, | 7 |
| atp5meb | atp5meb,atp5pb,atp5md, | 7 |
| atp5mf | atp5meb,atp5pb,atp5md, | 7 |
| atp5pb | atp5meb,atp5pb,atp5md, | 7 |
| atp5pd | atp5meb,atp5pb,atp5md, | 7 |
| atp5pf | atp5meb,atp5pb,atp5md, | 7 |
| atp5po | atp5meb,atp5pb,atp5md, | 7 |
| atp6v0b | atp6v0b,atp6v0ca | 2 |

| | | |
|---|---|---|
| atp6v0ca | atp6v0b,atp6v0ca | 2 |
| atp6v1d | atp6v1d,atp6v1f,atp6v1h | 3 |
| atp6v1f | atp6v1d,atp6v1f,atp6v1h | 3 |
| atp6v1h | atp6v1d,atp6v1f,atp6v1h | 3 |
| atp7a | atp7a,atp7b | 2 |
| atp7b | atp7a,atp7b | 2 |
| b3gat1a | b3gat1b,b3gat1a | 2 |
| b3gat1b | b3gat1b,b3gat1a | 2 |
| baz2a | baz2a,baz2ba | 2 |
| baz2ba | baz2a,baz2ba | 2 |
| bcl6aa | bcl6b,bcl6aa | 2 |
| bcl6b | bcl6b,bcl6aa | 2 |
| bcl7a | bcl7a,bcl7bb | 2 |
| bcl7bb | bcl7a,bcl7bb | 2 |
| bmp2a | bmp2a,bmp2k | 2 |
| bmp2k | bmp2a,bmp2k | 2 |
| bmpr1aa | bmpr1bb,bmpr1aa | 2 |
| bmpr1bb | bmpr1bb,bmpr1aa | 2 |
| c1d | c1d,c1qbp,c1qc,c1r | 4 |
| c1qbp | c1d,c1qbp,c1qc,c1r | 4 |
| c1qc | c1d,c1qbp,c1qc,c1r | 4 |
| c1r | c1d,c1qbp,c1qc,c1r | 4 |
| ca4a | ca4a,ca4c | 2 |
| ca4c | ca4a,ca4c | 2 |
| cacna1ba | cacna1sb,cacna1da,cacna | 9 |
| cacna1c | cacna1sb,cacna1da,cacna | 9 |
| cacna1da | cacna1sb,cacna1da,cacna | 9 |
| cacna1db | cacna1sb,cacna1da,cacna | 9 |
| cacna1ea | cacna1sb,cacna1da,cacna | 9 |
| cacna1g | cacna1sb,cacna1da,cacna | 9 |
| cacna1hb | cacna1sb,cacna1da,cacna | 9 |
| cacna1ia | cacna1sb,cacna1da,cacna | 9 |
| cacna1sb | cacna1sb,cacna1da,cacna | 9 |
| camk1a | camk1ga,camk1a,camk1d | 3 |
| camk1da | camk1ga,camk1a,camk1d | 3 |
| camk1ga | camk1ga,camk1a,camk1d | 3 |
| ccdc28a | ccdc28a,ccdc28b | 2 |
| ccdc28b | ccdc28a,ccdc28b | 2 |
| ccdc3a | ccdc3b,ccdc3a | 2 |
| ccdc3b | ccdc3b,ccdc3a | 2 |
| ccdc85a | ccdc85cb,ccdc85a,ccdc85 | 3 |
| ccdc85b | ccdc85cb,ccdc85a,ccdc85 | 3 |
| ccdc85cb | ccdc85cb,ccdc85a,ccdc85 | 3 |
| ccdc88aa | ccdc88c,ccdc88aa | 2 |
| ccdc88c | ccdc88c,ccdc88aa | 2 |
| ccdc9b | ccdc9,ccdc9b | 2 |
| ccn2a | ccn2a,ccn2b | 2 |
| ccn2b | ccn2a,ccn2b | 2 |
| cd79a | cd79b,cd79a | 2 |
| cd79b | cd79b,cd79a | 2 |

| | | |
|---|---|---|
| cdc42bpab | cdc42bpab,cdc42bpb,cdc4 | 3 |
| cdc42bpb | cdc42bpab,cdc42bpb,cdc4 | 3 |
| cdca7a | cdca7b,cdca7a | 2 |
| cdca7b | cdca7b,cdca7a | 2 |
| cdkn1a | cdkn1a,cdkn1ba,cdkn1cb | 3 |
| cdkn1ba | cdkn1a,cdkn1ba,cdkn1cb | 3 |
| cdkn1cb | cdkn1a,cdkn1ba,cdkn1cb | 3 |
| cdkn2aip | cdkn2aip,cdkn2d,cdkn2c | 3 |
| cdkn2c | cdkn2aip,cdkn2d,cdkn2c | 3 |
| cdkn2d | cdkn2aip,cdkn2d,cdkn2c | 3 |
| cep170aa | cep170b,cep170aa | 2 |
| cep170b | cep170b,cep170aa | 2 |
| chaf1a | chaf1b,chaf1a | 2 |
| chaf1b | chaf1b,chaf1a | 2 |
| chmp1a | chmp1b,chmp1a | 2 |
| chmp1b | chmp1b,chmp1a | 2 |
| chmp4ba | chmp4c,chmp4ba | 2 |
| chmp4c | chmp4c,chmp4ba | 2 |
| ciao2a | ciao2a,ciao2b | 2 |
| ciao2b | ciao2a,ciao2b | 2 |
| coq8aa | coq8aa,coq8b | 2 |
| coq8b | coq8aa,coq8b | 2 |
| coro2a | coro2ba,coro2a | 2 |
| coro2ba | coro2ba,coro2a | 2 |
| cplx4b | cplx4b,cplx4c | 2 |
| cplx4c | cplx4b,cplx4c | 2 |
| cpt1aa | cpt1b,cpt1aa | 2 |
| cpt1b | cpt1b,cpt1aa | 2 |
| csf1b | csf1b,csf1ra | 2 |
| csf1ra | csf1b,csf1ra | 2 |
| csnk1db | csnk1db,csnk1e | 2 |
| csnk1e | csnk1db,csnk1e | 2 |
| cul4a | cul4b,cul4a | 2 |
| cul4b | cul4b,cul4a | 2 |
| cyb5a | cyb5a,cyb5b | 2 |
| cyb5b | cyb5a,cyb5b | 2 |
| cyth1a | cyth1b,cyth1a | 2 |
| cyth1b | cyth1b,cyth1a | 2 |
| dclre1a | dclre1b,dclre1a,dclre1c | 3 |
| dclre1b | dclre1b,dclre1a,dclre1c | 3 |
| dclre1c | dclre1b,dclre1a,dclre1c | 3 |
| dcp1a | dcp1a,dcp1b | 2 |
| dcp1b | dcp1a,dcp1b | 2 |
| dennd1a | dennd1b,dennd1a | 2 |
| dennd1b | dennd1b,dennd1a | 2 |
| dennd2c | dennd2c,dennd2db | 2 |
| dennd2db | dennd2c,dennd2db | 2 |
| dennd4a | dennd4a,dennd4c,dennd4 | 3 |
| dennd4b | dennd4a,dennd4c,dennd4 | 3 |
| dennd4c | dennd4a,dennd4c,dennd4 | 3 |

| | | |
|---|---|---|
| dennd6aa | dennd6aa,dennd6b | 2 |
| dennd6b | dennd6aa,dennd6b | 2 |
| dhrs7b | dhrs7,dhrs7ca,dhrs7b | 3 |
| dhrs7ca | dhrs7,dhrs7ca,dhrs7b | 3 |
| dip2a | dip2bb,dip2ca,dip2a | 3 |
| dip2bb | dip2bb,dip2ca,dip2a | 3 |
| dip2ca | dip2bb,dip2ca,dip2a | 3 |
| dipk1ab | dipk1c,dipk1ab,dipk1b | 3 |
| dipk1b | dipk1c,dipk1ab,dipk1b | 3 |
| dipk1c | dipk1c,dipk1ab,dipk1b | 3 |
| dipk2aa | dipk2aa,dipk2b | 2 |
| dipk2b | dipk2aa,dipk2b | 2 |
| dmrt2a | dmrt2a,dmrt2b | 2 |
| dmrt2b | dmrt2a,dmrt2b | 2 |
| dnajc3a | dnajc3b,dnajc3a | 2 |
| dnajc3b | dnajc3b,dnajc3a | 2 |
| dnajc5aa | dnajc5b,dnajc5gb,dnajc5a | 3 |
| dnajc5b | dnajc5b,dnajc5gb,dnajc5a | 3 |
| dnajc5gb | dnajc5b,dnajc5gb,dnajc5a | 3 |
| dnmt3ab | dnmt3ab,dnmt3ba | 2 |
| dnmt3ba | dnmt3ab,dnmt3ba | 2 |
| dop1a | dop1a,dop1b | 2 |
| dop1b | dop1a,dop1b | 2 |
| dusp22a | dusp22b,dusp22a | 2 |
| dusp22b | dusp22b,dusp22a | 2 |
| dyrk1ab | dyrk1ab,dyrk1b | 2 |
| dyrk1b | dyrk1ab,dyrk1b | 2 |
| ece2a | ece2a,ece2b | 2 |
| ece2b | ece2a,ece2b | 2 |
| eef1db | eef1g,eef1db | 2 |
| eef1g | eef1g,eef1db | 2 |
| eef2b | eef2kmt,eef2k,eef2b | 3 |
| eef2k | eef2kmt,eef2k,eef2b | 3 |
| eef2kmt | eef2kmt,eef2k,eef2b | 3 |
| efr3a | efr3ba,efr3a | 2 |
| efr3ba | efr3ba,efr3a | 2 |
| eif1ad | eif1axb,eif1b,eif1ad | 3 |
| eif1axb | eif1axb,eif1b,eif1ad | 3 |
| eif1b | eif1axb,eif1b,eif1ad | 3 |
| eif2a | eif2d,eif2a | 2 |
| eif2d | eif2d,eif2a | 2 |
| eif3ba | eif3f,eif3k,eif3ja,eif3g,eif3 | 8 |
| eif3ea | eif3f,eif3k,eif3ja,eif3g,eif3 | 8 |
| eif3f | eif3f,eif3k,eif3ja,eif3g,eif3 | 8 |
| eif3g | eif3f,eif3k,eif3ja,eif3g,eif3 | 8 |
| eif3ha | eif3f,eif3k,eif3ja,eif3g,eif3 | 8 |
| eif3ja | eif3f,eif3k,eif3ja,eif3g,eif3 | 8 |
| eif3k | eif3f,eif3k,eif3ja,eif3g,eif3 | 8 |
| eif3m | eif3f,eif3k,eif3ja,eif3g,eif3 | 8 |
| eif4ba | eif4ba,eif4eb,eif4h | 3 |

| | | |
|---|---|---|
| eif4e1b | eif4e1c,eif4e1b | 2 |
| eif4e1c | eif4e1c,eif4e1b | 2 |
| eif4eb | eif4ba,eif4eb,eif4h | 3 |
| eif4h | eif4ba,eif4eb,eif4h | 3 |
| eif5a | eif5,eif5a | 2 |
| epb41l4a | epb41l4a,epb41l4b | 2 |
| epb41l4b | epb41l4a,epb41l4b | 2 |
| ero1a | ero1a,ero1b | 2 |
| ero1b | ero1a,ero1b | 2 |
| exoc6b | exoc6b,exoc6 | 2 |
| ext1b | ext1c,ext1b | 2 |
| ext1c | ext1c,ext1b | 2 |
| faim2a | faim2b,faim2a | 2 |
| faim2b | faim2b,faim2a | 2 |
| fam102aa | fam102bb,fam102aa | 2 |
| fam102bb | fam102bb,fam102aa | 2 |
| fam120a | fam120c,fam120a,fam12( | 3 |
| fam120b | fam120c,fam120a,fam12( | 3 |
| fam120c | fam120c,fam120a,fam12( | 3 |
| fam13a | fam13a,fam13b | 2 |
| fam13b | fam13a,fam13b | 2 |
| fam151a | fam151b,fam151a | 2 |
| fam151b | fam151b,fam151a | 2 |
| fam155a | fam155b,fam155a | 2 |
| fam155b | fam155b,fam155a | 2 |
| fam161a | fam161b,fam161a | 2 |
| fam161b | fam161b,fam161a | 2 |
| fam167ab | fam167ab,fam167b | 2 |
| fam167b | fam167ab,fam167b | 2 |
| fam168a | fam168a,fam168b | 2 |
| fam168b | fam168a,fam168b | 2 |
| fam169aa | fam169aa,fam169b | 2 |
| fam169b | fam169aa,fam169b | 2 |
| fam184a | fam184a,fam184b | 2 |
| fam184b | fam184a,fam184b | 2 |
| fam193a | fam193a,fam193b | 2 |
| fam193b | fam193a,fam193b | 2 |
| fam20a | fam20b,fam20a,fam20cb | 3 |
| fam20b | fam20b,fam20a,fam20cb | 3 |
| fam20cb | fam20b,fam20a,fam20cb | 3 |
| fam210aa | fam210aa,fam210b | 2 |
| fam210b | fam210aa,fam210b | 2 |
| fam214a | fam214b,fam214a | 2 |
| fam214b | fam214b,fam214a | 2 |
| fam219ab | fam219ab,fam219b | 2 |
| fam219b | fam219ab,fam219b | 2 |
| fam43a | fam43a,fam43b | 2 |
| fam43b | fam43a,fam43b | 2 |
| fam49a | fam49a,fam49ba | 2 |
| fam49ba | fam49a,fam49ba | 2 |

| | | |
|---|---|---|
| fam83b | fam83hb,fam83fb,fam83l | 5 |
| fam83c | fam83hb,fam83fb,fam83l | 5 |
| fam83d | fam83hb,fam83fb,fam83l | 5 |
| fam83fb | fam83hb,fam83fb,fam83l | 5 |
| fam83hb | fam83hb,fam83fb,fam83l | 5 |
| fam98a | fam98b,fam98a | 2 |
| fam98b | fam98b,fam98a | 2 |
| fbn2a | fbn2a,fbn2b | 2 |
| fbn2b | fbn2a,fbn2b | 2 |
| fem1b | fem1c,fem1b | 2 |
| fem1c | fem1c,fem1b | 2 |
| fgfrl1a | fgfrl1a,fgfrl1b | 2 |
| fgfrl1b | fgfrl1a,fgfrl1b | 2 |
| fkbp1aa | fkbp1b,fkbp1aa | 2 |
| fkbp1b | fkbp1b,fkbp1aa | 2 |
| fli1a | fli1b,fli1a | 2 |
| fli1b | fli1b,fli1a | 2 |
| fndc3a | fndc3a,fndc3ba | 2 |
| fndc3ba | fndc3a,fndc3ba | 2 |
| foxj1a | foxj1a,foxj1b | 2 |
| foxj1b | foxj1a,foxj1b | 2 |
| frem1a | frem1a,frem1b | 2 |
| frem1b | frem1a,frem1b | 2 |
| fzr1a | fzr1b,fzr1a | 2 |
| fzr1b | fzr1b,fzr1a | 2 |
| gadd45aa | gadd45ga,gadd45aa,gadd | 3 |
| gadd45ba | gadd45ga,gadd45aa,gadd | 3 |
| gadd45ga | gadd45ga,gadd45aa,gadd | 3 |
| galr1a | galr1a,galr1b | 2 |
| galr1b | galr1a,galr1b | 2 |
| galr2a | galr2a,galr2b | 2 |
| galr2b | galr2a,galr2b | 2 |
| gask1a | gask1b,gask1a | 2 |
| gask1b | gask1b,gask1a | 2 |
| gatad2ab | gatad2ab,gatad2b | 2 |
| gatad2b | gatad2ab,gatad2b | 2 |
| gfi1ab | gfi1ab,gfi1b | 2 |
| gfi1b | gfi1ab,gfi1b | 2 |
| gig2e | gig2p,gig2e,gig2o | 3 |
| gig2o | gig2p,gig2e,gig2o | 3 |
| gig2p | gig2p,gig2e,gig2o | 3 |
| golt1a | golt1a,golt1bb | 2 |
| golt1bb | golt1a,golt1bb | 2 |
| gpc5a | gpc5c,gpc5a | 2 |
| gpc5c | gpc5c,gpc5a | 2 |
| gpd1b | gpd1b,gpd1c | 2 |
| gpd1c | gpd1b,gpd1c | 2 |
| gpm6aa | gpm6aa,gpm6bb | 2 |
| gpm6bb | gpm6aa,gpm6bb | 2 |
| gpr137bb | gpr137c,gpr137bb | 2 |

| | | |
|---|---|---|
| gpr137c | gpr137c,gpr137bb | 2 |
| gprc5bb | gprc5c,gprc5bb | 2 |
| gprc5c | gprc5c,gprc5bb | 2 |
| gramd1a | gramd1c,gramd1a,gramd | 3 |
| gramd1bb | gramd1c,gramd1a,gramd | 3 |
| gramd1c | gramd1c,gramd1a,gramd | 3 |
| grid2ipa | grid2,grid2ipa | 2 |
| grin2aa | grin2cb,grin2bb,grin2aa | 3 |
| grin2bb | grin2cb,grin2bb,grin2aa | 3 |
| grin2cb | grin2cb,grin2bb,grin2aa | 3 |
| grk1a | grk1b,grk1a | 2 |
| grk1b | grk1b,grk1a | 2 |
| guca1a | guca1c,guca1b,guca1e,gu | 4 |
| guca1b | guca1c,guca1b,guca1e,gu | 4 |
| guca1c | guca1c,guca1b,guca1e,gu | 4 |
| guca1e | guca1c,guca1b,guca1e,gu | 4 |
| gucy2f | gucy2f,gucy2g | 2 |
| gucy2g | gucy2f,gucy2g | 2 |
| h2afva | h2afva,h2afy | 2 |
| h2afy | h2afva,h2afy | 2 |
| heatr5a | heatr5a,heatr5b | 2 |
| heatr5b | heatr5a,heatr5b | 2 |
| hiat1a | hiat1b,hiat1a | 2 |
| hiat1b | hiat1b,hiat1a | 2 |
| hip1rb | hip1rb,hip1 | 2 |
| hmg20a | hmg20a,hmg20b | 2 |
| hmg20b | hmg20a,hmg20b | 2 |
| hnf1a | hnf1a,hnf1bb | 2 |
| hnf1bb | hnf1a,hnf1bb | 2 |
| hnf4a | hnf4g,hnf4a | 2 |
| hnf4g | hnf4g,hnf4a | 2 |
| hsf2bp | hsf2,hsf2bp | 2 |
| hspa12a | hspa12b,hspa12a | 2 |
| hspa12b | hspa12b,hspa12a | 2 |
| htr2aa | htr2b,htr2aa | 2 |
| htr2b | htr2b,htr2aa | 2 |
| htr3a | htr3b,htr3a | 2 |
| htr3b | htr3b,htr3a | 2 |
| htr7a | htr7a,htr7b | 2 |
| htr7b | htr7a,htr7b | 2 |
| idh3a | idh3g,idh3b,idh3a | 3 |
| idh3b | idh3g,idh3b,idh3a | 3 |
| idh3g | idh3g,idh3b,idh3a | 3 |
| igf1ra | igf1ra,igf1 | 2 |
| igf2b | igf2r,igf2b | 2 |
| igf2r | igf2r,igf2b | 2 |
| igsf9b | igsf9b,igsf9bb | 2 |
| igsf9bb | igsf9b,igsf9bb | 2 |
| il17d | il17rd,il17d | 2 |
| il17rd | il17rd,il17d | 2 |

| | | |
|---|---|---|
| ino80b | ino80b,ino80,ino80c,ino8 | 5 |
| ino80c | ino80b,ino80,ino80c,ino8 | 5 |
| ino80db | ino80b,ino80,ino80c,ino8 | 5 |
| ino80e | ino80b,ino80,ino80c,ino8 | 5 |
| inpp4ab | inpp4b,inpp4ab | 2 |
| inpp4b | inpp4b,inpp4ab | 2 |
| inpp5b | inpp5f,inpp5b,inpp5d,inp | 6 |
| inpp5d | inpp5f,inpp5b,inpp5d,inp | 6 |
| inpp5e | inpp5f,inpp5b,inpp5d,inp | 6 |
| inpp5f | inpp5f,inpp5b,inpp5d,inp | 6 |
| inpp5ja | inpp5f,inpp5b,inpp5d,inp | 6 |
| inpp5kb | inpp5f,inpp5b,inpp5d,inp | 6 |
| inppl1a | inppl1a,inppl1b | 2 |
| inppl1b | inppl1a,inppl1b | 2 |
| irf1a | irf1a,irf1b | 2 |
| irf1b | irf1a,irf1b | 2 |
| itm2bb | itm2ca,itm2bb | 2 |
| itm2ca | itm2ca,itm2bb | 2 |
| kat2a | kat2b,kat2a | 2 |
| kat2b | kat2b,kat2a | 2 |
| kat6a | kat6a,kat6b | 2 |
| kat6b | kat6a,kat6b | 2 |
| kdm2aa | kdm2aa,kdm2ba | 2 |
| kdm2ba | kdm2aa,kdm2ba | 2 |
| kdm5a | kdm5ba,kdm5a | 2 |
| kdm5ba | kdm5ba,kdm5a | 2 |
| keap1a | keap1b,keap1a | 2 |
| keap1b | keap1b,keap1a | 2 |
| kif1aa | kif1bp,kif1aa,kif1b | 3 |
| kif1b | kif1bp,kif1aa,kif1b | 3 |
| kif1bp | kif1bp,kif1aa,kif1b | 3 |
| kif21a | kif21b,kif21a | 2 |
| kif21b | kif21b,kif21a | 2 |
| kif26aa | kif26ba,kif26aa | 2 |
| kif26ba | kif26ba,kif26aa | 2 |
| kif3a | kif3ca,kif3a,kif3b | 3 |
| kif3b | kif3ca,kif3a,kif3b | 3 |
| kif3ca | kif3ca,kif3a,kif3b | 3 |
| kif5aa | kif5aa,kif5ba | 2 |
| kif5ba | kif5aa,kif5ba | 2 |
| klhdc8a | klhdc8a,klhdc8b | 2 |
| klhdc8b | klhdc8a,klhdc8b | 2 |
| kmt2ba | kmt2d,kmt2e,kmt2ca,km | 4 |
| kmt2ca | kmt2d,kmt2e,kmt2ca,km | 4 |
| kmt2d | kmt2d,kmt2e,kmt2ca,km | 4 |
| kmt2e | kmt2d,kmt2e,kmt2ca,km | 4 |
| kmt5ab | kmt5ab,kmt5b | 2 |
| kmt5b | kmt5ab,kmt5b | 2 |
| l3mbtl1a | l3mbtl1b,l3mbtl1a | 2 |
| l3mbtl1b | l3mbtl1b,l3mbtl1a | 2 |

| | | |
|---|---|---|
| laptm4a | laptm4a,laptm4b | 2 |
| laptm4b | laptm4a,laptm4b | 2 |
| larp1b | larp1b,larp1 | 2 |
| larp4ab | larp4ab,larp4b | 2 |
| larp4b | larp4ab,larp4b | 2 |
| ldb3a | ldb3b,ldb3a | 2 |
| ldb3b | ldb3b,ldb3a | 2 |
| lhfpl4a | lhfpl4b,lhfpl4a | 2 |
| lhfpl4b | lhfpl4b,lhfpl4a | 2 |
| lin28a | lin28b,lin28a | 2 |
| lin28b | lin28b,lin28a | 2 |
| lin7a | lin7c,lin7a | 2 |
| lin7c | lin7c,lin7a | 2 |
| lmo4a | lmo4b,lmo4a | 2 |
| lmo4b | lmo4b,lmo4a | 2 |
| lnx2a | lnx2a,lnx2b | 2 |
| lnx2b | lnx2a,lnx2b | 2 |
| lpar5a | lpar5a,lpar5b | 2 |
| lpar5b | lpar5a,lpar5b | 2 |
| lrp1ab | lrp1ab,lrp1ba | 2 |
| lrp1ba | lrp1ab,lrp1ba | 2 |
| lrp2a | lrp2bp,lrp2a | 2 |
| lrp2bp | lrp2bp,lrp2a | 2 |
| lrrc3b | lrrc3b,lrrc3,lrrc3ca | 3 |
| lrrc3ca | lrrc3b,lrrc3,lrrc3ca | 3 |
| lrrc4bb | lrrc4bb,lrrc4ca | 2 |
| lrrc4ca | lrrc4bb,lrrc4ca | 2 |
| lrrc74a | lrrc74a,lrrc74b | 2 |
| lrrc74b | lrrc74a,lrrc74b | 2 |
| lrrc8ab | lrrc8da,lrrc8c,lrrc8ab | 3 |
| lrrc8c | lrrc8da,lrrc8c,lrrc8ab | 3 |
| lrrc8da | lrrc8da,lrrc8c,lrrc8ab | 3 |
| lsm14ab | lsm14b,lsm14ab | 2 |
| lsm14b | lsm14b,lsm14ab | 2 |
| lypd6b | lypd6b,lypd6 | 2 |
| mad2l1bp | mad2l1bp,mad2l1 | 2 |
| map1aa | map1b,map1aa,map1sb | 3 |
| map1b | map1b,map1aa,map1sb | 3 |
| map1lc3b | map1lc3c,map1lc3b | 2 |
| map1lc3c | map1lc3c,map1lc3b | 2 |
| map1sb | map1b,map1aa,map1sb | 3 |
| mat2ab | mat2ab,mat2b | 2 |
| mat2b | mat2ab,mat2b | 2 |
| mbd3a | mbd3a,mbd3b | 2 |
| mbd3b | mbd3a,mbd3b | 2 |
| mcm3ap | mcm3,mcm3ap | 2 |
| mdh1ab | mdh1b,mdh1ab | 2 |
| mdh1b | mdh1b,mdh1ab | 2 |
| mef2aa | mef2ca,mef2aa,mef2b,m[illegible] | 4 |
| mef2b | mef2ca,mef2aa,mef2b,m[illegible] | 4 |

| | | |
|---|---|---|
| mef2ca | mef2ca,mef2aa,mef2b,m | 4 |
| mef2d | mef2ca,mef2aa,mef2b,m | 4 |
| metap1d | metap1,metap1d | 2 |
| mfsd2aa | mfsd2b,mfsd2aa | 2 |
| mfsd2b | mfsd2b,mfsd2aa | 2 |
| mgat4a | mgat4b,mgat4c,mgat4a | 3 |
| mgat4b | mgat4b,mgat4c,mgat4a | 3 |
| mgat4c | mgat4b,mgat4c,mgat4a | 3 |
| mgst3a | mgst3b,mgst3a | 2 |
| mgst3b | mgst3b,mgst3a | 2 |
| mmp13a | mmp13b,mmp13a | 2 |
| mmp13b | mmp13b,mmp13a | 2 |
| mmp20a | mmp20b,mmp20a | 2 |
| mmp20b | mmp20b,mmp20a | 2 |
| mob3a | mob3c,mob3a | 2 |
| mob3c | mob3c,mob3a | 2 |
| mon1a | mon1bb,mon1a | 2 |
| mon1bb | mon1bb,mon1a | 2 |
| mrps18a | mrps18a,mrps18c | 2 |
| mrps18c | mrps18a,mrps18c | 2 |
| msrb1b | msrb1b,msrb1b | 2 |
| msrb1b | msrb1b,msrb1b | 2 |
| mst1rb | mst1rb,mst1 | 2 |
| mtnr1bb | mtnr1bb,mtnr1c | 2 |
| mtnr1c | mtnr1bb,mtnr1c | 2 |
| mul1a | mul1a,mul1b | 2 |
| mul1b | mul1a,mul1b | 2 |
| myo1b | myo1ea,myo1g,myo1b,m | 7 |
| myo1cb | myo1ea,myo1g,myo1b,m | 7 |
| myo1ea | myo1ea,myo1g,myo1b,m | 7 |
| myo1eb | myo1ea,myo1g,myo1b,m | 7 |
| myo1f | myo1ea,myo1g,myo1b,m | 7 |
| myo1g | myo1ea,myo1g,myo1b,m | 7 |
| myo1hb | myo1ea,myo1g,myo1b,m | 7 |
| myo3a | myo3a,myo3b | 2 |
| myo3b | myo3a,myo3b | 2 |
| myo5ab | myo5b,myo5c,myo5ab | 3 |
| myo5b | myo5b,myo5c,myo5ab | 3 |
| myo5c | myo5b,myo5c,myo5ab | 3 |
| myo7aa | myo7aa,myo7bb | 2 |
| myo7bb | myo7aa,myo7bb | 2 |
| myo9aa | myo9aa,myo9b | 2 |
| myo9b | myo9aa,myo9b | 2 |
| neurl1aa | neurl1aa,neurl1b | 2 |
| neurl1b | neurl1aa,neurl1b | 2 |
| nf2a | nf2a,nf2b | 2 |
| nf2b | nf2a,nf2b | 2 |
| nos1apa | nos1apa,nos1 | 2 |
| npm1a | npm1b,npm1a | 2 |
| npm1b | npm1b,npm1a | 2 |

| | | |
|---|---|---|
| nr2c2ap | nr2c2,nr2c2ap | 2 |
| nt5c1aa | nt5c1bb,nt5c1aa | 2 |
| nt5c1bb | nt5c1bb,nt5c1aa | 2 |
| oaz1a | oaz1b,oaz1a | 2 |
| oaz1b | oaz1b,oaz1a | 2 |
| opn4a | opn4xa,opn4a | 2 |
| opn4xa | opn4xa,opn4a | 2 |
| opn6a | opn6b,opn6a | 2 |
| opn6b | opn6b,opn6a | 2 |
| opn7a | opn7a,opn7b | 2 |
| opn7b | opn7a,opn7b | 2 |
| opn8a | opn8a,opn8c,opn8b | 3 |
| opn8b | opn8a,opn8c,opn8b | 3 |
| opn8c | opn8a,opn8c,opn8b | 3 |
| oscp1a | oscp1a,oscp1a | 2 |
| oscp1a | oscp1a,oscp1a | 2 |
| p4hb | p4htm,p4hb | 2 |
| p4htm | p4htm,p4hb | 2 |
| paqr7a | paqr7b,paqr7a | 2 |
| paqr7b | paqr7b,paqr7a | 2 |
| pard3ab | pard3ab,pard3bb | 2 |
| pard3bb | pard3ab,pard3bb | 2 |
| pard6a | pard6gb,pard6b,pard6a | 3 |
| pard6b | pard6gb,pard6b,pard6a | 3 |
| pard6gb | pard6gb,pard6b,pard6a | 3 |
| parp12a | parp12b,parp12a | 2 |
| parp12b | parp12b,parp12a | 2 |
| pcyt1ab | pcyt1bb,pcyt1ab | 2 |
| pcyt1bb | pcyt1bb,pcyt1ab | 2 |
| pdap1a | pdap1a,pdap1b | 2 |
| pdap1b | pdap1a,pdap1b | 2 |
| pdcd4a | pdcd4a,pdcd4b | 2 |
| pdcd4b | pdcd4a,pdcd4b | 2 |
| pdcd6ip | pdcd6,pdcd6ip | 2 |
| pde3a | pde3a,pde3b | 2 |
| pde3b | pde3a,pde3b | 2 |
| pde4a | pde4d,pde4a,pde4ba,pde | 4 |
| pde4ba | pde4d,pde4a,pde4ba,pde | 4 |
| pde4ca | pde4d,pde4a,pde4ba,pde | 4 |
| pde4d | pde4d,pde4a,pde4ba,pde | 4 |
| pde6a | pde6b,pde6c,pde6a,pde6 | 5 |
| pde6b | pde6b,pde6c,pde6a,pde6 | 5 |
| pde6c | pde6b,pde6c,pde6a,pde6 | 5 |
| pde6d | pde6b,pde6c,pde6a,pde6 | 5 |
| pde6gb | pde6b,pde6c,pde6a,pde6 | 5 |
| pde8a | pde8b,pde8a | 2 |
| pde8b | pde8b,pde8a | 2 |
| pds5a | pds5b,pds5a | 2 |
| pds5b | pds5b,pds5a | 2 |
| pex11a | pex11a,pex11g,pex11b | 3 |

| | | |
|---|---|---|
| pex11b | pex11a,pex11g,pex11b | 3 |
| pex11g | pex11a,pex11g,pex11b | 3 |
| phox2a | phox2a,phox2bb | 2 |
| phox2bb | phox2a,phox2bb | 2 |
| pi4k2a | pi4k2b,pi4k2a | 2 |
| pi4k2b | pi4k2b,pi4k2a | 2 |
| pi4kab | pi4kab,pi4kb | 2 |
| pi4kb | pi4kab,pi4kb | 2 |
| pik3c2a | pik3c2b,pik3c2a | 2 |
| pik3c2b | pik3c2b,pik3c2a | 2 |
| pik3ca | pik3cg,pik3ca,pik3cb,pik3 | 4 |
| pik3cb | pik3cg,pik3ca,pik3cb,pik3 | 4 |
| pik3cd | pik3cg,pik3ca,pik3cb,pik3 | 4 |
| pik3cg | pik3cg,pik3ca,pik3cb,pik3 | 4 |
| pip4k2ab | pip4k2cb,pip4k2ab | 2 |
| pip4k2cb | pip4k2cb,pip4k2ab | 2 |
| pip5k1ab | pip5k1bb,pip5k1ca,pip5k1 | 3 |
| pip5k1bb | pip5k1bb,pip5k1ca,pip5k1 | 3 |
| pip5k1ca | pip5k1bb,pip5k1ca,pip5k1 | 3 |
| pla2g12a | pla2g12a,pla2g12b | 2 |
| pla2g12b | pla2g12a,pla2g12b | 2 |
| pnp4a | pnp4b,pnp4a | 2 |
| pnp4b | pnp4b,pnp4a | 2 |
| polr1a | polr1a,polr1e,polr1d,polr | 5 |
| polr1b | polr1a,polr1e,polr1d,polr | 5 |
| polr1c | polr1a,polr1e,polr1d,polr | 5 |
| polr1d | polr1a,polr1e,polr1d,polr | 5 |
| polr1e | polr1a,polr1e,polr1d,polr | 5 |
| polr2a | polr2a,polr2i,polr2b,polr2 | 9 |
| polr2b | polr2a,polr2i,polr2b,polr2 | 9 |
| polr2c | polr2a,polr2i,polr2b,polr2 | 9 |
| polr2d | polr2a,polr2i,polr2b,polr2 | 9 |
| polr2f | polr2a,polr2i,polr2b,polr2 | 9 |
| polr2h | polr2a,polr2i,polr2b,polr2 | 9 |
| polr2i | polr2a,polr2i,polr2b,polr2 | 9 |
| polr2j | polr2a,polr2i,polr2b,polr2 | 9 |
| polr2k | polr2a,polr2i,polr2b,polr2 | 9 |
| polr3a | polr3d,polr3g,polr3a,polr3 | 9 |
| polr3b | polr3d,polr3g,polr3a,polr3 | 9 |
| polr3c | polr3d,polr3g,polr3a,polr3 | 9 |
| polr3d | polr3d,polr3g,polr3a,polr3 | 9 |
| polr3e | polr3d,polr3g,polr3a,polr3 | 9 |
| polr3f | polr3d,polr3g,polr3a,polr3 | 9 |
| polr3g | polr3d,polr3g,polr3a,polr3 | 9 |
| polr3h | polr3d,polr3g,polr3a,polr3 | 9 |
| polr3k | polr3d,polr3g,polr3a,polr3 | 9 |
| ppargc1a | ppargc1a,ppargc1b | 2 |
| ppargc1b | ppargc1a,ppargc1b | 2 |
| ppm1aa | ppm1nb,ppm1j,ppm1aa,p | 8 |
| ppm1bb | ppm1nb,ppm1j,ppm1aa,p | 8 |

| | | |
|---|---|---|
| ppm1da | ppm1nb,ppm1j,ppm1aa,p | 8 |
| ppm1e | ppm1nb,ppm1j,ppm1aa,p | 8 |
| ppm1f | ppm1nb,ppm1j,ppm1aa,p | 8 |
| ppm1g | ppm1nb,ppm1j,ppm1aa,p | 8 |
| ppm1j | ppm1nb,ppm1j,ppm1aa,p | 8 |
| ppm1nb | ppm1nb,ppm1j,ppm1aa,p | 8 |
| ppp1r14bb | ppp1r14c,ppp1r14bb | 2 |
| ppp1r14c | ppp1r14c,ppp1r14bb | 2 |
| ppp1r1b | ppp1r1c,ppp1r1b | 2 |
| ppp1r1c | ppp1r1c,ppp1r1b | 2 |
| ppp1r3b | ppp1r3b,ppp1r3ca,ppp1r | 3 |
| ppp1r3ca | ppp1r3b,ppp1r3ca,ppp1r | 3 |
| ppp1r3db | ppp1r3b,ppp1r3ca,ppp1r | 3 |
| ppp2r2aa | ppp2r2aa,ppp2r2d,ppp2r | 3 |
| ppp2r2bb | ppp2r2aa,ppp2r2d,ppp2r | 3 |
| ppp2r2d | ppp2r2aa,ppp2r2d,ppp2r | 3 |
| ppp2r3a | ppp2r3c,ppp2r3a | 2 |
| ppp2r3c | ppp2r3c,ppp2r3a | 2 |
| ppp2r5a | ppp2r5a,ppp2r5eb,ppp2r | 3 |
| ppp2r5d | ppp2r5a,ppp2r5eb,ppp2r | 3 |
| ppp2r5eb | ppp2r5a,ppp2r5eb,ppp2r | 3 |
| ppp3ca | ppp3ccb,ppp3ca | 2 |
| ppp3ccb | ppp3ccb,ppp3ca | 2 |
| prdm1a | prdm1a,prdm1b | 2 |
| prdm1b | prdm1a,prdm1b | 2 |
| prdm8b | prdm8b,prdm8 | 2 |
| prelid1a | prelid1b,prelid1a | 2 |
| prelid1b | prelid1b,prelid1a | 2 |
| prelid3a | prelid3a,prelid3b | 2 |
| prelid3b | prelid3a,prelid3b | 2 |
| prkar1ab | prkar1ab,prkar1b | 2 |
| prkar1b | prkar1ab,prkar1b | 2 |
| prpf4ba | prpf4ba,prpf4 | 2 |
| prxl2b | prxl2c,prxl2b | 2 |
| prxl2c | prxl2c,prxl2b | 2 |
| psmc3ip | psmc3ip,psmc3 | 2 |
| ptger1b | ptger1b,ptger1c | 2 |
| ptger1c | ptger1b,ptger1c | 2 |
| ptger4a | ptger4a,ptger4c | 2 |
| ptger4c | ptger4a,ptger4c | 2 |
| pth1ra | pth1ra,pth1rb | 2 |
| pth1rb | pth1ra,pth1rb | 2 |
| pth2r | pth2,pth2r | 2 |
| ptk2aa | ptk2bb,ptk2aa | 2 |
| ptk2bb | ptk2bb,ptk2aa | 2 |
| ptpn11a | ptpn11a,ptpn11b | 2 |
| ptpn11b | ptpn11a,ptpn11b | 2 |
| ptpn9a | ptpn9a,ptpn9b | 2 |
| ptpn9b | ptpn9a,ptpn9b | 2 |
| pwwp2a | pwwp2b,pwwp2a | 2 |

| | | |
|---|---|---|
| pwwp2b | pwwp2b,pwwp2a | 2 |
| rab11a | rab11a,rab11ba | 2 |
| rab11ba | rab11a,rab11ba | 2 |
| rab27a | rab27b,rab27a | 2 |
| rab27b | rab27b,rab27a | 2 |
| rab33a | rab33ba,rab33a | 2 |
| rab33ba | rab33ba,rab33a | 2 |
| rab3ab | rab3c,rab3db,rab3ip,rab3 | 5 |
| rab3b | rab3c,rab3db,rab3ip,rab3 | 5 |
| rab3c | rab3c,rab3db,rab3ip,rab3 | 5 |
| rab3db | rab3c,rab3db,rab3ip,rab3 | 5 |
| rab3ip | rab3c,rab3db,rab3ip,rab3 | 5 |
| rab40b | rab40b,rab40c | 2 |
| rab40c | rab40b,rab40c | 2 |
| rab4a | rab4b,rab4a | 2 |
| rab4b | rab4b,rab4a | 2 |
| rab5ab | rab5ab,rab5b,rab5c,rab5i | 4 |
| rab5b | rab5ab,rab5b,rab5c,rab5i | 4 |
| rab5c | rab5ab,rab5b,rab5c,rab5i | 4 |
| rab5if | rab5ab,rab5b,rab5c,rab5i | 4 |
| rab8a | rab8b,rab8a | 2 |
| rab8b | rab8b,rab8a | 2 |
| rad23aa | rad23b,rad23aa,rad23ab | 3 |
| rad23ab | rad23b,rad23aa,rad23ab | 3 |
| rad23b | rad23b,rad23aa,rad23ab | 3 |
| rad51b | rad51,rad51b,rad51c,rad5 | 4 |
| rad51c | rad51,rad51b,rad51c,rad5 | 4 |
| rad51d | rad51,rad51b,rad51c,rad5 | 4 |
| rap1ab | rap1ab,rap1b,rap1gap | 3 |
| rap1b | rap1ab,rap1b,rap1gap | 3 |
| rap1gap | rap1ab,rap1b,rap1gap | 3 |
| rap2b | rap2c,rap2b | 2 |
| rap2c | rap2c,rap2b | 2 |
| rasl11a | rasl11b,rasl11a | 2 |
| rasl11b | rasl11b,rasl11a | 2 |
| rbm12b | rbm12b,rbm12 | 2 |
| rbm15b | rbm15,rbm15b | 2 |
| rgs5a | rgs5a,rgs5b | 2 |
| rgs5b | rgs5a,rgs5b | 2 |
| rgs7a | rgs7bpa,rgs7a | 2 |
| rgs7bpa | rgs7bpa,rgs7a | 2 |
| rgs9b | rgs9b,rgs9bp | 2 |
| rgs9bp | rgs9b,rgs9bp | 2 |
| ric8a | ric8b,ric8a | 2 |
| ric8b | ric8b,ric8a | 2 |
| rnaseh2a | rnaseh2a,rnaseh2b | 2 |
| rnaseh2b | rnaseh2a,rnaseh2b | 2 |
| rnf144aa | rnf144aa,rnf144b | 2 |
| rnf144b | rnf144aa,rnf144b | 2 |
| rnf19a | rnf19b,rnf19a | 2 |

| | | |
|---|---|---|
| rnf19b | rnf19b,rnf19a | 2 |
| rnf213a | rnf213a,rnf213b | 2 |
| rnf213b | rnf213a,rnf213b | 2 |
| rpl10a | rpl10a,rpl10 | 2 |
| rpl13a | rpl13a,rpl13 | 2 |
| rpl18a | rpl18,rpl18a | 2 |
| rpl23a | rpl23,rpl23a | 2 |
| rpl35a | rpl35a,rpl35 | 2 |
| rpl36a | rpl36a,rpl36 | 2 |
| rprd1a | rprd1a,rprd1b | 2 |
| rprd1b | rprd1a,rprd1b | 2 |
| rps15a | rps15a,rps15 | 2 |
| rps3a | rps3,rps3a | 2 |
| rrm2b | rrm2,rrm2b | 2 |
| rtkn2a | rtkn2b,rtkn2a | 2 |
| rtkn2b | rtkn2b,rtkn2a | 2 |
| rtn4a | rtn4a,rtn4r | 2 |
| rtn4r | rtn4a,rtn4r | 2 |
| rundc3ab | rundc3b,rundc3ab | 2 |
| rundc3b | rundc3b,rundc3ab | 2 |
| s100b | s100z,s100b | 2 |
| s100z | s100z,s100b | 2 |
| scarb2a | scarb2c,scarb2a | 2 |
| scarb2c | scarb2c,scarb2a | 2 |
| sec22a | sec22c,sec22ba,sec22a | 3 |
| sec22ba | sec22c,sec22ba,sec22a | 3 |
| sec22c | sec22c,sec22ba,sec22a | 3 |
| sec23a | sec23ip,sec23a,sec23b | 3 |
| sec23b | sec23ip,sec23a,sec23b | 3 |
| sec23ip | sec23ip,sec23a,sec23b | 3 |
| sec24a | sec24b,sec24d,sec24c,sec | 4 |
| sec24b | sec24b,sec24d,sec24c,sec | 4 |
| sec24c | sec24b,sec24d,sec24c,sec | 4 |
| sec24d | sec24b,sec24d,sec24c,sec | 4 |
| sec31a | sec31a,sec31b | 2 |
| sec31b | sec31a,sec31b | 2 |
| sec61b | sec61g,sec61b | 2 |
| sec61g | sec61g,sec61b | 2 |
| sema3ab | sema3ga,sema3fb,sema3 | 7 |
| sema3b | sema3ga,sema3fb,sema3 | 7 |
| sema3c | sema3ga,sema3fb,sema3 | 7 |
| sema3d | sema3ga,sema3fb,sema3 | 7 |
| sema3e | sema3ga,sema3fb,sema3 | 7 |
| sema3fb | sema3ga,sema3fb,sema3 | 7 |
| sema3ga | sema3ga,sema3fb,sema3 | 7 |
| sema4ab | sema4ab,sema4c,sema4b | 4 |
| sema4ba | sema4ab,sema4c,sema4b | 4 |
| sema4c | sema4ab,sema4c,sema4b | 4 |
| sema4gb | sema4ab,sema4c,sema4b | 4 |
| sema5a | sema5a,sema5ba | 2 |

| | | |
|---|---|---|
| sema5ba | sema5a,sema5ba | 2 |
| sema6a | sema6a,sema6ba,sema6c | 3 |
| sema6ba | sema6a,sema6ba,sema6c | 3 |
| sema6d | sema6a,sema6ba,sema6c | 3 |
| sh3pxd2aa | sh3pxd2aa,sh3pxd2b | 2 |
| sh3pxd2b | sh3pxd2aa,sh3pxd2b | 2 |
| shisa2a | shisa2b,shisa2a | 2 |
| shisa2b | shisa2b,shisa2a | 2 |
| sin3aa | sin3aa,sin3b | 2 |
| sin3b | sin3aa,sin3b | 2 |
| slc19a3a | slc19a3b,slc19a3a | 2 |
| slc19a3b | slc19a3b,slc19a3a | 2 |
| slc25a1a | slc25a1b,slc25a1a | 2 |
| slc25a1b | slc25a1b,slc25a1a | 2 |
| slc2a11a | slc2a11b,slc2a11a | 2 |
| slc2a11b | slc2a11b,slc2a11a | 2 |
| slc4a1a | slc4a1ap,slc4a1a | 2 |
| slc4a1ap | slc4a1ap,slc4a1a | 2 |
| slc6a11a | slc6a11a,slc6a11b | 2 |
| slc6a11b | slc6a11a,slc6a11b | 2 |
| smpdl3a | smpdl3a,smpdl3b | 2 |
| smpdl3b | smpdl3a,smpdl3b | 2 |
| spon2a | spon2b,spon2a | 2 |
| spon2b | spon2b,spon2a | 2 |
| srsf5a | srsf5b,srsf5a | 2 |
| srsf5b | srsf5b,srsf5a | 2 |
| srsf7a | srsf7a,srsf7b | 2 |
| srsf7b | srsf7a,srsf7b | 2 |
| sstr1a | sstr1a,sstr1b | 2 |
| sstr1b | sstr1a,sstr1b | 2 |
| stk11ip | stk11ip,stk11 | 2 |
| stk17a | stk17a,stk17b | 2 |
| stk17b | stk17a,stk17b | 2 |
| stt3a | stt3b,stt3a | 2 |
| stt3b | stt3b,stt3a | 2 |
| sv2a | sv2ca,sv2bb,sv2,sv2a | 4 |
| sv2bb | sv2ca,sv2bb,sv2,sv2a | 4 |
| sv2ca | sv2ca,sv2bb,sv2,sv2a | 4 |
| tada2a | tada2b,tada2a | 2 |
| tada2b | tada2b,tada2a | 2 |
| taf1a | taf1a,taf1b,taf1 | 3 |
| taf1b | taf1a,taf1b,taf1 | 3 |
| tagln3a | tagln3a,tagln3b | 2 |
| tagln3b | tagln3a,tagln3b | 2 |
| tbc1d22a | tbc1d22b,tbc1d22a | 2 |
| tbc1d22b | tbc1d22b,tbc1d22a | 2 |
| tbc1d2b | tbc1d2b,tbc1d2 | 2 |
| tent4a | tent4a,tent4b | 2 |
| tent4b | tent4a,tent4b | 2 |
| tent5bb | tent5bb,tent5d,tent5c | 3 |

| | | |
|---|---|---|
| tent5c | tent5bb,tent5d,tent5c | 3 |
| tent5d | tent5bb,tent5d,tent5c | 3 |
| tfap2a | tfap2d,tfap2b,tfap2e,tfap | 5 |
| tfap2b | tfap2d,tfap2b,tfap2e,tfap | 5 |
| tfap2c | tfap2d,tfap2b,tfap2e,tfap | 5 |
| tfap2d | tfap2d,tfap2b,tfap2e,tfap | 5 |
| tfap2e | tfap2d,tfap2b,tfap2e,tfap | 5 |
| thsd7ab | thsd7ab,thsd7ba | 2 |
| thsd7ba | thsd7ab,thsd7ba | 2 |
| timm17a | timm17b,timm17a | 2 |
| timm17b | timm17b,timm17a | 2 |
| timm8a | timm8a,timm8b | 2 |
| timm8b | timm8a,timm8b | 2 |
| tlcd4a | tlcd4b,tlcd4a | 2 |
| tlcd4b | tlcd4b,tlcd4a | 2 |
| tmem106a | tmem106c,tmem106bb,tı | 3 |
| tmem106b | tmem106c,tmem106bb,tı | 3 |
| tmem106c | tmem106c,tmem106bb,tı | 3 |
| tmem121a | tmem121ab,tmem121b | 2 |
| tmem121b | tmem121ab,tmem121b | 2 |
| tmem150a | tmem150aa,tmem150c | 2 |
| tmem150c | tmem150aa,tmem150c | 2 |
| tmem151a | tmem151bb,tmem151a | 2 |
| tmem151b | tmem151bb,tmem151a | 2 |
| tmem161a | tmem161b,tmem161a | 2 |
| tmem161b | tmem161b,tmem161a | 2 |
| tmem167a | tmem167a,tmem167b | 2 |
| tmem167b | tmem167a,tmem167b | 2 |
| tmem170a | tmem170b,tmem170a | 2 |
| tmem170b | tmem170b,tmem170a | 2 |
| tmem184a | tmem184c,tmem184bb,tı | 3 |
| tmem184b | tmem184c,tmem184bb,tı | 3 |
| tmem184c | tmem184c,tmem184bb,tı | 3 |
| tmem200a | tmem200a,tmem200b | 2 |
| tmem200b | tmem200a,tmem200b | 2 |
| tmem30ab | tmem30ab,tmem30b,tme | 3 |
| tmem30b | tmem30ab,tmem30b,tme | 3 |
| tmem30c | tmem30ab,tmem30b,tme | 3 |
| tmem39a | tmem39b,tmem39a | 2 |
| tmem39b | tmem39b,tmem39a | 2 |
| tmem41ab | tmem41ab,tmem41b | 2 |
| tmem41b | tmem41ab,tmem41b | 2 |
| tmem45a | tmem45a,tmem45b | 2 |
| tmem45b | tmem45a,tmem45b | 2 |
| tmem63a | tmem63a,tmem63c,tmen | 3 |
| tmem63ba | tmem63a,tmem63c,tmen | 3 |
| tmem63c | tmem63a,tmem63c,tmen | 3 |
| tmem9b | tmem9,tmem9b | 2 |
| tnnc1a | tnnc1b,tnnc1a | 2 |
| tnnc1b | tnnc1b,tnnc1a | 2 |

| | | |
|---|---|---|
| tnnt2b | tnnt2e,tnnt2b,tnnt2c | 3 |
| tnnt2c | tnnt2e,tnnt2b,tnnt2c | 3 |
| tnnt2e | tnnt2e,tnnt2b,tnnt2c | 3 |
| tomm20a | tomm20a,tomm20b | 2 |
| tomm20b | tomm20a,tomm20b | 2 |
| top2a | top2b,top2a | 2 |
| top2b | top2b,top2a | 2 |
| top3a | top3a,top3b | 2 |
| top3b | top3a,top3b | 2 |
| tp53rk | tp53,tp53rk | 2 |
| trmt10a | trmt10a,trmt10b,trmt10c | 3 |
| trmt10b | trmt10a,trmt10b,trmt10c | 3 |
| trmt10c | trmt10a,trmt10b,trmt10c | 3 |
| trnau1apa | trnau1apa,trnau1apb | 2 |
| trnau1apb | trnau1apa,trnau1apb | 2 |
| trpc4apa | trpc4b,trpc4apa | 2 |
| trpc4b | trpc4b,trpc4apa | 2 |
| tspan33a | tspan33a,tspan33b | 2 |
| tspan33b | tspan33a,tspan33b | 2 |
| tspan4a | tspan4a,tspan4b | 2 |
| tspan4b | tspan4a,tspan4b | 2 |
| tspan7b | tspan7,tspan7b | 2 |
| ttc39a | ttc39b,ttc39c,ttc39a | 3 |
| ttc39b | ttc39b,ttc39c,ttc39a | 3 |
| ttc39c | ttc39b,ttc39c,ttc39a | 3 |
| ttc7a | ttc7b,ttc7a | 2 |
| ttc7b | ttc7b,ttc7a | 2 |
| ttc9b | ttc9b,ttc9c | 2 |
| ttc9c | ttc9b,ttc9c | 2 |
| tubb2b | tubb2,tubb2b | 2 |
| txnl4a | txnl4a,txnl4b | 2 |
| txnl4b | txnl4a,txnl4b | 2 |
| ube2a | ube2t,ube2kb,ube2b,ube2 | 11 |
| ube2b | ube2t,ube2kb,ube2b,ube2 | 11 |
| ube2c | ube2t,ube2kb,ube2b,ube2 | 11 |
| ube2f | ube2t,ube2kb,ube2b,ube2 | 11 |
| ube2h | ube2t,ube2kb,ube2b,ube2 | 11 |
| ube2ia | ube2t,ube2kb,ube2b,ube2 | 11 |
| ube2kb | ube2t,ube2kb,ube2b,ube2 | 11 |
| ube2na | ube2t,ube2kb,ube2b,ube2 | 11 |
| ube2t | ube2t,ube2kb,ube2b,ube2 | 11 |
| ube2w | ube2t,ube2kb,ube2b,ube2 | 11 |
| ube2z | ube2t,ube2kb,ube2b,ube2 | 11 |
| ube3a | ube3d,ube3c,ube3a,ube3 | 4 |
| ube3b | ube3d,ube3c,ube3a,ube3 | 4 |
| ube3c | ube3d,ube3c,ube3a,ube3 | 4 |
| ube3d | ube3d,ube3c,ube3a,ube3 | 4 |
| ube4a | ube4b,ube4a | 2 |
| ube4b | ube4b,ube4a | 2 |
| unc45a | unc45b,unc45a | 2 |

| | | |
|---|---|---|
| unc45b | unc45b,unc45a | 2 |
| unc5a | unc5db,unc5c,unc5b,unc[illegible] | 4 |
| unc5b | unc5db,unc5c,unc5b,unc[illegible] | 4 |
| unc5c | unc5db,unc5c,unc5b,unc[illegible] | 4 |
| unc5db | unc5db,unc5c,unc5b,unc[illegible] | 4 |
| upf3a | upf3b,upf3a | 2 |
| upf3b | upf3b,upf3a | 2 |
| ush1c | ush1gb,ush1c | 2 |
| ush1gb | ush1gb,ush1c | 2 |
| usp12a | usp12a,usp12b | 2 |
| usp12b | usp12a,usp12b | 2 |
| vav3b | vav3b,vav3b | 2 |
| vav3b | vav3b,vav3b | 2 |
| vgll2a | vgll2a,vgll2b | 2 |
| vgll2b | vgll2a,vgll2b | 2 |
| vps13a | vps13a,vps13c,vps13d | 3 |
| vps13c | vps13a,vps13c,vps13d | 3 |
| vps13d | vps13a,vps13c,vps13d | 3 |
| vps26a | vps26c,vps26a,vps26b | 3 |
| vps26b | vps26c,vps26a,vps26b | 3 |
| vps26c | vps26c,vps26a,vps26b | 3 |
| vps33a | vps33b,vps33a | 2 |
| vps33b | vps33b,vps33a | 2 |
| vps37a | vps37a,vps37c | 2 |
| vps37c | vps37a,vps37c | 2 |
| vps4a | vps4b,vps4a | 2 |
| vps4b | vps4b,vps4a | 2 |
| vti1a | vti1a,vti1b | 2 |
| vti1b | vti1a,vti1b | 2 |
| wdr45b | wdr45,wdr45b | 2 |
| wdr83os | wdr83,wdr83os | 2 |
| wnt10a | wnt10b,wnt10a | 2 |
| wnt10b | wnt10b,wnt10a | 2 |
| wnt11r | wnt11r,wnt11 | 2 |
| wnt2ba | wnt2ba,wnt2 | 2 |
| wnt3a | wnt3a,wnt3 | 2 |
| wnt4b | wnt4,wnt4b | 2 |
| wnt5a | wnt5a,wnt5b | 2 |
| wnt5b | wnt5a,wnt5b | 2 |
| wnt7aa | wnt7aa,wnt7bb | 2 |
| wnt7bb | wnt7aa,wnt7bb | 2 |
| wnt8a | wnt8b,wnt8a | 2 |
| wnt8b | wnt8b,wnt8a | 2 |
| wnt9a | wnt9a,wnt9b | 2 |
| wnt9b | wnt9a,wnt9b | 2 |
| yif1a | yif1b,yif1a | 2 |
| yif1b | yif1b,yif1a | 2 |
| zbtb7a | zbtb7c,zbtb7a | 2 |
| zbtb7c | zbtb7c,zbtb7a | 2 |
| zbtb8a | zbtb8b,zbtb8a,zbtb8os | 3 |

| | | |
|---|---|---|
| zbtb8b | zbtb8b,zbtb8a,zbtb8os | 3 |
| zbtb8os | zbtb8b,zbtb8a,zbtb8os | 3 |
| zc3h7a | zc3h7ba,zc3h7a | 2 |
| zc3h7ba | zc3h7ba,zc3h7a | 2 |
| znf385a | znf385a,znf385b,znf385c | 3 |
| znf385b | znf385a,znf385b,znf385c | 3 |
| znf385c | znf385a,znf385b,znf385c | 3 |
