## Supplemental file 4 for "Evolutionary context can clarify teleosts gene names"

single copy genes named as duplicated genes
a1cf
aak1b
abca1a
abca3b
abcb11b
abcb6b
abcd3a
abcf2b
abcg4a
abhd14a
abhd15a
abhd2a
abhd6a
abhd8b
abi2b
ablim1a
abtb2a
acap3a
acbd5a
ackr3b
ackr4a
acot11a
acsl1b
acsl3b
acsl4a
acta1b
actn2b
actn3a
actr2a
acvr1bb
ada2a
adam10b
adam17b
adam23a
adarb1a
adcy1b
adcy2a
adcy3b
adcy6a
adcyap1a
adcyap1r1a
add3a
adgrb1a
adgrf3b
adgrl1a
adgrl2a
adh8b
adipor1a
adm2b

adnp2b
adora1a
adrb2a
adrb3a
afap1l1a
agfg1b
agtr1b
ahr1a
ahsa1a
aimp1a
ak7b
akap12b
akap17a
akap1b
akt3a
aldh3a2b
alkal2b
als2b
alx4b
ambra1a
amotl2b
ampd2b
ampd3a
angptl1b
angptl2b
ank1b
ank2a
ank3a
ankef1b
ankib1b
ankmy2a
ankrd10a
ankrd1a
ankrd28b
ankrd34ba
ankrd46a
ankrd52a
ankrd6b
anks1b
anks4b
ano2b
ano5b
ano8b
ano9b
antxr2a
anxa11b
anxa1a
anxa2a
anxa5b
ap1ar

ap1s3a
ap2m1a
ap3b1a
apba1a
apba2b
apbb2b
aph1b
apoa1a
apobec2b
aqp10b
aqp3b
aqp9b
arf2b
arf6b
arfip2b
arglu1a
arhgap11a
arhgap12b
arhgap17b
arhgap21b
arhgap23a
arhgap32b
arhgap35b
arhgap42a
arhgap45a
arhgap4a
arhgef12b
arhgef18b
arhgef1b
arhgef28a
arhgef7a
arhgef9b
arid4a
arid5b
arl14ep
arl15b
arl5a
arl6ip5b
arntl1a
arpc5b
arrb2a
arrdc1b
arrdc3a
as3mt
asah1b
asap1b
asap2a
asb12a
asb14a
asb15b

asf1ba
asic4a
atad2b
atad5a
atf4b
atf5a
atp1b1b
atp1b2a
atp1b3b
atp2a2b
atp2b3b
atp6ap1a
atp6v0a1b
atp6v0a2a
atp6v1c1b
atp6v1e1a
atp8b5a
atp9b
atrnl1b
atxn1a
atxn7l2a
auts2a
avpr1ab
azin1b
b3galt1a
b3gnt2a
b3gnt5b
b4galnt1b
b4galnt3b
b4galnt4a
bach1b
bach2a
bahcc1a
baiap2b
baiap2l1b
baiap2l2a
barhl1b
baz1b
bcdin3d
bcl11aa
bcl2b
bco2b
bicc1a
bicd1a
bin1b
bin2a
birc5a
bmi1b
bmp1b
bmp7b

bmp8a
bmpr2a
bnip1a
brd1a
brf1b
bri3bp
brpf3a
brsk2b
btbd10a
btbd11b
btbd17b
btbd2a
btbd3b
btbd6b
bzw1a
c1galt1a
c1ql3b
c1ql4b
c1qtnf6a
c2cd4a
c7b
c8b
ca10a
ca5a
cables2b
cabp1a
cabp7b
cacna2d1a
cacna2d4a
cacnb3a
cacnb4a
cacng1b
cacng3b
cacng4a
cacng5a
cacng6b
cacng7a
cadm1b
cadm2a
calb2a
calcoco1b
cald1b
calml4b
calr3b
camk2n1a
camkk1a
camsap1b
camsap2a
cant1a
capn1a

capn2a
capn3a
capn5b
capns1b
caprin1b
capza1b
casp3a
casp6a
casq1b
cavin1b
cavin2b
cavin4b
cbln2a
cbx1b
cbx3a
cbx8a
cc2d1b
cc2d2a
ccdc102a
ccdc127a
ccdc136b
ccdc149b
ccdc6b
ccdc90b
ccl25b
ccn4a
ccnd2b
ccnl1a
ccnt2a
ccr12a
ccr6a
ccr9b
ccser2b
cct6a
cd2ap
cd3eap
cd74a
cd81b
cd82b
cd8b
cd9a
cdc14ab
cdc25d
cdc34a
cdc42ep4a
cdcp1a
cdh10a
cdh7a
cdhr1a
cdk11b

cdk5r1b
cdk5r2b
cdx1a
celf3b
celf5a
celsr1a
cept1b
cers2a
cers3a
cers4a
cert1a
ch25h
chchd3a
chchd4b
chd4a
chl1b
chmp2bb
chmp5a
chmp6b
chordc1a
chrm1a
chrm2a
chrm3b
chrm4a
chrm5b
chrna2a
chrna4b
chrnb3b
chst12a
chst2b
chst3a
cip2a
cited4b
ckmt2b
clasp1a
clcn1a
clcn2b
clcn5b
cldn10b
cldn11b
cldn15a
cldn23a
cldn5a
cldn7a
cldnd1b
clec11a
clec16a
clec19a
clec3bb
clic5b

clint1a
clip1a
clk2a
clk4a
cln6a
clns1a
cmtm8b
cnga1b
cnga2a
cnga3b
cnksr2b
cnn1b
cnn3a
cnnm2b
cnnm4b
cnot3b
cnot4a
cnot6b
cnrip1a
cntn1b
cntn3b
cntnap2b
cntnap5b
coa3a
col10a1a
col11a1a
col12a1b
col14a1a
col15a1b
col1a1b
col27a1b
col28a1b
col28a2a
col2a1b
col5a2a
col5a3a
col6a4a
col8a1b
col9a1b
cops7a
coq10b
coro1cb
cox5aa
cox6c
cox7a2a
cox7c
cpeb1b
cpeb4a
cplx3a
cpne4b

cpne5a
cpxm1b
crabp1b
crabp2b
cracr2ab
crb2b
creb1b
creb3l3a
creb5b
crispld1b
crlf1b
crtac1a
crtc1a
cry1b
cry3a
cryba1a
cryba2b
csdc2a
csf3r
csgalnact1a
csmd3a
csnk1g2a
csnk2a2b
csnk2b
cspg5a
cspp1a
csrnp1b
csrp1a
ctbp2a
ctdnep1b
ctdspl2a
cthrc1a
ctnnd2b
ctps1b
cuedc1b
cul1a
cul3a
cul5b
cux1a
cxcl12b
cxcr4a
cxxc1a
cxxc5b
cyb561a3b
cyp19a1a
cyp1a
cyth4a
d2hgdh
daam1a
dab1b

dab2ipa
dact3a
dapk2a
dbx1b
dchs1a
dclk1a
dclk2a
dctn1a
dcun1d2a
ddhd1b
ddr2a
ddx39aa
ddx3xa
def6a
dennd3a
dennd5a
depdc1a
depdc7a
desi1b
dgat1b
dhrs11a
dhrs3a
dhx32b
diras1b
dixdc1a
dkk1b
dkk3a
dlg4a
dlgap1a
dlgap2b
dlgap4b
dlx1a
dlx2a
dlx3b
dlx4b
dlx5a
dlx6a
dmbx1b
dmrt3a
dnai2b
dnaja2b
dnaja3b
dnajb12b
dnajb1b
dnajb9a
dnajc11a
dnase2b
dnm1a
dnm2a
doc2b

dock4b
dock9b
dok1b
dpp6b
dpysl2b
dpysl5a
dram2a
drd1b
drd4a
drd6b
dtnbp1a
dusp13a
dusp19a
dusp23b
dusp8a
dvl1a
dvl3a
dync1i2a
dynll2a
ebf1b
ebf3a
ecrg4a
edil3a
edn3b
efemp2a
efna2a
efna3b
efnb2a
efnb3a
egln1a
egr2b
ehd1b
ehd2a
ehmt1b
eif2s1a
eif3s6ip
eif4a1b
eif4g2a
eif4g3b
elavl1b
elf2b
elfn1b
elfn2a
elmsan1a
elovl1a
elovl4b
elovl7b
elovl8a
emilin1b
emilin2a

emilin3a
emp3b
en1b
en2b
eno1a
entpd2b
entpd5b
ep300a
epas1b
epb41a
epb41l3a
epc1a
epha2b
epha4b
ephb2b
ephb4b
epm2a
epn3b
eps15l1a
eps8a
eps8l3b
erap1b
erbb3b
erbb4b
erc1a
errfi1a
esr2a
esyt1a
esyt2a
etf1b
etv5a
eva1bb
evi5b
ewsr1b
exoc3l2b
f13a1b
f3a
f9b
fa2h
faah2b
fabp10a
fabp11b
fabp1a
fabp7a
fam107b
fam110a
fam117bb
fam118b
fam122b
fam124b

fam126a
fam131c
fam135a
fam136a
fam149a
fam160a1a
fam162a
fam163ba
fam166b
fam171a2b
fam172a
fam174b
fam183a
fam185a
fam192a
fam199x
fam204a
fam207a
fam221a
fam222ba
fam234a
fam241a
fam32a
fam3c
fam50a
fam53b
fam76b
fam78bb
fam89a
fam8a1b
fat1a
fat3a
fbp1b
fbxl14a
fbxl3a
fbxo11a
fbxo30a
fbxo36b
fdx1b
fermt3b
fgd4a
fgd5a
fgf10b
fgf11a
fgf12a
fgf18b
fgf20a
fgf6b
fgf8b
fgfbp1b

fgfbp2b
fgfr1b
fgl2a
fhl2b
fhl3a
fhod3b
filip1b
fip1l1b
fkbp10b
flot2a
flvcr2a
fmn2a
fmnl1a
fmnl2a
fn1a
fn3krp
fnbp1b
fndc4a
fndc5a
fndc7a
fosl1a
foxb1a
foxc1a
foxf2a
foxg1a
foxl2b
foxn2b
foxo1b
foxo6a
foxp1b
foxp3b
foxq1a
frem2b
frmd4bb
frmpd1b
frrs1a
frs2a
fscn1a
fscn2b
fstl1a
fut8a
fyco1a
fzd7a
fzd8a
fzd9b
g6pd
gabpb2b
gabra6a
gabrb2a
gabrr2a

gabrr3b
gad1a
gal3st1a
galnt18a
gas1a
gas2a
gas7a
gata1a
gata2a
gatd3a
gbe1b
gcnt4a
gdf10b
gdf6a
gdpd4a
gdpd5b
gfra1b
gfra2b
gfra4a
gga3a
ggt1a
gid8a
gigyf1b
git2a
gja5b
gja8a
gjd1a
gjd2b
glg1b
gli2b
glis1b
glis2a
glra4a
gls2b
glud1a
gm2a
gna11a
gna12a
gna13b
gna14a
gnai2b
gnao1b
gnb1b
gnb3a
gnb4b
gnb5b
gngt2a
golga7ba
gorasp1a
got2a

gp1bb
gpc1a
gpc6b
gpr132b
gpr155a
gpr158b
gpr183b
gpr185b
gpr22b
gpr34b
gpr37b
gpr37l1b
gpr55a
gpr78b
gprc6a
gpsm1b
gpx1b
gpx4b
gramd2aa
gramd4b
grap2a
grb2b
grem1a
grem2a
grhl2b
gria1b
gria2b
gria3a
gria4a
grid1b
grik1a
grin1b
grin3bb
grip2b
grk7b
grm1b
grm2b
grm5b
grm6a
grm8a
grtp1b
grxcr1a
gsg1l2b
gsk3bb
gtf2b
gtf2f2a
gtf3ab
guk1b
gulp1a
gxylt1b

gyg1b
h6pd
hapln1b
hcn2b
hdac7b
hecw1b
hecw2a
hephl1b
her8a
hid1b
hif1an
higd2a
hipk1a
hipk3b
hivep2a
hivep3b
hmbox1a
hmga1a
hmgb1a
hmgb2a
hmgb3a
hmox2b
hmx3a
homer1b
homer3b
hoxa10b
hoxa11a
hoxa13b
hoxa1a
hoxa4a
hoxa5a
hoxb10a
hoxb13a
hoxb1b
hoxb3a
hoxb5b
hoxb6a
hoxb7a
hoxb8a
hoxb9a
hoxc11a
hoxc12b
hoxc13a
hoxc4a
hoxc5a
hoxc6a
hoxd10a
hoxd11a
hoxd12a
hoxd3a

hoxd4a
hrh2a
hs2st1b
hs6st1b
hs6st3a
hsbp1b
hsd17b12a
hspa4b
htr1fa
htr5ab
htra1b
htra3a
hyal2b
id2b
iffo1a
igf2bp2b
igfbp1a
igfbp2b
igfbp5b
igfbp6b
igsf21a
il12ba
il2rb
ildr1a
ilf3b
impdh1a
impg1b
ing5a
inka1a
insl5a
insm1b
ip6k2a
iqsec1b
iqsec2b
iqsec3a
irf2bp2b
irf4a
irs2b
irs4a
irx1a
irx2a
irx3a
irx4a
irx5b
irx6a
isl2a
ism2a
itga11a
itga2b
itga3a

itga6b
itgb1b
itgb3b
itih3b
itpk1b
itpr1b
itsn2b
ivns1abpb
jag2b
jagn1b
jak2b
jam2a
jam3a
jarid2b
jazf1b
jdp2a
jmjd1cb
jph1a
jpt1b
kank1b
kansl1b
kat7b
kcna1b
kcna2a
kcnab1a
kcnab2a
kcnc1b
kcnc3b
kcnf1b
kcng4a
kcnh1b
kcnh2a
kcnh4b
kcnh5a
kcnh6a
kcnip1b
kcnj12b
kcnj19a
kcnj2a
kcnj3a
kcnk10a
kcnk13a
kcnk1a
kcnk2a
kcnk3b
kcnk5a
kcnma1a
kcnn1a
kcnq2a
kcnq5b

kcns3a
kcnv2a
kctd12b
kctd15a
kctd16b
kctd5a
kctd6a
kctd9b
kdelr2b
kdf1a
kdm1a
kdm3b
kdm4b
kdm6a
kdm7ab
khdrbs1a
kidins220b
kif13ba
kif16ba
kif20bb
kif2c
kifap3b
kirrel1b
kirrel3a
klc1a
klf11a
klf12b
klf2a
klf5b
klf6a
klhdc7a
klhl24a
klhl38b
klhl40b
klhl41b
krt18b
ksr1b
l1cama
l2hgdh
l3hypdh
lactbl1b
lamb1a
lamp1a
larp6a
lbx1a
ldb1a
ldb2a
ldlrad4b
ldlrap1a
lgals3a

lgals8b
lgi1b
lgi2b
lhfpl2b
lhfpl5a
lhx1a
lhx2b
lhx8b
lima1a
limch1b
limk1b
lingo1a
lingo2b
lingo3a
lmbrd2b
lmf2a
lmo7a
lmod1b
lmx1bb
loxhd1b
loxl2b
loxl3b
loxl5b
lpar6a
lrfn2b
lrfn4b
lrfn5a
lrit1a
lrit3b
lrrc14b
lrrc18a
lrrc30a
lrrc38b
lrrc58a
lrrc75bb
lrrfip1b
lrrn3b
lrtm2a
lsm12b
lta4h
ltc4s
lyrm5a
lzts2b
lzts3b
m1ap
maco1b
magi1a
magi2a
magi3b
mamdc2a

man1b1b
map2k2a
map2k4b
map3k14a
map4k3a
map6a
map7d1a
map7d2b
mapk12b
mapk14b
mapk8b
mapkapk2a
mapre1a
mapre3b
marcksl1a
mark2b
mark3a
marveld2a
mast1b
mast3a
mat1a
matn3b
mboat2a
mc1r
mc2r
mc3r
mc4r
mc5ra
mcf2b
mchr1a
mcl1b
mcoln1a
mctp1b
mctp2a
mdga2a
med13b
med19a
megf6b
meis1b
meis2b
meox2a
mep1b
metap2b
mettl11b
mettl21a
mettl2a
mex3b
mfge8b
mfn1b
mfsd12a

mfsd13a
mfsd4ab
mfsd6a
mgat3b
mgrn1a
mical2a
mical3a
micall2a
micu3b
mid1ip1a
mier1a
mier3b
mif4gda
mindy4b
minpp1b
mir196b
mknk2a
mllt1b
mmd2b
mmp11a
mmp15b
mmp16b
mmp17a
mmrn2a
mn1a
mnx2b
mob1bb
mob2a
mogat3b
mpdu1b
mpp2a
mpp3b
mpp4a
mpp5a
mpp7a
mpped2a
mrap2a
mrc1a
mre11a
msi2b
msl1a
msl2b
mthfd1b
mtmr1b
mtmr7a
mtx1b
muc13b
mustn1b
mvb12bb
mxra5b

mxra8b
mybbp1a
mybl2b
myct1b
myh7ba
myh9a
myl2a
myl9a
mylk4b
myo15ab
myo18ab
myo6a
myom1a
myom2a
myoz1b
myoz2b
myoz3a
myt1a
mzt2b
naa15b
nab1a
nabp1a
nap1l4b
nav1b
nav2a
nbr1b
ncam1a
nck1a
nck2b
ndel1b
ndrg1a
ndrg3b
ndst1a
ndst2a
ndufa4l2a
ndufa9a
ndufab1b
nectin1b
nectin3b
nedd4a
nell2a
neo1b
neto2b
neurod6a
nf1b
nfat5a
nfatc2a
nfatc3a
nfe2l1a
nfe2l2a

nhsl1b
niban1a
niban2a
nid1b
nid2a
nim1k
nipsnap3a
nkd2a
nkx2.2b
nkx2.4a
nlgn2b
nlgn3a
nlgn4xb
nmt1a
nmur1b
nos2a
notch1a
notum1a
noxo1b
npas4a
npbwr2a
npdc1b
npffr2b
npm2b
npr1a
nptx2a
npy1r
npy2r
npy7r
npy8br
nr0b2b
nr1d2a
nr1d4b
nr2f1a
nr2f6a
nr4a2b
nr5a1a
nr6a1a
nrbf2b
nrbp2a
nrd1b
nrg2b
nrip1b
nrn1a
nrp1a
nrp2a
nrxn1b
nrxn3a
nsfl1c
nsmce4a

nt5c2b
nt5c3a
nt5e
ntn1b
ntng2a
ntrk2a
ntrk3a
nuak1b
nucb2b
nucks1a
nudt3b
nudt4a
nupr1a
nxph2a
nyap2a
oaz2b
obsl1a
odf2a
odf3l2a
olfm1a
olfm2a
olfm3a
olfml2a
olfml3a
onecut3a
oprd1a
orai1b
osbpl10b
osbpl1a
osbpl2b
osbpl3b
otol1b
otud5a
otud6b
otud7b
otx2b
ovol1a
oxct1b
oxr1b
oxsr1b
p2rx4b
p4ha1b
pa2g4a
pacsin1a
pafah1b1b
paip2b
pak2b
pak6a
pald1b
palm1b

pank1b
panx1a
papss2b
paqr3b
paqr5a
parp6b
pax1a
pax2b
pax3b
pax6a
pax7b
pcdh10b
pcdh15b
pcdh18b
pcdh1a
pcdh7b
pcgf5b
pcmtd2a
pcolce2a
pcp4a
pcsk5b
pdcd10b
pde10a
pde11a
pde1a
pde5ab
pde7a
pde9a
pdha1b
pdk2a
pdk3a
pdlim3b
pdlim5b
pdpk1a
pdzd3b
pdzd7a
pdzrn3a
peli1b
per1a
pfkfb2b
pfkfb4a
pggt1b
phactr3b
phactr4b
phc2a
phf12a
phf20a
phf21ab
phf23b
phkg1b

phldb1b
phldb2a
pi15a
pias1b
pias4a
pik3r3b
pik3r6b
pip4p1a
pkd1b
pkn1b
pkp1b
pla2g4aa
plcd1a
plcd3b
plcd4a
plch2a
pld1a
plekha1b
plekha7b
plekhg5a
plekho1b
plk2b
plod1a
plp1a
plpp2b
plpp7a
plppr2a
plppr3b
plppr4a
plppr5a
plscr3b
plxna1a
plxnb1a
plxnb2b
pmp22b
pnp5a
pnpla7b
poc1a
pof1b
pou2f1b
pou2f2a
pou3f2b
pou3f3a
ppa1a
ppap2d
ppfibp1a
ppfibp2a
ppip5k1b
ppp1r12a
ppp1r13bb

ppp1r15b
ppp1r16a
ppp1r27b
ppp1r8b
ppp1r9a
ppp2cb
ppp2r1bb
ppp3r1a
ppp4r2b
ppp4r3b
ppp5c
ppp6c
ppp6r2a
pptc7b
prc1a
prdm12b
prdm2b
prg4a
prickle1a
prickle2b
prkab1b
prkag2b
prkag3b
prkar2ab
prkg1a
prlhr2b
prmt8b
prokr1b
prom1b
prox1a
prpf38a
prpf40a
prph2b
prps1b
prr5a
prrc2c
psma6a
psmc1a
psmd11b
psme4b
pstpip1b
ptbp1b
ptbp2a
ptdss1a
ptf1a
ptger2a
ptges3b
ptgs2b
ptk6a
ptk7a

ptp4a2b
ptp4a3b
ptpdc1b
ptpn23a
ptpn2a
ptpn4a
ptprz1a
ptx3b
puf60b
pwp2h
pxdc1b
pycr1a
rab11fip4b
rab18a
rab1aa
rab22a
rab25a
rab2a
rab32b
rab34a
rab35b
rab38b
rab39bb
rab42a
rab6a
rab9a
rabl6b
rac1b
rad21a
rad54b
rad9b
raf1b
ranbp3b
rangap1a
rap1gap2b
rapgef1a
rapgef5a
raph1a
rasa1b
rasgef1ba
rasgrf2b
rassf2b
rassf4a
rassf7a
rassf8a
rbfox3a
rbm24b
rbm25b
rbm39a
rbm8a

rbms1a
rbms2a
rbp2a
rbp7b
rbpms2b
rc3h1a
rcan1a
rdh10b
rdh14b
rdh8a
reep3a
rex1bd
rftn1a
rfx1b
rfx7a
rgl3a
rgs12b
rgs14a
rgs3b
rhbdf1b
rhobtb2a
rhot1a
rims1a
rlbp1a
rmnd5b
rnd1a
rnd3b
rnf113a
rnf11b
rnf128a
rnf145b
rnf150a
rnf165a
rnf207b
rnf220b
rnf34b
rock2b
rom1a
rp1l1a
rpe65b
rph3ab
rpl5b
rpp25b
rprd2a
rps27a
rps4x
rps6ka3a
rps6kb1b
rps8b
rrbp1b

rreb1b
rsph4a
rtn1b
rtn4rl1a
rtn4rl2b
runx2a
rwdd2b
rxfp2a
rxfp3.2b
ryr1b
ryr2b
s100a10b
s1pr3a
s1pr5a
sall1b
sall3a
samd10a
samd1b
samd4a
samsn1b
sap130a
sap30bp
sar1b
sash1b
sat1b
sat2a
sbno2a
sc5d
scaf4a
scamp5b
scg2b
scn12aa
scn1bb
scn2b
scn3b
scn4ba
scn8aa
scrt1b
sdk1a
sdk2b
sdr16c5b
sec11a
sec16b
selenot1a
selenou1a
sema7a
senp3a
sept7b
sept8a
sept9a

serbp1b
serpina10a
serpinf2b
serpinh1a
sertad2b
setd1ba
sez6a
sfrp1a
sft2d2a
sfxn5b
sgip1b
sgk2a
sgms2b
sgsm1a
sh2d1ab
sh2d3ca
sh2d4ba
sh3bgr
sh3bp5a
sh3gl1b
sh3gl3a
sh3glb1a
sh3glb2b
shank3a
shisa8b
shisa9a
shisal1a
shrprbck1r
sik2b
sim1a
six1b
six2a
six3a
six6a
skor1a
sla1a
slain1a
slc12a5a
slc13a5a
slc15a1b
slc16a12a
slc16a1a
slc16a5b
slc16a6a
slc16a9a
slc17a6b
slc17a9a
slc18a3b
slc1a2a
slc1a3b

slc1a7b
slc1a8b
slc20a1b
slc22a13b
slc22a7a
slc24a4a
slc24a6a
slc25a15a
slc25a22a
slc25a23a
slc25a25b
slc25a32a
slc25a36b
slc25a38b
slc25a3b
slc25a44b
slc25a47a
slc25a51b
slc25a55b
slc27a1a
slc27a2a
slc29a1a
slc2a13b
slc2a15a
slc2a1b
slc2a3b
slc30a1a
slc34a1a
slc34a2b
slc35a3a
slc35d1a
slc35f3a
slc35g2a
slc37a4a
slc38a3b
slc38a5a
slc38a8a
slc41a2b
slc43a1b
slc43a2a
slc43a3a
slc44a1b
slc44a5a
slc48a1b
slc4a10a
slc4a2b
slc4a4b
slc51a
slc5a3a
slc5a6b

slc5a7a
slc6a1a
slc6a4b
slc6a6b
slc7a10b
slc7a14a
slc7a1a
slc7a3a
slc8a1b
slc8a4b
slc9a1b
slc9a3r1a
slco5a1a
slit1a
slitrk3a
slitrk5a
slx1b
smad3b
smad4a
smad6a
smarca4a
smarcad1a
smarcb1b
smarcc1a
smarcd3b
smc1b
smcr8a
smdt1b
smpd2b
smu1a
smyd1b
smyd2a
snai1b
snap25b
snap91a
snapc1a
snu13a
snx10b
snx18b
snx19a
snx1b
snx27b
snx8b
snx9a
socs1a
socs3b
socs6b
sod3b
soga3a
sorbs2a

sort1a
sostdc1a
sox11a
sox19a
sox1b
sox21b
sox4b
sox8a
sox9a
sp3b
sp5a
sp8b
spag1a
spag9a
spi1b
spint1b
spire1b
spon1a
spred2b
spryd7b
spsb3a
spsb4a
sptbn4b
sptlc2a
srd5a2b
srgap1a
srpk1b
srsf1b
srsf2a
srsf3a
ssh1b
ssh2b
ssrp1a
sstr2a
ssx2ipa
st14b
st3gal3b
st6gal2b
st6galnac5a
stag1a
stag2b
stard13a
stat1a
stat5a
stc2b
stim1a
stim2a
stk25a
stk38a
stmn1b

stoml3a
stox2b
stx11a
stx1b
stx2b
stx5a
stxbp1a
stxbp5a
sub1a
sulf2b
sumo2b
sumo3a
supt3h
supt6h
suv39h1b
suz12b
swap70a
syn2a
syne1b
syngr2b
syngr3b
sypl2a
syt11b
syt14a
syt1a
syt2a
syt6b
syt9b
taar13b
taar1b
tacr1a
tacr3a
taf4a
tafa5a
tanc1b
tanc2a
taok1a
taok3a
tapt1b
tax1bp1b
tbc1d10ab
tbc1d12b
tbl1x
tbl1xr1a
tbr1a
tbx2b
tbx3a
tbx5a
tbxa2r
tcf3a

tcf7l1b
tcirg1b
tdo2b
tdp2b
tdrd7a
tead1b
tead3b
tecpr1a
tecrl2a
tex264a
tfb1m
tfb2m
tfdp1a
tfe3a
tfr1b
tgfb1a
tgfbr1a
tgfbr2b
tgm2a
thap12b
thbs1b
thbs2a
thbs3a
thbs4b
tiam1a
tiam2a
timm23a
timp2b
tjp1a
tjp2a
tlcd3a
tlcd5a
tle2a
tle3a
tlk1a
tln2a
tmbim1a
tmc2b
tmc6a
tmcc1b
tmed1a
tmeff1b
tmeff2b
tmem119b
tmem120b
tmem125b
tmem126a
tmem132e
tmem144a
tmem14ca

tmem163b
tmem168a
tmem169a
tmem179aa
tmem182b
tmem183a
tmem196b
tmem198b
tmem222a
tmem229b
tmem230b
tmem235b
tmem237b
tmem240b
tmem243a
tmem255a
tmem26a
tmem38a
tmem50a
tmem51b
tmem54a
tmem74b
tmem86a
tmem8a
tmprss3b
tmtc2a
tmtops2a
tmtops3a
tmx2b
tmx3a
tnfaip2a
tnfaip8l2b
tnfrsf9b
tnfsf13b
tnk2b
tnni1a
tnni3k
tnnt3a
tnrc6a
tns1b
tns2a
tob1a
tomm70a
tor2a
tor3a
tor4aa
tp53bp2a
tp53i11b
tpd52l2a
tph1a

tpi1b
tpm4a
tra2a
traf2b
traf4a
trak1a
trappc6b
trarg1a
trim2b
trim3a
trim46b
trim55b
trim8a
trip10b
trmt2a
trmt61a
trmt9b
trpa1a
trpc2b
trpc5a
trpc6a
trpc7b
trpm1b
trpm4a
tsc1a
tsg101a
tshz3b
tspan13b
tspan18b
tspan2a
tspan3a
tspan5b
tspan9b
ttbk1b
ttbk2b
ttc21b
ttyh3a
tulp1a
tulp4a
tusc2a
tvp23b
twf1b
twf2a
tyrp1a
u2af2b
u2surp
ubald1b
ubap2a
ubash3ba
ube2d1b

ube2g1a
ube2l3a
ubl3a
ubl7a
ubtd1b
ubxn2a
uck2a
uckl1b
ugp2a
ulk1a
unc119b
unc13d
unc93a
upk1a
uqcrc2b
ush2a
usp43b
usp53b
usp54b
uts2d
vcam1b
vezf1a
vgll4a
vipr1b
vps72a
vsig8b
vsnl1a
vstm4a
wasf3a
wdr20a
wdr26a
wdr47b
wdr48b
wfs1b
wipf1a
wnk1b
wnk4b
wscd1b
wt1a
xirp2a
xkr5a
xkr6a
xpo1a
xpr1a
ypel2a
yy1b
zbtb16a
zbtb2b
zc2hc1a
zc3h11a

zc3h12b
zdhhc12a
zdhhc15a
zdhhc16a
zdhhc18b
zdhhc20a
zdhhc3b
zdhhc5b
zdhhc8b
zeb1b
zeb2b
zfand2a
zfand5a
zfp36l1a
zfpm2a
zfyve9a
zhx2a
zic2a
zmat4b
zmiz1a
znf143b
znf207a
znf280d
znf281b
znf292a
znf362a
znf395b
znf513b
znf609a
znf644a
znf687b
znf710b
znf740b
znf800a
znf804a
znrf2b
zpld1b
zranb1b
